# Nucleosomal DNA has topological memory

**DOI:** 10.1101/2023.05.21.541612

**Authors:** Joana Segura, Christoforos Nikolaou, Joaquim Roca

## Abstract

One fundamental yet elusive aspect of the chromosome architecture is the constrained topolome, which refers to how chromatin elements restrain DNA topology. Nucleosomes stabilise negative DNA supercoils, with most nucleosomes typically restraining a DNA linking number difference (ΔLk) of about -1.26. However, whether this capacity is uniform across the genome is unknown. Here, we calculated the ΔLk restrained by over 4000 nucleosomes in yeast cells. To achieve this, we placed each nucleosome in a circular minichromosome and performed Topo-seq, a novel high-throughput procedure to inspect the topology of circular DNA libraries in a single gel electrophoresis. We found that nucleosomes inherently restrain distinct ΔLk values depending on their genomic origin. Nucleosome DNA topologies differ significantly at gene bodies (ΔLk=-1.29), intergenic regions (ΔLk=-1.23), rDNA genes (ΔLk=-1.24) and telomeric regions (ΔLk=-1.07). Nucleosomes nearby the transcription start and termination sites also exhibit singular DNA topologies. These findings demonstrate that nucleosome DNA topology is imprinted by its native chromatin context and persists even when the nucleosome is relocated. This imprinting contributes to nucleosome functional roles and chromatin folding architectures.

## INTRODUCTION

High-throughput analyses increasingly improve our knowledge of chromatin structure and function genome-wide ^1,2^. However, a fundamental trait that remains elusive to current technologies is the topology of the chromatinized DNA. We do not know how the DNA double helix is twisted and bent along chromatin. We define “constrained topolome” as DNA deformations stabilized by chromatin, and “unconstrained topolome” as those resulting from the mechanical stress the DNA undergoes during genome activities ^3-5^.

The principal actor of the constrained topolome in eukaryotic cells is the nucleosome, the DNA packaging unit of chromatin, in which about 147 base pairs (bp) of DNA make nearly 1.6 left-handed superhelical turns around a histone octamer ^6^. However, this canonical configuration is neither uniform nor static ^7-9^. The DNA nucleotide sequence, histone composition and modifications affect nucleosome conformation and stability ^10-12^. So far, numerous studies have mapped nucleosome structure and position in model organisms, such as budding yeast ^13-17^. However, no study has yet addressed how nucleosomes constrain the topology of DNA throughout the genome.

In a previous study, we developed a strategy to measure the DNA linking number difference (ΔLk) restrained by nucleosomes *in vivo* ^18^. We compared by DNA electrophoresis the ΔLk constrained by yeast circular minichromosomes before and after inserting an additional nucleosome. By doing so, we found that not all nucleosomes restrain ΔLk (DNA supercoils) to the same extent, being the average value of ΔLk=−1.26 ^18^. Our next goal was to determine the ΔLk constrained by all individual nucleosomes. However, since it was unfeasible to perform thousands of electrophoretic analyses (one for each nucleosome), we needed another strategy to circumvent this problem.

Here, we developed a novel high-throughput procedure termed “Topo-seq” to analyse the topology of large libraries of circular DNA molecules. We applied Topo-seq to a library of 4000 nucleosomes hosted in yeast circular minichromosomes *in vivo*. Our results uncovered that nucleosomes inherently restrain distinctive ΔLk values depending on their genomic origin. Therefore, nucleosome DNA topology is an intrinsic trait imprinted by the native chromatin context, which persists when allocated somewhere else. We discuss the implications of this new nucleosomal feature and the potential of Topo-seq to disclose further DNA topology genome-wide.

## RESULTS

### Construction of a nucleosome library within circular minichromosomes

As in our previous study ^18^, we digested budding yeast chromatin with Micrococcal nuclease and purified mono-nucleosomal DNA fragments (≈150 bp) obtained with increasing digestion rates (**Fig. 1a**). After repairing the DNA ends, we added adapters to insert the nucleosome DNA library in the YCp1.3 (1341 bp) yeast circular minichromosome (**Extended Data Fig. 1**). Next, we introduced these minichromosomes, each containing one nucleosome of the library, into yeast cells and obtained about ten thousand transformants. We pooled all these colonies and extracted their DNA (**Fig. 1a**). Parallel DNA sequencing of the minichromosomes revealed a library of 8369 mono-nucleosome DNA fragments of length 144±21 bp (mean ± SD). Nearly all (>91%) of these DNA fragments overlapped (>50 bp) with the genomic coordinates of 4276 previously referenced nucleosomes ^13^ **(Supplementary Table 1)**. The chromosomal distribution (**Fig. 1b**), genomic allocation (**Fig. 1c**) and stability (**Fig. 1d**) of the identified nucleosomes were comparable to that of the bulk of referenced nucleosomes ^13^, signifying the nucleosome library was representative.

**Fig. 1.**
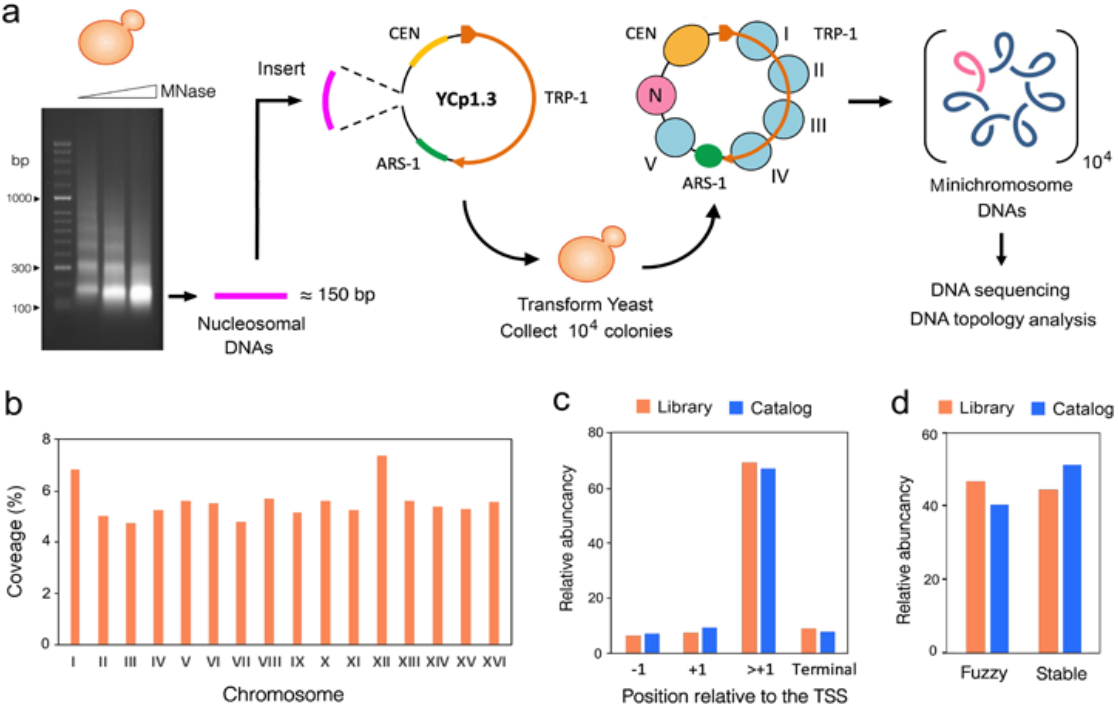
Mono-nucleosome library construction. **a**, The gel shows yeast chromatin digested with increasing amounts of Micrococcal nuclease. Nucleosome DNA fragments of about 150 bp (pink) were inserted into the YCp1.3 minichromosome DNA (1341 bp) to transform yeast cells (see **Extended Data Fig. 1**). About 10^4^ transformants were collected and pooled for sequencing and topology analysis of the minichromosome DNAs. The allocation of the nucleosome library (N) into the chromatin structure of YCp1.3 is illustrated (for more details, see Supplementary Fig. 1). **b**, Coverage (%) of the nucleosome library across the 16 yeast chromosomes (I-XVI) normalized per kb. **c**, Relative abundance of nucleosomes from different gene-body positions (−1, +1 >+1, terminal). **d**, Relative abundance of fuzzy and stable nucleosomes. In **c** and **d**, the nucleosome library (n=4276, pink) is compared to the full catalogue of yeast nucleosomes (n=61110, blue).

### Conceptualization leading to Topo-seq

Due to DNA thermodynamics, covalently closed circular DNA molecules present a Gaussian distribution of DNA linking number (Lk) topoisomers, visualized as a ladder of DNA bands by agarose gel electrophoresis ^19^. When the mean Lk value of a DNA molecule relaxed in vitro (Lk^0^) is compared to that of the DNA in a circular minichromosome (Lk^CH^), the Lk difference is the ΔLk constrained by the minichromosome chromatin (Lk^0^ - Lk^CH^ = ΔLk^CH^). Upon adding a new nucleosome to such minichromosome, ΔLk^CH^ changes to ΔLk^CH+nuc^ and the resulting difference (ΔLk^CH+nuc^ - ΔLk^CH^ = ΔLk^nuc^) equals the ΔLk restrained by the new nucleosome (**Extended Data Fig. 2**). Following this strategy, we previously showed that most nucleosomes restrain ΔLk values of about -1.26 *in vivo* ^18^. However, this strategy was unfeasible for determining the ΔLk of each nucleosome of our library, as it would require running thousands of electrophoresis. To circumvent this problem, we ran instead all minichromosome DNAs containing the library in a single electrophoresis lane (**Fig 2a**, lane 1). As a result, the pool of Lk distributions overlapped and produced a smear rather than a ladder of Lk topoisomers. This smear occurred because the nucleosome sequences of the library were not of equal length (144±21 bp) and, since these length differences were not multiples of the helical repeat of DNA (10.5 bp), most ladders of Lk topoisomers had different phasing. We evidenced this misalignment by running, in the same gel, the Lk distributions of individual minichromosomes that carried nucleosome DNA sequences differing by only 1 to 5 bp in length (**Fig 2a**, lanes 2-7). Note that Lk distributions can have equal or distinct Lk mean (red lines) irrespectively of whether their Lk topoisomers are aligned. This gel electrophoresis also evidenced that the mean Lk of the pool of Lk distributions produced by the nucleosome library (**Fig. 2b**, lane 1) virtually matched with that produced by individual nucleosomes that restrained ΔLk -1.26 (**Fig. 2b**, lanes 2-5), which corroborated that the average ΔLk constrained by nucleosomes is -1.26.

**Fig. 2.**
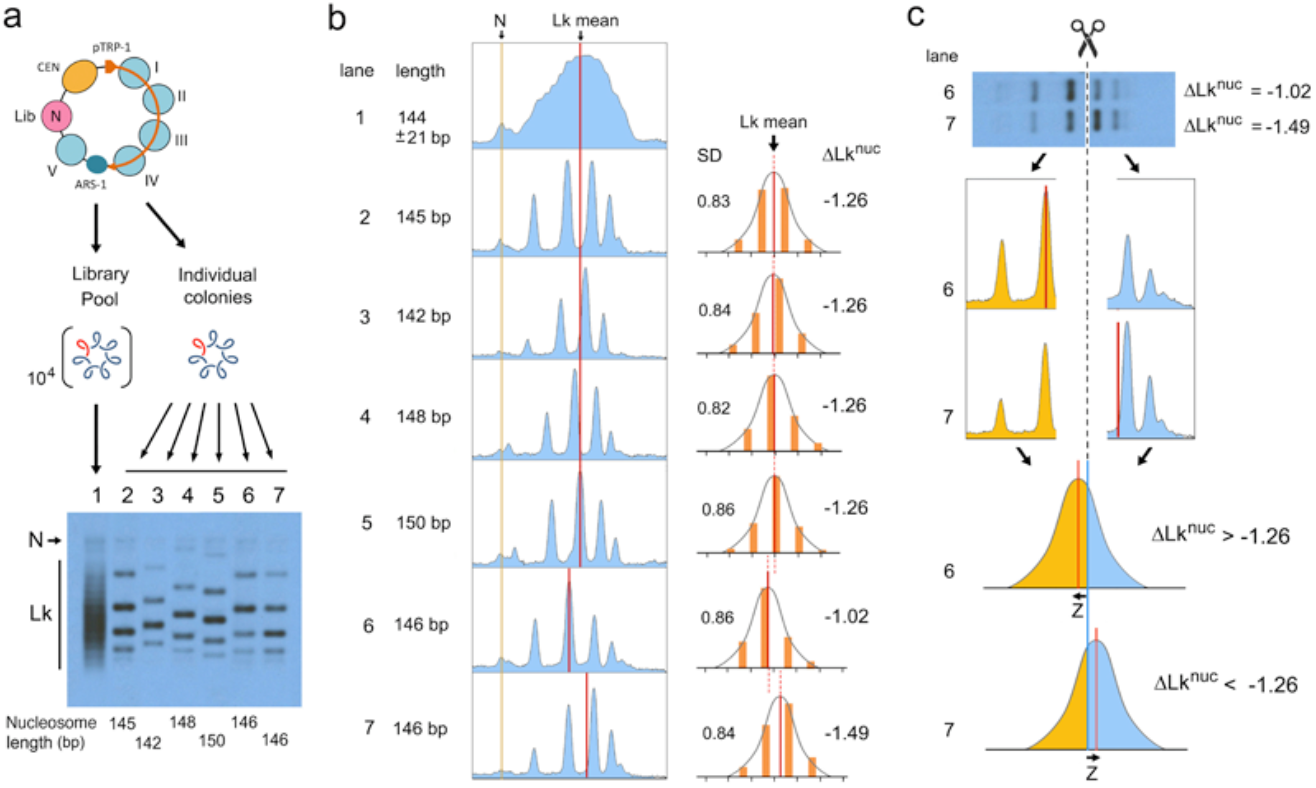
Topo-seq conceptualization. **a**, Gel electrophoresis of Lk distributions. Lane 1, pool of 10^4^ minichromosome DNAs hosting the nucleosome library. Lanes 2-7, individual minichromosome DNAs hosting a nucleosome with the indicated DNA length (bp). Electrophoresis conditions are described in Methods. N, nicked circles. Lk, topoisomers. **b**, Gel densitometry (left, in blue) and relative topoisomer intensities (right, in orange) of the previous Lk distributions (lanes 2-7), indicating their Lk mean position (red lines), standard deviation (SD) and the ΔLk restrained by the hosted nucleosome (ΔLk^nuc^) calculated as described in **Extended Data Fig 2. c**, Cutting the Lk distributions of sample 6 (ΔLk^nuc^ =-1.02) and 7 (ΔLk^nuc^ =-1.49) at the global Lk mean position (ΔLk^nuc^ =-1.26) produces unequal topoisomer DNA abundances at each side of the cut (orange and blue). As Lk distributions are Gaussian, the distinct partition probabilities of DNA abundance translate into Z-scores (Z) of a normal probability distribution. These Z-scores reflect how far the Lk mean of each distribution is from that of the global average (the cut site), where ΔLk^nuc^ =-1.26.

At this point, we realized that within the library pool of Lk distributions (**Fig. 2b**, lane 1), those from nucleosomes restraining ΔLk <-1.26 and >-1.26 should respectively have their Lk mean slightly ahead and behind the global average (for example, samples 6 and 7 in **Fig. 2b**). Accordingly, upon cutting the smear of Lk distributions at the level of the global mean, nucleosome DNA sequences restraining ΔLk <-1.26 and >-1.26 would differently enrich at each side of the cut (**Fig. 2c**, top). The partition probability of each nucleosome DNA sequence would reflect how far its ΔLk^nuc^ deviates from the global mean (ΔLk^nuc^=-1.26, at the cut site); and, since Lk distributions are Gaussian, this deviation would correlate to the Z-score of a normal distribution (**Fig. 2c**, bottom). In this way, we could classify all nucleosomes of the library by their capacity to restrain ΔLk. We termed this procedure “Topo-seq” since it relies on sequencing DNA topoisomers eluted from different sections of an electrophoresis gel.

### Calculation of the ΔLk restrained by individual nucleosomes via Topo-seq

We cut into two sections the gel slab comprising the overlapping Lk distributions of our nucleosome library (**Fig. 3a**), such that the top section (A) enriched nucleosome DNA sequences restraining ΔLk >-1.26 and the bottom section (B) those restraining ΔLk <-1.26. We eluted the DNA from both sections, amplified the nucleosome DNAs by PCR and sequenced them. As expected, all DNA sequences of the nucleosome library were present in both sections but with distinct partition probabilities (**Fig. 3b**). Although section B slightly enriched in short nucleosome DNA sequences, the partition probabilities within each length group denoted main differences in nucleosome DNA topology (**Fig. 3c**).

**Fig. 3.**
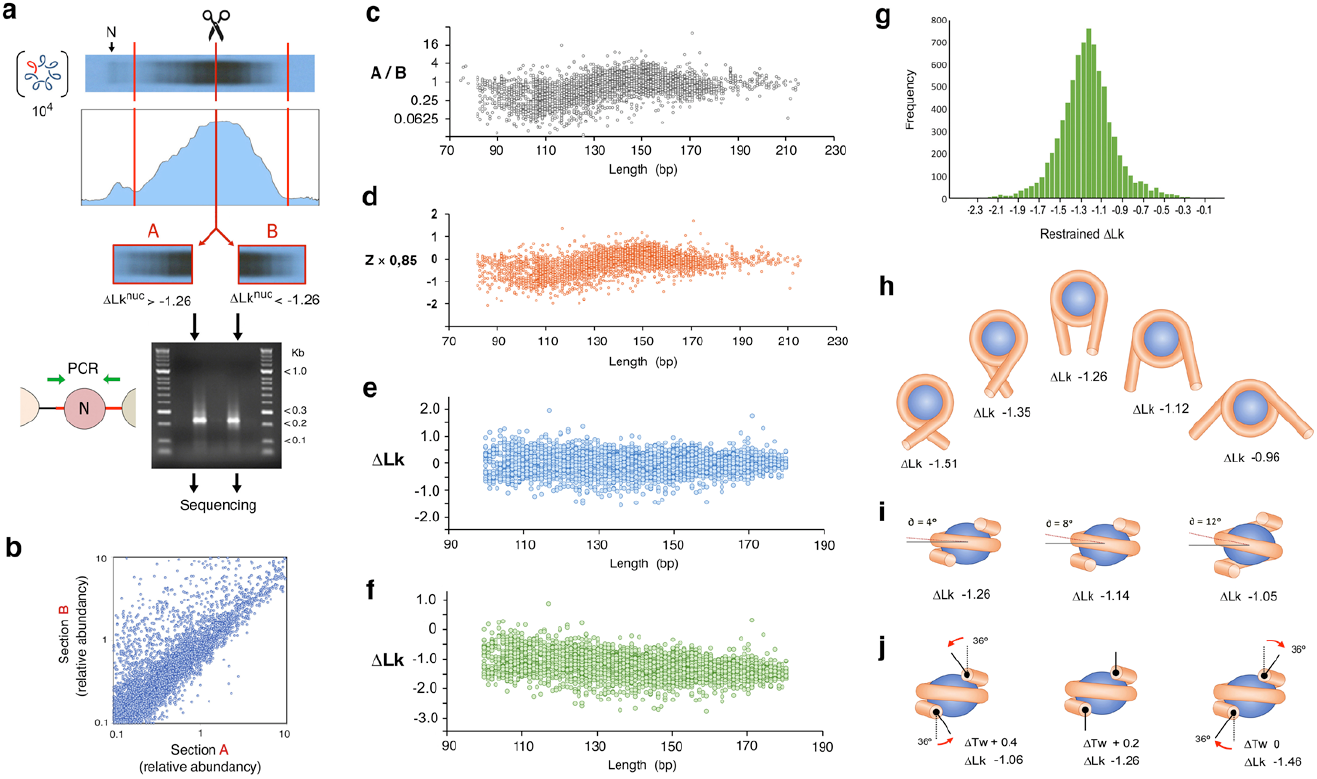
Calculation of the ΔLk restrained by individual nucleosomes via Topo-seq. **a**, Splitting of the Lk distributions of lane 1 in Fig. 2a at the level of the global Lk mean (ΔLk^nuc^=-1.26); and PCR products of the nucleosome sequences eluted from section A and B. **b**, Plot of the abundance of individual nucleosome DNA sequences in sections A and B. **c**, Plot of the partition probability of each nucleosome DNA sequence (ratio A/B) vs the sequence length (bp). **d**, Plot of the partition probabilities converted into Z-scores of a normal distribution and multiplied by 0.85 (SD of Lk distributions). **e**, Corrected ΔLk scores after subtracting the average ΔLk of all the sequences of the same length. **f**, Plot of ΔLk values after adding or subtracting 0.007 units for each bp difference from 144 bp and finally adding -1.26. See **Supplementary Table 2** for calculated values. **g**, Histogram of ΔLk values restrained by the nucleosome library (mean = -1.26, SD = 0.33). **h**, Models of restrained ΔLk as a function of the number of DNA super-helical turns (N), considering ΔTw=+0.2 and ΔWr=N(1–sin 4°). **i**, Models of restrained ΔLk as a function of the pitch angle of DNA superhelical turns (∂), considering ΔTw=+0.2 and ΔWr=1.56(1–sin ∂). **j**, Twisting angle (red arrows) at the entry and exit of DNA to produce ΔTw of 0, +0.2, +0.4 and the indicated ΔLk, considering ΔWr=1.56(1–sin 4°). See **Extended Data Fig.5** for extended correlations of ΔWr and ΔTw on ΔLk.

To calculate the ΔLk restrained by each nucleosome, we converted the partition probability of each nucleosome DNA sequence into a Z-score of a normal distribution of mean zero and a standard deviation one. Next, we multiplied these Z-scores by 0.85, which is the standard deviation of the Lk distributions of the DNAs hosting the nucleosome library (**Fig. 2b)**. The resulting values indicated how far (ΔLk units) the mean of each Lk distribution was from the global Lk average (ΔLk^nuc^ of -1.26 at the cut site) (**Fig. 3d**). Next, we adjusted these ΔLk scores by taking into account that the nucleosome DNA sequences had different lengths. Firstly, to correct the effect of misaligned Lk ladders (**Extended Data Fig. 3**), we subtracted from each ΔLk score the average ΔLk of all the sequences of the same length (i.e., aligned ladders). These corrected ΔLk values now reflected how the topology of each nucleosome compared to those of equal length (**Fig. 3e**). Secondly, we calculated that the overall electrophoretic velocity of the Lk distributions shifted the equivalent of 0.007 Lk units for each bp difference in length (**Extended Data Fig. 4**). Therefore, we adjusted the ΔLk values by adding or subtracting 0.007 for each bp difference from 144 bp. Finally, as the current ΔLk values were relative to the global average (ΔLk^nuc^ = -1.26), we added -1.26 to obtain the actual ΔLk restrained by each nucleosome (**Fig. 3f, Supplementary Table 2**).

**Fig. 4.**
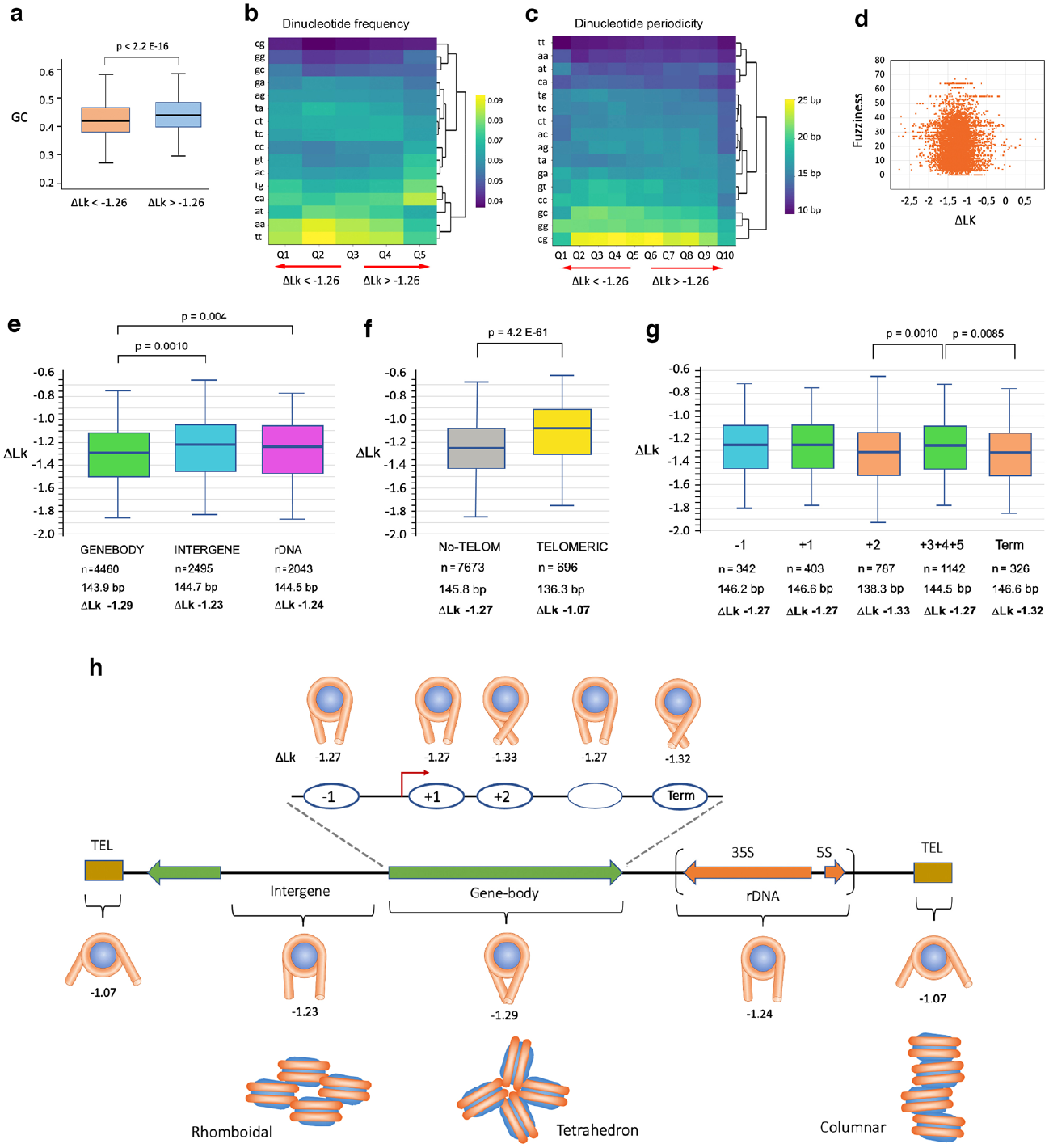
ΔLk correlation with DNA composition and genomic allocation of nucleosomes. **a**, Boxplots of GC content for ΔLk values above and below the mean. *p*-value determined by a Wilcoxon test. **b**, Heatmap of dinucleotide frequencies for ΔLk values in 5 quantiles. **c**, Heatmap of dinucleotide periodicities for ΔLk values in 10 quantiles. **d**, Scatter plot of ΔLk values against the native positional fuzziness of nucleosomes. **e**, Boxplot of ΔLk restrained by gene body, intergenic and rDNA nucleosomes. *p*-values determined by ANOVA with Tukey’s test. **f**, Boxplot of ΔLk restrained by non-telomeric and telomeric nucleosomes. *p*-value determined by unpaired two-tailed *t*-test. **g**, Boxplot of ΔLk restrained nucleosomes at positions - 1,+1,+2, 3 to 5 relative to the TSS and the TTS (Term). *p*-values determined by ANOVA with Tukey’s test. In **e-f**, the bottom rows indicate the number (n), average sequence length (bp) and mean ΔLk of nucleosomes in each class. The ΔLk and genomic allocation of individual nucleosomes are sorted in **Supplementary Table 3. h**, Nucleosome DNA topology depending on their genomic allocation. Restrained ΔLk values are modelled as different DNA wrapping degrees (**Extended Data Fig 5a**). Proposed folding motives of tetranucleosomes in gene bodies (tetrahedron) and intergenic regions (rhomboidal) ^16^ and the columnar architecture of telomeric chromatin ^47^ are illustrated.

The ΔLk values calculated via Topo-seq (mean -1.26, SD 0.33) denoted a substantial heterogeneity in the nucleosome DNA topology (**Fig. 3g**). Since ΔLk=ΔTw+ΔWr, nucleosomes could differ in restraining ΔTw (helical twist) but, more likely, by their capacity to constrain ΔWr (super-helical turns). We then modelled how ΔWr values would translate into ΔLk by considering ΔTw=+0.2 (as in the cannon nucleosome) and ΔWr=N(1–sin ∂)^18^, where N is the number of wrapped super-helical turns (**Fig. 3h, Extended Data Fig. 5a**) and ∂ their pitch angle (**Fig 3i, Extended Data Fig. 5b**). We also considered how ΔTw values would translate into ΔLk, being ΔWr=-1.46 (**Fig. 3j, Extended Data Fig. 5c**). For simplicity, we depicted only symmetric shapes, but asymmetric changes of ΔWr and ΔTw are likely to occur and combine producing a broad spectrum of possible conformations even for nucleosomes restraining similar ΔLk values.

### Correlation of ΔLk with structural and functional traits of nucleosomes

We examined whether the nucleosome capacity to restrain ΔLk would depend on the DNA nucleotide composition, which affects DNA flexibility and curvature ^20^. We found that GC content was significantly higher (p<2e-16) for nucleosomes restraining ΔLk>-1.26 (GC 44%) compared to those of ΔLk<-1.26 (GC 42%) (**Fig. 4a**). Regarding dinucleotide frequencies, nucleosomes of the library were enriched in AA/TT/TA dinucleotides, being CG/GG/GC dinucleotides the less abundant. However, this general trend faded in the nucleosomes restraining less negative ΔLk values, which exhibited an abundance of other dinucleotides such as CA/TG (**Fig. 4b**). As expected, AA/TT/TA dinucleotides had a periodical spacing close to the DNA helical repeat (10-11 nts), though nucleosomes restraining less negative ΔLk values also presented other dinucleotides with short periodic patterns (**Fig. 4c**). Since DNA composition affects the position stability of nucleosomes ^21,22^, we examined whether the restrained ΔLk correlated to nucleosome stability. We found that the positional fuzziness of nucleosomes, measured in their natural loci ^13^, had no relation with their intrinsic capacity to constrain ΔLk (**Fig. 4d**).

Next, we examined whether the nucleosome capacity to restrain ΔLk would depend on their genomic origin. Surprisingly, several unexpected correlations emerged (**Fig. 4e-g, Supplementary Table 3**). The mean ΔLk of -1.29 constrained by gene body nucleosomes was distinct (p<0.001) from the mean ΔLk of -1.23 constrained by the intergenic ones (**Fig. 4e)**. The ΔLk restrained by gene body nucleosomes (transcribed by Pol-II) was also different (p<0.004) from the rDNA genes (transcribed by Pol-I and -III), whose nucleosomes restrained a mean ΔLk of -1.24 (**Fig. 4e)**. However, the most marked difference was in the nucleosomes from the telomeric regions, which exhibited a mean ΔLk of -1.07, highly deviated (p<E-60) from the mean ΔLk of -1.27 of non-telomeric nucleosomes (**Fig. 4f)**. Within gene body nucleosomes, there are also significant differences depending on the nucleosome position relative to the transcription start site (TSS) and termination site (TTS) (**Fig. 4g**). Nucleosomes at position -1 from TSS restrained a mean ΔLk of - 1.27, a value more similar to gene body than intergenic nucleosomes. Nucleosomes at position +1, stably positioned downstream of the TSS, also restrained a mean ΔLk of -1.27, like the bulk of gene body nucleosomes. However, two unexpected gene body nucleosomes constrained ΔLk values significantly more-negative than the others. Nucleosomes at position +2 restrained a mean ΔLk of - 1.33 (p<0.001), and terminal nucleosomes at the TTS restrained a mean ΔLk of -1.32 (p<0.008). As discussed below, these distinctive capacities to constrain DNA topology might have been imprinted by specific nucleosome roles or by distinct folding architectures of chromatin.

## DISCUSSION

Our study provides two advances toward the unravelling of intracellular DNA topology. First, it shows the feasibility of Topo-seq, a new high-throughput procedure for examining topolomes. Second, it demonstrates that DNA retains a topological memory of its deformation by chromatin.

To date, gel electrophoresis of circular DNA molecules is the only procedure to quantitatively characterize changes in the linking number (supercoiling) and the degree of entanglement (knotting or catenation) of DNA. Therefore, to examine local DNA topology within cellular chromosomes, one strategy has been popping out DNA rings at specific chromatin regions ^23-25^. However, this procedure would be extremely tedious, if not impracticable, for conducting genome-wide analyses of DNA topology. Topo-seq relies on the sequencing of DNA topoisomers eluted from distinct sections of one gel electrophoresis and, therefore, permits inspecting the topology of multiple DNA molecules at once. We show that DNA amounts as little as 0.1 ng, comprising a library of thousands of DNA constructs, can be eluted from an agarose gel and processed for high-throughput sequencing. In the present study, since the circular DNAs containing the nucleosome library were of similar length (about 1.5 kb), we resolved the pool of ΔLk distributions in a one-dimensional gel, cut it into two halves, and applied corrections for the little length differences. However, for future developments of Topo-seq, the analysis of libraries containing multiple DNA lengths (1 to 15 kb) could be done by cutting the gel into many small sections to determine the position and abundance of any DNA topoisomer irrespective of its length. Another possibility is running the DNA library in a two-dimensional gel. In this case, Lk distributions would resolve into a diagonal of arched ladders, such that Lk topoisomers from one DNA length do not overlay with those of another.

After calculating the ΔLk values constrained by our nucleosome library, we found structural and functional correlations that validated the reliability of Topo-seq. Importantly, since we interrogated the nucleosomes outside their native genomic locus, the restrained ΔLk values had to rely on the intrinsic traits of their DNA sequences. In this respect, since high GC content makes the DNA stiffer ^26,27^, the increased GC content of nucleosomes restraining less negative ΔLk values might reflect a reduced capacity of DNA to completely or stably wrap around histone cores. As expected, the nucleosome library exhibited a 10-11 bp periodicity of AA/TT/TA dinucleotides, which is known to favour nucleosome positioning and stability ^21,22^. But, curiously, this dinucleotide pattern did not correlate with the capacity to restrain ΔLk. Only the nucleosomes constraining less negative ΔLk values presented atypical dinucleotide periodicities, which emerged from the repetitive DNA sequences of telomeric regions, as discussed below. Therefore, nucleosome DNA topology might not only depend on dinucleotide parameters, which are also weak predictors of the intrinsic cyclability of DNA ^28^. Moreover, the no correlation between the nucleosome capacity to restrain ΔLk and their native genomic stability also denoted these are independent structural traits. Conceivably, positional stability depends on the central turn of DNA around the H3-H4 tetramer; and DNA topology mainly relies on interactions with the H2A-H2B dimers configuring the geometry of nucleosome entry and exit DNA segments.

In contrast to the nucleotide composition, the genomic origin of nucleosomes presented striking correlations to their intrinsic capacity to restrain ΔLk (**Fig 4e-g**). The gene-body nucleosomes (mean ΔLk of -1.29) are distinct from the intergenic ones (mean ΔLk of -1.23). If this difference relies on the extent of DNA wrapping (ΔWr) and the subsequent orientation of DNA entry and exit segments, nucleosome arrays in genic and intergenic regions must adopt distinct architectures. Interestingly, this inference might relate to the distinctive folding motifs described in yeast chromatin, where the gene-body nucleosomes tend to present tetrahedron folds and the intergenic ones rhomboidal folds ^16^ (**Fig 4h**). The distinct DNA topology of gene-body and intergenic nucleosomes could also reflect that the firsts are suited to confront the torque generated during DNA transcription ^29^ and to facilitate DNA tearing by transcribing RNA polymerases ^30^. The topology of gene-body nucleosome is not conserved in the rDNA genes (mean ΔLk of -1.24). Since rDNA nucleosomes are evicted by the transcribing trains of RNA polymerase I ^31,32^, their distinctive DNA topology probably reflects this unique biophysical context, quite different from genes transcribed by RNA polymerase II.

Within gene-body nucleosomes, we expected nucleosomes at positions -1 and +1 relative to the TSS would have singular DNA topologies since they delimit a nucleosome-free region (NFR) in most gene promoters ^17,33,34^. Their allocation relies on transcription factors and chromatin remodelling activities that push them out of the NFR ^35-38^. The +1 nucleosomes also interact with the transcription preinitiation complex (PIC) ^39,40^. However, the intrinsic capacity of the -1 and +1 nucleosomes to restrain ΔLk is similar to that of most gene-body ones. In contrast, Topo-seq uncovered that nucleosomes at positions +2 and TTS have singular DNA topologies. The case of the +2 nucleosomes (ΔLk = -1.33) is striking because they presented DNA sequences much shorter (138,3 bp) than the global average and still restrained more negative ΔLk values than any other nucleosome. The +1 and +2 nucleosomes have similar positional stability ^34^ and are recognized as a dinucleosome by remodelling complexes ^35,41^. This feature and a plausible role in the promoter-proximal pausing of RNA Polymerase II ^42,43^ might explain the unexpected topology of +2 nucleosomes. Regarding the TTS nucleosomes, their high capacity to restrain DNA topology (ΔLk = -1.32) likely relates to their role as transcription elongation barriers. Interestingly, their capacity to mark transcription termination depends on a proper configuration of their DNA entry-exit site, thus, their DNA topology ^44^.

Lastly, Topo-seq uncovered nucleosomes from telomeric regions ^45^ as those with the most extreme DNA topology (ΔLk=-1.07). Remarkably, these nucleosomes presented shorter DNA sequences (136,3 bp) than the non-telomeric ones; and exhibited unusual patterns of dinucleotides since they host abundant short DNA sequence repeats ^46^. Therefore, their reduced capacity to restrain ΔLk likely reflects an incomplete or distinctive wrapping of the DNA (**Fig 4 b-c**). Their peculiar DNA topology might relate to the recently reported columnar architecture of telomeric chromatin ^47^ that shows nucleosomes stacked on each other with a repeat length of about 132 bp, letting the DNA wound as a continuous superhelix (**Fig. 4h**).

In conclusion, using Topo-seq, we uncovered several nucleosome classes based on their capacity to restrain ΔLk. Since this structural trait correlates to the genomic origin of the nucleosomes, the intrinsic topology of nucleosome DNA emerges as a new determinant of chromatin architecture and functions. We foresee that applying Topo-seq to more extensive libraries of chromatin elements or regions will provide unprecedented insights into the implications of DNA topology genome-wide. Ultimately, the need for circular minichromosomes to host these libraries could be dispensable if, for instance, the bulk of intracellular chromatin is fragmented and circularised in situ. In this case, upon running an entire genomic pool of DNA rings in a single gel electrophoresis, Topo-seq could readily disclose the topolome of any cell type.

## METHODS

### Construction of the nucleosome library

*Saccharomyces cerevisiae* FY251 cells (*MATa his3-Δ200 leu2-D1 trp1-Δ63 ura3–52*) were grown at 28ºC in 250 ml of rich medium until OD 1.0. Cells were collected, washed with water, and incubated with 80 ml of 1M Sorbitol, 30 mM DTT for 15 min at 28°C. Next, 625 U of Lyticase (Sigma-Aldrich L2524) and 10 μL of 4M NaOH were added to the cell’s suspension, and the incubation continued until >80% of cells converted into spheroplasts. Spheroplasts were washed with 1M Sorbitol and resuspended in 1.5 ml of hypotonic lysis buffer (1 mM CaCl_2_ 5 mM KH_2_PO_4_ 1 mM PMSF) at 24°C. After adding 30 units of micrococcal nuclease (Sigma-Aldrich N3755), the lysate was incubated at 24°C. Aliquots of 300 μl were quenched with 20 mM EDTA 1% SDS at different incubation times (3 to 30 min). Gel electrophoresis of digested chromatin was done in 1% agarose in TBE buffer, at 80 V for 3 h. Mono-nucleosome DNA fragments (about 150 bp in length) produced at successive digestion times were gel-eluted and pooled. The DNA ends produced by micrococcal nuclease were repaired by removing terminal 3’-phosphates with T4-polynucleotide kinase and filled with Klenow and T4-DNA polymerase activities. The resulting A-tailed DNA fragments were ligated to T-tailed adaptors that provided degenerated *Asc1* or *BamH1* ligation ends (see Supplementary Fig. 1). The adapted DNA fragments were inserted between the *Asc1* and *BamH1* sites of the YCp1.3 DNA (1341 bp), which was hosted in a bacterial plasmid. After amplifying these plasmids in *E. coli* cells, they were digested with *Not1* to release DNA segments of about 1.55 Kb that comprised YCp1.3 with the inserted library of mono-nucleosome DNAs (≈200 bp). These DNA segments were gel purified and circularised by ligating their *Not1* ends.

### Extraction of yeast minichromosomes hosting the nucleosome library

Monomeric circles of YCp1.3 hosting the nucleosome DNA library were gel-purified and used to transform FY251 via electroporation. About ten thousand colonies were collected from agar plates containing yeast synthetic media (TRP dropout). These colonies were pooled, washed with water, resuspended in 200 ml of TRP dropout and incubated at 28°C for 2 hours. The cells were then fixed by mixing the suspension with one cold volume (−20°C) of ET solution (Ethanol 95%, Toluene 28 mM, Tris HCl pH 8.8 20 mM, EDTA 5 mM). The cells were sedimented at room temperature, washed twice with water, resuspended in 400 μl of TE, and transferred to a 1.5-ml microfuge tube containing 400 μl of phenol and 400 μl of acid-washed glass beads (425–600 μm, Sigma). Mechanic lysis of >80% cells was achieved by shaking the tubes in a FastPrep® apparatus for 10 sec at power 5. The aqueous phase of the lysate was collected, extracted with chloroform, precipitated with ethanol, and dissolved in 100 μl of TE containing RNAse-A. After a 15-min incubation at 37°C, the samples were extracted with phenol and chloroform, the DNA precipitated with ethanol and dissolved in 50 μl of TE.

### Electrophoresis of *Lk* distributions

The DNA of minichromosomes that hosted the nucleosome library was loaded onto 1.4% (w/v) agarose gels. Electrophoreses were carried out at 2.5 V/cm for 18 h in TBE buffer (89 mM Tris-borate and 2 mM EDTA) containing 0.55 μg/ml chloroquine. Gels were blot-transferred to a nylon membrane and probed at 60°C with the YCp1.3 DNA labelled with AlkPhos Direct (GE Healthcare®). Chemiluminescent signals of increasing exposure periods were recorded on X-ray films. Non-saturated signals of individual Lk topoisomers and bins of pooled Lk distributions were quantified with ImageJ. The Lk mean of the Lk distributions was determined as previously described ^18^ and illustrated in Supplementary Fig. 2.

### Topo-seq

About 500 ng of yeast DNA, including that of minichromosomes hosting the nucleosome library (about 0.5 ng), were loaded in two adjacent lanes of a gel containing 1.4% (w/v) agarose (Ultrapure grade, nzytech® MB05202) and electrophoresed at 2.5 V/cm for 18 h in TBE buffer. The gel slab of the first lane was kept at 4°C in TBE, while the second lane was blot-transferred and probed to determine the position of the Lk mean of the overlapping Lk distributions. The gel slab of the first lane was then cut at the level of this Lk mean in two sections (A and B) of about 25×10×4 mm each, such that section A contained the top half of the overlapping Lk distributions; and section B, the bottom half. DNAs from both gel sections were eluted and captured using the Geneclean II kit®. DNA fragments of the nucleosome library present in each section were amplified via PCR (NEB Taq Polymerase) by using as primers the sequences of the *Asc1* and *BamH1* adapters described in Supplementary Fig. 1. The obtained amplicons of about 200 bp were sequenced (Illumina MiSeq v2 2×150pb, paired-end reads) and the resulting FASTQ data subjected to QC using Cutadapt (1.12). The library sequences were then mapped to the *Saccharomyces cerevisiae* reference genome (R64-1-1 from Ensembl Genomes) using bowtie (v1.1.2). Once nucleosome coordinates were established, alignment and size statistics were calculated using SAMTools and Picard. Subsequent analyses were performed by integrating published nucleosome data sets ^13,48^ and using BEDTools (v2.27) and Galaxy. The calculation of ΔLk values using the Topo-seq DNA sequencing data is described in the results section and Supplementary Table 2. This process includes cunning the relative abundancies of nucleosomal DNA sequences in the gel sections A and B, the conversion of these partition probabilities into Z scores of a normal distribution, the transformation of Z scores into Lk values, and the adjustment of these ΔLk values to take into account the length differences within the library of nucleosomal DNA sequences.

### ΔLk modelling of nucleosomes

To model restrained ΔLk values as a function of ΔWr, the value of ΔTw was fixed to +0.2 and restrained ΔWr values were calculated from ΔWr=N(1–sin ∂). The number of wrapped super-helical turns (N) was depicted as the length of the arcs in contact with the cylindrical histone core, leaving the entry and exit segments of DNA as detached tangents and with a pitch angle (∂) of 4°. As super-helical turns are left-handed, N<0. The pitch angle (∂) of the super-helical turns was calculated from the height at the entry and exit points of 1.56 super-helical turns. To model the restrained ΔLk as a function of ΔTw, the nucleosome ΔWr was fixed to -1.46 (=-1.56 (1–sin 4°) and restrained ΔTw values of 0 and +0.4 were illustrated as the twisting angle observed at the entry and exit segments of DNA relative to ΔTw in the cannon nucleosome (ΔTw=+0.2).

### ΔLk correlation with nucleosome structural traits and genomic allocation

Sequence properties of nucleosome DNAs including GC content, dinucleotide frequency and periodicity were analyzed in R using the seqinR package. Mean dinucleotide frequencies were calculated for nucleosome ΔLk values split into five quantiles. Dinucleotide periodicities were measured as the mean distance (in bps) of consecutive dinucleotides of the same type and calculated for nucleosome ΔLk values split into ten quantiles. The native allocation of identified nucleosomes within a gene body, intergenic region and rDNA genes was defined by Jiang and Pugh (2009). Positional fuzziness of nucleosomes was determined as the standard deviation in bp of the nucleosome position from the 6 datasets compiled by Jian & Puhg (2009) ^13^. Nucleosomes of telomeric regions were as defined in The *Saccharomyces* Genome Database (SGD) ^45^. Statistic tests for the significance of ΔLk correlations are described in the figure legends.

## Data Availability

The data discussed in this publication have been deposited in NCBI’s Gene Expression Omnibus ^49^ and are accessible through GEO Series accession number **<u>GSE228623</u>**.

## Acknowledgements

We acknowledge the CNAG Sequencing Service and the CRG Bioinformatics Core Facility for assistance.

## Author’s contributions

J.R. conceived Topo-seq and supervised the study.

J.S. designed and performed all the experiments.

J.S. and C.N. performed the bioinformatics analyses.

J.R. produced nucleosome models and wrote the manuscript.

