## Supplemental Data for "Nucleosomal DNA has topological memory"

**Extended Data Fig. 1**

**Extended Data Fig. 2**

**Extended Data Fig. 3**

**Extended Data Fig. 4**

**Extended Data Fig. 5**

**Supplementary Table 1**

**Supplementary Table 2**

**Supplementary Table 3**

**a**

Adaptor with BamH1 cohesive-end ligatable but not recutable

```
5′      *GGACGATTACAGCTACGTG        3′
3′      TCCTGCTAATGTCGATGCACCTAG    5′
```

Adaptor with Asc1 cohesive-end ligatable but not recutable

```
5′      *CGTTCGGATCCGTTTAAACGTGAAGAGGTAACTCAT        3′
3′      TGCAAGCCTAGGCAAATTTGCACTTCTCCATTGAGTAGCGC    5′
```

**b**

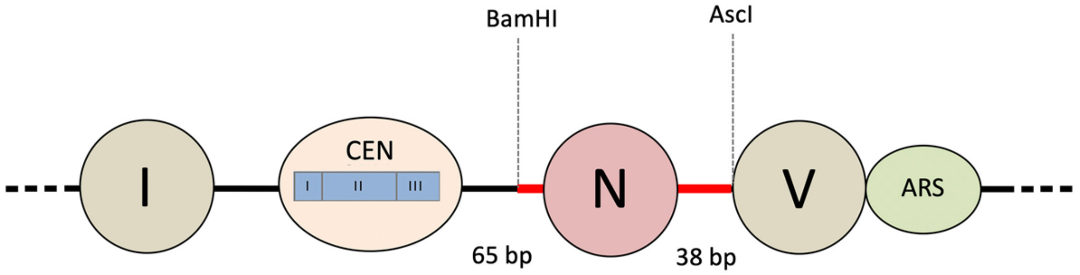


**Extended Data Fig. 1. Insertion of the nucleosome library into YCp1.3. a**, DNA adaptors ligated to A-tailed nucleosome DNA fragments to generate *BamH1* and *Asc1* cohesive ends. **b,** Position of the nucleosome library between nucleosome V and CEN2 of YCp1.3. The lengths (bp) of the resulting linker DNA segments are indicated.

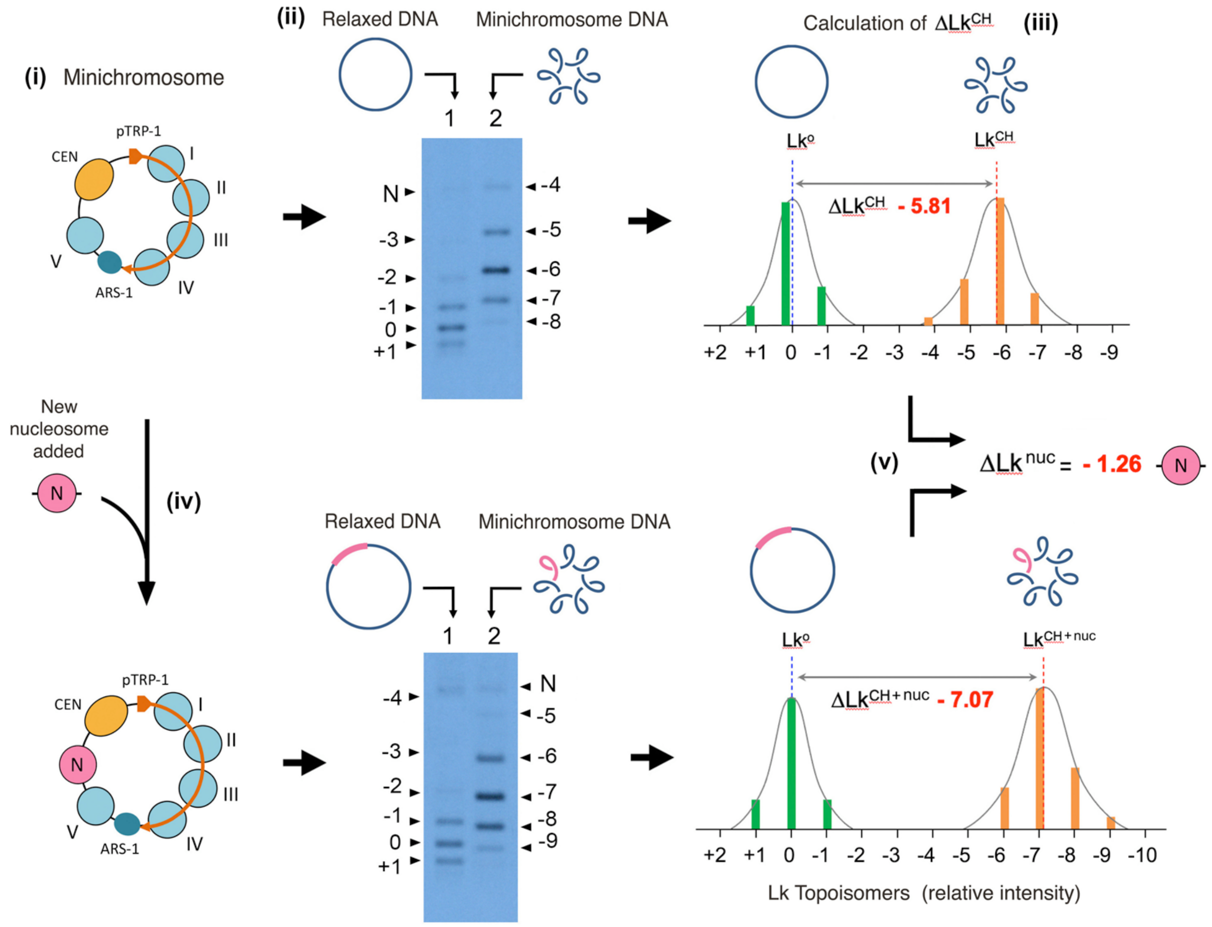


**Extended Data Fig. 2. Experimental layout to calculate the ΔLk restrained by nucleosomes *in vivo*.** In a circular minichromosome (i), the DNA linking number difference ($\Delta Lk^{CH}$) constrained by chromatin elements is calculated by comparing, via gel electrophoresis (ii), the distribution of Lk topoisomers of the DNA relaxed *in vitro* (lane 1) with that of the minichromosome DNA fixed *in vivo* (lane 2). The Lk mean of the relaxed DNA and the minichromosome DNA distributions ($Lk^0$ and $Lk^{CH}$) is determined by plotting the intensity of their corresponding Lk topoisomers along a scale of ΔLk units (iii). $\Delta Lk^{CH}$ is the distance in Lk units between the two means ($Lk^0 - Lk^{CH} = \Delta Lk^{CH}$). After adding a new nucleosome into the minichromosome (iv), repeating the same analysis produces $\Delta Lk^{CH+nuc}$. The difference between $\Delta Lk^{CH}$ and $\Delta Lk^{CH+nuc}$ indicates the ΔLk restrained by the new nucleosome ($\Delta Lk^{nuc}$) (v). $\Delta Lk^{nuc}$ is about -1.26 for most nucleosomes, as previously reported by Segura *et al* (2018).

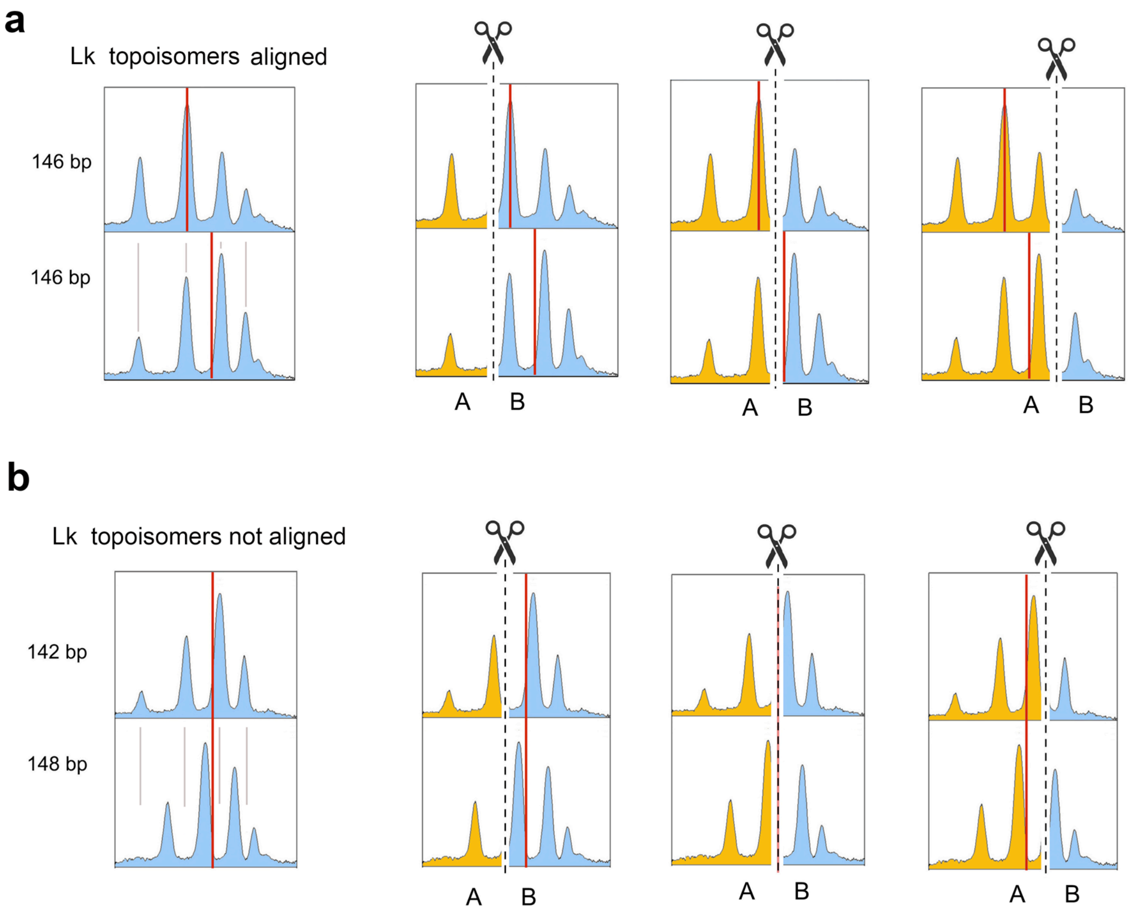


**Extended Data Fig. 3. Effect of aligned and non-aligned Lk distributions when conducting Topo-seq. a,** Example of two Lk distributions with different Lk mean (red lines), but with aligned topoisomers, since the DNA fragments of the nucleosome library have equal length (146 bp). Note that, whatever the position of a splitting cut (3 examples depicted), the partition of DNA intensities (coloured areas A and B) always denotes that the two distributions have different Lk mean. **b**, Example of two Lk distributions with equal Lk mean (red lines) but with not aligned topoisomers because the DNA fragments of the nucleosome library have different lengths (142 and 148 bp). In this case, depending on the position of the splitting cut, the partition of DNA intensities does not always denote that the two distributions have equal Lk mean.

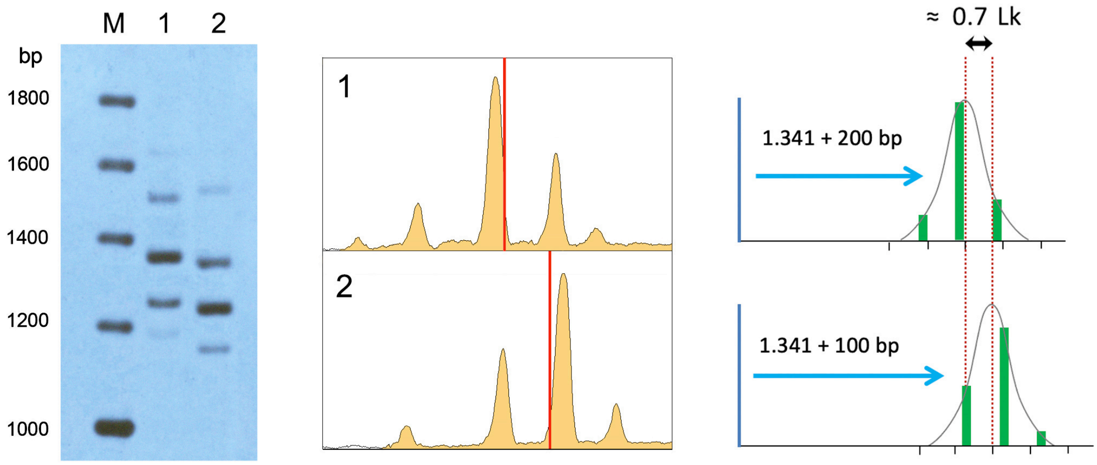


**Extended Data Fig. 4. Correlation of DNA length with the electrophoretic velocity of Lk distributions.** Left, the gel shows a marker of linear DNAs (M) and the Lk distributions of two relaxed DNA molecules differing in 100 bp in length. These molecules are the YCp1.3 minichromosome DNA (1.341 bp) with an insert of 100 bp (lane 1) and 200 bp (lane 2). Note that the size of these inserts encloses the length variability of the nucleosome library (144±33bp). Middle, densitometry of lanes 1 and 2 indicating the position of the Lk means (red lines). Right, the distance of the Lk means of these DNAs differing in 100 bp was equivalent to 0.7 Lk units, thus 0.007 Lk/bp.

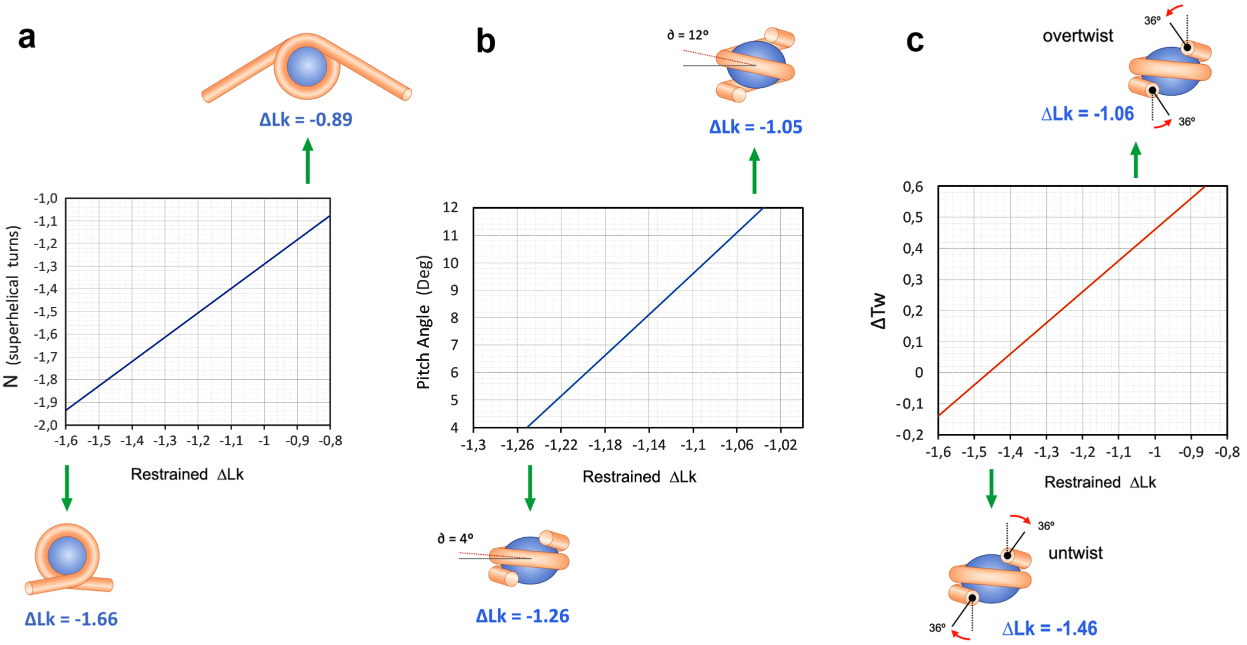


**Extended Data Fig. 5. Effect of nucleosome DNA wrapping (ΔWr) and twisting (ΔTw) on the restrained ΔLk. a,** Plot of restrained ΔLk as a function of the number of DNA super-helical turns (N) wrapped in the nucleosome. ΔLk values are calculated as ΔTw+ΔWr, where ΔTw=+0.2 and ΔWr=N(1–sin 4º). **b,** Plot of restrained ΔLk as a function of the pitch angle (∂) of the super-helical turns wrapped in the nucleosome. ΔLk values are calculated considering ΔTw=+0.2 and ΔWr=1.56(1–sin ∂). **c,** Plot of restrained ΔLk as a function of the DNA twist (ΔTw) restrained by the nucleosome. ΔLk values are calculated considering ΔWr=1.56(1–sin 4º). Nucleosome DNA geometries are modelled for the indicated ΔLk.

**Supplementary Table 1. Coordinates of the nucleosome DNA library and their corresponding nucleosome ID.** The four first columns indicate the chromosomal coordinates of the nucleosome library. The last column (nucleosome ID) tags those library sequences that overlapped >50 bp with the previously referenced nucleosomes compiled by Jiang and Pugh (2009). 91% of the library sequences matched with 4276 previously referenced nucleosomes.

**Supplementary Table 2. Calculation of the ΔLk restrained by individual nucleosomes.** The four first columns indicate the chromosomal coordinates of the nucleosome library. The next two columns indicate the relative abundance (number of reads relative to total reads) of each nucleosome sequence in sections A and B of the gel electrophoresis. Column A/A+B indicates the partition probability of each sequence in section A. Column Z indicates the corresponding Z-scores of the previous partition probabilities in a normal distribution of mean 0 and standard deviation 1. The next column, multiplies Z-scores by 0.85 (mean standard deviation of the Lk distributions of the library) to yield approximate ΔLk deviations within a normal distribution of mean 0. The next column, Phase correction, subtracts from the previous ΔLk values the average ΔLk of all the sequences of the same length. The next column, Length correction, adjusts the previous ΔLk value by adding or subtracting 0.007 for each bp difference from 144 bp (the average length of the library of nucleosome DNA sequences). The last column adds -1.26 to obtain the ΔLk restrained by each nucleosome.

**Supplementary Table 3. ΔLk values and genomic context of nucleosomes.** The first five columns indicate the chromosomal coordinates, length (bp) and the ΔLk restrained by the nucleosomes of the library. The next four columns are parameters defined in Jiang and Pugh (2009) that indicate the nucleosome IDs, the positional fuzziness, the gene ID, and the allocation within a gene body or intergenic region. Coordinates without a nucleosome ID are those that overlapped by <50 bp with referenced nucleosomes. The next two columns tag the gene terminal nucleosomes and rDNA nucleosomes. The last column tags the nucleosomes of telomeric regions, which are defined from The *Saccharomyces* Genome Database (SGD) (Cherry and Wong 2012) as those overlapping with chromosomal coordinates that include an X element core sequence, X element combinatorial repeats, short or long Y' elements or a terminal stretch of telomeric repeats. Depending on the chromosomal arm, the length of the telomeric regions spans from 430 to 13896 bp.

**Table 1. Coordinates of the nucleosome DNA library and their corresponding nucleosome ID.**

The four first columns indicate the chromosomal coordinates of the nucleosome library. The last column (nucleosome ID) tags those library sequences that overlapped >50 bp with the previously referenced nucleosomes compiled by Jiang and Pugh (2009). 91% of the library sequences matched with 4276 previously referenced nucleosomes.

| CHR | Start | End | Length (bp) | Nucleosome_ID |
|---|---|---|---|---|
| chrI | 375 | 485 | 110 | N1:382 |
| chrI | 1992 | 2143 | 151 | N1:2081 |
| chrI | 3900 | 4046 | 146 | |
| chrI | 7310 | 7448 | 138 | N1:7314 |
| chrI | 19698 | 19849 | 151 | N1:19743 |
| chrI | 24106 | 24256 | 150 | |
| chrI | 24106 | 24267 | 161 | |
| chrI | 24156 | 24264 | 108 | |
| chrI | 24156 | 24265 | 109 | |
| chrI | 24156 | 24266 | 110 | |
| chrI | 24156 | 24267 | 111 | |
| chrI | 25658 | 25790 | 132 | |
| chrI | 25658 | 25797 | 139 | |
| chrI | 25671 | 25797 | 126 | |
| chrI | 25885 | 26057 | 172 | |
| chrI | 26200 | 26329 | 129 | |
| chrI | 26200 | 26330 | 130 | |
| chrI | 26200 | 26337 | 137 | |
| chrI | 26204 | 26330 | 126 | |
| chrI | 26211 | 26330 | 119 | |
| chrI | 26268 | 26418 | 150 | N1:26380 |
| chrI | 26290 | 26465 | 175 | N1:26380 |
| chrI | 26425 | 26573 | 148 | N1:26590 |
| chrI | 26560 | 26708 | 148 | N1:26590 |
| chrI | 26695 | 26843 | 148 | N1:26787 |
| chrI | 27347 | 27495 | 148 | N1:27437 |
| chrI | 28545 | 28653 | 108 | N1:28676 |
| chrI | 28558 | 28705 | 147 | N1:28676 |
| chrI | 28865 | 28996 | 131 | N1:28942 |
| chrI | 31670 | 31815 | 145 | N1:31784 |
| chrI | 32825 | 32968 | 143 | N1:32915 |
| chrI | 38266 | 38455 | 189 | N1:38289 |
| chrI | 38266 | 38455 | 189 | N1:38459 |
| chrI | 39356 | 39504 | 148 | N1:39417 |
| chrI | 44014 | 44169 | 155 | N1:44128 |
| chrI | 48679 | 48833 | 154 | N1:48747 |
| chrI | 52107 | 52211 | 104 | N1:52114 |
| chrI | 52108 | 52200 | 92 | N1:52114 |
| chrI | 52108 | 52202 | 94 | N1:52114 |
| chrI | 52108 | 52203 | 95 | N1:52114 |
| chrI | 52108 | 52204 | 96 | N1:52114 |
| chrI | 52108 | 52205 | 97 | N1:52114 |
| chrI | 52108 | 52206 | 98 | N1:52114 |
| chrI | 52108 | 52207 | 99 | N1:52114 |
| chrI | 52108 | 52208 | 100 | N1:52114 |
| chrI | 52108 | 52209 | 101 | N1:52114 |
| chrI | 52108 | 52210 | 102 | N1:52114 |
| chrI | 52108 | 52211 | 103 | N1:52114 |
| chrI | 52112 | 52211 | 99 | N1:52114 |
| chrI | 54178 | 54332 | 154 | N1:54248 |
| chrI | 56067 | 56218 | 151 | N1:56143 |
| chrI | 57573 | 57723 | 150 | N1:57726 |
| chrI | 57573 | 57723 | 150 | N1:57555 |
| chrI | 57985 | 58142 | 157 | N1:58076 |
| chrI | 58357 | 58529 | 172 | N1:58432 |
| chrI | 59835 | 59985 | 150 | N1:59903 |
| chrI | 60956 | 61114 | 158 | N1:61025 |
| chrI | 61139 | 61253 | 114 | |
| chrI | 62731 | 62859 | 128 | N1:62815 |
| chrI | 73815 | 73966 | 151 | N1:73878 |
| chrI | 75929 | 76089 | 160 | N1:75990 |
| chrI | 77820 | 77969 | 149 | N1:77885 |
| chrI | 78867 | 79013 | 146 | N1:78863 |
| chrI | 79075 | 79223 | 148 | N1:79100 |
| chrI | 79820 | 79919 | 99 | N1:79895 |
| chrI | 80712 | 80877 | 165 | N1:80770 |
| chrI | 83877 | 84031 | 154 | N1:83953 |
| chrI | 87561 | 87716 | 155 | N1:87641 |
| chrI | 88852 | 89013 | 161 | N1:89028 |
| chrI | 88852 | 89013 | 161 | N1:88849 |
| chrI | 88889 | 89053 | 164 | N1:89028 |
| chrI | 91036 | 91130 | 94 | N1:91042 |
| chrI | 94271 | 94416 | 145 | |
| chrI | 97984 | 98151 | 167 | N1:98096 |
| chrI | 100299 | 100401 | 102 | N1:100335 |
| chrI | 100430 | 100592 | 162 | N1:100502 |
| chrI | 100744 | 100901 | 157 | N1:100813 |
| chrI | 100903 | 101028 | 125 | N1:100972 |
| chrI | 101774 | 101913 | 139 | N1:101784 |
| chrI | 102759 | 102918 | 159 | N1:102829 |
| chrI | 110352 | 110498 | 146 | N1:110405 |
| chrI | 110959 | 111107 | 148 | N1:110988 |
| chrI | 111829 | 111987 | 158 | N1:111914 |
| chrI | 111829 | 111989 | 160 | N1:111914 |
| chrI | 111835 | 111987 | 152 | N1:111914 |
| chrI | 111835 | 111989 | 154 | N1:111914 |
| chrI | 113198 | 113345 | 147 | N1:113359 |
| chrI | 113198 | 113345 | 147 | N1:113192 |
| chrI | 113272 | 113431 | 159 | N1:113359 |
| chrI | 114916 | 115064 | 148 | N1:115021 |
| chrI | 116310 | 116474 | 164 | N1:116396 |
| chrI | 119258 | 119408 | 150 | N1:119265 |
| chrI | 121555 | 121709 | 154 | N1:121654 |

| CHR | Start | End | Length (bp) | Nucleosome_ID |
|---|---|---|---|---|
| chrI | 128610 | 128759 | 149 | |
| chrI | 128749 | 128913 | 164 | N1:128845 |
| chrI | 132160 | 132339 | 179 | N1:132234 |
| chrI | 132322 | 132449 | 127 | N1:132398 |
| chrI | 133311 | 133456 | 145 | N1:133382 |
| chrI | 133444 | 133595 | 151 | N1:133558 |
| chrI | 134936 | 135082 | 146 | |
| chrI | 139478 | 139611 | 133 | N1:139570 |
| chrI | 144452 | 144595 | 143 | N1:144447 |
| chrI | 144452 | 144595 | 143 | N1:144596 |
| chrI | 145995 | 146131 | 136 | N1:145995 |
| chrI | 145995 | 146131 | 136 | N1:146151 |
| chrI | 157171 | 157325 | 154 | N1:157317 |
| chrI | 158225 | 158386 | 161 | N1:158305 |
| chrI | 158476 | 158631 | 155 | N1:158473 |
| chrI | 158476 | 158631 | 155 | N1:158641 |
| chrI | 159246 | 159353 | 107 | N1:159237 |
| chrI | 161173 | 161287 | 114 | N1:161301 |
| chrI | 163265 | 163422 | 157 | N1:163329 |
| chrI | 164304 | 164458 | 154 | N1:164322 |
| chrI | 164304 | 164458 | 154 | N1:164474 |
| chrI | 164952 | 165092 | 140 | N1:165025 |
| chrI | 164958 | 165092 | 134 | N1:165025 |
| chrI | 165370 | 165525 | 155 | N1:165400 |
| chrI | 165501 | 165609 | 108 | N1:165556 |
| chrI | 165531 | 165649 | 118 | N1:165556 |
| chrI | 165650 | 165799 | 149 | N1:165722 |
| chrI | 170185 | 170316 | 131 | |
| chrI | 170746 | 170892 | 146 | N1:170868 |
| chrI | 173251 | 173422 | 171 | |
| chrI | 174403 | 174559 | 156 | N1:174467 |
| chrI | 177062 | 177157 | 95 | |
| chrI | 178178 | 178314 | 136 | N1:178222 |
| chrI | 181632 | 181781 | 149 | N1:181719 |
| chrI | 184868 | 185010 | 142 | N1:184881 |
| chrI | 188940 | 189093 | 153 | N1:189025 |
| chrI | 189954 | 190061 | 107 | N1:190031 |
| chrI | 189954 | 190062 | 108 | N1:190031 |
| chrI | 189954 | 190063 | 109 | N1:190031 |
| chrI | 189954 | 190077 | 123 | N1:190031 |
| chrI | 191415 | 191573 | 158 | N1:191409 |
| chrI | 191722 | 191874 | 152 | N1:191782 |
| chrI | 192889 | 193041 | 152 | N1:192956 |
| chrI | 193467 | 193622 | 155 | N1:193586 |
| chrI | 194126 | 194279 | 153 | |
| chrI | 194141 | 194279 | 138 | |
| chrI | 194290 | 194442 | 152 | N1:194358 |
| chrI | 204527 | 204675 | 148 | N1:204558 |
| chrI | 204662 | 204810 | 148 | N1:204754 |
| chrI | 204797 | 204945 | 148 | |
| chrI | 205040 | 205170 | 130 | |
| chrI | 205573 | 205699 | 126 | N1:205618 |
| chrI | 205573 | 205705 | 132 | N1:205618 |
| chrI | 205573 | 205712 | 139 | N1:205618 |
| chrI | 205580 | 205712 | 132 | N1:205618 |
| chrI | 206390 | 206543 | 153 | N1:206455 |
| chrI | 207406 | 207531 | 125 | N1:207477 |
| chrI | 215442 | 215555 | 113 | N1:215495 |
| chrI | 217878 | 218015 | 137 | N1:217966 |
| chrI | 217916 | 218076 | 160 | N1:217966 |
| chrI | 218711 | 218856 | 145 | N1:218780 |
| chrI | 218738 | 218886 | 148 | N1:218780 |
| chrI | 222387 | 222554 | 167 | N1:222504 |
| chrI | 222751 | 222855 | 104 | N1:222820 |
| chrI | 223257 | 223420 | 163 | N1:223335 |
| chrI | 224065 | 224173 | 108 | N1:224083 |
| chrI | 224877 | 224997 | 120 | N1:224931 |
| chrI | 226540 | 226687 | 147 | N1:226556 |
| chrI | 229861 | 230005 | 144 | N1:229980 |
| chrI | 230119 | 230218 | 99 | N1:230156 |
| chrI | 230124 | 230218 | 94 | N1:230156 |
| chrII | 163 | 311 | 148 | N2:255 |
| chrII | 526 | 677 | 151 | N2:589 |
| chrII | 1089 | 1240 | 151 | N2:1150 |
| chrII | 1727 | 1858 | 131 | N2:1853 |
| chrII | 1727 | 1870 | 143 | N2:1853 |
| chrII | 1727 | 1914 | 187 | N2:1853 |
| chrII | 1931 | 2076 | 145 | |
| chrII | 1936 | 2043 | 107 | |
| chrII | 1937 | 2034 | 97 | |
| chrII | 1937 | 2042 | 105 | |
| chrII | 1937 | 2046 | 109 | |
| chrII | 1937 | 2051 | 114 | |
| chrII | 1937 | 2054 | 117 | |
| chrII | 1937 | 2057 | 120 | |
| chrII | 1937 | 2062 | 125 | |
| chrII | 1937 | 2071 | 134 | |
| chrII | 1937 | 2072 | 135 | |
| chrII | 1937 | 2073 | 136 | |
| chrII | 1937 | 2075 | 138 | |
| chrII | 1937 | 2076 | 139 | |
| chrII | 1937 | 2082 | 145 | |
| chrII | 1937 | 2086 | 149 | |
| chrII | 1938 | 2076 | 138 | |
| chrII | 1939 | 2076 | 137 | |
| chrII | 1940 | 2076 | 136 | |
| chrII | 1941 | 2076 | 135 | |
| chrII | 1942 | 2076 | 134 | |
| chrII | 1944 | 2076 | 132 | |
| chrII | 1947 | 2076 | 129 | |
| chrII | 1948 | 2076 | 128 | |
| chrII | 1952 | 2076 | 124 | |
| chrII | 1953 | 2076 | 123 | |

| CHR | Start | End | Length (bp) | Nucleosome_ID |
|---|---|---|---|---|
| chrII | 1964 | 2076 | 112 | |
| chrII | 1966 | 2076 | 110 | |
| chrII | 2124 | 2286 | 162 | N2:2264 |
| chrII | 2124 | 2286 | 162 | N2:2110 |
| chrII | 2191 | 2286 | 95 | N2:2264 |
| chrII | 2267 | 2374 | 107 | N2:2264 |
| chrII | 2267 | 2414 | 147 | N2:2264 |
| chrII | 2522 | 2620 | 98 | |
| chrII | 2522 | 2673 | 151 | |
| chrII | 3354 | 3498 | 144 | N2:3449 |
| chrII | 3419 | 3571 | 152 | N2:3449 |
| chrII | 3701 | 3844 | 143 | N2:3787 |
| chrII | 3992 | 4129 | 137 | N2:4071 |
| chrII | 7980 | 8135 | 155 | |
| chrII | 10712 | 10874 | 162 | N2:10765 |
| chrII | 20981 | 21152 | 171 | N2:21084 |
| chrII | 21442 | 21596 | 154 | N2:21498 |
| chrII | 24248 | 24399 | 151 | N2:24353 |
| chrII | 24299 | 24437 | 138 | N2:24353 |
| chrII | 25280 | 25416 | 136 | N2:25382 |
| chrII | 26126 | 26261 | 135 | N2:26202 |
| chrII | 30232 | 30334 | 102 | |
| chrII | 31655 | 31803 | 148 | |
| chrII | 32970 | 33121 | 151 | N2:32989 |
| chrII | 35988 | 36136 | 148 | N2:36067 |
| chrII | 48768 | 48912 | 144 | N2:48843 |
| chrII | 54394 | 54539 | 145 | N2:54546 |
| chrII | 54394 | 54539 | 145 | N2:54380 |
| chrII | 54551 | 54716 | 165 | N2:54732 |
| chrII | 54551 | 54716 | 165 | N2:54546 |
| chrII | 62389 | 62546 | 157 | N2:62463 |
| chrII | 62885 | 63033 | 148 | N2:62915 |
| chrII | 64104 | 64264 | 160 | |
| chrII | 64653 | 64794 | 141 | |
| chrII | 65899 | 66050 | 151 | N2:65948 |
| chrII | 67731 | 67881 | 150 | N2:67884 |
| chrII | 69118 | 69263 | 145 | N2:69156 |
| chrII | 79393 | 79540 | 147 | N2:79471 |
| chrII | 81527 | 81678 | 151 | N2:81676 |
| chrII | 81527 | 81678 | 151 | N2:81506 |
| chrII | 86475 | 86605 | 130 | N2:86602 |
| chrII | 87052 | 87211 | 159 | N2:87114 |
| chrII | 94866 | 95023 | 157 | N2:94979 |
| chrII | 97063 | 97217 | 154 | N2:97213 |
| chrII | 97108 | 97247 | 139 | N2:97213 |
| chrII | 98068 | 98229 | 161 | N2:98203 |
| chrII | 98405 | 98563 | 158 | N2:98532 |
| chrII | 102188 | 102336 | 148 | N2:102248 |
| chrII | 105450 | 105577 | 127 | N2:105509 |
| chrII | 105532 | 105684 | 152 | N2:105680 |
| chrII | 105532 | 105684 | 152 | N2:105509 |
| chrII | 105754 | 105910 | 156 | N2:105836 |
| chrII | 106088 | 106233 | 145 | |
| chrII | 112631 | 112737 | 106 | N2:112714 |
| chrII | 112631 | 112749 | 118 | N2:112714 |
| chrII | 117607 | 117737 | 130 | N2:117637 |
| chrII | 118185 | 118336 | 151 | N2:118300 |
| chrII | 118430 | 118588 | 158 | N2:118451 |
| chrII | 118430 | 118588 | 158 | N2:118611 |
| chrII | 118604 | 118737 | 133 | N2:118611 |
| chrII | 118662 | 118820 | 158 | N2:118790 |
| chrII | 120690 | 120830 | 140 | N2:120678 |
| chrII | 130330 | 130481 | 151 | N2:130370 |
| chrII | 133172 | 133320 | 148 | N2:133293 |
| chrII | 134044 | 134208 | 164 | N2:134212 |
| chrII | 134044 | 134208 | 164 | N2:134052 |
| chrII | 146223 | 146376 | 153 | N2:146325 |
| chrII | 147729 | 147892 | 163 | |
| chrII | 157306 | 157457 | 151 | N2:157445 |
| chrII | 158794 | 158975 | 181 | N2:158865 |
| chrII | 159579 | 159667 | 88 | N2:159667 |
| chrII | 159579 | 159734 | 155 | N2:159667 |
| chrII | 161150 | 161286 | 136 | N2:161276 |
| chrII | 161650 | 161786 | 136 | N2:161723 |
| chrII | 164812 | 164963 | 151 | N2:164880 |
| chrII | 169155 | 169305 | 150 | N2:169306 |
| chrII | 171571 | 171720 | 149 | N2:171641 |
| chrII | 171661 | 171810 | 149 | N2:171641 |
| chrII | 171661 | 171810 | 149 | N2:171803 |
| chrII | 174305 | 174447 | 142 | N2:174374 |
| chrII | 186048 | 186192 | 144 | N2:186122 |
| chrII | 187191 | 187351 | 160 | N2:187294 |
| chrII | 188175 | 188332 | 157 | N2:188328 |
| chrII | 189372 | 189539 | 167 | |
| chrII | 192490 | 192638 | 148 | N2:192564 |
| chrII | 195265 | 195410 | 145 | N2:195403 |
| chrII | 203726 | 203877 | 151 | N2:203822 |
| chrII | 204265 | 204408 | 143 | N2:204305 |
| chrII | 213654 | 213809 | 155 | N2:213693 |
| chrII | 214589 | 214739 | 150 | N2:214660 |
| chrII | 214869 | 215017 | 148 | N2:214983 |
| chrII | 215320 | 215464 | 144 | N2:215386 |
| chrII | 216639 | 216771 | 132 | N2:216655 |
| chrII | 217546 | 217694 | 148 | N2:217574 |
| chrII | 222615 | 222772 | 157 | N2:222719 |
| chrII | 222619 | 222772 | 153 | N2:222719 |
| chrII | 223146 | 223281 | 135 | N2:223185 |
| chrII | 223772 | 223929 | 157 | N2:223866 |
| chrII | 223872 | 224016 | 144 | N2:223866 |
| chrII | 223872 | 224016 | 144 | N2:224033 |
| chrII | 224435 | 224582 | 147 | N2:224426 |
| chrII | 224435 | 224582 | 147 | N2:224599 |
| chrII | 225175 | 225325 | 150 | N2:225274 |

| CHR | Start | End | Length (bp) | Nucleosome_ID |
|---|---|---|---|---|
| chrII | 226374 | 226481 | 107 | N2:226487 |
| chrII | 226832 | 226946 | 114 | N2:226921 |
| chrII | 228528 | 228693 | 165 | |
| chrII | 233542 | 233703 | 161 | N2:233579 |
| chrII | 234707 | 234798 | 91 | N2:234705 |
| chrII | 239108 | 239251 | 143 | N2:239202 |
| chrII | 249586 | 249752 | 166 | N2:249650 |
| chrII | 252919 | 253070 | 151 | |
| chrII | 253969 | 254127 | 158 | N2:254057 |
| chrII | 255566 | 255720 | 154 | N2:255613 |
| chrII | 260109 | 260232 | 123 | |
| chrII | 260209 | 260364 | 155 | N2:260319 |
| chrII | 260642 | 260776 | 134 | N2:260707 |
| chrII | 260642 | 260782 | 140 | N2:260707 |
| chrII | 261685 | 261820 | 135 | N2:261719 |
| chrII | 262132 | 262282 | 150 | N2:262199 |
| chrII | 262311 | 262468 | 157 | N2:262403 |
| chrII | 262974 | 263121 | 147 | N2:262967 |
| chrII | 263717 | 263867 | 150 | N2:263815 |
| chrII | 264916 | 265023 | 107 | |
| chrII | 269730 | 269883 | 153 | N2:269845 |
| chrII | 269730 | 269885 | 155 | N2:269845 |
| chrII | 269781 | 269930 | 149 | N2:269845 |
| chrII | 272917 | 273071 | 154 | N2:273057 |
| chrII | 279942 | 280075 | 133 | N2:280032 |
| chrII | 282915 | 283035 | 120 | N2:282943 |
| chrII | 288899 | 289053 | 154 | N2:288983 |
| chrII | 289452 | 289559 | 107 | N2:289558 |
| chrII | 289452 | 289573 | 121 | N2:289558 |
| chrII | 292636 | 292792 | 156 | N2:292683 |
| chrII | 295986 | 296116 | 130 | N2:296031 |
| chrII | 296940 | 297047 | 107 | N2:296987 |
| chrII | 296940 | 297056 | 116 | N2:296987 |
| chrII | 298361 | 298510 | 149 | N2:298428 |
| chrII | 298515 | 298656 | 141 | N2:298593 |
| chrII | 300341 | 300475 | 134 | N2:300360 |
| chrII | 303103 | 303264 | 161 | N2:303172 |
| chrII | 304167 | 304319 | 152 | N2:304247 |
| chrII | 306192 | 306322 | 130 | N2:306304 |
| chrII | 310844 | 311002 | 158 | N2:310950 |
| chrII | 314561 | 314691 | 130 | N2:314625 |
| chrII | 316179 | 316338 | 159 | N2:316235 |
| chrII | 320829 | 320984 | 155 | N2:320918 |
| chrII | 324261 | 324383 | 122 | N2:324329 |
| chrII | 324593 | 324738 | 145 | N2:324676 |
| chrII | 333410 | 333578 | 168 | N2:333441 |
| chrII | 334171 | 334335 | 164 | N2:334243 |
| chrII | 341568 | 341673 | 105 | |
| chrII | 344002 | 344150 | 148 | N2:344104 |
| chrII | 345574 | 345727 | 153 | N2:345699 |
| chrII | 348428 | 348599 | 171 | N2:348504 |
| chrII | 348488 | 348624 | 136 | N2:348504 |
| chrII | 352601 | 352731 | 130 | N2:352692 |
| chrII | 354955 | 355107 | 152 | N2:355010 |
| chrII | 360789 | 360938 | 149 | N2:360839 |
| chrII | 369124 | 369272 | 148 | N2:369186 |
| chrII | 369614 | 369763 | 149 | N2:369688 |
| chrII | 372782 | 372926 | 144 | N2:372886 |
| chrII | 379733 | 379878 | 145 | N2:379800 |
| chrII | 380438 | 380593 | 155 | N2:380439 |
| chrII | 380438 | 380593 | 155 | N2:380588 |
| chrII | 383232 | 383381 | 149 | N2:383330 |
| chrII | 384056 | 384165 | 109 | |
| chrII | 384888 | 385039 | 151 | N2:384899 |
| chrII | 385956 | 386120 | 164 | N2:386082 |
| chrII | 386724 | 386859 | 135 | N2:386790 |
| chrII | 402057 | 402202 | 145 | N2:402105 |
| chrII | 405294 | 405447 | 153 | N2:405333 |
| chrII | 411086 | 411218 | 132 | N2:411221 |
| chrII | 412922 | 413067 | 145 | N2:413068 |
| chrII | 417276 | 417442 | 166 | N2:417375 |
| chrII | 418844 | 418980 | 136 | N2:418877 |
| chrII | 420492 | 420630 | 138 | N2:420545 |
| chrII | 432763 | 432911 | 148 | N2:432818 |
| chrII | 435614 | 435764 | 150 | N2:435690 |
| chrII | 441246 | 441413 | 167 | N2:441290 |
| chrII | 442313 | 442462 | 149 | |
| chrII | 447274 | 447421 | 147 | N2:447332 |
| chrII | 448183 | 448311 | 128 | N2:448253 |
| chrII | 448687 | 448853 | 166 | |
| chrII | 450522 | 450683 | 161 | N2:450575 |
| chrII | 456967 | 457110 | 143 | N2:456997 |
| chrII | 459507 | 459619 | 112 | N2:459525 |
| chrII | 463510 | 463659 | 149 | N2:463671 |
| chrII | 464402 | 464557 | 155 | N2:464478 |
| chrII | 467973 | 468117 | 144 | N2:468080 |
| chrII | 469937 | 470099 | 162 | N2:469962 |
| chrII | 478544 | 478678 | 134 | N2:478563 |
| chrII | 479993 | 480154 | 161 | N2:480057 |
| chrII | 480073 | 480226 | 153 | N2:480057 |
| chrII | 483614 | 483775 | 161 | N2:483752 |
| chrII | 490237 | 490391 | 154 | N2:490316 |
| chrII | 496459 | 496596 | 137 | N2:496548 |
| chrII | 498698 | 498864 | 166 | N2:498780 |
| chrII | 499848 | 499990 | 142 | N2:499949 |
| chrII | 501137 | 501288 | 151 | N2:501227 |
| chrII | 503704 | 503871 | 167 | N2:503875 |
| chrII | 503704 | 503871 | 167 | N2:503696 |
| chrII | 505974 | 506148 | 174 | N2:506017 |
| chrII | 506025 | 506148 | 123 | N2:506017 |
| chrII | 506025 | 506179 | 154 | N2:506017 |
| chrII | 506025 | 506179 | 154 | N2:506177 |
| chrII | 509772 | 509912 | 140 | N2:509886 |

| CHR | Start | End | Length (bp) | Nucleosome_ID |
|---|---|---|---|---|
| chrII | 511646 | 511795 | 149 | N2:511651 |
| chrII | 512273 | 512423 | 150 | N2:512342 |
| chrII | 519148 | 519306 | 158 | N2:519220 |
| chrII | 523260 | 523407 | 147 | N2:523344 |
| chrII | 523774 | 523925 | 151 | |
| chrII | 525378 | 525528 | 150 | N2:525423 |
| chrII | 527770 | 527902 | 132 | N2:527832 |
| chrII | 528914 | 529061 | 147 | N2:528946 |
| chrII | 529231 | 529384 | 153 | N2:529279 |
| chrII | 533755 | 533907 | 152 | N2:533826 |
| chrII | 543934 | 544097 | 163 | N2:543989 |
| chrII | 545512 | 545645 | 133 | |
| chrII | 550054 | 550200 | 146 | N2:550101 |
| chrII | 551769 | 551916 | 147 | N2:551862 |
| chrII | 551821 | 551946 | 125 | N2:551862 |
| chrII | 552584 | 552756 | 172 | N2:552697 |
| chrII | 560316 | 560442 | 126 | N2:560343 |
| chrII | 568857 | 569009 | 152 | N2:568953 |
| chrII | 570513 | 570675 | 162 | N2:570686 |
| chrII | 570513 | 570675 | 162 | N2:570530 |
| chrII | 575360 | 575489 | 129 | N2:575420 |
| chrII | 575363 | 575489 | 126 | N2:575420 |
| chrII | 575635 | 575789 | 154 | N2:575736 |
| chrII | 577662 | 577794 | 132 | N2:577744 |
| chrII | 581371 | 581525 | 154 | N2:581425 |
| chrII | 583028 | 583186 | 158 | N2:583091 |
| chrII | 583277 | 583426 | 149 | |
| chrII | 589093 | 589249 | 156 | N2:589146 |
| chrII | 598510 | 598657 | 147 | N2:598568 |
| chrII | 599865 | 600024 | 159 | N2:599938 |
| chrII | 601519 | 601677 | 158 | N2:601690 |
| chrII | 601519 | 601677 | 158 | N2:601539 |
| chrII | 608166 | 608308 | 142 | N2:608176 |
| chrII | 608166 | 608308 | 142 | N2:608330 |
| chrII | 610440 | 610604 | 164 | N2:610482 |
| chrII | 615980 | 616137 | 157 | N2:616096 |
| chrII | 618802 | 618957 | 155 | N2:618851 |
| chrII | 621368 | 621503 | 135 | N2:621487 |
| chrII | 624538 | 624687 | 149 | N2:624638 |
| chrII | 626717 | 626881 | 164 | N2:626851 |
| chrII | 633744 | 633889 | 145 | N2:633802 |
| chrII | 633747 | 633889 | 142 | N2:633802 |
| chrII | 634590 | 634750 | 160 | N2:634645 |
| chrII | 635204 | 635350 | 146 | N2:635289 |
| chrII | 636694 | 636853 | 159 | N2:636766 |
| chrII | 636695 | 636853 | 158 | N2:636766 |
| chrII | 636699 | 636853 | 154 | N2:636766 |
| chrII | 641881 | 642011 | 130 | N2:641909 |
| chrII | 643000 | 643147 | 147 | N2:643066 |
| chrII | 643625 | 643773 | 148 | N2:643703 |
| chrII | 645958 | 646066 | 108 | N2:646057 |
| chrII | 652842 | 653006 | 164 | |
| chrII | 667311 | 667462 | 151 | N2:667376 |
| chrII | 670468 | 670600 | 132 | N2:670509 |
| chrII | 672951 | 673112 | 161 | N2:673046 |
| chrII | 673886 | 673964 | 78 | N2:673878 |
| chrII | 673886 | 673989 | 103 | N2:673878 |
| chrII | 674406 | 674512 | 106 | N2:674454 |
| chrII | 674406 | 674523 | 117 | N2:674454 |
| chrII | 678764 | 678912 | 148 | N2:678851 |
| chrII | 682036 | 682166 | 130 | N2:682093 |
| chrII | 683947 | 684082 | 135 | N2:684039 |
| chrII | 684506 | 684665 | 159 | N2:684541 |
| chrII | 685114 | 685267 | 153 | N2:685155 |
| chrII | 688568 | 688714 | 146 | N2:688658 |
| chrII | 693669 | 693820 | 151 | N2:693836 |
| chrII | 696303 | 696437 | 134 | |
| chrII | 698169 | 698308 | 139 | N2:698202 |
| chrII | 698728 | 698854 | 126 | N2:698800 |
| chrII | 698730 | 698836 | 106 | N2:698800 |
| chrII | 699474 | 699629 | 155 | N2:699534 |
| chrII | 699802 | 699945 | 143 | N2:699870 |
| chrII | 699802 | 699953 | 151 | N2:699870 |
| chrII | 699834 | 699945 | 111 | N2:699870 |
| chrII | 699834 | 699953 | 119 | N2:699870 |
| chrII | 702899 | 703031 | 132 | N2:702974 |
| chrII | 706525 | 706696 | 171 | N2:706600 |
| chrII | 708493 | 708657 | 164 | N2:708624 |
| chrII | 708881 | 709054 | 173 | N2:708943 |
| chrII | 716866 | 717023 | 157 | N2:716955 |
| chrII | 716874 | 717023 | 149 | N2:716955 |
| chrII | 718549 | 718695 | 146 | N2:718618 |
| chrII | 721490 | 721639 | 149 | N2:721551 |
| chrII | 723357 | 723507 | 150 | N2:723426 |
| chrII | 727858 | 728011 | 153 | N2:727936 |
| chrII | 733512 | 733676 | 164 | N2:733639 |
| chrII | 735258 | 735391 | 133 | N2:735332 |
| chrII | 736781 | 736927 | 146 | N2:736768 |
| chrII | 736912 | 737033 | 121 | |
| chrII | 740252 | 740411 | 159 | N2:740348 |
| chrII | 743678 | 743781 | 103 | N2:743723 |
| chrII | 746216 | 746373 | 157 | N2:746234 |
| chrII | 747192 | 747321 | 129 | N2:747277 |
| chrII | 748930 | 749090 | 160 | N2:749008 |
| chrII | 749254 | 749401 | 147 | N2:749334 |
| chrII | 750045 | 750206 | 161 | N2:750083 |
| chrII | 760513 | 760620 | 107 | |
| chrII | 760513 | 760634 | 121 | |
| chrII | 771359 | 771518 | 159 | |
| chrII | 771799 | 771953 | 154 | |
| chrII | 774249 | 774405 | 156 | N2:774294 |
| chrII | 775701 | 775861 | 160 | |
| chrII | 777331 | 777484 | 153 | N2:777349 |

| CHR | Start | End | Length (bp) | Nucleosome_ID |
|---|---|---|---|---|
| chrII | 779629 | 779798 | 169 | N2:779707 |
| chrII | 780145 | 780251 | 106 | N2:780192 |
| chrII | 780145 | 780260 | 115 | N2:780192 |
| chrII | 780582 | 780757 | 175 | N2:780666 |
| chrII | 781915 | 782083 | 168 | N2:782014 |
| chrII | 786436 | 786542 | 106 | N2:786464 |
| chrII | 786436 | 786560 | 124 | N2:786464 |
| chrII | 788442 | 788582 | 140 | N2:788514 |
| chrII | 788604 | 788759 | 155 | N2:788689 |
| chrII | 789546 | 789709 | 163 | N2:789650 |
| chrII | 791029 | 791155 | 126 | N2:791088 |
| chrII | 791029 | 791156 | 127 | N2:791088 |
| chrII | 791031 | 791136 | 105 | N2:791088 |
| chrII | 793238 | 793400 | 162 | N2:793321 |
| chrII | 795412 | 795559 | 147 | N2:795519 |
| chrII | 809074 | 809203 | 129 | N2:809118 |
| chrII | 812955 | 813098 | 143 | N2:812953 |
| chrIII | 2 | 145 | 143 | |
| chrIII | 2 | 176 | 174 | |
| chrIII | 4 | 145 | 141 | |
| chrIII | 10 | 145 | 135 | |
| chrIII | 10 | 146 | 136 | |
| chrIII | 14 | 145 | 131 | |
| chrIII | 19 | 145 | 126 | |
| chrIII | 26 | 145 | 119 | |
| chrIII | 37 | 145 | 108 | |
| chrIII | 41 | 145 | 104 | |
| chrIII | 50 | 145 | 95 | |
| chrIII | 50 | 176 | 126 | |
| chrIII | 50 | 185 | 135 | |
| chrIII | 50 | 201 | 151 | |
| chrIII | 50 | 220 | 170 | |
| chrIII | 73 | 263 | 190 | |
| chrIII | 88 | 231 | 143 | |
| chrIII | 97 | 212 | 115 | |
| chrIII | 110 | 263 | 153 | |
| chrIII | 110 | 280 | 170 | N3:298 |
| chrIII | 110 | 283 | 173 | N3:298 |
| chrIII | 115 | 280 | 165 | N3:298 |
| chrIII | 115 | 283 | 168 | N3:298 |
| chrIII | 118 | 284 | 166 | N3:298 |
| chrIII | 129 | 252 | 123 | |
| chrIII | 129 | 260 | 131 | |
| chrIII | 129 | 263 | 134 | |
| chrIII | 129 | 280 | 151 | N3:298 |
| chrIII | 129 | 283 | 154 | N3:298 |
| chrIII | 129 | 284 | 155 | N3:298 |
| chrIII | 129 | 301 | 172 | N3:298 |
| chrIII | 129 | 309 | 180 | N3:298 |
| chrIII | 129 | 326 | 197 | N3:298 |
| chrIII | 143 | 284 | 141 | N3:298 |
| chrIII | 164 | 284 | 120 | N3:298 |
| chrIII | 164 | 341 | 177 | N3:298 |
| chrIII | 186 | 284 | 98 | N3:298 |
| chrIII | 186 | 320 | 134 | N3:298 |
| chrIII | 186 | 337 | 151 | N3:298 |
| chrIII | 186 | 341 | 155 | N3:298 |
| chrIII | 186 | 352 | 166 | N3:298 |
| chrIII | 186 | 353 | 167 | N3:298 |
| chrIII | 186 | 355 | 169 | N3:298 |
| chrIII | 192 | 329 | 137 | N3:298 |
| chrIII | 246 | 341 | 95 | N3:298 |
| chrIII | 246 | 352 | 106 | N3:298 |
| chrIII | 5166 | 5314 | 148 | N3:5206 |
| chrIII | 5882 | 5994 | 112 | N3:5973 |
| chrIII | 7560 | 7714 | 154 | N3:7557 |
| chrIII | 9830 | 9940 | 110 | N3:9914 |
| chrIII | 9831 | 9940 | 109 | N3:9914 |
| chrIII | 9831 | 9941 | 110 | N3:9914 |
| chrIII | 9832 | 9955 | 123 | N3:9914 |
| chrIII | 11462 | 11613 | 151 | N3:11616 |
| chrIII | 17946 | 18093 | 147 | N3:18008 |
| chrIII | 23277 | 23427 | 150 | N3:23351 |
| chrIII | 27235 | 27381 | 146 | N3:27325 |
| chrIII | 32929 | 33083 | 154 | N3:33061 |
| chrIII | 34797 | 34957 | 160 | N3:34875 |
| chrIII | 36971 | 37125 | 154 | |
| chrIII | 37274 | 37421 | 147 | N3:37360 |
| chrIII | 41105 | 41261 | 156 | N3:41160 |
| chrIII | 42270 | 42419 | 149 | N3:42308 |
| chrIII | 56437 | 56565 | 128 | N3:56555 |
| chrIII | 62417 | 62568 | 151 | N3:62518 |
| chrIII | 62420 | 62568 | 148 | N3:62518 |
| chrIII | 62779 | 62946 | 167 | N3:62871 |
| chrIII | 62988 | 63143 | 155 | N3:63052 |
| chrIII | 68315 | 68492 | 177 | N3:68474 |
| chrIII | 68580 | 68746 | 166 | N3:68637 |
| chrIII | 74254 | 74383 | 129 | |
| chrIII | 75106 | 75269 | 163 | N3:75209 |
| chrIII | 82256 | 82435 | 179 | N3:82326 |
| chrIII | 86659 | 86788 | 129 | |
| chrIII | 86884 | 87013 | 129 | N3:86885 |
| chrIII | 88349 | 88492 | 143 | N3:88396 |
| chrIII | 89624 | 89790 | 166 | |
| chrIII | 90428 | 90577 | 149 | N3:90522 |
| chrIII | 90432 | 90582 | 150 | N3:90522 |
| chrIII | 92227 | 92379 | 152 | N3:92268 |
| chrIII | 93544 | 93697 | 153 | N3:93669 |
| chrIII | 94513 | 94671 | 158 | N3:94604 |
| chrIII | 95867 | 96028 | 161 | N3:95947 |
| chrIII | 95874 | 96028 | 154 | N3:95947 |
| chrIII | 97469 | 97602 | 133 | N3:97480 |
| chrIII | 100878 | 101025 | 147 | N3:100936 |

| CHR | Start | End | Length (bp) | Nucleosome_ID |
|---|---|---|---|---|
| chrIII | 100878 | 101048 | 170 | N3:100936 |
| chrIII | 101513 | 101652 | 139 | N3:101599 |
| chrIII | 101547 | 101689 | 142 | N3:101599 |
| chrIII | 101696 | 101853 | 157 | N3:101768 |
| chrIII | 103033 | 103165 | 132 | N3:103166 |
| chrIII | 103103 | 103249 | 146 | N3:103166 |
| chrIII | 103103 | 103251 | 148 | N3:103166 |
| chrIII | 104725 | 104877 | 152 | N3:104791 |
| chrIII | 106067 | 106230 | 163 | N3:106142 |
| chrIII | 116514 | 116644 | 130 | N3:116605 |
| chrIII | 117818 | 117976 | 158 | N3:117863 |
| chrIII | 132816 | 132923 | 107 | |
| chrIII | 132817 | 132923 | 106 | |
| chrIII | 133268 | 133417 | 149 | N3:133250 |
| chrIII | 134878 | 135010 | 132 | N3:134986 |
| chrIII | 136592 | 136744 | 152 | N3:136655 |
| chrIII | 137946 | 138097 | 151 | N3:138059 |
| chrIII | 138189 | 138339 | 150 | N3:138298 |
| chrIII | 146706 | 146857 | 151 | N3:146686 |
| chrIII | 154254 | 154394 | 140 | N3:154383 |
| chrIII | 154256 | 154394 | 138 | N3:154383 |
| chrIII | 156400 | 156550 | 150 | N3:156457 |
| chrIII | 156437 | 156550 | 113 | N3:156457 |
| chrIII | 156484 | 156645 | 161 | N3:156660 |
| chrIII | 158189 | 158338 | 149 | |
| chrIII | 159911 | 160084 | 173 | N3:160082 |
| chrIII | 159911 | 160084 | 173 | N3:159912 |
| chrIII | 161801 | 161938 | 137 | N3:161850 |
| chrIII | 175934 | 176084 | 150 | N3:176001 |
| chrIII | 186417 | 186569 | 152 | N3:186495 |
| chrIII | 188100 | 188260 | 160 | N3:188165 |
| chrIII | 191522 | 191646 | 124 | N3:191559 |
| chrIII | 195429 | 195578 | 149 | |
| chrIII | 201078 | 201230 | 152 | |
| chrIII | 208738 | 208879 | 141 | N3:208806 |
| chrIII | 209585 | 209734 | 149 | N3:209637 |
| chrIII | 210916 | 211072 | 156 | N3:210951 |
| chrIII | 213468 | 213602 | 134 | N3:213471 |
| chrIII | 215653 | 215790 | 137 | N3:215679 |
| chrIII | 217960 | 218102 | 142 | N3:218083 |
| chrIII | 228591 | 228741 | 150 | N3:228733 |
| chrIII | 231011 | 231166 | 155 | N3:231100 |
| chrIII | 234251 | 234402 | 151 | N3:234286 |
| chrIII | 234921 | 235052 | 131 | |
| chrIII | 236307 | 236469 | 162 | N3:236392 |
| chrIII | 238120 | 238291 | 171 | N3:238307 |
| chrIII | 238120 | 238291 | 171 | N3:238147 |
| chrIII | 240298 | 240452 | 154 | N3:240369 |
| chrIII | 240452 | 240614 | 162 | N3:240532 |
| chrIII | 242569 | 242719 | 150 | N3:242631 |
| chrIII | 246007 | 246188 | 181 | N3:246094 |
| chrIII | 246926 | 247057 | 131 | N3:246993 |
| chrIII | 246926 | 247058 | 132 | N3:246993 |
| chrIII | 248934 | 249085 | 151 | N3:249001 |
| chrIII | 251728 | 251875 | 147 | |
| chrIII | 252213 | 252361 | 148 | N3:252282 |
| chrIII | 252238 | 252397 | 159 | N3:252282 |
| chrIII | 254663 | 254821 | 158 | N3:254734 |
| chrIII | 257083 | 257228 | 145 | N3:257179 |
| chrIII | 257806 | 257969 | 163 | N3:257802 |
| chrIII | 258183 | 258325 | 142 | N3:258264 |
| chrIII | 259144 | 259292 | 148 | N3:259206 |
| chrIII | 262204 | 262356 | 152 | N3:262264 |
| chrIII | 269289 | 269448 | 159 | N3:269317 |
| chrIII | 272784 | 272929 | 145 | N3:272830 |
| chrIII | 276672 | 276825 | 153 | N3:276740 |
| chrIII | 280552 | 280680 | 128 | N3:280622 |
| chrIII | 281034 | 281170 | 136 | N3:281080 |
| chrIII | 286830 | 286938 | 108 | N3:286909 |
| chrIII | 286830 | 286939 | 109 | N3:286909 |
| chrIII | 286830 | 286940 | 110 | N3:286909 |
| chrIII | 286830 | 286941 | 111 | N3:286909 |
| chrIII | 286832 | 286938 | 106 | N3:286909 |
| chrIII | 286832 | 286939 | 107 | N3:286909 |
| chrIII | 286832 | 286955 | 123 | N3:286909 |
| chrIII | 288517 | 288666 | 149 | N3:288593 |
| chrIII | 303804 | 303963 | 159 | N3:303873 |
| chrIII | 307204 | 307345 | 141 | N3:307293 |
| chrIII | 307633 | 307788 | 155 | |
| chrIII | 308042 | 308191 | 149 | N3:308087 |
| chrIII | 310321 | 310472 | 151 | N3:310383 |
| chrIII | 310654 | 310803 | 149 | N3:310728 |
| chrIII | 311798 | 311947 | 149 | N3:311793 |
| chrIII | 314554 | 314707 | 153 | N3:314702 |
| chrIV | 0 | 126 | 126 | |
| chrIV | 0 | 145 | 145 | |
| chrIV | 0 | 155 | 155 | |
| chrIV | 5 | 126 | 121 | |
| chrIV | 14 | 126 | 112 | |
| chrIV | 14 | 134 | 120 | |
| chrIV | 14 | 155 | 141 | |
| chrIV | 44 | 126 | 82 | |
| chrIV | 44 | 134 | 90 | |
| chrIV | 44 | 145 | 101 | |
| chrIV | 44 | 155 | 111 | |
| chrIV | 57 | 155 | 98 | |
| chrIV | 1765 | 1923 | 158 | N4:1847 |
| chrIV | 6558 | 6652 | 94 | N4:6650 |
| chrIV | 8786 | 8917 | 131 | N4:8900 |
| chrIV | 10468 | 10622 | 154 | N4:10539 |
| chrIV | 11706 | 11865 | 159 | N4:11735 |
| chrIV | 12342 | 12498 | 156 | N4:12435 |
| chrIV | 12922 | 13062 | 140 | |

| CHR | Start | End | Length (bp) | Nucleosome_ID |
|---|---|---|---|---|
| chrIV | 13187 | 13340 | 153 | N4:13269 |
| chrIV | 15846 | 16000 | 154 | N4:15932 |
| chrIV | 16215 | 16372 | 157 | N4:16282 |
| chrIV | 16517 | 16676 | 159 | N4:16612 |
| chrIV | 22332 | 22485 | 153 | N4:22409 |
| chrIV | 22780 | 22921 | 141 | N4:22851 |
| chrIV | 24904 | 25025 | 121 | N4:25029 |
| chrIV | 26118 | 26260 | 142 | N4:26130 |
| chrIV | 28132 | 28284 | 152 | N4:28244 |
| chrIV | 28565 | 28649 | 84 | |
| chrIV | 28606 | 28762 | 156 | N4:28689 |
| chrIV | 29713 | 29864 | 151 | N4:29784 |
| chrIV | 30032 | 30191 | 159 | N4:30113 |
| chrIV | 31789 | 31954 | 165 | N4:31846 |
| chrIV | 32229 | 32385 | 156 | N4:32309 |
| chrIV | 33584 | 33743 | 159 | N4:33663 |
| chrIV | 34942 | 35098 | 156 | N4:35008 |
| chrIV | 43249 | 43399 | 150 | N4:43358 |
| chrIV | 43986 | 44132 | 146 | N4:44075 |
| chrIV | 51938 | 52044 | 106 | N4:51987 |
| chrIV | 55412 | 55541 | 129 | |
| chrIV | 56586 | 56757 | 171 | N4:56663 |
| chrIV | 60333 | 60467 | 134 | N4:60396 |
| chrIV | 61937 | 62077 | 140 | N4:62069 |
| chrIV | 77124 | 77282 | 158 | N4:77237 |
| chrIV | 78906 | 79062 | 156 | N4:78931 |
| chrIV | 86969 | 87132 | 163 | N4:87040 |
| chrIV | 90699 | 90847 | 148 | N4:90745 |
| chrIV | 94580 | 94722 | 142 | N4:94705 |
| chrIV | 97197 | 97345 | 148 | N4:97289 |
| chrIV | 101729 | 101892 | 163 | N4:101825 |
| chrIV | 103149 | 103279 | 130 | N4:103192 |
| chrIV | 109515 | 109652 | 137 | N4:109593 |
| chrIV | 109515 | 109656 | 141 | N4:109593 |
| chrIV | 109515 | 109666 | 151 | N4:109593 |
| chrIV | 109518 | 109656 | 138 | N4:109593 |
| chrIV | 110479 | 110614 | 135 | N4:110476 |
| chrIV | 113056 | 113240 | 184 | N4:113034 |
| chrIV | 118783 | 118936 | 153 | N4:118787 |
| chrIV | 121491 | 121639 | 148 | N4:121562 |
| chrIV | 122619 | 122777 | 158 | N4:122683 |
| chrIV | 124114 | 124238 | 124 | N4:124182 |
| chrIV | 125012 | 125174 | 162 | N4:125123 |
| chrIV | 126907 | 127064 | 157 | N4:126899 |
| chrIV | 126907 | 127064 | 157 | N4:127056 |
| chrIV | 127133 | 127254 | 121 | N4:127204 |
| chrIV | 130391 | 130540 | 149 | N4:130459 |
| chrIV | 134454 | 134627 | 173 | N4:134518 |
| chrIV | 137744 | 137906 | 162 | N4:137857 |
| chrIV | 147424 | 147558 | 134 | |
| chrIV | 148138 | 148259 | 121 | N4:148212 |
| chrIV | 148138 | 148287 | 149 | N4:148212 |
| chrIV | 148147 | 148256 | 109 | N4:148212 |
| chrIV | 148147 | 148259 | 112 | N4:148212 |
| chrIV | 148147 | 148287 | 140 | N4:148212 |
| chrIV | 148150 | 148259 | 109 | N4:148212 |
| chrIV | 148662 | 148839 | 177 | N4:148802 |
| chrIV | 150319 | 150438 | 119 | |
| chrIV | 150791 | 150954 | 163 | N4:150914 |
| chrIV | 151146 | 151295 | 149 | N4:151233 |
| chrIV | 154011 | 154153 | 142 | N4:154052 |
| chrIV | 156456 | 156616 | 160 | N4:156533 |
| chrIV | 156781 | 156930 | 149 | |
| chrIV | 158096 | 158254 | 158 | N4:158248 |
| chrIV | 158096 | 158254 | 158 | N4:158093 |
| chrIV | 158496 | 158646 | 150 | N4:158568 |
| chrIV | 162837 | 162965 | 128 | N4:162831 |
| chrIV | 163802 | 163934 | 132 | N4:163873 |
| chrIV | 163887 | 164037 | 150 | N4:163873 |
| chrIV | 163887 | 164037 | 150 | N4:164037 |
| chrIV | 164671 | 164820 | 149 | N4:164814 |
| chrIV | 168971 | 169122 | 151 | N4:169057 |
| chrIV | 180947 | 181107 | 160 | N4:181019 |
| chrIV | 185030 | 185179 | 149 | N4:185100 |
| chrIV | 185042 | 185156 | 114 | N4:185100 |
| chrIV | 196778 | 196946 | 168 | N4:196824 |
| chrIV | 198969 | 199115 | 146 | N4:199010 |
| chrIV | 200831 | 200925 | 94 | N4:200946 |
| chrIV | 202938 | 203097 | 159 | N4:203012 |
| chrIV | 204891 | 205041 | 150 | N4:204992 |
| chrIV | 205789 | 205917 | 128 | N4:205848 |
| chrIV | 214844 | 215003 | 159 | |
| chrIV | 224519 | 224676 | 157 | N4:224539 |
| chrIV | 224519 | 224676 | 157 | N4:224699 |
| chrIV | 228430 | 228579 | 149 | N4:228507 |
| chrIV | 230300 | 230458 | 158 | N4:230409 |
| chrIV | 235048 | 235188 | 140 | N4:235136 |
| chrIV | 235556 | 235704 | 148 | N4:235550 |
| chrIV | 235556 | 235704 | 148 | N4:235716 |
| chrIV | 240743 | 240895 | 152 | N4:240819 |
| chrIV | 241083 | 241236 | 153 | N4:241147 |
| chrIV | 243252 | 243404 | 152 | N4:243309 |
| chrIV | 247954 | 248116 | 162 | N4:247969 |
| chrIV | 249148 | 249302 | 154 | |
| chrIV | 249297 | 249444 | 147 | |
| chrIV | 253799 | 253940 | 141 | |
| chrIV | 262881 | 263029 | 148 | N4:262904 |
| chrIV | 268349 | 268491 | 142 | N4:268409 |
| chrIV | 269754 | 269886 | 132 | N4:269794 |
| chrIV | 275399 | 275562 | 163 | N4:275425 |
| chrIV | 277085 | 277203 | 118 | N4:277148 |
| chrIV | 281092 | 281244 | 152 | N4:281168 |
| chrIV | 282751 | 282862 | 111 | N4:282817 |

TABLE 1

| CHR | Start | End | Length (bp) | Nucleosome_ID |
|---|---|---|---|---|
| chrIV | 287334 | 287498 | 164 | N4:287398 |
| chrIV | 290503 | 290642 | 139 | N4:290580 |
| chrIV | 291728 | 291875 | 147 | N4:291860 |
| chrIV | 294716 | 294818 | 102 | N4:294818 |
| chrIV | 303242 | 303392 | 150 | N4:303310 |
| chrIV | 305394 | 305543 | 149 | N4:305450 |
| chrIV | 307871 | 308025 | 154 | N4:307996 |
| chrIV | 308997 | 309152 | 155 | N4:309056 |
| chrIV | 311358 | 311498 | 140 | N4:311442 |
| chrIV | 313115 | 313272 | 157 | N4:313232 |
| chrIV | 315132 | 315251 | 119 | N4:315205 |
| chrIV | 316257 | 316381 | 124 | N4:316392 |
| chrIV | 317932 | 318025 | 93 | |
| chrIV | 320773 | 320934 | 161 | N4:320798 |
| chrIV | 324041 | 324187 | 146 | N4:324118 |
| chrIV | 326087 | 326232 | 145 | N4:326162 |
| chrIV | 331066 | 331232 | 166 | N4:331089 |
| chrIV | 331066 | 331232 | 166 | N4:331253 |
| chrIV | 336500 | 336644 | 144 | N4:336586 |
| chrIV | 339003 | 339155 | 152 | N4:339069 |
| chrIV | 339626 | 339796 | 170 | N4:339709 |
| chrIV | 343754 | 343879 | 125 | N4:343821 |
| chrIV | 344625 | 344793 | 168 | N4:344682 |
| chrIV | 344904 | 345057 | 153 | N4:345036 |
| chrIV | 346054 | 346184 | 130 | N4:346140 |
| chrIV | 346695 | 346849 | 154 | |
| chrIV | 354236 | 354370 | 134 | N4:354255 |
| chrIV | 357373 | 357527 | 154 | N4:357441 |
| chrIV | 357519 | 357616 | 97 | N4:357598 |
| chrIV | 367261 | 367414 | 153 | N4:367365 |
| chrIV | 369734 | 369884 | 150 | N4:369786 |
| chrIV | 372733 | 372859 | 126 | N4:372737 |
| chrIV | 377302 | 377457 | 155 | N4:377352 |
| chrIV | 377961 | 378102 | 141 | N4:378096 |
| chrIV | 379551 | 379707 | 156 | N4:379708 |
| chrIV | 380490 | 380611 | 121 | N4:380618 |
| chrIV | 380895 | 381031 | 136 | N4:380935 |
| chrIV | 383984 | 384136 | 152 | N4:384003 |
| chrIV | 384406 | 384547 | 141 | |
| chrIV | 386414 | 386562 | 148 | N4:386473 |
| chrIV | 389563 | 389664 | 101 | N4:389614 |
| chrIV | 394898 | 395059 | 161 | N4:395016 |
| chrIV | 395659 | 395798 | 139 | N4:395816 |
| chrIV | 405838 | 405992 | 154 | N4:405924 |
| chrIV | 406138 | 406284 | 146 | |
| chrIV | 407097 | 407258 | 161 | N4:407198 |
| chrIV | 407639 | 407774 | 135 | |
| chrIV | 410001 | 410147 | 146 | N4:410035 |
| chrIV | 412510 | 412666 | 156 | N4:412554 |
| chrIV | 414505 | 414611 | 106 | |
| chrIV | 414505 | 414627 | 122 | |
| chrIV | 417857 | 418015 | 158 | N4:418030 |
| chrIV | 418210 | 418363 | 153 | N4:418242 |
| chrIV | 418328 | 418478 | 150 | |
| chrIV | 419977 | 420127 | 150 | |
| chrIV | 430775 | 430925 | 150 | |
| chrIV | 431218 | 431366 | 148 | N4:431307 |
| chrIV | 433624 | 433778 | 154 | N4:433704 |
| chrIV | 444592 | 444742 | 150 | N4:444681 |
| chrIV | 447153 | 447303 | 150 | N4:447172 |
| chrIV | 452119 | 452270 | 151 | N4:452201 |
| chrIV | 456027 | 456181 | 154 | N4:456128 |
| chrIV | 464196 | 464345 | 149 | N4:464243 |
| chrIV | 465523 | 465668 | 145 | N4:465537 |
| chrIV | 472028 | 472162 | 134 | N4:472039 |
| chrIV | 480768 | 480918 | 150 | N4:480858 |
| chrIV | 481714 | 481864 | 150 | N4:481812 |
| chrIV | 481903 | 482061 | 158 | N4:481994 |
| chrIV | 487395 | 487546 | 151 | N4:487410 |
| chrIV | 488401 | 488552 | 151 | N4:488452 |
| chrIV | 488508 | 488615 | 107 | N4:488621 |
| chrIV | 488508 | 488632 | 124 | N4:488621 |
| chrIV | 491144 | 491308 | 164 | N4:491134 |
| chrIV | 491144 | 491308 | 164 | N4:491316 |
| chrIV | 492176 | 492332 | 156 | N4:492275 |
| chrIV | 494125 | 494275 | 150 | N4:494221 |
| chrIV | 499197 | 499355 | 158 | N4:499232 |
| chrIV | 499814 | 499967 | 153 | N4:499892 |
| chrIV | 503596 | 503736 | 140 | N4:503693 |
| chrIV | 503768 | 503924 | 156 | |
| chrIV | 508653 | 508795 | 142 | N4:508667 |
| chrIV | 513880 | 514029 | 149 | N4:513900 |
| chrIV | 517444 | 517573 | 129 | |
| chrIV | 518819 | 518929 | 110 | N4:518890 |
| chrIV | 521357 | 521482 | 125 | N4:521409 |
| chrIV | 522483 | 522574 | 91 | N4:522532 |
| chrIV | 522691 | 522836 | 145 | N4:522707 |
| chrIV | 524988 | 525151 | 163 | N4:525064 |
| chrIV | 525488 | 525645 | 157 | |
| chrIV | 527030 | 527163 | 133 | N4:527071 |
| chrIV | 527878 | 528014 | 136 | N4:527977 |
| chrIV | 528421 | 528544 | 123 | N4:528503 |
| chrIV | 528421 | 528548 | 127 | N4:528503 |
| chrIV | 529656 | 529804 | 148 | |
| chrIV | 529731 | 529886 | 155 | |
| chrIV | 530359 | 530519 | 160 | N4:530430 |
| chrIV | 531763 | 531899 | 136 | N4:531865 |
| chrIV | 532306 | 532429 | 123 | N4:532381 |
| chrIV | 532306 | 532433 | 127 | N4:532381 |
| chrIV | 533541 | 533689 | 148 | |
| chrIV | 533616 | 533771 | 155 | |
| chrIV | 534244 | 534404 | 160 | N4:534319 |
| chrIV | 535648 | 535784 | 136 | N4:535758 |

| CHR | Start | End | Length (bp) | Nucleosome_ID |
|---|---|---|---|---|
| chrIV | 536109 | 536229 | 120 | |
| chrIV | 537426 | 537574 | 148 | N4:537413 |
| chrIV | 537501 | 537656 | 155 | |
| chrIV | 540906 | 541052 | 146 | N4:541006 |
| chrIV | 541731 | 541872 | 141 | N4:541799 |
| chrIV | 545183 | 545329 | 146 | N4:545271 |
| chrIV | 549706 | 549856 | 150 | N4:549849 |
| chrIV | 549706 | 549856 | 150 | N4:549694 |
| chrIV | 552005 | 552154 | 149 | N4:552080 |
| chrIV | 557304 | 557456 | 152 | N4:557386 |
| chrIV | 560465 | 560600 | 135 | N4:560548 |
| chrIV | 570638 | 570778 | 140 | N4:570621 |
| chrIV | 570638 | 570778 | 140 | N4:570783 |
| chrIV | 574209 | 574359 | 150 | N4:574313 |
| chrIV | 575376 | 575525 | 149 | N4:575491 |
| chrIV | 585684 | 585843 | 159 | N4:585698 |
| chrIV | 587983 | 588137 | 154 | N4:588040 |
| chrIV | 601658 | 601795 | 137 | N4:601720 |
| chrIV | 607967 | 608120 | 153 | N4:608009 |
| chrIV | 615631 | 615766 | 135 | N4:615640 |
| chrIV | 618044 | 618203 | 159 | N4:618139 |
| chrIV | 625810 | 625961 | 151 | N4:625892 |
| chrIV | 640754 | 640908 | 154 | N4:640848 |
| chrIV | 643481 | 643635 | 154 | N4:643559 |
| chrIV | 645695 | 645844 | 149 | N4:645718 |
| chrIV | 646491 | 646637 | 146 | N4:646571 |
| chrIV | 647130 | 647280 | 150 | N4:647179 |
| chrIV | 648526 | 648683 | 157 | N4:648585 |
| chrIV | 648994 | 649155 | 161 | N4:649119 |
| chrIV | 649174 | 649309 | 135 | N4:649272 |
| chrIV | 650212 | 650352 | 140 | N4:650289 |
| chrIV | 650218 | 650352 | 134 | N4:650289 |
| chrIV | 650630 | 650785 | 155 | N4:650651 |
| chrIV | 650761 | 650869 | 108 | N4:650859 |
| chrIV | 654067 | 654214 | 147 | N4:654056 |
| chrIV | 654067 | 654214 | 147 | N4:654214 |
| chrIV | 656839 | 656998 | 159 | N4:656949 |
| chrIV | 658487 | 658641 | 154 | N4:658664 |
| chrIV | 658600 | 658747 | 147 | N4:658664 |
| chrIV | 662245 | 662401 | 156 | N4:662311 |
| chrIV | 665084 | 665248 | 164 | N4:665151 |
| chrIV | 665597 | 665732 | 135 | N4:665678 |
| chrIV | 666941 | 667094 | 153 | N4:667026 |
| chrIV | 666948 | 667109 | 161 | N4:667026 |
| chrIV | 669497 | 669656 | 159 | N4:669574 |
| chrIV | 669741 | 669906 | 165 | N4:669781 |
| chrIV | 677555 | 677687 | 132 | N4:677564 |
| chrIV | 682440 | 682595 | 155 | N4:682517 |
| chrIV | 682928 | 683076 | 148 | N4:683045 |
| chrIV | 684905 | 685053 | 148 | N4:684976 |
| chrIV | 685367 | 685521 | 154 | N4:685462 |
| chrIV | 685409 | 685539 | 130 | N4:685462 |
| chrIV | 692976 | 693125 | 149 | N4:693039 |
| chrIV | 699427 | 699543 | 116 | N4:699492 |
| chrIV | 699429 | 699572 | 143 | N4:699492 |
| chrIV | 701924 | 702080 | 156 | N4:702005 |
| chrIV | 705337 | 705438 | 101 | N4:705421 |
| chrIV | 705533 | 705696 | 163 | |
| chrIV | 709396 | 709543 | 147 | N4:709492 |
| chrIV | 710394 | 710553 | 159 | N4:710486 |
| chrIV | 710648 | 710758 | 110 | N4:710780 |
| chrIV | 711680 | 711827 | 147 | N4:711712 |
| chrIV | 712887 | 713032 | 145 | N4:712928 |
| chrIV | 717311 | 717466 | 155 | N4:717376 |
| chrIV | 723685 | 723835 | 150 | |
| chrIV | 724692 | 724830 | 138 | N4:724752 |
| chrIV | 727870 | 728018 | 148 | N4:727949 |
| chrIV | 729848 | 729954 | 106 | |
| chrIV | 729848 | 729970 | 122 | N4:729980 |
| chrIV | 730903 | 730997 | 94 | N4:730925 |
| chrIV | 734743 | 734893 | 150 | N4:734827 |
| chrIV | 736895 | 737045 | 150 | |
| chrIV | 737420 | 737548 | 128 | N4:737479 |
| chrIV | 737638 | 737793 | 155 | N4:737705 |
| chrIV | 745330 | 745443 | 113 | N4:745385 |
| chrIV | 747582 | 747734 | 152 | N4:747675 |
| chrIV | 750682 | 750850 | 168 | N4:750748 |
| chrIV | 760658 | 760811 | 153 | N4:760826 |
| chrIV | 761833 | 761982 | 149 | N4:761970 |
| chrIV | 762005 | 762157 | 152 | N4:762143 |
| chrIV | 764493 | 764644 | 151 | N4:764585 |
| chrIV | 766598 | 766749 | 151 | N4:766690 |
| chrIV | 768253 | 768411 | 158 | N4:768405 |
| chrIV | 771149 | 771299 | 150 | |
| chrIV | 773973 | 774101 | 128 | |
| chrIV | 779473 | 779647 | 174 | N4:779537 |
| chrIV | 779988 | 780110 | 122 | N4:779996 |
| chrIV | 785291 | 785441 | 150 | N4:785366 |
| chrIV | 787217 | 787366 | 149 | N4:787358 |
| chrIV | 787217 | 787366 | 149 | N4:787195 |
| chrIV | 789145 | 789305 | 160 | N4:789230 |
| chrIV | 800500 | 800661 | 161 | N4:800519 |
| chrIV | 803067 | 803219 | 152 | N4:803114 |
| chrIV | 805417 | 805524 | 107 | N4:805529 |
| chrIV | 805417 | 805530 | 113 | N4:805529 |
| chrIV | 810559 | 810694 | 135 | N4:810661 |
| chrIV | 815236 | 815393 | 157 | N4:815317 |
| chrIV | 815520 | 815678 | 158 | N4:815624 |
| chrIV | 826418 | 826583 | 165 | N4:826491 |
| chrIV | 827506 | 827652 | 146 | N4:827588 |
| chrIV | 835382 | 835525 | 143 | N4:835504 |
| chrIV | 835382 | 835528 | 146 | N4:835504 |
| chrIV | 835383 | 835528 | 145 | N4:835504 |

| CHR | Start | End | Length (bp) | Nucleosome_ID |
|---|---|---|---|---|
| chrIV | 835384 | 835528 | 144 | N4:835504 |
| chrIV | 835385 | 835528 | 143 | N4:835504 |
| chrIV | 835386 | 835528 | 142 | N4:835504 |
| chrIV | 835932 | 836065 | 133 | N4:835972 |
| chrIV | 836520 | 836671 | 151 | N4:836563 |
| chrIV | 837322 | 837477 | 155 | N4:837482 |
| chrIV | 838477 | 838608 | 131 | N4:838553 |
| chrIV | 843960 | 844136 | 176 | N4:844047 |
| chrIV | 844482 | 844623 | 141 | N4:844544 |
| chrIV | 844482 | 844649 | 167 | N4:844544 |
| chrIV | 850466 | 850631 | 165 | N4:850636 |
| chrIV | 850466 | 850631 | 165 | N4:850477 |
| chrIV | 857181 | 857346 | 165 | |
| chrIV | 861657 | 861796 | 139 | N4:861762 |
| chrIV | 863651 | 863788 | 137 | N4:863721 |
| chrIV | 866143 | 866295 | 152 | N4:866178 |
| chrIV | 867460 | 867598 | 138 | N4:867531 |
| chrIV | 870210 | 870347 | 137 | N4:870239 |
| chrIV | 872409 | 872511 | 102 | N4:872520 |
| chrIV | 873832 | 873980 | 148 | |
| chrIV | 874621 | 874762 | 141 | |
| chrIV | 875359 | 875502 | 143 | N4:875420 |
| chrIV | 876645 | 876791 | 146 | N4:876701 |
| chrIV | 877438 | 877587 | 149 | N4:877571 |
| chrIV | 878298 | 878451 | 153 | N4:878280 |
| chrIV | 878496 | 878645 | 149 | N4:878510 |
| chrIV | 879292 | 879438 | 146 | N4:879370 |
| chrIV | 881327 | 881484 | 157 | N4:881391 |
| chrIV | 881795 | 881956 | 161 | |
| chrIV | 881975 | 882110 | 135 | N4:882066 |
| chrIV | 883013 | 883153 | 140 | N4:883091 |
| chrIV | 883019 | 883153 | 134 | N4:883091 |
| chrIV | 883431 | 883586 | 155 | N4:883463 |
| chrIV | 883562 | 883670 | 108 | |
| chrIV | 883696 | 883848 | 152 | N4:883720 |
| chrIV | 884819 | 884969 | 150 | N4:884882 |
| chrIV | 888390 | 888567 | 177 | N4:888453 |
| chrIV | 889668 | 889827 | 159 | N4:889734 |
| chrIV | 890834 | 890988 | 154 | N4:890967 |
| chrIV | 891389 | 891533 | 144 | N4:891510 |
| chrIV | 891986 | 892153 | 167 | N4:892055 |
| chrIV | 896334 | 896482 | 148 | N4:896422 |
| chrIV | 897629 | 897776 | 147 | N4:897659 |
| chrIV | 899028 | 899177 | 149 | N4:899049 |
| chrIV | 906466 | 906593 | 127 | N4:906545 |
| chrIV | 907937 | 908075 | 138 | |
| chrIV | 908538 | 908667 | 129 | N4:908599 |
| chrIV | 909620 | 909773 | 153 | N4:909630 |
| chrIV | 915796 | 915980 | 184 | N4:915793 |
| chrIV | 917617 | 917765 | 148 | N4:917653 |
| chrIV | 924934 | 925100 | 166 | N4:925073 |
| chrIV | 926425 | 926570 | 145 | N4:926519 |
| chrIV | 927161 | 927289 | 128 | N4:927242 |
| chrIV | 928961 | 929115 | 154 | N4:929074 |
| chrIV | 930238 | 930383 | 145 | N4:930295 |
| chrIV | 931111 | 931260 | 149 | N4:931170 |
| chrIV | 933440 | 933600 | 160 | N4:933517 |
| chrIV | 944365 | 944524 | 159 | N4:944467 |
| chrIV | 944403 | 944557 | 154 | N4:944467 |
| chrIV | 948002 | 948171 | 169 | N4:948157 |
| chrIV | 949233 | 949387 | 154 | N4:949302 |
| chrIV | 955015 | 955163 | 148 | N4:955072 |
| chrIV | 957192 | 957317 | 125 | N4:957237 |
| chrIV | 957192 | 957321 | 129 | N4:957237 |
| chrIV | 957196 | 957298 | 102 | N4:957237 |
| chrIV | 962120 | 962274 | 154 | N4:962102 |
| chrIV | 962406 | 962555 | 149 | N4:962480 |
| chrIV | 965842 | 965991 | 149 | N4:965942 |
| chrIV | 968475 | 968604 | 129 | N4:968597 |
| chrIV | 971192 | 971329 | 137 | N4:971253 |
| chrIV | 975387 | 975508 | 121 | N4:975430 |
| chrIV | 979606 | 979759 | 153 | N4:979673 |
| chrIV | 980738 | 980889 | 151 | N4:980809 |
| chrIV | 981759 | 981861 | 102 | N4:981871 |
| chrIV | 983182 | 983330 | 148 | |
| chrIV | 984709 | 984852 | 143 | N4:984756 |
| chrIV | 985995 | 986141 | 146 | |
| chrIV | 986788 | 986937 | 149 | N4:986912 |
| chrIV | 987343 | 987492 | 149 | N4:987353 |
| chrIV | 987657 | 987771 | 114 | N4:987788 |
| chrIV | 988353 | 988503 | 150 | N4:988403 |
| chrIV | 989662 | 989806 | 144 | N4:989808 |
| chrIV | 989749 | 989906 | 157 | N4:989808 |
| chrIV | 990217 | 990378 | 161 | N4:990348 |
| chrIV | 990397 | 990532 | 135 | N4:990499 |
| chrIV | 991435 | 991575 | 140 | N4:991512 |
| chrIV | 991441 | 991575 | 134 | N4:991512 |
| chrIV | 991853 | 992008 | 155 | N4:991879 |
| chrIV | 991984 | 992092 | 108 | N4:992082 |
| chrIV | 992133 | 992282 | 149 | |
| chrIV | 994735 | 994883 | 148 | N4:994783 |
| chrIV | 1001383 | 1001529 | 146 | N4:1001542 |
| chrIV | 1001383 | 1001529 | 146 | N4:1001379 |
| chrIV | 1005517 | 1005664 | 147 | N4:1005589 |
| chrIV | 1005876 | 1006022 | 146 | N4:1005930 |
| chrIV | 1012282 | 1012417 | 135 | N4:1012368 |
| chrIV | 1016514 | 1016662 | 148 | N4:1016582 |
| chrIV | 1019255 | 1019404 | 149 | N4:1019326 |
| chrIV | 1021134 | 1021284 | 150 | N4:1021147 |
| chrIV | 1021134 | 1021284 | 150 | N4:1021302 |
| chrIV | 1025226 | 1025374 | 148 | N4:1025251 |
| chrIV | 1028806 | 1028950 | 144 | N4:1028803 |
| chrIV | 1039896 | 1040063 | 167 | N4:1039922 |

TABLE 1

| CHR | Start | End | Length (bp) | Nucleosome_ID |
|---|---|---|---|---|
| chrIV | 1042109 | 1042252 | 143 | N4:1042173 |
| chrIV | 1052004 | 1052154 | 150 | N4:1052080 |
| chrIV | 1055603 | 1055756 | 153 | N4:1055666 |
| chrIV | 1058574 | 1058755 | 181 | N4:1058759 |
| chrIV | 1058574 | 1058755 | 181 | N4:1058587 |
| chrIV | 1063264 | 1063420 | 156 | N4:1063318 |
| chrIV | 1066695 | 1066823 | 128 | N4:1066763 |
| chrIV | 1075797 | 1075953 | 156 | N4:1075869 |
| chrIV | 1079066 | 1079219 | 153 | N4:1079155 |
| chrIV | 1079249 | 1079403 | 154 | N4:1079306 |
| chrIV | 1080074 | 1080246 | 172 | N4:1080115 |
| chrIV | 1090868 | 1091040 | 172 | N4:1090903 |
| chrIV | 1091309 | 1091450 | 141 | |
| chrIV | 1096304 | 1096427 | 123 | |
| chrIV | 1096404 | 1096559 | 155 | N4:1096525 |
| chrIV | 1096837 | 1096971 | 134 | N4:1096900 |
| chrIV | 1096837 | 1096977 | 140 | N4:1096900 |
| chrIV | 1097880 | 1098015 | 135 | N4:1097914 |
| chrIV | 1098034 | 1098195 | 161 | N4:1098134 |
| chrIV | 1098506 | 1098663 | 157 | N4:1098582 |
| chrIV | 1098997 | 1099126 | 129 | |
| chrIV | 1099909 | 1100059 | 150 | N4:1100005 |
| chrIV | 1100669 | 1100827 | 158 | N4:1100752 |
| chrIV | 1102545 | 1102694 | 149 | N4:1102665 |
| chrIV | 1102953 | 1103100 | 147 | N4:1102998 |
| chrIV | 1107156 | 1107303 | 147 | N4:1107223 |
| chrIV | 1109551 | 1109723 | 172 | N4:1109646 |
| chrIV | 1112644 | 1112796 | 152 | N4:1112718 |
| chrIV | 1117237 | 1117387 | 150 | N4:1117306 |
| chrIV | 1121058 | 1121205 | 147 | N4:1121186 |
| chrIV | 1121801 | 1121948 | 147 | N4:1121812 |
| chrIV | 1124473 | 1124641 | 168 | N4:1124564 |
| chrIV | 1128248 | 1128395 | 147 | N4:1128331 |
| chrIV | 1137826 | 1137974 | 148 | N4:1137842 |
| chrIV | 1140412 | 1140528 | 116 | N4:1140541 |
| chrIV | 1144636 | 1144783 | 147 | N4:1144674 |
| chrIV | 1154940 | 1155065 | 125 | N4:1155020 |
| chrIV | 1155027 | 1155183 | 156 | N4:1155020 |
| chrIV | 1155195 | 1155331 | 136 | N4:1155235 |
| chrIV | 1158507 | 1158654 | 147 | N4:1158572 |
| chrIV | 1160332 | 1160457 | 125 | N4:1160332 |
| chrIV | 1160419 | 1160575 | 156 | N4:1160485 |
| chrIV | 1160587 | 1160723 | 136 | N4:1160638 |
| chrIV | 1164215 | 1164366 | 151 | N4:1164258 |
| chrIV | 1166734 | 1166868 | 134 | N4:1166854 |
| chrIV | 1169216 | 1169370 | 154 | N4:1169282 |
| chrIV | 1170479 | 1170639 | 160 | N4:1170579 |
| chrIV | 1170874 | 1171037 | 163 | N4:1171049 |
| chrIV | 1170874 | 1171037 | 163 | N4:1170900 |
| chrIV | 1171736 | 1171886 | 150 | N4:1171844 |
| chrIV | 1175589 | 1175742 | 153 | N4:1175671 |
| chrIV | 1176500 | 1176647 | 147 | N4:1176576 |
| chrIV | 1179339 | 1179509 | 170 | N4:1179429 |
| chrIV | 1185766 | 1185913 | 147 | N4:1185835 |
| chrIV | 1186635 | 1186765 | 130 | N4:1186749 |
| chrIV | 1187842 | 1187997 | 155 | N4:1187892 |
| chrIV | 1199637 | 1199781 | 144 | N4:1199716 |
| chrIV | 1199946 | 1200093 | 147 | N4:1200030 |
| chrIV | 1202429 | 1202587 | 158 | N4:1202473 |
| chrIV | 1205211 | 1205359 | 148 | N4:1205237 |
| chrIV | 1207063 | 1207212 | 149 | N4:1207202 |
| chrIV | 1207237 | 1207360 | 123 | |
| chrIV | 1207337 | 1207492 | 155 | N4:1207459 |
| chrIV | 1207770 | 1207904 | 134 | N4:1207827 |
| chrIV | 1207770 | 1207910 | 140 | N4:1207827 |
| chrIV | 1207966 | 1208117 | 151 | N4:1208017 |
| chrIV | 1207976 | 1208121 | 145 | N4:1208017 |
| chrIV | 1207994 | 1208145 | 151 | N4:1208017 |
| chrIV | 1207996 | 1208145 | 149 | N4:1208017 |
| chrIV | 1208248 | 1208402 | 154 | N4:1208380 |
| chrIV | 1208282 | 1208426 | 144 | N4:1208380 |
| chrIV | 1208813 | 1208948 | 135 | N4:1208846 |
| chrIV | 1208967 | 1209128 | 161 | N4:1209000 |
| chrIV | 1209439 | 1209596 | 157 | N4:1209520 |
| chrIV | 1209930 | 1210059 | 129 | N4:1210072 |
| chrIV | 1210842 | 1210992 | 150 | N4:1210937 |
| chrIV | 1211574 | 1211688 | 114 | N4:1211673 |
| chrIV | 1215599 | 1215745 | 146 | N4:1215661 |
| chrIV | 1216176 | 1216327 | 151 | |
| chrIV | 1217346 | 1217492 | 146 | N4:1217374 |
| chrIV | 1218508 | 1218653 | 145 | N4:1218588 |
| chrIV | 1221088 | 1221239 | 151 | N4:1221154 |
| chrIV | 1225710 | 1225871 | 161 | N4:1225704 |
| chrIV | 1225924 | 1226071 | 147 | N4:1225999 |
| chrIV | 1227651 | 1227806 | 155 | N4:1227756 |
| chrIV | 1238397 | 1238557 | 160 | N4:1238463 |
| chrIV | 1240257 | 1240406 | 149 | N4:1240306 |
| chrIV | 1240385 | 1240526 | 141 | N4:1240462 |
| chrIV | 1244524 | 1244687 | 163 | N4:1244612 |
| chrIV | 1245862 | 1245987 | 125 | N4:1245846 |
| chrIV | 1251539 | 1251695 | 156 | N4:1251612 |
| chrIV | 1259546 | 1259668 | 122 | N4:1259621 |
| chrIV | 1265859 | 1266020 | 161 | N4:1265947 |
| chrIV | 1266398 | 1266553 | 155 | N4:1266476 |
| chrIV | 1267550 | 1267699 | 149 | N4:1267615 |
| chrIV | 1269091 | 1269198 | 107 | N4:1269188 |
| chrIV | 1269091 | 1269209 | 118 | N4:1269188 |
| chrIV | 1272800 | 1272981 | 181 | N4:1272869 |
| chrIV | 1277856 | 1277982 | 126 | N4:1277949 |
| chrIV | 1278160 | 1278320 | 160 | N4:1278264 |
| chrIV | 1287754 | 1287911 | 157 | N4:1287812 |
| chrIV | 1288372 | 1288504 | 132 | |
| chrIV | 1291651 | 1291799 | 148 | N4:1291819 |

| CHR | Start | End | Length (bp) | Nucleosome_ID |
|---|---|---|---|---|
| chrIV | 1293635 | 1293774 | 139 | N4:1293718 |
| chrIV | 1298059 | 1298223 | 164 | N4:1298152 |
| chrIV | 1300809 | 1300966 | 157 | N4:1300924 |
| chrIV | 1300809 | 1300970 | 161 | N4:1300924 |
| chrIV | 1300843 | 1301002 | 159 | N4:1300924 |
| chrIV | 1300844 | 1300970 | 126 | N4:1300924 |
| chrIV | 1300844 | 1300987 | 143 | N4:1300924 |
| chrIV | 1302267 | 1302373 | 106 | N4:1302245 |
| chrIV | 1302267 | 1302383 | 116 | N4:1302245 |
| chrIV | 1308150 | 1308260 | 110 | N4:1308222 |
| chrIV | 1308150 | 1308309 | 159 | N4:1308222 |
| chrIV | 1308153 | 1308309 | 156 | N4:1308222 |
| chrIV | 1308185 | 1308314 | 129 | N4:1308222 |
| chrIV | 1308191 | 1308309 | 118 | N4:1308222 |
| chrIV | 1308196 | 1308309 | 113 | N4:1308222 |
| chrIV | 1308234 | 1308344 | 110 | N4:1308222 |
| chrIV | 1308280 | 1308442 | 162 | N4:1308370 |
| chrIV | 1308280 | 1308477 | 197 | N4:1308370 |
| chrIV | 1308478 | 1308605 | 127 | N4:1308546 |
| chrIV | 1308858 | 1308982 | 124 | N4:1308860 |
| chrIV | 1312653 | 1312780 | 127 | N4:1312676 |
| chrIV | 1312778 | 1312924 | 146 | N4:1312823 |
| chrIV | 1324502 | 1324680 | 178 | N4:1324612 |
| chrIV | 1324730 | 1324882 | 152 | N4:1324791 |
| chrIV | 1329033 | 1329189 | 156 | |
| chrIV | 1331269 | 1331431 | 162 | N4:1331328 |
| chrIV | 1332261 | 1332409 | 148 | N4:1332317 |
| chrIV | 1334150 | 1334293 | 143 | N4:1334177 |
| chrIV | 1334858 | 1335004 | 146 | N4:1334915 |
| chrIV | 1335220 | 1335376 | 156 | N4:1335248 |
| chrIV | 1335220 | 1335376 | 156 | N4:1335399 |
| chrIV | 1337771 | 1337882 | 111 | N4:1337823 |
| chrIV | 1341919 | 1342069 | 150 | N4:1342003 |
| chrIV | 1354730 | 1354858 | 128 | N4:1354868 |
| chrIV | 1356736 | 1356889 | 153 | |
| chrIV | 1359819 | 1359977 | 158 | |
| chrIV | 1360674 | 1360822 | 148 | N4:1360796 |
| chrIV | 1361305 | 1361412 | 107 | N4:1361296 |
| chrIV | 1361305 | 1361420 | 115 | N4:1361443 |
| chrIV | 1361305 | 1361420 | 115 | N4:1361296 |
| chrIV | 1364336 | 1364488 | 152 | N4:1364372 |
| chrIV | 1364512 | 1364671 | 159 | N4:1364657 |
| chrIV | 1365728 | 1365875 | 147 | N4:1365799 |
| chrIV | 1365746 | 1365909 | 163 | N4:1365799 |
| chrIV | 1367419 | 1367567 | 148 | N4:1367482 |
| chrIV | 1367927 | 1368075 | 148 | N4:1367994 |
| chrIV | 1372245 | 1372378 | 133 | N4:1372326 |
| chrIV | 1376748 | 1376898 | 150 | N4:1376802 |
| chrIV | 1382369 | 1382521 | 152 | N4:1382446 |
| chrIV | 1387683 | 1387833 | 150 | N4:1387776 |
| chrIV | 1390839 | 1391009 | 170 | N4:1390850 |
| chrIV | 1391435 | 1391580 | 145 | N4:1391591 |
| chrIV | 1391940 | 1392060 | 120 | |
| chrIV | 1397500 | 1397655 | 155 | |
| chrIV | 1398087 | 1398256 | 169 | N4:1398124 |
| chrIV | 1406033 | 1406179 | 146 | N4:1406110 |
| chrIV | 1407803 | 1407943 | 140 | N4:1407944 |
| chrIV | 1407803 | 1407943 | 140 | N4:1407780 |
| chrIV | 1408060 | 1408202 | 142 | N4:1408111 |
| chrIV | 1408065 | 1408226 | 161 | N4:1408111 |
| chrIV | 1408340 | 1408489 | 149 | |
| chrIV | 1409678 | 1409825 | 147 | N4:1409695 |
| chrIV | 1413171 | 1413310 | 139 | N4:1413232 |
| chrIV | 1420451 | 1420608 | 157 | N4:1420526 |
| chrIV | 1420630 | 1420804 | 174 | N4:1420827 |
| chrIV | 1420630 | 1420804 | 174 | N4:1420679 |
| chrIV | 1420634 | 1420804 | 170 | N4:1420827 |
| chrIV | 1420634 | 1420804 | 170 | N4:1420679 |
| chrIV | 1424831 | 1424973 | 142 | N4:1424916 |
| chrIV | 1428254 | 1428414 | 160 | N4:1428344 |
| chrIV | 1430994 | 1431130 | 136 | N4:1431036 |
| chrIV | 1433662 | 1433813 | 151 | N4:1433787 |
| chrIV | 1447757 | 1447907 | 150 | N4:1447835 |
| chrIV | 1454001 | 1454153 | 152 | N4:1454073 |
| chrIV | 1460987 | 1461146 | 159 | N4:1461050 |
| chrIV | 1461681 | 1461828 | 147 | N4:1461801 |
| chrIV | 1462856 | 1462994 | 138 | |
| chrIV | 1462967 | 1463131 | 164 | N4:1463035 |
| chrIV | 1464532 | 1464669 | 137 | N4:1464622 |
| chrIV | 1465805 | 1465959 | 154 | N4:1465878 |
| chrIV | 1466131 | 1466277 | 146 | N4:1466195 |
| chrIV | 1473970 | 1474117 | 147 | N4:1474065 |
| chrIV | 1477488 | 1477647 | 159 | N4:1477526 |
| chrIV | 1478568 | 1478697 | 129 | N4:1478612 |
| chrIV | 1479228 | 1479388 | 160 | N4:1479283 |
| chrIV | 1479727 | 1479884 | 157 | N4:1479790 |
| chrIV | 1481305 | 1481467 | 162 | N4:1481450 |
| chrIV | 1489675 | 1489832 | 157 | N4:1489795 |
| chrIV | 1495628 | 1495753 | 125 | N4:1495663 |
| chrIV | 1496435 | 1496575 | 140 | N4:1496536 |
| chrIV | 1497627 | 1497776 | 149 | N4:1497727 |
| chrIV | 1498743 | 1498886 | 143 | N4:1498864 |
| chrIV | 1499148 | 1499307 | 159 | N4:1499221 |
| chrIV | 1503439 | 1503593 | 154 | N4:1503589 |
| chrIV | 1505128 | 1505252 | 124 | N4:1505200 |
| chrIV | 1507157 | 1507304 | 147 | N4:1507240 |
| chrIV | 1512418 | 1512571 | 153 | N4:1512551 |
| chrIV | 1512829 | 1512960 | 131 | N4:1512870 |
| chrIV | 1526039 | 1526194 | 155 | N4:1526079 |
| chrIV | 1527054 | 1527225 | 171 | N4:1527142 |
| chrIV | 1527204 | 1527351 | 147 | |
| chrIV | 1527360 | 1527490 | 130 | N4:1527407 |
| chrIV | 1527741 | 1527877 | 136 | N4:1527764 |

| CHR | Start | End | Length (bp) | Nucleosome_ID |
|---|---|---|---|---|
| chrIV | 1528122 | 1528284 | 162 | N4:1528239 |
| chrIV | 1528223 | 1528378 | 155 | N4:1528239 |
| chrIV | 1528357 | 1528494 | 137 | N4:1528410 |
| chrIV | 1528652 | 1528795 | 143 | N4:1528689 |
| chrIV | 1528700 | 1528825 | 125 | N4:1528689 |
| chrIV | 1528925 | 1529077 | 152 | N4:1529021 |
| chrIV | 1528998 | 1529142 | 144 | N4:1529021 |
| chrIV | 1529095 | 1529230 | 135 | N4:1529177 |
| chrIV | 1529095 | 1529282 | 187 | N4:1529177 |
| chrIV | 1529138 | 1529230 | 92 | N4:1529177 |
| chrIV | 1529138 | 1529282 | 144 | N4:1529177 |
| chrIV | 1529477 | 1529629 | 152 | N4:1529532 |
| chrIV | 1529778 | 1529908 | 130 | N4:1529917 |
| chrIV | 1529916 | 1530077 | 161 | N4:1529917 |
| chrIV | 1530973 | 1531125 | 152 | N4:1531041 |
| chrIV | 1531313 | 1531464 | 151 | N4:1531365 |
| chrIV | 1531313 | 1531484 | 171 | N4:1531365 |
| chrIX | 1597 | 1749 | 152 | N9:1629 |
| chrIX | 1686 | 1802 | 116 | N9:1790 |
| chrIX | 1686 | 1821 | 135 | N9:1790 |
| chrIX | 1686 | 1838 | 152 | N9:1790 |
| chrIX | 1710 | 1802 | 92 | N9:1790 |
| chrIX | 1710 | 1811 | 101 | N9:1790 |
| chrIX | 1710 | 1821 | 111 | N9:1790 |
| chrIX | 1710 | 1838 | 128 | N9:1790 |
| chrIX | 1710 | 1857 | 147 | N9:1790 |
| chrIX | 1746 | 1838 | 92 | N9:1790 |
| chrIX | 2215 | 2352 | 137 | N9:2320 |
| chrIX | 2382 | 2529 | 147 | N9:2535 |
| chrIX | 2382 | 2530 | 148 | N9:2535 |
| chrIX | 2382 | 2592 | 210 | N9:2535 |
| chrIX | 2383 | 2530 | 147 | N9:2535 |
| chrIX | 2385 | 2530 | 145 | N9:2535 |
| chrIX | 2397 | 2530 | 133 | N9:2535 |
| chrIX | 2437 | 2530 | 93 | N9:2535 |
| chrIX | 2437 | 2592 | 155 | N9:2535 |
| chrIX | 2558 | 2688 | 130 | N9:2535 |
| chrIX | 2879 | 3032 | 153 | N9:2921 |
| chrIX | 3324 | 3468 | 144 | N9:3423 |
| chrIX | 3531 | 3658 | 127 | N9:3591 |
| chrIX | 4088 | 4243 | 155 | N9:4203 |
| chrIX | 4805 | 4968 | 163 | N9:4855 |
| chrIX | 5216 | 5367 | 151 | |
| chrIX | 5429 | 5588 | 159 | N9:5453 |
| chrIX | 9290 | 9441 | 151 | N9:9376 |
| chrIX | 10228 | 10371 | 143 | N9:10309 |
| chrIX | 11460 | 11611 | 151 | N9:11554 |
| chrIX | 12345 | 12490 | 145 | |
| chrIX | 12910 | 13059 | 149 | |
| chrIX | 14208 | 14357 | 149 | N9:14236 |
| chrIX | 16078 | 16185 | 107 | N9:16064 |
| chrIX | 25656 | 25806 | 150 | N9:25676 |
| chrIX | 32330 | 32492 | 162 | N9:32393 |
| chrIX | 33763 | 33917 | 154 | N9:33762 |
| chrIX | 35803 | 35935 | 132 | N9:35809 |
| chrIX | 41284 | 41431 | 147 | N9:41356 |
| chrIX | 48410 | 48570 | 160 | N9:48477 |
| chrIX | 55336 | 55486 | 150 | N9:55401 |
| chrIX | 55497 | 55655 | 158 | N9:55554 |
| chrIX | 55498 | 55655 | 157 | N9:55554 |
| chrIX | 55500 | 55655 | 155 | N9:55554 |
| chrIX | 56537 | 56665 | 128 | N9:56598 |
| chrIX | 56786 | 56893 | 107 | N9:56789 |
| chrIX | 56786 | 56905 | 119 | N9:56789 |
| chrIX | 56791 | 56905 | 114 | N9:56789 |
| chrIX | 59003 | 59166 | 163 | N9:59158 |
| chrIX | 59003 | 59166 | 163 | N9:58990 |
| chrIX | 59591 | 59741 | 150 | N9:59671 |
| chrIX | 60731 | 60881 | 150 | N9:60802 |
| chrIX | 62677 | 62823 | 146 | N9:62723 |
| chrIX | 70968 | 71118 | 150 | N9:71141 |
| chrIX | 77046 | 77204 | 158 | N9:77127 |
| chrIX | 78123 | 78268 | 145 | N9:78189 |
| chrIX | 80413 | 80548 | 135 | N9:80482 |
| chrIX | 89142 | 89281 | 139 | N9:89221 |
| chrIX | 91580 | 91733 | 153 | N9:91642 |
| chrIX | 93328 | 93477 | 149 | N9:93388 |
| chrIX | 96377 | 96489 | 112 | N9:96391 |
| chrIX | 97100 | 97255 | 155 | N9:97083 |
| chrIX | 101159 | 101314 | 155 | N9:101310 |
| chrIX | 101159 | 101314 | 155 | N9:101155 |
| chrIX | 109382 | 109534 | 152 | N9:109439 |
| chrIX | 122918 | 123072 | 154 | N9:122984 |
| chrIX | 123420 | 123580 | 160 | N9:123402 |
| chrIX | 123420 | 123580 | 160 | N9:123567 |
| chrIX | 123668 | 123818 | 150 | N9:123750 |
| chrIX | 127761 | 127867 | 106 | N9:127744 |
| chrIX | 127761 | 127880 | 119 | N9:127744 |
| chrIX | 129250 | 129396 | 146 | N9:129359 |
| chrIX | 132115 | 132265 | 150 | N9:132195 |
| chrIX | 140470 | 140605 | 135 | N9:140507 |
| chrIX | 140794 | 140945 | 151 | N9:140849 |
| chrIX | 145476 | 145609 | 133 | |
| chrIX | 145755 | 145902 | 147 | N9:145791 |
| chrIX | 149396 | 149568 | 172 | N9:149494 |
| chrIX | 150158 | 150305 | 147 | N9:150234 |
| chrIX | 154400 | 154561 | 161 | N9:154551 |
| chrIX | 157942 | 158056 | 114 | N9:157996 |
| chrIX | 158500 | 158651 | 151 | |
| chrIX | 159528 | 159680 | 152 | N9:159539 |
| chrIX | 160370 | 160515 | 145 | N9:160446 |
| chrIX | 164945 | 165092 | 147 | N9:165040 |
| chrIX | 167654 | 167783 | 129 | |

| CHR | Start | End | Length (bp) | Nucleosome_ID |
|---|---|---|---|---|
| chrIX | 170117 | 170264 | 147 | N9:170186 |
| chrIX | 175448 | 175623 | 175 | N9:175501 |
| chrIX | 175932 | 176081 | 149 | N9:175976 |
| chrIX | 176147 | 176286 | 139 | N9:176155 |
| chrIX | 178557 | 178730 | 173 | N9:178691 |
| chrIX | 182864 | 183017 | 153 | N9:182946 |
| chrIX | 190132 | 190312 | 180 | N9:190192 |
| chrIX | 190978 | 191126 | 148 | N9:191107 |
| chrIX | 192456 | 192591 | 135 | N9:192542 |
| chrIX | 192983 | 193089 | 106 | N9:193029 |
| chrIX | 192983 | 193098 | 115 | N9:193029 |
| chrIX | 193702 | 193858 | 156 | N9:193815 |
| chrIX | 196790 | 196938 | 148 | N9:196869 |
| chrIX | 200251 | 200397 | 146 | N9:200295 |
| chrIX | 205441 | 205561 | 120 | |
| chrIX | 208655 | 208802 | 147 | N9:208742 |
| chrIX | 212863 | 213023 | 160 | N9:212982 |
| chrIX | 213578 | 213684 | 106 | |
| chrIX | 213578 | 213686 | 108 | |
| chrIX | 213578 | 213701 | 123 | |
| chrIX | 213866 | 214017 | 151 | N9:213871 |
| chrIX | 213866 | 214017 | 151 | N9:214029 |
| chrIX | 217876 | 218011 | 135 | N9:217888 |
| chrIX | 218740 | 218900 | 160 | N9:218850 |
| chrIX | 223366 | 223481 | 115 | N9:223420 |
| chrIX | 224421 | 224561 | 140 | N9:224566 |
| chrIX | 236168 | 236319 | 151 | N9:236230 |
| chrIX | 240288 | 240448 | 160 | N9:240358 |
| chrIX | 241031 | 241216 | 185 | N9:241153 |
| chrIX | 241349 | 241481 | 132 | |
| chrIX | 244655 | 244803 | 148 | N9:244726 |
| chrIX | 244820 | 244976 | 156 | N9:244898 |
| chrIX | 244822 | 244983 | 161 | N9:244898 |
| chrIX | 247146 | 247297 | 151 | N9:247208 |
| chrIX | 249643 | 249773 | 130 | N9:249728 |
| chrIX | 254780 | 254919 | 139 | N9:254874 |
| chrIX | 255472 | 255626 | 154 | N9:255553 |
| chrIX | 255629 | 255772 | 143 | N9:255778 |
| chrIX | 255629 | 255776 | 147 | N9:255778 |
| chrIX | 258137 | 258291 | 154 | N9:258298 |
| chrIX | 258137 | 258291 | 154 | N9:258138 |
| chrIX | 263024 | 263196 | 172 | N9:263062 |
| chrIX | 265122 | 265273 | 151 | N9:265235 |
| chrIX | 265238 | 265355 | 117 | N9:265235 |
| chrIX | 270073 | 270226 | 153 | N9:270153 |
| chrIX | 271691 | 271837 | 146 | N9:271766 |
| chrIX | 280995 | 281156 | 161 | N9:281073 |
| chrIX | 282582 | 282731 | 149 | N9:282678 |
| chrIX | 283375 | 283539 | 164 | N9:283458 |
| chrIX | 285840 | 285993 | 153 | N9:285843 |
| chrIX | 285840 | 285993 | 153 | N9:286010 |
| chrIX | 286233 | 286373 | 140 | |
| chrIX | 287987 | 288121 | 134 | N9:288045 |
| chrIX | 289697 | 289831 | 134 | N9:289737 |
| chrIX | 289822 | 289971 | 149 | N9:289899 |
| chrIX | 292476 | 292604 | 128 | N9:292512 |
| chrIX | 296896 | 297052 | 156 | N9:297004 |
| chrIX | 299139 | 299300 | 161 | N9:299210 |
| chrIX | 301522 | 301672 | 150 | N9:301595 |
| chrIX | 301692 | 301851 | 159 | N9:301782 |
| chrIX | 301709 | 301857 | 148 | N9:301782 |
| chrIX | 306823 | 306944 | 121 | N9:306927 |
| chrIX | 309339 | 309491 | 152 | N9:309417 |
| chrIX | 309860 | 310002 | 142 | N9:309904 |
| chrIX | 314424 | 314578 | 154 | |
| chrIX | 320445 | 320583 | 138 | N9:320481 |
| chrIX | 331176 | 331345 | 169 | N9:331307 |
| chrIX | 332149 | 332296 | 147 | N9:332195 |
| chrIX | 333705 | 333871 | 166 | N9:333780 |
| chrIX | 338938 | 339077 | 139 | N9:338970 |
| chrIX | 339456 | 339607 | 151 | N9:339523 |
| chrIX | 343521 | 343669 | 148 | N9:343647 |
| chrIX | 345865 | 346018 | 153 | N9:345939 |
| chrIX | 346400 | 346566 | 166 | N9:346455 |
| chrIX | 352082 | 352233 | 151 | N9:352150 |
| chrIX | 360987 | 361138 | 151 | N9:361056 |
| chrIX | 361485 | 361638 | 153 | N9:361563 |
| chrIX | 364237 | 364386 | 149 | N9:364329 |
| chrIX | 365450 | 365589 | 139 | N9:365504 |
| chrIX | 371073 | 371217 | 144 | N9:371134 |
| chrIX | 377621 | 377774 | 153 | N9:377728 |
| chrIX | 378406 | 378536 | 130 | N9:378506 |
| chrIX | 380201 | 380307 | 106 | N9:380285 |
| chrIX | 380201 | 380310 | 109 | N9:380285 |
| chrIX | 381638 | 381781 | 143 | N9:381710 |
| chrIX | 382286 | 382451 | 165 | N9:382299 |
| chrIX | 386066 | 386219 | 153 | N9:386093 |
| chrIX | 386767 | 386910 | 143 | N9:386861 |
| chrIX | 387904 | 388014 | 110 | N9:387935 |
| chrIX | 388900 | 389051 | 151 | N9:389018 |
| chrIX | 390177 | 390342 | 165 | N9:390256 |
| chrIX | 390386 | 390532 | 146 | N9:390414 |
| chrIX | 392788 | 392939 | 151 | N9:392855 |
| chrIX | 396135 | 396302 | 167 | N9:396197 |
| chrIX | 396795 | 396922 | 127 | N9:396865 |
| chrIX | 405278 | 405390 | 112 | N9:405412 |
| chrIX | 407378 | 407518 | 140 | N9:407408 |
| chrIX | 407520 | 407679 | 159 | N9:407581 |
| chrIX | 409036 | 409181 | 145 | N9:409099 |
| chrIX | 420519 | 420625 | 106 | N9:420590 |
| chrIX | 425922 | 426069 | 147 | N9:426001 |
| chrIX | 433552 | 433722 | 170 | N9:433692 |
| chrIX | 438504 | 438651 | 147 | N9:438544 |

| CHR | Start | End | Length (bp) | Nucleosome_ID |
|---|---|---|---|---|
| chrIX | 438570 | 438711 | 141 | |
| chrIX | 439579 | 439688 | 109 | N9:439641 |
| chrIX | 439796 | 439887 | 91 | N9:439845 |
| chrV | 1165 | 1320 | 155 | |
| chrV | 1854 | 1981 | 127 | N5:1841 |
| chrV | 3000 | 3144 | 144 | N5:3094 |
| chrV | 3648 | 3785 | 137 | N5:3709 |
| chrV | 3764 | 3919 | 155 | N5:3872 |
| chrV | 4024 | 4184 | 160 | N5:4031 |
| chrV | 4831 | 4982 | 151 | N5:4948 |
| chrV | 7115 | 7272 | 157 | N5:7242 |
| chrV | 9760 | 9930 | 170 | |
| chrV | 11061 | 11170 | 109 | N5:11123 |
| chrV | 23046 | 23205 | 159 | N5:23195 |
| chrV | 26697 | 26869 | 172 | N5:26684 |
| chrV | 26697 | 26869 | 172 | N5:26878 |
| chrV | 27053 | 27184 | 131 | N5:27040 |
| chrV | 28903 | 29024 | 121 | N5:29008 |
| chrV | 32832 | 32992 | 160 | N5:32875 |
| chrV | 41648 | 41754 | 106 | N5:41658 |
| chrV | 41648 | 41774 | 126 | N5:41658 |
| chrV | 41648 | 41782 | 134 | N5:41658 |
| chrV | 41648 | 41785 | 137 | N5:41658 |
| chrV | 41648 | 41795 | 147 | N5:41813 |
| chrV | 41648 | 41795 | 147 | N5:41658 |
| chrV | 41648 | 41797 | 149 | N5:41813 |
| chrV | 41648 | 41797 | 149 | N5:41658 |
| chrV | 41648 | 41798 | 150 | N5:41813 |
| chrV | 41648 | 41798 | 150 | N5:41658 |
| chrV | 41648 | 41799 | 151 | N5:41813 |
| chrV | 41648 | 41799 | 151 | N5:41658 |
| chrV | 41648 | 41800 | 152 | N5:41813 |
| chrV | 41648 | 41800 | 152 | N5:41658 |
| chrV | 41648 | 41801 | 153 | N5:41813 |
| chrV | 41648 | 41801 | 153 | N5:41658 |
| chrV | 41648 | 41802 | 154 | N5:41813 |
| chrV | 41648 | 41802 | 154 | N5:41658 |
| chrV | 41649 | 41802 | 153 | N5:41813 |
| chrV | 41649 | 41802 | 153 | N5:41658 |
| chrV | 41650 | 41802 | 152 | N5:41813 |
| chrV | 41650 | 41802 | 152 | N5:41658 |
| chrV | 41651 | 41802 | 151 | N5:41813 |
| chrV | 41651 | 41802 | 151 | N5:41658 |
| chrV | 41652 | 41802 | 150 | N5:41813 |
| chrV | 41652 | 41802 | 150 | N5:41658 |
| chrV | 41653 | 41802 | 149 | N5:41813 |
| chrV | 41653 | 41802 | 149 | N5:41658 |
| chrV | 41654 | 41802 | 148 | N5:41813 |
| chrV | 41654 | 41802 | 148 | N5:41658 |
| chrV | 41658 | 41802 | 144 | N5:41813 |
| chrV | 41658 | 41802 | 144 | N5:41658 |
| chrV | 42773 | 42909 | 136 | N5:42882 |
| chrV | 44013 | 44174 | 161 | N5:44097 |
| chrV | 51922 | 52070 | 148 | N5:52074 |
| chrV | 56549 | 56680 | 131 | N5:56569 |
| chrV | 58655 | 58795 | 140 | N5:58725 |
| chrV | 63504 | 63668 | 164 | N5:63566 |
| chrV | 64024 | 64155 | 131 | N5:64053 |
| chrV | 65475 | 65628 | 153 | N5:65581 |
| chrV | 69437 | 69601 | 164 | N5:69487 |
| chrV | 71549 | 71648 | 99 | N5:71615 |
| chrV | 72028 | 72188 | 160 | N5:72094 |
| chrV | 78159 | 78308 | 149 | N5:78154 |
| chrV | 78159 | 78308 | 149 | N5:78324 |
| chrV | 78447 | 78544 | 97 | N5:78484 |
| chrV | 78782 | 78922 | 140 | N5:78847 |
| chrV | 79095 | 79249 | 154 | N5:79182 |
| chrV | 83616 | 83770 | 154 | N5:83673 |
| chrV | 84397 | 84539 | 142 | N5:84467 |
| chrV | 87005 | 87158 | 153 | N5:87089 |
| chrV | 92533 | 92664 | 131 | N5:92614 |
| chrV | 94783 | 94926 | 143 | N5:94888 |
| chrV | 96167 | 96341 | 174 | N5:96189 |
| chrV | 105359 | 105516 | 157 | N5:105459 |
| chrV | 105504 | 105653 | 149 | N5:105613 |
| chrV | 106502 | 106638 | 136 | N5:106562 |
| chrV | 112707 | 112854 | 147 | N5:112767 |
| chrV | 114004 | 114167 | 163 | N5:114084 |
| chrV | 116119 | 116272 | 153 | N5:116132 |
| chrV | 116657 | 116785 | 128 | N5:116788 |
| chrV | 119860 | 120019 | 159 | N5:119994 |
| chrV | 121418 | 121572 | 154 | N5:121505 |
| chrV | 127589 | 127741 | 152 | N5:127639 |
| chrV | 129771 | 129919 | 148 | N5:129920 |
| chrV | 132213 | 132348 | 135 | N5:132209 |
| chrV | 134881 | 135033 | 152 | N5:134908 |
| chrV | 137057 | 137219 | 162 | N5:137135 |
| chrV | 137099 | 137245 | 146 | N5:137135 |
| chrV | 143216 | 143369 | 153 | N5:143330 |
| chrV | 146039 | 146192 | 153 | N5:146118 |
| chrV | 147368 | 147525 | 157 | N5:147443 |
| chrV | 147862 | 148011 | 149 | N5:147935 |
| chrV | 150905 | 151028 | 123 | N5:150967 |
| chrV | 154668 | 154823 | 155 | N5:154737 |
| chrV | 154964 | 155109 | 145 | N5:155071 |
| chrV | 156735 | 156886 | 151 | N5:156804 |
| chrV | 157741 | 157900 | 159 | N5:157806 |
| chrV | 158361 | 158517 | 156 | N5:158435 |
| chrV | 158999 | 159169 | 170 | N5:159105 |
| chrV | 161114 | 161267 | 153 | N5:161151 |
| chrV | 162828 | 162978 | 150 | N5:162902 |
| chrV | 162990 | 163135 | 145 | N5:163066 |
| chrV | 163162 | 163326 | 164 | N5:163230 |

TABLE 1

| CHR | Start | End | Length (bp) | Nucleosome_ID |
|---|---|---|---|---|
| chrV | 164105 | 164236 | 131 | N5:164174 |
| chrV | 165644 | 165751 | 107 | N5:165694 |
| chrV | 165645 | 165751 | 106 | N5:165694 |
| chrV | 165645 | 165753 | 108 | N5:165694 |
| chrV | 165645 | 165770 | 125 | N5:165694 |
| chrV | 169801 | 169967 | 166 | N5:169892 |
| chrV | 170735 | 170894 | 159 | N5:170840 |
| chrV | 175783 | 175962 | 179 | N5:175893 |
| chrV | 177369 | 177538 | 169 | N5:177535 |
| chrV | 177369 | 177538 | 169 | N5:177381 |
| chrV | 177766 | 177914 | 148 | N5:177821 |
| chrV | 179498 | 179607 | 109 | N5:179488 |
| chrV | 179498 | 179665 | 167 | N5:179488 |
| chrV | 179498 | 179665 | 167 | N5:179670 |
| chrV | 179498 | 179666 | 168 | N5:179488 |
| chrV | 179498 | 179666 | 168 | N5:179670 |
| chrV | 179498 | 179667 | 169 | N5:179488 |
| chrV | 179498 | 179667 | 169 | N5:179670 |
| chrV | 179498 | 179669 | 171 | N5:179488 |
| chrV | 179498 | 179669 | 171 | N5:179670 |
| chrV | 179498 | 179670 | 172 | N5:179488 |
| chrV | 179498 | 179670 | 172 | N5:179670 |
| chrV | 179499 | 179670 | 171 | N5:179488 |
| chrV | 179499 | 179670 | 171 | N5:179670 |
| chrV | 179502 | 179670 | 168 | N5:179488 |
| chrV | 179502 | 179670 | 168 | N5:179670 |
| chrV | 180478 | 180630 | 152 | N5:180475 |
| chrV | 180478 | 180630 | 152 | N5:180645 |
| chrV | 182865 | 183029 | 164 | N5:182948 |
| chrV | 186698 | 186836 | 138 | N5:186791 |
| chrV | 189568 | 189720 | 152 | N5:189634 |
| chrV | 190625 | 190773 | 148 | N5:190729 |
| chrV | 193660 | 193792 | 132 | N5:193731 |
| chrV | 194009 | 194135 | 126 | |
| chrV | 194030 | 194163 | 133 | N5:194161 |
| chrV | 195885 | 196040 | 155 | N5:195969 |
| chrV | 198265 | 198403 | 138 | N5:198244 |
| chrV | 201379 | 201531 | 152 | N5:201422 |
| chrV | 201380 | 201531 | 151 | N5:201422 |
| chrV | 201382 | 201531 | 149 | N5:201422 |
| chrV | 201383 | 201531 | 148 | N5:201422 |
| chrV | 206462 | 206572 | 110 | |
| chrV | 207496 | 207646 | 150 | N5:207609 |
| chrV | 212628 | 212767 | 139 | |
| chrV | 213547 | 213709 | 162 | N5:213639 |
| chrV | 215300 | 215450 | 150 | N5:215374 |
| chrV | 220704 | 220853 | 149 | N5:220767 |
| chrV | 226712 | 226861 | 149 | N5:226775 |
| chrV | 227455 | 227599 | 144 | N5:227578 |
| chrV | 234231 | 234385 | 154 | N5:234301 |
| chrV | 238011 | 238157 | 146 | N5:238087 |
| chrV | 239144 | 239298 | 154 | N5:239207 |
| chrV | 250567 | 250727 | 160 | N5:250698 |
| chrV | 250628 | 250757 | 129 | N5:250698 |
| chrV | 253010 | 253142 | 132 | N5:253162 |
| chrV | 255896 | 256035 | 139 | N5:256029 |
| chrV | 258267 | 258412 | 145 | N5:258341 |
| chrV | 263170 | 263306 | 136 | |
| chrV | 268138 | 268266 | 128 | N5:268214 |
| chrV | 271754 | 271916 | 162 | N5:271765 |
| chrV | 273268 | 273395 | 127 | N5:273323 |
| chrV | 274395 | 274545 | 150 | N5:274471 |
| chrV | 279045 | 279197 | 152 | N5:279105 |
| chrV | 282333 | 282495 | 162 | N5:282372 |
| chrV | 286305 | 286436 | 131 | N5:286328 |
| chrV | 293237 | 293387 | 150 | N5:293318 |
| chrV | 293855 | 294005 | 150 | N5:293929 |
| chrV | 298879 | 299018 | 139 | N5:298928 |
| chrV | 299205 | 299328 | 123 | N5:299273 |
| chrV | 299678 | 299831 | 153 | N5:299784 |
| chrV | 301505 | 301647 | 142 | N5:301602 |
| chrV | 314667 | 314817 | 150 | N5:314812 |
| chrV | 315905 | 316063 | 158 | N5:315980 |
| chrV | 320077 | 320234 | 157 | N5:320131 |
| chrV | 321656 | 321815 | 159 | N5:321825 |
| chrV | 321656 | 321815 | 159 | N5:321662 |
| chrV | 323513 | 323661 | 148 | N5:323606 |
| chrV | 324743 | 324895 | 152 | N5:324823 |
| chrV | 324904 | 325052 | 148 | N5:324975 |
| chrV | 325348 | 325499 | 151 | N5:325472 |
| chrV | 328966 | 329128 | 162 | N5:329104 |
| chrV | 331009 | 331174 | 165 | N5:331055 |
| chrV | 333088 | 333236 | 148 | N5:333122 |
| chrV | 340240 | 340377 | 137 | N5:340278 |
| chrV | 340798 | 340946 | 148 | N5:340808 |
| chrV | 341934 | 342080 | 146 | |
| chrV | 349486 | 349586 | 100 | |
| chrV | 351515 | 351672 | 157 | N5:351572 |
| chrV | 352323 | 352478 | 155 | N5:352405 |
| chrV | 360556 | 360703 | 147 | N5:360635 |
| chrV | 360556 | 360709 | 153 | N5:360635 |
| chrV | 360596 | 360757 | 161 | N5:360635 |
| chrV | 361101 | 361246 | 145 | N5:361124 |
| chrV | 364550 | 364707 | 157 | N5:364627 |
| chrV | 369752 | 369900 | 148 | |
| chrV | 370967 | 371076 | 109 | N5:371022 |
| chrV | 372915 | 373052 | 137 | N5:373029 |
| chrV | 373805 | 373958 | 153 | N5:373887 |
| chrV | 376017 | 376164 | 147 | N5:376149 |
| chrV | 376371 | 376519 | 148 | |
| chrV | 376832 | 376976 | 144 | N5:376938 |
| chrV | 377517 | 377651 | 134 | N5:377578 |
| chrV | 378722 | 378880 | 158 | N5:378804 |

| CHR | Start | End | Length (bp) | Nucleosome_ID |
|---|---|---|---|---|
| chrV | 389589 | 389739 | 150 | N5:389641 |
| chrV | 392068 | 392208 | 140 | N5:392146 |
| chrV | 408769 | 408926 | 157 | N5:408837 |
| chrV | 411385 | 411539 | 154 | N5:411472 |
| chrV | 421211 | 421362 | 151 | N5:421329 |
| chrV | 426798 | 426949 | 151 | |
| chrV | 426821 | 426975 | 154 | |
| chrV | 428076 | 428230 | 154 | N5:428133 |
| chrV | 429069 | 429212 | 143 | |
| chrV | 434847 | 434995 | 148 | N5:434827 |
| chrV | 440738 | 440898 | 160 | N5:440906 |
| chrV | 444332 | 444446 | 114 | N5:444468 |
| chrV | 445028 | 445178 | 150 | N5:445075 |
| chrV | 446200 | 446349 | 149 | N5:446313 |
| chrV | 446424 | 446581 | 157 | N5:446484 |
| chrV | 446610 | 446760 | 150 | N5:446692 |
| chrV | 446614 | 446760 | 146 | N5:446692 |
| chrV | 447072 | 447207 | 135 | N5:447121 |
| chrV | 447594 | 447738 | 144 | N5:447622 |
| chrV | 447618 | 447772 | 154 | N5:447622 |
| chrV | 447875 | 448026 | 151 | N5:447970 |
| chrV | 447899 | 448044 | 145 | N5:447970 |
| chrV | 447903 | 448054 | 151 | N5:447970 |
| chrV | 448110 | 448250 | 140 | N5:448190 |
| chrV | 448116 | 448250 | 134 | N5:448190 |
| chrV | 448528 | 448683 | 155 | N5:448518 |
| chrV | 448659 | 448767 | 108 | N5:448761 |
| chrV | 448808 | 448957 | 149 | |
| chrV | 450553 | 450702 | 149 | N5:450631 |
| chrV | 451474 | 451643 | 169 | N5:451553 |
| chrV | 456731 | 456859 | 128 | N5:456821 |
| chrV | 458282 | 458446 | 164 | |
| chrV | 458920 | 459063 | 143 | N5:458935 |
| chrV | 458920 | 459063 | 143 | N5:459085 |
| chrV | 460202 | 460309 | 107 | N5:460198 |
| chrV | 460202 | 460330 | 128 | N5:460198 |
| chrV | 460206 | 460309 | 103 | N5:460198 |
| chrV | 465104 | 465256 | 152 | N5:465150 |
| chrV | 466895 | 467059 | 164 | N5:466980 |
| chrV | 471982 | 472140 | 158 | N5:472119 |
| chrV | 472402 | 472551 | 149 | N5:472474 |
| chrV | 472672 | 472832 | 160 | N5:472741 |
| chrV | 472989 | 473145 | 156 | N5:473075 |
| chrV | 473919 | 474069 | 150 | N5:473967 |
| chrV | 481969 | 482118 | 149 | |
| chrV | 488812 | 488981 | 169 | N5:488949 |
| chrV | 489802 | 489965 | 163 | N5:489855 |
| chrV | 493537 | 493686 | 149 | N5:493561 |
| chrV | 495710 | 495860 | 150 | N5:495765 |
| chrV | 496172 | 496307 | 135 | N5:496222 |
| chrV | 497210 | 497350 | 140 | N5:497289 |
| chrV | 497216 | 497350 | 134 | N5:497289 |
| chrV | 497628 | 497783 | 155 | N5:497650 |
| chrV | 497759 | 497867 | 108 | N5:497866 |
| chrV | 497893 | 498045 | 152 | |
| chrV | 501813 | 501975 | 162 | N5:501881 |
| chrV | 502108 | 502246 | 138 | N5:502208 |
| chrV | 503589 | 503721 | 132 | N5:503607 |
| chrV | 504504 | 504656 | 152 | N5:504574 |
| chrV | 506128 | 506280 | 152 | N5:506233 |
| chrV | 508434 | 508572 | 138 | |
| chrV | 508812 | 508941 | 129 | N5:508869 |
| chrV | 517240 | 517402 | 162 | N5:517367 |
| chrV | 522575 | 522735 | 160 | N5:522650 |
| chrV | 525612 | 525757 | 145 | N5:525693 |
| chrV | 528789 | 528933 | 144 | N5:528845 |
| chrV | 536553 | 536715 | 162 | N5:536626 |
| chrV | 538015 | 538173 | 158 | N5:538060 |
| chrV | 540818 | 540968 | 150 | N5:540902 |
| chrV | 541050 | 541210 | 160 | N5:541230 |
| chrV | 541050 | 541210 | 160 | N5:541076 |
| chrV | 541616 | 541757 | 141 | N5:541694 |
| chrV | 541626 | 541757 | 131 | N5:541694 |
| chrV | 541626 | 541777 | 151 | N5:541694 |
| chrV | 542716 | 542874 | 158 | N5:542824 |
| chrV | 544584 | 544725 | 141 | N5:544632 |
| chrV | 560594 | 560752 | 158 | N5:560681 |
| chrV | 560787 | 560934 | 147 | N5:560849 |
| chrV | 561033 | 561174 | 141 | N5:561031 |
| chrV | 561033 | 561174 | 141 | N5:561188 |
| chrV | 571735 | 571876 | 141 | N5:571819 |
| chrV | 572939 | 573101 | 162 | N5:573065 |
| chrV | 573040 | 573195 | 155 | N5:573065 |
| chrV | 573174 | 573311 | 137 | N5:573237 |
| chrV | 573461 | 573612 | 151 | N5:573516 |
| chrV | 573469 | 573612 | 143 | N5:573516 |
| chrV | 573517 | 573642 | 125 | N5:573516 |
| chrV | 573742 | 573894 | 152 | N5:573853 |
| chrV | 573815 | 573959 | 144 | N5:573853 |
| chrV | 573912 | 574047 | 135 | N5:574037 |
| chrV | 573912 | 574099 | 187 | N5:574037 |
| chrV | 573955 | 574047 | 92 | N5:574037 |
| chrV | 573955 | 574099 | 144 | N5:574037 |
| chrV | 574294 | 574446 | 152 | N5:574369 |
| chrV | 574595 | 574725 | 130 | N5:574672 |
| chrV | 574965 | 575116 | 151 | N5:575085 |
| chrV | 574970 | 575104 | 134 | N5:575085 |
| chrV | 574983 | 575079 | 96 | N5:575085 |
| chrV | 575037 | 575152 | 115 | N5:575085 |
| chrV | 576130 | 576281 | 151 | N5:576205 |
| chrVI | 257 | 431 | 174 | N6:337 |
| chrVI | 377 | 525 | 148 | |
| chrVI | 2507 | 2632 | 125 | N6:2616 |

| CHR | Start | End | Length (bp) | Nucleosome_ID |
|---|---|---|---|---|
| chrVI | 2537 | 2680 | 143 | N6:2616 |
| chrVI | 2537 | 2688 | 151 | N6:2616 |
| chrVI | 2828 | 2965 | 137 | N6:2902 |
| chrVI | 3919 | 4067 | 148 | N6:4059 |
| chrVI | 4714 | 4814 | 100 | N6:4769 |
| chrVI | 6022 | 6182 | 160 | N6:6092 |
| chrVI | 13717 | 13867 | 150 | N6:13829 |
| chrVI | 15894 | 16044 | 150 | N6:15963 |
| chrVI | 24066 | 24172 | 106 | N6:24173 |
| chrVI | 24067 | 24172 | 105 | N6:24173 |
| chrVI | 29981 | 30130 | 149 | N6:30055 |
| chrVI | 36134 | 36286 | 152 | N6:36241 |
| chrVI | 36777 | 36932 | 155 | N6:36827 |
| chrVI | 41762 | 41909 | 147 | N6:41884 |
| chrVI | 48029 | 48199 | 170 | N6:48100 |
| chrVI | 49950 | 50105 | 155 | N6:50008 |
| chrVI | 55917 | 56077 | 160 | N6:55979 |
| chrVI | 60687 | 60850 | 163 | N6:60777 |
| chrVI | 65236 | 65388 | 152 | N6:65325 |
| chrVI | 65445 | 65606 | 161 | N6:65510 |
| chrVI | 68129 | 68276 | 147 | N6:68162 |
| chrVI | 68203 | 68367 | 164 | |
| chrVI | 71963 | 72118 | 155 | N6:72001 |
| chrVI | 72978 | 73128 | 150 | |
| chrVI | 76566 | 76713 | 147 | N6:76715 |
| chrVI | 77258 | 77379 | 121 | |
| chrVI | 79934 | 80100 | 166 | N6:80017 |
| chrVI | 81318 | 81445 | 127 | N6:81386 |
| chrVI | 82718 | 82885 | 167 | |
| chrVI | 84294 | 84451 | 157 | N6:84401 |
| chrVI | 91095 | 91248 | 153 | N6:91173 |
| chrVI | 94045 | 94152 | 107 | N6:94050 |
| chrVI | 94173 | 94319 | 146 | N6:94222 |
| chrVI | 94286 | 94435 | 149 | N6:94398 |
| chrVI | 97720 | 97865 | 145 | N6:97797 |
| chrVI | 97725 | 97874 | 149 | N6:97797 |
| chrVI | 100985 | 101138 | 153 | N6:101049 |
| chrVI | 108232 | 108382 | 150 | N6:108293 |
| chrVI | 108406 | 108531 | 125 | N6:108481 |
| chrVI | 110912 | 111064 | 152 | N6:110937 |
| chrVI | 110912 | 111064 | 152 | N6:111087 |
| chrVI | 111983 | 112114 | 131 | N6:112065 |
| chrVI | 115465 | 115634 | 169 | N6:115552 |
| chrVI | 123008 | 123146 | 138 | N6:123085 |
| chrVI | 130462 | 130613 | 151 | N6:130530 |
| chrVI | 131462 | 131615 | 153 | N6:131556 |
| chrVI | 139986 | 140115 | 129 | N6:139988 |
| chrVI | 140063 | 140227 | 164 | N6:140160 |
| chrVI | 141451 | 141594 | 143 | N6:141487 |
| chrVI | 142726 | 142892 | 166 | N6:142759 |
| chrVI | 143530 | 143679 | 149 | N6:143622 |
| chrVI | 148326 | 148498 | 172 | N6:148376 |
| chrVI | 151570 | 151741 | 171 | N6:151682 |
| chrVI | 151602 | 151755 | 153 | N6:151682 |
| chrVI | 151830 | 151995 | 165 | N6:151925 |
| chrVI | 152009 | 152167 | 158 | N6:152083 |
| chrVI | 152552 | 152696 | 144 | N6:152640 |
| chrVI | 155423 | 155586 | 163 | N6:155517 |
| chrVI | 162535 | 162694 | 159 | N6:162635 |
| chrVI | 174443 | 174598 | 155 | N6:174527 |
| chrVI | 175285 | 175426 | 141 | N6:175340 |
| chrVI | 176188 | 176314 | 126 | N6:176274 |
| chrVI | 177968 | 178139 | 171 | N6:178030 |
| chrVI | 180563 | 180715 | 152 | N6:180613 |
| chrVI | 182163 | 182304 | 141 | N6:182222 |
| chrVI | 183274 | 183405 | 131 | |
| chrVI | 183755 | 183882 | 127 | N6:183852 |
| chrVI | 185980 | 186131 | 151 | N6:186127 |
| chrVI | 186486 | 186602 | 116 | N6:186517 |
| chrVI | 192368 | 192528 | 160 | N6:192442 |
| chrVI | 193031 | 193186 | 155 | N6:193102 |
| chrVI | 195001 | 195152 | 151 | N6:195090 |
| chrVI | 197175 | 197337 | 162 | N6:197229 |
| chrVI | 197468 | 197595 | 127 | |
| chrVI | 198399 | 198562 | 163 | N6:198429 |
| chrVI | 202836 | 202999 | 163 | N6:202902 |
| chrVI | 204046 | 204202 | 156 | N6:204111 |
| chrVI | 208087 | 208238 | 151 | N6:208144 |
| chrVI | 208444 | 208571 | 127 | N6:208478 |
| chrVI | 208746 | 208898 | 152 | N6:208819 |
| chrVI | 210302 | 210398 | 96 | N6:210340 |
| chrVI | 210302 | 210400 | 98 | N6:210340 |
| chrVI | 210302 | 210402 | 100 | N6:210340 |
| chrVI | 210302 | 210403 | 101 | N6:210340 |
| chrVI | 210302 | 210404 | 102 | N6:210340 |
| chrVI | 210302 | 210405 | 103 | N6:210340 |
| chrVI | 210302 | 210406 | 104 | N6:210340 |
| chrVI | 210302 | 210407 | 105 | N6:210340 |
| chrVI | 210302 | 210409 | 107 | N6:210340 |
| chrVI | 214215 | 214366 | 151 | N6:214204 |
| chrVI | 214215 | 214366 | 151 | N6:214374 |
| chrVI | 224528 | 224665 | 137 | N6:224600 |
| chrVI | 225246 | 225383 | 137 | |
| chrVI | 225640 | 225800 | 160 | N6:225764 |
| chrVI | 228483 | 228624 | 141 | N6:228495 |
| chrVI | 232870 | 233015 | 145 | N6:232930 |
| chrVI | 236135 | 236288 | 153 | N6:236220 |
| chrVI | 236706 | 236866 | 160 | N6:236786 |
| chrVI | 237607 | 237756 | 149 | N6:237708 |
| chrVI | 237976 | 238116 | 140 | N6:238042 |
| chrVI | 239069 | 239219 | 150 | N6:239120 |
| chrVI | 242117 | 242275 | 158 | N6:242181 |
| chrVI | 242831 | 242985 | 154 | N6:242929 |

| CHR | Start | End | Length (bp) | Nucleosome_ID |
|---|---|---|---|---|
| chrVI | 242895 | 243045 | 150 | N6:242929 |
| chrVI | 244023 | 244165 | 142 | N6:244096 |
| chrVI | 244610 | 244760 | 150 | N6:244637 |
| chrVI | 247013 | 247149 | 136 | N6:247082 |
| chrVI | 258453 | 258604 | 151 | N6:258521 |
| chrVI | 260977 | 261105 | 128 | N6:261029 |
| chrVI | 264647 | 264774 | 127 | |
| chrVI | 265600 | 265745 | 145 | N6:265749 |
| chrVI | 267454 | 267607 | 153 | N6:267517 |
| chrVI | 269103 | 269252 | 149 | N6:269131 |
| chrVI | 270018 | 270158 | 140 | N6:270163 |
| chrVII | 6501 | 6604 | 103 | N7:6569 |
| chrVII | 7179 | 7332 | 153 | N7:7156 |
| chrVII | 7179 | 7332 | 153 | N7:7319 |
| chrVII | 8743 | 8898 | 155 | N7:8851 |
| chrVII | 8747 | 8898 | 151 | N7:8851 |
| chrVII | 9683 | 9793 | 110 | N7:9667 |
| chrVII | 15630 | 15761 | 131 | N7:15706 |
| chrVII | 15633 | 15761 | 128 | N7:15706 |
| chrVII | 15791 | 15935 | 144 | |
| chrVII | 20349 | 20487 | 138 | N7:20424 |
| chrVII | 21325 | 21484 | 159 | N7:21457 |
| chrVII | 22410 | 22546 | 136 | N7:22504 |
| chrVII | 23209 | 23365 | 156 | N7:23217 |
| chrVII | 32814 | 32961 | 147 | N7:32833 |
| chrVII | 33222 | 33355 | 133 | N7:33236 |
| chrVII | 36941 | 37083 | 142 | N7:36962 |
| chrVII | 41750 | 41885 | 135 | N7:41812 |
| chrVII | 52996 | 53149 | 153 | N7:53078 |
| chrVII | 53375 | 53486 | 111 | N7:53412 |
| chrVII | 62645 | 62823 | 178 | N7:62782 |
| chrVII | 69139 | 69303 | 164 | N7:69217 |
| chrVII | 72210 | 72368 | 158 | N7:72277 |
| chrVII | 73049 | 73197 | 148 | N7:73145 |
| chrVII | 78222 | 78356 | 134 | N7:78264 |
| chrVII | 83125 | 83232 | 107 | N7:83164 |
| chrVII | 83125 | 83247 | 122 | N7:83164 |
| chrVII | 84927 | 85059 | 132 | N7:84990 |
| chrVII | 88176 | 88368 | 192 | N7:88188 |
| chrVII | 88303 | 88443 | 140 | |
| chrVII | 88402 | 88555 | 153 | N7:88473 |
| chrVII | 89045 | 89177 | 132 | |
| chrVII | 90038 | 90161 | 123 | N7:90075 |
| chrVII | 91402 | 91556 | 154 | N7:91484 |
| chrVII | 92970 | 93138 | 168 | N7:93036 |
| chrVII | 97437 | 97599 | 162 | N7:97514 |
| chrVII | 102150 | 102299 | 149 | N7:102228 |
| chrVII | 102520 | 102673 | 153 | N7:102600 |
| chrVII | 103288 | 103440 | 152 | N7:103447 |
| chrVII | 103852 | 103984 | 132 | N7:103952 |
| chrVII | 108983 | 109147 | 164 | N7:109010 |
| chrVII | 116027 | 116173 | 146 | N7:116126 |
| chrVII | 119268 | 119414 | 146 | N7:119279 |
| chrVII | 125888 | 126050 | 162 | N7:125970 |
| chrVII | 138978 | 139125 | 147 | N7:139099 |
| chrVII | 147769 | 147920 | 151 | N7:147908 |
| chrVII | 152057 | 152208 | 151 | N7:152137 |
| chrVII | 156139 | 156292 | 153 | N7:156241 |
| chrVII | 167824 | 167963 | 139 | N7:167816 |
| chrVII | 167824 | 167963 | 139 | N7:167983 |
| chrVII | 168448 | 168554 | 106 | N7:168455 |
| chrVII | 168635 | 168789 | 154 | N7:168761 |
| chrVII | 171388 | 171548 | 160 | N7:171452 |
| chrVII | 174752 | 174849 | 97 | N7:174743 |
| chrVII | 174828 | 174980 | 152 | N7:174904 |
| chrVII | 178188 | 178320 | 132 | N7:178266 |
| chrVII | 183143 | 183279 | 136 | |
| chrVII | 184850 | 184994 | 144 | N7:184898 |
| chrVII | 196178 | 196334 | 156 | N7:196334 |
| chrVII | 196178 | 196334 | 156 | N7:196162 |
| chrVII | 204030 | 204183 | 153 | N7:204104 |
| chrVII | 204995 | 205129 | 134 | N7:205048 |
| chrVII | 209064 | 209201 | 137 | N7:209110 |
| chrVII | 214611 | 214774 | 163 | N7:214691 |
| chrVII | 217469 | 217614 | 145 | N7:217533 |
| chrVII | 221203 | 221378 | 175 | N7:221318 |
| chrVII | 223475 | 223638 | 163 | N7:223563 |
| chrVII | 226620 | 226777 | 157 | N7:226722 |
| chrVII | 226621 | 226770 | 149 | N7:226722 |
| chrVII | 233772 | 233924 | 152 | N7:233838 |
| chrVII | 238049 | 238197 | 148 | N7:238126 |
| chrVII | 243163 | 243328 | 165 | N7:243257 |
| chrVII | 249506 | 249649 | 143 | N7:249567 |
| chrVII | 251482 | 251633 | 151 | |
| chrVII | 266552 | 266693 | 141 | N7:266631 |
| chrVII | 267377 | 267528 | 151 | N7:267455 |
| chrVII | 268853 | 268959 | 106 | N7:268920 |
| chrVII | 268853 | 268977 | 124 | N7:268920 |
| chrVII | 270004 | 270159 | 155 | N7:270127 |
| chrVII | 273536 | 273656 | 120 | N7:273600 |
| chrVII | 275567 | 275723 | 156 | N7:275555 |
| chrVII | 275567 | 275723 | 156 | N7:275720 |
| chrVII | 279938 | 280099 | 161 | N7:280010 |
| chrVII | 281760 | 281908 | 148 | N7:281898 |
| chrVII | 281764 | 281908 | 144 | N7:281898 |
| chrVII | 284331 | 284485 | 154 | N7:284426 |
| chrVII | 289603 | 289760 | 157 | N7:289668 |
| chrVII | 293723 | 293871 | 148 | N7:293806 |
| chrVII | 294019 | 294183 | 164 | N7:294135 |
| chrVII | 297177 | 297313 | 136 | N7:297182 |
| chrVII | 299926 | 300073 | 147 | N7:300004 |
| chrVII | 300091 | 300244 | 153 | N7:300166 |
| chrVII | 300595 | 300755 | 160 | |

| CHR | Start | End | Length (bp) | Nucleosome_ID |
|---|---|---|---|---|
| chrVII | 313487 | 313646 | 159 | N7:313555 |
| chrVII | 314575 | 314719 | 144 | N7:314647 |
| chrVII | 320068 | 320221 | 153 | N7:320063 |
| chrVII | 320068 | 320221 | 153 | N7:320242 |
| chrVII | 323178 | 323326 | 148 | |
| chrVII | 324915 | 325051 | 136 | N7:325012 |
| chrVII | 327547 | 327717 | 170 | N7:327548 |
| chrVII | 327547 | 327717 | 170 | N7:327719 |
| chrVII | 328882 | 329034 | 152 | N7:329029 |
| chrVII | 329271 | 329388 | 117 | N7:329338 |
| chrVII | 330372 | 330517 | 145 | N7:330463 |
| chrVII | 331632 | 331780 | 148 | N7:331795 |
| chrVII | 331632 | 331780 | 148 | N7:331642 |
| chrVII | 341547 | 341699 | 152 | N7:341572 |
| chrVII | 342190 | 342339 | 149 | |
| chrVII | 342310 | 342453 | 143 | N7:342424 |
| chrVII | 343265 | 343403 | 138 | N7:343315 |
| chrVII | 352278 | 352440 | 162 | N7:352305 |
| chrVII | 354451 | 354624 | 173 | N7:354512 |
| chrVII | 356531 | 356676 | 145 | N7:356589 |
| chrVII | 364639 | 364801 | 162 | N7:364708 |
| chrVII | 367967 | 368120 | 153 | N7:368054 |
| chrVII | 370607 | 370760 | 153 | N7:370672 |
| chrVII | 382677 | 382828 | 151 | N7:382840 |
| chrVII | 382677 | 382828 | 151 | N7:382673 |
| chrVII | 383386 | 383517 | 131 | N7:383447 |
| chrVII | 383588 | 383739 | 151 | N7:383638 |
| chrVII | 384835 | 384978 | 143 | N7:384900 |
| chrVII | 384841 | 384978 | 137 | N7:384900 |
| chrVII | 386231 | 386378 | 147 | N7:386322 |
| chrVII | 386588 | 386752 | 164 | N7:386632 |
| chrVII | 398118 | 398266 | 148 | |
| chrVII | 398740 | 398893 | 153 | N7:398821 |
| chrVII | 399113 | 399273 | 160 | N7:399182 |
| chrVII | 402184 | 402330 | 146 | N7:402298 |
| chrVII | 406230 | 406375 | 145 | N7:406270 |
| chrVII | 412759 | 412909 | 150 | N7:412835 |
| chrVII | 413027 | 413197 | 170 | N7:413154 |
| chrVII | 416964 | 417114 | 150 | N7:417040 |
| chrVII | 419105 | 419256 | 151 | N7:419189 |
| chrVII | 422885 | 423015 | 130 | N7:422958 |
| chrVII | 424432 | 424587 | 155 | N7:424573 |
| chrVII | 426688 | 426828 | 140 | N7:426744 |
| chrVII | 430629 | 430785 | 156 | N7:430704 |
| chrVII | 438337 | 438504 | 167 | N7:438393 |
| chrVII | 445398 | 445550 | 152 | N7:445477 |
| chrVII | 445405 | 445562 | 157 | N7:445477 |
| chrVII | 448295 | 448446 | 151 | N7:448359 |
| chrVII | 449063 | 449214 | 151 | N7:449131 |
| chrVII | 449866 | 450017 | 151 | N7:449940 |
| chrVII | 450289 | 450475 | 186 | N7:450357 |
| chrVII | 450564 | 450727 | 163 | |
| chrVII | 451657 | 451811 | 154 | N7:451765 |
| chrVII | 452015 | 452160 | 145 | N7:452093 |
| chrVII | 454120 | 454220 | 100 | N7:454191 |
| chrVII | 457180 | 457330 | 150 | N7:457208 |
| chrVII | 457820 | 457954 | 134 | N7:457886 |
| chrVII | 458030 | 458181 | 151 | N7:458116 |
| chrVII | 464168 | 464326 | 158 | N7:464210 |
| chrVII | 468550 | 468701 | 151 | N7:468622 |
| chrVII | 470833 | 470993 | 160 | |
| chrVII | 471018 | 471094 | 76 | N7:471038 |
| chrVII | 471018 | 471123 | 105 | N7:471038 |
| chrVII | 471173 | 471322 | 149 | N7:471209 |
| chrVII | 475774 | 475931 | 157 | N7:475867 |
| chrVII | 476346 | 476454 | 108 | |
| chrVII | 476346 | 476465 | 119 | |
| chrVII | 478282 | 478437 | 155 | N7:478357 |
| chrVII | 482579 | 482701 | 122 | N7:482600 |
| chrVII | 483653 | 483754 | 101 | N7:483643 |
| chrVII | 495041 | 495196 | 155 | N7:495024 |
| chrVII | 501090 | 501258 | 168 | N7:501148 |
| chrVII | 502391 | 502522 | 131 | N7:502483 |
| chrVII | 502870 | 503027 | 157 | N7:502941 |
| chrVII | 504629 | 504778 | 149 | N7:504726 |
| chrVII | 505104 | 505255 | 151 | N7:505190 |
| chrVII | 509103 | 509251 | 148 | N7:509169 |
| chrVII | 509261 | 509410 | 149 | N7:509325 |
| chrVII | 510438 | 510590 | 152 | N7:510527 |
| chrVII | 512325 | 512500 | 175 | N7:512399 |
| chrVII | 515194 | 515319 | 125 | N7:515214 |
| chrVII | 516112 | 516262 | 150 | N7:516125 |
| chrVII | 516112 | 516262 | 150 | N7:516278 |
| chrVII | 517734 | 517882 | 148 | N7:517806 |
| chrVII | 517736 | 517882 | 146 | N7:517806 |
| chrVII | 518337 | 518484 | 147 | N7:518402 |
| chrVII | 519407 | 519563 | 156 | N7:519436 |
| chrVII | 519674 | 519834 | 160 | |
| chrVII | 519935 | 520102 | 167 | N7:520057 |
| chrVII | 520743 | 520902 | 159 | N7:520845 |
| chrVII | 520956 | 521107 | 151 | N7:521078 |
| chrVII | 526220 | 526384 | 164 | N7:526309 |
| chrVII | 529947 | 530056 | 109 | N7:529978 |
| chrVII | 535422 | 535536 | 114 | N7:535504 |
| chrVII | 536123 | 536272 | 149 | N7:536209 |
| chrVII | 536273 | 536379 | 106 | |
| chrVII | 536273 | 536391 | 118 | |
| chrVII | 536297 | 536420 | 123 | |
| chrVII | 536397 | 536552 | 155 | N7:536510 |
| chrVII | 536830 | 536964 | 134 | N7:536902 |
| chrVII | 536830 | 536970 | 140 | N7:536902 |
| chrVII | 537026 | 537177 | 151 | N7:537109 |
| chrVII | 537036 | 537181 | 145 | N7:537109 |

| CHR | Start | End | Length (bp) | Nucleosome_ID |
|---|---|---|---|---|
| chrVII | 537308 | 537462 | 154 | N7:537445 |
| chrVII | 537342 | 537486 | 144 | N7:537445 |
| chrVII | 537873 | 538008 | 135 | N7:537917 |
| chrVII | 538055 | 538187 | 132 | |
| chrVII | 538144 | 538282 | 138 | N7:538222 |
| chrVII | 538499 | 538656 | 157 | N7:538585 |
| chrVII | 538599 | 538743 | 144 | N7:538585 |
| chrVII | 538990 | 539119 | 129 | |
| chrVII | 539470 | 539601 | 131 | N7:539492 |
| chrVII | 539902 | 540052 | 150 | N7:539986 |
| chrVII | 540853 | 541004 | 151 | N7:540903 |
| chrVII | 542644 | 542795 | 151 | N7:542763 |
| chrVII | 544012 | 544140 | 128 | N7:544085 |
| chrVII | 545414 | 545580 | 166 | N7:545553 |
| chrVII | 546693 | 546840 | 147 | N7:546767 |
| chrVII | 548258 | 548409 | 151 | N7:548239 |
| chrVII | 554460 | 554622 | 162 | N7:554535 |
| chrVII | 554946 | 555089 | 143 | N7:555041 |
| chrVII | 562268 | 562415 | 147 | N7:562318 |
| chrVII | 562775 | 562889 | 114 | |
| chrVII | 563471 | 563621 | 150 | N7:563527 |
| chrVII | 564404 | 564533 | 129 | |
| chrVII | 564867 | 565024 | 157 | N7:564949 |
| chrVII | 565335 | 565496 | 161 | N7:565396 |
| chrVII | 565515 | 565650 | 135 | N7:565587 |
| chrVII | 566553 | 566693 | 140 | N7:566644 |
| chrVII | 566559 | 566693 | 134 | N7:566644 |
| chrVII | 566971 | 567126 | 155 | N7:567004 |
| chrVII | 567102 | 567210 | 108 | N7:567215 |
| chrVII | 568106 | 568272 | 166 | N7:568185 |
| chrVII | 568932 | 569081 | 149 | N7:568958 |
| chrVII | 569719 | 569885 | 166 | N7:569812 |
| chrVII | 571017 | 571160 | 143 | N7:571098 |
| chrVII | 572385 | 572549 | 164 | N7:572444 |
| chrVII | 572497 | 572626 | 129 | N7:572607 |
| chrVII | 573872 | 573982 | 110 | N7:573948 |
| chrVII | 575134 | 575270 | 136 | N7:575205 |
| chrVII | 575584 | 575763 | 179 | N7:575570 |
| chrVII | 575584 | 575763 | 179 | N7:575745 |
| chrVII | 578432 | 578606 | 174 | N7:578577 |
| chrVII | 578432 | 578606 | 174 | N7:578412 |
| chrVII | 578979 | 579129 | 150 | N7:579094 |
| chrVII | 579144 | 579294 | 150 | N7:579259 |
| chrVII | 583828 | 583997 | 169 | N7:583861 |
| chrVII | 584107 | 584258 | 151 | N7:584192 |
| chrVII | 584200 | 584336 | 136 | N7:584192 |
| chrVII | 584954 | 585099 | 145 | N7:584996 |
| chrVII | 594298 | 594445 | 147 | N7:594404 |
| chrVII | 595048 | 595213 | 165 | N7:595207 |
| chrVII | 595048 | 595213 | 165 | N7:595043 |
| chrVII | 595281 | 595432 | 151 | N7:595369 |
| chrVII | 610980 | 611160 | 180 | N7:610959 |
| chrVII | 610980 | 611160 | 180 | N7:611131 |
| chrVII | 613266 | 613414 | 148 | N7:613372 |
| chrVII | 637713 | 637869 | 156 | N7:637824 |
| chrVII | 640000 | 640148 | 148 | N7:640070 |
| chrVII | 640002 | 640148 | 146 | N7:640070 |
| chrVII | 640719 | 640865 | 146 | N7:640822 |
| chrVII | 652407 | 652570 | 163 | N7:652507 |
| chrVII | 652780 | 652925 | 145 | N7:652859 |
| chrVII | 654255 | 654401 | 146 | N7:654248 |
| chrVII | 654255 | 654401 | 146 | N7:654398 |
| chrVII | 668831 | 668991 | 160 | N7:668917 |
| chrVII | 671142 | 671294 | 152 | |
| chrVII | 671676 | 671849 | 173 | N7:671743 |
| chrVII | 674557 | 674713 | 156 | N7:674628 |
| chrVII | 676654 | 676817 | 163 | N7:676774 |
| chrVII | 680899 | 681051 | 152 | N7:680980 |
| chrVII | 681312 | 681478 | 166 | N7:681411 |
| chrVII | 682534 | 682696 | 162 | N7:682605 |
| chrVII | 687608 | 687758 | 150 | N7:687651 |
| chrVII | 689536 | 689689 | 153 | N7:689603 |
| chrVII | 692091 | 692197 | 106 | N7:692094 |
| chrVII | 692091 | 692218 | 127 | N7:692094 |
| chrVII | 692091 | 692219 | 128 | N7:692094 |
| chrVII | 692091 | 692220 | 129 | N7:692094 |
| chrVII | 692091 | 692221 | 130 | N7:692094 |
| chrVII | 692091 | 692222 | 131 | N7:692094 |
| chrVII | 692091 | 692223 | 132 | N7:692094 |
| chrVII | 692092 | 692223 | 131 | N7:692094 |
| chrVII | 692095 | 692223 | 128 | N7:692094 |
| chrVII | 692097 | 692197 | 100 | N7:692094 |
| chrVII | 693772 | 693921 | 149 | N7:693854 |
| chrVII | 697506 | 697652 | 146 | N7:697541 |
| chrVII | 697827 | 697977 | 150 | N7:697877 |
| chrVII | 701841 | 701992 | 151 | |
| chrVII | 707416 | 707536 | 120 | N7:707505 |
| chrVII | 709744 | 709890 | 146 | N7:709832 |
| chrVII | 712418 | 712569 | 151 | N7:712512 |
| chrVII | 714004 | 714140 | 136 | N7:714042 |
| chrVII | 716031 | 716189 | 158 | |
| chrVII | 716536 | 716689 | 153 | N7:716629 |
| chrVII | 726352 | 726496 | 144 | N7:726431 |
| chrVII | 731531 | 731680 | 149 | N7:731570 |
| chrVII | 740022 | 740127 | 105 | N7:740078 |
| chrVII | 746153 | 746303 | 150 | N7:746206 |
| chrVII | 747978 | 748102 | 124 | N7:748002 |
| chrVII | 748361 | 748501 | 140 | N7:748439 |
| chrVII | 751512 | 751624 | 112 | N7:751606 |
| chrVII | 758738 | 758894 | 156 | N7:758814 |
| chrVII | 760806 | 760962 | 156 | N7:760938 |
| chrVII | 760828 | 760978 | 150 | N7:760938 |
| chrVII | 762415 | 762565 | 150 | N7:762428 |

| CHR | Start | End | Length (bp) | Nucleosome_ID |
|---|---|---|---|---|
| chrVII | 766213 | 766368 | 155 | N7:766304 |
| chrVII | 769383 | 769497 | 114 | N7:769444 |
| chrVII | 773391 | 773557 | 166 | N7:773500 |
| chrVII | 776840 | 777001 | 161 | N7:777019 |
| chrVII | 776982 | 777149 | 167 | N7:777019 |
| chrVII | 780856 | 781004 | 148 | N7:780944 |
| chrVII | 781219 | 781380 | 161 | N7:781318 |
| chrVII | 782373 | 782522 | 149 | |
| chrVII | 782985 | 783136 | 151 | N7:783055 |
| chrVII | 811114 | 811251 | 137 | N7:811183 |
| chrVII | 813520 | 813649 | 129 | N7:813530 |
| chrVII | 813597 | 813761 | 164 | N7:813691 |
| chrVII | 814985 | 815128 | 143 | N7:815042 |
| chrVII | 816260 | 816426 | 166 | N7:816238 |
| chrVII | 817064 | 817213 | 149 | N7:817170 |
| chrVII | 817068 | 817218 | 150 | N7:817170 |
| chrVII | 817863 | 817968 | 105 | N7:817860 |
| chrVII | 817863 | 817970 | 107 | N7:817860 |
| chrVII | 819019 | 819169 | 150 | N7:819103 |
| chrVII | 819762 | 819909 | 147 | N7:819930 |
| chrVII | 820415 | 820572 | 157 | N7:820493 |
| chrVII | 820601 | 820751 | 150 | N7:820670 |
| chrVII | 820792 | 820937 | 145 | |
| chrVII | 821063 | 821198 | 135 | N7:821136 |
| chrVII | 821345 | 821513 | 168 | N7:821422 |
| chrVII | 822101 | 822241 | 140 | N7:822182 |
| chrVII | 822107 | 822241 | 134 | N7:822182 |
| chrVII | 822519 | 822674 | 155 | N7:822552 |
| chrVII | 822650 | 822758 | 108 | N7:822752 |
| chrVII | 825754 | 825897 | 143 | N7:825833 |
| chrVII | 831865 | 832015 | 150 | N7:831995 |
| chrVII | 841785 | 841939 | 154 | N7:841891 |
| chrVII | 842909 | 843085 | 176 | N7:843080 |
| chrVII | 842909 | 843085 | 176 | N7:842921 |
| chrVII | 843924 | 844090 | 166 | N7:844011 |
| chrVII | 844983 | 845124 | 141 | N7:845017 |
| chrVII | 847489 | 847638 | 149 | |
| chrVII | 853632 | 853775 | 143 | N7:853718 |
| chrVII | 855743 | 855883 | 140 | N7:855791 |
| chrVII | 855860 | 856013 | 153 | N7:855962 |
| chrVII | 861234 | 861372 | 138 | N7:861384 |
| chrVII | 869483 | 869631 | 148 | N7:869530 |
| chrVII | 873303 | 873436 | 133 | N7:873338 |
| chrVII | 873927 | 874079 | 152 | N7:873992 |
| chrVII | 878924 | 879054 | 130 | N7:878962 |
| chrVII | 879023 | 879172 | 149 | |
| chrVII | 879984 | 880088 | 104 | |
| chrVII | 882915 | 883023 | 108 | |
| chrVII | 882915 | 883031 | 116 | N7:883049 |
| chrVII | 887398 | 887557 | 159 | |
| chrVII | 889980 | 890139 | 159 | N7:890158 |
| chrVII | 889980 | 890139 | 159 | N7:890006 |
| chrVII | 892462 | 892609 | 147 | N7:892520 |
| chrVII | 895680 | 895828 | 148 | N7:895781 |
| chrVII | 911520 | 911682 | 162 | N7:911624 |
| chrVII | 914384 | 914518 | 134 | N7:914490 |
| chrVII | 920349 | 920433 | 84 | N7:920408 |
| chrVII | 922436 | 922538 | 102 | N7:922490 |
| chrVII | 925873 | 926024 | 151 | |
| chrVII | 928464 | 928607 | 143 | |
| chrVII | 928560 | 928710 | 150 | N7:928631 |
| chrVII | 929745 | 929899 | 154 | N7:929875 |
| chrVII | 938659 | 938812 | 153 | N7:938652 |
| chrVII | 943256 | 943409 | 153 | N7:943358 |
| chrVII | 943702 | 943854 | 152 | N7:943837 |
| chrVII | 948590 | 948746 | 156 | N7:948665 |
| chrVII | 949733 | 949881 | 148 | N7:949820 |
| chrVII | 960612 | 960754 | 142 | N7:960724 |
| chrVII | 961680 | 961814 | 134 | N7:961727 |
| chrVII | 961681 | 961788 | 107 | N7:961727 |
| chrVII | 961681 | 961791 | 110 | N7:961727 |
| chrVII | 961681 | 961794 | 113 | N7:961727 |
| chrVII | 961681 | 961795 | 114 | N7:961727 |
| chrVII | 961681 | 961796 | 115 | N7:961727 |
| chrVII | 961681 | 961797 | 116 | N7:961727 |
| chrVII | 961681 | 961798 | 117 | N7:961727 |
| chrVII | 961681 | 961799 | 118 | N7:961727 |
| chrVII | 961681 | 961800 | 119 | N7:961727 |
| chrVII | 961681 | 961801 | 120 | N7:961727 |
| chrVII | 961681 | 961802 | 121 | N7:961727 |
| chrVII | 961681 | 961803 | 122 | N7:961727 |
| chrVII | 961681 | 961804 | 123 | N7:961727 |
| chrVII | 961681 | 961805 | 124 | N7:961727 |
| chrVII | 961681 | 961806 | 125 | N7:961727 |
| chrVII | 961681 | 961807 | 126 | N7:961727 |
| chrVII | 961681 | 961808 | 127 | N7:961727 |
| chrVII | 961681 | 961809 | 128 | N7:961727 |
| chrVII | 961681 | 961810 | 129 | N7:961727 |
| chrVII | 961681 | 961811 | 130 | N7:961727 |
| chrVII | 961681 | 961812 | 131 | N7:961727 |
| chrVII | 961681 | 961813 | 132 | N7:961727 |
| chrVII | 961681 | 961814 | 133 | N7:961727 |
| chrVII | 961682 | 961814 | 132 | N7:961727 |
| chrVII | 961683 | 961814 | 131 | N7:961727 |
| chrVII | 961684 | 961814 | 130 | N7:961727 |
| chrVII | 961685 | 961814 | 129 | N7:961727 |
| chrVII | 961686 | 961814 | 128 | N7:961727 |
| chrVII | 961687 | 961814 | 127 | N7:961727 |
| chrVII | 961694 | 961814 | 120 | N7:961727 |
| chrVII | 961695 | 961814 | 119 | N7:961727 |
| chrVII | 961696 | 961814 | 118 | N7:961727 |
| chrVII | 961708 | 961806 | 98 | N7:961727 |
| chrVII | 962925 | 963072 | 147 | N7:963033 |

| CHR | Start | End | Length (bp) | Nucleosome_ID |
|---|---|---|---|---|
| chrVII | 966527 | 966637 | 110 | N7:966590 |
| chrVII | 974170 | 974327 | 157 | N7:974249 |
| chrVII | 975022 | 975129 | 107 | N7:975032 |
| chrVII | 975022 | 975130 | 108 | N7:975032 |
| chrVII | 975022 | 975131 | 109 | N7:975032 |
| chrVII | 975022 | 975142 | 120 | N7:975032 |
| chrVII | 979943 | 980105 | 162 | N7:980040 |
| chrVII | 982482 | 982630 | 148 | N7:982592 |
| chrVII | 987451 | 987604 | 153 | N7:987557 |
| chrVII | 987451 | 987606 | 155 | N7:987557 |
| chrVII | 987451 | 987608 | 157 | N7:987557 |
| chrVII | 987451 | 987609 | 158 | N7:987557 |
| chrVII | 988814 | 988967 | 153 | N7:988944 |
| chrVII | 988881 | 989019 | 138 | N7:988944 |
| chrVII | 989319 | 989475 | 156 | N7:989398 |
| chrVII | 989319 | 989477 | 158 | N7:989398 |
| chrVII | 991949 | 992103 | 154 | N7:992011 |
| chrVII | 999326 | 999473 | 147 | N7:999420 |
| chrVII | 1001929 | 1002078 | 149 | N7:1001976 |
| chrVII | 1012079 | 1012238 | 159 | N7:1012073 |
| chrVII | 1012603 | 1012726 | 123 | N7:1012698 |
| chrVII | 1016668 | 1016789 | 121 | N7:1016735 |
| chrVII | 1018529 | 1018686 | 157 | N7:1018544 |
| chrVII | 1018639 | 1018770 | 131 | N7:1018717 |
| chrVII | 1024687 | 1024847 | 160 | |
| chrVII | 1030700 | 1030865 | 165 | N7:1030777 |
| chrVII | 1030899 | 1031060 | 161 | N7:1030983 |
| chrVII | 1037731 | 1037891 | 160 | N7:1037843 |
| chrVII | 1039928 | 1040095 | 167 | N7:1040010 |
| chrVII | 1043906 | 1044058 | 152 | N7:1044003 |
| chrVII | 1044184 | 1044344 | 160 | N7:1044171 |
| chrVII | 1045755 | 1045901 | 146 | N7:1045865 |
| chrVII | 1049209 | 1049360 | 151 | N7:1049248 |
| chrVII | 1051642 | 1051789 | 147 | N7:1051693 |
| chrVII | 1052012 | 1052165 | 153 | N7:1052015 |
| chrVII | 1065087 | 1065242 | 155 | N7:1065159 |
| chrVII | 1065864 | 1066016 | 152 | N7:1065932 |
| chrVII | 1065866 | 1066016 | 150 | N7:1065932 |
| chrVII | 1065867 | 1066016 | 149 | N7:1065932 |
| chrVII | 1067431 | 1067584 | 153 | N7:1067518 |
| chrVII | 1071145 | 1071303 | 158 | N7:1071240 |
| chrVII | 1079049 | 1079187 | 138 | N7:1079141 |
| chrVII | 1079684 | 1079846 | 162 | N7:1079686 |
| chrVII | 1080323 | 1080452 | 129 | N7:1080409 |
| chrVII | 1085801 | 1085942 | 141 | N7:1085920 |
| chrVII | 1087005 | 1087167 | 162 | N7:1086993 |
| chrVII | 1087005 | 1087167 | 162 | N7:1087148 |
| chrVII | 1087106 | 1087261 | 155 | N7:1087148 |
| chrVII | 1087240 | 1087377 | 137 | N7:1087315 |
| chrVII | 1087527 | 1087678 | 151 | N7:1087597 |
| chrVII | 1087535 | 1087678 | 143 | N7:1087597 |
| chrVII | 1087583 | 1087708 | 125 | N7:1087597 |
| chrVII | 1087808 | 1087960 | 152 | N7:1087922 |
| chrVII | 1087881 | 1088025 | 144 | N7:1087922 |
| chrVII | 1087978 | 1088113 | 135 | N7:1088072 |
| chrVII | 1087978 | 1088165 | 187 | N7:1088072 |
| chrVII | 1088021 | 1088113 | 92 | N7:1088072 |
| chrVII | 1088021 | 1088165 | 144 | N7:1088072 |
| chrVII | 1088360 | 1088512 | 152 | N7:1088436 |
| chrVII | 1088661 | 1088791 | 130 | N7:1088758 |
| chrVII | 1089031 | 1089182 | 151 | N7:1089163 |
| chrVII | 1089036 | 1089170 | 134 | N7:1089163 |
| chrVII | 1089049 | 1089145 | 96 | N7:1089163 |
| chrVII | 1090196 | 1090347 | 151 | N7:1090283 |
| chrVIII | 691 | 842 | 151 | N8:754 |
| chrVIII | 1289 | 1405 | 116 | N8:1300 |
| chrVIII | 1346 | 1501 | 155 | |
| chrVIII | 1712 | 1899 | 187 | N8:1845 |
| chrVIII | 1733 | 1899 | 166 | N8:1845 |
| chrVIII | 1748 | 1846 | 98 | N8:1845 |
| chrVIII | 1748 | 1855 | 107 | N8:1845 |
| chrVIII | 1748 | 1878 | 130 | N8:1845 |
| chrVIII | 1748 | 1879 | 131 | N8:1845 |
| chrVIII | 1748 | 1880 | 132 | N8:1845 |
| chrVIII | 1748 | 1889 | 141 | N8:1845 |
| chrVIII | 1748 | 1891 | 143 | N8:1845 |
| chrVIII | 1748 | 1894 | 146 | N8:1845 |
| chrVIII | 1748 | 1895 | 147 | N8:1845 |
| chrVIII | 1748 | 1896 | 148 | N8:1845 |
| chrVIII | 1748 | 1897 | 149 | N8:1845 |
| chrVIII | 1748 | 1898 | 150 | N8:1845 |
| chrVIII | 1748 | 1899 | 151 | N8:1845 |
| chrVIII | 1748 | 1909 | 161 | N8:1845 |
| chrVIII | 1748 | 1932 | 184 | N8:1845 |
| chrVIII | 1749 | 1899 | 150 | N8:1845 |
| chrVIII | 1750 | 1899 | 149 | N8:1845 |
| chrVIII | 1751 | 1899 | 148 | N8:1845 |
| chrVIII | 1752 | 1899 | 147 | N8:1845 |
| chrVIII | 1753 | 1899 | 146 | N8:1845 |
| chrVIII | 1754 | 1899 | 145 | N8:1845 |
| chrVIII | 1755 | 1899 | 144 | N8:1845 |
| chrVIII | 1756 | 1899 | 143 | N8:1845 |
| chrVIII | 1757 | 1899 | 142 | N8:1845 |
| chrVIII | 1758 | 1899 | 141 | N8:1845 |
| chrVIII | 1759 | 1899 | 140 | N8:1845 |
| chrVIII | 1760 | 1899 | 139 | N8:1845 |
| chrVIII | 1761 | 1899 | 138 | N8:1845 |
| chrVIII | 1762 | 1899 | 137 | N8:1845 |
| chrVIII | 1764 | 1899 | 135 | N8:1845 |
| chrVIII | 1765 | 1899 | 134 | N8:1845 |
| chrVIII | 1766 | 1899 | 133 | N8:1845 |
| chrVIII | 1769 | 1899 | 130 | N8:1845 |
| chrVIII | 1771 | 1899 | 128 | N8:1845 |

| CHR | Start | End | Length (bp) | Nucleosome_ID |
|---|---|---|---|---|
| chrVIII | 1773 | 1899 | 126 | N8:1845 |
| chrVIII | 1784 | 1899 | 115 | N8:1845 |
| chrVIII | 1821 | 1968 | 147 | N8:1845 |
| chrVIII | 2051 | 2222 | 171 | |
| chrVIII | 2441 | 2594 | 153 | N8:2474 |
| chrVIII | 2951 | 3103 | 152 | N8:2994 |
| chrVIII | 3093 | 3220 | 127 | N8:3153 |
| chrVIII | 3233 | 3376 | 143 | N8:3313 |
| chrVIII | 3524 | 3661 | 137 | N8:3599 |
| chrVIII | 4667 | 4822 | 155 | N8:4741 |
| chrVIII | 9299 | 9453 | 154 | N8:9352 |
| chrVIII | 10577 | 10683 | 106 | N8:10600 |
| chrVIII | 10578 | 10683 | 105 | N8:10600 |
| chrVIII | 10864 | 11003 | 139 | N8:10940 |
| chrVIII | 12729 | 12877 | 148 | N8:12794 |
| chrVIII | 14994 | 15154 | 160 | N8:15064 |
| chrVIII | 15294 | 15466 | 172 | N8:15381 |
| chrVIII | 17876 | 18017 | 141 | N8:18009 |
| chrVIII | 18350 | 18522 | 172 | N8:18473 |
| chrVIII | 23920 | 24090 | 170 | N8:24051 |
| chrVIII | 26981 | 27134 | 153 | N8:27018 |
| chrVIII | 32520 | 32666 | 146 | N8:32599 |
| chrVIII | 33329 | 33481 | 152 | N8:33481 |
| chrVIII | 33329 | 33481 | 152 | N8:33328 |
| chrVIII | 33632 | 33782 | 150 | N8:33647 |
| chrVIII | 39986 | 40150 | 164 | N8:40093 |
| chrVIII | 46044 | 46208 | 164 | N8:46074 |
| chrVIII | 48223 | 48365 | 142 | N8:48283 |
| chrVIII | 49771 | 49922 | 151 | N8:49832 |
| chrVIII | 50265 | 50415 | 150 | N8:50342 |
| chrVIII | 51501 | 51665 | 164 | N8:51521 |
| chrVIII | 52096 | 52204 | 108 | N8:52145 |
| chrVIII | 52096 | 52213 | 117 | N8:52145 |
| chrVIII | 52629 | 52803 | 174 | N8:52786 |
| chrVIII | 53713 | 53863 | 150 | N8:53785 |
| chrVIII | 58605 | 58733 | 128 | N8:58676 |
| chrVIII | 59349 | 59502 | 153 | N8:59376 |
| chrVIII | 61087 | 61247 | 160 | N8:61111 |
| chrVIII | 62149 | 62261 | 112 | N8:62222 |
| chrVIII | 65254 | 65423 | 169 | N8:65391 |
| chrVIII | 66426 | 66576 | 150 | N8:66523 |
| chrVIII | 66660 | 66810 | 150 | N8:66683 |
| chrVIII | 66812 | 66936 | 124 | N8:66852 |
| chrVIII | 66813 | 66902 | 89 | N8:66852 |
| chrVIII | 66813 | 66904 | 91 | N8:66852 |
| chrVIII | 66813 | 66911 | 98 | N8:66852 |
| chrVIII | 66813 | 66912 | 99 | N8:66852 |
| chrVIII | 66813 | 66924 | 111 | N8:66852 |
| chrVIII | 66813 | 66925 | 112 | N8:66852 |
| chrVIII | 66813 | 66926 | 113 | N8:66852 |
| chrVIII | 66813 | 66928 | 115 | N8:66852 |
| chrVIII | 66813 | 66929 | 116 | N8:66852 |
| chrVIII | 66813 | 66930 | 117 | N8:66852 |
| chrVIII | 66813 | 66931 | 118 | N8:66852 |
| chrVIII | 66813 | 66932 | 119 | N8:66852 |
| chrVIII | 66813 | 66933 | 120 | N8:66852 |
| chrVIII | 66813 | 66934 | 121 | N8:66852 |
| chrVIII | 66813 | 66935 | 122 | N8:66852 |
| chrVIII | 66813 | 66936 | 123 | N8:66852 |
| chrVIII | 66815 | 66936 | 121 | N8:66852 |
| chrVIII | 66816 | 66936 | 120 | N8:66852 |
| chrVIII | 66817 | 66936 | 119 | N8:66852 |
| chrVIII | 66818 | 66936 | 118 | N8:66852 |
| chrVIII | 66820 | 66936 | 116 | N8:66852 |
| chrVIII | 66821 | 66936 | 115 | N8:66852 |
| chrVIII | 66822 | 66936 | 114 | N8:66852 |
| chrVIII | 66823 | 66936 | 113 | N8:66852 |
| chrVIII | 66830 | 66936 | 106 | N8:66852 |
| chrVIII | 68261 | 68399 | 138 | N8:68264 |
| chrVIII | 69975 | 70096 | 121 | N8:70043 |
| chrVIII | 69975 | 70122 | 147 | N8:70043 |
| chrVIII | 70402 | 70553 | 151 | N8:70464 |
| chrVIII | 72065 | 72199 | 134 | N8:72067 |
| chrVIII | 77481 | 77612 | 131 | N8:77523 |
| chrVIII | 86148 | 86310 | 162 | N8:86173 |
| chrVIII | 86176 | 86324 | 148 | N8:86173 |
| chrVIII | 86476 | 86623 | 147 | N8:86566 |
| chrVIII | 88963 | 89116 | 153 | N8:89119 |
| chrVIII | 91372 | 91534 | 162 | N8:91357 |
| chrVIII | 91372 | 91534 | 162 | N8:91547 |
| chrVIII | 94804 | 94953 | 149 | N8:94867 |
| chrVIII | 94985 | 95141 | 156 | N8:95070 |
| chrVIII | 96353 | 96506 | 153 | N8:96448 |
| chrVIII | 100385 | 100534 | 149 | N8:100544 |
| chrVIII | 107252 | 107412 | 160 | N8:107297 |
| chrVIII | 108345 | 108493 | 148 | N8:108419 |
| chrVIII | 109527 | 109701 | 174 | N8:109636 |
| chrVIII | 114779 | 114932 | 153 | N8:114850 |
| chrVIII | 115531 | 115709 | 178 | N8:115711 |
| chrVIII | 121476 | 121637 | 161 | N8:121495 |
| chrVIII | 122292 | 122451 | 159 | N8:122379 |
| chrVIII | 123820 | 123978 | 158 | N8:123911 |
| chrVIII | 125638 | 125758 | 120 | N8:125733 |
| chrVIII | 130844 | 131002 | 158 | N8:130922 |
| chrVIII | 131196 | 131345 | 149 | N8:131260 |
| chrVIII | 142630 | 142776 | 146 | N8:142760 |
| chrVIII | 144193 | 144334 | 141 | N8:144264 |
| chrVIII | 152201 | 152353 | 152 | N8:152258 |
| chrVIII | 153827 | 153949 | 122 | N8:153924 |
| chrVIII | 157804 | 157963 | 159 | N8:157803 |
| chrVIII | 158390 | 158542 | 152 | N8:158513 |
| chrVIII | 162439 | 162593 | 154 | N8:162524 |
| chrVIII | 162949 | 163087 | 138 | |

| CHR | Start | End | Length (bp) | Nucleosome_ID |
|---|---|---|---|---|
| chrVIII | 165932 | 166076 | 144 | N8:165989 |
| chrVIII | 167959 | 168110 | 151 | N8:168035 |
| chrVIII | 171020 | 171171 | 151 | N8:171084 |
| chrVIII | 171703 | 171855 | 152 | N8:171826 |
| chrVIII | 174446 | 174586 | 140 | |
| chrVIII | 174447 | 174533 | 86 | |
| chrVIII | 174447 | 174535 | 88 | |
| chrVIII | 174447 | 174538 | 91 | |
| chrVIII | 174447 | 174540 | 93 | |
| chrVIII | 174447 | 174541 | 94 | |
| chrVIII | 174447 | 174544 | 97 | |
| chrVIII | 174447 | 174545 | 98 | |
| chrVIII | 174447 | 174548 | 101 | |
| chrVIII | 174447 | 174550 | 103 | |
| chrVIII | 174447 | 174551 | 104 | |
| chrVIII | 174447 | 174552 | 105 | |
| chrVIII | 174447 | 174553 | 106 | |
| chrVIII | 174447 | 174555 | 108 | |
| chrVIII | 174447 | 174561 | 114 | |
| chrVIII | 174447 | 174565 | 118 | |
| chrVIII | 174447 | 174566 | 119 | |
| chrVIII | 174447 | 174567 | 120 | |
| chrVIII | 174447 | 174568 | 121 | |
| chrVIII | 174447 | 174569 | 122 | |
| chrVIII | 174447 | 174570 | 123 | |
| chrVIII | 174447 | 174571 | 124 | |
| chrVIII | 174447 | 174572 | 125 | |
| chrVIII | 174447 | 174573 | 126 | |
| chrVIII | 174447 | 174574 | 127 | |
| chrVIII | 174447 | 174575 | 128 | |
| chrVIII | 174447 | 174576 | 129 | |
| chrVIII | 174447 | 174577 | 130 | |
| chrVIII | 174447 | 174578 | 131 | |
| chrVIII | 174447 | 174579 | 132 | |
| chrVIII | 174447 | 174580 | 133 | |
| chrVIII | 174447 | 174581 | 134 | |
| chrVIII | 174447 | 174582 | 135 | |
| chrVIII | 174447 | 174583 | 136 | |
| chrVIII | 174447 | 174584 | 137 | |
| chrVIII | 174447 | 174585 | 138 | |
| chrVIII | 174447 | 174586 | 139 | |
| chrVIII | 174447 | 174587 | 140 | |
| chrVIII | 174448 | 174586 | 138 | |
| chrVIII | 174449 | 174586 | 137 | |
| chrVIII | 174450 | 174586 | 136 | |
| chrVIII | 174451 | 174586 | 135 | |
| chrVIII | 174452 | 174586 | 134 | |
| chrVIII | 174453 | 174586 | 133 | |
| chrVIII | 174454 | 174586 | 132 | |
| chrVIII | 174455 | 174586 | 131 | |
| chrVIII | 174456 | 174586 | 130 | |
| chrVIII | 174457 | 174586 | 129 | |
| chrVIII | 174458 | 174586 | 128 | |
| chrVIII | 174459 | 174586 | 127 | |
| chrVIII | 174460 | 174586 | 126 | |
| chrVIII | 174461 | 174586 | 125 | |
| chrVIII | 174462 | 174553 | 91 | |
| chrVIII | 174467 | 174586 | 119 | |
| chrVIII | 174928 | 175084 | 156 | N8:175011 |
| chrVIII | 176211 | 176366 | 155 | N8:176283 |
| chrVIII | 177719 | 177867 | 148 | N8:177804 |
| chrVIII | 177797 | 177947 | 150 | N8:177804 |
| chrVIII | 178837 | 178985 | 148 | N8:178874 |
| chrVIII | 180375 | 180509 | 134 | N8:180443 |
| chrVIII | 190177 | 190319 | 142 | N8:190199 |
| chrVIII | 194784 | 194941 | 157 | |
| chrVIII | 194987 | 195122 | 135 | N8:195028 |
| chrVIII | 205654 | 205804 | 150 | N8:205739 |
| chrVIII | 207083 | 207225 | 142 | N8:207213 |
| chrVIII | 212526 | 212683 | 157 | N8:212615 |
| chrVIII | 212729 | 212874 | 145 | N8:212883 |
| chrVIII | 212963 | 213069 | 106 | N8:213049 |
| chrVIII | 212963 | 213076 | 113 | N8:213049 |
| chrVIII | 213025 | 213187 | 162 | N8:213049 |
| chrVIII | 214079 | 214248 | 169 | N8:214186 |
| chrVIII | 214176 | 214329 | 153 | N8:214186 |
| chrVIII | 214524 | 214681 | 157 | N8:214628 |
| chrVIII | 214727 | 214872 | 145 | N8:214799 |
| chrVIII | 214961 | 215067 | 106 | N8:215040 |
| chrVIII | 214961 | 215074 | 113 | N8:215040 |
| chrVIII | 215023 | 215185 | 162 | N8:215040 |
| chrVIII | 216077 | 216246 | 169 | N8:216180 |
| chrVIII | 217808 | 217962 | 154 | N8:217835 |
| chrVIII | 218430 | 218569 | 139 | N8:218465 |
| chrVIII | 219279 | 219430 | 151 | N8:219365 |
| chrVIII | 223655 | 223760 | 105 | N8:223731 |
| chrVIII | 226108 | 226256 | 148 | N8:226185 |
| chrVIII | 227838 | 227999 | 161 | N8:228012 |
| chrVIII | 227838 | 227999 | 161 | N8:227836 |
| chrVIII | 228803 | 228946 | 143 | N8:228877 |
| chrVIII | 234383 | 234536 | 153 | N8:234466 |
| chrVIII | 234544 | 234693 | 149 | N8:234634 |
| chrVIII | 240775 | 240923 | 148 | N8:240835 |
| chrVIII | 250570 | 250724 | 154 | N8:250736 |
| chrVIII | 256594 | 256759 | 165 | N8:256675 |
| chrVIII | 258714 | 258856 | 142 | N8:258740 |
| chrVIII | 261335 | 261482 | 147 | N8:261454 |
| chrVIII | 264809 | 264956 | 147 | N8:264814 |
| chrVIII | 270516 | 270670 | 154 | N8:270646 |
| chrVIII | 272692 | 272812 | 120 | N8:272823 |
| chrVIII | 275159 | 275317 | 158 | |
| chrVIII | 281596 | 281745 | 149 | N8:281674 |
| chrVIII | 288944 | 289050 | 106 | N8:288960 |

| CHR | Start | End | Length (bp) | Nucleosome_ID |
|---|---|---|---|---|
| chrVIII | 291607 | 291756 | 149 | N8:291704 |
| chrVIII | 291938 | 292090 | 152 | N8:292062 |
| chrVIII | 294718 | 294863 | 145 | N8:294786 |
| chrVIII | 297474 | 297608 | 134 | |
| chrVIII | 297952 | 298079 | 127 | |
| chrVIII | 299058 | 299223 | 165 | N8:299169 |
| chrVIII | 299773 | 299910 | 137 | N8:299791 |
| chrVIII | 310121 | 310260 | 139 | N8:310259 |
| chrVIII | 310839 | 310998 | 159 | N8:310900 |
| chrVIII | 311969 | 312130 | 161 | N8:312117 |
| chrVIII | 333005 | 333157 | 152 | N8:333100 |
| chrVIII | 334511 | 334659 | 148 | N8:334641 |
| chrVIII | 341021 | 341173 | 152 | N8:341133 |
| chrVIII | 341021 | 341178 | 157 | N8:341133 |
| chrVIII | 342001 | 342158 | 157 | N8:342074 |
| chrVIII | 342217 | 342368 | 151 | N8:342346 |
| chrVIII | 343838 | 343999 | 161 | N8:344006 |
| chrVIII | 345951 | 346111 | 160 | N8:346037 |
| chrVIII | 346434 | 346582 | 148 | N8:346511 |
| chrVIII | 350191 | 350346 | 155 | N8:350266 |
| chrVIII | 353263 | 353415 | 152 | |
| chrVIII | 354917 | 355076 | 159 | N8:355010 |
| chrVIII | 355941 | 356089 | 148 | N8:356016 |
| chrVIII | 357549 | 357703 | 154 | N8:357581 |
| chrVIII | 360834 | 360982 | 148 | N8:360921 |
| chrVIII | 364130 | 364290 | 160 | N8:364200 |
| chrVIII | 370648 | 370829 | 181 | N8:370765 |
| chrVIII | 371736 | 371883 | 147 | N8:371818 |
| chrVIII | 374061 | 374221 | 160 | N8:374205 |
| chrVIII | 374061 | 374221 | 160 | N8:374049 |
| chrVIII | 374061 | 374255 | 194 | N8:374205 |
| chrVIII | 374061 | 374255 | 194 | N8:374049 |
| chrVIII | 374122 | 374209 | 87 | N8:374205 |
| chrVIII | 374122 | 374221 | 99 | N8:374205 |
| chrVIII | 374122 | 374255 | 133 | N8:374205 |
| chrVIII | 376459 | 376622 | 163 | N8:376456 |
| chrVIII | 376459 | 376622 | 163 | N8:376622 |
| chrVIII | 381490 | 381641 | 151 | N8:381642 |
| chrVIII | 382542 | 382702 | 160 | N8:382584 |
| chrVIII | 382976 | 383151 | 175 | N8:383155 |
| chrVIII | 399609 | 399757 | 148 | N8:399693 |
| chrVIII | 402529 | 402689 | 160 | N8:402659 |
| chrVIII | 402529 | 402689 | 160 | N8:402507 |
| chrVIII | 403584 | 403737 | 153 | N8:403661 |
| chrVIII | 404097 | 404230 | 133 | N8:404184 |
| chrVIII | 414157 | 414333 | 176 | N8:414190 |
| chrVIII | 414157 | 414333 | 176 | N8:414356 |
| chrVIII | 416537 | 416675 | 138 | N8:416611 |
| chrVIII | 424973 | 425127 | 154 | N8:425133 |
| chrVIII | 424973 | 425127 | 154 | N8:424954 |
| chrVIII | 427961 | 428122 | 161 | N8:428089 |
| chrVIII | 431042 | 431168 | 126 | N8:431037 |
| chrVIII | 433338 | 433488 | 150 | N8:433377 |
| chrVIII | 439467 | 439624 | 157 | N8:439505 |
| chrVIII | 444830 | 444988 | 158 | N8:444909 |
| chrVIII | 447873 | 448000 | 127 | N8:448007 |
| chrVIII | 456058 | 456162 | 104 | N8:456183 |
| chrVIII | 465568 | 465728 | 160 | N8:465665 |
| chrVIII | 469779 | 469934 | 155 | N8:469909 |
| chrVIII | 469928 | 470081 | 153 | N8:469909 |
| chrVIII | 469928 | 470081 | 153 | N8:470061 |
| chrVIII | 470995 | 471124 | 129 | N8:471041 |
| chrVIII | 482780 | 482943 | 163 | N8:482852 |
| chrVIII | 485160 | 485318 | 158 | N8:485166 |
| chrVIII | 485160 | 485318 | 158 | N8:485325 |
| chrVIII | 495830 | 495995 | 165 | N8:495933 |
| chrVIII | 497750 | 497909 | 159 | N8:497804 |
| chrVIII | 501432 | 501582 | 150 | N8:501512 |
| chrVIII | 503721 | 503876 | 155 | N8:503793 |
| chrVIII | 510274 | 510423 | 149 | N8:510320 |
| chrVIII | 518475 | 518638 | 163 | N8:518487 |
| chrVIII | 518475 | 518638 | 163 | N8:518641 |
| chrVIII | 521779 | 521913 | 134 | N8:521848 |
| chrVIII | 522442 | 522595 | 153 | N8:522525 |
| chrVIII | 523030 | 523182 | 152 | N8:523103 |
| chrVIII | 523197 | 523325 | 128 | N8:523263 |
| chrVIII | 525186 | 525342 | 156 | N8:525241 |
| chrVIII | 526030 | 526198 | 168 | N8:526058 |
| chrVIII | 526169 | 526320 | 151 | N8:526224 |
| chrVIII | 526293 | 526441 | 148 | N8:526381 |
| chrVIII | 526293 | 526481 | 188 | N8:526381 |
| chrVIII | 526600 | 526706 | 106 | N8:526683 |
| chrVIII | 526600 | 526743 | 143 | N8:526683 |
| chrVIII | 526600 | 526749 | 149 | N8:526683 |
| chrVIII | 526600 | 526751 | 151 | N8:526683 |
| chrVIII | 526624 | 526751 | 127 | N8:526683 |
| chrVIII | 526651 | 526751 | 100 | N8:526683 |
| chrVIII | 527864 | 528038 | 174 | N8:527985 |
| chrVIII | 536044 | 536157 | 113 | N8:536107 |
| chrVIII | 538480 | 538617 | 137 | N8:538580 |
| chrVIII | 539313 | 539458 | 145 | N8:539402 |
| chrVIII | 539340 | 539488 | 148 | N8:539402 |
| chrVIII | 542782 | 542926 | 144 | N8:542760 |
| chrVIII | 542782 | 542926 | 144 | N8:542922 |
| chrVIII | 542989 | 543156 | 167 | |
| chrVIII | 543353 | 543457 | 104 | N8:543426 |
| chrVIII | 543780 | 543928 | 148 | N8:543820 |
| chrVIII | 544540 | 544654 | 114 | |
| chrVIII | 546632 | 546789 | 157 | N8:546708 |
| chrVIII | 546818 | 546968 | 150 | N8:546888 |
| chrVIII | 546822 | 546968 | 146 | N8:546888 |
| chrVIII | 547280 | 547415 | 135 | N8:547333 |
| chrVIII | 548318 | 548458 | 140 | N8:548393 |

| CHR | Start | End | Length (bp) | Nucleosome_ID |
|---|---|---|---|---|
| chrVIII | 548324 | 548458 | 134 | N8:548393 |
| chrVIII | 548850 | 549005 | 155 | N8:548872 |
| chrVIII | 548981 | 549089 | 108 | N8:549081 |
| chrVIII | 549893 | 550056 | 163 | N8:549980 |
| chrVIII | 550702 | 550810 | 108 | N8:550729 |
| chrVIII | 551514 | 551634 | 120 | N8:551574 |
| chrVIII | 553179 | 553326 | 147 | N8:553215 |
| chrVIII | 553630 | 553778 | 148 | N8:553708 |
| chrVIII | 556401 | 556536 | 135 | N8:556476 |
| chrVIII | 556826 | 556854 | 28 | |
| chrVIII | 556826 | 556859 | 33 | |
| chrVIII | 556826 | 556865 | 39 | |
| chrVIII | 556826 | 556871 | 45 | |
| chrVIII | 556826 | 556873 | 47 | |
| chrVIII | 556826 | 556876 | 50 | |
| chrVIII | 556826 | 556884 | 58 | N8:556907 |
| chrVIII | 556826 | 556910 | 84 | N8:556907 |
| chrVIII | 556826 | 556914 | 88 | N8:556907 |
| chrVIII | 556826 | 556918 | 92 | N8:556907 |
| chrVIII | 556826 | 556919 | 93 | N8:556907 |
| chrVIII | 556826 | 556921 | 95 | N8:556907 |
| chrVIII | 556826 | 556923 | 97 | N8:556907 |
| chrVIII | 556826 | 556926 | 100 | N8:556907 |
| chrVIII | 556826 | 556930 | 104 | N8:556907 |
| chrVIII | 556826 | 556935 | 109 | N8:556907 |
| chrVIII | 556826 | 556936 | 110 | N8:556907 |
| chrVIII | 556826 | 556939 | 113 | N8:556907 |
| chrVIII | 556826 | 556942 | 116 | N8:556907 |
| chrVIII | 556826 | 556944 | 118 | N8:556907 |
| chrVIII | 556826 | 556950 | 124 | N8:556907 |
| chrVIII | 556826 | 556954 | 128 | N8:556907 |
| chrVIII | 556826 | 556956 | 130 | N8:556907 |
| chrVIII | 556826 | 556961 | 135 | N8:556907 |
| chrVIII | 556826 | 556963 | 137 | N8:556907 |
| chrVIII | 556826 | 556965 | 139 | N8:556907 |
| chrVIII | 556826 | 556967 | 141 | N8:556907 |
| chrVIII | 556826 | 556969 | 143 | N8:556907 |
| chrVIII | 556826 | 556970 | 144 | N8:556907 |
| chrVIII | 556826 | 556971 | 145 | N8:556907 |
| chrVIII | 556826 | 556972 | 146 | N8:556907 |
| chrVIII | 556826 | 556973 | 147 | N8:556907 |
| chrVIII | 556826 | 556975 | 149 | N8:556907 |
| chrVIII | 556826 | 556976 | 150 | N8:556907 |
| chrVIII | 556826 | 556977 | 151 | N8:556907 |
| chrVIII | 556826 | 556979 | 153 | N8:556907 |
| chrVIII | 556826 | 556981 | 155 | N8:556907 |
| chrVIII | 556826 | 556982 | 156 | N8:556907 |
| chrVIII | 556826 | 556983 | 157 | N8:556907 |
| chrVIII | 556826 | 556984 | 158 | N8:556907 |
| chrVIII | 556826 | 556985 | 159 | N8:556907 |
| chrVIII | 556826 | 556986 | 160 | N8:556907 |
| chrVIII | 556826 | 556987 | 161 | N8:556907 |
| chrVIII | 556826 | 556989 | 163 | N8:556907 |
| chrVIII | 556833 | 556986 | 153 | N8:556907 |
| chrVIII | 556840 | 556986 | 146 | N8:556907 |
| chrVIII | 556843 | 556989 | 146 | N8:556907 |
| chrVIII | 556853 | 556986 | 133 | N8:556907 |
| chrVIII | 556853 | 556989 | 136 | N8:556907 |
| chrVIII | 556858 | 556975 | 117 | N8:556907 |
| chrVIII | 556858 | 556989 | 131 | N8:556907 |
| chrVIII | 556859 | 556963 | 104 | N8:556907 |
| chrVIII | 556859 | 556989 | 130 | N8:556907 |
| chrVIII | 556863 | 556986 | 123 | N8:556907 |
| chrVIII | 556869 | 556986 | 117 | N8:556907 |
| chrVIII | 556878 | 556973 | 95 | N8:556907 |
| chrVIII | 556878 | 556975 | 97 | N8:556907 |
| chrVIII | 556878 | 556983 | 105 | N8:556907 |
| chrVIII | 556878 | 556989 | 111 | N8:556907 |
| chrVIII | 556883 | 556989 | 106 | N8:556907 |
| chrVIII | 556887 | 556986 | 99 | N8:556907 |
| chrVIII | 556893 | 556975 | 82 | N8:556907 |
| chrVIII | 556893 | 556989 | 96 | N8:556907 |
| chrVIII | 556901 | 556986 | 85 | N8:556907 |
| chrVIII | 556904 | 556989 | 85 | N8:556907 |
| chrVIII | 558701 | 558838 | 137 | N8:558769 |
| chrVIII | 558986 | 559129 | 143 | N8:559040 |
| chrVIII | 559259 | 559411 | 152 | N8:559370 |
| chrVIII | 559332 | 559476 | 144 | N8:559370 |
| chrVIII | 560157 | 560308 | 151 | |
| chrVIII | 560416 | 560563 | 147 | N8:560523 |
| chrVIII | 560448 | 560563 | 115 | N8:560523 |
| chrVIII | 560502 | 560671 | 169 | N8:560523 |
| chrVIII | 560544 | 560634 | 90 | N8:560523 |
| chrVIII | 560544 | 560639 | 95 | N8:560523 |
| chrVIII | 560784 | 560911 | 127 | N8:560813 |
| chrVIII | 561208 | 561356 | 148 | |
| chrVIII | 561302 | 561476 | 174 | N8:561386 |
| chrVIII | 561649 | 561800 | 151 | N8:561716 |
| chrVIII | 562455 | 562594 | 139 | N8:562514 |
| chrVIII | 562455 | 562640 | 185 | N8:562514 |
| chrVIII | 562469 | 562594 | 125 | N8:562514 |
| chrVIII | 562469 | 562614 | 145 | N8:562514 |
| chrVIII | 562469 | 562632 | 163 | N8:562514 |
| chrVIII | 562469 | 562636 | 167 | N8:562514 |
| chrVIII | 562469 | 562640 | 171 | N8:562514 |
| chrVIII | 562500 | 562636 | 136 | N8:562514 |
| chrVIII | 562506 | 562632 | 126 | N8:562514 |
| chrVIII | 562506 | 562636 | 130 | N8:562514 |
| chrVIII | 562506 | 562640 | 134 | N8:562514 |
| chrVIII | 562520 | 562621 | 101 | N8:562514 |
| chrVIII | 562520 | 562640 | 120 | N8:562514 |
| chrVIII | 562523 | 562640 | 117 | N8:562514 |
| chrVIII | 562552 | 562640 | 88 | |

| CHR | Start | End | Length (bp) | Nucleosome_ID |
|---|---|---|---|---|
| chrX | 1669 | 1785 | 116 | N10:1770 |
| chrX | 1669 | 1804 | 135 | N10:1770 |
| chrX | 1669 | 1821 | 152 | N10:1770 |
| chrX | 1693 | 1785 | 92 | N10:1770 |
| chrX | 1693 | 1794 | 101 | N10:1770 |
| chrX | 1693 | 1804 | 111 | N10:1770 |
| chrX | 1693 | 1821 | 128 | N10:1770 |
| chrX | 1698 | 1821 | 123 | N10:1770 |
| chrX | 1980 | 2120 | 140 | N10:2021 |
| chrX | 2198 | 2335 | 137 | N10:2294 |
| chrX | 2365 | 2509 | 144 | N10:2503 |
| chrX | 2365 | 2512 | 147 | N10:2503 |
| chrX | 2365 | 2513 | 148 | N10:2503 |
| chrX | 2365 | 2575 | 210 | N10:2503 |
| chrX | 2370 | 2513 | 143 | N10:2503 |
| chrX | 2380 | 2513 | 133 | N10:2503 |
| chrX | 2420 | 2513 | 93 | N10:2503 |
| chrX | 2420 | 2575 | 155 | N10:2503 |
| chrX | 2862 | 3015 | 153 | N10:2906 |
| chrX | 3307 | 3451 | 144 | N10:3406 |
| chrX | 3518 | 3609 | 91 | N10:3573 |
| chrX | 3518 | 3627 | 109 | N10:3573 |
| chrX | 3518 | 3628 | 110 | N10:3573 |
| chrX | 3518 | 3629 | 111 | N10:3573 |
| chrX | 3518 | 3644 | 126 | N10:3573 |
| chrX | 3520 | 3627 | 107 | N10:3573 |
| chrX | 3520 | 3644 | 124 | N10:3573 |
| chrX | 4071 | 4226 | 155 | N10:4185 |
| chrX | 4165 | 4327 | 162 | N10:4343 |
| chrX | 4165 | 4327 | 162 | N10:4185 |
| chrX | 4788 | 4951 | 163 | N10:4836 |
| chrX | 5199 | 5350 | 151 | |
| chrX | 5412 | 5571 | 159 | N10:5402 |
| chrX | 9273 | 9424 | 151 | N10:9358 |
| chrX | 10211 | 10354 | 143 | N10:10296 |
| chrX | 11443 | 11594 | 151 | N10:11535 |
| chrX | 12328 | 12473 | 145 | N10:12354 |
| chrX | 12893 | 13042 | 149 | |
| chrX | 14191 | 14340 | 149 | N10:14274 |
| chrX | 16003 | 16132 | 129 | N10:16047 |
| chrX | 16061 | 16168 | 107 | N10:16047 |
| chrX | 21416 | 21535 | 119 | N10:21469 |
| chrX | 28854 | 29005 | 151 | N10:28867 |
| chrX | 28854 | 29005 | 151 | N10:29020 |
| chrX | 30398 | 30549 | 151 | N10:30469 |
| chrX | 31626 | 31734 | 108 | N10:31614 |
| chrX | 32694 | 32839 | 145 | N10:32766 |
| chrX | 33933 | 34077 | 144 | N10:33990 |
| chrX | 37483 | 37615 | 132 | N10:37503 |
| chrX | 38720 | 38876 | 156 | N10:38782 |
| chrX | 40633 | 40803 | 170 | N10:40707 |
| chrX | 40783 | 40953 | 170 | N10:40870 |
| chrX | 46039 | 46190 | 151 | N10:46072 |
| chrX | 53460 | 53598 | 138 | N10:53518 |
| chrX | 55658 | 55819 | 161 | N10:55711 |
| chrX | 59858 | 59998 | 140 | N10:59955 |
| chrX | 64405 | 64545 | 140 | |
| chrX | 86256 | 86397 | 141 | N10:86335 |
| chrX | 88120 | 88277 | 157 | N10:88190 |
| chrX | 90912 | 91092 | 180 | N10:91046 |
| chrX | 94290 | 94441 | 151 | N10:94365 |
| chrX | 94366 | 94450 | 84 | N10:94365 |
| chrX | 94724 | 94862 | 138 | N10:94769 |
| chrX | 100058 | 100217 | 159 | N10:100203 |
| chrX | 100062 | 100217 | 155 | N10:100203 |
| chrX | 100066 | 100217 | 151 | N10:100203 |
| chrX | 105547 | 105698 | 151 | N10:105607 |
| chrX | 106971 | 107136 | 165 | |
| chrX | 108096 | 108187 | 91 | N10:108130 |
| chrX | 108096 | 108204 | 108 | N10:108130 |
| chrX | 108096 | 108205 | 109 | N10:108130 |
| chrX | 108096 | 108206 | 110 | N10:108130 |
| chrX | 108096 | 108207 | 111 | N10:108130 |
| chrX | 108096 | 108208 | 112 | N10:108130 |
| chrX | 108096 | 108220 | 124 | N10:108130 |
| chrX | 108097 | 108205 | 108 | N10:108130 |
| chrX | 108098 | 108200 | 102 | N10:108130 |
| chrX | 108098 | 108205 | 107 | N10:108130 |
| chrX | 108098 | 108206 | 108 | N10:108130 |
| chrX | 108098 | 108220 | 122 | N10:108130 |
| chrX | 108135 | 108220 | 85 | N10:108130 |
| chrX | 117567 | 117726 | 159 | |
| chrX | 118042 | 118180 | 138 | N10:118106 |
| chrX | 118045 | 118180 | 135 | N10:118106 |
| chrX | 118063 | 118213 | 150 | N10:118106 |
| chrX | 118208 | 118364 | 156 | N10:118386 |
| chrX | 122371 | 122478 | 107 | N10:122443 |
| chrX | 126969 | 127111 | 142 | N10:126996 |
| chrX | 127866 | 128026 | 160 | N10:127919 |
| chrX | 131244 | 131407 | 163 | N10:131345 |
| chrX | 139844 | 139996 | 152 | N10:139830 |
| chrX | 141168 | 141318 | 150 | N10:141239 |
| chrX | 142696 | 142846 | 150 | N10:142702 |
| chrX | 145224 | 145374 | 150 | N10:145222 |
| chrX | 145224 | 145374 | 150 | N10:145384 |
| chrX | 148124 | 148271 | 147 | N10:148257 |
| chrX | 149558 | 149726 | 168 | N10:149585 |
| chrX | 154966 | 155110 | 144 | N10:155017 |
| chrX | 162602 | 162770 | 168 | N10:162588 |
| chrX | 163009 | 163162 | 153 | N10:163000 |
| chrX | 169091 | 169251 | 160 | N10:169161 |
| chrX | 173679 | 173830 | 151 | N10:173775 |
| chrX | 173874 | 174016 | 142 | N10:173949 |

TABLE 1

| CHR | Start | End | Length (bp) | Nucleosome_ID |
|---|---|---|---|---|
| chrX | 178426 | 178580 | 154 | N10:178551 |
| chrX | 182961 | 183077 | 116 | N10:183051 |
| chrX | 182964 | 183077 | 113 | N10:183051 |
| chrX | 182966 | 183077 | 111 | N10:183051 |
| chrX | 185927 | 186084 | 157 | N10:186100 |
| chrX | 185927 | 186084 | 157 | N10:185938 |
| chrX | 189658 | 189821 | 163 | N10:189737 |
| chrX | 190658 | 190805 | 147 | N10:190730 |
| chrX | 191225 | 191359 | 134 | N10:191216 |
| chrX | 192006 | 192112 | 106 | |
| chrX | 192006 | 192128 | 122 | |
| chrX | 192132 | 192293 | 161 | N10:192220 |
| chrX | 192599 | 192736 | 137 | N10:192656 |
| chrX | 194181 | 194318 | 137 | N10:194236 |
| chrX | 197629 | 197777 | 148 | N10:197770 |
| chrX | 198154 | 198316 | 162 | N10:198291 |
| chrX | 198182 | 198330 | 148 | N10:198291 |
| chrX | 199707 | 199858 | 151 | N10:199867 |
| chrX | 203379 | 203541 | 162 | N10:203536 |
| chrX | 203482 | 203630 | 148 | N10:203536 |
| chrX | 206401 | 206560 | 159 | N10:206482 |
| chrX | 209002 | 209147 | 145 | N10:209025 |
| chrX | 210689 | 210857 | 168 | N10:210876 |
| chrX | 210689 | 210857 | 168 | N10:210724 |
| chrX | 212042 | 212183 | 141 | N10:212201 |
| chrX | 212042 | 212183 | 141 | N10:212021 |
| chrX | 215649 | 215789 | 140 | N10:215728 |
| chrX | 218927 | 219076 | 149 | |
| chrX | 219865 | 220016 | 151 | N10:219873 |
| chrX | 219865 | 220016 | 151 | N10:220031 |
| chrX | 220580 | 220715 | 135 | N10:220634 |
| chrX | 221358 | 221508 | 150 | N10:221437 |
| chrX | 225766 | 225921 | 155 | N10:225854 |
| chrX | 226052 | 226204 | 152 | N10:226103 |
| chrX | 226472 | 226633 | 161 | N10:226484 |
| chrX | 227406 | 227571 | 165 | N10:227419 |
| chrX | 227406 | 227571 | 165 | N10:227588 |
| chrX | 229005 | 229137 | 132 | N10:229146 |
| chrX | 229384 | 229535 | 151 | N10:229378 |
| chrX | 233516 | 233665 | 149 | |
| chrX | 235494 | 235644 | 150 | N10:235627 |
| chrX | 237835 | 237975 | 140 | N10:237857 |
| chrX | 240265 | 240414 | 149 | N10:240350 |
| chrX | 240554 | 240707 | 153 | N10:240699 |
| chrX | 240601 | 240747 | 146 | N10:240699 |
| chrX | 242702 | 242868 | 166 | |
| chrX | 243519 | 243619 | 100 | N10:243533 |
| chrX | 245287 | 245435 | 148 | N10:245271 |
| chrX | 247930 | 248081 | 151 | N10:247910 |
| chrX | 248270 | 248435 | 165 | N10:248400 |
| chrX | 251069 | 251224 | 155 | N10:251097 |
| chrX | 251916 | 252080 | 164 | N10:251970 |
| chrX | 253721 | 253885 | 164 | N10:253709 |
| chrX | 254829 | 255000 | 171 | N10:254961 |
| chrX | 255572 | 255735 | 163 | N10:255685 |
| chrX | 271510 | 271677 | 167 | N10:271629 |
| chrX | 271664 | 271795 | 131 | |
| chrX | 272130 | 272276 | 146 | N10:272201 |
| chrX | 273881 | 274032 | 151 | N10:273955 |
| chrX | 275971 | 276080 | 109 | N10:276085 |
| chrX | 276361 | 276455 | 94 | |
| chrX | 276361 | 276463 | 102 | |
| chrX | 276481 | 276614 | 133 | N10:276566 |
| chrX | 278918 | 279067 | 149 | N10:278918 |
| chrX | 283812 | 283977 | 165 | N10:283889 |
| chrX | 287746 | 287904 | 158 | N10:287749 |
| chrX | 292191 | 292322 | 131 | N10:292221 |
| chrX | 292971 | 293126 | 155 | |
| chrX | 293774 | 293933 | 159 | N10:293870 |
| chrX | 301109 | 301275 | 166 | N10:301192 |
| chrX | 303727 | 303887 | 160 | N10:303776 |
| chrX | 309182 | 309321 | 139 | N10:309163 |
| chrX | 310987 | 311146 | 159 | N10:310995 |
| chrX | 310987 | 311146 | 159 | N10:311155 |
| chrX | 321769 | 321919 | 150 | N10:321795 |
| chrX | 322337 | 322492 | 155 | N10:322482 |
| chrX | 325234 | 325383 | 149 | N10:325322 |
| chrX | 326985 | 327142 | 157 | N10:327050 |
| chrX | 327633 | 327785 | 152 | N10:327706 |
| chrX | 328569 | 328730 | 161 | |
| chrX | 329259 | 329407 | 148 | N10:329425 |
| chrX | 329259 | 329407 | 148 | N10:329251 |
| chrX | 331899 | 332052 | 153 | |
| chrX | 333305 | 333466 | 161 | N10:333312 |
| chrX | 333305 | 333466 | 161 | N10:333478 |
| chrX | 333361 | 333513 | 152 | N10:333478 |
| chrX | 337451 | 337612 | 161 | |
| chrX | 338776 | 338907 | 131 | N10:338772 |
| chrX | 343385 | 343533 | 148 | N10:343380 |
| chrX | 344740 | 344900 | 160 | |
| chrX | 350347 | 350517 | 170 | N10:350502 |
| chrX | 350347 | 350517 | 170 | N10:350327 |
| chrX | 357774 | 357934 | 160 | N10:357775 |
| chrX | 358769 | 358924 | 155 | N10:358940 |
| chrX | 359603 | 359757 | 154 | N10:359674 |
| chrX | 362162 | 362320 | 158 | N10:362260 |
| chrX | 365711 | 365851 | 140 | N10:365757 |
| chrX | 366070 | 366219 | 149 | N10:366182 |
| chrX | 366353 | 366502 | 149 | N10:366336 |
| chrX | 366353 | 366502 | 149 | N10:366486 |
| chrX | 366520 | 366671 | 151 | N10:366641 |
| chrX | 367696 | 367845 | 149 | N10:367708 |
| chrX | 367696 | 367845 | 149 | N10:367868 |

TABLE 1

| CHR | Start | End | Length (bp) | Nucleosome_ID |
|---|---|---|---|---|
| chrX | 369411 | 369539 | 128 | N10:369398 |
| chrX | 369776 | 369927 | 151 | N10:369846 |
| chrX | 374096 | 374246 | 150 | N10:374118 |
| chrX | 380580 | 380750 | 170 | N10:380752 |
| chrX | 380580 | 380750 | 170 | N10:380571 |
| chrX | 380923 | 381086 | 163 | N10:380993 |
| chrX | 385357 | 385485 | 128 | |
| chrX | 386997 | 387161 | 164 | N10:387073 |
| chrX | 392484 | 392621 | 137 | N10:392516 |
| chrX | 392991 | 393143 | 152 | N10:393070 |
| chrX | 394180 | 394326 | 146 | N10:394214 |
| chrX | 394276 | 394372 | 96 | N10:394374 |
| chrX | 398245 | 398394 | 149 | N10:398328 |
| chrX | 399541 | 399701 | 160 | N10:399582 |
| chrX | 407672 | 407824 | 152 | N10:407809 |
| chrX | 410284 | 410430 | 146 | N10:410379 |
| chrX | 410723 | 410889 | 166 | N10:410731 |
| chrX | 412052 | 412213 | 161 | |
| chrX | 425378 | 425524 | 146 | |
| chrX | 429176 | 429332 | 156 | N10:429178 |
| chrX | 429194 | 429332 | 138 | N10:429178 |
| chrX | 429194 | 429339 | 145 | N10:429178 |
| chrX | 435614 | 435721 | 107 | N10:435670 |
| chrX | 435614 | 435731 | 117 | N10:435670 |
| chrX | 439795 | 439950 | 155 | N10:439773 |
| chrX | 447409 | 447538 | 129 | |
| chrX | 448364 | 448517 | 153 | N10:448344 |
| chrX | 456180 | 456328 | 148 | N10:456302 |
| chrX | 459326 | 459480 | 154 | N10:459411 |
| chrX | 465469 | 465620 | 151 | N10:465481 |
| chrX | 466504 | 466659 | 155 | N10:466505 |
| chrX | 467197 | 467335 | 138 | N10:467195 |
| chrX | 467559 | 467705 | 146 | N10:467683 |
| chrX | 470836 | 470984 | 148 | N10:470847 |
| chrX | 472163 | 472315 | 152 | N10:472226 |
| chrX | 472826 | 472975 | 149 | N10:472972 |
| chrX | 473000 | 473123 | 123 | N10:473146 |
| chrX | 473100 | 473255 | 155 | N10:473146 |
| chrX | 473533 | 473667 | 134 | N10:473586 |
| chrX | 473533 | 473673 | 140 | N10:473586 |
| chrX | 473729 | 473880 | 151 | N10:473881 |
| chrX | 473739 | 473884 | 145 | N10:473881 |
| chrX | 473757 | 473908 | 151 | N10:473881 |
| chrX | 474011 | 474165 | 154 | N10:474051 |
| chrX | 474045 | 474189 | 144 | N10:474051 |
| chrX | 474045 | 474189 | 144 | N10:474202 |
| chrX | 474576 | 474711 | 135 | |
| chrX | 475023 | 475169 | 146 | |
| chrX | 475023 | 475173 | 150 | |
| chrX | 475693 | 475822 | 129 | N10:475768 |
| chrX | 477337 | 477451 | 114 | |
| chrX | 477657 | 477804 | 147 | N10:477753 |
| chrX | 478238 | 478390 | 152 | N10:478237 |
| chrX | 478410 | 478559 | 149 | N10:478482 |
| chrX | 478560 | 478666 | 106 | |
| chrX | 478560 | 478678 | 118 | |
| chrX | 478584 | 478707 | 123 | N10:478712 |
| chrX | 478684 | 478839 | 155 | N10:478712 |
| chrX | 479117 | 479251 | 134 | |
| chrX | 479117 | 479257 | 140 | N10:479275 |
| chrX | 480160 | 480295 | 135 | N10:480180 |
| chrX | 480607 | 480757 | 150 | N10:480589 |
| chrX | 480786 | 480943 | 157 | N10:480816 |
| chrX | 481277 | 481406 | 129 | N10:481374 |
| chrX | 481757 | 481888 | 131 | N10:481781 |
| chrX | 482189 | 482339 | 150 | N10:482287 |
| chrX | 483140 | 483291 | 151 | N10:483274 |
| chrX | 495866 | 496005 | 139 | N10:495880 |
| chrX | 496369 | 496517 | 148 | N10:496351 |
| chrX | 498437 | 498584 | 147 | N10:498554 |
| chrX | 503072 | 503238 | 166 | N10:503242 |
| chrX | 503072 | 503238 | 166 | N10:503073 |
| chrX | 503922 | 504069 | 147 | N10:504008 |
| chrX | 512701 | 512853 | 152 | N10:512703 |
| chrX | 516708 | 516897 | 189 | N10:516725 |
| chrX | 519161 | 519304 | 143 | N10:519222 |
| chrX | 519732 | 519838 | 106 | N10:519795 |
| chrX | 519732 | 519858 | 126 | N10:519795 |
| chrX | 523978 | 524108 | 130 | N10:523988 |
| chrX | 526346 | 526498 | 152 | N10:526454 |
| chrX | 527259 | 527387 | 128 | N10:527394 |
| chrX | 530647 | 530795 | 148 | N10:530763 |
| chrX | 531614 | 531753 | 139 | N10:531763 |
| chrX | 533693 | 533845 | 152 | N10:533743 |
| chrX | 537746 | 537899 | 153 | N10:537764 |
| chrX | 542575 | 542723 | 148 | |
| chrX | 544311 | 544443 | 132 | N10:544344 |
| chrX | 547133 | 547290 | 157 | N10:547288 |
| chrX | 554602 | 554737 | 135 | N10:554644 |
| chrX | 557805 | 557945 | 140 | N10:557963 |
| chrX | 557805 | 557945 | 140 | N10:557804 |
| chrX | 557934 | 558083 | 149 | N10:557963 |
| chrX | 558445 | 558574 | 129 | N10:558562 |
| chrX | 561483 | 561587 | 104 | |
| chrX | 561700 | 561836 | 136 | N10:561697 |
| chrX | 563670 | 563822 | 152 | N10:563737 |
| chrX | 567134 | 567269 | 135 | N10:567226 |
| chrX | 569137 | 569291 | 154 | N10:569134 |
| chrX | 572775 | 572903 | 128 | |
| chrX | 573799 | 573959 | 160 | N10:573951 |
| chrX | 582808 | 582958 | 150 | |
| chrX | 586031 | 586180 | 149 | N10:586162 |
| chrX | 590874 | 591029 | 155 | N10:591040 |

| CHR | Start | End | Length (bp) | Nucleosome_ID |
|---|---|---|---|---|
| chrX | 591957 | 592113 | 156 | N10:592071 |
| chrX | 594961 | 595112 | 151 | N10:594945 |
| chrX | 595496 | 595660 | 164 | N10:595504 |
| chrX | 598239 | 598403 | 164 | N10:598323 |
| chrX | 598789 | 598947 | 158 | N10:598769 |
| chrX | 600553 | 600699 | 146 | N10:600632 |
| chrX | 606604 | 606728 | 124 | N10:606660 |
| chrX | 608387 | 608544 | 157 | N10:608402 |
| chrX | 608709 | 608811 | 102 | N10:608690 |
| chrX | 610999 | 611149 | 150 | N10:610987 |
| chrX | 623512 | 623664 | 152 | N10:623582 |
| chrX | 624211 | 624342 | 131 | N10:624345 |
| chrX | 624949 | 625089 | 140 | N10:625080 |
| chrX | 632590 | 632738 | 148 | N10:632669 |
| chrX | 635827 | 635966 | 139 | N10:635846 |
| chrX | 636181 | 636337 | 156 | N10:636350 |
| chrX | 636181 | 636337 | 156 | N10:636186 |
| chrX | 636714 | 636864 | 150 | N10:636770 |
| chrX | 637154 | 637305 | 151 | N10:637147 |
| chrX | 637154 | 637305 | 151 | N10:637320 |
| chrX | 638977 | 639126 | 149 | N10:638974 |
| chrX | 653069 | 653214 | 145 | N10:653126 |
| chrX | 661255 | 661420 | 165 | N10:661267 |
| chrX | 662029 | 662161 | 132 | N10:662110 |
| chrX | 662034 | 662161 | 127 | N10:662110 |
| chrX | 662037 | 662135 | 98 | N10:662110 |
| chrX | 671104 | 671223 | 119 | |
| chrX | 671901 | 672050 | 149 | |
| chrX | 673361 | 673549 | 188 | |
| chrX | 673385 | 673539 | 154 | |
| chrX | 674527 | 674676 | 149 | N10:674605 |
| chrX | 684490 | 684642 | 152 | N10:684478 |
| chrX | 684490 | 684642 | 152 | N10:684641 |
| chrX | 685942 | 686069 | 127 | N10:686043 |
| chrX | 687031 | 687188 | 157 | N10:687180 |
| chrX | 687681 | 687842 | 161 | N10:687764 |
| chrX | 689548 | 689698 | 150 | N10:689678 |
| chrX | 695637 | 695789 | 152 | N10:695637 |
| chrX | 696260 | 696391 | 131 | N10:696342 |
| chrX | 697538 | 697692 | 154 | N10:697604 |
| chrX | 699421 | 699528 | 107 | N10:699480 |
| chrX | 699421 | 699559 | 138 | N10:699480 |
| chrX | 699421 | 699560 | 139 | N10:699480 |
| chrX | 699421 | 699561 | 140 | N10:699480 |
| chrX | 699421 | 699563 | 142 | N10:699480 |
| chrX | 699421 | 699564 | 143 | N10:699480 |
| chrX | 700845 | 700991 | 146 | N10:700894 |
| chrX | 701322 | 701470 | 148 | N10:701322 |
| chrX | 705757 | 705926 | 169 | N10:705760 |
| chrX | 705757 | 705926 | 169 | N10:705934 |
| chrX | 706905 | 707066 | 161 | N10:707000 |
| chrX | 712931 | 713070 | 139 | N10:712913 |
| chrX | 714988 | 715121 | 133 | N10:715105 |
| chrX | 720830 | 720969 | 139 | N10:720969 |
| chrX | 721017 | 721124 | 107 | |
| chrX | 721017 | 721148 | 131 | |
| chrX | 721017 | 721149 | 132 | |
| chrX | 721017 | 721153 | 136 | |
| chrX | 721018 | 721153 | 135 | |
| chrX | 721019 | 721153 | 134 | |
| chrX | 721020 | 721153 | 133 | |
| chrX | 723086 | 723238 | 152 | N10:723143 |
| chrX | 723301 | 723431 | 130 | N10:723416 |
| chrX | 723442 | 723594 | 152 | N10:723587 |
| chrX | 725588 | 725741 | 153 | N10:725606 |
| chrX | 729427 | 729584 | 157 | N10:729606 |
| chrX | 729427 | 729584 | 157 | N10:729424 |
| chrX | 729799 | 729953 | 154 | |
| chrX | 732737 | 732877 | 140 | N10:732736 |
| chrX | 735177 | 735331 | 154 | N10:735172 |
| chrX | 735177 | 735331 | 154 | N10:735346 |
| chrX | 736882 | 737013 | 131 | N10:736967 |
| chrX | 739148 | 739242 | 94 | N10:739259 |
| chrX | 745644 | 745739 | 95 | |
| chrX | 745644 | 745742 | 98 | |
| chrXI | 6938 | 7102 | 164 | N11:6990 |
| chrXI | 10271 | 10420 | 149 | N11:10428 |
| chrXI | 17984 | 18132 | 148 | N11:18117 |
| chrXI | 19266 | 19389 | 123 | N11:19336 |
| chrXI | 19267 | 19389 | 122 | N11:19336 |
| chrXI | 19267 | 19414 | 147 | N11:19336 |
| chrXI | 20703 | 20861 | 158 | N11:20761 |
| chrXI | 31510 | 31641 | 131 | |
| chrXI | 32617 | 32778 | 161 | N11:32717 |
| chrXI | 38219 | 38379 | 160 | |
| chrXI | 39078 | 39236 | 158 | N11:39170 |
| chrXI | 41766 | 41916 | 150 | |
| chrXI | 42476 | 42618 | 142 | N11:42545 |
| chrXI | 42479 | 42618 | 139 | N11:42545 |
| chrXI | 44129 | 44285 | 156 | N11:44262 |
| chrXI | 45474 | 45591 | 117 | |
| chrXI | 52845 | 53010 | 165 | N11:52908 |
| chrXI | 53086 | 53185 | 99 | N11:53095 |
| chrXI | 55429 | 55585 | 156 | N11:55522 |
| chrXI | 55463 | 55623 | 160 | N11:55522 |
| chrXI | 55924 | 56074 | 150 | N11:55980 |
| chrXI | 55924 | 56080 | 156 | N11:55980 |
| chrXI | 56846 | 56978 | 132 | N11:56861 |
| chrXI | 61848 | 62032 | 184 | N11:61896 |
| chrXI | 63877 | 64030 | 153 | N11:63947 |
| chrXI | 65154 | 65316 | 162 | N11:65282 |
| chrXI | 65672 | 65838 | 166 | N11:65746 |
| chrXI | 68614 | 68763 | 149 | N11:68734 |

TABLE 1

| CHR | Start | End | Length (bp) | Nucleosome_ID |
|---|---|---|---|---|
| chrXI | 73102 | 73261 | 159 | N11:73185 |
| chrXI | 86463 | 86611 | 148 | N11:86556 |
| chrXI | 86483 | 86645 | 162 | N11:86556 |
| chrXI | 88336 | 88484 | 148 | N11:88371 |
| chrXI | 89789 | 89957 | 168 | N11:89946 |
| chrXI | 89789 | 89957 | 168 | N11:89782 |
| chrXI | 91032 | 91184 | 152 | N11:91137 |
| chrXI | 97596 | 97707 | 111 | N11:97729 |
| chrXI | 97601 | 97707 | 106 | N11:97729 |
| chrXI | 99816 | 99946 | 130 | N11:99812 |
| chrXI | 99816 | 99946 | 130 | N11:99961 |
| chrXI | 103049 | 103197 | 148 | N11:103034 |
| chrXI | 103049 | 103197 | 148 | N11:103200 |
| chrXI | 103372 | 103526 | 154 | N11:103384 |
| chrXI | 105256 | 105405 | 149 | N11:105284 |
| chrXI | 105988 | 106134 | 146 | N11:106069 |
| chrXI | 110658 | 110811 | 153 | N11:110802 |
| chrXI | 115225 | 115382 | 157 | N11:115291 |
| chrXI | 116126 | 116262 | 136 | N11:116268 |
| chrXI | 116126 | 116262 | 136 | N11:116112 |
| chrXI | 116231 | 116308 | 77 | N11:116268 |
| chrXI | 116232 | 116308 | 76 | N11:116268 |
| chrXI | 116232 | 116357 | 125 | N11:116268 |
| chrXI | 117579 | 117716 | 137 | N11:117661 |
| chrXI | 120342 | 120499 | 157 | N11:120460 |
| chrXI | 120705 | 120857 | 152 | N11:120781 |
| chrXI | 121556 | 121713 | 157 | N11:121624 |
| chrXI | 130706 | 130855 | 149 | N11:130776 |
| chrXI | 130929 | 131079 | 150 | N11:130940 |
| chrXI | 132985 | 133151 | 166 | N11:133074 |
| chrXI | 134033 | 134163 | 130 | N11:134102 |
| chrXI | 136517 | 136676 | 159 | |
| chrXI | 140618 | 140768 | 150 | N11:140692 |
| chrXI | 142672 | 142804 | 132 | N11:142756 |
| chrXI | 145250 | 145428 | 178 | N11:145382 |
| chrXI | 146568 | 146719 | 151 | N11:146644 |
| chrXI | 147356 | 147520 | 164 | N11:147421 |
| chrXI | 149408 | 149568 | 160 | N11:149508 |
| chrXI | 149964 | 150108 | 144 | N11:149984 |
| chrXI | 151755 | 151890 | 135 | N11:151768 |
| chrXI | 161235 | 161384 | 149 | N11:161325 |
| chrXI | 161799 | 161928 | 129 | N11:161887 |
| chrXI | 162802 | 162952 | 150 | N11:162881 |
| chrXI | 170837 | 170984 | 147 | N11:170954 |
| chrXI | 171758 | 171891 | 133 | N11:171815 |
| chrXI | 172320 | 172456 | 136 | N11:172446 |
| chrXI | 178039 | 178227 | 188 | N11:178152 |
| chrXI | 178982 | 179138 | 156 | N11:179070 |
| chrXI | 179058 | 179209 | 151 | N11:179232 |
| chrXI | 179058 | 179209 | 151 | N11:179070 |
| chrXI | 182120 | 182298 | 178 | N11:182189 |
| chrXI | 183536 | 183690 | 154 | N11:183621 |
| chrXI | 183964 | 184081 | 117 | N11:183955 |
| chrXI | 187208 | 187352 | 144 | N11:187276 |
| chrXI | 188029 | 188179 | 150 | N11:188071 |
| chrXI | 190331 | 190479 | 148 | N11:190473 |
| chrXI | 191024 | 191171 | 147 | N11:191119 |
| chrXI | 195322 | 195449 | 127 | N11:195406 |
| chrXI | 196316 | 196481 | 165 | N11:196493 |
| chrXI | 196620 | 196752 | 132 | N11:196660 |
| chrXI | 201937 | 202099 | 162 | N11:202016 |
| chrXI | 204817 | 204983 | 166 | N11:204909 |
| chrXI | 207618 | 207770 | 152 | |
| chrXI | 209750 | 209876 | 126 | N11:209839 |
| chrXI | 214868 | 215018 | 150 | N11:214868 |
| chrXI | 214868 | 215019 | 151 | N11:214868 |
| chrXI | 214886 | 215018 | 132 | N11:214868 |
| chrXI | 214886 | 215019 | 133 | N11:214868 |
| chrXI | 216300 | 216452 | 152 | N11:216342 |
| chrXI | 216511 | 216657 | 146 | N11:216610 |
| chrXI | 226833 | 226983 | 150 | N11:226966 |
| chrXI | 226930 | 227062 | 132 | N11:226966 |
| chrXI | 227206 | 227326 | 120 | |
| chrXI | 229813 | 229973 | 160 | N11:229834 |
| chrXI | 233634 | 233784 | 150 | N11:233638 |
| chrXI | 233634 | 233784 | 150 | N11:233802 |
| chrXI | 236594 | 236764 | 170 | N11:236682 |
| chrXI | 241460 | 241610 | 150 | N11:241499 |
| chrXI | 245174 | 245347 | 173 | N11:245177 |
| chrXI | 245189 | 245339 | 150 | N11:245177 |
| chrXI | 245551 | 245725 | 174 | N11:245572 |
| chrXI | 247204 | 247369 | 165 | N11:247366 |
| chrXI | 247204 | 247369 | 165 | N11:247197 |
| chrXI | 249638 | 249773 | 135 | N11:249715 |
| chrXI | 251126 | 251263 | 137 | N11:251198 |
| chrXI | 253186 | 253299 | 113 | N11:253297 |
| chrXI | 253541 | 253688 | 147 | N11:253657 |
| chrXI | 257352 | 257460 | 108 | |
| chrXI | 257353 | 257460 | 107 | |
| chrXI | 257353 | 257478 | 125 | N11:257490 |
| chrXI | 257617 | 257767 | 150 | N11:257744 |
| chrXI | 258177 | 258336 | 159 | N11:258256 |
| chrXI | 261407 | 261543 | 136 | N11:261504 |
| chrXI | 269387 | 269539 | 152 | N11:269548 |
| chrXI | 269387 | 269539 | 152 | N11:269374 |
| chrXI | 270529 | 270668 | 139 | N11:270660 |
| chrXI | 275707 | 275869 | 162 | N11:275750 |
| chrXI | 275877 | 276030 | 153 | N11:276025 |
| chrXI | 277324 | 277474 | 150 | N11:277389 |
| chrXI | 278497 | 278637 | 140 | N11:278487 |
| chrXI | 278499 | 278637 | 138 | N11:278487 |
| chrXI | 278501 | 278637 | 136 | N11:278487 |
| chrXI | 278502 | 278637 | 135 | N11:278487 |

| CHR | Start | End | Length (bp) | Nucleosome_ID |
|---|---|---|---|---|
| chrXI | 282993 | 283156 | 163 | N11:283075 |
| chrXI | 286651 | 286824 | 173 | N11:286733 |
| chrXI | 287603 | 287693 | 90 | N11:287704 |
| chrXI | 288326 | 288474 | 148 | |
| chrXI | 290076 | 290228 | 152 | N11:290108 |
| chrXI | 292965 | 293112 | 147 | N11:293063 |
| chrXI | 297006 | 297125 | 119 | N11:297036 |
| chrXI | 297006 | 297156 | 150 | N11:297036 |
| chrXI | 298702 | 298832 | 130 | N11:298779 |
| chrXI | 298875 | 299007 | 132 | |
| chrXI | 300199 | 300305 | 106 | N11:300189 |
| chrXI | 300199 | 300317 | 118 | N11:300189 |
| chrXI | 303921 | 304068 | 147 | N11:304040 |
| chrXI | 305787 | 305949 | 162 | |
| chrXI | 318927 | 319072 | 145 | |
| chrXI | 319054 | 319202 | 148 | N11:319121 |
| chrXI | 322500 | 322670 | 170 | N11:322551 |
| chrXI | 329617 | 329749 | 132 | |
| chrXI | 329629 | 329749 | 120 | |
| chrXI | 329629 | 329775 | 146 | |
| chrXI | 337370 | 337525 | 155 | N11:337362 |
| chrXI | 337447 | 337600 | 153 | |
| chrXI | 337450 | 337600 | 150 | |
| chrXI | 337452 | 337600 | 148 | |
| chrXI | 337462 | 337600 | 138 | |
| chrXI | 337968 | 338118 | 150 | |
| chrXI | 340261 | 340401 | 140 | N11:340305 |
| chrXI | 341684 | 341830 | 146 | N11:341720 |
| chrXI | 347279 | 347450 | 171 | N11:347369 |
| chrXI | 347320 | 347467 | 147 | N11:347369 |
| chrXI | 347479 | 347608 | 129 | N11:347544 |
| chrXI | 351244 | 351421 | 177 | |
| chrXI | 351885 | 352030 | 145 | N11:351957 |
| chrXI | 352745 | 352882 | 137 | N11:352864 |
| chrXI | 353147 | 353287 | 140 | N11:353269 |
| chrXI | 355288 | 355448 | 160 | |
| chrXI | 357460 | 357614 | 154 | N11:357564 |
| chrXI | 357462 | 357614 | 152 | N11:357564 |
| chrXI | 362917 | 363070 | 153 | |
| chrXI | 364062 | 364210 | 148 | |
| chrXI | 364602 | 364723 | 121 | N11:364623 |
| chrXI | 368064 | 368220 | 156 | |
| chrXI | 368389 | 368522 | 133 | N11:368440 |
| chrXI | 368971 | 369078 | 107 | N11:368963 |
| chrXI | 368971 | 369121 | 150 | N11:368963 |
| chrXI | 368971 | 369122 | 151 | N11:368963 |
| chrXI | 372978 | 373087 | 109 | N11:372980 |
| chrXI | 373360 | 373511 | 151 | N11:373499 |
| chrXI | 377591 | 377751 | 160 | N11:377588 |
| chrXI | 379254 | 379407 | 153 | N11:379329 |
| chrXI | 385681 | 385837 | 156 | N11:385749 |
| chrXI | 394646 | 394790 | 144 | N11:394794 |
| chrXI | 397867 | 398011 | 144 | N11:398025 |
| chrXI | 397867 | 398011 | 144 | N11:397862 |
| chrXI | 399301 | 399460 | 159 | |
| chrXI | 400364 | 400506 | 142 | N11:400496 |
| chrXI | 401469 | 401625 | 156 | N11:401550 |
| chrXI | 401694 | 401852 | 158 | N11:401711 |
| chrXI | 403288 | 403463 | 175 | N11:403356 |
| chrXI | 406796 | 406955 | 159 | N11:406832 |
| chrXI | 407916 | 408065 | 149 | |
| chrXI | 416183 | 416327 | 144 | N11:416221 |
| chrXI | 417925 | 418084 | 159 | N11:417916 |
| chrXI | 417925 | 418084 | 159 | N11:418102 |
| chrXI | 434342 | 434496 | 154 | N11:434396 |
| chrXI | 434529 | 434684 | 155 | N11:434609 |
| chrXI | 434561 | 434692 | 131 | N11:434609 |
| chrXI | 434662 | 434792 | 130 | |
| chrXI | 439672 | 439806 | 134 | N11:439814 |
| chrXI | 439682 | 439780 | 98 | |
| chrXI | 447937 | 448094 | 157 | N11:447938 |
| chrXI | 448453 | 448602 | 149 | N11:448490 |
| chrXI | 450755 | 450913 | 158 | N11:450835 |
| chrXI | 454604 | 454760 | 156 | N11:454730 |
| chrXI | 455031 | 455138 | 107 | |
| chrXI | 455031 | 455148 | 117 | N11:455162 |
| chrXI | 456736 | 456888 | 152 | N11:456824 |
| chrXI | 459229 | 459336 | 107 | N11:459206 |
| chrXI | 459229 | 459341 | 112 | N11:459206 |
| chrXI | 462337 | 462501 | 164 | N11:462401 |
| chrXI | 468677 | 468825 | 148 | |
| chrXI | 470132 | 470274 | 142 | |
| chrXI | 473335 | 473488 | 153 | N11:473478 |
| chrXI | 479959 | 480086 | 127 | N11:479984 |
| chrXI | 483328 | 483476 | 148 | N11:483388 |
| chrXI | 484474 | 484623 | 149 | N11:484589 |
| chrXI | 488055 | 488207 | 152 | N11:488122 |
| chrXI | 488418 | 488598 | 180 | N11:488497 |
| chrXI | 496967 | 497107 | 140 | N11:497088 |
| chrXI | 501192 | 501346 | 154 | N11:501317 |
| chrXI | 506644 | 506790 | 146 | N11:506685 |
| chrXI | 509826 | 509992 | 166 | N11:509894 |
| chrXI | 514096 | 514200 | 104 | |
| chrXI | 516175 | 516336 | 161 | N11:516248 |
| chrXI | 517302 | 517453 | 151 | N11:517465 |
| chrXI | 517302 | 517453 | 151 | N11:517303 |
| chrXI | 517458 | 517572 | 114 | N11:517465 |
| chrXI | 519553 | 519695 | 142 | N11:519598 |
| chrXI | 527505 | 527640 | 135 | N11:527634 |
| chrXI | 527506 | 527600 | 94 | |
| chrXI | 527506 | 527630 | 124 | N11:527634 |
| chrXI | 527506 | 527632 | 126 | N11:527634 |
| chrXI | 527506 | 527635 | 129 | N11:527634 |

| CHR | Start | End | Length (bp) | Nucleosome_ID |
|---|---|---|---|---|
| chrXI | 527506 | 527636 | 130 | N11:527634 |
| chrXI | 527506 | 527637 | 131 | N11:527634 |
| chrXI | 527506 | 527638 | 132 | N11:527634 |
| chrXI | 527506 | 527639 | 133 | N11:527634 |
| chrXI | 527506 | 527640 | 134 | N11:527634 |
| chrXI | 527506 | 527641 | 135 | N11:527634 |
| chrXI | 527507 | 527640 | 133 | N11:527634 |
| chrXI | 527508 | 527640 | 132 | N11:527634 |
| chrXI | 527509 | 527640 | 131 | N11:527634 |
| chrXI | 527510 | 527640 | 130 | N11:527634 |
| chrXI | 527511 | 527640 | 129 | N11:527634 |
| chrXI | 527513 | 527640 | 127 | N11:527634 |
| chrXI | 527515 | 527640 | 125 | N11:527634 |
| chrXI | 527516 | 527640 | 124 | N11:527634 |
| chrXI | 529083 | 529220 | 137 | N11:529204 |
| chrXI | 529403 | 529549 | 146 | N11:529570 |
| chrXI | 532781 | 532933 | 152 | N11:532891 |
| chrXI | 533171 | 533320 | 149 | N11:533271 |
| chrXI | 540415 | 540570 | 155 | N11:540451 |
| chrXI | 540630 | 540787 | 157 | N11:540698 |
| chrXI | 540646 | 540797 | 151 | N11:540698 |
| chrXI | 541723 | 541874 | 151 | N11:541853 |
| chrXI | 542290 | 542416 | 126 | N11:542354 |
| chrXI | 542290 | 542440 | 150 | N11:542354 |
| chrXI | 544676 | 544823 | 147 | |
| chrXI | 545722 | 545853 | 131 | N11:545860 |
| chrXI | 547817 | 547975 | 158 | N11:547848 |
| chrXI | 548333 | 548481 | 148 | N11:548344 |
| chrXI | 548769 | 548931 | 162 | N11:548799 |
| chrXI | 548769 | 548931 | 162 | N11:548953 |
| chrXI | 549757 | 549907 | 150 | N11:549905 |
| chrXI | 550284 | 550422 | 138 | N11:550349 |
| chrXI | 554400 | 554551 | 151 | N11:554437 |
| chrXI | 558933 | 559085 | 152 | N11:558954 |
| chrXI | 562000 | 562173 | 173 | N11:562186 |
| chrXI | 571010 | 571152 | 142 | N11:571147 |
| chrXI | 573165 | 573306 | 141 | N11:573269 |
| chrXI | 574030 | 574178 | 148 | N11:574099 |
| chrXI | 574402 | 574555 | 153 | N11:574493 |
| chrXI | 575707 | 575877 | 170 | |
| chrXI | 577254 | 577388 | 134 | N11:577250 |
| chrXI | 577351 | 577483 | 132 | |
| chrXI | 577690 | 577817 | 127 | N11:577711 |
| chrXI | 578476 | 578613 | 137 | N11:578514 |
| chrXI | 579137 | 579295 | 158 | N11:579194 |
| chrXI | 590716 | 590822 | 106 | N11:590776 |
| chrXI | 590716 | 590835 | 119 | N11:590776 |
| chrXI | 600361 | 600514 | 153 | N11:600370 |
| chrXI | 600361 | 600514 | 153 | N11:600524 |
| chrXI | 601868 | 602039 | 171 | N11:601908 |
| chrXI | 602067 | 602222 | 155 | N11:602124 |
| chrXI | 603803 | 603944 | 141 | N11:603837 |
| chrXI | 604749 | 604856 | 107 | |
| chrXI | 604749 | 604865 | 116 | |
| chrXI | 607498 | 607646 | 148 | N11:607543 |
| chrXI | 607998 | 608143 | 145 | N11:608050 |
| chrXI | 613167 | 613294 | 127 | N11:613186 |
| chrXI | 614331 | 614461 | 130 | N11:614435 |
| chrXI | 618644 | 618800 | 156 | N11:618766 |
| chrXI | 621479 | 621632 | 153 | N11:621620 |
| chrXI | 624990 | 625141 | 151 | N11:625071 |
| chrXI | 632085 | 632240 | 155 | N11:632159 |
| chrXI | 632617 | 632753 | 136 | N11:632714 |
| chrXI | 638182 | 638313 | 131 | N11:638195 |
| chrXI | 638182 | 638316 | 134 | N11:638195 |
| chrXI | 643087 | 643244 | 157 | N11:643162 |
| chrXI | 646189 | 646315 | 126 | N11:646259 |
| chrXI | 647330 | 647482 | 152 | N11:647430 |
| chrXI | 648395 | 648512 | 117 | |
| chrXI | 650461 | 650612 | 151 | N11:650494 |
| chrXI | 651142 | 651292 | 150 | N11:651252 |
| chrXI | 654048 | 654197 | 149 | |
| chrXI | 655704 | 655870 | 166 | N11:655848 |
| chrXI | 655763 | 655920 | 157 | N11:655848 |
| chrXI | 657176 | 657328 | 152 | N11:657284 |
| chrXI | 658054 | 658203 | 149 | N11:658207 |
| chrXI | 659842 | 659965 | 123 | N11:659867 |
| chrXI | 661850 | 662006 | 156 | N11:662001 |
| chrXI | 663739 | 663846 | 107 | N11:663807 |
| chrXI | 663739 | 663851 | 112 | N11:663807 |
| chrXI | 664373 | 664519 | 146 | N11:664394 |
| chrXI | 666612 | 666748 | 136 | |
| chrXI | 666628 | 666815 | 187 | |
| chrXI | 666656 | 666815 | 159 | |
| chrXI | 666681 | 666771 | 90 | |
| chrXI | 666681 | 666810 | 129 | |
| chrXI | 666693 | 666810 | 117 | |
| chrXI | 666703 | 666728 | 25 | |
| chrXI | 666703 | 666733 | 30 | |
| chrXI | 666703 | 666737 | 34 | |
| chrXI | 666703 | 666739 | 36 | |
| chrXI | 666709 | 666792 | 83 | |
| chrXI | 666709 | 666810 | 101 | |
| chrXI | 666709 | 666815 | 106 | |
| chrXI | 666714 | 666815 | 101 | |
| chrXII | 723 | 874 | 151 | N12:805 |
| chrXII | 1167 | 1315 | 148 | |
| chrXII | 1996 | 2147 | 151 | N12:2014 |
| chrXII | 2008 | 2142 | 134 | N12:2014 |
| chrXII | 2033 | 2129 | 96 | N12:2014 |
| chrXII | 2033 | 2193 | 160 | N12:2177 |
| chrXII | 2033 | 2193 | 160 | N12:2014 |
| chrXII | 2211 | 2359 | 148 | N12:2350 |

TABLE 1

| CHR | Start | End | Length (bp) | Nucleosome_ID |
|---|---|---|---|---|
| chrXII | 2226 | 2355 | 129 | N12:2350 |
| chrXII | 2226 | 2359 | 133 | N12:2350 |
| chrXII | 2226 | 2421 | 195 | N12:2350 |
| chrXII | 2387 | 2517 | 130 | |
| chrXII | 2666 | 2818 | 152 | N12:2754 |
| chrXII | 3013 | 3157 | 144 | N12:3070 |
| chrXII | 3013 | 3200 | 187 | N12:3070 |
| chrXII | 3065 | 3157 | 92 | N12:3070 |
| chrXII | 3065 | 3200 | 135 | N12:3070 |
| chrXII | 3153 | 3297 | 144 | N12:3257 |
| chrXII | 3218 | 3370 | 152 | N12:3257 |
| chrXII | 3470 | 3595 | 125 | N12:3583 |
| chrXII | 3500 | 3643 | 143 | N12:3583 |
| chrXII | 3500 | 3651 | 151 | N12:3583 |
| chrXII | 3791 | 3928 | 137 | N12:3868 |
| chrXII | 6346 | 6517 | 171 | N12:6439 |
| chrXII | 6366 | 6517 | 151 | N12:6439 |
| chrXII | 6690 | 6864 | 174 | N12:6762 |
| chrXII | 6705 | 6857 | 152 | N12:6762 |
| chrXII | 7495 | 7682 | 187 | N12:7546 |
| chrXII | 7531 | 7682 | 151 | N12:7546 |
| chrXII | 7543 | 7677 | 134 | N12:7546 |
| chrXII | 7568 | 7664 | 96 | N12:7546 |
| chrXII | 7746 | 7894 | 148 | N12:7816 |
| chrXII | 7761 | 7894 | 133 | N12:7816 |
| chrXII | 7761 | 7956 | 195 | N12:7816 |
| chrXII | 7922 | 8052 | 130 | |
| chrXII | 8201 | 8353 | 152 | N12:8284 |
| chrXII | 8548 | 8692 | 144 | N12:8607 |
| chrXII | 8548 | 8735 | 187 | N12:8607 |
| chrXII | 8600 | 8692 | 92 | N12:8607 |
| chrXII | 8600 | 8735 | 135 | N12:8607 |
| chrXII | 8688 | 8832 | 144 | N12:8786 |
| chrXII | 8753 | 8905 | 152 | N12:8786 |
| chrXII | 9005 | 9130 | 125 | N12:9112 |
| chrXII | 9035 | 9178 | 143 | N12:9112 |
| chrXII | 9326 | 9463 | 137 | N12:9402 |
| chrXII | 11195 | 11357 | 162 | N12:11275 |
| chrXII | 11198 | 11354 | 156 | N12:11275 |
| chrXII | 11205 | 11354 | 149 | N12:11275 |
| chrXII | 11205 | 11357 | 152 | N12:11275 |
| chrXII | 11207 | 11357 | 150 | N12:11275 |
| chrXII | 11208 | 11354 | 146 | N12:11275 |
| chrXII | 11211 | 11357 | 146 | N12:11275 |
| chrXII | 11213 | 11310 | 97 | N12:11275 |
| chrXII | 11213 | 11346 | 133 | N12:11275 |
| chrXII | 11213 | 11352 | 139 | N12:11275 |
| chrXII | 11213 | 11355 | 142 | N12:11275 |
| chrXII | 11213 | 11357 | 144 | N12:11275 |
| chrXII | 11213 | 11358 | 145 | N12:11275 |
| chrXII | 11218 | 11354 | 136 | N12:11275 |
| chrXII | 11222 | 11354 | 132 | N12:11275 |
| chrXII | 11222 | 11357 | 135 | N12:11275 |
| chrXII | 11226 | 11354 | 128 | N12:11275 |
| chrXII | 11226 | 11357 | 131 | N12:11275 |
| chrXII | 11226 | 11358 | 132 | N12:11275 |
| chrXII | 11227 | 11357 | 130 | N12:11275 |
| chrXII | 11228 | 11357 | 129 | N12:11275 |
| chrXII | 11229 | 11357 | 128 | N12:11275 |
| chrXII | 11230 | 11357 | 127 | N12:11275 |
| chrXII | 11231 | 11354 | 123 | N12:11275 |
| chrXII | 11231 | 11357 | 126 | N12:11275 |
| chrXII | 11231 | 11358 | 127 | N12:11275 |
| chrXII | 11233 | 11357 | 124 | N12:11275 |
| chrXII | 11235 | 11357 | 122 | N12:11275 |
| chrXII | 11252 | 11354 | 102 | N12:11275 |
| chrXII | 11252 | 11357 | 105 | N12:11275 |
| chrXII | 11259 | 11357 | 98 | N12:11275 |
| chrXII | 11262 | 11357 | 95 | N12:11275 |
| chrXII | 11273 | 11357 | 84 | N12:11275 |
| chrXII | 13256 | 13420 | 164 | N12:13360 |
| chrXII | 13298 | 13427 | 129 | N12:13360 |
| chrXII | 13457 | 13612 | 155 | N12:13509 |
| chrXII | 15085 | 15257 | 172 | N12:15154 |
| chrXII | 15201 | 15361 | 160 | N12:15316 |
| chrXII | 16724 | 16865 | 141 | N12:16731 |
| chrXII | 18847 | 18996 | 149 | N12:18970 |
| chrXII | 25117 | 25257 | 140 | N12:25169 |
| chrXII | 27714 | 27865 | 151 | N12:27788 |
| chrXII | 35557 | 35688 | 131 | N12:35640 |
| chrXII | 39250 | 39399 | 149 | N12:39315 |
| chrXII | 48577 | 48723 | 146 | N12:48603 |
| chrXII | 51782 | 51932 | 150 | N12:51881 |
| chrXII | 56756 | 56881 | 125 | N12:56813 |
| chrXII | 57933 | 58079 | 146 | N12:58069 |
| chrXII | 60326 | 60478 | 152 | N12:60341 |
| chrXII | 62223 | 62381 | 158 | |
| chrXII | 67309 | 67460 | 151 | |
| chrXII | 70032 | 70180 | 148 | N12:70102 |
| chrXII | 70780 | 70917 | 137 | N12:70939 |
| chrXII | 70780 | 70917 | 137 | N12:70773 |
| chrXII | 71343 | 71501 | 158 | N12:71413 |
| chrXII | 73489 | 73643 | 154 | N12:73587 |
| chrXII | 79801 | 79962 | 161 | |
| chrXII | 99031 | 99175 | 144 | N12:99103 |
| chrXII | 104748 | 104900 | 152 | N12:104799 |
| chrXII | 104994 | 105164 | 170 | N12:104983 |
| chrXII | 106767 | 106921 | 154 | N12:106904 |
| chrXII | 107154 | 107242 | 88 | N12:107195 |
| chrXII | 107154 | 107250 | 96 | N12:107195 |
| chrXII | 107154 | 107252 | 98 | N12:107195 |
| chrXII | 107154 | 107253 | 99 | N12:107195 |
| chrXII | 107154 | 107255 | 101 | N12:107195 |

TABLE 1

| CHR | Start | End | Length (bp) | Nucleosome_ID |
|---|---|---|---|---|
| chrXII | 107154 | 107257 | 103 | N12:107195 |
| chrXII | 107154 | 107258 | 104 | N12:107195 |
| chrXII | 107154 | 107259 | 105 | N12:107195 |
| chrXII | 107154 | 107260 | 106 | N12:107195 |
| chrXII | 107154 | 107261 | 107 | N12:107195 |
| chrXII | 107154 | 107262 | 108 | N12:107195 |
| chrXII | 107154 | 107265 | 111 | N12:107195 |
| chrXII | 107154 | 107267 | 113 | N12:107195 |
| chrXII | 107154 | 107269 | 115 | N12:107195 |
| chrXII | 107154 | 107270 | 116 | N12:107195 |
| chrXII | 107154 | 107271 | 117 | N12:107195 |
| chrXII | 107154 | 107273 | 119 | N12:107195 |
| chrXII | 107154 | 107274 | 120 | N12:107195 |
| chrXII | 107154 | 107285 | 131 | N12:107195 |
| chrXII | 107271 | 107423 | 152 | N12:107365 |
| chrXII | 110633 | 110799 | 166 | N12:110737 |
| chrXII | 112506 | 112634 | 128 | N12:112551 |
| chrXII | 113051 | 113206 | 155 | N12:113139 |
| chrXII | 118248 | 118400 | 152 | N12:118377 |
| chrXII | 119597 | 119738 | 141 | N12:119672 |
| chrXII | 121717 | 121889 | 172 | N12:121848 |
| chrXII | 126647 | 126799 | 152 | N12:126629 |
| chrXII | 132281 | 132417 | 136 | |
| chrXII | 132310 | 132466 | 156 | |
| chrXII | 133301 | 133451 | 150 | N12:133308 |
| chrXII | 133762 | 133946 | 184 | N12:133881 |
| chrXII | 136653 | 136779 | 126 | N12:136749 |
| chrXII | 138426 | 138582 | 156 | N12:138554 |
| chrXII | 142345 | 142492 | 147 | N12:142429 |
| chrXII | 143973 | 144111 | 138 | N12:144010 |
| chrXII | 144765 | 144914 | 149 | N12:144832 |
| chrXII | 145430 | 145576 | 146 | N12:145533 |
| chrXII | 155097 | 155220 | 123 | N12:155205 |
| chrXII | 160945 | 161106 | 161 | N12:161017 |
| chrXII | 163470 | 163588 | 118 | N12:163522 |
| chrXII | 171732 | 171883 | 151 | N12:171897 |
| chrXII | 177830 | 177964 | 134 | N12:177819 |
| chrXII | 177830 | 177964 | 134 | N12:177986 |
| chrXII | 178588 | 178741 | 153 | |
| chrXII | 179076 | 179226 | 150 | N12:179114 |
| chrXII | 180676 | 180836 | 160 | N12:180728 |
| chrXII | 183315 | 183461 | 146 | N12:183389 |
| chrXII | 192705 | 192836 | 131 | N12:192762 |
| chrXII | 193019 | 193125 | 106 | N12:193086 |
| chrXII | 193019 | 193144 | 125 | N12:193086 |
| chrXII | 194389 | 194550 | 161 | N12:194429 |
| chrXII | 197350 | 197482 | 132 | N12:197391 |
| chrXII | 197802 | 197965 | 163 | |
| chrXII | 199062 | 199223 | 161 | N12:199130 |
| chrXII | 199271 | 199406 | 135 | N12:199299 |
| chrXII | 199721 | 199871 | 150 | N12:199806 |
| chrXII | 207917 | 208073 | 156 | N12:207961 |
| chrXII | 208414 | 208561 | 147 | N12:208481 |
| chrXII | 209818 | 209960 | 142 | N12:209848 |
| chrXII | 211059 | 211185 | 126 | N12:211093 |
| chrXII | 213217 | 213369 | 152 | N12:213269 |
| chrXII | 214356 | 214519 | 163 | N12:214440 |
| chrXII | 216121 | 216270 | 149 | N12:216153 |
| chrXII | 217884 | 218033 | 149 | N12:218006 |
| chrXII | 218108 | 218265 | 157 | N12:218186 |
| chrXII | 218294 | 218444 | 150 | N12:218356 |
| chrXII | 218298 | 218444 | 146 | N12:218356 |
| chrXII | 218756 | 218891 | 135 | N12:218862 |
| chrXII | 219794 | 219934 | 140 | N12:219876 |
| chrXII | 219800 | 219934 | 134 | N12:219876 |
| chrXII | 220212 | 220367 | 155 | N12:220242 |
| chrXII | 220343 | 220451 | 108 | N12:220455 |
| chrXII | 223444 | 223638 | 194 | N12:223614 |
| chrXII | 223444 | 223638 | 194 | N12:223443 |
| chrXII | 230481 | 230653 | 172 | N12:230555 |
| chrXII | 230739 | 230894 | 155 | N12:230860 |
| chrXII | 232813 | 232957 | 144 | N12:232966 |
| chrXII | 232813 | 232957 | 144 | N12:232814 |
| chrXII | 235469 | 235619 | 150 | N12:235506 |
| chrXII | 235483 | 235631 | 148 | N12:235506 |
| chrXII | 237484 | 237634 | 150 | N12:237551 |
| chrXII | 238295 | 238446 | 151 | N12:238404 |
| chrXII | 239615 | 239753 | 138 | N12:239689 |
| chrXII | 255888 | 256021 | 133 | |
| chrXII | 256324 | 256484 | 160 | N12:256416 |
| chrXII | 259071 | 259235 | 164 | N12:259139 |
| chrXII | 260642 | 260797 | 155 | N12:260719 |
| chrXII | 262571 | 262718 | 147 | N12:262667 |
| chrXII | 270658 | 270826 | 168 | N12:270763 |
| chrXII | 270673 | 270826 | 153 | N12:270763 |
| chrXII | 271752 | 271896 | 144 | N12:271899 |
| chrXII | 273309 | 273417 | 108 | N12:273383 |
| chrXII | 273309 | 273418 | 109 | N12:273383 |
| chrXII | 273309 | 273419 | 110 | N12:273383 |
| chrXII | 273309 | 273420 | 111 | N12:273383 |
| chrXII | 274415 | 274568 | 153 | N12:274467 |
| chrXII | 275945 | 276112 | 167 | N12:276130 |
| chrXII | 275945 | 276112 | 167 | N12:275964 |
| chrXII | 280170 | 280332 | 162 | N12:280290 |
| chrXII | 280378 | 280532 | 154 | N12:280450 |
| chrXII | 281608 | 281761 | 153 | N12:281710 |
| chrXII | 285125 | 285277 | 152 | N12:285193 |
| chrXII | 285128 | 285277 | 149 | N12:285193 |
| chrXII | 286283 | 286443 | 160 | N12:286398 |
| chrXII | 292173 | 292332 | 159 | N12:292269 |
| chrXII | 293330 | 293472 | 142 | N12:293400 |
| chrXII | 295117 | 295270 | 153 | N12:295096 |
| chrXII | 295472 | 295620 | 148 | N12:295545 |

TABLE 1

| CHR | Start | End | Length (bp) | Nucleosome_ID |
|---|---|---|---|---|
| chrXII | 296601 | 296740 | 139 | N12:296667 |
| chrXII | 300804 | 300959 | 155 | N12:300827 |
| chrXII | 301895 | 302037 | 142 | N12:301967 |
| chrXII | 302498 | 302646 | 148 | N12:302567 |
| chrXII | 304308 | 304455 | 147 | N12:304426 |
| chrXII | 304635 | 304774 | 139 | N12:304748 |
| chrXII | 310453 | 310583 | 130 | N12:310571 |
| chrXII | 311744 | 311903 | 159 | |
| chrXII | 312700 | 312835 | 135 | N12:312689 |
| chrXII | 314498 | 314655 | 157 | N12:314563 |
| chrXII | 319179 | 319325 | 146 | N12:319307 |
| chrXII | 320329 | 320487 | 158 | N12:320461 |
| chrXII | 320335 | 320487 | 152 | N12:320461 |
| chrXII | 320336 | 320487 | 151 | N12:320461 |
| chrXII | 321520 | 321655 | 135 | N12:321554 |
| chrXII | 322116 | 322262 | 146 | |
| chrXII | 327876 | 328044 | 168 | N12:327934 |
| chrXII | 338236 | 338387 | 151 | N12:338317 |
| chrXII | 340076 | 340209 | 133 | N12:340159 |
| chrXII | 340597 | 340761 | 164 | N12:340639 |
| chrXII | 340875 | 341034 | 159 | N12:340984 |
| chrXII | 347768 | 347916 | 148 | N12:347855 |
| chrXII | 347831 | 348011 | 180 | N12:348007 |
| chrXII | 347831 | 348011 | 180 | N12:347855 |
| chrXII | 348735 | 348938 | 203 | N12:348883 |
| chrXII | 348736 | 348938 | 202 | N12:348883 |
| chrXII | 350679 | 350830 | 151 | |
| chrXII | 355881 | 356007 | 126 | N12:355867 |
| chrXII | 355881 | 356007 | 126 | N12:356021 |
| chrXII | 358183 | 358323 | 140 | N12:358217 |
| chrXII | 359562 | 359714 | 152 | N12:359697 |
| chrXII | 360814 | 360966 | 152 | |
| chrXII | 367954 | 368092 | 138 | N12:368026 |
| chrXII | 372826 | 372968 | 142 | N12:372913 |
| chrXII | 375291 | 375434 | 143 | N12:375295 |
| chrXII | 375584 | 375750 | 166 | N12:375684 |
| chrXII | 388796 | 388957 | 161 | N12:388850 |
| chrXII | 388880 | 389024 | 144 | N12:389034 |
| chrXII | 391687 | 391826 | 139 | |
| chrXII | 395857 | 396006 | 149 | N12:396018 |
| chrXII | 395857 | 396006 | 149 | N12:395841 |
| chrXII | 397064 | 397220 | 156 | N12:397207 |
| chrXII | 398081 | 398231 | 150 | N12:398173 |
| chrXII | 400245 | 400397 | 152 | |
| chrXII | 401669 | 401819 | 150 | N12:401676 |
| chrXII | 401669 | 401819 | 150 | N12:401833 |
| chrXII | 405414 | 405520 | 106 | N12:405442 |
| chrXII | 405414 | 405538 | 124 | N12:405442 |
| chrXII | 406427 | 406580 | 153 | N12:406529 |
| chrXII | 406621 | 406768 | 147 | N12:406710 |
| chrXII | 408412 | 408522 | 110 | N12:408439 |
| chrXII | 408412 | 408523 | 111 | N12:408439 |
| chrXII | 408415 | 408537 | 122 | N12:408439 |
| chrXII | 411908 | 412040 | 132 | N12:411988 |
| chrXII | 412316 | 412427 | 111 | N12:412343 |
| chrXII | 412319 | 412427 | 108 | N12:412343 |
| chrXII | 412320 | 412427 | 107 | N12:412343 |
| chrXII | 419068 | 419220 | 152 | N12:419117 |
| chrXII | 419734 | 419893 | 159 | N12:419779 |
| chrXII | 421131 | 421283 | 152 | N12:421250 |
| chrXII | 423648 | 423793 | 145 | N12:423723 |
| chrXII | 424263 | 424441 | 178 | N12:424247 |
| chrXII | 425377 | 425494 | 117 | N12:425416 |
| chrXII | 428829 | 428982 | 153 | N12:428985 |
| chrXII | 433448 | 433586 | 138 | N12:433530 |
| chrXII | 433602 | 433749 | 147 | N12:433706 |
| chrXII | 434337 | 434490 | 153 | N12:434430 |
| chrXII | 443464 | 443629 | 165 | N12:443445 |
| chrXII | 446788 | 446962 | 174 | N12:446975 |
| chrXII | 446788 | 446962 | 174 | N12:446824 |
| chrXII | 447291 | 447454 | 163 | N12:447462 |
| chrXII | 447291 | 447454 | 163 | N12:447286 |
| chrXII | 450323 | 450477 | 154 | N12:450413 |
| chrXII | 451465 | 451626 | 161 | N12:451492 |
| chrXII | 451590 | 451738 | 148 | |
| chrXII | 451599 | 451733 | 134 | |
| chrXII | 451709 | 451842 | 133 | N12:451807 |
| chrXII | 451726 | 451885 | 159 | N12:451807 |
| chrXII | 451734 | 451869 | 135 | N12:451807 |
| chrXII | 451757 | 451842 | 85 | N12:451807 |
| chrXII | 451757 | 451908 | 151 | N12:451807 |
| chrXII | 451769 | 451932 | 163 | N12:451807 |
| chrXII | 451790 | 451908 | 118 | N12:451807 |
| chrXII | 451790 | 451932 | 142 | N12:451807 |
| chrXII | 451790 | 451960 | 170 | N12:451807 |
| chrXII | 451790 | 451960 | 170 | N12:451976 |
| chrXII | 451800 | 451934 | 134 | N12:451807 |
| chrXII | 451800 | 451961 | 161 | N12:451807 |
| chrXII | 451800 | 451961 | 161 | N12:451976 |
| chrXII | 451851 | 451934 | 83 | |
| chrXII | 451851 | 451958 | 107 | N12:451976 |
| chrXII | 451851 | 451959 | 108 | N12:451976 |
| chrXII | 451851 | 451960 | 109 | N12:451976 |
| chrXII | 451851 | 451961 | 110 | N12:451976 |
| chrXII | 451876 | 451960 | 84 | N12:451976 |
| chrXII | 451876 | 452009 | 133 | N12:451976 |
| chrXII | 451919 | 452075 | 156 | N12:451976 |
| chrXII | 451919 | 452077 | 158 | N12:451976 |
| chrXII | 451919 | 452078 | 159 | N12:451976 |
| chrXII | 451919 | 452092 | 173 | N12:451976 |
| chrXII | 451923 | 452075 | 152 | N12:451976 |
| chrXII | 451923 | 452077 | 154 | N12:451976 |
| chrXII | 451923 | 452092 | 169 | N12:451976 |

| CHR | Start | End | Length (bp) | Nucleosome_ID |
|---|---|---|---|---|
| chrXII | 451931 | 452062 | 131 | N12:451976 |
| chrXII | 451931 | 452077 | 146 | N12:451976 |
| chrXII | 451931 | 452109 | 178 | N12:451976 |
| chrXII | 451932 | 452062 | 130 | N12:451976 |
| chrXII | 451932 | 452075 | 143 | N12:451976 |
| chrXII | 451932 | 452077 | 145 | N12:451976 |
| chrXII | 451932 | 452109 | 177 | N12:451976 |
| chrXII | 451945 | 452051 | 106 | N12:451976 |
| chrXII | 451945 | 452062 | 117 | N12:451976 |
| chrXII | 451945 | 452075 | 130 | N12:451976 |
| chrXII | 451945 | 452077 | 132 | N12:451976 |
| chrXII | 451945 | 452078 | 133 | N12:451976 |
| chrXII | 451945 | 452092 | 147 | N12:451976 |
| chrXII | 451945 | 452109 | 164 | N12:451976 |
| chrXII | 451958 | 452109 | 151 | N12:451976 |
| chrXII | 452041 | 452196 | 155 | N12:452173 |
| chrXII | 452059 | 452215 | 156 | N12:452173 |
| chrXII | 452081 | 452227 | 146 | N12:452173 |
| chrXII | 452081 | 452236 | 155 | N12:452173 |
| chrXII | 452081 | 452247 | 166 | N12:452173 |
| chrXII | 452081 | 452251 | 170 | N12:452173 |
| chrXII | 452104 | 452247 | 143 | N12:452173 |
| chrXII | 452104 | 452251 | 147 | N12:452173 |
| chrXII | 452115 | 452271 | 156 | N12:452173 |
| chrXII | 452190 | 452344 | 154 | N12:452173 |
| chrXII | 452190 | 452344 | 154 | N12:452361 |
| chrXII | 452260 | 452414 | 154 | N12:452361 |
| chrXII | 452293 | 452400 | 107 | N12:452361 |
| chrXII | 452294 | 452455 | 161 | N12:452361 |
| chrXII | 452295 | 452455 | 160 | N12:452361 |
| chrXII | 452299 | 452455 | 156 | N12:452361 |
| chrXII | 452323 | 452455 | 132 | N12:452361 |
| chrXII | 452323 | 452485 | 162 | N12:452361 |
| chrXII | 452323 | 452486 | 163 | N12:452361 |
| chrXII | 452334 | 452441 | 107 | N12:452361 |
| chrXII | 452334 | 452455 | 121 | N12:452361 |
| chrXII | 452334 | 452485 | 151 | N12:452361 |
| chrXII | 452334 | 452486 | 152 | N12:452361 |
| chrXII | 452346 | 452496 | 150 | N12:452361 |
| chrXII | 452423 | 452584 | 161 | N12:452551 |
| chrXII | 452469 | 452584 | 115 | N12:452551 |
| chrXII | 452469 | 452620 | 151 | N12:452551 |
| chrXII | 452469 | 452621 | 152 | N12:452551 |
| chrXII | 452469 | 452629 | 160 | N12:452551 |
| chrXII | 452469 | 452630 | 161 | N12:452551 |
| chrXII | 452491 | 452621 | 130 | N12:452551 |
| chrXII | 452491 | 452629 | 138 | N12:452551 |
| chrXII | 452491 | 452663 | 172 | N12:452551 |
| chrXII | 452492 | 452584 | 92 | N12:452551 |
| chrXII | 452492 | 452620 | 128 | N12:452551 |
| chrXII | 452492 | 452621 | 129 | N12:452551 |
| chrXII | 452492 | 452629 | 137 | N12:452551 |
| chrXII | 452492 | 452663 | 171 | N12:452551 |
| chrXII | 452494 | 452621 | 127 | N12:452551 |
| chrXII | 452525 | 452695 | 170 | N12:452551 |
| chrXII | 452539 | 452673 | 134 | N12:452551 |
| chrXII | 452550 | 452672 | 122 | N12:452551 |
| chrXII | 452550 | 452702 | 152 | N12:452551 |
| chrXII | 452668 | 452796 | 128 | N12:452761 |
| chrXII | 452684 | 452796 | 112 | N12:452761 |
| chrXII | 452684 | 452835 | 151 | N12:452761 |
| chrXII | 452686 | 452814 | 128 | N12:452761 |
| chrXII | 452686 | 452884 | 198 | N12:452761 |
| chrXII | 452709 | 452864 | 155 | N12:452761 |
| chrXII | 452709 | 452884 | 175 | N12:452761 |
| chrXII | 452709 | 452900 | 191 | N12:452761 |
| chrXII | 452723 | 452814 | 91 | N12:452761 |
| chrXII | 452723 | 452864 | 141 | N12:452761 |
| chrXII | 452723 | 452884 | 161 | N12:452761 |
| chrXII | 452723 | 452900 | 177 | N12:452761 |
| chrXII | 452765 | 452864 | 99 | N12:452761 |
| chrXII | 452765 | 452871 | 106 | N12:452761 |
| chrXII | 452765 | 452884 | 119 | N12:452761 |
| chrXII | 452765 | 452900 | 135 | N12:452761 |
| chrXII | 452765 | 452925 | 160 | N12:452761 |
| chrXII | 452765 | 452925 | 160 | N12:452927 |
| chrXII | 452773 | 452864 | 91 | N12:452761 |
| chrXII | 452773 | 452879 | 106 | N12:452761 |
| chrXII | 452773 | 452884 | 111 | N12:452761 |
| chrXII | 452773 | 452900 | 127 | N12:452761 |
| chrXII | 452773 | 452925 | 152 | N12:452761 |
| chrXII | 452773 | 452925 | 152 | N12:452927 |
| chrXII | 452783 | 452962 | 179 | N12:452761 |
| chrXII | 452783 | 452962 | 179 | N12:452927 |
| chrXII | 452788 | 452923 | 135 | N12:452927 |
| chrXII | 452788 | 452953 | 165 | N12:452927 |
| chrXII | 452801 | 452923 | 122 | N12:452927 |
| chrXII | 452801 | 452953 | 152 | N12:452927 |
| chrXII | 452801 | 452999 | 198 | N12:452927 |
| chrXII | 452840 | 452953 | 113 | N12:452927 |
| chrXII | 452840 | 452999 | 159 | N12:452927 |
| chrXII | 452840 | 453023 | 183 | N12:452927 |
| chrXII | 452847 | 452953 | 106 | N12:452927 |
| chrXII | 452847 | 452996 | 149 | N12:452927 |
| chrXII | 452876 | 453034 | 158 | N12:452927 |
| chrXII | 452885 | 453034 | 149 | N12:452927 |
| chrXII | 452893 | 452999 | 106 | N12:452927 |
| chrXII | 452895 | 453023 | 128 | N12:452927 |
| chrXII | 452922 | 453029 | 107 | N12:452927 |
| chrXII | 452922 | 453031 | 109 | N12:452927 |
| chrXII | 452922 | 453034 | 112 | N12:452927 |
| chrXII | 452922 | 453056 | 134 | N12:452927 |
| chrXII | 452922 | 453056 | 134 | N12:453077 |

| CHR | Start | End | Length (bp) | Nucleosome_ID |
|---|---|---|---|---|
| chrXII | 452922 | 453075 | 153 | N12:452927 |
| chrXII | 452922 | 453075 | 153 | N12:453077 |
| chrXII | 452926 | 452997 | 71 | N12:452927 |
| chrXII | 452926 | 453033 | 107 | N12:452927 |
| chrXII | 452926 | 453034 | 108 | N12:452927 |
| chrXII | 452926 | 453056 | 130 | N12:452927 |
| chrXII | 452926 | 453056 | 130 | N12:453077 |
| chrXII | 452926 | 453070 | 144 | N12:452927 |
| chrXII | 452926 | 453070 | 144 | N12:453077 |
| chrXII | 452926 | 453075 | 149 | N12:452927 |
| chrXII | 452926 | 453075 | 149 | N12:453077 |
| chrXII | 452976 | 453138 | 162 | N12:453077 |
| chrXII | 452976 | 453163 | 187 | N12:453077 |
| chrXII | 452991 | 453141 | 150 | N12:453077 |
| chrXII | 453033 | 453163 | 130 | N12:453077 |
| chrXII | 453033 | 453187 | 154 | N12:453077 |
| chrXII | 453033 | 453193 | 160 | N12:453077 |
| chrXII | 453039 | 453163 | 124 | N12:453077 |
| chrXII | 453039 | 453187 | 148 | N12:453077 |
| chrXII | 453039 | 453193 | 154 | N12:453077 |
| chrXII | 453042 | 453141 | 99 | N12:453077 |
| chrXII | 453042 | 453212 | 170 | N12:453077 |
| chrXII | 453042 | 453218 | 176 | N12:453077 |
| chrXII | 453043 | 453163 | 120 | N12:453077 |
| chrXII | 453043 | 453187 | 144 | N12:453077 |
| chrXII | 453043 | 453190 | 147 | N12:453077 |
| chrXII | 453043 | 453193 | 150 | N12:453077 |
| chrXII | 453091 | 453199 | 108 | N12:453077 |
| chrXII | 453092 | 453199 | 107 | N12:453077 |
| chrXII | 453092 | 453212 | 120 | N12:453077 |
| chrXII | 453092 | 453236 | 144 | N12:453077 |
| chrXII | 453092 | 453245 | 153 | N12:453077 |
| chrXII | 453092 | 453270 | 178 | N12:453077 |
| chrXII | 453092 | 453282 | 190 | N12:453077 |
| chrXII | 453092 | 453286 | 194 | N12:453077 |
| chrXII | 453092 | 453288 | 196 | N12:453077 |
| chrXII | 453094 | 453236 | 142 | N12:453077 |
| chrXII | 453095 | 453201 | 106 | N12:453077 |
| chrXII | 453095 | 453212 | 117 | N12:453077 |
| chrXII | 453095 | 453218 | 123 | N12:453077 |
| chrXII | 453095 | 453236 | 141 | N12:453077 |
| chrXII | 453095 | 453245 | 150 | N12:453077 |
| chrXII | 453095 | 453270 | 175 | N12:453077 |
| chrXII | 453095 | 453286 | 191 | N12:453077 |
| chrXII | 453095 | 453288 | 193 | N12:453077 |
| chrXII | 453099 | 453236 | 137 | N12:453077 |
| chrXII | 453099 | 453287 | 188 | N12:453077 |
| chrXII | 453099 | 453288 | 189 | N12:453077 |
| chrXII | 453100 | 453238 | 138 | N12:453077 |
| chrXII | 453100 | 453245 | 145 | N12:453077 |
| chrXII | 453113 | 453270 | 157 | |
| chrXII | 453113 | 453286 | 173 | |
| chrXII | 453113 | 453288 | 175 | |
| chrXII | 453114 | 453236 | 122 | |
| chrXII | 453114 | 453269 | 155 | |
| chrXII | 453114 | 453270 | 156 | |
| chrXII | 453114 | 453286 | 172 | |
| chrXII | 453114 | 453287 | 173 | |
| chrXII | 453114 | 453288 | 174 | |
| chrXII | 453114 | 453302 | 188 | |
| chrXII | 453114 | 453309 | 195 | |
| chrXII | 453116 | 453269 | 153 | |
| chrXII | 453116 | 453270 | 154 | |
| chrXII | 453116 | 453288 | 172 | |
| chrXII | 453119 | 453236 | 117 | |
| chrXII | 453119 | 453245 | 126 | |
| chrXII | 453119 | 453269 | 150 | |
| chrXII | 453119 | 453270 | 151 | |
| chrXII | 453119 | 453282 | 163 | |
| chrXII | 453119 | 453286 | 167 | |
| chrXII | 453119 | 453287 | 168 | |
| chrXII | 453119 | 453288 | 169 | |
| chrXII | 453119 | 453302 | 183 | |
| chrXII | 453119 | 453309 | 190 | |
| chrXII | 453120 | 453228 | 108 | |
| chrXII | 453121 | 453270 | 149 | |
| chrXII | 453121 | 453282 | 161 | |
| chrXII | 453121 | 453286 | 165 | |
| chrXII | 453121 | 453288 | 167 | |
| chrXII | 453125 | 453231 | 106 | |
| chrXII | 453125 | 453236 | 111 | |
| chrXII | 453125 | 453270 | 145 | |
| chrXII | 453125 | 453286 | 161 | |
| chrXII | 453125 | 453287 | 162 | |
| chrXII | 453125 | 453288 | 163 | |
| chrXII | 453125 | 453302 | 177 | |
| chrXII | 453126 | 453248 | 122 | |
| chrXII | 453126 | 453270 | 144 | |
| chrXII | 453126 | 453286 | 160 | |
| chrXII | 453126 | 453287 | 161 | |
| chrXII | 453126 | 453288 | 162 | |
| chrXII | 453135 | 453270 | 135 | |
| chrXII | 453135 | 453282 | 147 | |
| chrXII | 453135 | 453286 | 151 | |
| chrXII | 453135 | 453288 | 153 | |
| chrXII | 453138 | 453286 | 148 | |
| chrXII | 453138 | 453288 | 150 | |
| chrXII | 453140 | 453245 | 105 | |
| chrXII | 453140 | 453246 | 106 | |
| chrXII | 453140 | 453251 | 111 | |
| chrXII | 453140 | 453270 | 130 | |
| chrXII | 453140 | 453282 | 142 | |
| chrXII | 453140 | 453286 | 146 | |

| CHR | Start | End | Length (bp) | Nucleosome_ID |
|---|---|---|---|---|
| chrXII | 453140 | 453287 | 147 | |
| chrXII | 453140 | 453288 | 148 | |
| chrXII | 453140 | 453302 | 162 | |
| chrXII | 453140 | 453309 | 169 | |
| chrXII | 453140 | 453323 | 183 | |
| chrXII | 453141 | 453236 | 95 | |
| chrXII | 453141 | 453245 | 104 | |
| chrXII | 453141 | 453251 | 110 | |
| chrXII | 453141 | 453269 | 128 | |
| chrXII | 453141 | 453270 | 129 | |
| chrXII | 453141 | 453281 | 140 | |
| chrXII | 453141 | 453282 | 141 | |
| chrXII | 453141 | 453286 | 145 | |
| chrXII | 453141 | 453287 | 146 | |
| chrXII | 453141 | 453288 | 147 | |
| chrXII | 453141 | 453302 | 161 | |
| chrXII | 453141 | 453309 | 168 | |
| chrXII | 453141 | 453323 | 182 | |
| chrXII | 453141 | 453324 | 183 | |
| chrXII | 453144 | 453288 | 144 | |
| chrXII | 453150 | 453286 | 136 | |
| chrXII | 453150 | 453287 | 137 | |
| chrXII | 453150 | 453288 | 138 | |
| chrXII | 453150 | 453295 | 145 | |
| chrXII | 453150 | 453302 | 152 | |
| chrXII | 453152 | 453240 | 88 | |
| chrXII | 453152 | 453270 | 118 | |
| chrXII | 453152 | 453286 | 134 | |
| chrXII | 453152 | 453287 | 135 | |
| chrXII | 453152 | 453288 | 136 | |
| chrXII | 453152 | 453302 | 150 | |
| chrXII | 453152 | 453309 | 157 | |
| chrXII | 453153 | 453286 | 133 | |
| chrXII | 453153 | 453288 | 135 | |
| chrXII | 453159 | 453270 | 111 | |
| chrXII | 453159 | 453286 | 127 | |
| chrXII | 453159 | 453287 | 128 | |
| chrXII | 453159 | 453288 | 129 | |
| chrXII | 453159 | 453302 | 143 | |
| chrXII | 453159 | 453309 | 150 | |
| chrXII | 453159 | 453335 | 176 | |
| chrXII | 453159 | 453342 | 183 | |
| chrXII | 453161 | 453245 | 84 | |
| chrXII | 453161 | 453268 | 107 | |
| chrXII | 453161 | 453270 | 109 | |
| chrXII | 453161 | 453282 | 121 | |
| chrXII | 453161 | 453286 | 125 | |
| chrXII | 453161 | 453288 | 127 | |
| chrXII | 453161 | 453309 | 148 | |
| chrXII | 453161 | 453323 | 162 | |
| chrXII | 453161 | 453324 | 163 | |
| chrXII | 453162 | 453269 | 107 | |
| chrXII | 453162 | 453286 | 124 | |
| chrXII | 453162 | 453288 | 126 | |
| chrXII | 453162 | 453323 | 161 | |
| chrXII | 453162 | 453324 | 162 | |
| chrXII | 453165 | 453286 | 121 | |
| chrXII | 453165 | 453287 | 122 | |
| chrXII | 453165 | 453288 | 123 | |
| chrXII | 453165 | 453302 | 137 | |
| chrXII | 453165 | 453342 | 177 | |
| chrXII | 453180 | 453286 | 106 | |
| chrXII | 453180 | 453288 | 108 | |
| chrXII | 453180 | 453340 | 160 | |
| chrXII | 453181 | 453270 | 89 | |
| chrXII | 453181 | 453282 | 101 | |
| chrXII | 453181 | 453286 | 105 | |
| chrXII | 453181 | 453288 | 107 | |
| chrXII | 453181 | 453323 | 142 | |
| chrXII | 453181 | 453324 | 143 | |
| chrXII | 453286 | 453432 | 146 | N12:453378 |
| chrXII | 453299 | 453425 | 126 | N12:453378 |
| chrXII | 453299 | 453440 | 141 | N12:453378 |
| chrXII | 453299 | 453460 | 161 | N12:453378 |
| chrXII | 453300 | 453410 | 110 | N12:453378 |
| chrXII | 453300 | 453414 | 114 | N12:453378 |
| chrXII | 453300 | 453440 | 140 | N12:453378 |
| chrXII | 453302 | 453440 | 138 | N12:453378 |
| chrXII | 453310 | 453440 | 130 | N12:453378 |
| chrXII | 453310 | 453460 | 150 | N12:453378 |
| chrXII | 453322 | 453481 | 159 | N12:453378 |
| chrXII | 453458 | 453595 | 137 | N12:453535 |
| chrXII | 453458 | 453610 | 152 | N12:453535 |
| chrXII | 453458 | 453612 | 154 | N12:453535 |
| chrXII | 453460 | 453612 | 152 | N12:453535 |
| chrXII | 453487 | 453595 | 108 | N12:453535 |
| chrXII | 453487 | 453610 | 123 | N12:453535 |
| chrXII | 453487 | 453612 | 125 | N12:453535 |
| chrXII | 453500 | 453652 | 152 | N12:453535 |
| chrXII | 453574 | 453727 | 153 | N12:453693 |
| chrXII | 453587 | 453737 | 150 | N12:453693 |
| chrXII | 453606 | 453774 | 168 | N12:453693 |
| chrXII | 453627 | 453776 | 149 | N12:453693 |
| chrXII | 453627 | 453779 | 152 | N12:453693 |
| chrXII | 453657 | 453814 | 157 | N12:453693 |
| chrXII | 453659 | 453776 | 117 | N12:453693 |
| chrXII | 453659 | 453779 | 120 | N12:453693 |
| chrXII | 453734 | 453853 | 119 | N12:453841 |
| chrXII | 453748 | 453910 | 162 | N12:453841 |
| chrXII | 453748 | 453913 | 165 | N12:453841 |
| chrXII | 453748 | 453919 | 171 | N12:453841 |
| chrXII | 453748 | 453922 | 174 | N12:453841 |
| chrXII | 453748 | 453962 | 214 | N12:453841 |

| CHR | Start | End | Length (bp) | Nucleosome_ID |
|---|---|---|---|---|
| chrXII | 453752 | 453853 | 101 | N12:453841 |
| chrXII | 453752 | 453859 | 107 | N12:453841 |
| chrXII | 453752 | 453908 | 156 | N12:453841 |
| chrXII | 453752 | 453911 | 159 | N12:453841 |
| chrXII | 453752 | 453914 | 162 | N12:453841 |
| chrXII | 453752 | 453916 | 164 | N12:453841 |
| chrXII | 453752 | 453922 | 170 | N12:453841 |
| chrXII | 453752 | 453924 | 172 | N12:453841 |
| chrXII | 453753 | 453913 | 160 | N12:453841 |
| chrXII | 453753 | 453919 | 166 | N12:453841 |
| chrXII | 453753 | 453922 | 169 | N12:453841 |
| chrXII | 453754 | 453910 | 156 | N12:453841 |
| chrXII | 453754 | 453924 | 170 | N12:453841 |
| chrXII | 453774 | 453910 | 136 | N12:453841 |
| chrXII | 453774 | 453913 | 139 | N12:453841 |
| chrXII | 453774 | 453914 | 140 | N12:453841 |
| chrXII | 453774 | 453916 | 142 | N12:453841 |
| chrXII | 453774 | 453919 | 145 | N12:453841 |
| chrXII | 453774 | 453922 | 148 | N12:453841 |
| chrXII | 453774 | 453925 | 151 | N12:453841 |
| chrXII | 453774 | 453962 | 188 | N12:453841 |
| chrXII | 453775 | 453908 | 133 | N12:453841 |
| chrXII | 453775 | 453911 | 136 | N12:453841 |
| chrXII | 453775 | 453914 | 139 | N12:453841 |
| chrXII | 453775 | 453916 | 141 | N12:453841 |
| chrXII | 453775 | 453919 | 144 | N12:453841 |
| chrXII | 453775 | 453922 | 147 | N12:453841 |
| chrXII | 453781 | 453908 | 127 | N12:453841 |
| chrXII | 453781 | 453911 | 130 | N12:453841 |
| chrXII | 453781 | 453914 | 133 | N12:453841 |
| chrXII | 453781 | 453916 | 135 | N12:453841 |
| chrXII | 453781 | 453922 | 141 | N12:453841 |
| chrXII | 453782 | 453908 | 126 | N12:453841 |
| chrXII | 453782 | 453911 | 129 | N12:453841 |
| chrXII | 453782 | 453913 | 131 | N12:453841 |
| chrXII | 453782 | 453914 | 132 | N12:453841 |
| chrXII | 453782 | 453919 | 137 | N12:453841 |
| chrXII | 453782 | 453922 | 140 | N12:453841 |
| chrXII | 453782 | 453925 | 143 | N12:453841 |
| chrXII | 453782 | 453929 | 147 | N12:453841 |
| chrXII | 453782 | 453937 | 155 | N12:453841 |
| chrXII | 453782 | 453962 | 180 | N12:453841 |
| chrXII | 453783 | 453911 | 128 | N12:453841 |
| chrXII | 453783 | 453913 | 130 | N12:453841 |
| chrXII | 453783 | 453914 | 131 | N12:453841 |
| chrXII | 453812 | 453913 | 101 | N12:453841 |
| chrXII | 453812 | 453919 | 107 | N12:453841 |
| chrXII | 453812 | 453962 | 150 | N12:453841 |
| chrXII | 453812 | 454001 | 189 | N12:453841 |
| chrXII | 453812 | 454001 | 189 | N12:454007 |
| chrXII | 453851 | 453958 | 107 | N12:453841 |
| chrXII | 453851 | 453961 | 110 | N12:453841 |
| chrXII | 453851 | 453962 | 111 | N12:453841 |
| chrXII | 453851 | 454001 | 150 | N12:453841 |
| chrXII | 453851 | 454001 | 150 | N12:454007 |
| chrXII | 453909 | 454072 | 163 | N12:454007 |
| chrXII | 453918 | 454080 | 162 | N12:454007 |
| chrXII | 453922 | 454079 | 157 | N12:454007 |
| chrXII | 453929 | 454079 | 150 | N12:454007 |
| chrXII | 453929 | 454097 | 168 | N12:454007 |
| chrXII | 453929 | 454108 | 179 | N12:454007 |
| chrXII | 453959 | 454097 | 138 | N12:454007 |
| chrXII | 453959 | 454108 | 149 | N12:454007 |
| chrXII | 453959 | 454112 | 153 | N12:454007 |
| chrXII | 453960 | 454116 | 156 | N12:454007 |
| chrXII | 453964 | 454097 | 133 | N12:454007 |
| chrXII | 453964 | 454108 | 144 | N12:454007 |
| chrXII | 453971 | 454123 | 152 | N12:454007 |
| chrXII | 454059 | 454216 | 157 | N12:454198 |
| chrXII | 454111 | 454268 | 157 | N12:454198 |
| chrXII | 454120 | 454267 | 147 | N12:454198 |
| chrXII | 454120 | 454277 | 157 | N12:454198 |
| chrXII | 454124 | 454277 | 153 | N12:454198 |
| chrXII | 454131 | 454277 | 146 | N12:454198 |
| chrXII | 454141 | 454277 | 136 | N12:454198 |
| chrXII | 454196 | 454346 | 150 | N12:454198 |
| chrXII | 454196 | 454346 | 150 | N12:454367 |
| chrXII | 454221 | 454392 | 171 | N12:454198 |
| chrXII | 454221 | 454392 | 171 | N12:454367 |
| chrXII | 454244 | 454392 | 148 | N12:454367 |
| chrXII | 454244 | 454402 | 158 | N12:454367 |
| chrXII | 454248 | 454374 | 126 | N12:454367 |
| chrXII | 454248 | 454392 | 144 | N12:454367 |
| chrXII | 454248 | 454398 | 150 | N12:454367 |
| chrXII | 454273 | 454436 | 163 | N12:454367 |
| chrXII | 454277 | 454398 | 121 | N12:454367 |
| chrXII | 454277 | 454402 | 125 | N12:454367 |
| chrXII | 454277 | 454438 | 161 | N12:454367 |
| chrXII | 454277 | 454446 | 169 | N12:454367 |
| chrXII | 454398 | 454551 | 153 | |
| chrXII | 454398 | 454552 | 154 | |
| chrXII | 454398 | 454555 | 157 | |
| chrXII | 454398 | 454567 | 169 | |
| chrXII | 454398 | 454573 | 175 | |
| chrXII | 454398 | 454575 | 177 | |
| chrXII | 454404 | 454552 | 148 | |
| chrXII | 454404 | 454555 | 151 | |
| chrXII | 454422 | 454552 | 130 | |
| chrXII | 454422 | 454555 | 133 | |
| chrXII | 454422 | 454567 | 145 | |
| chrXII | 454422 | 454573 | 151 | |
| chrXII | 454422 | 454575 | 153 | |
| chrXII | 454521 | 454678 | 157 | N12:454654 |

| CHR | Start | End | Length (bp) | Nucleosome_ID |
|---|---|---|---|---|
| chrXII | 454553 | 454712 | 159 | N12:454654 |
| chrXII | 454553 | 454715 | 162 | N12:454654 |
| chrXII | 454553 | 454733 | 180 | N12:454654 |
| chrXII | 454555 | 454703 | 148 | N12:454654 |
| chrXII | 454560 | 454678 | 118 | N12:454654 |
| chrXII | 454560 | 454712 | 152 | N12:454654 |
| chrXII | 454560 | 454715 | 155 | N12:454654 |
| chrXII | 454560 | 454724 | 164 | N12:454654 |
| chrXII | 454560 | 454733 | 173 | N12:454654 |
| chrXII | 454562 | 454712 | 150 | N12:454654 |
| chrXII | 454570 | 454712 | 142 | N12:454654 |
| chrXII | 454570 | 454715 | 145 | N12:454654 |
| chrXII | 454570 | 454724 | 154 | N12:454654 |
| chrXII | 454570 | 454733 | 163 | N12:454654 |
| chrXII | 454576 | 454712 | 136 | N12:454654 |
| chrXII | 454576 | 454715 | 139 | N12:454654 |
| chrXII | 454576 | 454724 | 148 | N12:454654 |
| chrXII | 454576 | 454733 | 157 | N12:454654 |
| chrXII | 454576 | 454772 | 196 | N12:454654 |
| chrXII | 454587 | 454746 | 159 | N12:454654 |
| chrXII | 454592 | 454742 | 150 | N12:454654 |
| chrXII | 454626 | 454724 | 98 | N12:454654 |
| chrXII | 454626 | 454733 | 107 | N12:454654 |
| chrXII | 454626 | 454772 | 146 | N12:454654 |
| chrXII | 454649 | 454741 | 92 | N12:454654 |
| chrXII | 454677 | 454828 | 151 | N12:454654 |
| chrXII | 454677 | 454828 | 151 | N12:454834 |
| chrXII | 454723 | 454854 | 131 | N12:454834 |
| chrXII | 454725 | 454863 | 138 | N12:454834 |
| chrXII | 454725 | 454886 | 161 | N12:454834 |
| chrXII | 454733 | 454863 | 130 | N12:454834 |
| chrXII | 454733 | 454886 | 153 | N12:454834 |
| chrXII | 454741 | 454896 | 155 | N12:454834 |
| chrXII | 454747 | 454853 | 106 | N12:454834 |
| chrXII | 454747 | 454886 | 139 | N12:454834 |
| chrXII | 454825 | 454972 | 147 | N12:454834 |
| chrXII | 454825 | 454972 | 147 | N12:454994 |
| chrXII | 454837 | 454991 | 154 | N12:454834 |
| chrXII | 454837 | 454991 | 154 | N12:454994 |
| chrXII | 454864 | 455021 | 157 | N12:454994 |
| chrXII | 454876 | 455023 | 147 | N12:454994 |
| chrXII | 454876 | 455029 | 153 | N12:454994 |
| chrXII | 454901 | 455029 | 128 | N12:454994 |
| chrXII | 454901 | 455074 | 173 | N12:454994 |
| chrXII | 454920 | 455087 | 167 | N12:454994 |
| chrXII | 454932 | 455073 | 141 | N12:454994 |
| chrXII | 454933 | 455023 | 90 | N12:454994 |
| chrXII | 454933 | 455024 | 91 | N12:454994 |
| chrXII | 454933 | 455025 | 92 | N12:454994 |
| chrXII | 454933 | 455027 | 94 | N12:454994 |
| chrXII | 454933 | 455029 | 96 | N12:454994 |
| chrXII | 454933 | 455073 | 140 | N12:454994 |
| chrXII | 454933 | 455074 | 141 | N12:454994 |
| chrXII | 454933 | 455076 | 143 | N12:454994 |
| chrXII | 454933 | 455132 | 199 | N12:454994 |
| chrXII | 454964 | 455132 | 168 | N12:454994 |
| chrXII | 454995 | 455164 | 169 | N12:454994 |
| chrXII | 455000 | 455132 | 132 | N12:454994 |
| chrXII | 455015 | 455166 | 151 | N12:454994 |
| chrXII | 455058 | 455208 | 150 | |
| chrXII | 455068 | 455223 | 155 | |
| chrXII | 455095 | 455269 | 174 | |
| chrXII | 455096 | 455223 | 127 | |
| chrXII | 455096 | 455240 | 144 | |
| chrXII | 455096 | 455243 | 147 | |
| chrXII | 455096 | 455245 | 149 | |
| chrXII | 455096 | 455249 | 153 | |
| chrXII | 455097 | 455240 | 143 | |
| chrXII | 455097 | 455243 | 146 | |
| chrXII | 455098 | 455243 | 145 | |
| chrXII | 455098 | 455249 | 151 | |
| chrXII | 455129 | 455293 | 164 | N12:455294 |
| chrXII | 455138 | 455289 | 151 | N12:455294 |
| chrXII | 455182 | 455338 | 156 | N12:455294 |
| chrXII | 455217 | 455366 | 149 | N12:455294 |
| chrXII | 455221 | 455395 | 174 | N12:455294 |
| chrXII | 455247 | 455381 | 134 | N12:455294 |
| chrXII | 455254 | 455401 | 147 | N12:455294 |
| chrXII | 455255 | 455400 | 145 | N12:455294 |
| chrXII | 455278 | 455423 | 145 | N12:455294 |
| chrXII | 455297 | 455447 | 150 | N12:455294 |
| chrXII | 455367 | 455512 | 145 | N12:455488 |
| chrXII | 455367 | 455531 | 164 | N12:455488 |
| chrXII | 455367 | 455549 | 182 | N12:455488 |
| chrXII | 455377 | 455512 | 135 | N12:455488 |
| chrXII | 455377 | 455531 | 154 | N12:455488 |
| chrXII | 455377 | 455549 | 172 | N12:455488 |
| chrXII | 455397 | 455503 | 106 | N12:455488 |
| chrXII | 455397 | 455512 | 115 | N12:455488 |
| chrXII | 455397 | 455531 | 134 | N12:455488 |
| chrXII | 455397 | 455549 | 152 | N12:455488 |
| chrXII | 455397 | 455568 | 171 | N12:455488 |
| chrXII | 455410 | 455512 | 102 | N12:455488 |
| chrXII | 455410 | 455549 | 139 | N12:455488 |
| chrXII | 455410 | 455560 | 150 | N12:455488 |
| chrXII | 455410 | 455568 | 158 | N12:455488 |
| chrXII | 455410 | 455569 | 159 | N12:455488 |
| chrXII | 455416 | 455571 | 155 | N12:455488 |
| chrXII | 455422 | 455569 | 147 | N12:455488 |
| chrXII | 455591 | 455751 | 160 | N12:455656 |
| chrXII | 455701 | 455870 | 169 | N12:455829 |
| chrXII | 455747 | 455895 | 148 | N12:455829 |
| chrXII | 455749 | 455889 | 140 | N12:455829 |

| CHR | Start | End | Length (bp) | Nucleosome_ID |
|---|---|---|---|---|
| chrXII | 455783 | 455958 | 175 | N12:455829 |
| chrXII | 455869 | 456021 | 152 | N12:456042 |
| chrXII | 455903 | 456067 | 164 | N12:456042 |
| chrXII | 455918 | 456039 | 121 | N12:456042 |
| chrXII | 455926 | 456086 | 160 | N12:456042 |
| chrXII | 456022 | 456173 | 151 | N12:456042 |
| chrXII | 456022 | 456205 | 183 | N12:456042 |
| chrXII | 456022 | 456205 | 183 | N12:456220 |
| chrXII | 456022 | 456211 | 189 | N12:456042 |
| chrXII | 456022 | 456211 | 189 | N12:456220 |
| chrXII | 456022 | 456220 | 198 | N12:456042 |
| chrXII | 456022 | 456220 | 198 | N12:456220 |
| chrXII | 456025 | 456173 | 148 | N12:456042 |
| chrXII | 456025 | 456182 | 157 | N12:456042 |
| chrXII | 456042 | 456173 | 131 | N12:456042 |
| chrXII | 456045 | 456173 | 128 | N12:456042 |
| chrXII | 456045 | 456175 | 130 | N12:456042 |
| chrXII | 456045 | 456205 | 160 | N12:456042 |
| chrXII | 456045 | 456205 | 160 | N12:456220 |
| chrXII | 456056 | 456173 | 117 | N12:456042 |
| chrXII | 456056 | 456175 | 119 | N12:456042 |
| chrXII | 456056 | 456205 | 149 | N12:456042 |
| chrXII | 456056 | 456205 | 149 | N12:456220 |
| chrXII | 456056 | 456211 | 155 | N12:456042 |
| chrXII | 456056 | 456211 | 155 | N12:456220 |
| chrXII | 456056 | 456220 | 164 | N12:456042 |
| chrXII | 456056 | 456220 | 164 | N12:456220 |
| chrXII | 456063 | 456215 | 152 | N12:456042 |
| chrXII | 456063 | 456215 | 152 | N12:456220 |
| chrXII | 456064 | 456214 | 150 | N12:456042 |
| chrXII | 456064 | 456214 | 150 | N12:456220 |
| chrXII | 456066 | 456173 | 107 | |
| chrXII | 456067 | 456205 | 138 | N12:456220 |
| chrXII | 456067 | 456211 | 144 | N12:456220 |
| chrXII | 456067 | 456215 | 148 | N12:456220 |
| chrXII | 456067 | 456220 | 153 | N12:456220 |
| chrXII | 456067 | 456266 | 199 | N12:456220 |
| chrXII | 456080 | 456173 | 93 | |
| chrXII | 456080 | 456205 | 125 | N12:456220 |
| chrXII | 456080 | 456211 | 131 | N12:456220 |
| chrXII | 456080 | 456220 | 140 | N12:456220 |
| chrXII | 456084 | 456215 | 131 | N12:456220 |
| chrXII | 456084 | 456243 | 159 | N12:456220 |
| chrXII | 456084 | 456266 | 182 | N12:456220 |
| chrXII | 456115 | 456215 | 100 | N12:456220 |
| chrXII | 456115 | 456243 | 128 | N12:456220 |
| chrXII | 456115 | 456266 | 151 | N12:456220 |
| chrXII | 456115 | 456281 | 166 | N12:456220 |
| chrXII | 456120 | 456290 | 170 | N12:456220 |
| chrXII | 456124 | 456215 | 91 | N12:456220 |
| chrXII | 456124 | 456266 | 142 | N12:456220 |
| chrXII | 456124 | 456281 | 157 | N12:456220 |
| chrXII | 456124 | 456333 | 209 | N12:456220 |
| chrXII | 456181 | 456266 | 85 | N12:456220 |
| chrXII | 456181 | 456281 | 100 | N12:456220 |
| chrXII | 456181 | 456287 | 106 | N12:456220 |
| chrXII | 456181 | 456306 | 125 | N12:456220 |
| chrXII | 456181 | 456310 | 129 | N12:456220 |
| chrXII | 456181 | 456329 | 148 | N12:456220 |
| chrXII | 456181 | 456332 | 151 | N12:456220 |
| chrXII | 456181 | 456333 | 152 | N12:456220 |
| chrXII | 456181 | 456336 | 155 | N12:456220 |
| chrXII | 456181 | 456349 | 168 | N12:456220 |
| chrXII | 456181 | 456387 | 206 | N12:456220 |
| chrXII | 456186 | 456333 | 147 | N12:456220 |
| chrXII | 456187 | 456333 | 146 | N12:456220 |
| chrXII | 456187 | 456349 | 162 | N12:456220 |
| chrXII | 456194 | 456333 | 139 | N12:456220 |
| chrXII | 456194 | 456336 | 142 | N12:456220 |
| chrXII | 456213 | 456319 | 106 | N12:456220 |
| chrXII | 456213 | 456320 | 107 | N12:456220 |
| chrXII | 456213 | 456321 | 108 | N12:456220 |
| chrXII | 456213 | 456333 | 120 | N12:456220 |
| chrXII | 456213 | 456336 | 123 | N12:456220 |
| chrXII | 456213 | 456349 | 136 | N12:456220 |
| chrXII | 456213 | 456352 | 139 | N12:456220 |
| chrXII | 456213 | 456387 | 174 | N12:456220 |
| chrXII | 456213 | 456419 | 206 | N12:456220 |
| chrXII | 456213 | 456419 | 206 | N12:456421 |
| chrXII | 456221 | 456386 | 165 | N12:456220 |
| chrXII | 456240 | 456333 | 93 | N12:456220 |
| chrXII | 456240 | 456386 | 146 | N12:456220 |
| chrXII | 456240 | 456387 | 147 | N12:456220 |
| chrXII | 456240 | 456419 | 179 | N12:456220 |
| chrXII | 456240 | 456419 | 179 | N12:456421 |
| chrXII | 456240 | 456447 | 207 | N12:456220 |
| chrXII | 456240 | 456447 | 207 | N12:456421 |
| chrXII | 456288 | 456387 | 99 | |
| chrXII | 456288 | 456395 | 107 | |
| chrXII | 456288 | 456419 | 131 | N12:456421 |
| chrXII | 456288 | 456447 | 159 | N12:456421 |
| chrXII | 456288 | 456457 | 169 | N12:456421 |
| chrXII | 456288 | 456471 | 183 | N12:456421 |
| chrXII | 456290 | 456419 | 129 | N12:456421 |
| chrXII | 456290 | 456447 | 157 | N12:456421 |
| chrXII | 456290 | 456471 | 181 | N12:456421 |
| chrXII | 456298 | 456447 | 149 | N12:456421 |
| chrXII | 456310 | 456417 | 107 | N12:456421 |
| chrXII | 456310 | 456419 | 109 | N12:456421 |
| chrXII | 456310 | 456447 | 137 | N12:456421 |
| chrXII | 456310 | 456471 | 161 | N12:456421 |
| chrXII | 456361 | 456447 | 86 | N12:456421 |
| chrXII | 456361 | 456466 | 105 | N12:456421 |

| CHR | Start | End | Length (bp) | Nucleosome_ID |
|---|---|---|---|---|
| chrXII | 456361 | 456468 | 107 | N12:456421 |
| chrXII | 456361 | 456470 | 109 | N12:456421 |
| chrXII | 456361 | 456471 | 110 | N12:456421 |
| chrXII | 456388 | 456471 | 83 | N12:456421 |
| chrXII | 456388 | 456551 | 163 | N12:456421 |
| chrXII | 456388 | 456551 | 163 | N12:456568 |
| chrXII | 456444 | 456594 | 150 | N12:456421 |
| chrXII | 456444 | 456594 | 150 | N12:456568 |
| chrXII | 456444 | 456595 | 151 | N12:456421 |
| chrXII | 456444 | 456595 | 151 | N12:456568 |
| chrXII | 456444 | 456619 | 175 | N12:456421 |
| chrXII | 456444 | 456619 | 175 | N12:456568 |
| chrXII | 456445 | 456595 | 150 | N12:456568 |
| chrXII | 456447 | 456594 | 147 | N12:456568 |
| chrXII | 456447 | 456595 | 148 | N12:456568 |
| chrXII | 456447 | 456608 | 161 | N12:456568 |
| chrXII | 456447 | 456619 | 172 | N12:456568 |
| chrXII | 456479 | 456585 | 106 | N12:456568 |
| chrXII | 456479 | 456594 | 115 | N12:456568 |
| chrXII | 456479 | 456608 | 129 | N12:456568 |
| chrXII | 456479 | 456609 | 130 | N12:456568 |
| chrXII | 456479 | 456653 | 174 | N12:456568 |
| chrXII | 456483 | 456594 | 111 | N12:456568 |
| chrXII | 456483 | 456608 | 125 | N12:456568 |
| chrXII | 456483 | 456609 | 126 | N12:456568 |
| chrXII | 456493 | 456599 | 106 | N12:456568 |
| chrXII | 456493 | 456608 | 115 | N12:456568 |
| chrXII | 456493 | 456653 | 160 | N12:456568 |
| chrXII | 456508 | 456594 | 86 | N12:456568 |
| chrXII | 456508 | 456608 | 100 | N12:456568 |
| chrXII | 456508 | 456642 | 134 | N12:456568 |
| chrXII | 456508 | 456653 | 145 | N12:456568 |
| chrXII | 456508 | 456670 | 162 | N12:456568 |
| chrXII | 456524 | 456695 | 171 | N12:456568 |
| chrXII | 456587 | 456738 | 151 | N12:456568 |
| chrXII | 456587 | 456738 | 151 | N12:456730 |
| chrXII | 456588 | 456751 | 163 | N12:456568 |
| chrXII | 456588 | 456751 | 163 | N12:456730 |
| chrXII | 456594 | 456751 | 157 | N12:456730 |
| chrXII | 456616 | 456772 | 156 | N12:456730 |
| chrXII | 456634 | 456751 | 117 | N12:456730 |
| chrXII | 456634 | 456762 | 128 | N12:456730 |
| chrXII | 456634 | 456781 | 147 | N12:456730 |
| chrXII | 456649 | 456803 | 154 | N12:456730 |
| chrXII | 456653 | 456762 | 109 | N12:456730 |
| chrXII | 456653 | 456803 | 150 | N12:456730 |
| chrXII | 456733 | 456869 | 136 | N12:456730 |
| chrXII | 456733 | 456869 | 136 | N12:456891 |
| chrXII | 456766 | 456920 | 154 | N12:456891 |
| chrXII | 456780 | 456887 | 107 | N12:456891 |
| chrXII | 456780 | 456890 | 110 | N12:456891 |
| chrXII | 456780 | 456920 | 140 | N12:456891 |
| chrXII | 456794 | 456903 | 109 | N12:456891 |
| chrXII | 456795 | 456903 | 108 | N12:456891 |
| chrXII | 456795 | 456904 | 109 | N12:456891 |
| chrXII | 456795 | 456905 | 110 | N12:456891 |
| chrXII | 456797 | 456890 | 93 | N12:456891 |
| chrXII | 456797 | 456920 | 123 | N12:456891 |
| chrXII | 456803 | 456959 | 156 | N12:456891 |
| chrXII | 456804 | 456911 | 107 | N12:456891 |
| chrXII | 456804 | 456920 | 116 | N12:456891 |
| chrXII | 456804 | 456960 | 156 | N12:456891 |
| chrXII | 456817 | 456920 | 103 | N12:456891 |
| chrXII | 456817 | 456971 | 154 | N12:456891 |
| chrXII | 456938 | 457104 | 166 | N12:457054 |
| chrXII | 456984 | 457118 | 134 | N12:457054 |
| chrXII | 456984 | 457127 | 143 | N12:457054 |
| chrXII | 456984 | 457130 | 146 | N12:457054 |
| chrXII | 456984 | 457140 | 156 | N12:457054 |
| chrXII | 456990 | 457127 | 137 | N12:457054 |
| chrXII | 456990 | 457140 | 150 | N12:457054 |
| chrXII | 457005 | 457166 | 161 | N12:457054 |
| chrXII | 457021 | 457127 | 106 | N12:457054 |
| chrXII | 457022 | 457205 | 183 | N12:457054 |
| chrXII | 457022 | 457205 | 183 | N12:457213 |
| chrXII | 457047 | 457207 | 160 | N12:457054 |
| chrXII | 457047 | 457207 | 160 | N12:457213 |
| chrXII | 457070 | 457200 | 130 | N12:457054 |
| chrXII | 457070 | 457200 | 130 | N12:457213 |
| chrXII | 457083 | 457217 | 134 | N12:457213 |
| chrXII | 457115 | 457248 | 133 | N12:457213 |
| chrXII | 457140 | 457305 | 165 | N12:457213 |
| chrXII | 457140 | 457312 | 172 | N12:457213 |
| chrXII | 457143 | 457248 | 105 | N12:457213 |
| chrXII | 457143 | 457279 | 136 | N12:457213 |
| chrXII | 457143 | 457316 | 173 | N12:457213 |
| chrXII | 457152 | 457305 | 153 | N12:457213 |
| chrXII | 457152 | 457312 | 160 | N12:457213 |
| chrXII | 457153 | 457248 | 95 | N12:457213 |
| chrXII | 457153 | 457279 | 126 | N12:457213 |
| chrXII | 457153 | 457312 | 159 | N12:457213 |
| chrXII | 457153 | 457316 | 163 | N12:457213 |
| chrXII | 457153 | 457317 | 164 | N12:457213 |
| chrXII | 457154 | 457312 | 158 | N12:457213 |
| chrXII | 457154 | 457316 | 162 | N12:457213 |
| chrXII | 457187 | 457316 | 129 | N12:457213 |
| chrXII | 457187 | 457317 | 130 | N12:457213 |
| chrXII | 457198 | 457346 | 148 | N12:457213 |
| chrXII | 457198 | 457365 | 167 | N12:457213 |
| chrXII | 457212 | 457365 | 153 | N12:457213 |
| chrXII | 457214 | 457363 | 149 | N12:457213 |
| chrXII | 457219 | 457312 | 93 | N12:457213 |
| chrXII | 457219 | 457346 | 127 | N12:457213 |

TABLE 1

| CHR | Start | End | Length (bp) | Nucleosome_ID |
|---|---|---|---|---|
| chrXII | 457219 | 457363 | 144 | N12:457213 |
| chrXII | 457219 | 457365 | 146 | N12:457213 |
| chrXII | 457219 | 457422 | 203 | N12:457213 |
| chrXII | 457227 | 457316 | 89 | N12:457213 |
| chrXII | 457227 | 457363 | 136 | N12:457213 |
| chrXII | 457227 | 457395 | 168 | N12:457213 |
| chrXII | 457230 | 457395 | 165 | N12:457213 |
| chrXII | 457230 | 457401 | 171 | N12:457213 |
| chrXII | 457248 | 457395 | 147 | |
| chrXII | 457248 | 457401 | 153 | |
| chrXII | 457255 | 457365 | 110 | |
| chrXII | 457255 | 457407 | 152 | |
| chrXII | 457255 | 457422 | 167 | |
| chrXII | 457266 | 457365 | 99 | |
| chrXII | 457266 | 457407 | 141 | |
| chrXII | 457266 | 457422 | 156 | |
| chrXII | 457295 | 457378 | 83 | |
| chrXII | 457295 | 457384 | 89 | |
| chrXII | 457295 | 457390 | 95 | |
| chrXII | 457295 | 457391 | 96 | |
| chrXII | 457295 | 457393 | 98 | |
| chrXII | 457295 | 457394 | 99 | |
| chrXII | 457295 | 457395 | 100 | |
| chrXII | 457295 | 457397 | 102 | |
| chrXII | 457295 | 457401 | 106 | |
| chrXII | 457295 | 457402 | 107 | |
| chrXII | 457337 | 457476 | 139 | N12:457494 |
| chrXII | 457337 | 457481 | 144 | N12:457494 |
| chrXII | 457360 | 457476 | 116 | N12:457494 |
| chrXII | 457360 | 457481 | 121 | N12:457494 |
| chrXII | 457360 | 457530 | 170 | N12:457494 |
| chrXII | 457370 | 457476 | 106 | N12:457494 |
| chrXII | 457372 | 457476 | 104 | N12:457494 |
| chrXII | 457420 | 457563 | 143 | N12:457494 |
| chrXII | 457420 | 457567 | 147 | N12:457494 |
| chrXII | 457420 | 457571 | 151 | N12:457494 |
| chrXII | 457420 | 457572 | 152 | N12:457494 |
| chrXII | 457437 | 457592 | 155 | N12:457494 |
| chrXII | 457440 | 457593 | 153 | N12:457494 |
| chrXII | 457540 | 457654 | 114 | |
| chrXII | 457553 | 457706 | 153 | N12:457682 |
| chrXII | 457569 | 457736 | 167 | N12:457682 |
| chrXII | 457569 | 457741 | 172 | N12:457682 |
| chrXII | 457600 | 457736 | 136 | N12:457682 |
| chrXII | 457600 | 457739 | 139 | N12:457682 |
| chrXII | 457600 | 457741 | 141 | N12:457682 |
| chrXII | 457606 | 457763 | 157 | N12:457682 |
| chrXII | 457620 | 457773 | 153 | N12:457682 |
| chrXII | 457620 | 457787 | 167 | N12:457682 |
| chrXII | 457622 | 457771 | 149 | N12:457682 |
| chrXII | 457634 | 457757 | 123 | N12:457682 |
| chrXII | 457650 | 457805 | 155 | N12:457682 |
| chrXII | 457667 | 457819 | 152 | N12:457682 |
| chrXII | 457803 | 457959 | 156 | N12:457885 |
| chrXII | 457816 | 457974 | 158 | N12:457885 |
| chrXII | 457820 | 457953 | 133 | N12:457885 |
| chrXII | 457872 | 458023 | 151 | N12:457885 |
| chrXII | 457945 | 458104 | 159 | |
| chrXII | 457949 | 458059 | 110 | |
| chrXII | 457949 | 458101 | 152 | |
| chrXII | 457950 | 458101 | 151 | |
| chrXII | 457950 | 458103 | 153 | |
| chrXII | 457959 | 458110 | 151 | |
| chrXII | 457976 | 458059 | 83 | |
| chrXII | 457976 | 458119 | 143 | |
| chrXII | 457983 | 458089 | 106 | |
| chrXII | 457983 | 458101 | 118 | |
| chrXII | 457983 | 458103 | 120 | |
| chrXII | 457983 | 458137 | 154 | N12:458145 |
| chrXII | 457986 | 458117 | 131 | |
| chrXII | 458025 | 458167 | 142 | N12:458145 |
| chrXII | 458055 | 458137 | 82 | N12:458145 |
| chrXII | 458055 | 458206 | 151 | N12:458145 |
| chrXII | 458079 | 458227 | 148 | N12:458145 |
| chrXII | 458087 | 458231 | 144 | N12:458145 |
| chrXII | 458127 | 458277 | 150 | N12:458145 |
| chrXII | 458130 | 458293 | 163 | N12:458145 |
| chrXII | 458134 | 458272 | 138 | N12:458145 |
| chrXII | 458134 | 458277 | 143 | N12:458145 |
| chrXII | 458136 | 458295 | 159 | N12:458145 |
| chrXII | 458143 | 458291 | 148 | N12:458145 |
| chrXII | 458166 | 458321 | 155 | N12:458145 |
| chrXII | 458166 | 458321 | 155 | N12:458325 |
| chrXII | 458166 | 458322 | 156 | N12:458145 |
| chrXII | 458166 | 458322 | 156 | N12:458325 |
| chrXII | 458167 | 458322 | 155 | N12:458145 |
| chrXII | 458167 | 458322 | 155 | N12:458325 |
| chrXII | 458177 | 458322 | 145 | N12:458325 |
| chrXII | 458177 | 458327 | 150 | N12:458325 |
| chrXII | 458177 | 458350 | 173 | N12:458325 |
| chrXII | 458179 | 458327 | 148 | N12:458325 |
| chrXII | 458191 | 458321 | 130 | N12:458325 |
| chrXII | 458191 | 458352 | 161 | N12:458325 |
| chrXII | 458195 | 458322 | 127 | N12:458325 |
| chrXII | 458195 | 458327 | 132 | N12:458325 |
| chrXII | 458195 | 458350 | 155 | N12:458325 |
| chrXII | 458204 | 458350 | 146 | N12:458325 |
| chrXII | 458204 | 458380 | 176 | N12:458325 |
| chrXII | 458291 | 458451 | 160 | N12:458325 |
| chrXII | 458297 | 458451 | 154 | N12:458325 |
| chrXII | 458298 | 458451 | 153 | N12:458325 |
| chrXII | 458319 | 458472 | 153 | N12:458325 |
| chrXII | 458338 | 458492 | 154 | N12:458325 |

| CHR | Start | End | Length (bp) | Nucleosome_ID |
|---|---|---|---|---|
| chrXII | 458356 | 458493 | 137 | |
| chrXII | 458356 | 458508 | 152 | N12:458524 |
| chrXII | 458359 | 458523 | 164 | N12:458524 |
| chrXII | 458365 | 458493 | 128 | |
| chrXII | 458373 | 458493 | 120 | |
| chrXII | 458373 | 458530 | 157 | N12:458524 |
| chrXII | 458374 | 458532 | 158 | N12:458524 |
| chrXII | 458395 | 458501 | 106 | N12:458524 |
| chrXII | 458395 | 458519 | 124 | N12:458524 |
| chrXII | 458419 | 458587 | 168 | N12:458524 |
| chrXII | 458508 | 458658 | 150 | N12:458524 |
| chrXII | 458508 | 458658 | 150 | N12:458681 |
| chrXII | 458508 | 458659 | 151 | N12:458524 |
| chrXII | 458508 | 458659 | 151 | N12:458681 |
| chrXII | 458746 | 458901 | 155 | N12:458830 |
| chrXII | 458746 | 458912 | 166 | N12:458830 |
| chrXII | 458750 | 458912 | 162 | N12:458830 |
| chrXII | 458768 | 458900 | 132 | N12:458830 |
| chrXII | 458768 | 458901 | 133 | N12:458830 |
| chrXII | 458768 | 458912 | 144 | N12:458830 |
| chrXII | 458769 | 458900 | 131 | N12:458830 |
| chrXII | 458769 | 458901 | 132 | N12:458830 |
| chrXII | 458769 | 458912 | 143 | N12:458830 |
| chrXII | 458769 | 458913 | 144 | N12:458830 |
| chrXII | 458770 | 458900 | 130 | N12:458830 |
| chrXII | 458770 | 458901 | 131 | N12:458830 |
| chrXII | 458770 | 458912 | 142 | N12:458830 |
| chrXII | 458846 | 459002 | 156 | N12:458830 |
| chrXII | 458846 | 459002 | 156 | N12:458985 |
| chrXII | 458902 | 459048 | 146 | N12:458985 |
| chrXII | 458908 | 459044 | 136 | N12:458985 |
| chrXII | 458908 | 459048 | 140 | N12:458985 |
| chrXII | 458910 | 459043 | 133 | N12:458985 |
| chrXII | 458910 | 459066 | 156 | N12:458985 |
| chrXII | 458913 | 459002 | 89 | N12:458985 |
| chrXII | 458913 | 459038 | 125 | N12:458985 |
| chrXII | 458913 | 459043 | 130 | N12:458985 |
| chrXII | 458913 | 459044 | 131 | N12:458985 |
| chrXII | 458913 | 459047 | 134 | N12:458985 |
| chrXII | 458913 | 459048 | 135 | N12:458985 |
| chrXII | 458913 | 459066 | 153 | N12:458985 |
| chrXII | 458932 | 459038 | 106 | N12:458985 |
| chrXII | 458933 | 459044 | 111 | N12:458985 |
| chrXII | 458933 | 459047 | 114 | N12:458985 |
| chrXII | 458933 | 459066 | 133 | N12:458985 |
| chrXII | 458957 | 459044 | 87 | N12:458985 |
| chrXII | 459050 | 459146 | 96 | |
| chrXII | 459089 | 459262 | 173 | N12:459173 |
| chrXII | 459089 | 459265 | 176 | N12:459173 |
| chrXII | 459095 | 459262 | 167 | N12:459173 |
| chrXII | 459095 | 459265 | 170 | N12:459173 |
| chrXII | 459098 | 459233 | 135 | N12:459173 |
| chrXII | 459098 | 459262 | 164 | N12:459173 |
| chrXII | 459166 | 459328 | 162 | N12:459173 |
| chrXII | 459185 | 459328 | 143 | N12:459173 |
| chrXII | 459204 | 459353 | 149 | |
| chrXII | 459204 | 459417 | 213 | |
| chrXII | 459204 | 459419 | 215 | |
| chrXII | 459227 | 459353 | 126 | |
| chrXII | 459227 | 459376 | 149 | |
| chrXII | 459227 | 459419 | 192 | |
| chrXII | 459227 | 459432 | 205 | |
| chrXII | 459235 | 459395 | 160 | |
| chrXII | 459236 | 459328 | 92 | |
| chrXII | 459236 | 459393 | 157 | |
| chrXII | 459236 | 459417 | 181 | |
| chrXII | 459236 | 459419 | 183 | |
| chrXII | 459264 | 459353 | 89 | |
| chrXII | 459264 | 459376 | 112 | |
| chrXII | 459264 | 459417 | 153 | |
| chrXII | 459264 | 459419 | 155 | |
| chrXII | 459264 | 459432 | 168 | |
| chrXII | 459264 | 459441 | 177 | |
| chrXII | 459265 | 459376 | 111 | |
| chrXII | 459265 | 459393 | 128 | |
| chrXII | 459265 | 459416 | 151 | |
| chrXII | 459265 | 459417 | 152 | |
| chrXII | 459265 | 459419 | 154 | |
| chrXII | 459265 | 459422 | 157 | |
| chrXII | 459265 | 459432 | 167 | |
| chrXII | 459265 | 459441 | 176 | |
| chrXII | 459265 | 459467 | 202 | N12:459485 |
| chrXII | 459265 | 459469 | 204 | N12:459485 |
| chrXII | 459267 | 459393 | 126 | |
| chrXII | 459267 | 459417 | 150 | |
| chrXII | 459267 | 459419 | 152 | |
| chrXII | 459267 | 459422 | 155 | |
| chrXII | 459267 | 459432 | 165 | |
| chrXII | 459267 | 459441 | 174 | |
| chrXII | 459267 | 459469 | 202 | N12:459485 |
| chrXII | 459268 | 459374 | 106 | |
| chrXII | 459268 | 459376 | 108 | |
| chrXII | 459268 | 459393 | 125 | |
| chrXII | 459268 | 459416 | 148 | |
| chrXII | 459268 | 459417 | 149 | |
| chrXII | 459268 | 459419 | 151 | |
| chrXII | 459268 | 459422 | 154 | |
| chrXII | 459268 | 459431 | 163 | |
| chrXII | 459268 | 459432 | 164 | |
| chrXII | 459268 | 459441 | 173 | |
| chrXII | 459268 | 459467 | 199 | N12:459485 |
| chrXII | 459268 | 459469 | 201 | N12:459485 |
| chrXII | 459269 | 459376 | 107 | |

| CHR | Start | End | Length (bp) | Nucleosome_ID |
|---|---|---|---|---|
| chrXII | 459269 | 459393 | 124 | |
| chrXII | 459269 | 459413 | 144 | |
| chrXII | 459269 | 459417 | 148 | |
| chrXII | 459269 | 459419 | 150 | |
| chrXII | 459269 | 459422 | 153 | |
| chrXII | 459269 | 459432 | 163 | |
| chrXII | 459269 | 459467 | 198 | N12:459485 |
| chrXII | 459269 | 459469 | 200 | N12:459485 |
| chrXII | 459271 | 459419 | 148 | |
| chrXII | 459308 | 459414 | 106 | |
| chrXII | 459308 | 459419 | 111 | |
| chrXII | 459308 | 459467 | 159 | N12:459485 |
| chrXII | 459308 | 459469 | 161 | N12:459485 |
| chrXII | 459325 | 459466 | 141 | N12:459485 |
| chrXII | 459326 | 459417 | 91 | |
| chrXII | 459326 | 459419 | 93 | |
| chrXII | 459326 | 459467 | 141 | N12:459485 |
| chrXII | 459326 | 459469 | 143 | N12:459485 |
| chrXII | 459408 | 459559 | 151 | N12:459485 |
| chrXII | 459429 | 459558 | 129 | N12:459485 |
| chrXII | 459429 | 459559 | 130 | N12:459485 |
| chrXII | 459429 | 459579 | 150 | N12:459485 |
| chrXII | 459430 | 459558 | 128 | N12:459485 |
| chrXII | 459430 | 459559 | 129 | N12:459485 |
| chrXII | 459430 | 459573 | 143 | N12:459485 |
| chrXII | 459430 | 459579 | 149 | N12:459485 |
| chrXII | 459519 | 459673 | 154 | |
| chrXII | 459521 | 459673 | 152 | |
| chrXII | 459602 | 459756 | 154 | N12:459720 |
| chrXII | 459602 | 459759 | 157 | N12:459720 |
| chrXII | 459604 | 459722 | 118 | N12:459720 |
| chrXII | 459604 | 459756 | 152 | N12:459720 |
| chrXII | 459604 | 459759 | 155 | N12:459720 |
| chrXII | 459604 | 459774 | 170 | N12:459720 |
| chrXII | 459604 | 459796 | 192 | N12:459720 |
| chrXII | 459604 | 459798 | 194 | N12:459720 |
| chrXII | 459604 | 459804 | 200 | N12:459720 |
| chrXII | 459604 | 459806 | 202 | N12:459720 |
| chrXII | 459618 | 459756 | 138 | N12:459720 |
| chrXII | 459618 | 459774 | 156 | N12:459720 |
| chrXII | 459618 | 459796 | 178 | N12:459720 |
| chrXII | 459621 | 459756 | 135 | N12:459720 |
| chrXII | 459621 | 459774 | 153 | N12:459720 |
| chrXII | 459625 | 459756 | 131 | N12:459720 |
| chrXII | 459625 | 459774 | 149 | N12:459720 |
| chrXII | 459625 | 459796 | 171 | N12:459720 |
| chrXII | 459625 | 459798 | 173 | N12:459720 |
| chrXII | 459625 | 459804 | 179 | N12:459720 |
| chrXII | 459625 | 459806 | 181 | N12:459720 |
| chrXII | 459628 | 459756 | 128 | N12:459720 |
| chrXII | 459628 | 459774 | 146 | N12:459720 |
| chrXII | 459628 | 459796 | 168 | N12:459720 |
| chrXII | 459628 | 459798 | 170 | N12:459720 |
| chrXII | 459628 | 459804 | 176 | N12:459720 |
| chrXII | 459628 | 459806 | 178 | N12:459720 |
| chrXII | 459633 | 459756 | 123 | N12:459720 |
| chrXII | 459633 | 459774 | 141 | N12:459720 |
| chrXII | 459633 | 459796 | 163 | N12:459720 |
| chrXII | 459633 | 459798 | 165 | N12:459720 |
| chrXII | 459633 | 459804 | 171 | N12:459720 |
| chrXII | 459633 | 459806 | 173 | N12:459720 |
| chrXII | 459648 | 459796 | 148 | N12:459720 |
| chrXII | 459673 | 459804 | 131 | N12:459720 |
| chrXII | 459673 | 459806 | 133 | N12:459720 |
| chrXII | 459686 | 459793 | 107 | N12:459720 |
| chrXII | 459686 | 459796 | 110 | N12:459720 |
| chrXII | 459686 | 459798 | 112 | N12:459720 |
| chrXII | 459686 | 459799 | 113 | N12:459720 |
| chrXII | 459686 | 459801 | 115 | N12:459720 |
| chrXII | 459686 | 459804 | 118 | N12:459720 |
| chrXII | 459686 | 459806 | 120 | N12:459720 |
| chrXII | 459686 | 459856 | 170 | N12:459720 |
| chrXII | 459688 | 459793 | 105 | N12:459720 |
| chrXII | 459688 | 459851 | 163 | N12:459720 |
| chrXII | 459698 | 459804 | 106 | N12:459720 |
| chrXII | 459698 | 459806 | 108 | N12:459720 |
| chrXII | 459698 | 459851 | 153 | N12:459720 |
| chrXII | 459698 | 459869 | 171 | N12:459720 |
| chrXII | 459818 | 459980 | 162 | N12:459899 |
| chrXII | 459834 | 459969 | 135 | N12:459899 |
| chrXII | 459846 | 459969 | 123 | N12:459899 |
| chrXII | 459846 | 460002 | 156 | N12:459899 |
| chrXII | 459899 | 460068 | 169 | N12:459899 |
| chrXII | 459899 | 460068 | 169 | N12:460071 |
| chrXII | 459930 | 460057 | 127 | N12:460071 |
| chrXII | 460017 | 460128 | 111 | N12:460071 |
| chrXII | 460017 | 460159 | 142 | N12:460071 |
| chrXII | 460032 | 460159 | 127 | N12:460071 |
| chrXII | 460102 | 460251 | 149 | |
| chrXII | 460102 | 460266 | 164 | |
| chrXII | 460112 | 460251 | 139 | |
| chrXII | 460112 | 460266 | 154 | |
| chrXII | 460188 | 460366 | 178 | N12:460347 |
| chrXII | 460243 | 460384 | 141 | N12:460347 |
| chrXII | 460256 | 460415 | 159 | N12:460347 |
| chrXII | 460262 | 460436 | 174 | N12:460347 |
| chrXII | 460302 | 460477 | 175 | N12:460347 |
| chrXII | 460314 | 460446 | 132 | N12:460347 |
| chrXII | 460402 | 460571 | 169 | |
| chrXII | 460461 | 460595 | 134 | N12:460603 |
| chrXII | 460487 | 460616 | 129 | N12:460603 |
| chrXII | 460493 | 460595 | 102 | N12:460603 |
| chrXII | 460493 | 460659 | 166 | N12:460603 |

| CHR | Start | End | Length (bp) | Nucleosome_ID |
|---|---|---|---|---|
| chrXII | 460512 | 460595 | 83 | N12:460603 |
| chrXII | 460512 | 460639 | 127 | N12:460603 |
| chrXII | 460521 | 460659 | 138 | N12:460603 |
| chrXII | 460602 | 460763 | 161 | N12:460603 |
| chrXII | 460846 | 460979 | 133 | N12:460826 |
| chrXII | 460863 | 461022 | 159 | |
| chrXII | 460871 | 461006 | 135 | |
| chrXII | 460894 | 460979 | 85 | |
| chrXII | 460894 | 461045 | 151 | |
| chrXII | 460894 | 461097 | 203 | |
| chrXII | 460906 | 461069 | 163 | |
| chrXII | 460927 | 461045 | 118 | |
| chrXII | 460927 | 461097 | 170 | |
| chrXII | 460937 | 461071 | 134 | |
| chrXII | 460937 | 461098 | 161 | |
| chrXII | 460988 | 461071 | 83 | |
| chrXII | 460988 | 461095 | 107 | |
| chrXII | 460988 | 461096 | 108 | |
| chrXII | 460988 | 461098 | 110 | |
| chrXII | 461013 | 461146 | 133 | N12:461125 |
| chrXII | 461056 | 461146 | 90 | N12:461125 |
| chrXII | 461056 | 461148 | 92 | N12:461125 |
| chrXII | 461056 | 461199 | 143 | N12:461125 |
| chrXII | 461056 | 461212 | 156 | N12:461125 |
| chrXII | 461056 | 461214 | 158 | N12:461125 |
| chrXII | 461056 | 461215 | 159 | N12:461125 |
| chrXII | 461056 | 461229 | 173 | N12:461125 |
| chrXII | 461060 | 461146 | 86 | N12:461125 |
| chrXII | 461060 | 461212 | 152 | N12:461125 |
| chrXII | 461060 | 461214 | 154 | N12:461125 |
| chrXII | 461060 | 461215 | 155 | N12:461125 |
| chrXII | 461060 | 461229 | 169 | N12:461125 |
| chrXII | 461068 | 461199 | 131 | N12:461125 |
| chrXII | 461068 | 461214 | 146 | N12:461125 |
| chrXII | 461068 | 461246 | 178 | N12:461125 |
| chrXII | 461069 | 461199 | 130 | N12:461125 |
| chrXII | 461069 | 461214 | 145 | N12:461125 |
| chrXII | 461069 | 461246 | 177 | N12:461125 |
| chrXII | 461082 | 461188 | 106 | N12:461125 |
| chrXII | 461082 | 461199 | 117 | N12:461125 |
| chrXII | 461082 | 461212 | 130 | N12:461125 |
| chrXII | 461082 | 461214 | 132 | N12:461125 |
| chrXII | 461082 | 461215 | 133 | N12:461125 |
| chrXII | 461082 | 461229 | 147 | N12:461125 |
| chrXII | 461082 | 461246 | 164 | N12:461125 |
| chrXII | 461095 | 461246 | 151 | N12:461125 |
| chrXII | 461178 | 461333 | 155 | N12:461308 |
| chrXII | 461196 | 461352 | 156 | N12:461308 |
| chrXII | 461218 | 461364 | 146 | N12:461308 |
| chrXII | 461218 | 461384 | 166 | N12:461308 |
| chrXII | 461218 | 461388 | 170 | N12:461308 |
| chrXII | 461241 | 461384 | 143 | N12:461308 |
| chrXII | 461241 | 461388 | 147 | N12:461308 |
| chrXII | 461252 | 461408 | 156 | N12:461308 |
| chrXII | 461260 | 461367 | 107 | N12:461308 |
| chrXII | 461327 | 461481 | 154 | N12:461308 |
| chrXII | 461397 | 461551 | 154 | N12:461507 |
| chrXII | 461430 | 461537 | 107 | N12:461507 |
| chrXII | 461431 | 461592 | 161 | N12:461507 |
| chrXII | 461432 | 461592 | 160 | N12:461507 |
| chrXII | 461436 | 461592 | 156 | N12:461507 |
| chrXII | 461460 | 461622 | 162 | N12:461507 |
| chrXII | 461460 | 461623 | 163 | N12:461507 |
| chrXII | 461471 | 461578 | 107 | N12:461507 |
| chrXII | 461471 | 461592 | 121 | N12:461507 |
| chrXII | 461471 | 461622 | 151 | N12:461507 |
| chrXII | 461471 | 461623 | 152 | N12:461507 |
| chrXII | 461483 | 461633 | 150 | N12:461507 |
| chrXII | 461536 | 461685 | 149 | N12:461692 |
| chrXII | 461560 | 461721 | 161 | N12:461692 |
| chrXII | 461606 | 461757 | 151 | N12:461692 |
| chrXII | 461606 | 461758 | 152 | N12:461692 |
| chrXII | 461606 | 461766 | 160 | N12:461692 |
| chrXII | 461606 | 461767 | 161 | N12:461692 |
| chrXII | 461628 | 461758 | 130 | N12:461692 |
| chrXII | 461628 | 461766 | 138 | N12:461692 |
| chrXII | 461628 | 461800 | 172 | N12:461692 |
| chrXII | 461629 | 461721 | 92 | N12:461692 |
| chrXII | 461629 | 461757 | 128 | N12:461692 |
| chrXII | 461629 | 461758 | 129 | N12:461692 |
| chrXII | 461629 | 461766 | 137 | N12:461692 |
| chrXII | 461629 | 461800 | 171 | N12:461692 |
| chrXII | 461662 | 461758 | 96 | N12:461692 |
| chrXII | 461662 | 461832 | 170 | N12:461692 |
| chrXII | 461676 | 461810 | 134 | N12:461692 |
| chrXII | 461687 | 461800 | 113 | N12:461692 |
| chrXII | 461687 | 461809 | 122 | N12:461692 |
| chrXII | 461687 | 461839 | 152 | N12:461692 |
| chrXII | 461805 | 461933 | 128 | N12:461891 |
| chrXII | 461805 | 461951 | 146 | N12:461891 |
| chrXII | 461821 | 461933 | 112 | N12:461891 |
| chrXII | 461821 | 461972 | 151 | N12:461891 |
| chrXII | 461823 | 461951 | 128 | N12:461891 |
| chrXII | 461823 | 462001 | 178 | N12:461891 |
| chrXII | 461846 | 461952 | 106 | N12:461891 |
| chrXII | 461846 | 462001 | 155 | N12:461891 |
| chrXII | 461846 | 462021 | 175 | N12:461891 |
| chrXII | 461846 | 462037 | 191 | N12:461891 |
| chrXII | 461860 | 461951 | 91 | N12:461891 |
| chrXII | 461860 | 462001 | 141 | N12:461891 |
| chrXII | 461860 | 462021 | 161 | N12:461891 |
| chrXII | 461860 | 462037 | 177 | N12:461891 |
| chrXII | 461902 | 462001 | 99 | N12:461891 |

| CHR | Start | End | Length (bp) | Nucleosome_ID |
|---|---|---|---|---|
| chrXII | 461902 | 462008 | 106 | N12:461891 |
| chrXII | 461902 | 462021 | 119 | N12:461891 |
| chrXII | 461902 | 462037 | 135 | N12:461891 |
| chrXII | 461902 | 462062 | 160 | N12:461891 |
| chrXII | 461902 | 462062 | 160 | N12:462072 |
| chrXII | 461910 | 462001 | 91 | N12:461891 |
| chrXII | 461910 | 462016 | 106 | N12:461891 |
| chrXII | 461910 | 462021 | 111 | N12:461891 |
| chrXII | 461910 | 462037 | 127 | N12:461891 |
| chrXII | 461910 | 462062 | 152 | N12:461891 |
| chrXII | 461910 | 462062 | 152 | N12:462072 |
| chrXII | 461920 | 462099 | 179 | N12:462072 |
| chrXII | 461925 | 462060 | 135 | N12:462072 |
| chrXII | 461925 | 462090 | 165 | N12:462072 |
| chrXII | 461938 | 462060 | 122 | N12:462072 |
| chrXII | 461938 | 462090 | 152 | N12:462072 |
| chrXII | 461938 | 462136 | 198 | N12:462072 |
| chrXII | 461977 | 462090 | 113 | N12:462072 |
| chrXII | 461977 | 462136 | 159 | N12:462072 |
| chrXII | 461977 | 462160 | 183 | N12:462072 |
| chrXII | 461984 | 462090 | 106 | N12:462072 |
| chrXII | 461984 | 462133 | 149 | N12:462072 |
| chrXII | 462013 | 462171 | 158 | N12:462072 |
| chrXII | 462022 | 462171 | 149 | N12:462072 |
| chrXII | 462030 | 462136 | 106 | N12:462072 |
| chrXII | 462032 | 462160 | 128 | N12:462072 |
| chrXII | 462059 | 462166 | 107 | N12:462072 |
| chrXII | 462059 | 462167 | 108 | N12:462072 |
| chrXII | 462059 | 462171 | 112 | N12:462072 |
| chrXII | 462059 | 462193 | 134 | N12:462072 |
| chrXII | 462059 | 462212 | 153 | N12:462072 |
| chrXII | 462063 | 462134 | 71 | N12:462072 |
| chrXII | 462063 | 462170 | 107 | N12:462072 |
| chrXII | 462063 | 462171 | 108 | N12:462072 |
| chrXII | 462063 | 462193 | 130 | N12:462072 |
| chrXII | 462063 | 462207 | 144 | N12:462072 |
| chrXII | 462063 | 462212 | 149 | N12:462072 |
| chrXII | 462113 | 462275 | 162 | N12:462253 |
| chrXII | 462128 | 462278 | 150 | N12:462253 |
| chrXII | 462168 | 462275 | 107 | N12:462253 |
| chrXII | 462170 | 462300 | 130 | N12:462253 |
| chrXII | 462170 | 462324 | 154 | N12:462253 |
| chrXII | 462170 | 462330 | 160 | N12:462253 |
| chrXII | 462176 | 462300 | 124 | N12:462253 |
| chrXII | 462176 | 462324 | 148 | N12:462253 |
| chrXII | 462176 | 462330 | 154 | N12:462253 |
| chrXII | 462179 | 462349 | 170 | N12:462253 |
| chrXII | 462179 | 462355 | 176 | N12:462253 |
| chrXII | 462180 | 462300 | 120 | N12:462253 |
| chrXII | 462180 | 462324 | 144 | N12:462253 |
| chrXII | 462180 | 462327 | 147 | N12:462253 |
| chrXII | 462180 | 462330 | 150 | N12:462253 |
| chrXII | 462228 | 462336 | 108 | N12:462253 |
| chrXII | 462229 | 462336 | 107 | N12:462253 |
| chrXII | 462229 | 462337 | 108 | N12:462253 |
| chrXII | 462229 | 462349 | 120 | N12:462253 |
| chrXII | 462229 | 462373 | 144 | N12:462253 |
| chrXII | 462229 | 462382 | 153 | N12:462253 |
| chrXII | 462229 | 462388 | 159 | N12:462253 |
| chrXII | 462229 | 462407 | 178 | N12:462253 |
| chrXII | 462229 | 462423 | 194 | N12:462253 |
| chrXII | 462229 | 462425 | 196 | N12:462253 |
| chrXII | 462232 | 462338 | 106 | N12:462253 |
| chrXII | 462232 | 462349 | 117 | N12:462253 |
| chrXII | 462232 | 462355 | 123 | N12:462253 |
| chrXII | 462232 | 462373 | 141 | N12:462253 |
| chrXII | 462232 | 462382 | 150 | N12:462253 |
| chrXII | 462232 | 462407 | 175 | N12:462253 |
| chrXII | 462232 | 462419 | 187 | N12:462253 |
| chrXII | 462232 | 462423 | 191 | N12:462253 |
| chrXII | 462232 | 462425 | 193 | N12:462253 |
| chrXII | 462236 | 462373 | 137 | N12:462253 |
| chrXII | 462236 | 462425 | 189 | N12:462253 |
| chrXII | 462237 | 462375 | 138 | N12:462253 |
| chrXII | 462237 | 462382 | 145 | N12:462253 |
| chrXII | 462250 | 462382 | 132 | N12:462253 |
| chrXII | 462250 | 462407 | 157 | N12:462253 |
| chrXII | 462250 | 462423 | 173 | N12:462253 |
| chrXII | 462250 | 462425 | 175 | N12:462253 |
| chrXII | 462251 | 462373 | 122 | N12:462253 |
| chrXII | 462251 | 462406 | 155 | N12:462253 |
| chrXII | 462251 | 462407 | 156 | N12:462253 |
| chrXII | 462251 | 462423 | 172 | N12:462253 |
| chrXII | 462251 | 462424 | 173 | N12:462253 |
| chrXII | 462251 | 462425 | 174 | N12:462253 |
| chrXII | 462251 | 462439 | 188 | N12:462253 |
| chrXII | 462253 | 462406 | 153 | N12:462253 |
| chrXII | 462253 | 462407 | 154 | N12:462253 |
| chrXII | 462253 | 462424 | 171 | N12:462253 |
| chrXII | 462253 | 462425 | 172 | N12:462253 |
| chrXII | 462256 | 462373 | 117 | N12:462253 |
| chrXII | 462256 | 462388 | 132 | N12:462253 |
| chrXII | 462256 | 462406 | 150 | N12:462253 |
| chrXII | 462256 | 462407 | 151 | N12:462253 |
| chrXII | 462256 | 462419 | 163 | N12:462253 |
| chrXII | 462256 | 462423 | 167 | N12:462253 |
| chrXII | 462256 | 462424 | 168 | N12:462253 |
| chrXII | 462256 | 462425 | 169 | N12:462253 |
| chrXII | 462256 | 462439 | 183 | N12:462253 |
| chrXII | 462256 | 462446 | 190 | N12:462253 |
| chrXII | 462257 | 462365 | 108 | N12:462253 |
| chrXII | 462258 | 462407 | 149 | N12:462253 |
| chrXII | 462258 | 462423 | 165 | N12:462253 |

| CHR | Start | End | Length (bp) | Nucleosome_ID |
|---|---|---|---|---|
| chrXII | 462258 | 462425 | 167 | N12:462253 |
| chrXII | 462262 | 462368 | 106 | N12:462253 |
| chrXII | 462262 | 462373 | 111 | N12:462253 |
| chrXII | 462262 | 462407 | 145 | N12:462253 |
| chrXII | 462262 | 462419 | 157 | N12:462253 |
| chrXII | 462262 | 462423 | 161 | N12:462253 |
| chrXII | 462262 | 462424 | 162 | N12:462253 |
| chrXII | 462262 | 462425 | 163 | N12:462253 |
| chrXII | 462262 | 462439 | 177 | N12:462253 |
| chrXII | 462263 | 462385 | 122 | N12:462253 |
| chrXII | 462263 | 462407 | 144 | N12:462253 |
| chrXII | 462263 | 462423 | 160 | N12:462253 |
| chrXII | 462263 | 462424 | 161 | N12:462253 |
| chrXII | 462263 | 462425 | 162 | N12:462253 |
| chrXII | 462272 | 462407 | 135 | N12:462253 |
| chrXII | 462272 | 462419 | 147 | N12:462253 |
| chrXII | 462272 | 462423 | 151 | N12:462253 |
| chrXII | 462272 | 462425 | 153 | N12:462253 |
| chrXII | 462275 | 462423 | 148 | N12:462253 |
| chrXII | 462275 | 462425 | 150 | N12:462253 |
| chrXII | 462277 | 462373 | 96 | |
| chrXII | 462277 | 462382 | 105 | |
| chrXII | 462277 | 462383 | 106 | |
| chrXII | 462277 | 462388 | 111 | |
| chrXII | 462277 | 462407 | 130 | |
| chrXII | 462277 | 462419 | 142 | |
| chrXII | 462277 | 462423 | 146 | |
| chrXII | 462277 | 462424 | 147 | |
| chrXII | 462277 | 462425 | 148 | |
| chrXII | 462277 | 462439 | 162 | |
| chrXII | 462277 | 462460 | 183 | |
| chrXII | 462278 | 462373 | 95 | |
| chrXII | 462278 | 462382 | 104 | |
| chrXII | 462278 | 462406 | 128 | |
| chrXII | 462278 | 462407 | 129 | |
| chrXII | 462278 | 462418 | 140 | |
| chrXII | 462278 | 462419 | 141 | |
| chrXII | 462278 | 462420 | 142 | |
| chrXII | 462278 | 462423 | 145 | |
| chrXII | 462278 | 462424 | 146 | |
| chrXII | 462278 | 462425 | 147 | |
| chrXII | 462278 | 462439 | 161 | |
| chrXII | 462278 | 462446 | 168 | |
| chrXII | 462278 | 462460 | 182 | |
| chrXII | 462278 | 462461 | 183 | |
| chrXII | 462278 | 462479 | 201 | |
| chrXII | 462281 | 462425 | 144 | |
| chrXII | 462287 | 462423 | 136 | |
| chrXII | 462287 | 462425 | 138 | |
| chrXII | 462287 | 462432 | 145 | |
| chrXII | 462287 | 462439 | 152 | |
| chrXII | 462289 | 462377 | 88 | |
| chrXII | 462289 | 462407 | 118 | |
| chrXII | 462289 | 462423 | 134 | |
| chrXII | 462289 | 462424 | 135 | |
| chrXII | 462289 | 462425 | 136 | |
| chrXII | 462289 | 462439 | 150 | |
| chrXII | 462290 | 462423 | 133 | |
| chrXII | 462290 | 462425 | 135 | |
| chrXII | 462296 | 462407 | 111 | |
| chrXII | 462296 | 462423 | 127 | |
| chrXII | 462296 | 462424 | 128 | |
| chrXII | 462296 | 462425 | 129 | |
| chrXII | 462296 | 462439 | 143 | |
| chrXII | 462296 | 462446 | 150 | |
| chrXII | 462296 | 462472 | 176 | |
| chrXII | 462296 | 462479 | 183 | |
| chrXII | 462298 | 462382 | 84 | |
| chrXII | 462298 | 462405 | 107 | |
| chrXII | 462298 | 462407 | 109 | |
| chrXII | 462298 | 462419 | 121 | |
| chrXII | 462298 | 462423 | 125 | |
| chrXII | 462298 | 462425 | 127 | |
| chrXII | 462298 | 462446 | 148 | |
| chrXII | 462298 | 462460 | 162 | |
| chrXII | 462298 | 462461 | 163 | |
| chrXII | 462299 | 462406 | 107 | |
| chrXII | 462299 | 462423 | 124 | |
| chrXII | 462299 | 462425 | 126 | |
| chrXII | 462299 | 462460 | 161 | |
| chrXII | 462299 | 462461 | 162 | |
| chrXII | 462302 | 462424 | 122 | |
| chrXII | 462302 | 462425 | 123 | |
| chrXII | 462302 | 462439 | 137 | |
| chrXII | 462302 | 462479 | 177 | |
| chrXII | 462308 | 462479 | 171 | |
| chrXII | 462317 | 462425 | 108 | |
| chrXII | 462317 | 462477 | 160 | |
| chrXII | 462318 | 462407 | 89 | |
| chrXII | 462318 | 462419 | 101 | |
| chrXII | 462318 | 462423 | 105 | |
| chrXII | 462318 | 462425 | 107 | |
| chrXII | 462318 | 462460 | 142 | |
| chrXII | 462318 | 462461 | 143 | |
| chrXII | 462423 | 462569 | 146 | N12:462515 |
| chrXII | 462436 | 462577 | 141 | N12:462515 |
| chrXII | 462436 | 462597 | 161 | N12:462515 |
| chrXII | 462437 | 462547 | 110 | N12:462515 |
| chrXII | 462437 | 462577 | 140 | N12:462515 |
| chrXII | 462439 | 462577 | 138 | N12:462515 |
| chrXII | 462440 | 462577 | 137 | N12:462515 |
| chrXII | 462447 | 462577 | 130 | N12:462515 |
| chrXII | 462447 | 462597 | 150 | N12:462515 |

| CHR | Start | End | Length (bp) | Nucleosome_ID |
|---|---|---|---|---|
| chrXII | 462459 | 462618 | 159 | N12:462515 |
| chrXII | 462595 | 462732 | 137 | N12:462669 |
| chrXII | 462595 | 462747 | 152 | N12:462669 |
| chrXII | 462595 | 462749 | 154 | N12:462669 |
| chrXII | 462597 | 462749 | 152 | N12:462669 |
| chrXII | 462624 | 462732 | 108 | N12:462669 |
| chrXII | 462624 | 462747 | 123 | N12:462669 |
| chrXII | 462624 | 462749 | 125 | N12:462669 |
| chrXII | 462637 | 462789 | 152 | N12:462669 |
| chrXII | 462711 | 462864 | 153 | N12:462833 |
| chrXII | 462724 | 462874 | 150 | N12:462833 |
| chrXII | 462764 | 462913 | 149 | N12:462833 |
| chrXII | 462764 | 462916 | 152 | N12:462833 |
| chrXII | 462794 | 462951 | 157 | N12:462833 |
| chrXII | 462796 | 462913 | 117 | N12:462833 |
| chrXII | 462796 | 462916 | 120 | N12:462833 |
| chrXII | 462871 | 462990 | 119 | N12:462990 |
| chrXII | 462885 | 463047 | 162 | N12:462990 |
| chrXII | 462885 | 463050 | 165 | N12:462990 |
| chrXII | 462885 | 463056 | 171 | N12:462990 |
| chrXII | 462885 | 463059 | 174 | N12:462990 |
| chrXII | 462885 | 463099 | 214 | N12:462990 |
| chrXII | 462889 | 462996 | 107 | N12:462990 |
| chrXII | 462889 | 463045 | 156 | N12:462990 |
| chrXII | 462889 | 463048 | 159 | N12:462990 |
| chrXII | 462889 | 463051 | 162 | N12:462990 |
| chrXII | 462889 | 463053 | 164 | N12:462990 |
| chrXII | 462889 | 463059 | 170 | N12:462990 |
| chrXII | 462889 | 463061 | 172 | N12:462990 |
| chrXII | 462889 | 463066 | 177 | N12:462990 |
| chrXII | 462890 | 463050 | 160 | N12:462990 |
| chrXII | 462890 | 463059 | 169 | N12:462990 |
| chrXII | 462891 | 463047 | 156 | N12:462990 |
| chrXII | 462891 | 463056 | 165 | N12:462990 |
| chrXII | 462891 | 463061 | 170 | N12:462990 |
| chrXII | 462911 | 463047 | 136 | N12:462990 |
| chrXII | 462911 | 463050 | 139 | N12:462990 |
| chrXII | 462911 | 463051 | 140 | N12:462990 |
| chrXII | 462911 | 463053 | 142 | N12:462990 |
| chrXII | 462911 | 463056 | 145 | N12:462990 |
| chrXII | 462911 | 463062 | 151 | N12:462990 |
| chrXII | 462911 | 463099 | 188 | N12:462990 |
| chrXII | 462912 | 463045 | 133 | N12:462990 |
| chrXII | 462912 | 463048 | 136 | N12:462990 |
| chrXII | 462912 | 463051 | 139 | N12:462990 |
| chrXII | 462912 | 463053 | 141 | N12:462990 |
| chrXII | 462912 | 463059 | 147 | N12:462990 |
| chrXII | 462918 | 463045 | 127 | N12:462990 |
| chrXII | 462918 | 463048 | 130 | N12:462990 |
| chrXII | 462918 | 463051 | 133 | N12:462990 |
| chrXII | 462918 | 463053 | 135 | N12:462990 |
| chrXII | 462918 | 463059 | 141 | N12:462990 |
| chrXII | 462919 | 463048 | 129 | N12:462990 |
| chrXII | 462919 | 463050 | 131 | N12:462990 |
| chrXII | 462919 | 463051 | 132 | N12:462990 |
| chrXII | 462919 | 463056 | 137 | N12:462990 |
| chrXII | 462919 | 463059 | 140 | N12:462990 |
| chrXII | 462919 | 463062 | 143 | N12:462990 |
| chrXII | 462919 | 463066 | 147 | N12:462990 |
| chrXII | 462919 | 463074 | 155 | N12:462990 |
| chrXII | 462919 | 463099 | 180 | N12:462990 |
| chrXII | 462920 | 463045 | 125 | N12:462990 |
| chrXII | 462920 | 463048 | 128 | N12:462990 |
| chrXII | 462920 | 463050 | 130 | N12:462990 |
| chrXII | 462920 | 463051 | 131 | N12:462990 |
| chrXII | 462949 | 463050 | 101 | N12:462990 |
| chrXII | 462949 | 463056 | 107 | N12:462990 |
| chrXII | 462949 | 463099 | 150 | N12:462990 |
| chrXII | 462949 | 463138 | 189 | N12:463152 |
| chrXII | 462949 | 463138 | 189 | N12:462990 |
| chrXII | 462988 | 463095 | 107 | N12:462990 |
| chrXII | 462988 | 463099 | 111 | N12:462990 |
| chrXII | 462988 | 463138 | 150 | N12:463152 |
| chrXII | 462988 | 463138 | 150 | N12:462990 |
| chrXII | 463046 | 463209 | 163 | N12:463152 |
| chrXII | 463055 | 463217 | 162 | N12:463152 |
| chrXII | 463059 | 463216 | 157 | N12:463152 |
| chrXII | 463059 | 463234 | 175 | N12:463152 |
| chrXII | 463066 | 463216 | 150 | N12:463152 |
| chrXII | 463066 | 463234 | 168 | N12:463152 |
| chrXII | 463066 | 463245 | 179 | N12:463152 |
| chrXII | 463096 | 463234 | 138 | N12:463152 |
| chrXII | 463096 | 463245 | 149 | N12:463152 |
| chrXII | 463096 | 463249 | 153 | N12:463152 |
| chrXII | 463097 | 463253 | 156 | N12:463152 |
| chrXII | 463101 | 463234 | 133 | N12:463152 |
| chrXII | 463101 | 463245 | 144 | N12:463152 |
| chrXII | 463108 | 463245 | 137 | N12:463152 |
| chrXII | 463108 | 463260 | 152 | N12:463152 |
| chrXII | 463196 | 463353 | 157 | N12:463325 |
| chrXII | 463248 | 463405 | 157 | N12:463325 |
| chrXII | 463257 | 463404 | 147 | N12:463325 |
| chrXII | 463257 | 463414 | 157 | N12:463325 |
| chrXII | 463261 | 463414 | 153 | N12:463325 |
| chrXII | 463268 | 463414 | 146 | N12:463325 |
| chrXII | 463278 | 463414 | 136 | N12:463325 |
| chrXII | 463333 | 463483 | 150 | N12:463325 |
| chrXII | 463333 | 463483 | 150 | N12:463504 |
| chrXII | 463358 | 463511 | 153 | N12:463504 |
| chrXII | 463358 | 463529 | 171 | N12:463504 |
| chrXII | 463381 | 463529 | 148 | N12:463504 |
| chrXII | 463381 | 463539 | 158 | N12:463504 |
| chrXII | 463385 | 463511 | 126 | N12:463504 |

| CHR | Start | End | Length (bp) | Nucleosome_ID |
|---|---|---|---|---|
| chrXII | 463385 | 463529 | 144 | N12:463504 |
| chrXII | 463385 | 463535 | 150 | N12:463504 |
| chrXII | 463385 | 463539 | 154 | N12:463504 |
| chrXII | 463410 | 463573 | 163 | N12:463504 |
| chrXII | 463414 | 463521 | 107 | N12:463504 |
| chrXII | 463414 | 463535 | 121 | N12:463504 |
| chrXII | 463414 | 463539 | 125 | N12:463504 |
| chrXII | 463414 | 463575 | 161 | N12:463504 |
| chrXII | 463414 | 463583 | 169 | N12:463504 |
| chrXII | 463463 | 463603 | 140 | N12:463504 |
| chrXII | 463471 | 463608 | 137 | N12:463504 |
| chrXII | 463535 | 463688 | 153 | |
| chrXII | 463535 | 463689 | 154 | |
| chrXII | 463535 | 463690 | 155 | |
| chrXII | 463535 | 463692 | 157 | |
| chrXII | 463535 | 463704 | 169 | |
| chrXII | 463535 | 463710 | 175 | |
| chrXII | 463535 | 463712 | 177 | |
| chrXII | 463541 | 463689 | 148 | |
| chrXII | 463541 | 463692 | 151 | |
| chrXII | 463559 | 463689 | 130 | |
| chrXII | 463559 | 463704 | 145 | |
| chrXII | 463559 | 463710 | 151 | |
| chrXII | 463559 | 463712 | 153 | |
| chrXII | 463658 | 463815 | 157 | N12:463786 |
| chrXII | 463658 | 463852 | 194 | N12:463786 |
| chrXII | 463658 | 463870 | 212 | N12:463786 |
| chrXII | 463690 | 463849 | 159 | N12:463786 |
| chrXII | 463690 | 463870 | 180 | N12:463786 |
| chrXII | 463692 | 463840 | 148 | N12:463786 |
| chrXII | 463697 | 463815 | 118 | N12:463786 |
| chrXII | 463697 | 463849 | 152 | N12:463786 |
| chrXII | 463697 | 463852 | 155 | N12:463786 |
| chrXII | 463697 | 463861 | 164 | N12:463786 |
| chrXII | 463697 | 463870 | 173 | N12:463786 |
| chrXII | 463699 | 463849 | 150 | N12:463786 |
| chrXII | 463707 | 463849 | 142 | N12:463786 |
| chrXII | 463707 | 463852 | 145 | N12:463786 |
| chrXII | 463707 | 463861 | 154 | N12:463786 |
| chrXII | 463707 | 463870 | 163 | N12:463786 |
| chrXII | 463713 | 463849 | 136 | N12:463786 |
| chrXII | 463713 | 463852 | 139 | N12:463786 |
| chrXII | 463713 | 463861 | 148 | N12:463786 |
| chrXII | 463713 | 463870 | 157 | N12:463786 |
| chrXII | 463713 | 463909 | 196 | N12:463786 |
| chrXII | 463724 | 463883 | 159 | N12:463786 |
| chrXII | 463729 | 463879 | 150 | N12:463786 |
| chrXII | 463763 | 463849 | 86 | N12:463786 |
| chrXII | 463763 | 463870 | 107 | N12:463786 |
| chrXII | 463763 | 463909 | 146 | N12:463786 |
| chrXII | 463860 | 463991 | 131 | N12:463954 |
| chrXII | 463862 | 464000 | 138 | N12:463954 |
| chrXII | 463862 | 464023 | 161 | N12:463954 |
| chrXII | 463870 | 464023 | 153 | N12:463954 |
| chrXII | 463878 | 464033 | 155 | N12:463954 |
| chrXII | 463884 | 464023 | 139 | N12:463954 |
| chrXII | 463974 | 464128 | 154 | N12:463954 |
| chrXII | 463974 | 464128 | 154 | N12:464114 |
| chrXII | 464001 | 464158 | 157 | N12:464114 |
| chrXII | 464013 | 464160 | 147 | N12:464114 |
| chrXII | 464013 | 464166 | 153 | N12:464114 |
| chrXII | 464038 | 464166 | 128 | N12:464114 |
| chrXII | 464038 | 464211 | 173 | N12:464114 |
| chrXII | 464057 | 464224 | 167 | N12:464114 |
| chrXII | 464069 | 464210 | 141 | N12:464114 |
| chrXII | 464070 | 464160 | 90 | N12:464114 |
| chrXII | 464070 | 464166 | 96 | N12:464114 |
| chrXII | 464070 | 464210 | 140 | N12:464114 |
| chrXII | 464070 | 464211 | 141 | N12:464114 |
| chrXII | 464070 | 464213 | 143 | N12:464114 |
| chrXII | 464101 | 464269 | 168 | N12:464114 |
| chrXII | 464101 | 464269 | 168 | N12:464279 |
| chrXII | 464132 | 464301 | 169 | N12:464114 |
| chrXII | 464132 | 464301 | 169 | N12:464279 |
| chrXII | 464137 | 464269 | 132 | N12:464114 |
| chrXII | 464137 | 464269 | 132 | N12:464279 |
| chrXII | 464152 | 464303 | 151 | N12:464279 |
| chrXII | 464195 | 464345 | 150 | N12:464279 |
| chrXII | 464205 | 464360 | 155 | N12:464279 |
| chrXII | 464232 | 464406 | 174 | N12:464279 |
| chrXII | 464232 | 464406 | 174 | N12:464427 |
| chrXII | 464233 | 464360 | 127 | N12:464279 |
| chrXII | 464233 | 464377 | 144 | N12:464279 |
| chrXII | 464233 | 464380 | 147 | N12:464279 |
| chrXII | 464233 | 464382 | 149 | N12:464279 |
| chrXII | 464233 | 464386 | 153 | N12:464279 |
| chrXII | 464234 | 464377 | 143 | N12:464279 |
| chrXII | 464234 | 464380 | 146 | N12:464279 |
| chrXII | 464234 | 464382 | 148 | N12:464279 |
| chrXII | 464235 | 464377 | 142 | N12:464279 |
| chrXII | 464235 | 464380 | 145 | N12:464279 |
| chrXII | 464235 | 464382 | 147 | N12:464279 |
| chrXII | 464235 | 464386 | 151 | N12:464279 |
| chrXII | 464266 | 464430 | 164 | N12:464279 |
| chrXII | 464266 | 464430 | 164 | N12:464427 |
| chrXII | 464275 | 464426 | 151 | N12:464279 |
| chrXII | 464275 | 464426 | 151 | N12:464427 |
| chrXII | 464319 | 464475 | 156 | N12:464427 |
| chrXII | 464354 | 464503 | 149 | N12:464427 |
| chrXII | 464358 | 464532 | 174 | N12:464427 |
| chrXII | 464384 | 464518 | 134 | N12:464427 |
| chrXII | 464391 | 464538 | 147 | N12:464427 |
| chrXII | 464392 | 464537 | 145 | N12:464427 |

| CHR | Start | End | Length (bp) | Nucleosome_ID |
|---|---|---|---|---|
| chrXII | 464415 | 464560 | 145 | N12:464427 |
| chrXII | 464434 | 464584 | 150 | N12:464427 |
| chrXII | 464504 | 464649 | 145 | N12:464616 |
| chrXII | 464504 | 464668 | 164 | N12:464616 |
| chrXII | 464504 | 464686 | 182 | N12:464616 |
| chrXII | 464514 | 464649 | 135 | N12:464616 |
| chrXII | 464514 | 464668 | 154 | N12:464616 |
| chrXII | 464514 | 464686 | 172 | N12:464616 |
| chrXII | 464534 | 464640 | 106 | N12:464616 |
| chrXII | 464534 | 464649 | 115 | N12:464616 |
| chrXII | 464534 | 464668 | 134 | N12:464616 |
| chrXII | 464534 | 464686 | 152 | N12:464616 |
| chrXII | 464547 | 464649 | 102 | N12:464616 |
| chrXII | 464547 | 464686 | 139 | N12:464616 |
| chrXII | 464547 | 464697 | 150 | N12:464616 |
| chrXII | 464547 | 464705 | 158 | N12:464616 |
| chrXII | 464547 | 464706 | 159 | N12:464616 |
| chrXII | 464553 | 464708 | 155 | N12:464616 |
| chrXII | 464559 | 464706 | 147 | N12:464616 |
| chrXII | 464728 | 464888 | 160 | N12:464796 |
| chrXII | 464838 | 465007 | 169 | N12:464962 |
| chrXII | 464884 | 465032 | 148 | N12:464962 |
| chrXII | 464886 | 465026 | 140 | N12:464962 |
| chrXII | 464920 | 465095 | 175 | N12:464962 |
| chrXII | 465006 | 465158 | 152 | N12:465169 |
| chrXII | 465040 | 465204 | 164 | N12:465169 |
| chrXII | 465055 | 465176 | 121 | N12:465169 |
| chrXII | 465063 | 465223 | 160 | N12:465169 |
| chrXII | 465159 | 465310 | 151 | N12:465169 |
| chrXII | 465159 | 465342 | 183 | N12:465169 |
| chrXII | 465159 | 465342 | 183 | N12:465356 |
| chrXII | 465159 | 465348 | 189 | N12:465169 |
| chrXII | 465159 | 465348 | 189 | N12:465356 |
| chrXII | 465159 | 465357 | 198 | N12:465169 |
| chrXII | 465159 | 465357 | 198 | N12:465356 |
| chrXII | 465179 | 465310 | 131 | N12:465169 |
| chrXII | 465182 | 465312 | 130 | N12:465169 |
| chrXII | 465193 | 465310 | 117 | |
| chrXII | 465193 | 465312 | 119 | |
| chrXII | 465193 | 465342 | 149 | N12:465356 |
| chrXII | 465193 | 465348 | 155 | N12:465356 |
| chrXII | 465193 | 465357 | 164 | N12:465356 |
| chrXII | 465200 | 465352 | 152 | N12:465356 |
| chrXII | 465201 | 465342 | 141 | N12:465356 |
| chrXII | 465201 | 465345 | 144 | N12:465356 |
| chrXII | 465201 | 465351 | 150 | N12:465356 |
| chrXII | 465203 | 465310 | 107 | |
| chrXII | 465204 | 465342 | 138 | N12:465356 |
| chrXII | 465204 | 465348 | 144 | N12:465356 |
| chrXII | 465204 | 465352 | 148 | N12:465356 |
| chrXII | 465204 | 465357 | 153 | N12:465356 |
| chrXII | 465204 | 465403 | 199 | N12:465356 |
| chrXII | 465204 | 465418 | 214 | N12:465356 |
| chrXII | 465217 | 465310 | 93 | |
| chrXII | 465217 | 465342 | 125 | N12:465356 |
| chrXII | 465217 | 465348 | 131 | N12:465356 |
| chrXII | 465217 | 465357 | 140 | N12:465356 |
| chrXII | 465221 | 465352 | 131 | N12:465356 |
| chrXII | 465221 | 465375 | 154 | N12:465356 |
| chrXII | 465221 | 465380 | 159 | N12:465356 |
| chrXII | 465221 | 465403 | 182 | N12:465356 |
| chrXII | 465252 | 465352 | 100 | N12:465356 |
| chrXII | 465252 | 465380 | 128 | N12:465356 |
| chrXII | 465252 | 465403 | 151 | N12:465356 |
| chrXII | 465252 | 465418 | 166 | N12:465356 |
| chrXII | 465257 | 465427 | 170 | N12:465356 |
| chrXII | 465261 | 465352 | 91 | N12:465356 |
| chrXII | 465261 | 465403 | 142 | N12:465356 |
| chrXII | 465261 | 465418 | 157 | N12:465356 |
| chrXII | 465261 | 465470 | 209 | N12:465356 |
| chrXII | 465318 | 465403 | 85 | N12:465356 |
| chrXII | 465318 | 465418 | 100 | N12:465356 |
| chrXII | 465318 | 465447 | 129 | N12:465356 |
| chrXII | 465318 | 465469 | 151 | N12:465356 |
| chrXII | 465318 | 465470 | 152 | N12:465356 |
| chrXII | 465318 | 465473 | 155 | N12:465356 |
| chrXII | 465318 | 465486 | 168 | N12:465356 |
| chrXII | 465318 | 465524 | 206 | N12:465356 |
| chrXII | 465318 | 465524 | 206 | N12:465541 |
| chrXII | 465323 | 465470 | 147 | N12:465356 |
| chrXII | 465324 | 465470 | 146 | N12:465356 |
| chrXII | 465324 | 465486 | 162 | N12:465356 |
| chrXII | 465331 | 465470 | 139 | N12:465356 |
| chrXII | 465331 | 465473 | 142 | N12:465356 |
| chrXII | 465350 | 465456 | 106 | N12:465356 |
| chrXII | 465350 | 465457 | 107 | N12:465356 |
| chrXII | 465350 | 465458 | 108 | N12:465356 |
| chrXII | 465350 | 465470 | 120 | N12:465356 |
| chrXII | 465350 | 465473 | 123 | N12:465356 |
| chrXII | 465350 | 465486 | 136 | N12:465356 |
| chrXII | 465350 | 465524 | 174 | N12:465356 |
| chrXII | 465350 | 465524 | 174 | N12:465541 |
| chrXII | 465350 | 465556 | 206 | N12:465356 |
| chrXII | 465350 | 465556 | 206 | N12:465541 |
| chrXII | 465358 | 465523 | 165 | N12:465356 |
| chrXII | 465358 | 465523 | 165 | N12:465541 |
| chrXII | 465377 | 465470 | 93 | N12:465356 |
| chrXII | 465377 | 465484 | 107 | N12:465356 |
| chrXII | 465377 | 465523 | 146 | N12:465356 |
| chrXII | 465377 | 465523 | 146 | N12:465541 |
| chrXII | 465377 | 465524 | 147 | N12:465356 |
| chrXII | 465377 | 465524 | 147 | N12:465541 |
| chrXII | 465377 | 465556 | 179 | N12:465356 |

| CHR | Start | End | Length (bp) | Nucleosome_ID |
|---|---|---|---|---|
| chrXII | 465377 | 465556 | 179 | N12:465541 |
| chrXII | 465377 | 465584 | 207 | N12:465356 |
| chrXII | 465377 | 465584 | 207 | N12:465541 |
| chrXII | 465425 | 465524 | 99 | N12:465541 |
| chrXII | 465425 | 465532 | 107 | N12:465541 |
| chrXII | 465425 | 465556 | 131 | N12:465541 |
| chrXII | 465425 | 465584 | 159 | N12:465541 |
| chrXII | 465425 | 465594 | 169 | N12:465541 |
| chrXII | 465425 | 465608 | 183 | N12:465541 |
| chrXII | 465427 | 465524 | 97 | N12:465541 |
| chrXII | 465427 | 465556 | 129 | N12:465541 |
| chrXII | 465427 | 465584 | 157 | N12:465541 |
| chrXII | 465427 | 465608 | 181 | N12:465541 |
| chrXII | 465435 | 465584 | 149 | N12:465541 |
| chrXII | 465447 | 465556 | 109 | N12:465541 |
| chrXII | 465447 | 465584 | 137 | N12:465541 |
| chrXII | 465447 | 465608 | 161 | N12:465541 |
| chrXII | 465498 | 465584 | 86 | N12:465541 |
| chrXII | 465498 | 465603 | 105 | N12:465541 |
| chrXII | 465498 | 465605 | 107 | N12:465541 |
| chrXII | 465498 | 465606 | 108 | N12:465541 |
| chrXII | 465498 | 465607 | 109 | N12:465541 |
| chrXII | 465498 | 465608 | 110 | N12:465541 |
| chrXII | 465525 | 465608 | 83 | N12:465541 |
| chrXII | 465525 | 465688 | 163 | N12:465541 |
| chrXII | 465525 | 465688 | 163 | N12:465711 |
| chrXII | 465562 | 465739 | 177 | N12:465541 |
| chrXII | 465562 | 465739 | 177 | N12:465711 |
| chrXII | 465581 | 465731 | 150 | N12:465711 |
| chrXII | 465581 | 465732 | 151 | N12:465711 |
| chrXII | 465581 | 465756 | 175 | N12:465711 |
| chrXII | 465582 | 465732 | 150 | N12:465711 |
| chrXII | 465584 | 465731 | 147 | N12:465711 |
| chrXII | 465584 | 465732 | 148 | N12:465711 |
| chrXII | 465584 | 465745 | 161 | N12:465711 |
| chrXII | 465584 | 465756 | 172 | N12:465711 |
| chrXII | 465616 | 465722 | 106 | N12:465711 |
| chrXII | 465616 | 465731 | 115 | N12:465711 |
| chrXII | 465616 | 465745 | 129 | N12:465711 |
| chrXII | 465616 | 465746 | 130 | N12:465711 |
| chrXII | 465616 | 465790 | 174 | N12:465711 |
| chrXII | 465620 | 465731 | 111 | N12:465711 |
| chrXII | 465620 | 465745 | 125 | N12:465711 |
| chrXII | 465620 | 465746 | 126 | N12:465711 |
| chrXII | 465630 | 465736 | 106 | N12:465711 |
| chrXII | 465630 | 465745 | 115 | N12:465711 |
| chrXII | 465630 | 465790 | 160 | N12:465711 |
| chrXII | 465645 | 465731 | 86 | N12:465711 |
| chrXII | 465645 | 465745 | 100 | N12:465711 |
| chrXII | 465645 | 465779 | 134 | N12:465711 |
| chrXII | 465645 | 465790 | 145 | N12:465711 |
| chrXII | 465645 | 465807 | 162 | N12:465711 |
| chrXII | 465661 | 465832 | 171 | N12:465711 |
| chrXII | 465724 | 465875 | 151 | N12:465711 |
| chrXII | 465724 | 465875 | 151 | N12:465875 |
| chrXII | 465725 | 465888 | 163 | N12:465711 |
| chrXII | 465725 | 465888 | 163 | N12:465875 |
| chrXII | 465731 | 465888 | 157 | N12:465711 |
| chrXII | 465731 | 465888 | 157 | N12:465875 |
| chrXII | 465753 | 465909 | 156 | N12:465875 |
| chrXII | 465771 | 465899 | 128 | N12:465875 |
| chrXII | 465771 | 465918 | 147 | N12:465875 |
| chrXII | 465786 | 465940 | 154 | N12:465875 |
| chrXII | 465790 | 465899 | 109 | N12:465875 |
| chrXII | 465790 | 465940 | 150 | N12:465875 |
| chrXII | 465903 | 466057 | 154 | N12:466028 |
| chrXII | 465917 | 466024 | 107 | N12:466028 |
| chrXII | 465917 | 466027 | 110 | N12:466028 |
| chrXII | 465917 | 466057 | 140 | N12:466028 |
| chrXII | 465931 | 466040 | 109 | N12:466028 |
| chrXII | 465932 | 466040 | 108 | N12:466028 |
| chrXII | 465932 | 466041 | 109 | N12:466028 |
| chrXII | 465934 | 466027 | 93 | N12:466028 |
| chrXII | 465934 | 466057 | 123 | N12:466028 |
| chrXII | 465940 | 466096 | 156 | N12:466028 |
| chrXII | 465941 | 466048 | 107 | N12:466028 |
| chrXII | 465941 | 466057 | 116 | N12:466028 |
| chrXII | 465941 | 466097 | 156 | N12:466028 |
| chrXII | 465954 | 466057 | 103 | N12:466028 |
| chrXII | 465954 | 466108 | 154 | N12:466028 |
| chrXII | 466075 | 466241 | 166 | N12:466190 |
| chrXII | 466105 | 466258 | 153 | N12:466190 |
| chrXII | 466121 | 466255 | 134 | N12:466190 |
| chrXII | 466121 | 466264 | 143 | N12:466190 |
| chrXII | 466121 | 466267 | 146 | N12:466190 |
| chrXII | 466121 | 466277 | 156 | N12:466190 |
| chrXII | 466127 | 466264 | 137 | N12:466190 |
| chrXII | 466127 | 466277 | 150 | N12:466190 |
| chrXII | 466142 | 466303 | 161 | N12:466190 |
| chrXII | 466158 | 466264 | 106 | N12:466190 |
| chrXII | 466159 | 466264 | 105 | N12:466190 |
| chrXII | 466159 | 466342 | 183 | N12:466190 |
| chrXII | 466159 | 466342 | 183 | N12:466349 |
| chrXII | 466184 | 466344 | 160 | N12:466190 |
| chrXII | 466184 | 466344 | 160 | N12:466349 |
| chrXII | 466207 | 466337 | 130 | N12:466190 |
| chrXII | 466207 | 466337 | 130 | N12:466349 |
| chrXII | 466220 | 466354 | 134 | N12:466349 |
| chrXII | 466252 | 466385 | 133 | N12:466349 |
| chrXII | 466277 | 466442 | 165 | N12:466349 |
| chrXII | 466277 | 466449 | 172 | N12:466349 |
| chrXII | 466280 | 466385 | 105 | N12:466349 |
| chrXII | 466280 | 466416 | 136 | N12:466349 |

| CHR | Start | End | Length (bp) | Nucleosome_ID |
|---|---|---|---|---|
| chrXII | 466280 | 466453 | 173 | N12:466349 |
| chrXII | 466289 | 466442 | 153 | N12:466349 |
| chrXII | 466289 | 466449 | 160 | N12:466349 |
| chrXII | 466290 | 466385 | 95 | N12:466349 |
| chrXII | 466290 | 466416 | 126 | N12:466349 |
| chrXII | 466290 | 466449 | 159 | N12:466349 |
| chrXII | 466290 | 466453 | 163 | N12:466349 |
| chrXII | 466290 | 466454 | 164 | N12:466349 |
| chrXII | 466291 | 466449 | 158 | N12:466349 |
| chrXII | 466291 | 466453 | 162 | N12:466349 |
| chrXII | 466324 | 466453 | 129 | N12:466349 |
| chrXII | 466324 | 466454 | 130 | N12:466349 |
| chrXII | 466335 | 466483 | 148 | N12:466349 |
| chrXII | 466349 | 466502 | 153 | N12:466349 |
| chrXII | 466351 | 466500 | 149 | N12:466349 |
| chrXII | 466356 | 466449 | 93 | N12:466349 |
| chrXII | 466356 | 466483 | 127 | N12:466349 |
| chrXII | 466356 | 466500 | 144 | N12:466349 |
| chrXII | 466356 | 466502 | 146 | N12:466349 |
| chrXII | 466364 | 466500 | 136 | N12:466349 |
| chrXII | 466367 | 466532 | 165 | N12:466349 |
| chrXII | 466367 | 466538 | 171 | N12:466349 |
| chrXII | 466385 | 466532 | 147 | |
| chrXII | 466385 | 466538 | 153 | |
| chrXII | 466392 | 466502 | 110 | |
| chrXII | 466392 | 466544 | 152 | |
| chrXII | 466403 | 466500 | 97 | |
| chrXII | 466403 | 466502 | 99 | |
| chrXII | 466403 | 466544 | 141 | |
| chrXII | 466403 | 466559 | 156 | |
| chrXII | 466432 | 466521 | 89 | |
| chrXII | 466432 | 466523 | 91 | |
| chrXII | 466432 | 466527 | 95 | |
| chrXII | 466432 | 466528 | 96 | |
| chrXII | 466432 | 466530 | 98 | |
| chrXII | 466432 | 466531 | 99 | |
| chrXII | 466432 | 466532 | 100 | |
| chrXII | 466432 | 466534 | 102 | |
| chrXII | 466432 | 466538 | 106 | |
| chrXII | 466432 | 466539 | 107 | |
| chrXII | 466474 | 466613 | 139 | N12:466630 |
| chrXII | 466474 | 466618 | 144 | N12:466630 |
| chrXII | 466497 | 466613 | 116 | N12:466630 |
| chrXII | 466497 | 466618 | 121 | N12:466630 |
| chrXII | 466497 | 466667 | 170 | N12:466630 |
| chrXII | 466507 | 466613 | 106 | N12:466630 |
| chrXII | 466509 | 466613 | 104 | N12:466630 |
| chrXII | 466509 | 466618 | 109 | N12:466630 |
| chrXII | 466557 | 466700 | 143 | N12:466630 |
| chrXII | 466557 | 466704 | 147 | N12:466630 |
| chrXII | 466557 | 466708 | 151 | N12:466630 |
| chrXII | 466557 | 466709 | 152 | N12:466630 |
| chrXII | 466574 | 466729 | 155 | N12:466630 |
| chrXII | 466577 | 466730 | 153 | N12:466630 |
| chrXII | 466611 | 466736 | 125 | N12:466630 |
| chrXII | 466690 | 466843 | 153 | N12:466825 |
| chrXII | 466706 | 466873 | 167 | N12:466825 |
| chrXII | 466706 | 466878 | 172 | N12:466825 |
| chrXII | 466737 | 466873 | 136 | N12:466825 |
| chrXII | 466737 | 466876 | 139 | N12:466825 |
| chrXII | 466737 | 466878 | 141 | N12:466825 |
| chrXII | 466743 | 466900 | 157 | N12:466825 |
| chrXII | 466757 | 466910 | 153 | N12:466825 |
| chrXII | 466757 | 466924 | 167 | N12:466825 |
| chrXII | 466771 | 466894 | 123 | N12:466825 |
| chrXII | 466804 | 466956 | 152 | N12:466825 |
| chrXII | 466940 | 467096 | 156 | N12:467036 |
| chrXII | 466947 | 467098 | 151 | N12:467036 |
| chrXII | 466953 | 467111 | 158 | N12:467036 |
| chrXII | 466957 | 467090 | 133 | N12:467036 |
| chrXII | 467009 | 467160 | 151 | N12:467036 |
| chrXII | 467063 | 467189 | 126 | |
| chrXII | 467082 | 467240 | 158 | |
| chrXII | 467082 | 467241 | 159 | |
| chrXII | 467086 | 467196 | 110 | |
| chrXII | 467086 | 467238 | 152 | |
| chrXII | 467087 | 467238 | 151 | |
| chrXII | 467087 | 467240 | 153 | |
| chrXII | 467096 | 467196 | 100 | |
| chrXII | 467096 | 467247 | 151 | |
| chrXII | 467113 | 467196 | 83 | |
| chrXII | 467113 | 467256 | 143 | |
| chrXII | 467120 | 467226 | 106 | |
| chrXII | 467120 | 467240 | 120 | |
| chrXII | 467120 | 467274 | 154 | |
| chrXII | 467123 | 467254 | 131 | |
| chrXII | 467162 | 467304 | 142 | N12:467327 |
| chrXII | 467192 | 467274 | 82 | |
| chrXII | 467192 | 467343 | 151 | N12:467327 |
| chrXII | 467216 | 467364 | 148 | N12:467327 |
| chrXII | 467224 | 467368 | 144 | N12:467327 |
| chrXII | 467264 | 467414 | 150 | N12:467327 |
| chrXII | 467267 | 467430 | 163 | N12:467327 |
| chrXII | 467271 | 467409 | 138 | N12:467327 |
| chrXII | 467271 | 467414 | 143 | N12:467327 |
| chrXII | 467273 | 467432 | 159 | N12:467327 |
| chrXII | 467280 | 467428 | 148 | N12:467327 |
| chrXII | 467303 | 467458 | 155 | N12:467327 |
| chrXII | 467303 | 467459 | 156 | N12:467327 |
| chrXII | 467304 | 467459 | 155 | N12:467327 |
| chrXII | 467314 | 467414 | 100 | N12:467327 |
| chrXII | 467314 | 467459 | 145 | N12:467327 |
| chrXII | 467314 | 467464 | 150 | N12:467327 |

TABLE 1

| CHR | Start | End | Length (bp) | Nucleosome_ID |
|---|---|---|---|---|
| chrXII | 467314 | 467487 | 173 | N12:467327 |
| chrXII | 467316 | 467464 | 148 | N12:467327 |
| chrXII | 467328 | 467458 | 130 | N12:467327 |
| chrXII | 467328 | 467489 | 161 | N12:467327 |
| chrXII | 467332 | 467459 | 127 | N12:467327 |
| chrXII | 467332 | 467487 | 155 | N12:467327 |
| chrXII | 467341 | 467487 | 146 | N12:467327 |
| chrXII | 467341 | 467517 | 176 | N12:467327 |
| chrXII | 467428 | 467588 | 160 | N12:467582 |
| chrXII | 467434 | 467588 | 154 | N12:467582 |
| chrXII | 467435 | 467598 | 163 | N12:467582 |
| chrXII | 467456 | 467609 | 153 | N12:467582 |
| chrXII | 467475 | 467629 | 154 | N12:467582 |
| chrXII | 467493 | 467630 | 137 | N12:467582 |
| chrXII | 467493 | 467645 | 152 | N12:467582 |
| chrXII | 467496 | 467660 | 164 | N12:467582 |
| chrXII | 467502 | 467630 | 128 | N12:467582 |
| chrXII | 467510 | 467630 | 120 | N12:467582 |
| chrXII | 467510 | 467645 | 135 | N12:467582 |
| chrXII | 467510 | 467667 | 157 | N12:467582 |
| chrXII | 467511 | 467669 | 158 | N12:467582 |
| chrXII | 467515 | 467649 | 134 | N12:467582 |
| chrXII | 467532 | 467638 | 106 | N12:467582 |
| chrXII | 467556 | 467724 | 168 | N12:467582 |
| chrXII | 467645 | 467795 | 150 | |
| chrXII | 467645 | 467796 | 151 | |
| chrXII | 467883 | 468037 | 154 | N12:467959 |
| chrXII | 467883 | 468038 | 155 | N12:467959 |
| chrXII | 467883 | 468049 | 166 | N12:467959 |
| chrXII | 467887 | 468049 | 162 | N12:467959 |
| chrXII | 467905 | 468037 | 132 | N12:467959 |
| chrXII | 467905 | 468038 | 133 | N12:467959 |
| chrXII | 467905 | 468049 | 144 | N12:467959 |
| chrXII | 467906 | 468037 | 131 | N12:467959 |
| chrXII | 467906 | 468038 | 132 | N12:467959 |
| chrXII | 467906 | 468049 | 143 | N12:467959 |
| chrXII | 467906 | 468050 | 144 | N12:467959 |
| chrXII | 467907 | 468037 | 130 | N12:467959 |
| chrXII | 467907 | 468038 | 131 | N12:467959 |
| chrXII | 467907 | 468049 | 142 | N12:467959 |
| chrXII | 467983 | 468139 | 156 | N12:468113 |
| chrXII | 468039 | 468185 | 146 | N12:468113 |
| chrXII | 468045 | 468181 | 136 | N12:468113 |
| chrXII | 468045 | 468185 | 140 | N12:468113 |
| chrXII | 468047 | 468203 | 156 | N12:468113 |
| chrXII | 468050 | 468139 | 89 | N12:468113 |
| chrXII | 468050 | 468175 | 125 | N12:468113 |
| chrXII | 468050 | 468176 | 126 | N12:468113 |
| chrXII | 468050 | 468180 | 130 | N12:468113 |
| chrXII | 468050 | 468181 | 131 | N12:468113 |
| chrXII | 468050 | 468184 | 134 | N12:468113 |
| chrXII | 468050 | 468185 | 135 | N12:468113 |
| chrXII | 468050 | 468203 | 153 | N12:468113 |
| chrXII | 468070 | 468181 | 111 | N12:468113 |
| chrXII | 468070 | 468184 | 114 | N12:468113 |
| chrXII | 468070 | 468203 | 133 | N12:468113 |
| chrXII | 468187 | 468283 | 96 | N12:468275 |
| chrXII | 468226 | 468399 | 173 | N12:468275 |
| chrXII | 468226 | 468402 | 176 | N12:468275 |
| chrXII | 468232 | 468399 | 167 | N12:468275 |
| chrXII | 468232 | 468402 | 170 | N12:468275 |
| chrXII | 468235 | 468370 | 135 | N12:468275 |
| chrXII | 468235 | 468399 | 164 | N12:468275 |
| chrXII | 468303 | 468465 | 162 | N12:468484 |
| chrXII | 468322 | 468465 | 143 | N12:468484 |
| chrXII | 468341 | 468490 | 149 | N12:468484 |
| chrXII | 468341 | 468513 | 172 | N12:468484 |
| chrXII | 468341 | 468556 | 215 | N12:468484 |
| chrXII | 468364 | 468513 | 149 | N12:468484 |
| chrXII | 468364 | 468554 | 190 | N12:468484 |
| chrXII | 468364 | 468556 | 192 | N12:468484 |
| chrXII | 468364 | 468569 | 205 | N12:468484 |
| chrXII | 468372 | 468532 | 160 | N12:468484 |
| chrXII | 468373 | 468530 | 157 | N12:468484 |
| chrXII | 468373 | 468554 | 181 | N12:468484 |
| chrXII | 468373 | 468556 | 183 | N12:468484 |
| chrXII | 468401 | 468490 | 89 | N12:468484 |
| chrXII | 468401 | 468513 | 112 | N12:468484 |
| chrXII | 468401 | 468532 | 131 | N12:468484 |
| chrXII | 468401 | 468554 | 153 | N12:468484 |
| chrXII | 468401 | 468556 | 155 | N12:468484 |
| chrXII | 468401 | 468569 | 168 | N12:468484 |
| chrXII | 468401 | 468578 | 177 | N12:468484 |
| chrXII | 468402 | 468513 | 111 | N12:468484 |
| chrXII | 468402 | 468530 | 128 | N12:468484 |
| chrXII | 468402 | 468550 | 148 | N12:468484 |
| chrXII | 468402 | 468553 | 151 | N12:468484 |
| chrXII | 468402 | 468554 | 152 | N12:468484 |
| chrXII | 468402 | 468556 | 154 | N12:468484 |
| chrXII | 468402 | 468559 | 157 | N12:468484 |
| chrXII | 468402 | 468569 | 167 | N12:468484 |
| chrXII | 468402 | 468578 | 176 | N12:468484 |
| chrXII | 468402 | 468604 | 202 | N12:468484 |
| chrXII | 468402 | 468606 | 204 | N12:468484 |
| chrXII | 468404 | 468530 | 126 | N12:468484 |
| chrXII | 468404 | 468554 | 150 | N12:468484 |
| chrXII | 468404 | 468556 | 152 | N12:468484 |
| chrXII | 468404 | 468559 | 155 | N12:468484 |
| chrXII | 468404 | 468569 | 165 | N12:468484 |
| chrXII | 468404 | 468578 | 174 | N12:468484 |
| chrXII | 468404 | 468606 | 202 | N12:468484 |
| chrXII | 468405 | 468513 | 108 | N12:468484 |
| chrXII | 468405 | 468530 | 125 | N12:468484 |

TABLE 1

| CHR | Start | End | Length (bp) | Nucleosome_ID |
|---|---|---|---|---|
| chrXII | 468405 | 468532 | 127 | N12:468484 |
| chrXII | 468405 | 468550 | 145 | N12:468484 |
| chrXII | 468405 | 468553 | 148 | N12:468484 |
| chrXII | 468405 | 468554 | 149 | N12:468484 |
| chrXII | 468405 | 468556 | 151 | N12:468484 |
| chrXII | 468405 | 468559 | 154 | N12:468484 |
| chrXII | 468405 | 468568 | 163 | N12:468484 |
| chrXII | 468405 | 468569 | 164 | N12:468484 |
| chrXII | 468405 | 468578 | 173 | N12:468484 |
| chrXII | 468405 | 468604 | 199 | N12:468484 |
| chrXII | 468405 | 468606 | 201 | N12:468484 |
| chrXII | 468406 | 468513 | 107 | N12:468484 |
| chrXII | 468406 | 468530 | 124 | N12:468484 |
| chrXII | 468406 | 468550 | 144 | N12:468484 |
| chrXII | 468406 | 468554 | 148 | N12:468484 |
| chrXII | 468406 | 468556 | 150 | N12:468484 |
| chrXII | 468406 | 468559 | 153 | N12:468484 |
| chrXII | 468406 | 468568 | 162 | N12:468484 |
| chrXII | 468406 | 468604 | 198 | N12:468484 |
| chrXII | 468406 | 468606 | 200 | N12:468484 |
| chrXII | 468445 | 468551 | 106 | N12:468484 |
| chrXII | 468445 | 468554 | 109 | N12:468484 |
| chrXII | 468445 | 468556 | 111 | N12:468484 |
| chrXII | 468445 | 468604 | 159 | N12:468484 |
| chrXII | 468445 | 468606 | 161 | N12:468484 |
| chrXII | 468462 | 468603 | 141 | N12:468484 |
| chrXII | 468463 | 468554 | 91 | N12:468484 |
| chrXII | 468463 | 468556 | 93 | N12:468484 |
| chrXII | 468463 | 468604 | 141 | N12:468484 |
| chrXII | 468463 | 468606 | 143 | N12:468484 |
| chrXII | 468545 | 468696 | 151 | N12:468638 |
| chrXII | 468566 | 468665 | 99 | N12:468638 |
| chrXII | 468566 | 468695 | 129 | N12:468638 |
| chrXII | 468566 | 468710 | 144 | N12:468638 |
| chrXII | 468566 | 468716 | 150 | N12:468638 |
| chrXII | 468567 | 468695 | 128 | N12:468638 |
| chrXII | 468567 | 468710 | 143 | N12:468638 |
| chrXII | 468567 | 468716 | 149 | N12:468638 |
| chrXII | 468656 | 468810 | 154 | N12:468638 |
| chrXII | 468658 | 468810 | 152 | N12:468638 |
| chrXII | 468703 | 468859 | 156 | N12:468850 |
| chrXII | 468739 | 468893 | 154 | N12:468850 |
| chrXII | 468739 | 468896 | 157 | N12:468850 |
| chrXII | 468741 | 468859 | 118 | N12:468850 |
| chrXII | 468741 | 468893 | 152 | N12:468850 |
| chrXII | 468741 | 468896 | 155 | N12:468850 |
| chrXII | 468741 | 468911 | 170 | N12:468850 |
| chrXII | 468755 | 468893 | 138 | N12:468850 |
| chrXII | 468755 | 468911 | 156 | N12:468850 |
| chrXII | 468758 | 468893 | 135 | N12:468850 |
| chrXII | 468762 | 468893 | 131 | N12:468850 |
| chrXII | 468762 | 468911 | 149 | N12:468850 |
| chrXII | 468765 | 468893 | 128 | N12:468850 |
| chrXII | 468765 | 468911 | 146 | N12:468850 |
| chrXII | 468770 | 468893 | 123 | N12:468850 |
| chrXII | 468770 | 468911 | 141 | N12:468850 |
| chrXII | 472060 | 472213 | 153 | N12:472134 |
| chrXII | 472461 | 472545 | 84 | N12:472515 |
| chrXII | 472461 | 472563 | 102 | N12:472515 |
| chrXII | 475711 | 475864 | 153 | N12:475777 |
| chrXII | 476909 | 477023 | 114 | |
| chrXII | 477605 | 477755 | 150 | N12:477669 |
| chrXII | 478538 | 478667 | 129 | N12:478520 |
| chrXII | 479001 | 479158 | 157 | N12:479064 |
| chrXII | 479187 | 479337 | 150 | N12:479289 |
| chrXII | 479469 | 479630 | 161 | N12:479454 |
| chrXII | 480195 | 480349 | 154 | N12:480208 |
| chrXII | 480476 | 480621 | 145 | N12:480542 |
| chrXII | 480480 | 480631 | 151 | N12:480542 |
| chrXII | 480687 | 480827 | 140 | N12:480768 |
| chrXII | 480693 | 480827 | 134 | N12:480768 |
| chrXII | 481105 | 481260 | 155 | N12:481138 |
| chrXII | 481236 | 481344 | 108 | |
| chrXII | 481385 | 481534 | 149 | N12:481399 |
| chrXII | 482041 | 482125 | 84 | N12:482076 |
| chrXII | 482041 | 482143 | 102 | N12:482076 |
| chrXII | 482885 | 483037 | 152 | N12:482962 |
| chrXII | 485292 | 485445 | 153 | N12:485368 |
| chrXII | 485693 | 485777 | 84 | N12:485732 |
| chrXII | 488944 | 489097 | 153 | N12:489014 |
| chrXII | 489345 | 489429 | 84 | N12:489399 |
| chrXII | 489345 | 489447 | 102 | N12:489399 |
| chrXII | 489345 | 489466 | 121 | N12:489399 |
| chrXII | 489345 | 489469 | 124 | N12:489399 |
| chrXII | 489345 | 489471 | 126 | N12:489399 |
| chrXII | 489359 | 489469 | 110 | N12:489399 |
| chrXII | 489359 | 489471 | 112 | N12:489399 |
| chrXII | 489361 | 489466 | 105 | N12:489399 |
| chrXII | 489556 | 489702 | 146 | N12:489601 |
| chrXII | 489926 | 490077 | 151 | N12:489932 |
| chrXII | 489933 | 490080 | 147 | N12:489932 |
| chrXII | 490009 | 490164 | 155 | N12:490150 |
| chrXII | 490027 | 490183 | 156 | N12:490150 |
| chrXII | 490049 | 490195 | 146 | N12:490150 |
| chrXII | 490049 | 490215 | 166 | N12:490150 |
| chrXII | 490049 | 490219 | 170 | N12:490150 |
| chrXII | 490072 | 490215 | 143 | N12:490150 |
| chrXII | 490072 | 490219 | 147 | N12:490150 |
| chrXII | 490083 | 490239 | 156 | N12:490150 |
| chrXII | 490158 | 490312 | 154 | N12:490324 |
| chrXII | 490158 | 490312 | 154 | N12:490150 |
| chrXII | 490228 | 490334 | 106 | N12:490324 |
| chrXII | 490228 | 490337 | 109 | N12:490324 |

| CHR | Start | End | Length (bp) | Nucleosome_ID |
|---|---|---|---|---|
| chrXII | 490228 | 490382 | 154 | N12:490324 |
| chrXII | 490261 | 490368 | 107 | N12:490324 |
| chrXII | 490262 | 490423 | 161 | N12:490324 |
| chrXII | 490263 | 490423 | 160 | N12:490324 |
| chrXII | 490267 | 490423 | 156 | N12:490324 |
| chrXII | 490291 | 490423 | 132 | N12:490324 |
| chrXII | 490302 | 490409 | 107 | N12:490324 |
| chrXII | 490302 | 490423 | 121 | N12:490324 |
| chrXII | 494931 | 495079 | 148 | N12:495031 |
| chrXII | 496357 | 496507 | 150 | |
| chrXII | 508585 | 508725 | 140 | N12:508668 |
| chrXII | 514529 | 514679 | 150 | N12:514599 |
| chrXII | 518369 | 518501 | 132 | N12:518444 |
| chrXII | 522434 | 522581 | 147 | N12:522432 |
| chrXII | 528715 | 528875 | 160 | N12:528770 |
| chrXII | 529249 | 529419 | 170 | N12:529380 |
| chrXII | 537872 | 537976 | 104 | N12:537943 |
| chrXII | 542120 | 542273 | 153 | N12:542188 |
| chrXII | 544133 | 544261 | 128 | N12:544269 |
| chrXII | 545204 | 545351 | 147 | N12:545241 |
| chrXII | 546037 | 546189 | 152 | N12:546112 |
| chrXII | 546038 | 546189 | 151 | N12:546112 |
| chrXII | 546040 | 546189 | 149 | N12:546112 |
| chrXII | 547890 | 548039 | 149 | N12:547946 |
| chrXII | 553542 | 553689 | 147 | N12:553631 |
| chrXII | 555694 | 555847 | 153 | N12:555721 |
| chrXII | 559139 | 559286 | 147 | N12:559216 |
| chrXII | 559606 | 559755 | 149 | N12:559682 |
| chrXII | 560049 | 560194 | 145 | N12:560179 |
| chrXII | 570252 | 570403 | 151 | N12:570424 |
| chrXII | 573533 | 573669 | 136 | N12:573595 |
| chrXII | 576803 | 576966 | 163 | N12:576902 |
| chrXII | 586445 | 586571 | 126 | N12:586538 |
| chrXII | 587477 | 587583 | 106 | N12:587489 |
| chrXII | 588088 | 588237 | 149 | N12:588110 |
| chrXII | 593332 | 593484 | 152 | N12:593448 |
| chrXII | 593504 | 593653 | 149 | N12:593653 |
| chrXII | 593678 | 593801 | 123 | |
| chrXII | 593778 | 593933 | 155 | N12:593897 |
| chrXII | 594211 | 594345 | 134 | N12:594281 |
| chrXII | 594211 | 594351 | 140 | N12:594281 |
| chrXII | 594407 | 594558 | 151 | N12:594490 |
| chrXII | 594417 | 594562 | 145 | N12:594490 |
| chrXII | 594435 | 594586 | 151 | N12:594490 |
| chrXII | 594689 | 594843 | 154 | N12:594825 |
| chrXII | 594723 | 594867 | 144 | N12:594825 |
| chrXII | 595254 | 595389 | 135 | N12:595293 |
| chrXII | 595436 | 595568 | 132 | N12:595587 |
| chrXII | 595525 | 595663 | 138 | N12:595587 |
| chrXII | 598015 | 598129 | 114 | N12:598127 |
| chrXII | 598489 | 598636 | 147 | N12:598658 |
| chrXII | 603237 | 603400 | 163 | N12:603419 |
| chrXII | 607039 | 607198 | 159 | N12:607110 |
| chrXII | 608189 | 608339 | 150 | N12:608276 |
| chrXII | 610208 | 610357 | 149 | N12:610297 |
| chrXII | 611103 | 611232 | 129 | N12:611216 |
| chrXII | 614828 | 614977 | 149 | N12:614901 |
| chrXII | 614828 | 614997 | 169 | N12:614901 |
| chrXII | 614830 | 614977 | 147 | N12:614901 |
| chrXII | 625233 | 625381 | 148 | N12:625336 |
| chrXII | 628912 | 629085 | 173 | N12:629078 |
| chrXII | 631988 | 632127 | 139 | N12:632062 |
| chrXII | 635940 | 636108 | 168 | N12:635980 |
| chrXII | 640543 | 640689 | 146 | N12:640632 |
| chrXII | 649877 | 650024 | 147 | N12:649854 |
| chrXII | 649877 | 650024 | 147 | N12:650003 |
| chrXII | 651183 | 651332 | 149 | N12:651337 |
| chrXII | 651357 | 651480 | 123 | N12:651337 |
| chrXII | 651457 | 651612 | 155 | N12:651585 |
| chrXII | 651890 | 652024 | 134 | N12:651960 |
| chrXII | 651890 | 652030 | 140 | N12:651960 |
| chrXII | 652086 | 652237 | 151 | N12:652161 |
| chrXII | 652096 | 652241 | 145 | N12:652161 |
| chrXII | 652114 | 652265 | 151 | N12:652161 |
| chrXII | 652116 | 652265 | 149 | N12:652161 |
| chrXII | 652368 | 652522 | 154 | N12:652504 |
| chrXII | 652618 | 652786 | 168 | N12:652671 |
| chrXII | 652933 | 653068 | 135 | N12:652970 |
| chrXII | 653087 | 653248 | 161 | N12:653246 |
| chrXII | 653194 | 653339 | 145 | N12:653246 |
| chrXII | 653561 | 653718 | 157 | N12:653642 |
| chrXII | 654052 | 654181 | 129 | |
| chrXII | 654964 | 655114 | 150 | N12:655066 |
| chrXII | 656400 | 656549 | 149 | N12:656524 |
| chrXII | 657729 | 657887 | 158 | N12:657828 |
| chrXII | 664370 | 664478 | 108 | N12:664374 |
| chrXII | 664370 | 664482 | 112 | N12:664374 |
| chrXII | 665850 | 666010 | 160 | N12:665933 |
| chrXII | 675650 | 675803 | 153 | N12:675721 |
| chrXII | 678419 | 678585 | 166 | N12:678473 |
| chrXII | 684089 | 684224 | 135 | N12:684178 |
| chrXII | 686656 | 686810 | 154 | N12:686728 |
| chrXII | 686971 | 687118 | 147 | N12:687063 |
| chrXII | 691050 | 691199 | 149 | N12:691100 |
| chrXII | 695440 | 695604 | 164 | N12:695602 |
| chrXII | 698814 | 698965 | 151 | N12:698805 |
| chrXII | 700732 | 700860 | 128 | N12:700798 |
| chrXII | 700732 | 700861 | 129 | N12:700798 |
| chrXII | 704633 | 704781 | 148 | N12:704801 |
| chrXII | 704633 | 704781 | 148 | N12:704613 |
| chrXII | 704633 | 704785 | 152 | N12:704801 |
| chrXII | 704633 | 704785 | 152 | N12:704613 |
| chrXII | 704637 | 704785 | 148 | N12:704801 |

| CHR | Start | End | Length (bp) | Nucleosome_ID |
|---|---|---|---|---|
| chrXII | 704738 | 704878 | 140 | N12:704801 |
| chrXII | 707321 | 707471 | 150 | N12:707390 |
| chrXII | 707855 | 708017 | 162 | N12:707899 |
| chrXII | 710806 | 710956 | 150 | N12:710883 |
| chrXII | 713802 | 713939 | 137 | N12:713872 |
| chrXII | 714204 | 714311 | 107 | N12:714214 |
| chrXII | 714204 | 714312 | 108 | N12:714214 |
| chrXII | 714794 | 714945 | 151 | N12:714880 |
| chrXII | 721307 | 721447 | 140 | N12:721455 |
| chrXII | 723297 | 723432 | 135 | N12:723451 |
| chrXII | 726306 | 726432 | 126 | N12:726432 |
| chrXII | 727018 | 727148 | 130 | N12:727116 |
| chrXII | 731448 | 731612 | 164 | N12:731474 |
| chrXII | 731448 | 731612 | 164 | N12:731633 |
| chrXII | 733723 | 733874 | 151 | N12:733718 |
| chrXII | 733723 | 733874 | 151 | N12:733875 |
| chrXII | 735532 | 735635 | 103 | N12:735568 |
| chrXII | 736027 | 736189 | 162 | N12:736034 |
| chrXII | 738785 | 738957 | 172 | N12:738797 |
| chrXII | 740597 | 740703 | 106 | N12:740592 |
| chrXII | 745846 | 745998 | 152 | N12:745926 |
| chrXII | 751027 | 751192 | 165 | N12:751096 |
| chrXII | 752922 | 753058 | 136 | N12:753013 |
| chrXII | 753617 | 753790 | 173 | N12:753609 |
| chrXII | 753617 | 753790 | 173 | N12:753760 |
| chrXII | 759427 | 759579 | 152 | N12:759504 |
| chrXII | 761633 | 761798 | 165 | |
| chrXII | 762105 | 762269 | 164 | N12:762185 |
| chrXII | 770786 | 770938 | 152 | N12:770870 |
| chrXII | 779526 | 779669 | 143 | N12:779529 |
| chrXII | 779740 | 779903 | 163 | N12:779869 |
| chrXII | 781130 | 781282 | 152 | N12:781211 |
| chrXII | 782565 | 782713 | 148 | N12:782593 |
| chrXII | 783914 | 784060 | 146 | N12:784073 |
| chrXII | 783914 | 784060 | 146 | N12:783921 |
| chrXII | 785535 | 785685 | 150 | N12:785589 |
| chrXII | 786502 | 786666 | 164 | N12:786585 |
| chrXII | 792530 | 792669 | 139 | N12:792622 |
| chrXII | 797667 | 797817 | 150 | N12:797755 |
| chrXII | 799286 | 799439 | 153 | N12:799357 |
| chrXII | 803374 | 803523 | 149 | N12:803394 |
| chrXII | 804062 | 804197 | 135 | N12:804104 |
| chrXII | 804515 | 804661 | 146 | N12:804614 |
| chrXII | 806849 | 807003 | 154 | N12:806937 |
| chrXII | 807887 | 808014 | 127 | N12:807864 |
| chrXII | 807887 | 808014 | 127 | N12:808034 |
| chrXII | 827684 | 827838 | 154 | N12:827706 |
| chrXII | 830101 | 830255 | 154 | N12:830181 |
| chrXII | 830655 | 830774 | 119 | N12:830722 |
| chrXII | 830666 | 830794 | 128 | N12:830722 |
| chrXII | 831008 | 831187 | 179 | N12:831140 |
| chrXII | 831542 | 831700 | 158 | N12:831639 |
| chrXII | 839642 | 839772 | 130 | N12:839733 |
| chrXII | 840366 | 840517 | 151 | |
| chrXII | 841812 | 841967 | 155 | N12:841835 |
| chrXII | 842219 | 842368 | 149 | N12:842300 |
| chrXII | 846202 | 846357 | 155 | N12:846233 |
| chrXII | 864284 | 864432 | 148 | N12:864280 |
| chrXII | 868597 | 868734 | 137 | N12:868681 |
| chrXII | 871320 | 871444 | 124 | N12:871354 |
| chrXII | 871320 | 871449 | 129 | N12:871354 |
| chrXII | 871322 | 871449 | 127 | N12:871354 |
| chrXII | 874679 | 874837 | 158 | N12:874753 |
| chrXII | 878567 | 878705 | 138 | N12:878691 |
| chrXII | 884555 | 884713 | 158 | N12:884629 |
| chrXII | 889517 | 889681 | 164 | N12:889511 |
| chrXII | 893166 | 893327 | 161 | N12:893245 |
| chrXII | 896277 | 896446 | 169 | N12:896387 |
| chrXII | 896668 | 896822 | 154 | N12:896735 |
| chrXII | 896811 | 896958 | 147 | N12:896895 |
| chrXII | 903630 | 903761 | 131 | N12:903719 |
| chrXII | 903784 | 903945 | 161 | N12:903890 |
| chrXII | 903961 | 904117 | 156 | N12:904050 |
| chrXII | 905180 | 905335 | 155 | |
| chrXII | 908649 | 908820 | 171 | N12:908748 |
| chrXII | 912499 | 912653 | 154 | N12:912578 |
| chrXII | 913527 | 913695 | 168 | N12:913636 |
| chrXII | 922406 | 922556 | 150 | N12:922523 |
| chrXII | 922509 | 922654 | 145 | N12:922523 |
| chrXII | 924375 | 924524 | 149 | N12:924433 |
| chrXII | 926149 | 926303 | 154 | N12:926312 |
| chrXII | 926358 | 926515 | 157 | N12:926512 |
| chrXII | 929019 | 929172 | 153 | N12:929097 |
| chrXII | 931573 | 931698 | 125 | |
| chrXII | 937960 | 938108 | 148 | N12:938021 |
| chrXII | 938740 | 938880 | 140 | N12:938790 |
| chrXII | 939726 | 939894 | 168 | N12:939769 |
| chrXII | 940022 | 940161 | 139 | N12:940088 |
| chrXII | 941909 | 942015 | 106 | N12:941949 |
| chrXII | 941909 | 942019 | 110 | N12:941949 |
| chrXII | 943265 | 943394 | 129 | N12:943276 |
| chrXII | 944730 | 944873 | 143 | N12:944794 |
| chrXII | 946105 | 946219 | 114 | N12:946210 |
| chrXII | 946809 | 946958 | 149 | N12:946920 |
| chrXII | 947096 | 947245 | 149 | N12:947106 |
| chrXII | 947588 | 947739 | 151 | N12:947709 |
| chrXII | 952012 | 952159 | 147 | N12:952105 |
| chrXII | 955881 | 956039 | 158 | N12:955940 |
| chrXII | 956586 | 956713 | 127 | N12:956629 |
| chrXII | 962105 | 962271 | 166 | N12:962110 |
| chrXII | 962105 | 962271 | 166 | N12:962293 |
| chrXII | 966042 | 966188 | 146 | N12:966156 |
| chrXII | 967180 | 967328 | 148 | N12:967232 |

| CHR | Start | End | Length (bp) | Nucleosome_ID |
|---|---|---|---|---|
| chrXII | 967209 | 967341 | 132 | N12:967232 |
| chrXII | 974145 | 974297 | 152 | N12:974223 |
| chrXII | 975018 | 975184 | 166 | N12:975031 |
| chrXII | 976449 | 976598 | 149 | N12:976465 |
| chrXII | 978018 | 978161 | 143 | N12:978103 |
| chrXII | 978601 | 978756 | 155 | N12:978687 |
| chrXII | 979540 | 979688 | 148 | |
| chrXII | 981009 | 981111 | 102 | |
| chrXII | 981009 | 981139 | 130 | |
| chrXII | 981073 | 981227 | 154 | N12:981187 |
| chrXII | 981080 | 981248 | 168 | N12:981187 |
| chrXII | 982426 | 982590 | 164 | N12:982511 |
| chrXII | 987388 | 987536 | 148 | N12:987440 |
| chrXII | 988804 | 988961 | 157 | N12:988850 |
| chrXII | 988850 | 988961 | 111 | N12:988850 |
| chrXII | 988957 | 989104 | 147 | N12:989040 |
| chrXII | 990093 | 990242 | 149 | N12:990189 |
| chrXII | 990292 | 990442 | 150 | N12:990360 |
| chrXII | 997620 | 997777 | 157 | N12:997691 |
| chrXII | 998925 | 999073 | 148 | N12:999015 |
| chrXII | 1001567 | 1001716 | 149 | N12:1001641 |
| chrXII | 1005509 | 1005668 | 159 | N12:1005574 |
| chrXII | 1005994 | 1006142 | 148 | N12:1006065 |
| chrXII | 1012419 | 1012577 | 158 | N12:1012484 |
| chrXII | 1012549 | 1012696 | 147 | N12:1012647 |
| chrXII | 1012550 | 1012632 | 82 | N12:1012647 |
| chrXII | 1012550 | 1012633 | 83 | N12:1012647 |
| chrXII | 1012550 | 1012634 | 84 | N12:1012647 |
| chrXII | 1012550 | 1012635 | 85 | N12:1012647 |
| chrXII | 1012550 | 1012636 | 86 | N12:1012647 |
| chrXII | 1012550 | 1012637 | 87 | N12:1012647 |
| chrXII | 1012550 | 1012638 | 88 | N12:1012647 |
| chrXII | 1012550 | 1012639 | 89 | N12:1012647 |
| chrXII | 1012550 | 1012640 | 90 | N12:1012647 |
| chrXII | 1012550 | 1012641 | 91 | N12:1012647 |
| chrXII | 1012550 | 1012642 | 92 | N12:1012647 |
| chrXII | 1012550 | 1012643 | 93 | N12:1012647 |
| chrXII | 1012550 | 1012644 | 94 | N12:1012647 |
| chrXII | 1012550 | 1012645 | 95 | N12:1012647 |
| chrXII | 1012550 | 1012646 | 96 | N12:1012647 |
| chrXII | 1012550 | 1012648 | 98 | N12:1012647 |
| chrXII | 1012550 | 1012649 | 99 | N12:1012647 |
| chrXII | 1012550 | 1012650 | 100 | N12:1012647 |
| chrXII | 1012550 | 1012651 | 101 | N12:1012647 |
| chrXII | 1012550 | 1012652 | 102 | N12:1012647 |
| chrXII | 1012550 | 1012653 | 103 | N12:1012647 |
| chrXII | 1012550 | 1012654 | 104 | N12:1012647 |
| chrXII | 1012550 | 1012655 | 105 | N12:1012647 |
| chrXII | 1012550 | 1012656 | 106 | N12:1012647 |
| chrXII | 1012550 | 1012657 | 107 | N12:1012647 |
| chrXII | 1012550 | 1012659 | 109 | N12:1012647 |
| chrXII | 1012550 | 1012660 | 110 | N12:1012647 |
| chrXII | 1012550 | 1012661 | 111 | N12:1012647 |
| chrXII | 1012550 | 1012662 | 112 | N12:1012647 |
| chrXII | 1012550 | 1012663 | 113 | N12:1012647 |
| chrXII | 1012550 | 1012664 | 114 | N12:1012647 |
| chrXII | 1012550 | 1012665 | 115 | N12:1012647 |
| chrXII | 1012550 | 1012666 | 116 | N12:1012647 |
| chrXII | 1012550 | 1012667 | 117 | N12:1012647 |
| chrXII | 1012550 | 1012668 | 118 | N12:1012647 |
| chrXII | 1012550 | 1012669 | 119 | N12:1012647 |
| chrXII | 1012550 | 1012670 | 120 | N12:1012647 |
| chrXII | 1012550 | 1012671 | 121 | N12:1012647 |
| chrXII | 1012550 | 1012672 | 122 | N12:1012647 |
| chrXII | 1012550 | 1012673 | 123 | N12:1012647 |
| chrXII | 1012550 | 1012674 | 124 | N12:1012647 |
| chrXII | 1012550 | 1012675 | 125 | N12:1012647 |
| chrXII | 1012550 | 1012676 | 126 | N12:1012647 |
| chrXII | 1012550 | 1012677 | 127 | N12:1012647 |
| chrXII | 1012550 | 1012678 | 128 | N12:1012647 |
| chrXII | 1012550 | 1012679 | 129 | N12:1012647 |
| chrXII | 1012550 | 1012680 | 130 | N12:1012647 |
| chrXII | 1012550 | 1012681 | 131 | N12:1012647 |
| chrXII | 1012550 | 1012682 | 132 | N12:1012647 |
| chrXII | 1012550 | 1012683 | 133 | N12:1012647 |
| chrXII | 1012550 | 1012684 | 134 | N12:1012647 |
| chrXII | 1012550 | 1012685 | 135 | N12:1012647 |
| chrXII | 1012550 | 1012686 | 136 | N12:1012647 |
| chrXII | 1012550 | 1012687 | 137 | N12:1012647 |
| chrXII | 1012550 | 1012688 | 138 | N12:1012647 |
| chrXII | 1012550 | 1012689 | 139 | N12:1012647 |
| chrXII | 1012550 | 1012690 | 140 | N12:1012647 |
| chrXII | 1012550 | 1012691 | 141 | N12:1012647 |
| chrXII | 1012550 | 1012692 | 142 | N12:1012647 |
| chrXII | 1012550 | 1012693 | 143 | N12:1012647 |
| chrXII | 1012550 | 1012694 | 144 | N12:1012647 |
| chrXII | 1012550 | 1012695 | 145 | N12:1012647 |
| chrXII | 1012550 | 1012696 | 146 | N12:1012647 |
| chrXII | 1012550 | 1012698 | 148 | N12:1012647 |
| chrXII | 1012551 | 1012693 | 142 | N12:1012647 |
| chrXII | 1012551 | 1012696 | 145 | N12:1012647 |
| chrXII | 1012552 | 1012696 | 144 | N12:1012647 |
| chrXII | 1012553 | 1012693 | 140 | N12:1012647 |
| chrXII | 1012553 | 1012696 | 143 | N12:1012647 |
| chrXII | 1012554 | 1012696 | 142 | N12:1012647 |
| chrXII | 1012555 | 1012696 | 141 | N12:1012647 |
| chrXII | 1012556 | 1012696 | 140 | N12:1012647 |
| chrXII | 1012557 | 1012696 | 139 | N12:1012647 |
| chrXII | 1012558 | 1012696 | 138 | N12:1012647 |
| chrXII | 1012559 | 1012696 | 137 | N12:1012647 |
| chrXII | 1012560 | 1012696 | 136 | N12:1012647 |
| chrXII | 1012561 | 1012696 | 135 | N12:1012647 |
| chrXII | 1012562 | 1012696 | 134 | N12:1012647 |

| CHR | Start | End | Length (bp) | Nucleosome_ID |
|---|---|---|---|---|
| chrXII | 1012563 | 1012696 | 133 | N12:1012647 |
| chrXII | 1012564 | 1012696 | 132 | N12:1012647 |
| chrXII | 1012565 | 1012696 | 131 | N12:1012647 |
| chrXII | 1012566 | 1012696 | 130 | N12:1012647 |
| chrXII | 1012567 | 1012696 | 129 | N12:1012647 |
| chrXII | 1012568 | 1012696 | 128 | N12:1012647 |
| chrXII | 1012569 | 1012696 | 127 | N12:1012647 |
| chrXII | 1012570 | 1012696 | 126 | N12:1012647 |
| chrXII | 1012571 | 1012696 | 125 | N12:1012647 |
| chrXII | 1012572 | 1012696 | 124 | N12:1012647 |
| chrXII | 1012573 | 1012696 | 123 | N12:1012647 |
| chrXII | 1012574 | 1012696 | 122 | N12:1012647 |
| chrXII | 1012575 | 1012696 | 121 | N12:1012647 |
| chrXII | 1012576 | 1012696 | 120 | N12:1012647 |
| chrXII | 1012578 | 1012696 | 118 | N12:1012647 |
| chrXII | 1012579 | 1012696 | 117 | N12:1012647 |
| chrXII | 1012580 | 1012696 | 116 | N12:1012647 |
| chrXII | 1012581 | 1012696 | 115 | N12:1012647 |
| chrXII | 1012583 | 1012696 | 113 | N12:1012647 |
| chrXII | 1012584 | 1012696 | 112 | N12:1012647 |
| chrXII | 1012586 | 1012696 | 110 | N12:1012647 |
| chrXII | 1012590 | 1012674 | 84 | N12:1012647 |
| chrXII | 1012610 | 1012696 | 86 | N12:1012647 |
| chrXII | 1013868 | 1014024 | 156 | N12:1013902 |
| chrXII | 1017667 | 1017816 | 149 | N12:1017757 |
| chrXII | 1023621 | 1023767 | 146 | N12:1023663 |
| chrXII | 1024132 | 1024289 | 157 | N12:1024222 |
| chrXII | 1031294 | 1031445 | 151 | N12:1031333 |
| chrXII | 1032438 | 1032586 | 148 | N12:1032481 |
| chrXII | 1050400 | 1050528 | 128 | N12:1050503 |
| chrXII | 1050732 | 1050875 | 143 | N12:1050868 |
| chrXII | 1059679 | 1059820 | 141 | N12:1059812 |
| chrXII | 1065029 | 1065154 | 125 | N12:1065098 |
| chrXII | 1065029 | 1065156 | 127 | N12:1065098 |
| chrXII | 1066703 | 1066854 | 151 | |
| chrXII | 1067646 | 1067808 | 162 | N12:1067769 |
| chrXII | 1067747 | 1067902 | 155 | N12:1067769 |
| chrXII | 1067881 | 1068018 | 137 | N12:1067950 |
| chrXII | 1068176 | 1068319 | 143 | N12:1068230 |
| chrXII | 1068224 | 1068349 | 125 | N12:1068230 |
| chrXII | 1068449 | 1068601 | 152 | |
| chrXII | 1068522 | 1068666 | 144 | |
| chrXII | 1068619 | 1068754 | 135 | N12:1068716 |
| chrXII | 1068619 | 1068806 | 187 | N12:1068716 |
| chrXII | 1068662 | 1068754 | 92 | N12:1068716 |
| chrXII | 1068662 | 1068806 | 144 | N12:1068716 |
| chrXII | 1069001 | 1069153 | 152 | N12:1069071 |
| chrXII | 1069302 | 1069432 | 130 | N12:1069283 |
| chrXII | 1069440 | 1069601 | 161 | |
| chrXII | 1070837 | 1070988 | 151 | N12:1070915 |
| chrXII | 1070837 | 1071008 | 171 | N12:1070915 |
| chrXII | 1072226 | 1072381 | 155 | N12:1072335 |
| chrXII | 1073391 | 1073538 | 147 | |
| chrXII | 1073547 | 1073677 | 130 | N12:1073613 |
| chrXII | 1073928 | 1074064 | 136 | N12:1073964 |
| chrXII | 1074309 | 1074471 | 162 | N12:1074439 |
| chrXII | 1074410 | 1074565 | 155 | N12:1074439 |
| chrXII | 1074544 | 1074681 | 137 | N12:1074612 |
| chrXII | 1074831 | 1074982 | 151 | N12:1074890 |
| chrXII | 1074839 | 1074982 | 143 | N12:1074890 |
| chrXII | 1074887 | 1075012 | 125 | N12:1074890 |
| chrXII | 1075112 | 1075264 | 152 | |
| chrXII | 1075185 | 1075329 | 144 | |
| chrXII | 1075282 | 1075417 | 135 | N12:1075369 |
| chrXII | 1075282 | 1075469 | 187 | N12:1075369 |
| chrXII | 1075325 | 1075417 | 92 | N12:1075369 |
| chrXII | 1075325 | 1075469 | 144 | N12:1075369 |
| chrXII | 1075664 | 1075816 | 152 | N12:1075734 |
| chrXII | 1075965 | 1076095 | 130 | N12:1076061 |
| chrXII | 1076103 | 1076264 | 161 | |
| chrXII | 1077160 | 1077312 | 152 | N12:1077229 |
| chrXII | 1077500 | 1077651 | 151 | N12:1077576 |
| chrXIII | 687 | 858 | 171 | N13:774 |
| chrXIII | 707 | 858 | 151 | N13:774 |
| chrXIII | 1046 | 1198 | 152 | N13:1107 |
| chrXIII | 1151 | 1299 | 148 | |
| chrXIII | 1836 | 2023 | 187 | N13:1892 |
| chrXIII | 1872 | 2023 | 151 | N13:1892 |
| chrXIII | 1884 | 2018 | 134 | N13:1892 |
| chrXIII | 1909 | 2005 | 96 | N13:1892 |
| chrXIII | 2542 | 2694 | 152 | N13:2630 |
| chrXIII | 2889 | 3033 | 144 | N13:2945 |
| chrXIII | 2889 | 3076 | 187 | N13:2945 |
| chrXIII | 2941 | 3033 | 92 | N13:2945 |
| chrXIII | 2941 | 3076 | 135 | N13:2945 |
| chrXIII | 3029 | 3173 | 144 | N13:3125 |
| chrXIII | 3094 | 3246 | 152 | N13:3125 |
| chrXIII | 3346 | 3471 | 125 | N13:3450 |
| chrXIII | 3376 | 3519 | 143 | N13:3450 |
| chrXIII | 3376 | 3527 | 151 | N13:3450 |
| chrXIII | 5624 | 5767 | 143 | N13:5763 |
| chrXIII | 9070 | 9207 | 137 | N13:9142 |
| chrXIII | 13123 | 13267 | 144 | N13:13226 |
| chrXIII | 14333 | 14505 | 172 | N13:14398 |
| chrXIII | 16025 | 16179 | 154 | N13:16099 |
| chrXIII | 18573 | 18744 | 171 | N13:18675 |
| chrXIII | 19858 | 19999 | 141 | N13:19950 |
| chrXIII | 20186 | 20333 | 147 | N13:20274 |
| chrXIII | 21593 | 21740 | 147 | N13:21676 |
| chrXIII | 23095 | 23242 | 147 | N13:23173 |
| chrXIII | 32413 | 32558 | 145 | N13:32440 |
| chrXIII | 33232 | 33384 | 152 | N13:33387 |
| chrXIII | 38992 | 39143 | 151 | N13:39050 |

| CHR | Start | End | Length (bp) | Nucleosome_ID |
|---|---|---|---|---|
| chrXIII | 40472 | 40624 | 152 | N13:40521 |
| chrXIII | 42972 | 43133 | 161 | N13:43054 |
| chrXIII | 47095 | 47254 | 159 | N13:47157 |
| chrXIII | 54675 | 54757 | 82 | |
| chrXIII | 54913 | 55020 | 107 | N13:54961 |
| chrXIII | 54913 | 55032 | 119 | N13:54961 |
| chrXIII | 59231 | 59396 | 165 | N13:59272 |
| chrXIII | 63883 | 64021 | 138 | N13:63895 |
| chrXIII | 64235 | 64393 | 158 | N13:64363 |
| chrXIII | 64871 | 65018 | 147 | N13:64980 |
| chrXIII | 66681 | 66829 | 148 | N13:66758 |
| chrXIII | 75788 | 75939 | 151 | N13:75859 |
| chrXIII | 76732 | 76885 | 153 | N13:76800 |
| chrXIII | 77591 | 77728 | 137 | N13:77626 |
| chrXIII | 83978 | 84137 | 159 | |
| chrXIII | 93745 | 93871 | 126 | N13:93877 |
| chrXIII | 98827 | 98976 | 149 | N13:98894 |
| chrXIII | 102493 | 102629 | 136 | N13:102572 |
| chrXIII | 104088 | 104231 | 143 | N13:104162 |
| chrXIII | 104601 | 104708 | 107 | N13:104679 |
| chrXIII | 104601 | 104720 | 119 | N13:104679 |
| chrXIII | 107860 | 108019 | 159 | N13:107958 |
| chrXIII | 109020 | 109176 | 156 | N13:109082 |
| chrXIII | 112712 | 112852 | 140 | N13:112775 |
| chrXIII | 113210 | 113365 | 155 | N13:113351 |
| chrXIII | 113210 | 113365 | 155 | N13:113196 |
| chrXIII | 116680 | 116830 | 150 | N13:116697 |
| chrXIII | 116680 | 116833 | 153 | N13:116697 |
| chrXIII | 116680 | 116835 | 155 | N13:116697 |
| chrXIII | 116684 | 116835 | 151 | N13:116697 |
| chrXIII | 116783 | 116928 | 145 | |
| chrXIII | 117349 | 117491 | 142 | N13:117410 |
| chrXIII | 117514 | 117618 | 104 | N13:117572 |
| chrXIII | 119216 | 119393 | 177 | N13:119273 |
| chrXIII | 120074 | 120243 | 169 | N13:120213 |
| chrXIII | 120074 | 120243 | 169 | N13:120063 |
| chrXIII | 123583 | 123746 | 163 | N13:123684 |
| chrXIII | 124934 | 125083 | 149 | N13:125011 |
| chrXIII | 125456 | 125612 | 156 | N13:125539 |
| chrXIII | 126339 | 126468 | 129 | |
| chrXIII | 129825 | 129930 | 105 | N13:129823 |
| chrXIII | 129825 | 129932 | 107 | N13:129823 |
| chrXIII | 129843 | 129930 | 87 | N13:129823 |
| chrXIII | 129843 | 130004 | 161 | N13:129987 |
| chrXIII | 129843 | 130004 | 161 | N13:129823 |
| chrXIII | 130886 | 131027 | 141 | N13:130956 |
| chrXIII | 131028 | 131190 | 162 | N13:131120 |
| chrXIII | 131602 | 131755 | 153 | N13:131595 |
| chrXIII | 133367 | 133536 | 169 | N13:133467 |
| chrXIII | 134348 | 134447 | 99 | N13:134415 |
| chrXIII | 135575 | 135725 | 150 | N13:135649 |
| chrXIII | 139632 | 139738 | 106 | N13:139678 |
| chrXIII | 139632 | 139741 | 109 | N13:139678 |
| chrXIII | 142450 | 142583 | 133 | N13:142508 |
| chrXIII | 147977 | 148129 | 152 | |
| chrXIII | 148322 | 148434 | 112 | |
| chrXIII | 154231 | 154388 | 157 | |
| chrXIII | 155621 | 155784 | 163 | |
| chrXIII | 163164 | 163259 | 95 | |
| chrXIII | 169237 | 169345 | 108 | N13:169314 |
| chrXIII | 170327 | 170483 | 156 | N13:170398 |
| chrXIII | 170574 | 170680 | 106 | N13:170557 |
| chrXIII | 170574 | 170696 | 122 | N13:170557 |
| chrXIII | 170574 | 170696 | 122 | N13:170707 |
| chrXIII | 171072 | 171220 | 148 | N13:171141 |
| chrXIII | 175762 | 175915 | 153 | N13:175792 |
| chrXIII | 179143 | 179321 | 178 | |
| chrXIII | 180747 | 180905 | 158 | N13:180820 |
| chrXIII | 181711 | 181851 | 140 | N13:181781 |
| chrXIII | 181848 | 181977 | 129 | |
| chrXIII | 184182 | 184330 | 148 | N13:184276 |
| chrXIII | 184701 | 184824 | 123 | |
| chrXIII | 184801 | 184956 | 155 | N13:184876 |
| chrXIII | 185234 | 185368 | 134 | N13:185305 |
| chrXIII | 185234 | 185374 | 140 | N13:185305 |
| chrXIII | 186277 | 186412 | 135 | N13:186311 |
| chrXIII | 186724 | 186874 | 150 | N13:186799 |
| chrXIII | 186903 | 187060 | 157 | N13:186999 |
| chrXIII | 187135 | 187284 | 149 | N13:187179 |
| chrXIII | 187394 | 187523 | 129 | |
| chrXIII | 188306 | 188456 | 150 | N13:188399 |
| chrXIII | 195171 | 195314 | 143 | N13:195288 |
| chrXIII | 196534 | 196661 | 127 | N13:196631 |
| chrXIII | 196694 | 196843 | 149 | N13:196842 |
| chrXIII | 196868 | 196991 | 123 | |
| chrXIII | 196968 | 197123 | 155 | N13:197081 |
| chrXIII | 197401 | 197535 | 134 | N13:197469 |
| chrXIII | 197401 | 197541 | 140 | N13:197469 |
| chrXIII | 197597 | 197748 | 151 | N13:197630 |
| chrXIII | 197607 | 197752 | 145 | N13:197630 |
| chrXIII | 197625 | 197776 | 151 | N13:197630 |
| chrXIII | 197879 | 198033 | 154 | N13:198018 |
| chrXIII | 197913 | 198057 | 144 | N13:198018 |
| chrXIII | 198891 | 199037 | 146 | N13:198960 |
| chrXIII | 198891 | 199041 | 150 | N13:198960 |
| chrXIII | 199070 | 199227 | 157 | N13:199146 |
| chrXIII | 199302 | 199451 | 149 | N13:199440 |
| chrXIII | 199561 | 199690 | 129 | N13:199711 |
| chrXIII | 200473 | 200623 | 150 | N13:200564 |
| chrXIII | 203778 | 203929 | 151 | N13:203861 |
| chrXIII | 204102 | 204256 | 154 | N13:204195 |
| chrXIII | 208554 | 208676 | 122 | N13:208664 |
| chrXIII | 211138 | 211313 | 175 | N13:211187 |

TABLE 1

| CHR | Start | End | Length (bp) | Nucleosome_ID |
|---|---|---|---|---|
| chrXIII | 218290 | 218401 | 111 | N13:218320 |
| chrXIII | 220837 | 220983 | 146 | N13:220846 |
| chrXIII | 228866 | 229000 | 134 | N13:228934 |
| chrXIII | 230938 | 231091 | 153 | N13:231007 |
| chrXIII | 231257 | 231408 | 151 | N13:231330 |
| chrXIII | 233569 | 233719 | 150 | N13:233649 |
| chrXIII | 240347 | 240506 | 159 | N13:240446 |
| chrXIII | 242945 | 243091 | 146 | N13:243011 |
| chrXIII | 243502 | 243662 | 160 | N13:243575 |
| chrXIII | 243825 | 243980 | 155 | N13:243892 |
| chrXIII | 244965 | 245130 | 165 | N13:244990 |
| chrXIII | 244965 | 245130 | 165 | N13:245141 |
| chrXIII | 245559 | 245710 | 151 | N13:245618 |
| chrXIII | 246032 | 246179 | 147 | N13:246115 |
| chrXIII | 246042 | 246200 | 158 | N13:246115 |
| chrXIII | 246354 | 246509 | 155 | N13:246437 |
| chrXIII | 246617 | 246737 | 120 | N13:246744 |
| chrXIII | 250435 | 250548 | 113 | N13:250485 |
| chrXIII | 250507 | 250631 | 124 | N13:250485 |
| chrXIII | 250507 | 250662 | 155 | N13:250485 |
| chrXIII | 250507 | 250664 | 157 | N13:250485 |
| chrXIII | 250507 | 250665 | 158 | N13:250485 |
| chrXIII | 250507 | 250666 | 159 | N13:250485 |
| chrXIII | 250507 | 250667 | 160 | N13:250485 |
| chrXIII | 250507 | 250668 | 161 | N13:250485 |
| chrXIII | 250507 | 250669 | 162 | N13:250485 |
| chrXIII | 250507 | 250670 | 163 | N13:250485 |
| chrXIII | 250507 | 250671 | 164 | N13:250485 |
| chrXIII | 250507 | 250673 | 166 | N13:250485 |
| chrXIII | 250507 | 250674 | 167 | N13:250485 |
| chrXIII | 250507 | 250675 | 168 | N13:250485 |
| chrXIII | 250507 | 250676 | 169 | N13:250485 |
| chrXIII | 250507 | 250677 | 170 | N13:250485 |
| chrXIII | 250507 | 250678 | 171 | N13:250485 |
| chrXIII | 250507 | 250718 | 211 | N13:250485 |
| chrXIII | 250507 | 250718 | 211 | N13:250740 |
| chrXIII | 250508 | 250678 | 170 | N13:250485 |
| chrXIII | 250509 | 250678 | 169 | |
| chrXIII | 250510 | 250678 | 168 | |
| chrXIII | 250511 | 250678 | 167 | |
| chrXIII | 250514 | 250678 | 164 | |
| chrXIII | 250515 | 250678 | 163 | |
| chrXIII | 250566 | 250718 | 152 | N13:250740 |
| chrXIII | 253372 | 253532 | 160 | N13:253539 |
| chrXIII | 256009 | 256161 | 152 | N13:256158 |
| chrXIII | 260961 | 261102 | 141 | N13:261093 |
| chrXIII | 261845 | 261977 | 132 | N13:261963 |
| chrXIII | 262094 | 262251 | 157 | N13:262151 |
| chrXIII | 266226 | 266377 | 151 | N13:266295 |
| chrXIII | 271419 | 271566 | 147 | N13:271513 |
| chrXIII | 273047 | 273197 | 150 | N13:273120 |
| chrXIII | 276628 | 276785 | 157 | N13:276642 |
| chrXIII | 276628 | 276785 | 157 | N13:276801 |
| chrXIII | 277879 | 278014 | 135 | N13:277912 |
| chrXIII | 279195 | 279369 | 174 | N13:279270 |
| chrXIII | 283987 | 284078 | 91 | N13:284101 |
| chrXIII | 283987 | 284079 | 92 | N13:284101 |
| chrXIII | 289854 | 289961 | 107 | N13:289883 |
| chrXIII | 289854 | 289977 | 123 | N13:289883 |
| chrXIII | 303791 | 303918 | 127 | N13:303847 |
| chrXIII | 304476 | 304622 | 146 | N13:304602 |
| chrXIII | 310201 | 310351 | 150 | N13:310222 |
| chrXIII | 313026 | 313180 | 154 | |
| chrXIII | 324504 | 324669 | 165 | N13:324559 |
| chrXIII | 325807 | 325958 | 151 | N13:325840 |
| chrXIII | 333017 | 333169 | 152 | N13:333098 |
| chrXIII | 335127 | 335272 | 145 | N13:335202 |
| chrXIII | 335472 | 335613 | 141 | N13:335545 |
| chrXIII | 336952 | 337113 | 161 | N13:337009 |
| chrXIII | 340438 | 340585 | 147 | N13:340517 |
| chrXIII | 342150 | 342302 | 152 | N13:342208 |
| chrXIII | 348582 | 348731 | 149 | N13:348736 |
| chrXIII | 351853 | 352022 | 169 | N13:351943 |
| chrXIII | 351863 | 352007 | 144 | N13:351943 |
| chrXIII | 357010 | 357124 | 114 | |
| chrXIII | 357475 | 357580 | 105 | N13:357470 |
| chrXIII | 358631 | 358781 | 150 | N13:358713 |
| chrXIII | 358907 | 359055 | 148 | N13:358983 |
| chrXIII | 360027 | 360184 | 157 | N13:360111 |
| chrXIII | 360404 | 360549 | 145 | N13:360557 |
| chrXIII | 360675 | 360810 | 135 | N13:360775 |
| chrXIII | 362078 | 362238 | 160 | N13:362147 |
| chrXIII | 362592 | 362706 | 114 | N13:362628 |
| chrXIII | 374780 | 374911 | 131 | N13:374845 |
| chrXIII | 375262 | 375391 | 129 | |
| chrXIII | 375501 | 375650 | 149 | N13:375616 |
| chrXIII | 375725 | 375882 | 157 | N13:375785 |
| chrXIII | 375911 | 376061 | 150 | N13:376003 |
| chrXIII | 376373 | 376508 | 135 | N13:376424 |
| chrXIII | 376763 | 376917 | 154 | N13:376925 |
| chrXIII | 377411 | 377551 | 140 | N13:377486 |
| chrXIII | 377417 | 377551 | 134 | N13:377486 |
| chrXIII | 377829 | 377984 | 155 | N13:377857 |
| chrXIII | 377960 | 378068 | 108 | |
| chrXIII | 379716 | 379822 | 106 | N13:379764 |
| chrXIII | 379716 | 379838 | 122 | N13:379764 |
| chrXIII | 381135 | 381283 | 148 | N13:381244 |
| chrXIII | 383859 | 384012 | 153 | N13:383963 |
| chrXIII | 384143 | 384283 | 140 | N13:384129 |
| chrXIII | 384143 | 384283 | 140 | N13:384300 |
| chrXIII | 384959 | 385107 | 148 | |
| chrXIII | 386345 | 386477 | 132 | N13:386478 |
| chrXIII | 386345 | 386477 | 132 | N13:386323 |

TABLE 1

| CHR | Start | End | Length (bp) | Nucleosome_ID |
|---|---|---|---|---|
| chrXIII | 387854 | 388024 | 170 | N13:387911 |
| chrXIII | 395268 | 395412 | 144 | N13:395361 |
| chrXIII | 400158 | 400292 | 134 | N13:400231 |
| chrXIII | 402458 | 402607 | 149 | N13:402522 |
| chrXIII | 406280 | 406447 | 167 | N13:406339 |
| chrXIII | 408429 | 408536 | 107 | N13:408500 |
| chrXIII | 411105 | 411252 | 147 | N13:411102 |
| chrXIII | 412475 | 412581 | 106 | N13:412535 |
| chrXIII | 412475 | 412589 | 114 | N13:412535 |
| chrXIII | 413837 | 414003 | 166 | N13:413996 |
| chrXIII | 422824 | 422985 | 161 | N13:422941 |
| chrXIII | 431294 | 431455 | 161 | N13:431429 |
| chrXIII | 431358 | 431515 | 157 | N13:431429 |
| chrXIII | 435010 | 435159 | 149 | N13:435081 |
| chrXIII | 436980 | 437142 | 162 | N13:437057 |
| chrXIII | 439283 | 439457 | 174 | N13:439388 |
| chrXIII | 443760 | 443869 | 109 | N13:443884 |
| chrXIII | 445137 | 445296 | 159 | N13:445171 |
| chrXIII | 448884 | 449018 | 134 | N13:448865 |
| chrXIII | 448987 | 449140 | 153 | N13:449042 |
| chrXIII | 452811 | 452953 | 142 | N13:452872 |
| chrXIII | 455777 | 455925 | 148 | N13:455844 |
| chrXIII | 456607 | 456760 | 153 | N13:456671 |
| chrXIII | 458537 | 458683 | 146 | |
| chrXIII | 460130 | 460281 | 151 | N13:460189 |
| chrXIII | 460228 | 460362 | 134 | N13:460349 |
| chrXIII | 460442 | 460567 | 125 | N13:460520 |
| chrXIII | 460442 | 460595 | 153 | N13:460520 |
| chrXIII | 460442 | 460598 | 156 | N13:460520 |
| chrXIII | 460442 | 460600 | 158 | N13:460520 |
| chrXIII | 461194 | 461345 | 151 | N13:461301 |
| chrXIII | 464378 | 464520 | 142 | N13:464473 |
| chrXIII | 467950 | 468100 | 150 | |
| chrXIII | 470902 | 471051 | 149 | N13:470916 |
| chrXIII | 473040 | 473146 | 106 | N13:473098 |
| chrXIII | 473040 | 473162 | 122 | N13:473098 |
| chrXIII | 473911 | 474081 | 170 | N13:474025 |
| chrXIII | 484473 | 484603 | 130 | N13:484551 |
| chrXIII | 488956 | 489063 | 107 | N13:489016 |
| chrXIII | 488956 | 489073 | 117 | N13:489016 |
| chrXIII | 491261 | 491409 | 148 | N13:491313 |
| chrXIII | 492970 | 493101 | 131 | N13:492992 |
| chrXIII | 495547 | 495700 | 153 | N13:495639 |
| chrXIII | 496801 | 496973 | 172 | N13:496955 |
| chrXIII | 496801 | 496973 | 172 | N13:496786 |
| chrXIII | 499783 | 499912 | 129 | N13:499808 |
| chrXIII | 503855 | 504003 | 148 | N13:503924 |
| chrXIII | 505412 | 505581 | 169 | N13:505521 |
| chrXIII | 512141 | 512286 | 145 | N13:512212 |
| chrXIII | 514521 | 514669 | 148 | N13:514629 |
| chrXIII | 514553 | 514703 | 150 | N13:514629 |
| chrXIII | 515323 | 515459 | 136 | N13:515420 |
| chrXIII | 520223 | 520378 | 155 | N13:520207 |
| chrXIII | 520223 | 520378 | 155 | N13:520395 |
| chrXIII | 520376 | 520508 | 132 | N13:520395 |
| chrXIII | 521730 | 521901 | 171 | N13:521801 |
| chrXIII | 522712 | 522817 | 105 | N13:522725 |
| chrXIII | 528716 | 528857 | 141 | N13:528767 |
| chrXIII | 533451 | 533610 | 159 | N13:533576 |
| chrXIII | 535081 | 535246 | 165 | N13:535146 |
| chrXIII | 536826 | 536976 | 150 | N13:536905 |
| chrXIII | 537368 | 537507 | 139 | N13:537514 |
| chrXIII | 539637 | 539760 | 123 | N13:539719 |
| chrXIII | 540611 | 540759 | 148 | N13:540749 |
| chrXIII | 541727 | 541880 | 153 | N13:541835 |
| chrXIII | 549153 | 549302 | 149 | N13:549269 |
| chrXIII | 549525 | 549679 | 154 | N13:549554 |
| chrXIII | 549713 | 549857 | 144 | N13:549747 |
| chrXIII | 555791 | 555929 | 138 | N13:555792 |
| chrXIII | 562267 | 562429 | 162 | N13:562366 |
| chrXIII | 562901 | 563062 | 161 | N13:562920 |
| chrXIII | 571585 | 571738 | 153 | N13:571681 |
| chrXIII | 571590 | 571738 | 148 | N13:571681 |
| chrXIII | 574736 | 574904 | 168 | N13:574834 |
| chrXIII | 580311 | 580467 | 156 | N13:580401 |
| chrXIII | 580644 | 580772 | 128 | N13:580704 |
| chrXIII | 581583 | 581719 | 136 | N13:581638 |
| chrXIII | 582011 | 582174 | 163 | N13:582099 |
| chrXIII | 582013 | 582174 | 161 | N13:582099 |
| chrXIII | 583110 | 583268 | 158 | |
| chrXIII | 588471 | 588586 | 115 | |
| chrXIII | 595003 | 595166 | 163 | N13:595084 |
| chrXIII | 596303 | 596450 | 147 | N13:596386 |
| chrXIII | 598087 | 598214 | 127 | N13:598137 |
| chrXIII | 598887 | 599027 | 140 | N13:598952 |
| chrXIII | 600432 | 600562 | 130 | N13:600483 |
| chrXIII | 600662 | 600821 | 159 | N13:600688 |
| chrXIII | 602872 | 603031 | 159 | N13:602904 |
| chrXIII | 608199 | 608341 | 142 | N13:608209 |
| chrXIII | 614091 | 614242 | 151 | N13:614214 |
| chrXIII | 617621 | 617773 | 152 | N13:617702 |
| chrXIII | 617962 | 618111 | 149 | N13:618058 |
| chrXIII | 622293 | 622443 | 150 | N13:622398 |
| chrXIII | 627780 | 627908 | 128 | N13:627847 |
| chrXIII | 627788 | 627919 | 131 | N13:627847 |
| chrXIII | 634568 | 634713 | 145 | N13:634666 |
| chrXIII | 637620 | 637765 | 145 | N13:637702 |
| chrXIII | 641481 | 641611 | 130 | N13:641533 |
| chrXIII | 642381 | 642525 | 144 | N13:642458 |
| chrXIII | 642389 | 642541 | 152 | N13:642458 |
| chrXIII | 649168 | 649320 | 152 | N13:649263 |
| chrXIII | 652798 | 652958 | 160 | N13:652939 |
| chrXIII | 657660 | 657789 | 129 | N13:657675 |

TABLE 1

| CHR | Start | End | Length (bp) | Nucleosome_ID |
|---|---|---|---|---|
| chrXIII | 658250 | 658404 | 154 | N13:658346 |
| chrXIII | 658827 | 658990 | 163 | N13:658863 |
| chrXIII | 658970 | 659105 | 135 | N13:659025 |
| chrXIII | 660778 | 660927 | 149 | N13:660944 |
| chrXIII | 662900 | 663052 | 152 | N13:662956 |
| chrXIII | 663803 | 663949 | 146 | N13:663870 |
| chrXIII | 663837 | 663944 | 107 | N13:663870 |
| chrXIII | 663837 | 663953 | 116 | N13:663870 |
| chrXIII | 667634 | 667789 | 155 | N13:667703 |
| chrXIII | 671983 | 672128 | 145 | N13:672090 |
| chrXIII | 675888 | 676037 | 149 | N13:675911 |
| chrXIII | 675888 | 676037 | 149 | N13:676060 |
| chrXIII | 675950 | 676037 | 87 | N13:676060 |
| chrXIII | 675950 | 676105 | 155 | N13:676060 |
| chrXIII | 677204 | 677344 | 140 | N13:677258 |
| chrXIII | 677920 | 678066 | 146 | N13:677969 |
| chrXIII | 678554 | 678723 | 169 | N13:678626 |
| chrXIII | 679422 | 679570 | 148 | |
| chrXIII | 680831 | 680955 | 124 | N13:680907 |
| chrXIII | 684362 | 684526 | 164 | N13:684432 |
| chrXIII | 686491 | 686656 | 165 | N13:686622 |
| chrXIII | 692898 | 693052 | 154 | N13:692964 |
| chrXIII | 693627 | 693771 | 144 | N13:693719 |
| chrXIII | 693831 | 693958 | 127 | N13:693886 |
| chrXIII | 694160 | 694314 | 154 | N13:694234 |
| chrXIII | 694285 | 694446 | 161 | |
| chrXIII | 694710 | 694842 | 132 | N13:694758 |
| chrXIII | 711631 | 711787 | 156 | N13:711674 |
| chrXIII | 713556 | 713705 | 149 | N13:713622 |
| chrXIII | 714522 | 714675 | 153 | N13:714616 |
| chrXIII | 715186 | 715332 | 146 | N13:715254 |
| chrXIII | 718847 | 718995 | 148 | N13:719006 |
| chrXIII | 718847 | 718995 | 148 | N13:718849 |
| chrXIII | 719890 | 720048 | 158 | N13:719976 |
| chrXIII | 723055 | 723205 | 150 | |
| chrXIII | 723972 | 724126 | 154 | N13:724061 |
| chrXIII | 730943 | 731091 | 148 | N13:731008 |
| chrXIII | 737903 | 738069 | 166 | N13:737964 |
| chrXIII | 748521 | 748674 | 153 | N13:748606 |
| chrXIII | 750522 | 750675 | 153 | N13:750581 |
| chrXIII | 757851 | 758017 | 166 | N13:757933 |
| chrXIII | 758839 | 758989 | 150 | N13:759009 |
| chrXIII | 758839 | 758989 | 150 | N13:758860 |
| chrXIII | 759376 | 759557 | 181 | N13:759498 |
| chrXIII | 765908 | 766066 | 158 | N13:766027 |
| chrXIII | 772941 | 773097 | 156 | N13:773019 |
| chrXIII | 773665 | 773803 | 138 | N13:773793 |
| chrXIII | 773665 | 773807 | 142 | N13:773793 |
| chrXIII | 774153 | 774312 | 159 | N13:774317 |
| chrXIII | 774153 | 774312 | 159 | N13:774158 |
| chrXIII | 778663 | 778795 | 132 | N13:778773 |
| chrXIII | 779035 | 779188 | 153 | N13:779109 |
| chrXIII | 780072 | 780218 | 146 | N13:780137 |
| chrXIII | 780801 | 780937 | 136 | N13:780898 |
| chrXIII | 790225 | 790394 | 169 | N13:790301 |
| chrXIII | 790368 | 790515 | 147 | N13:790486 |
| chrXIII | 790640 | 790805 | 165 | N13:790824 |
| chrXIII | 790640 | 790805 | 165 | N13:790646 |
| chrXIII | 791775 | 791919 | 144 | N13:791864 |
| chrXIII | 792265 | 792408 | 143 | N13:792303 |
| chrXIII | 792607 | 792777 | 170 | N13:792794 |
| chrXIII | 792607 | 792777 | 170 | N13:792641 |
| chrXIII | 792935 | 793098 | 163 | N13:793117 |
| chrXIII | 792935 | 793098 | 163 | N13:792956 |
| chrXIII | 793020 | 793160 | 140 | N13:793117 |
| chrXIII | 793713 | 793841 | 128 | N13:793740 |
| chrXIII | 795911 | 796064 | 153 | N13:795966 |
| chrXIII | 797737 | 797897 | 160 | N13:797874 |
| chrXIII | 797737 | 797940 | 203 | N13:797874 |
| chrXIII | 797778 | 797885 | 107 | N13:797874 |
| chrXIII | 797778 | 797897 | 119 | N13:797874 |
| chrXIII | 797778 | 797940 | 162 | N13:797874 |
| chrXIII | 799674 | 799836 | 162 | |
| chrXIII | 800670 | 800819 | 149 | |
| chrXIII | 801962 | 802127 | 165 | N13:802061 |
| chrXIII | 804104 | 804264 | 160 | N13:804203 |
| chrXIII | 806854 | 806962 | 108 | N13:806905 |
| chrXIII | 806855 | 806962 | 107 | N13:806905 |
| chrXIII | 808568 | 808720 | 152 | |
| chrXIII | 811849 | 811978 | 129 | N13:811868 |
| chrXIII | 816433 | 816597 | 164 | N13:816517 |
| chrXIII | 819869 | 819975 | 106 | N13:819933 |
| chrXIII | 819869 | 819988 | 119 | N13:819933 |
| chrXIII | 820608 | 820710 | 102 | N13:820698 |
| chrXIII | 820608 | 820711 | 103 | N13:820698 |
| chrXIII | 820608 | 820715 | 107 | N13:820698 |
| chrXIII | 820608 | 820717 | 109 | N13:820698 |
| chrXIII | 820608 | 820733 | 125 | N13:820698 |
| chrXIII | 820608 | 820737 | 129 | N13:820698 |
| chrXIII | 820608 | 820738 | 130 | N13:820698 |
| chrXIII | 820608 | 820740 | 132 | N13:820698 |
| chrXIII | 820608 | 820741 | 133 | N13:820698 |
| chrXIII | 820608 | 820742 | 134 | N13:820698 |
| chrXIII | 820608 | 820743 | 135 | N13:820698 |
| chrXIII | 820608 | 820744 | 136 | N13:820698 |
| chrXIII | 820608 | 820745 | 137 | N13:820698 |
| chrXIII | 820608 | 820746 | 138 | N13:820698 |
| chrXIII | 820608 | 820747 | 139 | N13:820698 |
| chrXIII | 820608 | 820748 | 140 | N13:820698 |
| chrXIII | 820608 | 820749 | 141 | N13:820698 |
| chrXIII | 820608 | 820750 | 142 | N13:820698 |
| chrXIII | 820608 | 820751 | 143 | N13:820698 |
| chrXIII | 820609 | 820751 | 142 | N13:820698 |

| CHR | Start | End | Length (bp) | Nucleosome_ID |
|---|---|---|---|---|
| chrXIII | 820610 | 820751 | 141 | N13:820698 |
| chrXIII | 820611 | 820751 | 140 | N13:820698 |
| chrXIII | 820612 | 820751 | 139 | N13:820698 |
| chrXIII | 820613 | 820751 | 138 | N13:820698 |
| chrXIII | 820614 | 820751 | 137 | N13:820698 |
| chrXIII | 820615 | 820751 | 136 | N13:820698 |
| chrXIII | 820616 | 820751 | 135 | N13:820698 |
| chrXIII | 820617 | 820751 | 134 | N13:820698 |
| chrXIII | 820619 | 820751 | 132 | N13:820698 |
| chrXIII | 820620 | 820751 | 131 | N13:820698 |
| chrXIII | 820621 | 820751 | 130 | N13:820698 |
| chrXIII | 820624 | 820751 | 127 | N13:820698 |
| chrXIII | 820626 | 820715 | 89 | N13:820698 |
| chrXIII | 828572 | 828720 | 148 | N13:828670 |
| chrXIII | 835990 | 836099 | 109 | |
| chrXIII | 836333 | 836463 | 130 | N13:836461 |
| chrXIII | 842098 | 842232 | 134 | N13:842139 |
| chrXIII | 843431 | 843583 | 152 | N13:843529 |
| chrXIII | 847888 | 848017 | 129 | N13:847969 |
| chrXIII | 847897 | 848050 | 153 | N13:847969 |
| chrXIII | 851484 | 851633 | 149 | N13:851582 |
| chrXIII | 864286 | 864433 | 147 | N13:864337 |
| chrXIII | 864644 | 864799 | 155 | N13:864820 |
| chrXIII | 864644 | 864799 | 155 | N13:864653 |
| chrXIII | 866005 | 866155 | 150 | |
| chrXIII | 868732 | 868890 | 158 | N13:868810 |
| chrXIII | 871472 | 871623 | 151 | N13:871631 |
| chrXIII | 871472 | 871623 | 151 | N13:871471 |
| chrXIII | 873640 | 873761 | 121 | N13:873682 |
| chrXIII | 873640 | 873789 | 149 | N13:873682 |
| chrXIII | 873640 | 873790 | 150 | N13:873682 |
| chrXIII | 873648 | 873761 | 113 | N13:873682 |
| chrXIII | 873648 | 873789 | 141 | N13:873682 |
| chrXIII | 873648 | 873790 | 142 | N13:873682 |
| chrXIII | 878188 | 878341 | 153 | N13:878290 |
| chrXIII | 879147 | 879302 | 155 | N13:879224 |
| chrXIII | 884345 | 884496 | 151 | N13:884479 |
| chrXIII | 886126 | 886284 | 158 | |
| chrXIII | 889335 | 889469 | 134 | N13:889367 |
| chrXIII | 890068 | 890170 | 102 | N13:890188 |
| chrXIII | 891024 | 891149 | 125 | N13:891052 |
| chrXIII | 891955 | 892105 | 150 | N13:892008 |
| chrXIII | 895348 | 895499 | 151 | N13:895429 |
| chrXIII | 902827 | 902965 | 138 | N13:902909 |
| chrXIII | 904201 | 904335 | 134 | N13:904278 |
| chrXIII | 904428 | 904582 | 154 | N13:904475 |
| chrXIII | 910695 | 910845 | 150 | N13:910757 |
| chrXIII | 911000 | 911154 | 154 | N13:911157 |
| chrXIII | 917204 | 917355 | 151 | N13:917298 |
| chrXIII | 921093 | 921253 | 160 | N13:921174 |
| chrXIII | 923830 | 923976 | 146 | N13:923958 |
| chrXIII | 924304 | 924426 | 122 | N13:924361 |
| chrXIII | 924311 | 924426 | 115 | N13:924361 |
| chrXIII | 924321 | 924426 | 105 | N13:924361 |
| chrXIV | 614 | 765 | 151 | N14:689 |
| chrXIV | 953 | 1105 | 152 | N14:1005 |
| chrXIV | 1779 | 1930 | 151 | N14:1793 |
| chrXIV | 1791 | 1925 | 134 | N14:1793 |
| chrXIV | 1816 | 1912 | 96 | N14:1793 |
| chrXIV | 2170 | 2300 | 130 | N14:2318 |
| chrXIV | 2449 | 2601 | 152 | N14:2530 |
| chrXIV | 2796 | 2940 | 144 | N14:2849 |
| chrXIV | 2796 | 2983 | 187 | N14:2849 |
| chrXIV | 2848 | 2940 | 92 | N14:2849 |
| chrXIV | 2848 | 2983 | 135 | N14:2849 |
| chrXIV | 2936 | 3080 | 144 | N14:3031 |
| chrXIV | 3001 | 3153 | 152 | N14:3031 |
| chrXIV | 3253 | 3378 | 125 | N14:3371 |
| chrXIV | 3283 | 3426 | 143 | N14:3371 |
| chrXIV | 3584 | 3721 | 137 | N14:3651 |
| chrXIV | 3700 | 3855 | 155 | N14:3808 |
| chrXIV | 3794 | 3956 | 162 | N14:3967 |
| chrXIV | 3794 | 3956 | 162 | N14:3808 |
| chrXIV | 5019 | 5160 | 141 | N14:5067 |
| chrXIV | 15620 | 15770 | 150 | N14:15736 |
| chrXIV | 18901 | 19054 | 153 | N14:18957 |
| chrXIV | 22418 | 22569 | 151 | N14:22459 |
| chrXIV | 24850 | 25019 | 169 | N14:24952 |
| chrXIV | 25836 | 25986 | 150 | N14:25940 |
| chrXIV | 26530 | 26680 | 150 | N14:26673 |
| chrXIV | 27268 | 27422 | 154 | |
| chrXIV | 30249 | 30400 | 151 | N14:30360 |
| chrXIV | 32059 | 32210 | 151 | N14:32102 |
| chrXIV | 33454 | 33599 | 145 | N14:33540 |
| chrXIV | 33454 | 33606 | 152 | N14:33540 |
| chrXIV | 33459 | 33599 | 140 | N14:33540 |
| chrXIV | 33461 | 33599 | 138 | N14:33540 |
| chrXIV | 33461 | 33603 | 142 | N14:33540 |
| chrXIV | 33461 | 33606 | 145 | N14:33540 |
| chrXIV | 33464 | 33606 | 142 | N14:33540 |
| chrXIV | 33466 | 33606 | 140 | N14:33540 |
| chrXIV | 41403 | 41548 | 145 | N14:41462 |
| chrXIV | 45261 | 45368 | 107 | N14:45343 |
| chrXIV | 45261 | 45374 | 113 | N14:45343 |
| chrXIV | 45783 | 45937 | 154 | N14:45851 |
| chrXIV | 49049 | 49199 | 150 | N14:49135 |
| chrXIV | 53101 | 53263 | 162 | N14:53189 |
| chrXIV | 55301 | 55450 | 149 | N14:55396 |
| chrXIV | 58407 | 58536 | 129 | N14:58525 |
| chrXIV | 60113 | 60210 | 97 | |
| chrXIV | 61184 | 61331 | 147 | N14:61254 |
| chrXIV | 63570 | 63733 | 163 | N14:63662 |
| chrXIV | 65407 | 65535 | 128 | N14:65439 |

| CHR | Start | End | Length (bp) | Nucleosome_ID |
|---|---|---|---|---|
| chrXIV | 73936 | 74081 | 145 | N14:73929 |
| chrXIV | 77741 | 77895 | 154 | N14:77801 |
| chrXIV | 79985 | 80135 | 150 | N14:80056 |
| chrXIV | 81420 | 81583 | 163 | N14:81459 |
| chrXIV | 91093 | 91242 | 149 | N14:91178 |
| chrXIV | 93267 | 93403 | 136 | N14:93374 |
| chrXIV | 97284 | 97435 | 151 | N14:97317 |
| chrXIV | 97941 | 98103 | 162 | N14:98029 |
| chrXIV | 99408 | 99557 | 149 | N14:99525 |
| chrXIV | 99632 | 99789 | 157 | N14:99713 |
| chrXIV | 99818 | 99968 | 150 | N14:99892 |
| chrXIV | 99822 | 99968 | 146 | N14:99892 |
| chrXIV | 100280 | 100415 | 135 | N14:100384 |
| chrXIV | 100932 | 101089 | 157 | N14:101020 |
| chrXIV | 102362 | 102510 | 148 | N14:102397 |
| chrXIV | 103973 | 104137 | 164 | N14:104026 |
| chrXIV | 107707 | 107834 | 127 | N14:107700 |
| chrXIV | 110079 | 110230 | 151 | N14:110172 |
| chrXIV | 111620 | 111775 | 155 | |
| chrXIV | 112009 | 112129 | 120 | N14:112032 |
| chrXIV | 116452 | 116578 | 126 | N14:116519 |
| chrXIV | 119252 | 119378 | 126 | N14:119284 |
| chrXIV | 121333 | 121464 | 131 | N14:121346 |
| chrXIV | 124849 | 124975 | 126 | N14:124938 |
| chrXIV | 129691 | 129843 | 152 | N14:129772 |
| chrXIV | 129719 | 129870 | 151 | N14:129772 |
| chrXIV | 131962 | 132135 | 173 | N14:131992 |
| chrXIV | 132312 | 132463 | 151 | N14:132390 |
| chrXIV | 133490 | 133654 | 164 | N14:133593 |
| chrXIV | 134176 | 134325 | 149 | N14:134291 |
| chrXIV | 135289 | 135440 | 151 | N14:135423 |
| chrXIV | 135829 | 135979 | 150 | N14:135901 |
| chrXIV | 139766 | 139921 | 155 | N14:139942 |
| chrXIV | 140110 | 140269 | 159 | N14:140187 |
| chrXIV | 141911 | 142060 | 149 | N14:142074 |
| chrXIV | 150518 | 150667 | 149 | N14:150605 |
| chrXIV | 153107 | 153271 | 164 | N14:153137 |
| chrXIV | 153107 | 153271 | 164 | N14:153294 |
| chrXIV | 155351 | 155502 | 151 | N14:155389 |
| chrXIV | 163862 | 164000 | 138 | N14:164012 |
| chrXIV | 163862 | 164000 | 138 | N14:163855 |
| chrXIV | 165154 | 165280 | 126 | N14:165235 |
| chrXIV | 167446 | 167585 | 139 | N14:167470 |
| chrXIV | 174863 | 175007 | 144 | N14:174931 |
| chrXIV | 177654 | 177807 | 153 | N14:177732 |
| chrXIV | 178329 | 178461 | 132 | N14:178391 |
| chrXIV | 183769 | 183934 | 165 | N14:183818 |
| chrXIV | 187776 | 187960 | 184 | N14:187755 |
| chrXIV | 188930 | 189078 | 148 | |
| chrXIV | 188930 | 189079 | 149 | |
| chrXIV | 188930 | 189080 | 150 | |
| chrXIV | 188930 | 189081 | 151 | |
| chrXIV | 188930 | 189083 | 153 | |
| chrXIV | 188931 | 189083 | 152 | |
| chrXIV | 188933 | 189083 | 150 | |
| chrXIV | 188934 | 189083 | 149 | |
| chrXIV | 188935 | 189083 | 148 | |
| chrXIV | 190891 | 191040 | 149 | N14:190930 |
| chrXIV | 192308 | 192455 | 147 | N14:192383 |
| chrXIV | 192813 | 192954 | 141 | N14:192825 |
| chrXIV | 193504 | 193657 | 153 | N14:193627 |
| chrXIV | 196760 | 196910 | 150 | N14:196922 |
| chrXIV | 202136 | 202292 | 156 | N14:202215 |
| chrXIV | 203542 | 203662 | 120 | N14:203649 |
| chrXIV | 206857 | 207023 | 166 | N14:206949 |
| chrXIV | 210139 | 210288 | 149 | N14:210220 |
| chrXIV | 216142 | 216308 | 166 | N14:216232 |
| chrXIV | 219506 | 219633 | 127 | N14:219552 |
| chrXIV | 224826 | 224979 | 153 | N14:224898 |
| chrXIV | 227320 | 227476 | 156 | N14:227397 |
| chrXIV | 235082 | 235232 | 150 | |
| chrXIV | 238200 | 238369 | 169 | N14:238254 |
| chrXIV | 246737 | 246897 | 160 | N14:246814 |
| chrXIV | 247839 | 247988 | 149 | N14:247931 |
| chrXIV | 254524 | 254675 | 151 | N14:254603 |
| chrXIV | 257454 | 257590 | 136 | N14:257486 |
| chrXIV | 257743 | 257893 | 150 | N14:257796 |
| chrXIV | 259132 | 259283 | 151 | N14:259110 |
| chrXIV | 259132 | 259283 | 151 | N14:259278 |
| chrXIV | 265572 | 265700 | 128 | N14:265642 |
| chrXIV | 271513 | 271667 | 154 | N14:271563 |
| chrXIV | 273060 | 273209 | 149 | N14:273068 |
| chrXIV | 273060 | 273209 | 149 | N14:273226 |
| chrXIV | 275916 | 276056 | 140 | N14:276038 |
| chrXIV | 281515 | 281622 | 107 | N14:281537 |
| chrXIV | 281515 | 281631 | 116 | N14:281537 |
| chrXIV | 281832 | 281976 | 144 | N14:281889 |
| chrXIV | 283804 | 283989 | 185 | N14:283836 |
| chrXIV | 287747 | 287882 | 135 | N14:287841 |
| chrXIV | 291077 | 291208 | 131 | N14:291175 |
| chrXIV | 292461 | 292633 | 172 | N14:292551 |
| chrXIV | 292477 | 292630 | 153 | N14:292551 |
| chrXIV | 296854 | 297025 | 171 | N14:296864 |
| chrXIV | 297410 | 297562 | 152 | N14:297470 |
| chrXIV | 298390 | 298549 | 159 | N14:298469 |
| chrXIV | 302946 | 303074 | 128 | N14:302995 |
| chrXIV | 303880 | 304030 | 150 | N14:303933 |
| chrXIV | 307042 | 307199 | 157 | N14:307110 |
| chrXIV | 308271 | 308446 | 175 | N14:308444 |
| chrXIV | 308271 | 308446 | 175 | N14:308284 |
| chrXIV | 309709 | 309860 | 151 | N14:309803 |
| chrXIV | 310999 | 311146 | 147 | N14:311066 |
| chrXIV | 314726 | 314877 | 151 | N14:314793 |

TABLE 1

| CHR | Start | End | Length (bp) | Nucleosome_ID |
|---|---|---|---|---|
| chrXIV | 317936 | 318050 | 114 | N14:318021 |
| chrXIV | 324620 | 324765 | 145 | N14:324681 |
| chrXIV | 324977 | 325117 | 140 | N14:325028 |
| chrXIV | 336864 | 337018 | 154 | N14:336949 |
| chrXIV | 339979 | 340132 | 153 | N14:340063 |
| chrXIV | 340143 | 340297 | 154 | |
| chrXIV | 343939 | 344095 | 156 | N14:344007 |
| chrXIV | 348424 | 348579 | 155 | N14:348518 |
| chrXIV | 349341 | 349497 | 156 | N14:349432 |
| chrXIV | 357765 | 357910 | 145 | N14:357862 |
| chrXIV | 357767 | 357910 | 143 | N14:357862 |
| chrXIV | 358288 | 358439 | 151 | N14:358379 |
| chrXIV | 360600 | 360759 | 159 | N14:360625 |
| chrXIV | 360762 | 360912 | 150 | N14:360803 |
| chrXIV | 361214 | 361352 | 138 | N14:361265 |
| chrXIV | 364273 | 364404 | 131 | N14:364350 |
| chrXIV | 365246 | 365404 | 158 | N14:365272 |
| chrXIV | 366145 | 366271 | 126 | |
| chrXIV | 368092 | 368240 | 148 | N14:368205 |
| chrXIV | 369049 | 369214 | 165 | N14:369138 |
| chrXIV | 376927 | 377089 | 162 | N14:377019 |
| chrXIV | 384387 | 384546 | 159 | N14:384456 |
| chrXIV | 388430 | 388576 | 146 | N14:388511 |
| chrXIV | 388719 | 388880 | 161 | N14:388701 |
| chrXIV | 388719 | 388880 | 161 | N14:388900 |
| chrXIV | 396710 | 396865 | 155 | N14:396752 |
| chrXIV | 403433 | 403582 | 149 | N14:403451 |
| chrXIV | 403995 | 404145 | 150 | |
| chrXIV | 410286 | 410440 | 154 | N14:410457 |
| chrXIV | 410286 | 410440 | 154 | N14:410304 |
| chrXIV | 410317 | 410470 | 153 | N14:410457 |
| chrXIV | 410317 | 410470 | 153 | N14:410304 |
| chrXIV | 410968 | 411126 | 158 | N14:411083 |
| chrXIV | 411008 | 411145 | 137 | N14:411083 |
| chrXIV | 416136 | 416292 | 156 | N14:416212 |
| chrXIV | 418446 | 418610 | 164 | N14:418509 |
| chrXIV | 419242 | 419381 | 139 | N14:419334 |
| chrXIV | 420269 | 420420 | 151 | N14:420290 |
| chrXIV | 421794 | 421931 | 137 | N14:421859 |
| chrXIV | 421911 | 422067 | 156 | N14:422017 |
| chrXIV | 423473 | 423612 | 139 | N14:423543 |
| chrXIV | 423497 | 423650 | 153 | N14:423543 |
| chrXIV | 426375 | 426492 | 117 | N14:426395 |
| chrXIV | 430126 | 430279 | 153 | N14:430203 |
| chrXIV | 430126 | 430286 | 160 | N14:430203 |
| chrXIV | 439717 | 439834 | 117 | N14:439718 |
| chrXIV | 459511 | 459626 | 115 | N14:459588 |
| chrXIV | 460164 | 460311 | 147 | N14:460238 |
| chrXIV | 464412 | 464538 | 126 | N14:464472 |
| chrXIV | 466515 | 466659 | 144 | N14:466599 |
| chrXIV | 467295 | 467445 | 150 | |
| chrXIV | 473999 | 474106 | 107 | N14:474082 |
| chrXIV | 475462 | 475606 | 144 | N14:475517 |
| chrXIV | 477594 | 477739 | 145 | N14:477658 |
| chrXIV | 478141 | 478275 | 134 | N14:478165 |
| chrXIV | 483192 | 483305 | 113 | |
| chrXIV | 483272 | 483422 | 150 | N14:483343 |
| chrXIV | 483529 | 483666 | 137 | N14:483509 |
| chrXIV | 483529 | 483666 | 137 | N14:483683 |
| chrXIV | 483752 | 483901 | 149 | N14:483855 |
| chrXIV | 484694 | 484843 | 149 | |
| chrXIV | 490422 | 490573 | 151 | N14:490508 |
| chrXIV | 490584 | 490699 | 115 | N14:490670 |
| chrXIV | 490584 | 490718 | 134 | N14:490670 |
| chrXIV | 490584 | 490722 | 138 | N14:490670 |
| chrXIV | 500789 | 500918 | 129 | N14:500830 |
| chrXIV | 502692 | 502828 | 136 | N14:502692 |
| chrXIV | 511107 | 511261 | 154 | |
| chrXIV | 515776 | 515877 | 101 | N14:515831 |
| chrXIV | 516941 | 517091 | 150 | N14:517027 |
| chrXIV | 517457 | 517589 | 132 | N14:517452 |
| chrXIV | 517457 | 517589 | 132 | N14:517609 |
| chrXIV | 520229 | 520363 | 134 | N14:520294 |
| chrXIV | 520229 | 520369 | 140 | N14:520294 |
| chrXIV | 521272 | 521407 | 135 | N14:521305 |
| chrXIV | 521719 | 521865 | 146 | N14:521781 |
| chrXIV | 521719 | 521869 | 150 | N14:521781 |
| chrXIV | 521898 | 522055 | 157 | N14:521995 |
| chrXIV | 524015 | 524129 | 114 | |
| chrXIV | 524719 | 524868 | 149 | N14:524853 |
| chrXIV | 524979 | 525110 | 131 | N14:525025 |
| chrXIV | 526013 | 526193 | 180 | N14:526078 |
| chrXIV | 526651 | 526800 | 149 | N14:526736 |
| chrXIV | 527097 | 527235 | 138 | N14:527172 |
| chrXIV | 527709 | 527871 | 162 | N14:527797 |
| chrXIV | 529282 | 529444 | 162 | N14:529327 |
| chrXIV | 529934 | 530079 | 145 | N14:530020 |
| chrXIV | 530913 | 531056 | 143 | |
| chrXIV | 532041 | 532181 | 140 | N14:532069 |
| chrXIV | 532139 | 532286 | 147 | N14:532293 |
| chrXIV | 536984 | 537144 | 160 | N14:537118 |
| chrXIV | 538071 | 538217 | 146 | N14:538181 |
| chrXIV | 538102 | 538265 | 163 | N14:538181 |
| chrXIV | 538210 | 538368 | 158 | N14:538353 |
| chrXIV | 540636 | 540777 | 141 | N14:540797 |
| chrXIV | 541287 | 541432 | 145 | N14:541365 |
| chrXIV | 541780 | 541943 | 163 | N14:541870 |
| chrXIV | 541920 | 542078 | 158 | |
| chrXIV | 542491 | 542640 | 149 | N14:542475 |
| chrXIV | 544884 | 545027 | 143 | N14:544952 |
| chrXIV | 544927 | 545097 | 170 | N14:544952 |
| chrXIV | 546338 | 546446 | 108 | N14:546423 |
| chrXIV | 546358 | 546520 | 162 | N14:546423 |

TABLE 1

| CHR | Start | End | Length (bp) | Nucleosome_ID |
|---|---|---|---|---|
| chrXIV | 551772 | 551902 | 130 | N14:551848 |
| chrXIV | 552571 | 552722 | 151 | N14:552590 |
| chrXIV | 553534 | 553684 | 150 | N14:553556 |
| chrXIV | 554534 | 554685 | 151 | N14:554609 |
| chrXIV | 555938 | 556085 | 147 | N14:555999 |
| chrXIV | 557972 | 558127 | 155 | N14:558022 |
| chrXIV | 560057 | 560198 | 141 | N14:560145 |
| chrXIV | 561823 | 561978 | 155 | N14:561825 |
| chrXIV | 562224 | 562373 | 149 | N14:562251 |
| chrXIV | 563011 | 563177 | 166 | N14:563098 |
| chrXIV | 564309 | 564452 | 143 | N14:564390 |
| chrXIV | 565788 | 565917 | 129 | N14:565899 |
| chrXIV | 567163 | 567273 | 110 | N14:567239 |
| chrXIV | 573304 | 573437 | 133 | N14:573369 |
| chrXIV | 575453 | 575613 | 160 | N14:575635 |
| chrXIV | 575453 | 575613 | 160 | N14:575487 |
| chrXIV | 576714 | 576844 | 130 | N14:576799 |
| chrXIV | 576714 | 576864 | 150 | N14:576799 |
| chrXIV | 578365 | 578517 | 152 | N14:578459 |
| chrXIV | 578706 | 578859 | 153 | N14:578787 |
| chrXIV | 585214 | 585366 | 152 | N14:585310 |
| chrXIV | 589016 | 589141 | 125 | N14:589151 |
| chrXIV | 589016 | 589141 | 125 | N14:589003 |
| chrXIV | 593245 | 593393 | 148 | N14:593327 |
| chrXIV | 597466 | 597618 | 152 | N14:597483 |
| chrXIV | 598791 | 598924 | 133 | N14:598861 |
| chrXIV | 602237 | 602346 | 109 | N14:602324 |
| chrXIV | 612323 | 612470 | 147 | N14:612426 |
| chrXIV | 615779 | 615947 | 168 | N14:615899 |
| chrXIV | 620267 | 620398 | 131 | N14:620245 |
| chrXIV | 620267 | 620398 | 131 | N14:620397 |
| chrXIV | 620785 | 620892 | 107 | N14:620770 |
| chrXIV | 620785 | 620946 | 161 | N14:620770 |
| chrXIV | 620785 | 620946 | 161 | N14:620937 |
| chrXIV | 625000 | 625108 | 108 | N14:625110 |
| chrXIV | 625453 | 625609 | 156 | N14:625501 |
| chrXIV | 630331 | 630483 | 152 | N14:630418 |
| chrXIV | 630745 | 630896 | 151 | N14:630914 |
| chrXIV | 630745 | 630896 | 151 | N14:630752 |
| chrXIV | 635632 | 635775 | 143 | N14:635681 |
| chrXIV | 640348 | 640505 | 157 | N14:640377 |
| chrXIV | 640382 | 640540 | 158 | N14:640377 |
| chrXIV | 640382 | 640540 | 158 | N14:640553 |
| chrXIV | 641911 | 642059 | 148 | N14:641943 |
| chrXIV | 645323 | 645472 | 149 | N14:645480 |
| chrXIV | 645323 | 645472 | 149 | N14:645332 |
| chrXIV | 646222 | 646370 | 148 | N14:646249 |
| chrXIV | 649139 | 649287 | 148 | N14:649302 |
| chrXIV | 649139 | 649287 | 148 | N14:649141 |
| chrXIV | 659033 | 659204 | 171 | N14:659107 |
| chrXIV | 660122 | 660273 | 151 | |
| chrXIV | 660694 | 660857 | 163 | N14:660797 |
| chrXIV | 669731 | 669882 | 151 | N14:669825 |
| chrXIV | 673304 | 673429 | 125 | N14:673428 |
| chrXIV | 673304 | 673441 | 137 | N14:673428 |
| chrXIV | 673304 | 673449 | 145 | N14:673428 |
| chrXIV | 673304 | 673450 | 146 | N14:673428 |
| chrXIV | 673304 | 673451 | 147 | N14:673428 |
| chrXIV | 673304 | 673452 | 148 | N14:673428 |
| chrXIV | 673304 | 673453 | 149 | N14:673428 |
| chrXIV | 673304 | 673454 | 150 | N14:673428 |
| chrXIV | 673304 | 673455 | 151 | N14:673428 |
| chrXIV | 673305 | 673455 | 150 | N14:673428 |
| chrXIV | 673306 | 673455 | 149 | N14:673428 |
| chrXIV | 673307 | 673455 | 148 | N14:673428 |
| chrXIV | 673308 | 673455 | 147 | N14:673428 |
| chrXIV | 673309 | 673455 | 146 | N14:673428 |
| chrXIV | 673310 | 673455 | 145 | N14:673428 |
| chrXIV | 673311 | 673455 | 144 | N14:673428 |
| chrXIV | 673313 | 673455 | 142 | N14:673428 |
| chrXIV | 673314 | 673455 | 141 | N14:673428 |
| chrXIV | 673321 | 673455 | 134 | N14:673428 |
| chrXIV | 673329 | 673455 | 126 | N14:673428 |
| chrXIV | 673357 | 673455 | 98 | N14:673428 |
| chrXIV | 675190 | 675330 | 140 | N14:675271 |
| chrXIV | 678978 | 679124 | 146 | N14:679095 |
| chrXIV | 679561 | 679717 | 156 | N14:679584 |
| chrXIV | 681968 | 682096 | 128 | N14:682040 |
| chrXIV | 681972 | 682075 | 103 | N14:682040 |
| chrXIV | 682856 | 682998 | 142 | N14:682935 |
| chrXIV | 689540 | 689646 | 106 | N14:689607 |
| chrXIV | 689540 | 689701 | 161 | N14:689607 |
| chrXIV | 689540 | 689704 | 164 | N14:689607 |
| chrXIV | 689540 | 689705 | 165 | N14:689607 |
| chrXIV | 689541 | 689705 | 164 | N14:689607 |
| chrXIV | 689543 | 689705 | 162 | N14:689607 |
| chrXIV | 689545 | 689705 | 160 | N14:689607 |
| chrXIV | 689550 | 689705 | 155 | N14:689607 |
| chrXIV | 689554 | 689705 | 151 | N14:689607 |
| chrXIV | 694543 | 694689 | 146 | N14:694619 |
| chrXIV | 694549 | 694699 | 150 | N14:694619 |
| chrXIV | 698880 | 699032 | 152 | N14:698920 |
| chrXIV | 704009 | 704157 | 148 | N14:704120 |
| chrXIV | 704660 | 704785 | 125 | N14:704768 |
| chrXIV | 705096 | 705262 | 166 | N14:705073 |
| chrXIV | 705096 | 705262 | 166 | N14:705244 |
| chrXIV | 705533 | 705682 | 149 | N14:705572 |
| chrXIV | 705820 | 705974 | 154 | N14:705901 |
| chrXIV | 706013 | 706167 | 154 | N14:706150 |
| chrXIV | 706675 | 706825 | 150 | N14:706652 |
| chrXIV | 706675 | 706825 | 150 | N14:706826 |
| chrXIV | 707449 | 707602 | 153 | N14:707511 |
| chrXIV | 723355 | 723513 | 158 | N14:723439 |

| CHR | Start | End | Length (bp) | Nucleosome_ID |
|---|---|---|---|---|
| chrXIV | 724785 | 724950 | 165 | N14:724864 |
| chrXIV | 727236 | 727386 | 150 | N14:727324 |
| chrXIV | 727633 | 727771 | 138 | N14:727696 |
| chrXIV | 731977 | 732139 | 162 | N14:732086 |
| chrXIV | 735524 | 735689 | 165 | N14:735604 |
| chrXIV | 735878 | 736030 | 152 | N14:735951 |
| chrXIV | 737786 | 737917 | 131 | N14:737841 |
| chrXIV | 749149 | 749259 | 110 | N14:749152 |
| chrXIV | 749725 | 749855 | 130 | N14:749877 |
| chrXIV | 749823 | 749962 | 139 | N14:749877 |
| chrXIV | 750208 | 750335 | 127 | |
| chrXIV | 753363 | 753527 | 164 | N14:753511 |
| chrXIV | 754387 | 754526 | 139 | N14:754466 |
| chrXIV | 761541 | 761643 | 102 | N14:761639 |
| chrXIV | 763360 | 763525 | 165 | N14:763380 |
| chrXIV | 763360 | 763525 | 165 | N14:763542 |
| chrXIV | 767084 | 767240 | 156 | N14:767184 |
| chrXIV | 773470 | 773620 | 150 | N14:773461 |
| chrXIV | 778474 | 778625 | 151 | N14:778523 |
| chrXIV | 779862 | 780020 | 158 | N14:779923 |
| chrXIV | 781645 | 781793 | 148 | N14:781689 |
| chrXIV | 781935 | 782064 | 129 | N14:781999 |
| chrXIV | 781942 | 782106 | 164 | N14:781999 |
| chrXIV | 783376 | 783501 | 125 | N14:783457 |
| chrXIV | 784039 | 784210 | 171 | N14:784097 |
| chrXIV | 784040 | 784199 | 159 | N14:784097 |
| chrXIV | 784044 | 784183 | 139 | N14:784097 |
| chrXIV | 784044 | 784199 | 155 | N14:784097 |
| chrXIV | 784049 | 784198 | 149 | N14:784097 |
| chrXIV | 784049 | 784203 | 154 | N14:784097 |
| chrXIV | 784063 | 784183 | 120 | N14:784097 |
| chrXIV | 784063 | 784195 | 132 | N14:784097 |
| chrXIV | 784063 | 784199 | 136 | N14:784097 |
| chrXIV | 784063 | 784202 | 139 | N14:784097 |
| chrXIV | 784063 | 784210 | 147 | N14:784097 |
| chrXIV | 784063 | 784233 | 170 | N14:784097 |
| chrXIV | 784063 | 784239 | 176 | N14:784097 |
| chrXIV | 784063 | 784239 | 176 | N14:784261 |
| chrXIV | 784063 | 784275 | 212 | N14:784097 |
| chrXIV | 784063 | 784275 | 212 | N14:784261 |
| chrXIV | 784064 | 784183 | 119 | N14:784097 |
| chrXIV | 784064 | 784199 | 135 | N14:784097 |
| chrXIV | 784101 | 784203 | 102 | N14:784097 |
| chrXIV | 784101 | 784210 | 109 | N14:784097 |
| chrXIV | 784106 | 784198 | 92 | N14:784097 |
| chrXIV | 784106 | 784203 | 97 | N14:784097 |
| chrXIV | 784106 | 784210 | 104 | N14:784097 |
| chrXIV | 784106 | 784238 | 132 | N14:784097 |
| chrXIV | 784106 | 784238 | 132 | N14:784261 |
| chrXIV | 784106 | 784265 | 159 | N14:784097 |
| chrXIV | 784106 | 784265 | 159 | N14:784261 |
| chrXIV | 784106 | 784274 | 168 | N14:784097 |
| chrXIV | 784106 | 784274 | 168 | N14:784261 |
| chrXIV | 784106 | 784275 | 169 | N14:784097 |
| chrXIV | 784106 | 784275 | 169 | N14:784261 |
| chrXIV | 784106 | 784277 | 171 | N14:784097 |
| chrXIV | 784106 | 784277 | 171 | N14:784261 |
| chrXIV | 784111 | 784233 | 122 | N14:784097 |
| chrXIV | 784111 | 784286 | 175 | N14:784097 |
| chrXIV | 784111 | 784286 | 175 | N14:784261 |
| chrXIV | 784130 | 784238 | 108 | N14:784261 |
| chrXIV | 784130 | 784274 | 144 | N14:784261 |
| chrXIV | 784130 | 784275 | 145 | N14:784261 |
| chrXIV | 784130 | 784277 | 147 | N14:784261 |
| chrXIV | 784130 | 784329 | 199 | N14:784261 |
| chrXIV | 784132 | 784275 | 143 | N14:784261 |
| chrXIV | 784135 | 784274 | 139 | N14:784261 |
| chrXIV | 784139 | 784275 | 136 | N14:784261 |
| chrXIV | 784139 | 784296 | 157 | N14:784261 |
| chrXIV | 784140 | 784275 | 135 | N14:784261 |
| chrXIV | 784145 | 784286 | 141 | N14:784261 |
| chrXIV | 784164 | 784275 | 111 | N14:784261 |
| chrXIV | 784169 | 784274 | 105 | N14:784261 |
| chrXIV | 784169 | 784275 | 106 | N14:784261 |
| chrXIV | 784169 | 784277 | 108 | N14:784261 |
| chrXIV | 784172 | 784265 | 93 | N14:784261 |
| chrXIV | 784172 | 784267 | 95 | N14:784261 |
| chrXIV | 784172 | 784271 | 99 | N14:784261 |
| chrXIV | 784172 | 784274 | 102 | N14:784261 |
| chrXIV | 784172 | 784275 | 103 | N14:784261 |
| chrXIV | 784172 | 784277 | 105 | N14:784261 |
| chrXIV | 784177 | 784271 | 94 | N14:784261 |
| chrXIV | 784177 | 784275 | 98 | N14:784261 |
| chrXV | 838 | 992 | 154 | N15:915 |
| chrXV | 6743 | 6894 | 151 | N15:6849 |
| chrXV | 8892 | 9048 | 156 | N15:8985 |
| chrXV | 10875 | 10982 | 107 | N15:10900 |
| chrXV | 23820 | 23972 | 152 | N15:23919 |
| chrXV | 29238 | 29397 | 159 | N15:29410 |
| chrXV | 29238 | 29397 | 159 | N15:29253 |
| chrXV | 36163 | 36311 | 148 | N15:36242 |
| chrXV | 39246 | 39406 | 160 | N15:39349 |
| chrXV | 43767 | 43929 | 162 | N15:43844 |
| chrXV | 44442 | 44591 | 149 | N15:44533 |
| chrXV | 49181 | 49338 | 157 | N15:49259 |
| chrXV | 51344 | 51469 | 125 | |
| chrXV | 53593 | 53699 | 106 | |
| chrXV | 53593 | 53708 | 115 | |
| chrXV | 53860 | 54011 | 151 | N15:53913 |
| chrXV | 53996 | 54153 | 157 | N15:54073 |
| chrXV | 63125 | 63255 | 130 | N15:63178 |
| chrXV | 64118 | 64281 | 163 | N15:64218 |
| chrXV | 66159 | 66329 | 170 | N15:66265 |

| CHR | Start | End | Length (bp) | Nucleosome_ID |
|---|---|---|---|---|
| chrXV | 72755 | 72904 | 149 | N15:72830 |
| chrXV | 74613 | 74750 | 137 | N15:74669 |
| chrXV | 74660 | 74806 | 146 | N15:74669 |
| chrXV | 74660 | 74806 | 146 | N15:74825 |
| chrXV | 75646 | 75807 | 161 | N15:75770 |
| chrXV | 80807 | 80978 | 171 | N15:80842 |
| chrXV | 101110 | 101258 | 148 | N15:101210 |
| chrXV | 108670 | 108830 | 160 | N15:108728 |
| chrXV | 110950 | 111091 | 141 | N15:111017 |
| chrXV | 112040 | 112197 | 157 | N15:112099 |
| chrXV | 113588 | 113739 | 151 | N15:113650 |
| chrXV | 116654 | 116808 | 154 | N15:116777 |
| chrXV | 118023 | 118177 | 154 | N15:118099 |
| chrXV | 118066 | 118215 | 149 | N15:118099 |
| chrXV | 118240 | 118363 | 123 | N15:118282 |
| chrXV | 118340 | 118495 | 155 | N15:118453 |
| chrXV | 118969 | 119120 | 151 | N15:119046 |
| chrXV | 118979 | 119124 | 145 | N15:119046 |
| chrXV | 118997 | 119148 | 151 | N15:119046 |
| chrXV | 119251 | 119405 | 154 | N15:119384 |
| chrXV | 119285 | 119429 | 144 | N15:119384 |
| chrXV | 119816 | 119951 | 135 | N15:119853 |
| chrXV | 120263 | 120409 | 146 | N15:120330 |
| chrXV | 120263 | 120413 | 150 | N15:120330 |
| chrXV | 120442 | 120599 | 157 | N15:120533 |
| chrXV | 120674 | 120823 | 149 | N15:120708 |
| chrXV | 120933 | 121062 | 129 | |
| chrXV | 121845 | 121995 | 150 | N15:121935 |
| chrXV | 122824 | 122966 | 142 | N15:122896 |
| chrXV | 132108 | 132259 | 151 | N15:132177 |
| chrXV | 132333 | 132482 | 149 | N15:132411 |
| chrXV | 138290 | 138437 | 147 | N15:138366 |
| chrXV | 141232 | 141379 | 147 | N15:141343 |
| chrXV | 142140 | 142290 | 150 | N15:142219 |
| chrXV | 143684 | 143828 | 144 | N15:143766 |
| chrXV | 147632 | 147784 | 152 | N15:147706 |
| chrXV | 152975 | 153131 | 156 | N15:153029 |
| chrXV | 160615 | 160744 | 129 | N15:160677 |
| chrXV | 162501 | 162638 | 137 | N15:162550 |
| chrXV | 163008 | 163166 | 158 | N15:163097 |
| chrXV | 163425 | 163564 | 139 | N15:163509 |
| chrXV | 164146 | 164288 | 142 | |
| chrXV | 169600 | 169748 | 148 | N15:169700 |
| chrXV | 171569 | 171743 | 174 | N15:171764 |
| chrXV | 173781 | 173933 | 152 | |
| chrXV | 177092 | 177238 | 146 | N15:177172 |
| chrXV | 177093 | 177211 | 118 | N15:177172 |
| chrXV | 177093 | 177214 | 121 | N15:177172 |
| chrXV | 177093 | 177217 | 124 | N15:177172 |
| chrXV | 177093 | 177220 | 127 | N15:177172 |
| chrXV | 177093 | 177221 | 128 | N15:177172 |
| chrXV | 177093 | 177222 | 129 | N15:177172 |
| chrXV | 177093 | 177223 | 130 | N15:177172 |
| chrXV | 177093 | 177224 | 131 | N15:177172 |
| chrXV | 177093 | 177225 | 132 | N15:177172 |
| chrXV | 177093 | 177226 | 133 | N15:177172 |
| chrXV | 177093 | 177227 | 134 | N15:177172 |
| chrXV | 177093 | 177228 | 135 | N15:177172 |
| chrXV | 177093 | 177229 | 136 | N15:177172 |
| chrXV | 177093 | 177230 | 137 | N15:177172 |
| chrXV | 177093 | 177231 | 138 | N15:177172 |
| chrXV | 177093 | 177232 | 139 | N15:177172 |
| chrXV | 177093 | 177233 | 140 | N15:177172 |
| chrXV | 177093 | 177234 | 141 | N15:177172 |
| chrXV | 177093 | 177235 | 142 | N15:177172 |
| chrXV | 177093 | 177236 | 143 | N15:177172 |
| chrXV | 177093 | 177237 | 144 | N15:177172 |
| chrXV | 177093 | 177238 | 145 | N15:177172 |
| chrXV | 177093 | 177239 | 146 | N15:177172 |
| chrXV | 177094 | 177238 | 144 | N15:177172 |
| chrXV | 177095 | 177238 | 143 | N15:177172 |
| chrXV | 177096 | 177238 | 142 | N15:177172 |
| chrXV | 177097 | 177238 | 141 | N15:177172 |
| chrXV | 177098 | 177238 | 140 | N15:177172 |
| chrXV | 177099 | 177238 | 139 | N15:177172 |
| chrXV | 177100 | 177238 | 138 | N15:177172 |
| chrXV | 177101 | 177238 | 137 | N15:177172 |
| chrXV | 177102 | 177238 | 136 | N15:177172 |
| chrXV | 177103 | 177238 | 135 | N15:177172 |
| chrXV | 177104 | 177238 | 134 | N15:177172 |
| chrXV | 177105 | 177238 | 133 | N15:177172 |
| chrXV | 177106 | 177238 | 132 | N15:177172 |
| chrXV | 177107 | 177238 | 131 | N15:177172 |
| chrXV | 177109 | 177238 | 129 | N15:177172 |
| chrXV | 177110 | 177238 | 128 | N15:177172 |
| chrXV | 177111 | 177238 | 127 | N15:177172 |
| chrXV | 177112 | 177238 | 126 | N15:177172 |
| chrXV | 177113 | 177238 | 125 | N15:177172 |
| chrXV | 177115 | 177238 | 123 | N15:177172 |
| chrXV | 177117 | 177238 | 121 | N15:177172 |
| chrXV | 177118 | 177238 | 120 | N15:177172 |
| chrXV | 177119 | 177238 | 119 | N15:177172 |
| chrXV | 177120 | 177238 | 118 | N15:177172 |
| chrXV | 177121 | 177238 | 117 | N15:177172 |
| chrXV | 177124 | 177238 | 114 | N15:177172 |
| chrXV | 177125 | 177238 | 113 | N15:177172 |
| chrXV | 177128 | 177238 | 110 | N15:177172 |
| chrXV | 177130 | 177238 | 108 | N15:177172 |
| chrXV | 177132 | 177217 | 85 | N15:177172 |
| chrXV | 177132 | 177238 | 106 | N15:177172 |
| chrXV | 177733 | 177893 | 160 | N15:177858 |
| chrXV | 178523 | 178704 | 181 | N15:178709 |
| chrXV | 185749 | 185889 | 140 | N15:185774 |

| CHR | Start | End | Length (bp) | Nucleosome_ID |
|---|---|---|---|---|
| chrXV | 187300 | 187456 | 156 | N15:187379 |
| chrXV | 189704 | 189862 | 158 | N15:189795 |
| chrXV | 192332 | 192491 | 159 | N15:192321 |
| chrXV | 196562 | 196707 | 145 | N15:196728 |
| chrXV | 196562 | 196707 | 145 | N15:196546 |
| chrXV | 198924 | 199072 | 148 | N15:198961 |
| chrXV | 202849 | 203002 | 153 | N15:202912 |
| chrXV | 205781 | 205911 | 130 | N15:205866 |
| chrXV | 208753 | 208900 | 147 | N15:208837 |
| chrXV | 209390 | 209535 | 145 | N15:209458 |
| chrXV | 212639 | 212808 | 169 | N15:212754 |
| chrXV | 214778 | 214916 | 138 | N15:214872 |
| chrXV | 215741 | 215894 | 153 | N15:215837 |
| chrXV | 218336 | 218507 | 171 | N15:218350 |
| chrXV | 220428 | 220557 | 129 | N15:220513 |
| chrXV | 222406 | 222534 | 128 | N15:222459 |
| chrXV | 225244 | 225389 | 145 | N15:225300 |
| chrXV | 227790 | 227919 | 129 | N15:227846 |
| chrXV | 229356 | 229535 | 179 | N15:229431 |
| chrXV | 229736 | 229903 | 167 | N15:229797 |
| chrXV | 233701 | 233860 | 159 | N15:233870 |
| chrXV | 233701 | 233860 | 159 | N15:233712 |
| chrXV | 237293 | 237436 | 143 | N15:237339 |
| chrXV | 242082 | 242240 | 158 | N15:242059 |
| chrXV | 242082 | 242240 | 158 | N15:242224 |
| chrXV | 243486 | 243636 | 150 | N15:243571 |
| chrXV | 244606 | 244758 | 152 | N15:244626 |
| chrXV | 244606 | 244758 | 152 | N15:244776 |
| chrXV | 248190 | 248334 | 144 | N15:248279 |
| chrXV | 248556 | 248722 | 166 | |
| chrXV | 255239 | 255421 | 182 | N15:255344 |
| chrXV | 256726 | 256872 | 146 | N15:256740 |
| chrXV | 260668 | 260810 | 142 | |
| chrXV | 261423 | 261595 | 172 | N15:261549 |
| chrXV | 263580 | 263731 | 151 | N15:263648 |
| chrXV | 265679 | 265840 | 161 | N15:265718 |
| chrXV | 266859 | 267005 | 146 | N15:266903 |
| chrXV | 268238 | 268379 | 141 | N15:268361 |
| chrXV | 270968 | 271134 | 166 | N15:271050 |
| chrXV | 271150 | 271308 | 158 | N15:271220 |
| chrXV | 273050 | 273212 | 162 | N15:273212 |
| chrXV | 276074 | 276211 | 137 | N15:276066 |
| chrXV | 276074 | 276211 | 137 | N15:276231 |
| chrXV | 276839 | 276983 | 144 | N15:276892 |
| chrXV | 279591 | 279744 | 153 | N15:279667 |
| chrXV | 280089 | 280217 | 128 | |
| chrXV | 282501 | 282650 | 149 | N15:282573 |
| chrXV | 284484 | 284582 | 98 | |
| chrXV | 284650 | 284789 | 139 | N15:284768 |
| chrXV | 284999 | 285154 | 155 | N15:285080 |
| chrXV | 285195 | 285326 | 131 | N15:285245 |
| chrXV | 292850 | 293005 | 155 | N15:292974 |
| chrXV | 292905 | 293050 | 145 | N15:292974 |
| chrXV | 293213 | 293372 | 159 | N15:293294 |
| chrXV | 294193 | 294362 | 169 | N15:294360 |
| chrXV | 298993 | 299142 | 149 | N15:299031 |
| chrXV | 306450 | 306602 | 152 | N15:306552 |
| chrXV | 311787 | 311943 | 156 | N15:311928 |
| chrXV | 319552 | 319692 | 140 | N15:319612 |
| chrXV | 320812 | 320963 | 151 | N15:320792 |
| chrXV | 321565 | 321713 | 148 | N15:321561 |
| chrXV | 327427 | 327560 | 133 | N15:327498 |
| chrXV | 328438 | 328583 | 145 | N15:328423 |
| chrXV | 328438 | 328583 | 145 | N15:328597 |
| chrXV | 329812 | 329980 | 168 | N15:329910 |
| chrXV | 338001 | 338161 | 160 | N15:338135 |
| chrXV | 341700 | 341803 | 103 | N15:341798 |
| chrXV | 348865 | 348972 | 107 | N15:348916 |
| chrXV | 349258 | 349411 | 153 | N15:349279 |
| chrXV | 351942 | 352092 | 150 | N15:351997 |
| chrXV | 353769 | 353922 | 153 | N15:353843 |
| chrXV | 356685 | 356815 | 130 | N15:356719 |
| chrXV | 357881 | 358029 | 148 | N15:357967 |
| chrXV | 359149 | 359277 | 128 | |
| chrXV | 359963 | 360110 | 147 | N15:360012 |
| chrXV | 363241 | 363404 | 163 | N15:363313 |
| chrXV | 363647 | 363809 | 162 | N15:363630 |
| chrXV | 365392 | 365542 | 150 | N15:365381 |
| chrXV | 365392 | 365542 | 150 | N15:365558 |
| chrXV | 369266 | 369425 | 159 | N15:369250 |
| chrXV | 369347 | 369504 | 157 | N15:369518 |
| chrXV | 371267 | 371398 | 131 | N15:371332 |
| chrXV | 371602 | 371749 | 147 | N15:371676 |
| chrXV | 377796 | 377936 | 140 | N15:377801 |
| chrXV | 384219 | 384368 | 149 | N15:384312 |
| chrXV | 387959 | 388110 | 151 | N15:388058 |
| chrXV | 389816 | 389942 | 126 | N15:389861 |
| chrXV | 392262 | 392373 | 111 | N15:392325 |
| chrXV | 392576 | 392737 | 161 | N15:392661 |
| chrXV | 401254 | 401398 | 144 | N15:401349 |
| chrXV | 403569 | 403709 | 140 | |
| chrXV | 405315 | 405460 | 145 | N15:405385 |
| chrXV | 405735 | 405887 | 152 | N15:405829 |
| chrXV | 408309 | 408462 | 153 | N15:408405 |
| chrXV | 413175 | 413325 | 150 | N15:413244 |
| chrXV | 413525 | 413650 | 125 | N15:413578 |
| chrXV | 419166 | 419317 | 151 | N15:419244 |
| chrXV | 420271 | 420429 | 158 | N15:420382 |
| chrXV | 423240 | 423391 | 151 | N15:423319 |
| chrXV | 423241 | 423391 | 150 | N15:423319 |
| chrXV | 423242 | 423391 | 149 | N15:423319 |
| chrXV | 423243 | 423391 | 148 | N15:423319 |
| chrXV | 423245 | 423391 | 146 | N15:423319 |

| CHR | Start | End | Length (bp) | Nucleosome_ID |
|---|---|---|---|---|
| chrXV | 433044 | 433202 | 158 | N15:433178 |
| chrXV | 433171 | 433320 | 149 | N15:433178 |
| chrXV | 433171 | 433320 | 149 | N15:433331 |
| chrXV | 435682 | 435849 | 167 | N15:435774 |
| chrXV | 441363 | 441514 | 151 | N15:441436 |
| chrXV | 443421 | 443578 | 157 | N15:443539 |
| chrXV | 445187 | 445337 | 150 | N15:445171 |
| chrXV | 445187 | 445337 | 150 | N15:445329 |
| chrXV | 446874 | 447012 | 138 | N15:446940 |
| chrXV | 450011 | 450165 | 154 | N15:450081 |
| chrXV | 456168 | 456329 | 161 | N15:456318 |
| chrXV | 457433 | 457588 | 155 | N15:457449 |
| chrXV | 457539 | 457691 | 152 | N15:457612 |
| chrXV | 459473 | 459604 | 131 | |
| chrXV | 465612 | 465776 | 164 | N15:465716 |
| chrXV | 469152 | 469262 | 110 | N15:469219 |
| chrXV | 469152 | 469277 | 125 | N15:469219 |
| chrXV | 480195 | 480360 | 165 | N15:480344 |
| chrXV | 480209 | 480363 | 154 | N15:480344 |
| chrXV | 480628 | 480755 | 127 | N15:480718 |
| chrXV | 480628 | 480787 | 159 | N15:480718 |
| chrXV | 480629 | 480736 | 107 | N15:480718 |
| chrXV | 480629 | 480737 | 108 | N15:480718 |
| chrXV | 480629 | 480738 | 109 | N15:480718 |
| chrXV | 480629 | 480739 | 110 | N15:480718 |
| chrXV | 480629 | 480741 | 112 | N15:480718 |
| chrXV | 480629 | 480742 | 113 | N15:480718 |
| chrXV | 480629 | 480743 | 114 | N15:480718 |
| chrXV | 480629 | 480744 | 115 | N15:480718 |
| chrXV | 480629 | 480745 | 116 | N15:480718 |
| chrXV | 480629 | 480746 | 117 | N15:480718 |
| chrXV | 480629 | 480747 | 118 | N15:480718 |
| chrXV | 480629 | 480748 | 119 | N15:480718 |
| chrXV | 480629 | 480749 | 120 | N15:480718 |
| chrXV | 480629 | 480750 | 121 | N15:480718 |
| chrXV | 480629 | 480751 | 122 | N15:480718 |
| chrXV | 480629 | 480752 | 123 | N15:480718 |
| chrXV | 480629 | 480753 | 124 | N15:480718 |
| chrXV | 480629 | 480754 | 125 | N15:480718 |
| chrXV | 480629 | 480755 | 126 | N15:480718 |
| chrXV | 480629 | 480756 | 127 | N15:480718 |
| chrXV | 480629 | 480757 | 128 | N15:480718 |
| chrXV | 480629 | 480758 | 129 | N15:480718 |
| chrXV | 480629 | 480759 | 130 | N15:480718 |
| chrXV | 480629 | 480760 | 131 | N15:480718 |
| chrXV | 480629 | 480761 | 132 | N15:480718 |
| chrXV | 480629 | 480762 | 133 | N15:480718 |
| chrXV | 480629 | 480763 | 134 | N15:480718 |
| chrXV | 480629 | 480764 | 135 | N15:480718 |
| chrXV | 480629 | 480765 | 136 | N15:480718 |
| chrXV | 480629 | 480766 | 137 | N15:480718 |
| chrXV | 480629 | 480767 | 138 | N15:480718 |
| chrXV | 480629 | 480768 | 139 | N15:480718 |
| chrXV | 480629 | 480769 | 140 | N15:480718 |
| chrXV | 480629 | 480770 | 141 | N15:480718 |
| chrXV | 480629 | 480771 | 142 | N15:480718 |
| chrXV | 480629 | 480772 | 143 | N15:480718 |
| chrXV | 480629 | 480773 | 144 | N15:480718 |
| chrXV | 480629 | 480774 | 145 | N15:480718 |
| chrXV | 480629 | 480775 | 146 | N15:480718 |
| chrXV | 480629 | 480776 | 147 | N15:480718 |
| chrXV | 480629 | 480777 | 148 | N15:480718 |
| chrXV | 480629 | 480778 | 149 | N15:480718 |
| chrXV | 480629 | 480779 | 150 | N15:480718 |
| chrXV | 480629 | 480780 | 151 | N15:480718 |
| chrXV | 480629 | 480781 | 152 | N15:480718 |
| chrXV | 480629 | 480782 | 153 | N15:480718 |
| chrXV | 480629 | 480783 | 154 | N15:480718 |
| chrXV | 480629 | 480784 | 155 | N15:480718 |
| chrXV | 480629 | 480785 | 156 | N15:480718 |
| chrXV | 480629 | 480786 | 157 | N15:480718 |
| chrXV | 480629 | 480787 | 158 | N15:480718 |
| chrXV | 480629 | 480788 | 159 | N15:480718 |
| chrXV | 480629 | 480789 | 160 | N15:480718 |
| chrXV | 480630 | 480787 | 157 | N15:480718 |
| chrXV | 480631 | 480787 | 156 | N15:480718 |
| chrXV | 480632 | 480787 | 155 | N15:480718 |
| chrXV | 480633 | 480787 | 154 | N15:480718 |
| chrXV | 480634 | 480787 | 153 | N15:480718 |
| chrXV | 480635 | 480787 | 152 | N15:480718 |
| chrXV | 480636 | 480787 | 151 | N15:480718 |
| chrXV | 480637 | 480787 | 150 | N15:480718 |
| chrXV | 480638 | 480787 | 149 | N15:480718 |
| chrXV | 480639 | 480787 | 148 | N15:480718 |
| chrXV | 480640 | 480787 | 147 | N15:480718 |
| chrXV | 480641 | 480787 | 146 | N15:480718 |
| chrXV | 480642 | 480787 | 145 | N15:480718 |
| chrXV | 480643 | 480787 | 144 | N15:480718 |
| chrXV | 480644 | 480787 | 143 | N15:480718 |
| chrXV | 480645 | 480787 | 142 | N15:480718 |
| chrXV | 480646 | 480787 | 141 | N15:480718 |
| chrXV | 480647 | 480787 | 140 | N15:480718 |
| chrXV | 480648 | 480787 | 139 | N15:480718 |
| chrXV | 480649 | 480787 | 138 | N15:480718 |
| chrXV | 480650 | 480787 | 137 | N15:480718 |
| chrXV | 480651 | 480787 | 136 | N15:480718 |
| chrXV | 480652 | 480787 | 135 | N15:480718 |
| chrXV | 480653 | 480787 | 134 | N15:480718 |
| chrXV | 480654 | 480787 | 133 | N15:480718 |
| chrXV | 480655 | 480787 | 132 | N15:480718 |
| chrXV | 480656 | 480787 | 131 | N15:480718 |
| chrXV | 480657 | 480787 | 130 | N15:480718 |
| chrXV | 480658 | 480787 | 129 | N15:480718 |

TABLE 1

| CHR | Start | End | Length (bp) | Nucleosome_ID |
|---|---|---|---|---|
| chrXV | 480659 | 480787 | 128 | N15:480718 |
| chrXV | 480660 | 480787 | 127 | N15:480718 |
| chrXV | 480661 | 480787 | 126 | N15:480718 |
| chrXV | 480662 | 480787 | 125 | N15:480718 |
| chrXV | 480663 | 480787 | 124 | N15:480718 |
| chrXV | 480664 | 480787 | 123 | N15:480718 |
| chrXV | 480665 | 480787 | 122 | N15:480718 |
| chrXV | 480666 | 480787 | 121 | N15:480718 |
| chrXV | 480667 | 480787 | 120 | N15:480718 |
| chrXV | 480668 | 480787 | 119 | N15:480718 |
| chrXV | 480669 | 480787 | 118 | N15:480718 |
| chrXV | 480670 | 480787 | 117 | N15:480718 |
| chrXV | 480671 | 480787 | 116 | N15:480718 |
| chrXV | 480672 | 480787 | 115 | N15:480718 |
| chrXV | 480673 | 480787 | 114 | N15:480718 |
| chrXV | 480674 | 480787 | 113 | N15:480718 |
| chrXV | 480675 | 480787 | 112 | N15:480718 |
| chrXV | 480676 | 480787 | 111 | N15:480718 |
| chrXV | 480677 | 480787 | 110 | N15:480718 |
| chrXV | 480678 | 480787 | 109 | N15:480718 |
| chrXV | 480679 | 480787 | 108 | N15:480718 |
| chrXV | 480680 | 480755 | 75 | N15:480718 |
| chrXV | 480680 | 480787 | 107 | N15:480718 |
| chrXV | 480681 | 480787 | 106 | N15:480718 |
| chrXV | 480682 | 480787 | 105 | N15:480718 |
| chrXV | 480683 | 480787 | 104 | N15:480718 |
| chrXV | 480684 | 480787 | 103 | N15:480718 |
| chrXV | 480685 | 480787 | 102 | N15:480718 |
| chrXV | 480686 | 480787 | 101 | N15:480718 |
| chrXV | 480687 | 480787 | 100 | N15:480718 |
| chrXV | 480688 | 480787 | 99 | N15:480718 |
| chrXV | 480689 | 480787 | 98 | N15:480718 |
| chrXV | 480690 | 480787 | 97 | N15:480718 |
| chrXV | 480691 | 480787 | 96 | N15:480718 |
| chrXV | 480692 | 480787 | 95 | N15:480718 |
| chrXV | 480693 | 480787 | 94 | N15:480718 |
| chrXV | 480695 | 480787 | 92 | N15:480718 |
| chrXV | 480696 | 480787 | 91 | N15:480718 |
| chrXV | 480697 | 480787 | 90 | N15:480718 |
| chrXV | 480698 | 480787 | 89 | N15:480718 |
| chrXV | 480700 | 480787 | 87 | N15:480718 |
| chrXV | 480701 | 480787 | 86 | N15:480718 |
| chrXV | 480703 | 480787 | 84 | N15:480718 |
| chrXV | 480704 | 480787 | 83 | N15:480718 |
| chrXV | 482098 | 482249 | 151 | N15:482205 |
| chrXV | 483506 | 483656 | 150 | N15:483559 |
| chrXV | 490437 | 490586 | 149 | |
| chrXV | 492152 | 492296 | 144 | |
| chrXV | 492636 | 492766 | 130 | N15:492727 |
| chrXV | 499981 | 500137 | 156 | N15:499985 |
| chrXV | 500251 | 500392 | 141 | N15:500266 |
| chrXV | 500870 | 501031 | 161 | N15:500992 |
| chrXV | 504656 | 504829 | 173 | N15:504757 |
| chrXV | 504839 | 504995 | 156 | N15:504924 |
| chrXV | 505651 | 505827 | 176 | N15:505793 |
| chrXV | 508541 | 508687 | 146 | N15:508522 |
| chrXV | 512017 | 512169 | 152 | N15:512077 |
| chrXV | 520467 | 520627 | 160 | N15:520577 |
| chrXV | 521328 | 521487 | 159 | N15:521406 |
| chrXV | 524434 | 524584 | 150 | N15:524552 |
| chrXV | 526545 | 526704 | 159 | N15:526589 |
| chrXV | 527282 | 527433 | 151 | N15:527364 |
| chrXV | 528345 | 528493 | 148 | N15:528465 |
| chrXV | 529088 | 529208 | 120 | |
| chrXV | 534111 | 534263 | 152 | N15:534188 |
| chrXV | 535122 | 535279 | 157 | |
| chrXV | 539793 | 539942 | 149 | N15:539889 |
| chrXV | 540123 | 540277 | 154 | N15:540228 |
| chrXV | 542672 | 542801 | 129 | N15:542768 |
| chrXV | 545231 | 545391 | 160 | N15:545321 |
| chrXV | 545588 | 545715 | 127 | N15:545625 |
| chrXV | 550006 | 550155 | 149 | N15:550070 |
| chrXV | 550413 | 550558 | 145 | N15:550502 |
| chrXV | 550604 | 550752 | 148 | N15:550670 |
| chrXV | 559497 | 559585 | 88 | N15:559589 |
| chrXV | 559497 | 559598 | 101 | N15:559589 |
| chrXV | 559497 | 559601 | 104 | N15:559589 |
| chrXV | 559497 | 559654 | 157 | N15:559589 |
| chrXV | 559919 | 560068 | 149 | N15:559990 |
| chrXV | 559929 | 560086 | 157 | N15:559990 |
| chrXV | 565053 | 565199 | 146 | N15:565222 |
| chrXV | 565053 | 565201 | 148 | N15:565222 |
| chrXV | 565053 | 565202 | 149 | N15:565222 |
| chrXV | 566775 | 566926 | 151 | N15:566870 |
| chrXV | 576392 | 576555 | 163 | N15:576489 |
| chrXV | 590114 | 590263 | 149 | N15:590176 |
| chrXV | 590721 | 590873 | 152 | N15:590856 |
| chrXV | 592017 | 592170 | 153 | N15:592096 |
| chrXV | 592017 | 592179 | 162 | N15:592096 |
| chrXV | 592023 | 592179 | 156 | N15:592096 |
| chrXV | 592044 | 592126 | 82 | N15:592096 |
| chrXV | 595352 | 595475 | 123 | |
| chrXV | 595452 | 595607 | 155 | N15:595551 |
| chrXV | 595885 | 596019 | 134 | N15:595950 |
| chrXV | 595885 | 596025 | 140 | N15:595950 |
| chrXV | 596928 | 597063 | 135 | N15:596968 |
| chrXV | 597189 | 597334 | 145 | N15:597178 |
| chrXV | 597375 | 597525 | 150 | N15:597445 |
| chrXV | 597554 | 597711 | 157 | N15:597645 |
| chrXV | 598045 | 598174 | 129 | |
| chrXV | 598957 | 599107 | 150 | N15:599055 |
| chrXV | 600156 | 600263 | 107 | |
| chrXV | 601179 | 601328 | 149 | N15:601265 |

| CHR | Start | End | Length (bp) | Nucleosome_ID |
|---|---|---|---|---|
| chrXV | 603463 | 603616 | 153 | N15:603547 |
| chrXV | 605449 | 605600 | 151 | N15:605543 |
| chrXV | 606728 | 606884 | 156 | N15:606775 |
| chrXV | 608095 | 608251 | 156 | N15:608159 |
| chrXV | 610131 | 610285 | 154 | N15:610149 |
| chrXV | 610432 | 610587 | 155 | N15:610582 |
| chrXV | 610432 | 610587 | 155 | N15:610423 |
| chrXV | 613403 | 613549 | 146 | N15:613529 |
| chrXV | 617632 | 617793 | 161 | N15:617725 |
| chrXV | 621001 | 621151 | 150 | N15:621077 |
| chrXV | 622311 | 622441 | 130 | |
| chrXV | 624028 | 624159 | 131 | N15:624066 |
| chrXV | 625834 | 625959 | 125 | N15:625885 |
| chrXV | 626417 | 626570 | 153 | N15:626508 |
| chrXV | 632144 | 632251 | 107 | N15:632173 |
| chrXV | 632144 | 632253 | 109 | N15:632173 |
| chrXV | 633485 | 633638 | 153 | N15:633575 |
| chrXV | 635429 | 635578 | 149 | N15:635550 |
| chrXV | 637512 | 637669 | 157 | N15:637581 |
| chrXV | 641886 | 642041 | 155 | N15:641956 |
| chrXV | 646176 | 646338 | 162 | N15:646189 |
| chrXV | 647481 | 647636 | 155 | N15:647630 |
| chrXV | 650796 | 650922 | 126 | |
| chrXV | 650923 | 651080 | 157 | N15:650997 |
| chrXV | 651588 | 651740 | 152 | N15:651687 |
| chrXV | 671862 | 672011 | 149 | N15:671896 |
| chrXV | 674697 | 674849 | 152 | N15:674786 |
| chrXV | 674884 | 675018 | 134 | N15:674983 |
| chrXV | 677635 | 677796 | 161 | N15:677669 |
| chrXV | 679372 | 679525 | 153 | N15:679447 |
| chrXV | 680555 | 680689 | 134 | N15:680616 |
| chrXV | 682390 | 682545 | 155 | N15:682487 |
| chrXV | 682421 | 682570 | 149 | N15:682487 |
| chrXV | 682600 | 682746 | 146 | N15:682669 |
| chrXV | 683178 | 683329 | 151 | N15:683179 |
| chrXV | 683178 | 683329 | 151 | N15:683348 |
| chrXV | 683542 | 683705 | 163 | N15:683602 |
| chrXV | 698388 | 698544 | 156 | N15:698448 |
| chrXV | 699187 | 699320 | 133 | N15:699226 |
| chrXV | 704256 | 704405 | 149 | N15:704277 |
| chrXV | 705052 | 705198 | 146 | N15:705134 |
| chrXV | 707863 | 708011 | 148 | |
| chrXV | 709269 | 709434 | 165 | N15:709269 |
| chrXV | 709269 | 709434 | 165 | N15:709445 |
| chrXV | 709269 | 709462 | 193 | N15:709269 |
| chrXV | 709269 | 709462 | 193 | N15:709445 |
| chrXV | 709332 | 709434 | 102 | N15:709445 |
| chrXV | 709332 | 709462 | 130 | N15:709445 |
| chrXV | 709396 | 709550 | 154 | N15:709445 |
| chrXV | 709396 | 709571 | 175 | N15:709445 |
| chrXV | 709403 | 709571 | 168 | N15:709445 |
| chrXV | 712148 | 712278 | 130 | N15:712206 |
| chrXV | 713754 | 713908 | 154 | |
| chrXV | 716082 | 716234 | 152 | N15:716164 |
| chrXV | 717294 | 717452 | 158 | N15:717288 |
| chrXV | 717294 | 717452 | 158 | N15:717452 |
| chrXV | 728130 | 728275 | 145 | N15:728119 |
| chrXV | 728130 | 728275 | 145 | N15:728288 |
| chrXV | 729241 | 729409 | 168 | N15:729229 |
| chrXV | 730369 | 730527 | 158 | N15:730492 |
| chrXV | 730626 | 730772 | 146 | N15:730666 |
| chrXV | 732904 | 733026 | 122 | N15:732962 |
| chrXV | 736114 | 736267 | 153 | N15:736157 |
| chrXV | 749719 | 749841 | 122 | N15:749778 |
| chrXV | 751696 | 751803 | 107 | N15:751682 |
| chrXV | 751696 | 751804 | 108 | N15:751682 |
| chrXV | 751696 | 751805 | 109 | N15:751682 |
| chrXV | 751696 | 751808 | 112 | N15:751682 |
| chrXV | 759323 | 759474 | 151 | N15:759342 |
| chrXV | 759886 | 760039 | 153 | N15:759976 |
| chrXV | 762029 | 762161 | 132 | N15:762082 |
| chrXV | 762283 | 762425 | 142 | N15:762373 |
| chrXV | 763377 | 763524 | 147 | N15:763477 |
| chrXV | 765404 | 765530 | 126 | N15:765450 |
| chrXV | 766078 | 766228 | 150 | N15:766247 |
| chrXV | 767013 | 767155 | 142 | N15:767164 |
| chrXV | 767013 | 767155 | 142 | N15:767010 |
| chrXV | 768735 | 768888 | 153 | N15:768713 |
| chrXV | 770268 | 770418 | 150 | |
| chrXV | 773124 | 773277 | 153 | N15:773165 |
| chrXV | 775176 | 775297 | 121 | N15:775265 |
| chrXV | 777968 | 778119 | 151 | N15:778110 |
| chrXV | 783192 | 783285 | 93 | N15:783252 |
| chrXV | 789013 | 789147 | 134 | N15:789066 |
| chrXV | 790833 | 790987 | 154 | N15:790822 |
| chrXV | 794701 | 794852 | 151 | |
| chrXV | 796032 | 796168 | 136 | N15:796058 |
| chrXV | 801709 | 801857 | 148 | N15:801792 |
| chrXV | 802168 | 802313 | 145 | N15:802287 |
| chrXV | 804681 | 804839 | 158 | N15:804764 |
| chrXV | 805157 | 805300 | 143 | N15:805287 |
| chrXV | 806800 | 806942 | 142 | N15:806847 |
| chrXV | 806939 | 807090 | 151 | N15:807004 |
| chrXV | 810091 | 810222 | 131 | N15:810121 |
| chrXV | 815641 | 815767 | 126 | N15:815726 |
| chrXV | 816418 | 816570 | 152 | N15:816506 |
| chrXV | 816891 | 817023 | 132 | N15:817009 |
| chrXV | 823212 | 823372 | 160 | N15:823302 |
| chrXV | 823715 | 823880 | 165 | N15:823900 |
| chrXV | 824770 | 824918 | 148 | N15:824827 |
| chrXV | 824898 | 825045 | 147 | N15:824984 |
| chrXV | 826750 | 826899 | 149 | N15:826822 |
| chrXV | 828443 | 828590 | 147 | N15:828514 |

TABLE 1

| CHR | Start | End | Length (bp) | Nucleosome_ID |
|---|---|---|---|---|
| chrXV | 829595 | 829749 | 154 | N15:829756 |
| chrXV | 832011 | 832158 | 147 | N15:832060 |
| chrXV | 838047 | 838199 | 152 | N15:838035 |
| chrXV | 838047 | 838199 | 152 | N15:838195 |
| chrXV | 838749 | 838897 | 148 | N15:838866 |
| chrXV | 840989 | 841143 | 154 | N15:841051 |
| chrXV | 845931 | 846091 | 160 | N15:845986 |
| chrXV | 851758 | 851911 | 153 | N15:851815 |
| chrXV | 855604 | 855755 | 151 | N15:855699 |
| chrXV | 856174 | 856339 | 165 | N15:856207 |
| chrXV | 856222 | 856364 | 142 | N15:856207 |
| chrXV | 860271 | 860423 | 152 | N15:860352 |
| chrXV | 863117 | 863234 | 117 | N15:863177 |
| chrXV | 866318 | 866474 | 156 | N15:866417 |
| chrXV | 866636 | 866779 | 143 | |
| chrXV | 873282 | 873420 | 138 | N15:873312 |
| chrXV | 873426 | 873558 | 132 | N15:873537 |
| chrXV | 873491 | 873643 | 152 | N15:873537 |
| chrXV | 883077 | 883230 | 153 | N15:883079 |
| chrXV | 883077 | 883230 | 153 | N15:883239 |
| chrXV | 884043 | 884200 | 157 | N15:884168 |
| chrXV | 895782 | 895933 | 151 | N15:895865 |
| chrXV | 900646 | 900802 | 156 | N15:900717 |
| chrXV | 902699 | 902828 | 129 | N15:902765 |
| chrXV | 904931 | 905079 | 148 | N15:904922 |
| chrXV | 917190 | 917343 | 153 | N15:917323 |
| chrXV | 920557 | 920665 | 108 | N15:920598 |
| chrXV | 928849 | 929014 | 165 | N15:928971 |
| chrXV | 932648 | 932807 | 159 | N15:932684 |
| chrXV | 932960 | 933112 | 152 | N15:933085 |
| chrXV | 937286 | 937429 | 143 | N15:937309 |
| chrXV | 941867 | 942022 | 155 | N15:941903 |
| chrXV | 943142 | 943285 | 143 | N15:943207 |
| chrXV | 956493 | 956644 | 151 | N15:956543 |
| chrXV | 958715 | 958860 | 145 | N15:958840 |
| chrXV | 961158 | 961313 | 155 | N15:961224 |
| chrXV | 963324 | 963474 | 150 | N15:963309 |
| chrXV | 967141 | 967307 | 166 | N15:967161 |
| chrXV | 967454 | 967605 | 151 | N15:967556 |
| chrXV | 970738 | 970906 | 168 | N15:970789 |
| chrXV | 970759 | 970913 | 154 | N15:970789 |
| chrXV | 970847 | 970977 | 130 | |
| chrXV | 970875 | 970977 | 102 | |
| chrXV | 972298 | 972446 | 148 | |
| chrXV | 975111 | 975257 | 146 | |
| chrXV | 975228 | 975386 | 158 | N15:975305 |
| chrXV | 975904 | 976053 | 149 | N15:975989 |
| chrXV | 975908 | 976058 | 150 | N15:975989 |
| chrXV | 977285 | 977436 | 151 | N15:977376 |
| chrXV | 977616 | 977761 | 145 | |
| chrXV | 981987 | 982146 | 159 | N15:982019 |
| chrXV | 986417 | 986558 | 141 | N15:986505 |
| chrXV | 986699 | 986864 | 165 | N15:986780 |
| chrXV | 987920 | 988058 | 138 | N15:987920 |
| chrXV | 988983 | 989114 | 131 | N15:989106 |
| chrXV | 989889 | 990033 | 144 | N15:989965 |
| chrXV | 996318 | 996470 | 152 | N15:996407 |
| chrXV | 997102 | 997264 | 162 | N15:997152 |
| chrXV | 1000108 | 1000248 | 140 | |
| chrXV | 1000780 | 1000917 | 137 | N15:1000816 |
| chrXV | 1001402 | 1001553 | 151 | N15:1001454 |
| chrXV | 1002082 | 1002260 | 178 | N15:1002190 |
| chrXV | 1004460 | 1004618 | 158 | N15:1004637 |
| chrXV | 1004460 | 1004618 | 158 | N15:1004454 |
| chrXV | 1005856 | 1006001 | 145 | |
| chrXV | 1008258 | 1008393 | 135 | N15:1008325 |
| chrXV | 1019659 | 1019809 | 150 | N15:1019734 |
| chrXV | 1019667 | 1019830 | 163 | N15:1019734 |
| chrXV | 1023929 | 1024082 | 153 | N15:1024013 |
| chrXV | 1035603 | 1035747 | 144 | N15:1035709 |
| chrXV | 1036929 | 1037060 | 131 | N15:1036972 |
| chrXV | 1041309 | 1041484 | 175 | N15:1041351 |
| chrXV | 1042260 | 1042368 | 108 | N15:1042335 |
| chrXV | 1042260 | 1042387 | 127 | N15:1042335 |
| chrXV | 1042260 | 1042389 | 129 | N15:1042335 |
| chrXV | 1042260 | 1042392 | 132 | N15:1042335 |
| chrXV | 1042265 | 1042392 | 127 | N15:1042335 |
| chrXV | 1042267 | 1042368 | 101 | N15:1042335 |
| chrXV | 1043663 | 1043770 | 107 | |
| chrXV | 1043663 | 1043771 | 108 | |
| chrXV | 1043663 | 1043772 | 109 | |
| chrXV | 1047470 | 1047621 | 151 | N15:1047478 |
| chrXV | 1047470 | 1047621 | 151 | N15:1047635 |
| chrXV | 1050132 | 1050280 | 148 | N15:1050219 |
| chrXV | 1050137 | 1050280 | 143 | N15:1050219 |
| chrXV | 1050608 | 1050773 | 165 | N15:1050703 |
| chrXV | 1050861 | 1050999 | 138 | N15:1050887 |
| chrXV | 1052062 | 1052210 | 148 | N15:1052087 |
| chrXV | 1056315 | 1056470 | 155 | N15:1056388 |
| chrXV | 1059723 | 1059876 | 153 | N15:1059763 |
| chrXV | 1060927 | 1061054 | 127 | N15:1060979 |
| chrXV | 1061685 | 1061838 | 153 | N15:1061757 |
| chrXV | 1062032 | 1062164 | 132 | N15:1062167 |
| chrXV | 1062744 | 1062896 | 152 | N15:1062785 |
| chrXV | 1071888 | 1072059 | 171 | N15:1071953 |
| chrXV | 1073330 | 1073480 | 150 | N15:1073396 |
| chrXV | 1080435 | 1080579 | 144 | N15:1080493 |
| chrXV | 1085191 | 1085346 | 155 | N15:1085246 |
| chrXV | 1086206 | 1086377 | 171 | N15:1086303 |
| chrXV | 1086356 | 1086503 | 147 | |
| chrXV | 1086512 | 1086642 | 130 | N15:1086572 |
| chrXV | 1086893 | 1087029 | 136 | N15:1086930 |
| chrXV | 1087274 | 1087436 | 162 | N15:1087405 |

| CHR | Start | End | Length (bp) | Nucleosome_ID |
|---|---|---|---|---|
| chrXV | 1087375 | 1087530 | 155 | N15:1087405 |
| chrXV | 1087509 | 1087646 | 137 | N15:1087575 |
| chrXV | 1087804 | 1087947 | 143 | N15:1087853 |
| chrXV | 1087852 | 1087977 | 125 | N15:1087853 |
| chrXV | 1088077 | 1088229 | 152 | N15:1088185 |
| chrXV | 1088150 | 1088294 | 144 | N15:1088185 |
| chrXV | 1088247 | 1088382 | 135 | N15:1088344 |
| chrXV | 1088247 | 1088434 | 187 | N15:1088344 |
| chrXV | 1088290 | 1088382 | 92 | N15:1088344 |
| chrXV | 1088290 | 1088434 | 144 | N15:1088344 |
| chrXV | 1088629 | 1088781 | 152 | N15:1088692 |
| chrXV | 1088930 | 1089060 | 130 | N15:1088919 |
| chrXV | 1089068 | 1089229 | 161 | N15:1089094 |
| chrXV | 1090465 | 1090616 | 151 | N15:1090543 |
| chrXVI | 523 | 674 | 151 | N16:600 |
| chrXVI | 847 | 1021 | 174 | N16:925 |
| chrXVI | 1688 | 1839 | 151 | N16:1728 |
| chrXVI | 1700 | 1834 | 134 | N16:1728 |
| chrXVI | 1725 | 1821 | 96 | N16:1728 |
| chrXVI | 1725 | 1834 | 109 | N16:1728 |
| chrXVI | 2079 | 2209 | 130 | N16:2222 |
| chrXVI | 2079 | 2209 | 130 | N16:2056 |
| chrXVI | 2358 | 2510 | 152 | N16:2443 |
| chrXVI | 2705 | 2849 | 144 | N16:2758 |
| chrXVI | 2705 | 2892 | 187 | N16:2758 |
| chrXVI | 2757 | 2849 | 92 | N16:2758 |
| chrXVI | 2757 | 2892 | 135 | N16:2758 |
| chrXVI | 2845 | 2989 | 144 | N16:2943 |
| chrXVI | 2910 | 3062 | 152 | N16:2943 |
| chrXVI | 3162 | 3287 | 125 | N16:3271 |
| chrXVI | 3192 | 3335 | 143 | N16:3271 |
| chrXVI | 3192 | 3343 | 151 | N16:3271 |
| chrXVI | 3493 | 3630 | 137 | N16:3559 |
| chrXVI | 3609 | 3764 | 155 | N16:3722 |
| chrXVI | 3703 | 3865 | 162 | N16:3880 |
| chrXVI | 3703 | 3865 | 162 | N16:3722 |
| chrXVI | 4928 | 5069 | 141 | N16:4975 |
| chrXVI | 8461 | 8628 | 167 | N16:8544 |
| chrXVI | 10566 | 10710 | 144 | N16:10653 |
| chrXVI | 23283 | 23434 | 151 | N16:23371 |
| chrXVI | 26164 | 26341 | 177 | N16:26333 |
| chrXVI | 26440 | 26591 | 151 | |
| chrXVI | 30175 | 30330 | 155 | N16:30253 |
| chrXVI | 30411 | 30566 | 155 | N16:30492 |
| chrXVI | 37895 | 38053 | 158 | N16:37928 |
| chrXVI | 41612 | 41762 | 150 | N16:41620 |
| chrXVI | 41938 | 42057 | 119 | |
| chrXVI | 44256 | 44374 | 118 | N16:44326 |
| chrXVI | 54075 | 54252 | 177 | N16:54126 |
| chrXVI | 55410 | 55560 | 150 | N16:55405 |
| chrXVI | 56814 | 56963 | 149 | N16:56959 |
| chrXVI | 56988 | 57111 | 123 | |
| chrXVI | 57088 | 57243 | 155 | N16:57189 |
| chrXVI | 57521 | 57655 | 134 | N16:57591 |
| chrXVI | 57521 | 57661 | 140 | N16:57591 |
| chrXVI | 57717 | 57868 | 151 | N16:57746 |
| chrXVI | 57727 | 57872 | 145 | N16:57746 |
| chrXVI | 57747 | 57896 | 149 | N16:57746 |
| chrXVI | 57999 | 58153 | 154 | N16:58141 |
| chrXVI | 58564 | 58699 | 135 | N16:58596 |
| chrXVI | 58718 | 58879 | 161 | N16:58834 |
| chrXVI | 59011 | 59161 | 150 | N16:59079 |
| chrXVI | 59190 | 59347 | 157 | N16:59296 |
| chrXVI | 59681 | 59810 | 129 | N16:59829 |
| chrXVI | 60593 | 60743 | 150 | N16:60686 |
| chrXVI | 61572 | 61714 | 142 | N16:61694 |
| chrXVI | 66888 | 67040 | 152 | N16:66990 |
| chrXVI | 67654 | 67814 | 160 | N16:67736 |
| chrXVI | 69456 | 69617 | 161 | N16:69502 |
| chrXVI | 73482 | 73631 | 149 | N16:73588 |
| chrXVI | 74222 | 74371 | 149 | N16:74321 |
| chrXVI | 85235 | 85390 | 155 | N16:85298 |
| chrXVI | 86229 | 86396 | 167 | N16:86219 |
| chrXVI | 93799 | 93933 | 134 | N16:93856 |
| chrXVI | 97906 | 98054 | 148 | N16:97910 |
| chrXVI | 100090 | 100196 | 106 | N16:100168 |
| chrXVI | 101988 | 102144 | 156 | N16:102072 |
| chrXVI | 111563 | 111694 | 131 | N16:111614 |
| chrXVI | 111839 | 111996 | 157 | N16:111919 |
| chrXVI | 112816 | 112964 | 148 | N16:112902 |
| chrXVI | 113851 | 114002 | 151 | N16:113884 |
| chrXVI | 115900 | 116007 | 107 | N16:115995 |
| chrXVI | 115900 | 116008 | 108 | N16:115995 |
| chrXVI | 115900 | 116009 | 109 | N16:115995 |
| chrXVI | 115900 | 116013 | 113 | N16:115995 |
| chrXVI | 117285 | 117438 | 153 | |
| chrXVI | 119500 | 119650 | 150 | N16:119550 |
| chrXVI | 119632 | 119770 | 138 | N16:119717 |
| chrXVI | 124878 | 125016 | 138 | N16:124969 |
| chrXVI | 127061 | 127209 | 148 | N16:127222 |
| chrXVI | 127061 | 127209 | 148 | N16:127054 |
| chrXVI | 127345 | 127506 | 161 | N16:127392 |
| chrXVI | 128019 | 128185 | 166 | N16:128056 |
| chrXVI | 128210 | 128316 | 106 | N16:128260 |
| chrXVI | 128210 | 128332 | 122 | N16:128260 |
| chrXVI | 130172 | 130324 | 152 | N16:130248 |
| chrXVI | 131123 | 131279 | 156 | N16:131253 |
| chrXVI | 136763 | 136914 | 151 | N16:136878 |
| chrXVI | 136805 | 136963 | 158 | N16:136878 |
| chrXVI | 139377 | 139534 | 157 | N16:139490 |
| chrXVI | 142443 | 142561 | 118 | N16:142521 |
| chrXVI | 144178 | 144336 | 158 | N16:144285 |
| chrXVI | 149028 | 149196 | 168 | N16:149123 |

| CHR | Start | End | Length (bp) | Nucleosome_ID |
|---|---|---|---|---|
| chrXVI | 152572 | 152722 | 150 | N16:152635 |
| chrXVI | 161247 | 161414 | 167 | N16:161357 |
| chrXVI | 164847 | 164957 | 110 | N16:164935 |
| chrXVI | 165348 | 165502 | 154 | N16:165511 |
| chrXVI | 167030 | 167184 | 154 | N16:167113 |
| chrXVI | 169266 | 169418 | 152 | N16:169376 |
| chrXVI | 173412 | 173549 | 137 | N16:173465 |
| chrXVI | 174194 | 174363 | 169 | N16:174232 |
| chrXVI | 175111 | 175256 | 145 | N16:175175 |
| chrXVI | 177466 | 177621 | 155 | N16:177552 |
| chrXVI | 177625 | 177797 | 172 | N16:177780 |
| chrXVI | 189167 | 189337 | 170 | N16:189180 |
| chrXVI | 190409 | 190560 | 151 | N16:190473 |
| chrXVI | 191083 | 191228 | 145 | N16:191166 |
| chrXVI | 193727 | 193886 | 159 | N16:193851 |
| chrXVI | 194794 | 194946 | 152 | N16:194885 |
| chrXVI | 198714 | 198866 | 152 | N16:198810 |
| chrXVI | 200785 | 200926 | 141 | |
| chrXVI | 203076 | 203237 | 161 | N16:203255 |
| chrXVI | 203882 | 204036 | 154 | N16:203940 |
| chrXVI | 205158 | 205313 | 155 | N16:205222 |
| chrXVI | 211347 | 211529 | 182 | N16:211407 |
| chrXVI | 212567 | 212715 | 148 | N16:212620 |
| chrXVI | 213085 | 213234 | 149 | N16:213180 |
| chrXVI | 220298 | 220444 | 146 | N16:220437 |
| chrXVI | 227791 | 227944 | 153 | |
| chrXVI | 228023 | 228173 | 150 | N16:228073 |
| chrXVI | 235024 | 235161 | 137 | N16:235068 |
| chrXVI | 238735 | 238888 | 153 | N16:238814 |
| chrXVI | 239149 | 239288 | 139 | N16:239177 |
| chrXVI | 240071 | 240209 | 138 | N16:240084 |
| chrXVI | 240947 | 241098 | 151 | N16:241005 |
| chrXVI | 242739 | 242891 | 152 | N16:242817 |
| chrXVI | 242995 | 243131 | 136 | N16:243075 |
| chrXVI | 244964 | 245114 | 150 | N16:245006 |
| chrXVI | 246169 | 246331 | 162 | N16:246246 |
| chrXVI | 247500 | 247655 | 155 | |
| chrXVI | 247870 | 248025 | 155 | N16:248033 |
| chrXVI | 248518 | 248667 | 149 | N16:248596 |
| chrXVI | 251350 | 251499 | 149 | N16:251383 |
| chrXVI | 251647 | 251802 | 155 | N16:251733 |
| chrXVI | 254749 | 254901 | 152 | N16:254752 |
| chrXVI | 254749 | 254901 | 152 | N16:254920 |
| chrXVI | 257303 | 257455 | 152 | N16:257371 |
| chrXVI | 258470 | 258614 | 144 | N16:258522 |
| chrXVI | 269441 | 269592 | 151 | N16:269499 |
| chrXVI | 272088 | 272225 | 137 | N16:272113 |
| chrXVI | 273849 | 274002 | 153 | N16:273925 |
| chrXVI | 275518 | 275683 | 165 | N16:275553 |
| chrXVI | 278767 | 278916 | 149 | N16:278752 |
| chrXVI | 284968 | 285121 | 153 | N16:285013 |
| chrXVI | 287278 | 287435 | 157 | N16:287376 |
| chrXVI | 287516 | 287643 | 127 | N16:287547 |
| chrXVI | 295119 | 295280 | 161 | N16:295214 |
| chrXVI | 296343 | 296490 | 147 | N16:296416 |
| chrXVI | 301254 | 301382 | 128 | N16:301333 |
| chrXVI | 305851 | 306015 | 164 | N16:305962 |
| chrXVI | 306387 | 306542 | 155 | N16:306463 |
| chrXVI | 317763 | 317907 | 144 | N16:317793 |
| chrXVI | 318699 | 318846 | 147 | |
| chrXVI | 322025 | 322158 | 133 | N16:322082 |
| chrXVI | 323166 | 323321 | 155 | N16:323256 |
| chrXVI | 332594 | 332745 | 151 | |
| chrXVI | 332948 | 333099 | 151 | N16:333032 |
| chrXVI | 336537 | 336643 | 106 | N16:336620 |
| chrXVI | 336537 | 336650 | 113 | N16:336620 |
| chrXVI | 341289 | 341447 | 158 | N16:341297 |
| chrXVI | 344751 | 344913 | 162 | N16:344736 |
| chrXVI | 344751 | 344913 | 162 | N16:344907 |
| chrXVI | 350439 | 350593 | 154 | N16:350549 |
| chrXVI | 351311 | 351471 | 160 | N16:351332 |
| chrXVI | 354968 | 355128 | 160 | N16:355047 |
| chrXVI | 361977 | 362073 | 96 | N16:362047 |
| chrXVI | 364336 | 364488 | 152 | N16:364416 |
| chrXVI | 366545 | 366652 | 107 | N16:366569 |
| chrXVI | 366545 | 366653 | 108 | N16:366569 |
| chrXVI | 367044 | 367180 | 136 | N16:367128 |
| chrXVI | 370599 | 370734 | 135 | N16:370630 |
| chrXVI | 370905 | 371054 | 149 | N16:370964 |
| chrXVI | 373474 | 373616 | 142 | N16:373635 |
| chrXVI | 379313 | 379470 | 157 | N16:379337 |
| chrXVI | 379794 | 379947 | 153 | |
| chrXVI | 379982 | 380135 | 153 | N16:379982 |
| chrXVI | 379982 | 380135 | 153 | N16:380145 |
| chrXVI | 383644 | 383787 | 143 | N16:383784 |
| chrXVI | 385233 | 385386 | 153 | N16:385330 |
| chrXVI | 389294 | 389391 | 97 | N16:389336 |
| chrXVI | 390081 | 390232 | 151 | N16:390091 |
| chrXVI | 390081 | 390232 | 151 | N16:390239 |
| chrXVI | 390545 | 390691 | 146 | N16:390526 |
| chrXVI | 392582 | 392731 | 149 | N16:392651 |
| chrXVI | 392582 | 392736 | 154 | N16:392651 |
| chrXVI | 394420 | 394573 | 153 | N16:394533 |
| chrXVI | 407155 | 407307 | 152 | N16:407273 |
| chrXVI | 407160 | 407307 | 147 | N16:407273 |
| chrXVI | 411818 | 411982 | 164 | N16:411890 |
| chrXVI | 411838 | 411991 | 153 | N16:411890 |
| chrXVI | 412957 | 413111 | 154 | |
| chrXVI | 415881 | 416027 | 146 | N16:415956 |
| chrXVI | 417433 | 417565 | 132 | N16:417568 |
| chrXVI | 418493 | 418641 | 148 | |
| chrXVI | 418678 | 418846 | 168 | N16:418685 |
| chrXVI | 418678 | 418846 | 168 | N16:418866 |

| CHR | Start | End | Length (bp) | Nucleosome_ID |
|---|---|---|---|---|
| chrXVI | 432250 | 432440 | 190 | N16:432391 |
| chrXVI | 432594 | 432737 | 143 | N16:432613 |
| chrXVI | 432908 | 433068 | 160 | N16:433087 |
| chrXVI | 433481 | 433638 | 157 | N16:433601 |
| chrXVI | 437116 | 437278 | 162 | N16:437280 |
| chrXVI | 439535 | 439688 | 153 | |
| chrXVI | 442027 | 442174 | 147 | N16:442080 |
| chrXVI | 442326 | 442474 | 148 | N16:442468 |
| chrXVI | 442340 | 442502 | 162 | N16:442468 |
| chrXVI | 446281 | 446447 | 166 | N16:446378 |
| chrXVI | 448877 | 449010 | 133 | N16:448925 |
| chrXVI | 454839 | 454945 | 106 | N16:454945 |
| chrXVI | 462929 | 463059 | 130 | N16:462975 |
| chrXVI | 465016 | 465172 | 156 | N16:465100 |
| chrXVI | 466157 | 466315 | 158 | N16:466243 |
| chrXVI | 474538 | 474690 | 152 | N16:474587 |
| chrXVI | 484126 | 484288 | 162 | N16:484123 |
| chrXVI | 484126 | 484288 | 162 | N16:484296 |
| chrXVI | 484807 | 484963 | 156 | |
| chrXVI | 487013 | 487184 | 171 | N16:487139 |
| chrXVI | 493217 | 493377 | 160 | N16:493271 |
| chrXVI | 497274 | 497428 | 154 | N16:497348 |
| chrXVI | 501076 | 501229 | 153 | N16:501190 |
| chrXVI | 501841 | 501999 | 158 | N16:501884 |
| chrXVI | 505233 | 505380 | 147 | N16:505303 |
| chrXVI | 506320 | 506478 | 158 | N16:506395 |
| chrXVI | 506594 | 506750 | 156 | N16:506676 |
| chrXVI | 509864 | 510024 | 160 | N16:509922 |
| chrXVI | 511391 | 511535 | 144 | N16:511439 |
| chrXVI | 517443 | 517595 | 152 | N16:517508 |
| chrXVI | 519049 | 519173 | 124 | N16:519058 |
| chrXVI | 520082 | 520209 | 127 | N16:520178 |
| chrXVI | 522508 | 522654 | 146 | N16:522677 |
| chrXVI | 522574 | 522734 | 160 | N16:522677 |
| chrXVI | 525954 | 526108 | 154 | N16:526036 |
| chrXVI | 536923 | 537076 | 153 | N16:537021 |
| chrXVI | 538001 | 538163 | 162 | N16:538105 |
| chrXVI | 541695 | 541831 | 136 | N16:541839 |
| chrXVI | 547021 | 547160 | 139 | N16:547005 |
| chrXVI | 547636 | 547771 | 135 | N16:547723 |
| chrXVI | 547851 | 548005 | 154 | N16:547889 |
| chrXVI | 550617 | 550788 | 171 | N16:550762 |
| chrXVI | 553457 | 553564 | 107 | N16:553491 |
| chrXVI | 553457 | 553565 | 108 | N16:553491 |
| chrXVI | 560771 | 560926 | 155 | N16:560761 |
| chrXVI | 560771 | 560926 | 155 | N16:560933 |
| chrXVI | 564313 | 564461 | 148 | N16:564375 |
| chrXVI | 564317 | 564438 | 121 | N16:564375 |
| chrXVI | 566468 | 566620 | 152 | N16:566631 |
| chrXVI | 566468 | 566620 | 152 | N16:566462 |
| chrXVI | 568077 | 568211 | 134 | N16:568157 |
| chrXVI | 573444 | 573609 | 165 | N16:573552 |
| chrXVI | 573688 | 573839 | 151 | N16:573733 |
| chrXVI | 578898 | 579048 | 150 | N16:579014 |
| chrXVI | 580106 | 580256 | 150 | N16:580137 |
| chrXVI | 581758 | 581907 | 149 | N16:581737 |
| chrXVI | 581758 | 581907 | 149 | N16:581900 |
| chrXVI | 582967 | 583116 | 149 | N16:583058 |
| chrXVI | 584868 | 585014 | 146 | N16:584868 |
| chrXVI | 591560 | 591713 | 153 | N16:591636 |
| chrXVI | 592210 | 592352 | 142 | N16:592274 |
| chrXVI | 593528 | 593683 | 155 | N16:593589 |
| chrXVI | 594996 | 595138 | 142 | N16:595089 |
| chrXVI | 599046 | 599200 | 154 | N16:599073 |
| chrXVI | 600831 | 600997 | 166 | N16:600995 |
| chrXVI | 600831 | 600997 | 166 | N16:600844 |
| chrXVI | 601108 | 601231 | 123 | N16:601151 |
| chrXVI | 603139 | 603315 | 176 | N16:603211 |
| chrXVI | 604400 | 604533 | 133 | N16:604513 |
| chrXVI | 611112 | 611263 | 151 | |
| chrXVI | 611769 | 611929 | 160 | N16:611782 |
| chrXVI | 611769 | 611929 | 160 | N16:611938 |
| chrXVI | 614264 | 614371 | 107 | N16:614344 |
| chrXVI | 614264 | 614381 | 117 | N16:614344 |
| chrXVI | 616962 | 617122 | 160 | N16:617008 |
| chrXVI | 617750 | 617867 | 117 | N16:617790 |
| chrXVI | 617888 | 618049 | 161 | N16:617954 |
| chrXVI | 623817 | 623987 | 170 | N16:623864 |
| chrXVI | 627166 | 627316 | 150 | N16:627249 |
| chrXVI | 630544 | 630672 | 128 | |
| chrXVI | 630787 | 630947 | 160 | N16:630889 |
| chrXVI | 639745 | 639881 | 136 | N16:639847 |
| chrXVI | 644095 | 644255 | 160 | N16:644162 |
| chrXVI | 647938 | 648089 | 151 | N16:647982 |
| chrXVI | 651006 | 651170 | 164 | N16:651112 |
| chrXVI | 651448 | 651577 | 129 | N16:651501 |
| chrXVI | 651814 | 651966 | 152 | |
| chrXVI | 655669 | 655818 | 149 | N16:655729 |
| chrXVI | 661336 | 661465 | 129 | N16:661468 |
| chrXVI | 664409 | 664571 | 162 | N16:664522 |
| chrXVI | 665020 | 665144 | 124 | N16:665145 |
| chrXVI | 670319 | 670426 | 107 | N16:670368 |
| chrXVI | 670319 | 670427 | 108 | N16:670368 |
| chrXVI | 670635 | 670782 | 147 | N16:670711 |
| chrXVI | 672000 | 672158 | 158 | N16:672118 |
| chrXVI | 678051 | 678201 | 150 | N16:678147 |
| chrXVI | 679094 | 679205 | 111 | N16:679146 |
| chrXVI | 679094 | 679206 | 112 | N16:679146 |
| chrXVI | 679094 | 679207 | 113 | N16:679146 |
| chrXVI | 679098 | 679205 | 107 | N16:679146 |
| chrXVI | 679098 | 679220 | 122 | N16:679146 |
| chrXVI | 681181 | 681332 | 151 | N16:681246 |
| chrXVI | 685561 | 685687 | 126 | N16:685673 |

TABLE 1

| CHR | Start | End | Length (bp) | Nucleosome_ID |
|---|---|---|---|---|
| chrXVI | 685866 | 686024 | 158 | N16:685893 |
| chrXVI | 689048 | 689183 | 135 | N16:689086 |
| chrXVI | 695989 | 696136 | 147 | N16:696027 |
| chrXVI | 697045 | 697195 | 150 | N16:697144 |
| chrXVI | 702075 | 702202 | 127 | N16:702218 |
| chrXVI | 706433 | 706576 | 143 | N16:706496 |
| chrXVI | 719549 | 719656 | 107 | N16:719618 |
| chrXVI | 719549 | 719694 | 145 | N16:719618 |
| chrXVI | 721766 | 721909 | 143 | N16:721871 |
| chrXVI | 722565 | 722701 | 136 | N16:722569 |
| chrXVI | 726383 | 726544 | 161 | N16:726563 |
| chrXVI | 726865 | 727012 | 147 | N16:726909 |
| chrXVI | 730093 | 730221 | 128 | N16:730129 |
| chrXVI | 732571 | 732734 | 163 | N16:732631 |
| chrXVI | 733908 | 734067 | 159 | N16:733903 |
| chrXVI | 734318 | 734467 | 149 | N16:734451 |
| chrXVI | 735641 | 735790 | 149 | N16:735692 |
| chrXVI | 735832 | 735978 | 146 | N16:735865 |
| chrXVI | 736889 | 737030 | 141 | N16:736965 |
| chrXVI | 737626 | 737736 | 110 | N16:737632 |
| chrXVI | 739761 | 739921 | 160 | N16:739810 |
| chrXVI | 740972 | 741124 | 152 | N16:741035 |
| chrXVI | 742321 | 742473 | 152 | N16:742396 |
| chrXVI | 745613 | 745766 | 153 | N16:745699 |
| chrXVI | 747443 | 747595 | 152 | N16:747452 |
| chrXVI | 748166 | 748325 | 159 | N16:748291 |
| chrXVI | 751513 | 751665 | 152 | N16:751585 |
| chrXVI | 753264 | 753415 | 151 | N16:753426 |
| chrXVI | 753264 | 753415 | 151 | N16:753263 |
| chrXVI | 753454 | 753608 | 154 | N16:753612 |
| chrXVI | 759400 | 759547 | 147 | N16:759472 |
| chrXVI | 761447 | 761589 | 142 | N16:761432 |
| chrXVI | 761447 | 761589 | 142 | N16:761585 |
| chrXVI | 763063 | 763215 | 152 | N16:763092 |
| chrXVI | 763430 | 763578 | 148 | N16:763564 |
| chrXVI | 764397 | 764544 | 147 | N16:764439 |
| chrXVI | 764502 | 764659 | 157 | N16:764652 |
| chrXVI | 770323 | 770455 | 132 | N16:770351 |
| chrXVI | 778559 | 778687 | 128 | N16:778565 |
| chrXVI | 786310 | 786475 | 165 | |
| chrXVI | 788541 | 788654 | 113 | |
| chrXVI | 788648 | 788776 | 128 | N16:788711 |
| chrXVI | 790762 | 790920 | 158 | N16:790928 |
| chrXVI | 791267 | 791414 | 147 | N16:791367 |
| chrXVI | 791438 | 791595 | 157 | N16:791540 |
| chrXVI | 791737 | 791890 | 153 | N16:791867 |
| chrXVI | 792774 | 792924 | 150 | N16:792865 |
| chrXVI | 794619 | 794743 | 124 | N16:794744 |
| chrXVI | 796842 | 796998 | 156 | N16:796924 |
| chrXVI | 798394 | 798554 | 160 | N16:798384 |
| chrXVI | 798628 | 798777 | 149 | N16:798655 |
| chrXVI | 805070 | 805217 | 147 | N16:805107 |
| chrXVI | 805577 | 805691 | 114 | |
| chrXVI | 807669 | 807826 | 157 | N16:807711 |
| chrXVI | 808317 | 808452 | 135 | N16:808366 |
| chrXVI | 809355 | 809495 | 140 | N16:809434 |
| chrXVI | 809361 | 809495 | 134 | N16:809434 |
| chrXVI | 809773 | 809928 | 155 | N16:809806 |
| chrXVI | 809904 | 810012 | 108 | N16:809957 |
| chrXVI | 809934 | 810052 | 118 | N16:809957 |
| chrXVI | 810053 | 810202 | 149 | N16:810107 |
| chrXVI | 813679 | 813806 | 127 | N16:813742 |
| chrXVI | 822438 | 822593 | 155 | |
| chrXVI | 822870 | 822974 | 104 | N16:822924 |
| chrXVI | 827045 | 827170 | 125 | N16:827139 |
| chrXVI | 827247 | 827389 | 142 | N16:827298 |
| chrXVI | 827869 | 828021 | 152 | N16:827957 |
| chrXVI | 830012 | 830156 | 144 | N16:830081 |
| chrXVI | 839224 | 839390 | 166 | N16:839304 |
| chrXVI | 841656 | 841805 | 149 | |
| chrXVI | 843827 | 843986 | 159 | N16:843902 |
| chrXVI | 844949 | 845072 | 123 | |
| chrXVI | 845049 | 845204 | 155 | N16:845158 |
| chrXVI | 845482 | 845616 | 134 | N16:845543 |
| chrXVI | 845482 | 845622 | 140 | N16:845543 |
| chrXVI | 846525 | 846660 | 135 | N16:846560 |
| chrXVI | 847154 | 847311 | 157 | N16:847244 |
| chrXVI | 848557 | 848707 | 150 | N16:848653 |
| chrXVI | 849289 | 849403 | 114 | N16:849394 |
| chrXVI | 851012 | 851164 | 152 | N16:851100 |
| chrXVI | 851565 | 851679 | 114 | N16:851699 |
| chrXVI | 853843 | 853993 | 150 | N16:853920 |
| chrXVI | 853847 | 853993 | 146 | N16:853920 |
| chrXVI | 854031 | 854169 | 138 | N16:854079 |
| chrXVI | 854126 | 854258 | 132 | N16:854239 |
| chrXVI | 854305 | 854440 | 135 | N16:854397 |
| chrXVI | 854851 | 855005 | 154 | N16:854861 |
| chrXVI | 855108 | 855259 | 151 | N16:855198 |
| chrXVI | 855132 | 855277 | 145 | N16:855198 |
| chrXVI | 855136 | 855287 | 151 | N16:855198 |
| chrXVI | 855343 | 855483 | 140 | N16:855418 |
| chrXVI | 855349 | 855483 | 134 | N16:855418 |
| chrXVI | 855892 | 856000 | 108 | N16:855993 |
| chrXVI | 856112 | 856214 | 102 | |
| chrXVI | 858667 | 858810 | 143 | N16:858736 |
| chrXVI | 863879 | 864033 | 154 | N16:863878 |
| chrXVI | 863879 | 864033 | 154 | N16:864033 |
| chrXVI | 864255 | 864385 | 130 | N16:864244 |
| chrXVI | 867145 | 867288 | 143 | |
| chrXVI | 868632 | 868780 | 148 | N16:868711 |
| chrXVI | 888008 | 888137 | 129 | N16:888074 |
| chrXVI | 891517 | 891679 | 162 | N16:891562 |
| chrXVI | 893449 | 893599 | 150 | N16:893574 |

TABLE 1

| CHR | Start | End | Length (bp) | Nucleosome_ID |
|---|---|---|---|---|
| chrXVI | 897488 | 897648 | 160 | N16:897619 |
| chrXVI | 897990 | 898142 | 152 | N16:898110 |
| chrXVI | 901200 | 901349 | 149 | N16:901238 |
| chrXVI | 905702 | 905842 | 140 | N16:905808 |
| chrXVI | 910317 | 910457 | 140 | N16:910477 |
| chrXVI | 910317 | 910457 | 140 | N16:910305 |
| chrXVI | 922519 | 922676 | 157 | N16:922581 |
| chrXVI | 922519 | 922681 | 162 | N16:922581 |
| chrXVI | 922523 | 922681 | 158 | N16:922581 |
| chrXVI | 927439 | 927574 | 135 | N16:927563 |
| chrXVI | 930833 | 931008 | 175 | N16:930869 |
| chrXVI | 931493 | 931642 | 149 | N16:931566 |
| chrXVI | 940363 | 940515 | 152 | N16:940386 |
| chrXVI | 943076 | 943229 | 153 | N16:943200 |
| chrXVI | 943085 | 943229 | 144 | N16:943200 |
| chrXVI | 944252 | 944389 | 137 | N16:944314 |
| chrXVI | 944537 | 944680 | 143 | N16:944589 |
| chrXVI | 944585 | 944710 | 125 | N16:944589 |
| chrXVI | 944810 | 944962 | 152 | N16:944922 |
| chrXVI | 944883 | 945027 | 144 | N16:944922 |
| chrXVI | 944980 | 945115 | 135 | N16:945080 |
| chrXVI | 944980 | 945167 | 187 | N16:945080 |
| chrXVI | 945023 | 945115 | 92 | N16:945080 |
| chrXVI | 945023 | 945167 | 144 | N16:945080 |
| chrXVI | 945362 | 945514 | 152 | N16:945422 |
| chrXVI | 945663 | 945793 | 130 | N16:945658 |
| chrXVI | 945663 | 945793 | 130 | N16:945816 |
| chrXVI | 945759 | 945954 | 195 | N16:945816 |
| chrXVI | 945821 | 945954 | 133 | N16:945816 |
| chrXVI | 945821 | 945969 | 148 | N16:945816 |
| chrXVI | 946033 | 946184 | 151 | N16:946078 |
| chrXVI | 946038 | 946172 | 134 | N16:946078 |
| chrXVI | 946051 | 946147 | 96 | N16:946078 |
| chrXVI | 946159 | 946281 | 122 | |
| chrXVI | 946966 | 947118 | 152 | N16:947053 |
| chrXVI | 947306 | 947457 | 151 | N16:947384 |

### TABLE 2. Calculation of the ΔLk restrained by individual nucleosomes

The four first columns indicate the chromosomal coordinates of the nucleosome library. The next two columns indicate the relative abundance (number of reads relative to total reads) of each nucleosome sequence in sections A and B of the gel electrophoresis. Column A/A+B indicates the partition probability of each sequence in section A. Column Z indicates the corresponding Z-scores of the previous partition probabilities in a normal distribution of mean 0 and standard deviation 1. The next column, multiplies Z-scores by 0.85 (mean standard deviation of the Lk distributions of the library) to yield approximate ΔLk deviations within a normal distribution of mean 0. The next column, Phase correction, subtracts from the previous ΔLk values the average ΔLk of all the sequences of the same length. The next column, Length correction, adjusts the previous ΔLk value by adding or subtracting 0.007 for each bp difference from 144 bp (the average length of the library of nucleosome DNA sequences). The last column adds -1.26 to obtain the ΔLk restrained by each nucleosome.

| Chrom | Start | End | Length | Section A | Section B | A/A+B | Z-score | Z * 0.85 | Phase correction | Length correction | ΔLK |
|---|---|---|---|---|---|---|---|---|---|---|---|
| chrI | 375 | 485 | 110 | 0,324849907 | 2,657108488 | 0,108938444 | -1,232193295 | -1,047364301 | -0,453273061 | -0,215273061 | **-1,475273061** |
| chrI | 1992 | 2143 | 151 | 1,929152515 | 0,952408482 | 0,669481756 | 0,438482589 | 0,372710201 | 0,163375138 | 0,114375138 | **-1,145624862** |
| chrI | 3900 | 4046 | 146 | 0,185221438 | 0,159352392 | 0,537537741 | 0,094232436 | 0,080097571 | -0,15491971 | -0,16891971 | **-1,42891971** |
| chrI | 7310 | 7448 | 138 | 0,088336378 | 0,059293913 | 0,598362147 | 0,249109985 | 0,211743487 | 0,118785994 | 0,160785994 | **-1,099214006** |
| chrI | 19698 | 19849 | 151 | 0,153876272 | 0,081529131 | 0,653664997 | 0,395234266 | 0,335949126 | 0,138279312 | 0,089279312 | **-1,170720688** |
| chrI | 24106 | 24256 | 150 | 0,373292436 | 0,133411305 | 0,73670748 | 0,633227577 | 0,53824344 | 0,299199857 | 0,257199857 | **-1,002800143** |
| chrI | 24106 | 24267 | 161 | 0,062690333 | 0,066705652 | 0,484484373 | -0,038901718 | -0,03306646 | 0,007591221 | -0,111408779 | **-1,371408779** |
| chrI | 24156 | 24264 | 108 | 3,769968652 | 21,72380747 | 0,147878001 | -1,045577803 | -0,888741132 | -0,21236164 | 0,03963836 | **-1,22036164** |
| chrI | 24156 | 24265 | 109 | 0,085486818 | 0,422469132 | 0,16829573 | -0,960921856 | -0,816783577 | -0,184955994 | 0,060044006 | **-1,199955994** |
| chrI | 24156 | 24266 | 110 | 0,051292091 | 0,604056741 | 0,078266853 | -1,416826344 | -1,204302393 | -0,642918922 | -0,404918922 | **-1,664918922** |
| chrI | 24156 | 24267 | 111 | 4,513703964 | 28,94284141 | 0,134912434 | -1,103465956 | -0,937946063 | -0,405214917 | -0,174214917 | **-1,434214917** |
| chrI | 25658 | 25790 | 132 | 0,330549028 | 0,233469783 | 0,586060289 | 0,217422085 | 0,184808773 | 0,27821694 | 0,36221694 | **-0,89778306** |
| chrI | 25658 | 25797 | 139 | 2,68428607 | 1,474936092 | 0,645381748 | 0,372881685 | 0,316949432 | 0,206470586 | 0,241470586 | **-1,018529414** |
| chrI | 25671 | 25797 | 126 | 0,094035499 | 0,114881957 | 0,450108387 | -0,125387512 | -0,106579385 | 0,227304304 | 0,353304304 | **-0,906695696** |
| chrI | 25885 | 26057 | 172 | 0,062690333 | 0,037058696 | 0,628480635 | 0,327831948 | 0,278657156 | 0,447151435 | 0,251151435 | **-1,008848565** |
| chrI | 26200 | 26329 | 129 | 0,028495606 | 0,100058479 | 0,221662392 | -0,766590907 | -0,651602271 | -0,419518143 | -0,314518143 | **-1,574518143** |
| chrI | 26200 | 26330 | 130 | 14,93454702 | 19,2594042 | 0,436759909 | -0,159189193 | -0,135310814 | 0,067390633 | 0,165390633 | **-1,094609367** |
| chrI | 26200 | 26337 | 137 | 0,772230918 | 0,61517435 | 0,556600826 | 0,142356591 | 0,121003103 | 0,041760198 | 0,090760198 | **-1,169239802** |
| chrI | 26204 | 26330 | 126 | 0,079787696 | 0,122293696 | 0,394829506 | -0,266753431 | -0,226740416 | 0,087246837 | 0,213246837 | **-1,046753163** |
| chrI | 26211 | 26330 | 119 | 0,225115286 | 0,285351958 | 0,440998495 | -0,148438156 | -0,126172432 | 0,30764548 | 0,48264548 | **-0,77735452** |
| chrI | 26268 | 26418 | 150 | 0,416035845 | 0,237175653 | 0,636908331 | 0,350207021 | 0,297675968 | 0,072283311 | 0,030283311 | **-1,229716689** |
| chrI | 26290 | 26465 | 175 | 0,034194727 | 0,092646739 | 0,269586342 | -0,614064532 | -0,521954852 | -0,350207551 | -0,567207551 | **-1,827207551** |
| chrI | 26425 | 26573 | 148 | 0,467327936 | 0,203822827 | 0,696308433 | 0,513812434 | 0,436740568 | 0,196075677 | 0,168075677 | **-1,091924323** |
| chrI | 26560 | 26708 | 148 | 0,3903898 | 0,222352175 | 0,637119401 | 0,35076961 | 0,298154169 | 0,036820934 | 0,008820934 | **-1,251179066** |
| chrI | 26695 | 26843 | 148 | 0,404637603 | 0,166764131 | 0,708149064 | 0,54798548 | 0,465787658 | 0,197216135 | 0,169216135 | **-1,090783865** |
| chrI | 27347 | 27495 | 148 | 0,447381012 | 0,307587175 | 0,592582601 | 0,234193452 | 0,199064434 | -0,069301342 | -0,097301342 | **-1,357301342** |
| chrI | 28545 | 28653 | 108 | 0,219416165 | 0,548468698 | 0,285740969 | -0,565870321 | -0,480989773 | 0,192229317 | 0,444229317 | **-0,815770683** |
| chrI | 28558 | 28705 | 147 | 0,29350474 | 0,329822392 | 0,470867903 | -0,073088357 | -0,062125103 | -0,304279852 | -0,325279852 | **-1,585279852** |
| chrI | 28865 | 28996 | 131 | 0,769381358 | 0,763409133 | 0,501948154 | 0,004883318 | 0,00415082 | 0,164044397 | 0,255044397 | **-1,004955603** |
| chrI | 31670 | 31815 | 145 | 0,572761677 | 0,478057176 | 0,54506224 | 0,113195553 | 0,09621622 | -0,115405277 | -0,122405277 | **-1,382405277** |
| chrI | 32825 | 32968 | 143 | 0,131079787 | 0,08894087 | 0,595761274 | 0,242390685 | 0,206032083 | 0,022965057 | 0,029965057 | **-1,230034943** |
| chrI | 38266 | 38455 | 189 | 0,447381012 | 0,70040935 | 0,389775892 | -0,279903186 | -0,237917708 | -0,058713958 | -0,373713958 | **-1,633713958** |
| chrI | 38266 | 38455 | 189 | 0,447381012 | 0,70040935 | 0,389775892 | -0,279903186 | -0,237917708 | -0,058713958 | -0,373713958 | **-1,633713958** |
| chrI | 39356 | 39504 | 148 | 0,068389454 | 0,040764565 | 0,626540868 | 0,322705529 | 0,274299699 | 0,00693267 | -0,02106733 | **-1,28106733** |
| chrI | 44014 | 44169 | 155 | 2,396480451 | 2,01969892 | 0,54265922 | 0,107135403 | 0,091065093 | 0,007886483 | -0,069113517 | **-1,329113517** |
| chrI | 48679 | 48833 | 154 | 1,493169746 | 1,3155837 | 0,531612964 | 0,079325064 | 0,067426304 | -0,034356893 | -0,104356893 | **-1,364356893** |
| chrI | 52107 | 52211 | 104 | 0,071239015 | 0,314998914 | 0,184443343 | -0,898560728 | -0,763776618 | -0,017609421 | 0,262390579 | **-0,997609421** |
| chrI | 52108 | 52200 | 92 | 0,039893848 | 0,314998914 | 0,112410994 | -1,213805546 | -1,031734714 | -0,298494905 | 0,065505095 | **-1,194494905** |
| chrI | 52108 | 52202 | 94 | 0,028495606 | 0,188999349 | 0,131017319 | -1,121595095 | -0,953355831 | -0,205050422 | 0,144949578 | **-1,115050422** |
| chrI | 52108 | 52203 | 95 | 0,042743409 | 0,326116523 | 0,115879783 | -1,195838559 | -1,016462775 | -0,286469923 | 0,056530077 | **-1,203469923** |
| chrI | 52108 | 52204 | 96 | 0,039893848 | 0,270528479 | 0,128514751 | -1,133440052 | -0,963424045 | -0,250333328 | 0,085666672 | **-1,174333328** |
| chrI | 52108 | 52205 | 97 | 0,042743409 | 0,203822827 | 0,173354671 | -0,940991251 | -0,799842563 | -0,077989827 | 0,251010173 | **-1,008989827** |
| chrI | 52108 | 52206 | 98 | 0,031345166 | 0,240881523 | 0,11514362 | -1,199619268 | -1,019676378 | -0,294433255 | 0,027566745 | **-1,232433255** |
| chrI | 52108 | 52207 | 99 | 0,037044288 | 0,555880437 | 0,062477219 | -1,53430581 | -1,304159938 | -0,588375011 | -0,273375011 | **-1,533375011** |
| chrI | 52108 | 52208 | 100 | 0,085486818 | 0,326116523 | 0,207692235 | -0,814454777 | -0,692286561 | 0,069901083 | 0,377901083 | **-0,882098917** |
| chrI | 52108 | 52209 | 101 | 0,136778908 | 0,874585221 | 0,135242001 | -1,101948639 | -0,936656343 | -0,168128482 | 0,132871518 | **-1,127128482** |
| chrI | 52108 | 52210 | 102 | 0,037044288 | 0,363175219 | 0,092559925 | -1,325154688 | -1,126381485 | -0,354587152 | -0,060587152 | **-1,320587152** |
| chrI | 52108 | 52211 | 103 | 35,59956037 | 163,3992015 | 0,178893376 | -0,919590578 | -0,781651991 | -0,03332145 | 0,25367855 | **-1,00632145** |
| chrI | 52112 | 52211 | 99 | 0,042743409 | 0,389116306 | 0,098975217 | -1,287412835 | -1,09430091 | -0,367463337 | -0,052463337 | **-1,312463337** |
| chrI | 54178 | 54332 | 154 | 0,29065518 | 0,181587609 | 0,61547828 | 0,293626349 | 0,249582396 | 0,150968964 | 0,080968964 | **-1,179031036** |
| chrI | 56067 | 56218 | 151 | 1,484621064 | 1,226642831 | 0,547575272 | 0,119537596 | 0,101606956 | -0,082434877 | -0,131434877 | **-1,391434877** |
| chrI | 57573 | 57723 | 150 | 0,262159574 | 0,133411305 | 0,662737294 | 0,419945303 | 0,356953508 | 0,125523607 | 0,083523607 | **-1,176476393** |
| chrI | 57573 | 57723 | 150 | 0,262159574 | 0,133411305 | 0,662737294 | 0,419945303 | 0,356953508 | 0,125523607 | 0,083523607 | **-1,176476393** |
| chrI | 57985 | 58142 | 157 | 0,108283302 | 0,133411305 | 0,448017039 | -0,130672891 | -0,111071958 | -0,140896053 | -0,231896053 | **-1,491896053** |
| chrI | 58357 | 58529 | 172 | 0,225115286 | 0,285351958 | 0,440998495 | -0,148438156 | -0,126172432 | 0,060582422 | -0,135417578 | **-1,395417578** |
| chrI | 59835 | 59985 | 150 | 0,267858695 | 0,26682261 | 0,500968881 | 0,002428627 | 0,002064333 | -0,223981569 | -0,265981569 | **-1,525981569** |
| chrI | 60956 | 61114 | 158 | 6,844644522 | 4,895453713 | 0,583014246 | 0,209610726 | 0,178169117 | 0,156739962 | 0,058739962 | **-1,201260038** |
| chrI | 61139 | 61253 | 114 | 0,253610892 | 1,775111528 | 0,125010149 | -1,15030008 | -0,977755068 | -0,464954291 | -0,254954291 | **-1,514954291** |
| chrI | 62731 | 62859 | 128 | 2,034586257 | 3,053636533 | 0,399861865 | -0,253704663 | -0,215648964 | 0,029545139 | 0,141545139 | **-1,118454861** |
| chrI | 73815 | 73966 | 151 | 0,142478029 | 0,133411305 | 0,516431814 | 0,041200102 | 0,035020087 | -0,147101359 | -0,196101359 | **-1,456101359** |
| chrI | 75929 | 76089 | 160 | 0,151026711 | 0,037058696 | 0,802968787 | 0,852273293 | 0,724432299 | 0,736809571 | 0,624809571 | **-0,635190429** |
| chrI | 77820 | 77969 | 149 | 0,353345512 | 0,237175653 | 0,598362147 | 0,249109985 | 0,211743487 | -0,034929447 | -0,069929447 | **-1,329929447** |
| chrI | 78867 | 79013 | 146 | 0,082637257 | 0,077823261 | 0,515000562 | 0,037609696 | 0,031968242 | -0,202014857 | -0,216014857 | **-1,476014857** |
| chrI | 79075 | 79223 | 148 | 0,270708255 | 0,222352175 | 0,549036667 | 0,123227859 | 0,104743681 | -0,157857802 | -0,185857802 | **-1,445857802** |
| chrI | 79820 | 79919 | 99 | 0,051292091 | 0,185293479 | 0,216801433 | -0,783041289 | -0,665585096 | 0,076654568 | 0,391654568 | **-0,868345432** |
| chrI | 80712 | 80877 | 165 | 0,074088575 | 0,051882174 | 0,588141101 | 0,222765786 | 0,189350918 | 0,304193553 | 0,157193553 | **-1,102806447** |
| chrI | 83877 | 84031 | 154 | 0,786478721 | 0,35205761 | 0,690780522 | 0,498063968 | 0,423354373 | 0,315267959 | 0,245267959 | **-1,014732041** |
| chrI | 87561 | 87716 | 155 | 0,102584181 | 0,140823044 | 0,421450846 | -0,198183249 | -0,168455762 | -0,231954094 | -0,308954094 | **-1,568954094** |
| chrI | 88852 | 89013 | 161 | 1,46467414 | 1,526818266 | 0,48961319 | -0,026038814 | -0,022132992 | 0,029070756 | -0,089929244 | **-1,349929244** |
| chrI | 88852 | 89013 | 161 | 1,46467414 | 1,526818266 | 0,48961319 | -0,026038814 | -0,022132992 | 0,029070756 | -0,089929244 | **-1,349929244** |
| chrI | 88889 | 89053 | 164 | 0,111132863 | 0,226058044 | 0,3295844 | -0,441061051 | -0,374901893 | -0,31832009 | -0,45832009 | **-1,71832009** |
| chrI | 91036 | 91130 | 94 | 0,028495606 | 0,055588044 | 0,338895921 | -0,415478239 | -0,353156503 | 0,380938839 | 0,730938839 | **-0,529061161** |
| chrI | 94271 | 94416 | 145 | 0,253610892 | 0,129705435 | 0,661622983 | 0,41689661 | 0,354362118 | 0,139529776 | 0,132529776 | **-1,127470224** |
| chrI | 97984 | 98151 | 167 | 0,156725832 | 0,203822827 | 0,434687048 | -0,164453569 | -0,139785534 | -0,018823481 | -0,179823481 | **-1,439823481** |
| chrI | 100299 | 100401 | 102 | 0,071239015 | 0,037058696 | 0,657807209 | 0,406485945 | 0,345513053 | 1,111125764 | 1,405125764 | **0,145125764** |
| chrI | 100430 | 100592 | 162 | 0,34194727 | 0,17417587 | 0,662530399 | 0,419378955 | 0,356472112 | 0,413429344 | 0,287429344 | **-0,972570656** |
| chrI | 100744 | 100901 | 157 | 0,219416165 | 0,151940653 | 0,590849971 | 0,229731961 | 0,195272167 | 0,173217738 | 0,082217738 | **-1,177782262** |
| chrI | 100903 | 101028 | 125 | 0,082637257 | 0,074117392 | 0,527175798 | 0,068172392 | 0,057946533 | 0,445541957 | 0,578541957 | **-0,681458043** |
| chrI | 101774 | 101913 | 139 | 0,424584527 | 0,363175219 | 0,538977181 | 0,097857261 | 0,083178671 | -0,043887199 | -0,008887199 | **-1,268887199** |
| chrI | 102759 | 102918 | 159 | 0,319150785 | 0,34835174 | 0,478126708 | -0,054855711 | -0,046627354 | -0,039046068 | -0,144046068 | **-1,404046068** |
| chrI | 110352 | 110498 | 146 | 4,00648218 | 2,497756096 | 0,615980229 | 0,294940225 | 0,250699191 | 0,011903264 | -0,002096736 | **-1,262096736** |
| chrI | 110959 | 111107 | 148 | 0,324849907 | 0,229763914 | 0,585722704 | 0,216555726 | 0,184072367 | -0,077708219 | -0,105708219 | **-1,365708219** |
| chrI | 111829 | 111987 | 158 | 0,034194727 | 0,214940436 | 0,137253717 | -1,092741227 | -0,928830043 | -0,946517965 | -1,044517965 | **-2,304517965** |

TABLE 2

| Chrom | Start | End | Length | Section A | Section B | A/A+B | Z-score | Z * 0.85 | Phase correction | Length correction | ΔLK |
|---|---|---|---|---|---|---|---|---|---|---|---|
| chrI | 111829 | 111989 | 160 | 7,66816753 | 5,996096978 | 0,561184067 | 0,153971933 | 0,130876143 | 0,151324568 | 0,039324568 | -1,220675432 |
| chrI | 111835 | 111987 | 152 | 4,339880769 | 3,146283272 | 0,579720234 | 0,201177822 | 0,171001149 | -0,005773661 | -0,061773661 | -1,321773661 |
| chrI | 111835 | 111989 | 154 | 0,091185939 | 0,218646305 | 0,294307453 | -0,540844298 | -0,459717653 | -0,573586461 | -0,643586461 | -1,903586461 |
| chrI | 113198 | 113345 | 147 | 0,09973462 | 0,066705652 | 0,599221683 | 0,251333038 | 0,213633083 | -0,032354286 | -0,053354286 | -1,313354286 |
| chrI | 113198 | 113345 | 147 | 0,09973462 | 0,066705652 | 0,599221683 | 0,251333038 | 0,213633083 | -0,032354286 | -0,053354286 | -1,313354286 |
| chrI | 113272 | 113431 | 159 | 1,877860425 | 1,663935441 | 0,530200072 | 0,075772802 | 0,064406881 | 0,069150553 | -0,035849447 | -1,295849447 |
| chrI | 114916 | 115064 | 148 | 0,179522317 | 0,062999783 | 0,740230754 | 0,644056972 | 0,547448426 | 0,287356404 | 0,259356404 | -1,000643596 |
| chrI | 116310 | 116474 | 164 | 2,185612968 | 1,697288267 | 0,562881422 | 0,158278745 | 0,134536933 | 0,213769775 | 0,073769775 | -1,186230225 |
| chrI | 119258 | 119408 | 150 | 2,34518836 | 0,229763914 | 0,910769642 | 1,345509621 | 1,143683178 | 0,925706228 | 0,883706228 | -0,376293772 |
| chrI | 121555 | 121709 | 154 | 0,555664314 | 0,396528045 | 0,583563089 | 0,211017235 | 0,17936465 | 0,082446566 | 0,012446566 | -1,247553434 |
| chrI | 128610 | 128759 | 149 | 0,116831984 | 0,081529131 | 0,588986325 | 0,224938202 | 0,191197471 | -0,043333574 | -0,078333574 | -1,338333574 |
| chrI | 128749 | 128913 | 164 | 0,752283994 | 0,956114351 | 0,440344605 | -0,150095577 | -0,127581241 | -0,02014932 | -0,16014932 | -1,42014932 |
| chrI | 132160 | 132339 | 179 | 0,843469933 | 1,463818484 | 0,365567619 | -0,343615804 | -0,292073433 | -0,120001105 | -0,365001105 | -1,625001105 |
| chrI | 132322 | 132449 | 127 | 0,045592969 | 0,074117392 | 0,380860679 | -0,303221116 | -0,257737948 | 0,01200283 | 0,13100283 | -1,12899717 |
| chrI | 133311 | 133456 | 145 | 4,274340875 | 2,90169588 | 0,595640884 | 0,242079926 | 0,205767937 | -0,012779418 | -0,019779418 | -1,279779418 |
| chrI | 133444 | 133595 | 151 | 0,775080479 | 0,559586306 | 0,580729578 | 0,203760266 | 0,173196226 | -0,003572395 | -0,052572395 | -1,312572395 |
| chrI | 134936 | 135082 | 146 | 0,612655525 | 0,363175219 | 0,627829702 | 0,326110708 | 0,277194102 | 0,040400686 | 0,026400686 | -1,233599314 |
| chrI | 139478 | 139611 | 133 | 13,67504124 | 8,486441335 | 0,617063466 | 0,297777395 | 0,253110785 | 0,245477013 | 0,322477013 | -0,937522987 |
| chrI | 144452 | 144595 | 143 | 0,302053422 | 0,192705218 | 0,610506614 | 0,280639687 | 0,238543734 | 0,049475346 | 0,056475346 | -1,203524654 |
| chrI | 144452 | 144595 | 143 | 0,302053422 | 0,192705218 | 0,610506614 | 0,280639687 | 0,238543734 | 0,049475346 | 0,056475346 | -1,203524654 |
| chrI | 145995 | 146131 | 136 | 0,507221784 | 0,226058044 | 0,691716538 | 0,500721804 | 0,425613534 | 0,379375359 | 0,435375359 | -0,824624641 |
| chrI | 145995 | 146131 | 136 | 0,507221784 | 0,226058044 | 0,691716538 | 0,500721804 | 0,425613534 | 0,379375359 | 0,435375359 | -0,824624641 |
| chrI | 157171 | 157325 | 154 | 0,09973462 | 0,100058479 | 0,499189516 | -0,002031583 | -0,001726846 | -0,090376368 | -0,160376368 | -1,420376368 |
| chrI | 158225 | 158386 | 161 | 0,085486818 | 0,062999783 | 0,575720754 | 0,190958009 | 0,162314307 | 0,216969523 | 0,097969523 | -1,162030477 |
| chrI | 158476 | 158631 | 155 | 0,265009134 | 0,307587175 | 0,462820193 | -0,093331276 | -0,079331585 | -0,129764872 | -0,206764872 | -1,466764872 |
| chrI | 158476 | 158631 | 155 | 0,265009134 | 0,307587175 | 0,462820193 | -0,093331276 | -0,079331585 | -0,129764872 | -0,206764872 | -1,466764872 |
| chrI | 159246 | 159353 | 107 | 0,48727486 | 4,006045014 | 0,108444285 | -1,234843997 | -1,049617397 | -0,32764883 | -0,06864883 | -1,32864883 |
| chrI | 161173 | 161287 | 114 | 0,068389454 | 0,389116306 | 0,149483263 | -1,038652184 | -0,882854356 | -0,39116562 | -0,18116562 | -1,44116562 |
| chrI | 163265 | 163422 | 157 | 0,059840772 | 0,059293913 | 0,502295129 | 0,005753068 | 0,004890107 | -0,009421984 | -0,100421984 | -1,360421984 |
| chrI | 164304 | 164458 | 154 | 0,04844253 | 0,048176305 | 0,50137771 | 0,003453413 | 0,002935401 | -0,076236959 | -0,146236959 | -1,406236959 |
| chrI | 164304 | 164458 | 154 | 0,04844253 | 0,048176305 | 0,50137771 | 0,003453413 | 0,002935401 | -0,076236959 | -0,146236959 | -1,406236959 |
| chrI | 164952 | 165092 | 140 | 0,159575393 | 0,122293696 | 0,566132999 | 0,166537466 | 0,141556846 | -0,007994795 | 0,020005205 | -1,239994795 |
| chrI | 164958 | 165092 | 134 | 0,136778908 | 0,137117174 | 0,499382491 | -0,001547865 | -0,001315685 | -0,033023256 | 0,036976744 | -1,223023256 |
| chrI | 165370 | 165525 | 155 | 0,102584181 | 0,081529131 | 0,557179598 | 0,143822286 | 0,122248943 | 0,089586931 | 0,012586931 | -1,247413069 |
| chrI | 165501 | 165609 | 108 | 0,045592969 | 0,155646522 | 0,226560746 | -0,750221201 | -0,637688021 | 0,033894185 | 0,285894185 | -0,974105815 |
| chrI | 165531 | 165649 | 118 | 1,704037229 | 4,873218496 | 0,259080276 | -0,646183457 | -0,549255938 | -0,08731138 | 0,09468862 | -1,16531138 |
| chrI | 165650 | 165799 | 149 | 0,085486818 | 0,044470435 | 0,657807209 | 0,406485945 | 0,345513053 | 0,110095556 | 0,075095556 | -1,184904444 |
| chrI | 170185 | 170316 | 131 | 0,367593315 | 0,333528262 | 0,524293257 | 0,060931847 | 0,05179207 | 0,177713203 | 0,268713203 | -0,991286797 |
| chrI | 170746 | 170892 | 146 | 0,296354301 | 0,251999131 | 0,540443961 | 0,101552255 | 0,086319417 | -0,145404324 | -0,159404324 | -1,419404324 |
| chrI | 173251 | 173422 | 171 | 0,170973635 | 0,159352392 | 0,517590565 | 0,044107306 | 0,03749121 | 0,201163341 | 0,012163341 | -1,247836659 |
| chrI | 174403 | 174559 | 156 | 1,020142689 | 0,837526525 | 0,54915196 | 0,123519062 | 0,104991202 | 0,075570974 | -0,008429026 | -1,268429026 |
| chrI | 177062 | 177157 | 95 | 0,723788388 | 3,016577837 | 0,193507359 | -0,865043858 | -0,735287279 | -0,008986621 | 0,334013379 | -0,925986621 |
| chrI | 178178 | 178314 | 136 | 0,265009134 | 0,218646305 | 0,547929606 | 0,120432196 | 0,102367367 | 0,046707401 | 0,102707401 | -1,157292599 |
| chrI | 181632 | 181781 | 149 | 0,119681545 | 0,144528914 | 0,452978074 | -0,118140734 | -0,100419624 | -0,3358092 | -0,3708092 | -1,6308092 |
| chrI | 184868 | 185010 | 142 | 1,048638295 | 0,685585872 | 0,604672865 | 0,265461103 | 0,225641937 | 0,065722461 | 0,079722461 | -1,180277539 |
| chrI | 188940 | 189093 | 153 | 0,085486818 | 0,051882174 | 0,622315244 | 0,311567137 | 0,264832067 | 0,11250467 | 0,04950467 | -1,21049533 |
| chrI | 189954 | 190061 | 107 | 1,894957788 | 4,335867407 | 0,304126296 | -0,512569358 | -0,435683954 | 0,261864949 | 0,520864949 | -0,739135051 |
| chrI | 189954 | 190062 | 108 | 0,04844253 | 0,100058479 | 0,326210107 | -0,450402539 | -0,382842158 | 0,269771927 | 0,521771927 | -0,738228073 |
| chrI | 189954 | 190063 | 109 | 0,045592969 | 0,107470218 | 0,297870247 | -0,530535801 | -0,450955431 | 0,164258858 | 0,409258858 | -0,850741142 |
| chrI | 189954 | 190077 | 123 | 1,85506394 | 5,666274586 | 0,246640134 | -0,685100876 | -0,582335745 | -0,172620828 | -0,025620828 | -1,285620828 |
| chrI | 191415 | 191573 | 158 | 0,062690333 | 0,085235 | 0,423797138 | -0,192188854 | -0,163360526 | -0,171273336 | -0,269273336 | -1,529273336 |
| chrI | 191722 | 191874 | 152 | 0,153876272 | 0,096352609 | 0,614942093 | 0,292223409 | 0,248389898 | 0,078993697 | 0,022993697 | -1,237006303 |
| chrI | 192889 | 193041 | 152 | 1,062886098 | 1,693582397 | 0,385597042 | -0,290813338 | -0,247191337 | -0,407783132 | -0,463783132 | -1,723783132 |
| chrI | 193467 | 193622 | 155 | 2,883755311 | 2,575579357 | 0,52822468 | 0,070807907 | 0,060186721 | 0,026701459 | -0,050298541 | -1,310298541 |
| chrI | 194126 | 194279 | 153 | 2,658640024 | 2,212404138 | 0,545804952 | 0,115069422 | 0,097809009 | -0,02265981 | -0,08565981 | -1,34565981 |
| chrI | 194141 | 194279 | 138 | 0,512920905 | 0,259410871 | 0,664119905 | 0,423733486 | 0,360173463 | 0,26050144 | 0,30250144 | -0,95749856 |
| chrI | 194290 | 194442 | 152 | 0,125380666 | 0,055588044 | 0,692830634 | 0,503889924 | 0,428306435 | 0,272098034 | 0,216098034 | -1,043901966 |
| chrI | 204527 | 204675 | 148 | 0,327699467 | 0,233469783 | 0,583958346 | 0,212030411 | 0,180225849 | -0,078287583 | -0,106287583 | -1,366287583 |
| chrI | 204662 | 204810 | 148 | 0,424584527 | 0,226058044 | 0,652561861 | 0,392246244 | 0,333409307 | 0,083481064 | 0,055481064 | -1,204518936 |
| chrI | 204797 | 204945 | 148 | 0,398938482 | 0,207528696 | 0,657807209 | 0,406485945 | 0,345513053 | 0,093736202 | 0,065736202 | -1,194263798 |
| chrI | 205040 | 205170 | 130 | 0,125380666 | 0,17417587 | 0,418554265 | -0,205593405 | -0,174754394 | 0,002376662 | 0,100376662 | -1,159623338 |
| chrI | 205573 | 205699 | 126 | 0,102584181 | 0,081529131 | 0,557179598 | 0,143822286 | 0,122248943 | 0,406093932 | 0,532093932 | -0,727906068 |
| chrI | 205573 | 205705 | 132 | 0,039893848 | 0,051882174 | 0,434687048 | -0,164453569 | -0,139785534 | -0,078685071 | 0,005314929 | -1,254685071 |
| chrI | 205573 | 205712 | 139 | 2,778321569 | 1,448995005 | 0,657230543 | 0,404916468 | 0,344178997 | 0,212715496 | 0,247715496 | -1,012284504 |
| chrI | 205580 | 205712 | 132 | 0,330549028 | 0,248293262 | 0,571051967 | 0,179053024 | 0,152195071 | 0,17803092 | 0,26203092 | -0,99796908 |
| chrI | 206390 | 206543 | 153 | 0,133929347 | 0,289057827 | 0,316627443 | -0,477150599 | -0,40557801 | -0,513820478 | -0,576820478 | -1,836820478 |
| chrI | 207406 | 207531 | 125 | 10,97365781 | 12,16636983 | 0,47422838 | -0,064644869 | -0,054948139 | 0,310469572 | 0,443469572 | -0,816530428 |
| chrI | 215442 | 215555 | 113 | 0,085486818 | 1,007996525 | 0,078178436 | -1,417431283 | -1,20481659 | -0,693345291 | -0,476345291 | -1,736345291 |
| chrI | 217878 | 218015 | 137 | 5,109262126 | 3,894868927 | 0,567435336 | 0,169848443 | 0,144371176 | 0,048411892 | 0,097411892 | -1,162588108 |
| chrI | 217916 | 218076 | 160 | 0,031345166 | 0,040764565 | 0,434687048 | -0,164453569 | -0,139785534 | -0,107307354 | -0,219307354 | -1,479307354 |
| chrI | 218711 | 218856 | 145 | 2,473418586 | 1,248878048 | 0,664487232 | 0,424740939 | 0,361029798 | 0,130349978 | 0,123349978 | -1,136650022 |
| chrI | 218738 | 218886 | 148 | 0,404637603 | 0,218646305 | 0,649202711 | 0,383168843 | 0,325693517 | 0,072440063 | 0,044440063 | -1,215559937 |
| chrI | 222387 | 222554 | 167 | 0,435982769 | 0,489174784 | 0,471252456 | -0,072121883 | -0,0613036 | 0,071391448 | -0,089608552 | -1,349608552 |
| chrI | 222751 | 222855 | 104 | 0,156725832 | 0,663350655 | 0,191111237 | -0,873808639 | -0,742737344 | -0,000770953 | 0,279229047 | -0,980770953 |
| chrI | 223257 | 223420 | 163 | 0,14817715 | 0,196411088 | 0,430012212 | -0,176343075 | -0,149891614 | -0,090544312 | -0,223544312 | -1,483544312 |
| chrI | 224065 | 224173 | 108 | 0,04844253 | 0,318704784 | 0,131943032 | -1,117253234 | -0,949665249 | -0,310042122 | -0,058042122 | -1,318042122 |
| chrI | 224877 | 224997 | 120 | 0,208017923 | 0,752291524 | 0,216615512 | -0,783674679 | -0,666123477 | -0,234308076 | -0,066308076 | -1,326308076 |
| chrI | 226540 | 226687 | 147 | 1,213912809 | 0,655938915 | 0,649202711 | 0,383168843 | 0,325693517 | 0,076075897 | 0,055075897 | -1,204924103 |
| chrI | 229861 | 230005 | 144 | 4,63908463 | 2,575579357 | 0,643007718 | 0,366509985 | 0,311533487 | 0,11349443 | 0,11349443 | -1,14650557 |
| chrI | 230119 | 230218 | 99 | 0,119681545 | 0,140823044 | 0,459422021 | -0,101889932 | -0,086606442 | 0,661749047 | 0,976749047 | -0,283250953 |
| chrI | 230124 | 230218 | 94 | 0,387540239 | 0,34835174 | 0,526626529 | 0,066792441 | 0,056773575 | 0,797592648 | 1,147592648 | -0,112407352 |
| chrII | 163 | 311 | 148 | 0,059840772 | 0,055588044 | 0,518421434 | 0,046192108 | 0,039263292 | -0,051618443 | -0,079618443 | -1,339618443 |
| chrII | 526 | 677 | 151 | 0,564212996 | 0,318704784 | 0,639032318 | 0,355873419 | 0,302492406 | 0,186299094 | 0,137299094 | -1,122700906 |
| chrII | 1089 | 1240 | 151 | 0,823523009 | 0,492880654 | 0,625585474 | 0,320183734 | 0,272156174 | 0,155357044 | 0,106357044 | -1,153642956 |
| chrII | 1727 | 1858 | 131 | 2,162816483 | 2,01969892 | 0,51710903 | 0,042899133 | 0,036464263 | 0,206368447 | 0,297368447 | -0,962631553 |
| chrII | 1727 | 1870 | 143 | 0,359044634 | 0,292763697 | 0,550843886 | 0,12779371 | 0,108624653 | 0,044015112 | 0,051015112 | -1,208984888 |
| chrII | 1727 | 1914 | 187 | 0,14532759 | 0,111176087 | 0,566571175 | 0,16765125 | 0,142503563 | 0,537148593 | 0,236148593 | -1,023851407 |
| chrII | 1931 | 2076 | 145 | 0,592708601 | 0,674468263 | 0,467739443 | -0,080953558 | -0,068810524 | -0,131920977 | -0,138920977 | -1,398920977 |
| chrII | 1936 | 2043 | 107 | 0,119681545 | 0,166764131 | 0,417815854 | -0,20748423 | -0,176361595 | 0,366147208 | 0,625147208 | -0,634852792 |
| chrII | 1937 | 2034 | 97 | 0,188070999 | 0,244587392 | 0,434687048 | -0,164453569 | -0,139785534 | 0,168480379 | 0,497480379 | -0,762519621 |
| chrII | 1937 | 2042 | 105 | 0,031345166 | 0,233469783 | 0,118366302 | -1,183193156 | -1,005714183 | -0,631967553 | -0,358967553 | -1,618967553 |
| chrII | 1937 | 2046 | 109 | 0,142478029 | 0,237175653 | 0,375284202 | -0,317889966 | -0,270206471 | 0,334933722 | 0,579933722 | -0,680066278 |
| chrII | 1937 | 2051 | 114 | 0,042743409 | 0,166764131 | 0,204018475 | -0,827353109 | -0,703250143 | -0,092580445 | 0,117419555 | -1,142580445 |
| chrII | 1937 | 2054 | 117 | 0,056991212 | 0,285351958 | 0,166473927 | -0,968193274 | -0,822964283 | -0,211598983 | -0,022598983 | -1,282598983 |
| chrII | 1937 | 2057 | 120 | 0,042743409 | 0,151940653 | 0,219552687 | -0,773704813 | -0,657649091 | -0,037549176 | 0,130450824 | -1,129549176 |
| chrII | 1937 | 2062 | 125 | 0,065539893 | 0,774526742 | 0,078017494 | -1,418533767 | -1,205753702 | -0,711064879 | -0,578064879 | -1,838064879 |
| chrII | 1937 | 2071 | 134 | 0,059840772 | 0,055588044 | 0,518421434 | 0,046192108 | 0,039263292 | 0,105040379 | 0,175040379 | -1,084959621 |
| chrII | 1937 | 2072 | 135 | 0,054141651 | 0,070411522 | 0,434687048 | -0,164453569 | -0,139785534 | -0,029959369 | 0,033040631 | -1,226959369 |
| chrII | 1937 | 2073 | 136 | 0,042743409 | 0,200116957 | 0,175999936 | -0,930717198 | -0,791109618 | -0,753126123 | -0,697126123 | -1,957126123 |
| chrII | 1937 | 2075 | 138 | 0,045592969 | 0,096352609 | 0,321200349 | -0,464344846 | -0,394693119 | -0,378758786 | -0,336758786 | -1,596758786 |
| chrII | 1937 | 2076 | 139 | 34,02375337 | 28,62413663 | 0,543094961 | 0,108233996 | 0,091998897 | 0,119744817 | 0,154744817 | -1,105255183 |

| Chrom | Start | End | Length | Section A | Section B | A/A+B | Z-score | Z * 0.85 | Phase correction | Length correction | ΔLK |
|---|---|---|---|---|---|---|---|---|---|---|---|
| chrII | 1937 | 2082 | 145 | 0,031345166 | 0,511410002 | 0,057751945 | -1,573928894 | -1,337839559 | -1,412580974 | -1,419580974 | -2,679580974 |
| chrII | 1937 | 2086 | 149 | 0,287805619 | 0,418763262 | 0,407328467 | -0,234422577 | -0,19925919 | -0,33674152 | -0,37174152 | -1,63174152 |
| chrII | 1938 | 2076 | 138 | 0,102584181 | 0,107470218 | 0,488369592 | -0,029157241 | -0,024783655 | -0,006450881 | 0,035549119 | -1,224450881 |
| chrII | 1939 | 2076 | 137 | 0,056991212 | 0,103764348 | 0,354520937 | -0,373143192 | -0,317171714 | -0,307387315 | -0,258387315 | -1,518387315 |
| chrII | 1940 | 2076 | 136 | 0,071239015 | 0,100058479 | 0,415878909 | -0,212447646 | -0,180580499 | -0,155600303 | -0,099600303 | -1,359600303 |
| chrII | 1941 | 2076 | 135 | 0,071239015 | 0,062999783 | 0,530688713 | 0,07700122 | 0,065451037 | 0,167857361 | 0,230857361 | -1,029142639 |
| chrII | 1942 | 2076 | 134 | 0,054141651 | 0,185293479 | 0,22612242 | -0,751677806 | -0,638926135 | -0,536837644 | -0,466837644 | -1,726837644 |
| chrII | 1944 | 2076 | 132 | 0,692443222 | 0,937585003 | 0,424804437 | -0,1896175 | -0,161174875 | -0,005981789 | 0,078018211 | -1,181981789 |
| chrII | 1947 | 2076 | 129 | 0,031345166 | 0,125999566 | 0,19921332 | -0,84443452 | -0,717769342 | -0,362897595 | -0,257897595 | -1,517897595 |
| chrII | 1948 | 2076 | 128 | 0,028495606 | 0,085235 | 0,250553539 | -0,67274886 | -0,571836531 | -0,207921231 | -0,095921231 | -1,355921231 |
| chrII | 1952 | 2076 | 124 | 0,042743409 | 0,08894087 | 0,324590066 | -0,454901459 | -0,386666241 | 0,137892569 | 0,277892569 | -0,982107431 |
| chrII | 1953 | 2076 | 123 | 0,076938136 | 0,081529131 | 0,485514374 | -0,036318061 | -0,030870352 | 0,528799631 | 0,675799631 | -0,584200369 |
| chrII | 1964 | 2076 | 112 | 0,09688506 | 0,233469783 | 0,293275736 | -0,543840157 | -0,462264134 | 0,138170855 | 0,362170855 | -0,897829145 |
| chrII | 1966 | 2076 | 110 | 0,102584181 | 0,166764131 | 0,380860679 | -0,303221116 | -0,257737948 | 0,358155933 | 0,596155933 | -0,663844067 |
| chrII | 2124 | 2286 | 162 | 0,037044288 | 0,048176305 | 0,434687048 | -0,164453569 | -0,139785534 | -0,103886698 | -0,229886698 | -1,489886698 |
| chrII | 2124 | 2286 | 162 | 0,037044288 | 0,048176305 | 0,434687048 | -0,164453569 | -0,139785534 | -0,103886698 | -0,229886698 | -1,489886698 |
| chrII | 2191 | 2286 | 95 | 0,039893848 | 0,062999783 | 0,387719315 | -0,285268256 | -0,242478018 | 0,053315126 | 0,396315126 | -0,863684874 |
| chrII | 2267 | 2374 | 107 | 0,071239015 | 0,133411305 | 0,348101165 | -0,390452017 | -0,331884215 | 0,215051727 | 0,474051727 | -0,785948273 |
| chrII | 2267 | 2414 | 147 | 0,079787696 | 0,114881957 | 0,409862015 | -0,227899939 | -0,193714948 | -0,278119201 | -0,299119201 | -1,559119201 |
| chrII | 2522 | 2620 | 98 | 0,031345166 | 0,037058696 | 0,458236793 | -0,104876779 | -0,089145262 | 0,218747595 | 0,540747595 | -0,719252405 |
| chrII | 2522 | 2673 | 151 | 0,795027403 | 0,415057393 | 0,657001398 | 0,404293094 | 0,34364913 | 0,232079266 | 0,183079266 | -1,076920734 |
| chrII | 3354 | 3498 | 144 | 0,128230226 | 0,059293913 | 0,683806504 | 0,478369846 | 0,406614369 | 0,307662419 | 0,307662419 | -0,952337581 |
| chrII | 3419 | 3571 | 152 | 0,116831984 | 0,085235 | 0,578184429 | 0,197250975 | 0,167663329 | 0,065281468 | 0,009281468 | -1,250718532 |
| chrII | 3701 | 3844 | 143 | 0,373292436 | 0,448410219 | 0,454291384 | -0,114826344 | -0,097602392 | -0,171193058 | -0,164193058 | -1,424193058 |
| chrII | 3992 | 4129 | 137 | 0,444531451 | 0,474351306 | 0,483773852 | -0,040684142 | -0,03458152 | -0,043485891 | 0,005514109 | -1,254485891 |
| chrII | 7980 | 8135 | 155 | 0,159575393 | 0,074117392 | 0,682842618 | 0,475662609 | 0,404313218 | 0,319117223 | 0,242117223 | -1,017882777 |
| chrII | 10712 | 10874 | 162 | 0,065539893 | 0,103764348 | 0,387113121 | -0,28685122 | -0,243823537 | -0,207400301 | -0,333400301 | -1,593400301 |
| chrII | 20981 | 21152 | 171 | 0,153876272 | 0,452116089 | 0,253924441 | -0,662190904 | -0,562862269 | -0,266983406 | -0,455983406 | -1,715983406 |
| chrII | 21442 | 21596 | 154 | 0,091185939 | 0,118587827 | 0,434687048 | -0,164453569 | -0,139785534 | -0,206203893 | -0,276203893 | -1,536203893 |
| chrII | 24248 | 24399 | 151 | 0,267858695 | 0,129705435 | 0,673749653 | 0,450290909 | 0,382747273 | 0,277943929 | 0,228943929 | -1,031056071 |
| chrII | 24299 | 24437 | 138 | 0,703841464 | 0,659644785 | 0,516207233 | 0,04063669 | 0,034541187 | 0,059825685 | 0,101825685 | -1,158174315 |
| chrII | 25280 | 25416 | 136 | 0,185221438 | 0,170470001 | 0,520736284 | 0,052001583 | 0,044201345 | 0,060620037 | 0,116620037 | -1,143379963 |
| chrII | 26126 | 26261 | 135 | 0,498673102 | 0,459527828 | 0,520426444 | 0,051223893 | 0,043540309 | 0,142174601 | 0,205174601 | -1,054825399 |
| chrII | 30232 | 30334 | 102 | 0,139628469 | 0,140823044 | 0,497870264 | -0,005338482 | -0,00453771 | 0,314168167 | 0,608168167 | -0,651831833 |
| chrII | 31655 | 31803 | 148 | 0,09688506 | 0,055588044 | 0,635423938 | 0,346253615 | 0,294315573 | 0,195989109 | 0,167989109 | -1,092010891 |
| chrII | 32970 | 33121 | 151 | 0,270708255 | 0,148234783 | 0,646169599 | 0,37499955 | 0,318749618 | 0,205206515 | 0,156206515 | -1,103793485 |
| chrII | 35988 | 36136 | 148 | 0,162424953 | 0,122293696 | 0,570475287 | 0,177584338 | 0,150946687 | 0,043798437 | 0,015798437 | -1,244201563 |
| chrII | 48768 | 48912 | 144 | 0,222265726 | 0,17788174 | 0,555459536 | 0,139467258 | 0,118547169 | 0,020446528 | 0,020446528 | -1,239553472 |
| chrII | 54394 | 54539 | 145 | 0,151026711 | 0,077823261 | 0,659937642 | 0,412292949 | 0,350449007 | 0,280783299 | 0,273783299 | -0,986216701 |
| chrII | 54394 | 54539 | 145 | 0,151026711 | 0,077823261 | 0,659937642 | 0,412292949 | 0,350449007 | 0,280783299 | 0,273783299 | -0,986216701 |
| chrII | 54551 | 54716 | 165 | 0,239363089 | 0,185293479 | 0,56366275 | 0,160262244 | 0,136222907 | 0,267326947 | 0,120326947 | -1,139673053 |
| chrII | 54551 | 54716 | 165 | 0,239363089 | 0,185293479 | 0,56366275 | 0,160262244 | 0,136222907 | 0,267326947 | 0,120326947 | -1,139673053 |
| chrII | 62389 | 62546 | 157 | 0,370442876 | 0,374292827 | 0,497415223 | -0,006479121 | -0,005507253 | -0,031861839 | -0,122861839 | -1,382861839 |
| chrII | 62885 | 63033 | 148 | 0,159575393 | 0,125999566 | 0,558786364 | 0,147892925 | 0,125708986 | 0,024882618 | -0,003117382 | -1,263117382 |
| chrII | 64104 | 64264 | 160 | 0,056991212 | 0,051882174 | 0,523463207 | 0,058847485 | 0,050020363 | 0,053194984 | -0,058805016 | -1,318805016 |
| chrII | 64653 | 64794 | 141 | 0,105433742 | 0,133411305 | 0,441431561 | -0,147340685 | -0,125239582 | -0,159025619 | -0,138025619 | -1,398025619 |
| chrII | 65899 | 66050 | 151 | 0,190920559 | 0,218646305 | 0,466152357 | -0,084945506 | -0,07220368 | -0,186715774 | -0,235715774 | -1,495715774 |
| chrII | 67731 | 67881 | 150 | 0,370442876 | 0,326116523 | 0,531818071 | 0,079840821 | 0,067864697 | -0,07433213 | -0,11633213 | -1,37633213 |
| chrII | 69118 | 69263 | 145 | 1,057186976 | 0,752291524 | 0,584249537 | 0,212776972 | 0,180860426 | 0,107846031 | 0,100846031 | -1,159153969 |
| chrII | 79393 | 79540 | 147 | 0,074088575 | 0,144528914 | 0,338895921 | -0,415478239 | -0,353156503 | -0,438609071 | -0,459609071 | -1,719609071 |
| chrII | 81527 | 81678 | 151 | 0,256460453 | 0,233469783 | 0,523463207 | 0,058847485 | 0,050020363 | -0,065821732 | -0,114821732 | -1,374821732 |
| chrII | 81527 | 81678 | 151 | 0,256460453 | 0,233469783 | 0,523463207 | 0,058847485 | 0,050020363 | -0,065821732 | -0,114821732 | -1,374821732 |
| chrII | 86475 | 86605 | 130 | 1,094231264 | 1,382289353 | 0,441842178 | -0,146300265 | -0,124355225 | 0,146709204 | 0,244709204 | -1,015290796 |
| chrII | 87052 | 87211 | 159 | 0,054141651 | 0,085235 | 0,388455674 | -0,283346352 | -0,2408444 | -0,248149953 | -0,353149953 | -1,613149953 |
| chrII | 94866 | 95023 | 157 | 0,076938136 | 0,096352609 | 0,443982948 | -0,140878539 | -0,119746758 | -0,145880406 | -0,236880406 | -1,496880406 |
| chrII | 97063 | 97217 | 154 | 0,823523009 | 0,655938915 | 0,556636839 | 0,142447782 | 0,121080615 | 0,056095479 | -0,013904521 | -1,273904521 |
| chrII | 97108 | 97247 | 139 | 0,122531105 | 0,070411522 | 0,635064977 | 0,345298405 | 0,293503645 | 0,312172713 | 0,347172713 | -0,912827287 |
| chrII | 98068 | 98229 | 161 | 0,14532759 | 0,17788174 | 0,449639217 | -0,126572917 | -0,10758698 | -0,084100968 | -0,203100968 | -1,463100968 |
| chrII | 98405 | 98563 | 158 | 0,644000692 | 0,689291742 | 0,483015335 | -0,04258711 | -0,036199044 | -0,042718636 | -0,140718636 | -1,400718636 |
| chrII | 102188 | 102336 | 148 | 0,04844253 | 0,059293913 | 0,449639217 | -0,126572917 | -0,10758698 | -0,19655441 | -0,22455441 | -1,48455441 |
| chrII | 105450 | 105577 | 127 | 1,262355338 | 2,308756747 | 0,35349082 | -0,375912914 | -0,319525977 | 0,072226215 | 0,191226215 | -1,068773785 |
| chrII | 105532 | 105684 | 152 | 0,133929347 | 0,140823044 | 0,487454711 | -0,031451562 | -0,026733828 | -0,12757567 | -0,18357567 | -1,44357567 |
| chrII | 105532 | 105684 | 152 | 0,133929347 | 0,140823044 | 0,487454711 | -0,031451562 | -0,026733828 | -0,12757567 | -0,18357567 | -1,44357567 |
| chrII | 105754 | 105910 | 156 | 0,125380666 | 0,040764565 | 0,754644987 | 0,68917996 | 0,585802966 | 0,528953837 | 0,444953837 | -0,815046163 |
| chrII | 106088 | 106233 | 145 | 0,444531451 | 0,292763697 | 0,602921981 | 0,260917618 | 0,221779975 | 0,155157764 | 0,148157764 | -1,111842236 |
| chrII | 112631 | 112737 | 106 | 1,820869213 | 5,54027502 | 0,247362252 | -0,682813808 | -0,580391737 | -0,144361425 | 0,121638575 | -1,138361425 |
| chrII | 112631 | 112749 | 118 | 1,877860425 | 7,263504374 | 0,205424514 | -0,82240044 | -0,699040374 | -0,087103148 | 0,094896852 | -1,165103148 |
| chrII | 117607 | 117737 | 130 | 1,225311051 | 1,211819352 | 0,502767948 | 0,006938271 | 0,005897531 | 0,265325358 | 0,363325358 | -0,896674642 |
| chrII | 118185 | 118336 | 151 | 0,230814407 | 0,196411088 | 0,540263655 | 0,101097969 | 0,085933273 | -0,02133991 | -0,07033991 | -1,33033991 |
| chrII | 118430 | 118588 | 158 | 0,102584181 | 0,070411522 | 0,592986873 | 0,235235114 | 0,199949847 | 0,162333021 | 0,064333021 | -1,195666979 |
| chrII | 118430 | 118588 | 158 | 0,102584181 | 0,070411522 | 0,592986873 | 0,235235114 | 0,199949847 | 0,162333021 | 0,064333021 | -1,195666979 |
| chrII | 118604 | 118737 | 133 | 11,30990596 | 7,600738506 | 0,59807089 | 0,248356974 | 0,211103428 | 0,317086836 | 0,394086836 | -0,865913164 |
| chrII | 118662 | 118820 | 158 | 1,250957096 | 1,445289136 | 0,463962483 | -0,090455864 | -0,076887484 | -0,10819699 | -0,20619699 | -1,46619699 |
| chrII | 120690 | 120830 | 140 | 0,256460453 | 0,207528696 | 0,552729419 | 0,13256026 | 0,112676221 | 0,12126134 | 0,14926134 | -1,11073866 |
| chrII | 130330 | 130481 | 151 | 2,789719811 | 2,471815009 | 0,530210273 | 0,075798443 | 0,064428677 | -0,045184127 | -0,094184127 | -1,354184127 |
| chrII | 133172 | 133320 | 148 | 0,054141651 | 0,081529131 | 0,399066404 | -0,255764342 | -0,217399691 | -0,320611205 | -0,348611205 | -1,608611205 |
| chrII | 134044 | 134208 | 164 | 0,606956404 | 0,629997828 | 0,490686226 | -0,02334829 | -0,019846047 | 0,056070417 | -0,083929583 | -1,343929583 |
| chrII | 134044 | 134208 | 164 | 0,606956404 | 0,629997828 | 0,490686226 | -0,02334829 | -0,019846047 | 0,056070417 | -0,083929583 | -1,343929583 |
| chrII | 146223 | 146376 | 153 | 0,108283302 | 0,096352609 | 0,529151025 | 0,07313593 | 0,062165541 | 0,001766017 | -0,061233983 | -1,321233983 |
| chrII | 147729 | 147892 | 163 | 0,159575393 | 0,289057827 | 0,355692324 | -0,369997104 | -0,314497538 | -0,25596304 | -0,38896304 | -1,64896304 |
| chrII | 157306 | 157457 | 151 | 0,122531105 | 0,107470218 | 0,532740871 | 0,082161538 | 0,069837308 | -0,03377469 | -0,08277469 | -1,34277469 |
| chrII | 158794 | 158975 | 181 | 0,353345512 | 0,70411522 | 0,33414528 | -0,428495293 | -0,364220999 | 0,029190346 | -0,229809654 | -1,489809654 |
| chrII | 159579 | 159667 | 88 | 0,039893848 | 0,044470435 | 0,472876041 | -0,06804215 | -0,057835827 | 0,195231863 | 0,587231863 | -0,672768137 |
| chrII | 159579 | 159734 | 155 | 4,015030862 | 3,965280449 | 0,503117072 | 0,007813421 | 0,006641408 | -0,07816889 | -0,15516889 | -1,41516889 |
| chrII | 161150 | 161286 | 136 | 0,729487509 | 0,863467612 | 0,457946052 | -0,105609604 | -0,089768163 | -0,073479554 | -0,017479554 | -1,277479554 |
| chrII | 161650 | 161786 | 136 | 2,718480797 | 2,664520227 | 0,505012127 | 0,01256387 | 0,010679289 | 0,02354348 | 0,07954348 | -1,18045652 |
| chrII | 164812 | 164963 | 151 | 0,153876272 | 0,111176087 | 0,58055047 | 0,203301913 | 0,172806626 | 0,071079698 | 0,022079698 | -1,237920302 |
| chrII | 169155 | 169305 | 150 | 0,695292782 | 0,537351089 | 0,564066231 | 0,161286776 | 0,137093759 | -0,00391113 | -0,04591113 | -1,30591113 |
| chrII | 171571 | 171720 | 149 | 0,806425645 | 0,489174784 | 0,622433913 | 0,311879406 | 0,265097495 | 0,133053285 | 0,098053285 | -1,161946715 |
| chrII | 171661 | 171810 | 149 | 0,63260245 | 0,444704349 | 0,58720733 | 0,220367006 | 0,187311955 | 0,054779123 | 0,019779123 | -1,240220877 |
| chrII | 171661 | 171810 | 149 | 0,63260245 | 0,444704349 | 0,58720733 | 0,220367006 | 0,187311955 | 0,054779123 | 0,019779123 | -1,240220877 |
| chrII | 174305 | 174447 | 142 | 0,273557816 | 0,203822827 | 0,573039188 | 0,18411706 | 0,156499501 | 0,154470237 | 0,168470237 | -1,091529763 |
| chrII | 186048 | 186192 | 144 | 0,558513874 | 0,326116523 | 0,631352796 | 0,335438395 | 0,285122636 | 0,197541935 | 0,197541935 | -1,062458065 |
| chrII | 187191 | 187351 | 160 | 0,108283302 | 0,17788174 | 0,378394585 | -0,309699913 | -0,263244926 | -0,255421457 | -0,367421457 | -1,627421457 |
| chrII | 188175 | 188332 | 157 | 0,424584527 | 0,392822175 | 0,519428732 | 0,048719876 | 0,041411895 | 0,007410682 | -0,083589318 | -1,343589318 |
| chrII | 189372 | 189539 | 167 | 0,131079787 | 0,218646305 | 0,374806999 | -0,319148382 | -0,271276125 | -0,099501284 | -0,260501284 | -1,520501284 |
| chrII | 192490 | 192638 | 148 | 1,894957788 | 1,14140783 | 0,624087487 | 0,316233837 | 0,268798762 | 0,159912897 | 0,131912897 | -1,128087103 |
| chrII | 195265 | 195410 | 145 | 0,957452356 | 0,70782109 | 0,574952035 | 0,188996027 | 0,160646623 | 0,095612534 | 0,088612534 | -1,171387466 |
| chrII | 203726 | 203877 | 151 | 1,470373261 | 1,082113917 | 0,576055102 | 0,191811584 | 0,163039847 | 0,058560082 | 0,009560082 | -1,250439918 |
| chrII | 204265 | 204408 | 143 | 0,376141997 | 0,589233263 | 0,389632936 | -0,280275857 | -0,238234478 | -0,318212415 | -0,311212415 | -1,571212415 |
| chrII | 213654 | 213809 | 155 | 0,262159574 | 0,059293913 | 0,815544346 | 0,898514518 | 0,763737341 | 0,678396211 | 0,601396211 | -0,658603789 |

TABLE 2

| Chrom | Start | End | Length | Section A | Section B | A/A+B | Z-score | Z * 0.85 | Phase correction | Length correction | ΔLK |
|---|---|---|---|---|---|---|---|---|---|---|---|
| chrII | 214589 | 214739 | 150 | 0,222265726 | 0,240881523 | 0,479902939 | -0,050397187 | -0,042837609 | -0,187830875 | -0,229830875 | -1,489830875 |
| chrII | 214869 | 215017 | 148 | 0,210867483 | 0,137117174 | 0,605967759 | 0,268824834 | 0,228501109 | 0,111380536 | 0,083380536 | -1,176619464 |
| chrII | 215320 | 215464 | 144 | 1,476072382 | 0,937585003 | 0,611550086 | 0,283361383 | 0,240857176 | 0,15608066 | 0,15608066 | -1,10391934 |
| chrII | 216639 | 216771 | 132 | 0,04844253 | 0,185293479 | 0,207253175 | -0,815989135 | -0,693590765 | -0,547397887 | -0,463397887 | -1,723397887 |
| chrII | 217546 | 217694 | 148 | 1,307948308 | 0,767115003 | 0,630317302 | 0,332693846 | 0,282789769 | 0,16136999 | 0,13336999 | -1,12663001 |
| chrII | 222615 | 222772 | 157 | 2,299595391 | 1,441583266 | 0,614671365 | 0,291515265 | 0,247787976 | 0,208974001 | 0,117974001 | -1,142025999 |
| chrII | 222619 | 222772 | 153 | 0,059840772 | 0,066705652 | 0,472876041 | -0,06804215 | -0,057835827 | -0,123041905 | -0,186041905 | -1,446041905 |
| chrII | 223146 | 223281 | 135 | 0,062690333 | 0,077823261 | 0,446151373 | -0,135390993 | -0,115082344 | -0,017836418 | 0,045163582 | -1,214836418 |
| chrII | 223772 | 223929 | 157 | 0,062690333 | 0,074117392 | 0,458236793 | -0,104876779 | -0,089145262 | -0,132106003 | -0,223106003 | -1,483106003 |
| chrII | 223872 | 224016 | 144 | 0,054141651 | 0,037058696 | 0,593656197 | 0,236960278 | 0,201416237 | 0,1242995 | 0,1242995 | -1,1357005 |
| chrII | 223872 | 224016 | 144 | 0,054141651 | 0,037058696 | 0,593656197 | 0,236960278 | 0,201416237 | 0,1242995 | 0,1242995 | -1,1357005 |
| chrII | 224435 | 224582 | 147 | 0,247911771 | 0,192705218 | 0,562646873 | 0,157683434 | 0,134030919 | 0,048101653 | 0,027101653 | -1,232898347 |
| chrII | 224435 | 224582 | 147 | 0,247911771 | 0,192705218 | 0,562646873 | 0,157683434 | 0,134030919 | 0,048101653 | 0,027101653 | -1,232898347 |
| chrII | 225175 | 225325 | 150 | 0,478726178 | 0,43729261 | 0,522616112 | 0,056720586 | 0,048212498 | -0,108601247 | -0,150601247 | -1,410601247 |
| chrII | 226374 | 226481 | 107 | 0,056991212 | 0,585527393 | 0,088699706 | -1,348805608 | -1,146484767 | -0,595265069 | -0,336265069 | -1,596265069 |
| chrII | 226832 | 226946 | 114 | 0,091185939 | 0,566998046 | 0,138541716 | -1,086894512 | -0,923860335 | -0,299868328 | -0,089868328 | -1,349868328 |
| chrII | 228528 | 228693 | 165 | 0,062690333 | 0,155646522 | 0,287126664 | -0,56179848 | -0,477528708 | -0,347544703 | -0,494544703 | -1,754544703 |
| chrII | 233542 | 233703 | 161 | 0,122531105 | 0,159352392 | 0,434687048 | -0,164453569 | -0,139785534 | -0,109641967 | -0,228641967 | -1,488641967 |
| chrII | 234707 | 234798 | 91 | 2,105825271 | 9,772378079 | 0,177284831 | -0,925762034 | -0,786897729 | -0,510796163 | -0,139796163 | -1,399796163 |
| chrII | 239108 | 239251 | 143 | 0,638301571 | 0,459527828 | 0,581421459 | 0,205531254 | 0,174701566 | 0,092067678 | 0,099067678 | -1,160932322 |
| chrII | 249586 | 249752 | 166 | 0,119681545 | 0,170470001 | 0,412479432 | -0,221171546 | -0,187995814 | -0,046240793 | -0,200240793 | -1,460240793 |
| chrII | 252919 | 253070 | 151 | 0,515770466 | 0,448410219 | 0,53493134 | 0,087672069 | 0,074521258 | -0,029288891 | -0,078288891 | -1,338288891 |
| chrII | 253969 | 254127 | 158 | 0,262159574 | 0,340940001 | 0,434687048 | -0,164453569 | -0,139785534 | -0,159313475 | -0,257313475 | -1,517313475 |
| chrII | 255566 | 255720 | 154 | 0,34194727 | 0,35205761 | 0,492715945 | -0,018259434 | -0,015520519 | -0,100021544 | -0,170021544 | -1,430021544 |
| chrII | 260109 | 260232 | 123 | 0,088336378 | 0,226058044 | 0,280973108 | -0,579953143 | -0,492960172 | 0,059807211 | 0,206807211 | -1,053192789 |
| chrII | 260209 | 260364 | 155 | 0,094035499 | 0,111176087 | 0,458236793 | -0,104876779 | -0,089145262 | -0,172131457 | -0,249131457 | -1,509131457 |
| chrII | 260642 | 260776 | 134 | 0,142478029 | 0,055588044 | 0,719345959 | 0,58089966 | 0,493764711 | 0,603929933 | 0,673929933 | -0,586070067 |
| chrII | 260642 | 260782 | 140 | 0,176672756 | 0,107470218 | 0,62177415 | 0,310143677 | 0,263622125 | 0,268242464 | 0,296242464 | -0,963757536 |
| chrII | 261685 | 261820 | 135 | 0,059840772 | 0,048176305 | 0,553993628 | 0,135757811 | 0,11539414 | 0,201273103 | 0,264273103 | -0,995726897 |
| chrII | 262132 | 262282 | 150 | 1,060036537 | 0,552174567 | 0,657504798 | 0,405662767 | 0,344813352 | 0,19221466 | 0,15021466 | -1,10978534 |
| chrII | 262311 | 262468 | 157 | 0,04844253 | 0,055588044 | 0,465656665 | -0,08619258 | -0,073263693 | -0,111281745 | -0,202281745 | -1,462281745 |
| chrII | 262974 | 263121 | 147 | 0,222265726 | 0,125999566 | 0,638208088 | 0,35367318 | 0,300622203 | 0,222279176 | 0,201279176 | -1,058720824 |
| chrII | 263717 | 263867 | 150 | 0,492973981 | 0,429880871 | 0,534183658 | 0,085790846 | 0,072922219 | -0,082266737 | -0,124266737 | -1,384266737 |
| chrII | 264916 | 265023 | 107 | 0,088336378 | 0,678174133 | 0,115244836 | -1,199098437 | -1,019233671 | -0,457520689 | -0,198520689 | -1,458520689 |
| chrII | 269730 | 269883 | 153 | 3,801313818 | 2,175345443 | 0,636026525 | 0,347857838 | 0,295679162 | 0,221615763 | 0,158615763 | -1,101384237 |
| chrII | 269730 | 269885 | 155 | 0,105433742 | 0,52623348 | 0,166913428 | -0,966434402 | -0,821469241 | -0,896425246 | -0,973425246 | -2,233425246 |
| chrII | 269781 | 269930 | 149 | 0,14817715 | 0,144528914 | 0,506231912 | 0,015621722 | 0,013278464 | -0,115364781 | -0,150364781 | -1,410364781 |
| chrII | 272917 | 273071 | 154 | 0,065539893 | 0,059293913 | 0,525017182 | 0,062749932 | 0,053337442 | -0,051071069 | -0,121071069 | -1,381071069 |
| chrII | 279942 | 280075 | 133 | 1,22246149 | 1,178466526 | 0,509162075 | 0,022967935 | 0,019522745 | 0,106456886 | 0,183456886 | -1,076543114 |
| chrII | 282915 | 283035 | 120 | 0,786478721 | 2,501461966 | 0,239201006 | -0,708875061 | -0,602543802 | -0,00349185 | 0,16450815 | -1,09549185 |
| chrII | 288899 | 289053 | 154 | 1,139824233 | 0,922761525 | 0,552619075 | 0,132281233 | 0,112439048 | -0,000754737 | -0,070754737 | -1,330754737 |
| chrII | 289452 | 289559 | 107 | 0,119681545 | 0,559586306 | 0,176191975 | -0,92997515 | -0,790478878 | -0,215533893 | 0,043466107 | -1,216533893 |
| chrII | 289452 | 289573 | 121 | 0,142478029 | 0,648527176 | 0,180122745 | -0,914897409 | -0,777662797 | -0,186147454 | -0,025147454 | -1,285147454 |
| chrII | 292636 | 292792 | 156 | 1,285151823 | 1,137701961 | 0,530428964 | 0,07634821 | 0,064895979 | 0,011153476 | -0,072846524 | -1,332846524 |
| chrII | 295986 | 296116 | 130 | 0,116831984 | 0,111176087 | 0,512402843 | 0,031094327 | 0,026430178 | 0,301059399 | 0,399059399 | -0,860940601 |
| chrII | 296940 | 297047 | 107 | 0,424584527 | 2,660814357 | 0,137610903 | -1,091116084 | -0,927448672 | -0,334508013 | -0,075508013 | -1,335508013 |
| chrII | 296940 | 297056 | 116 | 0,435982769 | 3,531693708 | 0,109883649 | -1,227147126 | -1,043075057 | -0,434188266 | -0,238188266 | -1,498188266 |
| chrII | 298361 | 298510 | 149 | 0,361894194 | 0,233469783 | 0,607853696 | 0,273729366 | 0,232669961 | 0,101731233 | 0,066731233 | -1,193268767 |
| chrII | 298515 | 298656 | 141 | 2,194161649 | 1,460112614 | 0,600437047 | 0,254478509 | 0,216306733 | 0,179726446 | 0,200726446 | -1,059273554 |
| chrII | 300341 | 300475 | 134 | 1,689789426 | 1,241466309 | 0,576472877 | 0,192878343 | 0,163946591 | 0,275637106 | 0,345637106 | -0,914362894 |
| chrII | 303103 | 303264 | 161 | 0,367593315 | 0,448410219 | 0,450480053 | -0,124448587 | -0,105781299 | -0,082900746 | -0,201900746 | -1,461900746 |
| chrII | 304167 | 304319 | 152 | 0,168124074 | 0,125999566 | 0,571610206 | 0,180475116 | 0,153403848 | 0,048789963 | -0,007210037 | -1,267210037 |
| chrII | 306192 | 306322 | 130 | 0,225115286 | 0,452116089 | 0,332405282 | -0,433281104 | -0,368288938 | -0,118327839 | -0,020327839 | -1,280327839 |
| chrII | 310844 | 311002 | 158 | 0,185221438 | 0,185293479 | 0,499902783 | -0,000243688 | -0,000207135 | -0,010252604 | -0,108252604 | -1,368252604 |
| chrII | 314561 | 314691 | 130 | 0,350495952 | 0,381704567 | 0,478688478 | -0,053445496 | -0,045428672 | 0,199479088 | 0,297479088 | -0,962520912 |
| chrII | 316179 | 316338 | 159 | 0,059840772 | 0,092646739 | 0,392430643 | -0,272989502 | -0,232041076 | -0,239946668 | -0,344946668 | -1,604946668 |
| chrII | 320829 | 320984 | 155 | 0,963151477 | 0,733762177 | 0,567590151 | 0,170242159 | 0,144705835 | 0,07510426 | -0,00189574 | -1,26189574 |
| chrII | 324261 | 324383 | 122 | 0,188070999 | 0,422469132 | 0,308040355 | -0,501412691 | -0,426200787 | 0,130540583 | 0,284540583 | -0,975459417 |
| chrII | 324593 | 324738 | 145 | 0,216566604 | 0,200116957 | 0,519738776 | 0,049497979 | 0,042073282 | -0,02086413 | -0,02786413 | -1,28786413 |
| chrII | 333410 | 333578 | 168 | 0,367593315 | 0,581821524 | 0,387178818 | -0,28667963 | -0,243677685 | 0,019902012 | -0,148097988 | -1,408097988 |
| chrII | 334171 | 334335 | 164 | 0,170973635 | 0,314998914 | 0,351817475 | -0,380418277 | -0,323355536 | -0,239948638 | -0,379948638 | -1,639948638 |
| chrII | 341568 | 341673 | 105 | 0,085486818 | 0,133411305 | 0,390532439 | -0,277931585 | -0,236241848 | 0,152424809 | 0,425424809 | -0,834575191 |
| chrII | 344002 | 344150 | 148 | 3,536304684 | 2,334697835 | 0,602334043 | 0,259393151 | 0,220484178 | 0,086011167 | 0,058011167 | -1,201988833 |
| chrII | 345574 | 345727 | 153 | 0,071239015 | 0,059293913 | 0,545755127 | 0,114943701 | 0,097702146 | 0,025621043 | -0,037378957 | -1,297378957 |
| chrII | 348428 | 348599 | 171 | 0,265009134 | 0,259410871 | 0,505337576 | 0,013379719 | 0,011372761 | 0,31746519 | 0,12846519 | -1,13153481 |
| chrII | 348488 | 348624 | 136 | 0,549965193 | 0,485468915 | 0,531144559 | 0,078147301 | 0,066425206 | 0,071222097 | 0,127222097 | -1,132777903 |
| chrII | 352601 | 352731 | 130 | 0,170973635 | 0,214940436 | 0,443035505 | -0,14327752 | -0,121785892 | 0,106689233 | 0,204689233 | -1,055310767 |
| chrII | 354955 | 355107 | 152 | 0,464478375 | 0,340940001 | 0,576692051 | 0,193438075 | 0,164422364 | 0,061765223 | 0,005765223 | -1,254234777 |
| chrII | 360789 | 360938 | 149 | 0,037044288 | 0,040764565 | 0,47609348 | -0,059960669 | -0,050966568 | -0,179616729 | -0,214616729 | -1,474616729 |
| chrII | 369124 | 369272 | 148 | 0,205168362 | 0,203822827 | 0,501644944 | 0,004123275 | 0,003504784 | -0,141493757 | -0,169493757 | -1,429493757 |
| chrII | 369614 | 369763 | 149 | 0,176672756 | 0,08894087 | 0,665149446 | 0,426558255 | 0,362574517 | 0,231627205 | 0,196627205 | -1,063372795 |
| chrII | 372782 | 372926 | 144 | 0,7807796 | 0,43729261 | 0,640996152 | 0,361122739 | 0,306954328 | 0,245257057 | 0,245257057 | -1,014742943 |
| chrII | 379733 | 379878 | 145 | 0,19661968 | 0,129705435 | 0,602526961 | 0,259893303 | 0,220909308 | 0,154047949 | 0,147047949 | -1,112952051 |
| chrII | 380438 | 380593 | 155 | 1,775276244 | 0,948702612 | 0,651721741 | 0,389972994 | 0,331477045 | 0,259634452 | 0,182634452 | -1,077365548 |
| chrII | 380438 | 380593 | 155 | 1,775276244 | 0,948702612 | 0,651721741 | 0,389972994 | 0,331477045 | 0,259634452 | 0,182634452 | -1,077365548 |
| chrII | 383232 | 383381 | 149 | 0,914708947 | 0,400233914 | 0,695626383 | 0,511862517 | 0,435083139 | 0,298313505 | 0,263313505 | -0,996686495 |
| chrII | 384056 | 384165 | 109 | 0,219416165 | 2,63487327 | 0,07687243 | -1,426427917 | -1,21246373 | -0,608780409 | -0,363780409 | -1,623780409 |
| chrII | 384888 | 385039 | 151 | 0,501522663 | 0,415057393 | 0,547167331 | 0,118507771 | 0,100731606 | -0,010140026 | -0,059140026 | -1,319140026 |
| chrII | 385956 | 386120 | 164 | 0,364743755 | 0,300175436 | 0,548553508 | 0,12200762 | 0,103706477 | 0,188306369 | 0,048306369 | -1,211693631 |
| chrII | 386724 | 386859 | 135 | 5,587988304 | 4,787983496 | 0,538550838 | 0,096783504 | 0,082265978 | 0,143856833 | 0,206856833 | -1,053143167 |
| chrII | 402057 | 402202 | 145 | 0,267858695 | 0,237175653 | 0,530377183 | 0,076218036 | 0,064785331 | -0,009781246 | -0,016781246 | -1,276781246 |
| chrII | 405294 | 405447 | 153 | 0,250761331 | 0,155646522 | 0,617018911 | 0,297660652 | 0,253011555 | 0,163447736 | 0,100447736 | -1,159552264 |
| chrII | 411086 | 411218 | 132 | 0,689593661 | 0,507704132 | 0,57595835 | 0,191564566 | 0,162829881 | 0,298959502 | 0,382959502 | -0,877040498 |
| chrII | 412922 | 413067 | 145 | 0,367593315 | 0,314998914 | 0,538525491 | 0,09671967 | 0,082211719 | 0,000707699 | -0,006292301 | -1,266292301 |
| chrII | 417276 | 417442 | 166 | 0,037044288 | 0,037058696 | 0,499902783 | -0,000243688 | -0,000207135 | 0,1552076 | 0,0012076 | -1,2587924 |
| chrII | 418844 | 418980 | 136 | 0,034194727 | 1,041349352 | 0,031792957 | -1,855072217 | -1,576811384 | -1,564708743 | -1,508708743 | -2,768708743 |
| chrII | 420492 | 420630 | 138 | 2,826764099 | 2,438462183 | 0,536874191 | 0,092561893 | 0,078677609 | 0,108551358 | 0,150551358 | -1,109448642 |
| chrII | 432763 | 432911 | 148 | 0,031345166 | 0,066705652 | 0,319682862 | -0,468585806 | -0,398297935 | -0,543373047 | -0,571373047 | -1,831373047 |
| chrII | 435614 | 435764 | 150 | 5,884342605 | 4,917688931 | 0,544744068 | 0,112392927 | 0,095533988 | -0,067178642 | -0,109178642 | -1,369178642 |
| chrII | 441246 | 441413 | 167 | 0,042743409 | 0,074117392 | 0,365763444 | -0,34309514 | -0,291630869 | -0,083828399 | -0,244828399 | -1,504828399 |
| chrII | 442313 | 442462 | 149 | 0,817823887 | 0,726350437 | 0,529618887 | 0,074311878 | 0,063165096 | -0,074837877 | -0,109837877 | -1,369837877 |
| chrII | 447274 | 447421 | 147 | 0,133929347 | 0,070411522 | 0,655421247 | 0,399998657 | 0,339998858 | 0,278840004 | 0,257840004 | -1,002159996 |
| chrII | 448183 | 448311 | 128 | 0,427434088 | 0,759703264 | 0,360054451 | -0,358313251 | -0,304566264 | 0,055054216 | 0,167054216 | -1,092945784 |
| chrII | 448687 | 448853 | 166 | 0,43883233 | 0,718938698 | 0,379032053 | -0,308023953 | -0,26182036 | -0,095032771 | -0,249032771 | -1,509032771 |
| chrII | 450522 | 450683 | 161 | 0,151026711 | 0,137117174 | 0,524136442 | 0,060538046 | 0,051457339 | 0,064698676 | -0,054301324 | -1,314301324 |
| chrII | 456967 | 457110 | 143 | 0,128230226 | 0,181587609 | 0,413889104 | -0,217551974 | -0,184919178 | -0,279475687 | -0,272475687 | -1,532475687 |
| chrII | 459507 | 459619 | 112 | 0,091185939 | 0,900526308 | 0,091947981 | -1,328854548 | -1,129526366 | -0,543094464 | -0,319094464 | -1,579094464 |
| chrII | 463510 | 463659 | 149 | 0,458779254 | 0,429880871 | 0,516259525 | 0,040767875 | 0,034652694 | -0,109145294 | -0,144145294 | -1,404145294 |
| chrII | 464402 | 464557 | 155 | 0,347646391 | 0,207528696 | 0,626192347 | 0,32178536 | 0,273517556 | 0,209587693 | 0,132587693 | -1,127412307 |
| chrII | 467973 | 468117 | 144 | 0,812124766 | 0,470645436 | 0,6331023 | 0,340081173 | 0,289068997 | 0,227761572 | 0,227761572 | -1,032238428 |
| chrII | 469937 | 470099 | 162 | 1,057186976 | 1,096937395 | 0,490773416 | -0,023129678 | -0,019660227 | 0,01445662 | -0,11154338 | -1,37154338 |

TABLE 2

| Chrom | Start | End | Length | Section A | Section B | A/A+B | Z-score | Z * 0.85 | Phase correction | Length correction | ΔLK |
|---|---|---|---|---|---|---|---|---|---|---|---|
| chrII | 478544 | 478678 | 134 | 0,111132863 | 0,066705652 | 0,624908854 | 0,318399007 | 0,270639156 | 0,374158766 | 0,444158766 | -0,815841234 |
| chrII | 479993 | 480154 | 161 | 0,071239015 | 0,059293913 | 0,545755127 | 0,114943701 | 0,097702146 | 0,119670466 | 0,000670466 | -1,259329534 |
| chrII | 480073 | 480226 | 153 | 1,818019652 | 1,300760222 | 0,582926569 | 0,209386076 | 0,177978165 | 0,09238891 | 0,02938891 | -1,23061109 |
| chrII | 483614 | 483775 | 161 | 3,71012788 | 2,834990228 | 0,566854229 | 0,168370847 | 0,14311522 | 0,164034151 | 0,045034151 | -1,214965849 |
| chrII | 490237 | 490391 | 154 | 0,219416165 | 0,092646739 | 0,703115179 | 0,533381326 | 0,453374127 | 0,348466602 | 0,278466602 | -0,981533398 |
| chrII | 496459 | 496596 | 137 | 0,219416165 | 0,237175653 | 0,480552117 | -0,048767936 | -0,041452746 | -0,035572939 | 0,013427061 | -1,246572939 |
| chrII | 498698 | 498864 | 166 | 0,176672756 | 0,229763914 | 0,434687048 | -0,164453569 | -0,139785534 | 0,034590256 | -0,119409744 | -1,379409744 |
| chrII | 499848 | 499990 | 142 | 1,165470279 | 0,622586089 | 0,651808466 | 0,390207566 | 0,331676431 | 0,318641738 | 0,332641738 | -0,927358262 |
| chrII | 501137 | 501288 | 151 | 1,929152515 | 1,004290656 | 0,657641005 | 0,406033495 | 0,345128471 | 0,228495722 | 0,179495722 | -1,080504278 |
| chrII | 503704 | 503871 | 167 | 1,601453048 | 2,645990879 | 0,377039244 | -0,313266119 | -0,266276202 | -0,031303465 | -0,192303465 | -1,452303465 |
| chrII | 503704 | 503871 | 167 | 1,601453048 | 2,645990879 | 0,377039244 | -0,313266119 | -0,266276202 | -0,031303465 | -0,192303465 | -1,452303465 |
| chrII | 505974 | 506148 | 174 | 0,401788042 | 0,659644785 | 0,37853365 | -0,309334224 | -0,26293409 | 0,075731346 | -0,134268654 | -1,394268654 |
| chrII | 506025 | 506148 | 123 | 0,065539893 | 0,070411522 | 0,4820832 | -0,044925865 | -0,038186985 | 0,48360805 | 0,63060805 | -0,62939195 |
| chrII | 506025 | 506179 | 154 | 0,715239706 | 0,496586524 | 0,590216393 | 0,228101654 | 0,193886406 | 0,088906439 | 0,018906439 | -1,241093561 |
| chrII | 506025 | 506179 | 154 | 0,715239706 | 0,496586524 | 0,590216393 | 0,228101654 | 0,193886406 | 0,088906439 | 0,018906439 | -1,241093561 |
| chrII | 509772 | 509912 | 140 | 0,726637949 | 0,548468698 | 0,569864451 | 0,176029078 | 0,149624716 | 0,149331902 | 0,177331902 | -1,082668098 |
| chrII | 511646 | 511795 | 149 | 0,233663968 | 0,229763914 | 0,504207833 | 0,010547668 | 0,008965518 | -0,139658486 | -0,174658486 | -1,434658486 |
| chrII | 512273 | 512423 | 150 | 0,407487163 | 0,415057393 | 0,495398286 | -0,011535042 | -0,009804786 | -0,14506331 | -0,18706331 | -1,44706331 |
| chrII | 519148 | 519306 | 158 | 0,464478375 | 0,389116306 | 0,544143943 | 0,110879232 | 0,094247347 | 0,086944464 | -0,011055536 | -1,271055536 |
| chrII | 523260 | 523407 | 147 | 0,307752543 | 0,300175436 | 0,506231912 | 0,015621722 | 0,013278464 | -0,056333998 | -0,077333998 | -1,337333998 |
| chrII | 523774 | 523925 | 151 | 0,453080133 | 0,374292827 | 0,547612932 | 0,119632674 | 0,101687773 | -0,007681204 | -0,056681204 | -1,316681204 |
| chrII | 525378 | 525528 | 150 | 0,766531797 | 0,589233263 | 0,565386895 | 0,164641441 | 0,139945225 | 0,014631912 | -0,027368088 | -1,287368088 |
| chrII | 527770 | 527902 | 132 | 0,236513528 | 0,240881523 | 0,495425179 | -0,011467628 | -0,009747484 | 0,109722164 | 0,193722164 | -1,066277836 |
| chrII | 528914 | 529061 | 147 | 0,076938136 | 0,044470435 | 0,633712557 | 0,341702373 | 0,290447017 | 0,181350619 | 0,160350619 | -1,099649381 |
| chrII | 529231 | 529384 | 153 | 0,045592969 | 0,037058696 | 0,551627959 | 0,129775456 | 0,110309137 | 0,049576779 | -0,013423221 | -1,273423221 |
| chrII | 533755 | 533907 | 152 | 0,105433742 | 0,048176305 | 0,686372697 | 0,485594627 | 0,412755433 | 0,326773409 | 0,270773409 | -0,989226591 |
| chrII | 543934 | 544097 | 163 | 0,153876272 | 0,159352392 | 0,49125859 | -0,021913219 | -0,018626236 | 0,04763648 | -0,08536352 | -1,34536352 |
| chrII | 545512 | 545645 | 133 | 2,581701889 | 2,538520661 | 0,504216733 | 0,01056998 | 0,008984483 | 0,094236569 | 0,171236569 | -1,088763431 |
| chrII | 550054 | 550200 | 146 | 0,225115286 | 0,085235 | 0,725358719 | 0,598835536 | 0,509010206 | 0,416119188 | 0,402119188 | -0,857880812 |
| chrII | 551769 | 551916 | 147 | 0,136778908 | 0,118587827 | 0,535617563 | 0,08939893 | 0,07598909 | -0,019364602 | -0,040364602 | -1,300364602 |
| chrII | 551821 | 551946 | 125 | 0,031345166 | 0,114881957 | 0,214359455 | -0,791385773 | -0,672677907 | -0,199670932 | -0,066670932 | -1,326670932 |
| chrII | 552584 | 552756 | 172 | 0,153876272 | 0,274234349 | 0,359431101 | -0,359979851 | -0,305982873 | 0,015374615 | -0,180625385 | -1,440625385 |
| chrII | 560316 | 560442 | 126 | 0,601257283 | 1,704700006 | 0,260740858 | -0,641063054 | -0,544903596 | -0,087672782 | 0,038327218 | -1,221672782 |
| chrII | 568857 | 569009 | 152 | 0,29350474 | 0,255705001 | 0,534412845 | 0,086367464 | 0,073412344 | 0,00253359 | -0,05346641 | -1,31346641 |
| chrII | 570513 | 570675 | 162 | 0,501522663 | 0,455821958 | 0,523868471 | 0,059865123 | 0,050885354 | 0,096410325 | -0,029589675 | -1,289589675 |
| chrII | 570513 | 570675 | 162 | 0,501522663 | 0,455821958 | 0,523868471 | 0,059865123 | 0,050885354 | 0,096410325 | -0,029589675 | -1,289589675 |
| chrII | 575360 | 575489 | 129 | 8,354911631 | 11,22507895 | 0,426706621 | -0,184765151 | -0,157050378 | 0,177480188 | 0,282480188 | -0,977519812 |
| chrII | 575363 | 575489 | 126 | 0,028495606 | 0,070411522 | 0,288104674 | -0,558930214 | -0,475090682 | -0,041867983 | 0,084132017 | -1,175867983 |
| chrII | 575635 | 575789 | 154 | 0,034194727 | 0,062999783 | 0,351817475 | -0,380418277 | -0,323355536 | -0,424362726 | -0,494362726 | -1,754362726 |
| chrII | 577662 | 577794 | 132 | 0,162424953 | 0,485468915 | 0,250696852 | -0,672298469 | -0,571453699 | -0,460094452 | -0,376094452 | -1,636094452 |
| chrII | 581371 | 581525 | 154 | 0,034194727 | 0,051882174 | 0,397257877 | -0,260451199 | -0,221383519 | -0,315286481 | -0,385286481 | -1,645286481 |
| chrII | 583028 | 583186 | 158 | 3,068976748 | 2,657108488 | 0,53596421 | 0,090271358 | 0,076730654 | 0,068094189 | -0,029905811 | -1,289905811 |
| chrII | 583277 | 583426 | 149 | 0,336248149 | 0,196411088 | 0,631263153 | 0,335200699 | 0,284920594 | 0,135412008 | 0,100412008 | -1,159587992 |
| chrII | 589093 | 589249 | 156 | 0,307752543 | 0,418763262 | 0,423600616 | -0,192690667 | -0,163787067 | -0,21610912 | -0,30010912 | -1,56010912 |
| chrII | 598510 | 598657 | 147 | 0,185221438 | 0,140823044 | 0,568086406 | 0,171504378 | 0,145778722 | 0,048319102 | 0,027319102 | -1,232680898 |
| chrII | 599865 | 600024 | 159 | 0,09973462 | 0,070411522 | 0,58617033 | 0,217704522 | 0,185048844 | 0,182642937 | 0,077642937 | -1,182357063 |
| chrII | 601519 | 601677 | 158 | 0,051292091 | 0,066705652 | 0,434687048 | -0,164453569 | -0,139785534 | -0,153310659 | -0,251310659 | -1,511310659 |
| chrII | 601519 | 601677 | 158 | 0,051292091 | 0,066705652 | 0,434687048 | -0,164453569 | -0,139785534 | -0,153310659 | -0,251310659 | -1,511310659 |
| chrII | 608166 | 608308 | 142 | 0,384690679 | 0,259410871 | 0,597251597 | 0,246239543 | 0,209303612 | 0,19201439 | 0,20601439 | -1,05398561 |
| chrII | 608166 | 608308 | 142 | 0,384690679 | 0,259410871 | 0,597251597 | 0,246239543 | 0,209303612 | 0,19201439 | 0,20601439 | -1,05398561 |
| chrII | 610440 | 610604 | 164 | 0,094035499 | 0,129705435 | 0,420287416 | -0,201158254 | -0,170984516 | -0,084272313 | -0,224272313 | -1,484272313 |
| chrII | 615980 | 616137 | 157 | 0,202318801 | 0,111176087 | 0,64536555 | 0,372838161 | 0,316912436 | 0,287751705 | 0,196751705 | -1,063248295 |
| chrII | 618802 | 618957 | 155 | 0,230814407 | 0,170470001 | 0,575189075 | 0,18960094 | 0,161160799 | 0,107692003 | 0,030692003 | -1,229307997 |
| chrII | 621368 | 621503 | 135 | 0,059840772 | 0,059293913 | 0,502295129 | 0,005753068 | 0,004890107 | 0,067734804 | 0,130734804 | -1,129265196 |
| chrII | 624538 | 624687 | 149 | 0,666797177 | 0,455821958 | 0,593965625 | 0,23775806 | 0,202094351 | 0,045900264 | 0,010900264 | -1,249099736 |
| chrII | 626717 | 626881 | 164 | 0,353345512 | 0,400233914 | 0,468889542 | -0,078061561 | -0,066352326 | 0,038569952 | -0,101430048 | -1,361430048 |
| chrII | 633744 | 633889 | 145 | 11,37829541 | 6,262919588 | 0,64498366 | 0,371812199 | 0,316040369 | 0,231545998 | 0,224545998 | -1,035454002 |
| chrII | 633747 | 633889 | 142 | 0,056991212 | 0,044470435 | 0,561702018 | 0,155285859 | 0,13199298 | 0,110819889 | 0,124819889 | -1,135180111 |
| chrII | 634590 | 634750 | 160 | 0,265009134 | 0,237175653 | 0,52771239 | 0,06952062 | 0,059092527 | 0,075334957 | -0,036665043 | -1,296665043 |
| chrII | 635204 | 635350 | 146 | 0,170973635 | 0,500292393 | 0,254703244 | -0,659762131 | -0,560797812 | -0,664915442 | -0,678915442 | -1,938915442 |
| chrII | 636694 | 636853 | 159 | 1,943400318 | 1,975228485 | 0,495938864 | -0,010179934 | -0,008652944 | -0,010705026 | -0,115705026 | -1,375705026 |
| chrII | 636695 | 636853 | 158 | 0,83207169 | 0,837526525 | 0,498366423 | -0,004094782 | -0,003480564 | -0,006353674 | -0,104353674 | -1,364353674 |
| chrII | 636699 | 636853 | 154 | 0,039893848 | 0,048176305 | 0,452978074 | -0,118140734 | -0,100419624 | -0,185038997 | -0,255038997 | -1,515038997 |
| chrII | 641881 | 642011 | 130 | 1,322196111 | 2,167933703 | 0,378838662 | -0,308532302 | -0,262252457 | -0,047245845 | 0,050754155 | -1,209245845 |
| chrII | 643000 | 643147 | 147 | 1,484621064 | 1,356348266 | 0,522575534 | 0,056618707 | 0,048125901 | -0,048472018 | -0,069472018 | -1,329472018 |
| chrII | 643625 | 643773 | 148 | 0,190920559 | 0,114881957 | 0,624326319 | 0,316863258 | 0,269333769 | 0,132845152 | 0,104845152 | -1,155154848 |
| chrII | 645958 | 646066 | 108 | 0,387540239 | 0,392822175 | 0,496615716 | -0,008483244 | -0,007210757 | 0,590769055 | 0,842769055 | -0,417230945 |
| chrII | 652842 | 653006 | 164 | 0,09973462 | 0,066705652 | 0,599221683 | 0,251333038 | 0,213633083 | 0,324402012 | 0,184402012 | -1,075597988 |
| chrII | 667311 | 667462 | 151 | 0,042743409 | 0,218646305 | 0,163523683 | -0,98007851 | -0,833066733 | -0,937118016 | -0,986118016 | -2,246118016 |
| chrII | 670468 | 670600 | 132 | 0,034194727 | 0,059293913 | 0,365763444 | -0,34309514 | -0,291630869 | -0,188029643 | -0,104029643 | -1,364029643 |
| chrII | 672951 | 673112 | 161 | 0,205168362 | 0,151940653 | 0,574525855 | 0,187908614 | 0,159722322 | 0,194497737 | 0,075497737 | -1,184502263 |
| chrII | 673886 | 673964 | 78 | 0,09688506 | 0,111176087 | 0,465656665 | -0,08619258 | -0,073263693 | | 0,462 | -0,798 |
| chrII | 673886 | 673989 | 103 | 0,079787696 | 0,08894087 | 0,472876041 | -0,06804215 | -0,057835827 | 0,300093445 | 0,587093445 | -0,672906555 |
| chrII | 674406 | 674512 | 106 | 0,131079787 | 1,063584569 | 0,109721016 | -1,228013149 | -1,043811177 | -0,588178995 | -0,322178995 | -1,582178995 |
| chrII | 674406 | 674523 | 117 | 0,185221438 | 1,348936527 | 0,12073166 | -1,171337061 | -0,995636502 | -0,383474226 | -0,194474226 | -1,454474226 |
| chrII | 678764 | 678912 | 148 | 0,116831984 | 0,074117392 | 0,61184795 | 0,284138687 | 0,241517884 | 0,112323926 | 0,084323926 | -1,175676074 |
| chrII | 682036 | 682166 | 130 | 0,763682236 | 0,874585221 | 0,466152357 | -0,084945506 | -0,07220368 | 0,116153727 | 0,214153727 | -1,045846273 |
| chrII | 683947 | 684082 | 135 | 0,413186285 | 0,377998697 | 0,52223727 | 0,055769465 | 0,047404045 | 0,103414501 | 0,166414501 | -1,093585499 |
| chrII | 684506 | 684665 | 159 | 0,24221265 | 0,255705001 | 0,486451222 | -0,033968281 | -0,028873039 | -0,035190924 | -0,140190924 | -1,400190924 |
| chrII | 685114 | 685267 | 153 | 1,501718428 | 0,956114351 | 0,61099292 | 0,281907864 | 0,239621685 | 0,179347677 | 0,116347677 | -1,143652323 |
| chrII | 688568 | 688714 | 146 | 0,082637257 | 0,074117392 | 0,527175798 | 0,068172392 | 0,057946533 | -0,039717319 | -0,053717319 | -1,313717319 |
| chrII | 693669 | 693820 | 151 | 0,09973462 | 0,148234783 | 0,402205349 | -0,247642923 | -0,210496484 | -0,315186433 | -0,364186433 | -1,624186433 |
| chrII | 696303 | 696437 | 134 | 0,53856695 | 0,452116089 | 0,543631948 | 0,10958803 | 0,093149826 | 0,205401047 | 0,275401047 | -0,984598953 |
| chrII | 698169 | 698308 | 139 | 0,413186285 | 0,244587392 | 0,628158741 | 0,326980653 | 0,277933555 | 0,290761886 | 0,325761886 | -0,934238114 |
| chrII | 698728 | 698854 | 126 | 7,762203029 | 13,34483635 | 0,367754231 | -0,33780723 | -0,287136145 | 0,137998008 | 0,263998008 | -0,996001992 |
| chrII | 698730 | 698836 | 106 | 0,059840772 | 0,085235 | 0,412479432 | -0,221171546 | -0,187995814 | 0,278221695 | 0,544221695 | -0,715778305 |
| chrII | 699474 | 699629 | 155 | 0,418885406 | 0,385410436 | 0,52081011 | 0,052186888 | 0,044358855 | -0,0077482 | -0,0847482 | -1,3447482 |
| chrII | 699802 | 699945 | 143 | 0,068389454 | 0,125999566 | 0,351817475 | -0,380418277 | -0,323355536 | -0,426432646 | -0,419432646 | -1,679432646 |
| chrII | 699802 | 699953 | 151 | 0,29350474 | 0,503998263 | 0,368029636 | -0,337076446 | -0,286514979 | -0,391062035 | -0,440062035 | -1,700062035 |
| chrII | 699834 | 699945 | 111 | 0,646850252 | 3,817045666 | 0,144907109 | -1,058529261 | -0,899749872 | -0,284347211 | -0,053347211 | -1,313347211 |
| chrII | 699834 | 699953 | 119 | 0,039893848 | 0,096352609 | 0,292806499 | -0,545204323 | -0,463423675 | 0,150677432 | 0,325677432 | -0,934322568 |
| chrII | 702899 | 703031 | 132 | 0,156725832 | 0,381704567 | 0,291079093 | -0,550235021 | -0,467699768 | -0,358616715 | -0,274616715 | -1,534616715 |
| chrII | 706525 | 706696 | 171 | 0,133929347 | 0,163058261 | 0,450959378 | -0,123237847 | -0,10475217 | 0,206150217 | 0,017150217 | -1,242849783 |
| chrII | 708493 | 708657 | 164 | 0,176672756 | 0,111176087 | 0,613769206 | 0,28915654 | 0,245783059 | 0,359630702 | 0,219630702 | -1,040369298 |
| chrII | 708881 | 709054 | 173 | 0,512920905 | 0,978349569 | 0,343948944 | -0,401709424 | -0,34145301 | -0,018569108 | -0,221569108 | -1,481569108 |
| chrII | 716866 | 717023 | 157 | 0,205168362 | 0,203822827 | 0,501644944 | 0,004123275 | 0,003504784 | -0,018908789 | -0,109908789 | -1,369908789 |
| chrII | 716874 | 717023 | 149 | 0,378991558 | 0,200116957 | 0,654439622 | 0,397334573 | 0,337734387 | 0,186111937 | 0,151111937 | -1,108888063 |
| chrII | 718549 | 718695 | 146 | 0,612655525 | 0,778232612 | 0,440477929 | -0,149757607 | -0,127293966 | -0,209357466 | -0,223357466 | -1,483357466 |
| chrII | 721490 | 721639 | 149 | 0,213717044 | 0,114881957 | 0,650388599 | 0,386369815 | 0,328414343 | 0,180893704 | 0,145893704 | -1,114106296 |
| chrII | 723357 | 723507 | 150 | 0,735186631 | 0,52993935 | 0,581117329 | 0,204752703 | 0,174039797 | 0,05622132 | 0,01422132 | -1,24577868 |
| chrII | 727858 | 728011 | 153 | 0,626903328 | 0,370586958 | 0,628480635 | 0,327831948 | 0,278657156 | 0,211959253 | 0,148959253 | -1,111040747 |

| Chrom | Start | End | Length | Section A | Section B | A/A+B | Z-score | Z * 0.85 | Phase correction | Length correction | ΔLK |
|---|---|---|---|---|---|---|---|---|---|---|---|
| chrII | 733512 | 733676 | 164 | 0,128230226 | 0,248293262 | 0,340563684 | -0,410925221 | -0,349286438 | -0,228574356 | -0,368574356 | -1,628574356 |
| chrII | 735258 | 735391 | 133 | 0,125380666 | 0,048176305 | 0,722417922 | 0,590039516 | 0,501533589 | 0,581374356 | 0,658374356 | -0,601625644 |
| chrII | 736781 | 736927 | 146 | 2,556055843 | 1,871464137 | 0,577310967 | 0,195019011 | 0,165766159 | 0,079946368 | 0,065946368 | -1,194053632 |
| chrII | 736912 | 737033 | 121 | 0,435982769 | 1,256289787 | 0,257631531 | -0,650664535 | -0,553064855 | 0,022614777 | 0,183614777 | -1,076385223 |
| chrII | 740252 | 740411 | 159 | 0,284956058 | 0,251999131 | 0,530688713 | 0,07700122 | 0,065451037 | 0,056331496 | -0,048668504 | -1,308668504 |
| chrII | 743678 | 743781 | 103 | 0,054141651 | 0,385410436 | 0,123174597 | -1,159262522 | -0,985373144 | -0,624747713 | -0,337747713 | -1,597747713 |
| chrII | 746216 | 746373 | 157 | 3,527756002 | 4,009750884 | 0,468026903 | -0,08023066 | -0,068196061 | -0,072993514 | -0,163993514 | -1,423993514 |
| chrII | 747192 | 747321 | 129 | 0,88051422 | 1,297054353 | 0,404356598 | -0,242086426 | -0,205773462 | 0,108827465 | 0,213827465 | -1,046172535 |
| chrII | 748930 | 749090 | 160 | 0,453080133 | 0,470645436 | 0,490492142 | -0,023834923 | -0,020259685 | 0,004154254 | -0,107845746 | -1,367845746 |
| chrII | 749254 | 749401 | 147 | 5,992625907 | 3,631752187 | 0,622650716 | 0,312449984 | 0,265582486 | 0,172668771 | 0,151668771 | -1,108331229 |
| chrII | 750045 | 750206 | 161 | 0,111132863 | 0,070411522 | 0,612152576 | 0,284933816 | 0,242193743 | 0,278174415 | 0,159174415 | -1,100825585 |
| chrII | 760513 | 760620 | 107 | 0,364743755 | 1,007996525 | 0,265704853 | -0,625855528 | -0,531977199 | 0,055117639 | 0,314117639 | -0,945882361 |
| chrII | 760513 | 760634 | 121 | 0,319150785 | 1,182172396 | 0,212579669 | -0,79750235 | -0,677876997 | -0,120239247 | 0,040760753 | -1,219239247 |
| chrII | 771359 | 771518 | 159 | 0,413186285 | 0,478057176 | 0,463606526 | -0,091351809 | -0,077649038 | -0,072711441 | -0,177711441 | -1,437711441 |
| chrII | 771799 | 771953 | 154 | 0,125380666 | 0,100058479 | 0,556161913 | 0,141245282 | 0,12005849 | 0,031079725 | -0,038920275 | -1,298920275 |
| chrII | 774249 | 774405 | 156 | 0,276407377 | 0,289057827 | 0,488814121 | -0,028042515 | -0,023836138 | -0,067326104 | -0,151326104 | -1,411326104 |
| chrII | 775701 | 775861 | 160 | 0,082637257 | 0,037058696 | 0,690393076 | 0,496964836 | 0,42242011 | 0,442536486 | 0,330536486 | -0,929463514 |
| chrII | 777331 | 777484 | 153 | 0,102584181 | 0,092646739 | 0,525450481 | 0,063838228 | 0,054262494 | -0,016755057 | -0,079755057 | -1,339755057 |
| chrII | 779629 | 779798 | 169 | 0,059840772 | 0,118587827 | 0,335376575 | -0,425114581 | -0,361347394 | -0,07207337 | -0,24707337 | -1,50707337 |
| chrII | 780145 | 780251 | 106 | 0,094035499 | 0,270528479 | 0,257939634 | -0,649710456 | -0,552253888 | -0,059022222 | 0,206977778 | -1,053022222 |
| chrII | 780145 | 780260 | 115 | 0,133929347 | 0,337234132 | 0,284252395 | -0,570254951 | -0,484716709 | 0,138388095 | 0,341388095 | -0,918611905 |
| chrII | 780582 | 780757 | 175 | 0,088336378 | 0,181587609 | 0,327263905 | -0,447480992 | -0,380358843 | -0,023281704 | -0,240281704 | -1,500281704 |
| chrII | 781915 | 782083 | 168 | 0,094035499 | 0,103764348 | 0,47540734 | -0,061683752 | -0,052431189 | 0,229523645 | 0,061523645 | -1,198476355 |
| chrII | 786436 | 786542 | 106 | 0,185221438 | 0,344645871 | 0,349561928 | -0,38650344 | -0,328527924 | 0,192244278 | 0,458244278 | -0,801755722 |
| chrII | 786436 | 786560 | 124 | 0,179522317 | 0,418763262 | 0,300061247 | -0,524224369 | -0,445590714 | 0,057610066 | 0,197610066 | -1,062389934 |
| chrII | 788442 | 788582 | 140 | 0,416035845 | 0,270528479 | 0,605967759 | 0,268824834 | 0,228501109 | 0,221719537 | 0,249719537 | -1,010280463 |
| chrII | 788604 | 788759 | 155 | 0,094035499 | 0,103764348 | 0,47540734 | -0,061683752 | -0,052431189 | -0,110389067 | -0,187389067 | -1,447389067 |
| chrII | 789546 | 789709 | 163 | 0,202318801 | 0,329822392 | 0,380197594 | -0,304961878 | -0,259217596 | -0,193585326 | -0,326585326 | -1,586585326 |
| chrII | 791029 | 791155 | 126 | 0,039893848 | 0,122293696 | 0,245973563 | -0,687215201 | -0,584132921 | -0,167735875 | -0,041735875 | -1,301735875 |
| chrII | 791029 | 791156 | 127 | 8,753850113 | 14,37136223 | 0,378541394 | -0,309313861 | -0,262916782 | 0,104188184 | 0,223188184 | -1,036811816 |
| chrII | 791031 | 791136 | 105 | 0,074088575 | 0,096352609 | 0,434687048 | -0,164453569 | -0,139785534 | 0,273929751 | 0,546929751 | -0,713070249 |
| chrII | 793238 | 793400 | 162 | 0,598407723 | 0,448410219 | 0,571644504 | 0,1805625 | 0,153478125 | 0,206077544 | 0,080077544 | -1,179922456 |
| chrII | 795412 | 795559 | 147 | 0,082637257 | 0,055588044 | 0,597844653 | 0,247772164 | 0,210606339 | 0,112927708 | 0,091927708 | -1,168072292 |
| chrII | 809074 | 809203 | 129 | 0,142478029 | 0,366881088 | 0,27972019 | -0,583672934 | -0,496121994 | -0,1955723 | -0,0905723 | -1,3505723 |
| chrII | 812955 | 813098 | 143 | 0,359044634 | 0,270528479 | 0,57029855 | 0,177134301 | 0,150564156 | 0,035625321 | 0,042625321 | -1,217374679 |
| chrIII | 2 | 145 | 143 | 0,344796831 | 0,159352392 | 0,683918204 | 0,478683799 | 0,406881229 | 0,298320548 | 0,305320548 | -0,954679452 |
| chrIII | 2 | 176 | 174 | 0,669646737 | 0,074117392 | 0,900348257 | 1,283538485 | 1,091007713 | 1,240915948 | 1,030915948 | -0,229084052 |
| chrIII | 4 | 145 | 141 | 0,210867483 | 0,270528479 | 0,438033344 | -0,155957289 | -0,132563695 | -0,167102945 | -0,146102945 | -1,406102945 |
| chrIII | 10 | 145 | 135 | 0,675345858 | 0,874585221 | 0,43572638 | -0,161813451 | -0,137541434 | -0,12229281 | -0,05929281 | -1,31929281 |
| chrIII | 10 | 146 | 136 | 0,074088575 | 0,051882174 | 0,588141101 | 0,222765786 | 0,189350918 | 0,208974256 | 0,264974256 | -0,995025744 |
| chrIII | 14 | 145 | 131 | 0,056991212 | 0,048176305 | 0,541908887 | 0,105243965 | 0,08945737 | 0,27623444 | 0,36723444 | -0,89276556 |
| chrIII | 19 | 145 | 126 | 0,065539893 | 0,051882174 | 0,558156527 | 0,146296985 | 0,124352437 | 0,457895969 | 0,583895969 | -0,676104031 |
| chrIII | 26 | 145 | 119 | 0,059840772 | 0,133411305 | 0,309651379 | -0,496838761 | -0,422312947 | -0,021431779 | 0,153568221 | -1,106431779 |
| chrIII | 37 | 145 | 108 | 0,074088575 | 0,133411305 | 0,357053581 | -0,366345661 | -0,311393812 | 0,096984747 | 0,348984747 | -0,911015253 |
| chrIII | 41 | 145 | 104 | 0,079787696 | 0,185293479 | 0,300993446 | -0,521545394 | -0,443313585 | -0,036883833 | 0,243116167 | -1,016883833 |
| chrIII | 50 | 145 | 95 | 0,199469241 | 0,370586958 | 0,349911537 | -0,385559308 | -0,327725412 | 0,076768639 | 0,419768639 | -0,840231361 |
| chrIII | 50 | 176 | 126 | 2,151418241 | 0,296469566 | 0,8788876 | 1,16944398 | 0,994027383 | 1,290676152 | 1,416676152 | 0,156676152 |
| chrIII | 50 | 185 | 135 | 0,14532759 | 0,062999783 | 0,69759239 | 0,517488464 | 0,439865194 | 0,450177698 | 0,513177698 | -0,746822302 |
| chrIII | 50 | 201 | 151 | 0,44168189 | 0,08894087 | 0,83238399 | 0,963628885 | 0,819084552 | 0,630061678 | 0,581061678 | -0,678938322 |
| chrIII | 50 | 220 | 170 | 0,606956404 | 0,148234783 | 0,803712245 | 0,854955993 | 0,726712594 | 0,738073111 | 0,556073111 | -0,703926889 |
| chrIII | 73 | 263 | 190 | 0,119681545 | 0,118587827 | 0,502295129 | 0,005753068 | 0,004890107 | 0,269206811 | -0,052793189 | -1,312793189 |
| chrIII | 88 | 231 | 143 | 0,04844253 | 0,040764565 | 0,543034495 | 0,108081541 | 0,09186931 | -0,014464587 | -0,007464587 | -1,267464587 |
| chrIII | 97 | 212 | 115 | 0,034194727 | 0,055588044 | 0,380860679 | -0,303221116 | -0,257737948 | 0,125668629 | 0,328668629 | -0,931331371 |
| chrIII | 110 | 263 | 153 | 0,367593315 | 0,26311674 | 0,582824568 | 0,209124736 | 0,177756026 | 0,053369477 | -0,009630523 | -1,269630523 |
| chrIII | 110 | 280 | 170 | 0,672496298 | 0,248293262 | 0,73034744 | 0,613864123 | 0,521784505 | 0,55339614 | 0,37139614 | -0,88860386 |
| chrIII | 110 | 283 | 173 | 0,455929693 | 0,289057827 | 0,611996417 | 0,284526191 | 0,241847262 | 0,346609158 | 0,143609158 | -1,116390842 |
| chrIII | 115 | 280 | 165 | 0,296354301 | 0,100058479 | 0,747590179 | 0,666925625 | 0,566886781 | 0,483721517 | 0,336721517 | -0,923278483 |
| chrIII | 115 | 283 | 168 | 0,108283302 | 0,062999783 | 0,632189116 | 0,337656888 | 0,287008354 | 0,280775739 | 0,112775739 | -1,147224261 |
| chrIII | 118 | 284 | 166 | 0,168124074 | 0,055588044 | 0,751519747 | 0,679279938 | 0,577387947 | 0,484847373 | 0,330847373 | -0,929152627 |
| chrIII | 129 | 252 | 123 | 0,076938136 | 0,111176087 | 0,408996909 | -0,230126057 | -0,195607148 | 0,193894175 | 0,340894175 | -0,919105825 |
| chrIII | 129 | 260 | 131 | 0,071239015 | 0,048176305 | 0,596565123 | 0,24446623 | 0,207796296 | 0,386187095 | 0,477187095 | -0,782812905 |
| chrIII | 129 | 263 | 134 | 3,336835443 | 1,641700223 | 0,670244358 | 0,440588012 | 0,37449981 | 0,46923419 | 0,53923419 | -0,72076581 |
| chrIII | 129 | 280 | 151 | 3,522056881 | 1,211819352 | 0,744011188 | 0,65576145 | 0,557397233 | 0,365105029 | 0,316105029 | -0,943894971 |
| chrIII | 129 | 283 | 154 | 2,769772887 | 1,130290221 | 0,710186684 | 0,553930177 | 0,470840651 | 0,374478112 | 0,304478112 | -0,955521888 |
| chrIII | 129 | 284 | 155 | 0,054141651 | 0,103764348 | 0,342872667 | -0,404635672 | -0,343940321 | -0,386242825 | -0,463242825 | -1,723242825 |
| chrIII | 129 | 301 | 172 | 0,076938136 | 0,144528914 | 0,34740218 | -0,39234359 | -0,333492051 | -0,239454046 | -0,435454046 | -1,695454046 |
| chrIII | 129 | 309 | 180 | 0,159575393 | 0,129705435 | 0,551627959 | 0,129775456 | 0,110309137 | 0,323534988 | 0,071534988 | -1,188465012 |
| chrIII | 129 | 326 | 197 | 0,760832676 | 0,396528045 | 0,657385949 | 0,405339328 | 0,344538429 | 0,626714483 | 0,255714483 | -1,004285517 |
| chrIII | 143 | 284 | 141 | 0,222265726 | 0,389116306 | 0,363546382 | -0,348995399 | -0,296646089 | -0,334492119 | -0,313492119 | -1,573492119 |
| chrIII | 164 | 284 | 120 | 0,105433742 | 0,133411305 | 0,441431561 | -0,147340685 | -0,125239582 | 0,263478412 | 0,431478412 | -0,828521588 |
| chrIII | 164 | 341 | 177 | 0,151026711 | 0,044470435 | 0,772526424 | 0,747192861 | 0,635113932 | 0,814217871 | 0,583217871 | -0,676782129 |
| chrIII | 186 | 284 | 98 | 0,675345858 | 1,007996525 | 0,401193402 | -0,250259331 | -0,212720432 | 0,18882432 | 0,51082432 | -0,74917568 |
| chrIII | 186 | 320 | 134 | 0,219416165 | 0,507704132 | 0,301760473 | -0,519343898 | -0,441442313 | -0,361770143 | -0,291770143 | -1,551770143 |
| chrIII | 186 | 337 | 151 | 0,034194727 | 0,048176305 | 0,415130494 | -0,214366854 | -0,182211826 | -0,362725041 | -0,411725041 | -1,671725041 |
| chrIII | 186 | 341 | 155 | 0,119681545 | 0,107470218 | 0,526879225 | 0,067427283 | 0,057313191 | 0,009551777 | -0,067448223 | -1,327448223 |
| chrIII | 186 | 352 | 166 | 1,114178188 | 1,011702395 | 0,524101964 | 0,060451462 | 0,051383743 | -0,012052459 | -0,166052459 | -1,426052459 |
| chrIII | 186 | 353 | 167 | 0,14532759 | 0,170470001 | 0,460192206 | -0,099949507 | -0,084957081 | -0,111908001 | -0,272908001 | -1,532908001 |
| chrIII | 186 | 355 | 169 | 0,202318801 | 0,055588044 | 0,784464644 | 0,787360758 | 0,669256644 | 0,662770011 | 0,487770011 | -0,772229989 |
| chrIII | 192 | 329 | 137 | 0,113982423 | 0,070411522 | 0,618146236 | 0,300615741 | 0,25552338 | 0,284763137 | 0,333763137 | -0,926236863 |
| chrIII | 246 | 341 | 95 | 0,156725832 | 0,326116523 | 0,324590066 | -0,454901459 | -0,386666241 | 0,016447417 | 0,359447417 | -0,900552583 |
| chrIII | 246 | 352 | 106 | 0,042743409 | 0,055588044 | 0,434687048 | -0,164453569 | -0,139785534 | 0,28829251 | 0,55429251 | -0,70570749 |
| chrIII | 5166 | 5314 | 148 | 3,171560929 | 2,167933703 | 0,593981481 | 0,237798943 | 0,202129102 | 0,055999786 | 0,027999786 | -1,232000214 |
| chrIII | 5882 | 5994 | 112 | 0,202318801 | 2,401403487 | 0,077703679 | -1,420688427 | -1,207585163 | -0,799019833 | -0,575019833 | -1,835019833 |
| chrIII | 7560 | 7714 | 154 | 0,062690333 | 0,08894087 | 0,413439528 | -0,218706025 | -0,185900121 | -0,259499231 | -0,329499231 | -1,589499231 |
| chrIII | 9830 | 9940 | 110 | 0,079787696 | 0,144528914 | 0,355692324 | -0,369997104 | -0,314497538 | 0,101456424 | 0,339456424 | -0,920543576 |
| chrIII | 9831 | 9940 | 109 | 2,641542661 | 5,959038282 | 0,307135376 | -0,503986659 | -0,42838866 | -0,016392141 | 0,228607859 | -1,031392141 |
| chrIII | 9831 | 9941 | 110 | 0,059840772 | 0,092646739 | 0,392430643 | -0,272989502 | -0,232041076 | 0,187696999 | 0,425696999 | -0,834303001 |
| chrIII | 9832 | 9955 | 123 | 0,045592969 | 0,448410219 | 0,092292865 | -1,326767123 | -1,127752055 | -0,752931237 | -0,605931237 | -1,865931237 |
| chrIII | 11462 | 11613 | 151 | 0,056991212 | 0,044470435 | 0,561702018 | 0,155285859 | 0,13199298 | -0,049056487 | -0,098056487 | -1,358056487 |
| chrIII | 17946 | 18093 | 147 | 0,205168362 | 0,200116957 | 0,506231912 | 0,015621722 | 0,013278464 | -0,100815646 | -0,121815646 | -1,381815646 |
| chrIII | 23277 | 23427 | 150 | 0,795027403 | 0,548468698 | 0,591760112 | 0,232074982 | 0,197263735 | -0,002688533 | -0,044688533 | -1,304688533 |
| chrIII | 27235 | 27381 | 146 | 0,165274514 | 0,137117174 | 0,546557727 | 0,116969093 | 0,099423729 | -0,013126281 | -0,027126281 | -1,287126281 |
| chrIII | 32929 | 33083 | 154 | 1,265204899 | 0,904232177 | 0,58319502 | 0,210073946 | 0,178562854 | 0,124326924 | 0,054326924 | -1,205673076 |
| chrIII | 34797 | 34957 | 160 | 0,458779254 | 0,544762828 | 0,457159956 | -0,107591283 | -0,091452591 | -0,182230317 | -0,294230317 | -1,554230317 |
| chrIII | 36971 | 37125 | 154 | 1,168319839 | 1,026525873 | 0,532301579 | 0,081056724 | 0,068898216 | 0,016647479 | -0,053352521 | -1,313352521 |
| chrIII | 37274 | 37421 | 147 | 0,29065518 | 0,163058261 | 0,640613994 | 0,360100456 | 0,306085387 | 0,193926136 | 0,172926136 | -1,087073864 |
| chrIII | 41105 | 41261 | 156 | 0,512920905 | 0,463233697 | 0,525450481 | 0,063838228 | 0,054262494 | -0,031512355 | -0,115512355 | -1,375512355 |
| chrIII | 42270 | 42419 | 149 | 0,199469241 | 0,096352609 | 0,674288397 | 0,45178593 | 0,38401804 | 0,174343281 | 0,139343281 | -1,120656719 |
| chrIII | 56437 | 56565 | 128 | 0,185221438 | 0,314998914 | 0,370279692 | -0,331112667 | -0,281445767 | -0,014300779 | 0,097699221 | -1,162300779 |
| chrIII | 62417 | 62568 | 151 | 13,98279378 | 10,67661026 | 0,567036971 | 0,168835471 | 0,14351015 | -0,029661321 | -0,078661321 | -1,338661321 |
| chrIII | 62420 | 62568 | 148 | 0,039893848 | 0,037058696 | 0,518421434 | 0,046192108 | 0,039263292 | -0,107111568 | -0,135111568 | -1,395111568 |

TABLE 2

| Chrom | Start | End | Length | Section A | Section B | A/A+B | Z-score | Z * 0.85 | Phase correction | Length correction | ΔLK |
|---|---|---|---|---|---|---|---|---|---|---|---|
| chrIII | 62779 | 62946 | 167 | 0,230814407 | 0,270528479 | 0,460392305 | -0,099445434 | -0,084528619 | -0,10512384 | -0,26612384 | -1,52612384 |
| chrIII | 62988 | 63143 | 155 | 0,159575393 | 0,181587609 | 0,467739443 | -0,080953558 | -0,068810524 | -0,123107083 | -0,200107083 | -1,460107083 |
| chrIII | 68315 | 68492 | 177 | 0,037044288 | 0,085235 | 0,302948179 | -0,51593994 | -0,438548949 | -0,235689326 | -0,466689326 | -1,726689326 |
| chrIII | 68580 | 68746 | 166 | 4,029278665 | 3,583575882 | 0,529273039 | 0,073442598 | 0,062426208 | 0,000224316 | -0,153775684 | -1,413775684 |
| chrIII | 74254 | 74383 | 129 | 1,005894886 | 1,167348917 | 0,462854137 | -0,093245819 | -0,079258946 | 0,145353208 | 0,250353208 | -1,009646792 |
| chrIII | 75106 | 75269 | 163 | 1,70688679 | 1,537935875 | 0,526033921 | 0,065303749 | 0,055508186 | -0,032221364 | -0,165221364 | -1,425221364 |
| chrIII | 82256 | 82435 | 179 | 0,028495606 | 0,118587827 | 0,1937377 | -0,864204794 | -0,734574075 | -0,522682426 | -0,767682426 | -2,027682426 |
| chrIII | 86659 | 86788 | 129 | 3,020534219 | 5,573627846 | 0,351463493 | -0,381372335 | -0,324166485 | -0,096349554 | 0,008650446 | -1,251349554 |
| chrIII | 86884 | 87013 | 129 | 0,176672756 | 0,366881088 | 0,325032668 | -0,453671427 | -0,385620713 | -0,176965765 | -0,071965765 | -1,331965765 |
| chrIII | 88349 | 88492 | 143 | 0,188070999 | 0,092646739 | 0,669964783 | 0,439815924 | 0,373843536 | 0,247127492 | 0,254127492 | -1,005872508 |
| chrIII | 89624 | 89790 | 166 | 0,042743409 | 0,074117392 | 0,365763444 | -0,34309514 | -0,291630869 | -0,357992996 | -0,511992996 | -1,771992996 |
| chrIII | 90428 | 90577 | 149 | 0,119681545 | 0,051882174 | 0,69759239 | 0,517488464 | 0,439865194 | 0,229648339 | 0,194648339 | -1,065351661 |
| chrIII | 90432 | 90582 | 150 | 0,056991212 | 0,040764565 | 0,582995843 | 0,209563572 | 0,178129036 | -0,018665557 | -0,060665557 | -1,320665557 |
| chrIII | 92227 | 92379 | 152 | 3,997933499 | 2,164227834 | 0,648787541 | 0,382049137 | 0,324741767 | 0,203568862 | 0,147568862 | -1,112431138 |
| chrIII | 93544 | 93697 | 153 | 1,681240744 | 1,267407396 | 0,570173403 | 0,176815652 | 0,150293304 | 0,029672552 | -0,033327448 | -1,293327448 |
| chrIII | 94513 | 94671 | 158 | 1,344992595 | 1,471230223 | 0,477587422 | -0,056209587 | -0,047778149 | -0,107807665 | -0,205807665 | -1,465807665 |
| chrIII | 95867 | 96028 | 161 | 0,076938136 | 0,125999566 | 0,379121943 | -0,307787693 | -0,261619539 | -0,331905784 | -0,450905784 | -1,710905784 |
| chrIII | 95874 | 96028 | 154 | 0,213717044 | 0,244587392 | 0,466321133 | -0,084520926 | -0,071842787 | -0,117836567 | -0,187836567 | -1,447836567 |
| chrIII | 97469 | 97602 | 133 | 0,028495606 | 0,062999783 | 0,311443082 | -0,491764035 | -0,41799943 | -0,312463613 | -0,235463613 | -1,495463613 |
| chrIII | 100878 | 101025 | 147 | 0,094035499 | 0,074117392 | 0,559226183 | 0,149007602 | 0,126656462 | -0,00886259 | -0,02986259 | -1,28986259 |
| chrIII | 100878 | 101048 | 170 | 0,085486818 | 0,077823261 | 0,523463207 | 0,058847485 | 0,050020363 | 0,106177271 | -0,075822729 | -1,335822729 |
| chrIII | 101513 | 101652 | 139 | 0,119681545 | 0,081529131 | 0,594807131 | 0,239928434 | 0,203939169 | 0,179601212 | 0,214601212 | -1,045398788 |
| chrIII | 101547 | 101689 | 142 | 1,045788734 | 0,755997394 | 0,580417796 | 0,202962418 | 0,172518055 | 0,10153868 | 0,11553868 | -1,14446132 |
| chrIII | 101696 | 101853 | 157 | 0,065539893 | 0,037058696 | 0,638799168 | 0,355250862 | 0,301963233 | 0,232088092 | 0,141088092 | -1,118911908 |
| chrIII | 103033 | 103165 | 132 | 10,5405246 | 6,470448284 | 0,619630909 | 0,304511568 | 0,258834833 | 0,401752809 | 0,485752809 | -0,774247191 |
| chrIII | 103103 | 103249 | 146 | 0,056991212 | 0,037058696 | 0,605967759 | 0,268824834 | 0,228501109 | 0,117216263 | 0,103216263 | -1,156783737 |
| chrIII | 103103 | 103251 | 148 | 17,08881482 | 8,983027859 | 0,655450979 | 0,400079394 | 0,340067485 | 0,185477499 | 0,157477499 | -1,102522501 |
| chrIII | 104725 | 104877 | 152 | 6,280431526 | 3,379753056 | 0,650135768 | 0,385687039 | 0,327833983 | 0,195910235 | 0,139910235 | -1,120089765 |
| chrIII | 106067 | 106230 | 163 | 0,227964847 | 0,159352392 | 0,588573975 | 0,223878237 | 0,190296502 | 0,078090595 | -0,054909405 | -1,314909405 |
| chrIII | 116514 | 116644 | 130 | 0,370442876 | 0,600350872 | 0,38158762 | -0,301313772 | -0,256116706 | -0,056378718 | 0,041621282 | -1,218378718 |
| chrIII | 117818 | 117976 | 158 | 0,125380666 | 0,103764348 | 0,547167331 | 0,118507771 | 0,100731606 | 0,049370399 | -0,048629601 | -1,308629601 |
| chrIII | 132816 | 132923 | 107 | 0,817823887 | 5,89233263 | 0,121878511 | -1,16564743 | -0,990800316 | -0,551858369 | -0,292858369 | -1,552858369 |
| chrIII | 132817 | 132923 | 106 | 0,04844253 | 1,082113917 | 0,042848396 | -1,718547538 | -1,460765408 | -1,011595077 | -0,745595077 | -2,005595077 |
| chrIII | 133268 | 133417 | 149 | 0,125380666 | 0,070411522 | 0,640376244 | 0,359464657 | 0,305544959 | 0,096053301 | 0,061053301 | -1,198946699 |
| chrIII | 134878 | 135010 | 132 | 0,037044288 | 0,107470218 | 0,25633612 | -0,65468243 | -0,556480065 | -0,426285663 | -0,342285663 | -1,602285663 |
| chrIII | 136592 | 136744 | 152 | 0,322000346 | 0,144528914 | 0,690203968 | 0,496428581 | 0,421964293 | 0,290980715 | 0,234980715 | -1,025019285 |
| chrIII | 137946 | 138097 | 151 | 0,29065518 | 0,192705218 | 0,601321873 | 0,256770098 | 0,218254584 | 0,057875812 | 0,008875812 | -1,251124188 |
| chrIII | 138189 | 138339 | 150 | 2,325241436 | 1,363760005 | 0,630317302 | 0,332693846 | 0,282789769 | 0,079782755 | 0,037782755 | -1,222217245 |
| chrIII | 146706 | 146857 | 151 | 0,128230226 | 0,17417587 | 0,424033205 | -0,191586128 | -0,162848208 | -0,308003251 | -0,357003251 | -1,617003251 |
| chrIII | 154254 | 154394 | 140 | 0,567062556 | 0,466939567 | 0,548415273 | 0,121658533 | 0,103409753 | 0,067615651 | 0,095615651 | -1,164384349 |
| chrIII | 154256 | 154394 | 138 | 0,153876272 | 0,207528696 | 0,42577243 | -0,18714766 | -0,159075511 | -0,163052849 | -0,121052849 | -1,381052849 |
| chrIII | 156400 | 156550 | 150 | 6,454254722 | 3,968986319 | 0,61921764 | 0,303426675 | 0,257912674 | 0,055700303 | 0,013700303 | -1,246299697 |
| chrIII | 156437 | 156550 | 113 | 0,97454972 | 2,075286964 | 0,319541609 | -0,468981 | -0,39863385 | 0,000349124 | 0,217349124 | -1,042650876 |
| chrIII | 156484 | 156645 | 161 | 0,359044634 | 0,448410219 | 0,444662178 | -0,13915919 | -0,118285311 | -0,175770183 | -0,294770183 | -1,554770183 |
| chrIII | 158189 | 158338 | 149 | 0,239363089 | 0,26682261 | 0,472876041 | -0,06804215 | -0,057835827 | -0,269439488 | -0,304439488 | -1,564439488 |
| chrIII | 159911 | 160084 | 173 | 0,239363089 | 0,541056958 | 0,306710585 | -0,505196015 | -0,429416612 | -0,306715977 | -0,509715977 | -1,769715977 |
| chrIII | 159911 | 160084 | 173 | 0,239363089 | 0,541056958 | 0,306710585 | -0,505196015 | -0,429416612 | -0,306715977 | -0,509715977 | -1,769715977 |
| chrIII | 161801 | 161938 | 137 | 0,028495606 | 0,100058479 | 0,221662392 | -0,766590907 | -0,651602271 | -0,634675426 | -0,585675426 | -1,845675426 |
| chrIII | 175934 | 176084 | 150 | 0,142478029 | 0,074117392 | 0,657807209 | 0,406485945 | 0,345513053 | 0,142023193 | 0,100023193 | -1,159976807 |
| chrIII | 186417 | 186569 | 152 | 0,296354301 | 0,244587392 | 0,54784888 | 0,120228376 | 0,10219412 | -0,029774075 | -0,085774075 | -1,345774075 |
| chrIII | 188100 | 188260 | 160 | 0,253610892 | 0,381704567 | 0,39918892 | -0,255447043 | -0,217129987 | -0,297387205 | -0,409387205 | -1,669387205 |
| chrIII | 191522 | 191646 | 124 | 2,045984499 | 3,457576317 | 0,371756499 | -0,327204791 | -0,278124073 | 0,088500058 | 0,228500058 | -1,031499942 |
| chrIII | 195429 | 195578 | 149 | 0,222265726 | 0,129705435 | 0,631488458 | 0,335798151 | 0,285428429 | 0,08556642 | 0,05056642 | -1,20943358 |
| chrIII | 201078 | 201230 | 152 | 0,139628469 | 0,085235 | 0,620947766 | 0,307970909 | 0,261775272 | 0,12718959 | 0,07118959 | -1,18881041 |
| chrIII | 208738 | 208879 | 141 | 0,236513528 | 0,718938698 | 0,247540925 | -0,68224847 | -0,579911199 | -0,643906559 | -0,622906559 | -1,882906559 |
| chrIII | 209585 | 209734 | 149 | 0,188070999 | 0,092646739 | 0,669964783 | 0,439815924 | 0,373843536 | 0,18593129 | 0,15093129 | -1,10906871 |
| chrIII | 210916 | 211072 | 156 | 0,227964847 | 0,163058261 | 0,582995843 | 0,209563572 | 0,178129036 | 0,099265444 | 0,015265444 | -1,244734556 |
| chrIII | 213468 | 213602 | 134 | 0,547115632 | 0,385410436 | 0,586702775 | 0,219071373 | 0,186210667 | 0,235392304 | 0,305392304 | -0,954607696 |
| chrIII | 215653 | 215790 | 137 | 5,562342259 | 2,935048706 | 0,654594132 | 0,397753717 | 0,338090659 | 0,345383895 | 0,394383895 | -0,865616105 |
| chrIII | 217960 | 218102 | 142 | 0,567062556 | 0,44099848 | 0,562527998 | 0,157381739 | 0,133774478 | 0,037886471 | 0,051886471 | -1,208113529 |
| chrIII | 228591 | 228741 | 150 | 1,307948308 | 0,874585221 | 0,599279824 | 0,251483454 | 0,213760936 | 0,017233845 | -0,024766155 | -1,284766155 |
| chrIII | 231011 | 231166 | 155 | 0,042743409 | 0,070411522 | 0,377742344 | -0,311415611 | -0,264703269 | -0,322060501 | -0,399060501 | -1,659060501 |
| chrIII | 234251 | 234402 | 151 | 0,094035499 | 0,107470218 | 0,466664175 | -0,083658001 | -0,071109301 | -0,209111731 | -0,258111731 | -1,518111731 |
| chrIII | 234921 | 235052 | 131 | 0,105433742 | 0,463233697 | 0,185404921 | -0,894957435 | -0,76071382 | -0,59644927 | -0,50544927 | -1,76544927 |
| chrIII | 236307 | 236469 | 162 | 0,247911771 | 0,222352175 | 0,527175798 | 0,068172392 | 0,057946533 | -0,003862065 | -0,129862065 | -1,389862065 |
| chrIII | 238120 | 238291 | 171 | 0,142478029 | 0,43729261 | 0,245748956 | -0,687928336 | -0,584739086 | -0,513163652 | -0,702163652 | -1,962163652 |
| chrIII | 238120 | 238291 | 171 | 0,142478029 | 0,43729261 | 0,245748956 | -0,687928336 | -0,584739086 | -0,513163652 | -0,702163652 | -1,962163652 |
| chrIII | 240298 | 240452 | 154 | 0,031345166 | 0,055588044 | 0,360566076 | -0,356946107 | -0,303404191 | -0,34725888 | -0,41725888 | -1,67725888 |
| chrIII | 240452 | 240614 | 162 | 0,661098055 | 0,492880654 | 0,572885834 | 0,183726101 | 0,156167186 | 0,103297933 | -0,022702067 | -1,282702067 |
| chrIII | 242569 | 242719 | 150 | 0,185221438 | 0,081529131 | 0,694361923 | 0,508252677 | 0,432014775 | 0,242769543 | 0,200769543 | -1,059230457 |
| chrIII | 246007 | 246188 | 181 | 0,04844253 | 0,166764131 | 0,225097726 | -0,755089219 | -0,641825836 | -0,410933178 | -0,669933178 | -1,929933178 |
| chrIII | 246926 | 247057 | 131 | 0,04844253 | 0,048176305 | 0,50137771 | 0,003453413 | 0,002935401 | 0,161545227 | 0,252545227 | -1,007454773 |
| chrIII | 246926 | 247058 | 132 | 11,87411895 | 8,808851988 | 0,574101225 | 0,186825376 | 0,158801569 | 0,278239571 | 0,362239571 | -0,897760429 |
| chrIII | 248934 | 249085 | 151 | 0,265009134 | 0,285351958 | 0,481518658 | -0,046342437 | -0,039391072 | -0,171690621 | -0,220690621 | -1,480690621 |
| chrIII | 251728 | 251875 | 147 | 0,393239361 | 0,17417587 | 0,693036315 | 0,504475364 | 0,428804059 | 0,295289457 | 0,274289457 | -0,985710543 |
| chrIII | 252213 | 252361 | 148 | 14,90605141 | 8,701381771 | 0,631413475 | 0,335579302 | 0,285259406 | 0,106878214 | 0,078878214 | -1,181121786 |
| chrIII | 252238 | 252397 | 159 | 1,843665698 | 1,712111745 | 0,51849862 | 0,046385793 | 0,039427924 | -0,005693883 | -0,110693883 | -1,370693883 |
| chrIII | 254663 | 254821 | 158 | 1,097080825 | 0,896820438 | 0,550218231 | 0,126212725 | 0,107280816 | 0,064890317 | -0,033109683 | -1,293109683 |
| chrIII | 257083 | 257228 | 145 | 0,108283302 | 0,125999566 | 0,462190442 | -0,094916836 | -0,080679311 | -0,196182799 | -0,203182799 | -1,463182799 |
| chrIII | 257806 | 257969 | 163 | 0,108283302 | 0,092646739 | 0,538910465 | 0,097689227 | 0,083035843 | -0,01677351 | -0,14977351 | -1,40977351 |
| chrIII | 258183 | 258325 | 142 | 1,296550065 | 0,726350437 | 0,640936153 | 0,360962213 | 0,306817881 | 0,20726129 | 0,22126129 | -1,03873871 |
| chrIII | 259144 | 259292 | 148 | 0,492973981 | 0,366881088 | 0,573322178 | 0,184838583 | 0,157112796 | -0,04261948 | -0,07061948 | -1,33061948 |
| chrIII | 262204 | 262356 | 152 | 0,68389454 | 0,407645654 | 0,626540868 | 0,322705529 | 0,274299699 | 0,145361696 | 0,089361696 | -1,170638304 |
| chrIII | 269289 | 269448 | 159 | 0,153876272 | 0,181587609 | 0,458696987 | -0,10371695 | -0,088159408 | -0,146255299 | -0,251255299 | -1,511255299 |
| chrIII | 272784 | 272929 | 145 | 0,062690333 | 0,100058479 | 0,38519687 | -0,291859905 | -0,248080919 | -0,369330826 | -0,376330826 | -1,636330826 |
| chrIII | 276672 | 276825 | 153 | 0,125380666 | 0,148234783 | 0,458236793 | -0,104876779 | -0,089145262 | -0,202113886 | -0,265113886 | -1,525113886 |
| chrIII | 280552 | 280680 | 128 | 0,606956404 | 1,511994788 | 0,286441899 | -0,563809481 | -0,479238059 | -0,240452975 | -0,128452975 | -1,388452975 |
| chrIII | 281034 | 281170 | 136 | 0,686744101 | 0,585527393 | 0,539777951 | 0,099874329 | 0,084893179 | 0,126881609 | 0,182881609 | -1,077118391 |
| chrIII | 286830 | 286938 | 108 | 7,719459621 | 20,97522181 | 0,269020572 | -0,615777856 | -0,523411177 | -0,104362447 | 0,147637553 | -1,112362447 |
| chrIII | 286830 | 286939 | 109 | 2,652940903 | 7,100446113 | 0,272002013 | -0,606769297 | -0,515753903 | -0,109724778 | 0,135275222 | -1,124724778 |
| chrIII | 286830 | 286940 | 110 | 0,165274514 | 0,637409568 | 0,205902319 | -0,820721997 | -0,697613698 | -0,287761628 | -0,049761628 | -1,309761628 |
| chrIII | 286830 | 286941 | 111 | 0,065539893 | 0,17788174 | 0,269244325 | -0,615100047 | -0,52283504 | -0,126714204 | 0,104285796 | -1,155714204 |
| chrIII | 286832 | 286938 | 106 | 0,176672756 | 1,40823044 | 0,111472269 | -1,218735659 | -1,03592531 | -0,586583781 | -0,320583781 | -1,580583781 |
| chrIII | 286832 | 286939 | 107 | 0,071239015 | 0,585527393 | 0,108469334 | -1,234709424 | -1,049503011 | -0,642068654 | -0,383068654 | -1,643068654 |
| chrIII | 286832 | 286955 | 123 | 0,404637603 | 2,868343054 | 0,123629696 | -1,157031743 | -0,983476982 | -0,608702946 | -0,461702946 | -1,721702946 |
| chrIII | 288517 | 288666 | 149 | 0,065539893 | 0,107470218 | 0,378821174 | -0,308578276 | -0,262291535 | -0,446922271 | -0,481922271 | -1,741922271 |
| chrIII | 303804 | 303963 | 159 | 0,359044634 | 0,314998914 | 0,532672755 | 0,081990221 | 0,069691688 | -0,00880065 | -0,11380065 | -1,37380065 |
| chrIII | 307204 | 307345 | 141 | 0,045592969 | 0,037058696 | 0,551627969 | 0,129775456 | 0,110309137 | 0,042459342 | 0,063459342 | -1,196540658 |
| chrIII | 307633 | 307788 | 155 | 0,190920559 | 0,070411522 | 0,730566865 | 0,614528311 | 0,522349064 | 0,44289037 | 0,36589037 | -0,89410963 |
| chrIII | 308042 | 308191 | 149 | 2,78117113 | 2,338403704 | 0,543242597 | 0,108606247 | 0,09231531 | -0,075040044 | -0,110040044 | -1,370040044 |
| chrIII | 310321 | 310472 | 151 | 0,056991212 | 0,103764348 | 0,354520937 | -0,373143192 | -0,317171714 | -0,438404911 | -0,487404911 | -1,747404911 |

| Chrom | Start | End | Length | Section A | Section B | A/A+B | Z-score | Z * 0.85 | Phase correction | Length correction | ΔLK |
|---|---|---|---|---|---|---|---|---|---|---|---|
| chrIII | 310654 | 310803 | 149 | 0,04844253 | 0,040764565 | 0,543034495 | 0,108081541 | 0,09186931 | -0,085621571 | -0,120621571 | -1,380621571 |
| chrIII | 311798 | 311947 | 149 | 0,601257283 | 0,463233697 | 0,564830792 | 0,16322864 | 0,138744344 | -0,05314441 | -0,08814441 | -1,34814441 |
| chrIII | 314554 | 314707 | 153 | 0,270708255 | 0,255705001 | 0,514250453 | 0,035728188 | 0,030368959 | -0,075051656 | -0,138051656 | -1,398051656 |
| chrIV | 0 | 126 | 126 | 0,170973635 | 0,292763697 | 0,368686373 | -0,335334533 | -0,285034353 | 0,03016491 | 0,15616491 | -1,10383509 |
| chrIV | 0 | 145 | 145 | 0,108283302 | 0,196411088 | 0,355383315 | -0,370826679 | -0,315202677 | -0,490917866 | -0,497917866 | -1,757917866 |
| chrIV | 0 | 155 | 155 | 0,119681545 | 0,044470435 | 0,729089865 | 0,610062706 | 0,5185533 | 0,407184851 | 0,330184851 | -0,929815149 |
| chrIV | 5 | 126 | 121 | 0,051292091 | 0,062999783 | 0,448781606 | -0,128740216 | -0,109429183 | 0,44863893 | 0,60963893 | -0,65036107 |
| chrIV | 14 | 126 | 112 | 0,056991212 | 0,044470435 | 0,561702018 | 0,155285859 | 0,13199298 | 0,694521463 | 0,918521463 | -0,341478537 |
| chrIV | 14 | 134 | 120 | 0,364743755 | 0,125999566 | 0,743247518 | 0,653389875 | 0,555381394 | 1,101244849 | 1,269244849 | 0,009244849 |
| chrIV | 14 | 155 | 141 | 0,624053768 | 0,070411522 | 0,898610452 | 1,273673656 | 1,082622608 | 0,918480777 | 0,939480777 | -0,320519223 |
| chrIV | 44 | 126 | 82 | 0,031345166 | 0,43729261 | 0,066885701 | -1,499394179 | -1,274485052 | -0,971422563 | -0,537422563 | -1,797422563 |
| chrIV | 44 | 134 | 90 | 0,071239015 | 0,125999566 | 0,361181948 | -0,355301282 | -0,302006089 | 0,015467902 | 0,393467902 | -0,866532098 |
| chrIV | 44 | 145 | 101 | 0,065539893 | 0,337234132 | 0,162721251 | -0,983335176 | -0,8358349 | -0,376249574 | -0,075249574 | -1,335249574 |
| chrIV | 44 | 155 | 111 | 0,059840772 | 0,077823261 | 0,434687048 | -0,164453569 | -0,139785534 | 0,401249498 | 0,632249498 | -0,627750502 |
| chrIV | 57 | 155 | 98 | 0,037044288 | 0,140823044 | 0,208269204 | -0,812441383 | -0,690575176 | -0,237481081 | 0,084518919 | -1,175481081 |
| chrIV | 1765 | 1923 | 158 | 0,19377012 | 0,200116957 | 0,491943329 | -0,020196453 | -0,017166985 | 0,00297484 | -0,09502516 | -1,35502516 |
| chrIV | 6558 | 6652 | 94 | 1,980444606 | 11,64754809 | 0,145321813 | -1,056710731 | -0,898204121 | -0,504337508 | -0,154337508 | -1,414337508 |
| chrIV | 8786 | 8917 | 131 | 0,219416165 | 0,311293045 | 0,413439528 | -0,218706025 | -0,185900121 | -0,076132351 | 0,014867649 | -1,245132351 |
| chrIV | 10468 | 10622 | 154 | 0,059840772 | 0,066705652 | 0,472876041 | -0,06804215 | -0,057835827 | -0,143432049 | -0,213432049 | -1,473432049 |
| chrIV | 11706 | 11865 | 159 | 0,054141651 | 0,037058696 | 0,593656197 | 0,236960278 | 0,201416237 | 0,214856707 | 0,109856707 | -1,150143293 |
| chrIV | 12342 | 12498 | 156 | 0,028495606 | 0,051882174 | 0,354520937 | -0,373143192 | -0,317171714 | -0,270260576 | -0,354260576 | -1,614260576 |
| chrIV | 12922 | 13062 | 140 | 2,191312089 | 1,378583483 | 0,613830866 | 0,289317699 | 0,245920045 | 0,134244554 | 0,162244554 | -1,097755446 |
| chrIV | 13187 | 13340 | 153 | 0,09688506 | 0,051882174 | 0,65125268 | 0,388704654 | 0,330398956 | 0,255320579 | 0,192320579 | -1,067679421 |
| chrIV | 15846 | 16000 | 154 | 0,524319147 | 0,452116089 | 0,536972784 | 0,092810095 | 0,07888858 | -0,012552837 | -0,082552837 | -1,342552837 |
| chrIV | 16215 | 16372 | 157 | 0,789328282 | 0,496586524 | 0,613826265 | 0,289305674 | 0,245909823 | 0,281892997 | 0,190892997 | -1,069107003 |
| chrIV | 16517 | 16676 | 159 | 4,827155628 | 3,60210523 | 0,572666537 | 0,183167078 | 0,155692016 | 0,172945287 | 0,067945287 | -1,192054713 |
| chrIV | 22332 | 22485 | 153 | 0,552814753 | 0,496586524 | 0,526790624 | 0,067204689 | 0,057123986 | -0,020674134 | -0,083674134 | -1,343674134 |
| chrIV | 22780 | 22921 | 141 | 0,216566604 | 0,103764348 | 0,676071427 | 0,456741101 | 0,388229936 | 0,222885276 | 0,243885276 | -1,016114724 |
| chrIV | 24904 | 25025 | 121 | 0,068389454 | 0,374292827 | 0,154488799 | -1,017369684 | -0,864764232 | -0,290808288 | -0,129808288 | -1,389808288 |
| chrIV | 26118 | 26260 | 142 | 0,113982423 | 0,096352609 | 0,541908887 | 0,105243965 | 0,08945737 | -0,068219095 | -0,054219095 | -1,314219095 |
| chrIV | 28132 | 28284 | 152 | 0,427434088 | 0,452116089 | 0,485968964 | -0,035177847 | -0,02990117 | -0,144268241 | -0,200268241 | -1,460268241 |
| chrIV | 28565 | 28649 | 84 | 0,205168362 | 0,188999349 | 0,520510322 | 0,051434423 | 0,043719259 | 0,494520691 | 0,914520691 | -0,345479309 |
| chrIV | 28606 | 28762 | 156 | 0,510071344 | 0,322410653 | 0,612711561 | 0,286393336 | 0,243434336 | 0,28800125 | 0,20400125 | -1,05599875 |
| chrIV | 29713 | 29864 | 151 | 0,062690333 | 0,074117392 | 0,458236793 | -0,104876779 | -0,089145262 | -0,150773062 | -0,199773062 | -1,459773062 |
| chrIV | 30032 | 30191 | 159 | 0,074088575 | 0,085235 | 0,465019536 | -0,087795679 | -0,074626327 | -0,057734494 | -0,162734494 | -1,422734494 |
| chrIV | 31789 | 31954 | 165 | 0,401788042 | 0,670762394 | 0,374609929 | -0,319668215 | -0,271717983 | -0,186088039 | -0,333088039 | -1,593088039 |
| chrIV | 32229 | 32385 | 156 | 0,108283302 | 0,192705218 | 0,359758911 | -0,359103289 | -0,305237796 | -0,26634737 | -0,35034737 | -1,61034737 |
| chrIV | 33584 | 33743 | 159 | 0,421734966 | 0,463233697 | 0,476553559 | -0,058805386 | -0,049984578 | -0,026758771 | -0,131758771 | -1,391758771 |
| chrIV | 34942 | 35098 | 156 | 0,247911771 | 0,289057827 | 0,461686791 | -0,096185077 | -0,081757315 | -0,046390195 | -0,130390195 | -1,390390195 |
| chrIV | 43249 | 43399 | 150 | 0,199469241 | 0,137117174 | 0,592624158 | 0,234300517 | 0,19915544 | 0,088461441 | 0,046461441 | -1,213538559 |
| chrIV | 43986 | 44132 | 146 | 0,173823196 | 0,148234783 | 0,539726407 | 0,09974448 | 0,084782808 | -0,111102157 | -0,125102157 | -1,385102157 |
| chrIV | 51938 | 52044 | 106 | 0,076938136 | 0,337234132 | 0,18576361 | -0,893616302 | -0,759573857 | -0,222172823 | 0,043827177 | -1,216172823 |
| chrIV | 55412 | 55541 | 129 | 2,060232302 | 3,472399795 | 0,37237833 | -0,325560819 | -0,276726696 | -0,041002235 | 0,063997765 | -1,196002235 |
| chrIV | 56586 | 56757 | 171 | 0,190920559 | 0,188999349 | 0,502528442 | 0,006337906 | 0,00538722 | 0,221133588 | 0,032133588 | -1,227866412 |
| chrIV | 60333 | 60467 | 134 | 0,547115632 | 0,515115871 | 0,515062517 | 0,037765106 | 0,03210034 | 0,092544827 | 0,162544827 | -1,097455173 |
| chrIV | 61937 | 62077 | 140 | 3,376729291 | 1,84922892 | 0,646145483 | 0,374934696 | 0,318694492 | 0,203565359 | 0,231565359 | -1,028434641 |
| chrIV | 77124 | 77282 | 158 | 0,054141651 | 0,081529131 | 0,399066404 | -0,255764342 | -0,217399691 | -0,193926047 | -0,291926047 | -1,551926047 |
| chrIV | 78906 | 79062 | 156 | 0,273557816 | 0,240881523 | 0,531759132 | 0,079692611 | 0,067738719 | 0,111806886 | 0,027806886 | -1,232193114 |
| chrIV | 86969 | 87132 | 163 | 0,273557816 | 0,185293479 | 0,596179675 | 0,243470869 | 0,206950239 | 0,234734919 | 0,101734919 | -1,158265081 |
| chrIV | 90699 | 90847 | 148 | 0,381841118 | 0,34835174 | 0,522931872 | 0,057513369 | 0,048886364 | -0,122873582 | -0,150873582 | -1,410873582 |
| chrIV | 94580 | 94722 | 142 | 0,353345512 | 0,363175219 | 0,493140669 | -0,017194641 | -0,014615445 | -0,17929725 | -0,16529725 | -1,42529725 |
| chrIV | 97197 | 97345 | 148 | 0,629752889 | 0,218646305 | 0,742283695 | 0,650401964 | 0,552841669 | 0,376156247 | 0,348156247 | -0,911843753 |
| chrIV | 101729 | 101892 | 163 | 0,102584181 | 0,070411522 | 0,592986873 | 0,235235114 | 0,199949847 | 0,224554779 | 0,091554779 | -1,168445221 |
| chrIV | 103149 | 103279 | 130 | 0,105433742 | 0,040764565 | 0,721169375 | 0,586318873 | 0,498371042 | 0,632787552 | 0,730787552 | -0,529212448 |
| chrIV | 109515 | 109652 | 137 | 0,042743409 | 0,062999783 | 0,404219015 | -0,242441566 | -0,206075331 | -0,232348703 | -0,183348703 | -1,443348703 |
| chrIV | 109515 | 109656 | 141 | 8,169690193 | 5,821921108 | 0,583899168 | 0,211878705 | 0,180096899 | 0,025168222 | 0,046168222 | -1,213831778 |
| chrIV | 109515 | 109666 | 151 | 1,929152515 | 0,989467177 | 0,660981121 | 0,415142268 | 0,352870928 | 0,293994135 | 0,244994135 | -1,015005865 |
| chrIV | 109518 | 109656 | 138 | 0,034194727 | 0,044470435 | 0,434687048 | -0,164453569 | -0,139785534 | -0,204156087 | -0,162156087 | -1,422156087 |
| chrIV | 110479 | 110614 | 135 | 1,242408414 | 1,215525222 | 0,505468657 | 0,01370832 | 0,011652072 | 0,008085083 | 0,071085083 | -1,188914917 |
| chrIV | 113056 | 113240 | 184 | 0,094035499 | 0,203822827 | 0,315705458 | -0,47974192 | -0,407780632 | -0,007311362 | -0,287311362 | -1,547311362 |
| chrIV | 118783 | 118936 | 153 | 0,544266071 | 0,455821958 | 0,544218164 | 0,111066427 | 0,094406463 | 0,031868623 | -0,031131377 | -1,291131377 |
| chrIV | 121491 | 121639 | 148 | 0,629752889 | 0,396528045 | 0,613626219 | 0,288782843 | 0,245465417 | 0,068811191 | 0,040811191 | -1,219188809 |
| chrIV | 122619 | 122777 | 158 | 0,356195073 | 0,407645654 | 0,466321133 | -0,084520926 | -0,071842787 | -0,080248176 | -0,178248176 | -1,438248176 |
| chrIV | 124114 | 124238 | 124 | 5,796006227 | 10,83966852 | 0,348408244 | -0,38962145 | -0,331178233 | 0,095008169 | 0,235008169 | -1,024991831 |
| chrIV | 125012 | 125174 | 162 | 0,122531105 | 0,151940653 | 0,446425184 | -0,134698361 | -0,114493607 | -0,106641519 | -0,232641519 | -1,492641519 |
| chrIV | 126907 | 127064 | 157 | 5,308731367 | 3,876339579 | 0,57797391 | 0,196712944 | 0,167206002 | 0,189707939 | 0,098707939 | -1,161292061 |
| chrIV | 126907 | 127064 | 157 | 5,308731367 | 3,876339579 | 0,57797391 | 0,196712944 | 0,167206002 | 0,189707939 | 0,098707939 | -1,161292061 |
| chrIV | 127133 | 127254 | 121 | 0,059840772 | 0,285351958 | 0,173354671 | -0,940991251 | -0,799842563 | -0,239539227 | -0,078539227 | -1,338539227 |
| chrIV | 130391 | 130540 | 149 | 0,139628469 | 0,114881957 | 0,54861591 | 0,122165207 | 0,103840426 | -0,076628326 | -0,111628326 | -1,371628326 |
| chrIV | 134454 | 134627 | 173 | 1,060036537 | 1,108055004 | 0,488926098 | -0,027761722 | -0,023597463 | 0,239164791 | 0,036164791 | -1,223835209 |
| chrIV | 137744 | 137906 | 162 | 0,512920905 | 0,533645219 | 0,490098899 | -0,024820929 | -0,021097789 | -0,012042036 | -0,138042036 | -1,398042036 |
| chrIV | 147424 | 147558 | 134 | 2,570303646 | 1,130290221 | 0,69456518 | 0,508832496 | 0,432507621 | 0,484370843 | 0,554370843 | -0,705629157 |
| chrIV | 148138 | 148259 | 121 | 0,837770812 | 1,360054135 | 0,381181774 | -0,302378494 | -0,25702172 | 0,285254248 | 0,446254248 | -0,813745752 |
| chrIV | 148138 | 148287 | 149 | 7,619725 | 4,57674893 | 0,624748189 | 0,317975367 | 0,270279062 | 0,086153719 | 0,051153719 | -1,208846281 |
| chrIV | 148147 | 148256 | 109 | 0,065539893 | 0,251999131 | 0,206399492 | -0,818977968 | -0,696131272 | -0,107393819 | 0,137606181 | -1,122393819 |
| chrIV | 148147 | 148259 | 112 | 11,95105709 | 41,96526711 | 0,221659345 | -0,766601154 | -0,651610981 | -0,105336115 | 0,118663885 | -1,141336115 |
| chrIV | 148147 | 148287 | 140 | 1,31934655 | 1,93075805 | 0,40593972 | -0,238002134 | -0,202301814 | -0,333186748 | -0,305186748 | -1,565186748 |
| chrIV | 148150 | 148259 | 109 | 0,028495606 | 0,08894087 | 0,242646977 | -0,69781331 | -0,593141313 | -0,009588753 | 0,235411247 | -1,024588753 |
| chrIV | 148662 | 148839 | 177 | 0,139628469 | 0,17788174 | 0,439760565 | -0,151576297 | -0,128839853 | 0,204776147 | -0,026223853 | -1,286223853 |
| chrIV | 150319 | 150438 | 119 | 2,288197149 | 11,37331374 | 0,167492246 | -0,964122541 | -0,81950416 | -0,268528407 | -0,093528407 | -1,353528407 |
| chrIV | 150791 | 150954 | 163 | 0,692443222 | 0,96352609 | 0,418149791 | -0,206629036 | -0,175634681 | -0,145821231 | -0,278821231 | -1,538821231 |
| chrIV | 151146 | 151295 | 149 | 1,305098747 | 0,70411522 | 0,649556876 | 0,384124399 | 0,326505739 | 0,145355341 | 0,110355341 | -1,149644659 |
| chrIV | 154011 | 154153 | 142 | 0,042743409 | 0,081529131 | 0,343948944 | -0,401709424 | -0,34145301 | -0,503524939 | -0,489524939 | -1,749524939 |
| chrIV | 156456 | 156616 | 160 | 0,125380666 | 0,151940653 | 0,452113333 | -0,120323781 | -0,102275214 | -0,100583844 | -0,212583844 | -1,472583844 |
| chrIV | 156781 | 156930 | 149 | 0,658248495 | 0,52623348 | 0,555726899 | 0,140144018 | 0,119122416 | -0,065907042 | -0,100907042 | -1,360907042 |
| chrIV | 158096 | 158254 | 158 | 0,567062556 | 0,544762828 | 0,510028431 | 0,025140196 | 0,021369167 | 0,014869717 | -0,083130283 | -1,343130283 |
| chrIV | 158096 | 158254 | 158 | 0,567062556 | 0,544762828 | 0,510028431 | 0,025140196 | 0,021369167 | 0,014869717 | -0,083130283 | -1,343130283 |
| chrIV | 158496 | 158646 | 150 | 0,105433742 | 0,059293913 | 0,640048823 | 0,358589298 | 0,304800903 | 0,195796353 | 0,153796353 | -1,106203647 |
| chrIV | 162837 | 162965 | 128 | 1,211063248 | 1,93816979 | 0,384558155 | -0,293530996 | -0,249501347 | 0,012581614 | 0,124581614 | -1,135418386 |
| chrIV | 163802 | 163934 | 132 | 1,236709293 | 1,14140783 | 0,520037168 | 0,050246866 | 0,042709836 | 0,141690088 | 0,225690088 | -1,034309912 |
| chrIV | 163887 | 164037 | 150 | 0,53571739 | 0,26682261 | 0,667527338 | 0,433095594 | 0,368131255 | 0,272521266 | 0,230521266 | -1,029478734 |
| chrIV | 163887 | 164037 | 150 | 0,53571739 | 0,26682261 | 0,667527338 | 0,433095594 | 0,368131255 | 0,272521266 | 0,230521266 | -1,029478734 |
| chrIV | 164671 | 164820 | 149 | 0,920408068 | 0,552174567 | 0,625029826 | 0,318718021 | 0,270910318 | 0,087536287 | 0,052536287 | -1,207463713 |
| chrIV | 168971 | 169122 | 151 | 0,450230572 | 0,229763914 | 0,662109151 | 0,418226261 | 0,355492322 | 0,303711889 | 0,254711889 | -1,005288111 |
| chrIV | 180947 | 181107 | 160 | 0,042743409 | 0,037058696 | 0,535617563 | 0,08939893 | 0,07598909 | 0,078506457 | -0,033493543 | -1,293493543 |
| chrIV | 185030 | 185179 | 149 | 3,453667427 | 2,327286095 | 0,59742176 | 0,246679232 | 0,209677347 | 0,031572531 | -0,003427469 | -1,263427469 |
| chrIV | 185042 | 185156 | 114 | 0,09973462 | 0,448410219 | 0,181949392 | -0,90796108 | -0,771766918 | -0,230167402 | -0,020167402 | -1,280167402 |
| chrIV | 196778 | 196946 | 168 | 0,065539893 | 0,044470435 | 0,595761274 | 0,242390685 | 0,206032083 | 0,364632468 | 0,196632468 | -1,063367532 |
| chrIV | 198969 | 199115 | 146 | 0,834921251 | 0,400233914 | 0,675964668 | 0,456444093 | 0,387977479 | 0,192389934 | 0,178389934 | -1,081610066 |
| chrIV | 200831 | 200925 | 94 | 0,159575393 | 0,96723196 | 0,14161728 | -1,073081478 | -0,912119256 | -0,491682972 | -0,141682972 | -1,401682972 |
| chrIV | 202938 | 203097 | 159 | 0,108283302 | 0,163058261 | 0,399066404 | -0,255764342 | -0,217399691 | -0,196079247 | -0,301079247 | -1,561079247 |

TABLE 2

| Chrom | Start | End | Length | Section A | Section B | A/A+B | Z-score | Z * 0.85 | Phase correction | Length correction | ΔLK |
|---|---|---|---|---|---|---|---|---|---|---|---|
| chrIV | 204891 | 205041 | 150 | 1,70688679 | 1,104349134 | 0,607165971 | 0,271940125 | 0,231149106 | 0,137165497 | 0,095165497 | **-1,164834503** |
| chrIV | 205789 | 205917 | 128 | 0,387540239 | 0,396528045 | 0,49426848 | -0,014367283 | -0,012212191 | 0,233289463 | 0,345289463 | **-0,914710537** |
| chrIV | 214844 | 215003 | 159 | 0,04844253 | 0,055588044 | 0,465656665 | -0,08619258 | -0,073263693 | -0,061304769 | -0,166304769 | **-1,426304769** |
| chrIV | 224519 | 224676 | 157 | 0,558513874 | 0,515115871 | 0,52021088 | 0,050682852 | 0,043080425 | 0,059131226 | -0,031868774 | **-1,291868774** |
| chrIV | 224519 | 224676 | 157 | 0,558513874 | 0,515115871 | 0,52021088 | 0,050682852 | 0,043080425 | 0,059131226 | -0,031868774 | **-1,291868774** |
| chrIV | 228430 | 228579 | 149 | 0,606956404 | 0,578115654 | 0,512168353 | 0,030506268 | 0,025930328 | -0,147284036 | -0,182284036 | **-1,442284036** |
| chrIV | 230300 | 230458 | 158 | 0,04844253 | 0,051882174 | 0,482857442 | -0,042983253 | -0,036535765 | -0,045873142 | -0,143873142 | **-1,403873142** |
| chrIV | 235048 | 235188 | 140 | 1,079983461 | 0,637409568 | 0,628850498 | 0,328810397 | 0,279488837 | 0,147889203 | 0,175889203 | **-1,084110797** |
| chrIV | 235556 | 235704 | 148 | 1,131275552 | 0,804173699 | 0,584502824 | 0,213426448 | 0,181412481 | 0,005979333 | -0,022020667 | **-1,282020667** |
| chrIV | 235556 | 235704 | 148 | 1,131275552 | 0,804173699 | 0,584502824 | 0,213426448 | 0,181412481 | 0,005979333 | -0,022020667 | **-1,282020667** |
| chrIV | 240743 | 240895 | 152 | 0,14532759 | 0,037058696 | 0,796812048 | 0,83028812 | 0,705744902 | 0,587609955 | 0,531609955 | **-0,728390045** |
| chrIV | 241083 | 241236 | 153 | 0,142478029 | 0,17788174 | 0,444743825 | -0,138952542 | -0,118109661 | -0,185388003 | -0,248388003 | **-1,508388003** |
| chrIV | 243252 | 243404 | 152 | 0,327699467 | 0,481763045 | 0,404835878 | -0,240849509 | -0,204722082 | -0,319774479 | -0,375774479 | **-1,635774479** |
| chrIV | 247954 | 248116 | 162 | 0,276407377 | 0,544762828 | 0,336601809 | -0,421755324 | -0,358492025 | -0,343018586 | -0,469018586 | **-1,729018586** |
| chrIV | 249148 | 249302 | 154 | 0,079787696 | 0,066705652 | 0,544650641 | 0,112157259 | 0,09533367 | 0,004636672 | -0,065363328 | **-1,325363328** |
| chrIV | 249297 | 249444 | 147 | 0,071239015 | 0,051882174 | 0,578608892 | 0,198335964 | 0,168585569 | -0,002329213 | -0,023329213 | **-1,283329213** |
| chrIV | 253799 | 253940 | 141 | 0,139628469 | 0,081529131 | 0,631352796 | 0,335438395 | 0,285122636 | 0,125831948 | 0,146831948 | **-1,113168052** |
| chrIV | 262881 | 263029 | 148 | 0,054141651 | 0,048176305 | 0,529151025 | 0,07313593 | 0,062165541 | -0,12841368 | -0,15641368 | **-1,41641368** |
| chrIV | 268349 | 268491 | 142 | 1,951949 | 0,893114568 | 0,686082737 | 0,484777019 | 0,412060466 | 0,248688773 | 0,262688773 | **-0,997311227** |
| chrIV | 269754 | 269886 | 132 | 0,567062556 | 0,503998263 | 0,529440108 | 0,073862515 | 0,062783138 | 0,176711988 | 0,260711988 | **-0,999288012** |
| chrIV | 275399 | 275562 | 163 | 0,606956404 | 0,681880002 | 0,470933628 | -0,072923171 | -0,061984696 | -0,029105141 | -0,162105141 | **-1,422105141** |
| chrIV | 277085 | 277203 | 118 | 0,042743409 | 0,322410653 | 0,117055822 | -1,189834 | -1,0113589 | -0,464351584 | -0,282351584 | **-1,542351584** |
| chrIV | 281092 | 281244 | 152 | 0,227964847 | 0,244587392 | 0,482411949 | -0,044100996 | -0,037485847 | -0,157127037 | -0,213127037 | **-1,473127037** |
| chrIV | 282751 | 282862 | 111 | 0,034194727 | 0,303881305 | 0,101145079 | -1,275053919 | -1,083795831 | -0,560919947 | -0,329919947 | **-1,589919947** |
| chrIV | 287334 | 287498 | 164 | 0,846319493 | 0,893114568 | 0,486548764 | -0,033723639 | -0,028665093 | 0,026443501 | -0,113556499 | **-1,373556499** |
| chrIV | 290503 | 290642 | 139 | 1,399134247 | 0,922761525 | 0,60258271 | 0,260037847 | 0,22103217 | 0,12500996 | 0,16000996 | **-1,09999004** |
| chrIV | 291728 | 291875 | 147 | 0,082637257 | 0,037058696 | 0,690393076 | 0,496964836 | 0,42242011 | 0,257577313 | 0,236577313 | **-1,023422687** |
| chrIV | 294716 | 294818 | 102 | 0,122531105 | 0,715232829 | 0,146259704 | -1,052610794 | -0,894719174 | -0,374369689 | -0,080369689 | **-1,340369689** |
| chrIV | 303242 | 303392 | 150 | 0,336248149 | 0,274234349 | 0,550790809 | 0,127659576 | 0,10851064 | 0,026362858 | -0,015637142 | **-1,275637142** |
| chrIV | 305394 | 305543 | 149 | 0,63545201 | 0,785644351 | 0,447156173 | -0,132849576 | -0,112922139 | -0,275453476 | -0,310453476 | **-1,570453476** |
| chrIV | 307871 | 308025 | 154 | 0,381841118 | 0,392822175 | 0,492912368 | -0,017766994 | -0,015101945 | -0,108235216 | -0,178235216 | **-1,438235216** |
| chrIV | 308997 | 309152 | 155 | 0,889062902 | 0,774526742 | 0,534424403 | 0,086396544 | 0,073437062 | -0,022995753 | -0,099995753 | **-1,359995753** |
| chrIV | 311358 | 311498 | 140 | 0,658248495 | 0,607762611 | 0,519938958 | 0,050000381 | 0,042500324 | -0,092086032 | -0,064086032 | **-1,324086032** |
| chrIV | 313115 | 313272 | 157 | 0,04844253 | 0,037058696 | 0,566571175 | 0,16765125 | 0,142503563 | 0,16365864 | 0,07265864 | **-1,18734136** |
| chrIV | 315132 | 315251 | 119 | 0,085486818 | 0,125999566 | 0,404219015 | -0,242441566 | -0,206075331 | 0,334562048 | 0,509562048 | **-0,750437952** |
| chrIV | 316257 | 316381 | 124 | 0,136778908 | 0,218646305 | 0,384831753 | -0,292815072 | -0,248892811 | 0,162674824 | 0,302674824 | **-0,957325176** |
| chrIV | 317932 | 318025 | 93 | 2,225506816 | 8,186265899 | 0,213749078 | -0,793480106 | -0,67445809 | -0,311586684 | 0,045413316 | **-1,214586684** |
| chrIV | 320773 | 320934 | 161 | 0,327699467 | 0,329822392 | 0,498385662 | -0,004046557 | -0,003439573 | 0,01191987 | -0,10708013 | **-1,36708013** |
| chrIV | 324041 | 324187 | 146 | 0,361894194 | 0,255705001 | 0,585969342 | 0,217188668 | 0,184610368 | -0,011155009 | -0,025155009 | **-1,285155009** |
| chrIV | 326087 | 326232 | 145 | 0,071239015 | 0,055588044 | 0,561702018 | 0,155285859 | 0,13199298 | -0,038473147 | -0,045473147 | **-1,305473147** |
| chrIV | 331066 | 331232 | 166 | 0,034194727 | 0,044470435 | 0,434687048 | -0,164453569 | -0,139785534 | -0,021411356 | -0,175411356 | **-1,435411356** |
| chrIV | 331066 | 331232 | 166 | 0,034194727 | 0,044470435 | 0,434687048 | -0,164453569 | -0,139785534 | -0,021411356 | -0,175411356 | **-1,435411356** |
| chrIV | 336500 | 336644 | 144 | 0,738036191 | 0,585527393 | 0,557612947 | 0,144919911 | 0,123181924 | -0,009088495 | -0,009088495 | **-1,269088495** |
| chrIV | 339003 | 339155 | 152 | 0,051292091 | 0,051882174 | 0,497140355 | -0,007168129 | -0,00609291 | -0,113959585 | -0,169959585 | **-1,429959585** |
| chrIV | 339626 | 339796 | 170 | 0,159575393 | 0,244587392 | 0,394829506 | -0,266753431 | -0,226740416 | -0,015911274 | -0,197911274 | **-1,457911274** |
| chrIV | 343754 | 343879 | 125 | 0,53856695 | 1,500877179 | 0,264075364 | -0,630831465 | -0,536206745 | -0,136432816 | -0,003432816 | **-1,263432816** |
| chrIV | 344625 | 344793 | 168 | 0,034194727 | 0,040764565 | 0,456177292 | -0,110069087 | -0,093558724 | 0,074357169 | -0,093642831 | **-1,353642831** |
| chrIV | 344904 | 345057 | 153 | 1,932002076 | 1,133996091 | 0,63013804 | 0,332218971 | 0,282386125 | 0,213629633 | 0,150629633 | **-1,109370367** |
| chrIV | 346054 | 346184 | 130 | 0,498673102 | 0,629997828 | 0,441823288 | -0,146348124 | -0,124395906 | 0,005861123 | 0,103861123 | **-1,156138877** |
| chrIV | 346695 | 346849 | 154 | 0,304902982 | 0,292763697 | 0,510155565 | 0,025458977 | 0,02164013 | -0,073788246 | -0,143788246 | **-1,403788246** |
| chrIV | 354236 | 354370 | 134 | 0,233663968 | 0,214940436 | 0,520868645 | 0,052333815 | 0,044483743 | 0,083318708 | 0,153318708 | **-1,106681292** |
| chrIV | 357373 | 357527 | 154 | 0,398938482 | 0,244587392 | 0,619926094 | 0,305286692 | 0,259493688 | 0,148148738 | 0,078148738 | **-1,181851262** |
| chrIV | 357519 | 357616 | 97 | 0,14532759 | 1,389701092 | 0,094674185 | -1,31250924 | -1,115632854 | -0,665323564 | -0,336323564 | **-1,596323564** |
| chrIV | 367261 | 367414 | 153 | 2,348037921 | 2,097522181 | 0,528175948 | 0,070685446 | 0,060082629 | -0,015313528 | -0,078313528 | **-1,338313528** |
| chrIV | 369734 | 369884 | 150 | 0,572761677 | 0,78935022 | 0,420495319 | -0,200626496 | -0,170532521 | -0,257769134 | -0,299769134 | **-1,559769134** |
| chrIV | 372733 | 372859 | 126 | 0,125380666 | 0,552174567 | 0,185048627 | -0,896291207 | -0,761847526 | -0,457174564 | -0,331174564 | **-1,591174564** |
| chrIV | 377302 | 377457 | 155 | 0,393239361 | 0,555880437 | 0,414320049 | -0,216446021 | -0,183979117 | -0,265995297 | -0,342995297 | **-1,602995297** |
| chrIV | 377961 | 378102 | 141 | 0,102584181 | 0,074117392 | 0,58055047 | 0,203301913 | 0,172806626 | 0,002616803 | 0,023616803 | **-1,236383197** |
| chrIV | 379551 | 379707 | 156 | 0,373292436 | 0,407645654 | 0,47800516 | -0,055160849 | -0,046886722 | -0,006401121 | -0,090401121 | **-1,350401121** |
| chrIV | 380490 | 380611 | 121 | 0,142478029 | 0,548468698 | 0,206206967 | -0,819653028 | -0,696705073 | -0,163833261 | -0,002833261 | **-1,262833261** |
| chrIV | 380895 | 381031 | 136 | 0,287805619 | 0,251999131 | 0,533166147 | 0,083231199 | 0,070746519 | 0,057012071 | 0,113012071 | **-1,146987929** |
| chrIV | 383984 | 384136 | 152 | 0,589859041 | 0,270528479 | 0,685573683 | 0,483342407 | 0,410841046 | 0,304802025 | 0,248802025 | **-1,011197975** |
| chrIV | 384406 | 384547 | 141 | 1,17401896 | 1,115466743 | 0,512787199 | 0,032058244 | 0,027249508 | -0,144858974 | -0,123858974 | **-1,383858974** |
| chrIV | 386414 | 386562 | 148 | 0,105433742 | 0,040764565 | 0,721169375 | 0,586318873 | 0,498371042 | 0,306521717 | 0,278521717 | **-0,981478283** |
| chrIV | 389563 | 389664 | 101 | 0,054141651 | 0,426175002 | 0,112720745 | -1,212185213 | -1,030357431 | -0,544448267 | -0,243448267 | **-1,503448267** |
| chrIV | 394898 | 395059 | 161 | 0,282106498 | 0,303881305 | 0,481420426 | -0,046588934 | -0,039600594 | -0,021132814 | -0,140132814 | **-1,400132814** |
| chrIV | 395659 | 395798 | 139 | 0,612655525 | 0,511410002 | 0,545035419 | 0,113127892 | 0,096158708 | -0,007800588 | 0,027199412 | **-1,232800588** |
| chrIV | 405838 | 405992 | 154 | 2,037435817 | 1,523112397 | 0,572225313 | 0,182042499 | 0,154736124 | 0,048614901 | -0,021385099 | **-1,281385099** |
| chrIV | 406138 | 406284 | 146 | 0,82922213 | 0,311293045 | 0,72705927 | 0,603943118 | 0,51335165 | 0,321241964 | 0,307241964 | **-0,952758036** |
| chrIV | 407097 | 407258 | 161 | 0,159575393 | 0,100058479 | 0,614617006 | 0,291373097 | 0,247667133 | 0,259288758 | 0,140288758 | **-1,119711242** |
| chrIV | 407639 | 407774 | 135 | 3,040481143 | 2,664520227 | 0,532950116 | 0,082687823 | 0,070284649 | 0,063855797 | 0,126855797 | **-1,133144203** |
| chrIV | 410001 | 410147 | 146 | 0,085486818 | 0,100058479 | 0,460732874 | -0,09858756 | -0,083799426 | -0,283684068 | -0,297684068 | **-1,557684068** |
| chrIV | 412510 | 412666 | 156 | 0,421734966 | 0,88199696 | 0,323482886 | -0,457981444 | -0,389284228 | -0,345284093 | -0,429284093 | **-1,689284093** |
| chrIV | 414505 | 414611 | 106 | 0,108283302 | 0,389116306 | 0,217698809 | -0,779988545 | -0,662990263 | -0,1449347 | 0,1210653 | **-1,1389347** |
| chrIV | 414505 | 414627 | 122 | 0,108283302 | 0,43729261 | 0,198475225 | -0,847080146 | -0,720018124 | -0,203091331 | -0,049091331 | **-1,309091331** |
| chrIV | 417857 | 418015 | 158 | 0,58700948 | 0,655938915 | 0,472271803 | -0,069560337 | -0,059126287 | -0,054643102 | -0,152643102 | **-1,412643102** |
| chrIV | 418210 | 418363 | 153 | 0,270708255 | 0,237175653 | 0,533012074 | 0,082843661 | 0,070417112 | 0,001423578 | -0,061576422 | **-1,321576422** |
| chrIV | 418328 | 418478 | 150 | 0,076938136 | 0,100058479 | 0,434687048 | -0,164453569 | -0,139785534 | -0,239852845 | -0,281852845 | **-1,541852845** |
| chrIV | 419977 | 420127 | 150 | 0,056991212 | 0,062999783 | 0,474962408 | -0,062801193 | -0,053381014 | -0,148435292 | -0,190435292 | **-1,450435292** |
| chrIV | 430775 | 430925 | 150 | 0,079787696 | 0,066705652 | 0,544650641 | 0,112157259 | 0,09533367 | -0,003814201 | -0,045814201 | **-1,305814201** |
| chrIV | 431218 | 431366 | 148 | 1,265204899 | 0,989467177 | 0,561148077 | 0,153880643 | 0,130798547 | -0,06736047 | -0,09536047 | **-1,35536047** |
| chrIV | 433624 | 433778 | 154 | 5,154855096 | 3,980103927 | 0,56429975 | 0,161879814 | 0,137597842 | 0,037578162 | -0,032421838 | **-1,292421838** |
| chrIV | 444592 | 444742 | 150 | 0,179522317 | 0,114881957 | 0,609781626 | 0,278749922 | 0,236937434 | 0,141476159 | 0,099476159 | **-1,160523841** |
| chrIV | 447153 | 447303 | 150 | 1,860763061 | 1,252583918 | 0,597672882 | 0,2473282 | 0,21022897 | 0,130187425 | 0,088187425 | **-1,171812575** |
| chrIV | 452119 | 452270 | 151 | 0,247911771 | 0,281646088 | 0,468148601 | -0,079924628 | -0,067935934 | -0,118081433 | -0,167081433 | **-1,427081433** |
| chrIV | 456027 | 456181 | 154 | 1,114178188 | 0,678174133 | 0,621629004 | 0,309761946 | 0,263297654 | 0,164999118 | 0,094999118 | **-1,165000882** |
| chrIV | 464196 | 464345 | 149 | 0,350495952 | 0,277940218 | 0,55772721 | 0,145209355 | 0,123427951 | -0,040150598 | -0,075150598 | **-1,335150598** |
| chrIV | 465523 | 465668 | 145 | 0,287805619 | 0,200116957 | 0,589859197 | 0,227182794 | 0,193105375 | 0,028778533 | 0,021778533 | **-1,238221467** |
| chrIV | 472028 | 472162 | 134 | 1,649895578 | 1,222936961 | 0,574309695 | 0,187357158 | 0,159253584 | 0,184583951 | 0,254583951 | **-1,005416049** |
| chrIV | 480768 | 480918 | 150 | 0,356195073 | 0,292763697 | 0,548871653 | 0,122811089 | 0,104389426 | 0,019671842 | -0,022328158 | **-1,282328158** |
| chrIV | 481714 | 481864 | 150 | 0,960301917 | 0,559586306 | 0,631824039 | 0,336688251 | 0,286185013 | 0,207850393 | 0,165850393 | **-1,094149607** |
| chrIV | 481903 | 482061 | 158 | 0,059840772 | 0,055588044 | 0,518421434 | 0,046192108 | 0,039263292 | 0,051683921 | -0,046316079 | **-1,306316079** |
| chrIV | 487395 | 487546 | 151 | 0,247911771 | 0,222352175 | 0,527175798 | 0,068172392 | 0,057946533 | 0,00069642 | -0,04830358 | **-1,30830358** |
| chrIV | 488401 | 488552 | 151 | 0,478726178 | 0,318704784 | 0,600335579 | 0,254215803 | 0,216083433 | 0,143503378 | 0,094503378 | **-1,165496622** |
| chrIV | 488508 | 488615 | 107 | 0,054141651 | 0,140823044 | 0,277699771 | -0,589688448 | -0,501235181 | 0,056980382 | 0,315980382 | **-0,944019618** |
| chrIV | 488508 | 488632 | 124 | 0,054141651 | 0,203822827 | 0,209880258 | -0,806836798 | -0,685811279 | -0,283800432 | -0,143800432 | **-1,403800432** |
| chrIV | 491144 | 491308 | 164 | 0,159575393 | 0,185293479 | 0,462713239 | -0,093600543 | -0,079560461 | -0,022663446 | -0,162663446 | **-1,422663446** |
| chrIV | 491144 | 491308 | 164 | 0,159575393 | 0,185293479 | 0,462713239 | -0,093600543 | -0,079560461 | -0,022663446 | -0,162663446 | **-1,422663446** |
| chrIV | 492176 | 492332 | 156 | 1,12272687 | 0,930173264 | 0,546897948 | 0,117827796 | 0,100153627 | 0,154909575 | 0,070909575 | **-1,189090425** |
| chrIV | 494125 | 494275 | 150 | 0,19661968 | 0,111176087 | 0,638799168 | 0,355250862 | 0,301963233 | 0,213833903 | 0,171833903 | **-1,088166097** |
| chrIV | 499197 | 499355 | 158 | 0,162424953 | 0,077823261 | 0,676071427 | 0,456741101 | 0,388229936 | 0,395689628 | 0,297689628 | **-0,962310372** |

TABLE 2

| Chrom | Start | End | Length | Section A | Section B | A/A+B | Z-score | Z * 0.85 | Phase correction | Length correction | ΔLK |
|---|---|---|---|---|---|---|---|---|---|---|---|
| chrIV | 499814 | 499967 | 153 | 0,102584181 | 0,118587827 | 0,463820816 | -0,090812431 | -0,077190567 | -0,142156544 | -0,205156544 | -1,465156544 |
| chrIV | 503596 | 503736 | 140 | 0,185221438 | 0,107470218 | 0,63282104 | 0,339334281 | 0,288434139 | 0,164015832 | 0,192015832 | -1,067984168 |
| chrIV | 503768 | 503924 | 156 | 0,034194727 | 0,040764565 | 0,456177292 | -0,110069087 | -0,093558724 | -0,042951835 | -0,126951835 | -1,386951835 |
| chrIV | 508653 | 508795 | 142 | 0,669646737 | 0,363175219 | 0,648366094 | 0,380912991 | 0,323776043 | 0,167412466 | 0,181412466 | -1,078587534 |
| chrIV | 513880 | 514029 | 149 | 0,091185939 | 0,051882174 | 0,637360324 | 0,351411905 | 0,298700119 | 0,141571865 | 0,106571865 | -1,153428135 |
| chrIV | 517444 | 517573 | 129 | 0,208017923 | 0,340940001 | 0,378932362 | -0,308285993 | -0,262043094 | -0,02776644 | 0,07723356 | -1,18276644 |
| chrIV | 518819 | 518929 | 110 | 0,151026711 | 0,737468046 | 0,169980419 | -0,954242636 | -0,81110624 | -0,219534086 | 0,018465914 | -1,241534086 |
| chrIV | 521357 | 521482 | 125 | 1,581506124 | 4,13204458 | 0,276799175 | -0,592376721 | -0,503520213 | -0,112691432 | 0,020308568 | -1,239691432 |
| chrIV | 522483 | 522574 | 91 | 0,407487163 | 1,004290656 | 0,28863406 | -0,557379572 | -0,473772636 | -0,145891693 | 0,225108307 | -1,034891693 |
| chrIV | 522691 | 522836 | 145 | 0,359044634 | 0,166764131 | 0,682842618 | 0,475662609 | 0,404313218 | 0,23553621 | 0,22853621 | -1,03146379 |
| chrIV | 524988 | 525151 | 163 | 0,09973462 | 0,062999783 | 0,61286746 | 0,286800501 | 0,243780426 | 0,28443055 | 0,15143055 | -1,10856945 |
| chrIV | 525488 | 525645 | 157 | 0,510071344 | 0,559586306 | 0,476854762 | -0,058049093 | -0,049341729 | -0,045523345 | -0,136523345 | -1,396523345 |
| chrIV | 527030 | 527163 | 133 | 1,450426337 | 1,371171744 | 0,514044274 | 0,035211049 | 0,029929391 | 0,123047456 | 0,200047456 | -1,059952544 |
| chrIV | 527878 | 528014 | 136 | 0,056991212 | 0,059293913 | 0,490098899 | -0,024820929 | -0,021097789 | -0,036376702 | 0,019623298 | -1,240376702 |
| chrIV | 528421 | 528544 | 123 | 0,039893848 | 0,044470435 | 0,472876041 | -0,06804215 | -0,057835827 | 0,429844903 | 0,576844903 | -0,683155097 |
| chrIV | 528421 | 528548 | 127 | 6,39726351 | 8,26408916 | 0,43633515 | -0,160267576 | -0,13622744 | 0,163004468 | 0,282004468 | -0,977995532 |
| chrIV | 529656 | 529804 | 148 | 0,227964847 | 0,159352392 | 0,588573975 | 0,223878237 | 0,190296502 | -0,015964923 | -0,043964923 | -1,303964923 |
| chrIV | 529731 | 529886 | 155 | 0,085486818 | 0,085235 | 0,500737507 | 0,001848658 | 0,001571359 | -0,082662276 | -0,159662276 | -1,419662276 |
| chrIV | 530359 | 530519 | 160 | 0,056991212 | 0,040764565 | 0,582995843 | 0,209563572 | 0,178129036 | 0,188408754 | 0,076408754 | -1,183591246 |
| chrIV | 531763 | 531899 | 136 | 0,071239015 | 0,066705652 | 0,516431814 | 0,041200102 | 0,035020087 | 0,0180789 | 0,0740789 | -1,1859211 |
| chrIV | 532306 | 532429 | 123 | 0,042743409 | 0,062999783 | 0,404219015 | -0,242441566 | -0,206075331 | 0,266195855 | 0,413195855 | -0,846804145 |
| chrIV | 532306 | 532433 | 127 | 6,414360873 | 8,293736117 | 0,436110863 | -0,160837072 | -0,136711511 | 0,166145066 | 0,285145066 | -0,974854934 |
| chrIV | 533541 | 533689 | 148 | 0,225115286 | 0,200116957 | 0,529393736 | 0,07374596 | 0,062684066 | -0,140567679 | -0,168567679 | -1,428567679 |
| chrIV | 533616 | 533771 | 155 | 0,136778908 | 0,077823261 | 0,637360324 | 0,351411905 | 0,298700119 | 0,225537375 | 0,148537375 | -1,111462625 |
| chrIV | 534244 | 534404 | 160 | 0,065539893 | 0,055588044 | 0,541079911 | 0,103154718 | 0,08768151 | 0,101896699 | -0,010103301 | -1,270103301 |
| chrIV | 535648 | 535784 | 136 | 0,065539893 | 0,051882174 | 0,558156527 | 0,146296985 | 0,124352437 | 0,107449203 | 0,163449203 | -1,096550797 |
| chrIV | 536109 | 536229 | 120 | 1,66129382 | 9,805730905 | 0,144875751 | -1,058666911 | -0,899866875 | -0,341892143 | -0,173892143 | -1,433892143 |
| chrIV | 537426 | 537574 | 148 | 0,205168362 | 0,129705435 | 0,612673681 | 0,286294411 | 0,24335025 | 0,040540711 | 0,012540711 | -1,247459289 |
| chrIV | 537501 | 537656 | 155 | 0,111132863 | 0,070411522 | 0,612152576 | 0,284933816 | 0,242193743 | 0,177657373 | 0,100657373 | -1,159342627 |
| chrIV | 540906 | 541052 | 146 | 0,051292091 | 0,066705652 | 0,434687048 | -0,164453569 | -0,139785534 | -0,337192237 | -0,351192237 | -1,611192237 |
| chrIV | 541731 | 541872 | 141 | 0,364743755 | 0,166764131 | 0,686243355 | 0,485229878 | 0,412445397 | 0,245874374 | 0,266874374 | -0,993125626 |
| chrIV | 545183 | 545329 | 146 | 3,61609238 | 1,830699572 | 0,663893979 | 0,423114063 | 0,359646953 | 0,162085357 | 0,148085357 | -1,111914643 |
| chrIV | 549706 | 549856 | 150 | 0,24506221 | 0,292763697 | 0,455653413 | -0,111390332 | -0,094681782 | -0,182719952 | -0,224719952 | -1,484719952 |
| chrIV | 549706 | 549856 | 150 | 0,24506221 | 0,292763697 | 0,455653413 | -0,111390332 | -0,094681782 | -0,182719952 | -0,224719952 | -1,484719952 |
| chrIV | 552005 | 552154 | 149 | 0,279256937 | 0,207528696 | 0,573675388 | 0,185739279 | 0,157878387 | -0,002012773 | -0,037012773 | -1,297012773 |
| chrIV | 557304 | 557456 | 152 | 0,136778908 | 0,077823261 | 0,637360324 | 0,351411905 | 0,298700119 | 0,189460635 | 0,133460635 | -1,126539365 |
| chrIV | 560465 | 560600 | 135 | 0,435982769 | 0,344645871 | 0,558502144 | 0,147172696 | 0,125096791 | 0,088646014 | 0,151646014 | -1,108353986 |
| chrIV | 570638 | 570778 | 140 | 0,131079787 | 0,100058479 | 0,567105523 | 0,169009772 | 0,143658307 | 0,019324095 | 0,047324095 | -1,212675905 |
| chrIV | 570638 | 570778 | 140 | 0,131079787 | 0,100058479 | 0,567105523 | 0,169009772 | 0,143658307 | 0,019324095 | 0,047324095 | -1,212675905 |
| chrIV | 574209 | 574359 | 150 | 0,170973635 | 0,107470218 | 0,614032715 | 0,289845327 | 0,246368528 | 0,16875484 | 0,12675484 | -1,13324516 |
| chrIV | 575376 | 575525 | 149 | 0,381841118 | 0,344645871 | 0,52559939 | 0,064212254 | 0,054580416 | -0,102404726 | -0,137404726 | -1,397404726 |
| chrIV | 585684 | 585843 | 159 | 0,19377012 | 0,100058479 | 0,659466508 | 0,41100757 | 0,349356434 | 0,35877657 | 0,25377657 | -1,00622343 |
| chrIV | 587983 | 588137 | 154 | 1,000195765 | 0,696703481 | 0,589425546 | 0,226067518 | 0,19215739 | 0,084461587 | 0,014461587 | -1,245538413 |
| chrIV | 601658 | 601795 | 137 | 0,444531451 | 0,17417587 | 0,718484226 | 0,578344513 | 0,491592836 | 0,460089359 | 0,509089359 | -0,750910641 |
| chrIV | 607967 | 608120 | 153 | 0,059840772 | 0,077823261 | 0,434687048 | -0,164453569 | -0,139785534 | -0,212272632 | -0,275272632 | -1,535272632 |
| chrIV | 615631 | 615766 | 135 | 0,034194727 | 0,092646739 | 0,269586342 | -0,614064532 | -0,521954852 | -0,570681315 | -0,507681315 | -1,767681315 |
| chrIV | 618044 | 618203 | 159 | 0,601257283 | 0,563292176 | 0,516300341 | 0,040870272 | 0,034739731 | 0,043236421 | -0,061763579 | -1,321763579 |
| chrIV | 625810 | 625961 | 151 | 0,136778908 | 0,140823044 | 0,492715945 | -0,018259434 | -0,015520519 | -0,084621247 | -0,133621247 | -1,393621247 |
| chrIV | 640754 | 640908 | 154 | 7,152397065 | 4,154279798 | 0,632581717 | 0,338698905 | 0,287894069 | 0,170648784 | 0,100648784 | -1,159351216 |
| chrIV | 643481 | 643635 | 154 | 2,672887827 | 2,012287181 | 0,570499036 | 0,177644815 | 0,150998092 | 0,03058229 | -0,03941771 | -1,29941771 |
| chrIV | 645695 | 645844 | 149 | 0,09973462 | 0,062999783 | 0,61286746 | 0,286800501 | 0,243780426 | 0,095451835 | 0,060451835 | -1,199548165 |
| chrIV | 646491 | 646637 | 146 | 0,074088575 | 0,062999783 | 0,540443961 | 0,101552255 | 0,086319417 | -0,102936315 | -0,116936315 | -1,376936315 |
| chrIV | 647130 | 647280 | 150 | 0,447381012 | 0,400233914 | 0,527811625 | 0,069769969 | 0,059304474 | -0,026296514 | -0,068296514 | -1,328296514 |
| chrIV | 648526 | 648683 | 157 | 0,076938136 | 0,037058696 | 0,674914686 | 0,453525163 | 0,385496389 | 0,393714289 | 0,302714289 | -0,957285711 |
| chrIV | 648994 | 649155 | 161 | 0,071239015 | 0,074117392 | 0,490098899 | -0,024820929 | -0,021097789 | -0,018407214 | -0,137407214 | -1,397407214 |
| chrIV | 649174 | 649309 | 135 | 0,105433742 | 0,066705652 | 0,612490489 | 0,28581604 | 0,242943634 | 0,192964386 | 0,255964386 | -1,004035614 |
| chrIV | 650212 | 650352 | 140 | 0,165274514 | 0,122293696 | 0,574731518 | 0,188433343 | 0,160168342 | 0,035811504 | 0,063811504 | -1,196188496 |
| chrIV | 650218 | 650352 | 134 | 0,125380666 | 0,118587827 | 0,51392155 | 0,034903236 | 0,029667751 | 0,044452902 | 0,114452902 | -1,145547098 |
| chrIV | 650630 | 650785 | 155 | 0,108283302 | 0,077823261 | 0,581834946 | 0,206589951 | 0,175601459 | 0,114175556 | 0,037175556 | -1,222824444 |
| chrIV | 650761 | 650869 | 108 | 0,088336378 | 0,166764131 | 0,346280681 | -0,395381497 | -0,336074272 | 0,261766643 | 0,513766643 | -0,746233357 |
| chrIV | 654067 | 654214 | 147 | 1,883559546 | 0,97093783 | 0,659856815 | 0,412072382 | 0,350261525 | 0,185407766 | 0,164407766 | -1,095592234 |
| chrIV | 654067 | 654214 | 147 | 1,883559546 | 0,97093783 | 0,659856815 | 0,412072382 | 0,350261525 | 0,185407766 | 0,164407766 | -1,095592234 |
| chrIV | 656839 | 656998 | 159 | 0,302053422 | 0,214940436 | 0,584249537 | 0,212776972 | 0,180860426 | 0,187677433 | 0,082677433 | -1,177322567 |
| chrIV | 658487 | 658641 | 154 | 0,333398588 | 0,389116306 | 0,461441821 | -0,096801992 | -0,082281694 | -0,209803086 | -0,279803086 | -1,539803086 |
| chrIV | 658600 | 658747 | 147 | 0,444531451 | 0,359469349 | 0,552899264 | 0,13298977 | 0,113041304 | -0,040267512 | -0,061267512 | -1,321267512 |
| chrIV | 662245 | 662401 | 156 | 0,34194727 | 0,281646088 | 0,548349763 | 0,121493107 | 0,103269141 | 0,142831041 | 0,058831041 | -1,201168959 |
| chrIV | 665084 | 665248 | 164 | 0,04844253 | 0,085235 | 0,36238349 | -0,352095051 | -0,299280793 | -0,237505983 | -0,377505983 | -1,637505983 |
| chrIV | 665597 | 665732 | 135 | 0,649699813 | 0,670762394 | 0,492024542 | -0,019992839 | -0,016993913 | -0,06501557 | -0,00201557 | -1,26201557 |
| chrIV | 666941 | 667094 | 153 | 1,22246149 | 1,115466743 | 0,522882385 | 0,057389119 | 0,048780751 | -0,021306476 | -0,084306476 | -1,344306476 |
| chrIV | 666948 | 667109 | 161 | 0,094035499 | 0,140823044 | 0,400392074 | -0,252332398 | -0,214482538 | -0,210752762 | -0,329752762 | -1,589752762 |
| chrIV | 669497 | 669656 | 159 | 0,492973981 | 0,52623348 | 0,483683646 | -0,040910443 | -0,034773876 | -0,028388218 | -0,133388218 | -1,393388218 |
| chrIV | 669741 | 669906 | 165 | 0,48727486 | 0,930173264 | 0,343769096 | -0,402198167 | -0,341868442 | -0,256900623 | -0,403900623 | -1,663900623 |
| chrIV | 677555 | 677687 | 132 | 0,581310359 | 1,167348917 | 0,332432033 | -0,43320745 | -0,368226333 | -0,264701538 | -0,180701538 | -1,440701538 |
| chrIV | 682440 | 682595 | 155 | 0,076938136 | 0,044470435 | 0,633712557 | 0,341702373 | 0,290447017 | 0,237390633 | 0,160390633 | -1,099609367 |
| chrIV | 682928 | 683076 | 148 | 0,646850252 | 0,452116089 | 0,588598784 | 0,223942004 | 0,190350703 | -0,011709395 | -0,039709395 | -1,299709395 |
| chrIV | 684905 | 685053 | 148 | 0,259310013 | 0,207528696 | 0,555459536 | 0,139467258 | 0,118547169 | -0,08843622 | -0,11643622 | -1,37643622 |
| chrIV | 685367 | 685521 | 154 | 0,253610892 | 0,122293696 | 0,674668254 | 0,452840646 | 0,384914549 | 0,25646396 | 0,18646396 | -1,07353604 |
| chrIV | 685409 | 685539 | 130 | 0,282106498 | 0,359469349 | 0,439708725 | -0,151707742 | -0,128951581 | -0,012834812 | 0,085165188 | -1,174834812 |
| chrIV | 692976 | 693125 | 149 | 0,062690333 | 0,037058696 | 0,628480635 | 0,327831948 | 0,278657156 | 0,133180597 | 0,098180597 | -1,161819403 |
| chrIV | 699427 | 699543 | 116 | 0,262159574 | 1,015408265 | 0,205202077 | -0,823182615 | -0,699705223 | -0,142881546 | 0,053118454 | -1,206881546 |
| chrIV | 699429 | 699572 | 143 | 0,14817715 | 0,070411522 | 0,677881195 | 0,461782069 | 0,392514759 | 0,235146202 | 0,242146202 | -1,017853798 |
| chrIV | 701924 | 702080 | 156 | 0,113982423 | 0,148234783 | 0,434687048 | -0,164453569 | -0,139785534 | -0,101347585 | -0,185347585 | -1,445347585 |
| chrIV | 705337 | 705438 | 101 | 0,14532759 | 1,096937395 | 0,116985982 | -1,190189384 | -1,011660976 | -0,535147377 | -0,234147377 | -1,494147377 |
| chrIV | 705533 | 705696 | 163 | 0,131079787 | 0,111176087 | 0,541079911 | 0,103154718 | 0,08768151 | 0,123220117 | -0,009779883 | -1,269779883 |
| chrIV | 709396 | 709543 | 147 | 0,407487163 | 0,389116306 | 0,511530742 | 0,028907309 | 0,024571212 | -0,133813573 | -0,154813573 | -1,414813573 |
| chrIV | 710394 | 710553 | 159 | 1,250957096 | 1,267407396 | 0,496733932 | -0,00818691 | -0,006958874 | -0,005386623 | -0,110386623 | -1,370386623 |
| chrIV | 710648 | 710758 | 110 | 0,418885406 | 0,463233697 | 0,47486264 | -0,063051771 | -0,053594005 | 0,534241165 | 0,772241165 | -0,487758835 |
| chrIV | 711680 | 711827 | 147 | 0,262159574 | 0,255705001 | 0,506231912 | 0,015621722 | 0,013278464 | -0,154110026 | -0,175110026 | -1,435110026 |
| chrIV | 712887 | 713032 | 145 | 2,159966922 | 1,337818918 | 0,617524063 | 0,298984497 | 0,254136822 | 0,068779988 | 0,061779988 | -1,198220012 |
| chrIV | 717311 | 717466 | 155 | 0,14532759 | 0,218646305 | 0,399280256 | -0,255210514 | -0,216928937 | -0,256478262 | -0,333478262 | -1,593478262 |
| chrIV | 723685 | 723835 | 150 | 0,094035499 | 0,074117392 | 0,559226183 | 0,149007602 | 0,126656462 | 0,034925894 | -0,007074106 | -1,267074106 |
| chrIV | 724692 | 724830 | 138 | 0,324849907 | 0,370586958 | 0,467116317 | -0,082520731 | -0,070142622 | -0,139829463 | -0,097829463 | -1,357829463 |
| chrIV | 727870 | 728018 | 148 | 0,87766466 | 0,663350655 | 0,569536624 | 0,175194569 | 0,148915383 | -0,056269525 | -0,084269525 | -1,344269525 |
| chrIV | 729848 | 729954 | 106 | 0,037044288 | 0,163058261 | 0,185126515 | -0,8959995 | -0,761599575 | -0,227131484 | 0,038868516 | -1,221131484 |
| chrIV | 729848 | 729970 | 122 | 0,071239015 | 0,270528479 | 0,208442921 | -0,811835827 | -0,690060453 | -0,19237542 | -0,03837542 | -1,29837542 |
| chrIV | 730903 | 730997 | 94 | 0,558513874 | 2,20128653 | 0,202374735 | -0,833168919 | -0,708193581 | -0,267649955 | 0,082350045 | -1,177649955 |
| chrIV | 734743 | 734893 | 150 | 0,079787696 | 0,144528914 | 0,355692324 | -0,369997104 | -0,314497538 | -0,401921558 | -0,443921558 | -1,703921558 |
| chrIV | 736895 | 737045 | 150 | 0,24221265 | 0,200116957 | 0,547584077 | 0,119559826 | 0,101625852 | 0,01731623 | -0,02468377 | -1,28468377 |
| chrIV | 737420 | 737548 | 128 | 0,359044634 | 0,522527611 | 0,407277606 | -0,234553618 | -0,199370575 | 0,035744442 | 0,147744442 | -1,112255558 |
| chrIV | 737638 | 737793 | 155 | 0,133929347 | 0,200116957 | 0,400930486 | -0,250939386 | -0,213298478 | -0,257408687 | -0,334408687 | -1,594408687 |
| chrIV | 745330 | 745443 | 113 | 2,020338454 | 17,69923311 | 0,102453466 | -1,267694886 | -1,077540653 | -0,526221438 | -0,309221438 | -1,569221438 |

TABLE 2

| Chrom | Start | End | Length | Section A | Section B | A/A+B | Z-score | Z * 0.85 | Phase correction | Length correction | ΔLK |
|---|---|---|---|---|---|---|---|---|---|---|---|
| chrIV | 747582 | 747734 | 152 | 0,749434433 | 0,804173699 | 0,482383181 | -0,044173177 | -0,0375472 | -0,147098449 | -0,203098449 | -1,463098449 |
| chrIV | 750682 | 750850 | 168 | 0,079787696 | 0,148234783 | 0,349911537 | -0,385559308 | -0,327725412 | -0,15250291 | -0,32050291 | -1,58050291 |
| chrIV | 760658 | 760811 | 153 | 0,284956058 | 0,281646088 | 0,502920895 | 0,007321663 | 0,006223413 | -0,074555805 | -0,137555805 | -1,397555805 |
| chrIV | 761833 | 761982 | 149 | 0,78362916 | 0,515115871 | 0,603374135 | 0,262090424 | 0,22277686 | 0,076154591 | 0,041154591 | -1,218845409 |
| chrIV | 762005 | 762157 | 152 | 1,461824579 | 0,848644133 | 0,632696115 | 0,339002603 | 0,288152213 | 0,185932814 | 0,129932814 | -1,130067186 |
| chrIV | 764493 | 764644 | 151 | 0,199469241 | 0,229763914 | 0,464710702 | -0,088572826 | -0,075286902 | -0,146314455 | -0,195314455 | -1,455314455 |
| chrIV | 766598 | 766749 | 151 | 0,182371877 | 0,129705435 | 0,58438044 | 0,213112623 | 0,181145729 | 0,105787675 | 0,056787675 | -1,203212325 |
| chrIV | 768253 | 768411 | 158 | 0,079787696 | 0,129705435 | 0,380860679 | -0,303221116 | -0,257737948 | -0,249347271 | -0,347347271 | -1,607347271 |
| chrIV | 771149 | 771299 | 150 | 0,424584527 | 0,285351958 | 0,598059877 | 0,248328506 | 0,21107923 | 0,123772928 | 0,081772928 | -1,178227072 |
| chrIV | 773973 | 774101 | 128 | 0,14817715 | 0,340940001 | 0,302948179 | -0,51593994 | -0,438548949 | -0,197804847 | -0,085804847 | -1,345804847 |
| chrIV | 779473 | 779647 | 174 | 0,034194727 | 0,125999566 | 0,213457836 | -0,794480647 | -0,67530855 | -0,379471933 | -0,589471933 | -1,849471933 |
| chrIV | 779988 | 780110 | 122 | 0,92895675 | 3,331576751 | 0,218037659 | -0,778837715 | -0,662012058 | -0,179242314 | -0,025242314 | -1,285242314 |
| chrIV | 785291 | 785441 | 150 | 0,153876272 | 0,277940218 | 0,356346446 | -0,368241871 | -0,31300559 | -0,394944994 | -0,436944994 | -1,696944994 |
| chrIV | 787217 | 787366 | 149 | 1,191116324 | 0,822703046 | 0,591471282 | 0,231331294 | 0,1966316 | 0,047899321 | 0,012899321 | -1,247100679 |
| chrIV | 787217 | 787366 | 149 | 1,191116324 | 0,822703046 | 0,591471282 | 0,231331294 | 0,1966316 | 0,047899321 | 0,012899321 | -1,247100679 |
| chrIV | 789145 | 789305 | 160 | 0,128230226 | 0,122293696 | 0,51184823 | 0,029703476 | 0,025247955 | 0,0383882 | -0,0736118 | -1,3336118 |
| chrIV | 800500 | 800661 | 161 | 0,051292091 | 0,059293913 | 0,463820816 | -0,090812431 | -0,077190567 | -0,066438319 | -0,185438319 | -1,445438319 |
| chrIV | 803067 | 803219 | 152 | 0,139628469 | 0,181587609 | 0,434687048 | -0,164453569 | -0,139785534 | -0,238898307 | -0,294898307 | -1,554898307 |
| chrIV | 805417 | 805524 | 107 | 0,173823196 | 0,489174784 | 0,262177564 | -0,636646501 | -0,541149525 | 0,021629282 | 0,280629282 | -0,979370718 |
| chrIV | 805417 | 805530 | 113 | 0,09973462 | 0,478057176 | 0,172613424 | -0,943888071 | -0,80230486 | -0,25002957 | -0,03302957 | -1,29302957 |
| chrIV | 810559 | 810694 | 135 | 0,227964847 | 0,277940218 | 0,450607955 | -0,124125498 | -0,105506674 | -0,16535258 | -0,10235258 | -1,36235258 |
| chrIV | 815236 | 815393 | 157 | 0,071239015 | 0,066705652 | 0,516431814 | 0,041200102 | 0,035020087 | 0,052672634 | -0,038327366 | -1,298327366 |
| chrIV | 815520 | 815678 | 158 | 0,284956058 | 0,511410002 | 0,357820445 | -0,364290771 | -0,309647155 | -0,301962576 | -0,399962576 | -1,659962576 |
| chrIV | 826418 | 826583 | 165 | 0,265009134 | 0,329822392 | 0,445519651 | -0,136989237 | -0,116440852 | -0,02680278 | -0,17380278 | -1,43380278 |
| chrIV | 827506 | 827652 | 146 | 0,97739928 | 0,400233914 | 0,709477155 | 0,551857937 | 0,469079246 | 0,282469943 | 0,268469943 | -0,991530057 |
| chrIV | 835382 | 835525 | 143 | 0,088336378 | 0,070411522 | 0,556456986 | 0,141992377 | 0,12069352 | -0,039362561 | -0,032362561 | -1,292362561 |
| chrIV | 835382 | 835528 | 146 | 21,76494374 | 15,62765201 | 0,582065602 | 0,207180626 | 0,176103532 | -0,0059835 | -0,0199835 | -1,2799835 |
| chrIV | 835383 | 835528 | 145 | 0,04844253 | 0,044470435 | 0,521375354 | 0,053605728 | 0,045564869 | -0,133809352 | -0,140809352 | -1,400809352 |
| chrIV | 835384 | 835528 | 144 | 0,105433742 | 0,077823261 | 0,575332675 | 0,189967434 | 0,161472318 | 0,024917168 | 0,024917168 | -1,235082832 |
| chrIV | 835385 | 835528 | 143 | 0,051292091 | 0,074117392 | 0,408996909 | -0,230126057 | -0,195607148 | -0,3334253 | -0,3264253 | -1,5864253 |
| chrIV | 835386 | 835528 | 142 | 0,028495606 | 0,070411522 | 0,288104674 | -0,558930214 | -0,475090682 | -0,634812245 | -0,620812245 | -1,880812245 |
| chrIV | 835932 | 836065 | 133 | 5,018076188 | 3,409400012 | 0,595442345 | 0,241567496 | 0,205332371 | 0,274766364 | 0,351766364 | -0,908233636 |
| chrIV | 836520 | 836671 | 151 | 0,111132863 | 0,040764565 | 0,731630971 | 0,617753167 | 0,525090192 | 0,443146629 | 0,394146629 | -0,865853371 |
| chrIV | 837322 | 837477 | 155 | 3,413773579 | 2,731225879 | 0,55553684 | 0,139662925 | 0,118713486 | 0,08656511 | 0,00956511 | -1,25043489 |
| chrIV | 838477 | 838608 | 131 | 0,302053422 | 5,870097412 | 0,048938114 | -1,655238008 | -1,406952307 | -1,302460103 | -1,211460103 | -2,471460103 |
| chrIV | 843960 | 844136 | 176 | 0,421734966 | 0,993173047 | 0,298065289 | -0,529973104 | -0,450477139 | -0,133694818 | -0,357694818 | -1,617694818 |
| chrIV | 844482 | 844623 | 141 | 9,451992456 | 5,707039151 | 0,623522181 | 0,314744522 | 0,267532844 | 0,10053779 | 0,12153779 | -1,13846221 |
| chrIV | 844482 | 844649 | 167 | 0,185221438 | 0,285351958 | 0,393607968 | -0,269927623 | -0,229438479 | -0,10155302 | -0,26255302 | -1,52255302 |
| chrIV | 850466 | 850631 | 165 | 0,470177496 | 0,463233697 | 0,503719582 | 0,009323745 | 0,007925183 | 0,113646055 | -0,033353945 | -1,293353945 |
| chrIV | 850466 | 850631 | 165 | 0,470177496 | 0,463233697 | 0,503719582 | 0,009323745 | 0,007925183 | 0,113646055 | -0,033353945 | -1,293353945 |
| chrIV | 857181 | 857346 | 165 | 0,085486818 | 0,081529131 | 0,51184823 | 0,029703476 | 0,025247955 | 0,146205924 | -0,000794076 | -1,260794076 |
| chrIV | 861657 | 861796 | 139 | 0,498673102 | 0,433586741 | 0,534907844 | 0,087612945 | 0,074471003 | -0,03451816 | 0,00048184 | -1,25951816 |
| chrIV | 863651 | 863788 | 137 | 3,476463912 | 2,920225228 | 0,54347864 | 0,109201437 | 0,092821221 | 0,063382656 | 0,112382656 | -1,147617344 |
| chrIV | 866143 | 866295 | 152 | 0,313451664 | 0,148234783 | 0,678927584 | 0,464702061 | 0,394996752 | 0,292782287 | 0,236782287 | -1,023217713 |
| chrIV | 867460 | 867598 | 138 | 0,062690333 | 0,122293696 | 0,338895921 | -0,415478239 | -0,353156503 | -0,425394345 | -0,383394345 | -1,643394345 |
| chrIV | 870210 | 870347 | 137 | 0,136778908 | 0,140823044 | 0,492715945 | -0,018259434 | -0,015520519 | -0,065218809 | -0,016218809 | -1,276218809 |
| chrIV | 872409 | 872511 | 102 | 0,094035499 | 0,129705435 | 0,420287416 | -0,201158254 | -0,170984516 | 0,377162022 | 0,671162022 | -0,588837978 |
| chrIV | 873832 | 873980 | 148 | 0,082637257 | 0,085235 | 0,492262738 | -0,019395657 | -0,016486308 | -0,228638041 | -0,256638041 | -1,516638041 |
| chrIV | 874621 | 874762 | 141 | 0,170973635 | 0,185293479 | 0,479902939 | -0,050397187 | -0,042837609 | -0,200702297 | -0,179702297 | -1,439702297 |
| chrIV | 875359 | 875502 | 143 | 0,185221438 | 0,111176087 | 0,624908854 | 0,318399007 | 0,270639156 | 0,132418923 | 0,139418923 | -1,120581077 |
| chrIV | 876645 | 876791 | 146 | 0,068389454 | 0,051882174 | 0,568624996 | 0,172874585 | 0,146943398 | -0,033156374 | -0,047156374 | -1,307156374 |
| chrIV | 877438 | 877587 | 149 | 0,085486818 | 0,077823261 | 0,523463207 | 0,058847485 | 0,050020363 | -0,102636453 | -0,137636453 | -1,397636453 |
| chrIV | 878298 | 878451 | 153 | 0,247911771 | 0,400233914 | 0,3824939 | -0,298937414 | -0,254096802 | -0,328776053 | -0,391776053 | -1,651776053 |
| chrIV | 878496 | 878645 | 149 | 0,065539893 | 0,100058479 | 0,395776194 | -0,264295292 | -0,224650998 | -0,380764605 | -0,415764605 | -1,675764605 |
| chrIV | 879292 | 879438 | 146 | 0,091185939 | 0,051882174 | 0,637360324 | 0,351411905 | 0,298700119 | 0,12084834 | 0,10684834 | -1,15315166 |
| chrIV | 881327 | 881484 | 157 | 0,045592969 | 0,059293913 | 0,434687048 | -0,164453569 | -0,139785534 | -0,124286737 | -0,215286737 | -1,475286737 |
| chrIV | 881795 | 881956 | 161 | 0,085486818 | 0,085235 | 0,500737507 | 0,001848658 | 0,001571359 | 0,013345779 | -0,105654221 | -1,365654221 |
| chrIV | 881975 | 882110 | 135 | 0,076938136 | 0,062999783 | 0,549801916 | 0,125160904 | 0,106386768 | 0,05340794 | 0,11640794 | -1,14359206 |
| chrIV | 883013 | 883153 | 140 | 0,136778908 | 0,077823261 | 0,637360324 | 0,351411905 | 0,298700119 | 0,185498198 | 0,213498198 | -1,046501802 |
| chrIV | 883019 | 883153 | 134 | 0,136778908 | 0,118587827 | 0,535617563 | 0,08939893 | 0,07598909 | 0,103760962 | 0,173760962 | -1,086239038 |
| chrIV | 883431 | 883586 | 155 | 0,079787696 | 0,08894087 | 0,472876041 | -0,06804215 | -0,057835827 | -0,07585025 | -0,15285025 | -1,41285025 |
| chrIV | 883562 | 883670 | 108 | 0,082637257 | 0,140823044 | 0,369807329 | -0,332363684 | -0,282509132 | 0,305213328 | 0,557213328 | -0,702786672 |
| chrIV | 883696 | 883848 | 152 | 0,039893848 | 0,037058696 | 0,518421434 | 0,046192108 | 0,039263292 | -0,058578115 | -0,114578115 | -1,374578115 |
| chrIV | 884819 | 884969 | 150 | 3,630340183 | 2,834990228 | 0,561508841 | 0,154795779 | 0,131576412 | 0,059139204 | 0,017139204 | -1,242860796 |
| chrIV | 888390 | 888567 | 177 | 0,416035845 | 0,61146848 | 0,404899361 | -0,2406857 | -0,204582845 | 0,162128822 | -0,068871178 | -1,328871178 |
| chrIV | 889668 | 889827 | 159 | 0,649699813 | 1,159937178 | 0,359022177 | -0,361073698 | -0,306912643 | -0,304720174 | -0,409720174 | -1,669720174 |
| chrIV | 890834 | 890988 | 154 | 0,14817715 | 0,040764565 | 0,784247935 | 0,786620371 | 0,668627316 | 0,538218417 | 0,468218417 | -0,791781583 |
| chrIV | 891389 | 891533 | 144 | 0,572761677 | 0,192705218 | 0,748251402 | 0,668997305 | 0,568647709 | 0,424514894 | 0,424514894 | -0,835485106 |
| chrIV | 891986 | 892153 | 167 | 0,125380666 | 0,237175653 | 0,345823971 | -0,396619676 | -0,337126724 | -0,187359805 | -0,348359805 | -1,608359805 |
| chrIV | 896334 | 896482 | 148 | 0,227964847 | 0,070411522 | 0,764017766 | 0,71928641 | 0,611393448 | 0,40732014 | 0,37932014 | -0,88067986 |
| chrIV | 897629 | 897776 | 147 | 0,113982423 | 0,085235 | 0,572150875 | 0,181852797 | 0,154574878 | -0,005044597 | -0,026044597 | -1,286044597 |
| chrIV | 899028 | 899177 | 149 | 0,199469241 | 0,26311674 | 0,431204682 | -0,173307964 | -0,147311769 | -0,297654491 | -0,332654491 | -1,592654491 |
| chrIV | 906466 | 906593 | 127 | 0,378991558 | 0,96352609 | 0,282299125 | -0,576025049 | -0,489621292 | -0,213777757 | -0,094777757 | -1,354777757 |
| chrIV | 907937 | 908075 | 138 | 0,056991212 | 0,125999566 | 0,311443082 | -0,491764035 | -0,41799943 | -0,49183835 | -0,44983835 | -1,70983835 |
| chrIV | 908538 | 908667 | 129 | 0,062690333 | 0,326116523 | 0,161237725 | -0,989383693 | -0,840976139 | -0,620390989 | -0,515390989 | -1,775390989 |
| chrIV | 909620 | 909773 | 153 | 0,042743409 | 0,074117392 | 0,365763444 | -0,34309514 | -0,291630869 | -0,363271496 | -0,426271496 | -1,686271496 |
| chrIV | 915796 | 915980 | 184 | 0,179522317 | 0,359469349 | 0,333070673 | -0,431449801 | -0,366732331 | 0,053933553 | -0,226066447 | -1,486066447 |
| chrIV | 917617 | 917765 | 148 | 0,082637257 | 0,048176305 | 0,631717813 | 0,336406466 | 0,285945496 | 0,07692569 | 0,04892569 | -1,21107431 |
| chrIV | 924934 | 925100 | 166 | 0,233663968 | 0,114881957 | 0,670396499 | 0,441008282 | 0,37485704 | 0,496978806 | 0,342978806 | -0,917021194 |
| chrIV | 926425 | 926570 | 145 | 0,393239361 | 0,240881523 | 0,620133118 | 0,305830423 | 0,259955859 | 0,08751488 | 0,08051488 | -1,17948512 |
| chrIV | 927161 | 927289 | 128 | 0,162424953 | 0,074117392 | 0,686663326 | 0,486414449 | 0,413452282 | 0,675965023 | 0,787965023 | -0,472034977 |
| chrIV | 928961 | 929115 | 154 | 0,054141651 | 0,048176305 | 0,529151025 | 0,07313593 | 0,062165541 | -0,070149499 | -0,140149499 | -1,400149499 |
| chrIV | 930238 | 930383 | 145 | 0,760832676 | 0,585527393 | 0,565103417 | 0,163921215 | 0,139333033 | -0,033991281 | -0,040991281 | -1,300991281 |
| chrIV | 931111 | 931260 | 149 | 0,923257629 | 0,478057176 | 0,658850978 | 0,409329261 | 0,347929872 | 0,213172003 | 0,178172003 | -1,081827997 |
| chrIV | 933440 | 933600 | 160 | 0,037044288 | 0,074117392 | 0,333246923 | -0,430964963 | -0,366320218 | -0,354468871 | -0,466468871 | -1,726468871 |
| chrIV | 944365 | 944524 | 159 | 0,142478029 | 0,188999349 | 0,429827309 | -0,17681384 | -0,150291764 | -0,148022674 | -0,253022674 | -1,513022674 |
| chrIV | 944403 | 944557 | 154 | 0,364743755 | 0,314998914 | 0,536590936 | 0,091848853 | 0,078071525 | -0,049890931 | -0,119890931 | -1,379890931 |
| chrIV | 948002 | 948171 | 169 | 0,034194727 | 0,100058479 | 0,254703244 | -0,659762131 | -0,560797812 | -0,379066687 | -0,554066687 | -1,814066687 |
| chrIV | 949233 | 949387 | 154 | 0,188070999 | 0,055588044 | 0,771861355 | 0,744990787 | 0,633242169 | 0,510635993 | 0,440635993 | -0,819364007 |
| chrIV | 955015 | 955163 | 148 | 1,188266763 | 0,685585872 | 0,634130316 | 0,342812703 | 0,291390798 | 0,088821498 | 0,060821498 | -1,199178502 |
| chrIV | 957192 | 957317 | 125 | 0,045592969 | 0,055588044 | 0,450607955 | -0,124125498 | -0,105506674 | 0,272250071 | 0,405250071 | -0,854749929 |
| chrIV | 957192 | 957321 | 129 | 9,614417409 | 9,609319817 | 0,500132586 | 0,000332343 | 0,000282492 | 0,212384766 | 0,317384766 | -0,942615234 |
| chrIV | 957196 | 957298 | 102 | 0,062690333 | 0,059293913 | 0,51392155 | 0,034903236 | 0,029667751 | 0,570345204 | 0,864345204 | -0,395654796 |
| chrIV | 962120 | 962274 | 154 | 0,347646391 | 0,381704567 | 0,476651724 | -0,0585589 | -0,049775065 | -0,173690862 | -0,243690862 | -1,503690862 |
| chrIV | 962406 | 962555 | 149 | 0,302053422 | 0,203822827 | 0,59708955 | 0,245820872 | 0,208947741 | 0,078621791 | 0,043621791 | -1,216378209 |
| chrIV | 965842 | 965991 | 149 | 0,065539893 | 0,051882174 | 0,558156527 | 0,146296985 | 0,124352437 | -0,002042977 | -0,037042977 | -1,297042977 |
| chrIV | 968475 | 968604 | 129 | 4,339880769 | 7,20791633 | 0,375818932 | -0,316480448 | -0,269008381 | -0,062294172 | 0,042705828 | -1,217294172 |
| chrIV | 971192 | 971329 | 137 | 0,108283302 | 0,08894087 | 0,549036667 | 0,123227859 | 0,104743681 | 0,057527524 | 0,106527524 | -1,153472476 |
| chrIV | 975387 | 975508 | 121 | 0,475876617 | 1,634288484 | 0,225516296 | -0,753694656 | -0,640640457 | -0,112154132 | 0,048845868 | -1,211154132 |
| chrIV | 979606 | 979759 | 153 | 0,284956058 | 0,26311674 | 0,519923739 | 0,049962185 | 0,042467857 | -0,038501555 | -0,101501555 | -1,361501555 |
| chrIV | 980738 | 980889 | 151 | 2,14856868 | 1,819581963 | 0,541453406 | 0,10409597 | 0,088481575 | -0,005028889 | -0,054028889 | -1,314028889 |

TABLE 2

| Chrom | Start | End | Length | Section A | Section B | A/A+B | Z-score | Z * 0.85 | Phase correction | Length correction | ΔLK |
|---|---|---|---|---|---|---|---|---|---|---|---|
| chrIV | 981759 | 981861 | 102 | 0,131079787 | 0,170470001 | 0,434687048 | -0,164453569 | -0,139785534 | 0,409622463 | 0,703622463 | -0,556377537 |
| chrIV | 983182 | 983330 | 148 | 0,074088575 | 0,074117392 | 0,499902783 | -0,000243688 | -0,000207135 | -0,201235912 | -0,229235912 | -1,489235912 |
| chrIV | 984709 | 984852 | 143 | 0,176672756 | 0,066705652 | 0,725917953 | 0,600513459 | 0,51043644 | 0,370365209 | 0,377365209 | -0,882634791 |
| chrIV | 985995 | 986141 | 146 | 0,108283302 | 0,044470435 | 0,708874979 | 0,550101085 | 0,467585923 | 0,292387819 | 0,278387819 | -0,981612181 |
| chrIV | 986788 | 986937 | 149 | 0,085486818 | 0,044470435 | 0,657807209 | 0,406485945 | 0,345513053 | 0,223491274 | 0,188491274 | -1,071508726 |
| chrIV | 987343 | 987492 | 149 | 0,094035499 | 0,085235 | 0,524545307 | 0,06156483 | 0,052330105 | -0,067452466 | -0,102452466 | -1,362452466 |
| chrIV | 987657 | 987771 | 114 | 0,088336378 | 0,403939784 | 0,179444761 | -0,917483145 | -0,779860673 | -0,235773884 | -0,025773884 | -1,285773884 |
| chrIV | 988353 | 988503 | 150 | 0,111132863 | 0,248293262 | 0,309195284 | -0,498132622 | -0,423412729 | -0,491175299 | -0,533175299 | -1,793175299 |
| chrIV | 989662 | 989806 | 144 | 0,039893848 | 0,037058696 | 0,518421434 | 0,046192108 | 0,039263292 | -0,107248348 | -0,107248348 | -1,367248348 |
| chrIV | 989749 | 989906 | 157 | 0,074088575 | 0,081529131 | 0,47609348 | -0,059960669 | -0,050966568 | -0,035005362 | -0,126005362 | -1,386005362 |
| chrIV | 990217 | 990378 | 161 | 0,059840772 | 0,096352609 | 0,38311977 | -0,297297305 | -0,252702709 | -0,244091229 | -0,363091229 | -1,623091229 |
| chrIV | 990397 | 990532 | 135 | 0,074088575 | 0,085235 | 0,465019536 | -0,087795679 | -0,074626327 | -0,130630588 | -0,067630588 | -1,327630588 |
| chrIV | 991435 | 991575 | 140 | 0,170973635 | 0,114881957 | 0,598111913 | 0,248463027 | 0,211193573 | 0,086786656 | 0,114786656 | -1,145213344 |
| chrIV | 991441 | 991575 | 134 | 0,119681545 | 0,081529131 | 0,594807131 | 0,239928434 | 0,203939169 | 0,227725641 | 0,297725641 | -0,962274359 |
| chrIV | 991853 | 992008 | 155 | 0,094035499 | 0,085235 | 0,524545307 | 0,06156483 | 0,052330105 | 0,03936446 | -0,03763554 | -1,29763554 |
| chrIV | 991984 | 992092 | 108 | 0,079787696 | 0,151940653 | 0,344315647 | -0,400713197 | -0,340606218 | 0,241805891 | 0,493805891 | -0,766194109 |
| chrIV | 992133 | 992282 | 149 | 0,062690333 | 0,070411522 | 0,470995186 | -0,072768458 | -0,061853189 | -0,183913425 | -0,218913425 | -1,478913425 |
| chrIV | 994735 | 994883 | 148 | 0,376141997 | 0,248293262 | 0,60237149 | 0,259490228 | 0,220566694 | 0,013359261 | -0,014640739 | -1,274640739 |
| chrIV | 1001383 | 1001529 | 146 | 0,113982423 | 0,107470218 | 0,514703382 | 0,036864261 | 0,031334622 | -0,136974783 | -0,150974783 | -1,410974783 |
| chrIV | 1001383 | 1001529 | 146 | 0,113982423 | 0,107470218 | 0,514703382 | 0,036864261 | 0,031334622 | -0,136974783 | -0,150974783 | -1,410974783 |
| chrIV | 1005517 | 1005664 | 147 | 0,53571739 | 0,322410653 | 0,624286077 | 0,316757196 | 0,269243616 | 0,097961085 | 0,076961085 | -1,183038915 |
| chrIV | 1005876 | 1006022 | 146 | 0,327699467 | 0,211234566 | 0,608051166 | 0,274243282 | 0,23310679 | 0,067203845 | 0,053203845 | -1,206796155 |
| chrIV | 1012282 | 1012417 | 135 | 0,450230572 | 0,329822392 | 0,57717949 | 0,194683129 | 0,16548066 | 0,118104347 | 0,181104347 | -1,078895653 |
| chrIV | 1016514 | 1016662 | 148 | 0,213717044 | 0,077823261 | 0,733061742 | 0,62209939 | 0,528784481 | 0,320611072 | 0,292611072 | -0,967388928 |
| chrIV | 1019255 | 1019404 | 149 | 0,233663968 | 0,34835174 | 0,401473645 | -0,249534587 | -0,212104399 | -0,330001612 | -0,365001612 | -1,625001612 |
| chrIV | 1021134 | 1021284 | 150 | 0,042743409 | 0,044470435 | 0,490098899 | -0,024820929 | -0,021097789 | -0,096208887 | -0,138208887 | -1,398208887 |
| chrIV | 1021134 | 1021284 | 150 | 0,042743409 | 0,044470435 | 0,490098899 | -0,024820929 | -0,021097789 | -0,096208887 | -0,138208887 | -1,398208887 |
| chrIV | 1025226 | 1025374 | 148 | 0,324849907 | 0,218646305 | 0,597704086 | 0,247408847 | 0,21029752 | 0,006268667 | -0,021731333 | -1,281731333 |
| chrIV | 1028806 | 1028950 | 144 | 0,453080133 | 0,303881305 | 0,598551141 | 0,249598683 | 0,21215888 | 0,052448063 | 0,052448063 | -1,207551937 |
| chrIV | 1039896 | 1040063 | 167 | 0,136778908 | 0,285351958 | 0,324020154 | -0,456486317 | -0,38801337 | -0,239125295 | -0,400125295 | -1,660125295 |
| chrIV | 1042109 | 1042252 | 143 | 1,017293128 | 0,785644351 | 0,564242044 | 0,16173326 | 0,137473271 | 0,003390984 | 0,010390984 | -1,249609016 |
| chrIV | 1052004 | 1052154 | 150 | 0,111132863 | 0,055588044 | 0,666580246 | 0,43048963 | 0,365916185 | 0,291305814 | 0,249305814 | -1,010694186 |
| chrIV | 1055603 | 1055756 | 153 | 2,402179572 | 1,93075805 | 0,554399759 | 0,136785326 | 0,116267527 | 0,043268067 | -0,019731933 | -1,279731933 |
| chrIV | 1058574 | 1058755 | 181 | 0,091185939 | 0,140823044 | 0,393027622 | -0,271436616 | -0,230721123 | 0,150923081 | -0,108076919 | -1,368076919 |
| chrIV | 1058574 | 1058755 | 181 | 0,091185939 | 0,140823044 | 0,393027622 | -0,271436616 | -0,230721123 | 0,150923081 | -0,108076919 | -1,368076919 |
| chrIV | 1063264 | 1063420 | 156 | 0,14817715 | 0,062999783 | 0,701672991 | 0,529218308 | 0,449835562 | 0,478301603 | 0,394301603 | -0,865698397 |
| chrIV | 1066695 | 1066823 | 128 | 0,14532759 | 0,125999566 | 0,535617563 | 0,08939893 | 0,07598909 | 0,325899691 | 0,437899691 | -0,822100309 |
| chrIV | 1075797 | 1075953 | 156 | 0,09973462 | 0,166764131 | 0,374240479 | -0,320642986 | -0,272546538 | -0,24149808 | -0,32549808 | -1,58549808 |
| chrIV | 1079066 | 1079219 | 153 | 0,122531105 | 0,074117392 | 0,623097085 | 0,313625053 | 0,266581295 | 0,197740224 | 0,134740224 | -1,125259776 |
| chrIV | 1079249 | 1079403 | 154 | 1,826568334 | 1,1451137 | 0,614658067 | 0,291480486 | 0,247758413 | 0,123957043 | 0,053957043 | -1,206042957 |
| chrIV | 1080074 | 1080246 | 172 | 0,168124074 | 0,281646088 | 0,373799973 | -0,321805635 | -0,273534789 | -0,043505391 | -0,239505391 | -1,499505391 |
| chrIV | 1090868 | 1091040 | 172 | 0,165274514 | 0,17417587 | 0,486888575 | -0,032871387 | -0,027940679 | 0,210997989 | 0,014997989 | -1,245002011 |
| chrIV | 1091309 | 1091450 | 141 | 0,119681545 | 0,140823044 | 0,459422021 | -0,101889932 | -0,086606442 | -0,24521628 | -0,22421628 | -1,48421628 |
| chrIV | 1096304 | 1096427 | 123 | 0,065539893 | 0,200116957 | 0,24670884 | -0,684883118 | -0,582150651 | -0,129179707 | 0,017820293 | -1,242179707 |
| chrIV | 1096404 | 1096559 | 155 | 0,09973462 | 0,103764348 | 0,490098899 | -0,024820929 | -0,021097789 | -0,043357536 | -0,120357536 | -1,380357536 |
| chrIV | 1096837 | 1096971 | 134 | 0,108283302 | 0,125999566 | 0,462190442 | -0,094916836 | -0,080679311 | -0,063225439 | 0,006774561 | -1,253225439 |
| chrIV | 1096837 | 1096977 | 140 | 0,168124074 | 0,129705435 | 0,564497704 | 0,162382575 | 0,138025189 | 0,009297538 | 0,037297538 | -1,222702462 |
| chrIV | 1097880 | 1098015 | 135 | 0,076938136 | 0,074117392 | 0,509336779 | 0,023405971 | 0,019895076 | -0,024031054 | 0,038968946 | -1,221031054 |
| chrIV | 1098034 | 1098195 | 161 | 0,062690333 | 0,096352609 | 0,394172367 | -0,268460684 | -0,228191582 | -0,219648121 | -0,338648121 | -1,598648121 |
| chrIV | 1098506 | 1098663 | 157 | 0,056991212 | 0,037058696 | 0,605967759 | 0,268824834 | 0,228501109 | 0,245984574 | 0,154984574 | -1,105015426 |
| chrIV | 1098997 | 1099126 | 129 | 0,068389454 | 0,125999566 | 0,351817475 | -0,380418277 | -0,323355536 | -0,125463803 | -0,020463803 | -1,280463803 |
| chrIV | 1099909 | 1100059 | 150 | 0,464478375 | 0,35205761 | 0,568840056 | 0,173421802 | 0,147408532 | 0,064320209 | 0,022320209 | -1,237679791 |
| chrIV | 1100669 | 1100827 | 158 | 0,039893848 | 0,037058696 | 0,518421434 | 0,046192108 | 0,039263292 | 0,05365371 | -0,04434629 | -1,30434629 |
| chrIV | 1102545 | 1102694 | 149 | 0,162424953 | 0,114881957 | 0,585722704 | 0,216555726 | 0,184072367 | 0,064262463 | 0,029262463 | -1,230737537 |
| chrIV | 1102953 | 1103100 | 147 | 0,068389454 | 0,144528914 | 0,321200349 | -0,464344846 | -0,394693119 | -0,560385305 | -0,581385305 | -1,841385305 |
| chrIV | 1107156 | 1107303 | 147 | 0,216566604 | 0,200116957 | 0,519738776 | 0,049497979 | 0,042073282 | -0,123751627 | -0,144751627 | -1,404751627 |
| chrIV | 1109551 | 1109723 | 172 | 1,376337762 | 1,941875659 | 0,414782772 | -0,215258806 | -0,182969985 | 0,065552071 | -0,130447929 | -1,390447929 |
| chrIV | 1112644 | 1112796 | 152 | 0,333398588 | 0,326116523 | 0,505520772 | 0,013838965 | 0,011763121 | -0,066598612 | -0,122598612 | -1,382598612 |
| chrIV | 1117237 | 1117387 | 150 | 0,307752543 | 0,151940653 | 0,669473784 | 0,43846059 | 0,372691501 | 0,288203525 | 0,246203525 | -1,013796475 |
| chrIV | 1121058 | 1121205 | 147 | 0,139628469 | 0,118587827 | 0,540742282 | 0,102303929 | 0,08695834 | -0,076747123 | -0,097747123 | -1,357747123 |
| chrIV | 1121801 | 1121948 | 147 | 0,467327936 | 0,366881088 | 0,560204843 | 0,151488587 | 0,128765299 | -0,039931937 | -0,060931937 | -1,320931937 |
| chrIV | 1124473 | 1124641 | 168 | 0,336248149 | 0,696703481 | 0,325521679 | -0,452313214 | -0,384466232 | -0,209973632 | -0,377973632 | -1,637973632 |
| chrIV | 1128248 | 1128395 | 147 | 0,398938482 | 0,296469566 | 0,573675388 | 0,185739279 | 0,157878387 | -0,002399164 | -0,023399164 | -1,283399164 |
| chrIV | 1137826 | 1137974 | 148 | 0,578460798 | 0,433586741 | 0,57157473 | 0,180384731 | 0,153327021 | -0,055574184 | -0,083574184 | -1,343574184 |
| chrIV | 1140412 | 1140528 | 116 | 2,459170784 | 16,59488397 | 0,129062859 | -1,130832215 | -0,961207383 | -0,394879409 | -0,198879409 | -1,458879409 |
| chrIV | 1144636 | 1144783 | 147 | 0,763682236 | 0,607762611 | 0,556845022 | 0,142974962 | 0,121528717 | -0,042722881 | -0,063722881 | -1,323722881 |
| chrIV | 1154940 | 1155065 | 125 | 1,347842156 | 2,645990879 | 0,337480847 | -0,419348174 | -0,356445948 | -0,002208036 | 0,130791964 | -1,129208036 |
| chrIV | 1155027 | 1155183 | 156 | 0,592708601 | 0,455821958 | 0,565275467 | 0,164358328 | 0,139704579 | 0,188745118 | 0,104745118 | -1,155254882 |
| chrIV | 1155195 | 1155331 | 136 | 0,58700948 | 0,389116306 | 0,601366636 | 0,256886065 | 0,218353155 | 0,205030227 | 0,261030227 | -0,998969773 |
| chrIV | 1158507 | 1158654 | 147 | 0,960301917 | 0,544762828 | 0,638046915 | 0,353243137 | 0,300256667 | 0,140265636 | 0,119265636 | -1,140734364 |
| chrIV | 1160332 | 1160457 | 125 | 1,242408414 | 2,790519793 | 0,308066088 | -0,50133955 | -0,426138618 | -0,087566306 | 0,045433694 | -1,214566306 |
| chrIV | 1160419 | 1160575 | 156 | 0,661098055 | 0,444704349 | 0,597844653 | 0,247772164 | 0,210606339 | 0,250750814 | 0,166750814 | -1,093249186 |
| chrIV | 1160587 | 1160723 | 136 | 0,615505086 | 0,366881088 | 0,626540868 | 0,322705529 | 0,274299699 | 0,257359042 | 0,313359042 | -0,946640958 |
| chrIV | 1164215 | 1164366 | 151 | 0,498673102 | 0,363175219 | 0,578608892 | 0,198335964 | 0,168585569 | 0,074253583 | 0,025253583 | -1,234746417 |
| chrIV | 1166734 | 1166868 | 134 | 0,549965193 | 0,592939133 | 0,481199677 | -0,047142876 | -0,040071445 | -0,040065171 | 0,029934829 | -1,230065171 |
| chrIV | 1169216 | 1169370 | 154 | 1,02584181 | 0,859761742 | 0,544038968 | 0,11061448 | 0,094022308 | -0,029826604 | -0,099826604 | -1,359826604 |
| chrIV | 1170479 | 1170639 | 160 | 0,350495952 | 0,244587392 | 0,588986325 | 0,224938202 | 0,191197471 | 0,199334089 | 0,087334089 | -1,172665911 |
| chrIV | 1170874 | 1171037 | 163 | 0,151026711 | 0,207528696 | 0,42120885 | -0,198801911 | -0,168981625 | -0,137421618 | -0,270421618 | -1,530421618 |
| chrIV | 1170874 | 1171037 | 163 | 0,151026711 | 0,207528696 | 0,42120885 | -0,198801911 | -0,168981625 | -0,137421618 | -0,270421618 | -1,530421618 |
| chrIV | 1171736 | 1171886 | 150 | 0,564212996 | 0,589233263 | 0,489154125 | -0,027189926 | -0,023111437 | -0,107341001 | -0,149341001 | -1,409341001 |
| chrIV | 1175589 | 1175742 | 153 | 0,461628815 | 0,52993935 | 0,465554292 | -0,086450148 | -0,073482626 | -0,143943524 | -0,206943524 | -1,466943524 |
| chrIV | 1176500 | 1176647 | 147 | 0,208017923 | 0,163058261 | 0,560580095 | 0,152440128 | 0,129574109 | -0,032180512 | -0,053180512 | -1,313180512 |
| chrIV | 1179339 | 1179509 | 170 | 0,111132863 | 0,226058044 | 0,3295844 | -0,441061051 | -0,374901893 | -0,156893454 | -0,338893454 | -1,598893454 |
| chrIV | 1185766 | 1185913 | 147 | 0,504372223 | 0,35205761 | 0,588924164 | 0,224778397 | 0,191061638 | 0,025305653 | 0,004305653 | -1,255694347 |
| chrIV | 1186635 | 1186765 | 130 | 0,136778908 | 0,340940001 | 0,286316713 | -0,564177371 | -0,479550765 | -0,354892523 | -0,256892523 | -1,516892523 |
| chrIV | 1187842 | 1187997 | 155 | 0,105433742 | 0,17417587 | 0,377074811 | -0,313172484 | -0,266196611 | -0,26581164 | -0,34281164 | -1,60281164 |
| chrIV | 1199637 | 1199781 | 144 | 0,299203861 | 0,289057827 | 0,508623742 | 0,021618199 | 0,01837547 | -0,147776687 | -0,147776687 | -1,407776687 |
| chrIV | 1199946 | 1200093 | 147 | 0,259310013 | 0,26311674 | 0,496356688 | -0,009132555 | -0,007762672 | -0,173907531 | -0,194907531 | -1,454907531 |
| chrIV | 1202429 | 1202587 | 158 | 0,051292091 | 0,111176087 | 0,315705458 | -0,47974192 | -0,407780632 | -0,39463873 | -0,49263873 | -1,75263873 |
| chrIV | 1205211 | 1205359 | 148 | 2,026037575 | 1,337818918 | 0,602296079 | 0,259294733 | 0,220400523 | 0,018353932 | -0,009646068 | -1,269646068 |
| chrIV | 1207063 | 1207212 | 149 | 0,082637257 | 0,044470435 | 0,650135768 | 0,385687039 | 0,327833983 | 0,201123255 | 0,166123255 | -1,093876745 |
| chrIV | 1207237 | 1207360 | 123 | 0,059840772 | 0,233469783 | 0,204018475 | -0,827353109 | -0,703250143 | -0,272145384 | -0,125145384 | -1,385145384 |
| chrIV | 1207337 | 1207492 | 155 | 0,091185939 | 0,111176087 | 0,450607955 | -0,124125498 | -0,105506674 | -0,109858693 | -0,186858693 | -1,446858693 |
| chrIV | 1207770 | 1207904 | 134 | 0,122531105 | 0,118587827 | 0,508177041 | 0,020498237 | 0,017423501 | 0,010008307 | 0,080008307 | -1,179991693 |
| chrIV | 1207770 | 1207910 | 140 | 0,159575393 | 0,096352609 | 0,623516738 | 0,314730183 | 0,267520656 | 0,147175634 | 0,175175634 | -1,084824366 |
| chrIV | 1207966 | 1208117 | 151 | 0,085486818 | 0,100058479 | 0,460732874 | -0,09858756 | -0,083799426 | -0,193669143 | -0,242669143 | -1,502669143 |
| chrIV | 1207976 | 1208121 | 145 | 0,09688506 | 0,092646739 | 0,511181027 | 0,028030349 | 0,023825797 | -0,154192634 | -0,161192634 | -1,421192634 |
| chrIV | 1207994 | 1208145 | 151 | 0,042743409 | 0,048176305 | 0,470122564 | -0,074961771 | -0,063717505 | -0,175614536 | -0,224614536 | -1,484614536 |
| chrIV | 1207996 | 1208145 | 149 | 0,051292091 | 0,048176305 | 0,515662191 | 0,039269381 | 0,033378974 | -0,093787762 | -0,128787762 | -1,388787762 |
| chrIV | 1208248 | 1208402 | 154 | 0,222265726 | 0,125999566 | 0,638208088 | 0,35367318 | 0,300622203 | 0,183411283 | 0,113411283 | -1,146588717 |
| chrIV | 1208282 | 1208426 | 144 | 0,071239015 | 0,037058696 | 0,657807209 | 0,406485945 | 0,345513053 | 0,193157046 | 0,193157046 | -1,066842954 |

TABLE 2

| Chrom | Start | End | Length | Section A | Section B | A/A+B | Z-score | Z * 0.85 | Phase correction | Length correction | ΔLK |
|---|---|---|---|---|---|---|---|---|---|---|---|
| chrIV | 1208813 | 1208948 | 135 | 0,076938136 | 0,074117392 | 0,509336779 | 0,023405971 | 0,019895076 | -0,014399633 | 0,048600367 | -1,211399633 |
| chrIV | 1208967 | 1209128 | 161 | 0,071239015 | 0,070411522 | 0,502920895 | 0,007321663 | 0,006223413 | 0,016480924 | -0,102519076 | -1,362519076 |
| chrIV | 1209439 | 1209596 | 157 | 0,065539893 | 0,048176305 | 0,576346155 | 0,192554744 | 0,163671533 | 0,173780624 | 0,082780624 | -1,177219376 |
| chrIV | 1209930 | 1210059 | 129 | 0,045592969 | 0,118587827 | 0,277699771 | -0,589688448 | -0,501235181 | -0,323192145 | -0,218192145 | -1,478192145 |
| chrIV | 1210842 | 1210992 | 150 | 0,131079787 | 0,218646305 | 0,374806999 | -0,319148382 | -0,271276125 | -0,347101236 | -0,389101236 | -1,649101236 |
| chrIV | 1211574 | 1211688 | 114 | 0,031345166 | 0,17788174 | 0,149814223 | -1,037230501 | -0,881645926 | -0,335958414 | -0,125958414 | -1,385958414 |
| chrIV | 1215599 | 1215745 | 146 | 0,076938136 | 0,055588044 | 0,58055047 | 0,203301913 | 0,172806626 | 0,005004586 | -0,008995414 | -1,268995414 |
| chrIV | 1216176 | 1216327 | 151 | 0,054141651 | 0,085235 | 0,388455674 | -0,283346352 | -0,2408444 | -0,343756105 | -0,392756105 | -1,652756105 |
| chrIV | 1217346 | 1217492 | 146 | 0,302053422 | 0,285351958 | 0,514216302 | 0,035642529 | 0,03029615 | -0,138513411 | -0,152513411 | -1,412513411 |
| chrIV | 1218508 | 1218653 | 145 | 0,735186631 | 0,552174567 | 0,571080309 | 0,179125217 | 0,152256434 | -0,021764769 | -0,028764769 | -1,288764769 |
| chrIV | 1221088 | 1221239 | 151 | 0,091185939 | 0,122293696 | 0,427141159 | -0,183657287 | -0,156108694 | -0,261973339 | -0,310973339 | -1,570973339 |
| chrIV | 1225710 | 1225871 | 161 | 0,253610892 | 0,311293045 | 0,448945167 | -0,12832683 | -0,109077806 | -0,102847672 | -0,221847672 | -1,481847672 |
| chrIV | 1225924 | 1226071 | 147 | 0,153876272 | 0,055588044 | 0,734618073 | 0,626840405 | 0,532814345 | 0,375707512 | 0,354707512 | -0,905292488 |
| chrIV | 1227651 | 1227806 | 155 | 0,034194727 | 0,062999783 | 0,351817475 | -0,380418277 | -0,323355536 | -0,326778754 | -0,403778754 | -1,663778754 |
| chrIV | 1238397 | 1238557 | 160 | 0,313451664 | 0,322410653 | 0,492955244 | -0,017659501 | -0,015010576 | -0,00584321 | -0,11784321 | -1,37784321 |
| chrIV | 1240257 | 1240406 | 149 | 0,250761331 | 0,203822827 | 0,551627959 | 0,129775456 | 0,110309137 | -0,019931113 | -0,054931113 | -1,314931113 |
| chrIV | 1240385 | 1240526 | 141 | 0,430283648 | 0,366881088 | 0,539767539 | 0,099848099 | 0,084870884 | -0,069831282 | -0,048831282 | -1,308831282 |
| chrIV | 1244524 | 1244687 | 163 | 0,173823196 | 0,155646522 | 0,52758474 | 0,069199879 | 0,058819897 | 0,09286825 | -0,04013175 | -1,30013175 |
| chrIV | 1245862 | 1245987 | 125 | 1,384886444 | 2,482932618 | 0,358053575 | -0,363666381 | -0,309116424 | 0,029948512 | 0,162948512 | -1,097051488 |
| chrIV | 1251539 | 1251695 | 156 | 0,940354993 | 0,685585872 | 0,578345137 | 0,197661741 | 0,16801248 | 0,206817998 | 0,122817998 | -1,137182002 |
| chrIV | 1259546 | 1259668 | 122 | 0,561363435 | 2,015993051 | 0,217805895 | -0,77962474 | -0,662681029 | -0,173201745 | -0,019201745 | -1,279201745 |
| chrIV | 1265859 | 1266020 | 161 | 0,09973462 | 0,100058479 | 0,499189516 | -0,002031583 | -0,001726846 | 0,001910915 | -0,117089085 | -1,377089085 |
| chrIV | 1266398 | 1266553 | 155 | 0,236513528 | 0,274234349 | 0,463072954 | -0,092694951 | -0,078790708 | -0,08406368 | -0,16106368 | -1,42106368 |
| chrIV | 1267550 | 1267699 | 149 | 0,302053422 | 0,248293262 | 0,548842086 | 0,122736414 | 0,104325951 | -0,019929408 | -0,054929408 | -1,314929408 |
| chrIV | 1269091 | 1269198 | 107 | 0,082637257 | 0,292763697 | 0,220130653 | -0,771752035 | -0,65598923 | -0,085760565 | 0,173239435 | -1,086760565 |
| chrIV | 1269091 | 1269209 | 118 | 0,139628469 | 0,52623348 | 0,209695822 | -0,807477133 | -0,686355563 | -0,139623014 | 0,042376986 | -1,217623014 |
| chrIV | 1272800 | 1272981 | 181 | 0,068389454 | 0,218646305 | 0,238261094 | -0,711907289 | -0,605121196 | -0,207560067 | -0,466560067 | -1,726560067 |
| chrIV | 1277856 | 1277982 | 126 | 0,273557816 | 0,915349786 | 0,230091738 | -0,738544765 | -0,62776305 | -0,331556501 | -0,205556501 | -1,465556501 |
| chrIV | 1278160 | 1278320 | 160 | 2,137170438 | 1,945581529 | 0,523463207 | 0,058847485 | 0,050020363 | 0,068449084 | -0,043550916 | -1,303550916 |
| chrIV | 1287754 | 1287911 | 157 | 0,826372569 | 0,433586741 | 0,655872426 | 0,401224087 | 0,341040474 | 0,349838993 | 0,258838993 | -1,001161007 |
| chrIV | 1288372 | 1288504 | 132 | 0,302053422 | 0,192705218 | 0,610506614 | 0,280639687 | 0,238543734 | 0,33557521 | 0,41957521 | -0,84042479 |
| chrIV | 1291651 | 1291799 | 148 | 1,108479067 | 0,744879785 | 0,598091981 | 0,248411498 | 0,211149773 | 0,009670395 | -0,018329605 | -1,278329605 |
| chrIV | 1293635 | 1293774 | 139 | 2,105825271 | 1,878875876 | 0,528477593 | 0,07144347 | 0,060726949 | -0,054514777 | -0,019514777 | -1,279514777 |
| chrIV | 1298059 | 1298223 | 164 | 0,065539893 | 0,048176305 | 0,576346155 | 0,192554744 | 0,163671533 | 0,234858175 | 0,094858175 | -1,165141825 |
| chrIV | 1300809 | 1300966 | 157 | 0,037044288 | 0,040764565 | 0,47609348 | -0,059960669 | -0,050966568 | -0,043797029 | -0,134797029 | -1,394797029 |
| chrIV | 1300809 | 1300970 | 161 | 2,818215417 | 2,99063675 | 0,485158743 | -0,037210099 | -0,031628584 | -0,02282097 | -0,14182097 | -1,40182097 |
| chrIV | 1300843 | 1301002 | 159 | 0,467327936 | 0,296469566 | 0,61184795 | 0,284138687 | 0,241517884 | 0,236117138 | 0,131117138 | -1,128882862 |
| chrIV | 1300844 | 1300970 | 126 | 0,059840772 | 0,114881957 | 0,342489913 | -0,405677162 | -0,344825587 | -0,052655185 | 0,073344815 | -1,186655185 |
| chrIV | 1300844 | 1300987 | 143 | 0,168124074 | 0,226058044 | 0,426513701 | -0,185257077 | -0,157468516 | -0,293729534 | -0,286729534 | -1,546729534 |
| chrIV | 1302267 | 1302373 | 106 | 0,182371877 | 1,030231743 | 0,150396943 | -1,034732432 | -0,879522567 | -0,309502499 | -0,043502499 | -1,303502499 |
| chrIV | 1302267 | 1302383 | 116 | 0,210867483 | 1,267407396 | 0,142644299 | -1,06851427 | -0,908237129 | -0,348978415 | -0,152978415 | -1,412978415 |
| chrIV | 1308150 | 1308260 | 110 | 0,071239015 | 0,255705001 | 0,217893619 | -0,779326791 | -0,662427772 | -0,099372724 | 0,138627276 | -1,121372724 |
| chrIV | 1308150 | 1308309 | 159 | 20,13784464 | 16,52447245 | 0,549279103 | 0,12384021 | 0,105264178 | 0,101372459 | -0,003627541 | -1,263627541 |
| chrIV | 1308153 | 1308309 | 156 | 0,031345166 | 0,285351958 | 0,098975217 | -1,287412835 | -1,09430091 | -1,063150106 | -1,147150106 | -2,407150106 |
| chrIV | 1308185 | 1308314 | 129 | 0,116831984 | 0,222352175 | 0,344450001 | -0,400348293 | -0,340296049 | -0,18875978 | -0,08375978 | -1,34375978 |
| chrIV | 1308191 | 1308309 | 118 | 0,082637257 | 0,08894087 | 0,481630488 | -0,046061821 | -0,039152548 | 0,504883598 | 0,686883598 | -0,573116402 |
| chrIV | 1308196 | 1308309 | 113 | 0,176672756 | 0,444704349 | 0,284324534 | -0,570042211 | -0,48453588 | 0,052176889 | 0,269176889 | -0,990823111 |
| chrIV | 1308234 | 1308344 | 110 | 0,091185939 | 0,226058044 | 0,287431578 | -0,560903746 | -0,476768184 | 0,081482944 | 0,319482944 | -0,940517056 |
| chrIV | 1308280 | 1308442 | 162 | 1,242408414 | 0,670762394 | 0,649397539 | 0,383694456 | 0,326140288 | 0,351410947 | 0,225410947 | -1,034589053 |
| chrIV | 1308280 | 1308477 | 197 | 0,042743409 | 0,066705652 | 0,390532439 | -0,277931585 | -0,236241848 | 0,197801403 | -0,173198597 | -1,433198597 |
| chrIV | 1308478 | 1308605 | 127 | 0,136778908 | 0,048176305 | 0,739524483 | 0,641880105 | 0,545598089 | 0,806970267 | 0,925970267 | -0,334029733 |
| chrIV | 1308858 | 1308982 | 124 | 4,741668811 | 15,75365158 | 0,23135373 | -0,734395942 | -0,624236551 | -0,221140054 | -0,081140054 | -1,341140054 |
| chrIV | 1312653 | 1312780 | 127 | 0,14532759 | 0,333528262 | 0,303489222 | -0,514391296 | -0,437232602 | -0,180916089 | -0,061916089 | -1,321916089 |
| chrIV | 1312778 | 1312924 | 146 | 0,287805619 | 0,237175653 | 0,548220735 | 0,121167293 | 0,102992199 | -0,065573388 | -0,079573388 | -1,339573388 |
| chrIV | 1324502 | 1324680 | 178 | 0,04844253 | 0,062999783 | 0,434687048 | -0,164453569 | -0,139785534 | 0,240834911 | 0,002834911 | -1,257165089 |
| chrIV | 1324730 | 1324882 | 152 | 0,265009134 | 0,037058696 | 0,877316642 | 1,161676939 | 0,987425398 | 0,918566477 | 0,862566477 | -0,397433523 |
| chrIV | 1329033 | 1329189 | 156 | 0,552814753 | 0,511410002 | 0,519453011 | 0,048780806 | 0,041463685 | 0,076079714 | -0,007920286 | -1,267920286 |
| chrIV | 1331269 | 1331431 | 162 | 0,076938136 | 0,040764565 | 0,653664997 | 0,395234266 | 0,335949126 | 0,359127163 | 0,233127163 | -1,026872837 |
| chrIV | 1332261 | 1332409 | 148 | 0,136778908 | 0,114881957 | 0,543504879 | 0,109267603 | 0,092877463 | -0,108982393 | -0,136982393 | -1,396982393 |
| chrIV | 1334150 | 1334293 | 143 | 0,766531797 | 0,466939567 | 0,621442718 | 0,309272085 | 0,262881272 | 0,123818166 | 0,130818166 | -1,129181834 |
| chrIV | 1334858 | 1335004 | 146 | 0,176672756 | 0,066705652 | 0,725917953 | 0,600513459 | 0,51043644 | 0,337689956 | 0,323689956 | -0,936310044 |
| chrIV | 1335220 | 1335376 | 156 | 0,159575393 | 0,200116957 | 0,443644111 | -0,141736401 | -0,120475941 | -0,082561302 | -0,166561302 | -1,426561302 |
| chrIV | 1335220 | 1335376 | 156 | 0,159575393 | 0,200116957 | 0,443644111 | -0,141736401 | -0,120475941 | -0,082561302 | -0,166561302 | -1,426561302 |
| chrIV | 1337771 | 1337882 | 111 | 0,299203861 | 0,096352609 | 0,756412507 | 0,694809033 | 0,590587678 | 1,136701928 | 1,367701928 | 0,107701928 |
| chrIV | 1341919 | 1342069 | 150 | 1,117027749 | 0,711526959 | 0,610880136 | 0,281613708 | 0,239371652 | 0,173372693 | 0,131372693 | -1,128627307 |
| chrIV | 1354730 | 1354858 | 128 | 0,481575739 | 0,667056524 | 0,419260153 | -0,203786546 | -0,173218564 | 0,072986624 | 0,184986624 | -1,075013376 |
| chrIV | 1356736 | 1356889 | 153 | 0,153876272 | 0,251999131 | 0,379121943 | -0,307787693 | -0,261619539 | -0,345536338 | -0,408536338 | -1,668536338 |
| chrIV | 1359819 | 1359977 | 158 | 0,239363089 | 0,222352175 | 0,518421434 | 0,046192108 | 0,039263292 | 0,051421046 | -0,046578954 | -1,306578954 |
| chrIV | 1360674 | 1360822 | 148 | 0,359044634 | 0,188999349 | 0,655138356 | 0,399230614 | 0,339346022 | 0,137789796 | 0,109789796 | -1,150210204 |
| chrIV | 1361305 | 1361412 | 107 | 0,29065518 | 0,867173481 | 0,251034708 | -0,671237229 | -0,570551645 | 0,022109575 | 0,281109575 | -0,978890425 |
| chrIV | 1361305 | 1361420 | 115 | 0,564212996 | 1,445289136 | 0,280772529 | -0,580548103 | -0,493465887 | 0,055209345 | 0,258209345 | -1,001790655 |
| chrIV | 1361305 | 1361420 | 115 | 0,564212996 | 1,445289136 | 0,280772529 | -0,580548103 | -0,493465887 | 0,055209345 | 0,258209345 | -1,001790655 |
| chrIV | 1364336 | 1364488 | 152 | 0,056991212 | 0,037058696 | 0,605967759 | 0,268824834 | 0,228501109 | 0,162795131 | 0,106795131 | -1,153204869 |
| chrIV | 1364512 | 1364671 | 159 | 0,612655525 | 0,637409568 | 0,490098899 | -0,024820929 | -0,021097789 | -0,009921696 | -0,114921696 | -1,374921696 |
| chrIV | 1365728 | 1365875 | 147 | 0,045592969 | 0,062999783 | 0,419852784 | -0,202270107 | -0,171929591 | -0,324996813 | -0,345996813 | -1,605996813 |
| chrIV | 1365746 | 1365909 | 163 | 0,176672756 | 0,125999566 | 0,583709654 | 0,211392905 | 0,179683969 | 0,223499563 | 0,090499563 | -1,169500437 |
| chrIV | 1367419 | 1367567 | 148 | 0,113982423 | 0,048176305 | 0,702906497 | 0,532778373 | 0,452861617 | 0,269380942 | 0,241380942 | -1,018619058 |
| chrIV | 1367927 | 1368075 | 148 | 0,170973635 | 0,159352392 | 0,517590565 | 0,044107306 | 0,03749121 | -0,139036708 | -0,167036708 | -1,427036708 |
| chrIV | 1372245 | 1372378 | 133 | 0,185221438 | 0,163058261 | 0,531818071 | 0,079840821 | 0,067864697 | 0,13917141 | 0,21617141 | -1,04382859 |
| chrIV | 1376748 | 1376898 | 150 | 0,042743409 | 0,077823261 | 0,354520937 | -0,373143192 | -0,317171714 | -0,374420309 | -0,416420309 | -1,676420309 |
| chrIV | 1382369 | 1382521 | 152 | 0,604106844 | 0,589233263 | 0,506231912 | 0,015621722 | 0,013278464 | -0,057429188 | -0,113429188 | -1,373429188 |
| chrIV | 1387683 | 1387833 | 150 | 0,530018269 | 0,377998697 | 0,583709654 | 0,211392905 | 0,179683969 | 0,120248556 | 0,078248556 | -1,181751444 |
| chrIV | 1390839 | 1391009 | 170 | 0,045592969 | 0,092646739 | 0,329810947 | -0,440435256 | -0,374369967 | -0,171104723 | -0,353104723 | -1,613104723 |
| chrIV | 1391435 | 1391580 | 145 | 0,074088575 | 0,051882174 | 0,588141101 | 0,222765786 | 0,189350918 | 0,0014058 | -0,0055942 | -1,2655942 |
| chrIV | 1391940 | 1392060 | 120 | 1,843665698 | 8,727322858 | 0,174408069 | -0,936888068 | -0,796354858 | -0,266456605 | -0,098456605 | -1,358456605 |
| chrIV | 1397500 | 1397655 | 155 | 0,282106498 | 0,251999131 | 0,528184843 | 0,070707798 | 0,060101628 | 0,089701734 | 0,012701734 | -1,247298266 |
| chrIV | 1398087 | 1398256 | 169 | 0,04844253 | 0,055588044 | 0,465656665 | -0,08619258 | -0,073263693 | 0,128087561 | -0,046912439 | -1,306912439 |
| chrIV | 1406033 | 1406179 | 146 | 0,58415992 | 0,444704349 | 0,567771607 | 0,170703659 | 0,14509811 | -0,023647896 | -0,037647896 | -1,297647896 |
| chrIV | 1407803 | 1407943 | 140 | 0,672496298 | 0,574409785 | 0,539331957 | 0,09875086 | 0,083938231 | -0,041460181 | -0,013460181 | -1,273460181 |
| chrIV | 1407803 | 1407943 | 140 | 0,672496298 | 0,574409785 | 0,539331957 | 0,09875086 | 0,083938231 | -0,041460181 | -0,013460181 | -1,273460181 |
| chrIV | 1408060 | 1408202 | 142 | 0,071239015 | 0,055588044 | 0,561702018 | 0,155285859 | 0,13199298 | -0,030788755 | -0,016788755 | -1,276788755 |
| chrIV | 1408065 | 1408226 | 161 | 0,170973635 | 0,170470001 | 0,500737507 | 0,001848658 | 0,001571359 | 0,014560023 | -0,104439977 | -1,364439977 |
| chrIV | 1408340 | 1408489 | 149 | 0,470177496 | 0,333528262 | 0,585011979 | 0,214732295 | 0,182522451 | 0,059047639 | 0,024047639 | -1,235952361 |
| chrIV | 1409678 | 1409825 | 147 | 0,467327936 | 0,35576348 | 0,567771607 | 0,170703659 | 0,14509811 | -0,02047182 | -0,04147182 | -1,30147182 |
| chrIV | 1413171 | 1413310 | 139 | 0,125380666 | 0,111176087 | 0,530023616 | 0,075329227 | 0,064029843 | -0,052423238 | -0,017423238 | -1,277423238 |
| chrIV | 1420451 | 1420608 | 157 | 0,082637257 | 0,34835174 | 0,19173867 | -0,871507085 | -0,740781022 | -0,72111598 | -0,81211598 | -2,07211598 |
| chrIV | 1420630 | 1420804 | 174 | 13,03958923 | 16,98029441 | 0,434365082 | -0,165271659 | -0,14048091 | 0,159807867 | -0,050192133 | -1,310192133 |
| chrIV | 1420630 | 1420804 | 174 | 13,03958923 | 16,98029441 | 0,434365082 | -0,165271659 | -0,14048091 | 0,159807867 | -0,050192133 | -1,310192133 |
| chrIV | 1420634 | 1420804 | 170 | 0,028495606 | 0,059293913 | 0,324590066 | -0,454901459 | -0,386666241 | -0,173403162 | -0,355403162 | -1,615403162 |
| chrIV | 1420634 | 1420804 | 170 | 0,028495606 | 0,059293913 | 0,324590066 | -0,454901459 | -0,386666241 | -0,173403162 | -0,355403162 | -1,615403162 |
| chrIV | 1424831 | 1424973 | 142 | 0,777930039 | 0,374292827 | 0,675155876 | 0,454195325 | 0,386066026 | 0,218718947 | 0,232718947 | -1,027281053 |

TABLE 2

| Chrom | Start | End | Length | Section A | Section B | A/A+B | Z-score | Z * 0.85 | Phase correction | Length correction | ΔLK |
|---|---|---|---|---|---|---|---|---|---|---|---|
| chrIV | 1428254 | 1428414 | 160 | 0,165274514 | 0,240881523 | 0,406923692 | -0,235465589 | -0,200145751 | -0,171886236 | -0,283886236 | -1,543886236 |
| chrIV | 1430994 | 1431130 | 136 | 0,102584181 | 0,203822827 | 0,334797111 | -0,426704981 | -0,362699234 | -0,386789679 | -0,330789679 | -1,590789679 |
| chrIV | 1433662 | 1433813 | 151 | 0,062690333 | 0,081529131 | 0,434687048 | -0,164453569 | -0,139785534 | -0,253571203 | -0,302571203 | -1,562571203 |
| chrIV | 1447757 | 1447907 | 150 | 0,359044634 | 0,255705001 | 0,584050178 | 0,21226584 | 0,180425964 | 0,128892706 | 0,086892706 | -1,173107294 |
| chrIV | 1454001 | 1454153 | 152 | 1,541612276 | 1,289642613 | 0,544497877 | 0,111771928 | 0,095006139 | 0,016041017 | -0,039958983 | -1,299958983 |
| chrIV | 1460987 | 1461146 | 159 | 2,060232302 | 1,652817832 | 0,554862506 | 0,137956258 | 0,117262819 | 0,123925227 | 0,018925227 | -1,241074773 |
| chrIV | 1461681 | 1461828 | 147 | 0,09973462 | 0,070411522 | 0,58617033 | 0,217704522 | 0,185048844 | 0,016544915 | -0,004455085 | -1,264455085 |
| chrIV | 1462856 | 1462994 | 138 | 1,108479067 | 0,96723196 | 0,534023789 | 0,085388642 | 0,072580346 | -0,001373496 | 0,040626504 | -1,219373496 |
| chrIV | 1462967 | 1463131 | 164 | 1,105629506 | 0,844938264 | 0,566824451 | 0,16829514 | 0,143050869 | 0,221876497 | 0,081876497 | -1,178123503 |
| chrIV | 1464532 | 1464669 | 137 | 0,273557816 | 0,292763697 | 0,483043306 | -0,042516934 | -0,036139394 | -0,087784607 | -0,038784607 | -1,298784607 |
| chrIV | 1465805 | 1465959 | 154 | 0,113982423 | 0,085235 | 0,572150875 | 0,181852797 | 0,154574878 | 0,04130594 | -0,02869406 | -1,28869406 |
| chrIV | 1466131 | 1466277 | 146 | 0,310602104 | 0,251999131 | 0,552082158 | 0,130923676 | 0,111285125 | -0,066047467 | -0,080047467 | -1,340047467 |
| chrIV | 1473970 | 1474117 | 147 | 0,424584527 | 0,385410436 | 0,524181688 | 0,060651668 | 0,051553918 | -0,113674278 | -0,134674278 | -1,394674278 |
| chrIV | 1477488 | 1477647 | 159 | 0,852018614 | 0,726350437 | 0,539809504 | 0,099953814 | 0,084960742 | 0,083842688 | -0,021157312 | -1,281157312 |
| chrIV | 1478568 | 1478697 | 129 | 0,709540585 | 1,378583483 | 0,339798097 | -0,413014221 | -0,351062088 | -0,209258552 | -0,104258552 | -1,364258552 |
| chrIV | 1479228 | 1479388 | 160 | 0,088336378 | 0,070411522 | 0,556456986 | 0,141992377 | 0,12069952 | 0,145321039 | 0,033321039 | -1,226678961 |
| chrIV | 1479727 | 1479884 | 157 | 0,618354647 | 0,522527611 | 0,5419969 | 0,105465808 | 0,089645937 | 0,107932308 | 0,016932308 | -1,243067692 |
| chrIV | 1481305 | 1481467 | 162 | 0,179522317 | 0,229763914 | 0,438622908 | -0,154461562 | -0,131292328 | -0,105295083 | -0,231295083 | -1,491295083 |
| chrIV | 1489675 | 1489832 | 157 | 0,561363435 | 0,61888022 | 0,475633512 | -0,061115751 | -0,051948389 | -0,02833781 | -0,11933781 | -1,37933781 |
| chrIV | 1495628 | 1495753 | 125 | 0,082637257 | 0,148234783 | 0,357935317 | -0,363983093 | -0,309385629 | 0,018812758 | 0,151812758 | -1,108187242 |
| chrIV | 1496435 | 1496575 | 140 | 0,182371877 | 0,17417587 | 0,511493562 | 0,028814074 | 0,024491963 | -0,119387013 | -0,091387013 | -1,351387013 |
| chrIV | 1497627 | 1497776 | 149 | 0,190920559 | 0,159352392 | 0,54506224 | 0,113195553 | 0,09621622 | -0,018031642 | -0,053031642 | -1,313031642 |
| chrIV | 1498743 | 1498886 | 143 | 0,903310705 | 0,985761308 | 0,478176956 | -0,054729569 | -0,046520133 | -0,175867749 | -0,168867749 | -1,428867749 |
| chrIV | 1499148 | 1499307 | 159 | 0,079787696 | 0,096352609 | 0,452978074 | -0,118140734 | -0,100419624 | -0,094238528 | -0,199238528 | -1,459238528 |
| chrIV | 1503439 | 1503593 | 154 | 0,34194727 | 0,240881523 | 0,586702775 | 0,219071373 | 0,186210667 | 0,073822531 | 0,003822531 | -1,256177469 |
| chrIV | 1505128 | 1505252 | 124 | 0,059840772 | 0,055588044 | 0,518421434 | 0,046192108 | 0,039263292 | 0,456549629 | 0,596549629 | -0,663450371 |
| chrIV | 1507157 | 1507304 | 147 | 0,273557816 | 0,218646305 | 0,555781238 | 0,140281573 | 0,119239337 | -0,037227142 | -0,058227142 | -1,318227142 |
| chrIV | 1512418 | 1512571 | 153 | 0,530018269 | 0,648527176 | 0,449722385 | -0,126362771 | -0,107408355 | -0,188941655 | -0,251941655 | -1,511941655 |
| chrIV | 1512829 | 1512960 | 131 | 1,376337762 | 1,374877614 | 0,500265364 | 0,000665169 | 0,000565394 | 0,092336758 | 0,183336758 | -1,076663242 |
| chrIV | 1526039 | 1526194 | 155 | 0,3903898 | 0,433586741 | 0,473787518 | -0,065752297 | -0,055889452 | -0,023461567 | -0,100461567 | -1,360461567 |
| chrIV | 1527054 | 1527225 | 171 | 0,031345166 | 0,08894087 | 0,260588572 | -0,641531928 | -0,545302139 | -0,320058663 | -0,509058663 | -1,769058663 |
| chrIV | 1527204 | 1527351 | 147 | 1,649895578 | 1,089525656 | 0,602278889 | 0,259250173 | 0,220362647 | 0,061208033 | 0,040208033 | -1,219791967 |
| chrIV | 1527360 | 1527490 | 130 | 0,259310013 | 0,270528479 | 0,489413315 | -0,026539999 | -0,022558999 | 0,093185411 | 0,191185411 | -1,068814589 |
| chrIV | 1527741 | 1527877 | 136 | 0,495823542 | 0,566998046 | 0,46651625 | -0,084030099 | -0,071425584 | -0,103047804 | -0,047047804 | -1,307047804 |
| chrIV | 1528122 | 1528284 | 162 | 0,062690333 | 0,066705652 | 0,484484373 | -0,038901718 | -0,03306646 | -0,007786404 | -0,133786404 | -1,393786404 |
| chrIV | 1528223 | 1528378 | 155 | 0,062690333 | 0,188999349 | 0,249077882 | -0,677394378 | -0,575785221 | -0,53569017 | -0,61269017 | -1,87269017 |
| chrIV | 1528357 | 1528494 | 137 | 0,162424953 | 0,122293696 | 0,570475287 | 0,177584338 | 0,150946687 | 0,089196304 | 0,138196304 | -1,121803696 |
| chrIV | 1528652 | 1528795 | 143 | 0,313451664 | 0,433586741 | 0,419592436 | -0,202936235 | -0,1724958 | -0,298961505 | -0,291961505 | -1,551961505 |
| chrIV | 1528700 | 1528825 | 125 | 0,210867483 | 0,214940436 | 0,495217383 | -0,01198853 | -0,010190251 | 0,323016528 | 0,456016528 | -0,803983472 |
| chrIV | 1528925 | 1529077 | 152 | 0,09688506 | 0,08894087 | 0,521375354 | 0,053605728 | 0,045564869 | -0,037584297 | -0,093584297 | -1,353584297 |
| chrIV | 1528998 | 1529142 | 144 | 0,09688506 | 0,059293913 | 0,620346375 | 0,30639062 | 0,260432027 | 0,107179247 | 0,107179247 | -1,152820753 |
| chrIV | 1529095 | 1529230 | 135 | 1,279452702 | 0,87829109 | 0,592958583 | 0,235162213 | 0,199887881 | 0,186855848 | 0,249855848 | -1,010144152 |
| chrIV | 1529095 | 1529282 | 187 | 0,128230226 | 0,074117392 | 0,633712557 | 0,341702373 | 0,290447017 | 0,705715081 | 0,404715081 | -0,855284919 |
| chrIV | 1529138 | 1529230 | 92 | 0,170973635 | 0,137117174 | 0,554945587 | 0,138166505 | 0,117441529 | 0,456737995 | 0,820737995 | -0,439262005 |
| chrIV | 1529138 | 1529282 | 144 | 1,439028095 | 0,61146848 | 0,701794927 | 0,529569933 | 0,450134443 | 0,282846411 | 0,282846411 | -0,977153589 |
| chrIV | 1529477 | 1529629 | 152 | 0,165274514 | 0,140823044 | 0,539940648 | 0,100284197 | 0,085241567 | 0,004730172 | -0,051269828 | -1,311269828 |
| chrIV | 1529778 | 1529908 | 130 | 0,327699467 | 0,485468915 | 0,402990911 | -0,245613003 | -0,208771053 | -0,098262391 | -0,000262391 | -1,260262391 |
| chrIV | 1529916 | 1530077 | 161 | 0,182371877 | 0,207528696 | 0,467739443 | -0,080953558 | -0,068810524 | -0,047708642 | -0,166708642 | -1,426708642 |
| chrIV | 1530973 | 1531125 | 152 | 0,037044288 | 0,048176305 | 0,434687048 | -0,164453569 | -0,139785534 | -0,221106359 | -0,277106359 | -1,537106359 |
| chrIV | 1531313 | 1531464 | 151 | 0,515770466 | 0,359469349 | 0,589290452 | 0,225720135 | 0,191862115 | 0,081929697 | 0,032929697 | -1,227070303 |
| chrIV | 1531313 | 1531484 | 171 | 0,028495606 | 0,051882174 | 0,354520937 | -0,373143192 | -0,317171714 | -0,098343973 | -0,287343973 | -1,547343973 |
| chrIX | 1597 | 1749 | 152 | 0,062690333 | 0,040764565 | 0,605967759 | 0,268824834 | 0,228501109 | 0,12314995 | 0,06714995 | -1,19285005 |
| chrIX | 1686 | 1802 | 116 | 0,119681545 | 0,114881957 | 0,510230892 | 0,025647854 | 0,021800676 | 0,549404827 | 0,745404827 | -0,514595173 |
| chrIX | 1686 | 1821 | 135 | 2,322391876 | 1,222936961 | 0,655056832 | 0,399009321 | 0,339157923 | 0,416188243 | 0,479188243 | -0,780811757 |
| chrIX | 1686 | 1838 | 152 | 0,333398588 | 0,259410871 | 0,562404299 | 0,157067817 | 0,133507644 | 0,045231069 | -0,010768931 | -1,270768931 |
| chrIX | 1710 | 1802 | 92 | 0,116831984 | 0,137117174 | 0,460060528 | -0,100281233 | -0,085239048 | 0,384362519 | 0,748362519 | -0,511637481 |
| chrIX | 1710 | 1811 | 101 | 0,09973462 | 0,196411088 | 0,336775505 | -0,421279483 | -0,358087561 | 0,129205877 | 0,430205877 | -0,829794123 |
| chrIX | 1710 | 1821 | 111 | 0,410336724 | 0,418763262 | 0,494918262 | -0,012738373 | -0,010827617 | 0,556370722 | 0,787370722 | -0,472629278 |
| chrIX | 1710 | 1838 | 128 | 7,195140473 | 7,82309068 | 0,479093736 | -0,052428241 | -0,044564005 | 0,314144122 | 0,426144122 | -0,833855878 |
| chrIX | 1710 | 1857 | 147 | 0,071239015 | 0,044470435 | 0,615671537 | 0,294132147 | 0,250012325 | 0,077120251 | 0,056120251 | -1,203879749 |
| chrIX | 1746 | 1838 | 92 | 0,042743409 | 0,125999566 | 0,25330482 | -0,664126048 | -0,564507141 | -0,086758461 | 0,277241539 | -0,982758461 |
| chrIX | 2215 | 2352 | 137 | 0,128230226 | 0,100058479 | 0,561702018 | 0,155285859 | 0,13199298 | 0,11547547 | 0,16447547 | -1,09552453 |
| chrIX | 2382 | 2529 | 147 | 0,062690333 | 0,040764565 | 0,605967759 | 0,268824834 | 0,228501109 | 0,061943972 | 0,040943972 | -1,219056028 |
| chrIX | 2382 | 2530 | 148 | 17,53334627 | 11,31772569 | 0,607719058 | 0,273379012 | 0,23237216 | 0,053883019 | 0,025883019 | -1,234116981 |
| chrIX | 2382 | 2592 | 210 | 0,091185939 | 0,122293696 | 0,427141159 | -0,183657287 | -0,156108694 | 0,217349873 | -0,244650127 | -1,504650127 |
| chrIX | 2383 | 2530 | 147 | 0,051292091 | 0,08894087 | 0,365763444 | -0,34309514 | -0,291630869 | -0,463892777 | -0,484892777 | -1,744892777 |
| chrIX | 2385 | 2530 | 145 | 0,028495606 | 0,037058696 | 0,434687048 | -0,164453569 | -0,139785534 | -0,285370664 | -0,292370664 | -1,552370664 |
| chrIX | 2397 | 2530 | 133 | 1,162620718 | 1,078408047 | 0,518788842 | 0,047114067 | 0,040046957 | 0,208947731 | 0,285947731 | -0,974052269 |
| chrIX | 2437 | 2530 | 93 | 0,210867483 | 0,407645654 | 0,340926442 | -0,409936016 | -0,348445614 | 0,146231211 | 0,503231211 | -0,756768789 |
| chrIX | 2437 | 2592 | 155 | 0,900461144 | 0,718938698 | 0,55604621 | 0,140952356 | 0,119809503 | 0,082881379 | 0,005881379 | -1,254118621 |
| chrIX | 2558 | 2688 | 130 | 0,3903898 | 0,444704349 | 0,467479984 | -0,081606077 | -0,069365165 | 0,209137097 | 0,307137097 | -0,952862903 |
| chrIX | 2879 | 3032 | 153 | 0,823523009 | 0,685585872 | 0,545701519 | 0,114808437 | 0,097587171 | 0,029805292 | -0,033194708 | -1,293194708 |
| chrIX | 3324 | 3468 | 144 | 0,09973462 | 0,066705652 | 0,599221683 | 0,251333038 | 0,213633083 | 0,093621403 | 0,093621403 | -1,166378597 |
| chrIX | 3531 | 3658 | 127 | 0,567062556 | 1,137701961 | 0,332633951 | -0,432651594 | -0,367753855 | 0,03588107 | 0,15488107 | -1,10511893 |
| chrIX | 4088 | 4243 | 155 | 0,088336378 | 0,196411088 | 0,310227091 | -0,495206755 | -0,420925742 | -0,451390823 | -0,528390823 | -1,788390823 |
| chrIX | 4805 | 4968 | 163 | 0,213717044 | 0,329822392 | 0,393195101 | -0,271001078 | -0,230350917 | -0,158739927 | -0,291739927 | -1,551739927 |
| chrIX | 5216 | 5367 | 151 | 0,205168362 | 0,118587827 | 0,633712557 | 0,341702373 | 0,290447017 | 0,138095623 | 0,089095623 | -1,170904377 |
| chrIX | 5429 | 5588 | 159 | 0,530018269 | 0,470645436 | 0,529666726 | 0,074432125 | 0,063267306 | 0,088734685 | -0,016265315 | -1,276265315 |
| chrIX | 9290 | 9441 | 151 | 0,589859041 | 0,470645436 | 0,556206083 | 0,14135711 | 0,120153544 | -0,031792373 | -0,080792373 | -1,340792373 |
| chrIX | 10228 | 10371 | 143 | 0,102584181 | 0,059293913 | 0,633712557 | 0,341702373 | 0,290447017 | 0,1800709 | 0,1870709 | -1,0729291 |
| chrIX | 11460 | 11611 | 151 | 0,319150785 | 0,289057827 | 0,52473901 | 0,0620513 | 0,052743605 | -0,09506829 | -0,14406829 | -1,40406829 |
| chrIX | 12345 | 12490 | 145 | 0,236513528 | 0,137117174 | 0,633014168 | 0,339847115 | 0,288870048 | 0,140066543 | 0,133066543 | -1,126933457 |
| chrIX | 12910 | 13059 | 149 | 0,068389454 | 0,055588044 | 0,551627959 | 0,129775456 | 0,110309137 | -0,065544233 | -0,100544233 | -1,360544233 |
| chrIX | 14208 | 14357 | 149 | 0,122531105 | 0,055588044 | 0,687916521 | 0,489953281 | 0,416460289 | 0,249257829 | 0,214257829 | -1,045742171 |
| chrIX | 16078 | 16185 | 107 | 0,168124074 | 1,615759136 | 0,094246122 | -1,315052545 | -1,117794663 | -0,554620992 | -0,295620992 | -1,555620992 |
| chrIX | 25656 | 25806 | 150 | 1,467523701 | 1,363760005 | 0,518324496 | 0,045948864 | 0,039056534 | -0,132893897 | -0,174893897 | -1,434893897 |
| chrIX | 32330 | 32492 | 162 | 0,076938136 | 0,055588044 | 0,58055047 | 0,203301913 | 0,172806626 | 0,235027004 | 0,109027004 | -1,150972996 |
| chrIX | 33763 | 33917 | 154 | 0,188070999 | 0,211234566 | 0,470995186 | -0,072768458 | -0,061853189 | -0,117263707 | -0,187263707 | -1,447263707 |
| chrIX | 35803 | 35935 | 132 | 0,373292436 | 0,300175436 | 0,554283956 | 0,136492329 | 0,11601848 | 0,321260624 | 0,405260624 | -0,854739376 |
| chrIX | 41284 | 41431 | 147 | 0,723788388 | 0,485468915 | 0,598539605 | 0,24956885 | 0,212133523 | 0,037046296 | 0,016046296 | -1,243953704 |
| chrIX | 48410 | 48570 | 160 | 0,706691025 | 0,407645654 | 0,634180888 | 0,342947146 | 0,291505074 | 0,356834562 | 0,244834562 | -1,015165438 |
| chrIX | 55336 | 55486 | 150 | 0,199469241 | 0,074117392 | 0,729089865 | 0,610062706 | 0,5185533 | 0,35763216 | 0,31563216 | -0,94436784 |
| chrIX | 55497 | 55655 | 158 | 10,26981634 | 9,631555035 | 0,516035611 | 0,040206146 | 0,034175224 | 0,051419363 | -0,046580637 | -1,306580637 |
| chrIX | 55498 | 55655 | 157 | 0,031345166 | 0,037058696 | 0,458236793 | -0,104876779 | -0,089145262 | -0,07861383 | -0,16961383 | -1,42961383 |
| chrIX | 55500 | 55655 | 155 | 0,034194727 | 0,062999783 | 0,351817475 | -0,380418277 | -0,323355536 | -0,352362206 | -0,429362206 | -1,689362206 |
| chrIX | 56537 | 56665 | 128 | 0,173823196 | 0,622586089 | 0,218258625 | -0,778087807 | -0,661374636 | -0,310850015 | -0,198850015 | -1,458850015 |
| chrIX | 56786 | 56893 | 107 | 0,985947962 | 3,450164578 | 0,222254948 | -0,764599787 | -0,649909819 | -0,080763297 | 0,178236703 | -1,081763297 |
| chrIX | 56786 | 56905 | 119 | 1,191116324 | 4,754630669 | 0,200330812 | -0,840440188 | -0,71437416 | -0,198552102 | -0,023552102 | -1,283552102 |
| chrIX | 56791 | 56905 | 114 | 0,054141651 | 0,148234783 | 0,267529425 | -0,620302195 | -0,527256866 | 0,042055219 | 0,252055219 | -1,007944781 |
| chrIX | 59003 | 59166 | 163 | 0,059840772 | 0,085235 | 0,412479432 | -0,221171546 | -0,187995814 | -0,097248005 | -0,230248005 | -1,490248005 |
| chrIX | 59003 | 59166 | 163 | 0,059840772 | 0,085235 | 0,412479432 | -0,221171546 | -0,187995814 | -0,097248005 | -0,230248005 | -1,490248005 |

TABLE 2

| Chrom | Start | End | Length | Section A | Section B | A/A+B | Z-score | Z * 0.85 | Phase correction | Length correction | ΔLK |
|---|---|---|---|---|---|---|---|---|---|---|---|
| chrIX | 59591 | 59741 | 150 | 0,230814407 | 0,140823044 | 0,621074131 | 0,308303061 | 0,262057602 | 0,09893919 | 0,05693919 | -1,20306081 |
| chrIX | 60731 | 60881 | 150 | 0,29065518 | 0,274234349 | 0,51453455 | 0,036440779 | 0,030974662 | -0,129779436 | -0,171779436 | -1,431779436 |
| chrIX | 62677 | 62823 | 146 | 0,413186285 | 0,311293045 | 0,570321703 | 0,177193254 | 0,150614266 | -0,003875339 | -0,017875339 | -1,277875339 |
| chrIX | 70968 | 71118 | 150 | 0,162424953 | 0,17417587 | 0,482544729 | -0,043767845 | -0,037202668 | -0,192228197 | -0,234228197 | -1,494228197 |
| chrIX | 77046 | 77204 | 158 | 0,869115978 | 0,826408916 | 0,512594053 | 0,031573855 | 0,026837777 | 0,057249924 | -0,040750076 | -1,300750076 |
| chrIX | 78123 | 78268 | 145 | 2,427825617 | 1,560171093 | 0,608783255 | 0,276149178 | 0,234726801 | 0,091962418 | 0,084962418 | -1,175037582 |
| chrIX | 80413 | 80548 | 135 | 0,119681545 | 0,122293696 | 0,49460244 | -0,013530088 | -0,011500575 | 0,047001473 | 0,110001473 | -1,149998527 |
| chrIX | 89142 | 89281 | 139 | 0,202318801 | 0,218646305 | 0,480607058 | -0,048630057 | -0,041335549 | -0,089422241 | -0,054422241 | -1,314422241 |
| chrIX | 91580 | 91733 | 153 | 0,296354301 | 0,211234566 | 0,583847125 | 0,211745295 | 0,1799835 | 0,113408267 | 0,050408267 | -1,209591733 |
| chrIX | 93328 | 93477 | 149 | 0,530018269 | 0,222352175 | 0,704464501 | 0,537284661 | 0,456691962 | 0,284259101 | 0,249259101 | -1,010740899 |
| chrIX | 96377 | 96489 | 112 | 0,424584527 | 4,157985667 | 0,092652051 | -1,324599259 | -1,12590937 | -0,56614521 | -0,34214521 | -1,60214521 |
| chrIX | 97100 | 97255 | 155 | 0,042743409 | 0,074117392 | 0,365763444 | -0,34309514 | -0,291630869 | -0,320090971 | -0,397090971 | -1,657090971 |
| chrIX | 101159 | 101314 | 155 | 0,156725832 | 0,200116957 | 0,439201342 | -0,152994409 | -0,130045247 | -0,159745126 | -0,236745126 | -1,496745126 |
| chrIX | 101159 | 101314 | 155 | 0,156725832 | 0,200116957 | 0,439201342 | -0,152994409 | -0,130045247 | -0,159745126 | -0,236745126 | -1,496745126 |
| chrIX | 109382 | 109534 | 152 | 0,284956058 | 0,226058044 | 0,55762856 | 0,144959461 | 0,123215542 | 0,043797221 | -0,012202779 | -1,272202779 |
| chrIX | 122918 | 123072 | 154 | 0,122531105 | 0,159352392 | 0,434687048 | -0,164453569 | -0,139785534 | -0,182403055 | -0,252403055 | -1,512403055 |
| chrIX | 123420 | 123580 | 160 | 0,042743409 | 0,059293913 | 0,418899751 | -0,204708983 | -0,174002636 | -0,097104963 | -0,209104963 | -1,469104963 |
| chrIX | 123420 | 123580 | 160 | 0,042743409 | 0,059293913 | 0,418899751 | -0,204708983 | -0,174002636 | -0,097104963 | -0,209104963 | -1,469104963 |
| chrIX | 123668 | 123818 | 150 | 0,102584181 | 0,096352609 | 0,515662191 | 0,039269381 | 0,033378974 | -0,120071678 | -0,162071678 | -1,422071678 |
| chrIX | 127761 | 127867 | 106 | 0,299203861 | 1,182172396 | 0,201976952 | -0,834580573 | -0,709393487 | -0,187987067 | 0,078012933 | -1,181987067 |
| chrIX | 127761 | 127880 | 119 | 0,376141997 | 1,423053918 | 0,209061167 | -0,8096831 | -0,688230635 | -0,198622363 | -0,023622363 | -1,283622363 |
| chrIX | 129250 | 129396 | 146 | 0,581310359 | 0,26682261 | 0,685400026 | 0,482853237 | 0,410425251 | 0,250406189 | 0,236406189 | -1,023593811 |
| chrIX | 132115 | 132265 | 150 | 0,307752543 | 0,244587392 | 0,557179598 | 0,143822286 | 0,122248943 | -0,035879065 | -0,077879065 | -1,337879065 |
| chrIX | 140470 | 140605 | 135 | 0,074088575 | 0,137117174 | 0,350788628 | -0,383192206 | -0,325713375 | -0,28975792 | -0,22675792 | -1,48675792 |
| chrIX | 140794 | 140945 | 151 | 0,128230226 | 0,125999566 | 0,504387095 | 0,010997038 | 0,009347483 | -0,136811521 | -0,185811521 | -1,445811521 |
| chrIX | 145476 | 145609 | 133 | 1,347842156 | 0,993173047 | 0,575751133 | 0,19103556 | 0,162380226 | 0,304864899 | 0,381864899 | -0,878135101 |
| chrIX | 145755 | 145902 | 147 | 0,210867483 | 0,203822827 | 0,508493876 | 0,021292599 | 0,018098709 | -0,154251811 | -0,175251811 | -1,435251811 |
| chrIX | 149396 | 149568 | 172 | 0,663947616 | 1,096937395 | 0,377053363 | -0,313228948 | -0,266244606 | -0,024252043 | -0,220252043 | -1,480252043 |
| chrIX | 150158 | 150305 | 147 | 0,470177496 | 0,385410436 | 0,549537316 | 0,124492463 | 0,105818594 | -0,062166142 | -0,083166142 | -1,343166142 |
| chrIX | 154400 | 154561 | 161 | 0,128230226 | 0,140823044 | 0,47659791 | -0,058694023 | -0,04988992 | 0,015001562 | -0,103998438 | -1,363998438 |
| chrIX | 157942 | 158056 | 114 | 0,344796831 | 1,104349134 | 0,237931057 | -0,712973568 | -0,606027533 | -0,038522929 | 0,171477071 | -1,088522929 |
| chrIX | 158500 | 158651 | 151 | 0,478726178 | 0,329822392 | 0,592080916 | 0,232901146 | 0,197965974 | 0,057280364 | 0,008280364 | -1,251719636 |
| chrIX | 159528 | 159680 | 152 | 1,966196803 | 1,226642831 | 0,615814456 | 0,294506251 | 0,250330313 | 0,177272698 | 0,121272698 | -1,138727302 |
| chrIX | 160370 | 160515 | 145 | 0,074088575 | 0,051882174 | 0,588141101 | 0,222765786 | 0,189350918 | 0,049174709 | 0,042174709 | -1,217825291 |
| chrIX | 164945 | 165092 | 147 | 0,666797177 | 0,400233914 | 0,624908854 | 0,318399007 | 0,270639156 | 0,098542471 | 0,077542471 | -1,182457529 |
| chrIX | 167654 | 167783 | 129 | 1,615700851 | 2,445873922 | 0,397801577 | -0,259041586 | -0,220185348 | 0,090794051 | 0,195794051 | -1,064205949 |
| chrIX | 170117 | 170264 | 147 | 0,968850598 | 0,596645002 | 0,61887788 | 0,302535022 | 0,257154768 | 0,081088001 | 0,060088001 | -1,199911999 |
| chrIX | 175448 | 175623 | 175 | 0,971700159 | 1,045055221 | 0,481813595 | -0,045602359 | -0,038762005 | 0,29007498 | 0,07307498 | -1,18692502 |
| chrIX | 175932 | 176081 | 149 | 0,435982769 | 0,296469566 | 0,595237052 | 0,241037703 | 0,204882048 | 0,049475732 | 0,014475732 | -1,245524268 |
| chrIX | 176147 | 176286 | 139 | 1,225311051 | 0,978349569 | 0,556034373 | 0,14092239 | 0,119784031 | 0,064799404 | 0,099799404 | -1,160200596 |
| chrIX | 178557 | 178730 | 173 | 0,889062902 | 0,818997177 | 0,520510322 | 0,051434423 | 0,043719259 | 0,351253929 | 0,148253929 | -1,111746071 |
| chrIX | 182864 | 183017 | 153 | 0,108283302 | 0,077823261 | 0,581834946 | 0,206589951 | 0,175601459 | 0,116675414 | 0,053675414 | -1,206324586 |
| chrIX | 190132 | 190312 | 180 | 0,059840772 | 0,148234783 | 0,287591554 | -0,560434497 | -0,476369322 | -0,12557736 | -0,37757736 | -1,63757736 |
| chrIX | 190978 | 191126 | 148 | 0,316301225 | 0,218646305 | 0,591275232 | 0,230826573 | 0,196202587 | 0,018654704 | -0,009345296 | -1,269345296 |
| chrIX | 192456 | 192591 | 135 | 0,322000346 | 0,289057827 | 0,526955305 | 0,067618422 | 0,057475659 | 0,073133909 | 0,136133909 | -1,123866091 |
| chrIX | 192983 | 193089 | 106 | 0,068389454 | 0,448410219 | 0,132332619 | -1,115432247 | -0,94811741 | -0,413425886 | -0,147425886 | -1,407425886 |
| chrIX | 192983 | 193098 | 115 | 0,085486818 | 0,474351306 | 0,152699171 | -1,024925498 | -0,871186673 | -0,322278188 | -0,119278188 | -1,379278188 |
| chrIX | 193702 | 193858 | 156 | 1,12557643 | 0,978349569 | 0,534988603 | 0,087816159 | 0,074643735 | 0,056960887 | -0,027039113 | -1,287039113 |
| chrIX | 196790 | 196938 | 148 | 0,162424953 | 0,085235 | 0,655838584 | 0,40113215 | 0,340962327 | 0,161937104 | 0,133937104 | -1,126062896 |
| chrIX | 200251 | 200397 | 146 | 0,199469241 | 0,244587392 | 0,449197751 | -0,127688487 | -0,108535214 | -0,279507125 | -0,293507125 | -1,553507125 |
| chrIX | 205441 | 205561 | 120 | 0,190920559 | 0,607762611 | 0,239044175 | -0,709380558 | -0,602973475 | -0,136780623 | 0,031219377 | -1,228780623 |
| chrIX | 208655 | 208802 | 147 | 0,518620026 | 0,281646088 | 0,64805946 | 0,380086672 | 0,323073671 | 0,134711989 | 0,113711989 | -1,146288011 |
| chrIX | 212863 | 213023 | 160 | 0,897611584 | 0,837526525 | 0,517314201 | 0,043413899 | 0,036901814 | 0,102378238 | -0,009621762 | -1,269621762 |
| chrIX | 213578 | 213684 | 106 | 1,151222476 | 2,494050226 | 0,315812443 | -0,479441066 | -0,407524906 | 0,143782447 | 0,409782447 | -0,850217553 |
| chrIX | 213578 | 213686 | 108 | 0,031345166 | 0,062999783 | 0,332240005 | -0,433736208 | -0,368675777 | 0,207129567 | 0,459129567 | -0,800870433 |
| chrIX | 213578 | 213701 | 123 | 1,179718082 | 3,161106751 | 0,271772791 | -0,607460147 | -0,516341125 | -0,078552165 | 0,068447835 | -1,191552165 |
| chrIX | 213866 | 214017 | 151 | 0,14532759 | 0,203822827 | 0,416232039 | -0,211542365 | -0,17981101 | -0,333303779 | -0,382303779 | -1,642303779 |
| chrIX | 213866 | 214017 | 151 | 0,14532759 | 0,203822827 | 0,416232039 | -0,211542365 | -0,17981101 | -0,333303779 | -0,382303779 | -1,642303779 |
| chrIX | 217876 | 218011 | 135 | 0,105433742 | 0,059293913 | 0,640048823 | 0,358589298 | 0,304800903 | 0,308413673 | 0,371413673 | -0,888586327 |
| chrIX | 218740 | 218900 | 160 | 2,689985191 | 2,875754793 | 0,483311329 | -0,041844502 | -0,035567826 | 0,025320978 | -0,086679022 | -1,346679022 |
| chrIX | 223366 | 223481 | 115 | 0,068389454 | 0,496586524 | 0,121048428 | -1,16976176 | -0,994297496 | -0,448521161 | -0,245521161 | -1,505521161 |
| chrIX | 224421 | 224561 | 140 | 0,225115286 | 0,233469783 | 0,490891006 | -0,022834846 | -0,019409619 | -0,113669602 | -0,085669602 | -1,345669602 |
| chrIX | 236168 | 236319 | 151 | 0,230814407 | 0,096352609 | 0,705494123 | 0,540268681 | 0,459228379 | 0,300224443 | 0,251224443 | -1,008775557 |
| chrIX | 240288 | 240448 | 160 | 0,541416511 | 0,522527611 | 0,508876829 | 0,022252747 | 0,018914835 | 0,084427963 | -0,027572037 | -1,287572037 |
| chrIX | 241031 | 241216 | 185 | 0,068389454 | 0,055588044 | 0,551627959 | 0,129775456 | 0,110309137 | 0,484910274 | 0,197910274 | -1,062089726 |
| chrIX | 241349 | 241481 | 132 | 1,846515258 | 1,552759353 | 0,543208616 | 0,108520566 | 0,092242481 | 0,278590676 | 0,362590676 | -0,897409324 |
| chrIX | 244655 | 244803 | 148 | 1,322196111 | 0,904232177 | 0,593864225 | 0,23749661 | 0,201872118 | 0,023608413 | -0,004391587 | -1,264391587 |
| chrIX | 244820 | 244976 | 156 | 0,481575739 | 0,396528045 | 0,548426903 | 0,121687903 | 0,103434718 | 0,088756425 | 0,004756425 | -1,255243575 |
| chrIX | 244822 | 244983 | 161 | 0,435982769 | 0,370586958 | 0,540539466 | 0,101792891 | 0,086523957 | 0,154595562 | 0,035595562 | -1,224404438 |
| chrIX | 247146 | 247297 | 151 | 0,031345166 | 0,074117392 | 0,297216064 | -0,532424343 | -0,452560692 | -0,599432857 | -0,648432857 | -1,908432857 |
| chrIX | 249643 | 249773 | 130 | 0,510071344 | 0,659644785 | 0,436064214 | -0,160955528 | -0,136812198 | 0,138956314 | 0,236956314 | -1,023043686 |
| chrIX | 254780 | 254919 | 139 | 0,142478029 | 0,222352175 | 0,390532439 | -0,277931585 | -0,236241848 | -0,311526198 | -0,276526198 | -1,536526198 |
| chrIX | 255472 | 255626 | 154 | 1,473222822 | 1,119172613 | 0,568286305 | 0,172012896 | 0,146210962 | 0,111460975 | 0,041460975 | -1,218539025 |
| chrIX | 255629 | 255772 | 143 | 0,700991904 | 0,600350872 | 0,538668149 | 0,097078943 | 0,082517102 | -0,024251977 | -0,017251977 | -1,277251977 |
| chrIX | 255629 | 255776 | 147 | 1,627099093 | 0,818997177 | 0,665181953 | 0,4266475 | 0,362650375 | 0,17307808 | 0,15207808 | -1,10792192 |
| chrIX | 258137 | 258291 | 154 | 0,031345166 | 0,081529131 | 0,277699771 | -0,589688448 | -0,501235181 | -0,540358384 | -0,610358384 | -1,870358384 |
| chrIX | 258137 | 258291 | 154 | 0,031345166 | 0,081529131 | 0,277699771 | -0,589688448 | -0,501235181 | -0,540358384 | -0,610358384 | -1,870358384 |
| chrIX | 263024 | 263196 | 172 | 0,125380666 | 0,059293913 | 0,678927584 | 0,464702061 | 0,394996752 | 0,669741742 | 0,473741742 | -0,786258258 |
| chrIX | 265122 | 265273 | 151 | 1,607152169 | 0,848644133 | 0,654432197 | 0,397314432 | 0,337717267 | 0,197350332 | 0,148350332 | -1,111649668 |
| chrIX | 265238 | 265355 | 117 | 0,056991212 | 1,067290439 | 0,050691223 | -1,638188175 | -1,392459948 | -0,865674668 | -0,676674668 | -1,936674668 |
| chrIX | 270073 | 270226 | 153 | 3,630340183 | 2,81275501 | 0,563446616 | 0,159713497 | 0,135756473 | 0,082062898 | 0,019062898 | -1,240937102 |
| chrIX | 271691 | 271837 | 146 | 0,381841118 | 0,344645871 | 0,52559939 | 0,064212254 | 0,054580416 | -0,111880416 | -0,125880416 | -1,385880416 |
| chrIX | 280995 | 281156 | 161 | 0,34194727 | 0,248293262 | 0,57933546 | 0,200193718 | 0,17016466 | 0,245134221 | 0,126134221 | -1,133865779 |
| chrIX | 282582 | 282731 | 149 | 0,042743409 | 0,037058696 | 0,535617563 | 0,08939893 | 0,07598909 | -0,095150929 | -0,130150929 | -1,390150929 |
| chrIX | 283375 | 283539 | 164 | 0,113982423 | 0,207528696 | 0,354520937 | -0,373143192 | -0,317171714 | -0,210878615 | -0,350878615 | -1,610878615 |
| chrIX | 285840 | 285993 | 153 | 0,287805619 | 0,277940218 | 0,508718933 | 0,021856865 | 0,018578335 | -0,03465103 | -0,09765103 | -1,35765103 |
| chrIX | 285840 | 285993 | 153 | 0,287805619 | 0,277940218 | 0,508718933 | 0,021856865 | 0,018578335 | -0,03465103 | -0,09765103 | -1,35765103 |
| chrIX | 286233 | 286373 | 140 | 0,310602104 | 0,085235 | 0,784671524 | 0,788067966 | 0,669857771 | 0,568021848 | 0,596021848 | -0,663978152 |
| chrIX | 287987 | 288121 | 134 | 0,287805619 | 0,522527611 | 0,355169464 | -0,371400936 | -0,315690796 | -0,205324736 | -0,135324736 | -1,395324736 |
| chrIX | 289697 | 289831 | 134 | 1,538762715 | 1,32299544 | 0,537698377 | 0,094636892 | 0,080441358 | 0,189289507 | 0,259289507 | -1,000710493 |
| chrIX | 289822 | 289971 | 149 | 0,552814753 | 0,444704349 | 0,554189641 | 0,136253708 | 0,115815652 | -0,058811109 | -0,093811109 | -1,353811109 |
| chrIX | 292476 | 292604 | 128 | 0,347646391 | 0,429880871 | 0,447117944 | -0,132946251 | -0,113004313 | 0,208997126 | 0,320997126 | -0,939002874 |
| chrIX | 296896 | 297052 | 156 | 0,136778908 | 0,140823044 | 0,492715945 | -0,018259434 | -0,015520519 | -0,01982984 | -0,10382984 | -1,36382984 |
| chrIX | 299139 | 299300 | 161 | 0,681044979 | 0,552174567 | 0,552249582 | 0,131346972 | 0,111644926 | 0,175505743 | 0,056505743 | -1,203494257 |
| chrIX | 301522 | 301672 | 150 | 0,182371877 | 0,137117174 | 0,570823558 | 0,178471268 | 0,151700578 | -0,00402044 | -0,04602044 | -1,30602044 |
| chrIX | 301692 | 301851 | 159 | 0,757983115 | 0,900526308 | 0,457026716 | -0,107927211 | -0,091738129 | -0,044464246 | -0,149464246 | -1,409464246 |
| chrIX | 301709 | 301857 | 148 | 0,504372223 | 0,322410653 | 0,610041932 | 0,279428325 | 0,237514076 | 0,068192412 | 0,040192412 | -1,219807588 |
| chrIX | 306823 | 306944 | 121 | 0,726637949 | 2,401403487 | 0,232298057 | -0,731299698 | -0,621604744 | -0,15745718 | 0,00354282 | -1,25645718 |
| chrIX | 309339 | 309491 | 152 | 0,652549374 | 0,741173916 | 0,468205833 | -0,079780711 | -0,067813604 | -0,136785614 | -0,192785614 | -1,452785614 |
| chrIX | 309860 | 310002 | 142 | 3,693030516 | 2,605226314 | 0,586357561 | 0,218185122 | 0,185457353 | 0,0793834 | 0,0933834 | -1,1666166 |
| chrIX | 314424 | 314578 | 154 | 0,336248149 | 0,214940436 | 0,610041932 | 0,279428325 | 0,237514076 | 0,203602018 | 0,133602018 | -1,126397982 |

TABLE 2

| Chrom | Start | End | Length | Section A | Section B | A/A+B | Z-score | Z * 0.85 | Phase correction | Length correction | ΔLK |
|---|---|---|---|---|---|---|---|---|---|---|---|
| chrIX | 320445 | 320583 | 138 | 0,210867483 | 0,255705001 | 0,451950105 | -0,120735916 | -0,102625529 | -0,134205055 | -0,092205055 | -1,352205055 |
| chrIX | 331176 | 331345 | 169 | 0,122531105 | 0,185293479 | 0,398054968 | -0,258384812 | -0,21962709 | 0,007193209 | -0,167806791 | -1,427806791 |
| chrIX | 332149 | 332296 | 147 | 0,068389454 | 0,048176305 | 0,586702775 | 0,219071373 | 0,186210667 | 0,001086874 | -0,019913126 | -1,279913126 |
| chrIX | 333705 | 333871 | 166 | 0,213717044 | 0,448410219 | 0,322773364 | -0,459957499 | -0,390963874 | -0,230484936 | -0,384484936 | -1,644484936 |
| chrIX | 338938 | 339077 | 139 | 0,131079787 | 0,059293913 | 0,688539366 | 0,491714385 | 0,417957228 | 0,329587138 | 0,364587138 | -0,895412862 |
| chrIX | 339456 | 339607 | 151 | 0,367593315 | 0,188999349 | 0,660435071 | 0,413650807 | 0,351603186 | 0,217904632 | 0,168904632 | -1,091095368 |
| chrIX | 343521 | 343669 | 148 | 0,612655525 | 0,400233914 | 0,604859229 | 0,265945033 | 0,226053278 | 0,053784546 | 0,025784546 | -1,234215454 |
| chrIX | 345865 | 346018 | 153 | 0,170973635 | 0,040764565 | 0,807476566 | 0,868634885 | 0,738339652 | 0,68467303 | 0,62167303 | -0,63832697 |
| chrIX | 346400 | 346566 | 166 | 0,24506221 | 0,163058261 | 0,60046537 | 0,254551841 | 0,216369065 | 0,384382689 | 0,230382689 | -1,029617311 |
| chrIX | 352082 | 352233 | 151 | 0,09973462 | 0,148234783 | 0,402205349 | -0,247642923 | -0,210496484 | -0,338800673 | -0,387800673 | -1,647800673 |
| chrIX | 360987 | 361138 | 151 | 5,265987958 | 3,43163523 | 0,60545138 | 0,267483079 | 0,227360617 | 0,098296546 | 0,049296546 | -1,210703454 |
| chrIX | 361485 | 361638 | 153 | 1,191116324 | 0,637409568 | 0,651407962 | 0,389124465 | 0,330755795 | 0,280456639 | 0,217456639 | -1,042543361 |
| chrIX | 364237 | 364386 | 149 | 0,082637257 | 0,051882174 | 0,614314648 | 0,290582424 | 0,24699506 | 0,078083179 | 0,043083179 | -1,216916821 |
| chrIX | 365450 | 365589 | 139 | 1,27090402 | 1,05617283 | 0,546137537 | 0,115908669 | 0,098522369 | 0,007361026 | 0,042361026 | -1,217638974 |
| chrIX | 371073 | 371217 | 144 | 0,82922213 | 0,559586306 | 0,59707452 | 0,24578204 | 0,208914734 | 0,080538698 | 0,080538698 | -1,179461302 |
| chrIX | 377621 | 377774 | 153 | 0,068389454 | 0,077823261 | 0,467739443 | -0,080953558 | -0,068810524 | -0,121311502 | -0,184311502 | -1,444311502 |
| chrIX | 378406 | 378536 | 130 | 0,225115286 | 0,322410653 | 0,411149993 | -0,224587763 | -0,190899598 | 0,05720642 | 0,15520642 | -1,10479358 |
| chrIX | 380201 | 380307 | 106 | 0,037044288 | 0,270528479 | 0,120440727 | -1,172786449 | -0,996868482 | -0,447369505 | -0,181369505 | -1,441369505 |
| chrIX | 380201 | 380310 | 109 | 0,039893848 | 0,307587175 | 0,11480871 | -1,20134495 | -1,021143207 | -0,447718401 | -0,202718401 | -1,462718401 |
| chrIX | 381638 | 381781 | 143 | 0,076938136 | 0,096352609 | 0,443982948 | -0,140878539 | -0,119746758 | -0,237053572 | -0,230053572 | -1,490053572 |
| chrIX | 382286 | 382451 | 165 | 0,09973462 | 0,218646305 | 0,313255639 | -0,486643094 | -0,41364663 | -0,29619655 | -0,44319655 | -1,70319655 |
| chrIX | 386066 | 386219 | 153 | 0,273557816 | 0,188999349 | 0,591403262 | 0,231156174 | 0,196482748 | 0,159673874 | 0,096673874 | -1,163326126 |
| chrIX | 386767 | 386910 | 143 | 2,014639333 | 1,915934572 | 0,512556024 | 0,031478483 | 0,02675671 | -0,095305049 | -0,088305049 | -1,348305049 |
| chrIX | 387904 | 388014 | 110 | 0,373292436 | 2,116051529 | 0,14995615 | -1,036621478 | -0,881128257 | -0,309240035 | -0,071240035 | -1,331240035 |
| chrIX | 388900 | 389051 | 151 | 0,190920559 | 0,26311674 | 0,420495319 | -0,200626496 | -0,170532521 | -0,280890837 | -0,329890837 | -1,589890837 |
| chrIX | 390177 | 390342 | 165 | 3,171560929 | 3,776281101 | 0,456481439 | -0,109302103 | -0,092906788 | 0,055295516 | -0,091704484 | -1,351704484 |
| chrIX | 390386 | 390532 | 146 | 0,547115632 | 0,400233914 | 0,577522451 | 0,195559327 | 0,166225428 | -0,003423677 | -0,017423677 | -1,277423677 |
| chrIX | 392788 | 392939 | 151 | 3,17441049 | 1,652817832 | 0,657605209 | 0,40593606 | 0,345045651 | 0,229725897 | 0,180725897 | -1,079274103 |
| chrIX | 396135 | 396302 | 167 | 0,259310013 | 0,333528262 | 0,437404304 | -0,15755355 | -0,133920518 | 0,070293696 | -0,090706304 | -1,350706304 |
| chrIX | 396795 | 396922 | 127 | 0,074088575 | 0,181587609 | 0,289775034 | -0,554042049 | -0,470935742 | -0,098545508 | 0,020454492 | -1,239545508 |
| chrIX | 405278 | 405390 | 112 | 2,604498373 | 21,17533877 | 0,109525492 | -1,229055535 | -1,044697205 | -0,474251564 | -0,250251564 | -1,510251564 |
| chrIX | 407378 | 407518 | 140 | 1,139824233 | 0,741173916 | 0,605967759 | 0,268824834 | 0,228501109 | 0,115642388 | 0,143642388 | -1,116357612 |
| chrIX | 407520 | 407679 | 159 | 0,051292091 | 0,070411522 | 0,421450846 | -0,198183249 | -0,168455762 | -0,107422306 | -0,212422306 | -1,472422306 |
| chrIX | 409036 | 409181 | 145 | 2,641542661 | 1,708405876 | 0,607258371 | 0,272180469 | 0,231353399 | 0,082802247 | 0,075802247 | -1,184197753 |
| chrIX | 420519 | 420625 | 106 | 3,536304684 | 29,85819119 | 0,105894837 | -1,248659409 | -1,061360498 | -0,499833426 | -0,233833426 | -1,493833426 |
| chrIX | 425922 | 426069 | 147 | 0,763682236 | 0,678174133 | 0,529652088 | 0,07439533 | 0,063236031 | -0,124452334 | -0,145452334 | -1,405452334 |
| chrIX | 433552 | 433722 | 170 | 0,678195419 | 1,067290439 | 0,388542489 | -0,283119835 | -0,24065186 | -0,008340168 | -0,190340168 | -1,450340168 |
| chrIX | 438504 | 438651 | 147 | 0,473027057 | 0,448410219 | 0,513357848 | 0,033489418 | 0,028466005 | -0,152835083 | -0,173835083 | -1,433835083 |
| chrIX | 438570 | 438711 | 141 | 0,09688506 | 0,081529131 | 0,543034495 | 0,108081541 | 0,09186931 | -0,023801477 | -0,002801477 | -1,262801477 |
| chrIX | 439579 | 439688 | 109 | 1,988993287 | 7,356151114 | 0,212837084 | -0,796615851 | -0,677123473 | -0,09293233 | 0,15206767 | -1,10793233 |
| chrIX | 439796 | 439887 | 91 | 0,170973635 | 0,600350872 | 0,221662392 | -0,766590907 | -0,651602271 | -0,197286073 | 0,173713927 | -1,086286073 |
| chrV | 1165 | 1320 | 155 | 0,547115632 | 0,518821741 | 0,513271836 | 0,033273697 | 0,028282642 | -0,068729016 | -0,145729016 | -1,405729016 |
| chrV | 1854 | 1981 | 127 | 0,265009134 | 0,166764131 | 0,613769206 | 0,28915654 | 0,245783059 | 0,646234699 | 0,765234699 | -0,494765301 |
| chrV | 3000 | 3144 | 144 | 0,091185939 | 0,037058696 | 0,711031218 | 0,556399818 | 0,472939845 | 0,300529854 | 0,300529854 | -0,959470146 |
| chrV | 3648 | 3785 | 137 | 0,151026711 | 0,163058261 | 0,480846663 | -0,048028754 | -0,040824441 | -0,085180049 | -0,036180049 | -1,296180049 |
| chrV | 3764 | 3919 | 155 | 0,062690333 | 0,185293479 | 0,252800102 | -0,665704177 | -0,56584855 | -0,666651267 | -0,743651267 | -2,003651267 |
| chrV | 4024 | 4184 | 160 | 0,58700948 | 0,492880654 | 0,543582594 | 0,109463573 | 0,093044037 | 0,089543487 | -0,022456513 | -1,282456513 |
| chrV | 4831 | 4982 | 151 | 0,236513528 | 0,159352392 | 0,597458676 | 0,246774626 | 0,209758432 | 0,098465856 | 0,049465856 | -1,210534144 |
| chrV | 7115 | 7272 | 157 | 0,173823196 | 0,237175653 | 0,422928668 | -0,194406838 | -0,165245812 | -0,214806674 | -0,305806674 | -1,565806674 |
| chrV | 9760 | 9930 | 170 | 0,071239015 | 0,037058696 | 0,657807209 | 0,406485945 | 0,345513053 | 0,743886826 | 0,561886826 | -0,698113174 |
| chrV | 11061 | 11170 | 109 | 0,324849907 | 2,490344357 | 0,115391648 | -1,198343556 | -1,018592022 | -0,53523065 | -0,29023065 | -1,55023065 |
| chrV | 23046 | 23205 | 159 | 0,236513528 | 0,17788174 | 0,570743796 | 0,178268128 | 0,151527909 | 0,14142897 | 0,03642897 | -1,22357103 |
| chrV | 26697 | 26869 | 172 | 0,116831984 | 0,159352392 | 0,423021699 | -0,194169201 | -0,165043821 | 0,306567242 | 0,110567242 | -1,149432758 |
| chrV | 26697 | 26869 | 172 | 0,116831984 | 0,159352392 | 0,423021699 | -0,194169201 | -0,165043821 | 0,306567242 | 0,110567242 | -1,149432758 |
| chrV | 27053 | 27184 | 131 | 7,545636425 | 7,011505243 | 0,518346019 | 0,046002869 | 0,039102439 | 0,263834413 | 0,354834413 | -0,905165587 |
| chrV | 28903 | 29024 | 121 | 0,866266417 | 2,186463051 | 0,28376783 | -0,571684613 | -0,485931921 | 0,016509854 | 0,177509854 | -1,082490146 |
| chrV | 32832 | 32992 | 160 | 0,128230226 | 0,188999349 | 0,404219015 | -0,242441566 | -0,206075331 | -0,202941414 | -0,314941414 | -1,574941414 |
| chrV | 41648 | 41754 | 106 | 0,071239015 | 0,926467395 | 0,071402783 | -1,465422925 | -1,245609486 | -0,786909181 | -0,520909181 | -1,780909181 |
| chrV | 41648 | 41774 | 126 | 0,031345166 | 0,059293913 | 0,345823971 | -0,396619676 | -0,337126724 | 0,086350864 | 0,212350864 | -1,047649136 |
| chrV | 41648 | 41782 | 134 | 0,028495606 | 0,192705218 | 0,128822331 | -1,131975672 | -0,962179321 | -0,897017223 | -0,827017223 | -2,087017223 |
| chrV | 41648 | 41785 | 137 | 0,037044288 | 0,044470435 | 0,454449042 | -0,114428548 | -0,097264266 | -0,151538346 | -0,102538346 | -1,362538346 |
| chrV | 41648 | 41795 | 147 | 0,037044288 | 0,040764565 | 0,47609348 | -0,059960669 | -0,050966568 | -0,226225129 | -0,247225129 | -1,507225129 |
| chrV | 41648 | 41795 | 147 | 0,037044288 | 0,040764565 | 0,47609348 | -0,059960669 | -0,050966568 | -0,226225129 | -0,247225129 | -1,507225129 |
| chrV | 41648 | 41797 | 149 | 0,09973462 | 0,055588044 | 0,642112476 | 0,3641111 | 0,309494435 | 0,1538125 | 0,1188125 | -1,1411875 |
| chrV | 41648 | 41797 | 149 | 0,09973462 | 0,055588044 | 0,642112476 | 0,3641111 | 0,309494435 | 0,1538125 | 0,1188125 | -1,1411875 |
| chrV | 41648 | 41798 | 150 | 0,153876272 | 0,100058479 | 0,605967759 | 0,268824834 | 0,228501109 | 0,096458292 | 0,054458292 | -1,205541708 |
| chrV | 41648 | 41798 | 150 | 0,153876272 | 0,100058479 | 0,605967759 | 0,268824834 | 0,228501109 | 0,096458292 | 0,054458292 | -1,205541708 |
| chrV | 41648 | 41799 | 151 | 0,076938136 | 0,044470435 | 0,633712557 | 0,341702373 | 0,290447017 | 0,179985154 | 0,130985154 | -1,129014846 |
| chrV | 41648 | 41799 | 151 | 0,076938136 | 0,044470435 | 0,633712557 | 0,341702373 | 0,290447017 | 0,179985154 | 0,130985154 | -1,129014846 |
| chrV | 41648 | 41800 | 152 | 0,031345166 | 0,048176305 | 0,394172367 | -0,268460684 | -0,228191582 | -0,328917889 | -0,384917889 | -1,644917889 |
| chrV | 41648 | 41800 | 152 | 0,031345166 | 0,048176305 | 0,394172367 | -0,268460684 | -0,228191582 | -0,328917889 | -0,384917889 | -1,644917889 |
| chrV | 41648 | 41801 | 153 | 0,076938136 | 0,107470218 | 0,417216109 | -0,209020522 | -0,177667444 | -0,322272097 | -0,385272097 | -1,645272097 |
| chrV | 41648 | 41801 | 153 | 0,076938136 | 0,107470218 | 0,417216109 | -0,209020522 | -0,177667444 | -0,322272097 | -0,385272097 | -1,645272097 |
| chrV | 41648 | 41802 | 154 | 47,4537324 | 27,73472793 | 0,631130524 | 0,334849059 | 0,2846217 | 0,173558409 | 0,103558409 | -1,156441591 |
| chrV | 41648 | 41802 | 154 | 47,4537324 | 27,73472793 | 0,631130524 | 0,334849059 | 0,2846217 | 0,173558409 | 0,103558409 | -1,156441591 |
| chrV | 41649 | 41802 | 153 | 0,051292091 | 0,081529131 | 0,386173911 | -0,289305214 | -0,245909432 | -0,390590505 | -0,453590505 | -1,713590505 |
| chrV | 41649 | 41802 | 153 | 0,051292091 | 0,081529131 | 0,386173911 | -0,289305214 | -0,245909432 | -0,390590505 | -0,453590505 | -1,713590505 |
| chrV | 41650 | 41802 | 152 | 0,045592969 | 0,08894087 | 0,338895921 | -0,415478239 | -0,353156503 | -0,465225545 | -0,521225545 | -1,781225545 |
| chrV | 41650 | 41802 | 152 | 0,045592969 | 0,08894087 | 0,338895921 | -0,415478239 | -0,353156503 | -0,465225545 | -0,521225545 | -1,781225545 |
| chrV | 41651 | 41802 | 151 | 0,076938136 | 0,092646739 | 0,453685128 | -0,116356189 | -0,098902761 | -0,209929443 | -0,258929443 | -1,518929443 |
| chrV | 41651 | 41802 | 151 | 0,076938136 | 0,092646739 | 0,453685128 | -0,116356189 | -0,098902761 | -0,209929443 | -0,258929443 | -1,518929443 |
| chrV | 41652 | 41802 | 150 | 0,09688506 | 0,077823261 | 0,554553208 | 0,137173592 | 0,116597553 | -0,007938999 | -0,049938999 | -1,309938999 |
| chrV | 41652 | 41802 | 150 | 0,09688506 | 0,077823261 | 0,554553208 | 0,137173592 | 0,116597553 | -0,007938999 | -0,049938999 | -1,309938999 |
| chrV | 41653 | 41802 | 149 | 0,065539893 | 0,081529131 | 0,445640364 | -0,13668381 | -0,116181238 | -0,262833551 | -0,297833551 | -1,557833551 |
| chrV | 41653 | 41802 | 149 | 0,065539893 | 0,081529131 | 0,445640364 | -0,13668381 | -0,116181238 | -0,262833551 | -0,297833551 | -1,557833551 |
| chrV | 41654 | 41802 | 148 | 0,076938136 | 0,100058479 | 0,434687048 | -0,164453569 | -0,139785534 | -0,327267201 | -0,355267201 | -1,615267201 |
| chrV | 41654 | 41802 | 148 | 0,076938136 | 0,100058479 | 0,434687048 | -0,164453569 | -0,139785534 | -0,327267201 | -0,355267201 | -1,615267201 |
| chrV | 41658 | 41802 | 144 | 0,028495606 | 0,062999783 | 0,311443082 | -0,491764035 | -0,41799943 | -0,597621742 | -0,597621742 | -1,857621742 |
| chrV | 41658 | 41802 | 144 | 0,028495606 | 0,062999783 | 0,311443082 | -0,491764035 | -0,41799943 | -0,597621742 | -0,597621742 | -1,857621742 |
| chrV | 42773 | 42909 | 136 | 0,045592969 | 0,040764565 | 0,527955893 | 0,070132481 | 0,059612609 | 0,038546852 | 0,094546852 | -1,165453148 |
| chrV | 44013 | 44174 | 161 | 0,119681545 | 0,085235 | 0,584050178 | 0,21226584 | 0,180425964 | 0,191151844 | 0,072151844 | -1,187848156 |
| chrV | 51922 | 52070 | 148 | 0,233663968 | 0,114881957 | 0,670396499 | 0,441008282 | 0,37485704 | 0,191077972 | 0,163077972 | -1,096922028 |
| chrV | 56549 | 56680 | 131 | 2,741277281 | 3,261165229 | 0,456693634 | -0,108767039 | -0,092451983 | 0,110409905 | 0,201409905 | -1,058590095 |
| chrV | 58655 | 58795 | 140 | 1,000195765 | 0,600350872 | 0,624908854 | 0,318399007 | 0,270639156 | 0,163878157 | 0,191878157 | -1,068121843 |
| chrV | 63504 | 63668 | 164 | 0,094035499 | 0,118587827 | 0,442263326 | -0,145233329 | -0,12344833 | 0,021092963 | -0,118907037 | -1,378907037 |
| chrV | 64024 | 64155 | 131 | 0,162424953 | 0,314998914 | 0,340211213 | -0,411886758 | -0,350103745 | -0,16730923 | -0,07630923 | -1,33630923 |
| chrV | 65475 | 65628 | 153 | 0,530018269 | 0,34835174 | 0,603411163 | 0,262186483 | 0,22285851 | 0,085660374 | 0,022660374 | -1,237339626 |
| chrV | 69437 | 69601 | 164 | 1,347842156 | 1,133996091 | 0,543082192 | 0,108201802 | 0,091971532 | 0,240350819 | 0,100350819 | -1,159649181 |
| chrV | 71549 | 71648 | 99 | 0,367593315 | 2,149404356 | 0,146044361 | -1,05355059 | -0,895518001 | -0,434388459 | -0,119388459 | -1,379388459 |
| chrV | 72028 | 72188 | 160 | 0,09973462 | 0,107470218 | 0,481333454 | -0,046807179 | -0,039786102 | -0,037697122 | -0,149697122 | -1,409697122 |
| chrV | 78159 | 78308 | 149 | 0,076938136 | 0,040764565 | 0,653664997 | 0,395234266 | 0,335949126 | 0,190857919 | 0,155857919 | -1,104142081 |

| Chrom | Start | End | Length | Section A | Section B | A/A+B | Z-score | Z * 0.85 | Phase correction | Length correction | ΔLK |
|---|---|---|---|---|---|---|---|---|---|---|---|
| chrV | 78159 | 78308 | 149 | 0,076938136 | 0,040764565 | 0,653664997 | 0,395234266 | 0,335949126 | 0,190857919 | 0,155857919 | -1,104142081 |
| chrV | 78447 | 78544 | 97 | 0,133929347 | 0,485468915 | 0,216224932 | -0,785006317 | -0,667255369 | -0,222851563 | 0,106148437 | -1,153851563 |
| chrV | 78782 | 78922 | 140 | 0,125380666 | 0,081529131 | 0,605967759 | 0,268824834 | 0,228501109 | 0,124749033 | 0,152749033 | -1,107250967 |
| chrV | 79095 | 79249 | 154 | 0,524319147 | 0,322410653 | 0,619228409 | 0,30345494 | 0,257936699 | 0,153993948 | 0,083993948 | -1,176006052 |
| chrV | 83616 | 83770 | 154 | 2,205559892 | 1,356348266 | 0,619207401 | 0,303399799 | 0,257889829 | 0,160306321 | 0,090306321 | -1,169693679 |
| chrV | 84397 | 84539 | 142 | 0,624053768 | 0,318704784 | 0,661944425 | 0,417775659 | 0,35510931 | 0,20199085 | 0,21599085 | -1,04400915 |
| chrV | 87005 | 87158 | 153 | 0,270708255 | 0,118587827 | 0,695378834 | 0,511155276 | 0,434481985 | 0,320577967 | 0,257577967 | -1,002422033 |
| chrV | 92533 | 92664 | 131 | 2,023188014 | 2,701578923 | 0,428209061 | -0,180935604 | -0,153795263 | 0,007108844 | 0,098108844 | -1,161891156 |
| chrV | 94783 | 94926 | 143 | 1,097080825 | 0,592939133 | 0,649152585 | 0,38303363 | 0,325578585 | 0,161088409 | 0,168088409 | -1,091911591 |
| chrV | 96167 | 96341 | 174 | 0,262159574 | 0,251999131 | 0,509880648 | 0,024769644 | 0,021054198 | 0,504189345 | 0,294189345 | -0,965810655 |
| chrV | 105359 | 105516 | 157 | 0,09688506 | 0,166764131 | 0,367477175 | -0,338542574 | -0,287761188 | -0,325054623 | -0,416054623 | -1,676054623 |
| chrV | 105504 | 105653 | 149 | 0,190920559 | 0,059293913 | 0,763027643 | 0,716075528 | 0,608664198 | 0,450475331 | 0,415475331 | -0,844524669 |
| chrV | 106502 | 106638 | 136 | 0,054141651 | 0,066705652 | 0,448017039 | -0,130672891 | -0,111071958 | -0,135012408 | -0,079012408 | -1,339012408 |
| chrV | 112707 | 112854 | 147 | 0,09688506 | 0,055588044 | 0,635423938 | 0,346253615 | 0,294315573 | 0,12058787 | 0,09958787 | -1,16041213 |
| chrV | 114004 | 114167 | 163 | 1,179718082 | 1,478641962 | 0,443776638 | -0,14140086 | -0,120190731 | 0,012290536 | -0,120709464 | -1,380709464 |
| chrV | 116119 | 116272 | 153 | 6,59103363 | 5,247511323 | 0,556743557 | 0,142718019 | 0,121310316 | 0,019072973 | -0,043927027 | -1,303927027 |
| chrV | 116657 | 116785 | 128 | 0,287805619 | 0,870879351 | 0,248389879 | -0,679565285 | -0,577630492 | -0,196624347 | -0,084624347 | -1,344624347 |
| chrV | 119860 | 120019 | 159 | 0,045592969 | 0,044470435 | 0,506231912 | 0,015621722 | 0,013278464 | -0,000154346 | -0,105154346 | -1,365154346 |
| chrV | 121418 | 121572 | 154 | 0,492973981 | 0,329822392 | 0,599144572 | 0,251133552 | 0,213463519 | 0,118194078 | 0,048194078 | -1,211805922 |
| chrV | 127589 | 127741 | 152 | 0,09973462 | 0,062999783 | 0,61286746 | 0,286800501 | 0,243780426 | 0,120464269 | 0,064464269 | -1,195535731 |
| chrV | 129771 | 129919 | 148 | 0,284956058 | 0,166764131 | 0,630824269 | 0,334037235 | 0,283931649 | 0,09111919 | 0,06311919 | -1,19688081 |
| chrV | 132213 | 132348 | 135 | 0,43883233 | 0,600350872 | 0,422285819 | -0,196049223 | -0,16664184 | -0,158209347 | -0,095209347 | -1,355209347 |
| chrV | 134881 | 135033 | 152 | 0,24221265 | 0,274234349 | 0,468998078 | -0,077788674 | -0,066120373 | -0,182693151 | -0,238693151 | -1,498693151 |
| chrV | 137057 | 137219 | 162 | 0,492973981 | 0,470645436 | 0,511585769 | 0,0290453 | 0,024688505 | 0,074786651 | -0,051213349 | -1,311213349 |
| chrV | 137099 | 137245 | 146 | 0,236513528 | 0,151940653 | 0,608858238 | 0,276344444 | 0,234892777 | 0,036372228 | 0,022372228 | -1,237627772 |
| chrV | 143216 | 143369 | 153 | 0,310602104 | 0,118587827 | 0,723693828 | 0,59385015 | 0,504772627 | 0,400696281 | 0,337696281 | -0,922303719 |
| chrV | 146039 | 146192 | 153 | 0,159575393 | 0,103764348 | 0,605967759 | 0,268824834 | 0,228501109 | 0,120440237 | 0,057440237 | -1,202559763 |
| chrV | 147368 | 147525 | 157 | 0,447381012 | 0,511410002 | 0,466609517 | -0,08379549 | -0,071226167 | -0,101076532 | -0,192076532 | -1,452076532 |
| chrV | 147862 | 148011 | 149 | 0,125380666 | 0,096352609 | 0,56545715 | 0,193905826 | 0,164819952 | 0,140096959 | -0,0120106 | -1,3070106 |
| chrV | 150905 | 151028 | 123 | 0,142478029 | 0,770820872 | 0,156003723 | -1,011018769 | -0,859365954 | -0,366539424 | -0,219539424 | -1,479539424 |
| chrV | 154668 | 154823 | 155 | 0,227964847 | 0,274234349 | 0,453933118 | -0,11573037 | -0,098370815 | -0,192104692 | -0,269104692 | -1,529104692 |
| chrV | 154964 | 155109 | 145 | 0,284956058 | 0,370586958 | 0,434687048 | -0,164453569 | -0,139785534 | -0,338098809 | -0,345098809 | -1,605098809 |
| chrV | 156735 | 156886 | 151 | 0,068389454 | 0,114881957 | 0,373159423 | -0,323497043 | -0,274972487 | -0,384406271 | -0,433406271 | -1,693406271 |
| chrV | 157741 | 157900 | 159 | 0,125380666 | 0,044470435 | 0,738179884 | 0,637744162 | 0,542082538 | 0,544873574 | 0,439873574 | -0,820126426 |
| chrV | 158361 | 158517 | 156 | 0,113982423 | 0,125999566 | 0,474962408 | -0,062801193 | -0,053381014 | -0,113691927 | -0,197691927 | -1,457691927 |
| chrV | 158999 | 159169 | 170 | 0,074088575 | 0,159352392 | 0,317376063 | -0,47504889 | -0,403791556 | 0,008831741 | -0,173168259 | -1,433168259 |
| chrV | 161114 | 161267 | 153 | 1,091381703 | 0,885702829 | 0,552015701 | 0,130755661 | 0,111142312 | -0,02070096 | -0,08370096 | -1,34370096 |
| chrV | 162828 | 162978 | 150 | 0,190920559 | 0,17417587 | 0,522931822 | 0,057513369 | 0,048886364 | -0,085541174 | -0,127541174 | -1,387541174 |
| chrV | 162990 | 163135 | 145 | 7,750804787 | 4,154279798 | 0,651049955 | 0,38815668 | 0,329933178 | 0,134311071 | 0,127311071 | -1,132688929 |
| chrV | 163162 | 163326 | 164 | 0,743735312 | 0,685585872 | 0,520341628 | 0,051011015 | 0,043359362 | 0,190217642 | 0,050217642 | -1,209782358 |
| chrV | 164105 | 164236 | 131 | 0,09688506 | 0,062999783 | 0,605967759 | 0,268824834 | 0,228501109 | 0,390321148 | 0,481321148 | -0,778678852 |
| chrV | 165644 | 165751 | 107 | 0,094035499 | 0,155646522 | 0,376621026 | -0,314367347 | -0,267212245 | 0,180257713 | 0,439257713 | -0,820742287 |
| chrV | 165645 | 165751 | 106 | 1,228160611 | 3,735516535 | 0,247429592 | -0,682600712 | -0,580210605 | -0,137996043 | 0,128003957 | -1,131996043 |
| chrV | 165645 | 165753 | 108 | 0,031345166 | 0,163058261 | 0,161237725 | -0,989383693 | -0,840976139 | -0,373562961 | -0,121562961 | -1,381562961 |
| chrV | 165645 | 165770 | 125 | 1,598603487 | 4,117221102 | 0,279680292 | -0,583791522 | -0,496222793 | -0,021063114 | 0,111936886 | -1,148063114 |
| chrV | 169801 | 169967 | 166 | 0,051292091 | 0,155646522 | 0,247861382 | -0,681235065 | -0,579049806 | -0,38064542 | -0,53464542 | -1,79464542 |
| chrV | 170735 | 170894 | 159 | 0,039893848 | 0,048176305 | 0,452978074 | -0,118140734 | -0,100419624 | -0,091763496 | -0,196763496 | -1,456763496 |
| chrV | 175783 | 175962 | 179 | 0,68389454 | 1,738052832 | 0,282373824 | -0,575804033 | -0,489433428 | -0,011190621 | -0,256190621 | -1,516190621 |
| chrV | 177369 | 177538 | 169 | 0,079787696 | 0,133411305 | 0,374240479 | -0,320642986 | -0,272546538 | 0,028469708 | -0,146530292 | -1,406530292 |
| chrV | 177369 | 177538 | 169 | 0,079787696 | 0,133411305 | 0,374240479 | -0,320642986 | -0,272546538 | 0,028469708 | -0,146530292 | -1,406530292 |
| chrV | 177766 | 177914 | 148 | 0,159575393 | 0,159352392 | 0,50034961 | 0,000876343 | 0,000744891 | -0,194333537 | -0,222333537 | -1,482333537 |
| chrV | 179498 | 179607 | 109 | 0,034194727 | 0,815291307 | 0,04025343 | -1,747752689 | -1,485589785 | -1,000610091 | -0,755610091 | -2,015610091 |
| chrV | 179498 | 179665 | 167 | 0,034194727 | 0,166764131 | 0,170157849 | -0,953541662 | -0,810510413 | -0,583173751 | -0,744173751 | -2,004173751 |
| chrV | 179498 | 179665 | 167 | 0,034194727 | 0,166764131 | 0,170157849 | -0,953541662 | -0,810510413 | -0,583173751 | -0,744173751 | -2,004173751 |
| chrV | 179498 | 179666 | 168 | 0,039893848 | 0,096352609 | 0,292806499 | -0,545204323 | -0,463423675 | -0,21057675 | -0,37857675 | -1,63857675 |
| chrV | 179498 | 179666 | 168 | 0,039893848 | 0,096352609 | 0,292806499 | -0,545204323 | -0,463423675 | -0,21057675 | -0,37857675 | -1,63857675 |
| chrV | 179498 | 179667 | 169 | 0,085486818 | 0,255705001 | 0,250553539 | -0,67274886 | -0,571836531 | -0,245425085 | -0,420425085 | -1,680425085 |
| chrV | 179498 | 179667 | 169 | 0,085486818 | 0,255705001 | 0,250553539 | -0,67274886 | -0,571836531 | -0,245425085 | -0,420425085 | -1,680425085 |
| chrV | 179498 | 179669 | 171 | 0,034194727 | 0,144528914 | 0,191327386 | -0,873015237 | -0,742062952 | -0,31083774 | -0,49983774 | -1,75983774 |
| chrV | 179498 | 179669 | 171 | 0,034194727 | 0,144528914 | 0,191327386 | -0,873015237 | -0,742062952 | -0,31083774 | -0,49983774 | -1,75983774 |
| chrV | 179498 | 179670 | 172 | 15,48166265 | 30,57712989 | 0,336128279 | -0,423053049 | -0,359595091 | 0,113381321 | -0,082618679 | -1,342618679 |
| chrV | 179498 | 179670 | 172 | 15,48166265 | 30,57712989 | 0,336128279 | -0,423053049 | -0,359595091 | 0,113381321 | -0,082618679 | -1,342618679 |
| chrV | 179499 | 179670 | 171 | 0,059840772 | 0,140823044 | 0,298214065 | -0,529544001 | -0,450112401 | 0,003907434 | -0,185092566 | -1,445092566 |
| chrV | 179499 | 179670 | 171 | 0,059840772 | 0,140823044 | 0,298214065 | -0,529544001 | -0,450112401 | 0,003907434 | -0,185092566 | -1,445092566 |
| chrV | 179502 | 179670 | 168 | 0,031345166 | 0,118587827 | 0,209061167 | -0,8096831 | -0,688230635 | -0,413093399 | -0,581093399 | -1,841093399 |
| chrV | 179502 | 179670 | 168 | 0,031345166 | 0,118587827 | 0,209061167 | -0,8096831 | -0,688230635 | -0,413093399 | -0,581093399 | -1,841093399 |
| chrV | 180478 | 180630 | 152 | 1,022992249 | 0,785644351 | 0,565615143 | 0,16522141 | 0,140438199 | 0,026604359 | -0,029395641 | -1,289395641 |
| chrV | 180478 | 180630 | 152 | 1,022992249 | 0,785644351 | 0,565615143 | 0,16522141 | 0,140438199 | 0,026604359 | -0,029395641 | -1,289395641 |
| chrV | 182865 | 183029 | 164 | 0,284956058 | 0,478057176 | 0,373461489 | -0,322699306 | -0,27429441 | -0,109791068 | -0,249791068 | -1,509791068 |
| chrV | 186698 | 186836 | 138 | 9,155638155 | 5,996096978 | 0,604263345 | 0,264397925 | 0,224738237 | 0,167258922 | 0,209258922 | -1,050741078 |
| chrV | 189568 | 189720 | 152 | 0,19661968 | 0,096352609 | 0,671120401 | 0,443009037 | 0,376557681 | 0,268755198 | 0,212755198 | -1,047244802 |
| chrV | 190625 | 190773 | 148 | 1,014443568 | 0,767115003 | 0,569413537 | 0,174881273 | 0,148649082 | -0,033681522 | -0,061681522 | -1,321681522 |
| chrV | 193660 | 193792 | 132 | 0,219416165 | 0,081529131 | 0,729089865 | 0,610062706 | 0,5185533 | 0,623434496 | 0,707434496 | -0,552565504 |
| chrV | 194009 | 194135 | 126 | 0,068389454 | 0,092646739 | 0,424683747 | -0,189925521 | -0,161436693 | 0,248227019 | 0,374227019 | -0,885772981 |
| chrV | 194030 | 194163 | 133 | 9,192682442 | 8,760675684 | 0,512031364 | 0,030162729 | 0,02563832 | 0,088255212 | 0,165255212 | -1,094744788 |
| chrV | 195885 | 196040 | 155 | 0,316301225 | 0,17788174 | 0,640048823 | 0,358589298 | 0,304800903 | 0,23311419 | 0,15611419 | -1,10388581 |
| chrV | 198265 | 198403 | 138 | 0,094035499 | 0,044470435 | 0,678927584 | 0,464702061 | 0,394996752 | 0,327236775 | 0,369236775 | -0,890763225 |
| chrV | 201379 | 201531 | 152 | 10,68015307 | 8,53091177 | 0,555937589 | 0,140677372 | 0,119575766 | 0,00493926 | -0,05106074 | -1,31106074 |
| chrV | 201380 | 201531 | 151 | 0,056991212 | 0,051882174 | 0,523463207 | 0,058847485 | 0,050020363 | -0,056803429 | -0,105803429 | -1,365803429 |
| chrV | 201382 | 201531 | 149 | 0,088336378 | 0,066705652 | 0,569757619 | 0,175757115 | 0,149393547 | 0,003451775 | -0,031548225 | -1,291548225 |
| chrV | 201383 | 201531 | 148 | 0,045592969 | 0,040764565 | 0,527955893 | 0,070132481 | 0,059612609 | -0,114498629 | -0,142498629 | -1,402498629 |
| chrV | 206462 | 206572 | 110 | 0,088336378 | 0,537351089 | 0,141182911 | -1,075019892 | -0,913766908 | -0,416113353 | -0,178113353 | -1,438113353 |
| chrV | 207496 | 207646 | 150 | 0,105433742 | 0,122293696 | 0,462982163 | -0,092923514 | -0,078984987 | -0,21106086 | -0,25306086 | -1,51306086 |
| chrV | 212628 | 212767 | 139 | 0,113982423 | 0,081529131 | 0,582995843 | 0,209563572 | 0,178129036 | 0,087573978 | 0,122573978 | -1,137426022 |
| chrV | 213547 | 213709 | 162 | 2,54180804 | 3,031401315 | 0,456076181 | -0,110324078 | -0,093775466 | -0,03126739 | -0,15726739 | -1,41726739 |
| chrV | 215300 | 215450 | 150 | 0,065539893 | 0,070411522 | 0,4820832 | -0,044925865 | -0,038186985 | -0,162001951 | -0,204001951 | -1,464001951 |
| chrV | 220704 | 220853 | 149 | 0,088336378 | 0,122293696 | 0,419391098 | -0,203451441 | -0,172933725 | -0,324077269 | -0,359077269 | -1,619077269 |
| chrV | 226712 | 226861 | 149 | 0,173823196 | 0,122293696 | 0,587008713 | 0,219856939 | 0,186878398 | 0,037830098 | 0,002830098 | -1,257169902 |
| chrV | 227455 | 227599 | 144 | 0,085486818 | 0,059293913 | 0,590457149 | 0,22872109 | 0,194412926 | 0,010215692 | 0,010215692 | -1,249784308 |
| chrV | 234231 | 234385 | 154 | 0,997346204 | 0,541056958 | 0,648299632 | 0,380733867 | 0,323623787 | 0,218628317 | 0,148628317 | -1,111371683 |
| chrV | 238011 | 238157 | 146 | 0,424584527 | 0,274234349 | 0,607574497 | 0,273002874 | 0,232052443 | 0,04056343 | 0,02656343 | -1,23343657 |
| chrV | 239144 | 239298 | 154 | 0,068389454 | 0,048176305 | 0,586702775 | 0,219071373 | 0,186210667 | 0,082089419 | 0,012089419 | -1,247910581 |
| chrV | 250567 | 250727 | 160 | 0,122531105 | 0,081529131 | 0,60046537 | 0,254551841 | 0,216369065 | 0,208964727 | 0,096964727 | -1,163035273 |
| chrV | 250628 | 250757 | 129 | 1,473222822 | 2,245756965 | 0,396136281 | -0,263360721 | -0,223856613 | 0,082996262 | 0,187996262 | -1,072003738 |
| chrV | 253010 | 253142 | 132 | 0,054141651 | 0,055588044 | 0,493409293 | -0,016521205 | -0,014043024 | 0,075990049 | 0,159990049 | -1,100009951 |
| chrV | 255896 | 256035 | 139 | 0,549965193 | 0,581821524 | 0,485926531 | -0,035284276 | -0,029991635 | -0,127552539 | -0,092552539 | -1,352552539 |
| chrV | 258267 | 258412 | 145 | 0,09973462 | 0,062999783 | 0,61286746 | 0,286800501 | 0,243780426 | 0,041710092 | 0,034710092 | -1,225289908 |
| chrV | 263170 | 263306 | 136 | 0,119681545 | 0,133411305 | 0,472876041 | -0,06804215 | -0,057835827 | -0,088138703 | -0,032138703 | -1,292138703 |
| chrV | 268138 | 268266 | 128 | 0,159575393 | 0,337234132 | 0,321200349 | -0,464344846 | -0,394693119 | -0,022559319 | 0,089440681 | -1,170559319 |
| chrV | 271754 | 271916 | 162 | 1,296550065 | 1,208113483 | 0,517654384 | 0,044267431 | 0,037627316 | 0,114984481 | -0,011015519 | -1,271015519 |
| chrV | 273268 | 273395 | 127 | 0,353345512 | 0,626291959 | 0,360690074 | -0,356614863 | -0,303122634 | 0,090243252 | 0,209243252 | -1,050756748 |

TABLE 2

| Chrom | Start | End | Length | Section A | Section B | A/A+B | Z-score | Z * 0.85 | Phase correction | Length correction | ΔLK |
|---|---|---|---|---|---|---|---|---|---|---|---|
| chrV | 274395 | 274545 | 150 | 0,085486818 | 0,051882174 | 0,622315244 | 0,311567137 | 0,264832067 | 0,136318916 | 0,094318916 | -1,165681084 |
| chrV | 279045 | 279197 | 152 | 0,518620026 | 0,552174567 | 0,484331943 | -0,039284096 | -0,033391482 | -0,149899569 | -0,205899569 | -1,465899569 |
| chrV | 282333 | 282495 | 162 | 0,122531105 | 0,181587609 | 0,402905508 | -0,245833639 | -0,208958593 | -0,126285142 | -0,252285142 | -1,512285142 |
| chrV | 286305 | 286436 | 131 | 0,116831984 | 0,081529131 | 0,588986325 | 0,224938202 | 0,191197471 | 0,343300819 | 0,434300819 | -0,825699181 |
| chrV | 293237 | 293387 | 150 | 0,575611238 | 0,626291959 | 0,478916471 | -0,052873194 | -0,044942215 | -0,177895789 | -0,219895789 | -1,479895789 |
| chrV | 293855 | 294005 | 150 | 0,29065518 | 0,196411088 | 0,596746683 | 0,244935165 | 0,20819489 | 0,073830772 | 0,031830772 | -1,228169228 |
| chrV | 298879 | 299018 | 139 | 0,527168708 | 0,470645436 | 0,528323547 | 0,071056352 | 0,060397899 | -0,043345527 | -0,008345527 | -1,268345527 |
| chrV | 299205 | 299328 | 123 | 0,039893848 | 0,389116306 | 0,092990452 | -1,322562523 | -1,124178145 | -0,635545952 | -0,488545952 | -1,748545952 |
| chrV | 299678 | 299831 | 153 | 0,649699813 | 0,481763045 | 0,574212232 | 0,187108535 | 0,159042255 | 0,029346533 | -0,033653467 | -1,293653467 |
| chrV | 301505 | 301647 | 142 | 0,125380666 | 0,08894087 | 0,585011979 | 0,214732295 | 0,182522451 | 0,021100065 | 0,035100065 | -1,224899935 |
| chrV | 314667 | 314817 | 150 | 0,227964847 | 0,092646739 | 0,711031218 | 0,556399818 | 0,472939845 | 0,326463792 | 0,284463792 | -0,975536208 |
| chrV | 315905 | 316063 | 158 | 0,085486818 | 0,066705652 | 0,561702018 | 0,155285859 | 0,13199298 | 0,120302862 | 0,022302862 | -1,237697138 |
| chrV | 320077 | 320234 | 157 | 0,082637257 | 0,037058696 | 0,690393076 | 0,496964836 | 0,42242011 | 0,393607764 | 0,302607764 | -0,957392236 |
| chrV | 321656 | 321815 | 159 | 0,09688506 | 0,148234783 | 0,395255882 | -0,265646117 | -0,2257992 | -0,230969993 | -0,335969993 | -1,595969993 |
| chrV | 321656 | 321815 | 159 | 0,09688506 | 0,148234783 | 0,395255882 | -0,265646117 | -0,2257992 | -0,230969993 | -0,335969993 | -1,595969993 |
| chrV | 323513 | 323661 | 148 | 0,852018614 | 0,403939784 | 0,678381239 | 0,46317697 | 0,393700425 | 0,212650082 | 0,184650082 | -1,075349918 |
| chrV | 324743 | 324895 | 152 | 0,63260245 | 0,607762611 | 0,510013116 | 0,025101796 | 0,021336526 | -0,094358804 | -0,150358804 | -1,410358804 |
| chrV | 324904 | 325052 | 148 | 0,629752889 | 0,381704567 | 0,622619256 | 0,312367184 | 0,265512106 | 0,091073817 | 0,063073817 | -1,196926183 |
| chrV | 325348 | 325499 | 151 | 0,302053422 | 0,322410653 | 0,483700238 | -0,040868818 | -0,034738495 | -0,136431373 | -0,185431373 | -1,445431373 |
| chrV | 328966 | 329128 | 162 | 0,074088575 | 0,070411522 | 0,512723359 | 0,03189814 | 0,027113419 | 0,112897669 | -0,013102331 | -1,273102331 |
| chrV | 331009 | 331174 | 165 | 0,142478029 | 0,188999349 | 0,429827309 | -0,17681384 | -0,150291764 | 0,058186504 | -0,088813496 | -1,348813496 |
| chrV | 333088 | 333236 | 148 | 0,071239015 | 0,096352609 | 0,425075031 | -0,188926962 | -0,160587917 | -0,334644224 | -0,362644224 | -1,622644224 |
| chrV | 340240 | 340377 | 137 | 0,937505432 | 0,763409133 | 0,551177261 | 0,128636255 | 0,109340817 | 0,051901532 | 0,100901532 | -1,159098468 |
| chrV | 340798 | 340946 | 148 | 0,179522317 | 0,163058261 | 0,524029464 | 0,060269401 | 0,051228991 | -0,125410293 | -0,153410293 | -1,413410293 |
| chrV | 341934 | 342080 | 146 | 0,14817715 | 0,081529131 | 0,645072262 | 0,372050197 | 0,316242667 | 0,125567135 | 0,111567135 | -1,148432865 |
| chrV | 349486 | 349586 | 100 | 9,993408966 | 28,46107836 | 0,259876276 | -0,643726886 | -0,547167853 | -0,088302446 | 0,219697554 | -1,040302446 |
| chrV | 351515 | 351672 | 157 | 0,125380666 | 0,17417587 | 0,418554265 | -0,205593405 | -0,174754394 | -0,201705746 | -0,292705746 | -1,552705746 |
| chrV | 352323 | 352478 | 155 | 0,113982423 | 0,129705435 | 0,467739443 | -0,080953558 | -0,068810524 | -0,139046162 | -0,216046162 | -1,476046162 |
| chrV | 360556 | 360703 | 147 | 0,042743409 | 0,040764565 | 0,51184823 | 0,029703476 | 0,025247955 | -0,165553627 | -0,186553627 | -1,446553627 |
| chrV | 360556 | 360709 | 153 | 0,065539893 | 0,048176305 | 0,576346155 | 0,192554744 | 0,163671533 | 0,045315248 | -0,017684752 | -1,277684752 |
| chrV | 360596 | 360757 | 161 | 0,370442876 | 0,340940001 | 0,520736284 | 0,052001583 | 0,044201345 | 0,070736906 | -0,048263094 | -1,308263094 |
| chrV | 361101 | 361246 | 145 | 0,695292782 | 0,678174133 | 0,506231912 | 0,015621722 | 0,013278464 | -0,191577418 | -0,198577418 | -1,458577418 |
| chrV | 364550 | 364707 | 157 | 0,3903898 | 0,429880871 | 0,475928024 | -0,060376156 | -0,051319733 | -0,071709887 | -0,162709887 | -1,422709887 |
| chrV | 369752 | 369900 | 148 | 0,091185939 | 0,037058696 | 0,711031218 | 0,556399818 | 0,472939845 | 0,305120146 | 0,277120146 | -0,982879854 |
| chrV | 370967 | 371076 | 109 | 0,133929347 | 0,659644785 | 0,16876728 | -0,959048011 | -0,81519081 | -0,317652675 | -0,072652675 | -1,332652675 |
| chrV | 372915 | 373052 | 137 | 0,641151131 | 0,941290873 | 0,405165642 | -0,239998674 | -0,203998873 | -0,261221282 | -0,212221282 | -1,472221282 |
| chrV | 373805 | 373958 | 153 | 0,638301571 | 0,496586524 | 0,562435692 | 0,157147484 | 0,133575361 | 0,020064557 | -0,042935443 | -1,302935443 |
| chrV | 376017 | 376164 | 147 | 0,948903674 | 0,685585872 | 0,58055047 | 0,203301913 | 0,172806626 | -0,009673883 | -0,030673883 | -1,290673883 |
| chrV | 376371 | 376519 | 148 | 0,353345512 | 0,307587175 | 0,534616488 | 0,086879841 | 0,073847865 | -0,087841574 | -0,115841574 | -1,375841574 |
| chrV | 376832 | 376976 | 144 | 0,418885406 | 0,222352175 | 0,653245253 | 0,394096909 | 0,334982372 | 0,146428127 | 0,146428127 | -1,113571873 |
| chrV | 377517 | 377651 | 134 | 0,914708947 | 1,074702178 | 0,459788797 | -0,10096582 | -0,085820947 | -0,014259516 | 0,055740484 | -1,204259516 |
| chrV | 378722 | 378880 | 158 | 0,287805619 | 0,270528479 | 0,515472044 | 0,038792391 | 0,032973532 | 0,019199567 | -0,078800433 | -1,338800433 |
| chrV | 389589 | 389739 | 150 | 1,456125458 | 1,32299544 | 0,523951822 | 0,060074429 | 0,051063265 | -0,09279508 | -0,13479508 | -1,39479508 |
| chrV | 392068 | 392208 | 140 | 0,04844253 | 0,077823261 | 0,383655221 | -0,295894775 | -0,251510559 | -0,370776899 | -0,342776899 | -1,602776899 |
| chrV | 408769 | 408926 | 157 | 0,267858695 | 0,200116957 | 0,572377417 | 0,18243015 | 0,155065628 | 0,145743206 | 0,054743206 | -1,205256794 |
| chrV | 411385 | 411539 | 154 | 2,05168362 | 1,374877614 | 0,598758779 | 0,250135657 | 0,212615308 | 0,11768312 | 0,04768312 | -1,21231688 |
| chrV | 421211 | 421362 | 151 | 0,230814407 | 0,148234783 | 0,608929957 | 0,276531218 | 0,235051535 | 0,131165119 | 0,082165119 | -1,177834881 |
| chrV | 426798 | 426949 | 151 | 0,034194727 | 0,037058696 | 0,479902939 | -0,050397187 | -0,042837609 | -0,148116882 | -0,197116882 | -1,457116882 |
| chrV | 426821 | 426975 | 154 | 0,359044634 | 0,374292827 | 0,489603563 | -0,026062954 | -0,022153511 | -0,116654544 | -0,186654544 | -1,446654544 |
| chrV | 428076 | 428230 | 154 | 0,082637257 | 0,08894087 | 0,481630488 | -0,046061821 | -0,039152548 | -0,12927449 | -0,19927449 | -1,45927449 |
| chrV | 429069 | 429212 | 143 | 0,102584181 | 0,040764565 | 0,715626635 | 0,569898223 | 0,48441349 | 0,306917275 | 0,313917275 | -0,946082725 |
| chrV | 434847 | 434995 | 148 | 0,569912117 | 0,429880871 | 0,57003012 | 0,176450848 | 0,149983221 | -0,004879224 | -0,032879224 | -1,292879224 |
| chrV | 440738 | 440898 | 160 | 0,433133209 | 0,333528262 | 0,564960188 | 0,163557348 | 0,139023746 | 0,12875477 | 0,01675477 | -1,24324523 |
| chrV | 444332 | 444446 | 114 | 0,04844253 | 0,307587175 | 0,136063169 | -1,098178991 | -0,933452143 | -0,434456338 | -0,224456338 | -1,484456338 |
| chrV | 445028 | 445178 | 150 | 0,119681545 | 0,181587609 | 0,397257877 | -0,260451199 | -0,221383519 | -0,344320469 | -0,386320469 | -1,646320469 |
| chrV | 446200 | 446349 | 149 | 0,09688506 | 0,037058696 | 0,723326439 | 0,592752021 | 0,503839218 | 0,341173135 | 0,306173135 | -0,953826865 |
| chrV | 446424 | 446581 | 157 | 0,065539893 | 0,059293913 | 0,525017182 | 0,062749932 | 0,053337442 | 0,046711811 | -0,044288189 | -1,304288189 |
| chrV | 446610 | 446760 | 150 | 8,944770672 | 4,346985016 | 0,672956296 | 0,448091159 | 0,380877485 | 0,270271871 | 0,228271871 | -1,031728129 |
| chrV | 446614 | 446760 | 146 | 0,074088575 | 0,044470435 | 0,624908854 | 0,318399007 | 0,270639156 | 0,080365727 | 0,066365727 | -1,193634273 |
| chrV | 447072 | 447207 | 135 | 0,076938136 | 0,044470435 | 0,633712557 | 0,341702373 | 0,290447017 | 0,287223637 | 0,350223637 | -0,909776363 |
| chrV | 447594 | 447738 | 144 | 0,04844253 | 0,037058696 | 0,566571175 | 0,16765125 | 0,142503563 | -0,043305836 | -0,043305836 | -1,303305836 |
| chrV | 447618 | 447772 | 154 | 0,213717044 | 0,129705435 | 0,622315244 | 0,311567137 | 0,264832067 | 0,173086433 | 0,103086433 | -1,156913567 |
| chrV | 447875 | 448026 | 151 | 0,04844253 | 0,037058696 | 0,566571175 | 0,16765125 | 0,142503563 | 0,032733921 | -0,016266079 | -1,276266079 |
| chrV | 447899 | 448044 | 145 | 0,111132863 | 0,051882174 | 0,681733814 | 0,472552647 | 0,40166975 | 0,204879692 | 0,197879692 | -1,062120308 |
| chrV | 447903 | 448054 | 151 | 0,139628469 | 0,085235 | 0,620947766 | 0,307970909 | 0,261775272 | 0,148354874 | 0,099354874 | -1,160645126 |
| chrV | 448110 | 448250 | 140 | 0,162424953 | 0,118587827 | 0,577998458 | 0,196775678 | 0,167259327 | 0,044908485 | 0,072908485 | -1,187091515 |
| chrV | 448116 | 448250 | 134 | 0,131079787 | 0,111176087 | 0,541079911 | 0,103154718 | 0,08768151 | 0,147211649 | 0,217211649 | -1,042788351 |
| chrV | 448528 | 448683 | 155 | 0,116831984 | 0,037058696 | 0,759188173 | 0,703693523 | 0,598139495 | 0,536734974 | 0,459734974 | -0,800265026 |
| chrV | 448659 | 448767 | 108 | 0,074088575 | 0,133411305 | 0,357053581 | -0,366345661 | -0,311393812 | 0,171934774 | 0,423934774 | -0,836065226 |
| chrV | 448808 | 448957 | 149 | 0,074088575 | 0,055588044 | 0,571333335 | 0,179769754 | 0,152804291 | 0,003004272 | -0,031995728 | -1,291995728 |
| chrV | 450553 | 450702 | 149 | 0,136778908 | 0,151940653 | 0,473743129 | -0,065863805 | -0,055984234 | -0,203929923 | -0,238929923 | -1,498929923 |
| chrV | 451474 | 451643 | 169 | 0,336248149 | 0,511410002 | 0,396678954 | -0,2619527 | -0,222659795 | 0,129226717 | -0,045773283 | -1,305773283 |
| chrV | 456731 | 456859 | 128 | 0,082637257 | 0,370586958 | 0,182331955 | -0,906513904 | -0,770536818 | -0,434477679 | -0,322477679 | -1,582477679 |
| chrV | 458282 | 458446 | 164 | 0,170973635 | 0,148234783 | 0,535617563 | 0,08939893 | 0,07598909 | 0,23683996 | 0,09683996 | -1,16316004 |
| chrV | 458920 | 459063 | 143 | 4,391172859 | 2,067875225 | 0,6798483 | 0,467274637 | 0,397183441 | 0,215363801 | 0,222363801 | -1,037636199 |
| chrV | 458920 | 459063 | 143 | 4,391172859 | 2,067875225 | 0,6798483 | 0,467274637 | 0,397183441 | 0,215363801 | 0,222363801 | -1,037636199 |
| chrV | 460202 | 460309 | 107 | 0,028495606 | 0,574409785 | 0,04726381 | -1,671983276 | -1,421185784 | -0,96904916 | -0,71004916 | -1,97004916 |
| chrV | 460202 | 460330 | 128 | 8,209584041 | 12,84824983 | 0,389858904 | -0,279686797 | -0,237733777 | 0,072737897 | 0,184737897 | -1,075262103 |
| chrV | 460206 | 460309 | 103 | 0,04844253 | 0,092646739 | 0,343346664 | -0,403346515 | -0,342844538 | 0,119749143 | 0,406749143 | -0,853250857 |
| chrV | 465104 | 465256 | 152 | 0,051292091 | 0,048176305 | 0,515662191 | 0,039269381 | 0,033378974 | -0,091236171 | -0,147236171 | -1,407236171 |
| chrV | 466895 | 467059 | 164 | 0,056991212 | 0,074117392 | 0,434687048 | -0,164453569 | -0,139785534 | 0,050733324 | -0,089266676 | -1,349266676 |
| chrV | 471982 | 472140 | 158 | 0,971700159 | 1,159937178 | 0,455846847 | -0,110902461 | -0,094267092 | -0,102037468 | -0,200037468 | -1,460037468 |
| chrV | 472402 | 472551 | 149 | 0,421734966 | 0,359469349 | 0,539852325 | 0,100061691 | 0,085052437 | -0,070135805 | -0,105135805 | -1,365135805 |
| chrV | 472672 | 472832 | 160 | 2,473418586 | 2,204992399 | 0,528687752 | 0,071971616 | 0,061175874 | 0,068324761 | -0,043675239 | -1,303675239 |
| chrV | 472989 | 473145 | 156 | 0,376141997 | 0,222352175 | 0,628480635 | 0,327831948 | 0,278657156 | 0,228083011 | 0,144083011 | -1,115916989 |
| chrV | 473919 | 474069 | 150 | 0,854868175 | 0,515115871 | 0,623998635 | 0,315999708 | 0,268599752 | 0,155634466 | 0,113634466 | -1,146365534 |
| chrV | 481969 | 482118 | 149 | 0,350495952 | 0,270528479 | 0,564383516 | 0,162092556 | 0,137778672 | -0,011220774 | -0,046220774 | -1,306220774 |
| chrV | 488812 | 488981 | 169 | 0,082637257 | 0,114881957 | 0,418375789 | -0,206050355 | -0,175142801 | 0,201633446 | 0,026633446 | -1,233366554 |
| chrV | 489802 | 489965 | 163 | 0,056991212 | 0,055588044 | 0,506231912 | 0,015621722 | 0,013278464 | 0,15136756 | 0,01836756 | -1,24163244 |
| chrV | 493537 | 493686 | 149 | 0,071239015 | 0,062999783 | 0,530688713 | 0,07700122 | 0,065451037 | -0,075183782 | -0,110183782 | -1,370183782 |
| chrV | 495710 | 495860 | 150 | 9,138540791 | 4,239514798 | 0,683099329 | 0,476383282 | 0,40492579 | 0,28929017 | 0,24729017 | -1,01270983 |
| chrV | 496172 | 496307 | 135 | 0,045592969 | 0,077823261 | 0,36942442 | -0,33337817 | -0,283371444 | -0,285903475 | -0,222903475 | -1,482903475 |
| chrV | 497210 | 497350 | 140 | 0,133929347 | 0,103764348 | 0,563453511 | 0,159731003 | 0,135771353 | 0,009830864 | 0,037830864 | -1,222169136 |
| chrV | 497216 | 497350 | 134 | 0,119681545 | 0,114881957 | 0,510230892 | 0,025647854 | 0,021800676 | 0,067639843 | 0,137639843 | -1,122360157 |
| chrV | 497628 | 497783 | 155 | 0,09973462 | 0,074117392 | 0,573675388 | 0,185739279 | 0,157878387 | 0,101854433 | 0,024854433 | -1,235145567 |
| chrV | 497759 | 497867 | 108 | 0,062690333 | 0,181587609 | 0,256635258 | -0,653753674 | -0,555690623 | -0,071909369 | 0,180090631 | -1,079909369 |
| chrV | 497893 | 498045 | 152 | 0,034194727 | 0,044470435 | 0,434687048 | -0,164453569 | -0,139785534 | -0,276611579 | -0,332611579 | -1,592611579 |
| chrV | 501813 | 501975 | 162 | 1,769577122 | 1,567582832 | 0,5302644 | 0,075934513 | 0,064544336 | 0,157476764 | 0,031476764 | -1,228523236 |
| chrV | 502108 | 502246 | 138 | 0,054141651 | 0,08894087 | 0,378394585 | -0,309699913 | -0,263244926 | -0,329012948 | -0,287012948 | -1,547012948 |
| chrV | 503589 | 503721 | 132 | 2,10012615 | 2,078992834 | 0,502528442 | 0,006337906 | 0,00538722 | 0,086307199 | 0,170307199 | -1,089692801 |
| chrV | 504504 | 504656 | 152 | 0,074088575 | 0,040764565 | 0,645072262 | 0,372050197 | 0,316242667 | 0,172576537 | 0,116576537 | -1,143423463 |

TABLE 2

| Chrom | Start | End | Length | Section A | Section B | A/A+B | Z-score | Z * 0.85 | Phase correction | Length correction | ΔLK |
|---|---|---|---|---|---|---|---|---|---|---|---|
| chrV | 506128 | 506280 | 152 | 1,21961193 | 0,88199696 | 0,580322978 | 0,202719804 | 0,172311833 | 0,023121974 | -0,032878026 | -1,292878026 |
| chrV | 508434 | 508572 | 138 | 0,210867483 | 0,214940436 | 0,495217383 | -0,01198853 | -0,010190251 | -0,087140811 | -0,045140811 | -1,305140811 |
| chrV | 508812 | 508941 | 129 | 0,091185939 | 0,170470001 | 0,348495581 | -0,389385276 | -0,330977484 | -0,04527805 | 0,05972195 | -1,20027805 |
| chrV | 517240 | 517402 | 162 | 1,812320531 | 1,993757833 | 0,476164797 | -0,059781583 | -0,050814346 | 0,05028485 | -0,07571515 | -1,33571515 |
| chrV | 522575 | 522735 | 160 | 0,484425299 | 0,785644351 | 0,381416326 | -0,301763114 | -0,256498647 | -0,255618347 | -0,367618347 | -1,627618347 |
| chrV | 525612 | 525757 | 145 | 0,227964847 | 0,166764131 | 0,577522451 | 0,195559327 | 0,166225428 | -0,026545217 | -0,033545217 | -1,293545217 |
| chrV | 528789 | 528933 | 144 | 0,054141651 | 0,048176305 | 0,529151025 | 0,07313593 | 0,062165541 | -0,131505336 | -0,131505336 | -1,391505336 |
| chrV | 536553 | 536715 | 162 | 0,407487163 | 0,470645436 | 0,464038305 | -0,090265028 | -0,076725274 | 0,016068058 | -0,109931942 | -1,369931942 |
| chrV | 538015 | 538173 | 158 | 0,122531105 | 0,137117174 | 0,47191187 | -0,070464771 | -0,059895055 | -0,064613978 | -0,162613978 | -1,422613978 |
| chrV | 540818 | 540968 | 150 | 0,056991212 | 0,074117392 | 0,434687048 | -0,164453569 | -0,139785534 | -0,254260148 | -0,296260148 | -1,556260148 |
| chrV | 541050 | 541210 | 160 | 0,108283302 | 0,08894087 | 0,549036667 | 0,123227859 | 0,104743681 | 0,107835625 | -0,004164375 | -1,264164375 |
| chrV | 541050 | 541210 | 160 | 0,108283302 | 0,08894087 | 0,549036667 | 0,123227859 | 0,104743681 | 0,107835625 | -0,004164375 | -1,264164375 |
| chrV | 541616 | 541757 | 141 | 1,490320185 | 1,115466743 | 0,571927109 | 0,181282576 | 0,15409019 | 0,025348572 | 0,046348572 | -1,213651428 |
| chrV | 541626 | 541757 | 131 | 0,119681545 | 0,070411522 | 0,629594476 | 0,330779497 | 0,281162572 | 0,409048251 | 0,500048251 | -0,759951749 |
| chrV | 541626 | 541777 | 151 | 0,530018269 | 0,285351958 | 0,650033876 | 0,385411926 | 0,327600137 | 0,21766675 | 0,16866675 | -1,09133325 |
| chrV | 542716 | 542874 | 158 | 0,056991212 | 0,133411305 | 0,299319635 | -0,52635832 | -0,447404572 | -0,467210392 | -0,565210392 | -1,825210392 |
| chrV | 544584 | 544725 | 141 | 2,6386931 | 1,945581529 | 0,575596646 | 0,190641201 | 0,162045021 | 0,028699208 | 0,049699208 | -1,210300792 |
| chrV | 560594 | 560752 | 158 | 0,530018269 | 0,470645436 | 0,529666726 | 0,074432125 | 0,063267306 | 0,04622618 | -0,05177382 | -1,31177382 |
| chrV | 560787 | 560934 | 147 | 0,108283302 | 0,055588044 | 0,660782406 | 0,414599399 | 0,352409489 | 0,155744156 | 0,134744156 | -1,125255844 |
| chrV | 561033 | 561174 | 141 | 0,718089267 | 0,481763045 | 0,59848138 | 0,249418287 | 0,212005544 | 0,07555097 | 0,09655097 | -1,16344903 |
| chrV | 561033 | 561174 | 141 | 0,718089267 | 0,481763045 | 0,59848138 | 0,249418287 | 0,212005544 | 0,07555097 | 0,09655097 | -1,16344903 |
| chrV | 571735 | 571876 | 141 | 0,105433742 | 0,066705652 | 0,612490489 | 0,28581604 | 0,242943634 | 0,095001421 | 0,116001421 | -1,143998579 |
| chrV | 572939 | 573101 | 162 | 0,051292091 | 0,048176305 | 0,515662191 | 0,039269381 | 0,033378974 | 0,149819157 | 0,023819157 | -1,236180843 |
| chrV | 573040 | 573195 | 155 | 0,076938136 | 0,207528696 | 0,270464346 | -0,611409232 | -0,519697847 | -0,586495307 | -0,663495307 | -1,923495307 |
| chrV | 573174 | 573311 | 137 | 0,142478029 | 0,118587827 | 0,545755127 | 0,114943701 | 0,097702146 | 0,039369336 | 0,088369336 | -1,171630664 |
| chrV | 573461 | 573612 | 151 | 0,074088575 | 0,037058696 | 0,666580246 | 0,43048963 | 0,365916185 | 0,268751335 | 0,219751335 | -1,040248665 |
| chrV | 573469 | 573612 | 143 | 0,393239361 | 0,452116089 | 0,465176348 | -0,087401099 | -0,074290934 | -0,249217954 | -0,242217954 | -1,502217954 |
| chrV | 573517 | 573642 | 125 | 0,173823196 | 0,226058044 | 0,434687048 | -0,164453569 | -0,139785534 | 0,305728929 | 0,438728929 | -0,821271071 |
| chrV | 573742 | 573894 | 152 | 0,091185939 | 0,059293913 | 0,605967759 | 0,268824834 | 0,228501109 | 0,079872155 | 0,023872155 | -1,236127845 |
| chrV | 573815 | 573959 | 144 | 0,105433742 | 0,044470435 | 0,703340921 | 0,534033788 | 0,453928719 | 0,25992553 | 0,25992553 | -1,00007447 |
| chrV | 573912 | 574047 | 135 | 1,262355338 | 0,837526525 | 0,601155408 | 0,256338873 | 0,217888042 | 0,20290009 | 0,26590009 | -0,99409991 |
| chrV | 573912 | 574099 | 187 | 0,139628469 | 0,081529131 | 0,631352796 | 0,335438395 | 0,285122636 | 0,776387644 | 0,475387644 | -0,784612356 |
| chrV | 573955 | 574047 | 92 | 0,151026711 | 0,144528914 | 0,510992512 | 0,027557628 | 0,023423984 | 0,411193949 | 0,775193949 | -0,484806051 |
| chrV | 573955 | 574099 | 144 | 1,373488201 | 0,626291959 | 0,686819596 | 0,486855398 | 0,413827089 | 0,225517805 | 0,225517805 | -1,034482195 |
| chrV | 574294 | 574446 | 152 | 0,168124074 | 0,148234783 | 0,5314347 | 0,07887682 | 0,067045297 | -0,082100759 | -0,138100759 | -1,398100759 |
| chrV | 574595 | 574725 | 130 | 0,384690679 | 0,470645436 | 0,449753812 | -0,126283363 | -0,107340859 | 0,159978061 | 0,257978061 | -1,002021939 |
| chrV | 574965 | 575116 | 151 | 0,498673102 | 0,248293262 | 0,667597801 | 0,433289593 | 0,368296154 | 0,264436286 | 0,215436286 | -1,044563714 |
| chrV | 574970 | 575104 | 134 | 0,626903328 | 0,444704349 | 0,585011979 | 0,214732295 | 0,182522451 | 0,216984343 | 0,286984343 | -0,973015657 |
| chrV | 574983 | 575079 | 96 | 0,068389454 | 0,111176087 | 0,380860679 | -0,303221116 | -0,257737948 | 0,155770281 | 0,491770281 | -0,768229719 |
| chrV | 575037 | 575152 | 115 | 0,113982423 | 0,114881957 | 0,498034789 | -0,004926073 | -0,004187162 | 0,491652003 | 0,694652003 | -0,565347997 |
| chrV | 576130 | 576281 | 151 | 0,549965193 | 0,35576348 | 0,607207445 | 0,272048002 | 0,231240802 | 0,121723756 | 0,072723756 | -1,187276244 |
| chrVI | 257 | 431 | 174 | 0,037044288 | 0,040764565 | 0,47609348 | -0,059960669 | -0,050966568 | 0,269885654 | 0,059885654 | -1,200114346 |
| chrVI | 377 | 525 | 148 | 0,056991212 | 0,062999783 | 0,474962408 | -0,062801193 | -0,053381014 | -0,19273657 | -0,22073657 | -1,48073657 |
| chrVI | 2507 | 2632 | 125 | 0,202318801 | 0,218646305 | 0,480607058 | -0,048630057 | -0,041335549 | 0,368938868 | 0,501938868 | -0,758061132 |
| chrVI | 2537 | 2680 | 143 | 0,307752543 | 0,422469132 | 0,421450846 | -0,198183249 | -0,168455762 | -0,185728731 | -0,178728731 | -1,438728731 |
| chrVI | 2537 | 2688 | 151 | 0,039893848 | 0,037058696 | 0,518421434 | 0,046192108 | 0,039263292 | -0,077433233 | -0,126433233 | -1,386433233 |
| chrVI | 2828 | 2965 | 137 | 0,558513874 | 0,518821741 | 0,518421434 | 0,046192108 | 0,039263292 | 0,200811074 | 0,249811074 | -1,010188926 |
| chrVI | 3919 | 4067 | 148 | 0,14532759 | 0,077823261 | 0,65125268 | 0,388704654 | 0,330398956 | 0,189217026 | 0,161217026 | -1,098782974 |
| chrVI | 4714 | 4814 | 100 | 0,527168708 | 0,200116957 | 0,724844079 | 0,597292902 | 0,507698967 | 1,014255146 | 1,322255146 | 0,062255146 |
| chrVI | 6022 | 6182 | 160 | 0,478726178 | 0,329822392 | 0,592080916 | 0,232901146 | 0,197965974 | 0,162385959 | 0,050385959 | -1,209614041 |
| chrVI | 13717 | 13867 | 150 | 0,062690333 | 0,044470435 | 0,585011979 | 0,214732295 | 0,182522451 | 0,051743114 | 0,009743114 | -1,250256886 |
| chrVI | 15894 | 16044 | 150 | 3,439419624 | 1,93446392 | 0,640024964 | 0,358525523 | 0,304746694 | 0,17766544 | 0,13566544 | -1,12433456 |
| chrVI | 24066 | 24172 | 106 | 0,507221784 | 2,416226965 | 0,173501172 | -0,940419656 | -0,799356708 | -0,338111344 | -0,072111344 | -1,332111344 |
| chrVI | 24067 | 24172 | 105 | 0,034194727 | 0,426175002 | 0,074276663 | -1,444660304 | -1,227961258 | -0,74528138 | -0,47228138 | -1,73228138 |
| chrVI | 29981 | 30130 | 149 | 0,507221784 | 0,592939133 | 0,461043268 | -0,097805758 | -0,083134894 | -0,217408731 | -0,252408731 | -1,512408731 |
| chrVI | 36134 | 36286 | 152 | 0,04844253 | 0,048176305 | 0,50137771 | 0,003453413 | 0,002935401 | -0,098020516 | -0,154020516 | -1,414020516 |
| chrVI | 36777 | 36932 | 155 | 0,216566604 | 0,08894087 | 0,708874979 | 0,550101085 | 0,467585923 | 0,367772466 | 0,290772466 | -0,969227534 |
| chrVI | 41762 | 41909 | 147 | 0,091185939 | 0,040764565 | 0,691061693 | 0,498861988 | 0,42403269 | 0,329548098 | 0,308548098 | -0,951451902 |
| chrVI | 48029 | 48199 | 170 | 2,704232994 | 3,368635447 | 0,445297477 | -0,137551417 | -0,116918705 | 0,110350476 | -0,071649524 | -1,331649524 |
| chrVI | 49950 | 50105 | 155 | 0,14817715 | 0,037058696 | 0,799937774 | 0,841398987 | 0,715189139 | 0,620358902 | 0,543358902 | -0,716641098 |
| chrVI | 55917 | 56077 | 160 | 0,162424953 | 0,166764131 | 0,493409293 | -0,016521205 | -0,014043024 | -0,041934322 | -0,153934322 | -1,413934322 |
| chrVI | 60687 | 60850 | 163 | 1,538762715 | 1,593523919 | 0,49125859 | -0,021913219 | -0,018626236 | -0,021413713 | -0,154413713 | -1,414413713 |
| chrVI | 65236 | 65388 | 152 | 0,43883233 | 0,233469783 | 0,65273085 | 0,392703751 | 0,333798188 | 0,223734835 | 0,167734835 | -1,092265165 |
| chrVI | 65445 | 65606 | 161 | 0,353345512 | 0,218646305 | 0,617745747 | 0,299565626 | 0,254630782 | 0,240376832 | 0,121376832 | -1,138623168 |
| chrVI | 68129 | 68276 | 147 | 0,250761331 | 0,233469783 | 0,517854644 | 0,044769906 | 0,03805442 | -0,066365696 | -0,087365696 | -1,347365696 |
| chrVI | 68203 | 68367 | 164 | 0,119681545 | 0,103764348 | 0,535617563 | 0,08939893 | 0,07598909 | 0,141372297 | 0,001372297 | -1,258627703 |
| chrVI | 71963 | 72118 | 155 | 0,202318801 | 0,140823044 | 0,589606905 | 0,226533908 | 0,192553821 | 0,096805439 | 0,019805439 | -1,240194561 |
| chrVI | 72978 | 73128 | 150 | 0,230814407 | 0,17788174 | 0,564757972 | 0,163043664 | 0,138587115 | 0,010207619 | -0,031792381 | -1,291792381 |
| chrVI | 76566 | 76713 | 147 | 0,444531451 | 0,196411088 | 0,693558976 | 0,505963822 | 0,430069249 | 0,292919404 | 0,271919404 | -0,988080596 |
| chrVI | 77258 | 77379 | 121 | 0,039893848 | 0,200116957 | 0,166216884 | -0,969223341 | -0,82383984 | -0,399582826 | -0,238582826 | -1,498582826 |
| chrVI | 79934 | 80100 | 166 | 0,04844253 | 0,188999349 | 0,204018475 | -0,827353109 | -0,703250143 | -0,55583285 | -0,70983285 | -1,96983285 |
| chrVI | 81318 | 81445 | 127 | 0,282106498 | 0,70040935 | 0,287126664 | -0,56179848 | -0,477528708 | -0,093917645 | 0,025082355 | -1,234917645 |
| chrVI | 82718 | 82885 | 167 | 0,267858695 | 0,311293045 | 0,462501753 | -0,094133 | -0,08001305 | 0,090036949 | -0,070963051 | -1,330963051 |
| chrVI | 84294 | 84451 | 157 | 0,176672756 | 0,192705218 | 0,478298027 | -0,054425638 | -0,046261792 | -0,097113915 | -0,188113915 | -1,448113915 |
| chrVI | 91095 | 91248 | 153 | 0,655398934 | 0,589233263 | 0,526580411 | 0,066676582 | 0,056675095 | -0,060248162 | -0,123248162 | -1,383248162 |
| chrVI | 94045 | 94152 | 107 | 0,153876272 | 0,507704132 | 0,23258892 | -0,730347436 | -0,62079532 | -0,172735745 | 0,086264255 | -1,173735745 |
| chrVI | 94173 | 94319 | 146 | 3,188658293 | 2,694167184 | 0,542028368 | 0,105545127 | 0,089713358 | 0,010993383 | -0,003006617 | -1,263006617 |
| chrVI | 94286 | 94435 | 149 | 0,088336378 | 0,062999783 | 0,583709654 | 0,211392905 | 0,179683969 | 0,054143965 | 0,019143965 | -1,240856035 |
| chrVI | 97720 | 97865 | 145 | 0,079787696 | 0,062999783 | 0,558786364 | 0,147892925 | 0,125708986 | 0,072146015 | 0,065146015 | -1,194853985 |
| chrVI | 97725 | 97874 | 149 | 0,034194727 | 0,118587827 | 0,223813035 | -0,759378669 | -0,645471868 | -0,77690461 | -0,81190461 | -2,07190461 |
| chrVI | 100985 | 101138 | 153 | 2,14856868 | 1,797346746 | 0,544504493 | 0,111788617 | 0,095020325 | -0,018002939 | -0,081002939 | -1,341002939 |
| chrVI | 108232 | 108382 | 150 | 0,324849907 | 0,196411088 | 0,623200105 | 0,313896315 | 0,266811868 | 0,130671474 | 0,088671474 | -1,171328526 |
| chrVI | 108406 | 108531 | 125 | 2,065931423 | 5,684803934 | 0,266546505 | -0,623291457 | -0,529797739 | -0,129091324 | 0,003908676 | -1,256091324 |
| chrVI | 110912 | 111064 | 152 | 0,467327936 | 0,489174784 | 0,488579829 | -0,028630034 | -0,024335529 | -0,133062316 | -0,189062316 | -1,449062316 |
| chrVI | 110912 | 111064 | 152 | 0,467327936 | 0,489174784 | 0,488579829 | -0,028630034 | -0,024335529 | -0,133062316 | -0,189062316 | -1,449062316 |
| chrVI | 111983 | 112114 | 131 | 2,843861462 | 4,684219147 | 0,377767137 | -0,311350375 | -0,264647819 | -0,006445116 | 0,084554884 | -1,175445116 |
| chrVI | 115465 | 115634 | 169 | 0,054141651 | 0,044470435 | 0,549036667 | 0,123227859 | 0,104743681 | 0,298760557 | 0,123760557 | -1,136239443 |
| chrVI | 123008 | 123146 | 138 | 0,24506221 | 0,203822827 | 0,545935351 | 0,115398462 | 0,098088692 | 0,17989455 | 0,22189455 | -1,03810545 |
| chrVI | 130462 | 130613 | 151 | 0,772230918 | 0,533645219 | 0,591350815 | 0,231021151 | 0,196367979 | 0,06944386 | 0,02044386 | -1,23955614 |
| chrVI | 131462 | 131615 | 153 | 0,208017923 | 0,155646522 | 0,572005115 | 0,181481351 | 0,154259148 | 0,044607726 | -0,018392274 | -1,278392274 |
| chrVI | 139986 | 140115 | 129 | 0,162424953 | 0,370586958 | 0,304730438 | -0,510843172 | -0,434216696 | -0,149365929 | -0,044365929 | -1,304365929 |
| chrVI | 140063 | 140227 | 164 | 0,53856695 | 0,541056958 | 0,498846817 | -0,002890606 | -0,002457015 | 0,089193282 | -0,050806718 | -1,310806718 |
| chrVI | 141451 | 141594 | 143 | 0,202318801 | 0,114881957 | 0,637825718 | 0,352653046 | 0,299755089 | 0,267079225 | 0,274079225 | -0,985920775 |
| chrVI | 142726 | 142892 | 166 | 0,059840772 | 0,066705652 | 0,472876041 | -0,06804215 | -0,057835827 | 0,112685298 | -0,041314702 | -1,301314702 |
| chrVI | 143530 | 143679 | 149 | 0,09688506 | 0,070411522 | 0,579121574 | 0,199646764 | 0,16969975 | 0,040860939 | 0,005860939 | -1,254139061 |
| chrVI | 148326 | 148498 | 172 | 0,056991212 | 0,077823261 | 0,422738082 | -0,194893703 | -0,165659647 | 0,129779351 | -0,066220649 | -1,326220649 |
| chrVI | 151570 | 151741 | 171 | 0,695292782 | 1,278525005 | 0,352257836 | -0,37923189 | -0,322347107 | -0,058539224 | -0,247539224 | -1,507539224 |
| chrVI | 151602 | 151755 | 153 | 0,866266417 | 0,663350655 | 0,566328942 | 0,167035499 | 0,141980174 | 0,03358165 | -0,02941835 | -1,28941835 |
| chrVI | 151830 | 151995 | 165 | 0,797876963 | 1,1451137 | 0,410643745 | -0,225889337 | -0,192005937 | -0,066101052 | -0,213101052 | -1,473101052 |
| chrVI | 152009 | 152167 | 158 | 1,453275898 | 1,523112397 | 0,488268248 | -0,029411381 | -0,024999674 | -0,071043515 | -0,169043515 | -1,429043515 |
| chrVI | 152552 | 152696 | 144 | 0,09973462 | 0,051882174 | 0,657807209 | 0,406485945 | 0,345513053 | 0,306475138 | 0,306475138 | -0,953524862 |

TABLE 2

| Chrom | Start | End | Length | Section A | Section B | A/A+B | Z-score | Z * 0.85 | Phase correction | Length correction | ΔLK |
|---|---|---|---|---|---|---|---|---|---|---|---|
| chrVI | 155423 | 155586 | 163 | 0,515770466 | 0,644821307 | 0,444403 | -0,139815204 | -0,118842923 | -0,104475844 | -0,237475844 | -1,497475844 |
| chrVI | 162535 | 162694 | 159 | 0,347646391 | 0,274234349 | 0,559024213 | 0,148495706 | 0,12622135 | 0,074237703 | -0,030762297 | -1,290762297 |
| chrVI | 174443 | 174598 | 155 | 1,521665352 | 1,541641745 | 0,496739407 | -0,008173185 | -0,006947207 | -0,097724964 | -0,174724964 | -1,434724964 |
| chrVI | 175285 | 175426 | 141 | 0,541416511 | 0,44099848 | 0,551107746 | 0,128460559 | 0,109191475 | 0,140399459 | 0,161399459 | -1,098600541 |
| chrVI | 176188 | 176314 | 126 | 0,133929347 | 0,077823261 | 0,632480272 | 0,33842962 | 0,287665177 | 0,673308123 | 0,799308123 | -0,460691877 |
| chrVI | 177968 | 178139 | 171 | 0,142478029 | 0,26682261 | 0,348101165 | -0,390452017 | -0,331884215 | -0,053795237 | -0,242795237 | -1,502795237 |
| chrVI | 180563 | 180715 | 152 | 1,823718773 | 1,367465874 | 0,571486446 | 0,180159812 | 0,15313584 | 0,023323106 | -0,032676894 | -1,292676894 |
| chrVI | 182163 | 182304 | 141 | 4,399721541 | 2,920225228 | 0,601059226 | 0,256089736 | 0,217676275 | 0,222190028 | 0,243190028 | -1,016809972 |
| chrVI | 183274 | 183405 | 131 | 0,088336378 | 1,000584786 | 0,081122841 | -1,397558522 | -1,187924744 | -0,961995957 | -0,870995957 | -2,130995957 |
| chrVI | 183755 | 183882 | 127 | 1,065735658 | 1,041349352 | 0,505786674 | 0,014505715 | 0,012329858 | 0,387916479 | 0,506916479 | -0,753083521 |
| chrVI | 185980 | 186131 | 151 | 1,940550757 | 1,511994788 | 0,562063768 | 0,156203695 | 0,13277314 | 0,00327801 | -0,04572199 | -1,30572199 |
| chrVI | 186486 | 186602 | 116 | 0,085486818 | 0,481763045 | 0,15070399 | -1,033418742 | -0,87840593 | -0,436241078 | -0,240241078 | -1,500241078 |
| chrVI | 192368 | 192528 | 160 | 0,24506221 | 0,289057827 | 0,45881486 | -0,103419898 | -0,087906914 | -0,107828325 | -0,219828325 | -1,479828325 |
| chrVI | 193031 | 193186 | 155 | 1,216762369 | 1,222936961 | 0,498734559 | -0,003171996 | -0,002696196 | -0,085354477 | -0,162354477 | -1,422354477 |
| chrVI | 195001 | 195152 | 151 | 0,139628469 | 0,111176087 | 0,556722218 | 0,142663981 | 0,121264384 | 0,002543751 | -0,046456249 | -1,306456249 |
| chrVI | 197175 | 197337 | 162 | 0,447381012 | 0,492880654 | 0,475804798 | -0,060685606 | -0,051582765 | -0,062677025 | -0,188677025 | -1,448677025 |
| chrVI | 197468 | 197595 | 127 | 0,772230918 | 1,990051964 | 0,279562576 | -0,584141446 | -0,496520229 | -0,121299419 | -0,002299419 | -1,262299419 |
| chrVI | 198399 | 198562 | 163 | 2,692834751 | 1,752876311 | 0,605715197 | 0,268168519 | 0,227943241 | 0,250926202 | 0,117926202 | -1,142073798 |
| chrVI | 202836 | 202999 | 163 | 0,934655871 | 0,959820221 | 0,493358494 | -0,016648556 | -0,014151273 | 0,036852612 | -0,096147388 | -1,356147388 |
| chrVI | 204046 | 204202 | 156 | 0,168124074 | 0,137117174 | 0,550790809 | 0,127659576 | 0,10851064 | 0,046465223 | -0,037534777 | -1,297534777 |
| chrVI | 208087 | 208238 | 151 | 2,661489585 | 1,271113265 | 0,676775583 | 0,458701085 | 0,389895922 | 0,272194055 | 0,223194055 | -1,036805945 |
| chrVI | 208444 | 208571 | 127 | 0,105433742 | 0,255705001 | 0,291948022 | -0,54770272 | -0,465547312 | -0,127102919 | -0,008102919 | -1,268102919 |
| chrVI | 208746 | 208898 | 152 | 5,662076879 | 4,120926971 | 0,578766703 | 0,198739409 | 0,168928498 | 0,043709333 | -0,012290667 | -1,272290667 |
| chrVI | 210302 | 210398 | 96 | 0,059840772 | 0,585527393 | 0,092723465 | -1,324168984 | -1,125543636 | -0,641444235 | -0,305444235 | -1,565444235 |
| chrVI | 210302 | 210400 | 98 | 0,19661968 | 1,367465874 | 0,125709031 | -1,14691183 | -0,974875055 | -0,476462921 | -0,154462921 | -1,414462921 |
| chrVI | 210302 | 210402 | 100 | 0,398938482 | 2,786813923 | 0,125225828 | -1,149253041 | -0,976865085 | -0,467983737 | -0,159983737 | -1,419983737 |
| chrVI | 210302 | 210403 | 101 | 0,031345166 | 0,218646305 | 0,125384943 | -1,148481403 | -0,976209192 | -0,477289577 | -0,176289577 | -1,436289577 |
| chrVI | 210302 | 210404 | 102 | 0,675345858 | 3,713281318 | 0,153885448 | -1,019910529 | -0,866923949 | -0,377411677 | -0,083411677 | -1,343411677 |
| chrVI | 210302 | 210405 | 103 | 0,034194727 | 0,337234132 | 0,092062655 | -1,328159833 | -1,128935858 | -0,644392982 | -0,357392982 | -1,617392982 |
| chrVI | 210302 | 210406 | 104 | 0,176672756 | 1,237760439 | 0,124907105 | -1,150800763 | -0,978180649 | -0,48114542 | -0,20114542 | -1,46114542 |
| chrVI | 210302 | 210407 | 105 | 0,031345166 | 0,214940436 | 0,127271615 | -1,13938351 | -0,968475983 | -0,5010736 | -0,2280736 | -1,4880736 |
| chrVI | 210302 | 210409 | 107 | 1,56440876 | 7,797149593 | 0,167109866 | -0,965649231 | -0,820801847 | -0,366556663 | -0,107556663 | -1,367556663 |
| chrVI | 214215 | 214366 | 151 | 0,453080133 | 0,366881088 | 0,552562879 | 0,132139134 | 0,112318264 | 0,002655362 | -0,046344638 | -1,306344638 |
| chrVI | 214215 | 214366 | 151 | 0,453080133 | 0,366881088 | 0,552562879 | 0,132139134 | 0,112318264 | 0,002655362 | -0,046344638 | -1,306344638 |
| chrVI | 224528 | 224665 | 137 | 0,592708601 | 0,566998046 | 0,511084939 | 0,027789399 | 0,023620989 | 0,155804265 | 0,204804265 | -1,055195735 |
| chrVI | 225246 | 225383 | 137 | 11,46093267 | 5,884920891 | 0,660730395 | 0,41445733 | 0,35228873 | 0,452000133 | 0,501000133 | -0,758999867 |
| chrVI | 225640 | 225800 | 160 | 1,085682582 | 1,204407613 | 0,474078525 | -0,065021289 | -0,055268095 | -0,06969213 | -0,18169213 | -1,44169213 |
| chrVI | 228483 | 228624 | 141 | 0,210867483 | 0,133411305 | 0,612490489 | 0,28581604 | 0,242943634 | 0,243905708 | 0,264905708 | -0,995094292 |
| chrVI | 232870 | 233015 | 145 | 3,556251608 | 2,145698486 | 0,623690413 | 0,31518766 | 0,267909511 | 0,205397532 | 0,198397532 | -1,061602468 |
| chrVI | 236135 | 236288 | 153 | 0,53571739 | 0,70040935 | 0,433383869 | -0,167765533 | -0,142600703 | -0,246244981 | -0,309244981 | -1,569244981 |
| chrVI | 236706 | 236866 | 160 | 0,182371877 | 0,166764131 | 0,522351957 | 0,056057394 | 0,047648785 | 0,020579733 | -0,091420267 | -1,351420267 |
| chrVI | 237607 | 237756 | 149 | 0,592708601 | 0,374292827 | 0,612934566 | 0,286975777 | 0,243929411 | 0,110618469 | 0,075618469 | -1,184381531 |
| chrVI | 237976 | 238116 | 140 | 0,886213341 | 0,763409133 | 0,537221913 | 0,093437281 | 0,079421689 | 0,141215564 | 0,169215564 | -1,090784436 |
| chrVI | 239069 | 239219 | 150 | 0,629752889 | 0,537351089 | 0,539585933 | 0,099390615 | 0,084482023 | -0,040599476 | -0,082599476 | -1,342599476 |
| chrVI | 242117 | 242275 | 158 | 0,359044634 | 0,211234566 | 0,629594476 | 0,330779497 | 0,281162572 | 0,226182572 | 0,128182572 | -1,131817428 |
| chrVI | 242831 | 242985 | 154 | 0,179522317 | 0,163058261 | 0,524029464 | 0,060269401 | 0,051228991 | -0,04019621 | -0,11019621 | -1,37019621 |
| chrVI | 242895 | 243045 | 150 | 0,04844253 | 0,059293913 | 0,449639217 | -0,126572917 | -0,10758678 | -0,221153594 | -0,263153594 | -1,523153594 |
| chrVI | 244023 | 244165 | 142 | 0,678195419 | 0,559586306 | 0,547911966 | 0,120387658 | 0,10232951 | 0,085483579 | 0,099483579 | -1,160516421 |
| chrVI | 244610 | 244760 | 150 | 1,233859733 | 0,870879351 | 0,586229306 | 0,217855901 | 0,185177516 | 0,073472902 | 0,031472902 | -1,228527098 |
| chrVI | 247013 | 247149 | 136 | 0,504372223 | 0,285351958 | 0,638668836 | 0,35490291 | 0,301667474 | 0,498421504 | 0,554421504 | -0,705578496 |
| chrVI | 258453 | 258604 | 151 | 0,059840772 | 0,096352609 | 0,38311977 | -0,297297305 | -0,252702709 | -0,362543592 | -0,411543592 | -1,671543592 |
| chrVI | 260977 | 261105 | 128 | 0,498673102 | 0,70040935 | 0,415878909 | -0,212447646 | -0,180580499 | 0,136344029 | 0,248344029 | -1,011655971 |
| chrVI | 264647 | 264774 | 127 | 2,043134938 | 3,231518273 | 0,387349624 | -0,286233551 | -0,243298518 | 0,067631273 | 0,186631273 | -1,073368727 |
| chrVI | 265600 | 265745 | 145 | 0,14532759 | 0,081529131 | 0,640613994 | 0,360100456 | 0,306085387 | 0,232861507 | 0,225861507 | -1,034138493 |
| chrVI | 267454 | 267607 | 153 | 0,125380666 | 0,155646522 | 0,446151373 | -0,135390993 | -0,115082344 | -0,211590669 | -0,274590669 | -1,534590669 |
| chrVI | 269103 | 269252 | 149 | 0,094035499 | 0,077823261 | 0,547167331 | 0,118507771 | 0,100731606 | -0,034072761 | -0,069072761 | -1,329072761 |
| chrVI | 270018 | 270158 | 140 | 0,48727486 | 0,511410002 | 0,487916538 | -0,030293381 | -0,025749374 | 0,031661252 | 0,059661252 | -1,200338748 |
| chrVII | 6501 | 6604 | 103 | 0,122531105 | 0,674468263 | 0,15374053 | -1,020521792 | -0,867443523 | -0,371732916 | -0,084732916 | -1,344732916 |
| chrVII | 7179 | 7332 | 153 | 0,111132863 | 0,137117174 | 0,44766504 | -0,13156284 | -0,111828414 | -0,219523593 | -0,282523593 | -1,542523593 |
| chrVII | 7179 | 7332 | 153 | 0,111132863 | 0,137117174 | 0,44766504 | -0,13156284 | -0,111828414 | -0,219523593 | -0,282523593 | -1,542523593 |
| chrVII | 8743 | 8898 | 155 | 8,232380526 | 6,659447633 | 0,552811947 | 0,132768958 | 0,112853614 | 0,049528237 | -0,027471763 | -1,287471763 |
| chrVII | 8747 | 8898 | 151 | 0,054141651 | 0,044470435 | 0,549036667 | 0,123227859 | 0,104743681 | -0,016453083 | -0,065453083 | -1,325453083 |
| chrVII | 9683 | 9793 | 110 | 0,29350474 | 1,57870044 | 0,156769537 | -1,007823765 | -0,8566502 | -0,174384333 | 0,063615667 | -1,196384333 |
| chrVII | 15630 | 15761 | 131 | 9,825284892 | 9,331379599 | 0,512891213 | 0,032319105 | 0,027471239 | 0,226185837 | 0,317185837 | -0,942814163 |
| chrVII | 15633 | 15761 | 128 | 0,045592969 | 0,085235 | 0,348495581 | -0,389385276 | -0,330977484 | 0,07335771 | 0,18535771 | -1,07464229 |
| chrVII | 15791 | 15935 | 144 | 0,302053422 | 0,340940001 | 0,469761293 | -0,075869923 | -0,064489434 | -0,239304451 | -0,239304451 | -1,499304451 |
| chrVII | 20349 | 20487 | 138 | 3,829809424 | 3,012871967 | 0,559694249 | 0,150194073 | 0,127664962 | 0,022793831 | 0,064793831 | -1,195206169 |
| chrVII | 21325 | 21484 | 159 | 0,256460453 | 0,277940218 | 0,479902939 | -0,050397187 | -0,042837609 | -0,030544131 | -0,135544131 | -1,395544131 |
| chrVII | 22410 | 22546 | 136 | 0,364743755 | 0,244587392 | 0,598596931 | 0,249717095 | 0,21225953 | 0,15362908 | 0,20962908 | -1,05037092 |
| chrVII | 23209 | 23365 | 156 | 0,042743409 | 0,044470435 | 0,490098899 | -0,024820929 | -0,021097789 | -0,082702114 | -0,166702114 | -1,426702114 |
| chrVII | 32814 | 32961 | 147 | 0,125380666 | 0,122293696 | 0,506231912 | 0,015621722 | 0,013278464 | -0,175570695 | -0,196570695 | -1,456570695 |
| chrVII | 33222 | 33355 | 133 | 0,199469241 | 0,270528479 | 0,424404699 | -0,190637769 | -0,162042103 | -0,123549984 | -0,046549984 | -1,306549984 |
| chrVII | 36941 | 37083 | 142 | 0,381841118 | 0,311293045 | 0,550890634 | 0,127911851 | 0,108725073 | -0,013313104 | 0,000686896 | -1,259313104 |
| chrVII | 41750 | 41885 | 135 | 0,182371877 | 0,35205761 | 0,341245911 | -0,409065189 | -0,34770541 | -0,377749961 | -0,314749961 | -1,574749961 |
| chrVII | 52996 | 53149 | 153 | 4,473810116 | 3,023989576 | 0,596683067 | 0,24477085 | 0,208055222 | 0,094797651 | 0,031797651 | -1,228202349 |
| chrVII | 53375 | 53486 | 111 | 0,131079787 | 0,941290873 | 0,122233656 | -1,163893157 | -0,989309184 | -0,335857664 | -0,104857664 | -1,364857664 |
| chrVII | 62645 | 62823 | 178 | 0,04844253 | 0,188999349 | 0,204018475 | -0,827353109 | -0,703250143 | -0,263732769 | -0,501732769 | -1,761732769 |
| chrVII | 69139 | 69303 | 164 | 0,564212996 | 0,559586306 | 0,502058503 | 0,005159925 | 0,004385937 | 0,069410458 | -0,070589542 | -1,330589542 |
| chrVII | 72210 | 72368 | 158 | 0,43883233 | 0,481763045 | 0,476683179 | -0,058479919 | -0,049707931 | -0,046539502 | -0,144539502 | -1,404539502 |
| chrVII | 73049 | 73197 | 148 | 0,076938136 | 0,066705652 | 0,535617563 | 0,08939893 | 0,07598909 | -0,105409235 | -0,133409235 | -1,393409235 |
| chrVII | 78222 | 78356 | 134 | 0,302053422 | 0,233469783 | 0,564034236 | 0,161205527 | 0,137024698 | 0,14029456 | 0,21029456 | -1,04970544 |
| chrVII | 83125 | 83232 | 107 | 0,151026711 | 0,61888022 | 0,196162296 | -0,85540931 | -0,727097914 | -0,092800065 | 0,166199935 | -1,093800065 |
| chrVII | 83125 | 83247 | 122 | 0,185221438 | 0,715232829 | 0,205697774 | -0,821440246 | -0,698224209 | -0,208671883 | -0,054671883 | -1,314671883 |
| chrVII | 84927 | 85059 | 132 | 0,498673102 | 0,511410002 | 0,493695123 | -0,015804641 | -0,013433945 | 0,105423188 | 0,189423188 | -1,070576812 |
| chrVII | 88176 | 88368 | 192 | 0,136778908 | 0,196411088 | 0,41051325 | -0,226224907 | -0,192291171 | 0,270843567 | -0,065156433 | -1,325156433 |
| chrVII | 88303 | 88443 | 140 | 0,475876617 | 0,285351958 | 0,625142872 | 0,319016163 | 0,271163738 | 0,139169989 | 0,167169989 | -1,092830011 |
| chrVII | 88402 | 88555 | 153 | 0,24506221 | 0,137117174 | 0,641222996 | 0,361729731 | 0,307470272 | 0,203403199 | 0,140403199 | -1,119596801 |
| chrVII | 89045 | 89177 | 132 | 0,19377012 | 0,218646305 | 0,469840938 | -0,07566971 | -0,064319253 | 0,054802347 | 0,138802347 | -1,121197653 |
| chrVII | 90038 | 90161 | 123 | 0,037044288 | 0,103764348 | 0,263082498 | -0,633871027 | -0,538790373 | -0,062779921 | 0,084220079 | -1,175779921 |
| chrVII | 91402 | 91556 | 154 | 0,208017923 | 0,133411305 | 0,609256343 | 0,277381333 | 0,235774133 | 0,173593639 | 0,103593639 | -1,156406361 |
| chrVII | 92970 | 93138 | 168 | 2,843861462 | 3,212988925 | 0,469528101 | -0,076456145 | -0,064987724 | 0,161782507 | -0,006217493 | -1,266217493 |
| chrVII | 97437 | 97599 | 162 | 0,296354301 | 0,344645871 | 0,462331079 | -0,094562726 | -0,080378317 | -0,041513424 | -0,167513424 | -1,427513424 |
| chrVII | 102150 | 102299 | 149 | 0,076938136 | 0,066705652 | 0,535617563 | 0,08939893 | 0,07598909 | -0,110788269 | -0,145788269 | -1,405788269 |
| chrVII | 102520 | 102673 | 153 | 0,042743409 | 0,059293913 | 0,418899751 | -0,204708983 | -0,174002636 | -0,280581125 | -0,343581125 | -1,603581125 |
| chrVII | 103288 | 103440 | 152 | 0,233663968 | 0,129705435 | 0,643048 | 0,366617972 | 0,311625276 | 0,179971599 | 0,123971599 | -1,136028401 |
| chrVII | 103852 | 103984 | 132 | 0,581310359 | 0,678174133 | 0,461546262 | -0,096538972 | -0,082058126 | 0,026244755 | 0,110244755 | -1,149755245 |
| chrVII | 108983 | 109147 | 164 | 0,757983115 | 0,889408699 | 0,460111012 | -0,100154051 | -0,085130943 | -0,010889484 | -0,150889484 | -1,410889484 |
| chrVII | 116027 | 116173 | 146 | 0,376141997 | 0,207528696 | 0,644442151 | 0,370358087 | 0,314804374 | 0,153355084 | 0,139355084 | -1,120644916 |
| chrVII | 119268 | 119414 | 146 | 0,396088921 | 0,237175653 | 0,625471465 | 0,319882942 | 0,271900501 | 0,10447816 | 0,09047816 | -1,16952184 |
| chrVII | 125888 | 126050 | 162 | 0,78362916 | 0,837526525 | 0,48337687 | -0,041680073 | -0,035428062 | 0,001570111 | -0,124429889 | -1,384429889 |
| chrVII | 138978 | 139125 | 147 | 0,435982769 | 0,337234132 | 0,563855716 | 0,160752208 | 0,136639377 | -0,047937718 | -0,068937718 | -1,328937718 |

TABLE 2

| Chrom | Start | End | Length | Section A | Section B | A/A+B | Z-score | Z * 0.85 | Phase correction | Length correction | ΔLK |
|---|---|---|---|---|---|---|---|---|---|---|---|
| chrVII | 147769 | 147920 | 151 | 0,19377012 | 0,181587609 | 0,516227867 | 0,040688453 | 0,034585185 | -0,086131641 | -0,135131641 | -1,395131641 |
| chrVII | 152057 | 152208 | 151 | 0,094035499 | 0,125999566 | 0,427365971 | -0,183084213 | -0,155621581 | -0,282722759 | -0,331722759 | -1,591722759 |
| chrVII | 156139 | 156292 | 153 | 0,076938136 | 0,111176087 | 0,408996909 | -0,230126057 | -0,195607148 | -0,301930912 | -0,364930912 | -1,624930912 |
| chrVII | 167824 | 167963 | 139 | 0,521469587 | 0,277940218 | 0,652318227 | 0,391586796 | 0,332848776 | 0,20671779 | 0,24171779 | -1,01828221 |
| chrVII | 167824 | 167963 | 139 | 0,521469587 | 0,277940218 | 0,652318227 | 0,391586796 | 0,332848776 | 0,20671779 | 0,24171779 | -1,01828221 |
| chrVII | 168448 | 168554 | 106 | 0,051292091 | 0,570703915 | 0,082463698 | -1,38868882 | -1,180385497 | -0,602342667 | -0,336342667 | -1,596342667 |
| chrVII | 168635 | 168789 | 154 | 0,042743409 | 0,074117392 | 0,365763444 | -0,34309514 | -0,291630869 | -0,356079628 | -0,426079628 | -1,686079628 |
| chrVII | 171388 | 171548 | 160 | 0,125380666 | 0,125999566 | 0,498768996 | -0,003085674 | -0,002622823 | 0,042768107 | -0,069231893 | -1,329231893 |
| chrVII | 174752 | 174849 | 97 | 1,393435125 | 7,063387417 | 0,164770529 | -0,975038711 | -0,828782904 | -0,394810284 | -0,065810284 | -1,325810284 |
| chrVII | 174828 | 174980 | 152 | 0,273557816 | 0,163058261 | 0,626540868 | 0,322705529 | 0,274299699 | 0,139848201 | 0,083848201 | -1,176151799 |
| chrVII | 178188 | 178320 | 132 | 0,051292091 | 0,044470435 | 0,535617563 | 0,08939893 | 0,07598909 | 0,171276535 | 0,255276535 | -1,004723465 |
| chrVII | 183143 | 183279 | 136 | 0,757983115 | 0,926467395 | 0,449988356 | -0,125690766 | -0,106837151 | -0,168240676 | -0,112240676 | -1,372240676 |
| chrVII | 184850 | 184994 | 144 | 0,678195419 | 0,329822392 | 0,672801027 | 0,447660898 | 0,380511763 | 0,208784636 | 0,208784636 | -1,051215364 |
| chrVII | 196178 | 196334 | 156 | 2,140019998 | 1,812170224 | 0,541476973 | 0,104155366 | 0,088532061 | 0,041015225 | -0,042984775 | -1,302984775 |
| chrVII | 196178 | 196334 | 156 | 2,140019998 | 1,812170224 | 0,541476973 | 0,104155366 | 0,088532061 | 0,041015225 | -0,042984775 | -1,302984775 |
| chrVII | 204030 | 204183 | 153 | 0,48727486 | 0,337234132 | 0,590987927 | 0,230087027 | 0,195573973 | 0,089649878 | 0,026649878 | -1,233350122 |
| chrVII | 204995 | 205129 | 134 | 1,339293474 | 0,974643699 | 0,578794226 | 0,198809775 | 0,168988309 | 0,169336543 | 0,239336543 | -1,020663457 |
| chrVII | 209064 | 209201 | 137 | 0,353345512 | 0,337234132 | 0,511665114 | 0,029244273 | 0,024857632 | -0,088980566 | -0,039980566 | -1,299980566 |
| chrVII | 214611 | 214774 | 163 | 0,661098055 | 0,752291524 | 0,467739443 | -0,080953558 | -0,068810524 | -0,01160981 | -0,14460981 | -1,40460981 |
| chrVII | 217469 | 217614 | 145 | 0,236513528 | 0,148234783 | 0,61472272 | 0,291649583 | 0,247902146 | 0,122466618 | 0,115466618 | -1,144533382 |
| chrVII | 221203 | 221378 | 175 | 0,501522663 | 0,503998263 | 0,498768996 | -0,003085674 | -0,002622823 | 0,378803359 | 0,161803359 | -1,098196641 |
| chrVII | 223475 | 223638 | 163 | 0,843469933 | 0,874585221 | 0,490944619 | -0,022700423 | -0,01929536 | 0,048443371 | -0,084556629 | -1,344556629 |
| chrVII | 226620 | 226777 | 157 | 0,330549028 | 0,08894087 | 0,787978518 | 0,799426819 | 0,679512796 | 0,64924189 | 0,55824189 | -0,70175811 |
| chrVII | 226621 | 226770 | 149 | 4,439615389 | 2,972107402 | 0,598999115 | 0,250757281 | 0,213143689 | 0,034999523 | -4,77E-07 | -1,260000477 |
| chrVII | 233772 | 233924 | 152 | 0,219416165 | 0,17788174 | 0,552271135 | 0,131401466 | 0,111691246 | -0,025728485 | -0,081728485 | -1,341728485 |
| chrVII | 238049 | 238197 | 148 | 0,350495952 | 0,237175653 | 0,596414645 | 0,244077615 | 0,207465972 | 0,015443579 | -0,012556421 | -1,272556421 |
| chrVII | 243163 | 243328 | 165 | 0,73233707 | 0,61146848 | 0,5449725 | 0,112969166 | 0,096023791 | 0,201988332 | 0,054988332 | -1,205011668 |
| chrVII | 249506 | 249649 | 143 | 0,034194727 | 0,070411522 | 0,326889907 | -0,448517424 | -0,381239811 | -0,532763635 | -0,525763635 | -1,785763635 |
| chrVII | 251482 | 251633 | 151 | 0,09688506 | 0,044470435 | 0,685400026 | 0,482853237 | 0,410425251 | 0,28058505 | 0,23158505 | -1,02841495 |
| chrVII | 266552 | 266693 | 141 | 0,113982423 | 0,074117392 | 0,605967759 | 0,268824834 | 0,228501109 | 0,101885783 | 0,122885783 | -1,137114217 |
| chrVII | 267377 | 267528 | 151 | 1,478921943 | 0,889408699 | 0,624457547 | 0,317209152 | 0,269627779 | 0,137615653 | 0,088615653 | -1,171384347 |
| chrVII | 268853 | 268959 | 106 | 0,068389454 | 0,159352392 | 0,300293755 | -0,523555831 | -0,445022456 | 0,151624118 | 0,417624118 | -0,842375882 |
| chrVII | 268853 | 268977 | 124 | 0,071239015 | 0,214940436 | 0,248931272 | -0,677856719 | -0,576178211 | -0,117224761 | 0,022775239 | -1,237224761 |
| chrVII | 270004 | 270159 | 155 | 0,678195419 | 0,648527176 | 0,511181027 | 0,028030349 | 0,023825797 | -0,03256371 | -0,10956371 | -1,36956371 |
| chrVII | 273536 | 273656 | 120 | 0,076938136 | 0,251999131 | 0,233899115 | -0,726066131 | -0,617156212 | -0,029368652 | 0,138631348 | -1,121368652 |
| chrVII | 275567 | 275723 | 156 | 1,718285032 | 1,371171744 | 0,556177075 | 0,141283669 | 0,120091118 | 0,058662709 | -0,025337291 | -1,285337291 |
| chrVII | 275567 | 275723 | 156 | 1,718285032 | 1,371171744 | 0,556177075 | 0,141283669 | 0,120091118 | 0,058662709 | -0,025337291 | -1,285337291 |
| chrVII | 279938 | 280099 | 161 | 1,182567642 | 0,763409133 | 0,607698744 | 0,273326155 | 0,232327231 | 0,253661488 | 0,134661488 | -1,125338512 |
| chrVII | 281760 | 281908 | 148 | 13,45277551 | 8,697675901 | 0,607336404 | 0,272383457 | 0,231525938 | 0,034874121 | 0,006874121 | -1,253125879 |
| chrVII | 281764 | 281908 | 144 | 0,082637257 | 0,077823261 | 0,515000562 | 0,037609696 | 0,031968242 | -0,128605836 | -0,128605836 | -1,388605836 |
| chrVII | 284331 | 284485 | 154 | 0,44168189 | 0,237175653 | 0,650625296 | 0,387009186 | 0,328957808 | 0,267497128 | 0,197497128 | -1,062502872 |
| chrVII | 289603 | 289760 | 157 | 0,071239015 | 0,085235 | 0,455276965 | -0,11233987 | -0,09548889 | -0,111679938 | -0,202679938 | -1,462679938 |
| chrVII | 293723 | 293871 | 148 | 1,031540931 | 0,433586741 | 0,704062145 | 0,536119867 | 0,455701887 | 0,246833074 | 0,218833074 | -1,041166926 |
| chrVII | 294019 | 294183 | 164 | 0,068389454 | 0,163058261 | 0,295485544 | -0,537429332 | -0,456814932 | -0,380793194 | -0,520793194 | -1,780793194 |
| chrVII | 297177 | 297313 | 136 | 1,849364819 | 1,460112614 | 0,558808711 | 0,147949557 | 0,125757123 | 0,05325725 | 0,10925725 | -1,15074275 |
| chrVII | 299926 | 300073 | 147 | 0,265009134 | 0,151940653 | 0,635590046 | 0,346695744 | 0,294691382 | 0,110323283 | 0,089323283 | -1,170676717 |
| chrVII | 300091 | 300244 | 153 | 0,527168708 | 0,340940001 | 0,607261167 | 0,272187743 | 0,231359582 | 0,132294679 | 0,069294679 | -1,190705321 |
| chrVII | 300595 | 300755 | 160 | 0,076938136 | 0,096352609 | 0,443982948 | -0,140878539 | -0,119746758 | -0,079002598 | -0,191002598 | -1,451002598 |
| chrVII | 313487 | 313646 | 159 | 0,202318801 | 0,26311674 | 0,434687048 | -0,164453569 | -0,139785534 | -0,122474576 | -0,227474576 | -1,487474576 |
| chrVII | 314575 | 314719 | 144 | 0,142478029 | 0,144528914 | 0,496427117 | -0,008956009 | -0,007612608 | -0,152057745 | -0,152057745 | -1,412057745 |
| chrVII | 320068 | 320221 | 153 | 0,720938828 | 0,452116089 | 0,614582333 | 0,291282418 | 0,247590055 | 0,163719947 | 0,100719947 | -1,159280053 |
| chrVII | 320068 | 320221 | 153 | 0,720938828 | 0,452116089 | 0,614582333 | 0,291282418 | 0,247590055 | 0,163719947 | 0,100719947 | -1,159280053 |
| chrVII | 323178 | 323326 | 148 | 0,416035845 | 0,411351523 | 0,502830791 | 0,007095801 | 0,006031431 | -0,202382885 | -0,230382885 | -1,490382885 |
| chrVII | 324915 | 325051 | 136 | 0,284956058 | 0,359469349 | 0,442186256 | -0,145428565 | -0,123614281 | -0,207869503 | -0,151869503 | -1,411869503 |
| chrVII | 327547 | 327717 | 170 | 0,199469241 | 0,248293262 | 0,445480003 | -0,137089559 | -0,116526125 | 0,194020159 | 0,012020159 | -1,247979841 |
| chrVII | 327547 | 327717 | 170 | 0,199469241 | 0,248293262 | 0,445480003 | -0,137089559 | -0,116526125 | 0,194020159 | 0,012020159 | -1,247979841 |
| chrVII | 328882 | 329034 | 152 | 0,131079787 | 0,129705435 | 0,502635026 | 0,006605078 | 0,005614317 | -0,131105949 | -0,187105949 | -1,447105949 |
| chrVII | 329271 | 329388 | 117 | 0,028495606 | 0,400233914 | 0,066465229 | -1,502645598 | -1,277248758 | -0,707571749 | -0,518571749 | -1,778571749 |
| chrVII | 330372 | 330517 | 145 | 0,393239361 | 0,259410871 | 0,602526961 | 0,259893303 | 0,220909308 | 0,099839043 | 0,092839043 | -1,167160957 |
| chrVII | 331632 | 331780 | 148 | 1,048638295 | 0,678174133 | 0,607268212 | 0,272206067 | 0,231375157 | 0,012582381 | -0,015417619 | -1,275417619 |
| chrVII | 331632 | 331780 | 148 | 1,048638295 | 0,678174133 | 0,607268212 | 0,272206067 | 0,231375157 | 0,012582381 | -0,015417619 | -1,275417619 |
| chrVII | 341547 | 341699 | 152 | 0,208017923 | 0,237175653 | 0,46725275 | -0,082177582 | -0,069850944 | -0,212194936 | -0,268194936 | -1,528194936 |
| chrVII | 342190 | 342339 | 149 | 0,427434088 | 0,289057827 | 0,596565123 | 0,24446623 | 0,207796296 | 0,029450745 | -0,005549255 | -1,265549255 |
| chrVII | 342310 | 342453 | 143 | 2,111524392 | 1,208113483 | 0,636070702 | 0,347975482 | 0,29577916 | 0,142117775 | 0,149117775 | -1,110882225 |
| chrVII | 343265 | 343403 | 138 | 2,350887481 | 1,745464572 | 0,573897812 | 0,186306545 | 0,158360563 | 0,048613243 | 0,090613243 | -1,169386757 |
| chrVII | 352278 | 352440 | 162 | 0,29350474 | 0,363175219 | 0,446952486 | -0,133364686 | -0,113359983 | -0,080163356 | -0,206163356 | -1,466163356 |
| chrVII | 354451 | 354624 | 173 | 0,079787696 | 0,151940653 | 0,344315647 | -0,400713197 | -0,340606218 | 0,001447473 | -0,201552527 | -1,461552527 |
| chrVII | 356531 | 356676 | 145 | 0,04844253 | 0,037058696 | 0,566571175 | 0,16765125 | 0,142503563 | 0,004535019 | -0,002464981 | -1,262464981 |
| chrVII | 364639 | 364801 | 162 | 0,578640798 | 0,637409568 | 0,475758613 | -0,060801588 | -0,05168135 | -0,018407366 | -0,144407366 | -1,404407366 |
| chrVII | 367967 | 368120 | 153 | 0,133929347 | 0,062999783 | 0,680089062 | 0,46794786 | 0,397755681 | 0,311627336 | 0,248627336 | -1,011372664 |
| chrVII | 370607 | 370760 | 153 | 0,267858695 | 0,311293045 | 0,462501753 | -0,094133 | -0,08001305 | -0,168822377 | -0,231822377 | -1,491822377 |
| chrVII | 382677 | 382828 | 151 | 0,216566604 | 0,211234566 | 0,506231912 | 0,015621722 | 0,013278464 | -0,119412347 | -0,168412347 | -1,428412347 |
| chrVII | 382677 | 382828 | 151 | 0,216566604 | 0,211234566 | 0,506231912 | 0,015621722 | 0,013278464 | -0,119412347 | -0,168412347 | -1,428412347 |
| chrVII | 383386 | 383517 | 131 | 0,122531105 | 0,100058479 | 0,55047996 | 0,126874057 | 0,107842948 | 0,294857491 | 0,385857491 | -0,874142509 |
| chrVII | 383588 | 383739 | 151 | 0,185221438 | 0,051882174 | 0,781183536 | 0,776196573 | 0,659767087 | 0,51947176 | 0,47047176 | -0,78952824 |
| chrVII | 384835 | 384978 | 143 | 0,088336378 | 0,196411088 | 0,310227091 | -0,495206755 | -0,420925742 | -0,574958092 | -0,567958092 | -1,827958092 |
| chrVII | 384841 | 384978 | 137 | 0,076938136 | 0,107470218 | 0,417216109 | -0,209020522 | -0,177667444 | -0,290809551 | -0,241809551 | -1,501809551 |
| chrVII | 386231 | 386378 | 147 | 3,718676561 | 1,804758485 | 0,67325433 | 0,448917266 | 0,381579676 | 0,198113304 | 0,177113304 | -1,082886696 |
| chrVII | 386588 | 386752 | 164 | 0,387540239 | 0,589233263 | 0,39675548 | -0,26175419 | -0,222491061 | -0,134194436 | -0,274194436 | -1,534194436 |
| chrVII | 398118 | 398266 | 148 | 5,462607639 | 2,860931315 | 0,656284264 | 0,402343192 | 0,341991714 | 0,12232298 | 0,09432298 | -1,16567702 |
| chrVII | 398740 | 398893 | 153 | 0,24221265 | 0,188999349 | 0,561702018 | 0,155285859 | 0,13199298 | 0,040841734 | -0,022158266 | -1,282158266 |
| chrVII | 399113 | 399273 | 160 | 2,407878693 | 2,234639356 | 0,518657907 | 0,046785499 | 0,039767674 | 0,07442469 | -0,03757531 | -1,29757531 |
| chrVII | 402184 | 402330 | 146 | 0,088336378 | 0,081529131 | 0,520037168 | 0,050246866 | 0,042709836 | -0,128219053 | -0,142219053 | -1,402219053 |
| chrVII | 406230 | 406375 | 145 | 0,071239015 | 0,037058696 | 0,657807209 | 0,406485945 | 0,345513053 | 0,205399496 | 0,198399496 | -1,061600504 |
| chrVII | 412759 | 412909 | 150 | 0,843469933 | 0,741173916 | 0,53227729 | 0,080995639 | 0,068846293 | -0,071005527 | -0,113005527 | -1,373005527 |
| chrVII | 413027 | 413197 | 170 | 0,034194727 | 0,155646522 | 0,180122745 | -0,914897409 | -0,777662797 | -0,438692813 | -0,620692813 | -1,880692813 |
| chrVII | 416964 | 417114 | 150 | 7,659618848 | 6,900329155 | 0,526074602 | 0,065405938 | 0,055595047 | -0,083567264 | -0,125567264 | -1,385567264 |
| chrVII | 419105 | 419256 | 151 | 0,210867483 | 0,218646305 | 0,490944619 | -0,022700423 | -0,01929536 | -0,162173429 | -0,211173429 | -1,471173429 |
| chrVII | 422885 | 423015 | 130 | 0,481575739 | 0,748585655 | 0,391473624 | -0,275480315 | -0,234158267 | -0,000825096 | 0,097174904 | -1,162825096 |
| chrVII | 424432 | 424587 | 155 | 0,091185939 | 0,103764348 | 0,467739443 | -0,080953558 | -0,068810524 | -0,122672603 | -0,199672603 | -1,459672603 |
| chrVII | 426688 | 426828 | 140 | 0,547115632 | 0,44099848 | 0,55369681 | 0,13500695 | 0,114755908 | -0,009124711 | 0,018875289 | -1,241124711 |
| chrVII | 430629 | 430785 | 156 | 0,202318801 | 0,192705218 | 0,512168353 | 0,030506268 | 0,025930328 | -0,026031438 | -0,110031438 | -1,370031438 |
| chrVII | 438337 | 438504 | 167 | 3,1231184 | 2,98693088 | 0,511144552 | 0,027938883 | 0,02374805 | 0,195631873 | 0,034631873 | -1,225368127 |
| chrVII | 445398 | 445550 | 152 | 0,14532759 | 0,114881957 | 0,558502144 | 0,147172696 | 0,125096791 | -0,01424666 | -0,07024666 | -1,33024666 |
| chrVII | 445405 | 445562 | 157 | 0,082637257 | 0,048176305 | 0,631717813 | 0,336406466 | 0,285945496 | 0,284646424 | 0,193646424 | -1,066353576 |
| chrVII | 448295 | 448446 | 151 | 0,889062902 | 0,496586524 | 0,641621817 | 0,362797223 | 0,30837764 | 0,16622849 | 0,11722849 | -1,14277151 |
| chrVII | 449063 | 449214 | 151 | 7,762203029 | 6,633506546 | 0,53920253 | 0,098424855 | 0,083661127 | -0,063267789 | -0,112267789 | -1,372267789 |
| chrVII | 449866 | 450017 | 151 | 3,530605563 | 2,857225445 | 0,552708041 | 0,132506202 | 0,112630272 | -0,040950154 | -0,089950154 | -1,349950154 |
| chrVII | 450289 | 450475 | 186 | 0,059840772 | 0,059293913 | 0,502295129 | 0,005753068 | 0,004890107 | 0,435588056 | 0,141588056 | -1,118411944 |
| chrVII | 450564 | 450727 | 163 | 0,176672756 | 0,188999349 | 0,483145293 | -0,042261061 | -0,035921902 | 0,022609663 | -0,110390337 | -1,370390337 |
| chrVII | 451657 | 451811 | 154 | 1,362089959 | 0,952408482 | 0,588503295 | 0,223696577 | 0,19014209 | 0,133365889 | 0,063365889 | -1,196634111 |

TABLE 2

| Chrom | Start | End | Length | Section A | Section B | A/A+B | Z-score | Z * 0.85 | Phase correction | Length correction | ΔLK |
|---|---|---|---|---|---|---|---|---|---|---|---|
| chrVII | 452015 | 452160 | 145 | 0,131079787 | 0,140823044 | 0,4820832 | -0,044925865 | -0,038186985 | -0,191813534 | -0,198813534 | **-1,458813534** |
| chrVII | 454120 | 454220 | 100 | 0,074088575 | 0,637409568 | 0,104130385 | -1,258362274 | -1,069607933 | -0,611903427 | -0,303903427 | **-1,563903427** |
| chrVII | 457180 | 457330 | 150 | 0,119681545 | 0,144528914 | 0,452978074 | -0,118140734 | -0,100419624 | -0,237786381 | -0,279786381 | **-1,539786381** |
| chrVII | 457820 | 457954 | 134 | 0,367593315 | 0,374292827 | 0,495484811 | -0,011318142 | -0,009620421 | -0,007197554 | 0,062802446 | **-1,197197554** |
| chrVII | 458030 | 458181 | 151 | 0,966001038 | 0,574409785 | 0,627106109 | 0,324198469 | 0,275568699 | 0,120546827 | 0,071546827 | **-1,188453173** |
| chrVII | 464168 | 464326 | 158 | 0,116831984 | 0,17417587 | 0,401473645 | -0,249534587 | -0,212104399 | -0,205938962 | -0,303938962 | **-1,563938962** |
| chrVII | 468550 | 468701 | 151 | 0,410336724 | 0,296469566 | 0,58055047 | 0,203301913 | 0,172806626 | 0,011497835 | -0,037502165 | **-1,297502165** |
| chrVII | 470833 | 470993 | 160 | 0,122531105 | 0,111176087 | 0,524293257 | 0,060931847 | 0,05179207 | 0,082832195 | -0,029167805 | **-1,289167805** |
| chrVII | 471018 | 471094 | 76 | 0,119681545 | 0,040764565 | 0,745929861 | 0,661736237 | 0,562475801 | | 0,476 | **-0,784** |
| chrVII | 471018 | 471123 | 105 | 0,108283302 | 0,085235 | 0,559550703 | 0,149830185 | 0,127355657 | 0,646562227 | 0,919562227 | **-0,340437773** |
| chrVII | 471173 | 471322 | 149 | 0,239363089 | 0,125999566 | 0,655138356 | 0,399230614 | 0,339346022 | 0,155064634 | 0,120064634 | **-1,139935366** |
| chrVII | 475774 | 475931 | 157 | 0,188070999 | 0,200116957 | 0,484484373 | -0,038901718 | -0,03306646 | -0,022988807 | -0,113988807 | **-1,373988807** |
| chrVII | 476346 | 476454 | 108 | 0,088336378 | 0,581821524 | 0,131814275 | -1,117855882 | -0,950177499 | -0,297185286 | -0,045185286 | **-1,305185286** |
| chrVII | 476346 | 476465 | 119 | 0,071239015 | 0,741173916 | 0,087688184 | -1,35512929 | -1,151859897 | -0,535063938 | -0,360063938 | **-1,620063938** |
| chrVII | 478282 | 478437 | 155 | 0,475876617 | 0,185293479 | 0,719749154 | 0,582096486 | 0,494782013 | 0,439899592 | 0,362899592 | **-0,897100408** |
| chrVII | 482579 | 482701 | 122 | 0,122531105 | 0,400233914 | 0,234390406 | -0,724464178 | -0,615794551 | -0,135825791 | 0,018174209 | **-1,241825791** |
| chrVII | 483653 | 483754 | 101 | 0,056991212 | 0,107470218 | 0,346532387 | -0,394699362 | -0,335494458 | 0,135109524 | 0,436109524 | **-0,823890476** |
| chrVII | 495041 | 495196 | 155 | 0,156725832 | 0,081529131 | 0,657807209 | 0,406485945 | 0,345513053 | 0,278376586 | 0,201376586 | **-1,058623414** |
| chrVII | 501090 | 501258 | 168 | 0,421734966 | 0,781938481 | 0,350373241 | -0,38431299 | -0,326666042 | -0,071675317 | -0,239675317 | **-1,499675317** |
| chrVII | 502391 | 502522 | 131 | 0,837770812 | 1,259995657 | 0,399363239 | -0,254995625 | -0,216746281 | -0,040325931 | 0,050674069 | **-1,209325931** |
| chrVII | 502870 | 503027 | 157 | 0,430283648 | 0,489174784 | 0,467975096 | -0,08036094 | -0,068306799 | -0,055405847 | -0,146405847 | **-1,406405847** |
| chrVII | 504629 | 504778 | 149 | 1,079983461 | 0,88199696 | 0,550455779 | 0,126812955 | 0,107791012 | -0,066011428 | -0,101011428 | **-1,361011428** |
| chrVII | 505104 | 505255 | 151 | 0,407487163 | 0,377998697 | 0,518770845 | 0,047068905 | 0,040008569 | -0,115476987 | -0,164476987 | **-1,424476987** |
| chrVII | 509103 | 509251 | 148 | 0,376141997 | 0,229763914 | 0,620792751 | 0,307563498 | 0,261428973 | 0,043608396 | 0,015608396 | **-1,244391604** |
| chrVII | 509261 | 509410 | 149 | 0,116831984 | 0,129705435 | 0,473891486 | -0,065491126 | -0,055667457 | -0,228709292 | -0,263709292 | **-1,523709292** |
| chrVII | 510438 | 510590 | 152 | 0,119681545 | 0,111176087 | 0,518421434 | 0,046192108 | 0,039263292 | -0,110014425 | -0,166014425 | **-1,426014425** |
| chrVII | 512325 | 512500 | 175 | 0,83207169 | 1,293348483 | 0,39148574 | -0,275448768 | -0,234131453 | 0,193783244 | -0,023216756 | **-1,283216756** |
| chrVII | 515194 | 515319 | 125 | 0,541416511 | 1,026525873 | 0,34530383 | -0,398030558 | -0,338325975 | 0,09761383 | 0,23061383 | **-1,02938617** |
| chrVII | 516112 | 516262 | 150 | 0,581310359 | 0,596645002 | 0,493490991 | -0,016316389 | -0,013868931 | -0,157946635 | -0,199946635 | **-1,459946635** |
| chrVII | 516112 | 516262 | 150 | 0,581310359 | 0,596645002 | 0,493490991 | -0,016316389 | -0,013868931 | -0,157946635 | -0,199946635 | **-1,459946635** |
| chrVII | 517734 | 517882 | 148 | 10,99930385 | 7,004093504 | 0,610957123 | 0,281814497 | 0,239542323 | 0,022014422 | -0,005985578 | **-1,265985578** |
| chrVII | 517736 | 517882 | 146 | 0,028495606 | 0,051882174 | 0,354520937 | -0,373143192 | -0,317171714 | -0,499840711 | -0,513840711 | **-1,773840711** |
| chrVII | 518337 | 518484 | 147 | 2,895153553 | 2,312462617 | 0,555946033 | 0,140698749 | 0,119593937 | -0,06490264 | -0,08590264 | **-1,34590264** |
| chrVII | 519407 | 519563 | 156 | 0,364743755 | 0,296469566 | 0,551627959 | 0,129775456 | 0,110309137 | 0,068403533 | -0,015596467 | **-1,275596467** |
| chrVII | 519674 | 519834 | 160 | 3,596145456 | 3,231518273 | 0,526702193 | 0,066982525 | 0,056935146 | 0,087109296 | -0,024890704 | **-1,284890704** |
| chrVII | 519935 | 520102 | 167 | 0,524319147 | 0,600350872 | 0,466198208 | -0,08483016 | -0,072105636 | 0,09191371 | -0,06908629 | **-1,32908629** |
| chrVII | 520743 | 520902 | 159 | 0,133929347 | 0,111176087 | 0,54641525 | 0,116609512 | 0,099118085 | 0,112786991 | 0,007786991 | **-1,252213009** |
| chrVII | 520956 | 521107 | 151 | 0,116831984 | 0,192705218 | 0,377440848 | -0,312208996 | -0,265377647 | -0,413404401 | -0,462404401 | **-1,722404401** |
| chrVII | 526220 | 526384 | 164 | 0,119681545 | 0,255705001 | 0,318822147 | -0,470995015 | -0,400345763 | -0,308989585 | -0,448989585 | **-1,708989585** |
| chrVII | 529947 | 530056 | 109 | 0,170973635 | 0,17417587 | 0,49536109 | -0,011628286 | -0,009884043 | 0,665803302 | 0,910803302 | **-0,349196698** |
| chrVII | 535422 | 535536 | 114 | 0,085486818 | 0,570703915 | 0,130277392 | -1,125080828 | -0,956318704 | -0,339281652 | -0,129281652 | **-1,389281652** |
| chrVII | 536123 | 536272 | 149 | 0,062690333 | 0,040764565 | 0,605967759 | 0,268824834 | 0,228501109 | 0,055229877 | 0,020229877 | **-1,239770123** |
| chrVII | 536273 | 536379 | 106 | 0,840620372 | 2,164227834 | 0,279754688 | -0,583570406 | -0,496034845 | 0,101769946 | 0,367769946 | **-0,892230054** |
| chrVII | 536273 | 536391 | 118 | 0,837770812 | 2,479226748 | 0,252569017 | -0,666427276 | -0,566463185 | 0,014833424 | 0,196833424 | **-1,063166576** |
| chrVII | 536297 | 536420 | 123 | 0,076938136 | 0,155646522 | 0,330796263 | -0,437715508 | -0,372058182 | 0,124498511 | 0,271498511 | **-0,988501489** |
| chrVII | 536397 | 536552 | 155 | 0,125380666 | 0,08894087 | 0,585011979 | 0,214732295 | 0,182522451 | 0,110216743 | 0,033216743 | **-1,226783257** |
| chrVII | 536830 | 536964 | 134 | 0,159575393 | 0,155646522 | 0,506231912 | 0,015621722 | 0,013278464 | 0,014724359 | 0,084724359 | **-1,175275641** |
| chrVII | 536830 | 536970 | 140 | 0,136778908 | 0,100058479 | 0,577522451 | 0,195559327 | 0,166225428 | 0,052443065 | 0,080443065 | **-1,179556935** |
| chrVII | 537026 | 537177 | 151 | 0,151026711 | 0,085235 | 0,639234814 | 0,356414232 | 0,302952097 | 0,155744882 | 0,106744882 | **-1,153255118** |
| chrVII | 537036 | 537181 | 145 | 0,116831984 | 0,062999783 | 0,649673781 | 0,384439894 | 0,326773909 | 0,151391116 | 0,144391116 | **-1,115608884** |
| chrVII | 537308 | 537462 | 154 | 0,230814407 | 0,170470001 | 0,575189075 | 0,18960094 | 0,161160799 | 0,099386273 | 0,029386273 | **-1,230613727** |
| chrVII | 537342 | 537486 | 144 | 0,062690333 | 0,037058696 | 0,628480635 | 0,327831948 | 0,278657156 | 0,145485296 | 0,145485296 | **-1,114514704** |
| chrVII | 537873 | 538008 | 135 | 0,079787696 | 0,074117392 | 0,518421434 | 0,046192108 | 0,039263292 | -0,003992722 | 0,059007278 | **-1,200992722** |
| chrVII | 538055 | 538187 | 132 | 0,159575393 | 0,211234566 | 0,430342791 | -0,175501512 | -0,149176285 | -0,063847503 | 0,020152497 | **-1,239847503** |
| chrVII | 538144 | 538282 | 138 | 0,561363435 | 0,52623348 | 0,516150264 | 0,040493772 | 0,034419707 | -0,05356563 | -0,01156563 | **-1,27156563** |
| chrVII | 538499 | 538656 | 157 | 0,045592969 | 0,059293913 | 0,434687048 | -0,164453569 | -0,139785534 | -0,114812911 | -0,205812911 | **-1,465812911** |
| chrVII | 538599 | 538743 | 144 | 0,037044288 | 0,037058696 | 0,499902783 | -0,000243688 | -0,000207135 | -0,13422548 | -0,13422548 | **-1,39422548** |
| chrVII | 538990 | 539119 | 129 | 0,045592969 | 0,100058479 | 0,313027917 | -0,487285766 | -0,414192901 | -0,078675531 | 0,026324469 | **-1,233675531** |
| chrVII | 539470 | 539601 | 131 | 0,065539893 | 0,155646522 | 0,296310663 | -0,535041262 | -0,454785073 | -0,285414983 | -0,194414983 | **-1,454414983** |
| chrVII | 539902 | 540052 | 150 | 0,125380666 | 0,192705218 | 0,394172367 | -0,268460684 | -0,228191582 | -0,369066088 | -0,411066088 | **-1,671066088** |
| chrVII | 540853 | 541004 | 151 | 0,085486818 | 0,059293913 | 0,590457149 | 0,22872109 | 0,194412926 | 0,049038511 | 3,8511E-05 | **-1,259961489** |
| chrVII | 542644 | 542795 | 151 | 0,689593661 | 0,541056958 | 0,560348851 | 0,151853737 | 0,129075677 | -0,007033793 | -0,056033793 | **-1,316033793** |
| chrVII | 544012 | 544140 | 128 | 0,304902982 | 0,492880654 | 0,382187561 | -0,299740475 | -0,254779404 | 0,132788472 | 0,244788472 | **-1,015211528** |
| chrVII | 545414 | 545580 | 166 | 0,128230226 | 0,066705652 | 0,657807209 | 0,406485945 | 0,345513053 | 0,467670817 | 0,313670817 | **-0,946329183** |
| chrVII | 546693 | 546840 | 147 | 0,378991558 | 0,077823261 | 0,829639368 | 0,9527411 | 0,809829935 | 0,622368144 | 0,601368144 | **-0,658631856** |
| chrVII | 548258 | 548409 | 151 | 2,481967268 | 1,637994354 | 0,602424851 | 0,259628569 | 0,220684284 | 0,079659318 | 0,030659318 | **-1,229340682** |
| chrVII | 554460 | 554622 | 162 | 0,142478029 | 0,137117174 | 0,509586815 | 0,024032894 | 0,02042796 | 0,060820549 | -0,065179451 | **-1,325179451** |
| chrVII | 554946 | 555089 | 143 | 0,3903898 | 0,43729261 | 0,471666179 | -0,07108217 | -0,060419845 | -0,219131007 | -0,212131007 | **-1,472131007** |
| chrVII | 562268 | 562415 | 147 | 0,062690333 | 0,044470435 | 0,585011979 | 0,214732295 | 0,182522451 | -0,007448153 | -0,028448153 | **-1,288448153** |
| chrVII | 562775 | 562889 | 114 | 0,051292091 | 0,377998697 | 0,119480995 | -1,177585257 | -1,000947468 | -0,375104861 | -0,165104861 | **-1,425104861** |
| chrVII | 563471 | 563621 | 150 | 0,515770466 | 0,407645654 | 0,558546093 | 0,147284059 | 0,12519145 | -0,014115291 | -0,056115291 | **-1,316115291** |
| chrVII | 564404 | 564533 | 129 | 0,054141651 | 0,066705652 | 0,448017039 | -0,130672891 | -0,111071958 | 0,211288898 | 0,316288898 | **-0,943711102** |
| chrVII | 564867 | 565024 | 157 | 0,065539893 | 0,066705652 | 0,495592445 | -0,011048326 | -0,009391077 | 0,00943547 | -0,08156453 | **-1,34156453** |
| chrVII | 565335 | 565496 | 161 | 0,071239015 | 0,070411522 | 0,502920895 | 0,007321663 | 0,006223413 | 0,022145412 | -0,096854588 | **-1,356854588** |
| chrVII | 565515 | 565650 | 135 | 0,085486818 | 0,066705652 | 0,561702018 | 0,155285859 | 0,13199298 | 0,095212873 | 0,158212873 | **-1,101787127** |
| chrVII | 566553 | 566693 | 140 | 0,165274514 | 0,129705435 | 0,560290672 | 0,151706213 | 0,128950281 | 0,022086642 | 0,050086642 | **-1,209913358** |
| chrVII | 566559 | 566693 | 134 | 0,14817715 | 0,074117392 | 0,666580246 | 0,43048963 | 0,365916185 | 0,364031075 | 0,434031075 | **-0,825968925** |
| chrVII | 566971 | 567126 | 155 | 0,113982423 | 0,092646739 | 0,551627959 | 0,129775456 | 0,110309137 | 0,029917646 | -0,047082354 | **-1,307082354** |
| chrVII | 567102 | 567210 | 108 | 0,062690333 | 0,192705218 | 0,245463684 | -0,688834586 | -0,585509398 | 0,082988296 | 0,334988296 | **-0,925011704** |
| chrVII | 568106 | 568272 | 166 | 0,14532759 | 0,151940653 | 0,488876943 | -0,027884981 | -0,023702234 | 0,096483813 | -0,057516187 | **-1,317516187** |
| chrVII | 568932 | 569081 | 149 | 0,094035499 | 0,066705652 | 0,585011979 | 0,214732295 | 0,182522451 | 0,018534006 | -0,016465994 | **-1,276465994** |
| chrVII | 569719 | 569885 | 166 | 0,031345166 | 0,107470218 | 0,225804702 | -0,752734617 | -0,639824425 | -0,509964984 | -0,663964984 | **-1,923964984** |
| chrVII | 571017 | 571160 | 143 | 0,185221438 | 0,103764348 | 0,640936153 | 0,360962213 | 0,306817881 | 0,149921887 | 0,156921887 | **-1,103078113** |
| chrVII | 572385 | 572549 | 164 | 0,527168708 | 0,496586524 | 0,514936277 | 0,037448446 | 0,03183118 | 0,118336157 | -0,021663843 | **-1,281663843** |
| chrVII | 572497 | 572626 | 129 | 0,19377012 | 0,366881088 | 0,345616164 | -0,397183258 | -0,337605769 | -0,019394484 | 0,085605516 | **-1,174394484** |
| chrVII | 573872 | 573982 | 110 | 0,116831984 | 0,818997177 | 0,124843282 | -1,151111022 | -0,978444369 | -0,320307143 | -0,082307143 | **-1,342307143** |
| chrVII | 575134 | 575270 | 136 | 0,461628815 | 0,463233697 | 0,499132367 | -0,002174836 | -0,00184861 | -0,094923694 | -0,038923694 | **-1,298923694** |
| chrVII | 575584 | 575763 | 179 | 0,094035499 | 0,192705218 | 0,327946098 | -0,445591717 | -0,378752959 | 0,063727262 | -0,181272738 | **-1,441272738** |
| chrVII | 575584 | 575763 | 179 | 0,094035499 | 0,192705218 | 0,327946098 | -0,445591717 | -0,378752959 | 0,063727262 | -0,181272738 | **-1,441272738** |
| chrVII | 578432 | 578606 | 174 | 0,039893848 | 0,100058479 | 0,285053125 | -0,567895024 | -0,48271077 | -0,100721487 | -0,310721487 | **-1,570721487** |
| chrVII | 578432 | 578606 | 174 | 0,039893848 | 0,100058479 | 0,285053125 | -0,567895024 | -0,48271077 | -0,100721487 | -0,310721487 | **-1,570721487** |
| chrVII | 578979 | 579129 | 150 | 1,524514912 | 0,989467177 | 0,606414389 | 0,269985742 | 0,229487881 | 0,079015555 | 0,037015555 | **-1,222984445** |
| chrVII | 579144 | 579294 | 150 | 0,216566604 | 0,222352175 | 0,493409293 | -0,016521205 | -0,014043024 | -0,158838059 | -0,200838059 | **-1,460838059** |
| chrVII | 583828 | 583997 | 169 | 0,102584181 | 0,092646739 | 0,525450481 | 0,063838228 | 0,054262494 | 0,345712178 | 0,170712178 | **-1,089287822** |
| chrVII | 584107 | 584258 | 151 | 0,256460453 | 0,181587609 | 0,585461904 | 0,215886537 | 0,183503557 | 0,037173223 | -0,011826777 | **-1,271826777** |
| chrVII | 584200 | 584336 | 136 | 0,125380666 | 0,037058696 | 0,771861355 | 0,744990787 | 0,633242169 | 0,529479713 | 0,585479713 | **-0,674520287** |
| chrVII | 584954 | 585099 | 145 | 0,139628469 | 0,155646522 | 0,472876041 | -0,06804215 | -0,057835827 | -0,230111235 | -0,237111235 | **-1,497111235** |
| chrVII | 594298 | 594445 | 147 | 0,903310705 | 0,403939784 | 0,691000472 | 0,498688204 | 0,423884973 | 0,244178703 | 0,223178703 | **-1,036821297** |
| chrVII | 595048 | 595213 | 165 | 0,028495606 | 0,055588044 | 0,338895921 | -0,415478239 | -0,353156503 | -0,241510807 | -0,388510807 | **-1,648510807** |
| chrVII | 595048 | 595213 | 165 | 0,028495606 | 0,055588044 | 0,338895921 | -0,415478239 | -0,353156503 | -0,241510807 | -0,388510807 | **-1,648510807** |
| chrVII | 595281 | 595432 | 151 | 0,230814407 | 0,155646522 | 0,597251597 | 0,246239543 | 0,209303612 | 0,059859581 | 0,010859581 | **-1,249140419** |

TABLE 2

| Chrom | Start | End | Length | Section A | Section B | A/A+B | Z-score | Z * 0.85 | Phase correction | Length correction | ΔLK |
|---|---|---|---|---|---|---|---|---|---|---|---|
| chrVII | 610980 | 611160 | 180 | 0,324849907 | 0,452116089 | 0,418100546 | -0,206755141 | -0,17574187 | 0,257034573 | 0,005034573 | -1,254965427 |
| chrVII | 610980 | 611160 | 180 | 0,324849907 | 0,452116089 | 0,418100546 | -0,206755141 | -0,17574187 | 0,257034573 | 0,005034573 | -1,254965427 |
| chrVII | 613266 | 613414 | 148 | 3,804163379 | 2,686755444 | 0,586074712 | 0,217459102 | 0,184840237 | -0,038462944 | -0,066462944 | -1,326462944 |
| chrVII | 637713 | 637869 | 156 | 0,213717044 | 0,185293479 | 0,535617563 | 0,08939893 | 0,07598909 | 0,046151127 | -0,037848873 | -1,297848873 |
| chrVII | 640000 | 640148 | 148 | 12,27305743 | 8,686558292 | 0,585557369 | 0,216131479 | 0,183711757 | -0,039519137 | -0,067519137 | -1,327519137 |
| chrVII | 640002 | 640148 | 146 | 0,054141651 | 0,085235 | 0,388455674 | -0,283346352 | -0,2408444 | -0,41590033 | -0,42990033 | -1,68990033 |
| chrVII | 640719 | 640865 | 146 | 0,735186631 | 1,200701743 | 0,379767057 | -0,306092637 | -0,260178741 | -0,439622115 | -0,453622115 | -1,713622115 |
| chrVII | 652407 | 652570 | 163 | 0,076938136 | 0,114881957 | 0,40109529 | -0,250513093 | -0,212936129 | -0,145933903 | -0,278933903 | -1,538933903 |
| chrVII | 652780 | 652925 | 145 | 2,288197149 | 1,245172178 | 0,64759637 | 0,37883922 | 0,322013337 | 0,150137435 | 0,143137435 | -1,116862565 |
| chrVII | 654255 | 654401 | 146 | 0,63260245 | 0,433586741 | 0,593330391 | 0,236120435 | 0,20070237 | 0,015280533 | 0,001280533 | -1,258719467 |
| chrVII | 654255 | 654401 | 146 | 0,63260245 | 0,433586741 | 0,593330391 | 0,236120435 | 0,20070237 | 0,015280533 | 0,001280533 | -1,258719467 |
| chrVII | 668831 | 668991 | 160 | 0,202318801 | 0,26311674 | 0,434687048 | -0,164453569 | -0,139785534 | -0,113326114 | -0,225326114 | -1,485326114 |
| chrVII | 671142 | 671294 | 152 | 0,04844253 | 0,055588044 | 0,465656665 | -0,08619258 | -0,073263693 | -0,21970138 | -0,27570138 | -1,53570138 |
| chrVII | 671676 | 671849 | 173 | 0,14532759 | 0,137117174 | 0,51453455 | 0,036440779 | 0,030974662 | 0,399705709 | 0,196705709 | -1,063294291 |
| chrVII | 674557 | 674713 | 156 | 1,065735658 | 1,033937612 | 0,507572141 | 0,018981684 | 0,016134431 | -0,010891259 | -0,094891259 | -1,354891259 |
| chrVII | 676654 | 676817 | 163 | 0,24506221 | 0,144528914 | 0,629024111 | 0,329269787 | 0,279879319 | 0,342444717 | 0,209444717 | -1,050555283 |
| chrVII | 680899 | 681051 | 152 | 0,803576085 | 0,444704349 | 0,643746439 | 0,368491045 | 0,313217388 | 0,179196486 | 0,123196486 | -1,136803514 |
| chrVII | 681312 | 681478 | 166 | 0,039893848 | 0,074117392 | 0,349911537 | -0,385559308 | -0,327725412 | -0,185713766 | -0,339713766 | -1,599713766 |
| chrVII | 682534 | 682696 | 162 | 0,43883233 | 0,511410002 | 0,461810967 | -0,095872377 | -0,08149152 | -0,048187556 | -0,174187556 | -1,434187556 |
| chrVII | 687608 | 687758 | 150 | 0,14532759 | 0,133411305 | 0,521375354 | 0,053605728 | 0,045564869 | -0,098455956 | -0,140455956 | -1,400455956 |
| chrVII | 689536 | 689689 | 153 | 0,074088575 | 0,051882174 | 0,588141101 | 0,222765786 | 0,189350918 | 0,103047355 | 0,040047355 | -1,219952645 |
| chrVII | 692091 | 692197 | 106 | 0,102584181 | 0,807879568 | 0,11267245 | -1,212437641 | -1,030571995 | -0,417156342 | -0,151156342 | -1,411156342 |
| chrVII | 692091 | 692218 | 127 | 0,039893848 | 0,055588044 | 0,417815854 | -0,20748423 | -0,176361595 | 0,244765772 | 0,363765772 | -0,896234228 |
| chrVII | 692091 | 692219 | 128 | 0,051292091 | 0,08894087 | 0,365763444 | -0,34309514 | -0,291630869 | 0,072034507 | 0,184034507 | -1,075965493 |
| chrVII | 692091 | 692220 | 129 | 0,031345166 | 0,059293913 | 0,345823971 | -0,396619676 | -0,337126724 | -0,036786782 | 0,068213218 | -1,191786782 |
| chrVII | 692091 | 692221 | 130 | 0,028495606 | 0,085235 | 0,250553539 | -0,67274886 | -0,571836531 | -0,371516088 | -0,273516088 | -1,533516088 |
| chrVII | 692091 | 692222 | 131 | 0,056991212 | 0,111176087 | 0,338895921 | -0,415478239 | -0,353156503 | -0,189763356 | -0,098763356 | -1,358763356 |
| chrVII | 692091 | 692223 | 132 | 25,52351415 | 23,00603834 | 0,525937554 | 0,065061679 | 0,055302427 | 0,124318559 | 0,208318559 | -1,051681441 |
| chrVII | 692092 | 692223 | 131 | 0,039893848 | 0,051882174 | 0,434687048 | -0,164453569 | -0,139785534 | 0,007618017 | 0,098618017 | -1,161381983 |
| chrVII | 692095 | 692223 | 128 | 0,04844253 | 0,074117392 | 0,395255882 | -0,265646117 | -0,2257992 | 0,13385168 | 0,24585168 | -1,01414832 |
| chrVII | 692097 | 692197 | 100 | 0,04844253 | 0,059293913 | 0,449639217 | -0,126572917 | -0,10758698 | 0,346591594 | 0,654591594 | -0,605408406 |
| chrVII | 693772 | 693921 | 149 | 0,270708255 | 0,085235 | 0,760537673 | 0,708033178 | 0,601828201 | 0,44326871 | 0,40826871 | -0,85173129 |
| chrVII | 697506 | 697652 | 146 | 0,24221265 | 0,170470001 | 0,586922298 | 0,219635036 | 0,18668978 | -0,003158175 | -0,017158175 | -1,277158175 |
| chrVII | 697827 | 697977 | 150 | 0,752283994 | 0,500292393 | 0,600589315 | 0,254872767 | 0,216641852 | 0,073224274 | 0,031224274 | -1,228775726 |
| chrVII | 701841 | 701992 | 151 | 0,068389454 | 0,037058696 | 0,648560019 | 0,381435717 | 0,324220359 | 0,169255456 | 0,120255456 | -1,139744544 |
| chrVII | 707416 | 707536 | 120 | 0,168124074 | 0,641115437 | 0,207755642 | -0,814233348 | -0,692098346 | -0,104073719 | 0,063926281 | -1,196073719 |
| chrVII | 709744 | 709890 | 146 | 0,202318801 | 0,159352392 | 0,559399823 | 0,149447726 | 0,127030567 | -0,062973352 | -0,076973352 | -1,336973352 |
| chrVII | 712418 | 712569 | 151 | 0,156725832 | 0,114881957 | 0,577029962 | 0,194301161 | 0,165155987 | 0,003351975 | -0,045648025 | -1,305648025 |
| chrVII | 714004 | 714140 | 136 | 0,082637257 | 0,062999783 | 0,56741923 | 0,169807485 | 0,144336362 | 0,03758359 | 0,09358359 | -1,16641641 |
| chrVII | 716031 | 716189 | 158 | 0,04844253 | 0,08894087 | 0,352608321 | -0,378288023 | -0,32154482 | -0,320539305 | -0,418539305 | -1,678539305 |
| chrVII | 716536 | 716689 | 153 | 0,173823196 | 0,307587175 | 0,361070813 | -0,355598233 | -0,302258498 | -0,399950029 | -0,462950029 | -1,722950029 |
| chrVII | 726352 | 726496 | 144 | 0,270708255 | 0,166764131 | 0,618800783 | 0,302332726 | 0,256982817 | 0,124009755 | 0,124009755 | -1,135990245 |
| chrVII | 731531 | 731680 | 149 | 0,723788388 | 0,533645219 | 0,575607638 | 0,190669259 | 0,16206887 | -0,002052363 | -0,037052363 | -1,297052363 |
| chrVII | 740022 | 740127 | 105 | 4,798660023 | 35,6801123 | 0,118547568 | -1,182278695 | -1,00493689 | -0,477218768 | -0,204218768 | -1,464218768 |
| chrVII | 746153 | 746303 | 150 | 0,188070999 | 0,244587392 | 0,434687048 | -0,164453569 | -0,139785534 | -0,287410555 | -0,329410555 | -1,589410555 |
| chrVII | 747978 | 748102 | 124 | 0,675345858 | 2,116051529 | 0,241938271 | -0,700081285 | -0,595069093 | -0,140440085 | -0,000440085 | -1,260440085 |
| chrVII | 748361 | 748501 | 140 | 0,319150785 | 0,148234783 | 0,682842618 | 0,475662609 | 0,404313218 | 0,285265856 | 0,313265856 | -0,946734144 |
| chrVII | 751512 | 751624 | 112 | 2,846711023 | 15,45718201 | 0,155524894 | -1,013021717 | -0,86106846 | -0,234813929 | -0,010813929 | -1,270813929 |
| chrVII | 758738 | 758894 | 156 | 0,738036191 | 0,389116306 | 0,654779361 | 0,398256287 | 0,338517844 | 0,302455459 | 0,218455459 | -1,041544541 |
| chrVII | 760806 | 760962 | 156 | 0,427434088 | 0,340940001 | 0,556283839 | 0,141553976 | 0,120320879 | 0,086265624 | 0,002265624 | -1,257734376 |
| chrVII | 760828 | 760978 | 150 | 1,963347242 | 1,252583918 | 0,610506614 | 0,280639687 | 0,238543734 | 0,089742023 | 0,047742023 | -1,212257977 |
| chrVII | 762415 | 762565 | 150 | 0,205168362 | 0,133411305 | 0,605967759 | 0,268824834 | 0,228501109 | 0,079942294 | 0,037942294 | -1,222057706 |
| chrVII | 766213 | 766368 | 155 | 1,584355684 | 0,796761959 | 0,665383203 | 0,427200092 | 0,363120078 | 0,278020661 | 0,201020661 | -1,058979339 |
| chrVII | 769383 | 769497 | 114 | 0,065539893 | 0,211234566 | 0,236798921 | -0,71663743 | -0,609141816 | 0,000369645 | 0,210369645 | -1,049630355 |
| chrVII | 773391 | 773557 | 166 | 0,051292091 | 0,166764131 | 0,235224155 | -0,721749813 | -0,613487341 | -0,453772188 | -0,607772188 | -1,867772188 |
| chrVII | 776840 | 777001 | 161 | 0,176672756 | 0,129705435 | 0,576649256 | 0,193328781 | 0,164329464 | 0,174341115 | 0,055341115 | -1,204658885 |
| chrVII | 776982 | 777149 | 167 | 0,567062556 | 0,681880002 | 0,454034136 | -0,115475457 | -0,098154138 | 0,065349084 | -0,095650916 | -1,355650916 |
| chrVII | 780856 | 781004 | 148 | 1,048638295 | 0,433586741 | 0,707475768 | 0,546025419 | 0,464121606 | 0,242424011 | 0,214424011 | -1,045575989 |
| chrVII | 781219 | 781380 | 161 | 1,162620718 | 1,041349352 | 0,527512027 | 0,069017177 | 0,058664601 | 0,078215792 | -0,040784208 | -1,300784208 |
| chrVII | 782373 | 782522 | 149 | 0,113982423 | 0,085235 | 0,572150875 | 0,181852797 | 0,154574878 | -0,007708936 | -0,042708936 | -1,302708936 |
| chrVII | 782985 | 783136 | 151 | 0,202318801 | 0,207528696 | 0,493644105 | -0,015932539 | -0,013542658 | -0,184305693 | -0,233305693 | -1,493305693 |
| chrVII | 811114 | 811251 | 137 | 0,04844253 | 0,08894087 | 0,352608321 | -0,378288023 | -0,32154482 | -0,437645565 | -0,388645565 | -1,648645565 |
| chrVII | 813520 | 813649 | 129 | 0,188070999 | 0,396528045 | 0,321709385 | -0,462924104 | -0,393485489 | -0,113882154 | -0,008882154 | -1,268882154 |
| chrVII | 813597 | 813761 | 164 | 0,504372223 | 0,492880654 | 0,505761613 | 0,014442723 | 0,012276315 | 0,099684987 | -0,040315013 | -1,300315013 |
| chrVII | 814985 | 815128 | 143 | 0,239363089 | 0,103764348 | 0,69759239 | 0,517488464 | 0,439865194 | 0,273929894 | 0,280929894 | -0,979070106 |
| chrVII | 816260 | 816426 | 166 | 0,042743409 | 0,085235 | 0,333989218 | -0,428924135 | -0,364585514 | -0,194770339 | -0,348770339 | -1,608770339 |
| chrVII | 817064 | 817213 | 149 | 0,085486818 | 0,085235 | 0,500737507 | 0,001848658 | 0,001571359 | -0,154977044 | -0,189977044 | -1,449977044 |
| chrVII | 817068 | 817218 | 150 | 0,059840772 | 0,044470435 | 0,573675388 | 0,185739279 | 0,157878387 | 0,023365063 | -0,018634937 | -1,278634937 |
| chrVII | 817863 | 817968 | 105 | 0,028495606 | 0,485468915 | 0,055442749 | -1,594225676 | -1,355091824 | -0,804400316 | -0,531400316 | -1,791400316 |
| chrVII | 817863 | 817970 | 107 | 0,028495606 | 0,311293045 | 0,083862736 | -1,379549252 | -1,172616864 | -0,524364634 | -0,265364634 | -1,525364634 |
| chrVII | 819019 | 819169 | 150 | 0,53571739 | 0,433586741 | 0,552682458 | 0,13244151 | 0,112575284 | -0,011251878 | -0,053251878 | -1,313251878 |
| chrVII | 819762 | 819909 | 147 | 0,19661968 | 0,163058261 | 0,546654819 | 0,117214141 | 0,09963202 | -0,085069831 | -0,106069831 | -1,366069831 |
| chrVII | 820415 | 820572 | 157 | 0,054141651 | 0,081529131 | 0,399066404 | -0,255764342 | -0,217399691 | -0,206311394 | -0,297311394 | -1,557311394 |
| chrVII | 820601 | 820751 | 150 | 1,091381703 | 0,622586089 | 0,636757416 | 0,349804839 | 0,297334113 | 0,169797521 | 0,127797521 | -1,132202479 |
| chrVII | 820792 | 820937 | 145 | 0,113982423 | 0,051882174 | 0,68720164 | 0,487933817 | 0,414743744 | 0,247060269 | 0,240060269 | -1,019939731 |
| chrVII | 821063 | 821198 | 135 | 0,113982423 | 0,077823261 | 0,594259882 | 0,238516864 | 0,202739335 | 0,155586555 | 0,218586555 | -1,041413445 |
| chrVII | 821345 | 821513 | 168 | 0,042743409 | 0,048176305 | 0,470122564 | -0,074961771 | -0,063717505 | 0,207632583 | 0,039632583 | -1,220367417 |
| chrVII | 822101 | 822241 | 140 | 0,153876272 | 0,107470218 | 0,588782623 | 0,224414545 | 0,190752364 | 0,074049724 | 0,102049724 | -1,157950276 |
| chrVII | 822107 | 822241 | 134 | 0,125380666 | 0,103764348 | 0,547167331 | 0,118507771 | 0,100731606 | 0,08663241 | 0,15663241 | -1,10336759 |
| chrVII | 822519 | 822674 | 155 | 0,085486818 | 0,125999566 | 0,404219015 | -0,242441566 | -0,206075331 | -0,284234488 | -0,361234488 | -1,621234488 |
| chrVII | 822650 | 822758 | 108 | 0,062690333 | 0,163058261 | 0,277699771 | -0,589688448 | -0,501235181 | 0,178474159 | 0,430474159 | -0,829525841 |
| chrVII | 825754 | 825897 | 143 | 0,601257283 | 0,363175219 | 0,62343117 | 0,314504815 | 0,267329093 | 0,115715303 | 0,122715303 | -1,137284697 |
| chrVII | 831865 | 832015 | 150 | 0,29065518 | 0,233469783 | 0,554553208 | 0,137173592 | 0,116597553 | -0,015760078 | -0,057760078 | -1,317760078 |
| chrVII | 841785 | 841939 | 154 | 0,151026711 | 0,140823044 | 0,517481027 | 0,043832468 | 0,037257598 | -0,032164835 | -0,102164835 | -1,362164835 |
| chrVII | 842909 | 843085 | 176 | 0,170973635 | 0,307587175 | 0,357266269 | -0,365775588 | -0,310909249 | 0,11531991 | -0,10868009 | -1,36868009 |
| chrVII | 842909 | 843085 | 176 | 0,170973635 | 0,307587175 | 0,357266269 | -0,365775588 | -0,310909249 | 0,11531991 | -0,10868009 | -1,36868009 |
| chrVII | 843924 | 844090 | 166 | 0,091185939 | 0,059293913 | 0,605967759 | 0,268824834 | 0,228501109 | 0,400989568 | 0,246989568 | -1,013010432 |
| chrVII | 844983 | 845124 | 141 | 0,370442876 | 0,166764131 | 0,689571936 | 0,494637357 | 0,420441753 | 0,30966266 | 0,33066266 | -0,92933734 |
| chrVII | 847489 | 847638 | 149 | 0,170973635 | 0,085235 | 0,667321907 | 0,432530091 | 0,367650578 | 0,210204546 | 0,175204546 | -1,084795454 |
| chrVII | 853632 | 853775 | 143 | 0,504372223 | 0,251999131 | 0,666831471 | 0,431180602 | 0,366503512 | 0,198755499 | 0,205755499 | -1,054244501 |
| chrVII | 855743 | 855883 | 140 | 1,359240398 | 0,993173047 | 0,577806763 | 0,196285802 | 0,166842931 | 0,04724025 | 0,07524025 | -1,18475975 |
| chrVII | 855860 | 856013 | 153 | 0,113982423 | 0,122293696 | 0,482411949 | -0,044100996 | -0,037485847 | -0,145646796 | -0,208646796 | -1,468646796 |
| chrVII | 861234 | 861372 | 138 | 0,874815099 | 0,933879134 | 0,483672189 | -0,040939185 | -0,034798307 | -0,134465861 | -0,092465861 | -1,352465861 |
| chrVII | 869483 | 869631 | 148 | 0,054141651 | 0,048176305 | 0,529151025 | 0,07313593 | 0,062165541 | -0,162254733 | -0,190254733 | -1,450254733 |
| chrVII | 873303 | 873436 | 133 | 0,410336724 | 0,670762394 | 0,379555137 | -0,306649365 | -0,260651961 | -0,225824646 | -0,148824646 | -1,408824646 |
| chrVII | 873927 | 874079 | 152 | 0,142478029 | 0,077823261 | 0,646741692 | 0,376538476 | 0,320057705 | 0,190200841 | 0,134200841 | -1,125799159 |
| chrVII | 878924 | 879054 | 130 | 1,250957096 | 1,767699789 | 0,414408508 | -0,216219036 | -0,18378618 | 0,024604983 | 0,122604983 | -1,137395017 |
| chrVII | 879023 | 879172 | 149 | 0,552814753 | 0,318704784 | 0,634311372 | 0,343294051 | 0,291799943 | 0,137427531 | 0,102427531 | -1,157572469 |
| chrVII | 879984 | 880088 | 104 | 0,364743755 | 2,28281566 | 0,137766032 | -1,09041117 | -0,926849494 | -0,41452278 | -0,13452278 | -1,39452278 |
| chrVII | 882915 | 883023 | 108 | 0,09688506 | 0,048176305 | 0,667890174 | 0,43409473 | 0,36898052 | 1,042251293 | 1,294251293 | 0,034251293 |
| chrVII | 882915 | 883031 | 116 | 0,088336378 | 0,040764565 | 0,6842427 | 0,479596127 | 0,407656708 | 0,993893655 | 1,189893655 | -0,070106345 |

TABLE 2

| Chrom | Start | End | Length | Section A | Section B | A/A+B | Z-score | Z * 0.85 | Phase correction | Length correction | ΔLK |
|---|---|---|---|---|---|---|---|---|---|---|---|
| chrVII | 887398 | 887557 | 159 | 0,544266071 | 0,61888022 | 0,467925725 | -0,080485095 | -0,068412331 | -0,04758926 | -0,15258926 | -1,41258926 |
| chrVII | 889980 | 890139 | 159 | 0,159575393 | 0,255705001 | 0,384259395 | -0,294312935 | -0,250165995 | -0,21106307 | -0,31606307 | -1,57606307 |
| chrVII | 889980 | 890139 | 159 | 0,159575393 | 0,255705001 | 0,384259395 | -0,294312935 | -0,250165995 | -0,21106307 | -0,31606307 | -1,57606307 |
| chrVII | 892462 | 892609 | 147 | 0,53571739 | 0,429880871 | 0,554803598 | 0,137807188 | 0,11713611 | -0,0694593 | -0,0904593 | -1,3504593 |
| chrVII | 895680 | 895828 | 148 | 0,085486818 | 0,100058479 | 0,460732874 | -0,09858756 | -0,083799426 | -0,289695109 | -0,317695109 | -1,577695109 |
| chrVII | 911520 | 911682 | 162 | 0,074088575 | 0,055588044 | 0,571333335 | 0,179769754 | 0,152804291 | 0,170656766 | 0,044656766 | -1,215343234 |
| chrVII | 914384 | 914518 | 134 | 0,304902982 | 0,129705435 | 0,701557931 | 0,528886572 | 0,449553586 | 0,428768128 | 0,498768128 | -0,761231872 |
| chrVII | 920349 | 920433 | 84 | 0,208017923 | 0,459527828 | 0,311615979 | -0,491275002 | -0,417583751 | -0,052818035 | 0,367181965 | -0,892818035 |
| chrVII | 922436 | 922538 | 102 | 0,267858695 | 2,960989793 | 0,082957963 | -1,385446676 | -1,177629675 | -0,704878864 | -0,410878864 | -1,670878864 |
| chrVII | 925873 | 926024 | 151 | 0,170973635 | 0,166764131 | 0,506231912 | 0,015621722 | 0,013278464 | -0,148316922 | -0,197316922 | -1,457316922 |
| chrVII | 928464 | 928607 | 143 | 0,085486818 | 0,107470218 | 0,443035505 | -0,14327752 | -0,121785892 | -0,291581278 | -0,284581278 | -1,544581278 |
| chrVII | 928560 | 928710 | 150 | 0,24506221 | 0,200116957 | 0,55047996 | 0,126874057 | 0,107842948 | -0,01855031 | -0,06055031 | -1,32055031 |
| chrVII | 929745 | 929899 | 154 | 0,076938136 | 0,144528914 | 0,34740218 | -0,39234359 | -0,333492051 | -0,399924862 | -0,469924862 | -1,729924862 |
| chrVII | 938659 | 938812 | 153 | 0,034194727 | 0,051882174 | 0,397257877 | -0,260451199 | -0,221383519 | -0,326277095 | -0,389277095 | -1,649277095 |
| chrVII | 943256 | 943409 | 153 | 2,231205937 | 1,504583049 | 0,597251597 | 0,246239543 | 0,209303612 | 0,10965375 | 0,04665375 | -1,21334625 |
| chrVII | 943702 | 943854 | 152 | 0,83207169 | 0,61146848 | 0,576410485 | 0,192719017 | 0,163811164 | 0,045005996 | -0,010994004 | -1,270994004 |
| chrVII | 948590 | 948746 | 156 | 0,105433742 | 0,148234783 | 0,415635884 | -0,213070763 | -0,181110149 | -0,211273123 | -0,295273123 | -1,555273123 |
| chrVII | 949733 | 949881 | 148 | 2,3423388 | 1,793640876 | 0,56633228 | 0,167043985 | 0,141987387 | -0,06073511 | -0,08873511 | -1,34873511 |
| chrVII | 960612 | 960754 | 142 | 1,134125112 | 0,629997828 | 0,642883263 | 0,366176369 | 0,311249914 | 0,173962263 | 0,187962263 | -1,072037737 |
| chrVII | 961680 | 961814 | 134 | 0,159575393 | 0,43729261 | 0,267354577 | -0,620833537 | -0,527708506 | -0,552766225 | -0,482766225 | -1,742766225 |
| chrVII | 961681 | 961788 | 107 | 0,028495606 | 0,185293479 | 0,133288404 | -1,11098035 | -0,944333298 | -0,28136431 | -0,02236431 | -1,28236431 |
| chrVII | 961681 | 961791 | 110 | 0,031345166 | 0,066705652 | 0,319682862 | -0,468585806 | -0,398297935 | 0,244361751 | 0,482361751 | -0,777638249 |
| chrVII | 961681 | 961794 | 113 | 0,031345166 | 0,062999783 | 0,332240005 | -0,433736208 | -0,368675777 | 0,2434063 | 0,4604063 | -0,7995937 |
| chrVII | 961681 | 961795 | 114 | 0,074088575 | 0,077823261 | 0,487707719 | -0,030817056 | -0,026194498 | 0,58259512 | 0,79259512 | -0,46740488 |
| chrVII | 961681 | 961796 | 115 | 0,031345166 | 0,066705652 | 0,319682862 | -0,468585806 | -0,398297935 | 0,196571125 | 0,399571125 | -0,860428875 |
| chrVII | 961681 | 961797 | 116 | 0,045592969 | 0,222352175 | 0,170157849 | -0,953541662 | -0,810510413 | -0,226380421 | -0,030380421 | -1,290380421 |
| chrVII | 961681 | 961798 | 117 | 0,045592969 | 0,066705652 | 0,405997586 | -0,23785292 | -0,202174982 | 0,370580049 | 0,559580049 | -0,700419951 |
| chrVII | 961681 | 961799 | 118 | 0,111132863 | 0,43729261 | 0,202639863 | -0,83222895 | -0,707394607 | -0,102469031 | 0,079530969 | -1,180469031 |
| chrVII | 961681 | 961800 | 119 | 0,071239015 | 0,059293913 | 0,545755127 | 0,114943701 | 0,097702146 | 0,699935031 | 0,874935031 | -0,385064969 |
| chrVII | 961681 | 961801 | 120 | 0,125380666 | 0,207528696 | 0,376621026 | -0,314367347 | -0,267212245 | 0,300247709 | 0,468247709 | -0,791752291 |
| chrVII | 961681 | 961802 | 121 | 0,045592969 | 0,240881523 | 0,159151934 | -0,997949458 | -0,84825704 | -0,323433316 | -0,162433316 | -1,422433316 |
| chrVII | 961681 | 961803 | 122 | 0,039893848 | 0,125999566 | 0,240478794 | -0,704763239 | -0,599048753 | -0,112893163 | 0,041106837 | -1,218893163 |
| chrVII | 961681 | 961804 | 123 | 0,085486818 | 0,337234132 | 0,202229905 | -0,833682698 | -0,708630293 | -0,229559562 | -0,082559562 | -1,342559562 |
| chrVII | 961681 | 961805 | 124 | 0,045592969 | 0,133411305 | 0,254703244 | -0,659762131 | -0,560797812 | -0,114138058 | 0,025861942 | -1,234138058 |
| chrVII | 961681 | 961806 | 125 | 0,111132863 | 1,515700658 | 0,068312376 | -1,488478487 | -1,265206714 | -0,848143821 | -0,715143821 | -1,975143821 |
| chrVII | 961681 | 961807 | 126 | 0,068389454 | 0,148234783 | 0,315705458 | -0,47974192 | -0,407780632 | 0,024180985 | 0,150180985 | -1,109819015 |
| chrVII | 961681 | 961808 | 127 | 0,04844253 | 0,044470435 | 0,521375354 | 0,053605728 | 0,045564869 | 0,456144939 | 0,575144939 | -0,684855061 |
| chrVII | 961681 | 961809 | 128 | 0,14817715 | 0,248293262 | 0,373740753 | -0,321961969 | -0,273667674 | 0,079897227 | 0,191897227 | -1,068102773 |
| chrVII | 961681 | 961810 | 129 | 0,094035499 | 0,196411088 | 0,323761764 | -0,457205247 | -0,38862446 | -0,1218176 | -0,0168176 | -1,2768176 |
| chrVII | 961681 | 961811 | 130 | 0,071239015 | 0,118587827 | 0,375284202 | -0,317889966 | -0,270206471 | -0,070362305 | 0,027637695 | -1,232362305 |
| chrVII | 961681 | 961812 | 131 | 0,267858695 | 0,274234349 | 0,49411941 | -0,014740987 | -0,012529839 | 0,132543111 | 0,223543111 | -1,036456889 |
| chrVII | 961681 | 961813 | 132 | 0,094035499 | 0,170470001 | 0,355514344 | -0,370474883 | -0,31490365 | -0,257047964 | -0,173047964 | -1,433047964 |
| chrVII | 961681 | 961814 | 133 | 96,44907707 | 78,60519963 | 0,5509667 | 0,128104088 | 0,108888475 | 0,12810508 | 0,20510508 | -1,05489492 |
| chrVII | 961682 | 961814 | 132 | 0,373292436 | 0,422469132 | 0,46910086 | -0,077530259 | -0,06590072 | -0,019670018 | 0,064329982 | -1,195670018 |
| chrVII | 961683 | 961814 | 131 | 0,085486818 | 0,125999566 | 0,404219015 | -0,242441566 | -0,206075331 | -0,082596042 | 0,008403958 | -1,251596042 |
| chrVII | 961684 | 961814 | 130 | 0,065539893 | 0,218646305 | 0,230623069 | -0,736796456 | -0,626276987 | -0,426790809 | -0,328790809 | -1,588790809 |
| chrVII | 961685 | 961814 | 129 | 0,128230226 | 0,185293479 | 0,408996909 | -0,230126057 | -0,195607148 | 0,058374438 | 0,163374438 | -1,096625562 |
| chrVII | 961686 | 961814 | 128 | 0,435982769 | 0,429880871 | 0,503523591 | 0,008832447 | 0,00750758 | 0,35450653 | 0,46650653 | -0,79349347 |
| chrVII | 961687 | 961814 | 127 | 0,062690333 | 0,096352609 | 0,394172367 | -0,268460684 | -0,228191582 | 0,191365107 | 0,310365107 | -0,949634893 |
| chrVII | 961694 | 961814 | 120 | 0,039893848 | 0,129705435 | 0,235224155 | -0,721749813 | -0,613487341 | -0,071899238 | 0,096100762 | -1,163899238 |
| chrVII | 961695 | 961814 | 119 | 0,045592969 | 0,081529131 | 0,358654941 | -0,362056402 | -0,307747942 | 0,304698856 | 0,479698856 | -0,780301144 |
| chrVII | 961696 | 961814 | 118 | 0,051292091 | 0,170470001 | 0,231293321 | -0,734594249 | -0,624405111 | -0,021381997 | 0,160618003 | -1,099381997 |
| chrVII | 961708 | 961806 | 98 | 0,091185939 | 0,218646305 | 0,294307453 | -0,540844298 | -0,459717653 | -0,011572954 | 0,310427046 | -0,949572954 |
| chrVII | 962925 | 963072 | 147 | 0,133929347 | 0,107470218 | 0,554803598 | 0,137807188 | 0,11713611 | -0,060990433 | -0,081990433 | -1,341990433 |
| chrVII | 966527 | 966637 | 110 | 1,31934655 | 9,679731339 | 0,119950651 | -1,175233524 | -0,998948496 | -0,337676012 | -0,099676012 | -1,359676012 |
| chrVII | 974170 | 974327 | 157 | 0,054141651 | 0,062999783 | 0,462190442 | -0,094916836 | -0,080679311 | -0,068815787 | -0,159815787 | -1,419815787 |
| chrVII | 975022 | 975129 | 107 | 2,809666735 | 9,312850251 | 0,231772555 | -0,733021838 | -0,623068562 | 0,03941828 | 0,29841828 | -0,96158172 |
| chrVII | 975022 | 975130 | 108 | 0,059840772 | 0,218646305 | 0,214878093 | -0,789608938 | -0,671167597 | -0,010475442 | 0,241524558 | -1,018475442 |
| chrVII | 975022 | 975131 | 109 | 0,045592969 | 0,281646088 | 0,139326185 | -1,083351646 | -0,920848899 | -0,237938788 | 0,007061212 | -1,252938788 |
| chrVII | 975022 | 975142 | 120 | 3,604694138 | 12,44431004 | 0,224605471 | -0,756731162 | -0,643221488 | -0,117455307 | 0,050544693 | -1,209455307 |
| chrVII | 979943 | 980105 | 162 | 1,020142689 | 0,733762177 | 0,581640834 | 0,206092917 | 0,175178979 | 0,193558439 | 0,067558439 | -1,192441561 |
| chrVII | 982482 | 982630 | 148 | 0,133929347 | 0,062999783 | 0,680089062 | 0,46794786 | 0,397755681 | 0,208140798 | 0,180140798 | -1,079859202 |
| chrVII | 987451 | 987604 | 153 | 0,039893848 | 0,055588044 | 0,417815854 | -0,20748423 | -0,176361595 | -0,28099418 | -0,34399418 | -1,60399418 |
| chrVII | 987451 | 987606 | 155 | 0,034194727 | 0,055588044 | 0,380860679 | -0,303221116 | -0,257737948 | -0,336811507 | -0,413811507 | -1,673811507 |
| chrVII | 987451 | 987608 | 157 | 0,039893848 | 0,074117392 | 0,349911537 | -0,385559308 | -0,327725412 | -0,323417946 | -0,414417946 | -1,674417946 |
| chrVII | 987451 | 987609 | 158 | 10,96225956 | 12,62589765 | 0,464735734 | -0,088509834 | -0,075233359 | -0,071625702 | -0,169625702 | -1,429625702 |
| chrVII | 988814 | 988967 | 153 | 0,051292091 | 0,037058696 | 0,58055047 | 0,203301913 | 0,172806626 | 0,08146539 | 0,01846539 | -1,24153461 |
| chrVII | 988881 | 989019 | 138 | 1,020142689 | 0,381704567 | 0,727713155 | 0,605911247 | 0,51502456 | 0,414284484 | 0,456284484 | -0,803715516 |
| chrVII | 989319 | 989475 | 156 | 0,051292091 | 0,037058696 | 0,58055047 | 0,203301913 | 0,172806626 | 0,146733935 | 0,062733935 | -1,197266065 |
| chrVII | 989319 | 989477 | 158 | 5,932785135 | 5,62921589 | 0,513127885 | 0,032912669 | 0,027975769 | 0,029374917 | -0,068625083 | -1,328625083 |
| chrVII | 991949 | 992103 | 154 | 0,997346204 | 0,681880002 | 0,593932015 | 0,237671396 | 0,202020687 | 0,140968505 | 0,070968505 | -1,189031495 |
| chrVII | 999326 | 999473 | 147 | 0,116831984 | 0,118587827 | 0,496270827 | -0,009347788 | -0,007945619 | -0,183530646 | -0,204530646 | -1,464530646 |
| chrVII | 1001929 | 1002078 | 149 | 0,373292436 | 0,203822827 | 0,646824751 | 0,376761979 | 0,320247683 | 0,167800202 | 0,132800202 | -1,127199798 |
| chrVII | 1012079 | 1012238 | 159 | 0,09973462 | 0,166764131 | 0,374240479 | -0,320642986 | -0,272546538 | -0,236341534 | -0,341341534 | -1,601341534 |
| chrVII | 1012603 | 1012726 | 123 | 0,347646391 | 1,815876094 | 0,160685361 | -0,991645016 | -0,842898264 | -0,387318607 | -0,240318607 | -1,500318607 |
| chrVII | 1016668 | 1016789 | 121 | 0,059840772 | 0,122293696 | 0,328552705 | -0,443913104 | -0,377326138 | 0,129959103 | 0,290959103 | -0,969040897 |
| chrVII | 1018529 | 1018686 | 157 | 0,170973635 | 0,103764348 | 0,622315244 | 0,311567137 | 0,264832067 | 0,259547725 | 0,168547725 | -1,091452275 |
| chrVII | 1018639 | 1018770 | 131 | 0,142478029 | 0,140823044 | 0,502920895 | 0,007321663 | 0,006223413 | 0,140118615 | 0,231118615 | -1,028881385 |
| chrVII | 1024687 | 1024847 | 160 | 0,088336378 | 0,044470435 | 0,665149446 | 0,426558255 | 0,362574517 | 0,387499338 | 0,275499338 | -0,984500662 |
| chrVII | 1030700 | 1030865 | 165 | 0,102584181 | 0,096352609 | 0,515662191 | 0,039269381 | 0,033378974 | 0,14295827 | -0,00404173 | -1,26404173 |
| chrVII | 1030899 | 1031060 | 161 | 0,034194727 | 0,044470435 | 0,434687048 | -0,164453569 | -0,139785534 | -0,113972672 | -0,232972672 | -1,492972672 |
| chrVII | 1037731 | 1037891 | 160 | 0,037044288 | 0,059293913 | 0,384523349 | -0,293622086 | -0,249578773 | -0,23194904 | -0,34394904 | -1,60394904 |
| chrVII | 1039928 | 1040095 | 167 | 0,883363781 | 1,14140783 | 0,436278233 | -0,16041209 | -0,136350277 | 0,067492627 | -0,093507373 | -1,353507373 |
| chrVII | 1043906 | 1044058 | 152 | 0,219416165 | 0,118587827 | 0,649152585 | 0,38303363 | 0,325578585 | 0,206128415 | 0,150128415 | -1,109871585 |
| chrVII | 1044184 | 1044344 | 160 | 0,726637949 | 0,726350437 | 0,500098938 | 0,000248001 | 0,000210801 | 0,029784739 | -0,082215261 | -1,342215261 |
| chrVII | 1045755 | 1045901 | 146 | 1,094231264 | 0,711526959 | 0,605967759 | 0,268824834 | 0,228501109 | 0,039137381 | 0,025137381 | -1,234862619 |
| chrVII | 1049209 | 1049360 | 151 | 0,059840772 | 0,051882174 | 0,535617563 | 0,08939893 | 0,07598909 | -0,07606989 | -0,12506989 | -1,38506989 |
| chrVII | 1051642 | 1051789 | 147 | 0,735186631 | 0,340940001 | 0,683178549 | 0,47660573 | 0,40511487 | 0,231132449 | 0,210132449 | -1,049867551 |
| chrVII | 1052012 | 1052165 | 153 | 0,29065518 | 0,222352175 | 0,566571175 | 0,16765125 | 0,142503563 | 0,065309513 | 0,002309513 | -1,257690487 |
| chrVII | 1065087 | 1065242 | 155 | 0,384690679 | 0,326116523 | 0,541202562 | 0,103463804 | 0,087944233 | 0,013425617 | -0,063574383 | -1,323574383 |
| chrVII | 1065864 | 1066016 | 152 | 13,65509432 | 7,630385462 | 0,641521566 | 0,36252885 | 0,308149523 | 0,198340983 | 0,142340983 | -1,117659017 |
| chrVII | 1065866 | 1066016 | 150 | 0,039893848 | 0,037058696 | 0,518421434 | 0,046192108 | 0,039263292 | -0,085352075 | -0,127352075 | -1,387352075 |
| chrVII | 1065867 | 1066016 | 149 | 0,056991212 | 0,037058696 | 0,605967759 | 0,268824834 | 0,228501109 | 0,079664841 | 0,044664841 | -1,215335159 |
| chrVII | 1067431 | 1067584 | 153 | 0,179522317 | 0,155646522 | 0,535617563 | 0,08939893 | 0,07598909 | 0,001664918 | -0,061335082 | -1,321335082 |
| chrVII | 1071145 | 1071303 | 158 | 0,53571739 | 0,426175002 | 0,556941083 | 0,143218227 | 0,121735493 | 0,122071387 | 0,024071387 | -1,235928613 |
| chrVII | 1079049 | 1079187 | 138 | 0,034194727 | 0,062999783 | 0,351817475 | -0,380418277 | -0,323355536 | -0,443784877 | -0,401784877 | -1,661784877 |
| chrVII | 1079684 | 1079846 | 162 | 0,119681545 | 0,118587827 | 0,502295129 | 0,005753068 | 0,004890107 | 0,036271509 | -0,089728491 | -1,349728491 |
| chrVII | 1080323 | 1080452 | 129 | 0,162424953 | 0,344645871 | 0,320320053 | -0,466804011 | -0,396783409 | -0,158721496 | -0,053721496 | -1,313721496 |
| chrVII | 1085801 | 1085942 | 141 | 0,111132863 | 0,085235 | 0,565942212 | 0,166052574 | 0,141144688 | 0,027549433 | 0,048549433 | -1,211450567 |
| chrVII | 1087005 | 1087167 | 162 | 0,042743409 | 0,040764565 | 0,51184823 | 0,029703476 | 0,025247955 | 0,065816684 | -0,060183316 | -1,320183316 |
| chrVII | 1087005 | 1087167 | 162 | 0,042743409 | 0,040764565 | 0,51184823 | 0,029703476 | 0,025247955 | 0,065816684 | -0,060183316 | -1,320183316 |

TABLE 2

| Chrom | Start | End | Length | Section A | Section B | A/A+B | Z-score | Z * 0.85 | Phase correction | Length correction | ΔLK |
|---|---|---|---|---|---|---|---|---|---|---|---|
| chrVII | 1087106 | 1087261 | 155 | 0,091185939 | 0,211234566 | 0,301520357 | -0,520032799 | -0,442027879 | -0,507548914 | -0,584548914 | **-1,844548914** |
| chrVII | 1087240 | 1087377 | 137 | 0,14817715 | 0,114881957 | 0,563284624 | 0,159302245 | 0,135406908 | 0,011856877 | 0,060856877 | **-1,199143123** |
| chrVII | 1087527 | 1087678 | 151 | 0,062690333 | 0,040764565 | 0,605967759 | 0,268824834 | 0,228501109 | 0,07157271 | 0,02257271 | **-1,23742729** |
| chrVII | 1087535 | 1087678 | 143 | 0,356195073 | 0,470645436 | 0,430790544 | -0,174361858 | -0,148207579 | -0,319093981 | -0,312093981 | **-1,572093981** |
| chrVII | 1087583 | 1087708 | 125 | 0,188070999 | 0,185293479 | 0,503719582 | 0,009323745 | 0,007925183 | 0,433284586 | 0,566284586 | **-0,693715414** |
| chrVII | 1087808 | 1087960 | 152 | 0,113982423 | 0,077823261 | 0,594259882 | 0,238516864 | 0,202739335 | 0,096716138 | 0,040716138 | **-1,219283862** |
| chrVII | 1087881 | 1088025 | 144 | 0,088336378 | 0,044470435 | 0,665149446 | 0,426558255 | 0,362574517 | 0,236936735 | 0,236936735 | **-1,023063265** |
| chrVII | 1087978 | 1088113 | 135 | 1,188266763 | 0,733762177 | 0,618235625 | 0,30085017 | 0,255722644 | 0,203690499 | 0,266690499 | **-0,993309501** |
| chrVII | 1087978 | 1088165 | 187 | 0,125380666 | 0,074117392 | 0,628480635 | 0,327831948 | 0,278657156 | 0,722103776 | 0,421103776 | **-0,838896224** |
| chrVII | 1088021 | 1088113 | 92 | 0,105433742 | 0,100058479 | 0,513078994 | 0,032790051 | 0,027871543 | 0,417436812 | 0,781436812 | **-0,478563188** |
| chrVII | 1088021 | 1088165 | 144 | 1,347842156 | 0,592939133 | 0,694484311 | 0,508601784 | 0,432311517 | 0,304489824 | 0,304489824 | **-0,955510176** |
| chrVII | 1088360 | 1088512 | 152 | 0,136778908 | 0,125999566 | 0,520510322 | 0,051434423 | 0,043719259 | -0,061737626 | -0,117737626 | **-1,377737626** |
| chrVII | 1088661 | 1088791 | 130 | 0,444531451 | 0,448410219 | 0,497828096 | -0,005444184 | -0,004627556 | 0,202514043 | 0,300514043 | **-0,959485957** |
| chrVII | 1089031 | 1089182 | 151 | 0,416035845 | 0,285351958 | 0,593160935 | 0,235683687 | 0,200331134 | 0,052210155 | 0,003210155 | **-1,256789845** |
| chrVII | 1089036 | 1089170 | 134 | 0,700991904 | 0,426175002 | 0,621906038 | 0,310490576 | 0,263916989 | 0,232846895 | 0,302846895 | **-0,957153105** |
| chrVII | 1089049 | 1089145 | 96 | 0,085486818 | 0,122293696 | 0,411428464 | -0,223871969 | -0,190291174 | 0,221216578 | 0,557216578 | **-0,702783422** |
| chrVII | 1090196 | 1090347 | 151 | 0,558513874 | 0,400233914 | 0,58254515 | 0,20840891 | 0,177147573 | 0,032894276 | -0,016105724 | **-1,276105724** |
| chrVIII | 691 | 842 | 151 | 0,58415992 | 0,392822175 | 0,597922851 | 0,247974292 | 0,210778148 | 0,094444938 | 0,045444938 | **-1,214555062** |
| chrVIII | 1289 | 1405 | 116 | 0,156725832 | 0,229763914 | 0,405510971 | -0,239107867 | -0,203241687 | 0,269748559 | 0,465748559 | **-0,794251441** |
| chrVIII | 1346 | 1501 | 155 | 0,53856695 | 0,522527611 | 0,507557922 | 0,018946034 | 0,016104129 | 0,026227512 | -0,050772488 | **-1,310772488** |
| chrVIII | 1712 | 1899 | 187 | 0,076938136 | 0,122293696 | 0,386173911 | -0,289305214 | -0,245909432 | 0,052637638 | -0,248362362 | **-1,508362362** |
| chrVIII | 1733 | 1899 | 166 | 0,102584181 | 0,741173916 | 0,121580085 | -1,167124315 | -0,992055668 | -0,83701612 | -0,99101612 | **-2,25101612** |
| chrVIII | 1748 | 1846 | 98 | 0,028495606 | 0,077823261 | 0,268020217 | -0,618811668 | -0,525989918 | 0,110862216 | 0,432862216 | **-0,827137784** |
| chrVIII | 1748 | 1855 | 107 | 0,624053768 | 2,797931532 | 0,182365999 | -0,90638521 | -0,770427429 | -0,083236509 | 0,175763491 | **-1,084236509** |
| chrVIII | 1748 | 1878 | 130 | 0,042743409 | 0,233469783 | 0,15474789 | -1,016280608 | -0,863838517 | -0,406753508 | -0,308753508 | **-1,568753508** |
| chrVIII | 1748 | 1879 | 131 | 0,273557816 | 0,444704349 | 0,380860679 | -0,303221116 | -0,257737948 | 0,153635608 | 0,244635608 | **-1,015364392** |
| chrVIII | 1748 | 1880 | 132 | 0,059840772 | 0,122293696 | 0,328552705 | -0,443913104 | -0,377326138 | -0,009239046 | 0,074760954 | **-1,185239046** |
| chrVIII | 1748 | 1889 | 141 | 0,056991212 | 0,100058479 | 0,362886495 | -0,350753893 | -0,298140809 | -0,177915625 | -0,156915625 | **-1,416915625** |
| chrVIII | 1748 | 1891 | 143 | 0,259310013 | 0,366881088 | 0,414106832 | -0,216993175 | -0,184444199 | -0,098785569 | -0,091785569 | **-1,351785569** |
| chrVIII | 1748 | 1894 | 146 | 0,239363089 | 0,151940653 | 0,611706619 | 0,28376985 | 0,241204373 | 0,246443036 | 0,232443036 | **-1,027556964** |
| chrVIII | 1748 | 1895 | 147 | 0,24221265 | 0,277940218 | 0,465656665 | -0,08619258 | -0,073263693 | -0,110704254 | -0,131704254 | **-1,391704254** |
| chrVIII | 1748 | 1896 | 148 | 0,225115286 | 0,144528914 | 0,609005325 | 0,276727507 | 0,235218381 | 0,159351768 | 0,131351768 | **-1,128648232** |
| chrVIII | 1748 | 1897 | 149 | 0,199469241 | 0,196411088 | 0,503862472 | 0,009681932 | 0,008229642 | -0,120258873 | -0,155258873 | **-1,415258873** |
| chrVIII | 1748 | 1898 | 150 | 0,116831984 | 0,211234566 | 0,356122817 | -0,368841816 | -0,313515544 | -0,417025954 | -0,459025954 | **-1,719025954** |
| chrVIII | 1748 | 1899 | 151 | 133,1257713 | 80,91766225 | 0,621956811 | 0,310624134 | 0,264030513 | 0,15163894 | 0,10263894 | **-1,15736106** |
| chrVIII | 1748 | 1909 | 161 | 0,122531105 | 0,604056741 | 0,168639079 | -0,959557122 | -0,815623554 | -0,73715602 | -0,85615602 | **-2,11615602** |
| chrVIII | 1748 | 1932 | 184 | 0,151026711 | 0,537351089 | 0,219395092 | -0,77423779 | -0,658102121 | -0,364963662 | -0,644963662 | **-1,904963662** |
| chrVIII | 1749 | 1899 | 150 | 0,170973635 | 0,233469783 | 0,422738082 | -0,194893703 | -0,165659647 | -0,273487647 | -0,315487647 | **-1,575487647** |
| chrVIII | 1750 | 1899 | 149 | 0,319150785 | 0,422469132 | 0,430342791 | -0,175501512 | -0,149176285 | -0,279888675 | -0,314888675 | **-1,574888675** |
| chrVIII | 1751 | 1899 | 148 | 0,111132863 | 0,185293479 | 0,374908863 | -0,318879716 | -0,271047759 | -0,346679602 | -0,374679602 | **-1,634679602** |
| chrVIII | 1752 | 1899 | 147 | 1,003045325 | 0,867173481 | 0,53632512 | 0,091179756 | 0,077502792 | 0,017650706 | -0,003349294 | **-1,263349294** |
| chrVIII | 1753 | 1899 | 146 | 0,159575393 | 0,133411305 | 0,544650641 | 0,112157259 | 0,09533367 | 0,111183624 | 0,097183624 | **-1,162816376** |
| chrVIII | 1754 | 1899 | 145 | 0,190920559 | 0,214940436 | 0,47040874 | -0,074242436 | -0,063106071 | -0,042440533 | -0,049440533 | **-1,309440533** |
| chrVIII | 1755 | 1899 | 144 | 0,051292091 | 0,211234566 | 0,1953786 | -0,858244988 | -0,72950824 | -0,617474038 | -0,617474038 | **-1,877474038** |
| chrVIII | 1756 | 1899 | 143 | 0,051292091 | 0,133411305 | 0,277699771 | -0,589688448 | -0,501235181 | -0,413952 | -0,406952 | **-1,666952** |
| chrVIII | 1757 | 1899 | 142 | 0,14817715 | 0,207528696 | 0,416572153 | -0,210670615 | -0,179070023 | -0,062406001 | -0,048406001 | **-1,308406001** |
| chrVIII | 1758 | 1899 | 141 | 0,262159574 | 0,503998263 | 0,342174368 | -0,406536101 | -0,345555686 | -0,200200269 | -0,179200269 | **-1,439200269** |
| chrVIII | 1759 | 1899 | 140 | 0,04844253 | 0,062999783 | 0,434687048 | -0,164453569 | -0,139785534 | -0,004253829 | 0,023746171 | **-1,236253829** |
| chrVIII | 1760 | 1899 | 139 | 0,133929347 | 0,270528479 | 0,331133034 | -0,436786668 | -0,371268668 | -0,213682121 | -0,178682121 | **-1,438682121** |
| chrVIII | 1761 | 1899 | 138 | 0,04844253 | 0,248293262 | 0,163251388 | -0,981182451 | -0,834005084 | -0,667350855 | -0,625350855 | **-1,885350855** |
| chrVIII | 1762 | 1899 | 137 | 0,079787696 | 0,307587175 | 0,205970243 | -0,820483581 | -0,697411043 | -0,527108503 | -0,478108503 | **-1,738108503** |
| chrVIII | 1764 | 1899 | 135 | 0,111132863 | 0,192705218 | 0,365763444 | -0,34309514 | -0,291630869 | -0,0436654 | 0,0193346 | **-1,2406654** |
| chrVIII | 1765 | 1899 | 134 | 0,059840772 | 0,203822827 | 0,226958793 | -0,748899824 | -0,63656485 | -0,356702362 | -0,286702362 | **-1,546702362** |
| chrVIII | 1766 | 1899 | 133 | 0,042743409 | 0,055588044 | 0,434687048 | -0,164453569 | -0,139785534 | 0,206191624 | 0,283191624 | **-0,976808376** |
| chrVIII | 1769 | 1899 | 130 | 0,142478029 | 0,166764131 | 0,460732874 | -0,09858756 | -0,083799426 | 0,361700525 | 0,459700525 | **-0,800299475** |
| chrVIII | 1771 | 1899 | 128 | 0,042743409 | 0,259410871 | 0,141462199 | -1,073773081 | -0,912707119 | -0,401747623 | -0,289747623 | **-1,549747623** |
| chrVIII | 1773 | 1899 | 126 | 0,054141651 | 0,26682261 | 0,168684361 | -0,959377271 | -0,81547068 | -0,267502018 | -0,141502018 | **-1,401502018** |
| chrVIII | 1784 | 1899 | 115 | 0,136778908 | 0,170470001 | 0,445172966 | -0,137866493 | -0,117186519 | 0,367194985 | 0,570194985 | **-0,689805015** |
| chrVIII | 1821 | 1968 | 147 | 0,199469241 | 0,051882174 | 0,793587102 | 0,818930974 | 0,696091328 | 0,644736724 | 0,623736724 | **-0,636263276** |
| chrVIII | 2051 | 2222 | 171 | 0,108283302 | 0,055588044 | 0,660782406 | 0,414599399 | 0,352409489 | 0,545568832 | 0,356568832 | **-0,903431168** |
| chrVIII | 2441 | 2594 | 153 | 0,803576085 | 0,500292393 | 0,616301489 | 0,295781409 | 0,251414198 | 0,181389708 | 0,118389708 | **-1,141610292** |
| chrVIII | 2951 | 3103 | 152 | 0,091185939 | 0,081529131 | 0,527955893 | 0,070132481 | 0,059612609 | -0,007567952 | -0,063567952 | **-1,323567952** |
| chrVIII | 3093 | 3220 | 127 | 0,555664314 | 1,174760656 | 0,321114364 | -0,464584928 | -0,394897188 | 0,144799093 | 0,263799093 | **-0,996200907** |
| chrVIII | 3233 | 3376 | 143 | 0,373292436 | 0,392822175 | 0,487254036 | -0,031954832 | -0,027161607 | 0,056225957 | 0,063225957 | **-1,196774043** |
| chrVIII | 3524 | 3661 | 137 | 0,470177496 | 0,533645219 | 0,468386986 | -0,079325188 | -0,06742641 | 0,059508263 | 0,108508263 | **-1,151491737** |
| chrVIII | 4667 | 4822 | 155 | 0,316301225 | 0,500292393 | 0,387342269 | -0,286252759 | -0,243314845 | -0,224009173 | -0,301009173 | **-1,561009173** |
| chrVIII | 9299 | 9453 | 154 | 0,53571739 | 0,629997828 | 0,459561119 | -0,101539457 | -0,086308538 | -0,093690734 | -0,163690734 | **-1,423690734** |
| chrVIII | 10577 | 10683 | 106 | 0,418885406 | 2,697873053 | 0,134397776 | -1,105840522 | -0,939964444 | -0,316720016 | -0,050720016 | **-1,310720016** |
| chrVIII | 10578 | 10683 | 105 | 0,059840772 | 0,885702829 | 0,063287163 | -1,527751185 | -1,298588507 | -0,650540864 | -0,377540864 | **-1,637540864** |
| chrVIII | 10864 | 11003 | 139 | 0,310602104 | 0,229763914 | 0,574799476 | 0,188606742 | 0,160315731 | 0,316148488 | 0,351148488 | **-0,908851512** |
| chrVIII | 12729 | 12877 | 148 | 0,028495606 | 0,044470435 | 0,390532439 | -0,277931585 | -0,236241848 | -0,314577493 | -0,342577493 | **-1,602577493** |
| chrVIII | 14994 | 15154 | 160 | 2,19701121 | 2,02340479 | 0,520567454 | 0,05157782 | 0,043841147 | 0,137831864 | 0,025831864 | **-1,234168136** |
| chrVIII | 15294 | 15466 | 172 | 0,034194727 | 0,055588044 | 0,380860679 | -0,303221116 | -0,257737948 | -0,041801652 | -0,237801652 | **-1,497801652** |
| chrVIII | 17876 | 18017 | 141 | 0,065539893 | 0,085235 | 0,434687048 | -0,164453569 | -0,139785534 | -0,003839156 | 0,017160844 | **-1,242839156** |
| chrVIII | 18350 | 18522 | 172 | 0,322000346 | 0,500292393 | 0,391588458 | -0,275181347 | -0,233904145 | -0,013435581 | -0,209435581 | **-1,469435581** |
| chrVIII | 23920 | 24090 | 170 | 0,173823196 | 0,400233914 | 0,30279774 | -0,516370765 | -0,438915151 | -0,260762371 | -0,442762371 | **-1,702762371** |
| chrVIII | 26981 | 27134 | 153 | 0,698142343 | 0,433586741 | 0,616881154 | 0,297299724 | 0,252704765 | 0,190740619 | 0,127740619 | **-1,132259381** |
| chrVIII | 32520 | 32666 | 146 | 0,210867483 | 0,203822827 | 0,508493876 | 0,021292599 | 0,018098709 | 0,035243604 | 0,021243604 | **-1,238756396** |
| chrVIII | 33329 | 33481 | 152 | 2,097276589 | 1,645406093 | 0,560367193 | 0,151900247 | 0,12911521 | 0,062739327 | 0,006739327 | **-1,253260673** |
| chrVIII | 33329 | 33481 | 152 | 2,097276589 | 1,645406093 | 0,560367193 | 0,151900247 | 0,12911521 | 0,062739327 | 0,006739327 | **-1,253260673** |
| chrVIII | 33632 | 33782 | 150 | 0,302053422 | 0,196411088 | 0,605967759 | 0,268824834 | 0,228501109 | 0,129956915 | 0,087956915 | **-1,172043085** |
| chrVIII | 39986 | 40150 | 164 | 0,071239015 | 0,100058479 | 0,415878909 | -0,212447646 | -0,180580499 | -0,03506912 | -0,17506912 | **-1,43506912** |
| chrVIII | 46044 | 46208 | 164 | 0,943204553 | 0,926467395 | 0,504475961 | 0,011219806 | 0,009536836 | 0,162822848 | 0,022822848 | **-1,237177152** |
| chrVIII | 48223 | 48365 | 142 | 0,279256937 | 0,185293479 | 0,601133757 | 0,25628279 | 0,217840372 | 0,336995803 | 0,350995803 | **-0,909004197** |
| chrVIII | 49771 | 49922 | 151 | 0,44168189 | 0,314998914 | 0,583709654 | 0,211392905 | 0,179683969 | 0,078215958 | 0,029215958 | **-1,230784042** |
| chrVIII | 50265 | 50415 | 150 | 0,19377012 | 0,059293913 | 0,765696008 | 0,724745812 | 0,61603394 | 0,506953564 | 0,464953564 | **-0,795046436** |
| chrVIII | 51501 | 51665 | 164 | 0,604106844 | 0,563292176 | 0,517481027 | 0,043832468 | 0,037257598 | 0,199471971 | 0,059471971 | **-1,200528029** |
| chrVIII | 52096 | 52204 | 108 | 0,031345166 | 0,188999349 | 0,142255261 | -1,070241716 | -0,909705459 | -0,247071045 | 0,004928955 | **-1,255071045** |
| chrVIII | 52096 | 52213 | 117 | 0,04844253 | 0,26311674 | 0,155484155 | -1,013192315 | -0,861213468 | -0,340502374 | -0,151502374 | **-1,411502374** |
| chrVIII | 52629 | 52803 | 174 | 1,46467414 | 2,038228268 | 0,418131586 | -0,206675654 | -0,175674306 | 0,051816378 | -0,158183622 | **-1,418183622** |
| chrVIII | 53713 | 53863 | 150 | 0,504372223 | 0,307587175 | 0,621179118 | 0,308579045 | 0,262292188 | 0,153381421 | 0,111381421 | **-1,148618579** |
| chrVIII | 58605 | 58733 | 128 | 0,373292436 | 0,289057827 | 0,563587662 | 0,160071596 | 0,136060856 | 0,633409479 | 0,745409479 | **-0,514590521** |
| chrVIII | 59349 | 59502 | 153 | 6,804750673 | 5,340158063 | 0,560296567 | 0,151721162 | 0,128962988 | 0,077061847 | 0,014061847 | **-1,245938153** |
| chrVIII | 61087 | 61247 | 160 | 0,031345166 | 0,096352609 | 0,245463684 | -0,688834586 | -0,585509398 | -0,50021545 | -0,61221545 | **-1,87221545** |
| chrVIII | 62149 | 62261 | 112 | 1,382036883 | 7,400621548 | 0,157359744 | -1,005368404 | -0,854563143 | -0,254683303 | -0,030683303 | **-1,290683303** |
| chrVIII | 65254 | 65423 | 169 | 0,082637257 | 0,074117392 | 0,527175798 | 0,068172392 | 0,057946533 | 0,220764326 | 0,045764326 | **-1,214235674** |
| chrVIII | 66426 | 66576 | 150 | 6,129404815 | 2,783108054 | 0,687730263 | 0,489426929 | 0,41601289 | 0,324511449 | 0,282511449 | **-0,977488551** |
| chrVIII | 66660 | 66810 | 150 | 0,897611584 | 0,796761959 | 0,529760151 | 0,074666958 | 0,063466914 | -0,024633142 | -0,066633142 | **-1,326633142** |
| chrVIII | 66812 | 66936 | 124 | 0,042743409 | 0,240881523 | 0,15070399 | -1,033418742 | -0,87840593 | -0,31438553 | -0,17438553 | **-1,43438553** |
| chrVIII | 66813 | 66902 | 89 | 0,037044288 | 0,062999783 | 0,370279692 | -0,331112667 | -0,281445767 | 0,390429391 | 0,775429391 | **-0,484570609** |
| chrVIII | 66813 | 66904 | 91 | 0,028495606 | 0,125999566 | 0,184443343 | -0,898560728 | -0,763776618 | -0,086525697 | 0,284474303 | **-0,975525697** |

TABLE 2

| Chrom | Start | End | Length | Section A | Section B | A/A+B | Z-score | Z * 0.85 | Phase correction | Length correction | ΔLK |
|---|---|---|---|---|---|---|---|---|---|---|---|
| chrVIII | 66813 | 66911 | 98 | 0,034194727 | 0,125999566 | 0,213457836 | -0,794480647 | -0,67530855 | -0,013864323 | 0,308135677 | -0,951864323 |
| chrVIII | 66813 | 66912 | 99 | 0,045592969 | 0,137117174 | 0,249537154 | -0,67594698 | -0,574554933 | 0,105360341 | 0,420360341 | -0,839639659 |
| chrVIII | 66813 | 66924 | 111 | 0,059840772 | 0,062999783 | 0,487141826 | -0,032236246 | -0,027400809 | 0,564809056 | 0,795809056 | -0,464190944 |
| chrVIII | 66813 | 66925 | 112 | 0,065539893 | 0,092646739 | 0,414320049 | -0,216446021 | -0,183979117 | 0,407903066 | 0,631903066 | -0,628096934 |
| chrVIII | 66813 | 66926 | 113 | 0,028495606 | 0,085235 | 0,250553539 | -0,67274886 | -0,571836531 | -0,010373951 | 0,206626049 | -1,053373951 |
| chrVIII | 66813 | 66928 | 115 | 0,037044288 | 0,092646739 | 0,285634931 | -0,566182298 | -0,481254953 | 0,021266584 | 0,224266584 | -1,035733416 |
| chrVIII | 66813 | 66929 | 116 | 0,074088575 | 0,140823044 | 0,344739737 | -0,399561558 | -0,339627325 | 0,143939247 | 0,339939247 | -0,920060753 |
| chrVIII | 66813 | 66930 | 117 | 0,042743409 | 0,159352392 | 0,211500727 | -0,801224908 | -0,681041172 | -0,15646726 | 0,03253274 | -1,22746726 |
| chrVIII | 66813 | 66931 | 118 | 0,128230226 | 0,403939784 | 0,240957258 | -0,703226647 | -0,59774265 | -0,079170762 | 0,102829238 | -1,157170762 |
| chrVIII | 66813 | 66932 | 119 | 0,139628469 | 0,289057827 | 0,325712462 | -0,451783548 | -0,384016016 | 0,109174674 | 0,284174674 | -0,975825326 |
| chrVIII | 66813 | 66933 | 120 | 0,176672756 | 0,333528262 | 0,346280681 | -0,395381497 | -0,336074272 | 0,152539665 | 0,320539665 | -0,939460335 |
| chrVIII | 66813 | 66934 | 121 | 0,133929347 | 0,159352392 | 0,456657642 | -0,108857794 | -0,092529125 | 0,458432272 | 0,619432272 | -0,640567728 |
| chrVIII | 66813 | 66935 | 122 | 0,065539893 | 0,181587609 | 0,265206797 | -0,627374783 | -0,533268565 | 0,02761144 | 0,18161144 | -1,07838856 |
| chrVIII | 66813 | 66936 | 123 | 49,87870845 | 96,96407752 | 0,339674218 | -0,413352411 | -0,35134955 | 0,242906556 | 0,389906556 | -0,870093444 |
| chrVIII | 66815 | 66936 | 121 | 0,065539893 | 0,244587392 | 0,211332238 | -0,801807229 | -0,681536144 | -0,151420064 | 0,009579936 | -1,250420064 |
| chrVIII | 66816 | 66936 | 120 | 0,037044288 | 0,181587609 | 0,169436794 | -0,956393263 | -0,812934274 | -0,313470966 | -0,145470966 | -1,405470966 |
| chrVIII | 66817 | 66936 | 119 | 0,185221438 | 0,533645219 | 0,257657573 | -0,65058387 | -0,55299629 | -0,04502513 | 0,12997487 | -1,13002513 |
| chrVIII | 66818 | 66936 | 118 | 0,085486818 | 0,118587827 | 0,418899751 | -0,204708983 | -0,174002636 | 0,339622688 | 0,521622688 | -0,738377312 |
| chrVIII | 66820 | 66936 | 116 | 0,088336378 | 0,34835174 | 0,202287111 | -0,833479734 | -0,708457774 | -0,216546637 | -0,020546637 | -1,280546637 |
| chrVIII | 66821 | 66936 | 115 | 0,054141651 | 0,051882174 | 0,510655515 | 0,026712592 | 0,022705704 | 0,510886461 | 0,713886461 | -0,546113539 |
| chrVIII | 66822 | 66936 | 114 | 0,051292091 | 0,066705652 | 0,434687048 | -0,164453569 | -0,139785534 | 0,316777536 | 0,526777536 | -0,733222464 |
| chrVIII | 66823 | 66936 | 113 | 0,031345166 | 0,103764348 | 0,231998216 | -0,73228205 | -0,622439743 | -0,063581507 | 0,153418493 | -1,106581507 |
| chrVIII | 66830 | 66936 | 106 | 0,105433742 | 0,274234349 | 0,277699771 | -0,589688448 | -0,501235181 | 0,124420492 | 0,390420492 | -0,869579508 |
| chrVIII | 68261 | 68399 | 138 | 0,153876272 | 0,125999566 | 0,549801916 | 0,125160904 | 0,106386768 | 0,28068908 | 0,32268908 | -0,93731092 |
| chrVIII | 69975 | 70096 | 121 | 0,170973635 | 0,122293696 | 0,582995843 | 0,209563572 | 0,178129036 | 0,701091517 | 0,862091517 | -0,397908483 |
| chrVIII | 69975 | 70122 | 147 | 0,316301225 | 0,170470001 | 0,649794417 | 0,384765495 | 0,327050671 | 0,275247812 | 0,254247812 | -1,005752188 |
| chrVIII | 70402 | 70553 | 151 | 0,455929693 | 0,385410436 | 0,541908887 | 0,105243965 | 0,08945737 | -0,003551993 | -0,052551993 | -1,312551993 |
| chrVIII | 72065 | 72199 | 134 | 0,53856695 | 0,848644133 | 0,388237202 | -0,283916456 | -0,241328988 | 0,053873791 | 0,123873791 | -1,136126209 |
| chrVIII | 77481 | 77612 | 131 | 0,082637257 | 0,066705652 | 0,553339005 | 0,13410191 | 0,113986624 | 0,525574091 | 0,616574091 | -0,643425909 |
| chrVIII | 86148 | 86310 | 162 | 0,039893848 | 0,092646739 | 0,300993446 | -0,521545394 | -0,443313585 | -0,298499431 | -0,424499431 | -1,684499431 |
| chrVIII | 86176 | 86324 | 148 | 0,284956058 | 0,181587609 | 0,610781108 | 0,281355451 | 0,239152133 | 0,172357835 | 0,144357835 | -1,115642165 |
| chrVIII | 86476 | 86623 | 147 | 0,102584181 | 0,114881957 | 0,471724849 | -0,070934736 | -0,060294526 | -0,130195094 | -0,151195094 | -1,411195094 |
| chrVIII | 88963 | 89116 | 153 | 0,159575393 | 0,448410219 | 0,262465739 | -0,635762125 | -0,540397806 | -0,603932052 | -0,666932052 | -1,926932052 |
| chrVIII | 91372 | 91534 | 162 | 0,113982423 | 0,218646305 | 0,342671614 | -0,405182689 | -0,344405286 | -0,207772132 | -0,333772132 | -1,593772132 |
| chrVIII | 91372 | 91534 | 162 | 0,113982423 | 0,218646305 | 0,342671614 | -0,405182689 | -0,344405286 | -0,207772132 | -0,333772132 | -1,593772132 |
| chrVIII | 94804 | 94953 | 149 | 0,19377012 | 0,100058479 | 0,659466508 | 0,41100757 | 0,349356434 | 0,229059008 | 0,194059008 | -1,065940992 |
| chrVIII | 94985 | 95141 | 156 | 0,076938136 | 0,085235 | 0,474419732 | -0,064164224 | -0,05453959 | -0,053346819 | -0,137346819 | -1,397346819 |
| chrVIII | 96353 | 96506 | 153 | 0,752283994 | 0,466939567 | 0,617018911 | 0,297660652 | 0,253011555 | 0,186925542 | 0,123925542 | -1,136074458 |
| chrVIII | 100385 | 100534 | 149 | 0,037044288 | 0,074117392 | 0,333246923 | -0,430964963 | -0,366320218 | -0,503941269 | -0,538941269 | -1,798941269 |
| chrVIII | 107252 | 107412 | 160 | 0,028495606 | 0,044470435 | 0,390532439 | -0,277931585 | -0,236241848 | -0,146043361 | -0,258043361 | -1,518043361 |
| chrVIII | 108345 | 108493 | 148 | 0,09688506 | 0,140823044 | 0,407579962 | -0,23377466 | -0,198708461 | -0,274329229 | -0,302329229 | -1,562329229 |
| chrVIII | 109527 | 109701 | 174 | 0,24506221 | 0,470645436 | 0,342405466 | -0,405907004 | -0,345020953 | -0,096944703 | -0,306944703 | -1,566944703 |
| chrVIII | 114779 | 114932 | 153 | 1,638497336 | 1,093231526 | 0,59980233 | 0,252835492 | 0,214910168 | 0,16095662 | 0,09795662 | -1,16204338 |
| chrVIII | 115531 | 115709 | 178 | 0,082637257 | 0,129705435 | 0,3891693 | -0,28148478 | -0,239262063 | 0,043980934 | -0,194019066 | -1,454019066 |
| chrVIII | 121476 | 121637 | 161 | 0,812124766 | 0,570703915 | 0,587292394 | 0,220585474 | 0,187497653 | 0,278590182 | 0,159590182 | -1,100409818 |
| chrVIII | 122292 | 122451 | 159 | 0,091185939 | 0,037058696 | 0,711031218 | 0,556399818 | 0,472939845 | 0,573935886 | 0,468935886 | -0,791064114 |
| chrVIII | 123820 | 123978 | 158 | 0,453080133 | 0,563292176 | 0,445781658 | -0,136326323 | -0,115877374 | -0,052352592 | -0,150352592 | -1,410352592 |
| chrVIII | 125638 | 125758 | 120 | 0,247911771 | 0,374292827 | 0,398440917 | -0,257384672 | -0,218776971 | 0,314988238 | 0,482988238 | -0,777011762 |
| chrVIII | 130844 | 131002 | 158 | 2,122922635 | 2,045640007 | 0,509269697 | 0,023237776 | 0,01975211 | 0,082472461 | -0,015527539 | -1,275527539 |
| chrVIII | 131196 | 131345 | 149 | 0,681044979 | 0,337234132 | 0,668819553 | 0,436655929 | 0,37115754 | 0,252654129 | 0,217654129 | -1,042345871 |
| chrVIII | 142630 | 142776 | 146 | 0,869115978 | 0,667056524 | 0,565767176 | 0,165607749 | 0,140766587 | 0,139401799 | 0,125401799 | -1,134598201 |
| chrVIII | 144193 | 144334 | 141 | 2,564604525 | 1,801052615 | 0,587449825 | 0,220989833 | 0,187841358 | 0,326286935 | 0,347286935 | -0,912713065 |
| chrVIII | 152201 | 152353 | 152 | 0,082637257 | 0,096352609 | 0,461686791 | -0,096185077 | -0,081757315 | -0,146538567 | -0,202538567 | -1,462538567 |
| chrVIII | 153827 | 153949 | 122 | 0,387540239 | 2,334697835 | 0,142360892 | -1,069772367 | -0,909306512 | -0,310468334 | -0,156468334 | -1,416468334 |
| chrVIII | 157804 | 157963 | 159 | 0,162424953 | 0,107470218 | 0,601807556 | 0,258028531 | 0,219324251 | 0,325196433 | 0,220196433 | -1,039803567 |
| chrVIII | 158390 | 158542 | 152 | 0,250761331 | 0,163058261 | 0,605967759 | 0,268824834 | 0,228501109 | 0,172034719 | 0,116034719 | -1,143965281 |
| chrVIII | 162439 | 162593 | 154 | 1,647046017 | 1,100643265 | 0,599429502 | 0,251870714 | 0,214090107 | 0,207284304 | 0,137284304 | -1,122715696 |
| chrVIII | 162949 | 163087 | 138 | 0,401788042 | 0,240881523 | 0,625186043 | 0,319130028 | 0,271260524 | 0,445857169 | 0,487857169 | -0,772142831 |
| chrVIII | 165932 | 166076 | 144 | 0,453080133 | 0,381704567 | 0,542750883 | 0,107366495 | 0,091261521 | 0,170537298 | 0,170537298 | -1,089462702 |
| chrVIII | 167959 | 168110 | 151 | 1,159771157 | 1,111760874 | 0,51056782 | 0,026492695 | 0,022518791 | -0,067459633 | -0,116459633 | -1,376459633 |
| chrVIII | 171020 | 171171 | 151 | 0,105433742 | 0,055588044 | 0,654779361 | 0,398256287 | 0,338517844 | 0,244739505 | 0,195739505 | -1,064260495 |
| chrVIII | 171703 | 171855 | 152 | 0,393239361 | 0,333528262 | 0,541079911 | 0,103154718 | 0,08768151 | 0,036875132 | -0,019124868 | -1,279124868 |
| chrVIII | 174446 | 174586 | 140 | 0,108283302 | 1,982640225 | 0,051787309 | -1,627765505 | -1,383600679 | -1,251796618 | -1,223796618 | -2,483796618 |
| chrVIII | 174447 | 174533 | 86 | 0,037044288 | 0,952408482 | 0,037439167 | -1,781208876 | -1,514027544 | -0,903935898 | -0,497935898 | -1,757935898 |
| chrVIII | 174447 | 174535 | 88 | 0,045592969 | 0,496586524 | 0,084092021 | -1,378062339 | -1,171352988 | -0,535734994 | -0,143734994 | -1,403734994 |
| chrVIII | 174447 | 174538 | 91 | 0,09688506 | 0,607762611 | 0,137494331 | -1,091646154 | -0,927899231 | -0,246103149 | 0,124896851 | -1,135103149 |
| chrVIII | 174447 | 174540 | 93 | 0,045592969 | 0,396528045 | 0,10312328 | -1,263953936 | -1,074360846 | -0,431705523 | -0,074705523 | -1,334705523 |
| chrVIII | 174447 | 174541 | 94 | 0,04844253 | 0,97093783 | 0,047521545 | -1,669374972 | -1,418968726 | -0,778759971 | -0,428759971 | -1,688759971 |
| chrVIII | 174447 | 174544 | 97 | 0,031345166 | 0,87829109 | 0,034459012 | -1,818956785 | -1,546113267 | -0,917430106 | -0,588430106 | -1,848430106 |
| chrVIII | 174447 | 174545 | 98 | 0,039893848 | 1,271113265 | 0,030429925 | -1,874512288 | -1,593335445 | -0,948644163 | -0,626644163 | -1,886644163 |
| chrVIII | 174447 | 174548 | 101 | 0,034194727 | 0,314998914 | 0,097924827 | -1,293466814 | -1,099446792 | -0,408309311 | -0,107309311 | -1,367309311 |
| chrVIII | 174447 | 174550 | 103 | 0,031345166 | 0,270528479 | 0,103835386 | -1,259996094 | -1,07099668 | -0,364354548 | -0,077354548 | -1,337354548 |
| chrVIII | 174447 | 174551 | 104 | 0,034194727 | 0,359469349 | 0,086862706 | -1,360330337 | -1,156280786 | -0,449594893 | -0,169594893 | -1,429594893 |
| chrVIII | 174447 | 174552 | 105 | 0,059840772 | 0,44099848 | 0,119480995 | -1,177585257 | -1,000947468 | -0,35218531 | -0,07918531 | -1,33918531 |
| chrVIII | 174447 | 174553 | 106 | 0,142478029 | 9,661201991 | 0,014533117 | -2,182587053 | -1,855198995 | -1,218746809 | -0,952746809 | -2,212746809 |
| chrVIII | 174447 | 174555 | 108 | 0,039893848 | 0,34835174 | 0,102754157 | -1,266013318 | -1,076111321 | -0,385899438 | -0,133899438 | -1,393899438 |
| chrVIII | 174447 | 174561 | 114 | 0,042743409 | 0,444704349 | 0,087688184 | -1,35512929 | -1,151859897 | -0,700334161 | -0,490334161 | -1,750334161 |
| chrVIII | 174447 | 174565 | 118 | 0,051292091 | 0,570703915 | 0,082463698 | -1,38868882 | -1,180385497 | -0,677634194 | -0,495634194 | -1,755634194 |
| chrVIII | 174447 | 174566 | 119 | 0,056991212 | 0,418763262 | 0,119791226 | -1,176031092 | -0,999626428 | -0,498491617 | -0,323491617 | -1,583491617 |
| chrVIII | 174447 | 174567 | 120 | 0,085486818 | 0,574409785 | 0,129545776 | -1,128540904 | -0,959259768 | -0,427140194 | -0,259140194 | -1,519140194 |
| chrVIII | 174447 | 174568 | 121 | 0,065539893 | 0,43729261 | 0,130341402 | -1,124778745 | -0,956061933 | -0,40579122 | -0,24479122 | -1,50479122 |
| chrVIII | 174447 | 174569 | 122 | 0,088336378 | 0,318704784 | 0,21702075 | -0,782294532 | -0,664950353 | -0,066773356 | 0,087226644 | -1,172773356 |
| chrVIII | 174447 | 174570 | 123 | 0,085486818 | 0,377998697 | 0,184443343 | -0,898560728 | -0,763776618 | -0,192586702 | -0,045586702 | -1,305586702 |
| chrVIII | 174447 | 174571 | 124 | 0,182371877 | 0,507704132 | 0,264277956 | -0,630211965 | -0,53568017 | 0,033754057 | 0,173754057 | -1,086245943 |
| chrVIII | 174447 | 174572 | 125 | 0,111132863 | 0,678174133 | 0,140798021 | -1,076740885 | -0,915229752 | -0,389021409 | -0,256021409 | -1,516021409 |
| chrVIII | 174447 | 174573 | 126 | 0,065539893 | 0,270528479 | 0,195019522 | -0,85954656 | -0,730614576 | -0,205498373 | -0,079498373 | -1,339498373 |
| chrVIII | 174447 | 174574 | 127 | 0,094035499 | 0,61888022 | 0,131902687 | -1,117442025 | -0,949825721 | -0,40227693 | -0,28327693 | -1,54327693 |
| chrVIII | 174447 | 174575 | 128 | 0,09688506 | 0,433586741 | 0,182639416 | -0,905352197 | -0,769549367 | -0,281042701 | -0,169042701 | -1,429042701 |
| chrVIII | 174447 | 174576 | 129 | 0,250761331 | 0,733762177 | 0,254703244 | -0,659762131 | -0,560797812 | -0,078539464 | 0,026460536 | -1,233539464 |
| chrVIII | 174447 | 174577 | 130 | 0,444531451 | 1,096937395 | 0,288381729 | -0,558118516 | -0,474400738 | -0,034052957 | 0,063947043 | -1,196052957 |
| chrVIII | 174447 | 174578 | 131 | 0,239363089 | 0,604056741 | 0,283800642 | -0,571587769 | -0,485849604 | -0,089911953 | 0,001088047 | -1,258911953 |
| chrVIII | 174447 | 174579 | 132 | 0,273557816 | 0,641115437 | 0,299077091 | -0,527056746 | -0,447998234 | -0,066932972 | 0,017067028 | -1,242932972 |
| chrVIII | 174447 | 174580 | 133 | 0,284956058 | 0,778232612 | 0,268020217 | -0,618811668 | -0,525989918 | -0,189127814 | -0,112127814 | -1,372127814 |
| chrVIII | 174447 | 174581 | 134 | 0,521469587 | 1,304466092 | 0,285590338 | -0,566313514 | -0,481366487 | -0,196338066 | -0,126338066 | -1,386338066 |
| chrVIII | 174447 | 174582 | 135 | 0,834921251 | 1,40452457 | 0,372824939 | -0,324380637 | -0,275725541 | -0,041892414 | 0,021107586 | -1,238892414 |
| chrVIII | 174447 | 174583 | 136 | 0,481575739 | 0,959820221 | 0,334103711 | -0,428609513 | -0,364318086 | -0,174172993 | -0,118172993 | -1,378172993 |
| chrVIII | 174447 | 174584 | 137 | 0,624053768 | 1,58240631 | 0,282830301 | -0,574454027 | -0,488285923 | -0,363889836 | -0,314889836 | -1,574889836 |
| chrVIII | 174447 | 174585 | 138 | 0,367593315 | 1,430465657 | 0,204438965 | -0,825869842 | -0,701989358 | -0,529809058 | -0,487809058 | -1,747809058 |
| chrVIII | 174447 | 174586 | 139 | 409,06582 | 529,931938 | 0,435640891 | -0,162030568 | -0,137725983 | 0,01363651 | 0,04863651 | -1,21136349 |
| chrVIII | 174447 | 174587 | 140 | 0,128230226 | 0,496586524 | 0,205228535 | -0,823089551 | -0,699626119 | -0,561791499 | -0,533791499 | -1,793791499 |
| chrVIII | 174448 | 174586 | 138 | 0,435982769 | 1,356348266 | 0,243249021 | -0,695889487 | -0,591506064 | -0,434348415 | -0,392348415 | -1,652348415 |

TABLE 2

| Chrom | Start | End | Length | Section A | Section B | A/A+B | Z-score | Z * 0.85 | Phase correction | Length correction | ΔLK |
|---|---|---|---|---|---|---|---|---|---|---|---|
| chrVIII | 174449 | 174586 | 137 | 0,53571739 | 1,489759571 | 0,264489501 | -0,629565348 | -0,535130546 | -0,388178604 | -0,339178604 | -1,599178604 |
| chrVIII | 174450 | 174586 | 136 | 0,624053768 | 1,808464354 | 0,2565464 | -0,654029499 | -0,555925074 | -0,383297859 | -0,327297859 | -1,587297859 |
| chrVIII | 174451 | 174586 | 135 | 0,97739928 | 2,01969892 | 0,3261152 | -0,450665847 | -0,38306597 | -0,161499898 | -0,098499898 | -1,358499898 |
| chrVIII | 174452 | 174586 | 134 | 0,638301571 | 1,927052181 | 0,248816199 | -0,678219707 | -0,576486751 | -0,296654464 | -0,226654464 | -1,486654464 |
| chrVIII | 174453 | 174586 | 133 | 0,265009134 | 1,371171744 | 0,161968116 | -0,986401291 | -0,838441097 | -0,515119925 | -0,438119925 | -1,698119925 |
| chrVIII | 174454 | 174586 | 132 | 0,213717044 | 1,715817615 | 0,110760925 | -1,22249141 | -1,039117699 | -0,67614847 | -0,59214847 | -1,85214847 |
| chrVIII | 174455 | 174586 | 131 | 0,304902982 | 0,730056307 | 0,294603841 | -0,539984556 | -0,458986873 | -0,075822515 | 0,015177485 | -1,244822515 |
| chrVIII | 174456 | 174586 | 130 | 0,19661968 | 0,755997394 | 0,206399492 | -0,818977968 | -0,696131272 | -0,252295965 | -0,154295965 | -1,414295965 |
| chrVIII | 174457 | 174586 | 129 | 0,227964847 | 1,026525873 | 0,181719038 | -0,908833395 | -0,772508386 | -0,303023269 | -0,198023269 | -1,458023269 |
| chrVIII | 174458 | 174586 | 128 | 0,230814407 | 0,581821524 | 0,284031752 | -0,570905793 | -0,485269924 | -0,00752345 | 0,10447655 | -1,15552345 |
| chrVIII | 174459 | 174586 | 127 | 0,14817715 | 0,900526308 | 0,141295568 | -1,074516764 | -0,91333925 | -0,359069553 | -0,240069553 | -1,500069553 |
| chrVIII | 174460 | 174586 | 126 | 0,199469241 | 1,041349352 | 0,160756167 | -0,99135486 | -0,842651631 | -0,306462389 | -0,180462389 | -1,440462389 |
| chrVIII | 174461 | 174586 | 125 | 0,131079787 | 2,20128653 | 0,056200343 | -1,587494526 | -1,349370347 | -0,808941559 | -0,675941559 | -1,935941559 |
| chrVIII | 174462 | 174553 | 91 | 1,148372915 | 1,374877614 | 0,455116486 | -0,112744686 | -0,095832983 | 0,591042643 | 0,962042643 | -0,297957357 |
| chrVIII | 174467 | 174586 | 119 | 0,037044288 | 0,907938047 | 0,039201037 | -1,760033809 | -1,496028738 | -1,006322845 | -0,831322845 | -2,091322845 |
| chrVIII | 174928 | 175084 | 156 | 0,028495606 | 0,059293913 | 0,324590066 | -0,454901459 | -0,386666241 | -0,381251745 | -0,465251745 | -1,725251745 |
| chrVIII | 176211 | 176366 | 155 | 11,64330454 | 5,566216107 | 0,676561816 | 0,458105889 | 0,389390006 | 0,414519494 | 0,337519494 | -0,922480506 |
| chrVIII | 177719 | 177867 | 148 | 0,97454972 | 0,670762394 | 0,592319057 | 0,233514533 | 0,198487353 | 0,12204331 | 0,09404331 | -1,16595669 |
| chrVIII | 177797 | 177947 | 150 | 2,00039153 | 1,208113483 | 0,623465297 | 0,314594696 | 0,267405492 | 0,167756456 | 0,125756456 | -1,134243544 |
| chrVIII | 178837 | 178985 | 148 | 2,04883406 | 1,05246696 | 0,660636954 | 0,414202117 | 0,352071799 | 0,272900181 | 0,244900181 | -1,015099819 |
| chrVIII | 180375 | 180509 | 134 | 1,5615592 | 1,14140783 | 0,577720402 | 0,196065121 | 0,166655353 | 0,434974991 | 0,504974991 | -0,755025009 |
| chrVIII | 190177 | 190319 | 142 | 0,404637603 | 0,485468915 | 0,454594585 | -0,114061338 | -0,096952138 | -0,002374651 | 0,011625349 | -1,248374651 |
| chrVIII | 194784 | 194941 | 157 | 0,065539893 | 0,133411305 | 0,329426985 | -0,441495981 | -0,375271584 | -0,354812859 | -0,445812859 | -1,705812859 |
| chrVIII | 194987 | 195122 | 135 | 0,105433742 | 0,114881957 | 0,478557553 | -0,053774149 | -0,045708026 | 0,183974627 | 0,246974627 | -1,013025373 |
| chrVIII | 205654 | 205804 | 150 | 0,256460453 | 0,214940436 | 0,544038968 | 0,11061448 | 0,094022308 | -0,00516269 | -0,04716269 | -1,30716269 |
| chrVIII | 207083 | 207225 | 142 | 1,65844426 | 1,078408047 | 0,605967759 | 0,268824834 | 0,228501109 | 0,320200177 | 0,334200177 | -0,925799823 |
| chrVIII | 212526 | 212683 | 157 | 1,188266763 | 1,05246696 | 0,53030253 | 0,076030366 | 0,064625811 | 0,073793991 | -0,017206009 | -1,277206009 |
| chrVIII | 212729 | 212874 | 145 | 0,054141651 | 0,059293913 | 0,477289917 | -0,056956516 | -0,048413039 | -0,033209905 | -0,040209905 | -1,300209905 |
| chrVIII | 212963 | 213069 | 106 | 0,056991212 | 0,459527828 | 0,110337098 | -1,224737354 | -1,041026751 | -0,416910171 | -0,150910171 | -1,410910171 |
| chrVIII | 212963 | 213076 | 113 | 0,054141651 | 0,507704132 | 0,0963639 | -1,302551864 | -1,107169084 | -0,604861438 | -0,387861438 | -1,647861438 |
| chrVIII | 213025 | 213187 | 162 | 0,19661968 | 0,218646305 | 0,473478896 | -0,066527591 | -0,056548453 | 0,081124089 | -0,044875911 | -1,304875911 |
| chrVIII | 214079 | 214248 | 169 | 0,037044288 | 0,040764565 | 0,47609348 | -0,059960669 | -0,050966568 | 0,10129583 | -0,07370417 | -1,33370417 |
| chrVIII | 214176 | 214329 | 153 | 0,265009134 | 0,374292827 | 0,414528893 | -0,21591015 | -0,183523627 | -0,23838711 | -0,30138711 | -1,56138711 |
| chrVIII | 214524 | 214681 | 157 | 1,22246149 | 1,115466743 | 0,522882385 | 0,057389119 | 0,048780751 | 0,070428953 | -0,020571047 | -1,280571047 |
| chrVIII | 214727 | 214872 | 145 | 0,079787696 | 0,055588044 | 0,589379577 | 0,225949308 | 0,192050912 | 0,2059234 | 0,1989234 | -1,0610766 |
| chrVIII | 214961 | 215067 | 106 | 0,04844253 | 0,411351523 | 0,105357017 | -1,251604445 | -1,063863778 | -0,419393278 | -0,153393278 | -1,413393278 |
| chrVIII | 214961 | 215074 | 113 | 0,062690333 | 0,578115654 | 0,097830442 | -1,294013136 | -1,099911165 | -0,595190833 | -0,378190833 | -1,638190833 |
| chrVIII | 215023 | 215185 | 162 | 0,179522317 | 0,188999349 | 0,487141826 | -0,032236246 | -0,027400809 | 0,110499756 | -0,015500244 | -1,275500244 |
| chrVIII | 216077 | 216246 | 169 | 0,034194727 | 0,062999783 | 0,351817475 | -0,380418277 | -0,323355536 | -0,176889619 | -0,351889619 | -1,611889619 |
| chrVIII | 217808 | 217962 | 154 | 0,302053422 | 0,259410871 | 0,537974411 | 0,095331934 | 0,081032144 | 0,077450723 | 0,007450723 | -1,252549277 |
| chrVIII | 218430 | 218569 | 139 | 0,14817715 | 0,34835174 | 0,298426039 | -0,528932784 | -0,449592866 | -0,294807445 | -0,259807445 | -1,519807445 |
| chrVIII | 219279 | 219430 | 151 | 0,569912117 | 0,422469132 | 0,57428747 | 0,187300463 | 0,159205394 | 0,063875473 | 0,014875473 | -1,245124527 |
| chrVIII | 223655 | 223760 | 105 | 1,162620718 | 10,10961221 | 0,103140232 | -1,263859491 | -1,074280567 | -0,442807162 | -0,169807162 | -1,429807162 |
| chrVIII | 226108 | 226256 | 148 | 0,267858695 | 0,044470435 | 0,857616755 | 1,069673076 | 0,909222114 | 0,798670411 | 0,770670411 | -0,489329589 |
| chrVIII | 227838 | 227999 | 161 | 0,113982423 | 0,17417587 | 0,395554894 | -0,264869768 | -0,225139303 | -0,136272845 | -0,255272845 | -1,515272845 |
| chrVIII | 227838 | 227999 | 161 | 0,113982423 | 0,17417587 | 0,395554894 | -0,264869768 | -0,225139303 | -0,136272845 | -0,255272845 | -1,515272845 |
| chrVIII | 228803 | 228946 | 143 | 0,225115286 | 0,17788174 | 0,558602847 | 0,147427874 | 0,125313693 | 0,193942271 | 0,200942271 | -1,059057729 |
| chrVIII | 234383 | 234536 | 153 | 0,219416165 | 0,196411088 | 0,52766182 | 0,069393554 | 0,058984521 | -0,011466065 | -0,074466065 | -1,334466065 |
| chrVIII | 234544 | 234693 | 149 | 0,678195419 | 0,489174784 | 0,580960022 | 0,204350056 | 0,173697548 | 0,055903487 | 0,020903487 | -1,239096513 |
| chrVIII | 240775 | 240923 | 148 | 1,057186976 | 0,730056307 | 0,591518226 | 0,231452159 | 0,196734335 | 0,090268591 | 0,062268591 | -1,197731409 |
| chrVIII | 250570 | 250724 | 154 | 0,501522663 | 0,26311674 | 0,655894348 | 0,401283645 | 0,341091098 | 0,342704377 | 0,272704377 | -0,987295623 |
| chrVIII | 256594 | 256759 | 165 | 0,034194727 | 0,070411522 | 0,326889907 | -0,448517424 | -0,381239811 | -0,220989876 | -0,367989876 | -1,627989876 |
| chrVIII | 258714 | 258856 | 142 | 0,504372223 | 0,377998697 | 0,571610206 | 0,180475116 | 0,153403848 | 0,254374304 | 0,268374304 | -0,991625696 |
| chrVIII | 261335 | 261482 | 147 | 0,279256937 | 0,240881523 | 0,536889615 | 0,092600721 | 0,078710613 | 0,011523758 | -0,009476242 | -1,269476242 |
| chrVIII | 264809 | 264956 | 147 | 0,356195073 | 0,226058044 | 0,611752969 | 0,283890806 | 0,241307185 | 0,16766423 | 0,14666423 | -1,11333577 |
| chrVIII | 270516 | 270670 | 154 | 0,219416165 | 0,248293262 | 0,469129234 | -0,077458923 | -0,065840085 | -0,07332622 | -0,14332622 | -1,40332622 |
| chrVIII | 272692 | 272812 | 120 | 9,480488061 | 21,07898616 | 0,310230732 | -0,495196439 | -0,420916973 | 0,126508326 | 0,294508326 | -0,965491674 |
| chrVIII | 275159 | 275317 | 158 | 0,182371877 | 0,144528914 | 0,55788142 | 0,145600009 | 0,123760008 | 0,191190839 | 0,093190839 | -1,166809161 |
| chrVIII | 281596 | 281745 | 149 | 1,36778908 | 0,941290873 | 0,592352412 | 0,233600453 | 0,198560385 | 0,06919785 | 0,03419785 | -1,22580215 |
| chrVIII | 288944 | 289050 | 106 | 0,031345166 | 0,096352609 | 0,245463684 | -0,688834586 | -0,585509398 | 0,061082463 | 0,327082463 | -0,932917537 |
| chrVIII | 291607 | 291756 | 149 | 0,202318801 | 0,096352609 | 0,677395942 | 0,460429288 | 0,391364895 | 0,262472444 | 0,227472444 | -1,032527556 |
| chrVIII | 291938 | 292090 | 152 | 1,470373261 | 1,05246696 | 0,582824568 | 0,209124736 | 0,177756026 | 0,139654792 | 0,083654792 | -1,176345208 |
| chrVIII | 294718 | 294863 | 145 | 0,250761331 | 0,181587609 | 0,579997562 | 0,201887241 | 0,171604155 | 0,181380176 | 0,174380176 | -1,085619824 |
| chrVIII | 297474 | 297608 | 134 | 0,860567296 | 1,030231743 | 0,455134194 | -0,112700017 | -0,095795015 | 0,167780066 | 0,237780066 | -1,022219934 |
| chrVIII | 297952 | 298079 | 127 | 0,227964847 | 0,329822392 | 0,408694984 | -0,230903248 | -0,196267761 | 0,348948915 | 0,467948915 | -0,792051085 |
| chrVIII | 299058 | 299223 | 165 | 0,185221438 | 0,211234566 | 0,46719292 | -0,082328063 | -0,069978854 | 0,071191272 | -0,075808728 | -1,335808728 |
| chrVIII | 299773 | 299910 | 137 | 2,348037921 | 1,949287398 | 0,546395199 | 0,116558907 | 0,099075071 | 0,249105497 | 0,298105497 | -0,961894503 |
| chrVIII | 310121 | 310260 | 139 | 0,091185939 | 0,040764565 | 0,691061693 | 0,498861988 | 0,42403269 | 0,565599023 | 0,600599023 | -0,659400977 |
| chrVIII | 310839 | 310998 | 159 | 0,333398588 | 0,422469132 | 0,441080601 | -0,14823007 | -0,125995559 | -0,039089165 | -0,144089165 | -1,404089165 |
| chrVIII | 311969 | 312130 | 161 | 0,393239361 | 0,411351523 | 0,488744489 | -0,028217126 | -0,023984557 | 0,072509865 | -0,046490135 | -1,306490135 |
| chrVIII | 333005 | 333157 | 152 | 0,592708601 | 0,396528045 | 0,599157546 | 0,251167115 | 0,213492048 | 0,145014874 | 0,089014874 | -1,170985126 |
| chrVIII | 334511 | 334659 | 148 | 0,461628815 | 0,337234132 | 0,577857337 | 0,196415039 | 0,166952783 | 0,049415472 | 0,021415472 | -1,238584528 |
| chrVIII | 341021 | 341173 | 152 | 0,054141651 | 0,08894087 | 0,378394585 | -0,309699913 | -0,263242926 | -0,328478449 | -0,384478449 | -1,644478449 |
| chrVIII | 341021 | 341178 | 157 | 3,382428413 | 3,346400229 | 0,502677151 | 0,006710673 | 0,005704072 | 0,038610573 | -0,052389427 | -1,312389427 |
| chrVIII | 342001 | 342158 | 157 | 2,852410144 | 2,360638922 | 0,547167331 | 0,118507771 | 0,100731606 | 0,142700054 | 0,051700054 | -1,208299946 |
| chrVIII | 342217 | 342368 | 151 | 0,151026711 | 0,17417587 | 0,464408094 | -0,089334359 | -0,075934205 | -0,171422533 | -0,220422533 | -1,480422533 |
| chrVIII | 343838 | 343999 | 161 | 0,433133209 | 0,496586524 | 0,465875031 | -0,085643193 | -0,072796714 | 0,048423088 | -0,070576912 | -1,330576912 |
| chrVIII | 345951 | 346111 | 160 | 0,037044288 | 0,062999783 | 0,370279692 | -0,331112667 | -0,281445767 | -0,192221048 | -0,304221048 | -1,564221048 |
| chrVIII | 346434 | 346582 | 148 | 0,09973462 | 0,159352392 | 0,38494643 | -0,292515043 | -0,248637787 | -0,374165903 | -0,402165903 | -1,662165903 |
| chrVIII | 350191 | 350346 | 155 | 0,786478721 | 0,904232177 | 0,465176348 | -0,087401099 | -0,074290934 | -0,049031371 | -0,126031371 | -1,386031371 |
| chrVIII | 353263 | 353415 | 152 | 0,085486818 | 0,111176087 | 0,434687048 | -0,164453569 | -0,139785534 | -0,206945572 | -0,262945572 | -1,522945572 |
| chrVIII | 354917 | 355076 | 159 | 0,051292091 | 0,085235 | 0,375691668 | -0,31681585 | -0,269293473 | -0,183652808 | -0,288652808 | -1,548652808 |
| chrVIII | 355941 | 356089 | 148 | 3,915296242 | 2,712696532 | 0,59072126 | 0,229400715 | 0,194990608 | 0,07923795 | 0,05123795 | -1,20876205 |
| chrVIII | 357549 | 357703 | 154 | 2,040285378 | 1,1451137 | 0,6405117 | 0,35982688 | 0,305852848 | 0,290343432 | 0,220343432 | -1,039656568 |
| chrVIII | 360834 | 360982 | 148 | 0,453080133 | 0,522527611 | 0,464408094 | -0,089334359 | -0,075934205 | -0,170143089 | -0,198143089 | -1,458143089 |
| chrVIII | 364130 | 364290 | 160 | 0,045592969 | 0,044470435 | 0,506231912 | 0,015621722 | 0,013278464 | 0,100653733 | -0,011346267 | -1,271346267 |
| chrVIII | 370648 | 370829 | 181 | 0,074088575 | 0,107470218 | 0,408069331 | -0,232514197 | -0,197637067 | 0,095693482 | -0,163306518 | -1,423306518 |
| chrVIII | 371736 | 371883 | 147 | 0,031345166 | 0,040764565 | 0,434687048 | -0,164453569 | -0,139785534 | -0,223276641 | -0,244276641 | -1,504276641 |
| chrVIII | 374061 | 374221 | 160 | 6,132254376 | 5,506922194 | 0,526863248 | 0,067387144 | 0,057279073 | 0,147601807 | 0,035601807 | -1,224398193 |
| chrVIII | 374061 | 374221 | 160 | 6,132254376 | 5,506922194 | 0,526863248 | 0,067387144 | 0,057279073 | 0,147601807 | 0,035601807 | -1,224398193 |
| chrVIII | 374061 | 374255 | 194 | 0,056991212 | 0,070411522 | 0,447331152 | -0,132407093 | -0,112546029 | 0,237900122 | -0,112099878 | -1,372099878 |
| chrVIII | 374061 | 374255 | 194 | 0,056991212 | 0,070411522 | 0,447331152 | -0,132407093 | -0,112546029 | 0,237900122 | -0,112099878 | -1,372099878 |
| chrVIII | 374122 | 374209 | 87 | 0,068389454 | 0,062999783 | 0,520510322 | 0,051434423 | 0,043719259 | 0,672040791 | 1,071040791 | -0,188959209 |
| chrVIII | 374122 | 374221 | 99 | 0,262159574 | 0,274234349 | 0,488744489 | -0,028217126 | -0,023984557 | 0,67632298 | 0,99132298 | -0,26867702 |
| chrVIII | 374122 | 374255 | 133 | 1,396284686 | 1,652817832 | 0,457933007 | -0,105642485 | -0,089796113 | 0,228613066 | 0,305613066 | -0,954386934 |
| chrVIII | 376459 | 376622 | 163 | 0,185221438 | 0,237175653 | 0,438500742 | -0,154771469 | -0,131555749 | 0,025805197 | -0,107194803 | -1,367194803 |
| chrVIII | 376459 | 376622 | 163 | 0,185221438 | 0,237175653 | 0,438500742 | -0,154771469 | -0,131555749 | 0,025805197 | -0,107194803 | -1,367194803 |
| chrVIII | 381490 | 381641 | 151 | 0,039893848 | 0,055588044 | 0,417815854 | -0,20748423 | -0,176361595 | -0,282911886 | -0,331911886 | -1,591911886 |
| chrVIII | 382542 | 382702 | 160 | 0,128230226 | 0,170470001 | 0,429294037 | -0,178171777 | -0,15144601 | -0,064106299 | -0,176106299 | -1,436106299 |
| chrVIII | 382976 | 383151 | 175 | 0,094035499 | 0,103764348 | 0,47540734 | -0,061683752 | -0,052431189 | 0,230660011 | 0,013660011 | -1,246339989 |
| chrVIII | 399609 | 399757 | 148 | 0,19377012 | 0,118587827 | 0,620346375 | 0,30639062 | 0,260432027 | 0,168686882 | 0,140686882 | -1,119313118 |

| Chrom | Start | End | Length | Section A | Section B | A/A+B | Z-score | Z * 0.85 | Phase correction | Length correction | ΔLK |
|---|---|---|---|---|---|---|---|---|---|---|---|
| chrVIII | 402529 | 402689 | 160 | 0,173823196 | 0,185293479 | 0,484029865 | -0,040041889 | -0,034035606 | 0,053157064 | -0,058842936 | -1,318842936 |
| chrVIII | 402529 | 402689 | 160 | 0,173823196 | 0,185293479 | 0,484029865 | -0,040041889 | -0,034035606 | 0,053157064 | -0,058842936 | -1,318842936 |
| chrVIII | 403584 | 403737 | 153 | 0,421734966 | 0,389116306 | 0,520113837 | 0,050439292 | 0,042873398 | -0,002040433 | -0,065040433 | -1,325040433 |
| chrVIII | 404097 | 404230 | 133 | 0,302053422 | 0,377998697 | 0,444162166 | -0,140424842 | -0,119361116 | 0,19363348 | 0,27063348 | -0,98936652 |
| chrVIII | 414157 | 414333 | 176 | 0,085486818 | 0,166764131 | 0,338895921 | -0,415478239 | -0,353156503 | -0,075454807 | -0,299454807 | -1,559454807 |
| chrVIII | 414157 | 414333 | 176 | 0,085486818 | 0,166764131 | 0,338895921 | -0,415478239 | -0,353156503 | -0,075454807 | -0,299454807 | -1,559454807 |
| chrVIII | 416537 | 416675 | 138 | 5,864395681 | 3,794810448 | 0,607130193 | 0,271847066 | 0,231070006 | 0,389508758 | 0,431508758 | -0,828491242 |
| chrVIII | 424973 | 425127 | 154 | 0,182371877 | 0,133411305 | 0,577522451 | 0,195559327 | 0,166225428 | 0,159062132 | 0,089062132 | -1,170937868 |
| chrVIII | 424973 | 425127 | 154 | 0,182371877 | 0,133411305 | 0,577522451 | 0,195559327 | 0,166225428 | 0,159062132 | 0,089062132 | -1,170937868 |
| chrVIII | 427961 | 428122 | 161 | 0,056991212 | 0,062999783 | 0,474962408 | -0,062801193 | -0,053381014 | 0,081986683 | -0,037013317 | -1,297013317 |
| chrVIII | 431042 | 431168 | 126 | 0,082637257 | 0,100058479 | 0,452321762 | -0,11979755 | -0,101827917 | 0,422001132 | 0,548001132 | -0,711998868 |
| chrVIII | 433338 | 433488 | 150 | 0,068389454 | 0,059293913 | 0,535617563 | 0,08939893 | 0,07598909 | -0,026091826 | -0,068091826 | -1,328091826 |
| chrVIII | 439467 | 439624 | 157 | 0,364743755 | 0,259410871 | 0,58438044 | 0,213112623 | 0,181145729 | 0,231309493 | 0,140309493 | -1,119690507 |
| chrVIII | 444830 | 444988 | 158 | 1,003045325 | 1,352642396 | 0,425797238 | -0,187084379 | -0,159021722 | -0,061465552 | -0,159465552 | -1,419465552 |
| chrVIII | 447873 | 448000 | 127 | 0,344796831 | 1,030231743 | 0,25075612 | -0,672112246 | -0,571295409 | -0,032966999 | 0,086033001 | -1,173966999 |
| chrVIII | 456058 | 456162 | 104 | 0,094035499 | 1,371171744 | 0,064178975 | -1,520609146 | -1,292517774 | -0,579025913 | -0,299025913 | -1,559025913 |
| chrVIII | 465568 | 465728 | 160 | 0,746584873 | 0,681880002 | 0,522648394 | 0,056801634 | 0,048281389 | 0,130321915 | 0,018321915 | -1,241678085 |
| chrVIII | 469779 | 469934 | 155 | 0,062690333 | 0,081529131 | 0,434687048 | -0,164453569 | -0,139785534 | -0,124060373 | -0,201060373 | -1,461060373 |
| chrVIII | 469928 | 470081 | 153 | 0,225115286 | 0,144528914 | 0,609005325 | 0,276727507 | 0,235218381 | 0,197310974 | 0,134310974 | -1,125689026 |
| chrVIII | 469928 | 470081 | 153 | 0,225115286 | 0,144528914 | 0,609005325 | 0,276727507 | 0,235218381 | 0,197310974 | 0,134310974 | -1,125689026 |
| chrVIII | 470995 | 471124 | 129 | 1,433328974 | 1,560171093 | 0,478813744 | -0,053131056 | -0,045161397 | 0,42932635 | 0,53432635 | -0,72567365 |
| chrVIII | 482780 | 482943 | 163 | 2,023188014 | 1,715817615 | 0,541103228 | 0,103213476 | 0,087731455 | 0,241427544 | 0,108427544 | -1,151572456 |
| chrVIII | 485160 | 485318 | 158 | 0,652549374 | 0,418763262 | 0,609111992 | 0,277005326 | 0,235454527 | 0,326465982 | 0,228465982 | -1,031534018 |
| chrVIII | 485160 | 485318 | 158 | 0,652549374 | 0,418763262 | 0,609111992 | 0,277005326 | 0,235454527 | 0,326465982 | 0,228465982 | -1,031534018 |
| chrVIII | 495830 | 495995 | 165 | 0,253610892 | 0,289057827 | 0,467340171 | -0,081957711 | -0,069664054 | 0,085022425 | -0,061977575 | -1,321977575 |
| chrVIII | 497750 | 497909 | 159 | 0,131079787 | 0,133411305 | 0,495592445 | -0,011048326 | -0,009391077 | 0,077665771 | -0,027334229 | -1,287334229 |
| chrVIII | 501432 | 501582 | 150 | 0,139628469 | 0,196411088 | 0,415512001 | -0,213388432 | -0,181380167 | -0,281211409 | -0,323211409 | -1,583211409 |
| chrVIII | 503721 | 503876 | 155 | 0,92895675 | 0,537351089 | 0,6335346 | 0,341229522 | 0,290045094 | 0,30620918 | 0,22920918 | -1,03079082 |
| chrVIII | 510274 | 510423 | 149 | 0,14817715 | 0,111176087 | 0,571333335 | 0,179769754 | 0,152804291 | 0,030477201 | -0,004522799 | -1,264522799 |
| chrVIII | 518475 | 518638 | 163 | 2,883755311 | 2,482932618 | 0,537343581 | 0,093743597 | 0,079682057 | 0,218661863 | 0,085661863 | -1,174338137 |
| chrVIII | 518475 | 518638 | 163 | 2,883755311 | 2,482932618 | 0,537343581 | 0,093743597 | 0,079682057 | 0,218661863 | 0,085661863 | -1,174338137 |
| chrVIII | 521779 | 521913 | 134 | 1,134125112 | 1,66764131 | 0,404789315 | -0,240969661 | -0,204824212 | 0,038768173 | 0,108768173 | -1,151231827 |
| chrVIII | 522442 | 522595 | 153 | 0,333398588 | 0,314998914 | 0,514188576 | 0,035572986 | 0,030237038 | 0,003486603 | -0,059513397 | -1,319513397 |
| chrVIII | 523030 | 523182 | 152 | 0,48727486 | 0,292763697 | 0,62468048 | 0,31779685 | 0,270127323 | 0,209105386 | 0,153105386 | -1,106894614 |
| chrVIII | 523197 | 523325 | 128 | 0,917558508 | 1,560171093 | 0,370322293 | -0,330999866 | -0,281349886 | 0,208982526 | 0,320982526 | -0,939017474 |
| chrVIII | 525186 | 525342 | 156 | 0,142478029 | 0,133411305 | 0,516431814 | 0,041200102 | 0,035020087 | 0,053047378 | -0,030952622 | -1,290952622 |
| chrVIII | 526030 | 526198 | 168 | 0,168124074 | 0,118587827 | 0,586386801 | 0,218260181 | 0,185521153 | 0,342288972 | 0,174288972 | -1,085711028 |
| chrVIII | 526169 | 526320 | 151 | 2,849560584 | 1,482347831 | 0,657807209 | 0,406485945 | 0,345513053 | 0,248331216 | 0,199331216 | -1,060668784 |
| chrVIII | 526293 | 526441 | 148 | 5,200448065 | 3,60210523 | 0,590788592 | 0,229573995 | 0,195137895 | 0,086484539 | 0,058484539 | -1,201515461 |
| chrVIII | 526293 | 526481 | 188 | 0,042743409 | 0,044470435 | 0,490098899 | -0,024820929 | -0,021097789 | 0,313288137 | 0,005288137 | -1,254711863 |
| chrVIII | 526600 | 526706 | 106 | 0,054141651 | 0,092646739 | 0,368841506 | -0,334923211 | -0,284684729 | 0,356354326 | 0,622354326 | -0,637645674 |
| chrVIII | 526600 | 526743 | 143 | 0,076938136 | 0,096352609 | 0,443982948 | -0,140878539 | -0,119746758 | -0,042753797 | -0,035753797 | -1,295753797 |
| chrVIII | 526600 | 526749 | 149 | 0,074088575 | 0,051882174 | 0,588141101 | 0,222765786 | 0,189350918 | 0,089191411 | 0,054191411 | -1,205808589 |
| chrVIII | 526600 | 526751 | 151 | 4,830005189 | 2,99434262 | 0,61730451 | 0,298409053 | 0,253647695 | 0,155853097 | 0,106853097 | -1,153146903 |
| chrVIII | 526624 | 526751 | 127 | 0,09688506 | 0,100058479 | 0,491943329 | -0,020196453 | -0,017166985 | 0,495579096 | 0,614579096 | -0,645420904 |
| chrVIII | 526651 | 526751 | 100 | 0,111132863 | 0,092646739 | 0,54535813 | 0,113942039 | 0,096850733 | 0,824320311 | 1,132320311 | -0,127679689 |
| chrVIII | 527864 | 528038 | 174 | 0,49012442 | 0,87829109 | 0,358169296 | -0,3633565 | -0,308853025 | -0,048700927 | -0,258700927 | -1,518700927 |
| chrVIII | 536044 | 536157 | 113 | 0,054141651 | 1,004290656 | 0,051152682 | -1,633778522 | -1,388711744 | -0,92602132 | -0,70902132 | -1,96902132 |
| chrVIII | 538480 | 538617 | 137 | 5,089315202 | 3,457576317 | 0,595458032 | 0,241607982 | 0,205366785 | 0,358582012 | 0,407582012 | -0,852417988 |
| chrVIII | 539313 | 539458 | 145 | 2,305294512 | 1,356348266 | 0,629579304 | 0,33073933 | 0,28112843 | 0,27798331 | 0,27098331 | -0,98901669 |
| chrVIII | 539340 | 539488 | 148 | 0,427434088 | 0,222352175 | 0,657807209 | 0,406485945 | 0,345513053 | 0,232270157 | 0,204270157 | -1,055729843 |
| chrVIII | 542782 | 542926 | 144 | 1,923453394 | 1,378583483 | 0,582505122 | 0,208306373 | 0,177060417 | 0,236907971 | 0,236907971 | -1,023092029 |
| chrVIII | 542782 | 542926 | 144 | 1,923453394 | 1,378583483 | 0,582505122 | 0,208306373 | 0,177060417 | 0,236907971 | 0,236907971 | -1,023092029 |
| chrVIII | 542989 | 543156 | 167 | 0,473027057 | 0,563292176 | 0,456449173 | -0,109383467 | -0,092975947 | 0,066404914 | -0,094595086 | -1,354595086 |
| chrVIII | 543353 | 543457 | 104 | 0,142478029 | 0,715232829 | 0,166114289 | -0,969634765 | -0,82418955 | -0,132766739 | 0,147233261 | -1,112766739 |
| chrVIII | 543780 | 543928 | 148 | 0,028495606 | 0,044470435 | 0,390532439 | -0,277931585 | -0,236241848 | -0,353852386 | -0,381852386 | -1,641852386 |
| chrVIII | 544540 | 544654 | 114 | 0,065539893 | 0,474351306 | 0,121394632 | -1,168043388 | -0,99283688 | -0,534603356 | -0,324603356 | -1,584603356 |
| chrVIII | 546632 | 546789 | 157 | 0,042743409 | 0,048176305 | 0,470122564 | -0,074961771 | -0,063717505 | -0,01663172 | -0,10763172 | -1,36763172 |
| chrVIII | 546818 | 546968 | 150 | 8,981814959 | 4,031986102 | 0,690176138 | 0,496349676 | 0,421897225 | 0,312624047 | 0,270624047 | -0,989375953 |
| chrVIII | 546822 | 546968 | 146 | 0,056991212 | 0,048176305 | 0,541908887 | 0,105243965 | 0,08945737 | 0,092381253 | 0,078381253 | -1,181618747 |
| chrVIII | 547280 | 547415 | 135 | 0,085486818 | 0,059293913 | 0,590457149 | 0,22872109 | 0,194412926 | 0,427683275 | 0,490683275 | -0,769316725 |
| chrVIII | 548318 | 548458 | 140 | 0,139628469 | 0,129705435 | 0,518421434 | 0,046192108 | 0,039263292 | 0,16143811 | 0,18943811 | -1,07056189 |
| chrVIII | 548324 | 548458 | 134 | 0,139628469 | 0,125999566 | 0,525654112 | 0,064349707 | 0,054697251 | 0,292030714 | 0,362030714 | -0,897969286 |
| chrVIII | 548850 | 549005 | 155 | 0,102584181 | 0,081529131 | 0,557179598 | 0,143822286 | 0,122248943 | 0,143861795 | 0,066861795 | -1,193138205 |
| chrVIII | 548981 | 549089 | 108 | 0,065539893 | 0,192705218 | 0,253789483 | -0,662612181 | -0,563220354 | 0,09964897 | 0,35164897 | -0,90835103 |
| chrVIII | 549893 | 550056 | 163 | 0,19377012 | 0,200116957 | 0,491943329 | -0,020196453 | -0,017166985 | 0,113683695 | -0,019316305 | -1,279316305 |
| chrVIII | 550702 | 550810 | 108 | 0,09973462 | 0,277940218 | 0,264075364 | -0,630831465 | -0,536206745 | 0,096643465 | 0,348643465 | -0,911356535 |
| chrVIII | 551514 | 551634 | 120 | 0,24221265 | 0,733762177 | 0,2481751 | -0,680243648 | -0,578207101 | -0,021007121 | 0,146992879 | -1,113007121 |
| chrVIII | 553179 | 553326 | 147 | 1,333594353 | 0,70411522 | 0,65445752 | 0,397383121 | 0,337775653 | 0,264992015 | 0,243992015 | -1,016007985 |
| chrVIII | 553630 | 553778 | 148 | 0,347646391 | 0,300175436 | 0,536638898 | 0,091969585 | 0,078174148 | -0,044517702 | -0,072517702 | -1,332517702 |
| chrVIII | 556401 | 556536 | 135 | 0,840620372 | 0,718938698 | 0,539011563 | 0,097943858 | 0,083252279 | 0,284767935 | 0,347767935 | -0,912232065 |
| chrVIII | 556826 | 556854 | 28 | 0,034194727 | 0,552174567 | 0,058316026 | -1,569068185 | -1,333707957 | | 0,812 | -0,448 |
| chrVIII | 556826 | 556859 | 33 | 0,068389454 | 0,415057393 | 0,141462199 | -1,073773081 | -0,912707119 | | 0,777 | -0,483 |
| chrVIII | 556826 | 556865 | 39 | 0,042743409 | 0,515115871 | 0,076620414 | -1,428177319 | -1,213950721 | | 0,735 | -0,525 |
| chrVIII | 556826 | 556871 | 45 | 0,051292091 | 0,485468915 | 0,095558526 | -1,307281714 | -1,111189457 | | 0,693 | -0,567 |
| chrVIII | 556826 | 556873 | 47 | 0,045592969 | 0,35576348 | 0,113597201 | -1,207617557 | -1,026474924 | | 0,679 | -0,581 |
| chrVIII | 556826 | 556876 | 50 | 0,04844253 | 0,518821741 | 0,085396758 | -1,369658539 | -1,164209758 | | 0,658 | -0,602 |
| chrVIII | 556826 | 556884 | 58 | 0,04844253 | 0,452116089 | 0,096776937 | -1,300137421 | -1,105116808 | | 0,602 | -0,658 |
| chrVIII | 556826 | 556910 | 84 | 0,094035499 | 0,648527176 | 0,126636447 | -1,142435854 | -0,971070476 | -0,386387911 | 0,033612089 | -1,226387911 |
| chrVIII | 556826 | 556914 | 88 | 0,139628469 | 0,326116523 | 0,299795964 | -0,524987432 | -0,446239318 | 0,198488658 | 0,590488658 | -0,669511342 |
| chrVIII | 556826 | 556918 | 92 | 0,051292091 | 0,34835174 | 0,128344507 | -1,134251626 | -0,964113882 | -0,283094785 | 0,080905215 | -1,179094785 |
| chrVIII | 556826 | 556919 | 93 | 0,575611238 | 0,996878917 | 0,366050774 | -0,342331347 | -0,290981645 | 0,348099827 | 0,705099827 | -0,554900173 |
| chrVIII | 556826 | 556921 | 95 | 0,068389454 | 0,188999349 | 0,265704853 | -0,625855528 | -0,531977199 | 0,114882628 | 0,457882628 | -0,802117372 |
| chrVIII | 556826 | 556923 | 97 | 0,056991212 | 0,111176087 | 0,338895921 | -0,415478239 | -0,353156503 | 0,278463196 | 0,607463196 | -0,652536804 |
| chrVIII | 556826 | 556926 | 100 | 0,14532759 | 0,307587175 | 0,320871831 | -0,465262255 | -0,395472916 | 0,320019364 | 0,628019364 | -0,631980636 |
| chrVIII | 556826 | 556930 | 104 | 0,068389454 | 0,300175436 | 0,185556074 | -0,894392081 | -0,760233269 | -0,079931527 | 0,200068473 | -1,059931527 |
| chrVIII | 556826 | 556935 | 109 | 0,151026711 | 0,17788174 | 0,459175526 | -0,102511037 | -0,087134382 | 0,518483926 | 0,763483926 | -0,496516074 |
| chrVIII | 556826 | 556936 | 110 | 0,558513874 | 0,415057393 | 0,573675388 | 0,185739279 | 0,157878387 | 0,75138264 | 0,98938264 | -0,27061736 |
| chrVIII | 556826 | 556939 | 113 | 0,302053422 | 0,370586958 | 0,449056332 | -0,12804588 | -0,108838998 | 0,357335394 | 0,574335394 | -0,685664606 |
| chrVIII | 556826 | 556942 | 116 | 0,188070999 | 0,474351306 | 0,283914049 | -0,571253086 | -0,485565123 | 0,024748151 | 0,220748151 | -1,039251849 |
| chrVIII | 556826 | 556944 | 118 | 0,059840772 | 0,222352175 | 0,212056229 | -0,799306935 | -0,679410895 | -0,202593646 | -0,020593646 | -1,280593646 |
| chrVIII | 556826 | 556950 | 124 | 0,413186285 | 0,474351306 | 0,465542292 | -0,086480339 | -0,073508288 | 0,483067027 | 0,623067027 | -0,636932973 |
| chrVIII | 556826 | 556954 | 128 | 0,04844253 | 0,289057827 | 0,14353327 | -1,064578883 | -0,90489205 | -0,412902706 | -0,300902706 | -1,560902706 |
| chrVIII | 556826 | 556956 | 130 | 0,028495606 | 0,207528696 | 0,12073166 | -1,171337061 | -0,995636502 | -0,540452052 | -0,442452052 | -1,702452052 |
| chrVIII | 556826 | 556961 | 135 | 0,09688506 | 0,192705218 | 0,33455909 | -0,427358568 | -0,363254783 | -0,155952049 | -0,092952049 | -1,352952049 |
| chrVIII | 556826 | 556963 | 137 | 0,168124074 | 0,170470001 | 0,496535784 | -0,008683612 | -0,00738107 | 0,150892802 | 0,199892802 | -1,060107198 |
| chrVIII | 556826 | 556965 | 139 | 0,105433742 | 0,207528696 | 0,336889444 | -0,4209674 | -0,35782229 | -0,215694378 | -0,180694378 | -1,440694378 |
| chrVIII | 556826 | 556967 | 141 | 0,125380666 | 0,222352175 | 0,360566076 | -0,356946107 | -0,303404191 | -0,184598416 | -0,163598416 | -1,423598416 |
| chrVIII | 556826 | 556969 | 143 | 0,122531105 | 0,133411305 | 0,478744828 | -0,053304046 | -0,045308439 | 0,044334928 | 0,051334928 | -1,208665072 |
| chrVIII | 556826 | 556970 | 144 | 0,051292091 | 0,244587392 | 0,173354671 | -0,940991251 | -0,799842563 | -0,75641571 | -0,75641571 | -2,01641571 |
| chrVIII | 556826 | 556971 | 145 | 0,059840772 | 0,148234783 | 0,287591554 | -0,560434497 | -0,476369322 | -0,468863071 | -0,475863071 | -1,735863071 |

TABLE 2

| Chrom | Start | End | Length | Section A | Section B | A/A+B | Z-score | Z * 0.85 | Phase correction | Length correction | ΔLK |
|---|---|---|---|---|---|---|---|---|---|---|---|
| chrVIII | 556826 | 556972 | 146 | 0,042743409 | 0,233469783 | 0,15474789 | -1,016280608 | -0,863838517 | -0,862743975 | -0,876743975 | **-2,136743975** |
| chrVIII | 556826 | 556973 | 147 | 0,826372569 | 1,222936961 | 0,403244389 | -0,244958225 | -0,208214491 | -0,278662379 | -0,299662379 | **-1,559662379** |
| chrVIII | 556826 | 556975 | 149 | 1,02584181 | 0,644821307 | 0,614032715 | 0,289845327 | 0,246368528 | 0,142664434 | 0,107664434 | **-1,152335566** |
| chrVIII | 556826 | 556976 | 150 | 0,034194727 | 0,251999131 | 0,119480995 | -1,177585257 | -1,000947468 | -1,103678773 | -1,145678773 | **-2,405678773** |
| chrVIII | 556826 | 556977 | 151 | 0,059840772 | 0,155646522 | 0,277699771 | -0,589688448 | -0,501235181 | -0,589001685 | -0,638001685 | **-1,898001685** |
| chrVIII | 556826 | 556979 | 153 | 0,265009134 | 0,429880871 | 0,381368464 | -0,301888676 | -0,256605374 | -0,281003431 | -0,344003431 | **-1,604003431** |
| chrVIII | 556826 | 556981 | 155 | 0,059840772 | 0,292763697 | 0,16971076 | -0,955308878 | -0,812012546 | -0,806811282 | -0,883811282 | **-2,143811282** |
| chrVIII | 556826 | 556982 | 156 | 0,473027057 | 0,637409568 | 0,425982939 | -0,186610698 | -0,158619094 | -0,139601099 | -0,223601099 | **-1,483601099** |
| chrVIII | 556826 | 556983 | 157 | 0,455929693 | 0,948702612 | 0,324590066 | -0,454901459 | -0,386666241 | -0,328147984 | -0,419147984 | **-1,679147984** |
| chrVIII | 556826 | 556984 | 158 | 0,213717044 | 0,214940436 | 0,498572996 | -0,003576976 | -0,00304043 | 0,09010487 | -0,00789513 | **-1,26789513** |
| chrVIII | 556826 | 556985 | 159 | 0,14532759 | 0,26311674 | 0,355807583 | -0,369687743 | -0,314234581 | -0,217692098 | -0,322692098 | **-1,582692098** |
| chrVIII | 556826 | 556986 | 160 | 4,912642446 | 11,15466743 | 0,305753887 | -0,507922388 | -0,431734029 | -0,352828859 | -0,464828859 | **-1,724828859** |
| chrVIII | 556826 | 556987 | 161 | 1,592904366 | 1,604641528 | 0,498164661 | -0,004600529 | -0,00391045 | 0,148691623 | 0,029691623 | **-1,230308377** |
| chrVIII | 556826 | 556989 | 163 | 22,00430683 | 12,47766287 | 0,638139498 | 0,35349016 | 0,300466636 | 0,438092727 | 0,305092727 | **-0,954907273** |
| chrVIII | 556833 | 556986 | 153 | 0,042743409 | 0,092646739 | 0,315705458 | -0,47974192 | -0,407780632 | -0,430257145 | -0,493257145 | **-1,753257145** |
| chrVIII | 556840 | 556986 | 146 | 0,074088575 | 0,040764565 | 0,645072262 | 0,372050197 | 0,316242667 | 0,305541745 | 0,291541745 | **-0,968458255** |
| chrVIII | 556843 | 556989 | 146 | 0,071239015 | 0,114881957 | 0,382756516 | -0,298249123 | -0,253511755 | -0,288075941 | -0,302075941 | **-1,562075941** |
| chrVIII | 556853 | 556986 | 133 | 0,031345166 | 0,051882174 | 0,376621026 | -0,314367347 | -0,267212245 | 0,040958895 | 0,117958895 | **-1,142041105** |
| chrVIII | 556853 | 556989 | 136 | 0,259310013 | 0,196411088 | 0,569010328 | 0,173855096 | 0,147776831 | 0,329398965 | 0,385398965 | **-0,874601035** |
| chrVIII | 556858 | 556975 | 117 | 0,04844253 | 0,037058696 | 0,566571175 | 0,16765125 | 0,142503563 | 0,662751799 | 0,851751799 | **-0,408248201** |
| chrVIII | 556858 | 556989 | 131 | 0,105433742 | 0,159352392 | 0,398184528 | -0,258049045 | -0,219341688 | 0,164669993 | 0,255669993 | **-1,004330007** |
| chrVIII | 556859 | 556963 | 104 | 0,102584181 | 0,062999783 | 0,619529685 | 0,304245806 | 0,258608935 | 0,912651864 | 1,192651864 | **-0,067348136** |
| chrVIII | 556859 | 556989 | 130 | 0,09688506 | 0,114881957 | 0,45750779 | -0,106714373 | -0,090707217 | 0,369016972 | 0,467016972 | **-0,792983028** |
| chrVIII | 556863 | 556986 | 123 | 0,04844253 | 0,200116957 | 0,194893104 | -0,860005143 | -0,731004372 | -0,153982049 | -0,006982049 | **-1,266982049** |
| chrVIII | 556869 | 556986 | 117 | 0,059840772 | 0,151940653 | 0,282559116 | -0,575255914 | -0,488967527 | 0,022228005 | 0,211228005 | **-1,048771995** |
| chrVIII | 556878 | 556973 | 95 | 0,042743409 | 0,248293262 | 0,146866059 | -1,04996954 | -0,892474109 | -0,254114788 | 0,088885212 | **-1,171114788** |
| chrVIII | 556878 | 556975 | 97 | 0,031345166 | 0,129705435 | 0,194629303 | -0,860962664 | -0,731818264 | -0,071943975 | 0,257056025 | **-1,002943975** |
| chrVIII | 556878 | 556983 | 105 | 0,028495606 | 0,08894087 | 0,242646977 | -0,69781331 | -0,593141313 | 0,034102657 | 0,307102657 | **-0,952897343** |
| chrVIII | 556878 | 556989 | 111 | 0,641151131 | 0,570703915 | 0,529065859 | 0,072921881 | 0,061983599 | 0,672797816 | 0,903797816 | **-0,356202184** |
| chrVIII | 556883 | 556989 | 106 | 0,521469587 | 0,478057176 | 0,521716483 | 0,054462061 | 0,046292752 | 0,718549573 | 0,984549573 | **-0,275450427** |
| chrVIII | 556887 | 556986 | 99 | 0,168124074 | 0,333528262 | 0,335140619 | -0,425762057 | -0,361897749 | 0,357970238 | 0,672970238 | **-0,587029762** |
| chrVIII | 556893 | 556975 | 82 | 0,054141651 | 0,066705652 | 0,448017039 | -0,130672891 | -0,111071958 | 0,486371872 | 0,920371872 | **-0,339628128** |
| chrVIII | 556893 | 556989 | 96 | 0,105433742 | 0,08894087 | 0,542425478 | 0,106546147 | 0,090564225 | 0,711059436 | 1,047059436 | **-0,212940564** |
| chrVIII | 556901 | 556986 | 85 | 0,059840772 | 0,340940001 | 0,149310486 | -1,039395206 | -0,883485925 | -0,30958262 | 0,10341738 | **-1,15658262** |
| chrVIII | 556904 | 556989 | 85 | 0,407487163 | 0,366881088 | 0,52621884 | 0,065768269 | 0,055903029 | 0,669233531 | 1,082233531 | **-0,177766469** |
| chrVIII | 558701 | 558838 | 137 | 0,504372223 | 0,470645436 | 0,517295475 | 0,043366916 | 0,036861878 | 0,198654037 | 0,247654037 | **-1,012345963** |
| chrVIII | 558986 | 559129 | 143 | 0,401788042 | 0,344645871 | 0,538276779 | 0,096093338 | 0,081679337 | 0,190870383 | 0,197870383 | **-1,062129617** |
| chrVIII | 559259 | 559411 | 152 | 0,105433742 | 0,059293913 | 0,640048823 | 0,358589298 | 0,304800903 | 0,23836908 | 0,18236908 | **-1,07763092** |
| chrVIII | 559332 | 559476 | 144 | 0,09688506 | 0,048176305 | 0,667890174 | 0,43409473 | 0,36898052 | 0,396777586 | 0,396777586 | **-0,863222414** |
| chrVIII | 560157 | 560308 | 151 | 0,698142343 | 0,470645436 | 0,597321734 | 0,246420767 | 0,209457652 | 0,132836544 | 0,083836544 | **-1,176163456** |
| chrVIII | 560416 | 560563 | 147 | 0,094035499 | 0,096352609 | 0,493914773 | -0,015253994 | -0,012965895 | -0,076717533 | -0,097717533 | **-1,357717533** |
| chrVIII | 560448 | 560563 | 115 | 0,116831984 | 0,107470218 | 0,520868645 | 0,052333815 | 0,044483743 | 0,526085074 | 0,729085074 | **-0,530914926** |
| chrVIII | 560502 | 560671 | 169 | 0,071239015 | 0,296469566 | 0,1937377 | -0,864204794 | -0,734574075 | -0,581890925 | -0,756890925 | **-2,016890925** |
| chrVIII | 560544 | 560634 | 90 | 0,111132863 | 0,188999349 | 0,370279692 | -0,331112667 | -0,281445767 | 0,386635458 | 0,764635458 | **-0,495364542** |
| chrVIII | 560544 | 560639 | 95 | 0,062690333 | 0,103764348 | 0,376621026 | -0,314367347 | -0,267212245 | 0,362524987 | 0,705524987 | **-0,554475013** |
| chrVIII | 560784 | 560911 | 127 | 0,034194727 | 0,055588044 | 0,380860679 | -0,303221116 | -0,257737948 | 0,250530614 | 0,369530614 | **-0,890469386** |
| chrVIII | 561208 | 561356 | 148 | 0,056991212 | 0,055588044 | 0,506231912 | 0,015621722 | 0,013278464 | -0,107654132 | -0,135654132 | **-1,395654132** |
| chrVIII | 561302 | 561476 | 174 | 0,028495606 | 0,055588044 | 0,338895921 | -0,415478239 | -0,353156503 | -0,075644246 | -0,285644246 | **-1,545644246** |
| chrVIII | 561649 | 561800 | 151 | 0,53856695 | 0,35205761 | 0,60470705 | 0,265549866 | 0,225717386 | 0,158021988 | 0,109021988 | **-1,150978012** |
| chrVIII | 562455 | 562594 | 139 | 0,612655525 | 0,066705652 | 0,901811209 | 1,29194099 | 1,098149841 | 1,249984111 | 1,284984111 | **0,024984111** |
| chrVIII | 562455 | 562640 | 185 | 0,176672756 | 0,051882174 | 0,772999103 | 0,748760132 | 0,636446112 | 0,930183756 | 0,643183756 | **-0,616816244** |
| chrVIII | 562469 | 562594 | 125 | 0,136778908 | 0,111176087 | 0,551627959 | 0,129775456 | 0,110309137 | 0,644434514 | 0,777434514 | **-0,482565486** |
| chrVIII | 562469 | 562614 | 145 | 0,028495606 | 0,037058696 | 0,434687048 | -0,164453569 | -0,139785534 | -0,134378939 | -0,141378939 | **-1,401378939** |
| chrVIII | 562469 | 562632 | 163 | 0,039893848 | 0,044470435 | 0,472876041 | -0,06804215 | -0,057835827 | 0,087069844 | -0,045930156 | **-1,305930156** |
| chrVIII | 562469 | 562636 | 167 | 0,168124074 | 0,411351523 | 0,290131414 | -0,553000851 | -0,470050723 | -0,296037227 | -0,457037227 | **-1,717037227** |
| chrVIII | 562469 | 562640 | 171 | 0,136778908 | 0,125999566 | 0,520510322 | 0,051434423 | 0,043719259 | 0,255977844 | 0,066977844 | **-1,193022156** |
| chrVIII | 562500 | 562636 | 136 | 0,233663968 | 0,140823044 | 0,623957468 | 0,315891237 | 0,268507552 | 0,436544841 | 0,492544841 | **-0,767455159** |
| chrVIII | 562506 | 562632 | 126 | 0,059840772 | 0,062999783 | 0,487141826 | -0,032236246 | -0,027400809 | 0,497205419 | 0,623205419 | **-0,636794581** |
| chrVIII | 562506 | 562636 | 130 | 0,14817715 | 0,463233697 | 0,242352832 | -0,698754187 | -0,593941059 | -0,162128897 | -0,064128897 | **-1,324128897** |
| chrVIII | 562506 | 562640 | 134 | 0,256460453 | 0,100058479 | 0,719345959 | 0,58089966 | 0,493764711 | 0,733026293 | 0,803026293 | **-0,456973707** |
| chrVIII | 562520 | 562621 | 101 | 0,367593315 | 0,111176087 | 0,767787819 | 0,731580977 | 0,62184383 | 1,329846173 | 1,630846173 | **0,370846173** |
| chrVIII | 562520 | 562640 | 120 | 0,079787696 | 0,185293479 | 0,300993446 | -0,521545394 | -0,443313585 | 0,117241241 | 0,285241241 | **-0,974758759** |
| chrVIII | 562523 | 562640 | 117 | 0,581310359 | 0,037058696 | 0,940070261 | 1,555363688 | 1,322059135 | 1,842860303 | 2,031860303 | **0,771860303** |
| chrVIII | 562552 | 562640 | 88 | 0,162424953 | 0,248293262 | 0,395465668 | -0,265101417 | -0,225336204 | 0,441643904 | 0,833643904 | **-0,426356096** |
| chrX | 1669 | 1785 | 116 | 0,139628469 | 0,085235 | 0,620947766 | 0,307970909 | 0,261775272 | 0,976875124 | 1,172875124 | **-0,087124876** |
| chrX | 1669 | 1804 | 135 | 2,353737042 | 1,237760439 | 0,65536369 | 0,399842372 | 0,339866016 | 0,361212062 | 0,424212062 | **-0,835787938** |
| chrX | 1669 | 1821 | 152 | 0,304902982 | 0,285351958 | 0,516561509 | 0,041525478 | 0,035296656 | -0,073767415 | -0,129767415 | **-1,389767415** |
| chrX | 1693 | 1785 | 92 | 0,156725832 | 0,17788174 | 0,468386986 | -0,079325188 | -0,06742641 | 0,379492978 | 0,743492978 | **-0,516507022** |
| chrX | 1693 | 1794 | 101 | 0,079787696 | 0,196411088 | 0,288877797 | -0,556666083 | -0,47316617 | 0,109043226 | 0,410043226 | **-0,849956774** |
| chrX | 1693 | 1804 | 111 | 0,416035845 | 0,370586958 | 0,528888615 | 0,072476421 | 0,061604958 | 0,848664579 | 1,079664579 | **-0,180335421** |
| chrX | 1693 | 1821 | 128 | 7,292025533 | 8,378971117 | 0,465319832 | -0,087040066 | -0,073984056 | 0,361257021 | 0,473257021 | **-0,786742979** |
| chrX | 1698 | 1821 | 123 | 0,056991212 | 0,074117392 | 0,434687048 | -0,164453569 | -0,139785534 | 0,450048044 | 0,597048044 | **-0,662951956** |
| chrX | 1980 | 2120 | 140 | 0,068389454 | 0,040764565 | 0,626540868 | 0,322705529 | 0,274299699 | 0,131516735 | 0,159516735 | **-1,100483265** |
| chrX | 2198 | 2335 | 137 | 0,111132863 | 0,081529131 | 0,576828158 | 0,1937857 | 0,164717845 | 0,10435634 | 0,15335634 | **-1,10664366** |
| chrX | 2365 | 2509 | 144 | 0,074088575 | 0,048176305 | 0,605967759 | 0,268824834 | 0,228501109 | 0,053138325 | 0,053138325 | **-1,206861675** |
| chrX | 2365 | 2512 | 147 | 0,04844253 | 0,040764565 | 0,543034495 | 0,108081541 | 0,09186931 | -0,029235743 | -0,050235743 | **-1,310235743** |
| chrX | 2365 | 2513 | 148 | 17,40511604 | 11,19913787 | 0,608479987 | 0,275359539 | 0,234055608 | 0,138148361 | 0,110148361 | **-1,149851639** |
| chrX | 2365 | 2575 | 210 | 0,091185939 | 0,085235 | 0,516865737 | 0,042288735 | 0,035945424 | 0,273164822 | -0,188835178 | **-1,448835178** |
| chrX | 2370 | 2513 | 143 | 0,045592969 | 0,037058696 | 0,551627959 | 0,129775456 | 0,110309137 | -0,074709072 | -0,067709072 | **-1,327709072** |
| chrX | 2380 | 2513 | 133 | 1,088532143 | 1,171054787 | 0,48173944 | -0,04578843 | -0,038920166 | 0,011656548 | 0,088656548 | **-1,171343452** |
| chrX | 2420 | 2513 | 93 | 0,316301225 | 0,459527828 | 0,407694483 | -0,233479654 | -0,198457706 | 0,274149528 | 0,631149528 | **-0,628850472** |
| chrX | 2420 | 2575 | 155 | 0,900461144 | 0,807879568 | 0,52709693 | 0,06797424 | 0,057778104 | 0,004136618 | -0,072863382 | **-1,332863382** |
| chrX | 2862 | 3015 | 153 | 0,83207169 | 0,604056741 | 0,579385292 | 0,200321158 | 0,170272984 | 0,094062119 | 0,031062119 | **-1,228937881** |
| chrX | 3307 | 3451 | 144 | 0,091185939 | 0,051882174 | 0,637360324 | 0,351411905 | 0,298700119 | 0,12560838 | 0,12560838 | **-1,13439162** |
| chrX | 3518 | 3609 | 91 | 0,031345166 | 0,129705435 | 0,194629303 | -0,860962664 | -0,731818264 | -0,341099332 | 0,029900668 | **-1,230099332** |
| chrX | 3518 | 3627 | 109 | 14,86045844 | 43,32902711 | 0,255380475 | -0,657653281 | -0,559005289 | 0,186674252 | 0,431674252 | **-0,828325748** |
| chrX | 3518 | 3628 | 110 | 0,262159574 | 0,800467829 | 0,24670884 | -0,684883118 | -0,582150651 | 0,14610346 | 0,38410346 | **-0,87589654** |
| chrX | 3518 | 3629 | 111 | 0,168124074 | 0,885702829 | 0,159536707 | -0,9963638 | -0,84690923 | -0,076302234 | 0,154697766 | **-1,105302234** |
| chrX | 3518 | 3644 | 126 | 0,034194727 | 2,093816312 | 0,016068867 | -2,142693289 | -1,821289296 | -1,323596047 | -1,197596047 | **-2,457596047** |
| chrX | 3520 | 3627 | 107 | 0,265009134 | 2,427344574 | 0,098430282 | -1,290547693 | -1,096965539 | -0,414407206 | -0,155407206 | **-1,415407206** |
| chrX | 3520 | 3644 | 124 | 0,424584527 | 4,120926971 | 0,093407426 | -1,320060405 | -1,122051344 | -0,599221196 | -0,459221196 | **-1,719221196** |
| chrX | 4071 | 4226 | 155 | 0,074088575 | 0,200116957 | 0,270193583 | -0,612227627 | -0,520393483 | -0,573277336 | -0,650277336 | **-1,910277336** |
| chrX | 4165 | 4327 | 162 | 0,054141651 | 0,070411522 | 0,434687048 | -0,164453569 | -0,139785534 | -0,064790461 | -0,190790461 | **-1,450790461** |
| chrX | 4165 | 4327 | 162 | 0,054141651 | 0,070411522 | 0,434687048 | -0,164453569 | -0,139785534 | -0,064790461 | -0,190790461 | **-1,450790461** |
| chrX | 4788 | 4951 | 163 | 0,253610892 | 0,329822392 | 0,434687048 | -0,164453569 | -0,139785534 | -0,050979978 | -0,183979978 | **-1,443979978** |
| chrX | 5199 | 5350 | 151 | 0,233663968 | 0,125999566 | 0,649673781 | 0,384439894 | 0,326773909 | 0,186702566 | 0,137702566 | **-1,122297434** |
| chrX | 5412 | 5571 | 159 | 0,492973981 | 0,452116089 | 0,521615872 | 0,054209495 | 0,046078071 | 0,043569487 | -0,061430513 | **-1,321430513** |
| chrX | 9273 | 9424 | 151 | 0,492973981 | 0,466939567 | 0,513560812 | 0,033998463 | 0,028898694 | -0,090793187 | -0,139793187 | **-1,399793187** |
| chrX | 10211 | 10354 | 143 | 0,102584181 | 0,055588044 | 0,648560019 | 0,381435717 | 0,324220359 | 0,15281907 | 0,15981907 | **-1,10018093** |
| chrX | 11443 | 11594 | 151 | 0,359044634 | 0,255705001 | 0,584050178 | 0,21226584 | 0,180425964 | 0,057161732 | 0,008161732 | **-1,251838268** |
| chrX | 12328 | 12473 | 145 | 0,279256937 | 0,133411305 | 0,676710512 | 0,458519889 | 0,389741906 | 0,215339177 | 0,208339177 | **-1,051660823** |

TABLE 2

| Chrom | Start | End | Length | Section A | Section B | A/A+B | Z-score | Z * 0.85 | Phase correction | Length correction | ΔLK |
|---|---|---|---|---|---|---|---|---|---|---|---|
| chrX | 12893 | 13042 | 149 | 0,039893848 | 0,048176305 | 0,452978074 | -0,118140734 | -0,100419624 | -0,202918187 | -0,237918187 | **-1,497918187** |
| chrX | 14191 | 14340 | 149 | 0,136778908 | 0,055588044 | 0,711031218 | 0,556399818 | 0,472939845 | 0,368803942 | 0,333803942 | **-0,926196058** |
| chrX | 16003 | 16132 | 129 | 0,133929347 | 0,125999566 | 0,515253751 | 0,038244804 | 0,032508084 | 0,417341776 | 0,522341776 | **-0,737658224** |
| chrX | 16061 | 16168 | 107 | 0,142478029 | 1,708405876 | 0,076978372 | -1,425693812 | -1,211839741 | -0,533038136 | -0,274038136 | **-1,534038136** |
| chrX | 21416 | 21535 | 119 | 0,065539893 | 0,500292393 | 0,115829186 | -1,196097859 | -1,01668318 | -0,375940079 | -0,200940079 | **-1,460940079** |
| chrX | 28854 | 29005 | 151 | 0,330549028 | 0,292763697 | 0,530310091 | 0,076049373 | 0,064641967 | -0,049456578 | -0,098456578 | **-1,358456578** |
| chrX | 28854 | 29005 | 151 | 0,330549028 | 0,292763697 | 0,530310091 | 0,076049373 | 0,064641967 | -0,049456578 | -0,098456578 | **-1,358456578** |
| chrX | 30398 | 30549 | 151 | 0,541416511 | 0,455821958 | 0,54291579 | 0,107782253 | 0,091614915 | -0,027830824 | -0,076830824 | **-1,336830824** |
| chrX | 31626 | 31734 | 108 | 0,720938828 | 1,964110877 | 0,268501111 | -0,617352537 | -0,524749656 | 0,171907592 | 0,423907592 | **-0,836092408** |
| chrX | 32694 | 32839 | 145 | 1,353541277 | 1,030231743 | 0,567814664 | 0,170813172 | 0,145191196 | -0,023824729 | -0,030824729 | **-1,290824729** |
| chrX | 33933 | 34077 | 144 | 1,202514566 | 0,856055873 | 0,584150313 | 0,212522567 | 0,180644182 | 0,004374341 | 0,004374341 | **-1,255625659** |
| chrX | 37483 | 37615 | 132 | 0,92895675 | 1,693582397 | 0,354220356 | -0,373951081 | -0,317858419 | -0,224493138 | -0,140493138 | **-1,400493138** |
| chrX | 38720 | 38876 | 156 | 0,404637603 | 0,326116523 | 0,55372606 | 0,13508094 | 0,114818799 | 0,09035836 | 0,00635836 | **-1,25364164** |
| chrX | 40633 | 40803 | 170 | 0,250761331 | 0,500292393 | 0,333879353 | -0,42922608 | -0,364842168 | -0,264630918 | -0,446630918 | **-1,706630918** |
| chrX | 40783 | 40953 | 170 | 0,185221438 | 0,285351958 | 0,393607968 | -0,269927623 | -0,229438479 | -0,095780227 | -0,277780227 | **-1,537780227** |
| chrX | 46039 | 46190 | 151 | 0,284956058 | 0,218646305 | 0,565835427 | 0,165781194 | 0,140914015 | 0,028632183 | -0,020367817 | **-1,280367817** |
| chrX | 53460 | 53598 | 138 | 0,14532759 | 0,125999566 | 0,535617563 | 0,08939893 | 0,07598909 | -0,012023216 | 0,029976784 | **-1,230023216** |
| chrX | 55658 | 55819 | 161 | 0,82922213 | 0,796761959 | 0,5099817 | 0,025023022 | 0,021269569 | 0,080330804 | -0,038669196 | **-1,298669196** |
| chrX | 59858 | 59998 | 140 | 3,997933499 | 2,834990228 | 0,585098511 | 0,214954263 | 0,182711124 | 0,041734344 | 0,069734344 | **-1,190265656** |
| chrX | 64405 | 64545 | 140 | 0,09688506 | 0,081529131 | 0,543034495 | 0,108081541 | 0,09186931 | -0,053034458 | -0,025034458 | **-1,285034458** |
| chrX | 86256 | 86397 | 141 | 0,176672756 | 0,111176087 | 0,613769206 | 0,28915654 | 0,245783059 | 0,065295833 | 0,086295833 | **-1,173704167** |
| chrX | 88120 | 88277 | 157 | 1,46467414 | 1,530524136 | 0,489007406 | -0,027557834 | -0,023424159 | -0,0303057 | -0,1213057 | **-1,3813057** |
| chrX | 90912 | 91092 | 180 | 0,720938828 | 1,58240631 | 0,31299644 | -0,487374615 | -0,414268423 | -0,207804921 | -0,459804921 | **-1,719804921** |
| chrX | 94290 | 94441 | 151 | 0,213717044 | 0,055588044 | 0,793587102 | 0,818930974 | 0,696091328 | 0,591519909 | 0,542519909 | **-0,717480091** |
| chrX | 94366 | 94450 | 84 | 0,216566604 | 0,385410436 | 0,359758911 | -0,359103289 | -0,305237796 | 0,058599328 | 0,478599328 | **-0,781400672** |
| chrX | 94724 | 94862 | 138 | 0,333398588 | 0,363175219 | 0,478626364 | -0,053601417 | -0,045561205 | -0,134699576 | -0,092699576 | **-1,352699576** |
| chrX | 100058 | 100217 | 159 | 11,01070209 | 8,693970031 | 0,558786364 | 0,147892925 | 0,125708986 | 0,135727092 | 0,030727092 | **-1,229272908** |
| chrX | 100062 | 100217 | 155 | 0,039893848 | 0,059293913 | 0,402205349 | -0,247642923 | -0,210496484 | -0,249128037 | -0,326128037 | **-1,586128037** |
| chrX | 100066 | 100217 | 151 | 0,039893848 | 0,040764565 | 0,49460244 | -0,013530088 | -0,011500575 | -0,113028956 | -0,162028956 | **-1,422028956** |
| chrX | 105547 | 105698 | 151 | 1,062886098 | 0,607762611 | 0,636211606 | 0,348350745 | 0,296098133 | 0,18935592 | 0,14035592 | **-1,11964408** |
| chrX | 106971 | 107136 | 165 | 0,344796831 | 0,34835174 | 0,49743568 | -0,006427842 | -0,005463666 | 0,118780462 | -0,028219538 | **-1,288219538** |
| chrX | 108096 | 108187 | 91 | 0,088336378 | 0,326116523 | 0,213139727 | -0,795574396 | -0,676238236 | -0,266196189 | 0,104803811 | **-1,155196189** |
| chrX | 108096 | 108204 | 108 | 0,028495606 | 0,100058479 | 0,221662392 | -0,766590907 | -0,651602271 | 0,053881696 | 0,305881696 | **-0,954118304** |
| chrX | 108096 | 108205 | 109 | 36,9929955 | 115,334073 | 0,242852409 | -0,697156566 | -0,592583081 | 0,149300326 | 0,394300326 | **-0,865699674** |
| chrX | 108096 | 108206 | 110 | 0,512920905 | 1,893699355 | 0,213129139 | -0,795610814 | -0,676269192 | 0,077112619 | 0,315112619 | **-0,944887381** |
| chrX | 108096 | 108207 | 111 | 0,376141997 | 2,516285444 | 0,130043711 | -1,126184528 | -0,957256849 | -0,155729181 | 0,075270819 | **-1,184729181** |
| chrX | 108096 | 108208 | 112 | 0,028495606 | 0,166764131 | 0,145936926 | -1,054019799 | -0,895916829 | -0,117019389 | 0,106980611 | **-1,153019389** |
| chrX | 108096 | 108220 | 124 | 0,051292091 | 5,62551002 | 0,009035385 | -2,364164857 | -2,009540128 | -1,506057229 | -1,366057229 | **-2,626057229** |
| chrX | 108097 | 108205 | 108 | 0,034194727 | 0,103764348 | 0,247861382 | -0,681235065 | -0,579049806 | 0,128730357 | 0,380730357 | **-0,879269643** |
| chrX | 108098 | 108200 | 102 | 0,037044288 | 0,26311674 | 0,123414715 | -1,158084807 | -0,984372086 | -0,393329665 | -0,099329665 | **-1,359329665** |
| chrX | 108098 | 108205 | 107 | 0,823523009 | 8,00467829 | 0,093283216 | -1,32080488 | -1,122684148 | -0,442181936 | -0,183181936 | **-1,443181936** |
| chrX | 108098 | 108206 | 108 | 0,031345166 | 0,277940218 | 0,101347066 | -1,273913333 | -1,082826333 | -0,35277316 | -0,10077316 | **-1,36077316** |
| chrX | 108098 | 108220 | 122 | 1,233859733 | 10,79149221 | 0,102604875 | -1,266847707 | -1,076820551 | -0,453361863 | -0,299361863 | **-1,559361863** |
| chrX | 108135 | 108220 | 85 | 0,031345166 | 0,255705001 | 0,10919752 | -1,230807045 | -1,046185988 | -0,664526842 | -0,251526842 | **-1,511526842** |
| chrX | 117567 | 117726 | 159 | 0,253610892 | 0,207528696 | 0,54996556 | 0,125574335 | 0,106738185 | 0,122510802 | 0,017510802 | **-1,242489198** |
| chrX | 118042 | 118180 | 138 | 8,870682096 | 5,518039803 | 0,616502436 | 0,29630767 | 0,25186152 | 0,150146802 | 0,192146802 | **-1,067853198** |
| chrX | 118045 | 118180 | 135 | 0,056991212 | 0,037058696 | 0,605967759 | 0,268824834 | 0,228501109 | 0,255711578 | 0,318711578 | **-0,941288422** |
| chrX | 118063 | 118213 | 150 | 4,69037672 | 2,979519141 | 0,611530692 | 0,283310777 | 0,24081416 | 0,106275804 | 0,064275804 | **-1,195724196** |
| chrX | 118208 | 118364 | 156 | 0,29065518 | 0,392822175 | 0,425259414 | -0,18845648 | -0,160188008 | -0,179681402 | -0,263681402 | **-1,523681402** |
| chrX | 122371 | 122478 | 107 | 0,062690333 | 0,444704349 | 0,12355339 | -1,157405375 | -0,983794569 | -0,298979864 | -0,039979864 | **-1,299979864** |
| chrX | 126969 | 127111 | 142 | 0,980248841 | 0,79305609 | 0,552780756 | 0,132690081 | 0,112786569 | -0,07384166 | -0,05984166 | **-1,31984166** |
| chrX | 127866 | 128026 | 160 | 0,239363089 | 0,203822827 | 0,540096335 | 0,100676421 | 0,085574958 | 0,123873309 | 0,011873309 | **-1,248126691** |
| chrX | 131244 | 131407 | 163 | 0,649699813 | 1,004290656 | 0,392807471 | -0,272009205 | -0,231207824 | -0,132964617 | -0,265964617 | **-1,525964617** |
| chrX | 139844 | 139996 | 152 | 0,059840772 | 0,070411522 | 0,459422021 | -0,101889932 | -0,086606442 | -0,196267132 | -0,252267132 | **-1,512267132** |
| chrX | 141168 | 141318 | 150 | 0,062690333 | 0,048176305 | 0,56545715 | 0,164819952 | 0,140096959 | 0,005226772 | -0,036773228 | **-1,296773228** |
| chrX | 142696 | 142846 | 150 | 0,73233707 | 0,544762828 | 0,573437576 | 0,185132835 | 0,15736291 | 0,008443327 | -0,033556673 | **-1,293556673** |
| chrX | 145224 | 145374 | 150 | 0,424584527 | 0,444704349 | 0,488427424 | -0,029012215 | -0,024660382 | -0,175802941 | -0,217802941 | **-1,477802941** |
| chrX | 145224 | 145374 | 150 | 0,424584527 | 0,444704349 | 0,488427424 | -0,029012215 | -0,024660382 | -0,175802941 | -0,217802941 | **-1,477802941** |
| chrX | 148124 | 148271 | 147 | 0,156725832 | 0,103764348 | 0,601657352 | 0,2576393 | 0,218993405 | 0,106111371 | 0,085111371 | **-1,174888629** |
| chrX | 149558 | 149726 | 168 | 0,988797522 | 1,693582397 | 0,368626948 | -0,33549211 | -0,285168294 | -0,189893786 | -0,357893786 | **-1,617893786** |
| chrX | 154966 | 155110 | 144 | 0,324849907 | 0,392822175 | 0,452643923 | -0,118984235 | -0,1011366 | -0,279786272 | -0,279786272 | **-1,539786272** |
| chrX | 162602 | 162770 | 168 | 0,48727486 | 0,652233046 | 0,427618674 | -0,182440113 | -0,155074096 | -0,05141696 | -0,21941696 | **-1,47941696** |
| chrX | 163009 | 163162 | 153 | 1,877860425 | 1,786229137 | 0,512503964 | 0,031347923 | 0,026645735 | -0,043731271 | -0,106731271 | **-1,366731271** |
| chrX | 169091 | 169251 | 160 | 0,094035499 | 0,148234783 | 0,388142938 | -0,284162468 | -0,241538098 | -0,19210159 | -0,30410159 | **-1,56410159** |
| chrX | 173679 | 173830 | 151 | 0,028495606 | 0,037058696 | 0,434687048 | -0,164453569 | -0,139785534 | -0,253499413 | -0,302499413 | **-1,562499413** |
| chrX | 173874 | 174016 | 142 | 0,960301917 | 0,711526959 | 0,57440204 | 0,187592737 | 0,159453827 | -0,026576999 | -0,012576999 | **-1,272576999** |
| chrX | 178426 | 178580 | 154 | 0,823523009 | 0,596645002 | 0,579877171 | 0,201579264 | 0,171342374 | 0,085839665 | 0,015839665 | **-1,244160335** |
| chrX | 182961 | 183077 | 116 | 8,728204067 | 26,16343922 | 0,250151705 | -0,674012431 | -0,572910567 | 0,130628926 | 0,326628926 | **-0,933371074** |
| chrX | 182964 | 183077 | 113 | 0,042743409 | 0,203822827 | 0,173354671 | -0,940991251 | -0,799842563 | -0,040078441 | 0,176921559 | **-1,083078441** |
| chrX | 182966 | 183077 | 111 | 0,039893848 | 0,096352609 | 0,292806499 | -0,545204323 | -0,463423675 | 0,336325301 | 0,567325301 | **-0,692674699** |
| chrX | 185927 | 186084 | 157 | 0,071239015 | 0,062999783 | 0,530688713 | 0,07700122 | 0,065451037 | 0,067124536 | -0,023875464 | **-1,283875464** |
| chrX | 185927 | 186084 | 157 | 0,071239015 | 0,062999783 | 0,530688713 | 0,07700122 | 0,065451037 | 0,067124536 | -0,023875464 | **-1,283875464** |
| chrX | 189658 | 189821 | 163 | 0,43883233 | 0,340940001 | 0,562769814 | 0,157995466 | 0,134296146 | 0,238555579 | 0,105555579 | **-1,154444421** |
| chrX | 190658 | 190805 | 147 | 1,188266763 | 0,737468046 | 0,617045897 | 0,297731361 | 0,253071657 | 0,135833629 | 0,114833629 | **-1,145166371** |
| chrX | 191225 | 191359 | 134 | 0,322000346 | 0,344645871 | 0,483015335 | -0,04258711 | -0,036199044 | -0,006156856 | 0,063843144 | **-1,196156856** |
| chrX | 192006 | 192112 | 106 | 0,262159574 | 0,804173699 | 0,245851443 | -0,687602893 | -0,584462459 | 0,052378171 | 0,318378171 | **-0,941621829** |
| chrX | 192006 | 192128 | 122 | 0,190920559 | 0,893114568 | 0,17612027 | -0,930252163 | -0,790714339 | -0,176933223 | -0,022933223 | **-1,282933223** |
| chrX | 192132 | 192293 | 161 | 0,53571739 | 0,470645436 | 0,532330265 | 0,081128865 | 0,068959535 | 0,114542894 | -0,004457106 | **-1,264457106** |
| chrX | 192599 | 192736 | 137 | 0,042743409 | 0,125999566 | 0,25330482 | -0,664126048 | -0,564507141 | -0,623692411 | -0,574692411 | **-1,834692411** |
| chrX | 194181 | 194318 | 137 | 5,303032246 | 4,065338928 | 0,566057018 | 0,166344351 | 0,141392699 | 0,071968938 | 0,120968938 | **-1,139031062** |
| chrX | 197629 | 197777 | 148 | 0,151026711 | 0,066705652 | 0,693634647 | 0,506179412 | 0,4302525 | 0,331527788 | 0,303527788 | **-0,956472212** |
| chrX | 198154 | 198316 | 162 | 0,037044288 | 0,077823261 | 0,322495674 | -0,460731366 | -0,391621661 | -0,309549094 | -0,435549094 | **-1,695549094** |
| chrX | 198182 | 198330 | 148 | 0,282106498 | 0,222352175 | 0,559226183 | 0,149007602 | 0,126656462 | 0,029934556 | 0,001934556 | **-1,258065444** |
| chrX | 199707 | 199858 | 151 | 0,051292091 | 0,070411522 | 0,421450846 | -0,198183249 | -0,168455762 | -0,283069346 | -0,332069346 | **-1,592069346** |
| chrX | 203379 | 203541 | 162 | 0,139628469 | 0,192705218 | 0,420145397 | -0,201521534 | -0,171293304 | -0,094224784 | -0,220224784 | **-1,480224784** |
| chrX | 203482 | 203630 | 148 | 0,136778908 | 0,08894087 | 0,605967759 | 0,268824834 | 0,228501109 | 0,127621958 | 0,099621958 | **-1,160378042** |
| chrX | 206401 | 206560 | 159 | 0,168124074 | 0,314998914 | 0,347994358 | -0,390740963 | -0,332129819 | -0,310180542 | -0,415180542 | **-1,675180542** |
| chrX | 209002 | 209147 | 145 | 0,578460798 | 0,444704349 | 0,565364056 | 0,164583414 | 0,139895902 | -0,03003576 | -0,03703576 | **-1,29703576** |
| chrX | 210689 | 210857 | 168 | 0,92610719 | 1,267407396 | 0,422202431 | -0,196262308 | -0,166822962 | -0,065451383 | -0,233451383 | **-1,493451383** |
| chrX | 210689 | 210857 | 168 | 0,92610719 | 1,267407396 | 0,422202431 | -0,196262308 | -0,166822962 | -0,065451383 | -0,233451383 | **-1,493451383** |
| chrX | 212042 | 212183 | 141 | 3,539154245 | 2,11234566 | 0,626232735 | 0,32189198 | 0,273608183 | 0,091915266 | 0,112915266 | **-1,147084734** |
| chrX | 212042 | 212183 | 141 | 3,539154245 | 2,11234566 | 0,626232735 | 0,32189198 | 0,273608183 | 0,091915266 | 0,112915266 | **-1,147084734** |
| chrX | 215649 | 215789 | 140 | 0,139628469 | 0,103764348 | 0,573675388 | 0,185739279 | 0,157878387 | 0,00753646 | 0,03553646 | **-1,22446354** |
| chrX | 218927 | 219076 | 149 | 0,037044288 | 0,055588044 | 0,399906675 | -0,253588671 | -0,21555037 | -0,313594929 | -0,348594929 | **-1,608594929** |
| chrX | 219865 | 220016 | 151 | 0,373292436 | 0,244587392 | 0,604150547 | 0,264105137 | 0,224489366 | 0,109423823 | 0,060423823 | **-1,199576177** |
| chrX | 219865 | 220016 | 151 | 0,373292436 | 0,244587392 | 0,604150547 | 0,264105137 | 0,224489366 | 0,109423823 | 0,060423823 | **-1,199576177** |
| chrX | 220580 | 220715 | 135 | 0,327699467 | 0,70411522 | 0,317595273 | -0,474433869 | -0,403268789 | -0,388892948 | -0,325892948 | **-1,585892948** |
| chrX | 221358 | 221508 | 150 | 0,265009134 | 0,203822827 | 0,565253985 | 0,164303749 | 0,139658187 | -0,007063329 | -0,049063329 | **-1,309063329** |
| chrX | 225766 | 225921 | 155 | 0,823523009 | 0,685585872 | 0,545701519 | 0,114808437 | 0,097587171 | 0,060791685 | -0,016208315 | **-1,276208315** |
| chrX | 226052 | 226204 | 152 | 0,179522317 | 0,203822827 | 0,468304659 | -0,079532203 | -0,067602373 | -0,172928758 | -0,228928758 | **-1,488928758** |
| chrX | 226472 | 226633 | 161 | 0,279256937 | 0,259410871 | 0,518421434 | 0,046192108 | 0,039263292 | 0,084475079 | -0,034524921 | **-1,294524921** |

TABLE 2

| Chrom | Start | End | Length | Section A | Section B | A/A+B | Z-score | Z * 0.85 | Phase correction | Length correction | ΔLK |
|---|---|---|---|---|---|---|---|---|---|---|---|
| chrX | 227406 | 227571 | 165 | 0,359044634 | 0,463233697 | 0,436646109 | -0,159478092 | -0,135556378 | -0,003733447 | -0,150733447 | -1,410733447 |
| chrX | 227406 | 227571 | 165 | 0,359044634 | 0,463233697 | 0,436646109 | -0,159478092 | -0,135556378 | -0,003733447 | -0,150733447 | -1,410733447 |
| chrX | 229005 | 229137 | 132 | 0,128230226 | 0,200116957 | 0,390532439 | -0,277931585 | -0,236241848 | -0,152275249 | -0,068275249 | -1,328275249 |
| chrX | 229384 | 229535 | 151 | 0,205168362 | 0,203822827 | 0,501644944 | 0,004123275 | 0,003504784 | -0,110721107 | -0,159721107 | -1,419721107 |
| chrX | 233516 | 233665 | 149 | 0,213717044 | 0,307587175 | 0,409966074 | -0,227632249 | -0,193487411 | -0,288696633 | -0,323696633 | -1,583696633 |
| chrX | 235494 | 235644 | 150 | 0,512920905 | 0,481763045 | 0,515662191 | 0,039269381 | 0,033378974 | -0,115236663 | -0,157236663 | -1,417236663 |
| chrX | 237835 | 237975 | 140 | 1,934851636 | 1,241466309 | 0,609149232 | 0,277102325 | 0,235536977 | 0,084796891 | 0,112796891 | -1,147203109 |
| chrX | 240265 | 240414 | 149 | 0,356195073 | 0,203822827 | 0,6360423 | 0,347899847 | 0,29571487 | 0,198447788 | 0,163447788 | -1,096552212 |
| chrX | 240554 | 240707 | 153 | 0,122531105 | 0,103764348 | 0,541465166 | 0,104125608 | 0,088506767 | 0,007812413 | -0,055187587 | -1,315187587 |
| chrX | 240601 | 240747 | 146 | 0,814974327 | 0,626291959 | 0,56545715 | 0,164819952 | 0,140096959 | -0,026899591 | -0,040899591 | -1,300899591 |
| chrX | 242702 | 242868 | 166 | 0,481575739 | 0,826408916 | 0,36818149 | -0,336673583 | -0,286172546 | -0,164505634 | -0,318505634 | -1,578505634 |
| chrX | 243519 | 243619 | 100 | 0,034194727 | 0,059293913 | 0,365763444 | -0,34309514 | -0,291630869 | 0,285432307 | 0,593432307 | -0,666567693 |
| chrX | 245287 | 245435 | 148 | 0,156725832 | 0,148234783 | 0,51392155 | 0,034903236 | 0,029667751 | -0,070801988 | -0,098801988 | -1,358801988 |
| chrX | 247930 | 248081 | 151 | 0,213717044 | 0,270528479 | 0,441340257 | -0,14757205 | -0,125436242 | -0,258695434 | -0,307695434 | -1,567695434 |
| chrX | 248270 | 248435 | 165 | 0,199469241 | 0,285351958 | 0,411428464 | -0,223871969 | -0,190291174 | -0,060112674 | -0,207112674 | -1,467112674 |
| chrX | 251069 | 251224 | 155 | 0,076938136 | 0,077823261 | 0,497140355 | -0,007168129 | -0,00609291 | -0,035474 | -0,112474 | -1,372474 |
| chrX | 251916 | 252080 | 164 | 0,037044288 | 0,044470435 | 0,454449042 | -0,114428548 | -0,097264266 | 0,020206835 | -0,119793165 | -1,379793165 |
| chrX | 253721 | 253885 | 164 | 1,835117016 | 2,419932835 | 0,431279792 | -0,173116845 | -0,147149318 | -0,026559303 | -0,166559303 | -1,426559303 |
| chrX | 254829 | 255000 | 171 | 0,176672756 | 0,251999131 | 0,412139824 | -0,222043979 | -0,188737382 | -0,01692918 | -0,20592918 | -1,46592918 |
| chrX | 255572 | 255735 | 163 | 0,142478029 | 0,303881305 | 0,31920029 | -0,469936228 | -0,399445793 | -0,297054238 | -0,430054238 | -1,690054238 |
| chrX | 271510 | 271677 | 167 | 0,045592969 | 0,074117392 | 0,380860679 | -0,303221116 | -0,257737948 | -0,177903556 | -0,338903556 | -1,598903556 |
| chrX | 271664 | 271795 | 131 | 0,313451664 | 0,259410871 | 0,547167331 | 0,118507771 | 0,100731606 | 0,370516535 | 0,461516535 | -0,798483465 |
| chrX | 272130 | 272276 | 146 | 3,533455124 | 2,64228501 | 0,572150875 | 0,181852797 | 0,154574878 | -0,000905833 | -0,014905833 | -1,274905833 |
| chrX | 273881 | 274032 | 151 | 0,302053422 | 0,237175653 | 0,560157892 | 0,15136954 | 0,128664109 | -0,008405679 | -0,057405679 | -1,317405679 |
| chrX | 275971 | 276080 | 109 | 1,723984153 | 15,91300397 | 0,097748218 | -1,29448938 | -1,100315973 | -0,357479318 | -0,112479318 | -1,372479318 |
| chrX | 276361 | 276455 | 94 | 0,24221265 | 0,233469783 | 0,509189814 | 0,023037485 | 0,019581862 | 0,505187217 | 0,855187217 | -0,404812783 |
| chrX | 276361 | 276463 | 102 | 0,270708255 | 0,337234132 | 0,445286036 | -0,137580369 | -0,116943314 | 0,49666608 | 0,79066608 | -0,46933392 |
| chrX | 276481 | 276614 | 133 | 0,262159574 | 0,544762828 | 0,324888209 | -0,454072817 | -0,385961895 | -0,338107296 | -0,261107296 | -1,521107296 |
| chrX | 278918 | 279067 | 149 | 0,322000346 | 0,333528262 | 0,49120716 | -0,022042166 | -0,018735841 | -0,113743397 | -0,148743397 | -1,408743397 |
| chrX | 283812 | 283977 | 165 | 0,347646391 | 0,318704784 | 0,521716483 | 0,054462061 | 0,046292752 | 0,173543533 | 0,026543533 | -1,233456467 |
| chrX | 287746 | 287904 | 158 | 0,250761331 | 0,148234783 | 0,628480635 | 0,327831948 | 0,278657156 | 0,277120717 | 0,179120717 | -1,080879283 |
| chrX | 292191 | 292322 | 131 | 0,63260245 | 0,44099848 | 0,58923426 | 0,225575652 | 0,191739305 | 0,456759888 | 0,547759888 | -0,712240112 |
| chrX | 292971 | 293126 | 155 | 0,34194727 | 0,196411088 | 0,635166642 | 0,345568907 | 0,293733571 | 0,272592108 | 0,195592108 | -1,064407892 |
| chrX | 293774 | 293933 | 159 | 0,088336378 | 0,059293913 | 0,598362147 | 0,249109985 | 0,211743487 | 0,243399097 | 0,138399097 | -1,121600903 |
| chrX | 301109 | 301275 | 166 | 0,24506221 | 0,240881523 | 0,504301617 | 0,010782764 | 0,009165349 | 0,12453526 | -0,02946474 | -1,28946474 |
| chrX | 303727 | 303887 | 160 | 0,256460453 | 0,233469783 | 0,523463207 | 0,058847485 | 0,050020363 | 0,098413314 | -0,013586686 | -1,273586686 |
| chrX | 309182 | 309321 | 139 | 0,102584181 | 0,111176087 | 0,479902939 | -0,050397187 | -0,042837609 | -0,166568691 | -0,131568691 | -1,391568691 |
| chrX | 310987 | 311146 | 159 | 1,176868521 | 0,989467177 | 0,543253071 | 0,108632656 | 0,092337758 | 0,131146171 | 0,026146171 | -1,233853829 |
| chrX | 310987 | 311146 | 159 | 1,176868521 | 0,989467177 | 0,543253071 | 0,108632656 | 0,092337758 | 0,131146171 | 0,026146171 | -1,233853829 |
| chrX | 321769 | 321919 | 150 | 0,153876272 | 0,144528914 | 0,515662191 | 0,039269381 | 0,033378974 | -0,115862454 | -0,157862454 | -1,417862454 |
| chrX | 322337 | 322492 | 155 | 0,076938136 | 0,092646739 | 0,453685128 | -0,116356189 | -0,098902761 | -0,124691398 | -0,201691398 | -1,461691398 |
| chrX | 325234 | 325383 | 149 | 0,367593315 | 0,226058044 | 0,619207401 | 0,303399799 | 0,257889829 | 0,171138877 | 0,136138877 | -1,123861123 |
| chrX | 326985 | 327142 | 157 | 1,718285032 | 1,571288701 | 0,522342763 | 0,05603431 | 0,047629164 | 0,042486558 | -0,048513442 | -1,308513442 |
| chrX | 327633 | 327785 | 152 | 0,09973462 | 0,096352609 | 0,508623742 | 0,021618199 | 0,01837547 | -0,089140593 | -0,145140593 | -1,405140593 |
| chrX | 328569 | 328730 | 161 | 0,170973635 | 0,237175653 | 0,418899751 | -0,204708983 | -0,174002636 | -0,13263706 | -0,25163706 | -1,51163706 |
| chrX | 329259 | 329407 | 148 | 0,430283648 | 0,644821307 | 0,400224784 | -0,252765321 | -0,214850523 | -0,308777234 | -0,336777234 | -1,596777234 |
| chrX | 329259 | 329407 | 148 | 0,430283648 | 0,644821307 | 0,400224784 | -0,252765321 | -0,214850523 | -0,308777234 | -0,336777234 | -1,596777234 |
| chrX | 331899 | 332052 | 153 | 0,082637257 | 0,096352609 | 0,461686791 | -0,096185077 | -0,081757315 | -0,159096683 | -0,222096683 | -1,482096683 |
| chrX | 333305 | 333466 | 161 | 1,473222822 | 1,715817615 | 0,461964296 | -0,095486273 | -0,081163332 | -0,031095529 | -0,150095529 | -1,410095529 |
| chrX | 333305 | 333466 | 161 | 1,473222822 | 1,715817615 | 0,461964296 | -0,095486273 | -0,081163332 | -0,031095529 | -0,150095529 | -1,410095529 |
| chrX | 333361 | 333513 | 152 | 0,111132863 | 0,055588044 | 0,666580246 | 0,43048963 | 0,365916185 | 0,283731699 | 0,227731699 | -1,032268301 |
| chrX | 337451 | 337612 | 161 | 0,085486818 | 0,062999783 | 0,575720754 | 0,190958009 | 0,162314307 | 0,224105819 | 0,105105819 | -1,154894181 |
| chrX | 338776 | 338907 | 131 | 0,575611238 | 1,037643482 | 0,356801211 | -0,367022249 | -0,311968912 | -0,083496263 | 0,007503737 | -1,252496263 |
| chrX | 343385 | 343533 | 148 | 0,547115632 | 0,389116306 | 0,58438044 | 0,213112623 | 0,181145729 | 0,084033425 | 0,056033425 | -1,203966575 |
| chrX | 344740 | 344900 | 160 | 0,071239015 | 0,096352609 | 0,425075031 | -0,188926962 | -0,160587917 | -0,105610994 | -0,217610994 | -1,477610994 |
| chrX | 350347 | 350517 | 170 | 0,401788042 | 0,566998046 | 0,414733497 | -0,215385217 | -0,183077435 | -0,046234514 | -0,228234514 | -1,488234514 |
| chrX | 350347 | 350517 | 170 | 0,401788042 | 0,566998046 | 0,414733497 | -0,215385217 | -0,183077435 | -0,046234514 | -0,228234514 | -1,488234514 |
| chrX | 357774 | 357934 | 160 | 0,49012442 | 0,596645002 | 0,450992097 | -0,123155209 | -0,104681928 | -0,052926248 | -0,164926248 | -1,424926248 |
| chrX | 358769 | 358924 | 155 | 0,116831984 | 0,107470218 | 0,520868645 | 0,052333815 | 0,044483743 | 0,013941346 | -0,063058654 | -1,323058654 |
| chrX | 359603 | 359757 | 154 | 4,080570756 | 2,557050009 | 0,61476407 | 0,291757738 | 0,247994077 | 0,166164504 | 0,096164504 | -1,163835496 |
| chrX | 362162 | 362320 | 158 | 4,681828039 | 4,547101973 | 0,507299116 | 0,018297192 | 0,015552613 | 0,013572689 | -0,084427311 | -1,344427311 |
| chrX | 365711 | 365851 | 140 | 1,715435471 | 1,686170658 | 0,504301617 | 0,010782764 | 0,009165349 | -0,153159 | -0,125159 | -1,385159 |
| chrX | 366070 | 366219 | 149 | 1,61285129 | 1,441583266 | 0,528035962 | 0,070333681 | 0,059783629 | -0,030857401 | -0,065857401 | -1,325857401 |
| chrX | 366353 | 366502 | 149 | 1,159771157 | 0,578115654 | 0,667345623 | 0,43259537 | 0,367706065 | 0,277746223 | 0,242746223 | -1,017253777 |
| chrX | 366353 | 366502 | 149 | 1,159771157 | 0,578115654 | 0,667345623 | 0,43259537 | 0,367706065 | 0,277746223 | 0,242746223 | -1,017253777 |
| chrX | 366520 | 366671 | 151 | 1,162620718 | 0,559586306 | 0,675076052 | 0,453973505 | 0,385877479 | 0,25754938 | 0,20854938 | -1,05145062 |
| chrX | 367696 | 367845 | 149 | 0,364743755 | 0,326116523 | 0,527955893 | 0,070132481 | 0,059612609 | -0,026155084 | -0,061155084 | -1,321155084 |
| chrX | 367696 | 367845 | 149 | 0,364743755 | 0,326116523 | 0,527955893 | 0,070132481 | 0,059612609 | -0,026155084 | -0,061155084 | -1,321155084 |
| chrX | 369411 | 369539 | 128 | 0,273557816 | 0,674468263 | 0,288555159 | -0,557610598 | -0,473969008 | -0,050777534 | 0,061222466 | -1,198777534 |
| chrX | 369776 | 369927 | 151 | 0,644000692 | 0,411351523 | 0,610223471 | 0,279901524 | 0,237916295 | 0,106330376 | 0,057330376 | -1,202669624 |
| chrX | 374096 | 374246 | 150 | 0,028495606 | 0,044470435 | 0,390532439 | -0,277931585 | -0,236241848 | -0,383702638 | -0,425702638 | -1,685702638 |
| chrX | 380580 | 380750 | 170 | 0,247911771 | 0,129705435 | 0,656516088 | 0,402973357 | 0,342527354 | 0,48555887 | 0,30355887 | -0,95644113 |
| chrX | 380580 | 380750 | 170 | 0,247911771 | 0,129705435 | 0,656516088 | 0,402973357 | 0,342527354 | 0,48555887 | 0,30355887 | -0,95644113 |
| chrX | 380923 | 381086 | 163 | 1,296550065 | 1,063584569 | 0,549354281 | 0,124030106 | 0,10542559 | 0,217790335 | 0,084790335 | -1,175209665 |
| chrX | 385357 | 385485 | 128 | 0,703841464 | 1,904816963 | 0,269809745 | -0,613388498 | -0,521380224 | -0,113501152 | -0,001501152 | -1,261501152 |
| chrX | 386997 | 387161 | 164 | 0,031245166 | 0,055588044 | 0,360566076 | -0,356946107 | -0,303404191 | -0,186430376 | -0,326430376 | -1,586430376 |
| chrX | 392484 | 392621 | 137 | 0,370442876 | 0,366881088 | 0,502415348 | 0,006054416 | 0,005146254 | -0,065479645 | -0,016479645 | -1,276479645 |
| chrX | 392991 | 393143 | 152 | 0,179522317 | 0,111176087 | 0,617555219 | 0,299066166 | 0,254206241 | 0,172978617 | 0,116978617 | -1,143021383 |
| chrX | 394180 | 394326 | 146 | 0,450230572 | 0,322410653 | 0,582716217 | 0,208847146 | 0,177520074 | 0,024350925 | 0,010350925 | -1,249649075 |
| chrX | 394276 | 394372 | 96 | 3,36818061 | 12,65554461 | 0,210199599 | -0,805728874 | -0,684869543 | -0,147782309 | 0,188217691 | -1,071782309 |
| chrX | 398245 | 398394 | 149 | 0,364743755 | 0,366881088 | 0,498539324 | -0,00366138 | -0,003112173 | -0,094163967 | -0,129163967 | -1,389163967 |
| chrX | 399541 | 399701 | 160 | 0,09973462 | 0,185293479 | 0,349911537 | -0,385559308 | -0,327725412 | -0,27190185 | -0,38390185 | -1,64390185 |
| chrX | 407672 | 407824 | 152 | 1,598603487 | 1,148819569 | 0,581855599 | 0,206642838 | 0,175646412 | 0,097537682 | 0,041537682 | -1,218462318 |
| chrX | 410284 | 410430 | 146 | 0,381841118 | 0,274234349 | 0,58200792 | 0,207032907 | 0,175977971 | 0,030037525 | 0,016037525 | -1,243962475 |
| chrX | 410723 | 410889 | 166 | 0,279256937 | 0,389116306 | 0,417815854 | -0,20748423 | -0,176361595 | -0,063927134 | -0,217927134 | -1,477927134 |
| chrX | 412052 | 412213 | 161 | 0,718089267 | 0,648527176 | 0,525450481 | 0,063838228 | 0,054262494 | 0,115285502 | -0,003714498 | -1,263714498 |
| chrX | 425378 | 425524 | 146 | 0,398938482 | 0,251999131 | 0,61286746 | 0,286800501 | 0,243780426 | 0,097607643 | 0,083607643 | -1,176392357 |
| chrX | 429176 | 429332 | 156 | 0,73233707 | 0,489174784 | 0,59953333 | 0,252139371 | 0,214318465 | 0,194852405 | 0,110852405 | -1,149147595 |
| chrX | 429194 | 429332 | 138 | 0,042743409 | 0,037058696 | 0,535617563 | 0,08939893 | 0,07598909 | -0,033419524 | 0,008580476 | -1,251419524 |
| chrX | 429194 | 429339 | 145 | 0,433133209 | 0,296469566 | 0,593656197 | 0,236960278 | 0,201416237 | 0,026624303 | 0,019624303 | -1,240375697 |
| chrX | 435614 | 435721 | 107 | 0,666797177 | 3,035107185 | 0,180122745 | -0,914897409 | -0,777662797 | -0,101078135 | 0,157921865 | -1,102078135 |
| chrX | 435614 | 435731 | 117 | 0,803576085 | 3,331576751 | 0,194328025 | -0,862057185 | -0,732748607 | -0,032064954 | 0,156935046 | -1,103064954 |
| chrX | 439795 | 439950 | 155 | 0,054141651 | 0,040764565 | 0,570475287 | 0,177584338 | 0,150946687 | 0,117971542 | 0,040971542 | -1,219028458 |
| chrX | 447409 | 447538 | 129 | 0,267858695 | 0,452116089 | 0,372038995 | -0,32645783 | -0,277489155 | 0,094383077 | 0,199383077 | -1,060616923 |
| chrX | 448364 | 448517 | 153 | 0,062690333 | 0,040764565 | 0,605967759 | 0,268824834 | 0,228501109 | 0,147884486 | 0,084884486 | -1,175115514 |
| chrX | 456180 | 456328 | 148 | 0,079787696 | 0,111176087 | 0,417815854 | -0,20748423 | -0,176361595 | -0,271992869 | -0,299992869 | -1,559992869 |
| chrX | 459326 | 459480 | 154 | 0,09688506 | 0,044470435 | 0,685400026 | 0,482853237 | 0,410425251 | 0,333071046 | 0,263071046 | -0,996928954 |
| chrX | 465469 | 465620 | 151 | 0,102584181 | 0,077823261 | 0,568624996 | 0,172874585 | 0,146943398 | 0,029981403 | -0,019018597 | -1,279018597 |
| chrX | 466504 | 466659 | 155 | 0,133929347 | 0,074117392 | 0,643746439 | 0,368491045 | 0,313217388 | 0,286107094 | 0,209107094 | -1,050892906 |
| chrX | 467197 | 467335 | 138 | 1,541612276 | 0,996878917 | 0,607294711 | 0,272275 | 0,23143375 | 0,118358531 | 0,160358531 | -1,099641469 |
| chrX | 467559 | 467705 | 146 | 0,302053422 | 0,982055438 | 0,235224155 | -0,721749813 | -0,613487341 | -0,766973872 | -0,780973872 | -2,040973872 |

TABLE 2

| Chrom | Start | End | Length | Section A | Section B | A/A+B | Z-score | Z * 0.85 | Phase correction | Length correction | ΔLK |
|---|---|---|---|---|---|---|---|---|---|---|---|
| chrX | 470836 | 470984 | 148 | 0,666797177 | 0,485468915 | 0,578683328 | 0,198526256 | 0,168747318 | 0,063492151 | 0,035492151 | -1,224507849 |
| chrX | 472163 | 472315 | 152 | 0,302053422 | 0,274234349 | 0,524136442 | 0,060538046 | 0,051457339 | -0,036345881 | -0,092345881 | -1,352345881 |
| chrX | 472826 | 472975 | 149 | 0,079787696 | 0,048176305 | 0,623516738 | 0,314730183 | 0,267520656 | 0,174960831 | 0,139960831 | -1,120039169 |
| chrX | 473000 | 473123 | 123 | 0,071239015 | 0,188999349 | 0,273745245 | -0,601524812 | -0,51129609 | 0,067607002 | 0,214607002 | -1,045392998 |
| chrX | 473100 | 473255 | 155 | 0,105433742 | 0,096352609 | 0,52250185 | 0,056433715 | 0,047968658 | 0,011777119 | -0,065222881 | -1,325222881 |
| chrX | 473533 | 473667 | 134 | 0,128230226 | 0,100058479 | 0,561702018 | 0,155285859 | 0,13199298 | 0,152615509 | 0,222615509 | -1,037384491 |
| chrX | 473533 | 473673 | 140 | 0,179522317 | 0,100058479 | 0,642112476 | 0,3641111 | 0,309494435 | 0,144406829 | 0,172406829 | -1,087593171 |
| chrX | 473729 | 473880 | 151 | 0,119681545 | 0,077823261 | 0,605967759 | 0,268824834 | 0,228501109 | 0,111840476 | 0,062840476 | -1,197159524 |
| chrX | 473739 | 473884 | 145 | 0,076938136 | 0,066705652 | 0,535617563 | 0,08939893 | 0,07598909 | -0,095180335 | -0,102180335 | -1,362180335 |
| chrX | 473757 | 473908 | 151 | 0,037044288 | 0,055588044 | 0,399906675 | -0,253588671 | -0,21555037 | -0,33497204 | -0,38397204 | -1,64397204 |
| chrX | 474011 | 474165 | 154 | 0,176672756 | 0,148234783 | 0,543763178 | 0,109918963 | 0,093431118 | 0,014492876 | -0,055507124 | -1,315507124 |
| chrX | 474045 | 474189 | 144 | 0,065539893 | 0,040764565 | 0,616530051 | 0,29638 | 0,251923 | 0,080385137 | 0,080385137 | -1,179614863 |
| chrX | 474045 | 474189 | 144 | 0,065539893 | 0,040764565 | 0,616530051 | 0,29638 | 0,251923 | 0,080385137 | 0,080385137 | -1,179614863 |
| chrX | 474576 | 474711 | 135 | 0,082637257 | 0,100058479 | 0,452321762 | -0,11979755 | -0,101827917 | -0,095907621 | -0,032907621 | -1,292907621 |
| chrX | 475023 | 475169 | 146 | 0,088336378 | 0,048176305 | 0,64709283 | 0,377483478 | 0,320860956 | 0,163898839 | 0,149898839 | -1,110101161 |
| chrX | 475023 | 475173 | 150 | 8,864982975 | 4,235808928 | 0,676675352 | 0,458421991 | 0,389658692 | 0,241641886 | 0,199641886 | -1,060358114 |
| chrX | 475693 | 475822 | 129 | 0,062690333 | 0,092646739 | 0,403576119 | -0,244101464 | -0,207486244 | 0,148617956 | 0,253617956 | -1,006382044 |
| chrX | 477337 | 477451 | 114 | 0,042743409 | 0,226058044 | 0,159014798 | -0,998515203 | -0,848737923 | -0,117420193 | 0,092579807 | -1,167420193 |
| chrX | 477657 | 477804 | 147 | 0,544266071 | 0,596645002 | 0,47704513 | -0,057571113 | -0,048935446 | -0,15626861 | -0,17726861 | -1,43726861 |
| chrX | 478238 | 478390 | 152 | 0,037044288 | 0,037058696 | 0,499902783 | -0,000243688 | -0,000207135 | -0,10248238 | -0,15848238 | -1,41848238 |
| chrX | 478410 | 478559 | 149 | 0,079787696 | 0,066705652 | 0,544650641 | 0,112157259 | 0,09533367 | -0,004871786 | -0,039871786 | -1,299871786 |
| chrX | 478560 | 478666 | 106 | 0,749434433 | 2,308756747 | 0,245058072 | -0,690124109 | -0,586605492 | 0,070880424 | 0,336880424 | -0,923119576 |
| chrX | 478560 | 478678 | 118 | 0,934655871 | 2,716402401 | 0,255995879 | -0,655739487 | -0,557378564 | 0,11621819 | 0,29821819 | -0,96178181 |
| chrX | 478584 | 478707 | 123 | 0,088336378 | 0,188999349 | 0,318517845 | -0,471847434 | -0,401070319 | 0,151786019 | 0,298786019 | -0,961213981 |
| chrX | 478684 | 478839 | 155 | 0,09973462 | 0,081529131 | 0,550218231 | 0,126212725 | 0,107280816 | 0,066702176 | -0,010297824 | -1,270297824 |
| chrX | 479117 | 479251 | 134 | 0,165274514 | 0,118587827 | 0,582234733 | 0,207613792 | 0,176471723 | 0,192149877 | 0,262149877 | -0,997850123 |
| chrX | 479117 | 479257 | 140 | 0,188070999 | 0,129705435 | 0,591834317 | 0,232266067 | 0,197426157 | 0,022723659 | 0,050723659 | -1,209276341 |
| chrX | 480160 | 480295 | 135 | 0,079787696 | 0,059293913 | 0,573675388 | 0,185739279 | 0,157878387 | 0,153648194 | 0,216648194 | -1,043351806 |
| chrX | 480607 | 480757 | 150 | 8,993213202 | 4,187632624 | 0,68229409 | 0,474123529 | 0,403005 | 0,264571345 | 0,222571345 | -1,037428655 |
| chrX | 480786 | 480943 | 157 | 0,031345166 | 0,062999783 | 0,332240005 | -0,433736208 | -0,368675777 | -0,364180794 | -0,455180794 | -1,715180794 |
| chrX | 481277 | 481406 | 129 | 0,082637257 | 0,118587827 | 0,410670755 | -0,225819886 | -0,191946903 | 0,140929774 | 0,245929774 | -1,014070226 |
| chrX | 481757 | 481888 | 131 | 0,094035499 | 0,107470218 | 0,466664175 | -0,083658001 | -0,071109301 | 0,101412317 | 0,192412317 | -1,067587683 |
| chrX | 482189 | 482339 | 150 | 0,168124074 | 0,181587609 | 0,480750522 | -0,048270025 | -0,041029521 | -0,181185188 | -0,223185188 | -1,483185188 |
| chrX | 483140 | 483291 | 151 | 0,059840772 | 0,096352609 | 0,38311977 | -0,297297305 | -0,252702709 | -0,36333072 | -0,41233072 | -1,67233072 |
| chrX | 495866 | 496005 | 139 | 0,054141651 | 0,037058696 | 0,593656197 | 0,236960278 | 0,201416237 | 0,079341601 | 0,114341601 | -1,145658399 |
| chrX | 496369 | 496517 | 148 | 0,495823542 | 0,340940001 | 0,592549169 | 0,23410732 | 0,198991222 | 0,09602632 | 0,06802632 | -1,19197368 |
| chrX | 498437 | 498584 | 147 | 0,182371877 | 0,144528914 | 0,55788142 | 0,145600009 | 0,123760008 | 0,018735614 | -0,002264386 | -1,262264386 |
| chrX | 503072 | 503238 | 166 | 0,97454972 | 1,248878048 | 0,438309593 | -0,155256401 | -0,131967941 | -0,027143302 | -0,181143302 | -1,441143302 |
| chrX | 503072 | 503238 | 166 | 0,97454972 | 1,248878048 | 0,438309593 | -0,155256401 | -0,131967941 | -0,027143302 | -0,181143302 | -1,441143302 |
| chrX | 503922 | 504069 | 147 | 0,384690679 | 0,322410653 | 0,544038968 | 0,11061448 | 0,094022308 | -0,012096882 | -0,033096882 | -1,293096882 |
| chrX | 512701 | 512853 | 152 | 0,111132863 | 0,040764565 | 0,731630971 | 0,617753167 | 0,525090192 | 0,426091403 | 0,370091403 | -0,889908597 |
| chrX | 516708 | 516897 | 189 | 0,14817715 | 0,181587609 | 0,449341981 | -0,127324005 | -0,108225404 | 0,129090868 | -0,185909132 | -1,445909132 |
| chrX | 519161 | 519304 | 143 | 0,530018269 | 0,400233914 | 0,569757619 | 0,175757115 | 0,149393547 | -0,026845374 | -0,019845374 | -1,279845374 |
| chrX | 519732 | 519838 | 106 | 0,042743409 | 0,340940001 | 0,111402807 | -1,219101651 | -1,036236403 | -0,357632717 | -0,091632717 | -1,351632717 |
| chrX | 519732 | 519858 | 126 | 1,90635603 | 5,09927654 | 0,272117616 | -0,606420994 | -0,515457845 | -0,033615602 | 0,092384398 | -1,167615602 |
| chrX | 523978 | 524108 | 130 | 0,085486818 | 0,107470218 | 0,443035505 | -0,14327752 | -0,121785892 | 0,171987695 | 0,269987695 | -0,990012305 |
| chrX | 526346 | 526498 | 152 | 0,814974327 | 0,289057827 | 0,738179884 | 0,637744162 | 0,542082538 | 0,434611459 | 0,378611459 | -0,881388541 |
| chrX | 527259 | 527387 | 128 | 0,182371877 | 0,655938915 | 0,217546856 | -0,780504955 | -0,663429212 | -0,245180911 | -0,133180911 | -1,393180911 |
| chrX | 530647 | 530795 | 148 | 0,367593315 | 0,229763914 | 0,615365978 | 0,29333246 | 0,249332591 | 0,146270521 | 0,118270521 | -1,141729479 |
| chrX | 531614 | 531753 | 139 | 0,282106498 | 0,170470001 | 0,623334395 | 0,314249947 | 0,267112455 | 0,147930683 | 0,182930683 | -1,077069317 |
| chrX | 533693 | 533845 | 152 | 0,327699667 | 0,296469566 | 0,525017182 | 0,062749932 | 0,053337442 | -0,059582729 | -0,115582729 | -1,375582729 |
| chrX | 537746 | 537899 | 153 | 0,028495606 | 0,051882174 | 0,354520937 | -0,373143192 | -0,317171714 | -0,403252063 | -0,466252063 | -1,726252063 |
| chrX | 542575 | 542723 | 148 | 0,156725832 | 1,133996091 | 0,12142494 | -1,167893119 | -0,992709151 | -1,106643628 | -1,134643628 | -2,394643628 |
| chrX | 544311 | 544443 | 132 | 1,017293128 | 1,356348266 | 0,428579115 | -0,179992788 | -0,15299387 | -0,082662035 | 0,001337965 | -1,258662035 |
| chrX | 547133 | 547290 | 157 | 0,062690333 | 0,077823261 | 0,446151373 | -0,135390993 | -0,115082344 | -0,122099822 | -0,213099822 | -1,473099822 |
| chrX | 554602 | 554737 | 135 | 0,133929347 | 0,151940653 | 0,468497385 | -0,079047591 | -0,067190452 | -0,070845257 | -0,007845257 | -1,267845257 |
| chrX | 557805 | 557945 | 140 | 0,618354647 | 0,396528045 | 0,609286819 | 0,277460721 | 0,235841613 | 0,059409062 | 0,087409062 | -1,172590938 |
| chrX | 557805 | 557945 | 140 | 0,618354647 | 0,396528045 | 0,609286819 | 0,277460721 | 0,235841613 | 0,059409062 | 0,087409062 | -1,172590938 |
| chrX | 557934 | 558083 | 149 | 0,119681545 | 0,040764565 | 0,745929861 | 0,661736237 | 0,562475801 | 0,459046321 | 0,424046321 | -0,835953679 |
| chrX | 558445 | 558574 | 129 | 2,271099785 | 2,872048923 | 0,441577701 | -0,146970376 | -0,12492482 | 0,18771165 | 0,29271165 | -0,96728835 |
| chrX | 561483 | 561587 | 104 | 0,14532759 | 1,901111094 | 0,071014876 | -1,468274214 | -1,248033082 | -0,603504592 | -0,323504592 | -1,583504592 |
| chrX | 561700 | 561836 | 136 | 0,165274514 | 0,144528914 | 0,533481877 | 0,084025388 | 0,071421579 | 0,027655478 | 0,083655478 | -1,176344522 |
| chrX | 563670 | 563822 | 152 | 3,348233686 | 2,831284358 | 0,541827641 | 0,105039183 | 0,089283305 | -0,019483118 | -0,075483118 | -1,335483118 |
| chrX | 567134 | 567269 | 135 | 0,464478375 | 0,370586958 | 0,556218007 | 0,1413873 | 0,120179205 | 0,111508917 | 0,174508917 | -1,085491083 |
| chrX | 569137 | 569291 | 154 | 3,561950729 | 1,864052398 | 0,656459395 | 0,402819234 | 0,342396349 | 0,267457629 | 0,197457629 | -1,062542371 |
| chrX | 572775 | 572903 | 128 | 0,068389454 | 0,059293913 | 0,535617563 | 0,08939893 | 0,07598909 | 0,474791432 | 0,586791432 | -0,673208568 |
| chrX | 573799 | 573959 | 160 | 0,122531105 | 0,137117174 | 0,47191187 | -0,070464771 | -0,059895055 | 0,000797969 | -0,111202031 | -1,371202031 |
| chrX | 582808 | 582958 | 150 | 0,128230226 | 0,055588044 | 0,69759239 | 0,517488464 | 0,439865194 | 0,299255242 | 0,257255242 | -1,002744758 |
| chrX | 586031 | 586180 | 149 | 0,182371877 | 0,155646522 | 0,539532397 | 0,099255756 | 0,084367393 | -0,031640475 | -0,066640475 | -1,326640475 |
| chrX | 590874 | 591029 | 155 | 0,478726178 | 0,481763045 | 0,498419104 | -0,003962729 | -0,00336832 | -0,042331855 | -0,119331855 | -1,379331855 |
| chrX | 591957 | 592113 | 156 | 0,604106844 | 0,507704132 | 0,543353912 | 0,108886928 | 0,092553889 | 0,079508363 | -0,004491637 | -1,264491637 |
| chrX | 594961 | 595112 | 151 | 2,422126496 | 1,57870044 | 0,605406466 | 0,267366398 | 0,227261438 | 0,117887219 | 0,068887219 | -1,191112781 |
| chrX | 595496 | 595660 | 164 | 0,091185939 | 0,040764565 | 0,691061693 | 0,498861988 | 0,42403269 | 0,530945945 | 0,390945945 | -0,869054055 |
| chrX | 598239 | 598403 | 164 | 0,433133209 | 0,43729261 | 0,497610709 | -0,005989101 | -0,005090736 | 0,107072693 | -0,032927307 | -1,292927307 |
| chrX | 598789 | 598947 | 158 | 0,128230226 | 0,129705435 | 0,497140355 | -0,007168129 | -0,00609291 | -0,011018598 | -0,109018598 | -1,369018598 |
| chrX | 600553 | 600699 | 146 | 0,153876272 | 0,066705652 | 0,69759239 | 0,517488464 | 0,439865194 | 0,315826314 | 0,301826314 | -0,958173686 |
| chrX | 606604 | 606728 | 124 | 0,037044288 | 0,114881957 | 0,243830733 | -0,694033076 | -0,589928115 | -0,095552723 | 0,044447277 | -1,215552723 |
| chrX | 608387 | 608544 | 157 | 0,663947616 | 0,689291742 | 0,490635757 | -0,023474833 | -0,019953608 | -0,032765238 | -0,123765238 | -1,383765238 |
| chrX | 608709 | 608811 | 102 | 0,071239015 | 0,255705001 | 0,217893619 | -0,779326791 | -0,662427772 | -0,050717165 | 0,243282835 | -1,016717165 |
| chrX | 610999 | 611149 | 150 | 0,219416165 | 0,185293479 | 0,542156996 | 0,105869357 | 0,089988953 | -0,055078591 | -0,097078591 | -1,357078591 |
| chrX | 623512 | 623664 | 152 | 0,567062556 | 0,744879785 | 0,432231309 | -0,170696241 | -0,145091805 | -0,248751633 | -0,304751633 | -1,564751633 |
| chrX | 624211 | 624342 | 131 | 5,063669157 | 7,55997394 | 0,401125817 | -0,250434134 | -0,212869014 | -0,045013879 | 0,045986121 | -1,214013879 |
| chrX | 624949 | 625089 | 140 | 1,088532143 | 0,652233046 | 0,625318193 | 0,319478604 | 0,271556814 | 0,090487835 | 0,118487835 | -1,141512165 |
| chrX | 632590 | 632738 | 148 | 0,692443222 | 0,544762828 | 0,559683023 | 0,150165614 | 0,127640772 | 0,016729073 | -0,011270927 | -1,271270927 |
| chrX | 635827 | 635966 | 139 | 4,322783405 | 3,005460228 | 0,58987987 | 0,22723597 | 0,193150575 | 0,059900737 | 0,094900737 | -1,165099263 |
| chrX | 636181 | 636337 | 156 | 0,410336724 | 0,52993935 | 0,436400261 | -0,160102259 | -0,13608692 | -0,140780806 | -0,224780806 | -1,484780806 |
| chrX | 636181 | 636337 | 156 | 0,410336724 | 0,52993935 | 0,436400261 | -0,160102259 | -0,13608692 | -0,140780806 | -0,224780806 | -1,484780806 |
| chrX | 636714 | 636864 | 150 | 0,307752543 | 0,259410871 | 0,542617058 | 0,107029113 | 0,090974746 | -0,057844187 | -0,099844187 | -1,359844187 |
| chrX | 637154 | 637305 | 151 | 0,444531451 | 0,555880437 | 0,444348429 | -0,139953338 | -0,118960337 | -0,231868913 | -0,280868913 | -1,540868913 |
| chrX | 637154 | 637305 | 151 | 0,444531451 | 0,555880437 | 0,444348429 | -0,139953338 | -0,118960337 | -0,231868913 | -0,280868913 | -1,540868913 |
| chrX | 638977 | 639126 | 149 | 0,319150785 | 0,200116957 | 0,614617006 | 0,291373097 | 0,247667133 | 0,135155481 | 0,100155481 | -1,159844519 |
| chrX | 653069 | 653214 | 145 | 0,615505086 | 0,377998697 | 0,619529685 | 0,304245806 | 0,258608935 | 0,088693411 | 0,081693411 | -1,178306589 |
| chrX | 661255 | 661420 | 165 | 0,19661968 | 0,244587392 | 0,445640364 | -0,13668381 | -0,116181238 | 0,007280614 | -0,139719386 | -1,399719386 |
| chrX | 662029 | 662161 | 132 | 13,16781946 | 8,953380902 | 0,595257908 | 0,241091522 | 0,204927794 | 0,260195779 | 0,344195779 | -0,915804221 |
| chrX | 662034 | 662161 | 127 | 0,045592969 | 0,066705652 | 0,405997586 | -0,23785292 | -0,202174982 | 0,256880714 | 0,375880714 | -0,884119286 |
| chrX | 662037 | 662135 | 98 | 0,031345166 | 0,044470435 | 0,413439528 | -0,218706025 | -0,185900121 | 0,359755844 | 0,681755844 | -0,578244156 |
| chrX | 671104 | 671223 | 119 | 0,618354647 | 2,809049141 | 0,180414881 | -0,913785129 | -0,776717359 | -0,142155771 | 0,032844229 | -1,227155771 |
| chrX | 671901 | 672050 | 149 | 0,225115286 | 0,314998914 | 0,416792015 | -0,210107168 | -0,178591093 | -0,290638614 | -0,325638614 | -1,585638614 |
| chrX | 673361 | 673549 | 188 | 0,09688506 | 0,159352392 | 0,378106554 | -0,310457456 | -0,263888837 | -0,040961129 | -0,348961129 | -1,608961129 |
| chrX | 673385 | 673539 | 154 | 0,418885406 | 0,35205761 | 0,543341592 | 0,108855862 | 0,092527483 | 0,025358056 | -0,044641944 | -1,304641944 |
| chrX | 674527 | 674676 | 149 | 0,307752543 | 0,211234566 | 0,592986873 | 0,235235114 | 0,199949847 | 0,092519592 | 0,057519592 | -1,202480408 |

TABLE 2

| Chrom | Start | End | Length | Section A | Section B | A/A+B | Z-score | Z * 0.85 | Phase correction | Length correction | ΔLK |
|---|---|---|---|---|---|---|---|---|---|---|---|
| chrX | 684490 | 684642 | 152 | 0,3903898 | 0,337234132 | 0,536526883 | 0,091687619 | 0,077934477 | -0,010344291 | -0,066344291 | -1,326344291 |
| chrX | 684490 | 684642 | 152 | 0,3903898 | 0,337234132 | 0,536526883 | 0,091687619 | 0,077934477 | -0,010344291 | -0,066344291 | -1,326344291 |
| chrX | 685942 | 686069 | 127 | 0,492973981 | 0,922761525 | 0,348210509 | -0,39015624 | -0,331632804 | 0,117668893 | 0,236668893 | -1,023331107 |
| chrX | 687031 | 687188 | 157 | 0,455929693 | 0,61517435 | 0,425663311 | -0,187426022 | -0,159312119 | -0,171408057 | -0,262408057 | -1,522408057 |
| chrX | 687681 | 687842 | 161 | 0,450230572 | 0,641115437 | 0,41254613 | -0,221000223 | -0,18785019 | -0,124074974 | -0,243074974 | -1,503074974 |
| chrX | 689548 | 689698 | 150 | 0,812124766 | 0,452116089 | 0,642381365 | 0,364831393 | 0,310106684 | 0,162263239 | 0,120263239 | -1,139736761 |
| chrX | 695637 | 695789 | 152 | 0,108283302 | 0,074117392 | 0,593656197 | 0,236960278 | 0,201416237 | 0,115366601 | 0,059366601 | -1,200633399 |
| chrX | 696260 | 696391 | 131 | 1,12557643 | 1,226642831 | 0,478516799 | -0,053876453 | -0,045794985 | 0,079073615 | 0,170073615 | -1,089926385 |
| chrX | 697538 | 697692 | 154 | 0,113982423 | 0,122293696 | 0,482411949 | -0,044100996 | -0,037485847 | -0,093417057 | -0,163417057 | -1,423417057 |
| chrX | 699421 | 699528 | 107 | 0,034194727 | 0,255705001 | 0,117953636 | -1,1852787 | -1,007486895 | -0,321540912 | -0,062540912 | -1,322540912 |
| chrX | 699421 | 699559 | 138 | 0,04844253 | 0,040764565 | 0,543034495 | 0,108081541 | 0,09186931 | -0,020454783 | 0,021545217 | -1,238454783 |
| chrX | 699421 | 699560 | 139 | 0,045592969 | 0,051882174 | 0,467739443 | -0,080953558 | -0,068810524 | -0,202267091 | -0,167267091 | -1,427267091 |
| chrX | 699421 | 699561 | 140 | 0,062690333 | 0,037058696 | 0,628480635 | 0,327831948 | 0,278657156 | 0,098849309 | 0,126849309 | -1,133150691 |
| chrX | 699421 | 699563 | 142 | 0,034194727 | 0,040764565 | 0,456177292 | -0,110069087 | -0,093558724 | -0,277311187 | -0,263311187 | -1,523311187 |
| chrX | 699421 | 699564 | 143 | 16,10286686 | 10,44684634 | 0,606517545 | 0,270253923 | 0,229715834 | 0,054434842 | 0,061434842 | -1,198565158 |
| chrX | 700845 | 700991 | 146 | 1,376337762 | 0,815291307 | 0,627997585 | 0,326554541 | 0,27757136 | 0,1540763 | 0,1400763 | -1,1199237 |
| chrX | 701322 | 701470 | 148 | 3,094622794 | 2,405109357 | 0,562686093 | 0,157782975 | 0,134115528 | 0,028180105 | 0,000180105 | -1,259819895 |
| chrX | 705757 | 705926 | 169 | 0,350495952 | 0,678174133 | 0,340727272 | -0,410479083 | -0,348907221 | -0,237219511 | -0,412219511 | -1,672219511 |
| chrX | 705757 | 705926 | 169 | 0,350495952 | 0,678174133 | 0,340727272 | -0,410479083 | -0,348907221 | -0,237219511 | -0,412219511 | -1,672219511 |
| chrX | 706905 | 707066 | 161 | 0,068389454 | 0,096352609 | 0,415130494 | -0,214366854 | -0,182211826 | -0,110130344 | -0,229130344 | -1,489130344 |
| chrX | 712931 | 713070 | 139 | 0,324849907 | 0,255705001 | 0,559550703 | 0,149830185 | 0,127355657 | -0,004911679 | 0,030088321 | -1,229911679 |
| chrX | 714988 | 715121 | 133 | 0,478726178 | 0,359469349 | 0,571139027 | 0,179274784 | 0,152383566 | 0,19317228 | 0,27017228 | -0,98982772 |
| chrX | 720830 | 720969 | 139 | 0,948903674 | 0,381704567 | 0,713135275 | 0,562567469 | 0,478182348 | 0,336129742 | 0,371129742 | -0,888870258 |
| chrX | 721017 | 721124 | 107 | 0,034194727 | 0,485468915 | 0,065801654 | -1,507809432 | -1,281638017 | -0,576883772 | -0,317883772 | -1,577883772 |
| chrX | 721017 | 721148 | 131 | 0,04844253 | 0,055588044 | 0,465656665 | -0,08619258 | -0,073263693 | 0,034771502 | 0,125771502 | -1,134228498 |
| chrX | 721017 | 721149 | 132 | 0,085486818 | 0,118587827 | 0,418899751 | -0,204708983 | -0,174002636 | -0,139438328 | -0,055438328 | -1,315438328 |
| chrX | 721017 | 721153 | 136 | 18,6589227 | 16,02047419 | 0,538040577 | 0,095498544 | 0,081173762 | 0,026933837 | 0,082933837 | -1,177066163 |
| chrX | 721018 | 721153 | 135 | 0,065539893 | 0,037058696 | 0,638799168 | 0,355250862 | 0,301963233 | 0,273001084 | 0,336001084 | -0,923998916 |
| chrX | 721019 | 721153 | 134 | 0,039893848 | 0,059293913 | 0,402205349 | -0,247642923 | -0,210496484 | -0,189371996 | -0,119371996 | -1,379371996 |
| chrX | 721020 | 721153 | 133 | 0,031345166 | 0,051882174 | 0,376621026 | -0,314367347 | -0,267212245 | -0,233510415 | -0,156510415 | -1,416510415 |
| chrX | 723086 | 723238 | 152 | 0,760832676 | 0,822703046 | 0,480464485 | -0,04898786 | -0,041639681 | -0,108966951 | -0,164966951 | -1,424966951 |
| chrX | 723301 | 723431 | 130 | 0,156725832 | 0,066705652 | 0,701449182 | 0,528573085 | 0,449287122 | 0,736266246 | 0,834266246 | -0,425733754 |
| chrX | 723442 | 723594 | 152 | 0,29350474 | 0,17417587 | 0,627575173 | 0,325437929 | 0,27662224 | 0,209168622 | 0,153168622 | -1,106831378 |
| chrX | 725588 | 725741 | 153 | 0,347646391 | 0,381704567 | 0,476651724 | -0,0585589 | -0,049775065 | -0,143226462 | -0,206226462 | -1,466226462 |
| chrX | 729427 | 729584 | 157 | 0,834921251 | 0,559586306 | 0,598721209 | 0,250038492 | 0,212532718 | 0,199302874 | 0,108302874 | -1,151697126 |
| chrX | 729427 | 729584 | 157 | 0,834921251 | 0,559586306 | 0,598721209 | 0,250038492 | 0,212532718 | 0,199302874 | 0,108302874 | -1,151697126 |
| chrX | 729799 | 729953 | 154 | 0,552814753 | 0,392822175 | 0,584595141 | 0,21366319 | 0,181613711 | 0,13041811 | 0,06041811 | -1,19958189 |
| chrX | 732737 | 732877 | 140 | 2,088727908 | 1,58611218 | 0,568386068 | 0,172266698 | 0,146426693 | -0,033381153 | -0,005381153 | -1,265381153 |
| chrX | 735177 | 735331 | 154 | 0,039893848 | 0,081529131 | 0,328552705 | -0,443913104 | -0,377326138 | -0,425704053 | -0,495704053 | -1,755704053 |
| chrX | 735177 | 735331 | 154 | 0,039893848 | 0,081529131 | 0,328552705 | -0,443913104 | -0,377326138 | -0,425704053 | -0,495704053 | -1,755704053 |
| chrX | 736882 | 737013 | 131 | 0,205168362 | 0,385410436 | 0,34740218 | -0,39234359 | -0,333492051 | -0,22883147 | -0,13783147 | -1,39783147 |
| chrX | 739148 | 739242 | 94 | 2,065931423 | 11,71425374 | 0,14992044 | -1,036774676 | -0,881258475 | -0,376313551 | -0,026313551 | -1,286313551 |
| chrX | 745644 | 745739 | 95 | 0,085486818 | 0,307587175 | 0,217482762 | -0,780722841 | -0,663614415 | -0,131020398 | 0,211979602 | -1,048020398 |
| chrX | 745644 | 745742 | 98 | 0,056991212 | 0,077823261 | 0,422738082 | -0,194893703 | -0,165659647 | 0,38564365 | 0,70764365 | -0,55235635 |
| chrXI | 6938 | 7102 | 164 | 3,259897308 | 2,886872402 | 0,530343166 | 0,07613252 | 0,064712642 | 0,122198787 | -0,017801213 | -1,277801213 |
| chrXI | 10271 | 10420 | 149 | 0,273557816 | 0,151940653 | 0,642911399 | 0,366251789 | 0,31131402 | 0,207069374 | 0,172069374 | -1,087930626 |
| chrXI | 17984 | 18132 | 148 | 0,213717044 | 0,129705435 | 0,622315244 | 0,311567137 | 0,264832067 | 0,144482326 | 0,116482326 | -1,143517674 |
| chrXI | 19266 | 19389 | 123 | 1,929152515 | 5,89233263 | 0,246647853 | -0,68507641 | -0,582314949 | 0,025296211 | 0,172296211 | -1,087703789 |
| chrXI | 19267 | 19389 | 122 | 0,065539893 | 0,207528696 | 0,240012568 | -0,706262136 | -0,600322816 | 0,028135062 | 0,182135062 | -1,077864938 |
| chrXI | 19267 | 19414 | 147 | 0,09973462 | 0,118587827 | 0,456822566 | -0,108441943 | -0,092175651 | -0,165101711 | -0,186101711 | -1,446101711 |
| chrXI | 20703 | 20861 | 158 | 0,447381012 | 0,52993935 | 0,457762909 | -0,106071255 | -0,090160566 | -0,067894316 | -0,165894316 | -1,425894316 |
| chrXI | 31510 | 31641 | 131 | 0,282106498 | 0,500292393 | 0,360566076 | -0,356946107 | -0,303404191 | -0,076794249 | 0,014205751 | -1,245794249 |
| chrXI | 32617 | 32778 | 161 | 0,14817715 | 0,188999349 | 0,439464645 | -0,152326671 | -0,12947767 | -0,123199352 | -0,242199352 | -1,502199352 |
| chrXI | 38219 | 38379 | 160 | 0,068389454 | 0,037058696 | 0,648560019 | 0,381435717 | 0,324220359 | 0,339421729 | 0,227421729 | -1,032578271 |
| chrXI | 39078 | 39236 | 158 | 0,396088921 | 0,61146848 | 0,393117971 | -0,271201653 | -0,230521405 | -0,210674681 | -0,308674681 | -1,568674681 |
| chrXI | 41766 | 41916 | 150 | 0,273557816 | 0,211234566 | 0,564278289 | 0,16182531 | 0,137551513 | 0,017444382 | -0,024555618 | -1,284555618 |
| chrXI | 42476 | 42618 | 142 | 8,534433948 | 5,770038934 | 0,596626944 | 0,244625896 | 0,207932012 | 0,102552613 | 0,116552613 | -1,143447387 |
| chrXI | 42479 | 42618 | 139 | 0,039893848 | 0,051882174 | 0,434687048 | -0,164453569 | -0,139785534 | -0,142993666 | -0,107993666 | -1,367993666 |
| chrXI | 44129 | 44285 | 156 | 0,153876272 | 0,211234566 | 0,421450846 | -0,198183249 | -0,168455762 | -0,179919203 | -0,263919203 | -1,523919203 |
| chrXI | 45474 | 45591 | 117 | 0,176672756 | 0,952408482 | 0,156474796 | -1,009052218 | -0,857694386 | -0,160710626 | 0,028289374 | -1,231710626 |
| chrXI | 52845 | 53010 | 165 | 1,344992595 | 1,66764131 | 0,446450726 | -0,134633754 | -0,114438691 | -0,033049486 | -0,180049486 | -1,440049486 |
| chrXI | 53086 | 53185 | 99 | 0,091185939 | 1,148819569 | 0,073536721 | -1,449946491 | -1,232454517 | -0,785695943 | -0,470695943 | -1,730695943 |
| chrXI | 55429 | 55585 | 156 | 0,307752543 | 0,244587392 | 0,557179598 | 0,143822286 | 0,12224843 | 0,097166489 | 0,013166489 | -1,246833511 |
| chrXI | 55463 | 55623 | 160 | 0,113982423 | 0,103764348 | 0,523463207 | 0,058847485 | 0,050020363 | 0,068429027 | -0,043570973 | -1,303570973 |
| chrXI | 55924 | 56074 | 150 | 0,863416857 | 0,800467829 | 0,518916283 | 0,047433872 | 0,040318791 | -0,08374672 | -0,12574672 | -1,38574672 |
| chrXI | 55924 | 56080 | 156 | 0,125380666 | 0,122293696 | 0,506231912 | 0,015621722 | 0,013278464 | -0,010459686 | -0,094459686 | -1,354459686 |
| chrXI | 56846 | 56978 | 132 | 0,156725832 | 0,170470001 | 0,47899703 | -0,052670982 | -0,044770335 | 0,136424479 | 0,220424479 | -1,039575521 |
| chrXI | 61848 | 62032 | 184 | 0,09688506 | 0,170470001 | 0,36238349 | -0,352095051 | -0,299280793 | 0,011385456 | -0,268614544 | -1,528614544 |
| chrXI | 63877 | 64030 | 153 | 0,076938136 | 0,118587827 | 0,393493196 | -0,270226 | -0,2296921 | -0,322514067 | -0,385514067 | -1,645514067 |
| chrXI | 65154 | 65316 | 162 | 0,210867483 | 0,155646522 | 0,575332675 | 0,189967434 | 0,161472318 | 0,169430117 | 0,043430117 | -1,216569883 |
| chrXI | 65672 | 65838 | 166 | 3,923844923 | 3,068460011 | 0,561166162 | 0,153926517 | 0,130837539 | 0,233631803 | 0,079631803 | -1,180368197 |
| chrXI | 68614 | 68763 | 149 | 1,085682582 | 0,885702829 | 0,550720613 | 0,127482183 | 0,108359856 | 0,001605651 | -0,033394349 | -1,293394349 |
| chrXI | 73102 | 73261 | 159 | 0,108283302 | 0,111176087 | 0,493409293 | -0,016521205 | -0,014043024 | 0,010190038 | -0,094809962 | -1,354809962 |
| chrXI | 86463 | 86611 | 148 | 0,153876272 | 0,125999566 | 0,549801916 | 0,125160904 | 0,106386768 | -0,006537371 | -0,034537371 | -1,294537371 |
| chrXI | 86483 | 86645 | 162 | 1,051487855 | 1,011702395 | 0,509641733 | 0,024170595 | 0,020545005 | 0,034350256 | -0,091649744 | -1,351649744 |
| chrXI | 88336 | 88484 | 148 | 0,09973462 | 0,092646739 | 0,518421434 | 0,046192108 | 0,039263292 | -0,072316734 | -0,100316734 | -1,360316734 |
| chrXI | 89789 | 89957 | 168 | 0,284956058 | 0,233469783 | 0,549656355 | 0,124793175 | 0,106074199 | 0,248118621 | 0,080118621 | -1,179881379 |
| chrXI | 89789 | 89957 | 168 | 0,284956058 | 0,233469783 | 0,549656355 | 0,124793175 | 0,106074199 | 0,248118621 | 0,080118621 | -1,179881379 |
| chrXI | 91032 | 91184 | 152 | 2,481967268 | 2,116051529 | 0,539790588 | 0,099906161 | 0,084920237 | 3,61E-05 | -0,055963927 | -1,315963927 |
| chrXI | 97596 | 97707 | 111 | 1,70973635 | 14,67153766 | 0,104371391 | -1,257029976 | -1,068475479 | -0,378182241 | -0,147182241 | -1,407182241 |
| chrXI | 97601 | 97707 | 106 | 0,042743409 | 0,289057827 | 0,128822331 | -1,131975672 | -0,962179321 | -0,441094211 | -0,175094211 | -1,435094211 |
| chrXI | 99816 | 99946 | 130 | 0,168124074 | 0,100058479 | 0,626901611 | 0,323658255 | 0,275109516 | 0,527325968 | 0,625325968 | -0,634674032 |
| chrXI | 99816 | 99946 | 130 | 0,168124074 | 0,100058479 | 0,626901611 | 0,323658255 | 0,275109516 | 0,527325968 | 0,625325968 | -0,634674032 |
| chrXI | 103049 | 103197 | 148 | 0,347646391 | 0,274234349 | 0,559024213 | 0,148495706 | 0,12622135 | 0,005110108 | -0,022889892 | -1,282889892 |
| chrXI | 103049 | 103197 | 148 | 0,347646391 | 0,274234349 | 0,559024213 | 0,148495706 | 0,12622135 | 0,005110108 | -0,022889892 | -1,282889892 |
| chrXI | 103372 | 103526 | 154 | 5,189049823 | 3,5205761 | 0,595783317 | 0,242447586 | 0,206080448 | 0,173625353 | 0,103625353 | -1,156374647 |
| chrXI | 105256 | 105405 | 149 | 0,339097709 | 0,192705218 | 0,637637914 | 0,352152134 | 0,299329314 | 0,185161926 | 0,150161926 | -1,109838074 |
| chrXI | 105988 | 106134 | 146 | 0,276407377 | 0,100058479 | 0,734216325 | 0,625615224 | 0,531772941 | 0,441727294 | 0,427727294 | -0,832272706 |
| chrXI | 110658 | 110811 | 153 | 0,128230226 | 0,125999566 | 0,504387095 | 0,010997038 | 0,009347483 | -0,075077383 | -0,138077383 | -1,398077383 |
| chrXI | 115225 | 115382 | 157 | 0,353345512 | 0,329822392 | 0,517216207 | 0,043168034 | 0,036692829 | 0,032830238 | -0,058169762 | -1,318169762 |
| chrXI | 116126 | 116262 | 136 | 0,031345166 | 0,070411522 | 0,308040355 | -0,501412691 | -0,426200787 | -0,320932821 | -0,264932821 | -1,524932821 |
| chrXI | 116126 | 116262 | 136 | 0,031345166 | 0,070411522 | 0,308040355 | -0,501412691 | -0,426200787 | -0,320932821 | -0,264932821 | -1,524932821 |
| chrXI | 116231 | 116308 | 77 | 0,131079787 | 0,040764565 | 0,762782048 | 0,715280226 | 0,607988192 | | 0,469 | -0,791 |
| chrXI | 116232 | 116308 | 76 | 1,242408414 | 0,911643916 | 0,576777266 | 0,193655717 | 0,16460736 | | 0,476 | -0,784 |
| chrXI | 116232 | 116357 | 125 | 1,70973635 | 1,545347614 | 0,525251075 | 0,06333738 | 0,053836773 | 0,572627202 | 0,705627202 | -0,554372798 |
| chrXI | 117579 | 117716 | 137 | 0,626903328 | 0,515115871 | 0,548942897 | 0,122991025 | 0,104542371 | 0,163653163 | 0,212653163 | -1,047346837 |
| chrXI | 120342 | 120499 | 157 | 0,44168189 | 0,363175219 | 0,548770565 | 0,122555783 | 0,104172416 | 0,097356327 | 0,006356327 | -1,253643673 |
| chrXI | 120705 | 120857 | 152 | 0,980248841 | 0,933879134 | 0,512112489 | 0,030366173 | 0,025811247 | -0,057594098 | -0,113594098 | -1,373594098 |
| chrXI | 121556 | 121713 | 157 | 0,037044288 | 0,092646739 | 0,285634931 | -0,566182298 | -0,481254953 | -0,489848996 | -0,580848996 | -1,840848996 |
| chrXI | 130706 | 130855 | 149 | 1,441877655 | 1,156231309 | 0,554971972 | 0,138233278 | 0,117498286 | 0,000210026 | -0,034789974 | -1,294789974 |
| chrXI | 130929 | 131079 | 150 | 0,102584181 | 0,074117392 | 0,58055047 | 0,203301913 | 0,172806626 | 0,058158787 | 0,016158787 | -1,243841213 |

TABLE 2

| Chrom | Start | End | Length | Section A | Section B | A/A+B | Z-score | Z * 0.85 | Phase correction | Length correction | ΔLK |
|---|---|---|---|---|---|---|---|---|---|---|---|
| chrXI | 132985 | 133151 | 166 | 1,179718082 | 1,100643265 | 0,517338221 | 0,043474165 | 0,03695304 | 0,148479833 | -0,005520167 | -1,265520167 |
| chrXI | 134033 | 134163 | 130 | 0,404637603 | 1,05246696 | 0,277699771 | -0,589688448 | -0,501235181 | -0,251200418 | -0,153200418 | -1,413200418 |
| chrXI | 136517 | 136676 | 159 | 0,162424953 | 0,281646088 | 0,365763444 | -0,34309514 | -0,291630869 | -0,275354503 | -0,380354503 | -1,640354503 |
| chrXI | 140618 | 140768 | 150 | 0,316301225 | 0,285351958 | 0,525720189 | 0,064515681 | 0,054838328 | -0,063274442 | -0,105274442 | -1,365274442 |
| chrXI | 142672 | 142804 | 132 | 1,695488547 | 1,285936744 | 0,568683895 | 0,173024449 | 0,147070782 | 0,302129605 | 0,386129605 | -0,873870395 |
| chrXI | 145250 | 145428 | 178 | 0,151026711 | 0,155646522 | 0,49246786 | -0,018881397 | -0,016049187 | 0,279449967 | 0,041449967 | -1,218550033 |
| chrXI | 146568 | 146719 | 151 | 0,473027057 | 0,337234132 | 0,583795773 | 0,211613657 | 0,179871608 | 0,101971149 | 0,052971149 | -1,207028851 |
| chrXI | 147356 | 147520 | 164 | 0,056991212 | 0,081529131 | 0,411428464 | -0,223871969 | -0,190291174 | -0,125104196 | -0,265104196 | -1,525104196 |
| chrXI | 149408 | 149568 | 160 | 0,108283302 | 0,125999566 | 0,462190442 | -0,094916836 | -0,080679311 | -0,062937275 | -0,174937275 | -1,434937275 |
| chrXI | 149964 | 150108 | 144 | 0,316301225 | 0,307587175 | 0,506983661 | 0,017506335 | 0,014880385 | -0,045308002 | -0,045308002 | -1,305308002 |
| chrXI | 151755 | 151890 | 135 | 0,054141651 | 0,137117174 | 0,283080537 | -0,573714411 | -0,48765725 | -0,37137928 | -0,30837928 | -1,56837928 |
| chrXI | 161235 | 161384 | 149 | 0,222265726 | 0,159352392 | 0,5824297 | 0,208113175 | 0,176896198 | 0,059452199 | 0,024452199 | -1,235547801 |
| chrXI | 161799 | 161928 | 129 | 1,8579135 | 2,616343922 | 0,415245107 | -0,214072894 | -0,18196196 | 0,112655679 | 0,217655679 | -1,042344321 |
| chrXI | 162802 | 162952 | 150 | 0,498673102 | 0,522527611 | 0,488320362 | -0,029280693 | -0,024888589 | -0,147849305 | -0,189849305 | -1,449849305 |
| chrXI | 170837 | 170984 | 147 | 0,082637257 | 0,26311674 | 0,239005934 | -0,709503843 | -0,603078266 | -0,675666349 | -0,696666349 | -1,956666349 |
| chrXI | 171758 | 171891 | 133 | 0,14817715 | 0,200116957 | 0,425436857 | -0,188003748 | -0,159803185 | -0,027196837 | 0,049803163 | -1,210196837 |
| chrXI | 172320 | 172456 | 136 | 1,134125112 | 1,178466526 | 0,490413047 | -0,02403324 | -0,020428254 | 0,075219961 | 0,131219961 | -1,128780039 |
| chrXI | 178039 | 178227 | 188 | 0,139628469 | 0,277940218 | 0,334384433 | -0,427838279 | -0,363662537 | -0,052970865 | -0,360970865 | -1,620970865 |
| chrXI | 178982 | 179138 | 156 | 0,119681545 | 0,133411305 | 0,472876041 | -0,06804215 | -0,057835827 | -0,07586694 | -0,15986694 | -1,41986694 |
| chrXI | 179058 | 179209 | 151 | 0,042743409 | 0,092646739 | 0,315705458 | -0,47974192 | -0,407780632 | -0,492591915 | -0,541591915 | -1,801591915 |
| chrXI | 179058 | 179209 | 151 | 0,042743409 | 0,092646739 | 0,315705458 | -0,47974192 | -0,407780632 | -0,492591915 | -0,541591915 | -1,801591915 |
| chrXI | 182120 | 182298 | 178 | 0,202318801 | 0,185293479 | 0,521961795 | 0,05507789 | 0,046816206 | 0,355868288 | 0,117868288 | -1,142131712 |
| chrXI | 183536 | 183690 | 154 | 0,284956058 | 0,296469566 | 0,490098899 | -0,024820929 | -0,021097789 | -0,053518065 | -0,123518065 | -1,383518065 |
| chrXI | 183964 | 184081 | 117 | 0,458779254 | 2,171639573 | 0,174413006 | -0,936868875 | -0,796338543 | -0,102982643 | 0,086017357 | -1,173982643 |
| chrXI | 187208 | 187352 | 144 | 1,208213687 | 0,844938264 | 0,588467739 | 0,223605195 | 0,190064415 | 0,117481947 | 0,117481947 | -1,142518053 |
| chrXI | 188029 | 188179 | 150 | 0,718089267 | 0,515115871 | 0,582295066 | 0,207768321 | 0,176603073 | 0,047590992 | 0,005590992 | -1,254409008 |
| chrXI | 190331 | 190479 | 148 | 0,364743755 | 0,322410653 | 0,530803194 | 0,077289038 | 0,065695682 | -0,054896657 | -0,082896657 | -1,342896657 |
| chrXI | 191024 | 191171 | 147 | 0,3903898 | 0,270528479 | 0,590677868 | 0,229289049 | 0,194895692 | 0,11557827 | 0,09457827 | -1,16542173 |
| chrXI | 195322 | 195449 | 127 | 0,128230226 | 0,370586958 | 0,257068582 | -0,652409303 | -0,554547908 | -0,166366987 | -0,047366987 | -1,307366987 |
| chrXI | 196316 | 196481 | 165 | 2,912250916 | 3,009166098 | 0,491816555 | -0,020514294 | -0,017437149 | 0,064685247 | -0,082314753 | -1,342314753 |
| chrXI | 196620 | 196752 | 132 | 0,527168708 | 0,678174133 | 0,437359969 | -0,157666069 | -0,134016159 | 0,003751455 | 0,087751455 | -1,172248545 |
| chrXI | 201937 | 202099 | 162 | 0,088336378 | 0,155646522 | 0,36205971 | -0,352958678 | -0,300014876 | -0,271529265 | -0,397529265 | -1,657529265 |
| chrXI | 204817 | 204983 | 166 | 0,133929347 | 0,188999349 | 0,414733497 | -0,215385217 | -0,183077435 | -0,071847362 | -0,225847362 | -1,485847362 |
| chrXI | 207618 | 207770 | 152 | 0,971700159 | 0,589233263 | 0,622512239 | 0,312085531 | 0,265272701 | 0,180054768 | 0,124054768 | -1,135945232 |
| chrXI | 209750 | 209876 | 126 | 0,595558162 | 1,84552305 | 0,243973105 | -0,693579092 | -0,589542228 | -0,123680546 | 0,002319454 | -1,257680546 |
| chrXI | 214868 | 215018 | 150 | 0,054141651 | 0,122293696 | 0,30686397 | -0,50475925 | -0,429045363 | -0,555026937 | -0,597026937 | -1,857026937 |
| chrXI | 214868 | 215019 | 151 | 0,658248495 | 0,607762611 | 0,519938958 | 0,050000381 | 0,042500324 | -0,040297145 | -0,089297145 | -1,349297145 |
| chrXI | 214886 | 215018 | 132 | 0,492973981 | 0,807879568 | 0,378961937 | -0,308208251 | -0,261977013 | -0,111451285 | -0,027451285 | -1,287451285 |
| chrXI | 214886 | 215019 | 133 | 0,153876272 | 0,181587609 | 0,458696987 | -0,10371695 | -0,088159408 | 0,037749368 | 0,114749368 | -1,145250632 |
| chrXI | 216300 | 216452 | 152 | 0,772230918 | 0,718938698 | 0,517869268 | 0,044806601 | 0,038085611 | -0,035795978 | -0,091795978 | -1,351795978 |
| chrXI | 216511 | 216657 | 146 | 1,131275552 | 0,633703698 | 0,640956857 | 0,361017604 | 0,306864964 | 0,218566995 | 0,204566995 | -1,055433005 |
| chrXI | 226833 | 226983 | 150 | 0,555664314 | 0,52623348 | 0,513601485 | 0,034100475 | 0,028985404 | -0,094144118 | -0,136144118 | -1,396144118 |
| chrXI | 226930 | 227062 | 132 | 7,286326412 | 8,786616771 | 0,453328698 | -0,117255741 | -0,09966738 | 0,061094685 | 0,145094685 | -1,114905315 |
| chrXI | 227206 | 227326 | 120 | 0,236513528 | 1,389701092 | 0,145438078 | -1,056201522 | -0,897771294 | -0,215397937 | -0,047397937 | -1,307397937 |
| chrXI | 229813 | 229973 | 160 | 0,3903898 | 0,381704567 | 0,505624464 | 0,014098907 | 0,011984071 | 0,016770907 | -0,095229093 | -1,355229093 |
| chrXI | 233634 | 233784 | 150 | 0,359044634 | 0,289057827 | 0,553993628 | 0,135757811 | 0,11539414 | -0,011879396 | -0,053879396 | -1,313879396 |
| chrXI | 233634 | 233784 | 150 | 0,359044634 | 0,289057827 | 0,553993628 | 0,135757811 | 0,11539414 | -0,011879396 | -0,053879396 | -1,313879396 |
| chrXI | 236594 | 236764 | 170 | 0,19377012 | 0,26311674 | 0,424109636 | -0,191390999 | -0,162682349 | -0,026111303 | -0,208111303 | -1,468111303 |
| chrXI | 241460 | 241610 | 150 | 0,310602104 | 0,17788174 | 0,635849287 | 0,347385899 | 0,295278014 | 0,173541198 | 0,131541198 | -1,128458802 |
| chrXI | 245174 | 245347 | 173 | 0,088336378 | 0,148234783 | 0,373402986 | -0,322853792 | -0,274425723 | -0,071240295 | -0,274240295 | -1,534240295 |
| chrXI | 245189 | 245339 | 150 | 0,609805965 | 0,463233697 | 0,568297693 | 0,172041869 | 0,146235588 | 0,01372297 | -0,02827703 | -1,28827703 |
| chrXI | 245551 | 245725 | 174 | 0,102584181 | 0,17788174 | 0,365763444 | -0,34309514 | -0,291630869 | -0,046690766 | -0,256690766 | -1,516690766 |
| chrXI | 247204 | 247369 | 165 | 0,492973981 | 0,585527393 | 0,457091658 | -0,107763477 | -0,091598955 | -0,008912496 | -0,155912496 | -1,415912496 |
| chrXI | 247204 | 247369 | 165 | 0,492973981 | 0,585527393 | 0,457091658 | -0,107763477 | -0,091598955 | -0,008912496 | -0,155912496 | -1,415912496 |
| chrXI | 249638 | 249773 | 135 | 0,045592969 | 0,092646739 | 0,329810947 | -0,440435256 | -0,374369967 | -0,257078261 | -0,194078261 | -1,454078261 |
| chrXI | 251126 | 251263 | 137 | 0,045592969 | 0,122293696 | 0,271569926 | -0,608071803 | -0,516861033 | -0,458708437 | -0,409708437 | -1,669708437 |
| chrXI | 253186 | 253299 | 113 | 0,330549028 | 2,208698269 | 0,130175989 | -1,125559597 | -0,956725658 | -0,261056574 | -0,044056574 | -1,304056574 |
| chrXI | 253541 | 253688 | 147 | 0,142478029 | 0,188999349 | 0,429827309 | -0,17681384 | -0,150291764 | -0,239689262 | -0,260689262 | -1,520689262 |
| chrXI | 257352 | 257460 | 108 | 0,039893848 | 0,048176305 | 0,452978074 | -0,118140734 | -0,100419624 | 0,566956687 | 0,818956687 | -0,441043313 |
| chrXI | 257353 | 257460 | 107 | 0,407487163 | 0,937585003 | 0,302948179 | -0,51593994 | -0,438548949 | 0,127014847 | 0,386014847 | -0,873985153 |
| chrXI | 257353 | 257478 | 125 | 0,755133555 | 1,682464789 | 0,309785883 | -0,496457358 | -0,421988754 | 0,071761483 | 0,204761483 | -1,055238517 |
| chrXI | 257617 | 257767 | 150 | 1,188266763 | 0,715232829 | 0,624253753 | 0,316672003 | 0,269171202 | 0,136265898 | 0,094265898 | -1,165734102 |
| chrXI | 258177 | 258336 | 159 | 0,324849907 | 0,314998914 | 0,507697906 | 0,019296987 | 0,016402439 | 0,032865901 | -0,072134099 | -1,332134099 |
| chrXI | 261407 | 261543 | 136 | 0,091185939 | 0,066705652 | 0,577522451 | 0,195559327 | 0,166225428 | 0,243684279 | 0,299684279 | -0,960315721 |
| chrXI | 269387 | 269539 | 152 | 0,347646391 | 0,340940001 | 0,504869679 | 0,012206779 | 0,010375762 | -0,068663016 | -0,124663016 | -1,384663016 |
| chrXI | 269387 | 269539 | 152 | 0,347646391 | 0,340940001 | 0,504869679 | 0,012206779 | 0,010375762 | -0,068663016 | -0,124663016 | -1,384663016 |
| chrXI | 270529 | 270668 | 139 | 0,646850252 | 0,544762828 | 0,542835811 | 0,10758061 | 0,091443518 | 0,083212133 | 0,118212133 | -1,141787867 |
| chrXI | 275707 | 275869 | 162 | 4,943987612 | 5,784862412 | 0,460812445 | -0,098387135 | -0,083629065 | -0,043772106 | -0,169772106 | -1,429772106 |
| chrXI | 275877 | 276030 | 153 | 0,102584181 | 0,059293913 | 0,633712557 | 0,341702373 | 0,290447017 | 0,205647948 | 0,142647948 | -1,117352052 |
| chrXI | 277324 | 277474 | 150 | 0,122531105 | 0,125999566 | 0,493022067 | -0,017491975 | -0,014868179 | -0,142413256 | -0,184413256 | -1,444413256 |
| chrXI | 278497 | 278637 | 140 | 20,10934904 | 14,40471505 | 0,582642165 | 0,208657437 | 0,177358821 | 0,143641681 | 0,171641681 | -1,088358319 |
| chrXI | 278499 | 278637 | 138 | 0,04844253 | 0,037058696 | 0,566571175 | 0,16765125 | 0,142503563 | 0,146950165 | 0,188950165 | -1,071049835 |
| chrXI | 278501 | 278637 | 136 | 0,056991212 | 0,051882174 | 0,523463207 | 0,058847485 | 0,050020363 | 0,12053464 | 0,17653464 | -1,08346536 |
| chrXI | 278502 | 278637 | 135 | 0,09688506 | 0,062999783 | 0,605967759 | 0,268824834 | 0,228501109 | 0,340239586 | 0,403239586 | -0,856760414 |
| chrXI | 282993 | 283156 | 163 | 0,14817715 | 0,181587609 | 0,449341981 | -0,127324005 | -0,108225404 | -0,048195826 | -0,181195826 | -1,441195826 |
| chrXI | 286651 | 286824 | 173 | 0,914708947 | 1,204407613 | 0,431646359 | -0,172184202 | -0,146356571 | 0,066114029 | -0,136885971 | -1,396885971 |
| chrXI | 287603 | 287693 | 90 | 0,028495606 | 0,070411522 | 0,288104674 | -0,558930214 | -0,475090682 | -0,110563336 | 0,267436664 | -0,992563336 |
| chrXI | 288326 | 288474 | 148 | 0,185221438 | 0,17788174 | 0,510106904 | 0,025336963 | 0,021536418 | -0,095497141 | -0,123497141 | -1,383497141 |
| chrXI | 290076 | 290228 | 152 | 3,66168535 | 3,650281535 | 0,500779805 | 0,001954682 | 0,00166148 | -0,074011365 | -0,130011365 | -1,390011365 |
| chrXI | 292965 | 293112 | 147 | 0,082637257 | 0,062999783 | 0,56741923 | 0,169807485 | 0,144336362 | 0,036694373 | 0,015694373 | -1,244305627 |
| chrXI | 297006 | 297125 | 119 | 2,131471316 | 8,834793075 | 0,194366216 | -0,861918383 | -0,732630625 | -0,072503768 | 0,102496232 | -1,157503768 |
| chrXI | 297006 | 297156 | 150 | 0,644000692 | 0,300175436 | 0,682076864 | 0,47351434 | 0,402487189 | 0,277234288 | 0,235234288 | -1,024765712 |
| chrXI | 298702 | 298832 | 130 | 0,276407377 | 0,418763262 | 0,397610833 | -0,259536057 | -0,220605649 | 0,010576411 | 0,108576411 | -1,151423589 |
| chrXI | 298875 | 299007 | 132 | 0,085486818 | 0,218646305 | 0,281083549 | -0,579625641 | -0,492681795 | -0,325964606 | -0,241964606 | -1,501964606 |
| chrXI | 300199 | 300305 | 106 | 0,378991558 | 1,363760005 | 0,217467346 | -0,780775251 | -0,663658963 | -0,128868376 | 0,137131624 | -1,122868376 |
| chrXI | 300199 | 300317 | 118 | 0,284956058 | 1,574994571 | 0,153206249 | -1,022778687 | -0,869361884 | -0,211205404 | -0,029205404 | -1,289205404 |
| chrXI | 303921 | 304068 | 147 | 0,170973635 | 0,17788174 | 0,490098899 | -0,024820929 | -0,021097789 | -0,139259895 | -0,160259895 | -1,420259895 |
| chrXI | 305787 | 305949 | 162 | 0,136778908 | 0,092646739 | 0,596179675 | 0,243470869 | 0,206950239 | 0,255257197 | 0,129257197 | -1,130742803 |
| chrXI | 318927 | 319072 | 145 | 0,113982423 | 0,166764131 | 0,405997586 | -0,23785292 | -0,202174982 | -0,280433426 | -0,287433426 | -1,547433426 |
| chrXI | 319054 | 319202 | 148 | 0,339097709 | 0,359469349 | 0,485418981 | -0,036557337 | -0,031073736 | -0,13825176 | -0,16625176 | -1,42625176 |
| chrXI | 322500 | 322670 | 170 | 0,225115286 | 0,544762828 | 0,2924038 | -0,54637586 | -0,464419481 | -0,32712977 | -0,50912977 | -1,76912977 |
| chrXI | 329617 | 329749 | 132 | 0,208017923 | 0,237175653 | 0,46725275 | -0,082177582 | -0,069850944 | 0,0872634 | 0,1712634 | -1,0887366 |
| chrXI | 329629 | 329749 | 120 | 0,037044288 | 0,062999783 | 0,370279692 | -0,331112667 | -0,281445767 | 0,390026352 | 0,558026352 | -0,701973648 |
| chrXI | 329629 | 329775 | 146 | 0,712390146 | 0,796761959 | 0,472046617 | -0,070126174 | -0,059607248 | -0,142694403 | -0,156694403 | -1,416694403 |
| chrXI | 337370 | 337525 | 155 | 1,923453394 | 2,004875442 | 0,489636554 | -0,025980229 | -0,022083195 | -0,048232608 | -0,125232608 | -1,385232608 |
| chrXI | 337447 | 337600 | 153 | 8,705407583 | 6,967034808 | 0,555459536 | 0,139467258 | 0,118547169 | 0,035492035 | -0,027507965 | -1,287507965 |
| chrXI | 337450 | 337600 | 150 | 0,037044288 | 0,037058696 | 0,499902783 | -0,000243688 | -0,000207135 | -0,127669448 | -0,169669448 | -1,429669448 |
| chrXI | 337452 | 337600 | 148 | 0,108283302 | 0,129705435 | 0,454993389 | -0,113055218 | -0,096096936 | -0,196611305 | -0,224611305 | -1,484611305 |
| chrXI | 337462 | 337600 | 138 | 0,039893848 | 0,037058696 | 0,518421434 | 0,046192108 | 0,039263292 | 0,036098418 | 0,078098418 | -1,181901582 |
| chrXI | 337968 | 338118 | 150 | 0,188070999 | 0,151940653 | 0,553131041 | 0,13357593 | 0,113539541 | -0,011410235 | -0,053410235 | -1,313410235 |
| chrXI | 340261 | 340401 | 140 | 0,227964847 | 0,181587609 | 0,556619411 | 0,142403651 | 0,121043103 | 0,07029509 | 0,09829509 | -1,16170491 |

TABLE 2

| Chrom | Start | End | Length | Section A | Section B | A/A+B | Z-score | Z * 0.85 | Phase correction | Length correction | ΔLK |
|---|---|---|---|---|---|---|---|---|---|---|---|
| chrXI | 341684 | 341830 | 146 | 0,860567296 | 0,674468263 | 0,560617173 | 0,152534155 | 0,129654032 | 0,051995378 | 0,037995378 | -1,222004622 |
| chrXI | 347279 | 347450 | 171 | 0,051292091 | 0,066705652 | 0,434687048 | -0,164453569 | -0,139785534 | 0,038800622 | -0,150199378 | -1,410199378 |
| chrXI | 347320 | 347467 | 147 | 0,558513874 | 0,689291742 | 0,447596859 | -0,131735231 | -0,111974946 | -0,2327025 | -0,2537025 | -1,5137025 |
| chrXI | 347479 | 347608 | 129 | 0,065539893 | 0,100058479 | 0,395776194 | -0,264295292 | -0,224650998 | 0,050872762 | 0,155872762 | -1,104127238 |
| chrXI | 351244 | 351421 | 177 | 0,176672756 | 0,322410653 | 0,353994448 | -0,374558427 | -0,318374663 | -0,05369598 | -0,28469598 | -1,54469598 |
| chrXI | 351885 | 352030 | 145 | 0,803576085 | 0,541056958 | 0,597617386 | 0,247184775 | 0,210107059 | 0,131848615 | 0,124848615 | -1,135151385 |
| chrXI | 352745 | 352882 | 137 | 0,208017923 | 0,181587609 | 0,533919325 | 0,085125837 | 0,072356961 | 0,127933562 | 0,176933562 | -1,083066438 |
| chrXI | 353147 | 353287 | 140 | 0,435982769 | 0,344645871 | 0,558502144 | 0,147172696 | 0,125096791 | 0,081551487 | 0,109551487 | -1,150448513 |
| chrXI | 355288 | 355448 | 160 | 0,552814753 | 0,463233697 | 0,544083063 | 0,11072569 | 0,094116836 | 0,098966096 | -0,013033904 | -1,273033904 |
| chrXI | 357460 | 357614 | 154 | 1,587205245 | 1,641700223 | 0,491561385 | -0,021154048 | -0,017980941 | -0,052746132 | -0,122746132 | -1,382746132 |
| chrXI | 357462 | 357614 | 152 | 0,031345166 | 0,055588044 | 0,360566076 | -0,356946107 | -0,303404191 | -0,376727786 | -0,432727786 | -1,692727786 |
| chrXI | 362917 | 363070 | 153 | 0,262159574 | 0,303881305 | 0,463146008 | -0,092511045 | -0,078634388 | -0,15123929 | -0,21423929 | -1,47423929 |
| chrXI | 364062 | 364210 | 148 | 0,364743755 | 0,466939567 | 0,438560862 | -0,154618956 | -0,131426113 | -0,23775083 | -0,26575083 | -1,52575083 |
| chrXI | 364602 | 364723 | 121 | 2,020338454 | 4,250632407 | 0,322173153 | -0,461630518 | -0,39238594 | 0,252526211 | 0,413526211 | -0,846473789 |
| chrXI | 368064 | 368220 | 156 | 0,361894194 | 0,43729261 | 0,45282804 | -0,118519456 | -0,100741538 | -0,115457895 | -0,199457895 | -1,459457895 |
| chrXI | 368389 | 368522 | 133 | 1,951949 | 2,594108705 | 0,429371804 | -0,177973727 | -0,151277668 | -0,022237908 | 0,054762092 | -1,205237908 |
| chrXI | 368971 | 369078 | 107 | 0,076938136 | 0,222352175 | 0,257068582 | -0,652409303 | -0,554547908 | 0,030230289 | 0,289230289 | -0,970769711 |
| chrXI | 368971 | 369121 | 150 | 0,059840772 | 0,051882174 | 0,535617563 | 0,08939893 | 0,07598909 | -0,04670179 | -0,08870179 | -1,34870179 |
| chrXI | 368971 | 369122 | 151 | 11,09333935 | 9,865024818 | 0,529303683 | 0,073519618 | 0,062491675 | -0,022130205 | -0,071130205 | -1,331130205 |
| chrXI | 372978 | 373087 | 109 | 1,66129382 | 15,76847506 | 0,095313589 | -1,308726012 | -1,11241711 | -0,415283221 | -0,170283221 | -1,430283221 |
| chrXI | 373360 | 373511 | 151 | 0,279256937 | 0,203822827 | 0,578076247 | 0,196974484 | 0,167428312 | 0,080158728 | 0,031158728 | -1,228841272 |
| chrXI | 377591 | 377751 | 160 | 0,059840772 | 0,066705652 | 0,472876041 | -0,06804215 | -0,057835827 | -0,051960104 | -0,163960104 | -1,423960104 |
| chrXI | 379254 | 379407 | 153 | 0,424584527 | 0,503998263 | 0,457239281 | -0,107391293 | -0,091282599 | -0,157716649 | -0,220716649 | -1,480716649 |
| chrXI | 385681 | 385837 | 156 | 0,276407377 | 0,285351958 | 0,492038779 | -0,019957148 | -0,016963575 | -0,026355021 | -0,110355021 | -1,370355021 |
| chrXI | 394646 | 394790 | 144 | 0,53571739 | 0,411351523 | 0,565658298 | 0,165331072 | 0,140531411 | 0,072557241 | 0,072557241 | -1,187442759 |
| chrXI | 397867 | 398011 | 144 | 0,133929347 | 0,062999783 | 0,680089062 | 0,46794786 | 0,397755681 | 0,321993033 | 0,321993033 | -0,938006967 |
| chrXI | 397867 | 398011 | 144 | 0,133929347 | 0,062999783 | 0,680089062 | 0,46794786 | 0,397755681 | 0,321993033 | 0,321993033 | -0,938006967 |
| chrXI | 399301 | 399460 | 159 | 0,404637603 | 0,403939784 | 0,50043151 | 0,001081636 | 0,000919391 | 0,015340948 | -0,089659052 | -1,349659052 |
| chrXI | 400364 | 400506 | 142 | 0,552814753 | 0,459527828 | 0,546074781 | 0,115750303 | 0,098387758 | -0,000531614 | 0,013468386 | -1,246531614 |
| chrXI | 401469 | 401625 | 156 | 0,812124766 | 0,722644568 | 0,529151025 | 0,07313593 | 0,062165541 | 0,050081969 | -0,033918031 | -1,293918031 |
| chrXI | 401694 | 401852 | 158 | 0,313451664 | 0,289057827 | 0,520243529 | 0,050764798 | 0,043150079 | 0,060076725 | -0,037923275 | -1,297923275 |
| chrXI | 403288 | 403463 | 175 | 0,373292436 | 0,696703481 | 0,348872767 | -0,38836555 | -0,330110718 | -0,069859788 | -0,286859788 | -1,546859788 |
| chrXI | 406796 | 406955 | 159 | 0,136778908 | 0,137117174 | 0,499382491 | -0,001547865 | -0,001315685 | 0,019417289 | -0,085582711 | -1,345582711 |
| chrXI | 407916 | 408065 | 149 | 0,076938136 | 0,059293913 | 0,564757972 | 0,163043664 | 0,138587115 | 0,008281562 | -0,026718438 | -1,286718438 |
| chrXI | 416183 | 416327 | 144 | 0,076938136 | 0,037058696 | 0,674914686 | 0,453525163 | 0,385496389 | 0,306335025 | 0,306335025 | -0,953664975 |
| chrXI | 417925 | 418084 | 159 | 1,094231264 | 1,067290439 | 0,506231912 | 0,015621722 | 0,013278464 | 0,028843489 | -0,076156511 | -1,336156511 |
| chrXI | 417925 | 418084 | 159 | 1,094231264 | 1,067290439 | 0,506231912 | 0,015621722 | 0,013278464 | 0,028843489 | -0,076156511 | -1,336156511 |
| chrXI | 434342 | 434496 | 154 | 0,450230572 | 0,448410219 | 0,501012837 | 0,002538808 | 0,002157987 | -0,020534294 | -0,090534294 | -1,350534294 |
| chrXI | 434529 | 434684 | 155 | 0,265009134 | 0,070411522 | 0,790079947 | 0,806698677 | 0,685693876 | 0,650303421 | 0,573303421 | -0,686696579 |
| chrXI | 434561 | 434692 | 131 | 0,122531105 | 0,070411522 | 0,635064977 | 0,345298405 | 0,293503645 | 0,528537142 | 0,619537142 | -0,640462858 |
| chrXI | 434662 | 434792 | 130 | 0,062690333 | 0,085235 | 0,423797138 | -0,192188854 | -0,163360526 | 0,073698243 | 0,171698243 | -1,088301757 |
| chrXI | 439672 | 439806 | 134 | 8,297920419 | 7,1263872 | 0,53797685 | 0,095338075 | 0,081037364 | 0,188557019 | 0,258557019 | -1,001442981 |
| chrXI | 439682 | 439780 | 98 | 0,045592969 | 0,040764565 | 0,527955893 | 0,070132481 | 0,059612609 | 0,491175723 | 0,813175723 | -0,446824277 |
| chrXI | 447937 | 448094 | 157 | 0,071239015 | 0,100058479 | 0,415878909 | -0,212447646 | -0,180580499 | -0,187607083 | -0,278607083 | -1,538607083 |
| chrXI | 448453 | 448602 | 149 | 0,188070999 | 0,155646522 | 0,547167331 | 0,118507771 | 0,100731606 | -0,018032092 | -0,053032092 | -1,313032092 |
| chrXI | 450755 | 450913 | 158 | 1,61285129 | 0,889408699 | 0,644557839 | 0,370668677 | 0,315068376 | 0,335707304 | 0,237707304 | -1,022292696 |
| chrXI | 454604 | 454760 | 156 | 0,068389454 | 0,070411522 | 0,492715945 | -0,018259434 | -0,015520519 | -0,022766752 | -0,106766752 | -1,366766752 |
| chrXI | 455031 | 455138 | 107 | 0,997346204 | 3,272282838 | 0,233590833 | -0,727072301 | -0,618011456 | -0,012180878 | 0,246819122 | -1,013180878 |
| chrXI | 455031 | 455148 | 117 | 1,333594353 | 5,577333716 | 0,192968924 | -0,867007596 | -0,736956456 | -0,066213638 | 0,122786362 | -1,137213638 |
| chrXI | 456736 | 456888 | 152 | 0,24221265 | 0,300175436 | 0,446567054 | -0,134339514 | -0,114188587 | -0,178743672 | -0,234743672 | -1,494743672 |
| chrXI | 459229 | 459336 | 107 | 0,045592969 | 0,655938915 | 0,064990587 | -1,514176126 | -1,287049707 | -0,655623344 | -0,396623344 | -1,656623344 |
| chrXI | 459229 | 459341 | 112 | 0,09973462 | 0,830114786 | 0,107258896 | -1,241238134 | -1,055052414 | -0,347589223 | -0,123589223 | -1,383589223 |
| chrXI | 462337 | 462501 | 164 | 0,233663968 | 0,163058261 | 0,588986325 | 0,224938202 | 0,191197471 | 0,264660202 | 0,124660202 | -1,135339798 |
| chrXI | 468677 | 468825 | 148 | 0,14817715 | 0,170470001 | 0,465019536 | -0,087795679 | -0,074626327 | -0,179080651 | -0,207080651 | -1,467080651 |
| chrXI | 470132 | 470274 | 142 | 0,165274514 | 0,196411088 | 0,456956299 | -0,108104753 | -0,09188904 | -0,174480946 | -0,160480946 | -1,420480946 |
| chrXI | 473335 | 473488 | 153 | 0,128230226 | 0,08894087 | 0,590457149 | 0,22872109 | 0,194412926 | 0,146961619 | 0,083961619 | -1,176038381 |
| chrXI | 479959 | 480086 | 127 | 0,09973462 | 0,429880871 | 0,188315149 | -0,88412211 | -0,751503794 | -0,403628482 | -0,284628482 | -1,544628482 |
| chrXI | 483328 | 483476 | 148 | 0,068389454 | 0,044470435 | 0,605967759 | 0,268824834 | 0,228501109 | 0,128183725 | 0,100183725 | -1,159816275 |
| chrXI | 484474 | 484623 | 149 | 1,117027749 | 0,596645002 | 0,651832591 | 0,390272821 | 0,331731898 | 0,216750513 | 0,181750513 | -1,078249487 |
| chrXI | 488055 | 488207 | 152 | 0,082637257 | 0,226058044 | 0,267698461 | -0,619788682 | -0,52682038 | -0,603810319 | -0,659810319 | -1,919810319 |
| chrXI | 488418 | 488598 | 180 | 0,350495952 | 0,874585221 | 0,286100186 | -0,564813869 | -0,480091788 | -0,164439491 | -0,416439491 | -1,676439491 |
| chrXI | 496967 | 497107 | 140 | 0,028495606 | 0,044470435 | 0,390532439 | -0,277931585 | -0,236241848 | -0,300335667 | -0,272335667 | -1,532335667 |
| chrXI | 501192 | 501346 | 154 | 0,190920559 | 0,085235 | 0,691351496 | 0,499684841 | 0,424732115 | 0,398969242 | 0,328969242 | -0,931030758 |
| chrXI | 506644 | 506790 | 146 | 0,473027057 | 0,363175219 | 0,565684967 | 0,165398841 | 0,140589015 | 0,069343434 | 0,055343434 | -1,204656566 |
| chrXI | 509826 | 509992 | 166 | 0,208017923 | 0,270528479 | 0,434687048 | -0,164453569 | -0,139785534 | -0,019164296 | -0,173164296 | -1,433164296 |
| chrXI | 514096 | 514200 | 104 | 0,082637257 | 1,815876094 | 0,043527351 | -1,711143087 | -1,454471624 | -0,961858161 | -0,681858161 | -1,941858161 |
| chrXI | 516175 | 516336 | 161 | 0,085486818 | 0,100058479 | 0,460732874 | -0,09858756 | -0,083799426 | -0,077654672 | -0,196654672 | -1,456654672 |
| chrXI | 517302 | 517453 | 151 | 0,199469241 | 0,163058261 | 0,550218231 | 0,126212725 | 0,107280816 | 0,025131321 | -0,023868679 | -1,283868679 |
| chrXI | 517302 | 517453 | 151 | 0,199469241 | 0,163058261 | 0,550218231 | 0,126212725 | 0,107280816 | 0,025131321 | -0,023868679 | -1,283868679 |
| chrXI | 517458 | 517572 | 114 | 0,068389454 | 0,281646088 | 0,1953786 | -0,858244988 | -0,72950824 | -0,021799122 | 0,188200878 | -1,071799122 |
| chrXI | 519553 | 519695 | 142 | 0,495823542 | 1,637994354 | 0,232364506 | -0,731082091 | -0,621419778 | -0,708964418 | -0,694964418 | -1,954964418 |
| chrXI | 527505 | 527640 | 135 | 0,042743409 | 0,163058261 | 0,207692235 | -0,814454777 | -0,692286561 | -0,579897392 | -0,516897392 | -1,776897392 |
| chrXI | 527506 | 527600 | 94 | 0,028495606 | 0,048176305 | 0,371656395 | -0,327469522 | -0,278349094 | 0,128260389 | 0,478260389 | -0,781739611 |
| chrXI | 527506 | 527630 | 124 | 0,059840772 | 0,637409568 | 0,085823941 | -1,366927974 | -1,161888778 | -0,608056172 | -0,468056172 | -1,728056172 |
| chrXI | 527506 | 527632 | 126 | 0,037044288 | 0,059293913 | 0,384523349 | -0,293622086 | -0,249578773 | 0,191353175 | 0,317353175 | -0,942646825 |
| chrXI | 527506 | 527635 | 129 | 0,059840772 | 0,085235 | 0,412479432 | -0,221171546 | -0,187995814 | 0,087041228 | 0,192041228 | -1,067958772 |
| chrXI | 527506 | 527636 | 130 | 0,037044288 | 0,077823261 | 0,322495674 | -0,460731366 | -0,391621661 | -0,149221893 | -0,051221893 | -1,311221893 |
| chrXI | 527506 | 527637 | 131 | 0,111132863 | 0,185293479 | 0,374908863 | -0,318879716 | -0,271047759 | -0,051788904 | 0,039211096 | -1,220788904 |
| chrXI | 527506 | 527638 | 132 | 0,042743409 | 0,096352609 | 0,307294266 | -0,503534497 | -0,428004322 | -0,258465716 | -0,174465716 | -1,434465716 |
| chrXI | 527506 | 527639 | 133 | 0,034194727 | 0,103764348 | 0,247861382 | -0,681235065 | -0,579049806 | -0,45629572 | -0,37929572 | -1,63929572 |
| chrXI | 527506 | 527640 | 134 | 36,15237512 | 35,83946469 | 0,502173235 | 0,00544752 | 0,004630392 | 0,115925049 | 0,185925049 | -1,074074951 |
| chrXI | 527506 | 527641 | 135 | 0,094035499 | 0,318704784 | 0,227832133 | -0,746005211 | -0,63410443 | -0,528193084 | -0,465193084 | -1,725193084 |
| chrXI | 527507 | 527640 | 133 | 0,054141651 | 0,166764131 | 0,245089334 | -0,690024678 | -0,586520976 | -0,46819413 | -0,39119413 | -1,65119413 |
| chrXI | 527508 | 527640 | 132 | 0,042743409 | 0,092646739 | 0,315705458 | -0,47974192 | -0,407780632 | -0,232614796 | -0,148614796 | -1,408614796 |
| chrXI | 527509 | 527640 | 131 | 0,136778908 | 0,08894087 | 0,605967759 | 0,268824834 | 0,228501109 | 0,437710722 | 0,528710722 | -0,731289278 |
| chrXI | 527510 | 527640 | 130 | 0,074088575 | 0,103764348 | 0,416572153 | -0,210670615 | -0,179070023 | 0,054984012 | 0,152984012 | -1,107015988 |
| chrXI | 527511 | 527640 | 129 | 0,091185939 | 0,196411088 | 0,31706148 | -0,47593181 | -0,404542039 | -0,136822961 | -0,031822961 | -1,291822961 |
| chrXI | 527513 | 527640 | 127 | 0,059840772 | 0,08894087 | 0,402205349 | -0,247642923 | -0,210496484 | 0,143233897 | 0,262233897 | -0,997766103 |
| chrXI | 527515 | 527640 | 125 | 0,037044288 | 0,085235 | 0,302948179 | -0,51593994 | -0,438548949 | 0,041073689 | 0,174073689 | -1,085926311 |
| chrXI | 527516 | 527640 | 124 | 0,037044288 | 0,037058696 | 0,499902783 | -0,000243688 | -0,000207135 | 0,539444463 | 0,679444463 | -0,580555537 |
| chrXI | 529083 | 529220 | 137 | 1,886409106 | 1,423053918 | 0,570004588 | 0,176385844 | 0,149927968 | 0,196971017 | 0,245971017 | -1,014028983 |
| chrXI | 529403 | 529549 | 146 | 1,396284686 | 0,78935022 | 0,63884626 | 0,355376596 | 0,302070107 | 0,226784138 | 0,212784138 | -1,047215862 |
| chrXI | 532781 | 532933 | 152 | 0,276407377 | 0,125999566 | 0,686885209 | 0,487040568 | 0,413984483 | 0,342205913 | 0,286205913 | -0,973794087 |
| chrXI | 533171 | 533320 | 149 | 0,581310359 | 0,415057393 | 0,58342952 | 0,210674902 | 0,179073667 | 0,064852224 | 0,029852224 | -1,230147776 |
| chrXI | 540415 | 540570 | 155 | 0,071239015 | 0,100058479 | 0,415878909 | -0,212447646 | -0,180580499 | -0,208139632 | -0,285139632 | -1,545139632 |
| chrXI | 540630 | 540787 | 157 | 0,344796831 | 0,415057393 | 0,453767078 | -0,116149378 | -0,098726971 | -0,105999207 | -0,196999207 | -1,456999207 |
| chrXI | 540646 | 540797 | 151 | 0,108283302 | 0,092646739 | 0,538910465 | 0,097689227 | 0,083035843 | 0,005067737 | -0,043932263 | -1,303932263 |
| chrXI | 541723 | 541874 | 151 | 0,102584181 | 0,051882174 | 0,664119905 | 0,423733486 | 0,360173463 | 0,264218048 | 0,215218048 | -1,044781952 |
| chrXI | 542290 | 542416 | 126 | 0,564212996 | 1,615759136 | 0,258816609 | -0,64699803 | -0,549948325 | -0,122873632 | 0,003126368 | -1,256873632 |
| chrXI | 542290 | 542440 | 150 | 0,202318801 | 0,103764348 | 0,660992942 | 0,415174565 | 0,35289838 | 0,238459261 | 0,196459261 | -1,063540739 |
| chrXI | 544676 | 544823 | 147 | 0,276407377 | 0,114881957 | 0,706401511 | 0,542902438 | 0,461467072 | 0,341068724 | 0,320068724 | -0,939931276 |

TABLE 2

| Chrom | Start | End | Length | Section A | Section B | A/A+B | Z-score | Z * 0.85 | Phase correction | Length correction | ΔLK |
|---|---|---|---|---|---|---|---|---|---|---|---|
| chrXI | 545722 | 545853 | 131 | 0,170973635 | 0,170470001 | 0,500737507 | 0,001848658 | 0,001571359 | 0,196877991 | 0,287877991 | -0,972122009 |
| chrXI | 547817 | 547975 | 158 | 0,954602795 | 1,037643482 | 0,479159031 | -0,052264347 | -0,044424695 | -0,020496788 | -0,118496788 | -1,378496788 |
| chrXI | 548333 | 548481 | 148 | 0,97739928 | 0,674468263 | 0,591693495 | 0,231903444 | 0,197117927 | 0,099937219 | 0,071937219 | -1,188062781 |
| chrXI | 548769 | 548931 | 162 | 0,14532759 | 0,08894087 | 0,620346375 | 0,30639062 | 0,260432027 | 0,317153766 | 0,191153766 | -1,068846234 |
| chrXI | 548769 | 548931 | 162 | 0,14532759 | 0,08894087 | 0,620346375 | 0,30639062 | 0,260432027 | 0,317153766 | 0,191153766 | -1,068846234 |
| chrXI | 549757 | 549907 | 150 | 0,168124074 | 0,192705218 | 0,465937988 | -0,085484805 | -0,072662084 | -0,175039257 | -0,217039257 | -1,477039257 |
| chrXI | 550284 | 550422 | 138 | 0,031345166 | 0,040764565 | 0,434687048 | -0,164453569 | -0,139785534 | -0,137780273 | -0,095780273 | -1,355780273 |
| chrXI | 554400 | 554551 | 151 | 0,253610892 | 0,192705218 | 0,568231543 | 0,171873585 | 0,146092547 | 0,047898088 | -0,001101912 | -1,261101912 |
| chrXI | 558933 | 559085 | 152 | 0,233663968 | 0,251999131 | 0,481123578 | -0,047333843 | -0,040233767 | -0,124657893 | -0,180657893 | -1,440657893 |
| chrXI | 562000 | 562173 | 173 | 0,344796831 | 0,492880654 | 0,411610479 | -0,223404172 | -0,189893546 | 0,048865595 | -0,154134405 | -1,414134405 |
| chrXI | 571010 | 571152 | 142 | 0,168124074 | 0,211234566 | 0,443179768 | -0,142912183 | -0,121475355 | -0,180668893 | -0,166668893 | -1,426668893 |
| chrXI | 573165 | 573306 | 141 | 0,418885406 | 0,244587392 | 0,631352796 | 0,335438395 | 0,285122636 | 0,20166148 | 0,22266148 | -1,03733852 |
| chrXI | 574030 | 574178 | 148 | 0,854868175 | 0,592939133 | 0,590457149 | 0,22872109 | 0,194412926 | 0,111159116 | 0,083159116 | -1,176840884 |
| chrXI | 574402 | 574555 | 153 | 5,117810808 | 3,446458708 | 0,597577038 | 0,2470805 | 0,210018425 | 0,164114661 | 0,101114661 | -1,158885339 |
| chrXI | 575707 | 575877 | 170 | 0,04844253 | 0,137117174 | 0,26106169 | -0,640075711 | -0,544064354 | -0,372578963 | -0,554578963 | -1,814578963 |
| chrXI | 577254 | 577388 | 134 | 0,170973635 | 0,170470001 | 0,500737507 | 0,001848658 | 0,001571359 | 0,10510824 | 0,17510824 | -1,08489176 |
| chrXI | 577351 | 577483 | 132 | 1,037240052 | 1,615759136 | 0,390968854 | -0,276794756 | -0,235275543 | -0,072560245 | 0,011439755 | -1,248560245 |
| chrXI | 577690 | 577817 | 127 | 0,44168189 | 8,871851771 | 0,047423664 | -1,670364202 | -1,419809571 | -1,091113062 | -0,972113062 | -2,232113062 |
| chrXI | 578476 | 578613 | 137 | 0,646850252 | 0,507704132 | 0,560259665 | 0,151627591 | 0,128883452 | 0,157936971 | 0,206936971 | -1,053063029 |
| chrXI | 579137 | 579295 | 158 | 0,162424953 | 0,144528914 | 0,529151025 | 0,07313593 | 0,062165541 | 0,091240397 | -0,006759603 | -1,266759603 |
| chrXI | 590716 | 590822 | 106 | 0,416035845 | 1,267407396 | 0,247133871 | -0,68353674 | -0,581006229 | -0,021979271 | 0,244020729 | -1,015979271 |
| chrXI | 590716 | 590835 | 119 | 0,521469587 | 1,430465657 | 0,267155167 | -0,621439735 | -0,528223775 | 0,156434597 | 0,331434597 | -0,928565403 |
| chrXI | 600361 | 600514 | 153 | 0,48727486 | 0,43729261 | 0,527030071 | 0,067806264 | 0,057635325 | 0,014242579 | -0,048757421 | -1,308757421 |
| chrXI | 600361 | 600514 | 153 | 0,48727486 | 0,43729261 | 0,527030071 | 0,067806264 | 0,057635325 | 0,014242579 | -0,048757421 | -1,308757421 |
| chrXI | 601868 | 602039 | 171 | 0,131079787 | 0,270528479 | 0,326387174 | -0,449911372 | -0,382424667 | -0,197568101 | -0,386568101 | -1,646568101 |
| chrXI | 602067 | 602222 | 155 | 0,282106498 | 0,259410871 | 0,520955586 | 0,052552043 | 0,044669236 | 0,018358273 | -0,058641727 | -1,318641727 |
| chrXI | 603803 | 603944 | 141 | 2,128621756 | 1,367465874 | 0,608858238 | 0,276344444 | 0,234892777 | 0,133224849 | 0,154224849 | -1,105775151 |
| chrXI | 604749 | 604856 | 107 | 0,128230226 | 0,674468263 | 0,159748932 | -0,995490292 | -0,846166748 | -0,206103495 | 0,052896505 | -1,207103495 |
| chrXI | 604749 | 604865 | 116 | 0,09973462 | 0,774526742 | 0,114078724 | -1,205118789 | -1,024350971 | -0,336489492 | -0,140489492 | -1,400489492 |
| chrXI | 607498 | 607646 | 148 | 0,720938828 | 0,359469349 | 0,667283757 | 0,432425091 | 0,367561327 | 0,281278491 | 0,253278491 | -1,006721509 |
| chrXI | 607998 | 608143 | 145 | 0,609805965 | 0,35205761 | 0,633983842 | 0,342423354 | 0,291059851 | 0,206151235 | 0,199151235 | -1,060848765 |
| chrXI | 613167 | 613294 | 127 | 0,812124766 | 1,960405007 | 0,292918321 | -0,544879143 | -0,463147271 | -0,151647269 | -0,032647269 | -1,292647269 |
| chrXI | 614331 | 614461 | 130 | 0,373292436 | 0,392822175 | 0,487254036 | -0,031954832 | -0,027161607 | 0,199710646 | 0,297710646 | -0,962289354 |
| chrXI | 618644 | 618800 | 156 | 0,319150785 | 0,300175436 | 0,515319349 | 0,038409356 | 0,032647953 | 0,030685073 | -0,053314927 | -1,313314927 |
| chrXI | 621479 | 621632 | 153 | 0,037044288 | 0,040764565 | 0,47609348 | -0,059960669 | -0,050966568 | -0,094530231 | -0,157530231 | -1,417530231 |
| chrXI | 624990 | 625141 | 151 | 0,188070999 | 0,044470435 | 0,808763392 | 0,873348384 | 0,742346126 | 0,64900238 | 0,60000238 | -0,65999762 |
| chrXI | 632085 | 632240 | 155 | 0,239363089 | 0,17417587 | 0,578816297 | 0,198866205 | 0,169036274 | 0,154632923 | 0,077632923 | -1,182367077 |
| chrXI | 632617 | 632753 | 136 | 0,151026711 | 0,151940653 | 0,498491683 | -0,0037808 | -0,00321368 | 0,054140796 | 0,110140796 | -1,149859204 |
| chrXI | 638182 | 638313 | 131 | 0,056991212 | 0,044470435 | 0,561702018 | 0,155285859 | 0,13199298 | 0,306258774 | 0,397258774 | -0,862741226 |
| chrXI | 638182 | 638316 | 134 | 6,246236799 | 5,27345241 | 0,542222684 | 0,106034937 | 0,090129697 | 0,202873744 | 0,272873744 | -0,987126256 |
| chrXI | 643087 | 643244 | 157 | 0,085486818 | 0,114881957 | 0,426647404 | -0,184916144 | -0,157178722 | -0,156185576 | -0,247185576 | -1,507185576 |
| chrXI | 646189 | 646315 | 126 | 0,085486818 | 0,181587609 | 0,320086122 | -0,467457987 | -0,397339289 | 0,002489349 | 0,128489349 | -1,131510651 |
| chrXI | 647330 | 647482 | 152 | 0,088336378 | 0,055588044 | 0,613769206 | 0,28915654 | 0,245783059 | 0,155678929 | 0,099678929 | -1,160321071 |
| chrXI | 648395 | 648512 | 117 | 0,034194727 | 0,626291959 | 0,051772016 | -1,627909715 | -1,383723258 | -0,72710144 | -0,53810144 | -1,79810144 |
| chrXI | 650461 | 650612 | 151 | 0,378991558 | 0,300175436 | 0,558024111 | 0,145961504 | 0,124067278 | 0,040387547 | -0,008612453 | -1,268612453 |
| chrXI | 651142 | 651292 | 150 | 0,720938828 | 0,507704132 | 0,58677651 | 0,219260691 | 0,186371588 | 0,091582902 | 0,049582902 | -1,210417098 |
| chrXI | 654048 | 654197 | 149 | 0,185221438 | 0,129705435 | 0,588141101 | 0,222765786 | 0,189350918 | 0,067288598 | 0,032288598 | -1,227711402 |
| chrXI | 655704 | 655870 | 166 | 0,131079787 | 0,181587609 | 0,419230749 | -0,203861798 | -0,173282528 | -0,041669477 | -0,195669477 | -1,455669477 |
| chrXI | 655763 | 655920 | 157 | 0,094035499 | 0,077823261 | 0,547167331 | 0,118507771 | 0,100731606 | 0,102618568 | 0,011618568 | -1,248381432 |
| chrXI | 657176 | 657328 | 152 | 0,396088921 | 0,292763697 | 0,574998063 | 0,189113484 | 0,160746461 | 0,070588109 | 0,014588109 | -1,245411891 |
| chrXI | 658054 | 658203 | 149 | 0,122531105 | 0,092646739 | 0,569441084 | 0,174951387 | 0,148708679 | 0,031618445 | -0,003381555 | -1,263381555 |
| chrXI | 659842 | 659965 | 123 | 2,898003113 | 7,748973289 | 0,272190245 | -0,606202204 | -0,515271873 | 0,073248898 | 0,220248898 | -1,039751102 |
| chrXI | 661850 | 662006 | 156 | 0,185221438 | 0,259410871 | 0,416572153 | -0,210670615 | -0,179070023 | -0,179923982 | -0,263923982 | -1,523923982 |
| chrXI | 663739 | 663846 | 107 | 0,082637257 | 1,05617283 | 0,072564564 | -1,456953824 | -1,23841075 | -0,590157694 | -0,331157694 | -1,591157694 |
| chrXI | 663739 | 663851 | 112 | 0,136778908 | 1,467524353 | 0,085257514 | -1,370550803 | -1,164968182 | -0,460994452 | -0,236994452 | -1,496994452 |
| chrXI | 664373 | 664519 | 146 | 1,26805446 | 0,996878917 | 0,559863912 | 0,150624192 | 0,128030563 | 0,054160132 | 0,040160132 | -1,219839868 |
| chrXI | 666612 | 666748 | 136 | 1,288001384 | 0,181587609 | 0,876436466 | 1,157355694 | 0,98375234 | 1,033209526 | 1,089209526 | -0,170790474 |
| chrXI | 666628 | 666815 | 187 | 0,045592969 | 0,051882174 | 0,467739443 | -0,080953558 | -0,068810524 | 0,243475613 | -0,057524387 | -1,317524387 |
| chrXI | 666656 | 666815 | 159 | 0,113982423 | 0,100058479 | 0,532526364 | 0,081622044 | 0,069378737 | 0,088620929 | -0,016379071 | -1,276379071 |
| chrXI | 666681 | 666771 | 90 | 0,031345166 | 0,040764565 | 0,434687048 | -0,164453569 | -0,139785534 | 0,244324492 | 0,622324492 | -0,637675508 |
| chrXI | 666681 | 666810 | 129 | 1,02584181 | 0,285351958 | 0,782372396 | 0,780230509 | 0,663195933 | 0,928223985 | 1,033223985 | -0,226776015 |
| chrXI | 666693 | 666810 | 117 | 0,208017923 | 0,133411305 | 0,609256343 | 0,277381333 | 0,235774133 | 0,889668174 | 1,078668174 | -0,181331826 |
| chrXI | 666703 | 666728 | 25 | 0,113982423 | 0,062999783 | 0,644033238 | 0,369260555 | 0,313871472 | | 0,805 | -0,455 |
| chrXI | 666703 | 666733 | 30 | 0,253610892 | 0,200116957 | 0,558949362 | 0,148306006 | 0,126060105 | | 0,798 | -0,462 |
| chrXI | 666703 | 666737 | 34 | 0,188070999 | 0,166764131 | 0,530023616 | 0,075329227 | 0,064029843 | | 0,77 | -0,49 |
| chrXI | 666703 | 666739 | 36 | 0,216566604 | 0,244587392 | 0,469618839 | -0,076228037 | -0,064793831 | | 0,756 | -0,504 |
| chrXI | 666709 | 666792 | 83 | 0,034194727 | 0,159352392 | 0,176673914 | -0,928115177 | -0,788897901 | -0,445737534 | -0,018737534 | -1,278737534 |
| chrXI | 666709 | 666810 | 101 | 0,786478721 | 1,200701743 | 0,395776194 | -0,264295292 | -0,224650998 | 0,231224229 | 0,532224229 | -0,727775771 |
| chrXI | 666709 | 666815 | 106 | 0,564212996 | 0,722644568 | 0,43844246 | -0,154919323 | -0,131681425 | 0,414853959 | 0,680853959 | -0,579146041 |
| chrXI | 666714 | 666815 | 101 | 0,045592969 | 0,096352609 | 0,321200349 | -0,464344846 | -0,394693119 | 0,081054919 | 0,382054919 | -0,877945081 |
| chrXII | 723 | 874 | 151 | 0,578460798 | 0,381704567 | 0,602459555 | 0,25971854 | 0,220760759 | 0,093970571 | 0,044970571 | -1,215029429 |
| chrXII | 1167 | 1315 | 148 | 0,045592969 | 0,051882174 | 0,467739443 | -0,080953558 | -0,068810524 | -0,162874716 | -0,190874716 | -1,450874716 |
| chrXII | 1996 | 2147 | 151 | 0,504372223 | 0,277940218 | 0,644719675 | 0,371103224 | 0,315437741 | 0,194715863 | 0,145715863 | -1,114284137 |
| chrXII | 2008 | 2142 | 134 | 0,592708601 | 0,418763262 | 0,585986247 | 0,217232053 | 0,184647245 | 0,171166216 | 0,241166216 | -1,018833784 |
| chrXII | 2033 | 2129 | 96 | 0,056991212 | 0,122293696 | 0,317880698 | -0,47363334 | -0,402588339 | 0,159797715 | 0,495797715 | -0,764202285 |
| chrXII | 2033 | 2193 | 160 | 0,037044288 | 0,044470435 | 0,454449042 | -0,114428548 | -0,097264266 | -0,106624355 | -0,218624355 | -1,478624355 |
| chrXII | 2033 | 2193 | 160 | 0,037044288 | 0,044470435 | 0,454449042 | -0,114428548 | -0,097264266 | -0,106624355 | -0,218624355 | -1,478624355 |
| chrXII | 2211 | 2359 | 148 | 0,775080479 | 0,522527611 | 0,597314771 | 0,246402775 | 0,209442359 | 0,114836564 | 0,086836564 | -1,173163436 |
| chrXII | 2226 | 2355 | 129 | 0,042743409 | 0,074117392 | 0,365763444 | -0,34309514 | -0,291630869 | -0,048857305 | 0,056142695 | -1,203857305 |
| chrXII | 2226 | 2359 | 133 | 13,34164265 | 9,831671992 | 0,57573303 | 0,190989346 | 0,162340944 | 0,232221032 | 0,309221032 | -0,950778968 |
| chrXII | 2226 | 2421 | 195 | 0,076938136 | 0,066705652 | 0,535617563 | 0,08939893 | 0,07598909 | 0,149474199 | -0,207525801 | -1,467525801 |
| chrXII | 2387 | 2517 | 130 | 0,350495952 | 0,463233697 | 0,430727763 | -0,174521638 | -0,148343393 | -0,018494239 | 0,079505761 | -1,180494239 |
| chrXII | 2666 | 2818 | 152 | 0,170973635 | 0,114881957 | 0,598111913 | 0,248463027 | 0,211193573 | 0,058948536 | 0,002948536 | -1,257051464 |
| chrXII | 3013 | 3157 | 144 | 1,299399626 | 0,592939133 | 0,686663326 | 0,486414449 | 0,413452282 | 0,280276974 | 0,280276974 | -0,979723026 |
| chrXII | 3013 | 3200 | 187 | 0,131079787 | 0,062999783 | 0,67539199 | 0,45485158 | 0,386623843 | 0,441936062 | 0,140936062 | -1,119063938 |
| chrXII | 3065 | 3157 | 92 | 0,153876272 | 0,114881957 | 0,572545341 | 0,182858153 | 0,15542943 | 0,533587426 | 0,897587426 | -0,362412574 |
| chrXII | 3065 | 3200 | 135 | 1,165470279 | 0,852350003 | 0,577588742 | 0,195728705 | 0,166369399 | 0,179887046 | 0,242887046 | -1,017112954 |
| chrXII | 3153 | 3297 | 144 | 0,105433742 | 0,037058696 | 0,739925174 | 0,64311474 | 0,546647529 | 0,410440407 | 0,410440407 | -0,849559593 |
| chrXII | 3218 | 3370 | 152 | 0,102584181 | 0,074117392 | 0,58055047 | 0,203301913 | 0,172806626 | 0,032133523 | -0,023866477 | -1,283866477 |
| chrXII | 3470 | 3595 | 125 | 0,230814407 | 0,237175653 | 0,493203653 | -0,01703674 | -0,014481229 | 0,385091451 | 0,518091451 | -0,741908549 |
| chrXII | 3500 | 3643 | 143 | 0,330549028 | 0,400233914 | 0,452321762 | -0,11979755 | -0,101827917 | -0,164970922 | -0,157970922 | -1,417970922 |
| chrXII | 3500 | 3651 | 151 | 0,045592969 | 0,037058696 | 0,551627959 | 0,129775456 | 0,110309137 | -0,018111307 | -0,067111307 | -1,327111307 |
| chrXII | 3791 | 3928 | 137 | 0,410336724 | 0,466939567 | 0,467739443 | -0,080953558 | -0,068810524 | -0,02038484 | 0,02861516 | -1,23138484 |
| chrXII | 6346 | 6517 | 171 | 0,034194727 | 0,059293913 | 0,365763444 | -0,34309514 | -0,291630869 | -0,13689495 | -0,32589495 | -1,58589495 |
| chrXII | 6366 | 6517 | 151 | 0,626903328 | 0,43729261 | 0,589086376 | 0,225195424 | 0,191416111 | 0,068977125 | 0,019977125 | -1,240022875 |
| chrXII | 6690 | 6864 | 174 | 0,028495606 | 0,048176305 | 0,371656395 | -0,327469522 | -0,278349094 | -0,072909836 | -0,282909836 | -1,542909836 |
| chrXII | 6705 | 6857 | 152 | 0,034194727 | 0,051882174 | 0,397257877 | -0,260451199 | -0,221383519 | -0,345251081 | -0,401251081 | -1,661251081 |
| chrXII | 7495 | 7682 | 187 | 0,065539893 | 0,037058696 | 0,638799168 | 0,355250862 | 0,301963233 | 0,353522145 | 0,052522145 | -1,207477855 |
| chrXII | 7531 | 7682 | 151 | 0,561363435 | 0,26311674 | 0,680869537 | 0,470131702 | 0,399611946 | 0,272527021 | 0,223527021 | -1,036472979 |
| chrXII | 7543 | 7677 | 134 | 0,652549374 | 0,444704349 | 0,594711469 | 0,23968165 | 0,203729402 | 0,210733109 | 0,280733109 | -0,979266891 |
| chrXII | 7568 | 7664 | 96 | 0,037044288 | 0,107470218 | 0,25633612 | -0,65468243 | -0,556480065 | 0,026047505 | 0,362047505 | -0,897952495 |

TABLE 2

| Chrom | Start | End | Length | Section A | Section B | A/A+B | Z-score | Z * 0.85 | Phase correction | Length correction | ΔLK |
|---|---|---|---|---|---|---|---|---|---|---|---|
| chrXII | 7746 | 7894 | 148 | 0,675345858 | 0,670762394 | 0,501702487 | 0,004267516 | 0,003627389 | -0,092037904 | -0,120037904 | **-1,380037904** |
| chrXII | 7761 | 7894 | 133 | 13,05383703 | 9,876142427 | 0,569291266 | 0,17457007 | 0,148384559 | 0,215679755 | 0,292679755 | **-0,967320245** |
| chrXII | 7761 | 7956 | 195 | 0,082637257 | 0,081529131 | 0,503375009 | 0,008459995 | 0,007190996 | 0,079047487 | -0,277952513 | **-1,537952513** |
| chrXII | 7922 | 8052 | 130 | 0,333398588 | 0,444704349 | 0,428476198 | -0,180254982 | -0,153216735 | -0,018881073 | 0,079118927 | **-1,180881073** |
| chrXII | 8201 | 8353 | 152 | 0,176672756 | 0,092646739 | 0,655996907 | 0,401562292 | 0,341327948 | 0,222993139 | 0,166993139 | **-1,093006861** |
| chrXII | 8548 | 8692 | 144 | 1,407682928 | 0,61517435 | 0,695888407 | 0,512611386 | 0,435719678 | 0,295105628 | 0,295105628 | **-0,964894372** |
| chrXII | 8548 | 8735 | 187 | 0,116831984 | 0,092646739 | 0,55772721 | 0,145209355 | 0,123427951 | 0,177653657 | -0,123346343 | **-1,383346343** |
| chrXII | 8600 | 8692 | 92 | 0,125380666 | 0,17417587 | 0,418554265 | -0,205593405 | -0,174754394 | 0,183007114 | 0,547007114 | **-0,712992886** |
| chrXII | 8600 | 8735 | 135 | 1,262355338 | 0,870879351 | 0,591756427 | 0,232065494 | 0,19725567 | 0,212034861 | 0,275034861 | **-0,984965139** |
| chrXII | 8688 | 8832 | 144 | 0,09688506 | 0,062999783 | 0,605967759 | 0,268824834 | 0,228501109 | 0,07798138 | 0,07798138 | **-1,18201862** |
| chrXII | 8753 | 8905 | 152 | 0,09688506 | 0,077823261 | 0,554553208 | 0,137173592 | 0,116597553 | 0,004423552 | -0,051576448 | **-1,311576448** |
| chrXII | 9005 | 9130 | 125 | 0,159575393 | 0,259410871 | 0,380860679 | -0,303221116 | -0,257737948 | 0,110673859 | 0,243673859 | **-1,016326141** |
| chrXII | 9035 | 9178 | 143 | 0,29350474 | 0,429880871 | 0,405737598 | -0,238523365 | -0,20274486 | -0,272598667 | -0,265598667 | **-1,525598667** |
| chrXII | 9326 | 9463 | 137 | 0,504372223 | 0,448410219 | 0,529367672 | 0,073680449 | 0,062628381 | 0,113587303 | 0,162587303 | **-1,097412697** |
| chrXII | 11195 | 11357 | 162 | 0,128230226 | 0,085235 | 0,600707798 | 0,255179578 | 0,216902641 | 0,213742641 | 0,087742641 | **-1,172257359** |
| chrXII | 11198 | 11354 | 156 | 0,185221438 | 0,077823261 | 0,704144347 | 0,536357779 | 0,455904112 | 0,461386243 | 0,377386243 | **-0,882613757** |
| chrXII | 11205 | 11354 | 149 | 0,039893848 | 0,059293913 | 0,402205349 | -0,247642923 | -0,210496484 | -0,353300921 | -0,388300921 | **-1,648300921** |
| chrXII | 11205 | 11357 | 152 | 0,128230226 | 0,037058696 | 0,775794437 | 0,75806658 | 0,644356593 | 0,527511964 | 0,471511964 | **-0,788488036** |
| chrXII | 11207 | 11357 | 150 | 0,125380666 | 0,081529131 | 0,605967759 | 0,268824834 | 0,228501109 | 0,131804571 | 0,089804571 | **-1,170195429** |
| chrXII | 11208 | 11354 | 146 | 0,34194727 | 0,181587609 | 0,653150885 | 0,393841275 | 0,334765084 | 0,185395834 | 0,171395834 | **-1,088604166** |
| chrXII | 11211 | 11357 | 146 | 0,088336378 | 0,051882174 | 0,629990659 | 0,331828607 | 0,282054316 | 0,136111871 | 0,122111871 | **-1,137888129** |
| chrXII | 11213 | 11310 | 97 | 0,094035499 | 0,077823261 | 0,547167331 | 0,118507771 | 0,100731606 | 0,655734131 | 0,984734131 | **-0,275265869** |
| chrXII | 11213 | 11346 | 133 | 0,108283302 | 0,037058696 | 0,745024175 | 0,658912981 | 0,560076034 | 0,624919884 | 0,701919884 | **-0,558080116** |
| chrXII | 11213 | 11352 | 139 | 0,034194727 | 0,100058479 | 0,254703244 | -0,659762131 | -0,560797812 | -0,590797997 | -0,555797997 | **-1,815797997** |
| chrXII | 11213 | 11355 | 142 | 0,031345166 | 0,103764348 | 0,231998216 | -0,73228205 | -0,622439743 | -0,657383612 | -0,643383612 | **-1,903383612** |
| chrXII | 11213 | 11357 | 144 | 3,724375683 | 5,658862847 | 0,396917937 | -0,2613328 | -0,22213288 | -0,378455941 | -0,378455941 | **-1,638455941** |
| chrXII | 11213 | 11358 | 145 | 0,034194727 | 0,111176087 | 0,235224155 | -0,721749813 | -0,613487341 | -0,807981651 | -0,814981651 | **-2,074981651** |
| chrXII | 11218 | 11354 | 136 | 0,056991212 | 0,040764565 | 0,582995843 | 0,209563572 | 0,178129036 | 0,144974149 | 0,200974149 | **-1,059025851** |
| chrXII | 11222 | 11354 | 132 | 0,042743409 | 0,040764565 | 0,51184823 | 0,029703476 | 0,025247955 | 0,146256695 | 0,230256695 | **-1,029743305** |
| chrXII | 11222 | 11357 | 135 | 0,056991212 | 0,074117392 | 0,434687048 | -0,164453569 | -0,139785534 | -0,134619259 | -0,071619259 | **-1,331619259** |
| chrXII | 11226 | 11354 | 128 | 0,718089267 | 0,448410219 | 0,615593299 | 0,293927372 | 0,249838266 | 0,498784127 | 0,610784127 | **-0,649215873** |
| chrXII | 11226 | 11357 | 131 | 0,934655871 | 0,641115437 | 0,593141826 | 0,235634438 | 0,200289273 | 0,272786581 | 0,363786581 | **-0,896213419** |
| chrXII | 11226 | 11358 | 132 | 0,119681545 | 0,114881957 | 0,510230892 | 0,025647854 | 0,021800676 | 0,14658318 | 0,23058318 | **-1,02941682** |
| chrXII | 11227 | 11357 | 130 | 0,09973462 | 0,08894087 | 0,528604008 | 0,071761159 | 0,060996985 | 0,202152468 | 0,300152468 | **-0,959847532** |
| chrXII | 11228 | 11357 | 129 | 0,039893848 | 0,118587827 | 0,251725307 | -0,669070333 | -0,568709783 | -0,340122644 | -0,235122644 | **-1,495122644** |
| chrXII | 11229 | 11357 | 128 | 0,934655871 | 0,896820438 | 0,510329217 | 0,025894402 | 0,022010241 | 0,26797266 | 0,37997266 | **-0,88002734** |
| chrXII | 11230 | 11357 | 127 | 0,031345166 | 0,085235 | 0,268872204 | -0,616227458 | -0,523793339 | -0,28934951 | -0,17034951 | **-1,43034951** |
| chrXII | 11231 | 11354 | 123 | 0,547115632 | 0,326116523 | 0,626540868 | 0,322705529 | 0,274299699 | 0,693320613 | 0,840320613 | **-0,419679387** |
| chrXII | 11231 | 11357 | 126 | 1,020142689 | 0,396528045 | 0,720098654 | 0,583134602 | 0,495664412 | 0,722730576 | 0,848730576 | **-0,411269424** |
| chrXII | 11231 | 11358 | 127 | 0,404637603 | 0,159352392 | 0,717455286 | 0,575298511 | 0,489003734 | 0,719559615 | 0,838559615 | **-0,421440385** |
| chrXII | 11233 | 11357 | 124 | 0,065539893 | 0,040764565 | 0,616530051 | 0,29638 | 0,251923 | 0,689774459 | 0,829774459 | **-0,430225541** |
| chrXII | 11235 | 11357 | 122 | 0,216566604 | 0,251999131 | 0,462190442 | -0,094916836 | -0,080679311 | 0,293098613 | 0,447098613 | **-0,812901387** |
| chrXII | 11252 | 11354 | 102 | 0,076938136 | 0,066705652 | 0,535617563 | 0,08939893 | 0,07598909 | 0,569578512 | 0,863578512 | **-0,396421488** |
| chrXII | 11252 | 11357 | 105 | 0,43883233 | 0,985761308 | 0,308040355 | -0,501412691 | -0,426200787 | 0,155973964 | 0,428973964 | **-0,831026036** |
| chrXII | 11259 | 11357 | 98 | 0,062690333 | 0,096352609 | 0,394172367 | -0,268460684 | -0,228191582 | 0,317686663 | 0,639686663 | **-0,620313337** |
| chrXII | 11262 | 11357 | 95 | 0,085486818 | 0,322410653 | 0,20957918 | -0,807882268 | -0,686699927 | -0,153259517 | 0,189740483 | **-1,070259517** |
| chrXII | 11273 | 11357 | 84 | 0,031345166 | 0,111176087 | 0,219933277 | -0,772418577 | -0,656555791 | -0,328520136 | 0,091479864 | **-1,168520136** |
| chrXII | 13256 | 13420 | 164 | 0,125380666 | 0,051882174 | 0,707315001 | 0,545557714 | 0,463724057 | 0,455770013 | 0,315770013 | **-0,944229987** |
| chrXII | 13298 | 13427 | 129 | 0,156725832 | 0,374292827 | 0,295141855 | -0,538424944 | -0,457661202 | -0,239486236 | -0,134486236 | **-1,394486236** |
| chrXII | 13457 | 13612 | 155 | 0,14532759 | 0,059293913 | 0,710226382 | 0,554046188 | 0,47093926 | 0,355620708 | 0,278620708 | **-0,981379292** |
| chrXII | 15085 | 15257 | 172 | 0,49012442 | 0,541056958 | 0,475303793 | -0,061943802 | -0,052652232 | 0,079247134 | -0,116752866 | **-1,376752866** |
| chrXII | 15201 | 15361 | 160 | 0,364743755 | 0,314998914 | 0,536590936 | 0,091848853 | 0,078071525 | 0,066694293 | -0,045305707 | **-1,305305707** |
| chrXII | 16724 | 16865 | 141 | 0,159575393 | 0,151940653 | 0,512254168 | 0,030721476 | 0,026113255 | 0,009913435 | 0,030913435 | **-1,229086565** |
| chrXII | 18847 | 18996 | 149 | 0,909009826 | 0,578115654 | 0,611252943 | 0,282586129 | 0,24019821 | 0,089176588 | 0,054176588 | **-1,205823412** |
| chrXII | 25117 | 25257 | 140 | 1,743931077 | 1,23405457 | 0,585607617 | 0,216260411 | 0,183821349 | 0,210367273 | 0,238367273 | **-1,021632727** |
| chrXII | 27714 | 27865 | 151 | 0,769381358 | 0,400233914 | 0,657807209 | 0,406485945 | 0,345513053 | 0,210723624 | 0,161723624 | **-1,098276376** |
| chrXII | 35557 | 35688 | 131 | 0,153876272 | 0,100058479 | 0,605967759 | 0,268824834 | 0,228501109 | 0,309415172 | 0,400415172 | **-0,859584828** |
| chrXII | 39250 | 39399 | 149 | 0,532867829 | 0,70782109 | 0,429493502 | -0,177663815 | -0,151014242 | -0,304332552 | -0,339332552 | **-1,599332552** |
| chrXII | 48577 | 48723 | 146 | 0,039893848 | 0,040764565 | 0,49460244 | -0,013530088 | -0,011500575 | -0,152204565 | -0,166204565 | **-1,426204565** |
| chrXII | 51782 | 51932 | 150 | 0,87766466 | 0,70782109 | 0,553561998 | 0,134665941 | 0,11446605 | 0,015408087 | -0,026591913 | **-1,286591913** |
| chrXII | 56756 | 56881 | 125 | 0,062690333 | 0,103764348 | 0,376621026 | -0,314367347 | -0,267212245 | 0,099125764 | 0,232125764 | **-1,027874236** |
| chrXII | 57933 | 58079 | 146 | 0,133929347 | 0,085235 | 0,611090941 | 0,282163534 | 0,239839004 | 0,10553226 | 0,09153226 | **-1,16846774** |
| chrXII | 60326 | 60478 | 152 | 0,074088575 | 0,066705652 | 0,52621884 | 0,065768269 | 0,055903029 | -0,0452987 | -0,1012987 | **-1,3612987** |
| chrXII | 62223 | 62381 | 158 | 0,250761331 | 0,248293262 | 0,502472745 | 0,006198292 | 0,005268549 | -0,046856717 | -0,144856717 | **-1,404856717** |
| chrXII | 67309 | 67460 | 151 | 0,088336378 | 0,166764131 | 0,346280681 | -0,395381497 | -0,336074272 | -0,466579442 | -0,515579442 | **-1,775579442** |
| chrXII | 70032 | 70180 | 148 | 0,188070999 | 0,151940653 | 0,553131041 | 0,13357593 | 0,113539541 | 0,025729856 | -0,002270144 | **-1,262270144** |
| chrXII | 70780 | 70917 | 137 | 1,581506124 | 1,374877614 | 0,534946159 | 0,087709357 | 0,074552953 | 0,13420713 | 0,18320713 | **-1,07679287** |
| chrXII | 70780 | 70917 | 137 | 1,581506124 | 1,374877614 | 0,534946159 | 0,087709357 | 0,074552953 | 0,13420713 | 0,18320713 | **-1,07679287** |
| chrXII | 71343 | 71501 | 158 | 0,299203861 | 0,429880871 | 0,410382838 | -0,226560288 | -0,192576245 | -0,233948363 | -0,331948363 | **-1,591948363** |
| chrXII | 73489 | 73643 | 154 | 0,045592969 | 0,040764565 | 0,527955893 | 0,070132481 | 0,059612609 | -0,121911524 | -0,191911524 | **-1,451911524** |
| chrXII | 79801 | 79962 | 161 | 0,131079787 | 0,085235 | 0,605967759 | 0,268824834 | 0,228501109 | 0,202542856 | 0,083542856 | **-1,176457144** |
| chrXII | 99031 | 99175 | 144 | 2,074480105 | 1,237760439 | 0,626307201 | 0,322088569 | 0,273775284 | 0,123978737 | 0,123978737 | **-1,136021263** |
| chrXII | 104748 | 104900 | 152 | 0,595558162 | 0,370586958 | 0,616427232 | 0,296110709 | 0,251694103 | 0,157276799 | 0,101276799 | **-1,158723201** |
| chrXII | 104994 | 105164 | 170 | 0,19377012 | 0,200116957 | 0,491943329 | -0,020196453 | -0,017166985 | 0,157926685 | -0,024073315 | **-1,284073315** |
| chrXII | 106767 | 106921 | 154 | 0,649699813 | 0,807879568 | 0,445738888 | -0,136434532 | -0,115969352 | -0,299492884 | -0,369492884 | **-1,629492884** |
| chrXII | 107154 | 107242 | 88 | 0,031345166 | 0,125999566 | 0,19921332 | -0,84443452 | -0,717769342 | -0,309940147 | 0,082059853 | **-1,177940147** |
| chrXII | 107154 | 107250 | 96 | 0,028495606 | 0,133411305 | 0,175999936 | -0,930717198 | -0,791109618 | -0,223943904 | 0,112056096 | **-1,147943904** |
| chrXII | 107154 | 107252 | 98 | 0,028495606 | 0,062999783 | 0,311443082 | -0,491764035 | -0,41799943 | 0,108189924 | 0,430189924 | **-0,829810076** |
| chrXII | 107154 | 107253 | 99 | 0,028495606 | 0,066705652 | 0,299319635 | -0,52635832 | -0,447404572 | 0,048964852 | 0,363964852 | **-0,896035148** |
| chrXII | 107154 | 107255 | 101 | 0,04844253 | 0,129705435 | 0,271923004 | -0,607007389 | -0,51595628 | -0,081529761 | 0,219470239 | **-1,040529761** |
| chrXII | 107154 | 107257 | 103 | 0,042743409 | 0,066705652 | 0,390532439 | -0,277931585 | -0,236241848 | 0,335249143 | 0,622249143 | **-0,637750857** |
| chrXII | 107154 | 107258 | 104 | 0,039893848 | 0,085235 | 0,318822147 | -0,470995015 | -0,400345763 | 0,18420368 | 0,46420368 | **-0,79579632** |
| chrXII | 107154 | 107259 | 105 | 0,037044288 | 0,103764348 | 0,263082498 | -0,633871027 | -0,538790373 | 0,027955547 | 0,300955547 | **-0,959044453** |
| chrXII | 107154 | 107260 | 106 | 10,13303743 | 23,328449 | 0,302826877 | -0,516287316 | -0,438844218 | 0,084082013 | 0,350082013 | **-0,909917987** |
| chrXII | 107154 | 107261 | 107 | 0,250761331 | 0,522527611 | 0,324278957 | -0,455766476 | -0,387401505 | 0,110046441 | 0,369046441 | **-0,890953559** |
| chrXII | 107154 | 107262 | 108 | 0,239363089 | 0,674468263 | 0,261933549 | -0,637395748 | -0,541786386 | -0,004194712 | 0,247805288 | **-1,012194712** |
| chrXII | 107154 | 107265 | 111 | 0,039893848 | 0,151940653 | 0,207959715 | -0,813520966 | -0,691492821 | -0,092650654 | 0,138349346 | **-1,121650654** |
| chrXII | 107154 | 107267 | 113 | 0,045592969 | 0,207528696 | 0,180122745 | -0,914897409 | -0,777662797 | -0,219695093 | -0,002695093 | **-1,262695093** |
| chrXII | 107154 | 107269 | 115 | 0,054141651 | 0,259410871 | 0,172671713 | -0,943659989 | -0,802110991 | -0,129868891 | 0,073131109 | **-1,186868891** |
| chrXII | 107154 | 107270 | 116 | 0,039893848 | 0,100058479 | 0,285053125 | -0,567895024 | -0,48271077 | 0,118324276 | 0,314324276 | **-0,945675724** |
| chrXII | 107154 | 107271 | 117 | 0,028495606 | 0,240881523 | 0,105783316 | -1,249269194 | -1,061878815 | -0,514444505 | -0,325444505 | **-1,585444505** |
| chrXII | 107154 | 107273 | 119 | 0,051292091 | 0,170470001 | 0,231293321 | -0,734594249 | -0,624405111 | -0,220611231 | -0,045611231 | **-1,305611231** |
| chrXII | 107154 | 107274 | 120 | 11,24721562 | 28,02749162 | 0,286372997 | -0,564011958 | -0,479410164 | -0,088463625 | 0,079536375 | **-1,180463625** |
| chrXII | 107154 | 107285 | 131 | 3,359631928 | 3,546517187 | 0,486469648 | -0,033922068 | -0,028833758 | 0,056012127 | 0,147012127 | **-1,112987873** |
| chrXII | 107271 | 107423 | 152 | 1,504567988 | 0,885702829 | 0,62945503 | 0,330410325 | 0,280848776 | 0,181151651 | 0,125151651 | **-1,134848349** |
| chrXII | 110633 | 110799 | 166 | 0,162424953 | 0,292763697 | 0,356829972 | -0,366945134 | -0,311903364 | -0,208628817 | -0,362628817 | **-1,622628817** |
| chrXII | 112506 | 112634 | 128 | 0,094035499 | 0,35205761 | 0,210797919 | -0,803655711 | -0,683107355 | -0,441031641 | -0,329031641 | **-1,589031641** |
| chrXII | 113051 | 113206 | 155 | 0,091185939 | 0,08894087 | 0,506231912 | 0,015621722 | 0,013278464 | -0,103164869 | -0,180164869 | **-1,440164869** |
| chrXII | 118248 | 118400 | 152 | 0,14817715 | 0,144528914 | 0,506231912 | 0,015621722 | 0,013278464 | -0,083982792 | -0,139982792 | **-1,399982792** |
| chrXII | 119597 | 119738 | 141 | 0,111132863 | 0,133411305 | 0,454449042 | -0,114428548 | -0,097264266 | -0,117877866 | -0,096877866 | **-1,356877866** |
| chrXII | 121717 | 121889 | 172 | 0,028495606 | 0,051882174 | 0,354520937 | -0,373143192 | -0,317171714 | -0,190744401 | -0,386744401 | **-1,646744401** |

TABLE 2

| Chrom | Start | End | Length | Section A | Section B | A/A+B | Z-score | Z * 0.85 | Phase correction | Length correction | ΔLK |
|---|---|---|---|---|---|---|---|---|---|---|---|
| chrXII | 126647 | 126799 | 152 | 0,122531105 | 0,107470218 | 0,532740871 | 0,082161538 | 0,069837308 | -0,036649175 | -0,092649175 | -1,352649175 |
| chrXII | 132281 | 132417 | 136 | 3,151614005 | 2,779402184 | 0,531378419 | 0,078735306 | 0,066925011 | 0,041156744 | 0,097156744 | -1,162843256 |
| chrXII | 132310 | 132466 | 156 | 0,111132863 | 0,118587827 | 0,483773852 | -0,040684142 | -0,03458152 | -0,026869368 | -0,110869368 | -1,370869368 |
| chrXII | 133301 | 133451 | 150 | 1,179718082 | 1,015408265 | 0,53742605 | 0,093951228 | 0,079858544 | -0,020944539 | -0,062944539 | -1,322944539 |
| chrXII | 133762 | 133946 | 184 | 0,059840772 | 0,103764348 | 0,365763444 | -0,34309514 | -0,291630869 | -0,246922939 | -0,526922939 | -1,786922939 |
| chrXII | 136653 | 136779 | 126 | 0,416035845 | 1,037643482 | 0,286195062 | -0,564534946 | -0,479854704 | -0,246410878 | -0,120410878 | -1,380410878 |
| chrXII | 138426 | 138582 | 156 | 0,404637603 | 0,474351306 | 0,460344378 | -0,099566165 | -0,08463124 | -0,101659153 | -0,185659153 | -1,445659153 |
| chrXII | 142345 | 142492 | 147 | 0,039893848 | 0,040764565 | 0,49460244 | -0,013530088 | -0,011500575 | -0,174631662 | -0,195631662 | -1,455631662 |
| chrXII | 143973 | 144111 | 138 | 0,424584527 | 0,659644785 | 0,391600303 | -0,275150512 | -0,233877935 | -0,155961249 | -0,113961249 | -1,373961249 |
| chrXII | 144765 | 144914 | 149 | 2,39363089 | 1,560171093 | 0,60539979 | 0,267349056 | 0,227246697 | 0,074078304 | 0,039078304 | -1,220921696 |
| chrXII | 145430 | 145576 | 146 | 0,330549028 | 0,289057827 | 0,533481877 | 0,084025388 | 0,071421579 | -0,053704722 | -0,067704722 | -1,327704722 |
| chrXII | 155097 | 155220 | 123 | 0,113982423 | 0,181587609 | 0,385635926 | -0,290711661 | -0,247104912 | 0,184641063 | 0,331641063 | -0,928358937 |
| chrXII | 160945 | 161106 | 161 | 0,071239015 | 0,062999783 | 0,530688713 | 0,07700122 | 0,065451037 | 0,049858502 | -0,069141498 | -1,329141498 |
| chrXII | 163470 | 163588 | 118 | 1,880709985 | 11,01384439 | 0,145853042 | -1,054386316 | -0,896228369 | -0,431741427 | -0,249741427 | -1,509741427 |
| chrXII | 171732 | 171883 | 151 | 0,09688506 | 0,118587827 | 0,449639217 | -0,126572917 | -0,10758698 | -0,239097036 | -0,288097036 | -1,548097036 |
| chrXII | 177830 | 177964 | 134 | 0,230814407 | 0,188999349 | 0,549801916 | 0,125160904 | 0,106386768 | 0,10180946 | 0,17180946 | -1,08819054 |
| chrXII | 177830 | 177964 | 134 | 0,230814407 | 0,188999349 | 0,549801916 | 0,125160904 | 0,106386768 | 0,10180946 | 0,17180946 | -1,08819054 |
| chrXII | 178588 | 178741 | 153 | 0,133929347 | 0,081529131 | 0,621601659 | 0,309690034 | 0,263236529 | 0,152146299 | 0,089146299 | -1,170853701 |
| chrXII | 179076 | 179226 | 150 | 0,04844253 | 0,111176087 | 0,303489222 | -0,514391296 | -0,437232602 | -0,515080258 | -0,557080258 | -1,817080258 |
| chrXII | 180676 | 180836 | 160 | 0,937505432 | 0,767115003 | 0,549978994 | 0,125608276 | 0,106767034 | 0,099076995 | -0,012923005 | -1,272923005 |
| chrXII | 183315 | 183461 | 146 | 1,393435125 | 0,767115003 | 0,644944594 | 0,37170727 | 0,315951179 | 0,184406624 | 0,170406624 | -1,089593376 |
| chrXII | 192705 | 192836 | 131 | 2,185612968 | 2,657108488 | 0,451319158 | -0,12232919 | -0,103979812 | -0,027488888 | 0,063511112 | -1,196488888 |
| chrXII | 193019 | 193125 | 106 | 0,153876272 | 0,340940001 | 0,310976579 | -0,493084111 | -0,419121495 | 0,125559079 | 0,391559079 | -0,868440921 |
| chrXII | 193019 | 193144 | 125 | 0,19661968 | 0,415057393 | 0,3214436 | -0,463665806 | -0,394115935 | -0,011759005 | 0,121240995 | -1,138759005 |
| chrXII | 194389 | 194550 | 161 | 0,091185939 | 0,125999566 | 0,419852784 | -0,202270107 | -0,171929591 | -0,171441442 | -0,290441442 | -1,550441442 |
| chrXII | 197350 | 197482 | 132 | 1,134125112 | 7,359856983 | 0,133521015 | -1,109900209 | -0,943415178 | -0,825958585 | -0,741958585 | -2,001958585 |
| chrXII | 197802 | 197965 | 163 | 9,605868727 | 10,40237591 | 0,480095526 | -0,049913836 | -0,042426761 | 0,030258379 | -0,102741621 | -1,362741621 |
| chrXII | 199062 | 199223 | 161 | 1,686939865 | 1,875170007 | 0,473578841 | -0,066276513 | -0,056335036 | -0,050848524 | -0,169848524 | -1,429848524 |
| chrXII | 199271 | 199406 | 135 | 0,450230572 | 0,370586958 | 0,548514811 | 0,121909896 | 0,103623411 | 0,114877818 | 0,177877818 | -1,082122182 |
| chrXII | 199721 | 199871 | 150 | 19,63632198 | 19,49287398 | 0,501833005 | 0,004594677 | 0,003905476 | -0,081895841 | -0,123895841 | -1,383895841 |
| chrXII | 207917 | 208073 | 156 | 0,131079787 | 0,255705001 | 0,338895921 | -0,415478239 | -0,353156503 | -0,364440981 | -0,448440981 | -1,708440981 |
| chrXII | 208414 | 208561 | 147 | 0,151026711 | 0,133411305 | 0,530965281 | 0,07769655 | 0,066042068 | -0,091723134 | -0,112723134 | -1,372723134 |
| chrXII | 209818 | 209960 | 142 | 0,444531451 | 0,233469783 | 0,655649914 | 0,400619655 | 0,340526707 | 0,311200063 | 0,325200063 | -0,934799937 |
| chrXII | 211059 | 211185 | 126 | 0,094035499 | 0,070411522 | 0,571828535 | 0,1810314 | 0,15387669 | 0,402331061 | 0,528331061 | -0,731668939 |
| chrXII | 213217 | 213369 | 152 | 0,156725832 | 0,096352609 | 0,619277689 | 0,303584289 | 0,258046646 | 0,152248929 | 0,096248929 | -1,163751071 |
| chrXII | 214356 | 214519 | 163 | 0,062690333 | 0,096352609 | 0,394172367 | -0,268460684 | -0,228191582 | -0,157549705 | -0,290549705 | -1,550549705 |
| chrXII | 216121 | 216270 | 149 | 0,076938136 | 0,044470435 | 0,633712557 | 0,341702373 | 0,290447017 | 0,136457726 | 0,101457726 | -1,158542274 |
| chrXII | 217884 | 218033 | 149 | 0,125380666 | 0,040764565 | 0,754644987 | 0,68917996 | 0,585802966 | 0,429462213 | 0,394462213 | -0,865537787 |
| chrXII | 218108 | 218265 | 157 | 0,051292091 | 0,062999783 | 0,448781606 | -0,128740216 | -0,109429183 | -0,194823073 | -0,285823073 | -1,545823073 |
| chrXII | 218294 | 218444 | 150 | 9,218328488 | 4,302514581 | 0,681786516 | 0,472700362 | 0,401795308 | 0,319207357 | 0,277207357 | -0,982792643 |
| chrXII | 218298 | 218444 | 146 | 0,09688506 | 0,051882174 | 0,65125268 | 0,388704654 | 0,330398956 | 0,193906975 | 0,179906975 | -1,080093025 |
| chrXII | 218756 | 218891 | 135 | 0,088336378 | 0,08894087 | 0,498295067 | -0,004273647 | -0,0036326 | 0,01444599 | 0,07744599 | -1,18255401 |
| chrXII | 219794 | 219934 | 140 | 0,14532759 | 0,125999566 | 0,535617563 | 0,08939893 | 0,07598909 | 0,133799211 | 0,161799211 | -1,098200789 |
| chrXII | 219800 | 219934 | 134 | 0,111132863 | 0,096352609 | 0,535617563 | 0,08939893 | 0,07598909 | 0,078651916 | 0,148651916 | -1,111348084 |
| chrXII | 220212 | 220367 | 155 | 0,116831984 | 0,037058696 | 0,759188173 | 0,703693523 | 0,598139495 | 0,4751371 | 0,3981371 | -0,8618629 |
| chrXII | 220343 | 220451 | 108 | 0,042743409 | 0,155646522 | 0,215451503 | -0,787647358 | -0,669500254 | -0,116492107 | 0,135507893 | -1,124492107 |
| chrXII | 223444 | 223638 | 194 | 0,04844253 | 0,185293479 | 0,207253175 | -0,815989135 | -0,693590765 | -0,622023883 | -0,972023883 | -2,232023883 |
| chrXII | 223444 | 223638 | 194 | 0,04844253 | 0,185293479 | 0,207253175 | -0,815989135 | -0,693590765 | -0,622023883 | -0,972023883 | -2,232023883 |
| chrXII | 230481 | 230653 | 172 | 0,213717044 | 0,478057176 | 0,308940457 | -0,498855884 | -0,424027501 | -0,305350347 | -0,501350347 | -1,761350347 |
| chrXII | 230739 | 230894 | 155 | 0,225115286 | 0,163058261 | 0,579934639 | 0,201726272 | 0,171467332 | 0,045097264 | -0,031902736 | -1,291902736 |
| chrXII | 232813 | 232957 | 144 | 1,054337416 | 0,985761308 | 0,516807056 | 0,042141512 | 0,035820285 | -0,116459762 | -0,116459762 | -1,376459762 |
| chrXII | 232813 | 232957 | 144 | 1,054337416 | 0,985761308 | 0,516807056 | 0,042141512 | 0,035820285 | -0,116459762 | -0,116459762 | -1,376459762 |
| chrXII | 235469 | 235619 | 150 | 0,116831984 | 0,044470435 | 0,72430398 | 0,595675482 | 0,506324159 | 0,420545047 | 0,378545047 | -0,881454953 |
| chrXII | 235483 | 235631 | 148 | 0,247911771 | 0,35576348 | 0,410670755 | -0,225819886 | -0,191946903 | -0,283333857 | -0,311333857 | -1,571333857 |
| chrXII | 237484 | 237634 | 150 | 0,319150785 | 0,326116523 | 0,49460244 | -0,013530088 | -0,011500575 | -0,112712146 | -0,154712146 | -1,414712146 |
| chrXII | 238295 | 238446 | 151 | 0,051292091 | 0,048176305 | 0,515662191 | 0,039269381 | 0,033378974 | -0,110905244 | -0,159905244 | -1,419905244 |
| chrXII | 239615 | 239753 | 138 | 0,34194727 | 0,392822175 | 0,465380361 | -0,086887768 | -0,073854602 | -0,001336787 | 0,040663213 | -1,219336787 |
| chrXII | 255888 | 256021 | 133 | 0,233663968 | 0,203822827 | 0,534105191 | 0,085593433 | 0,072754418 | 0,127162215 | 0,204162215 | -1,055837785 |
| chrXII | 256324 | 256484 | 160 | 0,094035499 | 0,055588044 | 0,628480635 | 0,327831948 | 0,278657156 | 0,25246996 | 0,14046996 | -1,11953004 |
| chrXII | 259071 | 259235 | 164 | 0,29065518 | 0,307587175 | 0,485848548 | -0,035479871 | -0,030157891 | -0,045486947 | -0,185486947 | -1,445486947 |
| chrXII | 260642 | 260797 | 155 | 0,173823196 | 0,081529131 | 0,68071906 | 0,46971048 | 0,399253908 | 0,275607702 | 0,198607702 | -1,061392298 |
| chrXII | 262571 | 262718 | 147 | 1,105629506 | 0,659644785 | 0,62632165 | 0,322126715 | 0,273807708 | 0,12498415 | 0,10398415 | -1,15601585 |
| chrXII | 270658 | 270826 | 168 | 0,236513528 | 0,366881088 | 0,391971559 | -0,274184138 | -0,233056517 | -0,098070813 | -0,266070813 | -1,526070813 |
| chrXII | 270673 | 270826 | 153 | 0,173823196 | 0,048176305 | 0,782989131 | 0,782328165 | 0,66497894 | 0,550453334 | 0,487453334 | -0,772546666 |
| chrXII | 271752 | 271896 | 144 | 4,49375704 | 3,231518273 | 0,581695391 | 0,206232607 | 0,175297716 | 0,023106802 | 0,023106802 | -1,236893198 |
| chrXII | 273309 | 273417 | 108 | 2,994888173 | 15,38306462 | 0,162960924 | -0,982361374 | -0,835007168 | -0,273740562 | -0,021740562 | -1,281740562 |
| chrXII | 273309 | 273418 | 109 | 0,068389454 | 0,292763697 | 0,189364135 | -0,880241917 | -0,74820563 | -0,109731612 | 0,135268388 | -1,124731612 |
| chrXII | 273309 | 273419 | 110 | 0,045592969 | 0,426175002 | 0,096642782 | -1,300920803 | -1,105782682 | -0,398347604 | -0,160347604 | -1,420347604 |
| chrXII | 273309 | 273420 | 111 | 2,826764099 | 19,84863746 | 0,124662141 | -1,151992193 | -0,979193364 | -0,39519264 | -0,16419264 | -1,42419264 |
| chrXII | 274415 | 274568 | 153 | 5,411315548 | 4,258044146 | 0,559635355 | 0,150044776 | 0,127538059 | 0,011215076 | -0,051784924 | -1,311784924 |
| chrXII | 275945 | 276112 | 167 | 0,233663968 | 0,35205761 | 0,398933515 | -0,256108537 | -0,217692257 | -0,054374505 | -0,215374505 | -1,475374505 |
| chrXII | 275945 | 276112 | 167 | 0,233663968 | 0,35205761 | 0,398933515 | -0,256108537 | -0,217692257 | -0,054374505 | -0,215374505 | -1,475374505 |
| chrXII | 280170 | 280332 | 162 | 0,256460453 | 0,148234783 | 0,633712557 | 0,341702373 | 0,290447017 | 0,295377389 | 0,169377389 | -1,090622611 |
| chrXII | 280378 | 280532 | 154 | 1,849364819 | 1,089525656 | 0,629273134 | 0,329928838 | 0,280439513 | 0,09784117 | 0,02784117 | -1,23215883 |
| chrXII | 281608 | 281761 | 153 | 0,071239015 | 0,125999566 | 0,361181948 | -0,355301282 | -0,302006089 | -0,423857777 | -0,486857777 | -1,746857777 |
| chrXII | 285125 | 285277 | 152 | 7,628273682 | 4,521160886 | 0,627870675 | 0,326219022 | 0,277286169 | 0,166931155 | 0,110931155 | -1,149068845 |
| chrXII | 285128 | 285277 | 149 | 0,051292091 | 0,048176305 | 0,515662191 | 0,039269381 | 0,033378974 | -0,123503979 | -0,158503979 | -1,418503979 |
| chrXII | 286283 | 286443 | 160 | 0,153876272 | 0,137117174 | 0,528796348 | 0,072244535 | 0,061407855 | 0,040409835 | -0,071590165 | -1,331590165 |
| chrXII | 292173 | 292332 | 159 | 0,088336378 | 0,107470218 | 0,451140973 | -0,122779201 | -0,104362321 | -0,139775814 | -0,244775814 | -1,504775814 |
| chrXII | 293330 | 293472 | 142 | 0,227964847 | 0,274234349 | 0,453933118 | -0,11573037 | -0,098370815 | -0,118315349 | -0,104315349 | -1,364315349 |
| chrXII | 295117 | 295270 | 153 | 0,29350474 | 0,270528479 | 0,520367826 | 0,051076768 | 0,043415253 | -0,081531963 | -0,144531963 | -1,404531963 |
| chrXII | 295472 | 295620 | 148 | 1,017293128 | 0,459527828 | 0,688839852 | 0,492564557 | 0,418679873 | 0,322988245 | 0,294988245 | -0,965011755 |
| chrXII | 296601 | 296740 | 139 | 0,262159574 | 0,170470001 | 0,605967759 | 0,268824834 | 0,228501109 | 0,202626761 | 0,237626761 | -1,022373239 |
| chrXII | 300804 | 300959 | 155 | 0,353345512 | 0,163058261 | 0,6842427 | 0,479596127 | 0,407656708 | 0,282738181 | 0,205738181 | -1,054261819 |
| chrXII | 301895 | 302037 | 142 | 0,056991212 | 0,085235 | 0,400708216 | -0,251514398 | -0,213787238 | -0,223906682 | -0,209906682 | -1,469906682 |
| chrXII | 302498 | 302646 | 148 | 0,903310705 | 0,61146848 | 0,596331606 | 0,243863179 | 0,207283702 | 0,104687712 | 0,076687712 | -1,183312288 |
| chrXII | 304308 | 304455 | 147 | 2,222657255 | 1,478641962 | 0,600507315 | 0,254660445 | 0,216461378 | 0,070027915 | 0,049027915 | -1,210972085 |
| chrXII | 304635 | 304774 | 139 | 0,182371877 | 0,114881957 | 0,613522371 | 0,288511463 | 0,245234744 | 0,206879322 | 0,241879322 | -1,018120678 |
| chrXII | 310453 | 310583 | 130 | 0,074088575 | 0,114881957 | 0,392064172 | -0,273943107 | -0,232851641 | -0,091342475 | 0,006657525 | -1,253342475 |
| chrXII | 311744 | 311903 | 159 | 0,031345166 | 0,055588044 | 0,360566076 | -0,356946107 | -0,303404191 | -0,341893012 | -0,446893012 | -1,706893012 |
| chrXII | 312700 | 312835 | 135 | 0,111132863 | 0,17417587 | 0,389517915 | -0,280575734 | -0,238489374 | -0,221469889 | -0,158469889 | -1,418469889 |
| chrXII | 314498 | 314655 | 157 | 0,430283648 | 0,429880871 | 0,500234128 | 0,000586872 | 0,000498841 | -0,088540057 | -0,179540057 | -1,439540057 |
| chrXII | 319179 | 319325 | 146 | 0,173823196 | 0,203822827 | 0,460280753 | -0,099726444 | -0,084767477 | -0,22371803 | -0,23771803 | -1,49771803 |
| chrXII | 320329 | 320487 | 158 | 0,44168189 | 0,633703698 | 0,410719555 | -0,225694405 | -0,191840244 | -0,233880671 | -0,331880671 | -1,591880671 |
| chrXII | 320335 | 320487 | 152 | 0,051292091 | 0,044470435 | 0,535617563 | 0,08939893 | 0,07598909 | -0,03533557 | -0,09133557 | -1,35133557 |
| chrXII | 320336 | 320487 | 151 | 0,094035499 | 0,129705435 | 0,420287416 | -0,201158254 | -0,170984516 | -0,306866232 | -0,355866232 | -1,615866232 |
| chrXII | 321520 | 321655 | 135 | 0,094035499 | 0,107470218 | 0,466664175 | -0,083658001 | -0,071109301 | -0,063570288 | -0,000570288 | -1,260570288 |
| chrXII | 322116 | 322262 | 146 | 0,051292091 | 0,037058696 | 0,58055047 | 0,203301913 | 0,172806626 | 0,030913728 | 0,016913728 | -1,243086272 |
| chrXII | 327876 | 328044 | 168 | 0,19377012 | 0,218646305 | 0,469840938 | -0,07566971 | -0,064319253 | 0,067556774 | -0,100443226 | -1,360443226 |
| chrXII | 338236 | 338387 | 151 | 0,105433742 | 0,081529131 | 0,563928764 | 0,160937697 | 0,136797043 | 0,010094553 | -0,038905447 | -1,298905447 |
| chrXII | 340076 | 340209 | 133 | 1,5615592 | 1,49346544 | 0,511144552 | 0,027938883 | 0,02374805 | 0,078747667 | 0,155747667 | -1,104252333 |

TABLE 2

| Chrom | Start | End | Length | Section A | Section B | A/A+B | Z-score | Z * 0.85 | Phase correction | Length correction | ΔLK |
|---|---|---|---|---|---|---|---|---|---|---|---|
| chrXII | 340597 | 340761 | 164 | 1,430479413 | 1,941875659 | 0,424178173 | -0,191216029 | -0,162533625 | -0,179950386 | -0,319950386 | -1,579950386 |
| chrXII | 340875 | 341034 | 159 | 0,082637257 | 0,055588044 | 0,597844653 | 0,247772164 | 0,210606339 | 0,176205254 | 0,071205254 | -1,188794746 |
| chrXII | 347768 | 347916 | 148 | 0,738036191 | 0,544762828 | 0,575332675 | 0,189967434 | 0,161472318 | 0,056600327 | 0,028600327 | -1,231399673 |
| chrXII | 347831 | 348011 | 180 | 0,202318801 | 0,452116089 | 0,30915039 | -0,498260025 | -0,423521021 | -0,267904971 | -0,519904971 | -1,779904971 |
| chrXII | 347831 | 348011 | 180 | 0,202318801 | 0,452116089 | 0,30915039 | -0,498260025 | -0,423521021 | -0,267904971 | -0,519904971 | -1,779904971 |
| chrXII | 348735 | 348938 | 203 | 0,088336378 | 0,125999566 | 0,412139824 | -0,222043979 | -0,188737382 | -0,152869978 | -0,565869978 | -1,825869978 |
| chrXII | 348736 | 348938 | 202 | 0,079787696 | 0,096352609 | 0,452978074 | -0,118140734 | -0,100419624 | -0,077917651 | -0,483917651 | -1,743917651 |
| chrXII | 350679 | 350830 | 151 | 3,453667427 | 2,190168921 | 0,61193614 | 0,284368859 | 0,241713531 | 0,135944322 | 0,086944322 | -1,173055678 |
| chrXII | 355881 | 356007 | 126 | 0,324849907 | 1,096937395 | 0,22847996 | -0,743862076 | -0,632282764 | -0,391486704 | -0,265486704 | -1,525486704 |
| chrXII | 355881 | 356007 | 126 | 0,324849907 | 1,096937395 | 0,22847996 | -0,743862076 | -0,632282764 | -0,391486704 | -0,265486704 | -1,525486704 |
| chrXII | 358183 | 358323 | 140 | 1,521665352 | 1,511994788 | 0,501593877 | 0,003995268 | 0,003395978 | 0,046140705 | 0,074140705 | -1,185859295 |
| chrXII | 359562 | 359714 | 152 | 0,304902982 | 0,259410871 | 0,540307456 | 0,101208326 | 0,086027077 | -0,021600487 | -0,077600487 | -1,337600487 |
| chrXII | 360814 | 360966 | 152 | 0,162424953 | 0,070411522 | 0,69759239 | 0,517488464 | 0,439865194 | 0,318432625 | 0,262432625 | -0,997567375 |
| chrXII | 367954 | 368092 | 138 | 0,054141651 | 0,100058479 | 0,351112876 | -0,382317666 | -0,324970016 | -0,268814766 | -0,226814766 | -1,486814766 |
| chrXII | 372826 | 372968 | 142 | 0,92610719 | 0,915349786 | 0,502920895 | 0,007321663 | 0,006223413 | -0,008508295 | 0,005491705 | -1,254508295 |
| chrXII | 375291 | 375434 | 143 | 8,0101148 | 4,495219799 | 0,640535824 | 0,359891395 | 0,305907686 | 0,2522374 | 0,2592374 | -1,0007626 |
| chrXII | 375584 | 375750 | 166 | 0,085486818 | 0,207528696 | 0,291748435 | -0,548284062 | -0,466041453 | -0,36642701 | -0,52042701 | -1,78042701 |
| chrXII | 388796 | 388957 | 161 | 2,741277281 | 2,612638053 | 0,512013566 | 0,030118096 | 0,025600382 | 0,02797669 | -0,09102331 | -1,35102331 |
| chrXII | 388880 | 389024 | 144 | 0,102584181 | 0,059293913 | 0,633712557 | 0,341702373 | 0,290447017 | 0,148105143 | 0,148105143 | -1,111894857 |
| chrXII | 391687 | 391826 | 139 | 0,170973635 | 0,103764348 | 0,622315244 | 0,311567137 | 0,264832067 | 0,228981162 | 0,263981162 | -0,996018838 |
| chrXII | 395857 | 396006 | 149 | 0,094035499 | 0,037058696 | 0,717312458 | 0,574876116 | 0,488644698 | 0,335033822 | 0,300033822 | -0,959966178 |
| chrXII | 395857 | 396006 | 149 | 0,094035499 | 0,037058696 | 0,717312458 | 0,574876116 | 0,488644698 | 0,335033822 | 0,300033822 | -0,959966178 |
| chrXII | 397064 | 397220 | 156 | 0,259310013 | 0,196411088 | 0,569010328 | 0,173855096 | 0,147776831 | 0,135868738 | 0,051868738 | -1,208131262 |
| chrXII | 398081 | 398231 | 150 | 0,34194727 | 0,188999349 | 0,644033238 | 0,369260555 | 0,313871472 | 0,214062848 | 0,172062848 | -1,087937152 |
| chrXII | 400245 | 400397 | 152 | 0,068389454 | 0,055588044 | 0,551627959 | 0,129775456 | 0,110309137 | -0,011418445 | -0,067418445 | -1,327418445 |
| chrXII | 401669 | 401819 | 150 | 0,259310013 | 0,200116957 | 0,564420528 | 0,162186559 | 0,137858575 | 0,040573111 | -0,001426889 | -1,261426889 |
| chrXII | 401669 | 401819 | 150 | 0,259310013 | 0,200116957 | 0,564420528 | 0,162186559 | 0,137858575 | 0,040573111 | -0,001426889 | -1,261426889 |
| chrXII | 405414 | 405520 | 106 | 0,205168362 | 0,826408916 | 0,19888802 | -0,845599796 | -0,718759827 | -0,20871227 | 0,05728773 | -1,20271227 |
| chrXII | 405414 | 405538 | 124 | 0,247911771 | 1,282230874 | 0,162018732 | -0,986194937 | -0,838265696 | -0,410378564 | -0,270378564 | -1,530378564 |
| chrXII | 406427 | 406580 | 153 | 0,809275206 | 0,837526525 | 0,491422368 | -0,021502592 | -0,018277203 | -0,1437777 | -0,2067777 | -1,4667777 |
| chrXII | 406621 | 406768 | 147 | 1,798072728 | 1,411936309 | 0,56014569 | 0,151338603 | 0,128637812 | -0,014054924 | -0,035054924 | -1,295054924 |
| chrXII | 408412 | 408522 | 110 | 1,057186976 | 2,90540175 | 0,266792001 | -0,622544332 | -0,529162682 | 0,165189926 | 0,403189926 | -0,856810074 |
| chrXII | 408412 | 408523 | 111 | 0,042743409 | 0,077823261 | 0,354520937 | -0,373143192 | -0,317171714 | 0,255088896 | 0,486088896 | -0,773911104 |
| chrXII | 408415 | 408537 | 122 | 0,028495606 | 0,300175436 | 0,086699472 | -1,361363185 | -1,157158707 | -0,765925096 | -0,611925096 | -1,871925096 |
| chrXII | 411908 | 412040 | 132 | 0,87766466 | 1,248878048 | 0,412719038 | -0,220556114 | -0,187472697 | -0,070599746 | 0,013400254 | -1,246599746 |
| chrXII | 412316 | 412427 | 111 | 6,140803057 | 36,49910948 | 0,144015376 | -1,062451526 | -0,903083797 | -0,348712832 | -0,117712832 | -1,377712832 |
| chrXII | 412319 | 412427 | 108 | 0,028495606 | 0,181587609 | 0,135639612 | -1,100121415 | -0,935103203 | -0,355948696 | -0,103948696 | -1,363948696 |
| chrXII | 412320 | 412427 | 107 | 0,054141651 | 0,340940001 | 0,137039143 | -1,093718891 | -0,929661058 | -0,409672298 | -0,150672298 | -1,410672298 |
| chrXII | 419068 | 419220 | 152 | 0,056991212 | 0,118587827 | 0,324590066 | -0,454901459 | -0,386666241 | -0,515397902 | -0,571397902 | -1,831397902 |
| chrXII | 419734 | 419893 | 159 | 2,154267801 | 1,482347831 | 0,592382594 | 0,233678201 | 0,198626471 | 0,167895595 | 0,062895595 | -1,197104405 |
| chrXII | 421131 | 421283 | 152 | 0,424584527 | 0,381704567 | 0,526590934 | 0,066703018 | 0,056697565 | -0,081536735 | -0,137536735 | -1,397536735 |
| chrXII | 423648 | 423793 | 145 | 0,176672756 | 0,037058696 | 0,826610939 | 0,940857046 | 0,799728489 | 0,587352627 | 0,580352627 | -0,679647373 |
| chrXII | 424263 | 424441 | 178 | 0,071239015 | 0,059293913 | 0,545755127 | 0,114943701 | 0,097702146 | 0,22770264 | -0,01029736 | -1,27029736 |
| chrXII | 425377 | 425494 | 117 | 0,712390146 | 5,547686759 | 0,113798945 | -1,206569731 | -1,025584272 | -0,493343258 | -0,304343258 | -1,564343258 |
| chrXII | 428829 | 428982 | 153 | 0,128230226 | 0,070411522 | 0,645535127 | 0,373293861 | 0,317299782 | 0,195225716 | 0,132225716 | -1,127774284 |
| chrXII | 433448 | 433586 | 138 | 8,517336584 | 5,729274369 | 0,597850016 | 0,247786025 | 0,210618121 | 0,268118424 | 0,310118424 | -0,949881576 |
| chrXII | 433602 | 433749 | 147 | 0,151026711 | 0,274234349 | 0,35513882 | -0,371483236 | -0,315760751 | -0,447650857 | -0,468650857 | -1,728650857 |
| chrXII | 434337 | 434490 | 153 | 2,778321569 | 1,815876094 | 0,604745763 | 0,265650389 | 0,22580283 | 0,100153744 | 0,037153744 | -1,222846256 |
| chrXII | 443464 | 443629 | 165 | 0,142478029 | 0,166764131 | 0,460732874 | -0,09858756 | -0,083799426 | -0,033060345 | -0,180060345 | -1,440060345 |
| chrXII | 446788 | 446962 | 174 | 0,108283302 | 0,17417587 | 0,383359129 | -0,29667027 | -0,252169729 | -0,042566972 | -0,252566972 | -1,512566972 |
| chrXII | 446788 | 446962 | 174 | 0,108283302 | 0,17417587 | 0,383359129 | -0,29667027 | -0,252169729 | -0,042566972 | -0,252566972 | -1,512566972 |
| chrXII | 447291 | 447454 | 163 | 0,125380666 | 0,196411088 | 0,389632936 | -0,280275857 | -0,238234478 | -0,151928781 | -0,284928781 | -1,544928781 |
| chrXII | 447291 | 447454 | 163 | 0,125380666 | 0,196411088 | 0,389632936 | -0,280275857 | -0,238234478 | -0,151928781 | -0,284928781 | -1,544928781 |
| chrXII | 450323 | 450477 | 154 | 0,840620372 | 0,444704349 | 0,654014008 | 0,396180352 | 0,336753299 | 0,166512966 | 0,096512966 | -1,163487034 |
| chrXII | 451465 | 451626 | 161 | 0,062690333 | 0,103764348 | 0,376621026 | -0,314367347 | -0,267212245 | -0,260740675 | -0,379740675 | -1,639740675 |
| chrXII | 451590 | 451738 | 148 | 0,04844253 | 0,059293913 | 0,449639217 | -0,126572917 | -0,10758698 | -0,206170043 | -0,234170043 | -1,494170043 |
| chrXII | 451599 | 451733 | 134 | 0,045592969 | 0,044470435 | 0,506231912 | 0,015621722 | 0,013278464 | 0,025931925 | 0,095931925 | -1,164068075 |
| chrXII | 451709 | 451842 | 133 | 0,384690679 | 0,685585872 | 0,359431101 | -0,359979851 | -0,305982873 | -0,250511471 | -0,173511471 | -1,433511471 |
| chrXII | 451726 | 451885 | 159 | 0,054141651 | 0,100058479 | 0,351112876 | -0,382317666 | -0,324970016 | -0,357956651 | -0,462956651 | -1,722956651 |
| chrXII | 451734 | 451869 | 135 | 0,045592969 | 0,092646739 | 0,329810947 | -0,440435256 | -0,374369967 | -0,372225321 | -0,309225321 | -1,569225321 |
| chrXII | 451757 | 451842 | 85 | 0,04844253 | 0,051882174 | 0,482857442 | -0,042983253 | -0,036535765 | 0,389672539 | 0,802672539 | -0,457327461 |
| chrXII | 451757 | 451908 | 151 | 0,464478375 | 0,511410002 | 0,475954408 | -0,060309901 | -0,051263415 | -0,153446946 | -0,202446946 | -1,462446946 |
| chrXII | 451769 | 451932 | 163 | 0,088336378 | 0,144528914 | 0,379345404 | -0,307200444 | -0,261120377 | -0,172028581 | -0,305028581 | -1,565028581 |
| chrXII | 451790 | 451908 | 118 | 0,054141651 | 0,155646522 | 0,258077709 | -0,649283084 | -0,551890622 | -0,087323576 | 0,094676424 | -1,165323576 |
| chrXII | 451790 | 451932 | 142 | 0,045592969 | 0,048176305 | 0,486225044 | -0,034535557 | -0,029355224 | -0,045093773 | -0,031093773 | -1,291093773 |
| chrXII | 451790 | 451960 | 170 | 1,399134247 | 1,508288918 | 0,481228279 | -0,047071103 | -0,040010437 | 0,139419652 | -0,042580348 | -1,302580348 |
| chrXII | 451790 | 451960 | 170 | 1,399134247 | 1,508288918 | 0,481228279 | -0,047071103 | -0,040010437 | 0,139419652 | -0,042580348 | -1,302580348 |
| chrXII | 451800 | 451934 | 134 | 1,21961193 | 0,974643699 | 0,555820349 | 0,140380579 | 0,119323492 | 0,125479691 | 0,195479691 | -1,064520309 |
| chrXII | 451800 | 451961 | 161 | 0,296354301 | 0,426175002 | 0,410162328 | -0,227127429 | -0,193058315 | -0,169953909 | -0,288953909 | -1,548953909 |
| chrXII | 451800 | 451961 | 161 | 0,296354301 | 0,426175002 | 0,410162328 | -0,227127429 | -0,193058315 | -0,169953909 | -0,288953909 | -1,548953909 |
| chrXII | 451851 | 451934 | 83 | 0,259310013 | 0,426175002 | 0,378286917 | -0,309983065 | -0,263485605 | 0,054184397 | 0,481184397 | -0,778815603 |
| chrXII | 451851 | 451958 | 107 | 2,288197149 | 12,388722 | 0,15590446 | -1,011433657 | -0,859718608 | -0,328802255 | -0,069802255 | -1,329802255 |
| chrXII | 451851 | 451959 | 108 | 0,065539893 | 0,34835174 | 0,15835037 | -1,001260798 | -0,851071678 | -0,248939933 | 0,003060067 | -1,256939933 |
| chrXII | 451851 | 451960 | 109 | 0,04844253 | 0,466939567 | 0,093993428 | -1,316557908 | -1,119074222 | -0,495613919 | -0,250613919 | -1,510613919 |
| chrXII | 451851 | 451961 | 110 | 2,507613313 | 16,39476702 | 0,132661245 | -1,113899065 | -0,946814205 | -0,263875966 | -0,025875966 | -1,285875966 |
| chrXII | 451876 | 451960 | 84 | 0,037044288 | 0,062999783 | 0,370279692 | -0,331112667 | -0,281445767 | 0,048417289 | 0,468417289 | -0,791582711 |
| chrXII | 451876 | 452009 | 133 | 0,897611584 | 1,011702395 | 0,470122564 | -0,074961771 | -0,063717505 | -0,010600939 | 0,066399061 | -1,193600939 |
| chrXII | 451919 | 452075 | 156 | 0,079787696 | 0,092646739 | 0,462713239 | -0,093600543 | -0,079560461 | -0,076369664 | -0,160369664 | -1,420369664 |
| chrXII | 451919 | 452077 | 158 | 0,113982423 | 0,081529131 | 0,582995843 | 0,209563572 | 0,178129036 | 0,136410237 | 0,038410237 | -1,221589763 |
| chrXII | 451919 | 452078 | 159 | 0,889062902 | 0,674468263 | 0,568624996 | 0,172874585 | 0,146943398 | 0,121453034 | 0,016453034 | -1,243546966 |
| chrXII | 451919 | 452092 | 173 | 0,09973462 | 0,096352609 | 0,508623742 | 0,021618199 | 0,01837547 | 0,15710684 | -0,04589316 | -1,30589316 |
| chrXII | 451923 | 452075 | 152 | 1,265204899 | 0,722644568 | 0,636469169 | 0,349036828 | 0,296681304 | 0,158237503 | 0,102237503 | -1,157762497 |
| chrXII | 451923 | 452077 | 154 | 0,054141651 | 0,070411522 | 0,434687048 | -0,164453569 | -0,139785534 | -0,309654166 | -0,379654166 | -1,639654166 |
| chrXII | 451923 | 452092 | 169 | 0,131079787 | 0,111176087 | 0,541079911 | 0,103154718 | 0,08768151 | 0,199492153 | 0,024492153 | -1,235507847 |
| chrXII | 451931 | 452062 | 131 | 0,364743755 | 0,485468915 | 0,429002963 | -0,178913112 | -0,152076146 | -0,085948784 | 0,005051216 | -1,254948784 |
| chrXII | 451931 | 452077 | 146 | 2,29389627 | 1,534230006 | 0,599221683 | 0,251333038 | 0,213633083 | 0,080212524 | 0,066212524 | -1,193787476 |
| chrXII | 451931 | 452109 | 178 | 0,091185939 | 0,103764348 | 0,467739443 | -0,080953558 | -0,068810524 | 0,060735308 | -0,177264692 | -1,437264692 |
| chrXII | 451932 | 452062 | 130 | 4,747367932 | 5,647745238 | 0,45669228 | -0,108770453 | -0,092454885 | 0,037504009 | 0,135504009 | -1,124495991 |
| chrXII | 451932 | 452075 | 143 | 0,051292091 | 0,048176305 | 0,515662191 | 0,039269381 | 0,033378974 | -0,029064137 | -0,022064137 | -1,282064137 |
| chrXII | 451932 | 452077 | 145 | 0,672496298 | 0,844938264 | 0,443179768 | -0,142912183 | -0,121475355 | -0,338633967 | -0,345633967 | -1,605633967 |
| chrXII | 451932 | 452109 | 177 | 0,125380666 | 0,125999566 | 0,498768996 | -0,003085674 | -0,002622833 | 0,159689408 | -0,071310592 | -1,331310592 |
| chrXII | 451945 | 452051 | 106 | 0,168124074 | 0,270528479 | 0,383273899 | -0,296893528 | -0,252359499 | 0,257845604 | 0,523845604 | -0,736154396 |
| chrXII | 451945 | 452062 | 117 | 0,14817715 | 0,26682261 | 0,357053581 | -0,366345661 | -0,311393812 | 0,202718613 | 0,391718613 | -0,868281387 |
| chrXII | 451945 | 452075 | 130 | 0,188070999 | 0,111176087 | 0,628480635 | 0,327831948 | 0,278657156 | 0,408185349 | 0,506185349 | -0,753814651 |
| chrXII | 451945 | 452077 | 132 | 3,342534564 | 2,879460663 | 0,537212653 | 0,093413969 | 0,079401874 | 0,193593672 | 0,277593672 | -0,982406328 |
| chrXII | 451945 | 452078 | 133 | 0,156725832 | 0,125999566 | 0,554339417 | 0,136632651 | 0,116137754 | 0,155615448 | 0,232615448 | -1,027384552 |
| chrXII | 451945 | 452092 | 147 | 3,160162687 | 1,534230006 | 0,673178171 | 0,448706136 | 0,381400215 | 0,241676572 | 0,220676572 | -1,039323428 |
| chrXII | 451945 | 452109 | 164 | 0,208017923 | 0,170470001 | 0,549602536 | 0,124657219 | 0,105958636 | 0,093266889 | -0,046733111 | -1,306733111 |
| chrXII | 451958 | 452109 | 151 | 0,478726178 | 0,251999131 | 0,655138356 | 0,399230614 | 0,339346022 | 0,220938216 | 0,171938216 | -1,088061784 |
| chrXII | 452041 | 452196 | 155 | 0,185221438 | 0,08894087 | 0,675590454 | 0,45540334 | 0,387092839 | 0,256637472 | 0,179637472 | -1,080362528 |
| chrXII | 452059 | 452215 | 156 | 0,253610892 | 0,229763914 | 0,524667171 | 0,06187088 | 0,052590248 | 0,049802559 | -0,034197441 | -1,294197441 |

TABLE 2

| Chrom | Start | End | Length | Section A | Section B | A/A+B | Z-score | Z * 0.85 | Phase correction | Length correction | ΔLK |
|---|---|---|---|---|---|---|---|---|---|---|---|
| chrXII | 452081 | 452227 | 146 | 0,68389454 | 0,377998697 | 0,644033238 | 0,369260555 | 0,313871472 | 0,168453549 | 0,154453549 | -1,105546451 |
| chrXII | 452081 | 452236 | 155 | 0,074088575 | 0,070411522 | 0,512723359 | 0,03189814 | 0,027113419 | -0,102692328 | -0,179692328 | -1,439692328 |
| chrXII | 452081 | 452247 | 166 | 1,461824579 | 1,174760656 | 0,554438582 | 0,136883557 | 0,116351023 | 0,218347231 | 0,064347231 | -1,195652769 |
| chrXII | 452081 | 452251 | 170 | 0,071239015 | 0,085235 | 0,455276965 | -0,11233987 | -0,09548889 | 0,078196631 | -0,103803369 | -1,363803369 |
| chrXII | 452104 | 452247 | 143 | 0,076938136 | 0,040764565 | 0,653664997 | 0,395234266 | 0,335949126 | 0,275132834 | 0,282132834 | -0,977867166 |
| chrXII | 452104 | 452251 | 147 | 0,313451664 | 0,163058261 | 0,657807209 | 0,406485945 | 0,345513053 | 0,20586995 | 0,18486995 | -1,07513005 |
| chrXII | 452115 | 452271 | 156 | 0,567062556 | 0,448410219 | 0,558422215 | 0,146970164 | 0,124924639 | 0,125004046 | 0,041004046 | -1,218995954 |
| chrXII | 452190 | 452344 | 154 | 0,618354647 | 0,43729261 | 0,585758777 | 0,216648296 | 0,184151051 | 0,01709395 | -0,05290605 | -1,31290605 |
| chrXII | 452190 | 452344 | 154 | 0,618354647 | 0,43729261 | 0,585758777 | 0,216648296 | 0,184151051 | 0,01709395 | -0,05290605 | -1,31290605 |
| chrXII | 452260 | 452414 | 154 | 0,373292436 | 0,285351958 | 0,566758694 | 0,168127962 | 0,142908767 | -0,024485193 | -0,094485193 | -1,354485193 |
| chrXII | 452293 | 452400 | 107 | 0,113982423 | 0,852350003 | 0,117953636 | -1,1852787 | -1,007486895 | -0,478635313 | -0,219635313 | -1,479635313 |
| chrXII | 452294 | 452455 | 161 | 0,558513874 | 0,541056958 | 0,507938059 | 0,019899076 | 0,016914214 | 0,035748986 | -0,083251014 | -1,343251014 |
| chrXII | 452295 | 452455 | 160 | 0,267858695 | 0,270528479 | 0,497520572 | -0,006215044 | -0,005282788 | -0,023266066 | -0,135266066 | -1,395266066 |
| chrXII | 452299 | 452455 | 156 | 0,153876272 | 0,196411088 | 0,439285825 | -0,152780153 | -0,12986313 | -0,127744026 | -0,211744026 | -1,471744026 |
| chrXII | 452323 | 452455 | 132 | 0,054141651 | 0,074117392 | 0,42212736 | -0,196454145 | -0,166986023 | -0,021849248 | 0,062150752 | -1,197849248 |
| chrXII | 452323 | 452485 | 162 | 0,071239015 | 0,059293913 | 0,545755127 | 0,114943701 | 0,097702146 | 0,098982299 | -0,027017701 | -1,287017701 |
| chrXII | 452323 | 452486 | 163 | 0,34194727 | 0,255705001 | 0,572150875 | 0,181852797 | 0,154574878 | 0,240182565 | 0,107182565 | -1,152817435 |
| chrXII | 452334 | 452441 | 107 | 0,031345166 | 0,070411522 | 0,308040355 | -0,501412691 | -0,426200787 | 0,104221579 | 0,363221579 | -0,896778421 |
| chrXII | 452334 | 452455 | 121 | 0,039893848 | 0,074117392 | 0,349911537 | -0,385559308 | -0,327725412 | 0,054617055 | 0,215617055 | -1,044382945 |
| chrXII | 452334 | 452485 | 151 | 0,624053768 | 0,400233914 | 0,609256343 | 0,277381333 | 0,235774133 | 0,109101522 | 0,060101522 | -1,199898478 |
| chrXII | 452334 | 452486 | 152 | 0,051292091 | 0,066705652 | 0,434687048 | -0,164453569 | -0,139785534 | -0,280333429 | -0,336333429 | -1,596333429 |
| chrXII | 452346 | 452496 | 150 | 0,182371877 | 0,114881957 | 0,613522371 | 0,288511463 | 0,245234744 | 0,132855305 | 0,090855305 | -1,169144695 |
| chrXII | 452423 | 452584 | 161 | 0,14817715 | 0,207528696 | 0,416572153 | -0,210670615 | -0,179070023 | -0,155443647 | -0,274443647 | -1,534443647 |
| chrXII | 452469 | 452584 | 115 | 0,028495606 | 0,151940653 | 0,157926162 | -1,003017694 | -0,852565039 | -0,177893602 | 0,025106398 | -1,234893602 |
| chrXII | 452469 | 452620 | 151 | 0,282106498 | 0,326116523 | 0,463820816 | -0,090812431 | -0,077190567 | -0,20661518 | -0,25561518 | -1,51561518 |
| chrXII | 452469 | 452621 | 152 | 0,330549028 | 0,392822175 | 0,456956299 | -0,108104753 | -0,09188904 | -0,236052632 | -0,292052632 | -1,552052632 |
| chrXII | 452469 | 452629 | 160 | 0,282106498 | 0,222352175 | 0,559226183 | 0,149007602 | 0,126656462 | 0,12092135 | 0,00892135 | -1,25107865 |
| chrXII | 452469 | 452630 | 161 | 0,225115286 | 0,274234349 | 0,450816963 | -0,123597557 | -0,105057923 | -0,091757401 | -0,210757401 | -1,470757401 |
| chrXII | 452491 | 452621 | 130 | 0,119681545 | 0,181587609 | 0,397257877 | -0,260451199 | -0,221383519 | -0,075878987 | 0,022121013 | -1,237878987 |
| chrXII | 452491 | 452629 | 138 | 0,034194727 | 0,044470435 | 0,434687048 | -0,164453569 | -0,139785534 | -0,095026015 | -0,053026015 | -1,313026015 |
| chrXII | 452491 | 452663 | 172 | 0,219416165 | 0,26682261 | 0,451251888 | -0,122499079 | -0,104124217 | 0,005679695 | -0,190320305 | -1,450320305 |
| chrXII | 452492 | 452584 | 92 | 0,034194727 | 0,107470218 | 0,241377477 | -0,701878473 | -0,596596702 | -0,242468787 | 0,121531213 | -1,138468787 |
| chrXII | 452492 | 452620 | 128 | 0,116831984 | 0,296469566 | 0,282679762 | -0,574899121 | -0,488664253 | -0,262493358 | -0,150493358 | -1,410493358 |
| chrXII | 452492 | 452621 | 129 | 1,46467414 | 3,742928274 | 0,281256905 | -0,579111693 | -0,492244939 | -0,273985011 | -0,168985011 | -1,428985011 |
| chrXII | 452492 | 452629 | 137 | 0,122531105 | 0,181587609 | 0,402905508 | -0,245833639 | -0,208958593 | -0,154106135 | -0,105106135 | -1,365106135 |
| chrXII | 452492 | 452663 | 171 | 0,039893848 | 0,133411305 | 0,230194241 | -0,738207309 | -0,627476213 | -0,467598169 | -0,656598169 | -1,916598169 |
| chrXII | 452494 | 452621 | 127 | 0,028495606 | 0,051882174 | 0,354520937 | -0,373143192 | -0,317171714 | -0,093550926 | 0,025449074 | -1,234550926 |
| chrXII | 452525 | 452695 | 170 | 0,04844253 | 0,085235 | 0,36238349 | -0,352095051 | -0,299280793 | -0,132892129 | -0,314892129 | -1,574892129 |
| chrXII | 452539 | 452673 | 134 | 0,273557816 | 0,370586958 | 0,424683747 | -0,189925521 | -0,161436693 | -0,15971257 | -0,08971257 | -1,34971257 |
| chrXII | 452550 | 452672 | 122 | 0,29350474 | 1,115466743 | 0,208311342 | -0,812294469 | -0,690450299 | -0,299173722 | -0,145173722 | -1,405173722 |
| chrXII | 452550 | 452702 | 152 | 0,165274514 | 0,163058261 | 0,503375009 | 0,008459995 | 0,007190996 | -0,137196622 | -0,193196622 | -1,453196622 |
| chrXII | 452668 | 452796 | 128 | 0,282106498 | 0,463233697 | 0,378493606 | -0,309439521 | -0,263023593 | -0,047558592 | 0,064441408 | -1,195558592 |
| chrXII | 452684 | 452796 | 112 | 0,034194727 | 0,044470435 | 0,434687048 | -0,164453569 | -0,139785534 | 0,351605931 | 0,575605931 | -0,684394069 |
| chrXII | 452684 | 452835 | 151 | 0,592708601 | 0,333528262 | 0,6399104 | 0,358219306 | 0,30448641 | 0,175959639 | 0,126959639 | -1,133040361 |
| chrXII | 452686 | 452814 | 128 | 0,73233707 | 1,185878265 | 0,381780427 | -0,30080807 | -0,25568686 | -0,040401963 | 0,071598037 | -1,188401963 |
| chrXII | 452686 | 452884 | 198 | 0,034194727 | 0,048176305 | 0,415130494 | -0,214366854 | -0,182211826 | -0,140198455 | -0,518198455 | -1,778198455 |
| chrXII | 452709 | 452864 | 155 | 0,871965539 | 0,692997611 | 0,557179598 | 0,143822286 | 0,122248943 | -0,003921092 | -0,080921092 | -1,340921092 |
| chrXII | 452709 | 452884 | 175 | 0,094035499 | 0,096352609 | 0,493914773 | -0,015253994 | -0,012965895 | 0,242883564 | 0,025883564 | -1,234116436 |
| chrXII | 452709 | 452900 | 191 | 0,094035499 | 0,062999783 | 0,598817654 | 0,250287928 | 0,212744739 | 0,255077275 | -0,073922725 | -1,333922725 |
| chrXII | 452723 | 452814 | 91 | 0,04844253 | 0,092646739 | 0,343346664 | -0,403346515 | -0,342844538 | 0,10127586 | 0,47227586 | -0,78772414 |
| chrXII | 452723 | 452864 | 141 | 0,076938136 | 0,155646522 | 0,330796263 | -0,437715508 | -0,372058182 | -0,398532571 | -0,377532571 | -1,637532571 |
| chrXII | 452723 | 452884 | 161 | 0,88051422 | 0,737468046 | 0,544205112 | 0,111033507 | 0,094378481 | 0,111132479 | -0,007867521 | -1,267867521 |
| chrXII | 452723 | 452900 | 177 | 0,125380666 | 0,122293696 | 0,506231912 | 0,015621722 | 0,013278464 | 0,163689555 | -0,067310445 | -1,327310445 |
| chrXII | 452765 | 452864 | 99 | 0,122531105 | 0,133411305 | 0,478744828 | -0,053304046 | -0,045308439 | 0,440643621 | 0,755643621 | -0,504356379 |
| chrXII | 452765 | 452871 | 106 | 0,091185939 | 0,188999349 | 0,325448704 | -0,452515846 | -0,384638469 | 0,093033171 | 0,359033171 | -0,900966829 |
| chrXII | 452765 | 452884 | 119 | 0,088336378 | 0,107470218 | 0,451140973 | -0,122779201 | -0,104362321 | 0,305468751 | 0,480468751 | -0,779531249 |
| chrXII | 452765 | 452900 | 135 | 2,405029132 | 2,02340479 | 0,543087957 | 0,108216337 | 0,091983887 | 0,102668836 | 0,165668836 | -1,094331164 |
| chrXII | 452765 | 452925 | 160 | 0,113982423 | 0,144528914 | 0,440918471 | -0,148640972 | -0,126344826 | -0,137037703 | -0,249037703 | -1,509037703 |
| chrXII | 452765 | 452925 | 160 | 0,113982423 | 0,144528914 | 0,440918471 | -0,148640972 | -0,126344826 | -0,137037703 | -0,249037703 | -1,509037703 |
| chrXII | 452773 | 452864 | 91 | 0,059840772 | 0,096352609 | 0,38311977 | -0,297297305 | -0,252702709 | 0,180943827 | 0,551943827 | -0,708056173 |
| chrXII | 452773 | 452879 | 106 | 0,042743409 | 0,129705435 | 0,247861382 | -0,681235065 | -0,579049806 | -0,10208273 | 0,16391727 | -1,09608273 |
| chrXII | 452773 | 452884 | 111 | 0,028495606 | 0,070411522 | 0,288104674 | -0,558930214 | -0,475090682 | 0,067993226 | 0,298993226 | -0,961006774 |
| chrXII | 452773 | 452900 | 127 | 0,170973635 | 0,259410871 | 0,397257877 | -0,260451199 | -0,221383519 | 0,008317586 | 0,127317586 | -1,132682414 |
| chrXII | 452773 | 452925 | 152 | 0,752283994 | 0,622586089 | 0,547167331 | 0,118507771 | 0,100731606 | -0,054232757 | -0,110232757 | -1,370232757 |
| chrXII | 452773 | 452925 | 152 | 0,752283994 | 0,622586089 | 0,547167331 | 0,118507771 | 0,100731606 | -0,054232757 | -0,110232757 | -1,370232757 |
| chrXII | 452783 | 452962 | 179 | 0,082637257 | 0,148234783 | 0,357935317 | -0,363983093 | -0,309385629 | -0,169350294 | -0,414350294 | -1,674350294 |
| chrXII | 452783 | 452962 | 179 | 0,082637257 | 0,148234783 | 0,357935317 | -0,363983093 | -0,309385629 | -0,169350294 | -0,414350294 | -1,674350294 |
| chrXII | 452788 | 452923 | 135 | 0,672496298 | 0,511410002 | 0,568031691 | 0,171365199 | 0,145660419 | 0,144369235 | 0,207369235 | -1,052630765 |
| chrXII | 452788 | 452953 | 165 | 0,088336378 | 0,055588044 | 0,613769206 | 0,28915654 | 0,245783059 | 0,302799342 | 0,155799342 | -1,104200658 |
| chrXII | 452801 | 452923 | 122 | 0,116831984 | 0,103764348 | 0,529618887 | 0,074311878 | 0,063165096 | 0,441038612 | 0,595038612 | -0,664961388 |
| chrXII | 452801 | 452953 | 152 | 2,063081862 | 1,415642179 | 0,593057063 | 0,235415993 | 0,200103594 | 0,047959902 | -0,008040098 | -1,268040098 |
| chrXII | 452801 | 452999 | 198 | 0,062690333 | 0,066705652 | 0,484484373 | -0,038901718 | -0,03306646 | 0,013493278 | -0,364506722 | -1,624506722 |
| chrXII | 452840 | 452953 | 113 | 0,116831984 | 0,170470001 | 0,406652199 | -0,236165307 | -0,200740511 | 0,351159875 | 0,568159875 | -0,691840125 |
| chrXII | 452840 | 452999 | 159 | 0,604106844 | 0,463233697 | 0,565992596 | 0,166180623 | 0,14125353 | 0,110269708 | 0,005269708 | -1,254730292 |
| chrXII | 452840 | 453023 | 183 | 0,065539893 | 0,096352609 | 0,404835878 | -0,240849509 | -0,204722082 | -0,025460647 | -0,298460647 | -1,558460647 |
| chrXII | 452847 | 452953 | 106 | 0,034194727 | 0,08894087 | 0,277699771 | -0,589688448 | -0,501235181 | -0,037668425 | 0,228331575 | -1,031668425 |
| chrXII | 452847 | 452996 | 149 | 0,170973635 | 0,200116957 | 0,460732874 | -0,09858756 | -0,083799426 | -0,232444518 | -0,267444518 | -1,527444518 |
| chrXII | 452876 | 453034 | 158 | 1,533063594 | 1,397112831 | 0,523198392 | 0,058182555 | 0,049455172 | 0,006910067 | -0,091089933 | -1,351089933 |
| chrXII | 452885 | 453034 | 149 | 0,324849907 | 0,144528914 | 0,692084714 | 0,501768218 | 0,426502985 | 0,277992086 | 0,242992086 | -1,017007914 |
| chrXII | 452893 | 452999 | 106 | 0,054141651 | 0,092646739 | 0,368841506 | -0,334923211 | -0,284684729 | 0,184618073 | 0,450618073 | -0,809381927 |
| chrXII | 452895 | 453023 | 128 | 1,478921943 | 1,700994137 | 0,465082067 | -0,087638333 | -0,074492583 | 0,141637186 | 0,253637186 | -1,006362814 |
| chrXII | 452922 | 453029 | 107 | 1,211063248 | 7,841620028 | 0,133779479 | -1,108701541 | -0,94239631 | -0,420338671 | -0,161338671 | -1,421338671 |
| chrXII | 452922 | 453031 | 109 | 0,031345166 | 0,163058261 | 0,161237725 | -0,989383693 | -0,840976139 | -0,210115667 | 0,034884333 | -1,225115667 |
| chrXII | 452922 | 453034 | 112 | 1,718285032 | 10,44684634 | 0,14124673 | -1,07473484 | -0,913524614 | -0,436328186 | -0,212328186 | -1,472328186 |
| chrXII | 452922 | 453056 | 134 | 0,085486818 | 0,137117174 | 0,384030928 | -0,294911018 | -0,250674365 | -0,251546027 | -0,181546027 | -1,441546027 |
| chrXII | 452922 | 453056 | 134 | 0,085486818 | 0,137117174 | 0,384030928 | -0,294911018 | -0,250674365 | -0,251546027 | -0,181546027 | -1,441546027 |
| chrXII | 452922 | 453075 | 153 | 0,031345166 | 0,077823261 | 0,287126664 | -0,56179848 | -0,477528708 | -0,595531986 | -0,658531986 | -1,918531986 |
| chrXII | 452922 | 453075 | 153 | 0,031345166 | 0,077823261 | 0,287126664 | -0,56179848 | -0,477528708 | -0,595531986 | -0,658531986 | -1,918531986 |
| chrXII | 452926 | 452997 | 71 | 0,065539893 | 0,125999566 | 0,342174368 | -0,406536101 | -0,345555686 | | 0,511 | -0,749 |
| chrXII | 452926 | 453033 | 107 | 0,159575393 | 0,289057827 | 0,355692324 | -0,369997104 | -0,314497538 | 0,223013604 | 0,482013604 | -0,777986396 |
| chrXII | 452926 | 453034 | 108 | 0,216566604 | 0,581821524 | 0,27125479 | -0,609022416 | -0,517669054 | 0,097481871 | 0,349481871 | -0,910518129 |
| chrXII | 452926 | 453056 | 130 | 0,58700948 | 0,503998263 | 0,538043368 | 0,095505572 | 0,081179736 | 0,221328974 | 0,319328974 | -0,940671026 |
| chrXII | 452926 | 453056 | 130 | 0,58700948 | 0,503998263 | 0,538043368 | 0,095505572 | 0,081179736 | 0,221328974 | 0,319328974 | -0,940671026 |
| chrXII | 452926 | 453070 | 144 | 0,076938136 | 0,044470435 | 0,633712557 | 0,341702373 | 0,290447017 | 0,154750695 | 0,154750695 | -1,105249305 |
| chrXII | 452926 | 453070 | 144 | 0,076938136 | 0,044470435 | 0,633712557 | 0,341702373 | 0,290447017 | 0,154750695 | 0,154750695 | -1,105249305 |
| chrXII | 452926 | 453075 | 149 | 0,401788042 | 0,26311674 | 0,604279068 | 0,26443874 | 0,224772929 | 0,071574536 | 0,036574536 | -1,223425464 |
| chrXII | 452926 | 453075 | 149 | 0,401788042 | 0,26311674 | 0,604279068 | 0,26443874 | 0,224772929 | 0,071574536 | 0,036574536 | -1,223425464 |
| chrXII | 452976 | 453138 | 162 | 0,239363089 | 0,35576348 | 0,402205349 | -0,247642923 | -0,210496484 | -0,204667628 | -0,330667628 | -1,590667628 |
| chrXII | 452976 | 453163 | 187 | 0,039893848 | 0,040764565 | 0,49460244 | -0,013530088 | -0,011500575 | 0,040879243 | -0,260120757 | -1,520120757 |
| chrXII | 452991 | 453141 | 150 | 0,775080479 | 0,641115437 | 0,547297496 | 0,118836352 | 0,101010899 | -0,021610662 | -0,063610662 | -1,323610662 |
| chrXII | 453033 | 453163 | 130 | 1,305098747 | 1,767699789 | 0,424726428 | -0,189816588 | -0,1613441 | -0,021362186 | 0,076637814 | -1,183362186 |

| Chrom | Start | End | Length | Section A | Section B | A/A+B | Z-score | Z * 0.85 | Phase correction | Length correction | ΔLK |
|---|---|---|---|---|---|---|---|---|---|---|---|
| chrXII | 453033 | 453187 | 154 | 0,102584181 | 0,08894087 | 0,535617563 | 0,08939893 | 0,07598909 | -0,081350238 | -0,151350238 | -1,411350238 |
| chrXII | 453033 | 453193 | 160 | 0,076938136 | 0,103764348 | 0,42577243 | -0,18714766 | -0,159075511 | -0,158822309 | -0,270822309 | -1,530822309 |
| chrXII | 453039 | 453163 | 124 | 0,102584181 | 0,137117174 | 0,427966629 | -0,181553355 | -0,154320352 | 0,279806752 | 0,419806752 | -0,840193248 |
| chrXII | 453039 | 453187 | 148 | 1,111328628 | 0,748585655 | 0,597516046 | 0,246922881 | 0,209884448 | 0,107966427 | 0,079966427 | -1,180033573 |
| chrXII | 453039 | 453193 | 154 | 0,094035499 | 0,096352609 | 0,493914773 | -0,015253994 | -0,012965895 | -0,173427977 | -0,243427977 | -1,503427977 |
| chrXII | 453042 | 453141 | 99 | 0,028495606 | 0,074117392 | 0,277699771 | -0,589688448 | -0,501235181 | -0,030505566 | 0,284494434 | -0,975505566 |
| chrXII | 453042 | 453212 | 170 | 0,062690333 | 0,107470218 | 0,368418723 | -0,336044318 | -0,28563767 | -0,134007874 | -0,316007874 | -1,576007874 |
| chrXII | 453042 | 453218 | 176 | 0,703841464 | 0,629997828 | 0,527680859 | 0,069441393 | 0,059025184 | 0,252519086 | 0,028519086 | -1,231480914 |
| chrXII | 453043 | 453163 | 120 | 0,122531105 | 0,233469783 | 0,344187638 | -0,401060914 | -0,340901777 | 0,066047091 | 0,234047091 | -1,025952909 |
| chrXII | 453043 | 453187 | 144 | 0,14532759 | 0,111176087 | 0,566571175 | 0,16765125 | 0,142503563 | 0,02833511 | 0,02833511 | -1,23166489 |
| chrXII | 453043 | 453190 | 147 | 0,279256937 | 0,244587392 | 0,533091458 | 0,083043332 | 0,070586832 | -0,078773731 | -0,099773731 | -1,359773731 |
| chrXII | 453043 | 453193 | 150 | 1,567258321 | 1,070996308 | 0,594051197 | 0,237978712 | 0,202281905 | 0,074416672 | 0,032416672 | -1,227583328 |
| chrXII | 453091 | 453199 | 108 | 0,065539893 | 0,140823044 | 0,317595273 | -0,474433869 | -0,403268789 | 0,222020228 | 0,474020228 | -0,785979772 |
| chrXII | 453092 | 453199 | 107 | 1,769577122 | 2,964695663 | 0,373780135 | -0,321858003 | -0,273579302 | 0,271312997 | 0,530312997 | -0,729687003 |
| chrXII | 453092 | 453212 | 120 | 1,393435125 | 3,357517838 | 0,293295921 | -0,543781498 | -0,462214274 | -0,064455905 | 0,103544095 | -1,156455905 |
| chrXII | 453092 | 453236 | 144 | 0,133929347 | 0,192705218 | 0,410028091 | -0,227472716 | -0,193351809 | -0,300901931 | -0,300901931 | -1,560901931 |
| chrXII | 453092 | 453245 | 153 | 0,210867483 | 0,240881523 | 0,466780182 | -0,083366199 | -0,070861269 | -0,18892762 | -0,25192762 | -1,51192762 |
| chrXII | 453092 | 453270 | 178 | 0,088336378 | 0,118587827 | 0,426902103 | -0,184266739 | -0,156626728 | -0,022025594 | -0,260025594 | -1,520025594 |
| chrXII | 453092 | 453282 | 190 | 0,039893848 | 0,111176087 | 0,264075364 | -0,630831465 | -0,536206745 | -0,515441483 | -0,837441483 | -2,097441483 |
| chrXII | 453092 | 453286 | 194 | 0,142478029 | 0,218646305 | 0,394540095 | -0,267505227 | -0,227379443 | -0,149000832 | -0,499000832 | -1,759000832 |
| chrXII | 453092 | 453288 | 196 | 0,139628469 | 0,151940653 | 0,478886337 | -0,052948834 | -0,045006509 | 0,01835609 | -0,34564391 | -1,60564391 |
| chrXII | 453094 | 453236 | 142 | 0,031345166 | 0,096352609 | 0,245463684 | -0,688834586 | -0,585509398 | -0,604329181 | -0,590329181 | -1,850329181 |
| chrXII | 453095 | 453201 | 106 | 0,108283302 | 0,237175653 | 0,313447663 | -0,486101327 | -0,413186128 | 0,052841431 | 0,318841431 | -0,941158569 |
| chrXII | 453095 | 453212 | 117 | 0,128230226 | 0,159352392 | 0,445890044 | -0,136052108 | -0,115644291 | 0,391670966 | 0,580670966 | -0,679329034 |
| chrXII | 453095 | 453218 | 123 | 0,079787696 | 0,051882174 | 0,605967759 | 0,268824834 | 0,228501109 | 0,654930591 | 0,801930591 | -0,458069409 |
| chrXII | 453095 | 453236 | 141 | 0,04844253 | 0,148234783 | 0,246304615 | -0,686164739 | -0,583240028 | -0,628564354 | -0,607564354 | -1,867564354 |
| chrXII | 453095 | 453245 | 150 | 2,476268147 | 1,40823044 | 0,637474333 | 0,351715902 | 0,298958517 | 0,177339301 | 0,135339301 | -1,124660699 |
| chrXII | 453095 | 453270 | 175 | 0,068389454 | 0,103764348 | 0,397257877 | -0,260451199 | -0,221383519 | 0,025067671 | -0,191932329 | -1,451932329 |
| chrXII | 453095 | 453286 | 191 | 0,179522317 | 0,185293479 | 0,492090307 | -0,01982796 | -0,016853766 | 0,026046178 | -0,302953822 | -1,562953822 |
| chrXII | 453095 | 453288 | 193 | 0,182371877 | 0,188999349 | 0,491077027 | -0,022368443 | -0,019013176 | 0,041561199 | -0,301438801 | -1,561438801 |
| chrXII | 453099 | 453236 | 137 | 1,339293474 | 0,570703915 | 0,70120173 | 0,527859957 | 0,448680963 | 0,504630723 | 0,553630723 | -0,706369277 |
| chrXII | 453099 | 453287 | 188 | 0,076938136 | 0,066705652 | 0,535617563 | 0,08939893 | 0,07598909 | 0,119094131 | -0,188905869 | -1,448905869 |
| chrXII | 453099 | 453288 | 189 | 0,139628469 | 0,137117174 | 0,504537188 | 0,01137329 | 0,009667296 | 0,045679934 | -0,269320066 | -1,529320066 |
| chrXII | 453100 | 453238 | 138 | 0,045592969 | 0,051882174 | 0,467739443 | -0,080953558 | -0,068810524 | -0,030023589 | 0,011976411 | -1,248023589 |
| chrXII | 453100 | 453245 | 145 | 0,190920559 | 0,040764565 | 0,824051866 | 0,930917448 | 0,791279831 | 0,565833646 | 0,558833646 | -0,701166354 |
| chrXII | 453113 | 453270 | 157 | 0,404637603 | 0,389116306 | 0,509777147 | 0,024510127 | 0,020833608 | -0,081507267 | -0,172507267 | -1,432507267 |
| chrXII | 453113 | 453286 | 173 | 0,059840772 | 0,100058479 | 0,374240479 | -0,320642986 | -0,272546538 | -0,129096458 | -0,332096458 | -1,592096458 |
| chrXII | 453113 | 453288 | 175 | 0,074088575 | 0,111176087 | 0,399906675 | -0,253588671 | -0,21555037 | 0,025066481 | -0,191933519 | -1,451933519 |
| chrXII | 453114 | 453236 | 122 | 0,131079787 | 0,185293479 | 0,414320049 | -0,216446021 | -0,183979117 | 0,191503816 | 0,345503816 | -0,914496184 |
| chrXII | 453114 | 453269 | 155 | 0,071239015 | 0,077823261 | 0,477914444 | -0,055388588 | -0,0470803 | -0,171105897 | -0,248105897 | -1,508105897 |
| chrXII | 453114 | 453270 | 156 | 3,194357414 | 1,875170007 | 0,630109505 | 0,332143387 | 0,282321879 | 0,294506455 | 0,210506455 | -1,049493545 |
| chrXII | 453114 | 453286 | 172 | 0,287805619 | 0,311293045 | 0,480397698 | -0,049155472 | -0,041782151 | 0,067273846 | -0,128726154 | -1,388726154 |
| chrXII | 453114 | 453287 | 173 | 0,396088921 | 0,444704349 | 0,471089547 | -0,072531304 | -0,061651608 | 0,072354329 | -0,130645671 | -1,390645671 |
| chrXII | 453114 | 453288 | 174 | 0,914708947 | 1,182172396 | 0,436223514 | -0,160551029 | -0,136468375 | 0,08363206 | -0,12636794 | -1,38636794 |
| chrXII | 453114 | 453302 | 188 | 0,113982423 | 0,111176087 | 0,506231912 | 0,015621722 | 0,013278464 | 0,045313105 | -0,262686895 | -1,522686895 |
| chrXII | 453114 | 453309 | 195 | 0,034194727 | 0,062999783 | 0,351817475 | -0,380418277 | -0,323355536 | -0,244987022 | -0,601987022 | -1,861987022 |
| chrXII | 453116 | 453269 | 153 | 0,589859041 | 0,196411088 | 0,75019897 | 0,675116015 | 0,573848613 | 0,453219102 | 0,390219102 | -0,869780898 |
| chrXII | 453116 | 453270 | 154 | 0,142478029 | 0,040764565 | 0,77753772 | 0,763903822 | 0,649318248 | 0,486762655 | 0,416762655 | -0,843237345 |
| chrXII | 453116 | 453288 | 172 | 0,168124074 | 0,151940653 | 0,525281483 | 0,063413755 | 0,053901691 | 0,163908041 | -0,032091959 | -1,292091959 |
| chrXII | 453119 | 453236 | 117 | 0,162424953 | 0,259410871 | 0,385043053 | -0,292262269 | -0,248422928 | 0,243045293 | 0,432045293 | -0,827954707 |
| chrXII | 453119 | 453245 | 126 | 0,071239015 | 0,100058479 | 0,415878909 | -0,212447646 | -0,180580499 | 0,065962342 | 0,191962342 | -1,068037658 |
| chrXII | 453119 | 453269 | 150 | 0,136778908 | 0,111176087 | 0,551627959 | 0,129775456 | 0,110309137 | -0,017742249 | -0,059742249 | -1,319742249 |
| chrXII | 453119 | 453270 | 151 | 1,188266763 | 0,985761308 | 0,54657379 | 0,117009634 | 0,099458189 | -0,026573212 | -0,075573212 | -1,335573212 |
| chrXII | 453119 | 453282 | 163 | 0,028495606 | 0,303881305 | 0,085732808 | -1,367509647 | -1,1623832 | -1,079034165 | -1,212034165 | -2,472034165 |
| chrXII | 453119 | 453286 | 167 | 0,720938828 | 0,893114568 | 0,446663555 | -0,134095435 | -0,113981119 | 0,046085585 | -0,114914415 | -1,374914415 |
| chrXII | 453119 | 453287 | 168 | 1,273753581 | 1,115466743 | 0,533125207 | 0,083128223 | 0,070658989 | 0,207152547 | 0,039152547 | -1,220847453 |
| chrXII | 453119 | 453288 | 169 | 7,651070167 | 7,534032853 | 0,503853688 | 0,009659915 | 0,008210928 | 0,128840312 | -0,046159688 | -1,306159688 |
| chrXII | 453119 | 453302 | 183 | 0,29350474 | 0,226058044 | 0,564907166 | 0,163422653 | 0,138909255 | 0,299436496 | 0,026436496 | -1,233563504 |
| chrXII | 453119 | 453309 | 190 | 0,079787696 | 0,170470001 | 0,318822147 | -0,470995015 | -0,400345763 | -0,375136229 | -0,697136229 | -1,957136229 |
| chrXII | 453120 | 453228 | 108 | 0,059840772 | 0,074117392 | 0,446712395 | -0,133971905 | -0,113876119 | 0,518112775 | 0,770112775 | -0,489887225 |
| chrXII | 453121 | 453270 | 149 | 1,382036883 | 0,763409133 | 0,644172295 | 0,369633736 | 0,314188676 | 0,160780882 | 0,125780882 | -1,134219118 |
| chrXII | 453121 | 453282 | 161 | 0,056991212 | 0,092646739 | 0,380860679 | -0,303221116 | -0,257737948 | -0,250300245 | -0,369300245 | -1,629300245 |
| chrXII | 453121 | 453286 | 165 | 0,250761331 | 0,300175436 | 0,455154468 | -0,112648873 | -0,095751542 | -0,039224619 | -0,186224619 | -1,446224619 |
| chrXII | 453121 | 453288 | 167 | 0,159575393 | 0,26311674 | 0,377521558 | -0,311996587 | -0,265197099 | -0,106854968 | -0,267854968 | -1,527854968 |
| chrXII | 453125 | 453231 | 106 | 0,296354301 | 0,870879351 | 0,253894583 | -0,6622841 | -0,562941485 | -0,089676986 | 0,176323014 | -1,083676986 |
| chrXII | 453125 | 453236 | 111 | 0,205168362 | 0,978349569 | 0,173354671 | -0,940991251 | -0,799842563 | -0,271868661 | -0,040868661 | -1,300868661 |
| chrXII | 453125 | 453270 | 145 | 0,424584527 | 0,35576348 | 0,544096382 | 0,11075928 | 0,094145388 | -0,122673767 | -0,129673767 | -1,389673767 |
| chrXII | 453125 | 453286 | 161 | 0,378991558 | 0,448410219 | 0,45805021 | -0,105347063 | -0,089545003 | -0,080005225 | -0,199005225 | -1,459005225 |
| chrXII | 453125 | 453287 | 162 | 2,78117113 | 2,193874791 | 0,559024213 | 0,148495706 | 0,12622135 | 0,14638526 | 0,02038526 | -1,23961474 |
| chrXII | 453125 | 453288 | 163 | 1,231010172 | 1,397112831 | 0,468398994 | -0,079294993 | -0,067400744 | 0,014033736 | -0,118966264 | -1,378966264 |
| chrXII | 453125 | 453302 | 177 | 0,14817715 | 0,148234783 | 0,499902783 | -0,000243688 | -0,000207135 | 0,143546357 | -0,087453643 | -1,347453643 |
| chrXII | 453126 | 453248 | 122 | 0,136778908 | 0,696703481 | 0,164105337 | -0,977724354 | -0,831065701 | -0,488624393 | -0,334624393 | -1,594624393 |
| chrXII | 453126 | 453270 | 144 | 0,168124074 | 0,170470001 | 0,496535784 | -0,008683612 | -0,00738107 | -0,110868156 | -0,110868156 | -1,370868156 |
| chrXII | 453126 | 453286 | 160 | 0,572761677 | 0,61888022 | 0,48064916 | -0,048524398 | -0,041245738 | -0,042253349 | -0,154253349 | -1,414253349 |
| chrXII | 453126 | 453287 | 161 | 0,151026711 | 0,140823044 | 0,517481027 | 0,043832468 | 0,037257598 | 0,044059254 | -0,074940746 | -1,334940746 |
| chrXII | 453126 | 453288 | 162 | 0,484425299 | 0,770820872 | 0,385920555 | -0,289967488 | -0,246472365 | -0,226408164 | -0,352408164 | -1,612408164 |
| chrXII | 453135 | 453270 | 135 | 0,119681545 | 0,144528914 | 0,452978074 | -0,118140734 | -0,100419624 | -0,091874124 | -0,028874124 | -1,288874124 |
| chrXII | 453135 | 453282 | 147 | 1,601453048 | 1,32299544 | 0,547608568 | 0,119621655 | 0,101678407 | -0,047952715 | -0,068952715 | -1,328952715 |
| chrXII | 453135 | 453286 | 151 | 0,430283648 | 0,61888022 | 0,410120536 | -0,227234924 | -0,193149685 | -0,306071406 | -0,355071406 | -1,615071406 |
| chrXII | 453135 | 453288 | 153 | 0,310602104 | 0,426175002 | 0,421568615 | -0,197882202 | -0,168199872 | -0,297874875 | -0,360874875 | -1,620874875 |
| chrXII | 453138 | 453286 | 148 | 0,068389454 | 0,055588044 | 0,551627959 | 0,129775456 | 0,110309137 | 0,013630939 | -0,014369061 | -1,274369061 |
| chrXII | 453138 | 453288 | 150 | 0,156725832 | 0,292763697 | 0,348675157 | -0,38889974 | -0,330564779 | -0,452737165 | -0,494737165 | -1,754737165 |
| chrXII | 453140 | 453245 | 105 | 0,108283302 | 0,077823261 | 0,581834946 | 0,206589951 | 0,175601459 | 0,748393557 | 1,021393557 | -0,238606443 |
| chrXII | 453140 | 453246 | 106 | 0,085486818 | 0,17788174 | 0,324590066 | -0,454901459 | -0,386666241 | 0,078767339 | 0,344767339 | -0,915232661 |
| chrXII | 453140 | 453251 | 111 | 0,122531105 | 0,70040935 | 0,148894254 | -1,041187561 | -0,885009427 | -0,384838737 | -0,153838737 | -1,413838737 |
| chrXII | 453140 | 453270 | 130 | 0,387540239 | 0,429880871 | 0,47410109 | -0,064964608 | -0,055219916 | 0,077825013 | 0,175825013 | -1,084174987 |
| chrXII | 453140 | 453282 | 142 | 0,259310013 | 0,389116306 | 0,399906675 | -0,253588671 | -0,21555037 | -0,232152831 | -0,218152831 | -1,478152831 |
| chrXII | 453140 | 453286 | 146 | 1,242408414 | 1,108055004 | 0,528580196 | 0,071701317 | 0,06094612 | -0,084675116 | -0,098675116 | -1,358675116 |
| chrXII | 453140 | 453287 | 147 | 1,288001384 | 1,167348917 | 0,524569298 | 0,06162508 | 0,052381318 | -0,099617363 | -0,120617363 | -1,380617363 |
| chrXII | 453140 | 453288 | 148 | 5,106412566 | 3,561340665 | 0,589127589 | 0,225301386 | 0,191506178 | 0,093680306 | 0,065680306 | -1,194319694 |
| chrXII | 453140 | 453302 | 162 | 0,065539893 | 0,114881957 | 0,36325918 | -0,349760614 | -0,297296522 | -0,280411185 | -0,406411185 | -1,666411185 |
| chrXII | 453140 | 453309 | 169 | 0,034194727 | 0,092646739 | 0,269586342 | -0,614064532 | -0,521954852 | -0,408043158 | -0,583043158 | -1,843043158 |
| chrXII | 453140 | 453323 | 183 | 0,074088575 | 0,077823261 | 0,487707719 | -0,030817056 | -0,026194498 | 0,124288316 | -0,148711684 | -1,408711684 |
| chrXII | 453141 | 453236 | 95 | 0,131079787 | 0,211234566 | 0,382922264 | -0,297814788 | -0,25314257 | 0,273132047 | 0,616132047 | -0,643867953 |
| chrXII | 453141 | 453245 | 104 | 0,199469241 | 0,211234566 | 0,48567663 | -0,035911081 | -0,030524419 | 0,552406369 | 0,832406369 | -0,427593631 |
| chrXII | 453141 | 453251 | 110 | 0,065539893 | 0,200116957 | 0,24670884 | -0,684883118 | -0,582150651 | 0,090098053 | 0,328098053 | -0,931901947 |
| chrXII | 453141 | 453269 | 128 | 0,116831984 | 0,103764348 | 0,529618887 | 0,074311878 | 0,063165096 | 0,2756474 | 0,3876474 | -0,8723526 |
| chrXII | 453141 | 453270 | 129 | 0,988797522 | 1,011702395 | 0,494275213 | -0,014350406 | -0,012197845 | 0,209114833 | 0,314114833 | -0,945885167 |
| chrXII | 453141 | 453281 | 140 | 0,054141651 | 0,062999783 | 0,462190442 | -0,094916836 | -0,080679311 | -0,04334751 | -0,01534751 | -1,27534751 |
| chrXII | 453141 | 453282 | 141 | 0,330549028 | 0,392822175 | 0,456956299 | -0,108104753 | -0,09188904 | -0,143524872 | -0,122524872 | -1,382524872 |
| chrXII | 453141 | 453286 | 145 | 6,491299009 | 4,139456319 | 0,610615032 | 0,280922379 | 0,238784023 | 0,032681886 | 0,025681886 | -1,234318114 |

TABLE 2

| Chrom | Start | End | Length | Section A | Section B | A/A+B | Z-score | Z * 0.85 | Phase correction | Length correction | ΔLK |
|---|---|---|---|---|---|---|---|---|---|---|---|
| chrXII | 453141 | 453287 | 146 | 1,062886098 | 1,022820004 | 0,509604923 | 0,024078299 | 0,020466554 | -0,124220625 | -0,138220625 | **-1,398220625** |
| chrXII | 453141 | 453288 | 147 | 7,349016745 | 5,429098933 | 0,575125232 | 0,18943801 | 0,161022309 | 0,009564057 | -0,011435943 | **-1,271435943** |
| chrXII | 453141 | 453302 | 161 | 0,398938482 | 0,35576348 | 0,528604008 | 0,071761159 | 0,060996985 | 0,067177375 | -0,051822625 | **-1,311822625** |
| chrXII | 453141 | 453309 | 168 | 0,746584873 | 0,863467612 | 0,463702196 | -0,091111 | -0,07744435 | 0,056858772 | -0,111141228 | **-1,371141228** |
| chrXII | 453141 | 453323 | 182 | 0,065539893 | 0,081529131 | 0,445640364 | -0,13668381 | -0,116181238 | 0,066974078 | -0,199025922 | **-1,459025922** |
| chrXII | 453141 | 453324 | 183 | 0,042743409 | 0,048176305 | 0,470122564 | -0,074961771 | -0,063717505 | 0,087250953 | -0,185749047 | **-1,445749047** |
| chrXII | 453144 | 453288 | 144 | 0,037044288 | 0,096352609 | 0,277699771 | -0,589688448 | -0,501235181 | -0,609085848 | -0,609085848 | **-1,869085848** |
| chrXII | 453150 | 453286 | 136 | 0,131079787 | 0,114881957 | 0,532927539 | 0,082631037 | 0,070236381 | 0,043906171 | 0,099906171 | **-1,160093829** |
| chrXII | 453150 | 453287 | 137 | 0,039893848 | 0,051882174 | 0,434687048 | -0,164453569 | -0,139785534 | -0,077786282 | -0,028786282 | **-1,288786282** |
| chrXII | 453150 | 453288 | 138 | 0,102584181 | 0,122293696 | 0,456177292 | -0,110069087 | -0,093558724 | -0,066642731 | -0,024642731 | **-1,284642731** |
| chrXII | 453150 | 453295 | 145 | 0,094035499 | 0,044470435 | 0,678927584 | 0,464702061 | 0,394996752 | 0,204011487 | 0,197011487 | **-1,062988513** |
| chrXII | 453150 | 453302 | 152 | 0,09973462 | 0,203822827 | 0,328552705 | -0,443913104 | -0,377326138 | -0,53351621 | -0,58951621 | **-1,84951621** |
| chrXII | 453152 | 453240 | 88 | 0,102584181 | 0,137117174 | 0,427966629 | -0,181553355 | -0,154320352 | 0,274284332 | 0,666284332 | **-0,593715668** |
| chrXII | 453152 | 453270 | 118 | 0,190920559 | 0,170470001 | 0,528294262 | 0,070982761 | 0,060335347 | 0,513510439 | 0,695510439 | **-0,564489561** |
| chrXII | 453152 | 453286 | 134 | 1,894957788 | 1,608347397 | 0,540905713 | 0,102715748 | 0,087308386 | 0,076732939 | 0,146732939 | **-1,113267061** |
| chrXII | 453152 | 453287 | 135 | 0,29350474 | 0,303881305 | 0,491315025 | -0,021771723 | -0,018505964 | -0,013451589 | 0,049548411 | **-1,210451589** |
| chrXII | 453152 | 453288 | 136 | 0,703841464 | 0,770820872 | 0,477289917 | -0,056956516 | -0,048413039 | -0,083494979 | -0,027494979 | **-1,287494979** |
| chrXII | 453152 | 453302 | 150 | 0,182371877 | 0,192705218 | 0,486225044 | -0,034535557 | -0,029355224 | -0,156554267 | -0,198554267 | **-1,458554267** |
| chrXII | 453152 | 453309 | 157 | 0,04844253 | 0,044470435 | 0,521375354 | 0,053605728 | 0,045564869 | -0,052027613 | -0,143027613 | **-1,403027613** |
| chrXII | 453153 | 453286 | 133 | 0,054141651 | 0,107470218 | 0,335010364 | -0,426119559 | -0,362201625 | -0,33187596 | -0,25487596 | **-1,51487596** |
| chrXII | 453153 | 453288 | 135 | 0,088336378 | 0,077823261 | 0,531635591 | 0,07938196 | 0,067474666 | 0,067082055 | 0,130082055 | **-1,129917945** |
| chrXII | 453159 | 453270 | 111 | 0,225115286 | 0,281646088 | 0,44422345 | -0,140269704 | -0,119229248 | 0,381265429 | 0,612265429 | **-0,647734571** |
| chrXII | 453159 | 453286 | 127 | 0,478726178 | 0,655938915 | 0,421909673 | -0,197010469 | -0,167458899 | 0,070134939 | 0,189134939 | **-1,070865061** |
| chrXII | 453159 | 453287 | 128 | 0,544266071 | 0,648527176 | 0,456295399 | -0,109771242 | -0,093305555 | 0,116665585 | 0,228665585 | **-1,031334415** |
| chrXII | 453159 | 453288 | 129 | 1,134125112 | 1,356348266 | 0,455385359 | -0,112066452 | -0,095256484 | 0,129126482 | 0,234126482 | **-1,025873518** |
| chrXII | 453159 | 453302 | 143 | 2,165666043 | 1,523112397 | 0,587095722 | 0,22008038 | 0,187068323 | 0,128321638 | 0,135321638 | **-1,124678362** |
| chrXII | 453159 | 453309 | 150 | 0,105433742 | 0,137117174 | 0,434687048 | -0,164453569 | -0,139785534 | -0,259989909 | -0,301989909 | **-1,561989909** |
| chrXII | 453159 | 453335 | 176 | 0,045592969 | 0,096352609 | 0,321200349 | -0,464344846 | -0,394693119 | -0,204153227 | -0,428153227 | **-1,688153227** |
| chrXII | 453159 | 453342 | 183 | 0,028495606 | 0,048176305 | 0,371656395 | -0,327469522 | -0,278349094 | -0,133339773 | -0,406339773 | **-1,666339773** |
| chrXII | 453161 | 453245 | 84 | 0,122531105 | 0,100058479 | 0,55047996 | 0,126874057 | 0,107842948 | 0,44136194 | 0,86136194 | **-0,39863806** |
| chrXII | 453161 | 453268 | 107 | 0,265009134 | 0,581821524 | 0,3129423 | -0,487527441 | -0,414398325 | 0,146960567 | 0,405960567 | **-0,854039433** |
| chrXII | 453161 | 453270 | 109 | 0,079787696 | 0,185293479 | 0,300993446 | -0,521545394 | -0,443313585 | 0,187194449 | 0,432194449 | **-0,827805551** |
| chrXII | 453161 | 453282 | 121 | 0,131079787 | 0,163058261 | 0,445640364 | -0,13668381 | -0,116181238 | 0,27406708 | 0,43506708 | **-0,82493292** |
| chrXII | 453161 | 453286 | 125 | 3,90674756 | 5,907156108 | 0,398082933 | -0,258312336 | -0,219565485 | 0,184542265 | 0,317542265 | **-0,942457735** |
| chrXII | 453161 | 453288 | 127 | 0,666797177 | 1,015408265 | 0,396382725 | -0,262721233 | -0,223313048 | 0,027655605 | 0,146655605 | **-1,113344395** |
| chrXII | 453161 | 453309 | 148 | 0,339097709 | 0,366881088 | 0,480322795 | -0,049343454 | -0,041941936 | -0,139578397 | -0,167578397 | **-1,427578397** |
| chrXII | 453161 | 453323 | 162 | 0,222265726 | 0,140823044 | 0,612152576 | 0,284933816 | 0,242193743 | 0,280547015 | 0,154547015 | **-1,105452985** |
| chrXII | 453161 | 453324 | 163 | 0,09973462 | 0,066705652 | 0,599221683 | 0,251333038 | 0,213633083 | 0,291452564 | 0,158452564 | **-1,101547436** |
| chrXII | 453162 | 453269 | 107 | 0,173823196 | 0,233469783 | 0,426776803 | -0,184586205 | -0,156898275 | 0,406945026 | 0,665945026 | **-0,594054974** |
| chrXII | 453162 | 453286 | 124 | 0,111132863 | 0,185293479 | 0,374908863 | -0,318879716 | -0,271047759 | 0,16264587 | 0,30264587 | **-0,95735413** |
| chrXII | 453162 | 453288 | 126 | 0,14532759 | 0,329822392 | 0,305856246 | -0,507630508 | -0,431485931 | -0,156809458 | -0,030809458 | **-1,290809458** |
| chrXII | 453162 | 453323 | 161 | 0,222265726 | 0,214940436 | 0,508377386 | 0,021000537 | 0,017850457 | 0,022278271 | -0,096721729 | **-1,356721729** |
| chrXII | 453162 | 453324 | 162 | 0,373292436 | 0,374292827 | 0,499330918 | -0,001677139 | -0,001425569 | 0,043632421 | -0,082367579 | **-1,342367579** |
| chrXII | 453165 | 453286 | 121 | 0,059840772 | 0,107470218 | 0,357661934 | -0,364715392 | -0,310008083 | 0,088920118 | 0,249920118 | **-1,010079882** |
| chrXII | 453165 | 453287 | 122 | 0,045592969 | 0,074117392 | 0,380860679 | -0,303221116 | -0,257737948 | 0,085000907 | 0,239000907 | **-1,020999093** |
| chrXII | 453165 | 453288 | 123 | 0,156725832 | 0,292763697 | 0,348675157 | -0,38889974 | -0,330564779 | 0,111176365 | 0,258176365 | **-1,001823635** |
| chrXII | 453165 | 453302 | 137 | 0,045592969 | 0,059293913 | 0,434687048 | -0,164453569 | -0,139785534 | -0,071037093 | -0,022037093 | **-1,282037093** |
| chrXII | 453165 | 453342 | 177 | 0,256460453 | 0,233469783 | 0,523463207 | 0,058847485 | 0,050020363 | 0,203049028 | -0,027950972 | **-1,287950972** |
| chrXII | 453180 | 453286 | 106 | 0,031345166 | 0,166764131 | 0,158221582 | -1,001793859 | -0,85152478 | -0,406973842 | -0,140973842 | **-1,400973842** |
| chrXII | 453180 | 453288 | 108 | 0,062690333 | 0,214940436 | 0,225804702 | -0,752734617 | -0,639824425 | 0,013653156 | 0,265653156 | **-0,994346844** |
| chrXII | 453180 | 453340 | 160 | 0,111132863 | 0,125999566 | 0,468653164 | -0,078655895 | -0,066857511 | -0,068407218 | -0,180407218 | **-1,440407218** |
| chrXII | 453181 | 453270 | 89 | 0,04844253 | 0,277940218 | 0,148422459 | -1,04322323 | -0,886739745 | -0,462550529 | -0,077550529 | **-1,337550529** |
| chrXII | 453181 | 453282 | 101 | 0,113982423 | 1,856640659 | 0,057840804 | -1,573160721 | -1,337186613 | -0,895233757 | -0,594233757 | **-1,854233757** |
| chrXII | 453181 | 453286 | 105 | 0,279256937 | 1,045055221 | 0,21086942 | -0,803408194 | -0,682896965 | -0,130474245 | 0,142525755 | **-1,117474245** |
| chrXII | 453181 | 453288 | 107 | 0,227964847 | 0,96723196 | 0,19073415 | -0,875194106 | -0,74391499 | -0,191141437 | 0,067858563 | **-1,192141437** |
| chrXII | 453181 | 453323 | 142 | 0,410336724 | 0,418763262 | 0,494918262 | -0,012738373 | -0,010827617 | -0,020947266 | -0,006947266 | **-1,266947266** |
| chrXII | 453181 | 453324 | 143 | 0,045592969 | 0,100058479 | 0,313027917 | -0,487285766 | -0,414192901 | -0,469976 | -0,462976 | **-1,722976** |
| chrXII | 453286 | 453432 | 146 | 0,53571739 | 0,214940436 | 0,713663898 | 0,564120388 | 0,47950233 | 0,329999522 | 0,315999522 | **-0,944000478** |
| chrXII | 453299 | 453425 | 126 | 0,037044288 | 0,133411305 | 0,217325152 | -0,781258786 | -0,664069968 | -0,381497291 | -0,255497291 | **-1,515497291** |
| chrXII | 453299 | 453440 | 141 | 13,33594353 | 8,542029379 | 0,609560291 | 0,278173185 | 0,236447207 | 0,171490712 | 0,192490712 | **-1,067509288** |
| chrXII | 453299 | 453460 | 161 | 0,079787696 | 0,055588044 | 0,589379577 | 0,225949308 | 0,192056912 | 0,198988413 | 0,079988413 | **-1,180011587** |
| chrXII | 453300 | 453410 | 110 | 0,028495606 | 0,218646305 | 0,115300581 | -1,198811728 | -1,018989969 | -0,343285834 | -0,105285834 | **-1,365285834** |
| chrXII | 453300 | 453414 | 114 | 0,031345166 | 0,255705001 | 0,10919752 | -1,230807045 | -1,046185988 | -0,449902166 | -0,239902166 | **-1,499902166** |
| chrXII | 453300 | 453440 | 140 | 0,031345166 | 0,144528914 | 0,178225048 | -0,922150465 | -0,783827895 | -0,746581283 | -0,718581283 | **-1,978581283** |
| chrXII | 453302 | 453440 | 138 | 0,042743409 | 0,040764565 | 0,51184823 | 0,029703476 | 0,025247955 | 0,053743977 | 0,095743977 | **-1,164256023** |
| chrXII | 453310 | 453440 | 130 | 0,116831984 | 0,077823261 | 0,600199516 | 0,25386356 | 0,215784026 | 0,344227911 | 0,442227911 | **-0,817772089** |
| chrXII | 453310 | 453460 | 150 | 0,119681545 | 0,085235 | 0,584050178 | 0,21226584 | 0,180425964 | 0,065115525 | 0,023115525 | **-1,236884475** |
| chrXII | 453322 | 453481 | 159 | 0,176672756 | 0,148234783 | 0,543763178 | 0,109918963 | 0,093431118 | 0,054934533 | -0,050065467 | **-1,310065467** |
| chrXII | 453458 | 453595 | 137 | 0,094035499 | 0,237175653 | 0,283914049 | -0,571253086 | -0,485565123 | -0,422667366 | -0,373667366 | **-1,633667366** |
| chrXII | 453458 | 453610 | 152 | 0,344796831 | 0,281646088 | 0,550404227 | 0,126682692 | 0,107680288 | -0,038015096 | -0,094015096 | **-1,354015096** |
| chrXII | 453458 | 453612 | 154 | 0,225115286 | 0,255705001 | 0,468190075 | -0,079820337 | -0,067847286 | -0,228180698 | -0,298180698 | **-1,558180698** |
| chrXII | 453460 | 453612 | 152 | 0,179522317 | 0,166764131 | 0,518421434 | 0,046192108 | 0,039263292 | -0,107694186 | -0,163694186 | **-1,423694186** |
| chrXII | 453487 | 453595 | 108 | 0,381841118 | 3,772575231 | 0,091912097 | -1,32907207 | -1,129711259 | -0,463679521 | -0,211679521 | **-1,471679521** |
| chrXII | 453487 | 453610 | 123 | 0,04844253 | 0,188999349 | 0,204018475 | -0,827353109 | -0,703250143 | -0,264331296 | -0,117331296 | **-1,377331296** |
| chrXII | 453487 | 453612 | 125 | 0,085486818 | 0,170470001 | 0,333989218 | -0,428924135 | -0,364585514 | 0,01586215 | 0,14886215 | **-1,11113785** |
| chrXII | 453500 | 453652 | 152 | 0,347646391 | 0,251999131 | 0,579753168 | 0,201262063 | 0,171072754 | 0,033024016 | -0,022975984 | **-1,282975984** |
| chrXII | 453574 | 453727 | 153 | 0,113982423 | 0,044470435 | 0,719345959 | 0,58089966 | 0,493764711 | 0,364325495 | 0,301325495 | **-0,958674505** |
| chrXII | 453587 | 453737 | 150 | 0,054141651 | 0,085235 | 0,388455674 | -0,283346352 | -0,2408444 | -0,362370448 | -0,404370448 | **-1,664370448** |
| chrXII | 453606 | 453774 | 168 | 0,062690333 | 0,044470435 | 0,585011979 | 0,214732295 | 0,182522451 | 0,313676152 | 0,145676152 | **-1,114323848** |
| chrXII | 453627 | 453776 | 149 | 0,074088575 | 0,133411305 | 0,357053581 | -0,366345661 | -0,311393812 | -0,447013422 | -0,482013422 | **-1,742013422** |
| chrXII | 453627 | 453779 | 152 | 1,376337762 | 0,667056524 | 0,673554669 | 0,449750072 | 0,382287561 | 0,246202221 | 0,190202221 | **-1,069797779** |
| chrXII | 453657 | 453814 | 157 | 0,247911771 | 0,096352609 | 0,720120307 | 0,583198937 | 0,495719096 | 0,406659632 | 0,315659632 | **-0,944340368** |
| chrXII | 453659 | 453776 | 117 | 1,057186976 | 5,392040237 | 0,163924598 | -0,978455287 | -0,831686994 | -0,337011648 | -0,148011648 | **-1,408011648** |
| chrXII | 453659 | 453779 | 120 | 0,082637257 | 0,200116957 | 0,292258268 | -0,546799428 | -0,464779514 | -0,065080791 | 0,102919209 | **-1,157080791** |
| chrXII | 453734 | 453853 | 119 | 0,410336724 | 0,848644133 | 0,325927691 | -0,451186162 | -0,383508238 | 0,031158634 | 0,206158634 | **-1,053841366** |
| chrXII | 453748 | 453910 | 162 | 0,039893848 | 0,111176087 | 0,264075364 | -0,630831465 | -0,536206745 | -0,490176937 | -0,616176937 | **-1,876176937** |
| chrXII | 453748 | 453913 | 165 | 3,182959172 | 3,290812186 | 0,491670001 | -0,020881728 | -0,017749469 | 0,039893353 | -0,107106647 | **-1,367106647** |
| chrXII | 453748 | 453919 | 171 | 0,624053768 | 0,741173916 | 0,457106002 | -0,107727312 | -0,091568216 | 0,069043188 | -0,119956812 | **-1,379956812** |
| chrXII | 453748 | 453922 | 174 | 0,031345166 | 0,077823261 | 0,287126664 | -0,56179848 | -0,477528708 | -0,248968275 | -0,458968275 | **-1,718968275** |
| chrXII | 453748 | 453962 | 214 | 0,119681545 | 0,188999349 | 0,387719315 | -0,285268256 | -0,242478018 | 0,018914742 | -0,471085258 | **-1,731085258** |
| chrXII | 453752 | 453853 | 101 | 0,056991212 | 0,163058261 | 0,25899272 | -0,646453904 | -0,549485818 | -0,066030532 | 0,234969468 | **-1,025030532** |
| chrXII | 453752 | 453859 | 107 | 0,202318801 | 0,237175653 | 0,460344378 | -0,099566165 | -0,08463124 | 0,46589067 | 0,72489067 | **-0,53510933** |
| chrXII | 453752 | 453908 | 156 | 0,678195419 | 0,959820221 | 0,414034764 | -0,21717813 | -0,18460141 | -0,178404847 | -0,262404847 | **-1,522404847** |
| chrXII | 453752 | 453911 | 159 | 0,63260245 | 0,667056524 | 0,486744956 | -0,033231584 | -0,028246847 | -0,062007064 | -0,167007064 | **-1,427007064** |
| chrXII | 453752 | 453914 | 162 | 1,07428434 | 1,05617283 | 0,504250616 | 0,010654916 | 0,009056678 | 0,044899598 | -0,081100402 | **-1,341100402** |
| chrXII | 453752 | 453916 | 164 | 2,011789772 | 1,912228703 | 0,512686111 | 0,031804726 | 0,027034017 | 0,011069017 | -0,128930983 | **-1,388930983** |
| chrXII | 453752 | 453922 | 170 | 2,63584354 | 3,25004762 | 0,447824037 | -0,13116084 | -0,111486714 | 0,054166177 | -0,127833823 | **-1,387833823** |
| chrXII | 453752 | 453924 | 172 | 0,330549028 | 0,329822392 | 0,500550172 | 0,001379076 | 0,001172215 | 0,106453431 | -0,089546569 | **-1,349546569** |
| chrXII | 453753 | 453913 | 160 | 0,037044288 | 0,062999783 | 0,370279692 | -0,331112667 | -0,281445767 | -0,273741933 | -0,385741933 | **-1,645741933** |
| chrXII | 453753 | 453919 | 166 | 0,085486818 | 0,062999783 | 0,575720754 | 0,190958009 | 0,162314307 | 0,279053599 | 0,125053599 | **-1,134946401** |
| chrXII | 453753 | 453922 | 169 | 0,102584181 | 0,122293696 | 0,456177292 | -0,110069087 | -0,093558724 | 0,021229332 | -0,153770668 | **-1,413770668** |

| Chrom | Start | End | Length | Section A | Section B | A/A+B | Z-score | Z * 0.85 | Phase correction | Length correction | ΔLK |
|---|---|---|---|---|---|---|---|---|---|---|---|
| chrXII | 453754 | 453910 | 156 | 0,182371877 | 0,333528262 | 0,353502284 | -0,375882075 | -0,319499764 | -0,306313296 | -0,390313296 | -1,650313296 |
| chrXII | 453754 | 453924 | 170 | 0,045592969 | 0,096352609 | 0,321200349 | -0,464344846 | -0,394693119 | -0,231342306 | -0,413342306 | -1,673342306 |
| chrXII | 453774 | 453910 | 136 | 0,09688506 | 0,077823261 | 0,554553208 | 0,137173592 | 0,116597553 | 0,07479581 | 0,13079581 | -1,12920419 |
| chrXII | 453774 | 453913 | 139 | 0,797876963 | 0,733762177 | 0,520930122 | 0,052488125 | 0,044614907 | 0,004857857 | 0,039857857 | -1,220142143 |
| chrXII | 453774 | 453914 | 140 | 0,091185939 | 0,17417587 | 0,34362872 | -0,402579708 | -0,342192752 | -0,310051489 | -0,282051489 | -1,542051489 |
| chrXII | 453774 | 453916 | 142 | 0,076938136 | 0,081529131 | 0,485514374 | -0,036318061 | -0,030870352 | -0,051713418 | -0,037713418 | -1,297713418 |
| chrXII | 453774 | 453919 | 145 | 9,870877861 | 5,45504002 | 0,644064384 | 0,369344136 | 0,313942516 | 0,127412679 | 0,120412679 | -1,139587321 |
| chrXII | 453774 | 453922 | 148 | 0,04844253 | 0,155646522 | 0,237359767 | -0,714821199 | -0,607598019 | -0,706673623 | -0,734673623 | -1,994673623 |
| chrXII | 453774 | 453925 | 151 | 0,085486818 | 0,081529131 | 0,51184823 | 0,029703476 | 0,025247955 | -0,099783162 | -0,148783162 | -1,408783162 |
| chrXII | 453774 | 453962 | 188 | 0,504372223 | 0,326116523 | 0,607319757 | 0,272340151 | 0,231489128 | 0,251958584 | -0,056041416 | -1,316041416 |
| chrXII | 453775 | 453908 | 133 | 0,703841464 | 0,930173264 | 0,430743647 | -0,174481212 | -0,14830903 | -0,120308731 | -0,043308731 | -1,303308731 |
| chrXII | 453775 | 453911 | 136 | 0,176672756 | 0,155646522 | 0,531635591 | 0,07938196 | 0,067474666 | 0,034226704 | 0,090226704 | -1,169773296 |
| chrXII | 453775 | 453914 | 139 | 7,81349512 | 5,27345241 | 0,597044888 | 0,245705487 | 0,208849664 | 0,16854541 | 0,20354541 | -1,05645459 |
| chrXII | 453775 | 453916 | 141 | 0,276407377 | 0,34835174 | 0,442422318 | -0,144830582 | -0,123105995 | -0,198427921 | -0,177427921 | -1,437427921 |
| chrXII | 453775 | 453919 | 144 | 0,037044288 | 0,181587609 | 0,169436794 | -0,956393263 | -0,812934274 | -0,918844761 | -0,918844761 | -2,178844761 |
| chrXII | 453775 | 453922 | 147 | 0,447381012 | 0,503998263 | 0,470244648 | -0,074654896 | -0,063456662 | -0,220005416 | -0,241005416 | -1,501005416 |
| chrXII | 453781 | 453908 | 127 | 6,308927132 | 6,407448501 | 0,496126201 | -0,009710327 | -0,008253778 | 0,268710526 | 0,387710526 | -0,872289474 |
| chrXII | 453781 | 453911 | 130 | 0,094035499 | 0,08894087 | 0,51392155 | 0,034903236 | 0,029667751 | 0,161371498 | 0,259371498 | -1,000628502 |
| chrXII | 453781 | 453914 | 133 | 0,777930039 | 0,796761959 | 0,494020443 | -0,014989087 | -0,012740724 | 0,00089945 | 0,07789945 | -1,18210055 |
| chrXII | 453781 | 453916 | 135 | 0,24506221 | 0,200116957 | 0,55047996 | 0,126874057 | 0,107842948 | 0,109813167 | 0,172813167 | -1,087186833 |
| chrXII | 453781 | 453922 | 141 | 0,473027057 | 0,507704132 | 0,482320805 | -0,044329684 | -0,037680231 | -0,106406678 | -0,085406678 | -1,345406678 |
| chrXII | 453782 | 453908 | 126 | 0,082637257 | 0,096352609 | 0,461686791 | -0,096185077 | -0,081757315 | 0,172933344 | 0,298933344 | -0,961066656 |
| chrXII | 453782 | 453911 | 129 | 0,085486818 | 0,17788174 | 0,324590066 | -0,454901459 | -0,386666241 | -0,157732098 | -0,052732098 | -1,312732098 |
| chrXII | 453782 | 453913 | 131 | 0,373292436 | 0,226058044 | 0,622828292 | 0,312917403 | 0,265979793 | 0,330479689 | 0,421479689 | -0,838520311 |
| chrXII | 453782 | 453914 | 132 | 0,088336378 | 0,096352609 | 0,478298027 | -0,054425638 | -0,046261792 | 0,085987547 | 0,169987547 | -1,090012453 |
| chrXII | 453782 | 453919 | 137 | 0,173823196 | 0,196411088 | 0,469495137 | -0,076539017 | -0,065058164 | -0,006429412 | 0,042570588 | -1,217429412 |
| chrXII | 453782 | 453922 | 140 | 0,068389454 | 0,062999783 | 0,520510322 | 0,051434423 | 0,043719259 | 0,085002832 | 0,113002832 | -1,146997168 |
| chrXII | 453782 | 453925 | 143 | 0,213717044 | 0,151940653 | 0,584472981 | 0,213349921 | 0,181347433 | 0,120070199 | 0,127070199 | -1,132929801 |
| chrXII | 453782 | 453929 | 147 | 0,190920559 | 0,151940653 | 0,556845022 | 0,142974962 | 0,121528717 | -0,029697855 | -0,050697855 | -1,310697855 |
| chrXII | 453782 | 453937 | 155 | 0,085486818 | 0,100058479 | 0,460732874 | -0,09858756 | -0,083799426 | -0,20462973 | -0,28162973 | -1,54162973 |
| chrXII | 453782 | 453962 | 180 | 0,059840772 | 0,074117392 | 0,446712395 | -0,133971905 | -0,113876119 | 0,049196035 | -0,202803965 | -1,462803965 |
| chrXII | 453783 | 453911 | 128 | 0,059840772 | 0,070411522 | 0,459422021 | -0,101889932 | -0,086606442 | 0,121313982 | 0,233313982 | -1,026686018 |
| chrXII | 453783 | 453913 | 130 | 0,059840772 | 0,059293913 | 0,502295129 | 0,005753068 | 0,004890107 | 0,137629418 | 0,235629418 | -1,024370582 |
| chrXII | 453783 | 453914 | 131 | 0,028495606 | 0,122293696 | 0,18897631 | -0,881674934 | -0,749423694 | -0,68682847 | -0,59582847 | -1,85582847 |
| chrXII | 453812 | 453913 | 101 | 0,156725832 | 0,222352175 | 0,413439528 | -0,218706025 | -0,185900121 | 0,286590897 | 0,587590897 | -0,672409103 |
| chrXII | 453812 | 453919 | 107 | 0,413186285 | 0,411351523 | 0,5011126 | 0,002788878 | 0,002370546 | 0,540754032 | 0,799754032 | -0,460245968 |
| chrXII | 453812 | 453962 | 150 | 2,68428607 | 1,571288701 | 0,630769335 | 0,333891639 | 0,283807893 | 0,154712147 | 0,112712147 | -1,147287853 |
| chrXII | 453812 | 454001 | 189 | 0,111132863 | 0,100058479 | 0,52621884 | 0,065768269 | 0,055903029 | 0,101547112 | -0,213452888 | -1,473452888 |
| chrXII | 453812 | 454001 | 189 | 0,111132863 | 0,100058479 | 0,52621884 | 0,065768269 | 0,055903029 | 0,101547112 | -0,213452888 | -1,473452888 |
| chrXII | 453851 | 453958 | 107 | 0,179522317 | 0,218646305 | 0,450870076 | -0,123463404 | -0,104943894 | 0,441364284 | 0,700364284 | -0,559635716 |
| chrXII | 453851 | 453961 | 110 | 0,028495606 | 0,133411305 | 0,175999936 | -0,930717198 | -0,791109618 | -0,118559031 | 0,119440969 | -1,140559031 |
| chrXII | 453851 | 453962 | 111 | 0,102584181 | 0,133411305 | 0,434687048 | -0,164453569 | -0,139785534 | 0,37049467 | 0,60149467 | -0,65850533 |
| chrXII | 453851 | 454001 | 150 | 1,421930731 | 0,770820872 | 0,648468677 | 0,381189491 | 0,324011068 | 0,202557613 | 0,160557613 | -1,099442387 |
| chrXII | 453851 | 454001 | 150 | 1,421930731 | 0,770820872 | 0,648468677 | 0,381189491 | 0,324011068 | 0,202557613 | 0,160557613 | -1,099442387 |
| chrXII | 453909 | 454072 | 163 | 0,094035499 | 0,129705435 | 0,420287416 | -0,201158254 | -0,170984516 | -0,107128358 | -0,240128358 | -1,500128358 |
| chrXII | 453918 | 454080 | 162 | 0,282106498 | 0,218646305 | 0,56336479 | 0,159505761 | 0,135579897 | 0,151011692 | 0,025011692 | -1,234988308 |
| chrXII | 453922 | 454079 | 157 | 0,666797177 | 0,548468698 | 0,548684194 | 0,122337654 | 0,103987006 | 0,004428209 | -0,086571791 | -1,346571791 |
| chrXII | 453929 | 454079 | 150 | 0,051292091 | 0,111176087 | 0,315705458 | -0,47974192 | -0,407780632 | -0,531161087 | -0,573161087 | -1,833161087 |
| chrXII | 453929 | 454097 | 168 | 0,652549374 | 1,119172613 | 0,36831364 | -0,336323037 | -0,285874581 | -0,159762446 | -0,327762446 | -1,587762446 |
| chrXII | 453929 | 454108 | 179 | 0,108283302 | 0,148234783 | 0,42212736 | -0,196454145 | -0,166986023 | -0,022671331 | -0,267671331 | -1,527671331 |
| chrXII | 453959 | 454097 | 138 | 0,108283302 | 0,111176087 | 0,493409293 | -0,016521205 | -0,014043024 | 0,015511747 | 0,057511747 | -1,202488253 |
| chrXII | 453959 | 454108 | 149 | 0,48727486 | 0,300175436 | 0,618800783 | 0,302332726 | 0,256982817 | 0,118025928 | 0,083025928 | -1,176974072 |
| chrXII | 453959 | 454112 | 153 | 0,156725832 | 0,077823261 | 0,668200546 | 0,434949741 | 0,36970728 | 0,238955318 | 0,175955318 | -1,084044682 |
| chrXII | 453960 | 454116 | 156 | 0,068389454 | 0,055588044 | 0,551627959 | 0,129775456 | 0,110309137 | 0,131896953 | 0,047896953 | -1,212103047 |
| chrXII | 453964 | 454097 | 133 | 0,076938136 | 0,107470218 | 0,417216109 | -0,209020522 | -0,177667444 | -0,160610196 | -0,083610196 | -1,343610196 |
| chrXII | 453964 | 454108 | 144 | 0,581310359 | 0,363175219 | 0,61547828 | 0,293626349 | 0,249582396 | 0,147478582 | 0,147478582 | -1,112521418 |
| chrXII | 453971 | 454123 | 152 | 0,108283302 | 0,077823261 | 0,581834946 | 0,206589951 | 0,175601459 | 0,035613079 | -0,020386921 | -1,280386921 |
| chrXII | 454059 | 454216 | 157 | 0,045592969 | 0,040764565 | 0,527955893 | 0,070132481 | 0,059612609 | -0,046576609 | -0,137576609 | -1,397576609 |
| chrXII | 454111 | 454268 | 157 | 0,108283302 | 0,122293696 | 0,469618839 | -0,076228037 | -0,064793831 | -0,180530374 | -0,271530374 | -1,531530374 |
| chrXII | 454120 | 454267 | 147 | 0,059840772 | 0,081529131 | 0,423292165 | -0,193478389 | -0,164456631 | -0,309901998 | -0,330901998 | -1,590901998 |
| chrXII | 454120 | 454277 | 157 | 0,09688506 | 0,08894087 | 0,521375354 | 0,053605728 | 0,045564869 | -0,073361653 | -0,164361653 | -1,424361653 |
| chrXII | 454124 | 454277 | 153 | 0,199469241 | 0,17788174 | 0,528604008 | 0,071761159 | 0,060996985 | -0,076493395 | -0,139493395 | -1,399493395 |
| chrXII | 454131 | 454277 | 146 | 0,14532759 | 0,148234783 | 0,495048423 | -0,012412082 | -0,010550269 | -0,151230991 | -0,165230991 | -1,425230991 |
| chrXII | 454141 | 454277 | 136 | 0,213717044 | 0,229763914 | 0,481908051 | -0,045365348 | -0,038560545 | -0,076053521 | -0,020053521 | -1,280053521 |
| chrXII | 454196 | 454346 | 150 | 0,105433742 | 0,037058696 | 0,739925174 | 0,64311474 | 0,546647529 | 0,420291519 | 0,378291519 | -0,881708481 |
| chrXII | 454196 | 454346 | 150 | 0,105433742 | 0,037058696 | 0,739925174 | 0,64311474 | 0,546647529 | 0,420291519 | 0,378291519 | -0,881708481 |
| chrXII | 454221 | 454392 | 171 | 0,501522663 | 0,511410002 | 0,495119449 | -0,012234032 | -0,010398928 | 0,152528067 | -0,036471933 | -1,296471933 |
| chrXII | 454221 | 454392 | 171 | 0,501522663 | 0,511410002 | 0,495119449 | -0,012234032 | -0,010398928 | 0,152528067 | -0,036471933 | -1,296471933 |
| chrXII | 454244 | 454392 | 148 | 0,04844253 | 0,066705652 | 0,420697304 | -0,200109928 | -0,170093439 | -0,27818022 | -0,30618022 | -1,56618022 |
| chrXII | 454244 | 454402 | 158 | 0,410336724 | 0,244587392 | 0,626540868 | 0,322705529 | 0,274299699 | 0,237017651 | 0,139017651 | -1,120982349 |
| chrXII | 454248 | 454374 | 126 | 0,433133209 | 0,633703698 | 0,405997586 | -0,23785292 | -0,202174982 | 0,055687691 | 0,181687691 | -1,078312309 |
| chrXII | 454248 | 454392 | 144 | 0,04844253 | 0,051882174 | 0,482857442 | -0,042983253 | -0,036535765 | -0,138153008 | -0,138153008 | -1,398153008 |
| chrXII | 454248 | 454398 | 150 | 0,444531451 | 0,307587175 | 0,591039014 | 0,230218519 | 0,195685741 | 0,05403379 | 0,01203379 | -1,24796621 |
| chrXII | 454273 | 454436 | 163 | 0,416035845 | 0,35205761 | 0,541647429 | 0,104584969 | 0,088897224 | 0,147307966 | 0,014307966 | -1,245692034 |
| chrXII | 454277 | 454398 | 121 | 0,054141651 | 0,048176305 | 0,529151025 | 0,07313593 | 0,062165541 | 0,464726559 | 0,625726559 | -0,634273441 |
| chrXII | 454277 | 454402 | 125 | 0,037044288 | 0,133411305 | 0,217325152 | -0,781258786 | -0,664069968 | -0,295699207 | -0,162699207 | -1,422699207 |
| chrXII | 454277 | 454438 | 161 | 0,373292436 | 0,248293262 | 0,600548625 | 0,254767408 | 0,216552296 | 0,230628115 | 0,111628115 | -1,148371885 |
| chrXII | 454277 | 454446 | 169 | 0,954602795 | 1,600935658 | 0,373542724 | -0,322484804 | -0,274112083 | -0,151449775 | -0,326449775 | -1,586449775 |
| chrXII | 454398 | 454551 | 153 | 0,091185939 | 0,085235 | 0,516865737 | 0,042288735 | 0,035945424 | -0,099643255 | -0,162643255 | -1,422643255 |
| chrXII | 454398 | 454552 | 154 | 5,220394989 | 2,894284141 | 0,643327346 | 0,367366967 | 0,312261922 | 0,148413881 | 0,078413881 | -1,181586119 |
| chrXII | 454398 | 454555 | 157 | 0,208017923 | 0,129705435 | 0,615941769 | 0,294839538 | 0,250613607 | 0,135507463 | 0,044507463 | -1,215492537 |
| chrXII | 454398 | 454567 | 169 | 0,176672756 | 0,125999566 | 0,583709654 | 0,211392905 | 0,179683969 | 0,297796923 | 0,122796923 | -1,137203077 |
| chrXII | 454398 | 454573 | 175 | 1,159771157 | 1,782523267 | 0,394172367 | -0,268460684 | -0,228191582 | 0,013821164 | -0,203178836 | -1,463178836 |
| chrXII | 454398 | 454575 | 177 | 0,062690333 | 0,037058696 | 0,628480635 | 0,327831948 | 0,278657156 | 0,439752423 | 0,208752423 | -1,051247577 |
| chrXII | 454404 | 454552 | 148 | 0,336248149 | 0,211234566 | 0,61417126 | 0,290207525 | 0,246676396 | 0,143569962 | 0,115569962 | -1,144430038 |
| chrXII | 454404 | 454555 | 151 | 0,960301917 | 0,507704132 | 0,654153924 | 0,396559731 | 0,337075771 | 0,208942116 | 0,159942116 | -1,100057884 |
| chrXII | 454422 | 454552 | 130 | 0,3903898 | 0,311293045 | 0,556362184 | 0,141752339 | 0,120489488 | 0,241473258 | 0,339473258 | -0,920526742 |
| chrXII | 454422 | 454555 | 133 | 0,088336378 | 0,044470435 | 0,665149446 | 0,426558255 | 0,362574517 | 0,38480568 | 0,46180568 | -0,79819432 |
| chrXII | 454422 | 454567 | 145 | 1,259505778 | 0,692997611 | 0,645072262 | 0,372050197 | 0,316242667 | 0,12828311 | 0,12128311 | -1,13871689 |
| chrXII | 454422 | 454573 | 151 | 0,116831984 | 0,070411522 | 0,623957468 | 0,315891237 | 0,268507552 | 0,147845437 | 0,098845437 | -1,161154563 |
| chrXII | 454422 | 454575 | 153 | 0,703841464 | 0,452116089 | 0,608881756 | 0,276405688 | 0,234944834 | 0,112068247 | 0,049068247 | -1,210931753 |
| chrXII | 454521 | 454678 | 157 | 0,561363435 | 0,522527611 | 0,517915004 | 0,044921358 | 0,038183155 | -0,075562991 | -0,166562991 | -1,426562991 |
| chrXII | 454553 | 454712 | 159 | 0,718089267 | 0,52623348 | 0,577092453 | 0,194460792 | 0,165291673 | 0,133082602 | 0,028082602 | -1,231917398 |
| chrXII | 454553 | 454715 | 162 | 0,065539893 | 0,048176305 | 0,576346155 | 0,192554744 | 0,163671533 | 0,188931362 | 0,062931362 | -1,197068638 |
| chrXII | 454553 | 454733 | 180 | 0,074088575 | 0,125999566 | 0,370279692 | -0,331112667 | -0,281445767 | -0,120580235 | -0,372580235 | -1,632580235 |
| chrXII | 454555 | 454703 | 148 | 0,766531797 | 0,507704132 | 0,601561908 | 0,257391992 | 0,218783194 | 0,112447845 | 0,084447845 | -1,175552155 |
| chrXII | 454560 | 454678 | 118 | 0,051292091 | 0,08894087 | 0,365763444 | -0,34309514 | -0,291630869 | 0,158964129 | 0,340964129 | -0,919035871 |
| chrXII | 454560 | 454712 | 152 | 0,381841118 | 0,366881088 | 0,509990374 | 0,025044772 | 0,021288056 | -0,122878957 | -0,178878957 | -1,438878957 |
| chrXII | 454560 | 454715 | 155 | 1,117027749 | 0,919055655 | 0,54861591 | 0,122165207 | 0,103840426 | -0,007898696 | -0,084898696 | -1,344898696 |
| chrXII | 454560 | 454724 | 164 | 0,142478029 | 0,074117392 | 0,657807209 | 0,406485945 | 0,345513053 | 0,331403884 | 0,191403884 | -1,068596116 |
| chrXII | 454560 | 454733 | 173 | 0,216566604 | 0,237175653 | 0,477289917 | -0,056956516 | -0,048413039 | 0,08994852 | -0,11305148 | -1,37305148 |

TABLE 2

| Chrom | Start | End | Length | Section A | Section B | A/A+B | Z-score | Z * 0.85 | Phase correction | Length correction | ΔLK |
|---|---|---|---|---|---|---|---|---|---|---|---|
| chrXII | 454562 | 454712 | 150 | 0,168124074 | 0,085235 | 0,663580235 | 0,422254138 | 0,358916017 | 0,212326297 | 0,170326297 | -1,089673703 |
| chrXII | 454570 | 454712 | 142 | 0,082637257 | 0,051882174 | 0,614314648 | 0,290582424 | 0,24699506 | 0,234180798 | 0,248180798 | -1,011819202 |
| chrXII | 454570 | 454715 | 145 | 0,059840772 | 0,048176305 | 0,553993628 | 0,135757811 | 0,11539414 | -0,066321638 | -0,073321638 | -1,333321638 |
| chrXII | 454570 | 454724 | 154 | 0,797876963 | 0,43729261 | 0,645965526 | 0,374450809 | 0,318283187 | 0,156046954 | 0,086046954 | -1,173953046 |
| chrXII | 454570 | 454733 | 163 | 0,113982423 | 0,129705435 | 0,467739443 | -0,080953558 | -0,068810524 | -0,013490175 | -0,146490175 | -1,406490175 |
| chrXII | 454576 | 454712 | 136 | 0,094035499 | 0,144528914 | 0,394172367 | -0,268460684 | -0,228191582 | -0,272847827 | -0,216847827 | -1,476847827 |
| chrXII | 454576 | 454715 | 139 | 0,116831984 | 0,122293696 | 0,488579829 | -0,028630034 | -0,024335529 | -0,066344565 | -0,031344565 | -1,291344565 |
| chrXII | 454576 | 454724 | 148 | 0,133929347 | 0,129705435 | 0,508010916 | 0,020081738 | 0,017069477 | -0,08348199 | -0,11148199 | -1,37148199 |
| chrXII | 454576 | 454733 | 157 | 1,789524046 | 1,619465006 | 0,524942738 | 0,062562961 | 0,053178517 | -0,061750071 | -0,152750071 | -1,412750071 |
| chrXII | 454576 | 454772 | 196 | 0,059840772 | 0,051882174 | 0,535617563 | 0,08939893 | 0,07598909 | 0,130076926 | -0,233923074 | -1,493923074 |
| chrXII | 454587 | 454746 | 159 | 0,085486818 | 0,085235 | 0,500737507 | 0,001848658 | 0,001571359 | -0,030125026 | -0,135125026 | -1,395125026 |
| chrXII | 454592 | 454742 | 150 | 0,14532759 | 0,107470218 | 0,574876781 | 0,188803997 | 0,160483398 | 0,019809661 | -0,022190339 | -1,282190339 |
| chrXII | 454626 | 454724 | 98 | 0,051292091 | 0,048176305 | 0,515662191 | 0,039269381 | 0,033378974 | 0,581862749 | 0,903862749 | -0,356137251 |
| chrXII | 454626 | 454733 | 107 | 0,128230226 | 0,163058261 | 0,440217282 | -0,150418352 | -0,127855599 | 0,425385357 | 0,684385357 | -0,575614643 |
| chrXII | 454626 | 454772 | 146 | 1,12557643 | 0,70411522 | 0,615172742 | 0,292826836 | 0,24890281 | 0,106705574 | 0,092705574 | -1,167294426 |
| chrXII | 454649 | 454741 | 92 | 0,031345166 | 0,037058696 | 0,458236793 | -0,104876779 | -0,089145262 | 0,277034387 | 0,641034387 | -0,618965613 |
| chrXII | 454677 | 454828 | 151 | 0,04844253 | 0,048176305 | 0,50137771 | 0,003453413 | 0,002935401 | -0,120219436 | -0,169219436 | -1,429219436 |
| chrXII | 454677 | 454828 | 151 | 0,04844253 | 0,048176305 | 0,50137771 | 0,003453413 | 0,002935401 | -0,120219436 | -0,169219436 | -1,429219436 |
| chrXII | 454723 | 454854 | 131 | 0,262159574 | 0,259410871 | 0,502635026 | 0,006605078 | 0,005614317 | 0,051885394 | 0,142885394 | -1,117114606 |
| chrXII | 454725 | 454863 | 138 | 0,92895675 | 0,752291524 | 0,552539898 | 0,132081025 | 0,112268872 | 0,135889795 | 0,177889795 | -1,082110205 |
| chrXII | 454725 | 454886 | 161 | 0,062690333 | 0,055588044 | 0,530023616 | 0,075329227 | 0,064029843 | 0,081234846 | -0,037765154 | -1,297765154 |
| chrXII | 454733 | 454863 | 130 | 0,056991212 | 0,077823261 | 0,422738082 | -0,194893703 | -0,165659647 | -0,037206879 | 0,060793121 | -1,199206879 |
| chrXII | 454733 | 454886 | 153 | 0,170973635 | 0,26311674 | 0,39386645 | -0,269255726 | -0,228867367 | -0,350674328 | -0,413674328 | -1,673674328 |
| chrXII | 454741 | 454896 | 155 | 0,039893848 | 0,044470435 | 0,472876041 | -0,06804215 | -0,057835827 | -0,168428713 | -0,245428713 | -1,505428713 |
| chrXII | 454747 | 454853 | 106 | 0,045592969 | 0,037058696 | 0,551627959 | 0,129775456 | 0,110309137 | 0,550377411 | 0,816377411 | -0,443622589 |
| chrXII | 454747 | 454886 | 139 | 0,356195073 | 0,255705001 | 0,582113139 | 0,207302373 | 0,176207017 | 0,147166905 | 0,182166905 | -1,077833095 |
| chrXII | 454825 | 454972 | 147 | 0,034194727 | 0,055588044 | 0,380860679 | -0,303221116 | -0,257737948 | -0,399443806 | -0,420443806 | -1,680443806 |
| chrXII | 454825 | 454972 | 147 | 0,034194727 | 0,055588044 | 0,380860679 | -0,303221116 | -0,257737948 | -0,399443806 | -0,420443806 | -1,680443806 |
| chrXII | 454837 | 454991 | 154 | 0,307752543 | 0,111176087 | 0,734618073 | 0,626840405 | 0,532814345 | 0,371795248 | 0,301795248 | -0,958204752 |
| chrXII | 454837 | 454991 | 154 | 0,307752543 | 0,111176087 | 0,734618073 | 0,626840405 | 0,532814345 | 0,371795248 | 0,301795248 | -0,958204752 |
| chrXII | 454864 | 455021 | 157 | 0,173823196 | 0,081529131 | 0,68071906 | 0,46971048 | 0,399253908 | 0,296990657 | 0,205990657 | -1,054009343 |
| chrXII | 454876 | 455023 | 147 | 0,812124766 | 0,411351523 | 0,663784638 | 0,422814341 | 0,35939219 | 0,215351213 | 0,194351213 | -1,065648787 |
| chrXII | 454876 | 455029 | 153 | 0,125380666 | 0,170470001 | 0,423797138 | -0,192188854 | -0,163360526 | -0,293796046 | -0,356796046 | -1,616796046 |
| chrXII | 454901 | 455029 | 128 | 0,094035499 | 0,248293262 | 0,27469354 | -0,598678825 | -0,508877001 | -0,303662519 | -0,191662519 | -1,451662519 |
| chrXII | 454901 | 455074 | 173 | 0,210867483 | 0,255705001 | 0,451950105 | -0,120735916 | -0,102625529 | 0,040995719 | -0,162004281 | -1,422004281 |
| chrXII | 454920 | 455087 | 167 | 0,09973462 | 0,103764348 | 0,490098899 | -0,024820929 | -0,021097789 | 0,147009799 | -0,013990201 | -1,273990201 |
| chrXII | 454932 | 455073 | 141 | 0,689593661 | 0,370586958 | 0,650449224 | 0,386533562 | 0,328553528 | 0,260644273 | 0,281644273 | -0,978355727 |
| chrXII | 454933 | 455023 | 90 | 0,190920559 | 0,407645654 | 0,318963141 | -0,470600177 | -0,40001015 | 0,034285231 | 0,412285231 | -0,847714769 |
| chrXII | 454933 | 455024 | 91 | 0,031345166 | 0,140823044 | 0,18206129 | -0,907537593 | -0,771406954 | -0,376537125 | -0,005537125 | -1,265537125 |
| chrXII | 454933 | 455025 | 92 | 0,031345166 | 0,044470435 | 0,413439528 | -0,218706025 | -0,185900121 | 0,20438963 | 0,56838963 | -0,69161037 |
| chrXII | 454933 | 455027 | 94 | 0,028495606 | 0,08894087 | 0,242646977 | -0,69781331 | -0,593141313 | -0,084638715 | 0,265361285 | -0,994638715 |
| chrXII | 454933 | 455029 | 96 | 5,867245241 | 28,82054771 | 0,16914438 | -0,957551906 | -0,81391912 | -0,257792976 | 0,078207024 | -1,181792976 |
| chrXII | 454933 | 455073 | 140 | 0,074088575 | 0,037058696 | 0,666580246 | 0,43048963 | 0,365916185 | 0,401637802 | 0,429637802 | -0,830362198 |
| chrXII | 454933 | 455074 | 141 | 0,24506221 | 0,237175653 | 0,508177041 | 0,020498237 | 0,017423501 | -0,087709506 | -0,066709506 | -1,326709506 |
| chrXII | 454933 | 455076 | 143 | 0,669646737 | 0,481763045 | 0,581588542 | 0,205959029 | 0,175065175 | 0,107599857 | 0,114599857 | -1,145400143 |
| chrXII | 454933 | 455132 | 199 | 0,031345166 | 0,096352609 | 0,245463684 | -0,688834586 | -0,585509398 | -0,535476381 | -0,920476381 | -2,180476381 |
| chrXII | 454964 | 455132 | 168 | 0,190920559 | 0,192705218 | 0,497679358 | -0,005830555 | -0,004955971 | 0,114320963 | -0,053679037 | -1,313679037 |
| chrXII | 454995 | 455164 | 169 | 0,125380666 | 0,259410871 | 0,325840498 | -0,451428154 | -0,383713931 | -0,262455115 | -0,437455115 | -1,697455115 |
| chrXII | 455000 | 455132 | 132 | 0,085486818 | 0,163058261 | 0,343948944 | -0,401709424 | -0,34145301 | -0,213881427 | -0,129881427 | -1,389881427 |
| chrXII | 455015 | 455166 | 151 | 0,481575739 | 0,318704784 | 0,601758665 | 0,257901833 | 0,219216558 | 0,093046579 | 0,044046579 | -1,215953421 |
| chrXII | 455058 | 455208 | 150 | 0,042743409 | 0,044470435 | 0,490098899 | -0,024820929 | -0,021097789 | -0,137311083 | -0,179311083 | -1,439311083 |
| chrXII | 455068 | 455223 | 155 | 0,162424953 | 0,118587827 | 0,577998458 | 0,196775678 | 0,167259327 | 0,055866116 | -0,021133884 | -1,281133884 |
| chrXII | 455095 | 455269 | 174 | 0,14817715 | 0,222352175 | 0,399906675 | -0,253588671 | -0,21555037 | 0,022525549 | -0,187474451 | -1,447474451 |
| chrXII | 455096 | 455223 | 127 | 0,054141651 | 0,096352609 | 0,359758911 | -0,359103289 | -0,305237796 | -0,048315108 | 0,070684892 | -1,189315108 |
| chrXII | 455096 | 455240 | 144 | 0,849169054 | 0,70782109 | 0,545391413 | 0,11402601 | 0,096922109 | -0,016225788 | -0,016225788 | -1,276225788 |
| chrXII | 455096 | 455243 | 147 | 0,14817715 | 0,111176087 | 0,571333335 | 0,179769754 | 0,152804291 | 0,023903006 | 0,002903006 | -1,257096994 |
| chrXII | 455096 | 455245 | 149 | 0,971700159 | 0,61146848 | 0,613769206 | 0,28915654 | 0,245783059 | 0,095716462 | 0,060716462 | -1,199283538 |
| chrXII | 455096 | 455249 | 153 | 0,056991212 | 0,037058696 | 0,605967759 | 0,268824834 | 0,228501109 | 0,111772271 | 0,048772271 | -1,211227729 |
| chrXII | 455097 | 455240 | 143 | 0,04844253 | 0,044470435 | 0,521375354 | 0,053605728 | 0,045564869 | -0,033935047 | -0,026935047 | -1,286935047 |
| chrXII | 455097 | 455243 | 146 | 0,527168708 | 0,318704784 | 0,623224056 | 0,313959381 | 0,266865474 | 0,131253075 | 0,117253075 | -1,142746925 |
| chrXII | 455098 | 455243 | 145 | 0,051292091 | 0,040764565 | 0,557179598 | 0,143822286 | 0,122248943 | -0,059139823 | -0,066139823 | -1,326139823 |
| chrXII | 455098 | 455249 | 151 | 0,151026711 | 0,151940653 | 0,498491683 | -0,0037808 | -0,00321368 | -0,120443897 | -0,169443897 | -1,429443897 |
| chrXII | 455129 | 455293 | 164 | 0,059840772 | 0,044470435 | 0,573675388 | 0,185739279 | 0,157878387 | 0,120678445 | -0,019321555 | -1,279321555 |
| chrXII | 455138 | 455289 | 151 | 0,091185939 | 0,081529131 | 0,527955893 | 0,070132481 | 0,059612609 | -0,054275748 | -0,103275748 | -1,363275748 |
| chrXII | 455182 | 455338 | 156 | 0,034194727 | 0,044470435 | 0,434687048 | -0,164453569 | -0,139785534 | -0,112895455 | -0,196895455 | -1,456895455 |
| chrXII | 455217 | 455366 | 149 | 0,059840772 | 0,051882174 | 0,535617563 | 0,08939893 | 0,07598909 | -0,076046834 | -0,111046834 | -1,371046834 |
| chrXII | 455221 | 455395 | 174 | 0,054141651 | 0,218646305 | 0,198475225 | -0,847080146 | -0,720018124 | -0,487052919 | -0,697052919 | -1,957052919 |
| chrXII | 455247 | 455381 | 134 | 0,085486818 | 0,08894087 | 0,490098899 | -0,024820929 | -0,021097789 | -0,033894193 | 0,036105807 | -1,223894193 |
| chrXII | 455254 | 455401 | 147 | 0,133929347 | 0,103764348 | 0,563453511 | 0,159731003 | 0,135771353 | 0,015886157 | -0,005113843 | -1,265113843 |
| chrXII | 455255 | 455400 | 145 | 0,102584181 | 0,074117392 | 0,58055047 | 0,203301913 | 0,172806626 | -0,000794559 | -0,007794559 | -1,267794559 |
| chrXII | 455278 | 455423 | 145 | 0,068389454 | 0,08894087 | 0,434687048 | -0,164453569 | -0,139785534 | -0,306385181 | -0,313385181 | -1,573385181 |
| chrXII | 455297 | 455447 | 150 | 0,265009134 | 0,26311674 | 0,501791613 | 0,004490922 | 0,003817284 | -0,123154852 | -0,165154852 | -1,425154852 |
| chrXII | 455367 | 455512 | 145 | 1,410532489 | 0,763409133 | 0,648836415 | 0,382180925 | 0,324853786 | 0,155291229 | 0,148291229 | -1,111708771 |
| chrXII | 455367 | 455531 | 164 | 0,068389454 | 0,085235 | 0,445172966 | -0,137866493 | -0,117186519 | -0,155598298 | -0,295598298 | -1,555598298 |
| chrXII | 455367 | 455549 | 182 | 0,219416165 | 0,222352175 | 0,49667698 | -0,008329671 | -0,007080221 | 0,166341286 | -0,099658714 | -1,359658714 |
| chrXII | 455377 | 455512 | 135 | 0,062690333 | 0,037058696 | 0,628480635 | 0,327831948 | 0,278657156 | 0,283073674 | 0,346073674 | -0,913926326 |
| chrXII | 455377 | 455531 | 154 | 0,567062556 | 0,233469783 | 0,708356837 | 0,548590764 | 0,466302149 | 0,324069805 | 0,254069805 | -1,005930195 |
| chrXII | 455377 | 455549 | 172 | 0,102584181 | 0,077823261 | 0,568624996 | 0,172874585 | 0,146943398 | 0,247276298 | 0,051276298 | -1,208723702 |
| chrXII | 455397 | 455503 | 106 | 0,085486818 | 0,103764348 | 0,451710916 | -0,121339882 | -0,103138899 | 0,317438125 | 0,583438125 | -0,676561875 |
| chrXII | 455397 | 455512 | 115 | 0,079787696 | 0,081529131 | 0,49460244 | -0,013530088 | -0,011500575 | 0,648722201 | 0,851722201 | -0,408277799 |
| chrXII | 455397 | 455531 | 134 | 0,105433742 | 0,100058479 | 0,513078994 | 0,032790051 | 0,027871543 | 0,014470214 | 0,084470214 | -1,175529786 |
| chrXII | 455397 | 455549 | 152 | 1,761028441 | 1,334113048 | 0,568965408 | 0,173740784 | 0,147679667 | 0,021277837 | -0,034722163 | -1,294722163 |
| chrXII | 455397 | 455568 | 171 | 0,034194727 | 0,059293913 | 0,365763444 | -0,34309514 | -0,291630869 | -0,132457993 | -0,321457993 | -1,581457993 |
| chrXII | 455410 | 455512 | 102 | 0,062690333 | 0,062999783 | 0,498768996 | -0,003085674 | -0,002622823 | 0,504219582 | 0,798219582 | -0,461780418 |
| chrXII | 455410 | 455549 | 139 | 0,128230226 | 0,196411088 | 0,394990473 | -0,266335355 | -0,226385052 | -0,270215213 | -0,235215213 | -1,495215213 |
| chrXII | 455410 | 455560 | 150 | 0,156725832 | 0,037058696 | 0,808763392 | 0,873348384 | 0,742346126 | 0,624840622 | 0,582840622 | -0,677159378 |
| chrXII | 455410 | 455568 | 158 | 0,524319147 | 0,344645871 | 0,603383492 | 0,262114698 | 0,222797493 | 0,199292228 | 0,101292228 | -1,158707772 |
| chrXII | 455410 | 455569 | 159 | 0,29065518 | 0,237175653 | 0,550659722 | 0,127328308 | 0,108229062 | 0,083393586 | -0,021606414 | -1,281606414 |
| chrXII | 455416 | 455571 | 155 | 0,088336378 | 0,118587827 | 0,426902103 | -0,184266739 | -0,156626728 | -0,270926045 | -0,347926045 | -1,607926045 |
| chrXII | 455422 | 455569 | 147 | 1,003045325 | 0,537351089 | 0,651160517 | 0,388455518 | 0,33018719 | 0,218737231 | 0,197737231 | -1,062262769 |
| chrXII | 455591 | 455751 | 160 | 0,045592969 | 0,077823261 | 0,36942442 | -0,33337817 | -0,283371444 | -0,273431743 | -0,385431743 | -1,645431743 |
| chrXII | 455701 | 455870 | 169 | 0,079787696 | 0,137117174 | 0,367846494 | -0,337562389 | -0,286928031 | -0,160963139 | -0,335963139 | -1,595963139 |
| chrXII | 455747 | 455895 | 148 | 0,960301917 | 0,544762828 | 0,638046915 | 0,353243137 | 0,300256667 | 0,200387699 | 0,172387699 | -1,087612301 |
| chrXII | 455749 | 455889 | 140 | 0,336248149 | 0,26682261 | 0,557560028 | 0,144785863 | 0,123067983 | 0,156871321 | 0,184871321 | -1,075128679 |
| chrXII | 455783 | 455958 | 175 | 0,045592969 | 0,111176087 | 0,290828881 | -0,550964862 | -0,468320133 | -0,227204164 | -0,444204164 | -1,704204164 |
| chrXII | 455869 | 456021 | 152 | 0,045592969 | 0,040764565 | 0,527955893 | 0,070132481 | 0,059612609 | -0,061701939 | -0,117701939 | -1,377701939 |
| chrXII | 455903 | 456067 | 164 | 0,085486818 | 0,077823261 | 0,523463207 | 0,058847485 | 0,050020363 | 0,010280425 | -0,129719575 | -1,389719575 |
| chrXII | 455918 | 456039 | 121 | 0,287805619 | 0,874585221 | 0,247597976 | -0,682068004 | -0,579757803 | -0,181252299 | -0,020252299 | -1,280252299 |
| chrXII | 455926 | 456086 | 160 | 0,262159574 | 0,259410871 | 0,502635026 | 0,006605078 | 0,005614317 | 0,007330386 | -0,104669614 | -1,364669614 |
| chrXII | 456022 | 456173 | 151 | 2,191312089 | 1,330407179 | 0,62222793 | 0,311337397 | 0,264636787 | 0,158908992 | 0,109908992 | -1,150091008 |
| chrXII | 456022 | 456205 | 183 | 0,210867483 | 0,26311674 | 0,444882916 | -0,138600522 | -0,117810443 | 0,005903191 | -0,267096809 | -1,527096809 |

TABLE 2

| Chrom | Start | End | Length | Section A | Section B | A/A+B | Z-score | Z * 0.85 | Phase correction | Length correction | ΔLK |
|---|---|---|---|---|---|---|---|---|---|---|---|
| chrXII | 456022 | 456205 | 183 | 0,210867483 | 0,26311674 | 0,444882916 | -0,138600522 | -0,117810443 | 0,005903191 | -0,267096809 | -1,527096809 |
| chrXII | 456022 | 456211 | 189 | 0,111132863 | 0,070411522 | 0,612152576 | 0,284933816 | 0,242193743 | 0,291546318 | -0,023453682 | -1,283453682 |
| chrXII | 456022 | 456211 | 189 | 0,111132863 | 0,070411522 | 0,612152576 | 0,284933816 | 0,242193743 | 0,291546318 | -0,023453682 | -1,283453682 |
| chrXII | 456022 | 456220 | 198 | 0,062690333 | 0,055588044 | 0,530023616 | 0,075329227 | 0,064029843 | 0,118059114 | -0,259940886 | -1,519940886 |
| chrXII | 456022 | 456220 | 198 | 0,062690333 | 0,055588044 | 0,530023616 | 0,075329227 | 0,064029843 | 0,118059114 | -0,259940886 | -1,519940886 |
| chrXII | 456025 | 456173 | 148 | 0,085486818 | 0,037058696 | 0,69759239 | 0,517488464 | 0,439865194 | 0,306358185 | 0,278358185 | -0,981641815 |
| chrXII | 456025 | 456182 | 157 | 0,071239015 | 0,037058696 | 0,657807209 | 0,406485945 | 0,345513053 | 0,229767352 | 0,138767352 | -1,121232648 |
| chrXII | 456042 | 456173 | 131 | 0,111132863 | 0,037058696 | 0,74992708 | 0,674260298 | 0,573121253 | 0,606712094 | 0,697712094 | -0,562287906 |
| chrXII | 456045 | 456173 | 128 | 0,091185939 | 0,081529131 | 0,527955893 | 0,070132481 | 0,059612609 | 0,254983651 | 0,366983651 | -0,893016349 |
| chrXII | 456045 | 456175 | 130 | 0,396088921 | 0,396528045 | 0,499722991 | -0,000694358 | -0,000590204 | 0,134288824 | 0,232288824 | -1,027711176 |
| chrXII | 456045 | 456205 | 160 | 0,088336378 | 0,081529131 | 0,520037168 | 0,050246866 | 0,042709836 | 0,035943381 | -0,076056619 | -1,336056619 |
| chrXII | 456045 | 456205 | 160 | 0,088336378 | 0,081529131 | 0,520037168 | 0,050246866 | 0,042709836 | 0,035943381 | -0,076056619 | -1,336056619 |
| chrXII | 456056 | 456173 | 117 | 0,424584527 | 0,570703915 | 0,426594451 | -0,185051167 | -0,157293492 | 0,332357375 | 0,521357375 | -0,738642625 |
| chrXII | 456056 | 456175 | 119 | 0,059840772 | 0,074117392 | 0,446712395 | -0,133971905 | -0,113876119 | 0,303114708 | 0,478114708 | -0,781885292 |
| chrXII | 456056 | 456205 | 149 | 4,453863192 | 3,146283272 | 0,586023337 | 0,217327245 | 0,184728158 | 0,033871174 | -0,001128826 | -1,261128826 |
| chrXII | 456056 | 456205 | 149 | 4,453863192 | 3,146283272 | 0,586023337 | 0,217327245 | 0,184728158 | 0,033871174 | -0,001128826 | -1,261128826 |
| chrXII | 456056 | 456211 | 155 | 0,250761331 | 0,26311674 | 0,487978268 | -0,030138575 | -0,025617789 | -0,140494129 | -0,217494129 | -1,477494129 |
| chrXII | 456056 | 456211 | 155 | 0,250761331 | 0,26311674 | 0,487978268 | -0,030138575 | -0,025617789 | -0,140494129 | -0,217494129 | -1,477494129 |
| chrXII | 456056 | 456220 | 164 | 0,262159574 | 0,248293262 | 0,513582363 | 0,034052516 | 0,028944639 | -0,010412803 | -0,150412803 | -1,410412803 |
| chrXII | 456056 | 456220 | 164 | 0,262159574 | 0,248293262 | 0,513582363 | 0,034052516 | 0,028944639 | -0,010412803 | -0,150412803 | -1,410412803 |
| chrXII | 456063 | 456215 | 152 | 0,361894194 | 0,170470001 | 0,679786878 | 0,467102923 | 0,397037484 | 0,269587555 | 0,213587555 | -1,046412445 |
| chrXII | 456063 | 456215 | 152 | 0,361894194 | 0,170470001 | 0,679786878 | 0,467102923 | 0,397037484 | 0,269587555 | 0,213587555 | -1,046412445 |
| chrXII | 456064 | 456214 | 150 | 0,122531105 | 0,070411522 | 0,635064977 | 0,345298405 | 0,293503645 | 0,176412996 | 0,134412996 | -1,125587004 |
| chrXII | 456064 | 456214 | 150 | 0,122531105 | 0,070411522 | 0,635064977 | 0,345298405 | 0,293503645 | 0,176412996 | 0,134412996 | -1,125587004 |
| chrXII | 456066 | 456173 | 107 | 0,165274514 | 0,155646522 | 0,515000562 | 0,037609696 | 0,031968242 | 0,585141421 | 0,844141421 | -0,415858579 |
| chrXII | 456067 | 456205 | 138 | 0,373292436 | 0,333528262 | 0,528128898 | 0,070567214 | 0,059982132 | 0,080974611 | 0,122974611 | -1,137025389 |
| chrXII | 456067 | 456211 | 144 | 1,880709985 | 0,852350003 | 0,688133445 | 0,490566465 | 0,416981495 | 0,2889066 | 0,2889066 | -0,9710934 |
| chrXII | 456067 | 456215 | 148 | 1,553010518 | 1,048761091 | 0,596905014 | 0,245344146 | 0,208542524 | 0,074794215 | 0,046794215 | -1,213205785 |
| chrXII | 456067 | 456220 | 153 | 0,14817715 | 0,074117392 | 0,666580246 | 0,43048963 | 0,365916185 | 0,241334896 | 0,178334896 | -1,081665104 |
| chrXII | 456067 | 456266 | 199 | 0,170973635 | 0,214940436 | 0,443035505 | -0,14327752 | -0,121785892 | -0,074861336 | -0,459861336 | -1,719861336 |
| chrXII | 456080 | 456173 | 93 | 0,205168362 | 0,196411088 | 0,510903539 | 0,027334523 | 0,023234344 | 0,422860872 | 0,779860872 | -0,480139128 |
| chrXII | 456080 | 456205 | 125 | 0,527168708 | 0,396528045 | 0,570716208 | 0,178197868 | 0,151468187 | 0,492116465 | 0,625116465 | -0,634883535 |
| chrXII | 456080 | 456211 | 131 | 0,199469241 | 0,092646739 | 0,682842618 | 0,475662609 | 0,404313218 | 0,44614102 | 0,53714102 | -0,72285898 |
| chrXII | 456080 | 456220 | 140 | 2,137170438 | 1,367465874 | 0,609812331 | 0,278829937 | 0,237005447 | 0,275744062 | 0,303744062 | -0,956255938 |
| chrXII | 456084 | 456215 | 131 | 0,068389454 | 0,070411522 | 0,492715945 | -0,018259434 | -0,015520519 | 0,030076617 | 0,121076617 | -1,138923383 |
| chrXII | 456084 | 456243 | 159 | 0,316301225 | 0,277940218 | 0,53227729 | 0,080995639 | 0,068846293 | 0,055517876 | -0,049482124 | -1,309482124 |
| chrXII | 456084 | 456266 | 182 | 0,082637257 | 0,103764348 | 0,443329106 | -0,142534017 | -0,121153915 | 0,053155818 | -0,212844182 | -1,472844182 |
| chrXII | 456115 | 456215 | 100 | 0,173823196 | 0,233469783 | 0,426776803 | -0,184586205 | -0,156898275 | 0,258010743 | 0,566010743 | -0,693989257 |
| chrXII | 456115 | 456243 | 128 | 0,062690333 | 0,103764348 | 0,376621026 | -0,314367347 | -0,267212245 | -0,067431524 | 0,044568476 | -1,215431524 |
| chrXII | 456115 | 456266 | 151 | 7,235034322 | 4,346985016 | 0,624678142 | 0,317790687 | 0,270122084 | 0,15948707 | 0,11048707 | -1,14951293 |
| chrXII | 456115 | 456281 | 166 | 0,373292436 | 0,548468698 | 0,40497741 | -0,240484318 | -0,20441167 | -0,083622718 | -0,237622718 | -1,497622718 |
| chrXII | 456120 | 456290 | 170 | 0,039893848 | 0,040764565 | 0,49460244 | -0,013530088 | -0,011500575 | 0,160363943 | -0,021636057 | -1,281636057 |
| chrXII | 456124 | 456215 | 91 | 0,059840772 | 0,100058479 | 0,374240479 | -0,320642986 | -0,272546538 | 0,103798198 | 0,474798198 | -0,785201802 |
| chrXII | 456124 | 456266 | 142 | 0,427434088 | 0,448410219 | 0,488025194 | -0,030020896 | -0,025517762 | -0,033910624 | -0,019910624 | -1,279910624 |
| chrXII | 456124 | 456281 | 157 | 1,478921943 | 1,382289353 | 0,516886657 | 0,042341221 | 0,035990038 | -0,078016872 | -0,169016872 | -1,429016872 |
| chrXII | 456124 | 456333 | 209 | 0,250761331 | 0,296469566 | 0,458236793 | -0,104876779 | -0,089145262 | 0,082135938 | -0,372864062 | -1,632864062 |
| chrXII | 456181 | 456266 | 85 | 1,071434779 | 1,385995222 | 0,435998087 | -0,161123447 | -0,13695493 | 0,256336774 | 0,669336774 | -0,590663226 |
| chrXII | 456181 | 456281 | 100 | 0,578460798 | 0,867173481 | 0,40014325 | -0,252976337 | -0,215029886 | 0,222190456 | 0,530190456 | -0,729809544 |
| chrXII | 456181 | 456287 | 106 | 0,09973462 | 0,344645871 | 0,224435191 | -0,757299614 | -0,643704672 | -0,208175501 | 0,057824499 | -1,202175501 |
| chrXII | 456181 | 456306 | 125 | 0,056991212 | 0,059293913 | 0,490098899 | -0,024820929 | -0,021097789 | 0,313456953 | 0,446456953 | -0,813543047 |
| chrXII | 456181 | 456310 | 129 | 1,56440876 | 1,763993919 | 0,470017877 | -0,075224925 | -0,063941186 | 0,18665456 | 0,29165456 | -0,96834544 |
| chrXII | 456181 | 456329 | 148 | 0,042743409 | 0,037058696 | 0,535617563 | 0,08939893 | 0,07598909 | -0,06181526 | -0,08981526 | -1,34981526 |
| chrXII | 456181 | 456332 | 151 | 0,037044288 | 0,081529131 | 0,31241646 | -0,489012398 | -0,415660538 | -0,531742954 | -0,580742954 | -1,840742954 |
| chrXII | 456181 | 456333 | 152 | 14,91460009 | 11,08425591 | 0,573663706 | 0,185709488 | 0,157853065 | 0,018413455 | -0,037586545 | -1,297586545 |
| chrXII | 456181 | 456336 | 155 | 0,250761331 | 0,248293262 | 0,502472745 | 0,006198292 | 0,005268549 | -0,11091953 | -0,18791953 | -1,44791953 |
| chrXII | 456181 | 456349 | 168 | 0,330549028 | 0,218646305 | 0,6018788 | 0,258213162 | 0,219481187 | 0,351279549 | 0,183279549 | -1,076720451 |
| chrXII | 456181 | 456387 | 206 | 0,239363089 | 0,163058261 | 0,594807131 | 0,239928434 | 0,203939169 | 0,280599079 | -0,153400921 | -1,413400921 |
| chrXII | 456186 | 456333 | 147 | 0,045592969 | 0,070411522 | 0,393027622 | -0,271436616 | -0,230721123 | -0,346256815 | -0,367256815 | -1,627256815 |
| chrXII | 456187 | 456333 | 146 | 0,458779254 | 0,652233046 | 0,412938051 | -0,219993647 | -0,1869946 | -0,32371293 | -0,33771293 | -1,59771293 |
| chrXII | 456187 | 456349 | 162 | 0,495823542 | 0,459527828 | 0,518996002 | 0,047633924 | 0,040488835 | 0,069862664 | -0,056137336 | -1,316137336 |
| chrXII | 456194 | 456333 | 139 | 0,3903898 | 0,381704567 | 0,505624464 | 0,014098907 | 0,011984071 | -0,03389814 | 0,00110186 | -1,25889814 |
| chrXII | 456194 | 456336 | 142 | 0,572761677 | 0,366881088 | 0,609552586 | 0,278153109 | 0,236430142 | 0,232499925 | 0,246499925 | -1,013500075 |
| chrXII | 456213 | 456319 | 106 | 3,975137014 | 8,638381988 | 0,31514893 | -0,481307652 | -0,409111504 | 0,024846884 | 0,290846884 | -0,969153116 |
| chrXII | 456213 | 456320 | 107 | 0,09688506 | 0,192705218 | 0,33455909 | -0,427358568 | -0,363254783 | 0,185371853 | 0,444371853 | -0,815628147 |
| chrXII | 456213 | 456321 | 108 | 0,065539893 | 0,229763914 | 0,221940564 | -0,76565581 | -0,650807439 | 0,023574618 | 0,275574618 | -0,984425382 |
| chrXII | 456213 | 456333 | 120 | 4,331332087 | 10,28749395 | 0,296284536 | -0,535116832 | -0,454849307 | -0,055941248 | 0,112058752 | -1,147941248 |
| chrXII | 456213 | 456336 | 123 | 0,068389454 | 0,125999566 | 0,351817475 | -0,380418277 | -0,323355536 | 0,103082503 | 0,250082503 | -1,009917497 |
| chrXII | 456213 | 456349 | 136 | 0,122531105 | 0,222352175 | 0,355282823 | -0,371096517 | -0,31543204 | -0,365253966 | -0,309253966 | -1,569253966 |
| chrXII | 456213 | 456352 | 139 | 0,054141651 | 0,051882174 | 0,510655515 | 0,026712592 | 0,022705704 | -0,029217269 | 0,005782731 | -1,254217269 |
| chrXII | 456213 | 456387 | 174 | 0,173823196 | 0,196411088 | 0,469495137 | -0,076539017 | -0,065058164 | 0,160254557 | -0,049745443 | -1,309745443 |
| chrXII | 456213 | 456419 | 206 | 0,133929347 | 0,159352392 | 0,456657642 | -0,108857794 | -0,092529125 | -0,0088479 | -0,4428479 | -1,7028479 |
| chrXII | 456213 | 456419 | 206 | 0,133929347 | 0,159352392 | 0,456657642 | -0,108857794 | -0,092529125 | -0,0088479 | -0,4428479 | -1,7028479 |
| chrXII | 456221 | 456386 | 165 | 0,615505086 | 0,377998697 | 0,619529685 | 0,304245806 | 0,258608935 | 0,321764652 | 0,174764652 | -1,085235348 |
| chrXII | 456240 | 456333 | 93 | 0,416035845 | 0,485468915 | 0,461490459 | -0,096679504 | -0,082177578 | 0,32754207 | 0,68454207 | -0,57545793 |
| chrXII | 456240 | 456386 | 146 | 0,253610892 | 0,114881957 | 0,6882383 | 0,490862927 | 0,417233488 | 0,286336317 | 0,272336317 | -0,987663683 |
| chrXII | 456240 | 456387 | 147 | 2,305294512 | 1,552759353 | 0,597527819 | 0,246953305 | 0,20991031 | 0,099791893 | 0,078791893 | -1,181208107 |
| chrXII | 456240 | 456419 | 179 | 0,29065518 | 0,311293045 | 0,482857442 | -0,042983253 | -0,036535765 | 0,116910517 | -0,128089483 | -1,388089483 |
| chrXII | 456240 | 456419 | 179 | 0,29065518 | 0,311293045 | 0,482857442 | -0,042983253 | -0,036535765 | 0,116910517 | -0,128089483 | -1,388089483 |
| chrXII | 456240 | 456447 | 207 | 0,165274514 | 0,129705435 | 0,560290672 | 0,151706213 | 0,128950281 | 0,261007412 | -0,179992588 | -1,439992588 |
| chrXII | 456240 | 456447 | 207 | 0,165274514 | 0,129705435 | 0,560290672 | 0,151706213 | 0,128950281 | 0,261007412 | -0,179992588 | -1,439992588 |
| chrXII | 456288 | 456387 | 99 | 0,330549028 | 0,403939784 | 0,450039568 | -0,125561379 | -0,106727172 | 0,355074839 | 0,670074839 | -0,589925161 |
| chrXII | 456288 | 456395 | 107 | 0,04844253 | 0,274234349 | 0,150127056 | -1,035888609 | -0,880505318 | -0,354476424 | -0,095476424 | -1,355476424 |
| chrXII | 456288 | 456419 | 131 | 6,274732405 | 6,296272414 | 0,499143266 | -0,002147515 | -0,001825388 | 0,041872249 | 0,132872249 | -1,127127751 |
| chrXII | 456288 | 456447 | 159 | 2,251152861 | 1,923346311 | 0,539262979 | 0,098577115 | 0,083790548 | 0,068411309 | -0,036588691 | -1,296588691 |
| chrXII | 456288 | 456457 | 169 | 0,165274514 | 0,188999349 | 0,46651625 | -0,084030099 | -0,071425584 | 0,066001998 | -0,108998002 | -1,368998002 |
| chrXII | 456288 | 456471 | 183 | 0,413186285 | 1,030231743 | 0,286255455 | -0,564357421 | -0,479703807 | -0,369593388 | -0,642593388 | -1,902593388 |
| chrXII | 456290 | 456419 | 129 | 0,179522317 | 0,329822392 | 0,352457409 | -0,378694391 | -0,321890232 | -0,080364681 | 0,024635319 | -1,235364681 |
| chrXII | 456290 | 456447 | 157 | 0,606956404 | 0,904232177 | 0,401641735 | -0,249099947 | -0,211734955 | -0,312206413 | -0,403206413 | -1,663206413 |
| chrXII | 456290 | 456471 | 181 | 0,074088575 | 0,17417587 | 0,298426039 | -0,528932784 | -0,449592866 | -0,257796652 | -0,516796652 | -1,776796652 |
| chrXII | 456298 | 456447 | 149 | 0,931806311 | 0,563292176 | 0,623240756 | 0,314003358 | 0,266902854 | 0,101352444 | 0,066352444 | -1,193647556 |
| chrXII | 456310 | 456417 | 107 | 0,071239015 | 0,055588044 | 0,561702018 | 0,155285859 | 0,13199298 | 0,63140811 | 0,89040811 | -0,36959189 |
| chrXII | 456310 | 456419 | 109 | 0,065539893 | 0,122293696 | 0,34892531 | -0,388223533 | -0,329990003 | 0,30713624 | 0,55213624 | -0,70786376 |
| chrXII | 456310 | 456447 | 137 | 0,082637257 | 0,118587827 | 0,410670755 | -0,225819886 | -0,191946903 | -0,136510747 | -0,087510747 | -1,347510747 |
| chrXII | 456310 | 456471 | 161 | 0,578460798 | 0,578115654 | 0,500149209 | 0,000374013 | 0,000317911 | 0,024638021 | -0,094361979 | -1,354361979 |
| chrXII | 456361 | 456447 | 86 | 0,128230226 | 0,70411522 | 0,154058903 | -1,019179398 | -0,866302488 | -0,474979226 | -0,068979226 | -1,328979226 |
| chrXII | 456361 | 456466 | 105 | 0,028495606 | 0,418763262 | 0,063711662 | -1,524341906 | -1,29569062 | -0,729721808 | -0,456721808 | -1,716721808 |
| chrXII | 456361 | 456468 | 107 | 1,17401896 | 8,734734597 | 0,118483011 | -1,182604262 | -1,005213622 | -0,521622032 | -0,262622032 | -1,522622032 |
| chrXII | 456361 | 456470 | 109 | 0,042743409 | 0,34835174 | 0,109291585 | -1,230304312 | -1,045758665 | -0,406743856 | -0,161743856 | -1,421743856 |
| chrXII | 456361 | 456471 | 110 | 1,256656217 | 14,53812636 | 0,079561476 | -1,408027415 | -1,196823303 | -0,50994537 | -0,27194537 | -1,53194537 |
| chrXII | 456388 | 456471 | 83 | 0,09688506 | 0,277940218 | 0,258480592 | -0,648036746 | -0,550831234 | -0,243338967 | 0,183661033 | -1,076338967 |
| chrXII | 456388 | 456551 | 163 | 0,322000346 | 0,281646088 | 0,533425409 | 0,083883345 | 0,071300843 | 0,121990496 | -0,011009504 | -1,271009504 |

TABLE 2

| Chrom | Start | End | Length | Section A | Section B | A/A+B | Z-score | Z * 0.85 | Phase correction | Length correction | ΔLK |
|---|---|---|---|---|---|---|---|---|---|---|---|
| chrXII | 456388 | 456551 | 163 | 0,322000346 | 0,281646088 | 0,533425409 | 0,083883345 | 0,071300843 | 0,121990496 | -0,011009504 | **-1,271009504** |
| chrXII | 456444 | 456594 | 150 | 0,473027057 | 0,281646088 | 0,626797257 | 0,323382625 | 0,274875231 | 0,164067773 | 0,122067773 | **-1,137932227** |
| chrXII | 456444 | 456594 | 150 | 0,473027057 | 0,281646088 | 0,626797257 | 0,323382625 | 0,274875231 | 0,164067773 | 0,122067773 | **-1,137932227** |
| chrXII | 456444 | 456595 | 151 | 0,199469241 | 0,188999349 | 0,513475854 | 0,033785384 | 0,028717576 | -0,095318252 | -0,144318252 | **-1,404318252** |
| chrXII | 456444 | 456595 | 151 | 0,199469241 | 0,188999349 | 0,513475854 | 0,033785384 | 0,028717576 | -0,095318252 | -0,144318252 | **-1,404318252** |
| chrXII | 456444 | 456619 | 175 | 0,051292091 | 0,118587827 | 0,301931455 | -0,518853495 | -0,441025471 | -0,190758771 | -0,407758771 | **-1,667758771** |
| chrXII | 456444 | 456619 | 175 | 0,051292091 | 0,118587827 | 0,301931455 | -0,518853495 | -0,441025471 | -0,190758771 | -0,407758771 | **-1,667758771** |
| chrXII | 456445 | 456595 | 150 | 0,071239015 | 0,103764348 | 0,407072261 | -0,235082731 | -0,199820321 | -0,316067859 | -0,358067859 | **-1,618067859** |
| chrXII | 456447 | 456594 | 147 | 1,142673794 | 0,926467395 | 0,552245444 | 0,131336508 | 0,111636032 | -0,001503292 | -0,022503292 | **-1,282503292** |
| chrXII | 456447 | 456595 | 148 | 0,074088575 | 0,096352609 | 0,434687048 | -0,164453569 | -0,139785534 | -0,276730455 | -0,304730455 | **-1,564730455** |
| chrXII | 456447 | 456608 | 161 | 0,085486818 | 0,222352175 | 0,277699771 | -0,589688448 | -0,501235181 | -0,466538213 | -0,585538213 | **-1,845538213** |
| chrXII | 456447 | 456619 | 172 | 0,58700948 | 0,944996743 | 0,383163901 | -0,297181688 | -0,252604435 | -0,146807876 | -0,342807876 | **-1,602807876** |
| chrXII | 456479 | 456585 | 106 | 0,113982423 | 0,17788174 | 0,390532439 | -0,277931585 | -0,236241848 | 0,179795974 | 0,445795974 | **-0,814204026** |
| chrXII | 456479 | 456594 | 115 | 0,068389454 | 0,125999566 | 0,351817475 | -0,380418277 | -0,323355536 | 0,320601485 | 0,523601485 | **-0,736398515** |
| chrXII | 456479 | 456608 | 129 | 1,800922289 | 1,912228703 | 0,485011865 | -0,037578525 | -0,031941746 | 0,220386243 | 0,325386243 | **-0,934613757** |
| chrXII | 456479 | 456609 | 130 | 0,088336378 | 0,100058479 | 0,468889542 | -0,078061561 | -0,066352326 | 0,051295948 | 0,149295948 | **-1,110704052** |
| chrXII | 456479 | 456653 | 174 | 0,056991212 | 0,059293913 | 0,490098899 | -0,024820929 | -0,021097789 | 0,196798578 | -0,013201422 | **-1,273201422** |
| chrXII | 456483 | 456594 | 111 | 0,031345166 | 0,111176087 | 0,219933277 | -0,772418577 | -0,656555791 | -0,142021328 | 0,088978672 | **-1,171021328** |
| chrXII | 456483 | 456608 | 125 | 0,270708255 | 0,489174784 | 0,356249898 | -0,368500871 | -0,31322574 | 0,012368728 | 0,145368728 | **-1,114631272** |
| chrXII | 456483 | 456609 | 126 | 0,247911771 | 0,392822175 | 0,386918427 | -0,287359779 | -0,244255812 | 0,002324501 | 0,128324501 | **-1,131675499** |
| chrXII | 456493 | 456599 | 106 | 0,065539893 | 0,062999783 | 0,509880648 | 0,024769644 | 0,021054198 | 0,424854448 | 0,690854448 | **-0,569145552** |
| chrXII | 456493 | 456608 | 115 | 0,054141651 | 0,100058479 | 0,351112876 | -0,382317666 | -0,324970016 | 0,329839263 | 0,532839263 | **-0,727160737** |
| chrXII | 456493 | 456653 | 160 | 0,284956058 | 0,340940001 | 0,455276965 | -0,11233987 | -0,09548889 | -0,110686853 | -0,222686853 | **-1,482686853** |
| chrXII | 456508 | 456594 | 86 | 0,039893848 | 0,055588044 | 0,417815854 | -0,20748423 | -0,176361595 | 0,220105638 | 0,626105638 | **-0,633894362** |
| chrXII | 456508 | 456608 | 100 | 0,153876272 | 0,185293479 | 0,453685128 | -0,116356189 | -0,098902761 | 0,359659711 | 0,667659711 | **-0,592340289** |
| chrXII | 456508 | 456642 | 134 | 0,227964847 | 0,26311674 | 0,464209722 | -0,089833605 | -0,076358564 | -0,091826631 | -0,021826631 | **-1,281826631** |
| chrXII | 456508 | 456653 | 145 | 0,09688506 | 0,055588044 | 0,635423938 | 0,346253615 | 0,294315573 | 0,12102827 | 0,11402827 | **-1,14597173** |
| chrXII | 456508 | 456670 | 162 | 0,159575393 | 0,133411305 | 0,544650641 | 0,112157259 | 0,09533367 | 0,118697713 | -0,007302287 | **-1,267302287** |
| chrXII | 456524 | 456695 | 171 | 0,199469241 | 0,359469349 | 0,356871478 | -0,36683385 | -0,311808773 | -0,160925187 | -0,349925187 | **-1,609925187** |
| chrXII | 456587 | 456738 | 151 | 0,051292091 | 0,096352609 | 0,34740218 | -0,39234359 | -0,333492051 | -0,461478345 | -0,510478345 | **-1,770478345** |
| chrXII | 456587 | 456738 | 151 | 0,051292091 | 0,096352609 | 0,34740218 | -0,39234359 | -0,333492051 | -0,461478345 | -0,510478345 | **-1,770478345** |
| chrXII | 456588 | 456751 | 163 | 0,068389454 | 0,037058696 | 0,648560019 | 0,381435717 | 0,324220359 | 0,375194338 | 0,242194338 | **-1,017805662** |
| chrXII | 456588 | 456751 | 163 | 0,068389454 | 0,037058696 | 0,648560019 | 0,381435717 | 0,324220359 | 0,375194338 | 0,242194338 | **-1,017805662** |
| chrXII | 456594 | 456751 | 157 | 0,102584181 | 0,122293696 | 0,456177292 | -0,110069087 | -0,093558724 | -0,190495057 | -0,281495057 | **-1,541495057** |
| chrXII | 456616 | 456772 | 156 | 0,307752543 | 0,218646305 | 0,584637569 | 0,213771999 | 0,181706199 | 0,211014995 | 0,127014995 | **-1,132985005** |
| chrXII | 456634 | 456751 | 117 | 0,039893848 | 0,059293913 | 0,402205349 | -0,247642923 | -0,210496484 | 0,28589714 | 0,47489714 | **-0,78510286** |
| chrXII | 456634 | 456762 | 128 | 0,738036191 | 1,045055221 | 0,413908219 | -0,217502911 | -0,184877475 | 0,024891251 | 0,136891251 | **-1,123108749** |
| chrXII | 456634 | 456781 | 147 | 0,270708255 | 0,229763914 | 0,540905713 | 0,102715748 | 0,087308386 | -0,019825975 | -0,040825975 | **-1,300825975** |
| chrXII | 456649 | 456803 | 154 | 0,250761331 | 0,207528696 | 0,547167331 | 0,118507771 | 0,100731606 | -0,036873488 | -0,106873488 | **-1,366873488** |
| chrXII | 456653 | 456762 | 109 | 0,031345166 | 0,044470435 | 0,413439528 | -0,218706025 | -0,185900121 | 0,468337565 | 0,713337565 | **-0,546662435** |
| chrXII | 456653 | 456803 | 150 | 0,19377012 | 0,081529131 | 0,703852696 | 0,535513815 | 0,455186742 | 0,336987848 | 0,294987848 | **-0,965012152** |
| chrXII | 456733 | 456869 | 136 | 0,045592969 | 0,037058696 | 0,551627959 | 0,129775456 | 0,110309137 | 0,066186264 | 0,122186264 | **-1,137813736** |
| chrXII | 456733 | 456869 | 136 | 0,045592969 | 0,037058696 | 0,551627959 | 0,129775456 | 0,110309137 | 0,066186264 | 0,122186264 | **-1,137813736** |
| chrXII | 456766 | 456920 | 154 | 0,062690333 | 0,048176305 | 0,56545715 | 0,164819952 | 0,140096959 | -0,00740753 | -0,07740753 | **-1,33740753** |
| chrXII | 456780 | 456887 | 107 | 0,039893848 | 0,470645436 | 0,078140604 | -1,417690283 | -1,20503674 | -0,744208602 | -0,485208602 | **-1,745208602** |
| chrXII | 456780 | 456890 | 110 | 0,037044288 | 0,489174784 | 0,070397083 | -1,472840067 | -1,251914057 | -0,579302824 | -0,341302824 | **-1,601302824** |
| chrXII | 456780 | 456920 | 140 | 0,045592969 | 0,314998914 | 0,126439256 | -1,143385642 | -0,971877795 | -0,941661016 | -0,913661016 | **-2,173661016** |
| chrXII | 456794 | 456903 | 109 | 0,076938136 | 0,200116957 | 0,277699771 | -0,589688448 | -0,501235181 | 0,15961202 | 0,40461202 | **-0,85538798** |
| chrXII | 456795 | 456903 | 108 | 2,815365857 | 8,842204815 | 0,241505365 | -0,701468431 | -0,596248166 | 0,0864811 | 0,3384811 | **-0,9215189** |
| chrXII | 456795 | 456904 | 109 | 0,037044288 | 0,137117174 | 0,21270083 | -0,797085011 | -0,67752226 | -0,000424336 | 0,244575664 | **-1,015424336** |
| chrXII | 456795 | 456905 | 110 | 0,031345166 | 0,185293479 | 0,144688711 | -1,059488371 | -0,900565115 | -0,212605321 | 0,025394679 | **-1,234605321** |
| chrXII | 456797 | 456890 | 93 | 0,056991212 | 0,481763045 | 0,105783316 | -1,249269194 | -1,061878815 | -0,648908122 | -0,291908122 | **-1,551908122** |
| chrXII | 456797 | 456920 | 123 | 0,051292091 | 0,596645002 | 0,079162146 | -1,410729832 | -1,199120357 | -0,772943385 | -0,625943385 | **-1,885943385** |
| chrXII | 456803 | 456959 | 156 | 0,085486818 | 0,103764348 | 0,451710916 | -0,121339881 | -0,103138899 | -0,074588796 | -0,158588796 | **-1,418588796** |
| chrXII | 456804 | 456911 | 107 | 0,04844253 | 0,166764131 | 0,225097726 | -0,755089219 | -0,641825836 | -0,187188098 | 0,071811902 | **-1,188188098** |
| chrXII | 456804 | 456920 | 116 | 0,042743409 | 0,218646305 | 0,163523683 | -0,98007851 | -0,833066733 | -0,235414435 | -0,039414435 | **-1,299414435** |
| chrXII | 456804 | 456960 | 156 | 0,082637257 | 0,203822827 | 0,288477389 | -0,557838343 | -0,474162592 | -0,45160905 | -0,53560905 | **-1,79560905** |
| chrXII | 456817 | 456920 | 103 | 0,051292091 | 0,192705218 | 0,210215804 | -0,805672675 | -0,684821774 | -0,100304825 | 0,186695175 | **-1,073304825** |
| chrXII | 456817 | 456971 | 154 | 0,051292091 | 0,044470435 | 0,535617563 | 0,08939893 | 0,07598909 | -0,084587486 | -0,154587486 | **-1,414587486** |
| chrXII | 456938 | 457104 | 166 | 0,062690333 | 0,077823261 | 0,446151373 | -0,135390993 | -0,115082344 | 0,019739117 | -0,134260883 | **-1,394260883** |
| chrXII | 456984 | 457118 | 134 | 0,09688506 | 0,074117392 | 0,566571175 | 0,16765125 | 0,142503563 | 0,143635748 | 0,213635748 | **-1,046364252** |
| chrXII | 456984 | 457127 | 143 | 0,492973981 | 0,559586306 | 0,468357002 | -0,079400583 | -0,067490496 | -0,154129479 | -0,147129479 | **-1,407129479** |
| chrXII | 456984 | 457130 | 146 | 0,051292091 | 0,077823261 | 0,397257877 | -0,260451199 | -0,221383519 | -0,353737478 | -0,367737478 | **-1,627737478** |
| chrXII | 456984 | 457140 | 156 | 0,037044288 | 0,066705652 | 0,357053581 | -0,366345661 | -0,311393812 | -0,283988076 | -0,367988076 | **-1,627988076** |
| chrXII | 456990 | 457127 | 137 | 0,105433742 | 0,155646522 | 0,403836506 | -0,243429088 | -0,206914725 | -0,146231234 | -0,097231234 | **-1,357231234** |
| chrXII | 456990 | 457140 | 150 | 0,552814753 | 0,429880871 | 0,562549318 | 0,157435847 | 0,13382047 | 0,036565297 | -0,005434703 | **-1,265434703** |
| chrXII | 457005 | 457166 | 161 | 0,091185939 | 0,181587609 | 0,33429172 | -0,428092961 | -0,363879017 | -0,328329845 | -0,447329845 | **-1,707329845** |
| chrXII | 457021 | 457127 | 106 | 0,065539893 | 0,670762394 | 0,089012209 | -1,346862817 | -1,144833394 | -0,735888188 | -0,469888188 | **-1,729888188** |
| chrXII | 457022 | 457205 | 183 | 0,071239015 | 0,255705001 | 0,217893619 | -0,779326791 | -0,662427772 | -0,561253224 | -0,834253224 | **-2,094253224** |
| chrXII | 457022 | 457205 | 183 | 0,071239015 | 0,255705001 | 0,217893619 | -0,779326791 | -0,662427772 | -0,561253224 | -0,834253224 | **-2,094253224** |
| chrXII | 457047 | 457207 | 160 | 0,094035499 | 0,103764348 | 0,47540734 | -0,061683752 | -0,052431189 | -0,068694378 | -0,180694378 | **-1,440694378** |
| chrXII | 457047 | 457207 | 160 | 0,094035499 | 0,103764348 | 0,47540734 | -0,061683752 | -0,052431189 | -0,068694378 | -0,180694378 | **-1,440694378** |
| chrXII | 457070 | 457200 | 130 | 0,267858695 | 0,537351089 | 0,33265703 | -0,432588068 | -0,367699858 | -0,253691373 | -0,155691373 | **-1,415691373** |
| chrXII | 457070 | 457200 | 130 | 0,267858695 | 0,537351089 | 0,33265703 | -0,432588068 | -0,367699858 | -0,253691373 | -0,155691373 | **-1,415691373** |
| chrXII | 457083 | 457217 | 134 | 0,051292091 | 0,085235 | 0,375691668 | -0,31681585 | -0,269293473 | -0,271019882 | -0,201019882 | **-1,461019882** |
| chrXII | 457115 | 457248 | 133 | 1,065735658 | 0,919055655 | 0,536950989 | 0,092755225 | 0,078841941 | 0,091402548 | 0,168402548 | **-1,091597452** |
| chrXII | 457140 | 457305 | 165 | 0,074088575 | 0,066705652 | 0,52621884 | 0,065768269 | 0,055903029 | 0,123287051 | -0,023712949 | **-1,283712949** |
| chrXII | 457140 | 457312 | 172 | 0,777930039 | 1,397112831 | 0,357661934 | -0,364715392 | -0,310008083 | -0,204956141 | -0,400956141 | **-1,660956141** |
| chrXII | 457143 | 457248 | 105 | 0,039893848 | 0,100058479 | 0,285053125 | -0,567895024 | -0,48271077 | 0,089865288 | 0,362865288 | **-0,897134712** |
| chrXII | 457143 | 457279 | 136 | 0,558513874 | 0,433586741 | 0,562960919 | 0,15848053 | 0,134708451 | 0,097578637 | 0,153578637 | **-1,106421363** |
| chrXII | 457143 | 457316 | 173 | 0,059840772 | 0,074117392 | 0,446712395 | -0,133971905 | -0,113876119 | 0,034504381 | -0,168495619 | **-1,428495619** |
| chrXII | 457152 | 457305 | 153 | 0,552814753 | 0,300175436 | 0,6480904 | 0,380170037 | 0,323144532 | 0,197570353 | 0,134570353 | **-1,125429647** |
| chrXII | 457152 | 457312 | 160 | 0,059840772 | 0,070411522 | 0,459422021 | -0,101889932 | -0,086606442 | -0,099274216 | -0,211274216 | **-1,471274216** |
| chrXII | 457153 | 457248 | 95 | 0,091185939 | 0,207528696 | 0,305261035 | -0,509328378 | -0,432929121 | 0,094245116 | 0,437245116 | **-0,822754884** |
| chrXII | 457153 | 457279 | 126 | 0,079787696 | 0,096352609 | 0,452978074 | -0,118140734 | -0,100419624 | 0,151398994 | 0,277398994 | **-0,982601006** |
| chrXII | 457153 | 457312 | 159 | 0,139628469 | 0,166764131 | 0,455717497 | -0,111228699 | -0,094544395 | -0,117818599 | -0,222818599 | **-1,482818599** |
| chrXII | 457153 | 457316 | 163 | 1,735382395 | 1,589818049 | 0,521888056 | 0,054892775 | 0,046658858 | 0,092452593 | -0,040547407 | **-1,300547407** |
| chrXII | 457153 | 457317 | 164 | 0,074088575 | 0,166764131 | 0,307609478 | -0,502637789 | -0,427242121 | -0,458981961 | -0,598981961 | **-1,858981961** |
| chrXII | 457154 | 457312 | 158 | 0,284956058 | 0,311293045 | 0,477914444 | -0,055388588 | -0,0470803 | -0,074625836 | -0,172625836 | **-1,432625836** |
| chrXII | 457154 | 457316 | 162 | 0,079787696 | 0,081529131 | 0,49460244 | -0,013530088 | -0,011500575 | 0,015251926 | -0,110748074 | **-1,370748074** |
| chrXII | 457187 | 457316 | 129 | 0,088336378 | 0,111176087 | 0,442761197 | -0,143972235 | -0,122376399 | 0,120739353 | 0,225739353 | **-1,034260647** |
| chrXII | 457187 | 457317 | 130 | 0,407487163 | 0,366881088 | 0,52621884 | 0,065768269 | 0,055903029 | 0,159829042 | 0,257829042 | **-1,002170958** |
| chrXII | 457198 | 457346 | 148 | 0,421734966 | 0,255705001 | 0,622542198 | 0,312164377 | 0,26533972 | 0,130367657 | 0,102367657 | **-1,157632343** |
| chrXII | 457198 | 457365 | 167 | 0,054141651 | 0,037058696 | 0,593656197 | 0,236960278 | 0,201416237 | 0,360214571 | 0,199214571 | **-1,060785429** |
| chrXII | 457212 | 457365 | 153 | 0,094035499 | 0,048176305 | 0,661235543 | 0,415837502 | 0,353461877 | 0,249050769 | 0,186050769 | **-1,073949231** |
| chrXII | 457214 | 457363 | 149 | 0,262159574 | 0,103764348 | 0,716431908 | 0,572274263 | 0,486433123 | 0,320715917 | 0,285715917 | **-0,974284083** |
| chrXII | 457219 | 457312 | 93 | 0,051292091 | 0,129705435 | 0,283385589 | -0,572813203 | -0,486891222 | -0,067833668 | 0,289166332 | **-0,970833668** |
| chrXII | 457219 | 457346 | 127 | 0,054141651 | 0,085235 | 0,388455674 | -0,283346352 | -0,2408444 | 0,001903039 | 0,120903039 | **-1,139096961** |
| chrXII | 457219 | 457363 | 144 | 0,045592969 | 0,044470435 | 0,506231912 | 0,015621722 | 0,013278464 | -0,118376444 | -0,118376444 | **-1,378376444** |
| chrXII | 457219 | 457365 | 146 | 0,869115978 | 0,681880002 | 0,560359916 | 0,151881795 | 0,129099526 | -0,012451175 | -0,026451175 | **-1,286451175** |
| chrXII | 457219 | 457422 | 203 | 0,028495606 | 0,048176305 | 0,371656395 | -0,327469522 | -0,278349094 | -0,242949793 | -0,655949793 | **-1,915949793** |

TABLE 2

| Chrom | Start | End | Length | Section A | Section B | A/A+B | Z-score | Z * 0.85 | Phase correction | Length correction | ΔLK |
|---|---|---|---|---|---|---|---|---|---|---|---|
| chrXII | 457227 | 457316 | 89 | 0,045592969 | 0,037058696 | 0,551627959 | 0,129775456 | 0,110309137 | 0,533144053 | 0,918144053 | -0,341855947 |
| chrXII | 457227 | 457363 | 136 | 0,473027057 | 0,311293045 | 0,603104595 | 0,261391241 | 0,222182555 | 0,189736683 | 0,245736683 | -1,014263317 |
| chrXII | 457227 | 457395 | 168 | 0,037044288 | 0,092646739 | 0,285634931 | -0,566182298 | -0,481254953 | -0,342082218 | -0,510082218 | -1,770082218 |
| chrXII | 457230 | 457395 | 165 | 0,054141651 | 0,192705218 | 0,219332946 | -0,774448026 | -0,658280822 | -0,589545303 | -0,736545303 | -1,996545303 |
| chrXII | 457230 | 457401 | 171 | 0,247911771 | 0,233469783 | 0,515000562 | 0,037609696 | 0,031968242 | 0,180262495 | -0,008737505 | -1,268737505 |
| chrXII | 457248 | 457395 | 147 | 0,059840772 | 0,100058479 | 0,374240479 | -0,320642986 | -0,272546538 | -0,378630334 | -0,399630334 | -1,659630334 |
| chrXII | 457248 | 457401 | 153 | 0,210867483 | 0,092646739 | 0,694753219 | 0,509369056 | 0,432963697 | 0,341698489 | 0,278698489 | -0,981301511 |
| chrXII | 457255 | 457365 | 110 | 0,062690333 | 0,059293913 | 0,51392155 | 0,034903236 | 0,029667751 | 0,727221901 | 0,965221901 | -0,294778099 |
| chrXII | 457255 | 457407 | 152 | 0,512920905 | 0,26311674 | 0,660948484 | 0,415053099 | 0,352795134 | 0,213627948 | 0,157627948 | -1,102372052 |
| chrXII | 457255 | 457422 | 167 | 0,037044288 | 0,051882174 | 0,416572153 | -0,210670615 | -0,179070023 | -0,033556303 | -0,194556303 | -1,454556303 |
| chrXII | 457266 | 457365 | 99 | 0,088336378 | 0,096352609 | 0,478298027 | -0,054425638 | -0,046261792 | 0,400334672 | 0,715334672 | -0,544665328 |
| chrXII | 457266 | 457407 | 141 | 0,116831984 | 0,062999783 | 0,649673781 | 0,384439894 | 0,326773909 | 0,222597822 | 0,243597822 | -1,016402178 |
| chrXII | 457266 | 457422 | 156 | 1,159771157 | 0,804173699 | 0,590531427 | 0,228912214 | 0,194575382 | 0,214166428 | 0,130166428 | -1,129833572 |
| chrXII | 457295 | 457378 | 83 | 0,028495606 | 0,140823044 | 0,16829573 | -0,960921856 | -0,816783577 | -0,504960809 | -0,077960809 | -1,337960809 |
| chrXII | 457295 | 457384 | 89 | 0,076938136 | 0,229763914 | 0,250856282 | -0,67179759 | -0,571027951 | -0,14434915 | 0,24065085 | -1,01934915 |
| chrXII | 457295 | 457390 | 95 | 0,062690333 | 0,359469349 | 0,1484991 | -1,042892249 | -0,886458411 | -0,376055282 | -0,033055282 | -1,293055282 |
| chrXII | 457295 | 457391 | 96 | 0,091185939 | 0,889408699 | 0,092990452 | -1,322562523 | -1,124178145 | -0,555094384 | -0,219094384 | -1,479094384 |
| chrXII | 457295 | 457393 | 98 | 0,037044288 | 0,344645871 | 0,097053295 | -1,298526168 | -1,103747243 | -0,560740471 | -0,238740471 | -1,498740471 |
| chrXII | 457295 | 457394 | 99 | 0,031345166 | 0,381704567 | 0,075887148 | -1,433292386 | -1,218298528 | -0,760260463 | -0,445260463 | -1,705260463 |
| chrXII | 457295 | 457395 | 100 | 14,8490602 | 81,00289725 | 0,154916609 | -1,015572055 | -0,863236247 | -0,409307624 | -0,101307624 | -1,361307624 |
| chrXII | 457295 | 457397 | 102 | 0,037044288 | 0,481763045 | 0,071402783 | -1,465422925 | -1,245609486 | -0,733109196 | -0,439109196 | -1,699109196 |
| chrXII | 457295 | 457401 | 106 | 0,136778908 | 0,318704784 | 0,300293755 | -0,523555831 | -0,445022456 | -0,046004191 | 0,219995809 | -1,040004191 |
| chrXII | 457295 | 457402 | 107 | 0,133929347 | 0,296469566 | 0,31117492 | -0,492522755 | -0,418644342 | 0,009189281 | 0,268189281 | -0,991810719 |
| chrXII | 457337 | 457476 | 139 | 0,071239015 | 0,085235 | 0,455276965 | -0,11233987 | -0,09548889 | -0,150582036 | -0,115582036 | -1,375582036 |
| chrXII | 457337 | 457481 | 144 | 0,649699813 | 0,277940218 | 0,700379232 | 0,525491536 | 0,446667806 | 0,300064179 | 0,300064179 | -0,959935821 |
| chrXII | 457360 | 457476 | 116 | 0,045592969 | 0,17788174 | 0,204018475 | -0,827353109 | -0,703250143 | -0,110417981 | 0,085582019 | -1,174417981 |
| chrXII | 457360 | 457481 | 121 | 0,042743409 | 0,037058696 | 0,535617563 | 0,08939893 | 0,07598909 | 0,459366615 | 0,620366615 | -0,639633385 |
| chrXII | 457360 | 457530 | 170 | 0,165274514 | 0,237175653 | 0,410670755 | -0,225819886 | -0,191946903 | -0,019578819 | -0,201578819 | -1,461578819 |
| chrXII | 457370 | 457476 | 106 | 0,792177842 | 4,684219147 | 0,144653107 | -1,05964482 | -0,900698097 | -0,502843022 | -0,236843022 | -1,496843022 |
| chrXII | 457372 | 457476 | 104 | 0,04844253 | 0,896820438 | 0,051247676 | -1,632874705 | -1,387943499 | -0,8246605 | -0,5446605 | -1,8046605 |
| chrXII | 457420 | 457563 | 143 | 0,105433742 | 0,092646739 | 0,53227729 | 0,080995639 | 0,068846293 | -0,025611526 | -0,018611526 | -1,278611526 |
| chrXII | 457420 | 457567 | 147 | 0,076938136 | 0,059293913 | 0,564757972 | 0,163043664 | 0,138587115 | 0,03266304 | 0,01166304 | -1,24833696 |
| chrXII | 457420 | 457571 | 151 | 0,085486818 | 0,051882174 | 0,622315244 | 0,311567137 | 0,264832067 | 0,13385782 | 0,08485782 | -1,17514218 |
| chrXII | 457420 | 457572 | 152 | 0,512920905 | 0,17417587 | 0,746504602 | 0,663530589 | 0,564001001 | 0,419854182 | 0,363854182 | -0,896145818 |
| chrXII | 457437 | 457592 | 155 | 0,045592969 | 0,037058696 | 0,551627959 | 0,129775456 | 0,110309137 | -0,012422117 | -0,089422117 | -1,349422117 |
| chrXII | 457440 | 457593 | 153 | 0,045592969 | 0,044470435 | 0,506231912 | 0,015621722 | 0,013278464 | -0,082659838 | -0,145659838 | -1,405659838 |
| chrXII | 457540 | 457654 | 114 | 0,04844253 | 0,415057393 | 0,104514645 | -1,256239112 | -1,067803245 | -0,459183439 | -0,249183439 | -1,509183439 |
| chrXII | 457553 | 457706 | 153 | 0,190920559 | 0,159352392 | 0,54506224 | 0,113195553 | 0,09621622 | -0,005591869 | -0,068591869 | -1,328591869 |
| chrXII | 457569 | 457736 | 167 | 0,054141651 | 0,103764348 | 0,342872667 | -0,404635672 | -0,343940321 | -0,198046263 | -0,359046263 | -1,619046263 |
| chrXII | 457569 | 457741 | 172 | 0,336248149 | 0,559586306 | 0,375346301 | -0,317726245 | -0,270067308 | -0,14254435 | -0,33854435 | -1,59854435 |
| chrXII | 457600 | 457736 | 136 | 0,769381358 | 0,755997394 | 0,504387095 | 0,010997038 | 0,009347483 | -0,025052112 | 0,030947888 | -1,229052112 |
| chrXII | 457600 | 457739 | 139 | 0,142478029 | 0,037058696 | 0,793587102 | 0,818930974 | 0,696091328 | 0,652400744 | 0,687400744 | -0,572599256 |
| chrXII | 457600 | 457741 | 141 | 0,076938136 | 0,125999566 | 0,379121943 | -0,307787693 | -0,261619539 | -0,357682821 | -0,336682821 | -1,596682821 |
| chrXII | 457606 | 457763 | 157 | 0,122531105 | 0,044470435 | 0,733712426 | 0,624079848 | 0,53046787 | 0,427229536 | 0,336229536 | -0,923770464 |
| chrXII | 457620 | 457773 | 153 | 0,507221784 | 0,470645436 | 0,518702103 | 0,046896405 | 0,039861944 | -0,061555884 | -0,124555884 | -1,384555884 |
| chrXII | 457620 | 457787 | 167 | 0,079787696 | 0,129705435 | 0,380860679 | -0,303221116 | -0,257737948 | -0,104940035 | -0,265940035 | -1,525940035 |
| chrXII | 457622 | 457771 | 149 | 0,054141651 | 0,059293913 | 0,477289917 | -0,056956516 | -0,048413039 | -0,226147421 | -0,261147421 | -1,521147421 |
| chrXII | 457634 | 457757 | 123 | 0,159575393 | 0,43729261 | 0,267354577 | -0,620833537 | -0,527708506 | -0,090859799 | 0,056140201 | -1,203859799 |
| chrXII | 457650 | 457805 | 155 | 0,039893848 | 0,074117392 | 0,349911537 | -0,385559308 | -0,327725412 | -0,449824643 | -0,526824643 | -1,786824643 |
| chrXII | 457667 | 457819 | 152 | 0,045592969 | 0,051882174 | 0,467739443 | -0,080953558 | -0,068810524 | -0,225599981 | -0,281599981 | -1,541599981 |
| chrXII | 457803 | 457959 | 156 | 0,239363089 | 0,159352392 | 0,600335579 | 0,254215803 | 0,216083433 | 0,227279109 | 0,143279109 | -1,116720891 |
| chrXII | 457816 | 457974 | 158 | 0,09973462 | 0,059293913 | 0,627149217 | 0,324312357 | 0,275665503 | 0,237309414 | 0,139309414 | -1,120690586 |
| chrXII | 457820 | 457953 | 133 | 0,116831984 | 0,070411522 | 0,623957468 | 0,315891237 | 0,268507552 | 0,276763448 | 0,353763448 | -0,906236552 |
| chrXII | 457872 | 458023 | 151 | 0,276407377 | 0,214940436 | 0,562549318 | 0,157435847 | 0,13382047 | 0,000251234 | -0,048748766 | -1,308748766 |
| chrXII | 457945 | 458104 | 159 | 0,350495952 | 0,385410436 | 0,476277904 | -0,059497562 | -0,050572928 | -0,07431108 | -0,17931108 | -1,43931108 |
| chrXII | 457949 | 458059 | 110 | 0,04844253 | 0,103764348 | 0,31826768 | -0,472548462 | -0,401666193 | 0,281930632 | 0,519930632 | -0,740069368 |
| chrXII | 457949 | 458101 | 152 | 0,34194727 | 0,207528696 | 0,622315244 | 0,311567137 | 0,264832067 | 0,105765228 | 0,049765228 | -1,210234772 |
| chrXII | 457950 | 458101 | 151 | 0,159575393 | 0,08894087 | 0,642112476 | 0,3641111 | 0,309494435 | 0,170696655 | 0,121696655 | -1,138303345 |
| chrXII | 457950 | 458103 | 153 | 0,527168708 | 0,370586958 | 0,58720733 | 0,220367006 | 0,187311955 | 0,099732668 | 0,036732668 | -1,223267332 |
| chrXII | 457959 | 458110 | 151 | 0,151026711 | 0,092646739 | 0,619791408 | 0,304932997 | 0,259193047 | 0,12199394 | 0,07299394 | -1,18700606 |
| chrXII | 457976 | 458059 | 83 | 0,059840772 | 0,081529131 | 0,423292165 | -0,193478389 | -0,164456631 | 0,141697137 | 0,568697137 | -0,691302863 |
| chrXII | 457976 | 458119 | 143 | 0,139628469 | 0,074117392 | 0,653245253 | 0,394096909 | 0,334982372 | 0,236524301 | 0,243524301 | -1,016475699 |
| chrXII | 457983 | 458089 | 106 | 0,062690333 | 0,062999783 | 0,498768996 | -0,003085674 | -0,002622823 | 0,417907765 | 0,683907765 | -0,576092235 |
| chrXII | 457983 | 458101 | 118 | 0,031345166 | 0,040764565 | 0,434687048 | -0,164453569 | -0,139785534 | 0,299375305 | 0,481375305 | -0,778624695 |
| chrXII | 457983 | 458103 | 120 | 0,039893848 | 0,070411522 | 0,361667325 | -0,354005648 | -0,300904801 | 0,089769079 | 0,257769079 | -1,002230921 |
| chrXII | 457983 | 458137 | 154 | 0,743735312 | 0,678174133 | 0,523053922 | 0,057819814 | 0,049146842 | -0,121691684 | -0,191691684 | -1,451691684 |
| chrXII | 457986 | 458117 | 131 | 0,091185939 | 0,040764565 | 0,691061693 | 0,498861988 | 0,42403269 | 0,475856577 | 0,566856577 | -0,693143423 |
| chrXII | 458025 | 458167 | 142 | 0,094035499 | 0,111176087 | 0,458236793 | -0,104876779 | -0,089145262 | -0,082913323 | -0,068913323 | -1,328913323 |
| chrXII | 458055 | 458137 | 82 | 0,056991212 | 0,048176305 | 0,541908887 | 0,105243965 | 0,08945737 | | 0,434 | -0,826 |
| chrXII | 458055 | 458206 | 151 | 0,931806311 | 0,52623348 | 0,639081537 | 0,35600486 | 0,302604131 | 0,164892351 | 0,115892351 | -1,144107649 |
| chrXII | 458079 | 458227 | 148 | 0,079787696 | 0,048176305 | 0,623516738 | 0,314730183 | 0,267520656 | 0,128403373 | 0,100403373 | -1,159596627 |
| chrXII | 458087 | 458231 | 144 | 0,350495952 | 0,181587609 | 0,658723511 | 0,408981855 | 0,347634577 | 0,205192381 | 0,205192381 | -1,054807619 |
| chrXII | 458127 | 458277 | 150 | 0,079787696 | 0,085235 | 0,483495289 | -0,041382985 | -0,035175537 | -0,135385509 | -0,177385509 | -1,437385509 |
| chrXII | 458130 | 458293 | 163 | 0,139628469 | 0,118587827 | 0,540742282 | 0,102303929 | 0,08695834 | 0,133112844 | 0,000112844 | -1,259887156 |
| chrXII | 458134 | 458272 | 138 | 0,142478029 | 0,08894087 | 0,615671537 | 0,294132147 | 0,250012325 | 0,253683895 | 0,295683895 | -0,964316105 |
| chrXII | 458134 | 458277 | 143 | 0,336248149 | 0,233469783 | 0,590201097 | 0,228062304 | 0,193852958 | 0,086485005 | 0,093485005 | -1,166514995 |
| chrXII | 458136 | 458295 | 159 | 0,094035499 | 0,081529131 | 0,535617563 | 0,08939893 | 0,07598909 | 0,064857272 | -0,040142728 | -1,300142728 |
| chrXII | 458143 | 458291 | 148 | 0,168124074 | 0,092646739 | 0,644719675 | 0,371103224 | 0,315437741 | 0,169967647 | 0,141967647 | -1,118032353 |
| chrXII | 458166 | 458321 | 155 | 0,481575739 | 0,507704132 | 0,486794236 | -0,03310799 | -0,028141791 | -0,139247261 | -0,216247261 | -1,476247261 |
| chrXII | 458166 | 458321 | 155 | 0,481575739 | 0,507704132 | 0,486794236 | -0,03310799 | -0,028141791 | -0,139247261 | -0,216247261 | -1,476247261 |
| chrXII | 458166 | 458322 | 156 | 0,056991212 | 0,048176305 | 0,541908887 | 0,105243965 | 0,08945737 | 0,079166027 | -0,004833973 | -1,264833973 |
| chrXII | 458166 | 458322 | 156 | 0,056991212 | 0,048176305 | 0,541908887 | 0,105243965 | 0,08945737 | 0,079166027 | -0,004833973 | -1,264833973 |
| chrXII | 458167 | 458322 | 155 | 0,170973635 | 0,103764348 | 0,622315244 | 0,311567137 | 0,264832067 | 0,156000788 | 0,079000788 | -1,180999212 |
| chrXII | 458167 | 458322 | 155 | 0,170973635 | 0,103764348 | 0,622315244 | 0,311567137 | 0,264832067 | 0,156000788 | 0,079000788 | -1,180999212 |
| chrXII | 458177 | 458322 | 145 | 0,823523009 | 0,541056958 | 0,603499266 | 0,262415054 | 0,223052796 | 0,039020365 | 0,032020365 | -1,227979635 |
| chrXII | 458177 | 458327 | 150 | 0,037044288 | 0,044470435 | 0,454449042 | -0,114428548 | -0,097264266 | -0,209203035 | -0,251203035 | -1,511203035 |
| chrXII | 458177 | 458350 | 173 | 0,555664314 | 0,78935022 | 0,413128855 | -0,219503686 | -0,186578133 | -0,02623011 | -0,22923011 | -1,48923011 |
| chrXII | 458179 | 458327 | 148 | 0,304902982 | 0,118587827 | 0,719975442 | 0,582768555 | 0,495353272 | 0,358663308 | 0,330663308 | -0,929336692 |
| chrXII | 458191 | 458321 | 130 | 0,054141651 | 0,048176305 | 0,529151025 | 0,07313593 | 0,062165541 | 0,173347052 | 0,271347052 | -0,988652948 |
| chrXII | 458191 | 458352 | 161 | 0,626903328 | 0,444704349 | 0,585011979 | 0,214732295 | 0,182522451 | 0,216507294 | 0,097507294 | -1,162492706 |
| chrXII | 458195 | 458322 | 127 | 0,062690333 | 0,103764348 | 0,376621026 | -0,314367347 | -0,267212245 | -0,033457517 | 0,085542483 | -1,174457517 |
| chrXII | 458195 | 458327 | 132 | 0,045592969 | 0,051882174 | 0,467739443 | -0,080953558 | -0,068810524 | 0,049619481 | 0,133619481 | -1,126380519 |
| chrXII | 458195 | 458350 | 155 | 0,891912463 | 0,492880654 | 0,644076326 | 0,369376183 | 0,313969755 | 0,211489392 | 0,134489392 | -1,125510608 |
| chrXII | 458204 | 458350 | 146 | 0,062690333 | 0,066705652 | 0,484484373 | -0,038901718 | -0,03306646 | -0,171133047 | -0,185133047 | -1,445133047 |
| chrXII | 458204 | 458380 | 176 | 0,199469241 | 0,281646088 | 0,414597559 | -0,215733974 | -0,183373878 | 0,006105358 | -0,217894642 | -1,477894642 |
| chrXII | 458291 | 458451 | 160 | 0,433133209 | 0,255705001 | 0,628788012 | 0,328645073 | 0,279348312 | 0,267588812 | 0,155588812 | -1,104411188 |
| chrXII | 458297 | 458451 | 154 | 1,27090402 | 0,733762177 | 0,633972889 | 0,342394241 | 0,291035105 | 0,116915377 | 0,046915377 | -1,213084623 |
| chrXII | 458298 | 458451 | 153 | 0,059840772 | 0,040764565 | 0,594807131 | 0,239928434 | 0,203939169 | 0,116906473 | 0,053906473 | -1,206093527 |
| chrXII | 458319 | 458472 | 153 | 0,208017923 | 0,107470218 | 0,65935259 | 0,410696874 | 0,349092343 | 0,240116559 | 0,177116559 | -1,082883441 |
| chrXII | 458338 | 458492 | 154 | 0,276407377 | 0,166764131 | 0,62370295 | 0,315220686 | 0,267937583 | 0,089711611 | 0,019711611 | -1,240288389 |
| chrXII | 458356 | 458493 | 137 | 0,051292091 | 0,051882174 | 0,497140355 | -0,007168129 | -0,00609291 | 0,05135137 | 0,10035137 | -1,15964863 |

TABLE 2

| Chrom | Start | End | Length | Section A | Section B | A/A+B | Z-score | Z * 0.85 | Phase correction | Length correction | ΔLK |
|---|---|---|---|---|---|---|---|---|---|---|---|
| chrXII | 458356 | 458508 | 152 | 0,29065518 | 0,148234783 | 0,662250688 | 0,418613496 | 0,355821471 | 0,202040626 | 0,146040626 | -1,113959374 |
| chrXII | 458359 | 458523 | 164 | 0,071239015 | 0,092646739 | 0,434687048 | -0,164453569 | -0,139785534 | -0,159855743 | -0,299855743 | -1,559855743 |
| chrXII | 458365 | 458493 | 128 | 0,720938828 | 1,074702178 | 0,401493854 | -0,249482329 | -0,21205998 | 0,020852559 | 0,132852559 | -1,127147441 |
| chrXII | 458373 | 458493 | 120 | 0,042743409 | 0,059293913 | 0,418899751 | -0,204708983 | -0,174002636 | 0,220005997 | 0,388005997 | -0,871994003 |
| chrXII | 458373 | 458530 | 157 | 0,219416165 | 0,17788174 | 0,552271135 | 0,131401466 | 0,111691246 | 0,014875251 | -0,076124749 | -1,336124749 |
| chrXII | 458374 | 458532 | 158 | 0,051292091 | 0,062999783 | 0,448781606 | -0,128740216 | -0,109429183 | -0,13867944 | -0,23667944 | -1,49667944 |
| chrXII | 458395 | 458501 | 106 | 0,051292091 | 0,055588044 | 0,479902939 | -0,050397187 | -0,042837609 | 0,380376981 | 0,646376981 | -0,613623019 |
| chrXII | 458395 | 458519 | 124 | 0,04844253 | 0,074117392 | 0,395255882 | -0,265646117 | -0,2257992 | 0,197359159 | 0,337359159 | -0,922640841 |
| chrXII | 458419 | 458587 | 168 | 0,074088575 | 0,100058479 | 0,425436857 | -0,188003748 | -0,159803185 | -0,018254399 | -0,186254399 | -1,446254399 |
| chrXII | 458508 | 458658 | 150 | 0,062690333 | 0,059293913 | 0,51392155 | 0,034903236 | 0,029667751 | -0,091642725 | -0,133642725 | -1,393642725 |
| chrXII | 458508 | 458658 | 150 | 0,062690333 | 0,059293913 | 0,51392155 | 0,034903236 | 0,029667751 | -0,091642725 | -0,133642725 | -1,393642725 |
| chrXII | 458508 | 458659 | 151 | 0,116831984 | 0,081529131 | 0,588986325 | 0,224938202 | 0,191197471 | 0,058288264 | 0,009288264 | -1,250711736 |
| chrXII | 458508 | 458659 | 151 | 0,116831984 | 0,081529131 | 0,588986325 | 0,224938202 | 0,191197471 | 0,058288264 | 0,009288264 | -1,250711736 |
| chrXII | 458746 | 458901 | 155 | 0,09688506 | 0,081529131 | 0,543034495 | 0,108081541 | 0,09186931 | -0,019935675 | -0,096935675 | -1,356935675 |
| chrXII | 458746 | 458912 | 166 | 0,239363089 | 0,200116957 | 0,544650641 | 0,112157259 | 0,09533367 | 0,243211807 | 0,089211807 | -1,170788193 |
| chrXII | 458750 | 458912 | 162 | 0,037044288 | 0,055588044 | 0,399906675 | -0,253588671 | -0,21555037 | -0,188860532 | -0,314860532 | -1,574860532 |
| chrXII | 458768 | 458900 | 132 | 0,14817715 | 0,129705435 | 0,533236547 | 0,083408278 | 0,070897036 | 0,192103715 | 0,276103715 | -0,983896285 |
| chrXII | 458768 | 458901 | 133 | 0,182371877 | 0,133411305 | 0,577522451 | 0,195559327 | 0,166225428 | 0,173072164 | 0,250072164 | -1,009927836 |
| chrXII | 458768 | 458912 | 144 | 0,712390146 | 0,466939567 | 0,604063595 | 0,263879451 | 0,224297533 | 0,09403121 | 0,09403121 | -1,16596879 |
| chrXII | 458769 | 458900 | 131 | 0,626903328 | 0,689291742 | 0,476299709 | -0,059442808 | -0,050526387 | -0,016463585 | 0,074536415 | -1,185463585 |
| chrXII | 458769 | 458901 | 132 | 1,587205245 | 1,426759788 | 0,526617007 | 0,066768519 | 0,056753241 | 0,185562068 | 0,269562068 | -0,990437932 |
| chrXII | 458769 | 458912 | 143 | 0,803576085 | 0,61517435 | 0,566397067 | 0,167208665 | 0,142127365 | 0,027433769 | 0,034433769 | -1,225566231 |
| chrXII | 458769 | 458913 | 144 | 0,233663968 | 0,155646522 | 0,600199516 | 0,25386356 | 0,215784026 | 0,092090529 | 0,092090529 | -1,167909471 |
| chrXII | 458770 | 458900 | 130 | 0,082637257 | 0,163058261 | 0,336340107 | -0,422472439 | -0,359101573 | -0,273229287 | -0,175229287 | -1,435229287 |
| chrXII | 458770 | 458901 | 131 | 0,740885752 | 1,156231309 | 0,390532439 | -0,277931585 | -0,236241848 | -0,210998286 | -0,119998286 | -1,379998286 |
| chrXII | 458770 | 458912 | 142 | 0,14817715 | 0,122293696 | 0,54784888 | 0,120228376 | 0,10219412 | 0,096050443 | 0,110050443 | -1,149949557 |
| chrXII | 458846 | 459002 | 156 | 0,176672756 | 0,051882174 | 0,772999103 | 0,748760132 | 0,636446112 | 0,636984729 | 0,552984729 | -0,707015271 |
| chrXII | 458846 | 459002 | 156 | 0,176672756 | 0,051882174 | 0,772999103 | 0,748760132 | 0,636446112 | 0,636984729 | 0,552984729 | -0,707015271 |
| chrXII | 458902 | 459048 | 146 | 0,094035499 | 0,085235 | 0,524545307 | 0,06156483 | 0,052330105 | -0,086067074 | -0,100067074 | -1,360067074 |
| chrXII | 458908 | 459044 | 136 | 0,142478029 | 0,085235 | 0,625691158 | 0,320462592 | 0,272393203 | 0,239976702 | 0,295976702 | -0,964023298 |
| chrXII | 458908 | 459048 | 140 | 0,136778908 | 0,140823044 | 0,492715945 | -0,018259434 | -0,015520519 | 0,010627022 | 0,038627022 | -1,221372978 |
| chrXII | 458910 | 459043 | 133 | 0,039893848 | 0,040764565 | 0,49460244 | -0,013530088 | -0,011500575 | -0,014184511 | 0,062815489 | -1,197184511 |
| chrXII | 458910 | 459066 | 156 | 0,074088575 | 0,107470218 | 0,408069331 | -0,232514197 | -0,197637067 | -0,213282324 | -0,297282324 | -1,557282324 |
| chrXII | 458913 | 459002 | 89 | 0,085486818 | 0,159352392 | 0,349154932 | -0,387602979 | -0,329462532 | 0,113233966 | 0,498233966 | -0,761766034 |
| chrXII | 458913 | 459038 | 125 | 1,117027749 | 1,901111094 | 0,370104825 | -0,331575727 | -0,281839368 | 0,036050174 | 0,169050174 | -1,090949826 |
| chrXII | 458913 | 459043 | 130 | 0,803576085 | 1,026525873 | 0,439088151 | -0,153281484 | -0,130289261 | -0,029761209 | 0,068238791 | -1,191761209 |
| chrXII | 458913 | 459044 | 131 | 3,701579198 | 4,947335887 | 0,42798191 | -0,181514414 | -0,154287252 | -0,111686726 | -0,020686726 | -1,280686726 |
| chrXII | 458913 | 459047 | 134 | 1,749630198 | 1,852934789 | 0,485662356 | -0,035946885 | -0,030554852 | -0,033438986 | 0,036561014 | -1,223438986 |
| chrXII | 458913 | 459048 | 135 | 0,19661968 | 0,192705218 | 0,505027244 | 0,012601764 | 0,0107115 | 0,01129021 | 0,07429021 | -1,18570979 |
| chrXII | 458913 | 459066 | 153 | 0,159575393 | 0,129705435 | 0,551627959 | 0,129775456 | 0,110309137 | 0,001993428 | -0,061006572 | -1,321006572 |
| chrXII | 458932 | 459038 | 106 | 0,039893848 | 0,044470435 | 0,472876041 | -0,06804215 | -0,057835827 | 0,370968565 | 0,636968565 | -0,623031435 |
| chrXII | 458933 | 459044 | 111 | 0,136778908 | 0,218646305 | 0,384831753 | -0,292815072 | -0,248892811 | 0,25839996 | 0,48939996 | -0,77060004 |
| chrXII | 458933 | 459047 | 114 | 0,059840772 | 0,092646739 | 0,392430643 | -0,272989502 | -0,232041076 | 0,379349407 | 0,589349407 | -0,670650593 |
| chrXII | 458933 | 459066 | 133 | 0,330549028 | 0,26311674 | 0,556793141 | 0,142843581 | 0,121417044 | 0,115936515 | 0,192936515 | -1,067063485 |
| chrXII | 458957 | 459044 | 87 | 0,09688506 | 0,159352392 | 0,378106554 | -0,310457456 | -0,263888837 | 0,124882335 | 0,523882335 | -0,736117665 |
| chrXII | 459050 | 459146 | 96 | 0,182371877 | 1,030231743 | 0,150396943 | -1,034732432 | -0,879522567 | -0,305102172 | 0,030897828 | -1,229102172 |
| chrXII | 459089 | 459262 | 173 | 0,313451664 | 0,544762828 | 0,365236974 | -0,344495143 | -0,292820872 | -0,123410505 | -0,326410505 | -1,586410505 |
| chrXII | 459089 | 459265 | 176 | 0,042743409 | 0,155646522 | 0,215451503 | -0,787647358 | -0,669500254 | -0,494183737 | -0,718183737 | -1,978183737 |
| chrXII | 459095 | 459262 | 167 | 0,068389454 | 0,207528696 | 0,247861382 | -0,681235065 | -0,579049806 | -0,424112943 | -0,585112943 | -1,845112943 |
| chrXII | 459095 | 459265 | 170 | 0,678195419 | 1,1451137 | 0,371958552 | -0,326670515 | -0,277669938 | -0,111855544 | -0,293855544 | -1,553855544 |
| chrXII | 459098 | 459233 | 135 | 1,880709985 | 1,504583049 | 0,555553084 | 0,139704043 | 0,118748436 | 0,122672455 | 0,185672455 | -1,074327545 |
| chrXII | 459098 | 459262 | 164 | 0,168124074 | 0,251999131 | 0,400178024 | -0,252886336 | -0,214953386 | -0,237625524 | -0,377625524 | -1,637625524 |
| chrXII | 459166 | 459328 | 162 | 0,068389454 | 0,062999783 | 0,520510322 | 0,051434423 | 0,043719259 | 0,07276317 | -0,05323683 | -1,31323683 |
| chrXII | 459185 | 459328 | 143 | 0,834921251 | 0,52993935 | 0,611726392 | 0,28382145 | 0,241248232 | 0,126029327 | 0,133029327 | -1,126970673 |
| chrXII | 459204 | 459353 | 149 | 0,886213341 | 0,422469132 | 0,677179804 | 0,459827015 | 0,390852962 | 0,211245938 | 0,176245938 | -1,083754062 |
| chrXII | 459204 | 459417 | 213 | 0,042743409 | 0,074117392 | 0,365763444 | -0,34309514 | -0,291630869 | -0,051668548 | -0,534668548 | -1,794668548 |
| chrXII | 459204 | 459419 | 215 | 0,076938136 | 0,085235 | 0,474419732 | -0,064164224 | -0,05453959 | 0,248559708 | -0,248440292 | -1,508440292 |
| chrXII | 459227 | 459353 | 126 | 0,045592969 | 0,066705652 | 0,405997586 | -0,23785292 | -0,202174982 | 0,050053594 | 0,176053594 | -1,083946406 |
| chrXII | 459227 | 459376 | 149 | 0,775080479 | 0,574409785 | 0,574350553 | 0,187461389 | 0,15934218 | -0,032600072 | -0,067600072 | -1,327600072 |
| chrXII | 459227 | 459419 | 192 | 0,065539893 | 0,085235 | 0,434687048 | -0,164453569 | -0,139785534 | -0,083369252 | -0,419369252 | -1,679369252 |
| chrXII | 459227 | 459432 | 205 | 0,031345166 | 0,055588044 | 0,360566076 | -0,356946107 | -0,303404191 | -0,239623666 | -0,666623666 | -1,926623666 |
| chrXII | 459235 | 459395 | 160 | 0,14532759 | 0,070411522 | 0,673626532 | 0,449949388 | 0,38245698 | 0,3644137 | 0,2524137 | -1,0075863 |
| chrXII | 459236 | 459328 | 92 | 0,028495606 | 0,051882174 | 0,354520937 | -0,373143192 | -0,317171714 | 0,07019594 | 0,43419594 | -0,82580406 |
| chrXII | 459236 | 459393 | 157 | 0,501522663 | 0,470645436 | 0,5158806 | 0,039817281 | 0,033844689 | -0,057383138 | -0,148383138 | -1,408383138 |
| chrXII | 459236 | 459417 | 181 | 0,105433742 | 0,08894087 | 0,542425478 | 0,106546147 | 0,090564225 | 0,279099634 | 0,020099634 | -1,239900366 |
| chrXII | 459236 | 459419 | 183 | 0,19377012 | 0,259410871 | 0,427577775 | -0,182544352 | -0,155162699 | -0,050105474 | -0,323105474 | -1,583105474 |
| chrXII | 459264 | 459353 | 89 | 0,082637257 | 0,085235 | 0,492262738 | -0,019395657 | -0,016486308 | 0,421551971 | 0,806551971 | -0,453448029 |
| chrXII | 459264 | 459376 | 112 | 0,113982423 | 0,226058044 | 0,335202525 | -0,425592166 | -0,361753341 | 0,100840471 | 0,324840471 | -0,935159529 |
| chrXII | 459264 | 459417 | 153 | 2,097276589 | 1,567582832 | 0,572266586 | 0,182147684 | 0,154825532 | 0,056156998 | -0,006843002 | -1,266843002 |
| chrXII | 459264 | 459419 | 155 | 2,00039153 | 1,567582832 | 0,560651879 | 0,152622168 | 0,129728843 | 0,03403389 | -0,04296611 | -1,30296611 |
| chrXII | 459264 | 459432 | 168 | 0,988797522 | 0,856055873 | 0,535976206 | 0,090301549 | 0,076756317 | 0,205214496 | 0,037214496 | -1,222785504 |
| chrXII | 459264 | 459441 | 177 | 0,73233707 | 1,019114134 | 0,418131586 | -0,206675654 | -0,175674306 | -0,018443008 | -0,249443008 | -1,509443008 |
| chrXII | 459265 | 459376 | 111 | 0,045592969 | 0,059293913 | 0,434687048 | -0,164453569 | -0,139785534 | 0,36628747 | 0,59728747 | -0,66271253 |
| chrXII | 459265 | 459393 | 128 | 0,188070999 | 0,233469783 | 0,446151373 | -0,135390993 | -0,115082344 | 0,110401581 | 0,222401581 | -1,037598419 |
| chrXII | 459265 | 459416 | 151 | 0,054141651 | 0,044470435 | 0,549036667 | 0,123227859 | 0,104743681 | -0,02405579 | -0,07305579 | -1,33305579 |
| chrXII | 459265 | 459417 | 152 | 1,664143381 | 1,182172396 | 0,584665761 | 0,213844301 | 0,181767656 | 0,019242725 | -0,036757275 | -1,296757275 |
| chrXII | 459265 | 459419 | 154 | 11,28710947 | 6,959623069 | 0,618582502 | 0,301760039 | 0,256496033 | 0,070696014 | 0,000696014 | -1,259303986 |
| chrXII | 459265 | 459422 | 157 | 0,236513528 | 0,226058044 | 0,511301477 | 0,028332392 | 0,024082533 | -0,062848697 | -0,153848697 | -1,413848697 |
| chrXII | 459265 | 459432 | 167 | 0,151026711 | 0,248293262 | 0,37820976 | -0,310185996 | -0,263658096 | -0,115426888 | -0,276426888 | -1,536426888 |
| chrXII | 459265 | 459441 | 176 | 0,039893848 | 0,140823044 | 0,220753288 | -0,769651619 | -0,654203876 | -0,479536502 | -0,703536502 | -1,963536502 |
| chrXII | 459265 | 459467 | 202 | 0,128230226 | 0,096352609 | 0,570970734 | 0,178846117 | 0,1520192 | 0,166797533 | -0,239202467 | -1,499202467 |
| chrXII | 459265 | 459469 | 204 | 0,162424953 | 0,170470001 | 0,487916538 | -0,030293381 | -0,025749374 | 0,034716958 | -0,385283042 | -1,645283042 |
| chrXII | 459267 | 459393 | 126 | 0,042743409 | 0,077823261 | 0,354520937 | -0,373143192 | -0,317171714 | -0,082318047 | 0,043681953 | -1,216318047 |
| chrXII | 459267 | 459417 | 150 | 0,53571739 | 0,400233914 | 0,572377417 | 0,18243015 | 0,155065628 | 0,028609461 | -0,013390539 | -1,273390539 |
| chrXII | 459267 | 459419 | 152 | 2,789719811 | 1,908522833 | 0,593779424 | 0,237277969 | 0,201686274 | 0,032628061 | -0,023371939 | -1,283371939 |
| chrXII | 459267 | 459422 | 155 | 0,681044979 | 0,363175219 | 0,652204373 | 0,391278682 | 0,33258688 | 0,238652723 | 0,161652723 | -1,098347277 |
| chrXII | 459267 | 459432 | 165 | 0,153876272 | 0,200116957 | 0,434687048 | -0,164453569 | -0,139785534 | -0,048804089 | -0,195804089 | -1,455804089 |
| chrXII | 459267 | 459441 | 174 | 0,065539893 | 0,051882174 | 0,558156527 | 0,146296985 | 0,124352437 | 0,344176444 | 0,134176444 | -1,125823556 |
| chrXII | 459267 | 459469 | 202 | 0,04844253 | 0,070411522 | 0,407579962 | -0,23377466 | -0,198708461 | -0,181406659 | -0,587406659 | -1,847406659 |
| chrXII | 459268 | 459374 | 106 | 0,04844253 | 0,248293262 | 0,163251388 | -0,981182451 | -0,834005084 | -0,411548963 | -0,145548963 | -1,405548963 |
| chrXII | 459268 | 459376 | 108 | 0,042743409 | 0,185293479 | 0,187440765 | -0,887366654 | -0,754261656 | -0,075596376 | 0,176403624 | -1,083596376 |
| chrXII | 459268 | 459393 | 125 | 0,24221265 | 0,200116957 | 0,547584077 | 0,119559826 | 0,101625852 | 0,428546269 | 0,561546269 | -0,698453731 |
| chrXII | 459268 | 459416 | 148 | 0,054141651 | 0,055588044 | 0,493409293 | -0,016521205 | -0,014043024 | -0,153000592 | -0,181000592 | -1,441000592 |
| chrXII | 459268 | 459417 | 149 | 2,179913846 | 1,256289787 | 0,634396002 | 0,343519071 | 0,29199121 | 0,103573735 | 0,068573735 | -1,191426265 |
| chrXII | 459268 | 459419 | 151 | 11,13893232 | 6,099861326 | 0,646154977 | 0,374960229 | 0,318716195 | 0,193474119 | 0,144474119 | -1,115525881 |
| chrXII | 459268 | 459422 | 154 | 0,310602104 | 0,214940436 | 0,591012298 | 0,230149754 | 0,195627291 | 0,007455482 | -0,062544518 | -1,322544518 |
| chrXII | 459268 | 459431 | 163 | 0,088336378 | 0,103764348 | 0,459844061 | -0,100826586 | -0,085702598 | -0,045496428 | -0,178496428 | -1,438496428 |
| chrXII | 459268 | 459432 | 164 | 1,635647775 | 1,363760005 | 0,545323576 | 0,11385486 | 0,096776631 | 0,080652704 | -0,059347296 | -1,319347296 |
| chrXII | 459268 | 459441 | 173 | 0,059840772 | 0,196411088 | 0,23352327 | -0,727292911 | -0,618198974 | -0,437103265 | -0,640103265 | -1,900103265 |
| chrXII | 459268 | 459467 | 199 | 0,156725832 | 0,100058479 | 0,610340373 | 0,280206274 | 0,238175333 | 0,268028998 | -0,116971002 | -1,376971002 |
| chrXII | 459268 | 459469 | 201 | 0,188070999 | 0,166764131 | 0,530023616 | 0,075329227 | 0,064029843 | 0,090624871 | -0,308375129 | -1,568375129 |

TABLE 2

| Chrom | Start | End | Length | Section A | Section B | A/A+B | Z-score | Z * 0.85 | Phase correction | Length correction | ΔLK |
|---|---|---|---|---|---|---|---|---|---|---|---|
| chrXII | 459269 | 459376 | 107 | 0,068389454 | 0,26682261 | 0,204018475 | -0,827353109 | -0,703250143 | -0,287326132 | -0,028326132 | -1,288326132 |
| chrXII | 459269 | 459393 | 124 | 0,113982423 | 0,300175436 | 0,275214923 | -0,597116135 | -0,507548715 | -0,095014891 | 0,044985109 | -1,215014891 |
| chrXII | 459269 | 459413 | 144 | 0,065539893 | 0,048176305 | 0,576346155 | 0,192554744 | 0,163671533 | 0,045439062 | 0,045439062 | -1,214560938 |
| chrXII | 459269 | 459417 | 148 | 4,963934536 | 2,542226531 | 0,661314684 | 0,416053806 | 0,353645735 | 0,214404809 | 0,186404809 | -1,073595191 |
| chrXII | 459269 | 459419 | 150 | 3,388127534 | 2,190168921 | 0,607376743 | 0,272488395 | 0,231615136 | 0,115909088 | 0,073909088 | -1,186090912 |
| chrXII | 459269 | 459422 | 153 | 0,108283302 | 0,144528914 | 0,42831515 | -0,180665296 | -0,153565501 | -0,263776623 | -0,326776623 | -1,586776623 |
| chrXII | 459269 | 459432 | 163 | 0,028495606 | 0,08894087 | 0,242646977 | -0,69781331 | -0,593141313 | -0,549235017 | -0,682235017 | -1,942235017 |
| chrXII | 459269 | 459467 | 198 | 0,102584181 | 0,085235 | 0,54618586 | 0,116030612 | 0,09862602 | 0,146674733 | -0,231325267 | -1,491325267 |
| chrXII | 459269 | 459469 | 200 | 0,136778908 | 0,133411305 | 0,506231912 | 0,015621722 | 0,013278464 | 0,034276504 | -0,357723496 | -1,617723496 |
| chrXII | 459271 | 459419 | 148 | 0,045592969 | 0,040764565 | 0,527955893 | 0,070132481 | 0,059612609 | -0,080597729 | -0,108597729 | -1,368597729 |
| chrXII | 459308 | 459414 | 106 | 0,182371877 | 0,285351958 | 0,389913585 | -0,279544268 | -0,237612628 | 0,179479867 | 0,445479867 | -0,814520133 |
| chrXII | 459308 | 459419 | 111 | 0,24506221 | 0,329822392 | 0,426280699 | -0,185851269 | -0,157973579 | 0,329440195 | 0,560440195 | -0,699559805 |
| chrXII | 459308 | 459467 | 159 | 0,088336378 | 0,077823261 | 0,531635591 | 0,07938196 | 0,067474666 | 0,046243575 | -0,058756425 | -1,318756425 |
| chrXII | 459308 | 459469 | 161 | 0,988797522 | 0,581821524 | 0,629559106 | 0,330685854 | 0,281082976 | 0,315178619 | 0,196178619 | -1,063821381 |
| chrXII | 459325 | 459466 | 141 | 0,222265726 | 0,08894087 | 0,714206347 | 0,565715338 | 0,480858037 | 0,384192582 | 0,405192582 | -0,854807418 |
| chrXII | 459326 | 459417 | 91 | 0,102584181 | 0,185293479 | 0,356346446 | -0,368241871 | -0,31300559 | 0,061535577 | 0,432535577 | -0,827464423 |
| chrXII | 459326 | 459419 | 93 | 0,401788042 | 0,389116306 | 0,508010916 | 0,020081738 | 0,017069477 | 0,452410134 | 0,809410134 | -0,450589866 |
| chrXII | 459326 | 459467 | 141 | 1,373488201 | 0,659644785 | 0,675552564 | 0,455297989 | 0,387003291 | 0,295791692 | 0,316791692 | -0,943208308 |
| chrXII | 459326 | 459469 | 143 | 0,19661968 | 0,107470218 | 0,646584058 | 0,376114353 | 0,3196972 | 0,18507962 | 0,19207962 | -1,06792038 |
| chrXII | 459408 | 459559 | 151 | 0,230814407 | 0,092646739 | 0,713576915 | 0,563864767 | 0,479285052 | 0,342247805 | 0,293247805 | -0,966752195 |
| chrXII | 459429 | 459558 | 129 | 1,131275552 | 1,58611218 | 0,41630995 | -0,211342655 | -0,179641257 | 0,088905802 | 0,193905802 | -1,066094198 |
| chrXII | 459429 | 459559 | 130 | 0,105433742 | 0,140823044 | 0,428145528 | -0,181097487 | -0,153932864 | -0,039653976 | 0,058346024 | -1,201653976 |
| chrXII | 459429 | 459579 | 150 | 0,076938136 | 0,08894087 | 0,463820816 | -0,090812431 | -0,077190567 | -0,202459762 | -0,244459762 | -1,504459762 |
| chrXII | 459430 | 459558 | 128 | 0,162424953 | 0,233469783 | 0,410273081 | -0,226842569 | -0,192816183 | 0,028038129 | 0,140038129 | -1,119961871 |
| chrXII | 459430 | 459559 | 129 | 0,04844253 | 0,037058696 | 0,566571175 | 0,16765125 | 0,142503563 | 0,416198501 | 0,521198501 | -0,738801499 |
| chrXII | 459430 | 459573 | 143 | 0,361894194 | 0,218646305 | 0,623374587 | 0,314355796 | 0,267202426 | 0,120743519 | 0,127743519 | -1,132256481 |
| chrXII | 459430 | 459579 | 149 | 0,644000692 | 0,43729261 | 0,595583724 | 0,24193239 | 0,205642532 | 0,015042638 | -0,019957362 | -1,279957362 |
| chrXII | 459519 | 459673 | 154 | 0,199469241 | 0,133411305 | 0,599221683 | 0,251333038 | 0,213633083 | 0,032105848 | -0,037894152 | -1,297894152 |
| chrXII | 459521 | 459673 | 152 | 0,19377012 | 0,125999566 | 0,605967759 | 0,268824834 | 0,228501109 | 0,051836236 | -0,004163764 | -1,264163764 |
| chrXII | 459602 | 459756 | 154 | 0,056991212 | 0,037058696 | 0,605967759 | 0,268824834 | 0,228501109 | 0,049896685 | -0,020103315 | -1,280103315 |
| chrXII | 459602 | 459759 | 157 | 0,076938136 | 0,051882174 | 0,597251597 | 0,246239543 | 0,209303612 | 0,115316486 | 0,024316486 | -1,235683514 |
| chrXII | 459604 | 459722 | 118 | 0,054141651 | 0,111176087 | 0,327500555 | -0,446825428 | -0,379801614 | 0,034575182 | 0,216575182 | -1,043424818 |
| chrXII | 459604 | 459756 | 152 | 5,852997439 | 3,461282186 | 0,628389706 | 0,32759145 | 0,278452733 | 0,091988041 | 0,035988041 | -1,224011959 |
| chrXII | 459604 | 459759 | 155 | 0,088336378 | 0,037058696 | 0,704464501 | 0,537284661 | 0,456691962 | 0,3653848 | 0,2883848 | -0,9716152 |
| chrXII | 459604 | 459774 | 170 | 0,29350474 | 0,35576348 | 0,452054684 | -0,120471861 | -0,102401082 | 0,060746365 | -0,121253365 | -1,381253365 |
| chrXII | 459604 | 459796 | 192 | 0,615505086 | 0,659644785 | 0,482692349 | -0,043397467 | -0,036887847 | 0,027819097 | -0,308180903 | -1,568180903 |
| chrXII | 459604 | 459798 | 194 | 0,139628469 | 0,214940436 | 0,39379784 | -0,269434058 | -0,22901895 | -0,15635304 | -0,50635304 | -1,76635304 |
| chrXII | 459604 | 459804 | 200 | 0,319150785 | 0,200116957 | 0,614617006 | 0,291373097 | 0,247667133 | 0,266221122 | -0,125778878 | -1,385778878 |
| chrXII | 459604 | 459806 | 202 | 0,168124074 | 0,274234349 | 0,380063011 | -0,305315302 | -0,259518007 | -0,239480729 | -0,645480729 | -1,905480729 |
| chrXII | 459618 | 459756 | 138 | 0,162424953 | 0,144528914 | 0,529151025 | 0,07313593 | 0,062165541 | 0,065556212 | 0,107556212 | -1,152443788 |
| chrXII | 459618 | 459774 | 156 | 0,227964847 | 0,144528914 | 0,611996417 | 0,284526191 | 0,241847262 | 0,231376996 | 0,147376996 | -1,112623004 |
| chrXII | 459618 | 459796 | 178 | 0,09688506 | 0,081529131 | 0,543034495 | 0,108081541 | 0,09186931 | 0,230896603 | -0,007103397 | -1,267103397 |
| chrXII | 459621 | 459756 | 135 | 0,062690333 | 0,066705652 | 0,484484373 | -0,038901718 | -0,03306646 | -0,031611434 | 0,031388566 | -1,228611434 |
| chrXII | 459621 | 459774 | 153 | 0,04844253 | 0,040764565 | 0,543034495 | 0,108081541 | 0,09186931 | -0,023246386 | -0,086246386 | -1,346246386 |
| chrXII | 459625 | 459756 | 131 | 0,074088575 | 0,155646522 | 0,322495674 | -0,460731366 | -0,391621661 | -0,352083941 | -0,261083941 | -1,521083941 |
| chrXII | 459625 | 459774 | 149 | 0,042743409 | 0,040764565 | 0,51184823 | 0,029703476 | 0,025247955 | -0,169308181 | -0,204308181 | -1,464308181 |
| chrXII | 459625 | 459796 | 171 | 0,116831984 | 0,214940436 | 0,352144956 | -0,379535953 | -0,32260556 | -0,183843104 | -0,372843104 | -1,632843104 |
| chrXII | 459625 | 459798 | 173 | 0,675345858 | 0,759703264 | 0,470608182 | -0,073741139 | -0,062679968 | 0,112291993 | -0,090708007 | -1,350708007 |
| chrXII | 459625 | 459804 | 179 | 0,062690333 | 0,08894087 | 0,413439528 | -0,218706025 | -0,185900121 | -0,037441342 | -0,282441342 | -1,542441342 |
| chrXII | 459625 | 459806 | 181 | 0,056991212 | 0,125999566 | 0,311443082 | -0,491764035 | -0,41799943 | -0,225677107 | -0,484677107 | -1,744677107 |
| chrXII | 459628 | 459756 | 128 | 0,339097709 | 0,344645871 | 0,495942806 | -0,010170054 | -0,008644546 | 0,225212864 | 0,337212864 | -0,922787136 |
| chrXII | 459628 | 459774 | 146 | 0,176672756 | 0,188999349 | 0,483145293 | -0,042261061 | -0,035921902 | -0,179679542 | -0,193679542 | -1,453679542 |
| chrXII | 459628 | 459796 | 168 | 3,755720849 | 3,331576751 | 0,529922837 | 0,075075895 | 0,063814511 | 0,19542209 | 0,02742209 | -1,23257791 |
| chrXII | 459628 | 459798 | 170 | 0,131079787 | 0,203822827 | 0,391396727 | -0,275680526 | -0,234328447 | -0,05370819 | -0,23570819 | -1,49570819 |
| chrXII | 459628 | 459804 | 176 | 0,250761331 | 0,188999349 | 0,570222266 | 0,176940065 | 0,150399055 | 0,332024167 | 0,108024167 | -1,151975833 |
| chrXII | 459628 | 459806 | 178 | 0,173823196 | 0,374292827 | 0,31712847 | -0,475743762 | -0,404382198 | -0,27017145 | -0,50817145 | -1,76817145 |
| chrXII | 459633 | 459756 | 123 | 0,310602104 | 0,17788174 | 0,635849287 | 0,347385899 | 0,295278014 | 0,704687935 | 0,851687935 | -0,408312065 |
| chrXII | 459633 | 459774 | 141 | 1,940550757 | 0,907938047 | 0,681256235 | 0,471214548 | 0,400532366 | 0,333141734 | 0,354141734 | -0,905858266 |
| chrXII | 459633 | 459796 | 163 | 0,284956058 | 0,26311674 | 0,519923739 | 0,049962185 | 0,042467857 | 0,075175677 | -0,057824323 | -1,317824323 |
| chrXII | 459633 | 459798 | 165 | 0,065539893 | 0,092646739 | 0,414320049 | -0,216446021 | -0,183979117 | -0,085815425 | -0,232815425 | -1,492815425 |
| chrXII | 459633 | 459804 | 171 | 0,151026711 | 0,08894087 | 0,629362977 | 0,330166647 | 0,28064165 | 0,413188338 | 0,224188338 | -1,035811662 |
| chrXII | 459633 | 459806 | 173 | 0,108283302 | 0,140823044 | 0,434687048 | -0,164453569 | -0,139785534 | 0,028962189 | -0,174037811 | -1,434037811 |
| chrXII | 459648 | 459796 | 148 | 0,262159574 | 0,17417587 | 0,600821174 | 0,255473185 | 0,217152207 | 0,074104443 | 0,046104443 | -1,213895557 |
| chrXII | 459673 | 459804 | 131 | 3,938092726 | 2,820166749 | 0,582708128 | 0,208826422 | 0,177502459 | 0,203789038 | 0,294789038 | -0,965210962 |
| chrXII | 459673 | 459806 | 133 | 0,740885752 | 1,030231743 | 0,418315416 | -0,206204936 | -0,175274196 | -0,178716608 | -0,101716608 | -1,361716608 |
| chrXII | 459686 | 459793 | 107 | 0,393239361 | 0,52623348 | 0,427679147 | -0,182285988 | -0,154943089 | 0,246148572 | 0,505148572 | -0,754851428 |
| chrXII | 459686 | 459796 | 110 | 0,210867483 | 0,715232829 | 0,227693999 | -0,746462631 | -0,634493236 | 0,043516857 | 0,281516857 | -0,978483143 |
| chrXII | 459686 | 459798 | 112 | 0,056991212 | 0,170470001 | 0,250553539 | -0,67274886 | -0,571836531 | -0,087389994 | 0,136610006 | -1,123389994 |
| chrXII | 459686 | 459799 | 113 | 0,43883233 | 0,767115003 | 0,363890129 | -0,348079796 | -0,295867827 | 0,250356662 | 0,467356662 | -0,792643338 |
| chrXII | 459686 | 459801 | 115 | 0,028495606 | 0,096352609 | 0,228241997 | -0,744648908 | -0,632951572 | -0,012719075 | 0,190280925 | -1,069719075 |
| chrXII | 459686 | 459804 | 118 | 0,692443222 | 1,31928957 | 0,344202383 | -0,401020859 | -0,34086773 | 0,058581584 | 0,240581584 | -1,019418416 |
| chrXII | 459686 | 459806 | 120 | 3,009135976 | 10,98419743 | 0,215040683 | -0,789052425 | -0,670694561 | -0,283134601 | -0,115134601 | -1,375134601 |
| chrXII | 459686 | 459856 | 170 | 0,051292091 | 0,133411305 | 0,277699771 | -0,589688448 | -0,501235181 | -0,321960877 | -0,503960877 | -1,763960877 |
| chrXII | 459688 | 459793 | 105 | 0,074088575 | 0,52623348 | 0,123414715 | -1,158084807 | -0,984372086 | -0,391871398 | -0,118871398 | -1,378871398 |
| chrXII | 459688 | 459851 | 163 | 0,051292091 | 0,048176305 | 0,515662191 | 0,039269381 | 0,033378974 | 0,067957472 | -0,065042528 | -1,325042528 |
| chrXII | 459698 | 459804 | 106 | 0,051292091 | 0,181587609 | 0,220251445 | -0,771344283 | -0,655642641 | -0,248933518 | 0,017066482 | -1,242933518 |
| chrXII | 459698 | 459806 | 108 | 0,037044288 | 0,148234783 | 0,199937788 | -0,84184347 | -0,715566949 | -0,057499049 | 0,194500951 | -1,065499049 |
| chrXII | 459698 | 459851 | 153 | 0,116831984 | 0,151940653 | 0,434687048 | -0,164453569 | -0,139785534 | -0,251371076 | -0,314371076 | -1,574371076 |
| chrXII | 459698 | 459869 | 171 | 0,062690333 | 0,08894087 | 0,413439528 | -0,218706025 | -0,185900121 | -0,048642081 | -0,237642081 | -1,497642081 |
| chrXII | 459818 | 459980 | 162 | 0,04844253 | 0,077823261 | 0,383655221 | -0,295894775 | -0,251510559 | -0,224125296 | -0,350125296 | -1,610125296 |
| chrXII | 459834 | 459969 | 135 | 0,555664314 | 0,726350437 | 0,433430515 | -0,167646954 | -0,142499911 | -0,137389082 | -0,074389082 | -1,334389082 |
| chrXII | 459846 | 459969 | 123 | 0,054141651 | 0,103764348 | 0,342872667 | -0,404635672 | -0,343940321 | 0,056182004 | 0,203182004 | -1,056817996 |
| chrXII | 459846 | 460002 | 156 | 0,233663968 | 0,35205761 | 0,398933515 | -0,256108537 | -0,217692257 | -0,239955114 | -0,323955114 | -1,583955114 |
| chrXII | 459899 | 460068 | 169 | 0,119681545 | 0,200116957 | 0,374240479 | -0,320642986 | -0,272546538 | -0,137256926 | -0,312256926 | -1,572256926 |
| chrXII | 459899 | 460068 | 169 | 0,119681545 | 0,200116957 | 0,374240479 | -0,320642986 | -0,272546538 | -0,137256926 | -0,312256926 | -1,572256926 |
| chrXII | 459930 | 460057 | 127 | 0,188070999 | 0,503998263 | 0,2717517 | -0,607523728 | -0,516395169 | -0,270320036 | -0,151320036 | -1,411320036 |
| chrXII | 460017 | 460128 | 111 | 0,054141651 | 0,070411522 | 0,434687048 | -0,164453569 | -0,139785534 | 0,336307004 | 0,567307004 | -0,692692996 |
| chrXII | 460017 | 460159 | 142 | 2,00039153 | 1,360054135 | 0,595275666 | 0,241137346 | 0,204966744 | 0,20019254 | 0,21419254 | -1,04580746 |
| chrXII | 460032 | 460159 | 127 | 0,63545201 | 1,196995874 | 0,346777671 | -0,394034805 | -0,334929585 | -0,078835518 | 0,040164482 | -1,219835518 |
| chrXII | 460102 | 460251 | 149 | 1,672692063 | 0,870879351 | 0,65761553 | 0,405964152 | 0,345069529 | 0,153040795 | 0,118040795 | -1,141959205 |
| chrXII | 460102 | 460266 | 164 | 0,136778908 | 0,092646739 | 0,596179675 | 0,243470869 | 0,206950239 | 0,185863164 | 0,045863164 | -1,214136836 |
| chrXII | 460112 | 460251 | 139 | 0,424584527 | 0,370586958 | 0,533953411 | 0,085211589 | 0,072429851 | 0,041041141 | 0,076041141 | -1,183958859 |
| chrXII | 460112 | 460266 | 154 | 1,934851636 | 1,623170875 | 0,543799717 | 0,110011108 | 0,093509442 | -0,089879802 | -0,159879802 | -1,419879802 |
| chrXII | 460188 | 460366 | 178 | 0,227964847 | 0,381704567 | 0,373915505 | -0,321500662 | -0,273275563 | -0,13467678 | -0,37267678 | -1,63267678 |
| chrXII | 460243 | 460384 | 141 | 0,504372223 | 0,307587175 | 0,621179118 | 0,308579045 | 0,262292188 | 0,193761091 | 0,214761091 | -1,045238909 |
| chrXII | 460256 | 460415 | 159 | 0,091185939 | 0,151940653 | 0,37505539 | -0,318493294 | -0,2707193 | -0,293160485 | -0,398160485 | -1,658160485 |
| chrXII | 460262 | 460436 | 174 | 0,034194727 | 0,111176087 | 0,235224155 | -0,721749813 | -0,613487341 | -0,390156586 | -0,600156586 | -1,860156586 |
| chrXII | 460302 | 460477 | 175 | 0,108283302 | 0,277940218 | 0,280364339 | -0,581759513 | -0,494495586 | -0,247575025 | -0,464575025 | -1,724575025 |
| chrXII | 460314 | 460446 | 132 | 0,051292091 | 0,107470218 | 0,323074734 | -0,459117949 | -0,390250257 | -0,25320196 | -0,16920196 | -1,42920196 |
| chrXII | 460402 | 460571 | 169 | 0,062690333 | 0,133411305 | 0,319682862 | -0,468585806 | -0,398297935 | -0,258998877 | -0,433998877 | -1,693998877 |
| chrXII | 460461 | 460595 | 134 | 0,575611238 | 0,463233697 | 0,554087736 | 0,135995892 | 0,115596508 | 0,120089627 | 0,190089627 | -1,069910373 |

TABLE 2

| Chrom | Start | End | Length | Section A | Section B | A/A+B | Z-score | Z * 0.85 | Phase correction | Length correction | ΔLK |
|---|---|---|---|---|---|---|---|---|---|---|---|
| chrXII | 460487 | 460616 | 129 | 0,065539893 | 0,085235 | 0,434687048 | -0,164453569 | -0,139785534 | 0,137388747 | 0,242388747 | -1,017611253 |
| chrXII | 460493 | 460595 | 102 | 2,236905058 | 21,52739638 | 0,094128795 | -1,315751121 | -1,118388453 | -0,590469984 | -0,296469984 | -1,556469984 |
| chrXII | 460493 | 460659 | 166 | 0,039893848 | 0,170470001 | 0,189642129 | -0,879215839 | -0,747333463 | -0,594001199 | -0,748001199 | -2,008001199 |
| chrXII | 460512 | 460595 | 83 | 0,042743409 | 0,200116957 | 0,175999936 | -0,930717198 | -0,791109618 | -0,477340248 | -0,050340248 | -1,310340248 |
| chrXII | 460512 | 460639 | 127 | 0,812124766 | 1,57870044 | 0,339683873 | -0,41332605 | -0,351327143 | -0,088662665 | 0,030337335 | -1,229662665 |
| chrXII | 460521 | 460659 | 138 | 0,470177496 | 0,518821741 | 0,47540734 | -0,061683752 | -0,052431189 | -0,057402451 | -0,015402451 | -1,275402451 |
| chrXII | 460602 | 460763 | 161 | 0,065539893 | 0,103764348 | 0,387113121 | -0,28685122 | -0,243823537 | -0,211887061 | -0,330887061 | -1,590887061 |
| chrXII | 460846 | 460979 | 133 | 0,401788042 | 0,70411522 | 0,363312105 | -0,349619586 | -0,297176648 | -0,296111747 | -0,219111747 | -1,479111747 |
| chrXII | 460863 | 461022 | 159 | 0,056991212 | 0,048176305 | 0,541908887 | 0,105243965 | 0,08945737 | 0,069176183 | -0,035823817 | -1,295823817 |
| chrXII | 460871 | 461006 | 135 | 0,051292091 | 0,08894087 | 0,365763444 | -0,34309514 | -0,291630869 | -0,286873741 | -0,223873741 | -1,483873741 |
| chrXII | 460894 | 460979 | 85 | 0,051292091 | 0,092646739 | 0,356346446 | -0,368241871 | -0,31300559 | 0,087974672 | 0,500974672 | -0,759025328 |
| chrXII | 460894 | 461045 | 151 | 0,416035845 | 0,481763045 | 0,463395366 | -0,091883336 | -0,078100835 | -0,214354561 | -0,263354561 | -1,523354561 |
| chrXII | 460894 | 461097 | 203 | 0,039893848 | 0,051882174 | 0,434687048 | -0,164453569 | -0,139785534 | -0,092695745 | -0,505695745 | -1,765695745 |
| chrXII | 460906 | 461069 | 163 | 0,108283302 | 0,118587827 | 0,477289917 | -0,056956516 | -0,048413039 | -0,006714976 | -0,139714976 | -1,399714976 |
| chrXII | 460927 | 461045 | 118 | 0,04844253 | 0,133411305 | 0,266381679 | -0,623793272 | -0,530224281 | -0,145043342 | 0,036956658 | -1,223043342 |
| chrXII | 460927 | 461097 | 170 | 1,350691717 | 1,697288267 | 0,443143237 | -0,143004694 | -0,12155399 | 0,064906065 | -0,117093935 | -1,377093935 |
| chrXII | 460937 | 461071 | 134 | 1,191116324 | 0,815291307 | 0,593656197 | 0,236960278 | 0,201416237 | 0,212795203 | 0,282795203 | -0,977204797 |
| chrXII | 460937 | 461098 | 161 | 0,279256937 | 0,403939784 | 0,408750406 | -0,230760574 | -0,196146488 | -0,163901964 | -0,282901964 | -1,542901964 |
| chrXII | 460988 | 461071 | 83 | 0,213717044 | 0,418763262 | 0,33790308 | -0,418192798 | -0,355463878 | -0,02369586 | 0,40330414 | -0,85669586 |
| chrXII | 460988 | 461095 | 107 | 2,276798906 | 12,01813504 | 0,159273132 | -0,997449727 | -0,847832268 | -0,43776598 | -0,17876598 | -1,43876598 |
| chrXII | 460988 | 461096 | 108 | 0,068389454 | 0,211234566 | 0,244576464 | -0,691656728 | -0,587908219 | 0,062926921 | 0,314926921 | -0,945073079 |
| chrXII | 460988 | 461098 | 110 | 2,447772541 | 16,56894289 | 0,128716894 | -1,132477383 | -0,962605776 | -0,277774635 | -0,039774635 | -1,299774635 |
| chrXII | 461013 | 461146 | 133 | 0,917558508 | 1,033937612 | 0,470182082 | -0,074812165 | -0,06359034 | -0,084856788 | -0,007856788 | -1,267856788 |
| chrXII | 461056 | 461146 | 90 | 0,037044288 | 0,062999783 | 0,370279692 | -0,331112667 | -0,281445767 | 0,143986723 | 0,521986723 | -0,738013277 |
| chrXII | 461056 | 461148 | 92 | 0,059840772 | 0,381704567 | 0,13552577 | -1,100644199 | -0,935547569 | -0,551851605 | -0,187851605 | -1,447851605 |
| chrXII | 461056 | 461199 | 143 | 0,056991212 | 0,085235 | 0,400708216 | -0,251514398 | -0,213787238 | -0,36288963 | -0,35588963 | -1,61588963 |
| chrXII | 461056 | 461212 | 156 | 0,122531105 | 0,085235 | 0,589755027 | 0,226914861 | 0,192877632 | 0,166618636 | 0,082618636 | -1,177381364 |
| chrXII | 461056 | 461214 | 158 | 0,088336378 | 0,08894087 | 0,498295067 | -0,004273647 | -0,0036326 | -0,024805765 | -0,122805765 | -1,382805765 |
| chrXII | 461056 | 461215 | 159 | 0,891912463 | 0,644821307 | 0,580394913 | 0,202903865 | 0,172468285 | 0,147797059 | 0,042797059 | -1,217202941 |
| chrXII | 461056 | 461229 | 173 | 0,085486818 | 0,077823261 | 0,523463207 | 0,058847485 | 0,050020363 | 0,210306733 | 0,007306733 | -1,252693267 |
| chrXII | 461060 | 461146 | 86 | 0,045592969 | 0,062999783 | 0,419852784 | -0,202270107 | -0,171929591 | 0,218427844 | 0,624427844 | -0,635572156 |
| chrXII | 461060 | 461212 | 152 | 1,142673794 | 0,748585655 | 0,604186694 | 0,264198961 | 0,224569117 | 0,040143336 | -0,015856664 | -1,275856664 |
| chrXII | 461060 | 461214 | 154 | 0,054141651 | 0,085235 | 0,388455674 | -0,283346352 | -0,2408444 | -0,418558927 | -0,488558927 | -1,748558927 |
| chrXII | 461060 | 461215 | 155 | 0,056991212 | 0,055588044 | 0,506231912 | 0,015621722 | 0,013278464 | -0,082763076 | -0,159763076 | -1,419763076 |
| chrXII | 461060 | 461229 | 169 | 0,185221438 | 0,08894087 | 0,675590454 | 0,45540334 | 0,387092839 | 0,528669881 | 0,353669881 | -0,906330119 |
| chrXII | 461068 | 461199 | 131 | 0,401788042 | 0,474351306 | 0,458589199 | -0,103988596 | -0,088390306 | -0,039852873 | 0,051147127 | -1,208852873 |
| chrXII | 461068 | 461214 | 146 | 2,239754619 | 1,608347397 | 0,582041383 | 0,207118603 | 0,176050812 | 0,033841175 | 0,019841175 | -1,240158825 |
| chrXII | 461068 | 461246 | 178 | 0,071239015 | 0,066705652 | 0,516431814 | 0,041200102 | 0,035020087 | 0,175386258 | -0,062613742 | -1,322613742 |
| chrXII | 461069 | 461199 | 130 | 4,938288491 | 5,677392194 | 0,465188115 | -0,08737149 | -0,074265766 | 0,042806057 | 0,140806057 | -1,119193943 |
| chrXII | 461069 | 461214 | 145 | 0,698142343 | 0,833820655 | 0,455717497 | -0,111228699 | -0,094544395 | -0,287933122 | -0,294933122 | -1,554933122 |
| chrXII | 461069 | 461246 | 177 | 0,094035499 | 0,103764348 | 0,47540734 | -0,061683752 | -0,052431189 | 0,102218864 | -0,128781136 | -1,388781136 |
| chrXII | 461082 | 461188 | 106 | 0,170973635 | 0,370586958 | 0,315705458 | -0,47974192 | -0,407780632 | 0,010167577 | 0,276167577 | -0,983832423 |
| chrXII | 461082 | 461199 | 117 | 0,179522317 | 0,292763697 | 0,380113558 | -0,305182559 | -0,259405175 | 0,261048571 | 0,450048571 | -0,809951429 |
| chrXII | 461082 | 461212 | 130 | 0,173823196 | 0,159352392 | 0,521716483 | 0,054462061 | 0,046292752 | 0,15997489 | 0,25797489 | -1,00202511 |
| chrXII | 461082 | 461214 | 132 | 3,345384125 | 3,264871099 | 0,506090009 | 0,015265981 | 0,012976084 | 0,154760044 | 0,238760044 | -1,021239956 |
| chrXII | 461082 | 461215 | 133 | 0,122531105 | 0,111176087 | 0,524293257 | 0,060931847 | 0,05179207 | 0,032401436 | 0,109401436 | -1,150598564 |
| chrXII | 461082 | 461229 | 147 | 2,766923327 | 1,348936527 | 0,672258878 | 0,446159211 | 0,37923533 | 0,263033894 | 0,242033894 | -1,017966106 |
| chrXII | 461082 | 461246 | 164 | 0,179522317 | 0,17417587 | 0,507557922 | 0,018946034 | 0,016104129 | 0,004248782 | -0,135751218 | -1,395751218 |
| chrXII | 461095 | 461246 | 151 | 0,521469587 | 0,237175653 | 0,687369484 | 0,488407779 | 0,415146613 | 0,283029678 | 0,234029678 | -1,025970322 |
| chrXII | 461178 | 461333 | 155 | 0,153876272 | 0,144528914 | 0,515662191 | 0,039269381 | 0,033378974 | -0,052661098 | -0,129661098 | -1,389661098 |
| chrXII | 461196 | 461352 | 156 | 0,256460453 | 0,26311674 | 0,493594515 | -0,016056859 | -0,01364833 | -0,046413046 | -0,130413046 | -1,390413046 |
| chrXII | 461218 | 461364 | 146 | 0,558513874 | 0,366881088 | 0,603541079 | 0,262523537 | 0,223145006 | 0,060113714 | 0,046113714 | -1,213886286 |
| chrXII | 461218 | 461384 | 166 | 1,225311051 | 1,297054353 | 0,485778567 | -0,0356554 | -0,03030709 | 0,112102742 | -0,041897258 | -1,301897258 |
| chrXII | 461218 | 461388 | 170 | 0,105433742 | 0,077823261 | 0,575332675 | 0,189967434 | 0,161472318 | 0,335294378 | 0,153294378 | -1,106705622 |
| chrXII | 461241 | 461384 | 143 | 0,062690333 | 0,040764565 | 0,605967759 | 0,268824834 | 0,228501109 | 0,079641393 | 0,086641393 | -1,173358607 |
| chrXII | 461241 | 461388 | 147 | 0,339097709 | 0,114881957 | 0,746944708 | 0,664906054 | 0,565170146 | 0,457522133 | 0,436522133 | -0,823477867 |
| chrXII | 461252 | 461408 | 156 | 0,558513874 | 0,374292827 | 0,598745564 | 0,250101478 | 0,212586256 | 0,175687621 | 0,091687621 | -1,168312379 |
| chrXII | 461260 | 461367 | 107 | 0,042743409 | 0,155646522 | 0,215451503 | -0,787647358 | -0,669500254 | -0,257905485 | 0,001094515 | -1,258905485 |
| chrXII | 461327 | 461481 | 154 | 0,592708601 | 0,433586741 | 0,577522451 | 0,195559327 | 0,166225428 | -0,013883463 | -0,083883463 | -1,343883463 |
| chrXII | 461397 | 461551 | 154 | 0,353345512 | 0,281646088 | 0,556456986 | 0,141992377 | 0,12069352 | -0,064557994 | -0,134557994 | -1,394557994 |
| chrXII | 461430 | 461537 | 107 | 0,113982423 | 0,893114568 | 0,113179192 | -1,209792869 | -1,028323939 | -0,592911389 | -0,333911389 | -1,593911389 |
| chrXII | 461431 | 461592 | 161 | 0,572761677 | 0,518821741 | 0,524707198 | 0,061971406 | 0,052675695 | 0,074769109 | -0,044230891 | -1,304230891 |
| chrXII | 461432 | 461592 | 160 | 0,313451664 | 0,233469783 | 0,573120081 | 0,184323297 | 0,156674802 | 0,132888108 | 0,020888108 | -1,239111892 |
| chrXII | 461436 | 461592 | 156 | 0,213717044 | 0,148234783 | 0,590457149 | 0,22872109 | 0,194412926 | 0,157208737 | 0,073208737 | -1,186791263 |
| chrXII | 461460 | 461622 | 162 | 0,042743409 | 0,048176305 | 0,470122564 | -0,074961771 | -0,063717505 | -0,033288148 | -0,159288148 | -1,419288148 |
| chrXII | 461460 | 461623 | 163 | 0,322000346 | 0,311293045 | 0,508453666 | 0,021191784 | 0,018013016 | 0,059358846 | -0,073641154 | -1,333641154 |
| chrXII | 461471 | 461578 | 107 | 0,042743409 | 0,059293913 | 0,418899751 | -0,204708983 | -0,174002636 | 0,270052943 | 0,529052943 | -0,730947057 |
| chrXII | 461471 | 461592 | 121 | 0,054141651 | 0,081529131 | 0,399066404 | -0,255764342 | -0,217399691 | 0,176353223 | 0,337353223 | -0,922646777 |
| chrXII | 461471 | 461622 | 151 | 0,624053768 | 0,403939784 | 0,607060002 | 0,271664505 | 0,23091483 | 0,099281351 | 0,050281351 | -1,209718649 |
| chrXII | 461471 | 461623 | 152 | 0,076938136 | 0,062999783 | 0,549801916 | 0,125160904 | 0,106386768 | -0,093184694 | -0,149184694 | -1,409184694 |
| chrXII | 461483 | 461633 | 150 | 0,19661968 | 0,137117174 | 0,589145842 | 0,225348315 | 0,191546068 | 0,053970159 | 0,011970159 | -1,248029841 |
| chrXII | 461536 | 461685 | 149 | 0,054141651 | 0,044470435 | 0,549036667 | 0,123227859 | 0,104743681 | -0,078995135 | -0,113995135 | -1,373995135 |
| chrXII | 461560 | 461721 | 161 | 0,14532759 | 0,188999349 | 0,434687048 | -0,164453569 | -0,139785534 | -0,123916154 | -0,242916154 | -1,502916154 |
| chrXII | 461606 | 461757 | 151 | 0,279256937 | 0,381704567 | 0,422501062 | -0,195499251 | -0,166174364 | -0,308421207 | -0,357421207 | -1,617421207 |
| chrXII | 461606 | 461758 | 152 | 0,356195073 | 0,433586741 | 0,4510044 | -0,123124135 | -0,104655515 | -0,307816206 | -0,363816206 | -1,623816206 |
| chrXII | 461606 | 461766 | 160 | 0,267858695 | 0,311293045 | 0,462501753 | -0,094133 | -0,08001305 | -0,104822976 | -0,216822976 | -1,476822976 |
| chrXII | 461606 | 461767 | 161 | 0,225115286 | 0,211234566 | 0,515905494 | 0,03987973 | 0,03389777 | 0,044849179 | -0,074150821 | -1,334150821 |
| chrXII | 461628 | 461758 | 130 | 0,074088575 | 0,270528479 | 0,214988127 | -0,789232287 | -0,670847444 | -0,546770553 | -0,448770553 | -1,708770553 |
| chrXII | 461628 | 461766 | 138 | 0,034194727 | 0,074117392 | 0,315705458 | -0,47974192 | -0,407780632 | -0,4071081 | -0,3651081 | -1,6251081 |
| chrXII | 461628 | 461800 | 172 | 0,208017923 | 0,311293045 | 0,400565241 | -0,251884318 | -0,21410167 | -0,089659216 | -0,285659216 | -1,545659216 |
| chrXII | 461629 | 461721 | 92 | 0,028495606 | 0,103764348 | 0,215451503 | -0,787647358 | -0,669500254 | -0,282017207 | 0,081982793 | -1,178017207 |
| chrXII | 461629 | 461757 | 128 | 0,085486818 | 0,289057827 | 0,228241997 | -0,744648908 | -0,632951572 | -0,406312406 | -0,294312406 | -1,554312406 |
| chrXII | 461629 | 461758 | 129 | 1,373488201 | 3,828163275 | 0,264048487 | -0,630913669 | -0,536276619 | -0,266107881 | -0,161107881 | -1,421107881 |
| chrXII | 461629 | 461766 | 137 | 0,082637257 | 0,17788174 | 0,317202423 | -0,475536189 | -0,40420576 | -0,335194707 | -0,286194707 | -1,546194707 |
| chrXII | 461629 | 461800 | 171 | 0,039893848 | 0,148234783 | 0,212056229 | -0,799306935 | -0,679410895 | -0,531364364 | -0,720364364 | -1,980364364 |
| chrXII | 461662 | 461758 | 96 | 0,045592969 | 0,092646739 | 0,329810947 | -0,440435256 | -0,374369967 | 0,197806774 | 0,533806774 | -0,726193226 |
| chrXII | 461662 | 461832 | 170 | 0,045592969 | 0,077823261 | 0,36942442 | -0,33337817 | -0,283371444 | -0,104249051 | -0,286249051 | -1,546249051 |
| chrXII | 461676 | 461810 | 134 | 0,19661968 | 0,389116306 | 0,3356797 | -0,424283048 | -0,360640591 | -0,352380032 | -0,282380032 | -1,542380032 |
| chrXII | 461687 | 461800 | 113 | 0,054141651 | 0,137117174 | 0,283080537 | -0,573714411 | -0,48765725 | 0,062123162 | 0,279123162 | -0,980876838 |
| chrXII | 461687 | 461809 | 122 | 0,24506221 | 1,304466092 | 0,158152781 | -1,002078746 | -0,851766934 | -0,496595426 | -0,342595426 | -1,602595426 |
| chrXII | 461687 | 461839 | 152 | 0,168124074 | 0,114881957 | 0,594065341 | 0,238015182 | 0,202312905 | -0,003174046 | -0,059174046 | -1,319174046 |
| chrXII | 461805 | 461933 | 128 | 0,239363089 | 0,537351089 | 0,308173966 | -0,501032951 | -0,425878008 | -0,220454211 | -0,108454211 | -1,368454211 |
| chrXII | 461805 | 461951 | 146 | 0,037044288 | 0,044470435 | 0,454449042 | -0,114428548 | -0,097264266 | -0,259184236 | -0,273184236 | -1,533184236 |
| chrXII | 461821 | 461933 | 112 | 0,059840772 | 0,114881957 | 0,342489913 | -0,405677162 | -0,344825587 | 0,16215198 | 0,38615198 | -0,87384802 |
| chrXII | 461821 | 461972 | 151 | 0,581310359 | 0,35205761 | 0,622809415 | 0,312867712 | 0,265937555 | 0,123803557 | 0,074803557 | -1,185196443 |
| chrXII | 461823 | 461951 | 128 | 0,752283994 | 1,067290439 | 0,413439528 | -0,218706025 | -0,185900121 | 0,048728662 | 0,160728662 | -1,099271338 |
| chrXII | 461823 | 462001 | 178 | 0,031345166 | 0,048176305 | 0,394172367 | -0,268460684 | -0,228191582 | -0,091564566 | -0,329564566 | -1,589564566 |
| chrXII | 461846 | 461952 | 106 | 0,059840772 | 0,074117392 | 0,446712395 | -0,133971905 | -0,113876119 | 0,318516123 | 0,584516123 | -0,675483877 |
| chrXII | 461846 | 462001 | 155 | 0,806425645 | 0,722644568 | 0,527396086 | 0,068725866 | 0,058416986 | -0,030892209 | -0,107892209 | -1,367892209 |
| chrXII | 461846 | 462021 | 175 | 0,111132863 | 0,107470218 | 0,508377386 | 0,021000537 | 0,017850457 | 0,252500839 | 0,035500839 | -1,224499161 |
| chrXII | 461846 | 462037 | 191 | 0,079787696 | 0,074117392 | 0,518421434 | 0,046192108 | 0,039263292 | 0,086431018 | -0,242568982 | -1,502568982 |

TABLE 2

| Chrom | Start | End | Length | Section A | Section B | A/A+B | Z-score | Z * 0.85 | Phase correction | Length correction | ΔLK |
|---|---|---|---|---|---|---|---|---|---|---|---|
| chrXII | 461860 | 461951 | 91 | 0,039893848 | 0,051882174 | 0,434687048 | -0,164453569 | -0,139785534 | 0,252830473 | 0,623830473 | -0,636169527 |
| chrXII | 461860 | 462001 | 141 | 0,128230226 | 0,159352392 | 0,445890044 | -0,136052108 | -0,115644291 | -0,180207735 | -0,159207735 | -1,419207735 |
| chrXII | 461860 | 462021 | 161 | 0,971700159 | 0,752291524 | 0,563633902 | 0,160188997 | 0,136160647 | 0,140268931 | 0,021268931 | -1,238731069 |
| chrXII | 461860 | 462037 | 177 | 0,131079787 | 0,129705435 | 0,502635026 | 0,006605078 | 0,005614317 | 0,157910486 | -0,073089514 | -1,333089514 |
| chrXII | 461902 | 462001 | 99 | 0,09688506 | 0,148234783 | 0,395255882 | -0,265646117 | -0,2257992 | 0,236616762 | 0,551616762 | -0,708383238 |
| chrXII | 461902 | 462008 | 106 | 0,088336378 | 0,17788174 | 0,331819558 | -0,434894351 | -0,369660198 | 0,078436835 | 0,344436835 | -0,915563165 |
| chrXII | 461902 | 462021 | 119 | 0,105433742 | 0,122293696 | 0,462982163 | -0,092923514 | -0,078984987 | 0,329290222 | 0,504290222 | -0,755709778 |
| chrXII | 461902 | 462037 | 135 | 2,396480451 | 2,015993051 | 0,543114978 | 0,108284468 | 0,092041798 | 0,071954108 | 0,134954108 | -1,125045892 |
| chrXII | 461902 | 462062 | 160 | 0,153876272 | 0,062999783 | 0,7095125 | 0,551961108 | 0,469166941 | 0,445668795 | 0,333668795 | -0,926331205 |
| chrXII | 461902 | 462062 | 160 | 0,153876272 | 0,062999783 | 0,7095125 | 0,551961108 | 0,469166941 | 0,445668795 | 0,333668795 | -0,926331205 |
| chrXII | 461910 | 462001 | 91 | 0,04844253 | 0,096352609 | 0,33455909 | -0,427358568 | -0,363254783 | 0,020030635 | 0,391030635 | -0,868969365 |
| chrXII | 461910 | 462016 | 106 | 0,037044288 | 0,08894087 | 0,294036919 | -0,541629394 | -0,460384985 | -0,008198979 | 0,257801021 | -1,002198979 |
| chrXII | 461910 | 462021 | 111 | 0,056991212 | 0,092646739 | 0,380860679 | -0,303221116 | -0,257737948 | 0,210131886 | 0,441131886 | -0,818868114 |
| chrXII | 461910 | 462037 | 127 | 0,185221438 | 0,233469783 | 0,442381948 | -0,144932843 | -0,123192917 | 0,147417871 | 0,266417871 | -0,993582129 |
| chrXII | 461910 | 462062 | 152 | 0,795027403 | 0,578115654 | 0,578983667 | 0,199294137 | 0,169400016 | -0,039538681 | -0,095538681 | -1,355538681 |
| chrXII | 461910 | 462062 | 152 | 0,795027403 | 0,578115654 | 0,578983667 | 0,199294137 | 0,169400016 | -0,039538681 | -0,095538681 | -1,355538681 |
| chrXII | 461920 | 462099 | 179 | 0,056991212 | 0,133411305 | 0,299319635 | -0,52635832 | -0,447404572 | -0,290874783 | -0,535874783 | -1,795874783 |
| chrXII | 461925 | 462060 | 135 | 0,681044979 | 0,548468698 | 0,553914114 | 0,135556657 | 0,115223159 | 0,094841478 | 0,157841478 | -1,102158522 |
| chrXII | 461925 | 462090 | 165 | 0,094035499 | 0,100058479 | 0,484484373 | -0,038901718 | -0,03306646 | 0,061915785 | -0,085084215 | -1,345084215 |
| chrXII | 461938 | 462060 | 122 | 0,125380666 | 0,163058261 | 0,434687048 | -0,164453569 | -0,139785534 | 0,220891379 | 0,374891379 | -0,885108621 |
| chrXII | 461938 | 462090 | 152 | 2,142869559 | 1,371171744 | 0,609802041 | 0,278803123 | 0,236982654 | 0,037822466 | -0,018177534 | -1,278177534 |
| chrXII | 461938 | 462136 | 198 | 0,04844253 | 0,055588044 | 0,465656665 | -0,08619258 | -0,073263693 | -0,032009443 | -0,410009443 | -1,670009443 |
| chrXII | 461977 | 462090 | 113 | 0,153876272 | 0,229763914 | 0,40109529 | -0,250513093 | -0,212936129 | 0,339026421 | 0,556026421 | -0,703973579 |
| chrXII | 461977 | 462136 | 159 | 0,629752889 | 0,511410002 | 0,551851882 | 0,130341515 | 0,110790288 | 0,081974808 | -0,023025192 | -1,283025192 |
| chrXII | 461977 | 462160 | 183 | 0,074088575 | 0,077823261 | 0,487707719 | -0,030817056 | -0,026194498 | 0,068079991 | -0,204920009 | -1,464920009 |
| chrXII | 461984 | 462090 | 106 | 0,031345166 | 0,055588044 | 0,360566076 | -0,356946107 | -0,303404191 | 0,167267369 | 0,433267369 | -0,826732631 |
| chrXII | 461984 | 462133 | 149 | 0,210867483 | 0,203822827 | 0,508493876 | 0,021292599 | 0,018098709 | -0,177566543 | -0,212566543 | -1,472566543 |
| chrXII | 462013 | 462171 | 158 | 1,590054806 | 1,330407179 | 0,544453177 | 0,111659181 | 0,094910303 | 0,067212132 | -0,030787868 | -1,290787868 |
| chrXII | 462022 | 462171 | 149 | 0,276407377 | 0,155646522 | 0,639752071 | 0,357796169 | 0,304126743 | 0,114965081 | 0,079965081 | -1,180034919 |
| chrXII | 462030 | 462136 | 106 | 0,065539893 | 0,096352609 | 0,404835878 | -0,240849509 | -0,204722082 | 0,254346396 | 0,520346396 | -0,739653604 |
| chrXII | 462032 | 462160 | 128 | 1,493169746 | 1,752876311 | 0,459996476 | -0,100442599 | -0,08537621 | 0,16694006 | 0,27894006 | -0,98105994 |
| chrXII | 462059 | 462166 | 107 | 1,339293474 | 7,960207855 | 0,144017773 | -1,062440962 | -0,903074817 | -0,463163252 | -0,204163252 | -1,464163252 |
| chrXII | 462059 | 462167 | 108 | 0,034194727 | 0,244587392 | 0,122657533 | -1,161804056 | -0,987533448 | -0,316770286 | -0,064770286 | -1,324770286 |
| chrXII | 462059 | 462171 | 112 | 1,689789426 | 10,28008221 | 0,141170221 | -1,075076584 | -0,913815096 | -0,398563087 | -0,174563087 | -1,434563087 |
| chrXII | 462059 | 462193 | 134 | 0,076938136 | 0,100058479 | 0,434687048 | -0,164453569 | -0,139785534 | -0,123044451 | -0,053044451 | -1,313044451 |
| chrXII | 462059 | 462212 | 153 | 0,042743409 | 0,070411522 | 0,377742344 | -0,311415611 | -0,264703269 | -0,383287945 | -0,446287945 | -1,706287945 |
| chrXII | 462063 | 462134 | 71 | 0,056991212 | 0,140823044 | 0,288104674 | -0,558930214 | -0,475090682 | | 0,511 | -0,749 |
| chrXII | 462063 | 462170 | 107 | 0,119681545 | 0,274234349 | 0,303825122 | -0,513430458 | -0,436415889 | 0,021477307 | 0,280477307 | -0,979522693 |
| chrXII | 462063 | 462171 | 108 | 0,29350474 | 0,470645436 | 0,384093008 | -0,294748492 | -0,250536218 | 0,402970359 | 0,654970359 | -0,605029641 |
| chrXII | 462063 | 462193 | 130 | 0,578460798 | 0,592939133 | 0,493820072 | -0,015491401 | -0,013167691 | 0,119210153 | 0,217210153 | -1,042789847 |
| chrXII | 462063 | 462207 | 144 | 0,074088575 | 0,040764565 | 0,645072262 | 0,372050197 | 0,316242667 | 0,197359946 | 0,197359946 | -1,062640054 |
| chrXII | 462063 | 462212 | 149 | 0,455929693 | 0,370586958 | 0,551627959 | 0,129775456 | 0,110309137 | -0,089053019 | -0,124053019 | -1,384053019 |
| chrXII | 462113 | 462275 | 162 | 0,24221265 | 0,344645871 | 0,412727499 | -0,220534382 | -0,187454224 | -0,153597824 | -0,279597824 | -1,539597824 |
| chrXII | 462128 | 462278 | 150 | 0,889062902 | 0,61146848 | 0,592498706 | 0,233977317 | 0,198880719 | 0,059440052 | 0,017440052 | -1,242559948 |
| chrXII | 462168 | 462275 | 107 | 0,034194727 | 0,081529131 | 0,295485544 | -0,537429332 | -0,456814932 | 0,007584165 | 0,266584165 | -0,993415835 |
| chrXII | 462170 | 462300 | 130 | 1,313647429 | 1,700994137 | 0,435755761 | -0,161738835 | -0,137478009 | -0,011539009 | 0,086460991 | -1,173539009 |
| chrXII | 462170 | 462324 | 154 | 0,14817715 | 0,118587827 | 0,555459536 | 0,139467258 | 0,118547169 | -0,072908952 | -0,142908952 | -1,402908952 |
| chrXII | 462170 | 462330 | 160 | 0,071239015 | 0,08894087 | 0,444743825 | -0,138952542 | -0,118109661 | -0,132258326 | -0,244258326 | -1,504258326 |
| chrXII | 462176 | 462300 | 124 | 0,142478029 | 0,159352392 | 0,472046617 | -0,070126174 | -0,059607248 | 0,328147196 | 0,468147196 | -0,791852804 |
| chrXII | 462176 | 462324 | 148 | 1,151222476 | 0,61888022 | 0,650370444 | 0,386320781 | 0,328372663 | 0,182191959 | 0,154191959 | -1,105808041 |
| chrXII | 462176 | 462330 | 154 | 0,113982423 | 0,066705652 | 0,630824269 | 0,334037235 | 0,283931649 | 0,107726902 | 0,037726902 | -1,222273098 |
| chrXII | 462179 | 462349 | 170 | 0,04844253 | 0,103764348 | 0,31826768 | -0,472548462 | -0,401666193 | -0,214822088 | -0,396822088 | -1,656822088 |
| chrXII | 462179 | 462355 | 176 | 0,644000692 | 0,641115437 | 0,501122566 | 0,002813859 | 0,00239178 | 0,173083655 | -0,050916345 | -1,310916345 |
| chrXII | 462180 | 462300 | 120 | 0,162424953 | 0,289057827 | 0,359758911 | -0,359103289 | -0,305237796 | 0,084472506 | 0,252472506 | -1,007527494 |
| chrXII | 462180 | 462324 | 144 | 0,165274514 | 0,111176087 | 0,597844653 | 0,247772164 | 0,210606339 | 0,093207335 | 0,093207335 | -1,166792665 |
| chrXII | 462180 | 462327 | 147 | 0,322000346 | 0,207528696 | 0,608088169 | 0,274339592 | 0,233188653 | 0,124070637 | 0,103070637 | -1,156929363 |
| chrXII | 462180 | 462330 | 150 | 1,51311667 | 1,089525656 | 0,581377108 | 0,205417712 | 0,174605055 | 0,020891488 | -0,021108512 | -1,281108512 |
| chrXII | 462228 | 462336 | 108 | 0,133929347 | 0,166764131 | 0,44540157 | -0,137288019 | -0,116694816 | 0,524238864 | 0,776238864 | -0,483761136 |
| chrXII | 462229 | 462336 | 107 | 1,578656563 | 2,864637184 | 0,355289714 | -0,371078012 | -0,315416311 | 0,159397885 | 0,418397885 | -0,841602115 |
| chrXII | 462229 | 462337 | 108 | 0,042743409 | 0,092646739 | 0,315705458 | -0,47974192 | -0,407780632 | 0,237560225 | 0,489560225 | -0,770439775 |
| chrXII | 462229 | 462349 | 120 | 1,407682928 | 3,313047403 | 0,298191769 | -0,529608301 | -0,450167056 | -0,056835858 | 0,111164142 | -1,148835858 |
| chrXII | 462229 | 462373 | 144 | 0,09973462 | 0,148234783 | 0,402205349 | -0,247642923 | -0,210496484 | -0,33916081 | -0,33916081 | -1,59916081 |
| chrXII | 462229 | 462382 | 153 | 0,247911771 | 0,222352175 | 0,527175798 | 0,068172392 | 0,057946533 | -0,061740384 | -0,124740384 | -1,384740384 |
| chrXII | 462229 | 462388 | 159 | 0,042743409 | 0,037058696 | 0,535617563 | 0,08939893 | 0,07598909 | 0,039720575 | -0,065279425 | -1,325279425 |
| chrXII | 462229 | 462407 | 178 | 0,09688506 | 0,188999349 | 0,338895921 | -0,415478239 | -0,353156503 | -0,213292995 | -0,451292995 | -1,711292995 |
| chrXII | 462229 | 462423 | 194 | 0,159575393 | 0,222352175 | 0,417815854 | -0,20748423 | -0,176361595 | -0,10060805 | -0,45060805 | -1,71060805 |
| chrXII | 462229 | 462425 | 196 | 0,119681545 | 0,218646305 | 0,353744289 | -0,375231134 | -0,318946464 | -0,273176877 | -0,637176877 | -1,897176877 |
| chrXII | 462232 | 462338 | 106 | 0,09688506 | 0,251999131 | 0,277699771 | -0,589688448 | -0,501235181 | -0,016167448 | 0,249832552 | -1,010167448 |
| chrXII | 462232 | 462349 | 117 | 0,122531105 | 0,229763914 | 0,347808225 | -0,391244593 | -0,332557904 | 0,203462798 | 0,392462798 | -0,867537202 |
| chrXII | 462232 | 462355 | 123 | 0,042743409 | 0,048176305 | 0,470122564 | -0,074961771 | -0,063717505 | 0,349751273 | 0,496751273 | -0,763248727 |
| chrXII | 462232 | 462373 | 141 | 0,054141651 | 0,092646739 | 0,368841506 | -0,334923211 | -0,284684729 | -0,344144002 | -0,323144002 | -1,583144002 |
| chrXII | 462232 | 462382 | 150 | 2,382232648 | 1,630582615 | 0,593656197 | 0,236960278 | 0,201416237 | 0,059277754 | 0,017277754 | -1,242722246 |
| chrXII | 462232 | 462407 | 175 | 0,091185939 | 0,08894087 | 0,506231912 | 0,015621722 | 0,013278464 | 0,242545265 | 0,025545265 | -1,234454735 |
| chrXII | 462232 | 462419 | 187 | 0,059840772 | 0,062999783 | 0,487141826 | -0,032236246 | -0,027400809 | 0,024745491 | -0,276254509 | -1,536254509 |
| chrXII | 462232 | 462423 | 191 | 0,182371877 | 0,181587609 | 0,501077411 | 0,002700672 | 0,002295572 | 0,049604939 | -0,279395061 | -1,539395061 |
| chrXII | 462232 | 462425 | 193 | 0,14532759 | 0,192705218 | 0,429921153 | -0,176573888 | -0,150087805 | -0,090854267 | -0,433854267 | -1,693854267 |
| chrXII | 462236 | 462373 | 137 | 1,31934655 | 0,763409133 | 0,633461985 | 0,341036599 | 0,289881109 | 0,355526437 | 0,404526437 | -0,855473563 |
| chrXII | 462236 | 462425 | 189 | 0,156725832 | 0,111176087 | 0,585011979 | 0,214732295 | 0,182522451 | 0,227455153 | -0,087544847 | -1,347544847 |
| chrXII | 462237 | 462375 | 138 | 0,051292091 | 0,051882174 | 0,497140355 | -0,007168129 | -0,00609291 | -0,004113025 | 0,037886975 | -1,222113025 |
| chrXII | 462237 | 462382 | 145 | 0,176672756 | 0,077823261 | 0,694206369 | 0,507809051 | 0,431637694 | 0,237608696 | 0,230608696 | -1,029391304 |
| chrXII | 462250 | 462382 | 132 | 0,068389454 | 0,103764348 | 0,397257877 | -0,260451199 | -0,221383519 | -0,082446751 | 0,001553249 | -1,258446751 |
| chrXII | 462250 | 462407 | 157 | 0,404637603 | 0,426175002 | 0,487038353 | -0,032495748 | -0,027621386 | -0,120150145 | -0,211150145 | -1,471150145 |
| chrXII | 462250 | 462423 | 173 | 0,074088575 | 0,144528914 | 0,338895921 | -0,415478239 | -0,353156503 | -0,174996903 | -0,377996903 | -1,637996903 |
| chrXII | 462250 | 462425 | 175 | 0,04844253 | 0,08894087 | 0,352608321 | -0,378288023 | -0,32154482 | -0,111528981 | -0,328528981 | -1,588528981 |
| chrXII | 462251 | 462373 | 122 | 0,159575393 | 0,125999566 | 0,558786364 | 0,147892925 | 0,125708986 | 0,522424323 | 0,676424323 | -0,583575677 |
| chrXII | 462251 | 462406 | 155 | 0,076938136 | 0,059293913 | 0,564757972 | 0,163043664 | 0,138587115 | 0,052002181 | -0,024997819 | -1,284997819 |
| chrXII | 462251 | 462407 | 156 | 3,083224551 | 2,012287181 | 0,605086341 | 0,266534858 | 0,226554629 | 0,180591704 | 0,096591704 | -1,163408296 |
| chrXII | 462251 | 462423 | 172 | 0,304902982 | 0,251999131 | 0,547498339 | 0,119343374 | 0,101441868 | 0,212393688 | 0,016393688 | -1,243606312 |
| chrXII | 462251 | 462424 | 173 | 0,3903898 | 0,444704349 | 0,467479984 | -0,081606077 | -0,069365165 | 0,117633691 | -0,085366309 | -1,345366309 |
| chrXII | 462251 | 462425 | 174 | 0,983098401 | 1,171054787 | 0,456373487 | -0,109574325 | -0,093138177 | 0,144537921 | -0,065462079 | -1,325462079 |
| chrXII | 462251 | 462439 | 188 | 0,09973462 | 0,118587827 | 0,456822566 | -0,108441943 | -0,092175651 | -0,075109934 | -0,383109934 | -1,643109934 |
| chrXII | 462253 | 462406 | 153 | 0,49012442 | 0,188999349 | 0,721701173 | 0,587902625 | 0,499717231 | 0,381870413 | 0,318870413 | -0,941129587 |
| chrXII | 462253 | 462407 | 154 | 0,102584181 | 0,066705652 | 0,605967759 | 0,268824834 | 0,228501109 | 0,046129994 | -0,023870006 | -1,283870006 |
| chrXII | 462253 | 462424 | 171 | 0,056991212 | 0,051882174 | 0,523463207 | 0,058847485 | 0,050020363 | 0,196335149 | 0,007335149 | -1,252664851 |
| chrXII | 462253 | 462425 | 172 | 0,14532759 | 0,08894087 | 0,620346375 | 0,30639062 | 0,260432027 | 0,371383847 | 0,175383847 | -1,084616153 |
| chrXII | 462256 | 462373 | 117 | 0,185221438 | 0,244587392 | 0,430939117 | -0,173983748 | -0,147886185 | 0,382935317 | 0,571935317 | -0,688064683 |
| chrXII | 462256 | 462388 | 132 | 0,054141651 | 0,133411305 | 0,288673942 | -0,557262806 | -0,473673385 | -0,330836606 | -0,246836606 | -1,506836606 |
| chrXII | 462256 | 462406 | 150 | 0,156725832 | 0,114881957 | 0,577029962 | 0,194301161 | 0,165155987 | 0,013414443 | -0,028585557 | -1,288585557 |
| chrXII | 462256 | 462407 | 151 | 1,225311051 | 1,182172396 | 0,508959284 | 0,022459482 | 0,01909056 | -0,120895149 | -0,169895149 | -1,429895149 |
| chrXII | 462256 | 462419 | 163 | 0,051292091 | 0,251999131 | 0,169118282 | -0,957655378 | -0,814007071 | -0,774397761 | -0,907397761 | -2,167397761 |
| chrXII | 462256 | 462423 | 167 | 0,658248495 | 0,893114568 | 0,424303318 | -0,19089656 | -0,162262076 | -0,014482441 | -0,175482441 | -1,435482441 |

TABLE 2

| Chrom | Start | End | Length | Section A | Section B | A/A+B | Z-score | Z * 0.85 | Phase correction | Length correction | ΔLK |
|---|---|---|---|---|---|---|---|---|---|---|---|
| chrXII | 462256 | 462424 | 168 | 1,031540931 | 1,237760439 | 0,454563217 | -0,114140479 | -0,097019408 | 0,027285127 | -0,140714873 | -1,400714873 |
| chrXII | 462256 | 462425 | 169 | 7,366114108 | 7,674855897 | 0,48973664 | -0,025729268 | -0,021869877 | 0,1235177 | -0,0514823 | -1,3114823 |
| chrXII | 462256 | 462439 | 183 | 0,284956058 | 0,318704784 | 0,472046617 | -0,070126174 | -0,059607248 | 0,025731811 | -0,247268189 | -1,507268189 |
| chrXII | 462256 | 462446 | 190 | 0,113982423 | 0,17417587 | 0,395554894 | -0,264869768 | -0,225139303 | -0,206095378 | -0,528095378 | -1,788095378 |
| chrXII | 462257 | 462365 | 108 | 0,079787696 | 0,074117392 | 0,518421434 | 0,046192108 | 0,039263292 | 0,671846555 | 0,923846555 | -0,336153445 |
| chrXII | 462258 | 462407 | 149 | 1,168319839 | 0,726350437 | 0,616634912 | 0,296654659 | 0,25215646 | 0,057675139 | 0,022675139 | -1,237324861 |
| chrXII | 462258 | 462423 | 165 | 0,250761331 | 0,289057827 | 0,464528403 | -0,089031589 | -0,075676851 | 0,02879259 | -0,11820741 | -1,37820741 |
| chrXII | 462258 | 462425 | 167 | 0,153876272 | 0,233469783 | 0,397257877 | -0,260451199 | -0,221383519 | -0,070431317 | -0,231431317 | -1,491431317 |
| chrXII | 462262 | 462368 | 106 | 0,24506221 | 0,841232394 | 0,225594612 | -0,753433889 | -0,640418806 | -0,13643658 | 0,12956342 | -1,13043658 |
| chrXII | 462262 | 462373 | 111 | 0,202318801 | 0,830114786 | 0,195963018 | -0,856129712 | -0,727710255 | -0,25956716 | -0,02856716 | -1,28856716 |
| chrXII | 462262 | 462407 | 145 | 0,461628815 | 0,43729261 | 0,513536336 | 0,033937075 | 0,028846514 | -0,153251734 | -0,160251734 | -1,420251734 |
| chrXII | 462262 | 462419 | 157 | 0,042743409 | 0,144528914 | 0,228241997 | -0,744648908 | -0,632951572 | -0,725701185 | -0,816701185 | -2,076701185 |
| chrXII | 462262 | 462423 | 161 | 0,401788042 | 0,418763262 | 0,48965621 | -0,025930941 | -0,0220413 | -0,025982031 | -0,144982031 | -1,404982031 |
| chrXII | 462262 | 462424 | 162 | 2,806817175 | 2,223521747 | 0,557977746 | 0,145844042 | 0,123967435 | 0,168951192 | 0,042951192 | -1,217048808 |
| chrXII | 462262 | 462425 | 163 | 1,339293474 | 1,308171961 | 0,505877605 | 0,014733504 | 0,012523479 | 0,037790173 | -0,095209827 | -1,355209827 |
| chrXII | 462262 | 462439 | 177 | 0,128230226 | 0,144528914 | 0,470122564 | -0,074961771 | -0,063717505 | 0,086668945 | -0,144331055 | -1,404331055 |
| chrXII | 462263 | 462385 | 122 | 0,165274514 | 0,718938698 | 0,186917037 | -0,889314512 | -0,755917335 | -0,359249066 | -0,205249066 | -1,465249066 |
| chrXII | 462263 | 462407 | 144 | 0,159575393 | 0,17417587 | 0,478126708 | -0,054855711 | -0,046627354 | -0,172089506 | -0,172089506 | -1,432089506 |
| chrXII | 462263 | 462423 | 160 | 0,58415992 | 0,52993935 | 0,524333813 | 0,061033699 | 0,051878644 | 0,049522866 | -0,062477134 | -1,322477134 |
| chrXII | 462263 | 462424 | 161 | 0,139628469 | 0,155646522 | 0,472876041 | -0,06804215 | -0,057835827 | -0,066678272 | -0,185678272 | -1,445678272 |
| chrXII | 462263 | 462425 | 162 | 0,492973981 | 0,774526742 | 0,388933885 | -0,282098777 | -0,239783961 | -0,181583167 | -0,307583167 | -1,567583167 |
| chrXII | 462272 | 462407 | 135 | 0,108283302 | 0,181587609 | 0,37355698 | -0,322447164 | -0,274080089 | -0,291203503 | -0,228203503 | -1,488203503 |
| chrXII | 462272 | 462419 | 147 | 1,678391184 | 1,385995222 | 0,547708729 | 0,119874529 | 0,10189335 | -0,011397848 | -0,032397848 | -1,292397848 |
| chrXII | 462272 | 462423 | 151 | 0,384690679 | 0,500292393 | 0,434687048 | -0,164453569 | -0,139785534 | -0,270990156 | -0,319990156 | -1,579990156 |
| chrXII | 462272 | 462425 | 153 | 0,262159574 | 0,452116089 | 0,367028568 | -0,339733626 | -0,288773582 | -0,413374966 | -0,476374966 | -1,736374966 |
| chrXII | 462275 | 462423 | 148 | 0,062690333 | 0,055588044 | 0,530023616 | 0,075329227 | 0,064029843 | -0,082781459 | -0,110781459 | -1,370781459 |
| chrXII | 462275 | 462425 | 150 | 0,102584181 | 0,251999131 | 0,289309106 | -0,555404213 | -0,472093581 | -0,613876347 | -0,655876347 | -1,915876347 |
| chrXII | 462277 | 462373 | 96 | 0,068389454 | 0,103764348 | 0,397257877 | -0,260451199 | -0,221383519 | 0,356878809 | 0,692878809 | -0,567121191 |
| chrXII | 462277 | 462382 | 105 | 0,111132863 | 0,092646739 | 0,54535813 | 0,113942039 | 0,096850733 | 0,684741492 | 0,957741492 | -0,302258508 |
| chrXII | 462277 | 462383 | 106 | 0,062690333 | 0,159352392 | 0,28233455 | -0,575920231 | -0,489532196 | 0,023544586 | 0,289544586 | -0,970455414 |
| chrXII | 462277 | 462388 | 111 | 0,14817715 | 0,830114786 | 0,151465166 | -1,03016974 | -0,875644279 | -0,37354736 | -0,14254736 | -1,40254736 |
| chrXII | 462277 | 462407 | 130 | 0,413186285 | 0,426175002 | 0,492262738 | -0,019395657 | -0,016486308 | 0,111542353 | 0,209542353 | -1,050457647 |
| chrXII | 462277 | 462419 | 142 | 0,276407377 | 0,292763697 | 0,485631455 | -0,036024391 | -0,030620732 | -0,014675382 | -0,000675382 | -1,260675382 |
| chrXII | 462277 | 462423 | 146 | 1,547311397 | 1,074702178 | 0,590123336 | 0,227862253 | 0,193682915 | 0,031090305 | 0,017090305 | -1,242909695 |
| chrXII | 462277 | 462424 | 147 | 1,296550065 | 1,189584135 | 0,521512501 | 0,053950004 | 0,045857503 | -0,070397919 | -0,091397919 | -1,351397919 |
| chrXII | 462277 | 462425 | 148 | 5,095014323 | 3,683634361 | 0,580387085 | 0,202883835 | 0,17245126 | 0,016854049 | -0,011145951 | -1,271145951 |
| chrXII | 462277 | 462439 | 162 | 0,071239015 | 0,133411305 | 0,348101165 | -0,390452017 | -0,331884215 | -0,264712858 | -0,390712858 | -1,650712858 |
| chrXII | 462277 | 462460 | 183 | 0,079787696 | 0,059293913 | 0,573675388 | 0,185739279 | 0,157878387 | 0,241714469 | -0,031285531 | -1,291285531 |
| chrXII | 462278 | 462373 | 95 | 0,168124074 | 0,159352392 | 0,513392844 | 0,03357719 | 0,028540612 | 0,527911024 | 0,870911024 | -0,389088976 |
| chrXII | 462278 | 462382 | 104 | 0,19377012 | 0,207528696 | 0,482857442 | -0,042983253 | -0,036535765 | 0,530633819 | 0,810633819 | -0,449366181 |
| chrXII | 462278 | 462406 | 128 | 0,09688506 | 0,048176305 | 0,667890174 | 0,43409473 | 0,36898052 | 0,634153169 | 0,746153169 | -0,513846831 |
| chrXII | 462278 | 462407 | 129 | 0,943204553 | 0,893114568 | 0,513638693 | 0,034193795 | 0,029064726 | 0,296966478 | 0,401966478 | -0,858033522 |
| chrXII | 462278 | 462418 | 140 | 0,042743409 | 0,051882174 | 0,451710916 | -0,121339881 | -0,103138899 | -0,080192965 | -0,052192965 | -1,312192965 |
| chrXII | 462278 | 462419 | 141 | 0,336248149 | 0,359469349 | 0,483311329 | -0,041844502 | -0,035567826 | -0,091812732 | -0,070812732 | -1,330812732 |
| chrXII | 462278 | 462420 | 142 | 0,031345166 | 0,051882174 | 0,376621026 | -0,314367347 | -0,267212245 | -0,247351734 | -0,233351734 | -1,493351734 |
| chrXII | 462278 | 462423 | 145 | 6,39726351 | 3,87263371 | 0,622914073 | 0,313143221 | 0,266171738 | 0,095348549 | 0,088348549 | -1,171651451 |
| chrXII | 462278 | 462424 | 146 | 0,988797522 | 1,019114134 | 0,492450711 | -0,018924391 | -0,016085733 | -0,186693145 | -0,200693145 | -1,460693145 |
| chrXII | 462278 | 462425 | 147 | 7,528539062 | 5,492098715 | 0,578200484 | 0,197292009 | 0,167698208 | 0,046022179 | 0,025022179 | -1,234977821 |
| chrXII | 462278 | 462439 | 161 | 0,467327936 | 0,296469566 | 0,61184795 | 0,284138687 | 0,241517884 | 0,223789437 | 0,104789437 | -1,155210563 |
| chrXII | 462278 | 462446 | 168 | 0,789328282 | 0,837526525 | 0,48518668 | -0,037140024 | -0,031569021 | 0,087929701 | -0,080070299 | -1,340070299 |
| chrXII | 462278 | 462460 | 182 | 0,059840772 | 0,070411522 | 0,459422021 | -0,101889932 | -0,086606442 | 0,101125749 | -0,164874251 | -1,424874251 |
| chrXII | 462278 | 462461 | 183 | 0,054141651 | 0,062999783 | 0,462190442 | -0,094916836 | -0,080679311 | -0,007534897 | -0,280534897 | -1,540534897 |
| chrXII | 462278 | 462479 | 201 | 0,045592969 | 0,048176305 | 0,486225044 | -0,034535557 | -0,029355224 | -0,014067841 | -0,413067841 | -1,673067841 |
| chrXII | 462281 | 462425 | 144 | 0,037044288 | 0,103764348 | 0,263082498 | -0,633871027 | -0,538790373 | -0,667457252 | -0,667457252 | -1,927457252 |
| chrXII | 462287 | 462423 | 136 | 0,14817715 | 0,122293696 | 0,54784888 | 0,120228376 | 0,10219412 | 0,080022787 | 0,136022787 | -1,123977213 |
| chrXII | 462287 | 462425 | 138 | 0,125380666 | 0,196411088 | 0,389632936 | -0,280275857 | -0,238234478 | -0,234089606 | -0,192089606 | -1,452089606 |
| chrXII | 462287 | 462432 | 145 | 0,082637257 | 0,059293913 | 0,582234733 | 0,207613792 | 0,176471723 | 0,007745309 | 0,000745309 | -1,259254691 |
| chrXII | 462287 | 462439 | 152 | 0,094035499 | 0,222352175 | 0,297216064 | -0,532424343 | -0,452560692 | -0,653162681 | -0,709162681 | -1,969162681 |
| chrXII | 462289 | 462377 | 88 | 0,105433742 | 0,200116957 | 0,345061366 | -0,398688514 | -0,338885237 | 0,096442296 | 0,488442296 | -0,771557704 |
| chrXII | 462289 | 462407 | 118 | 0,159575393 | 0,240881523 | 0,398483299 | -0,257274861 | -0,218683631 | 0,166235083 | 0,348235083 | -0,911764917 |
| chrXII | 462289 | 462423 | 134 | 1,877860425 | 1,793640876 | 0,511469361 | 0,028753386 | 0,024440378 | 0,032786959 | 0,102786959 | -1,157213041 |
| chrXII | 462289 | 462424 | 135 | 0,253610892 | 0,340940001 | 0,42655876 | -0,185142177 | -0,157370851 | -0,179744813 | -0,116744813 | -1,376744813 |
| chrXII | 462289 | 462425 | 136 | 0,720938828 | 0,726350437 | 0,498130433 | -0,004686328 | -0,003983379 | -0,035319998 | 0,020680002 | -1,239319998 |
| chrXII | 462289 | 462439 | 150 | 0,205168362 | 0,092646739 | 0,688911882 | 0,492768407 | 0,418853146 | 0,275027621 | 0,233027621 | -1,026972379 |
| chrXII | 462290 | 462423 | 133 | 0,085486818 | 0,074117392 | 0,535617563 | 0,08939893 | 0,07598909 | 0,060835515 | 0,137835515 | -1,122164485 |
| chrXII | 462290 | 462425 | 135 | 0,09973462 | 0,062999783 | 0,61286746 | 0,286800501 | 0,243780426 | 0,225207375 | 0,288207375 | -0,971792625 |
| chrXII | 462296 | 462407 | 111 | 0,202318801 | 0,311293045 | 0,393913814 | -0,26913262 | -0,228762727 | 0,277333998 | 0,508333998 | -0,751666002 |
| chrXII | 462296 | 462423 | 127 | 0,421734966 | 0,696703481 | 0,377074811 | -0,313172484 | -0,266196611 | 0,002781853 | 0,121781853 | -1,138218147 |
| chrXII | 462296 | 462424 | 128 | 0,461628815 | 0,544762828 | 0,458696987 | -0,10371695 | -0,088159408 | 0,160240504 | 0,272240504 | -0,987759496 |
| chrXII | 462296 | 462425 | 129 | 1,193965884 | 1,267407396 | 0,485081192 | -0,037404627 | -0,031793933 | 0,208376178 | 0,313376178 | -0,946623822 |
| chrXII | 462296 | 462439 | 143 | 2,174214725 | 1,389701092 | 0,610063435 | 0,279484371 | 0,237561716 | 0,104254767 | 0,111254767 | -1,148745233 |
| chrXII | 462296 | 462446 | 150 | 0,108283302 | 0,122293696 | 0,469618839 | -0,076228037 | -0,064793831 | -0,209981275 | -0,251981275 | -1,511981275 |
| chrXII | 462296 | 462472 | 176 | 0,039893848 | 0,066705652 | 0,374240479 | -0,320642986 | -0,272546538 | -0,100458547 | -0,324458547 | -1,584458547 |
| chrXII | 462296 | 462479 | 183 | 0,028495606 | 0,055588044 | 0,338895921 | -0,415478239 | -0,353156503 | -0,298632027 | -0,571632027 | -1,831632027 |
| chrXII | 462298 | 462382 | 84 | 0,151026711 | 0,133411305 | 0,530965281 | 0,07769655 | 0,066042068 | 0,417229998 | 0,837229998 | -0,422770002 |
| chrXII | 462298 | 462405 | 107 | 0,256460453 | 0,600350872 | 0,299319633 | -0,52635832 | -0,447404572 | 0,049307852 | 0,308307852 | -0,951692148 |
| chrXII | 462298 | 462407 | 109 | 0,09688506 | 0,192705218 | 0,33455909 | -0,427358568 | -0,363254783 | 0,310184077 | 0,555184077 | -0,704815923 |
| chrXII | 462298 | 462419 | 121 | 0,09973462 | 0,192705218 | 0,341043207 | -0,409617695 | -0,348175041 | 0,041438669 | 0,202438669 | -1,057561331 |
| chrXII | 462298 | 462423 | 125 | 4,066322953 | 6,014626326 | 0,403367068 | -0,244641361 | -0,207945156 | 0,117844793 | 0,250844793 | -1,009155207 |
| chrXII | 462298 | 462425 | 127 | 0,763682236 | 1,119172613 | 0,40559804 | -0,238883294 | -0,2030508 | 0,060173886 | 0,179173886 | -1,080826114 |
| chrXII | 462298 | 462446 | 148 | 0,270708255 | 0,322410653 | 0,456414812 | -0,109470114 | -0,093049597 | -0,261906994 | -0,289906994 | -1,549906994 |
| chrXII | 462298 | 462460 | 162 | 0,190920559 | 0,207528696 | 0,479159031 | -0,052264347 | -0,044424695 | 0,013619878 | -0,112380122 | -1,372380122 |
| chrXII | 462298 | 462461 | 163 | 0,079787696 | 0,096352609 | 0,452978074 | -0,118140734 | -0,100419624 | -0,085055751 | -0,218055751 | -1,478055751 |
| chrXII | 462299 | 462406 | 107 | 0,139628469 | 0,188999349 | 0,424883291 | -0,189416258 | -0,161003819 | 0,350349386 | 0,609349386 | -0,650650614 |
| chrXII | 462299 | 462423 | 124 | 0,119681545 | 0,218646305 | 0,353744289 | -0,375231134 | -0,318946464 | 0,071272192 | 0,211272192 | -1,048727808 |
| chrXII | 462299 | 462425 | 126 | 0,162424953 | 0,307587175 | 0,345576089 | -0,397291956 | -0,337698162 | -0,101184027 | 0,024815973 | -1,235184027 |
| chrXII | 462299 | 462460 | 161 | 0,216566604 | 0,237175653 | 0,477289917 | -0,056956516 | -0,048413039 | -0,063117698 | -0,182117698 | -1,442117698 |
| chrXII | 462299 | 462461 | 162 | 0,410336724 | 0,400233914 | 0,506231912 | 0,015621722 | 0,013278464 | 0,10353815 | -0,02246185 | -1,28246185 |
| chrXII | 462302 | 462424 | 122 | 0,074088575 | 0,129705435 | 0,363546382 | -0,348995399 | -0,296646089 | 0,098622083 | 0,252622083 | -1,007377917 |
| chrXII | 462302 | 462425 | 123 | 0,14817715 | 0,281646088 | 0,344739737 | -0,399561558 | -0,339627325 | 0,070830501 | 0,217830501 | -1,042169499 |
| chrXII | 462302 | 462439 | 137 | 0,028495606 | 0,055588044 | 0,338895921 | -0,415478239 | -0,353156503 | -0,281231006 | -0,232231006 | -1,492231006 |
| chrXII | 462302 | 462479 | 177 | 0,262159574 | 0,259410871 | 0,502635026 | 0,006605078 | 0,005614317 | 0,154766359 | -0,076233641 | -1,336233641 |
| chrXII | 462308 | 462479 | 171 | 0,071239015 | 0,085235 | 0,455276965 | -0,11233987 | -0,09548889 | 0,038248195 | -0,150751805 | -1,410751805 |
| chrXII | 462317 | 462425 | 108 | 0,062690333 | 0,285351958 | 0,180122745 | -0,914897409 | -0,777662797 | -0,152751066 | 0,099248934 | -1,160751066 |
| chrXII | 462317 | 462477 | 160 | 0,074088575 | 0,151940653 | 0,327783163 | -0,446042806 | -0,379136385 | -0,380379822 | -0,492379822 | -1,752379822 |
| chrXII | 462318 | 462407 | 89 | 0,051292091 | 0,244587392 | 0,173354671 | -0,940991251 | -0,799842563 | -0,343961854 | 0,041038146 | -1,218961854 |
| chrXII | 462318 | 462419 | 101 | 0,14532759 | 1,797346746 | 0,074808004 | -1,440889127 | -1,224755758 | -0,747901076 | -0,446901076 | -1,706901076 |
| chrXII | 462318 | 462423 | 105 | 0,270708255 | 1,05617283 | 0,204018475 | -0,827353109 | -0,703250143 | -0,146388482 | 0,126611518 | -1,133388482 |
| chrXII | 462318 | 462425 | 107 | 0,247911771 | 1,108055004 | 0,182830269 | -0,904631694 | -0,76893694 | -0,247235465 | 0,011764535 | -1,248235465 |
| chrXII | 462318 | 462460 | 142 | 0,416035845 | 0,503998263 | 0,452196111 | -0,120114784 | -0,102097566 | -0,089013962 | -0,075013962 | -1,335013962 |
| chrXII | 462318 | 462461 | 143 | 0,039893848 | 0,085235 | 0,318822147 | -0,470995015 | -0,400345763 | -0,537923092 | -0,530923092 | -1,790923092 |

TABLE 2

| Chrom | Start | End | Length | Section A | Section B | A/A+B | Z-score | Z * 0.85 | Phase correction | Length correction | ΔLK |
|---|---|---|---|---|---|---|---|---|---|---|---|
| chrXII | 462423 | 462569 | 146 | 0,53856695 | 0,181587609 | 0,747849115 | 0,667736558 | 0,567576075 | 0,401576342 | 0,387576342 | -0,872423658 |
| chrXII | 462436 | 462577 | 141 | 13,86881136 | 8,549441118 | 0,618639271 | 0,301908968 | 0,256622623 | 0,210383747 | 0,231383747 | -1,028616253 |
| chrXII | 462436 | 462597 | 161 | 0,059840772 | 0,074117392 | 0,446712395 | -0,133971905 | -0,113876119 | -0,130804872 | -0,249804872 | -1,509804872 |
| chrXII | 462437 | 462547 | 110 | 0,031345166 | 0,211234566 | 0,129215933 | -1,130105279 | -0,960589487 | -0,294457738 | -0,056457738 | -1,316457738 |
| chrXII | 462437 | 462577 | 140 | 0,039893848 | 0,070411522 | 0,361667325 | -0,354005648 | -0,300904801 | -0,280851225 | -0,252851225 | -1,512851225 |
| chrXII | 462439 | 462577 | 138 | 0,062690333 | 0,092646739 | 0,403576119 | -0,244101464 | -0,207486244 | -0,210489747 | -0,168489747 | -1,428489747 |
| chrXII | 462440 | 462577 | 137 | 0,039893848 | 0,066705652 | 0,374240479 | -0,320642986 | -0,272546538 | -0,201360047 | -0,152360047 | -1,412360047 |
| chrXII | 462447 | 462577 | 130 | 0,14817715 | 0,118587827 | 0,555459536 | 0,139467258 | 0,118547169 | 0,237534996 | 0,335534996 | -0,924465004 |
| chrXII | 462447 | 462597 | 150 | 0,139628469 | 0,111176087 | 0,556722218 | 0,142663981 | 0,121264384 | -0,016042782 | -0,058042782 | -1,318042782 |
| chrXII | 462459 | 462618 | 159 | 0,156725832 | 0,144528914 | 0,520243529 | 0,050764798 | 0,043150079 | 0,008980311 | -0,096019689 | -1,356019689 |
| chrXII | 462595 | 462732 | 137 | 0,082637257 | 0,222352175 | 0,270951215 | -0,609938679 | -0,518447877 | -0,440212947 | -0,391212947 | -1,651212947 |
| chrXII | 462595 | 462747 | 152 | 0,344796831 | 0,300175436 | 0,534591716 | 0,086817511 | 0,073794884 | -0,123696431 | -0,179696431 | -1,439696431 |
| chrXII | 462595 | 462749 | 154 | 0,233663968 | 0,196411088 | 0,543309743 | 0,108775554 | 0,092459221 | -0,085710209 | -0,155710209 | -1,415710209 |
| chrXII | 462597 | 462749 | 152 | 0,210867483 | 0,114881957 | 0,647330301 | 0,378122765 | 0,32140435 | 0,115276419 | 0,059276419 | -1,200723581 |
| chrXII | 462624 | 462732 | 108 | 0,424584527 | 3,909692405 | 0,097959714 | -1,293264983 | -1,099275236 | -0,462986452 | -0,210986452 | -1,470986452 |
| chrXII | 462624 | 462747 | 123 | 0,051292091 | 0,166764131 | 0,235224215 | -0,721749813 | -0,613487341 | -0,194820989 | -0,047820989 | -1,307820989 |
| chrXII | 462624 | 462749 | 125 | 0,065539893 | 0,218646305 | 0,230623069 | -0,736796456 | -0,626276987 | -0,302388607 | -0,169388607 | -1,429388607 |
| chrXII | 462637 | 462789 | 152 | 0,416035845 | 0,140823044 | 0,747111797 | 0,665428585 | 0,565614298 | 0,355941276 | 0,299941276 | -0,960058724 |
| chrXII | 462711 | 462864 | 153 | 0,119681545 | 0,062999783 | 0,655138356 | 0,399230614 | 0,339346022 | 0,209736798 | 0,146736798 | -1,113263202 |
| chrXII | 462724 | 462874 | 150 | 0,051292091 | 0,051882174 | 0,497140355 | -0,007168129 | -0,00609291 | -0,130615997 | -0,172615997 | -1,432615997 |
| chrXII | 462764 | 462913 | 149 | 0,056991212 | 0,107470218 | 0,346532387 | -0,394699362 | -0,335494458 | -0,512936083 | -0,547936083 | -1,807936083 |
| chrXII | 462764 | 462916 | 152 | 1,330744792 | 0,589233263 | 0,693104168 | 0,504668535 | 0,428968255 | 0,223057022 | 0,167057022 | -1,092942978 |
| chrXII | 462794 | 462951 | 157 | 0,230814407 | 0,077823261 | 0,747849115 | 0,667736558 | 0,567576075 | 0,474221815 | 0,383221815 | -0,876778185 |
| chrXII | 462796 | 462913 | 117 | 1,114178188 | 5,18451154 | 0,17689047 | -0,927280453 | -0,788188385 | -0,25478172 | -0,06578172 | -1,32578172 |
| chrXII | 462796 | 462916 | 120 | 0,059840772 | 0,240881523 | 0,198990142 | -0,845233583 | -0,718448775 | -0,320727582 | -0,152727582 | -1,412727582 |
| chrXII | 462871 | 462990 | 119 | 0,367593315 | 0,982055438 | 0,272362209 | -0,605684292 | -0,514831648 | -0,074231333 | 0,100768667 | -1,159231333 |
| chrXII | 462885 | 463047 | 162 | 0,04844253 | 0,155646522 | 0,237359767 | -0,714821199 | -0,607598019 | -0,521815123 | -0,647815123 | -1,907815123 |
| chrXII | 462885 | 463050 | 165 | 3,214304338 | 3,524281969 | 0,476999803 | -0,057684921 | -0,049032183 | 0,046955711 | -0,100044289 | -1,360044289 |
| chrXII | 462885 | 463056 | 171 | 0,675345858 | 0,781938481 | 0,463427651 | -0,091802067 | -0,078031757 | 0,046291972 | -0,142708028 | -1,402708028 |
| chrXII | 462885 | 463059 | 174 | 0,056991212 | 0,070411522 | 0,447331152 | -0,132407093 | -0,112546029 | 0,124494291 | -0,085505709 | -1,345505709 |
| chrXII | 462885 | 463099 | 214 | 0,153876272 | 0,133411305 | 0,535617563 | 0,08939893 | 0,07598909 | 0,361860225 | -0,128139775 | -1,388139775 |
| chrXII | 462889 | 462996 | 107 | 0,276407377 | 0,255705001 | 0,519453011 | 0,048780806 | 0,041463685 | 0,580701873 | 0,839701873 | -0,420298127 |
| chrXII | 462889 | 463045 | 156 | 0,766531797 | 0,96352609 | 0,443067138 | -0,143197409 | -0,121717797 | -0,168219097 | -0,252219097 | -1,512219097 |
| chrXII | 462889 | 463048 | 159 | 0,612655525 | 0,641115437 | 0,488650275 | -0,028453381 | -0,024185374 | -0,050230859 | -0,155230859 | -1,415230859 |
| chrXII | 462889 | 463051 | 162 | 1,062886098 | 1,1451137 | 0,481379617 | -0,046691337 | -0,039687637 | 0,033322019 | -0,092677981 | -1,352677981 |
| chrXII | 462889 | 463053 | 164 | 1,997541969 | 1,804758485 | 0,525350901 | 0,063588112 | 0,054049895 | 0,051474831 | -0,088525169 | -1,348525169 |
| chrXII | 462889 | 463059 | 170 | 2,613047055 | 3,25004762 | 0,445677104 | -0,136590852 | -0,116102224 | 0,060058218 | -0,121941782 | -1,381941782 |
| chrXII | 462889 | 463061 | 172 | 0,34194727 | 0,292763697 | 0,538744859 | 0,097272137 | 0,082681316 | 0,200074826 | 0,004074826 | -1,255925174 |
| chrXII | 462889 | 463066 | 177 | 0,031345166 | 0,040764565 | 0,434687048 | -0,164453569 | -0,139785534 | 0,010644009 | -0,220355991 | -1,480355991 |
| chrXII | 462890 | 463050 | 160 | 0,068389454 | 0,040764565 | 0,626540868 | 0,322705529 | 0,274299699 | 0,277400943 | 0,165400943 | -1,094599057 |
| chrXII | 462890 | 463059 | 169 | 0,079787696 | 0,085235 | 0,483495289 | -0,041382985 | -0,035175537 | 0,116437209 | -0,058562791 | -1,318562791 |
| chrXII | 462891 | 463047 | 156 | 0,151026711 | 0,277940218 | 0,352070756 | -0,379735842 | -0,322775466 | -0,371156071 | -0,455156071 | -1,715156071 |
| chrXII | 462891 | 463056 | 165 | 0,039893848 | 0,08894087 | 0,309651379 | -0,496838761 | -0,422312947 | -0,323680222 | -0,470680222 | -1,730680222 |
| chrXII | 462891 | 463061 | 170 | 0,045592969 | 0,070411522 | 0,393027622 | -0,271436616 | -0,230721123 | -0,054767484 | -0,236767484 | -1,496767484 |
| chrXII | 462911 | 463047 | 136 | 0,09688506 | 0,107470218 | 0,47410109 | -0,064964608 | -0,055219916 | -0,084132838 | -0,028132838 | -1,288132838 |
| chrXII | 462911 | 463050 | 139 | 0,729487509 | 0,752291524 | 0,49230519 | -0,019289223 | -0,01639584 | -0,048277334 | -0,013277334 | -1,273277334 |
| chrXII | 462911 | 463051 | 140 | 0,062690333 | 0,17417587 | 0,264665588 | -0,629027313 | -0,534673216 | -0,515682671 | -0,487682671 | -1,747682671 |
| chrXII | 462911 | 463053 | 142 | 0,068389454 | 0,081529131 | 0,456177292 | -0,110069087 | -0,093558724 | -0,080359236 | -0,066359236 | -1,326359236 |
| chrXII | 462911 | 463056 | 145 | 9,736948514 | 5,440216541 | 0,641552522 | 0,362611717 | 0,308219959 | 0,117105004 | 0,110105004 | -1,149894996 |
| chrXII | 462911 | 463062 | 151 | 0,113982423 | 0,092646739 | 0,551627959 | 0,129775456 | 0,110309137 | -0,02235686 | -0,07135686 | -1,33135686 |
| chrXII | 462911 | 463099 | 188 | 0,521469587 | 0,392822175 | 0,570353588 | 0,177274443 | 0,150683276 | 0,170846327 | -0,137153673 | -1,397153673 |
| chrXII | 462912 | 463045 | 133 | 0,769381358 | 0,915349786 | 0,456679014 | -0,108803903 | -0,092483317 | -0,09433178 | -0,01733178 | -1,27733178 |
| chrXII | 462912 | 463048 | 136 | 0,14817715 | 0,192705218 | 0,434687048 | -0,164453569 | -0,139785534 | -0,162103625 | -0,106103625 | -1,366103625 |
| chrXII | 462912 | 463051 | 139 | 7,719459621 | 5,421687193 | 0,587426633 | 0,220930261 | 0,187790722 | 0,157951021 | 0,192951021 | -1,067048979 |
| chrXII | 462912 | 463053 | 141 | 0,29350474 | 0,255705001 | 0,534412845 | 0,086367464 | 0,073412344 | 0,040365307 | 0,061365307 | -1,198634693 |
| chrXII | 462912 | 463059 | 147 | 0,527168708 | 0,544762828 | 0,491793263 | -0,02057269 | -0,017486786 | -0,139418388 | -0,160418388 | -1,420418388 |
| chrXII | 462918 | 463045 | 127 | 5,964130301 | 6,559389154 | 0,476234362 | -0,059606898 | -0,050665863 | 0,212300505 | 0,331300505 | -0,928699495 |
| chrXII | 462918 | 463048 | 130 | 0,085486818 | 0,111176087 | 0,434687048 | -0,164453569 | -0,139785534 | -0,027890733 | 0,070109267 | -1,189890733 |
| chrXII | 462918 | 463051 | 133 | 0,866266417 | 0,785644351 | 0,52440267 | 0,061206617 | 0,052025624 | 0,053904986 | 0,130904986 | -1,129095014 |
| chrXII | 462918 | 463053 | 135 | 0,287805619 | 0,277940218 | 0,508718933 | 0,021856865 | 0,018578335 | -0,003000469 | 0,059999531 | -1,200000469 |
| chrXII | 462918 | 463059 | 141 | 0,464478375 | 0,52623348 | 0,468832964 | -0,078203815 | -0,066473243 | -0,093478689 | -0,072478689 | -1,332478689 |
| chrXII | 462919 | 463048 | 129 | 0,142478029 | 0,218646305 | 0,394540095 | -0,267505227 | -0,227379443 | 0,003586197 | 0,108586197 | -1,151413803 |
| chrXII | 462919 | 463050 | 131 | 0,344796831 | 0,289057827 | 0,543968284 | 0,110436217 | 0,093870785 | 0,147999292 | 0,238999292 | -1,021000708 |
| chrXII | 462919 | 463051 | 132 | 0,071239015 | 0,118587827 | 0,375284202 | -0,317889966 | -0,270206471 | -0,127583622 | -0,043583622 | -1,303583622 |
| chrXII | 462919 | 463056 | 137 | 0,156725832 | 0,159352392 | 0,495845079 | -0,010415029 | -0,008852775 | 0,077813954 | 0,126813954 | -1,133186046 |
| chrXII | 462919 | 463059 | 140 | 0,056991212 | 0,096352609 | 0,371656395 | -0,327469522 | -0,278349094 | -0,237927096 | -0,209927096 | -1,469927096 |
| chrXII | 462919 | 463062 | 143 | 0,253610892 | 0,137117174 | 0,649072626 | 0,382817954 | 0,325395261 | 0,183052955 | 0,190052955 | -1,069947045 |
| chrXII | 462919 | 463066 | 147 | 0,188070999 | 0,166764131 | 0,530023616 | 0,075329227 | 0,064029843 | -0,067886903 | -0,088886903 | -1,348886903 |
| chrXII | 462919 | 463074 | 155 | 0,068389454 | 0,100058479 | 0,405997586 | -0,23785292 | -0,202174982 | -0,299297045 | -0,376297045 | -1,636297045 |
| chrXII | 462919 | 463099 | 180 | 0,071239015 | 0,051882174 | 0,578608892 | 0,198335964 | 0,168585569 | 0,339219731 | 0,087219731 | -1,172780269 |
| chrXII | 462920 | 463045 | 125 | 0,028495606 | 0,040764565 | 0,411428464 | -0,223871969 | -0,190291174 | 0,148708774 | 0,281708774 | -0,978291226 |
| chrXII | 462920 | 463048 | 128 | 0,059840772 | 0,111176087 | 0,349911537 | -0,385559308 | -0,327725412 | -0,089160695 | 0,022839305 | -1,237160695 |
| chrXII | 462920 | 463050 | 130 | 0,045592969 | 0,070411522 | 0,393027622 | -0,271436616 | -0,230721123 | -0,112710878 | -0,014710878 | -1,274710878 |
| chrXII | 462920 | 463051 | 131 | 0,062690333 | 0,122293696 | 0,338895921 | -0,415478239 | -0,353156503 | -0,30107321 | -0,21007321 | -1,47007321 |
| chrXII | 462949 | 463050 | 101 | 0,151026711 | 0,248293262 | 0,37820976 | -0,310185996 | -0,263658096 | 0,20979647 | 0,51079647 | -0,74920353 |
| chrXII | 462949 | 463056 | 107 | 0,464478375 | 0,444704349 | 0,510874616 | 0,027261996 | 0,023172697 | 0,574204694 | 0,833204694 | -0,426795306 |
| chrXII | 462949 | 463099 | 150 | 2,746976403 | 1,504583049 | 0,646110312 | 0,374840117 | 0,3186141 | 0,19581757 | 0,15381757 | -1,10618243 |
| chrXII | 462949 | 463138 | 189 | 0,125380666 | 0,08894087 | 0,585011979 | 0,214732295 | 0,182522451 | 0,236241452 | -0,078758548 | -1,338758548 |
| chrXII | 462949 | 463138 | 189 | 0,125380666 | 0,08894087 | 0,585011979 | 0,214732295 | 0,182522451 | 0,236241452 | -0,078758548 | -1,338758548 |
| chrXII | 462988 | 463095 | 107 | 0,170973635 | 0,203822827 | 0,456177292 | -0,110069087 | -0,093558724 | 0,45883822 | 0,71783822 | -0,54216178 |
| chrXII | 462988 | 463099 | 111 | 0,09688506 | 0,133411305 | 0,420697304 | -0,200109928 | -0,170093439 | 0,315641974 | 0,546641974 | -0,713358026 |
| chrXII | 462988 | 463138 | 150 | 1,427629852 | 0,841232394 | 0,629227206 | 0,329807278 | 0,280336186 | 0,170432647 | 0,128432647 | -1,131567353 |
| chrXII | 462988 | 463138 | 150 | 1,427629852 | 0,841232394 | 0,629227206 | 0,329807278 | 0,280336186 | 0,170432647 | 0,128432647 | -1,131567353 |
| chrXII | 463046 | 463209 | 163 | 0,094035499 | 0,148234783 | 0,388142938 | -0,284162468 | -0,241538098 | -0,229772572 | -0,362772572 | -1,622772572 |
| chrXII | 463055 | 463217 | 162 | 0,227964847 | 0,26311674 | 0,464209722 | -0,089833605 | -0,076358564 | 0,006980548 | -0,119019452 | -1,379019452 |
| chrXII | 463059 | 463216 | 157 | 0,681044979 | 0,496586524 | 0,578317562 | 0,197591256 | 0,167952568 | 0,085930515 | -0,005069485 | -1,265069485 |
| chrXII | 463059 | 463234 | 175 | 0,039893848 | 0,051882174 | 0,434687048 | -0,164453569 | -0,139785534 | 0,061637422 | -0,155362578 | -1,415362578 |
| chrXII | 463066 | 463216 | 150 | 0,037044288 | 0,103764348 | 0,263082498 | -0,633871027 | -0,538790373 | -0,651700408 | -0,693700408 | -1,953700408 |
| chrXII | 463066 | 463234 | 168 | 0,772230918 | 1,096937395 | 0,413141456 | -0,219471329 | -0,18655063 | -0,064294796 | -0,232294796 | -1,492294796 |
| chrXII | 463066 | 463245 | 179 | 0,085486818 | 0,144528914 | 0,371656395 | -0,327469522 | -0,278349094 | -0,116560895 | -0,361560895 | -1,621560895 |
| chrXII | 463096 | 463234 | 138 | 0,059840772 | 0,111176087 | 0,349911537 | -0,385559308 | -0,327725412 | -0,326826372 | -0,284826372 | -1,544826372 |
| chrXII | 463096 | 463245 | 149 | 0,467327936 | 0,277940218 | 0,627060117 | 0,324076964 | 0,27546542 | 0,105003009 | 0,070003009 | -1,189996991 |
| chrXII | 463096 | 463249 | 153 | 0,151026711 | 0,085235 | 0,639234814 | 0,356414232 | 0,302952097 | 0,173869539 | 0,110869539 | -1,149130461 |
| chrXII | 463097 | 463253 | 156 | 0,085486818 | 0,062999783 | 0,575720754 | 0,190958009 | 0,162314307 | 0,105933737 | 0,021933737 | -1,238066263 |
| chrXII | 463101 | 463234 | 133 | 0,108283302 | 0,081529131 | 0,570475287 | 0,177584338 | 0,150946687 | 0,149857161 | 0,226857161 | -1,033142839 |
| chrXII | 463101 | 463245 | 144 | 0,461628815 | 0,370586958 | 0,554698469 | 0,137541158 | 0,116909984 | 0,010841464 | 0,010841464 | -1,249158536 |
| chrXII | 463108 | 463245 | 137 | 0,062690333 | 0,048176305 | 0,56545715 | 0,164819952 | 0,140096959 | 0,224351152 | 0,273351152 | -0,986648848 |
| chrXII | 463108 | 463260 | 152 | 0,139628469 | 0,077823261 | 0,642112476 | 0,3641111 | 0,309494435 | 0,113817293 | 0,057817293 | -1,202182707 |
| chrXII | 463196 | 463353 | 157 | 0,062690333 | 0,044470435 | 0,585011979 | 0,214732295 | 0,182522451 | 0,104754633 | 0,013754633 | -1,246245367 |
| chrXII | 463248 | 463405 | 157 | 0,094035499 | 0,107470218 | 0,466664175 | -0,083658001 | -0,071109301 | -0,14445714 | -0,23545714 | -1,49545714 |
| chrXII | 463257 | 463404 | 147 | 0,065539893 | 0,059293913 | 0,525017182 | 0,062749932 | 0,053337442 | -0,075702689 | -0,096702689 | -1,356702689 |

TABLE 2

| Chrom | Start | End | Length | Section A | Section B | A/A+B | Z-score | Z * 0.85 | Phase correction | Length correction | ΔLK |
|---|---|---|---|---|---|---|---|---|---|---|---|
| chrXII | 463257 | 463414 | 157 | 0,09688506 | 0,096352609 | 0,50137771 | 0,003453413 | 0,002935401 | -0,072163998 | -0,163163998 | -1,423163998 |
| chrXII | 463261 | 463414 | 153 | 0,173823196 | 0,244587392 | 0,415436895 | -0,213581034 | -0,181543879 | -0,305062193 | -0,368062193 | -1,628062193 |
| chrXII | 463268 | 463414 | 146 | 0,208017923 | 0,140823044 | 0,596311622 | 0,243811574 | 0,207239838 | 0,044929881 | 0,030929881 | -1,229070119 |
| chrXII | 463278 | 463414 | 136 | 0,233663968 | 0,281646088 | 0,453443447 | -0,11696613 | -0,099421211 | -0,111356249 | -0,055356249 | -1,315356249 |
| chrXII | 463333 | 463483 | 150 | 0,119681545 | 0,077823261 | 0,605967759 | 0,268824834 | 0,228501109 | 0,117073577 | 0,075073577 | -1,184926423 |
| chrXII | 463333 | 463483 | 150 | 0,119681545 | 0,077823261 | 0,605967759 | 0,268824834 | 0,228501109 | 0,117073577 | 0,075073577 | -1,184926423 |
| chrXII | 463358 | 463511 | 153 | 0,031345166 | 0,066705652 | 0,319682862 | -0,468585806 | -0,398297935 | -0,521061972 | -0,584061972 | -1,844061972 |
| chrXII | 463358 | 463529 | 171 | 0,492973981 | 0,426175002 | 0,536337406 | 0,091210681 | 0,077529079 | 0,194017747 | 0,005017747 | -1,254982253 |
| chrXII | 463381 | 463529 | 148 | 0,065539893 | 0,037058696 | 0,638799168 | 0,355250862 | 0,301963233 | 0,122758989 | 0,094758989 | -1,165241011 |
| chrXII | 463381 | 463539 | 158 | 0,364743755 | 0,311293045 | 0,539532397 | 0,099255756 | 0,084367393 | 0,054257432 | -0,043742568 | -1,303742568 |
| chrXII | 463385 | 463511 | 126 | 0,455929693 | 0,652233046 | 0,411428464 | -0,223871969 | -0,190291174 | 0,048920659 | 0,174920659 | -1,085079341 |
| chrXII | 463385 | 463529 | 144 | 0,037044288 | 0,037058696 | 0,499902783 | -0,000243688 | -0,000207135 | -0,122706278 | -0,122706278 | -1,382706278 |
| chrXII | 463385 | 463535 | 150 | 0,384690679 | 0,296469566 | 0,564757972 | 0,163043664 | 0,138587115 | 0,017909114 | -0,024090886 | -1,284090886 |
| chrXII | 463385 | 463539 | 154 | 0,028495606 | 0,070411522 | 0,288104674 | -0,558930214 | -0,475090682 | -0,651963002 | -0,721963002 | -1,981963002 |
| chrXII | 463410 | 463573 | 163 | 0,381841118 | 0,359469349 | 0,515089338 | 0,037832385 | 0,032157528 | 0,046536916 | -0,086463084 | -1,346463084 |
| chrXII | 463414 | 463521 | 107 | 0,037044288 | 0,055588044 | 0,399906675 | -0,253588671 | -0,21555037 | 0,361423191 | 0,620423191 | -0,639576809 |
| chrXII | 463414 | 463535 | 121 | 0,039893848 | 0,040764565 | 0,49460244 | -0,013530088 | -0,011500575 | 0,382865359 | 0,543865359 | -0,716134641 |
| chrXII | 463414 | 463539 | 125 | 0,031345166 | 0,114881957 | 0,214359455 | -0,791385773 | -0,672677907 | -0,339607688 | -0,206607688 | -1,466607688 |
| chrXII | 463414 | 463575 | 161 | 0,339097709 | 0,211234566 | 0,616169039 | 0,295434579 | 0,251119392 | 0,241877373 | 0,122877373 | -1,137122627 |
| chrXII | 463414 | 463583 | 169 | 0,980248841 | 1,49717131 | 0,395673233 | -0,264562559 | -0,224878175 | -0,064229188 | -0,239229188 | -1,499229188 |
| chrXII | 463463 | 463603 | 140 | 0,054141651 | 0,059293913 | 0,477289917 | -0,056956516 | -0,048413039 | 0,006892165 | 0,034892165 | -1,225107835 |
| chrXII | 463471 | 463608 | 137 | 0,128230226 | 0,289057827 | 0,307294266 | -0,503534497 | -0,428004322 | -0,351700146 | -0,302700146 | -1,562700146 |
| chrXII | 463535 | 463688 | 153 | 0,062690333 | 0,077823261 | 0,446151373 | -0,135390993 | -0,115082344 | -0,232538451 | -0,295538451 | -1,555538451 |
| chrXII | 463535 | 463689 | 154 | 5,189049823 | 3,098106968 | 0,626155623 | 0,321688416 | 0,273435154 | 0,105846384 | 0,035846384 | -1,224153616 |
| chrXII | 463535 | 463690 | 155 | 0,054141651 | 0,062999783 | 0,462190442 | -0,094916836 | -0,080679311 | -0,175024304 | -0,252024304 | -1,512024304 |
| chrXII | 463535 | 463692 | 157 | 0,19661968 | 0,133411305 | 0,595761274 | 0,242390685 | 0,206032083 | 0,136886211 | 0,045886211 | -1,214113789 |
| chrXII | 463535 | 463704 | 169 | 0,159575393 | 0,151940653 | 0,512254168 | 0,030721476 | 0,026113255 | 0,191529162 | 0,016529162 | -1,243470838 |
| chrXII | 463535 | 463710 | 175 | 1,077133901 | 1,826993702 | 0,37089758 | -0,32947702 | -0,280055467 | -0,074768098 | -0,291768098 | -1,551768098 |
| chrXII | 463535 | 463712 | 177 | 0,076938136 | 0,074117392 | 0,509336779 | 0,023405971 | 0,019895076 | 0,167101448 | -0,063898552 | -1,323898552 |
| chrXII | 463541 | 463689 | 148 | 0,322000346 | 0,203822827 | 0,612373822 | 0,28551142 | 0,242684707 | 0,060246552 | 0,032246552 | -1,227753448 |
| chrXII | 463541 | 463692 | 151 | 0,994496644 | 0,511410002 | 0,660397274 | 0,413547605 | 0,351515464 | 0,211094203 | 0,162094203 | -1,097905797 |
| chrXII | 463559 | 463689 | 130 | 0,407487163 | 0,248293262 | 0,621377442 | 0,309100452 | 0,262735385 | 0,384086819 | 0,482086819 | -0,777913181 |
| chrXII | 463559 | 463704 | 145 | 1,185417203 | 0,633703698 | 0,651642891 | 0,389759738 | 0,331295778 | 0,160461549 | 0,153461549 | -1,106538451 |
| chrXII | 463559 | 463710 | 151 | 0,094035499 | 0,096352609 | 0,493914773 | -0,015253994 | -0,012965895 | -0,169491197 | -0,218491197 | -1,478491197 |
| chrXII | 463559 | 463712 | 153 | 0,663947616 | 0,392822175 | 0,628280276 | 0,327302042 | 0,278206735 | 0,164665235 | 0,101665235 | -1,158334765 |
| chrXII | 463658 | 463815 | 157 | 0,629752889 | 0,552174567 | 0,532818563 | 0,082356944 | 0,070003402 | 0,011918877 | -0,079081123 | -1,339081123 |
| chrXII | 463658 | 463852 | 194 | 0,028495606 | 0,040764565 | 0,411428464 | -0,223871969 | -0,190291174 | -0,121037165 | -0,471037165 | -1,731037165 |
| chrXII | 463658 | 463870 | 212 | 0,037044288 | 0,037058696 | 0,499902783 | -0,000243688 | -0,000207135 | 0,20878783 | -0,26721217 | -1,52721217 |
| chrXII | 463690 | 463849 | 159 | 0,615505086 | 0,500292393 | 0,551627959 | 0,129775456 | 0,110309137 | 0,080116289 | -0,024883711 | -1,284883711 |
| chrXII | 463690 | 463870 | 180 | 0,071239015 | 0,103764348 | 0,407072261 | -0,235082731 | -0,199820321 | -0,010182911 | -0,262182911 | -1,522182911 |
| chrXII | 463692 | 463840 | 148 | 0,760832676 | 0,463233697 | 0,621561618 | 0,309584739 | 0,263147028 | 0,082629095 | 0,054629095 | -1,205370905 |
| chrXII | 463697 | 463815 | 118 | 0,04844253 | 0,062999783 | 0,434687048 | -0,164453569 | -0,139785534 | 0,246592139 | 0,428592139 | -0,831407861 |
| chrXII | 463697 | 463849 | 152 | 0,384690679 | 0,307587175 | 0,555688264 | 0,14004622 | 0,119039287 | -0,071924403 | -0,127924403 | -1,387924403 |
| chrXII | 463697 | 463852 | 155 | 1,108479067 | 0,896820438 | 0,552774817 | 0,132675062 | 0,112773803 | 0,018093444 | -0,058906556 | -1,318906556 |
| chrXII | 463697 | 463861 | 164 | 0,088336378 | 0,066705652 | 0,569757619 | 0,175757115 | 0,149393547 | 0,140565794 | 0,000565794 | -1,259434206 |
| chrXII | 463697 | 463870 | 173 | 0,170973635 | 0,192705218 | 0,470122564 | -0,074961771 | -0,063717505 | 0,128162034 | -0,074837966 | -1,334837966 |
| chrXII | 463699 | 463849 | 150 | 0,199469241 | 0,103764348 | 0,657807209 | 0,406485945 | 0,345513053 | 0,213749184 | 0,171749184 | -1,088250816 |
| chrXII | 463707 | 463849 | 142 | 0,085486818 | 0,062999783 | 0,575720754 | 0,190958009 | 0,162314307 | 0,174465328 | 0,188465328 | -1,071534672 |
| chrXII | 463707 | 463852 | 145 | 0,054141651 | 0,062999783 | 0,462190442 | -0,094916836 | -0,080679311 | -0,260546399 | -0,267546399 | -1,527546399 |
| chrXII | 463707 | 463861 | 154 | 0,800726524 | 0,489174784 | 0,620765727 | 0,307492481 | 0,261368609 | 0,106308139 | 0,036308139 | -1,223691861 |
| chrXII | 463707 | 463870 | 163 | 0,139628469 | 0,103764348 | 0,573675388 | 0,185739279 | 0,157878387 | 0,14247458 | 0,00947458 | -1,25052542 |
| chrXII | 463713 | 463849 | 136 | 0,131079787 | 0,137117174 | 0,488744489 | -0,028217126 | -0,023984557 | -0,029972975 | 0,026027025 | -1,233972975 |
| chrXII | 463713 | 463852 | 139 | 0,085486818 | 0,125999566 | 0,404219015 | -0,242441566 | -0,206075331 | -0,240489107 | -0,205489107 | -1,465489107 |
| chrXII | 463713 | 463861 | 148 | 0,133929347 | 0,111176087 | 0,54641525 | 0,116609512 | 0,099118085 | -0,079426781 | -0,107426781 | -1,367426781 |
| chrXII | 463713 | 463870 | 157 | 1,963347242 | 1,608347397 | 0,549696276 | 0,124894026 | 0,106159922 | 0,04457675 | -0,04642325 | -1,30642325 |
| chrXII | 463713 | 463909 | 196 | 0,062690333 | 0,051882174 | 0,547167331 | 0,118507771 | 0,100731606 | 0,141226838 | -0,222773162 | -1,482773162 |
| chrXII | 463724 | 463883 | 159 | 0,116831984 | 0,066705652 | 0,636556002 | 0,349268164 | 0,296877939 | 0,271389819 | 0,166389819 | -1,093610181 |
| chrXII | 463729 | 463879 | 150 | 0,151026711 | 0,133411305 | 0,530965281 | 0,07769655 | 0,066042068 | -0,055274644 | -0,097274644 | -1,357274644 |
| chrXII | 463763 | 463849 | 86 | 0,039893848 | 0,037058696 | 0,518421434 | 0,046192108 | 0,039263292 | 0,412842017 | 0,818842017 | -0,441157983 |
| chrXII | 463763 | 463870 | 107 | 0,113982423 | 0,148234783 | 0,434687048 | -0,164453569 | -0,139785534 | 0,424999413 | 0,683999413 | -0,576000587 |
| chrXII | 463763 | 463909 | 146 | 1,077133901 | 0,689291742 | 0,609781626 | 0,278749922 | 0,236937434 | 0,075629071 | 0,061629071 | -1,198370929 |
| chrXII | 463860 | 463991 | 131 | 0,296354301 | 0,318704784 | 0,481830621 | -0,045559635 | -0,03872569 | 0,022539486 | 0,113539486 | -1,146460514 |
| chrXII | 463862 | 464000 | 138 | 1,031540931 | 0,778232612 | 0,56998343 | 0,176331979 | 0,149882182 | 0,153942811 | 0,195942811 | -1,064057189 |
| chrXII | 463862 | 464023 | 161 | 0,068389454 | 0,044470435 | 0,605967759 | 0,268824834 | 0,228501109 | 0,227137764 | 0,108137764 | -1,151862236 |
| chrXII | 463870 | 464023 | 153 | 0,205168362 | 0,218646305 | 0,484099249 | -0,039867831 | -0,033887657 | -0,158560325 | -0,221560325 | -1,481560325 |
| chrXII | 463878 | 464033 | 155 | 0,054141651 | 0,044470435 | 0,549036667 | 0,123227859 | 0,104743681 | 0,017045594 | -0,059954406 | -1,319954406 |
| chrXII | 463884 | 464023 | 139 | 0,361894194 | 0,244587392 | 0,596710934 | 0,244842828 | 0,208116404 | 0,170413131 | 0,205413131 | -1,054586869 |
| chrXII | 463974 | 464128 | 154 | 0,24506221 | 0,070411522 | 0,776807021 | 0,761453978 | 0,647235881 | 0,492674682 | 0,422674682 | -0,837325318 |
| chrXII | 463974 | 464128 | 154 | 0,24506221 | 0,070411522 | 0,776807021 | 0,761453978 | 0,647235881 | 0,492674682 | 0,422674682 | -0,837325318 |
| chrXII | 464001 | 464158 | 157 | 0,159575393 | 0,100058479 | 0,614617006 | 0,291373097 | 0,247667133 | 0,186177044 | 0,095177044 | -1,164822956 |
| chrXII | 464013 | 464160 | 147 | 0,729487509 | 0,43729261 | 0,625214209 | 0,319204319 | 0,271323671 | 0,142872019 | 0,121872019 | -1,138127981 |
| chrXII | 464013 | 464166 | 153 | 0,128230226 | 0,200116957 | 0,390532439 | -0,277931585 | -0,236241848 | -0,355143929 | -0,418143929 | -1,678143929 |
| chrXII | 464038 | 464166 | 128 | 0,119681545 | 0,248293262 | 0,325243855 | -0,453084759 | -0,385122045 | -0,143466261 | -0,031466261 | -1,291466261 |
| chrXII | 464038 | 464211 | 173 | 0,202318801 | 0,292763697 | 0,408656743 | -0,231001693 | -0,196351439 | 0,003503322 | -0,199496678 | -1,459496678 |
| chrXII | 464057 | 464224 | 167 | 0,091185939 | 0,129705435 | 0,412808961 | -0,220325165 | -0,18727639 | -0,050343759 | -0,211343759 | -1,471343759 |
| chrXII | 464069 | 464210 | 141 | 0,649699813 | 0,429880871 | 0,601807556 | 0,258028531 | 0,219324251 | 0,190157889 | 0,211157889 | -1,048842111 |
| chrXII | 464070 | 464160 | 90 | 0,185221438 | 0,359469349 | 0,340048781 | -0,412330002 | -0,350480502 | 0,096112046 | 0,474112046 | -0,785887954 |
| chrXII | 464070 | 464166 | 96 | 5,821652272 | 28,08678554 | 0,171687422 | -0,947518087 | -0,805390374 | -0,234013946 | 0,101986054 | -1,158013946 |
| chrXII | 464070 | 464210 | 140 | 0,042743409 | 0,040764565 | 0,51184823 | 0,029703476 | 0,025247955 | 0,088256082 | 0,116256082 | -1,143743918 |
| chrXII | 464070 | 464211 | 141 | 0,236513528 | 0,248293262 | 0,487851105 | -0,030457471 | -0,025888851 | -0,063584909 | -0,042584909 | -1,302584909 |
| chrXII | 464070 | 464213 | 143 | 0,624053768 | 0,452116089 | 0,579884081 | 0,201596939 | 0,171357398 | 0,026326478 | 0,033326478 | -1,226673522 |
| chrXII | 464101 | 464269 | 168 | 0,227964847 | 0,207528696 | 0,523463207 | 0,058847485 | 0,050020363 | 0,176699056 | 0,008699056 | -1,251300944 |
| chrXII | 464101 | 464269 | 168 | 0,227964847 | 0,207528696 | 0,523463207 | 0,058847485 | 0,050020363 | 0,176699056 | 0,008699056 | -1,251300944 |
| chrXII | 464132 | 464301 | 169 | 0,139628469 | 0,281646088 | 0,331442919 | -0,435932317 | -0,370542469 | -0,214425781 | -0,389425781 | -1,649425781 |
| chrXII | 464132 | 464301 | 169 | 0,139628469 | 0,281646088 | 0,331442919 | -0,435932317 | -0,370542469 | -0,214425781 | -0,389425781 | -1,649425781 |
| chrXII | 464137 | 464269 | 132 | 0,111132863 | 0,118587827 | 0,483773852 | -0,040684142 | -0,03458152 | 0,108072332 | 0,192072332 | -1,067927668 |
| chrXII | 464137 | 464269 | 132 | 0,111132863 | 0,118587827 | 0,483773852 | -0,040684142 | -0,03458152 | 0,108072332 | 0,192072332 | -1,067927668 |
| chrXII | 464152 | 464303 | 151 | 0,498673102 | 0,307587175 | 0,61850139 | 0,301547257 | 0,256315168 | 0,091828708 | 0,042828708 | -1,217171292 |
| chrXII | 464195 | 464345 | 150 | 0,037044288 | 0,059293913 | 0,384523349 | -0,293622086 | -0,249578773 | -0,369626346 | -0,411626346 | -1,671626346 |
| chrXII | 464205 | 464360 | 155 | 0,153876272 | 0,111176087 | 0,58055047 | 0,203301913 | 0,172806626 | 0,089422092 | 0,012422092 | -1,247577908 |
| chrXII | 464232 | 464406 | 174 | 0,102584181 | 0,144528914 | 0,415130494 | -0,214366854 | -0,182211826 | 0,051618736 | -0,158381264 | -1,418381264 |
| chrXII | 464232 | 464406 | 174 | 0,102584181 | 0,144528914 | 0,415130494 | -0,214366854 | -0,182211826 | 0,051618736 | -0,158381264 | -1,418381264 |
| chrXII | 464233 | 464360 | 127 | 0,034194727 | 0,074117392 | 0,315705458 | -0,47974192 | -0,407780632 | -0,158187093 | -0,039187093 | -1,299187093 |
| chrXII | 464233 | 464377 | 144 | 0,994496644 | 0,604056741 | 0,622122885 | 0,311061025 | 0,264401871 | 0,140105817 | 0,140105817 | -1,119894183 |
| chrXII | 464233 | 464380 | 147 | 0,136778908 | 0,077823261 | 0,637360324 | 0,351411905 | 0,298700119 | 0,161150912 | 0,140150912 | -1,119849088 |
| chrXII | 464233 | 464382 | 149 | 0,994496644 | 0,481763045 | 0,67365969 | 0,450041356 | 0,382535153 | 0,201669305 | 0,166669305 | -1,093330695 |
| chrXII | 464233 | 464386 | 153 | 0,059840772 | 0,040764565 | 0,594807131 | 0,239928434 | 0,203939169 | 0,082924825 | 0,019924825 | -1,240075175 |
| chrXII | 464234 | 464377 | 143 | 0,045592969 | 0,062999783 | 0,419852784 | -0,202270107 | -0,171929591 | -0,329792225 | -0,322792225 | -1,582792225 |
| chrXII | 464234 | 464380 | 146 | 0,450230572 | 0,333528262 | 0,574450395 | 0,187716099 | 0,159558684 | -0,00563756 | -0,01963756 | -1,27963756 |
| chrXII | 464234 | 464382 | 148 | 0,065539893 | 0,044470435 | 0,595761274 | 0,242390685 | 0,206032083 | 0,036630998 | 0,008630998 | -1,251369002 |
| chrXII | 464235 | 464377 | 142 | 0,042743409 | 0,040764565 | 0,51184823 | 0,029703476 | 0,025247955 | 0,020116824 | 0,034116824 | -1,225883176 |

TABLE 2

| Chrom | Start | End | Length | Section A | Section B | A/A+B | Z-score | Z * 0.85 | Phase correction | Length correction | ΔLK |
|---|---|---|---|---|---|---|---|---|---|---|---|
| chrXII | 464235 | 464380 | 145 | 0,042743409 | 0,040764565 | 0,51184823 | 0,029703476 | 0,025247955 | -0,136622677 | -0,143622677 | -1,403622677 |
| chrXII | 464235 | 464382 | 147 | 0,031345166 | 0,037058696 | 0,458236793 | -0,104876779 | -0,089145262 | -0,22202983 | -0,24302983 | -1,50302983 |
| chrXII | 464235 | 464386 | 151 | 0,227964847 | 0,118587827 | 0,657807209 | 0,406485945 | 0,345513053 | 0,172357329 | 0,123357329 | -1,136642671 |
| chrXII | 464266 | 464430 | 164 | 0,04844253 | 0,037058696 | 0,566571175 | 0,16765125 | 0,142503563 | 0,131004069 | -0,008995931 | -1,268995931 |
| chrXII | 464266 | 464430 | 164 | 0,04844253 | 0,037058696 | 0,566571175 | 0,16765125 | 0,142503563 | 0,131004069 | -0,008995931 | -1,268995931 |
| chrXII | 464275 | 464426 | 151 | 0,079787696 | 0,059293913 | 0,573675388 | 0,185739279 | 0,157878387 | -0,014866282 | -0,063866282 | -1,323866282 |
| chrXII | 464275 | 464426 | 151 | 0,079787696 | 0,059293913 | 0,573675388 | 0,185739279 | 0,157878387 | -0,014866282 | -0,063866282 | -1,323866282 |
| chrXII | 464319 | 464475 | 156 | 0,034194727 | 0,059293913 | 0,365763444 | -0,34309514 | -0,291630869 | -0,345774551 | -0,429774551 | -1,689774551 |
| chrXII | 464354 | 464503 | 149 | 0,045592969 | 0,044470435 | 0,506231912 | 0,015621722 | 0,013278464 | -0,164627491 | -0,199627491 | -1,459627491 |
| chrXII | 464358 | 464532 | 174 | 0,045592969 | 0,129705435 | 0,26008776 | -0,64307487 | -0,54661364 | -0,304508688 | -0,514508688 | -1,774508688 |
| chrXII | 464384 | 464518 | 134 | 0,068389454 | 0,163058261 | 0,295485544 | -0,537429332 | -0,456814932 | -0,445147555 | -0,375147555 | -1,635147555 |
| chrXII | 464391 | 464538 | 147 | 0,153876272 | 0,148234783 | 0,509336779 | 0,023405971 | 0,019895076 | -0,121020385 | -0,142020385 | -1,402020385 |
| chrXII | 464392 | 464537 | 145 | 0,051292091 | 0,074117392 | 0,408996909 | -0,230126057 | -0,195607148 | -0,363022925 | -0,370022925 | -1,630022925 |
| chrXII | 464415 | 464560 | 145 | 0,059840772 | 0,051882174 | 0,535617563 | 0,08939893 | 0,07598909 | -0,093717059 | -0,100717059 | -1,360717059 |
| chrXII | 464434 | 464584 | 150 | 0,367593315 | 0,281646088 | 0,566190705 | 0,166684133 | 0,141681513 | 0,021782757 | -0,020217243 | -1,280217243 |
| chrXII | 464504 | 464649 | 145 | 1,473222822 | 0,800467829 | 0,647943387 | 0,379773944 | 0,322807852 | 0,165095809 | 0,158095809 | -1,101904191 |
| chrXII | 464504 | 464668 | 164 | 0,108283302 | 0,066705652 | 0,618800783 | 0,302332726 | 0,256982817 | 0,269489425 | 0,129489425 | -1,130510575 |
| chrXII | 464504 | 464686 | 182 | 0,19377012 | 0,185293479 | 0,511181027 | 0,028030349 | 0,023825797 | 0,211022954 | -0,054977046 | -1,314977046 |
| chrXII | 464514 | 464649 | 135 | 0,076938136 | 0,066705652 | 0,535617563 | 0,08939893 | 0,07598909 | 0,046761185 | 0,109761185 | -1,150238815 |
| chrXII | 464514 | 464668 | 154 | 0,547115632 | 0,307587175 | 0,640123827 | 0,358789798 | 0,304971328 | 0,154200067 | 0,084200067 | -1,175799933 |
| chrXII | 464514 | 464686 | 172 | 0,105433742 | 0,081529131 | 0,563928764 | 0,160937697 | 0,136797043 | 0,253973888 | 0,057973888 | -1,202026112 |
| chrXII | 464534 | 464640 | 106 | 0,076938136 | 0,118587827 | 0,393493196 | -0,270226 | -0,2296921 | 0,283811914 | 0,549811914 | -0,710188086 |
| chrXII | 464534 | 464649 | 115 | 0,09973462 | 0,100058479 | 0,499189516 | -0,002031583 | -0,001726846 | 0,604326493 | 0,807326493 | -0,452673507 |
| chrXII | 464534 | 464668 | 134 | 0,125380666 | 0,044470435 | 0,738179884 | 0,637744162 | 0,542082538 | 0,556066499 | 0,626066499 | -0,633933501 |
| chrXII | 464534 | 464686 | 152 | 1,780975365 | 1,348936527 | 0,569017732 | 0,173873936 | 0,147792846 | -0,023918936 | -0,079918936 | -1,339918936 |
| chrXII | 464547 | 464649 | 102 | 0,068389454 | 0,074117392 | 0,479902939 | -0,050397187 | -0,042837609 | 0,490360132 | 0,784360132 | -0,475639868 |
| chrXII | 464547 | 464686 | 139 | 0,125380666 | 0,17788174 | 0,413439528 | -0,218706025 | -0,185900121 | -0,186086837 | -0,151086837 | -1,411086837 |
| chrXII | 464547 | 464697 | 150 | 0,168124074 | 0,037058696 | 0,819386902 | 0,913031079 | 0,776076417 | 0,653505093 | 0,611505093 | -0,648494907 |
| chrXII | 464547 | 464705 | 158 | 0,530018269 | 0,366881088 | 0,590945087 | 0,229976764 | 0,195480249 | 0,169376284 | 0,071376284 | -1,188623716 |
| chrXII | 464547 | 464706 | 159 | 0,279256937 | 0,333528262 | 0,455717497 | -0,111228699 | -0,094544395 | -0,114829738 | -0,219829738 | -1,479829738 |
| chrXII | 464553 | 464708 | 155 | 0,131079787 | 0,100058479 | 0,567105523 | 0,169009772 | 0,143658307 | 0,069140507 | -0,007859493 | -1,267859493 |
| chrXII | 464559 | 464706 | 147 | 1,042939174 | 0,489174784 | 0,68071906 | 0,46971048 | 0,399253908 | 0,254471556 | 0,233471556 | -1,026528444 |
| chrXII | 464728 | 464888 | 160 | 0,045592969 | 0,096352609 | 0,321200349 | -0,464344846 | -0,394693119 | -0,385799016 | -0,497799016 | -1,757799016 |
| chrXII | 464838 | 465007 | 169 | 0,059840772 | 0,122293696 | 0,328552705 | -0,443913104 | -0,377326138 | -0,21329069 | -0,38829069 | -1,64829069 |
| chrXII | 464884 | 465032 | 148 | 0,795027403 | 0,667056524 | 0,543763178 | 0,109918963 | 0,093431118 | -0,070902718 | -0,098902718 | -1,358902718 |
| chrXII | 464886 | 465026 | 140 | 0,324849907 | 0,26682261 | 0,549036667 | 0,123227859 | 0,104743681 | 0,16863814 | 0,19663814 | -1,06336186 |
| chrXII | 464920 | 465095 | 175 | 0,031345166 | 0,085235 | 0,268872204 | -0,616227458 | -0,523793339 | -0,31408638 | -0,53108638 | -1,79108638 |
| chrXII | 465006 | 465158 | 152 | 0,082637257 | 0,070411522 | 0,539940648 | 0,100284197 | 0,085241567 | -0,083748791 | -0,139748791 | -1,399748791 |
| chrXII | 465040 | 465204 | 164 | 0,056991212 | 0,100058479 | 0,362886495 | -0,350753893 | -0,298140809 | -0,288477357 | -0,428477357 | -1,688477357 |
| chrXII | 465055 | 465176 | 121 | 0,256460453 | 0,922761525 | 0,217482762 | -0,780722841 | -0,663614415 | -0,251886964 | -0,090886964 | -1,350886964 |
| chrXII | 465063 | 465223 | 160 | 0,276407377 | 0,26311674 | 0,512316999 | 0,030879046 | 0,026247189 | 0,02962321 | -0,08237679 | -1,34237679 |
| chrXII | 465159 | 465310 | 151 | 2,105825271 | 1,252583918 | 0,627030583 | 0,323998943 | 0,275399102 | 0,096523512 | 0,047523512 | -1,212476488 |
| chrXII | 465159 | 465342 | 183 | 0,230814407 | 0,170470001 | 0,575189075 | 0,18960094 | 0,161160799 | 0,213304016 | -0,059695984 | -1,319695984 |
| chrXII | 465159 | 465342 | 183 | 0,230814407 | 0,170470001 | 0,575189075 | 0,18960094 | 0,161160799 | 0,213304016 | -0,059695984 | -1,319695984 |
| chrXII | 465159 | 465348 | 189 | 0,091185939 | 0,070411522 | 0,564278289 | 0,16182531 | 0,137551513 | 0,179653649 | -0,135346351 | -1,395346351 |
| chrXII | 465159 | 465348 | 189 | 0,091185939 | 0,070411522 | 0,564278289 | 0,16182531 | 0,137551513 | 0,179653649 | -0,135346351 | -1,395346351 |
| chrXII | 465159 | 465357 | 198 | 0,082637257 | 0,044470435 | 0,650135768 | 0,385687039 | 0,327833983 | 0,372884471 | -0,005115529 | -1,265115529 |
| chrXII | 465159 | 465357 | 198 | 0,082637257 | 0,044470435 | 0,650135768 | 0,385687039 | 0,327833983 | 0,372884471 | -0,005115529 | -1,265115529 |
| chrXII | 465179 | 465310 | 131 | 0,085486818 | 0,059293913 | 0,590457149 | 0,22872109 | 0,194412926 | 0,262547175 | 0,353547175 | -0,906452825 |
| chrXII | 465182 | 465312 | 130 | 0,339097709 | 0,374292827 | 0,475332503 | -0,061871698 | -0,052590944 | 0,072684647 | 0,170684647 | -1,089315353 |
| chrXII | 465193 | 465310 | 117 | 0,467327936 | 0,555880437 | 0,456728022 | -0,108680331 | -0,092378281 | 0,42473058 | 0,61373058 | -0,64626942 |
| chrXII | 465193 | 465312 | 119 | 0,113982423 | 0,096352609 | 0,541908887 | 0,105243965 | 0,08945737 | 0,528840803 | 0,703840803 | -0,556159197 |
| chrXII | 465193 | 465342 | 149 | 4,525102207 | 3,16481262 | 0,588446336 | 0,223550187 | 0,190017659 | 0,021160812 | -0,013839188 | -1,273839188 |
| chrXII | 465193 | 465348 | 155 | 0,302053422 | 0,226058044 | 0,57195013 | 0,181341238 | 0,154140052 | 0,083483414 | 0,006483414 | -1,253516586 |
| chrXII | 465193 | 465357 | 164 | 0,333398588 | 0,26311674 | 0,558910346 | 0,148207125 | 0,125976056 | 0,151832088 | 0,011832088 | -1,248167912 |
| chrXII | 465200 | 465352 | 152 | 0,319150785 | 0,214940436 | 0,597558568 | 0,24703277 | 0,209977854 | 0,049201235 | -0,006798765 | -1,266798765 |
| chrXII | 465201 | 465342 | 141 | 0,102584181 | 0,040764565 | 0,715626635 | 0,569898223 | 0,48441349 | 0,425961554 | 0,446961554 | -0,813038446 |
| chrXII | 465201 | 465345 | 144 | 0,105433742 | 0,044470435 | 0,703340921 | 0,534033788 | 0,453928719 | 0,309666143 | 0,309666143 | -0,950333857 |
| chrXII | 465201 | 465351 | 150 | 0,102584181 | 0,048176305 | 0,680444751 | 0,468942833 | 0,398601408 | 0,266825013 | 0,224825013 | -1,035174987 |
| chrXII | 465203 | 465310 | 107 | 0,14532759 | 0,111176087 | 0,566571175 | 0,16765125 | 0,142503563 | 0,687244311 | 0,946244311 | -0,313755689 |
| chrXII | 465204 | 465342 | 138 | 0,333398588 | 0,318704784 | 0,511266469 | 0,028244604 | 0,024007913 | 0,011373325 | 0,053373325 | -1,206626675 |
| chrXII | 465204 | 465348 | 144 | 1,803771849 | 0,830114786 | 0,684832758 | 0,481256115 | 0,409067697 | 0,249920606 | 0,249920606 | -1,010079394 |
| chrXII | 465204 | 465352 | 148 | 1,607152169 | 1,156231309 | 0,581588542 | 0,205959029 | 0,175065175 | 0,026139418 | -0,001860582 | -1,261860582 |
| chrXII | 465204 | 465357 | 153 | 0,142478029 | 0,066705652 | 0,681114454 | 0,470817466 | 0,400194846 | 0,286500968 | 0,223500968 | -1,036499032 |
| chrXII | 465204 | 465403 | 199 | 0,179522317 | 0,203822827 | 0,468304659 | -0,079532203 | -0,067602373 | -0,03584809 | -0,42084809 | -1,68084809 |
| chrXII | 465204 | 465418 | 214 | 0,031345166 | 0,059293913 | 0,345823971 | -0,396619276 | -0,337126724 | -0,02160931 | -0,51160931 | -1,77160931 |
| chrXII | 465217 | 465310 | 93 | 0,253610892 | 0,207528696 | 0,54996556 | 0,125574335 | 0,106738185 | 0,55334545 | 0,91034545 | -0,34965455 |
| chrXII | 465217 | 465342 | 125 | 0,433133209 | 0,400233914 | 0,519738776 | 0,049497979 | 0,042073282 | 0,36843055 | 0,50143055 | -0,75856945 |
| chrXII | 465217 | 465348 | 131 | 0,19377012 | 0,137117174 | 0,585607617 | 0,216260411 | 0,183821349 | 0,259771079 | 0,350771079 | -0,909228921 |
| chrXII | 465217 | 465357 | 140 | 2,313843194 | 1,378583483 | 0,626645671 | 0,322982286 | 0,274534943 | 0,344765973 | 0,372765973 | -0,887234027 |
| chrXII | 465221 | 465352 | 131 | 0,082637257 | 0,044470435 | 0,650135768 | 0,385687039 | 0,327833983 | 0,40290448 | 0,49390448 | -0,76609552 |
| chrXII | 465221 | 465375 | 154 | 0,071239015 | 0,055588044 | 0,561702018 | 0,155285859 | 0,13199298 | -0,020621696 | -0,090621696 | -1,350621696 |
| chrXII | 465221 | 465380 | 159 | 0,282106498 | 0,270528479 | 0,510475286 | 0,026260665 | 0,022321566 | 0,011408144 | -0,093591856 | -1,353591856 |
| chrXII | 465221 | 465403 | 182 | 0,065539893 | 0,133411305 | 0,329426985 | -0,441495981 | -0,375271584 | -0,18169705 | -0,44769705 | -1,70769705 |
| chrXII | 465252 | 465352 | 100 | 0,199469241 | 0,240881523 | 0,452978074 | -0,118140734 | -0,100419624 | 0,373758045 | 0,681758045 | -0,578241955 |
| chrXII | 465252 | 465380 | 128 | 0,108283302 | 0,103764348 | 0,510655515 | 0,026712592 | 0,022705704 | 0,259567845 | 0,371567845 | -0,888432155 |
| chrXII | 465252 | 465403 | 151 | 6,878839249 | 4,243220667 | 0,618486081 | 0,301507098 | 0,256281033 | 0,089441044 | 0,040441044 | -1,219558956 |
| chrXII | 465252 | 465418 | 166 | 0,424584527 | 0,585527393 | 0,420334142 | -0,201038739 | -0,170882928 | -0,027285825 | -0,181285825 | -1,441285825 |
| chrXII | 465257 | 465427 | 170 | 0,034194727 | 0,037058696 | 0,479902939 | -0,050397187 | -0,042837609 | 0,125044255 | -0,056955745 | -1,316955745 |
| chrXII | 465261 | 465352 | 91 | 0,056991212 | 0,081529131 | 0,411428464 | -0,223871969 | -0,190291174 | 0,170399014 | 0,541399014 | -0,718600986 |
| chrXII | 465261 | 465403 | 142 | 0,504372223 | 0,644821307 | 0,438892328 | -0,153778159 | -0,130711435 | -0,135703957 | -0,121703957 | -1,381703957 |
| chrXII | 465261 | 465418 | 157 | 1,478921943 | 1,348936527 | 0,522983013 | 0,057641774 | 0,048995508 | -0,009370725 | -0,100370725 | -1,360370725 |
| chrXII | 465261 | 465470 | 209 | 0,276407377 | 0,281646088 | 0,495306264 | -0,011765723 | -0,010000865 | 0,192675678 | -0,262324322 | -1,522324322 |
| chrXII | 465318 | 465403 | 85 | 1,031540931 | 1,382289353 | 0,427346089 | -0,183134893 | -0,155664659 | 0,239781386 | 0,652781386 | -0,607218614 |
| chrXII | 465318 | 465418 | 100 | 0,552814753 | 0,830114786 | 0,399741807 | -0,25401546 | -0,215913141 | 0,259626447 | 0,567626447 | -0,692373553 |
| chrXII | 465318 | 465447 | 129 | 1,373488201 | 1,801052615 | 0,432657282 | -0,169612916 | -0,144170979 | 0,088834611 | 0,193834611 | -1,066165389 |
| chrXII | 465318 | 465469 | 151 | 0,051292091 | 0,085235 | 0,375691668 | -0,31681585 | -0,269293473 | -0,436366324 | -0,485366324 | -1,745366324 |
| chrXII | 465318 | 465470 | 152 | 15,1169189 | 10,99160917 | 0,579003108 | 0,199343845 | 0,169442268 | 0,011466675 | -0,044533325 | -1,304533325 |
| chrXII | 465318 | 465473 | 155 | 0,259310013 | 0,185293479 | 0,583238813 | 0,21018617 | 0,178658244 | 0,123109539 | 0,046109539 | -1,213890461 |
| chrXII | 465318 | 465486 | 168 | 0,361894194 | 0,35205761 | 0,506888829 | 0,017268592 | 0,014678304 | 0,126997767 | -0,041002233 | -1,301002233 |
| chrXII | 465318 | 465524 | 206 | 0,202318801 | 0,203822827 | 0,498148398 | -0,004641295 | -0,003945101 | 0,094123566 | -0,339876434 | -1,599876434 |
| chrXII | 465318 | 465524 | 206 | 0,202318801 | 0,203822827 | 0,498148398 | -0,004641295 | -0,003945101 | 0,094123566 | -0,339876434 | -1,599876434 |
| chrXII | 465323 | 465470 | 147 | 0,031345166 | 0,048176305 | 0,394172367 | -0,268460684 | -0,228191582 | -0,333559164 | -0,354559164 | -1,614559164 |
| chrXII | 465324 | 465470 | 146 | 0,501522663 | 0,696703481 | 0,418554265 | -0,205593405 | -0,174754394 | -0,34265492 | -0,35665492 | -1,61665492 |
| chrXII | 465324 | 465486 | 162 | 0,498673102 | 0,422469132 | 0,541363846 | 0,103870259 | 0,088289721 | 0,169370763 | 0,043370763 | -1,216629237 |
| chrXII | 465331 | 465470 | 139 | 0,347646391 | 0,411351523 | 0,458033395 | -0,105389448 | -0,08958103 | -0,086337704 | -0,051337704 | -1,311337704 |
| chrXII | 465331 | 465473 | 142 | 0,421734966 | 0,370586958 | 0,53227729 | 0,080995639 | 0,068846293 | 0,056548177 | 0,070548177 | -1,189451823 |
| chrXII | 465350 | 465456 | 106 | 4,080570756 | 8,290030247 | 0,329860348 | -0,440298818 | -0,374253995 | 0,15690388 | 0,42290388 | -0,83709612 |
| chrXII | 465350 | 465457 | 107 | 0,111132863 | 0,229763914 | 0,326001507 | -0,450981319 | -0,383334121 | 0,148207884 | 0,407207884 | -0,852792116 |
| chrXII | 465350 | 465458 | 108 | 0,065539893 | 0,240881523 | 0,213888097 | -0,793002797 | -0,674052378 | -0,042623833 | 0,209376167 | -1,050623833 |
| chrXII | 465350 | 465470 | 120 | 4,234447027 | 11,00643265 | 0,277834818 | -0,589285701 | -0,500892846 | -0,137149517 | 0,030850483 | -1,229149517 |

TABLE 2

| Chrom | Start | End | Length | Section A | Section B | A/A+B | Z-score | Z * 0.85 | Phase correction | Length correction | ΔLK |
|---|---|---|---|---|---|---|---|---|---|---|---|
| chrXII | 465350 | 465473 | 123 | 0,04844253 | 0,133411305 | 0,266381679 | -0,623793272 | -0,530224281 | -0,080061194 | 0,066938806 | -1,193061194 |
| chrXII | 465350 | 465486 | 136 | 0,133929347 | 0,196411088 | 0,405428259 | -0,239321213 | -0,203423031 | -0,198680295 | -0,142680295 | -1,402680295 |
| chrXII | 465350 | 465524 | 174 | 0,119681545 | 0,185293479 | 0,392430643 | -0,272989502 | -0,232041076 | 0,009739437 | -0,200260563 | -1,460260563 |
| chrXII | 465350 | 465524 | 174 | 0,119681545 | 0,185293479 | 0,392430643 | -0,272989502 | -0,232041076 | 0,009739437 | -0,200260563 | -1,460260563 |
| chrXII | 465350 | 465556 | 206 | 0,14532759 | 0,226058044 | 0,391311824 | -0,275901597 | -0,234516357 | -0,111916871 | -0,545916871 | -1,805916871 |
| chrXII | 465350 | 465556 | 206 | 0,14532759 | 0,226058044 | 0,391311824 | -0,275901597 | -0,234516357 | -0,111916871 | -0,545916871 | -1,805916871 |
| chrXII | 465358 | 465523 | 165 | 0,646850252 | 0,366881088 | 0,638088443 | 0,353353936 | 0,300350846 | 0,399853267 | 0,252853267 | -1,007146733 |
| chrXII | 465358 | 465523 | 165 | 0,646850252 | 0,366881088 | 0,638088443 | 0,353353936 | 0,300350846 | 0,399853267 | 0,252853267 | -1,007146733 |
| chrXII | 465377 | 465470 | 93 | 0,401788042 | 0,518821741 | 0,436436859 | -0,160009338 | -0,136007937 | 0,333946502 | 0,690946502 | -0,569053498 |
| chrXII | 465377 | 465484 | 107 | 0,045592969 | 0,100058479 | 0,313027917 | -0,487285766 | -0,414192901 | 0,122878606 | 0,381878606 | -0,878121394 |
| chrXII | 465377 | 465523 | 146 | 0,29065518 | 0,140823044 | 0,673626532 | 0,449949388 | 0,38245698 | 0,234438031 | 0,220438031 | -1,039561969 |
| chrXII | 465377 | 465523 | 146 | 0,29065518 | 0,140823044 | 0,673626532 | 0,449949388 | 0,38245698 | 0,234438031 | 0,220438031 | -1,039561969 |
| chrXII | 465377 | 465524 | 147 | 2,259701543 | 1,589818049 | 0,587008713 | 0,219856939 | 0,186878398 | 0,070833286 | 0,049833286 | -1,210166714 |
| chrXII | 465377 | 465524 | 147 | 2,259701543 | 1,589818049 | 0,587008713 | 0,219856939 | 0,186878398 | 0,070833286 | 0,049833286 | -1,210166714 |
| chrXII | 465377 | 465556 | 179 | 0,282106498 | 0,277940218 | 0,503719582 | 0,009323745 | 0,007925183 | 0,164647713 | -0,080352287 | -1,340352287 |
| chrXII | 465377 | 465556 | 179 | 0,282106498 | 0,277940218 | 0,503719582 | 0,009323745 | 0,007925183 | 0,164647713 | -0,080352287 | -1,340352287 |
| chrXII | 465377 | 465584 | 207 | 0,102584181 | 0,140823044 | 0,421450846 | -0,198183249 | -0,168455762 | -0,011269214 | -0,452269214 | -1,712269214 |
| chrXII | 465377 | 465584 | 207 | 0,102584181 | 0,140823044 | 0,421450846 | -0,198183249 | -0,168455762 | -0,011269214 | -0,452269214 | -1,712269214 |
| chrXII | 465425 | 465524 | 99 | 0,330549028 | 0,452116089 | 0,422337754 | -0,195916517 | -0,16652904 | 0,294999837 | 0,609999837 | -0,650000163 |
| chrXII | 465425 | 465532 | 107 | 0,051292091 | 0,237175653 | 0,177808756 | -0,923748045 | -0,785185838 | -0,237452803 | 0,021547197 | -1,238452803 |
| chrXII | 465425 | 465556 | 131 | 6,337422738 | 6,062802631 | 0,51107319 | 0,027759937 | 0,023595946 | 0,093092114 | 0,184092114 | -1,075907886 |
| chrXII | 465425 | 465584 | 159 | 2,513312435 | 1,808464354 | 0,5815461 | 0,205850361 | 0,174972807 | 0,1704375 | 0,0654375 | -1,1945625 |
| chrXII | 465425 | 465594 | 169 | 0,173823196 | 0,211234566 | 0,451421093 | -0,122071761 | -0,103760997 | 0,063978312 | -0,111021688 | -1,371021688 |
| chrXII | 465425 | 465608 | 183 | 0,455929693 | 0,952408482 | 0,323735947 | -0,45727709 | -0,388685526 | -0,344009901 | -0,617009901 | -1,877009901 |
| chrXII | 465427 | 465524 | 97 | 0,039893848 | 0,077823261 | 0,338895921 | -0,415478239 | -0,353156503 | 0,20057533 | 0,52957533 | -0,73042467 |
| chrXII | 465427 | 465556 | 129 | 0,159575393 | 0,277940218 | 0,36473074 | -0,345841972 | -0,293965676 | -0,088939783 | 0,016060217 | -1,243939783 |
| chrXII | 465427 | 465584 | 157 | 0,615505086 | 0,956114351 | 0,391637486 | -0,275053712 | -0,233795655 | -0,289094 | -0,380094 | -1,640094 |
| chrXII | 465427 | 465608 | 181 | 0,065539893 | 0,248293262 | 0,208836741 | -0,810464113 | -0,688894496 | -0,510628688 | -0,769628688 | -2,029628688 |
| chrXII | 465435 | 465584 | 149 | 0,97739928 | 0,696703481 | 0,583834698 | 0,211713438 | 0,179956422 | 0,011371093 | -0,023628907 | -1,283628907 |
| chrXII | 465447 | 465556 | 109 | 0,062690333 | 0,129705435 | 0,325840498 | -0,451428154 | -0,383713931 | 0,289572067 | 0,534572067 | -0,725427933 |
| chrXII | 465447 | 465584 | 137 | 0,102584181 | 0,096352609 | 0,515662191 | 0,039269381 | 0,033378974 | 0,104905919 | 0,153905919 | -1,106094081 |
| chrXII | 465447 | 465608 | 161 | 0,569912117 | 0,61146848 | 0,482411949 | -0,044100996 | -0,037485847 | -0,040102613 | -0,159102613 | -1,419102613 |
| chrXII | 465498 | 465584 | 86 | 0,182371877 | 0,667056524 | 0,214699529 | -0,790220412 | -0,67168735 | -0,287204061 | 0,118795939 | -1,141204061 |
| chrXII | 465498 | 465603 | 105 | 0,028495606 | 0,337234132 | 0,07791438 | -1,419241021 | -1,206354868 | -0,639306636 | -0,366306636 | -1,626306636 |
| chrXII | 465498 | 465605 | 107 | 1,099930385 | 8,152913073 | 0,118874851 | -1,180630098 | -1,003535583 | -0,44340594 | -0,18440594 | -1,44440594 |
| chrXII | 465498 | 465606 | 108 | 0,028495606 | 0,166764131 | 0,145936926 | -1,054019799 | -0,895916829 | -0,274178349 | -0,022178349 | -1,282178349 |
| chrXII | 465498 | 465607 | 109 | 0,037044288 | 0,270528479 | 0,120440727 | -1,172786449 | -0,996868482 | -0,30673857 | -0,06173857 | -1,32173857 |
| chrXII | 465498 | 465608 | 110 | 1,384886444 | 14,22683331 | 0,088708129 | -1,348753182 | -1,146440205 | -0,488691085 | -0,250691085 | -1,510691085 |
| chrXII | 465525 | 465608 | 83 | 0,074088575 | 0,211234566 | 0,259665497 | -0,644377003 | -0,547720452 | -0,216290733 | 0,210709267 | -1,049290733 |
| chrXII | 465525 | 465688 | 163 | 0,29065518 | 0,318704784 | 0,476984372 | -0,057723664 | -0,049065114 | -0,055472887 | -0,188472887 | -1,448472887 |
| chrXII | 465525 | 465688 | 163 | 0,29065518 | 0,318704784 | 0,476984372 | -0,057723664 | -0,049065114 | -0,055472887 | -0,188472887 | -1,448472887 |
| chrXII | 465562 | 465739 | 177 | 0,042743409 | 0,062999783 | 0,404219015 | -0,242441566 | -0,206075331 | -0,05286528 | -0,28386528 | -1,54386528 |
| chrXII | 465562 | 465739 | 177 | 0,042743409 | 0,062999783 | 0,404219015 | -0,242441566 | -0,206075331 | -0,05286528 | -0,28386528 | -1,54386528 |
| chrXII | 465581 | 465731 | 150 | 0,433133209 | 0,318704784 | 0,576099124 | 0,191923981 | 0,163135384 | 0,020419368 | -0,021580632 | -1,281580632 |
| chrXII | 465581 | 465732 | 151 | 0,188070999 | 0,144528914 | 0,56545715 | 0,164819952 | 0,140096959 | -0,019742751 | -0,068742751 | -1,328742751 |
| chrXII | 465581 | 465756 | 175 | 0,059840772 | 0,111176087 | 0,349911537 | -0,385559308 | -0,327725412 | -0,133495994 | -0,350495994 | -1,610495994 |
| chrXII | 465582 | 465732 | 150 | 0,09973462 | 0,085235 | 0,539194599 | 0,098404877 | 0,083644145 | -0,049958361 | -0,091958361 | -1,351958361 |
| chrXII | 465584 | 465731 | 147 | 1,014443568 | 0,952408482 | 0,515770146 | 0,039540193 | 0,033609164 | -0,075467928 | -0,096467928 | -1,356467928 |
| chrXII | 465584 | 465732 | 148 | 0,09973462 | 0,114881957 | 0,464710702 | -0,088572826 | -0,075286902 | -0,228457509 | -0,256457509 | -1,516457509 |
| chrXII | 465584 | 465745 | 161 | 0,105433742 | 0,192705218 | 0,353639597 | -0,375512712 | -0,319185805 | -0,316353124 | -0,435353124 | -1,695353124 |
| chrXII | 465584 | 465756 | 172 | 0,749434433 | 0,841232394 | 0,471144818 | -0,072392395 | -0,061533536 | 0,059174474 | -0,136825526 | -1,396825526 |
| chrXII | 465616 | 465722 | 106 | 0,088336378 | 0,17788174 | 0,331819558 | -0,434894351 | -0,369660198 | 0,16988647 | 0,43588647 | -0,82411353 |
| chrXII | 465616 | 465731 | 115 | 0,088336378 | 0,159352392 | 0,356642645 | -0,36744744 | -0,312330324 | 0,295189631 | 0,498189631 | -0,761810369 |
| chrXII | 465616 | 465745 | 129 | 1,769577122 | 2,001169572 | 0,469290903 | -0,077052467 | -0,065494597 | 0,13130193 | 0,23630193 | -1,02369807 |
| chrXII | 465616 | 465746 | 130 | 0,054141651 | 0,148234783 | 0,267529425 | -0,620302195 | -0,527256866 | -0,405618533 | -0,307618533 | -1,567618533 |
| chrXII | 465616 | 465790 | 174 | 0,074088575 | 0,051882174 | 0,588141101 | 0,222765786 | 0,189350918 | 0,443694287 | 0,233694287 | -1,026305713 |
| chrXII | 465620 | 465731 | 111 | 0,054141651 | 0,077823261 | 0,410273081 | -0,226842569 | -0,192816183 | 0,291260884 | 0,522260884 | -0,737739116 |
| chrXII | 465620 | 465745 | 125 | 0,316301225 | 0,466939567 | 0,403836506 | -0,243429088 | -0,206914725 | 0,123960413 | 0,256960413 | -1,003039587 |
| chrXII | 465620 | 465746 | 126 | 0,24506221 | 0,289057827 | 0,45881486 | -0,103419898 | -0,087906914 | 0,145612146 | 0,271612146 | -0,988387854 |
| chrXII | 465630 | 465736 | 106 | 0,074088575 | 0,085235 | 0,465019536 | -0,087795679 | -0,074626327 | 0,443138989 | 0,709138989 | -0,550861011 |
| chrXII | 465630 | 465745 | 115 | 0,065539893 | 0,111176087 | 0,370877003 | -0,329531476 | -0,280101754 | 0,328335453 | 0,531335453 | -0,728664547 |
| chrXII | 465630 | 465790 | 160 | 0,327699467 | 0,255705001 | 0,561702018 | 0,155285859 | 0,13199298 | 0,137835561 | 0,025835561 | -1,234164439 |
| chrXII | 465645 | 465731 | 86 | 0,031345166 | 0,037058696 | 0,458236793 | -0,104876779 | -0,089145262 | 0,279297746 | 0,685297746 | -0,574702254 |
| chrXII | 465645 | 465745 | 100 | 0,162424953 | 0,137117174 | 0,542244106 | 0,106088938 | 0,090175597 | 0,541180914 | 0,849180914 | -0,410819086 |
| chrXII | 465645 | 465779 | 134 | 0,299203861 | 0,311293045 | 0,490098899 | -0,024820929 | -0,021097789 | -0,005662104 | 0,064337896 | -1,195662104 |
| chrXII | 465645 | 465790 | 145 | 0,065539893 | 0,055588044 | 0,541079911 | 0,103154718 | 0,08768151 | -0,066502136 | -0,073502136 | -1,333502136 |
| chrXII | 465645 | 465807 | 162 | 0,273557816 | 0,137117174 | 0,666117544 | 0,429217551 | 0,364834918 | 0,434231055 | 0,308231055 | -0,951768945 |
| chrXII | 465661 | 465832 | 171 | 0,262159574 | 0,311293045 | 0,457159956 | -0,107591283 | -0,091452591 | 0,020545211 | -0,168454789 | -1,428454789 |
| chrXII | 465724 | 465875 | 151 | 0,068389454 | 0,074117392 | 0,479902939 | -0,050397187 | -0,042837609 | -0,202976185 | -0,251976185 | -1,511976185 |
| chrXII | 465724 | 465875 | 151 | 0,068389454 | 0,074117392 | 0,479902939 | -0,050397187 | -0,042837609 | -0,202976185 | -0,251976185 | -1,511976185 |
| chrXII | 465725 | 465888 | 163 | 0,056991212 | 0,040764565 | 0,582995843 | 0,209563572 | 0,178129036 | 0,180556729 | 0,047556729 | -1,212443271 |
| chrXII | 465725 | 465888 | 163 | 0,056991212 | 0,040764565 | 0,582995843 | 0,209563572 | 0,178129036 | 0,180556729 | 0,047556729 | -1,212443271 |
| chrXII | 465731 | 465888 | 157 | 0,139628469 | 0,129705435 | 0,518421434 | 0,046192108 | 0,039263292 | -0,013958294 | -0,104958294 | -1,364958294 |
| chrXII | 465731 | 465888 | 157 | 0,139628469 | 0,129705435 | 0,518421434 | 0,046192108 | 0,039263292 | -0,013958294 | -0,104958294 | -1,364958294 |
| chrXII | 465753 | 465909 | 156 | 0,327699467 | 0,248293262 | 0,568929868 | 0,173650344 | 0,147602792 | 0,088825351 | 0,004825351 | -1,255174649 |
| chrXII | 465771 | 465899 | 128 | 0,795027403 | 1,085819787 | 0,422696436 | -0,195000098 | -0,165750083 | 0,077297 | 0,189297 | -1,070703 |
| chrXII | 465771 | 465918 | 147 | 0,370442876 | 0,26682261 | 0,581300705 | 0,205222117 | 0,174438799 | 0,062916549 | 0,041916549 | -1,218083451 |
| chrXII | 465786 | 465940 | 154 | 0,302053422 | 0,17788174 | 0,629362977 | 0,330166647 | 0,28064165 | 0,126975925 | 0,056975925 | -1,203024075 |
| chrXII | 465790 | 465899 | 109 | 0,028495606 | 0,070411522 | 0,288104674 | -0,558930214 | -0,475090682 | 0,219662805 | 0,464662805 | -0,795337195 |
| chrXII | 465790 | 465940 | 150 | 0,236513528 | 0,077823261 | 0,752420767 | 0,682127291 | 0,579808197 | 0,441691229 | 0,399691229 | -0,860308771 |
| chrXII | 465903 | 466057 | 154 | 0,088336378 | 0,044470435 | 0,665149446 | 0,426558255 | 0,362574517 | 0,218874345 | 0,148874345 | -1,111125655 |
| chrXII | 465917 | 466024 | 107 | 0,037044288 | 0,507704132 | 0,068002561 | -1,490833852 | -1,267208774 | -0,713368733 | -0,454368733 | -1,714368733 |
| chrXII | 465917 | 466027 | 110 | 0,034194727 | 0,426175002 | 0,074276663 | -1,444660304 | -1,227961258 | -0,577447913 | -0,339447913 | -1,599447913 |
| chrXII | 465917 | 466057 | 140 | 0,045592969 | 0,396528045 | 0,10312328 | -1,263953936 | -1,074360846 | -1,017467514 | -0,989467514 | -2,249467514 |
| chrXII | 465931 | 466040 | 109 | 0,113982423 | 0,26682261 | 0,299319635 | -0,52635832 | -0,447404572 | 0,243881771 | 0,488881771 | -0,771118229 |
| chrXII | 465932 | 466040 | 108 | 2,701383433 | 8,79773438 | 0,234920928 | -0,722736385 | -0,614325927 | 0,004138527 | 0,256138527 | -1,003861473 |
| chrXII | 465932 | 466041 | 109 | 0,028495606 | 0,137117174 | 0,172061636 | -0,946049634 | -0,804142189 | -0,101046151 | 0,143953849 | -1,116046151 |
| chrXII | 465934 | 466027 | 93 | 0,065539893 | 0,455821958 | 0,125709031 | -1,14691183 | -0,974875055 | -0,502374122 | -0,145374122 | -1,405374122 |
| chrXII | 465934 | 466057 | 123 | 0,039893848 | 0,722644568 | 0,052317165 | -1,622789879 | -1,379371397 | -0,931151178 | -0,784151178 | -2,044151178 |
| chrXII | 465940 | 466096 | 156 | 0,074088575 | 0,08894087 | 0,454449042 | -0,114428548 | -0,097264266 | -0,16389203 | -0,24789203 | -1,50789203 |
| chrXII | 465941 | 466048 | 107 | 0,04844253 | 0,159352392 | 0,23312663 | -0,728588752 | -0,61930044 | -0,063341363 | 0,195658637 | -1,064341363 |
| chrXII | 465941 | 466057 | 116 | 0,039893848 | 0,17417587 | 0,186359138 | -0,891393179 | -0,757684202 | -0,169141412 | 0,026858588 | -1,233141412 |
| chrXII | 465941 | 466097 | 156 | 0,119681545 | 0,17417587 | 0,407277606 | -0,234553618 | -0,199370575 | -0,271176487 | -0,355176487 | -1,615176487 |
| chrXII | 465954 | 466057 | 103 | 0,068389454 | 0,148234783 | 0,315705458 | -0,47974192 | -0,407780632 | 0,195474815 | 0,482474815 | -0,777525185 |
| chrXII | 465954 | 466108 | 154 | 0,042743409 | 0,044470435 | 0,490098899 | -0,024820929 | -0,021097789 | -0,163198002 | -0,233198002 | -1,493198002 |
| chrXII | 466075 | 466241 | 166 | 0,071239015 | 0,114881957 | 0,382756516 | -0,298249123 | -0,253511755 | -0,102575912 | -0,256575912 | -1,516575912 |
| chrXII | 466105 | 466258 | 153 | 0,037044288 | 0,044470435 | 0,454449042 | -0,114428548 | -0,097264266 | -0,205878287 | -0,268878287 | -1,528878287 |
| chrXII | 466121 | 466255 | 134 | 0,088336378 | 0,137117174 | 0,391816306 | -0,274588228 | -0,233399994 | -0,212794148 | -0,142794148 | -1,402794148 |
| chrXII | 466121 | 466264 | 143 | 0,481575739 | 0,496586524 | 0,492327048 | -0,019234425 | -0,016349262 | -0,176706159 | -0,169706159 | -1,429706159 |
| chrXII | 466121 | 466267 | 146 | 0,082637257 | 0,059293913 | 0,582234733 | 0,207613792 | 0,176471723 | 0,018654012 | 0,004654012 | -1,255345988 |
| chrXII | 466121 | 466277 | 156 | 0,04844253 | 0,070411522 | 0,407579962 | -0,23377466 | -0,198708461 | -0,283858752 | -0,367858752 | -1,627858752 |

TABLE 2

| Chrom | Start | End | Length | Section A | Section B | A/A+B | Z-score | Z * 0.85 | Phase correction | Length correction | ΔLK |
|---|---|---|---|---|---|---|---|---|---|---|---|
| chrXII | 466127 | 466264 | 137 | 0,108283302 | 0,137117174 | 0,441251393 | -0,147797242 | -0,125627655 | -0,062071282 | -0,013071282 | -1,273071282 |
| chrXII | 466127 | 466277 | 150 | 0,447381012 | 0,481763045 | 0,481498007 | -0,046394257 | -0,039435118 | -0,1780357 | -0,2200357 | -1,4800357 |
| chrXII | 466142 | 466303 | 161 | 0,076938136 | 0,181587609 | 0,297603381 | -0,531305981 | -0,451610084 | -0,433771488 | -0,552771488 | -1,812771488 |
| chrXII | 466158 | 466264 | 106 | 0,028495606 | 0,474351306 | 0,056668551 | -1,58337024 | -1,345864704 | -0,828864991 | -0,562864991 | -1,822864991 |
| chrXII | 466159 | 466264 | 105 | 0,028495606 | 0,114881957 | 0,198745224 | -0,846111675 | -0,719194924 | -0,15903945 | 0,11396055 | -1,14603945 |
| chrXII | 466159 | 466342 | 183 | 0,062690333 | 0,329822392 | 0,159715415 | -0,995628195 | -0,846283966 | -0,806348407 | -1,079348407 | -2,339348407 |
| chrXII | 466159 | 466342 | 183 | 0,062690333 | 0,329822392 | 0,159715415 | -0,995628195 | -0,846283966 | -0,806348407 | -1,079348407 | -2,339348407 |
| chrXII | 466184 | 466344 | 160 | 0,09688506 | 0,111176087 | 0,465656665 | -0,08619258 | -0,073263693 | -0,066213629 | -0,178213629 | -1,438213629 |
| chrXII | 466184 | 466344 | 160 | 0,09688506 | 0,111176087 | 0,465656665 | -0,08619258 | -0,073263693 | -0,066213629 | -0,178213629 | -1,438213629 |
| chrXII | 466207 | 466337 | 130 | 0,19377012 | 0,507704132 | 0,27623269 | -0,594069873 | -0,504959392 | -0,368726957 | -0,270726957 | -1,530726957 |
| chrXII | 466207 | 466337 | 130 | 0,19377012 | 0,507704132 | 0,27623269 | -0,594069873 | -0,504959392 | -0,368726957 | -0,270726957 | -1,530726957 |
| chrXII | 466220 | 466354 | 134 | 0,076938136 | 0,048176305 | 0,614942093 | 0,292223409 | 0,248389898 | 0,27135812 | 0,34135812 | -0,91864188 |
| chrXII | 466252 | 466385 | 133 | 1,1711694 | 0,874585221 | 0,572487721 | 0,182711288 | 0,155304595 | 0,153297833 | 0,230297833 | -1,029702167 |
| chrXII | 466277 | 466442 | 165 | 0,04844253 | 0,070411522 | 0,407579962 | -0,23377466 | -0,198708461 | -0,09474518 | -0,24174518 | -1,50174518 |
| chrXII | 466277 | 466449 | 172 | 0,78362916 | 1,500877179 | 0,343019035 | -0,404237514 | -0,343601887 | -0,220271405 | -0,416271405 | -1,676271405 |
| chrXII | 466280 | 466385 | 105 | 0,051292091 | 0,070411522 | 0,421450846 | -0,198183249 | -0,168455762 | 0,387411243 | 0,660411243 | -0,599588757 |
| chrXII | 466280 | 466416 | 136 | 0,532867829 | 0,411351523 | 0,564347498 | 0,162001079 | 0,137700917 | 0,152069748 | 0,208069748 | -1,051930252 |
| chrXII | 466280 | 466453 | 173 | 0,054141651 | 0,074117392 | 0,42212736 | -0,196454145 | -0,166986023 | 0,044995741 | -0,158004259 | -1,418004259 |
| chrXII | 466289 | 466442 | 153 | 0,464478375 | 0,326116523 | 0,587504898 | 0,221131295 | 0,187961601 | 0,085165091 | 0,022165091 | -1,237834909 |
| chrXII | 466289 | 466449 | 160 | 0,065539893 | 0,092646739 | 0,414320049 | -0,216446021 | -0,183979117 | -0,179531932 | -0,291531932 | -1,551531932 |
| chrXII | 466290 | 466385 | 95 | 0,091185939 | 0,166764131 | 0,353502284 | -0,375882075 | -0,319499764 | 0,177373527 | 0,520373527 | -0,739626473 |
| chrXII | 466290 | 466416 | 126 | 0,111132863 | 0,111176087 | 0,499902783 | -0,000243688 | -0,000207135 | 0,233502708 | 0,359502708 | -0,900497292 |
| chrXII | 466290 | 466449 | 159 | 0,111132863 | 0,17788174 | 0,384523349 | -0,293622086 | -0,249578773 | -0,250122146 | -0,355122146 | -1,615122146 |
| chrXII | 466290 | 466453 | 163 | 1,721134592 | 1,574994571 | 0,522168431 | 0,055596643 | 0,047257147 | 0,050784061 | -0,082215939 | -1,342215939 |
| chrXII | 466290 | 466454 | 164 | 0,094035499 | 0,125999566 | 0,427365971 | -0,183084213 | -0,155621581 | -0,129692835 | -0,269692835 | -1,529692835 |
| chrXII | 466291 | 466449 | 158 | 0,353345512 | 0,337234132 | 0,511665114 | 0,029244273 | 0,024857632 | -0,001058459 | -0,099058459 | -1,359058459 |
| chrXII | 466291 | 466453 | 162 | 0,04844253 | 0,070411522 | 0,407579962 | -0,23377466 | -0,198708461 | -0,130868428 | -0,256868428 | -1,516868428 |
| chrXII | 466324 | 466453 | 129 | 0,065539893 | 0,066705652 | 0,495592445 | -0,011048326 | -0,009391077 | 0,181495176 | 0,286495176 | -0,973504824 |
| chrXII | 466324 | 466454 | 130 | 0,401788042 | 0,318704784 | 0,557657242 | 0,145032115 | 0,123277298 | 0,259354619 | 0,357354619 | -0,902645381 |
| chrXII | 466335 | 466483 | 148 | 0,433133209 | 0,255705001 | 0,628788012 | 0,328645073 | 0,279348312 | 0,125193558 | 0,097193558 | -1,162806442 |
| chrXII | 466349 | 466502 | 153 | 0,122531105 | 0,070411522 | 0,635064977 | 0,345298405 | 0,293503645 | 0,191974548 | 0,128974548 | -1,131025452 |
| chrXII | 466351 | 466500 | 149 | 0,256460453 | 0,159352392 | 0,616768953 | 0,297005782 | 0,252454915 | 0,08723728 | 0,05223728 | -1,20776272 |
| chrXII | 466356 | 466449 | 93 | 0,051292091 | 0,133411305 | 0,277699771 | -0,589688448 | -0,501235181 | -0,022541919 | 0,334458081 | -0,925541919 |
| chrXII | 466356 | 466483 | 127 | 0,068389454 | 0,074117392 | 0,479902939 | -0,050397187 | -0,042837609 | 0,219305522 | 0,338305522 | -0,921694478 |
| chrXII | 466356 | 466500 | 144 | 0,059840772 | 0,037058696 | 0,617555219 | 0,299066166 | 0,254206241 | 0,068766193 | 0,068766193 | -1,191233807 |
| chrXII | 466356 | 466502 | 146 | 0,97454972 | 0,641115437 | 0,603187929 | 0,261607395 | 0,222366286 | 0,062133318 | 0,048133318 | -1,211866682 |
| chrXII | 466364 | 466500 | 136 | 0,384690679 | 0,333528262 | 0,535617563 | 0,08939893 | 0,07598909 | 0,092183369 | 0,148183369 | -1,111816631 |
| chrXII | 466367 | 466532 | 165 | 0,125380666 | 0,185293479 | 0,403576119 | -0,244101464 | -0,207486244 | -0,097521289 | -0,244521289 | -1,504521289 |
| chrXII | 466367 | 466538 | 171 | 0,279256937 | 0,251999131 | 0,525654112 | 0,064349707 | 0,054697251 | 0,164102245 | -0,024897755 | -1,284897755 |
| chrXII | 466385 | 466532 | 147 | 0,051292091 | 0,040764565 | 0,557179598 | 0,143822286 | 0,122248943 | 0,012818718 | -0,008181282 | -1,268181282 |
| chrXII | 466385 | 466538 | 153 | 0,225115286 | 0,059293913 | 0,791519004 | 0,811703139 | 0,689947668 | 0,595188444 | 0,532188444 | -0,727811556 |
| chrXII | 466392 | 466502 | 110 | 0,042743409 | 0,08894087 | 0,324590066 | -0,454901459 | -0,386666241 | 0,247238188 | 0,485238188 | -0,774761812 |
| chrXII | 466392 | 466544 | 152 | 0,507221784 | 0,281646088 | 0,642974321 | 0,366420455 | 0,311457386 | 0,155507694 | 0,099507694 | -1,160492306 |
| chrXII | 466403 | 466500 | 97 | 0,042743409 | 0,070411522 | 0,377742344 | -0,311415611 | -0,264703269 | 0,286622456 | 0,615622456 | -0,644377544 |
| chrXII | 466403 | 466502 | 99 | 0,128230226 | 0,051882174 | 0,711945574 | 0,559077467 | 0,475215847 | 0,947228858 | 1,262228858 | 0,002228858 |
| chrXII | 466403 | 466544 | 141 | 0,094035499 | 0,040764565 | 0,69759239 | 0,517488464 | 0,439865194 | 0,379753977 | 0,400753977 | -0,859246023 |
| chrXII | 466403 | 466559 | 156 | 1,062886098 | 0,79305609 | 0,572693538 | 0,183235904 | 0,155750518 | 0,064435227 | -0,019564773 | -1,279564773 |
| chrXII | 466432 | 466521 | 89 | 0,037044288 | 0,270528479 | 0,120440727 | -1,172786449 | -0,996868482 | -0,559349753 | -0,174349753 | -1,434349753 |
| chrXII | 466432 | 466523 | 91 | 0,028495606 | 0,129705435 | 0,180122745 | -0,914897409 | -0,777662797 | -0,416883336 | -0,045883336 | -1,305883336 |
| chrXII | 466432 | 466527 | 95 | 0,028495606 | 0,389116306 | 0,068234658 | -1,489068562 | -1,265708277 | -0,76579282 | -0,42279282 | -1,68279282 |
| chrXII | 466432 | 466528 | 96 | 0,102584181 | 0,852350003 | 0,107425394 | -1,240336958 | -1,054286414 | -0,478346336 | -0,142346336 | -1,402346336 |
| chrXII | 466432 | 466530 | 98 | 0,031345166 | 0,344645871 | 0,083366792 | -1,382775925 | -1,175359536 | -0,658597643 | -0,336597643 | -1,596597643 |
| chrXII | 466432 | 466531 | 99 | 0,042743409 | 0,389116306 | 0,098975217 | -1,287412835 | -1,09430091 | -0,642053606 | -0,327053606 | -1,587053606 |
| chrXII | 466432 | 466532 | 100 | 15,31068902 | 81,57360116 | 0,158030667 | -1,002584589 | -0,852196901 | -0,429212083 | -0,121212083 | -1,381212083 |
| chrXII | 466432 | 466534 | 102 | 0,034194727 | 0,426175002 | 0,074276663 | -1,444660304 | -1,227961258 | -0,683766327 | -0,389766327 | -1,649766327 |
| chrXII | 466432 | 466538 | 106 | 0,139628469 | 0,381704567 | 0,267829697 | -0,619390113 | -0,526481596 | -0,028076878 | 0,237923122 | -1,022076878 |
| chrXII | 466432 | 466539 | 107 | 0,199469241 | 0,318704784 | 0,38494643 | -0,292515043 | -0,248637787 | 0,299502365 | 0,558502365 | -0,701497635 |
| chrXII | 466474 | 466613 | 139 | 0,074088575 | 0,096352609 | 0,434687048 | -0,164453569 | -0,139785534 | -0,142357428 | -0,107357428 | -1,367357428 |
| chrXII | 466474 | 466618 | 144 | 0,544266071 | 0,329822392 | 0,622667034 | 0,312492935 | 0,265618995 | 0,076543587 | 0,076543587 | -1,183456413 |
| chrXII | 466497 | 466613 | 116 | 0,056991212 | 0,17417587 | 0,246536882 | -0,685428183 | -0,582613956 | 0,013925073 | 0,209925073 | -1,050074927 |
| chrXII | 466497 | 466618 | 121 | 0,039893848 | 0,037058696 | 0,518421434 | 0,046192108 | 0,039263292 | 0,43661063 | 0,59761063 | -0,66238937 |
| chrXII | 466497 | 466667 | 170 | 0,173823196 | 0,237175653 | 0,422928668 | -0,194406838 | -0,165245812 | -0,003557855 | -0,185557855 | -1,445557855 |
| chrXII | 466507 | 466613 | 106 | 0,789528282 | 5,110394149 | 0,133790749 | -1,108649309 | -0,942351913 | -0,425414891 | -0,159414891 | -1,419414891 |
| chrXII | 466509 | 466613 | 104 | 0,04844253 | 0,833820655 | 0,054907119 | -1,599028642 | -1,359174346 | -0,778094708 | -0,498094708 | -1,758094708 |
| chrXII | 466509 | 466618 | 109 | 0,034194727 | 0,077823261 | 0,305261035 | -0,509328378 | -0,432929121 | 0,278468028 | 0,523468028 | -0,736531972 |
| chrXII | 466557 | 466700 | 143 | 0,190920559 | 0,085235 | 0,691351496 | 0,499684841 | 0,424732115 | 0,263339397 | 0,270339397 | -0,989660603 |
| chrXII | 466557 | 466704 | 147 | 0,079787696 | 0,066705652 | 0,544650641 | 0,112157259 | 0,09533367 | -0,023748707 | -0,044748707 | -1,304748707 |
| chrXII | 466557 | 466708 | 151 | 0,094035499 | 0,055588044 | 0,628480635 | 0,327831948 | 0,278657156 | 0,125162267 | 0,076162267 | -1,183837733 |
| chrXII | 466557 | 466709 | 152 | 0,464478375 | 0,181587609 | 0,718933339 | 0,579675715 | 0,492724358 | 0,337952451 | 0,281952451 | -0,978047549 |
| chrXII | 466574 | 466729 | 155 | 0,039893848 | 0,077823261 | 0,338895921 | -0,415478239 | -0,353156503 | -0,417345084 | -0,494345084 | -1,754345084 |
| chrXII | 466577 | 466730 | 153 | 0,062690333 | 0,040764565 | 0,605967759 | 0,268824834 | 0,228501109 | 0,122891759 | 0,059891759 | -1,200108241 |
| chrXII | 466611 | 466736 | 125 | 0,028495606 | 0,111176087 | 0,204018475 | -0,827353109 | -0,703250143 | -0,398849683 | -0,265849683 | -1,525849683 |
| chrXII | 466690 | 466843 | 153 | 0,162424953 | 0,085235 | 0,655838584 | 0,40113215 | 0,340962327 | 0,229230877 | 0,166230877 | -1,093769123 |
| chrXII | 466706 | 466873 | 167 | 0,031345166 | 0,077823261 | 0,287126664 | -0,56179848 | -0,477528708 | -0,342604263 | -0,503604263 | -1,763604263 |
| chrXII | 466706 | 466878 | 172 | 0,284956058 | 0,566998046 | 0,334473485 | -0,427593677 | -0,363454625 | -0,234582062 | -0,430582062 | -1,690582062 |
| chrXII | 466737 | 466873 | 136 | 0,817823887 | 0,663350655 | 0,55214552 | 0,131083871 | 0,11142129 | 0,139476622 | 0,195476622 | -1,064523378 |
| chrXII | 466737 | 466876 | 139 | 0,168124074 | 0,066705652 | 0,71594034 | 0,570823457 | 0,485199939 | 0,476130719 | 0,511130719 | -0,748869281 |
| chrXII | 466737 | 466878 | 141 | 0,076938136 | 0,103764348 | 0,42577243 | -0,18714766 | -0,159075511 | -0,219186302 | -0,198186302 | -1,458186302 |
| chrXII | 466743 | 466900 | 157 | 0,09973462 | 0,040764565 | 0,709859063 | 0,552973037 | 0,470027081 | 0,404620484 | 0,313620484 | -0,946379516 |
| chrXII | 466757 | 466910 | 153 | 0,564212996 | 0,485468915 | 0,537508544 | 0,094158927 | 0,080035088 | -0,031696362 | -0,094696362 | -1,354696362 |
| chrXII | 466757 | 466924 | 167 | 0,059840772 | 0,114881957 | 0,342489913 | -0,405677162 | -0,344825587 | -0,214540173 | -0,375540173 | -1,635540173 |
| chrXII | 466771 | 466894 | 123 | 0,173823196 | 0,52623348 | 0,248298747 | -0,679853078 | -0,577875116 | -0,132787505 | 0,014212495 | -1,245787505 |
| chrXII | 466804 | 466956 | 152 | 0,074088575 | 0,059293913 | 0,555459536 | 0,139467258 | 0,118547169 | -0,036999552 | -0,092999552 | -1,352999552 |
| chrXII | 466940 | 467096 | 156 | 0,19377012 | 0,118587827 | 0,620346375 | 0,30639062 | 0,260432027 | 0,172841998 | 0,088841998 | -1,171158002 |
| chrXII | 466947 | 467098 | 151 | 0,051292091 | 0,051882174 | 0,497140355 | -0,007168129 | -0,00609291 | -0,154342238 | -0,203342238 | -1,463342238 |
| chrXII | 466953 | 467111 | 158 | 0,122531105 | 0,059293913 | 0,673895739 | 0,450696199 | 0,383091769 | 0,355690498 | 0,257690498 | -1,002309502 |
| chrXII | 466957 | 467090 | 133 | 0,113982423 | 0,055588044 | 0,672183225 | 0,445949741 | 0,37905728 | 0,376221453 | 0,453221453 | -0,806778547 |
| chrXII | 467009 | 467160 | 151 | 0,250761331 | 0,181587609 | 0,579997562 | 0,201887241 | 0,171604155 | 0,030149541 | -0,018850459 | -1,278850459 |
| chrXII | 467063 | 467189 | 126 | 0,054141651 | 0,048176305 | 0,529151025 | 0,07313593 | 0,062165541 | 0,270447543 | 0,396447543 | -0,863552457 |
| chrXII | 467082 | 467240 | 158 | 0,042743409 | 0,040764565 | 0,51184823 | 0,029703476 | 0,025247955 | -0,004117349 | -0,102117349 | -1,362117349 |
| chrXII | 467082 | 467241 | 159 | 0,307752543 | 0,35576348 | 0,463820816 | -0,090812431 | -0,077190567 | -0,078762402 | -0,183762402 | -1,443762402 |
| chrXII | 467086 | 467196 | 110 | 0,034194727 | 0,048176305 | 0,415130494 | -0,214366854 | -0,182211826 | 0,447312824 | 0,685312824 | -0,574687176 |
| chrXII | 467086 | 467238 | 152 | 0,330549028 | 0,214940436 | 0,605967759 | 0,268824834 | 0,228501109 | 0,079284507 | 0,023284507 | -1,236715493 |
| chrXII | 467087 | 467238 | 151 | 0,14817715 | 0,092646739 | 0,615292571 | 0,293140374 | 0,249169318 | 0,109641251 | 0,060641251 | -1,199358749 |
| chrXII | 467087 | 467240 | 153 | 0,547115632 | 0,403939784 | 0,575272085 | 0,189812794 | 0,161340875 | 0,038388067 | -0,024611933 | -1,284611933 |
| chrXII | 467096 | 467196 | 100 | 0,034194727 | 0,044470435 | 0,434687048 | -0,164453569 | -0,139785534 | 0,262855529 | 0,570855529 | -0,689144471 |
| chrXII | 467096 | 467247 | 151 | 0,116831984 | 0,051882174 | 0,692484764 | 0,502905845 | 0,427469968 | 0,294669253 | 0,245669253 | -1,014330747 |
| chrXII | 467113 | 467196 | 83 | 0,068389454 | 0,137117174 | 0,332784663 | -0,432236787 | -0,367401269 | -0,038964081 | 0,388035919 | -0,871964081 |
| chrXII | 467113 | 467256 | 143 | 0,128230226 | 0,085235 | 0,600707798 | 0,255179578 | 0,216902641 | 0,050708423 | 0,057708423 | -1,202291577 |
| chrXII | 467120 | 467226 | 106 | 0,085486818 | 0,085235 | 0,500737507 | 0,001848658 | 0,001571359 | 0,519553222 | 0,785553222 | -0,474446778 |
| chrXII | 467120 | 467240 | 120 | 0,028495606 | 0,077823261 | 0,268020217 | -0,618811668 | -0,525989918 | -0,175839735 | -0,007839735 | -1,267839735 |

TABLE 2

| Chrom | Start | End | Length | Section A | Section B | A/A+B | Z-score | Z * 0.85 | Phase correction | Length correction | ΔLK |
|---|---|---|---|---|---|---|---|---|---|---|---|
| chrXII | 467120 | 467274 | 154 | 0,735186631 | 0,715232829 | 0,506878631 | 0,017243027 | 0,014656573 | -0,123196088 | -0,193196088 | -1,453196088 |
| chrXII | 467123 | 467254 | 131 | 0,079787696 | 0,062999783 | 0,558786364 | 0,147892925 | 0,125708986 | 0,200435818 | 0,291435818 | -0,968564182 |
| chrXII | 467162 | 467304 | 142 | 0,091185939 | 0,144528914 | 0,386848507 | -0,287542436 | -0,24441107 | -0,265607052 | -0,251607052 | -1,511607052 |
| chrXII | 467192 | 467274 | 82 | 0,056991212 | 0,070411522 | 0,447331152 | -0,132407093 | -0,112546029 | 0,209452455 | 0,643452455 | -0,616547545 |
| chrXII | 467192 | 467343 | 151 | 0,860567296 | 0,574409785 | 0,599708043 | 0,252591479 | 0,214702758 | 0,085859114 | 0,036859114 | -1,223140886 |
| chrXII | 467216 | 467364 | 148 | 0,054141651 | 0,048176305 | 0,529151025 | 0,07313593 | 0,062165541 | -0,085866419 | -0,113866419 | -1,373866419 |
| chrXII | 467224 | 467368 | 144 | 0,364743755 | 0,192705218 | 0,654308775 | 0,396979674 | 0,337432723 | 0,139595359 | 0,139595359 | -1,120404641 |
| chrXII | 467264 | 467414 | 150 | 0,111132863 | 0,070411522 | 0,612152576 | 0,284933816 | 0,242193743 | 0,099836118 | 0,057836118 | -1,202163882 |
| chrXII | 467267 | 467430 | 163 | 0,173823196 | 0,103764348 | 0,626192347 | 0,32178536 | 0,273517556 | 0,276847486 | 0,143847486 | -1,116152514 |
| chrXII | 467271 | 467409 | 138 | 0,190920559 | 0,081529131 | 0,700755282 | 0,526574021 | 0,447587917 | 0,427295635 | 0,469295635 | -0,790704365 |
| chrXII | 467271 | 467414 | 143 | 0,310602104 | 0,207528696 | 0,59946659 | 0,251966677 | 0,214171675 | 0,045666761 | 0,052666761 | -1,207333239 |
| chrXII | 467273 | 467432 | 159 | 0,09973462 | 0,081529131 | 0,550218231 | 0,126212725 | 0,107280816 | 0,11080193 | 0,00580193 | -1,25419807 |
| chrXII | 467280 | 467428 | 148 | 0,088336378 | 0,077823261 | 0,531635591 | 0,07938196 | 0,067474666 | -0,068606866 | -0,096606866 | -1,356606866 |
| chrXII | 467303 | 467458 | 155 | 0,547115632 | 0,515115871 | 0,515062517 | 0,037765106 | 0,03210034 | -0,039416837 | -0,116416837 | -1,376416837 |
| chrXII | 467303 | 467459 | 156 | 0,054141651 | 0,051882174 | 0,510655515 | 0,026712592 | 0,022705704 | -0,065747579 | -0,149747579 | -1,409747579 |
| chrXII | 467304 | 467459 | 155 | 0,128230226 | 0,085235 | 0,600707798 | 0,255179578 | 0,216902641 | 0,1467977 | 0,0697977 | -1,1902023 |
| chrXII | 467314 | 467414 | 100 | 0,034194727 | 0,048176305 | 0,415130494 | -0,214366854 | -0,182211826 | 0,242851694 | 0,550851694 | -0,709148306 |
| chrXII | 467314 | 467459 | 145 | 0,846319493 | 0,581821524 | 0,592602189 | 0,234243918 | 0,19910733 | 0,047488923 | 0,040488923 | -1,219511077 |
| chrXII | 467314 | 467464 | 150 | 0,051292091 | 0,048176305 | 0,515662191 | 0,039269381 | 0,033378974 | -0,12353872 | -0,16553872 | -1,42553872 |
| chrXII | 467314 | 467487 | 173 | 0,561363435 | 0,833820655 | 0,402357968 | -0,247248471 | -0,2101612 | -0,000968457 | -0,203968457 | -1,463968457 |
| chrXII | 467316 | 467464 | 148 | 0,299203861 | 0,170470001 | 0,637046013 | 0,350573989 | 0,297987891 | 0,163787347 | 0,135787347 | -1,124212653 |
| chrXII | 467328 | 467458 | 130 | 0,051292091 | 0,051882174 | 0,497140355 | -0,007168129 | -0,00609291 | 0,113997572 | 0,211997572 | -1,048002428 |
| chrXII | 467328 | 467489 | 161 | 0,598407723 | 0,415057393 | 0,590457149 | 0,22872109 | 0,194412926 | 0,212033052 | 0,093033052 | -1,166966948 |
| chrXII | 467332 | 467459 | 127 | 0,076938136 | 0,066705652 | 0,535617563 | 0,08939893 | 0,07598909 | 0,323849141 | 0,442849141 | -0,817150859 |
| chrXII | 467332 | 467487 | 155 | 0,963151477 | 0,463233697 | 0,675239405 | 0,454427464 | 0,386263344 | 0,331608117 | 0,254608117 | -1,005391883 |
| chrXII | 467341 | 467487 | 146 | 0,082637257 | 0,081529131 | 0,503375009 | 0,008459995 | 0,007190996 | -0,148135006 | -0,162135006 | -1,422135006 |
| chrXII | 467341 | 467517 | 176 | 0,208017923 | 0,251999131 | 0,452196111 | -0,120114784 | -0,102097566 | 0,068371505 | -0,155628495 | -1,415628495 |
| chrXII | 467428 | 467588 | 160 | 0,518620026 | 0,26682261 | 0,660290138 | 0,413255097 | 0,351266833 | 0,347152949 | 0,235152949 | -1,024847051 |
| chrXII | 467434 | 467588 | 154 | 1,205364127 | 0,796761959 | 0,602042067 | 0,258636308 | 0,219840862 | 0,083802184 | 0,013802184 | -1,246197816 |
| chrXII | 467435 | 467598 | 163 | 0,094035499 | 0,048176305 | 0,661235543 | 0,415837502 | 0,353461877 | 0,349897787 | 0,216897787 | -1,043102213 |
| chrXII | 467456 | 467609 | 153 | 0,14817715 | 0,137117174 | 0,519383449 | 0,048606234 | 0,041315299 | -0,083761565 | -0,146761565 | -1,406761565 |
| chrXII | 467475 | 467629 | 154 | 0,273557816 | 0,248293262 | 0,524206671 | 0,060714406 | 0,051607245 | -0,090864087 | -0,160864087 | -1,420864087 |
| chrXII | 467493 | 467630 | 137 | 0,054141651 | 0,037058696 | 0,593656197 | 0,236960278 | 0,201416237 | 0,264984181 | 0,313984181 | -0,946015819 |
| chrXII | 467493 | 467645 | 152 | 0,202318801 | 0,125999566 | 0,616227484 | 0,295587617 | 0,251249475 | 0,101066187 | 0,045066187 | -1,214933813 |
| chrXII | 467496 | 467660 | 164 | 0,116831984 | 0,100058479 | 0,538668149 | 0,097078943 | 0,082517102 | 0,106274428 | -0,033725572 | -1,293725572 |
| chrXII | 467502 | 467630 | 128 | 0,820673448 | 1,030231743 | 0,443390322 | -0,142379007 | -0,121022156 | 0,124402598 | 0,236402598 | -1,023597402 |
| chrXII | 467510 | 467630 | 120 | 0,054141651 | 0,085235 | 0,388455674 | -0,283346352 | -0,2408444 | 0,135625693 | 0,303625693 | -0,956374307 |
| chrXII | 467510 | 467645 | 135 | 0,028495606 | 0,051882174 | 0,354520937 | -0,373143192 | -0,317171714 | -0,347213338 | -0,284213338 | -1,544213338 |
| chrXII | 467510 | 467667 | 157 | 0,153876272 | 0,192705218 | 0,443982948 | -0,140878539 | -0,119746758 | -0,184756594 | -0,275756594 | -1,535756594 |
| chrXII | 467511 | 467669 | 158 | 0,074088575 | 0,081529131 | 0,47609348 | -0,059960669 | -0,050966568 | -0,08030926 | -0,17830926 | -1,43830926 |
| chrXII | 467515 | 467649 | 134 | 0,091185939 | 0,051882174 | 0,637360324 | 0,351411905 | 0,298700119 | 0,320313436 | 0,390313436 | -0,869686564 |
| chrXII | 467532 | 467638 | 106 | 0,04844253 | 0,066705652 | 0,420697304 | -0,200109928 | -0,170093439 | 0,346383264 | 0,612383264 | -0,647616736 |
| chrXII | 467556 | 467724 | 168 | 0,065539893 | 0,085235 | 0,434687048 | -0,164453569 | -0,139785534 | -0,038857542 | -0,206857542 | -1,466857542 |
| chrXII | 467645 | 467795 | 150 | 0,037044288 | 0,066705652 | 0,357053581 | -0,366345661 | -0,311393812 | -0,46262558 | -0,50462558 | -1,76462558 |
| chrXII | 467645 | 467796 | 151 | 0,094035499 | 0,111176087 | 0,458236793 | -0,104876779 | -0,089145262 | -0,226740651 | -0,275740651 | -1,535740651 |
| chrXII | 467883 | 468037 | 154 | 0,034194727 | 0,037058696 | 0,479902939 | -0,050397187 | -0,042837609 | -0,179928375 | -0,249928375 | -1,509928375 |
| chrXII | 467883 | 468038 | 155 | 0,088336378 | 0,096352609 | 0,478298027 | -0,054425638 | -0,046261792 | -0,096762196 | -0,173762196 | -1,433762196 |
| chrXII | 467883 | 468049 | 166 | 0,225115286 | 0,159352392 | 0,585524607 | 0,216047418 | 0,183640306 | 0,342370126 | 0,188370126 | -1,071629874 |
| chrXII | 467887 | 468049 | 162 | 0,04844253 | 0,051882174 | 0,482857442 | -0,042983253 | -0,036535765 | 0,026588256 | -0,099411744 | -1,359411744 |
| chrXII | 467905 | 468037 | 132 | 0,113982423 | 0,166764131 | 0,405997586 | -0,23785292 | -0,202174982 | -0,066353329 | 0,017646671 | -1,242353329 |
| chrXII | 467905 | 468038 | 133 | 0,133929347 | 0,170470001 | 0,439979088 | -0,151022236 | -0,128368901 | -0,131191197 | -0,054191197 | -1,314191197 |
| chrXII | 467905 | 468049 | 144 | 0,63545201 | 0,426175002 | 0,598564282 | 0,249632664 | 0,212187765 | 0,008867386 | 0,008867386 | -1,251132614 |
| chrXII | 467906 | 468037 | 131 | 0,681044979 | 0,678174133 | 0,501056065 | 0,002647165 | 0,00225009 | 0,067997586 | 0,158997586 | -1,101002414 |
| chrXII | 467906 | 468038 | 132 | 1,521665352 | 1,434171527 | 0,514800178 | 0,037107059 | 0,031541 | 0,157695023 | 0,241695023 | -1,018304977 |
| chrXII | 467906 | 468049 | 143 | 0,800726524 | 0,566998046 | 0,585444279 | 0,215841315 | 0,183465118 | 0,018441326 | 0,025441326 | -1,234558674 |
| chrXII | 467906 | 468050 | 144 | 0,19377012 | 0,062999783 | 0,754644987 | 0,68917996 | 0,585802966 | 0,390022271 | 0,390022271 | -0,869977729 |
| chrXII | 467907 | 468037 | 130 | 0,091185939 | 0,107470218 | 0,459013908 | -0,102918298 | -0,087480553 | 0,013309602 | 0,111309602 | -1,148690398 |
| chrXII | 467907 | 468038 | 131 | 0,703841464 | 1,037643482 | 0,40416167 | -0,242589598 | -0,206201158 | -0,140637707 | -0,049637707 | -1,309637707 |
| chrXII | 467907 | 468049 | 142 | 0,131079787 | 0,111176087 | 0,541079911 | 0,103154718 | 0,08768151 | 0,060609908 | 0,074609908 | -1,185390092 |
| chrXII | 467983 | 468139 | 156 | 0,216566604 | 0,066705652 | 0,764517524 | 0,720909899 | 0,612773414 | 0,525601987 | 0,441601987 | -0,818398013 |
| chrXII | 468039 | 468185 | 146 | 0,088336378 | 0,077823261 | 0,531635591 | 0,07938196 | 0,067474666 | -0,09506816 | -0,10906816 | -1,36906816 |
| chrXII | 468045 | 468181 | 136 | 0,113982423 | 0,137117174 | 0,453933118 | -0,11573037 | -0,098370815 | -0,078772836 | -0,022772836 | -1,282772836 |
| chrXII | 468045 | 468185 | 140 | 0,173823196 | 0,085235 | 0,670981263 | 0,442624345 | 0,376230693 | 0,42950572 | 0,45750572 | -0,80249428 |
| chrXII | 468047 | 468203 | 156 | 0,051292091 | 0,08894087 | 0,365763444 | -0,34309514 | -0,291630869 | -0,364220607 | -0,448220607 | -1,708220607 |
| chrXII | 468050 | 468139 | 89 | 0,108283302 | 0,133411305 | 0,448017039 | -0,130672891 | -0,111071958 | 0,309158069 | 0,694158069 | -0,565841931 |
| chrXII | 468050 | 468175 | 125 | 1,162620718 | 1,823287833 | 0,389369165 | -0,280963586 | -0,238819048 | 0,032695602 | 0,165695602 | -1,094304398 |
| chrXII | 468050 | 468176 | 126 | 0,028495606 | 0,037058696 | 0,434687048 | -0,164453569 | -0,139785534 | 0,09038191 | 0,21638191 | -1,04361809 |
| chrXII | 468050 | 468180 | 130 | 0,883363781 | 1,119172613 | 0,44112246 | -0,148123985 | -0,125905388 | -0,023329644 | 0,074670356 | -1,185329644 |
| chrXII | 468050 | 468181 | 131 | 3,744322607 | 5,028865018 | 0,42679158 | -0,18454853 | -0,15686625 | -0,071440179 | 0,019559821 | -1,240440179 |
| chrXII | 468050 | 468184 | 134 | 1,723984153 | 1,971522616 | 0,466508184 | -0,084050389 | -0,071442831 | -0,0495425 | 0,0204575 | -1,2395425 |
| chrXII | 468050 | 468185 | 135 | 0,227964847 | 0,218646305 | 0,5104325 | 0,02615338 | 0,022230373 | -0,011465711 | 0,051534289 | -1,208465711 |
| chrXII | 468050 | 468203 | 153 | 0,131079787 | 0,122293696 | 0,517338221 | 0,043474165 | 0,03695304 | -0,07753062 | -0,14053062 | -1,40053062 |
| chrXII | 468070 | 468181 | 111 | 0,119681545 | 0,233469783 | 0,338895921 | -0,415478239 | -0,353156503 | 0,135821224 | 0,366821224 | -0,893178776 |
| chrXII | 468070 | 468184 | 114 | 0,039893848 | 0,070411522 | 0,361667325 | -0,354005648 | -0,300904801 | 0,325933148 | 0,535933148 | -0,724066852 |
| chrXII | 468070 | 468203 | 133 | 0,313451664 | 0,259410871 | 0,547167331 | 0,118507771 | 0,100731606 | 0,083818863 | 0,160818863 | -1,099181137 |
| chrXII | 468187 | 468283 | 96 | 0,151026711 | 0,930173264 | 0,139684346 | -1,081738612 | -0,91947782 | -0,376706257 | -0,040706257 | -1,300706257 |
| chrXII | 468226 | 468399 | 173 | 0,353345512 | 0,548468698 | 0,391816306 | -0,274588228 | -0,233399994 | -0,038027388 | -0,241027388 | -1,501027388 |
| chrXII | 468226 | 468402 | 176 | 0,039893848 | 0,062999783 | 0,387719315 | -0,285268256 | -0,242478018 | -0,079694439 | -0,303694439 | -1,563694439 |
| chrXII | 468232 | 468399 | 167 | 0,091185939 | 0,125999566 | 0,419852784 | -0,202270107 | -0,171929591 | -0,044487332 | -0,205487332 | -1,465487332 |
| chrXII | 468232 | 468402 | 170 | 0,689593661 | 1,196995874 | 0,365523951 | -0,343731921 | -0,292172133 | -0,128627036 | -0,310627036 | -1,570627036 |
| chrXII | 468235 | 468370 | 135 | 1,940550757 | 1,467524353 | 0,569397884 | 0,174841432 | 0,148615217 | 0,125104532 | 0,188104532 | -1,071895468 |
| chrXII | 468235 | 468399 | 164 | 0,156725832 | 0,17417587 | 0,473632595 | -0,066141475 | -0,056220254 | -0,028588157 | -0,168588157 | -1,428588157 |
| chrXII | 468303 | 468465 | 162 | 0,062690333 | 0,055588044 | 0,530023616 | 0,075329227 | 0,064029843 | 0,130079181 | 0,004079181 | -1,255920819 |
| chrXII | 468322 | 468465 | 143 | 0,800726524 | 0,52993935 | 0,601748748 | 0,257876135 | 0,219194715 | 0,050927602 | 0,057927602 | -1,202072398 |
| chrXII | 468341 | 468490 | 149 | 0,814974327 | 0,396528045 | 0,672697261 | 0,4473734 | 0,38026739 | 0,216774829 | 0,181774829 | -1,078225171 |
| chrXII | 468341 | 468513 | 172 | 0,062690333 | 0,059293913 | 0,51392155 | 0,034903236 | 0,029667751 | 0,153402831 | -0,042597169 | -1,302597169 |
| chrXII | 468341 | 468556 | 215 | 0,04844253 | 0,062999783 | 0,434687048 | -0,164453569 | -0,139785534 | 0,185853124 | -0,311146876 | -1,571146876 |
| chrXII | 468364 | 468513 | 149 | 0,757983115 | 0,633703698 | 0,544650641 | 0,112157259 | 0,09533367 | -0,065222575 | -0,100222575 | -1,360222575 |
| chrXII | 468364 | 468554 | 190 | 0,037044288 | 0,066705652 | 0,357053581 | -0,366345661 | -0,311393812 | -0,284584018 | -0,606584018 | -1,866584018 |
| chrXII | 468364 | 468556 | 192 | 0,071239015 | 0,103764348 | 0,407072261 | -0,235082731 | -0,199820321 | -0,131382058 | -0,467382058 | -1,727382058 |
| chrXII | 468364 | 468569 | 205 | 0,037044288 | 0,051882174 | 0,416572153 | -0,210670615 | -0,179070023 | -0,113549754 | -0,540549754 | -1,800549754 |
| chrXII | 468372 | 468532 | 160 | 0,168124074 | 0,062999783 | 0,727419819 | 0,605028046 | 0,514273839 | 0,502629408 | 0,390629408 | -0,869370592 |
| chrXII | 468373 | 468530 | 157 | 0,427434088 | 0,429880871 | 0,498572996 | -0,003576976 | -0,00304043 | -0,054697754 | -0,145697754 | -1,405697754 |
| chrXII | 468373 | 468554 | 181 | 0,136778908 | 0,103764348 | 0,568624996 | 0,172874585 | 0,146943398 | 0,320394299 | 0,061394299 | -1,198605701 |
| chrXII | 468373 | 468556 | 183 | 0,259310013 | 0,340940001 | 0,432003343 | -0,171276081 | -0,145584669 | -0,112116746 | -0,385116746 | -1,645116746 |
| chrXII | 468401 | 468490 | 89 | 0,116831984 | 0,092646739 | 0,55772721 | 0,145209355 | 0,123427951 | 0,554163856 | 0,939163856 | -0,320836144 |
| chrXII | 468401 | 468513 | 112 | 0,151026711 | 0,226058044 | 0,400511314 | -0,252023851 | -0,214220273 | 0,321459698 | 0,545459698 | -0,714540302 |
| chrXII | 468401 | 468532 | 131 | 0,039893848 | 0,040764565 | 0,49460244 | -0,013530088 | -0,011500575 | 0,095875161 | 0,186875161 | -1,073124839 |
| chrXII | 468401 | 468554 | 153 | 2,239754619 | 1,515700658 | 0,596400291 | 0,244040545 | 0,207434463 | 0,086055611 | 0,023055611 | -1,236944389 |
| chrXII | 468401 | 468556 | 155 | 2,137170438 | 1,767699789 | 0,547308954 | 0,118865275 | 0,101035484 | 0,054820825 | -0,022179175 | -1,282179175 |
| chrXII | 468401 | 468569 | 168 | 0,87766466 | 0,770820872 | 0,53240665 | 0,081320967 | 0,069122822 | 0,162309563 | -0,005690437 | -1,265690437 |

TABLE 2

| Chrom | Start | End | Length | Section A | Section B | A/A+B | Z-score | Z * 0.85 | Phase correction | Length correction | ΔLK |
|---|---|---|---|---|---|---|---|---|---|---|---|
| chrXII | 468401 | 468578 | 177 | 0,812124766 | 0,959820221 | 0,458323916 | -0,10465719 | -0,088958611 | 0,050360836 | -0,180639164 | -1,440639164 |
| chrXII | 468402 | 468513 | 111 | 0,056991212 | 0,070411522 | 0,447331152 | -0,132407093 | -0,112546029 | 0,364467781 | 0,595467781 | -0,664532219 |
| chrXII | 468402 | 468530 | 128 | 0,202318801 | 0,181587609 | 0,52700032 | 0,067731518 | 0,05757179 | 0,303723315 | 0,415723315 | -0,844276685 |
| chrXII | 468402 | 468550 | 148 | 0,051292091 | 0,037058696 | 0,58055047 | 0,203301913 | 0,172806626 | 0,054765271 | 0,026765271 | -1,233234729 |
| chrXII | 468402 | 468553 | 151 | 0,042743409 | 0,074117392 | 0,365763444 | -0,34309514 | -0,291630869 | -0,420648055 | -0,469648055 | -1,729648055 |
| chrXII | 468402 | 468554 | 152 | 1,572957442 | 1,226642831 | 0,561850725 | 0,155663143 | 0,132313672 | -0,03183442 | -0,08783442 | -1,34783442 |
| chrXII | 468402 | 468556 | 154 | 11,17312705 | 7,1263872 | 0,610569597 | 0,280803909 | 0,238683323 | 0,088976806 | 0,018976806 | -1,241023194 |
| chrXII | 468402 | 468559 | 157 | 0,199469241 | 0,155646522 | 0,561702018 | 0,155285859 | 0,13199298 | 0,080755179 | -0,010244821 | -1,270244821 |
| chrXII | 468402 | 468569 | 167 | 0,168124074 | 0,259410871 | 0,393240544 | -0,270882911 | -0,230250475 | -0,099945279 | -0,260945279 | -1,520945279 |
| chrXII | 468402 | 468578 | 176 | 0,059840772 | 0,196411088 | 0,23352327 | -0,727292911 | -0,618198974 | -0,45101063 | -0,67501063 | -1,93501063 |
| chrXII | 468402 | 468604 | 202 | 0,116831984 | 0,074117392 | 0,61184795 | 0,284138687 | 0,241517884 | 0,26679045 | -0,13920955 | -1,39920955 |
| chrXII | 468402 | 468606 | 204 | 0,185221438 | 0,188999349 | 0,494952297 | -0,012653052 | -0,010755094 | 0,051015704 | -0,368984296 | -1,628984296 |
| chrXII | 468404 | 468530 | 126 | 0,054141651 | 0,074117392 | 0,42212736 | -0,196454145 | -0,166986023 | 0,062766605 | 0,188766605 | -1,071233395 |
| chrXII | 468404 | 468554 | 150 | 0,606956404 | 0,392822175 | 0,607090827 | 0,271744676 | 0,230982975 | 0,068141063 | 0,026141063 | -1,233858937 |
| chrXII | 468404 | 468556 | 152 | 2,823914538 | 1,574994571 | 0,641957919 | 0,363697164 | 0,309142589 | 0,139189658 | 0,083189658 | -1,176810342 |
| chrXII | 468404 | 468559 | 155 | 0,703841464 | 0,403939784 | 0,635361417 | 0,346087223 | 0,29417414 | 0,249887945 | 0,172887945 | -1,087112055 |
| chrXII | 468404 | 468569 | 165 | 0,188070999 | 0,151940653 | 0,553131041 | 0,13357593 | 0,113539541 | 0,22776447 | 0,08076447 | -1,17923553 |
| chrXII | 468404 | 468578 | 174 | 0,037044288 | 0,070411522 | 0,344739737 | -0,399561558 | -0,339627325 | -0,074182889 | -0,284182889 | -1,544182889 |
| chrXII | 468404 | 468606 | 202 | 0,056991212 | 0,044470435 | 0,561702018 | 0,155285859 | 0,13199298 | 0,156966565 | -0,249033435 | -1,509033435 |
| chrXII | 468405 | 468513 | 108 | 0,037044288 | 0,122293696 | 0,232488743 | -0,730675332 | -0,621074032 | -0,013063141 | 0,238936859 | -1,021063141 |
| chrXII | 468405 | 468530 | 125 | 0,222265726 | 0,285351958 | 0,437860486 | -0,156395895 | -0,132936511 | 0,127156126 | 0,260156126 | -0,999843874 |
| chrXII | 468405 | 468532 | 127 | 0,04844253 | 0,081529131 | 0,372716096 | -0,324668217 | -0,275967985 | -0,047020613 | 0,071979387 | -1,188020613 |
| chrXII | 468405 | 468550 | 145 | 0,068389454 | 0,040764565 | 0,626540868 | 0,322705529 | 0,274299699 | 0,117719359 | 0,110719359 | -1,149280641 |
| chrXII | 468405 | 468553 | 148 | 0,068389454 | 0,077823261 | 0,467739443 | -0,080953558 | -0,068810524 | -0,192884123 | -0,220884123 | -1,480884123 |
| chrXII | 468405 | 468554 | 149 | 2,117223514 | 1,241466309 | 0,630371849 | 0,332838358 | 0,282912604 | 0,127052079 | 0,092052079 | -1,167947921 |
| chrXII | 468405 | 468556 | 151 | 10,80268417 | 5,866391543 | 0,648067377 | 0,380108002 | 0,323091801 | 0,190035171 | 0,141035171 | -1,118964829 |
| chrXII | 468405 | 468559 | 154 | 0,296354301 | 0,17417587 | 0,629830602 | 0,331404725 | 0,281694016 | 0,121679706 | 0,051679706 | -1,208320294 |
| chrXII | 468405 | 468568 | 163 | 0,105433742 | 0,103764348 | 0,503989982 | 0,010001569 | 0,008501334 | 0,006317162 | -0,126682838 | -1,386682838 |
| chrXII | 468405 | 468569 | 164 | 1,766727562 | 1,437877396 | 0,551309002 | 0,128969231 | 0,109623846 | 0,148489617 | 0,008489617 | -1,251510383 |
| chrXII | 468405 | 468578 | 173 | 0,071239015 | 0,144528914 | 0,330164984 | -0,439457641 | -0,373538995 | -0,168934013 | -0,371934013 | -1,631934013 |
| chrXII | 468405 | 468604 | 199 | 0,173823196 | 0,066705652 | 0,722670886 | 0,590794338 | 0,502175188 | 0,532680747 | 0,147680747 | -1,112319253 |
| chrXII | 468405 | 468606 | 201 | 0,222265726 | 0,181587609 | 0,550362486 | 0,126577221 | 0,107590638 | 0,125964251 | -0,273035749 | -1,533035749 |
| chrXII | 468406 | 468513 | 107 | 0,045592969 | 0,255705001 | 0,15132186 | -1,030780594 | -0,876163505 | -0,317675714 | -0,058675714 | -1,318675714 |
| chrXII | 468406 | 468530 | 124 | 0,122531105 | 0,300175436 | 0,289872744 | -0,55375652 | -0,470693042 | -0,068335156 | 0,071664844 | -1,188335156 |
| chrXII | 468406 | 468550 | 144 | 0,082637257 | 0,051882174 | 0,614314648 | 0,290582424 | 0,24699506 | 0,048490104 | 0,048490104 | -1,211509896 |
| chrXII | 468406 | 468554 | 148 | 4,963934536 | 2,694167184 | 0,648193863 | 0,380448829 | 0,323381505 | 0,200887874 | 0,172887874 | -1,087112126 |
| chrXII | 468406 | 468556 | 150 | 3,31973808 | 2,175345443 | 0,604128776 | 0,264048627 | 0,224441333 | 0,055993204 | 0,013993204 | -1,246006796 |
| chrXII | 468406 | 468559 | 153 | 0,179522317 | 0,137117174 | 0,566961234 | 0,168642904 | 0,143346468 | 0,014218311 | -0,048781689 | -1,308781689 |
| chrXII | 468406 | 468568 | 162 | 0,045592969 | 0,055588044 | 0,450607955 | -0,124125498 | -0,105506674 | -0,040619073 | -0,166619073 | -1,426619073 |
| chrXII | 468406 | 468604 | 198 | 0,139628469 | 0,081529131 | 0,631352796 | 0,335438395 | 0,285122636 | 0,332136344 | -0,045863656 | -1,305863656 |
| chrXII | 468406 | 468606 | 200 | 0,091185939 | 0,114881957 | 0,442504343 | -0,144622812 | -0,12292939 | -0,102377046 | -0,494377046 | -1,754377046 |
| chrXII | 468445 | 468551 | 106 | 0,185221438 | 0,251999131 | 0,423633861 | -0,192605774 | -0,163714908 | 0,334535265 | 0,600535265 | -0,659464735 |
| chrXII | 468445 | 468554 | 109 | 0,076938136 | 0,08894087 | 0,463820816 | -0,090812431 | -0,077190567 | 0,62851509 | 0,87351509 | -0,38648491 |
| chrXII | 468445 | 468556 | 111 | 0,188070999 | 0,333528262 | 0,360566076 | -0,356946107 | -0,303404191 | 0,164728432 | 0,395728432 | -0,864271568 |
| chrXII | 468445 | 468604 | 159 | 0,085486818 | 0,051882174 | 0,622315244 | 0,311567137 | 0,264832067 | 0,271385118 | 0,166385118 | -1,093614882 |
| chrXII | 468445 | 468606 | 161 | 0,886213341 | 0,696703481 | 0,559860966 | 0,150616725 | 0,128024216 | 0,129723965 | 0,010723965 | -1,249276035 |
| chrXII | 468462 | 468603 | 141 | 0,227964847 | 0,144528914 | 0,611996417 | 0,284526191 | 0,241847262 | 0,180887453 | 0,201887453 | -1,058112547 |
| chrXII | 468463 | 468554 | 91 | 0,116831984 | 0,148234783 | 0,44076436 | -0,149031573 | -0,126676837 | 0,223227124 | 0,594227124 | -0,665772876 |
| chrXII | 468463 | 468556 | 93 | 0,370442876 | 0,359469349 | 0,507517018 | 0,018843485 | 0,016016962 | 0,478510765 | 0,835510765 | -0,424489235 |
| chrXII | 468463 | 468604 | 141 | 1,359240398 | 0,61888022 | 0,687137269 | 0,487752073 | 0,414589262 | 0,350132594 | 0,371132594 | -0,888867406 |
| chrXII | 468463 | 468606 | 143 | 0,156725832 | 0,085235 | 0,647732241 | 0,379205162 | 0,322324387 | 0,145108836 | 0,152108836 | -1,107891164 |
| chrXII | 468545 | 468696 | 151 | 0,259310013 | 0,085235 | 0,752615777 | 0,682744278 | 0,580332637 | 0,438501529 | 0,389501529 | -0,870498471 |
| chrXII | 468566 | 468665 | 99 | 0,028495606 | 0,144528914 | 0,164691143 | -0,975358855 | -0,829055027 | -0,395735727 | -0,080735727 | -1,340735727 |
| chrXII | 468566 | 468695 | 129 | 1,105629506 | 1,57870044 | 0,411882864 | -0,222704206 | -0,189298575 | -0,018024667 | 0,086975333 | -1,173024667 |
| chrXII | 468566 | 468710 | 144 | 0,045592969 | 0,040764565 | 0,527955893 | 0,070132481 | 0,059612609 | -0,132045818 | -0,132045818 | -1,392045818 |
| chrXII | 468566 | 468716 | 150 | 0,054141651 | 0,085235 | 0,388455674 | -0,283346352 | -0,2408444 | -0,401042994 | -0,443042994 | -1,703042994 |
| chrXII | 468567 | 468695 | 128 | 0,216566604 | 0,222352175 | 0,493409293 | -0,016521205 | -0,014043024 | 0,234434371 | 0,346434371 | -0,913565629 |
| chrXII | 468567 | 468710 | 143 | 0,381841118 | 0,185293479 | 0,673281299 | 0,448992035 | 0,38164323 | 0,198887661 | 0,205887661 | -1,054112339 |
| chrXII | 468567 | 468716 | 149 | 0,567062556 | 0,470645436 | 0,546456768 | 0,116714293 | 0,099207149 | -0,051445114 | -0,086445114 | -1,346445114 |
| chrXII | 468656 | 468810 | 154 | 0,225115286 | 0,144528914 | 0,609005325 | 0,276727507 | 0,235218381 | 0,069378358 | -0,000621642 | -1,260621642 |
| chrXII | 468658 | 468810 | 152 | 0,156725832 | 0,159352392 | 0,495845079 | -0,010415029 | -0,008852775 | -0,166197731 | -0,222197731 | -1,482197731 |
| chrXII | 468703 | 468859 | 156 | 0,074088575 | 0,059293913 | 0,555459536 | 0,139467258 | 0,118547169 | 0,031035157 | -0,052964843 | -1,312964843 |
| chrXII | 468739 | 468893 | 154 | 0,039893848 | 0,037058696 | 0,518421434 | 0,046192108 | 0,039263292 | -0,13375081 | -0,20375081 | -1,46375081 |
| chrXII | 468739 | 468896 | 157 | 0,051292091 | 0,037058696 | 0,58055047 | 0,203301913 | 0,172806626 | 0,120305758 | 0,029305758 | -1,230694242 |
| chrXII | 468741 | 468859 | 118 | 0,088336378 | 0,111176087 | 0,442761197 | -0,143972235 | -0,122376399 | 0,274558064 | 0,456558064 | -0,803441936 |
| chrXII | 468741 | 468893 | 152 | 5,693422046 | 3,668810883 | 0,608126511 | 0,274439387 | 0,233273479 | 0,079078142 | 0,023078142 | -1,236921858 |
| chrXII | 468741 | 468896 | 155 | 0,128230226 | 0,040764565 | 0,75878212 | 0,702390353 | 0,5970318 | 0,557937161 | 0,480937161 | -0,779062839 |
| chrXII | 468741 | 468911 | 170 | 0,387540239 | 0,26682261 | 0,592240589 | 0,233312412 | 0,19831555 | 0,359492955 | 0,177492955 | -1,082507045 |
| chrXII | 468755 | 468893 | 138 | 0,122531105 | 0,137117174 | 0,47191187 | -0,070464771 | -0,059895055 | -0,081623807 | -0,039623807 | -1,299623807 |
| chrXII | 468755 | 468911 | 156 | 0,265009134 | 0,148234783 | 0,641289861 | 0,361908673 | 0,307622372 | 0,217518715 | 0,133518715 | -1,126481285 |
| chrXII | 468758 | 468893 | 135 | 0,074088575 | 0,059293913 | 0,555459536 | 0,139467258 | 0,118547169 | 0,095920266 | 0,158920266 | -1,101079734 |
| chrXII | 468762 | 468893 | 131 | 0,045592969 | 0,107470218 | 0,297870247 | -0,530535801 | -0,450955431 | -0,337212546 | -0,246212546 | -1,506212546 |
| chrXII | 468762 | 468911 | 149 | 0,042743409 | 0,048176305 | 0,470122564 | -0,074961771 | -0,063717505 | -0,216021843 | -0,251021843 | -1,511021843 |
| chrXII | 468765 | 468893 | 128 | 0,299203861 | 0,452116089 | 0,39823761 | -0,257911486 | -0,219224763 | 0,012968118 | 0,124968118 | -1,135031882 |
| chrXII | 468765 | 468911 | 146 | 0,176672756 | 0,188999349 | 0,483145293 | -0,042261061 | -0,035921902 | -0,189343424 | -0,203343424 | -1,463343424 |
| chrXII | 468770 | 468893 | 123 | 0,327699467 | 0,255705001 | 0,561702018 | 0,155285859 | 0,13199298 | 0,586666765 | 0,733666765 | -0,526333235 |
| chrXII | 468770 | 468911 | 141 | 1,755329319 | 0,926467395 | 0,654534816 | 0,397592802 | 0,337953881 | 0,276586883 | 0,297586883 | -0,962413117 |
| chrXII | 472060 | 472213 | 153 | 0,14817715 | 0,085235 | 0,634830492 | 0,344674598 | 0,292973409 | 0,173115235 | 0,110115235 | -1,149884765 |
| chrXII | 472461 | 472545 | 84 | 0,094035499 | 0,077823261 | 0,547167331 | 0,118507771 | 0,100731606 | 0,451034737 | 0,871034737 | -0,388965263 |
| chrXII | 472461 | 472563 | 102 | 0,071239015 | 0,074117392 | 0,490098899 | -0,024820929 | -0,021097789 | 0,497126199 | 0,791126199 | -0,468873801 |
| chrXII | 475711 | 475864 | 153 | 0,111132863 | 0,085235 | 0,565942212 | 0,166052574 | 0,141144688 | 0,00894137 | -0,05405863 | -1,31405863 |
| chrXII | 476909 | 477023 | 114 | 0,059840772 | 0,433586741 | 0,121275711 | -1,168633262 | -0,993338272 | -0,374773089 | -0,164773089 | -1,424773089 |
| chrXII | 477605 | 477755 | 150 | 0,159575393 | 0,185293479 | 0,462713239 | -0,093600543 | -0,079560461 | -0,229104029 | -0,271104029 | -1,531104029 |
| chrXII | 478538 | 478667 | 129 | 0,062690333 | 0,092646739 | 0,403576119 | -0,244101464 | -0,207486244 | -0,04839556 | 0,05660444 | -1,20339556 |
| chrXII | 479001 | 479158 | 157 | 0,059840772 | 0,066705652 | 0,472876041 | -0,06804215 | -0,057835827 | -0,114621638 | -0,205621638 | -1,465621638 |
| chrXII | 479187 | 479337 | 150 | 8,705407583 | 4,109809363 | 0,679302397 | 0,465748954 | 0,395886611 | 0,252516974 | 0,210516974 | -1,049483026 |
| chrXII | 479469 | 479630 | 161 | 0,079787696 | 0,111176087 | 0,417815854 | -0,20748423 | -0,176361595 | -0,187850609 | -0,306850609 | -1,566850609 |
| chrXII | 480195 | 480349 | 154 | 0,202318801 | 0,144528914 | 0,583307292 | 0,210361658 | 0,178807409 | -0,000920335 | -0,070920335 | -1,330920335 |
| chrXII | 480476 | 480621 | 145 | 0,062690333 | 0,066705652 | 0,484484373 | -0,038901718 | -0,03306646 | -0,18811423 | -0,19511423 | -1,45511423 |
| chrXII | 480480 | 480631 | 151 | 0,125380666 | 0,081529131 | 0,605967759 | 0,268824834 | 0,228501109 | 0,085107564 | 0,036107564 | -1,223892436 |
| chrXII | 480687 | 480827 | 140 | 0,170973635 | 0,111176087 | 0,605967759 | 0,268824834 | 0,228501109 | 0,283240276 | 0,311240276 | -0,948759724 |
| chrXII | 480693 | 480827 | 134 | 0,153876272 | 0,129705435 | 0,542617058 | 0,107029113 | 0,090974746 | 0,10187273 | 0,17187273 | -1,08812727 |
| chrXII | 481105 | 481260 | 155 | 0,094035499 | 0,096352609 | 0,493914773 | -0,015253994 | -0,012965895 | -0,037396656 | -0,114396656 | -1,374396656 |
| chrXII | 481236 | 481344 | 108 | 0,068389454 | 0,148234783 | 0,315705458 | -0,47974192 | -0,407780632 | 0,189218296 | 0,441218296 | -0,818781704 |
| chrXII | 481385 | 481534 | 149 | 0,076938136 | 0,037058696 | 0,674914686 | 0,453525163 | 0,385496389 | 0,242830799 | 0,207830799 | -1,052169201 |
| chrXII | 482041 | 482125 | 84 | 0,091185939 | 0,081529131 | 0,527955893 | 0,070132481 | 0,059612609 | 0,442060116 | 0,862060116 | -0,397939884 |
| chrXII | 482041 | 482143 | 102 | 0,088336378 | 0,111176087 | 0,442761197 | -0,143972235 | -0,122376399 | 0,410409522 | 0,704409522 | -0,555590478 |
| chrXII | 482885 | 483037 | 152 | 0,042743409 | 0,051882174 | 0,451710916 | -0,121339881 | -0,103138899 | -0,250952739 | -0,306952739 | -1,566952739 |
| chrXII | 485292 | 485445 | 153 | 0,131079787 | 0,077823261 | 0,627467086 | 0,325152274 | 0,276379433 | 0,126261587 | 0,063261587 | -1,196738413 |
| chrXII | 485693 | 485777 | 84 | 0,088336378 | 0,08894087 | 0,498295067 | -0,004273647 | -0,0036326 | 0,387667577 | 0,807667577 | -0,452332423 |
| chrXII | 488944 | 489097 | 153 | 0,159575393 | 0,066705652 | 0,705208837 | 0,539441393 | 0,458525184 | 0,292209921 | 0,229209921 | -1,030790079 |

TABLE 2

| Chrom | Start | End | Length | Section A | Section B | A/A+B | Z-score | Z * 0.85 | Phase correction | Length correction | ΔLK |
|---|---|---|---|---|---|---|---|---|---|---|---|
| chrXII | 489345 | 489429 | 84 | 0,105433742 | 0,133411305 | 0,441431561 | -0,147340685 | -0,125239582 | 0,261279712 | 0,681279712 | -0,578720288 |
| chrXII | 489345 | 489447 | 102 | 0,076938136 | 0,085235 | 0,474419732 | -0,064164224 | -0,05453959 | 0,505240769 | 0,799240769 | -0,460759231 |
| chrXII | 489345 | 489466 | 121 | 0,04844253 | 0,074117392 | 0,395255882 | -0,265646117 | -0,2257992 | 0,173375695 | 0,334375695 | -0,925624305 |
| chrXII | 489345 | 489469 | 124 | 0,661098055 | 0,744879785 | 0,470205174 | -0,074754118 | -0,063541 | 0,347035325 | 0,487035325 | -0,772964675 |
| chrXII | 489345 | 489471 | 126 | 0,216566604 | 0,26311674 | 0,451478266 | -0,12192738 | -0,103638273 | 0,131176332 | 0,257176332 | -1,002823668 |
| chrXII | 489359 | 489469 | 110 | 0,202318801 | 0,663350655 | 0,233713688 | -0,726671238 | -0,617670553 | 0,028117089 | 0,266117089 | -0,993882911 |
| chrXII | 489359 | 489471 | 112 | 0,04844253 | 0,185293479 | 0,207253175 | -0,815989135 | -0,693590765 | -0,141352657 | 0,082647343 | -1,177352657 |
| chrXII | 489361 | 489466 | 105 | 0,108283302 | 0,481763045 | 0,183516605 | -0,902044538 | -0,766737857 | -0,254056183 | 0,018943817 | -1,241056183 |
| chrXII | 489556 | 489702 | 146 | 0,572761677 | 0,396528045 | 0,590908646 | 0,229882972 | 0,195400527 | 0,032418859 | 0,018418859 | -1,241581141 |
| chrXII | 489926 | 490077 | 151 | 0,450230572 | 0,233469783 | 0,658520313 | 0,408428145 | 0,347163923 | 0,218092853 | 0,169092853 | -1,090907147 |
| chrXII | 489933 | 490080 | 147 | 0,039893848 | 0,037058696 | 0,518421434 | 0,046192108 | 0,039263292 | -0,071555735 | -0,092555735 | -1,352555735 |
| chrXII | 490009 | 490164 | 155 | 0,199469241 | 0,148234783 | 0,573675388 | 0,185739279 | 0,157878387 | 0,12672154 | 0,04972154 | -1,21027846 |
| chrXII | 490027 | 490183 | 156 | 0,270708255 | 0,192705218 | 0,584161383 | 0,212550948 | 0,180668306 | 0,084222592 | 0,000222592 | -1,259777408 |
| chrXII | 490049 | 490195 | 146 | 0,621204207 | 0,35576348 | 0,635849287 | 0,347385899 | 0,295278014 | 0,137110251 | 0,123110251 | -1,136889749 |
| chrXII | 490049 | 490215 | 166 | 1,216762369 | 1,271113265 | 0,489076846 | -0,027383708 | -0,023276152 | 0,137859987 | -0,016140013 | -1,276140013 |
| chrXII | 490049 | 490219 | 170 | 0,074088575 | 0,040764565 | 0,645072262 | 0,372050197 | 0,316242667 | 0,485424364 | 0,303424364 | -0,956575636 |
| chrXII | 490072 | 490215 | 143 | 0,085486818 | 0,062999783 | 0,575720754 | 0,190958009 | 0,162314307 | -0,015629292 | -0,008629292 | -1,268629292 |
| chrXII | 490072 | 490219 | 147 | 0,279256937 | 0,129705435 | 0,682842618 | 0,475662609 | 0,404313218 | 0,292508077 | 0,271508077 | -0,988491923 |
| chrXII | 490083 | 490239 | 156 | 0,564212996 | 0,374292827 | 0,6011822 | 0,256408274 | 0,217947033 | 0,131616633 | 0,047616633 | -1,212383367 |
| chrXII | 490158 | 490312 | 154 | 0,626903328 | 0,478057176 | 0,567353608 | 0,169640608 | 0,144194517 | -0,029112771 | -0,099112771 | -1,359112771 |
| chrXII | 490158 | 490312 | 154 | 0,626903328 | 0,478057176 | 0,567353608 | 0,169640608 | 0,144194517 | -0,029112771 | -0,099112771 | -1,359112771 |
| chrXII | 490228 | 490334 | 106 | 0,039893848 | 0,26311674 | 0,131658264 | -1,118586632 | -0,950798637 | -0,45529239 | -0,18929239 | -1,44929239 |
| chrXII | 490228 | 490337 | 109 | 0,04844253 | 0,389116306 | 0,110710894 | -1,222756213 | -1,039342781 | -0,34460868 | -0,09960868 | -1,35960868 |
| chrXII | 490228 | 490382 | 154 | 0,410336724 | 0,248293262 | 0,623015552 | 0,313410384 | 0,266398826 | 0,095747842 | 0,025747842 | -1,234252158 |
| chrXII | 490261 | 490368 | 107 | 0,116831984 | 0,867173481 | 0,118731032 | -1,181354153 | -1,00415103 | -0,443551773 | -0,184551773 | -1,444551773 |
| chrXII | 490262 | 490423 | 161 | 0,521469587 | 0,492880654 | 0,514092239 | 0,035331355 | 0,030031652 | 0,022093478 | -0,096906522 | -1,356906522 |
| chrXII | 490263 | 490423 | 160 | 0,327699467 | 0,277940218 | 0,541079911 | 0,103154718 | 0,08768151 | 0,071376015 | -0,040623985 | -1,300623985 |
| chrXII | 490267 | 490423 | 156 | 0,19661968 | 0,151940653 | 0,564090809 | 0,161349191 | 0,137146813 | 0,052814173 | -0,031185827 | -1,291185827 |
| chrXII | 490291 | 490423 | 132 | 0,056991212 | 0,051882174 | 0,523463207 | 0,058847485 | 0,050020363 | 0,165441325 | 0,249441325 | -1,010558675 |
| chrXII | 490302 | 490409 | 107 | 0,037044288 | 0,096352609 | 0,277699771 | -0,589688448 | -0,501235181 | 0,054717084 | 0,313717084 | -0,946282916 |
| chrXII | 490302 | 490423 | 121 | 0,031345166 | 0,051882174 | 0,376621026 | -0,314367347 | -0,267212245 | 0,109482758 | 0,270482758 | -0,989517242 |
| chrXII | 494931 | 495079 | 148 | 0,849169054 | 0,544762828 | 0,609189778 | 0,277207939 | 0,235626748 | 0,099978359 | 0,071978359 | -1,188021641 |
| chrXII | 496357 | 496507 | 150 | 0,29065518 | 0,111176087 | 0,723326439 | 0,592752021 | 0,503839218 | 0,351274434 | 0,309274434 | -0,950725566 |
| chrXII | 508585 | 508725 | 140 | 0,159575393 | 0,151940653 | 0,512254168 | 0,030721476 | 0,026113255 | 0,072767973 | 0,100767973 | -1,159232027 |
| chrXII | 514529 | 514679 | 150 | 6,665122205 | 3,072165881 | 0,684494712 | 0,480304941 | 0,4082592 | 0,257987018 | 0,215987018 | -1,044012982 |
| chrXII | 518369 | 518501 | 132 | 0,108283302 | 0,192705218 | 0,359758911 | -0,359103289 | -0,305237796 | -0,215154497 | -0,131154497 | -1,391154497 |
| chrXII | 522434 | 522581 | 147 | 0,336248149 | 0,214940436 | 0,610041932 | 0,279428325 | 0,237514076 | 0,134656097 | 0,113656097 | -1,146343903 |
| chrXII | 528715 | 528875 | 160 | 0,333398588 | 0,35576348 | 0,483773852 | -0,040684142 | -0,03458152 | -0,055233078 | -0,167233078 | -1,427233078 |
| chrXII | 529249 | 529419 | 170 | 0,102584181 | 0,211234566 | 0,326889907 | -0,448517424 | -0,381239811 | -0,220679256 | -0,402679256 | -1,662679256 |
| chrXII | 537872 | 537976 | 104 | 0,153876272 | 0,837526525 | 0,155210649 | -1,014338418 | -0,862187655 | -0,272899303 | 0,007100697 | -1,252899303 |
| chrXII | 542120 | 542273 | 153 | 0,29350474 | 0,377998697 | 0,437085983 | -0,158361478 | -0,134607257 | -0,305624905 | -0,368624905 | -1,628624905 |
| chrXII | 544133 | 544261 | 128 | 0,498673102 | 0,919055655 | 0,351740839 | -0,380624799 | -0,323531079 | -0,086353244 | 0,025646756 | -1,234353244 |
| chrXII | 545204 | 545351 | 147 | 2,530409798 | 1,67134718 | 0,602226595 | 0,259114613 | 0,220247421 | 0,119182351 | 0,098182351 | -1,161817649 |
| chrXII | 546037 | 546189 | 152 | 17,92088651 | 7,000387634 | 0,719099931 | 0,580169768 | 0,493144303 | 0,326083951 | 0,270083951 | -0,989916049 |
| chrXII | 546038 | 546189 | 151 | 0,045592969 | 0,044470435 | 0,506231912 | 0,015621722 | 0,013278464 | -0,124649953 | -0,173649953 | -1,433649953 |
| chrXII | 546040 | 546189 | 149 | 0,14817715 | 0,103764348 | 0,588141101 | 0,222765786 | 0,189350918 | 0,053605712 | 0,018605712 | -1,241394288 |
| chrXII | 547890 | 548039 | 149 | 0,059840772 | 0,037058696 | 0,617555219 | 0,299066166 | 0,254206241 | 0,113379706 | 0,078379706 | -1,181620294 |
| chrXII | 553542 | 553689 | 147 | 2,259701543 | 1,360054135 | 0,624269079 | 0,316712396 | 0,269205536 | 0,173377474 | 0,152377474 | -1,107622526 |
| chrXII | 555694 | 555847 | 153 | 0,458779254 | 0,363175219 | 0,558156527 | 0,146296985 | 0,124352437 | -0,050030693 | -0,113030693 | -1,373030693 |
| chrXII | 559139 | 559286 | 147 | 0,772230918 | 0,559586306 | 0,579832505 | 0,201465007 | 0,171245256 | 0,071316519 | 0,050316519 | -1,209683481 |
| chrXII | 559606 | 559755 | 149 | 0,176672756 | 0,066705652 | 0,725917953 | 0,600513459 | 0,51043644 | 0,375374992 | 0,340374992 | -0,919625008 |
| chrXII | 560049 | 560194 | 145 | 0,056991212 | 0,040764565 | 0,582995843 | 0,209563572 | 0,178129036 | 0,026889769 | 0,019889769 | -1,240110231 |
| chrXII | 570252 | 570403 | 151 | 0,076938136 | 0,048176305 | 0,614942093 | 0,292223409 | 0,248389898 | 0,113954422 | 0,064954422 | -1,195045578 |
| chrXII | 573533 | 573669 | 136 | 11,95390665 | 8,386382856 | 0,587695993 | 0,221622182 | 0,188378855 | 0,219720685 | 0,275720685 | -0,984279315 |
| chrXII | 576803 | 576966 | 163 | 0,481575739 | 0,52993935 | 0,47609348 | -0,059960669 | -0,050966568 | -0,046396479 | -0,179396479 | -1,439396479 |
| chrXII | 586445 | 586571 | 126 | 0,769381358 | 2,542226531 | 0,23232864 | -0,731199541 | -0,62151961 | -0,369237124 | -0,243237124 | -1,503237124 |
| chrXII | 587477 | 587583 | 106 | 0,091185939 | 0,496586524 | 0,155138161 | -1,014562398 | -0,862446038 | -0,384130062 | -0,118130062 | -1,378130062 |
| chrXII | 588088 | 588237 | 149 | 0,153876272 | 0,100058479 | 0,605967759 | 0,268824834 | 0,228501109 | 0,094061789 | 0,059061789 | -1,200938211 |
| chrXII | 593332 | 593484 | 152 | 0,054141651 | 0,037058696 | 0,593656197 | 0,236960278 | 0,201416237 | 0,030555685 | -0,025444315 | -1,285444315 |
| chrXII | 593504 | 593653 | 149 | 0,102584181 | 0,055588044 | 0,648560019 | 0,381435717 | 0,324220359 | 0,201164322 | 0,166164322 | -1,093835678 |
| chrXII | 593678 | 593801 | 123 | 0,076938136 | 0,192705218 | 0,285332958 | -0,567071042 | -0,482010386 | -0,039554228 | 0,107445772 | -1,152554228 |
| chrXII | 593778 | 593933 | 155 | 0,079787696 | 0,077823261 | 0,506231912 | 0,015621722 | 0,013278464 | -0,012428874 | -0,089428874 | -1,349428874 |
| chrXII | 594211 | 594345 | 134 | 0,128230226 | 0,122293696 | 0,51184823 | 0,029703476 | 0,025247955 | 0,029611292 | 0,099611292 | -1,160388708 |
| chrXII | 594211 | 594351 | 140 | 0,190920559 | 0,155646522 | 0,550890634 | 0,127911851 | 0,108725073 | 0,159903297 | 0,187903297 | -1,072096703 |
| chrXII | 594407 | 594558 | 151 | 0,136778908 | 0,066705652 | 0,672183225 | 0,445949741 | 0,37905728 | 0,237150716 | 0,188150716 | -1,071849284 |
| chrXII | 594417 | 594562 | 145 | 0,122531105 | 0,066705652 | 0,64750161 | 0,378584033 | 0,321796428 | 0,155074964 | 0,148074964 | -1,111925036 |
| chrXII | 594435 | 594586 | 151 | 0,045592969 | 0,062999783 | 0,419852784 | -0,202270107 | -0,171929591 | -0,311286771 | -0,360286771 | -1,620286771 |
| chrXII | 594689 | 594843 | 154 | 0,210867483 | 0,155646522 | 0,575332675 | 0,189967434 | 0,161472318 | -0,005690177 | -0,075690177 | -1,335690177 |
| chrXII | 594723 | 594867 | 144 | 0,051292091 | 0,037058696 | 0,58055047 | 0,203301913 | 0,172806626 | -0,006666298 | -0,006666298 | -1,266666298 |
| chrXII | 595254 | 595389 | 135 | 0,068389454 | 0,070411522 | 0,492715945 | -0,018259434 | -0,015520519 | -0,044025624 | 0,018974376 | -1,241025624 |
| chrXII | 595436 | 595568 | 132 | 0,190920559 | 0,229763914 | 0,453833149 | -0,115982643 | -0,098585246 | -0,012026385 | 0,071973615 | -1,188026385 |
| chrXII | 595525 | 595663 | 138 | 0,555664314 | 0,455821958 | 0,549354281 | 0,124030106 | 0,10542559 | 0,07087782 | 0,11287782 | -1,14712218 |
| chrXII | 598015 | 598129 | 114 | 0,028495606 | 0,229763914 | 0,110337098 | -1,224737354 | -1,041026751 | -0,411548698 | -0,201548698 | -1,461548698 |
| chrXII | 598489 | 598636 | 147 | 0,076938136 | 0,040764565 | 0,653664997 | 0,395234266 | 0,335949126 | 0,237455117 | 0,216455117 | -1,043544883 |
| chrXII | 603237 | 603400 | 163 | 0,131079787 | 0,074117392 | 0,638799168 | 0,355250862 | 0,301963233 | 0,303964955 | 0,170964955 | -1,089035045 |
| chrXII | 607039 | 607198 | 159 | 0,094035499 | 0,133411305 | 0,413439528 | -0,218706025 | -0,185900121 | -0,175087145 | -0,280087145 | -1,540087145 |
| chrXII | 608189 | 608339 | 150 | 0,051292091 | 0,137117174 | 0,272237623 | -0,606059495 | -0,515150571 | -0,668000913 | -0,710000913 | -1,970000913 |
| chrXII | 610208 | 610357 | 149 | 1,632798214 | 1,05617283 | 0,607220453 | 0,272081837 | 0,231269561 | 0,115719056 | 0,080719056 | -1,179280944 |
| chrXII | 611103 | 611232 | 129 | 0,259310013 | 0,163058261 | 0,613942923 | 0,289610604 | 0,246169013 | 0,392831321 | 0,497831321 | -0,762168679 |
| chrXII | 614828 | 614977 | 149 | 0,051292091 | 0,122293696 | 0,295485544 | -0,537429332 | -0,456814932 | -0,579652591 | -0,614652591 | -1,874652591 |
| chrXII | 614828 | 614997 | 169 | 0,344796831 | 0,737468046 | 0,318588211 | -0,471650294 | -0,40090275 | -0,2325472 | -0,4075472 | -1,6675472 |
| chrXII | 614830 | 614977 | 147 | 0,663947616 | 0,655938915 | 0,503033859 | 0,007604831 | 0,006464107 | -0,097254727 | -0,118254727 | -1,378254727 |
| chrXII | 625233 | 625381 | 148 | 0,378991558 | 0,251999131 | 0,600629398 | 0,254976559 | 0,216730075 | 0,085676016 | 0,057676016 | -1,202323984 |
| chrXII | 628912 | 629085 | 173 | 0,153876272 | 0,403939784 | 0,275854863 | -0,595200097 | -0,505920082 | -0,305049895 | -0,508049895 | -1,768049895 |
| chrXII | 631988 | 632127 | 139 | 0,849169054 | 0,830114786 | 0,505673332 | 0,014221413 | 0,012088201 | 0,02209834 | 0,05709834 | -1,20290166 |
| chrXII | 635940 | 636108 | 168 | 0,299203861 | 0,444704349 | 0,402205349 | -0,247642923 | -0,210496484 | -0,115986029 | -0,283986029 | -1,543986029 |
| chrXII | 640543 | 640689 | 146 | 0,803576085 | 0,548468698 | 0,594341323 | 0,238726903 | 0,202917868 | 0,040885225 | 0,026885225 | -1,233114775 |
| chrXII | 649877 | 650024 | 147 | 0,059840772 | 0,085235 | 0,412479432 | -0,221171546 | -0,187995814 | -0,289065353 | -0,310065353 | -1,570065353 |
| chrXII | 649877 | 650024 | 147 | 0,059840772 | 0,085235 | 0,412479432 | -0,221171546 | -0,187995814 | -0,289065353 | -0,310065353 | -1,570065353 |
| chrXII | 651183 | 651332 | 149 | 0,071239015 | 0,066705652 | 0,516431814 | 0,041200102 | 0,035020087 | -0,094171765 | -0,129171765 | -1,389171765 |
| chrXII | 651357 | 651480 | 123 | 0,079787696 | 0,17788174 | 0,309651379 | -0,496838761 | -0,422312947 | 0,022579957 | 0,169579957 | -1,090420043 |
| chrXII | 651457 | 651612 | 155 | 0,105433742 | 0,051882174 | 0,670203909 | 0,440476288 | 0,374404845 | 0,358145426 | 0,281145426 | -0,978854574 |
| chrXII | 651890 | 652024 | 134 | 0,136778908 | 0,118587827 | 0,535617563 | 0,08939893 | 0,07598909 | 0,079566427 | 0,149566427 | -1,110433573 |
| chrXII | 651890 | 652030 | 140 | 0,19377012 | 0,103764348 | 0,65125268 | 0,388704654 | 0,330398956 | 0,349960031 | 0,377960031 | -0,882039969 |
| chrXII | 652086 | 652237 | 151 | 0,116831984 | 0,070411522 | 0,623957468 | 0,315891237 | 0,268507552 | 0,129742996 | 0,080742996 | -1,179257004 |
| chrXII | 652096 | 652241 | 145 | 0,119681545 | 0,051882174 | 0,69759239 | 0,517488464 | 0,439865194 | 0,281528831 | 0,274528831 | -0,985471169 |
| chrXII | 652114 | 652265 | 151 | 0,045592969 | 0,044470435 | 0,506231912 | 0,015621722 | 0,013278464 | -0,126100285 | -0,175100285 | -1,435100285 |
| chrXII | 652116 | 652265 | 149 | 0,054141651 | 0,044470435 | 0,549036667 | 0,123227859 | 0,104743681 | -0,032548019 | -0,067548019 | -1,327548019 |
| chrXII | 652368 | 652522 | 154 | 0,190920559 | 0,203822827 | 0,483657398 | -0,040976291 | -0,034829847 | -0,211774625 | -0,281774625 | -1,541774625 |
| chrXII | 652618 | 652786 | 168 | 0,051292091 | 0,048176305 | 0,515662191 | 0,039269381 | 0,033378974 | 0,121480336 | -0,046519664 | -1,306519664 |
| chrXII | 652933 | 653068 | 135 | 0,094035499 | 0,070411522 | 0,571828535 | 0,1810314 | 0,15387669 | 0,119354371 | 0,182354371 | -1,077645629 |

TABLE 2

| Chrom | Start | End | Length | Section A | Section B | A/A+B | Z-score | Z * 0.85 | Phase correction | Length correction | ΔLK |
|---|---|---|---|---|---|---|---|---|---|---|---|
| chrXII | 653087 | 653248 | 161 | 0,068389454 | 0,066705652 | 0,506231912 | 0,015621722 | 0,013278464 | 0,009984022 | -0,109015978 | -1,369015978 |
| chrXII | 653194 | 653339 | 145 | 0,113982423 | 0,081529131 | 0,582995843 | 0,209563572 | 0,178129036 | 0,020927992 | 0,013927992 | -1,246072008 |
| chrXII | 653561 | 653718 | 157 | 0,062690333 | 0,059293913 | 0,51392155 | 0,034903236 | 0,029667751 | -0,02250691 | -0,11350691 | -1,37350691 |
| chrXII | 654052 | 654181 | 129 | 0,071239015 | 0,118587827 | 0,375284202 | -0,317889966 | -0,270206471 | -0,126861347 | -0,021861347 | -1,281861347 |
| chrXII | 654964 | 655114 | 150 | 0,156725832 | 0,163058261 | 0,490098899 | -0,024820929 | -0,021097789 | -0,16402872 | -0,20602872 | -1,46602872 |
| chrXII | 656400 | 656549 | 149 | 0,076938136 | 0,059293913 | 0,564757972 | 0,163043664 | 0,138587115 | 0,000335264 | -0,034664736 | -1,294664736 |
| chrXII | 657729 | 657887 | 158 | 0,219416165 | 0,322410653 | 0,404956266 | -0,240538873 | -0,204458042 | -0,234834728 | -0,332834728 | -1,592834728 |
| chrXII | 664370 | 664478 | 108 | 0,561363435 | 2,253168704 | 0,199451776 | -0,843581058 | -0,717043899 | -0,104143121 | 0,147856879 | -1,112143121 |
| chrXII | 664370 | 664482 | 112 | 0,418885406 | 2,468109139 | 0,145093937 | -1,057709561 | -0,899053127 | -0,329450233 | -0,105450233 | -1,365450233 |
| chrXII | 665850 | 666010 | 160 | 0,04844253 | 0,051882174 | 0,482857442 | -0,042983253 | -0,036535765 | -0,061175291 | -0,173175291 | -1,433175291 |
| chrXII | 675650 | 675803 | 153 | 0,136778908 | 0,166764131 | 0,450607955 | -0,124125498 | -0,105506674 | -0,289298274 | -0,352298274 | -1,612298274 |
| chrXII | 678419 | 678585 | 166 | 0,162424953 | 0,192705218 | 0,457367372 | -0,107068364 | -0,09100811 | 0,074658487 | -0,079341513 | -1,339341513 |
| chrXII | 684089 | 684224 | 135 | 1,325045671 | 0,907938047 | 0,593396925 | 0,236291928 | 0,200848138 | 0,167778999 | 0,230778999 | -1,029221001 |
| chrXII | 686656 | 686810 | 154 | 0,159575393 | 0,137117174 | 0,537847625 | 0,095012688 | 0,080760785 | -0,091944125 | -0,161944125 | -1,421944125 |
| chrXII | 686971 | 687118 | 147 | 0,800726524 | 0,655938915 | 0,549698306 | 0,124899153 | 0,10616428 | 0,021618128 | 0,000618128 | -1,259381872 |
| chrXII | 691050 | 691199 | 149 | 0,19661968 | 0,244587392 | 0,445640364 | -0,13668381 | -0,116181238 | -0,248316174 | -0,283316174 | -1,543316174 |
| chrXII | 695440 | 695604 | 164 | 0,299203861 | 0,26682261 | 0,528604008 | 0,071761159 | 0,060996985 | 0,100991811 | -0,039008189 | -1,299008189 |
| chrXII | 698814 | 698965 | 151 | 0,347646391 | 0,237175653 | 0,594448165 | 0,239002467 | 0,203152097 | 0,05670905 | 0,00770905 | -1,25229095 |
| chrXII | 700732 | 700860 | 128 | 0,031345166 | 0,366881088 | 0,078711953 | -1,413788855 | -1,201720527 | -0,957816082 | -0,845816082 | -2,105816082 |
| chrXII | 700732 | 700861 | 129 | 4,10051768 | 47,14236691 | 0,080021211 | -1,404928899 | -1,194189564 | -1,049084361 | -0,944084361 | -2,204084361 |
| chrXII | 704633 | 704781 | 148 | 0,119681545 | 0,066705652 | 0,642112476 | 0,3641111 | 0,309494435 | 0,174433576 | 0,146433576 | -1,113566424 |
| chrXII | 704633 | 704781 | 148 | 0,119681545 | 0,066705652 | 0,642112476 | 0,3641111 | 0,309494435 | 0,174433576 | 0,146433576 | -1,113566424 |
| chrXII | 704633 | 704785 | 152 | 15,72957442 | 9,883554166 | 0,614121558 | 0,290077583 | 0,246565946 | 0,095678721 | 0,039678721 | -1,220321279 |
| chrXII | 704633 | 704785 | 152 | 15,72957442 | 9,883554166 | 0,614121558 | 0,290077583 | 0,246565946 | 0,095678721 | 0,039678721 | -1,220321279 |
| chrXII | 704637 | 704785 | 148 | 0,051292091 | 0,059293913 | 0,463820816 | -0,090812431 | -0,077190567 | -0,208555695 | -0,236555695 | -1,496555695 |
| chrXII | 704738 | 704878 | 140 | 2,017488893 | 1,156231309 | 0,635685809 | 0,346950665 | 0,294908066 | 0,29623924 | 0,32423924 | -0,93576076 |
| chrXII | 707321 | 707471 | 150 | 0,598407723 | 0,452116089 | 0,569627948 | 0,175427028 | 0,149112974 | -0,000650556 | -0,042650556 | -1,302650556 |
| chrXII | 707855 | 708017 | 162 | 0,404637603 | 0,392822175 | 0,507408165 | 0,018570584 | 0,015784996 | 0,085198503 | -0,040801497 | -1,300801497 |
| chrXII | 710806 | 710956 | 150 | 0,210867483 | 0,111176087 | 0,654779361 | 0,398256287 | 0,338517844 | 0,176620492 | 0,134620492 | -1,125379508 |
| chrXII | 713802 | 713939 | 137 | 0,068389454 | 0,044470435 | 0,605967759 | 0,268824834 | 0,228501109 | 0,295243657 | 0,344243657 | -0,915756343 |
| chrXII | 714204 | 714311 | 107 | 2,68713563 | 14,98653658 | 0,152041726 | -1,027715972 | -0,873558576 | -0,326109314 | -0,067109314 | -1,327109314 |
| chrXII | 714204 | 714312 | 108 | 0,028495606 | 0,163058261 | 0,148760274 | -1,041765208 | -0,885500426 | -0,258675365 | -0,006675365 | -1,266675365 |
| chrXII | 714794 | 714945 | 151 | 0,139628469 | 0,188999349 | 0,424883291 | -0,189416258 | -0,161003819 | -0,305690376 | -0,354690376 | -1,614690376 |
| chrXII | 721307 | 721447 | 140 | 0,381841118 | 0,340940001 | 0,528294262 | 0,070982761 | 0,060335347 | 0,052746194 | 0,080746194 | -1,179253806 |
| chrXII | 723297 | 723432 | 135 | 1,880709985 | 1,612053267 | 0,538459051 | 0,09655235 | 0,082069498 | 0,044886603 | 0,107886603 | -1,152113397 |
| chrXII | 726306 | 726432 | 126 | 0,270708255 | 0,274234349 | 0,496764711 | -0,008109755 | -0,006893292 | 0,234659473 | 0,360659473 | -0,899340527 |
| chrXII | 727018 | 727148 | 130 | 0,142478029 | 0,592939133 | 0,1937377 | -0,864204794 | -0,734574075 | -0,630398558 | -0,532398558 | -1,792398558 |
| chrXII | 731448 | 731612 | 164 | 0,339097709 | 0,370586958 | 0,47781462 | -0,055639195 | -0,047293316 | 0,001616181 | -0,138383819 | -1,398383819 |
| chrXII | 731448 | 731612 | 164 | 0,339097709 | 0,370586958 | 0,47781462 | -0,055639195 | -0,047293316 | 0,001616181 | -0,138383819 | -1,398383819 |
| chrXII | 733723 | 733874 | 151 | 0,213717044 | 0,17417587 | 0,550969188 | 0,128110377 | 0,10889382 | -0,043314842 | -0,092314842 | -1,352314842 |
| chrXII | 733723 | 733874 | 151 | 0,213717044 | 0,17417587 | 0,550969188 | 0,128110377 | 0,10889382 | -0,043314842 | -0,092314842 | -1,352314842 |
| chrXII | 735532 | 735635 | 103 | 0,031345166 | 1,371171744 | 0,022349225 | -2,007481199 | -1,706359019 | -1,098022605 | -0,811022605 | -2,071022605 |
| chrXII | 736027 | 736189 | 162 | 0,846319493 | 0,778232612 | 0,520955586 | 0,052552043 | 0,044669236 | 0,128623762 | 0,002623762 | -1,257376238 |
| chrXII | 738785 | 738957 | 172 | 0,065539893 | 0,08894087 | 0,424259254 | -0,191009043 | -0,162357687 | -0,031648485 | -0,227648485 | -1,487648485 |
| chrXII | 740597 | 740703 | 106 | 0,347646391 | 2,805343271 | 0,110259287 | -1,225150361 | -1,041377806 | -0,564174977 | -0,298174977 | -1,558174977 |
| chrXII | 745846 | 745998 | 152 | 0,04844253 | 0,077823261 | 0,383655221 | -0,295894775 | -0,251510559 | -0,386485895 | -0,442485895 | -1,702485895 |
| chrXII | 751027 | 751192 | 165 | 0,162424953 | 0,133411305 | 0,549036667 | 0,123227859 | 0,104743681 | 0,21539719 | 0,06839719 | -1,19160281 |
| chrXII | 752922 | 753058 | 136 | 2,259701543 | 1,137701961 | 0,665126041 | 0,426494002 | 0,362519902 | 0,402362262 | 0,458362262 | -0,801637738 |
| chrXII | 753617 | 753790 | 173 | 0,088336378 | 0,096352609 | 0,478298027 | -0,054425638 | -0,046261792 | 0,16060237 | -0,04239763 | -1,30239763 |
| chrXII | 753617 | 753790 | 173 | 0,088336378 | 0,096352609 | 0,478298027 | -0,054425638 | -0,046261792 | 0,16060237 | -0,04239763 | -1,30239763 |
| chrXII | 759427 | 759579 | 152 | 0,595558162 | 0,43729261 | 0,576615885 | 0,193243554 | 0,164257021 | 0,025659677 | -0,030340323 | -1,290340323 |
| chrXII | 761633 | 761798 | 165 | 0,039893848 | 0,081529131 | 0,328552705 | -0,443913104 | -0,377326138 | -0,266856365 | -0,413856365 | -1,673856365 |
| chrXII | 762105 | 762269 | 164 | 0,333398588 | 0,541056958 | 0,381264193 | -0,302162243 | -0,256837906 | -0,205670916 | -0,345670916 | -1,605670916 |
| chrXII | 770786 | 770938 | 152 | 0,128230226 | 0,188999349 | 0,404219015 | -0,242441566 | -0,206075331 | -0,332730318 | -0,388730318 | -1,648730318 |
| chrXII | 779526 | 779669 | 143 | 1,162620718 | 0,982055438 | 0,542096164 | 0,105716018 | 0,089858615 | -0,078404865 | -0,071404865 | -1,331404865 |
| chrXII | 779740 | 779903 | 163 | 0,284956058 | 0,459527828 | 0,382756516 | -0,298249123 | -0,253511755 | -0,252898663 | -0,385898663 | -1,645898663 |
| chrXII | 781130 | 781282 | 152 | 0,076938136 | 0,107470218 | 0,417216109 | -0,209020522 | -0,177667444 | -0,313690566 | -0,369690566 | -1,629690566 |
| chrXII | 782565 | 782713 | 148 | 0,09973462 | 0,062999783 | 0,61286746 | 0,286800501 | 0,243780426 | 0,106064005 | 0,078064005 | -1,181935995 |
| chrXII | 783914 | 784060 | 146 | 0,652549374 | 0,466939567 | 0,582899348 | 0,209316331 | 0,177918881 | 0,021305907 | 0,007305907 | -1,252694093 |
| chrXII | 783914 | 784060 | 146 | 0,652549374 | 0,466939567 | 0,582899348 | 0,209316331 | 0,177918881 | 0,021305907 | 0,007305907 | -1,252694093 |
| chrXII | 785535 | 785685 | 150 | 0,661098055 | 0,381704567 | 0,633962786 | 0,342367387 | 0,291012279 | 0,134586729 | 0,092586729 | -1,167413271 |
| chrXII | 786502 | 786666 | 164 | 0,179522317 | 0,300175436 | 0,374240479 | -0,320642986 | -0,272546538 | -0,222967527 | -0,362967527 | -1,622967527 |
| chrXII | 792530 | 792669 | 139 | 1,937701197 | 1,289642613 | 0,600401231 | 0,254385777 | 0,216227911 | 0,239327645 | 0,274327645 | -0,985672355 |
| chrXII | 797667 | 797817 | 150 | 0,136778908 | 0,040764565 | 0,770396711 | 0,740153964 | 0,62913087 | 0,48449507 | 0,44249507 | -0,81750493 |
| chrXII | 799286 | 799439 | 153 | 0,159575393 | 0,159352392 | 0,50034961 | 0,000876343 | 0,000744891 | -0,179847957 | -0,242847957 | -1,502847957 |
| chrXII | 803374 | 803523 | 149 | 0,82922213 | 0,733762177 | 0,53053772 | 0,076621618 | 0,065128376 | -0,072805467 | -0,107805467 | -1,367805467 |
| chrXII | 804062 | 804197 | 135 | 0,051292091 | 0,055588044 | 0,479902939 | -0,050397187 | -0,042837609 | -0,075608929 | -0,012608929 | -1,272608929 |
| chrXII | 804515 | 804661 | 146 | 0,265009134 | 0,044470435 | 0,856305749 | 1,063867998 | 0,904287798 | 0,753220225 | 0,739220225 | -0,520779775 |
| chrXII | 806849 | 807003 | 154 | 0,239363089 | 0,244587392 | 0,49460244 | -0,013530088 | -0,011500575 | -0,176225373 | -0,246225373 | -1,506225373 |
| chrXII | 807887 | 808014 | 127 | 0,128230226 | 0,26682261 | 0,324590066 | -0,454901459 | -0,386666241 | -0,166191765 | -0,047191765 | -1,307191765 |
| chrXII | 807887 | 808014 | 127 | 0,128230226 | 0,26682261 | 0,324590066 | -0,454901459 | -0,386666241 | -0,166191765 | -0,047191765 | -1,307191765 |
| chrXII | 827684 | 827838 | 154 | 6,069564043 | 4,580454799 | 0,569911108 | 0,176147858 | 0,14972568 | -0,002999705 | -0,072999705 | -1,332999705 |
| chrXII | 830101 | 830255 | 154 | 0,202318801 | 0,170470001 | 0,542716949 | 0,107280945 | 0,091188803 | -0,04999663 | -0,11999663 | -1,37999663 |
| chrXII | 830655 | 830774 | 119 | 0,863416857 | 3,390870664 | 0,202952164 | -0,831122683 | -0,70645428 | -0,28550083 | -0,11050083 | -1,37050083 |
| chrXII | 830666 | 830794 | 128 | 1,61000173 | 1,904816963 | 0,458061104 | -0,105319605 | -0,089521664 | 0,149034225 | 0,261034225 | -0,998965775 |
| chrXII | 831008 | 831187 | 179 | 0,074088575 | 0,122293696 | 0,377267126 | -0,312666232 | -0,265766297 | -0,108248547 | -0,353248547 | -1,613248547 |
| chrXII | 831542 | 831700 | 158 | 0,116831984 | 0,096352609 | 0,548032024 | 0,120690792 | 0,102587173 | 0,072919538 | -0,025080462 | -1,285080462 |
| chrXII | 839642 | 839772 | 130 | 0,185221438 | 0,722644568 | 0,204018475 | -0,827353109 | -0,703250143 | -0,618652983 | -0,520652983 | -1,780652983 |
| chrXII | 840366 | 840517 | 151 | 0,772230918 | 0,522527611 | 0,596428524 | 0,244113455 | 0,207496437 | 0,055322049 | 0,006322049 | -1,253677951 |
| chrXII | 841812 | 841967 | 155 | 0,208017923 | 0,300175436 | 0,409328298 | -0,229273181 | -0,194882204 | -0,210434673 | -0,287434673 | -1,547434673 |
| chrXII | 842219 | 842368 | 149 | 0,227964847 | 0,207528696 | 0,523463207 | 0,058847485 | 0,050020363 | -0,085688311 | -0,120688311 | -1,380688311 |
| chrXII | 846202 | 846357 | 155 | 0,43883233 | 0,35205761 | 0,55485891 | 0,137947158 | 0,117255084 | 0,100142879 | 0,023142879 | -1,236857121 |
| chrXII | 864284 | 864432 | 148 | 0,324849907 | 0,159352392 | 0,670897077 | 0,442391614 | 0,376032872 | 0,230370887 | 0,202370887 | -1,057629113 |
| chrXII | 868597 | 868734 | 137 | 0,039893848 | 0,051882174 | 0,434687048 | -0,164453569 | -0,139785534 | -0,068072435 | -0,019072435 | -1,279072435 |
| chrXII | 871320 | 871444 | 124 | 0,039893848 | 0,422469132 | 0,086282531 | -1,364007955 | -1,159406762 | -0,751475206 | -0,611475206 | -1,871475206 |
| chrXII | 871320 | 871449 | 129 | 15,88914981 | 16,56894289 | 0,489528142 | -0,02625207 | -0,02231426 | 0,113139043 | 0,218139043 | -1,041860957 |
| chrXII | 871322 | 871449 | 127 | 0,054141651 | 0,100058479 | 0,351112876 | -0,382317666 | -0,324970016 | -0,086968948 | 0,032031052 | -1,227968948 |
| chrXII | 874679 | 874837 | 158 | 0,042743409 | 0,111176087 | 0,277699771 | -0,589688448 | -0,501235181 | -0,532568156 | -0,630568156 | -1,890568156 |
| chrXII | 878567 | 878705 | 138 | 0,142478029 | 0,051882174 | 0,733061742 | 0,62209939 | 0,528784481 | 0,504654047 | 0,546654047 | -0,713345953 |
| chrXII | 884555 | 884713 | 158 | 0,156725832 | 0,148234783 | 0,51392155 | 0,034903236 | 0,029667751 | -0,003433238 | -0,101433238 | -1,361433238 |
| chrXII | 889517 | 889681 | 164 | 0,233663968 | 0,418763262 | 0,35814564 | -0,363419845 | -0,308906868 | -0,261222833 | -0,401222833 | -1,661222833 |
| chrXII | 893166 | 893327 | 161 | 1,929152515 | 1,486053701 | 0,564871458 | 0,163331944 | 0,138832153 | 0,129729637 | 0,010729637 | -1,249270363 |
| chrXII | 896277 | 896446 | 169 | 0,626903328 | 0,863467612 | 0,42063577 | -0,200267293 | -0,170227199 | 0,001603494 | -0,173396506 | -1,433396506 |
| chrXII | 896668 | 896822 | 154 | 0,09688506 | 0,118587827 | 0,449639217 | -0,126572917 | -0,10758698 | -0,244832143 | -0,314832143 | -1,574832143 |
| chrXII | 896811 | 896958 | 147 | 0,307752543 | 0,222352175 | 0,58055047 | 0,203301913 | 0,172806626 | 0,099295902 | 0,078295902 | -1,181704098 |
| chrXII | 903630 | 903761 | 131 | 1,12272687 | 1,174760656 | 0,488675937 | -0,028389029 | -0,024130675 | 0,090431113 | 0,181431113 | -1,078568887 |
| chrXII | 903784 | 903945 | 161 | 0,074088575 | 0,066705652 | 0,52621884 | 0,065768269 | 0,055903029 | 0,04728561 | -0,07171439 | -1,33171439 |
| chrXII | 903961 | 904117 | 156 | 1,117027749 | 0,737468046 | 0,602335013 | 0,259395665 | 0,220486315 | 0,128206635 | 0,044206635 | -1,215793365 |
| chrXII | 905180 | 905335 | 155 | 0,031345166 | 0,059293913 | 0,345823971 | -0,396619676 | -0,337126724 | -0,360798567 | -0,437798567 | -1,697798567 |
| chrXII | 908649 | 908820 | 171 | 0,037044288 | 0,081529131 | 0,31241646 | -0,489012398 | -0,415660538 | -0,3049698 | -0,4939698 | -1,7539698 |
| chrXII | 912499 | 912653 | 154 | 0,239363089 | 0,163058261 | 0,594807131 | 0,239928434 | 0,203939169 | 0,073578333 | 0,003578333 | -1,256421667 |

TABLE 2

| Chrom | Start | End | Length | Section A | Section B | A/A+B | Z-score | Z * 0.85 | Phase correction | Length correction | ΔLK |
|---|---|---|---|---|---|---|---|---|---|---|---|
| chrXII | 913527 | 913695 | 168 | 0,071239015 | 0,114881957 | 0,382756516 | -0,298249123 | -0,253511755 | -0,161973244 | -0,329973244 | -1,589973244 |
| chrXII | 922406 | 922556 | 150 | 0,056991212 | 0,077823261 | 0,422738082 | -0,194893703 | -0,165659647 | -0,302816867 | -0,344816867 | -1,604816867 |
| chrXII | 922509 | 922654 | 145 | 0,658248495 | 0,418763262 | 0,611180417 | 0,282396933 | 0,240037393 | 0,081741032 | 0,074741032 | -1,185258968 |
| chrXII | 924375 | 924524 | 149 | 2,818215417 | 1,941875659 | 0,592050734 | 0,232823413 | 0,197899901 | 0,06163309 | 0,02663309 | -1,23336691 |
| chrXII | 926149 | 926303 | 154 | 0,649699813 | 0,552174567 | 0,540572146 | 0,101875234 | 0,086593949 | -0,037460252 | -0,107460252 | -1,367460252 |
| chrXII | 926358 | 926515 | 157 | 1,812320531 | 1,589818049 | 0,532700385 | 0,082059712 | 0,069750755 | 0,007308265 | -0,083691735 | -1,343691735 |
| chrXII | 929019 | 929172 | 153 | 1,051487855 | 0,844938264 | 0,55445759 | 0,136931651 | 0,116391904 | -0,055424745 | -0,118424745 | -1,378424745 |
| chrXII | 931573 | 931698 | 125 | 1,040089613 | 2,671931966 | 0,280194926 | -0,582262541 | -0,49492316 | -0,226750099 | -0,093750099 | -1,353750099 |
| chrXII | 937960 | 938108 | 148 | 1,427629852 | 1,093231526 | 0,566326203 | 0,167028538 | 0,141974257 | -0,003993363 | -0,031993363 | -1,291993363 |
| chrXII | 938740 | 938880 | 140 | 0,367593315 | 0,374292827 | 0,495484811 | -0,011318142 | -0,009620421 | -0,014618974 | 0,013381026 | -1,246618974 |
| chrXII | 939726 | 939894 | 168 | 0,065539893 | 0,077823261 | 0,457159956 | -0,107591283 | -0,091452591 | 0,007911093 | -0,160088907 | -1,420088907 |
| chrXII | 940022 | 940161 | 139 | 0,396088921 | 0,237175653 | 0,625471465 | 0,319882942 | 0,271900501 | 0,302831363 | 0,337831363 | -0,922168637 |
| chrXII | 941909 | 942015 | 106 | 0,09688506 | 0,396528045 | 0,196356884 | -0,854706289 | -0,726500345 | -0,230131786 | 0,035868214 | -1,224131786 |
| chrXII | 941909 | 942019 | 110 | 0,056991212 | 0,433586741 | 0,116171572 | -1,194344741 | -1,01519303 | -0,373497851 | -0,135497851 | -1,395497851 |
| chrXII | 943265 | 943394 | 129 | 0,182371877 | 0,396528045 | 0,315031788 | -0,481637369 | -0,409391764 | -0,273878182 | -0,168878182 | -1,428878182 |
| chrXII | 944730 | 944873 | 143 | 0,225115286 | 0,081529131 | 0,734124849 | 0,625336388 | 0,53153593 | 0,367010161 | 0,374010161 | -0,885989839 |
| chrXII | 946105 | 946219 | 114 | 0,034194727 | 0,244587392 | 0,122657533 | -1,161804056 | -0,987533448 | -0,336011908 | -0,126011908 | -1,386011908 |
| chrXII | 946809 | 946958 | 149 | 0,09688506 | 0,048176305 | 0,667890174 | 0,43409473 | 0,36898052 | 0,231551119 | 0,196551119 | -1,063448881 |
| chrXII | 947096 | 947245 | 149 | 0,09973462 | 0,155646522 | 0,390532439 | -0,277931585 | -0,236241848 | -0,366922292 | -0,401922292 | -1,661922292 |
| chrXII | 947588 | 947739 | 151 | 0,09688506 | 0,129705435 | 0,427577775 | -0,182544352 | -0,155162699 | -0,309389137 | -0,358389137 | -1,618389137 |
| chrXII | 952012 | 952159 | 147 | 0,265009134 | 0,107470218 | 0,711473355 | 0,557694096 | 0,474039982 | 0,40182985 | 0,38082985 | -0,87917015 |
| chrXII | 955881 | 956039 | 158 | 0,453080133 | 0,314998914 | 0,589887375 | 0,227255273 | 0,193166982 | 0,153564721 | 0,055564721 | -1,204435279 |
| chrXII | 956586 | 956713 | 127 | 0,085486818 | 0,229763914 | 0,271170878 | -0,609275633 | -0,517884288 | -0,284935455 | -0,165935455 | -1,425935455 |
| chrXII | 962105 | 962271 | 166 | 0,136778908 | 0,155646522 | 0,467739443 | -0,080953558 | -0,068810524 | 0,092124662 | -0,061875338 | -1,321875338 |
| chrXII | 962105 | 962271 | 166 | 0,136778908 | 0,155646522 | 0,467739443 | -0,080953558 | -0,068810524 | 0,092124662 | -0,061875338 | -1,321875338 |
| chrXII | 966042 | 966188 | 146 | 0,581310359 | 0,34835174 | 0,625292092 | 0,319409753 | 0,27149829 | 0,115847183 | 0,101847183 | -1,158152817 |
| chrXII | 967180 | 967328 | 148 | 0,062690333 | 0,070411522 | 0,470995186 | -0,072768458 | -0,061853189 | -0,193968846 | -0,221968846 | -1,481968846 |
| chrXII | 967209 | 967341 | 132 | 0,265009134 | 0,181587609 | 0,593396925 | 0,236291928 | 0,200848138 | 0,288239532 | 0,372239532 | -0,887760468 |
| chrXII | 974145 | 974297 | 152 | 0,085486818 | 0,096352609 | 0,470122564 | -0,074961771 | -0,063717505 | -0,205288488 | -0,261288488 | -1,521288488 |
| chrXII | 975018 | 975184 | 166 | 0,222265726 | 0,311293045 | 0,416572153 | -0,210670615 | -0,179070023 | -0,009018085 | -0,163018085 | -1,423018085 |
| chrXII | 976449 | 976598 | 149 | 0,094035499 | 0,066705652 | 0,585011979 | 0,214732295 | 0,182522451 | 0,062489467 | 0,027489467 | -1,232510533 |
| chrXII | 978018 | 978161 | 143 | 0,227964847 | 0,077823261 | 0,745499386 | 0,660393679 | 0,561334627 | 0,415370943 | 0,422370943 | -0,837629057 |
| chrXII | 978601 | 978756 | 155 | 0,267858695 | 0,270528479 | 0,497520572 | -0,006215044 | -0,005282788 | -0,023556813 | -0,100556813 | -1,360556813 |
| chrXII | 979540 | 979688 | 148 | 0,065539893 | 0,074117392 | 0,469290903 | -0,077052467 | -0,065494597 | -0,197456149 | -0,225456149 | -1,485456149 |
| chrXII | 981009 | 981111 | 102 | 0,122531105 | 0,140823044 | 0,465271216 | -0,087162393 | -0,074088034 | 0,500014484 | 0,794014484 | -0,465985516 |
| chrXII | 981009 | 981139 | 130 | 1,681240744 | 1,84922892 | 0,476208806 | -0,059671071 | -0,05072041 | 0,031882678 | 0,129882678 | -1,130117322 |
| chrXII | 981073 | 981227 | 154 | 0,168124074 | 0,133411305 | 0,557560028 | 0,144785863 | 0,123067983 | 0,006679472 | -0,063320528 | -1,323320528 |
| chrXII | 981080 | 981248 | 168 | 0,823523009 | 0,978349569 | 0,457037317 | -0,107900483 | -0,09171541 | 0,012008773 | -0,155991227 | -1,415991227 |
| chrXII | 982426 | 982590 | 164 | 0,567062556 | 0,511410002 | 0,525801563 | 0,06472008 | 0,055012068 | 0,101448203 | -0,038551797 | -1,298551797 |
| chrXII | 987388 | 987536 | 148 | 1,22246149 | 0,781938481 | 0,609888998 | 0,279029737 | 0,237175277 | 0,1015471 | 0,0735471 | -1,1864529 |
| chrXII | 988804 | 988961 | 157 | 1,729683274 | 1,597229788 | 0,519906364 | 0,049918579 | 0,042430792 | -0,036466686 | -0,127466686 | -1,387466686 |
| chrXII | 988850 | 988961 | 111 | 0,037044288 | 0,070411522 | 0,344739737 | -0,399561558 | -0,339627325 | 0,133368771 | 0,364368771 | -0,895631229 |
| chrXII | 988957 | 989104 | 147 | 0,128230226 | 0,074117392 | 0,633712557 | 0,341702373 | 0,290447017 | 0,210538674 | 0,189538674 | -1,070461326 |
| chrXII | 990093 | 990242 | 149 | 0,48727486 | 0,318704784 | 0,604574649 | 0,265206091 | 0,225425178 | 0,11319889 | 0,07819889 | -1,18180111 |
| chrXII | 990292 | 990442 | 150 | 0,464478375 | 0,237175653 | 0,661976354 | 0,417862991 | 0,355183542 | 0,226933449 | 0,184933449 | -1,075066551 |
| chrXII | 997620 | 997777 | 157 | 0,330549028 | 0,374292827 | 0,468969068 | -0,077861613 | -0,066182371 | -0,129616283 | -0,220616283 | -1,480616283 |
| chrXII | 998925 | 999073 | 148 | 0,085486818 | 0,185293479 | 0,315705458 | -0,47974192 | -0,407780632 | -0,540157734 | -0,568157734 | -1,828157734 |
| chrXII | 1001567 | 1001716 | 149 | 0,205168362 | 0,244587392 | 0,456177292 | -0,110069087 | -0,093558724 | -0,199810195 | -0,234810195 | -1,494810195 |
| chrXII | 1005509 | 1005668 | 159 | 0,04844253 | 0,103764348 | 0,31826768 | -0,472548462 | -0,401666193 | -0,400200535 | -0,505200535 | -1,765200535 |
| chrXII | 1005994 | 1006142 | 148 | 0,088336378 | 0,055588044 | 0,613769206 | 0,28915654 | 0,245783059 | 0,111123532 | 0,083123532 | -1,176876468 |
| chrXII | 1012419 | 1012577 | 158 | 0,373292436 | 0,307587175 | 0,548250278 | 0,121241892 | 0,103055609 | 0,065425324 | -0,032574676 | -1,292574676 |
| chrXII | 1012549 | 1012696 | 147 | 0,14817715 | 1,93075805 | 0,071275502 | -1,466357184 | -1,246403606 | -1,32624131 | -1,34724131 | -2,60724131 |
| chrXII | 1012550 | 1012632 | 82 | 0,028495606 | 0,337234132 | 0,07791438 | -1,419241021 | -1,206354868 | -0,877987772 | -0,443987772 | -1,703987772 |
| chrXII | 1012550 | 1012633 | 83 | 0,068389454 | 0,552174567 | 0,110205316 | -1,225436951 | -1,041621408 | -0,722418571 | -0,295418571 | -1,555418571 |
| chrXII | 1012550 | 1012634 | 84 | 0,042743409 | 0,192705218 | 0,181540276 | -0,909510814 | -0,773084192 | -0,389970034 | 0,030029966 | -1,229970034 |
| chrXII | 1012550 | 1012635 | 85 | 0,051292091 | 0,155646522 | 0,247861382 | -0,681235065 | -0,579049806 | -0,203611192 | 0,209388808 | -1,050611192 |
| chrXII | 1012550 | 1012636 | 86 | 0,116831984 | 0,140823044 | 0,453443447 | -0,11696613 | -0,099421211 | 0,259644047 | 0,665644047 | -0,594355953 |
| chrXII | 1012550 | 1012637 | 87 | 0,034194727 | 0,581821524 | 0,055509456 | -1,593630085 | -1,354585572 | -0,955592235 | -0,556592235 | -1,816592235 |
| chrXII | 1012550 | 1012638 | 88 | 0,034194727 | 0,185293479 | 0,155793004 | -1,011899709 | -0,860114752 | -0,42009307 | -0,02809307 | -1,28809307 |
| chrXII | 1012550 | 1012639 | 89 | 0,065539893 | 0,314998914 | 0,172229197 | -0,945392764 | -0,80358385 | -0,374917554 | 0,010082446 | -1,249917554 |
| chrXII | 1012550 | 1012640 | 90 | 0,028495606 | 0,214940436 | 0,117055822 | -1,189834 | -1,0113589 | -0,550514376 | -0,172514376 | -1,432514376 |
| chrXII | 1012550 | 1012641 | 91 | 0,091185939 | 0,503998263 | 0,153206249 | -1,022778687 | -0,869361884 | -0,50332261 | -0,13232261 | -1,39232261 |
| chrXII | 1012550 | 1012642 | 92 | 0,128230226 | 0,344645871 | 0,271170878 | -0,609275633 | -0,517884288 | -0,117001793 | 0,246998207 | -1,013001793 |
| chrXII | 1012550 | 1012643 | 93 | 0,062690333 | 0,333528262 | 0,158221582 | -1,001793859 | -0,85152478 | -0,365010483 | -0,008010483 | -1,268010483 |
| chrXII | 1012550 | 1012644 | 94 | 0,079787696 | 0,415057393 | 0,161237725 | -0,989383693 | -0,840976139 | -0,324401513 | 0,025598487 | -1,234401513 |
| chrXII | 1012550 | 1012645 | 95 | 0,056991212 | 0,626291959 | 0,083407896 | -1,382507947 | -1,175131755 | -0,642710548 | -0,299710548 | -1,559710548 |
| chrXII | 1012550 | 1012646 | 96 | 0,042743409 | 0,188999349 | 0,184443343 | -0,898560728 | -0,763776618 | -0,208476066 | 0,127523934 | -1,132476066 |
| chrXII | 1012550 | 1012648 | 98 | 0,128230226 | 0,737468046 | 0,148123463 | -1,044515554 | -0,887838221 | -0,402709012 | -0,080709012 | -1,340709012 |
| chrXII | 1012550 | 1012649 | 99 | 0,074088575 | 0,251999131 | 0,227204441 | -0,748085011 | -0,635872259 | -0,223547102 | 0,091452898 | -1,168547102 |
| chrXII | 1012550 | 1012650 | 100 | 0,102584181 | 0,244587392 | 0,295485544 | -0,537429332 | -0,456814932 | -0,026978077 | 0,281021923 | -0,978978077 |
| chrXII | 1012550 | 1012651 | 101 | 0,065539893 | 0,08894087 | 0,424259254 | -0,191009043 | -0,162357687 | 0,357675864 | 0,658675864 | -0,601324136 |
| chrXII | 1012550 | 1012652 | 102 | 0,056991212 | 0,944996743 | 0,056878141 | -1,581532732 | -1,344302822 | -0,766895924 | -0,472895924 | -1,732895924 |
| chrXII | 1012550 | 1012653 | 103 | 0,094035499 | 0,429880871 | 0,179485705 | -0,917326814 | -0,779727792 | -0,20060965 | 0,08639035 | -1,17360965 |
| chrXII | 1012550 | 1012654 | 104 | 0,04844253 | 0,218646305 | 0,181372351 | -0,910147548 | -0,773625416 | -0,174786672 | 0,105213328 | -1,154786672 |
| chrXII | 1012550 | 1012655 | 105 | 0,082637257 | 0,103764348 | 0,443329106 | -0,142534017 | -0,121153915 | 0,4006554 | 0,6736554 | -0,5863446 |
| chrXII | 1012550 | 1012656 | 106 | 0,042743409 | 0,418763262 | 0,092617098 | -1,324809941 | -1,12608845 | -0,64226122 | -0,37626122 | -1,63626122 |
| chrXII | 1012550 | 1012657 | 107 | 0,037044288 | 0,140823044 | 0,208269204 | -0,812441383 | -0,690575176 | -0,140816806 | 0,118183194 | -1,141816806 |
| chrXII | 1012550 | 1012659 | 109 | 0,074088575 | 0,270528479 | 0,214988127 | -0,789232287 | -0,670847444 | 0,029133906 | 0,274133906 | -0,985866094 |
| chrXII | 1012550 | 1012660 | 110 | 0,054141651 | 0,107470218 | 0,335010364 | -0,426119559 | -0,362201625 | 0,268441436 | 0,506441436 | -0,753558564 |
| chrXII | 1012550 | 1012661 | 111 | 0,165274514 | 0,515115871 | 0,242911301 | -0,696968348 | -0,592423096 | -0,115849445 | 0,115150555 | -1,144849445 |
| chrXII | 1012550 | 1012662 | 112 | 0,116831984 | 0,248293262 | 0,31997783 | -0,467760794 | -0,397596675 | 0,168799096 | 0,392799096 | -0,867200904 |
| chrXII | 1012550 | 1012663 | 113 | 0,133929347 | 0,444704349 | 0,231457912 | -0,734054005 | -0,623945904 | -0,061667559 | 0,155332441 | -1,104667559 |
| chrXII | 1012550 | 1012664 | 114 | 0,162424953 | 1,385995222 | 0,104897208 | -1,25413094 | -1,066011299 | -0,396563429 | -0,186563429 | -1,446563429 |
| chrXII | 1012550 | 1012665 | 115 | 0,199469241 | 0,885702829 | 0,183813467 | -0,900927376 | -0,765788269 | -0,165845338 | 0,037154662 | -1,222845338 |
| chrXII | 1012550 | 1012666 | 116 | 0,156725832 | 0,744879785 | 0,173829698 | -0,939138975 | -0,798268129 | -0,210760103 | -0,014760103 | -1,274760103 |
| chrXII | 1012550 | 1012667 | 117 | 0,14532759 | 0,296469566 | 0,328946413 | -0,442824299 | -0,376400654 | 0,110346368 | 0,299346368 | -0,960653632 |
| chrXII | 1012550 | 1012668 | 118 | 0,076938136 | 0,240881523 | 0,24208111 | -0,699623888 | -0,594680304 | -0,210907071 | -0,028907071 | -1,288907071 |
| chrXII | 1012550 | 1012669 | 119 | 0,213717044 | 0,377998697 | 0,361181948 | -0,355301282 | -0,302006089 | 0,104291854 | 0,279291854 | -0,980708146 |
| chrXII | 1012550 | 1012670 | 120 | 0,062690333 | 0,144528914 | 0,30253142 | -0,517133686 | -0,439563633 | -0,055419989 | 0,112580011 | -1,147419989 |
| chrXII | 1012550 | 1012671 | 121 | 0,159575393 | 0,715232829 | 0,182411858 | -0,906211882 | -0,7702801 | -0,392951922 | -0,231951922 | -1,491951922 |
| chrXII | 1012550 | 1012672 | 122 | 0,111132863 | 0,392822175 | 0,220521385 | -0,770433532 | -0,654868502 | -0,252985062 | -0,098985062 | -1,358985062 |
| chrXII | 1012550 | 1012673 | 123 | 0,119681545 | 0,396528045 | 0,231846806 | -0,732778376 | -0,62286162 | -0,188851291 | -0,041851291 | -1,301851291 |
| chrXII | 1012550 | 1012674 | 124 | 0,208017923 | 5,399451976 | 0,037096574 | -1,785420424 | -1,517607361 | -1,11299362 | -0,97299362 | -2,23299362 |
| chrXII | 1012550 | 1012675 | 125 | 0,190920559 | 0,503998263 | 0,274737931 | -0,598545721 | -0,508763863 | -0,242276971 | -0,109276971 | -1,369276971 |
| chrXII | 1012550 | 1012676 | 126 | 0,205168362 | 0,544762828 | 0,273582916 | -0,602012476 | -0,511710605 | -0,256116407 | -0,130116407 | -1,390116407 |
| chrXII | 1012550 | 1012677 | 127 | 0,122531105 | 0,559586306 | 0,179633452 | -0,916762889 | -0,779248455 | -0,547013805 | -0,428013805 | -1,688013805 |
| chrXII | 1012550 | 1012678 | 128 | 0,125380666 | 0,163058261 | 0,434687048 | -0,164453569 | -0,139785534 | 0,094263704 | 0,206263704 | -1,053736296 |
| chrXII | 1012550 | 1012679 | 129 | 0,227964847 | 0,155646522 | 0,594259882 | 0,238516864 | 0,202739335 | 0,339398156 | 0,444398156 | -0,815601844 |
| chrXII | 1012550 | 1012680 | 130 | 0,185221438 | 0,222352175 | 0,454449042 | -0,114428548 | -0,097264266 | -0,012603453 | 0,085396547 | -1,174603453 |
| chrXII | 1012550 | 1012681 | 131 | 0,185221438 | 0,359469349 | 0,340048781 | -0,412330002 | -0,350480502 | -0,220263522 | -0,129263522 | -1,389263522 |
| chrXII | 1012550 | 1012682 | 132 | 0,364743755 | 0,35205761 | 0,508849135 | 0,022183312 | 0,018855816 | 0,079626403 | 0,163626403 | -1,096373597 |

TABLE 2

| Chrom | Start | End | Length | Section A | Section B | A/A+B | Z-score | Z * 0.85 | Phase correction | Length correction | ΔLK |
|---|---|---|---|---|---|---|---|---|---|---|---|
| chrXII | 1012550 | 1012683 | 133 | 0,330549028 | 0,715232829 | 0,316078373 | -0,478693422 | -0,406889409 | -0,418493863 | -0,341493863 | -1,601493863 |
| chrXII | 1012550 | 1012684 | 134 | 0,484425299 | 0,359469349 | 0,574035278 | 0,186657164 | 0,15865859 | 0,162535143 | 0,232535143 | -1,027464857 |
| chrXII | 1012550 | 1012685 | 135 | 0,626903328 | 0,496586524 | 0,557996432 | 0,14589138 | 0,124007673 | 0,091603725 | 0,154603725 | -1,105396275 |
| chrXII | 1012550 | 1012686 | 136 | 0,319150785 | 0,392822175 | 0,448262509 | -0,13005234 | -0,110544489 | -0,058704946 | -0,002704946 | -1,262704946 |
| chrXII | 1012550 | 1012687 | 137 | 0,236513528 | 0,396528045 | 0,373614528 | -0,322295218 | -0,273950935 | -0,19086849 | -0,14186849 | -1,40186849 |
| chrXII | 1012550 | 1012688 | 138 | 0,968850598 | 0,770820872 | 0,55691584 | 0,143154302 | 0,121681156 | 0,088853174 | 0,130853174 | -1,129146826 |
| chrXII | 1012550 | 1012689 | 139 | 0,307752543 | 0,240881523 | 0,56094319 | 0,153360973 | 0,130356827 | 0,161771863 | 0,196771863 | -1,063228137 |
| chrXII | 1012550 | 1012690 | 140 | 0,720938828 | 0,559586306 | 0,563002481 | 0,158586028 | 0,134798124 | 0,135767377 | 0,163767377 | -1,096232623 |
| chrXII | 1012550 | 1012691 | 141 | 0,504372223 | 0,604056741 | 0,455033421 | -0,112954231 | -0,096011096 | -0,156176989 | -0,135176989 | -1,395176989 |
| chrXII | 1012550 | 1012692 | 142 | 0,492973981 | 0,359469349 | 0,578307042 | 0,197564368 | 0,167929713 | 0,132131801 | 0,146131801 | -1,113868199 |
| chrXII | 1012550 | 1012693 | 143 | 3,881101515 | 2,316168487 | 0,626259872 | 0,321963618 | 0,273669075 | 0,156021178 | 0,163021178 | -1,096978822 |
| chrXII | 1012550 | 1012694 | 144 | 0,626903328 | 0,61517435 | 0,504721516 | 0,011835361 | 0,010060057 | -0,171926774 | -0,171926774 | -1,431926774 |
| chrXII | 1012550 | 1012695 | 145 | 0,618354647 | 0,607762611 | 0,50431934 | 0,010827192 | 0,009203114 | -0,146781992 | -0,153781992 | -1,413781992 |
| chrXII | 1012550 | 1012696 | 146 | 799,1792126 | 406,4227167 | 0,662888133 | 0,420358291 | 0,357304547 | 0,203411957 | 0,189411957 | -1,070588043 |
| chrXII | 1012550 | 1012698 | 148 | 0,239363089 | 0,289057827 | 0,452978074 | -0,118140734 | -0,100419624 | -0,237375093 | -0,265375093 | -1,525375093 |
| chrXII | 1012551 | 1012693 | 142 | 0,031345166 | 0,044470435 | 0,413439528 | -0,218706025 | -0,185900121 | -0,243015829 | -0,229015829 | -1,489015829 |
| chrXII | 1012551 | 1012696 | 145 | 0,666797177 | 1,434171527 | 0,317376063 | -0,47504889 | -0,403791556 | -0,559416557 | -0,566416557 | -1,826416557 |
| chrXII | 1012552 | 1012696 | 144 | 0,58415992 | 0,844938264 | 0,408761222 | -0,230732731 | -0,196122822 | -0,380072941 | -0,380072941 | -1,640072941 |
| chrXII | 1012553 | 1012693 | 140 | 0,179522317 | 0,08894087 | 0,668703665 | 0,436336407 | 0,370885946 | 0,384897455 | 0,412897455 | -0,847102545 |
| chrXII | 1012553 | 1012696 | 143 | 1,820869213 | 2,631167401 | 0,408996909 | -0,230126057 | -0,195607148 | -0,312814544 | -0,305814544 | -1,565814544 |
| chrXII | 1012554 | 1012696 | 142 | 1,111328628 | 0,559586306 | 0,665101858 | 0,426427614 | 0,362463472 | 0,305303686 | 0,319303686 | -0,940696314 |
| chrXII | 1012555 | 1012696 | 141 | 1,205364127 | 1,096937395 | 0,523547466 | 0,059059058 | 0,050200199 | 0,00488396 | 0,02588396 | -1,23411604 |
| chrXII | 1012556 | 1012696 | 140 | 0,259310013 | 0,289057827 | 0,472876041 | -0,06804215 | -0,057835827 | -0,067232068 | -0,039232068 | -1,299232068 |
| chrXII | 1012557 | 1012696 | 139 | 0,287805619 | 0,859761742 | 0,250796275 | -0,671986094 | -0,57118818 | -0,537170494 | -0,502170494 | -1,762170494 |
| chrXII | 1012558 | 1012696 | 138 | 0,404637603 | 0,763409133 | 0,346422442 | -0,394997295 | -0,335747701 | -0,365648443 | -0,323648443 | -1,583648443 |
| chrXII | 1012559 | 1012696 | 137 | 0,606956404 | 0,433586741 | 0,583307292 | 0,210361658 | 0,178807409 | 0,264351079 | 0,313351079 | -0,946648921 |
| chrXII | 1012560 | 1012696 | 136 | 0,515770466 | 0,759703264 | 0,404375609 | -0,242037356 | -0,205731753 | -0,15937568 | -0,10337568 | -1,36337568 |
| chrXII | 1012561 | 1012696 | 135 | 0,302053422 | 0,344645871 | 0,467069356 | -0,082638847 | -0,07024302 | -0,098297759 | -0,035297759 | -1,295297759 |
| chrXII | 1012562 | 1012696 | 134 | 0,122531105 | 0,533645219 | 0,186735029 | -0,88999223 | -0,756493395 | -0,748357556 | -0,678357556 | -1,938357556 |
| chrXII | 1012563 | 1012696 | 133 | 0,094035499 | 0,148234783 | 0,388142938 | -0,284162468 | -0,241538098 | -0,249966431 | -0,172966431 | -1,432966431 |
| chrXII | 1012564 | 1012696 | 132 | 0,09688506 | 0,989467177 | 0,089183836 | -1,345797995 | -1,143928296 | -1,083462671 | -0,999462671 | -2,259462671 |
| chrXII | 1012565 | 1012696 | 131 | 0,179522317 | 0,763409133 | 0,190387453 | -0,876469399 | -0,744998989 | -0,616833694 | -0,525833694 | -1,785833694 |
| chrXII | 1012566 | 1012696 | 130 | 0,102584181 | 0,270528479 | 0,274941571 | -0,597935244 | -0,508244957 | -0,421583607 | -0,323583607 | -1,583583607 |
| chrXII | 1012567 | 1012696 | 129 | 0,079787696 | 0,804173699 | 0,090261517 | -1,339146351 | -1,138274398 | -1,003811696 | -0,898811696 | -2,158811696 |
| chrXII | 1012568 | 1012696 | 128 | 0,054141651 | 0,433586741 | 0,11100779 | -1,221186061 | -1,038008152 | -0,801151486 | -0,689151486 | -1,949151486 |
| chrXII | 1012569 | 1012696 | 127 | 0,039893848 | 0,403939784 | 0,089884689 | -1,341465454 | -1,140245636 | -0,911981344 | -0,792981344 | -2,052981344 |
| chrXII | 1012570 | 1012696 | 126 | 0,076938136 | 0,596645002 | 0,114222182 | -1,204375793 | -1,023719424 | -0,782861355 | -0,656861355 | -1,916861355 |
| chrXII | 1012571 | 1012696 | 125 | 0,056991212 | 0,96352609 | 0,055845414 | -1,590639059 | -1,3520432 | -1,090038942 | -0,957038942 | -2,217038942 |
| chrXII | 1012572 | 1012696 | 124 | 0,054141651 | 0,244587392 | 0,181239998 | -0,910649661 | -0,774052212 | -0,367194401 | -0,227194401 | -1,487194401 |
| chrXII | 1012573 | 1012696 | 123 | 0,056991212 | 0,622586089 | 0,083862736 | -1,379549252 | -1,172616864 | -0,758224647 | -0,611224647 | -1,871224647 |
| chrXII | 1012574 | 1012696 | 122 | 0,059840772 | 0,61888022 | 0,088166968 | -1,352129359 | -1,149309955 | -0,745573068 | -0,591573068 | -1,851573068 |
| chrXII | 1012575 | 1012696 | 121 | 0,071239015 | 0,111176087 | 0,390532439 | -0,277931585 | -0,236241848 | 0,147413407 | 0,308413407 | -0,951586593 |
| chrXII | 1012576 | 1012696 | 120 | 0,039893848 | 0,733762177 | 0,051565356 | -1,629861787 | -1,385382519 | -1,005664905 | -0,837664905 | -2,097664905 |
| chrXII | 1012578 | 1012696 | 118 | 0,031345166 | 1,004290656 | 0,030266592 | -1,876889917 | -1,59535643 | -1,204261357 | -1,022261357 | -2,282261357 |
| chrXII | 1012579 | 1012696 | 117 | 0,079787696 | 0,426175002 | 0,15769482 | -1,003977126 | -0,853380557 | -0,365830513 | -0,176830513 | -1,436830513 |
| chrXII | 1012580 | 1012696 | 116 | 0,079787696 | 1,749170441 | 0,043624671 | -1,710089407 | -1,453575996 | -0,883610656 | -0,687610656 | -1,947610656 |
| chrXII | 1012581 | 1012696 | 115 | 0,059840772 | 0,874585221 | 0,064040141 | -1,5217159 | -1,293458515 | -0,679134278 | -0,476134278 | -1,736134278 |
| chrXII | 1012583 | 1012696 | 113 | 0,068389454 | 0,181587609 | 0,273582916 | -0,602012476 | -0,511710605 | 0,080090552 | 0,297090552 | -0,962909448 |
| chrXII | 1012584 | 1012696 | 112 | 0,039893848 | 1,100643265 | 0,034978124 | -1,812193861 | -1,540364782 | -0,972328572 | -0,748328572 | -2,008328572 |
| chrXII | 1012586 | 1012696 | 110 | 0,042743409 | 0,074117392 | 0,365763444 | -0,34309514 | -0,291630869 | 0,315619961 | 0,553619961 | -0,706380039 |
| chrXII | 1012590 | 1012674 | 84 | 0,151026711 | 0,333528262 | 0,311681273 | -0,49109035 | -0,417426797 | -0,010573336 | 0,409426664 | -0,850573336 |
| chrXII | 1012610 | 1012696 | 86 | 0,045592969 | 0,515115871 | 0,081313092 | -1,396293324 | -1,186849326 | -0,8260582 | -0,4200582 | -1,6800582 |
| chrXII | 1013868 | 1014024 | 156 | 3,946641408 | 2,223521747 | 0,639633233 | 0,357478614 | 0,303856822 | 0,213645184 | 0,129645184 | -1,130354816 |
| chrXII | 1017667 | 1017816 | 149 | 0,823523009 | 0,941290873 | 0,46663448 | -0,083732696 | -0,071172792 | -0,183527849 | -0,218527849 | -1,478527849 |
| chrXII | 1023621 | 1023767 | 146 | 2,479117708 | 1,589818049 | 0,609279147 | 0,277440736 | 0,235824626 | 0,076106146 | 0,062106146 | -1,197893854 |
| chrXII | 1024132 | 1024289 | 157 | 0,136778908 | 0,170470001 | 0,445172966 | -0,137866493 | -0,117186519 | -0,171310166 | -0,262310166 | -1,522310166 |
| chrXII | 1031294 | 1031445 | 151 | 0,726637949 | 0,552174567 | 0,568213041 | 0,171826517 | 0,14605254 | -0,006034471 | -0,055034471 | -1,315034471 |
| chrXII | 1032438 | 1032586 | 148 | 0,569912117 | 0,548468698 | 0,509586815 | 0,024032894 | 0,02042796 | -0,13057051 | -0,15857051 | -1,41857051 |
| chrXII | 1050400 | 1050528 | 128 | 0,205168362 | 0,522527611 | 0,281942418 | -0,577080856 | -0,490518728 | -0,254878192 | -0,142878192 | -1,402878192 |
| chrXII | 1050732 | 1050875 | 143 | 0,470177496 | 0,767115003 | 0,380005129 | -0,305467316 | -0,259647219 | -0,381285902 | -0,374285902 | -1,634285902 |
| chrXII | 1059679 | 1059820 | 141 | 0,051292091 | 0,062999783 | 0,448781606 | -0,128740216 | -0,109429183 | -0,158733471 | -0,137733471 | -1,397733471 |
| chrXII | 1065029 | 1065154 | 125 | 0,262159574 | 0,096352609 | 0,731243138 | 0,61657707 | 0,524090509 | 0,776247047 | 0,909247047 | -0,350752953 |
| chrXII | 1065029 | 1065156 | 127 | 0,527168708 | 0,166764131 | 0,759682607 | 0,705281957 | 0,599489663 | 0,837994918 | 0,956994918 | -0,303005082 |
| chrXII | 1066703 | 1066854 | 151 | 0,225115286 | 0,196411088 | 0,534047927 | 0,085449369 | 0,072631964 | -0,084222444 | -0,133222444 | -1,393222444 |
| chrXII | 1067646 | 1067808 | 162 | 0,051292091 | 0,040764565 | 0,557179598 | 0,143822286 | 0,122248943 | 0,199753013 | 0,073753013 | -1,186246987 |
| chrXII | 1067747 | 1067902 | 155 | 0,071239015 | 0,188999349 | 0,273745245 | -0,601524812 | -0,51129609 | -0,515097459 | -0,592097459 | -1,852097459 |
| chrXII | 1067881 | 1068018 | 137 | 0,14532759 | 0,137117174 | 0,51453455 | 0,036440779 | 0,030974662 | 0,118210849 | 0,167210849 | -1,092789151 |
| chrXII | 1068176 | 1068319 | 143 | 0,430283648 | 0,377998697 | 0,532343247 | 0,081161514 | 0,068987287 | -0,049361921 | -0,042361921 | -1,302361921 |
| chrXII | 1068224 | 1068349 | 125 | 0,210867483 | 0,274234349 | 0,434687048 | -0,164453569 | -0,139785534 | 0,105327729 | 0,238327729 | -1,021672271 |
| chrXII | 1068449 | 1068601 | 152 | 0,09973462 | 0,062999783 | 0,61286746 | 0,286800501 | 0,243780426 | 0,102815557 | 0,046815557 | -1,213184443 |
| chrXII | 1068522 | 1068666 | 144 | 0,105433742 | 0,070411522 | 0,599582493 | 0,252266586 | 0,214426598 | 0,028991948 | 0,028991948 | -1,231008052 |
| chrXII | 1068619 | 1068754 | 135 | 1,077133901 | 0,778232612 | 0,58055047 | 0,203301913 | 0,172806626 | 0,147419865 | 0,210419865 | -1,049580135 |
| chrXII | 1068619 | 1068806 | 187 | 0,125380666 | 0,055588044 | 0,692830634 | 0,503889924 | 0,428306435 | 0,477878216 | 0,176878216 | -1,083121784 |
| chrXII | 1068662 | 1068754 | 92 | 0,133929347 | 0,100058479 | 0,572377417 | 0,18243015 | 0,155065628 | 0,529950635 | 0,893950635 | -0,366049365 |
| chrXII | 1068662 | 1068806 | 144 | 1,310797868 | 0,518821741 | 0,716431908 | 0,572274263 | 0,486433123 | 0,311268149 | 0,311268149 | -0,948731851 |
| chrXII | 1069001 | 1069153 | 152 | 0,156725832 | 0,129705435 | 0,547167331 | 0,118507771 | 0,100731606 | -0,035882452 | -0,091882452 | -1,351882452 |
| chrXII | 1069302 | 1069432 | 130 | 0,398938482 | 0,44099848 | 0,474962408 | -0,062801193 | -0,053381014 | 0,044228196 | 0,142228196 | -1,117771804 |
| chrXII | 1069440 | 1069601 | 161 | 0,199469241 | 0,133411305 | 0,599221683 | 0,251333038 | 0,213633083 | 0,205239575 | 0,086239575 | -1,173760425 |
| chrXII | 1070837 | 1070988 | 151 | 0,549965193 | 0,389116306 | 0,585641601 | 0,216347614 | 0,183895472 | 0,034970085 | -0,014029915 | -1,274029915 |
| chrXII | 1070837 | 1071008 | 171 | 0,04844253 | 0,044470435 | 0,521375354 | 0,053605728 | 0,045564869 | 0,170299862 | -0,018700138 | -1,278700138 |
| chrXII | 1072226 | 1072381 | 155 | 0,324849907 | 0,455821958 | 0,416115812 | -0,211840305 | -0,180064259 | -0,171760617 | -0,248760617 | -1,508760617 |
| chrXII | 1073391 | 1073538 | 147 | 1,726833714 | 1,193290004 | 0,591356354 | 0,23103541 | 0,196380098 | 0,119017326 | 0,098017326 | -1,161982674 |
| chrXII | 1073547 | 1073677 | 130 | 0,29350474 | 0,274234349 | 0,516971168 | 0,042553249 | 0,036170262 | 0,112570725 | 0,210570725 | -1,049429275 |
| chrXII | 1073928 | 1074064 | 136 | 0,527168708 | 0,674468263 | 0,438708795 | -0,154243695 | -0,131107141 | -0,096139293 | -0,040139293 | -1,300139293 |
| chrXII | 1074309 | 1074471 | 162 | 0,04844253 | 0,040764565 | 0,543034495 | 0,108081541 | 0,09186931 | 0,169631887 | 0,043631887 | -1,216368113 |
| chrXII | 1074410 | 1074565 | 155 | 0,059840772 | 0,188999349 | 0,240478794 | -0,704763239 | -0,599048753 | -0,592018038 | -0,669018038 | -1,929018038 |
| chrXII | 1074544 | 1074681 | 137 | 0,136778908 | 0,129705435 | 0,513271836 | 0,033273697 | 0,028282642 | 0,111572837 | 0,160572837 | -1,099427163 |
| chrXII | 1074831 | 1074982 | 151 | 0,051292091 | 0,059293913 | 0,463820816 | -0,090812431 | -0,077190567 | -0,211081729 | -0,260081729 | -1,520081729 |
| chrXII | 1074839 | 1074982 | 143 | 0,427434088 | 0,43729261 | 0,494299631 | -0,014289193 | -0,012145814 | -0,134980516 | -0,127980516 | -1,387980516 |
| chrXII | 1074887 | 1075012 | 125 | 0,256460453 | 0,289057827 | 0,470122564 | -0,074961771 | -0,063717505 | 0,175775758 | 0,308775758 | -0,951224242 |
| chrXII | 1075112 | 1075264 | 152 | 0,113982423 | 0,085235 | 0,572150875 | 0,181852797 | 0,154574878 | 0,016323256 | -0,039676744 | -1,299676744 |
| chrXII | 1075185 | 1075329 | 144 | 0,076938136 | 0,070411522 | 0,522146688 | 0,055542058 | 0,047210749 | -0,133480009 | -0,133480009 | -1,393480009 |
| chrXII | 1075282 | 1075417 | 135 | 1,205364127 | 0,770820872 | 0,609944984 | 0,279175646 | 0,237299299 | 0,210835569 | 0,273835569 | -0,986164431 |
| chrXII | 1075282 | 1075469 | 187 | 0,151026711 | 0,066705652 | 0,693634647 | 0,506179412 | 0,4302525 | 0,472808164 | 0,171808164 | -1,088191836 |
| chrXII | 1075325 | 1075417 | 92 | 0,182371877 | 0,125999566 | 0,591403262 | 0,231156174 | 0,196482748 | 0,570784137 | 0,934784137 | -0,325215863 |
| chrXII | 1075325 | 1075469 | 144 | 1,353541277 | 0,61146848 | 0,688821657 | 0,492513068 | 0,418636108 | 0,231961775 | 0,231961775 | -1,028038225 |
| chrXII | 1075664 | 1075816 | 152 | 0,199469241 | 0,122293696 | 0,619926094 | 0,305286692 | 0,259493688 | 0,134750185 | 0,078750185 | -1,181249815 |
| chrXII | 1075965 | 1076095 | 130 | 0,413186285 | 0,415057393 | 0,498870436 | -0,0028314 | -0,00240669 | 0,075874338 | 0,173874338 | -1,086125662 |
| chrXII | 1076103 | 1076264 | 161 | 0,247911771 | 0,207528696 | 0,544334087 | 0,111358804 | 0,094654983 | 0,089054602 | -0,029945398 | -1,289945398 |
| chrXII | 1077160 | 1077312 | 152 | 0,042743409 | 0,040764565 | 0,51184823 | 0,029703476 | 0,025247955 | -0,083093427 | -0,139093427 | -1,399093427 |
| chrXII | 1077500 | 1077651 | 151 | 0,541416511 | 0,385410436 | 0,584161383 | 0,212550948 | 0,180668306 | 0,043131073 | -0,005868927 | -1,265868927 |

TABLE 2

| Chrom | Start | End | Length | Section A | Section B | A/A+B | Z-score | Z * 0.85 | Phase correction | Length correction | ΔLK |
|---|---|---|---|---|---|---|---|---|---|---|---|
| chrXIII | 687 | 858 | 171 | 0,045592969 | 0,048176305 | 0,486225044 | -0,034535557 | -0,029355224 | 0,265977604 | 0,076977604 | -1,183022396 |
| chrXIII | 707 | 858 | 151 | 0,58700948 | 0,340940001 | 0,632587756 | 0,338714936 | 0,287907696 | 0,143290712 | 0,094290712 | -1,165709288 |
| chrXIII | 1046 | 1198 | 152 | 0,045592969 | 0,051882174 | 0,467739443 | -0,080953558 | -0,068810524 | -0,164095132 | -0,220095132 | -1,480095132 |
| chrXIII | 1151 | 1299 | 148 | 0,062690333 | 0,059293913 | 0,51392155 | 0,034903236 | 0,029667751 | -0,151779997 | -0,179779997 | -1,439779997 |
| chrXIII | 1836 | 2023 | 187 | 0,071239015 | 0,040764565 | 0,6360423 | 0,347899847 | 0,29571487 | 0,650428694 | 0,349428694 | -0,910571306 |
| chrXIII | 1872 | 2023 | 151 | 0,473027057 | 0,289057827 | 0,620701272 | 0,307323098 | 0,261224633 | 0,12022764 | 0,07122764 | -1,18877236 |
| chrXIII | 1884 | 2018 | 134 | 0,572761677 | 0,489174784 | 0,539355882 | 0,098811125 | 0,083989456 | 0,15113177 | 0,22113177 | -1,03886823 |
| chrXIII | 1909 | 2005 | 96 | 0,039893848 | 0,103764348 | 0,277699771 | -0,589688448 | -0,501235181 | -0,033176331 | 0,302823669 | -0,957176331 |
| chrXIII | 2542 | 2694 | 152 | 0,165274514 | 0,107470218 | 0,605967759 | 0,268824834 | 0,228501109 | 0,139978606 | 0,083978606 | -1,176021394 |
| chrXIII | 2889 | 3033 | 144 | 1,342143035 | 0,626291959 | 0,681832542 | 0,472829373 | 0,401904967 | 0,256471872 | 0,256471872 | -1,003528128 |
| chrXIII | 2889 | 3076 | 187 | 0,119681545 | 0,081529131 | 0,594807131 | 0,239928434 | 0,203939169 | 0,557059861 | 0,256059861 | -1,003940139 |
| chrXIII | 2941 | 3033 | 92 | 0,208017923 | 0,137117174 | 0,602714486 | 0,260379534 | 0,221322604 | 0,624703275 | 0,988703275 | -0,271296725 |
| chrXIII | 2941 | 3076 | 135 | 1,199665006 | 0,711526959 | 0,627705133 | 0,325781423 | 0,27691421 | 0,337671475 | 0,400671475 | -0,859328525 |
| chrXIII | 3029 | 3173 | 144 | 0,059840772 | 0,048176305 | 0,553993628 | 0,135757811 | 0,11539414 | -0,034580427 | -0,034580427 | -1,294580427 |
| chrXIII | 3094 | 3246 | 152 | 0,125380666 | 0,085235 | 0,595305506 | 0,241214353 | 0,2050322 | 0,12159943 | 0,06559943 | -1,19440057 |
| chrXIII | 3346 | 3471 | 125 | 0,225115286 | 0,303881305 | 0,425551487 | -0,187711297 | -0,159554603 | 0,299345477 | 0,432345477 | -0,827654523 |
| chrXIII | 3376 | 3519 | 143 | 0,376141997 | 0,43729261 | 0,46241209 | -0,094358752 | -0,080204939 | -0,196081815 | -0,189081815 | -1,449081815 |
| chrXIII | 3376 | 3527 | 151 | 0,054141651 | 0,066705652 | 0,448017039 | -0,130672891 | -0,111071958 | -0,247775214 | -0,296775214 | -1,556775214 |
| chrXIII | 5624 | 5767 | 143 | 0,296354301 | 0,207528696 | 0,588141101 | 0,222765786 | 0,189350918 | 0,05900657 | 0,06600657 | -1,19399343 |
| chrXIII | 9070 | 9207 | 137 | 0,105433742 | 0,092646739 | 0,53227729 | 0,080995639 | 0,068846293 | 0,074698835 | 0,123698835 | -1,136301165 |
| chrXIII | 13123 | 13267 | 144 | 0,498673102 | 0,370586958 | 0,573675388 | 0,185739279 | 0,157878387 | 0,001693055 | 0,001693055 | -1,258306945 |
| chrXIII | 14333 | 14505 | 172 | 0,074088575 | 0,188999349 | 0,281611463 | -0,578061014 | -0,491351862 | -0,188543528 | -0,384543528 | -1,644543528 |
| chrXIII | 16025 | 16179 | 154 | 1,008744447 | 0,781938481 | 0,56332946 | 0,159416068 | 0,135503658 | 0,046243023 | -0,023756977 | -1,283756977 |
| chrXIII | 18573 | 18744 | 171 | 0,598407723 | 1,096937395 | 0,352971036 | -0,377311574 | -0,320714838 | -0,035644976 | -0,224644976 | -1,484644976 |
| chrXIII | 19858 | 19999 | 141 | 1,629948654 | 1,133996091 | 0,589718248 | 0,226820268 | 0,192797228 | 0,121333724 | 0,142333724 | -1,117666276 |
| chrXIII | 20186 | 20333 | 147 | 0,806425645 | 0,61888022 | 0,565791291 | 0,165669032 | 0,140818677 | -0,039687818 | -0,060687818 | -1,320687818 |
| chrXIII | 21593 | 21740 | 147 | 0,871965539 | 0,566998046 | 0,605967759 | 0,268824834 | 0,228501109 | 0,042672996 | 0,021672996 | -1,238327004 |
| chrXIII | 23095 | 23242 | 147 | 0,230814407 | 0,144528914 | 0,614942093 | 0,292223409 | 0,248389898 | 0,059233146 | 0,038233146 | -1,221766854 |
| chrXIII | 32413 | 32558 | 145 | 0,24221265 | 0,181587609 | 0,571525487 | 0,180259276 | 0,153220385 | -0,008997143 | -0,015997143 | -1,275997143 |
| chrXIII | 33232 | 33384 | 152 | 0,09688506 | 0,070411522 | 0,579121574 | 0,199646764 | 0,16969975 | 0,072463432 | 0,016463432 | -1,243536568 |
| chrXIII | 38992 | 39143 | 151 | 0,125380666 | 0,100058479 | 0,556161913 | 0,141245282 | 0,12005849 | -0,019742332 | -0,068742332 | -1,328742332 |
| chrXIII | 40472 | 40624 | 152 | 1,045788734 | 0,926467395 | 0,53024996 | 0,075898212 | 0,06451348 | -0,027492743 | -0,083492743 | -1,343492743 |
| chrXIII | 42972 | 43133 | 161 | 0,63260245 | 0,637409568 | 0,498107452 | -0,004743933 | -0,004032343 | 0,069202662 | -0,049797338 | -1,309797338 |
| chrXIII | 47095 | 47254 | 159 | 0,909009826 | 0,685585872 | 0,570056615 | 0,176518304 | 0,150040559 | 0,237685581 | 0,132685581 | -1,127314419 |
| chrXIII | 54675 | 54757 | 82 | 1,820869213 | 2,916519358 | 0,384361381 | -0,294045989 | -0,249939091 | 0,072235478 | 0,506235478 | -0,753764522 |
| chrXIII | 54913 | 55020 | 107 | 0,074088575 | 0,403939784 | 0,154987824 | -1,015273133 | -0,862982163 | -0,172871325 | 0,086128675 | -1,173871325 |
| chrXIII | 54913 | 55032 | 119 | 0,082637257 | 0,455821958 | 0,153469854 | -1,021664524 | -0,868414845 | -0,239745926 | -0,064745926 | -1,324745926 |
| chrXIII | 59231 | 59396 | 165 | 0,162424953 | 0,155646522 | 0,510655515 | 0,026712592 | 0,022705704 | 0,263807945 | 0,116807945 | -1,143192055 |
| chrXIII | 63883 | 64021 | 138 | 0,393239361 | 0,303881305 | 0,564090809 | 0,161349191 | 0,137146813 | 0,158086866 | 0,200086866 | -1,059913134 |
| chrXIII | 64235 | 64393 | 158 | 0,042743409 | 0,096352609 | 0,307294266 | -0,503534497 | -0,428004322 | -0,347670368 | -0,445670368 | -1,705670368 |
| chrXIII | 64871 | 65018 | 147 | 0,407487163 | 0,303881305 | 0,572821514 | 0,183562133 | 0,156027813 | -0,027217953 | -0,048217953 | -1,308217953 |
| chrXIII | 66681 | 66829 | 148 | 0,116831984 | 0,074117392 | 0,61184795 | 0,284138687 | 0,241517884 | 0,070515368 | 0,042515368 | -1,217484632 |
| chrXIII | 75788 | 75939 | 151 | 0,267858695 | 0,211234566 | 0,559095101 | 0,148675368 | 0,126374063 | -0,008183113 | -0,057183113 | -1,317183113 |
| chrXIII | 76732 | 76885 | 153 | 2,19701121 | 2,093816312 | 0,512025058 | 0,030146916 | 0,025624878 | -0,068942299 | -0,131942299 | -1,391942299 |
| chrXIII | 77591 | 77728 | 137 | 0,068389454 | 0,070411522 | 0,492715945 | -0,018259434 | -0,015520519 | 0,00337707 | 0,05237707 | -1,20762293 |
| chrXIII | 83978 | 84137 | 159 | 0,626903328 | 0,578115654 | 0,520243529 | 0,050764798 | 0,043150079 | 0,149088721 | 0,044088721 | -1,215911279 |
| chrXIII | 93745 | 93871 | 126 | 0,068389454 | 0,214940436 | 0,241377477 | -0,701878473 | -0,596596702 | -0,148405246 | -0,022405246 | -1,282405246 |
| chrXIII | 98827 | 98976 | 149 | 0,350495952 | 0,118587827 | 0,747192651 | 0,665681504 | 0,565829278 | 0,431325885 | 0,396325885 | -0,863674115 |
| chrXIII | 102493 | 102629 | 136 | 0,598407723 | 0,422469132 | 0,58617033 | 0,217704522 | 0,185048844 | 0,20021136 | 0,25621136 | -1,00378864 |
| chrXIII | 104088 | 104231 | 143 | 1,421930731 | 0,926467395 | 0,605489638 | 0,267582473 | 0,227445102 | 0,103169044 | 0,110169044 | -1,149830956 |
| chrXIII | 104601 | 104708 | 107 | 0,125380666 | 0,692997611 | 0,153206249 | -1,022778687 | -0,869361884 | -0,190806582 | 0,068193418 | -1,191806582 |
| chrXIII | 104601 | 104720 | 119 | 0,24221265 | 1,130290221 | 0,17647515 | -0,92888189 | -0,789549606 | -0,177640637 | -0,002640637 | -1,262640637 |
| chrXIII | 107860 | 108019 | 159 | 0,504372223 | 0,422469132 | 0,544183986 | 0,110980226 | 0,094333192 | 0,193405712 | 0,088405712 | -1,171594288 |
| chrXIII | 109020 | 109176 | 156 | 1,12272687 | 1,030231743 | 0,521480935 | 0,053870763 | 0,045790149 | 0,090124794 | 0,006124794 | -1,253875206 |
| chrXIII | 112712 | 112852 | 140 | 0,911859387 | 0,937585003 | 0,493045042 | -0,017434378 | -0,014819221 | -0,012036558 | 0,015963442 | -1,244036558 |
| chrXIII | 113210 | 113365 | 155 | 0,068389454 | 0,044470435 | 0,605967759 | 0,268824834 | 0,228501109 | 0,164884161 | 0,087884161 | -1,172115839 |
| chrXIII | 113210 | 113365 | 155 | 0,068389454 | 0,044470435 | 0,605967759 | 0,268824834 | 0,228501109 | 0,164884161 | 0,087884161 | -1,172115839 |
| chrXIII | 116680 | 116830 | 150 | 0,034194727 | 0,051882174 | 0,397257877 | -0,260451199 | -0,221383519 | -0,374344701 | -0,416344701 | -1,676344701 |
| chrXIII | 116680 | 116833 | 153 | 0,079787696 | 0,062999783 | 0,558786364 | 0,147892925 | 0,125708986 | 0,03145303 | -0,03154697 | -1,29154697 |
| chrXIII | 116680 | 116835 | 155 | 16,6784781 | 10,31714091 | 0,617821658 | 0,299764647 | 0,25479995 | 0,195429563 | 0,118429563 | -1,141570437 |
| chrXIII | 116684 | 116835 | 151 | 0,065539893 | 0,059293913 | 0,525017182 | 0,062749932 | 0,053337442 | -0,085129796 | -0,134129796 | -1,394129796 |
| chrXIII | 116783 | 116928 | 145 | 0,091185939 | 0,048176305 | 0,654308775 | 0,396979674 | 0,337432723 | 0,172376419 | 0,165376419 | -1,094623581 |
| chrXIII | 117349 | 117491 | 142 | 1,780975365 | 1,304466092 | 0,577218978 | 0,194784006 | 0,165566405 | 0,073633234 | 0,087633234 | -1,172366766 |
| chrXIII | 117514 | 117618 | 104 | 4,505155283 | 40,30133167 | 0,100546943 | -1,278441255 | -1,086675066 | -0,552456473 | -0,272456473 | -1,532456473 |
| chrXIII | 119216 | 119393 | 177 | 0,24221265 | 0,455821958 | 0,346992322 | -0,393453388 | -0,33443538 | 0,03299322 | -0,19800678 | -1,45800678 |
| chrXIII | 120074 | 120243 | 169 | 0,470177496 | 1,026525873 | 0,314142071 | -0,484143344 | -0,411521842 | -0,118893484 | -0,293893484 | -1,553893484 |
| chrXIII | 120074 | 120243 | 169 | 0,470177496 | 1,026525873 | 0,314142071 | -0,484143344 | -0,411521842 | -0,118893484 | -0,293893484 | -1,553893484 |
| chrXIII | 123583 | 123746 | 163 | 0,652549374 | 0,904232177 | 0,419165665 | -0,20402837 | -0,173424114 | -0,014894304 | -0,147894304 | -1,407894304 |
| chrXIII | 124934 | 125083 | 149 | 0,273557816 | 0,200116957 | 0,577522451 | 0,195559327 | 0,166225428 | 0,030863675 | -0,004136325 | -1,264136325 |
| chrXIII | 125456 | 125612 | 156 | 1,689789426 | 1,393406962 | 0,548064156 | 0,120771925 | 0,102656136 | 0,142102089 | 0,058102089 | -1,201897911 |
| chrXIII | 126339 | 126468 | 129 | 0,045592969 | 0,037058696 | 0,551627959 | 0,129775456 | 0,110309137 | 0,452252663 | 0,557252663 | -0,702747337 |
| chrXIII | 129825 | 129930 | 105 | 2,154267801 | 17,96605572 | 0,107069243 | -1,242265865 | -1,055925985 | -0,489714441 | -0,216714441 | -1,476714441 |
| chrXIII | 129825 | 129932 | 107 | 1,307948308 | 9,509261339 | 0,120913651 | -1,17043166 | -0,994866911 | -0,28524102 | -0,02624102 | -1,28624102 |
| chrXIII | 129843 | 129930 | 87 | 0,068389454 | 0,233469783 | 0,226560746 | -0,750221201 | -0,637688021 | -0,305659195 | 0,093340805 | -1,166659195 |
| chrXIII | 129843 | 130004 | 161 | 0,817823887 | 0,629997828 | 0,564865051 | 0,163315669 | 0,138818318 | 0,216212545 | 0,097212545 | -1,162787455 |
| chrXIII | 129843 | 130004 | 161 | 0,817823887 | 0,629997828 | 0,564865051 | 0,163315669 | 0,138818318 | 0,216212545 | 0,097212545 | -1,162787455 |
| chrXIII | 130886 | 131027 | 141 | 6,288980208 | 3,761457622 | 0,625741914 | 0,320596525 | 0,272507046 | 0,203352203 | 0,224352203 | -1,035647797 |
| chrXIII | 131028 | 131190 | 162 | 0,125380666 | 0,081529131 | 0,605967759 | 0,268824834 | 0,228501109 | 0,333154984 | 0,207154984 | -1,052845016 |
| chrXIII | 131602 | 131755 | 153 | 0,359044634 | 0,326116523 | 0,524029464 | 0,060269401 | 0,051228991 | -0,047976245 | -0,110976245 | -1,370976245 |
| chrXIII | 133367 | 133536 | 169 | 0,176672756 | 0,300175436 | 0,370501051 | -0,330526594 | -0,280947605 | 0,016410295 | -0,158589705 | -1,418589705 |
| chrXIII | 134348 | 134447 | 99 | 0,327699467 | 2,116051529 | 0,134096914 | -1,107231555 | -0,941146822 | -0,484663408 | -0,169663408 | -1,429663408 |
| chrXIII | 135575 | 135725 | 150 | 0,042743409 | 0,037058696 | 0,535617563 | 0,08939893 | 0,07598909 | -0,079659658 | -0,121659658 | -1,381659658 |
| chrXIII | 139632 | 139738 | 106 | 0,133929347 | 1,208113483 | 0,099795137 | -1,282719764 | -1,0903118 | -0,450777998 | -0,184777998 | -1,444777998 |
| chrXIII | 139632 | 139741 | 109 | 0,088336378 | 1,456406744 | 0,057185157 | -1,578850666 | -1,342023066 | -0,633734967 | -0,388734967 | -1,648734967 |
| chrXIII | 142450 | 142583 | 133 | 1,134125112 | 10,75813939 | 0,095366624 | -1,308413054 | -1,112151096 | -1,032482142 | -0,955482142 | -2,215482142 |
| chrXIII | 147977 | 148129 | 152 | 0,265009134 | 0,196411088 | 0,574333594 | 0,187418125 | 0,159305406 | 0,061820708 | 0,005820708 | -1,254179292 |
| chrXIII | 148322 | 148434 | 112 | 1,336443914 | 10,60619873 | 0,111905208 | -1,216458222 | -1,033989488 | -0,345900204 | -0,121900204 | -1,381900204 |
| chrXIII | 154231 | 154388 | 157 | 0,085486818 | 0,118587827 | 0,418899751 | -0,204708983 | -0,174002636 | -0,1377113 | -0,2287113 | -1,4887113 |
| chrXIII | 155621 | 155784 | 163 | 0,037044288 | 0,037058696 | 0,499902783 | -0,000243688 | -0,000207135 | 0,157913263 | 0,024913263 | -1,235086737 |
| chrXIII | 163164 | 163259 | 95 | 0,427434088 | 1,775111528 | 0,194063671 | -0,863018416 | -0,733565653 | -0,280794011 | 0,062205989 | -1,197794011 |
| chrXIII | 169237 | 169345 | 108 | 0,065539893 | 0,733762177 | 0,081996401 | -1,391767538 | -1,183002408 | -0,458604463 | -0,206604463 | -1,466604463 |
| chrXIII | 170327 | 170483 | 156 | 0,065539893 | 0,08894087 | 0,424259254 | -0,191009043 | -0,162357687 | -0,125530991 | -0,209530991 | -1,469530991 |
| chrXIII | 170574 | 170680 | 106 | 0,082637257 | 0,107470218 | 0,434687048 | -0,164453569 | -0,139785534 | 0,514602502 | 0,780602502 | -0,479397498 |
| chrXIII | 170574 | 170696 | 122 | 0,068389454 | 0,081529131 | 0,456177292 | -0,110069087 | -0,093558724 | 0,467199798 | 0,621199798 | -0,638800202 |
| chrXIII | 170574 | 170696 | 122 | 0,068389454 | 0,081529131 | 0,456177292 | -0,110069087 | -0,093558724 | 0,467199798 | 0,621199798 | -0,638800202 |
| chrXIII | 171072 | 171220 | 148 | 0,592708601 | 0,478057176 | 0,553537117 | 0,134603005 | 0,114412554 | -0,059544467 | -0,087544467 | -1,347544467 |
| chrXIII | 175762 | 175915 | 153 | 1,145523355 | 0,896820438 | 0,560886643 | 0,153217556 | 0,130234923 | 0,033738532 | -0,029261468 | -1,289261468 |
| chrXIII | 179143 | 179321 | 178 | 1,128425991 | 2,160521964 | 0,343096335 | -0,404027263 | -0,343423174 | 0,012339021 | -0,225660979 | -1,485660979 |
| chrXIII | 180747 | 180905 | 158 | 0,131079787 | 0,081529131 | 0,616530051 | 0,29638 | 0,251923 | 0,342010068 | 0,244010068 | -1,015989932 |
| chrXIII | 181711 | 181851 | 140 | 2,356586603 | 1,712111745 | 0,579199145 | 0,199845122 | 0,169868354 | 0,16196119 | 0,18996119 | -1,07003881 |
| chrXIII | 181848 | 181977 | 129 | 0,04844253 | 0,092646739 | 0,343346664 | -0,403346515 | -0,342844538 | 0,030213249 | 0,135213249 | -1,124786751 |

| Chrom | Start | End | Length | Section A | Section B | A/A+B | Z-score | Z * 0.85 | Phase correction | Length correction | ΔLK |
|---|---|---|---|---|---|---|---|---|---|---|---|
| chrXIII | 184182 | 184330 | 148 | 0,051292091 | 0,040764565 | 0,557179598 | 0,143822286 | 0,122248943 | -0,052931298 | -0,080931298 | -1,340931298 |
| chrXIII | 184701 | 184824 | 123 | 0,088336378 | 0,185293479 | 0,322831649 | -0,459795102 | -0,390825837 | 0,125433804 | 0,272433804 | -0,987566196 |
| chrXIII | 184801 | 184956 | 155 | 0,076938136 | 0,100058479 | 0,434687048 | -0,164453569 | -0,139785534 | -0,201963216 | -0,278963216 | -1,538963216 |
| chrXIII | 185234 | 185368 | 134 | 0,153876272 | 0,081529131 | 0,653664997 | 0,395234266 | 0,335949126 | 0,404001973 | 0,474001973 | -0,785998027 |
| chrXIII | 185234 | 185374 | 140 | 0,168124074 | 0,125999566 | 0,571610206 | 0,180475116 | 0,153403848 | 0,147504313 | 0,175504313 | -1,084495687 |
| chrXIII | 186277 | 186412 | 135 | 0,082637257 | 0,055588044 | 0,597844653 | 0,247772164 | 0,210606339 | 0,28176275 | 0,34476275 | -0,91523725 |
| chrXIII | 186724 | 186874 | 150 | 8,810841324 | 4,124632841 | 0,68113787 | 0,470883041 | 0,400250585 | 0,240242633 | 0,198242633 | -1,061757367 |
| chrXIII | 186903 | 187060 | 157 | 0,054141651 | 0,081529131 | 0,399066404 | -0,255764342 | -0,217399691 | -0,176474579 | -0,267474579 | -1,527474579 |
| chrXIII | 187135 | 187284 | 149 | 0,079787696 | 0,055588044 | 0,589379577 | 0,225949308 | 0,192056912 | 0,057636305 | 0,022636305 | -1,237363695 |
| chrXIII | 187394 | 187523 | 129 | 0,034194727 | 0,066705652 | 0,338895921 | -0,415478239 | -0,353156503 | 0,008667179 | 0,113667179 | -1,146332821 |
| chrXIII | 188306 | 188456 | 150 | 0,444531451 | 0,381704567 | 0,538019938 | 0,095446574 | 0,081129588 | -0,071263247 | -0,113263247 | -1,373263247 |
| chrXIII | 195171 | 195314 | 143 | 0,210867483 | 0,148234783 | 0,58720733 | 0,220367006 | 0,187311955 | 0,046659925 | 0,053659925 | -1,206340075 |
| chrXIII | 196534 | 196661 | 127 | 0,031345166 | 0,107470218 | 0,225804702 | -0,752734617 | -0,639824425 | -0,208408121 | -0,089408121 | -1,349408121 |
| chrXIII | 196694 | 196843 | 149 | 0,071239015 | 0,070411522 | 0,502920895 | 0,007321663 | 0,006223413 | -0,141177973 | -0,176177973 | -1,436177973 |
| chrXIII | 196868 | 196991 | 123 | 0,076938136 | 0,233469783 | 0,247861382 | -0,681235065 | -0,579049806 | -0,054746496 | 0,092253504 | -1,167746496 |
| chrXIII | 196968 | 197123 | 155 | 0,062690333 | 0,092646739 | 0,403576119 | -0,244101464 | -0,207486244 | -0,271269175 | -0,348269175 | -1,608269175 |
| chrXIII | 197401 | 197535 | 134 | 0,14817715 | 0,085235 | 0,634830492 | 0,344674598 | 0,292973409 | 0,323602132 | 0,393602132 | -0,866397868 |
| chrXIII | 197401 | 197541 | 140 | 0,179522317 | 0,081529131 | 0,687689413 | 0,48931151 | 0,415914784 | 0,402413702 | 0,430413702 | -0,829586298 |
| chrXIII | 197597 | 197748 | 151 | 0,113982423 | 0,066705652 | 0,630824269 | 0,334037235 | 0,283931649 | 0,14708664 | 0,09808664 | -1,16191336 |
| chrXIII | 197607 | 197752 | 145 | 0,113982423 | 0,077823261 | 0,594259882 | 0,238516864 | 0,202739335 | 0,016292599 | 0,009292599 | -1,250707401 |
| chrXIII | 197625 | 197776 | 151 | 0,059840772 | 0,059293913 | 0,502295129 | 0,005753068 | 0,004890107 | -0,138382794 | -0,187382794 | -1,447382794 |
| chrXIII | 197879 | 198033 | 154 | 0,225115286 | 0,159352392 | 0,585524607 | 0,216047418 | 0,183640306 | 0,095573711 | 0,025573711 | -1,234426289 |
| chrXIII | 197913 | 198057 | 144 | 0,062690333 | 0,048176305 | 0,56545715 | 0,164819952 | 0,140096959 | -0,013608373 | -0,013608373 | -1,273608373 |
| chrXIII | 198891 | 199037 | 146 | 0,079787696 | 0,037058696 | 0,682842618 | 0,475662609 | 0,404313218 | 0,208799295 | 0,194799295 | -1,065200705 |
| chrXIII | 198891 | 199041 | 150 | 8,882080339 | 4,328455668 | 0,672348218 | 0,446406602 | 0,379445612 | 0,233985211 | 0,191985211 | -1,068014789 |
| chrXIII | 199070 | 199227 | 157 | 0,054141651 | 0,040764565 | 0,570475287 | 0,177584338 | 0,150946687 | 0,195883468 | 0,104883468 | -1,155116532 |
| chrXIII | 199302 | 199451 | 149 | 0,09973462 | 0,051882174 | 0,657807209 | 0,406485945 | 0,345513053 | 0,201150587 | 0,166150587 | -1,093849413 |
| chrXIII | 199561 | 199690 | 129 | 0,04844253 | 0,085235 | 0,36238349 | -0,352095051 | -0,299280793 | 0,038738439 | 0,143738439 | -1,116261561 |
| chrXIII | 200473 | 200623 | 150 | 0,165274514 | 0,196411088 | 0,456956299 | -0,108104753 | -0,09188904 | -0,235549481 | -0,277549481 | -1,537549481 |
| chrXIII | 203778 | 203929 | 151 | 0,031345166 | 0,059293913 | 0,345823971 | -0,396619676 | -0,337126724 | -0,477825043 | -0,526825043 | -1,786825043 |
| chrXIII | 204102 | 204256 | 154 | 0,823523009 | 0,61517435 | 0,572408786 | 0,182510101 | 0,155133586 | 0,062922211 | -0,007077789 | -1,267077789 |
| chrXIII | 208554 | 208676 | 122 | 0,085486818 | 0,062999783 | 0,575720754 | 0,190958009 | 0,162314307 | 0,720592252 | 0,874592252 | -0,385407748 |
| chrXIII | 211138 | 211313 | 175 | 4,134712407 | 3,642869796 | 0,531619249 | 0,079340866 | 0,067439736 | 0,413215776 | 0,196215776 | -1,063784224 |
| chrXIII | 218290 | 218401 | 111 | 0,24221265 | 1,841817181 | 0,116223216 | -1,194080622 | -1,014968529 | -0,318884584 | -0,087884584 | -1,347884584 |
| chrXIII | 220837 | 220983 | 146 | 0,091185939 | 0,040764565 | 0,691061693 | 0,498861988 | 0,42403269 | 0,237761285 | 0,223761285 | -1,036238715 |
| chrXIII | 228866 | 229000 | 134 | 0,350495952 | 0,314998914 | 0,526669655 | 0,066900784 | 0,056865666 | 0,089086098 | 0,159086098 | -1,100913902 |
| chrXIII | 230938 | 231091 | 153 | 0,122531105 | 0,118587827 | 0,508177041 | 0,020498237 | 0,017423501 | -0,078564521 | -0,141564521 | -1,401564521 |
| chrXIII | 231257 | 231408 | 151 | 0,190920559 | 0,17417587 | 0,522931872 | 0,057513369 | 0,048886364 | -0,097040535 | -0,146040535 | -1,406040535 |
| chrXIII | 233569 | 233719 | 150 | 0,695292782 | 0,455821958 | 0,604016922 | 0,263758316 | 0,224194569 | 0,077530362 | 0,035530362 | -1,224469638 |
| chrXIII | 240347 | 240506 | 159 | 0,071239015 | 0,185293479 | 0,277699771 | -0,589688448 | -0,501235181 | -0,410706464 | -0,515706464 | -1,775706464 |
| chrXIII | 242945 | 243091 | 146 | 1,285151823 | 0,837526525 | 0,605438796 | 0,267450388 | 0,22733283 | 0,050367355 | 0,036367355 | -1,223632645 |
| chrXIII | 243502 | 243662 | 160 | 0,04844253 | 0,070411522 | 0,407579962 | -0,23377466 | -0,198708461 | -0,139479903 | -0,251479903 | -1,511479903 |
| chrXIII | 243825 | 243980 | 155 | 2,843861462 | 2,382874139 | 0,544098971 | 0,11076581 | 0,094150939 | 0,064556371 | -0,012443629 | -1,272443629 |
| chrXIII | 244965 | 245130 | 165 | 0,122531105 | 0,203822827 | 0,375454662 | -0,317440575 | -0,269824489 | -0,012863435 | -0,159863435 | -1,419863435 |
| chrXIII | 244965 | 245130 | 165 | 0,122531105 | 0,203822827 | 0,375454662 | -0,317440575 | -0,269824489 | -0,012863435 | -0,159863435 | -1,419863435 |
| chrXIII | 245559 | 245710 | 151 | 0,356195073 | 0,170470001 | 0,676321805 | 0,457437813 | 0,388822141 | 0,248959311 | 0,199959311 | -1,060040689 |
| chrXIII | 246029 | 246179 | 147 | 0,142478029 | 0,17788174 | 0,444743825 | -0,138952542 | -0,118109661 | -0,300607109 | -0,321607109 | -1,581607109 |
| chrXIII | 246042 | 246200 | 158 | 0,501522663 | 0,503998263 | 0,498768996 | -0,003085674 | -0,002622823 | 0,094767343 | -0,003232657 | -1,263232657 |
| chrXIII | 246354 | 246509 | 155 | 0,29350474 | 0,100058479 | 0,745762628 | 0,661214529 | 0,56203235 | 0,52102062 | 0,44402062 | -0,81597938 |
| chrXIII | 246617 | 246737 | 120 | 0,322000346 | 8,071383942 | 0,038363589 | -1,769999775 | -1,504499809 | -0,922758173 | -0,754758173 | -2,014758173 |
| chrXIII | 250435 | 250548 | 113 | 0,185221438 | 0,192705218 | 0,490098899 | -0,024820929 | -0,021097789 | 0,628743846 | 0,845743846 | -0,414256154 |
| chrXIII | 250507 | 250631 | 124 | 0,037044288 | 1,093231526 | 0,032774556 | -1,841494599 | -1,565270409 | -1,064232096 | -0,924232096 | -2,184232096 |
| chrXIII | 250507 | 250662 | 155 | 0,039893848 | 0,114881957 | 0,257752484 | -0,65028992 | -0,552746432 | -0,585632252 | -0,662632252 | -1,922632252 |
| chrXIII | 250507 | 250664 | 157 | 0,028495606 | 0,251999131 | 0,101590519 | -1,2725408 | -1,08165968 | -1,027686658 | -1,118686658 | -2,378686658 |
| chrXIII | 250507 | 250665 | 158 | 0,042743409 | 0,188999349 | 0,184443343 | -0,898560728 | -0,763776618 | -0,671478431 | -0,769478431 | -2,029478431 |
| chrXIII | 250507 | 250666 | 159 | 0,04844253 | 0,307587175 | 0,136063169 | -1,098178991 | -0,933452143 | -0,83565512 | -0,94065512 | -2,20065512 |
| chrXIII | 250507 | 250667 | 160 | 0,059840772 | 0,111176087 | 0,349911537 | -0,385559308 | -0,327725412 | -0,280154651 | -0,392154651 | -1,652154651 |
| chrXIII | 250507 | 250668 | 161 | 0,054141651 | 0,188999349 | 0,222675942 | -0,763186995 | -0,648708946 | -0,567263119 | -0,686263119 | -1,946263119 |
| chrXIII | 250507 | 250669 | 162 | 0,059840772 | 0,144528914 | 0,292806499 | -0,545204323 | -0,463423675 | -0,353441696 | -0,479441696 | -1,739441696 |
| chrXIII | 250507 | 250670 | 163 | 0,037044288 | 0,26311674 | 0,123414715 | -1,158084807 | -0,984372086 | -0,815768783 | -0,948768783 | -2,208768783 |
| chrXIII | 250507 | 250671 | 164 | 0,054141651 | 0,251999131 | 0,176852135 | -0,927428169 | -0,788313944 | -0,572065587 | -0,712065587 | -1,972065587 |
| chrXIII | 250507 | 250673 | 166 | 0,082637257 | 0,385410436 | 0,176557343 | -0,928564773 | -0,789280057 | -0,496430155 | -0,650430155 | -1,910430155 |
| chrXIII | 250507 | 250674 | 167 | 0,065539893 | 0,125999566 | 0,342174368 | -0,406536101 | -0,345555686 | -0,061753129 | -0,222753129 | -1,482753129 |
| chrXIII | 250507 | 250675 | 168 | 0,074088575 | 0,255705001 | 0,224651359 | -0,756578012 | -0,64309131 | -0,346771304 | -0,514771304 | -1,774771304 |
| chrXIII | 250507 | 250676 | 169 | 0,031345166 | 0,144528914 | 0,178225048 | -0,922150465 | -0,783827895 | -0,497912245 | -0,672912245 | -1,932912245 |
| chrXIII | 250507 | 250677 | 170 | 0,031345166 | 0,218646305 | 0,125384943 | -1,148481403 | -0,976209192 | -0,706264149 | -0,888264149 | -2,148264149 |
| chrXIII | 250507 | 250678 | 171 | 44,76944633 | 62,04366848 | 0,419138103 | -0,20409891 | -0,173484073 | 0,113141412 | -0,075858588 | -1,335858588 |
| chrXIII | 250507 | 250718 | 211 | 0,042743409 | 0,678174133 | 0,059290288 | -1,560759205 | -1,326645324 | -0,992214476 | -1,461214476 | -2,721214476 |
| chrXIII | 250507 | 250718 | 211 | 0,042743409 | 0,678174133 | 0,059290288 | -1,560759205 | -1,326645324 | -0,992214476 | -1,461214476 | -2,721214476 |
| chrXIII | 250508 | 250678 | 170 | 0,116831984 | 0,096352609 | 0,548032024 | 0,120690792 | 0,102587173 | 0,405013329 | 0,223013329 | -1,036986671 |
| chrXIII | 250509 | 250678 | 169 | 0,037044288 | 0,122293696 | 0,232488743 | -0,730675332 | -0,621074032 | -0,33337862 | -0,50837862 | -1,76837862 |
| chrXIII | 250510 | 250678 | 168 | 0,054141651 | 0,122293696 | 0,30686397 | -0,50475925 | -0,429045363 | -0,119810362 | -0,287810362 | -1,547810362 |
| chrXIII | 250511 | 250678 | 167 | 0,14817715 | 0,466939567 | 0,240892738 | -0,703433756 | -0,597918692 | -0,316106909 | -0,477106909 | -1,737106909 |
| chrXIII | 250514 | 250678 | 164 | 0,085486818 | 0,181587609 | 0,320086122 | -0,467457987 | -0,397339289 | -0,183573622 | -0,323573622 | -1,583573622 |
| chrXIII | 250515 | 250678 | 163 | 0,039893848 | 0,085235 | 0,318822147 | -0,470995015 | -0,400345763 | -0,22930085 | -0,36230085 | -1,62230085 |
| chrXIII | 250566 | 250718 | 152 | 6,109457891 | 3,668810883 | 0,624799546 | 0,318110779 | 0,270394162 | 0,167575663 | 0,111575663 | -1,148424337 |
| chrXIII | 253372 | 253532 | 160 | 0,09973462 | 0,044470435 | 0,691616672 | 0,500438064 | 0,425372355 | 0,476347675 | 0,364347675 | -0,895652325 |
| chrXIII | 256009 | 256161 | 152 | 0,071239015 | 0,077823261 | 0,477914444 | -0,055388588 | -0,0470803 | -0,143542201 | -0,199542201 | -1,459542201 |
| chrXIII | 260961 | 261102 | 141 | 0,43883233 | 0,277940218 | 0,612233729 | 0,285145669 | 0,242373819 | 0,167502451 | 0,188502451 | -1,071497549 |
| chrXIII | 261845 | 261977 | 132 | 0,205168362 | 0,474351306 | 0,301931455 | -0,518853495 | -0,441025471 | -0,290420039 | -0,206420039 | -1,466420039 |
| chrXIII | 262094 | 262251 | 157 | 0,079787696 | 0,062999783 | 0,558786364 | 0,147892925 | 0,125708986 | 0,197831156 | 0,106831156 | -1,153168844 |
| chrXIII | 266226 | 266377 | 151 | 0,094035499 | 0,08894087 | 0,51392155 | 0,034903236 | 0,029667751 | -0,120313389 | -0,169313389 | -1,429313389 |
| chrXIII | 271419 | 271566 | 147 | 0,236513528 | 0,122293696 | 0,659166015 | 0,4101881 | 0,348659885 | 0,172854195 | 0,151854195 | -1,108145805 |
| chrXIII | 273047 | 273197 | 150 | 0,136778908 | 0,133411305 | 0,506231912 | 0,015621722 | 0,013278464 | -0,126081353 | -0,168081353 | -1,428081353 |
| chrXIII | 276628 | 276785 | 157 | 0,131079787 | 0,185293479 | 0,414320049 | -0,216446021 | -0,183979117 | -0,100674262 | -0,191674262 | -1,451674262 |
| chrXIII | 276628 | 276785 | 157 | 0,131079787 | 0,185293479 | 0,414320049 | -0,216446021 | -0,183979117 | -0,100674262 | -0,191674262 | -1,451674262 |
| chrXIII | 277879 | 278014 | 135 | 2,348037921 | 1,686170658 | 0,582031859 | 0,207094214 | 0,176030082 | 0,24984962 | 0,31284962 | -0,94715038 |
| chrXIII | 279195 | 279369 | 174 | 0,726637949 | 0,844938264 | 0,462362527 | -0,094483544 | -0,080311013 | 0,246884606 | 0,036884606 | -1,223115394 |
| chrXIII | 283987 | 284078 | 91 | 0,059840772 | 0,159352392 | 0,273004738 | -0,603750588 | -0,513188 | -0,126905132 | 0,244094868 | -1,015905132 |
| chrXIII | 283987 | 284079 | 92 | 3,690180956 | 10,82855091 | 0,254166892 | -0,66143438 | -0,562219223 | -0,132006025 | 0,231993975 | -1,028006025 |
| chrXIII | 289854 | 289961 | 107 | 0,581310359 | 1,897405224 | 0,234520799 | -0,724039316 | -0,615433419 | 0,105027903 | 0,364027903 | -0,895972097 |
| chrXIII | 289854 | 289977 | 123 | 0,507221784 | 2,008581312 | 0,201614262 | -0,835869143 | -0,710488771 | -0,178630713 | -0,031630713 | -1,291630713 |
| chrXIII | 303791 | 303918 | 127 | 0,795027403 | 1,775111528 | 0,309332462 | -0,497743386 | -0,423081878 | -0,015799892 | 0,103200108 | -1,156799892 |
| chrXIII | 304476 | 304622 | 146 | 0,407487163 | 0,329822392 | 0,552667682 | 0,132404146 | 0,112543524 | -0,067575043 | -0,081575043 | -1,341575043 |
| chrXIII | 310201 | 310351 | 150 | 0,074088575 | 0,066705652 | 0,52621884 | 0,065768269 | 0,055903029 | -0,086518271 | -0,128518271 | -1,388518271 |
| chrXIII | 313026 | 313180 | 154 | 0,119681545 | 0,085235 | 0,584050178 | 0,21226584 | 0,180425964 | 0,092247352 | 0,022247352 | -1,237752648 |
| chrXIII | 324504 | 324669 | 165 | 0,054141651 | 0,096352609 | 0,359758911 | -0,359103289 | -0,305237796 | -0,049600636 | -0,196600636 | -1,456600636 |
| chrXIII | 325807 | 325958 | 151 | 0,205168362 | 0,137117174 | 0,599407045 | 0,251812611 | 0,214040719 | 0,055786457 | 0,006786457 | -1,253213543 |
| chrXIII | 333017 | 333169 | 152 | 1,182567642 | 0,61888022 | 0,656453993 | 0,40280455 | 0,342383867 | 0,246147724 | 0,190147724 | -1,069852276 |
| chrXIII | 335127 | 335272 | 145 | 0,076938136 | 0,055588044 | 0,58055047 | 0,203301913 | 0,172806626 | -0,015986666 | -0,022986666 | -1,282986666 |
| chrXIII | 335472 | 335613 | 141 | 0,156725832 | 0,163058261 | 0,490098899 | -0,024820929 | -0,021097789 | -0,107087722 | -0,086087722 | -1,346087722 |

TABLE 2

| Chrom | Start | End | Length | Section A | Section B | A/A+B | Z-score | Z * 0.85 | Phase correction | Length correction | ΔLK |
|---|---|---|---|---|---|---|---|---|---|---|---|
| chrXIII | 336952 | 337113 | 161 | 2,422126496 | 2,004875442 | 0,547125691 | 0,118402661 | 0,100642262 | 0,188959843 | 0,069959843 | **-1,190040157** |
| chrXIII | 340438 | 340585 | 147 | 0,105433742 | 0,118587827 | 0,47064103 | -0,073658577 | -0,06260979 | -0,229910328 | -0,250910328 | **-1,510910328** |
| chrXIII | 342150 | 342302 | 152 | 0,361894194 | 0,237175653 | 0,604093489 | 0,26395704 | 0,224363484 | 0,127941461 | 0,071941461 | **-1,188058539** |
| chrXIII | 348582 | 348731 | 149 | 0,527168708 | 0,389116306 | 0,575332675 | 0,189967434 | 0,161472318 | 0,017542935 | -0,017457065 | **-1,277457065** |
| chrXIII | 351853 | 352022 | 169 | 0,14532759 | 0,159352392 | 0,476984372 | -0,057723664 | -0,049065114 | 0,24778352 | 0,07278352 | **-1,18721648** |
| chrXIII | 351863 | 352007 | 144 | 1,604302609 | 1,612053267 | 0,498795118 | -0,003020196 | -0,002567166 | -0,163567584 | -0,163567584 | **-1,423567584** |
| chrXIII | 357010 | 357124 | 114 | 0,042743409 | 0,329822392 | 0,11472714 | -1,2017658 | -1,02150093 | -0,388113404 | -0,178113404 | **-1,438113404** |
| chrXIII | 357475 | 357580 | 105 | 0,034194727 | 0,43729261 | 0,072525229 | -1,457238859 | -1,23865303 | -0,650157197 | -0,377157197 | **-1,637157197** |
| chrXIII | 358631 | 358781 | 150 | 0,404637603 | 0,314998914 | 0,562280531 | 0,156753732 | 0,133240672 | -0,000298543 | -0,042298543 | **-1,302298543** |
| chrXIII | 358907 | 359055 | 148 | 1,114178188 | 0,896820438 | 0,554042242 | 0,135880798 | 0,115498679 | -0,058182852 | -0,086182852 | **-1,346182852** |
| chrXIII | 360027 | 360184 | 157 | 0,04844253 | 0,081529131 | 0,372716096 | -0,324668217 | -0,275967985 | -0,184050948 | -0,275050948 | **-1,535050948** |
| chrXIII | 360404 | 360549 | 145 | 0,122531105 | 0,059293913 | 0,673895739 | 0,450696199 | 0,383091769 | 0,202268778 | 0,195268778 | **-1,064731222** |
| chrXIII | 360675 | 360810 | 135 | 0,102584181 | 0,059293913 | 0,633712557 | 0,341702373 | 0,290447017 | 0,348176929 | 0,411176929 | **-0,848823071** |
| chrXIII | 362078 | 362238 | 160 | 0,091185939 | 0,081529131 | 0,527955893 | 0,070132481 | 0,059612609 | 0,102262385 | -0,009737615 | **-1,269737615** |
| chrXIII | 362592 | 362706 | 114 | 0,042743409 | 0,311293045 | 0,12073166 | -1,171337061 | -0,995636502 | -0,366220804 | -0,156220804 | **-1,416220804** |
| chrXIII | 374780 | 374911 | 131 | 0,076938136 | 0,100058479 | 0,434687048 | -0,164453569 | -0,139785534 | 0,047066863 | 0,138066863 | **-1,121933137** |
| chrXIII | 375262 | 375391 | 129 | 0,045592969 | 0,114881957 | 0,28411273 | -0,570666901 | -0,485066866 | -0,16465933 | -0,05965933 | **-1,31965933** |
| chrXIII | 375501 | 375650 | 149 | 0,102584181 | 0,051882174 | 0,664119905 | 0,423733486 | 0,360173463 | 0,218772432 | 0,183772432 | **-1,076227568** |
| chrXIII | 375725 | 375882 | 157 | 0,031345166 | 0,062999783 | 0,332240005 | -0,433736208 | -0,368675777 | -0,28827456 | -0,37927456 | **-1,63927456** |
| chrXIII | 375911 | 376061 | 150 | 9,201231124 | 3,78369284 | 0,70860878 | 0,54932499 | 0,466926241 | 0,349628586 | 0,307628586 | **-0,952371414** |
| chrXIII | 376373 | 376508 | 135 | 0,074088575 | 0,051882174 | 0,588141101 | 0,222765786 | 0,189350918 | 0,245031815 | 0,308031815 | **-0,951968185** |
| chrXIII | 376763 | 376917 | 154 | 0,031345166 | 0,037058696 | 0,458236793 | -0,104876779 | -0,089145262 | -0,18158548 | -0,25158548 | **-1,51158548** |
| chrXIII | 377411 | 377551 | 140 | 0,153876272 | 0,159352392 | 0,49125859 | -0,021913219 | -0,018626236 | -0,038495708 | -0,010495708 | **-1,270495708** |
| chrXIII | 377417 | 377551 | 134 | 0,153876272 | 0,114881957 | 0,572545341 | 0,182858153 | 0,15542943 | 0,208079393 | 0,278079393 | **-0,981920607** |
| chrXIII | 377829 | 377984 | 155 | 0,09973462 | 0,074117392 | 0,573675388 | 0,185739279 | 0,157878387 | 0,142738506 | 0,065738506 | **-1,194261494** |
| chrXIII | 377960 | 378068 | 108 | 0,065539893 | 0,140823044 | 0,317595273 | -0,474433869 | -0,403268789 | 0,30193043 | 0,55393043 | **-0,70606957** |
| chrXIII | 379716 | 379822 | 106 | 0,128230226 | 0,35576348 | 0,264941929 | -0,628183315 | -0,533955818 | 0,129468459 | 0,395468459 | **-0,864531541** |
| chrXIII | 379716 | 379838 | 122 | 0,088336378 | 0,426175002 | 0,171689843 | -0,94750858 | -0,805382293 | -0,250189495 | -0,096189495 | **-1,356189495** |
| chrXIII | 381135 | 381283 | 148 | 0,119681545 | 0,062999783 | 0,655138356 | 0,399230614 | 0,339346022 | 0,163368206 | 0,135368206 | **-1,124631794** |
| chrXIII | 383859 | 384012 | 153 | 0,48727486 | 0,43729261 | 0,527030071 | 0,067806264 | 0,057635325 | -0,040344256 | -0,103344256 | **-1,363344256** |
| chrXIII | 384143 | 384283 | 140 | 0,136778908 | 0,140823044 | 0,492715945 | -0,018259434 | -0,015520519 | -0,024686057 | 0,003313943 | **-1,256686057** |
| chrXIII | 384143 | 384283 | 140 | 0,136778908 | 0,140823044 | 0,492715945 | -0,018259434 | -0,015520519 | -0,024686057 | 0,003313943 | **-1,256686057** |
| chrXIII | 384959 | 385107 | 148 | 0,24221265 | 0,155646522 | 0,60878991 | 0,276166509 | 0,234741533 | 0,051523716 | 0,023523716 | **-1,236476284** |
| chrXIII | 386345 | 386477 | 132 | 0,128230226 | 0,114881957 | 0,527452901 | 0,068868618 | 0,058538325 | 0,210376561 | 0,294376561 | **-0,965623439** |
| chrXIII | 386345 | 386477 | 132 | 0,128230226 | 0,114881957 | 0,527452901 | 0,068868618 | 0,058538325 | 0,210376561 | 0,294376561 | **-0,965623439** |
| chrXIII | 387854 | 388024 | 170 | 0,475876617 | 0,648527176 | 0,423225731 | -0,193648063 | -0,164600853 | 0,128647487 | -0,053352513 | **-1,313352513** |
| chrXIII | 395268 | 395412 | 144 | 1,393435125 | 1,067290439 | 0,566270024 | 0,166885743 | 0,141852882 | -0,021700963 | -0,021700963 | **-1,281700963** |
| chrXIII | 400158 | 400292 | 134 | 0,045592969 | 0,114881957 | 0,28411273 | -0,570666901 | -0,485066866 | -0,442416398 | -0,372416398 | **-1,632416398** |
| chrXIII | 402458 | 402607 | 149 | 0,327699467 | 0,181587609 | 0,643447443 | 0,36768904 | 0,312535684 | 0,1727933 | 0,1377933 | **-1,1222067** |
| chrXIII | 406280 | 406447 | 167 | 0,188070999 | 0,292763697 | 0,391134418 | -0,276363569 | -0,234909033 | 0,052631556 | -0,108368444 | **-1,368368444** |
| chrXIII | 408429 | 408536 | 107 | 0,227964847 | 0,148234783 | 0,605967759 | 0,268824834 | 0,228501109 | 0,96074776 | 1,21974776 | **-0,04025224** |
| chrXIII | 411105 | 411252 | 147 | 1,439028095 | 0,692997611 | 0,674958135 | 0,453645875 | 0,385598993 | 0,214500101 | 0,193500101 | **-1,066499899** |
| chrXIII | 412475 | 412581 | 106 | 0,190920559 | 1,463818484 | 0,115378047 | -1,198413462 | -1,018651443 | -0,363510065 | -0,097510065 | **-1,357510065** |
| chrXIII | 412475 | 412589 | 114 | 0,162424953 | 1,915934572 | 0,078150556 | -1,41762214 | -1,204978819 | -0,584297997 | -0,374297997 | **-1,634297997** |
| chrXIII | 413837 | 414003 | 166 | 0,051292091 | 0,044470435 | 0,535617563 | 0,08939893 | 0,07598909 | 0,381213987 | 0,227213987 | **-1,032786013** |
| chrXIII | 422824 | 422985 | 161 | 0,131079787 | 0,103764348 | 0,558156527 | 0,146296985 | 0,124352437 | 0,233456619 | 0,114456619 | **-1,145543381** |
| chrXIII | 431294 | 431455 | 161 | 0,034194727 | 0,044470435 | 0,434687048 | -0,164453569 | -0,139785534 | -0,01838954 | -0,13738954 | **-1,39738954** |
| chrXIII | 431358 | 431515 | 157 | 0,04844253 | 0,092646739 | 0,343346664 | -0,403346515 | -0,342844538 | -0,256674928 | -0,347674928 | **-1,607674928** |
| chrXIII | 435010 | 435159 | 149 | 1,478921943 | 0,804173699 | 0,647770473 | 0,379308141 | 0,32241192 | 0,182225987 | 0,147225987 | **-1,112774013** |
| chrXIII | 436980 | 437142 | 162 | 0,153876272 | 0,125999566 | 0,549801916 | 0,125160904 | 0,106386768 | 0,224068124 | 0,098068124 | **-1,161931876** |
| chrXIII | 439283 | 439457 | 174 | 0,054141651 | 0,166764131 | 0,245089334 | -0,690024678 | -0,586520976 | -0,258658448 | -0,468658448 | **-1,728658448** |
| chrXIII | 443760 | 443869 | 109 | 0,054141651 | 0,188999349 | 0,222675942 | -0,763186995 | -0,648708946 | 0,07306066 | 0,31806066 | **-0,94193934** |
| chrXIII | 445137 | 445296 | 159 | 0,113982423 | 0,222352175 | 0,338895921 | -0,415478239 | -0,353156503 | -0,273215942 | -0,378215942 | **-1,638215942** |
| chrXIII | 448884 | 449018 | 134 | 4,225898345 | 4,161691537 | 0,503827489 | 0,009594238 | 0,008155102 | 0,06471085 | 0,13471085 | **-1,12528915** |
| chrXIII | 448987 | 449140 | 153 | 1,216762369 | 0,52623348 | 0,698086785 | 0,518905805 | 0,441069934 | 0,349055763 | 0,286055763 | **-0,973944237** |
| chrXIII | 452811 | 452953 | 142 | 0,723788388 | 0,581821524 | 0,554368025 | 0,136705034 | 0,116199279 | 0,021969308 | 0,035969308 | **-1,224030692** |
| chrXIII | 455777 | 455925 | 148 | 0,356195073 | 0,303881305 | 0,539627056 | 0,099494207 | 0,084570076 | -0,087167806 | -0,115167806 | **-1,375167806** |
| chrXIII | 456607 | 456760 | 153 | 0,094035499 | 0,111176087 | 0,458236793 | -0,104876779 | -0,089145262 | -0,17253899 | -0,23553899 | **-1,49553899** |
| chrXIII | 458537 | 458683 | 146 | 0,510071344 | 0,229763914 | 0,689439086 | 0,494261052 | 0,420121894 | 0,241089972 | 0,227089972 | **-1,032910028** |
| chrXIII | 460130 | 460281 | 151 | 0,492973981 | 0,518821741 | 0,487226789 | -0,032023164 | -0,027219689 | -0,173220402 | -0,222220402 | **-1,482220402** |
| chrXIII | 460228 | 460362 | 134 | 0,453080133 | 0,244587392 | 0,649421274 | 0,383758496 | 0,326194722 | 0,388423293 | 0,458423293 | **-0,801576707** |
| chrXIII | 460442 | 460567 | 125 | 0,031345166 | 0,051882174 | 0,376621026 | -0,314367347 | -0,267212245 | 0,207676006 | 0,340676006 | **-0,919323994** |
| chrXIII | 460442 | 460595 | 153 | 0,085486818 | 0,044470435 | 0,657807209 | 0,406485945 | 0,345513053 | 0,260649291 | 0,197649291 | **-1,062350709** |
| chrXIII | 460442 | 460598 | 156 | 0,028495606 | 0,059293913 | 0,324590066 | -0,454901459 | -0,386666241 | -0,354620617 | -0,438620617 | **-1,698620617** |
| chrXIII | 460442 | 460600 | 158 | 8,198185799 | 5,558804368 | 0,595928739 | 0,242822993 | 0,206399544 | 0,296429082 | 0,198429082 | **-1,061570918** |
| chrXIII | 461194 | 461345 | 151 | 0,108283302 | 0,077823261 | 0,581834946 | 0,206589951 | 0,175601459 | 0,046242148 | -0,002757852 | **-1,262757852** |
| chrXIII | 464378 | 464520 | 142 | 0,216566604 | 0,17788174 | 0,549036667 | 0,123227859 | 0,104743681 | 0,00187161 | 0,01587161 | **-1,24412839** |
| chrXIII | 467950 | 468100 | 150 | 0,119681545 | 0,144528914 | 0,452978074 | -0,118140734 | -0,100419624 | -0,21087914 | -0,25287914 | **-1,51287914** |
| chrXIII | 470902 | 471051 | 149 | 0,282106498 | 0,340940001 | 0,452785624 | -0,118626528 | -0,100832548 | -0,244207987 | -0,279207987 | **-1,539207987** |
| chrXIII | 473040 | 473146 | 106 | 0,740885752 | 3,031401315 | 0,196402272 | -0,854542368 | -0,726361013 | -0,061787622 | 0,204212378 | **-1,055787622** |
| chrXIII | 473040 | 473162 | 122 | 0,954602795 | 3,6910461 | 0,205483199 | -0,822194168 | -0,698865043 | -0,134734 | 0,019266 | **-1,240734** |
| chrXIII | 473911 | 474081 | 170 | 0,082637257 | 0,070411522 | 0,539940648 | 0,100284197 | 0,085241567 | 0,380076496 | 0,198076496 | **-1,061923504** |
| chrXIII | 484473 | 484603 | 130 | 0,059840772 | 0,915349786 | 0,061363158 | -1,543430674 | -1,311916073 | -1,035076681 | -0,937076681 | **-2,197076681** |
| chrXIII | 488956 | 489063 | 107 | 0,108283302 | 1,159937178 | 0,08538208 | -1,369752547 | -1,164289665 | -0,421093102 | -0,162093102 | **-1,422093102** |
| chrXIII | 488956 | 489073 | 117 | 0,131079787 | 1,397112831 | 0,085774388 | -1,367244197 | -1,162157567 | -0,548386548 | -0,359386548 | **-1,619386548** |
| chrXIII | 491261 | 491409 | 148 | 0,447381012 | 0,344645871 | 0,564855842 | 0,163292275 | 0,138798434 | -0,039601077 | -0,067601077 | **-1,327601077** |
| chrXIII | 492970 | 493101 | 131 | 1,627099093 | 1,986346094 | 0,450290238 | -0,124928095 | -0,106188881 | 0,058771503 | 0,149771503 | **-1,110228497** |
| chrXIII | 495547 | 495700 | 153 | 0,692443222 | 0,418763262 | 0,623145411 | 0,313752298 | 0,266689453 | 0,184144699 | 0,121144699 | **-1,138855301** |
| chrXIII | 496801 | 496973 | 172 | 0,715239706 | 0,982055438 | 0,421399725 | -0,198313936 | -0,168566845 | 0,144107123 | -0,051892877 | **-1,311892877** |
| chrXIII | 496801 | 496973 | 172 | 0,715239706 | 0,982055438 | 0,421399725 | -0,198313936 | -0,168566845 | 0,144107123 | -0,051892877 | **-1,311892877** |
| chrXIII | 499783 | 499912 | 129 | 1,042939174 | 1,467524353 | 0,415436895 | -0,213581034 | -0,181543879 | 0,132195684 | 0,237195684 | **-1,022804316** |
| chrXIII | 503855 | 504003 | 148 | 0,19377012 | 0,100058479 | 0,659466508 | 0,41100757 | 0,349356434 | 0,188809862 | 0,160809862 | **-1,099190138** |
| chrXIII | 505412 | 505581 | 169 | 1,134125112 | 1,971522616 | 0,365181505 | -0,344642686 | -0,292946283 | 0,007764389 | -0,167235611 | **-1,427235611** |
| chrXIII | 512141 | 512286 | 145 | 0,039893848 | 0,037058696 | 0,518421434 | 0,046192108 | 0,039263292 | -0,15228132 | -0,15928132 | **-1,41928132** |
| chrXIII | 514521 | 514669 | 148 | 0,273557816 | 0,159352392 | 0,631904286 | 0,336901138 | 0,286365968 | 0,126358879 | 0,098358879 | **-1,161641121** |
| chrXIII | 514553 | 514703 | 150 | 0,062690333 | 0,085235 | 0,423797138 | -0,192188854 | -0,163360526 | -0,28606141 | -0,32806141 | **-1,58806141** |
| chrXIII | 515323 | 515459 | 136 | 0,034194727 | 0,048176305 | 0,415130494 | -0,214366854 | -0,182211826 | -0,16289729 | -0,10689729 | **-1,36689729** |
| chrXIII | 520223 | 520378 | 155 | 0,353345512 | 0,259410871 | 0,576649256 | 0,193328781 | 0,164329464 | 0,16239202 | 0,08539202 | **-1,17460798** |
| chrXIII | 520223 | 520378 | 155 | 0,353345512 | 0,259410871 | 0,576649256 | 0,193328781 | 0,164329464 | 0,16239202 | 0,08539202 | **-1,17460798** |
| chrXIII | 520376 | 520508 | 132 | 0,082637257 | 0,040764565 | 0,66965994 | 0,438974353 | 0,3731282 | 0,521285501 | 0,605285501 | **-0,654714499** |
| chrXIII | 521730 | 521901 | 171 | 0,170973635 | 0,415057393 | 0,291748435 | -0,548284062 | -0,466041453 | -0,168508894 | -0,357508894 | **-1,617508894** |
| chrXIII | 522712 | 522817 | 105 | 0,031345166 | 0,203822827 | 0,133288404 | -1,11098035 | -0,944333298 | -0,351817696 | -0,078817696 | **-1,338817696** |
| chrXIII | 528716 | 528857 | 141 | 0,088336378 | 0,059293913 | 0,598362147 | 0,249109985 | 0,211743487 | 0,12005505 | 0,14105505 | **-1,11894495** |
| chrXIII | 533451 | 533610 | 159 | 0,407487163 | 0,270528479 | 0,600999649 | 0,255935423 | 0,21754511 | 0,300296435 | 0,195296435 | **-1,064703565** |
| chrXIII | 535081 | 535246 | 165 | 0,190920559 | 0,392822175 | 0,327062844 | -0,448038116 | -0,380832398 | -0,112159054 | -0,259159054 | **-1,519159054** |
| chrXIII | 536826 | 536976 | 150 | 1,496019306 | 1,093231526 | 0,577780757 | 0,196219347 | 0,166786445 | 0,041577329 | -0,000422671 | **-1,260422671** |
| chrXIII | 537368 | 537507 | 139 | 0,418885406 | 0,426175002 | 0,495686938 | -0,010811454 | -0,009189736 | 0,006568947 | 0,041568947 | **-1,218431053** |
| chrXIII | 539637 | 539760 | 123 | 0,059840772 | 0,207528696 | 0,223813035 | -0,759378669 | -0,645471868 | -0,131669314 | 0,015330686 | **-1,244669314** |
| chrXIII | 540611 | 540759 | 148 | 2,481967268 | 2,053051747 | 0,547289275 | 0,118815599 | 0,100993259 | -0,059502544 | -0,087502544 | **-1,347502544** |
| chrXIII | 541727 | 541880 | 153 | 0,188070999 | 0,17788174 | 0,51392155 | 0,034903236 | 0,029667751 | -0,052830119 | -0,115830119 | **-1,375830119** |
| chrXIII | 549153 | 549302 | 149 | 0,116831984 | 0,133411305 | 0,466873595 | -0,083131234 | -0,070661549 | -0,205657959 | -0,240657959 | **-1,500657959** |

TABLE 2

| Chrom | Start | End | Length | Section A | Section B | A/A+B | Z-score | Z * 0.85 | Phase correction | Length correction | ΔLK |
|---|---|---|---|---|---|---|---|---|---|---|---|
| chrXIII | 549525 | 549679 | 154 | 0,208017923 | 0,203822827 | 0,505093104 | 0,012766866 | 0,010851836 | -0,077554174 | -0,147554174 | -1,407554174 |
| chrXIII | 549713 | 549857 | 144 | 0,83207169 | 0,674468263 | 0,552306421 | 0,131490682 | 0,11176708 | -0,04361384 | -0,04361384 | -1,30361384 |
| chrXIII | 555791 | 555929 | 138 | 0,416035845 | 0,415057393 | 0,500588654 | 0,001475537 | 0,001254206 | 0,018899254 | 0,060899254 | -1,199100746 |
| chrXIII | 562267 | 562429 | 162 | 2,855259705 | 2,657108488 | 0,517973329 | 0,045067707 | 0,038307551 | 0,168592054 | 0,042592054 | -1,217407946 |
| chrXIII | 562901 | 563062 | 161 | 0,068389454 | 0,137117174 | 0,332784663 | -0,432236787 | -0,367401269 | -0,248779898 | -0,367779898 | -1,627779898 |
| chrXIII | 571585 | 571738 | 153 | 8,343513389 | 8,408618074 | 0,498056824 | -0,00487084 | -0,004140214 | -0,088203072 | -0,151203072 | -1,411203072 |
| chrXIII | 571590 | 571738 | 148 | 0,051292091 | 0,037058696 | 0,58055047 | 0,203301913 | 0,172806626 | 0,01298275 | -0,01501725 | -1,27501725 |
| chrXIII | 574736 | 574904 | 168 | 0,102584181 | 0,070411522 | 0,592986873 | 0,235235114 | 0,199949847 | 0,510278279 | 0,342278279 | -0,917721721 |
| chrXIII | 580311 | 580467 | 156 | 0,814974327 | 0,559586306 | 0,592898056 | 0,235006242 | 0,199755306 | 0,234213633 | 0,150213633 | -1,109786367 |
| chrXIII | 580644 | 580772 | 128 | 0,800726524 | 1,637994354 | 0,328338733 | -0,444505066 | -0,377829306 | -0,00637328 | 0,10562672 | -1,15437328 |
| chrXIII | 581583 | 581719 | 136 | 0,128230226 | 0,166764131 | 0,434687048 | -0,164453569 | -0,139785534 | -0,116981759 | -0,060981759 | -1,320981759 |
| chrXIII | 582011 | 582174 | 163 | 3,587596775 | 3,231518273 | 0,52610885 | 0,065491969 | 0,055668174 | 0,232853812 | 0,099853812 | -1,160146188 |
| chrXIII | 582013 | 582174 | 161 | 0,079787696 | 0,077823261 | 0,506231912 | 0,015621722 | 0,013278464 | 0,115144654 | -0,003855346 | -1,263855346 |
| chrXIII | 583110 | 583268 | 158 | 0,247911771 | 0,207528696 | 0,544334087 | 0,111358804 | 0,094654983 | 0,189323913 | 0,091323913 | -1,168676087 |
| chrXIII | 588471 | 588586 | 115 | 0,051292091 | 0,329822392 | 0,134584469 | -1,104978423 | -0,939231659 | -0,294804913 | -0,091804913 | -1,351804913 |
| chrXIII | 595003 | 595166 | 163 | 0,994496644 | 1,23405457 | 0,446252542 | -0,135135069 | -0,114864808 | 0,08155548 | -0,05144452 | -1,31144452 |
| chrXIII | 596303 | 596450 | 147 | 0,074088575 | 0,037058696 | 0,666580246 | 0,43048963 | 0,365916185 | 0,193878531 | 0,172878531 | -1,087121469 |
| chrXIII | 598087 | 598214 | 127 | 0,230814407 | 1,511994788 | 0,132438139 | -1,114939662 | -0,947698713 | -0,545991525 | -0,426991525 | -1,686991525 |
| chrXIII | 598887 | 599027 | 140 | 0,475876617 | 0,448410219 | 0,514858157 | 0,037252492 | 0,031664618 | 0,005968723 | 0,033968723 | -1,226031277 |
| chrXIII | 600432 | 600562 | 130 | 0,435982769 | 0,544762828 | 0,444542163 | -0,139462957 | -0,118543513 | 0,143068905 | 0,241068905 | -1,018931095 |
| chrXIII | 600662 | 600821 | 159 | 0,763682236 | 0,907938047 | 0,456851502 | -0,108368984 | -0,092113637 | -0,011411267 | -0,116411267 | -1,376411267 |
| chrXIII | 602872 | 603031 | 159 | 0,746584873 | 0,856055873 | 0,465846681 | -0,085714515 | -0,072857338 | -0,004766695 | -0,109766695 | -1,369766695 |
| chrXIII | 608199 | 608341 | 142 | 0,344796831 | 0,274234349 | 0,556994287 | 0,143352968 | 0,121850023 | 0,016869505 | 0,030869505 | -1,229130495 |
| chrXIII | 614091 | 614242 | 151 | 0,755133555 | 0,563292176 | 0,572753957 | 0,183389919 | 0,155881431 | 0,031790775 | -0,017209225 | -1,277209225 |
| chrXIII | 617621 | 617773 | 152 | 0,190920559 | 0,107470218 | 0,639833982 | 0,358015071 | 0,30431281 | 0,206944527 | 0,150944527 | -1,109055473 |
| chrXIII | 617962 | 618111 | 149 | 0,347646391 | 0,415057393 | 0,455807875 | -0,111000752 | -0,094350639 | -0,223148784 | -0,258148784 | -1,518148784 |
| chrXIII | 622293 | 622443 | 150 | 2,020338454 | 1,352642396 | 0,598977149 | 0,250700464 | 0,213095395 | 0,080176494 | 0,038176494 | -1,221823506 |
| chrXIII | 627780 | 627908 | 128 | 0,367593315 | 0,61888022 | 0,372633732 | -0,324885852 | -0,276152974 | 0,102015632 | 0,214015632 | -1,045984368 |
| chrXIII | 627788 | 627919 | 131 | 0,946054114 | 0,993173047 | 0,487851105 | -0,030457471 | -0,025888851 | 0,13564691 | 0,22664691 | -1,03335309 |
| chrXIII | 634568 | 634713 | 145 | 0,29065518 | 0,251999131 | 0,535617563 | 0,08939893 | 0,07598909 | -0,109256504 | -0,116256504 | -1,376256504 |
| chrXIII | 637620 | 637765 | 145 | 0,051292091 | 0,051882174 | 0,497140355 | -0,007168129 | -0,00609291 | -0,195292351 | -0,202292351 | -1,462292351 |
| chrXIII | 641481 | 641611 | 130 | 0,219416165 | 0,059293913 | 0,787255941 | 0,796936145 | 0,677395723 | 0,916768888 | 1,014768888 | -0,245231112 |
| chrXIII | 642381 | 642525 | 144 | 3,650287107 | 1,952993268 | 0,651455373 | 0,389252658 | 0,330864759 | 0,170663877 | 0,170663877 | -1,089336123 |
| chrXIII | 642389 | 642541 | 152 | 4,641934191 | 3,387164795 | 0,578138867 | 0,197134526 | 0,167564347 | 0,072769746 | 0,016769746 | -1,243230254 |
| chrXIII | 649168 | 649320 | 152 | 0,034194727 | 0,062999783 | 0,351817475 | -0,380418277 | -0,323355536 | -0,409134006 | -0,465134006 | -1,725134006 |
| chrXIII | 652798 | 652958 | 160 | 0,059840772 | 0,077823261 | 0,434687048 | -0,164453569 | -0,139785534 | -0,111894571 | -0,223894571 | -1,483894571 |
| chrXIII | 657660 | 657789 | 129 | 1,045788734 | 1,763993919 | 0,372195598 | -0,326043828 | -0,277137254 | -0,006476411 | 0,098523589 | -1,161476411 |
| chrXIII | 658250 | 658404 | 154 | 0,698142343 | 0,711526959 | 0,495252569 | -0,011900326 | -0,010115277 | -0,099230454 | -0,169230454 | -1,429230454 |
| chrXIII | 658827 | 658990 | 163 | 0,042743409 | 0,040764565 | 0,51184823 | 0,029703476 | 0,025247955 | 0,22744016 | 0,09444016 | -1,16555984 |
| chrXIII | 658970 | 659105 | 135 | 3,601844578 | 3,350106099 | 0,5181056 | 0,0453996 | 0,03858966 | 0,088965713 | 0,151965713 | -1,108034287 |
| chrXIII | 660778 | 660927 | 149 | 0,547115632 | 0,43729261 | 0,555787238 | 0,140281573 | 0,119239337 | -0,010258508 | -0,045258508 | -1,305258508 |
| chrXIII | 662900 | 663052 | 152 | 0,062690333 | 0,148234783 | 0,297216064 | -0,532424343 | -0,452560692 | -0,536514575 | -0,592514575 | -1,852514575 |
| chrXIII | 663803 | 663949 | 146 | 0,190920559 | 0,125999566 | 0,602424851 | 0,259628569 | 0,220684284 | 0,042098562 | 0,028098562 | -1,231901438 |
| chrXIII | 663837 | 663944 | 107 | 0,202318801 | 1,545347614 | 0,115765114 | -1,19642634 | -1,016962389 | -0,309336811 | -0,050336811 | -1,310336811 |
| chrXIII | 663837 | 663953 | 116 | 0,190920559 | 1,508288918 | 0,112358459 | -1,214080675 | -1,031968574 | -0,399453801 | -0,203453801 | -1,463453801 |
| chrXIII | 667634 | 667789 | 155 | 0,313451664 | 0,218646305 | 0,589086376 | 0,225195424 | 0,191416111 | 0,201437372 | 0,124437372 | -1,135562628 |
| chrXIII | 671983 | 672128 | 145 | 3,573348972 | 2,697873053 | 0,569801062 | 0,175867708 | 0,149487552 | -0,044176996 | -0,051176996 | -1,311176996 |
| chrXIII | 675888 | 676037 | 149 | 1,017293128 | 0,637409568 | 0,614789068 | 0,291823124 | 0,248049656 | 0,121958337 | 0,086958337 | -1,173041663 |
| chrXIII | 675888 | 676037 | 149 | 1,017293128 | 0,637409568 | 0,614789068 | 0,291823124 | 0,248049656 | 0,121958337 | 0,086958337 | -1,173041663 |
| chrXIII | 675950 | 676037 | 87 | 0,045592969 | 0,066705652 | 0,405997586 | -0,23785292 | -0,202174982 | 0,154676396 | 0,553676396 | -0,706323604 |
| chrXIII | 675950 | 676105 | 155 | 0,464478375 | 0,496586524 | 0,483295536 | -0,041884125 | -0,035601506 | -0,01028913 | -0,08728913 | -1,34728913 |
| chrXIII | 677204 | 677344 | 140 | 0,076938136 | 0,111176087 | 0,408996909 | -0,230126057 | -0,195607148 | -0,234891716 | -0,206891716 | -1,466891716 |
| chrXIII | 677920 | 678066 | 146 | 1,028691371 | 0,96723196 | 0,515396235 | 0,038602224 | 0,03281189 | -0,148725477 | -0,162725477 | -1,422725477 |
| chrXIII | 678554 | 678723 | 169 | 1,401983807 | 2,538520661 | 0,355787899 | -0,369740572 | -0,314279486 | -0,02859263 | -0,20359263 | -1,46359263 |
| chrXIII | 679422 | 679570 | 148 | 0,105433742 | 0,125999566 | 0,455568573 | -0,11160432 | -0,094863672 | -0,247292214 | -0,275292214 | -1,535292214 |
| chrXIII | 680831 | 680955 | 124 | 0,273557816 | 0,481763045 | 0,362174316 | -0,352652957 | -0,299755013 | 0,175697427 | 0,315697427 | -0,944302573 |
| chrXIII | 684362 | 684526 | 164 | 0,051292091 | 0,107470218 | 0,323074734 | -0,459117949 | -0,390250257 | -0,163087733 | -0,303087733 | -1,563087733 |
| chrXIII | 686491 | 686656 | 165 | 0,233663968 | 0,285351958 | 0,450205777 | -0,125141469 | -0,106370249 | 0,15857449 | 0,01157449 | -1,24842551 |
| chrXIII | 692898 | 693052 | 154 | 0,094035499 | 0,122293696 | 0,434687048 | -0,164453569 | -0,139785534 | -0,230026231 | -0,300026231 | -1,560026231 |
| chrXIII | 693627 | 693771 | 144 | 0,359044634 | 0,222352175 | 0,617555219 | 0,299066166 | 0,254206241 | 0,10533851 | 0,10533851 | -1,15466149 |
| chrXIII | 693831 | 693958 | 127 | 2,04883406 | 4,009750884 | 0,338170394 | -0,417461624 | -0,35484238 | 0,010323253 | 0,129323253 | -1,130676747 |
| chrXIII | 694160 | 694314 | 154 | 0,09688506 | 0,062999783 | 0,605967759 | 0,268824834 | 0,228501109 | 0,153346103 | 0,083346103 | -1,176653897 |
| chrXIII | 694285 | 694446 | 161 | 0,29065518 | 0,35205761 | 0,452231828 | -0,120024607 | -0,102020916 | -0,009551281 | -0,128551281 | -1,388551281 |
| chrXIII | 694710 | 694842 | 132 | 3,011985537 | 3,046224794 | 0,497174144 | -0,007083429 | -0,006020915 | 0,126384619 | 0,210384619 | -1,049615381 |
| chrXIII | 711631 | 711787 | 156 | 0,567062556 | 0,392822175 | 0,590761096 | 0,229503234 | 0,195077748 | 0,227542303 | 0,143542303 | -1,116457697 |
| chrXIII | 713556 | 713705 | 149 | 0,213717044 | 0,140823044 | 0,602800787 | 0,260603323 | 0,221512824 | 0,097966863 | 0,062966863 | -1,197033137 |
| chrXIII | 714522 | 714675 | 153 | 0,151026711 | 0,281646088 | 0,349055248 | -0,387872358 | -0,329691504 | -0,420643898 | -0,483643898 | -1,743643898 |
| chrXIII | 715186 | 715332 | 146 | 0,116831984 | 0,059293913 | 0,663343584 | 0,421605719 | 0,358364861 | 0,171890165 | 0,157890165 | -1,102109835 |
| chrXIII | 718847 | 718995 | 148 | 1,176868521 | 1,085819787 | 0,520119593 | 0,050453738 | 0,042885677 | -0,106105952 | -0,134105952 | -1,394105952 |
| chrXIII | 718847 | 718995 | 148 | 1,176868521 | 1,085819787 | 0,520119593 | 0,050453738 | 0,042885677 | -0,106105952 | -0,134105952 | -1,394105952 |
| chrXIII | 719890 | 720048 | 158 | 0,151026711 | 0,137117174 | 0,524136442 | 0,060538046 | 0,051457339 | 0,148115299 | 0,050115299 | -1,209884701 |
| chrXIII | 723055 | 723205 | 150 | 2,234055497 | 1,57870044 | 0,585942435 | 0,217119612 | 0,18455167 | 0,055294245 | 0,013294245 | -1,246705755 |
| chrXIII | 723972 | 724126 | 154 | 0,316301225 | 0,333528262 | 0,486744956 | -0,033231584 | -0,028246847 | -0,095963828 | -0,165963828 | -1,425963828 |
| chrXIII | 730943 | 731091 | 148 | 0,116831984 | 0,081529131 | 0,588986325 | 0,224938202 | 0,191197471 | 0,044984487 | 0,016984487 | -1,243015513 |
| chrXIII | 737903 | 738069 | 166 | 0,094035499 | 0,274234349 | 0,255344009 | -0,657766759 | -0,559101745 | -0,26034349 | -0,41434349 | -1,67434349 |
| chrXIII | 748521 | 748674 | 153 | 0,213717044 | 0,181587609 | 0,540638826 | 0,102043248 | 0,086736761 | -0,002026035 | -0,065026035 | -1,325026035 |
| chrXIII | 750522 | 750675 | 153 | 1,627099093 | 0,948702612 | 0,631686473 | 0,336323335 | 0,285874835 | 0,206215764 | 0,143215764 | -1,116784236 |
| chrXIII | 757851 | 758017 | 166 | 1,159771157 | 1,334113048 | 0,465046114 | -0,087728802 | -0,074569482 | 0,233108496 | 0,079108496 | -1,180891504 |
| chrXIII | 758839 | 758989 | 150 | 2,718480797 | 1,897405224 | 0,588940191 | 0,224819601 | 0,191096661 | 0,063650441 | 0,021650441 | -1,238349559 |
| chrXIII | 758839 | 758989 | 150 | 2,718480797 | 1,897405224 | 0,588940191 | 0,224819601 | 0,191096661 | 0,063650441 | 0,021650441 | -1,238349559 |
| chrXIII | 759376 | 759557 | 181 | 0,122531105 | 0,274234349 | 0,308825035 | -0,499183568 | -0,424306033 | -0,074404033 | -0,333404033 | -1,593404033 |
| chrXIII | 765908 | 766066 | 158 | 0,09973462 | 0,077823261 | 0,561702018 | 0,155285859 | 0,13199298 | 0,227090165 | 0,129090165 | -1,130909835 |
| chrXIII | 772941 | 773097 | 156 | 0,757983115 | 0,648527176 | 0,538910465 | 0,097689227 | 0,083035843 | 0,11475335 | 0,03075335 | -1,22924665 |
| chrXIII | 773665 | 773803 | 138 | 0,034194727 | 0,037058696 | 0,479902939 | -0,050397187 | -0,042837609 | -0,036895417 | 0,005104583 | -1,254895417 |
| chrXIII | 773665 | 773807 | 142 | 4,456712753 | 3,590987622 | 0,553787113 | 0,13523538 | 0,114950073 | 0,010392358 | 0,024392358 | -1,235607642 |
| chrXIII | 774153 | 774312 | 159 | 0,267858695 | 0,125999566 | 0,680089062 | 0,46794786 | 0,397755681 | 0,468215021 | 0,363215021 | -0,896784979 |
| chrXIII | 774153 | 774312 | 159 | 0,267858695 | 0,125999566 | 0,680089062 | 0,46794786 | 0,397755681 | 0,468215021 | 0,363215021 | -0,896784979 |
| chrXIII | 778663 | 778795 | 132 | 0,094035499 | 0,192705218 | 0,327946098 | -0,445591717 | -0,378752959 | -0,259094604 | -0,175094604 | -1,435094604 |
| chrXIII | 779035 | 779188 | 153 | 0,182371877 | 0,340940001 | 0,348495581 | -0,389385276 | -0,330977484 | -0,397841578 | -0,460841578 | -1,720841578 |
| chrXIII | 780072 | 780218 | 146 | 0,031345166 | 0,059293913 | 0,345823971 | -0,396619676 | -0,337126724 | -0,526675782 | -0,540675782 | -1,800675782 |
| chrXIII | 780801 | 780937 | 136 | 0,037044288 | 0,051882174 | 0,416572153 | -0,210670615 | -0,179070023 | -0,165242476 | -0,109242476 | -1,369242476 |
| chrXIII | 790225 | 790394 | 169 | 0,222265726 | 0,214940436 | 0,508377386 | 0,021000537 | 0,017850457 | 0,288918023 | 0,113918023 | -1,146081977 |
| chrXIII | 790368 | 790515 | 147 | 0,82922213 | 0,663350655 | 0,555565624 | 0,139735783 | 0,118775416 | -0,067560293 | -0,088560293 | -1,348560293 |
| chrXIII | 790640 | 790805 | 165 | 0,059840772 | 0,062999783 | 0,487141826 | -0,032236246 | -0,027400809 | 0,239757338 | 0,092757338 | -1,167242662 |
| chrXIII | 790640 | 790805 | 165 | 0,059840772 | 0,062999783 | 0,487141826 | -0,032236246 | -0,027400809 | 0,239757338 | 0,092757338 | -1,167242662 |
| chrXIII | 791775 | 791919 | 144 | 0,675345858 | 0,522527611 | 0,563787308 | 0,160578507 | 0,136491731 | -0,02525158 | -0,02525158 | -1,28525158 |
| chrXIII | 792265 | 792408 | 143 | 0,165274514 | 0,151940653 | 0,521017061 | 0,052706352 | 0,044800399 | -0,098420283 | -0,091420283 | -1,351420283 |
| chrXIII | 792607 | 792777 | 170 | 0,512920905 | 1,007996525 | 0,337244412 | -0,419995386 | -0,356996078 | -0,059527541 | -0,241527541 | -1,501527541 |
| chrXIII | 792607 | 792777 | 170 | 0,512920905 | 1,007996525 | 0,337244412 | -0,419995386 | -0,356996078 | -0,059527541 | -0,241527541 | -1,501527541 |
| chrXIII | 792935 | 793098 | 163 | 0,102584181 | 0,129705435 | 0,441621897 | -0,146858392 | -0,124829633 | 0,096910313 | -0,036089687 | -1,296089687 |
| chrXIII | 792935 | 793098 | 163 | 0,102584181 | 0,129705435 | 0,441621897 | -0,146858392 | -0,124829633 | 0,096910313 | -0,036089687 | -1,296089687 |

TABLE 2

| Chrom | Start | End | Length | Section A | Section B | A/A+B | Z-score | Z * 0.85 | Phase correction | Length correction | ΔLK |
|---|---|---|---|---|---|---|---|---|---|---|---|
| chrXIII | 793020 | 793160 | 140 | 1,798072728 | 1,05617283 | 0,629964273 | 0,331758724 | 0,281994916 | 0,246436629 | 0,274436629 | -0,985563371 |
| chrXIII | 793713 | 793841 | 128 | 0,091185939 | 0,144528914 | 0,386848507 | -0,287542436 | -0,24441107 | 0,1142702 | 0,2262702 | -1,0337298 |
| chrXIII | 795911 | 796064 | 153 | 7,517140819 | 5,766333064 | 0,565901728 | 0,165949687 | 0,141057234 | 0,076028475 | 0,013028475 | -1,246971525 |
| chrXIII | 797737 | 797897 | 160 | 3,901048439 | 3,735516535 | 0,510838113 | 0,027170462 | 0,023094893 | 0,056151023 | -0,055848977 | -1,315848977 |
| chrXIII | 797737 | 797940 | 203 | 0,208017923 | 0,233469783 | 0,471174893 | -0,072316812 | -0,06146929 | 0,272227782 | -0,140772218 | -1,400772218 |
| chrXIII | 797778 | 797885 | 107 | 0,279256937 | 0,35205761 | 0,442341996 | -0,145034044 | -0,123278937 | 0,595317323 | 0,854317323 | -0,405682677 |
| chrXIII | 797778 | 797897 | 119 | 0,307752543 | 0,52993935 | 0,367381546 | -0,338796427 | -0,287976963 | 0,306848795 | 0,481848795 | -0,778151205 |
| chrXIII | 797778 | 797940 | 162 | 3,02623334 | 2,434756313 | 0,554154747 | 0,136165427 | 0,115740613 | 0,241742425 | 0,115742425 | -1,144257575 |
| chrXIII | 799674 | 799836 | 162 | 0,09688506 | 0,051882174 | 0,65125268 | 0,388704654 | 0,330398956 | 0,458562455 | 0,332562455 | -0,927437545 |
| chrXIII | 800670 | 800819 | 149 | 0,208017923 | 0,270528479 | 0,434687048 | -0,164453569 | -0,139785534 | -0,265584081 | -0,300584081 | -1,560584081 |
| chrXIII | 801962 | 802127 | 165 | 0,091185939 | 0,200116957 | 0,313027917 | -0,487285766 | -0,414192901 | -0,121671835 | -0,268671835 | -1,528671835 |
| chrXIII | 804104 | 804264 | 160 | 0,236513528 | 0,285351958 | 0,453207838 | -0,117560789 | -0,099926671 | -0,039894864 | -0,151894864 | -1,411894864 |
| chrXIII | 806854 | 806962 | 108 | 0,034194727 | 0,163058261 | 0,173354671 | -0,940991251 | -0,799842563 | -0,105558627 | 0,146441373 | -1,113558627 |
| chrXIII | 806855 | 806962 | 107 | 1,615700851 | 10,48761091 | 0,133492459 | -1,11003274 | -0,943527829 | -0,224583207 | 0,034416793 | -1,225583207 |
| chrXIII | 808568 | 808720 | 152 | 0,074088575 | 0,077823261 | 0,487707719 | -0,030817056 | -0,026194498 | -0,121675613 | -0,177675613 | -1,437675613 |
| chrXIII | 811849 | 811978 | 129 | 3,046180264 | 4,098691754 | 0,426344972 | -0,185687355 | -0,157834252 | 0,110867304 | 0,215867304 | -1,044132696 |
| chrXIII | 816433 | 816597 | 164 | 0,162424953 | 0,26682261 | 0,378394585 | -0,309699913 | -0,263244926 | -0,03255335 | -0,17255335 | -1,43255335 |
| chrXIII | 819869 | 819975 | 106 | 0,250761331 | 1,111760874 | 0,184042014 | -0,900068066 | -0,765057856 | -0,079689103 | 0,186310897 | -1,073689103 |
| chrXIII | 819869 | 819988 | 119 | 0,279256937 | 1,337818918 | 0,17269254 | -0,943578505 | -0,80204173 | -0,220795421 | -0,045795421 | -1,305795421 |
| chrXIII | 820608 | 820710 | 102 | 0,028495606 | 0,08894087 | 0,242646977 | -0,69781331 | -0,593141313 | -0,104326901 | 0,189673099 | -1,070326901 |
| chrXIII | 820608 | 820711 | 103 | 0,028495606 | 0,055588044 | 0,338895921 | -0,415478239 | -0,353156503 | 0,157723916 | 0,444723916 | -0,815276084 |
| chrXIII | 820608 | 820715 | 107 | 0,088336378 | 1,248878048 | 0,066059995 | -1,50579429 | -1,279925146 | -0,548099066 | -0,289099066 | -1,549099066 |
| chrXIII | 820608 | 820717 | 109 | 0,034194727 | 0,08894087 | 0,277699771 | -0,589688448 | -0,501235181 | 0,195615598 | 0,440615598 | -0,819384402 |
| chrXIII | 820608 | 820733 | 125 | 0,028495606 | 0,114881957 | 0,198745224 | -0,846111675 | -0,719194924 | -0,259911372 | -0,126911372 | -1,386911372 |
| chrXIII | 820608 | 820737 | 129 | 0,034194727 | 0,074117392 | 0,315705458 | -0,47974192 | -0,407780632 | -0,115264164 | -0,010264164 | -1,270264164 |
| chrXIII | 820608 | 820738 | 130 | 0,042743409 | 0,129705435 | 0,247861382 | -0,681235065 | -0,579049806 | -0,357093893 | -0,259093893 | -1,519093893 |
| chrXIII | 820608 | 820740 | 132 | 0,059840772 | 0,044470435 | 0,573675388 | 0,185739279 | 0,157878387 | 0,261009318 | 0,345009318 | -0,914990682 |
| chrXIII | 820608 | 820741 | 133 | 0,039893848 | 0,092646739 | 0,300993446 | -0,521545394 | -0,443313585 | -0,368555228 | -0,291555228 | -1,551555228 |
| chrXIII | 820608 | 820742 | 134 | 0,054141651 | 0,085235 | 0,388455674 | -0,283346352 | -0,2408444 | -0,183831624 | -0,113831624 | -1,373831624 |
| chrXIII | 820608 | 820743 | 135 | 0,042743409 | 0,051882174 | 0,451710916 | -0,121339881 | -0,103138899 | -0,078866383 | -0,015866383 | -1,275866383 |
| chrXIII | 820608 | 820744 | 136 | 0,039893848 | 0,125999566 | 0,240478794 | -0,704763239 | -0,599048753 | -0,580869815 | -0,524869815 | -1,784869815 |
| chrXIII | 820608 | 820745 | 137 | 0,039893848 | 0,096352609 | 0,292806499 | -0,545204323 | -0,463423675 | -0,457597069 | -0,408597069 | -1,668597069 |
| chrXIII | 820608 | 820746 | 138 | 0,056991212 | 0,070411522 | 0,447331152 | -0,132407093 | -0,112546029 | -0,105740327 | -0,063740327 | -1,323740327 |
| chrXIII | 820608 | 820747 | 139 | 0,270708255 | 0,307587175 | 0,468114118 | -0,080011341 | -0,06800964 | -0,050480863 | -0,015480863 | -1,275480863 |
| chrXIII | 820608 | 820748 | 140 | 0,051292091 | 0,155646522 | 0,247861382 | -0,681235065 | -0,579049806 | -0,619729879 | -0,591729879 | -1,851729879 |
| chrXIII | 820608 | 820749 | 141 | 0,136778908 | 0,107470218 | 0,559997534 | 0,150962962 | 0,128318518 | 0,039303923 | 0,060303923 | -1,199696077 |
| chrXIII | 820608 | 820750 | 142 | 0,091185939 | 0,292763697 | 0,237494531 | -0,714385133 | -0,607227363 | -0,705217718 | -0,691217718 | -1,951217718 |
| chrXIII | 820608 | 820751 | 143 | 71,30455448 | 46,2418406 | 0,606607752 | 0,270488457 | 0,229915189 | 0,087777248 | 0,094777248 | -1,165222752 |
| chrXIII | 820609 | 820751 | 142 | 0,108283302 | 0,096352609 | 0,529151025 | 0,07313593 | 0,062165541 | -0,044726288 | -0,030726288 | -1,290726288 |
| chrXIII | 820610 | 820751 | 141 | 0,14817715 | 0,118587827 | 0,555459536 | 0,139467258 | 0,118547169 | 0,032026126 | 0,053026126 | -1,206973874 |
| chrXIII | 820611 | 820751 | 140 | 0,151026711 | 0,185293479 | 0,449056332 | -0,12804588 | -0,108838998 | -0,166790791 | -0,138790791 | -1,398790791 |
| chrXIII | 820612 | 820751 | 139 | 0,076938136 | 0,074117392 | 0,509336779 | 0,023405971 | 0,019895076 | 0,029395372 | 0,064395372 | -1,195604628 |
| chrXIII | 820613 | 820751 | 138 | 0,128230226 | 0,077823261 | 0,622315244 | 0,311567137 | 0,264832067 | 0,277430372 | 0,319430372 | -0,940569628 |
| chrXIII | 820614 | 820751 | 137 | 0,028495606 | 0,055588044 | 0,338895921 | -0,415478239 | -0,353156503 | -0,335491319 | -0,286491319 | -1,546491319 |
| chrXIII | 820615 | 820751 | 136 | 0,04844253 | 0,092646739 | 0,343346664 | -0,403346515 | -0,342844538 | -0,336404258 | -0,280404258 | -1,540404258 |
| chrXIII | 820616 | 820751 | 135 | 0,037044288 | 0,066705652 | 0,357053581 | -0,366345661 | -0,311393812 | -0,298701512 | -0,235701512 | -1,495701512 |
| chrXIII | 820617 | 820751 | 134 | 0,045592969 | 0,055588044 | 0,450607995 | -0,124125498 | -0,105506674 | -0,050760607 | 0,019239393 | -1,240760607 |
| chrXIII | 820619 | 820751 | 132 | 0,068389454 | 0,103764348 | 0,397257877 | -0,260451199 | -0,221383519 | -0,13005236 | -0,04605236 | -1,30605236 |
| chrXIII | 820620 | 820751 | 131 | 0,045592969 | 0,048176305 | 0,486225044 | -0,034535557 | -0,029355224 | 0,12791438 | 0,21891438 | -1,04108562 |
| chrXIII | 820621 | 820751 | 130 | 0,034194727 | 0,037058696 | 0,479902939 | -0,050397187 | -0,042837609 | 0,164655521 | 0,262655521 | -0,997344479 |
| chrXIII | 820624 | 820751 | 127 | 0,028495606 | 0,081529131 | 0,25899272 | -0,646453904 | -0,549485818 | -0,179185462 | -0,060185462 | -1,320185462 |
| chrXIII | 820626 | 820715 | 89 | 0,071239015 | 0,051882174 | 0,578608892 | 0,198335964 | 0,168585569 | 0,543327095 | 0,928327095 | -0,331672905 |
| chrXIII | 828572 | 828720 | 148 | 0,188070999 | 0,151940653 | 0,553131041 | 0,13357593 | 0,113539541 | -0,0287041 | -0,0567041 | -1,3167041 |
| chrXIII | 835990 | 836099 | 109 | 1,131275552 | 0,97093783 | 0,538135454 | 0,095737455 | 0,081376837 | 0,77067512 | 1,01567512 | -0,24432488 |
| chrXIII | 836333 | 836463 | 130 | 1,102779946 | 1,371171744 | 0,445756459 | -0,136390076 | -0,115931565 | 0,079995769 | 0,177995769 | -1,082004231 |
| chrXIII | 842098 | 842232 | 134 | 4,137561967 | 3,968986319 | 0,510397499 | 0,026065615 | 0,022155773 | 0,08784021 | 0,15784021 | -1,10215979 |
| chrXIII | 843431 | 843583 | 152 | 0,270708255 | 0,181587609 | 0,598520297 | 0,249518923 | 0,212091084 | 0,117014463 | 0,061014463 | -1,198985537 |
| chrXIII | 847888 | 848017 | 129 | 0,615505086 | 1,033937612 | 0,373159423 | -0,323497043 | -0,274972487 | 0,021946515 | 0,126946515 | -1,133053485 |
| chrXIII | 847897 | 848050 | 153 | 0,971700159 | 0,426175002 | 0,695126565 | 0,510434818 | 0,433869595 | 0,346706139 | 0,283706139 | -0,976293861 |
| chrXIII | 851484 | 851633 | 149 | 0,139628469 | 0,226058044 | 0,38182559 | -0,300689626 | -0,255586182 | -0,391947621 | -0,426947621 | -1,686947621 |
| chrXIII | 864286 | 864433 | 147 | 0,062690333 | 0,081529131 | 0,434687048 | -0,164453569 | -0,139785534 | -0,326539689 | -0,347539689 | -1,607539689 |
| chrXIII | 864644 | 864799 | 155 | 0,219416165 | 0,144528914 | 0,602882627 | 0,260815558 | 0,221693224 | 0,244585865 | 0,167585865 | -1,092414135 |
| chrXIII | 864644 | 864799 | 155 | 0,219416165 | 0,144528914 | 0,602882627 | 0,260815558 | 0,221693224 | 0,244585865 | 0,167585865 | -1,092414135 |
| chrXIII | 866005 | 866155 | 150 | 0,151026711 | 0,048176305 | 0,758154743 | 0,700379213 | 0,595322331 | 0,469516896 | 0,427516896 | -0,832483104 |
| chrXIII | 868732 | 868890 | 158 | 0,87766466 | 0,796761959 | 0,52415833 | 0,06059301 | 0,051504058 | 0,132647904 | 0,034647904 | -1,225352096 |
| chrXIII | 871472 | 871623 | 151 | 0,225115286 | 0,155646522 | 0,591223387 | 0,230693111 | 0,196089145 | 0,071473628 | 0,022473628 | -1,237526372 |
| chrXIII | 871472 | 871623 | 151 | 0,225115286 | 0,155646522 | 0,591223387 | 0,230693111 | 0,196089145 | 0,071473628 | 0,022473628 | -1,237526372 |
| chrXIII | 873640 | 873761 | 121 | 0,170973635 | 0,144528914 | 0,541908887 | 0,105243965 | 0,08945737 | 0,67489541 | 0,83589541 | -0,42410459 |
| chrXIII | 873640 | 873789 | 149 | 0,336248149 | 0,214940436 | 0,610041932 | 0,279428325 | 0,237514076 | 0,089806041 | 0,054806041 | -1,205193959 |
| chrXIII | 873640 | 873790 | 150 | 3,063277627 | 1,552759353 | 0,663616353 | 0,422353116 | 0,359000149 | 0,224797847 | 0,182797847 | -1,077202153 |
| chrXIII | 873648 | 873761 | 113 | 0,074088575 | 0,081529131 | 0,47609348 | -0,059960669 | -0,050966568 | 0,589519266 | 0,806519266 | -0,453480734 |
| chrXIII | 873648 | 873789 | 141 | 1,90350647 | 0,96352609 | 0,663929143 | 0,423210462 | 0,359728892 | 0,273042285 | 0,294042285 | -0,965957715 |
| chrXIII | 873648 | 873790 | 142 | 0,210867483 | 0,144528914 | 0,593330391 | 0,236120435 | 0,20070237 | 0,088354045 | 0,102354045 | -1,157645955 |
| chrXIII | 878188 | 878341 | 153 | 0,49012442 | 0,363175219 | 0,574387235 | 0,187554968 | 0,159421723 | 0,07476291 | 0,01176291 | -1,24823709 |
| chrXIII | 879147 | 879302 | 155 | 0,262159574 | 0,277940218 | 0,485390992 | -0,036627541 | -0,03113341 | -0,003664549 | -0,080664549 | -1,340664549 |
| chrXIII | 884345 | 884496 | 151 | 0,068389454 | 0,074117392 | 0,479902939 | -0,050397187 | -0,042837609 | -0,15321069 | -0,20221069 | -1,46221069 |
| chrXIII | 886126 | 886284 | 158 | 0,105433742 | 0,074117392 | 0,58720733 | 0,220367006 | 0,187311955 | 0,273550492 | 0,175550492 | -1,084449508 |
| chrXIII | 889335 | 889469 | 134 | 0,199469241 | 0,17788174 | 0,528604008 | 0,071761159 | 0,060996985 | 0,134776454 | 0,204776454 | -1,055223546 |
| chrXIII | 890068 | 890170 | 102 | 2,310993633 | 23,839859 | 0,088371636 | -1,350850672 | -1,148223071 | -0,638195994 | -0,344195994 | -1,604195994 |
| chrXIII | 891024 | 891149 | 125 | 1,994692408 | 3,33898849 | 0,373980455 | -0,321329227 | -0,273129843 | 0,163974082 | 0,296974082 | -0,963025918 |
| chrXIII | 891955 | 892105 | 150 | 0,125380666 | 0,077823261 | 0,617018911 | 0,297660652 | 0,253011555 | 0,113490538 | 0,071490538 | -1,188509462 |
| chrXIII | 895348 | 895499 | 151 | 0,447381012 | 0,259410871 | 0,632974179 | 0,339740923 | 0,288779784 | 0,184238983 | 0,135238983 | -1,124761017 |
| chrXIII | 902827 | 902965 | 138 | 0,133929347 | 0,085235 | 0,611090941 | 0,282163534 | 0,239839004 | 0,251544474 | 0,293544474 | -0,966455526 |
| chrXIII | 904201 | 904335 | 134 | 0,076938136 | 0,111176087 | 0,408996909 | -0,230126057 | -0,195607148 | -0,136281986 | -0,066281986 | -1,326281986 |
| chrXIII | 904428 | 904582 | 154 | 1,684090305 | 1,252583918 | 0,573468549 | 0,185211815 | 0,157430043 | 0,093357005 | 0,023357005 | -1,236642995 |
| chrXIII | 910695 | 910845 | 150 | 0,227964847 | 0,129705435 | 0,637360324 | 0,351411905 | 0,298700119 | 0,15789746 | 0,11589746 | -1,14410254 |
| chrXIII | 911000 | 911154 | 154 | 0,410336724 | 0,218646305 | 0,652381233 | 0,391757318 | 0,332993721 | 0,270710975 | 0,200710975 | -1,059289025 |
| chrXIII | 917204 | 917355 | 151 | 0,091185939 | 0,103764348 | 0,467739443 | -0,080953558 | -0,068810524 | -0,168306761 | -0,217306761 | -1,477306761 |
| chrXIII | 921093 | 921253 | 160 | 0,450230572 | 0,326116523 | 0,579934639 | 0,201726272 | 0,171467332 | 0,242794216 | 0,130794216 | -1,129205784 |
| chrXIII | 923830 | 923976 | 146 | 1,90350647 | 0,578115654 | 0,767041223 | 0,729137507 | 0,619766881 | 0,436375191 | 0,422375191 | -0,837624809 |
| chrXIII | 924304 | 924426 | 122 | 0,641151131 | 0,133411305 | 0,827759134 | 0,945347037 | 0,803544981 | 1,348412087 | 1,502412087 | 0,242412087 |
| chrXIII | 924311 | 924426 | 115 | 0,871965539 | 0,34835174 | 0,714540025 | 0,56669716 | 0,481692586 | 1,139325735 | 1,342325735 | 0,082325735 |
| chrXIII | 924321 | 924426 | 105 | 0,310602104 | 0,092646739 | 0,770249212 | 0,739667825 | 0,628717651 | 1,243129044 | 1,516129044 | 0,256129044 |
| chrXIV | 614 | 765 | 151 | 0,615505086 | 0,418763262 | 0,595111595 | 0,24071397 | 0,204606874 | 0,04785356 | -0,00114644 | -1,26114644 |
| chrXIV | 953 | 1105 | 152 | 0,062690333 | 0,040764565 | 0,605967759 | 0,268824834 | 0,228501109 | 0,148994944 | 0,092994944 | -1,167005056 |
| chrXIV | 1779 | 1930 | 151 | 0,455929693 | 0,300175436 | 0,602997752 | 0,26111413 | 0,22194701 | 0,07466169 | 0,02566169 | -1,23433831 |
| chrXIV | 1791 | 1925 | 134 | 0,681044979 | 0,418763262 | 0,619239749 | 0,303484704 | 0,257961999 | 0,28675712 | 0,35675712 | -0,90324288 |
| chrXIV | 1816 | 1912 | 96 | 0,051292091 | 0,111176087 | 0,315705458 | -0,47974192 | -0,407780632 | 0,014235878 | 0,350235878 | -0,909764122 |
| chrXIV | 2170 | 2300 | 130 | 0,333398588 | 0,433586741 | 0,434687048 | -0,164453569 | -0,139785534 | 0,003166341 | 0,101166341 | -1,158833659 |
| chrXIV | 2449 | 2601 | 152 | 0,239363089 | 0,100058479 | 0,705208837 | 0,539441393 | 0,458525184 | 0,382258698 | 0,326258698 | -0,933741302 |
| chrXIV | 2796 | 2940 | 144 | 1,379187322 | 0,548468698 | 0,715473771 | 0,569447547 | 0,484030415 | 0,315162396 | 0,315162396 | -0,944837604 |

TABLE 2

| Chrom | Start | End | Length | Section A | Section B | A/A+B | Z-score | Z * 0.85 | Phase correction | Length correction | ΔLK |
|---|---|---|---|---|---|---|---|---|---|---|---|
| chrXIV | 2796 | 2983 | 187 | 0,111132863 | 0,040764565 | 0,731630971 | 0,617753167 | 0,525090192 | 0,907509045 | 0,606509045 | -0,653490955 |
| chrXIV | 2848 | 2940 | 92 | 0,142478029 | 0,155646522 | 0,477914444 | -0,055388588 | -0,0470803 | 0,256539471 | 0,620539471 | -0,639460529 |
| chrXIV | 2848 | 2983 | 135 | 1,26805446 | 0,830114786 | 0,604362333 | 0,264654886 | 0,224956653 | 0,22780873 | 0,29080873 | -0,96919127 |
| chrXIV | 2936 | 3080 | 144 | 0,125380666 | 0,040764565 | 0,754644987 | 0,68917996 | 0,585802966 | 0,419727808 | 0,419727808 | -0,840272192 |
| chrXIV | 3001 | 3153 | 152 | 0,09973462 | 0,081529131 | 0,550218231 | 0,126212725 | 0,107280816 | 0,027135882 | -0,028864118 | -1,288864118 |
| chrXIV | 3253 | 3378 | 125 | 0,225115286 | 0,255705001 | 0,468190075 | -0,079820337 | -0,067847286 | 0,288699987 | 0,421699987 | -0,838300013 |
| chrXIV | 3283 | 3426 | 143 | 0,376141997 | 0,466939567 | 0,446151373 | -0,135390993 | -0,115082344 | -0,35052687 | -0,34352687 | -1,60352687 |
| chrXIV | 3584 | 3721 | 137 | 0,182371877 | 0,125999566 | 0,591403262 | 0,231156174 | 0,196482748 | 0,089827528 | 0,138827528 | -1,121172472 |
| chrXIV | 3700 | 3855 | 155 | 0,09688506 | 0,17417587 | 0,357429084 | -0,365339271 | -0,31053838 | -0,352110747 | -0,429110747 | -1,689110747 |
| chrXIV | 3794 | 3956 | 162 | 0,068389454 | 0,051882174 | 0,568624996 | 0,172874585 | 0,146943398 | 0,179118727 | 0,053118727 | -1,206881273 |
| chrXIV | 3794 | 3956 | 162 | 0,068389454 | 0,051882174 | 0,568624996 | 0,172874585 | 0,146943398 | 0,179118727 | 0,053118727 | -1,206881273 |
| chrXIV | 5019 | 5160 | 141 | 0,156725832 | 0,081529131 | 0,657807209 | 0,406485945 | 0,345513053 | 0,13555847 | 0,15655847 | -1,10344153 |
| chrXIV | 15620 | 15770 | 150 | 0,04844253 | 0,066705652 | 0,420697304 | -0,200109928 | -0,170093439 | -0,314250702 | -0,356250702 | -1,616250702 |
| chrXIV | 18901 | 19054 | 153 | 0,128230226 | 0,081529131 | 0,611320649 | 0,282762761 | 0,240348347 | 0,185485875 | 0,122485875 | -1,137514125 |
| chrXIV | 22418 | 22569 | 151 | 0,814974327 | 0,433586741 | 0,65273085 | 0,392703751 | 0,333798188 | 0,180539646 | 0,131539646 | -1,128460354 |
| chrXIV | 24850 | 25019 | 169 | 0,270708255 | 0,292763697 | 0,48042898 | -0,049076965 | -0,04171542 | 0,000251513 | -0,174748487 | -1,434748487 |
| chrXIV | 25836 | 25986 | 150 | 0,638301571 | 0,185293479 | 0,775018707 | 0,755477404 | 0,642155793 | 0,50357883 | 0,46157883 | -0,79842117 |
| chrXIV | 26530 | 26680 | 150 | 1,772426683 | 1,623170875 | 0,521977841 | 0,055118174 | 0,046850448 | -0,082838721 | -0,124838721 | -1,384838721 |
| chrXIV | 27268 | 27422 | 154 | 0,119681545 | 0,144528914 | 0,452978074 | -0,118140734 | -0,100419624 | -0,147611654 | -0,217611654 | -1,477611654 |
| chrXIV | 30249 | 30400 | 151 | 0,136778908 | 0,085235 | 0,61608261 | 0,295208275 | 0,250927034 | 0,103186702 | 0,054186702 | -1,205813298 |
| chrXIV | 32059 | 32210 | 151 | 0,339097709 | 0,296469566 | 0,533535508 | 0,084160298 | 0,071536253 | -0,078798737 | -0,127798737 | -1,387798737 |
| chrXIV | 33454 | 33599 | 145 | 7,950274028 | 5,429098933 | 0,594218731 | 0,238410741 | 0,20264913 | 0,040115665 | 0,033115665 | -1,226884335 |
| chrXIV | 33454 | 33606 | 152 | 2,29389627 | 2,041934138 | 0,52905581 | 0,072896625 | 0,061962132 | 0,002171909 | -0,053828091 | -1,313828091 |
| chrXIV | 33459 | 33599 | 140 | 0,045592969 | 0,037058696 | 0,551627959 | 0,129775456 | 0,110309137 | -0,038636694 | -0,010636694 | -1,270636694 |
| chrXIV | 33461 | 33599 | 138 | 2,131471316 | 2,093816312 | 0,50445591 | 0,011169543 | 0,009494111 | -0,106245062 | -0,064245062 | -1,324245062 |
| chrXIV | 33461 | 33603 | 142 | 0,056991212 | 0,040764565 | 0,582995843 | 0,209563572 | 0,178129036 | -0,065005072 | -0,051005072 | -1,311005072 |
| chrXIV | 33461 | 33606 | 145 | 26,10767407 | 14,02671635 | 0,650506306 | 0,386687748 | 0,328684586 | 0,176506802 | 0,169506802 | -1,090493198 |
| chrXIV | 33464 | 33606 | 142 | 0,042743409 | 0,048176305 | 0,470122564 | -0,074961771 | -0,063717505 | -0,314819366 | -0,300819366 | -1,560819366 |
| chrXIV | 33466 | 33606 | 140 | 0,142478029 | 0,114881957 | 0,553613758 | 0,134796867 | 0,114577337 | -0,033952191 | -0,005952191 | -1,265952191 |
| chrXIV | 41403 | 41548 | 145 | 1,336443914 | 1,026525873 | 0,565578079 | 0,165127229 | 0,140358145 | -0,007830872 | -0,014830872 | -1,274830872 |
| chrXIV | 45261 | 45368 | 107 | 0,521469587 | 3,542811317 | 0,128305498 | -1,13443769 | -0,964272037 | -0,442529939 | -0,183529939 | -1,443529939 |
| chrXIV | 45261 | 45374 | 113 | 0,424584527 | 4,187632624 | 0,092056491 | -1,328197161 | -1,128967587 | -0,706246433 | -0,489246433 | -1,749246433 |
| chrXIV | 45783 | 45937 | 154 | 0,911859387 | 0,585527393 | 0,608967168 | 0,276628128 | 0,235133909 | 0,167035336 | 0,097035336 | -1,162964664 |
| chrXIV | 49049 | 49199 | 150 | 0,567062556 | 0,392822175 | 0,590761096 | 0,229503234 | 0,195077748 | 0,077008882 | 0,035008882 | -1,224991118 |
| chrXIV | 53101 | 53263 | 162 | 0,205168362 | 0,359469349 | 0,36336284 | -0,349484402 | -0,297061742 | -0,251356206 | -0,377356206 | -1,637356206 |
| chrXIV | 55301 | 55450 | 149 | 0,247911771 | 0,140823044 | 0,637740077 | 0,352424614 | 0,299560922 | 0,206835369 | 0,171835369 | -1,088164631 |
| chrXIV | 58407 | 58536 | 129 | 0,455929693 | 0,581821524 | 0,439343925 | -0,152632808 | -0,129737887 | 0,049954033 | 0,154954033 | -1,105045967 |
| chrXIV | 60113 | 60210 | 97 | 0,216566604 | 1,719523484 | 0,11185771 | -1,216707773 | -1,034201607 | -0,597967024 | -0,268967024 | -1,528967024 |
| chrXIV | 61184 | 61331 | 147 | 0,179522317 | 0,114881957 | 0,609781626 | 0,278749922 | 0,236937434 | 0,161243621 | 0,140243621 | -1,119756379 |
| chrXIV | 63570 | 63733 | 163 | 0,433133209 | 0,452116089 | 0,489278229 | -0,02687873 | -0,02284692 | 0,056291692 | -0,076708308 | -1,336708308 |
| chrXIV | 65407 | 65535 | 128 | 2,151418241 | 3,498340882 | 0,380798224 | -0,303385036 | -0,257877281 | -0,065811438 | 0,046188562 | -1,213811438 |
| chrXIV | 73936 | 74081 | 145 | 0,162424953 | 0,240881523 | 0,402733313 | -0,246278533 | -0,209336753 | -0,36042696 | -0,36742696 | -1,62742696 |
| chrXIV | 77741 | 77895 | 154 | 0,173823196 | 0,237175653 | 0,422928668 | -0,194406838 | -0,165245812 | -0,225767322 | -0,295767322 | -1,555767322 |
| chrXIV | 79985 | 80135 | 150 | 0,14532759 | 0,125999566 | 0,535617563 | 0,08939893 | 0,07598909 | -0,057485803 | -0,099485803 | -1,359485803 |
| chrXIV | 81420 | 81583 | 163 | 0,304902982 | 0,466939567 | 0,395032617 | -0,266225903 | -0,226292018 | -0,144239866 | -0,277239866 | -1,537239866 |
| chrXIV | 91093 | 91242 | 149 | 1,382036883 | 0,726350437 | 0,65549478 | 0,400198336 | 0,340168586 | 0,247708432 | 0,212708432 | -1,047291568 |
| chrXIV | 93267 | 93403 | 136 | 0,119681545 | 0,107470218 | 0,526879225 | 0,067427283 | 0,057313191 | 0,034733336 | 0,090733336 | -1,169266664 |
| chrXIV | 97284 | 97435 | 151 | 0,04844253 | 0,051882174 | 0,482857442 | -0,042983253 | -0,036535765 | -0,182140008 | -0,231140008 | -1,491140008 |
| chrXIV | 97941 | 98103 | 162 | 0,34194727 | 0,214940436 | 0,614032715 | 0,289845327 | 0,246368528 | 0,284495878 | 0,158495878 | -1,101504122 |
| chrXIV | 99408 | 99557 | 149 | 0,108283302 | 0,059293913 | 0,646169599 | 0,37499955 | 0,318749618 | 0,239439385 | 0,204439385 | -1,055560615 |
| chrXIV | 99632 | 99789 | 157 | 0,062690333 | 0,044470435 | 0,585011979 | 0,214732295 | 0,182522451 | 0,096231744 | 0,005231744 | -1,254768256 |
| chrXIV | 99818 | 99968 | 150 | 8,930522869 | 4,376631972 | 0,671106858 | 0,442971589 | 0,376525851 | 0,237488158 | 0,195488158 | -1,064511842 |
| chrXIV | 99822 | 99968 | 146 | 0,071239015 | 0,051882174 | 0,578608892 | 0,198335964 | 0,168585569 | 0,029420429 | 0,015420429 | -1,244579571 |
| chrXIV | 100280 | 100415 | 135 | 0,04844253 | 0,048176305 | 0,50137771 | 0,003453413 | 0,002935401 | -0,004088959 | 0,058911041 | -1,201088959 |
| chrXIV | 100932 | 101089 | 157 | 0,153876272 | 0,08894087 | 0,633712557 | 0,341702373 | 0,290447017 | 0,203750247 | 0,112750247 | -1,147249753 |
| chrXIV | 102362 | 102510 | 148 | 0,034194727 | 0,037058696 | 0,479902939 | -0,050397187 | -0,042837609 | -0,128490541 | -0,156490541 | -1,416490541 |
| chrXIV | 103973 | 104137 | 164 | 0,29065518 | 0,307587175 | 0,485848548 | -0,035479871 | -0,030157891 | 0,044352063 | -0,095647937 | -1,355647937 |
| chrXIV | 107707 | 107834 | 127 | 2,216958134 | 2,749755227 | 0,446363213 | -0,134855118 | -0,11462685 | 0,121218978 | 0,240218978 | -1,019781022 |
| chrXIV | 110079 | 110230 | 151 | 1,020142689 | 0,982055438 | 0,509511359 | 0,023843701 | 0,020267146 | -0,120310439 | -0,169310439 | -1,429310439 |
| chrXIV | 111620 | 111775 | 155 | 0,165274514 | 0,188999349 | 0,46651625 | -0,084030099 | -0,071425584 | -0,127656367 | -0,204656367 | -1,464656367 |
| chrXIV | 112009 | 112129 | 120 | 0,265009134 | 0,874585221 | 0,2325469 | -0,730484965 | -0,620912221 | -0,235286399 | -0,067286399 | -1,327286399 |
| chrXIV | 116452 | 116578 | 126 | 0,054141651 | 0,192705218 | 0,219332946 | -0,774448026 | -0,658280822 | -0,295161036 | -0,169161036 | -1,429161036 |
| chrXIV | 119252 | 119378 | 126 | 0,29350474 | 0,455821958 | 0,391691289 | -0,274913653 | -0,233676605 | 0,116580392 | 0,242580392 | -1,017419608 |
| chrXIV | 121333 | 121464 | 131 | 0,065539893 | 0,044470435 | 0,595761274 | 0,242390685 | 0,206032083 | 0,331380734 | 0,422380734 | -0,837619266 |
| chrXIV | 124849 | 124975 | 126 | 0,105433742 | 0,151940653 | 0,409651247 | -0,228442189 | -0,194175861 | 0,140007787 | 0,266007787 | -0,993992213 |
| chrXIV | 129691 | 129843 | 152 | 0,042743409 | 0,066705652 | 0,390532439 | -0,277931585 | -0,236241848 | -0,293822261 | -0,349822261 | -1,609822261 |
| chrXIV | 129719 | 129870 | 151 | 0,350495952 | 0,192705218 | 0,645241526 | 0,372504922 | 0,316629184 | 0,179026794 | 0,130026794 | -1,129973206 |
| chrXIV | 131962 | 132135 | 173 | 0,113982423 | 0,237175653 | 0,324590066 | -0,454901459 | -0,386666241 | -0,168223871 | -0,371223871 | -1,631223871 |
| chrXIV | 132312 | 132463 | 151 | 0,709540585 | 0,837526525 | 0,458635945 | -0,103870788 | -0,08829017 | -0,224553729 | -0,273553729 | -1,533553729 |
| chrXIV | 133490 | 133654 | 164 | 0,151026711 | 0,196411088 | 0,434687048 | -0,164453569 | -0,139785534 | -0,065443021 | -0,205443021 | -1,465443021 |
| chrXIV | 134176 | 134325 | 149 | 0,276407377 | 0,203822827 | 0,575572662 | 0,190579981 | 0,161992984 | 0,083704125 | 0,048704125 | -1,211295875 |
| chrXIV | 135289 | 135440 | 151 | 0,222265726 | 0,148234783 | 0,599906668 | 0,253105531 | 0,215139702 | 0,068235154 | 0,019235154 | -1,240764846 |
| chrXIV | 135829 | 135979 | 150 | 1,154072036 | 0,715232829 | 0,61738032 | 0,29860774 | 0,253816579 | 0,108476634 | 0,066476634 | -1,193523366 |
| chrXIV | 139766 | 139921 | 155 | 0,190920559 | 0,044470435 | 0,811078435 | 0,881877363 | 0,749595759 | 0,689633657 | 0,612633657 | -0,647366343 |
| chrXIV | 140110 | 140269 | 159 | 0,210867483 | 0,159352392 | 0,569573644 | 0,175288799 | 0,148995479 | 0,085297053 | -0,019702947 | -1,279702947 |
| chrXIV | 141911 | 142060 | 149 | 0,612655525 | 0,292763697 | 0,676653986 | 0,4583625 | 0,389608125 | 0,30684807 | 0,27184807 | -0,98815193 |
| chrXIV | 150518 | 150667 | 149 | 1,248107536 | 0,711526959 | 0,636908331 | 0,350207021 | 0,297675968 | 0,21061933 | 0,17561933 | -1,08438067 |
| chrXIV | 153107 | 153271 | 164 | 0,247911771 | 0,377998697 | 0,396081842 | -0,263501999 | -0,223976699 | -0,147672375 | -0,287672375 | -1,547672375 |
| chrXIV | 153107 | 153271 | 164 | 0,247911771 | 0,377998697 | 0,396081842 | -0,263501999 | -0,223976699 | -0,147672375 | -0,287672375 | -1,547672375 |
| chrXIV | 155351 | 155502 | 151 | 0,059840772 | 0,037058696 | 0,617555219 | 0,299066166 | 0,254206241 | 0,103664999 | 0,054664999 | -1,205335001 |
| chrXIV | 163862 | 164000 | 138 | 0,299203861 | 0,233469783 | 0,561702018 | 0,155285859 | 0,13199298 | -0,001547147 | 0,040452853 | -1,219547147 |
| chrXIV | 163862 | 164000 | 138 | 0,299203861 | 0,233469783 | 0,561702018 | 0,155285859 | 0,13199298 | -0,001547147 | 0,040452853 | -1,219547147 |
| chrXIV | 165154 | 165280 | 126 | 2,131471316 | 6,985564156 | 0,233789955 | -0,726422322 | -0,617458974 | -0,312327677 | -0,186327677 | -1,446327677 |
| chrXIV | 167446 | 167585 | 139 | 1,715435471 | 1,619465006 | 0,514388805 | 0,036075209 | 0,030663927 | -0,114871234 | -0,079871234 | -1,339871234 |
| chrXIV | 174863 | 175007 | 144 | 0,322000346 | 0,285351958 | 0,530170618 | 0,075698758 | 0,064343944 | -0,094411165 | -0,094411165 | -1,354411165 |
| chrXIV | 177654 | 177807 | 153 | 0,49012442 | 0,541056958 | 0,475303793 | -0,061943802 | -0,052652232 | -0,105142246 | -0,168142246 | -1,428142246 |
| chrXIV | 178329 | 178461 | 132 | 0,669646737 | 1,182172396 | 0,361615627 | -0,354143619 | -0,301022076 | -0,234790425 | -0,150790425 | -1,410790425 |
| chrXIV | 183769 | 183934 | 165 | 0,316301225 | 0,370586958 | 0,460484301 | -0,099213694 | -0,08433164 | -0,061010197 | -0,208010197 | -1,468010197 |
| chrXIV | 187776 | 187960 | 184 | 0,071239015 | 0,17788174 | 0,285961781 | -0,565220842 | -0,480437716 | -0,15099612 | -0,43099612 | -1,69099612 |
| chrXIV | 188930 | 189078 | 148 | 0,051292091 | 0,037058696 | 0,58055047 | 0,203301913 | 0,172806626 | 0,078881341 | 0,050881341 | -1,209118659 |
| chrXIV | 188930 | 189079 | 149 | 0,039893848 | 0,059293913 | 0,402205349 | -0,247642923 | -0,210496484 | -0,321432157 | -0,356432157 | -1,616432157 |
| chrXIV | 188930 | 189080 | 150 | 0,091185939 | 0,037058696 | 0,711031218 | 0,556399818 | 0,472939845 | 0,321819881 | 0,279819881 | -0,980180119 |
| chrXIV | 188930 | 189081 | 151 | 0,051292091 | 0,048176305 | 0,515662191 | 0,039269381 | 0,033378974 | -0,105159259 | -0,154159259 | -1,414159259 |
| chrXIV | 188930 | 189083 | 153 | 28,49275627 | 21,14569182 | 0,574005783 | 0,186581933 | 0,158594643 | 0,109623367 | 0,046623367 | -1,213376633 |
| chrXIV | 188931 | 189083 | 152 | 0,031345166 | 0,062999783 | 0,332240005 | -0,433736208 | -0,368675777 | -0,422335508 | -0,478335508 | -1,738335508 |
| chrXIV | 188933 | 189083 | 150 | 0,108283302 | 0,107470218 | 0,50188429 | 0,004723232 | 0,004014747 | -0,148303976 | -0,190303976 | -1,450303976 |
| chrXIV | 188934 | 189083 | 149 | 0,074088575 | 0,074117392 | 0,499902783 | -0,000243688 | -0,000207135 | -0,105315724 | -0,140315724 | -1,400315724 |
| chrXIV | 188935 | 189083 | 148 | 0,085486818 | 0,122293696 | 0,411428464 | -0,223871969 | -0,190291174 | -0,283354733 | -0,311354733 | -1,571354733 |
| chrXIV | 190891 | 191040 | 149 | 0,09973462 | 0,092646739 | 0,518421434 | 0,046192108 | 0,039263292 | -0,056077167 | -0,091077167 | -1,351077167 |
| chrXIV | 192308 | 192455 | 147 | 0,287805619 | 0,277940218 | 0,508718933 | 0,021856865 | 0,018578335 | -0,029095422 | -0,050095422 | -1,310095422 |
| chrXIV | 192813 | 192954 | 141 | 0,344796831 | 0,17417587 | 0,664383368 | 0,424456031 | 0,360787627 | 0,139751378 | 0,160751378 | -1,099248622 |

TABLE 2

| Chrom | Start | End | Length | Section A | Section B | A/A+B | Z-score | Z * 0.85 | Phase correction | Length correction | ΔLK |
|---|---|---|---|---|---|---|---|---|---|---|---|
| chrXIV | 193504 | 193657 | 153 | 0,04844253 | 0,144528914 | 0,251034708 | -0,671237229 | -0,570551645 | -0,626134012 | -0,689134012 | -1,949134012 |
| chrXIV | 196760 | 196910 | 150 | 4,160358452 | 2,720108271 | 0,604662245 | 0,265433528 | 0,225618498 | 0,065971311 | 0,023971311 | -1,236028689 |
| chrXIV | 202136 | 202292 | 156 | 0,108283302 | 0,044470435 | 0,708874979 | 0,550101085 | 0,467585923 | 0,361169228 | 0,277169228 | -0,982830772 |
| chrXIV | 203542 | 203662 | 120 | 0,116831984 | 0,040764565 | 0,741335927 | 0,647469471 | 0,550349051 | 0,913212574 | 1,081212574 | -0,178787426 |
| chrXIV | 206857 | 207023 | 166 | 0,042743409 | 0,077823261 | 0,354520937 | -0,373143192 | -0,317171714 | -0,287452663 | -0,441452663 | -1,701452663 |
| chrXIV | 210139 | 210288 | 149 | 2,379383087 | 1,84922892 | 0,562686547 | 0,157784127 | 0,134116508 | 0,035805382 | 0,000805382 | -1,259194618 |
| chrXIV | 216142 | 216308 | 166 | 0,219416165 | 0,203822827 | 0,518421434 | 0,046192108 | 0,039263292 | 0,073692751 | -0,080307249 | -1,340307249 |
| chrXIV | 219506 | 219633 | 127 | 0,076938136 | 0,077823261 | 0,497140355 | -0,007168129 | -0,00609291 | 0,213472079 | 0,332472079 | -0,927527921 |
| chrXIV | 224826 | 224979 | 153 | 0,04844253 | 0,055588044 | 0,465656665 | -0,08619258 | -0,073263693 | -0,113847002 | -0,176847002 | -1,436847002 |
| chrXIV | 227320 | 227476 | 156 | 0,039893848 | 0,037058696 | 0,518421434 | 0,046192108 | 0,039263292 | -0,077656323 | -0,161656323 | -1,421656323 |
| chrXIV | 235082 | 235232 | 150 | 0,404637603 | 0,285351958 | 0,586440181 | 0,218397212 | 0,18563763 | 0,022894934 | -0,019105066 | -1,279105066 |
| chrXIV | 238200 | 238369 | 169 | 0,065539893 | 0,092646739 | 0,414320049 | -0,216446021 | -0,183979117 | -0,144285701 | -0,319285701 | -1,579285701 |
| chrXIV | 246737 | 246897 | 160 | 0,102584181 | 0,188999349 | 0,351817475 | -0,380418277 | -0,323355536 | -0,370123121 | -0,482123121 | -1,742123121 |
| chrXIV | 247839 | 247988 | 149 | 0,188070999 | 0,148234783 | 0,559226183 | 0,149007602 | 0,126656462 | 0,023252355 | -0,011747645 | -1,271747645 |
| chrXIV | 254524 | 254675 | 151 | 1,818019652 | 1,289642613 | 0,585011979 | 0,214732295 | 0,182522451 | 0,059930533 | 0,010930533 | -1,249069467 |
| chrXIV | 257454 | 257590 | 136 | 0,250761331 | 0,196411088 | 0,560771015 | 0,152924304 | 0,129985658 | 0,097921838 | 0,153921838 | -1,106078162 |
| chrXIV | 257743 | 257893 | 150 | 0,208017923 | 0,148234783 | 0,583905523 | 0,211894998 | 0,180110748 | 0,014233189 | -0,027766811 | -1,287766811 |
| chrXIV | 259132 | 259283 | 151 | 0,102584181 | 0,040764565 | 0,715626635 | 0,569898223 | 0,48441349 | 0,375331357 | 0,326331357 | -0,933668643 |
| chrXIV | 259132 | 259283 | 151 | 0,102584181 | 0,040764565 | 0,715626635 | 0,569898223 | 0,48441349 | 0,375331357 | 0,326331357 | -0,933668643 |
| chrXIV | 265572 | 265700 | 128 | 0,131079787 | 0,125999566 | 0,509880648 | 0,024769644 | 0,021054198 | 0,209083889 | 0,321083889 | -0,938916111 |
| chrXIV | 271513 | 271667 | 154 | 0,983098401 | 0,844938264 | 0,537789214 | 0,094865611 | 0,080635769 | 0,02724365 | -0,04275635 | -1,30275635 |
| chrXIV | 273060 | 273209 | 149 | 5,248890595 | 3,909692405 | 0,573111648 | 0,184301797 | 0,156656527 | 0,042149727 | 0,007149727 | -1,252850273 |
| chrXIV | 273060 | 273209 | 149 | 5,248890595 | 3,909692405 | 0,573111648 | 0,184301797 | 0,156656527 | 0,042149727 | 0,007149727 | -1,252850273 |
| chrXIV | 275916 | 276056 | 140 | 1,971895924 | 0,937585003 | 0,677748359 | 0,461411668 | 0,392199918 | 0,239678132 | 0,267678132 | -0,992321868 |
| chrXIV | 281515 | 281622 | 107 | 0,510071344 | 1,723229354 | 0,228393492 | -0,744147933 | -0,632525743 | -0,090421709 | 0,168578291 | -1,091421709 |
| chrXIV | 281515 | 281631 | 116 | 0,410336724 | 2,605226314 | 0,136073005 | -1,098133933 | -0,933413843 | -0,483908334 | -0,287908334 | -1,547908334 |
| chrXIV | 281832 | 281976 | 144 | 0,133929347 | 0,044470435 | 0,750725957 | 0,676776004 | 0,575259604 | 0,426794196 | 0,426794196 | -0,833205804 |
| chrXIV | 283804 | 283989 | 185 | 0,037044288 | 0,066705652 | 0,357053581 | -0,366345661 | -0,311393812 | 0,053634083 | -0,233365917 | -1,493365917 |
| chrXIV | 287747 | 287882 | 135 | 0,236513528 | 0,118587827 | 0,666045131 | 0,429018532 | 0,364665752 | 0,364786064 | 0,427786064 | -0,832213936 |
| chrXIV | 291077 | 291208 | 131 | 0,718089267 | 0,61517435 | 0,538595112 | 0,096895004 | 0,082360754 | 0,200195339 | 0,291195339 | -0,968804661 |
| chrXIV | 292461 | 292633 | 172 | 0,108283302 | 0,155646522 | 0,410273081 | -0,226842569 | -0,192816183 | -0,010114481 | -0,206114481 | -1,466114481 |
| chrXIV | 292477 | 292630 | 153 | 0,313451664 | 0,248293262 | 0,557996432 | 0,14589138 | 0,124007673 | 0,084982024 | 0,021982024 | -1,238017976 |
| chrXIV | 296854 | 297025 | 171 | 0,247911771 | 0,251999131 | 0,495911911 | -0,010247499 | -0,008710374 | 0,060048161 | -0,128951839 | -1,388951839 |
| chrXIV | 297410 | 297562 | 152 | 0,233663968 | 0,122293696 | 0,656437525 | 0,402759782 | 0,342345815 | 0,295759636 | 0,239759636 | -1,020240364 |
| chrXIV | 298390 | 298549 | 159 | 0,284956058 | 0,255705001 | 0,527051197 | 0,06785934 | 0,057680439 | -0,010091251 | -0,115091251 | -1,375091251 |
| chrXIV | 302946 | 303074 | 128 | 4,78441222 | 7,348739374 | 0,394325595 | -0,268062527 | -0,227853148 | -0,070272057 | 0,041727943 | -1,218272057 |
| chrXIV | 303880 | 304030 | 150 | 0,079787696 | 0,085235 | 0,483495289 | -0,041382985 | -0,035175537 | -0,199278913 | -0,241278913 | -1,501278913 |
| chrXIV | 307042 | 307199 | 157 | 0,404637603 | 0,392822175 | 0,507408165 | 0,018570584 | 0,015784996 | -0,062968104 | -0,153968104 | -1,413968104 |
| chrXIV | 308271 | 308446 | 175 | 0,108283302 | 0,181587609 | 0,37355698 | -0,322447164 | -0,274080089 | -0,019761126 | -0,236761126 | -1,496761126 |
| chrXIV | 308271 | 308446 | 175 | 0,108283302 | 0,181587609 | 0,37355698 | -0,322447164 | -0,274080089 | -0,019761126 | -0,236761126 | -1,496761126 |
| chrXIV | 309709 | 309860 | 151 | 2,00324109 | 1,801052615 | 0,52657372 | 0,066659774 | 0,056660808 | -0,056820068 | -0,105820068 | -1,365820068 |
| chrXIV | 310999 | 311146 | 147 | 0,282106498 | 0,255705001 | 0,524545307 | 0,06156483 | 0,052330105 | 0,004283084 | -0,016716916 | -1,276716916 |
| chrXIV | 314726 | 314877 | 151 | 0,162424953 | 0,111176087 | 0,593656197 | 0,236960278 | 0,201416237 | 0,080915047 | 0,031915047 | -1,228084953 |
| chrXIV | 317936 | 318050 | 114 | 0,037044288 | 0,604056741 | 0,057782293 | -1,573666432 | -1,337616467 | -0,915967979 | -0,705967979 | -1,965967979 |
| chrXIV | 324620 | 324765 | 145 | 0,515770466 | 0,426175002 | 0,547558732 | 0,11949584 | 0,101571464 | -0,067124924 | -0,074124924 | -1,334124924 |
| chrXIV | 324977 | 325117 | 140 | 1,473222822 | 0,841232394 | 0,636531142 | 0,349201932 | 0,296821642 | 0,148087413 | 0,176087413 | -1,083912587 |
| chrXIV | 336864 | 337018 | 154 | 0,119681545 | 0,037058696 | 0,763566167 | 0,717820996 | 0,610147846 | 0,55897416 | 0,48897416 | -0,77102584 |
| chrXIV | 339979 | 340132 | 153 | 0,176672756 | 0,289057827 | 0,379345404 | -0,307200444 | -0,261120377 | -0,324791967 | -0,387791967 | -1,647791967 |
| chrXIV | 340143 | 340297 | 154 | 0,062690333 | 0,055588044 | 0,530023616 | 0,075329227 | 0,064029843 | 0,017904282 | -0,052095718 | -1,312095718 |
| chrXIV | 343939 | 344095 | 156 | 0,230814407 | 0,259410871 | 0,470833345 | -0,073175214 | -0,062198932 | -0,173207307 | -0,257207307 | -1,517207307 |
| chrXIV | 348424 | 348579 | 155 | 0,450230572 | 0,303881305 | 0,597034188 | 0,245677844 | 0,208826167 | 0,14669575 | 0,06969575 | -1,19030425 |
| chrXIV | 349341 | 349497 | 156 | 1,444727216 | 1,537935875 | 0,484374927 | -0,03917627 | -0,033299829 | -0,152164237 | -0,236164237 | -1,496164237 |
| chrXIV | 357765 | 357910 | 145 | 13,52401453 | 8,049148725 | 0,626890659 | 0,323629327 | 0,275084928 | 0,102323921 | 0,095323921 | -1,164676079 |
| chrXIV | 357767 | 357910 | 143 | 0,054141651 | 0,037058696 | 0,593656197 | 0,236960278 | 0,201416237 | -0,029193499 | -0,022193499 | -1,282193499 |
| chrXIV | 358288 | 358439 | 151 | 0,056991212 | 0,040764565 | 0,582995843 | 0,209563572 | 0,178129036 | 0,039359196 | -0,009640804 | -1,269640804 |
| chrXIV | 360600 | 360759 | 159 | 0,037044288 | 0,044470435 | 0,454449042 | -0,114428548 | -0,097264266 | -0,162373813 | -0,267373813 | -1,527373813 |
| chrXIV | 360762 | 360912 | 150 | 0,24506221 | 0,289057827 | 0,45881486 | -0,103419898 | -0,087906914 | -0,247346224 | -0,289346224 | -1,549346224 |
| chrXIV | 361214 | 361352 | 138 | 2,128621756 | 1,993757833 | 0,516357533 | 0,041013751 | 0,034861688 | -0,108493272 | -0,066493272 | -1,326493272 |
| chrXIV | 364273 | 364404 | 131 | 0,34194727 | 0,459527828 | 0,426647404 | -0,184916144 | -0,157178722 | -0,061929249 | 0,029070751 | -1,230929249 |
| chrXIV | 365246 | 365404 | 158 | 0,265009134 | 0,237175653 | 0,52771239 | 0,06952062 | 0,059092527 | 0,003919977 | -0,094080023 | -1,354080023 |
| chrXIV | 366145 | 366271 | 126 | 1,966196803 | 4,939924148 | 0,284703499 | -0,568925058 | -0,4835863 | -0,213954433 | -0,087954433 | -1,347954433 |
| chrXIV | 368092 | 368240 | 148 | 0,119681545 | 0,074117392 | 0,617555219 | 0,299066166 | 0,254206241 | 0,155881285 | 0,127881285 | -1,132118715 |
| chrXIV | 369049 | 369214 | 165 | 0,094035499 | 0,17417587 | 0,350602211 | -0,38369513 | -0,32614086 | -0,308226837 | -0,455226837 | -1,715226837 |
| chrXIV | 376927 | 377089 | 162 | 1,547311397 | 1,356348266 | 0,532883181 | 0,082519467 | 0,070141547 | 0,11777006 | -0,00822994 | -1,26822994 |
| chrXIV | 384387 | 384546 | 159 | 0,909009826 | 0,678174133 | 0,572718632 | 0,183299871 | 0,15580489 | 0,097723837 | -0,007276163 | -1,267276163 |
| chrXIV | 388430 | 388576 | 146 | 0,056991212 | 0,066705652 | 0,460732874 | -0,09858756 | -0,083799426 | -0,215183971 | -0,229183971 | -1,489183971 |
| chrXIV | 388719 | 388880 | 161 | 0,159575393 | 0,140823044 | 0,531212461 | 0,078318026 | 0,066570322 | 0,046245796 | -0,072754204 | -1,332754204 |
| chrXIV | 388719 | 388880 | 161 | 0,159575393 | 0,140823044 | 0,531212461 | 0,078318026 | 0,066570322 | 0,046245796 | -0,072754204 | -1,332754204 |
| chrXIV | 396710 | 396865 | 155 | 0,265009134 | 0,196411088 | 0,574333594 | 0,187418125 | 0,159305406 | 0,076318949 | -0,000681051 | -1,260681051 |
| chrXIV | 403433 | 403582 | 149 | 0,63545201 | 0,726350437 | 0,466625692 | -0,083754801 | -0,071191581 | -0,191166722 | -0,226166722 | -1,486166722 |
| chrXIV | 403995 | 404145 | 150 | 0,088336378 | 0,051882174 | 0,629990659 | 0,331828607 | 0,282054316 | 0,129055182 | 0,087055182 | -1,172944818 |
| chrXIV | 410286 | 410440 | 154 | 0,58415992 | 0,403939784 | 0,591195319 | 0,230620858 | 0,19602773 | 0,149937439 | 0,079937439 | -1,180062561 |
| chrXIV | 410286 | 410440 | 154 | 0,58415992 | 0,403939784 | 0,591195319 | 0,230620858 | 0,19602773 | 0,149937439 | 0,079937439 | -1,180062561 |
| chrXIV | 410317 | 410470 | 153 | 0,279256937 | 0,196411088 | 0,587083686 | 0,220049468 | 0,187042048 | 0,112976937 | 0,049976937 | -1,210023063 |
| chrXIV | 410317 | 410470 | 153 | 0,279256937 | 0,196411088 | 0,587083686 | 0,220049468 | 0,187042048 | 0,112976937 | 0,049976937 | -1,210023063 |
| chrXIV | 410968 | 411126 | 158 | 0,569912117 | 0,396528045 | 0,589702435 | 0,226779597 | 0,192762657 | 0,143438401 | 0,045438401 | -1,214561599 |
| chrXIV | 411008 | 411145 | 137 | 2,131471316 | 1,778817398 | 0,545093079 | 0,113273355 | 0,096282351 | -0,001747297 | 0,047252703 | -1,212747297 |
| chrXIV | 416136 | 416292 | 156 | 5,704820288 | 4,302514581 | 0,570063894 | 0,176536837 | 0,150056312 | 0,031765787 | -0,052234213 | -1,312234213 |
| chrXIV | 418446 | 418610 | 164 | 0,039893848 | 0,08894087 | 0,309651379 | -0,496838761 | -0,422312947 | -0,355455222 | -0,495455222 | -1,755455222 |
| chrXIV | 419242 | 419381 | 139 | 1,068585219 | 1,048761091 | 0,504681362 | 0,011734703 | 0,009974498 | -0,142723137 | -0,107723137 | -1,367723137 |
| chrXIV | 420269 | 420420 | 151 | 0,056991212 | 0,066705652 | 0,460732874 | -0,09858756 | -0,083799426 | -0,21596795 | -0,26496795 | -1,52496795 |
| chrXIV | 421794 | 421931 | 137 | 0,897611584 | 0,741173916 | 0,547729757 | 0,119927619 | 0,101938476 | 0,009029625 | 0,058029625 | -1,201970375 |
| chrXIV | 421911 | 422067 | 156 | 0,222265726 | 0,233469783 | 0,487707719 | -0,030817056 | -0,026194498 | -0,12679972 | -0,21079972 | -1,47079972 |
| chrXIV | 423473 | 423612 | 139 | 0,051292091 | 0,044470435 | 0,535617563 | 0,08939893 | 0,07598909 | -0,078502101 | -0,043502101 | -1,303502101 |
| chrXIV | 423497 | 423650 | 153 | 0,575611238 | 0,507704132 | 0,531342261 | 0,07864439 | 0,066847732 | -0,005487487 | -0,068487487 | -1,328487487 |
| chrXIV | 426375 | 426492 | 117 | 0,116831984 | 0,61146848 | 0,160417286 | -0,992744317 | -0,843832669 | -0,415910803 | -0,226910803 | -1,486910803 |
| chrXIV | 430126 | 430279 | 153 | 0,039893848 | 0,125999566 | 0,240478794 | -0,704763239 | -0,599048753 | -0,663508783 | -0,726508783 | -1,986508783 |
| chrXIV | 430126 | 430286 | 160 | 8,560079993 | 6,852152851 | 0,555408167 | 0,139337237 | 0,118436651 | 0,074584108 | -0,037415892 | -1,297415892 |
| chrXIV | 439717 | 439834 | 117 | 0,262159574 | 0,893114568 | 0,226924125 | -0,749014859 | -0,63666263 | -0,232695242 | -0,043695242 | -1,303695242 |
| chrXIV | 459511 | 459626 | 115 | 0,091185939 | 0,885702829 | 0,093343215 | -1,32044517 | -1,122378395 | -0,661924657 | -0,458924657 | -1,718924657 |
| chrXIV | 460164 | 460311 | 147 | 0,524319147 | 0,552174567 | 0,487061968 | -0,032436523 | -0,027571045 | -0,078910805 | -0,099910805 | -1,359910805 |
| chrXIV | 464412 | 464538 | 126 | 2,211259013 | 4,643454582 | 0,322589556 | -0,460469706 | -0,39139925 | -0,143312664 | -0,017312664 | -1,277312664 |
| chrXIV | 466515 | 466659 | 144 | 1,065735658 | 0,648527176 | 0,621687431 | 0,309915603 | 0,263428262 | 0,117888154 | 0,117888154 | -1,142111846 |
| chrXIV | 467295 | 467445 | 150 | 0,236513528 | 0,222352175 | 0,515430826 | 0,038688995 | 0,032885645 | -0,129666583 | -0,171666583 | -1,431666583 |
| chrXIV | 473999 | 474106 | 107 | 0,296354301 | 2,449579791 | 0,107924768 | -1,237640106 | -1,05199409 | -0,505815325 | -0,246815325 | -1,506815325 |
| chrXIV | 475462 | 475606 | 144 | 0,378991558 | 0,200116957 | 0,654439622 | 0,397334573 | 0,337734387 | 0,200882007 | 0,200882007 | -1,059117993 |
| chrXIV | 477594 | 477739 | 145 | 0,763682236 | 0,392822175 | 0,660336639 | 0,413382053 | 0,351374745 | 0,168638388 | 0,161638388 | -1,098361612 |
| chrXIV | 478141 | 478275 | 134 | 1,008474447 | 1,304466092 | 0,436079825 | -0,160915886 | -0,136778503 | -0,107732191 | -0,037732191 | -1,297732191 |
| chrXIV | 483192 | 483305 | 113 | 0,065539893 | 0,43729261 | 0,130341402 | -1,124778745 | -0,956061933 | -0,533537317 | -0,316537317 | -1,576537317 |
| chrXIV | 483272 | 483422 | 150 | 0,065539893 | 0,070411522 | 0,4820832 | -0,044925865 | -0,038186985 | -0,194162295 | -0,236162295 | -1,496162295 |
| chrXIV | 483529 | 483666 | 137 | 0,037044288 | 0,100058479 | 0,270193583 | -0,612227627 | -0,520393483 | -0,623991947 | -0,574991947 | -1,834991947 |

TABLE 2

| Chrom | Start | End | Length | Section A | Section B | A/A+B | Z-score | Z * 0.85 | Phase correction | Length correction | ΔLK |
|---|---|---|---|---|---|---|---|---|---|---|---|
| chrXIV | 483529 | 483666 | 137 | 0,037044288 | 0,100058479 | 0,270193583 | -0,612227627 | -0,520393483 | -0,623991947 | -0,574991947 | -1,834991947 |
| chrXIV | 483752 | 483901 | 149 | 0,159575393 | 0,107470218 | 0,597558568 | 0,24703277 | 0,209977854 | 0,093960203 | 0,058960203 | -1,201039797 |
| chrXIV | 484694 | 484843 | 149 | 1,008744447 | 0,796761959 | 0,558704441 | 0,147685318 | 0,125532521 | 0,004009087 | -0,030990913 | -1,290990913 |
| chrXIV | 490422 | 490573 | 151 | 0,609805965 | 0,600350872 | 0,503906557 | 0,009792443 | 0,008323576 | -0,116680975 | -0,165680975 | -1,425680975 |
| chrXIV | 490584 | 490699 | 115 | 0,042743409 | 0,044470435 | 0,490098899 | -0,024820929 | -0,021097789 | 0,455546777 | 0,658546777 | -0,601453223 |
| chrXIV | 490584 | 490718 | 134 | 0,071239015 | 0,077823261 | 0,477914444 | -0,055388588 | -0,0470803 | -0,013006182 | 0,056993818 | -1,203006182 |
| chrXIV | 490584 | 490722 | 138 | 14,74362646 | 9,32026199 | 0,612686785 | 0,286328634 | 0,243379339 | 0,095219327 | 0,137219327 | -1,122780673 |
| chrXIV | 500789 | 500918 | 129 | 0,082637257 | 0,055588044 | 0,597844653 | 0,247772164 | 0,210606339 | 0,40210454 | 0,50710454 | -0,75289546 |
| chrXIV | 502692 | 502828 | 136 | 6,120856133 | 4,117221102 | 0,597852115 | 0,247791452 | 0,210622734 | 0,156326381 | 0,212326381 | -1,047673619 |
| chrXIV | 511107 | 511261 | 154 | 0,142478029 | 0,096352609 | 0,596565123 | 0,24446623 | 0,207796296 | 0,163045148 | 0,093045148 | -1,166954852 |
| chrXIV | 515776 | 515877 | 101 | 1,991842848 | 19,60405007 | 0,092232484 | -1,327132162 | -1,128062338 | -0,710420801 | -0,409420801 | -1,669420801 |
| chrXIV | 516941 | 517091 | 150 | 0,250761331 | 0,08894087 | 0,738179884 | 0,637744162 | 0,542082538 | 0,379774061 | 0,337774061 | -0,922225939 |
| chrXIV | 517457 | 517589 | 132 | 3,428021382 | 3,980103927 | 0,462738039 | -0,093538105 | -0,079507389 | -0,015756178 | 0,068243822 | -1,191756178 |
| chrXIV | 517457 | 517589 | 132 | 3,428021382 | 3,980103927 | 0,462738039 | -0,093538105 | -0,079507389 | -0,015756178 | 0,068243822 | -1,191756178 |
| chrXIV | 520229 | 520363 | 134 | 0,142478029 | 0,103764348 | 0,578608892 | 0,198335964 | 0,168585569 | 0,18908234 | 0,25908234 | -1,00091766 |
| chrXIV | 520229 | 520369 | 140 | 0,182371877 | 0,107470218 | 0,629211148 | 0,329764776 | 0,28030006 | 0,114967032 | 0,142967032 | -1,117032968 |
| chrXIV | 521272 | 521407 | 135 | 0,076938136 | 0,062999783 | 0,549801916 | 0,125160904 | 0,106386768 | 0,093589642 | 0,156589642 | -1,103410358 |
| chrXIV | 521719 | 521865 | 146 | 0,065539893 | 0,048176305 | 0,576346155 | 0,192554744 | 0,163671533 | 0,051689341 | 0,037689341 | -1,222310659 |
| chrXIV | 521719 | 521869 | 150 | 8,913425505 | 4,161691537 | 0,681709041 | 0,472483219 | 0,401610736 | 0,222315472 | 0,180315472 | -1,079684528 |
| chrXIV | 521898 | 522055 | 157 | 0,068389454 | 0,059293913 | 0,535617563 | 0,08939893 | 0,07598909 | -0,007889802 | -0,098889802 | -1,358889802 |
| chrXIV | 524015 | 524129 | 114 | 0,028495606 | 0,218646305 | 0,115300581 | -1,198811728 | -1,018989969 | -0,570338374 | -0,360338374 | -1,620338374 |
| chrXIV | 524719 | 524868 | 149 | 0,09688506 | 0,074117392 | 0,566571175 | 0,16765125 | 0,142503563 | 0,019901778 | -0,015098222 | -1,275098222 |
| chrXIV | 524979 | 525110 | 131 | 0,857717736 | 1,330407179 | 0,391987555 | -0,274142505 | -0,233021129 | -0,13452413 | -0,04352413 | -1,30352413 |
| chrXIV | 526013 | 526193 | 180 | 0,182371877 | 0,333528262 | 0,353502284 | -0,375882075 | -0,319499764 | -0,016319167 | -0,268319167 | -1,528319167 |
| chrXIV | 526651 | 526800 | 149 | 1,193965884 | 0,900526308 | 0,570050291 | 0,176502204 | 0,150026874 | 0,021648155 | -0,013351845 | -1,273351845 |
| chrXIV | 527097 | 527235 | 138 | 3,903897999 | 3,135165663 | 0,55460473 | 0,13730396 | 0,116708366 | -0,052138369 | -0,010138369 | -1,270138369 |
| chrXIV | 527709 | 527871 | 162 | 0,638301571 | 0,492880654 | 0,564278289 | 0,16182531 | 0,137551513 | 0,197370356 | 0,071370356 | -1,188629644 |
| chrXIV | 529282 | 529444 | 162 | 0,589859041 | 0,433586741 | 0,576346155 | 0,192554744 | 0,163671533 | 0,22846521 | 0,10246521 | -1,15753479 |
| chrXIV | 529934 | 530079 | 145 | 0,894762023 | 0,689291742 | 0,564855842 | 0,163292275 | 0,138798434 | -0,043222683 | -0,050222683 | -1,310222683 |
| chrXIV | 530913 | 531056 | 143 | 0,339097709 | 0,085235 | 0,799131676 | 0,83852369 | 0,712745137 | 0,486781727 | 0,493781727 | -0,766218273 |
| chrXIV | 532041 | 532181 | 140 | 2,499064632 | 1,686170658 | 0,597114489 | 0,245885302 | 0,209002507 | 0,018558882 | 0,046558882 | -1,213441118 |
| chrXIV | 532139 | 532286 | 147 | 0,094035499 | 0,077823261 | 0,547167331 | 0,118507771 | 0,100731606 | 0,036669403 | 0,015669403 | -1,244330597 |
| chrXIV | 536984 | 537144 | 160 | 0,054141651 | 0,059293913 | 0,477289917 | -0,056956516 | -0,048413039 | -0,088141096 | -0,200141096 | -1,460141096 |
| chrXIV | 538071 | 538217 | 146 | 0,079787696 | 0,051882174 | 0,605967759 | 0,268824834 | 0,228501109 | 0,11177236 | 0,09777236 | -1,16222764 |
| chrXIV | 538102 | 538265 | 163 | 0,262159574 | 0,363175219 | 0,419230749 | -0,203861798 | -0,173282528 | -0,088478864 | -0,221478864 | -1,481478864 |
| chrXIV | 538210 | 538368 | 158 | 0,296354301 | 0,340940001 | 0,465019536 | -0,087795679 | -0,074626327 | -0,124036228 | -0,222036228 | -1,482036228 |
| chrXIV | 540636 | 540777 | 141 | 0,210867483 | 0,122293696 | 0,632929334 | 0,339621835 | 0,28867856 | 0,061928147 | 0,082928147 | -1,177071853 |
| chrXIV | 541287 | 541432 | 145 | 0,091185939 | 0,044470435 | 0,672183225 | 0,445949741 | 0,37905728 | 0,223403711 | 0,216403711 | -1,043596289 |
| chrXIV | 541780 | 541943 | 163 | 1,875010864 | 2,078992834 | 0,474205643 | -0,064701982 | -0,054996684 | 0,043282864 | -0,089717136 | -1,349717136 |
| chrXIV | 541920 | 542078 | 158 | 0,213717044 | 0,122293696 | 0,6360423 | 0,347899847 | 0,29571487 | 0,246037814 | 0,148037814 | -1,111962186 |
| chrXIV | 542491 | 542640 | 149 | 0,381841118 | 0,226058044 | 0,628132331 | 0,32691082 | 0,277874197 | 0,145873261 | 0,110873261 | -1,149126739 |
| chrXIV | 544884 | 545027 | 143 | 0,116831984 | 0,096352609 | 0,548032024 | 0,120690792 | 0,102587173 | -0,107243496 | -0,100243496 | -1,360243496 |
| chrXIV | 544927 | 545097 | 170 | 0,208017923 | 0,163058261 | 0,560580095 | 0,152440128 | 0,129574109 | 0,189822255 | 0,007822255 | -1,252177745 |
| chrXIV | 546338 | 546446 | 108 | 0,250761331 | 0,570703915 | 0,305261035 | -0,509328378 | -0,432929121 | 0,119522807 | 0,371522807 | -0,888477193 |
| chrXIV | 546358 | 546520 | 162 | 0,344796831 | 0,285351958 | 0,547167331 | 0,118507771 | 0,100731606 | 0,163634961 | 0,037634961 | -1,222365039 |
| chrXIV | 551772 | 551902 | 130 | 0,085486818 | 0,144528914 | 0,371656395 | -0,327469522 | -0,278349094 | -0,115782883 | -0,017782883 | -1,277782883 |
| chrXIV | 552571 | 552722 | 151 | 0,094035499 | 0,085235 | 0,524545307 | 0,06156483 | 0,052330105 | -0,066558203 | -0,115558203 | -1,375558203 |
| chrXIV | 553534 | 553684 | 150 | 0,276407377 | 0,200116957 | 0,580048818 | 0,202018369 | 0,171715614 | -0,002337208 | -0,044337208 | -1,304337208 |
| chrXIV | 554534 | 554685 | 151 | 2,114373953 | 1,771405659 | 0,544131208 | 0,110847113 | 0,094220046 | -0,022815719 | -0,071815719 | -1,331815719 |
| chrXIV | 555938 | 556085 | 147 | 0,227964847 | 0,133411305 | 0,630824269 | 0,334037235 | 0,283931649 | 0,213904519 | 0,192904519 | -1,067095481 |
| chrXIV | 557972 | 558127 | 155 | 1,193965884 | 0,911643916 | 0,567040429 | 0,168844262 | 0,143517622 | 0,063729903 | -0,013270097 | -1,273270097 |
| chrXIV | 560057 | 560198 | 141 | 0,569912117 | 0,507704132 | 0,5288637 | 0,072413804 | 0,061551733 | -0,167557556 | -0,146557556 | -1,406557556 |
| chrXIV | 561823 | 561978 | 155 | 0,045592969 | 0,081529131 | 0,358654941 | -0,362056402 | -0,307747942 | -0,390683534 | -0,467683534 | -1,727683534 |
| chrXIV | 562224 | 562373 | 149 | 0,065539893 | 0,074117392 | 0,469290903 | -0,077052467 | -0,065494597 | -0,218251688 | -0,253251688 | -1,513251688 |
| chrXIV | 563011 | 563177 | 166 | 0,034194727 | 0,092646739 | 0,269586342 | -0,614064532 | -0,521954852 | -0,476085624 | -0,630085624 | -1,890085624 |
| chrXIV | 564309 | 564452 | 143 | 0,210867483 | 0,107470218 | 0,662401854 | 0,419027147 | 0,356173075 | 0,139623702 | 0,146623702 | -1,113376298 |
| chrXIV | 565788 | 565917 | 129 | 0,176672756 | 0,385410436 | 0,314317807 | -0,483648127 | -0,411100908 | -0,217879427 | -0,112879427 | -1,372879427 |
| chrXIV | 567163 | 567273 | 110 | 0,122531105 | 0,737468046 | 0,14247817 | -1,069251542 | -0,90886381 | -0,450287582 | -0,212287582 | -1,472287582 |
| chrXIV | 573304 | 573437 | 133 | 0,507221784 | 0,648527176 | 0,438868475 | -0,15383866 | -0,130762861 | -0,100556835 | -0,023556835 | -1,283556835 |
| chrXIV | 575453 | 575613 | 160 | 0,045592969 | 0,055588044 | 0,450607955 | -0,124125498 | -0,105506674 | -0,137219608 | -0,249219608 | -1,509219608 |
| chrXIV | 575453 | 575613 | 160 | 0,045592969 | 0,055588044 | 0,450607955 | -0,124125498 | -0,105506674 | -0,137219608 | -0,249219608 | -1,509219608 |
| chrXIV | 576714 | 576844 | 130 | 0,068389454 | 0,037058696 | 0,648560019 | 0,381435717 | 0,324220359 | 0,481057975 | 0,579057975 | -0,680942025 |
| chrXIV | 576714 | 576864 | 150 | 0,983098401 | 0,718938698 | 0,577601041 | 0,195760131 | 0,166396111 | -0,00342841 | -0,04542841 | -1,30542841 |
| chrXIV | 578365 | 578517 | 152 | 0,270708255 | 0,17788174 | 0,603464764 | 0,262325541 | 0,22297671 | 0,18262252 | 0,12662252 | -1,13337748 |
| chrXIV | 578706 | 578859 | 153 | 1,629948654 | 1,193290004 | 0,577332933 | 0,195075126 | 0,165813857 | 0,111729546 | 0,048729546 | -1,211270454 |
| chrXIV | 585214 | 585366 | 152 | 0,361894194 | 0,326116523 | 0,52600081 | 0,065220573 | 0,055437487 | 0,020630963 | -0,035369037 | -1,295369037 |
| chrXIV | 589016 | 589141 | 125 | 0,111132863 | 0,244587392 | 0,31241646 | -0,489012398 | -0,415660538 | -0,066717391 | 0,066282609 | -1,193717391 |
| chrXIV | 589016 | 589141 | 125 | 0,111132863 | 0,244587392 | 0,31241646 | -0,489012398 | -0,415660538 | -0,066717391 | 0,066282609 | -1,193717391 |
| chrXIV | 593245 | 593393 | 148 | 0,170973635 | 0,125999566 | 0,575720754 | 0,190958009 | 0,162314307 | 0,064164726 | 0,036164726 | -1,223835274 |
| chrXIV | 597466 | 597618 | 152 | 0,461628815 | 0,237175653 | 0,660597973 | 0,414095655 | 0,351981307 | 0,320194295 | 0,264194295 | -0,995805705 |
| chrXIV | 598791 | 598924 | 133 | 0,435982769 | 0,70411522 | 0,382408156 | -0,299162171 | -0,254287846 | -0,232326611 | -0,155326611 | -1,415326611 |
| chrXIV | 602237 | 602346 | 109 | 0,094035499 | 0,051882174 | 0,644442151 | 0,370358087 | 0,314804374 | 0,75001835 | 0,99501835 | -0,26498165 |
| chrXIV | 612323 | 612470 | 147 | 0,282106498 | 0,203822827 | 0,58055047 | 0,203301913 | 0,172806626 | 0,101687879 | 0,080687879 | -1,179312121 |
| chrXIV | 615779 | 615947 | 168 | 0,068389454 | 0,114881957 | 0,373159423 | -0,323497043 | -0,274972487 | -0,244435031 | -0,412435031 | -1,672435031 |
| chrXIV | 620267 | 620398 | 131 | 0,823523009 | 1,830699572 | 0,310269009 | -0,495087981 | -0,420824784 | -0,333545596 | -0,242545596 | -1,502545596 |
| chrXIV | 620267 | 620398 | 131 | 0,823523009 | 1,830699572 | 0,310269009 | -0,495087981 | -0,420824784 | -0,333545596 | -0,242545596 | -1,502545596 |
| chrXIV | 620785 | 620892 | 107 | 0,045592969 | 0,26311674 | 0,147688809 | -1,046397345 | -0,889437743 | -0,342314104 | -0,083314104 | -1,343314104 |
| chrXIV | 620785 | 620946 | 161 | 9,184133761 | 8,571676335 | 0,517246676 | 0,043244482 | 0,036757809 | 0,032259613 | -0,086740387 | -1,346740387 |
| chrXIV | 620785 | 620946 | 161 | 9,184133761 | 8,571676335 | 0,517246676 | 0,043244482 | 0,036757809 | 0,032259613 | -0,086740387 | -1,346740387 |
| chrXIV | 625000 | 625108 | 108 | 0,809275206 | 2,101228051 | 0,278053359 | -0,588634154 | -0,500339031 | 0,012499131 | 0,264499131 | -0,995500869 |
| chrXIV | 625453 | 625609 | 156 | 2,937896962 | 2,490344357 | 0,541224457 | 0,103518981 | 0,087991134 | -0,014908464 | -0,098908464 | -1,358908464 |
| chrXIV | 630331 | 630483 | 152 | 0,102584181 | 0,107470218 | 0,488369592 | -0,029157241 | -0,024783655 | -0,067371137 | -0,123371137 | -1,383371137 |
| chrXIV | 630745 | 630896 | 151 | 0,210867483 | 0,133411305 | 0,612490489 | 0,28581604 | 0,242943634 | 0,125014332 | 0,076014332 | -1,183985668 |
| chrXIV | 630745 | 630896 | 151 | 0,210867483 | 0,133411305 | 0,612490489 | 0,28581604 | 0,242943634 | 0,125014332 | 0,076014332 | -1,183985668 |
| chrXIV | 635632 | 635775 | 143 | 0,09688506 | 0,107470218 | 0,47410109 | -0,064964608 | -0,055219916 | -0,261614084 | -0,254614084 | -1,514614084 |
| chrXIV | 640348 | 640505 | 157 | 0,749434433 | 0,503998263 | 0,597905604 | 0,247929711 | 0,210740254 | 0,13419036 | 0,04319036 | -1,21680964 |
| chrXIV | 640382 | 640540 | 158 | 0,210867483 | 0,155646522 | 0,575332675 | 0,189967434 | 0,161472318 | 0,09939658 | 0,00139658 | -1,25860342 |
| chrXIV | 640382 | 640540 | 158 | 0,210867483 | 0,155646522 | 0,575332675 | 0,189967434 | 0,161472318 | 0,09939658 | 0,00139658 | -1,25860342 |
| chrXIV | 641911 | 642059 | 148 | 0,190920559 | 0,188999349 | 0,502528442 | 0,006337906 | 0,00538722 | -0,085270044 | -0,113270044 | -1,373270044 |
| chrXIV | 645323 | 645472 | 149 | 2,291046709 | 1,649111962 | 0,58146052 | 0,205631256 | 0,174786568 | 0,023568498 | -0,011431502 | -1,271431502 |
| chrXIV | 645323 | 645472 | 149 | 2,291046709 | 1,649111962 | 0,58146052 | 0,205631256 | 0,174786568 | 0,023568498 | -0,011431502 | -1,271431502 |
| chrXIV | 646222 | 646370 | 148 | 0,29065518 | 0,222352175 | 0,566571175 | 0,16765125 | 0,142503563 | 0,046928814 | 0,018928814 | -1,241071186 |
| chrXIV | 649139 | 649287 | 148 | 0,259310013 | 0,285351958 | 0,47609348 | -0,059960669 | -0,050966568 | -0,152489374 | -0,180489374 | -1,440489374 |
| chrXIV | 649139 | 649287 | 148 | 0,259310013 | 0,285351958 | 0,47609348 | -0,059960669 | -0,050966568 | -0,152489374 | -0,180489374 | -1,440489374 |
| chrXIV | 659033 | 659204 | 171 | 0,256460453 | 0,281646088 | 0,47659791 | -0,058694023 | -0,04988992 | 0,063217926 | -0,125782074 | -1,385782074 |
| chrXIV | 660122 | 660273 | 151 | 0,059840772 | 0,059293913 | 0,502295129 | 0,005753068 | 0,004890107 | -0,106174214 | -0,155174214 | -1,415174214 |
| chrXIV | 660694 | 660857 | 163 | 4,764465296 | 4,610101756 | 0,508233102 | 0,020638792 | 0,017542973 | 0,114612946 | -0,018387054 | -1,278387054 |
| chrXIV | 669731 | 669882 | 151 | 0,364743755 | 0,211234566 | 0,63325952 | 0,340498754 | 0,289423941 | 0,182739708 | 0,133739708 | -1,126260292 |
| chrXIV | 673304 | 673429 | 125 | 0,04844253 | 1,119172613 | 0,041488439 | -1,733668726 | -1,473618417 | -1,106527638 | -0,973527638 | -2,233527638 |
| chrXIV | 673304 | 673441 | 137 | 0,031345166 | 0,040764565 | 0,434687048 | -0,164453569 | -0,139785534 | -0,256748695 | -0,207748695 | -1,467748695 |
| chrXIV | 673304 | 673449 | 145 | 0,034194727 | 0,059293913 | 0,365763444 | -0,34309514 | -0,291630869 | -0,436894619 | -0,443894619 | -1,703894619 |

TABLE 2

| Chrom | Start | End | Length | Section A | Section B | A/A+B | Z-score | Z * 0.85 | Phase correction | Length correction | ΔLK |
|---|---|---|---|---|---|---|---|---|---|---|---|
| chrXIV | 673304 | 673450 | 146 | 0,068389454 | 0,066705652 | 0,506231912 | 0,015621722 | 0,013278464 | -0,093126653 | -0,107126653 | -1,367126653 |
| chrXIV | 673304 | 673451 | 147 | 0,133929347 | 0,255705001 | 0,343730854 | -0,402302101 | -0,341956786 | -0,408340989 | -0,429340989 | -1,689340989 |
| chrXIV | 673304 | 673452 | 148 | 0,108283302 | 0,151940653 | 0,416115812 | -0,211840305 | -0,180064259 | -0,278576302 | -0,306576302 | -1,566576302 |
| chrXIV | 673304 | 673453 | 149 | 0,065539893 | 0,070411522 | 0,4820832 | -0,044925865 | -0,038186985 | -0,183425504 | -0,218425504 | -1,478425504 |
| chrXIV | 673304 | 673454 | 150 | 0,119681545 | 0,133411305 | 0,472876041 | -0,06804215 | -0,057835827 | -0,244023022 | -0,286023022 | -1,546023022 |
| chrXIV | 673304 | 673455 | 151 | 50,87035554 | 51,7487628 | 0,49572006 | -0,010728423 | -0,00911916 | -0,095266426 | -0,144266426 | -1,404266426 |
| chrXIV | 673305 | 673455 | 150 | 0,051292091 | 0,188999349 | 0,213457836 | -0,794480647 | -0,67530855 | -0,846858369 | -0,888858369 | -2,148858369 |
| chrXIV | 673306 | 673455 | 149 | 0,082637257 | 0,185293479 | 0,308427686 | -0,500312047 | -0,42526524 | -0,562060061 | -0,597060061 | -1,857060061 |
| chrXIV | 673307 | 673455 | 148 | 0,051292091 | 0,314998914 | 0,140030986 | -1,080180137 | -0,918153116 | -1,008895662 | -1,036895662 | -2,296895662 |
| chrXIV | 673308 | 673455 | 147 | 0,102584181 | 0,185293479 | 0,356346446 | -0,368241871 | -0,31300559 | -0,385660036 | -0,406660036 | -1,666660036 |
| chrXIV | 673309 | 673455 | 146 | 0,510071344 | 0,629997828 | 0,447403856 | -0,132223249 | -0,112389761 | -0,212343852 | -0,226343852 | -1,486343852 |
| chrXIV | 673310 | 673455 | 145 | 0,037044288 | 0,100058479 | 0,270193583 | -0,612227627 | -0,520393483 | -0,663751251 | -0,670751251 | -1,930751251 |
| chrXIV | 673311 | 673455 | 144 | 0,042743409 | 0,066705652 | 0,390532439 | -0,277931585 | -0,236241848 | -0,363067519 | -0,363067519 | -1,623067519 |
| chrXIV | 673313 | 673455 | 142 | 0,031345166 | 0,129705435 | 0,194629303 | -0,860962664 | -0,731818264 | -0,984529351 | -0,970529351 | -2,230529351 |
| chrXIV | 673314 | 673455 | 141 | 0,034194727 | 0,151940653 | 0,183708906 | -0,901320735 | -0,766122625 | -0,986408159 | -0,965408159 | -2,225408159 |
| chrXIV | 673321 | 673455 | 134 | 0,037044288 | 0,103764348 | 0,263082498 | -0,633871027 | -0,538790373 | -0,526359174 | -0,456359174 | -1,716359174 |
| chrXIV | 673329 | 673455 | 126 | 0,042743409 | 0,08894087 | 0,324590066 | -0,454901459 | -0,386666241 | -0,160813587 | -0,034813587 | -1,294813587 |
| chrXIV | 673357 | 673455 | 98 | 0,037044288 | 0,296469566 | 0,11107271 | -1,22084313 | -1,037716661 | -0,594138512 | -0,272138512 | -1,532138512 |
| chrXIV | 675190 | 675330 | 140 | 0,133929347 | 0,044470435 | 0,750725957 | 0,676776004 | 0,575259604 | 0,366676266 | 0,394676266 | -0,865323734 |
| chrXIV | 678978 | 679124 | 146 | 0,450230572 | 0,274234349 | 0,621466353 | 0,309334231 | 0,262934096 | 0,167860777 | 0,153860777 | -1,106139223 |
| chrXIV | 679561 | 679717 | 156 | 0,852018614 | 0,904232177 | 0,485134935 | -0,037269817 | -0,031679345 | -0,11938089 | -0,20338089 | -1,46338089 |
| chrXIV | 681968 | 682096 | 128 | 13,28180188 | 21,15680942 | 0,385666012 | -0,290632993 | -0,247038044 | -0,072429156 | 0,039570844 | -1,220429156 |
| chrXIV | 681972 | 682075 | 103 | 0,094035499 | 0,148234783 | 0,388142938 | -0,284162468 | -0,241538098 | 0,172555612 | 0,459555612 | -0,800444388 |
| chrXIV | 682856 | 682998 | 142 | 0,125380666 | 0,140823044 | 0,470995186 | -0,072768458 | -0,061853189 | -0,298416865 | -0,284416865 | -1,544416865 |
| chrXIV | 689540 | 689646 | 106 | 0,028495606 | 0,655938915 | 0,041633794 | -1,732033557 | -1,472228523 | -0,95185362 | -0,68585362 | -1,94585362 |
| chrXIV | 689540 | 689701 | 161 | 0,074088575 | 0,085235 | 0,465019536 | -0,087795679 | -0,074626327 | -0,052265612 | -0,171265612 | -1,431265612 |
| chrXIV | 689540 | 689704 | 164 | 0,039893848 | 0,103764348 | 0,277699771 | -0,589688448 | -0,501235181 | -0,435952551 | -0,575952551 | -1,835952551 |
| chrXIV | 689540 | 689705 | 165 | 23,29800733 | 18,57011246 | 0,556461753 | 0,142004446 | 0,120703779 | 0,145191745 | -0,001808255 | -1,261808255 |
| chrXIV | 689541 | 689705 | 164 | 0,051292091 | 0,096352609 | 0,34740218 | -0,39234359 | -0,333492051 | -0,264238324 | -0,404238324 | -1,664238324 |
| chrXIV | 689543 | 689705 | 162 | 0,076938136 | 0,100058479 | 0,434687048 | -0,164453569 | -0,139785534 | -0,06765219 | -0,19365219 | -1,45365219 |
| chrXIV | 689545 | 689705 | 160 | 0,042743409 | 0,085235 | 0,333989218 | -0,428924135 | -0,364585514 | -0,392409185 | -0,504409185 | -1,764409185 |
| chrXIV | 689550 | 689705 | 155 | 0,037044288 | 0,059293913 | 0,384523349 | -0,293622086 | -0,249578773 | -0,345624926 | -0,422624926 | -1,682624926 |
| chrXIV | 689554 | 689705 | 151 | 0,028495606 | 0,044470435 | 0,390532439 | -0,277931585 | -0,236241848 | -0,318769115 | -0,367769115 | -1,627769115 |
| chrXIV | 694543 | 694689 | 146 | 0,142478029 | 0,059293913 | 0,706134002 | 0,542125583 | 0,460806745 | 0,361101543 | 0,347101543 | -0,912898457 |
| chrXIV | 694549 | 694699 | 150 | 0,14532759 | 0,044470435 | 0,765696008 | 0,724745812 | 0,61603394 | 0,440619976 | 0,398619976 | -0,861380024 |
| chrXIV | 698880 | 699032 | 152 | 1,276603141 | 0,978349569 | 0,566132999 | 0,166537466 | 0,141556846 | 0,095273632 | 0,039273632 | -1,220726368 |
| chrXIV | 704009 | 704157 | 148 | 2,7327286 | 1,426759788 | 0,656986712 | 0,404253146 | 0,343615174 | 0,25184921 | 0,22384921 | -1,03615079 |
| chrXIV | 704660 | 704785 | 125 | 0,088336378 | 0,055588044 | 0,613769206 | 0,28915654 | 0,245783059 | 0,634147349 | 0,767147349 | -0,492852651 |
| chrXIV | 705096 | 705262 | 166 | 0,911859387 | 1,030231743 | 0,46952451 | -0,076465173 | -0,064995397 | -0,016998624 | -0,170998624 | -1,430998624 |
| chrXIV | 705096 | 705262 | 166 | 0,911859387 | 1,030231743 | 0,46952451 | -0,076465173 | -0,064995397 | -0,016998624 | -0,170998624 | -1,430998624 |
| chrXIV | 705533 | 705682 | 149 | 0,626903328 | 0,396528045 | 0,612550431 | 0,285972559 | 0,243076675 | 0,101234006 | 0,066234006 | -1,193765994 |
| chrXIV | 705820 | 705974 | 154 | 0,322000346 | 0,322410653 | 0,499681642 | -0,000798006 | -0,000678305 | -0,03754358 | -0,10754358 | -1,36754358 |
| chrXIV | 706013 | 706167 | 154 | 0,740885752 | 0,61146848 | 0,54784888 | 0,120228376 | 0,10219412 | 0,06979743 | -0,00020257 | -1,26020257 |
| chrXIV | 706675 | 706825 | 150 | 0,279256937 | 0,170470001 | 0,620947766 | 0,307970909 | 0,261775272 | 0,086785026 | 0,044785026 | -1,215214974 |
| chrXIV | 706675 | 706825 | 150 | 0,279256937 | 0,170470001 | 0,620947766 | 0,307970909 | 0,261775272 | 0,086785026 | 0,044785026 | -1,215214974 |
| chrXIV | 707449 | 707602 | 153 | 0,094035499 | 0,118587827 | 0,442263326 | -0,145233329 | -0,12344833 | -0,169744118 | -0,232744118 | -1,492744118 |
| chrXIV | 723355 | 723513 | 158 | 0,19661968 | 0,107470218 | 0,646584058 | 0,376114353 | 0,3196972 | 0,25085812 | 0,15285812 | -1,10714188 |
| chrXIV | 724785 | 724950 | 165 | 0,447381012 | 0,389116306 | 0,534826595 | 0,087408503 | 0,074297228 | 0,111890638 | -0,035109362 | -1,295109362 |
| chrXIV | 727236 | 727386 | 150 | 0,208017923 | 0,100058479 | 0,675215374 | 0,454360676 | 0,386206574 | 0,215211041 | 0,173211041 | -1,086788959 |
| chrXIV | 727633 | 727771 | 138 | 1,461824579 | 1,360054135 | 0,518032392 | 0,045215905 | 0,038433519 | -0,126328093 | -0,084328093 | -1,344328093 |
| chrXIV | 731977 | 732139 | 162 | 0,700991904 | 0,896820438 | 0,438719795 | -0,154215792 | -0,131083423 | -0,057442498 | -0,183442498 | -1,443442498 |
| chrXIV | 735524 | 735689 | 165 | 0,364743755 | 0,552174567 | 0,397793071 | -0,259063637 | -0,220204091 | -0,17166689 | -0,31866689 | -1,57866689 |
| chrXIV | 735878 | 736030 | 152 | 1,886409106 | 1,104349134 | 0,630746103 | 0,333830067 | 0,283755557 | 0,232779739 | 0,176779739 | -1,083220261 |
| chrXIV | 737786 | 737917 | 131 | 0,598407723 | 1,019114134 | 0,369953407 | -0,33197675 | -0,282180238 | -0,207817347 | -0,116817347 | -1,376817347 |
| chrXIV | 749149 | 749259 | 110 | 0,319150785 | 0,859761742 | 0,270716256 | -0,610648191 | -0,519050963 | -0,09529556 | 0,14270444 | -1,11729556 |
| chrXIV | 749725 | 749855 | 130 | 0,04844253 | 0,122293696 | 0,283727309 | -0,571804221 | -0,486033588 | -0,345726378 | -0,247726378 | -1,507726378 |
| chrXIV | 749823 | 749962 | 139 | 0,44168189 | 0,192705218 | 0,696234026 | 0,513599615 | 0,436559673 | 0,290810057 | 0,325810057 | -0,934189943 |
| chrXIV | 750208 | 750335 | 127 | 0,045592969 | 0,17417587 | 0,207458753 | -0,815270475 | -0,692979903 | -0,48698518 | -0,36798518 | -1,62798518 |
| chrXIV | 753363 | 753527 | 164 | 0,105433742 | 0,203822827 | 0,340926442 | -0,409936016 | -0,348445614 | -0,280839786 | -0,420839786 | -1,680839786 |
| chrXIV | 754387 | 754526 | 139 | 1,068585219 | 0,811585438 | 0,568344802 | 0,172161713 | 0,146337456 | 0,003546114 | 0,038546114 | -1,221453886 |
| chrXIV | 761541 | 761643 | 102 | 0,190920559 | 1,964110877 | 0,088592935 | -1,349470563 | -1,147049979 | -0,758150947 | -0,464150947 | -1,724150947 |
| chrXIV | 763360 | 763525 | 165 | 0,062690333 | 0,062999783 | 0,498768996 | -0,003085674 | -0,002622823 | 0,037983738 | -0,109016262 | -1,369016262 |
| chrXIV | 763360 | 763525 | 165 | 0,062690333 | 0,062999783 | 0,498768996 | -0,003085674 | -0,002622823 | 0,037983738 | -0,109016262 | -1,369016262 |
| chrXIV | 767084 | 767240 | 156 | 0,208017923 | 0,185293479 | 0,528888615 | 0,072476421 | 0,061604958 | -0,032752679 | -0,116752679 | -1,376752679 |
| chrXIV | 773470 | 773620 | 150 | 0,364743755 | 0,185293479 | 0,663125571 | 0,421008524 | 0,357857245 | 0,196066768 | 0,154066768 | -1,105933232 |
| chrXIV | 778474 | 778625 | 151 | 0,105433742 | 0,155646522 | 0,403836506 | -0,243429088 | -0,206914725 | -0,293455578 | -0,342455578 | -1,602455578 |
| chrXIV | 779862 | 780020 | 158 | 0,136778908 | 0,137117174 | 0,499382491 | -0,001547865 | -0,001315685 | -0,061903906 | -0,159903906 | -1,419903906 |
| chrXIV | 781645 | 781793 | 148 | 1,399134247 | 0,837526525 | 0,625546021 | 0,320079643 | 0,272067696 | 0,182403555 | 0,154403555 | -1,105596445 |
| chrXIV | 781935 | 782064 | 129 | 0,19377012 | 0,333528262 | 0,367477175 | -0,338542574 | -0,287761188 | -0,103754213 | 0,001245787 | -1,258754213 |
| chrXIV | 781942 | 782106 | 164 | 0,125380666 | 0,081529131 | 0,605967759 | 0,268824834 | 0,228501109 | 0,30102777 | 0,16102777 | -1,09897223 |
| chrXIV | 783376 | 783501 | 125 | 0,113982423 | 0,114881957 | 0,498034789 | -0,004926073 | -0,004187162 | 0,36495017 | 0,49795017 | -0,76204983 |
| chrXIV | 784039 | 784210 | 171 | 2,19986077 | 0,055588044 | 0,975353888 | 1,966055295 | 1,671147001 | 1,807247291 | 1,618247291 | 0,358247291 |
| chrXIV | 784040 | 784199 | 159 | 0,071239015 | 0,155646522 | 0,313986584 | -0,484581602 | -0,411894361 | -0,470059105 | -0,575059105 | -1,835059105 |
| chrXIV | 784044 | 784183 | 139 | 1,36493952 | 0,559586306 | 0,709234192 | 0,551148886 | 0,468476553 | 0,32866437 | 0,36366437 | -0,89633563 |
| chrXIV | 784044 | 784199 | 155 | 0,094035499 | 0,137117174 | 0,406811212 | -0,235755471 | -0,20039215 | -0,299685829 | -0,376685829 | -1,636685829 |
| chrXIV | 784049 | 784198 | 149 | 1,701187668 | 0,874585221 | 0,660457168 | 0,413711146 | 0,351654474 | 0,207740276 | 0,172740276 | -1,087259724 |
| chrXIV | 784049 | 784203 | 154 | 0,190920559 | 0,114881957 | 0,624326319 | 0,316863258 | 0,269333769 | 0,231057712 | 0,161057712 | -1,098942288 |
| chrXIV | 784063 | 784183 | 120 | 0,111132863 | 0,122293696 | 0,47609348 | -0,059960669 | -0,050966568 | 0,306770056 | 0,474770056 | -0,785229944 |
| chrXIV | 784063 | 784195 | 132 | 0,151026711 | 0,062999783 | 0,705644933 | 0,540706158 | 0,459600234 | 0,503045964 | 0,587045964 | -0,672954036 |
| chrXIV | 784063 | 784199 | 136 | 1,977595045 | 1,660229571 | 0,543620227 | 0,109558474 | 0,093124703 | 0,018343149 | 0,074343149 | -1,185656851 |
| chrXIV | 784063 | 784202 | 139 | 0,19661968 | 0,070411522 | 0,736317249 | 0,632032712 | 0,537227806 | 0,403542322 | 0,438542322 | -0,821457678 |
| chrXIV | 784063 | 784210 | 147 | 0,703841464 | 0,415057393 | 0,629048336 | 0,329333894 | 0,27993381 | 0,209313897 | 0,188313897 | -1,071686103 |
| chrXIV | 784063 | 784233 | 170 | 0,173823196 | 0,218646305 | 0,44289606 | -0,14363067 | -0,12208607 | -0,044291294 | -0,226291294 | -1,486291294 |
| chrXIV | 784063 | 784239 | 176 | 0,071239015 | 0,040764565 | 0,6360423 | 0,347899847 | 0,29571487 | 0,591713451 | 0,367713451 | -0,892286549 |
| chrXIV | 784063 | 784239 | 176 | 0,071239015 | 0,040764565 | 0,6360423 | 0,347899847 | 0,29571487 | 0,591713451 | 0,367713451 | -0,892286549 |
| chrXIV | 784063 | 784275 | 212 | 0,028495606 | 0,103764348 | 0,215451503 | -0,787647358 | -0,669500254 | -0,227608692 | -0,703608692 | -1,963608692 |
| chrXIV | 784063 | 784275 | 212 | 0,028495606 | 0,103764348 | 0,215451503 | -0,787647358 | -0,669500254 | -0,227608692 | -0,703608692 | -1,963608692 |
| chrXIV | 784064 | 784183 | 119 | 0,3903898 | 0,222352175 | 0,637119401 | 0,35076961 | 0,298154169 | 0,703632911 | 0,878632911 | -0,381367089 |
| chrXIV | 784064 | 784199 | 135 | 0,082637257 | 0,048176305 | 0,631717813 | 0,336406466 | 0,285945496 | 0,278049809 | 0,341049809 | -0,918950191 |
| chrXIV | 784101 | 784203 | 102 | 0,028495606 | 0,051882174 | 0,354520937 | -0,373143192 | -0,317171714 | 0,096134907 | 0,390134907 | -0,869865093 |
| chrXIV | 784101 | 784210 | 109 | 0,185221438 | 0,062999783 | 0,746195017 | 0,662563791 | 0,563179222 | 1,013116549 | 1,258116549 | -0,001883451 |
| chrXIV | 784106 | 784198 | 92 | 0,364743755 | 1,945581529 | 0,157875498 | -1,003227731 | -0,852743571 | -0,509753408 | -0,145753408 | -1,405753408 |
| chrXIV | 784106 | 784203 | 97 | 0,111132863 | 0,259410871 | 0,299918344 | -0,524635379 | -0,445940072 | -0,003219363 | 0,325780637 | -0,934219363 |
| chrXIV | 784106 | 784210 | 104 | 0,589859041 | 0,244587392 | 0,706886647 | 0,544312122 | 0,462665304 | 0,925669973 | 1,205669973 | -0,054330027 |
| chrXIV | 784106 | 784238 | 132 | 0,384690679 | 0,122293696 | 0,75878212 | 0,702390353 | 0,5970318 | 0,629118529 | 0,713118529 | -0,546881471 |
| chrXIV | 784106 | 784238 | 132 | 0,384690679 | 0,122293696 | 0,75878212 | 0,702390353 | 0,5970318 | 0,629118529 | 0,713118529 | -0,546881471 |
| chrXIV | 784106 | 784265 | 159 | 0,265009134 | 0,107470218 | 0,711473355 | 0,557694096 | 0,474039982 | 0,423732317 | 0,318732317 | -0,941267683 |
| chrXIV | 784106 | 784265 | 159 | 0,265009134 | 0,107470218 | 0,711473355 | 0,557694096 | 0,474039982 | 0,423732317 | 0,318732317 | -0,941267683 |
| chrXIV | 784106 | 784274 | 168 | 2,826764099 | 0,366881088 | 0,885121525 | 1,200985181 | 1,020837404 | 1,045628256 | 0,877628256 | -0,382371744 |
| chrXIV | 784106 | 784274 | 168 | 2,826764099 | 0,366881088 | 0,885121525 | 1,200985181 | 1,020837404 | 1,045628256 | 0,877628256 | -0,382371744 |
| chrXIV | 784106 | 784275 | 169 | 0,079787696 | 0,077823261 | 0,506231912 | 0,015621722 | 0,013278464 | 0,045024016 | -0,129975984 | -1,389975984 |

TABLE 2

| Chrom | Start | End | Length | Section A | Section B | A/A+B | Z-score | Z * 0.85 | Phase correction | Length correction | ΔLK |
|---|---|---|---|---|---|---|---|---|---|---|---|
| chrXIV | 784106 | 784275 | 169 | 0,079787696 | 0,077823261 | 0,506231912 | 0,015621722 | 0,013278464 | 0,045024016 | -0,129975984 | -1,389975984 |
| chrXIV | 784106 | 784277 | 171 | 0,19661968 | 0,081529131 | 0,706886647 | 0,544312122 | 0,462665304 | 0,62594496 | 0,43694496 | -0,82305504 |
| chrXIV | 784106 | 784277 | 171 | 0,19661968 | 0,081529131 | 0,706886647 | 0,544312122 | 0,462665304 | 0,62594496 | 0,43694496 | -0,82305504 |
| chrXIV | 784111 | 784233 | 122 | 0,336248149 | 0,081529131 | 0,804850252 | 0,859074354 | 0,730213201 | 1,0816173 | 1,2356173 | -0,0243827 |
| chrXIV | 784111 | 784286 | 175 | 0,131079787 | 0,059293913 | 0,688539366 | 0,491714385 | 0,417957228 | 0,721360639 | 0,504360639 | -0,755639361 |
| chrXIV | 784111 | 784286 | 175 | 0,131079787 | 0,059293913 | 0,688539366 | 0,491714385 | 0,417957228 | 0,721360639 | 0,504360639 | -0,755639361 |
| chrXIV | 784130 | 784238 | 108 | 0,094035499 | 0,081529131 | 0,535617563 | 0,08939893 | 0,07598909 | 0,578952915 | 0,830952915 | -0,429047085 |
| chrXIV | 784130 | 784274 | 144 | 2,188462528 | 0,478057176 | 0,820718679 | 0,918107397 | 0,780391288 | 0,648531057 | 0,648531057 | -0,611468943 |
| chrXIV | 784130 | 784275 | 145 | 0,051292091 | 0,08894087 | 0,365763444 | -0,34309514 | -0,291630869 | -0,451014209 | -0,458014209 | -1,718014209 |
| chrXIV | 784130 | 784277 | 147 | 0,273557816 | 0,08894087 | 0,754644987 | 0,68917996 | 0,585802966 | 0,513154693 | 0,492154693 | -0,767845307 |
| chrXIV | 784130 | 784329 | 199 | 0,29350474 | 0,211234566 | 0,581497689 | 0,205726418 | 0,174867455 | 0,593221173 | 0,208221173 | -1,051778827 |
| chrXIV | 784132 | 784275 | 143 | 0,954602795 | 0,214940436 | 0,816218478 | 0,901047538 | 0,765890407 | 0,582311075 | 0,589311075 | -0,670688925 |
| chrXIV | 784135 | 784274 | 139 | 0,820673448 | 0,17788174 | 0,821860883 | 0,922480044 | 0,784108038 | 0,650652717 | 0,685652717 | -0,574347283 |
| chrXIV | 784139 | 784275 | 136 | 0,273557816 | 0,218646305 | 0,555781238 | 0,140281573 | 0,119239337 | 0,01732953 | 0,07332953 | -1,18667047 |
| chrXIV | 784139 | 784296 | 157 | 0,111132863 | 0,037058696 | 0,74992708 | 0,674260298 | 0,573121253 | 0,513646995 | 0,422646995 | -0,837353005 |
| chrXIV | 784140 | 784275 | 135 | 0,028495606 | 0,092646739 | 0,235224155 | -0,721749813 | -0,613487341 | -0,621223736 | -0,558223736 | -1,818223736 |
| chrXIV | 784145 | 784286 | 141 | 0,416035845 | 0,118587827 | 0,778184482 | 0,766076079 | 0,651164667 | 0,412330189 | 0,433330189 | -0,826669811 |
| chrXIV | 784164 | 784275 | 111 | 0,056991212 | 0,081529131 | 0,411428464 | -0,223871969 | -0,190291174 | 0,225639615 | 0,456639615 | -0,803360385 |
| chrXIV | 784169 | 784274 | 105 | 0,279256937 | 0,285351958 | 0,49460244 | -0,013530088 | -0,011500575 | 0,490042985 | 0,763042985 | -0,496957015 |
| chrXIV | 784169 | 784275 | 106 | 0,037044288 | 0,259410871 | 0,124957474 | -1,150555991 | -0,977972592 | -0,458247251 | -0,192247251 | -1,452247251 |
| chrXIV | 784169 | 784277 | 108 | 0,034194727 | 0,040764565 | 0,456177292 | -0,110069087 | -0,093558724 | 0,365206854 | 0,617206854 | -0,642793146 |
| chrXIV | 784172 | 784265 | 93 | 0,045592969 | 0,048176305 | 0,486225044 | -0,034535557 | -0,029355224 | 0,333969233 | 0,690969233 | -0,569030767 |
| chrXIV | 784172 | 784267 | 95 | 0,230814407 | 0,122293696 | 0,653664997 | 0,395234266 | 0,335949126 | 0,732399665 | 1,075399665 | -0,184600335 |
| chrXIV | 784172 | 784271 | 99 | 0,646850252 | 0,466939567 | 0,580765097 | 0,203851168 | 0,173273493 | 0,600792914 | 0,915792914 | -0,344207086 |
| chrXIV | 784172 | 784274 | 102 | 0,028495606 | 0,059293913 | 0,324590066 | -0,454901459 | -0,386666241 | 0,03125577 | 0,32525577 | -0,93474423 |
| chrXIV | 784172 | 784275 | 103 | 1,795223168 | 2,764578706 | 0,393706397 | -0,269671752 | -0,229220989 | 0,20248926 | 0,48948926 | -0,77051074 |
| chrXIV | 784172 | 784277 | 105 | 0,222265726 | 0,137117174 | 0,618464945 | 0,301451656 | 0,256233907 | 0,781715484 | 1,054715484 | -0,205284516 |
| chrXIV | 784177 | 784271 | 94 | 0,054141651 | 0,133411305 | 0,288673942 | -0,557262806 | -0,473673385 | -0,084111089 | 0,265888911 | -0,994111089 |
| chrXIV | 784177 | 784275 | 98 | 0,273557816 | 2,056757616 | 0,117390896 | -1,188131028 | -1,009911374 | -0,58051675 | -0,25851675 | -1,51851675 |
| chrXV | 838 | 992 | 154 | 1,51311667 | 1,452700875 | 0,510185353 | 0,025533667 | 0,021703617 | -0,050324046 | -0,120324046 | -1,380324046 |
| chrXV | 6743 | 6894 | 151 | 1,119877309 | 0,70040935 | 0,615220302 | 0,292951275 | 0,249008584 | 0,134861164 | 0,085861164 | -1,174138836 |
| chrXV | 8892 | 9048 | 156 | 0,039893848 | 0,055588044 | 0,417815854 | -0,20748423 | -0,176361595 | -0,154450814 | -0,238450814 | -1,498450814 |
| chrXV | 10875 | 10982 | 107 | 0,165274514 | 1,297054353 | 0,11302144 | -1,210615295 | -1,029023001 | -0,04672108 | 0,21227892 | -1,04772108 |
| chrXV | 23820 | 23972 | 152 | 0,387540239 | 0,26311674 | 0,595613744 | 0,242009873 | 0,205708392 | 0,137145589 | 0,081145589 | -1,178854411 |
| chrXV | 29238 | 29397 | 159 | 1,302249187 | 0,770820872 | 0,628174229 | 0,327021609 | 0,277968368 | 0,31276023 | 0,20776023 | -1,05223977 |
| chrXV | 29238 | 29397 | 159 | 1,302249187 | 0,770820872 | 0,628174229 | 0,327021609 | 0,277968368 | 0,31276023 | 0,20776023 | -1,05223977 |
| chrXV | 36163 | 36311 | 148 | 0,09973462 | 0,111176087 | 0,472876041 | -0,06804215 | -0,057835827 | -0,060033363 | -0,088033363 | -1,348033363 |
| chrXV | 39246 | 39406 | 160 | 0,302053422 | 0,255705001 | 0,541548831 | 0,104336469 | 0,088685998 | 0,138516678 | 0,026516678 | -1,233483322 |
| chrXV | 43767 | 43929 | 162 | 0,484425299 | 0,807879568 | 0,37485373 | -0,319025127 | -0,271171358 | -0,221068033 | -0,347068033 | -1,607068033 |
| chrXV | 44442 | 44591 | 149 | 1,917754273 | 1,082113917 | 0,639279512 | 0,356533624 | 0,303053581 | 0,272392175 | 0,237392175 | -1,022607825 |
| chrXV | 49181 | 49338 | 157 | 1,70688679 | 1,163643048 | 0,594624298 | 0,239456783 | 0,203538266 | 0,244524259 | 0,153524259 | -1,106475741 |
| chrXV | 51344 | 51469 | 125 | 0,085486818 | 0,144528914 | 0,371656395 | -0,327469522 | -0,278349094 | 0,45809714 | 0,59109714 | -0,66890286 |
| chrXV | 53593 | 53699 | 106 | 0,324849907 | 2,338403704 | 0,12197483 | -1,165171301 | -0,990395606 | 0,000651019 | 0,266651019 | -0,993348981 |
| chrXV | 53593 | 53708 | 115 | 0,339097709 | 2,946166315 | 0,103217795 | -1,263427488 | -1,073913365 | -0,090864622 | 0,112135378 | -1,147864622 |
| chrXV | 53860 | 54011 | 151 | 0,222265726 | 0,26311674 | 0,457918737 | -0,105678457 | -0,089826688 | -0,194230881 | -0,243230881 | -1,503230881 |
| chrXV | 53996 | 54153 | 157 | 0,763682236 | 0,663350655 | 0,535153914 | 0,08823214 | 0,074997319 | 0,115494397 | 0,024494397 | -1,235505603 |
| chrXV | 63125 | 63255 | 130 | 0,14817715 | 0,096352609 | 0,605967759 | 0,268824834 | 0,228501109 | 0,541492418 | 0,639492418 | -0,620507582 |
| chrXV | 64118 | 64281 | 163 | 0,678195419 | 0,948702612 | 0,416864122 | -0,209922394 | -0,178434035 | -0,104302814 | -0,237302814 | -1,497302814 |
| chrXV | 66159 | 66329 | 170 | 0,039893848 | 0,111176087 | 0,264075364 | -0,630831465 | -0,536206745 | -0,399108517 | -0,581108517 | -1,841108517 |
| chrXV | 72755 | 72904 | 149 | 0,378991558 | 0,318704784 | 0,543204164 | 0,108509339 | 0,092232938 | 0,055610787 | 0,020610787 | -1,239389213 |
| chrXV | 74613 | 74750 | 137 | 0,700991904 | 0,311293045 | 0,692484764 | 0,502905845 | 0,427469968 | 0,591494655 | 0,640494655 | -0,619505345 |
| chrXV | 74660 | 74806 | 146 | 1,473222822 | 0,96352609 | 0,604585402 | 0,265234012 | 0,22544891 | 0,248387375 | 0,234387375 | -1,025612625 |
| chrXV | 74660 | 74806 | 146 | 1,473222822 | 0,96352609 | 0,604585402 | 0,265234012 | 0,22544891 | 0,248387375 | 0,234387375 | -1,025612625 |
| chrXV | 75646 | 75807 | 161 | 0,108283302 | 0,196411088 | 0,355383315 | -0,370826679 | -0,315202677 | -0,257061107 | -0,376061107 | -1,636061107 |
| chrXV | 80807 | 80978 | 171 | 0,447381012 | 0,667056524 | 0,401441083 | -0,249618789 | -0,212175971 | -0,061279878 | -0,250279878 | -1,510279878 |
| chrXV | 101110 | 101258 | 148 | 0,549965193 | 0,418763262 | 0,567718631 | 0,170568921 | 0,144983583 | 0,141626669 | 0,113626669 | -1,146373331 |
| chrXV | 108670 | 108830 | 160 | 0,404637603 | 0,518821741 | 0,438175872 | -0,155595664 | -0,132256314 | -0,084753686 | -0,196753686 | -1,456753686 |
| chrXV | 110950 | 111091 | 141 | 0,481575739 | 0,385410436 | 0,555459536 | 0,139467258 | 0,118547169 | 0,197202826 | 0,218202826 | -1,041797174 |
| chrXV | 112040 | 112197 | 157 | 0,04844253 | 0,092646739 | 0,343346664 | -0,403346515 | -0,342844538 | -0,312471504 | -0,403471504 | -1,663471504 |
| chrXV | 113588 | 113739 | 151 | 0,116831984 | 0,044470435 | 0,72430398 | 0,595675482 | 0,506324159 | 0,407030525 | 0,358030525 | -0,901969475 |
| chrXV | 116654 | 116808 | 154 | 0,028495606 | 0,040764565 | 0,411428464 | -0,223871969 | -0,190291174 | -0,25598372 | -0,32598372 | -1,58598372 |
| chrXV | 118023 | 118177 | 154 | 0,484425299 | 0,370586958 | 0,566571175 | 0,16765125 | 0,142503563 | 0,07917029 | 0,00917029 | -1,25082971 |
| chrXV | 118066 | 118215 | 149 | 0,085486818 | 0,055588044 | 0,605967759 | 0,268824834 | 0,228501109 | 0,189276554 | 0,154276554 | -1,105723446 |
| chrXV | 118240 | 118363 | 123 | 0,09688506 | 0,277940218 | 0,258480592 | -0,648036746 | -0,550831234 | 0,196730019 | 0,343730019 | -0,916269981 |
| chrXV | 118340 | 118495 | 155 | 0,116831984 | 0,066705652 | 0,636556002 | 0,349268164 | 0,296877939 | 0,281221215 | 0,204221215 | -1,055778785 |
| chrXV | 118969 | 119120 | 151 | 0,119681545 | 0,122293696 | 0,49460244 | -0,013530088 | -0,011500575 | -0,105032477 | -0,154032477 | -1,414032477 |
| chrXV | 118979 | 119124 | 145 | 0,111132863 | 0,092646739 | 0,54535813 | 0,113942039 | 0,096850733 | 0,034332884 | 0,027332884 | -1,232667116 |
| chrXV | 118997 | 119148 | 151 | 0,056991212 | 0,037058696 | 0,605967759 | 0,268824834 | 0,228501109 | 0,134961158 | 0,085961158 | -1,174038842 |
| chrXV | 119251 | 119405 | 154 | 0,259310013 | 0,17417587 | 0,598197134 | 0,248683373 | 0,211380845 | 0,155347147 | 0,085347147 | -1,174652853 |
| chrXV | 119285 | 119429 | 144 | 0,062690333 | 0,048176305 | 0,56545715 | 0,164819952 | 0,140096959 | 0,107803736 | 0,107803736 | -1,152196264 |
| chrXV | 119816 | 119951 | 135 | 0,065539893 | 0,074117392 | 0,469290903 | -0,077052467 | -0,065494597 | 0,234993914 | 0,297993914 | -0,962006086 |
| chrXV | 120263 | 120409 | 146 | 0,071239015 | 0,051882174 | 0,578608892 | 0,198335964 | 0,168585569 | 0,192429279 | 0,178429279 | -1,081570721 |
| chrXV | 120263 | 120413 | 150 | 8,708257143 | 4,120926971 | 0,678784954 | 0,464303812 | 0,394658241 | 0,30160023 | 0,25960023 | -1,00039977 |
| chrXV | 120442 | 120599 | 157 | 0,062690333 | 0,055588044 | 0,530023616 | 0,075329227 | 0,064029843 | 0,091757309 | 0,000757309 | -1,259242691 |
| chrXV | 120674 | 120823 | 149 | 0,108283302 | 0,077823261 | 0,581834946 | 0,206589951 | 0,175601459 | 0,129503135 | 0,094503135 | -1,165496865 |
| chrXV | 120933 | 121062 | 129 | 0,056991212 | 0,103764348 | 0,354520937 | -0,373143192 | -0,317171714 | 0,136287842 | 0,241287842 | -1,018712158 |
| chrXV | 121845 | 121995 | 150 | 0,142478029 | 0,185293479 | 0,434687048 | -0,164453569 | -0,139785534 | -0,226664224 | -0,268664224 | -1,528664224 |
| chrXV | 122824 | 122966 | 142 | 0,042743409 | 0,044470435 | 0,490098899 | -0,024820929 | -0,021097789 | 0,030769416 | 0,044769416 | -1,215230584 |
| chrXV | 132108 | 132259 | 151 | 0,102584181 | 0,159352392 | 0,391637486 | -0,275053712 | -0,233795655 | -0,327024249 | -0,376024249 | -1,636024249 |
| chrXV | 132333 | 132482 | 149 | 0,122531105 | 0,133411305 | 0,478744828 | -0,053304046 | -0,045308439 | -0,08000934 | -0,11500934 | -1,37500934 |
| chrXV | 138290 | 138437 | 147 | 0,256460453 | 0,237175653 | 0,519533417 | 0,048982595 | 0,041635206 | 0,110909607 | 0,089909607 | -1,170090393 |
| chrXV | 141232 | 141379 | 147 | 0,370442876 | 0,333528262 | 0,52621884 | 0,065768269 | 0,055903029 | 0,128118531 | 0,107118531 | -1,152881469 |
| chrXV | 142140 | 142290 | 150 | 0,062690333 | 0,037058696 | 0,628480635 | 0,327831948 | 0,278657156 | 0,19267385 | 0,15067385 | -1,10932615 |
| chrXV | 143684 | 143828 | 144 | 3,168711369 | 2,393991748 | 0,569635176 | 0,175445428 | 0,149128613 | 0,12123166 | 0,12123166 | -1,13876834 |
| chrXV | 147632 | 147784 | 152 | 0,094035499 | 0,040764565 | 0,69759239 | 0,517488464 | 0,439865194 | 0,367015728 | 0,311015728 | -0,948984272 |
| chrXV | 152975 | 153131 | 156 | 0,29065518 | 0,277940218 | 0,511181027 | 0,028030349 | 0,023825797 | 0,048738125 | -0,035261875 | -1,295261875 |
| chrXV | 160615 | 160744 | 129 | 0,04844253 | 0,048176305 | 0,50137771 | 0,003453413 | 0,002935401 | 0,427041169 | 0,532041169 | -0,727958831 |
| chrXV | 162501 | 162638 | 137 | 0,304902982 | 0,166764131 | 0,646436806 | 0,375718225 | 0,319360491 | 0,471037129 | 0,520037129 | -0,739962871 |
| chrXV | 163008 | 163166 | 158 | 0,840620372 | 0,629997828 | 0,571610206 | 0,180475116 | 0,153403848 | 0,200525949 | 0,102525949 | -1,157474051 |
| chrXV | 163425 | 163564 | 139 | 0,692443222 | 0,459527828 | 0,601094291 | 0,256180562 | 0,217753478 | 0,280453925 | 0,315453925 | -0,944546075 |
| chrXV | 164146 | 164288 | 142 | 1,111328628 | 0,830114786 | 0,572423909 | 0,182548645 | 0,155166349 | 0,18415272 | 0,19815272 | -1,06184728 |
| chrXV | 169600 | 169748 | 148 | 0,444531451 | 0,515115871 | 0,46322377 | -0,09231529 | -0,078467996 | -0,081866628 | -0,109866628 | -1,369866628 |
| chrXV | 171569 | 171743 | 174 | 0,435982769 | 0,692997611 | 0,386173911 | -0,289305214 | -0,245909432 | 0,022479845 | -0,187520155 | -1,447520155 |
| chrXV | 173781 | 173933 | 152 | 1,168319839 | 0,874585221 | 0,571891402 | 0,18119159 | 0,154012851 | 0,072877569 | 0,016877569 | -1,243122431 |
| chrXV | 177092 | 177238 | 146 | 0,065539893 | 0,385410436 | 0,145337278 | -1,056642983 | -0,898146535 | -0,862246911 | -0,876246911 | -2,136246911 |
| chrXV | 177093 | 177211 | 118 | 0,042743409 | 0,17788174 | 0,1937377 | -0,864204794 | -0,734574075 | 0,244355211 | 0,426355211 | -0,833644789 |
| chrXV | 177093 | 177214 | 121 | 0,045592969 | 0,125999566 | 0,265704853 | -0,625855528 | -0,531977199 | 0,279754957 | 0,440754957 | -0,819245043 |
| chrXV | 177093 | 177217 | 124 | 0,131079787 | 1,889993485 | 0,064856524 | -1,515234426 | -1,287949263 | -0,581848134 | -0,441848134 | -1,701848134 |
| chrXV | 177093 | 177220 | 127 | 0,076938136 | 0,051882174 | 0,597251597 | 0,246239543 | 0,209303612 | 0,753197378 | 0,872197378 | -0,387802622 |
| chrXV | 177093 | 177221 | 128 | 0,111132863 | 0,144528914 | 0,434687048 | -0,164453569 | -0,139785534 | 0,401544342 | 0,513544342 | -0,746455658 |
| chrXV | 177093 | 177222 | 129 | 0,105433742 | 0,08894087 | 0,542425478 | 0,106546147 | 0,090564225 | 0,5092585 | 0,6142585 | -0,6457415 |

TABLE 2

| Chrom | Start | End | Length | Section A | Section B | A/A+B | Z-score | Z * 0.85 | Phase correction | Length correction | ΔLK |
|---|---|---|---|---|---|---|---|---|---|---|---|
| chrXV | 177093 | 177223 | 130 | 0,168124074 | 0,092646739 | 0,644719675 | 0,371103224 | 0,315437741 | 0,620739855 | 0,718739855 | -0,541260145 |
| chrXV | 177093 | 177224 | 131 | 0,14817715 | 0,129705435 | 0,533236547 | 0,083408278 | 0,070897036 | 0,301815019 | 0,392815019 | -0,867184981 |
| chrXV | 177093 | 177225 | 132 | 0,284956058 | 0,222352175 | 0,561702018 | 0,155285859 | 0,13199298 | 0,426511838 | 0,510511838 | -0,749488162 |
| chrXV | 177093 | 177226 | 133 | 0,09973462 | 0,274234349 | 0,266692236 | -0,62284791 | -0,529420724 | -0,247856435 | -0,170856435 | -1,430856435 |
| chrXV | 177093 | 177227 | 134 | 0,202318801 | 0,148234783 | 0,577140872 | 0,194584477 | 0,165396805 | 0,477938977 | 0,547938977 | -0,712061023 |
| chrXV | 177093 | 177228 | 135 | 0,202318801 | 0,148234783 | 0,577140872 | 0,194584477 | 0,165396805 | 0,453877291 | 0,516877291 | -0,743122709 |
| chrXV | 177093 | 177229 | 136 | 0,176672756 | 0,170470001 | 0,508934013 | 0,022396121 | 0,019036703 | 0,245519253 | 0,301519253 | -0,958480747 |
| chrXV | 177093 | 177230 | 137 | 0,208017923 | 0,240881523 | 0,463395366 | -0,091883336 | -0,078100835 | 0,052783278 | 0,101783278 | -1,158216722 |
| chrXV | 177093 | 177231 | 138 | 0,210867483 | 0,222352175 | 0,486744956 | -0,033231584 | -0,028246847 | 0,117737343 | 0,159737343 | -1,100262657 |
| chrXV | 177093 | 177232 | 139 | 0,119681545 | 0,096352609 | 0,553993628 | 0,135757811 | 0,11539414 | 0,196254366 | 0,231254366 | -1,028745634 |
| chrXV | 177093 | 177233 | 140 | 0,250761331 | 0,285351958 | 0,467739443 | -0,080953558 | -0,068810524 | 0,023305727 | 0,051305727 | -1,208694273 |
| chrXV | 177093 | 177234 | 141 | 0,250761331 | 0,222352175 | 0,530023616 | 0,075329227 | 0,064029843 | 0,131025469 | 0,152025469 | -1,107974531 |
| chrXV | 177093 | 177235 | 142 | 0,772230918 | 0,403939784 | 0,656563641 | 0,40310264 | 0,342637244 | 0,350797087 | 0,364797087 | -0,895202913 |
| chrXV | 177093 | 177236 | 143 | 0,259310013 | 0,163058261 | 0,613942923 | 0,289610604 | 0,246169013 | 0,285196127 | 0,292196127 | -0,967803873 |
| chrXV | 177093 | 177237 | 144 | 0,322000346 | 0,296469566 | 0,520640276 | 0,051760603 | 0,043996513 | 0,016616398 | 0,016616398 | -1,243383602 |
| chrXV | 177093 | 177238 | 145 | 249,0544446 | 103,5271725 | 0,706373879 | 0,542822178 | 0,461398851 | 0,436149214 | 0,429149214 | -0,830850786 |
| chrXV | 177093 | 177239 | 146 | 0,091185939 | 0,70411522 | 0,11465586 | -1,202133727 | -1,021813668 | -0,961173337 | -0,975173337 | -2,235173337 |
| chrXV | 177094 | 177238 | 144 | 0,279256937 | 0,296469566 | 0,485051384 | -0,037479397 | -0,031857487 | -0,070164434 | -0,070164434 | -1,330164434 |
| chrXV | 177095 | 177238 | 143 | 0,370442876 | 0,311293045 | 0,543381777 | 0,108957189 | 0,092613611 | 0,13029459 | 0,13729459 | -1,12270541 |
| chrXV | 177096 | 177238 | 142 | 1,040089613 | 0,61146848 | 0,629762657 | 0,331224804 | 0,281541083 | 0,28447824 | 0,29847824 | -0,96152176 |
| chrXV | 177097 | 177238 | 141 | 0,361894194 | 0,237175653 | 0,604093489 | 0,26395704 | 0,224363484 | 0,298607328 | 0,319607328 | -0,940392672 |
| chrXV | 177098 | 177238 | 140 | 0,58700948 | 0,426175002 | 0,579370777 | 0,200284037 | 0,170241431 | 0,264496599 | 0,292496599 | -0,967503401 |
| chrXV | 177099 | 177238 | 139 | 0,071239015 | 0,188999349 | 0,273745245 | -0,601524812 | -0,51129609 | -0,41743343 | -0,38243343 | -1,64243343 |
| chrXV | 177100 | 177238 | 138 | 0,091185939 | 0,107470218 | 0,459013908 | -0,102918298 | -0,087480553 | 0,024360604 | 0,066360604 | -1,193639396 |
| chrXV | 177101 | 177238 | 137 | 0,071239015 | 0,185293479 | 0,277699771 | -0,589688448 | -0,501235181 | -0,343361252 | -0,294361252 | -1,554361252 |
| chrXV | 177102 | 177238 | 136 | 0,088336378 | 0,240881523 | 0,268321917 | -0,617896095 | -0,525211681 | -0,289321503 | -0,233321503 | -1,493321503 |
| chrXV | 177103 | 177238 | 135 | 0,091185939 | 0,281646088 | 0,244576464 | -0,691656728 | -0,587908219 | -0,290576382 | -0,227576382 | -1,487576382 |
| chrXV | 177104 | 177238 | 134 | 0,068389454 | 0,326116523 | 0,173354671 | -0,940991251 | -0,799842563 | -0,503158055 | -0,433158055 | -1,693158055 |
| chrXV | 177105 | 177238 | 133 | 0,034194727 | 0,222352175 | 0,133288404 | -1,11098035 | -0,944333298 | -0,658043842 | -0,581043842 | -1,841043842 |
| chrXV | 177106 | 177238 | 132 | 0,062690333 | 0,074117392 | 0,458236793 | -0,104876779 | -0,089145262 | 0,188031908 | 0,272031908 | -0,987968092 |
| chrXV | 177107 | 177238 | 131 | 0,059840772 | 0,166764131 | 0,264075364 | -0,630831465 | -0,536206745 | -0,304584125 | -0,213584125 | -1,473584125 |
| chrXV | 177109 | 177238 | 129 | 0,094035499 | 0,326116523 | 0,223813035 | -0,759378669 | -0,645471868 | -0,24563726 | -0,14063726 | -1,40063726 |
| chrXV | 177110 | 177238 | 128 | 0,09688506 | 0,340940001 | 0,22128715 | -0,767853376 | -0,65267537 | -0,138237269 | -0,026237269 | -1,286237269 |
| chrXV | 177111 | 177238 | 127 | 0,037044288 | 0,085235 | 0,302948179 | -0,51593994 | -0,438548949 | 0,094569508 | 0,213569508 | -1,046430492 |
| chrXV | 177112 | 177238 | 126 | 0,051292091 | 0,233469783 | 0,180122745 | -0,914897409 | -0,777662797 | -0,17149581 | -0,04549581 | -1,30549581 |
| chrXV | 177113 | 177238 | 125 | 0,054141651 | 0,048176305 | 0,529151025 | 0,07313593 | 0,062165541 | 0,779487131 | 0,912487131 | -0,347512869 |
| chrXV | 177115 | 177238 | 123 | 0,031345166 | 0,118587827 | 0,209061167 | -0,8096831 | -0,688230635 | 0,0474975 | 0,1944975 | -1,0655025 |
| chrXV | 177117 | 177238 | 121 | 0,028495606 | 0,062999783 | 0,311443082 | -0,491764035 | -0,41800943 | 0,376491442 | 0,537491442 | -0,722508558 |
| chrXV | 177118 | 177238 | 120 | 0,054141651 | 0,048176305 | 0,529151025 | 0,07313593 | 0,062165541 | 0,961030763 | 1,129030763 | -0,130969237 |
| chrXV | 177119 | 177238 | 119 | 0,054141651 | 0,085235 | 0,388455674 | -0,283346352 | -0,2408444 | 0,692666629 | 0,867666629 | -0,392333371 |
| chrXV | 177120 | 177238 | 118 | 0,045592969 | 0,103764348 | 0,305261035 | -0,509328378 | -0,432929121 | 0,534930759 | 0,716930759 | -0,543069241 |
| chrXV | 177121 | 177238 | 117 | 0,039893848 | 0,114881957 | 0,257752484 | -0,65028992 | -0,552746432 | 0,40924629 | 0,59824629 | -0,66175371 |
| chrXV | 177124 | 177238 | 114 | 0,085486818 | 0,185293479 | 0,315705458 | -0,47974192 | -0,407780632 | 0,583965289 | 0,793965289 | -0,466034711 |
| chrXV | 177125 | 177238 | 113 | 0,054141651 | 0,203822827 | 0,209880258 | -0,806836798 | -0,685811279 | 0,322763491 | 0,539763491 | -0,720236509 |
| chrXV | 177128 | 177238 | 110 | 0,039893848 | 0,163058261 | 0,196567792 | -0,853944782 | -0,725853065 | 0,273640298 | 0,511640298 | -0,748359702 |
| chrXV | 177130 | 177238 | 108 | 0,028495606 | 0,059293913 | 0,324590066 | -0,454901459 | -0,386666241 | 0,56013891 | 0,81213891 | -0,44786109 |
| chrXV | 177132 | 177217 | 85 | 0,125380666 | 0,163058261 | 0,434687048 | -0,164453569 | -0,139785534 | 0,451462324 | 0,864462324 | -0,395537676 |
| chrXV | 177132 | 177238 | 106 | 0,031345166 | 0,570703915 | 0,052064138 | -1,625160916 | -1,381386779 | -0,375321852 | -0,109321852 | -1,369321852 |
| chrXV | 177733 | 177893 | 160 | 0,3903898 | 0,359469349 | 0,520617506 | 0,05170345 | 0,043947932 | 0,093220704 | -0,018779296 | -1,278779296 |
| chrXV | 178523 | 178704 | 181 | 0,068389454 | 0,163058261 | 0,295485544 | -0,537429332 | -0,456814932 | -0,0832499 | -0,3422499 | -1,6022499 |
| chrXV | 185749 | 185889 | 140 | 0,199469241 | 0,226058044 | 0,468757816 | -0,078392761 | -0,066633846 | -0,004430251 | 0,023569749 | -1,236430251 |
| chrXV | 187300 | 187456 | 156 | 0,316301225 | 0,363175219 | 0,465507271 | -0,086568454 | -0,073583186 | -0,043095032 | -0,127095032 | -1,387095032 |
| chrXV | 189704 | 189862 | 158 | 0,056991212 | 0,070411522 | 0,447331152 | -0,132407093 | -0,112546029 | -0,076006571 | -0,174006571 | -1,434006571 |
| chrXV | 192332 | 192491 | 159 | 0,162424953 | 0,129705435 | 0,556001565 | 0,140839333 | 0,119713433 | 0,15996104 | 0,05496104 | -1,20503896 |
| chrXV | 196562 | 196707 | 145 | 0,09688506 | 0,096352609 | 0,50137771 | 0,003453413 | 0,002935401 | -0,020089093 | -0,027089093 | -1,287089093 |
| chrXV | 196562 | 196707 | 145 | 0,09688506 | 0,096352609 | 0,50137771 | 0,003453413 | 0,002935401 | -0,020089093 | -0,027089093 | -1,287089093 |
| chrXV | 198924 | 199072 | 148 | 0,983098401 | 0,548468698 | 0,64189052 | 0,363516673 | 0,308989172 | 0,284769191 | 0,256769191 | -1,003230809 |
| chrXV | 202849 | 203002 | 153 | 0,065539893 | 0,066705652 | 0,495592445 | -0,011048326 | -0,009391077 | -0,089861553 | -0,152861553 | -1,412861553 |
| chrXV | 205781 | 205911 | 130 | 0,621204207 | 0,767115003 | 0,447450559 | -0,132105155 | -0,112289381 | 0,18829822 | 0,28629822 | -0,97370178 |
| chrXV | 208753 | 208900 | 147 | 0,621204207 | 0,455821958 | 0,576777266 | 0,193655717 | 0,16460736 | 0,243672574 | 0,222672574 | -1,037327426 |
| chrXV | 209390 | 209535 | 145 | 0,056991212 | 0,051882174 | 0,523463207 | 0,058847485 | 0,050020363 | 0,045755242 | 0,038755242 | -1,221244758 |
| chrXV | 212639 | 212808 | 169 | 0,176672756 | 0,26682261 | 0,398364379 | -0,25758299 | -0,218945542 | -0,075529614 | -0,250529614 | -1,510529614 |
| chrXV | 214778 | 214916 | 138 | 2,054533181 | 1,045055221 | 0,662840647 | 0,420228267 | 0,357194027 | 0,493612706 | 0,535612706 | -0,724387294 |
| chrXV | 215741 | 215894 | 153 | 0,547115632 | 0,500292393 | 0,522351957 | 0,056057394 | 0,047648785 | -0,030123569 | -0,093123569 | -1,353123569 |
| chrXV | 218336 | 218507 | 171 | 0,065539893 | 0,081529131 | 0,445640364 | -0,13668381 | -0,116181238 | 0,043004388 | -0,145995612 | -1,405995612 |
| chrXV | 220428 | 220557 | 129 | 2,627294858 | 5,080747192 | 0,340851132 | -0,410141347 | -0,348620145 | 0,029893116 | 0,134893116 | -1,125106884 |
| chrXV | 222406 | 222534 | 128 | 0,188070999 | 0,522527611 | 0,264665588 | -0,629027313 | -0,534673216 | -0,039436268 | 0,072563732 | -1,187436268 |
| chrXV | 225244 | 225389 | 145 | 10,69155131 | 6,285154805 | 0,629777722 | 0,331264697 | 0,281574992 | 0,279444375 | 0,272444375 | -0,987555625 |
| chrXV | 227790 | 227919 | 129 | 0,082637257 | 0,214940436 | 0,277699771 | -0,589688448 | -0,501235181 | -0,115952598 | -0,010952598 | -1,270952598 |
| chrXV | 229356 | 229535 | 179 | 0,042743409 | 0,077823261 | 0,354520937 | -0,373143192 | -0,317171714 | 0,042016297 | -0,202983703 | -1,462983703 |
| chrXV | 229736 | 229903 | 167 | 0,062690333 | 0,137117174 | 0,313753641 | -0,48523835 | -0,412452597 | -0,264697167 | -0,425697167 | -1,685697167 |
| chrXV | 233701 | 233860 | 159 | 0,173823196 | 0,129705435 | 0,572674792 | 0,18318812 | 0,155709902 | 0,200548692 | 0,095548692 | -1,164451308 |
| chrXV | 233701 | 233860 | 159 | 0,173823196 | 0,129705435 | 0,572674792 | 0,18318812 | 0,155709902 | 0,200548692 | 0,095548692 | -1,164451308 |
| chrXV | 237293 | 237436 | 143 | 0,133929347 | 0,155646522 | 0,462501753 | -0,094133 | -0,08001305 | -0,039865768 | -0,032865768 | -1,292865768 |
| chrXV | 242082 | 242240 | 158 | 0,125380666 | 0,140823044 | 0,470995186 | -0,072768458 | -0,061853189 | -0,032671127 | -0,130671127 | -1,390671127 |
| chrXV | 242082 | 242240 | 158 | 0,125380666 | 0,140823044 | 0,470995186 | -0,072768458 | -0,061853189 | -0,032671127 | -0,130671127 | -1,390671127 |
| chrXV | 243486 | 243636 | 150 | 0,510071344 | 0,248293262 | 0,672593816 | 0,44708683 | 0,380023805 | 0,29006655 | 0,24806655 | -1,01193345 |
| chrXV | 244606 | 244758 | 152 | 0,108283302 | 0,137117174 | 0,441251393 | -0,147797242 | -0,125627655 | -0,219663473 | -0,275663473 | -1,535663473 |
| chrXV | 244606 | 244758 | 152 | 0,108283302 | 0,137117174 | 0,441251393 | -0,147797242 | -0,125627655 | -0,219663473 | -0,275663473 | -1,535663473 |
| chrXV | 248190 | 248334 | 144 | 0,116831984 | 0,166764131 | 0,411966095 | -0,222490344 | -0,189116792 | -0,245492756 | -0,245492756 | -1,505492756 |
| chrXV | 248556 | 248722 | 166 | 0,883363781 | 1,263701526 | 0,411428464 | -0,223871969 | -0,190291174 | -0,061566659 | -0,215566659 | -1,475566659 |
| chrXV | 255239 | 255421 | 182 | 0,085486818 | 0,100058479 | 0,460732874 | -0,09858756 | -0,083799426 | 0,31879579 | 0,05279579 | -1,20720421 |
| chrXV | 256726 | 256872 | 146 | 1,336443914 | 1,196995874 | 0,527521483 | 0,069040937 | 0,058684797 | 0,123412242 | 0,109412242 | -1,150587758 |
| chrXV | 260668 | 260810 | 142 | 0,059840772 | 0,092646739 | 0,392430643 | -0,272989502 | -0,232041076 | -0,228576099 | -0,214576099 | -1,474576099 |
| chrXV | 261423 | 261595 | 172 | 0,53856695 | 0,796761959 | 0,403321568 | -0,244758877 | -0,208045045 | 0,020890716 | -0,175109284 | -1,435109284 |
| chrXV | 263580 | 263731 | 151 | 0,14817715 | 0,048176305 | 0,754644987 | 0,68917996 | 0,585802966 | 0,488547783 | 0,439547783 | -0,820452217 |
| chrXV | 265679 | 265840 | 161 | 0,151026711 | 0,159352392 | 0,486587884 | -0,033625524 | -0,028581696 | 0,040096818 | -0,078903182 | -1,338903182 |
| chrXV | 266859 | 267005 | 146 | 0,039893848 | 0,066705652 | 0,374240479 | -0,320642986 | -0,272546538 | -0,196278943 | -0,210278943 | -1,470278943 |
| chrXV | 268238 | 268379 | 141 | 0,661098055 | 0,533645219 | 0,553339005 | 0,13410191 | 0,113986624 | 0,1830404 | 0,2040404 | -1,0559596 |
| chrXV | 270968 | 271134 | 166 | 0,296354301 | 0,233469783 | 0,559344714 | 0,149308039 | 0,126911833 | 0,253677468 | 0,099677468 | -1,160322532 |
| chrXV | 271150 | 271308 | 158 | 0,079787696 | 0,085235 | 0,483495289 | -0,041382985 | -0,035175537 | 0,008695488 | -0,089304512 | -1,349304512 |
| chrXV | 273050 | 273212 | 162 | 0,076938136 | 0,118587827 | 0,393493196 | -0,270226 | -0,2296921 | -0,16451261 | -0,29051261 | -1,55051261 |
| chrXV | 276074 | 276211 | 137 | 0,869115978 | 0,385410436 | 0,69278412 | 0,503757552 | 0,428193919 | 0,606255311 | 0,655255311 | -0,604744689 |
| chrXV | 276074 | 276211 | 137 | 0,869115978 | 0,385410436 | 0,69278412 | 0,503757552 | 0,428193919 | 0,606255311 | 0,655255311 | -0,604744689 |
| chrXV | 276839 | 276983 | 144 | 0,051292091 | 0,062999783 | 0,448781606 | -0,128740216 | -0,109429183 | -0,159903964 | -0,159903964 | -1,419903964 |
| chrXV | 279591 | 279744 | 153 | 0,322000346 | 0,311293045 | 0,508453666 | 0,021191784 | 0,018013016 | -0,0644379 | -0,1274379 | -1,3874379 |
| chrXV | 280089 | 280217 | 128 | 0,492973981 | 0,622586089 | 0,441907159 | -0,146135633 | -0,124215288 | 0,358790379 | 0,470790379 | -0,789209621 |
| chrXV | 282501 | 282650 | 149 | 0,678195419 | 0,374292827 | 0,64437339 | 0,3701735 | 0,314647475 | 0,278839225 | 0,243839225 | -1,016160775 |
| chrXV | 284484 | 284582 | 98 | 0,555664314 | 6,737270894 | 0,076192136 | -1,431160321 | -1,216486272 | -0,304739685 | 0,017260315 | -1,242739685 |
| chrXV | 284650 | 284789 | 139 | 0,190920559 | 0,122293696 | 0,609552586 | 0,278153109 | 0,236430142 | 0,338870367 | 0,373870367 | -0,886129633 |
| chrXV | 284999 | 285154 | 155 | 0,165274514 | 0,100058479 | 0,622894697 | 0,313092211 | 0,266128379 | 0,26726287 | 0,19026287 | -1,06973713 |

TABLE 2

| Chrom | Start | End | Length | Section A | Section B | A/A+B | Z-score | Z * 0.85 | Phase correction | Length correction | ΔLK |
|---|---|---|---|---|---|---|---|---|---|---|---|
| chrXV | 285195 | 285326 | 131 | 1,544461836 | 1,704700006 | 0,475341615 | -0,061848815 | -0,052571493 | 0,189635695 | 0,280635695 | -0,979364305 |
| chrXV | 292850 | 293005 | 155 | 0,63545201 | 0,489174784 | 0,565033674 | 0,163744034 | 0,139182429 | 0,137803273 | 0,060803273 | -1,199196727 |
| chrXV | 292905 | 293050 | 145 | 0,299203861 | 0,218646305 | 0,577780757 | 0,196219347 | 0,166786445 | 0,203319709 | 0,196319709 | -1,063680291 |
| chrXV | 293213 | 293372 | 159 | 0,159575393 | 0,155646522 | 0,506231912 | 0,015621722 | 0,013278464 | 0,062555593 | -0,042444407 | -1,302444407 |
| chrXV | 294193 | 294362 | 169 | 0,202318801 | 0,181587609 | 0,52700032 | 0,067731518 | 0,05757179 | 0,193645257 | 0,018645257 | -1,241354743 |
| chrXV | 298993 | 299142 | 149 | 0,393239361 | 0,300175436 | 0,567105523 | 0,169009772 | 0,143658307 | 0,100227885 | 0,065227885 | -1,194772115 |
| chrXV | 306450 | 306602 | 152 | 0,225115286 | 0,192705218 | 0,538784679 | 0,097372425 | 0,082766561 | -0,01699903 | -0,07299903 | -1,33299903 |
| chrXV | 311787 | 311943 | 156 | 0,598407723 | 0,692997611 | 0,46337715 | -0,09192919 | -0,078139812 | -0,046667975 | -0,130667975 | -1,390667975 |
| chrXV | 319552 | 319692 | 140 | 3,499260397 | 1,84922892 | 0,654252106 | 0,396825983 | 0,337302085 | 0,3905154 | 0,4185154 | -0,8414846 |
| chrXV | 320812 | 320963 | 151 | 0,14532759 | 0,077823261 | 0,65125268 | 0,388704654 | 0,330398956 | 0,242395884 | 0,193395884 | -1,066604116 |
| chrXV | 321565 | 321713 | 148 | 0,111132863 | 0,092646739 | 0,54535813 | 0,113942039 | 0,096850733 | 0,060787195 | 0,032787195 | -1,227212805 |
| chrXV | 327427 | 327560 | 133 | 1,396284686 | 1,389701092 | 0,501181556 | 0,002961725 | 0,002517466 | 0,268839498 | 0,345839498 | -0,914160502 |
| chrXV | 328438 | 328583 | 145 | 1,290850944 | 0,78935022 | 0,620541401 | 0,306903009 | 0,260867558 | 0,294486284 | 0,287486284 | -0,972513716 |
| chrXV | 328438 | 328583 | 145 | 1,290850944 | 0,78935022 | 0,620541401 | 0,306903009 | 0,260867558 | 0,294486284 | 0,287486284 | -0,972513716 |
| chrXV | 329812 | 329980 | 168 | 0,074088575 | 0,066705652 | 0,52621884 | 0,065768269 | 0,055903029 | 0,222895876 | 0,054895876 | -1,205104124 |
| chrXV | 338001 | 338161 | 160 | 0,549965193 | 0,589233263 | 0,482765044 | -0,043215075 | -0,036732814 | 0,020786649 | -0,091213351 | -1,351213351 |
| chrXV | 341700 | 341803 | 103 | 0,398938482 | 5,651451107 | 0,065935999 | -1,506760733 | -1,280746623 | -0,298077672 | -0,011077672 | -1,271077672 |
| chrXV | 348865 | 348972 | 107 | 0,68389454 | 0,696703481 | 0,49536109 | -0,011628286 | -0,009884043 | 0,960124375 | 1,219124375 | -0,040875625 |
| chrXV | 349258 | 349411 | 153 | 0,287805619 | 0,229763914 | 0,556071408 | 0,14101615 | 0,119863728 | 0,03726466 | -0,02573534 | -1,28573534 |
| chrXV | 351942 | 352092 | 150 | 0,284956058 | 0,140823044 | 0,669257972 | 0,437865126 | 0,372185358 | 0,27339093 | 0,23139093 | -1,02860907 |
| chrXV | 353769 | 353922 | 153 | 0,028495606 | 0,040764565 | 0,411428464 | -0,223871969 | -0,190291174 | -0,287468578 | -0,350468578 | -1,610468578 |
| chrXV | 356685 | 356815 | 130 | 2,746976403 | 2,664520227 | 0,507618611 | 0,019098187 | 0,016233459 | 0,302677861 | 0,400677861 | -0,859322139 |
| chrXV | 357881 | 358029 | 148 | 1,114178188 | 0,941290873 | 0,542055441 | 0,105613368 | 0,089771362 | 0,06041723 | 0,03241723 | -1,22758277 |
| chrXV | 359149 | 359277 | 128 | 0,407487163 | 4,076456536 | 0,090876958 | -1,335374165 | -1,13506804 | -0,638539428 | -0,526539428 | -1,786539428 |
| chrXV | 359963 | 360110 | 147 | 0,031345166 | 0,074117392 | 0,297216064 | -0,532424343 | -0,452560692 | -0,371893437 | -0,392893437 | -1,652893437 |
| chrXV | 363241 | 363404 | 163 | 0,125380666 | 0,129705435 | 0,491522922 | -0,021250483 | -0,01806291 | 0,052851659 | -0,080148341 | -1,340148341 |
| chrXV | 363647 | 363809 | 162 | 0,179522317 | 0,188999349 | 0,487141826 | -0,032236246 | -0,027400809 | 0,036861631 | -0,089138369 | -1,349138369 |
| chrXV | 365392 | 365542 | 150 | 1,401983807 | 1,019114134 | 0,579069431 | 0,199513431 | 0,169586416 | 0,063091774 | 0,021091774 | -1,238908226 |
| chrXV | 365392 | 365542 | 150 | 1,401983807 | 1,019114134 | 0,579069431 | 0,199513431 | 0,169586416 | 0,063091774 | 0,021091774 | -1,238908226 |
| chrXV | 369266 | 369425 | 159 | 0,287805619 | 0,214940436 | 0,572467186 | 0,182658948 | 0,155260106 | 0,208237371 | 0,103237371 | -1,156762629 |
| chrXV | 369347 | 369504 | 157 | 0,039893848 | 0,077823261 | 0,338895921 | -0,415478239 | -0,353156503 | -0,321975616 | -0,412975616 | -1,672975616 |
| chrXV | 371267 | 371398 | 131 | 0,068389454 | 0,040764565 | 0,626540868 | 0,322705529 | 0,274299699 | 0,48437371 | 0,57537371 | -0,68462629 |
| chrXV | 371602 | 371749 | 147 | 0,418885406 | 0,229763914 | 0,645780999 | 0,373954724 | 0,317861515 | 0,424600295 | 0,403600295 | -0,856399705 |
| chrXV | 377796 | 377936 | 140 | 0,404637603 | 0,274234349 | 0,596044073 | 0,243120753 | 0,20665264 | 0,288888023 | 0,316888023 | -0,943111977 |
| chrXV | 384219 | 384368 | 149 | 0,983098401 | 0,830114786 | 0,542185777 | 0,105941906 | 0,09005062 | 0,050237089 | 0,015237089 | -1,244762911 |
| chrXV | 387959 | 388110 | 151 | 0,034194727 | 0,055588044 | 0,380860679 | -0,303221116 | -0,257737948 | -0,344310674 | -0,393310674 | -1,653310674 |
| chrXV | 389816 | 389942 | 126 | 0,128230226 | 0,226058044 | 0,361937543 | -0,353284604 | -0,300291913 | 0,316521175 | 0,442521175 | -0,817478825 |
| chrXV | 392262 | 392373 | 111 | 0,609805965 | 4,421102407 | 0,1212119 | -1,16894994 | -0,993607449 | 0,013359791 | 0,244359791 | -1,015640209 |
| chrXV | 392576 | 392737 | 161 | 0,122531105 | 0,17417587 | 0,412970086 | -0,219911382 | -0,186924674 | -0,114504576 | -0,233504576 | -1,493504576 |
| chrXV | 401254 | 401398 | 144 | 0,122531105 | 0,151940653 | 0,446425184 | -0,134698361 | -0,114493607 | -0,156581892 | -0,156581892 | -1,416581892 |
| chrXV | 403569 | 403709 | 140 | 0,461628815 | 0,240881523 | 0,657113198 | 0,404597219 | 0,343907636 | 0,440630931 | 0,468630931 | -0,791369069 |
| chrXV | 405315 | 405460 | 145 | 0,250761331 | 0,170470001 | 0,595305506 | 0,241214353 | 0,2050322 | 0,240115036 | 0,233115036 | -1,026884964 |
| chrXV | 405735 | 405887 | 152 | 1,798072728 | 1,426759788 | 0,557570888 | 0,144813371 | 0,123091365 | 0,021128945 | -0,034871055 | -1,294871055 |
| chrXV | 408309 | 408462 | 153 | 0,433133209 | 0,289057827 | 0,599748802 | 0,252696962 | 0,214792418 | 0,110242434 | 0,047242434 | -1,212757566 |
| chrXV | 413175 | 413325 | 150 | 0,755133555 | 0,670762394 | 0,529585315 | 0,074227492 | 0,063093368 | -0,053282683 | -0,095282683 | -1,355282683 |
| chrXV | 413525 | 413650 | 125 | 0,749434433 | 1,311877831 | 0,36357152 | -0,348928431 | -0,296589166 | 0,396959498 | 0,529959498 | -0,730040502 |
| chrXV | 419166 | 419317 | 151 | 0,168124074 | 0,122293696 | 0,578904225 | 0,199091015 | 0,169227363 | 0,075337828 | 0,026337828 | -1,233662172 |
| chrXV | 420271 | 420429 | 158 | 0,410336724 | 0,403939784 | 0,50392799 | 0,00984617 | 0,008369244 | 0,051837202 | -0,046162798 | -1,306162798 |
| chrXV | 423240 | 423391 | 151 | 19,55368472 | 10,35049373 | 0,653878011 | 0,395811655 | 0,336439906 | 0,248290539 | 0,199290539 | -1,060709461 |
| chrXV | 423241 | 423391 | 150 | 0,037044288 | 0,074117392 | 0,333246923 | -0,430964963 | -0,366320218 | -0,495723725 | -0,537723725 | -1,797723725 |
| chrXV | 423242 | 423391 | 149 | 0,039893848 | 0,092646739 | 0,300993446 | -0,521545394 | -0,443313585 | -0,500105226 | -0,535105226 | -1,795105226 |
| chrXV | 423243 | 423391 | 148 | 0,059840772 | 0,059293913 | 0,502295129 | 0,005753068 | 0,004890107 | -0,03134863 | -0,05934863 | -1,31934863 |
| chrXV | 423245 | 423391 | 146 | 0,065539893 | 0,066705652 | 0,495592445 | -0,011048326 | -0,009391077 | 0,063539942 | 0,049539942 | -1,210460058 |
| chrXV | 433044 | 433202 | 158 | 0,173823196 | 0,155646522 | 0,52758474 | 0,069199879 | 0,058819897 | 0,095857429 | -0,002142571 | -1,262142571 |
| chrXV | 433171 | 433320 | 149 | 0,19377012 | 0,133411305 | 0,592240589 | 0,233312412 | 0,19831555 | 0,131699167 | 0,096699167 | -1,163300833 |
| chrXV | 433171 | 433320 | 149 | 0,19377012 | 0,133411305 | 0,592240589 | 0,233312412 | 0,19831555 | 0,131699167 | 0,096699167 | -1,163300833 |
| chrXV | 435682 | 435849 | 167 | 0,128230226 | 0,125999566 | 0,504387095 | 0,010997038 | 0,009347483 | 0,166480869 | 0,005480869 | -1,254519131 |
| chrXV | 441363 | 441514 | 151 | 0,170973635 | 0,129705435 | 0,568624996 | 0,172874585 | 0,146943398 | 0,052939266 | 0,003939266 | -1,256060734 |
| chrXV | 443421 | 443578 | 157 | 0,04844253 | 0,051882174 | 0,482857442 | -0,042983253 | -0,036535765 | -0,007522809 | -0,098522809 | -1,358522809 |
| chrXV | 445187 | 445337 | 150 | 0,515770466 | 0,292763697 | 0,637908068 | 0,35287272 | 0,299941812 | 0,180660015 | 0,138660015 | -1,121339985 |
| chrXV | 445187 | 445337 | 150 | 0,515770466 | 0,292763697 | 0,637908068 | 0,35287272 | 0,299941812 | 0,180660015 | 0,138660015 | -1,121339985 |
| chrXV | 446874 | 447012 | 138 | 0,188070999 | 0,092646739 | 0,669964783 | 0,439815924 | 0,373843536 | 0,519779987 | 0,561779987 | -0,698220013 |
| chrXV | 450011 | 450165 | 154 | 0,225115286 | 0,192705218 | 0,538784679 | 0,097372425 | 0,082766561 | 0,017678533 | -0,052321467 | -1,312321467 |
| chrXV | 456168 | 456329 | 161 | 0,102584181 | 0,062999783 | 0,619529685 | 0,304245806 | 0,258608935 | 0,33213935 | 0,21313935 | -1,04686065 |
| chrXV | 457433 | 457588 | 155 | 8,978965399 | 6,303684153 | 0,587526748 | 0,221187421 | 0,188009308 | 0,180384098 | 0,103384098 | -1,156615902 |
| chrXV | 457539 | 457691 | 152 | 0,903310705 | 0,655938915 | 0,579323986 | 0,200164374 | 0,170139718 | 0,069901017 | 0,013901017 | -1,246098983 |
| chrXV | 459473 | 459604 | 131 | 0,262159574 | 0,563292176 | 0,317595273 | -0,474433869 | -0,403268789 | -0,193129338 | -0,102129338 | -1,362129338 |
| chrXV | 465612 | 465776 | 164 | 0,809275206 | 1,059878699 | 0,432963387 | -0,168834559 | -0,143509375 | -0,06017332 | -0,20017332 | -1,46017332 |
| chrXV | 469152 | 469262 | 110 | 0,079787696 | 0,207528696 | 0,277699771 | -0,589688448 | -0,501235181 | 0,486351268 | 0,724351268 | -0,535648732 |
| chrXV | 469152 | 469277 | 125 | 0,153876272 | 0,318704784 | 0,32560821 | -0,452072964 | -0,384262019 | 0,291475955 | 0,424475955 | -0,835524045 |
| chrXV | 480195 | 480360 | 165 | 5,100713444 | 5,395746106 | 0,485946087 | -0,035235226 | -0,029949942 | 0,050810446 | -0,096189554 | -1,356189554 |
| chrXV | 480209 | 480363 | 154 | 0,396088921 | 0,244587392 | 0,618235625 | 0,30085017 | 0,255722644 | 0,197348875 | 0,127348875 | -1,132651125 |
| chrXV | 480628 | 480755 | 127 | 0,045592969 | 0,96352609 | 0,045180961 | -1,69349165 | -1,439467903 | -0,906102363 | -0,787102363 | -2,047102363 |
| chrXV | 480628 | 480787 | 159 | 0,760832676 | 0,915349786 | 0,453908028 | -0,115793684 | -0,098424632 | -0,043184776 | -0,148184776 | -1,408184776 |
| chrXV | 480629 | 480736 | 107 | 0,04844253 | 1,511994788 | 0,031044201 | -1,865663908 | -1,585814322 | -0,640834775 | -0,381834775 | -1,641834775 |
| chrXV | 480629 | 480737 | 108 | 0,051292091 | 1,708405876 | 0,029148235 | -1,893461897 | -1,609442613 | -0,644642803 | -0,392642803 | -1,652642803 |
| chrXV | 480629 | 480738 | 109 | 0,111132863 | 1,541641745 | 0,06724018 | -1,496665334 | -1,272165534 | -0,258939347 | -0,013939347 | -1,273939347 |
| chrXV | 480629 | 480739 | 110 | 0,085486818 | 1,019114134 | 0,077391584 | -1,422837843 | -1,209412166 | -0,193774507 | 0,044225493 | -1,215774507 |
| chrXV | 480629 | 480741 | 112 | 0,059840772 | 2,927636967 | 0,020030533 | -2,053118702 | -1,745150897 | -0,751101329 | -0,527101329 | -1,787101329 |
| chrXV | 480629 | 480742 | 113 | 0,028495606 | 3,012871967 | 0,00936934 | -2,350689182 | -1,998085805 | -0,990901057 | -0,773901057 | -2,033901057 |
| chrXV | 480629 | 480743 | 114 | 0,042743409 | 1,560171093 | 0,026666057 | -1,932221982 | -1,642388685 | -0,652846269 | -0,442846269 | -1,702846269 |
| chrXV | 480629 | 480744 | 115 | 0,037044288 | 1,471230223 | 0,024560707 | -1,96753632 | -1,672405872 | -0,707861837 | -0,504861837 | -1,764861837 |
| chrXV | 480629 | 480745 | 116 | 0,045592969 | 0,300175436 | 0,131859848 | -1,117642532 | -0,949996152 | 0,005052415 | 0,201052415 | -1,058947585 |
| chrXV | 480629 | 480746 | 117 | 0,062690333 | 0,622586089 | 0,091481818 | -1,331685257 | -1,131932469 | -0,170355473 | 0,018644527 | -1,241355473 |
| chrXV | 480629 | 480747 | 118 | 0,039893848 | 0,444704349 | 0,082323559 | -1,389610724 | -1,181169116 | -0,219879372 | -0,037879372 | -1,297879372 |
| chrXV | 480629 | 480748 | 119 | 0,045592969 | 1,304466092 | 0,033771092 | -1,828049088 | -1,553841725 | -0,629200179 | -0,454200179 | -1,714200179 |
| chrXV | 480629 | 480749 | 120 | 0,128230226 | 1,274819135 | 0,091393952 | -1,332220006 | -1,132387005 | -0,255145544 | -0,087145544 | -1,347145544 |
| chrXV | 480629 | 480750 | 121 | 0,082637257 | 1,574994571 | 0,0498526 | -1,646284493 | -1,399341819 | -0,627655263 | -0,466655263 | -1,726655263 |
| chrXV | 480629 | 480751 | 122 | 0,076938136 | 0,303881305 | 0,202033109 | -0,834381181 | -0,709224004 | 0,050264393 | 0,204264393 | -1,055735607 |
| chrXV | 480629 | 480752 | 123 | 0,225115286 | 1,641700223 | 0,120587859 | -1,172053148 | -0,996245176 | -0,251898698 | -0,104898698 | -1,364898698 |
| chrXV | 480629 | 480753 | 124 | 0,208017923 | 0,415057393 | 0,333856787 | -0,429288104 | -0,364894888 | 0,338346895 | 0,478346895 | -0,781653105 |
| chrXV | 480629 | 480754 | 125 | 0,227964847 | 1,000584786 | 0,185556074 | -0,894392081 | -0,760233269 | -0,092940328 | 0,040059672 | -1,219940328 |
| chrXV | 480629 | 480755 | 126 | 0,344796831 | 19,20011029 | 0,017641262 | -2,105100918 | -1,78933578 | -1,197802401 | -1,071802401 | -2,331802401 |
| chrXV | 480629 | 480756 | 127 | 0,14817715 | 1,14140783 | 0,114902975 | -1,200858875 | -1,020730044 | -0,50653947 | -0,38753947 | -1,64753947 |
| chrXV | 480629 | 480757 | 128 | 0,188070999 | 0,359469349 | 0,34348336 | -0,402974858 | -0,342528629 | 0,155629347 | 0,267629347 | -0,992370653 |
| chrXV | 480629 | 480758 | 129 | 0,287805619 | 0,778232612 | 0,269976827 | -0,612883077 | -0,520950615 | -0,148404181 | -0,043404181 | -1,303404181 |
| chrXV | 480629 | 480759 | 130 | 0,256460453 | 0,88199696 | 0,225270133 | -0,754514627 | -0,641337433 | -0,364067322 | -0,266067322 | -1,526067322 |
| chrXV | 480629 | 480760 | 131 | 0,270708255 | 1,274819135 | 0,175155909 | -0,933984635 | -0,79388694 | -0,556088555 | -0,465088555 | -1,725088555 |
| chrXV | 480629 | 480761 | 132 | 0,532867829 | 1,019114134 | 0,343346664 | -0,403346515 | -0,342844538 | -0,084374692 | -0,000374692 | -1,260374692 |
| chrXV | 480629 | 480762 | 133 | 0,327699467 | 1,771405659 | 0,15611389 | -1,010558514 | -0,858974737 | -0,594442909 | -0,517442909 | -1,777442909 |
| chrXV | 480629 | 480763 | 134 | 0,367593315 | 2,734931749 | 0,118481981 | -1,18260946 | -1,005218041 | -0,702204097 | -0,632204097 | -1,892204097 |

TABLE 2

| Chrom | Start | End | Length | Section A | Section B | A/A+B | Z-score | Z * 0.85 | Phase correction | Length correction | ΔLK |
|---|---|---|---|---|---|---|---|---|---|---|---|
| chrXV | 480629 | 480764 | 135 | 0,227964847 | 1,76028805 | 0,11465586 | -1,202133727 | -1,021813668 | -0,741855951 | -0,678855951 | **-1,938855951** |
| chrXV | 480629 | 480765 | 136 | 0,216566604 | 1,637994354 | 0,116775134 | -1,19126322 | -1,012573737 | -0,803405819 | -0,747405819 | **-2,007405819** |
| chrXV | 480629 | 480766 | 137 | 0,176672756 | 1,300760222 | 0,119580894 | -1,177084483 | -1,000521811 | -0,823103766 | -0,774103766 | **-2,034103766** |
| chrXV | 480629 | 480767 | 138 | 0,44168189 | 0,863467612 | 0,338414787 | -0,416793341 | -0,35427434 | -0,215521314 | -0,173521314 | **-1,433521314** |
| chrXV | 480629 | 480768 | 139 | 0,19377012 | 1,004290656 | 0,161736469 | -0,987346222 | -0,839244289 | -0,729292923 | -0,694292923 | **-1,954292923** |
| chrXV | 480629 | 480769 | 140 | 0,44168189 | 1,952993268 | 0,184443343 | -0,898560728 | -0,763776618 | -0,675043766 | -0,647043766 | **-1,907043766** |
| chrXV | 480629 | 480770 | 141 | 0,29065518 | 0,870879351 | 0,250233783 | -0,673754249 | -0,572691112 | -0,489509808 | -0,468509808 | **-1,728509808** |
| chrXV | 480629 | 480771 | 142 | 0,19377012 | 2,012287181 | 0,087835488 | -1,354205026 | -1,151074272 | -1,150375054 | -1,136375054 | **-2,396375054** |
| chrXV | 480629 | 480772 | 143 | 0,24506221 | 1,137701961 | 0,177226323 | -0,925987172 | -0,787089096 | -0,750456916 | -0,743456916 | **-2,003456916** |
| chrXV | 480629 | 480773 | 144 | 0,267858695 | 1,070996308 | 0,200065499 | -0,841387298 | -0,715179204 | -0,753460411 | -0,753460411 | **-2,013460411** |
| chrXV | 480629 | 480774 | 145 | 0,182371877 | 0,411351523 | 0,307166396 | -0,503898375 | -0,428313619 | -0,377022048 | -0,384022048 | **-1,644022048** |
| chrXV | 480629 | 480775 | 146 | 0,202318801 | 4,613807625 | 0,042008615 | -1,727838236 | -1,4686625 | -1,397942306 | -1,411942306 | **-2,671942306** |
| chrXV | 480629 | 480776 | 147 | 0,324849907 | 0,359469349 | 0,474705196 | -0,063447213 | -0,053930131 | 0,053409465 | 0,032409465 | **-1,227590535** |
| chrXV | 480629 | 480777 | 148 | 0,527168708 | 1,126584352 | 0,318771116 | -0,471137942 | -0,400467251 | -0,438506313 | -0,466506313 | **-1,726506313** |
| chrXV | 480629 | 480778 | 149 | 0,287805619 | 0,622586089 | 0,316133832 | -0,478537538 | -0,406756908 | -0,481913413 | -0,516913413 | **-1,776913413** |
| chrXV | 480629 | 480779 | 150 | 0,624053768 | 0,518821741 | 0,546038272 | 0,115658174 | 0,098309448 | -0,030352265 | -0,072352265 | **-1,332352265** |
| chrXV | 480629 | 480780 | 151 | 0,544266071 | 0,796761959 | 0,405857341 | -0,238214562 | -0,202482378 | -0,29555997 | -0,34455997 | **-1,60455997** |
| chrXV | 480629 | 480781 | 152 | 0,755133555 | 2,212404138 | 0,254464688 | -0,660505676 | -0,561429825 | -0,655562384 | -0,711562384 | **-1,971562384** |
| chrXV | 480629 | 480782 | 153 | 2,989189052 | 4,358102624 | 0,406842301 | -0,235675346 | -0,200324044 | -0,308022204 | -0,371022204 | **-1,631022204** |
| chrXV | 480629 | 480783 | 154 | 3,07752543 | 4,120926971 | 0,427525982 | -0,182676363 | -0,155274908 | -0,207434115 | -0,277434115 | **-1,537434115** |
| chrXV | 480629 | 480784 | 155 | 2,923649159 | 3,757751753 | 0,437580262 | -0,157106997 | -0,133540948 | -0,133269874 | -0,210269874 | **-1,470269874** |
| chrXV | 480629 | 480785 | 156 | 1,812320531 | 4,072750667 | 0,307952184 | -0,501663322 | -0,426413824 | -0,395375556 | -0,479375556 | **-1,739375556** |
| chrXV | 480629 | 480786 | 157 | 1,843665698 | 3,009166098 | 0,379915434 | -0,305702896 | -0,259847462 | -0,23498926 | -0,32598926 | **-1,58598926** |
| chrXV | 480629 | 480787 | 158 | 2501,498156 | 1389,156329 | 0,642950477 | 0,366356538 | 0,311403057 | 0,352430347 | 0,254430347 | **-1,005569653** |
| chrXV | 480629 | 480788 | 159 | 0,051292091 | 1,167348917 | 0,042089582 | -1,726935968 | -1,467895573 | -1,414622115 | -1,519622115 | **-2,779622115** |
| chrXV | 480629 | 480789 | 160 | 0,726637949 | 1,159937178 | 0,385162477 | -0,291949869 | -0,248157388 | -0,181781909 | -0,293781909 | **-1,553781909** |
| chrXV | 480630 | 480787 | 157 | 3,901048439 | 4,680513278 | 0,454584907 | -0,114085754 | -0,096972891 | -0,070461779 | -0,161461779 | **-1,421461779** |
| chrXV | 480631 | 480787 | 156 | 3,778517334 | 3,279694577 | 0,53533634 | 0,088691209 | 0,075387528 | 0,103682384 | 0,019682384 | **-1,240317616** |
| chrXV | 480632 | 480787 | 155 | 2,644392221 | 4,524866755 | 0,368851541 | -0,334896607 | -0,284662116 | -0,289346486 | -0,366346486 | **-1,626346486** |
| chrXV | 480633 | 480787 | 154 | 3,456516988 | 3,920810014 | 0,468532436 | -0,078959457 | -0,067115538 | -0,116429767 | -0,186429767 | **-1,446429767** |
| chrXV | 480634 | 480787 | 153 | 3,413773579 | 2,204992399 | 0,607566429 | 0,272981882 | 0,2320346 | 0,128926838 | 0,065926838 | **-1,194073162** |
| chrXV | 480635 | 480787 | 152 | 0,735186631 | 1,852934789 | 0,284061878 | -0,570816915 | -0,485194378 | -0,582436973 | -0,638436973 | **-1,898436973** |
| chrXV | 480636 | 480787 | 151 | 1,057186976 | 1,967816746 | 0,349482868 | -0,386716991 | -0,328709443 | -0,430648531 | -0,479648531 | **-1,739648531** |
| chrXV | 480637 | 480787 | 150 | 1,604302609 | 2,056757616 | 0,438207107 | -0,155516416 | -0,132188953 | -0,270703578 | -0,312703578 | **-1,572703578** |
| chrXV | 480638 | 480787 | 149 | 0,772230918 | 1,341524787 | 0,365335936 | -0,34423193 | -0,292597141 | -0,367297273 | -0,402297273 | **-1,662297273** |
| chrXV | 480639 | 480787 | 148 | 0,940354993 | 0,96352609 | 0,493914773 | -0,015253994 | -0,012965895 | -0,052924261 | -0,080924261 | **-1,340924261** |
| chrXV | 480640 | 480787 | 147 | 0,618354647 | 1,823287833 | 0,253253559 | -0,664286254 | -0,564643316 | -0,45581727 | -0,47681727 | **-1,73681727** |
| chrXV | 480641 | 480787 | 146 | 0,547115632 | 0,337234132 | 0,618664305 | 0,301974647 | 0,25667845 | 0,32907 | 0,31507 | **-0,94493** |
| chrXV | 480642 | 480787 | 145 | 0,672496298 | 1,645406093 | 0,290131414 | -0,553000851 | -0,470050723 | -0,412102069 | -0,419102069 | **-1,679102069** |
| chrXV | 480643 | 480787 | 144 | 0,495823542 | 0,96723196 | 0,338895921 | -0,415478239 | -0,353156503 | -0,399589497 | -0,399589497 | **-1,659589497** |
| chrXV | 480644 | 480787 | 143 | 0,644000692 | 1,834405441 | 0,259844697 | -0,643824268 | -0,547250628 | -0,513861144 | -0,506861144 | **-1,766861144** |
| chrXV | 480645 | 480787 | 142 | 0,336248149 | 0,522527611 | 0,391543596 | -0,275298142 | -0,234003421 | -0,224075214 | -0,210075214 | **-1,470075214** |
| chrXV | 480646 | 480787 | 141 | 0,336248149 | 1,200701743 | 0,21877626 | -0,776332786 | -0,659882868 | -0,576668024 | -0,555668024 | **-1,815668024** |
| chrXV | 480647 | 480787 | 140 | 0,188070999 | 0,314998914 | 0,373846644 | -0,321682433 | -0,273430068 | -0,196032133 | -0,168032133 | **-1,428032133** |
| chrXV | 480648 | 480787 | 139 | 0,302053422 | 1,311877831 | 0,187153834 | -0,888433396 | -0,755168386 | -0,646642097 | -0,611642097 | **-1,871642097** |
| chrXV | 480649 | 480787 | 138 | 0,304902982 | 0,61888022 | 0,330059024 | -0,439750188 | -0,373787659 | -0,227458207 | -0,185458207 | **-1,445458207** |
| chrXV | 480650 | 480787 | 137 | 0,159575393 | 0,496586524 | 0,243195145 | -0,69606154 | -0,591652309 | -0,421534947 | -0,372534947 | **-1,632534947** |
| chrXV | 480651 | 480787 | 136 | 0,168124074 | 2,531108922 | 0,06228587 | -1,535864 | -1,3054844 | -1,104724396 | -1,048724396 | **-2,308724396** |
| chrXV | 480652 | 480787 | 135 | 0,162424953 | 0,107470218 | 0,601807556 | 0,258028531 | 0,219324251 | 0,499161478 | 0,562161478 | **-0,697838522** |
| chrXV | 480653 | 480787 | 134 | 0,333398588 | 1,045055221 | 0,241864171 | -0,700318625 | -0,595270832 | -0,301723761 | -0,231723761 | **-1,491723761** |
| chrXV | 480654 | 480787 | 133 | 0,128230226 | 2,056757616 | 0,058686929 | -1,565892193 | -1,331008364 | -1,046488752 | -0,969488752 | **-2,229488752** |
| chrXV | 480655 | 480787 | 132 | 0,122531105 | 1,059878699 | 0,103628289 | -1,261145089 | -1,071973325 | -0,815583892 | -0,731583892 | **-1,991583892** |
| chrXV | 480656 | 480787 | 131 | 0,359044634 | 0,896820438 | 0,285894275 | -0,565419375 | -0,480606469 | -0,23911072 | -0,14811072 | **-1,40811072** |
| chrXV | 480657 | 480787 | 130 | 0,179522317 | 0,937585003 | 0,160702838 | -0,991573388 | -0,84283738 | -0,571972476 | -0,473972476 | **-1,733972476** |
| chrXV | 480658 | 480787 | 129 | 0,219416165 | 3,124048055 | 0,065625396 | -1,509187816 | -1,282809644 | -0,932844739 | -0,827844739 | **-2,087844739** |
| chrXV | 480659 | 480787 | 128 | 0,173823196 | 0,822703046 | 0,17442912 | -0,93680623 | -0,796285295 | -0,300509558 | -0,188509558 | **-1,448509558** |
| chrXV | 480660 | 480787 | 127 | 0,133929347 | 1,523112397 | 0,080824365 | -1,399547947 | -1,189615755 | -0,652477312 | -0,533477312 | **-1,793477312** |
| chrXV | 480661 | 480787 | 126 | 0,151026711 | 1,66764131 | 0,083042484 | -1,384893716 | -1,177159658 | -0,57978471 | -0,45378471 | **-1,71378471** |
| chrXV | 480662 | 480787 | 125 | 0,316301225 | 1,789935006 | 0,15017367 | -1,035688819 | -0,880335497 | -0,219997903 | -0,086997903 | **-1,346997903** |
| chrXV | 480663 | 480787 | 124 | 0,247911771 | 0,578115654 | 0,300125351 | -0,524040025 | -0,445434021 | 0,259717681 | 0,399717681 | **-0,860282319** |
| chrXV | 480664 | 480787 | 123 | 0,492973981 | 2,253168704 | 0,179515064 | -0,917214731 | -0,779632522 | -0,044819208 | 0,102180792 | **-1,157819208** |
| chrXV | 480665 | 480787 | 122 | 0,208017923 | 3,531693708 | 0,055624054 | -1,592608219 | -1,353716987 | -0,588605681 | -0,434605681 | **-1,694605681** |
| chrXV | 480666 | 480787 | 121 | 0,19661968 | 2,935048706 | 0,062784323 | -1,53181276 | -1,302040846 | -0,51184054 | -0,35084054 | **-1,61084054** |
| chrXV | 480667 | 480787 | 120 | 0,139628469 | 1,57870044 | 0,08125829 | -1,396657535 | -1,187158905 | -0,313153232 | -0,145153232 | **-1,405153232** |
| chrXV | 480668 | 480787 | 119 | 0,182371877 | 1,185878265 | 0,133288404 | -1,11098035 | -0,944333298 | -0,004638303 | 0,170361697 | **-1,089638303** |
| chrXV | 480669 | 480787 | 118 | 0,113982423 | 1,1451137 | 0,090527182 | -1,337515693 | -1,136888339 | -0,194321239 | -0,012321239 | **-1,272321239** |
| chrXV | 480670 | 480787 | 117 | 0,125380666 | 3,16851849 | 0,038064513 | -1,773601889 | -1,507561605 | -0,524672565 | -0,335672565 | **-1,595672565** |
| chrXV | 480671 | 480787 | 116 | 0,151026711 | 0,804173699 | 0,158109973 | -1,002256045 | -0,851917638 | 0,11279301 | 0,30879301 | **-0,95120699** |
| chrXV | 480672 | 480787 | 115 | 0,133929347 | 0,17417587 | 0,434687048 | -0,164453569 | -0,139785534 | 0,802545376 | 1,005545376 | **-0,254454624** |
| chrXV | 480673 | 480787 | 114 | 0,111132863 | 0,985761308 | 0,101315939 | -1,274088996 | -1,082975647 | -0,103879603 | 0,106120397 | **-1,153879603** |
| chrXV | 480674 | 480787 | 113 | 0,131079787 | 1,997463703 | 0,061581916 | -1,541628761 | -1,310384447 | -0,299981955 | -0,082981955 | **-1,342981955** |
| chrXV | 480675 | 480787 | 112 | 0,185221438 | 1,882581746 | 0,089574017 | -1,343382859 | -1,14187543 | -0,13743774 | 0,08656226 | **-1,17343774** |
| chrXV | 480676 | 480787 | 111 | 0,153876272 | 1,015408265 | 0,131598654 | -1,118866003 | -0,951036102 | 0,048771881 | 0,279771881 | **-0,980228119** |
| chrXV | 480677 | 480787 | 110 | 0,108283302 | 0,185293479 | 0,368841506 | -0,334923211 | -0,284684729 | 0,728996613 | 0,966996613 | **-0,293003387** |
| chrXV | 480678 | 480787 | 109 | 0,094035499 | 1,419348049 | 0,062135933 | -1,537087569 | -1,306524434 | -0,286535234 | -0,041535234 | **-1,301535234** |
| chrXV | 480679 | 480787 | 108 | 0,122531105 | 0,604056741 | 0,168639079 | -0,959557122 | -0,815623554 | 0,13240042 | 0,38440042 | **-0,87559958** |
| chrXV | 480680 | 480755 | 75 | 0,589859041 | 0,52623348 | 0,528503712 | 0,071509109 | 0,060782743 | | 0,483 | **-0,777** |
| chrXV | 480680 | 480787 | 107 | 0,176672756 | 5,699627412 | 0,030065305 | -1,879834699 | -1,597859494 | -0,6530568 | -0,3940568 | **-1,6540568** |
| chrXV | 480681 | 480787 | 106 | 0,068389454 | 0,537351089 | 0,112902223 | -1,211237364 | -1,02955176 | -0,034788184 | 0,231211816 | **-1,028788184** |
| chrXV | 480682 | 480787 | 105 | 0,102584181 | 0,811585438 | 0,112215697 | -1,2148288 | -1,03260448 | -0,076120693 | 0,196879307 | **-1,063120693** |
| chrXV | 480683 | 480787 | 104 | 0,059840772 | 0,470645436 | 0,112803634 | -1,211752154 | -1,029989331 | -0,04210638 | 0,23789362 | **-1,02210638** |
| chrXV | 480684 | 480787 | 103 | 0,068389454 | 0,641115437 | 0,096390391 | -1,302396781 | -1,107037264 | -0,115385469 | 0,171614531 | **-1,088385469** |
| chrXV | 480685 | 480787 | 102 | 0,054141651 | 0,926467395 | 0,05521227 | -1,59628786 | -1,356844681 | -0,34604778 | -0,05204778 | **-1,31204778** |
| chrXV | 480686 | 480787 | 101 | 0,094035499 | 2,456991531 | 0,03686182 | -1,788324682 | -1,520075979 | -0,49302532 | -0,19202532 | **-1,45202532** |
| chrXV | 480687 | 480787 | 100 | 0,062690333 | 0,663350655 | 0,086345446 | -1,363608257 | -1,159067019 | -0,153051229 | 0,154948771 | **-1,105051229** |
| chrXV | 480688 | 480787 | 99 | 0,082637257 | 1,374877614 | 0,056697368 | -1,583117279 | -1,345649687 | -0,366677074 | -0,051677074 | **-1,311677074** |
| chrXV | 480689 | 480787 | 98 | 0,045592969 | 0,118587827 | 0,277699771 | -0,589688448 | -0,501235181 | 0,454528451 | 0,776528451 | **-0,483471549** |
| chrXV | 480690 | 480787 | 97 | 0,051292091 | 1,278525005 | 0,038570786 | -1,767517673 | -1,502390022 | -0,604017791 | -0,275017791 | **-1,535017791** |
| chrXV | 480691 | 480787 | 96 | 0,065539893 | 0,796761959 | 0,076005743 | -1,432462555 | -1,217593172 | -0,33716167 | -0,00116167 | **-1,26116167** |
| chrXV | 480692 | 480787 | 95 | 0,056991212 | 1,045055221 | 0,051713984 | -1,628457244 | -1,384188657 | -0,517485007 | -0,174485007 | **-1,434485007** |
| chrXV | 480693 | 480787 | 94 | 0,071239015 | 1,252583918 | 0,053813099 | -1,608954925 | -1,367611686 | -0,527455211 | -0,177455211 | **-1,437455211** |
| chrXV | 480695 | 480787 | 92 | 0,059840772 | 4,498925668 | 0,013126527 | -2,22244785 | -1,889080673 | -1,130573821 | -0,766573821 | **-2,026573821** |
| chrXV | 480696 | 480787 | 91 | 0,034194727 | 0,52623348 | 0,061015357 | -1,546305876 | -1,314359995 | -0,57193622 | -0,20093622 | **-1,46093622** |
| chrXV | 480697 | 480787 | 90 | 0,028495606 | 1,437877396 | 0,019432713 | -2,065608871 | -1,75576754 | -1,036199292 | -0,658199292 | **-1,918199292** |
| chrXV | 480698 | 480787 | 89 | 0,04844253 | 0,26311674 | 0,155484155 | -1,013192315 | -0,861213468 | -0,185230518 | 0,199769482 | **-1,060230518** |
| chrXV | 480700 | 480787 | 87 | 0,045592969 | 0,626291959 | 0,0678583 | -1,491933418 | -1,268143405 | -0,643085721 | -0,244085721 | **-1,504085721** |
| chrXV | 480701 | 480787 | 86 | 0,037044288 | 5,592157194 | 0,006580736 | -2,479369952 | -2,107464459 | -1,516110132 | -1,110110132 | **-2,370110132** |
| chrXV | 480703 | 480787 | 84 | 0,034194727 | 0,333528262 | 0,092990452 | -1,322562523 | -1,124178145 | -0,548925352 | -0,128925352 | **-1,388925352** |
| chrXV | 480704 | 480787 | 83 | 0,028495606 | 0,496586524 | 0,054268855 | -1,604800529 | -1,364080449 | -0,827161518 | -0,400161518 | **-1,660161518** |
| chrXV | 482098 | 482249 | 151 | 0,19377012 | 0,270528479 | 0,417339445 | -0,208704547 | -0,177398865 | -0,280170937 | -0,329170937 | **-1,589170937** |
| chrXV | 483506 | 483656 | 150 | 0,322000346 | 0,185293479 | 0,634741308 | 0,344437377 | 0,29277177 | 0,173185308 | 0,131185308 | **-1,128814692** |

TABLE 2

| Chrom | Start | End | Length | Section A | Section B | A/A+B | Z-score | Z * 0.85 | Phase correction | Length correction | ΔLK |
|---|---|---|---|---|---|---|---|---|---|---|---|
| chrXV | 490437 | 490586 | 149 | 0,347646391 | 0,251999131 | 0,579753168 | 0,201262063 | 0,171072754 | 0,067558079 | 0,032558079 | -1,227441921 |
| chrXV | 492152 | 492296 | 144 | 0,102584181 | 0,100058479 | 0,506231912 | 0,015621722 | 0,013278464 | -0,027616616 | -0,027616616 | -1,287616616 |
| chrXV | 492636 | 492766 | 130 | 2,892303992 | 4,254338276 | 0,404708097 | -0,241179248 | -0,205002361 | 0,039714474 | 0,137714474 | -1,122285526 |
| chrXV | 499981 | 500137 | 156 | 0,108283302 | 0,037058696 | 0,745024175 | 0,658912981 | 0,560076034 | 0,591350516 | 0,507350516 | -0,752649484 |
| chrXV | 500251 | 500392 | 141 | 0,182371877 | 0,155646522 | 0,539532397 | 0,099255756 | 0,084367393 | 0,15874422 | 0,17974422 | -1,08025578 |
| chrXV | 500870 | 501031 | 161 | 1,256656217 | 1,130290221 | 0,52647022 | 0,066399762 | 0,056439798 | 0,12899923 | 0,00999923 | -1,25000077 |
| chrXV | 504656 | 504829 | 173 | 0,29350474 | 0,418763262 | 0,412070652 | -0,222221698 | -0,188888443 | 0,069843956 | -0,133156044 | -1,393156044 |
| chrXV | 504839 | 504995 | 156 | 0,037044288 | 0,048176305 | 0,434687048 | -0,164453569 | -0,139785534 | -0,105906994 | -0,189906994 | -1,449906994 |
| chrXV | 505651 | 505827 | 176 | 0,270708255 | 0,570703915 | 0,321730853 | -0,462864205 | -0,393434574 | -0,073242507 | -0,297242507 | -1,557242507 |
| chrXV | 508541 | 508687 | 146 | 0,843469933 | 0,711526959 | 0,542425478 | 0,106546147 | 0,090564225 | 0,168571332 | 0,154571332 | -1,105428668 |
| chrXV | 512017 | 512169 | 152 | 0,615505086 | 0,596645002 | 0,507779599 | 0,0195018 | 0,01657653 | -0,072824829 | -0,128824829 | -1,388824829 |
| chrXV | 520467 | 520627 | 160 | 0,04844253 | 0,048176305 | 0,50137771 | 0,003453413 | 0,002935401 | 0,070129535 | -0,041870465 | -1,301870465 |
| chrXV | 521328 | 521487 | 159 | 8,802292642 | 6,259213718 | 0,584423127 | 0,213222081 | 0,181238769 | 0,226841157 | 0,121841157 | -1,138158843 |
| chrXV | 524434 | 524584 | 150 | 0,054141651 | 0,048176305 | 0,529151025 | 0,07313593 | 0,062165541 | -0,077271275 | -0,119271275 | -1,379271275 |
| chrXV | 526545 | 526704 | 159 | 0,105433742 | 0,048176305 | 0,686372697 | 0,485594627 | 0,412755433 | 0,44990439 | 0,34490439 | -0,91509561 |
| chrXV | 527282 | 527433 | 151 | 0,037044288 | 0,044470435 | 0,454449042 | -0,114428548 | -0,097264266 | -0,18965849 | -0,23865849 | -1,49865849 |
| chrXV | 528345 | 528493 | 148 | 0,83207169 | 0,785644351 | 0,514349657 | 0,035977015 | 0,030580463 | 0,002745817 | -0,025254183 | -1,285254183 |
| chrXV | 529088 | 529208 | 120 | 0,715239706 | 3,661399144 | 0,163422144 | -0,980490029 | -0,833416525 | -0,002024708 | 0,165975292 | -1,094024708 |
| chrXV | 534111 | 534263 | 152 | 0,094035499 | 0,044470435 | 0,678927584 | 0,464702061 | 0,394996752 | 0,297448784 | 0,241448784 | -1,018551216 |
| chrXV | 535122 | 535279 | 157 | 0,267858695 | 0,200116957 | 0,572377417 | 0,18243015 | 0,155065628 | 0,175223175 | 0,084223175 | -1,175776825 |
| chrXV | 539793 | 539942 | 149 | 0,082637257 | 0,055588044 | 0,597844653 | 0,247772164 | 0,210606339 | 0,121285076 | 0,086285076 | -1,173714924 |
| chrXV | 540123 | 540277 | 154 | 0,133929347 | 0,188999349 | 0,414733497 | -0,215385217 | -0,183077435 | -0,227816261 | -0,297816261 | -1,557816261 |
| chrXV | 542672 | 542801 | 129 | 0,085486818 | 0,459527828 | 0,156852331 | -1,007478964 | -0,856357119 | -0,530596178 | -0,425596178 | -1,685596178 |
| chrXV | 545231 | 545391 | 160 | 0,421734966 | 0,478057176 | 0,468702656 | -0,078531454 | -0,066751736 | -0,001952932 | -0,113952932 | -1,373952932 |
| chrXV | 545588 | 545715 | 127 | 1,914904712 | 3,839280883 | 0,332784663 | -0,432236787 | -0,367401269 | 0,180010251 | 0,299010251 | -0,960989749 |
| chrXV | 550006 | 550155 | 149 | 0,256460453 | 0,289057827 | 0,470122564 | -0,074961771 | -0,063717505 | -0,157809124 | -0,192809124 | -1,452809124 |
| chrXV | 550413 | 550558 | 145 | 0,24221265 | 0,26311674 | 0,479316372 | -0,051869417 | -0,044089004 | 0,003555036 | -0,003444964 | -1,263444964 |
| chrXV | 550604 | 550752 | 148 | 0,142478029 | 0,100058479 | 0,587449825 | 0,220989833 | 0,187841358 | 0,156446399 | 0,128446399 | -1,131553601 |
| chrXV | 559497 | 559585 | 88 | 0,037044288 | 0,200116957 | 0,15619874 | -1,010204174 | -0,858673548 | -0,231368473 | 0,160631527 | -1,099368473 |
| chrXV | 559497 | 559598 | 101 | 0,079787696 | 0,211234566 | 0,274163549 | -0,600268815 | -0,510228493 | 0,501861261 | 0,802861261 | -0,457138739 |
| chrXV | 559497 | 559601 | 104 | 9,67140862 | 46,46419278 | 0,172286542 | -0,94516806 | -0,803392851 | 0,168513814 | 0,448513814 | -0,811486186 |
| chrXV | 559497 | 559654 | 157 | 0,262159574 | 0,26682261 | 0,495592445 | -0,011048326 | -0,009391077 | 0,005934055 | -0,085065945 | -1,345065945 |
| chrXV | 559919 | 560068 | 149 | 0,518620026 | 0,200116957 | 0,721571365 | 0,587515907 | 0,499388521 | 0,394811242 | 0,359811242 | -0,900188758 |
| chrXV | 559929 | 560086 | 157 | 0,128230226 | 0,059293913 | 0,683806504 | 0,478369846 | 0,406614369 | 0,416093258 | 0,325093258 | -0,934906742 |
| chrXV | 565053 | 565199 | 146 | 0,031345166 | 0,100058479 | 0,238541073 | -0,711003374 | -0,604352868 | -0,523448662 | -0,537448662 | -1,797448662 |
| chrXV | 565053 | 565201 | 148 | 0,037044288 | 0,333528262 | 0,099965007 | -1,281750982 | -1,089488335 | -1,106599187 | -1,134599187 | -2,394599187 |
| chrXV | 565053 | 565202 | 149 | 0,644000692 | 2,141992616 | 0,231156582 | -0,735043233 | -0,624786748 | -0,719650298 | -0,754650298 | -2,014650298 |
| chrXV | 566775 | 566926 | 151 | 0,09688506 | 0,077823261 | 0,554553208 | 0,137173592 | 0,116597553 | 0,036515654 | -0,012484346 | -1,272484346 |
| chrXV | 576392 | 576555 | 163 | 0,139628469 | 0,196411088 | 0,415512001 | -0,213388432 | -0,181380167 | -0,107433993 | -0,240433993 | -1,500433993 |
| chrXV | 590114 | 590263 | 149 | 0,384690679 | 0,385410436 | 0,499532686 | -0,001171382 | -0,000995674 | -0,102404152 | -0,137404152 | -1,397404152 |
| chrXV | 590721 | 590873 | 152 | 3,051879385 | 1,952993268 | 0,609781626 | 0,278749922 | 0,236937434 | 0,148787604 | 0,092787604 | -1,167212396 |
| chrXV | 592017 | 592170 | 153 | 1,584355684 | 1,070996308 | 0,596665033 | 0,244724271 | 0,20801563 | 0,10876214 | 0,04576214 | -1,21423786 |
| chrXV | 592017 | 592179 | 162 | 0,082637257 | 0,074117392 | 0,527175798 | 0,068172392 | 0,057946533 | 0,123351908 | -0,002648092 | -1,262648092 |
| chrXV | 592023 | 592179 | 156 | 0,347646391 | 0,229763914 | 0,602078605 | 0,258731013 | 0,219921361 | 0,264716078 | 0,180716078 | -1,079283922 |
| chrXV | 592044 | 592126 | 82 | 0,056991212 | 0,040764565 | 0,582995843 | 0,209563572 | 0,178129036 | | 0,434 | -0,826 |
| chrXV | 595352 | 595475 | 123 | 0,074088575 | 0,188999349 | 0,281611463 | -0,578061014 | -0,491351862 | 0,209806653 | 0,356806653 | -0,903193347 |
| chrXV | 595452 | 595607 | 155 | 0,102584181 | 0,118587827 | 0,463820816 | -0,090812431 | -0,077190567 | -0,08636846 | -0,16336846 | -1,42336846 |
| chrXV | 595885 | 596019 | 134 | 0,128230226 | 0,081529131 | 0,611320649 | 0,282762761 | 0,240348347 | 0,52373772 | 0,59373772 | -0,66626228 |
| chrXV | 595885 | 596025 | 140 | 0,168124074 | 0,122293696 | 0,578904225 | 0,199091015 | 0,169227363 | 0,261702883 | 0,289702883 | -0,970297117 |
| chrXV | 596928 | 597063 | 135 | 0,09973462 | 0,081529131 | 0,550218231 | 0,126212725 | 0,107280816 | 0,387640454 | 0,450640454 | -0,809359546 |
| chrXV | 597189 | 597334 | 145 | 0,09973462 | 0,066705652 | 0,599221683 | 0,251333038 | 0,213633083 | 0,257559811 | 0,250559811 | -1,009440189 |
| chrXV | 597375 | 597525 | 150 | 0,960301917 | 0,607762611 | 0,612412244 | 0,285611739 | 0,242769978 | 0,094897274 | 0,052897274 | -1,207102726 |
| chrXV | 597554 | 597711 | 157 | 0,068389454 | 0,051882174 | 0,568624996 | 0,172874585 | 0,146943398 | 0,14542998 | 0,05442998 | -1,20557002 |
| chrXV | 598045 | 598174 | 129 | 0,065539893 | 0,100058479 | 0,395776194 | -0,264295292 | -0,224650998 | 0,064512057 | 0,169512057 | -1,090487943 |
| chrXV | 598957 | 599107 | 150 | 0,48727486 | 0,415057393 | 0,540017115 | 0,100476838 | 0,085405313 | -0,059578722 | -0,101578722 | -1,361578722 |
| chrXV | 600156 | 600263 | 107 | 0,076938136 | 0,592939133 | 0,114854078 | -1,201110978 | -1,020944331 | -0,081555398 | 0,177444602 | -1,082555398 |
| chrXV | 601179 | 601328 | 149 | 0,091185939 | 0,055588044 | 0,621267729 | 0,308811997 | 0,262490198 | 0,164261354 | 0,129261354 | -1,130738646 |
| chrXV | 603463 | 603616 | 153 | 0,3903898 | 0,292763697 | 0,571452539 | 0,180073428 | 0,153062414 | 0,054444922 | -0,008555078 | -1,268555078 |
| chrXV | 605449 | 605600 | 151 | 2,157117362 | 1,3155837 | 0,621164138 | 0,308539664 | 0,262258714 | 0,175177075 | 0,126177075 | -1,133822925 |
| chrXV | 606728 | 606884 | 156 | 0,581310359 | 0,592939133 | 0,495048423 | -0,012412082 | -0,010550269 | 0,036168336 | -0,047831664 | -1,307831664 |
| chrXV | 608095 | 608251 | 156 | 0,088336378 | 0,129705435 | 0,405135037 | -0,240077633 | -0,204065988 | -0,168299031 | -0,252299031 | -1,512299031 |
| chrXV | 610131 | 610285 | 154 | 0,988797522 | 0,748585655 | 0,569130365 | 0,174160574 | 0,148036488 | 0,113351294 | 0,043351294 | -1,216648706 |
| chrXV | 610432 | 610587 | 155 | 0,190920559 | 0,140823044 | 0,575506377 | 0,190410786 | 0,161849168 | 0,161335514 | 0,084335514 | -1,175664486 |
| chrXV | 610432 | 610587 | 155 | 0,190920559 | 0,140823044 | 0,575506377 | 0,190410786 | 0,161849168 | 0,161335514 | 0,084335514 | -1,175664486 |
| chrXV | 613403 | 613549 | 146 | 0,113982423 | 0,200116957 | 0,362886495 | -0,350753893 | -0,298140809 | -0,210149098 | -0,224149098 | -1,484149098 |
| chrXV | 617632 | 617793 | 161 | 0,541416511 | 0,43729261 | 0,553194508 | 0,133736447 | 0,11367598 | 0,189107239 | 0,070107239 | -1,189892761 |
| chrXV | 621001 | 621151 | 150 | 0,176672756 | 0,140823044 | 0,556456986 | 0,141992377 | 0,12069352 | -0,017977946 | -0,059977946 | -1,319977946 |
| chrXV | 622311 | 622441 | 130 | 0,219416165 | 0,26311674 | 0,454717518 | -0,113751185 | -0,096688507 | 0,148036225 | 0,246036225 | -1,013963775 |
| chrXV | 624028 | 624159 | 131 | 3,251348626 | 2,627461531 | 0,553062361 | 0,133402235 | 0,1133919 | 0,342714108 | 0,433714108 | -0,826285892 |
| chrXV | 625834 | 625959 | 125 | 0,185221438 | 0,455821958 | 0,28893744 | -0,556491534 | -0,473017804 | 0,168653128 | 0,301653128 | -0,958346872 |
| chrXV | 626417 | 626570 | 153 | 0,113982423 | 0,096352609 | 0,541908887 | 0,105243965 | 0,08945737 | 0,00511256 | -0,05788744 | -1,31788744 |
| chrXV | 632144 | 632251 | 107 | 0,14532759 | 0,522527611 | 0,217603441 | -0,780312627 | -0,663265733 | 0,267322144 | 0,526322144 | -0,733677856 |
| chrXV | 632144 | 632253 | 109 | 0,139628469 | 0,807879568 | 0,147363888 | -1,047806479 | -0,890635507 | 0,145348727 | 0,390348727 | -0,869651273 |
| chrXV | 633485 | 633638 | 153 | 0,88051422 | 0,663350655 | 0,570331144 | 0,177217294 | 0,1506347 | 0,071537658 | 0,008537658 | -1,251462342 |
| chrXV | 635429 | 635578 | 149 | 0,398938482 | 0,363175219 | 0,523463207 | 0,058847485 | 0,050020363 | -0,05159384 | -0,08659384 | -1,34659384 |
| chrXV | 637512 | 637669 | 157 | 0,071239015 | 0,066705652 | 0,516431814 | 0,041200102 | 0,035020087 | 0,031509855 | -0,059490145 | -1,319490145 |
| chrXV | 641886 | 642041 | 155 | 0,19377012 | 0,207528696 | 0,482857442 | -0,042983253 | -0,036535765 | -0,038090187 | -0,115090187 | -1,375090187 |
| chrXV | 646176 | 646338 | 162 | 0,547115632 | 0,61517435 | 0,470722144 | -0,073454704 | -0,062436499 | 0,015497341 | -0,110502659 | -1,370502659 |
| chrXV | 647481 | 647636 | 155 | 0,133929347 | 0,144528914 | 0,480967406 | -0,04772575 | -0,040566888 | -0,041866228 | -0,118866228 | -1,378866228 |
| chrXV | 650796 | 650922 | 126 | 1,786674486 | 2,923931098 | 0,379287642 | -0,30735223 | -0,261249395 | 0,324727741 | 0,450727741 | -0,809272259 |
| chrXV | 650923 | 651080 | 157 | 0,09973462 | 0,159352392 | 0,38494643 | -0,292515043 | -0,248637787 | -0,235685502 | -0,326685502 | -1,586685502 |
| chrXV | 651588 | 651740 | 152 | 0,119681545 | 0,070411522 | 0,629594476 | 0,330779497 | 0,281162572 | 0,190754779 | 0,134754779 | -1,125245221 |
| chrXV | 671862 | 672011 | 149 | 0,7807796 | 0,555880437 | 0,584127286 | 0,21246353 | 0,180594 | 0,090998247 | 0,055998247 | -1,204001753 |
| chrXV | 674697 | 674849 | 152 | 0,236513528 | 0,107470218 | 0,687571814 | 0,488979272 | 0,415632381 | 0,324271907 | 0,268271907 | -0,991728093 |
| chrXV | 674884 | 675018 | 134 | 0,376141997 | 0,340940001 | 0,524545307 | 0,06156483 | 0,052330105 | 0,33347946 | 0,40347946 | -0,85652054 |
| chrXV | 677635 | 677796 | 161 | 2,641542661 | 2,368050661 | 0,527296827 | 0,068476474 | 0,058205003 | 0,135862002 | 0,016862002 | -1,243137998 |
| chrXV | 679372 | 679525 | 153 | 1,829417895 | 1,482347831 | 0,552399549 | 0,13172615 | 0,111967227 | 0,048112048 | -0,014887952 | -1,274887952 |
| chrXV | 680555 | 680689 | 134 | 0,139628469 | 0,166764131 | 0,455717497 | -0,111228699 | -0,094544394 | 0,202435664 | 0,272435664 | -0,987564336 |
| chrXV | 682390 | 682545 | 155 | 0,259310013 | 0,240881523 | 0,518421434 | 0,046192108 | 0,039263292 | 0,054093852 | -0,022906148 | -1,282906148 |
| chrXV | 682421 | 682570 | 149 | 0,669646737 | 0,418763262 | 0,615252283 | 0,293034955 | 0,249079711 | 0,16404296 | 0,12904296 | -1,13095704 |
| chrXV | 682600 | 682746 | 146 | 1,376337762 | 0,904232177 | 0,603506053 | 0,262432662 | 0,223067762 | 0,292626903 | 0,278626903 | -0,981373097 |
| chrXV | 683178 | 683329 | 151 | 0,139628469 | 0,140823044 | 0,497870264 | -0,005338482 | -0,00453771 | -0,094272096 | -0,143272096 | -1,403272096 |
| chrXV | 683178 | 683329 | 151 | 0,139628469 | 0,140823044 | 0,497870264 | -0,005338482 | -0,00453771 | -0,094272096 | -0,143272096 | -1,403272096 |
| chrXV | 683542 | 683705 | 163 | 0,111132863 | 0,140823044 | 0,441080601 | -0,14823007 | -0,125995559 | -0,04824454 | -0,18124454 | -1,44124454 |
| chrXV | 698388 | 698544 | 156 | 0,125380666 | 0,077823261 | 0,617018911 | 0,297660652 | 0,253011555 | 0,289662398 | 0,205662398 | -1,054337602 |
| chrXV | 699187 | 699320 | 133 | 0,085486818 | 0,214940436 | 0,284550808 | -0,569375094 | -0,48396883 | -0,184216594 | -0,107216594 | -1,367216594 |
| chrXV | 704256 | 704405 | 149 | 0,091185939 | 0,062999783 | 0,591403262 | 0,231156174 | 0,196482748 | 0,106514664 | 0,071514664 | -1,188485336 |
| chrXV | 705052 | 705198 | 146 | 0,09973462 | 0,092646739 | 0,518421434 | 0,046192108 | 0,039263292 | 0,094649897 | 0,080649897 | -1,179350103 |
| chrXV | 707863 | 708011 | 148 | 0,108283302 | 0,074117392 | 0,593656197 | 0,236960278 | 0,201416237 | 0,180306295 | 0,152306295 | -1,107693705 |
| chrXV | 709269 | 709434 | 165 | 9,204080685 | 7,467327201 | 0,552087787 | 0,130937909 | 0,111297222 | 0,194247599 | 0,047247599 | -1,212752401 |
| chrXV | 709269 | 709434 | 165 | 9,204080685 | 7,467327201 | 0,552087787 | 0,130937909 | 0,111297222 | 0,194247599 | 0,047247599 | -1,212752401 |

TABLE 2

| Chrom | Start | End | Length | Section A | Section B | A/A+B | Z-score | Z * 0.85 | Phase correction | Length correction | ΔLK |
|---|---|---|---|---|---|---|---|---|---|---|---|
| chrXV | 709269 | 709462 | 193 | 0,706691025 | 0,79305609 | 0,471206791 | -0,072236647 | -0,06140115 | 0,383538613 | 0,040538613 | -1,219461387 |
| chrXV | 709269 | 709462 | 193 | 0,706691025 | 0,79305609 | 0,471206791 | -0,072236647 | -0,06140115 | 0,383538613 | 0,040538613 | -1,219461387 |
| chrXV | 709332 | 709434 | 102 | 0,113982423 | 0,148234783 | 0,434687048 | -0,164453569 | -0,139785534 | 0,85856082 | 1,15256082 | -0,10743918 |
| chrXV | 709332 | 709462 | 130 | 1,621399972 | 1,630582615 | 0,498588147 | -0,003538998 | -0,003008148 | 0,248608785 | 0,346608785 | -0,913391215 |
| chrXV | 709396 | 709550 | 154 | 0,185221438 | 0,140823044 | 0,568086406 | 0,171504378 | 0,145778722 | 0,113286189 | 0,043286189 | -1,216713811 |
| chrXV | 709396 | 709571 | 175 | 0,039893848 | 0,059293913 | 0,402205349 | -0,247642923 | -0,210496484 | 0,077063731 | -0,139936269 | -1,399936269 |
| chrXV | 709403 | 709571 | 168 | 0,852018614 | 0,915349786 | 0,4820832 | -0,044925865 | -0,038186985 | 0,124124654 | -0,043875346 | -1,303875346 |
| chrXV | 712148 | 712278 | 130 | 1,70688679 | 1,856640659 | 0,478987973 | -0,052693716 | -0,044789658 | 0,189801419 | 0,287801419 | -0,972198581 |
| chrXV | 713754 | 713908 | 154 | 0,059840772 | 0,059293913 | 0,502295129 | 0,005753068 | 0,004890107 | -0,032633769 | -0,102633769 | -1,362633769 |
| chrXV | 716082 | 716234 | 152 | 1,037240052 | 0,79305609 | 0,566706135 | 0,167994342 | 0,142795191 | 0,048874133 | -0,007125867 | -1,267125867 |
| chrXV | 717294 | 717452 | 158 | 1,273753581 | 1,67505305 | 0,431955615 | -0,171397488 | -0,145687865 | -0,101867322 | -0,199867322 | -1,459867322 |
| chrXV | 717294 | 717452 | 158 | 1,273753581 | 1,67505305 | 0,431955615 | -0,171397488 | -0,145687865 | -0,101867322 | -0,199867322 | -1,459867322 |
| chrXV | 728130 | 728275 | 145 | 1,641346896 | 1,411936309 | 0,537567853 | 0,094308252 | 0,080162014 | 0,11943527 | 0,11243527 | -1,14756473 |
| chrXV | 728130 | 728275 | 145 | 1,641346896 | 1,411936309 | 0,537567853 | 0,094308252 | 0,080162014 | 0,11943527 | 0,11243527 | -1,14756473 |
| chrXV | 729241 | 729409 | 168 | 0,227964847 | 0,285351958 | 0,444101664 | -0,140578003 | -0,119491302 | 0,030059749 | -0,137940251 | -1,397940251 |
| chrXV | 730369 | 730527 | 158 | 0,059840772 | 0,074117392 | 0,446712395 | -0,133971905 | -0,113876119 | -0,072422755 | -0,170422755 | -1,430422755 |
| chrXV | 730626 | 730772 | 146 | 0,347646391 | 0,303881305 | 0,533586512 | 0,0842886 | 0,07164531 | 0,123685406 | 0,109685406 | -1,150314594 |
| chrXV | 732904 | 733026 | 122 | 1,567258321 | 4,602690017 | 0,254014821 | -0,661908847 | -0,56262252 | 0,181697772 | 0,335697772 | -0,924302228 |
| chrXV | 736114 | 736267 | 153 | 0,119681545 | 0,196411088 | 0,378628074 | -0,309085948 | -0,262723056 | -0,323749186 | -0,386749186 | -1,646749186 |
| chrXV | 749719 | 749841 | 122 | 0,085486818 | 0,385410436 | 0,181540276 | -0,909510814 | -0,773084192 | -0,021719545 | 0,132280455 | -1,127719545 |
| chrXV | 751696 | 751803 | 107 | 3,137366202 | 13,92665788 | 0,183858519 | -0,900757933 | -0,765644243 | 0,159742862 | 0,418742862 | -0,841257138 |
| chrXV | 751696 | 751804 | 108 | 0,074088575 | 0,277940218 | 0,210461691 | -0,804820306 | -0,68409726 | 0,310384221 | 0,562384221 | -0,697615779 |
| chrXV | 751696 | 751805 | 109 | 0,065539893 | 0,400233914 | 0,140711848 | -1,077126632 | -0,915557637 | 0,099161345 | 0,344161345 | -0,915838655 |
| chrXV | 751696 | 751808 | 112 | 4,131862846 | 21,73121921 | 0,159759105 | -0,995448438 | -0,846131172 | 0,166089144 | 0,390089144 | -0,869910856 |
| chrXV | 759323 | 759474 | 151 | 0,869115978 | 0,689291742 | 0,557694862 | 0,145127411 | 0,1233583 | 0,049346526 | 0,000346526 | -1,259653474 |
| chrXV | 759886 | 760039 | 153 | 0,04844253 | 0,085235 | 0,36238349 | -0,352095051 | -0,299280793 | -0,36744603 | -0,43044603 | -1,69044603 |
| chrXV | 762029 | 762161 | 132 | 0,287805619 | 0,144528914 | 0,665701204 | 0,428073525 | 0,363862496 | 0,619472082 | 0,703472082 | -0,556527918 |
| chrXV | 762283 | 762425 | 142 | 0,404637603 | 0,233469783 | 0,634121484 | 0,342789226 | 0,291370842 | 0,311193285 | 0,325193285 | -0,934806715 |
| chrXV | 763377 | 763524 | 147 | 0,398938482 | 0,452116089 | 0,468757816 | -0,078392761 | -0,066633846 | 0,02776646 | 0,00676646 | -1,25323354 |
| chrXV | 765404 | 765530 | 126 | 0,045592969 | 0,207528696 | 0,180122745 | -0,914897409 | -0,777662797 | -0,205529827 | -0,079529827 | -1,339529827 |
| chrXV | 766078 | 766228 | 150 | 5,049421354 | 2,923931098 | 0,633287113 | 0,340572049 | 0,289486242 | 0,163547362 | 0,121547362 | -1,138452638 |
| chrXV | 767013 | 767155 | 142 | 0,481575739 | 0,240881523 | 0,666580246 | 0,43048963 | 0,365916185 | 0,397072049 | 0,411072049 | -0,848927951 |
| chrXV | 767013 | 767155 | 142 | 0,481575739 | 0,240881523 | 0,666580246 | 0,43048963 | 0,365916185 | 0,397072049 | 0,411072049 | -0,848927951 |
| chrXV | 768735 | 768888 | 153 | 0,102584181 | 0,062999783 | 0,619529685 | 0,304245806 | 0,258608935 | 0,187982478 | 0,124982478 | -1,135017522 |
| chrXV | 770268 | 770418 | 150 | 0,133929347 | 0,044470435 | 0,750725957 | 0,676776004 | 0,575259604 | 0,459982688 | 0,417982688 | -0,842017312 |
| chrXV | 773124 | 773277 | 153 | 0,427434088 | 0,429880871 | 0,498572996 | -0,003576976 | -0,00304043 | -0,064038035 | -0,127038035 | -1,387038035 |
| chrXV | 775176 | 775297 | 121 | 1,185417203 | 5,388334367 | 0,180325829 | -0,914124066 | -0,777005456 | 0,001270391 | 0,162270391 | -1,097729609 |
| chrXV | 777968 | 778119 | 151 | 0,162424953 | 0,200116957 | 0,448017039 | -0,130672891 | -0,111071958 | -0,170257799 | -0,219257799 | -1,479257799 |
| chrXV | 783192 | 783285 | 93 | 0,216566604 | 0,374292827 | 0,366528133 | -0,341062852 | -0,289903425 | 0,518100289 | 0,875100289 | -0,384899711 |
| chrXV | 789013 | 789147 | 134 | 0,34194727 | 0,429880871 | 0,443035505 | -0,14327752 | -0,121785892 | 0,162260687 | 0,232260687 | -1,027739313 |
| chrXV | 790833 | 790987 | 154 | 0,039893848 | 0,051882174 | 0,434687048 | -0,164453569 | -0,139785534 | -0,172707383 | -0,242707383 | -1,502707383 |
| chrXV | 794701 | 794852 | 151 | 0,324849907 | 0,233469783 | 0,581834946 | 0,206589951 | 0,175601459 | 0,122226419 | 0,073226419 | -1,186773581 |
| chrXV | 796032 | 796168 | 136 | 0,900461144 | 0,674468263 | 0,571746988 | 0,180823617 | 0,153700074 | 0,341914596 | 0,397914596 | -0,862085404 |
| chrXV | 801709 | 801857 | 148 | 1,792373607 | 1,460112614 | 0,551078002 | 0,128385385 | 0,109127577 | 0,091687355 | 0,063687355 | -1,196312645 |
| chrXV | 802168 | 802313 | 145 | 1,926302954 | 1,58240631 | 0,549006147 | 0,123150773 | 0,104678157 | 0,125739971 | 0,118739971 | -1,141260029 |
| chrXV | 804681 | 804839 | 158 | 0,042743409 | 0,040764565 | 0,51184823 | 0,029703476 | 0,025247955 | 0,06822868 | -0,02977132 | -1,28977132 |
| chrXV | 805157 | 805300 | 143 | 1,598603487 | 1,063584569 | 0,600484809 | 0,254602172 | 0,216411846 | 0,241943335 | 0,248943335 | -1,011056665 |
| chrXV | 806800 | 806942 | 142 | 0,051292091 | 0,081529131 | 0,386173911 | -0,289305214 | -0,245909432 | -0,230985644 | -0,216985644 | -1,476985644 |
| chrXV | 806939 | 807090 | 151 | 0,575611238 | 0,44099848 | 0,566206704 | 0,1667248 | 0,14171608 | 0,076220454 | 0,027220454 | -1,232779546 |
| chrXV | 810091 | 810222 | 131 | 0,353345512 | 0,322410653 | 0,522889069 | 0,057405902 | 0,048795016 | 0,303825816 | 0,394825816 | -0,865174184 |
| chrXV | 815641 | 815767 | 126 | 0,034194727 | 0,163058261 | 0,173354671 | -0,940991251 | -0,799842563 | -0,247179464 | -0,121179464 | -1,381179464 |
| chrXV | 816418 | 816570 | 152 | 1,199665006 | 0,533645219 | 0,692123654 | 0,501878922 | 0,426597084 | 0,325308175 | 0,269308175 | -0,990691825 |
| chrXV | 816891 | 817023 | 132 | 0,133929347 | 0,259410871 | 0,340492381 | -0,411119704 | -0,349451748 | -0,098052286 | -0,014052286 | -1,274052286 |
| chrXV | 823212 | 823372 | 160 | 1,282302263 | 1,634288484 | 0,439657934 | -0,151836531 | -0,129061052 | -0,059096545 | -0,171096545 | -1,431096545 |
| chrXV | 823715 | 823880 | 165 | 0,621204207 | 0,563292176 | 0,524445846 | 0,061315045 | 0,052117789 | 0,137108412 | -0,009891588 | -1,269891588 |
| chrXV | 824770 | 824918 | 148 | 1,450426337 | 2,660814357 | 0,352795286 | -0,377784657 | -0,321116959 | -0,345170593 | -0,373170593 | -1,633170593 |
| chrXV | 824898 | 825045 | 147 | 0,074088575 | 0,074117392 | 0,499902783 | -0,000243688 | -0,000207135 | 0,062238825 | 0,041238825 | -1,218761175 |
| chrXV | 826750 | 826899 | 149 | 0,210867483 | 0,277940218 | 0,431391491 | -0,172832638 | -0,146907742 | -0,245544788 | -0,280544788 | -1,540544788 |
| chrXV | 828443 | 828590 | 147 | 0,344796831 | 0,26682261 | 0,563744067 | 0,160468712 | 0,136398405 | 0,199537119 | 0,178537119 | -1,081462881 |
| chrXV | 829595 | 829749 | 154 | 0,253610892 | 0,200116957 | 0,558949362 | 0,148306006 | 0,126060105 | 0,10401252 | 0,03401252 | -1,22598748 |
| chrXV | 832011 | 832158 | 147 | 1,413382049 | 1,089525656 | 0,564696032 | 0,162886327 | 0,138453378 | 0,190044655 | 0,169044655 | -1,090955345 |
| chrXV | 838047 | 838199 | 152 | 1,467523701 | 0,896820438 | 0,62068955 | 0,307292296 | 0,261198452 | 0,154390109 | 0,098390109 | -1,161609891 |
| chrXV | 838047 | 838199 | 152 | 1,467523701 | 0,896820438 | 0,62068955 | 0,307292296 | 0,261198452 | 0,154390109 | 0,098390109 | -1,161609891 |
| chrXV | 838749 | 838897 | 148 | 1,604302609 | 0,989467177 | 0,618521589 | 0,301600243 | 0,256360206 | 0,219783427 | 0,191783427 | -1,068216573 |
| chrXV | 840989 | 841143 | 154 | 0,170973635 | 0,118587827 | 0,590457149 | 0,22872109 | 0,194412926 | 0,171693685 | 0,101693685 | -1,158306315 |
| chrXV | 845931 | 846091 | 160 | 0,225115286 | 0,285351958 | 0,440998495 | -0,148438156 | -0,126172432 | -0,06098733 | -0,17298733 | -1,43298733 |
| chrXV | 851758 | 851911 | 153 | 0,088336378 | 0,122293696 | 0,419391098 | -0,203451441 | -0,172933725 | -0,229122585 | -0,292122585 | -1,552122585 |
| chrXV | 855604 | 855755 | 151 | 1,142673794 | 0,830114786 | 0,579217563 | 0,199892222 | 0,169908389 | 0,111147431 | 0,062147431 | -1,197852569 |
| chrXV | 856174 | 856339 | 165 | 2,162816483 | 2,004875442 | 0,518948262 | 0,047514122 | 0,040387004 | 0,138878995 | -0,008121005 | -1,268121005 |
| chrXV | 856222 | 856364 | 142 | 0,151026711 | 0,081529131 | 0,649421274 | 0,383758496 | 0,326194722 | 0,352162238 | 0,366162238 | -0,893837762 |
| chrXV | 860271 | 860423 | 152 | 0,108283302 | 0,140823044 | 0,434687048 | -0,164453569 | -0,139785534 | -0,248365666 | -0,304365666 | -1,564365666 |
| chrXV | 863117 | 863234 | 117 | 0,578460798 | 9,238732859 | 0,058923234 | -1,563876953 | -1,32929541 | -0,362576111 | -0,173576111 | -1,433576111 |
| chrXV | 866318 | 866474 | 156 | 3,750021728 | 3,272282838 | 0,53401582 | 0,085368593 | 0,072563304 | 0,114806706 | 0,030806706 | -1,229193294 |
| chrXV | 866636 | 866779 | 143 | 2,379383087 | 0,974643699 | 0,709410878 | 0,551664489 | 0,468914816 | 0,469504403 | 0,476504403 | -0,783495597 |
| chrXV | 873282 | 873420 | 138 | 3,339685004 | 2,249462834 | 0,597530268 | 0,246959633 | 0,209915688 | 0,353757379 | 0,395757379 | -0,864242621 |
| chrXV | 873426 | 873558 | 132 | 0,094035499 | 0,100058479 | 0,484484373 | -0,038901718 | -0,03306646 | 0,231910966 | 0,315910966 | -0,944089034 |
| chrXV | 873491 | 873643 | 152 | 0,136778908 | 0,163058261 | 0,456177292 | -0,110069087 | -0,093558724 | -0,191558695 | -0,247558695 | -1,507558695 |
| chrXV | 883077 | 883230 | 153 | 0,170973635 | 0,096352609 | 0,639569211 | 0,357307551 | 0,303711418 | 0,245885992 | 0,182885992 | -1,077114008 |
| chrXV | 883077 | 883230 | 153 | 0,170973635 | 0,096352609 | 0,639569211 | 0,357307551 | 0,303711418 | 0,245885992 | 0,182885992 | -1,077114008 |
| chrXV | 884043 | 884200 | 157 | 0,039893848 | 0,059293913 | 0,402205349 | -0,247642923 | -0,210496484 | -0,164341449 | -0,255341449 | -1,515341449 |
| chrXV | 895782 | 895933 | 151 | 0,350495952 | 0,318704784 | 0,523753088 | 0,059575384 | 0,050639077 | -0,015438459 | -0,064438459 | -1,324438459 |
| chrXV | 900646 | 900802 | 156 | 0,065539893 | 0,137117174 | 0,323402949 | -0,458203983 | -0,389473386 | -0,347263989 | -0,431263989 | -1,691263989 |
| chrXV | 902699 | 902828 | 129 | 0,44168189 | 0,833820655 | 0,346280681 | -0,395381497 | -0,336074272 | -0,046014529 | 0,058985471 | -1,201014529 |
| chrXV | 904931 | 905079 | 148 | 0,14532759 | 0,137117174 | 0,51453455 | 0,036440779 | 0,030974662 | -0,019752677 | -0,047752677 | -1,307752677 |
| chrXV | 917190 | 917343 | 153 | 0,729487509 | 0,459527828 | 0,613522371 | 0,288511463 | 0,245234744 | 0,192760462 | 0,129760462 | -1,130239538 |
| chrXV | 920557 | 920665 | 108 | 0,082637257 | 1,011702395 | 0,075513353 | -1,435914377 | -1,220527221 | -0,200185328 | 0,051814672 | -1,208185328 |
| chrXV | 928849 | 929014 | 165 | 0,24221265 | 0,344645871 | 0,412727499 | -0,220534382 | -0,187454224 | -0,082350113 | -0,229350113 | -1,489350113 |
| chrXV | 932648 | 932807 | 159 | 1,510267109 | 1,274819135 | 0,542269422 | 0,106152755 | 0,090229842 | 0,137993179 | 0,032993179 | -1,227006821 |
| chrXV | 932960 | 933112 | 152 | 0,723788388 | 0,455821958 | 0,613582604 | 0,288668864 | 0,245368534 | 0,156429711 | 0,100429711 | -1,159570289 |
| chrXV | 937286 | 937429 | 143 | 2,313843194 | 1,752876311 | 0,568970442 | 0,173753594 | 0,147690555 | 0,129361802 | 0,136361802 | -1,123638198 |
| chrXV | 941867 | 942022 | 155 | 0,034194727 | 0,040764565 | 0,456177292 | -0,110069087 | -0,093558724 | -0,087698493 | -0,164698493 | -1,424698493 |
| chrXV | 943142 | 943285 | 143 | 0,119681545 | 0,048176305 | 0,712993436 | 0,562151022 | 0,477828368 | 0,450073829 | 0,457073829 | -0,802926171 |
| chrXV | 956493 | 956644 | 151 | 0,108283302 | 0,159352392 | 0,404592155 | -0,241478459 | -0,20525669 | -0,276012508 | -0,325012508 | -1,585012508 |
| chrXV | 958715 | 958860 | 145 | 0,803576085 | 0,452116089 | 0,639946718 | 0,358316375 | 0,304568919 | 0,31576094 | 0,30876094 | -0,95123906 |
| chrXV | 961158 | 961313 | 155 | 0,339097709 | 0,544762828 | 0,383655221 | -0,295894775 | -0,251510559 | -0,24920315 | -0,32620315 | -1,58620315 |
| chrXV | 963324 | 963474 | 150 | 0,173823196 | 0,233469783 | 0,426776803 | -0,184586205 | -0,156898275 | -0,264831515 | -0,306831515 | -1,566831515 |
| chrXV | 967141 | 967307 | 166 | 0,319150785 | 0,655938915 | 0,327304027 | -0,447369833 | -0,380264358 | -0,249405161 | -0,403405161 | -1,663405161 |
| chrXV | 967454 | 967605 | 151 | 0,162424953 | 0,185293479 | 0,467116317 | -0,082520731 | -0,070142622 | -0,150313211 | -0,199313211 | -1,459313211 |
| chrXV | 970738 | 970906 | 168 | 0,863416857 | 1,078408047 | 0,444641973 | -0,13921033 | -0,11832878 | 0,028296449 | -0,139703551 | -1,399703551 |
| chrXV | 970759 | 970913 | 154 | 0,133929347 | 0,140823044 | 0,487454711 | -0,031451562 | -0,026733828 | -0,051936832 | -0,121936832 | -1,381936832 |
| chrXV | 970847 | 970977 | 130 | 1,738231956 | 1,815876094 | 0,489076846 | -0,027383708 | -0,023276152 | 0,220124605 | 0,318124605 | -0,941875395 |

TABLE 2

| Chrom | Start | End | Length | Section A | Section B | A/A+B | Z-score | Z * 0.85 | Phase correction | Length correction | ΔLK |
|---|---|---|---|---|---|---|---|---|---|---|---|
| chrXV | 970875 | 970977 | 102 | 0,09688506 | 0,163058261 | 0,372716096 | -0,324668217 | -0,275967985 | 0,711065373 | 1,005065373 | **-0,254934627** |
| chrXV | 972298 | 972446 | 148 | 0,074088575 | 0,051882174 | 0,588141101 | 0,222765786 | 0,189350918 | 0,15423807 | 0,12623807 | **-1,13376193** |
| chrXV | 975111 | 975257 | 146 | 0,068389454 | 0,066705652 | 0,506231912 | 0,015621722 | 0,013278464 | 0,070537133 | 0,056537133 | **-1,203462867** |
| chrXV | 975228 | 975386 | 158 | 0,028495606 | 0,051882174 | 0,354520937 | -0,373143192 | -0,317171714 | -0,281351108 | -0,379351108 | **-1,639351108** |
| chrXV | 975904 | 976053 | 149 | 0,111132863 | 0,085235 | 0,565942212 | 0,166052574 | 0,141144688 | 0,050820135 | 0,015820135 | **-1,244179865** |
| chrXV | 975908 | 976058 | 150 | 0,062690333 | 0,055588044 | 0,530023616 | 0,075329227 | 0,064029843 | -0,05049103 | -0,09249103 | **-1,35249103** |
| chrXV | 977285 | 977436 | 151 | 1,493169746 | 1,382289353 | 0,519280468 | 0,048347795 | 0,041095625 | -0,035094817 | -0,084094817 | **-1,344094817** |
| chrXV | 977616 | 977761 | 145 | 0,065539893 | 0,040764565 | 0,616530051 | 0,29638 | 0,251923 | 0,262049406 | 0,255049406 | **-1,004950594** |
| chrXV | 981987 | 982146 | 159 | 0,108283302 | 0,203822827 | 0,346943851 | -0,39358467 | -0,33454697 | -0,28385498 | -0,38885498 | **-1,64885498** |
| chrXV | 986417 | 986558 | 141 | 0,541416511 | 0,522527611 | 0,508876829 | 0,022252747 | 0,018914835 | 0,076816996 | 0,097816996 | **-1,162183004** |
| chrXV | 986699 | 986864 | 165 | 0,031345166 | 0,051882174 | 0,376621026 | -0,314367347 | -0,267212245 | -0,154544023 | -0,301544023 | **-1,561544023** |
| chrXV | 987920 | 988058 | 138 | 0,04844253 | 0,35205761 | 0,120955089 | -1,17022564 | -0,994691794 | -0,865581141 | -0,823581141 | **-2,083581141** |
| chrXV | 988983 | 989114 | 131 | 3,040481143 | 3,579870013 | 0,45926282 | -0,102291073 | -0,086947412 | 0,201099782 | 0,292099782 | **-0,967900218** |
| chrXV | 989889 | 990033 | 144 | 0,153876272 | 0,148234783 | 0,509336779 | 0,023405971 | 0,019895076 | -0,022460001 | -0,022460001 | **-1,282460001** |
| chrXV | 996318 | 996470 | 152 | 2,191312089 | 1,200701743 | 0,646020977 | 0,374599901 | 0,318409916 | 0,230414771 | 0,174414771 | **-1,085585229** |
| chrXV | 997102 | 997264 | 162 | 0,111132863 | 0,185293479 | 0,374908863 | -0,318879716 | -0,271047759 | -0,196654014 | -0,322654014 | **-1,582654014** |
| chrXV | 1000108 | 1000248 | 140 | 0,133929347 | 0,144528914 | 0,480967406 | -0,04772575 | -0,040566888 | 0,053099853 | 0,081099853 | **-1,178900147** |
| chrXV | 1000780 | 1000917 | 137 | 0,29065518 | 0,285351958 | 0,504603434 | 0,011539355 | 0,009808451 | 0,179954295 | 0,228954295 | **-1,031045705** |
| chrXV | 1001402 | 1001553 | 151 | 0,14532759 | 0,107470218 | 0,574876781 | 0,188803997 | 0,160483398 | 0,086022968 | 0,037022968 | **-1,222977032** |
| chrXV | 1002082 | 1002260 | 178 | 0,079787696 | 0,311293045 | 0,204018475 | -0,827353109 | -0,703250143 | -0,358832932 | -0,596832932 | **-1,856832932** |
| chrXV | 1004460 | 1004618 | 158 | 0,039893848 | 0,062999783 | 0,387719315 | -0,285268256 | -0,242478018 | -0,20434428 | -0,30234428 | **-1,56234428** |
| chrXV | 1004460 | 1004618 | 158 | 0,039893848 | 0,062999783 | 0,387719315 | -0,285268256 | -0,242478018 | -0,20434428 | -0,30234428 | **-1,56234428** |
| chrXV | 1005856 | 1006001 | 145 | 0,159575393 | 0,17417587 | 0,478126708 | -0,054855711 | -0,046627354 | -0,040118755 | -0,047118755 | **-1,307118755** |
| chrXV | 1008258 | 1008393 | 135 | 0,213717044 | 0,26682261 | 0,444743825 | -0,138952542 | -0,118109661 | 0,14570597 | 0,20870597 | **-1,05129403** |
| chrXV | 1019659 | 1019809 | 150 | 0,088336378 | 0,077823261 | 0,531635591 | 0,07938196 | 0,067474666 | -0,050074057 | -0,092074057 | **-1,352074057** |
| chrXV | 1019667 | 1019830 | 163 | 0,09973462 | 0,092646739 | 0,518421434 | 0,046192108 | 0,039263292 | 0,11234849 | -0,02065151 | **-1,28065151** |
| chrXV | 1023929 | 1024082 | 153 | 0,279256937 | 0,292763697 | 0,488193818 | -0,029598031 | -0,025158326 | -0,083759754 | -0,146759754 | **-1,406759754** |
| chrXV | 1035603 | 1035747 | 144 | 0,985947962 | 0,670762394 | 0,59512392 | 0,240745775 | 0,204633908 | 0,158296097 | 0,158296097 | **-1,101703903** |
| chrXV | 1036929 | 1037060 | 131 | 0,105433742 | 0,144528914 | 0,421797974 | -0,19729595 | -0,167701557 | 0,132420173 | 0,223420173 | **-1,036579827** |
| chrXV | 1041309 | 1041484 | 175 | 0,378991558 | 0,544762828 | 0,410273081 | -0,226842569 | -0,192816183 | 0,10772199 | -0,10927801 | **-1,36927801** |
| chrXV | 1042260 | 1042368 | 108 | 0,028495606 | 0,937585003 | 0,029496095 | -1,888251531 | -1,605013801 | -0,606496058 | -0,354496058 | **-1,614496058** |
| chrXV | 1042260 | 1042387 | 127 | 0,056991212 | 0,08894087 | 0,390532439 | -0,277931585 | -0,236241848 | 0,30982722 | 0,42882722 | **-0,83117278** |
| chrXV | 1042260 | 1042389 | 129 | 0,04844253 | 0,08894087 | 0,352608321 | -0,378288023 | -0,32154482 | -0,017543949 | 0,087456051 | **-1,172543949** |
| chrXV | 1042260 | 1042392 | 132 | 23,98760099 | 27,18255336 | 0,468781095 | -0,078334229 | -0,066584095 | 0,223356769 | 0,307356769 | **-0,952643231** |
| chrXV | 1042265 | 1042392 | 127 | 0,327699467 | 0,411351523 | 0,443405761 | -0,142339911 | -0,120988925 | 0,435036139 | 0,554036139 | **-0,705963861** |
| chrXV | 1042267 | 1042368 | 101 | 0,113982423 | 0,144528914 | 0,440918471 | -0,148640972 | -0,126344826 | 0,878280057 | 1,179280057 | **-0,080719943** |
| chrXV | 1043663 | 1043770 | 107 | 2,282498027 | 11,49190156 | 0,165705809 | -0,971274471 | -0,8255833 | 0,105430677 | 0,364430677 | **-0,895569323** |
| chrXV | 1043663 | 1043771 | 108 | 0,068389454 | 0,251999131 | 0,213457836 | -0,794480647 | -0,67530855 | 0,336592979 | 0,588592979 | **-0,671407021** |
| chrXV | 1043663 | 1043772 | 109 | 2,707082554 | 15,49794658 | 0,14869971 | -1,042026441 | -0,885722475 | 0,118211741 | 0,363211741 | **-0,896788259** |
| chrXV | 1047470 | 1047621 | 151 | 0,233663968 | 0,207528696 | 0,529618887 | 0,074311878 | 0,063165096 | -0,014244143 | -0,063244143 | **-1,323244143** |
| chrXV | 1047470 | 1047621 | 151 | 0,233663968 | 0,207528696 | 0,529618887 | 0,074311878 | 0,063165096 | -0,014244143 | -0,063244143 | **-1,323244143** |
| chrXV | 1050132 | 1050280 | 148 | 3,681632274 | 3,009166098 | 0,550253059 | 0,126300727 | 0,107355618 | 0,075677621 | 0,047677621 | **-1,212322379** |
| chrXV | 1050137 | 1050280 | 143 | 0,042743409 | 0,037058696 | 0,535617563 | 0,08939893 | 0,07598909 | 0,048634037 | 0,055634037 | **-1,204365963** |
| chrXV | 1050608 | 1050773 | 165 | 1,818019652 | 1,556465223 | 0,538754719 | 0,097296971 | 0,082702426 | 0,202973509 | 0,055973509 | **-1,204026491** |
| chrXV | 1050861 | 1050999 | 138 | 0,111132863 | 0,107470218 | 0,508377386 | 0,021000537 | 0,017850457 | 0,116037679 | 0,158037679 | **-1,101962321** |
| chrXV | 1052062 | 1052210 | 148 | 0,142478029 | 0,048176305 | 0,747310729 | 0,66605094 | 0,566143299 | 0,531364382 | 0,503364382 | **-0,756635618** |
| chrXV | 1056315 | 1056470 | 155 | 0,139628469 | 0,133411305 | 0,511385088 | 0,028542059 | 0,02426075 | 0,030562337 | -0,046437663 | **-1,306437663** |
| chrXV | 1059723 | 1059876 | 153 | 1,344992595 | 0,748585655 | 0,642437222 | 0,364981045 | 0,310233888 | 0,244744473 | 0,181744473 | **-1,078255527** |
| chrXV | 1060927 | 1061054 | 127 | 0,105433742 | 0,151940653 | 0,409651247 | -0,228442189 | -0,194175861 | 0,362473094 | 0,481473094 | **-0,778526906** |
| chrXV | 1061685 | 1061838 | 153 | 1,393435125 | 1,000584786 | 0,582048261 | 0,207136216 | 0,176065784 | 0,108288028 | 0,045288028 | **-1,214711972** |
| chrXV | 1062032 | 1062164 | 132 | 0,170973635 | 0,303881305 | 0,360054451 | -0,358313251 | -0,304566264 | -0,018783223 | 0,065216777 | **-1,194783223** |
| chrXV | 1062744 | 1062896 | 152 | 0,165274514 | 0,129705435 | 0,560290672 | 0,151706213 | 0,128950281 | 0,046173857 | -0,009826143 | **-1,269826143** |
| chrXV | 1071888 | 1072059 | 171 | 1,373488201 | 2,279109791 | 0,376030487 | -0,315922974 | -0,268534528 | -0,094313612 | -0,283313612 | **-1,543313612** |
| chrXV | 1073330 | 1073480 | 150 | 0,122531105 | 0,100058479 | 0,55047996 | 0,126874057 | 0,107842948 | 0,000274124 | -0,041725876 | **-1,301725876** |
| chrXV | 1080435 | 1080579 | 144 | 1,310797868 | 0,670762394 | 0,661497858 | 0,416554516 | 0,354071338 | 0,306898884 | 0,306898884 | **-0,953101116** |
| chrXV | 1085191 | 1085346 | 155 | 0,416035845 | 0,466939567 | 0,471174893 | -0,072316812 | -0,06146929 | -0,053067537 | -0,130067537 | **-1,390067537** |
| chrXV | 1086206 | 1086377 | 171 | 0,037044288 | 0,062999783 | 0,370279692 | -0,331112667 | -0,281445767 | -0,078536511 | -0,267536511 | **-1,527536511** |
| chrXV | 1086356 | 1086503 | 147 | 1,615700851 | 1,078408047 | 0,599716237 | 0,252612686 | 0,214720783 | 0,263393393 | 0,242393393 | **-1,017606607** |
| chrXV | 1086512 | 1086642 | 130 | 0,29065518 | 0,26311674 | 0,524864424 | 0,062366274 | 0,053011333 | 0,299502856 | 0,397502856 | **-0,862497144** |
| chrXV | 1086893 | 1087029 | 136 | 0,530018269 | 0,689291742 | 0,434687048 | -0,164453569 | -0,139785534 | 0,047120071 | 0,103120071 | **-1,156879929** |
| chrXV | 1087274 | 1087436 | 162 | 0,051292091 | 0,055588044 | 0,479902939 | -0,050397187 | -0,042837609 | 0,031257258 | -0,094742742 | **-1,354742742** |
| chrXV | 1087375 | 1087530 | 155 | 0,082637257 | 0,207528696 | 0,284793085 | -0,568661074 | -0,483361913 | -0,462486779 | -0,539486779 | **-1,799486779** |
| chrXV | 1087509 | 1087646 | 137 | 0,185221438 | 0,096352609 | 0,657807209 | 0,406485945 | 0,345513053 | 0,511361264 | 0,560361264 | **-0,699638736** |
| chrXV | 1087804 | 1087947 | 143 | 0,398938482 | 0,344645871 | 0,536507365 | 0,09163849 | 0,077892716 | 0,048759709 | 0,055759709 | **-1,204240291** |
| chrXV | 1087852 | 1087977 | 125 | 0,159575393 | 0,192705218 | 0,452978074 | -0,118140734 | -0,100419624 | 0,522067118 | 0,655067118 | **-0,604932882** |
| chrXV | 1088077 | 1088229 | 152 | 0,105433742 | 0,074117392 | 0,58720733 | 0,220367006 | 0,187311955 | 0,119855504 | 0,063855504 | **-1,196144496** |
| chrXV | 1088150 | 1088294 | 144 | 0,125380666 | 0,055588044 | 0,692830634 | 0,503889924 | 0,428306435 | 0,355820531 | 0,355820531 | **-0,904179469** |
| chrXV | 1088247 | 1088382 | 135 | 1,185417203 | 0,722644568 | 0,621267729 | 0,308811997 | 0,262490198 | 0,503724222 | 0,566724222 | **-0,693275778** |
| chrXV | 1088247 | 1088434 | 187 | 0,162424953 | 0,066705652 | 0,708874979 | 0,550101085 | 0,467585923 | 0,892753984 | 0,591753984 | **-0,668246016** |
| chrXV | 1088290 | 1088382 | 92 | 0,162424953 | 0,140823044 | 0,535617563 | 0,08939893 | 0,07598909 | 0,847728761 | 1,211728761 | **-0,048271239** |
| chrXV | 1088290 | 1088434 | 144 | 1,262355338 | 0,563292176 | 0,691456225 | 0,499982287 | 0,424984944 | 0,334603135 | 0,334603135 | **-0,925396865** |
| chrXV | 1088629 | 1088781 | 152 | 0,122531105 | 0,111176087 | 0,524293257 | 0,060931847 | 0,05179207 | -0,019406845 | -0,075406845 | **-1,335406845** |
| chrXV | 1088930 | 1089060 | 130 | 0,416035845 | 0,466939567 | 0,471174893 | -0,072316812 | -0,06146929 | 0,170550039 | 0,268550039 | **-0,991449961** |
| chrXV | 1089068 | 1089229 | 161 | 0,230814407 | 0,181587609 | 0,559683023 | 0,150165614 | 0,127640772 | 0,175280688 | 0,056280688 | **-1,203719312** |
| chrXV | 1090465 | 1090616 | 151 | 0,63545201 | 0,363175219 | 0,636325539 | 0,348654215 | 0,296356083 | 0,225516495 | 0,176516495 | **-1,083483505** |
| chrXVI | 523 | 674 | 151 | 0,572761677 | 0,403939784 | 0,586424511 | 0,218356986 | 0,185603438 | 0,035128748 | -0,013871252 | **-1,273871252** |
| chrXVI | 847 | 1021 | 174 | 0,028495606 | 0,059293913 | 0,324590066 | -0,454901459 | -0,386666241 | -0,169074792 | -0,379074792 | **-1,639074792** |
| chrXVI | 1688 | 1839 | 151 | 0,416035845 | 0,240881523 | 0,633315339 | 0,340647026 | 0,289549972 | 0,14256344 | 0,09356344 | **-1,16643656** |
| chrXVI | 1700 | 1834 | 134 | 0,703841464 | 0,478057176 | 0,595517619 | 0,241761772 | 0,205497507 | 0,263119268 | 0,333119268 | **-0,926880732** |
| chrXVI | 1725 | 1821 | 96 | 0,054141651 | 0,074117392 | 0,42212736 | -0,196454145 | -0,166986023 | 0,343875265 | 0,679875265 | **-0,580124735** |
| chrXVI | 1725 | 1834 | 109 | 0,042743409 | 0,051882174 | 0,451710916 | -0,121339881 | -0,103138899 | 0,754581057 | 0,999581057 | **-0,260418943** |
| chrXVI | 2079 | 2209 | 130 | 0,361894194 | 0,407645654 | 0,470273495 | -0,074582384 | -0,063395026 | 0,141565089 | 0,239565089 | **-1,020434911** |
| chrXVI | 2079 | 2209 | 130 | 0,361894194 | 0,407645654 | 0,470273495 | -0,074582384 | -0,063395026 | 0,141565089 | 0,239565089 | **-1,020434911** |
| chrXVI | 2358 | 2510 | 152 | 0,168124074 | 0,125999566 | 0,571610206 | 0,180475116 | 0,153403848 | 0,059413024 | 0,003413024 | **-1,256586976** |
| chrXVI | 2705 | 2849 | 144 | 1,256656217 | 0,559586306 | 0,691898907 | 0,501240058 | 0,426054049 | 0,311657962 | 0,311657962 | **-0,948342038** |
| chrXVI | 2705 | 2892 | 187 | 0,125380666 | 0,096352609 | 0,56545715 | 0,164819952 | 0,140096959 | 0,333227232 | 0,032227232 | **-1,227772768** |
| chrXVI | 2757 | 2849 | 92 | 0,136778908 | 0,111176087 | 0,551627959 | 0,129775456 | 0,110309137 | 0,579086463 | 0,943086463 | **-0,316913537** |
| chrXVI | 2757 | 2892 | 135 | 1,276603141 | 0,774526742 | 0,622390202 | 0,311764379 | 0,264999722 | 0,262389421 | 0,325389421 | **-0,934610579** |
| chrXVI | 2845 | 2989 | 144 | 0,125380666 | 0,044470435 | 0,738179884 | 0,637744162 | 0,542082538 | 0,4307497 | 0,4307497 | **-0,8292503** |
| chrXVI | 2910 | 3062 | 152 | 0,088336378 | 0,066705652 | 0,569757619 | 0,175757115 | 0,149393547 | 0,059400239 | 0,003400239 | **-1,256599761** |
| chrXVI | 3162 | 3287 | 125 | 0,19377012 | 0,240881523 | 0,445805562 | -0,136265845 | -0,115825968 | 0,400110375 | 0,533110375 | **-0,726889625** |
| chrXVI | 3192 | 3335 | 143 | 0,350495952 | 0,400233914 | 0,466873595 | -0,083131234 | -0,070661549 | -0,16927426 | -0,16227426 | **-1,42227426** |
| chrXVI | 3192 | 3343 | 151 | 0,045592969 | 0,040764565 | 0,527955893 | 0,070132481 | 0,059612609 | -0,088479633 | -0,137479633 | **-1,397479633** |
| chrXVI | 3493 | 3630 | 137 | 0,159575393 | 0,103764348 | 0,605967759 | 0,268824834 | 0,228501109 | 0,162904284 | 0,211904284 | **-1,048095716** |
| chrXVI | 3609 | 3764 | 155 | 0,079787696 | 0,17417587 | 0,314169853 | -0,484065048 | -0,411455291 | -0,5158086 | -0,5928086 | **-1,8528086** |
| chrXVI | 3703 | 3865 | 162 | 0,059840772 | 0,055588044 | 0,518421434 | 0,046192108 | 0,039263292 | 0,104344522 | -0,021655478 | **-1,281655478** |
| chrXVI | 3703 | 3865 | 162 | 0,059840772 | 0,055588044 | 0,518421434 | 0,046192108 | 0,039263292 | 0,104344522 | -0,021655478 | **-1,281655478** |
| chrXVI | 4928 | 5069 | 141 | 0,125380666 | 0,096352609 | 0,56545715 | 0,164819952 | 0,140096959 | 0,030414419 | 0,051414419 | **-1,208585581** |
| chrXVI | 8461 | 8628 | 167 | 0,045592969 | 0,044470435 | 0,506231912 | 0,015621722 | 0,013278464 | 0,194044472 | 0,033044472 | **-1,226955528** |
| chrXVI | 10566 | 10710 | 144 | 1,250957096 | 0,822703046 | 0,603260424 | 0,261795443 | 0,222526126 | 0,103256195 | 0,103256195 | **-1,156743805** |
| chrXVI | 23283 | 23434 | 151 | 3,735773925 | 2,116051529 | 0,638394626 | 0,354170984 | 0,301045337 | 0,149497775 | 0,100497775 | **-1,159502225** |

TABLE 2

| Chrom | Start | End | Length | Section A | Section B | A/A+B | Z-score | Z * 0.85 | Phase correction | Length correction | ΔLK |
|---|---|---|---|---|---|---|---|---|---|---|---|
| chrXVI | 26164 | 26341 | 177 | 0,182371877 | 0,344645871 | 0,346045039 | -0,396020265 | -0,336617225 | -0,128423797 | -0,359423797 | -1,619423797 |
| chrXVI | 26440 | 26591 | 151 | 0,111132863 | 0,092646739 | 0,54535813 | 0,113942039 | 0,096850733 | -0,054898592 | -0,103898592 | -1,363898592 |
| chrXVI | 30175 | 30330 | 155 | 0,039893848 | 0,055588044 | 0,417815854 | -0,20748423 | -0,176361595 | -0,280591609 | -0,357591609 | -1,617591609 |
| chrXVI | 30411 | 30566 | 155 | 0,313451664 | 0,244587392 | 0,561702018 | 0,155285859 | 0,13199298 | 0,029291549 | -0,047708451 | -1,307708451 |
| chrXVI | 37895 | 38053 | 158 | 0,199469241 | 0,122293696 | 0,619926094 | 0,305286692 | 0,259493688 | 0,218325368 | 0,120325368 | -1,139674632 |
| chrXVI | 41612 | 41762 | 150 | 0,267858695 | 0,203822827 | 0,567880408 | 0,170980391 | 0,145333332 | -0,025971575 | -0,067971575 | -1,327971575 |
| chrXVI | 41938 | 42057 | 119 | 0,039893848 | 0,170470001 | 0,189642129 | -0,879215839 | -0,747333463 | -0,104522898 | 0,070477102 | -1,189522898 |
| chrXVI | 44256 | 44374 | 118 | 0,071239015 | 0,366881088 | 0,162601565 | -0,983821816 | -0,836248543 | -0,142080111 | 0,039919889 | -1,220080111 |
| chrXVI | 54075 | 54252 | 177 | 0,037044288 | 0,077823261 | 0,322495674 | -0,460731366 | -0,391621661 | -0,189003736 | -0,420003736 | -1,680003736 |
| chrXVI | 55410 | 55560 | 150 | 0,376141997 | 0,237175653 | 0,613290677 | 0,287906064 | 0,244720154 | 0,073759223 | 0,031759223 | -1,228240777 |
| chrXVI | 56814 | 56963 | 149 | 0,094035499 | 0,051882174 | 0,644442151 | 0,370358087 | 0,314804374 | 0,158809887 | 0,123809887 | -1,136190113 |
| chrXVI | 56988 | 57111 | 123 | 0,088336378 | 0,214940436 | 0,291273101 | -0,549669321 | -0,467218923 | 0,13827172 | 0,28527172 | -0,97472828 |
| chrXVI | 57088 | 57243 | 155 | 0,116831984 | 0,096352609 | 0,548032024 | 0,120690792 | 0,102587173 | 0,00098854 | -0,07601146 | -1,33601146 |
| chrXVI | 57521 | 57655 | 134 | 0,111132863 | 0,122293696 | 0,47609348 | -0,059960669 | -0,050966568 | -0,005236979 | 0,064763021 | -1,195236979 |
| chrXVI | 57521 | 57661 | 140 | 0,156725832 | 0,144528914 | 0,520243529 | 0,050764798 | 0,043150079 | -0,044093078 | -0,016093078 | -1,276093078 |
| chrXVI | 57717 | 57868 | 151 | 0,116831984 | 0,081529131 | 0,588986325 | 0,224938202 | 0,191197471 | 0,041831312 | -0,007168688 | -1,267168688 |
| chrXVI | 57727 | 57872 | 145 | 0,091185939 | 0,08894087 | 0,506231912 | 0,015621722 | 0,013278464 | -0,086081866 | -0,093081866 | -1,353081866 |
| chrXVI | 57747 | 57896 | 149 | 0,051292091 | 0,037058696 | 0,58055047 | 0,203301913 | 0,172806626 | 0,013173628 | -0,021826372 | -1,281826372 |
| chrXVI | 57999 | 58153 | 154 | 0,208017923 | 0,122293696 | 0,629762657 | 0,331224804 | 0,281541083 | 0,213862576 | 0,143862576 | -1,116137424 |
| chrXVI | 58564 | 58699 | 135 | 0,068389454 | 0,077823261 | 0,467739443 | -0,080953558 | -0,068810524 | -0,068287266 | -0,005287266 | -1,265287266 |
| chrXVI | 58718 | 58879 | 161 | 0,076938136 | 0,085235 | 0,474419732 | -0,064164224 | -0,05453959 | -0,025572534 | -0,144572534 | -1,404572534 |
| chrXVI | 59011 | 59161 | 150 | 8,842186491 | 4,22098545 | 0,676878979 | 0,458989034 | 0,390140678 | 0,221078645 | 0,179078645 | -1,080921355 |
| chrXVI | 59190 | 59347 | 157 | 0,068389454 | 0,070411522 | 0,492715945 | -0,018259434 | -0,015520519 | -0,089055979 | -0,180055979 | -1,440055979 |
| chrXVI | 59681 | 59810 | 129 | 0,037044288 | 0,103764348 | 0,263082498 | -0,633871027 | -0,538790373 | -0,289006599 | -0,184006599 | -1,444006599 |
| chrXVI | 60593 | 60743 | 150 | 0,14532759 | 0,181587609 | 0,444542163 | -0,139462957 | -0,118543513 | -0,277459688 | -0,319459688 | -1,579459688 |
| chrXVI | 61572 | 61714 | 142 | 0,04844253 | 0,037058696 | 0,566571175 | 0,16765125 | 0,142503563 | 0,036424799 | 0,050424799 | -1,209575201 |
| chrXVI | 66888 | 67040 | 152 | 2,242604179 | 1,656523702 | 0,575155329 | 0,189514819 | 0,161087596 | 0,074766204 | 0,018766204 | -1,241233796 |
| chrXVI | 67654 | 67814 | 160 | 0,273557816 | 0,270528479 | 0,502783875 | 0,006978197 | 0,005931468 | 0,004210129 | -0,107789871 | -1,367789871 |
| chrXVI | 69456 | 69617 | 161 | 0,455929693 | 0,452116089 | 0,502099897 | 0,005263685 | 0,004474132 | 0,027062486 | -0,091937514 | -1,351937514 |
| chrXVI | 73482 | 73631 | 149 | 0,128230226 | 0,040764565 | 0,75878212 | 0,702390353 | 0,5970318 | 0,42884919 | 0,39384919 | -0,86615081 |
| chrXVI | 74222 | 74371 | 149 | 3,507809078 | 1,511994788 | 0,698794051 | 0,520935222 | 0,442794938 | 0,264020507 | 0,229020507 | -1,030979493 |
| chrXVI | 85235 | 85390 | 155 | 0,071239015 | 0,077823261 | 0,477914444 | -0,055388588 | -0,0470803 | -0,153098199 | -0,230098199 | -1,490098199 |
| chrXVI | 86229 | 86396 | 167 | 0,113982423 | 0,166764131 | 0,405997586 | -0,23785292 | -0,202174982 | -0,016505095 | -0,177505095 | -1,437505095 |
| chrXVI | 93799 | 93933 | 134 | 0,213717044 | 0,218646305 | 0,494299631 | -0,014289193 | -0,012145814 | 0,037767312 | 0,107767312 | -1,152232688 |
| chrXVI | 97906 | 98054 | 148 | 0,361894194 | 0,188999349 | 0,656922193 | 0,404077658 | 0,343466009 | 0,191920125 | 0,163920125 | -1,096079875 |
| chrXVI | 100090 | 100196 | 106 | 0,398938482 | 4,146868058 | 0,08775967 | -1,354680605 | -1,151478514 | -0,562415834 | -0,296415834 | -1,556415834 |
| chrXVI | 101988 | 102144 | 156 | 0,376141997 | 0,17417587 | 0,683499518 | 0,477507244 | 0,405881157 | 0,329821332 | 0,245821332 | -1,014178668 |
| chrXVI | 111563 | 111694 | 131 | 1,279452702 | 1,271113265 | 0,501634821 | 0,004097899 | 0,003483215 | 0,147795452 | 0,238795452 | -1,021204548 |
| chrXVI | 111839 | 111996 | 157 | 0,789328282 | 0,70040935 | 0,529843823 | 0,074877279 | 0,063645687 | 0,001503818 | -0,089496182 | -1,349496182 |
| chrXVI | 112816 | 112964 | 148 | 0,091185939 | 0,040764565 | 0,691061693 | 0,498861988 | 0,42403269 | 0,278587346 | 0,250587346 | -1,009412654 |
| chrXVI | 113851 | 114002 | 151 | 2,675737388 | 1,634288484 | 0,620817013 | 0,307627261 | 0,261483172 | 0,11784334 | 0,06884334 | -1,19115666 |
| chrXVI | 115900 | 116007 | 107 | 1,376337762 | 9,605613948 | 0,125327246 | -1,148761129 | -0,97644696 | -0,335787112 | -0,076787112 | -1,336787112 |
| chrXVI | 115900 | 116008 | 108 | 0,034194727 | 0,296469566 | 0,103412215 | -1,262345669 | -1,072993819 | -0,306082644 | -0,054082644 | -1,314082644 |
| chrXVI | 115900 | 116009 | 109 | 0,037044288 | 0,270528479 | 0,120440727 | -1,172786449 | -0,996868482 | -0,136531721 | 0,108468279 | -1,151531721 |
| chrXVI | 115900 | 116013 | 113 | 1,327895232 | 12,17378157 | 0,098350394 | -1,291008333 | -1,097357083 | -0,302128504 | -0,085128504 | -1,345128504 |
| chrXVI | 117285 | 117438 | 153 | 0,222265726 | 0,059293913 | 0,789409045 | 0,804372445 | 0,683716578 | 0,628640195 | 0,565640195 | -0,694359805 |
| chrXVI | 119500 | 119650 | 150 | 1,102779946 | 1,007996525 | 0,522452264 | 0,056309222 | 0,047862839 | -0,106520542 | -0,148520542 | -1,408520542 |
| chrXVI | 119632 | 119770 | 138 | 0,088336378 | 0,125999566 | 0,412139824 | -0,222043979 | -0,188737382 | -0,269245546 | -0,227245546 | -1,487245546 |
| chrXVI | 124878 | 125016 | 138 | 0,954602795 | 0,637409568 | 0,599620215 | 0,252364199 | 0,214509569 | 0,147026653 | 0,189026653 | -1,070973347 |
| chrXVI | 127061 | 127209 | 148 | 0,455929693 | 0,26682261 | 0,630824269 | 0,334037235 | 0,283931649 | 0,139485645 | 0,111485645 | -1,148514355 |
| chrXVI | 127061 | 127209 | 148 | 0,455929693 | 0,26682261 | 0,630824269 | 0,334037235 | 0,283931649 | 0,139485645 | 0,111485645 | -1,148514355 |
| chrXVI | 127345 | 127506 | 161 | 0,333398588 | 0,359469349 | 0,481186342 | -0,047176341 | -0,04009989 | -0,006029955 | -0,125029955 | -1,385029955 |
| chrXVI | 128019 | 128185 | 166 | 0,037044288 | 0,100058479 | 0,270193583 | -0,612227627 | -0,520393483 | -0,34916196 | -0,50316196 | -1,76316196 |
| chrXVI | 128210 | 128316 | 106 | 0,655398934 | 1,967816746 | 0,249845615 | -0,674975659 | -0,573729311 | 0,038575618 | 0,304575618 | -0,955424382 |
| chrXVI | 128210 | 128332 | 122 | 0,530018269 | 2,212404138 | 0,193266459 | -0,865922035 | -0,736033729 | -0,112193357 | 0,041806643 | -1,218193357 |
| chrXVI | 130172 | 130324 | 152 | 0,094035499 | 0,218646305 | 0,300738636 | -0,522277301 | -0,443935706 | -0,521982522 | -0,577982522 | -1,837982522 |
| chrXVI | 131123 | 131279 | 156 | 0,92610719 | 0,604056741 | 0,605233969 | 0,266918306 | 0,22688056 | 0,151880866 | 0,067880866 | -1,192119134 |
| chrXVI | 136763 | 136914 | 151 | 1,188266763 | 0,667056524 | 0,640463455 | 0,359697864 | 0,305743184 | 0,170238157 | 0,121238157 | -1,138761843 |
| chrXVI | 136805 | 136963 | 158 | 0,168124074 | 0,125999566 | 0,571610206 | 0,180475116 | 0,153403848 | 0,118564759 | 0,020564759 | -1,239435241 |
| chrXVI | 139377 | 139534 | 157 | 0,24506221 | 0,274234349 | 0,47191187 | -0,070464771 | -0,059895055 | -0,119719293 | -0,210719293 | -1,470719293 |
| chrXVI | 142443 | 142561 | 118 | 0,153876272 | 1,285936744 | 0,106872399 | -1,24333395 | -1,056833857 | -0,375646725 | -0,193646725 | -1,453646725 |
| chrXVI | 144178 | 144336 | 158 | 0,082637257 | 0,077823261 | 0,515000562 | 0,037609696 | 0,031968242 | -0,003994703 | -0,101994703 | -1,361994703 |
| chrXVI | 149028 | 149196 | 168 | 0,113982423 | 0,107470218 | 0,514703382 | 0,036864261 | 0,031334622 | 0,223188971 | 0,055188971 | -1,204811029 |
| chrXVI | 152572 | 152722 | 150 | 1,826568334 | 0,915349786 | 0,666164435 | 0,429346432 | 0,364944468 | 0,205408307 | 0,163408307 | -1,096591693 |
| chrXVI | 161247 | 161414 | 167 | 0,871965539 | 0,915349786 | 0,487863292 | -0,030426909 | -0,025862873 | 0,161367021 | 0,000367021 | -1,259632979 |
| chrXVI | 164847 | 164957 | 110 | 0,230814407 | 1,84922892 | 0,110966154 | -1,221406078 | -1,038195166 | -0,184490994 | 0,053509006 | -1,206490994 |
| chrXVI | 165348 | 165502 | 154 | 0,262159574 | 0,377998697 | 0,40952306 | -0,228772014 | -0,194456212 | -0,264971934 | -0,334971934 | -1,594971934 |
| chrXVI | 167030 | 167184 | 154 | 0,304902982 | 0,137117174 | 0,689794295 | 0,495267357 | 0,420977253 | 0,349594429 | 0,279594429 | -0,980405571 |
| chrXVI | 169266 | 169418 | 152 | 0,270708255 | 0,196411088 | 0,579526966 | 0,200683491 | 0,170580968 | 0,094184225 | 0,038184225 | -1,221815775 |
| chrXVI | 173412 | 173549 | 137 | 0,424584527 | 0,240881523 | 0,638025828 | 0,353186878 | 0,300208846 | 0,229733126 | 0,278733126 | -0,981266874 |
| chrXVI | 174194 | 174363 | 169 | 0,752283994 | 1,041349352 | 0,41941905 | -0,203379911 | -0,172872925 | 0,024995935 | -0,150004065 | -1,410004065 |
| chrXVI | 175111 | 175256 | 145 | 0,210867483 | 0,207528696 | 0,503989982 | 0,010001569 | 0,008501334 | -0,090605346 | -0,097605346 | -1,357605346 |
| chrXVI | 177466 | 177621 | 155 | 0,259310013 | 0,211234566 | 0,551084902 | 0,128402824 | 0,1091424 | 0,0055767 | -0,0714233 | -1,3314233 |
| chrXVI | 177625 | 177797 | 172 | 0,039893848 | 0,066705652 | 0,374240479 | -0,320642986 | -0,272546538 | -0,05437597 | -0,25037597 | -1,51037597 |
| chrXVI | 189167 | 189337 | 170 | 0,037044288 | 0,062999783 | 0,370279692 | -0,331112667 | -0,281445767 | -0,085417555 | -0,267417555 | -1,527417555 |
| chrXVI | 190409 | 190560 | 151 | 0,447381012 | 0,415057393 | 0,518739668 | 0,04699067 | 0,039942069 | -0,098630912 | -0,147630912 | -1,407630912 |
| chrXVI | 191083 | 191228 | 145 | 0,515770466 | 0,459527828 | 0,528833557 | 0,072338048 | 0,061487341 | -0,04132992 | -0,04832992 | -1,30832992 |
| chrXVI | 193727 | 193886 | 159 | 0,265009134 | 0,137117174 | 0,659019638 | 0,409789016 | 0,348320663 | 0,344312842 | 0,239312842 | -1,020687158 |
| chrXVI | 194794 | 194946 | 152 | 0,450230572 | 0,455821958 | 0,496914425 | -0,007734467 | -0,006574297 | -0,077461999 | -0,133461999 | -1,393461999 |
| chrXVI | 198714 | 198866 | 152 | 0,222265726 | 0,237175653 | 0,483773852 | -0,040684142 | -0,03458152 | -0,108144256 | -0,164144256 | -1,424144256 |
| chrXVI | 200785 | 200926 | 141 | 0,04844253 | 0,040764565 | 0,543034495 | 0,108081541 | 0,09186931 | -0,011227923 | 0,009772077 | -1,250227923 |
| chrXVI | 203076 | 203237 | 161 | 0,14532759 | 0,144528914 | 0,50137771 | 0,003453413 | 0,002935401 | 0,054542265 | -0,064457735 | -1,324457735 |
| chrXVI | 203882 | 204036 | 154 | 0,467327936 | 0,385410436 | 0,548032024 | 0,120690792 | 0,102587173 | 0,037576264 | -0,032423736 | -1,292423736 |
| chrXVI | 205158 | 205313 | 155 | 0,339097709 | 0,207528696 | 0,620346375 | 0,30639062 | 0,260432027 | 0,155654951 | 0,078654951 | -1,181345049 |
| chrXVI | 211347 | 211529 | 182 | 0,210867483 | 0,329822392 | 0,389997099 | -0,279326594 | -0,237427605 | -0,031206804 | -0,297206804 | -1,557206804 |
| chrXVI | 212567 | 212715 | 148 | 1,427629852 | 1,011702395 | 0,585254368 | 0,215354086 | 0,183050973 | 0,033816915 | 0,005816915 | -1,254183085 |
| chrXVI | 213085 | 213234 | 149 | 0,589859041 | 0,489174784 | 0,546654819 | 0,117214141 | 0,09963202 | -0,068917648 | -0,103917648 | -1,363917648 |
| chrXVI | 220298 | 220444 | 146 | 0,435982769 | 0,270528479 | 0,617092467 | 0,297853387 | 0,253175379 | 0,148159146 | 0,134159146 | -1,125840854 |
| chrXVI | 227791 | 227944 | 153 | 3,27414511 | 2,501461966 | 0,566891942 | 0,168466728 | 0,143196719 | 0,077127488 | 0,014127488 | -1,245872512 |
| chrXVI | 228023 | 228173 | 150 | 0,78362916 | 0,718938698 | 0,521526636 | 0,053985485 | 0,045887663 | -0,113878485 | -0,155878485 | -1,415878485 |
| chrXVI | 235024 | 235161 | 137 | 0,14817715 | 0,188999349 | 0,439464645 | -0,152326671 | -0,12947767 | -0,202746474 | -0,153746474 | -1,413746474 |
| chrXVI | 238735 | 238888 | 153 | 0,521469587 | 0,511410002 | 0,504869679 | 0,012206779 | 0,010375762 | -0,057864364 | -0,120864364 | -1,380864364 |
| chrXVI | 239149 | 239288 | 139 | 0,227964847 | 0,270528479 | 0,457307721 | -0,107218749 | -0,091135937 | -0,170078608 | -0,135078608 | -1,395078608 |
| chrXVI | 240071 | 240209 | 138 | 0,156725832 | 0,114881957 | 0,577029962 | 0,194301161 | 0,165155987 | 0,097719347 | 0,139719347 | -1,120280653 |
| chrXVI | 240947 | 241098 | 151 | 0,125380666 | 0,070411522 | 0,640376244 | 0,359464657 | 0,305544959 | 0,176259947 | 0,127259947 | -1,132740053 |
| chrXVI | 242739 | 242891 | 152 | 0,453080133 | 0,374292827 | 0,547612932 | 0,119632674 | 0,101687773 | 0,032261237 | -0,023738763 | -1,283738763 |
| chrXVI | 242995 | 243131 | 136 | 0,530018269 | 0,663350655 | 0,444136141 | -0,140490722 | -0,119417114 | -0,17129551 | -0,11529551 | -1,37529551 |
| chrXVI | 244964 | 245114 | 150 | 4,325632966 | 3,090695229 | 0,583258029 | 0,210235413 | 0,178700101 | 0,010835994 | -0,031164006 | -1,291164006 |
| chrXVI | 246169 | 246331 | 162 | 0,339097709 | 0,34835174 | 0,493269301 | -0,016872161 | -0,014341337 | 0,052344659 | -0,073655341 | -1,333655341 |
| chrXVI | 247500 | 247655 | 155 | 0,612655525 | 0,44099848 | 0,581457976 | 0,205624744 | 0,174781032 | 0,073479895 | -0,003520105 | -1,263520105 |
| chrXVI | 247870 | 248025 | 155 | 0,142478029 | 0,155646522 | 0,477914444 | -0,055388588 | -0,0470803 | -0,152905312 | -0,229905312 | -1,489905312 |

TABLE 2

| Chrom | Start | End | Length | Section A | Section B | A/A+B | Z-score | Z * 0.85 | Phase correction | Length correction | ΔLK |
|---|---|---|---|---|---|---|---|---|---|---|---|
| chrXVI | 248518 | 248667 | 149 | 0,353345512 | 0,333528262 | 0,514425686 | 0,036167715 | 0,030742558 | -0,133423709 | -0,168423709 | -1,428423709 |
| chrXVI | 251350 | 251499 | 149 | 0,139628469 | 0,144528914 | 0,491377234 | -0,021615751 | -0,018373388 | -0,185166043 | -0,220166043 | -1,480166043 |
| chrXVI | 251647 | 251802 | 155 | 0,398938482 | 0,181587609 | 0,68720164 | 0,487933817 | 0,414743744 | 0,316276752 | 0,239276752 | -1,020723248 |
| chrXVI | 254749 | 254901 | 152 | 0,031345166 | 0,059293913 | 0,345823971 | -0,396619676 | -0,337126724 | -0,419333272 | -0,475333272 | -1,735333272 |
| chrXVI | 254749 | 254901 | 152 | 0,031345166 | 0,059293913 | 0,345823971 | -0,396619676 | -0,337126724 | -0,419333272 | -0,475333272 | -1,735333272 |
| chrXVI | 257303 | 257455 | 152 | 0,250761331 | 0,203822827 | 0,551627959 | 0,129775456 | 0,110309137 | 0,024207269 | -0,031792731 | -1,291792731 |
| chrXVI | 258470 | 258614 | 144 | 0,24506221 | 0,237175653 | 0,508177041 | 0,020498237 | 0,017423501 | -0,101375157 | -0,101375157 | -1,361375157 |
| chrXVI | 269441 | 269592 | 151 | 0,062690333 | 0,111176087 | 0,360566076 | -0,356946107 | -0,303404191 | -0,430677473 | -0,479677473 | -1,739677473 |
| chrXVI | 272088 | 272225 | 137 | 0,424584527 | 0,329822392 | 0,562805717 | 0,158086591 | 0,134373602 | 0,056416884 | 0,105416884 | -1,154583116 |
| chrXVI | 273849 | 274002 | 153 | 0,028495606 | 0,08894087 | 0,242646977 | -0,69781331 | -0,593141313 | -0,66119593 | -0,72419593 | -1,98419593 |
| chrXVI | 275518 | 275683 | 165 | 0,039893848 | 0,125999566 | 0,240478794 | -0,704763239 | -0,599048753 | -0,45657557 | -0,60357557 | -1,86357557 |
| chrXVI | 278767 | 278916 | 149 | 0,404637603 | 0,277940218 | 0,592808014 | 0,234774229 | 0,199558094 | 0,036683217 | 0,001683217 | -1,258316783 |
| chrXVI | 284968 | 285121 | 153 | 1,302249187 | 0,711526959 | 0,646670281 | 0,376346334 | 0,319894384 | 0,254644964 | 0,191644964 | -1,068355036 |
| chrXVI | 287278 | 287435 | 157 | 0,04844253 | 0,037058696 | 0,566571175 | 0,16765125 | 0,142503563 | 0,076699869 | -0,014300131 | -1,274300131 |
| chrXVI | 287516 | 287643 | 127 | 0,156725832 | 0,303881305 | 0,340259235 | -0,411755734 | -0,349992374 | 0,079661292 | 0,198661292 | -1,061338708 |
| chrXVI | 295119 | 295280 | 161 | 0,256460453 | 0,192705218 | 0,570970734 | 0,178846117 | 0,1520192 | 0,206771676 | 0,087771676 | -1,172228324 |
| chrXVI | 296343 | 296490 | 147 | 0,720938828 | 0,52993935 | 0,576346155 | 0,192554744 | 0,163671533 | 0,069383491 | 0,048383491 | -1,211616509 |
| chrXVI | 301254 | 301382 | 128 | 0,034194727 | 0,114881957 | 0,229376762 | -0,740900911 | -0,629765774 | -0,287572917 | -0,175572917 | -1,435572917 |
| chrXVI | 305851 | 306015 | 164 | 0,359044634 | 0,559586306 | 0,39084753 | -0,277110761 | -0,235544147 | -0,11799359 | -0,25799359 | -1,51799359 |
| chrXVI | 306387 | 306542 | 155 | 0,507221784 | 0,585527393 | 0,464170364 | -0,089932658 | -0,076442759 | -0,167147561 | -0,244147561 | -1,504147561 |
| chrXVI | 317763 | 317907 | 144 | 0,082637257 | 0,066705652 | 0,553339005 | 0,13410191 | 0,113986624 | 0,003337893 | 0,003337893 | -1,256662107 |
| chrXVI | 318699 | 318846 | 147 | 0,031345166 | 0,044470435 | 0,413439528 | -0,218706025 | -0,185900121 | -0,281030858 | -0,302030858 | -1,562030858 |
| chrXVI | 322025 | 322158 | 133 | 0,262159574 | 0,281646088 | 0,4820832 | -0,044925865 | -0,038186985 | 0,066638742 | 0,143638742 | -1,116361258 |
| chrXVI | 323166 | 323321 | 155 | 0,208017923 | 0,218646305 | 0,487544793 | -0,031225648 | -0,0265418 | -0,117678642 | -0,194678642 | -1,454678642 |
| chrXVI | 332594 | 332745 | 151 | 0,467327936 | 0,170470001 | 0,732720991 | 0,621063234 | 0,527903749 | 0,402555775 | 0,353555775 | -0,906444225 |
| chrXVI | 332948 | 333099 | 151 | 0,421734966 | 0,329822392 | 0,561148077 | 0,153880643 | 0,130798547 | 0,024875371 | -0,024124629 | -1,284124629 |
| chrXVI | 336537 | 336643 | 106 | 0,359044634 | 2,849813706 | 0,111891706 | -1,216529155 | -1,034049782 | -0,417968532 | -0,151968532 | -1,411968532 |
| chrXVI | 336537 | 336650 | 113 | 0,473027057 | 3,943045232 | 0,107114881 | -1,242018432 | -1,055715667 | -0,252826184 | -0,035826184 | -1,295826184 |
| chrXVI | 341289 | 341447 | 158 | 0,624053768 | 0,470645436 | 0,570068714 | 0,176549108 | 0,150066742 | 0,124506631 | 0,026506631 | -1,233493369 |
| chrXVI | 344751 | 344913 | 162 | 0,102584181 | 0,129705435 | 0,441621897 | -0,146858392 | -0,124829633 | -0,056602772 | -0,182602772 | -1,442602772 |
| chrXVI | 344751 | 344913 | 162 | 0,102584181 | 0,129705435 | 0,441621897 | -0,146858392 | -0,124829633 | -0,056602772 | -0,182602772 | -1,442602772 |
| chrXVI | 350439 | 350593 | 154 | 2,142869559 | 1,478641962 | 0,591705852 | 0,231935261 | 0,197144972 | 0,138858384 | 0,068858384 | -1,191141616 |
| chrXVI | 351311 | 351471 | 160 | 0,558513874 | 0,770820872 | 0,420145397 | -0,201521534 | -0,171293304 | -0,17899552 | -0,29099552 | -1,55099552 |
| chrXVI | 354968 | 355128 | 160 | 0,703841464 | 0,61146848 | 0,535114531 | 0,088133038 | 0,074913082 | 0,071589949 | -0,040410051 | -1,300410051 |
| chrXVI | 361977 | 362073 | 96 | 0,062690333 | 0,166764131 | 0,273214701 | -0,603119188 | -0,512651309 | 0,031229568 | 0,367229568 | -0,892770432 |
| chrXVI | 364336 | 364488 | 152 | 0,923257629 | 0,722644568 | 0,56094319 | 0,153360973 | 0,130356827 | 0,043945004 | -0,012054996 | -1,272054996 |
| chrXVI | 366545 | 366652 | 107 | 0,299203861 | 1,560171093 | 0,160916366 | -0,990698686 | -0,842093883 | -0,179861549 | 0,079138451 | -1,180861549 |
| chrXVI | 366545 | 366653 | 108 | 0,364743755 | 2,364344791 | 0,133650392 | -1,109300001 | -0,942905001 | -0,157589457 | 0,094410543 | -1,165589457 |
| chrXVI | 367044 | 367180 | 136 | 0,595558162 | 0,541056958 | 0,523975224 | 0,060133194 | 0,051113215 | -0,004608464 | 0,051391536 | -1,208608464 |
| chrXVI | 370599 | 370734 | 135 | 6,032519755 | 4,947335887 | 0,549417037 | 0,124188628 | 0,105560334 | 0,092826834 | 0,155826834 | -1,104173166 |
| chrXVI | 370905 | 371054 | 149 | 0,917558508 | 0,455821958 | 0,668102198 | 0,434678778 | 0,369476962 | 0,199444908 | 0,164444908 | -1,095555092 |
| chrXVI | 373474 | 373616 | 142 | 0,19377012 | 0,118587827 | 0,620346375 | 0,30639062 | 0,260432027 | 0,153772514 | 0,167772514 | -1,092227486 |
| chrXVI | 379313 | 379470 | 157 | 1,307948308 | 1,341524787 | 0,493663555 | -0,015883779 | -0,013501212 | -0,073126068 | -0,164126068 | -1,424126068 |
| chrXVI | 379794 | 379947 | 153 | 1,541612276 | 0,922761525 | 0,625559432 | 0,320115026 | 0,272097772 | 0,207517019 | 0,144517019 | -1,115482981 |
| chrXVI | 379982 | 380135 | 153 | 0,19661968 | 0,214940436 | 0,47774231 | -0,055820732 | -0,047447622 | -0,105326443 | -0,168326443 | -1,428326443 |
| chrXVI | 379982 | 380135 | 153 | 0,19661968 | 0,214940436 | 0,47774231 | -0,055820732 | -0,047447622 | -0,105326443 | -0,168326443 | -1,428326443 |
| chrXVI | 383644 | 383787 | 143 | 1,134125112 | 0,696703481 | 0,619460017 | 0,304062908 | 0,258453472 | 0,163666316 | 0,170666316 | -1,089333684 |
| chrXVI | 385233 | 385386 | 153 | 0,723788388 | 0,463233697 | 0,609751408 | 0,278671178 | 0,236870501 | 0,17357967 | 0,11057967 | -1,14942033 |
| chrXVI | 389294 | 389391 | 97 | 0,34194727 | 1,152525439 | 0,228807972 | -0,742778252 | -0,631361514 | -0,079099859 | 0,249900141 | -1,010099859 |
| chrXVI | 390081 | 390232 | 151 | 6,568237145 | 3,828163275 | 0,631779932 | 0,336571244 | 0,286085557 | 0,1806827 | 0,1316827 | -1,1283173 |
| chrXVI | 390081 | 390232 | 151 | 6,568237145 | 3,828163275 | 0,631779932 | 0,336571244 | 0,286085557 | 0,1806827 | 0,1316827 | -1,1283173 |
| chrXVI | 390545 | 390691 | 146 | 0,225115286 | 0,200116957 | 0,529393736 | 0,07374596 | 0,062684066 | -0,047041103 | -0,061041103 | -1,321041103 |
| chrXVI | 392582 | 392731 | 149 | 0,045592969 | 0,048176305 | 0,486225044 | -0,034535557 | -0,029355224 | -0,210449478 | -0,245449478 | -1,505449478 |
| chrXVI | 392582 | 392736 | 154 | 13,41858079 | 10,65066917 | 0,557498917 | 0,144631069 | 0,122936409 | 0,070140807 | 0,000140807 | -1,259859193 |
| chrXVI | 394420 | 394573 | 153 | 0,133929347 | 0,166764131 | 0,44540157 | -0,137288019 | -0,116694816 | -0,177244918 | -0,240244918 | -1,500244918 |
| chrXVI | 407155 | 407307 | 152 | 11,85417203 | 7,156034156 | 0,62356883 | 0,314867393 | 0,267637284 | 0,188118205 | 0,132118205 | -1,127881795 |
| chrXVI | 407160 | 407307 | 147 | 0,037044288 | 0,037058696 | 0,499902783 | -0,000243688 | -0,000207135 | -0,097323552 | -0,118323552 | -1,378323552 |
| chrXVI | 411818 | 411982 | 164 | 0,182371877 | 0,270528479 | 0,4026755 | -0,246427915 | -0,209463728 | -0,080884868 | -0,220884868 | -1,480884868 |
| chrXVI | 411838 | 411991 | 153 | 0,039893848 | 0,062999783 | 0,387719315 | -0,285268256 | -0,242478018 | -0,307554274 | -0,370554274 | -1,630554274 |
| chrXVI | 412957 | 413111 | 154 | 0,153876272 | 0,114881957 | 0,572545341 | 0,182858153 | 0,15542943 | 0,103556768 | 0,033556768 | -1,226443232 |
| chrXVI | 415881 | 416027 | 146 | 0,062690333 | 0,051882174 | 0,547167331 | 0,118507771 | 0,100731606 | -0,016705482 | -0,030705482 | -1,290705482 |
| chrXVI | 417433 | 417565 | 132 | 0,299203861 | 0,607762611 | 0,329895173 | -0,440202642 | -0,374172246 | -0,23443892 | -0,15043892 | -1,41043892 |
| chrXVI | 418493 | 418641 | 148 | 0,686744101 | 0,622586089 | 0,524500318 | 0,061451844 | 0,052234068 | -0,102537882 | -0,130537882 | -1,390537882 |
| chrXVI | 418678 | 418846 | 168 | 0,495823542 | 0,692997611 | 0,417071601 | -0,209390764 | -0,17798215 | 0,01831689 | -0,14968311 | -1,40968311 |
| chrXVI | 418678 | 418846 | 168 | 0,495823542 | 0,692997611 | 0,417071601 | -0,209390764 | -0,17798215 | 0,01831689 | -0,14968311 | -1,40968311 |
| chrXVI | 432250 | 432440 | 190 | 0,712390146 | 1,289642613 | 0,355833411 | -0,369618423 | -0,31417566 | -0,113972439 | -0,435972439 | -1,695972439 |
| chrXVI | 432594 | 432737 | 143 | 1,929152515 | 1,059878699 | 0,645410629 | 0,372959291 | 0,317015397 | 0,219382274 | 0,226382274 | -1,033617726 |
| chrXVI | 432908 | 433068 | 160 | 0,467327936 | 0,207528696 | 0,692484764 | 0,502905845 | 0,427469968 | 0,422488385 | 0,310488385 | -0,949511615 |
| chrXVI | 433481 | 433638 | 157 | 0,282106498 | 0,222352175 | 0,559226183 | 0,149007602 | 0,126656462 | 0,079414991 | -0,011585009 | -1,271585009 |
| chrXVI | 437116 | 437278 | 162 | 0,159575393 | 0,163058261 | 0,49460244 | -0,013530088 | -0,011500575 | 0,07246741 | -0,05353259 | -1,31353259 |
| chrXVI | 439535 | 439688 | 153 | 0,128230226 | 0,370586958 | 0,257068582 | -0,652409303 | -0,554547908 | -0,620673809 | -0,683673809 | -1,943673809 |
| chrXVI | 442027 | 442174 | 147 | 0,111132863 | 0,133411305 | 0,454449042 | -0,114428548 | -0,097264266 | -0,199401127 | -0,220401127 | -1,480401127 |
| chrXVI | 442326 | 442474 | 148 | 0,256460453 | 0,259410871 | 0,497140355 | -0,007168129 | -0,00609291 | -0,159406976 | -0,187406976 | -1,447406976 |
| chrXVI | 442340 | 442502 | 162 | 0,059840772 | 0,062999783 | 0,487141826 | -0,032236246 | -0,027400809 | 0,059165536 | -0,066834464 | -1,326834464 |
| chrXVI | 446281 | 446447 | 166 | 0,071239015 | 0,08894087 | 0,444743825 | -0,138952542 | -0,118109661 | 0,062858032 | -0,091141968 | -1,351141968 |
| chrXVI | 448877 | 449010 | 133 | 0,869115978 | 1,222936961 | 0,415436895 | -0,213581034 | -0,181543879 | -0,090597369 | -0,013597369 | -1,273597369 |
| chrXVI | 454839 | 454945 | 106 | 0,119681545 | 0,274234349 | 0,303825122 | -0,513430458 | -0,436415889 | 0,182072443 | 0,448072443 | -0,811927557 |
| chrXVI | 462929 | 463059 | 130 | 0,151026711 | 0,274234349 | 0,35513882 | -0,371483236 | -0,315760751 | -0,13599693 | -0,03799693 | -1,29799693 |
| chrXVI | 465016 | 465172 | 156 | 0,327699467 | 0,26311674 | 0,554655514 | 0,137432464 | 0,116817594 | 0,046743756 | -0,037256244 | -1,297256244 |
| chrXVI | 466157 | 466315 | 158 | 1,134125112 | 1,096937395 | 0,508334082 | 0,020891966 | 0,017758171 | 0,005085613 | -0,092914387 | -1,352914387 |
| chrXVI | 474538 | 474690 | 152 | 0,914708947 | 0,555880437 | 0,622001598 | 0,31074195 | 0,264130657 | 0,206252996 | 0,150252996 | -1,109747004 |
| chrXVI | 484126 | 484288 | 162 | 0,062690333 | 0,081529131 | 0,434687048 | -0,164453569 | -0,139785534 | -0,06068286 | -0,18668286 | -1,44668286 |
| chrXVI | 484126 | 484288 | 162 | 0,062690333 | 0,081529131 | 0,434687048 | -0,164453569 | -0,139785534 | -0,06068286 | -0,18668286 | -1,44668286 |
| chrXVI | 484807 | 484963 | 156 | 0,612655525 | 0,596645002 | 0,506619745 | 0,016594003 | 0,014104902 | -0,047719747 | -0,131719747 | -1,391719747 |
| chrXVI | 487013 | 487184 | 171 | 0,071239015 | 0,196411088 | 0,26616472 | -0,624454047 | -0,53078594 | -0,323409896 | -0,512409896 | -1,772409896 |
| chrXVI | 493217 | 493377 | 160 | 0,364743755 | 0,314998914 | 0,536590936 | 0,091848853 | 0,078071525 | 0,070304637 | -0,041695363 | -1,301695363 |
| chrXVI | 497274 | 497428 | 154 | 1,193965884 | 0,763409133 | 0,609983204 | 0,279275258 | 0,23738397 | 0,163330908 | 0,093330908 | -1,166669092 |
| chrXVI | 501076 | 501229 | 153 | 0,088336378 | 0,085235 | 0,508934013 | 0,022396121 | 0,019036703 | -0,061027529 | -0,124027529 | -1,384027529 |
| chrXVI | 501841 | 501999 | 158 | 0,216566604 | 0,203822827 | 0,51515711 | 0,038002385 | 0,032302027 | 0,02105665 | -0,07694335 | -1,33694335 |
| chrXVI | 505233 | 505380 | 147 | 1,085682582 | 1,156231309 | 0,48426596 | -0,03944962 | -0,033532177 | -0,152088424 | -0,173088424 | -1,433088424 |
| chrXVI | 506320 | 506478 | 158 | 0,076938136 | 0,081529131 | 0,485514374 | -0,036318061 | -0,030870352 | -0,048132268 | -0,146132268 | -1,406132268 |
| chrXVI | 506594 | 506750 | 156 | 0,344796831 | 0,281646088 | 0,550404227 | 0,126682692 | 0,107680288 | 0,036341015 | -0,047658985 | -1,307658985 |
| chrXVI | 509864 | 510024 | 160 | 0,094035499 | 0,155646522 | 0,376621026 | -0,314367347 | -0,267212245 | -0,278989859 | -0,390989859 | -1,650989859 |
| chrXVI | 511391 | 511535 | 144 | 0,042743409 | 0,107470218 | 0,284550808 | -0,569375094 | -0,48396883 | -0,59231565 | -0,59231565 | -1,85231565 |
| chrXVI | 517443 | 517595 | 152 | 0,273557816 | 0,222352175 | 0,551627959 | 0,129775456 | 0,110309137 | 0,060769391 | 0,004769391 | -1,255230609 |
| chrXVI | 519049 | 519173 | 124 | 0,133929347 | 0,566998046 | 0,191074495 | -0,873943563 | -0,742852028 | -0,173611838 | -0,033611838 | -1,293611838 |
| chrXVI | 520082 | 520209 | 127 | 0,165274514 | 0,296469566 | 0,357935317 | -0,363983093 | -0,309385629 | 0,090349118 | 0,209349118 | -1,050650882 |
| chrXVI | 522508 | 522654 | 146 | 1,005894886 | 0,959820221 | 0,511719568 | 0,029380828 | 0,024973704 | -0,098160162 | -0,112160162 | -1,372160162 |
| chrXVI | 522574 | 522734 | 160 | 3,37103017 | 3,494635013 | 0,490998335 | -0,022565744 | -0,019180883 | -0,023786133 | -0,135786133 | -1,395786133 |
| chrXVI | 525954 | 526108 | 154 | 0,082637257 | 0,111176087 | 0,426375476 | -0,185609565 | -0,15776813 | -0,22572356 | -0,29572356 | -1,55572356 |
| chrXVI | 536923 | 537076 | 153 | 0,453080133 | 0,329822392 | 0,578718446 | 0,198616037 | 0,168823631 | 0,098347265 | 0,035347265 | -1,224652735 |

TABLE 2

| Chrom | Start | End | Length | Section A | Section B | A/A+B | Z-score | Z * 0.85 | Phase correction | Length correction | ΔLK |
|---|---|---|---|---|---|---|---|---|---|---|---|
| chrXVI | 538001 | 538163 | 162 | 0,131079787 | 0,214940436 | 0,378821174 | -0,308578276 | -0,262291535 | -0,17921883 | -0,30521883 | -1,56521883 |
| chrXVI | 541695 | 541831 | 136 | 0,963151477 | 0,566998046 | 0,629449255 | 0,330395039 | 0,280835783 | 0,224662772 | 0,280662772 | -0,979337228 |
| chrXVI | 547021 | 547160 | 139 | 2,866657947 | 2,056757616 | 0,582249845 | 0,207652497 | 0,176504623 | 0,086538654 | 0,121538654 | -1,138461346 |
| chrXVI | 547636 | 547771 | 135 | 0,837770812 | 0,822703046 | 0,504537188 | 0,01137329 | 0,009667296 | -0,013576363 | 0,049423637 | -1,210576363 |
| chrXVI | 547851 | 548005 | 154 | 0,681044979 | 0,485468915 | 0,583829291 | 0,211699576 | 0,17994464 | 0,11914273 | 0,04914273 | -1,21085727 |
| chrXVI | 550617 | 550788 | 171 | 0,039893848 | 0,051882174 | 0,434687048 | -0,164453569 | -0,139785534 | 0,071112288 | -0,117887712 | -1,377887712 |
| chrXVI | 553457 | 553564 | 107 | 0,854868175 | 5,8960385 | 0,126630128 | -1,142466277 | -0,971096336 | -0,303362682 | -0,044362682 | -1,304362682 |
| chrXVI | 553457 | 553565 | 108 | 0,983098401 | 8,61614677 | 0,102414136 | -1,267915097 | -1,077727832 | -0,268748346 | -0,016748346 | -1,276748346 |
| chrXVI | 560771 | 560926 | 155 | 0,413186285 | 0,237175653 | 0,635317445 | 0,345970201 | 0,294074671 | 0,208028222 | 0,131028222 | -1,128971778 |
| chrXVI | 560771 | 560926 | 155 | 0,413186285 | 0,237175653 | 0,635317445 | 0,345970201 | 0,294074671 | 0,208028222 | 0,131028222 | -1,128971778 |
| chrXVI | 564313 | 564461 | 148 | 0,068389454 | 0,107470218 | 0,38888651 | -0,282222352 | -0,239889 | -0,394215094 | -0,422215094 | -1,682215094 |
| chrXVI | 564317 | 564438 | 121 | 1,082833022 | 3,583575882 | 0,232048465 | -0,732117375 | -0,622299769 | 0,012372237 | 0,173372237 | -1,086627763 |
| chrXVI | 566468 | 566620 | 152 | 0,04844253 | 0,051882174 | 0,482857442 | -0,042983253 | -0,036535765 | -0,095216924 | -0,151216924 | -1,411216924 |
| chrXVI | 566468 | 566620 | 152 | 0,04844253 | 0,051882174 | 0,482857442 | -0,042983253 | -0,036535765 | -0,095216924 | -0,151216924 | -1,411216924 |
| chrXVI | 568077 | 568211 | 134 | 0,24506221 | 0,270528479 | 0,475303793 | -0,061943802 | -0,052652232 | -0,01877957 | 0,05122043 | -1,20877957 |
| chrXVI | 573444 | 573609 | 165 | 0,079787696 | 0,155646522 | 0,338895921 | -0,415478239 | -0,353156503 | -0,19920184 | -0,34620184 | -1,60620184 |
| chrXVI | 573688 | 573839 | 151 | 0,532867829 | 0,318704784 | 0,625745616 | 0,320606293 | 0,272515349 | 0,164946553 | 0,115946553 | -1,144053447 |
| chrXVI | 578898 | 579048 | 150 | 0,102584181 | 0,044470435 | 0,69759239 | 0,517488464 | 0,439865194 | 0,284575145 | 0,242575145 | -1,017424855 |
| chrXVI | 580106 | 580256 | 150 | 0,578460798 | 0,452116089 | 0,561298051 | 0,154261063 | 0,131121903 | -0,028128816 | -0,070128816 | -1,330128816 |
| chrXVI | 581758 | 581907 | 149 | 0,136778908 | 0,140823044 | 0,492715945 | -0,018259434 | -0,015520519 | -0,193594746 | -0,228594746 | -1,488594746 |
| chrXVI | 581758 | 581907 | 149 | 0,136778908 | 0,140823044 | 0,492715945 | -0,018259434 | -0,015520519 | -0,193594746 | -0,228594746 | -1,488594746 |
| chrXVI | 582967 | 583116 | 149 | 0,19377012 | 0,129705435 | 0,599025542 | 0,250825642 | 0,213201796 | 0,042621237 | 0,007621237 | -1,252378763 |
| chrXVI | 584868 | 585014 | 146 | 4,174606255 | 3,187047838 | 0,567074492 | 0,168930872 | 0,143591241 | 0,023436187 | 0,009436187 | -1,250563813 |
| chrXVI | 591560 | 591713 | 153 | 1,655594699 | 1,122878482 | 0,59586492 | 0,24265824 | 0,206259504 | 0,143720795 | 0,080720795 | -1,179279205 |
| chrXVI | 592210 | 592352 | 142 | 0,233663968 | 0,163058261 | 0,588986325 | 0,224938202 | 0,191197471 | 0,104031949 | 0,118031949 | -1,141968051 |
| chrXVI | 593528 | 593683 | 155 | 0,119681545 | 0,103764348 | 0,535617563 | 0,08939893 | 0,07598909 | -0,000265577 | -0,077265577 | -1,337265577 |
| chrXVI | 594996 | 595138 | 142 | 0,706691025 | 0,563292176 | 0,556456986 | 0,141992377 | 0,12069352 | 0,034375069 | 0,048375069 | -1,211624931 |
| chrXVI | 599046 | 599200 | 154 | 1,550160957 | 1,489759571 | 0,509934698 | 0,02490517 | 0,021169395 | -0,039215328 | -0,109215328 | -1,369215328 |
| chrXVI | 600831 | 600997 | 166 | 0,227964847 | 0,281646088 | 0,447331152 | -0,132407093 | -0,112546029 | 0,064655484 | -0,089344516 | -1,349344516 |
| chrXVI | 600831 | 600997 | 166 | 0,227964847 | 0,281646088 | 0,447331152 | -0,132407093 | -0,112546029 | 0,064655484 | -0,089344516 | -1,349344516 |
| chrXVI | 601108 | 601231 | 123 | 0,159575393 | 0,444704349 | 0,264075364 | -0,630831465 | -0,536206745 | 0,040828219 | 0,187828219 | -1,072171781 |
| chrXVI | 603139 | 603315 | 176 | 0,065539893 | 0,155646522 | 0,296310663 | -0,535041262 | -0,454785073 | -0,232632526 | -0,456632526 | -1,716632526 |
| chrXVI | 604400 | 604533 | 133 | 0,598407723 | 0,822703046 | 0,421084503 | -0,199119834 | -0,169251859 | -0,093190771 | -0,016190771 | -1,276190771 |
| chrXVI | 611112 | 611263 | 151 | 0,156725832 | 0,044470435 | 0,778969881 | 0,768718839 | 0,653411013 | 0,545270986 | 0,496270986 | -0,763729014 |
| chrXVI | 611769 | 611929 | 160 | 0,256460453 | 0,300175436 | 0,460732874 | -0,09858756 | -0,083799426 | -0,081074942 | -0,193074942 | -1,453074942 |
| chrXVI | 611769 | 611929 | 160 | 0,256460453 | 0,300175436 | 0,460732874 | -0,09858756 | -0,083799426 | -0,081074942 | -0,193074942 | -1,453074942 |
| chrXVI | 614264 | 614371 | 107 | 0,056991212 | 0,407645654 | 0,122657533 | -1,161804056 | -0,987533448 | -0,319799794 | -0,060799794 | -1,320799794 |
| chrXVI | 614264 | 614381 | 117 | 0,039893848 | 0,500292393 | 0,073852026 | -1,447688987 | -1,230535639 | -0,505300435 | -0,316300435 | -1,576300435 |
| chrXVI | 616962 | 617122 | 160 | 0,464478375 | 0,570703915 | 0,448692351 | -0,128965812 | -0,10962094 | -0,099539962 | -0,211539962 | -1,471539962 |
| chrXVI | 617750 | 617867 | 117 | 0,92610719 | 4,39516132 | 0,174038801 | -0,938324639 | -0,797575943 | -0,084317306 | 0,104682694 | -1,155317306 |
| chrXVI | 617888 | 618049 | 161 | 0,29065518 | 0,296469566 | 0,495048423 | -0,012412082 | -0,010550269 | 0,049198185 | -0,069801815 | -1,329801815 |
| chrXVI | 623817 | 623987 | 170 | 0,276407377 | 0,26311674 | 0,512316999 | 0,030879046 | 0,026247189 | 0,225770625 | 0,043770625 | -1,216229375 |
| chrXVI | 627166 | 627316 | 150 | 0,068389454 | 0,062999783 | 0,520510322 | 0,051434423 | 0,043719259 | -0,119535304 | -0,161535304 | -1,421535304 |
| chrXVI | 630544 | 630672 | 128 | 0,598407723 | 0,863467612 | 0,409342513 | -0,229236602 | -0,194851112 | 0,129090979 | 0,241090979 | -1,018909021 |
| chrXVI | 630787 | 630947 | 160 | 1,698338108 | 1,23405457 | 0,579164626 | 0,199756852 | 0,169793324 | 0,185555735 | 0,073555735 | -1,186444265 |
| chrXVI | 639745 | 639881 | 136 | 0,131079787 | 0,229763914 | 0,36325918 | -0,349760614 | -0,297296522 | -0,351834519 | -0,295834519 | -1,555834519 |
| chrXVI | 644095 | 644255 | 160 | 0,313451664 | 0,159352392 | 0,662963145 | 0,420563695 | 0,357479141 | 0,37219519 | 0,26019519 | -0,99980481 |
| chrXVI | 647938 | 648089 | 151 | 0,410336724 | 0,489174784 | 0,456177292 | -0,110069087 | -0,093558724 | -0,210657531 | -0,259657531 | -1,519657531 |
| chrXVI | 651006 | 651170 | 164 | 0,339097709 | 0,407645654 | 0,454102073 | -0,115304027 | -0,098008423 | 0,03365597 | -0,10634403 | -1,36634403 |
| chrXVI | 651448 | 651577 | 129 | 1,95479856 | 1,952993268 | 0,500230986 | 0,000578997 | 0,000492147 | 0,254196828 | 0,359196828 | -0,900803172 |
| chrXVI | 651814 | 651966 | 152 | 0,421734966 | 0,329822392 | 0,561148077 | 0,153880643 | 0,130798547 | 0,063436251 | 0,007436251 | -1,252563749 |
| chrXVI | 655669 | 655818 | 149 | 0,059840772 | 0,037058696 | 0,617555219 | 0,299066166 | 0,254206241 | 0,088106798 | 0,053106798 | -1,206893202 |
| chrXVI | 661336 | 661465 | 129 | 6,830396719 | 10,99531504 | 0,383176661 | -0,297148258 | -0,252576019 | -0,010643834 | 0,094356166 | -1,165643834 |
| chrXVI | 664409 | 664571 | 162 | 0,102584181 | 0,118587827 | 0,463820816 | -0,090812431 | -0,077190567 | 0,008108976 | -0,117891024 | -1,377891024 |
| chrXVI | 665020 | 665144 | 124 | 0,273557816 | 1,000584786 | 0,214699529 | -0,790220412 | -0,67168735 | -0,128296211 | 0,011703789 | -1,248296211 |
| chrXVI | 670319 | 670426 | 107 | 0,165274514 | 1,719523484 | 0,087688184 | -1,35512929 | -1,151859897 | -0,463598676 | -0,204598676 | -1,464598676 |
| chrXVI | 670319 | 670427 | 108 | 0,156725832 | 2,127169138 | 0,068622171 | -1,486131496 | -1,263211772 | -0,425003482 | -0,173003482 | -1,433003482 |
| chrXVI | 670635 | 670782 | 147 | 0,256460453 | 0,148234783 | 0,633712557 | 0,341702373 | 0,290447017 | 0,163258251 | 0,142258251 | -1,117741749 |
| chrXVI | 672000 | 672158 | 158 | 0,549965193 | 0,518821741 | 0,514569532 | 0,036528524 | 0,031049245 | 0,015474477 | -0,082525523 | -1,342525523 |
| chrXVI | 678051 | 678201 | 150 | 5,525297971 | 3,483517404 | 0,613321257 | 0,28798596 | 0,244788066 | 0,073993351 | 0,031993351 | -1,228006649 |
| chrXVI | 679094 | 679205 | 111 | 7,468698289 | 14,20459809 | 0,344603708 | -0,399930896 | -0,339941262 | 0,464294236 | 0,695294236 | -0,564705764 |
| chrXVI | 679094 | 679206 | 112 | 0,182371877 | 0,34835174 | 0,34362872 | -0,402579708 | -0,342192752 | 0,461104681 | 0,685104681 | -0,574895319 |
| chrXVI | 679094 | 679207 | 113 | 0,108283302 | 0,35205761 | 0,235224155 | -0,721749813 | -0,613487341 | 0,181783152 | 0,398783152 | -0,861216848 |
| chrXVI | 679098 | 679205 | 107 | 0,165274514 | 0,870879351 | 0,159507694 | -0,996483277 | -0,847010785 | -0,143856442 | 0,115143558 | -1,144856442 |
| chrXVI | 679098 | 679220 | 122 | 0,378991558 | 1,700994137 | 0,182208733 | -0,906979826 | -0,770932852 | -0,128831814 | 0,025168186 | -1,234831814 |
| chrXVI | 681181 | 681332 | 151 | 0,113982423 | 0,240881523 | 0,321200349 | -0,464344846 | -0,394693119 | -0,512606949 | -0,561606949 | -1,821606949 |
| chrXVI | 685561 | 685687 | 126 | 0,045592969 | 0,207528696 | 0,180122745 | -0,914897409 | -0,777662797 | -0,32376229 | -0,19776229 | -1,45776229 |
| chrXVI | 685866 | 686024 | 158 | 0,595558162 | 0,581821524 | 0,505833563 | 0,014623096 | 0,012429632 | -0,004667769 | -0,102667769 | -1,362667769 |
| chrXVI | 689048 | 689183 | 135 | 0,108283302 | 0,159352392 | 0,404592155 | -0,241478459 | -0,20525669 | -0,226976798 | -0,163976798 | -1,423976798 |
| chrXVI | 695989 | 696136 | 147 | 1,410532489 | 0,926467395 | 0,603565494 | 0,262586882 | 0,223198849 | 0,072951441 | 0,051951441 | -1,208048559 |
| chrXVI | 697045 | 697195 | 150 | 0,208017923 | 0,248293262 | 0,45586856 | -0,110847699 | -0,094220544 | -0,266498098 | -0,308498098 | -1,568498098 |
| chrXVI | 702075 | 702202 | 127 | 0,455929693 | 1,019114134 | 0,309095693 | -0,498415257 | -0,423652969 | -0,042902945 | 0,076097055 | -1,183902945 |
| chrXVI | 706433 | 706576 | 143 | 0,421734966 | 0,270528479 | 0,609211665 | 0,277264951 | 0,235675208 | 0,129757457 | 0,136757457 | -1,123242543 |
| chrXVI | 719549 | 719656 | 107 | 0,028495606 | 0,144528914 | 0,164691143 | -0,975358855 | -0,829055027 | -0,104596891 | 0,154403109 | -1,105596891 |
| chrXVI | 719549 | 719694 | 145 | 13,29889924 | 7,856443507 | 0,628630762 | 0,328229063 | 0,278994703 | 0,174476475 | 0,167476475 | -1,092523525 |
| chrXVI | 721766 | 721909 | 143 | 1,535913155 | 1,608347397 | 0,488481514 | -0,028876575 | -0,024545089 | -0,139070623 | -0,132070623 | -1,392070623 |
| chrXVI | 722565 | 722701 | 136 | 0,082637257 | 0,118587827 | 0,410670755 | -0,225819886 | -0,191946903 | -0,250055797 | -0,194055797 | -1,454055797 |
| chrXVI | 726383 | 726544 | 161 | 0,113982423 | 0,107470218 | 0,514703382 | 0,036864261 | 0,031334622 | 0,093453914 | -0,025546086 | -1,285546086 |
| chrXVI | 726865 | 727012 | 147 | 0,344796831 | 0,211234566 | 0,620103168 | 0,305751756 | 0,259888992 | 0,10306677 | 0,08206677 | -1,17793323 |
| chrXVI | 730093 | 730221 | 128 | 0,621204207 | 0,79305609 | 0,439243192 | -0,152888273 | -0,129955032 | 0,170166285 | 0,282166285 | -0,977833715 |
| chrXVI | 732571 | 732734 | 163 | 0,792177842 | 0,770820872 | 0,50683205 | 0,017126246 | 0,014557309 | 0,129947753 | -0,003052247 | -1,263052247 |
| chrXVI | 733908 | 734067 | 159 | 0,056991212 | 0,055588044 | 0,506231912 | 0,015621722 | 0,013278464 | 0,008775763 | -0,096224237 | -1,356224237 |
| chrXVI | 734318 | 734467 | 149 | 5,872944363 | 3,194459577 | 0,647698548 | 0,379114412 | 0,32224725 | 0,165601612 | 0,130601612 | -1,129398388 |
| chrXVI | 735641 | 735790 | 149 | 0,612655525 | 0,652233046 | 0,484355333 | -0,039225421 | -0,033341608 | -0,187353676 | -0,222353676 | -1,482353676 |
| chrXVI | 735832 | 735978 | 146 | 1,689789426 | 1,059878699 | 0,614543046 | 0,291179674 | 0,247502723 | 0,127043061 | 0,113043061 | -1,146956939 |
| chrXVI | 736889 | 737030 | 141 | 0,484425299 | 0,281646088 | 0,632350075 | 0,33808405 | 0,287371443 | 0,17399213 | 0,19499213 | -1,06500787 |
| chrXVI | 737626 | 737736 | 110 | 0,87766466 | 8,834793075 | 0,090364837 | -1,338511746 | -1,137734984 | -0,316392997 | -0,078392997 | -1,338392997 |
| chrXVI | 739761 | 739921 | 160 | 0,170973635 | 0,188999349 | 0,474962408 | -0,062801193 | -0,053381014 | -0,035490435 | -0,147490435 | -1,407490435 |
| chrXVI | 740972 | 741124 | 152 | 1,635647775 | 1,600935658 | 0,505362463 | 0,013442106 | 0,01142579 | -0,063536127 | -0,119536127 | -1,379536127 |
| chrXVI | 742321 | 742473 | 152 | 0,615505086 | 0,444704349 | 0,58055047 | 0,203301913 | 0,172806626 | 0,096587429 | 0,040587429 | -1,219412571 |
| chrXVI | 745613 | 745766 | 153 | 0,136778908 | 0,196411088 | 0,41051325 | -0,226224907 | -0,192291171 | -0,259380352 | -0,322380352 | -1,582380352 |
| chrXVI | 747443 | 747595 | 152 | 0,43883233 | 0,633703698 | 0,409153929 | -0,229721923 | -0,195263635 | -0,262364658 | -0,318364658 | -1,578364658 |
| chrXVI | 748166 | 748325 | 159 | 0,062690333 | 0,114881957 | 0,353041192 | -0,377122752 | -0,320554339 | -0,321567836 | -0,426567836 | -1,686567836 |
| chrXVI | 751513 | 751665 | 152 | 0,165274514 | 0,17417587 | 0,486888575 | -0,032871387 | -0,027940679 | -0,074822849 | -0,130822849 | -1,390822849 |
| chrXVI | 753264 | 753415 | 151 | 2,430675178 | 2,757166966 | 0,468532987 | -0,078958069 | -0,067114359 | -0,183754888 | -0,232754888 | -1,492754888 |
| chrXVI | 753264 | 753415 | 151 | 2,430675178 | 2,757166966 | 0,468532987 | -0,078958069 | -0,067114359 | -0,183754888 | -0,232754888 | -1,492754888 |
| chrXVI | 753454 | 753608 | 154 | 0,484425299 | 0,337234132 | 0,589569451 | 0,226437584 | 0,192471947 | 0,129701139 | 0,059701139 | -1,200298861 |
| chrXVI | 759400 | 759547 | 147 | 0,510071344 | 0,34835174 | 0,594195745 | 0,238351462 | 0,202598743 | 0,055028974 | 0,034028974 | -1,225971026 |
| chrXVI | 761447 | 761589 | 142 | 0,188070999 | 0,125999566 | 0,598817654 | 0,250287928 | 0,212744739 | 0,119764611 | 0,133764611 | -1,126235389 |
| chrXVI | 761447 | 761589 | 142 | 0,188070999 | 0,125999566 | 0,598817654 | 0,250287928 | 0,212744739 | 0,119764611 | 0,133764611 | -1,126235389 |
| chrXVI | 763063 | 763215 | 152 | 0,111132863 | 0,085235 | 0,565942212 | 0,166052574 | 0,141144688 | 0,098397692 | 0,042397692 | -1,217602308 |

TABLE 2

| Chrom | Start | End | Length | Section A | Section B | A/A+B | Z-score | Z * 0.85 | Phase correction | Length correction | ΔLK |
|---|---|---|---|---|---|---|---|---|---|---|---|
| chrXVI | 763430 | 763578 | 148 | 0,230814407 | 0,170470001 | 0,575189075 | 0,18960094 | 0,161160799 | 0,011122545 | -0,016877455 | **-1,276877455** |
| chrXVI | 764397 | 764544 | 147 | 0,04844253 | 0,059293913 | 0,449639217 | -0,126572917 | -0,10758698 | -0,254365453 | -0,275365453 | **-1,535365453** |
| chrXVI | 764502 | 764659 | 157 | 0,29065518 | 0,429880871 | 0,403387421 | -0,244588792 | -0,207900474 | -0,255737048 | -0,346737048 | **-1,606737048** |
| chrXVI | 770323 | 770455 | 132 | 1,07428434 | 1,007996525 | 0,515917117 | 0,039908886 | 0,033922553 | 0,168242995 | 0,252242995 | **-1,007757005** |
| chrXVI | 778559 | 778687 | 128 | 0,045592969 | 0,233469783 | 0,163378913 | -0,980665288 | -0,833565495 | -0,547727508 | -0,435727508 | **-1,695727508** |
| chrXVI | 786310 | 786475 | 165 | 0,056991212 | 0,070411522 | 0,447331152 | -0,132407093 | -0,112546029 | 0,048821568 | -0,098178432 | **-1,358178432** |
| chrXVI | 788541 | 788654 | 113 | 0,059840772 | 0,233469783 | 0,204018475 | -0,827353109 | -0,703250143 | 0,085571326 | 0,302571326 | **-0,957428674** |
| chrXVI | 788648 | 788776 | 128 | 1,256656217 | 1,334113048 | 0,485051384 | -0,037479397 | -0,031857487 | 0,235883302 | 0,347883302 | **-0,912116698** |
| chrXVI | 790762 | 790920 | 158 | 0,045592969 | 0,08894087 | 0,338895921 | -0,415478239 | -0,353156503 | -0,359619091 | -0,457619091 | **-1,717619091** |
| chrXVI | 791267 | 791414 | 147 | 0,062690333 | 0,062999783 | 0,498768996 | -0,003085674 | -0,002622823 | -0,154177715 | -0,175177715 | **-1,435177715** |
| chrXVI | 791438 | 791595 | 157 | 0,105433742 | 0,070411522 | 0,599582493 | 0,252266586 | 0,214426598 | 0,176810331 | 0,085810331 | **-1,174189669** |
| chrXVI | 791737 | 791890 | 153 | 1,376337762 | 1,015408265 | 0,57545314 | 0,190274903 | 0,161733667 | 0,088585168 | 0,025585168 | **-1,234414832** |
| chrXVI | 792774 | 792924 | 150 | 0,44168189 | 0,381704567 | 0,536421126 | 0,091421413 | 0,077708201 | -0,104549472 | -0,146549472 | **-1,406549472** |
| chrXVI | 794619 | 794743 | 124 | 0,039893848 | 0,122293696 | 0,245973563 | -0,687215201 | -0,584132921 | -0,059240655 | 0,080759345 | **-1,179240655** |
| chrXVI | 796842 | 796998 | 156 | 0,532867829 | 0,392822175 | 0,575643927 | 0,190761892 | 0,162147608 | 0,085623064 | 0,001623064 | **-1,258376936** |
| chrXVI | 798394 | 798554 | 160 | 0,142478029 | 0,103764348 | 0,578608892 | 0,198335964 | 0,168585569 | 0,185340456 | 0,073340456 | **-1,186659544** |
| chrXVI | 798628 | 798777 | 149 | 0,162424953 | 0,114881957 | 0,585722704 | 0,216555726 | 0,184072367 | 0,031006243 | -0,003993757 | **-1,263993757** |
| chrXVI | 805070 | 805217 | 147 | 0,065539893 | 0,040764565 | 0,616530051 | 0,29638 | 0,251923 | 0,092668367 | 0,071668367 | **-1,188331633** |
| chrXVI | 805577 | 805691 | 114 | 0,045592969 | 0,459527828 | 0,090261517 | -1,339146351 | -1,138274398 | -0,370834688 | -0,160834688 | **-1,420834688** |
| chrXVI | 807669 | 807826 | 157 | 0,054141651 | 0,062999783 | 0,462190442 | -0,094916836 | -0,080679311 | -0,115924315 | -0,206924315 | **-1,466924315** |
| chrXVI | 808317 | 808452 | 135 | 0,091185939 | 0,077823261 | 0,539532397 | 0,099255756 | 0,084367393 | 0,065012764 | 0,128012764 | **-1,131987236** |
| chrXVI | 809355 | 809495 | 140 | 0,151026711 | 0,125999566 | 0,545171067 | 0,113470102 | 0,096449587 | 0,0131766 | 0,0411766 | **-1,2188234** |
| chrXVI | 809361 | 809495 | 134 | 0,122531105 | 0,092646739 | 0,569441084 | 0,174951387 | 0,148708679 | 0,176072376 | 0,246072376 | **-1,013927624** |
| chrXVI | 809773 | 809928 | 155 | 0,108283302 | 0,111176087 | 0,493409293 | -0,016521205 | -0,014043024 | -0,080064852 | -0,157064852 | **-1,417064852** |
| chrXVI | 809904 | 810012 | 108 | 0,074088575 | 0,200116957 | 0,270193583 | -0,612227627 | -0,520393483 | 0,327779192 | 0,579779192 | **-0,680220808** |
| chrXVI | 809934 | 810052 | 118 | 1,595753927 | 4,713866104 | 0,252908086 | -0,665366397 | -0,565561437 | 0,093085278 | 0,275085278 | **-0,984914722** |
| chrXVI | 810053 | 810202 | 149 | 0,068389454 | 0,059293913 | 0,535617563 | 0,08939893 | 0,07598909 | -0,087210704 | -0,122210704 | **-1,382210704** |
| chrXVI | 813679 | 813806 | 127 | 1,205364127 | 1,678758919 | 0,417930895 | -0,207189598 | -0,176111158 | 0,188271248 | 0,307271248 | **-0,952728752** |
| chrXVI | 822438 | 822593 | 155 | 0,558513874 | 0,629997828 | 0,469927114 | -0,07545308 | -0,064135118 | -0,140588491 | -0,217588491 | **-1,477588491** |
| chrXVI | 822870 | 822974 | 104 | 0,082637257 | 0,592939133 | 0,122321115 | -1,163461699 | -0,988942444 | -0,418798499 | -0,138798499 | **-1,398798499** |
| chrXVI | 827045 | 827170 | 125 | 0,527168708 | 1,100643265 | 0,323851106 | -0,456956639 | -0,388413143 | 0,110990494 | 0,243990494 | **-1,016009506** |
| chrXVI | 827247 | 827389 | 142 | 0,173823196 | 0,281646088 | 0,381635385 | -0,301188484 | -0,256010211 | -0,349691954 | -0,335691954 | **-1,595691954** |
| chrXVI | 827869 | 828021 | 152 | 0,29350474 | 0,322410653 | 0,476534185 | -0,058854034 | -0,050025929 | -0,09315842 | -0,14915842 | **-1,40915842** |
| chrXVI | 830012 | 830156 | 144 | 2,841011902 | 1,767699789 | 0,616443833 | 0,296154188 | 0,25173106 | 0,153000132 | 0,153000132 | **-1,106999868** |
| chrXVI | 839224 | 839390 | 166 | 0,330549028 | 0,515115871 | 0,390874716 | -0,277039947 | -0,235483955 | -0,06355392 | -0,21755392 | **-1,47755392** |
| chrXVI | 841656 | 841805 | 149 | 1,256656217 | 0,767115003 | 0,620947766 | 0,307970909 | 0,261775272 | 0,09098338 | 0,05598338 | **-1,20401662** |
| chrXVI | 843827 | 843986 | 159 | 0,173823196 | 0,181587609 | 0,489076846 | -0,027383708 | -0,023276152 | -0,028427659 | -0,133427659 | **-1,393427659** |
| chrXVI | 844949 | 845072 | 123 | 0,056991212 | 0,226058044 | 0,201347329 | -0,836818391 | -0,711295633 | -0,136444799 | 0,010555201 | **-1,249444799** |
| chrXVI | 845049 | 845204 | 155 | 0,111132863 | 0,059293913 | 0,652085695 | 0,390957556 | 0,332313923 | 0,256524069 | 0,179524069 | **-1,080475931** |
| chrXVI | 845482 | 845616 | 134 | 0,156725832 | 0,155646522 | 0,501727601 | 0,004330468 | 0,003680898 | 0,010069614 | 0,080069614 | **-1,179930386** |
| chrXVI | 845482 | 845622 | 140 | 0,131079787 | 0,125999566 | 0,509880648 | 0,024769644 | 0,021054198 | -0,068049783 | -0,040049783 | **-1,300049783** |
| chrXVI | 846525 | 846660 | 135 | 0,082637257 | 0,066705652 | 0,553339005 | 0,13410191 | 0,113986624 | 0,080865073 | 0,143865073 | **-1,116134927** |
| chrXVI | 847154 | 847311 | 157 | 0,042743409 | 0,066705652 | 0,390532439 | -0,277931585 | -0,236241848 | -0,274023224 | -0,365023224 | **-1,625023224** |
| chrXVI | 848557 | 848707 | 150 | 0,453080133 | 0,44099848 | 0,506756482 | 0,0169368 | 0,01439628 | -0,157466244 | -0,199466244 | **-1,459466244** |
| chrXVI | 849289 | 849403 | 114 | 0,031345166 | 0,259410871 | 0,107805729 | -1,238282157 | -1,052539834 | -0,300078132 | -0,090078132 | **-1,350078132** |
| chrXVI | 851012 | 851164 | 152 | 0,028495606 | 0,037058696 | 0,434687048 | -0,164453569 | -0,139785534 | -0,184339674 | -0,240339674 | **-1,500339674** |
| chrXVI | 851565 | 851679 | 114 | 0,062690333 | 0,400233914 | 0,135422444 | -1,101118957 | -0,935951113 | -0,20898221 | 0,00101779 | **-1,25898221** |
| chrXVI | 853843 | 853993 | 150 | 8,69115978 | 4,076456536 | 0,68071906 | 0,46971048 | 0,399253908 | 0,236425172 | 0,194425172 | **-1,065574828** |
| chrXVI | 853847 | 853993 | 146 | 0,065539893 | 0,037058696 | 0,638799168 | 0,355250862 | 0,301963233 | 0,198152147 | 0,184152147 | **-1,075847853** |
| chrXVI | 854031 | 854169 | 138 | 0,424584527 | 0,515115871 | 0,451829676 | -0,12104 | -0,102884 | -0,174142968 | -0,132142968 | **-1,392142968** |
| chrXVI | 854126 | 854258 | 132 | 0,236513528 | 0,251999131 | 0,484150254 | -0,039739878 | -0,033778897 | 0,07825031 | 0,16225031 | **-1,09774969** |
| chrXVI | 854305 | 854440 | 135 | 0,062690333 | 0,070411522 | 0,470995186 | -0,072768458 | -0,061853189 | -0,095205428 | -0,032205428 | **-1,292205428** |
| chrXVI | 854851 | 855005 | 154 | 0,216566604 | 0,144528914 | 0,599748802 | 0,252696962 | 0,214792418 | 0,138957211 | 0,068957211 | **-1,191042789** |
| chrXVI | 855108 | 855259 | 151 | 0,054141651 | 0,074117392 | 0,42212736 | -0,196454145 | -0,166986023 | -0,276915168 | -0,325915168 | **-1,585915168** |
| chrXVI | 855132 | 855277 | 145 | 0,113982423 | 0,074117392 | 0,605967759 | 0,268824834 | 0,228501109 | 0,124146142 | 0,117146142 | **-1,142853858** |
| chrXVI | 855136 | 855287 | 151 | 0,102584181 | 0,081529131 | 0,557179598 | 0,143822286 | 0,122248943 | 0,013629026 | -0,035370974 | **-1,295370974** |
| chrXVI | 855343 | 855483 | 140 | 0,216566604 | 0,107470218 | 0,66833949 | 0,435332606 | 0,370032715 | 0,287530242 | 0,315530242 | **-0,944469758** |
| chrXVI | 855349 | 855483 | 134 | 0,14817715 | 0,129705435 | 0,533236547 | 0,083408278 | 0,070897036 | 0,080534998 | 0,150534998 | **-1,109465002** |
| chrXVI | 855892 | 856000 | 108 | 0,065539893 | 0,192705218 | 0,253789483 | -0,662612181 | -0,563220354 | 0,290819375 | 0,542819375 | **-0,717180625** |
| chrXVI | 856112 | 856214 | 102 | 0,68389454 | 5,45504002 | 0,111402807 | -1,219101651 | -1,036236403 | -0,484300807 | -0,190300807 | **-1,450300807** |
| chrXVI | 858667 | 858810 | 143 | 1,986143727 | 2,160521964 | 0,47897368 | -0,052729592 | -0,044820154 | -0,156500174 | -0,149500174 | **-1,409500174** |
| chrXVI | 863879 | 864033 | 154 | 0,404637603 | 0,203822827 | 0,665018764 | 0,426199513 | 0,362269586 | 0,274700415 | 0,204700415 | **-1,055299585** |
| chrXVI | 863879 | 864033 | 154 | 0,404637603 | 0,203822827 | 0,665018764 | 0,426199513 | 0,362269586 | 0,274700415 | 0,204700415 | **-1,055299585** |
| chrXVI | 864255 | 864385 | 130 | 2,416427375 | 3,909692405 | 0,381976229 | -0,300294593 | -0,255250404 | -0,090558504 | 0,007441496 | **-1,252558504** |
| chrXVI | 867145 | 867288 | 143 | 0,424584527 | 0,307587175 | 0,579897483 | 0,201631223 | 0,171386539 | 0,058266981 | 0,065266981 | **-1,194733019** |
| chrXVI | 868632 | 868780 | 148 | 0,222265726 | 0,137117174 | 0,618464945 | 0,301451656 | 0,256233907 | 0,107200352 | 0,079200352 | **-1,180799648** |
| chrXVI | 888008 | 888137 | 129 | 1,473222822 | 1,838111311 | 0,4449031 | -0,138549438 | -0,117767022 | 0,108987568 | 0,213987568 | **-1,046012432** |
| chrXVI | 891517 | 891679 | 162 | 0,091185939 | 0,081529131 | 0,527955893 | 0,070132481 | 0,059612609 | 0,149207772 | 0,023207772 | **-1,236792228** |
| chrXVI | 893449 | 893599 | 150 | 0,430283648 | 0,381704567 | 0,529913662 | 0,075052832 | 0,063794907 | -0,092754161 | -0,134754161 | **-1,394754161** |
| chrXVI | 897488 | 897648 | 160 | 0,054141651 | 0,08894087 | 0,378394585 | -0,309699913 | -0,263244926 | -0,239825204 | -0,351825204 | **-1,611825204** |
| chrXVI | 897990 | 898142 | 152 | 0,530018269 | 0,366881088 | 0,590945087 | 0,229976764 | 0,195480249 | 0,159760579 | 0,103760579 | **-1,156239421** |
| chrXVI | 901200 | 901349 | 149 | 0,319150785 | 0,214940436 | 0,597558568 | 0,24703277 | 0,209977854 | 0,038574896 | 0,003574896 | **-1,256425104** |
| chrXVI | 905702 | 905842 | 140 | 0,68389454 | 0,70411522 | 0,492715945 | -0,018259434 | -0,015520519 | -0,104380245 | -0,076380245 | **-1,336380245** |
| chrXVI | 910317 | 910457 | 140 | 0,344796831 | 0,244587392 | 0,585011979 | 0,214732295 | 0,182522451 | 0,077125233 | 0,105125233 | **-1,154874767** |
| chrXVI | 910317 | 910457 | 140 | 0,344796831 | 0,244587392 | 0,585011979 | 0,214732295 | 0,182522451 | 0,077125233 | 0,105125233 | **-1,154874767** |
| chrXVI | 922519 | 922676 | 157 | 0,045592969 | 0,062999783 | 0,419852784 | -0,202270107 | -0,171929591 | -0,213045849 | -0,304045849 | **-1,564045849** |
| chrXVI | 922519 | 922681 | 162 | 7,246432564 | 8,656911336 | 0,455654648 | -0,111387218 | -0,094679135 | -0,001495318 | -0,127495318 | **-1,387495318** |
| chrXVI | 922523 | 922681 | 158 | 0,045592969 | 0,059293913 | 0,434687048 | -0,164453569 | -0,139785534 | -0,145272644 | -0,243272644 | **-1,503272644** |
| chrXVI | 927439 | 927574 | 135 | 0,088336378 | 0,055588044 | 0,613769206 | 0,28915654 | 0,245783059 | 0,209175108 | 0,272175108 | **-0,987824892** |
| chrXVI | 930833 | 931008 | 175 | 0,051292091 | 0,129705435 | 0,283385589 | -0,572813203 | -0,486891222 | -0,269019507 | -0,486019507 | **-1,746019507** |
| chrXVI | 931493 | 931642 | 149 | 0,569912117 | 0,522527611 | 0,521687469 | 0,054389227 | 0,046230843 | -0,126005016 | -0,161005016 | **-1,421005016** |
| chrXVI | 940363 | 940515 | 152 | 0,569912117 | 0,485468915 | 0,540006026 | 0,100448902 | 0,085381566 | 0,034520162 | -0,021479838 | **-1,281479838** |
| chrXVI | 943076 | 943229 | 153 | 0,495823542 | 0,607762611 | 0,449283946 | -0,127470662 | -0,108350062 | -0,175434548 | -0,238434548 | **-1,498434548** |
| chrXVI | 943085 | 943229 | 144 | 0,034194727 | 0,085235 | 0,286316713 | -0,564177371 | -0,479550765 | -0,573880799 | -0,573880799 | **-1,833880799** |
| chrXVI | 944252 | 944389 | 137 | 0,544266071 | 0,559586306 | 0,49306056 | -0,017395474 | -0,014786153 | -0,093987783 | -0,044987783 | **-1,304987783** |
| chrXVI | 944537 | 944680 | 143 | 0,404637603 | 0,418763262 | 0,491422368 | -0,021502592 | -0,018277203 | -0,129609864 | -0,122609864 | **-1,382609864** |
| chrXVI | 944585 | 944710 | 125 | 0,199469241 | 0,255705001 | 0,438226118 | -0,155468183 | -0,132147955 | 0,338711742 | 0,471711742 | **-0,788288258** |
| chrXVI | 944810 | 944962 | 152 | 0,102584181 | 0,077823261 | 0,568624996 | 0,172874585 | 0,146943398 | 0,103055269 | 0,047055269 | **-1,212944731** |
| chrXVI | 944883 | 945027 | 144 | 0,062690333 | 0,085235 | 0,423797138 | -0,192188854 | -0,163360526 | -0,258440935 | -0,258440935 | **-1,518440935** |
| chrXVI | 944980 | 945115 | 135 | 1,156921597 | 0,837526525 | 0,58007104 | 0,202075221 | 0,171763938 | 0,133674957 | 0,196674957 | **-1,063325043** |
| chrXVI | 944980 | 945167 | 187 | 0,111132863 | 0,070411522 | 0,612152576 | 0,284933816 | 0,242193743 | 0,438585782 | 0,137585782 | **-1,122414218** |
| chrXVI | 945023 | 945115 | 92 | 0,151026711 | 0,122293696 | 0,552562879 | 0,132139134 | 0,112318264 | 0,601709995 | 0,965709995 | **-0,294290005** |
| chrXVI | 945023 | 945167 | 144 | 1,26805446 | 0,592939133 | 0,68138572 | 0,471577263 | 0,400840674 | 0,304960231 | 0,304960231 | **-0,955039769** |
| chrXVI | 945362 | 945514 | 152 | 0,188070999 | 0,144528914 | 0,56545715 | 0,164819952 | 0,140096959 | 0,089005946 | 0,033005946 | **-1,226994054** |
| chrXVI | 945663 | 945793 | 130 | 0,339097709 | 0,366881088 | 0,480322795 | -0,049343454 | -0,041941936 | 0,105507633 | 0,203507633 | **-1,056492367** |
| chrXVI | 945663 | 945793 | 130 | 0,339097709 | 0,366881088 | 0,480322795 | -0,049343454 | -0,041941936 | 0,105507633 | 0,203507633 | **-1,056492367** |
| chrXVI | 945759 | 945954 | 195 | 0,079787696 | 0,118587827 | 0,402205349 | -0,247642923 | -0,210496484 | -0,011973299 | -0,368973299 | **-1,628973299** |
| chrXVI | 945821 | 945954 | 133 | 12,97119978 | 9,520378947 | 0,57671362 | 0,193493162 | 0,164469188 | 0,239112924 | 0,316112924 | **-0,943887076** |
| chrXVI | 945821 | 945969 | 148 | 0,666797177 | 0,555880437 | 0,54535813 | 0,113942039 | 0,096850733 | -0,05334048 | -0,08134048 | **-1,34134048** |
| chrXVI | 946033 | 946184 | 151 | 0,478726178 | 0,296469566 | 0,617555219 | 0,299066166 | 0,254206241 | 0,145906245 | 0,096906245 | **-1,163093755** |
| chrXVI | 946038 | 946172 | 134 | 0,581310359 | 0,429880871 | 0,574876781 | 0,188803997 | 0,160483398 | 0,160447619 | 0,230447619 | **-1,029552381** |

TABLE 2

| Chrom | Start | End | Length | Section A | Section B | A/A+B | Z-score | Z * 0.85 | Phase correction | Length correction | ΔLK |
|---|---|---|---|---|---|---|---|---|---|---|---|
| chrXVI | 946051 | 946147 | 96 | 0,056991212 | 0,140823044 | 0,288104674 | -0,558930214 | -0,475090682 | 0,068394864 | 0,404394864 | **-0,855605136** |
| chrXVI | 946159 | 946281 | 122 | 0,125380666 | 0,070411522 | 0,640376244 | 0,359464657 | 0,305544959 | 0,923520722 | 1,077520722 | **-0,182479278** |
| chrXVI | 946966 | 947118 | 152 | 0,071239015 | 0,048176305 | 0,596565123 | 0,24446623 | 0,207796296 | 0,160702149 | 0,104702149 | **-1,155297851** |
| chrXVI | 947306 | 947457 | 151 | 0,524319147 | 0,370586958 | 0,585892916 | 0,216992528 | 0,184443648 | 0,088263992 | 0,039263992 | **-1,220736008** |

### Supplementary Table 3. ΔLk values and genomic context of nucleosomes

The first five columns indicate the chromosomal coordinates, length (bp) and the ΔLk restrained by the nucleosomes of the library. The next four columns are parameters defined in Jiang and Pugh (2009) that indicate the nucleosome IDs, the positional fuzziness, the gene ID, and the allocation within a gene body or intergenic region. Coordinates without a nucleosome ID are those that overlapped by <50 bp with referenced nucleosomes. The next two columns tag the gene terminal nucleosomes and rDNA nucleosomes. The last column tags the nucleosomes of telomeric regions, which are defined from The Saccharomyces Genome Database (SGD) (Cherry and Wong 2012) as those overlapping with chromosomal coordinates that include an X element core sequence, X element combinatorial repeats, short or long Y' elements or a terminal stretch of telomeric repeats. Depending on the chromosomal arm, the length of the telomeric regions spans from 430 to 13896 bp.

| CHR | START | END | L(bp) | ΔLK | Nuc ID | Fuzziness | Gene ID | Gene body postion or intergene | Terminal | rDNA | Telomeric |
|---|---|---|---|---|---|---|---|---|---|---|---|
| chrI | 375 | 485 | 110 | -1,475273061 | N1:382 | 21,916 | | Intergene | | | TEL LEFT |
| chrI | 1992 | 2143 | 151 | -1,145624862 | N1:2081 | 37,206 | +2:YAL068C; | 2 | | | |
| chrI | 3900 | 4046 | 146 | -1,42891971 | | | | Overlap <50 bp | | | |
| chrI | 7310 | 7448 | 138 | -1,099214006 | N1:7314 | 32,078 | +12:YAL067C; | 12 | | | |
| chrI | 19698 | 19849 | 151 | -1,170720688 | N1:19743 | 9,192 | | Intergene | | | |
| chrI | 24106 | 24256 | 150 | -1,002800143 | | | | Overlap <50 bp | | | |
| chrI | 24106 | 24267 | 161 | -1,371408779 | | | | Overlap <50 bp | | | |
| chrI | 24156 | 24264 | 108 | -1,22036164 | | | | Overlap <50 bp | | | |
| chrI | 24156 | 24265 | 109 | -1,199955994 | | | | Overlap <50 bp | | | |
| chrI | 24156 | 24266 | 110 | -1,664918922 | | | | Overlap <50 bp | | | |
| chrI | 24156 | 24267 | 111 | -1,434214917 | | | | Overlap <50 bp | | | |
| chrI | 25658 | 25790 | 132 | -0,89778306 | | | | Overlap <50 bp | | | |
| chrI | 25658 | 25797 | 139 | -1,018529414 | | | | Overlap <50 bp | | | |
| chrI | 25671 | 25797 | 126 | -0,906695696 | | | | Overlap <50 bp | | | |
| chrI | 25885 | 26057 | 172 | -1,008848565 | | | | Overlap <50 bp | | | |
| chrI | 26200 | 26329 | 129 | -1,574518143 | | | | Overlap <50 bp | | | |
| chrI | 26200 | 26330 | 130 | -1,094609367 | | | | Overlap <50 bp | | | |
| chrI | 26200 | 26337 | 137 | -1,169239802 | | | | Overlap <50 bp | | | |
| chrI | 26204 | 26330 | 126 | -1,046753163 | | | | Overlap <50 bp | | | |
| chrI | 26211 | 26330 | 119 | -0,77735452 | | | | Overlap <50 bp | | | |
| chrI | 26268 | 26418 | 150 | -1,229716689 | N1:26380 | 32,909 | +11:YAL063C; | 11 | | | |
| chrI | 26290 | 26465 | 175 | -1,827207551 | N1:26380 | 32,909 | +11:YAL063C; | 11 | | | |
| chrI | 26425 | 26573 | 148 | -1,091924323 | N1:26590 | 51,619 | +10:YAL063C; | 10 | | | |
| chrI | 26560 | 26708 | 148 | -1,251179066 | N1:26590 | 51,619 | +10:YAL063C; | 10 | | | |
| chrI | 26695 | 26843 | 148 | -1,090783865 | N1:26787 | 43,547 | +9:YAL063C; | 9 | | | |
| chrI | 27347 | 27495 | 148 | -1,357301342 | N1:27437 | 37,487 | +5:YAL063C; | 5 | | | |
| chrI | 28545 | 28653 | 108 | -0,815770683 | N1:28676 | 33,941 | +7:SUT433; | 7 | | | |
| chrI | 28558 | 28705 | 147 | -1,585279852 | N1:28676 | 33,941 | +7:SUT433; | 7 | | | |
| chrI | 28865 | 28996 | 131 | -1,004955603 | N1:28942 | 15,556 | +5:SUT433; | 5 | | | |
| chrI | 31670 | 31815 | 145 | -1,382405277 | N1:31784 | 18,209 | +6:SUT434; | 6 | | | |
| chrI | 32825 | 32968 | 143 | -1,230034943 | N1:32915 | 24,186 | -1:SUT434; | -1 | | | |
| chrI | 38266 | 38455 | 189 | -1,633713958 | N1:38289 | 16 | +6:YAL058W; | 6 | | | |
| chrI | 38266 | 38455 | 189 | -1,633713958 | N1:38459 | 37,157 | +7:YAL058W; | 7 | | | |
| chrI | 39356 | 39504 | 148 | -1,28106733 | N1:39417 | 8,503 | +2:YAL056W; | 2 | | | |
| chrI | 44014 | 44169 | 155 | -1,329113517 | N1:44128 | 16,158 | +6:YAL054C; +5:SUT002; | 6 | | | |
| chrI | 48679 | 48833 | 154 | -1,364356893 | N1:48747 | 8,256 | +2:YAL051W; | 2 | | | |
| chrI | 52107 | 52211 | 104 | -0,997609421 | N1:52114 | 21,119 | +4:YAL049C; | 4 | | | |
| chrI | 52108 | 52200 | 92 | -1,194494905 | N1:52114 | 21,119 | +4:YAL049C; | 4 | | | |
| chrI | 52108 | 52202 | 94 | -1,115050422 | N1:52114 | 21,119 | +4:YAL049C; | 4 | | | |
| chrI | 52108 | 52203 | 95 | -1,203469923 | N1:52114 | 21,119 | +4:YAL049C; | 4 | | | |
| chrI | 52108 | 52204 | 96 | -1,174333328 | N1:52114 | 21,119 | +4:YAL049C; | 4 | | | |
| chrI | 52108 | 52205 | 97 | -1,008989827 | N1:52114 | 21,119 | +4:YAL049C; | 4 | | | |
| chrI | 52108 | 52206 | 98 | -1,232433255 | N1:52114 | 21,119 | +4:YAL049C; | 4 | | | |
| chrI | 52108 | 52207 | 99 | -1,533375011 | N1:52114 | 21,119 | +4:YAL049C; | 4 | | | |
| chrI | 52108 | 52208 | 100 | -0,882098917 | N1:52114 | 21,119 | +4:YAL049C; | 4 | | | |
| chrI | 52108 | 52209 | 101 | -1,127128482 | N1:52114 | 21,119 | +4:YAL049C; | 4 | | | |
| chrI | 52108 | 52210 | 102 | -1,320587152 | N1:52114 | 21,119 | +4:YAL049C; | 4 | | | |
| chrI | 52108 | 52211 | 103 | -1,00632145 | N1:52114 | 21,119 | +4:YAL049C; | 4 | | | |
| chrI | 52112 | 52211 | 99 | -1,312463337 | N1:52114 | 21,119 | +4:YAL049C; | 4 | | | |
| chrI | 54178 | 54332 | 154 | -1,179031036 | N1:54248 | 6,306 | +4:YAL048C; | 4 | | | |
| chrI | 56067 | 56218 | 151 | -1,391434877 | N1:56143 | 16,79 | +6:YAL047C; | 6 | | | |
| chrI | 57573 | 57723 | 150 | -1,176476393 | N1:57555 | 3,899 | -1:YAL046C; +1:YAL044W-A; | -1 | | | |
| chrI | 57573 | 57723 | 150 | -1,176476393 | N1:57726 | 5,502 | +2:YAL044W-A; | 2 | | | |
| chrI | 57985 | 58142 | 157 | -1,491896053 | N1:58076 | 12,422 | +3:YAL044C; +4*:YAL044W-A; | 3 | | | |
| chrI | 58357 | 58529 | 172 | -1,395417578 | N1:58432 | 12,859 | +1:YAL044C; | 1 | | | |
| chrI | 59835 | 59985 | 150 | -1,525981569 | N1:59903 | 12,276 | +8:YAL043C; | 8 | | | |
| chrI | 60956 | 61114 | 158 | -1,201260038 | N1:61025 | 8,618 | -1:YAL042W; +1:YAL043C; | -1 | | | |
| chrI | 61139 | 61253 | 114 | -1,514954291 | | | | Overlap <50 bp | | | |
| chrI | 62731 | 62859 | 128 | -1,118454861 | N1:62815 | 15,092 | +1:YAL041W; | 1 | | | |
| chrI | 73815 | 73966 | 151 | -1,456101359 | N1:73878 | 12,462 | YAL037C-A; 0:YAL037W; 0:Unit002; | -1 | | | |
| chrI | 75929 | 76089 | 160 | -0,635190429 | N1:75990 | 4,207 | +2:YAL036C; | 2 | | | |
| chrI | 77820 | 77969 | 149 | -1,329929447 | N1:77885 | 28,324 | | Intergene | | | |
| chrI | 78867 | 79013 | 146 | -1,476014857 | N1:78863 | 32,79 | | Intergene | | | |
| chrI | 79075 | 79223 | 148 | -1,445857802 | N1:79100 | 26,734 | | Intergene | | | |
| chrI | 79820 | 79919 | 99 | -0,868345432 | N1:79895 | 4,29 | +2:YAL034W-A; | 2 | | | |
| chrI | 80712 | 80877 | 165 | -1,102806447 | N1:80770 | 14,428 | +8:YAL034C; | 8 | | | |
| chrI | 83877 | 84031 | 154 | -1,014732041 | N1:83953 | 6,494 | +4:YAL032C; | 4 | | | |
| chrI | 87561 | 87716 | 155 | -1,568954094 | N1:87641 | 9,368 | | Intergene | | | |
| chrI | 88852 | 89013 | 161 | -1,349929244 | N1:89028 | 31,911 | +22:YAL029C; | 22 | | | |
| chrI | 88852 | 89013 | 161 | -1,349929244 | N1:88849 | 19,502 | +23:YAL029C; | 23 | | | |
| chrI | 88889 | 89053 | 164 | -1,71832009 | N1:89028 | 31,911 | +22:YAL029C; | 22 | | | |
| chrI | 91036 | 91130 | 94 | -0,529061161 | N1:91042 | 21,515 | +10:YAL029C; | 10 | | | |
| chrI | 94271 | 94416 | 145 | -1,127470224 | | | | Overlap <50 bp | | | |
| chrI | 97984 | 98151 | 167 | -1,439823481 | N1:98096 | 12,437 | +11:YAL026C; | 11 | | | |
| chrI | 100299 | 100401 | 102 | 0,145125764 | N1:100335 | 20,156 | +6:YAL025C; | 6 | | | |
| chrI | 100430 | 100592 | 162 | -0,972570656 | N1:100502 | 12,449 | +5:YAL025C; | 5 | | | |
| chrI | 100744 | 100901 | 157 | -1,177782262 | N1:100813 | 8,781 | +3:YAL025C; | 3 | | | |
| chrI | 100903 | 101028 | 125 | -0,681458043 | N1:100972 | 4,401 | +2:YAL025C; | 2 | | | |
| chrI | 101774 | 101913 | 139 | -1,268887199 | N1:101784 | 18,177 | +26:YAL024C; | 26 | | | |
| chrI | 102759 | 102918 | 159 | -1,404046068 | N1:102829 | 19,565 | +20:YAL024C; | 20 | | | |
| chrI | 110352 | 110498 | 146 | -1,262096736 | N1:110405 | 10,033 | +1:YAL022C; | 1 | | | |
| chrI | 110959 | 111107 | 148 | -1,365708219 | N1:110988 | 7,005 | +16:YAL021C; | 16 | | | |
| chrI | 111829 | 111987 | 158 | -2,304517965 | N1:111914 | 12,91 | +10:YAL021C; | 10 | | | |

TABLE 3

| CHR | START | END | L(bp) | ΔLK | Nuc ID | Fuzziness | Gene ID | Gene body postion or intergene | Terminal | rDNA | Telomeric |
|---|---|---|---|---|---|---|---|---|---|---|---|
| chrI | 111829 | 111989 | 160 | -1,220675432 | N1:111914 | 12,91 | +10:YAL021C; | 10 | | | |
| chrI | 111835 | 111987 | 152 | -1,321773661 | N1:111914 | 12,91 | +10:YAL021C; | 10 | | | |
| chrI | 111835 | 111989 | 154 | -1,903586461 | N1:111914 | 12,91 | +10:YAL021C; | 10 | | | |
| chrI | 113198 | 113345 | 147 | -1,313354286 | N1:113359 | 19,466 | +1:YAL021C; | 1 | | | |
| chrI | 113198 | 113345 | 147 | -1,313354286 | N1:113192 | 8,824 | +2:YAL021C; | 2 | | | |
| chrI | 113272 | 113431 | 159 | -1,295849447 | N1:113359 | 19,466 | +1:YAL021C; | 1 | | | |
| chrI | 114916 | 115064 | 148 | -1,000643596 | N1:115021 | 8,649 | +2:YAL019W; | 2 | | | |
| chrI | 116310 | 116474 | 164 | -1,186230225 | N1:116396 | 42,101 | +11:YAL019W; | 11 | | | |
| chrI | 119258 | 119408 | 150 | -0,376293772 | N1:119265 | 14,519 | +3:YAL018C; | 3 | | | |
| chrI | 121555 | 121709 | 154 | -1,247553434 | N1:121654 | 14,905 | +12:YAL017W; | 12 | | | |
| chrI | 128610 | 128759 | 149 | -1,338333574 | | | | Overlap <50 bp | | | |
| chrI | 128749 | 128913 | 164 | -1,42014932 | N1:128845 | 11,978 | +2:YAL014C; | 2 | | | |
| chrI | 132160 | 132339 | 179 | -1,625001105 | N1:132234 | 8,735 | +1:YAL011W; | 1 | | | |
| chrI | 132322 | 132449 | 127 | -1,12899717 | N1:132398 | 10,521 | +2:YAL011W; | 2 | | | |
| chrI | 133311 | 133456 | 145 | -1,279779418 | N1:133382 | 11,127 | +8:YAL011W; | 8 | | | |
| chrI | 133444 | 133595 | 151 | -1,312572395 | N1:133558 | 12,813 | +9:YAL011W; | 9 | | | |
| chrI | 134936 | 135082 | 146 | -1,233599314 | | | | Overlap <50 bp | | | |
| chrI | 139478 | 139611 | 133 | -0,937522987 | N1:139570 | 10,426 | +13:YAL005C; | 13 | | | |
| chrI | 144452 | 144595 | 143 | -1,203524654 | N1:144447 | 8,18 | +6:YAL002W; | 6 | | | |
| chrI | 144452 | 144595 | 143 | -1,203524654 | N1:144596 | 18,794 | +7:YAL002W; | 7 | | | |
| chrI | 145995 | 146131 | 136 | -0,824624641 | N1:145995 | 30,089 | +15:YAL002W; | 15 | | | |
| chrI | 145995 | 146131 | 136 | -0,824624641 | N1:146151 | 24,542 | +16:YAL002W; | 16 | | | |
| chrI | 157171 | 157325 | 154 | -1,420376368 | N1:157317 | 31,92 | +9:YAR007C; | 9 | | | |
| chrI | 158225 | 158386 | 161 | -1,162030477 | N1:158305 | 6,164 | +3:YAR007C; | 3 | | | |
| chrI | 158476 | 158631 | 155 | -1,466764872 | N1:158641 | 1,751 | -1:YAR008W; +1:YAR007C; | -1 | | | |
| chrI | 158476 | 158631 | 155 | -1,466764872 | N1:158473 | 6,314 | +2:YAR007C; | 2 | | | |
| chrI | 159246 | 159353 | 107 | -1,32864883 | N1:159237 | 21,23 | +3:YAR008W; | 3 | | | |
| chrI | 161173 | 161287 | 114 | -1,44116562 | N1:161301 | 30,254 | | Intergene | | | |
| chrI | 163265 | 163422 | 157 | -1,360421984 | N1:163329 | 36,373 | | Intergene | | | |
| chrI | 164304 | 164458 | 154 | -1,406236959 | N1:164322 | 46,436 | | Intergene | | | |
| chrI | 164304 | 164458 | 154 | -1,406236959 | N1:164474 | 15,524 | | Intergene | | | |
| chrI | 164952 | 165092 | 140 | -1,239994795 | N1:165025 | 16,503 | | Intergene | | | |
| chrI | 164958 | 165092 | 134 | -1,223023256 | N1:165025 | 16,503 | | Intergene | | | |
| chrI | 165370 | 165525 | 155 | -1,247413069 | N1:165400 | 21,385 | | Intergene | | | |
| chrI | 165501 | 165609 | 108 | -0,974105815 | N1:165556 | 35,162 | | Intergene | | | |
| chrI | 165531 | 165649 | 118 | -1,16531138 | N1:165556 | 35,162 | | Intergene | | | |
| chrI | 165650 | 165799 | 149 | -1,184904444 | N1:165722 | 1,414 | | Intergene | | | |
| chrI | 170185 | 170316 | 131 | -0,991286797 | | | | Overlap <50 bp | | | |
| chrI | 170746 | 170892 | 146 | -1,419404324 | N1:170868 | 11,895 | +3:CUT004; +6:YAR018C; | 3 | | | |
| chrI | 173251 | 173422 | 171 | -1,247836659 | | | | Overlap <50 bp | | | |
| chrI | 174403 | 174559 | 156 | -1,268429026 | N1:174467 | 7,257 | +5:YAR019C; | 5 | | | |
| chrI | 177062 | 177157 | 95 | -0,925986621 | | | | Overlap <50 bp | | | |
| chrI | 178178 | 178314 | 136 | -1,157292599 | N1:178222 | 37,276 | | Intergene | | | |
| chrI | 181632 | 181781 | 149 | -1,6308092 | N1:181719 | 10,926 | | Intergene | | | |
| chrI | 184868 | 185010 | 142 | -1,180277539 | N1:184881 | 10,053 | +1:YAR028W; | 1 | | | |
| chrI | 188940 | 189093 | 153 | -1,21049533 | N1:189025 | 15,588 | +1:anti-YAR033W-2; +6*:YAR033W; | 1 | | | |
| chrI | 189954 | 190061 | 107 | -0,739135051 | N1:190031 | 20,354 | 0:YAR035W; +9:anti006; | 0 | | | |
| chrI | 189954 | 190062 | 108 | -0,738228073 | N1:190031 | 20,354 | 0:YAR035W; +9:anti006; | 0 | | | |
| chrI | 189954 | 190063 | 109 | -0,850741142 | N1:190031 | 20,354 | 0:YAR035W; +9:anti006; | 0 | | | |
| chrI | 189954 | 190077 | 123 | -1,285620828 | N1:190031 | 20,354 | 0:YAR035W; +9:anti006; | 0 | | | |
| chrI | 191415 | 191573 | 158 | -1,529273336 | N1:191409 | 39,245 | +9:YAR035W; 0:anti006; | 9 | | | |
| chrI | 191722 | 191874 | 152 | -1,237006303 | N1:191782 | 11,389 | +11:YAR035W; +3:SUT436; | 11 | | | |
| chrI | 192889 | 193041 | 152 | -1,723783132 | N1:192956 | 22,963 | +3:YAR042W; | 3 | | | |
| chrI | 193467 | 193622 | 155 | -1,310298541 | N1:193586 | 11 | +7:YAR042W; | 7 | | | |
| chrI | 194126 | 194279 | 153 | -1,34565981 | | | | Overlap <50 bp | | | |
| chrI | 194141 | 194279 | 138 | -0,95749856 | | | | Overlap <50 bp | | | |
| chrI | 194290 | 194442 | 152 | -1,043901966 | N1:194358 | 7,396 | +12:YAR042W; | 12 | | | |
| chrI | 204527 | 204675 | 148 | -1,366287583 | N1:204558 | 21,213 | +8:YAR050W; | 8 | | | |
| chrI | 204662 | 204810 | 148 | -1,204518936 | N1:204754 | 36,77 | +9:YAR050W; | 9 | | | |
| chrI | 204797 | 204945 | 148 | -1,194263798 | | | | Overlap <50 bp | | | |
| chrI | 205040 | 205170 | 130 | -1,159623338 | | | | Overlap <50 bp | | | |
| chrI | 205573 | 205699 | 126 | -0,727906068 | N1:205618 | 18,385 | +14:YAR050W; | 14 | | | |
| chrI | 205573 | 205705 | 132 | -1,254685071 | N1:205618 | 18,385 | +14:YAR050W; | 14 | | | |
| chrI | 205573 | 205712 | 139 | -1,012284504 | N1:205618 | 18,385 | +14:YAR050W; | 14 | | | |
| chrI | 205580 | 205712 | 132 | -0,99796908 | N1:205618 | 18,385 | +14:YAR050W; | 14 | | | |
| chrI | 206390 | 206543 | 153 | -1,836820478 | N1:206455 | 34,269 | +19:YAR050W; | 19 | | | |
| chrI | 207406 | 207531 | 125 | -0,816530428 | N1:207477 | 12,021 | +25:YAR050W; | 25 | | | |
| chrI | 215442 | 215555 | 113 | -1,736345291 | N1:215495 | 14,849 | | Intergene | | | |
| chrI | 217878 | 218015 | 137 | -1,162588108 | N1:217966 | 3,512 | | Intergene | | | |
| chrI | 217916 | 218076 | 160 | -1,479307354 | N1:217966 | 3,512 | | Intergene | | | |
| chrI | 218711 | 218856 | 145 | -1,136650022 | N1:218780 | 38,158 | | Intergene | | | |
| chrI | 218738 | 218886 | 148 | -1,215559937 | N1:218780 | 38,158 | | Intergene | | | |
| chrI | 222387 | 222554 | 167 | -1,349608552 | N1:222504 | 27,109 | +2:YAR068W; | 2 | | | |
| chrI | 222751 | 222855 | 104 | -0,980770953 | N1:222820 | 21,572 | +4:YAR068W; | 4 | | | |
| chrI | 223257 | 223420 | 163 | -1,483544312 | N1:223335 | 24,826 | | Intergene | | | |
| chrI | 224065 | 224173 | 108 | -1,318042122 | N1:224083 | 3,786 | | Intergene | | | |
| chrI | 224877 | 224997 | 120 | -1,326308076 | N1:224931 | 13,892 | | Intergene | | | |
| chrI | 226540 | 226687 | 147 | -1,204924103 | N1:226556 | 39,737 | +8:YAR071W; | 8 | | | |
| chrI | 229861 | 230005 | 144 | -1,14650557 | N1:229980 | 25,565 | | Intergene | | | TEL RIGHT |
| chrI | 230119 | 230218 | 99 | -0,283250953 | N1:230156 | 20,664 | | Intergene | | | TEL RIGHT |
| chrI | 230124 | 230218 | 94 | -0,112407352 | N1:230156 | 20,664 | | Intergene | | | TEL RIGHT |
| chrII | 163 | 311 | 148 | -1,339618443 | N2:255 | 24,749 | +17:YBL113C; | 17 | | | TEL LEFT |
| chrII | 526 | 677 | 151 | -1,122700906 | N2:589 | 17,214 | +15:YBL113C; | 15 | | | TEL LEFT |
| chrII | 1089 | 1240 | 151 | -1,153642956 | N2:1150 | 29,783 | +11:YBL113C; | 11 | | | TEL LEFT |
| chrII | 1727 | 1858 | 131 | -0,962631553 | N2:1853 | 30,854 | +7:YBL113C; | 7 | | | TEL LEFT |
| chrII | 1727 | 1870 | 143 | -1,208984888 | N2:1853 | 30,854 | +7:YBL113C; | 7 | | | TEL LEFT |
| chrII | 1727 | 1914 | 187 | -1,023851407 | N2:1853 | 30,854 | +7:YBL113C; | 7 | | | TEL LEFT |
| chrII | 1931 | 2076 | 145 | -1,398920977 | | | | Overlap <50 bp | | | TEL LEFT |
| chrII | 1936 | 2043 | 107 | -0,634852792 | | | | Overlap <50 bp | | | TEL LEFT |
| chrII | 1937 | 2034 | 97 | -0,762519621 | | | | Overlap <50 bp | | | TEL LEFT |
| chrII | 1937 | 2042 | 105 | -1,618967553 | | | | Overlap <50 bp | | | TEL LEFT |
| chrII | 1937 | 2046 | 109 | -0,680066278 | | | | Overlap <50 bp | | | TEL LEFT |
| chrII | 1937 | 2051 | 114 | -1,142580445 | | | | Overlap <50 bp | | | TEL LEFT |
| chrII | 1937 | 2054 | 117 | -1,282598983 | | | | Overlap <50 bp | | | TEL LEFT |
| chrII | 1937 | 2057 | 120 | -1,129549176 | | | | Overlap <50 bp | | | TEL LEFT |
| chrII | 1937 | 2062 | 125 | -1,838064879 | | | | Overlap <50 bp | | | TEL LEFT |
| chrII | 1937 | 2071 | 134 | -1,084959621 | | | | Overlap <50 bp | | | TEL LEFT |

| CHR | START | END | L(bp) | ΔLK | Nuc ID | Fuzziness | Gene ID | Gene body postion or intergene | Terminal | rDNA | Telomeric |
|---|---|---|---|---|---|---|---|---|---|---|---|
| chrII | 1937 | 2072 | 135 | -1,226959369 | | | | Overlap <50 bp | | | TEL LEFT |
| chrII | 1937 | 2073 | 136 | -1,957126123 | | | | Overlap <50 bp | | | TEL LEFT |
| chrII | 1937 | 2075 | 138 | -1,596758786 | | | | Overlap <50 bp | | | TEL LEFT |
| chrII | 1937 | 2076 | 139 | -1,105255183 | | | | Overlap <50 bp | | | TEL LEFT |
| chrII | 1937 | 2082 | 145 | -2,679580974 | | | | Overlap <50 bp | | | TEL LEFT |
| chrII | 1937 | 2086 | 149 | -1,63174152 | | | | Overlap <50 bp | | | TEL LEFT |
| chrII | 1938 | 2076 | 138 | -1,224450881 | | | | Overlap <50 bp | | | TEL LEFT |
| chrII | 1939 | 2076 | 137 | -1,518387315 | | | | Overlap <50 bp | | | TEL LEFT |
| chrII | 1940 | 2076 | 136 | -1,359600303 | | | | Overlap <50 bp | | | TEL LEFT |
| chrII | 1941 | 2076 | 135 | -1,029142639 | | | | Overlap <50 bp | | | TEL LEFT |
| chrII | 1942 | 2076 | 134 | -1,726837644 | | | | Overlap <50 bp | | | TEL LEFT |
| chrII | 1944 | 2076 | 132 | -1,181981789 | | | | Overlap <50 bp | | | TEL LEFT |
| chrII | 1947 | 2076 | 129 | -1,517897595 | | | | Overlap <50 bp | | | TEL LEFT |
| chrII | 1948 | 2076 | 128 | -1,355921231 | | | | Overlap <50 bp | | | TEL LEFT |
| chrII | 1952 | 2076 | 124 | -0,982107431 | | | | Overlap <50 bp | | | TEL LEFT |
| chrII | 1953 | 2076 | 123 | -0,584200369 | | | | Overlap <50 bp | | | TEL LEFT |
| chrII | 1964 | 2076 | 112 | -0,897829145 | | | | Overlap <50 bp | | | TEL LEFT |
| chrII | 1966 | 2076 | 110 | -0,663844067 | | | | Overlap <50 bp | | | TEL LEFT |
| chrII | 2124 | 2286 | 162 | -1,489886698 | N2:2264 | 49,166 | +4:YBL113C; | 4 | | | TEL LEFT |
| chrII | 2124 | 2286 | 162 | -1,489886698 | N2:2110 | 20,033 | +5:YBL113C; | 5 | | | TEL LEFT |
| chrII | 2191 | 2286 | 95 | -0,863684874 | N2:2264 | 49,166 | +4:YBL113C; | 4 | | | TEL LEFT |
| chrII | 2267 | 2374 | 107 | -0,785948273 | N2:2264 | 49,166 | +4:YBL113C; | 4 | | | TEL LEFT |
| chrII | 2267 | 2414 | 147 | -1,559119201 | N2:2264 | 49,166 | +4:YBL113C; | 4 | | | TEL LEFT |
| chrII | 2522 | 2620 | 98 | -0,719252405 | | | | Overlap <50 bp | | | TEL LEFT |
| chrII | 2522 | 2673 | 151 | -1,076920734 | | | | Overlap <50 bp | | | TEL LEFT |
| chrII | 3354 | 3498 | 144 | -0,952337581 | N2:3449 | 18,475 | +11:YBL111C; | 11 | | | TEL LEFT |
| chrII | 3419 | 3571 | 152 | -1,250718532 | N2:3449 | 18,475 | +11:YBL111C; | 11 | | | TEL LEFT |
| chrII | 3701 | 3844 | 143 | -1,424193058 | N2:3787 | 13,868 | +9:YBL111C; | 9 | | | TEL LEFT |
| chrII | 3992 | 4129 | 137 | -1,254485891 | N2:4071 | 14,731 | +7:YBL111C; | 7 | | | TEL LEFT |
| chrII | 7980 | 8135 | 155 | -1,017882777 | | | | Overlap <50 bp | | | |
| chrII | 10712 | 10874 | 162 | -1,593400301 | N2:10765 | 10,926 | +20:YBL106C; -1:YBL107C; | 20 | | | |
| chrII | 20981 | 21152 | 171 | -1,715983406 | N2:21084 | 6,706 | +2:YBL104C; | 2 | | | |
| chrII | 21442 | 21596 | 154 | -1,536203893 | N2:21498 | 18,577 | -1:YBL104C; | -1 | | | |
| chrII | 24248 | 24399 | 151 | -1,031056071 | N2:24353 | 4,583 | +3:YBL102W; | 3 | | | |
| chrII | 24299 | 24437 | 138 | -1,158174315 | N2:24353 | 4,583 | +3:YBL102W; | 3 | | | |
| chrII | 25280 | 25416 | 136 | -1,143379963 | N2:25382 | 6,348 | +20:YBL101C; | 20 | | | |
| chrII | 26126 | 26261 | 135 | -1,054825399 | N2:26202 | 33,963 | +15:YBL101C; | 15 | | | |
| chrII | 30232 | 30334 | 102 | -0,651831833 | | | | Overlap <50 bp | | | |
| chrII | 31655 | 31803 | 148 | -1,092010891 | | | | Overlap <50 bp | | | |
| chrII | 32970 | 33121 | 151 | -1,103793485 | N2:32989 | 9,866 | | Intergene | | | |
| chrII | 35988 | 36136 | 148 | -1,244201563 | N2:36067 | 5,859 | | Intergene | | | |
| chrII | 48768 | 48912 | 144 | -1,239553472 | N2:48843 | 7,74 | +1:YBL090W; -1:YBL091C; | 1 | | | |
| chrII | 54394 | 54539 | 145 | -0,986216701 | N2:54546 | 32,988 | +31:YBL088C; | 31 | | | |
| chrII | 54394 | 54539 | 145 | -0,986216701 | N2:54380 | 27,379 | +32:YBL088C; | 32 | | | |
| chrII | 54551 | 54716 | 165 | -1,139673053 | N2:54732 | 19,799 | +30:YBL088C; | 30 | | | |
| chrII | 54551 | 54716 | 165 | -1,139673053 | N2:54546 | 32,988 | +31:YBL088C; | 31 | | | |
| chrII | 62389 | 62546 | 157 | -1,382861839 | N2:62463 | 4,875 | +2:YBL086C; | 2 | | | |
| chrII | 62885 | 63033 | 148 | -1,263117382 | N2:62915 | 14,569 | -1:YBL086C; | -1 | | | |
| chrII | 64104 | 64264 | 160 | -1,318805016 | | | | Overlap <50 bp | | | |
| chrII | 64653 | 64794 | 141 | -1,398025619 | | | | Overlap <50 bp | | | |
| chrII | 65899 | 66050 | 151 | -1,495715774 | N2:65948 | 12,404 | +14:YBL085W; | 14 | | | |
| chrII | 67731 | 67881 | 150 | -1,37633213 | N2:67884 | 32,216 | | Intergene | | | |
| chrII | 69118 | 69263 | 145 | -1,159153969 | N2:69156 | 9,333 | +3:YBL084C; | 3 | | | |
| chrII | 79393 | 79540 | 147 | -1,719609071 | N2:79471 | 11,701 | +26:YBL079W; | 26 | | | |
| chrII | 81527 | 81678 | 151 | -1,374821732 | N2:81676 | 23,415 | +16:YBL076C; | 16 | | | |
| chrII | 81527 | 81678 | 151 | -1,374821732 | N2:81506 | 24,489 | +17:YBL076C; | 17 | | | |
| chrII | 86475 | 86605 | 130 | -1,015290796 | N2:86602 | 9,283 | 0:YBL075C; +8:YBL074C; | 0 | | | |
| chrII | 87052 | 87211 | 159 | -1,613149953 | N2:87114 | 15,611 | +5:YBL074C; | 5 | | | |
| chrII | 94866 | 95023 | 157 | -1,496880406 | N2:94979 | 10,407 | +8:YBL067C; | 8 | | | |
| chrII | 97063 | 97217 | 154 | -1,273904521 | N2:97213 | 41,347 | +21:YBL066C; | 21 | | | |
| chrII | 97108 | 97247 | 139 | -0,912827287 | N2:97213 | 41,347 | +21:YBL066C; | 21 | | | |
| chrII | 98068 | 98229 | 161 | -1,463100968 | N2:98203 | 16,861 | +15:YBL066C; | 15 | | | |
| chrII | 98405 | 98563 | 158 | -1,400718636 | N2:98532 | 27,178 | +13:YBL066C; | 13 | | | |
| chrII | 102188 | 102336 | 148 | -1,48455441 | N2:102248 | 6,686 | +3:YBL063W; | 3 | | | |
| chrII | 105450 | 105577 | 127 | -1,068773785 | N2:105509 | 25,398 | +12:YBL061C; | 12 | | | |
| chrII | 105532 | 105684 | 152 | -1,44357567 | N2:105680 | 26,153 | +11:YBL061C; | 11 | | | |
| chrII | 105532 | 105684 | 152 | -1,44357567 | N2:105509 | 25,398 | +12:YBL061C; | 12 | | | |
| chrII | 105754 | 105910 | 156 | -0,815046163 | N2:105836 | 34,085 | +10:YBL061C; | 10 | | | |
| chrII | 106088 | 106233 | 145 | -1,111842236 | | | | Overlap <50 bp | | | |
| chrII | 112631 | 112737 | 106 | -1,138361425 | N2:112714 | 14,148 | +5*:YBL057C; +9*:YBL058W; | 5 | TERM | | |
| chrII | 112631 | 112749 | 118 | -1,165103148 | N2:112714 | 14,148 | +5*:YBL057C; +9*:YBL058W; | 5 | TERM | | |
| chrII | 117607 | 117737 | 130 | -0,896674642 | N2:117637 | 15,007 | +2:YBL054W; | 2 | | | |
| chrII | 118185 | 118336 | 151 | -1,33033991 | N2:118300 | 12,57 | +6:YBL054W; | 6 | | | |
| chrII | 118430 | 118588 | 158 | -1,195666979 | N2:118451 | 26,306 | +7:YBL054W; | 7 | | | |
| chrII | 118430 | 118588 | 158 | -1,195666979 | N2:118611 | 18,099 | +8:YBL054W; | 8 | | | |
| chrII | 118604 | 118737 | 133 | -0,865913164 | N2:118611 | 18,099 | +8:YBL054W; | 8 | | | |
| chrII | 118662 | 118820 | 158 | -1,46619699 | N2:118790 | 26,216 | +9:YBL054W; | 9 | | | |
| chrII | 120690 | 120830 | 140 | -1,11073866 | N2:120678 | 17,053 | +10:YBL052C; | 10 | | | |
| chrII | 130330 | 130481 | 151 | -1,354184127 | N2:130370 | 38,561 | +11:YBL047C; | 11 | | | |
| chrII | 133172 | 133320 | 148 | -1,608611205 | N2:133293 | 14,293 | +6:YBL046W; | 6 | | | |
| chrII | 134044 | 134208 | 164 | -1,343929583 | N2:134212 | 38,539 | -1:Unit009; +10:YBL045C; | -1 | | | |
| chrII | 134044 | 134208 | 164 | -1,343929583 | N2:134052 | 32,455 | 0:Unit009; +11:YBL045C; | 0 | | | |
| chrII | 146223 | 146376 | 153 | -1,321233983 | N2:146325 | 8,017 | +2:YBL038W; | 2 | | | |
| chrII | 147729 | 147892 | 163 | -1,64896304 | | | | Overlap <50 bp | | | |
| chrII | 157306 | 157457 | 151 | -1,34277469 | N2:157445 | 18,162 | +9:YBL034C; | 9 | | | |
| chrII | 158794 | 158975 | 181 | -1,489809654 | N2:158865 | 12,624 | +6:YBL033C; -1:YBL034C; | 6 | | | |
| chrII | 159579 | 159667 | 88 | -0,672768137 | N2:159667 | 4,324 | +1:YBL033C; | 1 | | | |
| chrII | 159579 | 159734 | 155 | -1,41516889 | N2:159667 | 4,324 | +1:YBL033C; | 1 | | | |
| chrII | 161150 | 161286 | 136 | -1,277479554 | N2:161276 | 10,559 | +8*:YBL032W; | 8 | TERM | | |
| chrII | 161650 | 161786 | 136 | -1,18045652 | N2:161723 | 6,676 | +1:YBL031W; | 1 | | | |
| chrII | 164812 | 164963 | 151 | -1,237920302 | N2:164880 | 12,501 | +1:YBL029C-A; +3*:SUT006; | 1 | | | |
| chrII | 169155 | 169305 | 150 | -1,30591113 | N2:169306 | 20,56 | +7:YBL027W; | 7 | | | |
| chrII | 171571 | 171720 | 149 | -1,161946715 | N2:171641 | 19,308 | +2:YBL025W; | 2 | | | |
| chrII | 171661 | 171810 | 149 | -1,240220877 | N2:171641 | 19,308 | +2:YBL025W; | 2 | | | |
| chrII | 171661 | 171810 | 149 | -1,240220877 | N2:171803 | 10,807 | +3:YBL025W; | 3 | | | |
| chrII | 174305 | 174447 | 142 | -1,091529763 | N2:174374 | 30,192 | +13:YBL024W; | 13 | | | |
| chrII | 186048 | 186192 | 144 | -1,062458065 | N2:186122 | 13,126 | | Intergene | | | |
| chrII | 187191 | 187351 | 160 | -1,627421457 | N2:187294 | 16,294 | +28:YBL017C; | 28 | | | |

| CHR | START | END | L(bp) | ΔLK | Nuc ID | Fuzziness | Gene ID | Gene body postion or intergene | Terminal | rDNA | Telomeric |
|---|---|---|---|---|---|---|---|---|---|---|---|
| chrII | 188175 | 188332 | 157 | -1,343589318 | N2:188328 | 34,317 | +22:YBL017C; | 22 | | | |
| chrII | 189372 | 189539 | 167 | -1,520501284 | | | | Overlap <50 bp | | | |
| chrII | 192490 | 192638 | 148 | -1,128087103 | N2:192564 | 9,283 | +2:YBL016W; | 2 | | | |
| chrII | 195265 | 195410 | 145 | -1,171387466 | N2:195403 | 43,981 | +9:YBL015W; | 9 | | | |
| chrII | 203726 | 203877 | 151 | -1,250439918 | N2:203822 | 8,846 | +3:YBL011W; | 3 | | | |
| chrII | 204265 | 204408 | 143 | -1,571212415 | N2:204305 | 18,267 | +6:YBL011W; | 6 | | | |
| chrII | 213654 | 213809 | 155 | -0,658603789 | N2:213693 | 21,109 | +16:YBL007C; | 16 | | | |
| chrII | 214589 | 214739 | 150 | -1,489830875 | N2:214660 | 10,33 | +11:YBL007C; | 11 | | | |
| chrII | 214869 | 215017 | 148 | -1,176619464 | N2:214983 | 23,267 | +9:YBL007C; | 9 | | | |
| chrII | 215320 | 215464 | 144 | -1,10391934 | N2:215386 | 24,924 | +7:YBL007C; | 7 | | | |
| chrII | 216639 | 216771 | 132 | -1,723397887 | N2:216655 | 7,789 | +4:YBL006C; -1:YBL007C; | 4 | | | |
| chrII | 217546 | 217694 | 148 | -1,12663001 | N2:217574 | 15,84 | +2:YBL005W; | 2 | | | |
| chrII | 222615 | 222772 | 157 | -1,142025999 | N2:222719 | 20,075 | | Intergene | | | |
| chrII | 222619 | 222772 | 153 | -1,446041905 | N2:222719 | 20,075 | | Intergene | | | |
| chrII | 223146 | 223281 | 135 | -1,214836418 | N2:223185 | 24,269 | | Intergene | | | |
| chrII | 223772 | 223929 | 157 | -1,483106003 | N2:223866 | 14,572 | | Intergene | | | |
| chrII | 223872 | 224016 | 144 | -1,1357005 | N2:223866 | 14,572 | | Intergene | | | |
| chrII | 223872 | 224016 | 144 | -1,1357005 | N2:224033 | 28,711 | | Intergene | | | |
| chrII | 224435 | 224582 | 147 | -1,232898347 | N2:224426 | 19,502 | | Intergene | | | |
| chrII | 224435 | 224582 | 147 | -1,232898347 | N2:224599 | 21,92 | | Intergene | | | |
| chrII | 225175 | 225325 | 150 | -1,410601247 | N2:225274 | 23,824 | | Intergene | | | |
| chrII | 226374 | 226481 | 107 | -1,596265069 | N2:226487 | 21,92 | | Intergene | | | |
| chrII | 226832 | 226946 | 114 | -1,349868328 | N2:226921 | 17,037 | | Intergene | | | |
| chrII | 228528 | 228693 | 165 | -1,754544703 | | | | Overlap <50 bp | | | |
| chrII | 233542 | 233703 | 161 | -1,488641967 | N2:233579 | 26,011 | | Intergene | | | |
| chrII | 234707 | 234798 | 91 | -1,399796163 | N2:234705 | 33,041 | | Intergene | | | |
| chrII | 239108 | 239251 | 143 | -1,160932322 | N2:239202 | 30,612 | +13:YBR001C; | 13 | | | |
| chrII | 249586 | 249752 | 166 | -1,460240793 | N2:249650 | 13,428 | +9:YBR007C; | 9 | | | |
| chrII | 252919 | 253070 | 151 | -1,338288891 | | | | Overlap <50 bp | | | |
| chrII | 253969 | 254127 | 158 | -1,517313475 | N2:254057 | 17,173 | +4:YBR008C; | 4 | | | |
| chrII | 255566 | 255720 | 154 | -1,430021544 | N2:255613 | 29,262 | +1:YBR009C; | 1 | | | |
| chrII | 260109 | 260232 | 123 | -1,053192789 | | | | Overlap <50 bp | | | |
| chrII | 260209 | 260364 | 155 | -1,509131457 | N2:260319 | 31,723 | | Intergene | | | |
| chrII | 260642 | 260776 | 134 | -0,586070067 | N2:260707 | 28,618 | | Intergene | | | |
| chrII | 260642 | 260782 | 140 | -0,963757536 | N2:260707 | 28,618 | | Intergene | | | |
| chrII | 261685 | 261820 | 135 | -0,995726897 | N2:261719 | 2,646 | | Intergene | | | |
| chrII | 262132 | 262282 | 150 | -1,10978534 | N2:262199 | 21,197 | | Intergene | | | |
| chrII | 262311 | 262468 | 157 | -1,462281745 | N2:262403 | 32,047 | | Intergene | | | |
| chrII | 262974 | 263121 | 147 | -1,058720824 | N2:262967 | 21,197 | | Intergene | | | |
| chrII | 263717 | 263867 | 150 | -1,384266737 | N2:263815 | 64,023 | | Intergene | | | |
| chrII | 264916 | 265023 | 107 | -1,458520689 | | | | Overlap <50 bp | | | |
| chrII | 269730 | 269883 | 153 | -1,101384237 | N2:269845 | 8,62 | -1:YBR015C; | -1 | | | |
| chrII | 269730 | 269885 | 155 | -2,233425246 | N2:269845 | 8,62 | -1:YBR015C; | -1 | | | |
| chrII | 269781 | 269930 | 149 | -1,410364781 | N2:269845 | 8,62 | -1:YBR015C; | -1 | | | |
| chrII | 272917 | 273071 | 154 | -1,381071069 | N2:273057 | 9,695 | +5:YBR017C; | 5 | | | |
| chrII | 279942 | 280075 | 133 | -1,076543114 | N2:280032 | 11,349 | +7:YBR020W; | 7 | | | |
| chrII | 282915 | 283035 | 120 | -1,09549185 | N2:282943 | 35,064 | +11:YBR021W; | 11 | | | |
| chrII | 288899 | 289053 | 154 | -1,330754737 | N2:288983 | 4,899 | | Intergene | | | |
| chrII | 289452 | 289559 | 107 | -1,216533893 | N2:289558 | 6,616 | +2:YBR024W; | 2 | | | |
| chrII | 289452 | 289573 | 121 | -1,285147454 | N2:289558 | 6,616 | +2:YBR024W; | 2 | | | |
| chrII | 292636 | 292792 | 156 | -1,332846524 | N2:292683 | 7,759 | | Intergene | | | |
| chrII | 295986 | 296116 | 130 | -0,860940601 | N2:296031 | 8,479 | +1:YBR028C; -1:CUT010; | 1 | | | |
| chrII | 296940 | 297047 | 107 | -1,335508013 | N2:296987 | 24,977 | +6:YBR029C; | 6 | | | |
| chrII | 296940 | 297056 | 116 | -1,498188266 | N2:296987 | 24,977 | +6:YBR029C; | 6 | | | |
| chrII | 298361 | 298510 | 149 | -1,193268767 | N2:298428 | 10,52 | +2:YBR030W; | 2 | | | |
| chrII | 298515 | 298656 | 141 | -1,059273554 | N2:298593 | 8,539 | +3:YBR030W; | 3 | | | |
| chrII | 300341 | 300475 | 134 | -0,914362894 | N2:300360 | 24,322 | +2:YBR031W; | 2 | | | |
| chrII | 303103 | 303264 | 161 | -1,461900746 | N2:303172 | 49,379 | +9:YBR033W; +9:SUT445; | 9 | | | |
| chrII | 304167 | 304319 | 152 | -1,267210037 | N2:304247 | 16,721 | +15:YBR033W; +3:SUT445; | 15 | | | |
| chrII | 306192 | 306322 | 130 | -1,280327839 | N2:306304 | 12,317 | +5:YBR035C; | 5 | | | |
| chrII | 310844 | 311002 | 158 | -1,368252604 | N2:310950 | 6,408 | +4:YBR037C; +3*:CUT013; | 4 | | | |
| chrII | 314561 | 314691 | 130 | -0,962520912 | N2:314625 | 45,711 | +18*:YBR038W; | 18 | TERM | | |
| chrII | 316179 | 316338 | 159 | -1,604946668 | N2:316235 | 23,964 | +5:YBR039W; | 5 | | | |
| chrII | 320829 | 320984 | 155 | -1,26189574 | N2:320918 | 9,165 | +5:YBR042C; | 5 | | | |
| chrII | 324261 | 324383 | 122 | -0,975459417 | N2:324329 | 21,524 | 3R043C; +11*:YBR044C; -1:CUT014; | -1 | | | |
| chrII | 324593 | 324738 | 145 | -1,28786413 | N2:324676 | 5,718 | +9:YBR044C; +2:CUT014; | 9 | | | |
| chrII | 333410 | 333578 | 168 | -1,408097988 | N2:333441 | 40,576 | | Intergene | | | |
| chrII | 334171 | 334335 | 164 | -1,639948638 | N2:334243 | 18,608 | +16*:YBR049C; | 16 | TERM | | |
| chrII | 341568 | 341673 | 105 | -0,834575191 | | | | Overlap <50 bp | | | |
| chrII | 344002 | 344150 | 148 | -1,201988833 | N2:344104 | 18,285 | +8:YBR054W; | 8 | | | |
| chrII | 345574 | 345727 | 153 | -1,297378957 | N2:345699 | 34,213 | +11:YBR055C; | 11 | | | |
| chrII | 348428 | 348599 | 171 | -1,13153481 | N2:348504 | 6,131 | +5:YBR056W; | 5 | | | |
| chrII | 348488 | 348624 | 136 | -1,132777903 | N2:348504 | 6,131 | +5:YBR056W; | 5 | | | |
| chrII | 352601 | 352731 | 130 | -1,055310767 | N2:352692 | 22,232 | +5:YBR057C; | 5 | | | |
| chrII | 354955 | 355107 | 152 | -1,254234777 | N2:355010 | 14,844 | +7:YBR058C; | 7 | | | |
| chrII | 360789 | 360938 | 149 | -1,474616729 | N2:360839 | 5,797 | +11:YBR060C; +2:CUT018; | 11 | | | |
| chrII | 369124 | 369272 | 148 | -1,429493757 | N2:369186 | 29,645 | +4:YBR065C; | 4 | | | |
| chrII | 369614 | 369763 | 149 | -1,063372795 | N2:369688 | 6,683 | +1:YBR065C; | 1 | | | |
| chrII | 372782 | 372926 | 144 | -1,014742943 | N2:372886 | 30,028 | +11:SUT016; 0:YBR067C; | 11 | | | |
| chrII | 379733 | 379878 | 145 | -1,112952051 | N2:379800 | 14,16 | +2:YBR070C; | 2 | | | |
| chrII | 380438 | 380593 | 155 | -1,077365548 | N2:380439 | 10,198 | +1:YBR071W; | 1 | | | |
| chrII | 380438 | 380593 | 155 | -1,077365548 | N2:380588 | 15,868 | +2:YBR071W; | 2 | | | |
| chrII | 383232 | 383381 | 149 | -0,996686495 | N2:383330 | 10,633 | +2:YBR073W; -1:YBR072C-A; | 2 | | | |
| chrII | 384056 | 384165 | 109 | -1,623780409 | | | | Overlap <50 bp | | | |
| chrII | 384888 | 385039 | 151 | -1,319140026 | N2:384899 | 36,378 | +12:YBR073W; | 12 | | | |
| chrII | 385956 | 386120 | 164 | -1,211693631 | N2:386082 | 32,347 | -1:YBR074W; | -1 | | | |
| chrII | 386724 | 386859 | 135 | -1,053143167 | N2:386790 | 13,644 | +4:YBR074W; | 4 | | | |
| chrII | 402057 | 402202 | 145 | -1,276781246 | N2:402105 | 40,195 | +21:YBR081C; | 21 | | | |
| chrII | 405294 | 405447 | 153 | -1,159552264 | N2:405333 | 7,083 | +1:YBR081C; | 1 | | | |
| chrII | 411086 | 411218 | 132 | -0,877040498 | N2:411221 | 7,092 | +2:YBR084W; | 2 | | | |
| chrII | 412922 | 413067 | 145 | -1,266292301 | N2:413068 | 10,368 | +13:YBR084W; | 13 | | | |
| chrII | 417276 | 417442 | 166 | -1,2587924 | N2:417375 | 10,21 | +4:Unit017; | 4 | | | |
| chrII | 418844 | 418980 | 136 | -2,768708743 | N2:418877 | 4,021 | +3:YBR085C-A; | 3 | | | |
| chrII | 420492 | 420630 | 138 | -1,109448642 | N2:420545 | 31,738 | +17:YBR086C; | 17 | | | |
| chrII | 432763 | 432911 | 148 | -1,831373047 | N2:432818 | 16,227 | +6:YBR094W; | 6 | | | |
| chrII | 435614 | 435764 | 150 | -1,369178642 | N2:435690 | 11,118 | +1:YBR095C; -1:YBR096W; | 1 | | | |
| chrII | 441246 | 441413 | 167 | -1,504828399 | N2:441290 | 29,543 | +29:YBR097W; -1:YBR098W; | 29 | | | |
| chrII | 442313 | 442462 | 149 | -1,369837877 | | | | Overlap <50 bp | | | |

TABLE 3

| CHR | START | END | L(bp) | ΔLK | Nuc ID | Fuzziness | Gene ID | Gene body postion or intergene | Terminal | rDNA | Telomeric |
|---|---|---|---|---|---|---|---|---|---|---|---|
| chrII | 447274 | 447421 | 147 | -1,002159996 | N2:447332 | 3,656 | -1:YBR103W; +1:YBR102C; | -1 | | | |
| chrII | 448183 | 448311 | 128 | -1,092945784 | N2:448253 | 12,973 | +5:YBR103W; | 5 | | | |
| chrII | 448687 | 448853 | 166 | -1,509032771 | | | | Overlap <50 bp | | | |
| chrII | 450522 | 450683 | 161 | -1,314301324 | N2:450575 | 13,726 | +7*:YBR104W; | 7 | TERM | | |
| chrII | 456967 | 457110 | 143 | -1,532475687 | N2:456997 | 9,602 | +14:YBR108W; | 14 | | | |
| chrII | 459507 | 459619 | 112 | -1,579094464 | N2:459525 | 13,172 | +5:YBR110W; | 5 | | | |
| chrII | 463510 | 463659 | 149 | -1,404145294 | N2:463671 | 35,929 | | Intergene | | | |
| chrII | 464402 | 464557 | 155 | -1,127412307 | N2:464478 | 22,113 | +10:YBR112C; | 10 | | | |
| chrII | 467973 | 468117 | 144 | -1,032238428 | N2:468080 | 9,236 | +6:YBR114W; | 6 | | | |
| chrII | 469937 | 470099 | 162 | -1,37154338 | N2:469962 | 17,234 | +27:YBR115C; | 27 | | | |
| chrII | 478544 | 478678 | 134 | -0,815841234 | N2:478563 | 46,083 | +8:YBR118W; | 8 | | | |
| chrII | 479993 | 480154 | 161 | -1,259329534 | N2:480057 | 49,497 | | Intergene | | | |
| chrII | 480073 | 480226 | 153 | -1,23061109 | N2:480057 | 49,497 | | Intergene | | | |
| chrII | 483614 | 483775 | 161 | -1,214965849 | N2:483752 | 12,416 | +1:CUT026; -1:YBR121C; | 1 | | | |
| chrII | 490237 | 490391 | 154 | -0,981533398 | N2:490316 | 21,747 | +2:YBR126C; | 2 | | | |
| chrII | 496459 | 496596 | 137 | -1,246572939 | N2:496548 | 5,805 | +3:YBR130C; | 3 | | | |
| chrII | 498698 | 498864 | 166 | -1,379409744 | N2:498780 | 27,474 | +11:YBR131W; | 11 | | | |
| chrII | 499848 | 499990 | 142 | -0,927358262 | N2:499949 | 23,845 | +12:YBR132C; | 12 | | | |
| chrII | 501137 | 501288 | 151 | -1,080504278 | N2:501227 | 9,888 | +4:YBR132C; | 4 | | | |
| chrII | 503704 | 503871 | 167 | -1,452303465 | N2:503875 | 21,622 | +4:YBR133C; | 4 | | | |
| chrII | 503704 | 503871 | 167 | -1,452303465 | N2:503696 | 25,832 | +5:YBR133C; | 5 | | | |
| chrII | 505974 | 506148 | 174 | -1,394268654 | N2:506017 | 6,87 | +4:YBR136W; | 4 | | | |
| chrII | 506025 | 506148 | 123 | -0,62939195 | N2:506017 | 6,87 | +4:YBR136W; | 4 | | | |
| chrII | 506025 | 506179 | 154 | -1,241093561 | N2:506017 | 6,87 | +4:YBR136W; | 4 | | | |
| chrII | 506025 | 506179 | 154 | -1,241093561 | N2:506177 | 7,225 | +5:YBR136W; | 5 | | | |
| chrII | 509772 | 509912 | 140 | -1,082668098 | N2:509886 | 33,421 | +28:YBR136W; | 28 | | | |
| chrII | 511646 | 511795 | 149 | -1,434658486 | N2:511651 | 27,644 | +38:YBR136W; | 38 | | | |
| chrII | 512273 | 512423 | 150 | -1,44706331 | N2:512342 | 8,937 | +42:YBR136W; | 42 | | | |
| chrII | 519148 | 519306 | 158 | -1,271055536 | N2:519220 | 10,668 | +46:YBR140C; +54:YBR141C; | 46 | | | |
| chrII | 523260 | 523407 | 147 | -1,337333998 | N2:523344 | 0 | +21:YBR140C; +29:YBR141C; | 21 | | | |
| chrII | 523774 | 523925 | 151 | -1,316681204 | | | | Overlap <50 bp | | | |
| chrII | 525378 | 525528 | 150 | -1,287368088 | N2:525423 | 30,425 | +9:YBR140C; +17:YBR141C; | 9 | | | |
| chrII | 527770 | 527902 | 132 | -1,066277836 | N2:527832 | 3,204 | +2:YBR141C; | 2 | | | |
| chrII | 528914 | 529061 | 147 | -1,099649381 | N2:528946 | 34,173 | +5:YBR142W; | 5 | | | |
| chrII | 529231 | 529384 | 153 | -1,273423221 | N2:529279 | 12,75 | +7:YBR142W; | 7 | | | |
| chrII | 533755 | 533907 | 152 | -0,989226591 | N2:533826 | 29,992 | +1:YBR145W; | 1 | | | |
| chrII | 543934 | 544097 | 163 | -1,34536352 | N2:543989 | 13,867 | +4:YBR150C; | 4 | | | |
| chrII | 545512 | 545645 | 133 | -1,088763431 | | | | Overlap <50 bp | | | |
| chrII | 550054 | 550200 | 146 | -0,857880812 | N2:550101 | 17,509 | +3:YBR155W; | 3 | | | |
| chrII | 551769 | 551916 | 147 | -1,300364602 | N2:551862 | 37,91 | +9:YBR156C; | 9 | | | |
| chrII | 551821 | 551946 | 125 | -1,326670932 | N2:551862 | 37,91 | +9:YBR156C; | 9 | | | |
| chrII | 552584 | 552756 | 172 | -1,440625385 | N2:552697 | 13,633 | +4:YBR156C; | 4 | | | |
| chrII | 560316 | 560442 | 126 | -1,221672782 | N2:560343 | 13,038 | +3:YBR160W; | 3 | | | |
| chrII | 568857 | 569009 | 152 | -1,31346641 | N2:568953 | 8,085 | +2:YBR165W; | 2 | | | |
| chrII | 570513 | 570675 | 162 | -1,289589675 | N2:570686 | 8,733 | +4:YBR166C; | 4 | | | |
| chrII | 570513 | 570675 | 162 | -1,289589675 | N2:570530 | 24,785 | +5:YBR166C; | 5 | | | |
| chrII | 575360 | 575489 | 129 | -0,977519812 | N2:575420 | 12,49 | +4:YBR169C; | 4 | | | |
| chrII | 575363 | 575489 | 126 | -1,175867983 | N2:575420 | 12,49 | +4:YBR169C; | 4 | | | |
| chrII | 575635 | 575789 | 154 | -1,754362726 | N2:575736 | 6,775 | +2:YBR169C; | 2 | | | |
| chrII | 577662 | 577794 | 132 | -1,636094452 | N2:577744 | 7,321 | +3:YBR170C; | 3 | | | |
| chrII | 581371 | 581525 | 154 | -1,645286481 | N2:581425 | 9,182 | +1:YBR172C; | 1 | | | |
| chrII | 583028 | 583186 | 158 | -1,289905811 | N2:583091 | 13,088 | +5:YBR175W; | 5 | | | |
| chrII | 583277 | 583426 | 149 | -1,159587992 | | | | Overlap <50 bp | | | |
| chrII | 589093 | 589249 | 156 | -1,56010912 | N2:589146 | 13,038 | +1:YBR179C; | 1 | | | |
| chrII | 598510 | 598657 | 147 | -1,232680898 | N2:598568 | 32,623 | +8:YBR184W; | 8 | | | |
| chrII | 599865 | 600024 | 159 | -1,182357063 | N2:599938 | 9,094 | +1:YBR185C; | 1 | | | |
| chrII | 601519 | 601677 | 158 | -1,511310659 | N2:601539 | 8,526 | +5*:anti012; +7:YBR186W; | 5 | TERM | | |
| chrII | 601519 | 601677 | 158 | -1,511310659 | N2:601690 | 18,091 | +4:anti012; +8:YBR186W; | 4 | | | |
| chrII | 608166 | 608308 | 142 | -1,05398561 | N2:608176 | 2,191 | +5:YBR192W; | 5 | | | |
| chrII | 608166 | 608308 | 142 | -1,05398561 | N2:608330 | 8,025 | +6:YBR192W; | 6 | | | |
| chrII | 610440 | 610604 | 164 | -1,484272313 | N2:610482 | 28,991 | +4*:YBR194W; | 4 | TERM | | |
| chrII | 615980 | 616137 | 157 | -1,063248295 | N2:616096 | 25,129 | -1:YBR197C; | -1 | | | |
| chrII | 618802 | 618957 | 155 | -1,229307997 | N2:618851 | 6,772 | +1:YBR199W; -1:YBR198C; | 1 | | | |
| chrII | 621368 | 621503 | 135 | -1,129265196 | N2:621487 | 21,166 | +5:YBR200W; | 5 | | | |
| chrII | 624538 | 624687 | 149 | -1,249099736 | N2:624638 | 10,756 | +2:YBR201C-A; | 2 | | | |
| chrII | 626717 | 626881 | 164 | -1,361430048 | N2:626851 | 25,153 | +8:YBR202W; | 8 | | | |
| chrII | 633744 | 633889 | 145 | -1,035454002 | N2:633802 | 9,98 | +2:YBR205W; | 2 | | | |
| chrII | 633747 | 633889 | 142 | -1,135180111 | N2:633802 | 9,98 | +2:YBR205W; | 2 | | | |
| chrII | 634590 | 634750 | 160 | -1,296665043 | N2:634645 | 27,668 | +7:YBR205W; | 7 | | | |
| chrII | 635204 | 635350 | 146 | -1,938915442 | N2:635289 | 8,289 | +2:YBR207W; | 2 | | | |
| chrII | 636694 | 636853 | 159 | -1,375705026 | N2:636766 | 22,941 | | Intergene | | | |
| chrII | 636695 | 636853 | 158 | -1,364353674 | N2:636766 | 22,941 | | Intergene | | | |
| chrII | 636699 | 636853 | 154 | -1,515038997 | N2:636766 | 22,941 | | Intergene | | | |
| chrII | 641881 | 642011 | 130 | -1,209245845 | N2:641909 | 5,037 | +3:YBR208C; | 3 | | | |
| chrII | 643000 | 643147 | 147 | -1,329472018 | N2:643066 | 11,003 | | Intergene | | | |
| chrII | 643625 | 643773 | 148 | -1,155154848 | N2:643703 | 20,516 | | Intergene | | | |
| chrII | 645958 | 646066 | 108 | -0,417230945 | N2:646057 | 9,452 | +5*:YBR210W; | 5 | TERM | | |
| chrII | 652842 | 653006 | 164 | -1,075597988 | | | | Overlap <50 bp | | | |
| chrII | 667311 | 667462 | 151 | -2,246118016 | N2:667376 | 25,608 | +7:YBR222C; | 7 | | | |
| chrII | 670468 | 670600 | 132 | -1,364029643 | N2:670509 | 22,176 | -1:YBR223C; +1:YBR225W; | -1 | | | |
| chrII | 672951 | 673112 | 161 | -1,184502263 | N2:673046 | 8,585 | +16:YBR225W; | 16 | | | |
| chrII | 673886 | 673964 | 78 | -0,798 | N2:673878 | 30,222 | +8:YBR227C; | 8 | | | |
| chrII | 673886 | 673989 | 103 | -0,672906555 | N2:673878 | 30,222 | +8:YBR227C; | 8 | | | |
| chrII | 674406 | 674512 | 106 | -1,582178995 | N2:674454 | 9,668 | +5:YBR227C; | 5 | | | |
| chrII | 674406 | 674523 | 117 | -1,454474226 | N2:674454 | 9,668 | +5:YBR227C; | 5 | | | |
| chrII | 678764 | 678912 | 148 | -1,175676074 | N2:678851 | 8,477 | +3:YBR229C; | 3 | | | |
| chrII | 682036 | 682166 | 130 | -1,045846273 | N2:682093 | 41,012 | -1:CUT034; | -1 | | | |
| chrII | 683947 | 684082 | 135 | -1,093585499 | N2:684039 | 9,867 | +5:YBR233W; | 5 | | | |
| chrII | 684506 | 684665 | 159 | -1,400190924 | N2:684541 | 13,867 | +8:YBR233W; | 8 | | | |
| chrII | 685114 | 685267 | 153 | -1,143652323 | N2:685155 | 9,201 | +2:YBR233W-A; | 2 | | | |
| chrII | 688568 | 688714 | 146 | -1,313717319 | N2:688658 | 27,969 | +12:YBR235W; | 12 | | | |
| chrII | 693669 | 693820 | 151 | -1,624186433 | N2:693836 | 6,429 | +13*:YBR237W; | 13 | TERM | | |
| chrII | 696303 | 696437 | 134 | -0,984598953 | | | | Overlap <50 bp | | | |
| chrII | 698169 | 698308 | 139 | -0,934238114 | N2:698202 | 28,19 | +3:anti016; | 3 | | | |
| chrII | 698728 | 698854 | 126 | -0,996001992 | N2:698800 | 13,206 | +8:YBR239C; | 8 | | | |
| chrII | 698730 | 698836 | 106 | -0,715778305 | N2:698800 | 13,206 | +8:YBR239C; | 8 | | | |
| chrII | 699474 | 699629 | 155 | -1,3447482 | N2:699534 | 13,05 | +4:YBR239C; | 4 | | | |
| chrII | 699802 | 699945 | 143 | -1,679432646 | N2:699870 | 11,622 | +2:YBR239C; | 2 | | | |
| chrII | 699802 | 699953 | 151 | -1,700062035 | N2:699870 | 11,622 | +2:YBR239C; | 2 | | | |

| CHR | START | END | L(bp) | ΔLK | Nuc ID | Fuzziness | Gene ID | Gene body postion or intergene | Terminal | rDNA | Telomeric |
|---|---|---|---|---|---|---|---|---|---|---|---|
| chrII | 699834 | 699945 | 111 | -1,313347211 | N2:699870 | 11,622 | +2:YBR239C; | 2 | | | |
| chrII | 699834 | 699953 | 119 | -0,934322568 | N2:699870 | 11,622 | +2:YBR239C; | 2 | | | |
| chrII | 702899 | 703031 | 132 | -1,534616715 | N2:702974 | 14,422 | +7*:YBR241C; | 7 | TERM | | |
| chrII | 706525 | 706696 | 171 | -1,242849783 | N2:706600 | 3,125 | +2:YBR243C; | 2 | | | |
| chrII | 708493 | 708657 | 164 | -1,040369298 | N2:708624 | 31,291 | +18:YBR245C; | 18 | | | |
| chrII | 708881 | 709054 | 173 | -1,481569108 | N2:708943 | 34,96 | +16:YBR245C; | 16 | | | |
| chrII | 716866 | 717023 | 157 | -1,369908789 | N2:716955 | 10,727 | +8:YBR249C; | 8 | | | |
| chrII | 716874 | 717023 | 149 | -1,108888063 | N2:716955 | 10,727 | +8:YBR249C; | 8 | | | |
| chrII | 718549 | 718695 | 146 | -1,483357466 | N2:718618 | 16,956 | +2:Unit021; | 2 | | | |
| chrII | 721490 | 721639 | 149 | -1,114106296 | N2:721551 | 12,073 | +2:YBR251W; | 2 | | | |
| chrII | 723357 | 723507 | 150 | -1,24577868 | N2:723426 | 6,221 | +2:YBR253W; | 2 | | | |
| chrII | 727858 | 728011 | 153 | -1,111040747 | N2:727936 | 4,135 | +2:YBR256C; | 2 | | | |
| chrII | 733512 | 733676 | 164 | -1,628574356 | N2:733639 | 30,94 | +8:YBR260C; | 8 | | | |
| chrII | 735258 | 735391 | 133 | -0,601625644 | N2:735332 | 8,118 | +2:YBR261C; | 2 | | | |
| chrII | 736781 | 736927 | 146 | -1,194053632 | N2:736768 | 5,989 | +4:YBR263W; | 4 | | | |
| chrII | 736912 | 737033 | 121 | -1,076385223 | | | | Overlap <50 bp | | | |
| chrII | 740252 | 740411 | 159 | -1,308668504 | N2:740348 | 11,413 | +4:YBR267W; | 4 | | | |
| chrII | 743678 | 743781 | 103 | -1,597747713 | N2:743723 | 18,469 | +6:YBR270C; | 6 | | | |
| chrII | 746216 | 746373 | 157 | -1,423993514 | N2:746234 | 22,405 | +9*:YBR271W; +10*:YBR272C; | 9 | TERM | | |
| chrII | 747192 | 747321 | 129 | -1,046172535 | N2:747277 | 11,185 | +4:YBR272C; | 4 | | | |
| chrII | 748930 | 749090 | 160 | -1,367845746 | N2:749008 | 9,935 | +3:YBR273C; | 3 | | | |
| chrII | 749254 | 749401 | 147 | -1,108331229 | N2:749334 | 7,026 | -1:YBR274W; +1:YBR273C; | -1 | | | |
| chrII | 750045 | 750206 | 161 | -1,100825585 | N2:750083 | 17,143 | +4:YBR274W; | 4 | | | |
| chrII | 760513 | 760620 | 107 | -0,945882361 | | | | Overlap <50 bp | | | |
| chrII | 760513 | 760634 | 121 | -1,219239247 | | | | Overlap <50 bp | | | |
| chrII | 771359 | 771518 | 159 | -1,437711441 | | | | Overlap <50 bp | | | |
| chrII | 771799 | 771953 | 154 | -1,298920275 | | | | Overlap <50 bp | | | |
| chrII | 774249 | 774405 | 156 | -1,411326104 | N2:774294 | 5,404 | +3:YBR285W; | 3 | | | |
| chrII | 775701 | 775861 | 160 | -0,929463514 | | | | Overlap <50 bp | | | |
| chrII | 777331 | 777484 | 153 | -1,339755057 | N2:777349 | 9,607 | +5:YBR287W; | 5 | | | |
| chrII | 779629 | 779798 | 169 | -1,50707337 | N2:779707 | 7,899 | +1:YBR289W; -1:YBR288C; | 1 | | | |
| chrII | 780145 | 780251 | 106 | -1,053022222 | N2:780192 | 4,676 | +4:YBR289W; | 4 | | | |
| chrII | 780145 | 780260 | 115 | -0,918611905 | N2:780192 | 4,676 | +4:YBR289W; | 4 | | | |
| chrII | 780582 | 780757 | 175 | -1,500281704 | N2:780666 | 31,575 | | Intergene | | | |
| chrII | 781915 | 782083 | 168 | -1,198476355 | N2:782014 | 4 | | Intergene | | | |
| chrII | 786436 | 786542 | 106 | -0,801755722 | N2:786464 | 5,128 | :Unit023; +3*:SUT451; +7*:SUT023; | -1 | | | |
| chrII | 786436 | 786560 | 124 | -1,062389934 | N2:786464 | 5,128 | :Unit023; +3*:SUT451; +7*:SUT023; | -1 | | | |
| chrII | 788442 | 788582 | 140 | -1,010280463 | N2:788514 | 10,759 | +10*:YBR293W; | 10 | TERM | | |
| chrII | 788604 | 788759 | 155 | -1,447389067 | N2:788689 | 4,848 | | Intergene | | | |
| chrII | 789546 | 789709 | 163 | -1,586585326 | N2:789650 | 8,66 | +16:anti017; +4:YBR294W; | 16 | | | |
| chrII | 791029 | 791155 | 126 | -1,301735875 | N2:791088 | 39,659 | +7:anti017; +13:YBR294W; | 7 | | | |
| chrII | 791029 | 791156 | 127 | -1,036811816 | N2:791088 | 39,659 | +7:anti017; +13:YBR294W; | 7 | | | |
| chrII | 791031 | 791136 | 105 | -0,713070249 | N2:791088 | 39,659 | +7:anti017; +13:YBR294W; | 7 | | | |
| chrII | 793238 | 793400 | 162 | -1,179922456 | N2:793321 | 24,028 | +4:YBR295W; | 4 | | | |
| chrII | 795412 | 795559 | 147 | -1,168072292 | N2:795519 | 41,841 | +18:YBR295W; | 18 | | | |
| chrII | 809074 | 809203 | 129 | -1,3505723 | N2:809118 | 18,385 | +2:YBR301W; | 2 | | | |
| chrII | 812955 | 813098 | 143 | -1,217374679 | N2:812953 | 6,083 | | Intergene | | | TEL RIGHT |
| chrIII | 2 | 145 | 143 | -0,954679452 | | | | Overlap <50 bp | | | TEL LEFT |
| chrIII | 2 | 176 | 174 | -0,229084052 | | | | Overlap <50 bp | | | TEL LEFT |
| chrIII | 4 | 145 | 141 | -1,406102945 | | | | Overlap <50 bp | | | TEL LEFT |
| chrIII | 10 | 145 | 135 | -1,31929281 | | | | Overlap <50 bp | | | TEL LEFT |
| chrIII | 10 | 146 | 136 | -0,995025744 | | | | Overlap <50 bp | | | TEL LEFT |
| chrIII | 14 | 145 | 131 | -0,89276556 | | | | Overlap <50 bp | | | TEL LEFT |
| chrIII | 19 | 145 | 126 | -0,676104031 | | | | Overlap <50 bp | | | TEL LEFT |
| chrIII | 26 | 145 | 119 | -1,106431779 | | | | Overlap <50 bp | | | TEL LEFT |
| chrIII | 37 | 145 | 108 | -0,911015253 | | | | Overlap <50 bp | | | TEL LEFT |
| chrIII | 41 | 145 | 104 | -1,016883833 | | | | Overlap <50 bp | | | TEL LEFT |
| chrIII | 50 | 145 | 95 | -0,840231361 | | | | Overlap <50 bp | | | TEL LEFT |
| chrIII | 50 | 176 | 126 | 0,156676152 | | | | Overlap <50 bp | | | TEL LEFT |
| chrIII | 50 | 185 | 135 | -0,746822302 | | | | Overlap <50 bp | | | TEL LEFT |
| chrIII | 50 | 201 | 151 | -0,678938322 | | | | Overlap <50 bp | | | TEL LEFT |
| chrIII | 50 | 220 | 170 | -0,703926889 | | | | Overlap <50 bp | | | TEL LEFT |
| chrIII | 73 | 263 | 190 | -1,312793189 | | | | Overlap <50 bp | | | TEL LEFT |
| chrIII | 88 | 231 | 143 | -1,267464587 | | | | Overlap <50 bp | | | TEL LEFT |
| chrIII | 97 | 212 | 115 | -0,931331371 | | | | Overlap <50 bp | | | TEL LEFT |
| chrIII | 110 | 263 | 153 | -1,269630523 | | | | Overlap <50 bp | | | TEL LEFT |
| chrIII | 110 | 280 | 170 | -0,88860386 | N3:298 | 23,979 | | Intergene | | | TEL LEFT |
| chrIII | 110 | 283 | 173 | -1,116390842 | N3:298 | 23,979 | | Intergene | | | TEL LEFT |
| chrIII | 115 | 280 | 165 | -0,923278483 | N3:298 | 23,979 | | Intergene | | | TEL LEFT |
| chrIII | 115 | 283 | 168 | -1,147224261 | N3:298 | 23,979 | | Intergene | | | TEL LEFT |
| chrIII | 118 | 284 | 166 | -0,929152627 | N3:298 | 23,979 | | Intergene | | | TEL LEFT |
| chrIII | 129 | 252 | 123 | -0,919105825 | | | | Overlap <50 bp | | | TEL LEFT |
| chrIII | 129 | 260 | 131 | -0,782812905 | | | | Overlap <50 bp | | | TEL LEFT |
| chrIII | 129 | 263 | 134 | -0,72076581 | | | | Overlap <50 bp | | | TEL LEFT |
| chrIII | 129 | 280 | 151 | -0,943894971 | N3:298 | 23,979 | | Intergene | | | TEL LEFT |
| chrIII | 129 | 283 | 154 | -0,955521888 | N3:298 | 23,979 | | Intergene | | | TEL LEFT |
| chrIII | 129 | 284 | 155 | -1,723242825 | N3:298 | 23,979 | | Intergene | | | TEL LEFT |
| chrIII | 129 | 301 | 172 | -1,695454046 | N3:298 | 23,979 | | Intergene | | | TEL LEFT |
| chrIII | 129 | 309 | 180 | -1,188465012 | N3:298 | 23,979 | | Intergene | | | TEL LEFT |
| chrIII | 129 | 326 | 197 | -1,004285517 | N3:298 | 23,979 | | Intergene | | | TEL LEFT |
| chrIII | 143 | 284 | 141 | -1,573492119 | N3:298 | 23,979 | | Intergene | | | TEL LEFT |
| chrIII | 164 | 284 | 120 | -0,828521588 | N3:298 | 23,979 | | Intergene | | | TEL LEFT |
| chrIII | 164 | 341 | 177 | -0,676782129 | N3:298 | 23,979 | | Intergene | | | TEL LEFT |
| chrIII | 186 | 284 | 98 | -0,74917568 | N3:298 | 23,979 | | Intergene | | | TEL LEFT |
| chrIII | 186 | 320 | 134 | -1,551770143 | N3:298 | 23,979 | | Intergene | | | TEL LEFT |
| chrIII | 186 | 337 | 151 | -1,671725041 | N3:298 | 23,979 | | Intergene | | | TEL LEFT |
| chrIII | 186 | 341 | 155 | -1,327448223 | N3:298 | 23,979 | | Intergene | | | TEL LEFT |
| chrIII | 186 | 352 | 166 | -1,426052459 | N3:298 | 23,979 | | Intergene | | | TEL LEFT |
| chrIII | 186 | 353 | 167 | -1,532908001 | N3:298 | 23,979 | | Intergene | | | TEL LEFT |
| chrIII | 186 | 355 | 169 | -0,772229989 | N3:298 | 23,979 | | Intergene | | | TEL LEFT |
| chrIII | 192 | 329 | 137 | -0,926236863 | N3:298 | 23,979 | | Intergene | | | TEL LEFT |
| chrIII | 246 | 341 | 95 | -0,900552583 | N3:298 | 23,979 | | Intergene | | | TEL LEFT |
| chrIII | 246 | 352 | 106 | -0,70570749 | N3:298 | 23,979 | | Intergene | | | TEL LEFT |
| chrIII | 5166 | 5314 | 148 | -1,232000214 | N3:5206 | 29,597 | | Intergene | | | |
| chrIII | 5882 | 5994 | 112 | -1,835019833 | N3:5973 | 10,599 | | Intergene | | | |
| chrIII | 7560 | 7714 | 154 | -1,589499231 | N3:7557 | 35,473 | +6:YCL073C; | 6 | | | |

| CHR | START | END | L(bp) | ΔLK | Nuc ID | Fuzziness | Gene ID | Gene body postion or intergene | Terminal | rDNA | Telomeric |
|---|---|---|---|---|---|---|---|---|---|---|---|
| chrIII | 9830 | 9940 | 110 | -0,920543576 | N3:9914 | 23,714 | +2:YCL069W; | 2 | | | |
| chrIII | 9831 | 9940 | 109 | -1,031392141 | N3:9914 | 23,714 | +2:YCL069W; | 2 | | | |
| chrIII | 9831 | 9941 | 110 | -0,834303001 | N3:9914 | 23,714 | +2:YCL069W; | 2 | | | |
| chrIII | 9832 | 9955 | 123 | -1,865931237 | N3:9914 | 23,714 | +2:YCL069W; | 2 | | | |
| chrIII | 11462 | 11613 | 151 | -1,358056487 | N3:11616 | 29,676 | +5:YCL068C; | 5 | | | |
| chrIII | 17946 | 18093 | 147 | -1,381815646 | N3:18008 | 16,096 | +5:YCL063W; | 5 | | | |
| chrIII | 23277 | 23427 | 150 | -1,304688533 | N3:23351 | 10,386 | +1:YCL059C; -1:YCL058W-A; | 1 | | | |
| chrIII | 27235 | 27381 | 146 | -1,287126281 | N3:27325 | 26,748 | +2:YCL056C; | 2 | | | |
| chrIII | 32929 | 33083 | 154 | -1,205673076 | N3:33061 | 29,942 | +10:YCL054W; | 10 | | | |
| chrIII | 34797 | 34957 | 160 | -1,554230317 | N3:34875 | 5,68 | +4:YCL052C; | 4 | | | |
| chrIII | 36971 | 37125 | 154 | -1,313352521 | | | | Overlap <50 bp | | | |
| chrIII | 37274 | 37421 | 147 | -1,087073864 | N3:37360 | 22,565 | +12:YCL051W; | 12 | | | |
| chrIII | 41105 | 41261 | 156 | -1,375512355 | N3:41160 | 18,78 | -1:YCL048W-A; | -1 | | | |
| chrIII | 42270 | 42419 | 149 | -1,120656719 | N3:42308 | 15,28 | +2:YCL048W; | 2 | | | |
| chrIII | 56437 | 56565 | 128 | -1,162300779 | N3:56555 | 12,423 | +1:YCL038C; | 1 | | | |
| chrIII | 62417 | 62568 | 151 | -1,338661321 | N3:62518 | 44,392 | +6:YCL034W; | 6 | | | |
| chrIII | 62420 | 62568 | 148 | -1,395111568 | N3:62518 | 44,392 | +6:YCL034W; | 6 | | | |
| chrIII | 62779 | 62946 | 167 | -1,52612384 | N3:62871 | 14,007 | +3*:YCL033C; | 3 | TERM | | |
| chrIII | 62988 | 63143 | 155 | -1,460107083 | N3:63052 | 10,71 | +2:YCL033C; | 2 | | | |
| chrIII | 68315 | 68492 | 177 | -1,726689326 | N3:68474 | 39,598 | +10*:YCL029C; 0:YCL030C; | 10 | TERM | | |
| chrIII | 68580 | 68746 | 166 | -1,413775684 | N3:68637 | 37,978 | 9:YCL029C; -1:CUT041; -1:YCL030C; | 9 | | | |
| chrIII | 74254 | 74383 | 129 | -1,009646792 | | | | Overlap <50 bp | | | |
| chrIII | 75106 | 75269 | 163 | -1,425221364 | N3:75209 | 19,463 | +2:YCL026C-A; | 2 | | | |
| chrIII | 82256 | 82435 | 179 | -2,027682426 | N3:82326 | 31,277 | +21*:YCL024W; | 21 | TERM | | |
| chrIII | 86659 | 86788 | 129 | -1,251349554 | | | | Overlap <50 bp | | | |
| chrIII | 86884 | 87013 | 129 | -1,331965765 | N3:86885 | 15,588 | | Intergene | | | |
| chrIII | 88349 | 88492 | 143 | -1,005872508 | N3:88396 | 13,748 | | Intergene | | | |
| chrIII | 89624 | 89790 | 166 | -1,771992996 | | | | Overlap <50 bp | | | |
| chrIII | 90428 | 90577 | 149 | -1,065351661 | N3:90522 | 24,846 | | Intergene | | | |
| chrIII | 90432 | 90582 | 150 | -1,320665557 | N3:90522 | 24,846 | | Intergene | | | |
| chrIII | 92227 | 92379 | 152 | -1,112431138 | N3:92268 | 8,485 | +7:YCL018W; | 7 | | | |
| chrIII | 93544 | 93697 | 153 | -1,293327448 | N3:93669 | 7,463 | +5:YCL017C; | 5 | | | |
| chrIII | 94513 | 94671 | 158 | -1,465807665 | N3:94604 | 12,106 | +8*:YCL016C; -1:YCL017C; | 8 | TERM | | |
| chrIII | 95867 | 96028 | 161 | -1,710905784 | N3:95947 | 9,065 | -1:YCL014W; 0:YCL016C; | -1 | | | |
| chrIII | 95874 | 96028 | 154 | -1,447836567 | N3:95947 | 9,065 | -1:YCL014W; 0:YCL016C; | -1 | | | |
| chrIII | 97469 | 97602 | 133 | -1,495463613 | N3:97480 | 31,252 | +9:YCL014W; | 9 | | | |
| chrIII | 100878 | 101025 | 147 | -1,28986259 | N3:100936 | 15,083 | +29:YCL014W; | 29 | | | |
| chrIII | 100878 | 101048 | 170 | -1,335822729 | N3:100936 | 15,083 | +29:YCL014W; | 29 | | | |
| chrIII | 101513 | 101652 | 139 | -1,045398788 | N3:101599 | 8,55 | +2:YCL012C; | 2 | | | |
| chrIII | 101547 | 101689 | 142 | -1,14446132 | N3:101599 | 8,55 | +2:YCL012C; | 2 | | | |
| chrIII | 101696 | 101853 | 157 | -1,118911908 | N3:101768 | 5,854 | +1:YCL012C; | 1 | | | |
| chrIII | 103033 | 103165 | 132 | -0,774247191 | N3:103166 | 25,719 | +2:YCL011C; | 2 | | | |
| chrIII | 103103 | 103249 | 146 | -1,156783737 | N3:103166 | 25,719 | +2:YCL011C; | 2 | | | |
| chrIII | 103103 | 103251 | 148 | -1,102522501 | N3:103166 | 25,719 | +2:YCL011C; | 2 | | | |
| chrIII | 104725 | 104877 | 152 | -1,120089765 | N3:104791 | 9,066 | +6:YCL009C; | 6 | | | |
| chrIII | 106067 | 106230 | 163 | -1,314909405 | N3:106142 | 4,967 | +5:YCL008C; -1:SUT027; | 5 | | | |
| chrIII | 116514 | 116644 | 130 | -1,218378718 | N3:116605 | 17,682 | +5:SUT456; +2:Unit067; | 5 | | | |
| chrIII | 117818 | 117976 | 158 | -1,308629601 | N3:117863 | 14,865 | +4:YCR002C; | 4 | | | |
| chrIII | 132816 | 132923 | 107 | -1,552858369 | | | | Overlap <50 bp | | | |
| chrIII | 132817 | 132923 | 106 | -2,005595077 | | | | Overlap <50 bp | | | |
| chrIII | 133268 | 133417 | 149 | -1,198946699 | N3:133250 | 24,654 | 0:YCR010C; | 0 | | | |
| chrIII | 134878 | 135010 | 132 | -1,602285663 | N3:134986 | 45,665 | +13:YCR011C; | 13 | | | |
| chrIII | 136592 | 136744 | 152 | -1,025019285 | N3:136655 | 25,667 | +3:YCR011C; | 3 | | | |
| chrIII | 137946 | 138097 | 151 | -1,251124188 | N3:138059 | 33,63 | +3:YCR012W; | 3 | | | |
| chrIII | 138189 | 138339 | 150 | -1,222217245 | N3:138298 | 30,452 | +4:YCR012W; | 4 | | | |
| chrIII | 146706 | 146857 | 151 | -1,617003251 | N3:146686 | 11,149 | +7:YCR017C; | 7 | | | |
| chrIII | 154254 | 154394 | 140 | -1,164384349 | N3:154383 | 21,554 | | Intergene | | | |
| chrIII | 154256 | 154394 | 138 | -1,381052849 | N3:154383 | 21,554 | | Intergene | | | |
| chrIII | 156400 | 156550 | 150 | -1,246299697 | N3:156457 | 28,745 | +5:YCR021C; | 5 | | | |
| chrIII | 156437 | 156550 | 113 | -1,042650876 | N3:156457 | 28,745 | +5:YCR021C; | 5 | | | |
| chrIII | 156484 | 156645 | 161 | -1,554770183 | N3:156660 | 32,7 | +4:YCR021C; | 4 | | | |
| chrIII | 158189 | 158338 | 149 | -1,564439488 | | | | Overlap <50 bp | | | |
| chrIII | 159911 | 160084 | 173 | -1,769715977 | N3:160082 | 7,148 | +3:YCR023C; | 3 | | | |
| chrIII | 159911 | 160084 | 173 | -1,769715977 | N3:159912 | 18,454 | +4:YCR023C; | 4 | | | |
| chrIII | 161801 | 161938 | 137 | -1,845675426 | N3:161850 | 16,664 | +4:YCR024C; | 4 | | | |
| chrIII | 175934 | 176084 | 150 | -1,159976807 | N3:176001 | 18,243 | +4:YCR030C; | 4 | | | |
| chrIII | 186417 | 186569 | 152 | -1,345774075 | N3:186495 | 7,483 | +2:YCR033W; | 2 | | | |
| chrIII | 188100 | 188260 | 160 | -1,669387205 | N3:188165 | 17,349 | +13:YCR033W; | 13 | | | |
| chrIII | 191522 | 191646 | 124 | -1,031499942 | N3:191559 | 45,337 | +7:YCR034W; | 7 | | | |
| chrIII | 195429 | 195578 | 149 | -1,20943358 | | | | Overlap <50 bp | | | |
| chrIII | 201078 | 201230 | 152 | -1,18881041 | | | | Overlap <50 bp | | | |
| chrIII | 208738 | 208879 | 141 | -1,882906559 | N3:208806 | 20,744 | +6:YCR045C; | 6 | | | |
| chrIII | 209585 | 209734 | 149 | -1,10906871 | N3:209637 | 10,635 | +1:YCR045C; | 1 | | | |
| chrIII | 210916 | 211072 | 156 | -1,244734556 | N3:210951 | 13,817 | +5:YCR047C; | 5 | | | |
| chrIII | 213468 | 213602 | 134 | -0,954607696 | N3:213471 | 16,678 | +3:YCR050C; +10:YCR048W; | 3 | | | |
| chrIII | 215653 | 215790 | 137 | -0,865616105 | N3:215679 | 16,303 | +6:YCR052W; | 6 | | | |
| chrIII | 217960 | 218102 | 142 | -1,208113529 | N3:218083 | 30,45 | +10*:YCR053W; | 10 | TERM | | |
| chrIII | 228591 | 228741 | 150 | -1,284766155 | N3:228733 | 34,94 | +3:YCR063W; | 3 | | | |
| chrIII | 231011 | 231166 | 155 | -1,659060501 | N3:231100 | 18,298 | +12*:YCR065W; | 12 | TERM | | |
| chrIII | 234251 | 234402 | 151 | -1,518111731 | N3:234286 | 23,258 | +13:YCR067C; | 13 | | | |
| chrIII | 234921 | 235052 | 131 | -1,76544927 | | | | Overlap <50 bp | | | |
| chrIII | 236307 | 236469 | 162 | -1,389862065 | N3:236392 | 4,412 | +1:YCR067C; | 1 | | | |
| chrIII | 238120 | 238291 | 171 | -1,962163652 | N3:238307 | 2,646 | +3:CUT467; +8:YCR068W; | 3 | | | |
| chrIII | 238120 | 238291 | 171 | -1,962163652 | N3:238147 | 55,567 | +4:CUT467; +7:YCR068W; | 4 | | | |
| chrIII | 240298 | 240452 | 154 | -1,67725888 | N3:240369 | 4,622 | +2:YCR071C; | 2 | | | |
| chrIII | 240452 | 240614 | 162 | -1,282702067 | N3:240532 | 4,792 | +1:YCR071C; | 1 | | | |
| chrIII | 242569 | 242719 | 150 | -1,059230457 | N3:242631 | 10,431 | :YCR073C; +1:SUT038; -1:YCR072C; | 25 | | | |
| chrIII | 246007 | 246188 | 181 | -1,929933178 | N3:246094 | 9,786 | +4:YCR073C; | 4 | | | |
| chrIII | 246926 | 247057 | 131 | -1,007454773 | N3:246993 | 9,368 | -1:YCR073C; +1:YCR073W-A; | -1 | | | |
| chrIII | 246926 | 247058 | 132 | -0,897760429 | N3:246993 | 9,368 | -1:YCR073C; +1:YCR073W-A; | -1 | | | |
| chrIII | 248934 | 249085 | 151 | -1,480690621 | N3:249001 | 8,01 | -1:YCR075C; +1:YCR075W-A; | -1 | | | |
| chrIII | 251728 | 251875 | 147 | -0,985710543 | | | | Overlap <50 bp | | | |
| chrIII | 252213 | 252361 | 148 | -1,181121786 | N3:252282 | 2,608 | +3:YCR077C; | 3 | | | |
| chrIII | 252238 | 252397 | 159 | -1,370693883 | N3:252282 | 2,608 | +3:YCR077C; | 3 | | | |
| chrIII | 254663 | 254821 | 158 | -1,293109683 | N3:254734 | 3,937 | +4:YCR081W; | 4 | | | |
| chrIII | 257083 | 257228 | 145 | -1,463182799 | N3:257179 | 22,247 | +18:YCR081W; | 18 | | | |
| chrIII | 257806 | 257969 | 163 | -1,40977351 | N3:257802 | 27,308 | +22:YCR081W; | 22 | | | |
| chrIII | 258183 | 258325 | 142 | -1,03873871 | N3:258264 | 20,248 | +25:YCR081W; | 25 | | | |
| chrIII | 259144 | 259292 | 148 | -1,33061948 | N3:259206 | 9,131 | +3:YCR082W; | 3 | | | |

| CHR | START | END | L(bp) | ΔLK | Nuc ID | Fuzziness | Gene ID | Gene body postion or intergene | Terminal | rDNA | Telomeric |
|---|---|---|---|---|---|---|---|---|---|---|---|
| chrIII | 262204 | 262356 | 152 | -1,170638304 | N3:262264 | 7,778 | +3:YCR084C; | 3 | | | |
| chrIII | 269289 | 269448 | 159 | -1,511255299 | N3:269317 | 29,513 | +13:YCR089W; | 13 | | | |
| chrIII | 272784 | 272929 | 145 | -1,636330826 | N3:272830 | 30,64 | +1:YCR090C; | 1 | | | |
| chrIII | 276672 | 276825 | 153 | -1,525113886 | N3:276740 | 25,57 | +15:YCR091W; +21*:YCR092C; | 15 | | | |
| chrIII | 280552 | 280680 | 128 | -1,388452975 | N3:280622 | 24,214 | +4:YCR093W; | 4 | | | |
| chrIII | 281034 | 281170 | 136 | -1,077118391 | N3:281080 | 15,057 | +7:YCR093W; | 7 | | | |
| chrIII | 286830 | 286938 | 108 | -1,112362447 | N3:286909 | 6,882 | +2:YCR094W; | 2 | | | |
| chrIII | 286830 | 286939 | 109 | -1,124724778 | N3:286909 | 6,882 | +2:YCR094W; | 2 | | | |
| chrIII | 286830 | 286940 | 110 | -1,309761628 | N3:286909 | 6,882 | +2:YCR094W; | 2 | | | |
| chrIII | 286830 | 286941 | 111 | -1,155714204 | N3:286909 | 6,882 | +2:YCR094W; | 2 | | | |
| chrIII | 286832 | 286938 | 106 | -1,580583781 | N3:286909 | 6,882 | +2:YCR094W; | 2 | | | |
| chrIII | 286832 | 286939 | 107 | -1,643068654 | N3:286909 | 6,882 | +2:YCR094W; | 2 | | | |
| chrIII | 286832 | 286955 | 123 | -1,721702946 | N3:286909 | 6,882 | +2:YCR094W; | 2 | | | |
| chrIII | 288517 | 288666 | 149 | -1,741922271 | N3:288593 | 13,989 | +5:YCR095C; | 5 | | | |
| chrIII | 303804 | 303963 | 159 | -1,37380065 | N3:303873 | 17,951 | +1:anti-YCR102C; | 1 | | | |
| chrIII | 307204 | 307345 | 141 | -1,196540658 | N3:307293 | 5,354 | | Intergene | | | |
| chrIII | 307633 | 307788 | 155 | -0,89410963 | | | | Overlap <50 bp | | | |
| chrIII | 308042 | 308191 | 149 | -1,370040044 | N3:308087 | 19,807 | +3:YCR104W; | 3 | | | |
| chrIII | 310321 | 310472 | 151 | -1,747404911 | N3:310383 | 14,711 | | Intergene | | | |
| chrIII | 310654 | 310803 | 149 | -1,380621571 | N3:310728 | 17,914 | -1:YCR106W; | -1 | | | |
| chrIII | 311798 | 311947 | 149 | -1,34814441 | N3:311793 | 20,917 | +6:YCR106W; | 6 | | | |
| chrIII | 314554 | 314707 | 153 | -1,398051656 | N3:314702 | 50,43 | +6:YCR107W; | 6 | | | |
| chrIV | 0 | 126 | 126 | -1,10383509 | | | | Overlap <50 bp | | | TEL LEFT |
| chrIV | 0 | 145 | 145 | -1,757917866 | | | | Overlap <50 bp | | | TEL LEFT |
| chrIV | 0 | 155 | 155 | -0,929815149 | | | | Overlap <50 bp | | | TEL LEFT |
| chrIV | 5 | 126 | 121 | -0,65036107 | | | | Overlap <50 bp | | | TEL LEFT |
| chrIV | 14 | 126 | 112 | -0,341478537 | | | | Overlap <50 bp | | | TEL LEFT |
| chrIV | 14 | 134 | 120 | 0,009244849 | | | | Overlap <50 bp | | | TEL LEFT |
| chrIV | 14 | 155 | 141 | -0,320519223 | | | | Overlap <50 bp | | | TEL LEFT |
| chrIV | 44 | 126 | 82 | -1,797422563 | | | | Overlap <50 bp | | | TEL LEFT |
| chrIV | 44 | 134 | 90 | -0,866532098 | | | | Overlap <50 bp | | | TEL LEFT |
| chrIV | 44 | 145 | 101 | -1,335249574 | | | | Overlap <50 bp | | | TEL LEFT |
| chrIV | 44 | 155 | 111 | -0,627750502 | | | | Overlap <50 bp | | | TEL LEFT |
| chrIV | 57 | 155 | 98 | -1,175481081 | | | | Overlap <50 bp | | | TEL LEFT |
| chrIV | 1765 | 1923 | 158 | -1,35502516 | N4:1847 | 27,354 | +2:YDL248W; | 2 | | | |
| chrIV | 6558 | 6652 | 94 | -1,414337508 | N4:6650 | 21,932 | +5:YDL247W; | 5 | | | |
| chrIV | 8786 | 8917 | 131 | -1,245132351 | N4:8900 | 7,071 | +6:YDL246C; | 6 | | | |
| chrIV | 10468 | 10622 | 154 | -1,473432049 | N4:10539 | 3,464 | | Intergene | | | |
| chrIV | 11706 | 11865 | 159 | -1,150143293 | N4:11735 | 6,348 | +11:YDL245C; | 11 | | | |
| chrIV | 12342 | 12498 | 156 | -1,614260576 | N4:12435 | 26,163 | +7:YDL245C; | 7 | | | |
| chrIV | 12922 | 13062 | 140 | -1,097755446 | | | | Overlap <50 bp | | | |
| chrIV | 13187 | 13340 | 153 | -1,067679421 | N4:13269 | 4,583 | +2:YDL245C; | 2 | | | |
| chrIV | 15846 | 16000 | 154 | -1,342552837 | N4:15932 | 4,359 | -1:YDL244W; | -1 | | | |
| chrIV | 16215 | 16372 | 157 | -1,069107003 | N4:16282 | 7,024 | +2:YDL244W; | 2 | | | |
| chrIV | 16517 | 16676 | 159 | -1,192054713 | N4:16612 | 14,519 | +4:YDL244W; | 4 | | | |
| chrIV | 22332 | 22485 | 153 | -1,343674134 | N4:22409 | 3,882 | -1:YDL240W; +1:CUT468; | -1 | | | |
| chrIV | 22780 | 22921 | 141 | -1,016114724 | N4:22851 | 8,112 | +2:YDL240W; | 2 | | | |
| chrIV | 24904 | 25025 | 121 | -1,389808288 | N4:25029 | 29,024 | +15:YDL240W; | 15 | | | |
| chrIV | 26118 | 26260 | 142 | -1,314219095 | N4:26130 | 27,577 | | Intergene | | | |
| chrIV | 28132 | 28284 | 152 | -1,460268241 | N4:28244 | 31,106 | +15:YDL238C; +5:YDL239C; | 15 | | | |
| chrIV | 28565 | 28649 | 84 | -0,345479309 | | | | Overlap <50 bp | | | |
| chrIV | 28606 | 28762 | 156 | -1,05599875 | N4:28689 | 9,659 | +12:YDL238C; +2:YDL239C; | 12 | | | |
| chrIV | 29713 | 29864 | 151 | -1,459773062 | N4:29784 | 20,5 | +5:YDL238C; | 5 | | | |
| chrIV | 30032 | 30191 | 159 | -1,422734494 | N4:30113 | 8,159 | +3:YDL238C; | 3 | | | |
| chrIV | 31789 | 31954 | 165 | -1,593088039 | N4:31846 | 6,76 | +2:CUT469; +8*:YDL237W; | 2 | | | |
| chrIV | 32229 | 32385 | 156 | -1,61034737 | N4:32309 | 10,218 | +1:YDL236W; | 1 | | | |
| chrIV | 33584 | 33743 | 159 | -1,391758771 | N4:33663 | 4,858 | +3:YDL235C; | 3 | | | |
| chrIV | 34942 | 35098 | 156 | -1,390390195 | N4:35008 | 6,602 | +11:YDL234C; | 11 | | | |
| chrIV | 43249 | 43399 | 150 | -1,213538559 | N4:43358 | 10,483 | +5:YDL230W; | 5 | | | |
| chrIV | 43986 | 44132 | 146 | -1,385102157 | N4:44075 | 13,952 | +1:YDL229W; | 1 | | | |
| chrIV | 51938 | 52044 | 106 | -1,216172823 | N4:51987 | 7,12 | +2:YDL226C; | 2 | | | |
| chrIV | 55412 | 55541 | 129 | -1,196002235 | | | | Overlap <50 bp | | | |
| chrIV | 56586 | 56757 | 171 | -1,227866412 | N4:56663 | 7,23 | +1:YDL224C; | 1 | | | |
| chrIV | 60333 | 60467 | 134 | -1,097455173 | N4:60396 | 7,339 | +1:YDL223C; +8:SUT041; | 1 | | | |
| chrIV | 61937 | 62077 | 140 | -1,028434641 | N4:62069 | 6,221 | +18:YDL220C; -1:YDL222C; | 18 | | | |
| chrIV | 77124 | 77282 | 158 | -1,551926047 | N4:77237 | 42,658 | +5*:YDL213C; | 5 | TERM | | |
| chrIV | 78906 | 79062 | 156 | -1,232193114 | N4:78931 | 10,635 | +4:YDL212W; | 4 | | | |
| chrIV | 86969 | 87132 | 163 | -1,158265081 | N4:87040 | 6,047 | +2:YDL209C; | 2 | | | |
| chrIV | 90699 | 90847 | 148 | -1,410873582 | N4:90745 | 8,989 | +5:YDL206W; | 5 | | | |
| chrIV | 94580 | 94722 | 142 | -1,42529725 | N4:94705 | 27,009 | +4:YDL204W; | 4 | | | |
| chrIV | 97197 | 97345 | 148 | -0,911843753 | N4:97289 | 4,32 | +6:YDL203C; | 6 | | | |
| chrIV | 101729 | 101892 | 163 | -1,168445221 | N4:101825 | 20,245 | +12:YDL199C; | 12 | | | |
| chrIV | 103149 | 103279 | 130 | -0,529212448 | N4:103192 | 9,585 | +4:YDL199C; | 4 | | | |
| chrIV | 109515 | 109652 | 137 | -1,443348703 | N4:109593 | 10,599 | | Intergene | | | |
| chrIV | 109515 | 109656 | 141 | -1,213831778 | N4:109593 | 10,599 | | Intergene | | | |
| chrIV | 109515 | 109666 | 151 | -1,015005865 | N4:109593 | 10,599 | | Intergene | | | |
| chrIV | 109518 | 109656 | 138 | -1,422156087 | N4:109593 | 10,599 | | Intergene | | | |
| chrIV | 110479 | 110614 | 135 | -1,188914917 | N4:110476 | 39,939 | | Intergene | | | |
| chrIV | 113056 | 113240 | 184 | -1,547311362 | N4:113034 | 35,62 | +10:YDL194W; | 10 | | | |
| chrIV | 118783 | 118936 | 153 | -1,291131377 | N4:118787 | 40,577 | +19:YDL190C; | 19 | | | |
| chrIV | 121491 | 121639 | 148 | -1,219188809 | N4:121562 | 26,283 | +2:YDL190C; | 2 | | | |
| chrIV | 122619 | 122777 | 158 | -1,438248176 | N4:122683 | 13,914 | +4:YDL189W; | 4 | | | |
| chrIV | 124114 | 124238 | 124 | -1,024991831 | N4:124182 | 10,017 | +7:YDL188C; | 7 | | | |
| chrIV | 125012 | 125174 | 162 | -1,492641519 | N4:125123 | 7,95 | +1:YDL188C; | 1 | | | |
| chrIV | 126907 | 127064 | 157 | -1,161292061 | N4:126899 | 4,506 | +2:YDL185W; | 2 | | | |
| chrIV | 126907 | 127064 | 157 | -1,161292061 | N4:127056 | 1,826 | +3:YDL185W; | 3 | | | |
| chrIV | 127133 | 127254 | 121 | -1,338539227 | N4:127204 | 2,95 | +4:YDL185W; | 4 | | | |
| chrIV | 130391 | 130540 | 149 | -1,371628326 | N4:130459 | 10,834 | +1:YDL184C; | 1 | | | |
| chrIV | 134454 | 134627 | 173 | -1,223835209 | N4:134518 | 7,906 | +8:YDL182W; | 8 | | | |
| chrIV | 137744 | 137906 | 162 | -1,398042036 | N4:137857 | 32,549 | | Intergene | | | |
| chrIV | 147424 | 147558 | 134 | -0,705629157 | | | | Overlap <50 bp | | | |
| chrIV | 148138 | 148259 | 121 | -0,813745752 | N4:148212 | 3,082 | +1:YDL173W; | 1 | | | |
| chrIV | 148138 | 148287 | 149 | -1,208846281 | N4:148212 | 3,082 | +1:YDL173W; | 1 | | | |
| chrIV | 148147 | 148256 | 109 | -1,122393819 | N4:148212 | 3,082 | +1:YDL173W; | 1 | | | |
| chrIV | 148147 | 148259 | 112 | -1,141336115 | N4:148212 | 3,082 | +1:YDL173W; | 1 | | | |
| chrIV | 148147 | 148287 | 140 | -1,565186748 | N4:148212 | 3,082 | +1:YDL173W; | 1 | | | |
| chrIV | 148150 | 148259 | 109 | -1,024588753 | N4:148212 | 3,082 | +1:YDL173W; | 1 | | | |
| chrIV | 148662 | 148839 | 177 | -1,286223853 | N4:148802 | 33,352 | +4:YDL173W; | 4 | | | |

| CHR | START | END | L(bp) | ΔLK | Nuc ID | Fuzziness | Gene ID | Gene body postion or intergene | Terminal | rDNA | Telomeric |
|---|---|---|---|---|---|---|---|---|---|---|---|
| chrIV | 150319 | 150438 | 119 | -1,353528407 | | | | Overlap <50 bp | | | |
| chrIV | 150791 | 150954 | 163 | -1,538821231 | N4:150914 | 16,008 | +33:YDL171C; | 33 | | | |
| chrIV | 151146 | 151295 | 149 | -1,149644659 | N4:151233 | 24,993 | +31:YDL171C; | 31 | | | |
| chrIV | 154011 | 154153 | 142 | -1,749524939 | N4:154052 | 36,679 | +13:YDL171C; | 13 | | | |
| chrIV | 156456 | 156616 | 160 | -1,472583844 | N4:156533 | 4,622 | +2:YDL170W; | 2 | | | |
| chrIV | 156781 | 156930 | 149 | -1,360907042 | | | | Overlap <50 bp | | | |
| chrIV | 158096 | 158254 | 158 | -1,343130283 | N4:158248 | 11,576 | +4:YDL169C; | 4 | | | |
| chrIV | 158096 | 158254 | 158 | -1,343130283 | N4:158093 | 50,153 | +5:YDL169C; | 5 | | | |
| chrIV | 158496 | 158646 | 150 | -1,106203647 | N4:158568 | 5,601 | +2:YDL169C; | 2 | | | |
| chrIV | 162837 | 162965 | 128 | -1,135418386 | N4:162831 | 10,188 | +3:YDL167C; | 3 | | | |
| chrIV | 163802 | 163934 | 132 | -1,034309912 | N4:163873 | 11,718 | +2:YDL166C; | 2 | | | |
| chrIV | 163887 | 164037 | 150 | -1,029478734 | N4:164037 | 7,211 | -1:YDL165W; +1:YDL166C; | -1 | | | |
| chrIV | 163887 | 164037 | 150 | -1,029478734 | N4:163873 | 11,718 | +2:YDL166C; | 2 | | | |
| chrIV | 164671 | 164820 | 149 | -1,207463713 | N4:164814 | 27,338 | +4*:YDL165W; | 4 | TERM | | |
| chrIV | 168971 | 169122 | 151 | -1,005288111 | N4:169057 | 14,738 | +9*:YDL161W; | 9 | TERM | | |
| chrIV | 180947 | 181107 | 160 | -1,293493543 | N4:181019 | 33,306 | +18:YDL154W; | 18 | | | |
| chrIV | 185030 | 185179 | 149 | -1,263427469 | N4:185100 | 8,456 | +3:YDL149W; | 3 | | | |
| chrIV | 185042 | 185156 | 114 | -1,280167402 | N4:185100 | 8,456 | +3:YDL149W; | 3 | | | |
| chrIV | 196778 | 196946 | 168 | -1,063367532 | N4:196824 | 40,782 | +11:YDL145C; | 11 | | | |
| chrIV | 198969 | 199115 | 146 | -1,081610066 | N4:199010 | 15,975 | +5:YDL144C; | 5 | | | |
| chrIV | 200831 | 200925 | 94 | -1,401682972 | N4:200946 | 30,748 | +7:YDL143W; | 7 | | | |
| chrIV | 202938 | 203097 | 159 | -1,561079247 | N4:203012 | 6,686 | +2:YDL141W; | 2 | | | |
| chrIV | 204891 | 205041 | 150 | -1,164834503 | N4:204992 | 19,395 | +14:YDL141W; | 14 | | | |
| chrIV | 205789 | 205917 | 128 | -0,914710537 | N4:205848 | 37,034 | +31:YDL140C; | 31 | | | |
| chrIV | 214844 | 215003 | 159 | -1,426304769 | | | | Overlap <50 bp | | | |
| chrIV | 224519 | 224676 | 157 | -1,291868774 | N4:224539 | 22,057 | +4:YDL132W; | 4 | | | |
| chrIV | 224519 | 224676 | 157 | -1,291868774 | N4:224699 | 13,672 | +5:YDL132W; | 5 | | | |
| chrIV | 228430 | 228579 | 149 | -1,442284036 | N4:228507 | 10,922 | +7:YDL131W; | 7 | | | |
| chrIV | 230300 | 230458 | 158 | -1,403873142 | N4:230409 | 29,012 | | Intergene | | | |
| chrIV | 235048 | 235188 | 140 | -1,084110797 | N4:235136 | 30,684 | +2:YDL127W; | 2 | | | |
| chrIV | 235556 | 235704 | 148 | -1,282020667 | N4:235550 | 29,368 | +4:YDL127W; | 4 | | | |
| chrIV | 235556 | 235704 | 148 | -1,282020667 | N4:235716 | 16,932 | +5:YDL127W; | 5 | | | |
| chrIV | 240743 | 240895 | 152 | -0,728390045 | N4:240819 | 28,978 | +4:YDL124W; | 4 | | | |
| chrIV | 241083 | 241236 | 153 | -1,508388003 | N4:241147 | 7,25 | +6*:YDL124W; -1:YDL123W; | 6 | TERM | | |
| chrIV | 243252 | 243404 | 152 | -1,635774479 | N4:243309 | 7,906 | +7:YDL122W; | 7 | | | |
| chrIV | 247954 | 248116 | 162 | -1,729018586 | N4:247969 | 5,428 | -1:YDL117W; -1:YDL119C; | -1 | | | |
| chrIV | 249148 | 249302 | 154 | -1,325363328 | | | | Overlap <50 bp | | | |
| chrIV | 249297 | 249444 | 147 | -1,283329213 | | | | Overlap <50 bp | | | |
| chrIV | 253799 | 253940 | 141 | -1,113168052 | | | | Overlap <50 bp | | | |
| chrIV | 262881 | 263029 | 148 | -1,41641368 | N4:262904 | 24,803 | +25:YDL112W; | 25 | | | |
| chrIV | 268349 | 268491 | 142 | -0,997311227 | N4:268409 | 5,02 | +5:YDL108W; | 5 | | | |
| chrIV | 269754 | 269886 | 132 | -0,999288012 | N4:269794 | 15,672 | +6:YDL107W; | 6 | | | |
| chrIV | 275399 | 275562 | 163 | -1,422105141 | N4:275425 | 13,206 | +8:YDL103C; | 8 | | | |
| chrIV | 277085 | 277203 | 118 | -1,542351584 | N4:277148 | 6,091 | +3:YDL102W; | 3 | | | |
| chrIV | 281092 | 281244 | 152 | -1,473127037 | N4:281168 | 12,95 | +5:YDL101C; | 5 | | | |
| chrIV | 282751 | 282862 | 111 | -1,589919947 | N4:282817 | 7,301 | +3:YDL100C; | 3 | | | |
| chrIV | 287334 | 287498 | 164 | -1,373556499 | N4:287398 | 14,029 | +3:YDL095W; | 3 | | | |
| chrIV | 290503 | 290642 | 139 | -1,09999004 | N4:290580 | 6,025 | +5:YDL093W; | 5 | | | |
| chrIV | 291728 | 291875 | 147 | -1,023422687 | N4:291860 | 9,218 | +13:YDL093W; | 13 | | | |
| chrIV | 294716 | 294818 | 102 | -1,340369689 | N4:294818 | 7,376 | +1:YDL091C; +10:YDL090C; | 1 | | | |
| chrIV | 303242 | 303392 | 150 | -1,275637142 | N4:303310 | 15,604 | +2:YDL085W; | 2 | | | |
| chrIV | 305394 | 305543 | 149 | -1,570453476 | N4:305450 | 33,906 | +3:YDL084W; | 3 | | | |
| chrIV | 307871 | 308025 | 154 | -1,438235216 | N4:307996 | 0,577 | -1:YDL083C; | -1 | | | |
| chrIV | 308997 | 309152 | 155 | -1,359995753 | N4:309056 | 8,183 | | Intergene | | | |
| chrIV | 311358 | 311498 | 140 | -1,324086032 | N4:311442 | 8,808 | +7:YDL080C; | 7 | | | |
| chrIV | 313115 | 313272 | 157 | -1,18734136 | N4:313232 | 21,674 | +10:YDL079C; +3:SUT043; | 10 | | | |
| chrIV | 315132 | 315251 | 119 | -0,750437952 | N4:315205 | 6,186 | +2:SUT044; | 2 | | | |
| chrIV | 316257 | 316381 | 124 | -0,957325176 | N4:316392 | 11,454 | +1:YDL078C; +9:SUT044; | 1 | | | |
| chrIV | 317932 | 318025 | 93 | -1,214586684 | | | | Overlap <50 bp | | | |
| chrIV | 320773 | 320934 | 161 | -1,36708013 | N4:320798 | 13,517 | +3:anti030; +6:YDL076C; | 3 | | | |
| chrIV | 324041 | 324187 | 146 | -1,285155009 | N4:324118 | 13,545 | +14*:YDL074C; +3:SUT046; | 14 | TERM | | |
| chrIV | 326087 | 326232 | 145 | -1,305473147 | N4:326162 | 12,156 | +1:YDL074C; | 1 | | | |
| chrIV | 331066 | 331232 | 166 | -1,435411356 | N4:331089 | 11,127 | +1:YDL070W; +20*:YDL069C; | 1 | | | |
| chrIV | 331066 | 331232 | 166 | -1,435411356 | N4:331253 | 13,964 | +2:YDL070W; +19:YDL069C; | 2 | | | |
| chrIV | 336500 | 336644 | 144 | -1,269088495 | N4:336586 | 11,776 | +5:YDL065C; | 5 | | | |
| chrIV | 339003 | 339155 | 152 | -1,429959585 | N4:339069 | 33,411 | +8:YDL063C; | 8 | | | |
| chrIV | 339626 | 339796 | 170 | -1,457911274 | N4:339709 | 7,855 | +4:YDL063C; | 4 | | | |
| chrIV | 343754 | 343879 | 125 | -1,263432816 | N4:343821 | 19,281 | | Intergene | | | |
| chrIV | 344625 | 344793 | 168 | -1,353642831 | N4:344682 | 3,67 | +3:YDL059C; | 3 | | | |
| chrIV | 344904 | 345057 | 153 | -1,109370367 | N4:345036 | 4,844 | +1:YDL059C; | 1 | | | |
| chrIV | 346054 | 346184 | 130 | -1,156138877 | N4:346140 | 9,834 | +4:YDL058W; | 4 | | | |
| chrIV | 346695 | 346849 | 154 | -1,403788246 | | | | Overlap <50 bp | | | |
| chrIV | 354236 | 354370 | 134 | -1,106681292 | N4:354255 | 24,166 | +10:YDL056W; | 10 | | | |
| chrIV | 357373 | 357527 | 154 | -1,181851262 | N4:357441 | 26,552 | | Intergene | | | |
| chrIV | 357519 | 357616 | 97 | -1,596323564 | N4:357598 | 23,627 | | Intergene | | | |
| chrIV | 367261 | 367414 | 153 | -1,338313528 | N4:367365 | 15,497 | +6:YDL048C; | 6 | | | |
| chrIV | 369734 | 369884 | 150 | -1,559769134 | N4:369786 | 19,343 | +2:YDL047W; | 2 | | | |
| chrIV | 372733 | 372859 | 126 | -1,591174564 | N4:372737 | 18,759 | +7:YDL045C; | 7 | | | |
| chrIV | 377302 | 377457 | 155 | -1,602995297 | N4:377352 | 9,138 | +8:YDL042C; | 8 | | | |
| chrIV | 377961 | 378102 | 141 | -1,236383197 | N4:378096 | 13,382 | +3:YDL042C; | 3 | | | |
| chrIV | 379551 | 379707 | 156 | -1,350401121 | N4:379708 | 30,649 | +12:YDL040C; | 12 | | | |
| chrIV | 380490 | 380611 | 121 | -1,262833261 | N4:380618 | 5,357 | +6:YDL040C; | 6 | | | |
| chrIV | 380895 | 381031 | 136 | -1,146987929 | N4:380935 | 6,593 | +4:YDL040C; | 4 | | | |
| chrIV | 383984 | 384136 | 152 | -1,011197975 | N4:384003 | 46,392 | 'DL037C; +2:YDL038C; +15:SUT048; | 11 | | | |
| chrIV | 384406 | 384547 | 141 | -1,383858974 | | | | Overlap <50 bp | | | |
| chrIV | 386414 | 386562 | 148 | -0,981478283 | N4:386473 | 30,287 | nit123; +4*:Unit079; +29*:SUT048; | 2 | | | |
| chrIV | 389563 | 389664 | 101 | -1,503448267 | N4:389614 | 24,214 | +17:YDL035C; | 17 | | | |
| chrIV | 394898 | 395059 | 161 | -1,400132814 | N4:395016 | 28,599 | +6:YDL031W; | 6 | | | |
| chrIV | 395659 | 395798 | 139 | -1,232800588 | N4:395816 | 41,967 | +11:YDL031W; | 11 | | | |
| chrIV | 405838 | 405992 | 154 | -1,281385099 | N4:405924 | 18,349 | +10:YDL025C; | 10 | | | |
| chrIV | 406138 | 406284 | 146 | -0,952758036 | | | | Overlap <50 bp | | | |
| chrIV | 407097 | 407258 | 161 | -1,119711242 | N4:407198 | 16,597 | +2:YDL025C; | 2 | | | |
| chrIV | 407639 | 407774 | 135 | -1,133144203 | | | | Overlap <50 bp | | | |
| chrIV | 410001 | 410147 | 146 | -1,557684068 | N4:410035 | 14,72 | 0:YDL024C; | 0 | | | |
| chrIV | 412510 | 412666 | 156 | -1,689284093 | N4:412554 | 21,014 | +5:YDL022W; | 5 | | | |
| chrIV | 414505 | 414611 | 106 | -1,1389347 | | | | Overlap <50 bp | | | |
| chrIV | 414505 | 414627 | 122 | -1,309091331 | | | | Overlap <50 bp | | | |
| chrIV | 417857 | 418015 | 158 | -1,412643102 | N4:418030 | 16,739 | | Intergene | | | |
| chrIV | 418210 | 418363 | 153 | -1,321576422 | N4:418242 | 30,003 | | Intergene | | | |

TABLE 3

| CHR | START | END | L(bp) | ΔLK | Nuc ID | Fuzziness | Gene ID | Gene body postion or intergene | Terminal | rDNA | Telomeric |
|---|---|---|---|---|---|---|---|---|---|---|---|
| chrIV | 418328 | 418478 | 150 | -1,541852845 | | | | Overlap <50 bp | | | |
| chrIV | 419977 | 420127 | 150 | -1,450435292 | | | | Overlap <50 bp | | | |
| chrIV | 430775 | 430925 | 150 | -1,305814201 | | | | Overlap <50 bp | | | |
| chrIV | 431218 | 431366 | 148 | -1,35536047 | N4:431307 | 9,579 | +2:YDL012C; | 2 | | | |
| chrIV | 433624 | 433778 | 154 | -1,292421838 | N4:433704 | 20,216 | +2:YDL008W; +3:SUT464; | 2 | | | |
| chrIV | 444592 | 444742 | 150 | -1,160523841 | N4:444681 | 7,19 | +1:YDL003W; | 1 | | | |
| chrIV | 447153 | 447303 | 150 | -1,171812575 | N4:447172 | 10,905 | +4:YDL002C; | 4 | | | |
| chrIV | 452119 | 452270 | 151 | -1,427081433 | N4:452201 | 6,419 | +4:YDR001C; | 4 | | | |
| chrIV | 456027 | 456181 | 154 | -1,165000882 | N4:456128 | 24,884 | +6:YDR004W; | 6 | | | |
| chrIV | 464196 | 464345 | 149 | -1,335150598 | N4:464243 | 15,055 | +6:YDR009W; | 6 | | | |
| chrIV | 465523 | 465668 | 145 | -1,238221467 | N4:465537 | 27,213 | 0:YDR011W; | 0 | | | |
| chrIV | 472028 | 472162 | 134 | -1,005416049 | N4:472039 | 17,914 | +2:YDR012W; | 2 | | | |
| chrIV | 480768 | 480918 | 150 | -1,282328158 | N4:480858 | 16,51 | +10:YDR017C; | 10 | | | |
| chrIV | 481714 | 481864 | 150 | -1,094149607 | N4:481812 | 29,575 | +4:YDR017C; -1:anti-YDR017C; | 4 | | | |
| chrIV | 481903 | 482061 | 158 | -1,306316079 | N4:481994 | 19,073 | +3:YDR017C; +1:anti-YDR017C; | 3 | | | |
| chrIV | 487395 | 487546 | 151 | -1,30830358 | N4:487410 | 13,372 | +5:YDR021W; | 5 | | | |
| chrIV | 488401 | 488552 | 151 | -1,165496622 | N4:488452 | 17,773 | +2:YDR022C; | 2 | | | |
| chrIV | 488508 | 488615 | 107 | -0,944019618 | N4:488621 | 22,914 | +1:YDR022C; | 1 | | | |
| chrIV | 488508 | 488632 | 124 | -1,403800432 | N4:488621 | 22,914 | +1:YDR022C; | 1 | | | |
| chrIV | 491144 | 491308 | 164 | -1,422663446 | N4:491316 | 12,897 | -1:YDR025W; | -1 | | | |
| chrIV | 491144 | 491308 | 164 | -1,422663446 | N4:491134 | 19,092 | | Intergene | | | |
| chrIV | 492176 | 492332 | 156 | -1,189090425 | N4:492275 | 22,819 | | Intergene | | | |
| chrIV | 494125 | 494275 | 150 | -1,088166097 | N4:494221 | 17,648 | +2:YDR026C; | 2 | | | |
| chrIV | 499197 | 499355 | 158 | -0,962310372 | N4:499232 | 49,303 | +14:YDR028C; | 14 | | | |
| chrIV | 499814 | 499967 | 153 | -1,465156544 | N4:499892 | 31,785 | +10:YDR028C; | 10 | | | |
| chrIV | 503596 | 503736 | 140 | -1,067984168 | N4:503693 | 10,206 | +2:YDR031W; | 2 | | | |
| chrIV | 503768 | 503924 | 156 | -1,386951835 | | | | Overlap <50 bp | | | |
| chrIV | 508653 | 508795 | 142 | -1,078587534 | N4:508667 | 3,899 | +6:YDR033W; | 6 | | | |
| chrIV | 513880 | 514029 | 149 | -1,153428135 | N4:513900 | 18,175 | | Intergene | | | |
| chrIV | 517444 | 517573 | 129 | -1,18276644 | | | | Overlap <50 bp | | | |
| chrIV | 518819 | 518929 | 110 | -1,241534086 | N4:518890 | 20,075 | | Intergene | | | |
| chrIV | 521357 | 521482 | 125 | -1,239691432 | N4:521409 | 13,667 | +2:YDR034W-B; | 2 | | | |
| chrIV | 522483 | 522574 | 91 | -1,034891693 | N4:522532 | 38,701 | +6:YDR035W; | 6 | | | |
| chrIV | 522691 | 522836 | 145 | -1,03146379 | N4:522707 | 26,26 | +7:YDR035W; | 7 | | | |
| chrIV | 524988 | 525151 | 163 | -1,10856945 | N4:525064 | 20,449 | -1:YDR037W; | -1 | | | |
| chrIV | 525488 | 525645 | 157 | -1,396523345 | | | | Overlap <50 bp | | | |
| chrIV | 527030 | 527163 | 133 | -1,059952544 | N4:527071 | 27,833 | +12:YDR037W; | 12 | | | |
| chrIV | 527878 | 528014 | 136 | -1,240376702 | N4:527977 | 28,355 | +18:YDR038C; | 18 | | | |
| chrIV | 528421 | 528544 | 123 | -0,683155097 | N4:528503 | 3,536 | +15:YDR038C; | 15 | | | |
| chrIV | 528421 | 528548 | 127 | -0,977995532 | N4:528503 | 3,536 | +15:YDR038C; | 15 | | | |
| chrIV | 529656 | 529804 | 148 | -1,303964923 | | | | Overlap <50 bp | | | |
| chrIV | 529731 | 529886 | 155 | -1,419662276 | | | | Overlap <50 bp | | | |
| chrIV | 530359 | 530519 | 160 | -1,183591246 | N4:530430 | 7,81 | +3:YDR038C; | 3 | | | |
| chrIV | 531763 | 531899 | 136 | -1,1859211 | N4:531865 | 26,102 | +19:YDR039C; | 19 | | | |
| chrIV | 532306 | 532429 | 123 | -0,846804145 | N4:532381 | 19,732 | +15:YDR039C; | 15 | | | |
| chrIV | 532306 | 532433 | 127 | -0,974854934 | N4:532381 | 19,732 | +15:YDR039C; | 15 | | | |
| chrIV | 533541 | 533689 | 148 | -1,428567679 | | | | Overlap <50 bp | | | |
| chrIV | 533616 | 533771 | 155 | -1,111462625 | | | | Overlap <50 bp | | | |
| chrIV | 534244 | 534404 | 160 | -1,270103301 | N4:534319 | 7,572 | +3:YDR039C; | 3 | | | |
| chrIV | 535648 | 535784 | 136 | -1,096550797 | N4:535758 | 21,502 | +17:YDR040C; | 17 | | | |
| chrIV | 536109 | 536229 | 120 | -1,433892143 | | | | Overlap <50 bp | | | |
| chrIV | 537426 | 537574 | 148 | -1,247459289 | N4:537413 | 3,536 | +8:YDR040C; | 8 | | | |
| chrIV | 537501 | 537656 | 155 | -1,159342627 | | | | Overlap <50 bp | | | |
| chrIV | 540906 | 541052 | 146 | -1,611192237 | N4:541006 | 6,723 | +5:YDR042C; | 5 | | | |
| chrIV | 541731 | 541872 | 141 | -0,993125626 | N4:541799 | 28,657 | -1:YDR042C; | -1 | | | |
| chrIV | 545183 | 545329 | 146 | -1,111914643 | N4:545271 | 9,352 | +5:SUT469; +5:Unit130; | 5 | | | |
| chrIV | 549706 | 549856 | 150 | -1,484719952 | N4:549849 | 34,637 | +6:YDR046C; | 6 | | | |
| chrIV | 549706 | 549856 | 150 | -1,484719952 | N4:549694 | 0 | +7:YDR046C; | 7 | | | |
| chrIV | 552005 | 552154 | 149 | -1,297012773 | N4:552080 | 20,322 | +4:YDR047W; | 4 | | | |
| chrIV | 557304 | 557456 | 152 | -1,126539365 | N4:557386 | 11,635 | +5:YDR051C; | 5 | | | |
| chrIV | 560465 | 560600 | 135 | -1,108353986 | N4:560548 | 9,004 | +2:YDR052C; | 2 | | | |
| chrIV | 570638 | 570778 | 140 | -1,212675905 | N4:570621 | 8,335 | +1:YDR060W; | 1 | | | |
| chrIV | 570638 | 570778 | 140 | -1,212675905 | N4:570783 | 6,998 | +2:YDR060W; | 2 | | | |
| chrIV | 574209 | 574359 | 150 | -1,13324516 | N4:574313 | 43,29 | +2:YDR061W; | 2 | | | |
| chrIV | 575376 | 575525 | 149 | -1,397404726 | N4:575491 | 11,652 | +9:YDR061W; +3:SUT470; | 9 | | | |
| chrIV | 585684 | 585843 | 159 | -1,00622343 | N4:585698 | 22,72 | +13:YDR069C; | 13 | | | |
| chrIV | 587983 | 588137 | 154 | -1,245538413 | N4:588040 | 21,649 | DR069C; +4*:YDR070C; +1:CUT067; | -1 | | | |
| chrIV | 601658 | 601795 | 137 | -0,750910641 | N4:601720 | 42,687 | 'DR077W; +8:YDR078C; -1:Unit131; | 5 | | | |
| chrIV | 607967 | 608120 | 153 | -1,535272632 | N4:608009 | 45,5 | +14:YDR081C; | 14 | | | |
| chrIV | 615631 | 615766 | 135 | -1,767681315 | N4:615640 | 15,162 | +4:YDR085C; +9:SUT057; | 4 | | | |
| chrIV | 618044 | 618203 | 159 | -1,321763579 | N4:618139 | 5,413 | +2:YDR087C; | 2 | | | |
| chrIV | 625810 | 625961 | 151 | -1,393621247 | N4:625892 | 15,947 | +2:YDR090C; | 2 | | | |
| chrIV | 640754 | 640908 | 154 | -1,159351216 | N4:640848 | 33,065 | +19:YDR097C; | 19 | | | |
| chrIV | 643481 | 643635 | 154 | -1,29941771 | N4:643559 | 13,777 | +3:YDR097C; | 3 | | | |
| chrIV | 645695 | 645844 | 149 | -1,199548165 | N4:645718 | 10,693 | | Intergene | | | |
| chrIV | 646491 | 646637 | 146 | -1,376936315 | N4:646571 | 14,468 | | Intergene | | | |
| chrIV | 647130 | 647280 | 150 | -1,328296514 | N4:647179 | 16,803 | | Intergene | | | |
| chrIV | 648526 | 648683 | 157 | -0,957285711 | N4:648585 | 34,646 | | Intergene | | | |
| chrIV | 648994 | 649155 | 161 | -1,397407214 | N4:649119 | 45,962 | | Intergene | | | |
| chrIV | 649174 | 649309 | 135 | -1,004035614 | N4:649272 | 23,065 | | Intergene | | | |
| chrIV | 650212 | 650352 | 140 | -1,196188496 | N4:650289 | 52,532 | | Intergene | | | |
| chrIV | 650218 | 650352 | 134 | -1,145547098 | N4:650289 | 52,532 | | Intergene | | | |
| chrIV | 650630 | 650785 | 155 | -1,222824444 | N4:650651 | 15,044 | | Intergene | | | |
| chrIV | 650761 | 650869 | 108 | -0,746233357 | N4:650859 | 51,118 | | Intergene | | | |
| chrIV | 654067 | 654214 | 147 | -1,095592234 | N4:654056 | 31,161 | +4:YDR099W; | 4 | | | |
| chrIV | 654067 | 654214 | 147 | -1,095592234 | N4:654214 | 32,997 | +5:YDR099W; | 5 | | | |
| chrIV | 656839 | 656998 | 159 | -1,177322567 | N4:656949 | 6,419 | +4:YDR101C; | 4 | | | |
| chrIV | 658487 | 658641 | 154 | -1,539803086 | N4:658664 | 14,177 | +3:YDR103W; | 3 | | | |
| chrIV | 658600 | 658747 | 147 | -1,321267512 | N4:658664 | 14,177 | +3:YDR103W; | 3 | | | |
| chrIV | 662245 | 662401 | 156 | -1,201168959 | N4:662311 | 41,701 | +17:YDR104C; | 17 | | | |
| chrIV | 665084 | 665248 | 164 | -1,637505983 | N4:665151 | 29,838 | -1:YDR104C; | -1 | | | |
| chrIV | 665597 | 665732 | 135 | -1,26201557 | N4:665678 | 14,886 | +8:YDR105C; | 8 | | | |
| chrIV | 666941 | 667094 | 153 | -1,344306476 | N4:667026 | 8,532 | -1:YDR105C; +1:YDR106W; | -1 | | | |
| chrIV | 666948 | 667109 | 161 | -1,589752762 | N4:667026 | 8,532 | -1:YDR105C; +1:YDR106W; | -1 | | | |
| chrIV | 669497 | 669656 | 159 | -1,393388218 | N4:669574 | 13,11 | +9:YDR107C; | 9 | | | |
| chrIV | 669741 | 669906 | 165 | -1,663900623 | N4:669781 | 43,632 | +8:YDR107C; | 8 | | | |
| chrIV | 677555 | 677687 | 132 | -1,440701538 | N4:677564 | 10,672 | +10:YDR110W; | 10 | | | |
| chrIV | 682440 | 682595 | 155 | -1,099609367 | N4:682517 | 11,446 | -1:YDR114C; +3:YDR115W; | -1 | | | |

| CHR | START | END | L(bp) | ΔLK | Nuc ID | Fuzziness | Gene ID | Gene body postion or intergene | Terminal | rDNA | Telomeric |
|---|---|---|---|---|---|---|---|---|---|---|---|
| chrIV | 682928 | 683076 | 148 | -1,299709395 | N4:683045 | 23,424 | +4:YDR116C; | 4 | | | |
| chrIV | 684905 | 685053 | 148 | -1,37643622 | N4:684976 | 22,003 | +5:YDR117C; | 5 | | | |
| chrIV | 685367 | 685521 | 154 | -1,07353604 | N4:685462 | 7,659 | +2:YDR117C; | 2 | | | |
| chrIV | 685409 | 685539 | 130 | -1,174834812 | N4:685462 | 7,659 | +2:YDR117C; | 2 | | | |
| chrIV | 692976 | 693125 | 149 | -1,161819403 | N4:693039 | 17,201 | +2:YDR120C; | 2 | | | |
| chrIV | 699427 | 699543 | 116 | -1,206881546 | N4:699492 | 6,919 | +1:YDR123C; | 1 | | | |
| chrIV | 699429 | 699572 | 143 | -1,017853798 | N4:699492 | 6,919 | +1:YDR123C; | 1 | | | |
| chrIV | 701924 | 702080 | 156 | -1,445347585 | N4:702005 | 22,546 | +5:YDR125C; | 5 | | | |
| chrIV | 705337 | 705438 | 101 | -1,494147377 | N4:705421 | 25,314 | +7:YDR127W; | 7 | | | |
| chrIV | 705533 | 705696 | 163 | -1,269779883 | | | | Overlap <50 bp | | | |
| chrIV | 709396 | 709543 | 147 | -1,414813573 | N4:709492 | 29,663 | 0:YDR128W; | 0 | | | |
| chrIV | 710394 | 710553 | 159 | -1,370386623 | N4:710486 | 41,504 | +7:YDR128W; | 7 | | | |
| chrIV | 710648 | 710758 | 110 | -0,487758835 | N4:710780 | 38,698 | +9:YDR128W; | 9 | | | |
| chrIV | 711680 | 711827 | 147 | -1,435110026 | N4:711712 | 33,139 | | Intergene | | | |
| chrIV | 712887 | 713032 | 145 | -1,198220012 | N4:712928 | 16,376 | | Intergene | | | |
| chrIV | 717311 | 717466 | 155 | -1,593478262 | N4:717376 | 7,19 | +8:YDR131C; | 8 | | | |
| chrIV | 723685 | 723835 | 150 | -1,267074106 | | | | Overlap <50 bp | | | |
| chrIV | 724692 | 724830 | 138 | -1,357829463 | N4:724752 | 43,68 | +18:YDR135C; +16:SUT060; | 18 | | | |
| chrIV | 727870 | 728018 | 148 | -1,344269525 | N4:727949 | 17,366 | -1:YDR137W; | -1 | | | |
| chrIV | 729848 | 729970 | 122 | -1,29837542 | N4:729980 | 23,761 | +12:YDR137W; | 12 | | | |
| chrIV | 729848 | 729954 | 106 | -1,221131484 | | | | Overlap <50 bp | | | |
| chrIV | 730903 | 730997 | 94 | -1,177649955 | N4:730925 | 5,686 | +3:YDR138W; | 3 | | | |
| chrIV | 734743 | 734893 | 150 | -1,703921558 | N4:734827 | 15,305 | +5*:YDR140W; | 5 | TERM | | |
| chrIV | 736895 | 737045 | 150 | -1,28468377 | | | | Overlap <50 bp | | | |
| chrIV | 737420 | 737548 | 128 | -1,112255558 | N4:737479 | 34,438 | +16:YDR141C; | 16 | | | |
| chrIV | 737638 | 737793 | 155 | -1,594408687 | N4:737705 | 21,498 | +15:YDR141C; | 15 | | | |
| chrIV | 745330 | 745443 | 113 | -1,569221438 | N4:745385 | 10,747 | +6:YDR144C; | 6 | | | |
| chrIV | 747582 | 747734 | 152 | -1,463098449 | N4:747675 | 7,174 | +7:YDR145W; | 7 | | | |
| chrIV | 750682 | 750850 | 168 | -1,58050291 | N4:750748 | 27,362 | +2:YDR146C; | 2 | | | |
| chrIV | 760658 | 760811 | 153 | -1,397555805 | N4:760826 | 15,436 | | Intergene | | | |
| chrIV | 761833 | 761982 | 149 | -1,218845409 | N4:761970 | 22,612 | | Intergene | | | |
| chrIV | 762005 | 762157 | 152 | -1,130067186 | N4:762143 | 11,314 | | Intergene | | | |
| chrIV | 764493 | 764644 | 151 | -1,455314455 | N4:764585 | 8,06 | +5:YDR151C; | 5 | | | |
| chrIV | 766598 | 766749 | 151 | -1,203212325 | N4:766690 | 16,514 | +10*:YDR153C; | 10 | TERM | | |
| chrIV | 768253 | 768411 | 158 | -1,607347271 | N4:768405 | 14,08 | +4*:YDR155C; | 4 | TERM | | |
| chrIV | 771149 | 771299 | 150 | -1,178227072 | | | | Overlap <50 bp | | | |
| chrIV | 773973 | 774101 | 128 | -1,345804847 | | | | Overlap <50 bp | | | |
| chrIV | 779473 | 779647 | 174 | -1,849471933 | N4:779537 | 10,237 | +4:YDR161W; | 4 | | | |
| chrIV | 779988 | 780110 | 122 | -1,285242314 | N4:779996 | 18,284 | +7:YDR161W; | 7 | | | |
| chrIV | 785291 | 785441 | 150 | -1,696944994 | N4:785366 | 21,213 | +4:YDR165W; | 4 | | | |
| chrIV | 787217 | 787366 | 149 | -1,247100679 | N4:787358 | 22,528 | +13:YDR166C; | 13 | | | |
| chrIV | 787217 | 787366 | 149 | -1,247100679 | N4:787195 | 23,788 | +14:YDR166C; | 14 | | | |
| chrIV | 789145 | 789305 | 160 | -1,3336118 | N4:789230 | 19,692 | +1:YDR166C; -1:YDR167W; | 1 | | | |
| chrIV | 800500 | 800661 | 161 | -1,445438319 | N4:800519 | 14,064 | | Intergene | | | |
| chrIV | 803067 | 803219 | 152 | -1,554898307 | N4:803114 | 40,951 | | Intergene | | | |
| chrIV | 805417 | 805524 | 107 | -0,979370718 | N4:805529 | 17,01 | | Intergene | | | |
| chrIV | 805417 | 805530 | 113 | -1,29302957 | N4:805529 | 17,01 | | Intergene | | | |
| chrIV | 810559 | 810694 | 135 | -1,36235258 | N4:810661 | 7,668 | +7*:YDR173C; | 7 | TERM | | |
| chrIV | 815236 | 815393 | 157 | -1,298327366 | N4:815317 | 26,988 | +6:YDR176W; | 6 | | | |
| chrIV | 815520 | 815678 | 158 | -1,659962576 | N4:815624 | 44,294 | +8:YDR176W; | 8 | | | |
| chrIV | 826418 | 826583 | 165 | -1,43380278 | N4:826491 | 11,286 | +6:YDR181C; | 6 | | | |
| chrIV | 827506 | 827652 | 146 | -0,991530057 | N4:827588 | 9,411 | -1:YDR181C; +1:YDR182W; | -1 | | | |
| chrIV | 835382 | 835525 | 143 | -1,292362561 | N4:835504 | 19,705 | +1:YDR186C; | 1 | | | |
| chrIV | 835382 | 835528 | 146 | -1,2799835 | N4:835504 | 19,705 | +1:YDR186C; | 1 | | | |
| chrIV | 835383 | 835528 | 145 | -1,400809352 | N4:835504 | 19,705 | +1:YDR186C; | 1 | | | |
| chrIV | 835384 | 835528 | 144 | -1,235082832 | N4:835504 | 19,705 | +1:YDR186C; | 1 | | | |
| chrIV | 835385 | 835528 | 143 | -1,5864253 | N4:835504 | 19,705 | +1:YDR186C; | 1 | | | |
| chrIV | 835386 | 835528 | 142 | -1,880812245 | N4:835504 | 19,705 | +1:YDR186C; | 1 | | | |
| chrIV | 835932 | 836065 | 133 | -0,908233636 | N4:835972 | 14,114 | | Intergene | | | |
| chrIV | 836520 | 836671 | 151 | -0,865853371 | N4:836563 | 5,541 | +2:YDR188W; | 2 | | | |
| chrIV | 837322 | 837477 | 155 | -1,25043489 | N4:837482 | 29,372 | +8:YDR188W; | 8 | | | |
| chrIV | 838477 | 838608 | 131 | -2,471460103 | N4:838553 | 7,195 | +2:YDR189W; | 2 | | | |
| chrIV | 843960 | 844136 | 176 | -1,617694818 | N4:844047 | 23,356 | +6:YDR192C; | 6 | | | |
| chrIV | 844482 | 844623 | 141 | -1,13846221 | N4:844544 | 2,63 | +3:YDR192C; | 3 | | | |
| chrIV | 844482 | 844649 | 167 | -1,52255302 | N4:844544 | 2,63 | +3:YDR192C; | 3 | | | |
| chrIV | 850466 | 850631 | 165 | -1,293353945 | N4:850636 | 8,841 | +3:YDR196C; | 3 | | | |
| chrIV | 850466 | 850631 | 165 | -1,293353945 | N4:850477 | 5,745 | +4:YDR196C; | 4 | | | |
| chrIV | 857181 | 857346 | 165 | -1,260794076 | | | | Overlap <50 bp | | | |
| chrIV | 861657 | 861796 | 139 | -1,25951816 | N4:861762 | 4,879 | +1:YDR206W; | 1 | | | |
| chrIV | 863651 | 863788 | 137 | -1,147617344 | N4:863721 | 31,07 | +13:YDR206W; | 13 | | | |
| chrIV | 866143 | 866295 | 152 | -1,023217713 | N4:866178 | 6,841 | +9:YDR207C; | 9 | | | |
| chrIV | 867460 | 867598 | 138 | -1,643394345 | N4:867531 | 9,783 | +1:YDR207C; | 1 | | | |
| chrIV | 870210 | 870347 | 137 | -1,276218809 | N4:870239 | 20,312 | +15:YDR208W; | 15 | | | |
| chrIV | 872409 | 872511 | 102 | -0,588837978 | N4:872520 | 29,28 | | Intergene | | | |
| chrIV | 873832 | 873980 | 148 | -1,516638041 | | | | Overlap <50 bp | | | |
| chrIV | 874621 | 874762 | 141 | -1,439702297 | | | | Overlap <50 bp | | | |
| chrIV | 875359 | 875502 | 143 | -1,120581077 | N4:875420 | 19,858 | | Intergene | | | |
| chrIV | 876645 | 876791 | 146 | -1,307156374 | N4:876701 | 12,728 | | Intergene | | | |
| chrIV | 877438 | 877587 | 149 | -1,397636453 | N4:877571 | 0,707 | | Intergene | | | |
| chrIV | 878298 | 878451 | 153 | -1,651776053 | N4:878280 | 33,279 | | Intergene | | | |
| chrIV | 878496 | 878645 | 149 | -1,675764605 | N4:878510 | 27,429 | | Intergene | | | |
| chrIV | 879292 | 879438 | 146 | -1,15315166 | N4:879370 | 5,686 | | Intergene | | | |
| chrIV | 881327 | 881484 | 157 | -1,475286737 | N4:881391 | 30,551 | | Intergene | | | |
| chrIV | 881795 | 881956 | 161 | -1,365654221 | | | | Overlap <50 bp | | | |
| chrIV | 881975 | 882110 | 135 | -1,14359206 | N4:882066 | 27,301 | | Intergene | | | |
| chrIV | 883013 | 883153 | 140 | -1,046501802 | N4:883091 | 39,887 | | Intergene | | | |
| chrIV | 883019 | 883153 | 134 | -1,086239038 | N4:883091 | 39,887 | | Intergene | | | |
| chrIV | 883431 | 883586 | 155 | -1,41285025 | N4:883463 | 24,132 | | Intergene | | | |
| chrIV | 883562 | 883670 | 108 | -0,702786672 | | | | Overlap <50 bp | | | |
| chrIV | 883696 | 883848 | 152 | -1,374578115 | N4:883720 | 47,606 | | Intergene | | | |
| chrIV | 884819 | 884969 | 150 | -1,242860796 | N4:884882 | 7,861 | +2:YDR211W; | 2 | | | |
| chrIV | 888390 | 888567 | 177 | -1,328871178 | N4:888453 | 33,49 | +8:YDR212W; | 8 | | | |
| chrIV | 889668 | 889827 | 159 | -1,669720174 | N4:889734 | 12,903 | +2:YDR213W; | 2 | | | |
| chrIV | 890834 | 890988 | 154 | -0,791781583 | N4:890967 | 31,753 | +10:YDR213W; | 10 | | | |
| chrIV | 891389 | 891533 | 144 | -0,835485106 | N4:891510 | 38,291 | +13:YDR213W; | 13 | | | |
| chrIV | 891986 | 892153 | 167 | -1,608359805 | N4:892055 | 27,537 | +16:YDR213W; +3*:CUT515; | 16 | | | |
| chrIV | 896334 | 896482 | 148 | -0,88067986 | N4:896422 | 15,438 | +10:YDR216W; | 10 | | | |
| chrIV | 897629 | 897776 | 147 | -1,286044597 | N4:897659 | 36,527 | +17:YDR216W; | 17 | | | |

| CHR | START | END | L(bp) | ΔLK | Nuc ID | Fuzziness | Gene ID | Gene body postion or intergene | Terminal | rDNA | Telomeric |
|---|---|---|---|---|---|---|---|---|---|---|---|
| chrIV | 899028 | 899177 | 149 | -1,592654491 | N4:899049 | 18,159 | +25*:YDR216W; | 25 | TERM | | |
| chrIV | 906466 | 906593 | 127 | -1,354777757 | N4:906545 | 17,068 | +4:YDR219C; | 4 | | | |
| chrIV | 907937 | 908075 | 138 | -1,70983835 | | | | Overlap <50 bp | | | |
| chrIV | 908538 | 908667 | 129 | -1,775390989 | N4:908599 | 35,786 | +6*:SUT474; +9:YDR221W; | 6 | TERM | | |
| chrIV | 909620 | 909773 | 153 | -1,686271496 | N4:909630 | 14,849 | -1:SUT474; +15*:YDR221W; | -1 | | | |
| chrIV | 915796 | 915980 | 184 | -1,486066447 | N4:915793 | 10,472 | +3:YDR225W; | 3 | | | |
| chrIV | 917617 | 917765 | 148 | -1,21107431 | N4:917653 | 10,498 | +2:YDR227W; | 2 | | | |
| chrIV | 924934 | 925100 | 166 | -0,917021194 | N4:925073 | 25,319 | +3:YDR229W; | 3 | | | |
| chrIV | 926425 | 926570 | 145 | -1,17948512 | N4:926519 | 11,437 | +3:YDR231C; | 3 | | | |
| chrIV | 927161 | 927289 | 128 | -0,472034977 | N4:927242 | 9,326 | +1:YDR232W; | 1 | | | |
| chrIV | 928961 | 929115 | 154 | -1,400149499 | N4:929074 | 12,458 | +11:YDR232W; | 11 | | | |
| chrIV | 930238 | 930383 | 145 | -1,300991281 | N4:930295 | 36,425 | +1:YDR233C; | 1 | | | |
| chrIV | 931111 | 931260 | 149 | -1,081827997 | N4:931170 | 10,602 | +2:YDR234W; | 2 | | | |
| chrIV | 933440 | 933600 | 160 | -1,726468871 | N4:933517 | 5,225 | +1:YDR235W; | 1 | | | |
| chrIV | 944365 | 944524 | 159 | -1,513022674 | N4:944467 | 19,672 | +5:YDR240C; | 5 | | | |
| chrIV | 944403 | 944557 | 154 | -1,379890931 | N4:944467 | 19,672 | +5:YDR240C; | 5 | | | |
| chrIV | 948002 | 948171 | 169 | -1,814066687 | N4:948157 | 42,823 | +10:YDR242W; | 10 | | | |
| chrIV | 949233 | 949387 | 154 | -0,819364007 | N4:949302 | 16,661 | +7:YDR243C; | 7 | | | |
| chrIV | 955015 | 955163 | 148 | -1,199178502 | N4:955072 | 27,826 | +5*:YDR246W; +1:YDR246W-A; | 5 | TERM | | |
| chrIV | 957192 | 957317 | 125 | -0,854749929 | N4:957237 | 22,485 | +10:YDR247W; | 10 | | | |
| chrIV | 957192 | 957321 | 129 | -0,942615234 | N4:957237 | 22,485 | +10:YDR247W; | 10 | | | |
| chrIV | 957196 | 957298 | 102 | -0,395654796 | N4:957237 | 22,485 | +10:YDR247W; | 10 | | | |
| chrIV | 962120 | 962274 | 154 | -1,503690862 | N4:962102 | 30,697 | +12:YDR251W; | 12 | | | |
| chrIV | 962406 | 962555 | 149 | -1,216378209 | N4:962480 | 29,614 | +14:YDR251W; | 14 | | | |
| chrIV | 965842 | 965991 | 149 | -1,297042977 | N4:965942 | 7,211 | +6:YDR254W; | 6 | | | |
| chrIV | 968475 | 968604 | 129 | -1,217294172 | N4:968597 | 21,904 | +4:anti039; +8:YDR256C; | 4 | | | |
| chrIV | 971192 | 971329 | 137 | -1,153472476 | N4:971253 | 10,583 | +2:YDR257C; | 2 | | | |
| chrIV | 975387 | 975508 | 121 | -1,211154132 | N4:975430 | 23,039 | +3:YDR259C; | 3 | | | |
| chrIV | 979606 | 979759 | 153 | -1,361501555 | N4:979673 | 31,32 | +2:Unit141; | 2 | | | |
| chrIV | 980738 | 980889 | 151 | -1,314028889 | N4:980809 | 11,437 | +1:SUT479; | 1 | | | |
| chrIV | 981759 | 981861 | 102 | -0,556377537 | N4:981871 | 42,414 | | Intergene | | | |
| chrIV | 983182 | 983330 | 148 | -1,489235912 | | | | Overlap <50 bp | | | |
| chrIV | 984709 | 984852 | 143 | -0,882634791 | N4:984756 | 11,504 | | Intergene | | | |
| chrIV | 985995 | 986141 | 146 | -0,981612181 | | | | Overlap <50 bp | | | |
| chrIV | 986788 | 986937 | 149 | -1,071508726 | N4:986912 | 25,456 | | Intergene | | | |
| chrIV | 987343 | 987492 | 149 | -1,362452466 | N4:987353 | 12,503 | | Intergene | | | |
| chrIV | 987657 | 987771 | 114 | -1,285773884 | N4:987788 | 2,517 | | Intergene | | | |
| chrIV | 988353 | 988503 | 150 | -1,793175299 | N4:988403 | 10,017 | | Intergene | | | |
| chrIV | 989662 | 989806 | 144 | -1,367248348 | N4:989808 | 37 | | Intergene | | | |
| chrIV | 989749 | 989906 | 157 | -1,386005362 | N4:989808 | 37 | | Intergene | | | |
| chrIV | 990217 | 990378 | 161 | -1,623091229 | N4:990348 | 50,912 | | Intergene | | | |
| chrIV | 990397 | 990532 | 135 | -1,327630588 | N4:990499 | 32,593 | | Intergene | | | |
| chrIV | 991435 | 991575 | 140 | -1,145213344 | N4:991512 | 35,553 | | Intergene | | | |
| chrIV | 991441 | 991575 | 134 | -0,962274359 | N4:991512 | 35,553 | | Intergene | | | |
| chrIV | 991853 | 992008 | 155 | -1,29763554 | N4:991879 | 21,221 | | Intergene | | | |
| chrIV | 991984 | 992092 | 108 | -0,766194109 | N4:992082 | 47,648 | | Intergene | | | |
| chrIV | 992133 | 992282 | 149 | -1,478913425 | | | | Overlap <50 bp | | | |
| chrIV | 994735 | 994883 | 148 | -1,274640739 | N4:994783 | 34,313 | +7:YDR263C; | 7 | | | |
| chrIV | 1001383 | 1001529 | 146 | -1,410974783 | N4:1001542 | 5,315 | +5:YDR266C; | 5 | | | |
| chrIV | 1001383 | 1001529 | 146 | -1,410974783 | N4:1001379 | 8,802 | +6:YDR266C; | 6 | | | |
| chrIV | 1005517 | 1005664 | 147 | -1,183038915 | N4:1005589 | 6,994 | +1:YDR270W; | 1 | | | |
| chrIV | 1005876 | 1006022 | 146 | -1,206736155 | N4:1005930 | 12,922 | +3:YDR270W; | 3 | | | |
| chrIV | 1012282 | 1012417 | 135 | -1,078895653 | N4:1012368 | 27,109 | +2:YDR275W; | 2 | | | |
| chrIV | 1016514 | 1016662 | 148 | -0,967388928 | N4:1016582 | 30,801 | +14:SUT065; | 14 | | | |
| chrIV | 1019255 | 1019404 | 149 | -1,625001612 | N4:1019326 | 11,828 | +1:YDR279W; -1:SUT480; | 1 | | | |
| chrIV | 1021134 | 1021284 | 150 | -1,398208887 | N4:1021147 | 6,95 | +4:YDR280W; | 4 | | | |
| chrIV | 1021134 | 1021284 | 150 | -1,398208887 | N4:1021302 | 4,933 | +5:YDR280W; | 5 | | | |
| chrIV | 1025226 | 1025374 | 148 | -1,281731333 | N4:1025251 | 7,411 | | Intergene | | | |
| chrIV | 1028806 | 1028950 | 144 | -1,207551937 | N4:1028803 | 22,345 | | Intergene | | | |
| chrIV | 1039896 | 1040063 | 167 | -1,660125295 | N4:1039922 | 6,186 | +2:YDR291W; -1:YDR289C; | 2 | | | |
| chrIV | 1042109 | 1042252 | 143 | -1,249609016 | N4:1042173 | 23,619 | +16:YDR291W; | 16 | | | |
| chrIV | 1052004 | 1052154 | 150 | -1,010694186 | N4:1052080 | 8,55 | +2:YDR294C; | 2 | | | |
| chrIV | 1055603 | 1055756 | 153 | -1,279731933 | N4:1055666 | 5,707 | +4*:YDR296W; | 4 | TERM | | |
| chrIV | 1058574 | 1058755 | 181 | -1,368076919 | N4:1058759 | 6,861 | +2:YDR298C; | 2 | | | |
| chrIV | 1058574 | 1058755 | 181 | -1,368076919 | N4:1058587 | 19,194 | +3:YDR298C; | 3 | | | |
| chrIV | 1063264 | 1063420 | 156 | -0,865698397 | N4:1063318 | 5,215 | +1:YDR301W; | 1 | | | |
| chrIV | 1066695 | 1066823 | 128 | -0,822100309 | N4:1066763 | 26,306 | +21:YDR301W; | 21 | | | |
| chrIV | 1075797 | 1075953 | 156 | -1,58549808 | N4:1075869 | 5,269 | +1:YDR307W; | 1 | | | |
| chrIV | 1079066 | 1079219 | 153 | -1,125259776 | N4:1079155 | 10,607 | +7:YDR309C; | 7 | | | |
| chrIV | 1079249 | 1079403 | 154 | -1,206042957 | N4:1079306 | 6,025 | +6:YDR309C; | 6 | | | |
| chrIV | 1080074 | 1080246 | 172 | -1,499505391 | N4:1080115 | 14,195 | +1:YDR309C; | 1 | | | |
| chrIV | 1090868 | 1091040 | 172 | -1,245002011 | N4:1090903 | 21,687 | +12:YDR314C; +4:CUT079; | 12 | | | |
| chrIV | 1091309 | 1091450 | 141 | -1,48421628 | | | | Overlap <50 bp | | | |
| chrIV | 1096304 | 1096427 | 123 | -1,242179707 | | | | Overlap <50 bp | | | |
| chrIV | 1096404 | 1096559 | 155 | -1,380357536 | N4:1096525 | 34,962 | | Intergene | | | |
| chrIV | 1096837 | 1096971 | 134 | -1,253225439 | N4:1096900 | 24,214 | | Intergene | | | |
| chrIV | 1096837 | 1096977 | 140 | -1,222702462 | N4:1096900 | 24,214 | | Intergene | | | |
| chrIV | 1097880 | 1098015 | 135 | -1,221031054 | N4:1097914 | 19,502 | | Intergene | | | |
| chrIV | 1098034 | 1098195 | 161 | -1,598648121 | N4:1098134 | 26,458 | | Intergene | | | |
| chrIV | 1098506 | 1098663 | 157 | -1,105015426 | N4:1098582 | 22,053 | | Intergene | | | |
| chrIV | 1098997 | 1099126 | 129 | -1,280463803 | | | | Overlap <50 bp | | | |
| chrIV | 1099909 | 1100059 | 150 | -1,237679791 | N4:1100005 | 8,622 | | Intergene | | | |
| chrIV | 1100669 | 1100827 | 158 | -1,30434629 | N4:1100752 | 28,219 | | Intergene | | | |
| chrIV | 1102545 | 1102694 | 149 | -1,230737537 | N4:1102665 | 32,019 | +4:YDR317W; +6:SUT481; | 4 | | | |
| chrIV | 1102953 | 1103100 | 147 | -1,841385305 | N4:1102998 | 35,444 | +6:YDR317W; +4:SUT481; | 6 | | | |
| chrIV | 1107156 | 1107303 | 147 | -1,404751627 | N4:1107223 | 31,356 | +7:YDR320C; | 7 | | | |
| chrIV | 1109551 | 1109723 | 172 | -1,390447929 | N4:1109646 | 15,865 | +7:YDR321W; | 7 | | | |
| chrIV | 1112644 | 1112796 | 152 | -1,382598612 | N4:1112718 | 6,066 | +9:YDR323C; | 9 | | | |
| chrIV | 1117237 | 1117387 | 150 | -1,013796475 | N4:1117306 | 8,809 | +2:YDR325W; | 2 | | | |
| chrIV | 1121058 | 1121205 | 147 | -1,357747123 | N4:1121186 | 12,971 | | Intergene | | | |
| chrIV | 1121801 | 1121948 | 147 | -1,320931937 | N4:1121812 | 30,534 | +21*:YDR326C; | 21 | TERM | | |
| chrIV | 1124473 | 1124641 | 168 | -1,637973632 | N4:1124564 | 25,882 | +4:YDR326C; | 4 | | | |
| chrIV | 1128248 | 1128395 | 147 | -1,283399164 | N4:1128331 | 15,042 | +4:YDR330W; | 4 | | | |
| chrIV | 1137826 | 1137974 | 148 | -1,343574184 | N4:1137842 | 30,178 | +13:YDR334W; | 13 | | | |
| chrIV | 1140412 | 1140528 | 116 | -1,458879409 | N4:1140541 | 12,189 | [illegible]335W; +31*:YDR334W; -1:SUT067; | -1 | | | |
| chrIV | 1144636 | 1144783 | 147 | -1,323722881 | N4:1144674 | 21,217 | -1:YDR336W; | -1 | | | |
| chrIV | 1154940 | 1155065 | 125 | -1,129208036 | N4:1155020 | 40,204 | +7:YDR342C; | 7 | | | |
| chrIV | 1155027 | 1155183 | 156 | -1,155254882 | N4:1155020 | 40,204 | +7:YDR342C; | 7 | | | |

| CHR | START | END | L(bp) | ΔLK | Nuc ID | Fuzziness | Gene ID | Gene body postion or intergene | Terminal | rDNA | Telomeric |
|---|---|---|---|---|---|---|---|---|---|---|---|
| chrIV | 1155195 | 1155331 | 136 | -0,998969773 | N4:1155235 | 0,707 | +6:YDR342C; | 6 | | | |
| chrIV | 1158507 | 1158654 | 147 | -1,140734364 | N4:1158572 | 30,138 | +8:SUT068; | 8 | | | |
| chrIV | 1160332 | 1160457 | 125 | -1,214566306 | N4:1160332 | 0 | +8:YDR343C; | 8 | | | |
| chrIV | 1160419 | 1160575 | 156 | -1,093249186 | N4:1160485 | 38,371 | +7:YDR343C; | 7 | | | |
| chrIV | 1160587 | 1160723 | 136 | -0,946640958 | N4:1160638 | 19,092 | +6:YDR343C; | 6 | | | |
| chrIV | 1164215 | 1164366 | 151 | -1,234746417 | N4:1164258 | 25,502 | +3:YDR345C; +11:SUT069; | 3 | | | |
| chrIV | 1166734 | 1166868 | 134 | -1,230065171 | N4:1166854 | 16,183 | +2:Unit103; +2:SUT070; | 2 | | | |
| chrIV | 1169216 | 1169370 | 154 | -1,359826604 | N4:1169282 | 11,397 | +2:YDR347W; | 2 | | | |
| chrIV | 1170479 | 1170639 | 160 | -1,172665911 | N4:1170579 | 27,306 | +10:YDR348C; | 10 | | | |
| chrIV | 1170874 | 1171037 | 163 | -1,530421618 | N4:1171049 | 28,358 | +7:YDR348C; | 7 | | | |
| chrIV | 1170874 | 1171037 | 163 | -1,530421618 | N4:1170900 | 37,528 | +8:YDR348C; | 8 | | | |
| chrIV | 1171736 | 1171886 | 150 | -1,409341001 | N4:1171844 | 18,902 | +2:YDR348C; | 2 | | | |
| chrIV | 1175589 | 1175742 | 153 | -1,466943524 | N4:1175671 | 10,085 | | Intergene | | | |
| chrIV | 1176500 | 1176647 | 147 | -1,313180512 | N4:1176576 | 19,271 | +11:YDR350C; | 11 | | | |
| chrIV | 1179339 | 1179509 | 170 | -1,598893454 | N4:1179429 | 15,384 | +7:YDR351W; | 7 | | | |
| chrIV | 1185766 | 1185913 | 147 | -1,255694347 | N4:1185835 | 4,93 | +8*:YDR354W; +1:YDR356W; | 8 | TERM | | |
| chrIV | 1186635 | 1186765 | 130 | -1,516892523 | N4:1186749 | 15,681 | +7:YDR356W; | 7 | | | |
| chrIV | 1187842 | 1187997 | 155 | -1,60281164 | N4:1187892 | 47,06 | +14:YDR356W; | 14 | | | |
| chrIV | 1199637 | 1199781 | 144 | -1,407776687 | N4:1199716 | 15,283 | +5:YDR363W; | 5 | | | |
| chrIV | 1199946 | 1200093 | 147 | -1,454907531 | N4:1200030 | 33,208 | +7:YDR363W; | 7 | | | |
| chrIV | 1202429 | 1202587 | 158 | -1,75263873 | N4:1202473 | 7 | +11*:YDR364C; +3*:YDR363W-A; | 11 | TERM | | |
| chrIV | 1205211 | 1205359 | 148 | -1,269646068 | N4:1205237 | 15,962 | +8:YDR365C; | 8 | | | |
| chrIV | 1207063 | 1207212 | 149 | -1,093876745 | N4:1207202 | 34,598 | | Intergene | | | |
| chrIV | 1207237 | 1207360 | 123 | -1,385145384 | | | | Overlap <50 bp | | | |
| chrIV | 1207337 | 1207492 | 155 | -1,446858693 | N4:1207459 | 34,646 | | Intergene | | | |
| chrIV | 1207770 | 1207904 | 134 | -1,179991693 | N4:1207827 | 23,065 | | Intergene | | | |
| chrIV | 1207770 | 1207910 | 140 | -1,084824366 | N4:1207827 | 23,065 | | Intergene | | | |
| chrIV | 1207966 | 1208117 | 151 | -1,502669143 | N4:1208017 | 24,062 | | Intergene | | | |
| chrIV | 1207976 | 1208121 | 145 | -1,421192634 | N4:1208017 | 24,062 | | Intergene | | | |
| chrIV | 1207994 | 1208145 | 151 | -1,484614536 | N4:1208017 | 24,062 | | Intergene | | | |
| chrIV | 1207996 | 1208145 | 149 | -1,388787762 | N4:1208017 | 24,062 | | Intergene | | | |
| chrIV | 1208248 | 1208402 | 154 | -1,146588717 | N4:1208380 | 21,502 | | Intergene | | | |
| chrIV | 1208282 | 1208426 | 144 | -1,066842954 | N4:1208380 | 21,502 | | Intergene | | | |
| chrIV | 1208813 | 1208948 | 135 | -1,211399633 | N4:1208846 | 3,786 | | Intergene | | | |
| chrIV | 1208967 | 1209128 | 161 | -1,362519076 | N4:1209000 | 34,646 | | Intergene | | | |
| chrIV | 1209439 | 1209596 | 157 | -1,177219376 | N4:1209520 | 25,58 | | Intergene | | | |
| chrIV | 1209930 | 1210059 | 129 | -1,478192145 | N4:1210072 | 10,408 | | Intergene | | | |
| chrIV | 1210842 | 1210992 | 150 | -1,649101236 | N4:1210937 | 4,243 | | Intergene | | | |
| chrIV | 1211574 | 1211688 | 114 | -1,385958414 | N4:1211673 | 34,933 | | Intergene | | | |
| chrIV | 1215599 | 1215745 | 146 | -1,268995414 | N4:1215661 | 31,31 | +13:YDR369C; | 13 | | | |
| chrIV | 1216176 | 1216327 | 151 | -1,652756105 | | | | Overlap <50 bp | | | |
| chrIV | 1217346 | 1217492 | 146 | -1,412513411 | N4:1217374 | 18,661 | +3:YDR369C; | 3 | | | |
| chrIV | 1218508 | 1218653 | 145 | -1,288764769 | N4:1218588 | 5,727 | +4:YDR370C; | 4 | | | |
| chrIV | 1221088 | 1221239 | 151 | -1,570973339 | N4:1221154 | 32,036 | +9:YDR372C; | 9 | | | |
| chrIV | 1225710 | 1225871 | 161 | -1,481847672 | N4:1225704 | 18,619 | +6:YDR375C; | 6 | | | |
| chrIV | 1225924 | 1226071 | 147 | -0,905292488 | N4:1225999 | 7,95 | +4:YDR375C; | 4 | | | |
| chrIV | 1227651 | 1227806 | 155 | -1,663778754 | N4:1227756 | 6,099 | +7:YDR376W; | 7 | | | |
| chrIV | 1238397 | 1238557 | 160 | -1,37784321 | N4:1238463 | 11,535 | +4:YDR381C-A; | 4 | | | |
| chrIV | 1240257 | 1240406 | 149 | -1,314931113 | N4:1240306 | 6,121 | +2:CUT088; +5:YDR383C; | 2 | | | |
| chrIV | 1240385 | 1240526 | 141 | -1,308831282 | N4:1240462 | 2,805 | +3:CUT088; +4:YDR383C; | 3 | | | |
| chrIV | 1244524 | 1244687 | 163 | -1,30013175 | N4:1244612 | 39,4 | +9:YDR385W; | 9 | | | |
| chrIV | 1245862 | 1245987 | 125 | -1,097051488 | N4:1245846 | 27,677 | 0:YDR386W; +17*:YDR385W; | 0 | | | |
| chrIV | 1251539 | 1251695 | 156 | -1,137182002 | N4:1251612 | 11,384 | +10*:YDR388W; -1:SUT486; | 10 | TERM | | |
| chrIV | 1259546 | 1259668 | 122 | -1,279201745 | N4:1259621 | 5,857 | -1:YDR393W; +7:YDR392W; | -1 | | | |
| chrIV | 1265859 | 1266020 | 161 | -1,377089085 | N4:1265947 | 32,749 | +17*:YDR395W; | 17 | TERM | | |
| chrIV | 1266398 | 1266553 | 155 | -1,42106368 | N4:1266476 | 8,524 | | Intergene | | | |
| chrIV | 1267550 | 1267699 | 149 | -1,314929408 | N4:1267615 | 6,314 | +2:YDR398W; | 2 | | | |
| chrIV | 1269091 | 1269198 | 107 | -1,086760565 | N4:1269188 | 36,028 | +12:YDR398W; +4:CUT528; | 12 | | | |
| chrIV | 1269091 | 1269209 | 118 | -1,217623014 | N4:1269188 | 36,028 | +12:YDR398W; +4:CUT528; | 12 | | | |
| chrIV | 1272800 | 1272981 | 181 | -1,726560067 | N4:1272869 | 10,488 | +6:YDR402C; | 6 | | | |
| chrIV | 1277856 | 1277982 | 126 | -1,465556501 | N4:1277949 | 5,683 | +3:YDR405W; | 3 | | | |
| chrIV | 1278160 | 1278320 | 160 | -1,303550916 | N4:1278264 | 14,096 | +5:YDR405W; | 5 | | | |
| chrIV | 1287754 | 1287911 | 157 | -1,001161007 | N4:1287812 | 7,396 | +2:YDR407C; | 2 | | | |
| chrIV | 1288372 | 1288504 | 132 | -0,84042479 | | | | Overlap <50 bp | | | |
| chrIV | 1291651 | 1291799 | 148 | -1,278329605 | N4:1291819 | 23,549 | +3:SUT490; +15:YDR409W; | 3 | | | |
| chrIV | 1293635 | 1293774 | 139 | -1,279514777 | N4:1293718 | 2,658 | +5:YDR411C; | 5 | | | |
| chrIV | 1298059 | 1298223 | 164 | -1,165141825 | N4:1298152 | 26,529 | +1:YDR415C; -1:YDR416W; | 1 | | | |
| chrIV | 1300809 | 1300966 | 157 | -1,394797029 | N4:1300924 | 10,907 | +17*:YDR416W; +1:CUT532; | 17 | TERM | | |
| chrIV | 1300809 | 1300970 | 161 | -1,40182097 | N4:1300924 | 10,907 | +17*:YDR416W; +1:CUT532; | 17 | TERM | | |
| chrIV | 1300843 | 1301002 | 159 | -1,128882862 | N4:1300924 | 10,907 | +17*:YDR416W; +1:CUT532; | 17 | TERM | | |
| chrIV | 1300844 | 1300970 | 126 | -1,186655185 | N4:1300924 | 10,907 | +17*:YDR416W; +1:CUT532; | 17 | TERM | | |
| chrIV | 1300844 | 1300987 | 143 | -1,546729534 | N4:1300924 | 10,907 | +17*:YDR416W; +1:CUT532; | 17 | TERM | | |
| chrIV | 1302267 | 1302373 | 106 | -1,303502499 | N4:1302245 | 32,813 | +5*:YDR418W; | 5 | TERM | | |
| chrIV | 1302267 | 1302383 | 116 | -1,412978415 | N4:1302245 | 32,813 | +5*:YDR418W; | 5 | TERM | | |
| chrIV | 1308150 | 1308260 | 110 | -1,121372724 | N4:1308222 | 30,406 | | Intergene | | | |
| chrIV | 1308150 | 1308309 | 159 | -1,263627541 | N4:1308222 | 30,406 | | Intergene | | | |
| chrIV | 1308153 | 1308309 | 156 | -2,407150106 | N4:1308222 | 30,406 | | Intergene | | | |
| chrIV | 1308185 | 1308314 | 129 | -1,34375978 | N4:1308222 | 30,406 | | Intergene | | | |
| chrIV | 1308191 | 1308309 | 118 | -0,573116402 | N4:1308222 | 30,406 | | Intergene | | | |
| chrIV | 1308196 | 1308309 | 113 | -0,990823111 | N4:1308222 | 30,406 | | Intergene | | | |
| chrIV | 1308234 | 1308344 | 110 | -0,940517056 | N4:1308222 | 30,406 | | Intergene | | | |
| chrIV | 1308280 | 1308442 | 162 | -1,034589053 | N4:1308370 | 24,749 | | Intergene | | | |
| chrIV | 1308280 | 1308477 | 197 | -1,433198597 | N4:1308370 | 24,749 | | Intergene | | | |
| chrIV | 1308478 | 1308605 | 127 | -0,334029733 | N4:1308546 | 31,711 | | Intergene | | | |
| chrIV | 1308858 | 1308982 | 124 | -1,341140054 | N4:1308860 | 40,536 | | Intergene | | | |
| chrIV | 1312653 | 1312780 | 127 | -1,321916089 | N4:1312676 | 20,705 | +5:YDR421W; | 5 | | | |
| chrIV | 1312778 | 1312924 | 146 | -1,339573388 | N4:1312823 | 27,048 | +6:YDR421W; | 6 | | | |
| chrIV | 1324502 | 1324680 | 178 | -1,257165089 | N4:1324612 | 12,518 | 0:YDR428C; +5*:YDR429C; | 0 | | | |
| chrIV | 1324730 | 1324882 | 152 | -0,397433523 | N4:1324791 | 12,74 | -1:YDR428C; +4:YDR429C; | -1 | | | |
| chrIV | 1329033 | 1329189 | 156 | -1,267920286 | | | | Overlap <50 bp | | | |
| chrIV | 1331269 | 1331431 | 162 | -1,026872837 | N4:1331328 | 17,002 | +2:YDR434W; | 2 | | | |
| chrIV | 1332261 | 1332409 | 148 | -1,396982393 | N4:1332317 | 20,971 | +8:YDR434W; | 8 | | | |
| chrIV | 1334150 | 1334293 | 143 | -1,129181834 | N4:1334177 | 16,61 | -1:YDR435C; | -1 | | | |
| chrIV | 1334858 | 1335004 | 146 | -0,936310044 | N4:1334915 | 8,472 | +3:YDR436W; | 3 | | | |
| chrIV | 1335220 | 1335376 | 156 | -1,426561302 | N4:1335248 | 33,548 | +5:YDR436W; | 5 | | | |
| chrIV | 1335220 | 1335376 | 156 | -1,426561302 | N4:1335399 | 7,638 | +6:YDR436W; | 6 | | | |
| chrIV | 1337771 | 1337882 | 111 | 0,107701928 | N4:1337823 | 5,913 | +4:YDR437W; | 4 | | | |
| chrIV | 1341919 | 1342069 | 150 | -1,128627307 | N4:1342003 | 3,688 | +2:SUT492; | 2 | | | |

TABLE 3

| CHR | START | END | L(bp) | ΔLK | Nuc ID | Fuzziness | Gene ID | Gene body postion or intergene | Terminal | rDNA | Telomeric |
|---|---|---|---|---|---|---|---|---|---|---|---|
| chrIV | 1354730 | 1354858 | 128 | -1,075013376 | N4:1354868 | 44,223 | | Intergene | | | |
| chrIV | 1356736 | 1356889 | 153 | -1,668536338 | | | | Overlap <50 bp | | | |
| chrIV | 1359819 | 1359977 | 158 | -1,306578954 | | | | Overlap <50 bp | | | |
| chrIV | 1360674 | 1360822 | 148 | -1,150210204 | N4:1360796 | 29,792 | -1:CUT096; | -1 | | | |
| chrIV | 1361305 | 1361420 | 115 | -1,001790655 | N4:1361443 | 11,927 | +5:YDR451C; | 5 | | | |
| chrIV | 1361305 | 1361412 | 107 | -0,978890425 | N4:1361296 | 13,609 | +6:YDR451C; | 6 | | | |
| chrIV | 1361305 | 1361420 | 115 | -1,001790655 | N4:1361296 | 13,609 | +6:YDR451C; | 6 | | | |
| chrIV | 1364336 | 1364488 | 152 | -1,153204869 | N4:1364372 | 15,073 | +11:YDR452W; | 11 | | | |
| chrIV | 1364512 | 1364671 | 159 | -1,374921696 | N4:1364657 | 14,818 | +13*:YDR452W; | 13 | TERM | | |
| chrIV | 1365728 | 1365875 | 147 | -1,605996813 | N4:1365799 | 9,607 | 0:YDR453C; | 0 | | | |
| chrIV | 1365746 | 1365909 | 163 | -1,169500437 | N4:1365799 | 9,607 | 0:YDR453C; | 0 | | | |
| chrIV | 1367419 | 1367567 | 148 | -1,018619058 | N4:1367482 | 12,133 | +1:YDR456W; | 1 | | | |
| chrIV | 1367927 | 1368075 | 148 | -1,427036708 | N4:1367994 | 21,942 | +4:YDR456W; | 4 | | | |
| chrIV | 1372245 | 1372378 | 133 | -1,04382859 | N4:1372326 | 22,7 | | Intergene | | | |
| chrIV | 1376748 | 1376898 | 150 | -1,676420309 | N4:1376802 | 22,55 | | Intergene | | | |
| chrIV | 1382369 | 1382521 | 152 | -1,373429188 | N4:1382446 | 13,501 | -1:YDR458C; +7:YDR459C; | -1 | | | |
| chrIV | 1387683 | 1387833 | 150 | -1,181751444 | N4:1387776 | 33,429 | +7:YDR463W; | 7 | | | |
| chrIV | 1390839 | 1391009 | 170 | -1,613104723 | N4:1390850 | 23,902 | +14*:YDR464W; | 14 | TERM | | |
| chrIV | 1391435 | 1391580 | 145 | -1,2655942 | N4:1391591 | 9,68 | | Intergene | | | |
| chrIV | 1391940 | 1392060 | 120 | -1,358456605 | | | | Overlap <50 bp | | | |
| chrIV | 1397500 | 1397655 | 155 | -1,247298266 | | | | Overlap <50 bp | | | |
| chrIV | 1398087 | 1398256 | 169 | -1,306912439 | N4:1398124 | 27,844 | +5:YDR468C; | 5 | | | |
| chrIV | 1406033 | 1406179 | 146 | -1,297647896 | N4:1406110 | 15,237 | -1:YDR473C; +1:SUT075; | -1 | | | |
| chrIV | 1407803 | 1407943 | 140 | -1,273460181 | N4:1407944 | 22,59 | +13:YDR475C; +12:SUT075; | 13 | | | |
| chrIV | 1407803 | 1407943 | 140 | -1,273460181 | N4:1407780 | 16,765 | +14:YDR475C; +11:SUT075; | 14 | | | |
| chrIV | 1408060 | 1408202 | 142 | -1,276788755 | N4:1408111 | 21,788 | +12:YDR475C; +13*:SUT075; | 12 | | | |
| chrIV | 1408065 | 1408226 | 161 | -1,364439977 | N4:1408111 | 21,788 | +12:YDR475C; +13*:SUT075; | 12 | | | |
| chrIV | 1408340 | 1408489 | 149 | -1,235952361 | | | | Overlap <50 bp | | | |
| chrIV | 1409678 | 1409825 | 147 | -1,30147182 | N4:1409695 | 22,722 | +4:YDR475C; | 4 | | | |
| chrIV | 1413171 | 1413310 | 139 | -1,277423238 | N4:1413232 | 4,95 | +8:YDR477W; | 8 | | | |
| chrIV | 1420451 | 1420608 | 157 | -2,07211598 | N4:1420526 | 28,932 | -1:YDR481C; +3:YDR482C; | -1 | | | |
| chrIV | 1420630 | 1420804 | 174 | -1,310192133 | N4:1420827 | 38,771 | +1:YDR482C; | 1 | | | |
| chrIV | 1420630 | 1420804 | 174 | -1,310192133 | N4:1420679 | 4,967 | +2:YDR482C; | 2 | | | |
| chrIV | 1420634 | 1420804 | 170 | -1,615403162 | N4:1420827 | 38,771 | +1:YDR482C; | 1 | | | |
| chrIV | 1420634 | 1420804 | 170 | -1,615403162 | N4:1420679 | 4,967 | +2:YDR482C; | 2 | | | |
| chrIV | 1424831 | 1424973 | 142 | -1,027281053 | N4:1424916 | 22,382 | +17:YDR485C; | 17 | | | |
| chrIV | 1428254 | 1428414 | 160 | -1,543886236 | N4:1428344 | 13,05 | +5*:YDR487C; -1:YDR486C; | 5 | TERM | | |
| chrIV | 1430994 | 1431130 | 136 | -1,590789679 | N4:1431036 | 3,559 | +1:YDR489W; -1:YDR488C; | 1 | | | |
| chrIV | 1433662 | 1433813 | 151 | -1,562571203 | N4:1433787 | 23,266 | +4:YDR490C; | 4 | | | |
| chrIV | 1447757 | 1447907 | 150 | -1,173107294 | N4:1447835 | 10,405 | +1:YDR499W; | 1 | | | |
| chrIV | 1454001 | 1454153 | 152 | -1,299958983 | N4:1454073 | 16,281 | +3:YDR502C; | 3 | | | |
| chrIV | 1460987 | 1461146 | 159 | -1,241074773 | N4:1461050 | 8,39 | +4:YDR506C; | 4 | | | |
| chrIV | 1461681 | 1461828 | 147 | -1,264455085 | N4:1461801 | 44,636 | -1:YDR506C; | -1 | | | |
| chrIV | 1462856 | 1462994 | 138 | -1,219373496 | | | | Overlap <50 bp | | | |
| chrIV | 1462967 | 1463131 | 164 | -1,178123503 | N4:1463035 | 24,28 | +18:YDR507C; | 18 | | | |
| chrIV | 1464532 | 1464669 | 137 | -1,298784607 | N4:1464622 | 35,517 | +9:YDR507C; | 9 | | | |
| chrIV | 1465805 | 1465959 | 154 | -1,28869406 | N4:1465878 | 10,237 | +1:YDR507C; | 1 | | | |
| chrIV | 1466131 | 1466277 | 146 | -1,340047467 | N4:1466195 | 16,068 | 0:SUT080; | 0 | | | |
| chrIV | 1473970 | 1474117 | 147 | -1,394674278 | N4:1474065 | 31,316 | +6:YDR515W; | 6 | | | |
| chrIV | 1477488 | 1477647 | 159 | -1,281157312 | N4:1477526 | 8,981 | +3:YDR517W; | 3 | | | |
| chrIV | 1478568 | 1478697 | 129 | -1,364258552 | N4:1478612 | 15,796 | +1:YDR518W; | 1 | | | |
| chrIV | 1479228 | 1479388 | 160 | -1,226678961 | N4:1479283 | 15,192 | +5:YDR518W; | 5 | | | |
| chrIV | 1479727 | 1479884 | 157 | -1,243067692 | N4:1479790 | 8,746 | +8*:YDR518W; | 8 | TERM | | |
| chrIV | 1481305 | 1481467 | 162 | -1,491295083 | N4:1481450 | 14,184 | +13:YDR520C; | 13 | | | |
| chrIV | 1489675 | 1489832 | 157 | -1,37933781 | N4:1489795 | 32,828 | +1:YDR524C-B; | 1 | | | |
| chrIV | 1495628 | 1495753 | 125 | -1,108187242 | N4:1495663 | 31,925 | +10:YDR528W; | 10 | | | |
| chrIV | 1496435 | 1496575 | 140 | -1,351387013 | N4:1496536 | 27,154 | +9*:YDR530C; +1:YDR529C; | 9 | TERM | | |
| chrIV | 1497627 | 1497776 | 149 | -1,313031642 | N4:1497727 | 15,807 | +2:YDR530C; | 2 | | | |
| chrIV | 1498743 | 1498886 | 143 | -1,428867749 | N4:1498864 | 18,389 | +5:YDR531W; | 5 | | | |
| chrIV | 1499148 | 1499307 | 159 | -1,459238528 | N4:1499221 | 45,768 | +7*:YDR531W; | 7 | TERM | | |
| chrIV | 1503439 | 1503593 | 154 | -1,256177469 | N4:1503589 | 15,156 | +9:YDR534C; | 9 | | | |
| chrIV | 1505128 | 1505252 | 124 | -0,663450371 | N4:1505200 | 17,186 | -1:YDR534C; | -1 | | | |
| chrIV | 1507157 | 1507304 | 147 | -1,318227142 | N4:1507240 | 34,464 | | Intergene | | | |
| chrIV | 1512418 | 1512571 | 153 | -1,511941655 | N4:1512551 | 9,899 | +4:YDR539W; | 4 | | | |
| chrIV | 1512829 | 1512960 | 131 | -1,076663242 | N4:1512870 | 31,852 | +6:YDR539W; | 6 | | | |
| chrIV | 1526039 | 1526194 | 155 | -1,360461567 | N4:1526079 | 28,919 | -1:YDR545W; | -1 | | | TEL RIGHT |
| chrIV | 1527054 | 1527225 | 171 | -1,769058663 | N4:1527142 | 3,536 | +6:YDR545W; | 6 | | | TEL RIGHT |
| chrIV | 1527204 | 1527351 | 147 | -1,219791967 | | | | Overlap <50 bp | | | TEL RIGHT |
| chrIV | 1527360 | 1527490 | 130 | -1,068814589 | N4:1527407 | 23,065 | +8:YDR545W; | 8 | | | TEL RIGHT |
| chrIV | 1527741 | 1527877 | 136 | -1,307047804 | N4:1527764 | 0 | +10:YDR545W; | 10 | | | TEL RIGHT |
| chrIV | 1528122 | 1528284 | 162 | -1,393786404 | N4:1528239 | 10,97 | +13:YDR545W; | 13 | | | TEL RIGHT |
| chrIV | 1528223 | 1528378 | 155 | -1,87269017 | N4:1528239 | 10,97 | +13:YDR545W; | 13 | | | TEL RIGHT |
| chrIV | 1528357 | 1528494 | 137 | -1,121803696 | N4:1528410 | 10,693 | +14:YDR545W; | 14 | | | TEL RIGHT |
| chrIV | 1528652 | 1528795 | 143 | -1,551961505 | N4:1528689 | 11,358 | +16:YDR545W; | 16 | | | TEL RIGHT |
| chrIV | 1528700 | 1528825 | 125 | -0,803983472 | N4:1528689 | 11,358 | +16:YDR545W; | 16 | | | TEL RIGHT |
| chrIV | 1528925 | 1529077 | 152 | -1,353584297 | N4:1529021 | 12,423 | +18:YDR545W; | 18 | | | TEL RIGHT |
| chrIV | 1528998 | 1529142 | 144 | -1,152820753 | N4:1529021 | 12,423 | +18:YDR545W; | 18 | | | TEL RIGHT |
| chrIV | 1529095 | 1529230 | 135 | -1,010144152 | N4:1529177 | 23,388 | +19:YDR545W; | 19 | | | TEL RIGHT |
| chrIV | 1529095 | 1529282 | 187 | -0,855284919 | N4:1529177 | 23,388 | +19:YDR545W; | 19 | | | TEL RIGHT |
| chrIV | 1529138 | 1529230 | 92 | -0,439262005 | N4:1529177 | 23,388 | +19:YDR545W; | 19 | | | TEL RIGHT |
| chrIV | 1529138 | 1529282 | 144 | -0,977153589 | N4:1529177 | 23,388 | +19:YDR545W; | 19 | | | TEL RIGHT |
| chrIV | 1529477 | 1529629 | 152 | -1,311269828 | N4:1529532 | 30,172 | +21:YDR545W; | 21 | | | TEL RIGHT |
| chrIV | 1529778 | 1529908 | 130 | -1,260262391 | N4:1529917 | 35,341 | +23:YDR545W; | 23 | | | TEL RIGHT |
| chrIV | 1529916 | 1530077 | 161 | -1,426708642 | N4:1529917 | 35,341 | +23:YDR545W; | 23 | | | TEL RIGHT |
| chrIV | 1530973 | 1531125 | 152 | -1,537106359 | N4:1531041 | 9,899 | +30:YDR545W; | 30 | | | TEL RIGHT |
| chrIV | 1531313 | 1531464 | 151 | -1,227070303 | N4:1531365 | 42,454 | +32:YDR545W; | 32 | | | TEL RIGHT |
| chrIV | 1531313 | 1531484 | 171 | -1,547343973 | N4:1531365 | 42,454 | +32:YDR545W; | 32 | | | TEL RIGHT |
| chrIX | 1597 | 1749 | 152 | -1,19285005 | N9:1629 | 28,006 | +29:YIL177C; | 29 | | | TEL LEFT |
| chrIX | 1686 | 1802 | 116 | -0,514595173 | N9:1790 | 44,523 | +28:YIL177C; | 28 | | | TEL LEFT |
| chrIX | 1686 | 1821 | 135 | -0,780811757 | N9:1790 | 44,523 | +28:YIL177C; | 28 | | | TEL LEFT |
| chrIX | 1686 | 1838 | 152 | -1,270768931 | N9:1790 | 44,523 | +28:YIL177C; | 28 | | | TEL LEFT |
| chrIX | 1710 | 1802 | 92 | -0,511637481 | N9:1790 | 44,523 | +28:YIL177C; | 28 | | | TEL LEFT |
| chrIX | 1710 | 1811 | 101 | -0,829794123 | N9:1790 | 44,523 | +28:YIL177C; | 28 | | | TEL LEFT |
| chrIX | 1710 | 1821 | 111 | -0,472629278 | N9:1790 | 44,523 | +28:YIL177C; | 28 | | | TEL LEFT |
| chrIX | 1710 | 1838 | 128 | -0,833855878 | N9:1790 | 44,523 | +28:YIL177C; | 28 | | | TEL LEFT |
| chrIX | 1710 | 1857 | 147 | -1,203879749 | N9:1790 | 44,523 | +28:YIL177C; | 28 | | | TEL LEFT |
| chrIX | 1746 | 1838 | 92 | -0,982758461 | N9:1790 | 44,523 | +28:YIL177C; | 28 | | | TEL LEFT |
| chrIX | 2215 | 2352 | 137 | -1,09552453 | N9:2320 | 18,385 | +24:YIL177C; | 24 | | | TEL LEFT |
| chrIX | 2382 | 2529 | 147 | -1,219056028 | N9:2535 | 41,012 | +23:YIL177C; | 23 | | | TEL LEFT |

| CHR | START | END | L(bp) | ΔLK | Nuc ID | Fuzziness | Gene ID | Gene body postion or intergene | Terminal | rDNA | Telomeric |
|---|---|---|---|---|---|---|---|---|---|---|---|
| chrIX | 2382 | 2530 | 148 | -1,234116981 | N9:2535 | 41,012 | +23:YIL177C; | 23 | | | TEL LEFT |
| chrIX | 2382 | 2592 | 210 | -1,504650127 | N9:2535 | 41,012 | +23:YIL177C; | 23 | | | TEL LEFT |
| chrIX | 2383 | 2530 | 147 | -1,744892777 | N9:2535 | 41,012 | +23:YIL177C; | 23 | | | TEL LEFT |
| chrIX | 2385 | 2530 | 145 | -1,552370664 | N9:2535 | 41,012 | +23:YIL177C; | 23 | | | TEL LEFT |
| chrIX | 2397 | 2530 | 133 | -0,974052269 | N9:2535 | 41,012 | +23:YIL177C; | 23 | | | TEL LEFT |
| chrIX | 2437 | 2530 | 93 | -0,756768789 | N9:2535 | 41,012 | +23:YIL177C; | 23 | | | TEL LEFT |
| chrIX | 2437 | 2592 | 155 | -1,254118621 | N9:2535 | 41,012 | +23:YIL177C; | 23 | | | TEL LEFT |
| chrIX | 2558 | 2688 | 130 | -0,952862903 | N9:2535 | 41,012 | +23:YIL177C; | 23 | | | TEL LEFT |
| chrIX | 2879 | 3032 | 153 | -1,293194708 | N9:2921 | 22,301 | +21:YIL177C; | 21 | | | TEL LEFT |
| chrIX | 3324 | 3468 | 144 | -1,166378597 | N9:3423 | 11,533 | +18:YIL177C; | 18 | | | TEL LEFT |
| chrIX | 3531 | 3658 | 127 | -1,10511893 | N9:3591 | 6,028 | +17:YIL177C; | 17 | | | TEL LEFT |
| chrIX | 4088 | 4243 | 155 | -1,788390823 | N9:4203 | 6,807 | +13:YIL177C; | 13 | | | TEL LEFT |
| chrIX | 4805 | 4968 | 163 | -1,551739927 | N9:4855 | 10,504 | +9:YIL177C; | 9 | | | TEL LEFT |
| chrIX | 5216 | 5367 | 151 | -1,170904377 | | | | Overlap <50 bp | | | TEL LEFT |
| chrIX | 5429 | 5588 | 159 | -1,276265315 | N9:5453 | 17,243 | +5:YIL177C; | 5 | | | TEL LEFT |
| chrIX | 9290 | 9441 | 151 | -1,340792373 | N9:9376 | 3,055 | -1:YIL176C; | -1 | | | |
| chrIX | 10228 | 10371 | 143 | -1,0729291 | N9:10309 | 5,292 | | Intergene | | | |
| chrIX | 11460 | 11611 | 151 | -1,40406829 | N9:11554 | 17,898 | +2:YIL173W; | 2 | | | |
| chrIX | 12345 | 12490 | 145 | -1,126933457 | | | | Overlap <50 bp | | | |
| chrIX | 12910 | 13059 | 149 | -1,360544233 | | | | Overlap <50 bp | | | |
| chrIX | 14208 | 14357 | 149 | -1,045742171 | N9:14236 | 56,964 | +19:YIL173W; | 19 | | | |
| chrIX | 16078 | 16185 | 107 | -1,555620992 | N9:16064 | 4,041 | +30:YIL173W; | 30 | | | |
| chrIX | 25656 | 25806 | 150 | -1,434893897 | N9:25676 | 26,163 | +19:anti079; +3:YIL169C; | 19 | | | |
| chrIX | 32330 | 32492 | 162 | -1,150972996 | N9:32393 | 15,629 | +2:YIL166C; +8:anti080; | 2 | | | |
| chrIX | 33763 | 33917 | 154 | -1,447263707 | N9:33762 | 35,155 | +8*:YIL164C; +3:YIL165C; | 8 | TERM | | |
| chrIX | 35803 | 35935 | 132 | -0,854739376 | N9:35809 | 22,95 | +2:SUT602; +5:anti081; | 2 | | | |
| chrIX | 41284 | 41431 | 147 | -1,243953704 | N9:41356 | 6,94 | +2:YIL160C; +7:SUT177; | 2 | | | |
| chrIX | 48410 | 48570 | 160 | -1,015165438 | N9:48477 | 19,201 | +3:YIL156W; | 3 | | | |
| chrIX | 55336 | 55486 | 150 | -0,94436784 | N9:55401 | 7,95 | +2:YIL153W; | 2 | | | |
| chrIX | 55497 | 55655 | 158 | -1,306580637 | N9:55554 | 14,592 | +3:YIL153W; | 3 | | | |
| chrIX | 55498 | 55655 | 157 | -1,42961383 | N9:55554 | 14,592 | +3:YIL153W; | 3 | | | |
| chrIX | 55500 | 55655 | 155 | -1,689362206 | N9:55554 | 14,592 | +3:YIL153W; | 3 | | | |
| chrIX | 56537 | 56665 | 128 | -1,458850015 | N9:56598 | 12,734 | +1:YIL152W; | 1 | | | |
| chrIX | 56786 | 56893 | 107 | -1,081763297 | N9:56789 | 33,126 | +2:YIL152W; | 2 | | | |
| chrIX | 56786 | 56905 | 119 | -1,283552102 | N9:56789 | 33,126 | +2:YIL152W; | 2 | | | |
| chrIX | 56791 | 56905 | 114 | -1,007944781 | N9:56789 | 33,126 | +2:YIL152W; | 2 | | | |
| chrIX | 59003 | 59166 | 163 | -1,490248005 | N9:59158 | 26,885 | +11:YIL151C; | 11 | | | |
| chrIX | 59003 | 59166 | 163 | -1,490248005 | N9:58990 | 17,349 | +12:YIL151C; | 12 | | | |
| chrIX | 59591 | 59741 | 150 | -1,20306081 | N9:59671 | 26,95 | +8:YIL151C; | 8 | | | |
| chrIX | 60731 | 60881 | 150 | -1,431779436 | N9:60802 | 2,229 | +1:YIL151C; | 1 | | | |
| chrIX | 62677 | 62823 | 146 | -1,277875339 | N9:62723 | 10,226 | +1:YIL150C; | 1 | | | |
| chrIX | 70968 | 71118 | 150 | -1,494228197 | N9:71141 | 30,666 | +16:YIL147C; | 16 | | | |
| chrIX | 77046 | 77204 | 158 | -1,300750076 | N9:77127 | 9,75 | +2:YIL145C; | 2 | | | |
| chrIX | 78123 | 78268 | 145 | -1,175037582 | N9:78189 | 4,69 | +2:YIL144W; | 2 | | | |
| chrIX | 80413 | 80548 | 135 | -1,149998527 | N9:80482 | 19,858 | | Intergene | | | |
| chrIX | 89142 | 89281 | 139 | -1,314422241 | N9:89221 | 22,687 | +4:YIL138C; | 4 | | | |
| chrIX | 91580 | 91733 | 153 | -1,209591733 | N9:91642 | 18,061 | +8:YIL137C; | 8 | | | |
| chrIX | 93328 | 93477 | 149 | -1,010740899 | N9:93388 | 13,92 | -1:YIL136W; -1:Unit327; | -1 | | | |
| chrIX | 96377 | 96489 | 112 | -1,60214521 | N9:96391 | 20,798 | +1:YIL135C; +4*:YIL134C-A; | 1 | | | |
| chrIX | 97100 | 97255 | 155 | -1,657090971 | N9:97083 | 46,769 | -1:YIL134C-A; | -1 | | | |
| chrIX | 101159 | 101314 | 155 | -1,496745126 | N9:101310 | 9,055 | +7:YIL131C; | 7 | | | |
| chrIX | 101159 | 101314 | 155 | -1,496745126 | N9:101155 | 12,3 | +8:YIL131C; | 8 | | | |
| chrIX | 109382 | 109534 | 152 | -1,272202779 | N9:109439 | 6,5 | +23:YIL129C; | 23 | | | |
| chrIX | 122918 | 123072 | 154 | -1,512403055 | N9:122984 | 29,558 | +4:YIL125W; | 4 | | | |
| chrIX | 123420 | 123580 | 160 | -1,469104963 | N9:123402 | 18,464 | +7:YIL125W; | 7 | | | |
| chrIX | 123420 | 123580 | 160 | -1,469104963 | N9:123567 | 8,963 | +8:YIL125W; | 8 | | | |
| chrIX | 123668 | 123818 | 150 | -1,422071678 | N9:123750 | 15,526 | +9:YIL125W; | 9 | | | |
| chrIX | 127761 | 127867 | 106 | -1,181987067 | N9:127744 | 12,66 | | Intergene | | | |
| chrIX | 127761 | 127880 | 119 | -1,283622363 | N9:127744 | 12,66 | | Intergene | | | |
| chrIX | 129250 | 129396 | 146 | -1,023593811 | N9:129359 | 30,164 | +8:YIL123W; | 8 | | | |
| chrIX | 132115 | 132265 | 150 | -1,337879065 | N9:132195 | 23,529 | 0:YIL121W; | 0 | | | |
| chrIX | 140470 | 140605 | 135 | -1,48675792 | N9:140507 | 18,886 | +6*:YIL118W; | 6 | TERM | | |
| chrIX | 140794 | 140945 | 151 | -1,445811521 | N9:140849 | 18,093 | +6*:YIL117C; | 6 | TERM | | |
| chrIX | 145476 | 145609 | 133 | -0,878135101 | | | | Overlap <50 bp | | | |
| chrIX | 145755 | 145902 | 147 | -1,435251811 | N9:145791 | 22,42 | +19:YIL115C; | 19 | | | |
| chrIX | 149396 | 149568 | 172 | -1,480252043 | N9:149494 | 5,701 | +4:YIL114C; | 4 | | | |
| chrIX | 150158 | 150305 | 147 | -1,343166142 | N9:150234 | 4,733 | -1:YIL114C; -1:YIL113W; | -1 | | | |
| chrIX | 154400 | 154561 | 161 | -1,363998438 | N9:154551 | 18,708 | +3:anti083; | 3 | | | |
| chrIX | 157942 | 158056 | 114 | -1,088522929 | N9:157996 | 21,695 | | Intergene | | | |
| chrIX | 158500 | 158651 | 151 | -1,251719636 | | | | Overlap <50 bp | | | |
| chrIX | 159528 | 159680 | 152 | -1,138727302 | N9:159539 | 7,127 | +5:YIL109C; | 5 | | | |
| chrIX | 160370 | 160515 | 145 | -1,217825291 | N9:160446 | 9,899 | -1:YIL109C; -1:YIL108W; | -1 | | | |
| chrIX | 164945 | 165092 | 147 | -1,182457529 | N9:165040 | 31,488 | +6:YIL107C; | 6 | | | |
| chrIX | 167654 | 167783 | 129 | -1,064205949 | | | | Overlap <50 bp | | | |
| chrIX | 170117 | 170264 | 147 | -1,199911999 | N9:170186 | 8,017 | +9:YIL104C; | 9 | | | |
| chrIX | 175448 | 175623 | 175 | -1,18692502 | N9:175501 | 8,562 | +13:YIL101C; +2:anti085; | 13 | | | |
| chrIX | 175932 | 176081 | 149 | -1,245524268 | N9:175976 | 6,656 | +10:YIL101C; +5:anti085; | 10 | | | |
| chrIX | 176147 | 176286 | 139 | -1,160200596 | N9:176155 | 18,281 | +9:YIL101C; +6:anti085; | 9 | | | |
| chrIX | 178557 | 178730 | 173 | -1,111746071 | N9:178691 | 35,197 | +5:YIL099W; | 5 | | | |
| chrIX | 182864 | 183017 | 153 | -1,206324586 | N9:182946 | 10,559 | +2:YIL096C; | 2 | | | |
| chrIX | 190132 | 190312 | 180 | -1,63757736 | N9:190192 | 25,517 | +8:YIL092W; | 8 | | | |
| chrIX | 190978 | 191126 | 148 | -1,269345296 | N9:191107 | 40,441 | +13*:YIL092W; +14*:YIL091C; | 13 | TERM | | |
| chrIX | 192456 | 192591 | 135 | -1,123866091 | N9:192542 | 16,546 | +5:YIL091C; | 5 | | | |
| chrIX | 192983 | 193089 | 106 | -1,407425886 | N9:193029 | 7,026 | +2:YIL091C; | 2 | | | |
| chrIX | 192983 | 193098 | 115 | -1,379278188 | N9:193029 | 7,026 | +2:YIL091C; | 2 | | | |
| chrIX | 193702 | 193858 | 156 | -1,287039113 | N9:193815 | 11,619 | +3:YIL090W; | 3 | | | |
| chrIX | 196790 | 196938 | 148 | -1,126062896 | N9:196869 | 32,578 | +8*:YIL089W; | 8 | TERM | | |
| chrIX | 200251 | 200397 | 146 | -1,553507125 | N9:200295 | 31,754 | 0:YIL087C; | 0 | | | |
| chrIX | 205441 | 205561 | 120 | -1,228780623 | | | | Overlap <50 bp | | | |
| chrIX | 208655 | 208802 | 147 | -1,146288011 | N9:208742 | 9,394 | | Intergene | | | |
| chrIX | 212863 | 213023 | 160 | -1,269621762 | N9:212982 | 26,996 | +4:YIL078W; | 4 | | | |
| chrIX | 213578 | 213684 | 106 | -0,850217553 | | | | Overlap <50 bp | | | |
| chrIX | 213578 | 213686 | 108 | -0,800870433 | | | | Overlap <50 bp | | | |
| chrIX | 213578 | 213701 | 123 | -1,191552165 | | | | Overlap <50 bp | | | |
| chrIX | 213866 | 214017 | 151 | -1,642303779 | N9:213871 | 32,025 | +9:YIL078W; | 9 | | | |
| chrIX | 213866 | 214017 | 151 | -1,642303779 | N9:214029 | 15,819 | +10:YIL078W; | 10 | | | |
| chrIX | 217876 | 218011 | 135 | -0,888586327 | N9:217888 | 17,138 | +18*:YIL075C; | 18 | TERM | | |
| chrIX | 218740 | 218900 | 160 | -1,346679022 | N9:218850 | 30,111 | +12:YIL075C; | 12 | | | |
| chrIX | 223366 | 223481 | 115 | -1,505521161 | N9:223420 | 12,287 | +4:SUT183; +17:YIL073C; | 4 | | | |

TABLE 3

| CHR | START | END | L(bp) | ΔLK | Nuc ID | Fuzziness | Gene ID | Gene body postion or intergene | Terminal | rDNA | Telomeric |
|---|---|---|---|---|---|---|---|---|---|---|---|
| chrIX | 224421 | 224561 | 140 | -1,345669602 | N9:224566 | 36,238 | +10:YIL073C; | 10 | | | |
| chrIX | 236168 | 236319 | 151 | -1,008775557 | N9:236230 | 10,784 | +10:YIL067C; | 10 | | | |
| chrIX | 240288 | 240448 | 160 | -1,287572037 | N9:240358 | 6,633 | +3:YIL066C; | 3 | | | |
| chrIX | 241031 | 241216 | 185 | -1,062089726 | N9:241153 | 13,773 | | Intergene | | | |
| chrIX | 241349 | 241481 | 132 | -0,897409324 | | | | Overlap <50 bp | | | |
| chrIX | 244655 | 244803 | 148 | -1,264391587 | N9:244726 | 6,861 | -1:YIL062C; +6:YIL061C; | -1 | | | |
| chrIX | 244820 | 244976 | 156 | -1,255243575 | N9:244898 | 3,05 | +5:YIL061C; | 5 | | | |
| chrIX | 244822 | 244983 | 161 | -1,224404438 | N9:244898 | 3,05 | +5:YIL061C; | 5 | | | |
| chrIX | 247146 | 247297 | 151 | -1,908432857 | N9:247208 | 15,362 | | Intergene | | | |
| chrIX | 249643 | 249773 | 130 | -1,023043686 | N9:249728 | 29,438 | +1:Unit338; +1:YIL056W; | 1 | | | |
| chrIX | 254780 | 254919 | 139 | -1,536526198 | N9:254874 | 41,004 | -1:YIL053W; | -1 | | | |
| chrIX | 255472 | 255626 | 154 | -1,218539025 | N9:255553 | 51,733 | | Intergene | | | |
| chrIX | 255629 | 255772 | 143 | -1,277251977 | N9:255778 | 22,427 | -1:Unit339; | -1 | | | |
| chrIX | 255629 | 255776 | 147 | -1,10792192 | N9:255778 | 22,427 | -1:Unit339; | -1 | | | |
| chrIX | 258137 | 258291 | 154 | -1,870358384 | N9:258298 | 11,077 | +1:YIL051C; | 1 | | | |
| chrIX | 258137 | 258291 | 154 | -1,870358384 | N9:258138 | 25,962 | +2:YIL051C; | 2 | | | |
| chrIX | 263024 | 263196 | 172 | -0,786258258 | N9:263062 | 25,173 | | Intergene | | | |
| chrIX | 265122 | 265273 | 151 | -1,111649668 | N9:265235 | 27,545 | +17:YIL047C; | 17 | | | |
| chrIX | 265238 | 265355 | 117 | -1,936674668 | N9:265235 | 27,545 | +17:YIL047C; | 17 | | | |
| chrIX | 270073 | 270226 | 153 | -1,240937102 | N9:270153 | 16,633 | +11:YIL046W; | 11 | | | |
| chrIX | 271691 | 271837 | 146 | -1,385880416 | N9:271766 | 24,031 | +5:YIL045W; | 5 | | | |
| chrIX | 280995 | 281156 | 161 | -1,133865779 | N9:281073 | 36,927 | +12:YIL038C; 0:anti088; | 12 | | | |
| chrIX | 282582 | 282731 | 149 | -1,390150929 | N9:282678 | 15,398 | +2:YIL038C; | 2 | | | |
| chrIX | 283375 | 283539 | 164 | -1,610878615 | N9:283458 | 4,179 | +3:SUT186; +10:YIL037C; | 3 | | | |
| chrIX | 285840 | 285993 | 153 | -1,35765103 | N9:285843 | 8,198 | +3:YIL036W; | 3 | | | |
| chrIX | 285840 | 285993 | 153 | -1,35765103 | N9:286010 | 2,927 | +4:YIL036W; | 4 | | | |
| chrIX | 286233 | 286373 | 140 | -0,663978152 | | | | Overlap <50 bp | | | |
| chrIX | 287987 | 288121 | 134 | -1,395324736 | N9:288045 | 4,561 | +6:YIL035C; | 6 | | | |
| chrIX | 289697 | 289831 | 134 | -1,000710493 | N9:289737 | 11,367 | +4:SUT187; +3:YIL034C; | 4 | | | |
| chrIX | 289822 | 289971 | 149 | -1,353811109 | N9:289899 | 4,435 | +5:SUT187; +2:YIL034C; | 5 | | | |
| chrIX | 292476 | 292604 | 128 | -0,939002874 | N9:292512 | 11,983 | +1:YIL031W; | 1 | | | |
| chrIX | 296896 | 297052 | 156 | -1,36382984 | N9:297004 | 41,557 | +19:YIL030C; | 19 | | | |
| chrIX | 299139 | 299300 | 161 | -1,203494257 | N9:299210 | 11,236 | +6:YIL030C; | 6 | | | |
| chrIX | 301522 | 301672 | 150 | -1,30602044 | N9:301595 | 3,869 | -1:YIL029C; | -1 | | | |
| chrIX | 301692 | 301851 | 159 | -1,409464246 | N9:301782 | 5,888 | | Intergene | | | |
| chrIX | 301709 | 301857 | 148 | -1,219807588 | N9:301782 | 5,888 | | Intergene | | | |
| chrIX | 306823 | 306944 | 121 | -1,25645718 | N9:306927 | 20,499 | +7:YIL026C; | 7 | | | |
| chrIX | 309339 | 309491 | 152 | -1,452785614 | N9:309417 | 6,261 | +7*:YIL023C; 0:YIL024C; | 7 | TERM | | |
| chrIX | 309860 | 310002 | 142 | -1,1666166 | N9:309904 | 8,556 | +4:YIL023C; | 4 | | | |
| chrIX | 314424 | 314578 | 154 | -1,126397982 | | | | Overlap <50 bp | | | |
| chrIX | 320445 | 320583 | 138 | -1,352205055 | N9:320481 | 17,63 | +4:YIL017C; | 4 | | | |
| chrIX | 331176 | 331345 | 169 | -1,427806791 | N9:331307 | 44,998 | +20:SUT189; +8:YIL013C; | 20 | | | |
| chrIX | 332149 | 332296 | 147 | -1,279913126 | N9:332195 | 9,559 | +3:YIL013C; | 3 | | | |
| chrIX | 333705 | 333871 | 166 | -1,644484936 | N9:333780 | 13,301 | +6:SUT613; +1:YIL011W; | 6 | | | |
| chrIX | 338938 | 339077 | 139 | -0,895412862 | N9:338970 | 23,377 | -1:Unit331; -1:YIL009W; | -1 | | | |
| chrIX | 339456 | 339607 | 151 | -1,091095368 | N9:339523 | 8,112 | +2:YIL009W; | 2 | | | |
| chrIX | 343521 | 343669 | 148 | -1,234215454 | N9:343647 | 10,922 | +1:YIL007C; | 1 | | | |
| chrIX | 345865 | 346018 | 153 | -0,63832697 | N9:345939 | 13,426 | +3:YIL005W; | 3 | | | |
| chrIX | 346400 | 346566 | 166 | -1,029617311 | N9:346455 | 20,253 | +6:YIL005W; | 6 | | | |
| chrIX | 352082 | 352233 | 151 | -1,647800673 | N9:352150 | 27,725 | +9:YIL002C; | 9 | | | |
| chrIX | 360987 | 361138 | 151 | -1,210703454 | N9:361056 | 4,438 | +2:YIR003W; | 2 | | | |
| chrIX | 361485 | 361638 | 153 | -1,042543361 | N9:361563 | 10,515 | +5:YIR003W; | 5 | | | |
| chrIX | 364237 | 364386 | 149 | -1,216916821 | N9:364329 | 13,916 | +8:YIR004W; | 8 | | | |
| chrIX | 365450 | 365589 | 139 | -1,217638974 | N9:365504 | 8,687 | +28*:YIR006C; | 28 | TERM | | |
| chrIX | 371073 | 371217 | 144 | -1,179461302 | N9:371134 | 4,262 | +3:YIR007W; | 3 | | | |
| chrIX | 377621 | 377774 | 153 | -1,444311502 | N9:377728 | 9,663 | +4:YIR011C; | 4 | | | |
| chrIX | 378406 | 378536 | 130 | -1,10479358 | N9:378506 | 4,561 | +1:YIR012W; -1:YIR011C; | 1 | | | |
| chrIX | 380201 | 380307 | 106 | -1,441369505 | N9:380285 | 34,457 | +2:YIR013C; | 2 | | | |
| chrIX | 380201 | 380310 | 109 | -1,462718401 | N9:380285 | 34,457 | +2:YIR013C; | 2 | | | |
| chrIX | 381638 | 381781 | 143 | -1,490053572 | N9:381710 | 16,293 | CUT669; 0:YIR015W; +5*:YIR014W; | 4 | | | |
| chrIX | 382286 | 382451 | 165 | -1,70319655 | N9:382299 | 17,578 | :CUT669; +4:YIR015W; -1:YIR016W; | -1 | | | |
| chrIX | 386066 | 386219 | 153 | -1,163326126 | N9:386093 | 32,897 | | Intergene | | | |
| chrIX | 386767 | 386910 | 143 | -1,348305049 | N9:386861 | 26,153 | | Intergene | | | |
| chrIX | 387904 | 388014 | 110 | -1,331240035 | N9:387935 | 7,047 | +1:Unit342; | 1 | | | |
| chrIX | 388900 | 389051 | 151 | -1,589890837 | N9:389018 | 7,448 | -1:SUT194; | -1 | | | |
| chrIX | 390177 | 390342 | 165 | -1,351704484 | N9:390256 | 34,664 | +7:SUT194; | 7 | | | |
| chrIX | 390386 | 390532 | 146 | -1,277423677 | N9:390414 | 30,376 | +8:SUT194; | 8 | | | |
| chrIX | 392788 | 392939 | 151 | -1,079274103 | N9:392855 | 31,658 | +23:SUT194; +6*:YIR019C; | 23 | | | |
| chrIX | 396135 | 396302 | 167 | -1,350706304 | N9:396197 | 15,385 | | Intergene | | | |
| chrIX | 396795 | 396922 | 127 | -1,239545508 | N9:396865 | 11,996 | | Intergene | | | |
| chrIX | 405278 | 405390 | 112 | -1,510251564 | N9:405412 | 16,415 | +4:YIR026C; | 4 | | | |
| chrIX | 407378 | 407518 | 140 | -1,116357612 | N9:407408 | 5,958 | +3:YIR027C; +5:SUT195; | 3 | | | |
| chrIX | 407520 | 407679 | 159 | -1,472422306 | N9:407581 | 5,02 | +2:YIR027C; +6:SUT195; | 2 | | | |
| chrIX | 409036 | 409181 | 145 | -1,184197753 | N9:409099 | 16,245 | +7:SUT614; +5:YIR028W; | 7 | | | |
| chrIX | 420519 | 420625 | 106 | -1,493833426 | N9:420590 | 6,863 | +2:YIR034C; | 2 | | | |
| chrIX | 425922 | 426069 | 147 | -1,405452334 | N9:426001 | 7,441 | | Intergene | | | |
| chrIX | 433552 | 433722 | 170 | -1,450340168 | N9:433692 | 21,749 | -1:YIR041W; | -1 | | | |
| chrIX | 438504 | 438651 | 147 | -1,433835083 | N9:438544 | 28,569 | | Intergene | | | |
| chrIX | 438570 | 438711 | 141 | -1,262801477 | | | | Overlap <50 bp | | | |
| chrIX | 439579 | 439688 | 109 | -1,10793233 | N9:439641 | 36,222 | | Intergene | | | TEL RIGHT |
| chrIX | 439796 | 439887 | 91 | -1,086286073 | N9:439845 | 9,539 | | Intergene | | | TEL RIGHT |
| chrV | 1165 | 1320 | 155 | -1,405729016 | | | | Overlap <50 bp | | | TEL LEFT |
| chrV | 1854 | 1981 | 127 | -0,494765301 | N5:1841 | 19,33 | +16:YEL077C; | 16 | | | TEL LEFT |
| chrV | 3000 | 3144 | 144 | -0,959470146 | N5:3094 | 3,055 | +8:YEL077C; | 8 | | | TEL LEFT |
| chrV | 3648 | 3785 | 137 | -1,296180049 | N5:3709 | 8,66 | +4:YEL077C; | 4 | | | TEL LEFT |
| chrV | 3764 | 3919 | 155 | -2,003651267 | N5:3872 | 9,849 | +3:YEL077C; | 3 | | | TEL LEFT |
| chrV | 4024 | 4184 | 160 | -1,282456513 | N5:4031 | 6,557 | +8*:YEL076C-A; +2:YEL077C; | 8 | TERM | | TEL LEFT |
| chrV | 4831 | 4982 | 151 | -1,210534144 | N5:4948 | 15,885 | +2:YEL076C; +2:YEL076C-A; | 2 | | | TEL LEFT |
| chrV | 7115 | 7272 | 157 | -1,565806674 | N5:7242 | 34,646 | +4:YEL073C; | 4 | | | |
| chrV | 9760 | 9930 | 170 | -0,698113174 | | | | Overlap <50 bp | | | |
| chrV | 11061 | 11170 | 109 | -1,55023065 | N5:11123 | 2,858 | +2:SUT501; | 2 | | | |
| chrV | 23046 | 23205 | 159 | -1,22357103 | N5:23195 | 30,03 | +1*:YEL069C; | 1 | TERM | | |
| chrV | 26697 | 26869 | 172 | -1,149432758 | N5:26684 | 26,838 | +1:YEL066W; -1:YEL067C; | 1 | | | |
| chrV | 26697 | 26869 | 172 | -1,149432758 | N5:26878 | 19,396 | +2:YEL066W; | 2 | | | |
| chrV | 27053 | 27184 | 131 | -0,905165587 | N5:27040 | 24,698 | +3*:YEL066W; | 3 | TERM | | |
| chrV | 28903 | 29024 | 121 | -1,082490146 | N5:29008 | 21,777 | +9:YEL065W; | 9 | | | |
| chrV | 32832 | 32992 | 160 | -1,574941414 | N5:32875 | 40,033 | +5:YEL063C; | 5 | | | |
| chrV | 41648 | 41795 | 147 | -1,507225129 | N5:41813 | 32,66 | +2:YEL060C; | 2 | | | |
| chrV | 41648 | 41797 | 149 | -1,1411875 | N5:41813 | 32,66 | +2:YEL060C; | 2 | | | |

| CHR | START | END | L(bp) | ΔLK | Nuc ID | Fuzziness | Gene ID | Gene body postion or intergene | Terminal | rDNA | Telomeric |
|---|---|---|---|---|---|---|---|---|---|---|---|
| chrV | 41648 | 41798 | 150 | -1,205541708 | N5:41813 | 32,66 | +2:YEL060C; | 2 | | | |
| chrV | 41648 | 41799 | 151 | -1,129014846 | N5:41813 | 32,66 | +2:YEL060C; | 2 | | | |
| chrV | 41648 | 41800 | 152 | -1,644917889 | N5:41813 | 32,66 | +2:YEL060C; | 2 | | | |
| chrV | 41648 | 41801 | 153 | -1,645272097 | N5:41813 | 32,66 | +2:YEL060C; | 2 | | | |
| chrV | 41648 | 41802 | 154 | -1,156441591 | N5:41813 | 32,66 | +2:YEL060C; | 2 | | | |
| chrV | 41648 | 41754 | 106 | -1,780909181 | N5:41658 | 22,906 | +3:YEL060C; | 3 | | | |
| chrV | 41648 | 41774 | 126 | -1,047649136 | N5:41658 | 22,906 | +3:YEL060C; | 3 | | | |
| chrV | 41648 | 41782 | 134 | -2,087017223 | N5:41658 | 22,906 | +3:YEL060C; | 3 | | | |
| chrV | 41648 | 41785 | 137 | -1,362538346 | N5:41658 | 22,906 | +3:YEL060C; | 3 | | | |
| chrV | 41648 | 41795 | 147 | -1,507225129 | N5:41658 | 22,906 | +3:YEL060C; | 3 | | | |
| chrV | 41648 | 41797 | 149 | -1,1411875 | N5:41658 | 22,906 | +3:YEL060C; | 3 | | | |
| chrV | 41648 | 41798 | 150 | -1,205541708 | N5:41658 | 22,906 | +3:YEL060C; | 3 | | | |
| chrV | 41648 | 41799 | 151 | -1,129014846 | N5:41658 | 22,906 | +3:YEL060C; | 3 | | | |
| chrV | 41648 | 41800 | 152 | -1,644917889 | N5:41658 | 22,906 | +3:YEL060C; | 3 | | | |
| chrV | 41648 | 41801 | 153 | -1,645272097 | N5:41658 | 22,906 | +3:YEL060C; | 3 | | | |
| chrV | 41648 | 41802 | 154 | -1,156441591 | N5:41658 | 22,906 | +3:YEL060C; | 3 | | | |
| chrV | 41649 | 41802 | 153 | -1,713590505 | N5:41813 | 32,66 | +2:YEL060C; | 2 | | | |
| chrV | 41649 | 41802 | 153 | -1,713590505 | N5:41658 | 22,906 | +3:YEL060C; | 3 | | | |
| chrV | 41650 | 41802 | 152 | -1,781225545 | N5:41813 | 32,66 | +2:YEL060C; | 2 | | | |
| chrV | 41650 | 41802 | 152 | -1,781225545 | N5:41658 | 22,906 | +3:YEL060C; | 3 | | | |
| chrV | 41651 | 41802 | 151 | -1,518929443 | N5:41813 | 32,66 | +2:YEL060C; | 2 | | | |
| chrV | 41651 | 41802 | 151 | -1,518929443 | N5:41658 | 22,906 | +3:YEL060C; | 3 | | | |
| chrV | 41652 | 41802 | 150 | -1,309938999 | N5:41813 | 32,66 | +2:YEL060C; | 2 | | | |
| chrV | 41652 | 41802 | 150 | -1,309938999 | N5:41658 | 22,906 | +3:YEL060C; | 3 | | | |
| chrV | 41653 | 41802 | 149 | -1,557833551 | N5:41813 | 32,66 | +2:YEL060C; | 2 | | | |
| chrV | 41653 | 41802 | 149 | -1,557833551 | N5:41658 | 22,906 | +3:YEL060C; | 3 | | | |
| chrV | 41654 | 41802 | 148 | -1,615267201 | N5:41813 | 32,66 | +2:YEL060C; | 2 | | | |
| chrV | 41654 | 41802 | 148 | -1,615267201 | N5:41658 | 22,906 | +3:YEL060C; | 3 | | | |
| chrV | 41658 | 41802 | 144 | -1,857621742 | N5:41813 | 32,66 | +2:YEL060C; | 2 | | | |
| chrV | 41658 | 41802 | 144 | -1,857621742 | N5:41658 | 22,906 | +3:YEL060C; | 3 | | | |
| chrV | 42773 | 42909 | 136 | -1,165453148 | N5:42882 | 7,448 | -1:YEL059C-A; | -1 | | | |
| chrV | 44013 | 44174 | 161 | -1,187848156 | N5:44097 | 21,735 | +6:YEL058W; | 6 | | | |
| chrV | 51922 | 52070 | 148 | -1,096922028 | N5:52074 | 19,837 | 0:CUT543; | 0 | | | |
| chrV | 56549 | 56680 | 131 | -1,058590095 | N5:56569 | 10,147 | +1:YEL052W; | 1 | | | |
| chrV | 58655 | 58795 | 140 | -1,068121843 | N5:58725 | 20,849 | +3:YEL051W; | 3 | | | |
| chrV | 63504 | 63668 | 164 | -1,378907037 | N5:63566 | 28,732 | 0:YEL049W; +2:CUT546; | 0 | | | |
| chrV | 64024 | 64155 | 131 | -1,33630923 | N5:64053 | 24,166 | +3:YEL049W; | 3 | | | |
| chrV | 65475 | 65628 | 153 | -1,237339626 | N5:65581 | 5,119 | +8:YEL047C; | 8 | | | |
| chrV | 69437 | 69601 | 164 | -1,159649181 | N5:69487 | 41,142 | -1:YEL044W; | -1 | | | |
| chrV | 71549 | 71648 | 99 | -1,379388459 | N5:71615 | 9,968 | +9:YEL043W; | 9 | | | |
| chrV | 72028 | 72188 | 160 | -1,409697122 | N5:72094 | 8,562 | | Intergene | | | |
| chrV | 78159 | 78308 | 149 | -1,104142081 | N5:78154 | 26,666 | +1:YEL040W; | 1 | | | |
| chrV | 78159 | 78308 | 149 | -1,104142081 | N5:78324 | 39,084 | +2:YEL040W; | 2 | | | |
| chrV | 78447 | 78544 | 97 | -1,153851563 | N5:78484 | 28,766 | +3:YEL040W; | 3 | | | |
| chrV | 78782 | 78922 | 140 | -1,107250967 | N5:78847 | 12,641 | +5:YEL040W; | 5 | | | |
| chrV | 79095 | 79249 | 154 | -1,176006052 | N5:79182 | 9,43 | +7:YEL040W; | 7 | | | |
| chrV | 83616 | 83770 | 154 | -1,169693679 | N5:83673 | 9,737 | +7:YEL036C; | 7 | | | |
| chrV | 84397 | 84539 | 142 | -1,04400915 | N5:84467 | 1,472 | +2:YEL036C; | 2 | | | |
| chrV | 87005 | 87158 | 153 | -1,002422033 | N5:87089 | 11,777 | +2:YEL032W; | 2 | | | |
| chrV | 92533 | 92664 | 131 | -1,161891156 | N5:92614 | 23,554 | +16:YEL031W; | 16 | | | |
| chrV | 94783 | 94926 | 143 | -1,091911591 | N5:94888 | 17,871 | +3:YEL030W; | 3 | | | |
| chrV | 96167 | 96341 | 174 | -0,965810655 | N5:96189 | 45,96 | +11:YEL030W; | 11 | | | |
| chrV | 105359 | 105516 | 157 | -1,676054623 | N5:105459 | 20,025 | +5:YEL025C; | 5 | | | |
| chrV | 105504 | 105653 | 149 | -0,844524669 | N5:105613 | 9,011 | +4:YEL025C; | 4 | | | |
| chrV | 106502 | 106638 | 136 | -1,339012408 | N5:106562 | 16,709 | | Intergene | | | |
| chrV | 112707 | 112854 | 147 | -1,16041213 | N5:112767 | 39,414 | +10:YEL022W; | 10 | | | |
| chrV | 114004 | 114167 | 163 | -1,380709464 | N5:114084 | 25,367 | +18:YEL022W; | 18 | | | |
| chrV | 116119 | 116272 | 153 | -1,303927027 | N5:116132 | 21,032 | +1:YEL021W; | 1 | | | |
| chrV | 116657 | 116785 | 128 | -1,344624347 | N5:116788 | 20,551 | +5:YEL021W; | 5 | | | |
| chrV | 119860 | 120019 | 159 | -1,365154346 | N5:119994 | 9,849 | +3:YEL020C; +9:YEL019C; | 3 | | | |
| chrV | 121418 | 121572 | 154 | -1,211805922 | N5:121505 | 5,231 | +1:YEL018W; -1:YEL019C; | 1 | | | |
| chrV | 127589 | 127741 | 152 | -1,195535731 | N5:127639 | 34,888 | +7:YEL015W; | 7 | | | |
| chrV | 129771 | 129919 | 148 | -1,19688081 | N5:129920 | 19,722 | +9:YEL013W; | 9 | | | |
| chrV | 132213 | 132348 | 135 | -1,355209347 | N5:132209 | 5,086 | | Intergene | | | |
| chrV | 134881 | 135033 | 152 | -1,498693151 | N5:134908 | 8,562 | +4:SUT504; +12*:YEL011W; | 4 | | | |
| chrV | 137057 | 137219 | 162 | -1,311213349 | N5:137135 | 14,152 | | Intergene | | | |
| chrV | 137099 | 137245 | 146 | -1,237627772 | N5:137135 | 14,152 | | Intergene | | | |
| chrV | 143216 | 143369 | 153 | -0,922303719 | N5:143330 | 30,179 | +12:YEL007W; | 12 | | | |
| chrV | 146039 | 146192 | 153 | -1,202559763 | N5:146118 | 13,967 | +4:YEL005C; | 4 | | | |
| chrV | 147368 | 147525 | 157 | -1,452076532 | N5:147443 | 6,113 | +4:YEL004W; | 4 | | | |
| chrV | 147862 | 148011 | 149 | -1,3070106 | N5:147935 | 7,259 | +7:YEL004W; -1:YEL003W; | 7 | | | |
| chrV | 150905 | 151028 | 123 | -1,479539424 | N5:150967 | 7,468 | +5:CUT106; +1:YEL001C; | 5 | | | |
| chrV | 154668 | 154823 | 155 | -1,529104692 | N5:154737 | 8,756 | +8:YER001W; | 8 | | | |
| chrV | 154964 | 155109 | 145 | -1,605098809 | N5:155071 | 18,679 | +10:YER001W; | 10 | | | |
| chrV | 156735 | 156886 | 151 | -1,693406271 | N5:156804 | 11,274 | +1:YER002W; | 1 | | | |
| chrV | 157741 | 157900 | 159 | -0,820126426 | N5:157806 | 25,965 | | Intergene | | | |
| chrV | 158361 | 158517 | 156 | -1,457691927 | N5:158435 | 16,216 | | Intergene | | | |
| chrV | 158999 | 159169 | 170 | -1,433168259 | N5:159105 | 7,396 | +1*:YER003C; | 1 | TERM | | |
| chrV | 161114 | 161267 | 153 | -1,34370096 | N5:161151 | 18,777 | +4:YER005W; | 4 | | | |
| chrV | 162828 | 162978 | 150 | -1,387541174 | N5:162902 | 13,027 | +2:YER006W; | 2 | | | |
| chrV | 162990 | 163135 | 145 | -1,132688929 | N5:163066 | 4,461 | +3:YER006W; | 3 | | | |
| chrV | 163162 | 163326 | 164 | -1,209782358 | N5:163230 | 22,107 | +4:YER006W; | 4 | | | |
| chrV | 164105 | 164236 | 131 | -0,778678852 | N5:164174 | 27,019 | -1:YER007W; +10:YER006W; | -1 | | | |
| chrV | 165644 | 165751 | 107 | -0,820742287 | N5:165694 | 6,38 | +9:YER007W; | 9 | | | |
| chrV | 165645 | 165751 | 106 | -1,131996043 | N5:165694 | 6,38 | +9:YER007W; | 9 | | | |
| chrV | 165645 | 165753 | 108 | -1,381562961 | N5:165694 | 6,38 | +9:YER007W; | 9 | | | |
| chrV | 165645 | 165770 | 125 | -1,148063114 | N5:165694 | 6,38 | +9:YER007W; | 9 | | | |
| chrV | 169801 | 169967 | 166 | -1,79464542 | N5:169892 | 22,692 | +13:YER008C; | 13 | | | |
| chrV | 170735 | 170894 | 159 | -1,456763496 | N5:170840 | 12,383 | +7:YER008C; | 7 | | | |
| chrV | 175783 | 175962 | 179 | -1,516190621 | N5:175893 | 20,008 | +6:YER011W; | 6 | | | |
| chrV | 177369 | 177538 | 169 | -1,406530292 | N5:177535 | 9,311 | +1:SUT508; -1:YER012W; | 1 | | | |
| chrV | 177369 | 177538 | 169 | -1,406530292 | N5:177381 | 24,234 | +2:SUT508; | 2 | | | |
| chrV | 177766 | 177914 | 148 | -1,482333537 | N5:177821 | 12,858 | -1:SUT508; +1:YER012W; | -1 | | | |
| chrV | 179498 | 179607 | 109 | -2,015610091 | N5:179488 | 12,644 | +5:YER013W; | 5 | | | |
| chrV | 179498 | 179665 | 167 | -2,004173751 | N5:179488 | 12,644 | +5:YER013W; | 5 | | | |
| chrV | 179498 | 179666 | 168 | -1,63857675 | N5:179488 | 12,644 | +5:YER013W; | 5 | | | |
| chrV | 179498 | 179667 | 169 | -1,680425085 | N5:179488 | 12,644 | +5:YER013W; | 5 | | | |
| chrV | 179498 | 179669 | 171 | -1,75983774 | N5:179488 | 12,644 | +5:YER013W; | 5 | | | |
| chrV | 179498 | 179670 | 172 | -1,342618679 | N5:179488 | 12,644 | +5:YER013W; | 5 | | | |
| chrV | 179498 | 179665 | 167 | -2,004173751 | N5:179670 | 27,842 | +6:YER013W; | 6 | | | |

| CHR | START | END | L(bp) | ΔLK | Nuc ID | Fuzziness | Gene ID | Gene body postion or intergene | Terminal | rDNA | Telomeric |
|---|---|---|---|---|---|---|---|---|---|---|---|
| chrV | 179498 | 179666 | 168 | -1,63857675 | N5:179670 | 27,842 | +6:YER013W; | 6 | | | |
| chrV | 179498 | 179667 | 169 | -1,680425085 | N5:179670 | 27,842 | +6:YER013W; | 6 | | | |
| chrV | 179498 | 179669 | 171 | -1,75983774 | N5:179670 | 27,842 | +6:YER013W; | 6 | | | |
| chrV | 179498 | 179670 | 172 | -1,342618679 | N5:179670 | 27,842 | +6:YER013W; | 6 | | | |
| chrV | 179499 | 179670 | 171 | -1,445092566 | N5:179488 | 12,644 | +5:YER013W; | 5 | | | |
| chrV | 179499 | 179670 | 171 | -1,445092566 | N5:179670 | 27,842 | +6:YER013W; | 6 | | | |
| chrV | 179502 | 179670 | 168 | -1,841093399 | N5:179488 | 12,644 | +5:YER013W; | 5 | | | |
| chrV | 179502 | 179670 | 168 | -1,841093399 | N5:179670 | 27,842 | +6:YER013W; | 6 | | | |
| chrV | 180478 | 180630 | 152 | -1,289395641 | N5:180475 | 42,543 | +11:YER013W; | 11 | | | |
| chrV | 180478 | 180630 | 152 | -1,289395641 | N5:180645 | 23,945 | +12:YER013W; | 12 | | | |
| chrV | 182865 | 183029 | 164 | -1,509791068 | N5:182948 | 7,554 | +3:YER014W; | 3 | | | |
| chrV | 186698 | 186836 | 138 | -1,050741078 | N5:186791 | 10,083 | +13:YER015W; | 13 | | | |
| chrV | 189568 | 189720 | 152 | -1,047244802 | N5:189634 | 26,397 | +15:YER017C; | 15 | | | |
| chrV | 190625 | 190773 | 148 | -1,321681522 | N5:190729 | 28,404 | +8:YER017C; | 8 | | | |
| chrV | 193660 | 193792 | 132 | -0,552565504 | N5:193731 | 8,106 | +3:SUT510; | 3 | | | |
| chrV | 194009 | 194135 | 126 | -0,885772981 | | | | Overlap <50 bp | | | |
| chrV | 194030 | 194163 | 133 | -1,094744788 | N5:194161 | 6,156 | 0:SUT510; | 0 | | | |
| chrV | 195885 | 196040 | 155 | -1,10388581 | N5:195969 | 9,798 | +6:YER020W; | 6 | | | |
| chrV | 198265 | 198403 | 138 | -0,890763225 | N5:198244 | 16,025 | +9:YER021W; | 9 | | | |
| chrV | 201379 | 201531 | 152 | -1,31106074 | N5:201422 | 5,385 | +3:YER023W; | 3 | | | |
| chrV | 201380 | 201531 | 151 | -1,365803429 | N5:201422 | 5,385 | +3:YER023W; | 3 | | | |
| chrV | 201382 | 201531 | 149 | -1,291548225 | N5:201422 | 5,385 | +3:YER023W; | 3 | | | |
| chrV | 201383 | 201531 | 148 | -1,402498629 | N5:201422 | 5,385 | +3:YER023W; | 3 | | | |
| chrV | 206462 | 206572 | 110 | -1,438113353 | | | | Overlap <50 bp | | | |
| chrV | 207496 | 207646 | 150 | -1,51306086 | N5:207609 | 14,821 | +7*:YER026C; | 7 | TERM | | |
| chrV | 212628 | 212767 | 139 | -1,137426022 | | | | Overlap <50 bp | | | |
| chrV | 213547 | 213709 | 162 | -1,41726739 | N5:213639 | 13,761 | +2:YER030W; | 2 | | | |
| chrV | 215300 | 215450 | 150 | -1,464001951 | N5:215374 | 8,961 | +3:YER032W; | 3 | | | |
| chrV | 220704 | 220853 | 149 | -1,619077269 | N5:220767 | 15,329 | +4:YER033C; | 4 | | | |
| chrV | 226712 | 226861 | 149 | -1,257169902 | N5:226775 | 12,078 | R038C; +7:YER037W; +18:YER039C; | 10 | TERM | | |
| chrV | 227455 | 227599 | 144 | -1,249784308 | N5:227578 | 17,421 | +5:YER038C; +13:YER039C; | 5 | | | |
| chrV | 234231 | 234385 | 154 | -1,111371683 | N5:234301 | 5,574 | | Intergene | | | |
| chrV | 238011 | 238157 | 146 | -1,23343657 | N5:238087 | 10,45 | +1:YER044C; +8*:YER044C-A; | 1 | | | |
| chrV | 239144 | 239298 | 154 | -1,247910581 | N5:239207 | 7,887 | +1:YER044C-A; +4*:CUT109; | 1 | | | |
| chrV | 250567 | 250727 | 160 | -1,163035273 | N5:250698 | 7,688 | +1:YER048W-A; | 1 | | | |
| chrV | 250628 | 250757 | 129 | -1,072003738 | N5:250698 | 7,688 | +1:YER048W-A; | 1 | | | |
| chrV | 253010 | 253142 | 132 | -1,100009951 | N5:253162 | 32,519 | +10:YER049W; | 10 | | | |
| chrV | 255896 | 256035 | 139 | -1,352552539 | N5:256029 | 34,914 | +9:YER051W; | 9 | | | |
| chrV | 258267 | 258412 | 145 | -1,225289908 | N5:258341 | 18,228 | -1:YER052C; | -1 | | | |
| chrV | 263170 | 263306 | 136 | -1,292138703 | | | | Overlap <50 bp | | | |
| chrV | 268138 | 268266 | 128 | -1,170559319 | N5:268214 | 8,386 | 0:YER056C; | 0 | | | |
| chrV | 271754 | 271916 | 162 | -1,271015519 | N5:271765 | 7,679 | +1:YER058W; | 1 | | | |
| chrV | 273268 | 273395 | 127 | -1,050756748 | N5:273323 | 27,749 | +6:YER059W; | 6 | | | |
| chrV | 274395 | 274545 | 150 | -1,165681084 | N5:274471 | 20,294 | +22:SUT514; +1:YER060W; | 22 | | | |
| chrV | 279045 | 279197 | 152 | -1,465899569 | N5:279105 | 8,515 | +4:YER061C; | 4 | | | |
| chrV | 282333 | 282495 | 162 | -1,512285142 | N5:282372 | 25,461 | +5*:YER063W; +1:CUT557; | 5 | TERM | | |
| chrV | 286305 | 286436 | 131 | -0,825699181 | N5:286328 | 14,009 | +8:anti052; +6:YER065C; | 8 | | | |
| chrV | 293237 | 293387 | 150 | -1,479895789 | N5:293318 | 9,973 | +3:YER068W; | 3 | | | |
| chrV | 293855 | 294005 | 150 | -1,228169228 | N5:293929 | 9,292 | +7:YER068W; | 7 | | | |
| chrV | 298879 | 299018 | 139 | -1,268345527 | N5:298928 | 15,218 | +1:YER070W; | 1 | | | |
| chrV | 299205 | 299328 | 123 | -1,748545952 | N5:299273 | 9,183 | +3:YER070W; | 3 | | | |
| chrV | 299678 | 299831 | 153 | -1,293653467 | N5:299784 | 13,809 | +6:YER070W; | 6 | | | |
| chrV | 301505 | 301647 | 142 | -1,224899935 | N5:301602 | 18,777 | +17:YER070W; +1:CUT559; | 17 | | | |
| chrV | 314667 | 314817 | 150 | -0,975536208 | N5:314812 | 27,626 | | Intergene | | | |
| chrV | 315905 | 316063 | 158 | -1,237697138 | N5:315980 | 9,67 | +4:YER077C; | 4 | | | |
| chrV | 320077 | 320234 | 157 | -0,957392236 | N5:320131 | 14,134 | +2:YER080W; | 2 | | | |
| chrV | 321656 | 321815 | 159 | -1,595969993 | N5:321662 | 22,068 | +2*:CUT563; +12:YER080W; | 2 | TERM | | |
| chrV | 321656 | 321815 | 159 | -1,595969993 | N5:321825 | 38,767 | +1:CUT563; +13*:YER080W; | 1 | | | |
| chrV | 323513 | 323661 | 148 | -1,075349918 | N5:323606 | 6,557 | +7:YER081W; | 7 | | | |
| chrV | 324743 | 324895 | 152 | -1,410358804 | N5:324823 | 32,406 | +8:YER082C; | 8 | | | |
| chrV | 324904 | 325052 | 148 | -1,196926183 | N5:324975 | 31,489 | +7:YER082C; | 7 | | | |
| chrV | 325348 | 325499 | 151 | -1,445431373 | N5:325472 | 17,501 | +4:YER082C; | 4 | | | |
| chrV | 328966 | 329128 | 162 | -1,273102331 | N5:329104 | 18,91 | +5:YER086W; | 5 | | | |
| chrV | 331009 | 331174 | 165 | -1,348813496 | N5:331055 | 6,156 | +4:YER087W; | 4 | | | |
| chrV | 333088 | 333236 | 148 | -1,622644224 | N5:333122 | 14,975 | -1:YER087C-B; +1*:Unit175; | -1 | | | |
| chrV | 340240 | 340377 | 137 | -1,159098468 | N5:340278 | 28,039 | +12:YER091C; | 12 | | | |
| chrV | 340798 | 340946 | 148 | -1,413410293 | N5:340808 | 15,143 | +9:YER091C; | 9 | | | |
| chrV | 341934 | 342080 | 146 | -1,148432865 | | | | Overlap <50 bp | | | |
| chrV | 349486 | 349586 | 100 | -1,040302446 | | | | Overlap <50 bp | | | |
| chrV | 351515 | 351672 | 157 | -1,552705746 | N5:351572 | 9,783 | 0:YER096W; | 0 | | | |
| chrV | 352323 | 352478 | 155 | -1,476046162 | N5:352405 | 18,501 | +6:YER096W; | 6 | | | |
| chrV | 360556 | 360703 | 147 | -1,446553627 | N5:360635 | 27,381 | +8*:YER101C; | 8 | TERM | | |
| chrV | 360556 | 360709 | 153 | -1,277684752 | N5:360635 | 27,381 | +8*:YER101C; | 8 | TERM | | |
| chrV | 360596 | 360757 | 161 | -1,308263094 | N5:360635 | 27,381 | +8*:YER101C; | 8 | TERM | | |
| chrV | 361101 | 361246 | 145 | -1,458577418 | N5:361124 | 17,146 | +5:YER101C; | 5 | | | |
| chrV | 364550 | 364707 | 157 | -1,422709887 | N5:364627 | 11,269 | +2:YER103W; | 2 | | | |
| chrV | 369752 | 369900 | 148 | -0,982879854 | | | | Overlap <50 bp | | | |
| chrV | 370967 | 371076 | 109 | -1,332652675 | N5:371022 | 7,234 | +7:YER105C; | 7 | | | |
| chrV | 372915 | 373052 | 137 | -1,472221282 | N5:373029 | 29,3 | +5:YER106W; | 5 | | | |
| chrV | 373805 | 373958 | 153 | -1,302935443 | N5:373887 | 10,967 | +6:YER107C; | 6 | | | |
| chrV | 376017 | 376164 | 147 | -1,290673883 | N5:376149 | 19,149 | +11:YER109C; | 11 | | | |
| chrV | 376371 | 376519 | 148 | -1,375841574 | | | | Overlap <50 bp | | | |
| chrV | 376832 | 376976 | 144 | -1,113571873 | N5:376938 | 5,252 | +6:YER109C; | 6 | | | |
| chrV | 377517 | 377651 | 134 | -1,204259516 | N5:377578 | 5,854 | +2:YER109C; | 2 | | | |
| chrV | 378722 | 378880 | 158 | -1,338800433 | N5:378804 | 19,408 | +22:YER110C; | 22 | | | |
| chrV | 389589 | 389739 | 150 | -1,39479508 | N5:389641 | 11,967 | +4:YER113C; | 4 | | | |
| chrV | 392068 | 392208 | 140 | -1,602776899 | N5:392146 | 36,065 | +12:YER114C; | 12 | | | |
| chrV | 408769 | 408926 | 157 | -1,205256794 | N5:408837 | 22,819 | +2:YER124C; | 2 | | | |
| chrV | 411385 | 411539 | 154 | -1,21231688 | N5:411472 | 26,414 | +9:YER125W; | 9 | | | |
| chrV | 421211 | 421362 | 151 | -1,177834881 | N5:421329 | 21,066 | +8:YER130C; | 8 | | | |
| chrV | 426798 | 426949 | 151 | -1,457116882 | | | | Overlap <50 bp | | | |
| chrV | 426821 | 426975 | 154 | -1,446654544 | | | | Overlap <50 bp | | | |
| chrV | 428076 | 428230 | 154 | -1,45927449 | N5:428133 | 20,599 | +17:YER132C; | 17 | | | |
| chrV | 429069 | 429212 | 143 | -0,946082725 | | | | Overlap <50 bp | | | |
| chrV | 434847 | 434995 | 148 | -1,292879224 | N5:434827 | 11,15 | | Intergene | | | |
| chrV | 440738 | 440898 | 160 | -1,24324523 | N5:440906 | 46,309 | +9:YER136W; +6:YER137C; | 9 | | | |
| chrV | 444332 | 444446 | 114 | -1,484456338 | N5:444468 | 28,688 | | Intergene | | | |
| chrV | 445028 | 445178 | 150 | -1,646320469 | N5:445075 | 14,503 | | Intergene | | | |
| chrV | 446200 | 446349 | 149 | -0,953826865 | N5:446313 | 30,348 | | Intergene | | | |

TABLE 3

| CHR | START | END | L(bp) | ΔLK | Nuc ID | Fuzziness | Gene ID | Gene body postion or intergene | Terminal | rDNA | Telomeric |
|---|---|---|---|---|---|---|---|---|---|---|---|
| chrV | 446424 | 446581 | 157 | -1,304288189 | N5:446484 | 31,66 | | Intergene | | | |
| chrV | 446610 | 446760 | 150 | -1,031728129 | N5:446692 | 15,1 | | Intergene | | | |
| chrV | 446614 | 446760 | 146 | -1,193634273 | N5:446692 | 15,1 | | Intergene | | | |
| chrV | 447072 | 447207 | 135 | -0,909776363 | N5:447121 | 27,025 | | Intergene | | | |
| chrV | 447594 | 447738 | 144 | -1,303305836 | N5:447622 | 21,733 | | Intergene | | | |
| chrV | 447618 | 447772 | 154 | -1,156913567 | N5:447622 | 21,733 | | Intergene | | | |
| chrV | 447875 | 448026 | 151 | -1,276266079 | N5:447970 | 17,098 | | Intergene | | | |
| chrV | 447899 | 448044 | 145 | -1,062120308 | N5:447970 | 17,098 | | Intergene | | | |
| chrV | 447903 | 448054 | 151 | -1,160645126 | N5:447970 | 17,098 | | Intergene | | | |
| chrV | 448110 | 448250 | 140 | -1,187091515 | N5:448190 | 53,181 | | Intergene | | | |
| chrV | 448116 | 448250 | 134 | -1,042788351 | N5:448190 | 53,181 | | Intergene | | | |
| chrV | 448528 | 448683 | 155 | -0,800265026 | N5:448518 | 26,539 | | Intergene | | | |
| chrV | 448659 | 448767 | 108 | -0,836065226 | N5:448761 | 30,86 | | Intergene | | | |
| chrV | 448808 | 448957 | 149 | -1,291995728 | | | | Overlap <50 bp | | | |
| chrV | 450553 | 450702 | 149 | -1,498929923 | N5:450631 | 11,149 | +5:YER139C; | 5 | | | |
| chrV | 451474 | 451643 | 169 | -1,305773283 | N5:451553 | 3,777 | +1:YER140W; -1:YER139C; | 1 | | | |
| chrV | 456731 | 456859 | 128 | -1,582477679 | N5:456821 | 6,346 | +4:YER143W; | 4 | | | |
| chrV | 458282 | 458446 | 164 | -1,16316004 | | | | Overlap <50 bp | | | |
| chrV | 458920 | 459063 | 143 | -1,037636199 | N5:458935 | 35,138 | | Intergene | | | |
| chrV | 458920 | 459063 | 143 | -1,037636199 | N5:459085 | 25,589 | | Intergene | | | |
| chrV | 460202 | 460309 | 107 | -1,97004916 | N5:460198 | 15,639 | +2*:YER144C; | 2 | TERM | | |
| chrV | 460202 | 460330 | 128 | -1,075262103 | N5:460198 | 15,639 | +2*:YER144C; | 2 | TERM | | |
| chrV | 460206 | 460309 | 103 | -0,853250857 | N5:460198 | 15,639 | +2*:YER144C; | 2 | TERM | | |
| chrV | 465104 | 465256 | 152 | -1,407236171 | N5:465150 | 10,521 | +1:YER148W; -1:YER147C; | 1 | | | |
| chrV | 466895 | 467059 | 164 | -1,349266676 | N5:466980 | 8,421 | +4:YER149C; | 4 | | | |
| chrV | 471982 | 472140 | 158 | -1,460037468 | N5:472119 | 11,718 | +12:YER152C; +3:YER151C; | 12 | | | |
| chrV | 472402 | 472551 | 149 | -1,365135805 | N5:472474 | 17,601 | +10:YER152C; +1:YER151C; | 10 | | | |
| chrV | 472672 | 472832 | 160 | -1,303675239 | N5:472741 | 8,085 | +8:YER152C; -1:YER151C; | 8 | | | |
| chrV | 472989 | 473145 | 156 | -1,115916989 | N5:473075 | 7,047 | +6:YER152C; | 6 | | | |
| chrV | 473919 | 474069 | 150 | -1,146365534 | N5:473967 | 10,572 | +1:YER152C; | 1 | | | |
| chrV | 481969 | 482118 | 149 | -1,306220774 | | | | Overlap <50 bp | | | |
| chrV | 488812 | 488981 | 169 | -1,233366554 | N5:488949 | 4,278 | | Intergene | | | |
| chrV | 489802 | 489965 | 163 | -1,24163244 | N5:489855 | 28,19 | 0:SUT097; +6:YER158C; | 0 | | | |
| chrV | 493537 | 493686 | 149 | -1,370183782 | N5:493561 | 30,616 | | Intergene | | | |
| chrV | 495710 | 495860 | 150 | -1,01270983 | N5:495765 | 28,054 | | Intergene | | | |
| chrV | 496172 | 496307 | 135 | -1,482903475 | N5:496222 | 62,83 | | Intergene | | | |
| chrV | 497210 | 497350 | 140 | -1,222169136 | N5:497289 | 53,072 | | Intergene | | | |
| chrV | 497216 | 497350 | 134 | -1,122360157 | N5:497289 | 53,072 | | Intergene | | | |
| chrV | 497628 | 497783 | 155 | -1,235145567 | N5:497650 | 19,348 | | Intergene | | | |
| chrV | 497759 | 497867 | 108 | -1,079909369 | N5:497866 | 50,013 | | Intergene | | | |
| chrV | 497893 | 498045 | 152 | -1,592611579 | | | | Overlap <50 bp | | | |
| chrV | 501813 | 501975 | 162 | -1,228523236 | N5:501881 | 22,788 | +9:YER162C; | 9 | | | |
| chrV | 502108 | 502246 | 138 | -1,547012948 | N5:502208 | 10,183 | +7:YER162C; | 7 | | | |
| chrV | 503589 | 503721 | 132 | -1,089692801 | N5:503607 | 12,728 | +2:YER163C; | 2 | | | |
| chrV | 504504 | 504656 | 152 | -1,143423463 | N5:504574 | 15,604 | +4:SUT098; | 4 | | | |
| chrV | 506128 | 506280 | 152 | -1,292878026 | N5:506233 | 19,502 | +8:YER164W; | 8 | | | |
| chrV | 508434 | 508572 | 138 | -1,305140811 | | | | Overlap <50 bp | | | |
| chrV | 508812 | 508941 | 129 | -1,20027805 | N5:508869 | 22,167 | +24:YER164W; | 24 | | | |
| chrV | 517240 | 517402 | 162 | -1,33571515 | N5:517367 | 9,813 | | Intergene | | | |
| chrV | 522575 | 522735 | 160 | -1,627618347 | N5:522650 | 3,983 | +1:YER168C; | 1 | | | |
| chrV | 525612 | 525757 | 145 | -1,293545217 | N5:525693 | 10,443 | +17:YER169W; -1:YER170W; | 17 | | | |
| chrV | 528789 | 528933 | 144 | -1,391505336 | N5:528845 | 21,927 | +12:YER171W; +37:YER169W; | 12 | | | |
| chrV | 536553 | 536715 | 162 | -1,369931942 | N5:536626 | 4,792 | +3:YER173W; | 3 | | | |
| chrV | 538015 | 538173 | 158 | -1,422613978 | N5:538060 | 16,486 | +12:YER173W; | 12 | | | |
| chrV | 540818 | 540968 | 150 | -1,556260148 | N5:540902 | 24,345 | +3*:YER175W-A; +4:anti054; | 3 | TERM | | |
| chrV | 541050 | 541210 | 160 | -1,264164375 | N5:541230 | 5,568 | +2:anti054; | 2 | | | |
| chrV | 541050 | 541210 | 160 | -1,264164375 | N5:541076 | 6,427 | +3:anti054; | 3 | | | |
| chrV | 541616 | 541757 | 141 | -1,213651428 | N5:541694 | 3,869 | -1:anti054; +1:YER176W; | -1 | | | |
| chrV | 541626 | 541757 | 131 | -0,759951749 | N5:541694 | 3,869 | -1:anti054; +1:YER176W; | -1 | | | |
| chrV | 541626 | 541777 | 151 | -1,09133325 | N5:541694 | 3,869 | -1:anti054; +1:YER176W; | -1 | | | |
| chrV | 542716 | 542874 | 158 | -1,825210392 | N5:542824 | 35,576 | +8:YER176W; | 8 | | | |
| chrV | 544584 | 544725 | 141 | -1,210300792 | N5:544632 | 19,98 | +19:YER176W; +3*:CUT571; | 19 | | | |
| chrV | 560594 | 560752 | 158 | -1,31177382 | N5:560681 | 13,292 | +1:Unit200; +3:SUT522; | 1 | | | |
| chrV | 560787 | 560934 | 147 | -1,125255844 | N5:560849 | 6,911 | +2:Unit200; +2:SUT522; | 2 | | | |
| chrV | 561033 | 561174 | 141 | -1,16344903 | N5:561031 | 2,563 | +3*:Unit200; +1:SUT522; | 3 | TERM | | |
| chrV | 561033 | 561174 | 141 | -1,16344903 | N5:561188 | 9,879 | 0:SUT522; | 0 | | | |
| chrV | 571735 | 571876 | 141 | -1,143998579 | N5:571819 | 39,305 | +3:YER190W; | 3 | | | TEL RIGHT |
| chrV | 572939 | 573101 | 162 | -1,236180843 | N5:573065 | 8,718 | +11:YER190W; | 11 | | | TEL RIGHT |
| chrV | 573040 | 573195 | 155 | -1,923495307 | N5:573065 | 8,718 | +11:YER190W; | 11 | | | TEL RIGHT |
| chrV | 573174 | 573311 | 137 | -1,171630664 | N5:573237 | 10,817 | +12:YER190W; | 12 | | | TEL RIGHT |
| chrV | 573461 | 573612 | 151 | -1,040248665 | N5:573516 | 22,368 | +14:YER190W; | 14 | | | TEL RIGHT |
| chrV | 573469 | 573612 | 143 | -1,502217954 | N5:573516 | 22,368 | +14:YER190W; | 14 | | | TEL RIGHT |
| chrV | 573517 | 573642 | 125 | -0,821271071 | N5:573516 | 22,368 | +14:YER190W; | 14 | | | TEL RIGHT |
| chrV | 573742 | 573894 | 152 | -1,236127845 | N5:573853 | 12,014 | +16:YER190W; | 16 | | | TEL RIGHT |
| chrV | 573815 | 573959 | 144 | -1,00007447 | N5:573853 | 12,014 | +16:YER190W; | 16 | | | TEL RIGHT |
| chrV | 573912 | 574047 | 135 | -0,99409991 | N5:574037 | 55,842 | +17:YER190W; | 17 | | | TEL RIGHT |
| chrV | 573912 | 574099 | 187 | -0,784612356 | N5:574037 | 55,842 | +17:YER190W; | 17 | | | TEL RIGHT |
| chrV | 573955 | 574047 | 92 | -0,484806051 | N5:574037 | 55,842 | +17:YER190W; | 17 | | | TEL RIGHT |
| chrV | 573955 | 574099 | 144 | -1,034482195 | N5:574037 | 55,842 | +17:YER190W; | 17 | | | TEL RIGHT |
| chrV | 574294 | 574446 | 152 | -1,398100759 | N5:574369 | 12,53 | +19:YER190W; | 19 | | | TEL RIGHT |
| chrV | 574595 | 574725 | 130 | -1,002021939 | N5:574672 | 41,677 | +21:YER190W; | 21 | | | TEL RIGHT |
| chrV | 574965 | 575116 | 151 | -1,044563714 | N5:575085 | 20,306 | +24:YER190W; | 24 | | | TEL RIGHT |
| chrV | 574970 | 575104 | 134 | -0,973015657 | N5:575085 | 20,306 | +24:YER190W; | 24 | | | TEL RIGHT |
| chrV | 574983 | 575079 | 96 | -0,768229719 | N5:575085 | 20,306 | +24:YER190W; | 24 | | | TEL RIGHT |
| chrV | 575037 | 575152 | 115 | -0,565347997 | N5:575085 | 20,306 | +24:YER190W; | 24 | | | TEL RIGHT |
| chrV | 576130 | 576281 | 151 | -1,187276244 | N5:576205 | 0 | +31:YER190W; +1:YER190C-B; | 31 | | | TEL RIGHT |
| chrVI | 257 | 431 | 174 | -1,200114346 | N6:337 | 43,267 | +3:YFL068W; | 3 | | | TEL LEFT |
| chrVI | 377 | 525 | 148 | -1,48073657 | | | | Overlap <50 bp | | | TEL LEFT |
| chrVI | 2507 | 2632 | 125 | -0,758061132 | N6:2616 | 10,066 | +1:YFL066C; | 1 | | | TEL LEFT |
| chrVI | 2537 | 2680 | 143 | -1,438728731 | N6:2616 | 10,066 | +1:YFL066C; | 1 | | | TEL LEFT |
| chrVI | 2537 | 2688 | 151 | -1,386433233 | N6:2616 | 10,066 | +1:YFL066C; | 1 | | | TEL LEFT |
| chrVI | 2828 | 2965 | 137 | -1,010188926 | N6:2902 | 5,196 | -1:YFL066C; +4*:YFL065C; | -1 | | | TEL LEFT |
| chrVI | 3919 | 4067 | 148 | -1,098782974 | N6:4059 | 27,074 | -1:YFL064C; | -1 | | | TEL LEFT |
| chrVI | 4714 | 4814 | 100 | 0,062255146 | N6:4769 | 17,678 | | Intergene | | | TEL LEFT |
| chrVI | 6022 | 6182 | 160 | -1,209614041 | N6:6092 | 34,131 | -1:YFL062W; | -1 | | | |
| chrVI | 13717 | 13867 | 150 | -1,250256886 | N6:13829 | 5,657 | +7:YFL058W; | 7 | | | |
| chrVI | 15894 | 16044 | 150 | -1,12433456 | N6:15963 | 11,994 | | Intergene | | | |
| chrVI | 24066 | 24172 | 106 | -1,332111344 | N6:24173 | 13,416 | +6:YFL053W; | 6 | | | |

| CHR | START | END | L(bp) | ΔLK | Nuc ID | Fuzziness | Gene ID | Gene body postion or intergene | Terminal | rDNA | Telomeric |
|---|---|---|---|---|---|---|---|---|---|---|---|
| chrVI | 24067 | 24172 | 105 | -1,73228138 | N6:24173 | 13,416 | +6:YFL053W; | 6 | | | |
| chrVI | 29981 | 30130 | 149 | -1,512408731 | N6:30055 | 18,228 | +4:YFL051C; | 4 | | | |
| chrVI | 36134 | 36286 | 152 | -1,414020516 | N6:36241 | 4,764 | | Intergene | | | |
| chrVI | 36777 | 36932 | 155 | -0,969227534 | N6:36827 | 6,314 | +1:YFL049W; | 1 | | | |
| chrVI | 41762 | 41909 | 147 | -0,951451902 | N6:41884 | 23,166 | +10:YFL047W; | 10 | | | |
| chrVI | 48029 | 48199 | 170 | -1,331649524 | N6:48100 | 14,542 | | Intergene | | | |
| chrVI | 49950 | 50105 | 155 | -0,716641098 | N6:50008 | 9,579 | +6:YFL041W; | 6 | | | |
| chrVI | 55917 | 56077 | 160 | -1,413934322 | N6:55979 | 7,174 | +1:YFL038C; -1:YFL037W; | 1 | | | |
| chrVI | 60687 | 60850 | 163 | -1,414413713 | N6:60777 | 39,259 | +14:YFL036W; | 14 | | | |
| chrVI | 65236 | 65388 | 152 | -1,092265165 | N6:65325 | 8,232 | +1:YFL034W; | 1 | | | |
| chrVI | 65445 | 65606 | 161 | -1,138623168 | N6:65510 | 26,489 | +2:YFL034W; | 2 | | | |
| chrVI | 68129 | 68276 | 147 | -1,347365696 | N6:68162 | 46,167 | +19:YFL034W; | 19 | | | |
| chrVI | 68203 | 68367 | 164 | -1,258627703 | | | | Overlap <50 bp | | | |
| chrVI | 71963 | 72118 | 155 | -1,240194561 | N6:72001 | 12,021 | | Intergene | | | |
| chrVI | 72978 | 73128 | 150 | -1,291792381 | | | | Overlap <50 bp | | | |
| chrVI | 76566 | 76713 | 147 | -0,988080596 | N6:76715 | 22,509 | 0:YFL030W; +7*:anti055; | 0 | | | |
| chrVI | 77258 | 77379 | 121 | -1,498582826 | | | | Overlap <50 bp | | | |
| chrVI | 79934 | 80100 | 166 | -1,96983285 | N6:80017 | 11,635 | +2:YFL028C; | 2 | | | |
| chrVI | 81318 | 81445 | 127 | -1,234917645 | N6:81386 | 12,042 | +4:YFL027C; | 4 | | | |
| chrVI | 82718 | 82885 | 167 | -1,330963051 | | | | Overlap <50 bp | | | |
| chrVI | 84294 | 84451 | 157 | -1,448113915 | N6:84401 | 43,535 | +18:YFL025C; | 18 | | | |
| chrVI | 91095 | 91248 | 153 | -1,383248162 | N6:91173 | 6,356 | +2:YFL023W; | 2 | | | |
| chrVI | 94045 | 94152 | 107 | -1,173735745 | N6:94050 | 19,305 | +7*:YFL022C; | 7 | TERM | | |
| chrVI | 94173 | 94319 | 146 | -1,263006617 | N6:94222 | 13,892 | +6:YFL022C; | 6 | | | |
| chrVI | 94286 | 94435 | 149 | -1,240856035 | N6:94398 | 14,348 | +5:YFL022C; | 5 | | | |
| chrVI | 97720 | 97865 | 145 | -1,194853985 | N6:97797 | 30,798 | +2:Unit204; | 2 | | | |
| chrVI | 97725 | 97874 | 149 | -2,07190461 | N6:97797 | 30,798 | +2:Unit204; | 2 | | | |
| chrVI | 100985 | 101138 | 153 | -1,341002939 | N6:101049 | 8,361 | | Intergene | | | |
| chrVI | 108232 | 108382 | 150 | -1,171328526 | N6:108293 | 20,809 | +11:YFL013C; | 11 | | | |
| chrVI | 108406 | 108531 | 125 | -1,256091324 | N6:108481 | 12,832 | +10:YFL013C; | 10 | | | |
| chrVI | 110912 | 111064 | 152 | -1,449062316 | N6:110937 | 25,521 | +3:YFL012W; | 3 | | | |
| chrVI | 110912 | 111064 | 152 | -1,449062316 | N6:111087 | 19,217 | +4:YFL012W; | 4 | | | |
| chrVI | 111983 | 112114 | 131 | -1,175445116 | N6:112065 | 15,026 | :YFL011W; +1:Unit217; -1:Unit205; | -1 | | | |
| chrVI | 115465 | 115634 | 169 | -1,136239443 | N6:115552 | 11,76 | +2:YFL010C; | 2 | | | |
| chrVI | 123008 | 123146 | 138 | -1,03810545 | N6:123085 | 13,52 | -1:YFL007W; | -1 | | | |
| chrVI | 130462 | 130613 | 151 | -1,23955614 | N6:130530 | 3,098 | +3:YFL005W; | 3 | | | |
| chrVI | 131462 | 131615 | 153 | -1,278392274 | N6:131556 | 23,69 | -1:Unit206; 0:YFL004W; | -1 | | | |
| chrVI | 139986 | 140115 | 129 | -1,304365929 | N6:139988 | 18,248 | | Intergene | | | |
| chrVI | 140063 | 140227 | 164 | -1,310806718 | N6:140160 | 24,269 | | Intergene | | | |
| chrVI | 141451 | 141594 | 143 | -0,985920775 | N6:141487 | 30,006 | | Intergene | | | |
| chrVI | 142726 | 142892 | 166 | -1,301314702 | N6:142759 | 24,434 | | Intergene | | | |
| chrVI | 143530 | 143679 | 149 | -1,254139061 | N6:143622 | 24,194 | | Intergene | | | |
| chrVI | 148326 | 148498 | 172 | -1,326220649 | N6:148376 | 9,288 | +8*:YFL001W; | 8 | TERM | | |
| chrVI | 151570 | 151741 | 171 | -1,507539224 | N6:151682 | 32,887 | +11:YFR002W; | 11 | | | |
| chrVI | 151602 | 151755 | 153 | -1,28941835 | N6:151682 | 32,887 | +11:YFR002W; | 11 | | | |
| chrVI | 151830 | 151995 | 165 | -1,473101052 | N6:151925 | 16,258 | +12:YFR002W; | 12 | | | |
| chrVI | 152009 | 152167 | 158 | -1,429043515 | N6:152083 | 17,855 | +13:YFR002W; | 13 | | | |
| chrVI | 152552 | 152696 | 144 | -0,953524862 | N6:152640 | 4,708 | +4*:YFR003C; | 4 | TERM | | |
| chrVI | 155423 | 155586 | 163 | -1,497475844 | N6:155517 | 10,71 | +3:YFR005C; | 3 | | | |
| chrVI | 162535 | 162694 | 159 | -1,290762297 | N6:162635 | 6,309 | +2:YFR009W; | 2 | | | |
| chrVI | 174443 | 174598 | 155 | -1,434724964 | N6:174527 | 14,046 | +12:YFR015C; | 12 | | | |
| chrVI | 175285 | 175426 | 141 | -1,098600541 | N6:175340 | 22,993 | +7:YFR015C; | 7 | | | |
| chrVI | 176188 | 176314 | 126 | -0,460691877 | N6:176274 | 18,228 | +2:YFR015C; | 2 | | | |
| chrVI | 177968 | 178139 | 171 | -1,502795237 | N6:178030 | 38,026 | | Intergene | | | |
| chrVI | 180563 | 180715 | 152 | -1,292676894 | N6:180613 | 25,528 | +2:YFR016C; | 2 | | | |
| chrVI | 182163 | 182304 | 141 | -1,016809972 | N6:182222 | 6,058 | +5*:YFR017C; | 5 | TERM | | |
| chrVI | 183274 | 183405 | 131 | -2,130995957 | | | | Overlap <50 bp | | | |
| chrVI | 183755 | 183882 | 127 | -0,753083521 | N6:183852 | 9,96 | +3:YFR018C; | 3 | | | |
| chrVI | 185980 | 186131 | 151 | -1,30572199 | N6:186127 | 16,489 | +10:YFR019W; | 10 | | | |
| chrVI | 186486 | 186602 | 116 | -1,500241078 | N6:186517 | 20,777 | +12:YFR019W; | 12 | | | |
| chrVI | 192368 | 192528 | 160 | -1,479828325 | N6:192442 | 6,535 | | Intergene | | | |
| chrVI | 193031 | 193186 | 155 | -1,422354477 | N6:193102 | 1,915 | | Intergene | | | |
| chrVI | 195001 | 195152 | 151 | -1,306456249 | N6:195090 | 9,394 | +3:YFR021W; | 3 | | | |
| chrVI | 197175 | 197337 | 162 | -1,448677025 | N6:197229 | 24,205 | +4:YFR022W; | 4 | | | |
| chrVI | 197468 | 197595 | 127 | -1,262299419 | | | | Overlap <50 bp | | | |
| chrVI | 198399 | 198562 | 163 | -1,142073798 | N6:198429 | 7,789 | +11:YFR022W; | 11 | | | |
| chrVI | 202836 | 202999 | 163 | -1,356147388 | N6:202902 | 9,98 | | Intergene | | | |
| chrVI | 204046 | 204202 | 156 | -1,297534777 | N6:204111 | 1,643 | +5:YFR025C; | 5 | | | |
| chrVI | 208087 | 208238 | 151 | -1,036805945 | N6:208144 | 24,403 | +5:YFR027W; | 5 | | | |
| chrVI | 208444 | 208571 | 127 | -1,268102919 | N6:208478 | 17,871 | +11*:YFR028C; +7:YFR027W; | 11 | TERM | | |
| chrVI | 208746 | 208898 | 152 | -1,272290667 | N6:208819 | 26,853 | +9:YFR028C; +9:YFR027W; | 9 | | | |
| chrVI | 210302 | 210398 | 96 | -1,565444235 | N6:210340 | 4,98 | -1:YFR028C; | -1 | | | |
| chrVI | 210302 | 210400 | 98 | -1,414462921 | N6:210340 | 4,98 | -1:YFR028C; | -1 | | | |
| chrVI | 210302 | 210402 | 100 | -1,419983737 | N6:210340 | 4,98 | -1:YFR028C; | -1 | | | |
| chrVI | 210302 | 210403 | 101 | -1,436289577 | N6:210340 | 4,98 | -1:YFR028C; | -1 | | | |
| chrVI | 210302 | 210404 | 102 | -1,343411677 | N6:210340 | 4,98 | -1:YFR028C; | -1 | | | |
| chrVI | 210302 | 210405 | 103 | -1,617392982 | N6:210340 | 4,98 | -1:YFR028C; | -1 | | | |
| chrVI | 210302 | 210406 | 104 | -1,46114542 | N6:210340 | 4,98 | -1:YFR028C; | -1 | | | |
| chrVI | 210302 | 210407 | 105 | -1,4880736 | N6:210340 | 4,98 | -1:YFR028C; | -1 | | | |
| chrVI | 210302 | 210409 | 107 | -1,367556663 | N6:210340 | 4,98 | -1:YFR028C; | -1 | | | |
| chrVI | 214215 | 214366 | 151 | -1,306344638 | N6:214204 | 32,337 | +7:YFR030W; | 7 | | | |
| chrVI | 214215 | 214366 | 151 | -1,306344638 | N6:214374 | 16,008 | +8:YFR030W; | 8 | | | |
| chrVI | 224528 | 224665 | 137 | -1,055195735 | N6:224600 | 41,62 | +2:YFR033C; | 2 | | | |
| chrVI | 225246 | 225383 | 137 | -0,758999867 | | | | Overlap <50 bp | | | |
| chrVI | 225640 | 225800 | 160 | -1,44169213 | N6:225764 | 5,441 | +2:YFR034C; | 2 | | | |
| chrVI | 228483 | 228624 | 141 | -0,995094292 | N6:228495 | 14,751 | +5:YFR037C; | 5 | | | |
| chrVI | 232870 | 233015 | 145 | -1,061602468 | N6:232930 | 12,637 | +5:YFR039C; | 5 | | | |
| chrVI | 236135 | 236288 | 153 | -1,569244981 | N6:236220 | 42,965 | +15:YFR040W; | 15 | | | |
| chrVI | 236706 | 236866 | 160 | -1,351420267 | N6:236786 | 35,865 | +18:YFR040W; | 18 | | | |
| chrVI | 237607 | 237756 | 149 | -1,184381531 | N6:237708 | 6,504 | +4:YFR041C; | 4 | | | |
| chrVI | 237976 | 238116 | 140 | -1,090784436 | N6:238042 | 8,379 | +2:YFR041C; | 2 | | | |
| chrVI | 239069 | 239219 | 150 | -1,342599476 | N6:239120 | 13,831 | +5*:YFR042W; | 5 | TERM | | |
| chrVI | 242117 | 242275 | 158 | -1,131817428 | N6:242181 | 10,159 | +2:YFR045W; | 2 | | | |
| chrVI | 242831 | 242985 | 154 | -1,37019621 | N6:242929 | 45,045 | +6*:YFR045W; +9*:YFR046C; | 6 | TERM | | |
| chrVI | 242895 | 243045 | 150 | -1,523153594 | N6:242929 | 45,045 | +6*:YFR045W; +9*:YFR046C; | 6 | TERM | | |
| chrVI | 244023 | 244165 | 142 | -1,160516421 | N6:244096 | 10,309 | +2:YFR046C; | 2 | | | |
| chrVI | 244610 | 244760 | 150 | -1,228527098 | N6:244637 | 0 | +4:YFR047C; | 4 | | | |
| chrVI | 247013 | 247149 | 136 | -0,705578496 | N6:247082 | 23,398 | +7:YFR048W; | 7 | | | |
| chrVI | 258453 | 258604 | 151 | -1,671543592 | N6:258521 | 27,206 | | Intergene | | | |

TABLE 3

| CHR | START | END | L(bp) | ΔLK | Nuc ID | Fuzziness | Gene ID | Gene body postion or intergene | Terminal | rDNA | Telomeric |
|---|---|---|---|---|---|---|---|---|---|---|---|
| chrVI | 260977 | 261105 | 128 | -1,011655971 | N6:261029 | 34,994 | | Intergene | | | |
| chrVI | 264647 | 264774 | 127 | -1,073368727 | | | | Overlap <50 bp | | | |
| chrVI | 265600 | 265745 | 145 | -1,034138493 | N6:265749 | 18,434 | | Intergene | | | |
| chrVI | 267454 | 267607 | 153 | -1,534590669 | N6:267517 | 27,951 | | Intergene | | | |
| chrVI | 269103 | 269252 | 149 | -1,329072761 | N6:269131 | 22,098 | +2:YFR057W; | 2 | | | |
| chrVI | 270018 | 270158 | 140 | -1,200338748 | N6:270163 | 33,941 | | Intergene | | | TEL RIGHT |
| chrVII | 6501 | 6604 | 103 | -1,344732916 | N7:6569 | 11,504 | -1:YGL260W; +2:YGL261C; | -1 | | | |
| chrVII | 7179 | 7332 | 153 | -1,542523593 | N7:7319 | 7,805 | +4*:YGL260W; | 4 | TERM | | |
| chrVII | 7179 | 7332 | 153 | -1,542523593 | N7:7156 | 4,272 | +3:YGL260W; | 3 | | | |
| chrVII | 8743 | 8898 | 155 | -1,287471763 | N7:8851 | 28,687 | +4:YGL259W; -1:YGL258W-A; | 4 | | | |
| chrVII | 8747 | 8898 | 151 | -1,325453083 | N7:8851 | 28,687 | +4:YGL259W; -1:YGL258W-A; | 4 | | | |
| chrVII | 9683 | 9793 | 110 | -1,196384333 | N7:9667 | 21,339 | +9:YGL259W; | 9 | | | |
| chrVII | 15630 | 15761 | 131 | -0,942814163 | N7:15706 | 47,019 | +7:YGL256W; | 7 | | | |
| chrVII | 15633 | 15761 | 128 | -1,07464229 | N7:15706 | 47,019 | +7:YGL256W; | 7 | | | |
| chrVII | 15791 | 15935 | 144 | -1,499304451 | | | | Overlap <50 bp | | | |
| chrVII | 20349 | 20487 | 138 | -1,195206169 | N7:20424 | 5,899 | | Intergene | | | |
| chrVII | 21325 | 21484 | 159 | -1,395544131 | N7:21457 | 24,473 | +5:YGL255W; | 5 | | | |
| chrVII | 22410 | 22546 | 136 | -1,05037092 | N7:22504 | 5,879 | +5:CUT583; +2:YGL254W; | 5 | | | |
| chrVII | 23209 | 23365 | 156 | -1,426702114 | N7:23217 | 14,748 | +1:CUT583; +6:YGL254W; | 1 | | | |
| chrVII | 32814 | 32961 | 147 | -1,456570695 | N7:32833 | 24,46 | 'GL250W; +1:SUT533; -1:YGL249W; | 6 | TERM | | |
| chrVII | 33222 | 33355 | 133 | -1,306549984 | N7:33236 | 10,496 | +11:SUT534; +2:YGL249W; | 11 | | | |
| chrVII | 36941 | 37083 | 142 | -1,259313104 | N7:36962 | 8,687 | +1:YGL247W; | 1 | | | |
| chrVII | 41750 | 41885 | 135 | -1,574749961 | N7:41812 | 5,785 | +3:YGL244W; | 3 | | | |
| chrVII | 52996 | 53149 | 153 | -1,228202349 | N7:53078 | 32,752 | +4:YGL237C; | 4 | | | |
| chrVII | 53375 | 53486 | 111 | -1,364857664 | N7:53412 | 3,502 | +2:YGL237C; | 2 | | | |
| chrVII | 62645 | 62823 | 178 | -1,761732769 | N7:62782 | 30,625 | | Intergene | | | |
| chrVII | 69139 | 69303 | 164 | -1,330589542 | N7:69217 | 9,757 | +11:YGL228W; -1:YGL227W; | 11 | | | |
| chrVII | 72210 | 72368 | 158 | -1,404539502 | N7:72277 | 8,042 | | Intergene | | | |
| chrVII | 73049 | 73197 | 148 | -1,393409235 | N7:73145 | 16,622 | +1*:YGL226C-A; -1:YGL226W; | 1 | TERM | | |
| chrVII | 78222 | 78356 | 134 | -1,04970544 | N7:78264 | 9,585 | +14:YGL223C; +5:YGL224C; | 14 | | | |
| chrVII | 83125 | 83232 | 107 | -1,093800065 | N7:83164 | 17,63 | +8:YGL219C; | 8 | | | |
| chrVII | 83125 | 83247 | 122 | -1,314671883 | N7:83164 | 17,63 | +8:YGL219C; | 8 | | | |
| chrVII | 84927 | 85059 | 132 | -1,070576812 | N7:84990 | 14,085 | +3:YGL216W; | 3 | | | |
| chrVII | 88176 | 88368 | 192 | -1,325156433 | N7:88188 | 9,988 | +5:YGL215W; | 5 | | | |
| chrVII | 88303 | 88443 | 140 | -1,092830011 | | | | Overlap <50 bp | | | |
| chrVII | 88402 | 88555 | 153 | -1,119596801 | N7:88473 | 12,759 | +7:YGL215W; | 7 | | | |
| chrVII | 89045 | 89177 | 132 | -1,121197653 | | | | Overlap <50 bp | | | |
| chrVII | 90038 | 90161 | 123 | -1,175779921 | N7:90075 | 15,222 | +8*:YGL213C; | 8 | TERM | | |
| chrVII | 91402 | 91556 | 154 | -1,156406361 | N7:91484 | 22,807 | +1:YGL212W; -1:YGL213C; | 1 | | | |
| chrVII | 92970 | 93138 | 168 | -1,266217493 | N7:93036 | 24,978 | +4:YGL211W; | 4 | | | |
| chrVII | 97437 | 97599 | 162 | -1,427513424 | N7:97514 | 14,549 | +2:YGL208W; | 2 | | | |
| chrVII | 102150 | 102299 | 149 | -1,405788269 | N7:102228 | 18,52 | | Intergene | | | |
| chrVII | 102520 | 102673 | 153 | -1,603581125 | N7:102600 | 27,313 | +32:YGL206C; | 32 | | | |
| chrVII | 103288 | 103440 | 152 | -1,136028401 | N7:103447 | 49,516 | +27:YGL206C; | 27 | | | |
| chrVII | 103852 | 103984 | 132 | -1,149755245 | N7:103952 | 22,777 | +24:YGL206C; | 24 | | | |
| chrVII | 108983 | 109147 | 164 | -1,410889484 | N7:109010 | 40,278 | +8:SUT537; +6:YGL205W; | 8 | | | |
| chrVII | 116027 | 116173 | 146 | -1,120644916 | N7:116126 | 14,519 | +2:YGL202W; | 2 | | | |
| chrVII | 119268 | 119414 | 146 | -1,16952184 | N7:119279 | 40,009 | +11:YGL201C; | 11 | | | |
| chrVII | 125888 | 126050 | 162 | -1,384429889 | N7:125970 | 26,986 | +9:YGL197W; | 9 | | | |
| chrVII | 138978 | 139125 | 147 | -1,328937718 | N7:139099 | 25,612 | | Intergene | | | |
| chrVII | 147769 | 147920 | 151 | -1,395131641 | N7:147908 | 6,535 | 0:Unit261; | 0 | | | |
| chrVII | 152057 | 152208 | 151 | -1,591722759 | N7:152137 | 19,449 | +5:YGL186C; | 5 | | | |
| chrVII | 156139 | 156292 | 153 | -1,624930912 | N7:156241 | 20,868 | -1:YGL184C; | -1 | | | |
| chrVII | 167824 | 167963 | 139 | -1,01828221 | N7:167816 | 7,935 | | Intergene | | | |
| chrVII | 167824 | 167963 | 139 | -1,01828221 | N7:167983 | 28,836 | | Intergene | | | |
| chrVII | 168448 | 168554 | 106 | -1,596342667 | N7:168455 | 9,368 | | Intergene | | | |
| chrVII | 168635 | 168789 | 154 | -1,686079628 | N7:168761 | 37,278 | | Intergene | | | |
| chrVII | 171388 | 171548 | 160 | -1,329231893 | N7:171452 | 17,19 | +10*:YGL176C; | 10 | TERM | | |
| chrVII | 174752 | 174849 | 97 | -1,325810284 | N7:174743 | 2,422 | +2:YGL174W; | 2 | | | |
| chrVII | 174828 | 174980 | 152 | -1,176151799 | N7:174904 | 5,989 | +3:YGL174W; | 3 | | | |
| chrVII | 178188 | 178320 | 132 | -1,004723465 | N7:178266 | 27,142 | +15:YGL173C; | 15 | | | |
| chrVII | 183143 | 183279 | 136 | -1,372240676 | | | | Overlap <50 bp | | | |
| chrVII | 184850 | 184994 | 144 | -1,051215364 | N7:184898 | 19,765 | +4:YGL170C; | 4 | | | |
| chrVII | 196178 | 196334 | 156 | -1,302984775 | N7:196334 | 11,032 | +2:YGL163C; | 2 | | | |
| chrVII | 196178 | 196334 | 156 | -1,302984775 | N7:196162 | 11,619 | +3:YGL163C; | 3 | | | |
| chrVII | 204030 | 204183 | 153 | -1,233350122 | N7:204104 | 5,468 | +9*:YGL159W; | 9 | TERM | | |
| chrVII | 204995 | 205129 | 134 | -1,020663457 | N7:205048 | 8,021 | | Intergene | | | |
| chrVII | 209064 | 209201 | 137 | -1,299980566 | N7:209110 | 12,687 | +2:YGL157W; | 2 | | | |
| chrVII | 214611 | 214774 | 163 | -1,40460981 | N7:214691 | 10,944 | +5:YGL155W; | 5 | | | |
| chrVII | 217469 | 217614 | 145 | -1,144533382 | N7:217533 | 8,542 | +1:YGL151W; | 1 | | | |
| chrVII | 221203 | 221378 | 175 | -1,098196641 | N7:221318 | 17,398 | +28:YGL150C; | 28 | | | |
| chrVII | 223475 | 223638 | 163 | -1,344556629 | N7:223563 | 16,468 | +15:YGL150C; | 15 | | | |
| chrVII | 226620 | 226777 | 157 | -0,70175811 | N7:226722 | 16,053 | +3:YGL148W; | 3 | | | |
| chrVII | 226621 | 226770 | 149 | -1,260000477 | N7:226722 | 16,053 | +3:YGL148W; | 3 | | | |
| chrVII | 233772 | 233924 | 152 | -1,341728485 | N7:233838 | 7,47 | +6:YGL144C; | 6 | | | |
| chrVII | 238049 | 238197 | 148 | -1,272556421 | N7:238126 | 20,169 | -1:YGL141W; +1:YGL142C; | -1 | | | |
| chrVII | 243163 | 243328 | 165 | -1,205011668 | N7:243257 | 26,011 | +13:YGL140C; | 13 | | | |
| chrVII | 249506 | 249649 | 143 | -1,785763635 | N7:249567 | 4,933 | +1:YGL138C; | 1 | | | |
| chrVII | 251482 | 251633 | 151 | -1,02841495 | | | | Overlap <50 bp | | | |
| chrVII | 266552 | 266693 | 141 | -1,137114217 | N7:266631 | 15,972 | +4:YGL130W; | 4 | | | |
| chrVII | 267377 | 267528 | 151 | -1,171384347 | N7:267455 | 27,294 | +9:YGL130W; | 9 | | | |
| chrVII | 268853 | 268959 | 106 | -0,842375882 | N7:268920 | 11,524 | +2:YGL129C; | 2 | | | |
| chrVII | 268853 | 268977 | 124 | -1,237224761 | N7:268920 | 11,524 | +2:YGL129C; | 2 | | | |
| chrVII | 270004 | 270159 | 155 | -1,36956371 | N7:270127 | 17,201 | +1:YGL128C; | 1 | | | |
| chrVII | 273536 | 273656 | 120 | -1,121368652 | N7:273600 | 25,616 | | Intergene | | | |
| chrVII | 275567 | 275723 | 156 | -1,285337291 | N7:275555 | 16,263 | | Intergene | | | |
| chrVII | 275567 | 275723 | 156 | -1,285337291 | N7:275720 | 26,82 | | Intergene | | | |
| chrVII | 279938 | 280099 | 161 | -1,125338512 | N7:280010 | 22,5 | +4:YGL122C; | 4 | | | |
| chrVII | 281760 | 281908 | 148 | -1,253125879 | N7:281898 | 21,64 | +14:YGL120C; | 14 | | | |
| chrVII | 281764 | 281908 | 144 | -1,388605836 | N7:281898 | 21,64 | +14:YGL120C; | 14 | | | |
| chrVII | 284331 | 284485 | 154 | -1,062502872 | N7:284426 | 7,899 | +1:YGL119W; | 1 | | | |
| chrVII | 289603 | 289760 | 157 | -1,462679938 | N7:289668 | 37,131 | +1:YGL116W; | 1 | | | |
| chrVII | 293723 | 293871 | 148 | -1,041166926 | N7:293806 | 25,272 | +4:YGL114W; | 4 | | | |
| chrVII | 294019 | 294183 | 164 | -1,780793194 | N7:294135 | 15,498 | +6:YGL114W; | 6 | | | |
| chrVII | 297177 | 297313 | 136 | -1,15074275 | N7:297182 | 35,303 | +9:YGL113W; | 9 | | | |
| chrVII | 299926 | 300073 | 147 | -1,170676717 | N7:300004 | 6,735 | -1:YGL112C; +1:YGL111W; | -1 | | | |
| chrVII | 300091 | 300244 | 153 | -1,190705321 | N7:300166 | 6,465 | +2:YGL111W; | 2 | | | |
| chrVII | 300595 | 300755 | 160 | -1,451002598 | | | | Overlap <50 bp | | | |
| chrVII | 313487 | 313646 | 159 | -1,487474576 | N7:313555 | 8,204 | +3:YGL100W; | 3 | | | |

| CHR | START | END | L(bp) | ΔLK | Nuc ID | Fuzziness | Gene ID | Gene body postion or intergene | Terminal | rDNA | Telomeric |
|---|---|---|---|---|---|---|---|---|---|---|---|
| chrVII | 314575 | 314719 | 144 | -1,412057745 | N7:314647 | 5,01 | +1:YGL099W; -1:CUT595; | 1 | | | |
| chrVII | 320068 | 320221 | 153 | -1,159280053 | N7:320063 | 7,859 | +1:SUT114; | 1 | | | |
| chrVII | 320068 | 320221 | 153 | -1,159280053 | N7:320242 | 21,417 | +2:SUT114; | 2 | | | |
| chrVII | 323178 | 323326 | 148 | -1,490382885 | | | | Overlap <50 bp | | | |
| chrVII | 324915 | 325051 | 136 | -1,411869503 | N7:325012 | 11,082 | -1:YGL096W; | -1 | | | |
| chrVII | 327547 | 327717 | 170 | -1,247979841 | N7:327548 | 34,723 | | Intergene | | | |
| chrVII | 327547 | 327717 | 170 | -1,247979841 | N7:327719 | 24,372 | | Intergene | | | |
| chrVII | 328882 | 329034 | 152 | -1,447105949 | N7:329029 | 16,562 | +11:YGL095C; | 11 | | | |
| chrVII | 329271 | 329388 | 117 | -1,778571749 | N7:329338 | 20,498 | +9:YGL095C; | 9 | | | |
| chrVII | 330372 | 330517 | 145 | -1,167160957 | N7:330463 | 11,79 | +2:YGL095C; | 2 | | | |
| chrVII | 331632 | 331780 | 148 | -1,275417619 | N7:331795 | 31,694 | +18:YGL094C; | 18 | | | |
| chrVII | 331632 | 331780 | 148 | -1,275417619 | N7:331642 | 32,828 | +19:YGL094C; | 19 | | | |
| chrVII | 341547 | 341699 | 152 | -1,528194936 | N7:341572 | 4,546 | +24:YGL092W; | 24 | | | |
| chrVII | 342190 | 342339 | 149 | -1,265549255 | | | | Overlap <50 bp | | | |
| chrVII | 342310 | 342453 | 143 | -1,110882225 | N7:342424 | 27,427 | +5:YGL091C; | 5 | | | |
| chrVII | 343265 | 343403 | 138 | -1,169386757 | N7:343315 | 4,844 | +1:YGL090W; -1:YGL091C; | 1 | | | |
| chrVII | 352278 | 352440 | 162 | -1,466163356 | N7:352305 | 6,439 | +1:YGL084C; | 1 | | | |
| chrVII | 354451 | 354624 | 173 | -1,461552527 | N7:354512 | 51,432 | +10:YGL083W; | 10 | | | |
| chrVII | 356531 | 356676 | 145 | -1,262464981 | N7:356589 | 11,61 | +6:YGL082W; | 6 | | | |
| chrVII | 364639 | 364801 | 162 | -1,404407366 | N7:364708 | 1,414 | | Intergene | | | |
| chrVII | 367967 | 368120 | 153 | -1,011372664 | N7:368054 | 8,802 | +2:YGL075C; | 2 | | | |
| chrVII | 370607 | 370760 | 153 | -1,491822377 | N7:370672 | 4,97 | +12:YGL073W; | 12 | | | |
| chrVII | 382677 | 382828 | 151 | -1,428412347 | N7:382840 | 6,042 | +3:YGL064C; | 3 | | | |
| chrVII | 382677 | 382828 | 151 | -1,428412347 | N7:382673 | 17,697 | +4:YGL064C; | 4 | | | |
| chrVII | 383386 | 383517 | 131 | -0,874142509 | N7:383447 | 21,685 | +1:YGL063W; -1:YGL064C; | 1 | | | |
| chrVII | 383588 | 383739 | 151 | -0,78952824 | N7:383638 | 24,47 | +2:YGL063W; | 2 | | | |
| chrVII | 384835 | 384978 | 143 | -1,827958092 | N7:384900 | 12,22 | +9:YGL063W; -1:YGL062W; | 9 | | | |
| chrVII | 384841 | 384978 | 137 | -1,501809551 | N7:384900 | 12,22 | +9:YGL063W; -1:YGL062W; | 9 | | | |
| chrVII | 386231 | 386378 | 147 | -1,082886696 | N7:386322 | 34,798 | +8:YGL062W; | 8 | | | |
| chrVII | 386588 | 386752 | 164 | -1,534194436 | N7:386632 | 45,997 | +10:YGL062W; | 10 | | | |
| chrVII | 398118 | 398266 | 148 | -1,16567702 | | | | Overlap <50 bp | | | |
| chrVII | 398740 | 398893 | 153 | -1,282158266 | N7:398821 | 14,799 | +2:YGL055W; | 2 | | | |
| chrVII | 399113 | 399273 | 160 | -1,29757531 | N7:399182 | 11,256 | +4:YGL055W; | 4 | | | |
| chrVII | 402184 | 402330 | 146 | -1,402219053 | N7:402298 | 26,92 | -1:YGL053W; | -1 | | | |
| chrVII | 406230 | 406375 | 145 | -1,061600504 | N7:406270 | 19,026 | +4:YGL050W; | 4 | | | |
| chrVII | 412759 | 412909 | 150 | -1,373005527 | N7:412835 | 6,745 | +3:CUT600; | 3 | | | |
| chrVII | 413027 | 413197 | 170 | -1,880692813 | N7:413154 | 24,499 | +1:CUT600; | 1 | | | |
| chrVII | 416964 | 417114 | 150 | -1,385567264 | N7:417040 | 5,086 | +2:YGL044C; | 2 | | | |
| chrVII | 419105 | 419256 | 151 | -1,471173429 | N7:419189 | 16,861 | -1:YGL041C-B; +3:YGL041W-A; | -1 | | | |
| chrVII | 422885 | 423015 | 130 | -1,162825096 | N7:422958 | 7,071 | +5*:SUT118; | 5 | TERM | | |
| chrVII | 424432 | 424587 | 155 | -1,459672603 | N7:424573 | 47,995 | +5:YGL039W; | 5 | | | |
| chrVII | 426688 | 426828 | 140 | -1,241124711 | N7:426744 | 16,661 | +2:YGL038C; | 2 | | | |
| chrVII | 430629 | 430785 | 156 | -1,370031438 | N7:430704 | 34,387 | +14:YGL036W; | 14 | | | |
| chrVII | 438337 | 438504 | 167 | -1,225368127 | N7:438393 | 27,83 | | Intergene | | | |
| chrVII | 445398 | 445550 | 152 | -1,33024666 | N7:445477 | 5,762 | +5:YGL027C; | 5 | | | |
| chrVII | 445405 | 445562 | 157 | -1,066353576 | N7:445477 | 5,762 | +5:YGL027C; | 5 | | | |
| chrVII | 448295 | 448446 | 151 | -1,14277151 | N7:448359 | 5,612 | +2:YGL026C; | 2 | | | |
| chrVII | 449063 | 449214 | 151 | -1,372267789 | N7:449131 | 13,626 | +6:YGL025C; | 6 | | | |
| chrVII | 449866 | 450017 | 151 | -1,349950154 | N7:449940 | 9,933 | +1:YGL025C; | 1 | | | |
| chrVII | 450289 | 450475 | 186 | -1,118411944 | N7:450357 | 12,05 | +12*:YGL023C; | 12 | TERM | | |
| chrVII | 450564 | 450727 | 163 | -1,370390337 | | | | Overlap <50 bp | | | |
| chrVII | 451657 | 451811 | 154 | -1,196634111 | N7:451765 | 9,301 | +3:YGL023C; | 3 | | | |
| chrVII | 452015 | 452160 | 145 | -1,458813534 | N7:452093 | 5,788 | +1:YGL023C; -1:YGL022W; | 1 | | | |
| chrVII | 454120 | 454220 | 100 | -1,563903427 | N7:454191 | 22,142 | +12:YGL022W; | 12 | | | |
| chrVII | 457180 | 457330 | 150 | -1,539786381 | N7:457208 | 7,868 | +5*:YGL020C; | 5 | TERM | | |
| chrVII | 457820 | 457954 | 134 | -1,197197554 | N7:457886 | 3,983 | -1:YGL019W; +1:YGL020C; | -1 | | | |
| chrVII | 458030 | 458181 | 151 | -1,188453173 | N7:458116 | 14,538 | +1:YGL019W; -1:YGL020C; | 1 | | | |
| chrVII | 464168 | 464326 | 158 | -1,563938962 | N7:464210 | 35,227 | +16:YGL016W; | 16 | | | |
| chrVII | 468550 | 468701 | 151 | -1,297502165 | N7:468622 | 12,5 | +19:YGL014W; | 19 | | | |
| chrVII | 470833 | 470993 | 160 | -1,289167805 | | | | Overlap <50 bp | | | |
| chrVII | 471018 | 471094 | 76 | -0,784 | N7:471038 | 27,175 | +9:YGL013C; | 9 | | | |
| chrVII | 471018 | 471123 | 105 | -0,340437773 | N7:471038 | 27,175 | +9:YGL013C; | 9 | | | |
| chrVII | 471173 | 471322 | 149 | -1,139935366 | N7:471209 | 46,098 | +8:YGL013C; | 8 | | | |
| chrVII | 475774 | 475931 | 157 | -1,373988807 | N7:475867 | 10,889 | +3:YGL010W; | 3 | | | |
| chrVII | 476346 | 476454 | 108 | -1,305185286 | | | | Overlap <50 bp | | | |
| chrVII | 476346 | 476465 | 119 | -1,620063938 | | | | Overlap <50 bp | | | |
| chrVII | 478282 | 478437 | 155 | -0,897100408 | N7:478357 | 17,783 | +3:YGL009C; | 3 | | | |
| chrVII | 482579 | 482701 | 122 | -1,241825791 | N7:482600 | 20,659 | +3:YGL008C; | 3 | | | |
| chrVII | 483653 | 483754 | 101 | -0,823890476 | N7:483643 | 36,062 | +2:SUT548; | 2 | | | |
| chrVII | 495041 | 495196 | 155 | -1,058623414 | N7:495024 | 24,326 | +4:YGL002W; | 4 | | | |
| chrVII | 501090 | 501258 | 168 | -1,499675317 | N7:501148 | 9,873 | +7:YGR003W; | 7 | | | |
| chrVII | 502391 | 502522 | 131 | -1,209325931 | N7:502483 | 13,866 | +15*:YGR003W; | 15 | TERM | | |
| chrVII | 502870 | 503027 | 157 | -1,406405847 | N7:502941 | 6,976 | +1:YGR004W; | 1 | | | |
| chrVII | 504629 | 504778 | 149 | -1,361011428 | N7:504726 | 14,919 | +8:YGR005C; | 8 | | | |
| chrVII | 505104 | 505255 | 151 | -1,424476987 | N7:505190 | 8,62 | +5:YGR005C; | 5 | | | |
| chrVII | 509103 | 509251 | 148 | -1,244391604 | N7:509169 | 19,167 | | Intergene | | | |
| chrVII | 509261 | 509410 | 149 | -1,523709292 | N7:509325 | 12,936 | +12*:YGR009C; | 12 | TERM | | |
| chrVII | 510438 | 510590 | 152 | -1,426014425 | N7:510527 | 7,765 | +4:YGR009C; | 4 | | | |
| chrVII | 512325 | 512500 | 175 | -1,283216756 | N7:512399 | 15,62 | +6:YGR010W; | 6 | | | |
| chrVII | 515194 | 515319 | 125 | -1,02938617 | N7:515214 | 14,283 | +5:YGR013W; | 5 | | | |
| chrVII | 516112 | 516262 | 150 | -1,459946635 | N7:516278 | 20,731 | 0:CUT604; +12:YGR013W; | 0 | | | |
| chrVII | 516112 | 516262 | 150 | -1,459946635 | N7:516125 | 8,77 | +2:CUT604; +11:YGR013W; | 2 | | | |
| chrVII | 517734 | 517882 | 148 | -1,265985578 | N7:517806 | 10,64 | +8:YGR014W; | 8 | | | |
| chrVII | 517736 | 517882 | 146 | -1,773840711 | N7:517806 | 10,64 | +8:YGR014W; | 8 | | | |
| chrVII | 518337 | 518484 | 147 | -1,34590264 | N7:518402 | 6,058 | +12:YGR014W; | 12 | | | |
| chrVII | 519407 | 519563 | 156 | -1,275596467 | N7:519436 | 17,328 | +18:YGR014W; | 18 | | | |
| chrVII | 519674 | 519834 | 160 | -1,284890704 | | | | Overlap <50 bp | | | |
| chrVII | 519935 | 520102 | 167 | -1,32908629 | N7:520057 | 32,474 | +21:YGR014W; | 21 | | | |
| chrVII | 520743 | 520902 | 159 | -1,252213009 | N7:520845 | 32,141 | +25*:YGR014W; | 25 | TERM | | |
| chrVII | 520956 | 521107 | 151 | -1,722404401 | N7:521078 | 28,902 | +7*:YGR015C; | 7 | TERM | | |
| chrVII | 526220 | 526384 | 164 | -1,708989585 | N7:526309 | 8,06 | | Intergene | | | |
| chrVII | 529947 | 530056 | 109 | -0,349196698 | N7:529978 | 13,7 | +6:YGR023W; | 6 | | | |
| chrVII | 535422 | 535536 | 114 | -1,389281652 | N7:535504 | 49,636 | -1:CUT606; | -1 | | | |
| chrVII | 536123 | 536272 | 149 | -1,239770123 | N7:536209 | 2,828 | | Intergene | | | |
| chrVII | 536273 | 536379 | 106 | -0,892230054 | | | | Overlap <50 bp | | | |
| chrVII | 536273 | 536391 | 118 | -1,063166576 | | | | Overlap <50 bp | | | |
| chrVII | 536297 | 536420 | 123 | -0,988501489 | | | | Overlap <50 bp | | | |
| chrVII | 536397 | 536552 | 155 | -1,226783257 | N7:536510 | 31,048 | | Intergene | | | |
| chrVII | 536830 | 536964 | 134 | -1,175275641 | N7:536902 | 26,577 | | Intergene | | | |

| CHR | START | END | L(bp) | ΔLK | Nuc ID | Fuzziness | Gene ID | Gene body postion or intergene | Terminal | rDNA | Telomeric |
|---|---|---|---|---|---|---|---|---|---|---|---|
| chrVII | 536830 | 536970 | 140 | -1,179556935 | N7:536902 | 26,577 | | Intergene | | | |
| chrVII | 537026 | 537177 | 151 | -1,153255118 | N7:537109 | 26,458 | | Intergene | | | |
| chrVII | 537036 | 537181 | 145 | -1,115608884 | N7:537109 | 26,458 | | Intergene | | | |
| chrVII | 537308 | 537462 | 154 | -1,230613727 | N7:537445 | 10 | | Intergene | | | |
| chrVII | 537342 | 537486 | 144 | -1,114514704 | N7:537445 | 10 | | Intergene | | | |
| chrVII | 537873 | 538008 | 135 | -1,200992722 | N7:537917 | 2,887 | | Intergene | | | |
| chrVII | 538055 | 538187 | 132 | -1,239847503 | | | | Overlap <50 bp | | | |
| chrVII | 538144 | 538282 | 138 | -1,27156563 | N7:538222 | 17,678 | | Intergene | | | |
| chrVII | 538499 | 538656 | 157 | -1,465812911 | N7:538585 | 30 | | Intergene | | | |
| chrVII | 538599 | 538743 | 144 | -1,39422548 | N7:538585 | 30 | | Intergene | | | |
| chrVII | 538990 | 539119 | 129 | -1,233675531 | | | | Overlap <50 bp | | | |
| chrVII | 539470 | 539601 | 131 | -1,454414983 | N7:539492 | 52,747 | | Intergene | | | |
| chrVII | 539902 | 540052 | 150 | -1,671066088 | N7:539986 | 21,502 | | Intergene | | | |
| chrVII | 540853 | 541004 | 151 | -1,259961489 | N7:540903 | 16,971 | | Intergene | | | |
| chrVII | 542644 | 542795 | 151 | -1,316033793 | N7:542763 | 21,005 | +5:YGR028W; | 5 | | | |
| chrVII | 544012 | 544140 | 128 | -1,015211528 | N7:544085 | 10,257 | +3:SUT550; | 3 | | | |
| chrVII | 545414 | 545580 | 166 | -0,946329183 | N7:545553 | 40,847 | | Intergene | | | |
| chrVII | 546693 | 546840 | 147 | -0,658631856 | N7:546767 | 5,565 | +3:YGR031W; | 3 | | | |
| chrVII | 548258 | 548409 | 151 | -1,229340682 | N7:548239 | 22,448 | +1:YGR032W; | 1 | | | |
| chrVII | 554460 | 554622 | 162 | -1,325179451 | N7:554535 | 15,349 | +4:YGR033C; | 4 | | | |
| chrVII | 554946 | 555089 | 143 | -1,472131007 | N7:555041 | 5,845 | +1:YGR033C; | 1 | | | |
| chrVII | 562268 | 562415 | 147 | -1,288448153 | N7:562318 | 11,79 | | Intergene | | | |
| chrVII | 562775 | 562889 | 114 | -1,425104861 | | | | Overlap <50 bp | | | |
| chrVII | 563471 | 563621 | 150 | -1,316115291 | N7:563527 | 36,665 | | Intergene | | | |
| chrVII | 564404 | 564533 | 129 | -0,943711102 | | | | Overlap <50 bp | | | |
| chrVII | 564867 | 565024 | 157 | -1,34156453 | N7:564949 | 19,975 | | Intergene | | | |
| chrVII | 565335 | 565496 | 161 | -1,356854588 | N7:565396 | 7,778 | | Intergene | | | |
| chrVII | 565515 | 565650 | 135 | -1,101787127 | N7:565587 | 55,126 | | Intergene | | | |
| chrVII | 566553 | 566693 | 140 | -1,209913358 | N7:566644 | 34,598 | | Intergene | | | |
| chrVII | 566559 | 566693 | 134 | -0,825968925 | N7:566644 | 34,598 | | Intergene | | | |
| chrVII | 566971 | 567126 | 155 | -1,307082354 | N7:567004 | 20,809 | | Intergene | | | |
| chrVII | 567102 | 567210 | 108 | -0,925011704 | N7:567215 | 49,003 | | Intergene | | | |
| chrVII | 568106 | 568272 | 166 | -1,317516187 | N7:568185 | 17,729 | | Intergene | | | |
| chrVII | 568932 | 569081 | 149 | -1,276465994 | N7:568958 | 7,55 | | Intergene | | | |
| chrVII | 569719 | 569885 | 166 | -1,923964984 | N7:569812 | 18,148 | | Intergene | | | |
| chrVII | 571017 | 571160 | 143 | -1,103078113 | N7:571098 | 15,885 | | Intergene | | | |
| chrVII | 572385 | 572549 | 164 | -1,281663843 | N7:572444 | 21,455 | | Intergene | | | |
| chrVII | 572497 | 572626 | 129 | -1,174394484 | N7:572607 | 26,577 | | Intergene | | | |
| chrVII | 573872 | 573982 | 110 | -1,342307143 | N7:573948 | 17,214 | | Intergene | | | |
| chrVII | 575134 | 575270 | 136 | -1,298923694 | N7:575205 | 10,187 | +2:YGR040W; | 2 | | | |
| chrVII | 575584 | 575763 | 179 | -1,441272738 | N7:575570 | 24,546 | | Intergene | | | |
| chrVII | 575584 | 575763 | 179 | -1,441272738 | N7:575745 | 8,905 | | Intergene | | | |
| chrVII | 578432 | 578606 | 174 | -1,570721487 | N7:578577 | 30,004 | +5:SUT551; +9:YGR041W; | 5 | | | |
| chrVII | 578432 | 578606 | 174 | -1,570721487 | N7:578412 | 51,237 | +6:SUT551; +8:YGR041W; | 6 | | | |
| chrVII | 578979 | 579129 | 150 | -1,222984445 | N7:579094 | 11,234 | +2:SUT551; +12*:YGR041W; | 2 | | | |
| chrVII | 579144 | 579294 | 150 | -1,460838059 | N7:579259 | 11,786 | +1:SUT551; -1:YGR042W; | 1 | | | |
| chrVII | 583828 | 583997 | 169 | -1,089287822 | N7:583861 | 10,759 | +1:YGR044C; | 1 | | | |
| chrVII | 584107 | 584258 | 151 | -1,271826777 | N7:584192 | 28,369 | -1:YGR044C; | -1 | | | |
| chrVII | 584200 | 584336 | 136 | -0,674520287 | N7:584192 | 28,369 | -1:YGR044C; | -1 | | | |
| chrVII | 584954 | 585099 | 145 | -1,497111235 | N7:584996 | 7,859 | +2:YGR046W; | 2 | | | |
| chrVII | 594298 | 594445 | 147 | -1,036821297 | N7:594404 | 21,664 | +6:YGR052W; | 6 | | | |
| chrVII | 595048 | 595213 | 165 | -1,648510807 | N7:595207 | 9,416 | +5:YGR053C; | 5 | | | |
| chrVII | 595048 | 595213 | 165 | -1,648510807 | N7:595043 | 17,394 | +6:YGR053C; | 6 | | | |
| chrVII | 595281 | 595432 | 151 | -1,249140419 | N7:595369 | 5,955 | +4:YGR053C; | 4 | | | |
| chrVII | 610980 | 611160 | 180 | -1,254965427 | N7:610959 | 31,098 | +4:YGR060W; | 4 | | | |
| chrVII | 610980 | 611160 | 180 | -1,254965427 | N7:611131 | 11,845 | +5:YGR060W; | 5 | | | |
| chrVII | 613266 | 613414 | 148 | -1,326462944 | N7:613372 | 40,623 | +17:YGR061C; | 17 | | | |
| chrVII | 637713 | 637869 | 156 | -1,297848873 | N7:637824 | 8,14 | +11:YGR077C; -1:YGR076C; | 11 | | | |
| chrVII | 640000 | 640148 | 148 | -1,327519137 | N7:640070 | 9,883 | +3:YGR078C; | 3 | | | |
| chrVII | 640002 | 640148 | 146 | -1,68990033 | N7:640070 | 9,883 | +3:YGR078C; | 3 | | | |
| chrVII | 640719 | 640865 | 146 | -1,713622115 | N7:640822 | 24,427 | +1:YGR079W; | 1 | | | |
| chrVII | 652407 | 652570 | 163 | -1,538933903 | N7:652507 | 3,371 | +4:YGR087C; | 4 | | | |
| chrVII | 652780 | 652925 | 145 | -1,116862565 | N7:652859 | 7,563 | +2:YGR087C; | 2 | | | |
| chrVII | 654255 | 654401 | 146 | -1,258719467 | N7:654398 | 16,94 | -1:YGR088W; | -1 | | | |
| chrVII | 654255 | 654401 | 146 | -1,258719467 | N7:654248 | 17,349 | | Intergene | | | |
| chrVII | 668831 | 668991 | 160 | -1,485326114 | N7:668917 | 30,091 | +6:YGR092W; | 6 | | | |
| chrVII | 671142 | 671294 | 152 | -1,53570138 | | | | Overlap <50 bp | | | |
| chrVII | 671676 | 671849 | 173 | -1,063294291 | N7:671743 | 11,76 | +9:YGR093W; +2:CUT614; | 9 | | | |
| chrVII | 674557 | 674713 | 156 | -1,354891259 | N7:674628 | 18,992 | +16:YGR094W; | 16 | | | |
| chrVII | 676654 | 676817 | 163 | -1,050555283 | N7:676774 | 11,432 | +2:YGR096W; | 2 | | | |
| chrVII | 680899 | 681051 | 152 | -1,136803514 | N7:680980 | 14,61 | +16:YGR097W; | 16 | | | |
| chrVII | 681312 | 681478 | 166 | -1,599713766 | N7:681411 | 21,461 | | Intergene | | | |
| chrVII | 682534 | 682696 | 162 | -1,434187556 | N7:682605 | 12,75 | +32:YGR098C; | 32 | | | |
| chrVII | 687608 | 687758 | 150 | -1,400455956 | N7:687651 | 5,01 | -1:YGR099W; +1:YGR098C; | -1 | | | |
| chrVII | 689536 | 689689 | 153 | -1,219952645 | N7:689603 | 31,394 | +11:YGR099W; -1:YGR100W; | 11 | | | |
| chrVII | 692091 | 692197 | 106 | -1,411156342 | N7:692094 | 15,549 | +15:YGR100W; | 15 | | | |
| chrVII | 692091 | 692218 | 127 | -0,896234228 | N7:692094 | 15,549 | +15:YGR100W; | 15 | | | |
| chrVII | 692091 | 692219 | 128 | -1,075965493 | N7:692094 | 15,549 | +15:YGR100W; | 15 | | | |
| chrVII | 692091 | 692220 | 129 | -1,191786782 | N7:692094 | 15,549 | +15:YGR100W; | 15 | | | |
| chrVII | 692091 | 692221 | 130 | -1,533516088 | N7:692094 | 15,549 | +15:YGR100W; | 15 | | | |
| chrVII | 692091 | 692222 | 131 | -1,358763356 | N7:692094 | 15,549 | +15:YGR100W; | 15 | | | |
| chrVII | 692091 | 692223 | 132 | -1,051681441 | N7:692094 | 15,549 | +15:YGR100W; | 15 | | | |
| chrVII | 692092 | 692223 | 131 | -1,161381983 | N7:692094 | 15,549 | +15:YGR100W; | 15 | | | |
| chrVII | 692095 | 692223 | 128 | -1,01414832 | N7:692094 | 15,549 | +15:YGR100W; | 15 | | | |
| chrVII | 692097 | 692197 | 100 | -0,605408406 | N7:692094 | 15,549 | +15:YGR100W; | 15 | | | |
| chrVII | 693772 | 693921 | 149 | -0,85173129 | N7:693854 | 9,116 | +4:YGR101W; | 4 | | | |
| chrVII | 697506 | 697652 | 146 | -1,277158175 | N7:697541 | 26,334 | +6*:YGR104C; | 6 | TERM | | |
| chrVII | 697827 | 697977 | 150 | -1,228775726 | N7:697877 | 27,396 | +4:YGR104C; | 4 | | | |
| chrVII | 701841 | 701992 | 151 | -1,139744544 | | | | Overlap <50 bp | | | |
| chrVII | 707416 | 707536 | 120 | -1,196073719 | N7:707505 | 13,204 | | Intergene | | | |
| chrVII | 709744 | 709890 | 146 | -1,336973352 | N7:709832 | 3,055 | | Intergene | | | |
| chrVII | 712418 | 712569 | 151 | -1,305648025 | N7:712512 | 26,569 | | Intergene | | | |
| chrVII | 714004 | 714140 | 136 | -1,16641641 | N7:714042 | 30,224 | +3:YGR110W; | 3 | | | |
| chrVII | 716031 | 716189 | 158 | -1,678539305 | | | | Overlap <50 bp | | | |
| chrVII | 716536 | 716689 | 153 | -1,722950029 | N7:716629 | 4,924 | +7:YGR111W; | 7 | | | |
| chrVII | 726352 | 726496 | 144 | -1,135990245 | N7:726431 | 19,356 | +1:YGR117C; | 1 | | | |
| chrVII | 731531 | 731680 | 149 | -1,297052363 | N7:731570 | 10,178 | +2:SUT128; +8:YGR121C; | 2 | | | |

| CHR | START | END | L(bp) | ΔLK | Nuc ID | Fuzziness | Gene ID | Gene body postion or intergene | Terminal | rDNA | Telomeric |
|---|---|---|---|---|---|---|---|---|---|---|---|
| chrVII | 740022 | 740127 | 105 | -1,464218768 | N7:740078 | 19,476 | +2:YGR124W; | 2 | | | |
| chrVII | 746153 | 746303 | 150 | -1,589410555 | N7:746206 | 16,724 | GR126W; +3:CUT618; -1:YGR127W; | 5 | | | |
| chrVII | 747978 | 748102 | 124 | -1,260440085 | N7:748002 | 23,392 | +13:YGR128C; | 13 | | | |
| chrVII | 748361 | 748501 | 140 | -0,946734144 | N7:748439 | 49,872 | +11:YGR128C; | 11 | | | |
| chrVII | 751512 | 751624 | 112 | -1,270813929 | N7:751606 | 17,302 | +14:YGR130C; +2:SUT130; | 14 | | | |
| chrVII | 758738 | 758894 | 156 | -1,041544541 | N7:758814 | 16,395 | +8:YGR134W; +13:YGR133W; | 8 | | | |
| chrVII | 760806 | 760962 | 156 | -1,257734376 | N7:760938 | 6,351 | R134W; +2:CUT619; +25:YGR133W; | 20 | | | |
| chrVII | 760828 | 760978 | 150 | -1,212257977 | N7:760938 | 6,351 | R134W; +2:CUT619; +25:YGR133W; | 20 | | | |
| chrVII | 762415 | 762565 | 150 | -1,222057706 | N7:762428 | 16,741 | -1:CUT620; +1:YGR136W; | -1 | | | |
| chrVII | 766213 | 766368 | 155 | -1,058979339 | N7:766304 | 36,17 | +4:Unit239; | 4 | | | |
| chrVII | 769383 | 769497 | 114 | -1,049630355 | N7:769444 | 31,04 | +13:YGR140W; | 13 | | | |
| chrVII | 773391 | 773557 | 166 | -1,867772188 | N7:773500 | 35,612 | +7:YGR142W; | 7 | | | |
| chrVII | 776840 | 777001 | 161 | -1,204658885 | N7:777019 | 36,146 | +14:YGR143W; +5:SUT562; | 14 | | | |
| chrVII | 776982 | 777149 | 167 | -1,355650916 | N7:777019 | 36,146 | +14:YGR143W; +5:SUT562; | 14 | | | |
| chrVII | 780856 | 781004 | 148 | -1,045575989 | N7:780944 | 21,173 | +4:YGR144W; | 4 | | | |
| chrVII | 781219 | 781380 | 161 | -1,300784208 | N7:781318 | 18,534 | +6:YGR144W; | 6 | | | |
| chrVII | 782373 | 782522 | 149 | -1,302708936 | | | | Overlap <50 bp | | | |
| chrVII | 782985 | 783136 | 151 | -1,493305693 | N7:783055 | 19,553 | +9:YGR145W; | 9 | | | |
| chrVII | 811114 | 811251 | 137 | -1,648645565 | N7:811183 | 20,461 | | Intergene | | | |
| chrVII | 813520 | 813649 | 129 | -1,268882154 | N7:813530 | 18,175 | | Intergene | | | |
| chrVII | 813597 | 813761 | 164 | -1,300315013 | N7:813691 | 22,502 | | Intergene | | | |
| chrVII | 814985 | 815128 | 143 | -0,979070106 | N7:815042 | 14,189 | | Intergene | | | |
| chrVII | 816260 | 816426 | 166 | -1,608770339 | N7:816238 | 45,181 | | Intergene | | | |
| chrVII | 817064 | 817213 | 149 | -1,449977044 | N7:817170 | 21,572 | | Intergene | | | |
| chrVII | 817068 | 817218 | 150 | -1,278634937 | N7:817170 | 21,572 | | Intergene | | | |
| chrVII | 817863 | 817968 | 105 | -1,791400316 | N7:817860 | 38,837 | | Intergene | | | |
| chrVII | 817863 | 817970 | 107 | -1,525364634 | N7:817860 | 38,837 | | Intergene | | | |
| chrVII | 819019 | 819169 | 150 | -1,313251878 | N7:819103 | 24,007 | | Intergene | | | |
| chrVII | 819762 | 819909 | 147 | -1,366069831 | N7:819930 | 29,698 | | Intergene | | | |
| chrVII | 820415 | 820572 | 157 | -1,557311394 | N7:820493 | 15,948 | | Intergene | | | |
| chrVII | 820601 | 820751 | 150 | -1,132202479 | N7:820670 | 23,352 | | Intergene | | | |
| chrVII | 820792 | 820937 | 145 | -1,019939731 | | | | Overlap <50 bp | | | |
| chrVII | 821063 | 821198 | 135 | -1,041413445 | N7:821136 | 20,841 | | Intergene | | | |
| chrVII | 821345 | 821513 | 168 | -1,220367417 | N7:821422 | 24,96 | | Intergene | | | |
| chrVII | 822101 | 822241 | 140 | -1,157950276 | N7:822182 | 22,338 | | Intergene | | | |
| chrVII | 822107 | 822241 | 134 | -1,10336759 | N7:822182 | 22,338 | | Intergene | | | |
| chrVII | 822519 | 822674 | 155 | -1,621234488 | N7:822552 | 24,502 | | Intergene | | | |
| chrVII | 822650 | 822758 | 108 | -0,829525841 | N7:822752 | 38,083 | | Intergene | | | |
| chrVII | 825754 | 825897 | 143 | -1,137284697 | N7:825833 | 17,326 | +11:YGR162W; | 11 | | | |
| chrVII | 831865 | 832015 | 150 | -1,317760078 | N7:831995 | 7,176 | +10:YGR166W; +2:SUT565; | 10 | | | |
| chrVII | 841785 | 841939 | 154 | -1,362164835 | N7:841891 | 16,415 | +5:YGR171C; | 5 | | | |
| chrVII | 842909 | 843085 | 176 | -1,36868009 | N7:843080 | 3,886 | +4:YGR172C; | 4 | | | |
| chrVII | 842909 | 843085 | 176 | -1,36868009 | N7:842921 | 10,381 | +5:YGR172C; | 5 | | | |
| chrVII | 843924 | 844090 | 166 | -1,013010432 | N7:844011 | 8,477 | +2:YGR173W; | 2 | | | |
| chrVII | 844983 | 845124 | 141 | -0,92933734 | N7:845017 | 21,939 | | Intergene | | | |
| chrVII | 847489 | 847638 | 149 | -1,084795454 | | | | Overlap <50 bp | | | |
| chrVII | 853632 | 853775 | 143 | -1,054244501 | N7:853718 | 6,473 | +8*:YGR179C; | 8 | TERM | | |
| chrVII | 855743 | 855883 | 140 | -1,18475975 | N7:855791 | 22,301 | +5:YGR180C; | 5 | | | |
| chrVII | 855860 | 856013 | 153 | -1,468646796 | N7:855962 | 20,579 | +4:YGR180C; | 4 | | | |
| chrVII | 861234 | 861372 | 138 | -1,352465861 | N7:861384 | 22,728 | +28:YGR184C; | 28 | | | |
| chrVII | 869483 | 869631 | 148 | -1,450254733 | N7:869530 | 23,106 | +12:YGR186W; | 12 | | | |
| chrVII | 873303 | 873436 | 133 | -1,408824646 | N7:873338 | 16,958 | | Intergene | | | |
| chrVII | 873927 | 874079 | 152 | -1,125799159 | N7:873992 | 17,844 | | Intergene | | | |
| chrVII | 878924 | 879054 | 130 | -1,137395017 | N7:878962 | 4,69 | | Intergene | | | |
| chrVII | 879023 | 879172 | 149 | -1,157572469 | | | | Overlap <50 bp | | | |
| chrVII | 879984 | 880088 | 104 | -1,39452278 | | | | Overlap <50 bp | | | |
| chrVII | 882915 | 883031 | 116 | -0,070106345 | N7:883049 | 13,478 | +5:YGR192C; | 5 | | | |
| chrVII | 882915 | 883023 | 108 | 0,034251293 | | | | Overlap <50 bp | | | |
| chrVII | 887398 | 887557 | 159 | -1,41258926 | | | | Overlap <50 bp | | | |
| chrVII | 889980 | 890139 | 159 | -1,57606307 | N7:890158 | 32,249 | +13:YGR196C; | 13 | | | |
| chrVII | 889980 | 890139 | 159 | -1,57606307 | N7:890006 | 29,841 | +14:YGR196C; | 14 | | | |
| chrVII | 892462 | 892609 | 147 | -1,3504593 | N7:892520 | 10,991 | | Intergene | | | |
| chrVII | 895680 | 895828 | 148 | -1,577695109 | N7:895781 | 23,114 | +8:YGR198W; | 8 | | | |
| chrVII | 911520 | 911682 | 162 | -1,215343234 | N7:911624 | 11,489 | +1:YGR207C; -1:YGR208W; | 1 | | | |
| chrVII | 914384 | 914518 | 134 | -0,761231872 | N7:914490 | 8,246 | +3:YGR210C; | 3 | | | |
| chrVII | 920349 | 920433 | 84 | -0,892818035 | N7:920408 | 36,509 | -1:YGR214W; | -1 | | | |
| chrVII | 922436 | 922538 | 102 | -1,670878864 | N7:922490 | 5,345 | +3*:YGR215W; | 3 | TERM | | |
| chrVII | 925873 | 926024 | 151 | -1,457316922 | | | | Overlap <50 bp | | | |
| chrVII | 928464 | 928607 | 143 | -1,544581278 | | | | Overlap <50 bp | | | |
| chrVII | 928560 | 928710 | 150 | -1,32055031 | N7:928631 | 26,21 | +25:YGR217W; | 25 | | | |
| chrVII | 929745 | 929899 | 154 | -1,729924862 | N7:929875 | 25,703 | | Intergene | | | |
| chrVII | 938659 | 938812 | 153 | -1,649277095 | N7:938652 | 16,879 | +4:YGR221C; | 4 | | | |
| chrVII | 943256 | 943409 | 153 | -1,21334625 | N7:943358 | 25,948 | +10:SUT569; +5:YGR224W; | 10 | | | |
| chrVII | 943702 | 943854 | 152 | -1,270994004 | N7:943837 | 20,389 | +7:SUT569; +8:YGR224W; | 7 | | | |
| chrVII | 948590 | 948746 | 156 | -1,555273123 | N7:948665 | 11,219 | +9:YGR227W; | 9 | | | |
| chrVII | 949733 | 949881 | 148 | -1,34873511 | N7:949820 | 22,248 | +8:YGR229C; | 8 | | | |
| chrVII | 960612 | 960754 | 142 | -1,072037737 | N7:960724 | 28,562 | +6:YGR234W; | 6 | | | |
| chrVII | 961680 | 961814 | 134 | -1,742766225 | N7:961727 | 7,403 | +3:YGR235C; | 3 | | | |
| chrVII | 961681 | 961788 | 107 | -1,28236431 | N7:961727 | 7,403 | +3:YGR235C; | 3 | | | |
| chrVII | 961681 | 961791 | 110 | -0,777638249 | N7:961727 | 7,403 | +3:YGR235C; | 3 | | | |
| chrVII | 961681 | 961794 | 113 | -0,7995937 | N7:961727 | 7,403 | +3:YGR235C; | 3 | | | |
| chrVII | 961681 | 961795 | 114 | -0,46740488 | N7:961727 | 7,403 | +3:YGR235C; | 3 | | | |
| chrVII | 961681 | 961796 | 115 | -0,860428875 | N7:961727 | 7,403 | +3:YGR235C; | 3 | | | |
| chrVII | 961681 | 961797 | 116 | -1,290380421 | N7:961727 | 7,403 | +3:YGR235C; | 3 | | | |
| chrVII | 961681 | 961798 | 117 | -0,700419951 | N7:961727 | 7,403 | +3:YGR235C; | 3 | | | |
| chrVII | 961681 | 961799 | 118 | -1,180469031 | N7:961727 | 7,403 | +3:YGR235C; | 3 | | | |
| chrVII | 961681 | 961800 | 119 | -0,385064969 | N7:961727 | 7,403 | +3:YGR235C; | 3 | | | |
| chrVII | 961681 | 961801 | 120 | -0,791752291 | N7:961727 | 7,403 | +3:YGR235C; | 3 | | | |
| chrVII | 961681 | 961802 | 121 | -1,422433316 | N7:961727 | 7,403 | +3:YGR235C; | 3 | | | |
| chrVII | 961681 | 961803 | 122 | -1,218893163 | N7:961727 | 7,403 | +3:YGR235C; | 3 | | | |
| chrVII | 961681 | 961804 | 123 | -1,342559562 | N7:961727 | 7,403 | +3:YGR235C; | 3 | | | |
| chrVII | 961681 | 961805 | 124 | -1,234138058 | N7:961727 | 7,403 | +3:YGR235C; | 3 | | | |
| chrVII | 961681 | 961806 | 125 | -1,975143821 | N7:961727 | 7,403 | +3:YGR235C; | 3 | | | |
| chrVII | 961681 | 961807 | 126 | -1,109819015 | N7:961727 | 7,403 | +3:YGR235C; | 3 | | | |
| chrVII | 961681 | 961808 | 127 | -0,684855061 | N7:961727 | 7,403 | +3:YGR235C; | 3 | | | |
| chrVII | 961681 | 961809 | 128 | -1,068102773 | N7:961727 | 7,403 | +3:YGR235C; | 3 | | | |
| chrVII | 961681 | 961810 | 129 | -1,2768176 | N7:961727 | 7,403 | +3:YGR235C; | 3 | | | |
| chrVII | 961681 | 961811 | 130 | -1,232362305 | N7:961727 | 7,403 | +3:YGR235C; | 3 | | | |

TABLE 3

| CHR | START | END | L(bp) | ΔLK | Nuc ID | Fuzziness | Gene ID | Gene body postion or intergene | Terminal | rDNA | Telomeric |
|---|---|---|---|---|---|---|---|---|---|---|---|
| chrVII | 961681 | 961812 | 131 | -1,036456889 | N7:961727 | 7,403 | +3:YGR235C; | 3 | | | |
| chrVII | 961681 | 961813 | 132 | -1,433047964 | N7:961727 | 7,403 | +3:YGR235C; | 3 | | | |
| chrVII | 961681 | 961814 | 133 | -1,05489492 | N7:961727 | 7,403 | +3:YGR235C; | 3 | | | |
| chrVII | 961682 | 961814 | 132 | -1,195670018 | N7:961727 | 7,403 | +3:YGR235C; | 3 | | | |
| chrVII | 961683 | 961814 | 131 | -1,251596042 | N7:961727 | 7,403 | +3:YGR235C; | 3 | | | |
| chrVII | 961684 | 961814 | 130 | -1,588790809 | N7:961727 | 7,403 | +3:YGR235C; | 3 | | | |
| chrVII | 961685 | 961814 | 129 | -1,096625562 | N7:961727 | 7,403 | +3:YGR235C; | 3 | | | |
| chrVII | 961686 | 961814 | 128 | -0,79349347 | N7:961727 | 7,403 | +3:YGR235C; | 3 | | | |
| chrVII | 961687 | 961814 | 127 | -0,949634893 | N7:961727 | 7,403 | +3:YGR235C; | 3 | | | |
| chrVII | 961694 | 961814 | 120 | -1,163899238 | N7:961727 | 7,403 | +3:YGR235C; | 3 | | | |
| chrVII | 961695 | 961814 | 119 | -0,780301144 | N7:961727 | 7,403 | +3:YGR235C; | 3 | | | |
| chrVII | 961696 | 961814 | 118 | -1,099381997 | N7:961727 | 7,403 | +3:YGR235C; | 3 | | | |
| chrVII | 961708 | 961806 | 98 | -0,949572954 | N7:961727 | 7,403 | +3:YGR235C; | 3 | | | |
| chrVII | 962925 | 963072 | 147 | -1,341990433 | N7:963033 | 25,365 | 0:YGR236C; | 0 | | | |
| chrVII | 966527 | 966637 | 110 | -1,359676012 | N7:966590 | 31,713 | +14:YGR238C; | 14 | | | |
| chrVII | 974170 | 974327 | 157 | -1,419815787 | N7:974249 | 28,729 | -1:YGR240C; 0:SUT141; | -1 | | | |
| chrVII | 975022 | 975129 | 107 | -0,96158172 | N7:975032 | 21,194 | T164; -1:YGR240C-A; +12:YGR241C; | 1 | | | |
| chrVII | 975022 | 975130 | 108 | -1,018475442 | N7:975032 | 21,194 | T164; -1:YGR240C-A; +12:YGR241C; | 1 | | | |
| chrVII | 975022 | 975131 | 109 | -1,252938788 | N7:975032 | 21,194 | T164; -1:YGR240C-A; +12:YGR241C; | 1 | | | |
| chrVII | 975022 | 975142 | 120 | -1,209455307 | N7:975032 | 21,194 | T164; -1:YGR240C-A; +12:YGR241C; | 1 | | | |
| chrVII | 979943 | 980105 | 162 | -1,192441561 | N7:980040 | 9,68 | +14:YGR245C; | 14 | | | |
| chrVII | 982482 | 982630 | 148 | -1,079859202 | N7:982592 | 16,598 | +11:YGR246C; | 11 | | | |
| chrVII | 987451 | 987604 | 153 | -1,60399418 | N7:987557 | 29,569 | -1:YGR249W; | -1 | | | |
| chrVII | 987451 | 987606 | 155 | -1,673811507 | N7:987557 | 29,569 | -1:YGR249W; | -1 | | | |
| chrVII | 987451 | 987608 | 157 | -1,674417946 | N7:987557 | 29,569 | -1:YGR249W; | -1 | | | |
| chrVII | 987451 | 987609 | 158 | -1,429625702 | N7:987557 | 29,569 | -1:YGR249W; | -1 | | | |
| chrVII | 988814 | 988967 | 153 | -1,24153461 | N7:988944 | 20,516 | +8:YGR249W; | 8 | | | |
| chrVII | 988881 | 989019 | 138 | -0,803715516 | N7:988944 | 20,516 | +8:YGR249W; | 8 | | | |
| chrVII | 989319 | 989475 | 156 | -1,197266065 | N7:989398 | 12,219 | +11:YGR249W; | 11 | | | |
| chrVII | 989319 | 989477 | 158 | -1,328625083 | N7:989398 | 12,219 | +11:YGR249W; | 11 | | | |
| chrVII | 991949 | 992103 | 154 | -1,189031495 | N7:992011 | 23,264 | +12:YGR250C; | 12 | | | |
| chrVII | 999326 | 999473 | 147 | -1,464530646 | N7:999420 | 27,779 | -1:YGR253C; | -1 | | | |
| chrVII | 1001929 | 1002078 | 149 | -1,127199798 | N7:1001976 | 38,888 | +7:YGR254W; | 7 | | | |
| chrVII | 1012079 | 1012238 | 159 | -1,601341534 | N7:1012073 | 20,617 | -1:YGR260W; +2:CUT634; | -1 | | | |
| chrVII | 1012603 | 1012726 | 123 | -1,500318607 | N7:1012698 | 5,523 | | Intergene | | | |
| chrVII | 1016668 | 1016789 | 121 | -0,969040897 | N7:1016735 | 11,409 | +1:YGR261C; | 1 | | | |
| chrVII | 1018529 | 1018686 | 157 | -1,091452275 | N7:1018544 | 6,325 | +4:CUT167; +5:YGR263C; | 4 | | | |
| chrVII | 1018639 | 1018770 | 131 | -1,028881385 | N7:1018717 | 13,049 | +5:CUT167; +4:YGR263C; | 5 | | | |
| chrVII | 1024687 | 1024847 | 160 | -0,984500662 | | | | Overlap <50 bp | | | |
| chrVII | 1030700 | 1030865 | 165 | -1,26404173 | N7:1030777 | 23,824 | +22:YGR270W; | 22 | | | |
| chrVII | 1030899 | 1031060 | 161 | -1,492972672 | N7:1030983 | 49,859 | +23:YGR270W; | 23 | | | |
| chrVII | 1037731 | 1037891 | 160 | -1,60394904 | N7:1037843 | 23,104 | +2:YGR271C-A; | 2 | | | |
| chrVII | 1039928 | 1040095 | 167 | -1,353507373 | N7:1040010 | 30,636 | +20*:YGR274C; | 20 | TERM | | |
| chrVII | 1043906 | 1044058 | 152 | -1,109871585 | N7:1044003 | 17,964 | | Intergene | | | |
| chrVII | 1044184 | 1044344 | 160 | -1,342215261 | N7:1044171 | 23,02 | | Intergene | | | |
| chrVII | 1045755 | 1045901 | 146 | -1,234862619 | N7:1045865 | 12,75 | +5:YGR277C; -1:YGR276C; | 5 | | | |
| chrVII | 1049209 | 1049360 | 151 | -1,38506989 | N7:1049248 | 27,636 | -1:SUT144; +6:YGR279C; | -1 | | | |
| chrVII | 1051642 | 1051789 | 147 | -1,049867551 | N7:1051693 | 14,588 | +1:YGR280C; | 1 | | | |
| chrVII | 1052012 | 1052165 | 153 | -1,257690487 | N7:1052015 | 11,021 | 0:Unit253; -1:YGR280C; | 0 | | | |
| chrVII | 1065087 | 1065242 | 155 | -1,323574383 | N7:1065159 | 8,976 | 0:YGR286C; | 0 | | | |
| chrVII | 1065864 | 1066016 | 152 | -1,117659017 | N7:1065932 | 11,345 | +3*:CUT170; | 3 | TERM | | |
| chrVII | 1065866 | 1066016 | 150 | -1,387352075 | N7:1065932 | 11,345 | +3*:CUT170; | 3 | TERM | | |
| chrVII | 1065867 | 1066016 | 149 | -1,215335159 | N7:1065932 | 11,345 | +3*:CUT170; | 3 | TERM | | |
| chrVII | 1067431 | 1067584 | 153 | -1,321335082 | N7:1067518 | 36,979 | +11:YGR287C; | 11 | | | |
| chrVII | 1071145 | 1071303 | 158 | -1,235928613 | N7:1071240 | 35,836 | +7:YGR288W; | 7 | | | |
| chrVII | 1079049 | 1079187 | 138 | -1,661784877 | N7:1079141 | 4,792 | +4:Unit255; +4:Unit287; | 4 | | | |
| chrVII | 1079684 | 1079846 | 162 | -1,349728491 | N7:1079686 | 11,345 | +1:Unit255; | 1 | | | |
| chrVII | 1080323 | 1080452 | 129 | -1,313721496 | N7:1080409 | 41,191 | +2:YGR294W; | 2 | | | |
| chrVII | 1085801 | 1085942 | 141 | -1,211450567 | N7:1085920 | 22,113 | +7:YGR296W; | 7 | | | TEL RIGHT |
| chrVII | 1087005 | 1087167 | 162 | -1,320183316 | N7:1086993 | 15,567 | +13:YGR296W; | 13 | | | TEL RIGHT |
| chrVII | 1087005 | 1087167 | 162 | -1,320183316 | N7:1087148 | 18 | +14:YGR296W; | 14 | | | TEL RIGHT |
| chrVII | 1087106 | 1087261 | 155 | -1,844548914 | N7:1087148 | 18 | +14:YGR296W; | 14 | | | TEL RIGHT |
| chrVII | 1087240 | 1087377 | 137 | -1,199143123 | N7:1087315 | 9,815 | +15:YGR296W; | 15 | | | TEL RIGHT |
| chrVII | 1087527 | 1087678 | 151 | -1,23742729 | N7:1087597 | 17 | +17:YGR296W; | 17 | | | TEL RIGHT |
| chrVII | 1087535 | 1087678 | 143 | -1,572093981 | N7:1087597 | 17 | +17:YGR296W; | 17 | | | TEL RIGHT |
| chrVII | 1087583 | 1087708 | 125 | -0,693715414 | N7:1087597 | 17 | +17:YGR296W; | 17 | | | TEL RIGHT |
| chrVII | 1087808 | 1087960 | 152 | -1,219283862 | N7:1087922 | 14,224 | +19:YGR296W; | 19 | | | TEL RIGHT |
| chrVII | 1087881 | 1088025 | 144 | -1,023063265 | N7:1087922 | 14,224 | +19:YGR296W; | 19 | | | TEL RIGHT |
| chrVII | 1087978 | 1088113 | 135 | -0,993309501 | N7:1088072 | 37,041 | +20:YGR296W; | 20 | | | TEL RIGHT |
| chrVII | 1087978 | 1088165 | 187 | -0,838896224 | N7:1088072 | 37,041 | +20:YGR296W; | 20 | | | TEL RIGHT |
| chrVII | 1088021 | 1088113 | 92 | -0,478563188 | N7:1088072 | 37,041 | +20:YGR296W; | 20 | | | TEL RIGHT |
| chrVII | 1088021 | 1088165 | 144 | -0,955510176 | N7:1088072 | 37,041 | +20:YGR296W; | 20 | | | TEL RIGHT |
| chrVII | 1088360 | 1088512 | 152 | -1,377737626 | N7:1088436 | 17,673 | +22:YGR296W; | 22 | | | TEL RIGHT |
| chrVII | 1088661 | 1088791 | 130 | -0,959485957 | N7:1088758 | 37,242 | +24:YGR296W; | 24 | | | TEL RIGHT |
| chrVII | 1089031 | 1089182 | 151 | -1,256789845 | N7:1089163 | 38,734 | +26:YGR296W; | 26 | | | TEL RIGHT |
| chrVII | 1089036 | 1089170 | 134 | -0,957153105 | N7:1089163 | 38,734 | +26:YGR296W; | 26 | | | TEL RIGHT |
| chrVII | 1089049 | 1089145 | 96 | -0,702783422 | N7:1089163 | 38,734 | +26:YGR296W; | 26 | | | TEL RIGHT |
| chrVII | 1090196 | 1090347 | 151 | -1,276105724 | N7:1090283 | 36,77 | +32:YGR296W; | 32 | | | TEL RIGHT |
| chrVIII | 691 | 842 | 151 | -1,214555062 | N8:754 | 11,136 | +18:YHL050C; | 18 | | | TEL LEFT |
| chrVIII | 1289 | 1405 | 116 | -0,794251441 | N8:1300 | 28,74 | +14:YHL050C; | 14 | | | TEL LEFT |
| chrVIII | 1346 | 1501 | 155 | -1,310772488 | | | | Overlap <50 bp | | | TEL LEFT |
| chrVIII | 1712 | 1899 | 187 | -1,508362362 | N8:1845 | 18,083 | +10:YHL050C; | 10 | | | TEL LEFT |
| chrVIII | 1733 | 1899 | 166 | -2,25101612 | N8:1845 | 18,083 | +10:YHL050C; | 10 | | | TEL LEFT |
| chrVIII | 1748 | 1846 | 98 | -0,827137784 | N8:1845 | 18,083 | +10:YHL050C; | 10 | | | TEL LEFT |
| chrVIII | 1748 | 1855 | 107 | -1,084236509 | N8:1845 | 18,083 | +10:YHL050C; | 10 | | | TEL LEFT |
| chrVIII | 1748 | 1878 | 130 | -1,568753508 | N8:1845 | 18,083 | +10:YHL050C; | 10 | | | TEL LEFT |
| chrVIII | 1748 | 1879 | 131 | -1,015364392 | N8:1845 | 18,083 | +10:YHL050C; | 10 | | | TEL LEFT |
| chrVIII | 1748 | 1880 | 132 | -1,185239046 | N8:1845 | 18,083 | +10:YHL050C; | 10 | | | TEL LEFT |
| chrVIII | 1748 | 1889 | 141 | -1,416915625 | N8:1845 | 18,083 | +10:YHL050C; | 10 | | | TEL LEFT |
| chrVIII | 1748 | 1891 | 143 | -1,351785569 | N8:1845 | 18,083 | +10:YHL050C; | 10 | | | TEL LEFT |
| chrVIII | 1748 | 1894 | 146 | -1,027556964 | N8:1845 | 18,083 | +10:YHL050C; | 10 | | | TEL LEFT |
| chrVIII | 1748 | 1895 | 147 | -1,391704254 | N8:1845 | 18,083 | +10:YHL050C; | 10 | | | TEL LEFT |
| chrVIII | 1748 | 1896 | 148 | -1,128648232 | N8:1845 | 18,083 | +10:YHL050C; | 10 | | | TEL LEFT |
| chrVIII | 1748 | 1897 | 149 | -1,415258873 | N8:1845 | 18,083 | +10:YHL050C; | 10 | | | TEL LEFT |
| chrVIII | 1748 | 1898 | 150 | -1,719025954 | N8:1845 | 18,083 | +10:YHL050C; | 10 | | | TEL LEFT |
| chrVIII | 1748 | 1899 | 151 | -1,15736106 | N8:1845 | 18,083 | +10:YHL050C; | 10 | | | TEL LEFT |
| chrVIII | 1748 | 1909 | 161 | -2,11615602 | N8:1845 | 18,083 | +10:YHL050C; | 10 | | | TEL LEFT |
| chrVIII | 1748 | 1932 | 184 | -1,904963662 | N8:1845 | 18,083 | +10:YHL050C; | 10 | | | TEL LEFT |
| chrVIII | 1749 | 1899 | 150 | -1,575487647 | N8:1845 | 18,083 | +10:YHL050C; | 10 | | | TEL LEFT |
| chrVIII | 1750 | 1899 | 149 | -1,574888675 | N8:1845 | 18,083 | +10:YHL050C; | 10 | | | TEL LEFT |

| CHR | START | END | L(bp) | ΔLK | Nuc ID | Fuzziness | Gene ID | Gene body postion or intergene | Terminal | rDNA | Telomeric |
|---|---|---|---|---|---|---|---|---|---|---|---|
| chrVIII | 1751 | 1899 | 148 | -1,634679602 | N8:1845 | 18,083 | +10:YHL050C; | 10 | | | TEL LEFT |
| chrVIII | 1752 | 1899 | 147 | -1,263349294 | N8:1845 | 18,083 | +10:YHL050C; | 10 | | | TEL LEFT |
| chrVIII | 1753 | 1899 | 146 | -1,162816376 | N8:1845 | 18,083 | +10:YHL050C; | 10 | | | TEL LEFT |
| chrVIII | 1754 | 1899 | 145 | -1,309440533 | N8:1845 | 18,083 | +10:YHL050C; | 10 | | | TEL LEFT |
| chrVIII | 1755 | 1899 | 144 | -1,877474038 | N8:1845 | 18,083 | +10:YHL050C; | 10 | | | TEL LEFT |
| chrVIII | 1756 | 1899 | 143 | -1,666952 | N8:1845 | 18,083 | +10:YHL050C; | 10 | | | TEL LEFT |
| chrVIII | 1757 | 1899 | 142 | -1,308406001 | N8:1845 | 18,083 | +10:YHL050C; | 10 | | | TEL LEFT |
| chrVIII | 1758 | 1899 | 141 | -1,439200269 | N8:1845 | 18,083 | +10:YHL050C; | 10 | | | TEL LEFT |
| chrVIII | 1759 | 1899 | 140 | -1,236253829 | N8:1845 | 18,083 | +10:YHL050C; | 10 | | | TEL LEFT |
| chrVIII | 1760 | 1899 | 139 | -1,438682121 | N8:1845 | 18,083 | +10:YHL050C; | 10 | | | TEL LEFT |
| chrVIII | 1761 | 1899 | 138 | -1,885350855 | N8:1845 | 18,083 | +10:YHL050C; | 10 | | | TEL LEFT |
| chrVIII | 1762 | 1899 | 137 | -1,738108503 | N8:1845 | 18,083 | +10:YHL050C; | 10 | | | TEL LEFT |
| chrVIII | 1764 | 1899 | 135 | -1,2406654 | N8:1845 | 18,083 | +10:YHL050C; | 10 | | | TEL LEFT |
| chrVIII | 1765 | 1899 | 134 | -1,546702362 | N8:1845 | 18,083 | +10:YHL050C; | 10 | | | TEL LEFT |
| chrVIII | 1766 | 1899 | 133 | -0,976808376 | N8:1845 | 18,083 | +10:YHL050C; | 10 | | | TEL LEFT |
| chrVIII | 1769 | 1899 | 130 | -0,800299475 | N8:1845 | 18,083 | +10:YHL050C; | 10 | | | TEL LEFT |
| chrVIII | 1771 | 1899 | 128 | -1,549747623 | N8:1845 | 18,083 | +10:YHL050C; | 10 | | | TEL LEFT |
| chrVIII | 1773 | 1899 | 126 | -1,401502018 | N8:1845 | 18,083 | +10:YHL050C; | 10 | | | TEL LEFT |
| chrVIII | 1784 | 1899 | 115 | -0,689805015 | N8:1845 | 18,083 | +10:YHL050C; | 10 | | | TEL LEFT |
| chrVIII | 1821 | 1968 | 147 | -0,636263276 | N8:1845 | 18,083 | +10:YHL050C; | 10 | | | TEL LEFT |
| chrVIII | 2051 | 2222 | 171 | -0,903431168 | | | | Overlap <50 bp | | | TEL LEFT |
| chrVIII | 2441 | 2594 | 153 | -1,141610292 | N8:2474 | 10,408 | +6:YHL050C; | 6 | | | TEL LEFT |
| chrVIII | 2951 | 3103 | 152 | -1,323567952 | N8:2994 | 19,192 | +3:YHL050C; | 3 | | | TEL LEFT |
| chrVIII | 3093 | 3220 | 127 | -0,996200907 | N8:3153 | 6,325 | +2:YHL050C; | 2 | | | TEL LEFT |
| chrVIII | 3233 | 3376 | 143 | -1,196774043 | N8:3313 | 10,786 | +1:YHL050C; | 1 | | | TEL LEFT |
| chrVIII | 3524 | 3661 | 137 | -1,151491737 | N8:3599 | 7,095 | -1:YHL050C; +7*:YHL049C; | -1 | | | TEL LEFT |
| chrVIII | 4667 | 4822 | 155 | -1,561009173 | N8:4741 | 23,335 | 0:YHL049C; | 0 | | | TEL LEFT |
| chrVIII | 9299 | 9453 | 154 | -1,423690734 | N8:9352 | 24,533 | +6:YHL047C; | 6 | | | |
| chrVIII | 10577 | 10683 | 106 | -1,310720016 | N8:10600 | 5,292 | 0:SUT147; | 0 | | | |
| chrVIII | 10578 | 10683 | 105 | -1,637540864 | N8:10600 | 5,292 | 0:SUT147; | 0 | | | |
| chrVIII | 10864 | 11003 | 139 | -0,908851512 | N8:10940 | 16,869 | +2:SUT147; | 2 | | | |
| chrVIII | 12729 | 12877 | 148 | -1,602577493 | N8:12794 | 9,311 | | Intergene | | | |
| chrVIII | 14994 | 15154 | 160 | -1,234168136 | N8:15064 | 43,65 | +2:YHL043W; +6:SUT573; | 2 | | | |
| chrVIII | 15294 | 15466 | 172 | -1,497801652 | N8:15381 | 9,839 | +4:YHL043W; +4:SUT573; | 4 | | | |
| chrVIII | 17876 | 18017 | 141 | -1,242839156 | N8:18009 | 25,395 | | Intergene | | | |
| chrVIII | 18350 | 18522 | 172 | -1,469435581 | N8:18473 | 28,374 | | Intergene | | | |
| chrVIII | 23920 | 24090 | 170 | -1,702762371 | N8:24051 | 18,708 | +10:YHL038C; | 10 | | | |
| chrVIII | 26981 | 27134 | 153 | -1,132259381 | N8:27018 | 21,775 | +6:YHL036W; | 6 | | | |
| chrVIII | 32520 | 32666 | 146 | -1,238756396 | N8:32599 | 26,876 | +2:YHL035C; | 2 | | | |
| chrVIII | 33329 | 33481 | 152 | -1,253260673 | N8:33481 | 35,088 | +5:YHL034C; | 5 | | | |
| chrVIII | 33329 | 33481 | 152 | -1,253260673 | N8:33328 | 25,521 | +6:YHL034C; | 6 | | | |
| chrVIII | 33632 | 33782 | 150 | -1,172043085 | N8:33647 | 7,765 | +4:YHL034C; | 4 | | | |
| chrVIII | 39986 | 40150 | 164 | -1,43506912 | N8:40093 | 7,202 | +1:YHL030W; | 1 | | | |
| chrVIII | 46044 | 46208 | 164 | -1,237177152 | N8:46074 | 8,335 | +13:YHL029C; | 13 | | | |
| chrVIII | 48223 | 48365 | 142 | -0,909004197 | N8:48283 | 3,536 | -1:YHL029C; | -1 | | | |
| chrVIII | 49771 | 49922 | 151 | -1,230784042 | N8:49832 | 12,816 | +7:YHL028W; | 7 | | | |
| chrVIII | 50265 | 50415 | 150 | -0,795046436 | N8:50342 | 18,779 | +2:CUT637; +10:YHL028W; | 2 | | | |
| chrVIII | 51501 | 51665 | 164 | -1,200528029 | N8:51521 | 23,53 | +4:YHL027W; | 4 | | | |
| chrVIII | 52096 | 52204 | 108 | -1,255071045 | N8:52145 | 35,726 | +8:YHL027W; | 8 | | | |
| chrVIII | 52096 | 52213 | 117 | -1,411502374 | N8:52145 | 35,726 | +8:YHL027W; | 8 | | | |
| chrVIII | 52629 | 52803 | 174 | -1,418183622 | N8:52786 | 5,468 | +11:YHL027W; | 11 | | | |
| chrVIII | 53713 | 53863 | 150 | -1,148618579 | N8:53785 | 11,374 | +3:YHL026C; | 3 | | | |
| chrVIII | 58605 | 58733 | 128 | -0,514590521 | N8:58676 | 27,193 | +13:YHL024W; +2:SUT574; | 13 | | | |
| chrVIII | 59349 | 59502 | 153 | -1,245938153 | N8:59376 | 25,918 | +2:SUT148; | 2 | | | |
| chrVIII | 61087 | 61247 | 160 | -1,87221545 | N8:61111 | 11,354 | +10:YHL023C; | 10 | | | |
| chrVIII | 62149 | 62261 | 112 | -1,290683303 | N8:62222 | 5,404 | +3:YHL023C; | 3 | | | |
| chrVIII | 65254 | 65423 | 169 | -1,214235674 | N8:65391 | 17,011 | +4:YHL021C; | 4 | | | |
| chrVIII | 66426 | 66576 | 150 | -0,977488551 | N8:66523 | 12,069 | +7:YHL020C; | 7 | | | |
| chrVIII | 66660 | 66810 | 150 | -1,326633142 | N8:66683 | 8,044 | +6:YHL020C; | 6 | | | |
| chrVIII | 66812 | 66936 | 124 | -1,43438553 | N8:66852 | 3 | +5:YHL020C; | 5 | | | |
| chrVIII | 66813 | 66902 | 89 | -0,484570609 | N8:66852 | 3 | +5:YHL020C; | 5 | | | |
| chrVIII | 66813 | 66904 | 91 | -0,975525697 | N8:66852 | 3 | +5:YHL020C; | 5 | | | |
| chrVIII | 66813 | 66911 | 98 | -0,951864323 | N8:66852 | 3 | +5:YHL020C; | 5 | | | |
| chrVIII | 66813 | 66912 | 99 | -0,839639659 | N8:66852 | 3 | +5:YHL020C; | 5 | | | |
| chrVIII | 66813 | 66924 | 111 | -0,464190944 | N8:66852 | 3 | +5:YHL020C; | 5 | | | |
| chrVIII | 66813 | 66925 | 112 | -0,628096934 | N8:66852 | 3 | +5:YHL020C; | 5 | | | |
| chrVIII | 66813 | 66926 | 113 | -1,053373951 | N8:66852 | 3 | +5:YHL020C; | 5 | | | |
| chrVIII | 66813 | 66928 | 115 | -1,035733416 | N8:66852 | 3 | +5:YHL020C; | 5 | | | |
| chrVIII | 66813 | 66929 | 116 | -0,920060753 | N8:66852 | 3 | +5:YHL020C; | 5 | | | |
| chrVIII | 66813 | 66930 | 117 | -1,22746726 | N8:66852 | 3 | +5:YHL020C; | 5 | | | |
| chrVIII | 66813 | 66931 | 118 | -1,157170762 | N8:66852 | 3 | +5:YHL020C; | 5 | | | |
| chrVIII | 66813 | 66932 | 119 | -0,975825326 | N8:66852 | 3 | +5:YHL020C; | 5 | | | |
| chrVIII | 66813 | 66933 | 120 | -0,939460335 | N8:66852 | 3 | +5:YHL020C; | 5 | | | |
| chrVIII | 66813 | 66934 | 121 | -0,640567728 | N8:66852 | 3 | +5:YHL020C; | 5 | | | |
| chrVIII | 66813 | 66935 | 122 | -1,07838856 | N8:66852 | 3 | +5:YHL020C; | 5 | | | |
| chrVIII | 66813 | 66936 | 123 | -0,870093444 | N8:66852 | 3 | +5:YHL020C; | 5 | | | |
| chrVIII | 66815 | 66936 | 121 | -1,250420064 | N8:66852 | 3 | +5:YHL020C; | 5 | | | |
| chrVIII | 66816 | 66936 | 120 | -1,405470966 | N8:66852 | 3 | +5:YHL020C; | 5 | | | |
| chrVIII | 66817 | 66936 | 119 | -1,13002513 | N8:66852 | 3 | +5:YHL020C; | 5 | | | |
| chrVIII | 66818 | 66936 | 118 | -0,738377312 | N8:66852 | 3 | +5:YHL020C; | 5 | | | |
| chrVIII | 66820 | 66936 | 116 | -1,280546637 | N8:66852 | 3 | +5:YHL020C; | 5 | | | |
| chrVIII | 66821 | 66936 | 115 | -0,546113539 | N8:66852 | 3 | +5:YHL020C; | 5 | | | |
| chrVIII | 66822 | 66936 | 114 | -0,733222464 | N8:66852 | 3 | +5:YHL020C; | 5 | | | |
| chrVIII | 66823 | 66936 | 113 | -1,106581507 | N8:66852 | 3 | +5:YHL020C; | 5 | | | |
| chrVIII | 66830 | 66936 | 106 | -0,869579508 | N8:66852 | 3 | +5:YHL020C; | 5 | | | |
| chrVIII | 68261 | 68399 | 138 | -0,93731092 | N8:68264 | 38,448 | | Intergene | | | |
| chrVIII | 69975 | 70096 | 121 | -0,397908483 | N8:70043 | 7,603 | -1:YHL017W; +3:YHL018W; | -1 | | | |
| chrVIII | 69975 | 70122 | 147 | -1,005752188 | N8:70043 | 7,603 | -1:YHL017W; +3:YHL018W; | -1 | | | |
| chrVIII | 70402 | 70553 | 151 | -1,312551993 | N8:70464 | 9,475 | +2:YHL017W; | 2 | | | |
| chrVIII | 72065 | 72199 | 134 | -1,136126209 | N8:72067 | 24,511 | +14:YHL016C; | 14 | | | |
| chrVIII | 77481 | 77612 | 131 | -0,643425909 | N8:77523 | 20,918 | -1:YHL014C; | -1 | | | |
| chrVIII | 86148 | 86310 | 162 | -1,684499431 | N8:86173 | 7,024 | | Intergene | | | |
| chrVIII | 86176 | 86324 | 148 | -1,115642165 | N8:86173 | 7,024 | | Intergene | | | |
| chrVIII | 86476 | 86623 | 147 | -1,411195094 | N8:86566 | 12,858 | | Intergene | | | |
| chrVIII | 88963 | 89116 | 153 | -1,926932052 | N8:89119 | 23,847 | | Intergene | | | |
| chrVIII | 91372 | 91534 | 162 | -1,593772132 | N8:91357 | 31,113 | | Intergene | | | |
| chrVIII | 91372 | 91534 | 162 | -1,593772132 | N8:91547 | 35,557 | | Intergene | | | |
| chrVIII | 94804 | 94953 | 149 | -1,065940992 | N8:94867 | 15,145 | -1:YHL008C; | -1 | | | |
| chrVIII | 94985 | 95141 | 156 | -1,397346819 | N8:95070 | 19,956 | +19*:YHL007C; | 19 | TERM | | |

TABLE 3

| CHR | START | END | L(bp) | ΔLK | Nuc ID | Fuzziness | Gene ID | Gene body postion or intergene | Terminal | rDNA | Telomeric |
|---|---|---|---|---|---|---|---|---|---|---|---|
| chrVIII | 96353 | 96506 | 153 | -1,136074458 | N8:96448 | 34,452 | +11:YHL007C; +2*:SUT150; | 11 | | | |
| chrVIII | 100385 | 100534 | 149 | -1,798941269 | N8:100544 | 36,981 | +10*:YHL003C; +9*:YHL004W; | 10 | TERM | | |
| chrVIII | 107252 | 107412 | 160 | -1,518043361 | N8:107297 | 1,949 | +9*:YHR001W; | 9 | TERM | | |
| chrVIII | 108345 | 108493 | 148 | -1,562329229 | N8:108419 | 9,99 | -1:YHR002W; | -1 | | | |
| chrVIII | 109527 | 109701 | 174 | -1,566944703 | N8:109636 | 27,373 | +7*:YHR002W; | 7 | TERM | | |
| chrVIII | 114779 | 114932 | 153 | -1,16204338 | N8:114850 | 11,147 | +2:YHR005C; | 2 | | | |
| chrVIII | 115531 | 115709 | 178 | -1,454019066 | N8:115711 | 7,868 | +2*:CUT173; +2:YHR005C-A; | 2 | TERM | | |
| chrVIII | 121476 | 121637 | 161 | -1,100409818 | N8:121495 | 19,604 | +3:YHR007C; | 3 | | | |
| chrVIII | 122292 | 122451 | 159 | -0,791064114 | N8:122379 | 11,269 | +4*:YHR007C-A; +4:SUT153; | 4 | TERM | | |
| chrVIII | 123820 | 123978 | 158 | -1,410352592 | N8:123911 | 7,653 | -1:YHR008C; +14*:SUT153; | -1 | | | |
| chrVIII | 125638 | 125758 | 120 | -0,777011762 | N8:125733 | 19,601 | +2:YHR009C; | 2 | | | |
| chrVIII | 130844 | 131002 | 158 | -1,275527539 | N8:130922 | 7,731 | +4:YHR013C; | 4 | | | |
| chrVIII | 131196 | 131345 | 149 | -1,042345871 | N8:131260 | 6,802 | +2:YHR013C; | 2 | | | |
| chrVIII | 142630 | 142776 | 146 | -1,134598201 | N8:142760 | 8,438 | +6:YHR019C; | 6 | | | |
| chrVIII | 144193 | 144334 | 141 | -0,912713065 | N8:144264 | 7,085 | +3:YHR020W; | 3 | | | |
| chrVIII | 152201 | 152353 | 152 | -1,462538567 | N8:152258 | 6,274 | +5:YHR023W; | 5 | | | |
| chrVIII | 153827 | 153949 | 122 | -1,416468334 | N8:153924 | 14,572 | +15:YHR023W; | 15 | | | |
| chrVIII | 157804 | 157963 | 159 | -1,039803567 | N8:157803 | 35,25 | | Intergene | | | |
| chrVIII | 158390 | 158542 | 152 | -1,143965281 | N8:158513 | 30 | +5:YHR024C; | 5 | | | |
| chrVIII | 162439 | 162593 | 154 | -1,122715696 | N8:162524 | 24,161 | +14:YHR027C; | 14 | | | |
| chrVIII | 162949 | 163087 | 138 | -0,772142831 | | | | Overlap <50 bp | | | |
| chrVIII | 165932 | 166076 | 144 | -1,089462702 | N8:165989 | 4,868 | +10:YHR028C; +7:SUT155; | 10 | | | |
| chrVIII | 167959 | 168110 | 151 | -1,376459633 | N8:168035 | 6,47 | +4:YHR029C; | 4 | | | |
| chrVIII | 171020 | 171171 | 151 | -1,064260495 | N8:171084 | 8,214 | +3:CUT175; +13:YHR031C; | 3 | | | |
| chrVIII | 171703 | 171855 | 152 | -1,279124868 | N8:171826 | 49,31 | +8:YHR031C; | 8 | | | |
| chrVIII | 174446 | 174586 | 140 | -2,483796618 | | | | Overlap <50 bp | | | |
| chrVIII | 174447 | 174533 | 86 | -1,757935898 | | | | Overlap <50 bp | | | |
| chrVIII | 174447 | 174535 | 88 | -1,403734994 | | | | Overlap <50 bp | | | |
| chrVIII | 174447 | 174538 | 91 | -1,135103149 | | | | Overlap <50 bp | | | |
| chrVIII | 174447 | 174540 | 93 | -1,334705523 | | | | Overlap <50 bp | | | |
| chrVIII | 174447 | 174541 | 94 | -1,688759971 | | | | Overlap <50 bp | | | |
| chrVIII | 174447 | 174544 | 97 | -1,848430106 | | | | Overlap <50 bp | | | |
| chrVIII | 174447 | 174545 | 98 | -1,886644163 | | | | Overlap <50 bp | | | |
| chrVIII | 174447 | 174548 | 101 | -1,367309311 | | | | Overlap <50 bp | | | |
| chrVIII | 174447 | 174550 | 103 | -1,337354548 | | | | Overlap <50 bp | | | |
| chrVIII | 174447 | 174551 | 104 | -1,429594893 | | | | Overlap <50 bp | | | |
| chrVIII | 174447 | 174552 | 105 | -1,33918531 | | | | Overlap <50 bp | | | |
| chrVIII | 174447 | 174553 | 106 | -2,212746809 | | | | Overlap <50 bp | | | |
| chrVIII | 174447 | 174555 | 108 | -1,393899438 | | | | Overlap <50 bp | | | |
| chrVIII | 174447 | 174561 | 114 | -1,750334161 | | | | Overlap <50 bp | | | |
| chrVIII | 174447 | 174565 | 118 | -1,755634194 | | | | Overlap <50 bp | | | |
| chrVIII | 174447 | 174566 | 119 | -1,583491617 | | | | Overlap <50 bp | | | |
| chrVIII | 174447 | 174567 | 120 | -1,519140194 | | | | Overlap <50 bp | | | |
| chrVIII | 174447 | 174568 | 121 | -1,50479122 | | | | Overlap <50 bp | | | |
| chrVIII | 174447 | 174569 | 122 | -1,172773356 | | | | Overlap <50 bp | | | |
| chrVIII | 174447 | 174570 | 123 | -1,305586702 | | | | Overlap <50 bp | | | |
| chrVIII | 174447 | 174571 | 124 | -1,086245943 | | | | Overlap <50 bp | | | |
| chrVIII | 174447 | 174572 | 125 | -1,516021409 | | | | Overlap <50 bp | | | |
| chrVIII | 174447 | 174573 | 126 | -1,339498373 | | | | Overlap <50 bp | | | |
| chrVIII | 174447 | 174574 | 127 | -1,54327693 | | | | Overlap <50 bp | | | |
| chrVIII | 174447 | 174575 | 128 | -1,429042701 | | | | Overlap <50 bp | | | |
| chrVIII | 174447 | 174576 | 129 | -1,233539464 | | | | Overlap <50 bp | | | |
| chrVIII | 174447 | 174577 | 130 | -1,196052957 | | | | Overlap <50 bp | | | |
| chrVIII | 174447 | 174578 | 131 | -1,258911953 | | | | Overlap <50 bp | | | |
| chrVIII | 174447 | 174579 | 132 | -1,242932972 | | | | Overlap <50 bp | | | |
| chrVIII | 174447 | 174580 | 133 | -1,372127814 | | | | Overlap <50 bp | | | |
| chrVIII | 174447 | 174581 | 134 | -1,386338066 | | | | Overlap <50 bp | | | |
| chrVIII | 174447 | 174582 | 135 | -1,238892414 | | | | Overlap <50 bp | | | |
| chrVIII | 174447 | 174583 | 136 | -1,378172993 | | | | Overlap <50 bp | | | |
| chrVIII | 174447 | 174584 | 137 | -1,574889836 | | | | Overlap <50 bp | | | |
| chrVIII | 174447 | 174585 | 138 | -1,747809058 | | | | Overlap <50 bp | | | |
| chrVIII | 174447 | 174586 | 139 | -1,21136349 | | | | Overlap <50 bp | | | |
| chrVIII | 174447 | 174587 | 140 | -1,793791499 | | | | Overlap <50 bp | | | |
| chrVIII | 174448 | 174586 | 138 | -1,652348415 | | | | Overlap <50 bp | | | |
| chrVIII | 174449 | 174586 | 137 | -1,599178604 | | | | Overlap <50 bp | | | |
| chrVIII | 174450 | 174586 | 136 | -1,587297859 | | | | Overlap <50 bp | | | |
| chrVIII | 174451 | 174586 | 135 | -1,358499898 | | | | Overlap <50 bp | | | |
| chrVIII | 174452 | 174586 | 134 | -1,486654464 | | | | Overlap <50 bp | | | |
| chrVIII | 174453 | 174586 | 133 | -1,698119925 | | | | Overlap <50 bp | | | |
| chrVIII | 174454 | 174586 | 132 | -1,85214847 | | | | Overlap <50 bp | | | |
| chrVIII | 174455 | 174586 | 131 | -1,244822515 | | | | Overlap <50 bp | | | |
| chrVIII | 174456 | 174586 | 130 | -1,414295965 | | | | Overlap <50 bp | | | |
| chrVIII | 174457 | 174586 | 129 | -1,458023269 | | | | Overlap <50 bp | | | |
| chrVIII | 174458 | 174586 | 128 | -1,15552345 | | | | Overlap <50 bp | | | |
| chrVIII | 174459 | 174586 | 127 | -1,500069553 | | | | Overlap <50 bp | | | |
| chrVIII | 174460 | 174586 | 126 | -1,440462389 | | | | Overlap <50 bp | | | |
| chrVIII | 174461 | 174586 | 125 | -1,935941559 | | | | Overlap <50 bp | | | |
| chrVIII | 174462 | 174553 | 91 | -0,297957357 | | | | Overlap <50 bp | | | |
| chrVIII | 174467 | 174586 | 119 | -2,091322845 | | | | Overlap <50 bp | | | |
| chrVIII | 174928 | 175084 | 156 | -1,725251745 | N8:175011 | 18,389 | +11:YHR032W; | 11 | | | |
| chrVIII | 176211 | 176366 | 155 | -0,922480506 | N8:176283 | 9,287 | +6:YHR033W; | 6 | | | |
| chrVIII | 177719 | 177867 | 148 | -1,16595669 | N8:177804 | 4,875 | +2:YHR034C; | 2 | | | |
| chrVIII | 177797 | 177947 | 150 | -1,134243544 | N8:177804 | 4,875 | +2:YHR034C; | 2 | | | |
| chrVIII | 178837 | 178985 | 148 | -1,015099819 | N8:178874 | 14,152 | +5:YHR035W; | 5 | | | |
| chrVIII | 180375 | 180509 | 134 | -0,755025009 | N8:180443 | 11,415 | +2:YHR036W; | 2 | | | |
| chrVIII | 190177 | 190319 | 142 | -1,248374651 | N8:190199 | 11,93 | +7*:SUT156; | 7 | TERM | | |
| chrVIII | 194784 | 194941 | 157 | -1,705812859 | | | | Overlap <50 bp | | | |
| chrVIII | 194987 | 195122 | 135 | -1,013025373 | N8:195028 | 18,082 | -1:YHR044C; | -1 | | | |
| chrVIII | 205654 | 205804 | 150 | -1,30716269 | N8:205739 | 10,012 | +8:YHR048W; | 8 | | | |
| chrVIII | 207083 | 207225 | 142 | -0,925799823 | N8:207213 | 3,83 | -1:YHR050W; +6*:YHR049W; | -1 | | | |
| chrVIII | 212526 | 212683 | 157 | -1,277206009 | N8:212615 | 13,503 | +2:YHR053C; | 2 | | | |
| chrVIII | 212729 | 212874 | 145 | -1,300209905 | N8:212883 | 12,728 | 0:YHR053C; | 0 | | | |
| chrVIII | 212963 | 213069 | 106 | -1,410910171 | N8:213049 | 35,355 | +9*:YHR054C; -1:YHR053C; | 9 | TERM | | |
| chrVIII | 212963 | 213076 | 113 | -1,647861438 | N8:213049 | 35,355 | +9*:YHR054C; -1:YHR053C; | 9 | TERM | | |
| chrVIII | 213025 | 213187 | 162 | -1,304875911 | N8:213049 | 35,355 | +9*:YHR054C; -1:YHR053C; | 9 | TERM | | |
| chrVIII | 214079 | 214248 | 169 | -1,33370417 | N8:214186 | 30,414 | +2:YHR054C; | 2 | | | |

TABLE 3

| CHR | START | END | L(bp) | ΔLK | Nuc ID | Fuzziness | Gene ID | Gene body postion or intergene | Terminal | rDNA | Telomeric |
|---|---|---|---|---|---|---|---|---|---|---|---|
| chrVIII | 214176 | 214329 | 153 | -1,56138711 | N8:214186 | 30,414 | +2:YHR054C; | 2 | | | |
| chrVIII | 214524 | 214681 | 157 | -1,280571047 | N8:214628 | 15,631 | -1:YHR054C; +2:YHR055C; | -1 | | | |
| chrVIII | 214727 | 214872 | 145 | -1,0610766 | N8:214799 | 4,95 | +1:YHR055C; | 1 | | | |
| chrVIII | 214961 | 215067 | 106 | -1,413393278 | N8:215040 | 24,749 | -1:YHR055C; | -1 | | | |
| chrVIII | 214961 | 215074 | 113 | -1,638190833 | N8:215040 | 24,749 | -1:YHR055C; | -1 | | | |
| chrVIII | 215023 | 215185 | 162 | -1,275500244 | N8:215040 | 24,749 | -1:YHR055C; | -1 | | | |
| chrVIII | 216077 | 216246 | 169 | -1,611889619 | N8:216180 | 23,866 | +10*:YHR056C; | 10 | TERM | | |
| chrVIII | 217808 | 217962 | 154 | -1,252549277 | N8:217835 | 7,394 | +1:YHR056C; -1:anti-YHR057C; | 1 | | | |
| chrVIII | 218430 | 218569 | 139 | -1,519807445 | N8:218465 | 31,519 | +3:YHR057C; | 3 | | | |
| chrVIII | 219279 | 219430 | 151 | -1,245124527 | N8:219365 | 5,345 | +3:CUT179; +4:YHR058C; | 3 | | | |
| chrVIII | 223655 | 223760 | 105 | -1,429807162 | N8:223731 | 8,612 | +5*:SUT159; +1:YHR062C; | 5 | TERM | | |
| chrVIII | 226108 | 226256 | 148 | -0,489329589 | N8:226185 | 7,81 | +7:YHR064C; | 7 | | | |
| chrVIII | 227838 | 227999 | 161 | -1,515272845 | N8:228012 | 53,866 | +7:YHR065C; | 7 | | | |
| chrVIII | 227838 | 227999 | 161 | -1,515272845 | N8:227836 | 26,621 | +8:YHR065C; | 8 | | | |
| chrVIII | 228803 | 228946 | 143 | -1,059057729 | N8:228877 | 11,788 | +2:YHR065C; | 2 | | | |
| chrVIII | 234383 | 234536 | 153 | -1,334466065 | N8:234466 | 3,619 | +2:YHR069C; | 2 | | | |
| chrVIII | 234544 | 234693 | 149 | -1,239096513 | N8:234634 | 7,969 | -1:YHR070W; +1:YHR069C; | -1 | | | |
| chrVIII | 240775 | 240923 | 148 | -1,197731409 | N8:240835 | 30,507 | +12:YHR072W; | 12 | | | |
| chrVIII | 250570 | 250724 | 154 | -0,987295623 | N8:250736 | 14,77 | +2:SUT581; | 2 | | | |
| chrVIII | 256594 | 256759 | 165 | -1,627989876 | N8:256675 | 19,235 | +3:YHR078W; | 3 | | | |
| chrVIII | 258714 | 258856 | 142 | -0,991625696 | N8:258740 | 30,827 | +19:YHR079C; | 19 | | | |
| chrVIII | 261335 | 261482 | 147 | -1,269476242 | N8:261454 | 6,463 | +3:YHR079C; | 3 | | | |
| chrVIII | 264809 | 264956 | 147 | -1,11333577 | N8:264814 | 27,568 | +14:YHR080C; | 14 | | | |
| chrVIII | 270516 | 270670 | 154 | -1,40332622 | N8:270646 | 24,98 | +7:YHR082C; | 7 | | | |
| chrVIII | 272692 | 272812 | 120 | -0,965491674 | N8:272823 | 2,345 | +2:YHR083W; | 2 | | | |
| chrVIII | 275159 | 275317 | 158 | -1,166809161 | | | | Overlap <50 bp | | | |
| chrVIII | 281596 | 281745 | 149 | -1,22580215 | N8:281674 | 6,512 | +2:YHR088W; | 2 | | | |
| chrVIII | 288944 | 289050 | 106 | -0,932917537 | N8:288960 | 4,163 | 0:YHR092C; | 0 | | | |
| chrVIII | 291607 | 291756 | 149 | -1,032527556 | N8:291704 | 10 | +6:YHR094C; +5:SUT163; | 6 | | | |
| chrVIII | 291938 | 292090 | 152 | -1,176345208 | N8:292062 | 41,088 | +4:YHR094C; +7:SUT163; | 4 | | | |
| chrVIII | 294718 | 294863 | 145 | -1,085619824 | N8:294786 | 16,078 | +11:YHR096C; +5:SUT164; | 11 | | | |
| chrVIII | 297474 | 297608 | 134 | -1,022219934 | | | | Overlap <50 bp | | | |
| chrVIII | 297952 | 298079 | 127 | -0,792051085 | | | | Overlap <50 bp | | | |
| chrVIII | 299058 | 299223 | 165 | -1,335808728 | N8:299169 | 16,753 | +19*:YHR098C; | 19 | TERM | | |
| chrVIII | 299773 | 299910 | 137 | -0,961894503 | N8:299791 | 50,553 | +15:YHR098C; | 15 | | | |
| chrVIII | 310121 | 310260 | 139 | -0,659400977 | N8:310259 | 29,744 | +47:YHR099W; | 47 | | | |
| chrVIII | 310839 | 310998 | 159 | -1,404089165 | N8:310900 | 14,96 | +51:YHR099W; | 51 | | | |
| chrVIII | 311969 | 312130 | 161 | -1,306490135 | N8:312117 | 34,174 | +58:YHR099W; | 58 | | | |
| chrVIII | 333005 | 333157 | 152 | -1,170985126 | N8:333100 | 13,467 | +1:YHR111W; | 1 | | | |
| chrVIII | 334511 | 334659 | 148 | -1,238584528 | N8:334641 | 12,39 | +7:YHR112C; +2:SUT166; | 7 | | | |
| chrVIII | 341021 | 341173 | 152 | -1,644478449 | N8:341133 | 6,058 | +3:YHR115C; | 3 | | | |
| chrVIII | 341021 | 341178 | 157 | -1,312389427 | N8:341133 | 6,058 | +3:YHR115C; | 3 | | | |
| chrVIII | 342001 | 342158 | 157 | -1,208299946 | N8:342074 | 21,135 | +3:YHR116W; -1:YHR117W; | 3 | | | |
| chrVIII | 342217 | 342368 | 151 | -1,480422533 | N8:342346 | 17,487 | +5:YHR116W; +1:YHR117W; | 5 | | | |
| chrVIII | 343838 | 343999 | 161 | -1,330576912 | N8:344006 | 22,521 | +15:YHR116W; +11:YHR117W; | 15 | | | |
| chrVIII | 345951 | 346111 | 160 | -1,564221048 | N8:346037 | 30,244 | +2:YHR119W; | 2 | | | |
| chrVIII | 346434 | 346582 | 148 | -1,662165903 | N8:346511 | 21,836 | +5:YHR119W; | 5 | | | |
| chrVIII | 350191 | 350346 | 155 | -1,386031371 | N8:350266 | 8,612 | +5:YHR120W; | 5 | | | |
| chrVIII | 353263 | 353415 | 152 | -1,522945572 | | | | Overlap <50 bp | | | |
| chrVIII | 354917 | 355076 | 159 | -1,548652808 | N8:355010 | 8,542 | | Intergene | | | |
| chrVIII | 355941 | 356089 | 148 | -1,20876205 | N8:356016 | 8,914 | | Intergene | | | |
| chrVIII | 357549 | 357703 | 154 | -1,039656568 | N8:357581 | 28,314 | +7:YHR124W; +4:anti074; | 7 | | | |
| chrVIII | 360834 | 360982 | 148 | -1,458143089 | N8:360921 | 13,197 | +1:YHR127W; | 1 | | | |
| chrVIII | 364130 | 364290 | 160 | -1,271346267 | N8:364200 | 8,612 | -1:SUT167; +1:YHR129C; | -1 | | | |
| chrVIII | 370648 | 370829 | 181 | -1,423306518 | N8:370765 | 23,128 | +6:YHR133C; | 6 | | | |
| chrVIII | 371736 | 371883 | 147 | -1,504276641 | N8:371818 | 5,727 | -1:YHR133C; +1:YHR134W; | -1 | | | |
| chrVIII | 374061 | 374221 | 160 | -1,224398193 | N8:374205 | 4,658 | +2:YHR135C; | 2 | | | |
| chrVIII | 374061 | 374255 | 194 | -1,372099878 | N8:374205 | 4,658 | +2:YHR135C; | 2 | | | |
| chrVIII | 374061 | 374221 | 160 | -1,224398193 | N8:374049 | 6,033 | +3:YHR135C; | 3 | | | |
| chrVIII | 374061 | 374255 | 194 | -1,372099878 | N8:374049 | 6,033 | +3:YHR135C; | 3 | | | |
| chrVIII | 374122 | 374209 | 87 | -0,188959209 | N8:374205 | 4,658 | +2:YHR135C; | 2 | | | |
| chrVIII | 374122 | 374221 | 99 | -0,26867702 | N8:374205 | 4,658 | +2:YHR135C; | 2 | | | |
| chrVIII | 374122 | 374255 | 133 | -0,954386934 | N8:374205 | 4,658 | +2:YHR135C; | 2 | | | |
| chrVIII | 376459 | 376622 | 163 | -1,367194803 | N8:376456 | 32,253 | +6:YHR137W; | 6 | | | |
| chrVIII | 376459 | 376622 | 163 | -1,367194803 | N8:376622 | 10,63 | +7:YHR137W; | 7 | | | |
| chrVIII | 381490 | 381641 | 151 | -1,591911886 | N8:381642 | 25,373 | -1:CUT650; | -1 | | | |
| chrVIII | 382542 | 382702 | 160 | -1,436106299 | N8:382584 | 5,292 | +2:YHR141C; | 2 | | | |
| chrVIII | 382976 | 383151 | 175 | -1,246339989 | N8:383155 | 42,505 | -1:YHR142W; | -1 | | | |
| chrVIII | 399609 | 399757 | 148 | -1,119313118 | N8:399693 | 19,343 | +9:YHR151C; | 9 | | | |
| chrVIII | 402529 | 402689 | 160 | -1,318842936 | N8:402659 | 11,15 | +1:YHR153C; -1:YHR154W; | 1 | | | |
| chrVIII | 402529 | 402689 | 160 | -1,318842936 | N8:402507 | 13,837 | +2:YHR153C; | 2 | | | |
| chrVIII | 403584 | 403737 | 153 | -1,325040433 | N8:403661 | 23,944 | +5:YHR154W; | 5 | | | |
| chrVIII | 404097 | 404230 | 133 | -0,98936652 | N8:404184 | 31,659 | +8:YHR154W; | 8 | | | |
| chrVIII | 414157 | 414333 | 176 | -1,559454807 | N8:414190 | 28,902 | | Intergene | | | |
| chrVIII | 414157 | 414333 | 176 | -1,559454807 | N8:414356 | 19,07 | | Intergene | | | |
| chrVIII | 416537 | 416675 | 138 | -0,828491242 | N8:416611 | 6,38 | +5:YHR158C; | 5 | | | |
| chrVIII | 424973 | 425127 | 154 | -1,170937868 | N8:425133 | 16,664 | +27:YHR164C; | 27 | | | |
| chrVIII | 424973 | 425127 | 154 | -1,170937868 | N8:424954 | 27,398 | +28:YHR164C; | 28 | | | |
| chrVIII | 427961 | 428122 | 161 | -1,297013317 | N8:428089 | 14,29 | +8:YHR164C; | 8 | | | |
| chrVIII | 431042 | 431168 | 126 | -0,711998868 | N8:431037 | 19,435 | +38:YHR165C; | 38 | | | |
| chrVIII | 433338 | 433488 | 150 | -1,328091826 | N8:433377 | 10,626 | +23:YHR165C; | 23 | | | |
| chrVIII | 439467 | 439624 | 157 | -1,119690507 | N8:439505 | 10,962 | +2:YHR167W; | 2 | | | |
| chrVIII | 444830 | 444988 | 158 | -1,419465552 | N8:444909 | 11,502 | +8:YHR170W; | 8 | | | |
| chrVIII | 447873 | 448000 | 127 | -1,173966999 | N8:448007 | 29,228 | -1:YHR172W; | -1 | | | |
| chrVIII | 456058 | 456162 | 104 | -1,559025913 | N8:456183 | 19,411 | | Intergene | | | |
| chrVIII | 465568 | 465728 | 160 | -1,241678085 | N8:465665 | 14,935 | -1:SUT593; 0:Unit319; | -1 | | | |
| chrVIII | 469779 | 469934 | 155 | -1,461060373 | N8:469909 | 7,188 | | Intergene | | | |
| chrVIII | 469928 | 470081 | 153 | -1,125689026 | N8:469909 | 7,188 | | Intergene | | | |
| chrVIII | 469928 | 470081 | 153 | -1,125689026 | N8:470061 | 25,066 | | Intergene | | | |
| chrVIII | 470995 | 471124 | 129 | -0,72567365 | N8:471041 | 21,059 | -1:SUT594; +2:YHR183W; | -1 | | | |
| chrVIII | 482780 | 482943 | 163 | -1,151572456 | N8:482852 | 30,892 | +7:YHR188C; | 7 | | | |
| chrVIII | 485160 | 485318 | 158 | -1,031534018 | N8:485166 | 15,71 | +3:YHR190W; | 3 | | | |
| chrVIII | 485160 | 485318 | 158 | -1,031534018 | N8:485325 | 34,672 | +4:YHR190W; | 4 | | | |
| chrVIII | 495830 | 495995 | 165 | -1,321977575 | N8:495933 | 28,957 | +13:YHR197W; | 13 | | | |
| chrVIII | 497750 | 497909 | 159 | -1,287334229 | N8:497804 | 8,456 | +6:YHR199C; | 6 | | | |
| chrVIII | 501432 | 501582 | 150 | -1,583211409 | N8:501512 | 22,95 | +2:CUT183; | 2 | | | |
| chrVIII | 503721 | 503876 | 155 | -1,03079082 | N8:503793 | 30,696 | +9:YHR202W; | 9 | | | |
| chrVIII | 510274 | 510423 | 149 | -1,264522799 | N8:510320 | 41,677 | +9:YHR205W; | 9 | | | |
| chrVIII | 518475 | 518638 | 163 | -1,174338137 | N8:518487 | 33,259 | +7:YHR208W; | 7 | | | |

| CHR | START | END | L(bp) | ΔLK | Nuc ID | Fuzziness | Gene ID | Gene body postion or intergene | Terminal | rDNA | Telomeric |
|---|---|---|---|---|---|---|---|---|---|---|---|
| chrVIII | 518475 | 518638 | 163 | -1,174338137 | N8:518641 | 31,729 | +8:YHR208W; | 8 | | | |
| chrVIII | 521779 | 521913 | 134 | -1,151231827 | N8:521848 | 15,9 | 0:YHR210C; | 0 | | | |
| chrVIII | 522442 | 522595 | 153 | -1,319513397 | N8:522525 | 4,992 | +3*:SUT174; | 3 | TERM | | |
| chrVIII | 523030 | 523182 | 152 | -1,106894614 | N8:523103 | 10,265 | +15:SUT600; | 15 | | | |
| chrVIII | 523197 | 523325 | 128 | -0,939017474 | N8:523263 | 12,987 | +14:SUT600; | 14 | | | |
| chrVIII | 525186 | 525342 | 156 | -1,290952622 | N8:525241 | 18,638 | +3:YHR211W; +2:SUT600; | 3 | | | |
| chrVIII | 526030 | 526198 | 168 | -1,085711028 | N8:526058 | 12,288 | +8:YHR211W; | 8 | | | |
| chrVIII | 526169 | 526320 | 151 | -1,060668784 | N8:526224 | 23,742 | +9*:YHR211W; | 9 | TERM | | |
| chrVIII | 526293 | 526441 | 148 | -1,201515461 | N8:526381 | 27,465 | | Intergene | | | |
| chrVIII | 526293 | 526481 | 188 | -1,254711863 | N8:526381 | 27,465 | | Intergene | | | |
| chrVIII | 526600 | 526706 | 106 | -0,637645674 | N8:526683 | 30,347 | | Intergene | | | |
| chrVIII | 526600 | 526743 | 143 | -1,295753797 | N8:526683 | 30,347 | | Intergene | | | |
| chrVIII | 526600 | 526749 | 149 | -1,205808589 | N8:526683 | 30,347 | | Intergene | | | |
| chrVIII | 526600 | 526751 | 151 | -1,153146903 | N8:526683 | 30,347 | | Intergene | | | |
| chrVIII | 526624 | 526751 | 127 | -0,645420904 | N8:526683 | 30,347 | | Intergene | | | |
| chrVIII | 526651 | 526751 | 100 | -0,127679689 | N8:526683 | 30,347 | | Intergene | | | |
| chrVIII | 527864 | 528038 | 174 | -1,518700927 | N8:527985 | 32,393 | | Intergene | | | |
| chrVIII | 536044 | 536157 | 113 | -1,96902132 | N8:536107 | 8,888 | | Intergene | | | |
| chrVIII | 538480 | 538617 | 137 | -0,852417988 | N8:538580 | 7 | 0:YHR212W-A; | 0 | | | |
| chrVIII | 539313 | 539458 | 145 | -0,98901669 | N8:539402 | 29,687 | +3:YHR213W; | 3 | | | |
| chrVIII | 539340 | 539488 | 148 | -1,055729843 | N8:539402 | 29,687 | +3:YHR213W; | 3 | | | |
| chrVIII | 542782 | 542926 | 144 | -1,023092029 | N8:542760 | 3,512 | | Intergene | | | |
| chrVIII | 542782 | 542926 | 144 | -1,023092029 | N8:542922 | 7,136 | | Intergene | | | |
| chrVIII | 542989 | 543156 | 167 | -1,354595086 | | | | Overlap <50 bp | | | |
| chrVIII | 543353 | 543457 | 104 | -1,112766739 | N8:543426 | 31,749 | | Intergene | | | |
| chrVIII | 543780 | 543928 | 148 | -1,641852386 | N8:543820 | 4,726 | | Intergene | | | |
| chrVIII | 544540 | 544654 | 114 | -1,584603356 | | | | Overlap <50 bp | | | |
| chrVIII | 546632 | 546789 | 157 | -1,36763172 | N8:546708 | 28 | | Intergene | | | |
| chrVIII | 546818 | 546968 | 150 | -0,989375953 | N8:546888 | 18,52 | | Intergene | | | |
| chrVIII | 546822 | 546968 | 146 | -1,181618747 | N8:546888 | 18,52 | | Intergene | | | |
| chrVIII | 547280 | 547415 | 135 | -0,769316725 | N8:547333 | 41,016 | | Intergene | | | |
| chrVIII | 548318 | 548458 | 140 | -1,07056189 | N8:548393 | 32,97 | | Intergene | | | |
| chrVIII | 548324 | 548458 | 134 | -0,897969286 | N8:548393 | 32,97 | | Intergene | | | |
| chrVIII | 548850 | 549005 | 155 | -1,193138205 | N8:548872 | 26,23 | | Intergene | | | |
| chrVIII | 548981 | 549089 | 108 | -0,90835103 | N8:549081 | 34,675 | | Intergene | | | |
| chrVIII | 549893 | 550056 | 163 | -1,279316305 | N8:549980 | 23,798 | | Intergene | | | |
| chrVIII | 550702 | 550810 | 108 | -0,911356535 | N8:550729 | 5,859 | +3:YHR214C-D; | 3 | | | |
| chrVIII | 551514 | 551634 | 120 | -1,113007121 | N8:551574 | 15,62 | +1:YHR214C-E; | 1 | | | |
| chrVIII | 553179 | 553326 | 147 | -1,016007985 | N8:553215 | 37,358 | +8:YHR215W; | 8 | | | |
| chrVIII | 553630 | 553778 | 148 | -1,332517702 | N8:553708 | 12,222 | +11*:YHR215W; | 11 | TERM | | |
| chrVIII | 556401 | 556536 | 135 | -0,912232065 | N8:556476 | 25,401 | | Intergene | | | TEL RIGHT |
| chrVIII | 556826 | 556854 | 28 | -0,448 | | | | Overlap <50 bp | | | TEL RIGHT |
| chrVIII | 556826 | 556859 | 33 | -0,483 | | | | Overlap <50 bp | | | TEL RIGHT |
| chrVIII | 556826 | 556865 | 39 | -0,525 | | | | Overlap <50 bp | | | TEL RIGHT |
| chrVIII | 556826 | 556871 | 45 | -0,567 | | | | Overlap <50 bp | | | TEL RIGHT |
| chrVIII | 556826 | 556873 | 47 | -0,581 | | | | Overlap <50 bp | | | TEL RIGHT |
| chrVIII | 556826 | 556876 | 50 | -0,602 | | | | Overlap <50 bp | | | TEL RIGHT |
| chrVIII | 556826 | 556884 | 58 | -0,658 | N8:556907 | 34,186 | | Intergene | | | TEL RIGHT |
| chrVIII | 556826 | 556910 | 84 | -1,226387911 | N8:556907 | 34,186 | | Intergene | | | TEL RIGHT |
| chrVIII | 556826 | 556914 | 88 | -0,669511342 | N8:556907 | 34,186 | | Intergene | | | TEL RIGHT |
| chrVIII | 556826 | 556918 | 92 | -1,179094785 | N8:556907 | 34,186 | | Intergene | | | TEL RIGHT |
| chrVIII | 556826 | 556919 | 93 | -0,554900173 | N8:556907 | 34,186 | | Intergene | | | TEL RIGHT |
| chrVIII | 556826 | 556921 | 95 | -0,802117372 | N8:556907 | 34,186 | | Intergene | | | TEL RIGHT |
| chrVIII | 556826 | 556923 | 97 | -0,652536804 | N8:556907 | 34,186 | | Intergene | | | TEL RIGHT |
| chrVIII | 556826 | 556926 | 100 | -0,631980636 | N8:556907 | 34,186 | | Intergene | | | TEL RIGHT |
| chrVIII | 556826 | 556930 | 104 | -1,059931527 | N8:556907 | 34,186 | | Intergene | | | TEL RIGHT |
| chrVIII | 556826 | 556935 | 109 | -0,496516074 | N8:556907 | 34,186 | | Intergene | | | TEL RIGHT |
| chrVIII | 556826 | 556936 | 110 | -0,27061736 | N8:556907 | 34,186 | | Intergene | | | TEL RIGHT |
| chrVIII | 556826 | 556939 | 113 | -0,685664606 | N8:556907 | 34,186 | | Intergene | | | TEL RIGHT |
| chrVIII | 556826 | 556942 | 116 | -1,039251849 | N8:556907 | 34,186 | | Intergene | | | TEL RIGHT |
| chrVIII | 556826 | 556944 | 118 | -1,280593646 | N8:556907 | 34,186 | | Intergene | | | TEL RIGHT |
| chrVIII | 556826 | 556950 | 124 | -0,636932973 | N8:556907 | 34,186 | | Intergene | | | TEL RIGHT |
| chrVIII | 556826 | 556954 | 128 | -1,560902706 | N8:556907 | 34,186 | | Intergene | | | TEL RIGHT |
| chrVIII | 556826 | 556956 | 130 | -1,702452052 | N8:556907 | 34,186 | | Intergene | | | TEL RIGHT |
| chrVIII | 556826 | 556961 | 135 | -1,352952049 | N8:556907 | 34,186 | | Intergene | | | TEL RIGHT |
| chrVIII | 556826 | 556963 | 137 | -1,060107198 | N8:556907 | 34,186 | | Intergene | | | TEL RIGHT |
| chrVIII | 556826 | 556965 | 139 | -1,440694378 | N8:556907 | 34,186 | | Intergene | | | TEL RIGHT |
| chrVIII | 556826 | 556967 | 141 | -1,423598416 | N8:556907 | 34,186 | | Intergene | | | TEL RIGHT |
| chrVIII | 556826 | 556969 | 143 | -1,208665072 | N8:556907 | 34,186 | | Intergene | | | TEL RIGHT |
| chrVIII | 556826 | 556970 | 144 | -2,01641571 | N8:556907 | 34,186 | | Intergene | | | TEL RIGHT |
| chrVIII | 556826 | 556971 | 145 | -1,735863071 | N8:556907 | 34,186 | | Intergene | | | TEL RIGHT |
| chrVIII | 556826 | 556972 | 146 | -2,136743975 | N8:556907 | 34,186 | | Intergene | | | TEL RIGHT |
| chrVIII | 556826 | 556973 | 147 | -1,559662379 | N8:556907 | 34,186 | | Intergene | | | TEL RIGHT |
| chrVIII | 556826 | 556975 | 149 | -1,152335566 | N8:556907 | 34,186 | | Intergene | | | TEL RIGHT |
| chrVIII | 556826 | 556976 | 150 | -2,405678773 | N8:556907 | 34,186 | | Intergene | | | TEL RIGHT |
| chrVIII | 556826 | 556977 | 151 | -1,898001685 | N8:556907 | 34,186 | | Intergene | | | TEL RIGHT |
| chrVIII | 556826 | 556979 | 153 | -1,604003431 | N8:556907 | 34,186 | | Intergene | | | TEL RIGHT |
| chrVIII | 556826 | 556981 | 155 | -2,143811282 | N8:556907 | 34,186 | | Intergene | | | TEL RIGHT |
| chrVIII | 556826 | 556982 | 156 | -1,483601099 | N8:556907 | 34,186 | | Intergene | | | TEL RIGHT |
| chrVIII | 556826 | 556983 | 157 | -1,679147984 | N8:556907 | 34,186 | | Intergene | | | TEL RIGHT |
| chrVIII | 556826 | 556984 | 158 | -1,26789513 | N8:556907 | 34,186 | | Intergene | | | TEL RIGHT |
| chrVIII | 556826 | 556985 | 159 | -1,582692098 | N8:556907 | 34,186 | | Intergene | | | TEL RIGHT |
| chrVIII | 556826 | 556986 | 160 | -1,724828859 | N8:556907 | 34,186 | | Intergene | | | TEL RIGHT |
| chrVIII | 556826 | 556987 | 161 | -1,230308377 | N8:556907 | 34,186 | | Intergene | | | TEL RIGHT |
| chrVIII | 556826 | 556989 | 163 | -0,954907273 | N8:556907 | 34,186 | | Intergene | | | TEL RIGHT |
| chrVIII | 556833 | 556986 | 153 | -1,753257145 | N8:556907 | 34,186 | | Intergene | | | TEL RIGHT |
| chrVIII | 556840 | 556986 | 146 | -0,968458255 | N8:556907 | 34,186 | | Intergene | | | TEL RIGHT |
| chrVIII | 556843 | 556989 | 146 | -1,562075941 | N8:556907 | 34,186 | | Intergene | | | TEL RIGHT |
| chrVIII | 556853 | 556986 | 133 | -1,142041105 | N8:556907 | 34,186 | | Intergene | | | TEL RIGHT |
| chrVIII | 556853 | 556989 | 136 | -0,874601035 | N8:556907 | 34,186 | | Intergene | | | TEL RIGHT |
| chrVIII | 556858 | 556975 | 117 | -0,408248201 | N8:556907 | 34,186 | | Intergene | | | TEL RIGHT |
| chrVIII | 556858 | 556989 | 131 | -1,004330007 | N8:556907 | 34,186 | | Intergene | | | TEL RIGHT |
| chrVIII | 556859 | 556963 | 104 | -0,067348136 | N8:556907 | 34,186 | | Intergene | | | TEL RIGHT |
| chrVIII | 556859 | 556989 | 130 | -0,792983028 | N8:556907 | 34,186 | | Intergene | | | TEL RIGHT |
| chrVIII | 556863 | 556986 | 123 | -1,266982049 | N8:556907 | 34,186 | | Intergene | | | TEL RIGHT |

TABLE 3

| CHR | START | END | L(bp) | ΔLK | Nuc ID | Fuzziness | Gene ID | Gene body postion or intergene | Terminal | rDNA | Telomeric |
|---|---|---|---|---|---|---|---|---|---|---|---|
| chrVIII | 556869 | 556986 | 117 | -1,048771995 | N8:556907 | 34,186 | | Intergene | | | TEL RIGHT |
| chrVIII | 556878 | 556973 | 95 | -1,171114788 | N8:556907 | 34,186 | | Intergene | | | TEL RIGHT |
| chrVIII | 556878 | 556975 | 97 | -1,002943975 | N8:556907 | 34,186 | | Intergene | | | TEL RIGHT |
| chrVIII | 556878 | 556983 | 105 | -0,952897343 | N8:556907 | 34,186 | | Intergene | | | TEL RIGHT |
| chrVIII | 556878 | 556989 | 111 | -0,356202184 | N8:556907 | 34,186 | | Intergene | | | TEL RIGHT |
| chrVIII | 556883 | 556989 | 106 | -0,275450427 | N8:556907 | 34,186 | | Intergene | | | TEL RIGHT |
| chrVIII | 556887 | 556986 | 99 | -0,587029762 | N8:556907 | 34,186 | | Intergene | | | TEL RIGHT |
| chrVIII | 556893 | 556975 | 82 | -0,339628128 | N8:556907 | 34,186 | | Intergene | | | TEL RIGHT |
| chrVIII | 556893 | 556989 | 96 | -0,212940564 | N8:556907 | 34,186 | | Intergene | | | TEL RIGHT |
| chrVIII | 556901 | 556986 | 85 | -1,15658262 | N8:556907 | 34,186 | | Intergene | | | TEL RIGHT |
| chrVIII | 556904 | 556989 | 85 | -0,177766469 | N8:556907 | 34,186 | | Intergene | | | TEL RIGHT |
| chrVIII | 558701 | 558838 | 137 | -1,012345963 | N8:558769 | 8,185 | +5:YHR218W; | 5 | | | TEL RIGHT |
| chrVIII | 558986 | 559129 | 143 | -1,062129617 | N8:559040 | 9,165 | +7:YHR218W; | 7 | | | TEL RIGHT |
| chrVIII | 559259 | 559411 | 152 | -1,07763092 | N8:559370 | 13,317 | +9:YHR218W; | 9 | | | TEL RIGHT |
| chrVIII | 559332 | 559476 | 144 | -0,863222414 | N8:559370 | 13,317 | +9:YHR218W; | 9 | | | TEL RIGHT |
| chrVIII | 560157 | 560308 | 151 | -1,176163456 | | | | Overlap <50 bp | | | TEL RIGHT |
| chrVIII | 560416 | 560563 | 147 | -1,357717533 | N8:560523 | 23,245 | +4:YHR219W; | 4 | | | TEL RIGHT |
| chrVIII | 560448 | 560563 | 115 | -0,530914926 | N8:560523 | 23,245 | +4:YHR219W; | 4 | | | TEL RIGHT |
| chrVIII | 560502 | 560671 | 169 | -2,016890925 | N8:560523 | 23,245 | +4:YHR219W; | 4 | | | TEL RIGHT |
| chrVIII | 560544 | 560634 | 90 | -0,495364542 | N8:560523 | 23,245 | +4:YHR219W; | 4 | | | TEL RIGHT |
| chrVIII | 560544 | 560639 | 95 | -0,554475013 | N8:560523 | 23,245 | +4:YHR219W; | 4 | | | TEL RIGHT |
| chrVIII | 560784 | 560911 | 127 | -0,890469386 | N8:560813 | 40,104 | +6:YHR219W; | 6 | | | TEL RIGHT |
| chrVIII | 561208 | 561356 | 148 | -1,395654132 | | | | Overlap <50 bp | | | TEL RIGHT |
| chrVIII | 561302 | 561476 | 174 | -1,545644246 | N8:561386 | 33,234 | +9:YHR219W; | 9 | | | TEL RIGHT |
| chrVIII | 561649 | 561800 | 151 | -1,150978012 | N8:561716 | 37,643 | +11:YHR219W; | 11 | | | TEL RIGHT |
| chrVIII | 562455 | 562594 | 139 | 0,024984111 | N8:562514 | 24,98 | | Intergene | | | TEL RIGHT |
| chrVIII | 562455 | 562640 | 185 | -0,616816244 | N8:562514 | 24,98 | | Intergene | | | TEL RIGHT |
| chrVIII | 562469 | 562594 | 125 | -0,482565486 | N8:562514 | 24,98 | | Intergene | | | TEL RIGHT |
| chrVIII | 562469 | 562614 | 145 | -1,401378939 | N8:562514 | 24,98 | | Intergene | | | TEL RIGHT |
| chrVIII | 562469 | 562632 | 163 | -1,305930156 | N8:562514 | 24,98 | | Intergene | | | TEL RIGHT |
| chrVIII | 562469 | 562636 | 167 | -1,717037227 | N8:562514 | 24,98 | | Intergene | | | TEL RIGHT |
| chrVIII | 562469 | 562640 | 171 | -1,193022156 | N8:562514 | 24,98 | | Intergene | | | TEL RIGHT |
| chrVIII | 562500 | 562636 | 136 | -0,767455159 | N8:562514 | 24,98 | | Intergene | | | TEL RIGHT |
| chrVIII | 562506 | 562632 | 126 | -0,636794581 | N8:562514 | 24,98 | | Intergene | | | TEL RIGHT |
| chrVIII | 562506 | 562636 | 130 | -1,324128897 | N8:562514 | 24,98 | | Intergene | | | TEL RIGHT |
| chrVIII | 562506 | 562640 | 134 | -0,456973707 | N8:562514 | 24,98 | | Intergene | | | TEL RIGHT |
| chrVIII | 562520 | 562621 | 101 | 0,370846173 | N8:562514 | 24,98 | | Intergene | | | TEL RIGHT |
| chrVIII | 562520 | 562640 | 120 | -0,974758759 | N8:562514 | 24,98 | | Intergene | | | TEL RIGHT |
| chrVIII | 562523 | 562640 | 117 | 0,771860303 | N8:562514 | 24,98 | | Intergene | | | TEL RIGHT |
| chrVIII | 562552 | 562640 | 88 | -0,426356096 | | | | Overlap <50 bp | | | TEL RIGHT |
| chrX | 1669 | 1785 | 116 | -0,087124876 | N10:1770 | 35,355 | +29:YJL225C; | 29 | | | TEL LEFT |
| chrX | 1669 | 1804 | 135 | -0,835787938 | N10:1770 | 35,355 | +29:YJL225C; | 29 | | | TEL LEFT |
| chrX | 1669 | 1821 | 152 | -1,389767415 | N10:1770 | 35,355 | +29:YJL225C; | 29 | | | TEL LEFT |
| chrX | 1693 | 1785 | 92 | -0,516507022 | N10:1770 | 35,355 | +29:YJL225C; | 29 | | | TEL LEFT |
| chrX | 1693 | 1794 | 101 | -0,849956774 | N10:1770 | 35,355 | +29:YJL225C; | 29 | | | TEL LEFT |
| chrX | 1693 | 1804 | 111 | -0,180335421 | N10:1770 | 35,355 | +29:YJL225C; | 29 | | | TEL LEFT |
| chrX | 1693 | 1821 | 128 | -0,786742979 | N10:1770 | 35,355 | +29:YJL225C; | 29 | | | TEL LEFT |
| chrX | 1698 | 1821 | 123 | -0,662951956 | N10:1770 | 35,355 | +29:YJL225C; | 29 | | | TEL LEFT |
| chrX | 1980 | 2120 | 140 | -1,100483265 | N10:2021 | 28,284 | +27:YJL225C; | 27 | | | TEL LEFT |
| chrX | 2198 | 2335 | 137 | -1,10664366 | N10:2294 | 49,87 | +25:YJL225C; | 25 | | | TEL LEFT |
| chrX | 2365 | 2509 | 144 | -1,206861675 | N10:2503 | 42,123 | +24:YJL225C; | 24 | | | TEL LEFT |
| chrX | 2365 | 2512 | 147 | -1,310235743 | N10:2503 | 42,123 | +24:YJL225C; | 24 | | | TEL LEFT |
| chrX | 2365 | 2513 | 148 | -1,149851639 | N10:2503 | 42,123 | +24:YJL225C; | 24 | | | TEL LEFT |
| chrX | 2365 | 2575 | 210 | -1,448835178 | N10:2503 | 42,123 | +24:YJL225C; | 24 | | | TEL LEFT |
| chrX | 2370 | 2513 | 143 | -1,327709072 | N10:2503 | 42,123 | +24:YJL225C; | 24 | | | TEL LEFT |
| chrX | 2380 | 2513 | 133 | -1,171343452 | N10:2503 | 42,123 | +24:YJL225C; | 24 | | | TEL LEFT |
| chrX | 2420 | 2513 | 93 | -0,628850472 | N10:2503 | 42,123 | +24:YJL225C; | 24 | | | TEL LEFT |
| chrX | 2420 | 2575 | 155 | -1,332863382 | N10:2503 | 42,123 | +24:YJL225C; | 24 | | | TEL LEFT |
| chrX | 2862 | 3015 | 153 | -1,228937881 | N10:2906 | 19,858 | +22:YJL225C; | 22 | | | TEL LEFT |
| chrX | 3307 | 3451 | 144 | -1,13439162 | N10:3406 | 9,609 | +19:YJL225C; | 19 | | | TEL LEFT |
| chrX | 3518 | 3609 | 91 | -1,230099332 | N10:3573 | 3,512 | +18:YJL225C; | 18 | | | TEL LEFT |
| chrX | 3518 | 3627 | 109 | -0,828325748 | N10:3573 | 3,512 | +18:YJL225C; | 18 | | | TEL LEFT |
| chrX | 3518 | 3628 | 110 | -0,87589654 | N10:3573 | 3,512 | +18:YJL225C; | 18 | | | TEL LEFT |
| chrX | 3518 | 3629 | 111 | -1,105302234 | N10:3573 | 3,512 | +18:YJL225C; | 18 | | | TEL LEFT |
| chrX | 3518 | 3644 | 126 | -2,457596047 | N10:3573 | 3,512 | +18:YJL225C; | 18 | | | TEL LEFT |
| chrX | 3520 | 3627 | 107 | -1,415407206 | N10:3573 | 3,512 | +18:YJL225C; | 18 | | | TEL LEFT |
| chrX | 3520 | 3644 | 124 | -1,719221196 | N10:3573 | 3,512 | +18:YJL225C; | 18 | | | TEL LEFT |
| chrX | 4071 | 4226 | 155 | -1,910277336 | N10:4185 | 5,508 | +14:YJL225C; | 14 | | | TEL LEFT |
| chrX | 4165 | 4327 | 162 | -1,450790461 | N10:4343 | 6,429 | +13:YJL225C; | 13 | | | TEL LEFT |
| chrX | 4165 | 4327 | 162 | -1,450790461 | N10:4185 | 5,508 | +14:YJL225C; | 14 | | | TEL LEFT |
| chrX | 4788 | 4951 | 163 | -1,443979978 | N10:4836 | 12,055 | +10:YJL225C; | 10 | | | TEL LEFT |
| chrX | 5199 | 5350 | 151 | -1,122297434 | | | | Overlap <50 bp | | | TEL LEFT |
| chrX | 5412 | 5571 | 159 | -1,321430513 | N10:5402 | 32,563 | +6:YJL225C; | 6 | | | TEL LEFT |
| chrX | 9273 | 9424 | 151 | -1,399793187 | N10:9358 | 3,606 | -1:YJL223C; | -1 | | | |
| chrX | 10211 | 10354 | 143 | -1,10018093 | N10:10296 | 5,132 | | Intergene | | | |
| chrX | 11443 | 11594 | 151 | -1,251838268 | N10:11535 | 2 | +2:YJL222W; | 2 | | | |
| chrX | 12328 | 12473 | 145 | -1,051660823 | N10:12354 | 30,139 | +7:YJL222W; | 7 | | | |
| chrX | 12893 | 13042 | 149 | -1,497918187 | | | | Overlap <50 bp | | | |
| chrX | 14191 | 14340 | 149 | -0,926196058 | N10:14274 | 40,447 | +20:YJL222W; | 20 | | | |
| chrX | 16003 | 16132 | 129 | -0,737658224 | N10:16047 | 20,091 | +31:YJL222W; | 31 | | | |
| chrX | 16061 | 16168 | 107 | -1,534038136 | N10:16047 | 20,091 | +31:YJL222W; | 31 | | | |
| chrX | 21416 | 21535 | 119 | -1,460940079 | N10:21469 | 5,167 | | Intergene | | | |
| chrX | 28854 | 29005 | 151 | -1,358456578 | N10:28867 | 7,861 | | Intergene | | | |
| chrX | 28854 | 29005 | 151 | -1,358456578 | N10:29020 | 40,305 | | Intergene | | | |
| chrX | 30398 | 30549 | 151 | -1,336830824 | N10:30469 | 24,095 | +5:Unit347; | 5 | | | |
| chrX | 31626 | 31734 | 108 | -0,836092408 | N10:31614 | 28,374 | -1:YJL213W; | -1 | | | |
| chrX | 32694 | 32839 | 145 | -1,290824729 | N10:32766 | 15,684 | +6:YJL213W; | 6 | | | |
| chrX | 33933 | 34077 | 144 | -1,255625659 | N10:33990 | 7,294 | +15:YJL212C; | 15 | | | |
| chrX | 37483 | 37615 | 132 | -1,400493138 | N10:37503 | 40,631 | +5*:YJL210W; | 5 | TERM | | |
| chrX | 38720 | 38876 | 156 | -1,25364164 | N10:38782 | 8,167 | +6:YJL209W; | 6 | | | |
| chrX | 40633 | 40803 | 170 | -1,706630918 | N10:40707 | 8,573 | +4:YJL208C; | 4 | | | |
| chrX | 40783 | 40953 | 170 | -1,537780227 | N10:40870 | 5,138 | +3:YJL208C; | 3 | | | |
| chrX | 46039 | 46190 | 151 | -1,280367817 | N10:46072 | 47,91 | | Intergene | | | |
| chrX | 53460 | 53598 | 138 | -1,230023216 | N10:53518 | 4,472 | +2:YJL203W; | 2 | | | |
| chrX | 55658 | 55819 | 161 | -1,298669196 | N10:55711 | 12,723 | +9:YJL201W; | 9 | | | |
| chrX | 59858 | 59998 | 140 | -1,190265656 | N10:59955 | 7,668 | | Intergene | | | |
| chrX | 64405 | 64545 | 140 | -1,285034458 | | | | Overlap <50 bp | | | |
| chrX | 86256 | 86397 | 141 | -1,173704167 | N10:86335 | 2,875 | +5:YJL181W; | 5 | | | |

TABLE 3

| CHR | START | END | L(bp) | ΔLK | Nuc ID | Fuzziness | Gene ID | Gene body postion or intergene | Terminal | rDNA | Telomeric |
|---|---|---|---|---|---|---|---|---|---|---|---|
| chrX | 88120 | 88277 | 157 | -1,3813057 | N10:88190 | 2,401 | +3:YJL180C; | 3 | | | |
| chrX | 90912 | 91092 | 180 | -1,719804921 | N10:91046 | 29,945 | +2*:YJL177W; | 2 | TERM | | |
| chrX | 94290 | 94441 | 151 | -0,717480091 | N10:94365 | 4,98 | +2:YJL176C; | 2 | | | |
| chrX | 94366 | 94450 | 84 | -0,781400672 | N10:94365 | 4,98 | +2:YJL176C; | 2 | | | |
| chrX | 94724 | 94862 | 138 | -1,352699576 | N10:94769 | 7,148 | -1:YJL174W; -1:YJL176C; | -1 | | | |
| chrX | 100058 | 100217 | 159 | -1,229272908 | N10:100203 | 6,058 | +5:YJL171C; | 5 | | | |
| chrX | 100062 | 100217 | 155 | -1,586128037 | N10:100203 | 6,058 | +5:YJL171C; | 5 | | | |
| chrX | 100066 | 100217 | 151 | -1,422028956 | N10:100203 | 6,058 | +5:YJL171C; | 5 | | | |
| chrX | 105547 | 105698 | 151 | -1,11964408 | N10:105607 | 28,499 | +5:YJL167W; | 5 | | | |
| chrX | 106971 | 107136 | 165 | -1,288219538 | | | | Overlap <50 bp | | | |
| chrX | 108096 | 108187 | 91 | -1,155196189 | N10:108130 | 32,496 | +10:YJL165C; | 10 | | | |
| chrX | 108096 | 108204 | 108 | -0,954118304 | N10:108130 | 32,496 | +10:YJL165C; | 10 | | | |
| chrX | 108096 | 108205 | 109 | -0,865699674 | N10:108130 | 32,496 | +10:YJL165C; | 10 | | | |
| chrX | 108096 | 108206 | 110 | -0,944887381 | N10:108130 | 32,496 | +10:YJL165C; | 10 | | | |
| chrX | 108096 | 108207 | 111 | -1,184729181 | N10:108130 | 32,496 | +10:YJL165C; | 10 | | | |
| chrX | 108096 | 108208 | 112 | -1,153019389 | N10:108130 | 32,496 | +10:YJL165C; | 10 | | | |
| chrX | 108096 | 108220 | 124 | -2,626057229 | N10:108130 | 32,496 | +10:YJL165C; | 10 | | | |
| chrX | 108097 | 108205 | 108 | -0,879269643 | N10:108130 | 32,496 | +10:YJL165C; | 10 | | | |
| chrX | 108098 | 108200 | 102 | -1,359329665 | N10:108130 | 32,496 | +10:YJL165C; | 10 | | | |
| chrX | 108098 | 108205 | 107 | -1,443181936 | N10:108130 | 32,496 | +10:YJL165C; | 10 | | | |
| chrX | 108098 | 108206 | 108 | -1,36077316 | N10:108130 | 32,496 | +10:YJL165C; | 10 | | | |
| chrX | 108098 | 108220 | 122 | -1,559361863 | N10:108130 | 32,496 | +10:YJL165C; | 10 | | | |
| chrX | 108135 | 108220 | 85 | -1,511526842 | N10:108130 | 32,496 | +10:YJL165C; | 10 | | | |
| chrX | 117567 | 117726 | 159 | -1,242489198 | | | | Overlap <50 bp | | | |
| chrX | 118042 | 118180 | 138 | -1,067853198 | N10:118106 | 19,442 | +1:CUT199; +6:YJL160C; | 1 | | | |
| chrX | 118045 | 118180 | 135 | -0,941288422 | N10:118106 | 19,442 | +1:CUT199; +6:YJL160C; | 1 | | | |
| chrX | 118063 | 118213 | 150 | -1,195724196 | N10:118106 | 19,442 | +1:CUT199; +6:YJL160C; | 1 | | | |
| chrX | 118208 | 118364 | 156 | -1,523681402 | N10:118386 | 11,769 | +3:CUT199; +4:YJL160C; | 3 | | | |
| chrX | 122371 | 122478 | 107 | -1,299979864 | N10:122443 | 42,087 | +3:YJL158C; | 3 | | | |
| chrX | 126969 | 127111 | 142 | -1,31984166 | N10:126996 | 12,661 | +13:YJL156C; | 13 | | | |
| chrX | 127866 | 128026 | 160 | -1,248126691 | N10:127919 | 32,764 | +7:YJL156C; | 7 | | | |
| chrX | 131244 | 131407 | 163 | -1,525964617 | N10:131345 | 42,669 | +17:YJL154C; | 17 | | | |
| chrX | 139844 | 139996 | 152 | -1,512267132 | N10:139830 | 39,009 | | Intergene | | | |
| chrX | 141168 | 141318 | 150 | -1,296773228 | N10:141239 | 26,038 | +6*:YJL148W; +9:YJL147C; | 6 | TERM | | |
| chrX | 142696 | 142846 | 150 | -1,293556673 | N10:142702 | 38,514 | -1:YJL147C; | -1 | | | |
| chrX | 145224 | 145374 | 150 | -1,477802941 | N10:145222 | 49,666 | +2:YJL145W; | 2 | | | |
| chrX | 145224 | 145374 | 150 | -1,477802941 | N10:145384 | 34,029 | +3:YJL145W; | 3 | | | |
| chrX | 148124 | 148271 | 147 | -1,174888629 | N10:148257 | 26,077 | +14:YJL141C; | 14 | | | |
| chrX | 149558 | 149726 | 168 | -1,617893786 | N10:149585 | 44,825 | +6:YJL141C; | 6 | | | |
| chrX | 154966 | 155110 | 144 | -1,539786272 | N10:155017 | 51,915 | +7:YJL137C; +1:CUT204; | 7 | | | |
| chrX | 162602 | 162770 | 168 | -1,47941696 | N10:162588 | 28,464 | | Intergene | | | |
| chrX | 163009 | 163162 | 153 | -1,366731271 | N10:163000 | 32,042 | | Intergene | | | |
| chrX | 169091 | 169251 | 160 | -1,56410159 | N10:169161 | 33,306 | +19:YJL130C; | 19 | | | |
| chrX | 173679 | 173830 | 151 | -1,562499413 | N10:173775 | 23,452 | | Intergene | | | |
| chrX | 173874 | 174016 | 142 | -1,272576999 | N10:173949 | 3,055 | | Intergene | | | |
| chrX | 178426 | 178580 | 154 | -1,244160335 | N10:178551 | 28,717 | +12:YJL128C; | 12 | | | |
| chrX | 182961 | 183077 | 116 | -0,933371074 | N10:183051 | 43,298 | +5:CUT206; +8:YJL127C; | 5 | | | |
| chrX | 182964 | 183077 | 113 | -1,083078441 | N10:183051 | 43,298 | +5:CUT206; +8:YJL127C; | 5 | | | |
| chrX | 182966 | 183077 | 111 | -0,692674699 | N10:183051 | 43,298 | +5:CUT206; +8:YJL127C; | 5 | | | |
| chrX | 185927 | 186084 | 157 | -1,283875464 | N10:186100 | 34,903 | +4:YJL125C; | 4 | | | |
| chrX | 185927 | 186084 | 157 | -1,283875464 | N10:185938 | 24,838 | +5:YJL125C; | 5 | | | |
| chrX | 189658 | 189821 | 163 | -1,154444421 | N10:189737 | 28,077 | +1:YJL122W; | 1 | | | |
| chrX | 190658 | 190805 | 147 | -1,145166371 | N10:190730 | 48,552 | +3:YJL121C; | 3 | | | |
| chrX | 191225 | 191359 | 134 | -1,196156856 | N10:191216 | 36,482 | -1:YJL121C; | -1 | | | |
| chrX | 192006 | 192112 | 106 | -0,941621829 | | | | Overlap <50 bp | | | |
| chrX | 192006 | 192128 | 122 | -1,282933223 | | | | Overlap <50 bp | | | |
| chrX | 192132 | 192293 | 161 | -1,264457106 | N10:192220 | 1 | -1:YJL117W; +5*:YJL118W; | -1 | | | |
| chrX | 192599 | 192736 | 137 | -1,834692411 | N10:192656 | 36,725 | +2:YJL117W; | 2 | | | |
| chrX | 194181 | 194318 | 137 | -1,139031062 | N10:194236 | 34,21 | +4:YJL116C; +3:SUT206; | 4 | | | |
| chrX | 197629 | 197777 | 148 | -0,956472212 | N10:197770 | 6,364 | | Intergene | | | |
| chrX | 198154 | 198316 | 162 | -1,695549094 | N10:198291 | 45,177 | | Intergene | | | |
| chrX | 198182 | 198330 | 148 | -1,258065444 | N10:198291 | 45,177 | | Intergene | | | |
| chrX | 199707 | 199858 | 151 | -1,592069346 | N10:199867 | 44,529 | | Intergene | | | |
| chrX | 203379 | 203541 | 162 | -1,480224784 | N10:203536 | 13,435 | | Intergene | | | |
| chrX | 203482 | 203630 | 148 | -1,160378042 | N10:203536 | 13,435 | | Intergene | | | |
| chrX | 206401 | 206560 | 159 | -1,675180542 | N10:206482 | 30,659 | +9:YJL112W; | 9 | | | |
| chrX | 209002 | 209147 | 145 | -1,29703576 | N10:209025 | 33,537 | +9:YJL111W; | 9 | | | |
| chrX | 210689 | 210857 | 168 | -1,493451383 | N10:210876 | 33,935 | +6:YJL110C; | 6 | | | |
| chrX | 210689 | 210857 | 168 | -1,493451383 | N10:210724 | 33,777 | +7:YJL110C; | 7 | | | |
| chrX | 212042 | 212183 | 141 | -1,147084734 | N10:212201 | 36,483 | +30:YJL109C; | 30 | | | |
| chrX | 212042 | 212183 | 141 | -1,147084734 | N10:212021 | 34,303 | +31:YJL109C; | 31 | | | |
| chrX | 215649 | 215789 | 140 | -1,22446354 | N10:215728 | 45,002 | +10:YJL109C; | 10 | | | |
| chrX | 218927 | 219076 | 149 | -1,608594929 | | | | Overlap <50 bp | | | |
| chrX | 219865 | 220016 | 151 | -1,199576177 | N10:220031 | 40,698 | 0:YJL107C; +4*:Unit351; | 0 | | | |
| chrX | 219865 | 220016 | 151 | -1,199576177 | N10:219873 | 26,588 | +1:YJL107C; | 1 | | | |
| chrX | 220580 | 220715 | 135 | -1,585892948 | N10:220634 | 37,474 | +2:Unit372; 0:Unit351; | 2 | | | |
| chrX | 221358 | 221508 | 150 | -1,309063329 | N10:221437 | 47,182 | +2:YJL106W; | 2 | | | |
| chrX | 225766 | 225921 | 155 | -1,276208315 | N10:225854 | 32,578 | +6:YJL105W; | 6 | | | |
| chrX | 226052 | 226204 | 152 | -1,488928758 | N10:226103 | 51,122 | +8:YJL105W; | 8 | | | |
| chrX | 226472 | 226633 | 161 | -1,294524921 | N10:226484 | 33,16 | +10:YJL105W; +4:SUT628; | 10 | | | |
| chrX | 227406 | 227571 | 165 | -1,410733447 | N10:227419 | 41,205 | +2:YJL104W; | 2 | | | |
| chrX | 227406 | 227571 | 165 | -1,410733447 | N10:227588 | 34,326 | +3:YJL104W; | 3 | | | |
| chrX | 229005 | 229137 | 132 | -1,328275249 | N10:229146 | 23,742 | +11*:YJL103C; | 11 | TERM | | |
| chrX | 229384 | 229535 | 151 | -1,419721107 | N10:229378 | 30,578 | +10:YJL103C; | 10 | | | |
| chrX | 233516 | 233665 | 149 | -1,583696633 | | | | Overlap <50 bp | | | |
| chrX | 235494 | 235644 | 150 | -1,417236663 | N10:235627 | 35,36 | +6:YJL101C; | 6 | | | |
| chrX | 237835 | 237975 | 140 | -1,147203109 | N10:237857 | 8,185 | +8:YJL100W; | 8 | | | |
| chrX | 240265 | 240414 | 149 | -1,096552212 | N10:240350 | 34,189 | | Intergene | | | |
| chrX | 240554 | 240707 | 153 | -1,315187587 | N10:240699 | 27,785 | | Intergene | | | |
| chrX | 240601 | 240747 | 146 | -1,300899591 | N10:240699 | 27,785 | | Intergene | | | |
| chrX | 242702 | 242868 | 166 | -1,578505634 | | | | Overlap <50 bp | | | |
| chrX | 243519 | 243619 | 100 | -0,666567693 | N10:243533 | 34,634 | +12:YJL098W; | 12 | | | |
| chrX | 245287 | 245435 | 148 | -1,358801988 | N10:245271 | 29,24 | -1:YJL097W; | -1 | | | |
| chrX | 247930 | 248081 | 151 | -1,567695434 | N10:247910 | 35,901 | +8:YJL095W; | 8 | | | |
| chrX | 248270 | 248435 | 165 | -1,467112674 | N10:248400 | 28,162 | +11:YJL095W; | 11 | | | |
| chrX | 251069 | 251224 | 155 | -1,372474 | N10:251097 | 33,323 | +28:YJL095W; | 28 | | | |
| chrX | 251916 | 252080 | 164 | -1,379793165 | N10:251970 | 34,616 | +15:YJL094C; | 15 | | | |
| chrX | 253721 | 253885 | 164 | -1,426559303 | N10:253709 | 45,908 | +4:YJL094C; | 4 | | | |
| chrX | 254829 | 255000 | 171 | -1,46592918 | N10:254961 | 5,568 | +11:YJL093C; | 11 | | | |

TABLE 3

| CHR | START | END | L(bp) | ΔLK | Nuc ID | Fuzziness | Gene ID | Gene body postion or intergene | Terminal | rDNA | Telomeric |
|---|---|---|---|---|---|---|---|---|---|---|---|
| chrX | 255572 | 255735 | 163 | -1,690054238 | N10:255685 | 33,927 | +6:YJL093C; | 6 | | | |
| chrX | 271510 | 271677 | 167 | -1,598903556 | N10:271629 | 42,484 | +6:YJL087C; | 6 | | | |
| chrX | 271664 | 271795 | 131 | -0,798483465 | | | | Overlap <50 bp | | | |
| chrX | 272130 | 272276 | 146 | -1,274905833 | N10:272201 | 34,004 | +2:YJL087C; | 2 | | | |
| chrX | 273881 | 274032 | 151 | -1,317405679 | N10:273955 | 31,329 | +9:YJL085W; | 9 | | | |
| chrX | 275971 | 276080 | 109 | -1,372479318 | N10:276085 | 33,62 | +13:YJL084C; | 13 | | | |
| chrX | 276361 | 276455 | 94 | -0,404812783 | | | | Overlap <50 bp | | | |
| chrX | 276361 | 276463 | 102 | -0,46933392 | | | | Overlap <50 bp | | | |
| chrX | 276481 | 276614 | 133 | -1,521107296 | N10:276566 | 35,204 | +10:YJL084C; | 10 | | | |
| chrX | 278918 | 279067 | 149 | -1,408743397 | N10:278918 | 2,517 | +2:YJL083W; | 2 | | | |
| chrX | 283812 | 283977 | 165 | -1,233456467 | N10:283889 | 37,704 | +9:YJL081C; | 9 | | | |
| chrX | 287746 | 287904 | 158 | -1,080879283 | N10:287749 | 6,557 | +10:YJL080C; | 10 | | | |
| chrX | 292191 | 292322 | 131 | -0,712240112 | N10:292221 | 37,136 | +12:YJL078C; | 12 | | | |
| chrX | 292971 | 293126 | 155 | -1,064407892 | | | | Overlap <50 bp | | | |
| chrX | 293774 | 293933 | 159 | -1,121600903 | N10:293870 | 26,577 | +1:YJL078C; +1:YJL077W-B; | 1 | | | |
| chrX | 301109 | 301275 | 166 | -1,28946474 | N10:301192 | 25,305 | +10:YJL074C; | 10 | | | |
| chrX | 303727 | 303887 | 160 | -1,273586686 | N10:303776 | 36,009 | +6*:YJL073W; | 6 | TERM | | |
| chrX | 309182 | 309321 | 139 | -1,391568691 | N10:309163 | 36,678 | +8:YJL070C; | 8 | | | |
| chrX | 310987 | 311146 | 159 | -1,233853829 | N10:310995 | 48,895 | | Intergene | | | |
| chrX | 310987 | 311146 | 159 | -1,233853829 | N10:311155 | 12,014 | | Intergene | | | |
| chrX | 321769 | 321919 | 150 | -1,417862454 | N10:321795 | 49,992 | +12:YJL061W; | 12 | | | |
| chrX | 322337 | 322492 | 155 | -1,461691398 | N10:322482 | 48,304 | | Intergene | | | |
| chrX | 325234 | 325383 | 149 | -1,123861123 | N10:325322 | 46,647 | +4:YJL059W; | 4 | | | |
| chrX | 326985 | 327142 | 157 | -1,308513442 | N10:327050 | 19,772 | +5:YJL058C; | 5 | | | |
| chrX | 327633 | 327785 | 152 | -1,405140593 | N10:327706 | 35,566 | +1:YJL058C; | 1 | | | |
| chrX | 328569 | 328730 | 161 | -1,51163706 | | | | Overlap <50 bp | | | |
| chrX | 329259 | 329407 | 148 | -1,596777234 | N10:329251 | 33,381 | +6*:YJL057C; | 6 | TERM | | |
| chrX | 329259 | 329407 | 148 | -1,596777234 | N10:329425 | 36,325 | +5:YJL057C; | 5 | | | |
| chrX | 331899 | 332052 | 153 | -1,482096683 | | | | Overlap <50 bp | | | |
| chrX | 333305 | 333466 | 161 | -1,410095529 | N10:333312 | 35,767 | +1:YJL055W; -1:YJL056C; | 1 | | | |
| chrX | 333305 | 333466 | 161 | -1,410095529 | N10:333478 | 3,215 | +2:YJL055W; | 2 | | | |
| chrX | 333361 | 333513 | 152 | -1,032268301 | N10:333478 | 3,215 | +2:YJL055W; | 2 | | | |
| chrX | 337451 | 337612 | 161 | -1,154894181 | | | | Overlap <50 bp | | | |
| chrX | 338776 | 338907 | 131 | -1,252496263 | N10:338772 | 2,828 | +5:YJL052W; | 5 | | | |
| chrX | 343385 | 343533 | 148 | -1,203966575 | N10:343380 | 8,327 | +7:YJL050W; | 7 | | | |
| chrX | 344740 | 344900 | 160 | -1,477610994 | | | | Overlap <50 bp | | | |
| chrX | 350347 | 350517 | 170 | -1,488234514 | N10:350502 | 31,67 | +11:YJL047C; | 11 | | | |
| chrX | 350347 | 350517 | 170 | -1,488234514 | N10:350327 | 49,044 | +12:YJL047C; | 12 | | | |
| chrX | 357774 | 357934 | 160 | -1,424926248 | N10:357775 | 40,351 | +13:YJL045W; | 13 | | | |
| chrX | 358769 | 358924 | 155 | -1,323058654 | N10:358940 | 39,357 | +5:YJL044C; | 5 | | | |
| chrX | 359603 | 359757 | 154 | -1,163835496 | N10:359674 | 3,786 | 0:YJL044C; -1:YJL043W; | 0 | | | |
| chrX | 362162 | 362320 | 158 | -1,344427311 | N10:362260 | 26,5 | +8:YJL042W; | 8 | | | |
| chrX | 365711 | 365851 | 140 | -1,385159 | N10:365757 | 3,215 | | Intergene | | | |
| chrX | 366070 | 366219 | 149 | -1,325857401 | N10:366182 | 14,107 | | Intergene | | | |
| chrX | 366353 | 366502 | 149 | -1,017253777 | N10:366336 | 30,237 | | Intergene | | | |
| chrX | 366353 | 366502 | 149 | -1,017253777 | N10:366486 | 48,083 | | Intergene | | | |
| chrX | 366520 | 366671 | 151 | -1,05145062 | N10:366641 | 32,083 | | Intergene | | | |
| chrX | 367696 | 367845 | 149 | -1,321155084 | N10:367708 | 34,691 | | Intergene | | | |
| chrX | 367696 | 367845 | 149 | -1,321155084 | N10:367868 | 47,107 | | Intergene | | | |
| chrX | 369411 | 369539 | 128 | -1,198777534 | N10:369398 | 27,3 | +28:YJL039C; | 28 | | | |
| chrX | 369776 | 369927 | 151 | -1,202669624 | N10:369846 | 34,392 | +26:YJL039C; | 26 | | | |
| chrX | 374096 | 374246 | 150 | -1,685702638 | N10:374118 | 34,861 | +2*:Unit353; -1:YJL039C; | 2 | TERM | | |
| chrX | 380580 | 380750 | 170 | -0,95644113 | N10:380752 | 32,65 | +2:YJL035C; | 2 | | | |
| chrX | 380580 | 380750 | 170 | -0,95644113 | N10:380571 | 6,11 | +3:YJL035C; | 3 | | | |
| chrX | 380923 | 381086 | 163 | -1,175209665 | N10:380993 | 31,549 | -1:YJL034W; 0:YJL035C; | -1 | | | |
| chrX | 385357 | 385485 | 128 | -1,261501152 | | | | Overlap <50 bp | | | |
| chrX | 386997 | 387161 | 164 | -1,586430376 | N10:387073 | 39,461 | +3:YJL031C; | 3 | | | |
| chrX | 392484 | 392621 | 137 | -1,276479645 | N10:392516 | 34,593 | +2:YJL026W; -1:YJL027C; | 2 | | | |
| chrX | 392991 | 393143 | 152 | -1,143021383 | N10:393070 | 46,144 | +5:YJL026W; | 5 | | | |
| chrX | 394180 | 394326 | 146 | -1,249649075 | N10:394214 | 24,228 | +3:YJL025W; | 3 | | | |
| chrX | 394276 | 394372 | 96 | -1,071782309 | N10:394374 | 36,682 | +4:YJL025W; | 4 | | | |
| chrX | 398245 | 398394 | 149 | -1,389163967 | N10:398328 | 34,519 | -1:Unit375; +1:YJL023C; | -1 | | | |
| chrX | 399541 | 399701 | 160 | -1,64390185 | N10:399582 | 51,041 | +17:YJL020C; | 17 | | | |
| chrX | 407672 | 407824 | 152 | -1,218462318 | N10:407809 | 32,908 | +3:YJL014W; | 3 | | | |
| chrX | 410284 | 410430 | 146 | -1,243962475 | N10:410379 | 44,132 | +4:YJL013C; | 4 | | | |
| chrX | 410723 | 410889 | 166 | -1,477927134 | N10:410731 | 56,691 | +2:YJL013C; | 2 | | | |
| chrX | 412052 | 412213 | 161 | -1,263714498 | | | | Overlap <50 bp | | | |
| chrX | 425378 | 425524 | 146 | -1,176392357 | | | | Overlap <50 bp | | | |
| chrX | 429176 | 429332 | 156 | -1,149147595 | N10:429178 | 47,831 | +26:YJL005W; | 26 | | | |
| chrX | 429194 | 429332 | 138 | -1,251419524 | N10:429178 | 47,831 | +26:YJL005W; | 26 | | | |
| chrX | 429194 | 429339 | 145 | -1,240375697 | N10:429178 | 47,831 | +26:YJL005W; | 26 | | | |
| chrX | 435614 | 435721 | 107 | -1,102078135 | N10:435670 | 37,917 | | Intergene | | | |
| chrX | 435614 | 435731 | 117 | -1,103064954 | N10:435670 | 37,917 | | Intergene | | | |
| chrX | 439795 | 439950 | 155 | -1,219028458 | N10:439773 | 35,566 | +6:YJR002W; | 6 | | | |
| chrX | 447409 | 447538 | 129 | -1,060616923 | | | | Overlap <50 bp | | | |
| chrX | 448364 | 448517 | 153 | -1,175115514 | N10:448344 | 23,242 | +4:YJR005C-A; | 4 | | | |
| chrX | 456180 | 456328 | 148 | -1,559992869 | N10:456302 | 36,148 | +3:YJR010W; | 3 | | | |
| chrX | 459326 | 459480 | 154 | -0,996928954 | N10:459411 | 29,08 | 0:YJR011C; 0:SUT214; | 0 | | | |
| chrX | 465469 | 465620 | 151 | -1,279018597 | N10:465481 | 49,761 | +4:YJR016C; | 4 | | | |
| chrX | 466504 | 466659 | 155 | -1,050892906 | N10:466505 | 18,751 | +4:YJR017C; | 4 | | | |
| chrX | 467197 | 467335 | 138 | -1,099641469 | N10:467195 | 36,606 | +7*:YJR019C; -1:YJR017C; | 7 | TERM | | |
| chrX | 467559 | 467705 | 146 | -2,040973872 | N10:467683 | 46,753 | +4:YJR019C; | 4 | | | |
| chrX | 470836 | 470984 | 148 | -1,224507849 | N10:470847 | 7 | +1:YJR024C; | 1 | | | |
| chrX | 472163 | 472315 | 152 | -1,352345881 | N10:472226 | 51,389 | | Intergene | | | |
| chrX | 472826 | 472975 | 149 | -1,120039169 | N10:472972 | 43,981 | | Intergene | | | |
| chrX | 473000 | 473123 | 123 | -1,045392998 | N10:473146 | 67,089 | | Intergene | | | |
| chrX | 473100 | 473255 | 155 | -1,325222881 | N10:473146 | 67,089 | | Intergene | | | |
| chrX | 473533 | 473667 | 134 | -1,037384491 | N10:473586 | 11,314 | | Intergene | | | |
| chrX | 473533 | 473673 | 140 | -1,087593171 | N10:473586 | 11,314 | | Intergene | | | |
| chrX | 473729 | 473880 | 151 | -1,197159524 | N10:473881 | 1,414 | | Intergene | | | |
| chrX | 473739 | 473884 | 145 | -1,362180335 | N10:473881 | 1,414 | | Intergene | | | |
| chrX | 473757 | 473908 | 151 | -1,64397204 | N10:473881 | 1,414 | | Intergene | | | |
| chrX | 474011 | 474165 | 154 | -1,315507124 | N10:474051 | 12,728 | | Intergene | | | |
| chrX | 474045 | 474189 | 144 | -1,179614863 | N10:474051 | 12,728 | | Intergene | | | |
| chrX | 474045 | 474189 | 144 | -1,179614863 | N10:474202 | 38,891 | | Intergene | | | |
| chrX | 474576 | 474711 | 135 | -1,292907621 | | | | Overlap <50 bp | | | |
| chrX | 475023 | 475169 | 146 | -1,110101161 | | | | Overlap <50 bp | | | |

| CHR | START | END | L(bp) | ΔLK | Nuc ID | Fuzziness | Gene ID | Gene body postion or intergene | Terminal | rDNA | Telomeric |
|---|---|---|---|---|---|---|---|---|---|---|---|
| chrX | 475023 | 475173 | 150 | -1,060358114 | | | | Overlap <50 bp | | | |
| chrX | 475693 | 475822 | 129 | -1,006382044 | N10:475768 | 41,356 | | Intergene | | | |
| chrX | 477337 | 477451 | 114 | -1,167420193 | | | | Overlap <50 bp | | | |
| chrX | 477657 | 477804 | 147 | -1,43726861 | N10:477753 | 33,941 | | Intergene | | | |
| chrX | 478238 | 478390 | 152 | -1,41848238 | N10:478237 | 52,918 | | Intergene | | | |
| chrX | 478410 | 478559 | 149 | -1,299871786 | N10:478482 | 41,885 | | Intergene | | | |
| chrX | 478560 | 478666 | 106 | -0,923119576 | | | | Overlap <50 bp | | | |
| chrX | 478560 | 478678 | 118 | -0,96178181 | | | | Overlap <50 bp | | | |
| chrX | 478584 | 478707 | 123 | -0,961213981 | N10:478712 | 31,113 | | Intergene | | | |
| chrX | 478684 | 478839 | 155 | -1,270297824 | N10:478712 | 31,113 | | Intergene | | | |
| chrX | 479117 | 479251 | 134 | -0,997850123 | | | | Overlap <50 bp | | | |
| chrX | 479117 | 479257 | 140 | -1,209276341 | N10:479275 | 21,213 | | Intergene | | | |
| chrX | 480160 | 480295 | 135 | -1,043351806 | N10:480180 | 39,247 | | Intergene | | | |
| chrX | 480607 | 480757 | 150 | -1,037428655 | N10:480589 | 19,092 | | Intergene | | | |
| chrX | 480786 | 480943 | 157 | -1,715180794 | N10:480816 | 49,541 | | Intergene | | | |
| chrX | 481277 | 481406 | 129 | -1,014070226 | N10:481374 | 34,044 | | Intergene | | | |
| chrX | 481757 | 481888 | 131 | -1,067587683 | N10:481781 | 7,071 | | Intergene | | | |
| chrX | 482189 | 482339 | 150 | -1,483185188 | N10:482287 | 43,097 | | Intergene | | | |
| chrX | 483140 | 483291 | 151 | -1,67233072 | N10:483274 | 17,616 | | Intergene | | | |
| chrX | 495866 | 496005 | 139 | -1,145658399 | N10:495880 | 36,396 | +4:YJR033C; | 4 | | | |
| chrX | 496369 | 496517 | 148 | -1,19197368 | N10:496351 | 3,215 | +1:YJR033C; -1:YJR034W; | 1 | | | |
| chrX | 498437 | 498584 | 147 | -1,262264386 | N10:498554 | 27,129 | +11:YJR035W; | 11 | | | |
| chrX | 503072 | 503238 | 166 | -1,441143302 | N10:503242 | 34,881 | +3:YJR036C; | 3 | | | |
| chrX | 503072 | 503238 | 166 | -1,441143302 | N10:503073 | 34,49 | +4:YJR036C; | 4 | | | |
| chrX | 503922 | 504069 | 147 | -1,293096882 | N10:504008 | 42,922 | +2:YJR039W; | 2 | | | |
| chrX | 512701 | 512853 | 152 | -0,889908597 | N10:512703 | 34,322 | +6:YJR041C; | 6 | | | |
| chrX | 516708 | 516897 | 189 | -1,445909132 | N10:516725 | 28,289 | +6:YJR043C; | 6 | | | |
| chrX | 519161 | 519304 | 143 | -1,279845374 | N10:519222 | 47,376 | +1:YJR044C; | 1 | | | |
| chrX | 519732 | 519838 | 106 | -1,351632717 | N10:519795 | 34,961 | +12:YJR045C; | 12 | | | |
| chrX | 519732 | 519858 | 126 | -1,167615602 | N10:519795 | 34,961 | +12:YJR045C; | 12 | | | |
| chrX | 523978 | 524108 | 130 | -0,990012305 | N10:523988 | 36,396 | | Intergene | | | |
| chrX | 526346 | 526498 | 152 | -0,881388541 | N10:526454 | 29,134 | +2:SUT641; +2:YJR048W; | 2 | | | |
| chrX | 527259 | 527387 | 128 | -1,393180911 | N10:527394 | 37,757 | +6:YJR049C; | 6 | | | |
| chrX | 530647 | 530795 | 148 | -1,141729479 | N10:530763 | 18,771 | +7:YJR051W; | 7 | | | |
| chrX | 531614 | 531753 | 139 | -1,077069317 | N10:531763 | 9,539 | -1:YJR052W; | -1 | | | |
| chrX | 533693 | 533845 | 152 | -1,375582729 | N10:533743 | 6,245 | +2:YJR053W; | 2 | | | |
| chrX | 537746 | 537899 | 153 | -1,726252063 | N10:537764 | 48,32 | | Intergene | | | |
| chrX | 542575 | 542723 | 148 | -2,394643628 | | | | Overlap <50 bp | | | |
| chrX | 544311 | 544443 | 132 | -1,258662035 | N10:544344 | 10,708 | +3:YJR057W; | 3 | | | |
| chrX | 547133 | 547290 | 157 | -1,473099822 | N10:547288 | 17,671 | +11:YJR059W; | 11 | | | |
| chrX | 554602 | 554737 | 135 | -1,267845257 | N10:554644 | 7,371 | +1:YJR062C; | 1 | | | |
| chrX | 557805 | 557945 | 140 | -1,172590938 | N10:557963 | 22,716 | +8:YJR065C; | 8 | | | |
| chrX | 557805 | 557945 | 140 | -1,172590938 | N10:557804 | 31,858 | +9:YJR065C; | 9 | | | |
| chrX | 557934 | 558083 | 149 | -0,835953679 | N10:557963 | 22,716 | +8:YJR065C; | 8 | | | |
| chrX | 558445 | 558574 | 129 | -0,96728835 | N10:558562 | 35,584 | +4:YJR065C; | 4 | | | |
| chrX | 561483 | 561587 | 104 | -1,583504592 | | | | Overlap <50 bp | | | |
| chrX | 561700 | 561836 | 136 | -1,176344522 | N10:561697 | 40,28 | +15:YJR066W; | 15 | | | |
| chrX | 563670 | 563822 | 152 | -1,335483118 | N10:563737 | 34,256 | +28:YJR066W; | 28 | | | |
| chrX | 567134 | 567269 | 135 | -1,085491083 | N10:567226 | 34,364 | +2:YJR067C; -1:YJR068W; | 2 | | | |
| chrX | 569137 | 569291 | 154 | -1,062542371 | N10:569134 | 34,974 | +2:YJR069C; | 2 | | | |
| chrX | 572775 | 572903 | 128 | -0,673208568 | | | | Overlap <50 bp | | | |
| chrX | 573799 | 573959 | 160 | -1,371202031 | N10:573951 | 37,712 | +1:YJR075W; | 1 | | | |
| chrX | 582808 | 582958 | 150 | -1,002744758 | | | | Overlap <50 bp | | | |
| chrX | 586031 | 586180 | 149 | -1,326640475 | N10:586162 | 37,954 | +2*:YJR086W; | 2 | TERM | | |
| chrX | 590874 | 591029 | 155 | -1,379331855 | N10:591040 | 38,188 | | Intergene | | | |
| chrX | 591957 | 592113 | 156 | -1,264491637 | N10:592071 | 23,195 | | Intergene | | | |
| chrX | 594961 | 595112 | 151 | -1,191112781 | N10:594945 | 37,249 | +18:YJR091C; | 18 | | | |
| chrX | 595496 | 595660 | 164 | -0,869054055 | N10:595504 | 47,178 | +15:YJR091C; | 15 | | | |
| chrX | 598239 | 598403 | 164 | -1,292927307 | N10:598323 | 36,065 | 0:YJR091C; -1:YJR092W; | 0 | | | |
| chrX | 598789 | 598947 | 158 | -1,369018598 | N10:598769 | 34,496 | +2:YJR092W; | 2 | | | |
| chrX | 600553 | 600699 | 146 | -0,958173686 | N10:600632 | 32,685 | +14:YJR092W; | 14 | | | |
| chrX | 606604 | 606728 | 124 | -1,215552723 | N10:606660 | 14,107 | +5:SUT643; | 5 | | | |
| chrX | 608387 | 608544 | 157 | -1,383765238 | N10:608402 | 48,976 | +2*:YJR094W-A; | 2 | TERM | | |
| chrX | 608709 | 608811 | 102 | -1,016717165 | N10:608690 | 45,318 | | Intergene | | | |
| chrX | 610999 | 611149 | 150 | -1,357078591 | N10:610987 | 31,919 | +1:YJR096W; +7*:SUT644; | 1 | | | |
| chrX | 623512 | 623664 | 152 | -1,564751633 | N10:623582 | 26,986 | +2:YJR105W; | 2 | | | |
| chrX | 624211 | 624342 | 131 | -1,214013879 | N10:624345 | 23,796 | +7:YJR105W; | 7 | | | |
| chrX | 624949 | 625089 | 140 | -1,141512165 | N10:625080 | 32,388 | +3:YJR106W; | 3 | | | |
| chrX | 632590 | 632738 | 148 | -1,271270927 | N10:632669 | 44,076 | +2:YJR109C; | 2 | | | |
| chrX | 635827 | 635966 | 139 | -1,165099263 | N10:635846 | 26,87 | +6*:YJR111C; | 6 | TERM | | |
| chrX | 636181 | 636337 | 156 | -1,484780806 | N10:636350 | 33,776 | +3:YJR111C; | 3 | | | |
| chrX | 636181 | 636337 | 156 | -1,484780806 | N10:636186 | 30,397 | +4:YJR111C; | 4 | | | |
| chrX | 636714 | 636864 | 150 | -1,359844187 | N10:636770 | 35,584 | 0:YJR111C; -1:YJR112W; | 0 | | | |
| chrX | 637154 | 637305 | 151 | -1,540868913 | N10:637320 | 48,003 | -1:YJR112W-A; +3:YJR112W; | -1 | | | |
| chrX | 637154 | 637305 | 151 | -1,540868913 | N10:637147 | 3,215 | +2:YJR112W; | 2 | | | |
| chrX | 638977 | 639126 | 149 | -1,159844519 | N10:638974 | 18,903 | 0:YJR113C; | 0 | | | |
| chrX | 653069 | 653214 | 145 | -1,178306589 | N10:653126 | 33,955 | +7:YJR124C; | 7 | | | |
| chrX | 661255 | 661420 | 165 | -1,399719386 | N10:661267 | 27,789 | +9:YJR127C; | 9 | | | |
| chrX | 662029 | 662161 | 132 | -0,915804221 | N10:662110 | 43,279 | +4:YJR127C; | 4 | | | |
| chrX | 662034 | 662161 | 127 | -0,884119286 | N10:662110 | 43,279 | +4:YJR127C; | 4 | | | |
| chrX | 662037 | 662135 | 98 | -0,578244156 | N10:662110 | 43,279 | +4:YJR127C; | 4 | | | |
| chrX | 671104 | 671223 | 119 | -1,227155771 | | | | Overlap <50 bp | | | |
| chrX | 671901 | 672050 | 149 | -1,585638614 | | | | Overlap <50 bp | | | |
| chrX | 673361 | 673549 | 188 | -1,608961129 | | | | Overlap <50 bp | | | |
| chrX | 673385 | 673539 | 154 | -1,304641944 | | | | Overlap <50 bp | | | |
| chrX | 674527 | 674676 | 149 | -1,202480408 | N10:674605 | 35,449 | +12:YJR135C; +9:YJR134C; | 12 | | | |
| chrX | 684490 | 684642 | 152 | -1,326344291 | N10:684478 | 38,073 | +1:YJR138W; | 1 | | | |
| chrX | 684490 | 684642 | 152 | -1,326344291 | N10:684641 | 38,118 | +2:YJR138W; | 2 | | | |
| chrX | 685942 | 686069 | 127 | -1,023331107 | N10:686043 | 45,181 | +11:YJR138W; | 11 | | | |
| chrX | 687031 | 687188 | 157 | -1,522408057 | N10:687180 | 29,092 | +18:YJR138W; | 18 | | | |
| chrX | 687681 | 687842 | 161 | -1,503074974 | N10:687764 | 6,856 | +22*:YJR138W; | 22 | TERM | | |
| chrX | 689548 | 689698 | 150 | -1,139736761 | N10:689678 | 19,883 | +6:YJR139C; | 6 | | | |
| chrX | 695637 | 695789 | 152 | -1,200633399 | N10:695637 | 7,55 | +1:YJR140C; -1:YJR141W; | 1 | | | |
| chrX | 696260 | 696391 | 131 | -1,089926385 | N10:696342 | 35,763 | +4:YJR141W; | 4 | | | |
| chrX | 697538 | 697692 | 154 | -1,423417057 | N10:697604 | 34,568 | +4:YJR142W; | 4 | | | |
| chrX | 699421 | 699528 | 107 | -1,322540912 | N10:699480 | 35,259 | +8:YJR143C; | 8 | | | |
| chrX | 699421 | 699559 | 138 | -1,238454783 | N10:699480 | 35,259 | +8:YJR143C; | 8 | | | |
| chrX | 699421 | 699560 | 139 | -1,427267091 | N10:699480 | 35,259 | +8:YJR143C; | 8 | | | |

| CHR | START | END | L(bp) | ΔLK | Nuc ID | Fuzziness | Gene ID | Gene body postion or intergene | Terminal | rDNA | Telomeric |
|---|---|---|---|---|---|---|---|---|---|---|---|
| chrX | 699421 | 699561 | 140 | -1,133150691 | N10:699480 | 35,259 | +8:YJR143C; | 8 | | | |
| chrX | 699421 | 699563 | 142 | -1,523311187 | N10:699480 | 35,259 | +8:YJR143C; | 8 | | | |
| chrX | 699421 | 699564 | 143 | -1,198565158 | N10:699480 | 35,259 | +8:YJR143C; | 8 | | | |
| chrX | 700845 | 700991 | 146 | -1,1199237 | N10:700894 | 36,581 | -1:YJR143C; +2:YJR144W; | -1 | | | |
| chrX | 701322 | 701470 | 148 | -1,259819895 | N10:701322 | 32,342 | +5:YJR144W; | 5 | | | |
| chrX | 705757 | 705926 | 169 | -1,672219511 | N10:705760 | 32,609 | +2:YJR148W; | 2 | | | |
| chrX | 705757 | 705926 | 169 | -1,672219511 | N10:705934 | 32,725 | +3:YJR148W; | 3 | | | |
| chrX | 706905 | 707066 | 161 | -1,489130344 | N10:707000 | 42,289 | +2*:YJR149W; | 2 | TERM | | |
| chrX | 712931 | 713070 | 139 | -1,229911679 | N10:712913 | 47,376 | +17:YJR151C; | 17 | | | |
| chrX | 714988 | 715121 | 133 | -0,98982772 | N10:715105 | 7,572 | +5:YJR151C; | 5 | | | |
| chrX | 720830 | 720969 | 139 | -0,888870258 | N10:720969 | 46,456 | +9:YJR152W; | 9 | | | |
| chrX | 721017 | 721124 | 107 | -1,577883772 | | | | Overlap <50 bp | | | |
| chrX | 721017 | 721148 | 131 | -1,134228498 | | | | Overlap <50 bp | | | |
| chrX | 721017 | 721149 | 132 | -1,315438328 | | | | Overlap <50 bp | | | |
| chrX | 721017 | 721153 | 136 | -1,177066163 | | | | Overlap <50 bp | | | |
| chrX | 721018 | 721153 | 135 | -0,923998916 | | | | Overlap <50 bp | | | |
| chrX | 721019 | 721153 | 134 | -1,379371996 | | | | Overlap <50 bp | | | |
| chrX | 721020 | 721153 | 133 | -1,416510415 | | | | Overlap <50 bp | | | |
| chrX | 723086 | 723238 | 152 | -1,424966951 | N10:723143 | 40,454 | +3:YJR153W; +7:anti102; | 3 | | | |
| chrX | 723301 | 723431 | 130 | -0,425733754 | N10:723416 | 25,487 | +5:YJR153W; +5:anti102; | 5 | | | |
| chrX | 723442 | 723594 | 152 | -1,106831378 | N10:723587 | 8,505 | +6:YJR153W; +4:anti102; | 6 | | | |
| chrX | 725588 | 725741 | 153 | -1,466226462 | N10:725606 | 7,371 | +5*:Unit382; +2:YJR154W; | 5 | TERM | | |
| chrX | 729427 | 729584 | 157 | -1,151697126 | N10:729606 | 0,707 | +1:YJR156C; | 1 | | | |
| chrX | 729427 | 729584 | 157 | -1,151697126 | N10:729424 | 8,485 | +2:YJR156C; | 2 | | | |
| chrX | 729799 | 729953 | 154 | -1,19958189 | | | | Overlap <50 bp | | | |
| chrX | 732737 | 732877 | 140 | -1,265381153 | N10:732736 | 20,664 | +4:YJR158W; | 4 | | | |
| chrX | 735177 | 735331 | 154 | -1,755704053 | N10:735172 | 9,192 | | Intergene | | | |
| chrX | 735177 | 735331 | 154 | -1,755704053 | N10:735346 | 0,707 | | Intergene | | | |
| chrX | 736882 | 737013 | 131 | -1,39783147 | N10:736967 | 3,536 | +7:YJR159W; | 7 | | | |
| chrX | 739148 | 739242 | 94 | -1,286313551 | N10:739259 | 35,355 | +4:YJR160C; | 4 | | | |
| chrX | 745644 | 745739 | 95 | -1,048020398 | | | | Overlap <50 bp | | | TEL RIGHT |
| chrX | 745644 | 745742 | 98 | -0,55235635 | | | | Overlap <50 bp | | | TEL RIGHT |
| chrXI | 6938 | 7102 | 164 | -1,277801213 | N11:6990 | 9,888 | +6:YKL221W; +5:SUT652; | 6 | | | |
| chrXI | 10271 | 10420 | 149 | -1,087930626 | N11:10428 | 25,377 | +7:YKL220C; | 7 | | | |
| chrXI | 17984 | 18132 | 148 | -1,143517674 | N11:18117 | 31,079 | +3:YKL218C; | 3 | | | |
| chrXI | 19266 | 19389 | 123 | -1,087703789 | N11:19336 | 3,724 | +4:Unit402; | 4 | | | |
| chrXI | 19267 | 19389 | 122 | -1,077864938 | N11:19336 | 3,724 | +4:Unit402; | 4 | | | |
| chrXI | 19267 | 19414 | 147 | -1,446101711 | N11:19336 | 3,724 | +4:Unit402; | 4 | | | |
| chrXI | 20703 | 20861 | 158 | -1,425894316 | N11:20761 | 35,26 | +3:SUT656; | 3 | | | |
| chrXI | 31510 | 31641 | 131 | -1,245794249 | | | | Overlap <50 bp | | | |
| chrXI | 32617 | 32778 | 161 | -1,502199352 | N11:32717 | 20,549 | +10:YKL213C; | 10 | | | |
| chrXI | 38219 | 38379 | 160 | -1,032578271 | | | | Overlap <50 bp | | | |
| chrXI | 39078 | 39236 | 158 | -1,568674681 | N11:39170 | 9,099 | +1:YKL210W; | 1 | | | |
| chrXI | 41766 | 41916 | 150 | -1,284555618 | | | | Overlap <50 bp | | | |
| chrXI | 42476 | 42618 | 142 | -1,143447387 | N11:42545 | 29,925 | +25:YKL209C; | 25 | | | |
| chrXI | 42479 | 42618 | 139 | -1,367993666 | N11:42545 | 29,925 | +25:YKL209C; | 25 | | | |
| chrXI | 44129 | 44285 | 156 | -1,523919203 | N11:44262 | 39,703 | +14:YKL209C; | 14 | | | |
| chrXI | 45474 | 45591 | 117 | -1,231710626 | | | | Overlap <50 bp | | | |
| chrXI | 52845 | 53010 | 165 | -1,440049486 | N11:52908 | 27,244 | +18:YKL205W; | 18 | | | |
| chrXI | 53086 | 53185 | 99 | -1,730695943 | N11:53095 | 29,811 | +19:YKL205W; | 19 | | | |
| chrXI | 55429 | 55585 | 156 | -1,246833511 | N11:55522 | 19,892 | +12*:YKL204W; | 12 | TERM | | |
| chrXI | 55463 | 55623 | 160 | -1,303570973 | N11:55522 | 19,892 | +12*:YKL204W; | 12 | TERM | | |
| chrXI | 55924 | 56074 | 150 | -1,38574672 | N11:55980 | 9,894 | | Intergene | | | |
| chrXI | 55924 | 56080 | 156 | -1,354459686 | N11:55980 | 9,894 | | Intergene | | | |
| chrXI | 56846 | 56978 | 132 | -1,039575521 | N11:56861 | 21,047 | | Intergene | | | |
| chrXI | 61848 | 62032 | 184 | -1,528614544 | N11:61896 | 24,205 | | Intergene | | | |
| chrXI | 63877 | 64030 | 153 | -1,645514067 | N11:63947 | 7,823 | | Intergene | | | |
| chrXI | 65154 | 65316 | 162 | -1,216569883 | N11:65282 | 34,252 | | Intergene | | | |
| chrXI | 65672 | 65838 | 166 | -1,180368197 | N11:65746 | 17,971 | | Intergene | | | |
| chrXI | 68614 | 68763 | 149 | -1,293394349 | N11:68734 | 35,242 | +33:YKL197C; +11:YKL198C; | 33 | | | |
| chrXI | 73102 | 73261 | 159 | -1,354809962 | N11:73185 | 9,317 | +5:YKL197C; | 5 | | | |
| chrXI | 86463 | 86611 | 148 | -1,294537371 | N11:86556 | 25,561 | +13:YKL188C; | 13 | | | |
| chrXI | 86483 | 86645 | 162 | -1,351649744 | N11:86556 | 25,561 | +13:YKL188C; | 13 | | | |
| chrXI | 88336 | 88484 | 148 | -1,360316734 | N11:88371 | 8,75 | +2:YKL188C; | 2 | | | |
| chrXI | 89789 | 89957 | 168 | -1,179881379 | N11:89946 | 7,815 | +8:YKL187C; +8:SUT223; | 8 | | | |
| chrXI | 89789 | 89957 | 168 | -1,179881379 | N11:89782 | 11,075 | +9:YKL187C; +7:SUT223; | 9 | | | |
| chrXI | 91032 | 91184 | 152 | -1,315963927 | N11:91137 | 10,652 | +1:YKL187C; +15:SUT223; | 1 | | | |
| chrXI | 97596 | 97707 | 111 | -1,407182241 | N11:97729 | 17,103 | +8:YKL184W; | 8 | | | |
| chrXI | 97601 | 97707 | 106 | -1,435094211 | N11:97729 | 17,103 | +8:YKL184W; | 8 | | | |
| chrXI | 99816 | 99946 | 130 | -0,634674032 | N11:99812 | 0 | | Intergene | | | |
| chrXI | 99816 | 99946 | 130 | -0,634674032 | N11:99961 | 26,7 | | Intergene | | | |
| chrXI | 103049 | 103197 | 148 | -1,282889892 | N11:103034 | 27,998 | +15:YKL182W; | 15 | | | |
| chrXI | 103049 | 103197 | 148 | -1,282889892 | N11:103200 | 36,661 | +16:YKL182W; | 16 | | | |
| chrXI | 103372 | 103526 | 154 | -1,156374647 | N11:103384 | 31,835 | +17:YKL182W; | 17 | | | |
| chrXI | 105256 | 105405 | 149 | -1,109838074 | N11:105284 | 18,267 | +29:YKL182W; | 29 | | | |
| chrXI | 105988 | 106134 | 146 | -0,832272706 | N11:106069 | 15,457 | +34:YKL182W; | 34 | | | |
| chrXI | 110658 | 110811 | 153 | -1,398077383 | N11:110802 | 28,341 | +12:YKL179C; | 12 | | | |
| chrXI | 115225 | 115382 | 157 | -1,318169762 | N11:115291 | 36,875 | +2:anti-YKL176C-2; | 2 | | | |
| chrXI | 116126 | 116262 | 136 | -1,524932821 | N11:116268 | 37,211 | +14:YKL176C; | 14 | | | |
| chrXI | 116126 | 116262 | 136 | -1,524932821 | N11:116112 | 30,053 | +15:YKL176C; | 15 | | | |
| chrXI | 116231 | 116308 | 77 | -0,791 | N11:116268 | 37,211 | +14:YKL176C; | 14 | | | |
| chrXI | 116232 | 116308 | 76 | -0,784 | N11:116268 | 37,211 | +14:YKL176C; | 14 | | | |
| chrXI | 116232 | 116357 | 125 | -0,554372798 | N11:116268 | 37,211 | +14:YKL176C; | 14 | | | |
| chrXI | 117579 | 117716 | 137 | -1,047346837 | N11:117661 | 14,618 | +5:YKL176C; | 5 | | | |
| chrXI | 120342 | 120499 | 157 | -1,253643673 | N11:120460 | 22,418 | +12:YKL174C; | 12 | | | |
| chrXI | 120705 | 120857 | 152 | -1,373594098 | N11:120781 | 22,241 | +10:YKL174C; | 10 | | | |
| chrXI | 121556 | 121713 | 157 | -1,840848996 | N11:121624 | 28,284 | +5:YKL174C; | 5 | | | |
| chrXI | 130706 | 130855 | 149 | -1,294789974 | N11:130776 | 9,564 | +2:YKL170W; | 2 | | | |
| chrXI | 130929 | 131079 | 150 | -1,243841213 | N11:130940 | 7,333 | +3:YKL170W; | 3 | | | |
| chrXI | 132985 | 133151 | 166 | -1,265520167 | N11:133074 | 6,573 | +3:YKL168C; | 3 | | | |
| chrXI | 134033 | 134163 | 130 | -1,413200418 | N11:134102 | 15,895 | +1:YKL167C; | 1 | | | |
| chrXI | 136517 | 136676 | 159 | -1,640354503 | | | | Overlap <50 bp | | | |
| chrXI | 140618 | 140768 | 150 | -1,365274442 | N11:140692 | 25,563 | +1:YKL165C; | 1 | | | |
| chrXI | 142672 | 142804 | 132 | -0,873870395 | N11:142756 | 11,5 | +9*:SUT227; +2:YKL164C; | 9 | TERM | | |
| chrXI | 145250 | 145428 | 178 | -1,218550033 | N11:145382 | 18,7 | +7:YKL163W; | 7 | | | |
| chrXI | 146568 | 146719 | 151 | -1,207028851 | N11:146644 | 7,448 | | Intergene | | | |
| chrXI | 147356 | 147520 | 164 | -1,525104196 | N11:147421 | 6,274 | | Intergene | | | |
| chrXI | 149408 | 149568 | 160 | -1,434937275 | N11:149508 | 6,892 | +9:YKL161C; | 9 | | | |

| CHR | START | END | L(bp) | ΔLK | Nuc ID | Fuzziness | Gene ID | Gene body postion or intergene | Terminal | rDNA | Telomeric |
|---|---|---|---|---|---|---|---|---|---|---|---|
| chrXI | 149964 | 150108 | 144 | -1,305308002 | N11:149984 | 38,811 | +6:YKL161C; | 6 | | | |
| chrXI | 151755 | 151890 | 135 | -1,56837928 | N11:151768 | 26,098 | +2:Unit408; | 2 | | | |
| chrXI | 161235 | 161384 | 149 | -1,235547801 | N11:161325 | 6,573 | +1:YKL155C; -1:YKL154W; | 1 | | | |
| chrXI | 161799 | 161928 | 129 | -1,042344321 | N11:161887 | 12,317 | +3:YKL154W; | 3 | | | |
| chrXI | 162802 | 162952 | 150 | -1,449849305 | N11:162881 | 9,708 | | Intergene | | | |
| chrXI | 170837 | 170984 | 147 | -1,956666349 | N11:170954 | 14,012 | +3:YKL148C; | 3 | | | |
| chrXI | 171758 | 171891 | 133 | -1,210196837 | N11:171815 | 21,408 | +1:YKL146W; | 1 | | | |
| chrXI | 172320 | 172456 | 136 | -1,128780039 | N11:172446 | 19,442 | +5:YKL146W; | 5 | | | |
| chrXI | 178039 | 178227 | 188 | -1,620970865 | N11:178152 | 9,365 | +9*:YKL143W; | 9 | TERM | | |
| chrXI | 178982 | 179138 | 156 | -1,41986694 | N11:179070 | 28,487 | +4:YKL142W; | 4 | | | |
| chrXI | 179058 | 179209 | 151 | -1,801591915 | N11:179232 | 11,314 | +1*:Unit387; +5*:YKL142W; | 1 | TERM | | |
| chrXI | 179058 | 179209 | 151 | -1,801591915 | N11:179070 | 28,487 | +4:YKL142W; | 4 | | | |
| chrXI | 182120 | 182298 | 178 | -1,142131712 | N11:182189 | 5,128 | +11:YKL140W; | 11 | | | |
| chrXI | 183536 | 183690 | 154 | -1,383518065 | N11:183621 | 15,501 | +6:YKL139W; | 6 | | | |
| chrXI | 183964 | 184081 | 117 | -1,173982643 | N11:183955 | 25,587 | +8:YKL139W; | 8 | | | |
| chrXI | 187208 | 187352 | 144 | -1,142518053 | N11:187276 | 5,231 | +10:YKL135C; | 10 | | | |
| chrXI | 188029 | 188179 | 150 | -1,254409008 | N11:188071 | 21,144 | +5:YKL135C; | 5 | | | |
| chrXI | 190331 | 190479 | 148 | -1,342896657 | N11:190473 | 13,732 | +7:YKL134C; | 7 | | | |
| chrXI | 191024 | 191171 | 147 | -1,16542173 | N11:191119 | 8,448 | +3:YKL134C; | 3 | | | |
| chrXI | 195322 | 195449 | 127 | -1,307366987 | N11:195406 | 13,648 | +2:SUT231; | 2 | | | |
| chrXI | 196316 | 196481 | 165 | -1,342314753 | N11:196493 | 8,585 | +25:YKL129C; | 25 | | | |
| chrXI | 196620 | 196752 | 132 | -1,172248545 | N11:196660 | 22,379 | +24:YKL129C; | 24 | | | |
| chrXI | 201937 | 202099 | 162 | -1,657529265 | N11:202016 | 9,317 | +3:anti105; | 3 | | | |
| chrXI | 204817 | 204983 | 166 | -1,485847362 | N11:204909 | 7,439 | YKL127W; -1:YKL126W; +2:SUT663; | 12 | TERM | | |
| chrXI | 207618 | 207770 | 152 | -1,135945232 | | | | Overlap <50 bp | | | |
| chrXI | 209750 | 209876 | 126 | -1,257680546 | N11:209839 | 15,742 | -1:YKL124W; | -1 | | | |
| chrXI | 214868 | 215018 | 150 | -1,857026937 | N11:214868 | 21,197 | +9:YKL121W; | 9 | | | |
| chrXI | 214868 | 215019 | 151 | -1,349297145 | N11:214868 | 21,197 | +9:YKL121W; | 9 | | | |
| chrXI | 214886 | 215018 | 132 | -1,287451285 | N11:214868 | 21,197 | +9:YKL121W; | 9 | | | |
| chrXI | 214886 | 215019 | 133 | -1,145250632 | N11:214868 | 21,197 | +9:YKL121W; | 9 | | | |
| chrXI | 216300 | 216452 | 152 | -1,351795978 | N11:216342 | 6,387 | +18*:YKL121W; | 18 | TERM | | |
| chrXI | 216511 | 216657 | 146 | -1,055433005 | N11:216610 | 41,801 | -1:YKL120W; | -1 | | | |
| chrXI | 226833 | 226983 | 150 | -1,396144118 | N11:226966 | 16,021 | +6:YKL112W; | 6 | | | |
| chrXI | 226930 | 227062 | 132 | -1,114905315 | N11:226966 | 16,021 | +6:YKL112W; | 6 | | | |
| chrXI | 227206 | 227326 | 120 | -1,307397937 | | | | Overlap <50 bp | | | |
| chrXI | 229813 | 229973 | 160 | -1,355229093 | N11:229834 | 45,727 | -1:YKL110C; | -1 | | | |
| chrXI | 233634 | 233784 | 150 | -1,313879396 | N11:233802 | 9,899 | -1:YKL108W; | -1 | | | |
| chrXI | 233634 | 233784 | 150 | -1,313879396 | N11:233638 | 27,845 | | Intergene | | | |
| chrXI | 236594 | 236764 | 170 | -1,468111303 | N11:236682 | 12,661 | +2*:YKL106C-A; +7:YKL107W; | 2 | TERM | | |
| chrXI | 241460 | 241610 | 150 | -1,128458802 | N11:241499 | 30,545 | +6:YKL105C; | 6 | | | |
| chrXI | 245174 | 245347 | 173 | -1,534240295 | N11:245177 | 23,08 | +1:YKL104C; | 1 | | | |
| chrXI | 245189 | 245339 | 150 | -1,28827703 | N11:245177 | 23,08 | +1:YKL104C; | 1 | | | |
| chrXI | 245551 | 245725 | 174 | -1,516690766 | N11:245572 | 13,378 | -1:CUT229; +12*:YKL103C; | -1 | | | |
| chrXI | 247204 | 247369 | 165 | -1,415912496 | N11:247366 | 10,69 | +1:YKL103C; | 1 | | | |
| chrXI | 247204 | 247369 | 165 | -1,415912496 | N11:247197 | 5,715 | +2:YKL103C; | 2 | | | |
| chrXI | 249638 | 249773 | 135 | -1,454078261 | N11:249715 | 14,782 | +9*:YKL101W; | 9 | TERM | | |
| chrXI | 251126 | 251263 | 137 | -1,669708437 | N11:251198 | 23,215 | | Intergene | | | |
| chrXI | 253186 | 253299 | 113 | -1,304056574 | N11:253297 | 14,096 | +12:YKL100C; | 12 | | | |
| chrXI | 253541 | 253688 | 147 | -1,520689262 | N11:253657 | 9,529 | +10:YKL100C; | 10 | | | |
| chrXI | 257352 | 257460 | 108 | -0,441043313 | | | | Overlap <50 bp | | | |
| chrXI | 257353 | 257478 | 125 | -1,055238517 | N11:257490 | 10,685 | +4*:CUT708; +8*:YKL098W; | 4 | TERM | | |
| chrXI | 257353 | 257460 | 107 | -0,873985153 | | | | Overlap <50 bp | | | |
| chrXI | 257617 | 257767 | 150 | -1,165734102 | N11:257744 | 10,895 | +2:CUT708; | 2 | | | |
| chrXI | 258177 | 258336 | 159 | -1,332134099 | N11:258256 | 40,596 | -1:CUT708; | -1 | | | |
| chrXI | 261407 | 261543 | 136 | -0,960315721 | N11:261504 | 9,674 | -1:YKL095W; +5*:YKL096W; | -1 | | | |
| chrXI | 269387 | 269539 | 152 | -1,384663016 | N11:269374 | 12,178 | +6*:YKL091C; -1:YKL092C; | 6 | TERM | | |
| chrXI | 269387 | 269539 | 152 | -1,384663016 | N11:269548 | 11,255 | +5:YKL091C; | 5 | | | |
| chrXI | 270529 | 270668 | 139 | -1,141787867 | N11:270660 | 5,776 | +1:Unit413; +7:SUT666; | 1 | | | |
| chrXI | 275707 | 275869 | 162 | -1,429772106 | N11:275750 | 17,297 | +6:YKL088W; | 6 | | | |
| chrXI | 275877 | 276030 | 153 | -1,117352052 | N11:276025 | 28,839 | +8:YKL088W; | 8 | | | |
| chrXI | 277324 | 277474 | 150 | -1,444413256 | N11:277389 | 10,132 | +2:YKL087C; | 2 | | | |
| chrXI | 278497 | 278637 | 140 | -1,088358319 | N11:278487 | 22,78 | ıit414; +5*:YKL086W; -1:YKL085W; | 1 | | | |
| chrXI | 278499 | 278637 | 138 | -1,071049835 | N11:278487 | 22,78 | ıit414; +5*:YKL086W; -1:YKL085W; | 1 | | | |
| chrXI | 278501 | 278637 | 136 | -1,08346536 | N11:278487 | 22,78 | ıit414; +5*:YKL086W; -1:YKL085W; | 1 | | | |
| chrXI | 278502 | 278637 | 135 | -0,856760414 | N11:278487 | 22,78 | ıit414; +5*:YKL086W; -1:YKL085W; | 1 | | | |
| chrXI | 282993 | 283156 | 163 | -1,441195826 | N11:283075 | 28,618 | | Intergene | | | |
| chrXI | 286651 | 286824 | 173 | -1,396885971 | N11:286733 | 14,89 | +5:YKL079W; | 5 | | | |
| chrXI | 287603 | 287693 | 90 | -0,992563336 | N11:287704 | 46,226 | +11:YKL079W; | 11 | | | |
| chrXI | 288326 | 288474 | 148 | -1,383497141 | | | | Overlap <50 bp | | | |
| chrXI | 290076 | 290228 | 152 | -1,390011365 | N11:290108 | 19,357 | +11:YKL078W; | 11 | | | |
| chrXI | 292965 | 293112 | 147 | -1,244305627 | N11:293063 | 26,326 | +7:YKL075C; | 7 | | | |
| chrXI | 297006 | 297125 | 119 | -1,157503768 | N11:297036 | 15,007 | +7:YKL073W; | 7 | | | |
| chrXI | 297006 | 297156 | 150 | -1,024765712 | N11:297036 | 15,007 | +7:YKL073W; | 7 | | | |
| chrXI | 298702 | 298832 | 130 | -1,151423589 | N11:298779 | 28,827 | 'KL073W; +1:CUT713; -1:YKL072W; | 18 | TERM | | |
| chrXI | 298875 | 299007 | 132 | -1,501964606 | | | | Overlap <50 bp | | | |
| chrXI | 300199 | 300305 | 106 | -1,122868376 | N11:300189 | 14,849 | +11:SUT669; +8:YKL072W; | 11 | | | |
| chrXI | 300199 | 300317 | 118 | -1,289205404 | N11:300189 | 14,849 | +11:SUT669; +8:YKL072W; | 11 | | | |
| chrXI | 303921 | 304068 | 147 | -1,420259895 | N11:304040 | 12,728 | +3:Unit394; | 3 | | | |
| chrXI | 305787 | 305949 | 162 | -1,130742803 | | | | Overlap <50 bp | | | |
| chrXI | 318927 | 319072 | 145 | -1,547433426 | | | | Overlap <50 bp | | | |
| chrXI | 319054 | 319202 | 148 | -1,42625176 | N11:319121 | 18,148 | +13:YKL064W; | 13 | | | |
| chrXI | 322500 | 322670 | 170 | -1,76912977 | N11:322551 | 13,736 | -1:YKL062W; | -1 | | | |
| chrXI | 329617 | 329749 | 132 | -1,0887366 | | | | Overlap <50 bp | | | |
| chrXI | 329629 | 329749 | 120 | -0,701973648 | | | | Overlap <50 bp | | | |
| chrXI | 329629 | 329775 | 146 | -1,416694403 | | | | Overlap <50 bp | | | |
| chrXI | 337370 | 337525 | 155 | -1,385232608 | N11:337362 | 13,686 | +8:YKL054C; | 8 | | | |
| chrXI | 337447 | 337600 | 153 | -1,287507965 | | | | Overlap <50 bp | | | |
| chrXI | 337450 | 337600 | 150 | -1,429669448 | | | | Overlap <50 bp | | | |
| chrXI | 337452 | 337600 | 148 | -1,484611305 | | | | Overlap <50 bp | | | |
| chrXI | 337462 | 337600 | 138 | -1,181901582 | | | | Overlap <50 bp | | | |
| chrXI | 337968 | 338118 | 150 | -1,313410235 | | | | Overlap <50 bp | | | |
| chrXI | 340261 | 340401 | 140 | -1,16170491 | N11:340305 | 20,506 | 0:YKL052C; -1:YKL051W; | 0 | | | |
| chrXI | 341684 | 341830 | 146 | -1,222004622 | N11:341720 | 15,777 | +7:YKL051W; +1:anti108; | 7 | | | |
| chrXI | 347279 | 347450 | 171 | -1,410199378 | N11:347369 | 11,283 | +10:YKL048C; | 10 | | | |
| chrXI | 347320 | 347467 | 147 | -1,5137025 | N11:347369 | 11,283 | +10:YKL048C; | 10 | | | |
| chrXI | 347479 | 347608 | 129 | -1,104127238 | N11:347544 | 6,95 | +9:YKL048C; | 9 | | | |
| chrXI | 351244 | 351421 | 177 | -1,54469598 | | | | Overlap <50 bp | | | |
| chrXI | 351885 | 352030 | 145 | -1,135151385 | N11:351957 | 9,68 | +4:YKL046C; | 4 | | | |
| chrXI | 352745 | 352882 | 137 | -1,083066438 | N11:352864 | 15,553 | -1:YKL045W; | -1 | | | |

| CHR | START | END | L(bp) | ΔLK | Nuc ID | Fuzziness | Gene ID | Gene body postion or intergene | Terminal | rDNA | Telomeric |
|---|---|---|---|---|---|---|---|---|---|---|---|
| chrXI | 353147 | 353287 | 140 | -1,150448513 | N11:353269 | 12,7 | +2:YKL045W; | 2 | | | |
| chrXI | 355288 | 355448 | 160 | -1,273033904 | | | | Overlap <50 bp | | | |
| chrXI | 357460 | 357614 | 154 | -1,382746132 | N11:357564 | 11,325 | +9*:YKL043W; | 9 | TERM | | |
| chrXI | 357462 | 357614 | 152 | -1,692727786 | N11:357564 | 11,325 | +9*:YKL043W; | 9 | TERM | | |
| chrXI | 362917 | 363070 | 153 | -1,47423929 | | | | Overlap <50 bp | | | |
| chrXI | 364062 | 364210 | 148 | -1,52575083 | | | | Overlap <50 bp | | | |
| chrXI | 364602 | 364723 | 121 | -0,846473789 | N11:364623 | 6,535 | +2:anti109; | 2 | | | |
| chrXI | 368064 | 368220 | 156 | -1,459457895 | | | | Overlap <50 bp | | | |
| chrXI | 368389 | 368522 | 133 | -1,205237908 | N11:368440 | 13,136 | +22:YKL038W; | 22 | | | |
| chrXI | 368971 | 369078 | 107 | -0,970769711 | N11:368963 | 29,344 | +1:YKL037W; +1:anti-YKL037W; | 1 | | | |
| chrXI | 368971 | 369121 | 150 | -1,34870179 | N11:368963 | 29,344 | +1:YKL037W; +1:anti-YKL037W; | 1 | | | |
| chrXI | 368971 | 369122 | 151 | -1,331130205 | N11:368963 | 29,344 | +1:YKL037W; +1:anti-YKL037W; | 1 | | | |
| chrXI | 372978 | 373087 | 109 | -1,430283221 | N11:372980 | 13,682 | +10:YKL034W; | 10 | | | |
| chrXI | 373360 | 373511 | 151 | -1,228841272 | N11:373499 | 11,514 | +13*:YKL034W; | 13 | TERM | | |
| chrXI | 377591 | 377751 | 160 | -1,423960104 | N11:377588 | 14,078 | +16:YKL033W; | 16 | | | |
| chrXI | 379254 | 379407 | 153 | -1,480716649 | N11:379329 | 40,278 | | Intergene | | | |
| chrXI | 385681 | 385837 | 156 | -1,370355021 | N11:385749 | 5,431 | +3:YKL028W; | 3 | | | |
| chrXI | 394646 | 394790 | 144 | -1,187442759 | N11:394794 | 41,012 | +13:YKL022C; | 13 | | | |
| chrXI | 397867 | 398011 | 144 | -0,938006967 | N11:398025 | 2,066 | +2:YKL021C; | 2 | | | |
| chrXI | 397867 | 398011 | 144 | -0,938006967 | N11:397862 | 6,969 | +3:YKL021C; | 3 | | | |
| chrXI | 399301 | 399460 | 159 | -1,349659052 | | | | Overlap <50 bp | | | |
| chrXI | 400364 | 400506 | 142 | -1,246531614 | N11:400496 | 32,782 | +9:YKL020C; | 9 | | | |
| chrXI | 401469 | 401625 | 156 | -1,293918031 | N11:401550 | 1,817 | +2:YKL020C; | 2 | | | |
| chrXI | 401694 | 401852 | 158 | -1,297923275 | N11:401711 | 12,34 | +1:YKL020C; | 1 | | | |
| chrXI | 403288 | 403463 | 175 | -1,546859788 | N11:403356 | 7,627 | +2:YKL018C-A; | 2 | | | |
| chrXI | 406796 | 406955 | 159 | -1,345582711 | N11:406832 | 9,411 | +1:YKL017C; | 1 | | | |
| chrXI | 407916 | 408065 | 149 | -1,286718438 | | | | Overlap <50 bp | | | |
| chrXI | 416183 | 416327 | 144 | -0,953664975 | N11:416221 | 9,566 | +3:YKL014C; | 3 | | | |
| chrXI | 417925 | 418084 | 159 | -1,336156511 | N11:417916 | 9,772 | +1:YKL012W; -1:YKL013C; | 1 | | | |
| chrXI | 417925 | 418084 | 159 | -1,336156511 | N11:418102 | 12,287 | +2:YKL012W; | 2 | | | |
| chrXI | 434342 | 434496 | 154 | -1,350534294 | N11:434396 | 8,329 | +2:YKL005C; | 2 | | | |
| chrXI | 434529 | 434684 | 155 | -0,686696579 | N11:434609 | 3,114 | -1:YKL004W; +1:YKL005C; | -1 | | | |
| chrXI | 434561 | 434692 | 131 | -0,640462858 | N11:434609 | 3,114 | -1:YKL004W; +1:YKL005C; | -1 | | | |
| chrXI | 434662 | 434792 | 130 | -1,088301757 | | | | Overlap <50 bp | | | |
| chrXI | 439672 | 439806 | 134 | -1,001442981 | N11:439814 | 7,767 | | Intergene | | | |
| chrXI | 439682 | 439780 | 98 | -0,446824277 | | | | Overlap <50 bp | | | |
| chrXI | 447937 | 448094 | 157 | -1,538607083 | N11:447938 | 9,247 | 'KR005C; -1:YKR004C; +2*:Unit396; | 10 | TERM | | |
| chrXI | 448453 | 448602 | 149 | -1,313032092 | N11:448490 | 17,581 | +7:YKR005C; | 7 | | | |
| chrXI | 450755 | 450913 | 158 | -1,022292696 | N11:450835 | 17,815 | -1:YKR007W; +1:YKR006C; | -1 | | | |
| chrXI | 454604 | 454760 | 156 | -1,366766752 | N11:454730 | 21,986 | +13:YKR009C; | 13 | | | |
| chrXI | 455031 | 455148 | 117 | -1,137213638 | N11:455162 | 46,231 | +10:YKR009C; -1:anti110; | 10 | | | |
| chrXI | 455031 | 455138 | 107 | -1,013180878 | | | | Overlap <50 bp | | | |
| chrXI | 456736 | 456888 | 152 | -1,494743672 | N11:456824 | 11,843 | 0:YKR009C; | 0 | | | |
| chrXI | 459229 | 459336 | 107 | -1,656623344 | N11:459206 | 18,877 | +12:YKR010C; | 12 | | | |
| chrXI | 459229 | 459341 | 112 | -1,383589223 | N11:459206 | 18,877 | +12:YKR010C; | 12 | | | |
| chrXI | 462337 | 462501 | 164 | -1,135339798 | N11:462401 | 7,607 | +1:YKR011C; +8*:SUT237; | 1 | | | |
| chrXI | 468677 | 468825 | 148 | -1,467080651 | | | | Overlap <50 bp | | | |
| chrXI | 470132 | 470274 | 142 | -1,420480946 | | | | Overlap <50 bp | | | |
| chrXI | 473335 | 473488 | 153 | -1,176038381 | N11:473478 | 15,401 | -1:YKR017C; +14:YKR018C; | -1 | | | |
| chrXI | 479959 | 480086 | 127 | -1,544628482 | N11:479984 | 23,099 | +8:YKR021W; | 8 | | | |
| chrXI | 483328 | 483476 | 148 | -1,159816275 | N11:483388 | 19,159 | +3:YKR023W; | 3 | | | |
| chrXI | 484474 | 484623 | 149 | -1,078249487 | N11:484589 | 4,472 | +10*:YKR023W; | 10 | TERM | | |
| chrXI | 488055 | 488207 | 152 | -1,919810319 | N11:488122 | 34,518 | +5*:YKR025W; | 5 | TERM | | |
| chrXI | 488418 | 488598 | 180 | -1,676439491 | N11:488497 | 42,997 | +6:YKR026C; | 6 | | | |
| chrXI | 496967 | 497107 | 140 | -1,532335667 | N11:497088 | 24,042 | +23*:YKR028W; +16*:YKR029C; | 23 | TERM | | |
| chrXI | 501192 | 501346 | 154 | -0,931030758 | N11:501317 | 13,229 | +31:YKR031C; | 31 | | | |
| chrXI | 506644 | 506790 | 146 | -1,204656566 | N11:506685 | 8,775 | +2:YKR034W; | 2 | | | |
| chrXI | 509826 | 509992 | 166 | -1,433164296 | N11:509894 | 14,798 | +3:YKR036C; | 3 | | | |
| chrXI | 514096 | 514200 | 104 | -1,941858161 | | | | Overlap <50 bp | | | |
| chrXI | 516175 | 516336 | 161 | -1,456654672 | N11:516248 | 10,289 | +10:YKR039W; +2*:CUT722; | 10 | | | |
| chrXI | 517302 | 517453 | 151 | -1,283868679 | N11:517465 | 7,312 | +1*:CUT243; | 1 | TERM | | |
| chrXI | 517302 | 517453 | 151 | -1,283868679 | N11:517303 | 5,859 | 0:CUT243; | 0 | | | |
| chrXI | 517458 | 517572 | 114 | -1,071799122 | N11:517465 | 7,312 | +1*:CUT243; | 1 | TERM | | |
| chrXI | 519553 | 519695 | 142 | -1,954964418 | N11:519598 | 24,595 | -1:SUT680; +3:YKR042W; | -1 | | | |
| chrXI | 527505 | 527640 | 135 | -1,776897392 | N11:527634 | 3,421 | +2:YKR050W; | 2 | | | |
| chrXI | 527506 | 527630 | 124 | -1,728056172 | N11:527634 | 3,421 | +2:YKR050W; | 2 | | | |
| chrXI | 527506 | 527632 | 126 | -0,942646825 | N11:527634 | 3,421 | +2:YKR050W; | 2 | | | |
| chrXI | 527506 | 527635 | 129 | -1,067958772 | N11:527634 | 3,421 | +2:YKR050W; | 2 | | | |
| chrXI | 527506 | 527636 | 130 | -1,311221893 | N11:527634 | 3,421 | +2:YKR050W; | 2 | | | |
| chrXI | 527506 | 527637 | 131 | -1,220788904 | N11:527634 | 3,421 | +2:YKR050W; | 2 | | | |
| chrXI | 527506 | 527638 | 132 | -1,434465716 | N11:527634 | 3,421 | +2:YKR050W; | 2 | | | |
| chrXI | 527506 | 527639 | 133 | -1,63929572 | N11:527634 | 3,421 | +2:YKR050W; | 2 | | | |
| chrXI | 527506 | 527640 | 134 | -1,074074951 | N11:527634 | 3,421 | +2:YKR050W; | 2 | | | |
| chrXI | 527506 | 527641 | 135 | -1,725193084 | N11:527634 | 3,421 | +2:YKR050W; | 2 | | | |
| chrXI | 527506 | 527600 | 94 | -0,781739611 | | | | Overlap <50 bp | | | |
| chrXI | 527507 | 527640 | 133 | -1,65119413 | N11:527634 | 3,421 | +2:YKR050W; | 2 | | | |
| chrXI | 527508 | 527640 | 132 | -1,408614796 | N11:527634 | 3,421 | +2:YKR050W; | 2 | | | |
| chrXI | 527509 | 527640 | 131 | -0,731289278 | N11:527634 | 3,421 | +2:YKR050W; | 2 | | | |
| chrXI | 527510 | 527640 | 130 | -1,107015988 | N11:527634 | 3,421 | +2:YKR050W; | 2 | | | |
| chrXI | 527511 | 527640 | 129 | -1,291822961 | N11:527634 | 3,421 | +2:YKR050W; | 2 | | | |
| chrXI | 527513 | 527640 | 127 | -0,997766103 | N11:527634 | 3,421 | +2:YKR050W; | 2 | | | |
| chrXI | 527515 | 527640 | 125 | -1,085926311 | N11:527634 | 3,421 | +2:YKR050W; | 2 | | | |
| chrXI | 527516 | 527640 | 124 | -0,580555537 | N11:527634 | 3,421 | +2:YKR050W; | 2 | | | |
| chrXI | 529083 | 529220 | 137 | -1,014028983 | N11:529204 | 40,962 | +12:YKR050W; | 12 | | | |
| chrXI | 529403 | 529549 | 146 | -1,047215862 | N11:529570 | 19,88 | +14:YKR050W; | 14 | | | |
| chrXI | 532781 | 532933 | 152 | -0,973794087 | N11:532891 | 2,582 | +2:YKR052C; | 2 | | | |
| chrXI | 533171 | 533320 | 149 | -1,230147776 | N11:533271 | 26,626 | 0:YKR052C; | 0 | | | |
| chrXI | 540415 | 540570 | 155 | -1,545139632 | N11:540451 | 10,801 | | Intergene | | | |
| chrXI | 540630 | 540787 | 157 | -1,456999207 | N11:540698 | 31,054 | | Intergene | | | |
| chrXI | 540646 | 540797 | 151 | -1,303932263 | N11:540698 | 31,054 | | Intergene | | | |
| chrXI | 541723 | 541874 | 151 | -1,044781952 | N11:541853 | 6,238 | | Intergene | | | |
| chrXI | 542290 | 542416 | 126 | -1,256873632 | N11:542354 | 21,299 | | Intergene | | | |
| chrXI | 542290 | 542440 | 150 | -1,063540739 | N11:542354 | 21,299 | | Intergene | | | |
| chrXI | 544676 | 544823 | 147 | -0,939931276 | | | | Overlap <50 bp | | | |
| chrXI | 545722 | 545853 | 131 | -0,972122009 | N11:545860 | 29,126 | +12:YKR054C; | 12 | | | |
| chrXI | 547817 | 547975 | 158 | -1,378496788 | N11:547848 | 12,42 | -1:YKR054C; +1:YKR055W; | -1 | | | |
| chrXI | 548333 | 548481 | 148 | -1,188062781 | N11:548344 | 9,094 | +4:YKR055W; | 4 | | | |
| chrXI | 548769 | 548931 | 162 | -1,068846234 | N11:548953 | 10,607 | -1:YKR056W; +8*:YKR055W; | -1 | | | |
| chrXI | 548769 | 548931 | 162 | -1,068846234 | N11:548799 | 12,814 | +7:YKR055W; | 7 | | | |

| CHR | START | END | L(bp) | ΔLK | Nuc ID | Fuzziness | Gene ID | Gene body postion or intergene | Terminal | rDNA | Telomeric |
|---|---|---|---|---|---|---|---|---|---|---|---|
| chrXI | 549757 | 549907 | 150 | -1,477039257 | N11:549905 | 7,815 | +5:YKR056W; | 5 | | | |
| chrXI | 550284 | 550422 | 138 | -1,355780273 | N11:550349 | 33,534 | +8:YKR056W; | 8 | | | |
| chrXI | 554400 | 554551 | 151 | -1,261101912 | N11:554437 | 9,899 | 0:CUT723; 0:YKR059W; | 0 | | | |
| chrXI | 558933 | 559085 | 152 | -1,440657893 | N11:558954 | 20,841 | | Intergene | | | |
| chrXI | 562000 | 562173 | 173 | -1,414134405 | N11:562186 | 10,108 | +1:YKR064W; -1:YKR063C; | 1 | | | |
| chrXI | 571010 | 571152 | 142 | -1,426668893 | N11:571147 | 7,234 | +1:YKR069W; | 1 | | | |
| chrXI | 573165 | 573306 | 141 | -1,03733852 | N11:573269 | 4,457 | +1:YKR070W; | 1 | | | |
| chrXI | 574030 | 574178 | 148 | -1,176840884 | N11:574099 | 7,893 | | Intergene | | | |
| chrXI | 574402 | 574555 | 153 | -1,158885339 | N11:574493 | 31,675 | +8*:YKR071C; | 8 | TERM | | |
| chrXI | 575707 | 575877 | 170 | -1,814578963 | | | | Overlap <50 bp | | | |
| chrXI | 577254 | 577388 | 134 | -1,08489176 | N11:577250 | 28,862 | +5:YKR072C; | 5 | | | |
| chrXI | 577351 | 577483 | 132 | -1,248560245 | | | | Overlap <50 bp | | | |
| chrXI | 577690 | 577817 | 127 | -2,232113062 | N11:577711 | 8,571 | +2:YKR072C; | 2 | | | |
| chrXI | 578476 | 578613 | 137 | -1,053063029 | N11:578514 | 43,317 | | Intergene | | | |
| chrXI | 579137 | 579295 | 158 | -1,266759603 | N11:579194 | 5,514 | +3*:YKR074W; | 3 | TERM | | |
| chrXI | 590716 | 590822 | 106 | -1,015979271 | N11:590776 | 16,754 | | Intergene | | | |
| chrXI | 590716 | 590835 | 119 | -0,928565403 | N11:590776 | 16,754 | | Intergene | | | |
| chrXI | 600361 | 600514 | 153 | -1,308757421 | N11:600370 | 5,115 | +6:YKR086W; | 6 | | | |
| chrXI | 600361 | 600514 | 153 | -1,308757421 | N11:600524 | 8,544 | +7:YKR086W; | 7 | | | |
| chrXI | 601868 | 602039 | 171 | -1,646568101 | N11:601908 | 24,105 | +15:YKR086W; | 15 | | | |
| chrXI | 602067 | 602222 | 155 | -1,318641727 | N11:602124 | 10,496 | +16:YKR086W; | 16 | | | |
| chrXI | 603803 | 603944 | 141 | -1,105775151 | N11:603837 | 21,977 | +1:YKR087C; -1:CUT245; | 1 | | | |
| chrXI | 604749 | 604856 | 107 | -1,207103495 | | | | Overlap <50 bp | | | |
| chrXI | 604749 | 604865 | 116 | -1,400489492 | | | | Overlap <50 bp | | | |
| chrXI | 607498 | 607646 | 148 | -1,006721509 | N11:607543 | 11,017 | +15:SUT241; +4:YKR089C; | 15 | | | |
| chrXI | 607998 | 608143 | 145 | -1,060848765 | N11:608050 | 7,635 | +18*:SUT241; +1:YKR089C; | 18 | TERM | | |
| chrXI | 613167 | 613294 | 127 | -1,292647269 | N11:613186 | 6,348 | +3:YKR092C; | 3 | | | |
| chrXI | 614331 | 614461 | 130 | -0,962289354 | N11:614435 | 21,213 | +1:Unit424; | 1 | | | |
| chrXI | 618644 | 618800 | 156 | -1,313314927 | N11:618766 | 22,095 | +2:SUT242; | 2 | | | |
| chrXI | 621479 | 621632 | 153 | -1,417530231 | N11:621620 | 21,733 | +16:YKR095W; | 16 | | | |
| chrXI | 624990 | 625141 | 151 | -0,65999762 | N11:625071 | 10,095 | | Intergene | | | |
| chrXI | 632085 | 632240 | 155 | -1,182367077 | N11:632159 | 7,874 | +9:YKR097W; | 9 | | | |
| chrXI | 632617 | 632753 | 136 | -1,149859204 | N11:632714 | 21,925 | +15*:YKR098C; | 15 | TERM | | |
| chrXI | 638182 | 638313 | 131 | -0,862741226 | N11:638195 | 7,629 | 0:Unit399; | 0 | | | |
| chrXI | 638182 | 638316 | 134 | -0,987126256 | N11:638195 | 7,629 | 0:Unit399; | 0 | | | |
| chrXI | 643087 | 643244 | 157 | -1,507185576 | N11:643162 | 17,502 | +4:Unit426; | 4 | | | |
| chrXI | 646189 | 646315 | 126 | -1,131510651 | N11:646259 | 24,505 | +3:YKR102W; | 3 | | | |
| chrXI | 647330 | 647482 | 152 | -1,160321071 | N11:647430 | 61,175 | +10:YKR102W; | 10 | | | |
| chrXI | 648395 | 648512 | 117 | -1,79810144 | | | | Overlap <50 bp | | | |
| chrXI | 650461 | 650612 | 151 | -1,268612453 | N11:650494 | 16,068 | | Intergene | | | |
| chrXI | 651142 | 651292 | 150 | -1,210417098 | N11:651252 | 57,57 | | Intergene | | | |
| chrXI | 654048 | 654197 | 149 | -1,227711402 | | | | Overlap <50 bp | | | |
| chrXI | 655704 | 655870 | 166 | -1,455669477 | N11:655848 | 10,909 | +20:YKR103W; | 20 | | | |
| chrXI | 655763 | 655920 | 157 | -1,248381432 | N11:655848 | 10,909 | +20:YKR103W; | 20 | | | |
| chrXI | 657176 | 657328 | 152 | -1,245411891 | N11:657284 | 18,889 | +6:YKR104W; | 6 | | | |
| chrXI | 658054 | 658203 | 149 | -1,263381555 | N11:658207 | 19,992 | +13*:YKR105C; | 13 | TERM | | |
| chrXI | 659842 | 659965 | 123 | -1,039751102 | N11:659867 | 6,083 | +3:YKR105C; | 3 | | | |
| chrXI | 661850 | 662006 | 156 | -1,523923982 | N11:662001 | 30,265 | +6:YKR106W; | 6 | | | |
| chrXI | 663739 | 663846 | 107 | -1,591157694 | N11:663807 | 18,484 | | Intergene | | | |
| chrXI | 663739 | 663851 | 112 | -1,496994452 | N11:663807 | 18,484 | | Intergene | | | |
| chrXI | 664373 | 664519 | 146 | -1,219839868 | N11:664394 | 30,288 | | Intergene | | | |
| chrXI | 666612 | 666748 | 136 | -0,170790474 | | | | Overlap <50 bp | | | TEL RIGHT |
| chrXI | 666628 | 666815 | 187 | -1,317524387 | | | | Overlap <50 bp | | | TEL RIGHT |
| chrXI | 666656 | 666815 | 159 | -1,276379071 | | | | Overlap <50 bp | | | TEL RIGHT |
| chrXI | 666681 | 666771 | 90 | -0,637675508 | | | | Overlap <50 bp | | | TEL RIGHT |
| chrXI | 666681 | 666810 | 129 | -0,226776015 | | | | Overlap <50 bp | | | TEL RIGHT |
| chrXI | 666693 | 666810 | 117 | -0,181331826 | | | | Overlap <50 bp | | | TEL RIGHT |
| chrXI | 666703 | 666728 | 25 | -0,455 | | | | Overlap <50 bp | | | TEL RIGHT |
| chrXI | 666703 | 666733 | 30 | -0,462 | | | | Overlap <50 bp | | | TEL RIGHT |
| chrXI | 666703 | 666737 | 34 | -0,49 | | | | Overlap <50 bp | | | TEL RIGHT |
| chrXI | 666703 | 666739 | 36 | -0,504 | | | | Overlap <50 bp | | | TEL RIGHT |
| chrXI | 666709 | 666792 | 83 | -1,278737534 | | | | Overlap <50 bp | | | TEL RIGHT |
| chrXI | 666709 | 666810 | 101 | -0,727775771 | | | | Overlap <50 bp | | | TEL RIGHT |
| chrXI | 666709 | 666815 | 106 | -0,579146041 | | | | Overlap <50 bp | | | TEL RIGHT |
| chrXI | 666714 | 666815 | 101 | -0,877945081 | | | | Overlap <50 bp | | | TEL RIGHT |
| chrXII | 723 | 874 | 151 | -1,215029429 | N12:805 | 36,056 | +22:YLL067C; | 22 | | | TEL LEFT |
| chrXII | 1167 | 1315 | 148 | -1,450874716 | | | | Overlap <50 bp | | | TEL LEFT |
| chrXII | 1996 | 2147 | 151 | -1,114284137 | N12:2014 | 33,886 | +15:YLL067C; | 15 | | | TEL LEFT |
| chrXII | 2008 | 2142 | 134 | -1,018833784 | N12:2014 | 33,886 | +15:YLL067C; | 15 | | | TEL LEFT |
| chrXII | 2033 | 2193 | 160 | -1,478624355 | N12:2177 | 0 | +14:YLL067C; | 14 | | | TEL LEFT |
| chrXII | 2033 | 2129 | 96 | -0,764202285 | N12:2014 | 33,886 | +15:YLL067C; | 15 | | | TEL LEFT |
| chrXII | 2033 | 2193 | 160 | -1,478624355 | N12:2014 | 33,886 | +15:YLL067C; | 15 | | | TEL LEFT |
| chrXII | 2211 | 2359 | 148 | -1,173163436 | N12:2350 | 42,884 | +13:YLL067C; | 13 | | | TEL LEFT |
| chrXII | 2226 | 2355 | 129 | -1,203857305 | N12:2350 | 42,884 | +13:YLL067C; | 13 | | | TEL LEFT |
| chrXII | 2226 | 2359 | 133 | -0,950778968 | N12:2350 | 42,884 | +13:YLL067C; | 13 | | | TEL LEFT |
| chrXII | 2226 | 2421 | 195 | -1,467525801 | N12:2350 | 42,884 | +13:YLL067C; | 13 | | | TEL LEFT |
| chrXII | 2387 | 2517 | 130 | -1,180494239 | | | | Overlap <50 bp | | | TEL LEFT |
| chrXII | 2666 | 2818 | 152 | -1,257051464 | N12:2754 | 19,655 | +11:YLL067C; | 11 | | | TEL LEFT |
| chrXII | 3013 | 3157 | 144 | -0,979723026 | N12:3070 | 21,794 | +9:YLL067C; | 9 | | | TEL LEFT |
| chrXII | 3013 | 3200 | 187 | -1,119063938 | N12:3070 | 21,794 | +9:YLL067C; | 9 | | | TEL LEFT |
| chrXII | 3065 | 3157 | 92 | -0,362412574 | N12:3070 | 21,794 | +9:YLL067C; | 9 | | | TEL LEFT |
| chrXII | 3065 | 3200 | 135 | -1,017112954 | N12:3070 | 21,794 | +9:YLL067C; | 9 | | | TEL LEFT |
| chrXII | 3153 | 3297 | 144 | -0,849559593 | N12:3257 | 10,97 | +8:YLL067C; | 8 | | | TEL LEFT |
| chrXII | 3218 | 3370 | 152 | -1,283866477 | N12:3257 | 10,97 | +8:YLL067C; | 8 | | | TEL LEFT |
| chrXII | 3470 | 3595 | 125 | -0,741908549 | N12:3583 | 6,807 | +6:YLL067C; | 6 | | | TEL LEFT |
| chrXII | 3500 | 3643 | 143 | -1,417970922 | N12:3583 | 6,807 | +6:YLL067C; | 6 | | | TEL LEFT |
| chrXII | 3500 | 3651 | 151 | -1,327111307 | N12:3583 | 6,807 | +6:YLL067C; | 6 | | | TEL LEFT |
| chrXII | 3791 | 3928 | 137 | -1,23138484 | N12:3868 | 7,371 | +4:YLL067C; | 4 | | | TEL LEFT |
| chrXII | 6346 | 6517 | 171 | -1,58589495 | N12:6439 | 52,275 | +23:YLL066C; | 23 | | | TEL LEFT |
| chrXII | 6366 | 6517 | 151 | -1,240022875 | N12:6439 | 52,275 | +23:YLL066C; | 23 | | | TEL LEFT |
| chrXII | 6690 | 6864 | 174 | -1,542909836 | N12:6762 | 16,258 | +21:YLL066C; | 21 | | | TEL LEFT |
| chrXII | 6705 | 6857 | 152 | -1,661251081 | N12:6762 | 16,258 | +21:YLL066C; | 21 | | | TEL LEFT |
| chrXII | 7495 | 7682 | 187 | -1,207477855 | N12:7546 | 19,313 | +16:YLL066C; | 16 | | | TEL LEFT |
| chrXII | 7531 | 7682 | 151 | -1,036472979 | N12:7546 | 19,313 | +16:YLL066C; | 16 | | | TEL LEFT |
| chrXII | 7543 | 7677 | 134 | -0,979266891 | N12:7546 | 19,313 | +16:YLL066C; | 16 | | | TEL LEFT |
| chrXII | 7568 | 7664 | 96 | -0,897952495 | N12:7546 | 19,313 | +16:YLL066C; | 16 | | | TEL LEFT |
| chrXII | 7746 | 7894 | 148 | -1,380037904 | N12:7816 | 1,414 | +14:YLL066C; | 14 | | | TEL LEFT |
| chrXII | 7761 | 7894 | 133 | -0,967320245 | N12:7816 | 1,414 | +14:YLL066C; | 14 | | | TEL LEFT |

TABLE 3

| CHR | START | END | L(bp) | ΔLK | Nuc ID | Fuzziness | Gene ID | Gene body postion or intergene | Terminal | rDNA | Telomeric |
|---|---|---|---|---|---|---|---|---|---|---|---|
| chrXII | 7761 | 7956 | 195 | -1,537952513 | N12:7816 | 1,414 | +14:YLL066C; | 14 | | | TEL LEFT |
| chrXII | 7922 | 8052 | 130 | -1,180881073 | | | | Overlap <50 bp | | | TEL LEFT |
| chrXII | 8201 | 8353 | 152 | -1,093006861 | N12:8284 | 26,502 | +11:YLL066C; | 11 | | | TEL LEFT |
| chrXII | 8548 | 8692 | 144 | -0,964894372 | N12:8607 | 17,786 | +9:YLL066C; | 9 | | | TEL LEFT |
| chrXII | 8548 | 8735 | 187 | -1,383346343 | N12:8607 | 17,786 | +9:YLL066C; | 9 | | | TEL LEFT |
| chrXII | 8600 | 8692 | 92 | -0,712992886 | N12:8607 | 17,786 | +9:YLL066C; | 9 | | | TEL LEFT |
| chrXII | 8600 | 8735 | 135 | -0,984965139 | N12:8607 | 17,786 | +9:YLL066C; | 9 | | | TEL LEFT |
| chrXII | 8688 | 8832 | 144 | -1,18201862 | N12:8786 | 7,024 | +8:YLL066C; | 8 | | | TEL LEFT |
| chrXII | 8753 | 8905 | 152 | -1,311576448 | N12:8786 | 7,024 | +8:YLL066C; | 8 | | | TEL LEFT |
| chrXII | 9005 | 9130 | 125 | -1,016326141 | N12:9112 | 11,533 | +6:YLL066C; | 6 | | | TEL LEFT |
| chrXII | 9035 | 9178 | 143 | -1,525598667 | N12:9112 | 11,533 | +6:YLL066C; | 6 | | | TEL LEFT |
| chrXII | 9326 | 9463 | 137 | -1,097412697 | N12:9402 | 8,021 | +4:YLL066C; | 4 | | | TEL LEFT |
| chrXII | 11195 | 11357 | 162 | -1,172257359 | N12:11275 | 20,614 | | Intergene | | | TEL LEFT |
| chrXII | 11198 | 11354 | 156 | -0,882613757 | N12:11275 | 20,614 | | Intergene | | | TEL LEFT |
| chrXII | 11205 | 11354 | 149 | -1,648300921 | N12:11275 | 20,614 | | Intergene | | | TEL LEFT |
| chrXII | 11205 | 11357 | 152 | -0,788488036 | N12:11275 | 20,614 | | Intergene | | | TEL LEFT |
| chrXII | 11207 | 11357 | 150 | -1,170195429 | N12:11275 | 20,614 | | Intergene | | | TEL LEFT |
| chrXII | 11208 | 11354 | 146 | -1,088604166 | N12:11275 | 20,614 | | Intergene | | | TEL LEFT |
| chrXII | 11211 | 11357 | 146 | -1,137888129 | N12:11275 | 20,614 | | Intergene | | | TEL LEFT |
| chrXII | 11213 | 11310 | 97 | -0,275265869 | N12:11275 | 20,614 | | Intergene | | | TEL LEFT |
| chrXII | 11213 | 11346 | 133 | -0,558080116 | N12:11275 | 20,614 | | Intergene | | | TEL LEFT |
| chrXII | 11213 | 11352 | 139 | -1,815797997 | N12:11275 | 20,614 | | Intergene | | | TEL LEFT |
| chrXII | 11213 | 11355 | 142 | -1,903383612 | N12:11275 | 20,614 | | Intergene | | | TEL LEFT |
| chrXII | 11213 | 11357 | 144 | -1,638455941 | N12:11275 | 20,614 | | Intergene | | | TEL LEFT |
| chrXII | 11213 | 11358 | 145 | -2,074981651 | N12:11275 | 20,614 | | Intergene | | | TEL LEFT |
| chrXII | 11218 | 11354 | 136 | -1,059025851 | N12:11275 | 20,614 | | Intergene | | | TEL LEFT |
| chrXII | 11222 | 11354 | 132 | -1,029743305 | N12:11275 | 20,614 | | Intergene | | | TEL LEFT |
| chrXII | 11222 | 11357 | 135 | -1,331619259 | N12:11275 | 20,614 | | Intergene | | | TEL LEFT |
| chrXII | 11226 | 11354 | 128 | -0,649215873 | N12:11275 | 20,614 | | Intergene | | | TEL LEFT |
| chrXII | 11226 | 11357 | 131 | -0,896213419 | N12:11275 | 20,614 | | Intergene | | | TEL LEFT |
| chrXII | 11226 | 11358 | 132 | -1,02941682 | N12:11275 | 20,614 | | Intergene | | | TEL LEFT |
| chrXII | 11227 | 11357 | 130 | -0,959847532 | N12:11275 | 20,614 | | Intergene | | | TEL LEFT |
| chrXII | 11228 | 11357 | 129 | -1,495122644 | N12:11275 | 20,614 | | Intergene | | | TEL LEFT |
| chrXII | 11229 | 11357 | 128 | -0,88002734 | N12:11275 | 20,614 | | Intergene | | | TEL LEFT |
| chrXII | 11230 | 11357 | 127 | -1,43034951 | N12:11275 | 20,614 | | Intergene | | | TEL LEFT |
| chrXII | 11231 | 11354 | 123 | -0,419679387 | N12:11275 | 20,614 | | Intergene | | | TEL LEFT |
| chrXII | 11231 | 11357 | 126 | -0,411269424 | N12:11275 | 20,614 | | Intergene | | | TEL LEFT |
| chrXII | 11231 | 11358 | 127 | -0,421440385 | N12:11275 | 20,614 | | Intergene | | | TEL LEFT |
| chrXII | 11233 | 11357 | 124 | -0,430225541 | N12:11275 | 20,614 | | Intergene | | | TEL LEFT |
| chrXII | 11235 | 11357 | 122 | -0,812901387 | N12:11275 | 20,614 | | Intergene | | | TEL LEFT |
| chrXII | 11252 | 11354 | 102 | -0,396421488 | N12:11275 | 20,614 | | Intergene | | | TEL LEFT |
| chrXII | 11252 | 11357 | 105 | -0,831026036 | N12:11275 | 20,614 | | Intergene | | | TEL LEFT |
| chrXII | 11259 | 11357 | 98 | -0,620313337 | N12:11275 | 20,614 | | Intergene | | | TEL LEFT |
| chrXII | 11262 | 11357 | 95 | -1,070259517 | N12:11275 | 20,614 | | Intergene | | | TEL LEFT |
| chrXII | 11273 | 11357 | 84 | -1,168520136 | N12:11275 | 20,614 | | Intergene | | | TEL LEFT |
| chrXII | 13256 | 13420 | 164 | -0,944229987 | N12:13360 | 13,115 | +2:YLL064C; | 2 | | | |
| chrXII | 13298 | 13427 | 129 | -1,394486236 | N12:13360 | 13,115 | +2:YLL064C; | 2 | | | |
| chrXII | 13457 | 13612 | 155 | -0,981379292 | N12:13509 | 0,707 | +1:YLL064C; | 1 | | | |
| chrXII | 15085 | 15257 | 172 | -1,376752866 | N12:15154 | 16,622 | +6:YLL063C; | 6 | | | |
| chrXII | 15201 | 15361 | 160 | -1,305305707 | N12:15316 | 26,994 | +5:YLL063C; | 5 | | | |
| chrXII | 16724 | 16865 | 141 | -1,229086565 | N12:16731 | 5,128 | +7*:YLL062C; | 7 | TERM | | |
| chrXII | 18847 | 18996 | 149 | -1,205823412 | N12:18970 | 11,546 | +7:YLL061W; | 7 | | | |
| chrXII | 25117 | 25257 | 140 | -1,021632727 | N12:25169 | 33,344 | +11*:YLL058W; | 11 | TERM | | |
| chrXII | 27714 | 27865 | 151 | -1,098276376 | N12:27788 | 23,692 | +5:YLL056C; | 5 | | | |
| chrXII | 35557 | 35688 | 131 | -0,859584828 | N12:35640 | 5,505 | +3:YLL053C; | 3 | | | |
| chrXII | 39250 | 39399 | 149 | -1,599332552 | N12:39315 | 14,152 | +2:YLL051C; | 2 | | | |
| chrXII | 48577 | 48723 | 146 | -1,426204565 | N12:48603 | 33,129 | +1:YLL045C; | 1 | | | |
| chrXII | 51782 | 51932 | 150 | -1,286591913 | N12:51881 | 32,34 | | Intergene | | | |
| chrXII | 56756 | 56881 | 125 | -1,027874236 | N12:56813 | 23,372 | +43*:YLL040C; | 43 | TERM | | |
| chrXII | 57933 | 58079 | 146 | -1,16846774 | N12:58069 | 15,659 | +35:YLL040C; | 35 | | | |
| chrXII | 60326 | 60478 | 152 | -1,3612987 | N12:60341 | 25,056 | +21:YLL040C; | 21 | | | |
| chrXII | 62223 | 62381 | 158 | -1,404856717 | | | | Overlap <50 bp | | | |
| chrXII | 67309 | 67460 | 151 | -1,775579442 | | | | Overlap <50 bp | | | |
| chrXII | 70032 | 70180 | 148 | -1,262270144 | N12:70102 | 20,315 | +11:YLL035W; | 11 | | | |
| chrXII | 70780 | 70917 | 137 | -1,07679287 | N12:70939 | 17,869 | +15:YLL034C; | 15 | | | |
| chrXII | 70780 | 70917 | 137 | -1,07679287 | N12:70773 | 35,665 | +16:YLL034C; | 16 | | | |
| chrXII | 71343 | 71501 | 158 | -1,591948363 | N12:71413 | 23,137 | +12:YLL034C; | 12 | | | |
| chrXII | 73489 | 73643 | 154 | -1,451911524 | N12:73587 | 11,161 | +3:anti-YLL033W; +2:YLL033W; | 3 | | | |
| chrXII | 79801 | 79962 | 161 | -1,176457144 | | | | Overlap <50 bp | | | |
| chrXII | 99031 | 99175 | 144 | -1,136021263 | N12:99103 | 10,373 | :CUT246; +7*:YLL022C; -1:YLL023C; | 8 | TERM | | |
| chrXII | 104748 | 104900 | 152 | -1,158723201 | N12:104799 | 23,028 | | Intergene | | | |
| chrXII | 104994 | 105164 | 170 | -1,284073315 | N12:104983 | 15,052 | | Intergene | | | |
| chrXII | 106767 | 106921 | 154 | -1,629492884 | N12:106904 | 35,972 | +8:YLL019C; | 8 | | | |
| chrXII | 107154 | 107242 | 88 | -1,177940147 | N12:107195 | 27,508 | +6:YLL019C; | 6 | | | |
| chrXII | 107154 | 107250 | 96 | -1,147943904 | N12:107195 | 27,508 | +6:YLL019C; | 6 | | | |
| chrXII | 107154 | 107252 | 98 | -0,829810076 | N12:107195 | 27,508 | +6:YLL019C; | 6 | | | |
| chrXII | 107154 | 107253 | 99 | -0,896035148 | N12:107195 | 27,508 | +6:YLL019C; | 6 | | | |
| chrXII | 107154 | 107255 | 101 | -1,040529761 | N12:107195 | 27,508 | +6:YLL019C; | 6 | | | |
| chrXII | 107154 | 107257 | 103 | -0,637750857 | N12:107195 | 27,508 | +6:YLL019C; | 6 | | | |
| chrXII | 107154 | 107258 | 104 | -0,79579632 | N12:107195 | 27,508 | +6:YLL019C; | 6 | | | |
| chrXII | 107154 | 107259 | 105 | -0,959044453 | N12:107195 | 27,508 | +6:YLL019C; | 6 | | | |
| chrXII | 107154 | 107260 | 106 | -0,909917987 | N12:107195 | 27,508 | +6:YLL019C; | 6 | | | |
| chrXII | 107154 | 107261 | 107 | -0,890953559 | N12:107195 | 27,508 | +6:YLL019C; | 6 | | | |
| chrXII | 107154 | 107262 | 108 | -1,012194712 | N12:107195 | 27,508 | +6:YLL019C; | 6 | | | |
| chrXII | 107154 | 107265 | 111 | -1,121650654 | N12:107195 | 27,508 | +6:YLL019C; | 6 | | | |
| chrXII | 107154 | 107267 | 113 | -1,262695093 | N12:107195 | 27,508 | +6:YLL019C; | 6 | | | |
| chrXII | 107154 | 107269 | 115 | -1,186868891 | N12:107195 | 27,508 | +6:YLL019C; | 6 | | | |
| chrXII | 107154 | 107270 | 116 | -0,945675724 | N12:107195 | 27,508 | +6:YLL019C; | 6 | | | |
| chrXII | 107154 | 107271 | 117 | -1,585444505 | N12:107195 | 27,508 | +6:YLL019C; | 6 | | | |
| chrXII | 107154 | 107273 | 119 | -1,305611231 | N12:107195 | 27,508 | +6:YLL019C; | 6 | | | |
| chrXII | 107154 | 107274 | 120 | -1,180463625 | N12:107195 | 27,508 | +6:YLL019C; | 6 | | | |
| chrXII | 107154 | 107285 | 131 | -1,112987873 | N12:107195 | 27,508 | +6:YLL019C; | 6 | | | |
| chrXII | 107271 | 107423 | 152 | -1,134848349 | N12:107365 | 32,347 | +5:YLL019C; | 5 | | | |
| chrXII | 110633 | 110799 | 166 | -1,622628817 | N12:110737 | 24,954 | +6:YLL018C; | 6 | | | |
| chrXII | 112506 | 112634 | 128 | -1,589031641 | N12:112551 | 10,998 | +11:anti115; | 11 | | | |
| chrXII | 113051 | 113206 | 155 | -1,440164869 | N12:113139 | 42,716 | +8:anti115; | 8 | | | |
| chrXII | 118248 | 118400 | 152 | -1,399982792 | N12:118377 | 9,504 | +14:YLL015W; | 14 | | | |

TABLE 3

| CHR | START | END | L(bp) | ΔLK | Nuc ID | Fuzziness | Gene ID | Gene body postion or intergene | Terminal | rDNA | Telomeric |
|---|---|---|---|---|---|---|---|---|---|---|---|
| chrXII | 119597 | 119738 | 141 | -1,356877866 | N12:119672 | 27,683 | +21:YLL015W; | 21 | | | |
| chrXII | 121717 | 121889 | 172 | -1,646744401 | N12:121848 | 32,485 | | Intergene | | | |
| chrXII | 126647 | 126799 | 152 | -1,352649175 | N12:126629 | 5,657 | +8:YLL012W; | 8 | | | |
| chrXII | 132281 | 132417 | 136 | -1,162843256 | | | | Overlap <50 bp | | | |
| chrXII | 132310 | 132466 | 156 | -1,370869368 | | | | Overlap <50 bp | | | |
| chrXII | 133301 | 133451 | 150 | -1,322944539 | N12:133308 | 20,688 | +11:YLL008W; | 11 | | | |
| chrXII | 133762 | 133946 | 184 | -1,786922939 | N12:133881 | 17,521 | +15:YLL008W; | 15 | | | |
| chrXII | 136653 | 136779 | 126 | -1,380410878 | N12:136749 | 12,656 | +2:YLL006W; +3*:YLL006W-A; | 2 | | | |
| chrXII | 138426 | 138582 | 156 | -1,445659153 | N12:138554 | 20,664 | +13:YLL005C; +12*:YLL006W; | 13 | | | |
| chrXII | 142345 | 142492 | 147 | -1,455631662 | N12:142429 | 11,673 | +9:YLL004W; | 9 | | | |
| chrXII | 143973 | 144111 | 138 | -1,373961249 | N12:144010 | 17,904 | +6:YLL003W; | 6 | | | |
| chrXII | 144765 | 144914 | 149 | -1,220921696 | N12:144832 | 22,013 | +11:YLL003W; | 11 | | | |
| chrXII | 145430 | 145576 | 146 | -1,327704722 | N12:145533 | 19,524 | +15:YLL003W; | 15 | | | |
| chrXII | 155097 | 155220 | 123 | -0,928358937 | N12:155205 | 20,149 | +8:YLR002C; | 8 | | | |
| chrXII | 160945 | 161106 | 161 | -1,329141498 | N12:161017 | 21,909 | +6:YLR005W; | 6 | | | |
| chrXII | 163470 | 163588 | 118 | -1,509741427 | N12:163522 | 13,565 | +4:YLR006C; | 4 | | | |
| chrXII | 171732 | 171883 | 151 | -1,548097036 | N12:171897 | 23,169 | +4*:YLR013W; | 4 | TERM | | |
| chrXII | 177830 | 177964 | 134 | -1,08819054 | N12:177819 | 6,542 | +2:YLR017W; | 2 | | | |
| chrXII | 177830 | 177964 | 134 | -1,08819054 | N12:177986 | 6,686 | +3:YLR017W; | 3 | | | |
| chrXII | 178588 | 178741 | 153 | -1,170853701 | | | | Overlap <50 bp | | | |
| chrXII | 179076 | 179226 | 150 | -1,817080258 | N12:179114 | 7,808 | +4:YLR018C; | 4 | | | |
| chrXII | 180676 | 180836 | 160 | -1,272923005 | N12:180728 | 11,726 | +5:YLR019W; | 5 | | | |
| chrXII | 183315 | 183461 | 146 | -1,089593376 | N12:183389 | 4,037 | +1:YLR020C; -1:YLR021W; | 1 | | | |
| chrXII | 192705 | 192836 | 131 | -1,196488888 | N12:192762 | 22,418 | +5:YLR024C; | 5 | | | |
| chrXII | 193019 | 193125 | 106 | -0,868440921 | N12:193086 | 6,646 | +3:YLR024C; | 3 | | | |
| chrXII | 193019 | 193144 | 125 | -1,138759005 | N12:193086 | 6,646 | +3:YLR024C; | 3 | | | |
| chrXII | 194389 | 194550 | 161 | -1,550441442 | N12:194429 | 16,669 | +1:YLR025W; | 1 | | | |
| chrXII | 197350 | 197482 | 132 | -2,001958585 | N12:197391 | 23,618 | +4*:CUT252; +5:YLR027C; | 4 | TERM | | |
| chrXII | 197802 | 197965 | 163 | -1,362741621 | | | | Overlap <50 bp | | | |
| chrXII | 199062 | 199223 | 161 | -1,429848524 | N12:199130 | 20,257 | | Intergene | | | |
| chrXII | 199271 | 199406 | 135 | -1,082122182 | N12:199299 | 18,673 | | Intergene | | | |
| chrXII | 199721 | 199871 | 150 | -1,383895841 | N12:199806 | 27,862 | +10:YLR028C; | 10 | | | |
| chrXII | 207917 | 208073 | 156 | -1,708440981 | N12:207961 | 16,362 | +19:YLR032W; | 19 | | | |
| chrXII | 208414 | 208561 | 147 | -1,372723134 | N12:208481 | 8,746 | +22*:YLR032W; -1:YLR033W; | 22 | TERM | | |
| chrXII | 209818 | 209960 | 142 | -0,934799937 | N12:209848 | 25,883 | +8:YLR033W; | 8 | | | |
| chrXII | 211059 | 211185 | 126 | -0,731668939 | N12:211093 | 24,468 | +7:YLR034C; | 7 | | | |
| chrXII | 213217 | 213369 | 152 | -1,163751071 | N12:213269 | 5,269 | +8:YLR035C; | 8 | | | |
| chrXII | 214356 | 214519 | 163 | -1,550549705 | N12:214440 | 10,328 | +1:YLR035C; | 1 | | | |
| chrXII | 216121 | 216270 | 149 | -1,158542274 | N12:216153 | 39,703 | | Intergene | | | |
| chrXII | 217884 | 218033 | 149 | -0,865537787 | N12:218006 | 19,732 | | Intergene | | | |
| chrXII | 218108 | 218265 | 157 | -1,545823073 | N12:218186 | 18,877 | | Intergene | | | |
| chrXII | 218294 | 218444 | 150 | -0,982792643 | N12:218356 | 27,099 | | Intergene | | | |
| chrXII | 218298 | 218444 | 146 | -1,080093025 | N12:218356 | 27,099 | | Intergene | | | |
| chrXII | 218756 | 218891 | 135 | -1,18255401 | N12:218862 | 33,645 | | Intergene | | | |
| chrXII | 219794 | 219934 | 140 | -1,098200789 | N12:219876 | 37,634 | | Intergene | | | |
| chrXII | 219800 | 219934 | 134 | -1,111348084 | N12:219876 | 37,634 | | Intergene | | | |
| chrXII | 220212 | 220367 | 155 | -0,8618629 | N12:220242 | 19,925 | | Intergene | | | |
| chrXII | 220343 | 220451 | 108 | -1,124492107 | N12:220455 | 49,003 | | Intergene | | | |
| chrXII | 223444 | 223638 | 194 | -2,232023883 | N12:223443 | 6,524 | -1:YLR037C; +7:SUT688; | -1 | | | |
| chrXII | 223444 | 223638 | 194 | -2,232023883 | N12:223614 | 15,123 | +6:SUT688; | 6 | | | |
| chrXII | 230481 | 230653 | 172 | -1,761350347 | N12:230555 | 17,166 | -1:SUT251; 0:YLR042C; | -1 | | | |
| chrXII | 230739 | 230894 | 155 | -1,291902736 | N12:230860 | 10,967 | +1:SUT251; | 1 | | | |
| chrXII | 232813 | 232957 | 144 | -1,376459762 | N12:232966 | 14,154 | +7:YLR044C; | 7 | | | |
| chrXII | 232813 | 232957 | 144 | -1,376459762 | N12:232814 | 22,311 | +8:YLR044C; | 8 | | | |
| chrXII | 235469 | 235619 | 150 | -0,881454953 | N12:235506 | 16,538 | +14:YLR045C; +6:SUT252; | 14 | | | |
| chrXII | 235483 | 235631 | 148 | -1,571333857 | N12:235506 | 16,538 | +14:YLR045C; +6:SUT252; | 14 | | | |
| chrXII | 237484 | 237634 | 150 | -1,414712146 | N12:237551 | 10,87 | +2:YLR045C; | 2 | | | |
| chrXII | 238295 | 238446 | 151 | -1,419905244 | N12:238404 | 24,977 | +21*:YLR047C; +4:YLR046C; | 21 | TERM | | |
| chrXII | 239615 | 239753 | 138 | -1,219336787 | N12:239689 | 6,921 | +13:YLR047C; | 13 | | | |
| chrXII | 255888 | 256021 | 133 | -1,055837785 | | | | Overlap <50 bp | | | |
| chrXII | 256324 | 256484 | 160 | -1,11953004 | N12:256416 | 10,284 | +8:YLR057W; | 8 | | | |
| chrXII | 259071 | 259235 | 164 | -1,445486947 | N12:259139 | 18,69 | +3:YLR058C; | 3 | | | |
| chrXII | 260642 | 260797 | 155 | -1,061392298 | N12:260719 | 3,312 | 0:YLR059C; -1:YLR060W; | 0 | | | |
| chrXII | 262571 | 262718 | 147 | -1,15601585 | N12:262667 | 32,722 | | Intergene | | | |
| chrXII | 270658 | 270826 | 168 | -1,526070813 | N12:270763 | 5,465 | +1:YLR067C; -1:YLR068W; | 1 | | | |
| chrXII | 270673 | 270826 | 153 | -0,772546666 | N12:270763 | 5,465 | +1:YLR067C; -1:YLR068W; | 1 | | | |
| chrXII | 271752 | 271896 | 144 | -1,236893198 | N12:271899 | 25,981 | +13:YLR069C; | 13 | | | |
| chrXII | 273309 | 273417 | 108 | -1,281740562 | N12:273383 | 18,217 | +4:YLR069C; | 4 | | | |
| chrXII | 273309 | 273418 | 109 | -1,124731612 | N12:273383 | 18,217 | +4:YLR069C; | 4 | | | |
| chrXII | 273309 | 273419 | 110 | -1,420347604 | N12:273383 | 18,217 | +4:YLR069C; | 4 | | | |
| chrXII | 273309 | 273420 | 111 | -1,42419264 | N12:273383 | 18,217 | +4:YLR069C; | 4 | | | |
| chrXII | 274415 | 274568 | 153 | -1,311784924 | N12:274467 | 16,86 | +3:anti119; +5:YLR070C; | 3 | | | |
| chrXII | 275945 | 276112 | 167 | -1,475374505 | N12:276130 | 34,15 | +16:YLR071C; | 16 | | | |
| chrXII | 275945 | 276112 | 167 | -1,475374505 | N12:275964 | 18,771 | +17:YLR071C; | 17 | | | |
| chrXII | 280170 | 280332 | 162 | -1,090622611 | N12:280290 | 22,143 | +9:YLR072W; | 9 | | | |
| chrXII | 280378 | 280532 | 154 | -1,23215883 | N12:280450 | 34,574 | +10:YLR072W; | 10 | | | |
| chrXII | 281608 | 281761 | 153 | -1,746857777 | N12:281710 | 25,034 | +1:YLR073C; +4:SUT257; | 1 | | | |
| chrXII | 285125 | 285277 | 152 | -1,149068845 | N12:285193 | 27,083 | +9:YLR077W; | 9 | | | |
| chrXII | 285128 | 285277 | 149 | -1,418503979 | N12:285193 | 27,083 | +9:YLR077W; | 9 | | | |
| chrXII | 286283 | 286443 | 160 | -1,331590165 | N12:286398 | 3,724 | +1:YLR078C; | 1 | | | |
| chrXII | 292173 | 292332 | 159 | -1,504775814 | N12:292269 | 6,245 | +9*:YLR082C; | 9 | TERM | | |
| chrXII | 293330 | 293472 | 142 | -1,364315349 | N12:293400 | 19,892 | +2:YLR082C; | 2 | | | |
| chrXII | 295117 | 295270 | 153 | -1,404531963 | N12:295096 | 27,545 | +8:YLR083C; | 8 | | | |
| chrXII | 295472 | 295620 | 148 | -0,965011755 | N12:295545 | 10,863 | +5:YLR083C; | 5 | | | |
| chrXII | 296601 | 296740 | 139 | -1,022373239 | N12:296667 | 9,45 | +24*:YLR084C; | 24 | TERM | | |
| chrXII | 300804 | 300959 | 155 | -1,054261819 | N12:300827 | 33,018 | +8:YLR085C; | 8 | | | |
| chrXII | 301895 | 302037 | 142 | -1,469906682 | N12:301967 | 8,773 | -1:YLR086W; +1:YLR085C; | -1 | | | |
| chrXII | 302498 | 302646 | 148 | -1,183312288 | N12:302567 | 3,71 | +3:YLR086W; | 3 | | | |
| chrXII | 304308 | 304455 | 147 | -1,210972085 | N12:304426 | 22,884 | +14:YLR086W; | 14 | | | |
| chrXII | 304635 | 304774 | 139 | -1,018120678 | N12:304748 | 16,754 | +16:YLR086W; | 16 | | | |
| chrXII | 310453 | 310583 | 130 | -1,253342475 | N12:310571 | 35,549 | +32:YLR087C; | 32 | | | |
| chrXII | 311744 | 311903 | 159 | -1,706893012 | | | | Overlap <50 bp | | | |
| chrXII | 312700 | 312835 | 135 | -1,418469889 | N12:312689 | 30,859 | +19:YLR087C; | 19 | | | |
| chrXII | 314498 | 314655 | 157 | -1,439540057 | N12:314563 | 26,84 | +8:YLR087C; | 8 | | | |
| chrXII | 319179 | 319325 | 146 | -1,49771803 | N12:319307 | 38,105 | +6:YLR089C; | 6 | | | |
| chrXII | 320329 | 320487 | 158 | -1,591880671 | N12:320461 | 19,14 | LR089C; -1:YLR090W; +2*:Unit433; | -1 | | | |
| chrXII | 320335 | 320487 | 152 | -1,35133557 | N12:320461 | 19,14 | LR089C; -1:YLR090W; +2*:Unit433; | -1 | | | |
| chrXII | 320336 | 320487 | 151 | -1,615866232 | N12:320461 | 19,14 | LR089C; -1:YLR090W; +2*:Unit433; | -1 | | | |
| chrXII | 321520 | 321655 | 135 | -1,260570288 | N12:321554 | 31,642 | +6:YLR090W; | 6 | | | |

TABLE 3

| CHR | START | END | L(bp) | ΔLK | Nuc ID | Fuzziness | Gene ID | Gene body postion or intergene | Terminal | rDNA | Telomeric |
|---|---|---|---|---|---|---|---|---|---|---|---|
| chrXII | 322116 | 322262 | 146 | -1,243086272 | | | | Overlap <50 bp | | | |
| chrXII | 327876 | 328044 | 168 | -1,360443226 | N12:327934 | 21,847 | +9:YLR094C; | 9 | | | |
| chrXII | 338236 | 338387 | 151 | -1,298905447 | N12:338317 | 25,519 | +8:YLR098C; | 8 | | | |
| chrXII | 340076 | 340209 | 133 | -1,104252333 | N12:340159 | 23,969 | +5:SUT261; +5:YLR099C; | 5 | | | |
| chrXII | 340597 | 340761 | 164 | -1,579950386 | N12:340639 | 18,276 | +8:SUT261; +2:YLR099C; | 8 | | | |
| chrXII | 340875 | 341034 | 159 | -1,188794746 | N12:340984 | 11,389 | SUT261; -1:YLR099W-A; 0:YLR099C; | 10 | | | |
| chrXII | 347768 | 347916 | 148 | -1,231399673 | N12:347855 | 31,324 | +3:YLR105C; | 3 | | | |
| chrXII | 347831 | 348011 | 180 | -1,779904971 | N12:348007 | 34,01 | +2:YLR105C; | 2 | | | |
| chrXII | 347831 | 348011 | 180 | -1,779904971 | N12:347855 | 31,324 | +3:YLR105C; | 3 | | | |
| chrXII | 348735 | 348938 | 203 | -1,825869978 | N12:348883 | 16,971 | +1*:Unit434; | 1 | TERM | | |
| chrXII | 348736 | 348938 | 202 | -1,743917651 | N12:348883 | 16,971 | +1*:Unit434; | 1 | TERM | | |
| chrXII | 350679 | 350830 | 151 | -1,173055678 | | | | Overlap <50 bp | | | |
| chrXII | 355881 | 356007 | 126 | -1,525486704 | N12:355867 | 7,211 | | Intergene | | | |
| chrXII | 355881 | 356007 | 126 | -1,525486704 | N12:356021 | 30,128 | | Intergene | | | |
| chrXII | 358183 | 358323 | 140 | -1,185859295 | N12:358217 | 18,166 | | Intergene | | | |
| chrXII | 359562 | 359714 | 152 | -1,337600487 | N12:359697 | 24,334 | | Intergene | | | |
| chrXII | 360814 | 360966 | 152 | -0,997567375 | | | | Overlap <50 bp | | | |
| chrXII | 367954 | 368092 | 138 | -1,486814766 | N12:368026 | 16,087 | +2:YLR108C; | 2 | | | |
| chrXII | 372826 | 372968 | 142 | -1,254508295 | N12:372913 | 22,665 | +10*:YLR113W; | 10 | TERM | | |
| chrXII | 375291 | 375434 | 143 | -1,0007626 | N12:375295 | 9,539 | +13:YLR114C; | 13 | | | |
| chrXII | 375584 | 375750 | 166 | -1,78042701 | N12:375684 | 28,702 | +11:YLR114C; | 11 | | | |
| chrXII | 388796 | 388957 | 161 | -1,35102331 | N12:388850 | 23,676 | -1:Unit436; +8:YLR121C; 0:anti122; | -1 | | | |
| chrXII | 388880 | 389024 | 144 | -1,111894857 | N12:389034 | 42,293 | +7:YLR121C; +2:anti122; | 7 | | | |
| chrXII | 391687 | 391826 | 139 | -0,996018838 | | | | Overlap <50 bp | | | |
| chrXII | 395857 | 396006 | 149 | -0,959966178 | N12:395841 | 5,225 | YLR126C; +1:SUT263; +17:YLR127C; | -1 | | | |
| chrXII | 395857 | 396006 | 149 | -0,959966178 | N12:396018 | 7,685 | +2:SUT263; +16:YLR127C; | 2 | | | |
| chrXII | 397064 | 397220 | 156 | -1,208131262 | N12:397207 | 7,396 | +8:YLR127C; | 8 | | | |
| chrXII | 398081 | 398231 | 150 | -1,087937152 | N12:398173 | 2,757 | +2:YLR127C; | 2 | | | |
| chrXII | 400245 | 400397 | 152 | -1,327418445 | | | | Overlap <50 bp | | | |
| chrXII | 401669 | 401819 | 150 | -1,261426889 | N12:401676 | 20,371 | +13:YLR129W; | 13 | | | |
| chrXII | 401669 | 401819 | 150 | -1,261426889 | N12:401833 | 28,733 | +14:YLR129W; | 14 | | | |
| chrXII | 405414 | 405520 | 106 | -1,20271227 | N12:405442 | 26,857 | +18:YLR132C; +10:YLR131C; | 18 | | | |
| chrXII | 405414 | 405538 | 124 | -1,530378564 | N12:405442 | 26,857 | +18:YLR132C; +10:YLR131C; | 18 | | | |
| chrXII | 406427 | 406580 | 153 | -1,4667777 | N12:406529 | 22,876 | +11:YLR132C; +3:YLR131C; | 11 | | | |
| chrXII | 406621 | 406768 | 147 | -1,295054924 | N12:406710 | 17,175 | +10:YLR132C; +2:YLR131C; | 10 | | | |
| chrXII | 408412 | 408522 | 110 | -0,856810074 | N12:408439 | 9,564 | +1:YLR133W; -1:YLR132C; | 1 | | | |
| chrXII | 408412 | 408523 | 111 | -0,773911104 | N12:408439 | 9,564 | +1:YLR133W; -1:YLR132C; | 1 | | | |
| chrXII | 408415 | 408537 | 122 | -1,871925096 | N12:408439 | 9,564 | +1:YLR133W; -1:YLR132C; | 1 | | | |
| chrXII | 411908 | 412040 | 132 | -1,246599746 | N12:411988 | 27,15 | | Intergene | | | |
| chrXII | 412316 | 412427 | 111 | -1,377712832 | N12:412343 | 15,55 | | Intergene | | | |
| chrXII | 412319 | 412427 | 108 | -1,363948696 | N12:412343 | 15,55 | | Intergene | | | |
| chrXII | 412320 | 412427 | 107 | -1,410672298 | N12:412343 | 15,55 | | Intergene | | | |
| chrXII | 419068 | 419220 | 152 | -1,831397902 | N12:419117 | 3,564 | +6:YLR138W; | 6 | | | |
| chrXII | 419734 | 419893 | 159 | -1,197104405 | N12:419779 | 16,709 | | Intergene | | | |
| chrXII | 421131 | 421283 | 152 | -1,397536735 | N12:421250 | 7,603 | | Intergene | | | |
| chrXII | 423648 | 423793 | 145 | -0,679647373 | N12:423723 | 9,888 | -1:YLR139C; +1:YLR141W; | -1 | | | |
| chrXII | 424263 | 424441 | 178 | -1,27029736 | N12:424247 | 2,828 | +4:YLR141W; | 4 | | | |
| chrXII | 425377 | 425494 | 117 | -1,564343258 | N12:425416 | 10,685 | +2:YLR142W; | 2 | | | |
| chrXII | 428829 | 428982 | 153 | -1,127774284 | N12:428985 | 31,284 | +10:YLR143W; | 10 | | | |
| chrXII | 433448 | 433586 | 138 | -0,949881576 | N12:433530 | 10,431 | +2:YLR146C; -1:YLR146W-A; | 2 | | | |
| chrXII | 433602 | 433749 | 147 | -1,728650857 | N12:433706 | 8,802 | +1:YLR146C; 0:YLR146W-A; | 1 | | | |
| chrXII | 434337 | 434490 | 153 | -1,222846256 | N12:434430 | 3,559 | -1:YLR148W; +1:YLR147C; | -1 | | | |
| chrXII | 443464 | 443629 | 165 | -1,440060345 | N12:443445 | 27,112 | +8*:YLR152C; | 8 | TERM | | |
| chrXII | 446788 | 446962 | 174 | -1,512566972 | N12:446975 | 12,675 | +5:YLR153C; | 5 | | | |
| chrXII | 446788 | 446962 | 174 | -1,512566972 | N12:446824 | 24,183 | +6:YLR153C; | 6 | | | |
| chrXII | 447291 | 447454 | 163 | -1,544928781 | N12:447462 | 18,694 | +2:YLR153C; | 2 | | | |
| chrXII | 447291 | 447454 | 163 | -1,544928781 | N12:447286 | 3,271 | +3:YLR153C; | 3 | | | |
| chrXII | 450323 | 450477 | 154 | -1,163487034 | N12:450413 | 20,919 | +5:Unit457; | 5 | | | |
| chrXII | 451465 | 451626 | 161 | -1,639740675 | N12:451492 | 34,463 | | | | rDNA | |
| chrXII | 451590 | 451738 | 148 | -1,494170043 | | | | Overlap <50 bp | | rDNA | |
| chrXII | 451599 | 451733 | 134 | -1,164068075 | | | | Overlap <50 bp | | rDNA | |
| chrXII | 451709 | 451842 | 133 | -1,433511471 | N12:451807 | 35,838 | | | | rDNA | |
| chrXII | 451726 | 451885 | 159 | -1,722956651 | N12:451807 | 35,838 | | | | rDNA | |
| chrXII | 451734 | 451869 | 135 | -1,569225321 | N12:451807 | 35,838 | | | | rDNA | |
| chrXII | 451757 | 451842 | 85 | -0,457327461 | N12:451807 | 35,838 | | | | rDNA | |
| chrXII | 451757 | 451908 | 151 | -1,462446946 | N12:451807 | 35,838 | | | | rDNA | |
| chrXII | 451769 | 451932 | 163 | -1,565028581 | N12:451807 | 35,838 | | | | rDNA | |
| chrXII | 451790 | 451908 | 118 | -1,165323576 | N12:451807 | 35,838 | | | | rDNA | |
| chrXII | 451790 | 451932 | 142 | -1,291093773 | N12:451807 | 35,838 | | | | rDNA | |
| chrXII | 451790 | 451960 | 170 | -1,302580348 | N12:451807 | 35,838 | | | | rDNA | |
| chrXII | 451790 | 451960 | 170 | -1,302580348 | N12:451976 | 40,079 | | | | rDNA | |
| chrXII | 451800 | 451934 | 134 | -1,064520309 | N12:451807 | 35,838 | | | | rDNA | |
| chrXII | 451800 | 451961 | 161 | -1,548953909 | N12:451807 | 35,838 | | | | rDNA | |
| chrXII | 451800 | 451961 | 161 | -1,548953909 | N12:451976 | 40,079 | | | | rDNA | |
| chrXII | 451851 | 451934 | 83 | -0,778815603 | | | | Overlap <50 bp | | rDNA | |
| chrXII | 451851 | 451958 | 107 | -1,329802255 | N12:451976 | 40,079 | | | | rDNA | |
| chrXII | 451851 | 451959 | 108 | -1,256939933 | N12:451976 | 40,079 | | | | rDNA | |
| chrXII | 451851 | 451960 | 109 | -1,510613919 | N12:451976 | 40,079 | | | | rDNA | |
| chrXII | 451851 | 451961 | 110 | -1,285875966 | N12:451976 | 40,079 | | | | rDNA | |
| chrXII | 451876 | 451960 | 84 | -0,791582711 | N12:451976 | 40,079 | | | | rDNA | |
| chrXII | 451876 | 452009 | 133 | -1,193600939 | N12:451976 | 40,079 | | | | rDNA | |
| chrXII | 451919 | 452075 | 156 | -1,420369664 | N12:451976 | 40,079 | | | | rDNA | |
| chrXII | 451919 | 452077 | 158 | -1,221589763 | N12:451976 | 40,079 | | | | rDNA | |
| chrXII | 451919 | 452078 | 159 | -1,243546966 | N12:451976 | 40,079 | | | | rDNA | |
| chrXII | 451919 | 452092 | 173 | -1,30589316 | N12:451976 | 40,079 | | | | rDNA | |
| chrXII | 451923 | 452075 | 152 | -1,157762497 | N12:451976 | 40,079 | | | | rDNA | |
| chrXII | 451923 | 452077 | 154 | -1,639654166 | N12:451976 | 40,079 | | | | rDNA | |
| chrXII | 451923 | 452092 | 169 | -1,235507847 | N12:451976 | 40,079 | | | | rDNA | |
| chrXII | 451931 | 452062 | 131 | -1,254948784 | N12:451976 | 40,079 | | | | rDNA | |
| chrXII | 451931 | 452077 | 146 | -1,193787476 | N12:451976 | 40,079 | | | | rDNA | |
| chrXII | 451931 | 452109 | 178 | -1,437264692 | N12:451976 | 40,079 | | | | rDNA | |
| chrXII | 451932 | 452062 | 130 | -1,124495991 | N12:451976 | 40,079 | | | | rDNA | |
| chrXII | 451932 | 452075 | 143 | -1,282064137 | N12:451976 | 40,079 | | | | rDNA | |
| chrXII | 451932 | 452077 | 145 | -1,605633967 | N12:451976 | 40,079 | | | | rDNA | |
| chrXII | 451932 | 452109 | 177 | -1,331310592 | N12:451976 | 40,079 | | | | rDNA | |
| chrXII | 451945 | 452051 | 106 | -0,736154396 | N12:451976 | 40,079 | | | | rDNA | |
| chrXII | 451945 | 452062 | 117 | -0,868281387 | N12:451976 | 40,079 | | | | rDNA | |
| chrXII | 451945 | 452075 | 130 | -0,753814651 | N12:451976 | 40,079 | | | | rDNA | |
| chrXII | 451945 | 452077 | 132 | -0,982406328 | N12:451976 | 40,079 | | | | rDNA | |

| CHR | START | END | L(bp) | ΔLK | Nuc ID | Fuzziness | Gene ID | Gene body postion or intergene | Terminal | rDNA | Telomeric |
|---|---|---|---|---|---|---|---|---|---|---|---|
| chrXII | 451945 | 452078 | 133 | -1,027384552 | N12:451976 | 40,079 | | | | rDNA | |
| chrXII | 451945 | 452092 | 147 | -1,039323428 | N12:451976 | 40,079 | | | | rDNA | |
| chrXII | 451945 | 452109 | 164 | -1,306733111 | N12:451976 | 40,079 | | | | rDNA | |
| chrXII | 451958 | 452109 | 151 | -1,088061784 | N12:451976 | 40,079 | | | | rDNA | |
| chrXII | 452041 | 452196 | 155 | -1,080362528 | N12:452173 | 28,583 | | | | rDNA | |
| chrXII | 452059 | 452215 | 156 | -1,294197441 | N12:452173 | 28,583 | | | | rDNA | |
| chrXII | 452081 | 452227 | 146 | -1,105546451 | N12:452173 | 28,583 | | | | rDNA | |
| chrXII | 452081 | 452236 | 155 | -1,439692328 | N12:452173 | 28,583 | | | | rDNA | |
| chrXII | 452081 | 452247 | 166 | -1,195652769 | N12:452173 | 28,583 | | | | rDNA | |
| chrXII | 452081 | 452251 | 170 | -1,363803369 | N12:452173 | 28,583 | | | | rDNA | |
| chrXII | 452104 | 452247 | 143 | -0,977867166 | N12:452173 | 28,583 | | | | rDNA | |
| chrXII | 452104 | 452251 | 147 | -1,07513005 | N12:452173 | 28,583 | | | | rDNA | |
| chrXII | 452115 | 452271 | 156 | -1,218995954 | N12:452173 | 28,583 | | | | rDNA | |
| chrXII | 452190 | 452344 | 154 | -1,31290605 | N12:452173 | 28,583 | | | | rDNA | |
| chrXII | 452190 | 452344 | 154 | -1,31290605 | N12:452361 | 15,716 | | | | rDNA | |
| chrXII | 452260 | 452414 | 154 | -1,354485193 | N12:452361 | 15,716 | | | | rDNA | |
| chrXII | 452293 | 452400 | 107 | -1,479635313 | N12:452361 | 15,716 | | | | rDNA | |
| chrXII | 452294 | 452455 | 161 | -1,343251014 | N12:452361 | 15,716 | | | | rDNA | |
| chrXII | 452295 | 452455 | 160 | -1,395266066 | N12:452361 | 15,716 | | | | rDNA | |
| chrXII | 452299 | 452455 | 156 | -1,471744026 | N12:452361 | 15,716 | | | | rDNA | |
| chrXII | 452323 | 452455 | 132 | -1,197849248 | N12:452361 | 15,716 | | | | rDNA | |
| chrXII | 452323 | 452485 | 162 | -1,287017701 | N12:452361 | 15,716 | | | | rDNA | |
| chrXII | 452323 | 452486 | 163 | -1,152817435 | N12:452361 | 15,716 | | | | rDNA | |
| chrXII | 452334 | 452441 | 107 | -0,896778421 | N12:452361 | 15,716 | | | | rDNA | |
| chrXII | 452334 | 452455 | 121 | -1,044382945 | N12:452361 | 15,716 | | | | rDNA | |
| chrXII | 452334 | 452485 | 151 | -1,199898478 | N12:452361 | 15,716 | | | | rDNA | |
| chrXII | 452334 | 452486 | 152 | -1,596333429 | N12:452361 | 15,716 | | | | rDNA | |
| chrXII | 452346 | 452496 | 150 | -1,169144695 | N12:452361 | 15,716 | | | | rDNA | |
| chrXII | 452423 | 452584 | 161 | -1,534443647 | N12:452551 | 34,395 | | | | rDNA | |
| chrXII | 452469 | 452584 | 115 | -1,234893602 | N12:452551 | 34,395 | | | | rDNA | |
| chrXII | 452469 | 452620 | 151 | -1,51561518 | N12:452551 | 34,395 | | | | rDNA | |
| chrXII | 452469 | 452621 | 152 | -1,552052632 | N12:452551 | 34,395 | | | | rDNA | |
| chrXII | 452469 | 452629 | 160 | -1,25107865 | N12:452551 | 34,395 | | | | rDNA | |
| chrXII | 452469 | 452630 | 161 | -1,470757401 | N12:452551 | 34,395 | | | | rDNA | |
| chrXII | 452491 | 452621 | 130 | -1,237878987 | N12:452551 | 34,395 | | | | rDNA | |
| chrXII | 452491 | 452629 | 138 | -1,313026015 | N12:452551 | 34,395 | | | | rDNA | |
| chrXII | 452491 | 452663 | 172 | -1,450320305 | N12:452551 | 34,395 | | | | rDNA | |
| chrXII | 452492 | 452584 | 92 | -1,138468787 | N12:452551 | 34,395 | | | | rDNA | |
| chrXII | 452492 | 452620 | 128 | -1,410493358 | N12:452551 | 34,395 | | | | rDNA | |
| chrXII | 452492 | 452621 | 129 | -1,428985011 | N12:452551 | 34,395 | | | | rDNA | |
| chrXII | 452492 | 452629 | 137 | -1,365106135 | N12:452551 | 34,395 | | | | rDNA | |
| chrXII | 452492 | 452663 | 171 | -1,916598169 | N12:452551 | 34,395 | | | | rDNA | |
| chrXII | 452494 | 452621 | 127 | -1,234550926 | N12:452551 | 34,395 | | | | rDNA | |
| chrXII | 452525 | 452695 | 170 | -1,574892129 | N12:452551 | 34,395 | | | | rDNA | |
| chrXII | 452539 | 452673 | 134 | -1,34971257 | N12:452551 | 34,395 | | | | rDNA | |
| chrXII | 452550 | 452672 | 122 | -1,405173722 | N12:452551 | 34,395 | | | | rDNA | |
| chrXII | 452550 | 452702 | 152 | -1,453196622 | N12:452551 | 34,395 | | | | rDNA | |
| chrXII | 452668 | 452796 | 128 | -1,195558592 | N12:452761 | 20,502 | | | | rDNA | |
| chrXII | 452684 | 452796 | 112 | -0,684394069 | N12:452761 | 20,502 | | | | rDNA | |
| chrXII | 452684 | 452835 | 151 | -1,133040361 | N12:452761 | 20,502 | | | | rDNA | |
| chrXII | 452686 | 452814 | 128 | -1,188401963 | N12:452761 | 20,502 | | | | rDNA | |
| chrXII | 452686 | 452884 | 198 | -1,778198455 | N12:452761 | 20,502 | | | | rDNA | |
| chrXII | 452709 | 452864 | 155 | -1,340921092 | N12:452761 | 20,502 | | | | rDNA | |
| chrXII | 452709 | 452884 | 175 | -1,234116436 | N12:452761 | 20,502 | | | | rDNA | |
| chrXII | 452709 | 452900 | 191 | -1,333922725 | N12:452761 | 20,502 | | | | rDNA | |
| chrXII | 452723 | 452814 | 91 | -0,78772414 | N12:452761 | 20,502 | | | | rDNA | |
| chrXII | 452723 | 452864 | 141 | -1,637532571 | N12:452761 | 20,502 | | | | rDNA | |
| chrXII | 452723 | 452884 | 161 | -1,267867521 | N12:452761 | 20,502 | | | | rDNA | |
| chrXII | 452723 | 452900 | 177 | -1,327310445 | N12:452761 | 20,502 | | | | rDNA | |
| chrXII | 452765 | 452864 | 99 | -0,504356379 | N12:452761 | 20,502 | | | | rDNA | |
| chrXII | 452765 | 452871 | 106 | -0,900966829 | N12:452761 | 20,502 | | | | rDNA | |
| chrXII | 452765 | 452884 | 119 | -0,779531249 | N12:452761 | 20,502 | | | | rDNA | |
| chrXII | 452765 | 452900 | 135 | -1,094331164 | N12:452761 | 20,502 | | | | rDNA | |
| chrXII | 452765 | 452925 | 160 | -1,509037703 | N12:452761 | 20,502 | | | | rDNA | |
| chrXII | 452765 | 452925 | 160 | -1,509037703 | N12:452927 | 2,887 | | | | rDNA | |
| chrXII | 452773 | 452864 | 91 | -0,708056173 | N12:452761 | 20,502 | | | | rDNA | |
| chrXII | 452773 | 452879 | 106 | -1,09608273 | N12:452761 | 20,502 | | | | rDNA | |
| chrXII | 452773 | 452884 | 111 | -0,961006774 | N12:452761 | 20,502 | | | | rDNA | |
| chrXII | 452773 | 452900 | 127 | -1,132682414 | N12:452761 | 20,502 | | | | rDNA | |
| chrXII | 452773 | 452925 | 152 | -1,370232757 | N12:452761 | 20,502 | | | | rDNA | |
| chrXII | 452773 | 452925 | 152 | -1,370232757 | N12:452927 | 2,887 | | | | rDNA | |
| chrXII | 452783 | 452962 | 179 | -1,674350294 | N12:452761 | 20,502 | | | | rDNA | |
| chrXII | 452783 | 452962 | 179 | -1,674350294 | N12:452927 | 2,887 | | | | rDNA | |
| chrXII | 452788 | 452923 | 135 | -1,052630765 | N12:452927 | 2,887 | | | | rDNA | |
| chrXII | 452788 | 452953 | 165 | -1,104200658 | N12:452927 | 2,887 | | | | rDNA | |
| chrXII | 452801 | 452923 | 122 | -0,664961388 | N12:452927 | 2,887 | | | | rDNA | |
| chrXII | 452801 | 452953 | 152 | -1,268040098 | N12:452927 | 2,887 | | | | rDNA | |
| chrXII | 452801 | 452999 | 198 | -1,624506722 | N12:452927 | 2,887 | | | | rDNA | |
| chrXII | 452840 | 452953 | 113 | -0,691840125 | N12:452927 | 2,887 | | | | rDNA | |
| chrXII | 452840 | 452999 | 159 | -1,254730292 | N12:452927 | 2,887 | | | | rDNA | |
| chrXII | 452840 | 453023 | 183 | -1,558460647 | N12:452927 | 2,887 | | | | rDNA | |
| chrXII | 452847 | 452953 | 106 | -1,031668425 | N12:452927 | 2,887 | | | | rDNA | |
| chrXII | 452847 | 452996 | 149 | -1,527444518 | N12:452927 | 2,887 | | | | rDNA | |
| chrXII | 452876 | 453034 | 158 | -1,351089933 | N12:452927 | 2,887 | | | | rDNA | |
| chrXII | 452885 | 453034 | 149 | -1,017007914 | N12:452927 | 2,887 | | | | rDNA | |
| chrXII | 452893 | 452999 | 106 | -0,809381927 | N12:452927 | 2,887 | | | | rDNA | |
| chrXII | 452895 | 453023 | 128 | -1,006362814 | N12:452927 | 2,887 | | | | rDNA | |
| chrXII | 452922 | 453029 | 107 | -1,421338671 | N12:452927 | 2,887 | | | | rDNA | |
| chrXII | 452922 | 453031 | 109 | -1,225115667 | N12:452927 | 2,887 | | | | rDNA | |
| chrXII | 452922 | 453034 | 112 | -1,472328186 | N12:452927 | 2,887 | | | | rDNA | |
| chrXII | 452922 | 453056 | 134 | -1,441546027 | N12:452927 | 2,887 | | | | rDNA | |
| chrXII | 452922 | 453056 | 134 | -1,441546027 | N12:453077 | 11,314 | | | | rDNA | |
| chrXII | 452922 | 453075 | 153 | -1,918531986 | N12:452927 | 2,887 | | | | rDNA | |
| chrXII | 452922 | 453075 | 153 | -1,918531986 | N12:453077 | 11,314 | | | | rDNA | |
| chrXII | 452926 | 452997 | 71 | -0,749 | N12:452927 | 2,887 | | | | rDNA | |
| chrXII | 452926 | 453033 | 107 | -0,777986396 | N12:452927 | 2,887 | | | | rDNA | |
| chrXII | 452926 | 453034 | 108 | -0,910518129 | N12:452927 | 2,887 | | | | rDNA | |
| chrXII | 452926 | 453056 | 130 | -0,940671026 | N12:452927 | 2,887 | | | | rDNA | |
| chrXII | 452926 | 453056 | 130 | -0,940671026 | N12:453077 | 11,314 | | | | rDNA | |
| chrXII | 452926 | 453070 | 144 | -1,105249305 | N12:452927 | 2,887 | | | | rDNA | |

TABLE 3

| CHR | START | END | L(bp) | ΔLK | Nuc ID | Fuzziness | Gene ID | Gene body postion or intergene | Terminal | rDNA | Telomeric |
|---|---|---|---|---|---|---|---|---|---|---|---|
| chrXII | 452926 | 453070 | 144 | -1,105249305 | N12:453077 | 11,314 | | | | rDNA | |
| chrXII | 452926 | 453075 | 149 | -1,223425464 | N12:452927 | 2,887 | | | | rDNA | |
| chrXII | 452926 | 453075 | 149 | -1,223425464 | N12:453077 | 11,314 | | | | rDNA | |
| chrXII | 452976 | 453138 | 162 | -1,590667628 | N12:453077 | 11,314 | | | | rDNA | |
| chrXII | 452976 | 453163 | 187 | -1,520120757 | N12:453077 | 11,314 | | | | rDNA | |
| chrXII | 452991 | 453141 | 150 | -1,323610662 | N12:453077 | 11,314 | | | | rDNA | |
| chrXII | 453033 | 453163 | 130 | -1,183362186 | N12:453077 | 11,314 | | | | rDNA | |
| chrXII | 453033 | 453187 | 154 | -1,411350238 | N12:453077 | 11,314 | | | | rDNA | |
| chrXII | 453033 | 453193 | 160 | -1,530822309 | N12:453077 | 11,314 | | | | rDNA | |
| chrXII | 453039 | 453163 | 124 | -0,840193248 | N12:453077 | 11,314 | | | | rDNA | |
| chrXII | 453039 | 453187 | 148 | -1,180033573 | N12:453077 | 11,314 | | | | rDNA | |
| chrXII | 453039 | 453193 | 154 | -1,503427977 | N12:453077 | 11,314 | | | | rDNA | |
| chrXII | 453042 | 453141 | 99 | -0,975505566 | N12:453077 | 11,314 | | | | rDNA | |
| chrXII | 453042 | 453212 | 170 | -1,576007874 | N12:453077 | 11,314 | | | | rDNA | |
| chrXII | 453042 | 453218 | 176 | -1,231480914 | N12:453077 | 11,314 | | | | rDNA | |
| chrXII | 453043 | 453163 | 120 | -1,025952909 | N12:453077 | 11,314 | | | | rDNA | |
| chrXII | 453043 | 453187 | 144 | -1,23166489 | N12:453077 | 11,314 | | | | rDNA | |
| chrXII | 453043 | 453190 | 147 | -1,359773731 | N12:453077 | 11,314 | | | | rDNA | |
| chrXII | 453043 | 453193 | 150 | -1,227583328 | N12:453077 | 11,314 | | | | rDNA | |
| chrXII | 453091 | 453199 | 108 | -0,785979772 | N12:453077 | 11,314 | | | | rDNA | |
| chrXII | 453092 | 453199 | 107 | -0,729687003 | N12:453077 | 11,314 | | | | rDNA | |
| chrXII | 453092 | 453212 | 120 | -1,156455905 | N12:453077 | 11,314 | | | | rDNA | |
| chrXII | 453092 | 453236 | 144 | -1,560901931 | N12:453077 | 11,314 | | | | rDNA | |
| chrXII | 453092 | 453245 | 153 | -1,51192762 | N12:453077 | 11,314 | | | | rDNA | |
| chrXII | 453092 | 453270 | 178 | -1,520025594 | N12:453077 | 11,314 | | | | rDNA | |
| chrXII | 453092 | 453282 | 190 | -2,097441483 | N12:453077 | 11,314 | | | | rDNA | |
| chrXII | 453092 | 453286 | 194 | -1,759000832 | N12:453077 | 11,314 | | | | rDNA | |
| chrXII | 453092 | 453288 | 196 | -1,60564391 | N12:453077 | 11,314 | | | | rDNA | |
| chrXII | 453094 | 453236 | 142 | -1,850329181 | N12:453077 | 11,314 | | | | rDNA | |
| chrXII | 453095 | 453201 | 106 | -0,941158569 | N12:453077 | 11,314 | | | | rDNA | |
| chrXII | 453095 | 453212 | 117 | -0,679329034 | N12:453077 | 11,314 | | | | rDNA | |
| chrXII | 453095 | 453218 | 123 | -0,458069409 | N12:453077 | 11,314 | | | | rDNA | |
| chrXII | 453095 | 453236 | 141 | -1,867564354 | N12:453077 | 11,314 | | | | rDNA | |
| chrXII | 453095 | 453245 | 150 | -1,124660699 | N12:453077 | 11,314 | | | | rDNA | |
| chrXII | 453095 | 453270 | 175 | -1,451932329 | N12:453077 | 11,314 | | | | rDNA | |
| chrXII | 453095 | 453286 | 191 | -1,562953822 | N12:453077 | 11,314 | | | | rDNA | |
| chrXII | 453095 | 453288 | 193 | -1,561438801 | N12:453077 | 11,314 | | | | rDNA | |
| chrXII | 453099 | 453236 | 137 | -0,706369277 | N12:453077 | 11,314 | | | | rDNA | |
| chrXII | 453099 | 453287 | 188 | -1,448905869 | N12:453077 | 11,314 | | | | rDNA | |
| chrXII | 453099 | 453288 | 189 | -1,529320066 | N12:453077 | 11,314 | | | | rDNA | |
| chrXII | 453100 | 453238 | 138 | -1,248023589 | N12:453077 | 11,314 | | | | rDNA | |
| chrXII | 453100 | 453245 | 145 | -0,701166354 | N12:453077 | 11,314 | | | | rDNA | |
| chrXII | 453113 | 453270 | 157 | -1,432507267 | | | | Overlap <50 bp | | rDNA | |
| chrXII | 453113 | 453286 | 173 | -1,592096458 | | | | Overlap <50 bp | | rDNA | |
| chrXII | 453113 | 453288 | 175 | -1,451933519 | | | | Overlap <50 bp | | rDNA | |
| chrXII | 453114 | 453236 | 122 | -0,914496184 | | | | Overlap <50 bp | | rDNA | |
| chrXII | 453114 | 453269 | 155 | -1,508105897 | | | | Overlap <50 bp | | rDNA | |
| chrXII | 453114 | 453270 | 156 | -1,049493545 | | | | Overlap <50 bp | | rDNA | |
| chrXII | 453114 | 453286 | 172 | -1,388726154 | | | | Overlap <50 bp | | rDNA | |
| chrXII | 453114 | 453287 | 173 | -1,390645671 | | | | Overlap <50 bp | | rDNA | |
| chrXII | 453114 | 453288 | 174 | -1,38636794 | | | | Overlap <50 bp | | rDNA | |
| chrXII | 453114 | 453302 | 188 | -1,522686895 | | | | Overlap <50 bp | | rDNA | |
| chrXII | 453114 | 453309 | 195 | -1,861987022 | | | | Overlap <50 bp | | rDNA | |
| chrXII | 453116 | 453269 | 153 | -0,869780898 | | | | Overlap <50 bp | | rDNA | |
| chrXII | 453116 | 453270 | 154 | -0,843237345 | | | | Overlap <50 bp | | rDNA | |
| chrXII | 453116 | 453288 | 172 | -1,292091959 | | | | Overlap <50 bp | | rDNA | |
| chrXII | 453119 | 453236 | 117 | -0,827954707 | | | | Overlap <50 bp | | rDNA | |
| chrXII | 453119 | 453245 | 126 | -1,068037658 | | | | Overlap <50 bp | | rDNA | |
| chrXII | 453119 | 453269 | 150 | -1,319742249 | | | | Overlap <50 bp | | rDNA | |
| chrXII | 453119 | 453270 | 151 | -1,335573212 | | | | Overlap <50 bp | | rDNA | |
| chrXII | 453119 | 453282 | 163 | -2,472034165 | | | | Overlap <50 bp | | rDNA | |
| chrXII | 453119 | 453286 | 167 | -1,374914415 | | | | Overlap <50 bp | | rDNA | |
| chrXII | 453119 | 453287 | 168 | -1,220847453 | | | | Overlap <50 bp | | rDNA | |
| chrXII | 453119 | 453288 | 169 | -1,306159688 | | | | Overlap <50 bp | | rDNA | |
| chrXII | 453119 | 453302 | 183 | -1,233563504 | | | | Overlap <50 bp | | rDNA | |
| chrXII | 453119 | 453309 | 190 | -1,957136229 | | | | Overlap <50 bp | | rDNA | |
| chrXII | 453120 | 453228 | 108 | -0,489887225 | | | | Overlap <50 bp | | rDNA | |
| chrXII | 453121 | 453270 | 149 | -1,134219118 | | | | Overlap <50 bp | | rDNA | |
| chrXII | 453121 | 453282 | 161 | -1,629300245 | | | | Overlap <50 bp | | rDNA | |
| chrXII | 453121 | 453286 | 165 | -1,446224619 | | | | Overlap <50 bp | | rDNA | |
| chrXII | 453121 | 453288 | 167 | -1,527854968 | | | | Overlap <50 bp | | rDNA | |
| chrXII | 453125 | 453231 | 106 | -1,083676986 | | | | Overlap <50 bp | | rDNA | |
| chrXII | 453125 | 453236 | 111 | -1,300868661 | | | | Overlap <50 bp | | rDNA | |
| chrXII | 453125 | 453270 | 145 | -1,389673767 | | | | Overlap <50 bp | | rDNA | |
| chrXII | 453125 | 453286 | 161 | -1,459005225 | | | | Overlap <50 bp | | rDNA | |
| chrXII | 453125 | 453287 | 162 | -1,23961474 | | | | Overlap <50 bp | | rDNA | |
| chrXII | 453125 | 453288 | 163 | -1,378966264 | | | | Overlap <50 bp | | rDNA | |
| chrXII | 453125 | 453302 | 177 | -1,347453643 | | | | Overlap <50 bp | | rDNA | |
| chrXII | 453126 | 453248 | 122 | -1,594624393 | | | | Overlap <50 bp | | rDNA | |
| chrXII | 453126 | 453270 | 144 | -1,370868156 | | | | Overlap <50 bp | | rDNA | |
| chrXII | 453126 | 453286 | 160 | -1,414253349 | | | | Overlap <50 bp | | rDNA | |
| chrXII | 453126 | 453287 | 161 | -1,334940746 | | | | Overlap <50 bp | | rDNA | |
| chrXII | 453126 | 453288 | 162 | -1,612408164 | | | | Overlap <50 bp | | rDNA | |
| chrXII | 453135 | 453270 | 135 | -1,288874124 | | | | Overlap <50 bp | | rDNA | |
| chrXII | 453135 | 453282 | 147 | -1,328952715 | | | | Overlap <50 bp | | rDNA | |
| chrXII | 453135 | 453286 | 151 | -1,615071406 | | | | Overlap <50 bp | | rDNA | |
| chrXII | 453135 | 453288 | 153 | -1,620874875 | | | | Overlap <50 bp | | rDNA | |
| chrXII | 453138 | 453286 | 148 | -1,274369061 | | | | Overlap <50 bp | | rDNA | |
| chrXII | 453138 | 453288 | 150 | -1,754737165 | | | | Overlap <50 bp | | rDNA | |
| chrXII | 453140 | 453245 | 105 | -0,238606443 | | | | Overlap <50 bp | | rDNA | |
| chrXII | 453140 | 453246 | 106 | -0,915232661 | | | | Overlap <50 bp | | rDNA | |
| chrXII | 453140 | 453251 | 111 | -1,413838737 | | | | Overlap <50 bp | | rDNA | |
| chrXII | 453140 | 453270 | 130 | -1,084174987 | | | | Overlap <50 bp | | rDNA | |
| chrXII | 453140 | 453282 | 142 | -1,478152831 | | | | Overlap <50 bp | | rDNA | |
| chrXII | 453140 | 453286 | 146 | -1,358675116 | | | | Overlap <50 bp | | rDNA | |
| chrXII | 453140 | 453287 | 147 | -1,380617363 | | | | Overlap <50 bp | | rDNA | |

TABLE 3

| CHR | START | END | L(bp) | ΔLK | Nuc ID | Fuzziness | Gene ID | Gene body postion or intergene | Terminal | rDNA | Telomeric |
|---|---|---|---|---|---|---|---|---|---|---|---|
| chrXII | 453140 | 453288 | 148 | -1,194319694 | | | | Overlap <50 bp | | rDNA | |
| chrXII | 453140 | 453302 | 162 | -1,666411185 | | | | Overlap <50 bp | | rDNA | |
| chrXII | 453140 | 453309 | 169 | -1,843043158 | | | | Overlap <50 bp | | rDNA | |
| chrXII | 453140 | 453323 | 183 | -1,408711684 | | | | Overlap <50 bp | | rDNA | |
| chrXII | 453141 | 453236 | 95 | -0,643867953 | | | | Overlap <50 bp | | rDNA | |
| chrXII | 453141 | 453245 | 104 | -0,427593631 | | | | Overlap <50 bp | | rDNA | |
| chrXII | 453141 | 453251 | 110 | -0,931901947 | | | | Overlap <50 bp | | rDNA | |
| chrXII | 453141 | 453269 | 128 | -0,8723526 | | | | Overlap <50 bp | | rDNA | |
| chrXII | 453141 | 453270 | 129 | -0,945885167 | | | | Overlap <50 bp | | rDNA | |
| chrXII | 453141 | 453281 | 140 | -1,27534751 | | | | Overlap <50 bp | | rDNA | |
| chrXII | 453141 | 453282 | 141 | -1,382524872 | | | | Overlap <50 bp | | rDNA | |
| chrXII | 453141 | 453286 | 145 | -1,234318114 | | | | Overlap <50 bp | | rDNA | |
| chrXII | 453141 | 453287 | 146 | -1,398220625 | | | | Overlap <50 bp | | rDNA | |
| chrXII | 453141 | 453288 | 147 | -1,271435943 | | | | Overlap <50 bp | | rDNA | |
| chrXII | 453141 | 453302 | 161 | -1,311822625 | | | | Overlap <50 bp | | rDNA | |
| chrXII | 453141 | 453309 | 168 | -1,371141228 | | | | Overlap <50 bp | | rDNA | |
| chrXII | 453141 | 453323 | 182 | -1,459025922 | | | | Overlap <50 bp | | rDNA | |
| chrXII | 453141 | 453324 | 183 | -1,445749047 | | | | Overlap <50 bp | | rDNA | |
| chrXII | 453144 | 453288 | 144 | -1,869085848 | | | | Overlap <50 bp | | rDNA | |
| chrXII | 453150 | 453286 | 136 | -1,160093829 | | | | Overlap <50 bp | | rDNA | |
| chrXII | 453150 | 453287 | 137 | -1,288786282 | | | | Overlap <50 bp | | rDNA | |
| chrXII | 453150 | 453288 | 138 | -1,284642731 | | | | Overlap <50 bp | | rDNA | |
| chrXII | 453150 | 453295 | 145 | -1,062988513 | | | | Overlap <50 bp | | rDNA | |
| chrXII | 453150 | 453302 | 152 | -1,84951621 | | | | Overlap <50 bp | | rDNA | |
| chrXII | 453152 | 453240 | 88 | -0,593715668 | | | | Overlap <50 bp | | rDNA | |
| chrXII | 453152 | 453270 | 118 | -0,564489561 | | | | Overlap <50 bp | | rDNA | |
| chrXII | 453152 | 453286 | 134 | -1,113267061 | | | | Overlap <50 bp | | rDNA | |
| chrXII | 453152 | 453287 | 135 | -1,210451589 | | | | Overlap <50 bp | | rDNA | |
| chrXII | 453152 | 453288 | 136 | -1,287494979 | | | | Overlap <50 bp | | rDNA | |
| chrXII | 453152 | 453302 | 150 | -1,458554267 | | | | Overlap <50 bp | | rDNA | |
| chrXII | 453152 | 453309 | 157 | -1,403027613 | | | | Overlap <50 bp | | rDNA | |
| chrXII | 453153 | 453286 | 133 | -1,51487596 | | | | Overlap <50 bp | | rDNA | |
| chrXII | 453153 | 453288 | 135 | -1,129917945 | | | | Overlap <50 bp | | rDNA | |
| chrXII | 453159 | 453270 | 111 | -0,647734571 | | | | Overlap <50 bp | | rDNA | |
| chrXII | 453159 | 453286 | 127 | -1,070865061 | | | | Overlap <50 bp | | rDNA | |
| chrXII | 453159 | 453287 | 128 | -1,031334415 | | | | Overlap <50 bp | | rDNA | |
| chrXII | 453159 | 453288 | 129 | -1,025873518 | | | | Overlap <50 bp | | rDNA | |
| chrXII | 453159 | 453302 | 143 | -1,124678362 | | | | Overlap <50 bp | | rDNA | |
| chrXII | 453159 | 453309 | 150 | -1,561989909 | | | | Overlap <50 bp | | rDNA | |
| chrXII | 453159 | 453335 | 176 | -1,688153227 | | | | Overlap <50 bp | | rDNA | |
| chrXII | 453159 | 453342 | 183 | -1,666339773 | | | | Overlap <50 bp | | rDNA | |
| chrXII | 453161 | 453245 | 84 | -0,39863806 | | | | Overlap <50 bp | | rDNA | |
| chrXII | 453161 | 453268 | 107 | -0,854039433 | | | | Overlap <50 bp | | rDNA | |
| chrXII | 453161 | 453270 | 109 | -0,827805551 | | | | Overlap <50 bp | | rDNA | |
| chrXII | 453161 | 453282 | 121 | -0,82493292 | | | | Overlap <50 bp | | rDNA | |
| chrXII | 453161 | 453286 | 125 | -0,942457735 | | | | Overlap <50 bp | | rDNA | |
| chrXII | 453161 | 453288 | 127 | -1,113344395 | | | | Overlap <50 bp | | rDNA | |
| chrXII | 453161 | 453309 | 148 | -1,427578397 | | | | Overlap <50 bp | | rDNA | |
| chrXII | 453161 | 453323 | 162 | -1,105452985 | | | | Overlap <50 bp | | rDNA | |
| chrXII | 453161 | 453324 | 163 | -1,101547436 | | | | Overlap <50 bp | | rDNA | |
| chrXII | 453162 | 453269 | 107 | -0,594054974 | | | | Overlap <50 bp | | rDNA | |
| chrXII | 453162 | 453286 | 124 | -0,95735413 | | | | Overlap <50 bp | | rDNA | |
| chrXII | 453162 | 453288 | 126 | -1,290809458 | | | | Overlap <50 bp | | rDNA | |
| chrXII | 453162 | 453323 | 161 | -1,356721729 | | | | Overlap <50 bp | | rDNA | |
| chrXII | 453162 | 453324 | 162 | -1,342367579 | | | | Overlap <50 bp | | rDNA | |
| chrXII | 453165 | 453286 | 121 | -1,010079882 | | | | Overlap <50 bp | | rDNA | |
| chrXII | 453165 | 453287 | 122 | -1,020999093 | | | | Overlap <50 bp | | rDNA | |
| chrXII | 453165 | 453288 | 123 | -1,001823635 | | | | Overlap <50 bp | | rDNA | |
| chrXII | 453165 | 453302 | 137 | -1,282037093 | | | | Overlap <50 bp | | rDNA | |
| chrXII | 453165 | 453342 | 177 | -1,287950972 | | | | Overlap <50 bp | | rDNA | |
| chrXII | 453180 | 453286 | 106 | -1,400973842 | | | | Overlap <50 bp | | rDNA | |
| chrXII | 453180 | 453288 | 108 | -0,994346844 | | | | Overlap <50 bp | | rDNA | |
| chrXII | 453180 | 453340 | 160 | -1,440407218 | | | | Overlap <50 bp | | rDNA | |
| chrXII | 453181 | 453270 | 89 | -1,337550529 | | | | Overlap <50 bp | | rDNA | |
| chrXII | 453181 | 453282 | 101 | -1,854233757 | | | | Overlap <50 bp | | rDNA | |
| chrXII | 453181 | 453286 | 105 | -1,117474245 | | | | Overlap <50 bp | | rDNA | |
| chrXII | 453181 | 453288 | 107 | -1,192141437 | | | | Overlap <50 bp | | rDNA | |
| chrXII | 453181 | 453323 | 142 | -1,266947266 | | | | Overlap <50 bp | | rDNA | |
| chrXII | 453181 | 453324 | 143 | -1,722976 | | | | Overlap <50 bp | | rDNA | |
| chrXII | 453286 | 453432 | 146 | -0,944000478 | N12:453378 | 19,858 | | | | rDNA | |
| chrXII | 453299 | 453425 | 126 | -1,515497291 | N12:453378 | 19,858 | | | | rDNA | |
| chrXII | 453299 | 453440 | 141 | -1,067509288 | N12:453378 | 19,858 | | | | rDNA | |
| chrXII | 453299 | 453460 | 161 | -1,180011587 | N12:453378 | 19,858 | | | | rDNA | |
| chrXII | 453300 | 453410 | 110 | -1,365285834 | N12:453378 | 19,858 | | | | rDNA | |
| chrXII | 453300 | 453414 | 114 | -1,499902166 | N12:453378 | 19,858 | | | | rDNA | |
| chrXII | 453300 | 453440 | 140 | -1,978581283 | N12:453378 | 19,858 | | | | rDNA | |
| chrXII | 453302 | 453440 | 138 | -1,164256023 | N12:453378 | 19,858 | | | | rDNA | |
| chrXII | 453310 | 453440 | 130 | -0,817772089 | N12:453378 | 19,858 | | | | rDNA | |
| chrXII | 453310 | 453460 | 150 | -1,236884475 | N12:453378 | 19,858 | | | | rDNA | |
| chrXII | 453322 | 453481 | 159 | -1,310065467 | N12:453378 | 19,858 | | | | rDNA | |
| chrXII | 453458 | 453595 | 137 | -1,633667366 | N12:453535 | 22,121 | | | | rDNA | |
| chrXII | 453458 | 453610 | 152 | -1,354015096 | N12:453535 | 22,121 | | | | rDNA | |
| chrXII | 453458 | 453612 | 154 | -1,558180698 | N12:453535 | 22,121 | | | | rDNA | |
| chrXII | 453460 | 453612 | 152 | -1,423694186 | N12:453535 | 22,121 | | | | rDNA | |
| chrXII | 453487 | 453595 | 108 | -1,471679521 | N12:453535 | 22,121 | | | | rDNA | |
| chrXII | 453487 | 453610 | 123 | -1,377331296 | N12:453535 | 22,121 | | | | rDNA | |
| chrXII | 453487 | 453612 | 125 | -1,11113785 | N12:453535 | 22,121 | | | | rDNA | |
| chrXII | 453500 | 453652 | 152 | -1,282975984 | N12:453535 | 22,121 | | | | rDNA | |
| chrXII | 453574 | 453727 | 153 | -0,958674505 | N12:453693 | 25,423 | | | | rDNA | |
| chrXII | 453587 | 453737 | 150 | -1,664370448 | N12:453693 | 25,423 | | | | rDNA | |
| chrXII | 453606 | 453774 | 168 | -1,114323848 | N12:453693 | 25,423 | | | | rDNA | |
| chrXII | 453627 | 453776 | 149 | -1,742013422 | N12:453693 | 25,423 | | | | rDNA | |
| chrXII | 453627 | 453779 | 152 | -1,069797779 | N12:453693 | 25,423 | | | | rDNA | |
| chrXII | 453657 | 453814 | 157 | -0,944340368 | N12:453693 | 25,423 | | | | rDNA | |
| chrXII | 453659 | 453776 | 117 | -1,408011648 | N12:453693 | 25,423 | | | | rDNA | |

TABLE 3

| CHR | START | END | L(bp) | ΔLK | Nuc ID | Fuzziness | Gene ID | Gene body postion or intergene | Terminal | rDNA | Telomeric |
|---|---|---|---|---|---|---|---|---|---|---|---|
| chrXII | 453659 | 453779 | 120 | -1,157080791 | N12:453693 | 25,423 | | | | rDNA | |
| chrXII | 453734 | 453853 | 119 | -1,053841366 | N12:453841 | 6,364 | | | | rDNA | |
| chrXII | 453748 | 453910 | 162 | -1,876176937 | N12:453841 | 6,364 | | | | rDNA | |
| chrXII | 453748 | 453913 | 165 | -1,367106647 | N12:453841 | 6,364 | | | | rDNA | |
| chrXII | 453748 | 453919 | 171 | -1,379956812 | N12:453841 | 6,364 | | | | rDNA | |
| chrXII | 453748 | 453922 | 174 | -1,718968275 | N12:453841 | 6,364 | | | | rDNA | |
| chrXII | 453748 | 453962 | 214 | -1,731085258 | N12:453841 | 6,364 | | | | rDNA | |
| chrXII | 453752 | 453853 | 101 | -1,025030532 | N12:453841 | 6,364 | | | | rDNA | |
| chrXII | 453752 | 453859 | 107 | -0,53510933 | N12:453841 | 6,364 | | | | rDNA | |
| chrXII | 453752 | 453908 | 156 | -1,522404847 | N12:453841 | 6,364 | | | | rDNA | |
| chrXII | 453752 | 453911 | 159 | -1,427007064 | N12:453841 | 6,364 | | | | rDNA | |
| chrXII | 453752 | 453914 | 162 | -1,341100402 | N12:453841 | 6,364 | | | | rDNA | |
| chrXII | 453752 | 453916 | 164 | -1,388930983 | N12:453841 | 6,364 | | | | rDNA | |
| chrXII | 453752 | 453922 | 170 | -1,387833823 | N12:453841 | 6,364 | | | | rDNA | |
| chrXII | 453752 | 453924 | 172 | -1,349546569 | N12:453841 | 6,364 | | | | rDNA | |
| chrXII | 453753 | 453913 | 160 | -1,645741933 | N12:453841 | 6,364 | | | | rDNA | |
| chrXII | 453753 | 453919 | 166 | -1,134946401 | N12:453841 | 6,364 | | | | rDNA | |
| chrXII | 453753 | 453922 | 169 | -1,413770668 | N12:453841 | 6,364 | | | | rDNA | |
| chrXII | 453754 | 453910 | 156 | -1,650313296 | N12:453841 | 6,364 | | | | rDNA | |
| chrXII | 453754 | 453924 | 170 | -1,673342306 | N12:453841 | 6,364 | | | | rDNA | |
| chrXII | 453774 | 453910 | 136 | -1,12920419 | N12:453841 | 6,364 | | | | rDNA | |
| chrXII | 453774 | 453913 | 139 | -1,220142143 | N12:453841 | 6,364 | | | | rDNA | |
| chrXII | 453774 | 453914 | 140 | -1,542051489 | N12:453841 | 6,364 | | | | rDNA | |
| chrXII | 453774 | 453916 | 142 | -1,297713418 | N12:453841 | 6,364 | | | | rDNA | |
| chrXII | 453774 | 453919 | 145 | -1,139587321 | N12:453841 | 6,364 | | | | rDNA | |
| chrXII | 453774 | 453922 | 148 | -1,994673623 | N12:453841 | 6,364 | | | | rDNA | |
| chrXII | 453774 | 453925 | 151 | -1,408783162 | N12:453841 | 6,364 | | | | rDNA | |
| chrXII | 453774 | 453962 | 188 | -1,316041416 | N12:453841 | 6,364 | | | | rDNA | |
| chrXII | 453775 | 453908 | 133 | -1,303308731 | N12:453841 | 6,364 | | | | rDNA | |
| chrXII | 453775 | 453911 | 136 | -1,169773296 | N12:453841 | 6,364 | | | | rDNA | |
| chrXII | 453775 | 453914 | 139 | -1,05645459 | N12:453841 | 6,364 | | | | rDNA | |
| chrXII | 453775 | 453916 | 141 | -1,437427921 | N12:453841 | 6,364 | | | | rDNA | |
| chrXII | 453775 | 453919 | 144 | -2,178844761 | N12:453841 | 6,364 | | | | rDNA | |
| chrXII | 453775 | 453922 | 147 | -1,501005416 | N12:453841 | 6,364 | | | | rDNA | |
| chrXII | 453781 | 453908 | 127 | -0,872289474 | N12:453841 | 6,364 | | | | rDNA | |
| chrXII | 453781 | 453911 | 130 | -1,000628502 | N12:453841 | 6,364 | | | | rDNA | |
| chrXII | 453781 | 453914 | 133 | -1,18210055 | N12:453841 | 6,364 | | | | rDNA | |
| chrXII | 453781 | 453916 | 135 | -1,087186833 | N12:453841 | 6,364 | | | | rDNA | |
| chrXII | 453781 | 453922 | 141 | -1,345406678 | N12:453841 | 6,364 | | | | rDNA | |
| chrXII | 453782 | 453908 | 126 | -0,961066656 | N12:453841 | 6,364 | | | | rDNA | |
| chrXII | 453782 | 453911 | 129 | -1,312732098 | N12:453841 | 6,364 | | | | rDNA | |
| chrXII | 453782 | 453913 | 131 | -0,838520311 | N12:453841 | 6,364 | | | | rDNA | |
| chrXII | 453782 | 453914 | 132 | -1,090012453 | N12:453841 | 6,364 | | | | rDNA | |
| chrXII | 453782 | 453919 | 137 | -1,217429412 | N12:453841 | 6,364 | | | | rDNA | |
| chrXII | 453782 | 453922 | 140 | -1,146997168 | N12:453841 | 6,364 | | | | rDNA | |
| chrXII | 453782 | 453925 | 143 | -1,132929801 | N12:453841 | 6,364 | | | | rDNA | |
| chrXII | 453782 | 453929 | 147 | -1,310697855 | N12:453841 | 6,364 | | | | rDNA | |
| chrXII | 453782 | 453937 | 155 | -1,54162973 | N12:453841 | 6,364 | | | | rDNA | |
| chrXII | 453782 | 453962 | 180 | -1,462803965 | N12:453841 | 6,364 | | | | rDNA | |
| chrXII | 453783 | 453911 | 128 | -1,026686018 | N12:453841 | 6,364 | | | | rDNA | |
| chrXII | 453783 | 453913 | 130 | -1,024370582 | N12:453841 | 6,364 | | | | rDNA | |
| chrXII | 453783 | 453914 | 131 | -1,85582847 | N12:453841 | 6,364 | | | | rDNA | |
| chrXII | 453812 | 453913 | 101 | -0,672409103 | N12:453841 | 6,364 | | | | rDNA | |
| chrXII | 453812 | 453919 | 107 | -0,460245968 | N12:453841 | 6,364 | | | | rDNA | |
| chrXII | 453812 | 453962 | 150 | -1,147287853 | N12:453841 | 6,364 | | | | rDNA | |
| chrXII | 453812 | 454001 | 189 | -1,473452888 | N12:453841 | 6,364 | | | | rDNA | |
| chrXII | 453812 | 454001 | 189 | -1,473452888 | N12:454007 | 8,505 | | | | rDNA | |
| chrXII | 453851 | 453958 | 107 | -0,559635716 | N12:453841 | 6,364 | | | | rDNA | |
| chrXII | 453851 | 453961 | 110 | -1,140559031 | N12:453841 | 6,364 | | | | rDNA | |
| chrXII | 453851 | 453962 | 111 | -0,65850533 | N12:453841 | 6,364 | | | | rDNA | |
| chrXII | 453851 | 454001 | 150 | -1,099442387 | N12:453841 | 6,364 | | | | rDNA | |
| chrXII | 453851 | 454001 | 150 | -1,099442387 | N12:454007 | 8,505 | | | | rDNA | |
| chrXII | 453909 | 454072 | 163 | -1,500128358 | N12:454007 | 8,505 | | | | rDNA | |
| chrXII | 453918 | 454080 | 162 | -1,234988308 | N12:454007 | 8,505 | | | | rDNA | |
| chrXII | 453922 | 454079 | 157 | -1,346571791 | N12:454007 | 8,505 | | | | rDNA | |
| chrXII | 453929 | 454079 | 150 | -1,833161087 | N12:454007 | 8,505 | | | | rDNA | |
| chrXII | 453929 | 454097 | 168 | -1,587762446 | N12:454007 | 8,505 | | | | rDNA | |
| chrXII | 453929 | 454108 | 179 | -1,527671331 | N12:454007 | 8,505 | | | | rDNA | |
| chrXII | 453959 | 454097 | 138 | -1,202488253 | N12:454007 | 8,505 | | | | rDNA | |
| chrXII | 453959 | 454108 | 149 | -1,176974072 | N12:454007 | 8,505 | | | | rDNA | |
| chrXII | 453959 | 454112 | 153 | -1,084044682 | N12:454007 | 8,505 | | | | rDNA | |
| chrXII | 453960 | 454116 | 156 | -1,212103047 | N12:454007 | 8,505 | | | | rDNA | |
| chrXII | 453964 | 454097 | 133 | -1,343610196 | N12:454007 | 8,505 | | | | rDNA | |
| chrXII | 453964 | 454108 | 144 | -1,112521418 | N12:454007 | 8,505 | | | | rDNA | |
| chrXII | 453971 | 454123 | 152 | -1,280386921 | N12:454007 | 8,505 | | | | rDNA | |
| chrXII | 454059 | 454216 | 157 | -1,397576609 | N12:454198 | 24,269 | | | | rDNA | |
| chrXII | 454111 | 454268 | 157 | -1,531530374 | N12:454198 | 24,269 | | | | rDNA | |
| chrXII | 454120 | 454267 | 147 | -1,590901998 | N12:454198 | 24,269 | | | | rDNA | |
| chrXII | 454120 | 454277 | 157 | -1,424361653 | N12:454198 | 24,269 | | | | rDNA | |
| chrXII | 454124 | 454277 | 153 | -1,399493395 | N12:454198 | 24,269 | | | | rDNA | |
| chrXII | 454131 | 454277 | 146 | -1,425230991 | N12:454198 | 24,269 | | | | rDNA | |
| chrXII | 454141 | 454277 | 136 | -1,280053521 | N12:454198 | 24,269 | | | | rDNA | |
| chrXII | 454196 | 454346 | 150 | -0,881708481 | N12:454367 | 64,01 | -1:YLR154W-C; | -1 | | rDNA | |
| chrXII | 454196 | 454346 | 150 | -0,881708481 | N12:454198 | 24,269 | | | | rDNA | |
| chrXII | 454221 | 454392 | 171 | -1,296471933 | N12:454367 | 64,01 | -1:YLR154W-C; | -1 | | rDNA | |
| chrXII | 454221 | 454392 | 171 | -1,296471933 | N12:454198 | 24,269 | | | | rDNA | |
| chrXII | 454244 | 454392 | 148 | -1,56618022 | N12:454367 | 64,01 | -1:YLR154W-C; | -1 | | rDNA | |
| chrXII | 454244 | 454402 | 158 | -1,120982349 | N12:454367 | 64,01 | -1:YLR154W-C; | -1 | | rDNA | |
| chrXII | 454248 | 454374 | 126 | -1,078312309 | N12:454367 | 64,01 | -1:YLR154W-C; | -1 | | rDNA | |
| chrXII | 454248 | 454392 | 144 | -1,398153008 | N12:454367 | 64,01 | -1:YLR154W-C; | -1 | | rDNA | |
| chrXII | 454248 | 454398 | 150 | -1,24796621 | N12:454367 | 64,01 | -1:YLR154W-C; | -1 | | rDNA | |
| chrXII | 454273 | 454436 | 163 | -1,245692034 | N12:454367 | 64,01 | -1:YLR154W-C; | -1 | | rDNA | |
| chrXII | 454277 | 454398 | 121 | -0,634273441 | N12:454367 | 64,01 | -1:YLR154W-C; | -1 | | rDNA | |
| chrXII | 454277 | 454402 | 125 | -1,422699207 | N12:454367 | 64,01 | -1:YLR154W-C; | -1 | | rDNA | |
| chrXII | 454277 | 454438 | 161 | -1,148371885 | N12:454367 | 64,01 | -1:YLR154W-C; | -1 | | rDNA | |
| chrXII | 454277 | 454446 | 169 | -1,586449775 | N12:454367 | 64,01 | -1:YLR154W-C; | -1 | | rDNA | |
| chrXII | 454398 | 454551 | 153 | -1,422643255 | | | | Overlap <50 bp | | rDNA | |
| chrXII | 454398 | 454552 | 154 | -1,181586119 | | | | Overlap <50 bp | | rDNA | |
| chrXII | 454398 | 454555 | 157 | -1,215492537 | | | | Overlap <50 bp | | rDNA | |
| chrXII | 454398 | 454567 | 169 | -1,137203077 | | | | Overlap <50 bp | | rDNA | |

| CHR | START | END | L(bp) | ΔLK | Nuc ID | Fuzziness | Gene ID | Gene body postion or intergene | Terminal | rDNA | Telomeric |
|---|---|---|---|---|---|---|---|---|---|---|---|
| chrXII | 454398 | 454573 | 175 | -1,463178836 | | | | Overlap <50 bp | | rDNA | |
| chrXII | 454398 | 454575 | 177 | -1,051247577 | | | | Overlap <50 bp | | rDNA | |
| chrXII | 454404 | 454552 | 148 | -1,144430038 | | | | Overlap <50 bp | | rDNA | |
| chrXII | 454404 | 454555 | 151 | -1,100057884 | | | | Overlap <50 bp | | rDNA | |
| chrXII | 454422 | 454552 | 130 | -0,920526742 | | | | Overlap <50 bp | | rDNA | |
| chrXII | 454422 | 454555 | 133 | -0,79819432 | | | | Overlap <50 bp | | rDNA | |
| chrXII | 454422 | 454567 | 145 | -1,13871689 | | | | Overlap <50 bp | | rDNA | |
| chrXII | 454422 | 454573 | 151 | -1,161154563 | | | | Overlap <50 bp | | rDNA | |
| chrXII | 454422 | 454575 | 153 | -1,210931753 | | | | Overlap <50 bp | | rDNA | |
| chrXII | 454521 | 454678 | 157 | -1,426562991 | N12:454654 | 30,199 | +1:YLR154W-C; | 1 | | rDNA | |
| chrXII | 454553 | 454712 | 159 | -1,231917398 | N12:454654 | 30,199 | +1:YLR154W-C; | 1 | | rDNA | |
| chrXII | 454553 | 454715 | 162 | -1,197068638 | N12:454654 | 30,199 | +1:YLR154W-C; | 1 | | rDNA | |
| chrXII | 454553 | 454733 | 180 | -1,632580235 | N12:454654 | 30,199 | +1:YLR154W-C; | 1 | | rDNA | |
| chrXII | 454555 | 454703 | 148 | -1,175552155 | N12:454654 | 30,199 | +1:YLR154W-C; | 1 | | rDNA | |
| chrXII | 454560 | 454678 | 118 | -0,919035871 | N12:454654 | 30,199 | +1:YLR154W-C; | 1 | | rDNA | |
| chrXII | 454560 | 454712 | 152 | -1,438878957 | N12:454654 | 30,199 | +1:YLR154W-C; | 1 | | rDNA | |
| chrXII | 454560 | 454715 | 155 | -1,344898696 | N12:454654 | 30,199 | +1:YLR154W-C; | 1 | | rDNA | |
| chrXII | 454560 | 454724 | 164 | -1,068596116 | N12:454654 | 30,199 | +1:YLR154W-C; | 1 | | rDNA | |
| chrXII | 454560 | 454733 | 173 | -1,37305148 | N12:454654 | 30,199 | +1:YLR154W-C; | 1 | | rDNA | |
| chrXII | 454562 | 454712 | 150 | -1,089673703 | N12:454654 | 30,199 | +1:YLR154W-C; | 1 | | rDNA | |
| chrXII | 454570 | 454712 | 142 | -1,011819202 | N12:454654 | 30,199 | +1:YLR154W-C; | 1 | | rDNA | |
| chrXII | 454570 | 454715 | 145 | -1,333321638 | N12:454654 | 30,199 | +1:YLR154W-C; | 1 | | rDNA | |
| chrXII | 454570 | 454724 | 154 | -1,173953046 | N12:454654 | 30,199 | +1:YLR154W-C; | 1 | | rDNA | |
| chrXII | 454570 | 454733 | 163 | -1,406490175 | N12:454654 | 30,199 | +1:YLR154W-C; | 1 | | rDNA | |
| chrXII | 454576 | 454712 | 136 | -1,476847827 | N12:454654 | 30,199 | +1:YLR154W-C; | 1 | | rDNA | |
| chrXII | 454576 | 454715 | 139 | -1,291344565 | N12:454654 | 30,199 | +1:YLR154W-C; | 1 | | rDNA | |
| chrXII | 454576 | 454724 | 148 | -1,37148199 | N12:454654 | 30,199 | +1:YLR154W-C; | 1 | | rDNA | |
| chrXII | 454576 | 454733 | 157 | -1,412750071 | N12:454654 | 30,199 | +1:YLR154W-C; | 1 | | rDNA | |
| chrXII | 454576 | 454772 | 196 | -1,493923074 | N12:454654 | 30,199 | +1:YLR154W-C; | 1 | | rDNA | |
| chrXII | 454587 | 454746 | 159 | -1,395125026 | N12:454654 | 30,199 | +1:YLR154W-C; | 1 | | rDNA | |
| chrXII | 454592 | 454742 | 150 | -1,282190339 | N12:454654 | 30,199 | +1:YLR154W-C; | 1 | | rDNA | |
| chrXII | 454626 | 454724 | 98 | -0,356137251 | N12:454654 | 30,199 | +1:YLR154W-C; | 1 | | rDNA | |
| chrXII | 454626 | 454733 | 107 | -0,575614643 | N12:454654 | 30,199 | +1:YLR154W-C; | 1 | | rDNA | |
| chrXII | 454626 | 454772 | 146 | -1,167294426 | N12:454654 | 30,199 | +1:YLR154W-C; | 1 | | rDNA | |
| chrXII | 454649 | 454741 | 92 | -0,618965613 | N12:454654 | 30,199 | +1:YLR154W-C; | 1 | | rDNA | |
| chrXII | 454677 | 454828 | 151 | -1,429219436 | N12:454654 | 30,199 | +1:YLR154W-C; | 1 | | rDNA | |
| chrXII | 454677 | 454828 | 151 | -1,429219436 | N12:454834 | 31,88 | +2:YLR154W-C; | 2 | | rDNA | |
| chrXII | 454723 | 454854 | 131 | -1,117114606 | N12:454834 | 31,88 | +2:YLR154W-C; | 2 | | rDNA | |
| chrXII | 454725 | 454863 | 138 | -1,082110205 | N12:454834 | 31,88 | +2:YLR154W-C; | 2 | | rDNA | |
| chrXII | 454725 | 454886 | 161 | -1,297765154 | N12:454834 | 31,88 | +2:YLR154W-C; | 2 | | rDNA | |
| chrXII | 454733 | 454863 | 130 | -1,199206879 | N12:454834 | 31,88 | +2:YLR154W-C; | 2 | | rDNA | |
| chrXII | 454733 | 454886 | 153 | -1,673674328 | N12:454834 | 31,88 | +2:YLR154W-C; | 2 | | rDNA | |
| chrXII | 454741 | 454896 | 155 | -1,505428713 | N12:454834 | 31,88 | +2:YLR154W-C; | 2 | | rDNA | |
| chrXII | 454747 | 454853 | 106 | -0,443622589 | N12:454834 | 31,88 | +2:YLR154W-C; | 2 | | rDNA | |
| chrXII | 454747 | 454886 | 139 | -1,077833095 | N12:454834 | 31,88 | +2:YLR154W-C; | 2 | | rDNA | |
| chrXII | 454825 | 454972 | 147 | -1,680443806 | N12:454834 | 31,88 | +2:YLR154W-C; | 2 | | rDNA | |
| chrXII | 454825 | 454972 | 147 | -1,680443806 | N12:454994 | 15,556 | +3:YLR154W-C; | 3 | | rDNA | |
| chrXII | 454837 | 454991 | 154 | -0,958204752 | N12:454834 | 31,88 | +2:YLR154W-C; | 2 | | rDNA | |
| chrXII | 454837 | 454991 | 154 | -0,958204752 | N12:454994 | 15,556 | +3:YLR154W-C; | 3 | | rDNA | |
| chrXII | 454864 | 455021 | 157 | -1,054009343 | N12:454994 | 15,556 | +3:YLR154W-C; | 3 | | rDNA | |
| chrXII | 454876 | 455023 | 147 | -1,065648787 | N12:454994 | 15,556 | +3:YLR154W-C; | 3 | | rDNA | |
| chrXII | 454876 | 455029 | 153 | -1,616796046 | N12:454994 | 15,556 | +3:YLR154W-C; | 3 | | rDNA | |
| chrXII | 454901 | 455029 | 128 | -1,451662519 | N12:454994 | 15,556 | +3:YLR154W-C; | 3 | | rDNA | |
| chrXII | 454901 | 455074 | 173 | -1,422004281 | N12:454994 | 15,556 | +3:YLR154W-C; | 3 | | rDNA | |
| chrXII | 454920 | 455087 | 167 | -1,273990201 | N12:454994 | 15,556 | +3:YLR154W-C; | 3 | | rDNA | |
| chrXII | 454932 | 455073 | 141 | -0,978355727 | N12:454994 | 15,556 | +3:YLR154W-C; | 3 | | rDNA | |
| chrXII | 454933 | 455023 | 90 | -0,847714769 | N12:454994 | 15,556 | +3:YLR154W-C; | 3 | | rDNA | |
| chrXII | 454933 | 455024 | 91 | -1,265537125 | N12:454994 | 15,556 | +3:YLR154W-C; | 3 | | rDNA | |
| chrXII | 454933 | 455025 | 92 | -0,69161037 | N12:454994 | 15,556 | +3:YLR154W-C; | 3 | | rDNA | |
| chrXII | 454933 | 455027 | 94 | -0,994638715 | N12:454994 | 15,556 | +3:YLR154W-C; | 3 | | rDNA | |
| chrXII | 454933 | 455029 | 96 | -1,181792976 | N12:454994 | 15,556 | +3:YLR154W-C; | 3 | | rDNA | |
| chrXII | 454933 | 455073 | 140 | -0,830362198 | N12:454994 | 15,556 | +3:YLR154W-C; | 3 | | rDNA | |
| chrXII | 454933 | 455074 | 141 | -1,326709506 | N12:454994 | 15,556 | +3:YLR154W-C; | 3 | | rDNA | |
| chrXII | 454933 | 455076 | 143 | -1,145400143 | N12:454994 | 15,556 | +3:YLR154W-C; | 3 | | rDNA | |
| chrXII | 454933 | 455132 | 199 | -2,180476381 | N12:454994 | 15,556 | +3:YLR154W-C; | 3 | | rDNA | |
| chrXII | 454964 | 455132 | 168 | -1,313679037 | N12:454994 | 15,556 | +3:YLR154W-C; | 3 | | rDNA | |
| chrXII | 454995 | 455164 | 169 | -1,697455115 | N12:454994 | 15,556 | +3:YLR154W-C; | 3 | | rDNA | |
| chrXII | 455000 | 455132 | 132 | -1,389881427 | N12:454994 | 15,556 | +3:YLR154W-C; | 3 | | rDNA | |
| chrXII | 455015 | 455166 | 151 | -1,215953421 | N12:454994 | 15,556 | +3:YLR154W-C; | 3 | | rDNA | |
| chrXII | 455058 | 455208 | 150 | -1,439311083 | | | | Overlap <50 bp | | rDNA | |
| chrXII | 455068 | 455223 | 155 | -1,281133884 | | | | Overlap <50 bp | | rDNA | |
| chrXII | 455095 | 455269 | 174 | -1,447474451 | | | | Overlap <50 bp | | rDNA | |
| chrXII | 455096 | 455223 | 127 | -1,189315108 | | | | Overlap <50 bp | | rDNA | |
| chrXII | 455096 | 455240 | 144 | -1,276225788 | | | | Overlap <50 bp | | rDNA | |
| chrXII | 455096 | 455243 | 147 | -1,257096994 | | | | Overlap <50 bp | | rDNA | |
| chrXII | 455096 | 455245 | 149 | -1,199283538 | | | | Overlap <50 bp | | rDNA | |
| chrXII | 455096 | 455249 | 153 | -1,211227729 | | | | Overlap <50 bp | | rDNA | |
| chrXII | 455097 | 455240 | 143 | -1,286935047 | | | | Overlap <50 bp | | rDNA | |
| chrXII | 455097 | 455243 | 146 | -1,142746925 | | | | Overlap <50 bp | | rDNA | |
| chrXII | 455098 | 455243 | 145 | -1,326139823 | | | | Overlap <50 bp | | rDNA | |
| chrXII | 455098 | 455249 | 151 | -1,429443897 | | | | Overlap <50 bp | | rDNA | |
| chrXII | 455129 | 455293 | 164 | -1,279321555 | N12:455294 | 19,975 | +5*:YLR154W-C; | 5 | TERM | rDNA | |
| chrXII | 455138 | 455289 | 151 | -1,363275748 | N12:455294 | 19,975 | +5*:YLR154W-C; | 5 | TERM | rDNA | |
| chrXII | 455182 | 455338 | 156 | -1,456895455 | N12:455294 | 19,975 | +5*:YLR154W-C; | 5 | TERM | rDNA | |
| chrXII | 455217 | 455366 | 149 | -1,371046834 | N12:455294 | 19,975 | +5*:YLR154W-C; | 5 | TERM | rDNA | |
| chrXII | 455221 | 455395 | 174 | -1,957052919 | N12:455294 | 19,975 | +5*:YLR154W-C; | 5 | TERM | rDNA | |
| chrXII | 455247 | 455381 | 134 | -1,223894193 | N12:455294 | 19,975 | +5*:YLR154W-C; | 5 | TERM | rDNA | |
| chrXII | 455254 | 455401 | 147 | -1,265113843 | N12:455294 | 19,975 | +5*:YLR154W-C; | 5 | TERM | rDNA | |
| chrXII | 455255 | 455400 | 145 | -1,267794559 | N12:455294 | 19,975 | +5*:YLR154W-C; | 5 | TERM | rDNA | |
| chrXII | 455278 | 455423 | 145 | -1,573385181 | N12:455294 | 19,975 | +5*:YLR154W-C; | 5 | TERM | rDNA | |
| chrXII | 455297 | 455447 | 150 | -1,425154852 | N12:455294 | 19,975 | +5*:YLR154W-C; | 5 | TERM | rDNA | |
| chrXII | 455367 | 455512 | 145 | -1,111708771 | N12:455488 | 36,77 | | | | rDNA | |
| chrXII | 455367 | 455531 | 164 | -1,555598298 | N12:455488 | 36,77 | | | | rDNA | |
| chrXII | 455367 | 455549 | 182 | -1,359658714 | N12:455488 | 36,77 | | | | rDNA | |
| chrXII | 455377 | 455512 | 135 | -0,913926326 | N12:455488 | 36,77 | | | | rDNA | |
| chrXII | 455377 | 455531 | 154 | -1,005930195 | N12:455488 | 36,77 | | | | rDNA | |
| chrXII | 455377 | 455549 | 172 | -1,208723702 | N12:455488 | 36,77 | | | | rDNA | |
| chrXII | 455397 | 455503 | 106 | -0,676561875 | N12:455488 | 36,77 | | | | rDNA | |
| chrXII | 455397 | 455512 | 115 | -0,408277799 | N12:455488 | 36,77 | | | | rDNA | |

TABLE 3

| CHR | START | END | L(bp) | ΔLK | Nuc ID | Fuzziness | Gene ID | Gene body postion or intergene | Terminal | rDNA | Telomeric |
|---|---|---|---|---|---|---|---|---|---|---|---|
| chrXII | 455397 | 455531 | 134 | -1,175529786 | N12:455488 | 36,77 | | | | rDNA | |
| chrXII | 455397 | 455549 | 152 | -1,294722163 | N12:455488 | 36,77 | | | | rDNA | |
| chrXII | 455397 | 455568 | 171 | -1,581457993 | N12:455488 | 36,77 | | | | rDNA | |
| chrXII | 455410 | 455512 | 102 | -0,461780418 | N12:455488 | 36,77 | | | | rDNA | |
| chrXII | 455410 | 455549 | 139 | -1,495215213 | N12:455488 | 36,77 | | | | rDNA | |
| chrXII | 455410 | 455560 | 150 | -0,677159378 | N12:455488 | 36,77 | | | | rDNA | |
| chrXII | 455410 | 455568 | 158 | -1,158707772 | N12:455488 | 36,77 | | | | rDNA | |
| chrXII | 455410 | 455569 | 159 | -1,281606414 | N12:455488 | 36,77 | | | | rDNA | |
| chrXII | 455416 | 455571 | 155 | -1,607926045 | N12:455488 | 36,77 | | | | rDNA | |
| chrXII | 455422 | 455569 | 147 | -1,062262769 | N12:455488 | 36,77 | | | | rDNA | |
| chrXII | 455591 | 455751 | 160 | -1,645431743 | N12:455656 | 22,627 | | | | rDNA | |
| chrXII | 455701 | 455870 | 169 | -1,595963139 | N12:455829 | 32,808 | | | | rDNA | |
| chrXII | 455747 | 455895 | 148 | -1,087612301 | N12:455829 | 32,808 | | | | rDNA | |
| chrXII | 455749 | 455889 | 140 | -1,075128679 | N12:455829 | 32,808 | | | | rDNA | |
| chrXII | 455783 | 455958 | 175 | -1,704204164 | N12:455829 | 32,808 | | | | rDNA | |
| chrXII | 455869 | 456021 | 152 | -1,377701939 | N12:456042 | 22,48 | | | | rDNA | |
| chrXII | 455903 | 456067 | 164 | -1,389719575 | N12:456042 | 22,48 | | | | rDNA | |
| chrXII | 455918 | 456039 | 121 | -1,280252299 | N12:456042 | 22,48 | | | | rDNA | |
| chrXII | 455926 | 456086 | 160 | -1,364669614 | N12:456042 | 22,48 | | | | rDNA | |
| chrXII | 456022 | 456173 | 151 | -1,150091008 | N12:456042 | 22,48 | | | | rDNA | |
| chrXII | 456022 | 456205 | 183 | -1,527096809 | N12:456042 | 22,48 | | | | rDNA | |
| chrXII | 456022 | 456205 | 183 | -1,527096809 | N12:456220 | 28 | | | | rDNA | |
| chrXII | 456022 | 456211 | 189 | -1,283453682 | N12:456042 | 22,48 | | | | rDNA | |
| chrXII | 456022 | 456211 | 189 | -1,283453682 | N12:456220 | 28 | | | | rDNA | |
| chrXII | 456022 | 456220 | 198 | -1,519940886 | N12:456042 | 22,48 | | | | rDNA | |
| chrXII | 456022 | 456220 | 198 | -1,519940886 | N12:456220 | 28 | | | | rDNA | |
| chrXII | 456025 | 456173 | 148 | -0,981641815 | N12:456042 | 22,48 | | | | rDNA | |
| chrXII | 456025 | 456182 | 157 | -1,121232648 | N12:456042 | 22,48 | | | | rDNA | |
| chrXII | 456042 | 456173 | 131 | -0,562287906 | N12:456042 | 22,48 | | | | rDNA | |
| chrXII | 456045 | 456173 | 128 | -0,893016349 | N12:456042 | 22,48 | | | | rDNA | |
| chrXII | 456045 | 456175 | 130 | -1,027711176 | N12:456042 | 22,48 | | | | rDNA | |
| chrXII | 456045 | 456205 | 160 | -1,336056619 | N12:456042 | 22,48 | | | | rDNA | |
| chrXII | 456045 | 456205 | 160 | -1,336056619 | N12:456220 | 28 | | | | rDNA | |
| chrXII | 456056 | 456173 | 117 | -0,738642625 | N12:456042 | 22,48 | | | | rDNA | |
| chrXII | 456056 | 456175 | 119 | -0,781885292 | N12:456042 | 22,48 | | | | rDNA | |
| chrXII | 456056 | 456205 | 149 | -1,261128826 | N12:456042 | 22,48 | | | | rDNA | |
| chrXII | 456056 | 456205 | 149 | -1,261128826 | N12:456220 | 28 | | | | rDNA | |
| chrXII | 456056 | 456211 | 155 | -1,477494129 | N12:456042 | 22,48 | | | | rDNA | |
| chrXII | 456056 | 456211 | 155 | -1,477494129 | N12:456220 | 28 | | | | rDNA | |
| chrXII | 456056 | 456220 | 164 | -1,410412803 | N12:456042 | 22,48 | | | | rDNA | |
| chrXII | 456056 | 456220 | 164 | -1,410412803 | N12:456220 | 28 | | | | rDNA | |
| chrXII | 456063 | 456215 | 152 | -1,046412445 | N12:456042 | 22,48 | | | | rDNA | |
| chrXII | 456063 | 456215 | 152 | -1,046412445 | N12:456220 | 28 | | | | rDNA | |
| chrXII | 456064 | 456214 | 150 | -1,125587004 | N12:456042 | 22,48 | | | | rDNA | |
| chrXII | 456064 | 456214 | 150 | -1,125587004 | N12:456220 | 28 | | | | rDNA | |
| chrXII | 456066 | 456173 | 107 | -0,415858579 | | | | Overlap <50 bp | | rDNA | |
| chrXII | 456067 | 456205 | 138 | -1,137025389 | N12:456220 | 28 | | | | rDNA | |
| chrXII | 456067 | 456211 | 144 | -0,9710934 | N12:456220 | 28 | | | | rDNA | |
| chrXII | 456067 | 456215 | 148 | -1,213205785 | N12:456220 | 28 | | | | rDNA | |
| chrXII | 456067 | 456220 | 153 | -1,081665104 | N12:456220 | 28 | | | | rDNA | |
| chrXII | 456067 | 456266 | 199 | -1,719861336 | N12:456220 | 28 | | | | rDNA | |
| chrXII | 456080 | 456173 | 93 | -0,480139128 | | | | Overlap <50 bp | | rDNA | |
| chrXII | 456080 | 456205 | 125 | -0,634883535 | N12:456220 | 28 | | | | rDNA | |
| chrXII | 456080 | 456211 | 131 | -0,72285898 | N12:456220 | 28 | | | | rDNA | |
| chrXII | 456080 | 456220 | 140 | -0,956255938 | N12:456220 | 28 | | | | rDNA | |
| chrXII | 456084 | 456215 | 131 | -1,138923383 | N12:456220 | 28 | | | | rDNA | |
| chrXII | 456084 | 456243 | 159 | -1,309482124 | N12:456220 | 28 | | | | rDNA | |
| chrXII | 456084 | 456266 | 182 | -1,472844182 | N12:456220 | 28 | | | | rDNA | |
| chrXII | 456115 | 456215 | 100 | -0,693989257 | N12:456220 | 28 | | | | rDNA | |
| chrXII | 456115 | 456243 | 128 | -1,215431524 | N12:456220 | 28 | | | | rDNA | |
| chrXII | 456115 | 456266 | 151 | -1,14951293 | N12:456220 | 28 | | | | rDNA | |
| chrXII | 456115 | 456281 | 166 | -1,497622718 | N12:456220 | 28 | | | | rDNA | |
| chrXII | 456120 | 456290 | 170 | -1,281636057 | N12:456220 | 28 | | | | rDNA | |
| chrXII | 456124 | 456215 | 91 | -0,785201802 | N12:456220 | 28 | | | | rDNA | |
| chrXII | 456124 | 456266 | 142 | -1,279910624 | N12:456220 | 28 | | | | rDNA | |
| chrXII | 456124 | 456281 | 157 | -1,429016872 | N12:456220 | 28 | | | | rDNA | |
| chrXII | 456124 | 456333 | 209 | -1,632864062 | N12:456220 | 28 | | | | rDNA | |
| chrXII | 456181 | 456266 | 85 | -0,590663226 | N12:456220 | 28 | | | | rDNA | |
| chrXII | 456181 | 456281 | 100 | -0,729809544 | N12:456220 | 28 | | | | rDNA | |
| chrXII | 456181 | 456287 | 106 | -1,202175501 | N12:456220 | 28 | | | | rDNA | |
| chrXII | 456181 | 456306 | 125 | -0,813543047 | N12:456220 | 28 | | | | rDNA | |
| chrXII | 456181 | 456310 | 129 | -0,96834544 | N12:456220 | 28 | | | | rDNA | |
| chrXII | 456181 | 456329 | 148 | -1,34981526 | N12:456220 | 28 | | | | rDNA | |
| chrXII | 456181 | 456332 | 151 | -1,840742954 | N12:456220 | 28 | | | | rDNA | |
| chrXII | 456181 | 456333 | 152 | -1,297586545 | N12:456220 | 28 | | | | rDNA | |
| chrXII | 456181 | 456336 | 155 | -1,44791953 | N12:456220 | 28 | | | | rDNA | |
| chrXII | 456181 | 456349 | 168 | -1,076720451 | N12:456220 | 28 | | | | rDNA | |
| chrXII | 456181 | 456387 | 206 | -1,413400921 | N12:456220 | 28 | | | | rDNA | |
| chrXII | 456186 | 456333 | 147 | -1,627256815 | N12:456220 | 28 | | | | rDNA | |
| chrXII | 456187 | 456333 | 146 | -1,59771293 | N12:456220 | 28 | | | | rDNA | |
| chrXII | 456187 | 456349 | 162 | -1,316137336 | N12:456220 | 28 | | | | rDNA | |
| chrXII | 456194 | 456333 | 139 | -1,25889814 | N12:456220 | 28 | | | | rDNA | |
| chrXII | 456194 | 456336 | 142 | -1,013500075 | N12:456220 | 28 | | | | rDNA | |
| chrXII | 456213 | 456319 | 106 | -0,969153116 | N12:456220 | 28 | | | | rDNA | |
| chrXII | 456213 | 456320 | 107 | -0,815628147 | N12:456220 | 28 | | | | rDNA | |
| chrXII | 456213 | 456321 | 108 | -0,984425382 | N12:456220 | 28 | | | | rDNA | |
| chrXII | 456213 | 456333 | 120 | -1,147941248 | N12:456220 | 28 | | | | rDNA | |
| chrXII | 456213 | 456336 | 123 | -1,009917497 | N12:456220 | 28 | | | | rDNA | |
| chrXII | 456213 | 456349 | 136 | -1,569253966 | N12:456220 | 28 | | | | rDNA | |
| chrXII | 456213 | 456352 | 139 | -1,254217269 | N12:456220 | 28 | | | | rDNA | |
| chrXII | 456213 | 456387 | 174 | -1,309745443 | N12:456220 | 28 | | | | rDNA | |
| chrXII | 456213 | 456419 | 206 | -1,7028479 | N12:456220 | 28 | | | | rDNA | |
| chrXII | 456213 | 456419 | 206 | -1,7028479 | N12:456421 | 19,858 | | | | rDNA | |
| chrXII | 456221 | 456386 | 165 | -1,085235348 | N12:456220 | 28 | | | | rDNA | |
| chrXII | 456240 | 456333 | 93 | -0,57545793 | N12:456220 | 28 | | | | rDNA | |
| chrXII | 456240 | 456386 | 146 | -0,987663683 | N12:456220 | 28 | | | | rDNA | |
| chrXII | 456240 | 456387 | 147 | -1,181208107 | N12:456220 | 28 | | | | rDNA | |
| chrXII | 456240 | 456419 | 179 | -1,388089483 | N12:456220 | 28 | | | | rDNA | |
| chrXII | 456240 | 456419 | 179 | -1,388089483 | N12:456421 | 19,858 | | | | rDNA | |
| chrXII | 456240 | 456447 | 207 | -1,439992588 | N12:456220 | 28 | | | | rDNA | |

TABLE 3

| CHR | START | END | L(bp) | ΔLK | Nuc ID | Fuzziness | Gene ID | Gene body postion or intergene | Terminal | rDNA | Telomeric |
|---|---|---|---|---|---|---|---|---|---|---|---|
| chrXII | 456240 | 456447 | 207 | -1,439992588 | N12:456421 | 19,858 | | | | rDNA | |
| chrXII | 456288 | 456387 | 99 | -0,589925161 | | | | Overlap <50 bp | | rDNA | |
| chrXII | 456288 | 456395 | 107 | -1,355476424 | | | | Overlap <50 bp | | rDNA | |
| chrXII | 456288 | 456419 | 131 | -1,127127751 | N12:456421 | 19,858 | | | | rDNA | |
| chrXII | 456288 | 456447 | 159 | -1,296588691 | N12:456421 | 19,858 | | | | rDNA | |
| chrXII | 456288 | 456457 | 169 | -1,368998002 | N12:456421 | 19,858 | | | | rDNA | |
| chrXII | 456288 | 456471 | 183 | -1,902593388 | N12:456421 | 19,858 | | | | rDNA | |
| chrXII | 456290 | 456419 | 129 | -1,235364681 | N12:456421 | 19,858 | | | | rDNA | |
| chrXII | 456290 | 456447 | 157 | -1,663206413 | N12:456421 | 19,858 | | | | rDNA | |
| chrXII | 456290 | 456471 | 181 | -1,776796652 | N12:456421 | 19,858 | | | | rDNA | |
| chrXII | 456298 | 456447 | 149 | -1,193647556 | N12:456421 | 19,858 | | | | rDNA | |
| chrXII | 456310 | 456417 | 107 | -0,36959189 | N12:456421 | 19,858 | | | | rDNA | |
| chrXII | 456310 | 456419 | 109 | -0,70786376 | N12:456421 | 19,858 | | | | rDNA | |
| chrXII | 456310 | 456447 | 137 | -1,347510747 | N12:456421 | 19,858 | | | | rDNA | |
| chrXII | 456310 | 456471 | 161 | -1,354361979 | N12:456421 | 19,858 | | | | rDNA | |
| chrXII | 456361 | 456447 | 86 | -1,328979226 | N12:456421 | 19,858 | | | | rDNA | |
| chrXII | 456361 | 456466 | 105 | -1,716721808 | N12:456421 | 19,858 | | | | rDNA | |
| chrXII | 456361 | 456468 | 107 | -1,522622032 | N12:456421 | 19,858 | | | | rDNA | |
| chrXII | 456361 | 456470 | 109 | -1,421743856 | N12:456421 | 19,858 | | | | rDNA | |
| chrXII | 456361 | 456471 | 110 | -1,53194537 | N12:456421 | 19,858 | | | | rDNA | |
| chrXII | 456388 | 456471 | 83 | -1,076338967 | N12:456421 | 19,858 | | | | rDNA | |
| chrXII | 456388 | 456551 | 163 | -1,271009504 | N12:456421 | 19,858 | | | | rDNA | |
| chrXII | 456388 | 456551 | 163 | -1,271009504 | N12:456568 | 7,071 | | | | rDNA | |
| chrXII | 456444 | 456594 | 150 | -1,137932227 | N12:456421 | 19,858 | | | | rDNA | |
| chrXII | 456444 | 456594 | 150 | -1,137932227 | N12:456568 | 7,071 | | | | rDNA | |
| chrXII | 456444 | 456595 | 151 | -1,404318252 | N12:456421 | 19,858 | | | | rDNA | |
| chrXII | 456444 | 456595 | 151 | -1,404318252 | N12:456568 | 7,071 | | | | rDNA | |
| chrXII | 456444 | 456619 | 175 | -1,667758771 | N12:456421 | 19,858 | | | | rDNA | |
| chrXII | 456444 | 456619 | 175 | -1,667758771 | N12:456568 | 7,071 | | | | rDNA | |
| chrXII | 456445 | 456595 | 150 | -1,618067859 | N12:456568 | 7,071 | | | | rDNA | |
| chrXII | 456447 | 456594 | 147 | -1,282503292 | N12:456568 | 7,071 | | | | rDNA | |
| chrXII | 456447 | 456595 | 148 | -1,564730455 | N12:456568 | 7,071 | | | | rDNA | |
| chrXII | 456447 | 456608 | 161 | -1,845538213 | N12:456568 | 7,071 | | | | rDNA | |
| chrXII | 456447 | 456619 | 172 | -1,602807876 | N12:456568 | 7,071 | | | | rDNA | |
| chrXII | 456479 | 456585 | 106 | -0,814204026 | N12:456568 | 7,071 | | | | rDNA | |
| chrXII | 456479 | 456594 | 115 | -0,736398515 | N12:456568 | 7,071 | | | | rDNA | |
| chrXII | 456479 | 456608 | 129 | -0,934613757 | N12:456568 | 7,071 | | | | rDNA | |
| chrXII | 456479 | 456609 | 130 | -1,110704052 | N12:456568 | 7,071 | | | | rDNA | |
| chrXII | 456479 | 456653 | 174 | -1,273201422 | N12:456568 | 7,071 | | | | rDNA | |
| chrXII | 456483 | 456594 | 111 | -1,171021328 | N12:456568 | 7,071 | | | | rDNA | |
| chrXII | 456483 | 456608 | 125 | -1,114631272 | N12:456568 | 7,071 | | | | rDNA | |
| chrXII | 456483 | 456609 | 126 | -1,131675499 | N12:456568 | 7,071 | | | | rDNA | |
| chrXII | 456493 | 456599 | 106 | -0,569145552 | N12:456568 | 7,071 | | | | rDNA | |
| chrXII | 456493 | 456608 | 115 | -0,727160737 | N12:456568 | 7,071 | | | | rDNA | |
| chrXII | 456493 | 456653 | 160 | -1,482686853 | N12:456568 | 7,071 | | | | rDNA | |
| chrXII | 456508 | 456594 | 86 | -0,633894362 | N12:456568 | 7,071 | | | | rDNA | |
| chrXII | 456508 | 456608 | 100 | -0,592340289 | N12:456568 | 7,071 | | | | rDNA | |
| chrXII | 456508 | 456642 | 134 | -1,281826631 | N12:456568 | 7,071 | | | | rDNA | |
| chrXII | 456508 | 456653 | 145 | -1,14597173 | N12:456568 | 7,071 | | | | rDNA | |
| chrXII | 456508 | 456670 | 162 | -1,267302287 | N12:456568 | 7,071 | | | | rDNA | |
| chrXII | 456524 | 456695 | 171 | -1,609925187 | N12:456568 | 7,071 | | | | rDNA | |
| chrXII | 456587 | 456738 | 151 | -1,770478345 | N12:456568 | 7,071 | | | | rDNA | |
| chrXII | 456587 | 456738 | 151 | -1,770478345 | N12:456730 | 37,899 | | | | rDNA | |
| chrXII | 456588 | 456751 | 163 | -1,017805662 | N12:456568 | 7,071 | | | | rDNA | |
| chrXII | 456588 | 456751 | 163 | -1,017805662 | N12:456730 | 37,899 | | | | rDNA | |
| chrXII | 456594 | 456751 | 157 | -1,541495057 | N12:456730 | 37,899 | | | | rDNA | |
| chrXII | 456616 | 456772 | 156 | -1,132985005 | N12:456730 | 37,899 | | | | rDNA | |
| chrXII | 456634 | 456751 | 117 | -0,78510286 | N12:456730 | 37,899 | | | | rDNA | |
| chrXII | 456634 | 456762 | 128 | -1,123108749 | N12:456730 | 37,899 | | | | rDNA | |
| chrXII | 456634 | 456781 | 147 | -1,300825975 | N12:456730 | 37,899 | | | | rDNA | |
| chrXII | 456649 | 456803 | 154 | -1,366873488 | N12:456730 | 37,899 | | | | rDNA | |
| chrXII | 456653 | 456762 | 109 | -0,546662435 | N12:456730 | 37,899 | | | | rDNA | |
| chrXII | 456653 | 456803 | 150 | -0,965012152 | N12:456730 | 37,899 | | | | rDNA | |
| chrXII | 456733 | 456869 | 136 | -1,137813736 | N12:456730 | 37,899 | | | | rDNA | |
| chrXII | 456733 | 456869 | 136 | -1,137813736 | N12:456891 | 32,655 | | | | rDNA | |
| chrXII | 456766 | 456920 | 154 | -1,33740753 | N12:456891 | 32,655 | | | | rDNA | |
| chrXII | 456780 | 456887 | 107 | -1,745208602 | N12:456891 | 32,655 | | | | rDNA | |
| chrXII | 456780 | 456890 | 110 | -1,601302824 | N12:456891 | 32,655 | | | | rDNA | |
| chrXII | 456780 | 456920 | 140 | -2,173661016 | N12:456891 | 32,655 | | | | rDNA | |
| chrXII | 456794 | 456903 | 109 | -0,85538798 | N12:456891 | 32,655 | | | | rDNA | |
| chrXII | 456795 | 456903 | 108 | -0,9215189 | N12:456891 | 32,655 | | | | rDNA | |
| chrXII | 456795 | 456904 | 109 | -1,015424336 | N12:456891 | 32,655 | | | | rDNA | |
| chrXII | 456795 | 456905 | 110 | -1,234605321 | N12:456891 | 32,655 | | | | rDNA | |
| chrXII | 456797 | 456890 | 93 | -1,551908122 | N12:456891 | 32,655 | | | | rDNA | |
| chrXII | 456797 | 456920 | 123 | -1,885943385 | N12:456891 | 32,655 | | | | rDNA | |
| chrXII | 456803 | 456959 | 156 | -1,418588796 | N12:456891 | 32,655 | | | | rDNA | |
| chrXII | 456804 | 456911 | 107 | -1,188188098 | N12:456891 | 32,655 | | | | rDNA | |
| chrXII | 456804 | 456920 | 116 | -1,299414435 | N12:456891 | 32,655 | | | | rDNA | |
| chrXII | 456804 | 456960 | 156 | -1,79560905 | N12:456891 | 32,655 | | | | rDNA | |
| chrXII | 456817 | 456920 | 103 | -1,073304825 | N12:456891 | 32,655 | | | | rDNA | |
| chrXII | 456817 | 456971 | 154 | -1,414587486 | N12:456891 | 32,655 | | | | rDNA | |
| chrXII | 456938 | 457104 | 166 | -1,394260883 | N12:457054 | 34,078 | | | | rDNA | |
| chrXII | 456984 | 457118 | 134 | -1,046364252 | N12:457054 | 34,078 | | | | rDNA | |
| chrXII | 456984 | 457127 | 143 | -1,407129479 | N12:457054 | 34,078 | | | | rDNA | |
| chrXII | 456984 | 457130 | 146 | -1,627737478 | N12:457054 | 34,078 | | | | rDNA | |
| chrXII | 456984 | 457140 | 156 | -1,627988076 | N12:457054 | 34,078 | | | | rDNA | |
| chrXII | 456990 | 457127 | 137 | -1,357231234 | N12:457054 | 34,078 | | | | rDNA | |
| chrXII | 456990 | 457140 | 150 | -1,265434703 | N12:457054 | 34,078 | | | | rDNA | |
| chrXII | 457005 | 457166 | 161 | -1,707329845 | N12:457054 | 34,078 | | | | rDNA | |
| chrXII | 457021 | 457127 | 106 | -1,729888188 | N12:457054 | 34,078 | | | | rDNA | |
| chrXII | 457022 | 457205 | 183 | -2,094253224 | N12:457054 | 34,078 | | | | rDNA | |
| chrXII | 457022 | 457205 | 183 | -2,094253224 | N12:457213 | 27,713 | | | | rDNA | |
| chrXII | 457047 | 457207 | 160 | -1,440694378 | N12:457054 | 34,078 | | | | rDNA | |
| chrXII | 457047 | 457207 | 160 | -1,440694378 | N12:457213 | 27,713 | | | | rDNA | |
| chrXII | 457070 | 457200 | 130 | -1,415691373 | N12:457054 | 34,078 | | | | rDNA | |
| chrXII | 457070 | 457200 | 130 | -1,415691373 | N12:457213 | 27,713 | | | | rDNA | |
| chrXII | 457083 | 457217 | 134 | -1,461019882 | N12:457213 | 27,713 | | | | rDNA | |
| chrXII | 457115 | 457248 | 133 | -1,091597452 | N12:457213 | 27,713 | | | | rDNA | |
| chrXII | 457140 | 457305 | 165 | -1,283712949 | N12:457213 | 27,713 | | | | rDNA | |
| chrXII | 457140 | 457312 | 172 | -1,660956141 | N12:457213 | 27,713 | | | | rDNA | |

TABLE 3

| CHR | START | END | L(bp) | ΔLK | Nuc ID | Fuzziness | Gene ID | Gene body postion or intergene | Terminal | rDNA | Telomeric |
|---|---|---|---|---|---|---|---|---|---|---|---|
| chrXII | 457143 | 457248 | 105 | -0,897134712 | N12:457213 | 27,713 | | | | rDNA | |
| chrXII | 457143 | 457279 | 136 | -1,106421363 | N12:457213 | 27,713 | | | | rDNA | |
| chrXII | 457143 | 457316 | 173 | -1,428495619 | N12:457213 | 27,713 | | | | rDNA | |
| chrXII | 457152 | 457305 | 153 | -1,125429647 | N12:457213 | 27,713 | | | | rDNA | |
| chrXII | 457152 | 457312 | 160 | -1,471274216 | N12:457213 | 27,713 | | | | rDNA | |
| chrXII | 457153 | 457248 | 95 | -0,822754884 | N12:457213 | 27,713 | | | | rDNA | |
| chrXII | 457153 | 457279 | 126 | -0,982601006 | N12:457213 | 27,713 | | | | rDNA | |
| chrXII | 457153 | 457312 | 159 | -1,482818599 | N12:457213 | 27,713 | | | | rDNA | |
| chrXII | 457153 | 457316 | 163 | -1,300547407 | N12:457213 | 27,713 | | | | rDNA | |
| chrXII | 457153 | 457317 | 164 | -1,858981961 | N12:457213 | 27,713 | | | | rDNA | |
| chrXII | 457154 | 457312 | 158 | -1,432625836 | N12:457213 | 27,713 | | | | rDNA | |
| chrXII | 457154 | 457316 | 162 | -1,370748074 | N12:457213 | 27,713 | | | | rDNA | |
| chrXII | 457187 | 457316 | 129 | -1,034260647 | N12:457213 | 27,713 | | | | rDNA | |
| chrXII | 457187 | 457317 | 130 | -1,002170958 | N12:457213 | 27,713 | | | | rDNA | |
| chrXII | 457198 | 457346 | 148 | -1,157632343 | N12:457213 | 27,713 | | | | rDNA | |
| chrXII | 457198 | 457365 | 167 | -1,060785429 | N12:457213 | 27,713 | | | | rDNA | |
| chrXII | 457212 | 457365 | 153 | -1,073949231 | N12:457213 | 27,713 | | | | rDNA | |
| chrXII | 457214 | 457363 | 149 | -0,974284083 | N12:457213 | 27,713 | | | | rDNA | |
| chrXII | 457219 | 457312 | 93 | -0,970833668 | N12:457213 | 27,713 | | | | rDNA | |
| chrXII | 457219 | 457346 | 127 | -1,139096961 | N12:457213 | 27,713 | | | | rDNA | |
| chrXII | 457219 | 457363 | 144 | -1,378376444 | N12:457213 | 27,713 | | | | rDNA | |
| chrXII | 457219 | 457365 | 146 | -1,286451175 | N12:457213 | 27,713 | | | | rDNA | |
| chrXII | 457219 | 457422 | 203 | -1,915949793 | N12:457213 | 27,713 | | | | rDNA | |
| chrXII | 457227 | 457316 | 89 | -0,341855947 | N12:457213 | 27,713 | | | | rDNA | |
| chrXII | 457227 | 457363 | 136 | -1,014263317 | N12:457213 | 27,713 | | | | rDNA | |
| chrXII | 457227 | 457395 | 168 | -1,770082218 | N12:457213 | 27,713 | | | | rDNA | |
| chrXII | 457230 | 457395 | 165 | -1,996545303 | N12:457213 | 27,713 | | | | rDNA | |
| chrXII | 457230 | 457401 | 171 | -1,268737505 | N12:457213 | 27,713 | | | | rDNA | |
| chrXII | 457248 | 457395 | 147 | -1,659630334 | | | | Overlap <50 bp | | rDNA | |
| chrXII | 457248 | 457401 | 153 | -0,981301511 | | | | Overlap <50 bp | | rDNA | |
| chrXII | 457255 | 457365 | 110 | -0,294778099 | | | | Overlap <50 bp | | rDNA | |
| chrXII | 457255 | 457407 | 152 | -1,102372052 | | | | Overlap <50 bp | | rDNA | |
| chrXII | 457255 | 457422 | 167 | -1,454556303 | | | | Overlap <50 bp | | rDNA | |
| chrXII | 457266 | 457365 | 99 | -0,544665328 | | | | Overlap <50 bp | | rDNA | |
| chrXII | 457266 | 457407 | 141 | -1,016402178 | | | | Overlap <50 bp | | rDNA | |
| chrXII | 457266 | 457422 | 156 | -1,129833572 | | | | Overlap <50 bp | | rDNA | |
| chrXII | 457295 | 457378 | 83 | -1,337960809 | | | | Overlap <50 bp | | rDNA | |
| chrXII | 457295 | 457384 | 89 | -1,01934915 | | | | Overlap <50 bp | | rDNA | |
| chrXII | 457295 | 457390 | 95 | -1,293055282 | | | | Overlap <50 bp | | rDNA | |
| chrXII | 457295 | 457391 | 96 | -1,479094384 | | | | Overlap <50 bp | | rDNA | |
| chrXII | 457295 | 457393 | 98 | -1,498740471 | | | | Overlap <50 bp | | rDNA | |
| chrXII | 457295 | 457394 | 99 | -1,705260463 | | | | Overlap <50 bp | | rDNA | |
| chrXII | 457295 | 457395 | 100 | -1,361307624 | | | | Overlap <50 bp | | rDNA | |
| chrXII | 457295 | 457397 | 102 | -1,699109196 | | | | Overlap <50 bp | | rDNA | |
| chrXII | 457295 | 457401 | 106 | -1,040004191 | | | | Overlap <50 bp | | rDNA | |
| chrXII | 457295 | 457402 | 107 | -0,991810719 | | | | Overlap <50 bp | | rDNA | |
| chrXII | 457337 | 457476 | 139 | -1,375582036 | N12:457494 | 9,899 | | | | rDNA | |
| chrXII | 457337 | 457481 | 144 | -0,959935821 | N12:457494 | 9,899 | | | | rDNA | |
| chrXII | 457360 | 457476 | 116 | -1,174417981 | N12:457494 | 9,899 | | | | rDNA | |
| chrXII | 457360 | 457481 | 121 | -0,639633385 | N12:457494 | 9,899 | | | | rDNA | |
| chrXII | 457360 | 457530 | 170 | -1,461578819 | N12:457494 | 9,899 | | | | rDNA | |
| chrXII | 457370 | 457476 | 106 | -1,496843022 | N12:457494 | 9,899 | | | | rDNA | |
| chrXII | 457372 | 457476 | 104 | -1,8046605 | N12:457494 | 9,899 | | | | rDNA | |
| chrXII | 457420 | 457563 | 143 | -1,278611526 | N12:457494 | 9,899 | | | | rDNA | |
| chrXII | 457420 | 457567 | 147 | -1,24833696 | N12:457494 | 9,899 | | | | rDNA | |
| chrXII | 457420 | 457571 | 151 | -1,17514218 | N12:457494 | 9,899 | | | | rDNA | |
| chrXII | 457420 | 457572 | 152 | -0,896145818 | N12:457494 | 9,899 | | | | rDNA | |
| chrXII | 457437 | 457592 | 155 | -1,349422117 | N12:457494 | 9,899 | | | | rDNA | |
| chrXII | 457440 | 457593 | 153 | -1,405659838 | N12:457494 | 9,899 | | | | rDNA | |
| chrXII | 457540 | 457654 | 114 | -1,509183439 | | | | Overlap <50 bp | | rDNA | |
| chrXII | 457553 | 457706 | 153 | -1,328591869 | N12:457682 | 13,229 | | | | rDNA | |
| chrXII | 457569 | 457736 | 167 | -1,619046263 | N12:457682 | 13,229 | | | | rDNA | |
| chrXII | 457569 | 457741 | 172 | -1,59854435 | N12:457682 | 13,229 | | | | rDNA | |
| chrXII | 457600 | 457736 | 136 | -1,229052112 | N12:457682 | 13,229 | | | | rDNA | |
| chrXII | 457600 | 457739 | 139 | -0,572599256 | N12:457682 | 13,229 | | | | rDNA | |
| chrXII | 457600 | 457741 | 141 | -1,596682821 | N12:457682 | 13,229 | | | | rDNA | |
| chrXII | 457606 | 457763 | 157 | -0,923770464 | N12:457682 | 13,229 | | | | rDNA | |
| chrXII | 457620 | 457773 | 153 | -1,384555884 | N12:457682 | 13,229 | | | | rDNA | |
| chrXII | 457620 | 457787 | 167 | -1,525940035 | N12:457682 | 13,229 | | | | rDNA | |
| chrXII | 457622 | 457771 | 149 | -1,521147421 | N12:457682 | 13,229 | | | | rDNA | |
| chrXII | 457634 | 457757 | 123 | -1,203859799 | N12:457682 | 13,229 | | | | rDNA | |
| chrXII | 457650 | 457805 | 155 | -1,786824643 | N12:457682 | 13,229 | | | | rDNA | |
| chrXII | 457667 | 457819 | 152 | -1,541599981 | N12:457682 | 13,229 | | | | rDNA | |
| chrXII | 457803 | 457959 | 156 | -1,116720891 | N12:457885 | 36,199 | | | | rDNA | |
| chrXII | 457816 | 457974 | 158 | -1,120690586 | N12:457885 | 36,199 | | | | rDNA | |
| chrXII | 457820 | 457953 | 133 | -0,906236552 | N12:457885 | 36,199 | | | | rDNA | |
| chrXII | 457872 | 458023 | 151 | -1,308748766 | N12:457885 | 36,199 | | | | rDNA | |
| chrXII | 457945 | 458104 | 159 | -1,43931108 | | | | Overlap <50 bp | | rDNA | |
| chrXII | 457949 | 458059 | 110 | -0,740069368 | | | | Overlap <50 bp | | rDNA | |
| chrXII | 457949 | 458101 | 152 | -1,210234772 | | | | Overlap <50 bp | | rDNA | |
| chrXII | 457950 | 458101 | 151 | -1,138303345 | | | | Overlap <50 bp | | rDNA | |
| chrXII | 457950 | 458103 | 153 | -1,223267332 | | | | Overlap <50 bp | | rDNA | |
| chrXII | 457959 | 458110 | 151 | -1,18700606 | | | | Overlap <50 bp | | rDNA | |
| chrXII | 457976 | 458059 | 83 | -0,691302863 | | | | Overlap <50 bp | | rDNA | |
| chrXII | 457976 | 458119 | 143 | -1,016475699 | | | | Overlap <50 bp | | rDNA | |
| chrXII | 457983 | 458089 | 106 | -0,576092235 | | | | Overlap <50 bp | | rDNA | |
| chrXII | 457983 | 458101 | 118 | -0,778624695 | | | | Overlap <50 bp | | rDNA | |
| chrXII | 457983 | 458103 | 120 | -1,002230921 | | | | Overlap <50 bp | | rDNA | |
| chrXII | 457983 | 458137 | 154 | -1,451691684 | N12:458145 | 37,162 | | | | rDNA | |
| chrXII | 457986 | 458117 | 131 | -0,693143423 | | | | Overlap <50 bp | | rDNA | |
| chrXII | 458025 | 458167 | 142 | -1,328913323 | N12:458145 | 37,162 | | | | rDNA | |
| chrXII | 458055 | 458137 | 82 | -0,826 | N12:458145 | 37,162 | | | | rDNA | |
| chrXII | 458055 | 458206 | 151 | -1,144107649 | N12:458145 | 37,162 | | | | rDNA | |
| chrXII | 458079 | 458227 | 148 | -1,159596627 | N12:458145 | 37,162 | | | | rDNA | |
| chrXII | 458087 | 458231 | 144 | -1,054807619 | N12:458145 | 37,162 | | | | rDNA | |
| chrXII | 458127 | 458277 | 150 | -1,437385509 | N12:458145 | 37,162 | | | | rDNA | |
| chrXII | 458130 | 458293 | 163 | -1,259887156 | N12:458145 | 37,162 | | | | rDNA | |
| chrXII | 458134 | 458272 | 138 | -0,964316105 | N12:458145 | 37,162 | | | | rDNA | |

| CHR | START | END | L(bp) | ΔLK | Nuc ID | Fuzziness | Gene ID | Gene body postion or intergene | Terminal | rDNA | Telomeric |
|---|---|---|---|---|---|---|---|---|---|---|---|
| chrXII | 458134 | 458277 | 143 | -1,166514995 | N12:458145 | 37,162 | | | | rDNA | |
| chrXII | 458136 | 458295 | 159 | -1,300142728 | N12:458145 | 37,162 | | | | rDNA | |
| chrXII | 458143 | 458291 | 148 | -1,118032353 | N12:458145 | 37,162 | | | | rDNA | |
| chrXII | 458166 | 458321 | 155 | -1,476247261 | N12:458145 | 37,162 | | | | rDNA | |
| chrXII | 458166 | 458321 | 155 | -1,476247261 | N12:458325 | 64,042 | | | | rDNA | |
| chrXII | 458166 | 458322 | 156 | -1,264833973 | N12:458145 | 37,162 | | | | rDNA | |
| chrXII | 458166 | 458322 | 156 | -1,264833973 | N12:458325 | 64,042 | | | | rDNA | |
| chrXII | 458167 | 458322 | 155 | -1,180999212 | N12:458145 | 37,162 | | | | rDNA | |
| chrXII | 458167 | 458322 | 155 | -1,180999212 | N12:458325 | 64,042 | | | | rDNA | |
| chrXII | 458177 | 458322 | 145 | -1,227979635 | N12:458325 | 64,042 | | | | rDNA | |
| chrXII | 458177 | 458327 | 150 | -1,511203035 | N12:458325 | 64,042 | | | | rDNA | |
| chrXII | 458177 | 458350 | 173 | -1,48923011 | N12:458325 | 64,042 | | | | rDNA | |
| chrXII | 458179 | 458327 | 148 | -0,929336692 | N12:458325 | 64,042 | | | | rDNA | |
| chrXII | 458191 | 458321 | 130 | -0,988652948 | N12:458325 | 64,042 | | | | rDNA | |
| chrXII | 458191 | 458352 | 161 | -1,162492706 | N12:458325 | 64,042 | | | | rDNA | |
| chrXII | 458195 | 458322 | 127 | -1,174457517 | N12:458325 | 64,042 | | | | rDNA | |
| chrXII | 458195 | 458327 | 132 | -1,126380519 | N12:458325 | 64,042 | | | | rDNA | |
| chrXII | 458195 | 458350 | 155 | -1,125510608 | N12:458325 | 64,042 | | | | rDNA | |
| chrXII | 458204 | 458350 | 146 | -1,445133047 | N12:458325 | 64,042 | | | | rDNA | |
| chrXII | 458204 | 458380 | 176 | -1,477894642 | N12:458325 | 64,042 | | | | rDNA | |
| chrXII | 458291 | 458451 | 160 | -1,104411188 | N12:458325 | 64,042 | | | | rDNA | |
| chrXII | 458297 | 458451 | 154 | -1,213084623 | N12:458325 | 64,042 | | | | rDNA | |
| chrXII | 458298 | 458451 | 153 | -1,206093527 | N12:458325 | 64,042 | | | | rDNA | |
| chrXII | 458319 | 458472 | 153 | -1,082883441 | N12:458325 | 64,042 | | | | rDNA | |
| chrXII | 458338 | 458492 | 154 | -1,240288389 | N12:458325 | 64,042 | | | | rDNA | |
| chrXII | 458356 | 458493 | 137 | -1,15964863 | | | | Overlap <50 bp | | rDNA | |
| chrXII | 458356 | 458508 | 152 | -1,113959374 | N12:458524 | 15,556 | | | | rDNA | |
| chrXII | 458359 | 458523 | 164 | -1,559855743 | N12:458524 | 15,556 | | | | rDNA | |
| chrXII | 458365 | 458493 | 128 | -1,127147441 | | | | Overlap <50 bp | | rDNA | |
| chrXII | 458373 | 458493 | 120 | -0,871994003 | | | | Overlap <50 bp | | rDNA | |
| chrXII | 458373 | 458530 | 157 | -1,336124749 | N12:458524 | 15,556 | | | | rDNA | |
| chrXII | 458374 | 458532 | 158 | -1,49667944 | N12:458524 | 15,556 | | | | rDNA | |
| chrXII | 458395 | 458501 | 106 | -0,613623019 | N12:458524 | 15,556 | | | | rDNA | |
| chrXII | 458395 | 458519 | 124 | -0,922640841 | N12:458524 | 15,556 | | | | rDNA | |
| chrXII | 458419 | 458587 | 168 | -1,446254399 | N12:458524 | 15,556 | | | | rDNA | |
| chrXII | 458508 | 458658 | 150 | -1,393642725 | N12:458524 | 15,556 | | | | rDNA | |
| chrXII | 458508 | 458658 | 150 | -1,393642725 | N12:458681 | 0 | | | | rDNA | |
| chrXII | 458508 | 458659 | 151 | -1,250711736 | N12:458524 | 15,556 | | | | rDNA | |
| chrXII | 458508 | 458659 | 151 | -1,250711736 | N12:458681 | 0 | | | | rDNA | |
| chrXII | 458746 | 458901 | 155 | -1,356935675 | N12:458830 | 17,205 | | | | rDNA | |
| chrXII | 458746 | 458912 | 166 | -1,170788193 | N12:458830 | 17,205 | | | | rDNA | |
| chrXII | 458750 | 458912 | 162 | -1,574860532 | N12:458830 | 17,205 | | | | rDNA | |
| chrXII | 458768 | 458900 | 132 | -0,983896285 | N12:458830 | 17,205 | | | | rDNA | |
| chrXII | 458768 | 458901 | 133 | -1,009927836 | N12:458830 | 17,205 | | | | rDNA | |
| chrXII | 458768 | 458912 | 144 | -1,16596879 | N12:458830 | 17,205 | | | | rDNA | |
| chrXII | 458769 | 458900 | 131 | -1,185463585 | N12:458830 | 17,205 | | | | rDNA | |
| chrXII | 458769 | 458901 | 132 | -0,990437932 | N12:458830 | 17,205 | | | | rDNA | |
| chrXII | 458769 | 458912 | 143 | -1,225566231 | N12:458830 | 17,205 | | | | rDNA | |
| chrXII | 458769 | 458913 | 144 | -1,167909471 | N12:458830 | 17,205 | | | | rDNA | |
| chrXII | 458770 | 458900 | 130 | -1,435229287 | N12:458830 | 17,205 | | | | rDNA | |
| chrXII | 458770 | 458901 | 131 | -1,379998286 | N12:458830 | 17,205 | | | | rDNA | |
| chrXII | 458770 | 458912 | 142 | -1,149949557 | N12:458830 | 17,205 | | | | rDNA | |
| chrXII | 458846 | 459002 | 156 | -0,707015271 | N12:458830 | 17,205 | | | | rDNA | |
| chrXII | 458846 | 459002 | 156 | -0,707015271 | N12:458985 | 0 | | | | rDNA | |
| chrXII | 458902 | 459048 | 146 | -1,360067074 | N12:458985 | 0 | | | | rDNA | |
| chrXII | 458908 | 459044 | 136 | -0,964023298 | N12:458985 | 0 | | | | rDNA | |
| chrXII | 458908 | 459048 | 140 | -1,221372978 | N12:458985 | 0 | | | | rDNA | |
| chrXII | 458910 | 459043 | 133 | -1,197184511 | N12:458985 | 0 | | | | rDNA | |
| chrXII | 458910 | 459066 | 156 | -1,557282324 | N12:458985 | 0 | | | | rDNA | |
| chrXII | 458913 | 459002 | 89 | -0,761766034 | N12:458985 | 0 | | | | rDNA | |
| chrXII | 458913 | 459038 | 125 | -1,090949826 | N12:458985 | 0 | | | | rDNA | |
| chrXII | 458913 | 459043 | 130 | -1,191761209 | N12:458985 | 0 | | | | rDNA | |
| chrXII | 458913 | 459044 | 131 | -1,280686726 | N12:458985 | 0 | | | | rDNA | |
| chrXII | 458913 | 459047 | 134 | -1,223438986 | N12:458985 | 0 | | | | rDNA | |
| chrXII | 458913 | 459048 | 135 | -1,18570979 | N12:458985 | 0 | | | | rDNA | |
| chrXII | 458913 | 459066 | 153 | -1,321006572 | N12:458985 | 0 | | | | rDNA | |
| chrXII | 458932 | 459038 | 106 | -0,623031435 | N12:458985 | 0 | | | | rDNA | |
| chrXII | 458933 | 459044 | 111 | -0,77060004 | N12:458985 | 0 | | | | rDNA | |
| chrXII | 458933 | 459047 | 114 | -0,670650593 | N12:458985 | 0 | | | | rDNA | |
| chrXII | 458933 | 459066 | 133 | -1,067063485 | N12:458985 | 0 | | | | rDNA | |
| chrXII | 458957 | 459044 | 87 | -0,736117665 | N12:458985 | 0 | | | | rDNA | |
| chrXII | 459050 | 459146 | 96 | -1,229102172 | | | | Overlap <50 bp | | rDNA | |
| chrXII | 459089 | 459262 | 173 | -1,586410505 | N12:459173 | 25,456 | | | | rDNA | |
| chrXII | 459089 | 459265 | 176 | -1,978183737 | N12:459173 | 25,456 | | | | rDNA | |
| chrXII | 459095 | 459262 | 167 | -1,845112943 | N12:459173 | 25,456 | | | | rDNA | |
| chrXII | 459095 | 459265 | 170 | -1,553855544 | N12:459173 | 25,456 | | | | rDNA | |
| chrXII | 459098 | 459233 | 135 | -1,074327545 | N12:459173 | 25,456 | | | | rDNA | |
| chrXII | 459098 | 459262 | 164 | -1,637625524 | N12:459173 | 25,456 | | | | rDNA | |
| chrXII | 459166 | 459328 | 162 | -1,31323683 | N12:459173 | 25,456 | | | | rDNA | |
| chrXII | 459185 | 459328 | 143 | -1,126970673 | N12:459173 | 25,456 | | | | rDNA | |
| chrXII | 459204 | 459353 | 149 | -1,083754062 | | | | Overlap <50 bp | | rDNA | |
| chrXII | 459204 | 459417 | 213 | -1,794668548 | | | | Overlap <50 bp | | rDNA | |
| chrXII | 459204 | 459419 | 215 | -1,508440292 | | | | Overlap <50 bp | | rDNA | |
| chrXII | 459227 | 459353 | 126 | -1,083946406 | | | | Overlap <50 bp | | rDNA | |
| chrXII | 459227 | 459376 | 149 | -1,327600072 | | | | Overlap <50 bp | | rDNA | |
| chrXII | 459227 | 459419 | 192 | -1,679369252 | | | | Overlap <50 bp | | rDNA | |
| chrXII | 459227 | 459432 | 205 | -1,926623666 | | | | Overlap <50 bp | | rDNA | |
| chrXII | 459235 | 459395 | 160 | -1,0075863 | | | | Overlap <50 bp | | rDNA | |
| chrXII | 459236 | 459328 | 92 | -0,82580406 | | | | Overlap <50 bp | | rDNA | |
| chrXII | 459236 | 459393 | 157 | -1,408383138 | | | | Overlap <50 bp | | rDNA | |
| chrXII | 459236 | 459417 | 181 | -1,239900366 | | | | Overlap <50 bp | | rDNA | |
| chrXII | 459236 | 459419 | 183 | -1,583105474 | | | | Overlap <50 bp | | rDNA | |
| chrXII | 459264 | 459353 | 89 | -0,453448029 | | | | Overlap <50 bp | | rDNA | |
| chrXII | 459264 | 459376 | 112 | -0,935159529 | | | | Overlap <50 bp | | rDNA | |
| chrXII | 459264 | 459417 | 153 | -1,266843002 | | | | Overlap <50 bp | | rDNA | |
| chrXII | 459264 | 459419 | 155 | -1,30296611 | | | | Overlap <50 bp | | rDNA | |
| chrXII | 459264 | 459432 | 168 | -1,222785504 | | | | Overlap <50 bp | | rDNA | |
| chrXII | 459264 | 459441 | 177 | -1,509443008 | | | | Overlap <50 bp | | rDNA | |
| chrXII | 459265 | 459376 | 111 | -0,66271253 | | | | Overlap <50 bp | | rDNA | |

TABLE 3

| CHR | START | END | L(bp) | ΔLK | Nuc ID | Fuzziness | Gene ID | Gene body postion or intergene | Terminal | rDNA | Telomeric |
|---|---|---|---|---|---|---|---|---|---|---|---|
| chrXII | 459265 | 459393 | 128 | -1,037598419 | | | | Overlap <50 bp | | rDNA | |
| chrXII | 459265 | 459416 | 151 | -1,33305579 | | | | Overlap <50 bp | | rDNA | |
| chrXII | 459265 | 459417 | 152 | -1,296757275 | | | | Overlap <50 bp | | rDNA | |
| chrXII | 459265 | 459419 | 154 | -1,259303986 | | | | Overlap <50 bp | | rDNA | |
| chrXII | 459265 | 459422 | 157 | -1,413848697 | | | | Overlap <50 bp | | rDNA | |
| chrXII | 459265 | 459432 | 167 | -1,536426888 | | | | Overlap <50 bp | | rDNA | |
| chrXII | 459265 | 459441 | 176 | -1,963536502 | | | | Overlap <50 bp | | rDNA | |
| chrXII | 459265 | 459467 | 202 | -1,499202467 | N12:459485 | 42,477 | | | | rDNA | |
| chrXII | 459265 | 459469 | 204 | -1,645283042 | N12:459485 | 42,477 | | | | rDNA | |
| chrXII | 459267 | 459393 | 126 | -1,216318047 | | | | Overlap <50 bp | | rDNA | |
| chrXII | 459267 | 459417 | 150 | -1,273390539 | | | | Overlap <50 bp | | rDNA | |
| chrXII | 459267 | 459419 | 152 | -1,283371939 | | | | Overlap <50 bp | | rDNA | |
| chrXII | 459267 | 459422 | 155 | -1,098347277 | | | | Overlap <50 bp | | rDNA | |
| chrXII | 459267 | 459432 | 165 | -1,455804089 | | | | Overlap <50 bp | | rDNA | |
| chrXII | 459267 | 459441 | 174 | -1,125823556 | | | | Overlap <50 bp | | rDNA | |
| chrXII | 459267 | 459469 | 202 | -1,847406659 | N12:459485 | 42,477 | | | | rDNA | |
| chrXII | 459268 | 459374 | 106 | -1,405548963 | | | | Overlap <50 bp | | rDNA | |
| chrXII | 459268 | 459376 | 108 | -1,083596376 | | | | Overlap <50 bp | | rDNA | |
| chrXII | 459268 | 459393 | 125 | -0,698453731 | | | | Overlap <50 bp | | rDNA | |
| chrXII | 459268 | 459416 | 148 | -1,441000592 | | | | Overlap <50 bp | | rDNA | |
| chrXII | 459268 | 459417 | 149 | -1,191426265 | | | | Overlap <50 bp | | rDNA | |
| chrXII | 459268 | 459419 | 151 | -1,115525881 | | | | Overlap <50 bp | | rDNA | |
| chrXII | 459268 | 459422 | 154 | -1,322544518 | | | | Overlap <50 bp | | rDNA | |
| chrXII | 459268 | 459431 | 163 | -1,438496428 | | | | Overlap <50 bp | | rDNA | |
| chrXII | 459268 | 459432 | 164 | -1,319347296 | | | | Overlap <50 bp | | rDNA | |
| chrXII | 459268 | 459441 | 173 | -1,900103265 | | | | Overlap <50 bp | | rDNA | |
| chrXII | 459268 | 459467 | 199 | -1,376971002 | N12:459485 | 42,477 | | | | rDNA | |
| chrXII | 459268 | 459469 | 201 | -1,568375129 | N12:459485 | 42,477 | | | | rDNA | |
| chrXII | 459269 | 459376 | 107 | -1,288326132 | | | | Overlap <50 bp | | rDNA | |
| chrXII | 459269 | 459393 | 124 | -1,215014891 | | | | Overlap <50 bp | | rDNA | |
| chrXII | 459269 | 459413 | 144 | -1,214560938 | | | | Overlap <50 bp | | rDNA | |
| chrXII | 459269 | 459417 | 148 | -1,073595191 | | | | Overlap <50 bp | | rDNA | |
| chrXII | 459269 | 459419 | 150 | -1,186090912 | | | | Overlap <50 bp | | rDNA | |
| chrXII | 459269 | 459422 | 153 | -1,586776623 | | | | Overlap <50 bp | | rDNA | |
| chrXII | 459269 | 459432 | 163 | -1,942235017 | | | | Overlap <50 bp | | rDNA | |
| chrXII | 459269 | 459467 | 198 | -1,491325267 | N12:459485 | 42,477 | | | | rDNA | |
| chrXII | 459269 | 459469 | 200 | -1,617723496 | N12:459485 | 42,477 | | | | rDNA | |
| chrXII | 459271 | 459419 | 148 | -1,368597729 | | | | Overlap <50 bp | | rDNA | |
| chrXII | 459308 | 459414 | 106 | -0,814520133 | | | | Overlap <50 bp | | rDNA | |
| chrXII | 459308 | 459419 | 111 | -0,699559805 | | | | Overlap <50 bp | | rDNA | |
| chrXII | 459308 | 459467 | 159 | -1,318756425 | N12:459485 | 42,477 | | | | rDNA | |
| chrXII | 459308 | 459469 | 161 | -1,063821381 | N12:459485 | 42,477 | | | | rDNA | |
| chrXII | 459325 | 459466 | 141 | -0,854807418 | N12:459485 | 42,477 | | | | rDNA | |
| chrXII | 459326 | 459417 | 91 | -0,827464423 | | | | Overlap <50 bp | | rDNA | |
| chrXII | 459326 | 459419 | 93 | -0,450589866 | | | | Overlap <50 bp | | rDNA | |
| chrXII | 459326 | 459467 | 141 | -0,943208308 | N12:459485 | 42,477 | | | | rDNA | |
| chrXII | 459326 | 459469 | 143 | -1,06792038 | N12:459485 | 42,477 | | | | rDNA | |
| chrXII | 459408 | 459559 | 151 | -0,966752195 | N12:459485 | 42,477 | | | | rDNA | |
| chrXII | 459429 | 459558 | 129 | -1,066094198 | N12:459485 | 42,477 | | | | rDNA | |
| chrXII | 459429 | 459559 | 130 | -1,201653976 | N12:459485 | 42,477 | | | | rDNA | |
| chrXII | 459429 | 459579 | 150 | -1,504459762 | N12:459485 | 42,477 | | | | rDNA | |
| chrXII | 459430 | 459558 | 128 | -1,119961871 | N12:459485 | 42,477 | | | | rDNA | |
| chrXII | 459430 | 459559 | 129 | -0,738801499 | N12:459485 | 42,477 | | | | rDNA | |
| chrXII | 459430 | 459573 | 143 | -1,132256481 | N12:459485 | 42,477 | | | | rDNA | |
| chrXII | 459430 | 459579 | 149 | -1,279957362 | N12:459485 | 42,477 | | | | rDNA | |
| chrXII | 459519 | 459673 | 154 | -1,297894152 | | | | Overlap <50 bp | | rDNA | |
| chrXII | 459521 | 459673 | 152 | -1,264163764 | | | | Overlap <50 bp | | rDNA | |
| chrXII | 459602 | 459756 | 154 | -1,280103315 | N12:459720 | 38,118 | | | | rDNA | |
| chrXII | 459602 | 459759 | 157 | -1,235683514 | N12:459720 | 38,118 | | | | rDNA | |
| chrXII | 459604 | 459722 | 118 | -1,043424818 | N12:459720 | 38,118 | | | | rDNA | |
| chrXII | 459604 | 459756 | 152 | -1,224011959 | N12:459720 | 38,118 | | | | rDNA | |
| chrXII | 459604 | 459759 | 155 | -0,9716152 | N12:459720 | 38,118 | | | | rDNA | |
| chrXII | 459604 | 459774 | 170 | -1,381253365 | N12:459720 | 38,118 | | | | rDNA | |
| chrXII | 459604 | 459796 | 192 | -1,568180903 | N12:459720 | 38,118 | | | | rDNA | |
| chrXII | 459604 | 459798 | 194 | -1,76635304 | N12:459720 | 38,118 | | | | rDNA | |
| chrXII | 459604 | 459804 | 200 | -1,385778878 | N12:459720 | 38,118 | | | | rDNA | |
| chrXII | 459604 | 459806 | 202 | -1,905480729 | N12:459720 | 38,118 | | | | rDNA | |
| chrXII | 459618 | 459756 | 138 | -1,152443788 | N12:459720 | 38,118 | | | | rDNA | |
| chrXII | 459618 | 459774 | 156 | -1,112623004 | N12:459720 | 38,118 | | | | rDNA | |
| chrXII | 459618 | 459796 | 178 | -1,267103397 | N12:459720 | 38,118 | | | | rDNA | |
| chrXII | 459621 | 459756 | 135 | -1,228611434 | N12:459720 | 38,118 | | | | rDNA | |
| chrXII | 459621 | 459774 | 153 | -1,346246386 | N12:459720 | 38,118 | | | | rDNA | |
| chrXII | 459625 | 459756 | 131 | -1,521083941 | N12:459720 | 38,118 | | | | rDNA | |
| chrXII | 459625 | 459774 | 149 | -1,464308181 | N12:459720 | 38,118 | | | | rDNA | |
| chrXII | 459625 | 459796 | 171 | -1,632843104 | N12:459720 | 38,118 | | | | rDNA | |
| chrXII | 459625 | 459798 | 173 | -1,350708007 | N12:459720 | 38,118 | | | | rDNA | |
| chrXII | 459625 | 459804 | 179 | -1,542441342 | N12:459720 | 38,118 | | | | rDNA | |
| chrXII | 459625 | 459806 | 181 | -1,744677107 | N12:459720 | 38,118 | | | | rDNA | |
| chrXII | 459628 | 459756 | 128 | -0,922787136 | N12:459720 | 38,118 | | | | rDNA | |
| chrXII | 459628 | 459774 | 146 | -1,453679542 | N12:459720 | 38,118 | | | | rDNA | |
| chrXII | 459628 | 459796 | 168 | -1,23257791 | N12:459720 | 38,118 | | | | rDNA | |
| chrXII | 459628 | 459798 | 170 | -1,49570819 | N12:459720 | 38,118 | | | | rDNA | |
| chrXII | 459628 | 459804 | 176 | -1,151975833 | N12:459720 | 38,118 | | | | rDNA | |
| chrXII | 459628 | 459806 | 178 | -1,76817145 | N12:459720 | 38,118 | | | | rDNA | |
| chrXII | 459633 | 459756 | 123 | -0,408312065 | N12:459720 | 38,118 | | | | rDNA | |
| chrXII | 459633 | 459774 | 141 | -0,905858266 | N12:459720 | 38,118 | | | | rDNA | |
| chrXII | 459633 | 459796 | 163 | -1,317824323 | N12:459720 | 38,118 | | | | rDNA | |
| chrXII | 459633 | 459798 | 165 | -1,492815425 | N12:459720 | 38,118 | | | | rDNA | |
| chrXII | 459633 | 459804 | 171 | -1,035811662 | N12:459720 | 38,118 | | | | rDNA | |
| chrXII | 459633 | 459806 | 173 | -1,434037811 | N12:459720 | 38,118 | | | | rDNA | |
| chrXII | 459648 | 459796 | 148 | -1,213895557 | N12:459720 | 38,118 | | | | rDNA | |
| chrXII | 459673 | 459804 | 131 | -0,965210962 | N12:459720 | 38,118 | | | | rDNA | |
| chrXII | 459673 | 459806 | 133 | -1,361716608 | N12:459720 | 38,118 | | | | rDNA | |
| chrXII | 459686 | 459793 | 107 | -0,754851428 | N12:459720 | 38,118 | | | | rDNA | |
| chrXII | 459686 | 459796 | 110 | -0,978483143 | N12:459720 | 38,118 | | | | rDNA | |
| chrXII | 459686 | 459798 | 112 | -1,123389994 | N12:459720 | 38,118 | | | | rDNA | |
| chrXII | 459686 | 459799 | 113 | -0,792643338 | N12:459720 | 38,118 | | | | rDNA | |
| chrXII | 459686 | 459801 | 115 | -1,069719075 | N12:459720 | 38,118 | | | | rDNA | |

TABLE 3

| CHR | START | END | L(bp) | ΔLK | Nuc ID | Fuzziness | Gene ID | Gene body postion or intergene | Terminal | rDNA | Telomeric |
|---|---|---|---|---|---|---|---|---|---|---|---|
| chrXII | 459686 | 459804 | 118 | -1,019418416 | N12:459720 | 38,118 | | | | rDNA | |
| chrXII | 459686 | 459806 | 120 | -1,375134601 | N12:459720 | 38,118 | | | | rDNA | |
| chrXII | 459686 | 459856 | 170 | -1,763960877 | N12:459720 | 38,118 | | | | rDNA | |
| chrXII | 459688 | 459793 | 105 | -1,378871398 | N12:459720 | 38,118 | | | | rDNA | |
| chrXII | 459688 | 459851 | 163 | -1,325042528 | N12:459720 | 38,118 | | | | rDNA | |
| chrXII | 459698 | 459804 | 106 | -1,242933518 | N12:459720 | 38,118 | | | | rDNA | |
| chrXII | 459698 | 459806 | 108 | -1,065499049 | N12:459720 | 38,118 | | | | rDNA | |
| chrXII | 459698 | 459851 | 153 | -1,574371076 | N12:459720 | 38,118 | | | | rDNA | |
| chrXII | 459698 | 459869 | 171 | -1,497642081 | N12:459720 | 38,118 | | | | rDNA | |
| chrXII | 459818 | 459980 | 162 | -1,610125296 | N12:459899 | 7,43 | | | | rDNA | |
| chrXII | 459834 | 459969 | 135 | -1,334389082 | N12:459899 | 7,43 | | | | rDNA | |
| chrXII | 459846 | 459969 | 123 | -1,056817996 | N12:459899 | 7,43 | | | | rDNA | |
| chrXII | 459846 | 460002 | 156 | -1,583955114 | N12:459899 | 7,43 | | | | rDNA | |
| chrXII | 459899 | 460068 | 169 | -1,572256926 | N12:459899 | 7,43 | | | | rDNA | |
| chrXII | 459899 | 460068 | 169 | -1,572256926 | N12:460071 | 11,206 | | | | rDNA | |
| chrXII | 459930 | 460057 | 127 | -1,411320036 | N12:460071 | 11,206 | | | | rDNA | |
| chrXII | 460017 | 460128 | 111 | -0,692692996 | N12:460071 | 11,206 | | | | rDNA | |
| chrXII | 460017 | 460159 | 142 | -1,04580746 | N12:460071 | 11,206 | | | | rDNA | |
| chrXII | 460032 | 460159 | 127 | -1,219835518 | N12:460071 | 11,206 | | | | rDNA | |
| chrXII | 460102 | 460251 | 149 | -1,141959205 | | | | Overlap <50 bp | | rDNA | |
| chrXII | 460102 | 460266 | 164 | -1,214136836 | | | | Overlap <50 bp | | rDNA | |
| chrXII | 460112 | 460251 | 139 | -1,183958859 | | | | Overlap <50 bp | | rDNA | |
| chrXII | 460112 | 460266 | 154 | -1,419879802 | | | | Overlap <50 bp | | rDNA | |
| chrXII | 460188 | 460366 | 178 | -1,63267678 | N12:460347 | 14,446 | | | | rDNA | |
| chrXII | 460243 | 460384 | 141 | -1,045238909 | N12:460347 | 14,446 | | | | rDNA | |
| chrXII | 460256 | 460415 | 159 | -1,658160485 | N12:460347 | 14,446 | | | | rDNA | |
| chrXII | 460262 | 460436 | 174 | -1,860156586 | N12:460347 | 14,446 | | | | rDNA | |
| chrXII | 460302 | 460477 | 175 | -1,724575025 | N12:460347 | 14,446 | | | | rDNA | |
| chrXII | 460314 | 460446 | 132 | -1,42920196 | N12:460347 | 14,446 | | | | rDNA | |
| chrXII | 460402 | 460571 | 169 | -1,693998877 | | | | Overlap <50 bp | | rDNA | |
| chrXII | 460461 | 460595 | 134 | -1,069910373 | N12:460603 | 33,387 | | | | rDNA | |
| chrXII | 460487 | 460616 | 129 | -1,017611253 | N12:460603 | 33,387 | | | | rDNA | |
| chrXII | 460493 | 460595 | 102 | -1,556469984 | N12:460603 | 33,387 | | | | rDNA | |
| chrXII | 460493 | 460659 | 166 | -2,008001199 | N12:460603 | 33,387 | | | | rDNA | |
| chrXII | 460512 | 460595 | 83 | -1,310340248 | N12:460603 | 33,387 | | | | rDNA | |
| chrXII | 460512 | 460639 | 127 | -1,229662665 | N12:460603 | 33,387 | | | | rDNA | |
| chrXII | 460521 | 460659 | 138 | -1,275402451 | N12:460603 | 33,387 | | | | rDNA | |
| chrXII | 460602 | 460763 | 161 | -1,590887061 | N12:460603 | 33,387 | | | | rDNA | |
| chrXII | 460846 | 460979 | 133 | -1,479111747 | N12:460826 | 15,556 | | | | rDNA | |
| chrXII | 460863 | 461022 | 159 | -1,295823817 | | | | Overlap <50 bp | | rDNA | |
| chrXII | 460871 | 461006 | 135 | -1,483873741 | | | | Overlap <50 bp | | rDNA | |
| chrXII | 460894 | 460979 | 85 | -0,759025328 | | | | Overlap <50 bp | | rDNA | |
| chrXII | 460894 | 461045 | 151 | -1,523354561 | | | | Overlap <50 bp | | rDNA | |
| chrXII | 460894 | 461097 | 203 | -1,765695745 | | | | Overlap <50 bp | | rDNA | |
| chrXII | 460906 | 461069 | 163 | -1,399714976 | | | | Overlap <50 bp | | rDNA | |
| chrXII | 460927 | 461045 | 118 | -1,223043342 | | | | Overlap <50 bp | | rDNA | |
| chrXII | 460927 | 461097 | 170 | -1,377093935 | | | | Overlap <50 bp | | rDNA | |
| chrXII | 460937 | 461071 | 134 | -0,977204797 | | | | Overlap <50 bp | | rDNA | |
| chrXII | 460937 | 461098 | 161 | -1,542901964 | | | | Overlap <50 bp | | rDNA | |
| chrXII | 460988 | 461071 | 83 | -0,85669586 | | | | Overlap <50 bp | | rDNA | |
| chrXII | 460988 | 461095 | 107 | -1,43876598 | | | | Overlap <50 bp | | rDNA | |
| chrXII | 460988 | 461096 | 108 | -0,945073079 | | | | Overlap <50 bp | | rDNA | |
| chrXII | 460988 | 461098 | 110 | -1,299774635 | | | | Overlap <50 bp | | rDNA | |
| chrXII | 461013 | 461146 | 133 | -1,267856788 | N12:461125 | 34,641 | | | | rDNA | |
| chrXII | 461056 | 461146 | 90 | -0,738013277 | N12:461125 | 34,641 | | | | rDNA | |
| chrXII | 461056 | 461148 | 92 | -1,447851605 | N12:461125 | 34,641 | | | | rDNA | |
| chrXII | 461056 | 461199 | 143 | -1,61588963 | N12:461125 | 34,641 | | | | rDNA | |
| chrXII | 461056 | 461212 | 156 | -1,177381364 | N12:461125 | 34,641 | | | | rDNA | |
| chrXII | 461056 | 461214 | 158 | -1,382805765 | N12:461125 | 34,641 | | | | rDNA | |
| chrXII | 461056 | 461215 | 159 | -1,217202941 | N12:461125 | 34,641 | | | | rDNA | |
| chrXII | 461056 | 461229 | 173 | -1,252693267 | N12:461125 | 34,641 | | | | rDNA | |
| chrXII | 461060 | 461146 | 86 | -0,635572156 | N12:461125 | 34,641 | | | | rDNA | |
| chrXII | 461060 | 461212 | 152 | -1,275856664 | N12:461125 | 34,641 | | | | rDNA | |
| chrXII | 461060 | 461214 | 154 | -1,748558927 | N12:461125 | 34,641 | | | | rDNA | |
| chrXII | 461060 | 461215 | 155 | -1,419763076 | N12:461125 | 34,641 | | | | rDNA | |
| chrXII | 461060 | 461229 | 169 | -0,906330119 | N12:461125 | 34,641 | | | | rDNA | |
| chrXII | 461068 | 461199 | 131 | -1,208852873 | N12:461125 | 34,641 | | | | rDNA | |
| chrXII | 461068 | 461214 | 146 | -1,240158825 | N12:461125 | 34,641 | | | | rDNA | |
| chrXII | 461068 | 461246 | 178 | -1,322613742 | N12:461125 | 34,641 | | | | rDNA | |
| chrXII | 461069 | 461199 | 130 | -1,119193943 | N12:461125 | 34,641 | | | | rDNA | |
| chrXII | 461069 | 461214 | 145 | -1,554933122 | N12:461125 | 34,641 | | | | rDNA | |
| chrXII | 461069 | 461246 | 177 | -1,388781136 | N12:461125 | 34,641 | | | | rDNA | |
| chrXII | 461082 | 461188 | 106 | -0,983832423 | N12:461125 | 34,641 | | | | rDNA | |
| chrXII | 461082 | 461199 | 117 | -0,809951429 | N12:461125 | 34,641 | | | | rDNA | |
| chrXII | 461082 | 461212 | 130 | -1,00202511 | N12:461125 | 34,641 | | | | rDNA | |
| chrXII | 461082 | 461214 | 132 | -1,021239956 | N12:461125 | 34,641 | | | | rDNA | |
| chrXII | 461082 | 461215 | 133 | -1,150598564 | N12:461125 | 34,641 | | | | rDNA | |
| chrXII | 461082 | 461229 | 147 | -1,017966106 | N12:461125 | 34,641 | | | | rDNA | |
| chrXII | 461082 | 461246 | 164 | -1,395751218 | N12:461125 | 34,641 | | | | rDNA | |
| chrXII | 461095 | 461246 | 151 | -1,025970322 | N12:461125 | 34,641 | | | | rDNA | |
| chrXII | 461178 | 461333 | 155 | -1,389661098 | N12:461308 | 61,052 | | | | rDNA | |
| chrXII | 461196 | 461352 | 156 | -1,390413046 | N12:461308 | 61,052 | | | | rDNA | |
| chrXII | 461218 | 461364 | 146 | -1,213886286 | N12:461308 | 61,052 | | | | rDNA | |
| chrXII | 461218 | 461384 | 166 | -1,301897258 | N12:461308 | 61,052 | | | | rDNA | |
| chrXII | 461218 | 461388 | 170 | -1,106705622 | N12:461308 | 61,052 | | | | rDNA | |
| chrXII | 461241 | 461384 | 143 | -1,173358607 | N12:461308 | 61,052 | | | | rDNA | |
| chrXII | 461241 | 461388 | 147 | -0,823477867 | N12:461308 | 61,052 | | | | rDNA | |
| chrXII | 461252 | 461408 | 156 | -1,168312379 | N12:461308 | 61,052 | | | | rDNA | |
| chrXII | 461260 | 461367 | 107 | -1,258905485 | N12:461308 | 61,052 | | | | rDNA | |
| chrXII | 461327 | 461481 | 154 | -1,343883463 | N12:461308 | 61,052 | | | | rDNA | |
| chrXII | 461397 | 461551 | 154 | -1,394557994 | N12:461507 | 14,422 | | | | rDNA | |
| chrXII | 461430 | 461537 | 107 | -1,593911389 | N12:461507 | 14,422 | | | | rDNA | |
| chrXII | 461431 | 461592 | 161 | -1,304230891 | N12:461507 | 14,422 | | | | rDNA | |
| chrXII | 461432 | 461592 | 160 | -1,239111892 | N12:461507 | 14,422 | | | | rDNA | |
| chrXII | 461436 | 461592 | 156 | -1,186791263 | N12:461507 | 14,422 | | | | rDNA | |
| chrXII | 461460 | 461622 | 162 | -1,419288148 | N12:461507 | 14,422 | | | | rDNA | |
| chrXII | 461460 | 461623 | 163 | -1,333641154 | N12:461507 | 14,422 | | | | rDNA | |
| chrXII | 461471 | 461578 | 107 | -0,730947057 | N12:461507 | 14,422 | | | | rDNA | |
| chrXII | 461471 | 461592 | 121 | -0,922646777 | N12:461507 | 14,422 | | | | rDNA | |

TABLE 3

| CHR | START | END | L(bp) | ΔLK | Nuc ID | Fuzziness | Gene ID | Gene body postion or intergene | Terminal | rDNA | Telomeric |
|---|---|---|---|---|---|---|---|---|---|---|---|
| chrXII | 461471 | 461622 | 151 | -1,209718649 | N12:461507 | 14,422 | | | | rDNA | |
| chrXII | 461471 | 461623 | 152 | -1,409184694 | N12:461507 | 14,422 | | | | rDNA | |
| chrXII | 461483 | 461633 | 150 | -1,248029841 | N12:461507 | 14,422 | | | | rDNA | |
| chrXII | 461536 | 461685 | 149 | -1,373995135 | N12:461692 | 5,774 | | | | rDNA | |
| chrXII | 461560 | 461721 | 161 | -1,502916154 | N12:461692 | 5,774 | | | | rDNA | |
| chrXII | 461606 | 461757 | 151 | -1,617421207 | N12:461692 | 5,774 | | | | rDNA | |
| chrXII | 461606 | 461758 | 152 | -1,623816206 | N12:461692 | 5,774 | | | | rDNA | |
| chrXII | 461606 | 461766 | 160 | -1,476822976 | N12:461692 | 5,774 | | | | rDNA | |
| chrXII | 461606 | 461767 | 161 | -1,334150821 | N12:461692 | 5,774 | | | | rDNA | |
| chrXII | 461628 | 461758 | 130 | -1,708770553 | N12:461692 | 5,774 | | | | rDNA | |
| chrXII | 461628 | 461766 | 138 | -1,6251081 | N12:461692 | 5,774 | | | | rDNA | |
| chrXII | 461628 | 461800 | 172 | -1,545659216 | N12:461692 | 5,774 | | | | rDNA | |
| chrXII | 461629 | 461721 | 92 | -1,178017207 | N12:461692 | 5,774 | | | | rDNA | |
| chrXII | 461629 | 461757 | 128 | -1,554312406 | N12:461692 | 5,774 | | | | rDNA | |
| chrXII | 461629 | 461758 | 129 | -1,421107881 | N12:461692 | 5,774 | | | | rDNA | |
| chrXII | 461629 | 461766 | 137 | -1,546194707 | N12:461692 | 5,774 | | | | rDNA | |
| chrXII | 461629 | 461800 | 171 | -1,980364364 | N12:461692 | 5,774 | | | | rDNA | |
| chrXII | 461662 | 461758 | 96 | -0,726193226 | N12:461692 | 5,774 | | | | rDNA | |
| chrXII | 461662 | 461832 | 170 | -1,546249051 | N12:461692 | 5,774 | | | | rDNA | |
| chrXII | 461676 | 461810 | 134 | -1,542380032 | N12:461692 | 5,774 | | | | rDNA | |
| chrXII | 461687 | 461800 | 113 | -0,980876838 | N12:461692 | 5,774 | | | | rDNA | |
| chrXII | 461687 | 461809 | 122 | -1,602595426 | N12:461692 | 5,774 | | | | rDNA | |
| chrXII | 461687 | 461839 | 152 | -1,319174046 | N12:461692 | 5,774 | | | | rDNA | |
| chrXII | 461805 | 461933 | 128 | -1,368454211 | N12:461891 | 27,622 | | | | rDNA | |
| chrXII | 461805 | 461951 | 146 | -1,533184236 | N12:461891 | 27,622 | | | | rDNA | |
| chrXII | 461821 | 461933 | 112 | -0,87384802 | N12:461891 | 27,622 | | | | rDNA | |
| chrXII | 461821 | 461972 | 151 | -1,185196443 | N12:461891 | 27,622 | | | | rDNA | |
| chrXII | 461823 | 461951 | 128 | -1,099271338 | N12:461891 | 27,622 | | | | rDNA | |
| chrXII | 461823 | 462001 | 178 | -1,589564566 | N12:461891 | 27,622 | | | | rDNA | |
| chrXII | 461846 | 461952 | 106 | -0,675483877 | N12:461891 | 27,622 | | | | rDNA | |
| chrXII | 461846 | 462001 | 155 | -1,367892209 | N12:461891 | 27,622 | | | | rDNA | |
| chrXII | 461846 | 462021 | 175 | -1,224499161 | N12:461891 | 27,622 | | | | rDNA | |
| chrXII | 461846 | 462037 | 191 | -1,502568982 | N12:461891 | 27,622 | | | | rDNA | |
| chrXII | 461860 | 461951 | 91 | -0,636169527 | N12:461891 | 27,622 | | | | rDNA | |
| chrXII | 461860 | 462001 | 141 | -1,419207735 | N12:461891 | 27,622 | | | | rDNA | |
| chrXII | 461860 | 462021 | 161 | -1,238731069 | N12:461891 | 27,622 | | | | rDNA | |
| chrXII | 461860 | 462037 | 177 | -1,333089514 | N12:461891 | 27,622 | | | | rDNA | |
| chrXII | 461902 | 462001 | 99 | -0,708383238 | N12:461891 | 27,622 | | | | rDNA | |
| chrXII | 461902 | 462008 | 106 | -0,915563165 | N12:461891 | 27,622 | | | | rDNA | |
| chrXII | 461902 | 462021 | 119 | -0,755709778 | N12:461891 | 27,622 | | | | rDNA | |
| chrXII | 461902 | 462037 | 135 | -1,125045892 | N12:461891 | 27,622 | | | | rDNA | |
| chrXII | 461902 | 462062 | 160 | -0,926331205 | N12:461891 | 27,622 | | | | rDNA | |
| chrXII | 461902 | 462062 | 160 | -0,926331205 | N12:462072 | 28,042 | | | | rDNA | |
| chrXII | 461910 | 462001 | 91 | -0,868969365 | N12:461891 | 27,622 | | | | rDNA | |
| chrXII | 461910 | 462016 | 106 | -1,002198979 | N12:461891 | 27,622 | | | | rDNA | |
| chrXII | 461910 | 462021 | 111 | -0,818868114 | N12:461891 | 27,622 | | | | rDNA | |
| chrXII | 461910 | 462037 | 127 | -0,993582129 | N12:461891 | 27,622 | | | | rDNA | |
| chrXII | 461910 | 462062 | 152 | -1,355538681 | N12:461891 | 27,622 | | | | rDNA | |
| chrXII | 461910 | 462062 | 152 | -1,355538681 | N12:462072 | 28,042 | | | | rDNA | |
| chrXII | 461920 | 462099 | 179 | -1,795874783 | N12:462072 | 28,042 | | | | rDNA | |
| chrXII | 461925 | 462060 | 135 | -1,102158522 | N12:462072 | 28,042 | | | | rDNA | |
| chrXII | 461925 | 462090 | 165 | -1,345084215 | N12:462072 | 28,042 | | | | rDNA | |
| chrXII | 461938 | 462060 | 122 | -0,885108621 | N12:462072 | 28,042 | | | | rDNA | |
| chrXII | 461938 | 462090 | 152 | -1,278177534 | N12:462072 | 28,042 | | | | rDNA | |
| chrXII | 461938 | 462136 | 198 | -1,670009443 | N12:462072 | 28,042 | | | | rDNA | |
| chrXII | 461977 | 462090 | 113 | -0,703973579 | N12:462072 | 28,042 | | | | rDNA | |
| chrXII | 461977 | 462136 | 159 | -1,283025192 | N12:462072 | 28,042 | | | | rDNA | |
| chrXII | 461977 | 462160 | 183 | -1,464920009 | N12:462072 | 28,042 | | | | rDNA | |
| chrXII | 461984 | 462090 | 106 | -0,826732631 | N12:462072 | 28,042 | | | | rDNA | |
| chrXII | 461984 | 462133 | 149 | -1,472566543 | N12:462072 | 28,042 | | | | rDNA | |
| chrXII | 462013 | 462171 | 158 | -1,290787868 | N12:462072 | 28,042 | | | | rDNA | |
| chrXII | 462022 | 462171 | 149 | -1,180034919 | N12:462072 | 28,042 | | | | rDNA | |
| chrXII | 462030 | 462136 | 106 | -0,739653604 | N12:462072 | 28,042 | | | | rDNA | |
| chrXII | 462032 | 462160 | 128 | -0,98105994 | N12:462072 | 28,042 | | | | rDNA | |
| chrXII | 462059 | 462166 | 107 | -1,464163252 | N12:462072 | 28,042 | | | | rDNA | |
| chrXII | 462059 | 462167 | 108 | -1,324770286 | N12:462072 | 28,042 | | | | rDNA | |
| chrXII | 462059 | 462171 | 112 | -1,434563087 | N12:462072 | 28,042 | | | | rDNA | |
| chrXII | 462059 | 462193 | 134 | -1,313044451 | N12:462072 | 28,042 | | | | rDNA | |
| chrXII | 462059 | 462212 | 153 | -1,706287945 | N12:462072 | 28,042 | | | | rDNA | |
| chrXII | 462063 | 462134 | 71 | -0,749 | N12:462072 | 28,042 | | | | rDNA | |
| chrXII | 462063 | 462170 | 107 | -0,979522693 | N12:462072 | 28,042 | | | | rDNA | |
| chrXII | 462063 | 462171 | 108 | -0,605029641 | N12:462072 | 28,042 | | | | rDNA | |
| chrXII | 462063 | 462193 | 130 | -1,042789847 | N12:462072 | 28,042 | | | | rDNA | |
| chrXII | 462063 | 462207 | 144 | -1,062640054 | N12:462072 | 28,042 | | | | rDNA | |
| chrXII | 462063 | 462212 | 149 | -1,384053019 | N12:462072 | 28,042 | | | | rDNA | |
| chrXII | 462113 | 462275 | 162 | -1,539597824 | N12:462253 | 25,456 | | | | rDNA | |
| chrXII | 462128 | 462278 | 150 | -1,242559948 | N12:462253 | 25,456 | | | | rDNA | |
| chrXII | 462168 | 462275 | 107 | -0,993415835 | N12:462253 | 25,456 | | | | rDNA | |
| chrXII | 462170 | 462300 | 130 | -1,173539009 | N12:462253 | 25,456 | | | | rDNA | |
| chrXII | 462170 | 462324 | 154 | -1,402908952 | N12:462253 | 25,456 | | | | rDNA | |
| chrXII | 462170 | 462330 | 160 | -1,504258326 | N12:462253 | 25,456 | | | | rDNA | |
| chrXII | 462176 | 462300 | 124 | -0,791852804 | N12:462253 | 25,456 | | | | rDNA | |
| chrXII | 462176 | 462324 | 148 | -1,105808041 | N12:462253 | 25,456 | | | | rDNA | |
| chrXII | 462176 | 462330 | 154 | -1,222273098 | N12:462253 | 25,456 | | | | rDNA | |
| chrXII | 462179 | 462349 | 170 | -1,656822088 | N12:462253 | 25,456 | | | | rDNA | |
| chrXII | 462179 | 462355 | 176 | -1,310916345 | N12:462253 | 25,456 | | | | rDNA | |
| chrXII | 462180 | 462300 | 120 | -1,007527494 | N12:462253 | 25,456 | | | | rDNA | |
| chrXII | 462180 | 462324 | 144 | -1,166792665 | N12:462253 | 25,456 | | | | rDNA | |
| chrXII | 462180 | 462327 | 147 | -1,156929363 | N12:462253 | 25,456 | | | | rDNA | |
| chrXII | 462180 | 462330 | 150 | -1,281108512 | N12:462253 | 25,456 | | | | rDNA | |
| chrXII | 462228 | 462336 | 108 | -0,483761136 | N12:462253 | 25,456 | | | | rDNA | |
| chrXII | 462229 | 462336 | 107 | -0,841602115 | N12:462253 | 25,456 | | | | rDNA | |
| chrXII | 462229 | 462337 | 108 | -0,770439775 | N12:462253 | 25,456 | | | | rDNA | |
| chrXII | 462229 | 462349 | 120 | -1,148835858 | N12:462253 | 25,456 | | | | rDNA | |
| chrXII | 462229 | 462373 | 144 | -1,59916081 | N12:462253 | 25,456 | | | | rDNA | |
| chrXII | 462229 | 462382 | 153 | -1,384740384 | N12:462253 | 25,456 | | | | rDNA | |
| chrXII | 462229 | 462388 | 159 | -1,325279425 | N12:462253 | 25,456 | | | | rDNA | |
| chrXII | 462229 | 462407 | 178 | -1,711292995 | N12:462253 | 25,456 | | | | rDNA | |
| chrXII | 462229 | 462423 | 194 | -1,71060805 | N12:462253 | 25,456 | | | | rDNA | |
| chrXII | 462229 | 462425 | 196 | -1,897176877 | N12:462253 | 25,456 | | | | rDNA | |

TABLE 3

| CHR | START | END | L(bp) | ΔLK | Nuc ID | Fuzziness | Gene ID | Gene body postion or intergene | Terminal | rDNA | Telomeric |
|---|---|---|---|---|---|---|---|---|---|---|---|
| chrXII | 462232 | 462338 | 106 | -1,010167448 | N12:462253 | 25,456 | | | | rDNA | |
| chrXII | 462232 | 462349 | 117 | -0,867537202 | N12:462253 | 25,456 | | | | rDNA | |
| chrXII | 462232 | 462355 | 123 | -0,763248727 | N12:462253 | 25,456 | | | | rDNA | |
| chrXII | 462232 | 462373 | 141 | -1,583144002 | N12:462253 | 25,456 | | | | rDNA | |
| chrXII | 462232 | 462382 | 150 | -1,242722246 | N12:462253 | 25,456 | | | | rDNA | |
| chrXII | 462232 | 462407 | 175 | -1,234454735 | N12:462253 | 25,456 | | | | rDNA | |
| chrXII | 462232 | 462419 | 187 | -1,536254509 | N12:462253 | 25,456 | | | | rDNA | |
| chrXII | 462232 | 462423 | 191 | -1,539395061 | N12:462253 | 25,456 | | | | rDNA | |
| chrXII | 462232 | 462425 | 193 | -1,693854267 | N12:462253 | 25,456 | | | | rDNA | |
| chrXII | 462236 | 462373 | 137 | -0,855473563 | N12:462253 | 25,456 | | | | rDNA | |
| chrXII | 462236 | 462425 | 189 | -1,347544847 | N12:462253 | 25,456 | | | | rDNA | |
| chrXII | 462237 | 462375 | 138 | -1,222113025 | N12:462253 | 25,456 | | | | rDNA | |
| chrXII | 462237 | 462382 | 145 | -1,029391304 | N12:462253 | 25,456 | | | | rDNA | |
| chrXII | 462250 | 462382 | 132 | -1,258446751 | N12:462253 | 25,456 | | | | rDNA | |
| chrXII | 462250 | 462407 | 157 | -1,471150145 | N12:462253 | 25,456 | | | | rDNA | |
| chrXII | 462250 | 462423 | 173 | -1,637996903 | N12:462253 | 25,456 | | | | rDNA | |
| chrXII | 462250 | 462425 | 175 | -1,588528981 | N12:462253 | 25,456 | | | | rDNA | |
| chrXII | 462251 | 462373 | 122 | -0,583575677 | N12:462253 | 25,456 | | | | rDNA | |
| chrXII | 462251 | 462406 | 155 | -1,284997819 | N12:462253 | 25,456 | | | | rDNA | |
| chrXII | 462251 | 462407 | 156 | -1,163408296 | N12:462253 | 25,456 | | | | rDNA | |
| chrXII | 462251 | 462423 | 172 | -1,243606312 | N12:462253 | 25,456 | | | | rDNA | |
| chrXII | 462251 | 462424 | 173 | -1,345366309 | N12:462253 | 25,456 | | | | rDNA | |
| chrXII | 462251 | 462425 | 174 | -1,325462079 | N12:462253 | 25,456 | | | | rDNA | |
| chrXII | 462251 | 462439 | 188 | -1,643109934 | N12:462253 | 25,456 | | | | rDNA | |
| chrXII | 462253 | 462406 | 153 | -0,941129587 | N12:462253 | 25,456 | | | | rDNA | |
| chrXII | 462253 | 462407 | 154 | -1,283870006 | N12:462253 | 25,456 | | | | rDNA | |
| chrXII | 462253 | 462424 | 171 | -1,252664851 | N12:462253 | 25,456 | | | | rDNA | |
| chrXII | 462253 | 462425 | 172 | -1,084616153 | N12:462253 | 25,456 | | | | rDNA | |
| chrXII | 462256 | 462373 | 117 | -0,688064683 | N12:462253 | 25,456 | | | | rDNA | |
| chrXII | 462256 | 462388 | 132 | -1,506836606 | N12:462253 | 25,456 | | | | rDNA | |
| chrXII | 462256 | 462406 | 150 | -1,288585557 | N12:462253 | 25,456 | | | | rDNA | |
| chrXII | 462256 | 462407 | 151 | -1,429895149 | N12:462253 | 25,456 | | | | rDNA | |
| chrXII | 462256 | 462419 | 163 | -2,167397761 | N12:462253 | 25,456 | | | | rDNA | |
| chrXII | 462256 | 462423 | 167 | -1,435482441 | N12:462253 | 25,456 | | | | rDNA | |
| chrXII | 462256 | 462424 | 168 | -1,400714873 | N12:462253 | 25,456 | | | | rDNA | |
| chrXII | 462256 | 462425 | 169 | -1,3114823 | N12:462253 | 25,456 | | | | rDNA | |
| chrXII | 462256 | 462439 | 183 | -1,507268189 | N12:462253 | 25,456 | | | | rDNA | |
| chrXII | 462256 | 462446 | 190 | -1,788095378 | N12:462253 | 25,456 | | | | rDNA | |
| chrXII | 462257 | 462365 | 108 | -0,336153445 | N12:462253 | 25,456 | | | | rDNA | |
| chrXII | 462258 | 462407 | 149 | -1,237324861 | N12:462253 | 25,456 | | | | rDNA | |
| chrXII | 462258 | 462423 | 165 | -1,37820741 | N12:462253 | 25,456 | | | | rDNA | |
| chrXII | 462258 | 462425 | 167 | -1,491431317 | N12:462253 | 25,456 | | | | rDNA | |
| chrXII | 462262 | 462368 | 106 | -1,13043658 | N12:462253 | 25,456 | | | | rDNA | |
| chrXII | 462262 | 462373 | 111 | -1,28856716 | N12:462253 | 25,456 | | | | rDNA | |
| chrXII | 462262 | 462407 | 145 | -1,420251734 | N12:462253 | 25,456 | | | | rDNA | |
| chrXII | 462262 | 462419 | 157 | -2,076701185 | N12:462253 | 25,456 | | | | rDNA | |
| chrXII | 462262 | 462423 | 161 | -1,404982031 | N12:462253 | 25,456 | | | | rDNA | |
| chrXII | 462262 | 462424 | 162 | -1,217048808 | N12:462253 | 25,456 | | | | rDNA | |
| chrXII | 462262 | 462425 | 163 | -1,355209827 | N12:462253 | 25,456 | | | | rDNA | |
| chrXII | 462262 | 462439 | 177 | -1,404331055 | N12:462253 | 25,456 | | | | rDNA | |
| chrXII | 462263 | 462385 | 122 | -1,465249066 | N12:462253 | 25,456 | | | | rDNA | |
| chrXII | 462263 | 462407 | 144 | -1,432089506 | N12:462253 | 25,456 | | | | rDNA | |
| chrXII | 462263 | 462423 | 160 | -1,322477134 | N12:462253 | 25,456 | | | | rDNA | |
| chrXII | 462263 | 462424 | 161 | -1,445678272 | N12:462253 | 25,456 | | | | rDNA | |
| chrXII | 462263 | 462425 | 162 | -1,567583167 | N12:462253 | 25,456 | | | | rDNA | |
| chrXII | 462272 | 462407 | 135 | -1,488203503 | N12:462253 | 25,456 | | | | rDNA | |
| chrXII | 462272 | 462419 | 147 | -1,292397848 | N12:462253 | 25,456 | | | | rDNA | |
| chrXII | 462272 | 462423 | 151 | -1,579990156 | N12:462253 | 25,456 | | | | rDNA | |
| chrXII | 462272 | 462425 | 153 | -1,736374966 | N12:462253 | 25,456 | | | | rDNA | |
| chrXII | 462275 | 462423 | 148 | -1,370781459 | N12:462253 | 25,456 | | | | rDNA | |
| chrXII | 462275 | 462425 | 150 | -1,915876347 | N12:462253 | 25,456 | | | | rDNA | |
| chrXII | 462277 | 462373 | 96 | -0,567121191 | | | | Overlap <50 bp | | rDNA | |
| chrXII | 462277 | 462382 | 105 | -0,302258508 | | | | Overlap <50 bp | | rDNA | |
| chrXII | 462277 | 462383 | 106 | -0,970455414 | | | | Overlap <50 bp | | rDNA | |
| chrXII | 462277 | 462388 | 111 | -1,40254736 | | | | Overlap <50 bp | | rDNA | |
| chrXII | 462277 | 462407 | 130 | -1,050457647 | | | | Overlap <50 bp | | rDNA | |
| chrXII | 462277 | 462419 | 142 | -1,260675382 | | | | Overlap <50 bp | | rDNA | |
| chrXII | 462277 | 462423 | 146 | -1,242909695 | | | | Overlap <50 bp | | rDNA | |
| chrXII | 462277 | 462424 | 147 | -1,351397919 | | | | Overlap <50 bp | | rDNA | |
| chrXII | 462277 | 462425 | 148 | -1,271145951 | | | | Overlap <50 bp | | rDNA | |
| chrXII | 462277 | 462439 | 162 | -1,650712858 | | | | Overlap <50 bp | | rDNA | |
| chrXII | 462277 | 462460 | 183 | -1,291285531 | | | | Overlap <50 bp | | rDNA | |
| chrXII | 462278 | 462373 | 95 | -0,389088976 | | | | Overlap <50 bp | | rDNA | |
| chrXII | 462278 | 462382 | 104 | -0,449366181 | | | | Overlap <50 bp | | rDNA | |
| chrXII | 462278 | 462406 | 128 | -0,513846831 | | | | Overlap <50 bp | | rDNA | |
| chrXII | 462278 | 462407 | 129 | -0,858033522 | | | | Overlap <50 bp | | rDNA | |
| chrXII | 462278 | 462418 | 140 | -1,312192965 | | | | Overlap <50 bp | | rDNA | |
| chrXII | 462278 | 462419 | 141 | -1,330812732 | | | | Overlap <50 bp | | rDNA | |
| chrXII | 462278 | 462420 | 142 | -1,493351734 | | | | Overlap <50 bp | | rDNA | |
| chrXII | 462278 | 462423 | 145 | -1,171651451 | | | | Overlap <50 bp | | rDNA | |
| chrXII | 462278 | 462424 | 146 | -1,460693145 | | | | Overlap <50 bp | | rDNA | |
| chrXII | 462278 | 462425 | 147 | -1,234977821 | | | | Overlap <50 bp | | rDNA | |
| chrXII | 462278 | 462439 | 161 | -1,155210563 | | | | Overlap <50 bp | | rDNA | |
| chrXII | 462278 | 462446 | 168 | -1,340070299 | | | | Overlap <50 bp | | rDNA | |
| chrXII | 462278 | 462460 | 182 | -1,424874251 | | | | Overlap <50 bp | | rDNA | |
| chrXII | 462278 | 462461 | 183 | -1,540534897 | | | | Overlap <50 bp | | rDNA | |
| chrXII | 462278 | 462479 | 201 | -1,673067841 | | | | Overlap <50 bp | | rDNA | |
| chrXII | 462281 | 462425 | 144 | -1,927457252 | | | | Overlap <50 bp | | rDNA | |
| chrXII | 462287 | 462423 | 136 | -1,123977213 | | | | Overlap <50 bp | | rDNA | |
| chrXII | 462287 | 462425 | 138 | -1,452089606 | | | | Overlap <50 bp | | rDNA | |
| chrXII | 462287 | 462432 | 145 | -1,259254691 | | | | Overlap <50 bp | | rDNA | |
| chrXII | 462287 | 462439 | 152 | -1,969162681 | | | | Overlap <50 bp | | rDNA | |
| chrXII | 462289 | 462377 | 88 | -0,771557704 | | | | Overlap <50 bp | | rDNA | |
| chrXII | 462289 | 462407 | 118 | -0,911764917 | | | | Overlap <50 bp | | rDNA | |
| chrXII | 462289 | 462423 | 134 | -1,157213041 | | | | Overlap <50 bp | | rDNA | |
| chrXII | 462289 | 462424 | 135 | -1,376744813 | | | | Overlap <50 bp | | rDNA | |
| chrXII | 462289 | 462425 | 136 | -1,239319998 | | | | Overlap <50 bp | | rDNA | |
| chrXII | 462289 | 462439 | 150 | -1,026972379 | | | | Overlap <50 bp | | rDNA | |

TABLE 3

| CHR | START | END | L(bp) | ΔLK | Nuc ID | Fuzziness | Gene ID | Gene body postion or intergene | Terminal | rDNA | Telomeric |
|---|---|---|---|---|---|---|---|---|---|---|---|
| chrXII | 462290 | 462423 | 133 | -1,122164485 | | | | Overlap <50 bp | | rDNA | |
| chrXII | 462290 | 462425 | 135 | -0,971792625 | | | | Overlap <50 bp | | rDNA | |
| chrXII | 462296 | 462407 | 111 | -0,751666002 | | | | Overlap <50 bp | | rDNA | |
| chrXII | 462296 | 462423 | 127 | -1,138218147 | | | | Overlap <50 bp | | rDNA | |
| chrXII | 462296 | 462424 | 128 | -0,987759496 | | | | Overlap <50 bp | | rDNA | |
| chrXII | 462296 | 462425 | 129 | -0,946623822 | | | | Overlap <50 bp | | rDNA | |
| chrXII | 462296 | 462439 | 143 | -1,148745233 | | | | Overlap <50 bp | | rDNA | |
| chrXII | 462296 | 462446 | 150 | -1,511981275 | | | | Overlap <50 bp | | rDNA | |
| chrXII | 462296 | 462472 | 176 | -1,584458547 | | | | Overlap <50 bp | | rDNA | |
| chrXII | 462296 | 462479 | 183 | -1,831632027 | | | | Overlap <50 bp | | rDNA | |
| chrXII | 462298 | 462382 | 84 | -0,422770002 | | | | Overlap <50 bp | | rDNA | |
| chrXII | 462298 | 462405 | 107 | -0,951692148 | | | | Overlap <50 bp | | rDNA | |
| chrXII | 462298 | 462407 | 109 | -0,704815923 | | | | Overlap <50 bp | | rDNA | |
| chrXII | 462298 | 462419 | 121 | -1,057561331 | | | | Overlap <50 bp | | rDNA | |
| chrXII | 462298 | 462423 | 125 | -1,009155207 | | | | Overlap <50 bp | | rDNA | |
| chrXII | 462298 | 462425 | 127 | -1,080826114 | | | | Overlap <50 bp | | rDNA | |
| chrXII | 462298 | 462446 | 148 | -1,549906994 | | | | Overlap <50 bp | | rDNA | |
| chrXII | 462298 | 462460 | 162 | -1,372380122 | | | | Overlap <50 bp | | rDNA | |
| chrXII | 462298 | 462461 | 163 | -1,478055751 | | | | Overlap <50 bp | | rDNA | |
| chrXII | 462299 | 462406 | 107 | -0,650650614 | | | | Overlap <50 bp | | rDNA | |
| chrXII | 462299 | 462423 | 124 | -1,048727808 | | | | Overlap <50 bp | | rDNA | |
| chrXII | 462299 | 462425 | 126 | -1,235184027 | | | | Overlap <50 bp | | rDNA | |
| chrXII | 462299 | 462460 | 161 | -1,442117698 | | | | Overlap <50 bp | | rDNA | |
| chrXII | 462299 | 462461 | 162 | -1,28246185 | | | | Overlap <50 bp | | rDNA | |
| chrXII | 462302 | 462424 | 122 | -1,007377917 | | | | Overlap <50 bp | | rDNA | |
| chrXII | 462302 | 462425 | 123 | -1,042169499 | | | | Overlap <50 bp | | rDNA | |
| chrXII | 462302 | 462439 | 137 | -1,492231006 | | | | Overlap <50 bp | | rDNA | |
| chrXII | 462302 | 462479 | 177 | -1,336233641 | | | | Overlap <50 bp | | rDNA | |
| chrXII | 462308 | 462479 | 171 | -1,410751805 | | | | Overlap <50 bp | | rDNA | |
| chrXII | 462317 | 462425 | 108 | -1,160751066 | | | | Overlap <50 bp | | rDNA | |
| chrXII | 462317 | 462477 | 160 | -1,752379822 | | | | Overlap <50 bp | | rDNA | |
| chrXII | 462318 | 462407 | 89 | -1,218961854 | | | | Overlap <50 bp | | rDNA | |
| chrXII | 462318 | 462419 | 101 | -1,706901076 | | | | Overlap <50 bp | | rDNA | |
| chrXII | 462318 | 462423 | 105 | -1,133388482 | | | | Overlap <50 bp | | rDNA | |
| chrXII | 462318 | 462425 | 107 | -1,248235465 | | | | Overlap <50 bp | | rDNA | |
| chrXII | 462318 | 462460 | 142 | -1,335013962 | | | | Overlap <50 bp | | rDNA | |
| chrXII | 462318 | 462461 | 143 | -1,790923092 | | | | Overlap <50 bp | | rDNA | |
| chrXII | 462423 | 462569 | 146 | -0,872423658 | N12:462515 | 21,517 | +2:YLR154C-G; | 2 | | rDNA | |
| chrXII | 462436 | 462577 | 141 | -1,028616253 | N12:462515 | 21,517 | +2:YLR154C-G; | 2 | | rDNA | |
| chrXII | 462436 | 462597 | 161 | -1,509804872 | N12:462515 | 21,517 | +2:YLR154C-G; | 2 | | rDNA | |
| chrXII | 462437 | 462547 | 110 | -1,316457738 | N12:462515 | 21,517 | +2:YLR154C-G; | 2 | | rDNA | |
| chrXII | 462437 | 462577 | 140 | -1,512851225 | N12:462515 | 21,517 | +2:YLR154C-G; | 2 | | rDNA | |
| chrXII | 462439 | 462577 | 138 | -1,428489747 | N12:462515 | 21,517 | +2:YLR154C-G; | 2 | | rDNA | |
| chrXII | 462440 | 462577 | 137 | -1,412360047 | N12:462515 | 21,517 | +2:YLR154C-G; | 2 | | rDNA | |
| chrXII | 462447 | 462577 | 130 | -0,924465004 | N12:462515 | 21,517 | +2:YLR154C-G; | 2 | | rDNA | |
| chrXII | 462447 | 462597 | 150 | -1,318042782 | N12:462515 | 21,517 | +2:YLR154C-G; | 2 | | rDNA | |
| chrXII | 462459 | 462618 | 159 | -1,356019689 | N12:462515 | 21,517 | +2:YLR154C-G; | 2 | | rDNA | |
| chrXII | 462595 | 462732 | 137 | -1,651212947 | N12:462669 | 19,296 | +1:YLR154C-G; | 1 | | rDNA | |
| chrXII | 462595 | 462747 | 152 | -1,439696431 | N12:462669 | 19,296 | +1:YLR154C-G; | 1 | | rDNA | |
| chrXII | 462595 | 462749 | 154 | -1,415710209 | N12:462669 | 19,296 | +1:YLR154C-G; | 1 | | rDNA | |
| chrXII | 462597 | 462749 | 152 | -1,200723581 | N12:462669 | 19,296 | +1:YLR154C-G; | 1 | | rDNA | |
| chrXII | 462624 | 462732 | 108 | -1,470986452 | N12:462669 | 19,296 | +1:YLR154C-G; | 1 | | rDNA | |
| chrXII | 462624 | 462747 | 123 | -1,307820989 | N12:462669 | 19,296 | +1:YLR154C-G; | 1 | | rDNA | |
| chrXII | 462624 | 462749 | 125 | -1,429388607 | N12:462669 | 19,296 | +1:YLR154C-G; | 1 | | rDNA | |
| chrXII | 462637 | 462789 | 152 | -0,960058724 | N12:462669 | 19,296 | +1:YLR154C-G; | 1 | | rDNA | |
| chrXII | 462711 | 462864 | 153 | -1,113263202 | N12:462833 | 22,942 | 0:YLR154C-G; | 0 | | rDNA | |
| chrXII | 462724 | 462874 | 150 | -1,432615997 | N12:462833 | 22,942 | 0:YLR154C-G; | 0 | | rDNA | |
| chrXII | 462764 | 462913 | 149 | -1,807936083 | N12:462833 | 22,942 | 0:YLR154C-G; | 0 | | rDNA | |
| chrXII | 462764 | 462916 | 152 | -1,092942978 | N12:462833 | 22,942 | 0:YLR154C-G; | 0 | | rDNA | |
| chrXII | 462794 | 462951 | 157 | -0,876778185 | N12:462833 | 22,942 | 0:YLR154C-G; | 0 | | rDNA | |
| chrXII | 462796 | 462913 | 117 | -1,32578172 | N12:462833 | 22,942 | 0:YLR154C-G; | 0 | | rDNA | |
| chrXII | 462796 | 462916 | 120 | -1,412727582 | N12:462833 | 22,942 | 0:YLR154C-G; | 0 | | rDNA | |
| chrXII | 462871 | 462990 | 119 | -1,159231333 | N12:462990 | 9,899 | -1:YLR154C-G; | -1 | | rDNA | |
| chrXII | 462885 | 463047 | 162 | -1,907815123 | N12:462990 | 9,899 | -1:YLR154C-G; | -1 | | rDNA | |
| chrXII | 462885 | 463050 | 165 | -1,360044289 | N12:462990 | 9,899 | -1:YLR154C-G; | -1 | | rDNA | |
| chrXII | 462885 | 463056 | 171 | -1,402708028 | N12:462990 | 9,899 | -1:YLR154C-G; | -1 | | rDNA | |
| chrXII | 462885 | 463059 | 174 | -1,345505709 | N12:462990 | 9,899 | -1:YLR154C-G; | -1 | | rDNA | |
| chrXII | 462885 | 463099 | 214 | -1,388139775 | N12:462990 | 9,899 | -1:YLR154C-G; | -1 | | rDNA | |
| chrXII | 462889 | 462996 | 107 | -0,420298127 | N12:462990 | 9,899 | -1:YLR154C-G; | -1 | | rDNA | |
| chrXII | 462889 | 463045 | 156 | -1,512219097 | N12:462990 | 9,899 | -1:YLR154C-G; | -1 | | rDNA | |
| chrXII | 462889 | 463048 | 159 | -1,415230859 | N12:462990 | 9,899 | -1:YLR154C-G; | -1 | | rDNA | |
| chrXII | 462889 | 463051 | 162 | -1,352677981 | N12:462990 | 9,899 | -1:YLR154C-G; | -1 | | rDNA | |
| chrXII | 462889 | 463053 | 164 | -1,348525169 | N12:462990 | 9,899 | -1:YLR154C-G; | -1 | | rDNA | |
| chrXII | 462889 | 463059 | 170 | -1,381941782 | N12:462990 | 9,899 | -1:YLR154C-G; | -1 | | rDNA | |
| chrXII | 462889 | 463061 | 172 | -1,255925174 | N12:462990 | 9,899 | -1:YLR154C-G; | -1 | | rDNA | |
| chrXII | 462889 | 463066 | 177 | -1,480355991 | N12:462990 | 9,899 | -1:YLR154C-G; | -1 | | rDNA | |
| chrXII | 462890 | 463050 | 160 | -1,094599057 | N12:462990 | 9,899 | -1:YLR154C-G; | -1 | | rDNA | |
| chrXII | 462890 | 463059 | 169 | -1,318562791 | N12:462990 | 9,899 | -1:YLR154C-G; | -1 | | rDNA | |
| chrXII | 462891 | 463047 | 156 | -1,715156071 | N12:462990 | 9,899 | -1:YLR154C-G; | -1 | | rDNA | |
| chrXII | 462891 | 463056 | 165 | -1,730680222 | N12:462990 | 9,899 | -1:YLR154C-G; | -1 | | rDNA | |
| chrXII | 462891 | 463061 | 170 | -1,496767484 | N12:462990 | 9,899 | -1:YLR154C-G; | -1 | | rDNA | |
| chrXII | 462911 | 463047 | 136 | -1,288132838 | N12:462990 | 9,899 | -1:YLR154C-G; | -1 | | rDNA | |
| chrXII | 462911 | 463050 | 139 | -1,273277334 | N12:462990 | 9,899 | -1:YLR154C-G; | -1 | | rDNA | |
| chrXII | 462911 | 463051 | 140 | -1,747682671 | N12:462990 | 9,899 | -1:YLR154C-G; | -1 | | rDNA | |
| chrXII | 462911 | 463053 | 142 | -1,326359236 | N12:462990 | 9,899 | -1:YLR154C-G; | -1 | | rDNA | |
| chrXII | 462911 | 463056 | 145 | -1,149894996 | N12:462990 | 9,899 | -1:YLR154C-G; | -1 | | rDNA | |
| chrXII | 462911 | 463062 | 151 | -1,33135686 | N12:462990 | 9,899 | -1:YLR154C-G; | -1 | | rDNA | |
| chrXII | 462911 | 463099 | 188 | -1,397153673 | N12:462990 | 9,899 | -1:YLR154C-G; | -1 | | rDNA | |
| chrXII | 462912 | 463045 | 133 | -1,27733178 | N12:462990 | 9,899 | -1:YLR154C-G; | -1 | | rDNA | |
| chrXII | 462912 | 463048 | 136 | -1,366103625 | N12:462990 | 9,899 | -1:YLR154C-G; | -1 | | rDNA | |
| chrXII | 462912 | 463051 | 139 | -1,067048979 | N12:462990 | 9,899 | -1:YLR154C-G; | -1 | | rDNA | |
| chrXII | 462912 | 463053 | 141 | -1,198634693 | N12:462990 | 9,899 | -1:YLR154C-G; | -1 | | rDNA | |
| chrXII | 462912 | 463059 | 147 | -1,420418388 | N12:462990 | 9,899 | -1:YLR154C-G; | -1 | | rDNA | |
| chrXII | 462918 | 463045 | 127 | -0,928699495 | N12:462990 | 9,899 | -1:YLR154C-G; | -1 | | rDNA | |
| chrXII | 462918 | 463048 | 130 | -1,189890733 | N12:462990 | 9,899 | -1:YLR154C-G; | -1 | | rDNA | |
| chrXII | 462918 | 463051 | 133 | -1,129095014 | N12:462990 | 9,899 | -1:YLR154C-G; | -1 | | rDNA | |
| chrXII | 462918 | 463053 | 135 | -1,200000469 | N12:462990 | 9,899 | -1:YLR154C-G; | -1 | | rDNA | |
| chrXII | 462918 | 463059 | 141 | -1,332478689 | N12:462990 | 9,899 | -1:YLR154C-G; | -1 | | rDNA | |

| CHR | START | END | L(bp) | ΔLK | Nuc ID | Fuzziness | Gene ID | Gene body postion or intergene | Terminal | rDNA | Telomeric |
|---|---|---|---|---|---|---|---|---|---|---|---|
| chrXII | 462919 | 463048 | 129 | -1,151413803 | N12:462990 | 9,899 | -1:YLR154C-G; | -1 | | rDNA | |
| chrXII | 462919 | 463050 | 131 | -1,021000708 | N12:462990 | 9,899 | -1:YLR154C-G; | -1 | | rDNA | |
| chrXII | 462919 | 463051 | 132 | -1,303583622 | N12:462990 | 9,899 | -1:YLR154C-G; | -1 | | rDNA | |
| chrXII | 462919 | 463056 | 137 | -1,133186046 | N12:462990 | 9,899 | -1:YLR154C-G; | -1 | | rDNA | |
| chrXII | 462919 | 463059 | 140 | -1,469927096 | N12:462990 | 9,899 | -1:YLR154C-G; | -1 | | rDNA | |
| chrXII | 462919 | 463062 | 143 | -1,069947045 | N12:462990 | 9,899 | -1:YLR154C-G; | -1 | | rDNA | |
| chrXII | 462919 | 463066 | 147 | -1,348886903 | N12:462990 | 9,899 | -1:YLR154C-G; | -1 | | rDNA | |
| chrXII | 462919 | 463074 | 155 | -1,636297045 | N12:462990 | 9,899 | -1:YLR154C-G; | -1 | | rDNA | |
| chrXII | 462919 | 463099 | 180 | -1,172780269 | N12:462990 | 9,899 | -1:YLR154C-G; | -1 | | rDNA | |
| chrXII | 462920 | 463045 | 125 | -0,978291226 | N12:462990 | 9,899 | -1:YLR154C-G; | -1 | | rDNA | |
| chrXII | 462920 | 463048 | 128 | -1,237160695 | N12:462990 | 9,899 | -1:YLR154C-G; | -1 | | rDNA | |
| chrXII | 462920 | 463050 | 130 | -1,274710878 | N12:462990 | 9,899 | -1:YLR154C-G; | -1 | | rDNA | |
| chrXII | 462920 | 463051 | 131 | -1,47007321 | N12:462990 | 9,899 | -1:YLR154C-G; | -1 | | rDNA | |
| chrXII | 462949 | 463050 | 101 | -0,74920353 | N12:462990 | 9,899 | -1:YLR154C-G; | -1 | | rDNA | |
| chrXII | 462949 | 463056 | 107 | -0,426795306 | N12:462990 | 9,899 | -1:YLR154C-G; | -1 | | rDNA | |
| chrXII | 462949 | 463099 | 150 | -1,10618243 | N12:462990 | 9,899 | -1:YLR154C-G; | -1 | | rDNA | |
| chrXII | 462949 | 463138 | 189 | -1,338758548 | N12:462990 | 9,899 | -1:YLR154C-G; | -1 | | rDNA | |
| chrXII | 462949 | 463138 | 189 | -1,338758548 | N12:463152 | 4,041 | | | | rDNA | |
| chrXII | 462988 | 463095 | 107 | -0,54216178 | N12:462990 | 9,899 | -1:YLR154C-G; | -1 | | rDNA | |
| chrXII | 462988 | 463099 | 111 | -0,713358026 | N12:462990 | 9,899 | -1:YLR154C-G; | -1 | | rDNA | |
| chrXII | 462988 | 463138 | 150 | -1,131567353 | N12:462990 | 9,899 | -1:YLR154C-G; | -1 | | rDNA | |
| chrXII | 462988 | 463138 | 150 | -1,131567353 | N12:463152 | 4,041 | | | | rDNA | |
| chrXII | 463046 | 463209 | 163 | -1,622772572 | N12:463152 | 4,041 | | | | rDNA | |
| chrXII | 463055 | 463217 | 162 | -1,379019452 | N12:463152 | 4,041 | | | | rDNA | |
| chrXII | 463059 | 463216 | 157 | -1,265069485 | N12:463152 | 4,041 | | | | rDNA | |
| chrXII | 463059 | 463234 | 175 | -1,415362578 | N12:463152 | 4,041 | | | | rDNA | |
| chrXII | 463066 | 463216 | 150 | -1,953700408 | N12:463152 | 4,041 | | | | rDNA | |
| chrXII | 463066 | 463234 | 168 | -1,492294796 | N12:463152 | 4,041 | | | | rDNA | |
| chrXII | 463066 | 463245 | 179 | -1,621560895 | N12:463152 | 4,041 | | | | rDNA | |
| chrXII | 463096 | 463234 | 138 | -1,544826372 | N12:463152 | 4,041 | | | | rDNA | |
| chrXII | 463096 | 463245 | 149 | -1,189996991 | N12:463152 | 4,041 | | | | rDNA | |
| chrXII | 463096 | 463249 | 153 | -1,149130461 | N12:463152 | 4,041 | | | | rDNA | |
| chrXII | 463097 | 463253 | 156 | -1,238066263 | N12:463152 | 4,041 | | | | rDNA | |
| chrXII | 463101 | 463234 | 133 | -1,033142839 | N12:463152 | 4,041 | | | | rDNA | |
| chrXII | 463101 | 463245 | 144 | -1,249158536 | N12:463152 | 4,041 | | | | rDNA | |
| chrXII | 463108 | 463245 | 137 | -0,986648848 | N12:463152 | 4,041 | | | | rDNA | |
| chrXII | 463108 | 463260 | 152 | -1,202182707 | N12:463152 | 4,041 | | | | rDNA | |
| chrXII | 463196 | 463353 | 157 | -1,246245367 | N12:463325 | 17,474 | | | | rDNA | |
| chrXII | 463248 | 463405 | 157 | -1,49545714 | N12:463325 | 17,474 | | | | rDNA | |
| chrXII | 463257 | 463404 | 147 | -1,356702689 | N12:463325 | 17,474 | | | | rDNA | |
| chrXII | 463257 | 463414 | 157 | -1,423163998 | N12:463325 | 17,474 | | | | rDNA | |
| chrXII | 463261 | 463414 | 153 | -1,628062193 | N12:463325 | 17,474 | | | | rDNA | |
| chrXII | 463268 | 463414 | 146 | -1,229070119 | N12:463325 | 17,474 | | | | rDNA | |
| chrXII | 463278 | 463414 | 136 | -1,315356249 | N12:463325 | 17,474 | | | | rDNA | |
| chrXII | 463333 | 463483 | 150 | -1,184926423 | N12:463325 | 17,474 | | | | rDNA | |
| chrXII | 463333 | 463483 | 150 | -1,184926423 | N12:463504 | 37,477 | | | | rDNA | |
| chrXII | 463358 | 463511 | 153 | -1,844061972 | N12:463504 | 37,477 | | | | rDNA | |
| chrXII | 463358 | 463529 | 171 | -1,254982253 | N12:463504 | 37,477 | | | | rDNA | |
| chrXII | 463381 | 463529 | 148 | -1,165241011 | N12:463504 | 37,477 | | | | rDNA | |
| chrXII | 463381 | 463539 | 158 | -1,303742568 | N12:463504 | 37,477 | | | | rDNA | |
| chrXII | 463385 | 463511 | 126 | -1,085079341 | N12:463504 | 37,477 | | | | rDNA | |
| chrXII | 463385 | 463529 | 144 | -1,382706278 | N12:463504 | 37,477 | | | | rDNA | |
| chrXII | 463385 | 463535 | 150 | -1,284090886 | N12:463504 | 37,477 | | | | rDNA | |
| chrXII | 463385 | 463539 | 154 | -1,981963002 | N12:463504 | 37,477 | | | | rDNA | |
| chrXII | 463410 | 463573 | 163 | -1,346463084 | N12:463504 | 37,477 | | | | rDNA | |
| chrXII | 463414 | 463521 | 107 | -0,639576809 | N12:463504 | 37,477 | | | | rDNA | |
| chrXII | 463414 | 463535 | 121 | -0,716134641 | N12:463504 | 37,477 | | | | rDNA | |
| chrXII | 463414 | 463539 | 125 | -1,466607688 | N12:463504 | 37,477 | | | | rDNA | |
| chrXII | 463414 | 463575 | 161 | -1,137122627 | N12:463504 | 37,477 | | | | rDNA | |
| chrXII | 463414 | 463583 | 169 | -1,499229188 | N12:463504 | 37,477 | | | | rDNA | |
| chrXII | 463463 | 463603 | 140 | -1,225107835 | N12:463504 | 37,477 | | | | rDNA | |
| chrXII | 463471 | 463608 | 137 | -1,562700146 | N12:463504 | 37,477 | | | | rDNA | |
| chrXII | 463535 | 463688 | 153 | -1,555538451 | | | | Overlap <50 bp | | rDNA | |
| chrXII | 463535 | 463689 | 154 | -1,224153616 | | | | Overlap <50 bp | | rDNA | |
| chrXII | 463535 | 463690 | 155 | -1,512024304 | | | | Overlap <50 bp | | rDNA | |
| chrXII | 463535 | 463692 | 157 | -1,214113789 | | | | Overlap <50 bp | | rDNA | |
| chrXII | 463535 | 463704 | 169 | -1,243470838 | | | | Overlap <50 bp | | rDNA | |
| chrXII | 463535 | 463710 | 175 | -1,551768098 | | | | Overlap <50 bp | | rDNA | |
| chrXII | 463535 | 463712 | 177 | -1,323898552 | | | | Overlap <50 bp | | rDNA | |
| chrXII | 463541 | 463689 | 148 | -1,227753448 | | | | Overlap <50 bp | | rDNA | |
| chrXII | 463541 | 463692 | 151 | -1,097905797 | | | | Overlap <50 bp | | rDNA | |
| chrXII | 463559 | 463689 | 130 | -0,777913181 | | | | Overlap <50 bp | | rDNA | |
| chrXII | 463559 | 463704 | 145 | -1,106538451 | | | | Overlap <50 bp | | rDNA | |
| chrXII | 463559 | 463710 | 151 | -1,478491197 | | | | Overlap <50 bp | | rDNA | |
| chrXII | 463559 | 463712 | 153 | -1,158334765 | | | | Overlap <50 bp | | rDNA | |
| chrXII | 463658 | 463815 | 157 | -1,339081123 | N12:463786 | 6,658 | | | | rDNA | |
| chrXII | 463658 | 463852 | 194 | -1,731037165 | N12:463786 | 6,658 | | | | rDNA | |
| chrXII | 463658 | 463870 | 212 | -1,52721217 | N12:463786 | 6,658 | | | | rDNA | |
| chrXII | 463690 | 463849 | 159 | -1,284883711 | N12:463786 | 6,658 | | | | rDNA | |
| chrXII | 463690 | 463870 | 180 | -1,522182911 | N12:463786 | 6,658 | | | | rDNA | |
| chrXII | 463692 | 463840 | 148 | -1,205370905 | N12:463786 | 6,658 | | | | rDNA | |
| chrXII | 463697 | 463815 | 118 | -0,831407861 | N12:463786 | 6,658 | | | | rDNA | |
| chrXII | 463697 | 463849 | 152 | -1,387924403 | N12:463786 | 6,658 | | | | rDNA | |
| chrXII | 463697 | 463852 | 155 | -1,318906556 | N12:463786 | 6,658 | | | | rDNA | |
| chrXII | 463697 | 463861 | 164 | -1,259434206 | N12:463786 | 6,658 | | | | rDNA | |
| chrXII | 463697 | 463870 | 173 | -1,334837966 | N12:463786 | 6,658 | | | | rDNA | |
| chrXII | 463699 | 463849 | 150 | -1,088250816 | N12:463786 | 6,658 | | | | rDNA | |
| chrXII | 463707 | 463849 | 142 | -1,071534672 | N12:463786 | 6,658 | | | | rDNA | |
| chrXII | 463707 | 463852 | 145 | -1,527546399 | N12:463786 | 6,658 | | | | rDNA | |
| chrXII | 463707 | 463861 | 154 | -1,223691861 | N12:463786 | 6,658 | | | | rDNA | |
| chrXII | 463707 | 463870 | 163 | -1,25052542 | N12:463786 | 6,658 | | | | rDNA | |
| chrXII | 463713 | 463849 | 136 | -1,233972975 | N12:463786 | 6,658 | | | | rDNA | |
| chrXII | 463713 | 463852 | 139 | -1,465489107 | N12:463786 | 6,658 | | | | rDNA | |
| chrXII | 463713 | 463861 | 148 | -1,367426781 | N12:463786 | 6,658 | | | | rDNA | |
| chrXII | 463713 | 463870 | 157 | -1,30642325 | N12:463786 | 6,658 | | | | rDNA | |
| chrXII | 463713 | 463909 | 196 | -1,482773162 | N12:463786 | 6,658 | | | | rDNA | |
| chrXII | 463724 | 463883 | 159 | -1,093610181 | N12:463786 | 6,658 | | | | rDNA | |
| chrXII | 463729 | 463879 | 150 | -1,357274644 | N12:463786 | 6,658 | | | | rDNA | |
| chrXII | 463763 | 463849 | 86 | -0,441157983 | N12:463786 | 6,658 | | | | rDNA | |

| CHR | START | END | L(bp) | ΔLK | Nuc ID | Fuzziness | Gene ID | Gene body postion or intergene | Terminal | rDNA | Telomeric |
|---|---|---|---|---|---|---|---|---|---|---|---|
| chrXII | 463763 | 463870 | 107 | -0,576000587 | N12:463786 | 6,658 | | | | rDNA | |
| chrXII | 463763 | 463909 | 146 | -1,198370929 | N12:463786 | 6,658 | | | | rDNA | |
| chrXII | 463860 | 463991 | 131 | -1,146460514 | N12:463954 | 7,071 | | | | rDNA | |
| chrXII | 463862 | 464000 | 138 | -1,064057189 | N12:463954 | 7,071 | | | | rDNA | |
| chrXII | 463862 | 464023 | 161 | -1,151862236 | N12:463954 | 7,071 | | | | rDNA | |
| chrXII | 463870 | 464023 | 153 | -1,481560325 | N12:463954 | 7,071 | | | | rDNA | |
| chrXII | 463878 | 464033 | 155 | -1,319954406 | N12:463954 | 7,071 | | | | rDNA | |
| chrXII | 463884 | 464023 | 139 | -1,054586869 | N12:463954 | 7,071 | | | | rDNA | |
| chrXII | 463974 | 464128 | 154 | -0,837325318 | N12:463954 | 7,071 | | | | rDNA | |
| chrXII | 463974 | 464128 | 154 | -0,837325318 | N12:464114 | 7,778 | | | | rDNA | |
| chrXII | 464001 | 464158 | 157 | -1,164822956 | N12:464114 | 7,778 | | | | rDNA | |
| chrXII | 464013 | 464160 | 147 | -1,138127981 | N12:464114 | 7,778 | | | | rDNA | |
| chrXII | 464013 | 464166 | 153 | -1,678143929 | N12:464114 | 7,778 | | | | rDNA | |
| chrXII | 464038 | 464166 | 128 | -1,291466261 | N12:464114 | 7,778 | | | | rDNA | |
| chrXII | 464038 | 464211 | 173 | -1,459496678 | N12:464114 | 7,778 | | | | rDNA | |
| chrXII | 464057 | 464224 | 167 | -1,471343759 | N12:464114 | 7,778 | | | | rDNA | |
| chrXII | 464069 | 464210 | 141 | -1,048842111 | N12:464114 | 7,778 | | | | rDNA | |
| chrXII | 464070 | 464160 | 90 | -0,785887954 | N12:464114 | 7,778 | | | | rDNA | |
| chrXII | 464070 | 464166 | 96 | -1,158013946 | N12:464114 | 7,778 | | | | rDNA | |
| chrXII | 464070 | 464210 | 140 | -1,143743918 | N12:464114 | 7,778 | | | | rDNA | |
| chrXII | 464070 | 464211 | 141 | -1,302584909 | N12:464114 | 7,778 | | | | rDNA | |
| chrXII | 464070 | 464213 | 143 | -1,226673522 | N12:464114 | 7,778 | | | | rDNA | |
| chrXII | 464101 | 464269 | 168 | -1,251300944 | N12:464114 | 7,778 | | | | rDNA | |
| chrXII | 464101 | 464269 | 168 | -1,251300944 | N12:464279 | 14,142 | | | | rDNA | |
| chrXII | 464132 | 464301 | 169 | -1,649425781 | N12:464114 | 7,778 | | | | rDNA | |
| chrXII | 464132 | 464301 | 169 | -1,649425781 | N12:464279 | 14,142 | | | | rDNA | |
| chrXII | 464137 | 464269 | 132 | -1,067927668 | N12:464114 | 7,778 | | | | rDNA | |
| chrXII | 464137 | 464269 | 132 | -1,067927668 | N12:464279 | 14,142 | | | | rDNA | |
| chrXII | 464152 | 464303 | 151 | -1,217171292 | N12:464279 | 14,142 | | | | rDNA | |
| chrXII | 464195 | 464345 | 150 | -1,671626346 | N12:464279 | 14,142 | | | | rDNA | |
| chrXII | 464205 | 464360 | 155 | -1,247577908 | N12:464279 | 14,142 | | | | rDNA | |
| chrXII | 464232 | 464406 | 174 | -1,418381264 | N12:464279 | 14,142 | | | | rDNA | |
| chrXII | 464232 | 464406 | 174 | -1,418381264 | N12:464427 | 22,679 | | | | rDNA | |
| chrXII | 464233 | 464360 | 127 | -1,299187093 | N12:464279 | 14,142 | | | | rDNA | |
| chrXII | 464233 | 464377 | 144 | -1,119894183 | N12:464279 | 14,142 | | | | rDNA | |
| chrXII | 464233 | 464380 | 147 | -1,119849088 | N12:464279 | 14,142 | | | | rDNA | |
| chrXII | 464233 | 464382 | 149 | -1,093330695 | N12:464279 | 14,142 | | | | rDNA | |
| chrXII | 464233 | 464386 | 153 | -1,240075175 | N12:464279 | 14,142 | | | | rDNA | |
| chrXII | 464234 | 464377 | 143 | -1,582792225 | N12:464279 | 14,142 | | | | rDNA | |
| chrXII | 464234 | 464380 | 146 | -1,27963756 | N12:464279 | 14,142 | | | | rDNA | |
| chrXII | 464234 | 464382 | 148 | -1,251369002 | N12:464279 | 14,142 | | | | rDNA | |
| chrXII | 464235 | 464377 | 142 | -1,225883176 | N12:464279 | 14,142 | | | | rDNA | |
| chrXII | 464235 | 464380 | 145 | -1,403622677 | N12:464279 | 14,142 | | | | rDNA | |
| chrXII | 464235 | 464382 | 147 | -1,50302983 | N12:464279 | 14,142 | | | | rDNA | |
| chrXII | 464235 | 464386 | 151 | -1,136642671 | N12:464279 | 14,142 | | | | rDNA | |
| chrXII | 464266 | 464430 | 164 | -1,268995931 | N12:464279 | 14,142 | | | | rDNA | |
| chrXII | 464266 | 464430 | 164 | -1,268995931 | N12:464427 | 22,679 | | | | rDNA | |
| chrXII | 464275 | 464426 | 151 | -1,323866282 | N12:464279 | 14,142 | | | | rDNA | |
| chrXII | 464275 | 464426 | 151 | -1,323866282 | N12:464427 | 22,679 | | | | rDNA | |
| chrXII | 464319 | 464475 | 156 | -1,689774551 | N12:464427 | 22,679 | | | | rDNA | |
| chrXII | 464354 | 464503 | 149 | -1,459627491 | N12:464427 | 22,679 | | | | rDNA | |
| chrXII | 464358 | 464532 | 174 | -1,774508688 | N12:464427 | 22,679 | | | | rDNA | |
| chrXII | 464384 | 464518 | 134 | -1,635147555 | N12:464427 | 22,679 | | | | rDNA | |
| chrXII | 464391 | 464538 | 147 | -1,402020385 | N12:464427 | 22,679 | | | | rDNA | |
| chrXII | 464392 | 464537 | 145 | -1,630022925 | N12:464427 | 22,679 | | | | rDNA | |
| chrXII | 464415 | 464560 | 145 | -1,360717059 | N12:464427 | 22,679 | | | | rDNA | |
| chrXII | 464434 | 464584 | 150 | -1,280217243 | N12:464427 | 22,679 | | | | rDNA | |
| chrXII | 464504 | 464649 | 145 | -1,101904191 | N12:464616 | 19 | | | | rDNA | |
| chrXII | 464504 | 464668 | 164 | -1,130510575 | N12:464616 | 19 | | | | rDNA | |
| chrXII | 464504 | 464686 | 182 | -1,314977046 | N12:464616 | 19 | | | | rDNA | |
| chrXII | 464514 | 464649 | 135 | -1,150238815 | N12:464616 | 19 | | | | rDNA | |
| chrXII | 464514 | 464668 | 154 | -1,175799933 | N12:464616 | 19 | | | | rDNA | |
| chrXII | 464514 | 464686 | 172 | -1,202026112 | N12:464616 | 19 | | | | rDNA | |
| chrXII | 464534 | 464640 | 106 | -0,710188086 | N12:464616 | 19 | | | | rDNA | |
| chrXII | 464534 | 464649 | 115 | -0,452673507 | N12:464616 | 19 | | | | rDNA | |
| chrXII | 464534 | 464668 | 134 | -0,633933501 | N12:464616 | 19 | | | | rDNA | |
| chrXII | 464534 | 464686 | 152 | -1,339918936 | N12:464616 | 19 | | | | rDNA | |
| chrXII | 464547 | 464649 | 102 | -0,475639868 | N12:464616 | 19 | | | | rDNA | |
| chrXII | 464547 | 464686 | 139 | -1,411086837 | N12:464616 | 19 | | | | rDNA | |
| chrXII | 464547 | 464697 | 150 | -0,648494907 | N12:464616 | 19 | | | | rDNA | |
| chrXII | 464547 | 464705 | 158 | -1,188623716 | N12:464616 | 19 | | | | rDNA | |
| chrXII | 464547 | 464706 | 159 | -1,479829738 | N12:464616 | 19 | | | | rDNA | |
| chrXII | 464553 | 464708 | 155 | -1,267859493 | N12:464616 | 19 | | | | rDNA | |
| chrXII | 464559 | 464706 | 147 | -1,026528444 | N12:464616 | 19 | | | | rDNA | |
| chrXII | 464728 | 464888 | 160 | -1,757799016 | N12:464796 | 30,199 | | | | rDNA | |
| chrXII | 464838 | 465007 | 169 | -1,64829069 | N12:464962 | 52,647 | | | | rDNA | |
| chrXII | 464884 | 465032 | 148 | -1,358902718 | N12:464962 | 52,647 | | | | rDNA | |
| chrXII | 464886 | 465026 | 140 | -1,06336186 | N12:464962 | 52,647 | | | | rDNA | |
| chrXII | 464920 | 465095 | 175 | -1,79108638 | N12:464962 | 52,647 | | | | rDNA | |
| chrXII | 465006 | 465158 | 152 | -1,399748791 | N12:465169 | 29,501 | | | | rDNA | |
| chrXII | 465040 | 465204 | 164 | -1,688477357 | N12:465169 | 29,501 | | | | rDNA | |
| chrXII | 465055 | 465176 | 121 | -1,350886964 | N12:465169 | 29,501 | | | | rDNA | |
| chrXII | 465063 | 465223 | 160 | -1,34237679 | N12:465169 | 29,501 | | | | rDNA | |
| chrXII | 465159 | 465310 | 151 | -1,212476488 | N12:465169 | 29,501 | | | | rDNA | |
| chrXII | 465159 | 465342 | 183 | -1,319695984 | N12:465169 | 29,501 | | | | rDNA | |
| chrXII | 465159 | 465342 | 183 | -1,319695984 | N12:465356 | 20,664 | | | | rDNA | |
| chrXII | 465159 | 465348 | 189 | -1,395346351 | N12:465169 | 29,501 | | | | rDNA | |
| chrXII | 465159 | 465348 | 189 | -1,395346351 | N12:465356 | 20,664 | | | | rDNA | |
| chrXII | 465159 | 465357 | 198 | -1,265115529 | N12:465169 | 29,501 | | | | rDNA | |
| chrXII | 465159 | 465357 | 198 | -1,265115529 | N12:465356 | 20,664 | | | | rDNA | |
| chrXII | 465179 | 465310 | 131 | -0,906452825 | N12:465169 | 29,501 | | | | rDNA | |
| chrXII | 465182 | 465312 | 130 | -1,089315353 | N12:465169 | 29,501 | | | | rDNA | |
| chrXII | 465193 | 465310 | 117 | -0,64626942 | | | | Overlap <50 bp | | rDNA | |
| chrXII | 465193 | 465312 | 119 | -0,556159197 | | | | Overlap <50 bp | | rDNA | |
| chrXII | 465193 | 465342 | 149 | -1,273839188 | N12:465356 | 20,664 | | | | rDNA | |
| chrXII | 465193 | 465348 | 155 | -1,253516586 | N12:465356 | 20,664 | | | | rDNA | |
| chrXII | 465193 | 465357 | 164 | -1,248167912 | N12:465356 | 20,664 | | | | rDNA | |
| chrXII | 465200 | 465352 | 152 | -1,266798765 | N12:465356 | 20,664 | | | | rDNA | |
| chrXII | 465201 | 465342 | 141 | -0,813038446 | N12:465356 | 20,664 | | | | rDNA | |
| chrXII | 465201 | 465345 | 144 | -0,950333857 | N12:465356 | 20,664 | | | | rDNA | |

| CHR | START | END | L(bp) | ΔLK | Nuc ID | Fuzziness | Gene ID | Gene body postion or intergene | Terminal | rDNA | Telomeric |
|---|---|---|---|---|---|---|---|---|---|---|---|
| chrXII | 465201 | 465351 | 150 | -1,035174987 | N12:465356 | 20,664 | | | | rDNA | |
| chrXII | 465203 | 465310 | 107 | -0,313755689 | | | | Overlap <50 bp | | rDNA | |
| chrXII | 465204 | 465342 | 138 | -1,206626675 | N12:465356 | 20,664 | | | | rDNA | |
| chrXII | 465204 | 465348 | 144 | -1,010079394 | N12:465356 | 20,664 | | | | rDNA | |
| chrXII | 465204 | 465352 | 148 | -1,261860582 | N12:465356 | 20,664 | | | | rDNA | |
| chrXII | 465204 | 465357 | 153 | -1,036499032 | N12:465356 | 20,664 | | | | rDNA | |
| chrXII | 465204 | 465403 | 199 | -1,68084809 | N12:465356 | 20,664 | | | | rDNA | |
| chrXII | 465204 | 465418 | 214 | -1,77160931 | N12:465356 | 20,664 | | | | rDNA | |
| chrXII | 465217 | 465310 | 93 | -0,34965455 | | | | Overlap <50 bp | | rDNA | |
| chrXII | 465217 | 465342 | 125 | -0,75856945 | N12:465356 | 20,664 | | | | rDNA | |
| chrXII | 465217 | 465348 | 131 | -0,909228921 | N12:465356 | 20,664 | | | | rDNA | |
| chrXII | 465217 | 465357 | 140 | -0,887234027 | N12:465356 | 20,664 | | | | rDNA | |
| chrXII | 465221 | 465352 | 131 | -0,76609552 | N12:465356 | 20,664 | | | | rDNA | |
| chrXII | 465221 | 465375 | 154 | -1,350621696 | N12:465356 | 20,664 | | | | rDNA | |
| chrXII | 465221 | 465380 | 159 | -1,353591856 | N12:465356 | 20,664 | | | | rDNA | |
| chrXII | 465221 | 465403 | 182 | -1,70769705 | N12:465356 | 20,664 | | | | rDNA | |
| chrXII | 465252 | 465352 | 100 | -0,578241955 | N12:465356 | 20,664 | | | | rDNA | |
| chrXII | 465252 | 465380 | 128 | -0,888432155 | N12:465356 | 20,664 | | | | rDNA | |
| chrXII | 465252 | 465403 | 151 | -1,219558956 | N12:465356 | 20,664 | | | | rDNA | |
| chrXII | 465252 | 465418 | 166 | -1,441285825 | N12:465356 | 20,664 | | | | rDNA | |
| chrXII | 465257 | 465427 | 170 | -1,316955745 | N12:465356 | 20,664 | | | | rDNA | |
| chrXII | 465261 | 465352 | 91 | -0,718600986 | N12:465356 | 20,664 | | | | rDNA | |
| chrXII | 465261 | 465403 | 142 | -1,381703957 | N12:465356 | 20,664 | | | | rDNA | |
| chrXII | 465261 | 465418 | 157 | -1,360370725 | N12:465356 | 20,664 | | | | rDNA | |
| chrXII | 465261 | 465470 | 209 | -1,522324322 | N12:465356 | 20,664 | | | | rDNA | |
| chrXII | 465318 | 465403 | 85 | -0,607218614 | N12:465356 | 20,664 | | | | rDNA | |
| chrXII | 465318 | 465418 | 100 | -0,692373553 | N12:465356 | 20,664 | | | | rDNA | |
| chrXII | 465318 | 465447 | 129 | -1,066165389 | N12:465356 | 20,664 | | | | rDNA | |
| chrXII | 465318 | 465469 | 151 | -1,745366324 | N12:465356 | 20,664 | | | | rDNA | |
| chrXII | 465318 | 465470 | 152 | -1,304533325 | N12:465356 | 20,664 | | | | rDNA | |
| chrXII | 465318 | 465473 | 155 | -1,213890461 | N12:465356 | 20,664 | | | | rDNA | |
| chrXII | 465318 | 465486 | 168 | -1,301002233 | N12:465356 | 20,664 | | | | rDNA | |
| chrXII | 465318 | 465524 | 206 | -1,599876434 | N12:465356 | 20,664 | | | | rDNA | |
| chrXII | 465318 | 465524 | 206 | -1,599876434 | N12:465541 | 35,303 | | | | rDNA | |
| chrXII | 465323 | 465470 | 147 | -1,614559164 | N12:465356 | 20,664 | | | | rDNA | |
| chrXII | 465324 | 465470 | 146 | -1,61665492 | N12:465356 | 20,664 | | | | rDNA | |
| chrXII | 465324 | 465486 | 162 | -1,216629237 | N12:465356 | 20,664 | | | | rDNA | |
| chrXII | 465331 | 465470 | 139 | -1,311337704 | N12:465356 | 20,664 | | | | rDNA | |
| chrXII | 465331 | 465473 | 142 | -1,189451823 | N12:465356 | 20,664 | | | | rDNA | |
| chrXII | 465350 | 465456 | 106 | -0,83709612 | N12:465356 | 20,664 | | | | rDNA | |
| chrXII | 465350 | 465457 | 107 | -0,852792116 | N12:465356 | 20,664 | | | | rDNA | |
| chrXII | 465350 | 465458 | 108 | -1,050623833 | N12:465356 | 20,664 | | | | rDNA | |
| chrXII | 465350 | 465470 | 120 | -1,229149517 | N12:465356 | 20,664 | | | | rDNA | |
| chrXII | 465350 | 465473 | 123 | -1,193061194 | N12:465356 | 20,664 | | | | rDNA | |
| chrXII | 465350 | 465486 | 136 | -1,402680295 | N12:465356 | 20,664 | | | | rDNA | |
| chrXII | 465350 | 465524 | 174 | -1,460260563 | N12:465356 | 20,664 | | | | rDNA | |
| chrXII | 465350 | 465524 | 174 | -1,460260563 | N12:465541 | 35,303 | | | | rDNA | |
| chrXII | 465350 | 465556 | 206 | -1,805916871 | N12:465356 | 20,664 | | | | rDNA | |
| chrXII | 465350 | 465556 | 206 | -1,805916871 | N12:465541 | 35,303 | | | | rDNA | |
| chrXII | 465358 | 465523 | 165 | -1,007146733 | N12:465356 | 20,664 | | | | rDNA | |
| chrXII | 465358 | 465523 | 165 | -1,007146733 | N12:465541 | 35,303 | | | | rDNA | |
| chrXII | 465377 | 465470 | 93 | -0,569053498 | N12:465356 | 20,664 | | | | rDNA | |
| chrXII | 465377 | 465484 | 107 | -0,878121394 | N12:465356 | 20,664 | | | | rDNA | |
| chrXII | 465377 | 465523 | 146 | -1,039561969 | N12:465356 | 20,664 | | | | rDNA | |
| chrXII | 465377 | 465523 | 146 | -1,039561969 | N12:465541 | 35,303 | | | | rDNA | |
| chrXII | 465377 | 465524 | 147 | -1,210166714 | N12:465356 | 20,664 | | | | rDNA | |
| chrXII | 465377 | 465524 | 147 | -1,210166714 | N12:465541 | 35,303 | | | | rDNA | |
| chrXII | 465377 | 465556 | 179 | -1,340352287 | N12:465356 | 20,664 | | | | rDNA | |
| chrXII | 465377 | 465556 | 179 | -1,340352287 | N12:465541 | 35,303 | | | | rDNA | |
| chrXII | 465377 | 465584 | 207 | -1,712269214 | N12:465356 | 20,664 | | | | rDNA | |
| chrXII | 465377 | 465584 | 207 | -1,712269214 | N12:465541 | 35,303 | | | | rDNA | |
| chrXII | 465425 | 465524 | 99 | -0,650000163 | N12:465541 | 35,303 | | | | rDNA | |
| chrXII | 465425 | 465532 | 107 | -1,238452803 | N12:465541 | 35,303 | | | | rDNA | |
| chrXII | 465425 | 465556 | 131 | -1,075907886 | N12:465541 | 35,303 | | | | rDNA | |
| chrXII | 465425 | 465584 | 159 | -1,1945625 | N12:465541 | 35,303 | | | | rDNA | |
| chrXII | 465425 | 465594 | 169 | -1,371021688 | N12:465541 | 35,303 | | | | rDNA | |
| chrXII | 465425 | 465608 | 183 | -1,877009901 | N12:465541 | 35,303 | | | | rDNA | |
| chrXII | 465427 | 465524 | 97 | -0,73042467 | N12:465541 | 35,303 | | | | rDNA | |
| chrXII | 465427 | 465556 | 129 | -1,243939783 | N12:465541 | 35,303 | | | | rDNA | |
| chrXII | 465427 | 465584 | 157 | -1,640094 | N12:465541 | 35,303 | | | | rDNA | |
| chrXII | 465427 | 465608 | 181 | -2,029628688 | N12:465541 | 35,303 | | | | rDNA | |
| chrXII | 465435 | 465584 | 149 | -1,283628907 | N12:465541 | 35,303 | | | | rDNA | |
| chrXII | 465447 | 465556 | 109 | -0,725427933 | N12:465541 | 35,303 | | | | rDNA | |
| chrXII | 465447 | 465584 | 137 | -1,106094081 | N12:465541 | 35,303 | | | | rDNA | |
| chrXII | 465447 | 465608 | 161 | -1,419102613 | N12:465541 | 35,303 | | | | rDNA | |
| chrXII | 465498 | 465584 | 86 | -1,141204061 | N12:465541 | 35,303 | | | | rDNA | |
| chrXII | 465498 | 465603 | 105 | -1,626306636 | N12:465541 | 35,303 | | | | rDNA | |
| chrXII | 465498 | 465605 | 107 | -1,44440594 | N12:465541 | 35,303 | | | | rDNA | |
| chrXII | 465498 | 465606 | 108 | -1,282178349 | N12:465541 | 35,303 | | | | rDNA | |
| chrXII | 465498 | 465607 | 109 | -1,32173857 | N12:465541 | 35,303 | | | | rDNA | |
| chrXII | 465498 | 465608 | 110 | -1,510691085 | N12:465541 | 35,303 | | | | rDNA | |
| chrXII | 465525 | 465608 | 83 | -1,049290733 | N12:465541 | 35,303 | | | | rDNA | |
| chrXII | 465525 | 465688 | 163 | -1,448472887 | N12:465541 | 35,303 | | | | rDNA | |
| chrXII | 465525 | 465688 | 163 | -1,448472887 | N12:465711 | 28,919 | | | | rDNA | |
| chrXII | 465562 | 465739 | 177 | -1,54386528 | N12:465541 | 35,303 | | | | rDNA | |
| chrXII | 465562 | 465739 | 177 | -1,54386528 | N12:465711 | 28,919 | | | | rDNA | |
| chrXII | 465581 | 465731 | 150 | -1,281580632 | N12:465711 | 28,919 | | | | rDNA | |
| chrXII | 465581 | 465732 | 151 | -1,328742751 | N12:465711 | 28,919 | | | | rDNA | |
| chrXII | 465581 | 465756 | 175 | -1,610495994 | N12:465711 | 28,919 | | | | rDNA | |
| chrXII | 465582 | 465732 | 150 | -1,351958361 | N12:465711 | 28,919 | | | | rDNA | |
| chrXII | 465584 | 465731 | 147 | -1,356467928 | N12:465711 | 28,919 | | | | rDNA | |
| chrXII | 465584 | 465732 | 148 | -1,516457509 | N12:465711 | 28,919 | | | | rDNA | |
| chrXII | 465584 | 465745 | 161 | -1,695353124 | N12:465711 | 28,919 | | | | rDNA | |
| chrXII | 465584 | 465756 | 172 | -1,396825526 | N12:465711 | 28,919 | | | | rDNA | |
| chrXII | 465616 | 465722 | 106 | -0,82411353 | N12:465711 | 28,919 | | | | rDNA | |
| chrXII | 465616 | 465731 | 115 | -0,761810369 | N12:465711 | 28,919 | | | | rDNA | |
| chrXII | 465616 | 465745 | 129 | -1,02369807 | N12:465711 | 28,919 | | | | rDNA | |
| chrXII | 465616 | 465746 | 130 | -1,567618533 | N12:465711 | 28,919 | | | | rDNA | |
| chrXII | 465616 | 465790 | 174 | -1,026305713 | N12:465711 | 28,919 | | | | rDNA | |
| chrXII | 465620 | 465731 | 111 | -0,737739116 | N12:465711 | 28,919 | | | | rDNA | |

| CHR | START | END | L(bp) | ΔLK | Nuc ID | Fuzziness | Gene ID | Gene body postion or intergene | Terminal | rDNA | Telomeric |
|---|---|---|---|---|---|---|---|---|---|---|---|
| chrXII | 465620 | 465745 | 125 | -1,003039587 | N12:465711 | 28,919 | | | | rDNA | |
| chrXII | 465620 | 465746 | 126 | -0,988387854 | N12:465711 | 28,919 | | | | rDNA | |
| chrXII | 465630 | 465736 | 106 | -0,550861011 | N12:465711 | 28,919 | | | | rDNA | |
| chrXII | 465630 | 465745 | 115 | -0,728664547 | N12:465711 | 28,919 | | | | rDNA | |
| chrXII | 465630 | 465790 | 160 | -1,234164439 | N12:465711 | 28,919 | | | | rDNA | |
| chrXII | 465645 | 465731 | 86 | -0,574702254 | N12:465711 | 28,919 | | | | rDNA | |
| chrXII | 465645 | 465745 | 100 | -0,410819086 | N12:465711 | 28,919 | | | | rDNA | |
| chrXII | 465645 | 465779 | 134 | -1,195662104 | N12:465711 | 28,919 | | | | rDNA | |
| chrXII | 465645 | 465790 | 145 | -1,333502136 | N12:465711 | 28,919 | | | | rDNA | |
| chrXII | 465645 | 465807 | 162 | -0,951768945 | N12:465711 | 28,919 | | | | rDNA | |
| chrXII | 465661 | 465832 | 171 | -1,428454789 | N12:465711 | 28,919 | | | | rDNA | |
| chrXII | 465724 | 465875 | 151 | -1,511976185 | N12:465711 | 28,919 | | | | rDNA | |
| chrXII | 465724 | 465875 | 151 | -1,511976185 | N12:465875 | 14,142 | | | | rDNA | |
| chrXII | 465725 | 465888 | 163 | -1,212443271 | N12:465711 | 28,919 | | | | rDNA | |
| chrXII | 465725 | 465888 | 163 | -1,212443271 | N12:465875 | 14,142 | | | | rDNA | |
| chrXII | 465731 | 465888 | 157 | -1,364958294 | N12:465711 | 28,919 | | | | rDNA | |
| chrXII | 465731 | 465888 | 157 | -1,364958294 | N12:465875 | 14,142 | | | | rDNA | |
| chrXII | 465753 | 465909 | 156 | -1,255174649 | N12:465875 | 14,142 | | | | rDNA | |
| chrXII | 465771 | 465899 | 128 | -1,070703 | N12:465875 | 14,142 | | | | rDNA | |
| chrXII | 465771 | 465918 | 147 | -1,218083451 | N12:465875 | 14,142 | | | | rDNA | |
| chrXII | 465786 | 465940 | 154 | -1,203024075 | N12:465875 | 14,142 | | | | rDNA | |
| chrXII | 465790 | 465899 | 109 | -0,795337195 | N12:465875 | 14,142 | | | | rDNA | |
| chrXII | 465790 | 465940 | 150 | -0,860308771 | N12:465875 | 14,142 | | | | rDNA | |
| chrXII | 465903 | 466057 | 154 | -1,111125655 | N12:466028 | 33,501 | | | | rDNA | |
| chrXII | 465917 | 466024 | 107 | -1,714368733 | N12:466028 | 33,501 | | | | rDNA | |
| chrXII | 465917 | 466027 | 110 | -1,599447913 | N12:466028 | 33,501 | | | | rDNA | |
| chrXII | 465917 | 466057 | 140 | -2,249467514 | N12:466028 | 33,501 | | | | rDNA | |
| chrXII | 465931 | 466040 | 109 | -0,771118229 | N12:466028 | 33,501 | | | | rDNA | |
| chrXII | 465932 | 466040 | 108 | -1,003861473 | N12:466028 | 33,501 | | | | rDNA | |
| chrXII | 465932 | 466041 | 109 | -1,116046151 | N12:466028 | 33,501 | | | | rDNA | |
| chrXII | 465934 | 466027 | 93 | -1,405374122 | N12:466028 | 33,501 | | | | rDNA | |
| chrXII | 465934 | 466057 | 123 | -2,044151178 | N12:466028 | 33,501 | | | | rDNA | |
| chrXII | 465940 | 466096 | 156 | -1,50789203 | N12:466028 | 33,501 | | | | rDNA | |
| chrXII | 465941 | 466048 | 107 | -1,064341363 | N12:466028 | 33,501 | | | | rDNA | |
| chrXII | 465941 | 466057 | 116 | -1,233141412 | N12:466028 | 33,501 | | | | rDNA | |
| chrXII | 465941 | 466097 | 156 | -1,615176487 | N12:466028 | 33,501 | | | | rDNA | |
| chrXII | 465954 | 466057 | 103 | -0,777525185 | N12:466028 | 33,501 | | | | rDNA | |
| chrXII | 465954 | 466108 | 154 | -1,493198002 | N12:466028 | 33,501 | | | | rDNA | |
| chrXII | 466075 | 466241 | 166 | -1,516575912 | N12:466190 | 42,899 | | | | rDNA | |
| chrXII | 466105 | 466258 | 153 | -1,528878287 | N12:466190 | 42,899 | | | | rDNA | |
| chrXII | 466121 | 466255 | 134 | -1,402794148 | N12:466190 | 42,899 | | | | rDNA | |
| chrXII | 466121 | 466264 | 143 | -1,429706159 | N12:466190 | 42,899 | | | | rDNA | |
| chrXII | 466121 | 466267 | 146 | -1,255345988 | N12:466190 | 42,899 | | | | rDNA | |
| chrXII | 466121 | 466277 | 156 | -1,627858752 | N12:466190 | 42,899 | | | | rDNA | |
| chrXII | 466127 | 466264 | 137 | -1,273071282 | N12:466190 | 42,899 | | | | rDNA | |
| chrXII | 466127 | 466277 | 150 | -1,4800357 | N12:466190 | 42,899 | | | | rDNA | |
| chrXII | 466142 | 466303 | 161 | -1,812771488 | N12:466190 | 42,899 | | | | rDNA | |
| chrXII | 466158 | 466264 | 106 | -1,822864991 | N12:466190 | 42,899 | | | | rDNA | |
| chrXII | 466159 | 466264 | 105 | -1,14603945 | N12:466190 | 42,899 | | | | rDNA | |
| chrXII | 466159 | 466342 | 183 | -2,339348407 | N12:466190 | 42,899 | | | | rDNA | |
| chrXII | 466159 | 466342 | 183 | -2,339348407 | N12:466349 | 34,356 | | | | rDNA | |
| chrXII | 466184 | 466344 | 160 | -1,438213629 | N12:466190 | 42,899 | | | | rDNA | |
| chrXII | 466184 | 466344 | 160 | -1,438213629 | N12:466349 | 34,356 | | | | rDNA | |
| chrXII | 466207 | 466337 | 130 | -1,530726957 | N12:466190 | 42,899 | | | | rDNA | |
| chrXII | 466207 | 466337 | 130 | -1,530726957 | N12:466349 | 34,356 | | | | rDNA | |
| chrXII | 466220 | 466354 | 134 | -0,91864188 | N12:466349 | 34,356 | | | | rDNA | |
| chrXII | 466252 | 466385 | 133 | -1,029702167 | N12:466349 | 34,356 | | | | rDNA | |
| chrXII | 466277 | 466442 | 165 | -1,50174518 | N12:466349 | 34,356 | | | | rDNA | |
| chrXII | 466277 | 466449 | 172 | -1,676271405 | N12:466349 | 34,356 | | | | rDNA | |
| chrXII | 466280 | 466385 | 105 | -0,599588757 | N12:466349 | 34,356 | | | | rDNA | |
| chrXII | 466280 | 466416 | 136 | -1,051930252 | N12:466349 | 34,356 | | | | rDNA | |
| chrXII | 466280 | 466453 | 173 | -1,418004259 | N12:466349 | 34,356 | | | | rDNA | |
| chrXII | 466289 | 466442 | 153 | -1,237834909 | N12:466349 | 34,356 | | | | rDNA | |
| chrXII | 466289 | 466449 | 160 | -1,551531932 | N12:466349 | 34,356 | | | | rDNA | |
| chrXII | 466290 | 466385 | 95 | -0,739626473 | N12:466349 | 34,356 | | | | rDNA | |
| chrXII | 466290 | 466416 | 126 | -0,900497292 | N12:466349 | 34,356 | | | | rDNA | |
| chrXII | 466290 | 466449 | 159 | -1,615122146 | N12:466349 | 34,356 | | | | rDNA | |
| chrXII | 466290 | 466453 | 163 | -1,342215939 | N12:466349 | 34,356 | | | | rDNA | |
| chrXII | 466290 | 466454 | 164 | -1,529692835 | N12:466349 | 34,356 | | | | rDNA | |
| chrXII | 466291 | 466449 | 158 | -1,359058459 | N12:466349 | 34,356 | | | | rDNA | |
| chrXII | 466291 | 466453 | 162 | -1,516868428 | N12:466349 | 34,356 | | | | rDNA | |
| chrXII | 466324 | 466453 | 129 | -0,973504824 | N12:466349 | 34,356 | | | | rDNA | |
| chrXII | 466324 | 466454 | 130 | -0,902645381 | N12:466349 | 34,356 | | | | rDNA | |
| chrXII | 466335 | 466483 | 148 | -1,162806442 | N12:466349 | 34,356 | | | | rDNA | |
| chrXII | 466349 | 466502 | 153 | -1,131025452 | N12:466349 | 34,356 | | | | rDNA | |
| chrXII | 466351 | 466500 | 149 | -1,20776272 | N12:466349 | 34,356 | | | | rDNA | |
| chrXII | 466356 | 466449 | 93 | -0,925541919 | N12:466349 | 34,356 | | | | rDNA | |
| chrXII | 466356 | 466483 | 127 | -0,921694478 | N12:466349 | 34,356 | | | | rDNA | |
| chrXII | 466356 | 466500 | 144 | -1,191233807 | N12:466349 | 34,356 | | | | rDNA | |
| chrXII | 466356 | 466502 | 146 | -1,211866682 | N12:466349 | 34,356 | | | | rDNA | |
| chrXII | 466364 | 466500 | 136 | -1,111816631 | N12:466349 | 34,356 | | | | rDNA | |
| chrXII | 466367 | 466532 | 165 | -1,504521289 | N12:466349 | 34,356 | | | | rDNA | |
| chrXII | 466367 | 466538 | 171 | -1,284897755 | N12:466349 | 34,356 | | | | rDNA | |
| chrXII | 466385 | 466532 | 147 | -1,268181282 | | | | Overlap <50 bp | | rDNA | |
| chrXII | 466385 | 466538 | 153 | -0,727811556 | | | | Overlap <50 bp | | rDNA | |
| chrXII | 466392 | 466502 | 110 | -0,774761812 | | | | Overlap <50 bp | | rDNA | |
| chrXII | 466392 | 466544 | 152 | -1,160492306 | | | | Overlap <50 bp | | rDNA | |
| chrXII | 466403 | 466500 | 97 | -0,644377544 | | | | Overlap <50 bp | | rDNA | |
| chrXII | 466403 | 466502 | 99 | 0,002228858 | | | | Overlap <50 bp | | rDNA | |
| chrXII | 466403 | 466544 | 141 | -0,859246023 | | | | Overlap <50 bp | | rDNA | |
| chrXII | 466403 | 466559 | 156 | -1,279564773 | | | | Overlap <50 bp | | rDNA | |
| chrXII | 466432 | 466521 | 89 | -1,434349753 | | | | Overlap <50 bp | | rDNA | |
| chrXII | 466432 | 466523 | 91 | -1,305883336 | | | | Overlap <50 bp | | rDNA | |
| chrXII | 466432 | 466527 | 95 | -1,68279282 | | | | Overlap <50 bp | | rDNA | |
| chrXII | 466432 | 466528 | 96 | -1,402346336 | | | | Overlap <50 bp | | rDNA | |
| chrXII | 466432 | 466530 | 98 | -1,596597643 | | | | Overlap <50 bp | | rDNA | |
| chrXII | 466432 | 466531 | 99 | -1,587053606 | | | | Overlap <50 bp | | rDNA | |
| chrXII | 466432 | 466532 | 100 | -1,381212083 | | | | Overlap <50 bp | | rDNA | |
| chrXII | 466432 | 466534 | 102 | -1,649766327 | | | | Overlap <50 bp | | rDNA | |

TABLE 3

| CHR | START | END | L(bp) | ΔLK | Nuc ID | Fuzziness | Gene ID | Gene body postion or intergene | Terminal | rDNA | Telomeric |
|---|---|---|---|---|---|---|---|---|---|---|---|
| chrXII | 466432 | 466538 | 106 | -1,022076878 | | | | Overlap <50 bp | | rDNA | |
| chrXII | 466432 | 466539 | 107 | -0,701497635 | | | | Overlap <50 bp | | rDNA | |
| chrXII | 466474 | 466613 | 139 | -1,367357428 | N12:466630 | 9,192 | | | | rDNA | |
| chrXII | 466474 | 466618 | 144 | -1,183456413 | N12:466630 | 9,192 | | | | rDNA | |
| chrXII | 466497 | 466613 | 116 | -1,050074927 | N12:466630 | 9,192 | | | | rDNA | |
| chrXII | 466497 | 466618 | 121 | -0,66238937 | N12:466630 | 9,192 | | | | rDNA | |
| chrXII | 466497 | 466667 | 170 | -1,445557855 | N12:466630 | 9,192 | | | | rDNA | |
| chrXII | 466507 | 466613 | 106 | -1,419414891 | N12:466630 | 9,192 | | | | rDNA | |
| chrXII | 466509 | 466613 | 104 | -1,758094708 | N12:466630 | 9,192 | | | | rDNA | |
| chrXII | 466509 | 466618 | 109 | -0,736531972 | N12:466630 | 9,192 | | | | rDNA | |
| chrXII | 466557 | 466700 | 143 | -0,989660603 | N12:466630 | 9,192 | | | | rDNA | |
| chrXII | 466557 | 466704 | 147 | -1,304748707 | N12:466630 | 9,192 | | | | rDNA | |
| chrXII | 466557 | 466708 | 151 | -1,183837733 | N12:466630 | 9,192 | | | | rDNA | |
| chrXII | 466557 | 466709 | 152 | -0,978047549 | N12:466630 | 9,192 | | | | rDNA | |
| chrXII | 466574 | 466729 | 155 | -1,754345084 | N12:466630 | 9,192 | | | | rDNA | |
| chrXII | 466577 | 466730 | 153 | -1,200108241 | N12:466630 | 9,192 | | | | rDNA | |
| chrXII | 466611 | 466736 | 125 | -1,525849683 | N12:466630 | 9,192 | | | | rDNA | |
| chrXII | 466690 | 466843 | 153 | -1,093769123 | N12:466825 | 26 | | | | rDNA | |
| chrXII | 466706 | 466873 | 167 | -1,763604263 | N12:466825 | 26 | | | | rDNA | |
| chrXII | 466706 | 466878 | 172 | -1,690582062 | N12:466825 | 26 | | | | rDNA | |
| chrXII | 466737 | 466873 | 136 | -1,064523378 | N12:466825 | 26 | | | | rDNA | |
| chrXII | 466737 | 466876 | 139 | -0,748869281 | N12:466825 | 26 | | | | rDNA | |
| chrXII | 466737 | 466878 | 141 | -1,458186302 | N12:466825 | 26 | | | | rDNA | |
| chrXII | 466743 | 466900 | 157 | -0,946379516 | N12:466825 | 26 | | | | rDNA | |
| chrXII | 466757 | 466910 | 153 | -1,354696362 | N12:466825 | 26 | | | | rDNA | |
| chrXII | 466757 | 466924 | 167 | -1,635540173 | N12:466825 | 26 | | | | rDNA | |
| chrXII | 466771 | 466894 | 123 | -1,245787505 | N12:466825 | 26 | | | | rDNA | |
| chrXII | 466804 | 466956 | 152 | -1,352999552 | N12:466825 | 26 | | | | rDNA | |
| chrXII | 466940 | 467096 | 156 | -1,171158002 | N12:467036 | 20,033 | | | | rDNA | |
| chrXII | 466947 | 467098 | 151 | -1,463342238 | N12:467036 | 20,033 | | | | rDNA | |
| chrXII | 466953 | 467111 | 158 | -1,002309502 | N12:467036 | 20,033 | | | | rDNA | |
| chrXII | 466957 | 467090 | 133 | -0,806778547 | N12:467036 | 20,033 | | | | rDNA | |
| chrXII | 467009 | 467160 | 151 | -1,278850459 | N12:467036 | 20,033 | | | | rDNA | |
| chrXII | 467063 | 467189 | 126 | -0,863552457 | | | | Overlap <50 bp | | rDNA | |
| chrXII | 467082 | 467240 | 158 | -1,362117349 | | | | Overlap <50 bp | | rDNA | |
| chrXII | 467082 | 467241 | 159 | -1,443762402 | | | | Overlap <50 bp | | rDNA | |
| chrXII | 467086 | 467196 | 110 | -0,574687176 | | | | Overlap <50 bp | | rDNA | |
| chrXII | 467086 | 467238 | 152 | -1,236715493 | | | | Overlap <50 bp | | rDNA | |
| chrXII | 467087 | 467238 | 151 | -1,199358749 | | | | Overlap <50 bp | | rDNA | |
| chrXII | 467087 | 467240 | 153 | -1,284611933 | | | | Overlap <50 bp | | rDNA | |
| chrXII | 467096 | 467196 | 100 | -0,689144471 | | | | Overlap <50 bp | | rDNA | |
| chrXII | 467096 | 467247 | 151 | -1,014330747 | | | | Overlap <50 bp | | rDNA | |
| chrXII | 467113 | 467196 | 83 | -0,871964081 | | | | Overlap <50 bp | | rDNA | |
| chrXII | 467113 | 467256 | 143 | -1,202291577 | | | | Overlap <50 bp | | rDNA | |
| chrXII | 467120 | 467226 | 106 | -0,474446778 | | | | Overlap <50 bp | | rDNA | |
| chrXII | 467120 | 467240 | 120 | -1,267839735 | | | | Overlap <50 bp | | rDNA | |
| chrXII | 467120 | 467274 | 154 | -1,453196088 | | | | Overlap <50 bp | | rDNA | |
| chrXII | 467123 | 467254 | 131 | -0,968564182 | | | | Overlap <50 bp | | rDNA | |
| chrXII | 467162 | 467304 | 142 | -1,511607052 | N12:467327 | 20,207 | | | | rDNA | |
| chrXII | 467192 | 467274 | 82 | -0,616547545 | | | | Overlap <50 bp | | rDNA | |
| chrXII | 467192 | 467343 | 151 | -1,223140886 | N12:467327 | 20,207 | | | | rDNA | |
| chrXII | 467216 | 467364 | 148 | -1,373866419 | N12:467327 | 20,207 | | | | rDNA | |
| chrXII | 467224 | 467368 | 144 | -1,120404641 | N12:467327 | 20,207 | | | | rDNA | |
| chrXII | 467264 | 467414 | 150 | -1,202163882 | N12:467327 | 20,207 | | | | rDNA | |
| chrXII | 467267 | 467430 | 163 | -1,116152514 | N12:467327 | 20,207 | | | | rDNA | |
| chrXII | 467271 | 467409 | 138 | -0,790704365 | N12:467327 | 20,207 | | | | rDNA | |
| chrXII | 467271 | 467414 | 143 | -1,207333239 | N12:467327 | 20,207 | | | | rDNA | |
| chrXII | 467273 | 467432 | 159 | -1,25419807 | N12:467327 | 20,207 | | | | rDNA | |
| chrXII | 467280 | 467428 | 148 | -1,356606866 | N12:467327 | 20,207 | | | | rDNA | |
| chrXII | 467303 | 467458 | 155 | -1,376416837 | N12:467327 | 20,207 | | | | rDNA | |
| chrXII | 467303 | 467459 | 156 | -1,409747579 | N12:467327 | 20,207 | | | | rDNA | |
| chrXII | 467304 | 467459 | 155 | -1,1902023 | N12:467327 | 20,207 | | | | rDNA | |
| chrXII | 467314 | 467414 | 100 | -0,709148306 | N12:467327 | 20,207 | | | | rDNA | |
| chrXII | 467314 | 467459 | 145 | -1,219511077 | N12:467327 | 20,207 | | | | rDNA | |
| chrXII | 467314 | 467464 | 150 | -1,42553872 | N12:467327 | 20,207 | | | | rDNA | |
| chrXII | 467314 | 467487 | 173 | -1,463968457 | N12:467327 | 20,207 | | | | rDNA | |
| chrXII | 467316 | 467464 | 148 | -1,124212653 | N12:467327 | 20,207 | | | | rDNA | |
| chrXII | 467328 | 467458 | 130 | -1,048002428 | N12:467327 | 20,207 | | | | rDNA | |
| chrXII | 467328 | 467489 | 161 | -1,166966948 | N12:467327 | 20,207 | | | | rDNA | |
| chrXII | 467332 | 467459 | 127 | -0,817150859 | N12:467327 | 20,207 | | | | rDNA | |
| chrXII | 467332 | 467487 | 155 | -1,005391883 | N12:467327 | 20,207 | | | | rDNA | |
| chrXII | 467341 | 467487 | 146 | -1,422135006 | N12:467327 | 20,207 | | | | rDNA | |
| chrXII | 467341 | 467517 | 176 | -1,415628495 | N12:467327 | 20,207 | | | | rDNA | |
| chrXII | 467428 | 467588 | 160 | -1,024847051 | N12:467582 | 21,213 | | | | rDNA | |
| chrXII | 467434 | 467588 | 154 | -1,246197816 | N12:467582 | 21,213 | | | | rDNA | |
| chrXII | 467435 | 467598 | 163 | -1,043102213 | N12:467582 | 21,213 | | | | rDNA | |
| chrXII | 467456 | 467609 | 153 | -1,406761565 | N12:467582 | 21,213 | | | | rDNA | |
| chrXII | 467475 | 467629 | 154 | -1,420864087 | N12:467582 | 21,213 | | | | rDNA | |
| chrXII | 467493 | 467630 | 137 | -0,946015819 | N12:467582 | 21,213 | | | | rDNA | |
| chrXII | 467493 | 467645 | 152 | -1,214933813 | N12:467582 | 21,213 | | | | rDNA | |
| chrXII | 467496 | 467660 | 164 | -1,293725572 | N12:467582 | 21,213 | | | | rDNA | |
| chrXII | 467502 | 467630 | 128 | -1,023597402 | N12:467582 | 21,213 | | | | rDNA | |
| chrXII | 467510 | 467630 | 120 | -0,956374307 | N12:467582 | 21,213 | | | | rDNA | |
| chrXII | 467510 | 467645 | 135 | -1,544213338 | N12:467582 | 21,213 | | | | rDNA | |
| chrXII | 467510 | 467667 | 157 | -1,535756594 | N12:467582 | 21,213 | | | | rDNA | |
| chrXII | 467511 | 467669 | 158 | -1,43830926 | N12:467582 | 21,213 | | | | rDNA | |
| chrXII | 467515 | 467649 | 134 | -0,869686564 | N12:467582 | 21,213 | | | | rDNA | |
| chrXII | 467532 | 467638 | 106 | -0,647616736 | N12:467582 | 21,213 | | | | rDNA | |
| chrXII | 467556 | 467724 | 168 | -1,466857542 | N12:467582 | 21,213 | | | | rDNA | |
| chrXII | 467645 | 467795 | 150 | -1,76462558 | | | | Overlap <50 bp | | rDNA | |
| chrXII | 467645 | 467796 | 151 | -1,535740651 | | | | Overlap <50 bp | | rDNA | |
| chrXII | 467883 | 468037 | 154 | -1,509928375 | N12:467959 | 20,928 | | | | rDNA | |
| chrXII | 467883 | 468038 | 155 | -1,433762196 | N12:467959 | 20,928 | | | | rDNA | |
| chrXII | 467883 | 468049 | 166 | -1,071629874 | N12:467959 | 20,928 | | | | rDNA | |
| chrXII | 467887 | 468049 | 162 | -1,359411744 | N12:467959 | 20,928 | | | | rDNA | |
| chrXII | 467905 | 468037 | 132 | -1,242353329 | N12:467959 | 20,928 | | | | rDNA | |
| chrXII | 467905 | 468038 | 133 | -1,314191197 | N12:467959 | 20,928 | | | | rDNA | |
| chrXII | 467905 | 468049 | 144 | -1,251132614 | N12:467959 | 20,928 | | | | rDNA | |
| chrXII | 467906 | 468037 | 131 | -1,101002414 | N12:467959 | 20,928 | | | | rDNA | |

| CHR | START | END | L(bp) | ΔLK | Nuc ID | Fuzziness | Gene ID | Gene body postion or intergene | Terminal | rDNA | Telomeric |
|---|---|---|---|---|---|---|---|---|---|---|---|
| chrXII | 467906 | 468038 | 132 | -1,018304977 | N12:467959 | 20,928 | | | | rDNA | |
| chrXII | 467906 | 468049 | 143 | -1,234558674 | N12:467959 | 20,928 | | | | rDNA | |
| chrXII | 467906 | 468050 | 144 | -0,869977729 | N12:467959 | 20,928 | | | | rDNA | |
| chrXII | 467907 | 468037 | 130 | -1,148690398 | N12:467959 | 20,928 | | | | rDNA | |
| chrXII | 467907 | 468038 | 131 | -1,309637707 | N12:467959 | 20,928 | | | | rDNA | |
| chrXII | 467907 | 468049 | 142 | -1,185390092 | N12:467959 | 20,928 | | | | rDNA | |
| chrXII | 467983 | 468139 | 156 | -0,818398013 | N12:468113 | 0 | | | | rDNA | |
| chrXII | 468039 | 468185 | 146 | -1,36906816 | N12:468113 | 0 | | | | rDNA | |
| chrXII | 468045 | 468181 | 136 | -1,282772836 | N12:468113 | 0 | | | | rDNA | |
| chrXII | 468045 | 468185 | 140 | -0,80249428 | N12:468113 | 0 | | | | rDNA | |
| chrXII | 468047 | 468203 | 156 | -1,708220607 | N12:468113 | 0 | | | | rDNA | |
| chrXII | 468050 | 468139 | 89 | -0,565841931 | N12:468113 | 0 | | | | rDNA | |
| chrXII | 468050 | 468175 | 125 | -1,094304398 | N12:468113 | 0 | | | | rDNA | |
| chrXII | 468050 | 468176 | 126 | -1,04361809 | N12:468113 | 0 | | | | rDNA | |
| chrXII | 468050 | 468180 | 130 | -1,185329644 | N12:468113 | 0 | | | | rDNA | |
| chrXII | 468050 | 468181 | 131 | -1,240440179 | N12:468113 | 0 | | | | rDNA | |
| chrXII | 468050 | 468184 | 134 | -1,2395425 | N12:468113 | 0 | | | | rDNA | |
| chrXII | 468050 | 468185 | 135 | -1,208465711 | N12:468113 | 0 | | | | rDNA | |
| chrXII | 468050 | 468203 | 153 | -1,40053062 | N12:468113 | 0 | | | | rDNA | |
| chrXII | 468070 | 468181 | 111 | -0,893178776 | N12:468113 | 0 | | | | rDNA | |
| chrXII | 468070 | 468184 | 114 | -0,724066852 | N12:468113 | 0 | | | | rDNA | |
| chrXII | 468070 | 468203 | 133 | -1,099181137 | N12:468113 | 0 | | | | rDNA | |
| chrXII | 468187 | 468283 | 96 | -1,300706257 | N12:468275 | 25,423 | | | | rDNA | |
| chrXII | 468226 | 468399 | 173 | -1,501027388 | N12:468275 | 25,423 | | | | rDNA | |
| chrXII | 468226 | 468402 | 176 | -1,563694439 | N12:468275 | 25,423 | | | | rDNA | |
| chrXII | 468232 | 468399 | 167 | -1,465487332 | N12:468275 | 25,423 | | | | rDNA | |
| chrXII | 468232 | 468402 | 170 | -1,570627036 | N12:468275 | 25,423 | | | | rDNA | |
| chrXII | 468235 | 468370 | 135 | -1,071895468 | N12:468275 | 25,423 | | | | rDNA | |
| chrXII | 468235 | 468399 | 164 | -1,428588157 | N12:468275 | 25,423 | | | | rDNA | |
| chrXII | 468303 | 468465 | 162 | -1,255920819 | N12:468484 | 36,77 | | | | rDNA | |
| chrXII | 468322 | 468465 | 143 | -1,202072398 | N12:468484 | 36,77 | | | | rDNA | |
| chrXII | 468341 | 468490 | 149 | -1,078225171 | N12:468484 | 36,77 | | | | rDNA | |
| chrXII | 468341 | 468513 | 172 | -1,302597169 | N12:468484 | 36,77 | | | | rDNA | |
| chrXII | 468341 | 468556 | 215 | -1,571146876 | N12:468484 | 36,77 | | | | rDNA | |
| chrXII | 468364 | 468513 | 149 | -1,360222575 | N12:468484 | 36,77 | | | | rDNA | |
| chrXII | 468364 | 468554 | 190 | -1,866584018 | N12:468484 | 36,77 | | | | rDNA | |
| chrXII | 468364 | 468556 | 192 | -1,727382058 | N12:468484 | 36,77 | | | | rDNA | |
| chrXII | 468364 | 468569 | 205 | -1,800549754 | N12:468484 | 36,77 | | | | rDNA | |
| chrXII | 468372 | 468532 | 160 | -0,869370592 | N12:468484 | 36,77 | | | | rDNA | |
| chrXII | 468373 | 468530 | 157 | -1,405697754 | N12:468484 | 36,77 | | | | rDNA | |
| chrXII | 468373 | 468554 | 181 | -1,198605701 | N12:468484 | 36,77 | | | | rDNA | |
| chrXII | 468373 | 468556 | 183 | -1,645116746 | N12:468484 | 36,77 | | | | rDNA | |
| chrXII | 468401 | 468490 | 89 | -0,320836144 | N12:468484 | 36,77 | | | | rDNA | |
| chrXII | 468401 | 468513 | 112 | -0,714540302 | N12:468484 | 36,77 | | | | rDNA | |
| chrXII | 468401 | 468532 | 131 | -1,073124839 | N12:468484 | 36,77 | | | | rDNA | |
| chrXII | 468401 | 468554 | 153 | -1,236944389 | N12:468484 | 36,77 | | | | rDNA | |
| chrXII | 468401 | 468556 | 155 | -1,282179175 | N12:468484 | 36,77 | | | | rDNA | |
| chrXII | 468401 | 468569 | 168 | -1,265690437 | N12:468484 | 36,77 | | | | rDNA | |
| chrXII | 468401 | 468578 | 177 | -1,440639164 | N12:468484 | 36,77 | | | | rDNA | |
| chrXII | 468402 | 468513 | 111 | -0,664532219 | N12:468484 | 36,77 | | | | rDNA | |
| chrXII | 468402 | 468530 | 128 | -0,844276685 | N12:468484 | 36,77 | | | | rDNA | |
| chrXII | 468402 | 468550 | 148 | -1,233234729 | N12:468484 | 36,77 | | | | rDNA | |
| chrXII | 468402 | 468553 | 151 | -1,729648055 | N12:468484 | 36,77 | | | | rDNA | |
| chrXII | 468402 | 468554 | 152 | -1,34783442 | N12:468484 | 36,77 | | | | rDNA | |
| chrXII | 468402 | 468556 | 154 | -1,241023194 | N12:468484 | 36,77 | | | | rDNA | |
| chrXII | 468402 | 468559 | 157 | -1,270244821 | N12:468484 | 36,77 | | | | rDNA | |
| chrXII | 468402 | 468569 | 167 | -1,520945279 | N12:468484 | 36,77 | | | | rDNA | |
| chrXII | 468402 | 468578 | 176 | -1,93501063 | N12:468484 | 36,77 | | | | rDNA | |
| chrXII | 468402 | 468604 | 202 | -1,39920955 | N12:468484 | 36,77 | | | | rDNA | |
| chrXII | 468402 | 468606 | 204 | -1,628984296 | N12:468484 | 36,77 | | | | rDNA | |
| chrXII | 468404 | 468530 | 126 | -1,071233395 | N12:468484 | 36,77 | | | | rDNA | |
| chrXII | 468404 | 468554 | 150 | -1,233858937 | N12:468484 | 36,77 | | | | rDNA | |
| chrXII | 468404 | 468556 | 152 | -1,176810342 | N12:468484 | 36,77 | | | | rDNA | |
| chrXII | 468404 | 468559 | 155 | -1,087112055 | N12:468484 | 36,77 | | | | rDNA | |
| chrXII | 468404 | 468569 | 165 | -1,17923553 | N12:468484 | 36,77 | | | | rDNA | |
| chrXII | 468404 | 468578 | 174 | -1,544182889 | N12:468484 | 36,77 | | | | rDNA | |
| chrXII | 468404 | 468606 | 202 | -1,509033435 | N12:468484 | 36,77 | | | | rDNA | |
| chrXII | 468405 | 468513 | 108 | -1,021063141 | N12:468484 | 36,77 | | | | rDNA | |
| chrXII | 468405 | 468530 | 125 | -0,999843874 | N12:468484 | 36,77 | | | | rDNA | |
| chrXII | 468405 | 468532 | 127 | -1,188020613 | N12:468484 | 36,77 | | | | rDNA | |
| chrXII | 468405 | 468550 | 145 | -1,149280641 | N12:468484 | 36,77 | | | | rDNA | |
| chrXII | 468405 | 468553 | 148 | -1,480884123 | N12:468484 | 36,77 | | | | rDNA | |
| chrXII | 468405 | 468554 | 149 | -1,167947921 | N12:468484 | 36,77 | | | | rDNA | |
| chrXII | 468405 | 468556 | 151 | -1,118964829 | N12:468484 | 36,77 | | | | rDNA | |
| chrXII | 468405 | 468559 | 154 | -1,208320294 | N12:468484 | 36,77 | | | | rDNA | |
| chrXII | 468405 | 468568 | 163 | -1,386682838 | N12:468484 | 36,77 | | | | rDNA | |
| chrXII | 468405 | 468569 | 164 | -1,251510383 | N12:468484 | 36,77 | | | | rDNA | |
| chrXII | 468405 | 468578 | 173 | -1,631934013 | N12:468484 | 36,77 | | | | rDNA | |
| chrXII | 468405 | 468604 | 199 | -1,112319253 | N12:468484 | 36,77 | | | | rDNA | |
| chrXII | 468405 | 468606 | 201 | -1,533035749 | N12:468484 | 36,77 | | | | rDNA | |
| chrXII | 468406 | 468513 | 107 | -1,318675714 | N12:468484 | 36,77 | | | | rDNA | |
| chrXII | 468406 | 468530 | 124 | -1,188335156 | N12:468484 | 36,77 | | | | rDNA | |
| chrXII | 468406 | 468550 | 144 | -1,211509896 | N12:468484 | 36,77 | | | | rDNA | |
| chrXII | 468406 | 468554 | 148 | -1,087112126 | N12:468484 | 36,77 | | | | rDNA | |
| chrXII | 468406 | 468556 | 150 | -1,246006796 | N12:468484 | 36,77 | | | | rDNA | |
| chrXII | 468406 | 468559 | 153 | -1,308781689 | N12:468484 | 36,77 | | | | rDNA | |
| chrXII | 468406 | 468568 | 162 | -1,426619073 | N12:468484 | 36,77 | | | | rDNA | |
| chrXII | 468406 | 468604 | 198 | -1,305863656 | N12:468484 | 36,77 | | | | rDNA | |
| chrXII | 468406 | 468606 | 200 | -1,754377046 | N12:468484 | 36,77 | | | | rDNA | |
| chrXII | 468445 | 468551 | 106 | -0,659464735 | N12:468484 | 36,77 | | | | rDNA | |
| chrXII | 468445 | 468554 | 109 | -0,38648491 | N12:468484 | 36,77 | | | | rDNA | |
| chrXII | 468445 | 468556 | 111 | -0,864271568 | N12:468484 | 36,77 | | | | rDNA | |
| chrXII | 468445 | 468604 | 159 | -1,093614882 | N12:468484 | 36,77 | | | | rDNA | |
| chrXII | 468445 | 468606 | 161 | -1,249276035 | N12:468484 | 36,77 | | | | rDNA | |
| chrXII | 468462 | 468603 | 141 | -1,058112547 | N12:468484 | 36,77 | | | | rDNA | |
| chrXII | 468463 | 468554 | 91 | -0,665772876 | N12:468484 | 36,77 | | | | rDNA | |
| chrXII | 468463 | 468556 | 93 | -0,424489235 | N12:468484 | 36,77 | | | | rDNA | |
| chrXII | 468463 | 468604 | 141 | -0,888867406 | N12:468484 | 36,77 | | | | rDNA | |
| chrXII | 468463 | 468606 | 143 | -1,107891164 | N12:468484 | 36,77 | | | | rDNA | |
| chrXII | 468545 | 468696 | 151 | -0,870498471 | N12:468638 | 64,023 | +3*:YLR154C-H; | 3 | TERM | rDNA | |

TABLE 3

| CHR | START | END | L(bp) | ΔLK | Nuc ID | Fuzziness | Gene ID | Gene body postion or intergene | Terminal | rDNA | Telomeric |
|---|---|---|---|---|---|---|---|---|---|---|---|
| chrXII | 468566 | 468665 | 99 | -1,340735727 | N12:468638 | 64,023 | +3*:YLR154C-H; | 3 | TERM | rDNA | |
| chrXII | 468566 | 468695 | 129 | -1,173024667 | N12:468638 | 64,023 | +3*:YLR154C-H; | 3 | TERM | rDNA | |
| chrXII | 468566 | 468710 | 144 | -1,392045818 | N12:468638 | 64,023 | +3*:YLR154C-H; | 3 | TERM | rDNA | |
| chrXII | 468566 | 468716 | 150 | -1,703042994 | N12:468638 | 64,023 | +3*:YLR154C-H; | 3 | TERM | rDNA | |
| chrXII | 468567 | 468695 | 128 | -0,913565629 | N12:468638 | 64,023 | +3*:YLR154C-H; | 3 | TERM | rDNA | |
| chrXII | 468567 | 468710 | 143 | -1,054112339 | N12:468638 | 64,023 | +3*:YLR154C-H; | 3 | TERM | rDNA | |
| chrXII | 468567 | 468716 | 149 | -1,346445114 | N12:468638 | 64,023 | +3*:YLR154C-H; | 3 | TERM | rDNA | |
| chrXII | 468656 | 468810 | 154 | -1,260621642 | N12:468638 | 64,023 | +3*:YLR154C-H; | 3 | TERM | rDNA | |
| chrXII | 468658 | 468810 | 152 | -1,482197731 | N12:468638 | 64,023 | +3*:YLR154C-H; | 3 | TERM | rDNA | |
| chrXII | 468703 | 468859 | 156 | -1,312964843 | N12:468850 | 34,655 | +2:YLR154C-H; | 2 | | rDNA | |
| chrXII | 468739 | 468893 | 154 | -1,46375081 | N12:468850 | 34,655 | +2:YLR154C-H; | 2 | | rDNA | |
| chrXII | 468739 | 468896 | 157 | -1,230694242 | N12:468850 | 34,655 | +2:YLR154C-H; | 2 | | rDNA | |
| chrXII | 468741 | 468859 | 118 | -0,803441936 | N12:468850 | 34,655 | +2:YLR154C-H; | 2 | | rDNA | |
| chrXII | 468741 | 468893 | 152 | -1,236921858 | N12:468850 | 34,655 | +2:YLR154C-H; | 2 | | rDNA | |
| chrXII | 468741 | 468896 | 155 | -0,779062839 | N12:468850 | 34,655 | +2:YLR154C-H; | 2 | | rDNA | |
| chrXII | 468741 | 468911 | 170 | -1,082507045 | N12:468850 | 34,655 | +2:YLR154C-H; | 2 | | rDNA | |
| chrXII | 468755 | 468893 | 138 | -1,299623807 | N12:468850 | 34,655 | +2:YLR154C-H; | 2 | | rDNA | |
| chrXII | 468755 | 468911 | 156 | -1,126481285 | N12:468850 | 34,655 | +2:YLR154C-H; | 2 | | rDNA | |
| chrXII | 468758 | 468893 | 135 | -1,101079734 | N12:468850 | 34,655 | +2:YLR154C-H; | 2 | | rDNA | |
| chrXII | 468762 | 468893 | 131 | -1,506212546 | N12:468850 | 34,655 | +2:YLR154C-H; | 2 | | rDNA | |
| chrXII | 468762 | 468911 | 149 | -1,511021843 | N12:468850 | 34,655 | +2:YLR154C-H; | 2 | | rDNA | |
| chrXII | 468765 | 468893 | 128 | -1,135031882 | N12:468850 | 34,655 | +2:YLR154C-H; | 2 | | rDNA | |
| chrXII | 468765 | 468911 | 146 | -1,463343424 | N12:468850 | 34,655 | +2:YLR154C-H; | 2 | | rDNA | |
| chrXII | 468770 | 468893 | 123 | -0,526333235 | N12:468850 | 34,655 | +2:YLR154C-H; | 2 | | rDNA | |
| chrXII | 468770 | 468911 | 141 | -0,962413117 | N12:468850 | 34,655 | +2:YLR154C-H; | 2 | | rDNA | |
| chrXII | 472060 | 472213 | 153 | -1,149884765 | N12:472134 | 7,81 | +1:YLR156W; | 1 | | | |
| chrXII | 472461 | 472545 | 84 | -0,388965263 | N12:472515 | 24,637 | +2:YLR156C-A; +3*:YLR156W; | 2 | | | |
| chrXII | 472461 | 472563 | 102 | -0,468873801 | N12:472515 | 24,637 | +2:YLR156C-A; +3*:YLR156W; | 2 | | | |
| chrXII | 475711 | 475864 | 153 | -1,31405863 | N12:475777 | 13,077 | +1:YLR157W-D; | 1 | | | |
| chrXII | 476909 | 477023 | 114 | -1,424773089 | | | | Overlap <50 bp | | | |
| chrXII | 477605 | 477755 | 150 | -1,531104029 | N12:477669 | 18,23 | | Intergene | | | |
| chrXII | 478538 | 478667 | 129 | -1,20339556 | N12:478520 | 6,364 | | Intergene | | | |
| chrXII | 479001 | 479158 | 157 | -1,465621638 | N12:479064 | 36,828 | | Intergene | | | |
| chrXII | 479187 | 479337 | 150 | -1,049483026 | N12:479289 | 19,975 | | Intergene | | | |
| chrXII | 479469 | 479630 | 161 | -1,566850609 | N12:479454 | 1,414 | | Intergene | | | |
| chrXII | 480195 | 480349 | 154 | -1,330920335 | N12:480208 | 13 | | Intergene | | | |
| chrXII | 480476 | 480621 | 145 | -1,45511423 | N12:480542 | 14,107 | | Intergene | | | |
| chrXII | 480480 | 480631 | 151 | -1,223892436 | N12:480542 | 14,107 | | Intergene | | | |
| chrXII | 480687 | 480827 | 140 | -0,948759724 | N12:480768 | 39,311 | | Intergene | | | |
| chrXII | 480693 | 480827 | 134 | -1,08812727 | N12:480768 | 39,311 | | Intergene | | | |
| chrXII | 481105 | 481260 | 155 | -1,374396656 | N12:481138 | 19,858 | | Intergene | | | |
| chrXII | 481236 | 481344 | 108 | -0,818781704 | | | | Overlap <50 bp | | | |
| chrXII | 481385 | 481534 | 149 | -1,052169201 | N12:481399 | 46,09 | | Intergene | | | |
| chrXII | 482041 | 482125 | 84 | -0,397939884 | N12:482076 | 14,572 | +2:YLR157C-C; +3:YLR157W-E; | 2 | | | |
| chrXII | 482041 | 482143 | 102 | -0,555590478 | N12:482076 | 14,572 | +2:YLR157C-C; +3:YLR157W-E; | 2 | | | |
| chrXII | 482885 | 483037 | 152 | -1,566952739 | N12:482962 | 13,229 | +6:YLR158C; | 6 | | | |
| chrXII | 485292 | 485445 | 153 | -1,196738413 | N12:485368 | 9,292 | +1:YLR159W; | 1 | | | |
| chrXII | 485693 | 485777 | 84 | -0,452332423 | N12:485732 | 28,827 | +3*:YLR159W; +2:YLR159C-A; | 3 | TERM | | |
| chrXII | 488944 | 489097 | 153 | -1,030790079 | N12:489014 | 10,116 | +1:YLR161W; | 1 | | | |
| chrXII | 489345 | 489429 | 84 | -0,578720288 | N12:489399 | 19,975 | +3*:YLR161W; 0:YLR162W; | 3 | TERM | | |
| chrXII | 489345 | 489447 | 102 | -0,460759231 | N12:489399 | 19,975 | +3*:YLR161W; 0:YLR162W; | 3 | TERM | | |
| chrXII | 489345 | 489466 | 121 | -0,925624305 | N12:489399 | 19,975 | +3*:YLR161W; 0:YLR162W; | 3 | TERM | | |
| chrXII | 489345 | 489469 | 124 | -0,772964675 | N12:489399 | 19,975 | +3*:YLR161W; 0:YLR162W; | 3 | TERM | | |
| chrXII | 489345 | 489471 | 126 | -1,002823668 | N12:489399 | 19,975 | +3*:YLR161W; 0:YLR162W; | 3 | TERM | | |
| chrXII | 489359 | 489469 | 110 | -0,993882911 | N12:489399 | 19,975 | +3*:YLR161W; 0:YLR162W; | 3 | TERM | | |
| chrXII | 489359 | 489471 | 112 | -1,177352657 | N12:489399 | 19,975 | +3*:YLR161W; 0:YLR162W; | 3 | TERM | | |
| chrXII | 489361 | 489466 | 105 | -1,241056183 | N12:489399 | 19,975 | +3*:YLR161W; 0:YLR162W; | 3 | TERM | | |
| chrXII | 489556 | 489702 | 146 | -1,241581141 | N12:489601 | 15,011 | +1:YLR162W; | 1 | | | |
| chrXII | 489926 | 490077 | 151 | -1,090907147 | N12:489932 | 48,407 | +3:YLR162W; | 3 | | | |
| chrXII | 489933 | 490080 | 147 | -1,352555735 | N12:489932 | 48,407 | +3:YLR162W; | 3 | | | |
| chrXII | 490009 | 490164 | 155 | -1,21027846 | N12:490150 | 25,865 | -1:YLR162W-A; +4*:YLR162W; | -1 | | | |
| chrXII | 490027 | 490183 | 156 | -1,259777408 | N12:490150 | 25,865 | -1:YLR162W-A; +4*:YLR162W; | -1 | | | |
| chrXII | 490049 | 490195 | 146 | -1,136889749 | N12:490150 | 25,865 | -1:YLR162W-A; +4*:YLR162W; | -1 | | | |
| chrXII | 490049 | 490215 | 166 | -1,276140013 | N12:490150 | 25,865 | -1:YLR162W-A; +4*:YLR162W; | -1 | | | |
| chrXII | 490049 | 490219 | 170 | -0,956575636 | N12:490150 | 25,865 | -1:YLR162W-A; +4*:YLR162W; | -1 | | | |
| chrXII | 490072 | 490215 | 143 | -1,268629292 | N12:490150 | 25,865 | -1:YLR162W-A; +4*:YLR162W; | -1 | | | |
| chrXII | 490072 | 490219 | 147 | -0,988491923 | N12:490150 | 25,865 | -1:YLR162W-A; +4*:YLR162W; | -1 | | | |
| chrXII | 490083 | 490239 | 156 | -1,212383367 | N12:490150 | 25,865 | -1:YLR162W-A; +4*:YLR162W; | -1 | | | |
| chrXII | 490158 | 490312 | 154 | -1,359112771 | N12:490150 | 25,865 | -1:YLR162W-A; +4*:YLR162W; | -1 | | | |
| chrXII | 490158 | 490312 | 154 | -1,359112771 | N12:490324 | 25,146 | +1:YLR162W-A; | 1 | | | |
| chrXII | 490228 | 490334 | 106 | -1,44929239 | N12:490324 | 25,146 | +1:YLR162W-A; | 1 | | | |
| chrXII | 490228 | 490337 | 109 | -1,35960868 | N12:490324 | 25,146 | +1:YLR162W-A; | 1 | | | |
| chrXII | 490228 | 490382 | 154 | -1,234252158 | N12:490324 | 25,146 | +1:YLR162W-A; | 1 | | | |
| chrXII | 490261 | 490368 | 107 | -1,444551773 | N12:490324 | 25,146 | +1:YLR162W-A; | 1 | | | |
| chrXII | 490262 | 490423 | 161 | -1,356906522 | N12:490324 | 25,146 | +1:YLR162W-A; | 1 | | | |
| chrXII | 490263 | 490423 | 160 | -1,300623985 | N12:490324 | 25,146 | +1:YLR162W-A; | 1 | | | |
| chrXII | 490267 | 490423 | 156 | -1,291185827 | N12:490324 | 25,146 | +1:YLR162W-A; | 1 | | | |
| chrXII | 490291 | 490423 | 132 | -1,010558675 | N12:490324 | 25,146 | +1:YLR162W-A; | 1 | | | |
| chrXII | 490302 | 490409 | 107 | -0,946282916 | N12:490324 | 25,146 | +1:YLR162W-A; | 1 | | | |
| chrXII | 490302 | 490423 | 121 | -0,989517242 | N12:490324 | 25,146 | +1:YLR162W-A; | 1 | | | |
| chrXII | 494931 | 495079 | 148 | -1,188021641 | N12:495031 | 11,669 | +2:YLR165C; | 2 | | | |
| chrXII | 496357 | 496507 | 150 | -0,950725566 | | | | Overlap <50 bp | | | |
| chrXII | 508585 | 508725 | 140 | -1,159232027 | N12:508668 | 26,883 | +11:YLR176C; | 11 | | | |
| chrXII | 514529 | 514679 | 150 | -1,044012982 | N12:514599 | 7,376 | +2:YLR179C; | 2 | | | |
| chrXII | 518369 | 518501 | 132 | -1,391154497 | N12:518444 | 10,237 | +4:YLR182W; | 4 | | | |
| chrXII | 522434 | 522581 | 147 | -1,146343903 | N12:522432 | 4,243 | -1:YLR185W; | -1 | | | |
| chrXII | 528715 | 528875 | 160 | -1,427233078 | N12:528770 | 19,139 | +4:YLR188W; | 4 | | | |
| chrXII | 529249 | 529419 | 170 | -1,662679256 | N12:529380 | 8,185 | +8:YLR188W; | 8 | | | |
| chrXII | 537872 | 537976 | 104 | -1,252899303 | N12:537943 | 7,136 | +5:YLR191W; | 5 | | | |
| chrXII | 542120 | 542273 | 153 | -1,628624905 | N12:542188 | 12,738 | +8:YLR195C; +2:CUT265; | 8 | | | |
| chrXII | 544133 | 544261 | 128 | -1,234353244 | N12:544269 | 4,32 | +3:YLR196W; | 3 | | | |
| chrXII | 545204 | 545351 | 147 | -1,161817649 | N12:545241 | 12,755 | +9:YLR196W; | 9 | | | |
| chrXII | 546037 | 546189 | 152 | -0,989916049 | N12:546112 | 8,981 | +1:YLR197W; | 1 | | | |
| chrXII | 546038 | 546189 | 151 | -1,433649953 | N12:546112 | 8,981 | +1:YLR197W; | 1 | | | |
| chrXII | 546040 | 546189 | 149 | -1,241394288 | N12:546112 | 8,981 | +1:YLR197W; | 1 | | | |
| chrXII | 547890 | 548039 | 149 | -1,181620294 | N12:547946 | 15,655 | | Intergene | | | |
| chrXII | 553542 | 553689 | 147 | -1,107622526 | N12:553631 | 7,642 | +2:YLR205C; | 2 | | | |
| chrXII | 555694 | 555847 | 153 | -1,373030693 | N12:555721 | 8,963 | +9:YLR206W; | 9 | | | |
| chrXII | 559139 | 559286 | 147 | -1,209683481 | N12:559216 | 3,189 | +15:YLR207W; -1:YLR208W; | 15 | | | |

| CHR | START | END | L(bp) | ΔLK | Nuc ID | Fuzziness | Gene ID | Gene body postion or intergene | Terminal | rDNA | Telomeric |
|---|---|---|---|---|---|---|---|---|---|---|---|
| chrXII | 559606 | 559755 | 149 | -0,919625008 | N12:559682 | 6,242 | +2:YLR208W; | 2 | | | |
| chrXII | 560049 | 560194 | 145 | -1,240110231 | N12:560179 | 9,68 | +5:YLR208W; | 5 | | | |
| chrXII | 570252 | 570403 | 151 | -1,195045578 | N12:570424 | 17,024 | +14:YLR214W; | 14 | | | |
| chrXII | 573533 | 573669 | 136 | -0,984279315 | N12:573595 | 15,659 | +2:YLR218C; | 2 | | | |
| chrXII | 576803 | 576966 | 163 | -1,439396479 | N12:576902 | 26,235 | +2:YLR220W; | 2 | | | |
| chrXII | 586445 | 586571 | 126 | -1,503237124 | N12:586538 | 12,565 | +2:YLR224W; | 2 | | | |
| chrXII | 587477 | 587583 | 106 | -1,378130062 | N12:587489 | 9,381 | +8*:YLR224W; | 8 | TERM | | |
| chrXII | 588088 | 588237 | 149 | -1,200938211 | N12:588110 | 13,435 | +6:YLR225C; | 6 | | | |
| chrXII | 593332 | 593484 | 152 | -1,285444315 | N12:593448 | 24,021 | | Intergene | | | |
| chrXII | 593504 | 593653 | 149 | -1,093835678 | N12:593653 | 9,899 | | Intergene | | | |
| chrXII | 593678 | 593801 | 123 | -1,152554228 | | | | Overlap <50 bp | | | |
| chrXII | 593778 | 593933 | 155 | -1,349428874 | N12:593897 | 31,48 | | Intergene | | | |
| chrXII | 594211 | 594345 | 134 | -1,160388708 | N12:594281 | 32,624 | | Intergene | | | |
| chrXII | 594211 | 594351 | 140 | -1,072096703 | N12:594281 | 32,624 | | Intergene | | | |
| chrXII | 594407 | 594558 | 151 | -1,071849284 | N12:594490 | 27,062 | | Intergene | | | |
| chrXII | 594417 | 594562 | 145 | -1,111925036 | N12:594490 | 27,062 | | Intergene | | | |
| chrXII | 594435 | 594586 | 151 | -1,620286771 | N12:594490 | 27,062 | | Intergene | | | |
| chrXII | 594689 | 594843 | 154 | -1,335690177 | N12:594825 | 8,505 | | Intergene | | | |
| chrXII | 594723 | 594867 | 144 | -1,266666298 | N12:594825 | 8,505 | | Intergene | | | |
| chrXII | 595254 | 595389 | 135 | -1,241025624 | N12:595293 | 18,717 | | Intergene | | | |
| chrXII | 595436 | 595568 | 132 | -1,188026385 | N12:595587 | 3,536 | | Intergene | | | |
| chrXII | 595525 | 595663 | 138 | -1,14712218 | N12:595587 | 3,536 | | Intergene | | | |
| chrXII | 598015 | 598129 | 114 | -1,461548698 | N12:598127 | 33,181 | | Intergene | | | |
| chrXII | 598489 | 598636 | 147 | -1,043544883 | N12:598658 | 29,771 | | Intergene | | | |
| chrXII | 603237 | 603400 | 163 | -1,089035045 | N12:603419 | 20,203 | 0:Unit459; +3*:SUT698; 0:Unit440; | 0 | | | |
| chrXII | 607039 | 607198 | 159 | -1,540087145 | N12:607110 | 2,658 | *:YLR233C; +1:YLR231C; -1:SUT276; | 16 | TERM | | |
| chrXII | 608189 | 608339 | 150 | -1,970000913 | N12:608276 | 38,05 | +9:YLR233C; +6:SUT276; | 9 | | | |
| chrXII | 610208 | 610357 | 149 | -1,179280944 | N12:610297 | 9,121 | +4:YLR234W; | 4 | | | |
| chrXII | 611103 | 611232 | 129 | -0,762168679 | N12:611216 | 20,535 | +10:YLR234W; | 10 | | | |
| chrXII | 614828 | 614977 | 149 | -1,874652591 | N12:614901 | 6,998 | +2:YLR238W; | 2 | | | |
| chrXII | 614828 | 614997 | 169 | -1,6675472 | N12:614901 | 6,998 | +2:YLR238W; | 2 | | | |
| chrXII | 614830 | 614977 | 147 | -1,378254727 | N12:614901 | 6,998 | +2:YLR238W; | 2 | | | |
| chrXII | 625233 | 625381 | 148 | -1,202323984 | N12:625336 | 34,788 | | Intergene | | | |
| chrXII | 628912 | 629085 | 173 | -1,768049895 | N12:629078 | 27,506 | +25:YLR247C; | 25 | | | |
| chrXII | 631988 | 632127 | 139 | -1,20290166 | N12:632062 | 13,809 | +9:YLR247C; | 9 | | | |
| chrXII | 635940 | 636108 | 168 | -1,543986029 | N12:635980 | 10,498 | +12:YLR248W; | 12 | | | |
| chrXII | 640543 | 640689 | 146 | -1,233114775 | N12:640632 | 11 | +3:YLR250W; | 3 | | | |
| chrXII | 649877 | 650024 | 147 | -1,570065353 | N12:649854 | 31,564 | +23:YLR256W; | 23 | | | |
| chrXII | 649877 | 650024 | 147 | -1,570065353 | N12:650003 | 40,336 | +24:YLR256W; | 24 | | | |
| chrXII | 651183 | 651332 | 149 | -1,389171765 | N12:651337 | 43,684 | | Intergene | | | |
| chrXII | 651357 | 651480 | 123 | -1,090420043 | N12:651337 | 43,684 | | Intergene | | | |
| chrXII | 651457 | 651612 | 155 | -0,978854574 | N12:651585 | 34,641 | | Intergene | | | |
| chrXII | 651890 | 652024 | 134 | -1,110433573 | N12:651960 | 52,412 | | Intergene | | | |
| chrXII | 651890 | 652030 | 140 | -0,882039969 | N12:651960 | 52,412 | | Intergene | | | |
| chrXII | 652086 | 652237 | 151 | -1,179257004 | N12:652161 | 24,269 | | Intergene | | | |
| chrXII | 652096 | 652241 | 145 | -0,985471169 | N12:652161 | 24,269 | | Intergene | | | |
| chrXII | 652114 | 652265 | 151 | -1,435100285 | N12:652161 | 24,269 | | Intergene | | | |
| chrXII | 652116 | 652265 | 149 | -1,327548019 | N12:652161 | 24,269 | | Intergene | | | |
| chrXII | 652368 | 652522 | 154 | -1,541774625 | N12:652504 | 16,653 | | Intergene | | | |
| chrXII | 652618 | 652786 | 168 | -1,306519664 | N12:652671 | 19,348 | | Intergene | | | |
| chrXII | 652933 | 653068 | 135 | -1,077645629 | N12:652970 | 7,572 | | Intergene | | | |
| chrXII | 653087 | 653248 | 161 | -1,369015978 | N12:653246 | 53,119 | | Intergene | | | |
| chrXII | 653194 | 653339 | 145 | -1,246072008 | N12:653246 | 53,119 | | Intergene | | | |
| chrXII | 653561 | 653718 | 157 | -1,37350691 | N12:653642 | 21,633 | | Intergene | | | |
| chrXII | 654052 | 654181 | 129 | -1,281861347 | | | | Overlap <50 bp | | | |
| chrXII | 654964 | 655114 | 150 | -1,46602872 | N12:655066 | 6,506 | | Intergene | | | |
| chrXII | 656400 | 656549 | 149 | -1,294664736 | N12:656524 | 53,569 | | Intergene | | | |
| chrXII | 657729 | 657887 | 158 | -1,592834728 | N12:657828 | 21,638 | +1:anti126; | 1 | | | |
| chrXII | 664370 | 664478 | 108 | -1,112143121 | N12:664374 | 19,891 | +5:YLR259C; | 5 | | | |
| chrXII | 664370 | 664482 | 112 | -1,365450233 | N12:664374 | 19,891 | +5:YLR259C; | 5 | | | |
| chrXII | 665850 | 666010 | 160 | -1,433175291 | N12:665933 | 10,445 | +2:YLR260W; | 2 | | | |
| chrXII | 675650 | 675803 | 153 | -1,612298274 | N12:675721 | 5,508 | .R265C; +1:CUT268; +13*:YLR266C; | -1 | | | |
| chrXII | 678419 | 678585 | 166 | -1,339341513 | N12:678473 | 28,407 | +3:YLR267W; | 3 | | | |
| chrXII | 684089 | 684224 | 135 | -1,029221001 | N12:684178 | 29,296 | +19:YLR272C; | 19 | | | |
| chrXII | 686656 | 686810 | 154 | -1,421944125 | N12:686728 | 6,346 | +4:YLR272C; | 4 | | | |
| chrXII | 686971 | 687118 | 147 | -1,259381872 | N12:687063 | 8,704 | +2:YLR272C; | 2 | | | |
| chrXII | 691050 | 691199 | 149 | -1,543316174 | N12:691100 | 8,86 | +2:YLR273C; | 2 | | | |
| chrXII | 695440 | 695604 | 164 | -1,299008189 | N12:695602 | 37,758 | +9:YLR276C; | 9 | | | |
| chrXII | 698814 | 698965 | 151 | -1,25229095 | N12:698805 | 4,868 | +6:YLR277C; | 6 | | | |
| chrXII | 700732 | 700860 | 128 | -2,105816082 | N12:700798 | 27,391 | +21:YLR278C; +7:anti128; | 21 | | | |
| chrXII | 700732 | 700861 | 129 | -2,204084361 | N12:700798 | 27,391 | +21:YLR278C; +7:anti128; | 21 | | | |
| chrXII | 704633 | 704781 | 148 | -1,113566424 | N12:704801 | 35,242 | +2:YLR281C; | 2 | | | |
| chrXII | 704633 | 704785 | 152 | -1,220321279 | N12:704801 | 35,242 | +2:YLR281C; | 2 | | | |
| chrXII | 704633 | 704781 | 148 | -1,113566424 | N12:704613 | 0 | +3:YLR281C; | 3 | | | |
| chrXII | 704633 | 704785 | 152 | -1,220321279 | N12:704613 | 0 | +3:YLR281C; | 3 | | | |
| chrXII | 704637 | 704785 | 148 | -1,496555695 | N12:704801 | 35,242 | +2:YLR281C; | 2 | | | |
| chrXII | 704738 | 704878 | 140 | -0,93576076 | N12:704801 | 35,242 | +2:YLR281C; | 2 | | | |
| chrXII | 707321 | 707471 | 150 | -1,302650556 | N12:707390 | 3,742 | +1:YLR285W; -1:YLR284C; | 1 | | | |
| chrXII | 707855 | 708017 | 162 | -1,300801497 | N12:707899 | 24,295 | +4:YLR285W; | 4 | | | |
| chrXII | 710806 | 710956 | 150 | -1,125379508 | N12:710883 | 25,079 | +8*:YLR287C; | 8 | TERM | | |
| chrXII | 713802 | 713939 | 137 | -0,915756343 | N12:713872 | 34,268 | +7:YLR288C; | 7 | | | |
| chrXII | 714204 | 714311 | 107 | -1,327109314 | N12:714214 | 7,083 | +5:YLR288C; | 5 | | | |
| chrXII | 714204 | 714312 | 108 | -1,266675365 | N12:714214 | 7,083 | +5:YLR288C; | 5 | | | |
| chrXII | 714794 | 714945 | 151 | -1,614690376 | N12:714880 | 17,952 | -1:YLR289W; +1:YLR288C; | -1 | | | |
| chrXII | 721307 | 721447 | 140 | -1,179253806 | N12:721455 | 23,215 | +1:YLR293C; | 1 | | | |
| chrXII | 723297 | 723432 | 135 | -1,152113397 | N12:723451 | 34,025 | | Intergene | | | |
| chrXII | 726306 | 726432 | 126 | -0,899340527 | N12:726432 | 8,548 | +3:YLR299W; | 3 | | | |
| chrXII | 727018 | 727148 | 130 | -1,792398558 | N12:727116 | 23 | +7:YLR299W; | 7 | | | |
| chrXII | 731448 | 731612 | 164 | -1,398383819 | N12:731633 | 17,01 | +6*:YLR301W; | 6 | TERM | | |
| chrXII | 731448 | 731612 | 164 | -1,398383819 | N12:731474 | 4,619 | +5:YLR301W; | 5 | | | |
| chrXII | 733723 | 733874 | 151 | -1,352314842 | N12:733875 | 29,092 | +9*:YLR303W; | 9 | TERM | | |
| chrXII | 733723 | 733874 | 151 | -1,352314842 | N12:733718 | 4,243 | +8:YLR303W; | 8 | | | |
| chrXII | 735532 | 735635 | 103 | -2,071022605 | N12:735568 | 9,395 | +13:YLR304C; | 13 | | | |
| chrXII | 736027 | 736189 | 162 | -1,257376238 | N12:736034 | 34,677 | +10:YLR304C; | 10 | | | |
| chrXII | 738785 | 738957 | 172 | -1,487648485 | N12:738797 | 4,792 | +32:YLR305C; | 32 | | | |
| chrXII | 740597 | 740703 | 106 | -1,558174977 | N12:740592 | 23,78 | +21:YLR305C; | 21 | | | |
| chrXII | 745846 | 745998 | 152 | -1,702485895 | N12:745926 | 16,947 | +3:YLR307W; | 3 | | | |
| chrXII | 751027 | 751192 | 165 | -1,19160281 | N12:751096 | 26,661 | +5:YLR309C; | 5 | | | |

TABLE 3

| CHR | START | END | L(bp) | ΔLK | Nuc ID | Fuzziness | Gene ID | Gene body postion or intergene | Terminal | rDNA | Telomeric |
|---|---|---|---|---|---|---|---|---|---|---|---|
| chrXII | 752922 | 753058 | 136 | -0,801637738 | N12:753013 | 14,67 | | Intergene | | | |
| chrXII | 753617 | 753790 | 173 | -1,30239763 | N12:753609 | 33,231 | | Intergene | | | |
| chrXII | 753617 | 753790 | 173 | -1,30239763 | N12:753760 | 30,729 | | Intergene | | | |
| chrXII | 759427 | 759579 | 152 | -1,290340323 | N12:759504 | 11,143 | +2:YLR312W-A; | 2 | | | |
| chrXII | 761633 | 761798 | 165 | -1,673856365 | | | | Overlap <50 bp | | | |
| chrXII | 762105 | 762269 | 164 | -1,605670916 | N12:762185 | 8,01 | +2:YLR313C; | 2 | | | |
| chrXII | 770786 | 770938 | 152 | -1,648730318 | N12:770870 | 8,886 | +6:YLR319C; | 6 | | | |
| chrXII | 779526 | 779669 | 143 | -1,331404865 | N12:779529 | 6,025 | +3:YLR324W; | 3 | | | |
| chrXII | 779740 | 779903 | 163 | -1,645898663 | N12:779869 | 8,28 | +5:YLR324W; | 5 | | | |
| chrXII | 781130 | 781282 | 152 | -1,629690566 | N12:781211 | 16,145 | +2:YLR325C; | 2 | | | |
| chrXII | 782565 | 782713 | 148 | -1,181935995 | N12:782593 | 19,449 | +4:YLR326W; | 4 | | | |
| chrXII | 783914 | 784060 | 146 | -1,252694093 | N12:783921 | 31,634 | -1:YLR327C; 0:Unit466; | -1 | | | |
| chrXII | 783914 | 784060 | 146 | -1,252694093 | N12:784073 | 31,129 | +1:Unit466; | 1 | | | |
| chrXII | 785535 | 785685 | 150 | -1,167413271 | N12:785589 | 36,322 | +6:YLR328W; | 6 | | | |
| chrXII | 786502 | 786666 | 164 | -1,622967527 | N12:786585 | 5,419 | +2:YLR329W; +6:CUT746; | 2 | | | |
| chrXII | 792530 | 792669 | 139 | -0,985672355 | N12:792622 | 18,052 | -1:Unit447; +3*:Unit448; | -1 | | | |
| chrXII | 797667 | 797817 | 150 | -0,81750493 | N12:797755 | 9,574 | +3:YLR335W; | 3 | | | |
| chrXII | 799286 | 799439 | 153 | -1,502847957 | N12:799357 | 5,273 | +13:YLR335W; | 13 | | | |
| chrXII | 803374 | 803523 | 149 | -1,367805467 | N12:803394 | 37,532 | +12:YLR337C; | 12 | | | |
| chrXII | 804062 | 804197 | 135 | -1,272608929 | N12:804104 | 25,936 | +7:YLR337C; | 7 | | | |
| chrXII | 804515 | 804661 | 146 | -0,520779775 | N12:804614 | 5,03 | +4:YLR337C; | 4 | | | |
| chrXII | 806849 | 807003 | 154 | -1,506225373 | N12:806937 | 21,954 | +12*:SUT707; | 12 | TERM | | |
| chrXII | 807887 | 808014 | 127 | -1,307191765 | N12:807864 | 17,291 | +4:YLR341W; +7:SUT707; | 4 | | | |
| chrXII | 807887 | 808014 | 127 | -1,307191765 | N12:808034 | 40,873 | +5:YLR341W; +6:SUT707; | 5 | | | |
| chrXII | 827684 | 827838 | 154 | -1,332999705 | N12:827706 | 9,708 | +2:YLR348C; | 2 | | | |
| chrXII | 830101 | 830255 | 154 | -1,37999663 | N12:830181 | 13,243 | +2:YLR351C; | 2 | | | |
| chrXII | 830655 | 830774 | 119 | -1,37050083 | N12:830722 | 5,845 | -1:YLR352W; | -1 | | | |
| chrXII | 830666 | 830794 | 128 | -0,998965775 | N12:830722 | 5,845 | -1:YLR352W; | -1 | | | |
| chrXII | 831008 | 831187 | 179 | -1,613248547 | N12:831140 | 8,841 | +2:YLR352W; | 2 | | | |
| chrXII | 831542 | 831700 | 158 | -1,285080462 | N12:831639 | 25,066 | +5:YLR352W; | 5 | | | |
| chrXII | 839642 | 839772 | 130 | -1,780652983 | N12:839733 | 26,666 | | Intergene | | | |
| chrXII | 840366 | 840517 | 151 | -1,253677951 | | | | Overlap <50 bp | | | |
| chrXII | 841812 | 841967 | 155 | -1,547434673 | N12:841835 | 12,073 | +4:YLR357W; | 4 | | | |
| chrXII | 842219 | 842368 | 149 | -1,380688311 | N12:842300 | 15,834 | +7:YLR357W; | 7 | | | |
| chrXII | 846202 | 846357 | 155 | -1,236857121 | N12:846233 | 7,457 | +2:YLR360W; | 2 | | | |
| chrXII | 864284 | 864432 | 148 | -1,057629113 | N12:864280 | 9,912 | +11:YLR371W; | 11 | | | |
| chrXII | 868597 | 868734 | 137 | -1,279072435 | N12:868681 | 17,493 | +17*:YLR373C; | 17 | TERM | | |
| chrXII | 871320 | 871444 | 124 | -1,871475206 | N12:871354 | 10,035 | +1:YLR373C; | 1 | | | |
| chrXII | 871320 | 871449 | 129 | -1,041860957 | N12:871354 | 10,035 | +1:YLR373C; | 1 | | | |
| chrXII | 871322 | 871449 | 127 | -1,227968948 | N12:871354 | 10,035 | +1:YLR373C; | 1 | | | |
| chrXII | 874679 | 874837 | 158 | -1,890568156 | N12:874753 | 6,055 | +1:YLR377C; +7:SUT281; | 1 | | | |
| chrXII | 878567 | 878705 | 138 | -0,713345953 | N12:878691 | 35,912 | +4:YLR380W; | 4 | | | |
| chrXII | 884555 | 884713 | 158 | -1,361433238 | N12:884629 | 14,796 | +2:YLR382C; | 2 | | | |
| chrXII | 889517 | 889681 | 164 | -1,661222833 | N12:889511 | 15,355 | +22:YLR384C; | 22 | | | |
| chrXII | 893166 | 893327 | 161 | -1,249270363 | N12:893245 | 1,708 | -1:YLR384C; +2:YLR385C; | -1 | | | |
| chrXII | 896277 | 896446 | 169 | -1,433396506 | N12:896387 | 20,801 | +18*:YLR386W; +9*:YLR387C; | 18 | TERM | | |
| chrXII | 896668 | 896822 | 154 | -1,574832143 | N12:896735 | 18,488 | +7:YLR387C; | 7 | | | |
| chrXII | 896811 | 896958 | 147 | -1,181704098 | N12:896895 | 4,719 | +6:YLR387C; | 6 | | | |
| chrXII | 903630 | 903761 | 131 | -1,078568887 | N12:903719 | 11,432 | +1:YLR390W-A; +18*:YLR392C; | 1 | | | |
| chrXII | 903784 | 903945 | 161 | -1,33171439 | N12:903890 | 5,076 | +2:YLR390W-A; +17:YLR392C; | 2 | | | |
| chrXII | 903961 | 904117 | 156 | -1,215793365 | N12:904050 | 10,035 | +3:YLR390W-A; +16:YLR392C; | 3 | | | |
| chrXII | 905180 | 905335 | 155 | -1,697798567 | | | | Overlap <50 bp | | | |
| chrXII | 908649 | 908820 | 171 | -1,7539698 | N12:908748 | 20,169 | +6:YLR394W; +11:YLR393W; | 6 | | | |
| chrXII | 912499 | 912653 | 154 | -1,256421667 | N12:912578 | 15,845 | -1:YLR396C; +14*:YLR397C; | -1 | | | |
| chrXII | 913527 | 913695 | 168 | -1,589973244 | N12:913636 | 55,94 | +8:YLR397C; | 8 | | | |
| chrXII | 922406 | 922556 | 150 | -1,604816867 | N12:922523 | 20,657 | +12*:YLR401C; | 12 | TERM | | |
| chrXII | 922509 | 922654 | 145 | -1,185258968 | N12:922523 | 20,657 | +12*:YLR401C; | 12 | TERM | | |
| chrXII | 924375 | 924524 | 149 | -1,23336691 | N12:924433 | 6,338 | +1:YLR401C; | 1 | | | |
| chrXII | 926149 | 926303 | 154 | -1,367460252 | N12:926312 | 33,277 | +5:YLR403W; | 5 | | | |
| chrXII | 926358 | 926515 | 157 | -1,343691735 | N12:926512 | 47,794 | +6:YLR403W; | 6 | | | |
| chrXII | 929019 | 929172 | 153 | -1,378424745 | N12:929097 | 9,538 | +3:YLR404W; +4:CUT753; | 3 | | | |
| chrXII | 931573 | 931698 | 125 | -1,353750099 | | | | Overlap <50 bp | | | |
| chrXII | 937960 | 938108 | 148 | -1,291993363 | N12:938021 | 7,033 | +4:YLR410W; | 4 | | | |
| chrXII | 938740 | 938880 | 140 | -1,246618974 | N12:938790 | 9,535 | +8:YLR410W; | 8 | | | |
| chrXII | 939726 | 939894 | 168 | -1,420088907 | N12:939769 | 15,955 | | Intergene | | | |
| chrXII | 940022 | 940161 | 139 | -0,922168637 | N12:940088 | 15,176 | | Intergene | | | |
| chrXII | 941909 | 942015 | 106 | -1,224131786 | N12:941949 | 36,042 | | Intergene | | | |
| chrXII | 941909 | 942019 | 110 | -1,395497851 | N12:941949 | 36,042 | | Intergene | | | |
| chrXII | 943265 | 943394 | 129 | -1,428878182 | N12:943276 | 16,263 | | Intergene | | | |
| chrXII | 944730 | 944873 | 143 | -0,885989839 | N12:944794 | 21,656 | | Intergene | | | |
| chrXII | 946105 | 946219 | 114 | -1,386011908 | N12:946210 | 33,292 | | Intergene | | | |
| chrXII | 946809 | 946958 | 149 | -1,063448881 | N12:946920 | 31,507 | -1:YLR411W; | -1 | | | |
| chrXII | 947096 | 947245 | 149 | -1,661922292 | N12:947106 | 31,07 | 0:YLR411W; | 0 | | | |
| chrXII | 947588 | 947739 | 151 | -1,618389137 | N12:947709 | 32,591 | +4:YLR411W; | 4 | | | |
| chrXII | 952012 | 952159 | 147 | -0,87917015 | N12:952105 | 30,279 | +7:YLR413W; | 7 | | | |
| chrXII | 955881 | 956039 | 158 | -1,204435279 | N12:955940 | 15,369 | +7:YLR417W; | 7 | | | |
| chrXII | 956586 | 956713 | 127 | -1,425935455 | N12:956629 | 44,319 | +10*:YLR418C; +12:YLR417W; | 10 | TERM | | |
| chrXII | 962105 | 962271 | 166 | -1,321875338 | N12:962110 | 20,942 | +23:YLR419W; | 23 | | | |
| chrXII | 962105 | 962271 | 166 | -1,321875338 | N12:962293 | 8,864 | +24:YLR419W; | 24 | | | |
| chrXII | 966042 | 966188 | 146 | -1,158152817 | N12:966156 | 18,446 | +3:YLR422W; | 3 | | | |
| chrXII | 967180 | 967328 | 148 | -1,481968846 | N12:967232 | 33,921 | +10:YLR422W; | 10 | | | |
| chrXII | 967209 | 967341 | 132 | -0,887760468 | N12:967232 | 33,921 | +10:YLR422W; | 10 | | | |
| chrXII | 974145 | 974297 | 152 | -1,521288488 | N12:974223 | 4,243 | +6:YLR424W; | 6 | | | |
| chrXII | 975018 | 975184 | 166 | -1,423018085 | N12:975031 | 34,852 | +11:YLR424W; | 11 | | | |
| chrXII | 976449 | 976598 | 149 | -1,232510533 | N12:976465 | 29,866 | | Intergene | | | |
| chrXII | 978018 | 978161 | 143 | -0,837629057 | N12:978103 | 25,697 | | Intergene | | | |
| chrXII | 978601 | 978756 | 155 | -1,360556813 | N12:978687 | 47,658 | | Intergene | | | |
| chrXII | 979540 | 979688 | 148 | -1,485456149 | | | | Overlap <50 bp | | | |
| chrXII | 981009 | 981111 | 102 | -0,465985516 | | | | Overlap <50 bp | | | |
| chrXII | 981009 | 981139 | 130 | -1,130117322 | | | | Overlap <50 bp | | | |
| chrXII | 981073 | 981227 | 154 | -1,323320528 | N12:981187 | 32,868 | | Intergene | | | |
| chrXII | 981080 | 981248 | 168 | -1,415991227 | N12:981187 | 32,868 | | Intergene | | | |
| chrXII | 982426 | 982590 | 164 | -1,298551797 | N12:982511 | 15,007 | -1:YLR425W; | -1 | | | |
| chrXII | 987388 | 987536 | 148 | -1,1864529 | N12:987440 | 6,178 | +3:YLR426W; | 3 | | | |
| chrXII | 988804 | 988961 | 157 | -1,387466686 | N12:988850 | 10,114 | +4:YLR427W; | 4 | | | |
| chrXII | 988850 | 988961 | 111 | -0,895631229 | N12:988850 | 10,114 | +4:YLR427W; | 4 | | | |
| chrXII | 988957 | 989104 | 147 | -1,070461326 | N12:989040 | 19,887 | +5:YLR427W; | 5 | | | |
| chrXII | 990093 | 990242 | 149 | -1,18180111 | N12:990189 | 11,541 | +12:YLR427W; | 12 | | | |
| chrXII | 990292 | 990442 | 150 | -1,075066551 | N12:990360 | 11,978 | +13*:YLR427W; | 13 | TERM | | |

TABLE 3

| CHR | START | END | L(bp) | ΔLK | Nuc ID | Fuzziness | Gene ID | Gene body postion or intergene | Terminal | rDNA | Telomeric |
|---|---|---|---|---|---|---|---|---|---|---|---|
| chrXII | 997620 | 997777 | 157 | -1,480616283 | N12:997691 | 37,877 | | Intergene | | | |
| chrXII | 998925 | 999073 | 148 | -1,828157734 | N12:999015 | 28,169 | | Intergene | | | |
| chrXII | 1001567 | 1001716 | 149 | -1,494810195 | N12:1001641 | 7,731 | +3:YLR431C; | 3 | | | |
| chrXII | 1005509 | 1005668 | 159 | -1,765200535 | N12:1005574 | 10,474 | +4:YLR433C; | 4 | | | |
| chrXII | 1005994 | 1006142 | 148 | -1,176876468 | N12:1006065 | 4,761 | +1:YLR433C; -1:YLR435W; | 1 | | | |
| chrXII | 1012419 | 1012577 | 158 | -1,292574676 | N12:1012484 | 16,355 | +1:YLR438W; | 1 | | | |
| chrXII | 1012549 | 1012696 | 147 | -2,60724131 | N12:1012647 | 29,704 | +2:YLR438W; | 2 | | | |
| chrXII | 1012550 | 1012632 | 82 | -1,703987772 | N12:1012647 | 29,704 | +2:YLR438W; | 2 | | | |
| chrXII | 1012550 | 1012633 | 83 | -1,555418571 | N12:1012647 | 29,704 | +2:YLR438W; | 2 | | | |
| chrXII | 1012550 | 1012634 | 84 | -1,229970034 | N12:1012647 | 29,704 | +2:YLR438W; | 2 | | | |
| chrXII | 1012550 | 1012635 | 85 | -1,050611192 | N12:1012647 | 29,704 | +2:YLR438W; | 2 | | | |
| chrXII | 1012550 | 1012636 | 86 | -0,594355953 | N12:1012647 | 29,704 | +2:YLR438W; | 2 | | | |
| chrXII | 1012550 | 1012637 | 87 | -1,816592235 | N12:1012647 | 29,704 | +2:YLR438W; | 2 | | | |
| chrXII | 1012550 | 1012638 | 88 | -1,28809307 | N12:1012647 | 29,704 | +2:YLR438W; | 2 | | | |
| chrXII | 1012550 | 1012639 | 89 | -1,249917554 | N12:1012647 | 29,704 | +2:YLR438W; | 2 | | | |
| chrXII | 1012550 | 1012640 | 90 | -1,432514376 | N12:1012647 | 29,704 | +2:YLR438W; | 2 | | | |
| chrXII | 1012550 | 1012641 | 91 | -1,39232261 | N12:1012647 | 29,704 | +2:YLR438W; | 2 | | | |
| chrXII | 1012550 | 1012642 | 92 | -1,013001793 | N12:1012647 | 29,704 | +2:YLR438W; | 2 | | | |
| chrXII | 1012550 | 1012643 | 93 | -1,268010483 | N12:1012647 | 29,704 | +2:YLR438W; | 2 | | | |
| chrXII | 1012550 | 1012644 | 94 | -1,234401513 | N12:1012647 | 29,704 | +2:YLR438W; | 2 | | | |
| chrXII | 1012550 | 1012645 | 95 | -1,559710548 | N12:1012647 | 29,704 | +2:YLR438W; | 2 | | | |
| chrXII | 1012550 | 1012646 | 96 | -1,132476066 | N12:1012647 | 29,704 | +2:YLR438W; | 2 | | | |
| chrXII | 1012550 | 1012648 | 98 | -1,340709012 | N12:1012647 | 29,704 | +2:YLR438W; | 2 | | | |
| chrXII | 1012550 | 1012649 | 99 | -1,168547102 | N12:1012647 | 29,704 | +2:YLR438W; | 2 | | | |
| chrXII | 1012550 | 1012650 | 100 | -0,978978077 | N12:1012647 | 29,704 | +2:YLR438W; | 2 | | | |
| chrXII | 1012550 | 1012651 | 101 | -0,601324136 | N12:1012647 | 29,704 | +2:YLR438W; | 2 | | | |
| chrXII | 1012550 | 1012652 | 102 | -1,732895924 | N12:1012647 | 29,704 | +2:YLR438W; | 2 | | | |
| chrXII | 1012550 | 1012653 | 103 | -1,17360965 | N12:1012647 | 29,704 | +2:YLR438W; | 2 | | | |
| chrXII | 1012550 | 1012654 | 104 | -1,154786672 | N12:1012647 | 29,704 | +2:YLR438W; | 2 | | | |
| chrXII | 1012550 | 1012655 | 105 | -0,5863446 | N12:1012647 | 29,704 | +2:YLR438W; | 2 | | | |
| chrXII | 1012550 | 1012656 | 106 | -1,63626122 | N12:1012647 | 29,704 | +2:YLR438W; | 2 | | | |
| chrXII | 1012550 | 1012657 | 107 | -1,141816806 | N12:1012647 | 29,704 | +2:YLR438W; | 2 | | | |
| chrXII | 1012550 | 1012659 | 109 | -0,985866094 | N12:1012647 | 29,704 | +2:YLR438W; | 2 | | | |
| chrXII | 1012550 | 1012660 | 110 | -0,753558564 | N12:1012647 | 29,704 | +2:YLR438W; | 2 | | | |
| chrXII | 1012550 | 1012661 | 111 | -1,144849445 | N12:1012647 | 29,704 | +2:YLR438W; | 2 | | | |
| chrXII | 1012550 | 1012662 | 112 | -0,867200904 | N12:1012647 | 29,704 | +2:YLR438W; | 2 | | | |
| chrXII | 1012550 | 1012663 | 113 | -1,104667559 | N12:1012647 | 29,704 | +2:YLR438W; | 2 | | | |
| chrXII | 1012550 | 1012664 | 114 | -1,446563429 | N12:1012647 | 29,704 | +2:YLR438W; | 2 | | | |
| chrXII | 1012550 | 1012665 | 115 | -1,222845338 | N12:1012647 | 29,704 | +2:YLR438W; | 2 | | | |
| chrXII | 1012550 | 1012666 | 116 | -1,274760103 | N12:1012647 | 29,704 | +2:YLR438W; | 2 | | | |
| chrXII | 1012550 | 1012667 | 117 | -0,960653632 | N12:1012647 | 29,704 | +2:YLR438W; | 2 | | | |
| chrXII | 1012550 | 1012668 | 118 | -1,288907071 | N12:1012647 | 29,704 | +2:YLR438W; | 2 | | | |
| chrXII | 1012550 | 1012669 | 119 | -0,980708146 | N12:1012647 | 29,704 | +2:YLR438W; | 2 | | | |
| chrXII | 1012550 | 1012670 | 120 | -1,147419989 | N12:1012647 | 29,704 | +2:YLR438W; | 2 | | | |
| chrXII | 1012550 | 1012671 | 121 | -1,491951922 | N12:1012647 | 29,704 | +2:YLR438W; | 2 | | | |
| chrXII | 1012550 | 1012672 | 122 | -1,358985062 | N12:1012647 | 29,704 | +2:YLR438W; | 2 | | | |
| chrXII | 1012550 | 1012673 | 123 | -1,301851291 | N12:1012647 | 29,704 | +2:YLR438W; | 2 | | | |
| chrXII | 1012550 | 1012674 | 124 | -2,23299362 | N12:1012647 | 29,704 | +2:YLR438W; | 2 | | | |
| chrXII | 1012550 | 1012675 | 125 | -1,369276971 | N12:1012647 | 29,704 | +2:YLR438W; | 2 | | | |
| chrXII | 1012550 | 1012676 | 126 | -1,390116407 | N12:1012647 | 29,704 | +2:YLR438W; | 2 | | | |
| chrXII | 1012550 | 1012677 | 127 | -1,688013805 | N12:1012647 | 29,704 | +2:YLR438W; | 2 | | | |
| chrXII | 1012550 | 1012678 | 128 | -1,053736296 | N12:1012647 | 29,704 | +2:YLR438W; | 2 | | | |
| chrXII | 1012550 | 1012679 | 129 | -0,815601844 | N12:1012647 | 29,704 | +2:YLR438W; | 2 | | | |
| chrXII | 1012550 | 1012680 | 130 | -1,174603453 | N12:1012647 | 29,704 | +2:YLR438W; | 2 | | | |
| chrXII | 1012550 | 1012681 | 131 | -1,389263522 | N12:1012647 | 29,704 | +2:YLR438W; | 2 | | | |
| chrXII | 1012550 | 1012682 | 132 | -1,096373597 | N12:1012647 | 29,704 | +2:YLR438W; | 2 | | | |
| chrXII | 1012550 | 1012683 | 133 | -1,601493863 | N12:1012647 | 29,704 | +2:YLR438W; | 2 | | | |
| chrXII | 1012550 | 1012684 | 134 | -1,027464857 | N12:1012647 | 29,704 | +2:YLR438W; | 2 | | | |
| chrXII | 1012550 | 1012685 | 135 | -1,105396275 | N12:1012647 | 29,704 | +2:YLR438W; | 2 | | | |
| chrXII | 1012550 | 1012686 | 136 | -1,262704946 | N12:1012647 | 29,704 | +2:YLR438W; | 2 | | | |
| chrXII | 1012550 | 1012687 | 137 | -1,40186849 | N12:1012647 | 29,704 | +2:YLR438W; | 2 | | | |
| chrXII | 1012550 | 1012688 | 138 | -1,129146826 | N12:1012647 | 29,704 | +2:YLR438W; | 2 | | | |
| chrXII | 1012550 | 1012689 | 139 | -1,063228137 | N12:1012647 | 29,704 | +2:YLR438W; | 2 | | | |
| chrXII | 1012550 | 1012690 | 140 | -1,096232623 | N12:1012647 | 29,704 | +2:YLR438W; | 2 | | | |
| chrXII | 1012550 | 1012691 | 141 | -1,395176989 | N12:1012647 | 29,704 | +2:YLR438W; | 2 | | | |
| chrXII | 1012550 | 1012692 | 142 | -1,113868199 | N12:1012647 | 29,704 | +2:YLR438W; | 2 | | | |
| chrXII | 1012550 | 1012693 | 143 | -1,096978822 | N12:1012647 | 29,704 | +2:YLR438W; | 2 | | | |
| chrXII | 1012550 | 1012694 | 144 | -1,431926774 | N12:1012647 | 29,704 | +2:YLR438W; | 2 | | | |
| chrXII | 1012550 | 1012695 | 145 | -1,413781992 | N12:1012647 | 29,704 | +2:YLR438W; | 2 | | | |
| chrXII | 1012550 | 1012696 | 146 | -1,070588043 | N12:1012647 | 29,704 | +2:YLR438W; | 2 | | | |
| chrXII | 1012550 | 1012698 | 148 | -1,525375093 | N12:1012647 | 29,704 | +2:YLR438W; | 2 | | | |
| chrXII | 1012551 | 1012693 | 142 | -1,489015829 | N12:1012647 | 29,704 | +2:YLR438W; | 2 | | | |
| chrXII | 1012551 | 1012696 | 145 | -1,826416557 | N12:1012647 | 29,704 | +2:YLR438W; | 2 | | | |
| chrXII | 1012552 | 1012696 | 144 | -1,640072941 | N12:1012647 | 29,704 | +2:YLR438W; | 2 | | | |
| chrXII | 1012553 | 1012693 | 140 | -0,847102545 | N12:1012647 | 29,704 | +2:YLR438W; | 2 | | | |
| chrXII | 1012553 | 1012696 | 143 | -1,565814544 | N12:1012647 | 29,704 | +2:YLR438W; | 2 | | | |
| chrXII | 1012554 | 1012696 | 142 | -0,940696314 | N12:1012647 | 29,704 | +2:YLR438W; | 2 | | | |
| chrXII | 1012555 | 1012696 | 141 | -1,23411604 | N12:1012647 | 29,704 | +2:YLR438W; | 2 | | | |
| chrXII | 1012556 | 1012696 | 140 | -1,299232068 | N12:1012647 | 29,704 | +2:YLR438W; | 2 | | | |
| chrXII | 1012557 | 1012696 | 139 | -1,762170494 | N12:1012647 | 29,704 | +2:YLR438W; | 2 | | | |
| chrXII | 1012558 | 1012696 | 138 | -1,583648443 | N12:1012647 | 29,704 | +2:YLR438W; | 2 | | | |
| chrXII | 1012559 | 1012696 | 137 | -0,946648921 | N12:1012647 | 29,704 | +2:YLR438W; | 2 | | | |
| chrXII | 1012560 | 1012696 | 136 | -1,36337568 | N12:1012647 | 29,704 | +2:YLR438W; | 2 | | | |
| chrXII | 1012561 | 1012696 | 135 | -1,295297759 | N12:1012647 | 29,704 | +2:YLR438W; | 2 | | | |
| chrXII | 1012562 | 1012696 | 134 | -1,938357556 | N12:1012647 | 29,704 | +2:YLR438W; | 2 | | | |
| chrXII | 1012563 | 1012696 | 133 | -1,432966431 | N12:1012647 | 29,704 | +2:YLR438W; | 2 | | | |
| chrXII | 1012564 | 1012696 | 132 | -2,259462671 | N12:1012647 | 29,704 | +2:YLR438W; | 2 | | | |
| chrXII | 1012565 | 1012696 | 131 | -1,785833694 | N12:1012647 | 29,704 | +2:YLR438W; | 2 | | | |
| chrXII | 1012566 | 1012696 | 130 | -1,583583607 | N12:1012647 | 29,704 | +2:YLR438W; | 2 | | | |
| chrXII | 1012567 | 1012696 | 129 | -2,158811696 | N12:1012647 | 29,704 | +2:YLR438W; | 2 | | | |
| chrXII | 1012568 | 1012696 | 128 | -1,949151486 | N12:1012647 | 29,704 | +2:YLR438W; | 2 | | | |
| chrXII | 1012569 | 1012696 | 127 | -2,052981344 | N12:1012647 | 29,704 | +2:YLR438W; | 2 | | | |
| chrXII | 1012570 | 1012696 | 126 | -1,916861355 | N12:1012647 | 29,704 | +2:YLR438W; | 2 | | | |
| chrXII | 1012571 | 1012696 | 125 | -2,217038942 | N12:1012647 | 29,704 | +2:YLR438W; | 2 | | | |
| chrXII | 1012572 | 1012696 | 124 | -1,487194401 | N12:1012647 | 29,704 | +2:YLR438W; | 2 | | | |
| chrXII | 1012573 | 1012696 | 123 | -1,871224647 | N12:1012647 | 29,704 | +2:YLR438W; | 2 | | | |
| chrXII | 1012574 | 1012696 | 122 | -1,851573068 | N12:1012647 | 29,704 | +2:YLR438W; | 2 | | | |
| chrXII | 1012575 | 1012696 | 121 | -0,951586593 | N12:1012647 | 29,704 | +2:YLR438W; | 2 | | | |
| chrXII | 1012576 | 1012696 | 120 | -2,097664905 | N12:1012647 | 29,704 | +2:YLR438W; | 2 | | | |
| chrXII | 1012578 | 1012696 | 118 | -2,282261357 | N12:1012647 | 29,704 | +2:YLR438W; | 2 | | | |

| CHR | START | END | L(bp) | ΔLK | Nuc ID | Fuzziness | Gene ID | Gene body postion or intergene | Terminal | rDNA | Telomeric |
|---|---|---|---|---|---|---|---|---|---|---|---|
| chrXII | 1012579 | 1012696 | 117 | -1,436830513 | N12:1012647 | 29,704 | +2:YLR438W; | 2 | | | |
| chrXII | 1012580 | 1012696 | 116 | -1,947610656 | N12:1012647 | 29,704 | +2:YLR438W; | 2 | | | |
| chrXII | 1012581 | 1012696 | 115 | -1,736134278 | N12:1012647 | 29,704 | +2:YLR438W; | 2 | | | |
| chrXII | 1012583 | 1012696 | 113 | -0,962909448 | N12:1012647 | 29,704 | +2:YLR438W; | 2 | | | |
| chrXII | 1012584 | 1012696 | 112 | -2,008328572 | N12:1012647 | 29,704 | +2:YLR438W; | 2 | | | |
| chrXII | 1012586 | 1012696 | 110 | -0,706380039 | N12:1012647 | 29,704 | +2:YLR438W; | 2 | | | |
| chrXII | 1012590 | 1012674 | 84 | -0,850573336 | N12:1012647 | 29,704 | +2:YLR438W; | 2 | | | |
| chrXII | 1012610 | 1012696 | 86 | -1,6800582 | N12:1012647 | 29,704 | +2:YLR438W; | 2 | | | |
| chrXII | 1013868 | 1014024 | 156 | -1,130354816 | N12:1013902 | 23,483 | +3:YLR438C-A; | 3 | | | |
| chrXII | 1017667 | 1017816 | 149 | -1,478527849 | N12:1017757 | 36,682 | +1:YLR440C; | 1 | | | |
| chrXII | 1023621 | 1023767 | 146 | -1,197893854 | N12:1023663 | 7,563 | +8:YLR443W; | 8 | | | |
| chrXII | 1024132 | 1024289 | 157 | -1,522310166 | N12:1024222 | 10,035 | +1:YLR445W; | 1 | | | |
| chrXII | 1031294 | 1031445 | 151 | -1,315034471 | N12:1031333 | 26,332 | +4:YLR449W; | 4 | | | |
| chrXII | 1032438 | 1032586 | 148 | -1,41857051 | N12:1032481 | 9,508 | +1:YLR450W; | 1 | | | |
| chrXII | 1050400 | 1050528 | 128 | -1,402878192 | N12:1050503 | 12,161 | | Intergene | | | |
| chrXII | 1050732 | 1050875 | 143 | -1,634285902 | N12:1050868 | 32,723 | | Intergene | | | |
| chrXII | 1059679 | 1059820 | 141 | -1,397733471 | N12:1059812 | 9,854 | +8*:YLR460C; | 8 | TERM | | |
| chrXII | 1065029 | 1065154 | 125 | -0,350752953 | N12:1065098 | 32,501 | | Intergene | | | TEL RIGHT |
| chrXII | 1065029 | 1065156 | 127 | -0,303005082 | N12:1065098 | 32,501 | | Intergene | | | TEL RIGHT |
| chrXII | 1066703 | 1066854 | 151 | -1,393222444 | | | | Overlap <50 bp | | | TEL RIGHT |
| chrXII | 1067646 | 1067808 | 162 | -1,186246987 | N12:1067769 | 9,539 | +5:YLR466W; | 5 | | | TEL RIGHT |
| chrXII | 1067747 | 1067902 | 155 | -1,852097459 | N12:1067769 | 9,539 | +5:YLR466W; | 5 | | | TEL RIGHT |
| chrXII | 1067881 | 1068018 | 137 | -1,092789151 | N12:1067950 | 18,583 | +6:YLR466W; | 6 | | | TEL RIGHT |
| chrXII | 1068176 | 1068319 | 143 | -1,302361921 | N12:1068230 | 14,503 | +8:YLR466W; | 8 | | | TEL RIGHT |
| chrXII | 1068224 | 1068349 | 125 | -1,021672271 | N12:1068230 | 14,503 | +8:YLR466W; | 8 | | | TEL RIGHT |
| chrXII | 1068449 | 1068601 | 152 | -1,213184443 | | | | Overlap <50 bp | | | TEL RIGHT |
| chrXII | 1068522 | 1068666 | 144 | -1,231008052 | | | | Overlap <50 bp | | | TEL RIGHT |
| chrXII | 1068619 | 1068754 | 135 | -1,049580135 | N12:1068716 | 23,302 | +11:YLR466W; | 11 | | | TEL RIGHT |
| chrXII | 1068619 | 1068806 | 187 | -1,083121784 | N12:1068716 | 23,302 | +11:YLR466W; | 11 | | | TEL RIGHT |
| chrXII | 1068662 | 1068754 | 92 | -0,366049365 | N12:1068716 | 23,302 | +11:YLR466W; | 11 | | | TEL RIGHT |
| chrXII | 1068662 | 1068806 | 144 | -0,948731851 | N12:1068716 | 23,302 | +11:YLR466W; | 11 | | | TEL RIGHT |
| chrXII | 1069001 | 1069153 | 152 | -1,351882452 | N12:1069071 | 40,377 | +13:YLR466W; | 13 | | | TEL RIGHT |
| chrXII | 1069302 | 1069432 | 130 | -1,117771804 | N12:1069283 | 42,438 | +14:YLR466W; | 14 | | | TEL RIGHT |
| chrXII | 1069440 | 1069601 | 161 | -1,173760425 | | | | Overlap <50 bp | | | TEL RIGHT |
| chrXII | 1070837 | 1070988 | 151 | -1,274029915 | N12:1070915 | 64,003 | +23:YLR466W; | 23 | | | TEL RIGHT |
| chrXII | 1070837 | 1071008 | 171 | -1,278700138 | N12:1070915 | 64,003 | +23:YLR466W; | 23 | | | TEL RIGHT |
| chrXII | 1072226 | 1072381 | 155 | -1,508760617 | N12:1072335 | 30,271 | 0:YLR467W; | 0 | | | TEL RIGHT |
| chrXII | 1073391 | 1073538 | 147 | -1,161982674 | | | | Overlap <50 bp | | | TEL RIGHT |
| chrXII | 1073547 | 1073677 | 130 | -1,049429275 | N12:1073613 | 19,63 | +8:YLR467W; | 8 | | | TEL RIGHT |
| chrXII | 1073928 | 1074064 | 136 | -1,300139293 | N12:1073964 | 0 | +10:YLR467W; | 10 | | | TEL RIGHT |
| chrXII | 1074309 | 1074471 | 162 | -1,216368113 | N12:1074439 | 10,97 | +13:YLR467W; | 13 | | | TEL RIGHT |
| chrXII | 1074410 | 1074565 | 155 | -1,929018038 | N12:1074439 | 10,97 | +13:YLR467W; | 13 | | | TEL RIGHT |
| chrXII | 1074544 | 1074681 | 137 | -1,099427163 | N12:1074612 | 9,713 | +14:YLR467W; | 14 | | | TEL RIGHT |
| chrXII | 1074831 | 1074982 | 151 | -1,520081729 | N12:1074890 | 4,359 | +16:YLR467W; | 16 | | | TEL RIGHT |
| chrXII | 1074839 | 1074982 | 143 | -1,387980516 | N12:1074890 | 4,359 | +16:YLR467W; | 16 | | | TEL RIGHT |
| chrXII | 1074887 | 1075012 | 125 | -0,951224242 | N12:1074890 | 4,359 | +16:YLR467W; | 16 | | | TEL RIGHT |
| chrXII | 1075112 | 1075264 | 152 | -1,299676744 | | | | Overlap <50 bp | | | TEL RIGHT |
| chrXII | 1075185 | 1075329 | 144 | -1,393480009 | | | | Overlap <50 bp | | | TEL RIGHT |
| chrXII | 1075282 | 1075417 | 135 | -0,986164431 | N12:1075369 | 30,199 | +19:YLR467W; | 19 | | | TEL RIGHT |
| chrXII | 1075282 | 1075469 | 187 | -1,088191836 | N12:1075369 | 30,199 | +19:YLR467W; | 19 | | | TEL RIGHT |
| chrXII | 1075325 | 1075417 | 92 | -0,325215863 | N12:1075369 | 30,199 | +19:YLR467W; | 19 | | | TEL RIGHT |
| chrXII | 1075325 | 1075469 | 144 | -1,028038225 | N12:1075369 | 30,199 | +19:YLR467W; | 19 | | | TEL RIGHT |
| chrXII | 1075664 | 1075816 | 152 | -1,181249815 | N12:1075734 | 7,778 | +21:YLR467W; | 21 | | | TEL RIGHT |
| chrXII | 1075965 | 1076095 | 130 | -1,086125662 | N12:1076061 | 36,638 | +23:YLR467W; | 23 | | | TEL RIGHT |
| chrXII | 1076103 | 1076264 | 161 | -1,289945398 | | | | Overlap <50 bp | | | TEL RIGHT |
| chrXII | 1077160 | 1077312 | 152 | -1,399093427 | N12:1077229 | 52,624 | +30:YLR467W; | 30 | | | TEL RIGHT |
| chrXII | 1077500 | 1077651 | 151 | -1,265868927 | N12:1077576 | 34,962 | +32:YLR467W; | 32 | | | TEL RIGHT |
| chrXIII | 687 | 858 | 171 | -1,183022396 | N13:774 | 35,791 | +25:YML133C; | 25 | | | TEL LEFT |
| chrXIII | 707 | 858 | 151 | -1,165709288 | N13:774 | 35,791 | +25:YML133C; | 25 | | | TEL LEFT |
| chrXIII | 1046 | 1198 | 152 | -1,480095132 | N13:1107 | 43,92 | +23:YML133C; | 23 | | | TEL LEFT |
| chrXIII | 1151 | 1299 | 148 | -1,439779997 | | | | Overlap <50 bp | | | TEL LEFT |
| chrXIII | 1836 | 2023 | 187 | -0,910571306 | N13:1892 | 9,452 | +18:YML133C; | 18 | | | TEL LEFT |
| chrXIII | 1872 | 2023 | 151 | -1,18877236 | N13:1892 | 9,452 | +18:YML133C; | 18 | | | TEL LEFT |
| chrXIII | 1884 | 2018 | 134 | -1,03886823 | N13:1892 | 9,452 | +18:YML133C; | 18 | | | TEL LEFT |
| chrXIII | 1909 | 2005 | 96 | -0,957176331 | N13:1892 | 9,452 | +18:YML133C; | 18 | | | TEL LEFT |
| chrXIII | 2542 | 2694 | 152 | -1,176021394 | N13:2630 | 14,978 | +14:YML133C; | 14 | | | TEL LEFT |
| chrXIII | 2889 | 3033 | 144 | -1,003528128 | N13:2945 | 16,462 | +12:YML133C; | 12 | | | TEL LEFT |
| chrXIII | 2889 | 3076 | 187 | -1,003940139 | N13:2945 | 16,462 | +12:YML133C; | 12 | | | TEL LEFT |
| chrXIII | 2941 | 3033 | 92 | -0,271296725 | N13:2945 | 16,462 | +12:YML133C; | 12 | | | TEL LEFT |
| chrXIII | 2941 | 3076 | 135 | -0,859328525 | N13:2945 | 16,462 | +12:YML133C; | 12 | | | TEL LEFT |
| chrXIII | 3029 | 3173 | 144 | -1,294580427 | N13:3125 | 4,509 | +11:YML133C; | 11 | | | TEL LEFT |
| chrXIII | 3094 | 3246 | 152 | -1,19440057 | N13:3125 | 4,509 | +11:YML133C; | 11 | | | TEL LEFT |
| chrXIII | 3346 | 3471 | 125 | -0,827654523 | N13:3450 | 12,124 | +9:YML133C; | 9 | | | TEL LEFT |
| chrXIII | 3376 | 3519 | 143 | -1,449081815 | N13:3450 | 12,124 | +9:YML133C; | 9 | | | TEL LEFT |
| chrXIII | 3376 | 3527 | 151 | -1,556775214 | N13:3450 | 12,124 | +9:YML133C; | 9 | | | TEL LEFT |
| chrXIII | 5624 | 5767 | 143 | -1,19399343 | N13:5763 | 5,686 | | Intergene | | | TEL LEFT |
| chrXIII | 9070 | 9207 | 137 | -1,136301165 | N13:9142 | 17,711 | 0:Unit471; | 0 | | | |
| chrXIII | 13123 | 13267 | 144 | -1,258306945 | N13:13226 | 17,268 | -1:SUT285; +1:YML130C; | -1 | | | |
| chrXIII | 14333 | 14505 | 172 | -1,644543528 | N13:14398 | 2,881 | +3*:YML129C; | 3 | TERM | | |
| chrXIII | 16025 | 16179 | 154 | -1,283756977 | N13:16099 | 9,452 | +5:YML128C; | 5 | | | |
| chrXIII | 18573 | 18744 | 171 | -1,484644976 | N13:18675 | 25,185 | +11:YML127W; | 11 | | | |
| chrXIII | 19858 | 19999 | 141 | -1,117666276 | N13:19950 | 6,768 | +4:YML126C; | 4 | | | |
| chrXIII | 20186 | 20333 | 147 | -1,320687818 | N13:20274 | 13,808 | +2:YML126C; | 2 | | | |
| chrXIII | 21593 | 21740 | 147 | -1,238327004 | N13:21676 | 8,718 | +1:YML125C; -1:anti-YML124C; | 1 | | | |
| chrXIII | 23095 | 23242 | 147 | -1,221766854 | N13:23173 | 7,662 | +4:YML124C; | 4 | | | |
| chrXIII | 32413 | 32558 | 145 | -1,275997143 | N13:32440 | 18,802 | +2:YML118W; | 2 | | | |
| chrXIII | 33232 | 33384 | 152 | -1,243536568 | N13:33387 | 20,253 | +8:YML118W; | 8 | | | |
| chrXIII | 38992 | 39143 | 151 | -1,328742332 | N13:39050 | 7,071 | +7:YML116W; | 7 | | | |
| chrXIII | 40472 | 40624 | 152 | -1,343492743 | N13:40521 | 24,185 | +9:YML115C; | 9 | | | |
| chrXIII | 42972 | 43133 | 161 | -1,309797338 | N13:43054 | 4,848 | +4:YML114C; | 4 | | | |
| chrXIII | 47095 | 47254 | 159 | -1,127314419 | N13:47157 | 13,074 | +3:YML111W; | 3 | | | |
| chrXIII | 54675 | 54757 | 82 | -0,753764522 | | | | Overlap <50 bp | | | |
| chrXIII | 54913 | 55020 | 107 | -1,173871325 | N13:54961 | 12,817 | +2:YML108W; +2:CUT759; | 2 | | | |
| chrXIII | 54913 | 55032 | 119 | -1,324745926 | N13:54961 | 12,817 | +2:YML108W; +2:CUT759; | 2 | | | |
| chrXIII | 59231 | 59396 | 165 | -1,143192055 | N13:59272 | 19,191 | +20:YML104C; +3*:CUT283; | 20 | | | |
| chrXIII | 63883 | 64021 | 138 | -1,059913134 | N13:63895 | 39,306 | | Intergene | | | |
| chrXIII | 64235 | 64393 | 158 | -1,705670368 | N13:64363 | 38,475 | | Intergene | | | |
| chrXIII | 64871 | 65018 | 147 | -1,308217953 | N13:64980 | 24,9 | | Intergene | | | |
| chrXIII | 66681 | 66829 | 148 | -1,217484632 | N13:66758 | 26,701 | +6:YML103C; | 6 | | | |

| CHR | START | END | L(bp) | ΔLK | Nuc ID | Fuzziness | Gene ID | Gene body postion or intergene | Terminal | rDNA | Telomeric |
|---|---|---|---|---|---|---|---|---|---|---|---|
| chrXIII | 75788 | 75939 | 151 | -1,317183113 | N13:75859 | 21,265 | +7:YML099C; | 7 | | | |
| chrXIII | 76732 | 76885 | 153 | -1,391942299 | N13:76800 | 13,794 | +2:YML099C; | 2 | | | |
| chrXIII | 77591 | 77728 | 137 | -1,20762293 | N13:77626 | 5,568 | +3:YML098W; | 3 | | | |
| chrXIII | 83978 | 84137 | 159 | -1,215911279 | | | | Overlap <50 bp | | | |
| chrXIII | 93745 | 93871 | 126 | -1,282405246 | N13:93877 | 21,352 | +11:YML088W; | 11 | | | |
| chrXIII | 98827 | 98976 | 149 | -0,863674115 | N13:98894 | 8,385 | | Intergene | | | |
| chrXIII | 102493 | 102629 | 136 | -1,00378864 | N13:102572 | 20,94 | +5:YML082W; | 5 | | | |
| chrXIII | 104088 | 104231 | 143 | -1,149830956 | N13:104162 | 13,029 | +1:YML081C-A; | 1 | | | |
| chrXIII | 104601 | 104708 | 107 | -1,191806582 | N13:104679 | 5,007 | +1:YML081W; | 1 | | | |
| chrXIII | 104601 | 104720 | 119 | -1,262640637 | N13:104679 | 5,007 | +1:YML081W; | 1 | | | |
| chrXIII | 107860 | 108019 | 159 | -1,171594288 | N13:107958 | 44,191 | +22:YML081W; | 22 | | | |
| chrXIII | 109020 | 109176 | 156 | -1,253875206 | N13:109082 | 7,007 | +3:YML080W; | 3 | | | |
| chrXIII | 112712 | 112852 | 140 | -1,244036558 | N13:112775 | 5,167 | +17:YML076C; | 17 | | | |
| chrXIII | 113210 | 113365 | 155 | -1,172115839 | N13:113351 | 22,543 | +13:YML076C; | 13 | | | |
| chrXIII | 113210 | 113365 | 155 | -1,172115839 | N13:113196 | 37,179 | +14:YML076C; | 14 | | | |
| chrXIII | 116680 | 116830 | 150 | -1,676344701 | N13:116697 | 12,723 | +16:YML075C; | 16 | | | |
| chrXIII | 116680 | 116833 | 153 | -1,29154697 | N13:116697 | 12,723 | +16:YML075C; | 16 | | | |
| chrXIII | 116680 | 116835 | 155 | -1,141570437 | N13:116697 | 12,723 | +16:YML075C; | 16 | | | |
| chrXIII | 116684 | 116835 | 151 | -1,394129796 | N13:116697 | 12,723 | +16:YML075C; | 16 | | | |
| chrXIII | 116783 | 116928 | 145 | -1,094623581 | | | | Overlap <50 bp | | | |
| chrXIII | 117349 | 117491 | 142 | -1,172366766 | N13:117410 | 14,328 | +11:YML075C; | 11 | | | |
| chrXIII | 117514 | 117618 | 104 | -1,532456473 | N13:117572 | 7,594 | +10:YML075C; | 10 | | | |
| chrXIII | 119216 | 119393 | 177 | -1,45800678 | N13:119273 | 28,583 | -1:SUT288; -1:YML075C; | -1 | | | |
| chrXIII | 120074 | 120243 | 169 | -1,553893484 | N13:120063 | 19,092 | +4*:SUT288; | 4 | TERM | | |
| chrXIII | 120074 | 120243 | 169 | -1,553893484 | N13:120213 | 12,925 | | Intergene | | | |
| chrXIII | 123583 | 123746 | 163 | -1,407894304 | N13:123684 | 9,065 | | Intergene | | | |
| chrXIII | 124934 | 125083 | 149 | -1,264136325 | N13:125011 | 35,084 | +29:YML072C; | 29 | | | |
| chrXIII | 125456 | 125612 | 156 | -1,201897911 | N13:125539 | 48,813 | +26:YML072C; | 26 | | | |
| chrXIII | 126339 | 126468 | 129 | -0,702747337 | | | | Overlap <50 bp | | | |
| chrXIII | 129825 | 129930 | 105 | -1,476714441 | N13:129823 | 11,396 | -1:YML072C; +1:CUT285; | -1 | | | |
| chrXIII | 129825 | 129932 | 107 | -1,28624102 | N13:129823 | 11,396 | -1:YML072C; +1:CUT285; | -1 | | | |
| chrXIII | 129843 | 130004 | 161 | -1,162787455 | N13:129987 | 10,863 | +2*:CUT285; | 2 | TERM | | |
| chrXIII | 129843 | 129930 | 87 | -1,166659195 | N13:129823 | 11,396 | -1:YML072C; +1:CUT285; | -1 | | | |
| chrXIII | 129843 | 130004 | 161 | -1,162787455 | N13:129823 | 11,396 | -1:YML072C; +1:CUT285; | -1 | | | |
| chrXIII | 130886 | 131027 | 141 | -1,035647797 | N13:130956 | 24,214 | +5:YML071C; | 5 | | | |
| chrXIII | 131028 | 131190 | 162 | -1,052845016 | N13:131120 | 9,973 | +4:YML071C; | 4 | | | |
| chrXIII | 131602 | 131755 | 153 | -1,370976245 | N13:131595 | 39,185 | +1:YML071C; | 1 | | | |
| chrXIII | 133367 | 133536 | 169 | -1,418589705 | N13:133467 | 19,94 | +1:YML070W; | 1 | | | |
| chrXIII | 134348 | 134447 | 99 | -1,429663408 | N13:134415 | 29,538 | +7:YML070W; | 7 | | | |
| chrXIII | 135575 | 135725 | 150 | -1,381659658 | N13:135649 | 9,158 | +2:YML069W; | 2 | | | |
| chrXIII | 139632 | 139738 | 106 | -1,444777998 | N13:139678 | 7,439 | | Intergene | | | |
| chrXIII | 139632 | 139741 | 109 | -1,648734967 | N13:139678 | 7,439 | | Intergene | | | |
| chrXIII | 142450 | 142583 | 133 | -2,215482142 | N13:142508 | 7,43 | +3:YML065W; | 3 | | | |
| chrXIII | 147977 | 148129 | 152 | -1,254179292 | | | | Overlap <50 bp | | | |
| chrXIII | 148322 | 148434 | 112 | -1,381900204 | | | | Overlap <50 bp | | | |
| chrXIII | 154231 | 154388 | 157 | -1,4887113 | | | | Overlap <50 bp | | | |
| chrXIII | 155621 | 155784 | 163 | -1,235086737 | | | | Overlap <50 bp | | | |
| chrXIII | 163164 | 163259 | 95 | -1,197794011 | | | | Overlap <50 bp | | | |
| chrXIII | 169237 | 169345 | 108 | -1,466604463 | N13:169314 | 1,5 | +4:YML053C; | 4 | | | |
| chrXIII | 170327 | 170483 | 156 | -1,469530991 | N13:170398 | 9,6 | +1:YML052W; | 1 | | | |
| chrXIII | 170574 | 170680 | 106 | -0,479397498 | N13:170557 | 5,167 | +2:YML052W; | 2 | | | |
| chrXIII | 170574 | 170696 | 122 | -0,638800202 | N13:170557 | 5,167 | +2:YML052W; | 2 | | | |
| chrXIII | 170574 | 170696 | 122 | -0,638800202 | N13:170707 | 9,192 | +3:YML052W; | 3 | | | |
| chrXIII | 171072 | 171220 | 148 | -1,347544467 | N13:171141 | 18,841 | +6:YML052W; | 6 | | | |
| chrXIII | 175762 | 175915 | 153 | -1,289261468 | N13:175792 | 10,04 | | Intergene | | | |
| chrXIII | 179143 | 179321 | 178 | -1,485660979 | | | | Overlap <50 bp | | | |
| chrXIII | 180747 | 180905 | 158 | -1,015989932 | N13:180820 | 9,048 | +3:YML047C; | 3 | | | |
| chrXIII | 181711 | 181851 | 140 | -1,07003881 | N13:181781 | 7,789 | +3:YML046W; | 3 | | | |
| chrXIII | 181848 | 181977 | 129 | -1,124786751 | | | | Overlap <50 bp | | | |
| chrXIII | 184182 | 184330 | 148 | -1,340931298 | N13:184276 | 24,173 | | Intergene | | | |
| chrXIII | 184701 | 184824 | 123 | -0,987566196 | | | | Overlap <50 bp | | | |
| chrXIII | 184801 | 184956 | 155 | -1,538963216 | N13:184876 | 30,238 | | Intergene | | | |
| chrXIII | 185234 | 185368 | 134 | -0,785998027 | N13:185305 | 24,007 | | Intergene | | | |
| chrXIII | 185234 | 185374 | 140 | -1,084495687 | N13:185305 | 24,007 | | Intergene | | | |
| chrXIII | 186277 | 186412 | 135 | -0,91523725 | N13:186311 | 22,942 | | Intergene | | | |
| chrXIII | 186724 | 186874 | 150 | -1,061757367 | N13:186799 | 16,643 | | Intergene | | | |
| chrXIII | 186903 | 187060 | 157 | -1,527474579 | N13:186999 | 35,119 | | Intergene | | | |
| chrXIII | 187135 | 187284 | 149 | -1,237363695 | N13:187179 | 14,849 | | Intergene | | | |
| chrXIII | 187394 | 187523 | 129 | -1,146332821 | | | | Overlap <50 bp | | | |
| chrXIII | 188306 | 188456 | 150 | -1,373263247 | N13:188399 | 8,185 | | Intergene | | | |
| chrXIII | 195171 | 195314 | 143 | -1,206340075 | N13:195288 | 11,554 | | Intergene | | | |
| chrXIII | 196534 | 196661 | 127 | -1,349408121 | N13:196631 | 7,767 | | Intergene | | | |
| chrXIII | 196694 | 196843 | 149 | -1,436177973 | N13:196842 | 59,152 | | Intergene | | | |
| chrXIII | 196868 | 196991 | 123 | -1,167746496 | | | | Overlap <50 bp | | | |
| chrXIII | 196968 | 197123 | 155 | -1,608269175 | N13:197081 | 27,221 | | Intergene | | | |
| chrXIII | 197401 | 197535 | 134 | -0,866397868 | N13:197469 | 22,03 | | Intergene | | | |
| chrXIII | 197401 | 197541 | 140 | -0,829586298 | N13:197469 | 22,03 | | Intergene | | | |
| chrXIII | 197597 | 197748 | 151 | -1,16191336 | N13:197630 | 25,146 | | Intergene | | | |
| chrXIII | 197607 | 197752 | 145 | -1,250707401 | N13:197630 | 25,146 | | Intergene | | | |
| chrXIII | 197625 | 197776 | 151 | -1,447382794 | N13:197630 | 25,146 | | Intergene | | | |
| chrXIII | 197879 | 198033 | 154 | -1,234426289 | N13:198018 | 3,606 | | Intergene | | | |
| chrXIII | 197913 | 198057 | 144 | -1,273608373 | N13:198018 | 3,606 | | Intergene | | | |
| chrXIII | 198891 | 199037 | 146 | -1,065200705 | N13:198960 | 14,048 | | Intergene | | | |
| chrXIII | 198891 | 199041 | 150 | -1,068014789 | N13:198960 | 14,048 | | Intergene | | | |
| chrXIII | 199070 | 199227 | 157 | -1,155116532 | N13:199146 | 32,005 | | Intergene | | | |
| chrXIII | 199302 | 199451 | 149 | -1,093849413 | N13:199440 | 39,038 | | Intergene | | | |
| chrXIII | 199561 | 199690 | 129 | -1,116261561 | N13:199711 | 11,93 | | Intergene | | | |
| chrXIII | 200473 | 200623 | 150 | -1,537549481 | N13:200564 | 22,605 | | Intergene | | | |
| chrXIII | 203778 | 203929 | 151 | -1,786825043 | N13:203861 | 11,197 | +3:YML038C; | 3 | | | |
| chrXIII | 204102 | 204256 | 154 | -1,267077789 | N13:204195 | 15,664 | ML038C; +9*:YML037C; -1:CUT288; | 1 | | | |
| chrXIII | 208554 | 208676 | 122 | -0,385407748 | N13:208664 | 11,295 | +3:YML035C; | 3 | | | |
| chrXIII | 211138 | 211313 | 175 | -1,063784224 | N13:211187 | 46,082 | +11:YML034W; | 11 | | | |
| chrXIII | 218290 | 218401 | 111 | -1,347884584 | N13:218320 | 22,821 | +7:YML029W; | 7 | | | |
| chrXIII | 220837 | 220983 | 146 | -1,036238715 | N13:220846 | 28,247 | +5*:YML028W; | 5 | TERM | | |
| chrXIII | 228866 | 229000 | 134 | -1,100913902 | N13:228934 | 16,112 | +1:YML022W; -1:YML023C; | 1 | | | |
| chrXIII | 230938 | 231091 | 153 | -1,401564521 | N13:231007 | 13,186 | +1:YML020W; -1:YML021C; | 1 | | | |
| chrXIII | 231257 | 231408 | 151 | -1,406040535 | N13:231330 | 5,909 | +3:YML020W; | 3 | | | |

TABLE 3

| CHR | START | END | L(bp) | ΔLK | Nuc ID | Fuzziness | Gene ID | Gene body postion or intergene | Terminal | rDNA | Telomeric |
|---|---|---|---|---|---|---|---|---|---|---|---|
| chrXIII | 233569 | 233719 | 150 | -1,224469638 | N13:233649 | 10,354 | +2:YML019W; | 2 | | | |
| chrXIII | 240347 | 240506 | 159 | -1,775706464 | N13:240446 | 17,877 | | Intergene | | | |
| chrXIII | 242945 | 243091 | 146 | -1,223632645 | N13:243011 | 11,771 | +1:YML015C; -1:YML014W; | 1 | | | |
| chrXIII | 243502 | 243662 | 160 | -1,511479903 | N13:243575 | 1,924 | +3:YML014W; | 3 | | | |
| chrXIII | 243825 | 243980 | 155 | -1,272443629 | N13:243892 | 4,856 | 0:YML013W; +5*:YML014W; | 0 | | | |
| chrXIII | 244965 | 245130 | 165 | -1,419863435 | N13:244990 | 8,066 | +7:YML013W; | 7 | | | |
| chrXIII | 244965 | 245130 | 165 | -1,419863435 | N13:245141 | 2,608 | +8:YML013W; | 8 | | | |
| chrXIII | 245559 | 245710 | 151 | -1,060040689 | N13:245618 | 9,543 | +11:YML013W; | 11 | | | |
| chrXIII | 246032 | 246179 | 147 | -1,581607109 | N13:246115 | 24,238 | +14*:YML013W; +1:YML012W; | 14 | TERM | | |
| chrXIII | 246042 | 246200 | 158 | -1,263232657 | N13:246115 | 24,238 | +14*:YML013W; +1:YML012W; | 14 | TERM | | |
| chrXIII | 246354 | 246509 | 155 | -0,81597938 | N13:246437 | 11,356 | +3:YML012W; | 3 | | | |
| chrXIII | 246617 | 246737 | 120 | -2,014758173 | N13:246744 | 9,745 | +5:YML012W; | 5 | | | |
| chrXIII | 250435 | 250548 | 113 | -0,414256154 | N13:250485 | 35,796 | +19:YML010W; | 19 | | | |
| chrXIII | 250507 | 250631 | 124 | -2,184232096 | N13:250485 | 35,796 | +19:YML010W; | 19 | | | |
| chrXIII | 250507 | 250662 | 155 | -1,922632252 | N13:250485 | 35,796 | +19:YML010W; | 19 | | | |
| chrXIII | 250507 | 250664 | 157 | -2,378686658 | N13:250485 | 35,796 | +19:YML010W; | 19 | | | |
| chrXIII | 250507 | 250665 | 158 | -2,029478431 | N13:250485 | 35,796 | +19:YML010W; | 19 | | | |
| chrXIII | 250507 | 250666 | 159 | -2,20065512 | N13:250485 | 35,796 | +19:YML010W; | 19 | | | |
| chrXIII | 250507 | 250667 | 160 | -1,652154651 | N13:250485 | 35,796 | +19:YML010W; | 19 | | | |
| chrXIII | 250507 | 250668 | 161 | -1,946263119 | N13:250485 | 35,796 | +19:YML010W; | 19 | | | |
| chrXIII | 250507 | 250669 | 162 | -1,739441696 | N13:250485 | 35,796 | +19:YML010W; | 19 | | | |
| chrXIII | 250507 | 250670 | 163 | -2,208768783 | N13:250485 | 35,796 | +19:YML010W; | 19 | | | |
| chrXIII | 250507 | 250671 | 164 | -1,972065587 | N13:250485 | 35,796 | +19:YML010W; | 19 | | | |
| chrXIII | 250507 | 250673 | 166 | -1,910430155 | N13:250485 | 35,796 | +19:YML010W; | 19 | | | |
| chrXIII | 250507 | 250674 | 167 | -1,482753129 | N13:250485 | 35,796 | +19:YML010W; | 19 | | | |
| chrXIII | 250507 | 250675 | 168 | -1,774771304 | N13:250485 | 35,796 | +19:YML010W; | 19 | | | |
| chrXIII | 250507 | 250676 | 169 | -1,932912245 | N13:250485 | 35,796 | +19:YML010W; | 19 | | | |
| chrXIII | 250507 | 250677 | 170 | -2,148264149 | N13:250485 | 35,796 | +19:YML010W; | 19 | | | |
| chrXIII | 250507 | 250678 | 171 | -1,335858588 | N13:250485 | 35,796 | +19:YML010W; | 19 | | | |
| chrXIII | 250507 | 250718 | 211 | -2,721214476 | N13:250485 | 35,796 | +19:YML010W; | 19 | | | |
| chrXIII | 250507 | 250718 | 211 | -2,721214476 | N13:250740 | 40,82 | +21:YML010W; | 21 | | | |
| chrXIII | 250508 | 250678 | 170 | -1,036986671 | N13:250485 | 35,796 | +19:YML010W; | 19 | | | |
| chrXIII | 250509 | 250678 | 169 | -1,76837862 | | | | Overlap <50 bp | | | |
| chrXIII | 250510 | 250678 | 168 | -1,547810362 | | | | Overlap <50 bp | | | |
| chrXIII | 250511 | 250678 | 167 | -1,737106909 | | | | Overlap <50 bp | | | |
| chrXIII | 250514 | 250678 | 164 | -1,583573622 | | | | Overlap <50 bp | | | |
| chrXIII | 250515 | 250678 | 163 | -1,62230085 | | | | Overlap <50 bp | | | |
| chrXIII | 250566 | 250718 | 152 | -1,148424337 | N13:250740 | 40,82 | +21:YML010W; | 21 | | | |
| chrXIII | 253372 | 253532 | 160 | -0,895652325 | N13:253539 | 55,194 | -1:YML007C-A; -1:YML007W; | -1 | | | |
| chrXIII | 256009 | 256161 | 152 | -1,459542201 | N13:256158 | 23,422 | +15:YML006C; | 15 | | | |
| chrXIII | 260961 | 261102 | 141 | -1,071497549 | N13:261093 | 16,056 | +6*:YML005W; | 6 | TERM | | |
| chrXIII | 261845 | 261977 | 132 | -1,466420039 | N13:261963 | 32,192 | +5:YML004C; | 5 | | | |
| chrXIII | 262094 | 262251 | 157 | -1,153168844 | N13:262151 | 9,888 | +4:YML004C; | 4 | | | |
| chrXIII | 266226 | 266377 | 151 | -1,429313389 | N13:266295 | 10,932 | L003W; +12:YML002W; +8:SUT724; | 18 | | | |
| chrXIII | 271419 | 271566 | 147 | -1,108145805 | N13:271513 | 12,629 | )1C-A; +1:YMR001C; +2:YMR001C-A; | 1 | | | |
| chrXIII | 273047 | 273197 | 150 | -1,428081353 | N13:273120 | 6,285 | +1:YMR003W; | 1 | | | |
| chrXIII | 276628 | 276785 | 157 | -1,451674262 | N13:276642 | 8,155 | +5:YMR005W; | 5 | | | |
| chrXIII | 276628 | 276785 | 157 | -1,451674262 | N13:276801 | 18,019 | +6:YMR005W; | 6 | | | |
| chrXIII | 277879 | 278014 | 135 | -0,94715038 | N13:277912 | 47,171 | +11:YMR006C; | 11 | | | |
| chrXIII | 279195 | 279369 | 174 | -1,223115394 | N13:279270 | 29,816 | +3:YMR006C; | 3 | | | |
| chrXIII | 283987 | 284078 | 91 | -1,015905132 | N13:284101 | 37,356 | +1:YMR009W; | 1 | | | |
| chrXIII | 283987 | 284079 | 92 | -1,028006025 | N13:284101 | 37,356 | +1:YMR009W; | 1 | | | |
| chrXIII | 289854 | 289961 | 107 | -0,895972097 | N13:289883 | 27,96 | -1:SUT727; 0:SUT297; | -1 | | | |
| chrXIII | 289854 | 289977 | 123 | -1,291630713 | N13:289883 | 27,96 | -1:SUT727; 0:SUT297; | -1 | | | |
| chrXIII | 303791 | 303918 | 127 | -1,156799892 | N13:303847 | 4,231 | +13:YMR016C; | 13 | | | |
| chrXIII | 304476 | 304622 | 146 | -1,341575043 | N13:304602 | 8,385 | +8:YMR016C; | 8 | | | |
| chrXIII | 310201 | 310351 | 150 | -1,388518271 | N13:310222 | 13,471 | +1:YMR018W; | 1 | | | |
| chrXIII | 313026 | 313180 | 154 | -1,237752648 | | | | Overlap <50 bp | | | |
| chrXIII | 324504 | 324669 | 165 | -1,456600636 | N13:324559 | 20,268 | +7:YMR026C; | 7 | | | |
| chrXIII | 325807 | 325958 | 151 | -1,253213543 | N13:325840 | 12,145 | -1:YMR026C; +1:YMR027W; | -1 | | | |
| chrXIII | 333017 | 333169 | 152 | -1,069852276 | N13:333098 | 27,704 | +12:YMR031C; | 12 | | | |
| chrXIII | 335127 | 335272 | 145 | -1,282986666 | N13:335202 | 14,516 | +1:YMR032W; | 1 | | | |
| chrXIII | 335472 | 335613 | 141 | -1,346087722 | N13:335545 | 8,287 | +3:YMR032W; | 3 | | | |
| chrXIII | 336952 | 337113 | 161 | -1,190040157 | N13:337009 | 16,583 | -1:YMR030W-A; +12:YMR032W; | -1 | | | |
| chrXIII | 340438 | 340585 | 147 | -1,510910328 | N13:340517 | 13,823 | +3:YMR034C; | 3 | | | |
| chrXIII | 342150 | 342302 | 152 | -1,188058539 | N13:342208 | 18,435 | +9:YMR036C; | 9 | | | |
| chrXIII | 348582 | 348731 | 149 | -1,277457065 | N13:348736 | 26,216 | +6*:YMR039C; +1*:CUT294; | 6 | TERM | | |
| chrXIII | 351853 | 352022 | 169 | -1,18721648 | N13:351943 | 10,685 | +1:YMR041C; | 1 | | | |
| chrXIII | 351863 | 352007 | 144 | -1,423567584 | N13:351943 | 10,685 | +1:YMR041C; | 1 | | | |
| chrXIII | 357010 | 357124 | 114 | -1,438113404 | | | | Overlap <50 bp | | | |
| chrXIII | 357475 | 357580 | 105 | -1,637157197 | N13:357470 | 40,501 | | Intergene | | | |
| chrXIII | 358631 | 358781 | 150 | -1,302298543 | N13:358713 | 20,232 | | Intergene | | | |
| chrXIII | 358907 | 359055 | 148 | -1,346182852 | N13:358983 | 27,635 | | Intergene | | | |
| chrXIII | 360027 | 360184 | 157 | -1,535050948 | N13:360111 | 27,622 | | Intergene | | | |
| chrXIII | 360404 | 360549 | 145 | -1,064731222 | N13:360557 | 0 | | Intergene | | | |
| chrXIII | 360675 | 360810 | 135 | -0,848823071 | N13:360775 | 51,856 | | Intergene | | | |
| chrXIII | 362078 | 362238 | 160 | -1,269737615 | N13:362147 | 10,714 | | Intergene | | | |
| chrXIII | 362592 | 362706 | 114 | -1,416220804 | N13:362628 | 0,707 | | Intergene | | | |
| chrXIII | 374780 | 374911 | 131 | -1,121933137 | N13:374845 | 40 | | Intergene | | | |
| chrXIII | 375262 | 375391 | 129 | -1,31965933 | | | | Overlap <50 bp | | | |
| chrXIII | 375501 | 375650 | 149 | -1,076227568 | N13:375616 | 40,129 | | Intergene | | | |
| chrXIII | 375725 | 375882 | 157 | -1,63927456 | N13:375785 | 37,749 | | Intergene | | | |
| chrXIII | 375911 | 376061 | 150 | -0,952371414 | N13:376003 | 16,643 | | Intergene | | | |
| chrXIII | 376373 | 376508 | 135 | -0,951968185 | N13:376424 | 32,532 | | Intergene | | | |
| chrXIII | 376763 | 376917 | 154 | -1,51158548 | N13:376925 | 25,534 | | Intergene | | | |
| chrXIII | 377411 | 377551 | 140 | -1,270495708 | N13:377486 | 52,468 | | Intergene | | | |
| chrXIII | 377417 | 377551 | 134 | -0,981920607 | N13:377486 | 52,468 | | Intergene | | | |
| chrXIII | 377829 | 377984 | 155 | -1,194261494 | N13:377857 | 20,232 | | Intergene | | | |
| chrXIII | 377960 | 378068 | 108 | -0,70606957 | | | | Overlap <50 bp | | | |
| chrXIII | 379716 | 379822 | 106 | -0,864531541 | N13:379764 | 20,096 | +2:YMR052W; | 2 | | | |
| chrXIII | 379716 | 379838 | 122 | -1,356189495 | N13:379764 | 20,096 | +2:YMR052W; | 2 | | | |
| chrXIII | 381135 | 381283 | 148 | -1,124631794 | N13:381244 | 27,959 | +11:YMR053C; | 11 | | | |
| chrXIII | 383859 | 384012 | 153 | -1,363344256 | N13:383963 | 25,28 | +6:YMR054W; | 6 | | | |
| chrXIII | 384143 | 384283 | 140 | -1,256686057 | N13:384129 | 22,14 | +7:YMR054W; | 7 | | | |
| chrXIII | 384143 | 384283 | 140 | -1,256686057 | N13:384300 | 9,933 | +8:YMR054W; | 8 | | | |
| chrXIII | 384959 | 385107 | 148 | -1,236476284 | | | | Overlap <50 bp | | | |
| chrXIII | 386345 | 386477 | 132 | -0,965623439 | N13:386478 | 11,76 | +4:YMR055C; | 4 | | | |
| chrXIII | 386345 | 386477 | 132 | -0,965623439 | N13:386323 | 7,937 | +5:YMR055C; | 5 | | | |

| CHR | START | END | L(bp) | ΔLK | Nuc ID | Fuzziness | Gene ID | Gene body postion or intergene | Terminal | rDNA | Telomeric |
|---|---|---|---|---|---|---|---|---|---|---|---|
| chrXIII | 387854 | 388024 | 170 | -1,313352513 | N13:387911 | 15,308 | +2:YMR056C; | 2 | | | |
| chrXIII | 395268 | 395412 | 144 | -1,281700963 | N13:395361 | 16,979 | +7:YMR062C; | 7 | | | |
| chrXIII | 400158 | 400292 | 134 | -1,632416398 | N13:400231 | 13,293 | +3:YMR065W; | 3 | | | |
| chrXIII | 402458 | 402607 | 149 | -1,1222067 | N13:402522 | 14,17 | +7:YMR066W; | 7 | | | |
| chrXIII | 406280 | 406447 | 167 | -1,368368444 | N13:406339 | 11,709 | +1:YMR068W; | 1 | | | |
| chrXIII | 408429 | 408536 | 107 | -0,04025224 | N13:408500 | 7,071 | +6:YMR069W; | 6 | | | |
| chrXIII | 411105 | 411252 | 147 | -1,066499899 | N13:411102 | 3,061 | +2:YMR071C; | 2 | | | |
| chrXIII | 412475 | 412581 | 106 | -1,357510065 | N13:412535 | 12,153 | +5:YMR073C; | 5 | | | |
| chrXIII | 412475 | 412589 | 114 | -1,634297997 | N13:412535 | 12,153 | +5:YMR073C; | 5 | | | |
| chrXIII | 413837 | 414003 | 166 | -1,032786013 | N13:413996 | 41,884 | +1:YMR075W; | 1 | | | |
| chrXIII | 422824 | 422985 | 161 | -1,145543381 | N13:422941 | 8,764 | +12:YMR078C; | 12 | | | |
| chrXIII | 431294 | 431455 | 161 | -1,39738954 | N13:431429 | 30,875 | -1:YMR081C; | -1 | | | |
| chrXIII | 431358 | 431515 | 157 | -1,607674928 | N13:431429 | 30,875 | -1:YMR081C; | -1 | | | |
| chrXIII | 435010 | 435159 | 149 | -1,112774013 | N13:435081 | 17,972 | +3:YMR083W; | 3 | | | |
| chrXIII | 436980 | 437142 | 162 | -1,161931876 | N13:437057 | 3,5 | +4:YMR084W; | 4 | | | |
| chrXIII | 439283 | 439457 | 174 | -1,728658448 | N13:439388 | 11,149 | +2:YMR086W; | 2 | | | |
| chrXIII | 443760 | 443869 | 109 | -0,94193934 | N13:443884 | 7,024 | +9:YMR088C; | 9 | | | |
| chrXIII | 445137 | 445296 | 159 | -1,638215942 | N13:445171 | 29,416 | +1:YMR088C; | 1 | | | |
| chrXIII | 448884 | 449018 | 134 | -1,12528915 | N13:448865 | 8,222 | -1:YMR090W; | -1 | | | |
| chrXIII | 448987 | 449140 | 153 | -0,973944237 | N13:449042 | 17,251 | 0:YMR090W; | 0 | | | |
| chrXIII | 452811 | 452953 | 142 | -1,224030692 | N13:452872 | 8,276 | +5:YMR092C; | 5 | | | |
| chrXIII | 455777 | 455925 | 148 | -1,375167806 | N13:455844 | 17,963 | +1:YMR094W; | 1 | | | |
| chrXIII | 456607 | 456760 | 153 | -1,49553899 | N13:456671 | 16,196 | +6:YMR094W; | 6 | | | |
| chrXIII | 458537 | 458683 | 146 | -1,032910028 | | | | Overlap <50 bp | | | |
| chrXIII | 460130 | 460281 | 151 | -1,482220402 | N13:460189 | 8,583 | +6:YMR097C; | 6 | | | |
| chrXIII | 460228 | 460362 | 134 | -0,801576707 | N13:460349 | 11,389 | +5:YMR097C; | 5 | | | |
| chrXIII | 460442 | 460567 | 125 | -0,919323994 | N13:460520 | 10,813 | +4:YMR097C; | 4 | | | |
| chrXIII | 460442 | 460595 | 153 | -1,062350709 | N13:460520 | 10,813 | +4:YMR097C; | 4 | | | |
| chrXIII | 460442 | 460598 | 156 | -1,698620617 | N13:460520 | 10,813 | +4:YMR097C; | 4 | | | |
| chrXIII | 460442 | 460600 | 158 | -1,061570918 | N13:460520 | 10,813 | +4:YMR097C; | 4 | | | |
| chrXIII | 461194 | 461345 | 151 | -1,262757852 | N13:461301 | 12,021 | +9:YMR098C; | 9 | | | |
| chrXIII | 464378 | 464520 | 142 | -1,24412839 | N13:464473 | 7,501 | +3:YMR099C; | 3 | | | |
| chrXIII | 467950 | 468100 | 150 | -1,51287914 | | | | Overlap <50 bp | | | |
| chrXIII | 470902 | 471051 | 149 | -1,539207987 | N13:470916 | 10,017 | +10:YMR102C; | 10 | | | |
| chrXIII | 473040 | 473146 | 106 | -1,055787622 | N13:473098 | 31,984 | +13:YMR104C; | 13 | | | |
| chrXIII | 473040 | 473162 | 122 | -1,240734 | N13:473098 | 31,984 | +13:YMR104C; | 13 | | | |
| chrXIII | 473911 | 474081 | 170 | -1,061923504 | N13:474025 | 7,925 | +8:YMR104C; | 8 | | | |
| chrXIII | 484473 | 484603 | 130 | -2,197076681 | N13:484551 | 12,657 | +5:YMR108W; | 5 | | | |
| chrXIII | 488956 | 489063 | 107 | -1,422093102 | N13:489016 | 37,901 | +15:YMR109W; | 15 | | | |
| chrXIII | 488956 | 489073 | 117 | -1,619386548 | N13:489016 | 37,901 | +15:YMR109W; | 15 | | | |
| chrXIII | 491261 | 491409 | 148 | -1,327601077 | N13:491313 | 8,886 | +5:YMR110C; | 5 | | | |
| chrXIII | 492970 | 493101 | 131 | -1,110228497 | N13:492992 | 35,329 | +8:YMR111C; | 8 | | | |
| chrXIII | 495547 | 495700 | 153 | -1,138855301 | N13:495639 | 20,75 | +5:YMR113W; | 5 | | | |
| chrXIII | 496801 | 496973 | 172 | -1,311892877 | N13:496786 | 17,487 | +5*:YMR114C; | 5 | TERM | | |
| chrXIII | 496801 | 496973 | 172 | -1,311892877 | N13:496955 | 23,034 | +4:YMR114C; | 4 | | | |
| chrXIII | 499783 | 499912 | 129 | -1,022804316 | N13:499808 | 15,947 | | Intergene | | | |
| chrXIII | 503855 | 504003 | 148 | -1,099190138 | N13:503924 | 27,185 | | Intergene | | | |
| chrXIII | 505412 | 505581 | 169 | -1,427235611 | N13:505521 | 21,658 | +3:YMR119W; | 3 | | | |
| chrXIII | 512141 | 512286 | 145 | -1,41928132 | N13:512212 | 22,879 | +8:CUT775; | 8 | | | |
| chrXIII | 514521 | 514669 | 148 | -1,161641121 | N13:514629 | 8,424 | +3:YMR124W; | 3 | | | |
| chrXIII | 514553 | 514703 | 150 | -1,58806141 | N13:514629 | 8,424 | +3:YMR124W; | 3 | | | |
| chrXIII | 515323 | 515459 | 136 | -1,36689729 | N13:515420 | 23,756 | +8:YMR124W; | 8 | | | |
| chrXIII | 520223 | 520378 | 155 | -1,17460798 | N13:520207 | 21,349 | | Intergene | | | |
| chrXIII | 520223 | 520378 | 155 | -1,17460798 | N13:520395 | 28,077 | | Intergene | | | |
| chrXIII | 520376 | 520508 | 132 | -0,654714499 | N13:520395 | 28,077 | | Intergene | | | |
| chrXIII | 521730 | 521901 | 171 | -1,617508894 | N13:521801 | 10,685 | +2:YMR126C; | 2 | | | |
| chrXIII | 522712 | 522817 | 105 | -1,338817696 | N13:522725 | 7,789 | +5:YMR127C; +4*:anti141; | 5 | | | |
| chrXIII | 528716 | 528857 | 141 | -1,11894495 | N13:528767 | 2,828 | +7:YMR129W; | 7 | | | |
| chrXIII | 533451 | 533610 | 159 | -1,064703565 | N13:533576 | 31,143 | +8:YMR131C; | 8 | | | |
| chrXIII | 535081 | 535246 | 165 | -1,519159054 | N13:535146 | 4,803 | +2:CUT301; +4:YMR132C; | 2 | | | |
| chrXIII | 536826 | 536976 | 150 | -1,260422671 | N13:536905 | 10,445 | +5:YMR133W; +2:SUT732; | 5 | | | |
| chrXIII | 537368 | 537507 | 139 | -1,218431053 | N13:537514 | 25,573 | +8:YMR133W; -1:YMR134W; | 8 | | | |
| chrXIII | 539637 | 539760 | 123 | -1,244669314 | N13:539719 | 8,937 | +3:YMR135C; | 3 | | | |
| chrXIII | 540611 | 540759 | 148 | -1,347502544 | N13:540749 | 18,556 | | Intergene | | | |
| chrXIII | 541727 | 541880 | 153 | -1,375830119 | N13:541835 | 29,208 | +4:YMR136W; | 4 | | | |
| chrXIII | 549153 | 549302 | 149 | -1,500657959 | N13:549269 | 21,03 | +11:YMR140W; | 11 | | | |
| chrXIII | 549525 | 549679 | 154 | -1,407554174 | N13:549554 | 28,294 | +13*:YMR140W; | 13 | TERM | | |
| chrXIII | 549713 | 549857 | 144 | -1,30361384 | N13:549747 | 27,727 | | Intergene | | | |
| chrXIII | 555791 | 555929 | 138 | -1,199100746 | N13:555792 | 26,291 | +6:YMR145C; | 6 | | | |
| chrXIII | 562267 | 562429 | 162 | -1,217407946 | N13:562366 | 29,778 | +2:YMR150C; | 2 | | | |
| chrXIII | 562901 | 563062 | 161 | -1,627779898 | N13:562920 | 22,133 | 0:YMR152W; | 0 | | | |
| chrXIII | 571585 | 571738 | 153 | -1,411203072 | N13:571681 | 19,23 | -1:YMR156C; +3:YMR157C; | -1 | | | |
| chrXIII | 571590 | 571738 | 148 | -1,27501725 | N13:571681 | 19,23 | -1:YMR156C; +3:YMR157C; | -1 | | | |
| chrXIII | 574736 | 574904 | 168 | -0,917721721 | N13:574834 | 9,867 | -1:YMR160W; +1:YMR159C; | -1 | | | |
| chrXIII | 580311 | 580467 | 156 | -1,109786367 | N13:580401 | 13,467 | +23:YMR162C; | 23 | | | |
| chrXIII | 580644 | 580772 | 128 | -1,15437328 | N13:580704 | 28,693 | +21:YMR162C; | 21 | | | |
| chrXIII | 581583 | 581719 | 136 | -1,320981759 | N13:581638 | 19,344 | +16:YMR162C; | 16 | | | |
| chrXIII | 582011 | 582174 | 163 | -1,160146188 | N13:582099 | 17,761 | +13:YMR162C; | 13 | | | |
| chrXIII | 582013 | 582174 | 161 | -1,263855346 | N13:582099 | 17,761 | +13:YMR162C; | 13 | | | |
| chrXIII | 583110 | 583268 | 158 | -1,168676087 | | | | Overlap <50 bp | | | |
| chrXIII | 588471 | 588586 | 115 | -1,351804913 | | | | Overlap <50 bp | | | |
| chrXIII | 595003 | 595166 | 163 | -1,31144452 | N13:595084 | 4,183 | +2:YMR167W; | 2 | | | |
| chrXIII | 596303 | 596450 | 147 | -1,087121469 | N13:596386 | 17,111 | +10:YMR167W; | 10 | | | |
| chrXIII | 598087 | 598214 | 127 | -1,686991525 | N13:598137 | 11,929 | +7:YMR168C; | 7 | | | |
| chrXIII | 598887 | 599027 | 140 | -1,226031277 | N13:598952 | 3,435 | +2:YMR168C; | 2 | | | |
| chrXIII | 600432 | 600562 | 130 | -1,018931095 | N13:600483 | 40,294 | +3:YMR169C; | 3 | | | |
| chrXIII | 600662 | 600821 | 159 | -1,376411267 | N13:600688 | 6,658 | +2:YMR169C; | 2 | | | |
| chrXIII | 602872 | 603031 | 159 | -1,369766695 | N13:602904 | 37,477 | +2:YMR170C; | 2 | | | |
| chrXIII | 608199 | 608341 | 142 | -1,229130495 | N13:608209 | 24,222 | +14*:YMR172W; | 14 | TERM | | |
| chrXIII | 614091 | 614242 | 151 | -1,277209225 | N13:614214 | 20,03 | +17:YMR176W; | 17 | | | |
| chrXIII | 617621 | 617773 | 152 | -1,109055473 | N13:617702 | 23,192 | +8:YMR177W; | 8 | | | |
| chrXIII | 617962 | 618111 | 149 | -1,518148784 | N13:618058 | 15,485 | +10:YMR177W; | 10 | | | |
| chrXIII | 622293 | 622443 | 150 | -1,221823506 | N13:622398 | 23,499 | +8:YMR180C; | 8 | | | |
| chrXIII | 627780 | 627908 | 128 | -1,045984368 | N13:627847 | 7,733 | +1:YMR183C; | 1 | | | |
| chrXIII | 627788 | 627919 | 131 | -1,03335309 | N13:627847 | 7,733 | +1:YMR183C; | 1 | | | |
| chrXIII | 634568 | 634713 | 145 | -1,376256504 | N13:634666 | 37,098 | +9*:YMR187C; | 9 | TERM | | |
| chrXIII | 637620 | 637765 | 145 | -1,462292351 | N13:637702 | 7,287 | +3:YMR189W; | 3 | | | |
| chrXIII | 641481 | 641611 | 130 | -0,245231112 | N13:641533 | 33,088 | +24:YMR190C; | 24 | | | |
| chrXIII | 642381 | 642525 | 144 | -1,089336123 | N13:642458 | 32,517 | +18:YMR190C; | 18 | | | |

TABLE 3

| CHR | START | END | L(bp) | ΔLK | Nuc ID | Fuzziness | Gene ID | Gene body postion or intergene | Terminal | rDNA | Telomeric |
|---|---|---|---|---|---|---|---|---|---|---|---|
| chrXIII | 642389 | 642541 | 152 | -1,243230254 | N13:642458 | 32,517 | +18:YMR190C; | 18 | | | |
| chrXIII | 649168 | 649320 | 152 | -1,725134006 | N13:649263 | 12,554 | | Intergene | | | |
| chrXIII | 652798 | 652958 | 160 | -1,483894571 | N13:652939 | 37,727 | +1:YMR194C-B; | 1 | | | |
| chrXIII | 657660 | 657789 | 129 | -1,161476411 | N13:657675 | 16,832 | +5:CUT784; | 5 | | | |
| chrXIII | 658250 | 658404 | 154 | -1,429230454 | N13:658346 | 11,692 | +1:CUT784; | 1 | | | |
| chrXIII | 658827 | 658990 | 163 | -1,16555984 | N13:658863 | 7,967 | +3:YMR197C; | 3 | | | |
| chrXIII | 658970 | 659105 | 135 | -1,108034287 | N13:659025 | 9,606 | +2:YMR197C; | 2 | | | |
| chrXIII | 660778 | 660927 | 149 | -1,305258508 | N13:660944 | 33,887 | +10:YMR198W; | 10 | | | |
| chrXIII | 662900 | 663052 | 152 | -1,852514575 | N13:662956 | 20,659 | +4:YMR199W; | 4 | | | |
| chrXIII | 663803 | 663949 | 146 | -1,231901438 | N13:663870 | 9,762 | +9:YMR199W; | 9 | | | |
| chrXIII | 663837 | 663944 | 107 | -1,310336811 | N13:663870 | 9,762 | +9:YMR199W; | 9 | | | |
| chrXIII | 663837 | 663953 | 116 | -1,463453801 | N13:663870 | 9,762 | +9:YMR199W; | 9 | | | |
| chrXIII | 667634 | 667789 | 155 | -1,135562628 | N13:667703 | 8,124 | +2:YMR202W; | 2 | | | |
| chrXIII | 671983 | 672128 | 145 | -1,311176996 | N13:672090 | 21,407 | +17:YMR205C; | 17 | | | |
| chrXIII | 675888 | 676037 | 149 | -1,173041663 | N13:675911 | 24,481 | +1:YMR206W; | 1 | | | |
| chrXIII | 675888 | 676037 | 149 | -1,173041663 | N13:676060 | 8,145 | +2:YMR206W; | 2 | | | |
| chrXIII | 675950 | 676037 | 87 | -0,706323604 | N13:676060 | 8,145 | +2:YMR206W; | 2 | | | |
| chrXIII | 675950 | 676105 | 155 | -1,34728913 | N13:676060 | 8,145 | +2:YMR206W; | 2 | | | |
| chrXIII | 677204 | 677344 | 140 | -1,466891716 | N13:677258 | 15,741 | +40:YMR207C; | 40 | | | |
| chrXIII | 677920 | 678066 | 146 | -1,422725477 | N13:677969 | 32,254 | +36:YMR207C; | 36 | | | |
| chrXIII | 678554 | 678723 | 169 | -1,46359263 | N13:678626 | 13,527 | +32:YMR207C; | 32 | | | |
| chrXIII | 679422 | 679570 | 148 | -1,535292214 | | | | Overlap <50 bp | | | |
| chrXIII | 680831 | 680955 | 124 | -0,944302573 | N13:680907 | 26 | +18:YMR207C; | 18 | | | |
| chrXIII | 684362 | 684526 | 164 | -1,563087733 | N13:684432 | 13,442 | 0:YMR208W; | 0 | | | |
| chrXIII | 686491 | 686656 | 165 | -1,24842551 | N13:686622 | 3,715 | +5:YMR209C; | 5 | | | |
| chrXIII | 692898 | 693052 | 154 | -1,560026231 | N13:692964 | 12,972 | +2:YMR212C; | 2 | | | |
| chrXIII | 693627 | 693771 | 144 | -1,15466149 | N13:693719 | 4,561 | +3:YMR213W; | 3 | | | |
| chrXIII | 693831 | 693958 | 127 | -1,130676747 | N13:693886 | 3,271 | +4:YMR213W; | 4 | | | |
| chrXIII | 694160 | 694314 | 154 | -1,176653897 | N13:694234 | 5,07 | +6:YMR213W; | 6 | | | |
| chrXIII | 694285 | 694446 | 161 | -1,388551281 | | | | Overlap <50 bp | | | |
| chrXIII | 694710 | 694842 | 132 | -1,049615381 | N13:694758 | 28,877 | +9:YMR213W; | 9 | | | |
| chrXIII | 711631 | 711787 | 156 | -1,116457697 | N13:711674 | 23,923 | | Intergene | | | |
| chrXIII | 713556 | 713705 | 149 | -1,197033137 | N13:713622 | 32,896 | +9:YMR220W; | 9 | | | |
| chrXIII | 714522 | 714675 | 153 | -1,743643898 | N13:714616 | 14,48 | +6:YMR221C; | 6 | | | |
| chrXIII | 715186 | 715332 | 146 | -1,102109835 | N13:715254 | 5,727 | +2:YMR221C; | 2 | | | |
| chrXIII | 718847 | 718995 | 148 | -1,394105952 | N13:719006 | 38,532 | +11:YMR224C; | 11 | | | |
| chrXIII | 718847 | 718995 | 148 | -1,394105952 | N13:718849 | 32,887 | +12:YMR224C; | 12 | | | |
| chrXIII | 719890 | 720048 | 158 | -1,209884701 | N13:719976 | 9,453 | +5:YMR224C; | 5 | | | |
| chrXIII | 723055 | 723205 | 150 | -1,246705755 | | | | Overlap <50 bp | | | |
| chrXIII | 723972 | 724126 | 154 | -1,425963828 | N13:724061 | 7,23 | +3:YMR227C; | 3 | | | |
| chrXIII | 730943 | 731091 | 148 | -1,243015513 | N13:731008 | 3,92 | +2:YMR229C; | 2 | | | |
| chrXIII | 737903 | 738069 | 166 | -1,67434349 | N13:737964 | 20,992 | +8:YMR232W; | 8 | | | |
| chrXIII | 748521 | 748674 | 153 | -1,325026035 | N13:748606 | 19,524 | +8:YMR239C; | 8 | | | |
| chrXIII | 750522 | 750675 | 153 | -1,116784236 | N13:750581 | 5,177 | +6:YMR240C; | 6 | | | |
| chrXIII | 757851 | 758017 | 166 | -1,180891504 | N13:757933 | 6,76 | +6:YMR244W; | 6 | | | |
| chrXIII | 758839 | 758989 | 150 | -1,238349559 | N13:759009 | 35,487 | +1:YMR244C-A; | 1 | | | |
| chrXIII | 758839 | 758989 | 150 | -1,238349559 | N13:758860 | 52,169 | +2:YMR244C-A; | 2 | | | |
| chrXIII | 759376 | 759557 | 181 | -1,593404033 | N13:759498 | 23,295 | 0:YMR246W; | 0 | | | |
| chrXIII | 765908 | 766066 | 158 | -1,130909835 | N13:766027 | 28,492 | +14:YMR247C; | 14 | | | |
| chrXIII | 772941 | 773097 | 156 | -1,22924665 | N13:773019 | 9,783 | +2:YMR251W; +4:SUT738; | 2 | | | |
| chrXIII | 773665 | 773803 | 138 | -1,254895417 | N13:773793 | 7,842 | +7:YMR251W; -1:SUT738; | 7 | | | |
| chrXIII | 773665 | 773807 | 142 | -1,235607642 | N13:773793 | 7,842 | +7:YMR251W; -1:SUT738; | 7 | | | |
| chrXIII | 774153 | 774312 | 159 | -0,896784979 | N13:774158 | 12,361 | +9*:YMR251W; | 9 | TERM | | |
| chrXIII | 774153 | 774312 | 159 | -0,896784979 | N13:774317 | 12,276 | | Intergene | | | |
| chrXIII | 778663 | 778795 | 132 | -1,435094604 | N13:778773 | 23,569 | +6*:YMR255W; +3*:YMR256C; | 6 | TERM | | |
| chrXIII | 779035 | 779188 | 153 | -1,720841578 | N13:779109 | 3,033 | +1:YMR256C; | 1 | | | |
| chrXIII | 780072 | 780218 | 146 | -1,800675782 | N13:780137 | 27,495 | +14:YMR257C; | 14 | | | |
| chrXIII | 780801 | 780937 | 136 | -1,369242476 | N13:780898 | 64,519 | +10:YMR257C; | 10 | | | |
| chrXIII | 790225 | 790394 | 169 | -1,146081977 | N13:790301 | 17,725 | +20:YMR261C; | 20 | | | |
| chrXIII | 790368 | 790515 | 147 | -1,348560293 | N13:790486 | 14,46 | +19:YMR261C; | 19 | | | |
| chrXIII | 790640 | 790805 | 165 | -1,167242662 | N13:790824 | 29,813 | +17:YMR261C; | 17 | | | |
| chrXIII | 790640 | 790805 | 165 | -1,167242662 | N13:790646 | 7,228 | +18:YMR261C; | 18 | | | |
| chrXIII | 791775 | 791919 | 144 | -1,28525158 | N13:791864 | 4,573 | +11:YMR261C; | 11 | | | |
| chrXIII | 792265 | 792408 | 143 | -1,351420283 | N13:792303 | 22,621 | +8:YMR261C; | 8 | | | |
| chrXIII | 792607 | 792777 | 170 | -1,501527541 | N13:792794 | 10,661 | +5:YMR261C; | 5 | | | |
| chrXIII | 792607 | 792777 | 170 | -1,501527541 | N13:792641 | 13,077 | +6:YMR261C; | 6 | | | |
| chrXIII | 792935 | 793098 | 163 | -1,296089687 | N13:793117 | 6,481 | +3:YMR261C; | 3 | | | |
| chrXIII | 792935 | 793098 | 163 | -1,296089687 | N13:792956 | 12,872 | +4:YMR261C; | 4 | | | |
| chrXIII | 793020 | 793160 | 140 | -0,985563371 | N13:793117 | 6,481 | +3:YMR261C; | 3 | | | |
| chrXIII | 793713 | 793841 | 128 | -1,0337298 | N13:793740 | 8,847 | -1:YMR261C; +1:YMR262W; | -1 | | | |
| chrXIII | 795911 | 796064 | 153 | -1,246971525 | N13:795966 | 4,336 | +2:YMR264W; | 2 | | | |
| chrXIII | 797737 | 797897 | 160 | -1,315848977 | N13:797874 | 16,582 | +1:YMR265C; | 1 | | | |
| chrXIII | 797737 | 797940 | 203 | -1,400772218 | N13:797874 | 16,582 | +1:YMR265C; | 1 | | | |
| chrXIII | 797778 | 797885 | 107 | -0,405682677 | N13:797874 | 16,582 | +1:YMR265C; | 1 | | | |
| chrXIII | 797778 | 797897 | 119 | -0,778151205 | N13:797874 | 16,582 | +1:YMR265C; | 1 | | | |
| chrXIII | 797778 | 797940 | 162 | -1,144257575 | N13:797874 | 16,582 | +1:YMR265C; | 1 | | | |
| chrXIII | 799674 | 799836 | 162 | -0,927437545 | | | | Overlap <50 bp | | | |
| chrXIII | 800670 | 800819 | 149 | -1,560584081 | | | | Overlap <50 bp | | | |
| chrXIII | 801962 | 802127 | 165 | -1,528671835 | N13:802061 | 7,791 | +3:YMR267W; | 3 | | | |
| chrXIII | 804104 | 804264 | 160 | -1,411894864 | N13:804203 | 7,055 | +1:YMR268C; -1:YMR269W; | 1 | | | |
| chrXIII | 806854 | 806962 | 108 | -1,113558627 | N13:806905 | 9,555 | +6*:YMR271C; | 6 | TERM | | |
| chrXIII | 806855 | 806962 | 107 | -1,225583207 | N13:806905 | 9,555 | +6*:YMR271C; | 6 | TERM | | |
| chrXIII | 808568 | 808720 | 152 | -1,437675613 | | | | Overlap <50 bp | | | |
| chrXIII | 811849 | 811978 | 129 | -1,044132696 | N13:811868 | 21,483 | +15:YMR273C; | 15 | | | |
| chrXIII | 816433 | 816597 | 164 | -1,43255335 | N13:816517 | 21,716 | +14:YMR275C; | 14 | | | |
| chrXIII | 819869 | 819975 | 106 | -1,073689103 | N13:819933 | 11,431 | +8*:YMR276W; -1:YMR277W; | 8 | TERM | | |
| chrXIII | 819869 | 819988 | 119 | -1,305795421 | N13:819933 | 11,431 | +8*:YMR276W; -1:YMR277W; | 8 | TERM | | |
| chrXIII | 820608 | 820710 | 102 | -1,070326901 | N13:820698 | 7,314 | +4:YMR277W; | 4 | | | |
| chrXIII | 820608 | 820711 | 103 | -0,815276084 | N13:820698 | 7,314 | +4:YMR277W; | 4 | | | |
| chrXIII | 820608 | 820715 | 107 | -1,549099066 | N13:820698 | 7,314 | +4:YMR277W; | 4 | | | |
| chrXIII | 820608 | 820717 | 109 | -0,819384402 | N13:820698 | 7,314 | +4:YMR277W; | 4 | | | |
| chrXIII | 820608 | 820733 | 125 | -1,386911372 | N13:820698 | 7,314 | +4:YMR277W; | 4 | | | |
| chrXIII | 820608 | 820737 | 129 | -1,270264164 | N13:820698 | 7,314 | +4:YMR277W; | 4 | | | |
| chrXIII | 820608 | 820738 | 130 | -1,519093893 | N13:820698 | 7,314 | +4:YMR277W; | 4 | | | |
| chrXIII | 820608 | 820740 | 132 | -0,914990682 | N13:820698 | 7,314 | +4:YMR277W; | 4 | | | |
| chrXIII | 820608 | 820741 | 133 | -1,551555228 | N13:820698 | 7,314 | +4:YMR277W; | 4 | | | |
| chrXIII | 820608 | 820742 | 134 | -1,373831624 | N13:820698 | 7,314 | +4:YMR277W; | 4 | | | |
| chrXIII | 820608 | 820743 | 135 | -1,275866383 | N13:820698 | 7,314 | +4:YMR277W; | 4 | | | |
| chrXIII | 820608 | 820744 | 136 | -1,784869815 | N13:820698 | 7,314 | +4:YMR277W; | 4 | | | |
| chrXIII | 820608 | 820745 | 137 | -1,668597069 | N13:820698 | 7,314 | +4:YMR277W; | 4 | | | |

TABLE 3

| CHR | START | END | L(bp) | ΔLK | Nuc ID | Fuzziness | Gene ID | Gene body postion or intergene | Terminal | rDNA | Telomeric |
|---|---|---|---|---|---|---|---|---|---|---|---|
| chrXIII | 820608 | 820746 | 138 | -1,323740327 | N13:820698 | 7,314 | +4:YMR277W; | 4 | | | |
| chrXIII | 820608 | 820747 | 139 | -1,275480863 | N13:820698 | 7,314 | +4:YMR277W; | 4 | | | |
| chrXIII | 820608 | 820748 | 140 | -1,851729879 | N13:820698 | 7,314 | +4:YMR277W; | 4 | | | |
| chrXIII | 820608 | 820749 | 141 | -1,199696077 | N13:820698 | 7,314 | +4:YMR277W; | 4 | | | |
| chrXIII | 820608 | 820750 | 142 | -1,951217718 | N13:820698 | 7,314 | +4:YMR277W; | 4 | | | |
| chrXIII | 820608 | 820751 | 143 | -1,165222752 | N13:820698 | 7,314 | +4:YMR277W; | 4 | | | |
| chrXIII | 820609 | 820751 | 142 | -1,290726288 | N13:820698 | 7,314 | +4:YMR277W; | 4 | | | |
| chrXIII | 820610 | 820751 | 141 | -1,206973874 | N13:820698 | 7,314 | +4:YMR277W; | 4 | | | |
| chrXIII | 820611 | 820751 | 140 | -1,398790791 | N13:820698 | 7,314 | +4:YMR277W; | 4 | | | |
| chrXIII | 820612 | 820751 | 139 | -1,195604628 | N13:820698 | 7,314 | +4:YMR277W; | 4 | | | |
| chrXIII | 820613 | 820751 | 138 | -0,940569628 | N13:820698 | 7,314 | +4:YMR277W; | 4 | | | |
| chrXIII | 820614 | 820751 | 137 | -1,546491319 | N13:820698 | 7,314 | +4:YMR277W; | 4 | | | |
| chrXIII | 820615 | 820751 | 136 | -1,540404258 | N13:820698 | 7,314 | +4:YMR277W; | 4 | | | |
| chrXIII | 820616 | 820751 | 135 | -1,495701512 | N13:820698 | 7,314 | +4:YMR277W; | 4 | | | |
| chrXIII | 820617 | 820751 | 134 | -1,240760607 | N13:820698 | 7,314 | +4:YMR277W; | 4 | | | |
| chrXIII | 820619 | 820751 | 132 | -1,30605236 | N13:820698 | 7,314 | +4:YMR277W; | 4 | | | |
| chrXIII | 820620 | 820751 | 131 | -1,04108562 | N13:820698 | 7,314 | +4:YMR277W; | 4 | | | |
| chrXIII | 820621 | 820751 | 130 | -0,997344479 | N13:820698 | 7,314 | +4:YMR277W; | 4 | | | |
| chrXIII | 820624 | 820751 | 127 | -1,320185462 | N13:820698 | 7,314 | +4:YMR277W; | 4 | | | |
| chrXIII | 820626 | 820715 | 89 | -0,331672905 | N13:820698 | 7,314 | +4:YMR277W; | 4 | | | |
| chrXIII | 828572 | 828720 | 148 | -1,3167041 | N13:828670 | 12,4 | +17:YMR280C; | 17 | | | |
| chrXIII | 835990 | 836099 | 109 | -0,24432488 | | | | Overlap <50 bp | | | |
| chrXIII | 836333 | 836463 | 130 | -1,082004231 | N13:836461 | 20,877 | +3:YMR283C; | 3 | | | |
| chrXIII | 842098 | 842232 | 134 | -1,10215979 | N13:842139 | 11,743 | +2:YMR286W; | 2 | | | |
| chrXIII | 843431 | 843583 | 152 | -1,198985537 | N13:843529 | 44,584 | | Intergene | | | |
| chrXIII | 847888 | 848017 | 129 | -1,133053485 | N13:847969 | 12,983 | +15:YMR288W; | 15 | | | |
| chrXIII | 847897 | 848050 | 153 | -0,976293861 | N13:847969 | 12,983 | +15:YMR288W; | 15 | | | |
| chrXIII | 851484 | 851633 | 149 | -1,686947621 | N13:851582 | 14,261 | +1:YMR290C; | 1 | | | |
| chrXIII | 864286 | 864433 | 147 | -1,607539689 | N13:864337 | 12,285 | +7*:YMR299C; | 7 | TERM | | |
| chrXIII | 864644 | 864799 | 155 | -1,092414135 | N13:864820 | 7,548 | +4:YMR299C; | 4 | | | |
| chrXIII | 864644 | 864799 | 155 | -1,092414135 | N13:864653 | 29,305 | +5:YMR299C; | 5 | | | |
| chrXIII | 866005 | 866155 | 150 | -0,832483104 | | | | Overlap <50 bp | | | |
| chrXIII | 868732 | 868890 | 158 | -1,225352096 | N13:868810 | 10,644 | +6:YMR301C; | 6 | | | |
| chrXIII | 871472 | 871623 | 151 | -1,237526372 | N13:871631 | 10,741 | +7:YMR302C; | 7 | | | |
| chrXIII | 871472 | 871623 | 151 | -1,237526372 | N13:871471 | 13,502 | +8:YMR302C; | 8 | | | |
| chrXIII | 873640 | 873761 | 121 | -0,42410459 | N13:873682 | 27,883 | | Intergene | | | |
| chrXIII | 873640 | 873789 | 149 | -1,205193959 | N13:873682 | 27,883 | | Intergene | | | |
| chrXIII | 873640 | 873790 | 150 | -1,077202153 | N13:873682 | 27,883 | | Intergene | | | |
| chrXIII | 873648 | 873761 | 113 | -0,453480734 | N13:873682 | 27,883 | | Intergene | | | |
| chrXIII | 873648 | 873789 | 141 | -0,965957715 | N13:873682 | 27,883 | | Intergene | | | |
| chrXIII | 873648 | 873790 | 142 | -1,157645955 | N13:873682 | 27,883 | | Intergene | | | |
| chrXIII | 878188 | 878341 | 153 | -1,24823709 | N13:878290 | 25,795 | | Intergene | | | |
| chrXIII | 879147 | 879302 | 155 | -1,340664549 | N13:879224 | 28,107 | +8:YMR305C; | 8 | | | |
| chrXIII | 884345 | 884496 | 151 | -1,46221069 | N13:884479 | 35,302 | +7:CUT792; | 7 | | | |
| chrXIII | 886126 | 886284 | 158 | -1,084449508 | | | | Overlap <50 bp | | | |
| chrXIII | 889335 | 889469 | 134 | -1,055223546 | N13:889367 | 19,917 | +19:YMR308C; | 19 | | | |
| chrXIII | 890068 | 890170 | 102 | -1,604195994 | N13:890188 | 29,579 | +14:YMR308C; | 14 | | | |
| chrXIII | 891024 | 891149 | 125 | -0,963025918 | N13:891052 | 13,096 | +9:YMR308C; | 9 | | | |
| chrXIII | 891955 | 892105 | 150 | -1,188509462 | N13:892008 | 3,782 | +3:YMR308C; | 3 | | | |
| chrXIII | 895348 | 895499 | 151 | -1,124761017 | N13:895429 | 10,616 | +1:YMR309C; | 1 | | | |
| chrXIII | 902827 | 902965 | 138 | -0,966455526 | N13:902909 | 40,773 | +2:YMR315W; | 2 | | | |
| chrXIII | 904201 | 904335 | 134 | -1,326281986 | N13:904278 | 13,693 | +1:YMR315W-A; | 1 | | | |
| chrXIII | 904428 | 904582 | 154 | -1,236642995 | N13:904475 | 14,106 | +2*:YMR315W-A; -1:YMR316W; | 2 | TERM | | |
| chrXIII | 910695 | 910845 | 150 | -1,14410254 | N13:910757 | 14,404 | +22:YMR317W; -1:SUT321; | 22 | | | |
| chrXIII | 911000 | 911154 | 154 | -1,059289025 | N13:911157 | 31,729 | +7*:YMR318C; +2:SUT321; | 7 | TERM | | |
| chrXIII | 917204 | 917355 | 151 | -1,477306761 | N13:917298 | 10,572 | | Intergene | | | |
| chrXIII | 921093 | 921253 | 160 | -1,129205784 | N13:921174 | 15,275 | +8:YMR323W; | 8 | | | |
| chrXIII | 923830 | 923976 | 146 | -0,837624809 | N13:923958 | 13,026 | | Intergene | | | TEL RIGHT |
| chrXIII | 924304 | 924426 | 122 | 0,242412087 | N13:924361 | 19,218 | | Intergene | | | TEL RIGHT |
| chrXIII | 924311 | 924426 | 115 | 0,082325735 | N13:924361 | 19,218 | | Intergene | | | TEL RIGHT |
| chrXIII | 924321 | 924426 | 105 | 0,256129044 | N13:924361 | 19,218 | | Intergene | | | TEL RIGHT |
| chrXIV | 614 | 765 | 151 | -1,26114644 | N14:689 | 23,756 | +34:YNL339C; | 34 | | | TEL LEFT |
| chrXIV | 953 | 1105 | 152 | -1,167005056 | N14:1005 | 17,039 | +32:YNL339C; | 32 | | | TEL LEFT |
| chrXIV | 1779 | 1930 | 151 | -1,23433831 | N14:1793 | 10,97 | +28:YNL339C; | 28 | | | TEL LEFT |
| chrXIV | 1791 | 1925 | 134 | -0,90324288 | N14:1793 | 10,97 | +28:YNL339C; | 28 | | | TEL LEFT |
| chrXIV | 1816 | 1912 | 96 | -0,909764122 | N14:1793 | 10,97 | +28:YNL339C; | 28 | | | TEL LEFT |
| chrXIV | 2170 | 2300 | 130 | -1,158833659 | N14:2318 | 20,075 | +24:YNL339C; | 24 | | | TEL LEFT |
| chrXIV | 2449 | 2601 | 152 | -0,933741302 | N14:2530 | 42,442 | +23:YNL339C; | 23 | | | TEL LEFT |
| chrXIV | 2796 | 2940 | 144 | -0,944837604 | N14:2849 | 32,347 | +21:YNL339C; | 21 | | | TEL LEFT |
| chrXIV | 2796 | 2983 | 187 | -0,653490955 | N14:2849 | 32,347 | +21:YNL339C; | 21 | | | TEL LEFT |
| chrXIV | 2848 | 2940 | 92 | -0,639460529 | N14:2849 | 32,347 | +21:YNL339C; | 21 | | | TEL LEFT |
| chrXIV | 2848 | 2983 | 135 | -0,96919127 | N14:2849 | 32,347 | +21:YNL339C; | 21 | | | TEL LEFT |
| chrXIV | 2936 | 3080 | 144 | -0,840272192 | N14:3031 | 21,127 | +20:YNL339C; | 20 | | | TEL LEFT |
| chrXIV | 3001 | 3153 | 152 | -1,288864118 | N14:3031 | 21,127 | +20:YNL339C; | 20 | | | TEL LEFT |
| chrXIV | 3253 | 3378 | 125 | -0,838300013 | N14:3371 | 17,616 | +18:YNL339C; | 18 | | | TEL LEFT |
| chrXIV | 3283 | 3426 | 143 | -1,60352687 | N14:3371 | 17,616 | +18:YNL339C; | 18 | | | TEL LEFT |
| chrXIV | 3584 | 3721 | 137 | -1,121172472 | N14:3651 | 8,718 | +16:YNL339C; | 16 | | | TEL LEFT |
| chrXIV | 3700 | 3855 | 155 | -1,689110747 | N14:3808 | 10,017 | +15:YNL339C; | 15 | | | TEL LEFT |
| chrXIV | 3794 | 3956 | 162 | -1,206881273 | N14:3967 | 7,211 | +14:YNL339C; | 14 | | | TEL LEFT |
| chrXIV | 3794 | 3956 | 162 | -1,206881273 | N14:3808 | 10,017 | +15:YNL339C; | 15 | | | TEL LEFT |
| chrXIV | 5019 | 5160 | 141 | -1,10344153 | N14:5067 | 19,672 | +7:YNL339C; | 7 | | | TEL LEFT |
| chrXIV | 15620 | 15770 | 150 | -1,616250702 | N14:15736 | 19,425 | +7:YNL332W; | 7 | | | |
| chrXIV | 18901 | 19054 | 153 | -1,137514125 | N14:18957 | 7,223 | +3:YNL330C; | 3 | | | |
| chrXIV | 22418 | 22569 | 151 | -1,128460354 | N14:22459 | 29,615 | +6:YNL328C; +2:YNL329C; | 6 | | | |
| chrXIV | 24850 | 25019 | 169 | -1,434748487 | N14:24952 | 26,405 | +6:YNL327W; | 6 | | | |
| chrXIV | 25836 | 25986 | 150 | -0,79842117 | N14:25940 | 33,589 | +12:YNL327W; | 12 | | | |
| chrXIV | 26530 | 26680 | 150 | -1,384838721 | N14:26673 | 28,253 | +16:YNL327W; | 16 | | | |
| chrXIV | 27268 | 27422 | 154 | -1,477611654 | | | | Overlap <50 bp | | | |
| chrXIV | 30249 | 30400 | 151 | -1,205813298 | N14:30360 | 10,854 | +7:YNL325C; | 7 | | | |
| chrXIV | 32059 | 32210 | 151 | -1,387798737 | N14:32102 | 7,649 | +2:YNL323W; | 2 | | | |
| chrXIV | 33454 | 33599 | 145 | -1,226884335 | N14:33540 | 30,632 | +5:YNL322C; | 5 | | | |
| chrXIV | 33454 | 33606 | 152 | -1,313828091 | N14:33540 | 30,632 | +5:YNL322C; | 5 | | | |
| chrXIV | 33459 | 33599 | 140 | -1,270636694 | N14:33540 | 30,632 | +5:YNL322C; | 5 | | | |
| chrXIV | 33461 | 33599 | 138 | -1,324245062 | N14:33540 | 30,632 | +5:YNL322C; | 5 | | | |
| chrXIV | 33461 | 33603 | 142 | -1,311005072 | N14:33540 | 30,632 | +5:YNL322C; | 5 | | | |
| chrXIV | 33461 | 33606 | 145 | -1,090493198 | N14:33540 | 30,632 | +5:YNL322C; | 5 | | | |
| chrXIV | 33464 | 33606 | 142 | -1,560819366 | N14:33540 | 30,632 | +5:YNL322C; | 5 | | | |
| chrXIV | 33466 | 33606 | 140 | -1,265952191 | N14:33540 | 30,632 | +5:YNL322C; | 5 | | | |
| chrXIV | 41403 | 41548 | 145 | -1,274830872 | N14:41462 | 15,123 | +6:YNL317W; | 6 | | | |

TABLE 3

| CHR | START | END | L(bp) | ΔLK | Nuc ID | Fuzziness | Gene ID | Gene body postion or intergene | Terminal | rDNA | Telomeric |
|---|---|---|---|---|---|---|---|---|---|---|---|
| chrXIV | 45261 | 45368 | 107 | -1,443529939 | N14:45343 | 13,171 | +17*:YNL313C; | 17 | TERM | | |
| chrXIV | 45261 | 45374 | 113 | -1,749246433 | N14:45343 | 13,171 | +17*:YNL313C; | 17 | TERM | | |
| chrXIV | 45783 | 45937 | 154 | -1,162964664 | N14:45851 | 24,025 | +14:YNL313C; | 14 | | | |
| chrXIV | 49049 | 49199 | 150 | -1,224991118 | N14:49135 | 9,96 | +6:YNL312W; | 6 | | | |
| chrXIV | 53101 | 53263 | 162 | -1,637356206 | N14:53189 | 11,462 | +4:YNL309W; | 4 | | | |
| chrXIV | 55301 | 55450 | 149 | -1,088164631 | N14:55396 | 16,82 | +4:YNL308C; | 4 | | | |
| chrXIV | 58407 | 58536 | 129 | -1,105045967 | N14:58525 | 21,102 | +3:YNL306W; | 3 | | | |
| chrXIV | 60113 | 60210 | 97 | -1,528967024 | | | | Overlap <50 bp | | | |
| chrXIV | 61184 | 61331 | 147 | -1,119756379 | N14:61254 | 14,37 | +7:YNL304W; | 7 | | | |
| chrXIV | 63570 | 63733 | 163 | -1,336708308 | N14:63662 | 16,41 | | Intergene | | | |
| chrXIV | 65407 | 65535 | 128 | -1,213811438 | N14:65439 | 5,981 | -1:Unit516; -1:YNL300W; | -1 | | | |
| chrXIV | 73936 | 74081 | 145 | -1,62742696 | N14:73929 | 21,37 | +17:YNL297C; | 17 | | | |
| chrXIV | 77741 | 77895 | 154 | -1,555767322 | N14:77801 | 39,107 | +7:YNL295W; | 7 | | | |
| chrXIV | 79985 | 80135 | 150 | -1,359485803 | N14:80056 | 4,367 | +3:YNL294C; | 3 | | | |
| chrXIV | 81420 | 81583 | 163 | -1,537239866 | N14:81459 | 11,295 | +6:YNL293W; | 6 | | | |
| chrXIV | 91093 | 91242 | 149 | -1,047291568 | N14:91178 | 3,435 | +7:YNL288W; | 7 | | | |
| chrXIV | 93267 | 93403 | 136 | -1,169266664 | N14:93374 | 29,579 | +11:YNL287W; | 11 | | | |
| chrXIV | 97284 | 97435 | 151 | -1,491140008 | N14:97317 | 31,82 | | Intergene | | | |
| chrXIV | 97941 | 98103 | 162 | -1,101504122 | N14:98029 | 22,332 | | Intergene | | | |
| chrXIV | 99408 | 99557 | 149 | -1,055560615 | N14:99525 | 52,354 | | Intergene | | | |
| chrXIV | 99632 | 99789 | 157 | -1,254768256 | N14:99713 | 23,643 | | Intergene | | | |
| chrXIV | 99818 | 99968 | 150 | -1,064511842 | N14:99892 | 16,166 | | Intergene | | | |
| chrXIV | 99822 | 99968 | 146 | -1,244579571 | N14:99892 | 16,166 | | Intergene | | | |
| chrXIV | 100280 | 100415 | 135 | -1,201088959 | N14:100384 | 42,618 | | Intergene | | | |
| chrXIV | 100932 | 101089 | 157 | -1,147249753 | N14:101020 | 32,14 | | Intergene | | | |
| chrXIV | 102362 | 102510 | 148 | -1,416490541 | N14:102397 | 6,658 | | Intergene | | | |
| chrXIV | 103973 | 104137 | 164 | -1,355647937 | N14:104026 | 5,167 | +1:YNL284C; -1:CUT315; | 1 | | | |
| chrXIV | 107707 | 107834 | 127 | -1,019781022 | N14:107700 | 12,287 | +1:YNL282W; | 1 | | | |
| chrXIV | 110079 | 110230 | 151 | -1,429310439 | N14:110172 | 5,465 | +2:YNL280C; | 2 | | | |
| chrXIV | 111620 | 111775 | 155 | -1,464656367 | | | | Overlap <50 bp | | | |
| chrXIV | 112009 | 112129 | 120 | -1,327286399 | N14:112032 | 11,846 | +7:YNL279W; +6:SUT743; | 7 | | | |
| chrXIV | 116452 | 116578 | 126 | -1,429161036 | N14:116519 | 22,941 | IL278W; +1:CUT802; 0:YNL277W-A; | 22 | TERM | | |
| chrXIV | 119252 | 119378 | 126 | -1,017419608 | N14:119284 | 9,973 | +13*:YNL277W; +1:YNL275W; | 13 | TERM | | |
| chrXIV | 121333 | 121464 | 131 | -0,837619266 | N14:121346 | 34,219 | +6:YNL274C; | 6 | | | |
| chrXIV | 124849 | 124975 | 126 | -0,993992213 | N14:124938 | 13,279 | +14:YNL273W; | 14 | | | |
| chrXIV | 129691 | 129843 | 152 | -1,609822261 | N14:129772 | 22,828 | | Intergene | | | |
| chrXIV | 129719 | 129870 | 151 | -1,129973206 | N14:129772 | 22,828 | | Intergene | | | |
| chrXIV | 131962 | 132135 | 173 | -1,631223871 | N14:131992 | 36,936 | +22:YNL271C; | 22 | | | |
| chrXIV | 132312 | 132463 | 151 | -1,533553729 | N14:132390 | 22,05 | +20:YNL271C; | 20 | | | |
| chrXIV | 133490 | 133654 | 164 | -1,465443021 | N14:133593 | 30,406 | +13:YNL271C; | 13 | | | |
| chrXIV | 134176 | 134325 | 149 | -1,211295875 | N14:134291 | 25,981 | +9:YNL271C; | 9 | | | |
| chrXIV | 135289 | 135440 | 151 | -1,240764846 | N14:135423 | 10,304 | +2:YNL271C; | 2 | | | |
| chrXIV | 135829 | 135979 | 150 | -1,193523366 | N14:135901 | 10,551 | -1:YNL271C; +11:YNL270C; | -1 | | | |
| chrXIV | 139766 | 139921 | 155 | -0,647366343 | N14:139942 | 37,76 | +10:YNL268W; | 10 | | | |
| chrXIV | 140110 | 140269 | 159 | -1,279702947 | N14:140187 | 32,047 | +11:YNL268W; | 11 | | | |
| chrXIV | 141911 | 142060 | 149 | -0,98815193 | N14:142074 | 10,905 | +9:YNL267W; | 9 | | | |
| chrXIV | 150518 | 150667 | 149 | -1,08438067 | N14:150605 | 14,011 | +14:YNL262W; | 14 | | | |
| chrXIV | 153107 | 153271 | 164 | -1,547672375 | N14:153137 | 34,072 | +29:YNL262W; | 29 | | | |
| chrXIV | 153107 | 153271 | 164 | -1,547672375 | N14:153294 | 12,565 | +30:YNL262W; | 30 | | | |
| chrXIV | 155351 | 155502 | 151 | -1,205335001 | N14:155389 | 10,616 | +3:YNL261W; | 3 | | | |
| chrXIV | 163862 | 164000 | 138 | -1,219547147 | N14:164012 | 26,711 | +3:YNL257C; | 3 | | | |
| chrXIV | 163862 | 164000 | 138 | -1,219547147 | N14:163855 | 2,582 | +4:YNL257C; | 4 | | | |
| chrXIV | 165154 | 165280 | 126 | -1,446327677 | N14:165235 | 7,956 | +5:YNL256W; | 5 | | | |
| chrXIV | 167446 | 167585 | 139 | -1,339871234 | N14:167470 | 13,15 | +3:YNL255C; | 3 | | | |
| chrXIV | 174863 | 175007 | 144 | -1,354411165 | N14:174931 | 7,414 | +1:CUT320; | 1 | | | |
| chrXIV | 177654 | 177807 | 153 | -1,428142246 | N14:177732 | 36,891 | +15:YNL250W; | 15 | | | |
| chrXIV | 178329 | 178461 | 132 | -1,410790425 | N14:178391 | 25,861 | +19:YNL250W; | 19 | | | |
| chrXIV | 183769 | 183934 | 165 | -1,468010197 | N14:183818 | 8,963 | +7:YNL247W; | 7 | | | |
| chrXIV | 187776 | 187960 | 184 | -1,69099612 | N14:187755 | 32,449 | -1:YNL243W; -1:YNL244C; | -1 | | | |
| chrXIV | 188930 | 189078 | 148 | -1,209118659 | | | | Overlap <50 bp | | | |
| chrXIV | 188930 | 189079 | 149 | -1,616432157 | | | | Overlap <50 bp | | | |
| chrXIV | 188930 | 189080 | 150 | -0,980180119 | | | | Overlap <50 bp | | | |
| chrXIV | 188930 | 189081 | 151 | -1,414159259 | | | | Overlap <50 bp | | | |
| chrXIV | 188930 | 189083 | 153 | -1,213376633 | | | | Overlap <50 bp | | | |
| chrXIV | 188931 | 189083 | 152 | -1,738335508 | | | | Overlap <50 bp | | | |
| chrXIV | 188933 | 189083 | 150 | -1,450303976 | | | | Overlap <50 bp | | | |
| chrXIV | 188934 | 189083 | 149 | -1,400315724 | | | | Overlap <50 bp | | | |
| chrXIV | 188935 | 189083 | 148 | -1,571354733 | | | | Overlap <50 bp | | | |
| chrXIV | 190891 | 191040 | 149 | -1,351077167 | N14:190930 | 12,014 | +18*:YNL243W; 0:YNL242W; | 18 | TERM | | |
| chrXIV | 192308 | 192455 | 147 | -1,310095422 | N14:192383 | 27,389 | +9:YNL242W; | 9 | | | |
| chrXIV | 192813 | 192954 | 141 | -1,099248622 | N14:192825 | 36,813 | +11:YNL242W; | 11 | | | |
| chrXIV | 193504 | 193657 | 153 | -1,949134012 | N14:193627 | 36,56 | +16:YNL242W; | 16 | | | |
| chrXIV | 196760 | 196910 | 150 | -1,236028689 | N14:196922 | 9,678 | +7:YNL241C; +36:YNL242W; | 7 | | | |
| chrXIV | 202136 | 202292 | 156 | -0,982830772 | N14:202215 | 4,528 | +1:YNL238W; -1:SUT744; | 1 | | | |
| chrXIV | 203542 | 203662 | 120 | -0,178787426 | N14:203649 | 41,856 | +10:YNL238W; | 10 | | | |
| chrXIV | 206857 | 207023 | 166 | -1,701452663 | N14:206949 | 4,844 | -1:anti149; +1:YNL236W; | -1 | | | |
| chrXIV | 210139 | 210288 | 149 | -1,259194618 | N14:210220 | 10,597 | +1:YNL234W; | 1 | | | |
| chrXIV | 216142 | 216308 | 166 | -1,340307249 | N14:216232 | 10,113 | +7:YNL231C; | 7 | | | |
| chrXIV | 219506 | 219633 | 127 | -0,927527921 | N14:219552 | 28,527 | +6:YNL229C; | 6 | | | |
| chrXIV | 224826 | 224979 | 153 | -1,436847002 | N14:224898 | 10,342 | +15*:YNL224C; | 15 | TERM | | |
| chrXIV | 227320 | 227476 | 156 | -1,421656323 | N14:227397 | 6,229 | -1:YNL224C; +1:YNL223W; | -1 | | | |
| chrXIV | 235082 | 235232 | 150 | -1,279105066 | | | | Overlap <50 bp | | | |
| chrXIV | 238200 | 238369 | 169 | -1,579285701 | N14:238254 | 4,546 | +1:YNL218W; | 1 | | | |
| chrXIV | 246737 | 246897 | 160 | -1,742123121 | N14:246814 | 5,888 | +3:YNL213C; | 3 | | | |
| chrXIV | 247839 | 247988 | 149 | -1,271747645 | N14:247931 | 11,584 | +4:YNL212W; | 4 | | | |
| chrXIV | 254524 | 254675 | 151 | -1,249069467 | N14:254603 | 21,237 | +2:YNL208W; | 2 | | | |
| chrXIV | 257454 | 257590 | 136 | -1,106078162 | N14:257486 | 20,957 | +5:YNL206C; | 5 | | | |
| chrXIV | 257743 | 257893 | 150 | -1,287766811 | N14:257796 | 10,281 | +3:YNL206C; | 3 | | | |
| chrXIV | 259132 | 259283 | 151 | -0,933668643 | N14:259278 | 34,57 | -1:YNL202W; +1:YNL204C; | -1 | | | |
| chrXIV | 259132 | 259283 | 151 | -0,933668643 | N14:259110 | 26,195 | +2:YNL204C; | 2 | | | |
| chrXIV | 265572 | 265700 | 128 | -0,938916111 | N14:265642 | 18,97 | +7:YNL199C; | 7 | | | |
| chrXIV | 271513 | 271667 | 154 | -1,30275635 | N14:271563 | 21,615 | +3*:YNL195C; | 3 | TERM | | |
| chrXIV | 273060 | 273209 | 149 | -1,252850273 | N14:273068 | 42,111 | +6:SUT326; +5:YNL194C; | 6 | | | |
| chrXIV | 273060 | 273209 | 149 | -1,252850273 | N14:273226 | 15,642 | +7:SUT326; +4:YNL194C; | 7 | | | |
| chrXIV | 275916 | 276056 | 140 | -0,992321868 | N14:276038 | 27,532 | +11*:YNL193W; +1:CUT806; | 11 | TERM | | |
| chrXIV | 281515 | 281622 | 107 | -1,091421709 | N14:281537 | 13,653 | +8*:YNL191W; | 8 | TERM | | |
| chrXIV | 281515 | 281631 | 116 | -1,547908334 | N14:281537 | 13,653 | +8*:YNL191W; | 8 | TERM | | |
| chrXIV | 281832 | 281976 | 144 | -0,833205804 | N14:281889 | 14,048 | -1:YNL190W; | -1 | | | |
| chrXIV | 283804 | 283989 | 185 | -1,493365917 | N14:283836 | 11,916 | -1:YNL189W; | -1 | | | |

TABLE 3

| CHR | START | END | L(bp) | ΔLK | Nuc ID | Fuzziness | Gene ID | Gene body postion or intergene | Terminal | rDNA | Telomeric |
|---|---|---|---|---|---|---|---|---|---|---|---|
| chrXIV | 287747 | 287882 | 135 | -0,832213936 | N14:287841 | 8,264 | +1:YNL187W; | 1 | | | |
| chrXIV | 291077 | 291208 | 131 | -0,968804661 | N14:291175 | 15,438 | +12:YNL186W; | 12 | | | |
| chrXIV | 292461 | 292633 | 172 | -1,466114481 | N14:292551 | 7,468 | +2:YNL185C; | 2 | | | |
| chrXIV | 292477 | 292630 | 153 | -1,238017976 | N14:292551 | 7,468 | +2:YNL185C; | 2 | | | |
| chrXIV | 296854 | 297025 | 171 | -1,388951839 | N14:296864 | 42,64 | +6:YNL182C; | 6 | | | |
| chrXIV | 297410 | 297562 | 152 | -1,020240364 | N14:297470 | 12,422 | +2:YNL182C; | 2 | | | |
| chrXIV | 298390 | 298549 | 159 | -1,375091251 | N14:298469 | 10,677 | +2:YNL181W; | 2 | | | |
| chrXIV | 302946 | 303074 | 128 | -1,218272057 | N14:302995 | 12,05 | +3:YNL178W; | 3 | | | |
| chrXIV | 303880 | 304030 | 150 | -1,501278913 | N14:303933 | 10,068 | +5:YNL177C; | 5 | | | |
| chrXIV | 307042 | 307199 | 157 | -1,413968104 | N14:307110 | 8,998 | +1:YNL176C; | 1 | | | |
| chrXIV | 308271 | 308446 | 175 | -1,496761126 | N14:308444 | 10,52 | +2:YNL175C; | 2 | | | |
| chrXIV | 308271 | 308446 | 175 | -1,496761126 | N14:308284 | 6,325 | +3:YNL175C; | 3 | | | |
| chrXIV | 309709 | 309860 | 151 | -1,365820068 | N14:309803 | 2,872 | +2:YNL173C; | 2 | | | |
| chrXIV | 310999 | 311146 | 147 | -1,276716916 | N14:311066 | 8,302 | +3:YNL172W; | 3 | | | |
| chrXIV | 314726 | 314877 | 151 | -1,228084953 | N14:314793 | 16,072 | +27:YNL172W; | 27 | | | |
| chrXIV | 317936 | 318050 | 114 | -1,965967979 | N14:318021 | 5,241 | -1:YNL169C; +6:YNL168C; | -1 | | | |
| chrXIV | 324620 | 324765 | 145 | -1,334124924 | N14:324681 | 26,049 | +6:YNL165W; | 6 | | | |
| chrXIV | 324977 | 325117 | 140 | -1,083912587 | N14:325028 | 13,091 | +8*:YNL165W; | 8 | TERM | | |
| chrXIV | 336864 | 337018 | 154 | -0,77102584 | N14:336949 | 6,626 | +5:SUT747; +4:YNL160W; | 5 | | | |
| chrXIV | 339979 | 340132 | 153 | -1,647791967 | N14:340063 | 7,369 | +4:YNL158W; -1:YNL157W; | 4 | | | |
| chrXIV | 340143 | 340297 | 154 | -1,312095718 | | | | Overlap <50 bp | | | |
| chrXIV | 343939 | 344095 | 156 | -1,517207307 | N14:344007 | 20,224 | +9:YNL154C; +14:YNL153C; | 9 | | | |
| chrXIV | 348424 | 348579 | 155 | -1,19030425 | N14:348518 | 11,514 | +1:YNL151C; -1:CUT332; | 1 | | | |
| chrXIV | 349341 | 349497 | 156 | -1,496164237 | N14:349432 | 8,622 | +3*:YNL149C; | 3 | TERM | | |
| chrXIV | 357765 | 357910 | 145 | -1,164676079 | N14:357862 | 10,7 | +5:YNL142W; | 5 | | | |
| chrXIV | 357767 | 357910 | 143 | -1,282193499 | N14:357862 | 10,7 | +5:YNL142W; | 5 | | | |
| chrXIV | 358288 | 358439 | 151 | -1,269640804 | N14:358379 | 41,494 | +8:YNL142W; | 8 | | | |
| chrXIV | 360600 | 360759 | 159 | -1,527373813 | N14:360625 | 33,342 | | Intergene | | | |
| chrXIV | 360762 | 360912 | 150 | -1,549346224 | N14:360803 | 10,21 | +33*:YNL139C; | 33 | TERM | | |
| chrXIV | 361214 | 361352 | 138 | -1,326493272 | N14:361265 | 29,938 | +30:YNL139C; | 30 | | | |
| chrXIV | 364273 | 364404 | 131 | -1,230929249 | N14:364350 | 8,386 | +10:YNL139C; | 10 | | | |
| chrXIV | 365246 | 365404 | 158 | -1,354080023 | N14:365272 | 14,66 | +4:YNL139C; | 4 | | | |
| chrXIV | 366145 | 366271 | 126 | -1,347954433 | | | | Overlap <50 bp | | | |
| chrXIV | 368092 | 368240 | 148 | -1,132118715 | N14:368205 | 22,678 | +10:YNL138W; | 10 | | | |
| chrXIV | 369049 | 369214 | 165 | -1,715226837 | N14:369138 | 22,052 | +7:YNL137C; | 7 | | | |
| chrXIV | 376927 | 377089 | 162 | -1,26822994 | N14:377019 | 12,755 | +12:YNL132W; | 12 | | | |
| chrXIV | 384387 | 384546 | 159 | -1,267276163 | N14:384456 | 9,642 | +4:YNL127W; | 4 | | | |
| chrXIV | 388430 | 388576 | 146 | -1,489183971 | N14:388511 | 8 | +9:YNL126W; | 9 | | | |
| chrXIV | 388719 | 388880 | 161 | -1,332754204 | N14:388701 | 22,067 | +10:YNL126W; | 10 | | | |
| chrXIV | 388719 | 388880 | 161 | -1,332754204 | N14:388900 | 32,085 | +11:YNL126W; | 11 | | | |
| chrXIV | 396710 | 396865 | 155 | -1,260681051 | N14:396752 | 40,296 | +14:YNL123W; | 14 | | | |
| chrXIV | 403433 | 403582 | 149 | -1,486166722 | N14:403451 | 27,415 | +14:YNL118C; | 14 | | | |
| chrXIV | 403995 | 404145 | 150 | -1,172944818 | | | | Overlap <50 bp | | | |
| chrXIV | 410286 | 410440 | 154 | -1,180062561 | N14:410457 | 49,662 | +10:YNL115C; | 10 | | | |
| chrXIV | 410286 | 410440 | 154 | -1,180062561 | N14:410304 | 41,071 | +11:YNL115C; | 11 | | | |
| chrXIV | 410317 | 410470 | 153 | -1,210023063 | N14:410457 | 49,662 | +10:YNL115C; | 10 | | | |
| chrXIV | 410317 | 410470 | 153 | -1,210023063 | N14:410304 | 41,071 | +11:YNL115C; | 11 | | | |
| chrXIV | 410968 | 411126 | 158 | -1,214561599 | N14:411083 | 29,918 | +7:YNL115C; | 7 | | | |
| chrXIV | 411008 | 411145 | 137 | -1,212747297 | N14:411083 | 29,918 | +7:YNL115C; | 7 | | | |
| chrXIV | 416136 | 416292 | 156 | -1,312234213 | N14:416212 | 15,789 | | Intergene | | | |
| chrXIV | 418446 | 418610 | 164 | -1,755455222 | N14:418509 | 14,55 | +1:YNL110C; | 1 | | | |
| chrXIV | 419242 | 419381 | 139 | -1,367723137 | N14:419334 | 12,612 | +4:YNL108C; | 4 | | | |
| chrXIV | 420269 | 420420 | 151 | -1,52496795 | N14:420290 | 10,257 | +2:YNL107W; | 2 | | | |
| chrXIV | 421794 | 421931 | 137 | -1,201970375 | N14:421859 | 28,993 | +18:YNL106C; | 18 | | | |
| chrXIV | 421911 | 422067 | 156 | -1,47079972 | N14:422017 | 26,329 | +17:YNL106C; | 17 | | | |
| chrXIV | 423473 | 423612 | 139 | -1,303502101 | N14:423543 | 28,726 | +8:YNL106C; | 8 | | | |
| chrXIV | 423497 | 423650 | 153 | -1,328487487 | N14:423543 | 28,726 | +8:YNL106C; | 8 | | | |
| chrXIV | 426375 | 426492 | 117 | -1,486910803 | N14:426395 | 26,397 | +4:YNL104C; | 4 | | | |
| chrXIV | 430126 | 430279 | 153 | -1,986508783 | N14:430203 | 18,608 | +2:YNL102W; | 2 | | | |
| chrXIV | 430126 | 430286 | 160 | -1,297415892 | N14:430203 | 18,608 | +2:YNL102W; | 2 | | | |
| chrXIV | 439717 | 439834 | 117 | -1,303695242 | N14:439718 | 28,219 | +6:YNL098C; | 6 | | | |
| chrXIV | 459511 | 459626 | 115 | -1,718924657 | N14:459588 | 4,359 | -1:CUT813; +15:YNL088W; | -1 | | | |
| chrXIV | 460164 | 460311 | 147 | -1,359910805 | N14:460238 | 28,077 | +19:YNL088W; | 19 | | | |
| chrXIV | 464412 | 464538 | 126 | -1,277312664 | N14:464472 | 47,776 | +14:YNL087W; | 14 | | | |
| chrXIV | 466515 | 466659 | 144 | -1,142111846 | N14:466599 | 7,537 | UT748; -1:YNL085W; +4:YNL086W; | 1 | | | |
| chrXIV | 467295 | 467445 | 150 | -1,431666583 | | | | Overlap <50 bp | | | |
| chrXIV | 473999 | 474106 | 107 | -1,506815325 | N14:474082 | 8,989 | +18:YNL083W; +5:YNL082W; | 18 | | | |
| chrXIV | 475462 | 475606 | 144 | -1,059117993 | N14:475517 | 36,679 | +26:YNL083W; +13:YNL082W; | 26 | | | |
| chrXIV | 477594 | 477739 | 145 | -1,098361612 | N14:477658 | 22,863 | +4:YNL080C; | 4 | | | |
| chrXIV | 478141 | 478275 | 134 | -1,297732191 | N14:478165 | 17,994 | +1:YNL080C; | 1 | | | |
| chrXIV | 483192 | 483305 | 113 | -1,576537317 | | | | Overlap <50 bp | | | |
| chrXIV | 483272 | 483422 | 150 | -1,496162295 | N14:483343 | 6,463 | +1:YNL076W; | 1 | | | |
| chrXIV | 483529 | 483666 | 137 | -1,834991947 | N14:483509 | 3,189 | +2:YNL076W; | 2 | | | |
| chrXIV | 483529 | 483666 | 137 | -1,834991947 | N14:483683 | 5,82 | +3:YNL076W; +11*:CUT815; | 3 | | | |
| chrXIV | 483752 | 483901 | 149 | -1,201039797 | N14:483855 | 18,801 | +4:YNL076W; +10:CUT815; | 4 | | | |
| chrXIV | 484694 | 484843 | 149 | -1,290990913 | | | | Overlap <50 bp | | | |
| chrXIV | 490422 | 490573 | 151 | -1,425680975 | N14:490508 | 4,708 | +2:YNL072W; | 2 | | | |
| chrXIV | 490584 | 490699 | 115 | -0,601453223 | N14:490670 | 9,039 | +3:YNL072W; | 3 | | | |
| chrXIV | 490584 | 490718 | 134 | -1,203006182 | N14:490670 | 9,039 | +3:YNL072W; | 3 | | | |
| chrXIV | 490584 | 490722 | 138 | -1,122780673 | N14:490670 | 9,039 | +3:YNL072W; | 3 | | | |
| chrXIV | 500789 | 500918 | 129 | -0,75289546 | N14:500830 | 31,204 | | Intergene | | | |
| chrXIV | 502692 | 502828 | 136 | -1,047673619 | N14:502692 | 35,576 | +7:YNL066W; | 7 | | | |
| chrXIV | 511107 | 511261 | 154 | -1,166954852 | | | | Overlap <50 bp | | | |
| chrXIV | 515776 | 515877 | 101 | -1,669420801 | N14:515831 | 8,689 | | Intergene | | | |
| chrXIV | 516941 | 517091 | 150 | -0,922225939 | N14:517027 | 28,81 | YNL058C; 0:YNL056W; +1:Unit525; | -1 | | | |
| chrXIV | 517457 | 517589 | 132 | -1,191756178 | N14:517452 | 9,67 | +2:YNL056W; | 2 | | | |
| chrXIV | 517457 | 517589 | 132 | -1,191756178 | N14:517609 | 6,812 | +3:YNL056W; | 3 | | | |
| chrXIV | 520229 | 520363 | 134 | -1,00091766 | N14:520294 | 52,37 | | Intergene | | | |
| chrXIV | 520229 | 520369 | 140 | -1,117032968 | N14:520294 | 52,37 | | Intergene | | | |
| chrXIV | 521272 | 521407 | 135 | -1,103410358 | N14:521305 | 5,508 | | Intergene | | | |
| chrXIV | 521719 | 521865 | 146 | -1,222310659 | N14:521781 | 16,371 | | Intergene | | | |
| chrXIV | 521719 | 521869 | 150 | -1,079684528 | N14:521781 | 16,371 | | Intergene | | | |
| chrXIV | 521898 | 522055 | 157 | -1,358889802 | N14:521995 | 35,171 | | Intergene | | | |
| chrXIV | 524015 | 524129 | 114 | -1,620338374 | | | | Overlap <50 bp | | | |
| chrXIV | 524719 | 524868 | 149 | -1,275098222 | N14:524853 | 25,482 | | Intergene | | | |
| chrXIV | 524979 | 525110 | 131 | -1,30352413 | N14:525025 | 32,441 | | Intergene | | | |
| chrXIV | 526013 | 526193 | 180 | -1,528319167 | N14:526078 | 8,931 | -1:CUT816; +1:YNL054W; | -1 | | | |
| chrXIV | 526651 | 526800 | 149 | -1,273351845 | N14:526736 | 8,983 | +5:YNL054W; | 5 | | | |
| chrXIV | 527097 | 527235 | 138 | -1,270138369 | N14:527172 | 40,714 | +8:YNL054W; | 8 | | | |

| CHR | START | END | L(bp) | ΔLK | Nuc ID | Fuzziness | Gene ID | Gene body postion or intergene | Terminal | rDNA | Telomeric |
|---|---|---|---|---|---|---|---|---|---|---|---|
| chrXIV | 527709 | 527871 | 162 | -1,188629644 | N14:527797 | 30,415 | +12:YNL054W; | 12 | | | |
| chrXIV | 529282 | 529444 | 162 | -1,15753479 | N14:529327 | 11,692 | +20:YNL054W; +2:CUT817; | 20 | | | |
| chrXIV | 529934 | 530079 | 145 | -1,310222683 | N14:530020 | 33,33 | NL054W; +2:YNL053W; +9:SUT749; | 24 | | | |
| chrXIV | 530913 | 531056 | 143 | -0,766218273 | | | | Overlap <50 bp | | | |
| chrXIV | 532041 | 532181 | 140 | -1,213441118 | N14:532069 | 19,07 | +3:YNL052W; | 3 | | | |
| chrXIV | 532139 | 532286 | 147 | -1,244330597 | N14:532293 | 15,769 | +4*:YNL052W; -1:YNL051W; | 4 | TERM | | |
| chrXIV | 536984 | 537144 | 160 | -1,460141096 | N14:537118 | 6,238 | +6:YNL049C; | 6 | | | |
| chrXIV | 538071 | 538217 | 146 | -1,16222764 | N14:538181 | 6,919 | +1:YNL048W; -1:YNL049C; | 1 | | | |
| chrXIV | 538102 | 538265 | 163 | -1,481478864 | N14:538181 | 6,919 | +1:YNL048W; -1:YNL049C; | 1 | | | |
| chrXIV | 538210 | 538368 | 158 | -1,482036228 | N14:538353 | 9,503 | +2:YNL048W; | 2 | | | |
| chrXIV | 540636 | 540777 | 141 | -1,177071853 | N14:540797 | 32,015 | +8:YNL047C; | 8 | | | |
| chrXIV | 541287 | 541432 | 145 | -1,043596289 | N14:541365 | 16,092 | +4:YNL047C; | 4 | | | |
| chrXIV | 541780 | 541943 | 163 | -1,349717136 | N14:541870 | 13,258 | +1:YNL047C; | 1 | | | |
| chrXIV | 541920 | 542078 | 158 | -1,111962186 | | | | Overlap <50 bp | | | |
| chrXIV | 542491 | 542640 | 149 | -1,149126739 | N14:542475 | 11,547 | 'NL046W; -1:YNL045W; +4:SUT750; | 2 | TERM | | |
| chrXIV | 544884 | 545027 | 143 | -1,360243496 | N14:544952 | 4,082 | -1:YNL044W; | -1 | | | |
| chrXIV | 544927 | 545097 | 170 | -1,252177745 | N14:544952 | 4,082 | -1:YNL044W; | -1 | | | |
| chrXIV | 546338 | 546446 | 108 | -0,888477193 | N14:546423 | 10,727 | | Intergene | | | |
| chrXIV | 546358 | 546520 | 162 | -1,222365039 | N14:546423 | 10,727 | | Intergene | | | |
| chrXIV | 551772 | 551902 | 130 | -1,277782883 | N14:551848 | 6,797 | +2:YNL041C; | 2 | | | |
| chrXIV | 552571 | 552722 | 151 | -1,375558203 | N14:552590 | 10,708 | | Intergene | | | |
| chrXIV | 553534 | 553684 | 150 | -1,304337208 | N14:553556 | 9,18 | +2:YNL040W; | 2 | | | |
| chrXIV | 554534 | 554685 | 151 | -1,331815719 | N14:554609 | 11,149 | +9:YNL040W; | 9 | | | |
| chrXIV | 555938 | 556085 | 147 | -1,067095481 | N14:555999 | 12,58 | +18:YNL040W; +7:YNL039W; | 18 | | | |
| chrXIV | 557972 | 558127 | 155 | -1,273270097 | N14:558022 | 18,912 | +7:YNL037C; | 7 | | | |
| chrXIV | 560057 | 560198 | 141 | -1,406557556 | N14:560145 | 19,856 | +4*:YNL036W; | 4 | TERM | | |
| chrXIV | 561823 | 561978 | 155 | -1,727683534 | N14:561825 | 41,073 | | Intergene | | | |
| chrXIV | 562224 | 562373 | 149 | -1,513251688 | N14:562251 | 14,64 | | Intergene | | | |
| chrXIV | 563011 | 563177 | 166 | -1,890085624 | N14:563098 | 21,939 | | Intergene | | | |
| chrXIV | 564309 | 564452 | 143 | -1,113376298 | N14:564390 | 19,502 | | Intergene | | | |
| chrXIV | 565788 | 565917 | 129 | -1,372879427 | N14:565899 | 24,664 | | Intergene | | | |
| chrXIV | 567163 | 567273 | 110 | -1,472287582 | N14:567239 | 17,301 | | Intergene | | | |
| chrXIV | 573304 | 573437 | 133 | -1,283556835 | N14:573369 | 27,074 | +4:YNL033W; | 4 | | | |
| chrXIV | 575453 | 575613 | 160 | -1,509219608 | N14:575487 | 22,93 | +4*:YNL031C; | 4 | TERM | | |
| chrXIV | 575453 | 575613 | 160 | -1,509219608 | N14:575635 | 37,233 | +3:YNL031C; | 3 | | | |
| chrXIV | 576714 | 576844 | 130 | -0,680942025 | N14:576799 | 11,846 | +1:YNL030W; | 1 | | | |
| chrXIV | 576714 | 576864 | 150 | -1,30542841 | N14:576799 | 11,846 | +1:YNL030W; | 1 | | | |
| chrXIV | 578365 | 578517 | 152 | -1,13337748 | N14:578459 | 6,022 | +3:YNL029C; | 3 | | | |
| chrXIV | 578706 | 578859 | 153 | -1,211270454 | N14:578787 | 6,979 | +1:YNL029C; | 1 | | | |
| chrXIV | 585214 | 585366 | 152 | -1,295369037 | N14:585310 | 8,258 | +1:YNL025C; | 1 | | | |
| chrXIV | 589016 | 589141 | 125 | -1,193717391 | N14:589151 | 10,986 | +13:YNL023C; | 13 | | | |
| chrXIV | 589016 | 589141 | 125 | -1,193717391 | N14:589003 | 31,456 | +14:YNL023C; | 14 | | | |
| chrXIV | 593245 | 593393 | 148 | -1,223835274 | N14:593327 | 8,343 | +2:YNL021W; | 2 | | | |
| chrXIV | 597466 | 597618 | 152 | -0,995805705 | N14:597483 | 10,815 | +2:YNL020C; | 2 | | | |
| chrXIV | 598791 | 598924 | 133 | -1,415326611 | N14:598861 | 23,352 | +3:YNL019C; | 3 | | | |
| chrXIV | 602237 | 602346 | 109 | -0,26498165 | N14:602324 | 11 | | Intergene | | | |
| chrXIV | 612323 | 612470 | 147 | -1,179312121 | N14:612426 | 45,358 | +6:YNL011C; | 6 | | | |
| chrXIV | 615779 | 615947 | 168 | -1,672435031 | N14:615899 | 17,367 | +7:YNL009W; | 7 | | | |
| chrXIV | 620267 | 620398 | 131 | -1,502545596 | N14:620245 | 11,389 | +2:YNL006W; | 2 | | | |
| chrXIV | 620267 | 620398 | 131 | -1,502545596 | N14:620397 | 6,501 | +3:YNL006W; | 3 | | | |
| chrXIV | 620785 | 620892 | 107 | -1,343314104 | N14:620770 | 28,87 | +5:YNL006W; | 5 | | | |
| chrXIV | 620785 | 620946 | 161 | -1,346740387 | N14:620770 | 28,87 | +5:YNL006W; | 5 | | | |
| chrXIV | 620785 | 620946 | 161 | -1,346740387 | N14:620937 | 39,069 | +6:YNL006W; | 6 | | | |
| chrXIV | 625000 | 625108 | 108 | -0,995500869 | N14:625110 | 27,682 | +5*:YNL003C; | 5 | TERM | | |
| chrXIV | 625453 | 625609 | 156 | -1,358908464 | N14:625501 | 15,253 | +3:YNL003C; | 3 | | | |
| chrXIV | 630331 | 630483 | 152 | -1,383371137 | N14:630418 | 30,1 | +6:YNR001C; | 6 | | | |
| chrXIV | 630745 | 630896 | 151 | -1,183985668 | N14:630914 | 23,664 | +3:YNR001C; | 3 | | | |
| chrXIV | 630745 | 630896 | 151 | -1,183985668 | N14:630752 | 21,175 | +4:YNR001C; | 4 | | | |
| chrXIV | 635632 | 635775 | 143 | -1,514614084 | N14:635681 | 12,708 | Unit535; +2:YNR004W; +8:SUT753; | 2 | | | |
| chrXIV | 640348 | 640505 | 157 | -1,21680964 | N14:640377 | 24,696 | +1:YNR008W; -1:YNR007C; | 1 | | | |
| chrXIV | 640382 | 640540 | 158 | -1,25860342 | N14:640377 | 24,696 | +1:YNR008W; -1:YNR007C; | 1 | | | |
| chrXIV | 640382 | 640540 | 158 | -1,25860342 | N14:640553 | 22,991 | +2:YNR008W; | 2 | | | |
| chrXIV | 641911 | 642059 | 148 | -1,373270044 | N14:641943 | 22,14 | +10:YNR008W; | 10 | | | |
| chrXIV | 645323 | 645472 | 149 | -1,271431502 | N14:645480 | 38,301 | +10:YNR011C; | 10 | | | |
| chrXIV | 645323 | 645472 | 149 | -1,271431502 | N14:645332 | 11,923 | +11:YNR011C; | 11 | | | |
| chrXIV | 646222 | 646370 | 148 | -1,241071186 | N14:646249 | 13,442 | +5:YNR011C; | 5 | | | |
| chrXIV | 649139 | 649287 | 148 | -1,440489374 | N14:649302 | 9,309 | +16:YNR013C; | 16 | | | |
| chrXIV | 649139 | 649287 | 148 | -1,440489374 | N14:649141 | 26,986 | +17:YNR013C; | 17 | | | |
| chrXIV | 659033 | 659204 | 171 | -1,385782074 | N14:659107 | 12,871 | +15:YNR016C; | 15 | | | |
| chrXIV | 660122 | 660273 | 151 | -1,415174214 | | | | Overlap <50 bp | | | |
| chrXIV | 660694 | 660857 | 163 | -1,278387054 | N14:660797 | 11,719 | +5:YNR016C; | 5 | | | |
| chrXIV | 669731 | 669882 | 151 | -1,126260292 | N14:669825 | 8,28 | +3*:YNR022C; | 3 | TERM | | |
| chrXIV | 673304 | 673429 | 125 | -2,233527638 | N14:673428 | 50,012 | +9:YNR026C; | 9 | | | |
| chrXIV | 673304 | 673441 | 137 | -1,467748695 | N14:673428 | 50,012 | +9:YNR026C; | 9 | | | |
| chrXIV | 673304 | 673449 | 145 | -1,703894619 | N14:673428 | 50,012 | +9:YNR026C; | 9 | | | |
| chrXIV | 673304 | 673450 | 146 | -1,367126653 | N14:673428 | 50,012 | +9:YNR026C; | 9 | | | |
| chrXIV | 673304 | 673451 | 147 | -1,689340989 | N14:673428 | 50,012 | +9:YNR026C; | 9 | | | |
| chrXIV | 673304 | 673452 | 148 | -1,566576302 | N14:673428 | 50,012 | +9:YNR026C; | 9 | | | |
| chrXIV | 673304 | 673453 | 149 | -1,478425504 | N14:673428 | 50,012 | +9:YNR026C; | 9 | | | |
| chrXIV | 673304 | 673454 | 150 | -1,546023022 | N14:673428 | 50,012 | +9:YNR026C; | 9 | | | |
| chrXIV | 673304 | 673455 | 151 | -1,404266426 | N14:673428 | 50,012 | +9:YNR026C; | 9 | | | |
| chrXIV | 673305 | 673455 | 150 | -2,148858369 | N14:673428 | 50,012 | +9:YNR026C; | 9 | | | |
| chrXIV | 673306 | 673455 | 149 | -1,857060061 | N14:673428 | 50,012 | +9:YNR026C; | 9 | | | |
| chrXIV | 673307 | 673455 | 148 | -2,296895662 | N14:673428 | 50,012 | +9:YNR026C; | 9 | | | |
| chrXIV | 673308 | 673455 | 147 | -1,666660036 | N14:673428 | 50,012 | +9:YNR026C; | 9 | | | |
| chrXIV | 673309 | 673455 | 146 | -1,486343852 | N14:673428 | 50,012 | +9:YNR026C; | 9 | | | |
| chrXIV | 673310 | 673455 | 145 | -1,930751251 | N14:673428 | 50,012 | +9:YNR026C; | 9 | | | |
| chrXIV | 673311 | 673455 | 144 | -1,623067519 | N14:673428 | 50,012 | +9:YNR026C; | 9 | | | |
| chrXIV | 673313 | 673455 | 142 | -2,230529351 | N14:673428 | 50,012 | +9:YNR026C; | 9 | | | |
| chrXIV | 673314 | 673455 | 141 | -2,225408159 | N14:673428 | 50,012 | +9:YNR026C; | 9 | | | |
| chrXIV | 673321 | 673455 | 134 | -1,716359174 | N14:673428 | 50,012 | +9:YNR026C; | 9 | | | |
| chrXIV | 673329 | 673455 | 126 | -1,294813587 | N14:673428 | 50,012 | +9:YNR026C; | 9 | | | |
| chrXIV | 673357 | 673455 | 98 | -1,532138512 | N14:673428 | 50,012 | +9:YNR026C; | 9 | | | |
| chrXIV | 675190 | 675330 | 140 | -0,865323734 | N14:675271 | 10,289 | +3:YNR027W; | 3 | | | |
| chrXIV | 678978 | 679124 | 146 | -1,106139223 | N14:679095 | 14,675 | +3:YNR030W; | 3 | | | |
| chrXIV | 679561 | 679717 | 156 | -1,46338089 | N14:679584 | 16,502 | +6:YNR030W; +36*:YNR031C; | 6 | | | |
| chrXIV | 681968 | 682096 | 128 | -1,220429156 | N14:682040 | 42,039 | +22:YNR031C; | 22 | | | |
| chrXIV | 681972 | 682075 | 103 | -0,800444388 | N14:682040 | 42,039 | +22:YNR031C; | 22 | | | |
| chrXIV | 682856 | 682998 | 142 | -1,544416865 | N14:682935 | 12,55 | +17:YNR031C; | 17 | | | |

| CHR | START | END | L(bp) | ΔLK | Nuc ID | Fuzziness | Gene ID | Gene body postion or intergene | Terminal | rDNA | Telomeric |
|---|---|---|---|---|---|---|---|---|---|---|---|
| chrXIV | 689540 | 689646 | 106 | -1,94585362 | N14:689607 | 12,133 | +12:YNR033W; | 12 | | | |
| chrXIV | 689540 | 689701 | 161 | -1,431265612 | N14:689607 | 12,133 | +12:YNR033W; | 12 | | | |
| chrXIV | 689540 | 689704 | 164 | -1,835952551 | N14:689607 | 12,133 | +12:YNR033W; | 12 | | | |
| chrXIV | 689540 | 689705 | 165 | -1,261808255 | N14:689607 | 12,133 | +12:YNR033W; | 12 | | | |
| chrXIV | 689541 | 689705 | 164 | -1,664238324 | N14:689607 | 12,133 | +12:YNR033W; | 12 | | | |
| chrXIV | 689543 | 689705 | 162 | -1,45365219 | N14:689607 | 12,133 | +12:YNR033W; | 12 | | | |
| chrXIV | 689545 | 689705 | 160 | -1,764409185 | N14:689607 | 12,133 | +12:YNR033W; | 12 | | | |
| chrXIV | 689550 | 689705 | 155 | -1,682624926 | N14:689607 | 12,133 | +12:YNR033W; | 12 | | | |
| chrXIV | 689554 | 689705 | 151 | -1,627769115 | N14:689607 | 12,133 | +12:YNR033W; | 12 | | | |
| chrXIV | 694543 | 694689 | 146 | -0,912898457 | N14:694619 | 4,528 | +2:YNR036C; | 2 | | | |
| chrXIV | 694549 | 694699 | 150 | -0,861380024 | N14:694619 | 4,528 | +2:YNR036C; | 2 | | | |
| chrXIV | 698880 | 699032 | 152 | -1,220726368 | N14:698920 | 23,309 | +4:YNR039C; | 4 | | | |
| chrXIV | 704009 | 704157 | 148 | -1,03615079 | N14:704120 | 18,735 | +3:YNR044W; | 3 | | | |
| chrXIV | 704660 | 704785 | 125 | -0,492852651 | N14:704768 | 7,396 | +7:YNR044W; | 7 | | | |
| chrXIV | 705096 | 705262 | 166 | -1,430998624 | N14:705073 | 15,822 | +9:YNR044W; | 9 | | | |
| chrXIV | 705096 | 705262 | 166 | -1,430998624 | N14:705244 | 36,116 | +10:YNR044W; | 10 | | | |
| chrXIV | 705533 | 705682 | 149 | -1,193765994 | N14:705572 | 14,825 | +12*:YNR044W; +3:SUT755; | 12 | TERM | | |
| chrXIV | 705820 | 705974 | 154 | -1,36754358 | N14:705901 | 36,918 | -1:YNR045W; +1:SUT755; | -1 | | | |
| chrXIV | 706013 | 706167 | 154 | -1,26020257 | N14:706150 | 8,524 | +1:YNR045W; -1:SUT755; | 1 | | | |
| chrXIV | 706675 | 706825 | 150 | -1,215214974 | N14:706652 | 7,815 | +4:YNR045W; | 4 | | | |
| chrXIV | 706675 | 706825 | 150 | -1,215214974 | N14:706826 | 11,431 | +5:YNR045W; | 5 | | | |
| chrXIV | 707449 | 707602 | 153 | -1,492744118 | N14:707511 | 10,262 | NR045W; +1:CUT819; -1:YNR046W; | 9 | | | |
| chrXIV | 723355 | 723513 | 158 | -1,10714188 | N14:723439 | 9,011 | +6:YNR054C; +1:CUT351; | 6 | | | |
| chrXIV | 724785 | 724950 | 165 | -1,295109362 | N14:724864 | 13,759 | +3:SUT339; | 3 | | | |
| chrXIV | 727236 | 727386 | 150 | -1,086788959 | N14:727324 | 23,881 | | Intergene | | | |
| chrXIV | 727633 | 727771 | 138 | -1,344328093 | N14:727696 | 16,802 | -1:SUT340; | -1 | | | |
| chrXIV | 731977 | 732139 | 162 | -1,443442498 | N14:732086 | 8,287 | +8:YNR056C; +5:SUT342; | 8 | | | |
| chrXIV | 735524 | 735689 | 165 | -1,57866689 | N14:735604 | 19,149 | +8*:YNR058W; | 8 | TERM | | |
| chrXIV | 735878 | 736030 | 152 | -1,083220261 | N14:735951 | 20,216 | +2*:SUT756; | 2 | TERM | | |
| chrXIV | 737786 | 737917 | 131 | -1,376817347 | N14:737841 | 21,267 | +8:YNR059W; | 8 | | | |
| chrXIV | 749149 | 749259 | 110 | -1,11729556 | N14:749152 | 22,888 | +15:YNR063W; +6:YNR064C; | 15 | | | |
| chrXIV | 749725 | 749855 | 130 | -1,507726378 | N14:749877 | 26,846 | +2:YNR064C; | 2 | | | |
| chrXIV | 749823 | 749962 | 139 | -0,934189943 | N14:749877 | 26,846 | +2:YNR064C; | 2 | | | |
| chrXIV | 750208 | 750335 | 127 | -1,62798518 | | | | Overlap <50 bp | | | |
| chrXIV | 753363 | 753527 | 164 | -1,680839786 | N14:753511 | 24,417 | +10:YNR066C; +3:YNR065C; | 10 | | | |
| chrXIV | 754387 | 754526 | 139 | -1,221453886 | N14:754466 | 21,424 | +4:YNR066C; | 4 | | | |
| chrXIV | 761541 | 761643 | 102 | -1,724150947 | N14:761639 | 45,249 | +9:YNR069C; | 9 | | | |
| chrXIV | 763360 | 763525 | 165 | -1,369016262 | N14:763380 | 19,259 | +3:CUT353; | 3 | | | |
| chrXIV | 763360 | 763525 | 165 | -1,369016262 | N14:763542 | 13,737 | +4:CUT353; | 4 | | | |
| chrXIV | 767084 | 767240 | 156 | -1,376752679 | N14:767184 | 24,921 | +11:YNR070W; +14:SUT760; | 11 | | | |
| chrXIV | 773470 | 773620 | 150 | -1,105933232 | N14:773461 | 25,007 | +6:YNR072W; +7:anti162; | 6 | | | |
| chrXIV | 778474 | 778625 | 151 | -1,602455578 | N14:778523 | 7,5 | +3:YNR074C; | 3 | | | |
| chrXIV | 779862 | 780020 | 158 | -1,419903906 | N14:779923 | 12,127 | +1:YNR075W; | 1 | | | |
| chrXIV | 781645 | 781793 | 148 | -1,105596445 | N14:781689 | 14,224 | +1:YNR075C-A; -1:YNR076W; | 1 | | | |
| chrXIV | 781935 | 782064 | 129 | -1,258754213 | N14:781999 | 13,856 | +2:YNR076W; | 2 | | | |
| chrXIV | 781942 | 782106 | 164 | -1,09897223 | N14:781999 | 13,856 | +2:YNR076W; | 2 | | | |
| chrXIV | 783376 | 783501 | 125 | -0,76204983 | N14:783457 | 20,599 | | Intergene | | | TEL RIGHT |
| chrXIV | 784039 | 784210 | 171 | 0,358247291 | N14:784097 | 20,984 | | Intergene | | | TEL RIGHT |
| chrXIV | 784040 | 784199 | 159 | -1,835059105 | N14:784097 | 20,984 | | Intergene | | | TEL RIGHT |
| chrXIV | 784044 | 784183 | 139 | -0,89633563 | N14:784097 | 20,984 | | Intergene | | | TEL RIGHT |
| chrXIV | 784044 | 784199 | 155 | -1,636685829 | N14:784097 | 20,984 | | Intergene | | | TEL RIGHT |
| chrXIV | 784049 | 784198 | 149 | -1,087259724 | N14:784097 | 20,984 | | Intergene | | | TEL RIGHT |
| chrXIV | 784049 | 784203 | 154 | -1,098942288 | N14:784097 | 20,984 | | Intergene | | | TEL RIGHT |
| chrXIV | 784063 | 784183 | 120 | -0,785229944 | N14:784097 | 20,984 | | Intergene | | | TEL RIGHT |
| chrXIV | 784063 | 784195 | 132 | -0,672954036 | N14:784097 | 20,984 | | Intergene | | | TEL RIGHT |
| chrXIV | 784063 | 784199 | 136 | -1,185656851 | N14:784097 | 20,984 | | Intergene | | | TEL RIGHT |
| chrXIV | 784063 | 784202 | 139 | -0,821457678 | N14:784097 | 20,984 | | Intergene | | | TEL RIGHT |
| chrXIV | 784063 | 784210 | 147 | -1,071686103 | N14:784097 | 20,984 | | Intergene | | | TEL RIGHT |
| chrXIV | 784063 | 784233 | 170 | -1,486291294 | N14:784097 | 20,984 | | Intergene | | | TEL RIGHT |
| chrXIV | 784063 | 784239 | 176 | -0,892286549 | N14:784097 | 20,984 | | Intergene | | | TEL RIGHT |
| chrXIV | 784063 | 784239 | 176 | -0,892286549 | N14:784261 | 54,976 | | Intergene | | | TEL RIGHT |
| chrXIV | 784063 | 784275 | 212 | -1,963608692 | N14:784097 | 20,984 | | Intergene | | | TEL RIGHT |
| chrXIV | 784063 | 784275 | 212 | -1,963608692 | N14:784261 | 54,976 | | Intergene | | | TEL RIGHT |
| chrXIV | 784064 | 784183 | 119 | -0,381367089 | N14:784097 | 20,984 | | Intergene | | | TEL RIGHT |
| chrXIV | 784064 | 784199 | 135 | -0,918950191 | N14:784097 | 20,984 | | Intergene | | | TEL RIGHT |
| chrXIV | 784101 | 784203 | 102 | -0,869865093 | N14:784097 | 20,984 | | Intergene | | | TEL RIGHT |
| chrXIV | 784101 | 784210 | 109 | -0,001883451 | N14:784097 | 20,984 | | Intergene | | | TEL RIGHT |
| chrXIV | 784106 | 784198 | 92 | -1,405753408 | N14:784097 | 20,984 | | Intergene | | | TEL RIGHT |
| chrXIV | 784106 | 784203 | 97 | -0,934219363 | N14:784097 | 20,984 | | Intergene | | | TEL RIGHT |
| chrXIV | 784106 | 784210 | 104 | -0,054330027 | N14:784097 | 20,984 | | Intergene | | | TEL RIGHT |
| chrXIV | 784106 | 784238 | 132 | -0,546881471 | N14:784097 | 20,984 | | Intergene | | | TEL RIGHT |
| chrXIV | 784106 | 784238 | 132 | -0,546881471 | N14:784261 | 54,976 | | Intergene | | | TEL RIGHT |
| chrXIV | 784106 | 784265 | 159 | -0,941267683 | N14:784097 | 20,984 | | Intergene | | | TEL RIGHT |
| chrXIV | 784106 | 784265 | 159 | -0,941267683 | N14:784261 | 54,976 | | Intergene | | | TEL RIGHT |
| chrXIV | 784106 | 784274 | 168 | -0,382371744 | N14:784097 | 20,984 | | Intergene | | | TEL RIGHT |
| chrXIV | 784106 | 784274 | 168 | -0,382371744 | N14:784261 | 54,976 | | Intergene | | | TEL RIGHT |
| chrXIV | 784106 | 784275 | 169 | -1,389975984 | N14:784097 | 20,984 | | Intergene | | | TEL RIGHT |
| chrXIV | 784106 | 784275 | 169 | -1,389975984 | N14:784261 | 54,976 | | Intergene | | | TEL RIGHT |
| chrXIV | 784106 | 784277 | 171 | -0,82305504 | N14:784097 | 20,984 | | Intergene | | | TEL RIGHT |
| chrXIV | 784106 | 784277 | 171 | -0,82305504 | N14:784261 | 54,976 | | Intergene | | | TEL RIGHT |
| chrXIV | 784111 | 784233 | 122 | -0,0243827 | N14:784097 | 20,984 | | Intergene | | | TEL RIGHT |
| chrXIV | 784111 | 784286 | 175 | -0,755639361 | N14:784097 | 20,984 | | Intergene | | | TEL RIGHT |
| chrXIV | 784111 | 784286 | 175 | -0,755639361 | N14:784261 | 54,976 | | Intergene | | | TEL RIGHT |
| chrXIV | 784130 | 784238 | 108 | -0,429047085 | N14:784261 | 54,976 | | Intergene | | | TEL RIGHT |
| chrXIV | 784130 | 784274 | 144 | -0,611468943 | N14:784261 | 54,976 | | Intergene | | | TEL RIGHT |
| chrXIV | 784130 | 784275 | 145 | -1,718014209 | N14:784261 | 54,976 | | Intergene | | | TEL RIGHT |
| chrXIV | 784130 | 784277 | 147 | -0,767845307 | N14:784261 | 54,976 | | Intergene | | | TEL RIGHT |
| chrXIV | 784130 | 784329 | 199 | -1,051778827 | N14:784261 | 54,976 | | Intergene | | | TEL RIGHT |
| chrXIV | 784132 | 784275 | 143 | -0,670688925 | N14:784261 | 54,976 | | Intergene | | | TEL RIGHT |
| chrXIV | 784135 | 784274 | 139 | -0,574347283 | N14:784261 | 54,976 | | Intergene | | | TEL RIGHT |
| chrXIV | 784139 | 784275 | 136 | -1,18667047 | N14:784261 | 54,976 | | Intergene | | | TEL RIGHT |
| chrXIV | 784139 | 784296 | 157 | -0,837353005 | N14:784261 | 54,976 | | Intergene | | | TEL RIGHT |
| chrXIV | 784140 | 784275 | 135 | -1,818223736 | N14:784261 | 54,976 | | Intergene | | | TEL RIGHT |
| chrXIV | 784145 | 784286 | 141 | -0,826669811 | N14:784261 | 54,976 | | Intergene | | | TEL RIGHT |
| chrXIV | 784164 | 784275 | 111 | -0,803360385 | N14:784261 | 54,976 | | Intergene | | | TEL RIGHT |
| chrXIV | 784169 | 784274 | 105 | -0,496957015 | N14:784261 | 54,976 | | Intergene | | | TEL RIGHT |

TABLE 3

| CHR | START | END | L(bp) | ΔLK | Nuc ID | Fuzziness | Gene ID | Gene body postion or intergene | Terminal | rDNA | Telomeric |
|---|---|---|---|---|---|---|---|---|---|---|---|
| chrXIV | 784169 | 784275 | 106 | -1,452247251 | N14:784261 | 54,976 | | Intergene | | | TEL RIGHT |
| chrXIV | 784169 | 784277 | 108 | -0,642793146 | N14:784261 | 54,976 | | Intergene | | | TEL RIGHT |
| chrXIV | 784172 | 784265 | 93 | -0,569030767 | N14:784261 | 54,976 | | Intergene | | | TEL RIGHT |
| chrXIV | 784172 | 784267 | 95 | -0,184600335 | N14:784261 | 54,976 | | Intergene | | | TEL RIGHT |
| chrXIV | 784172 | 784271 | 99 | -0,344207086 | N14:784261 | 54,976 | | Intergene | | | TEL RIGHT |
| chrXIV | 784172 | 784274 | 102 | -0,93474423 | N14:784261 | 54,976 | | Intergene | | | TEL RIGHT |
| chrXIV | 784172 | 784275 | 103 | -0,77051074 | N14:784261 | 54,976 | | Intergene | | | TEL RIGHT |
| chrXIV | 784172 | 784277 | 105 | -0,205284516 | N14:784261 | 54,976 | | Intergene | | | TEL RIGHT |
| chrXIV | 784177 | 784271 | 94 | -0,994111089 | N14:784261 | 54,976 | | Intergene | | | TEL RIGHT |
| chrXIV | 784177 | 784275 | 98 | -1,51851675 | N14:784261 | 54,976 | | Intergene | | | TEL RIGHT |
| chrXV | 838 | 992 | 154 | -1,380324046 | N15:915 | 15,086 | +3*:YOL166W-A; | 3 | TERM | | TEL LEFT |
| chrXV | 6743 | 6894 | 151 | -1,174138836 | N15:6849 | 19,188 | +7:YOL164W; +1:SUT761; | 7 | | | |
| chrXV | 8892 | 9048 | 156 | -1,498450814 | N15:8985 | 11,411 | +3*:SUT347; | 3 | TERM | | |
| chrXV | 10875 | 10982 | 107 | -1,04772108 | N15:10900 | 18,446 | +6*:YOL162W; | 6 | TERM | | |
| chrXV | 23820 | 23972 | 152 | -1,178854411 | N15:23919 | 14,142 | +3:YOL157C; | 3 | | | |
| chrXV | 29238 | 29397 | 159 | -1,05223977 | N15:29410 | 42,658 | +15:YOL155C; +7:anti164; | 15 | | | |
| chrXV | 29238 | 29397 | 159 | -1,05223977 | N15:29253 | 42,426 | +16:YOL155C; +6:anti164; | 16 | | | |
| chrXV | 36163 | 36311 | 148 | -1,348033363 | N15:36242 | 12,069 | +3:SUT352; | 3 | | | |
| chrXV | 39246 | 39406 | 160 | -1,233483322 | N15:39349 | 23,908 | +4:Unit583; +6:SUT764; | 4 | | | |
| chrXV | 43767 | 43929 | 162 | -1,607068033 | N15:43844 | 3,869 | +2:YOL151W; | 2 | | | |
| chrXV | 44442 | 44591 | 149 | -1,022607825 | N15:44533 | 7,521 | | Intergene | | | |
| chrXV | 49181 | 49338 | 157 | -1,106475741 | N15:49259 | 27,242 | +3:YOL146W; | 3 | | | |
| chrXV | 51344 | 51469 | 125 | -0,66890286 | | | | Overlap <50 bp | | | |
| chrXV | 53593 | 53699 | 106 | -0,993348981 | | | | Overlap <50 bp | | | |
| chrXV | 53593 | 53708 | 115 | -1,147864622 | | | | Overlap <50 bp | | | |
| chrXV | 53860 | 54011 | 151 | -1,503230881 | N15:53913 | 9,743 | +6*:YOL144W; | 6 | TERM | | |
| chrXV | 53996 | 54153 | 157 | -1,235505603 | N15:54073 | 31,044 | | Intergene | | | |
| chrXV | 63125 | 63255 | 130 | -0,620507582 | N15:63178 | 14,886 | +15:YOL138C; | 15 | | | |
| chrXV | 64118 | 64281 | 163 | -1,497302814 | N15:64218 | 28,312 | +8:YOL138C; | 8 | | | |
| chrXV | 66159 | 66329 | 170 | -1,841108517 | N15:66265 | 21,442 | +5:YOL137W; | 5 | | | |
| chrXV | 72755 | 72904 | 149 | -1,239389213 | N15:72830 | 16,477 | 0:YOL131W; +10*:YOL132W; | 0 | | | |
| chrXV | 74613 | 74750 | 137 | -0,619505345 | N15:74669 | 4,309 | +4:YOL130W; | 4 | | | |
| chrXV | 74660 | 74806 | 146 | -1,025612625 | N15:74669 | 4,309 | +4:YOL130W; | 4 | | | |
| chrXV | 74660 | 74806 | 146 | -1,025612625 | N15:74825 | 13,401 | +5:YOL130W; | 5 | | | |
| chrXV | 75646 | 75807 | 161 | -1,636061107 | N15:75770 | 20,057 | +11:YOL130W; | 11 | | | |
| chrXV | 80807 | 80978 | 171 | -1,510279878 | N15:80842 | 39,144 | | Intergene | | | |
| chrXV | 101110 | 101258 | 148 | -1,146373331 | N15:101210 | 10,443 | 0:YOL115W; +1:SUT767; | 0 | | | |
| chrXV | 108670 | 108830 | 160 | -1,456753686 | N15:108728 | 3,189 | +2:YOL111C; | 2 | | | |
| chrXV | 110950 | 111091 | 141 | -1,041797174 | N15:111017 | 31,321 | | Intergene | | | |
| chrXV | 112040 | 112197 | 157 | -1,663471504 | N15:112099 | 3,033 | -1:YOL108C; +1:YOL107W; | -1 | | | |
| chrXV | 113588 | 113739 | 151 | -0,901969475 | N15:113650 | 9,368 | | Intergene | | | |
| chrXV | 116654 | 116808 | 154 | -1,58598372 | N15:116777 | 43,911 | +6:YOL104C; | 6 | | | |
| chrXV | 118023 | 118177 | 154 | -1,25082971 | N15:118099 | 16,442 | | Intergene | | | |
| chrXV | 118066 | 118215 | 149 | -1,105723446 | N15:118099 | 16,442 | | Intergene | | | |
| chrXV | 118240 | 118363 | 123 | -0,916269981 | N15:118282 | 47,376 | | Intergene | | | |
| chrXV | 118340 | 118495 | 155 | -1,055778785 | N15:118453 | 30,73 | | Intergene | | | |
| chrXV | 118969 | 119120 | 151 | -1,414032477 | N15:119046 | 27,301 | | Intergene | | | |
| chrXV | 118979 | 119124 | 145 | -1,232667116 | N15:119046 | 27,301 | | Intergene | | | |
| chrXV | 118997 | 119148 | 151 | -1,174038842 | N15:119046 | 27,301 | | Intergene | | | |
| chrXV | 119251 | 119405 | 154 | -1,174652853 | N15:119384 | 12,124 | | Intergene | | | |
| chrXV | 119285 | 119429 | 144 | -1,152196264 | N15:119384 | 12,124 | | Intergene | | | |
| chrXV | 119816 | 119951 | 135 | -0,962006086 | N15:119853 | 2,646 | | Intergene | | | |
| chrXV | 120263 | 120409 | 146 | -1,081570721 | N15:120330 | 17,559 | | Intergene | | | |
| chrXV | 120263 | 120413 | 150 | -1,00039977 | N15:120330 | 17,559 | | Intergene | | | |
| chrXV | 120442 | 120599 | 157 | -1,259242691 | N15:120533 | 32,512 | | Intergene | | | |
| chrXV | 120674 | 120823 | 149 | -1,165496865 | N15:120708 | 41,012 | | Intergene | | | |
| chrXV | 120933 | 121062 | 129 | -1,018712158 | | | | Overlap <50 bp | | | |
| chrXV | 121845 | 121995 | 150 | -1,528664224 | N15:121935 | 24,007 | | Intergene | | | |
| chrXV | 122824 | 122966 | 142 | -1,215230584 | N15:122896 | 37,581 | | Intergene | | | |
| chrXV | 132108 | 132259 | 151 | -1,636024249 | N15:132177 | 18,73 | +20:YOL100W; | 20 | | | |
| chrXV | 132333 | 132482 | 149 | -1,37500934 | N15:132411 | 23,803 | +21*:YOL100W; | 21 | TERM | | |
| chrXV | 138290 | 138437 | 147 | -1,170090393 | N15:138366 | 8,206 | +3*:CUT358; +5:YOL096C; | 3 | TERM | | |
| chrXV | 141232 | 141379 | 147 | -1,152881469 | N15:141343 | 14,096 | +1:YOL095C; | 1 | | | |
| chrXV | 142140 | 142290 | 150 | -1,10932615 | N15:142219 | 4,05 | +3:YOL094C; | 3 | | | |
| chrXV | 143684 | 143828 | 144 | -1,13876834 | N15:143766 | 17,672 | +7*:YOL093W; | 7 | TERM | | |
| chrXV | 147632 | 147784 | 152 | -0,948984272 | N15:147706 | 6,401 | +3:YOL090W; | 3 | | | |
| chrXV | 152975 | 153131 | 156 | -1,295261875 | N15:153029 | 5,132 | +4:YOL089C; | 4 | | | |
| chrXV | 160615 | 160744 | 129 | -0,727958831 | N15:160677 | 39,887 | 0:YOL086C; | 0 | | | |
| chrXV | 162501 | 162638 | 137 | -0,739962871 | N15:162550 | 6,14 | +2:YOL084W; | 2 | | | |
| chrXV | 163008 | 163166 | 158 | -1,157474051 | N15:163097 | 9,288 | +5:YOL084W; | 5 | | | |
| chrXV | 163425 | 163564 | 139 | -0,944546075 | N15:163509 | 7,967 | +7:YOL084W; | 7 | | | |
| chrXV | 164146 | 164288 | 142 | -1,06184728 | | | | Overlap <50 bp | | | |
| chrXV | 169600 | 169748 | 148 | -1,369866628 | N15:169700 | 15,65 | +8:YOL082W; | 8 | | | |
| chrXV | 171569 | 171743 | 174 | -1,447520155 | N15:171764 | 24,647 | +7:YOL081W; | 7 | | | |
| chrXV | 173781 | 173933 | 152 | -1,243122431 | | | | Overlap <50 bp | | | |
| chrXV | 177092 | 177238 | 146 | -2,136246911 | N15:177172 | 13,52 | +41:YOL081W; | 41 | | | |
| chrXV | 177093 | 177211 | 118 | -0,833644789 | N15:177172 | 13,52 | +41:YOL081W; | 41 | | | |
| chrXV | 177093 | 177214 | 121 | -0,819245043 | N15:177172 | 13,52 | +41:YOL081W; | 41 | | | |
| chrXV | 177093 | 177217 | 124 | -1,701848134 | N15:177172 | 13,52 | +41:YOL081W; | 41 | | | |
| chrXV | 177093 | 177220 | 127 | -0,387802622 | N15:177172 | 13,52 | +41:YOL081W; | 41 | | | |
| chrXV | 177093 | 177221 | 128 | -0,746455658 | N15:177172 | 13,52 | +41:YOL081W; | 41 | | | |
| chrXV | 177093 | 177222 | 129 | -0,6457415 | N15:177172 | 13,52 | +41:YOL081W; | 41 | | | |
| chrXV | 177093 | 177223 | 130 | -0,541260145 | N15:177172 | 13,52 | +41:YOL081W; | 41 | | | |
| chrXV | 177093 | 177224 | 131 | -0,867184981 | N15:177172 | 13,52 | +41:YOL081W; | 41 | | | |
| chrXV | 177093 | 177225 | 132 | -0,749488162 | N15:177172 | 13,52 | +41:YOL081W; | 41 | | | |
| chrXV | 177093 | 177226 | 133 | -1,430856435 | N15:177172 | 13,52 | +41:YOL081W; | 41 | | | |
| chrXV | 177093 | 177227 | 134 | -0,712061023 | N15:177172 | 13,52 | +41:YOL081W; | 41 | | | |
| chrXV | 177093 | 177228 | 135 | -0,743122709 | N15:177172 | 13,52 | +41:YOL081W; | 41 | | | |
| chrXV | 177093 | 177229 | 136 | -0,958480747 | N15:177172 | 13,52 | +41:YOL081W; | 41 | | | |
| chrXV | 177093 | 177230 | 137 | -1,158216722 | N15:177172 | 13,52 | +41:YOL081W; | 41 | | | |
| chrXV | 177093 | 177231 | 138 | -1,100262657 | N15:177172 | 13,52 | +41:YOL081W; | 41 | | | |
| chrXV | 177093 | 177232 | 139 | -1,028745634 | N15:177172 | 13,52 | +41:YOL081W; | 41 | | | |
| chrXV | 177093 | 177233 | 140 | -1,208694273 | N15:177172 | 13,52 | +41:YOL081W; | 41 | | | |
| chrXV | 177093 | 177234 | 141 | -1,107974531 | N15:177172 | 13,52 | +41:YOL081W; | 41 | | | |
| chrXV | 177093 | 177235 | 142 | -0,895202913 | N15:177172 | 13,52 | +41:YOL081W; | 41 | | | |
| chrXV | 177093 | 177236 | 143 | -0,967803873 | N15:177172 | 13,52 | +41:YOL081W; | 41 | | | |
| chrXV | 177093 | 177237 | 144 | -1,243383602 | N15:177172 | 13,52 | +41:YOL081W; | 41 | | | |
| chrXV | 177093 | 177238 | 145 | -0,830850786 | N15:177172 | 13,52 | +41:YOL081W; | 41 | | | |

| CHR | START | END | L(bp) | ΔLK | Nuc ID | Fuzziness | Gene ID | Gene body postion or intergene | Terminal | rDNA | Telomeric |
|---|---|---|---|---|---|---|---|---|---|---|---|
| chrXV | 177093 | 177239 | 146 | -2,235173337 | N15:177172 | 13,52 | +41:YOL081W; | 41 | | | |
| chrXV | 177094 | 177238 | 144 | -1,330164434 | N15:177172 | 13,52 | +41:YOL081W; | 41 | | | |
| chrXV | 177095 | 177238 | 143 | -1,12270541 | N15:177172 | 13,52 | +41:YOL081W; | 41 | | | |
| chrXV | 177096 | 177238 | 142 | -0,96152176 | N15:177172 | 13,52 | +41:YOL081W; | 41 | | | |
| chrXV | 177097 | 177238 | 141 | -0,940392672 | N15:177172 | 13,52 | +41:YOL081W; | 41 | | | |
| chrXV | 177098 | 177238 | 140 | -0,967503401 | N15:177172 | 13,52 | +41:YOL081W; | 41 | | | |
| chrXV | 177099 | 177238 | 139 | -1,64243343 | N15:177172 | 13,52 | +41:YOL081W; | 41 | | | |
| chrXV | 177100 | 177238 | 138 | -1,193639396 | N15:177172 | 13,52 | +41:YOL081W; | 41 | | | |
| chrXV | 177101 | 177238 | 137 | -1,554361252 | N15:177172 | 13,52 | +41:YOL081W; | 41 | | | |
| chrXV | 177102 | 177238 | 136 | -1,493321503 | N15:177172 | 13,52 | +41:YOL081W; | 41 | | | |
| chrXV | 177103 | 177238 | 135 | -1,487576382 | N15:177172 | 13,52 | +41:YOL081W; | 41 | | | |
| chrXV | 177104 | 177238 | 134 | -1,693158055 | N15:177172 | 13,52 | +41:YOL081W; | 41 | | | |
| chrXV | 177105 | 177238 | 133 | -1,841043842 | N15:177172 | 13,52 | +41:YOL081W; | 41 | | | |
| chrXV | 177106 | 177238 | 132 | -0,987968092 | N15:177172 | 13,52 | +41:YOL081W; | 41 | | | |
| chrXV | 177107 | 177238 | 131 | -1,473584125 | N15:177172 | 13,52 | +41:YOL081W; | 41 | | | |
| chrXV | 177109 | 177238 | 129 | -1,40063726 | N15:177172 | 13,52 | +41:YOL081W; | 41 | | | |
| chrXV | 177110 | 177238 | 128 | -1,286237269 | N15:177172 | 13,52 | +41:YOL081W; | 41 | | | |
| chrXV | 177111 | 177238 | 127 | -1,046430492 | N15:177172 | 13,52 | +41:YOL081W; | 41 | | | |
| chrXV | 177112 | 177238 | 126 | -1,30549581 | N15:177172 | 13,52 | +41:YOL081W; | 41 | | | |
| chrXV | 177113 | 177238 | 125 | -0,347512869 | N15:177172 | 13,52 | +41:YOL081W; | 41 | | | |
| chrXV | 177115 | 177238 | 123 | -1,0655025 | N15:177172 | 13,52 | +41:YOL081W; | 41 | | | |
| chrXV | 177117 | 177238 | 121 | -0,722508558 | N15:177172 | 13,52 | +41:YOL081W; | 41 | | | |
| chrXV | 177118 | 177238 | 120 | -0,130969237 | N15:177172 | 13,52 | +41:YOL081W; | 41 | | | |
| chrXV | 177119 | 177238 | 119 | -0,392333371 | N15:177172 | 13,52 | +41:YOL081W; | 41 | | | |
| chrXV | 177120 | 177238 | 118 | -0,543069241 | N15:177172 | 13,52 | +41:YOL081W; | 41 | | | |
| chrXV | 177121 | 177238 | 117 | -0,66175371 | N15:177172 | 13,52 | +41:YOL081W; | 41 | | | |
| chrXV | 177124 | 177238 | 114 | -0,466034711 | N15:177172 | 13,52 | +41:YOL081W; | 41 | | | |
| chrXV | 177125 | 177238 | 113 | -0,720236509 | N15:177172 | 13,52 | +41:YOL081W; | 41 | | | |
| chrXV | 177128 | 177238 | 110 | -0,748359702 | N15:177172 | 13,52 | +41:YOL081W; | 41 | | | |
| chrXV | 177130 | 177238 | 108 | -0,44786109 | N15:177172 | 13,52 | +41:YOL081W; | 41 | | | |
| chrXV | 177132 | 177217 | 85 | -0,395537676 | N15:177172 | 13,52 | +41:YOL081W; | 41 | | | |
| chrXV | 177132 | 177238 | 106 | -1,369321852 | N15:177172 | 13,52 | +41:YOL081W; | 41 | | | |
| chrXV | 177733 | 177893 | 160 | -1,278779296 | N15:177858 | 30,517 | +45:YOL081W; | 45 | | | |
| chrXV | 178523 | 178704 | 181 | -1,6022499 | N15:178709 | 11,015 | +51:YOL081W; | 51 | | | |
| chrXV | 185749 | 185889 | 140 | -1,236430251 | N15:185774 | 30,288 | +3*:YOL077W-A; +7*:YOL077C; | 3 | TERM | | |
| chrXV | 187300 | 187456 | 156 | -1,387095032 | N15:187379 | 9,813 | +3:YOL076W; | 3 | | | |
| chrXV | 189704 | 189862 | 158 | -1,434006571 | N15:189795 | 17,838 | +23:YOL075C; | 23 | | | |
| chrXV | 192332 | 192491 | 159 | -1,20503896 | N15:192321 | 30,993 | +9:YOL075C; | 9 | | | |
| chrXV | 196562 | 196707 | 145 | -1,287089093 | N15:196546 | 19,26 | +10*:YOL072W; +1:YOL071W; | 10 | TERM | | |
| chrXV | 196562 | 196707 | 145 | -1,287089093 | N15:196728 | 15,172 | +2:YOL071W; | 2 | | | |
| chrXV | 198924 | 199072 | 148 | -1,003230809 | N15:198961 | 11,219 | -1:YOL070C; +1:YOL069W; | -1 | | | |
| chrXV | 202849 | 203002 | 153 | -1,412861553 | N15:202912 | 19,627 | +10:YOL066C; +18:YOL065C; | 10 | | | |
| chrXV | 205781 | 205911 | 130 | -0,97370178 | N15:205866 | 13,943 | +1:YOL065C; | 1 | | | |
| chrXV | 208753 | 208900 | 147 | -1,037327426 | N15:208837 | 10,61 | +10:YOL063C; | 10 | | | |
| chrXV | 209390 | 209535 | 145 | -1,221244758 | N15:209458 | 29,068 | +7:YOL063C; | 7 | | | |
| chrXV | 212639 | 212808 | 169 | -1,510529614 | N15:212754 | 23,37 | +4:YOL061W; | 4 | | | |
| chrXV | 214778 | 214916 | 138 | -0,724387294 | N15:214872 | 17,056 | +9:YOL060C; | 9 | | | |
| chrXV | 215741 | 215894 | 153 | -1,353123569 | N15:215837 | 20,917 | +3:YOL060C; | 3 | | | |
| chrXV | 218336 | 218507 | 171 | -1,405995612 | N15:218350 | 26,499 | +9:YOL059W; | 9 | | | |
| chrXV | 220428 | 220557 | 129 | -1,125106884 | N15:220513 | 10,61 | 'OL058W; -1:YOL057W; +1:anti167; | 8 | TERM | | |
| chrXV | 222406 | 222534 | 128 | -1,187436268 | N15:222459 | 11,161 | +11:YOL057W; | 11 | | | |
| chrXV | 225244 | 225389 | 145 | -0,987555625 | N15:225300 | 6,419 | | Intergene | | | |
| chrXV | 227790 | 227919 | 129 | -1,270952598 | N15:227846 | 36,592 | | Intergene | | | |
| chrXV | 229356 | 229535 | 179 | -1,462983703 | N15:229431 | 34,896 | +6:YOL054W; | 6 | | | |
| chrXV | 229736 | 229903 | 167 | -1,685697167 | N15:229797 | 15,293 | +8*:YOL054W; | 8 | TERM | | |
| chrXV | 233701 | 233860 | 159 | -1,164451308 | N15:233870 | 7,778 | -1:YOL052C; | -1 | | | |
| chrXV | 233701 | 233860 | 159 | -1,164451308 | N15:233712 | 24,612 | 0:YOL052C; | 0 | | | |
| chrXV | 237293 | 237436 | 143 | -1,292865768 | N15:237339 | 5,802 | +16:YOL051W; | 16 | | | |
| chrXV | 242082 | 242240 | 158 | -1,390671127 | N15:242059 | 10,747 | +3:SUT357; +5:YOL047C; | 3 | | | |
| chrXV | 242082 | 242240 | 158 | -1,390671127 | N15:242224 | 12,971 | +4:SUT357; +4:YOL047C; | 4 | | | |
| chrXV | 243486 | 243636 | 150 | -1,01193345 | N15:243571 | 6,285 | +4:YOL045W; | 4 | | | |
| chrXV | 244606 | 244758 | 152 | -1,535663473 | N15:244626 | 13,846 | +11:YOL045W; | 11 | | | |
| chrXV | 244606 | 244758 | 152 | -1,535663473 | N15:244776 | 39,569 | +12:YOL045W; | 12 | | | |
| chrXV | 248190 | 248334 | 144 | -1,505492756 | N15:248279 | 17,462 | +8*:YOL044W; | 8 | TERM | | |
| chrXV | 248556 | 248722 | 166 | -1,475566659 | | | | Overlap <50 bp | | | |
| chrXV | 255239 | 255421 | 182 | -1,20720421 | N15:255344 | 8,618 | -1:YOL038C-A; +1:YOL038W; | -1 | | | |
| chrXV | 256726 | 256872 | 146 | -1,150587758 | N15:256740 | 13,546 | +3:YOL036W; | 3 | | | |
| chrXV | 260668 | 260810 | 142 | -1,474576099 | | | | Overlap <50 bp | | | |
| chrXV | 261423 | 261595 | 172 | -1,435109284 | N15:261549 | 31,264 | +11:YOL034W; | 11 | | | |
| chrXV | 263580 | 263731 | 151 | -0,820452217 | N15:263648 | 8,497 | 0:anti-YOL033W; +2:YOL033W; | 0 | | | |
| chrXV | 265679 | 265840 | 161 | -1,338903182 | N15:265718 | 8,727 | +3:YOL032W; | 3 | | | |
| chrXV | 266859 | 267005 | 146 | -1,470278943 | N15:266903 | 10,559 | +5:YOL031C; | 5 | | | |
| chrXV | 268238 | 268379 | 141 | -1,0559596 | N15:268361 | 4,98 | +2:YOL030W; | 2 | | | |
| chrXV | 270968 | 271134 | 166 | -1,160322532 | N15:271050 | 9,948 | +4:YOL028C; | 4 | | | |
| chrXV | 271150 | 271308 | 158 | -1,349304512 | N15:271220 | 6,723 | +3:YOL028C; | 3 | | | |
| chrXV | 273050 | 273212 | 162 | -1,55051261 | N15:273212 | 30,589 | +4:YOL027C; | 4 | | | |
| chrXV | 276074 | 276211 | 137 | -0,604744689 | N15:276066 | 24,854 | +8:YOL025W; | 8 | | | |
| chrXV | 276074 | 276211 | 137 | -0,604744689 | N15:276231 | 22,633 | +9:YOL025W; | 9 | | | |
| chrXV | 276839 | 276983 | 144 | -1,419903964 | N15:276892 | 10,614 | +13*:YOL025W; 0:YOL024W; | 13 | TERM | | |
| chrXV | 279591 | 279744 | 153 | -1,3874379 | N15:279667 | 22,007 | +11:YOL023W; | 11 | | | |
| chrXV | 280089 | 280217 | 128 | -0,789209621 | | | | Overlap <50 bp | | | |
| chrXV | 282501 | 282650 | 149 | -1,016160775 | N15:282573 | 19,856 | +19:YOL021C; | 19 | | | |
| chrXV | 284484 | 284582 | 98 | -1,242739685 | | | | Overlap <50 bp | | | |
| chrXV | 284650 | 284789 | 139 | -0,886129633 | N15:284768 | 27,722 | +5:YOL021C; | 5 | | | |
| chrXV | 284999 | 285154 | 155 | -1,06973713 | N15:285080 | 11,548 | +3:YOL021C; | 3 | | | |
| chrXV | 285195 | 285326 | 131 | -0,979364305 | N15:285245 | 12,071 | +2:YOL021C; | 2 | | | |
| chrXV | 292850 | 293005 | 155 | -1,199196727 | N15:292974 | 3,651 | +4:YOL017W; | 4 | | | |
| chrXV | 292905 | 293050 | 145 | -1,063680291 | N15:292974 | 3,651 | +4:YOL017W; | 4 | | | |
| chrXV | 293213 | 293372 | 159 | -1,302444407 | N15:293294 | 6,907 | +6:YOL017W; | 6 | | | |
| chrXV | 294193 | 294362 | 169 | -1,241354743 | N15:294360 | 12,576 | +12:YOL017W; | 12 | | | |
| chrXV | 298993 | 299142 | 149 | -1,194772115 | N15:299031 | 18,702 | +13*:YOL015W; -1:Unit592; | 13 | TERM | | |
| chrXV | 306450 | 306602 | 152 | -1,33299903 | N15:306552 | 11,59 | +8:YOL011W; | 8 | | | |
| chrXV | 311787 | 311943 | 156 | -1,390667975 | N15:311928 | 17,315 | +4:YOL007C; | 4 | | | |
| chrXV | 319552 | 319692 | 140 | -0,8414846 | N15:319612 | 19,675 | +19:YOL004W; | 19 | | | |
| chrXV | 320812 | 320963 | 151 | -1,066604116 | N15:320792 | 45,768 | +26:YOL004W; | 26 | | | |
| chrXV | 321565 | 321713 | 148 | -1,227212805 | N15:321561 | 10,149 | +30*:YOL004W; | 30 | TERM | | |
| chrXV | 327427 | 327560 | 133 | -0,914160502 | N15:327498 | 11,537 | +5:YOR001W; | 5 | | | |
| chrXV | 328438 | 328583 | 145 | -0,972513716 | N15:328423 | 33,596 | +10:YOR001W; | 10 | | | |
| chrXV | 328438 | 328583 | 145 | -0,972513716 | N15:328597 | 24,648 | +11:YOR001W; | 11 | | | |
| chrXV | 329812 | 329980 | 168 | -1,205104124 | N15:329910 | 8,424 | +4:YOR002W; | 4 | | | |

| CHR | START | END | L(bp) | ΔLK | Nuc ID | Fuzziness | Gene ID | Gene body postion or intergene | Terminal | rDNA | Telomeric |
|---|---|---|---|---|---|---|---|---|---|---|---|
| chrXV | 338001 | 338161 | 160 | -1,351213351 | N15:338135 | 10,053 | +4:YOR006C; +4:SUT360; | 4 | | | |
| chrXV | 341700 | 341803 | 103 | -1,271077672 | N15:341798 | 15,84 | +6:YOR008C; | 6 | | | |
| chrXV | 348865 | 348972 | 107 | -0,040875625 | N15:348916 | 16,293 | +2*:SUT361; | 2 | TERM | | |
| chrXV | 349258 | 349411 | 153 | -1,28573534 | N15:349279 | 15,864 | +4:SUT770; | 4 | | | |
| chrXV | 351942 | 352092 | 150 | -1,02860907 | N15:351997 | 40,035 | +13:YOR011W; | 13 | | | |
| chrXV | 353769 | 353922 | 153 | -1,610468578 | N15:353843 | 3,536 | +24*:YOR011W; | 24 | TERM | | |
| chrXV | 356685 | 356815 | 130 | -0,859322139 | N15:356719 | 8,585 | +2:YOR012W; +4:SUT772; | 2 | | | |
| chrXV | 357881 | 358029 | 148 | -1,22758277 | N15:357967 | 9,209 | +4:YOR014W; | 4 | | | |
| chrXV | 359149 | 359277 | 128 | -1,786539428 | | | | Overlap <50 bp | | | |
| chrXV | 359963 | 360110 | 147 | -1,652893437 | N15:360012 | 28,378 | +17*:YOR014W; | 17 | TERM | | |
| chrXV | 363241 | 363404 | 163 | -1,340148341 | N15:363313 | 23,231 | +13:YOR017W; | 13 | | | |
| chrXV | 363647 | 363809 | 162 | -1,349138369 | N15:363630 | 9,503 | +4*:CUT834; +15:YOR017W; | 4 | TERM | | |
| chrXV | 365392 | 365542 | 150 | -1,238908226 | N15:365381 | 33,966 | +7:YOR018W; | 7 | | | |
| chrXV | 365392 | 365542 | 150 | -1,238908226 | N15:365558 | 50,406 | +8:YOR018W; | 8 | | | |
| chrXV | 369266 | 369425 | 159 | -1,156762629 | N15:369250 | 26,615 | +7:YOR019W; +2:anti170; | 7 | | | |
| chrXV | 369347 | 369504 | 157 | -1,672975616 | N15:369518 | 12,775 | +9:YOR019W; 0:anti170; | 9 | | | |
| chrXV | 371267 | 371398 | 131 | -0,68462629 | N15:371332 | 16,558 | +1:SUT774; -1:YOR020W-A; | 1 | | | |
| chrXV | 371602 | 371749 | 147 | -0,856399705 | N15:371676 | 16,485 | +5*:SUT775; +1:YOR020W-A; | 5 | TERM | | |
| chrXV | 377796 | 377936 | 140 | -0,943111977 | N15:377801 | 3,775 | +1:YOR023C; | 1 | | | |
| chrXV | 384219 | 384368 | 149 | -1,244762911 | N15:384312 | 13,906 | +3:YOR028C; | 3 | | | |
| chrXV | 387959 | 388110 | 151 | -1,653310674 | N15:388058 | 9,95 | +9:YOR030W; | 9 | | | |
| chrXV | 389816 | 389942 | 126 | -0,817478825 | N15:389861 | 7,234 | +8:YOR032C; | 8 | | | |
| chrXV | 392262 | 392373 | 111 | -1,015640209 | N15:392325 | 26,655 | R033C; +2:YOR032W-A; -1:CUT371; | 14 | TERM | | |
| chrXV | 392576 | 392737 | 161 | -1,493504576 | N15:392661 | 28,648 | OR033C; +29:YOR034C; +1:CUT371; | 12 | | | |
| chrXV | 401254 | 401398 | 144 | -1,416581892 | N15:401349 | 6,348 | -1:YOR037W; | -1 | | | |
| chrXV | 403569 | 403709 | 140 | -0,791369069 | | | | Overlap <50 bp | | | |
| chrXV | 405315 | 405460 | 145 | -1,026884964 | N15:405385 | 4,412 | -1:YOR039W; +1:YOR038C; | -1 | | | |
| chrXV | 405735 | 405887 | 152 | -1,294871055 | N15:405829 | 6,033 | +2:YOR039W; | 2 | | | |
| chrXV | 408309 | 408462 | 153 | -1,212757566 | N15:408405 | 5,565 | +1:YOR042W; | 1 | | | |
| chrXV | 413175 | 413325 | 150 | -1,355282683 | N15:413244 | 9,576 | +4:YOR044W; | 4 | | | |
| chrXV | 413525 | 413650 | 125 | -0,730040502 | N15:413578 | 9,136 | +6*:YOR044W; -1:YOR045W; | 6 | TERM | | |
| chrXV | 419166 | 419317 | 151 | -1,233662172 | N15:419244 | 21,394 | +16:YOR048C; | 16 | | | |
| chrXV | 420271 | 420429 | 158 | -1,306162798 | N15:420382 | 6,397 | +9:YOR048C; | 9 | | | |
| chrXV | 423240 | 423391 | 151 | -1,060709461 | N15:423319 | 4,278 | +6:anti172; +5:YOR049C; | 6 | | | |
| chrXV | 423241 | 423391 | 150 | -1,797723725 | N15:423319 | 4,278 | +6:anti172; +5:YOR049C; | 6 | | | |
| chrXV | 423242 | 423391 | 149 | -1,795105226 | N15:423319 | 4,278 | +6:anti172; +5:YOR049C; | 6 | | | |
| chrXV | 423243 | 423391 | 148 | -1,31934863 | N15:423319 | 4,278 | +6:anti172; +5:YOR049C; | 6 | | | |
| chrXV | 423245 | 423391 | 146 | -1,210460058 | N15:423319 | 4,278 | +6:anti172; +5:YOR049C; | 6 | | | |
| chrXV | 433044 | 433202 | 158 | -1,262142571 | N15:433178 | 22,196 | +7:YOR057W; | 7 | | | |
| chrXV | 433171 | 433320 | 149 | -1,163300833 | N15:433331 | 22,264 | +8*:YOR057W; | 8 | TERM | | |
| chrXV | 433171 | 433320 | 149 | -1,163300833 | N15:433178 | 22,196 | +7:YOR057W; | 7 | | | |
| chrXV | 435682 | 435849 | 167 | -1,254519131 | N15:435774 | 9,543 | +5:YOR058C; | 5 | | | |
| chrXV | 441363 | 441514 | 151 | -1,256060734 | N15:441436 | 4,494 | +1:YOR061W; -1:YOR060C; | 1 | | | |
| chrXV | 443421 | 443578 | 157 | -1,358522809 | N15:443539 | 18,119 | +2:YOR062C; | 2 | | | |
| chrXV | 445187 | 445337 | 150 | -1,121339985 | N15:445171 | 30,447 | +3:YOR063W; | 3 | | | |
| chrXV | 445187 | 445337 | 150 | -1,121339985 | N15:445329 | 15,953 | +4:YOR063W; | 4 | | | |
| chrXV | 446874 | 447012 | 138 | -0,698220013 | N15:446940 | 5 | -1:YOR064C; -1:YOR065W; | -1 | | | |
| chrXV | 450011 | 450165 | 154 | -1,312321467 | N15:450081 | 16,563 | +5:YOR066W; | 5 | | | |
| chrXV | 456168 | 456329 | 161 | -1,04686065 | N15:456318 | 17,474 | +11:YOR070C; | 11 | | | |
| chrXV | 457433 | 457588 | 155 | -1,156615902 | N15:457449 | 2,708 | +4:YOR070C; | 4 | | | |
| chrXV | 457539 | 457691 | 152 | -1,246098983 | N15:457612 | 2,317 | +3:YOR070C; | 3 | | | |
| chrXV | 459473 | 459604 | 131 | -1,362129338 | | | | Overlap <50 bp | | | |
| chrXV | 465612 | 465776 | 164 | -1,46017332 | N15:465716 | 10,577 | +7:YOR073W; | 7 | | | |
| chrXV | 469152 | 469262 | 110 | -0,535648732 | N15:469219 | 14,637 | +7*:YOR075W; | 7 | TERM | | |
| chrXV | 469152 | 469277 | 125 | -0,835524045 | N15:469219 | 14,637 | +7*:YOR075W; | 7 | TERM | | |
| chrXV | 480195 | 480360 | 165 | -1,356189554 | N15:480344 | 20,72 | -1:YOR084W; +6:YOR083W; | -1 | | | |
| chrXV | 480209 | 480363 | 154 | -1,132651125 | N15:480344 | 20,72 | -1:YOR084W; +6:YOR083W; | -1 | | | |
| chrXV | 480628 | 480755 | 127 | -2,047102363 | N15:480718 | 26,92 | +2:YOR084W; +8:YOR083W; | 2 | | | |
| chrXV | 480628 | 480787 | 159 | -1,408184776 | N15:480718 | 26,92 | +2:YOR084W; +8:YOR083W; | 2 | | | |
| chrXV | 480629 | 480736 | 107 | -1,641834775 | N15:480718 | 26,92 | +2:YOR084W; +8:YOR083W; | 2 | | | |
| chrXV | 480629 | 480737 | 108 | -1,652642803 | N15:480718 | 26,92 | +2:YOR084W; +8:YOR083W; | 2 | | | |
| chrXV | 480629 | 480738 | 109 | -1,273939347 | N15:480718 | 26,92 | +2:YOR084W; +8:YOR083W; | 2 | | | |
| chrXV | 480629 | 480739 | 110 | -1,215774507 | N15:480718 | 26,92 | +2:YOR084W; +8:YOR083W; | 2 | | | |
| chrXV | 480629 | 480741 | 112 | -1,787101329 | N15:480718 | 26,92 | +2:YOR084W; +8:YOR083W; | 2 | | | |
| chrXV | 480629 | 480742 | 113 | -2,033901057 | N15:480718 | 26,92 | +2:YOR084W; +8:YOR083W; | 2 | | | |
| chrXV | 480629 | 480743 | 114 | -1,702846269 | N15:480718 | 26,92 | +2:YOR084W; +8:YOR083W; | 2 | | | |
| chrXV | 480629 | 480744 | 115 | -1,764861837 | N15:480718 | 26,92 | +2:YOR084W; +8:YOR083W; | 2 | | | |
| chrXV | 480629 | 480745 | 116 | -1,058947585 | N15:480718 | 26,92 | +2:YOR084W; +8:YOR083W; | 2 | | | |
| chrXV | 480629 | 480746 | 117 | -1,241355473 | N15:480718 | 26,92 | +2:YOR084W; +8:YOR083W; | 2 | | | |
| chrXV | 480629 | 480747 | 118 | -1,297879372 | N15:480718 | 26,92 | +2:YOR084W; +8:YOR083W; | 2 | | | |
| chrXV | 480629 | 480748 | 119 | -1,714200179 | N15:480718 | 26,92 | +2:YOR084W; +8:YOR083W; | 2 | | | |
| chrXV | 480629 | 480749 | 120 | -1,347145544 | N15:480718 | 26,92 | +2:YOR084W; +8:YOR083W; | 2 | | | |
| chrXV | 480629 | 480750 | 121 | -1,726655263 | N15:480718 | 26,92 | +2:YOR084W; +8:YOR083W; | 2 | | | |
| chrXV | 480629 | 480751 | 122 | -1,055735607 | N15:480718 | 26,92 | +2:YOR084W; +8:YOR083W; | 2 | | | |
| chrXV | 480629 | 480752 | 123 | -1,364898698 | N15:480718 | 26,92 | +2:YOR084W; +8:YOR083W; | 2 | | | |
| chrXV | 480629 | 480753 | 124 | -0,781653105 | N15:480718 | 26,92 | +2:YOR084W; +8:YOR083W; | 2 | | | |
| chrXV | 480629 | 480754 | 125 | -1,219940328 | N15:480718 | 26,92 | +2:YOR084W; +8:YOR083W; | 2 | | | |
| chrXV | 480629 | 480755 | 126 | -2,331802401 | N15:480718 | 26,92 | +2:YOR084W; +8:YOR083W; | 2 | | | |
| chrXV | 480629 | 480756 | 127 | -1,64753947 | N15:480718 | 26,92 | +2:YOR084W; +8:YOR083W; | 2 | | | |
| chrXV | 480629 | 480757 | 128 | -0,992370653 | N15:480718 | 26,92 | +2:YOR084W; +8:YOR083W; | 2 | | | |
| chrXV | 480629 | 480758 | 129 | -1,303404181 | N15:480718 | 26,92 | +2:YOR084W; +8:YOR083W; | 2 | | | |
| chrXV | 480629 | 480759 | 130 | -1,526067322 | N15:480718 | 26,92 | +2:YOR084W; +8:YOR083W; | 2 | | | |
| chrXV | 480629 | 480760 | 131 | -1,725088555 | N15:480718 | 26,92 | +2:YOR084W; +8:YOR083W; | 2 | | | |
| chrXV | 480629 | 480761 | 132 | -1,260374692 | N15:480718 | 26,92 | +2:YOR084W; +8:YOR083W; | 2 | | | |
| chrXV | 480629 | 480762 | 133 | -1,777442909 | N15:480718 | 26,92 | +2:YOR084W; +8:YOR083W; | 2 | | | |
| chrXV | 480629 | 480763 | 134 | -1,892204097 | N15:480718 | 26,92 | +2:YOR084W; +8:YOR083W; | 2 | | | |
| chrXV | 480629 | 480764 | 135 | -1,938855951 | N15:480718 | 26,92 | +2:YOR084W; +8:YOR083W; | 2 | | | |
| chrXV | 480629 | 480765 | 136 | -2,007405819 | N15:480718 | 26,92 | +2:YOR084W; +8:YOR083W; | 2 | | | |
| chrXV | 480629 | 480766 | 137 | -2,034103766 | N15:480718 | 26,92 | +2:YOR084W; +8:YOR083W; | 2 | | | |
| chrXV | 480629 | 480767 | 138 | -1,433521314 | N15:480718 | 26,92 | +2:YOR084W; +8:YOR083W; | 2 | | | |
| chrXV | 480629 | 480768 | 139 | -1,954292923 | N15:480718 | 26,92 | +2:YOR084W; +8:YOR083W; | 2 | | | |
| chrXV | 480629 | 480769 | 140 | -1,907043766 | N15:480718 | 26,92 | +2:YOR084W; +8:YOR083W; | 2 | | | |
| chrXV | 480629 | 480770 | 141 | -1,728509808 | N15:480718 | 26,92 | +2:YOR084W; +8:YOR083W; | 2 | | | |
| chrXV | 480629 | 480771 | 142 | -2,396375054 | N15:480718 | 26,92 | +2:YOR084W; +8:YOR083W; | 2 | | | |
| chrXV | 480629 | 480772 | 143 | -2,003456916 | N15:480718 | 26,92 | +2:YOR084W; +8:YOR083W; | 2 | | | |
| chrXV | 480629 | 480773 | 144 | -2,013460411 | N15:480718 | 26,92 | +2:YOR084W; +8:YOR083W; | 2 | | | |
| chrXV | 480629 | 480774 | 145 | -1,644022048 | N15:480718 | 26,92 | +2:YOR084W; +8:YOR083W; | 2 | | | |
| chrXV | 480629 | 480775 | 146 | -2,671942306 | N15:480718 | 26,92 | +2:YOR084W; +8:YOR083W; | 2 | | | |
| chrXV | 480629 | 480776 | 147 | -1,227590535 | N15:480718 | 26,92 | +2:YOR084W; +8:YOR083W; | 2 | | | |
| chrXV | 480629 | 480777 | 148 | -1,726506313 | N15:480718 | 26,92 | +2:YOR084W; +8:YOR083W; | 2 | | | |

| CHR | START | END | L(bp) | ΔLK | Nuc ID | Fuzziness | Gene ID | Gene body postion or intergene | Terminal | rDNA | Telomeric |
|---|---|---|---|---|---|---|---|---|---|---|---|
| chrXV | 480629 | 480778 | 149 | -1,776913413 | N15:480718 | 26,92 | +2:YOR084W; +8:YOR083W; | 2 | | | |
| chrXV | 480629 | 480779 | 150 | -1,332352265 | N15:480718 | 26,92 | +2:YOR084W; +8:YOR083W; | 2 | | | |
| chrXV | 480629 | 480780 | 151 | -1,60455997 | N15:480718 | 26,92 | +2:YOR084W; +8:YOR083W; | 2 | | | |
| chrXV | 480629 | 480781 | 152 | -1,971562384 | N15:480718 | 26,92 | +2:YOR084W; +8:YOR083W; | 2 | | | |
| chrXV | 480629 | 480782 | 153 | -1,631022204 | N15:480718 | 26,92 | +2:YOR084W; +8:YOR083W; | 2 | | | |
| chrXV | 480629 | 480783 | 154 | -1,537434115 | N15:480718 | 26,92 | +2:YOR084W; +8:YOR083W; | 2 | | | |
| chrXV | 480629 | 480784 | 155 | -1,470269874 | N15:480718 | 26,92 | +2:YOR084W; +8:YOR083W; | 2 | | | |
| chrXV | 480629 | 480785 | 156 | -1,739375556 | N15:480718 | 26,92 | +2:YOR084W; +8:YOR083W; | 2 | | | |
| chrXV | 480629 | 480786 | 157 | -1,58598926 | N15:480718 | 26,92 | +2:YOR084W; +8:YOR083W; | 2 | | | |
| chrXV | 480629 | 480787 | 158 | -1,005569653 | N15:480718 | 26,92 | +2:YOR084W; +8:YOR083W; | 2 | | | |
| chrXV | 480629 | 480788 | 159 | -2,779622115 | N15:480718 | 26,92 | +2:YOR084W; +8:YOR083W; | 2 | | | |
| chrXV | 480629 | 480789 | 160 | -1,553781909 | N15:480718 | 26,92 | +2:YOR084W; +8:YOR083W; | 2 | | | |
| chrXV | 480630 | 480787 | 157 | -1,421461779 | N15:480718 | 26,92 | +2:YOR084W; +8:YOR083W; | 2 | | | |
| chrXV | 480631 | 480787 | 156 | -1,240317616 | N15:480718 | 26,92 | +2:YOR084W; +8:YOR083W; | 2 | | | |
| chrXV | 480632 | 480787 | 155 | -1,626346486 | N15:480718 | 26,92 | +2:YOR084W; +8:YOR083W; | 2 | | | |
| chrXV | 480633 | 480787 | 154 | -1,446429767 | N15:480718 | 26,92 | +2:YOR084W; +8:YOR083W; | 2 | | | |
| chrXV | 480634 | 480787 | 153 | -1,194073162 | N15:480718 | 26,92 | +2:YOR084W; +8:YOR083W; | 2 | | | |
| chrXV | 480635 | 480787 | 152 | -1,898436973 | N15:480718 | 26,92 | +2:YOR084W; +8:YOR083W; | 2 | | | |
| chrXV | 480636 | 480787 | 151 | -1,739648531 | N15:480718 | 26,92 | +2:YOR084W; +8:YOR083W; | 2 | | | |
| chrXV | 480637 | 480787 | 150 | -1,572703578 | N15:480718 | 26,92 | +2:YOR084W; +8:YOR083W; | 2 | | | |
| chrXV | 480638 | 480787 | 149 | -1,662297273 | N15:480718 | 26,92 | +2:YOR084W; +8:YOR083W; | 2 | | | |
| chrXV | 480639 | 480787 | 148 | -1,340924261 | N15:480718 | 26,92 | +2:YOR084W; +8:YOR083W; | 2 | | | |
| chrXV | 480640 | 480787 | 147 | -1,73681727 | N15:480718 | 26,92 | +2:YOR084W; +8:YOR083W; | 2 | | | |
| chrXV | 480641 | 480787 | 146 | -0,94493 | N15:480718 | 26,92 | +2:YOR084W; +8:YOR083W; | 2 | | | |
| chrXV | 480642 | 480787 | 145 | -1,679102069 | N15:480718 | 26,92 | +2:YOR084W; +8:YOR083W; | 2 | | | |
| chrXV | 480643 | 480787 | 144 | -1,659589497 | N15:480718 | 26,92 | +2:YOR084W; +8:YOR083W; | 2 | | | |
| chrXV | 480644 | 480787 | 143 | -1,766861144 | N15:480718 | 26,92 | +2:YOR084W; +8:YOR083W; | 2 | | | |
| chrXV | 480645 | 480787 | 142 | -1,470075214 | N15:480718 | 26,92 | +2:YOR084W; +8:YOR083W; | 2 | | | |
| chrXV | 480646 | 480787 | 141 | -1,815668024 | N15:480718 | 26,92 | +2:YOR084W; +8:YOR083W; | 2 | | | |
| chrXV | 480647 | 480787 | 140 | -1,428032133 | N15:480718 | 26,92 | +2:YOR084W; +8:YOR083W; | 2 | | | |
| chrXV | 480648 | 480787 | 139 | -1,871642097 | N15:480718 | 26,92 | +2:YOR084W; +8:YOR083W; | 2 | | | |
| chrXV | 480649 | 480787 | 138 | -1,445458207 | N15:480718 | 26,92 | +2:YOR084W; +8:YOR083W; | 2 | | | |
| chrXV | 480650 | 480787 | 137 | -1,632534947 | N15:480718 | 26,92 | +2:YOR084W; +8:YOR083W; | 2 | | | |
| chrXV | 480651 | 480787 | 136 | -2,308724396 | N15:480718 | 26,92 | +2:YOR084W; +8:YOR083W; | 2 | | | |
| chrXV | 480652 | 480787 | 135 | -0,697838522 | N15:480718 | 26,92 | +2:YOR084W; +8:YOR083W; | 2 | | | |
| chrXV | 480653 | 480787 | 134 | -1,491723761 | N15:480718 | 26,92 | +2:YOR084W; +8:YOR083W; | 2 | | | |
| chrXV | 480654 | 480787 | 133 | -2,229488752 | N15:480718 | 26,92 | +2:YOR084W; +8:YOR083W; | 2 | | | |
| chrXV | 480655 | 480787 | 132 | -1,991583892 | N15:480718 | 26,92 | +2:YOR084W; +8:YOR083W; | 2 | | | |
| chrXV | 480656 | 480787 | 131 | -1,40811072 | N15:480718 | 26,92 | +2:YOR084W; +8:YOR083W; | 2 | | | |
| chrXV | 480657 | 480787 | 130 | -1,733972476 | N15:480718 | 26,92 | +2:YOR084W; +8:YOR083W; | 2 | | | |
| chrXV | 480658 | 480787 | 129 | -2,087844739 | N15:480718 | 26,92 | +2:YOR084W; +8:YOR083W; | 2 | | | |
| chrXV | 480659 | 480787 | 128 | -1,448509558 | N15:480718 | 26,92 | +2:YOR084W; +8:YOR083W; | 2 | | | |
| chrXV | 480660 | 480787 | 127 | -1,793477312 | N15:480718 | 26,92 | +2:YOR084W; +8:YOR083W; | 2 | | | |
| chrXV | 480661 | 480787 | 126 | -1,71378471 | N15:480718 | 26,92 | +2:YOR084W; +8:YOR083W; | 2 | | | |
| chrXV | 480662 | 480787 | 125 | -1,346997903 | N15:480718 | 26,92 | +2:YOR084W; +8:YOR083W; | 2 | | | |
| chrXV | 480663 | 480787 | 124 | -0,860282319 | N15:480718 | 26,92 | +2:YOR084W; +8:YOR083W; | 2 | | | |
| chrXV | 480664 | 480787 | 123 | -1,157819208 | N15:480718 | 26,92 | +2:YOR084W; +8:YOR083W; | 2 | | | |
| chrXV | 480665 | 480787 | 122 | -1,694605681 | N15:480718 | 26,92 | +2:YOR084W; +8:YOR083W; | 2 | | | |
| chrXV | 480666 | 480787 | 121 | -1,61084054 | N15:480718 | 26,92 | +2:YOR084W; +8:YOR083W; | 2 | | | |
| chrXV | 480667 | 480787 | 120 | -1,405153232 | N15:480718 | 26,92 | +2:YOR084W; +8:YOR083W; | 2 | | | |
| chrXV | 480668 | 480787 | 119 | -1,089638303 | N15:480718 | 26,92 | +2:YOR084W; +8:YOR083W; | 2 | | | |
| chrXV | 480669 | 480787 | 118 | -1,272321239 | N15:480718 | 26,92 | +2:YOR084W; +8:YOR083W; | 2 | | | |
| chrXV | 480670 | 480787 | 117 | -1,595672565 | N15:480718 | 26,92 | +2:YOR084W; +8:YOR083W; | 2 | | | |
| chrXV | 480671 | 480787 | 116 | -0,95120699 | N15:480718 | 26,92 | +2:YOR084W; +8:YOR083W; | 2 | | | |
| chrXV | 480672 | 480787 | 115 | -0,254454624 | N15:480718 | 26,92 | +2:YOR084W; +8:YOR083W; | 2 | | | |
| chrXV | 480673 | 480787 | 114 | -1,153879603 | N15:480718 | 26,92 | +2:YOR084W; +8:YOR083W; | 2 | | | |
| chrXV | 480674 | 480787 | 113 | -1,342981955 | N15:480718 | 26,92 | +2:YOR084W; +8:YOR083W; | 2 | | | |
| chrXV | 480675 | 480787 | 112 | -1,17343774 | N15:480718 | 26,92 | +2:YOR084W; +8:YOR083W; | 2 | | | |
| chrXV | 480676 | 480787 | 111 | -0,980228119 | N15:480718 | 26,92 | +2:YOR084W; +8:YOR083W; | 2 | | | |
| chrXV | 480677 | 480787 | 110 | -0,293003387 | N15:480718 | 26,92 | +2:YOR084W; +8:YOR083W; | 2 | | | |
| chrXV | 480678 | 480787 | 109 | -1,301535234 | N15:480718 | 26,92 | +2:YOR084W; +8:YOR083W; | 2 | | | |
| chrXV | 480679 | 480787 | 108 | -0,87559958 | N15:480718 | 26,92 | +2:YOR084W; +8:YOR083W; | 2 | | | |
| chrXV | 480680 | 480755 | 75 | -0,777 | N15:480718 | 26,92 | +2:YOR084W; +8:YOR083W; | 2 | | | |
| chrXV | 480680 | 480787 | 107 | -1,6540568 | N15:480718 | 26,92 | +2:YOR084W; +8:YOR083W; | 2 | | | |
| chrXV | 480681 | 480787 | 106 | -1,028788184 | N15:480718 | 26,92 | +2:YOR084W; +8:YOR083W; | 2 | | | |
| chrXV | 480682 | 480787 | 105 | -1,063120693 | N15:480718 | 26,92 | +2:YOR084W; +8:YOR083W; | 2 | | | |
| chrXV | 480683 | 480787 | 104 | -1,02210638 | N15:480718 | 26,92 | +2:YOR084W; +8:YOR083W; | 2 | | | |
| chrXV | 480684 | 480787 | 103 | -1,088385469 | N15:480718 | 26,92 | +2:YOR084W; +8:YOR083W; | 2 | | | |
| chrXV | 480685 | 480787 | 102 | -1,31204778 | N15:480718 | 26,92 | +2:YOR084W; +8:YOR083W; | 2 | | | |
| chrXV | 480686 | 480787 | 101 | -1,45202532 | N15:480718 | 26,92 | +2:YOR084W; +8:YOR083W; | 2 | | | |
| chrXV | 480687 | 480787 | 100 | -1,105051229 | N15:480718 | 26,92 | +2:YOR084W; +8:YOR083W; | 2 | | | |
| chrXV | 480688 | 480787 | 99 | -1,311677074 | N15:480718 | 26,92 | +2:YOR084W; +8:YOR083W; | 2 | | | |
| chrXV | 480689 | 480787 | 98 | -0,483471549 | N15:480718 | 26,92 | +2:YOR084W; +8:YOR083W; | 2 | | | |
| chrXV | 480690 | 480787 | 97 | -1,535017791 | N15:480718 | 26,92 | +2:YOR084W; +8:YOR083W; | 2 | | | |
| chrXV | 480691 | 480787 | 96 | -1,26116167 | N15:480718 | 26,92 | +2:YOR084W; +8:YOR083W; | 2 | | | |
| chrXV | 480692 | 480787 | 95 | -1,434485007 | N15:480718 | 26,92 | +2:YOR084W; +8:YOR083W; | 2 | | | |
| chrXV | 480693 | 480787 | 94 | -1,437455211 | N15:480718 | 26,92 | +2:YOR084W; +8:YOR083W; | 2 | | | |
| chrXV | 480695 | 480787 | 92 | -2,026573821 | N15:480718 | 26,92 | +2:YOR084W; +8:YOR083W; | 2 | | | |
| chrXV | 480696 | 480787 | 91 | -1,46093622 | N15:480718 | 26,92 | +2:YOR084W; +8:YOR083W; | 2 | | | |
| chrXV | 480697 | 480787 | 90 | -1,918199292 | N15:480718 | 26,92 | +2:YOR084W; +8:YOR083W; | 2 | | | |
| chrXV | 480698 | 480787 | 89 | -1,060230518 | N15:480718 | 26,92 | +2:YOR084W; +8:YOR083W; | 2 | | | |
| chrXV | 480700 | 480787 | 87 | -1,504085721 | N15:480718 | 26,92 | +2:YOR084W; +8:YOR083W; | 2 | | | |
| chrXV | 480701 | 480787 | 86 | -2,370110132 | N15:480718 | 26,92 | +2:YOR084W; +8:YOR083W; | 2 | | | |
| chrXV | 480703 | 480787 | 84 | -1,388925352 | N15:480718 | 26,92 | +2:YOR084W; +8:YOR083W; | 2 | | | |
| chrXV | 480704 | 480787 | 83 | -1,660161518 | N15:480718 | 26,92 | +2:YOR084W; +8:YOR083W; | 2 | | | |
| chrXV | 482098 | 482249 | 151 | -1,589170937 | N15:482205 | 8,05 | +2:YOR085W; | 2 | | | |
| chrXV | 483506 | 483656 | 150 | -1,128814692 | N15:483559 | 37,401 | +22:YOR086C; | 22 | | | |
| chrXV | 490437 | 490586 | 149 | -1,227441921 | | | | Overlap <50 bp | | | |
| chrXV | 492152 | 492296 | 144 | -1,287616616 | | | | Overlap <50 bp | | | |
| chrXV | 492636 | 492766 | 130 | -1,122285526 | N15:492727 | 11,535 | +2:YOR090C; | 2 | | | |
| chrXV | 499981 | 500137 | 156 | -0,752649484 | N15:499985 | 29,69 | | Intergene | | | |
| chrXV | 500251 | 500392 | 141 | -1,08025578 | N15:500266 | 37,264 | | Intergene | | | |
| chrXV | 500870 | 501031 | 161 | -1,25000077 | N15:500992 | 38,972 | | Intergene | | | |
| chrXV | 504656 | 504829 | 173 | -1,393156044 | N15:504757 | 7,985 | -1:YOR095C; | -1 | | | |
| chrXV | 504839 | 504995 | 156 | -1,449906994 | N15:504924 | 6,573 | | Intergene | | | |
| chrXV | 505651 | 505827 | 176 | -1,557242507 | N15:505793 | 30,433 | +1:YOR096W; | 1 | | | |
| chrXV | 508541 | 508687 | 146 | -1,105428668 | N15:508522 | 35,529 | +18:YOR098C; | 18 | | | |
| chrXV | 512017 | 512169 | 152 | -1,388824829 | N15:512077 | 11,221 | +3:YOR099W; | 3 | | | |
| chrXV | 520467 | 520627 | 160 | -1,301870465 | N15:520577 | 0,707 | -1:CUT843; | -1 | | | |
| chrXV | 521328 | 521487 | 159 | -1,138158843 | N15:521406 | 17,512 | +3:YOR107W; | 3 | | | |

TABLE 3

| CHR | START | END | L(bp) | ΔLK | Nuc ID | Fuzziness | Gene ID | Gene body postion or intergene | Terminal | rDNA | Telomeric |
|---|---|---|---|---|---|---|---|---|---|---|---|
| chrXV | 524434 | 524584 | 150 | -1,379271275 | N15:524552 | 11,303 | +11:YOR108W; | 11 | | | |
| chrXV | 526545 | 526704 | 159 | -0,91509561 | N15:526589 | 22,811 | +10:YOR109W; | 10 | | | |
| chrXV | 527282 | 527433 | 151 | -1,49865849 | N15:527364 | 13,18 | +15:YOR109W; | 15 | | | |
| chrXV | 528345 | 528493 | 148 | -1,285254183 | N15:528465 | 16,461 | +22:YOR109W; | 22 | | | |
| chrXV | 529088 | 529208 | 120 | -1,094024708 | | | | Overlap <50 bp | | | |
| chrXV | 534111 | 534263 | 152 | -1,018551216 | N15:534188 | 4,494 | +2:YOR113W; | 2 | | | |
| chrXV | 535122 | 535279 | 157 | -1,175776825 | | | | Overlap <50 bp | | | |
| chrXV | 539793 | 539942 | 149 | -1,173714924 | N15:539889 | 19,429 | | Intergene | | | |
| chrXV | 540123 | 540277 | 154 | -1,557816261 | N15:540228 | 21,037 | +26:YOR116C; | 26 | | | |
| chrXV | 542672 | 542801 | 129 | -1,685596178 | N15:542768 | 19,242 | +11:YOR116C; | 11 | | | |
| chrXV | 545231 | 545391 | 160 | -1,373952932 | N15:545321 | 11,589 | +3:YOR117W; | 3 | | | |
| chrXV | 545588 | 545715 | 127 | -0,960989749 | N15:545625 | 8,302 | +5:YOR117W; | 5 | | | |
| chrXV | 550006 | 550155 | 149 | -1,452809124 | N15:550070 | 5,776 | +2:YOR119C; | 2 | | | |
| chrXV | 550413 | 550558 | 145 | -1,263444964 | N15:550502 | 8,472 | -1:YOR119C; | -1 | | | |
| chrXV | 550604 | 550752 | 148 | -1,131553601 | N15:550670 | 15,946 | | Intergene | | | |
| chrXV | 559497 | 559585 | 88 | -1,099368473 | N15:559589 | 8,764 | +2:YOR125C; | 2 | | | |
| chrXV | 559497 | 559598 | 101 | -0,457138739 | N15:559589 | 8,764 | +2:YOR125C; | 2 | | | |
| chrXV | 559497 | 559601 | 104 | -0,811486186 | N15:559589 | 8,764 | +2:YOR125C; | 2 | | | |
| chrXV | 559497 | 559654 | 157 | -1,345065945 | N15:559589 | 8,764 | +2:YOR125C; | 2 | | | |
| chrXV | 559919 | 560068 | 149 | -0,900188758 | N15:559990 | 9,136 | +5*:YOR126C; -1:YOR125C; | 5 | TERM | | |
| chrXV | 559929 | 560086 | 157 | -0,934906742 | N15:559990 | 9,136 | +5*:YOR126C; -1:YOR125C; | 5 | TERM | | |
| chrXV | 565053 | 565199 | 146 | -1,797448662 | N15:565222 | 25,393 | +7*:YOR128C; | 7 | TERM | | |
| chrXV | 565053 | 565201 | 148 | -2,394599187 | N15:565222 | 25,393 | +7*:YOR128C; | 7 | TERM | | |
| chrXV | 565053 | 565202 | 149 | -2,014650298 | N15:565222 | 25,393 | +7*:YOR128C; | 7 | TERM | | |
| chrXV | 566775 | 566926 | 151 | -1,272484346 | N15:566870 | 23,51 | +18*:YOR129C; | 18 | TERM | | |
| chrXV | 576392 | 576555 | 163 | -1,500433993 | N15:576489 | 38,592 | +9:YOR133W; | 9 | | | |
| chrXV | 590114 | 590263 | 149 | -1,397404152 | N15:590176 | 16,104 | +16:YOR141C; | 16 | | | |
| chrXV | 590721 | 590873 | 152 | -1,167212396 | N15:590856 | 36,321 | +11:YOR141C; | 11 | | | |
| chrXV | 592017 | 592170 | 153 | -1,21423786 | N15:592096 | 28,268 | +4:YOR141C; | 4 | | | |
| chrXV | 592017 | 592179 | 162 | -1,262648092 | N15:592096 | 28,268 | +4:YOR141C; | 4 | | | |
| chrXV | 592023 | 592179 | 156 | -1,079283922 | N15:592096 | 28,268 | +4:YOR141C; | 4 | | | |
| chrXV | 592044 | 592126 | 82 | -0,826 | N15:592096 | 28,268 | +4:YOR141C; | 4 | | | |
| chrXV | 595352 | 595475 | 123 | -0,903193347 | | | | Overlap <50 bp | | | |
| chrXV | 595452 | 595607 | 155 | -1,42336846 | N15:595551 | 22,068 | | Intergene | | | |
| chrXV | 595885 | 596019 | 134 | -0,66626228 | N15:595950 | 30,643 | | Intergene | | | |
| chrXV | 595885 | 596025 | 140 | -0,970297117 | N15:595950 | 30,643 | | Intergene | | | |
| chrXV | 596928 | 597063 | 135 | -0,809359546 | N15:596968 | 29,195 | | Intergene | | | |
| chrXV | 597189 | 597334 | 145 | -1,009440189 | N15:597178 | 5,657 | | Intergene | | | |
| chrXV | 597375 | 597525 | 150 | -1,207102726 | N15:597445 | 14,048 | | Intergene | | | |
| chrXV | 597554 | 597711 | 157 | -1,20557002 | N15:597645 | 29,513 | | Intergene | | | |
| chrXV | 598045 | 598174 | 129 | -1,090487943 | | | | Overlap <50 bp | | | |
| chrXV | 598957 | 599107 | 150 | -1,361578722 | N15:599055 | 14,572 | | Intergene | | | |
| chrXV | 600156 | 600263 | 107 | -1,082555398 | | | | Overlap <50 bp | | | |
| chrXV | 601179 | 601328 | 149 | -1,130738646 | N15:601265 | 12,561 | | Intergene | | | |
| chrXV | 603463 | 603616 | 153 | -1,268555078 | N15:603547 | 12,629 | +10:YOR144C; | 10 | | | |
| chrXV | 605449 | 605600 | 151 | -1,133822925 | N15:605543 | 14,944 | +2*:CUT384; +5:YOR145C; | 2 | TERM | | |
| chrXV | 606728 | 606884 | 156 | -1,307831664 | N15:606775 | 5,128 | +2:YOR147W; | 2 | | | |
| chrXV | 608095 | 608251 | 156 | -1,512299031 | N15:608159 | 32,269 | +11:YOR147W; | 11 | | | |
| chrXV | 610131 | 610285 | 154 | -1,216648706 | N15:610149 | 21,151 | +9:YOR149C; | 9 | | | |
| chrXV | 610432 | 610587 | 155 | -1,175664486 | N15:610582 | 10,45 | +6:YOR149C; | 6 | | | |
| chrXV | 610432 | 610587 | 155 | -1,175664486 | N15:610423 | 18,327 | +7:YOR149C; | 7 | | | |
| chrXV | 613403 | 613549 | 146 | -1,484149098 | N15:613529 | 27,017 | +21:YOR151C; | 21 | | | |
| chrXV | 617632 | 617793 | 161 | -1,189892761 | N15:617725 | 15,758 | +4:YOR152C; | 4 | | | |
| chrXV | 621001 | 621151 | 150 | -1,319977946 | N15:621077 | 32,654 | +10:YOR153W; | 10 | | | |
| chrXV | 622311 | 622441 | 130 | -1,013963775 | | | | Overlap <50 bp | | | |
| chrXV | 624028 | 624159 | 131 | -0,826285892 | N15:624066 | 28,715 | +26:YOR153W; | 26 | | | |
| chrXV | 625834 | 625959 | 125 | -0,958346872 | N15:625885 | 12,285 | +8:YOR154W; | 8 | | | |
| chrXV | 626417 | 626570 | 153 | -1,31788744 | N15:626508 | 32,039 | +12*:YOR154W; | 12 | TERM | | |
| chrXV | 632144 | 632251 | 107 | -0,733677856 | N15:632173 | 3,619 | +1:YOR158W; | 1 | | | |
| chrXV | 632144 | 632253 | 109 | -0,869651273 | N15:632173 | 3,619 | +1:YOR158W; | 1 | | | |
| chrXV | 633485 | 633638 | 153 | -1,251462342 | N15:633575 | 8,468 | -1:YOR160W; +1:YOR159C; | -1 | | | |
| chrXV | 635429 | 635578 | 149 | -1,34659384 | N15:635550 | 24,098 | +11:YOR160W; | 11 | | | |
| chrXV | 637512 | 637669 | 157 | -1,319490145 | N15:637581 | 32,836 | +7:YOR161C; | 7 | | | |
| chrXV | 641886 | 642041 | 155 | -1,375090187 | N15:641956 | 15,201 | +2:YOR162C; | 2 | | | |
| chrXV | 646176 | 646338 | 162 | -1,370502659 | N15:646189 | 23,384 | +11:YOR165W; | 11 | | | |
| chrXV | 647481 | 647636 | 155 | -1,378866228 | N15:647630 | 14,061 | +6:YOR166C; | 6 | | | |
| chrXV | 650796 | 650922 | 126 | -0,809272259 | | | | Overlap <50 bp | | | |
| chrXV | 650923 | 651080 | 157 | -1,586685502 | N15:650997 | 13,704 | +12:YOR168W; | 12 | | | |
| chrXV | 651588 | 651740 | 152 | -1,125245221 | N15:651687 | 17,587 | | Intergene | | | |
| chrXV | 671862 | 672011 | 149 | -1,204001753 | N15:671896 | 6,408 | +4:YOR179C; | 4 | | | |
| chrXV | 674697 | 674849 | 152 | -0,991728093 | N15:674786 | 33,65 | +3:YOR180C; | 3 | | | |
| chrXV | 674884 | 675018 | 134 | -0,85652054 | N15:674983 | 36,793 | +2:YOR180C; | 2 | | | |
| chrXV | 677635 | 677796 | 161 | -1,243137998 | N15:677669 | 31,321 | -1:anti176; +15:YOR181W; | -1 | | | |
| chrXV | 679372 | 679525 | 153 | -1,274887952 | N15:679447 | 6,314 | +2:YOR183W; +2:YOR184W; | 2 | | | |
| chrXV | 680555 | 680689 | 134 | -0,987564336 | N15:680616 | 11,06 | +9*:YOR183W; +9*:YOR184W; | 9 | TERM | | |
| chrXV | 682390 | 682545 | 155 | -1,282906148 | N15:682487 | 7,106 | | Intergene | | | |
| chrXV | 682421 | 682570 | 149 | -1,13095704 | N15:682487 | 7,106 | | Intergene | | | |
| chrXV | 682600 | 682746 | 146 | -0,981373097 | N15:682669 | 23,768 | | Intergene | | | |
| chrXV | 683178 | 683329 | 151 | -1,403272096 | N15:683179 | 19,274 | +2:YOR186W; | 2 | | | |
| chrXV | 683178 | 683329 | 151 | -1,403272096 | N15:683348 | 39,191 | +3:YOR186W; | 3 | | | |
| chrXV | 683542 | 683705 | 163 | -1,44124454 | N15:683602 | 15,188 | +5:YOR186W; | 5 | | | |
| chrXV | 698388 | 698544 | 156 | -1,054337602 | N15:698448 | 9,02 | | Intergene | | | |
| chrXV | 699187 | 699320 | 133 | -1,367216594 | N15:699226 | 40,166 | +9:YOR192C; | 9 | | | |
| chrXV | 704256 | 704405 | 149 | -1,188485336 | N15:704277 | 28,583 | +1:YOR192C-C; | 1 | | | |
| chrXV | 705052 | 705198 | 146 | -1,179350103 | N15:705134 | 6,429 | | Intergene | | | |
| chrXV | 707863 | 708011 | 148 | -1,107693705 | | | | Overlap <50 bp | | | |
| chrXV | 709269 | 709434 | 165 | -1,212752401 | N15:709269 | 19,95 | | Intergene | | | |
| chrXV | 709269 | 709434 | 165 | -1,212752401 | N15:709445 | 21,213 | | Intergene | | | |
| chrXV | 709269 | 709462 | 193 | -1,219461387 | N15:709269 | 19,95 | | Intergene | | | |
| chrXV | 709269 | 709462 | 193 | -1,219461387 | N15:709445 | 21,213 | | Intergene | | | |
| chrXV | 709332 | 709434 | 102 | -0,10743918 | N15:709445 | 21,213 | | Intergene | | | |
| chrXV | 709332 | 709462 | 130 | -0,913391215 | N15:709445 | 21,213 | | Intergene | | | |
| chrXV | 709396 | 709550 | 154 | -1,216713811 | N15:709445 | 21,213 | | Intergene | | | |
| chrXV | 709396 | 709571 | 175 | -1,399936269 | N15:709445 | 21,213 | | Intergene | | | |
| chrXV | 709403 | 709571 | 168 | -1,303875346 | N15:709445 | 21,213 | | Intergene | | | |
| chrXV | 712148 | 712278 | 130 | -0,972198581 | N15:712206 | 4,916 | +3:YOR194C; | 3 | | | |
| chrXV | 713754 | 713908 | 154 | -1,362633769 | | | | Overlap <50 bp | | | |
| chrXV | 716082 | 716234 | 152 | -1,267125867 | N15:716164 | 13,59 | +5:YOR196C; | 5 | | | |

| CHR | START | END | L(bp) | ΔLK | Nuc ID | Fuzziness | Gene ID | Gene body postion or intergene | Terminal | rDNA | Telomeric |
|---|---|---|---|---|---|---|---|---|---|---|---|
| chrXV | 717294 | 717452 | 158 | -1,459867322 | N15:717288 | 5,391 | +2:YOR197W; | 2 | | | |
| chrXV | 717294 | 717452 | 158 | -1,459867322 | N15:717452 | 4,412 | +3:YOR197W; | 3 | | | |
| chrXV | 728130 | 728275 | 145 | -1,14756473 | N15:728119 | 15,822 | +5:YOR206W; | 5 | | | |
| chrXV | 728130 | 728275 | 145 | -1,14756473 | N15:728288 | 8,735 | +6:YOR206W; | 6 | | | |
| chrXV | 729241 | 729409 | 168 | -1,397940251 | N15:729229 | 32,517 | +12:YOR206W; | 12 | | | |
| chrXV | 730369 | 730527 | 158 | -1,430422755 | N15:730492 | 43,783 | +18:YOR207C; | 18 | | | |
| chrXV | 730626 | 730772 | 146 | -1,150314594 | N15:730666 | 35,066 | +17:YOR207C; | 17 | | | |
| chrXV | 732904 | 733026 | 122 | -0,924302228 | N15:732962 | 10,61 | +4:YOR207C; | 4 | | | |
| chrXV | 736114 | 736267 | 153 | -1,646749186 | N15:736157 | 26,475 | +15*:YOR208W; | 15 | TERM | | |
| chrXV | 749719 | 749841 | 122 | -1,127719545 | N15:749778 | 6,555 | +4:YOR217W; | 4 | | | |
| chrXV | 751696 | 751803 | 107 | -0,841257138 | N15:751682 | 34,112 | +16:YOR217W; | 16 | | | |
| chrXV | 751696 | 751804 | 108 | -0,697615779 | N15:751682 | 34,112 | +16:YOR217W; | 16 | | | |
| chrXV | 751696 | 751805 | 109 | -0,915838655 | N15:751682 | 34,112 | +16:YOR217W; | 16 | | | |
| chrXV | 751696 | 751808 | 112 | -0,869910856 | N15:751682 | 34,112 | +16:YOR217W; | 16 | | | |
| chrXV | 759323 | 759474 | 151 | -1,259653474 | N15:759342 | 12,219 | | Intergene | | | |
| chrXV | 759886 | 760039 | 153 | -1,69044603 | N15:759976 | 11,29 | +2:YOR223W; | 2 | | | |
| chrXV | 762029 | 762161 | 132 | -0,556527918 | N15:762082 | 8,462 | +1:YOR226C; | 1 | | | |
| chrXV | 762283 | 762425 | 142 | -0,934806715 | N15:762373 | 18,815 | -1:YOR227W; -1:YOR226C; | -1 | | | |
| chrXV | 763377 | 763524 | 147 | -1,25323354 | N15:763477 | 13,164 | +6:YOR227W; | 6 | | | |
| chrXV | 765404 | 765530 | 126 | -1,339529827 | N15:765450 | 31,757 | +17:YOR227W; | 17 | | | |
| chrXV | 766078 | 766228 | 150 | -1,138452638 | N15:766247 | 28,768 | | Intergene | | | |
| chrXV | 767013 | 767155 | 142 | -0,848927951 | N15:767164 | 18,501 | +6:YOR228C; | 6 | | | |
| chrXV | 767013 | 767155 | 142 | -0,848927951 | N15:767010 | 10,689 | +7:YOR228C; | 7 | | | |
| chrXV | 768735 | 768888 | 153 | -1,135017522 | N15:768713 | 6,066 | +3:YOR229W; | 3 | | | |
| chrXV | 770268 | 770418 | 150 | -0,842017312 | | | | Overlap <50 bp | | | |
| chrXV | 773124 | 773277 | 153 | -1,387038035 | N15:773165 | 8,86 | +5:YOR231W; | 5 | | | |
| chrXV | 775176 | 775297 | 121 | -1,097729609 | N15:775265 | 23,329 | +6*:YOR232W; +2:CUT860; | 6 | TERM | | |
| chrXV | 777968 | 778119 | 151 | -1,479257799 | N15:778110 | 24,499 | +16*:YOR233W; | 16 | TERM | | |
| chrXV | 783192 | 783285 | 93 | -0,384899711 | N15:783252 | 18,833 | +10:YOR237W; | 10 | | | |
| chrXV | 789013 | 789147 | 134 | -1,027739313 | N15:789066 | 4,796 | +6:YOR242C; | 6 | | | |
| chrXV | 790833 | 790987 | 154 | -1,502707383 | N15:790822 | 41,429 | +10:YOR243C; | 10 | | | |
| chrXV | 794701 | 794852 | 151 | -1,186773581 | | | | Overlap <50 bp | | | |
| chrXV | 796032 | 796168 | 136 | -0,862085404 | N15:796058 | 27,477 | +6:YOR246C; | 6 | | | |
| chrXV | 801709 | 801857 | 148 | -1,196312645 | N15:801792 | 9,854 | +4:YOR250C; | 4 | | | |
| chrXV | 802168 | 802313 | 145 | -1,141260029 | N15:802287 | 9,121 | -1:anti-YOR251C; +1:YOR250C; | -1 | | | |
| chrXV | 804681 | 804839 | 158 | -1,28977132 | N15:804764 | 16,916 | +1:SUT788; +3*:YOR253W; | 1 | | | |
| chrXV | 805157 | 805300 | 143 | -1,011056665 | N15:805287 | 18,794 | +12:YOR254C; | 12 | | | |
| chrXV | 806800 | 806942 | 142 | -1,476985644 | N15:806847 | 6,309 | +2:YOR254C; | 2 | | | |
| chrXV | 806939 | 807090 | 151 | -1,232779546 | N15:807004 | 13,971 | -1:YOR255W; +1:YOR254C; | -1 | | | |
| chrXV | 810091 | 810222 | 131 | -0,865174184 | N15:810121 | 29,58 | +5:YOR256C; | 5 | | | |
| chrXV | 815641 | 815767 | 126 | -1,381179464 | N15:815726 | 31,945 | +12*:YOR260W; | 12 | TERM | | |
| chrXV | 816418 | 816570 | 152 | -0,990691825 | N15:816506 | 14,905 | +4:YOR261C; | 4 | | | |
| chrXV | 816891 | 817023 | 132 | -1,274052286 | N15:817009 | 12,738 | +1:YOR261C; -1:YOR262W; | 1 | | | |
| chrXV | 823212 | 823372 | 160 | -1,431096545 | N15:823302 | 8,699 | +13:YOR267C; | 13 | | | |
| chrXV | 823715 | 823880 | 165 | -1,269891588 | N15:823900 | 30,616 | +9:YOR267C; | 9 | | | |
| chrXV | 824770 | 824918 | 148 | -1,633170593 | N15:824827 | 14,414 | +3:YOR267C; | 3 | | | |
| chrXV | 824898 | 825045 | 147 | -1,218761175 | N15:824984 | 11,454 | +2:YOR267C; | 2 | | | |
| chrXV | 826750 | 826899 | 149 | -1,540544788 | N15:826822 | 8,819 | +4:YOR269W; | 4 | | | |
| chrXV | 828443 | 828590 | 147 | -1,081462881 | N15:828514 | 14,39 | +14:YOR270C; | 14 | | | |
| chrXV | 829595 | 829749 | 154 | -1,22598748 | N15:829756 | 17,773 | +7:YOR270C; | 7 | | | |
| chrXV | 832011 | 832158 | 147 | -1,090955345 | N15:832060 | 7,139 | +2*:SUT790; +1:YOR271C; | 2 | TERM | | |
| chrXV | 838047 | 838199 | 152 | -1,161609891 | N15:838035 | 12,703 | +4:YOR274W; | 4 | | | |
| chrXV | 838047 | 838199 | 152 | -1,161609891 | N15:838195 | 8,468 | +5:YOR274W; | 5 | | | |
| chrXV | 838749 | 838897 | 148 | -1,068216573 | N15:838866 | 26,12 | +9:YOR274W; | 9 | | | |
| chrXV | 840989 | 841143 | 154 | -1,158306315 | N15:841051 | 10,147 | -1:YOR276W; +1:YOR275C; | -1 | | | |
| chrXV | 845931 | 846091 | 160 | -1,43298733 | N15:845986 | 13,292 | -1:YOR280C; | -1 | | | |
| chrXV | 851758 | 851911 | 153 | -1,552122585 | N15:851815 | 9,524 | +1:YOR287C; | 1 | | | |
| chrXV | 855604 | 855755 | 151 | -1,197852569 | N15:855699 | 29,815 | +29:YOR290C; | 29 | | | |
| chrXV | 856174 | 856339 | 165 | -1,268121005 | N15:856207 | 14,44 | +26:YOR290C; | 26 | | | |
| chrXV | 856222 | 856364 | 142 | -0,893837762 | N15:856207 | 14,44 | +26:YOR290C; | 26 | | | |
| chrXV | 860271 | 860423 | 152 | -1,564365666 | N15:860352 | 4,561 | +1:YOR290C; | 1 | | | |
| chrXV | 863117 | 863234 | 117 | -1,433576111 | N15:863177 | 11,759 | +15:YOR291W; | 15 | | | |
| chrXV | 866318 | 866474 | 156 | -1,229193294 | N15:866417 | 12,28 | +2:YOR292C; | 2 | | | |
| chrXV | 866636 | 866779 | 143 | -0,783495597 | | | | Overlap <50 bp | | | |
| chrXV | 873282 | 873420 | 138 | -0,864242621 | N15:873312 | 10,858 | +20:YOR296W; | 20 | | | |
| chrXV | 873426 | 873558 | 132 | -0,944089034 | N15:873537 | 26,853 | +21:YOR296W; | 21 | | | |
| chrXV | 873491 | 873643 | 152 | -1,507558695 | N15:873537 | 26,853 | +21:YOR296W; | 21 | | | |
| chrXV | 883077 | 883230 | 153 | -1,077114008 | N15:883079 | 18,005 | +3:YOR303W; +3:YOR302W; | 3 | | | |
| chrXV | 883077 | 883230 | 153 | -1,077114008 | N15:883239 | 22,73 | +4:YOR303W; +4:YOR302W; | 4 | | | |
| chrXV | 884043 | 884200 | 157 | -1,515341449 | N15:884168 | 36,873 | 303W; +9*:YOR302W; -1:YOR304W; | 9 | TERM | | |
| chrXV | 895782 | 895933 | 151 | -1,324438459 | N15:895865 | 5,086 | +4:YOR308C; | 4 | | | |
| chrXV | 900646 | 900802 | 156 | -1,691263989 | N15:900717 | 12,137 | +4*:YOR312C; | 4 | TERM | | |
| chrXV | 902699 | 902828 | 129 | -1,201014529 | N15:902765 | 15,329 | +2:YOR313C; | 2 | | | |
| chrXV | 904931 | 905079 | 148 | -1,307752677 | N15:904922 | 20,404 | +2:YOR315W; | 2 | | | |
| chrXV | 917190 | 917343 | 153 | -1,130239538 | N15:917323 | 14,184 | +9:YOR321W; | 9 | | | |
| chrXV | 920557 | 920665 | 108 | -1,208185328 | N15:920598 | 9,731 | +5:YOR322C; | 5 | | | |
| chrXV | 928849 | 929014 | 165 | -1,489350113 | N15:928971 | 4,655 | +21:YOR326W; | 21 | | | |
| chrXV | 932648 | 932807 | 159 | -1,227006821 | N15:932684 | 31,149 | +7:YOR328W; | 7 | | | |
| chrXV | 932960 | 933112 | 152 | -1,159570289 | N15:933085 | 17,672 | +10:YOR328W; | 10 | | | |
| chrXV | 937286 | 937429 | 143 | -1,123638198 | N15:937309 | 27,898 | +14:YOR329C; +4:SUT383; | 14 | | | |
| chrXV | 941867 | 942022 | 155 | -1,424698493 | N15:941903 | 38,923 | +10:YOR330C; | 10 | | | |
| chrXV | 943142 | 943285 | 143 | -0,802926171 | N15:943207 | 14,3 | +2:YOR330C; | 2 | | | |
| chrXV | 956493 | 956644 | 151 | -1,585012508 | N15:956543 | 16,171 | +14:YOR337W; -1:YOR338W; | 14 | | | |
| chrXV | 958715 | 958860 | 145 | -0,95123906 | N15:958840 | 31,909 | *:SUT384; +1:YOR339C; -1:Unit611; | 4 | TERM | | |
| chrXV | 961158 | 961313 | 155 | -1,58620315 | N15:961224 | 14,445 | +3:YOR341W; | 3 | | | |
| chrXV | 963324 | 963474 | 150 | -1,566831515 | N15:963309 | 7,455 | +16:YOR341W; | 16 | | | |
| chrXV | 967141 | 967307 | 166 | -1,663405161 | N15:967161 | 27,577 | +4:YOR342C; | 4 | | | |
| chrXV | 967454 | 967605 | 151 | -1,459313211 | N15:967556 | 20,229 | +2:YOR342C; | 2 | | | |
| chrXV | 970738 | 970906 | 168 | -1,399703551 | N15:970789 | 9,713 | | Intergene | | | |
| chrXV | 970759 | 970913 | 154 | -1,381936832 | N15:970789 | 9,713 | | Intergene | | | |
| chrXV | 970847 | 970977 | 130 | -0,941875395 | | | | Overlap <50 bp | | | |
| chrXV | 970875 | 970977 | 102 | -0,254934627 | | | | Overlap <50 bp | | | |
| chrXV | 972298 | 972446 | 148 | -1,13376193 | | | | Overlap <50 bp | | | |
| chrXV | 975111 | 975257 | 146 | -1,203462867 | | | | Overlap <50 bp | | | |
| chrXV | 975228 | 975386 | 158 | -1,639351108 | N15:975305 | 32,512 | | Intergene | | | |
| chrXV | 975904 | 976053 | 149 | -1,244179865 | N15:975989 | 28,042 | | Intergene | | | |
| chrXV | 975908 | 976058 | 150 | -1,35249103 | N15:975989 | 28,042 | | Intergene | | | |
| chrXV | 977285 | 977436 | 151 | -1,344094817 | N15:977376 | 19,483 | +6:YOR344C; | 6 | | | |
| chrXV | 977616 | 977761 | 145 | -1,004950594 | | | | Overlap <50 bp | | | |

TABLE 3

| CHR | START | END | L(bp) | ΔLK | Nuc ID | Fuzziness | Gene ID | Gene body postion or intergene | Terminal | rDNA | Telomeric |
|---|---|---|---|---|---|---|---|---|---|---|---|
| chrXV | 981987 | 982146 | 159 | -1,64885498 | N15:982019 | 4,393 | +3:YOR346W; | 3 | | | |
| chrXV | 986417 | 986558 | 141 | -1,162183004 | N15:986505 | 5,076 | +1:YOR347C; | 1 | | | |
| chrXV | 986699 | 986864 | 165 | -1,561544023 | N15:986780 | 28,922 | -1:YOR347C; +13*:YOR348C; | -1 | | | |
| chrXV | 987920 | 988058 | 138 | -2,083581141 | N15:987920 | 30,887 | +6:YOR348C; | 6 | | | |
| chrXV | 988983 | 989114 | 131 | -0,967900218 | N15:989106 | 25,548 | -1:YOR348C; +3:SUT798; | -1 | | | |
| chrXV | 989889 | 990033 | 144 | -1,282460001 | N15:989965 | 12,545 | +2:YOR349W; | 2 | | | |
| chrXV | 996318 | 996470 | 152 | -1,085585229 | N15:996407 | 7,574 | +2:YOR351C; | 2 | | | |
| chrXV | 997102 | 997264 | 162 | -1,582654014 | N15:997152 | 7,797 | +1:YOR352W; | 1 | | | |
| chrXV | 1000108 | 1000248 | 140 | -1,178900147 | | | | Overlap <50 bp | | | |
| chrXV | 1000780 | 1000917 | 137 | -1,031045705 | N15:1000816 | 4,167 | +1:YOR353C; | 1 | | | |
| chrXV | 1001402 | 1001553 | 151 | -1,222977032 | N15:1001454 | 10,083 | +3*:CUT396; +11*:YOR354C; | 3 | TERM | | |
| chrXV | 1002082 | 1002260 | 178 | -1,856832932 | N15:1002190 | 48,149 | +7:YOR354C; | 7 | | | |
| chrXV | 1004460 | 1004618 | 158 | -1,56234428 | N15:1004454 | 50,205 | +4*:SUT388; | 4 | TERM | | |
| chrXV | 1004460 | 1004618 | 158 | -1,56234428 | N15:1004637 | 17,349 | | Intergene | | | |
| chrXV | 1005856 | 1006001 | 145 | -1,307118755 | | | | Overlap <50 bp | | | |
| chrXV | 1008258 | 1008393 | 135 | -1,05129403 | N15:1008325 | 5,508 | +8:YOR356W; | 8 | | | |
| chrXV | 1019659 | 1019809 | 150 | -1,352074057 | N15:1019734 | 9,883 | | Intergene | | | |
| chrXV | 1019667 | 1019830 | 163 | -1,28065151 | N15:1019734 | 9,883 | | Intergene | | | |
| chrXV | 1023929 | 1024082 | 153 | -1,406759754 | N15:1024013 | 16,86 | +10:YOR365C; | 10 | | | |
| chrXV | 1035603 | 1035747 | 144 | -1,101703903 | N15:1035709 | 23,887 | +6:YOR372C; | 6 | | | |
| chrXV | 1036929 | 1037060 | 131 | -1,036579827 | N15:1036972 | 5,565 | +2:YOR373W; | 2 | | | |
| chrXV | 1041309 | 1041484 | 175 | -1,36927801 | N15:1041351 | 34,665 | +11*:YOR374W; | 11 | TERM | | |
| chrXV | 1042260 | 1042368 | 108 | -1,614496058 | N15:1042335 | 3,962 | +5:YOR375C; | 5 | | | |
| chrXV | 1042260 | 1042387 | 127 | -0,83117278 | N15:1042335 | 3,962 | +5:YOR375C; | 5 | | | |
| chrXV | 1042260 | 1042389 | 129 | -1,172543949 | N15:1042335 | 3,962 | +5:YOR375C; | 5 | | | |
| chrXV | 1042260 | 1042392 | 132 | -0,952643231 | N15:1042335 | 3,962 | +5:YOR375C; | 5 | | | |
| chrXV | 1042265 | 1042392 | 127 | -0,705963861 | N15:1042335 | 3,962 | +5:YOR375C; | 5 | | | |
| chrXV | 1042267 | 1042368 | 101 | -0,080719943 | N15:1042335 | 3,962 | +5:YOR375C; | 5 | | | |
| chrXV | 1043663 | 1043770 | 107 | -0,895569323 | | | | Overlap <50 bp | | | |
| chrXV | 1043663 | 1043771 | 108 | -0,671407021 | | | | Overlap <50 bp | | | |
| chrXV | 1043663 | 1043772 | 109 | -0,896788259 | | | | Overlap <50 bp | | | |
| chrXV | 1047470 | 1047621 | 151 | -1,323244143 | N15:1047478 | 18,606 | +9:YOR377W; | 9 | | | |
| chrXV | 1047470 | 1047621 | 151 | -1,323244143 | N15:1047635 | 27,803 | +10:YOR377W; | 10 | | | |
| chrXV | 1050132 | 1050280 | 148 | -1,212322379 | N15:1050219 | 11,683 | +6:YOR378W; +4:anti179; | 6 | | | |
| chrXV | 1050137 | 1050280 | 143 | -1,204365963 | N15:1050219 | 11,683 | +6:YOR378W; +4:anti179; | 6 | | | |
| chrXV | 1050608 | 1050773 | 165 | -1,204026491 | N15:1050703 | 11,077 | +9:YOR378W; +1:anti179; | 9 | | | |
| chrXV | 1050861 | 1050999 | 138 | -1,101962321 | N15:1050887 | 27,595 | 'OR378W; -1:anti179; -1:YOR380W; | 10 | | | |
| chrXV | 1052062 | 1052210 | 148 | -0,756635618 | N15:1052087 | 22,623 | +7:YOR380W; | 7 | | | |
| chrXV | 1056315 | 1056470 | 155 | -1,306437663 | N15:1056388 | 5,263 | +6:YOR381W; | 6 | | | |
| chrXV | 1059723 | 1059876 | 153 | -1,078255527 | N15:1059763 | 9,158 | +4:YOR382W; | 4 | | | |
| chrXV | 1060927 | 1061054 | 127 | -0,778526906 | N15:1060979 | 35,816 | +2:YOR383C; | 2 | | | |
| chrXV | 1061685 | 1061838 | 153 | -1,214711972 | N15:1061757 | 11,136 | +2:YOR384W; | 2 | | | |
| chrXV | 1062032 | 1062164 | 132 | -1,194783223 | N15:1062167 | 32,908 | +4:YOR384W; | 4 | | | |
| chrXV | 1062744 | 1062896 | 152 | -1,269826143 | N15:1062785 | 34,965 | +8:YOR384W; | 8 | | | |
| chrXV | 1071888 | 1072059 | 171 | -1,543313612 | N15:1071953 | 16,442 | +7:YOR388C; | 7 | | | |
| chrXV | 1073330 | 1073480 | 150 | -1,301725876 | N15:1073396 | 14,261 | | Intergene | | | |
| chrXV | 1080435 | 1080579 | 144 | -0,953101116 | N15:1080493 | 2,646 | +3:YOR393W; | 3 | | | |
| chrXV | 1085191 | 1085346 | 155 | -1,390067537 | N15:1085246 | 25,325 | -1:YOR396W; | -1 | | | TEL RIGHT |
| chrXV | 1086206 | 1086377 | 171 | -1,527536511 | N15:1086303 | 16,971 | +6:YOR396W; | 6 | | | TEL RIGHT |
| chrXV | 1086356 | 1086503 | 147 | -1,017606607 | | | | Overlap <50 bp | | | TEL RIGHT |
| chrXV | 1086512 | 1086642 | 130 | -0,862497144 | N15:1086572 | 24,194 | +8:YOR396W; | 8 | | | TEL RIGHT |
| chrXV | 1086893 | 1087029 | 136 | -1,156879929 | N15:1086930 | 1,414 | +10:YOR396W; | 10 | | | TEL RIGHT |
| chrXV | 1087274 | 1087436 | 162 | -1,354742742 | N15:1087405 | 11,59 | +13:YOR396W; | 13 | | | TEL RIGHT |
| chrXV | 1087375 | 1087530 | 155 | -1,799486779 | N15:1087405 | 11,59 | +13:YOR396W; | 13 | | | TEL RIGHT |
| chrXV | 1087509 | 1087646 | 137 | -0,699638736 | N15:1087575 | 10,116 | +14:YOR396W; | 14 | | | TEL RIGHT |
| chrXV | 1087804 | 1087947 | 143 | -1,204240291 | N15:1087853 | 10,693 | +16:YOR396W; | 16 | | | TEL RIGHT |
| chrXV | 1087852 | 1087977 | 125 | -0,604932882 | N15:1087853 | 10,693 | +16:YOR396W; | 16 | | | TEL RIGHT |
| chrXV | 1088077 | 1088229 | 152 | -1,196144496 | N15:1088185 | 13 | +18:YOR396W; | 18 | | | TEL RIGHT |
| chrXV | 1088150 | 1088294 | 144 | -0,904179469 | N15:1088185 | 13 | +18:YOR396W; | 18 | | | TEL RIGHT |
| chrXV | 1088247 | 1088382 | 135 | -0,693275778 | N15:1088344 | 22,716 | +19:YOR396W; | 19 | | | TEL RIGHT |
| chrXV | 1088247 | 1088434 | 187 | -0,668246016 | N15:1088344 | 22,716 | +19:YOR396W; | 19 | | | TEL RIGHT |
| chrXV | 1088290 | 1088382 | 92 | -0,048271239 | N15:1088344 | 22,716 | +19:YOR396W; | 19 | | | TEL RIGHT |
| chrXV | 1088290 | 1088434 | 144 | -0,925396865 | N15:1088344 | 22,716 | +19:YOR396W; | 19 | | | TEL RIGHT |
| chrXV | 1088629 | 1088781 | 152 | -1,335406845 | N15:1088692 | 20,502 | +21:YOR396W; | 21 | | | TEL RIGHT |
| chrXV | 1088930 | 1089060 | 130 | -0,991449961 | N15:1088919 | 25,456 | +22:YOR396W; | 22 | | | TEL RIGHT |
| chrXV | 1089068 | 1089229 | 161 | -1,203719312 | N15:1089094 | 17,678 | +23:YOR396W; | 23 | | | TEL RIGHT |
| chrXV | 1090465 | 1090616 | 151 | -1,083483505 | N15:1090543 | 42,426 | +32:YOR396W; | 32 | | | TEL RIGHT |
| chrXVI | 523 | 674 | 151 | -1,273871252 | N16:600 | 36,715 | +34:YPL283C; | 34 | | | TEL LEFT |
| chrXVI | 847 | 1021 | 174 | -1,639074792 | N16:925 | 37,82 | +32:YPL283C; | 32 | | | TEL LEFT |
| chrXVI | 1688 | 1839 | 151 | -1,16643656 | N16:1728 | 11,24 | +27:YPL283C; | 27 | | | TEL LEFT |
| chrXVI | 1700 | 1834 | 134 | -0,926880732 | N16:1728 | 11,24 | +27:YPL283C; | 27 | | | TEL LEFT |
| chrXVI | 1725 | 1821 | 96 | -0,580124735 | N16:1728 | 11,24 | +27:YPL283C; | 27 | | | TEL LEFT |
| chrXVI | 1725 | 1834 | 109 | -0,260418943 | N16:1728 | 11,24 | +27:YPL283C; | 27 | | | TEL LEFT |
| chrXVI | 2079 | 2209 | 130 | -1,020434911 | N16:2222 | 27,006 | +24:YPL283C; | 24 | | | TEL LEFT |
| chrXVI | 2079 | 2209 | 130 | -1,020434911 | N16:2056 | 0,707 | +25:YPL283C; | 25 | | | TEL LEFT |
| chrXVI | 2358 | 2510 | 152 | -1,256586976 | N16:2443 | 19,858 | +23:YPL283C; | 23 | | | TEL LEFT |
| chrXVI | 2705 | 2849 | 144 | -0,948342038 | N16:2758 | 21,385 | +21:YPL283C; | 21 | | | TEL LEFT |
| chrXVI | 2705 | 2892 | 187 | -1,227772768 | N16:2758 | 21,385 | +21:YPL283C; | 21 | | | TEL LEFT |
| chrXVI | 2757 | 2849 | 92 | -0,316913537 | N16:2758 | 21,385 | +21:YPL283C; | 21 | | | TEL LEFT |
| chrXVI | 2757 | 2892 | 135 | -0,934610579 | N16:2758 | 21,385 | +21:YPL283C; | 21 | | | TEL LEFT |
| chrXVI | 2845 | 2989 | 144 | -0,8292503 | N16:2943 | 8,888 | +20:YPL283C; | 20 | | | TEL LEFT |
| chrXVI | 2910 | 3062 | 152 | -1,256599761 | N16:2943 | 8,888 | +20:YPL283C; | 20 | | | TEL LEFT |
| chrXVI | 3162 | 3287 | 125 | -0,726889625 | N16:3271 | 12,897 | +18:YPL283C; | 18 | | | TEL LEFT |
| chrXVI | 3192 | 3335 | 143 | -1,42227426 | N16:3271 | 12,897 | +18:YPL283C; | 18 | | | TEL LEFT |
| chrXVI | 3192 | 3343 | 151 | -1,397479633 | N16:3271 | 12,897 | +18:YPL283C; | 18 | | | TEL LEFT |
| chrXVI | 3493 | 3630 | 137 | -1,048095716 | N16:3559 | 7,767 | +16:YPL283C; | 16 | | | TEL LEFT |
| chrXVI | 3609 | 3764 | 155 | -1,8528086 | N16:3722 | 7,024 | +15:YPL283C; | 15 | | | TEL LEFT |
| chrXVI | 3703 | 3865 | 162 | -1,281655478 | N16:3880 | 4,583 | +14:YPL283C; | 14 | | | TEL LEFT |
| chrXVI | 3703 | 3865 | 162 | -1,281655478 | N16:3722 | 7,024 | +15:YPL283C; | 15 | | | TEL LEFT |
| chrXVI | 4928 | 5069 | 141 | -1,208585581 | N16:4975 | 20,075 | +7:YPL283C; | 7 | | | TEL LEFT |
| chrXVI | 8461 | 8628 | 167 | -1,226955528 | N16:8544 | 10,97 | 0:YPL282C; | 0 | | | |
| chrXVI | 10566 | 10710 | 144 | -1,156743805 | N16:10653 | 6,083 | +3:YPL281C; | 3 | | | |
| chrXVI | 23283 | 23434 | 151 | -1,159502225 | N16:23371 | 6,573 | +4:YPL274W; | 4 | | | |
| chrXVI | 26164 | 26341 | 177 | -1,619423797 | N16:26333 | 2,483 | +1:anti-YPL272C; | 1 | | | |
| chrXVI | 26440 | 26591 | 151 | -1,363898592 | | | | Overlap <50 bp | | | |
| chrXVI | 30175 | 30330 | 155 | -1,617591609 | N16:30253 | 10,629 | -1:YPL270W; +2*:YPL271W; | -1 | | | |
| chrXVI | 30411 | 30566 | 155 | -1,307708451 | N16:30492 | 11,147 | +1:YPL270W; | 1 | | | |
| chrXVI | 37895 | 38053 | 158 | -1,139674632 | N16:37928 | 29,233 | +17*:YPL268W; -1:YPL267W; | 17 | TERM | | |
| chrXVI | 41612 | 41762 | 150 | -1,327971575 | N16:41620 | 14,747 | +5:YPL265W; +18:YPL264C; | 5 | | | |

TABLE 3

| CHR | START | END | L(bp) | ΔLK | Nuc ID | Fuzziness | Gene ID | Gene body postion or intergene | Terminal | rDNA | Telomeric |
|---|---|---|---|---|---|---|---|---|---|---|---|
| chrXVI | 41938 | 42057 | 119 | -1,189522898 | | | | Overlap <50 bp | | | |
| chrXVI | 44256 | 44374 | 118 | -1,220080111 | N16:44326 | 28,272 | +3:YPL264C; | 3 | | | |
| chrXVI | 54075 | 54252 | 177 | -1,680003736 | N16:54126 | 22,106 | +9:YPL258C; | 9 | | | |
| chrXVI | 55410 | 55560 | 150 | -1,228240777 | N16:55405 | 35,216 | +1:YPL258C; | 1 | | | |
| chrXVI | 56814 | 56963 | 149 | -1,136190113 | N16:56959 | 36,501 | | Intergene | | | |
| chrXVI | 56988 | 57111 | 123 | -0,97472828 | | | | Overlap <50 bp | | | |
| chrXVI | 57088 | 57243 | 155 | -1,33601146 | N16:57189 | 29,501 | | Intergene | | | |
| chrXVI | 57521 | 57655 | 134 | -1,195236979 | N16:57591 | 17,436 | | Intergene | | | |
| chrXVI | 57521 | 57661 | 140 | -1,276093078 | N16:57591 | 17,436 | | Intergene | | | |
| chrXVI | 57717 | 57868 | 151 | -1,267168688 | N16:57746 | 27,135 | | Intergene | | | |
| chrXVI | 57727 | 57872 | 145 | -1,353081866 | N16:57746 | 27,135 | | Intergene | | | |
| chrXVI | 57747 | 57896 | 149 | -1,281826372 | N16:57746 | 27,135 | | Intergene | | | |
| chrXVI | 57999 | 58153 | 154 | -1,116137424 | N16:58141 | 5,132 | | Intergene | | | |
| chrXVI | 58564 | 58699 | 135 | -1,265287266 | N16:58596 | 13,892 | | Intergene | | | |
| chrXVI | 58718 | 58879 | 161 | -1,404572534 | N16:58834 | 41,405 | | Intergene | | | |
| chrXVI | 59011 | 59161 | 150 | -1,080921355 | N16:59079 | 42,771 | | Intergene | | | |
| chrXVI | 59190 | 59347 | 157 | -1,440055979 | N16:59296 | 28,501 | | Intergene | | | |
| chrXVI | 59681 | 59810 | 129 | -1,444006599 | N16:59829 | 14,731 | | Intergene | | | |
| chrXVI | 60593 | 60743 | 150 | -1,579459688 | N16:60686 | 10,786 | | Intergene | | | |
| chrXVI | 61572 | 61714 | 142 | -1,209575201 | N16:61694 | 32,254 | | Intergene | | | |
| chrXVI | 66888 | 67040 | 152 | -1,241233796 | N16:66990 | 12,156 | -1:YPL256C; | -1 | | | |
| chrXVI | 67654 | 67814 | 160 | -1,367789871 | N16:67736 | 6,899 | +1:YPL255W; | 1 | | | |
| chrXVI | 69456 | 69617 | 161 | -1,351937514 | N16:69502 | 6,824 | +2:YPL254W; | 2 | | | |
| chrXVI | 73482 | 73631 | 149 | -0,86615081 | N16:73588 | 12,92 | +3:YPL252C; | 3 | | | |
| chrXVI | 74222 | 74371 | 149 | -1,030979493 | N16:74321 | 17,044 | +2:CUT401; +3:YPL250C; | 2 | | | |
| chrXVI | 85235 | 85390 | 155 | -1,490098199 | N16:85298 | 8,456 | +1:YPL246C; -1:YPL245W; | 1 | | | |
| chrXVI | 86229 | 86396 | 167 | -1,437505095 | N16:86219 | 13,029 | +5:YPL245W; | 5 | | | |
| chrXVI | 93799 | 93933 | 134 | -1,152232688 | N16:93856 | 30,448 | +9:YPL242C; | 9 | | | |
| chrXVI | 97906 | 98054 | 148 | -1,096079875 | N16:97910 | 33,147 | +6:YPL240C; | 6 | | | |
| chrXVI | 100090 | 100196 | 106 | -1,556415834 | N16:100168 | 6,907 | -1:YPL237W; +5*:YPL239W; | -1 | | | |
| chrXVI | 101988 | 102144 | 156 | -1,014178668 | N16:102072 | 5,814 | +5:YPL236C; | 5 | | | |
| chrXVI | 111563 | 111694 | 131 | -1,021204548 | N16:111614 | 32,322 | +20:YPL231W; | 20 | | | |
| chrXVI | 111839 | 111996 | 157 | -1,349496182 | N16:111919 | 37,173 | | Intergene | | | |
| chrXVI | 112816 | 112964 | 148 | -1,009412654 | N16:112902 | 31,405 | | Intergene | | | |
| chrXVI | 113851 | 114002 | 151 | -1,19115666 | N16:113884 | 13,142 | | Intergene | | | |
| chrXVI | 115900 | 116007 | 107 | -1,336787112 | N16:115995 | 32,411 | +3:SUT807; +2:YPL230W; | 3 | | | |
| chrXVI | 115900 | 116008 | 108 | -1,314082644 | N16:115995 | 32,411 | +3:SUT807; +2:YPL230W; | 3 | | | |
| chrXVI | 115900 | 116009 | 109 | -1,151531721 | N16:115995 | 32,411 | +3:SUT807; +2:YPL230W; | 3 | | | |
| chrXVI | 115900 | 116013 | 113 | -1,345128504 | N16:115995 | 32,411 | +3:SUT807; +2:YPL230W; | 3 | | | |
| chrXVI | 117285 | 117438 | 153 | -0,694359805 | | | | Overlap <50 bp | | | |
| chrXVI | 119500 | 119650 | 150 | -1,408520542 | N16:119550 | 20,761 | +9:YPL228W; | 9 | | | |
| chrXVI | 119632 | 119770 | 138 | -1,487245546 | N16:119717 | 8,325 | +10:YPL228W; | 10 | | | |
| chrXVI | 124878 | 125016 | 138 | -1,070973347 | N16:124969 | 11,777 | +19:YPL226W; | 19 | | | |
| chrXVI | 127061 | 127209 | 148 | -1,148514355 | N16:127222 | 6,713 | +7:YPL224C; | 7 | | | |
| chrXVI | 127061 | 127209 | 148 | -1,148514355 | N16:127054 | 5,419 | +8:YPL224C; | 8 | | | |
| chrXVI | 127345 | 127506 | 161 | -1,385029955 | N16:127392 | 16,881 | +6:YPL224C; | 6 | | | |
| chrXVI | 128019 | 128185 | 166 | -1,76316196 | N16:128056 | 8,741 | -1:Unit649; +2:YPL224C; | -1 | | | |
| chrXVI | 128210 | 128316 | 106 | -0,955424382 | N16:128260 | 23,322 | +1:Unit649; +1:YPL224C; | 1 | | | |
| chrXVI | 128210 | 128332 | 122 | -1,218193357 | N16:128260 | 23,322 | +1:Unit649; +1:YPL224C; | 1 | | | |
| chrXVI | 130172 | 130324 | 152 | -1,837982522 | N16:130248 | 18,302 | +2:YPL222W; | 2 | | | |
| chrXVI | 131123 | 131279 | 156 | -1,192119134 | N16:131253 | 37,125 | +8:YPL222W; | 8 | | | |
| chrXVI | 136763 | 136914 | 151 | -1,138761843 | N16:136878 | 7,563 | +2:YPL219W; | 2 | | | |
| chrXVI | 136805 | 136963 | 158 | -1,239435241 | N16:136878 | 7,563 | +2:YPL219W; | 2 | | | |
| chrXVI | 139377 | 139534 | 157 | -1,470719293 | N16:139490 | 11,446 | | Intergene | | | |
| chrXVI | 142443 | 142561 | 118 | -1,453646725 | N16:142521 | 10,046 | +5:YPL217C; | 5 | | | |
| chrXVI | 144178 | 144336 | 158 | -1,361994703 | N16:144285 | 18,687 | +5:YPL216W; | 5 | | | |
| chrXVI | 149028 | 149196 | 168 | -1,204811029 | N16:149123 | 8,408 | +8:YPL214C; | 8 | | | |
| chrXVI | 152572 | 152722 | 150 | -1,096591693 | N16:152635 | 5,857 | +4:YPL212C; | 4 | | | |
| chrXVI | 161247 | 161414 | 167 | -1,259632979 | N16:161357 | 51,735 | +10:YPL207W; | 10 | | | |
| chrXVI | 164847 | 164957 | 110 | -1,206490994 | N16:164935 | 8,958 | +6:YPL204W; | 6 | | | |
| chrXVI | 165348 | 165502 | 154 | -1,594971934 | N16:165511 | 34,953 | +10:YPL204W; +1:SUT811; | 10 | | | |
| chrXVI | 167030 | 167184 | 154 | -0,980405571 | N16:167113 | 32,5 | +7:YPL203W; | 7 | | | |
| chrXVI | 169266 | 169418 | 152 | -1,221815775 | N16:169376 | 17,029 | +1:YPL202C; | 1 | | | |
| chrXVI | 173412 | 173549 | 137 | -0,981266874 | N16:173465 | 6,364 | | Intergene | | | |
| chrXVI | 174194 | 174363 | 169 | -1,410004065 | N16:174232 | 28,827 | | Intergene | | | |
| chrXVI | 175111 | 175256 | 145 | -1,357605346 | N16:175175 | 6,834 | +2:YPL196W; | 2 | | | |
| chrXVI | 177466 | 177621 | 155 | -1,3314233 | N16:177552 | 31,17 | +9:YPL195W; | 9 | | | |
| chrXVI | 177625 | 177797 | 172 | -1,51037597 | N16:177780 | 21,378 | +10:YPL195W; | 10 | | | |
| chrXVI | 189167 | 189337 | 170 | -1,527417555 | N16:189180 | 28,27 | +1:YPL189W; | 1 | | | |
| chrXVI | 190409 | 190560 | 151 | -1,407630912 | N16:190473 | 22,412 | +9:YPL189W; | 9 | | | |
| chrXVI | 191083 | 191228 | 145 | -1,30832992 | N16:191166 | 17,058 | YPL189W; +1:CUT888; -1:YPL188W; | 13 | TERM | | |
| chrXVI | 193727 | 193886 | 159 | -1,020687158 | N16:193851 | 10,402 | +2:YPL187W; | 2 | | | |
| chrXVI | 194794 | 194946 | 152 | -1,393461999 | N16:194885 | 7,823 | +3:YPL186C; | 3 | | | |
| chrXVI | 198714 | 198866 | 152 | -1,424144256 | N16:198810 | 5,203 | -1:YPL183W-A; | -1 | | | |
| chrXVI | 200785 | 200926 | 141 | -1,250227923 | | | | Overlap <50 bp | | | |
| chrXVI | 203076 | 203237 | 161 | -1,324457735 | N16:203255 | 19,502 | +1:YPL181W; | 1 | | | |
| chrXVI | 203882 | 204036 | 154 | -1,292423736 | N16:203940 | 13,722 | +5:YPL181W; | 5 | | | |
| chrXVI | 205158 | 205313 | 155 | -1,181345049 | N16:205222 | 6,293 | 0:YPL180W; -1:CUT890; | 0 | | | |
| chrXVI | 211347 | 211529 | 182 | -1,557206804 | N16:211407 | 47,917 | | Intergene | | | |
| chrXVI | 212567 | 212715 | 148 | -1,254183085 | N16:212620 | 11,769 | +4:YPL178W; | 4 | | | |
| chrXVI | 213085 | 213234 | 149 | -1,363917648 | N16:213180 | 30,105 | +7*:YPL177C; | 7 | TERM | | |
| chrXVI | 220298 | 220444 | 146 | -1,125840854 | N16:220437 | 26,974 | +16:YPL174C; | 16 | | | |
| chrXVI | 227791 | 227944 | 153 | -1,245872512 | | | | Overlap <50 bp | | | |
| chrXVI | 228023 | 228173 | 150 | -1,415878485 | N16:228073 | 4,95 | -1:YPL170W; | -1 | | | |
| chrXVI | 235024 | 235161 | 137 | -1,413746474 | N16:235068 | 23,858 | +14:YPL167C; | 14 | | | |
| chrXVI | 238735 | 238888 | 153 | -1,380864364 | N16:238814 | 17,992 | +9:YPL166W; +3:YPL165C; | 9 | | | |
| chrXVI | 239149 | 239288 | 139 | -1,395078608 | N16:239177 | 17,627 | 166W; +1:YPL165C; +15*:YPL164C; | 11 | TERM | | |
| chrXVI | 240071 | 240209 | 138 | -1,120280653 | N16:240084 | 21,307 | +10:YPL164C; | 10 | | | |
| chrXVI | 240947 | 241098 | 151 | -1,132740053 | N16:241005 | 8,826 | +4:YPL164C; | 4 | | | |
| chrXVI | 242739 | 242891 | 152 | -1,283738763 | N16:242817 | 22,634 | +9*:YPL162C; 0:YPL163C; | 9 | TERM | | |
| chrXVI | 242995 | 243131 | 136 | -1,37529551 | N16:243075 | 19,263 | +7:YPL162C; -1:anti182; | 7 | | | |
| chrXVI | 244964 | 245114 | 150 | -1,291164006 | N16:245006 | 15,132 | +9:YPL161C; | 9 | | | |
| chrXVI | 246169 | 246331 | 162 | -1,333655341 | N16:246246 | 14,621 | +1:YPL161C; | 1 | | | |
| chrXVI | 247500 | 247655 | 155 | -1,263520105 | | | | Overlap <50 bp | | | |
| chrXVI | 247870 | 248025 | 155 | -1,489905312 | N16:248033 | 42,382 | +8:YPL160W; | 8 | | | |
| chrXVI | 248518 | 248667 | 149 | -1,428423709 | N16:248596 | 31,886 | +11:YPL160W; | 11 | | | |
| chrXVI | 251350 | 251499 | 149 | -1,480166043 | N16:251383 | 22,257 | +3:YPL159C; | 3 | | | |
| chrXVI | 251647 | 251802 | 155 | -1,020723248 | N16:251733 | 5,797 | +1:YPL159C; | 1 | | | |

TABLE 3

| CHR | START | END | L(bp) | ΔLK | Nuc ID | Fuzziness | Gene ID | Gene body postion or intergene | Terminal | rDNA | Telomeric |
|---|---|---|---|---|---|---|---|---|---|---|---|
| chrXVI | 254749 | 254901 | 152 | -1,735333272 | N16:254752 | 5,046 | +1:YPL157W; | 1 | | | |
| chrXVI | 254749 | 254901 | 152 | -1,735333272 | N16:254920 | 8,813 | +2:YPL157W; | 2 | | | |
| chrXVI | 257303 | 257455 | 152 | -1,291792731 | N16:257371 | 12,319 | +13:YPL155C; | 13 | | | |
| chrXVI | 258470 | 258614 | 144 | -1,361375157 | N16:258522 | 9,933 | +6:YPL155C; | 6 | | | |
| chrXVI | 269441 | 269592 | 151 | -1,739677473 | N16:269499 | 29,331 | +10:YPL150W; | 10 | | | |
| chrXVI | 272088 | 272225 | 137 | -1,154583116 | N16:272113 | 33,171 | +6*:YPL148C; | 6 | TERM | | |
| chrXVI | 273849 | 274002 | 153 | -1,98419593 | N16:273925 | 33,991 | +5:YPL147W; | 5 | | | |
| chrXVI | 275518 | 275683 | 165 | -1,86357557 | N16:275553 | 22,26 | +14:YPL147W; | 14 | | | |
| chrXVI | 278767 | 278916 | 149 | -1,258316783 | N16:278752 | 21,479 | +8:YPL145C; | 8 | | | |
| chrXVI | 284968 | 285121 | 153 | -1,068355036 | N16:285013 | 29,392 | +8:YPL141C; | 8 | | | |
| chrXVI | 287278 | 287435 | 157 | -1,274300131 | N16:287376 | 11,3 | +12*:YPL140C; | 12 | TERM | | |
| chrXVI | 287516 | 287643 | 127 | -1,061338708 | N16:287547 | 21,811 | +11:YPL140C; | 11 | | | |
| chrXVI | 295119 | 295280 | 161 | -1,172228324 | N16:295214 | 31,102 | +11:YPL137C; | 11 | | | |
| chrXVI | 296343 | 296490 | 147 | -1,211616509 | N16:296416 | 17,992 | +4:YPL137C; | 4 | | | |
| chrXVI | 301254 | 301382 | 128 | -1,435572917 | N16:301333 | 19 | -1:YPL132W; +1:YPL133C; | -1 | | | |
| chrXVI | 305851 | 306015 | 164 | -1,51799359 | N16:305962 | 6,807 | | Intergene | | | |
| chrXVI | 306387 | 306542 | 155 | -1,504147561 | N16:306463 | 42,894 | +12*:YPL128C; | 12 | TERM | | |
| chrXVI | 317763 | 317907 | 144 | -1,256662107 | N16:317793 | 33,037 | | Intergene | | | |
| chrXVI | 318699 | 318846 | 147 | -1,562030858 | | | | Overlap <50 bp | | | |
| chrXVI | 322025 | 322158 | 133 | -1,116361258 | N16:322082 | 3,808 | +1:YPL120W; | 1 | | | |
| chrXVI | 323166 | 323321 | 155 | -1,454678642 | N16:323256 | 23,173 | +8:YPL120W; | 8 | | | |
| chrXVI | 332594 | 332745 | 151 | -0,906444225 | | | | Overlap <50 bp | | | |
| chrXVI | 332948 | 333099 | 151 | -1,284124629 | N16:333032 | 23,522 | +17:YPL115C; | 17 | | | |
| chrXVI | 336537 | 336643 | 106 | -1,411968532 | N16:336620 | 3,987 | +5:YPL113C; | 5 | | | |
| chrXVI | 336537 | 336650 | 113 | -1,295826184 | N16:336620 | 3,987 | +5:YPL113C; | 5 | | | |
| chrXVI | 341289 | 341447 | 158 | -1,233493369 | N16:341297 | 29,523 | +22*:YPL110C; | 22 | TERM | | |
| chrXVI | 344751 | 344913 | 162 | -1,442602772 | N16:344907 | 35,117 | -1:SUT402; 0:YPL110C; | -1 | | | |
| chrXVI | 344751 | 344913 | 162 | -1,442602772 | N16:344736 | 12,754 | +2:YPL110C; | 2 | | | |
| chrXVI | 350439 | 350593 | 154 | -1,191141616 | N16:350549 | 31,706 | +12:YPL106C; | 12 | | | |
| chrXVI | 351311 | 351471 | 160 | -1,55099552 | N16:351332 | 7,05 | +7:YPL106C; | 7 | | | |
| chrXVI | 354968 | 355128 | 160 | -1,300410051 | N16:355047 | 5,857 | +3:YPL105C; | 3 | | | |
| chrXVI | 361977 | 362073 | 96 | -0,892770432 | N16:362047 | 4,55 | +2:YPL100W; | 2 | | | |
| chrXVI | 364336 | 364488 | 152 | -1,272054996 | N16:364416 | 8,916 | +3:YPL098C; | 3 | | | |
| chrXVI | 366545 | 366652 | 107 | -1,180861549 | N16:366569 | 3,601 | +2:YPL096C-A; | 2 | | | |
| chrXVI | 366545 | 366653 | 108 | -1,165589457 | N16:366569 | 3,601 | +2:YPL096C-A; | 2 | | | |
| chrXVI | 367044 | 367180 | 136 | -1,208608464 | N16:367128 | 9,928 | +2:YPL096W; | 2 | | | |
| chrXVI | 370599 | 370734 | 135 | -1,104173166 | N16:370630 | 8,361 | +1:YPL094C; -1:YPL093W; | 1 | | | |
| chrXVI | 370905 | 371054 | 149 | -1,095555092 | N16:370964 | 15,531 | -1:YPL094C; +1:YPL093W; | -1 | | | |
| chrXVI | 373474 | 373616 | 142 | -1,092227486 | N16:373635 | 17,473 | +1:YPL092W; -1:CUT895; | 1 | | | |
| chrXVI | 379313 | 379470 | 157 | -1,424126068 | N16:379337 | 56,889 | +2:SUT406; | 2 | | | |
| chrXVI | 379794 | 379947 | 153 | -1,115482981 | | | | Overlap <50 bp | | | |
| chrXVI | 379982 | 380135 | 153 | -1,428326443 | N16:379982 | 17,445 | +6:SUT406; | 6 | | | |
| chrXVI | 379982 | 380135 | 153 | -1,428326443 | N16:380145 | 14,986 | +7:SUT406; | 7 | | | |
| chrXVI | 383644 | 383787 | 143 | -1,089333684 | N16:383784 | 18,111 | +4:YPL087W; | 4 | | | |
| chrXVI | 385233 | 385386 | 153 | -1,14942033 | N16:385330 | 17,297 | +8:YPL086C; | 8 | | | |
| chrXVI | 389294 | 389391 | 97 | -1,010099859 | N16:389336 | 40,087 | +16:YPL085W; | 16 | | | |
| chrXVI | 390081 | 390232 | 151 | -1,1283173 | N16:390091 | 25,843 | +21:YPL085W; | 21 | | | |
| chrXVI | 390081 | 390232 | 151 | -1,1283173 | N16:390239 | 0 | +22:YPL085W; | 22 | | | |
| chrXVI | 390545 | 390691 | 146 | -1,321041103 | N16:390526 | 34,385 | +24:YPL085W; | 24 | | | |
| chrXVI | 392582 | 392731 | 149 | -1,505449478 | N16:392651 | 35,2 | | Intergene | | | |
| chrXVI | 392582 | 392736 | 154 | -1,259859193 | N16:392651 | 35,2 | | Intergene | | | |
| chrXVI | 394420 | 394573 | 153 | -1,500244918 | N16:394533 | 3,962 | +4:YPL084W; | 4 | | | |
| chrXVI | 407155 | 407307 | 152 | -1,127881795 | N16:407273 | 15,834 | | Intergene | | | |
| chrXVI | 407160 | 407307 | 147 | -1,378323552 | N16:407273 | 15,834 | | Intergene | | | |
| chrXVI | 411818 | 411982 | 164 | -1,480884868 | N16:411890 | 10,252 | -1:YPL075W; -1:CUT897; | -1 | | | |
| chrXVI | 411838 | 411991 | 153 | -1,630554274 | N16:411890 | 10,252 | -1:YPL075W; -1:CUT897; | -1 | | | |
| chrXVI | 412957 | 413111 | 154 | -1,226443232 | | | | Overlap <50 bp | | | |
| chrXVI | 415881 | 416027 | 146 | -1,290705482 | N16:415956 | 10,991 | +2:YPL074W; | 2 | | | |
| chrXVI | 417433 | 417565 | 132 | -1,41043892 | N16:417568 | 26,063 | +12:YPL074W; | 12 | | | |
| chrXVI | 418493 | 418641 | 148 | -1,390537882 | | | | Overlap <50 bp | | | |
| chrXVI | 418678 | 418846 | 168 | -1,40968311 | N16:418685 | 22,013 | +3:YPL072W; | 3 | | | |
| chrXVI | 418678 | 418846 | 168 | -1,40968311 | N16:418866 | 7,301 | +4:YPL072W; | 4 | | | |
| chrXVI | 432250 | 432440 | 190 | -1,695972439 | N16:432391 | 28,898 | +1:Unit653; -1:YPL061W; | 1 | | | |
| chrXVI | 432594 | 432737 | 143 | -1,033617726 | N16:432613 | 20,408 | +1:YPL061W; | 1 | | | |
| chrXVI | 432908 | 433068 | 160 | -0,949511615 | N16:433087 | 53,51 | +4:YPL061W; | 4 | | | |
| chrXVI | 433481 | 433638 | 157 | -1,271585009 | N16:433601 | 26,912 | +7:YPL061W; | 7 | | | |
| chrXVI | 437116 | 437278 | 162 | -1,31353259 | N16:437280 | 12,702 | | Intergene | | | |
| chrXVI | 439535 | 439688 | 153 | -1,943673809 | | | | Overlap <50 bp | | | |
| chrXVI | 442027 | 442174 | 147 | -1,480401127 | N16:442080 | 10,017 | | Intergene | | | |
| chrXVI | 442326 | 442474 | 148 | -1,447406976 | N16:442468 | 19,553 | | Intergene | | | |
| chrXVI | 442340 | 442502 | 162 | -1,326834464 | N16:442468 | 19,553 | | Intergene | | | |
| chrXVI | 446281 | 446447 | 166 | -1,351141968 | N16:446378 | 7,182 | +25:YPL058C; | 25 | | | |
| chrXVI | 448877 | 449010 | 133 | -1,273597369 | N16:448925 | 17,257 | +11:YPL058C; | 11 | | | |
| chrXVI | 454839 | 454945 | 106 | -0,811927557 | N16:454945 | 4,604 | +1:YPL055C; | 1 | | | |
| chrXVI | 462929 | 463059 | 130 | -1,29799693 | N16:462975 | 23,302 | +7:YPL049C; | 7 | | | |
| chrXVI | 465016 | 465172 | 156 | -1,297256244 | N16:465100 | 13,985 | +6:YPL048W; | 6 | | | |
| chrXVI | 466157 | 466315 | 158 | -1,352914387 | N16:466243 | 4,926 | +3:YPL047W; | 3 | | | |
| chrXVI | 474538 | 474690 | 152 | -1,109747004 | N16:474587 | 12,49 | +3:YPL042C; | 3 | | | |
| chrXVI | 484126 | 484288 | 162 | -1,44668286 | N16:484123 | 4,722 | +9:YPL036W; | 9 | | | |
| chrXVI | 484126 | 484288 | 162 | -1,44668286 | N16:484296 | 18,118 | +10:YPL036W; | 10 | | | |
| chrXVI | 484807 | 484963 | 156 | -1,391719747 | | | | Overlap <50 bp | | | |
| chrXVI | 487013 | 487184 | 171 | -1,772409896 | N16:487139 | 30,232 | +5*:YPL034W; +7*:YPL033C; | 5 | TERM | | |
| chrXVI | 493217 | 493377 | 160 | -1,301695363 | N16:493271 | 33,361 | -1:YPL031C; -1:YPL030W; | -1 | | | |
| chrXVI | 497274 | 497428 | 154 | -1,166669092 | N16:497348 | 7,662 | +12:YPL029W; +3*:CUT903; | 12 | | | |
| chrXVI | 501076 | 501229 | 153 | -1,384027529 | N16:501190 | 23,255 | +7*:YPL026C; | 7 | TERM | | |
| chrXVI | 501841 | 501999 | 158 | -1,33694335 | N16:501884 | 19,243 | +3:YPL026C; | 3 | | | |
| chrXVI | 505233 | 505380 | 147 | -1,433088424 | N16:505303 | 36,928 | +8:YPL023C; | 8 | | | |
| chrXVI | 506320 | 506478 | 158 | -1,406132268 | N16:506395 | 2,881 | +1:YPL023C; -1:YPL022W; | 1 | | | |
| chrXVI | 506594 | 506750 | 156 | -1,307658985 | N16:506676 | 24,906 | -1:YPL023C; +1:YPL022W; | -1 | | | |
| chrXVI | 509864 | 510024 | 160 | -1,650989859 | N16:509922 | 26,571 | +21:YPL022W; | 21 | | | |
| chrXVI | 511391 | 511535 | 144 | -1,85231565 | N16:511439 | 13,342 | +3:YPL021W; | 3 | | | |
| chrXVI | 517443 | 517595 | 152 | -1,255230609 | N16:517508 | 19,453 | 0:YPL018W; | 0 | | | |
| chrXVI | 519049 | 519173 | 124 | -1,293611838 | N16:519058 | 35,379 | +9:YPL018W; +9:YPL017C; | 9 | | | |
| chrXVI | 520082 | 520209 | 127 | -1,050650882 | N16:520178 | 33,32 | +2:YPL017C; | 2 | | | |
| chrXVI | 522508 | 522654 | 146 | -1,372160162 | N16:522677 | 10,832 | +14:YPL016W; +1*:anti186; | 14 | | | |
| chrXVI | 522574 | 522734 | 160 | -1,395786133 | N16:522677 | 10,832 | +14:YPL016W; +1*:anti186; | 14 | | | |
| chrXVI | 525954 | 526108 | 154 | -1,55572356 | N16:526036 | 10,69 | +6:YPL015C; | 6 | | | |
| chrXVI | 536923 | 537076 | 153 | -1,224652735 | N16:537021 | 31,915 | +13:YPL009C; | 13 | | | |
| chrXVI | 538001 | 538163 | 162 | -1,56521883 | N16:538105 | 11,874 | +6:YPL009C; | 6 | | | |

TABLE 3

| CHR | START | END | L(bp) | ΔLK | Nuc ID | Fuzziness | Gene ID | Gene body postion or intergene | Terminal | rDNA | Telomeric |
|---|---|---|---|---|---|---|---|---|---|---|---|
| chrXVI | 541695 | 541831 | 136 | -0,979337228 | N16:541839 | 46,217 | +18:YPL008W; | 18 | | | |
| chrXVI | 547021 | 547160 | 139 | -1,138461346 | N16:547005 | 28,754 | +16:YPL006W; | 16 | | | |
| chrXVI | 547636 | 547771 | 135 | -1,210576363 | N16:547723 | 5,601 | +20:YPL006W; | 20 | | | |
| chrXVI | 547851 | 548005 | 154 | -1,21085727 | N16:547889 | 12,608 | +21*:YPL006W; | 21 | TERM | | |
| chrXVI | 550617 | 550788 | 171 | -1,377887712 | N16:550762 | 34,631 | +6*:YPL004C; | 6 | TERM | | |
| chrXVI | 553457 | 553564 | 107 | -1,304362682 | N16:553491 | 32,716 | +6:YPL002C; | 6 | | | |
| chrXVI | 553457 | 553565 | 108 | -1,276748346 | N16:553491 | 32,716 | +6:YPL002C; | 6 | | | |
| chrXVI | 560771 | 560926 | 155 | -1,128971778 | N16:560761 | 20,182 | | Intergene | | | |
| chrXVI | 560771 | 560926 | 155 | -1,128971778 | N16:560933 | 8,042 | | Intergene | | | |
| chrXVI | 564313 | 564461 | 148 | -1,682215094 | N16:564375 | 11,256 | +5:YPR004C; | 5 | | | |
| chrXVI | 564317 | 564438 | 121 | -1,086627763 | N16:564375 | 11,256 | +5:YPR004C; | 5 | | | |
| chrXVI | 566468 | 566620 | 152 | -1,411216924 | N16:566631 | 20,839 | +1:YPR005C; | 1 | | | |
| chrXVI | 566468 | 566620 | 152 | -1,411216924 | N16:566462 | 44,641 | +2:YPR005C; | 2 | | | |
| chrXVI | 568077 | 568211 | 134 | -1,20877957 | N16:568157 | 11,349 | +6:YPR006C; | 6 | | | |
| chrXVI | 573444 | 573609 | 165 | -1,60620184 | N16:573552 | 11,533 | +7:YPR008W; | 7 | | | |
| chrXVI | 573688 | 573839 | 151 | -1,144053447 | N16:573733 | 13,408 | +8:YPR008W; | 8 | | | |
| chrXVI | 578898 | 579048 | 150 | -1,017424855 | N16:579014 | 14,839 | +13:YPR010C; | 13 | | | |
| chrXVI | 580106 | 580256 | 150 | -1,330128816 | N16:580137 | 29,606 | +7:YPR010C; | 7 | | | |
| chrXVI | 581758 | 581907 | 149 | -1,488594746 | N16:581900 | 7,521 | +3*:CUT424; | 3 | TERM | | |
| chrXVI | 581758 | 581907 | 149 | -1,488594746 | N16:581737 | 3,899 | +2:CUT424; | 2 | | | |
| chrXVI | 582967 | 583116 | 149 | -1,252378763 | N16:583058 | 11,125 | SUT416; -1:Unit633; -1:YPR010C-A; | 1 | | | |
| chrXVI | 584868 | 585014 | 146 | -1,250563813 | N16:584868 | 19,99 | +8:YPR013C; | 8 | | | |
| chrXVI | 591560 | 591713 | 153 | -1,179279205 | N16:591636 | 21,79 | | Intergene | | | |
| chrXVI | 592210 | 592352 | 142 | -1,141968051 | N16:592274 | 34,238 | +6*:YPR016C; | 6 | TERM | | |
| chrXVI | 593528 | 593683 | 155 | -1,337265577 | N16:593589 | 7,767 | +3:YPR017C; | 3 | | | |
| chrXVI | 594996 | 595138 | 142 | -1,211624931 | N16:595089 | 43,597 | +5:YPR018W; | 5 | | | |
| chrXVI | 599046 | 599200 | 154 | -1,369215328 | N16:599073 | 8,562 | +16:YPR019W; | 16 | | | |
| chrXVI | 600831 | 600997 | 166 | -1,349344516 | N16:600995 | 6,928 | +17:YPR021C; | 17 | | | |
| chrXVI | 600831 | 600997 | 166 | -1,349344516 | N16:600844 | 4,879 | +18:YPR021C; | 18 | | | |
| chrXVI | 601108 | 601231 | 123 | -1,072171781 | N16:601151 | 14,846 | +16:YPR021C; | 16 | | | |
| chrXVI | 603139 | 603315 | 176 | -1,716632526 | N16:603211 | 4,59 | +3:YPR021C; | 3 | | | |
| chrXVI | 604400 | 604533 | 133 | -1,276190771 | N16:604513 | 17,952 | | Intergene | | | |
| chrXVI | 611112 | 611263 | 151 | -0,763729014 | | | | Overlap <50 bp | | | |
| chrXVI | 611769 | 611929 | 160 | -1,453074942 | N16:611782 | 8,655 | +10:YPR024W; | 10 | | | |
| chrXVI | 611769 | 611929 | 160 | -1,453074942 | N16:611938 | 33,548 | +11:YPR024W; | 11 | | | |
| chrXVI | 614264 | 614371 | 107 | -1,320799794 | N16:614344 | 5,958 | +2:YPR025C; | 2 | | | |
| chrXVI | 614264 | 614381 | 117 | -1,576300435 | N16:614344 | 5,958 | +2:YPR025C; | 2 | | | |
| chrXVI | 616962 | 617122 | 160 | -1,471539962 | N16:617008 | 22,426 | +11:YPR026W; | 11 | | | |
| chrXVI | 617750 | 617867 | 117 | -1,155317306 | N16:617790 | 18,584 | +16:YPR026W; | 16 | | | |
| chrXVI | 617888 | 618049 | 161 | -1,329801815 | N16:617954 | 26,435 | +17:YPR026W; | 17 | | | |
| chrXVI | 623817 | 623987 | 170 | -1,216229375 | N16:623864 | 10,677 | | Intergene | | | |
| chrXVI | 627166 | 627316 | 150 | -1,421535304 | N16:627249 | 13,008 | -1:YPR029C; | -1 | | | |
| chrXVI | 630544 | 630672 | 128 | -1,018909021 | | | | Overlap <50 bp | | | |
| chrXVI | 630787 | 630947 | 160 | -1,186444265 | N16:630889 | 9,311 | +19:YPR030W; | 19 | | | |
| chrXVI | 639745 | 639881 | 136 | -1,555834519 | N16:639847 | 6,156 | +3:YPR034W; | 3 | | | |
| chrXVI | 644095 | 644255 | 160 | -0,99980481 | N16:644162 | 26,136 | +6:SUT829; +3:YPR036W; | 6 | | | |
| chrXVI | 647938 | 648089 | 151 | -1,519657531 | N16:647982 | 8,548 | +5:YPR040W; | 5 | | | |
| chrXVI | 651006 | 651170 | 164 | -1,36634403 | N16:651112 | 4,243 | +16:YPR042C; | 16 | | | |
| chrXVI | 651448 | 651577 | 129 | -0,900803172 | N16:651501 | 45,675 | +14:YPR042C; | 14 | | | |
| chrXVI | 651814 | 651966 | 152 | -1,252563749 | | | | Overlap <50 bp | | | |
| chrXVI | 655669 | 655818 | 149 | -1,206893202 | N16:655729 | 25,381 | +6:YPR045C; | 6 | | | |
| chrXVI | 661336 | 661465 | 129 | -1,165643834 | N16:661468 | 46,144 | +21:YPR049C; | 21 | | | |
| chrXVI | 664409 | 664571 | 162 | -1,377891024 | N16:664522 | 7,057 | +2:YPR049C; | 2 | | | |
| chrXVI | 665020 | 665144 | 124 | -1,248296211 | N16:665145 | 12,656 | +2:YPR051W; | 2 | | | |
| chrXVI | 670319 | 670426 | 107 | -1,464598676 | N16:670368 | 6,758 | +18:YPR055W; | 18 | | | |
| chrXVI | 670319 | 670427 | 108 | -1,433003482 | N16:670368 | 6,758 | +18:YPR055W; | 18 | | | |
| chrXVI | 670635 | 670782 | 147 | -1,117741749 | N16:670711 | 15,362 | +20:YPR055W; | 20 | | | |
| chrXVI | 672000 | 672158 | 158 | -1,342525523 | N16:672118 | 25,399 | +1:YPR057W; +9:CUT908; | 1 | | | |
| chrXVI | 678051 | 678201 | 150 | -1,228006649 | N16:678147 | 4,535 | | Intergene | | | |
| chrXVI | 679094 | 679205 | 111 | -0,564705764 | N16:679146 | 16,523 | | Intergene | | | |
| chrXVI | 679094 | 679206 | 112 | -0,574895319 | N16:679146 | 16,523 | | Intergene | | | |
| chrXVI | 679094 | 679207 | 113 | -0,861216848 | N16:679146 | 16,523 | | Intergene | | | |
| chrXVI | 679098 | 679205 | 107 | -1,144856442 | N16:679146 | 16,523 | | Intergene | | | |
| chrXVI | 679098 | 679220 | 122 | -1,234831814 | N16:679146 | 16,523 | | Intergene | | | |
| chrXVI | 681181 | 681332 | 151 | -1,821606949 | N16:681246 | 15,293 | +2:YPR066W; +5*:SUT833; | 2 | | | |
| chrXVI | 685561 | 685687 | 126 | -1,45776229 | N16:685673 | 39,469 | -1:YPR070W; -1:YPR069C; | -1 | | | |
| chrXVI | 685866 | 686024 | 158 | -1,362667769 | N16:685893 | 14,109 | +1:YPR070W; | 1 | | | |
| chrXVI | 689048 | 689183 | 135 | -1,423976798 | N16:689086 | 28,7 | +7*:YPR071W; | 7 | TERM | | |
| chrXVI | 695989 | 696136 | 147 | -1,208048559 | N16:696027 | 10,445 | +6:YPR075C; | 6 | | | |
| chrXVI | 697045 | 697195 | 150 | -1,568498098 | N16:697144 | 20,341 | +8:YPR078C; +1:CUT431; | 8 | | | |
| chrXVI | 702075 | 702202 | 127 | -1,183902945 | N16:702218 | 18,022 | | Intergene | | | |
| chrXVI | 706433 | 706576 | 143 | -1,123242543 | N16:706496 | 2,16 | +2:CUT911; +11:YPR083W; | 2 | | | |
| chrXVI | 719549 | 719656 | 107 | -1,105596891 | N16:719618 | 6,831 | +6:YPR093C; | 6 | | | |
| chrXVI | 719549 | 719694 | 145 | -1,092523525 | N16:719618 | 6,831 | +6:YPR093C; | 6 | | | |
| chrXVI | 721766 | 721909 | 143 | -1,392070623 | N16:721871 | 22,506 | +20:YPR095C; +20:YPR096C; | 20 | | | |
| chrXVI | 722565 | 722701 | 136 | -1,454055797 | N16:722569 | 35,379 | +16:YPR095C; +16:YPR096C; | 16 | | | |
| chrXVI | 726383 | 726544 | 161 | -1,285546086 | N16:726563 | 37,57 | +10:YPR097W; | 10 | | | |
| chrXVI | 726865 | 727012 | 147 | -1,17793323 | N16:726909 | 24,315 | +12:YPR097W; | 12 | | | |
| chrXVI | 730093 | 730221 | 128 | -0,977833715 | N16:730129 | 46,934 | +3*:YPR100W; | 3 | TERM | | |
| chrXVI | 732571 | 732734 | 163 | -1,263052247 | N16:732631 | 8,846 | +3:YPR103W; | 3 | | | |
| chrXVI | 733908 | 734067 | 159 | -1,356224237 | N16:733903 | 31,934 | +18:YPR104C; | 18 | | | |
| chrXVI | 734318 | 734467 | 149 | -1,129398388 | N16:734451 | 40,054 | +15:YPR104C; | 15 | | | |
| chrXVI | 735641 | 735790 | 149 | -1,482353676 | N16:735692 | 19,149 | +7:YPR104C; | 7 | | | |
| chrXVI | 735832 | 735978 | 146 | -1,146956939 | N16:735865 | 9,072 | +6:YPR104C; | 6 | | | |
| chrXVI | 736889 | 737030 | 141 | -1,06500787 | N16:736965 | 7,925 | -1:YPR104C; +18*:YPR105C; | -1 | | | |
| chrXVI | 737626 | 737736 | 110 | -1,338392997 | N16:737632 | 27,399 | +14:YPR105C; | 14 | | | |
| chrXVI | 739761 | 739921 | 160 | -1,407490435 | N16:739810 | 18,995 | -1:YPR106W; 0:YPR105C; | -1 | | | |
| chrXVI | 740972 | 741124 | 152 | -1,379536127 | N16:741035 | 19,352 | +7:YPR106W; | 7 | | | |
| chrXVI | 742321 | 742473 | 152 | -1,219412571 | N16:742396 | 8,526 | +1:YPR108W; -1:YPR107C; | 1 | | | |
| chrXVI | 745613 | 745766 | 153 | -1,582380352 | N16:745699 | 19,502 | +8*:YPR110C; | 8 | TERM | | |
| chrXVI | 747443 | 747595 | 152 | -1,578364658 | N16:747452 | 9,854 | +2:YPR111W; | 2 | | | |
| chrXVI | 748166 | 748325 | 159 | -1,686567836 | N16:748291 | 22,076 | +7:YPR111W; | 7 | | | |
| chrXVI | 751513 | 751665 | 152 | -1,390822849 | N16:751585 | 9,988 | +3:YPR112C; | 3 | | | |
| chrXVI | 753264 | 753415 | 151 | -1,492754888 | N16:753263 | 10,895 | -1:CUT912; +3:YPR114W; | -1 | | | |
| chrXVI | 753264 | 753415 | 151 | -1,492754888 | N16:753426 | 8,701 | +4:YPR114W; | 4 | | | |
| chrXVI | 753454 | 753608 | 154 | -1,200298861 | N16:753612 | 19,753 | +5:YPR114W; | 5 | | | |
| chrXVI | 759400 | 759547 | 147 | -1,225971026 | N16:759472 | 10,284 | 0:CUT913; +6:YPR116W; | 0 | | | |
| chrXVI | 761447 | 761589 | 142 | -1,126235389 | N16:761432 | 27,999 | +17:YPR116W; +9:YPR117W; | 17 | | | |
| chrXVI | 761447 | 761589 | 142 | -1,126235389 | N16:761585 | 22,142 | +18:YPR116W; +10:YPR117W; | 18 | | | |

TABLE 3

| CHR | START | END | L(bp) | ΔLK | Nuc ID | Fuzziness | Gene ID | Gene body postion or intergene | Terminal | rDNA | Telomeric |
|---|---|---|---|---|---|---|---|---|---|---|---|
| chrXVI | 763063 | 763215 | 152 | -1,217602308 | N16:763092 | 14,986 | +27:YPR116W; +19:YPR117W; | 27 | | | |
| chrXVI | 763430 | 763578 | 148 | -1,276877455 | N16:763564 | 27,998 | +30:YPR116W; +22:YPR117W; | 30 | | | |
| chrXVI | 764397 | 764544 | 147 | -1,535365453 | N16:764439 | 30,391 | +35:YPR116W; +27:YPR117W; | 35 | | | |
| chrXVI | 764502 | 764659 | 157 | -1,606737048 | N16:764652 | 30,882 | +36:YPR116W; +28:YPR117W; | 36 | | | |
| chrXVI | 770323 | 770455 | 132 | -1,007757005 | N16:770351 | 5,357 | -1:Unit663; +3:SUT834; | -1 | | | |
| chrXVI | 778559 | 778687 | 128 | -1,695727508 | N16:778565 | 9,397 | +1:YPR121W; | 1 | | | |
| chrXVI | 786310 | 786475 | 165 | -1,358178432 | | | | Overlap <50 bp | | | |
| chrXVI | 788541 | 788654 | 113 | -0,957428674 | | | | Overlap <50 bp | | | |
| chrXVI | 788648 | 788776 | 128 | -0,912116698 | N16:788711 | 27,346 | +6:YPR125W; | 6 | | | |
| chrXVI | 790762 | 790920 | 158 | -1,717619091 | N16:790928 | 22,465 | -1:SUT839; +6:YPR127W; | -1 | | | |
| chrXVI | 791267 | 791414 | 147 | -1,435177715 | N16:791367 | 20,597 | +6:YPR128C; | 6 | | | |
| chrXVI | 791438 | 791595 | 157 | -1,174189669 | N16:791540 | 15,057 | +5:YPR128C; | 5 | | | |
| chrXVI | 791737 | 791890 | 153 | -1,234414832 | N16:791867 | 11,384 | +3:YPR128C; | 3 | | | |
| chrXVI | 792774 | 792924 | 150 | -1,406549472 | N16:792865 | 9,535 | +2:YPR129W; | 2 | | | |
| chrXVI | 794619 | 794743 | 124 | -1,179240655 | N16:794744 | 0 | -1:YPR132W; -1:YPR131C; | -1 | | | |
| chrXVI | 796842 | 796998 | 156 | -1,258376936 | N16:796924 | 5,167 | +3:YPR133C; | 3 | | | |
| chrXVI | 798394 | 798554 | 160 | -1,186659544 | N16:798384 | 26,696 | +5:SUT840; +3:YPR134W; | 5 | | | |
| chrXVI | 798628 | 798777 | 149 | -1,263993757 | N16:798655 | 37,61 | +3:SUT840; +5:YPR134W; | 3 | | | |
| chrXVI | 805070 | 805217 | 147 | -1,188331633 | N16:805107 | 12,021 | | Intergene | | | |
| chrXVI | 805577 | 805691 | 114 | -1,420834688 | | | | Overlap <50 bp | | | |
| chrXVI | 807669 | 807826 | 157 | -1,466924315 | N16:807711 | 23,065 | | Intergene | | | |
| chrXVI | 808317 | 808452 | 135 | -1,131987236 | N16:808366 | 33,843 | | Intergene | | | |
| chrXVI | 809355 | 809495 | 140 | -1,2188234 | N16:809434 | 35,921 | | Intergene | | | |
| chrXVI | 809361 | 809495 | 134 | -1,013927624 | N16:809434 | 35,921 | | Intergene | | | |
| chrXVI | 809773 | 809928 | 155 | -1,417064852 | N16:809806 | 26,539 | | Intergene | | | |
| chrXVI | 809904 | 810012 | 108 | -0,680220808 | N16:809957 | 38,136 | | Intergene | | | |
| chrXVI | 809934 | 810052 | 118 | -0,984914722 | N16:809957 | 38,136 | | Intergene | | | |
| chrXVI | 810053 | 810202 | 149 | -1,382210704 | N16:810107 | 16,971 | | Intergene | | | |
| chrXVI | 813679 | 813806 | 127 | -0,952728752 | N16:813742 | 6,369 | +4*:SUT424; +3:YPR139C; | 4 | TERM | | |
| chrXVI | 822438 | 822593 | 155 | -1,477588491 | | | | Overlap <50 bp | | | |
| chrXVI | 822870 | 822974 | 104 | -1,398798499 | N16:822924 | 21,198 | +3:YPR145W; | 3 | | | |
| chrXVI | 827045 | 827170 | 125 | -1,016009506 | N16:827139 | 12,522 | +7:YPR148C; | 7 | | | |
| chrXVI | 827247 | 827389 | 142 | -1,595691954 | N16:827298 | 19,325 | +6:YPR148C; | 6 | | | |
| chrXVI | 827869 | 828021 | 152 | -1,40915842 | N16:827957 | 7,57 | +2:YPR148C; | 2 | | | |
| chrXVI | 830012 | 830156 | 144 | -1,106999868 | N16:830081 | 24,766 | +3:YPR149W; | 3 | | | |
| chrXVI | 839224 | 839390 | 166 | -1,47755392 | N16:839304 | 25,12 | +6*:YPR156C; | 6 | TERM | | |
| chrXVI | 841656 | 841805 | 149 | -1,20401662 | | | | Overlap <50 bp | | | |
| chrXVI | 843827 | 843986 | 159 | -1,393427659 | N16:843902 | 35,884 | +5*:YPR158W; | 5 | TERM | | |
| chrXVI | 844949 | 845072 | 123 | -1,249444799 | | | | Overlap <50 bp | | | |
| chrXVI | 845049 | 845204 | 155 | -1,080475931 | N16:845158 | 30,73 | | Intergene | | | |
| chrXVI | 845482 | 845616 | 134 | -1,179930386 | N16:845543 | 30,172 | | Intergene | | | |
| chrXVI | 845482 | 845622 | 140 | -1,300049783 | N16:845543 | 30,172 | | Intergene | | | |
| chrXVI | 846525 | 846660 | 135 | -1,116134927 | N16:846560 | 5,033 | | Intergene | | | |
| chrXVI | 847154 | 847311 | 157 | -1,625023224 | N16:847244 | 32,604 | | Intergene | | | |
| chrXVI | 848557 | 848707 | 150 | -1,459466244 | N16:848653 | 13,503 | | Intergene | | | |
| chrXVI | 849289 | 849403 | 114 | -1,350078132 | N16:849394 | 31,565 | | Intergene | | | |
| chrXVI | 851012 | 851164 | 152 | -1,500339674 | N16:851100 | 18,009 | | Intergene | | | |
| chrXVI | 851565 | 851679 | 114 | -1,25898221 | N16:851699 | 5,132 | | Intergene | | | |
| chrXVI | 853843 | 853993 | 150 | -1,065574828 | N16:853920 | 16,258 | | Intergene | | | |
| chrXVI | 853847 | 853993 | 146 | -1,075847853 | N16:853920 | 16,258 | | Intergene | | | |
| chrXVI | 854031 | 854169 | 138 | -1,392142968 | N16:854079 | 0 | | Intergene | | | |
| chrXVI | 854126 | 854258 | 132 | -1,09774969 | N16:854239 | 38,891 | | Intergene | | | |
| chrXVI | 854305 | 854440 | 135 | -1,292205428 | N16:854397 | 22,517 | | Intergene | | | |
| chrXVI | 854851 | 855005 | 154 | -1,191042789 | N16:854861 | 22,03 | | Intergene | | | |
| chrXVI | 855108 | 855259 | 151 | -1,585915168 | N16:855198 | 12,055 | | Intergene | | | |
| chrXVI | 855132 | 855277 | 145 | -1,142853858 | N16:855198 | 12,055 | | Intergene | | | |
| chrXVI | 855136 | 855287 | 151 | -1,295370974 | N16:855198 | 12,055 | | Intergene | | | |
| chrXVI | 855343 | 855483 | 140 | -0,944469758 | N16:855418 | 34,962 | | Intergene | | | |
| chrXVI | 855349 | 855483 | 134 | -1,109465002 | N16:855418 | 34,962 | | Intergene | | | |
| chrXVI | 855892 | 856000 | 108 | -0,717180625 | N16:855993 | 49,602 | | Intergene | | | |
| chrXVI | 856112 | 856214 | 102 | -1,450300807 | | | | Overlap <50 bp | | | |
| chrXVI | 858667 | 858810 | 143 | -1,409500174 | N16:858736 | 38,96 | +10:YPR159W; | 10 | | | |
| chrXVI | 863879 | 864033 | 154 | -1,055299585 | N16:864033 | 22,755 | +17*:YPR160W; | 17 | TERM | | |
| chrXVI | 863879 | 864033 | 154 | -1,055299585 | N16:863878 | 41,094 | +16:YPR160W; | 16 | | | |
| chrXVI | 864255 | 864385 | 130 | -1,252558504 | N16:864244 | 23,845 | | Intergene | | | |
| chrXVI | 867145 | 867288 | 143 | -1,194733019 | | | | Overlap <50 bp | | | |
| chrXVI | 868632 | 868780 | 148 | -1,180799648 | N16:868711 | 11,606 | +9*:YPR163C; | 9 | TERM | | |
| chrXVI | 888008 | 888137 | 129 | -1,046012432 | N16:888074 | 10,323 | [illegible]*:YPR174C; 0:SUT427; -1:YPR173C; | 5 | TERM | | |
| chrXVI | 891517 | 891679 | 162 | -1,236792228 | N16:891562 | 6,501 | +4:YPR176C; | 4 | | | |
| chrXVI | 893449 | 893599 | 150 | -1,394754161 | N16:893574 | 34,979 | +8:YPR178W; | 8 | | | |
| chrXVI | 897488 | 897648 | 160 | -1,611825204 | N16:897619 | 22,599 | +14:YPR181C; | 14 | | | |
| chrXVI | 897990 | 898142 | 152 | -1,156239421 | N16:898110 | 26,338 | +11:YPR181C; | 11 | | | |
| chrXVI | 901200 | 901349 | 149 | -1,256425104 | N16:901238 | 6,397 | +4:YPR183W; +3:CUT923; | 4 | | | |
| chrXVI | 905702 | 905842 | 140 | -1,336380245 | N16:905808 | 23,973 | | Intergene | | | |
| chrXVI | 910317 | 910457 | 140 | -1,154874767 | N16:910477 | 3,017 | +4:YPR186C; | 4 | | | |
| chrXVI | 910317 | 910457 | 140 | -1,154874767 | N16:910305 | 13,424 | +5:YPR186C; | 5 | | | |
| chrXVI | 922519 | 922676 | 157 | -1,564045849 | N16:922581 | 2,828 | +2:anti192; +5:YPR192W; | 2 | | | |
| chrXVI | 922519 | 922681 | 162 | -1,387495318 | N16:922581 | 2,828 | +2:anti192; +5:YPR192W; | 2 | | | |
| chrXVI | 922523 | 922681 | 158 | -1,503272644 | N16:922581 | 2,828 | +2:anti192; +5:YPR192W; | 2 | | | |
| chrXVI | 927439 | 927574 | 135 | -0,987824892 | N16:927563 | 36,449 | 0:SUT429; | 0 | | | |
| chrXVI | 930833 | 931008 | 175 | -1,746019507 | N16:930869 | 24,595 | +3:SUT846; +14:SUT430; | 3 | | | |
| chrXVI | 931493 | 931642 | 149 | -1,421005016 | N16:931566 | 13,692 | +2:YPR196W; | 2 | | | |
| chrXVI | 940363 | 940515 | 152 | -1,281479838 | N16:940386 | 7,036 | +4:YPR201W; | 4 | | | |
| chrXVI | 943076 | 943229 | 153 | -1,498434548 | N16:943200 | 29,905 | +2:YPR202W; | 2 | | | TEL RIGHT |
| chrXVI | 943085 | 943229 | 144 | -1,833880799 | N16:943200 | 29,905 | +2:YPR202W; | 2 | | | TEL RIGHT |
| chrXVI | 944252 | 944389 | 137 | -1,304987783 | N16:944314 | 4,041 | -1:YPR204W; +4*:YPR203W; | -1 | | | TEL RIGHT |
| chrXVI | 944537 | 944680 | 143 | -1,382609864 | N16:944589 | 14 | +1:YPR204W; | 1 | | | TEL RIGHT |
| chrXVI | 944585 | 944710 | 125 | -0,788288258 | N16:944589 | 14 | +1:YPR204W; | 1 | | | TEL RIGHT |
| chrXVI | 944810 | 944962 | 152 | -1,212944731 | N16:944922 | 11,533 | +3:YPR204W; | 3 | | | TEL RIGHT |
| chrXVI | 944883 | 945027 | 144 | -1,518440935 | N16:944922 | 11,533 | +3:YPR204W; | 3 | | | TEL RIGHT |
| chrXVI | 944980 | 945115 | 135 | -1,063325043 | N16:945080 | 21,502 | +4:YPR204W; | 4 | | | TEL RIGHT |
| chrXVI | 944980 | 945167 | 187 | -1,122414218 | N16:945080 | 21,502 | +4:YPR204W; | 4 | | | TEL RIGHT |
| chrXVI | 945023 | 945115 | 92 | -0,294290005 | N16:945080 | 21,502 | +4:YPR204W; | 4 | | | TEL RIGHT |
| chrXVI | 945023 | 945167 | 144 | -0,955039769 | N16:945080 | 21,502 | +4:YPR204W; | 4 | | | TEL RIGHT |
| chrXVI | 945362 | 945514 | 152 | -1,226994054 | N16:945422 | 14,364 | +6:YPR204W; | 6 | | | TEL RIGHT |
| chrXVI | 945663 | 945793 | 130 | -1,056492367 | N16:945658 | 23,335 | +7:YPR204W; | 7 | | | TEL RIGHT |
| chrXVI | 945663 | 945793 | 130 | -1,056492367 | N16:945816 | 41,102 | +8:YPR204W; | 8 | | | TEL RIGHT |

| CHR | START | END | L(bp) | ΔLK | Nuc ID | Fuzziness | Gene ID | Gene body postion or intergene | Terminal | rDNA | Telomeric |
|---|---|---|---|---|---|---|---|---|---|---|---|
| chrXVI | 945759 | 945954 | 195 | **-1,628973299** | N16:945816 | 41,102 | +8:YPR204W; | 8 | | | TEL RIGHT |
| chrXVI | 945821 | 945954 | 133 | **-0,943887076** | N16:945816 | 41,102 | +8:YPR204W; | 8 | | | TEL RIGHT |
| chrXVI | 945821 | 945969 | 148 | **-1,34134048** | N16:945816 | 41,102 | +8:YPR204W; | 8 | | | TEL RIGHT |
| chrXVI | 946033 | 946184 | 151 | **-1,163093755** | N16:946078 | 43,134 | +10:YPR204W; | 10 | | | TEL RIGHT |
| chrXVI | 946038 | 946172 | 134 | **-1,029552381** | N16:946078 | 43,134 | +10:YPR204W; | 10 | | | TEL RIGHT |
| chrXVI | 946051 | 946147 | 96 | **-0,855605136** | N16:946078 | 43,134 | +10:YPR204W; | 10 | | | TEL RIGHT |
| chrXVI | 946159 | 946281 | 122 | **-0,182479278** | | | | Overlap <50 bp | | | TEL RIGHT |
| chrXVI | 946966 | 947118 | 152 | **-1,155297851** | N16:947053 | 20,506 | +16:YPR204W; | 16 | | | TEL RIGHT |
| chrXVI | 947306 | 947457 | 151 | **-1,220736008** | N16:947384 | 3,536 | +18:YPR204W; | 18 | | | TEL RIGHT |
